# TCRfp: a new fingerprint-based approach for TCR repertoire analysis

**DOI:** 10.1101/2023.12.19.572261

**Authors:** Francesca Mayol-Rullan, Marine Bugnon, Marta A. S. Perez, Vincent Zoete

## Abstract

The development of cancer immunotherapy has accelerated in recent years. Understanding the specificity of T cell receptors (TCR) for peptides presented by the major histocompatibility complex (pMHC) is a major step towards improving immunotherapy approaches, such as adoptive cell transfer and peptide vaccination. Despite recent computational advances, the unambiguous pairing of TCR with pMHC, from pools of thousands of candidates, remains out of reach. To tackle this challenge, we have developed a new tool that converts the 3D structure of TCR into individual one-dimensional structural fingerprints (TCRfp). We have modelled over 10’000 3D structures of paired TCR alpha and beta chains with known sequences and pMHC specificity and encoded them into 1D TCRfp. For future clinical needs, we have translated the TCR modelling process into a fast pipeline. Similarity measures between TCR FPs correlate with their ability to recognise similar or identical epitopes in the training set and in the external validation sets. TCRfp constitutes the first rapid approach for high-throughput TCR comparison and repertoire analysis based on molecular 3D structures, which is efficient enough to complement sequence-based approaches.

## 1. Introduction

Cancer immunotherapy development has been in the rise over the last years. One of the most promising research topics in this field involves the understanding and unambiguous prediction of T-cell receptor (TCR) recognition of a specific cancer epitope (i.e., peptide-MHC, pMHC). Predicting the TCR-pMHC specific interactions is essential for improving immunotherapies such as adoptive cell therapy or peptide vaccination [1–3]. Nevertheless, understanding the mechanisms underlying the TCR-pMHC specificity remains practically unresolved due to its complexity [4, 5].

Over the past years, TCR-pMHC specificity has been widely explored by several authors who developed numerous computational approaches [5, 6]. Those that predict TCR-pMHC specificity can be divided into sequence-based and structure-based methodologies. Ongoing improvements in sequencing technology [7–10] are yielding more numerous and reliable TCR-pMHC sequence pairs [11–13], accelerating the development of better, more accurate in silico approaches. As an example, biotech companies such as Adaptive Biotechnologies in partnership Microsoft Healthcare NExT initiative [14] are providing extensive TCR data and TCR mapping. Still, the number of TCR-pMHC pairs available corresponds to a tiny fraction of the overall possibilities. Typically, sequence-based approaches use machine learning techniques such as logistic regressions and deep neural networks that use sequences to train a function that attempts to correctly predict the epitope for a given TCR [15–18]. These models can achieve high performance for TCR classification, but they still require substantial sequence data sets to provide sufficient predictive power for their algorithms. Consequently, they show limited success in predicting TCR specificities for unseen epitopes or epitopes and alleles with very limited representation in the training data. Structure-based approaches provide a learning set-independent alternative. They use experimental structures or structural models of the TCR-pMHC complexes to determine the most likely ones based on estimates of the affinities between partners, using universal physics-based scoring functions [19–23]. Despite great successes, structure-based approaches are time-consuming and hardly tractable for large scale predictions. The increase of the number of TCR-pMHC structures (230 TCR-pMHC class I and 82 TCR-pMHC class II as of 5 July 2023) in the Protein Data Bank (PDB) [24] together with the appearance of more powerful tools to create 3D models out of sequences such as TCRmodel, AlphaFold and LYRA [25–27] are contributing to the development of new structure-based approaches [22].

Here, we present TCRfp, a totally new 3D-based approach that tackles TCR-pMHC specificities using the 5-dimensional ElectroShape approach (ES5D) [28]. Initially developed for computer-aided drug design application, ES5D extends classical shape-based methods by applying on top of the atomic 3D Cartesian coordinates, 4^th^ and 5^th^ dimensions corresponding to the atomic contributions of the charge and lipophilicity. TCRfp thus provides a rapid, tractable, and robust approach to quantify the similarity between TCRs, as a function of their 3D shapes, spatial distribution of charges and lipophilic groups.

TCRfp consists in 4 main steps. First, TCR sequences are converted into 3D structures using TCRmodel [26]. Second, the 3D structure of each TCR is converted into a simple fixed-length numerical representation, the fingerprint (FP), using a variant of the ES5D algorithm adapted to TCR structures. Third, TCR comparison is performed via the calculation of Manhattan distance between 1D-vectors. TCRfp is based on the similarity principle [29] according to which two structurally similar TCRs - that share close FPs - are more likely to bind to the same or to a similar pMHC. Fourth, TCR repertoires are analysed by clustering TCRs on the basis of the distance between their FPs (See figure 1).

**Figure 1.**
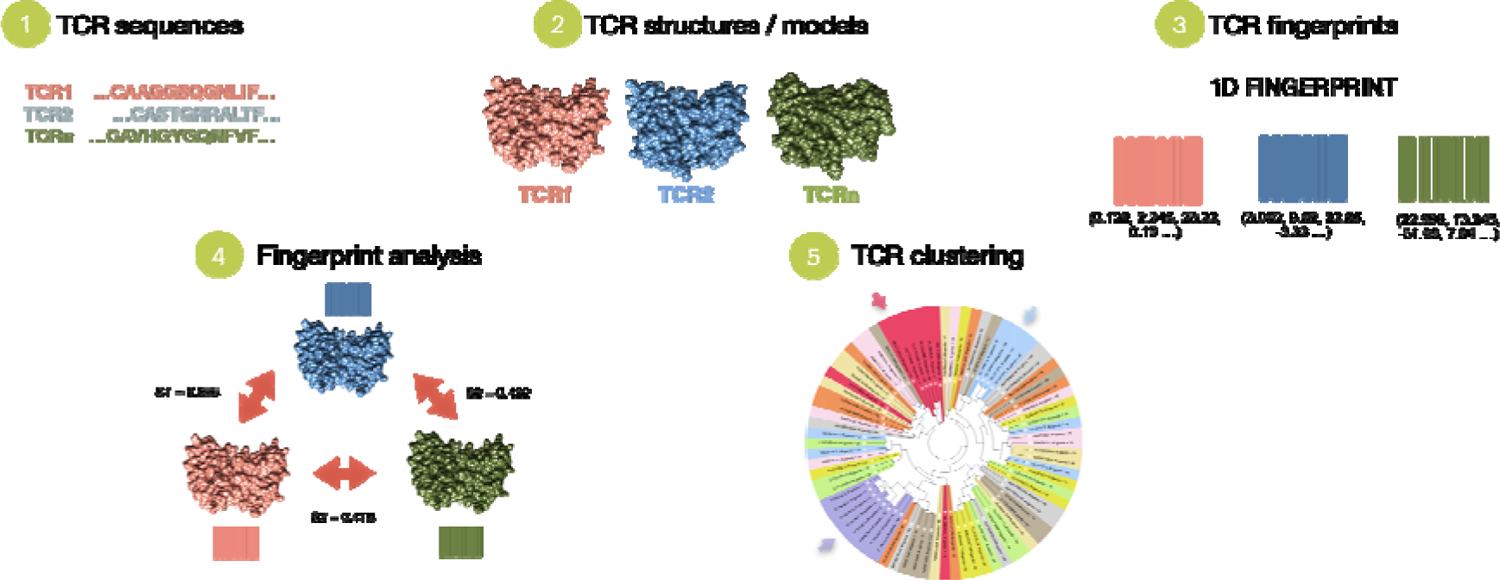
Schematic overview of the TCRfp approach. TCRfp is described in the main first three steps: 1. TCR sequences are retrieved and 2. converted into structures using homology modelling to be further 3. translated into a 1D numerical representation known as the FPs. With this vectorial representation, the TCR structures can be easily compared through the FPs (4) and even clustered (5) using different methods such as UPGMA hierarchical clustering.

In this paper we present a new approach (Figure 1) that demonstrates it is possible to cluster TCRs as a function of their specificity, when the latter is dominated by their shape and not by their sequence similarity. Our approach is capable of clustering rapidly a high number of TCRs. Since calculating the FP of a TCR structure is immediate, the time limiting step of our pipeline is the generation of each TCR model, which currently takes only 1.4 minutes on a 16-CPU machine. Our approach is thus much faster than the usual purely 3D-based approaches which rely on the modelling of the full TCR-pMHC complex and binding free energy estimation. Additionally, contrarily to sequence-based approaches, it does not require to be retrained for each pMHC. TCRfp introduces a new class of fast structure-based approaches for TCR analysis, clustering and potentially deorphanisation.

## 2. Results

### 2.1. Adaptation of the ES5D fingerprinting approach from small molecules to TCR experimental 3D structures (PDB)

The ES5D fingerprinting approach was initially developed for small drug-like molecules. We have adapted it for TCRs by changing the definition of the so-called centroids (See Methods). As a proof of concept, the ES5D-based TCR FPs were tested on a set of 74 3D-structures of TCR-pMHC (HLA-A*02) complexes from the PDB. Just the fraction of the 3D structures corresponding to the CDRs of the TCRs, which constitute the most variable part of the receptor and are responsible for pMHC binding, were considered. Then, the respective FPs were calculated using 6 centroids, each one placed on carbon alpha of the middle residue of the 6 CDR loops (Figure 2). The default weighting values of 25 and 4 were applied to the charge and lipophilicity, respectively. For each given TCR_ref_ in this set we identified the TCR with the highest Manhattan-based similarity according to their fingerprints (excluding TCR_ref_ itself), and compared the peptides they recognize. We obtained an average sequence identity of 76%, suggesting that the similarity calculated with this definition of the TCRfp strongly correlates with the pMHC these TCRs recognize. Interestingly, when a random TCR was chosen instead of the closest according to TCRfp, the average sequence identity dropped to 32%, showing that our approach is much better than random (p-value < 0.0001). Of note, 28 TCRs are singletons (no other TCR in the set binds the same epitope). Removing them led to an average sequence identity of 92%. Next, we extended the by adding 66 non-HLA-A*02 restricted TCRs, totalizing 140 TCRs. For this larger and more challenging set, we found that the closest TCR to a given TCR_ref_ according to TCRfp (excluding TCR_ref_ itself), was binding the same pMHC in 64% of the cases. This is significantly more than the average sequence recapitulation that could be obtained by random picking 12% (p<0.0001). The sequence recapitulation for this set, after removing 40 singletons, is 84%, showing once again that TCRfp strongly correlates with the pMHC these TCRs recognize.

**Figure 2.**
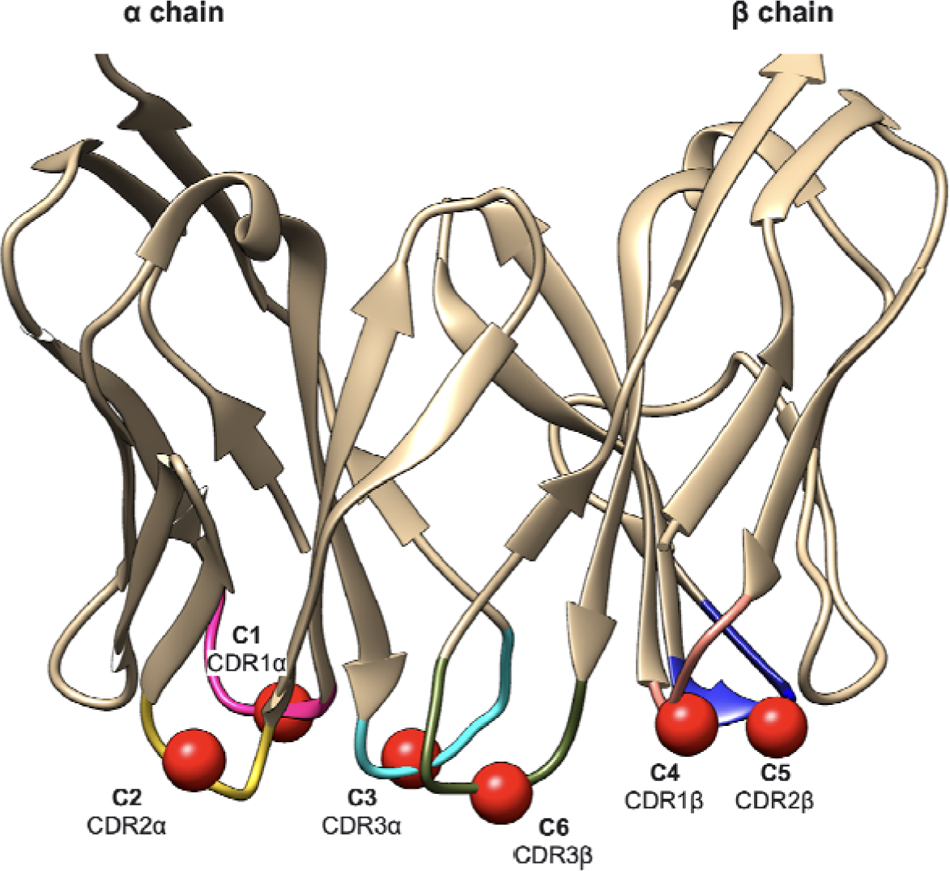
Positions of the 6 centroids (C1, C2, C3, C4, C5 and C6) in the TCR structure/model according to the first definition of the ES5D FP. Each loop is coloured differently and highlighted with a red sphere that represents the α carbon of the middle residue, which corresponds to the centroid. CDR1α is coloured in magenta, CDR2α in yellow, CDR3α in cyan, CDR1β in light red, CDR2β in dark blue and CDR3β in green.

Since CDR loops of TCR are particularly flexible and since only TCR 3D models will be available instead of X-ray structures in general applications, we have recalculated average sequence identity on TCRs after remodelling the CDR loops. This exercise is important to assess if the results obtained by TCRfp are sensitive to the imperfections inherent to structural models. Since only CDR residues are encoded in the TCRfp, this exercise could prove extremely challenging for our approach. Strikingly, the sequence recapitulation was maintained at 67% on the 74 individual HLA-A*02 restricted TCRs considered. This shows that X-ray structures can be replaced by structural homology models in our approach at a cost of a small reduction (9%) in the accuracy. These results proved that the similarity between TCR FPs calculated using this version of TCRfp strongly correlates with the pMHC they recognize, allowing to cluster TCRs showing the same pMHC specificity when using TCR models.

We therefore analysed the efficiency of TCRfp using much larger databases of TCR-pMHC sequences. We started by converting TCR sequences with known specificity into 3D structural models. Second, we calculated the TCRfp of each TCR. Third, we computed TCR similarities via the calculation of Manhattan distance between the TCRfp and finally we evaluated if TCRs with highest similarity share the same pMHC.

### 2.2. Development of a pipeline to obtain accurate TCR 3D structural models from their sequences

We have worked on the development and automatization of a pipeline based on the TCRmodel approach, which reads a large number of TCR sequences and constructs 3D models in a reasonable time. The ability of TCRmodel to accurately predict TCR structures was proved in its seminal paper [25]. Here, applying the approach to model thousands of TCRs sequences from 10xGenomics [30], we challenged the approach with many more sequences, and diversity of TCRs than in the original publication, and realized that the modeling of some TCRs could be particularly difficult due to the high flexibility of the CDR loops and the limited number of templates available. We have obtained several TCR models with erroneously disordered loops as can be seen in the example in Figure 3a. This example corresponds to TCR ID 10219 (TRAV19, TRAJ24, CDR3α: CALSEADDSWGKLQF, TRBV27, TRBJ2-2, CDR3 β: CASSLYGNLGTGELFF). Of note, all TCRs with their ID can be retrieved from the Supplementary Table 1. To automatically detect these problematic models, we have implemented distance-based filters. For this, we selected specific non-variable residues and measured distances between their Cα atoms (Figure 3) and checked if they fall within acceptable ranges. The latter were calculated using 92 experimental TCR class I structures available in the Protein Data Bank (PDB) [31], and were defined as the mean ± four standard deviations (STD) for each one of the five distances considered (Figure 3b). If at least one of the distances fell outside the acceptable ranges, the corresponding TCR model was considered erroneous and was therefore discarded. Since TCRmodel can provide somewhat different models if applied several times to the same sequence, various attempts were performed in order to find 3D models that fell within the accepted ranges. All distances are displayed in the Figure 3b.

**Figure 3.**
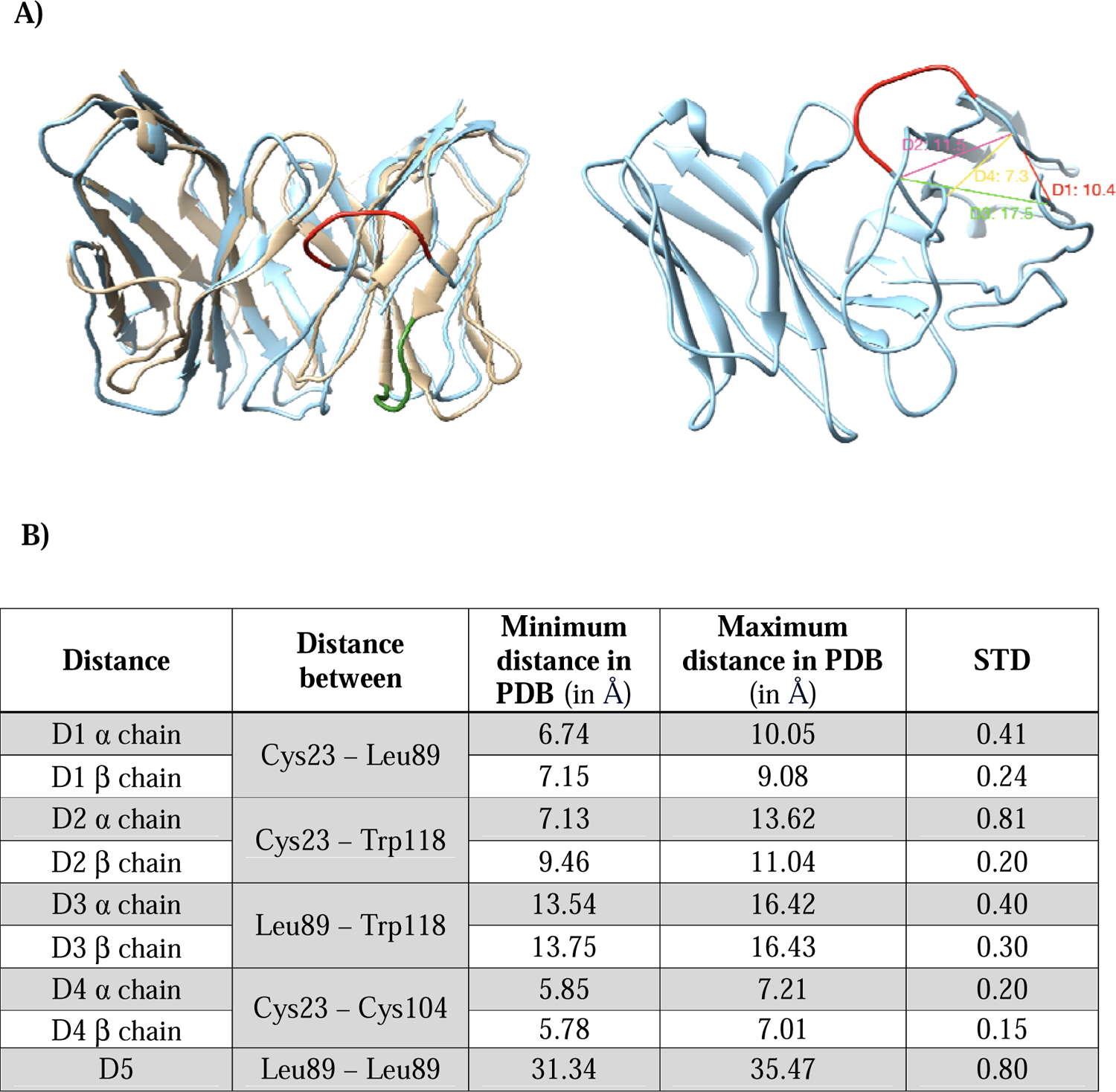
Distance-based validation filters for the TCR modeling. **A)** *<u>on the left side</u>*: Superimposition of a wrong TCR model (10X ID 10219: TRAV19, TRAJ24, CDR3α: CALSEADDSWGKLQF, TRBV27, TRBJ2-2, CDR3β: CASSLYGNLGTGELFF, binding the peptide KLGGALQAK) with a good TCR model (10X ID 1627: TRAV38-2DV8, TRAJ48, CDR3α: CAYRAPRGNEKLTF, TRBV2, TRBJ1-1, CDR3β: CASSDRMNTEAFF, binding the peptide AVFDRKSDAK), colored in light blue and sand, respectively. A fraction of the β chain badly modelled is colored in red and for comparison, the corresponding structural element of the correct model is displayed and colored in green. *On the right side*: Th model exceeded the distance thresholds for D1, D2, D3 and D4 of the β chain. Each distance is shown in the figure with a different color. The red representation corresponds to the same one shown in the left figure for a better visual comparison. B) Table detailing the distance ranges (in Å) and standard deviations (STD) used for the distance-based filters, calculated based on 92 experimental TCR structures. Accepted ranges of distances used to define satisfactory models are equal to the mean ± 4 standard deviations.

In brief, our modelling pipeline creates TCR sequences of both α and β chains using the TRAV, TRAJ, TRBV and TRBJ genes, as well as the CDR3α and CDR3β sequences as input. All the gene information and all the TCRs used for the modeling pipeline are given in the Supplementary Table 1, including the information regarding the specificity. To track down each TCR, an internal numbering system has been followed in which each TCR is given a unique ID. The latter are also given in the Supplementary Table 1. 10 attempts of conformations per TCR were constructed by homology modeling with the Rosetta TCRModel approach and for each attempt a quality check was performed with distance-based filters, that could be applied thanks to the renumbering of the residues using ANARCI and the IMGT scheme that provides the same numbers to conserved residues. Only the conformations that passed the quality checks were further considered and the lowest energy one according to Rosetta REF15 scoring function was used to calculate the TCFfp. If after 10 attempts, none of the models pass the quality check, we assumed that it was not possible to obtain a reliable model using this pipeline, due to a lack of relevant templates, and the corresponding TCR was excluded from the set. The accuracy of our pipeline was tested with a set of 187 TCRs with known experimental structures, targeting class I and class II pMHCs. Importantly, to model a given TCR, the experimental structures sharing the same genes and that of this TCR itself were removed from the list of possible templates, to make the benchmark modelling exercise more challenging and closer to the typical real application. Comparing the models and the experimental structures of these TCRs showed that our pipeline could predict TCRs structures with a satisfying accuracy, illustrated by a global RMSD (on the heavy atoms excluding CDR3s) of 2.1 Å between models and experimental structures. However, 78 TCRs out of 187 could not be modeled due to a lack of relevant templates. We attempted to increase modeling accuracy, especially for CDR3s, by exploring different options for loop refinement provided by TCRmodel (data not shown). However, none of the options achieved a better average accuracy, while computation time increased substantially. Consequently, these options were not applied to the final pipeline. Our modelling pipeline takes on average 1.40 minutes to model a TCR and can be distributed upon multiple cores, allowing 6171 TCRs to be modelled in ∼9 hours on a 16-cores machine. Using our modelling pipeline, 10’703 TCRs models were constructed for the 14’479 sequences in the 10X genomics database. As for now, 10X ID refers to the identification value given to the TCRs obtained from 10X database.

### 2.3. Exploration of an improved FP definition with heuristic search

In the initial definition of TCRfp, the centroids are placed on the Cα of the tip of the loop and are dependent on the TCR under investigation. This idea was inspired by the original version of ES5D developed for small drug-like compounds. Defining centroids as structural elements of the molecules under study has the advantage of making it possible to calculate the FP for these compounds without having to superimpose them all first. Indeed, FP values depend on the position of the molecule atoms around the centroids. If the latter are constant for all molecules and therefore fixed in space, then not only will the FP values depend on the shape of the compounds, as intended, but also on the overall absolute position and orientation of the molecule in space, which is to be avoided. To correct this, it would then be necessary to superimpose all the molecules on each other before calculating their FP, a task not easily achievable for large number of very diverse small molecules. Using a definition of centroids based on structural elements, which “follow” the position and global orientation of the molecules in space, makes it possible to correct this bias, and ensure that FP values are only a function of the shape of the compounds studied. One drawback of this approach, however, if that this does not allow to easily optimise the definition of the centroids to get the best similarity estimation. However, since all the TCRs share the same common global 3D structure, notably for the constant part, TCR superimposition is straightforward to perform and we therefore decided to explore the possibility of using universal centroid positions, where the same 6 centroids defined by constant 5D coordinates, can be used to describe all the TCRs. As there is an infinite number of combinations of 6 centroids in 5D, we have explored them systematically, making use of a heuristic search (MATCH) based on genetic algorithms. The algorithm changed centroids positions, charge, and lipophilicity while optimising the measure of similarity between a training set of TCRs binding the same pMHC. The explorations were done using 10 training sets comprising an equal number of TCRs binding the same peptide. Each training set was composed of groups of 9 TCRs each binding the same antigen, among 13 different ones, generating a total of 117 TCRs per set (as described in the Methods).

Several individual MATCH runs were performed using different combinations of parameters (mutation ranges, crossover types, random generations, and restraints of the movement). Supplementary Table 2 provides a detailed summary of the parameters used during the MATCH explorations. The genetic algorithm first defined a population of vectors made up of the collection of the six centroids coordinates, randomly chosen. During the evolution of that population mutations to a single entry or to multiple entries of the vector chosen at random were applied. The latter consisted in subtracting or adding a small, randomly chosen quantity to the corresponding entries. At the same time, the vectors underwent interchanges of their values with other vectors selected randomly by the algorithm – mimicking cross-over - to increase diversity and variability (see methods for the details and Supplementary Table 2 for specific values). At each generation, only the vectors corresponding to the highest values of the readout were allowed to generate offspring by mutation and crossover. As expected, starting from the first generation of vectors, the algorithm was progressively modifying them (so, the centroid coordinates) in order to increase the readout, i.e. the success rate for ensuring that the TCR closest to a given TCR, other than itself, is able to bind the same pMHC. Eventually, MATCH runs reached a plateau where none of the modifications provided a better readout and was stopped.

Starting from a totally random initial position of the centroids, our hypothesis was that the MATCH would move them towards the CDRs or at the vicinity of the place where the TCR-pMHC interaction takes place, due to the importance of this region for the TCR specificity. Interestingly, some centroids coordinates converge towards the same positions in different MATCH runs. These positions are far from the tip of the loop and sometimes even from the TCR surface, as seen in the Figure 4. We compared the best two solutions obtained by MATCH via random initial exploration (See Table 1).

**Figure 4.**
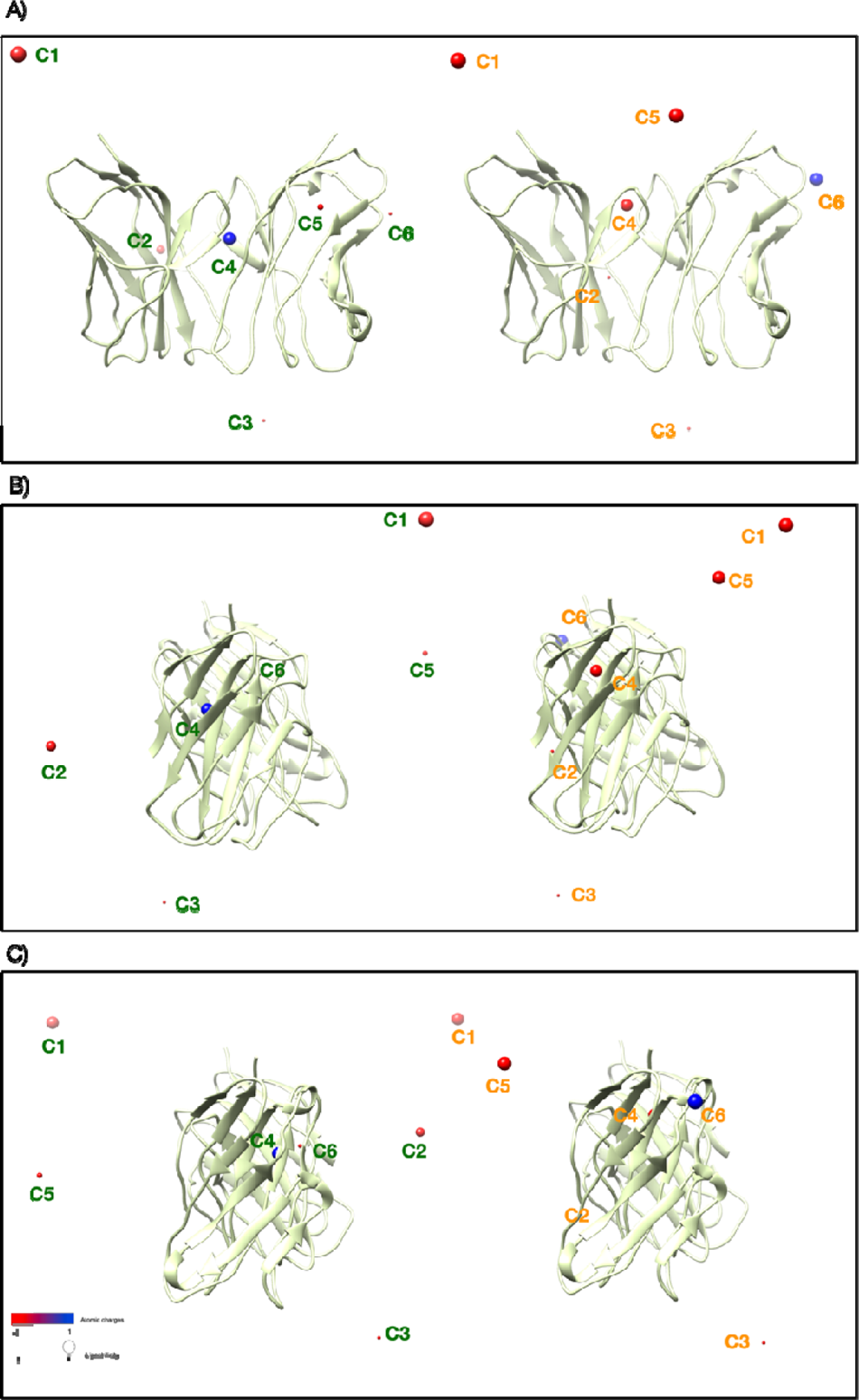
Comparison of the centroids obtained with the two best genomes from the algorithm starting from a random initial position. For each subfigure. The values for the charge range from −1 (in red) to 1 (in blue) and the diameter of each centroid correlates with the normalized charge and lipophilicity. See the scales in the bottom left of the last subfigure. The centroids of the best genome are coloured in green and the centroids of the second-best genome in orange. Different views of the two genome solutions represented with a TCR for reference: A. Front view of the TCR B) Side view rotated 90° to the right C) Side view rotated 90° to the left. The reference TCR used for the visualization is a model obtained with our pipeline corresponding to the ID 46 (encoded by the TRAV8-4, TRAJ20, TRBV19 and TRBJ1-1 genes, CDR3α: CAVSPNDYKLSF and CDR3β: CASSIRSTTEAFF, binding the peptide GILGFVFTL.

**Table 1.** Best 2 solutions obtained with the TCRfp MATCH GA-based algorithm. The genomes correspond to the FP values which are organized according to the weighted parameters (WP) and the 6 centroids (C1, C2, C3, C4, C5, C6). The numbers given to each centroid are just a reference, as they all originate from randomized values they hold no representation of their position in space for the GA-based algorithms, unlike the tip-of-the-loop original TCRfp.

| Genome | WP | C1 | C2 | C3 | C4 | C5 | C6 |
| --- | --- | --- | --- | --- | --- | --- | --- |
| Solution 1 | 43 5 | 25.816 | 1.444 | 5.886 | -23.905 | 13.805 - | -10.432 |
|  |  | -7.425 | -3.827 | 23.593 | -24.316 | 4.239 - | -4.425 |
|  |  | 7.769 | 3.993 | 17.028 | -31.71 | 29.239 | 25.446 - |
|  |  | -1.494 | -2.07 | 0.222 | -2.743 | 1.191 | 2.079 |
|  |  | -2.267 | -1.145 | -0.479 | 1.113 | 0.487 | -2.242 |
| Solution 2 | 37 1 | 27.147 | 6.103 | -4.875 | -1.594 | 6.838 | -21.546 |
|  |  | -12.329 | -17.662 | 1.829 | -8.229 | 25.258 | -23.554 |
|  |  | 8.209 | -19.221 | 10.115 | 1.465 | 15.477 | -29.691 |
|  |  | -5.032 | 0.647 | 0.685 | -0.405 | 1.289 | -1.42 |
|  |  | 0.519 | 0.201 | 0.164 | -1.499 | -0.053 | 3.258 |

As an illustration, Figure 4, shows the 6 centroids obtained as best solutions of two different MATCH runs, plotted in the cartesian space, along with a TCR model as a reference. The charge and lipophilicity values per each centroid are represented by colour and sphere radius, respectively. In both solutions, 4 of the 6 centroids, labelled as 1, 3, 4 and 6, are placed in equivalent regions (Figure 4 A, B and C). Strikingly, centroids 1 share almost identical charges (genome 1: 0.90 and genome 2: 1) and the same lipophilicity (1.0), like centroids 3 (charges 0.99 and 1, for genomes 1 and 2 respectively; and lipophilicity 1 for both). Centroids 3 (2.5 Å) are also closer among them than centroids 1 (3.2 Å). Centroids 6 are also quite close to each other (5.1 Å), as are centroids 4 which are only 5.9 Å apart. Interestingly, centroids 4 of both genomes are situated within the TCR, as well as the centroid 2 of the second genome. Both centroids 6 are placed also quite closely to the TCR. Since the initial positions of the centroids can be placed anywhere within a range of −30 and 30 Å in three dimensions and knowing that the averaged centre of all the TCRs is located at the origin of the Cartesian space, these centroid positions at the end of both match runs indicate that this region is of particular interest for describing TCRs shapes in a way that correlates with their binding specificity. Most importantly, the fact that our MATCH move the positions of the centroids – initially placed totally at random - towards similar positions at the end of the runs demonstrates the ability of the algorithm to converge towards optimal solutions. The huge sample space, the high number of parameters to calibrate combined with the relatively small training set make the search very difficult and time consuming. Still, we assume that we found near optimal solutions for 4 out of 6 centroids.

Since we obtained satisfactory results for TCR experimental structures by placing arbitrarily the centroids at the tips of the CDR loops, we decided to initiate the MATCH performed on homology models by placing the centroids closer to the tip of the loop, as described in section 2.1. Indeed, as previously hypothesized, the loop regions should bring the best descriptive ability into TCRfp definition. However, despite the use of different combinations of parameters, none of the GA MATCH runs starting closer to the CDRs succeeded to improve on the best readout obtained with the first MATCH runs starting from random positions.

When using the same algorithm parameters, the MATCH runs initiated by placing centroids at random position within a range of −30 to 30 Å for the cartesian coordinates and of −1 to 1 Å for the C and P values obtained a larger mean final *peptide identity* score (31.4% on average for 50 runs). On the contrary the MATCH runs started by placing the centroids close to the tip of the CDR loops, with a range of only −1 to 1 from their centre but with wider C and P initial range of −30 to 30 obtained an average *peptide identity* score of 30% over 60 runs. Of note, the best overall genome obtained with the TCRfp MATCH is the one displayed in the Figure 4 (on the left side in A, B, C). This MATCH run converged to a peptide identity score of 36.5% ± 5.0 (averaged over the 10 sets used in the training), which is significantly higher than the one obtained using random centroids, and C and P values, i.e. 7.7% ± 2.5 (p<0.0001).

The MATCH algorithm leading to the best readout used a population size of 400 genomes. The global weighting of the charges, C, and lipophilicity, P, in the encoding of TCRs by TCRfp (data not shown) were also part of the optimised parameters (See Methods). Their initial values could range from 0 to 50 for C and from 0 to 20 for P. For the X, Y and Z Cartesian coordinates, a random value between −30 and 30 Å was used to initiate the runs, whereas the charge, c, and lipophilicity, p, values of each centroid could range between −1 and 1, mimicking the values seen in initial arbitrary definition of TCRfp successfully applied to experimental structures. 200 new offspring genomes were generated at each generation, derived from a pool of the best 200 parent genomes from the total population. A low number of mutations per generation was applied (only 1 or 2 genes selected for mutation) leading to a steady and slow improve rate of the readout. For any gene selected for a mutation, a random value between 0 and 2 was either added or subtracted to the original value, to drastically limit the changes from one generation to another and ensure a slow but steady convergence. Finally, multiple centroid crossovers proved to be the more efficient for our algorithms than single crossovers and interchanging the whole centroid each time also performed better than making a crossover within the centroid itself. The rest of the parameters tested for the algorithms are explained in the Methods and the best algorithms values and best centroids are provided in the Supplementary Tables 2 and 3.

To assess the ability of the MATCH-optimised TCRfp to generate TCR clusters that correlate with their specificity, we built hierarchical trees for the training set. Even though only TCR structures were used to makes these trees, regardless of their actual specificity, each TCR branch in the tree was coloured afterward according to its cognate pMHC. As a measure of the quality of the clustering, we determined the number of times the colour changed between two successive nodes, starting from the upper node of the cluster and turning clockwise. The quality of the cluster was also determined by a more usual and quantitative metric, pMHC-distance. The pMHC-distance was defined as the average branch length distance between all possible pairs of TCR nodes that recognize the same pMHC. When comparing two trees, larger number of colour changes and pMHC-distance mean lower ability to cluster TCRs based on the pMHC they recognize. The ability of TCRfp MATCH to cluster 1 of the 10 sets of the training set is represented in a clustering tree in Figure 5 (the other 9 trees for the remaining 9 sets can be found in the Supplementary Figure 1). Our best MATCH-determined TCRfp version achieved 46% of *peptide identity* (with an average score of 36.5% ± 5.0 over the 10 sets used for the run). It also provides the lowest number of colour changes from node to node, with 83 changes compared to a mean of 88.60 ± 4.03 on the ten sets. This tree obtained a pMHC-distance a 0.79. The average pMHC-distance of this TCRfp version over all the ten sets of TCRs is 0.760 ± 0.03. Reversely, the set on which this TCRfp version achieved the lowest *peptide identity* score (29%) accounted for the highest number of colour changes from node to node (97 changes). We observe that TCRs binding to the same peptide tend to cluster together in the tree. This clustering is more pronounced for TCRs binding the GLCTLVAML, ELAGIGILTV or the FLYALALLL peptides. This is because of the small variance of the TCRs from the training set itself. For instance, the TCRs binding the peptide ELAGIGILTV and FLYALALLL both come from only 7 different TRAV genes, out of a pool of 38 TRAV genes represented in the entire TCR set used for this exercise. Moreover, if we group all the TRAV, TRBV, TRAJ and TRBJ genes together, the TCRs binding to the FLYALALLL peptide account for the smallest diversity of genes, with only 23 genes present from a total of 140 different genes. Also, only 26 TCRs of the whole training set bind to the FLYALALLL peptide, representing the peptide with less TCR diversity.

**Figure 5.**
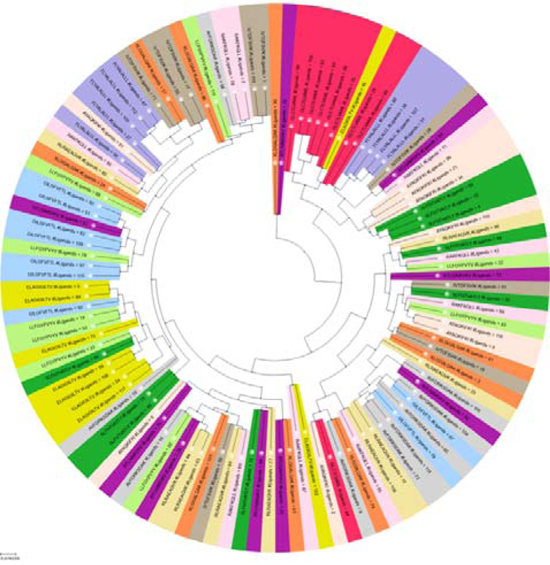
TCR hierarchical tree built with ES5D FPs for the TCRfp MATCH approach applied to a set of 117 TCRs. Importantly, the tree was obtained using only the ES5D-based similarities between TCRs, without any information regarding the pMHC they bind. It is only after the creation of this tree that the TCRs were coloured according to the pMHC they recognise. The sequence of the latter is also noted in the tree. The tree was obtained from the highest-ranked set of the best-ranked TCRfp parameters determined by GA MATCH, with a score of 45.0% according to the *peptide identity score* (average score of the 10 sets according to the *peptide identity score:* 36.5% ± 5.0). The values obtained with the pMHC distance, and the node colour change are 0.79 and 83, respectively.

Therefore, the majority of those TCRs appear multiple times within the training sets potentially causing an increased risk of data overfitting. The TCRs binding the peptide KLGGALQAK account for the highest diversity of each gene (31 TRAV, 30 TRBV, 40 TRAJ, 12 TRBJ) and the cluster of these TCRs is the worst. These tendencies are observed in the 9 remaining sets (Supplementary Figure 1). The fact that TCRs binding peptides GLCTLVAML, ELAGIGILTV and FLYALALLL, appear better clustered by TCRfp MATCH, could also be explained due to the lower variability of these peptides specifically. As we have seen, there is a large standard deviation of the *peptide identity* score (± 5.0) obtained by TCRfp on the ten TCR sets. This score ranges from 46% of the easiest set to only 29% on the most difficult one. This underlines a substantial dependency of TCRfp efficiency on the specific set of TCRs under investigation. This could be due to some overfitting of the centroids obtained by MATCH or due to the small size of TCR sets that are available – the latter itself a consequence of the fact that only a small number of pMHC have a large number of TCRs known to bind to them – or both. Overall, the algorithm succeeded to cluster together TCRs based on their peptide recognition.

To improve the ability of the heuristic search to find centroids, a new readout approach was introduced (MaxD). Contrarily to the previous MATCH, where we optimized centroids, so that every TCR close to a reference TCR was likely to bind the same pMHC, the goal of the MATCH based on the MaxD readout is to separate all TCRs as much as possible, hoping that the TCRs with closest distances will spontaneously correspond the TCRs binding the same antigens. One advantage of this approach is that it does not require sets of TCR with known specificities, with the same number of TCR per peptide, to train the centroids and C or P weighting parameters. All available TCRs can be used, whatever their specificities. Thus, we could constitute a much larger training set of 2831 TCRs. As for the previous MATCH runs, all information and parameters can be found in the Supplementary Table 1. In these new MATCH runs, the MaxD scoring function is thus the average distance between all possible TCRfp pairs in the training set and the objective is to converge to centroids that maximize as much as possible this value.

The parameters that provided the best results of the sequence identity-based MATCH were adapted to the new MaxD readout. Max D runs led to increased C and P values, compared to the initial MATCH runs. Obtained values of 118 and 57 for C and P, respectively, compared to 22 and 1 at the start of the GA runs. Visual inspection showed that centroids coordinates determined using the GA MaxD algorithm were situated further from the TCR than those obtained from MATCH runs (Figure 4). Even though the MaxD runs succeeded in maximizing distance scores, we decided to restrict the initial state to lower values of C and P to explore the performance within these limitations in line with the biological role of the CDRs. To reduce the noise induced by TCR pairs binding the same antigen we subtracted those pairs from the distance calculation of MaxD runs. These new MaxD runs obtained lower distances scores than those starting from higher C and P values, with the best result scoring 0.63. In the last attempt, we hypothesized that the bigger sample size could require a higher variability of the different values of the algorithm. Hence, we explored various number of mutations, range of mutations and we tested the different crossover parameters.

The genome from which we obtained a higher predictive ability for the TCRfp after a MaxD optimisation is the following: C=1, P=2, centroid 1 = (10.663, 16.277, −10.741, 2.462, −0.548), centroid 2 = (19.366, 16.652, −1.838, −0.223, −0.461), centroid 3 = (−1.249, 17.586, −13.000, 1.610, −0.533), centroid 4 = (−7.824, 11.080, 13.731, 0.799, 0.388), centroid 5 = (−19.457, 16.546, 2.955, −1.104, 1.109) and centroid 6 = (3.042, 17.347, 11.183, 1.265, 0.484).

This solution was obtained with an algorithm using a population size of 400 individuals, a reduced initial range for the C and P from 0 to 20 for both, and an even more reduced range for the rest of parameters, i.e. −1 to 1. Each generation created 200 new offspring genomes derived from a pool of the best 200 parent genomes of the total population. The reproduction parameters consisted of an slowed version of the best MATCH algorithm, allowing a small range of increments for the weighted C and P weighting parameters and the c and p coordinates of the centroids (random values between −2 and 2) and even smaller for the Cartesian coordinates (increment values ranging from −1 to 1), but forcing 4 genes to mutate each time to enhance a quicker variability within generations. We used a single centroid crossover, swapping the integrity of a genome between parents for the generation of a child genome.

We compared the performance all the best genomes - obtained from MaxD runs - using an external validation set (described in the Methods). The sigmoid curves, displaying the relationship between the TCR similarity (it was calculated using similarity = 1 - distance) and the probability of binding the same peptide, showed that the genomes with a better specificity prediction capacity were the ones having lower C and P values and coming from the fitness score that removed the noise from the TCRs binding the same peptide. Intriguingly, with a MaxD fitness of 0.6, the genome providing the best predictive ability (described previously) does not correspond to the genome with the best MaxD fitness value (best MaxD fitness score over all runs: 0.7).

For comparison, the same analysis of the sigmoid curve applied to the external validation set was performed for the best genomes from the MATCH runs, using different algorithms (data not shown). In this case, the genome which achieved the highest *peptide identity score* also obtained the best sigmoid score.

All the parameters tested for the MATCH and MaxD GA runs are explained in the Methods. The best algorithms values and best centroids are provided in the Supplementary Tables 2 and 3.

In conclusion, the GA exploration provides a good method to increase the predictive ability of TCRfp, but the complexity of the TCR interaction, the large number of parameters, and the high influence of the training sets on the algorithms’ outcome made it challenging to find an optimal set of parameters for TCRfp so far.

### 2.4. Analysis of the performance of the different centroids with an external validation set

The preliminary definition of TCRfp, with centroids positioned on the tips of the loop, provided an averaged *peptide identity score* of 29.8% (Rank 1; no threshold, p-value <0.0001 when compared with random) on the external test set. The best genomes of the MATCH and MaxD runs were led to averaged *peptide identity scores* of 28.2% and 29.6%, respectively (Rank 1; no threshold, p-value <0.0001 when compared with random).

For a better understanding of the different approaches influence over the FP vector, the mean and standard deviation as well as the minimum and maximum of the 18 values that describe the FP vector were calculated for the entire validation set and summarized in the following Table 2 and Table 3.

**Table 2.** Comparison of the mean and standard deviation of the FP vectors obtained from the TCRs of the external validation set with the three different approaches. The three approaches being compared are: TCRfp preliminary, TCRfp from MATCH runs and TCRfp from MaxD runs. Each vector is constructed with 18 values coming from three values per each centroid according to the ES5D protocol described in Methods. In the table we can see the mean obtained for each value for all the TCRs from the set according to each approach as well as the standard deviation given inside the brackets.

| Approach | C1 |  |  | C2 |  |  | C3 |  |  | C4 |  |  | C5 |  |  | C6 |  |  |
| --- | --- | --- | --- | --- | --- | --- | --- | --- | --- | --- | --- | --- | --- | --- | --- | --- | --- | --- |
| *TCRfp preliminary* | 18.81<br>(1.19) | 7.36<br>(0.59) | -0.84<br>(3.10) | 20.40<br>(0.99) | 8.24<br>(0.56) | -0.39<br>(3.32) | 16.70<br>(1.50) | 5.79<br>(0.63) | -1.01<br>(2.57) | 17.87<br>(0.96) | 7.53<br>(0.52) | 2.38<br>(2.54) | 19.24<br>(1.54) | 8.34<br>(0.59) | 4.28<br>(2.02) | 15.47<br>(1.18) | 5.42<br>(0.47) | 0.66<br>(2.74) |
| *TCRfp from MATCH runs* | 28.99<br>(0.71) | 9.47<br>(0.48) | -0.02<br>(3.85) | 16.96<br>(0.44) | 4.65<br>(0.27) | -0.13<br>(2.18) | 41.82<br>(0.41) | 4.93<br>(0.35) | -0.66<br>(2.14) | 43.77<br>(0.50) | 8.90<br>(0.50) | 1.33<br>(3.34) | 34.80<br>(0.49) | 6.29<br>(0.31) | 1.40<br>(2.4) | 32.58<br>(0.51) | 6.24<br>(0.27) | 2.07<br>(2.08) |
| *TCRfp from MaxD runs* | 33.14<br>(0.48) | 4.79<br>(0.27) | 1.41<br>(1.97) | 35.23<br>(0.58) | 6.16<br>(0.31) | 1.59<br>(2.56) | 33.66<br>(0.38) | 4.76<br>(0.32) | -1.46<br>(1.89) | 30.52<br>(0.45) | 4.93<br>(0.25) | -0.62<br>(2.22) | 36.58<br>(0.51) | 6.27<br>(0.28) | 2.85<br>(1.78) | 33.34<br>(0.40) | 4.44<br>(0.32) | -0.56<br>(2.00) |

**Table 3.**
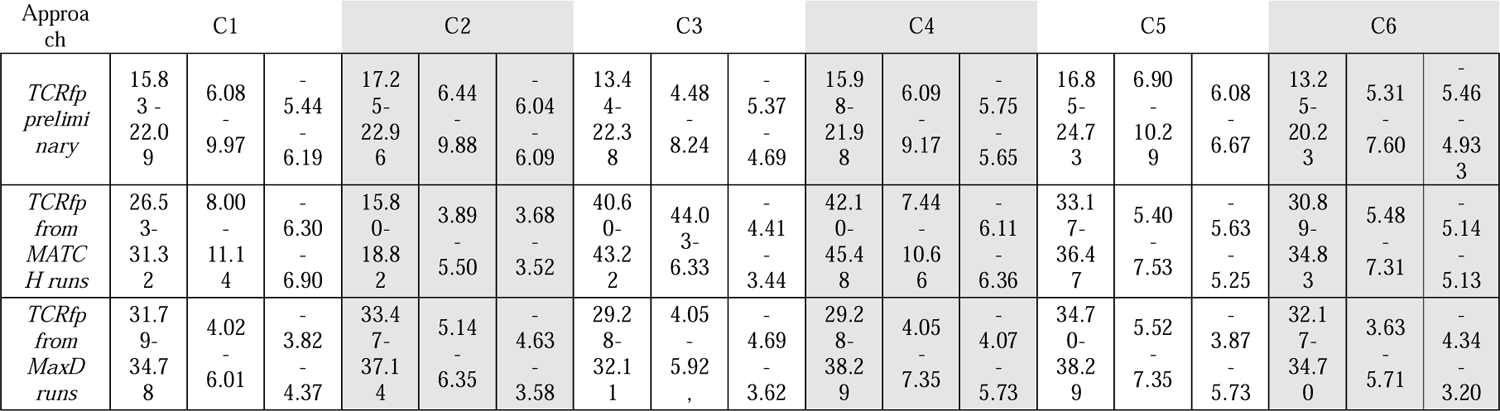
Comparison of the minimum and maximum values of the FP vectors obtained from the TCRs of the external validation set with the three different approaches. The three approaches being compared are: TCRfp preliminary, TCRfp from MATCH runs and TCRfp from MaxD runs. Each vector is constructed with 18 values coming from three values per each centroid according to the ES5D protocol described in Methods. In the table we can see the minimum and maximum numbers obtained for each value for all the TCRs from the set according to each approach.

The average standard deviation of the 18 values of the TCRfp vector is higher (4.31) compared to the heuristic searches (TCRfp MATCH: 1.18 and TCRfp MaxD: 2.40). Each TCRfp vector entry was more variable in the original tip-of-the-loop approach. The centroid with the highest variability was the one corresponding to the tip of CDR1α (4.9). The less variable one was the tip of CDR3β (3.22). Of note CDR3β is known to be the most important loop for pMHC binding.

These results allow us to conclude that using the initial tip of the loop approach provided more variability to the FP entry values, thus, making them more distinguishable. In addition, the preliminary definition of the centroids has the advantage of being independent from the position and the orientation of the molecule in space, making the ES5D vectors themselves independent from this, i.e. there is no need to align the TCR on the Cartesian centre and axes before calculating its FP. Also, it may indicate that an extensive tunning of the parameters translates into a potential overfitting of the algorithm, thus, the reduced variability and efficacy of the FPs generated using the parameters optimised by heuristic search approaches.

To assess if the similarity score given by TCRfp is able to cluster TCRs in a way it correlates with their peptide specificity, we calculated the relationship between the TCRfp similarity between pairs of TCRs and the probability of these two TCRs bind the same peptide (Figure 6). Interestingly, we find a clear sigmoid-like relationship between the similarity calculated by our approach and the probability of binding the same peptide. This relationship is very similar to the one found in the context of small drug-like molecules [32, 33], and supports the use of ES5D in the context of TCR repertoire analysis and specificity prediction. We observe that the inflection point of the sigmoid is obtained at lower similarity values for the tip-of-the-loop definition of TCRfp compared to the other two variants optimized by GA. At a similarity threshold of 0.6, the probability of binding the same peptide for two TCRs is 16.68% for the tip-of-the-loop TCRfp, 13.02% for TCRfp optimized by GA MATCH and 9.23% for TCRfp optimized by GA MaxD. These probabilities increase substantially at a similarity threshold of 0.8: 91.05% and 79.28% for the tip-of-the-loop TCRfp and the MATCH-optimised TCRfp, respectively, and 46.78% for the MaxD-optimised TCRfp. For the latter, a 80.5% probability of binding the same peptide requires a 0.85 similarity threshold. These values are in line with the previous analysis of the variability of the similarity values as a function of the TCRfp variant.

**Figure 6.**
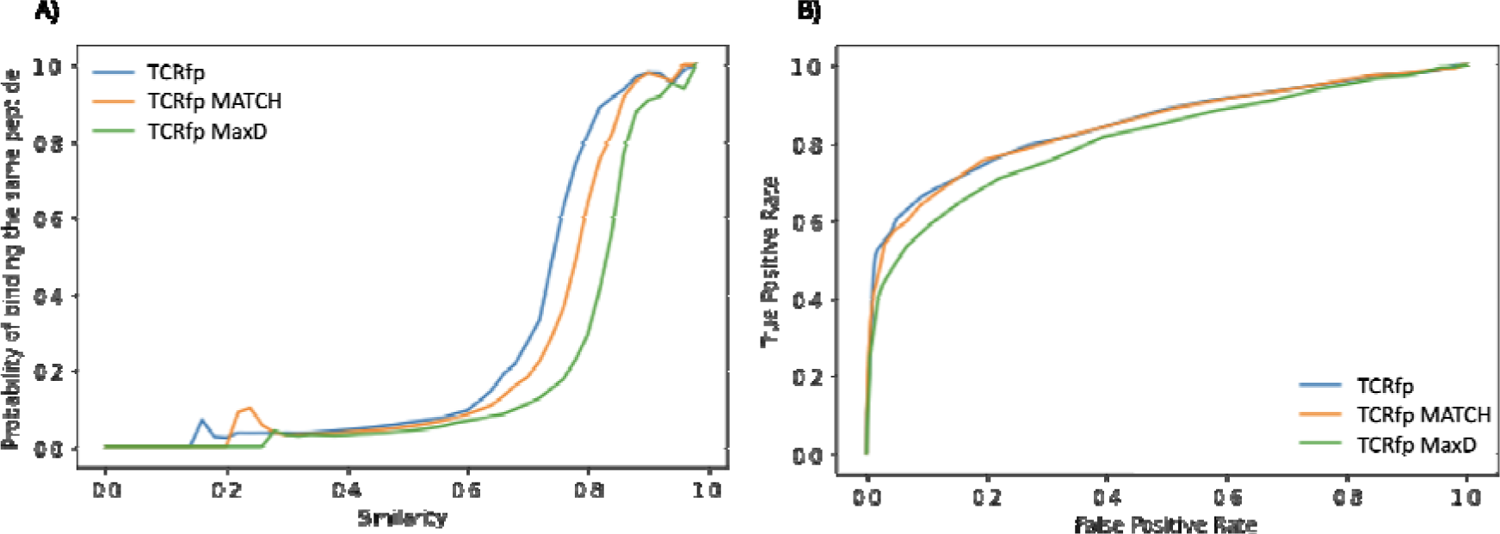
A) Relationship between TCR similarity (as calculated for an external validation set) and the probability of binding the same peptide. Comparison between the centroids placed in the tip of the loop approach (tip-of-the-loop TCRfp) and the best centroids position for the heuristic search (MATCH-optimised TCRfp) and the best centroids position obtained with the distance-based scoring (MaxD-optimised TCRfp). B) Relationship between the True Positive Rate and the False Positive Rate obtained with the three different approaches. The three approaches are much better than random.

To assess the ability of our approach to pair TCRs with the same specificity at different FP similarity values we have calculated how often the TCR with closest distance (Rank 1) the two TCRs with closest distances (Rank 2) and the 5 TCRs with closest distances (Rank 5) share the same specificity vs how often these pairs do not share the same specificity. The distribution of these frequencies can be seen in the Figure 7A. Again, the tip-of-the-loop TCRfp approach provides the best predictive ability, with a tipping point of positive and negative MATCHes increasing following this order: TCRfp - Rank 5; 0.74, Rank 2: 0.77; Rank 1: 0.85, TCRfp MATCH: Rank 5: 0.78; Rank 2: 0.82; Rank 1: 0.86, TCRfp MaxD: Rank 5: 0.82; Rank 2: 0.86; Rank 1: 0.88. After these tipping values, the frequency of positive pairs is higher than the frequency of negatives ones. This important finding supports again the idea that our tool can be used to predict specificities above these thesholds of similarity. Figure 7B shows how often the closest TCR share the same specificity at different thresholds (7B, left figure) and the number of TCRs considered at each threshold (7B, right figure).

**Figure 7.**
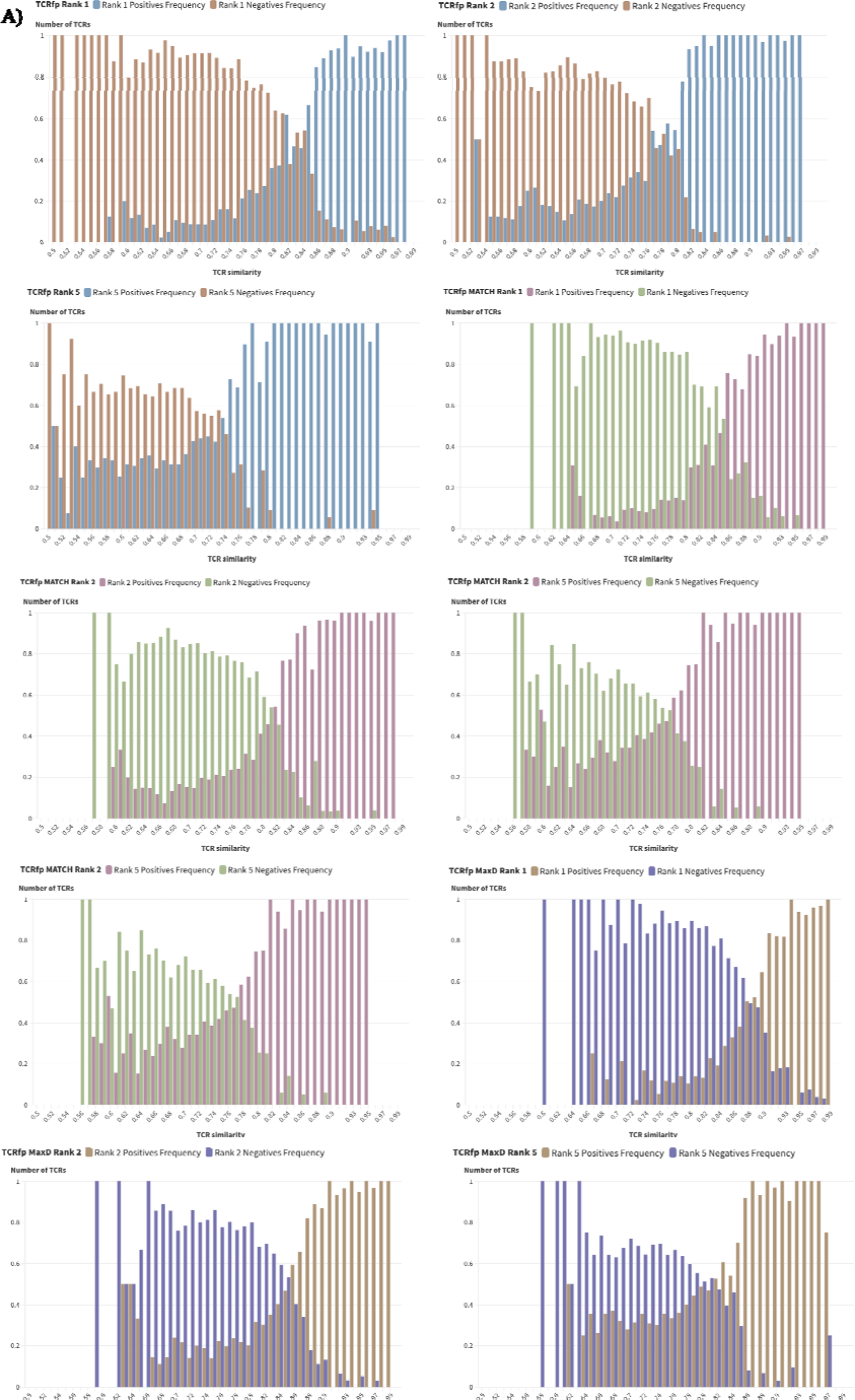
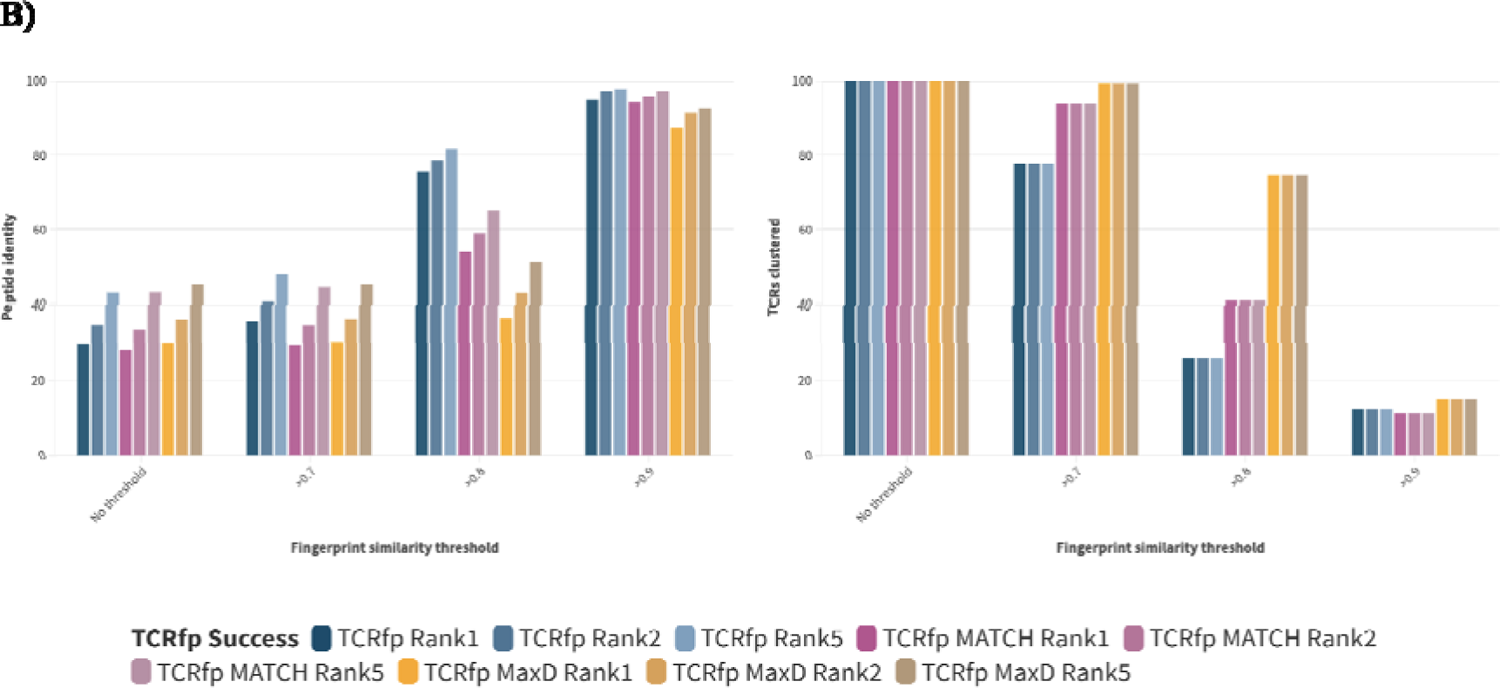
A) Distributions of the frequency with which the closest TCR (Rank1), at least one of the two closest (Rank2) or among the 5 closest TCRs (Rank 5) to a reference TCR bind the same pMHC as a function of the similarity calculated with TCRfp for the three different approaches: tip-of-the-loop TCRfp, MATCH-optimized TCRfp or MaxD-optimized TCRfp. B) Left: TCRfp success rates on the whole validation set as a function of the similarity thresholds (0.7, 0.8, 0.9). Right: number of TCRs considered at each threshold of similarity.

These results (Figure 7B) are in agreement with the better performance of the top-of-the-loop TCRfp original approach (Figure 6). When no similarity threshold was used, the three approaches gave similar *peptide identity scores* at Rank 1: tip-of-the-loop TCRfp: 29.75%, MATCH-optimized TCRfp: 28.17% and MaxD-optimized TCRfp: 29.97%. However, when considering higher ranks, and a similarity threshold of 0.8, thus close to the tipping value mentioned above, tip-of-the-loop TCRfp achieved a higher score (75.48%; Rank 5), substantially better than MATCH-optimized TCRfp (54.09%) and MaxD-optimized TCRfp (36.50%). The success rates of all TCRfp variants reach 90% and above when a similarity threshold of 0.9 was used. Of note, however, increasing the similarity threshold necessarily decreased the number of TCR that could be processed (Figure 7B). For instance, at a similarity threshold of 0.8, MaxD-optimised TCRfp MaxD was able to process 74,60% of the TCRs in the validation set, compared to 40,74% with MATCH-optimised TCRfp and only 26,02% with the tip-of-the-loop TCRfp. Hence, the amount of TCRs with the TCRfp MaxD was lower than for the rest of the approaches. This relationship between prediction accuracy and number of TCRs that could be processed, as seen in Figure 8, is shared by other TCR-clustering algorithms[15, 34], which can predict TCR specificity with 94% accuracy but only for 12% of the TCRs. In comparison, TCRfp could MATCH TCR specificities with an accuracy of 45% for 100% of the TCRs in the dataset (MaxD-optimised TCRfp at Rank 5, without similarity threshold). The best compromise between success rate and scope of application was obtained for tip-of-the-loop TCRfp at Rank 2, applied with a 0.7 similarity threshold, for which we were able to process 64.33% of the dataset with a success rate of 78.74%.

**Figure 8.**
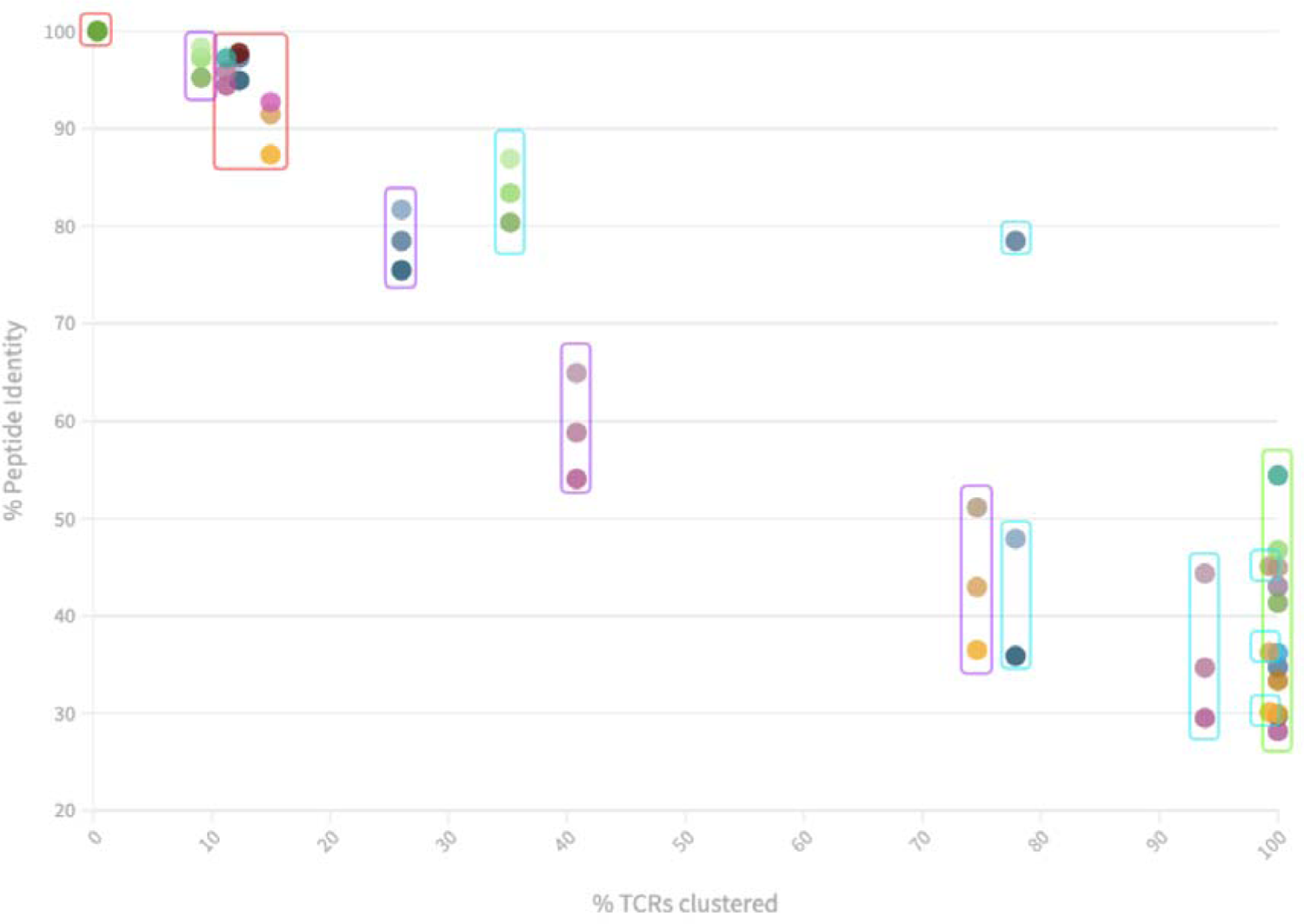
Relationship between the % peptide identity and the % of TCRs clustered by approach. Each TCRfp variant and rank are represented following the colour code of Figure 7B. The thresholds are represented with a square englobing the corresponding data points in different colours respective to their value: No threshold, cyan: 0.7, purple: 0.8 and red: 0.9. We observe higher % peptide identity for a reduced amount of clustered TCRs. All the approaches providing at least 70% peptide identity are only able to process 30% of the TCRs or less. Noticeably, the data point outside the linear tendency corresponds to the TCRfp approach for the 0.7 similarity threshold at Rank 2.

More efforts have been done to fine-tune the parameters of the MATCH GA. However, the MATCH optimisation presents a limitation over the MaxD optimisation, as it requires small datasets with equal numbers of TCR binding different pMHC to avoid bias. Such limited training datasets could lead to a risk of overfitting the parameters. On the contrary, MaxD optimisation, trained on a much larger amount of TCRs, has the advantage of avoiding the overfitting. However, since the fitness scored used in the MaxD approach is averaged over a higher number of data, it lacks the predictive ability shown by the MATCH optimisation. Consequently MATCH-optimized TCRfp performs better for higher similarity thresholds as it is more trained to discern among similar TCRs, while MaxD-optimised TCRfp performs better at smaller thresholds (all the values obtained are described in the Table 4). Further work is required to improve the MaxD approach, since a proper tunning of the fitness score could lead to better prediction of TCR specificity.

**Table 4.** Values obtained with tip-of-the-loop TCRfp, MATCH-optimised TCRfp, and MaxD-optimised TCRfp and the sequence-based approach, applying the different thresholds and ranks to the validation set. The value presented in the table is the data used for the Figure 7B visualization.

|  | Threshold | Rank1 | Paired TCRs | Rank2 | Paired TCRs | Rank5 | Paired TCRs | Total TCRs | TCRs clustered |
| --- | --- | --- | --- | --- | --- | --- | --- | --- | --- |
| TCRfp | no threshold | 29.75 | 956 | 34.77 | 1117 | 43.01 | 1382 | 3213 | 100.00% |
|  | >0.7 | 35.89 | 898 | 40.57 | 1015 | 47.92 | 1199 | 2502 | 77.87% |
|  | >0.8 | 75.48 | 631 | 78.47 | 656 | 81.70 | 683 | 836 | 26.02% |
|  | >0.9 | 94.95 | 276 | 97.22 | 385 | 97.73 | 387 | 396 | 12.32% |
| TCRfp MATCH | no threshold | 28.17 | 905 | 33.36 | 1072 | 43.08 | 1384 | 3213 | 100.00% |
|  | >0.7 | 29.51 | 890 | 34.71 | 1047 | 44.36 | 1338 | 3016 | 93.87% |
|  | >0.8 | 54.09 | 708 | 58.82 | 770 | 64.94 | 850 | 1309 | 40.74% |
|  | >0.9 | 94.43 | 339 | 95.82 | 344 | 97.21 | 349 | 359 | 11.17% |
| TCRfp MaxD | no threshold | 29.97 | 963 | 36.17 | 1162 | 45.00 | 1446 | 3213 | 100.00% |
|  | >0.7 | 30.12 | 961 | 36.26 | 1157 | 45.10 | 1439 | 3191 | 99.32% |
|  | >0.8 | 36.50 | 875 | 42.97 | 1030 | 51.15 | 1226 | 2397 | 74.60% |
|  | >0.9 | 87.32 | 420 | 91.48 | 440 | 92.72 | 446 | 481 | 14.97% |
| Sequence-based | no threshold | 41.36 | 1329 | 46.75 | 1502 | 54.44 | 1749 | 3213 | 100.00% |
|  | >0.7 | 80.37 | 909 | 83.38 | 943 | 86.91 | 983 | 1131 | 35.20% |
|  | >0.8 | 95.22 | 279 | 97.27 | 286 | 98.29 | 288 | 293 | 9.12% |
|  | >0.9 | 100.00 | 11 | 100.00 | 11 | 100.00 | 11 | 11 | 0.34% |

### 2.5. FPs can complement sequence-based approaches

To compare our structure-based approach with purely sequence-based approaches we applied a BLOSUM similarity score to our validation set. Briefly, we aligned the sequences of the 6 CDRs loops using pairwise alignment and compared their similarity using the BLOSUM62 matrix. An open gap penalty of −3 and an extension gap penalty of −1 were used [35]. The details of this sequence-based approach details can be found in the methods section. The distribution of the sequence-based score compared to TCRfp is also provided in Figure 9. The sequence-based approach consistently obtained higher *peptide identity score* than TCRfp variants. However, the amount of TCRs processed with the FP approaches, especially on the case of TCR MaxD, was higher than the sequence-based approach for all similarity thresholds. For instance, the sequence-based approach treated only 12.8% of the TCRs for a similarity threshold of 0.7 (2082 TCRs), while the MaxD-optimised TCRfp processed 99.3% of the data (Figure 9).

**Figure 9.**
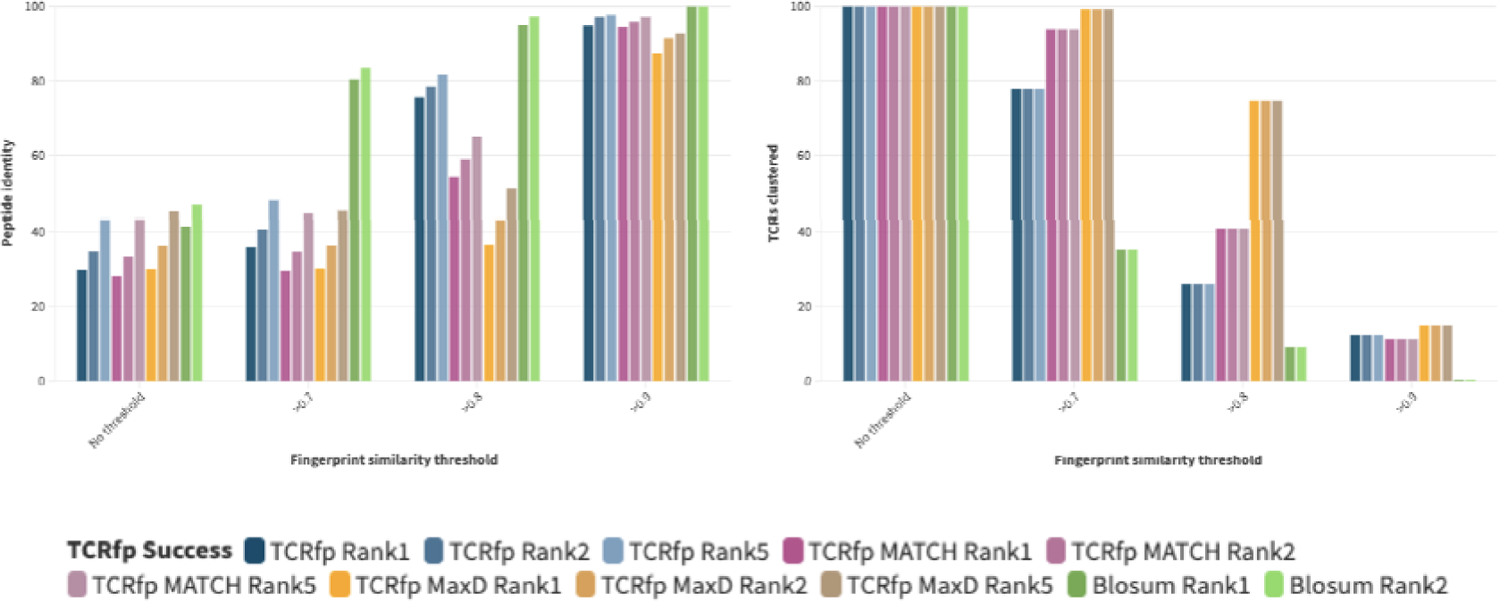
Comparison of the distribution of the *peptide identity score* and number of TCRs processed according to the TCRfp and sequence-based similarities, for matching at Rank 1, Rank 2 and Rank 5 and different thresholds of similarities (0.7, 0.8 and 0.9).

The sequence-based approach was able to MATCH correctly TCRs binding identical pMHC in 41.3% of the cases, while the success rate of the tip-of-the-loop TCRfp approach was only 29.8%, as previously discussed. Interestingly, as can been seen in Figure 10, 26.1% of the TCRs were correctly matched by both approaches, 15.2% only by the sequence-based approach and 3.4% only by TCRfp. Thus, TCRfp could provide additional information in some cases where a sequence-based approach did not work. Interestingly, when the length of the CDR3β of the reference TCR is 13, both approaches work better altogether and individually, with 59.6% of the TCRs properly matched, irrespective to the threshold (Figure 10).

**Figure 10.**
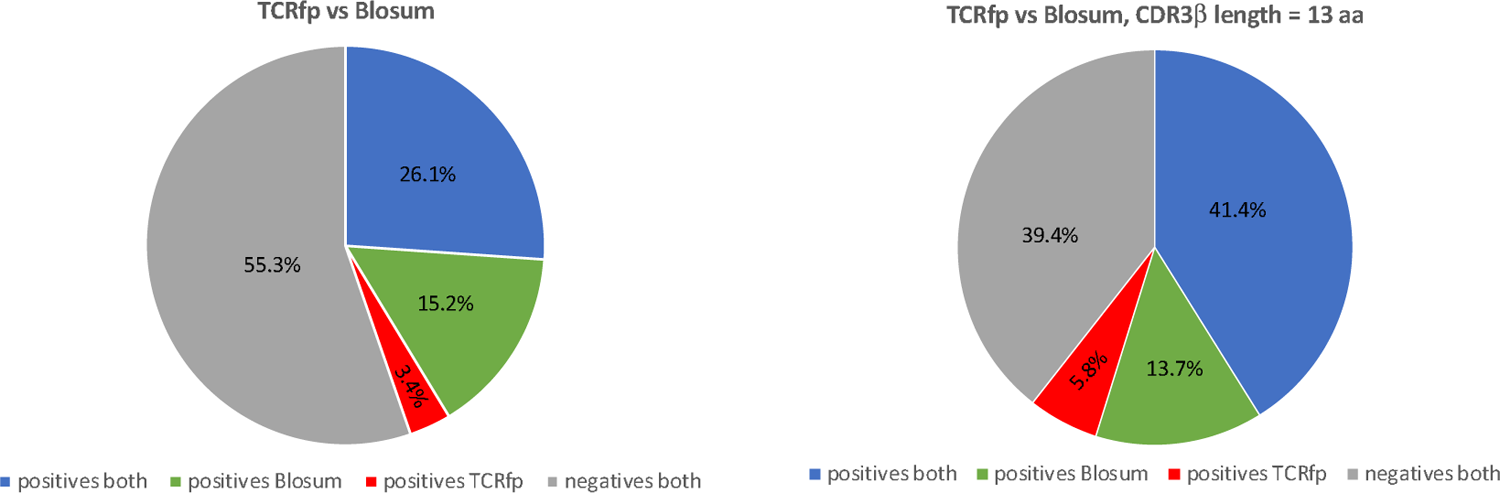
Success of the sequence-based approach and TCRfp for pairing two TCRs of the same specificity at Rank 1. The % of TCRs successfully paired by both approaches in blue, only by the sequence-based approach in green and only by TCRfp in red. The % of TCRs not paired, whatever the approach, is in grey.

Investigating the cases where TCRfp correctly matched TCRs sharing the same specificity while the sequence-based approach failed, we observed that despite the sequence dissimilarity, they shared a similar 3D shape. As for example 10X ID 00018 (formed by the genes: TRAV12-2, TRAJ45, TRBV28, TRBJ1-5, CDR3α: CAGGGGGADGLTF and CDR3β: CASTLTGLGQPQHF) and 10X ID 00017 (formed by the genes: TRAV12-2, TRAJ42, TRBV28, TRBJ2-3, CDR3α: CAVTHYGGSQGNLI and CDRβ: CASLRSAVWADTQYF), which both bind peptide ELAGIGILTV, are correctly matched by TCRfp (TCRfp similarity: 0.80). The root mean square deviation (RMSD) between their structures was 0.757 Å for the all the atoms. The same pair was scored 0.56 according to the sequence-based approach. Reversely, the 10X ID 00018 was erroneously paired by the sequence-based approach with the 10X ID 01495 (formed by the genes: TRAV12-2, CDRα: CAVISGGGADGLTF, TRAJ45, TRBV28, CDRβ: CASTIALGYEQYF), which binds the NLNCCSVPV peptide, with a score of 0.7 according to BLOSUM62, while the TCRfp similarity between these two TCRs of different specificities is only 0.44. Very interestingly, for these two TCRs of different specificities, the RMSD between the 3D structures is only 0.31 Å. The fact that these two TCRs displayed different specificities despite they exhibited a high sequence similarity and small structural RMSD, while 10X ID 00018 and 10X ID 00017 (correctly matched by TCRfp) displayed the same specificity despite large sequence and structure dissimilarities highlights the importance of the biophysiochemical properties encoded by TCRfp.

Figure 11, shows TCRs taken from the external test set, scored according to the TCRfp and BLOSUM62 values of the closest TCRs in the dataset according to each approach used separately. The colour coding indicates if the closest TCR according to each approach was sharing the same specificity. We observed that the majority of the TCRs correctly matched by both approaches are within the area comprised above a 0.8 TCRfp score and a 0.7 sequence-based score. Interestingly, some TCR that show a relative low sequence-based score are correctly paired using TCRfp, as it can be seen below the 0.6 threshold of the sequence-based score, illustrating again that our new TCRfp approach is able to correctly predict the specificity in some cases where the sequence-based approach would fail. All the data utilized to construct the Figure 29 can be found in the Supplementary Table 4.

**Figure 11.**
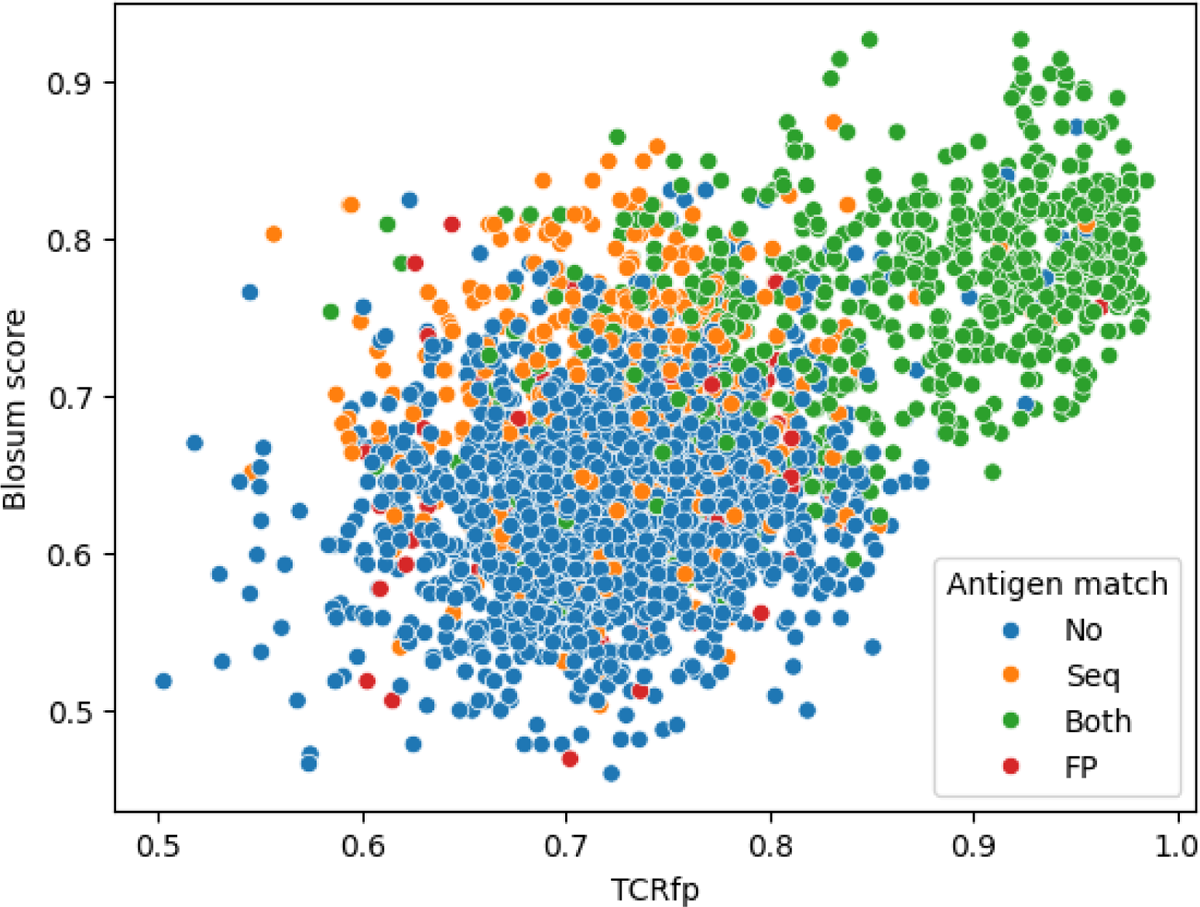
Relationship between the TCRfp and sequence-based scores obtained for each possible TCR, used as a reference, and its closest TCR according to each method, taken from the external validation set. The BLOSUM score is normalized for an easier comparison. Each dot represents a TCR according to the similarity of its closest TCR obtained with the TCRfp placed in the X axis and the sequence similarity of its closest TCR according to the sequence-based Blosum score placed in the Y axis. The colouring system represents the antigen pairing match according to the reference TCR and its closest TCR using the different approaches. No: reference TCR peptide did not match the peptide of the closest TCR of any approach, Seq: reference TCR peptide matched with the closest TCR peptide according to the sequence-based approach but not with the TCRfp, FP: reference TCR peptide matched with the closest TCR peptide according to the TCRfp approach but not with the sequence-based, Both: both approaches closest TCR binding peptide is the same as the reference TCR.

Table 5 shows the representation of each peptide in the subset of TCRs with specificity correctly predicted by: (i) both approaches, (ii) uniquely by sequence-based score and (iii) uniquely by TCRfp. The representation is calculated as the ratio between the number of TCRs correctly paired binding a given peptide and the total number of TCRs correctly paired, at rank 1. The 30 most frequent peptides in the validation set are described and their respective frequency in the validation set is also presented. We observe that GILGFVFTL is the most frequent peptide in the validation set, representing 14.6% of the TCRs. This is also largely the most frequent peptide in the subset of TCRs correctly predicted by both approaches, with 34.6 % of the TCRs correctly paired by both approaches recognizing this peptide. Interestingly, we observe that TCRs recognizing RLRAEAQVK are never correctly paired when using the sequence-based score indicating that shape has, more than a sequence, an effect in the recognition of the peptide. Interesting too is the fact that TCRs recognizing AVFDRKSDAK were never correctly paired by both approaches at the same time and are more frequent in the subset of TCRs correctly paired by the TCRfp (11.9% for TCRfp-uniq and 3.7% for SeqBased-uniq). The comparisons presented in the table 5 allowed us to understand peptides where shape-based approaches can be extremely relevant to find the TCR specificity.

**Table 5.**
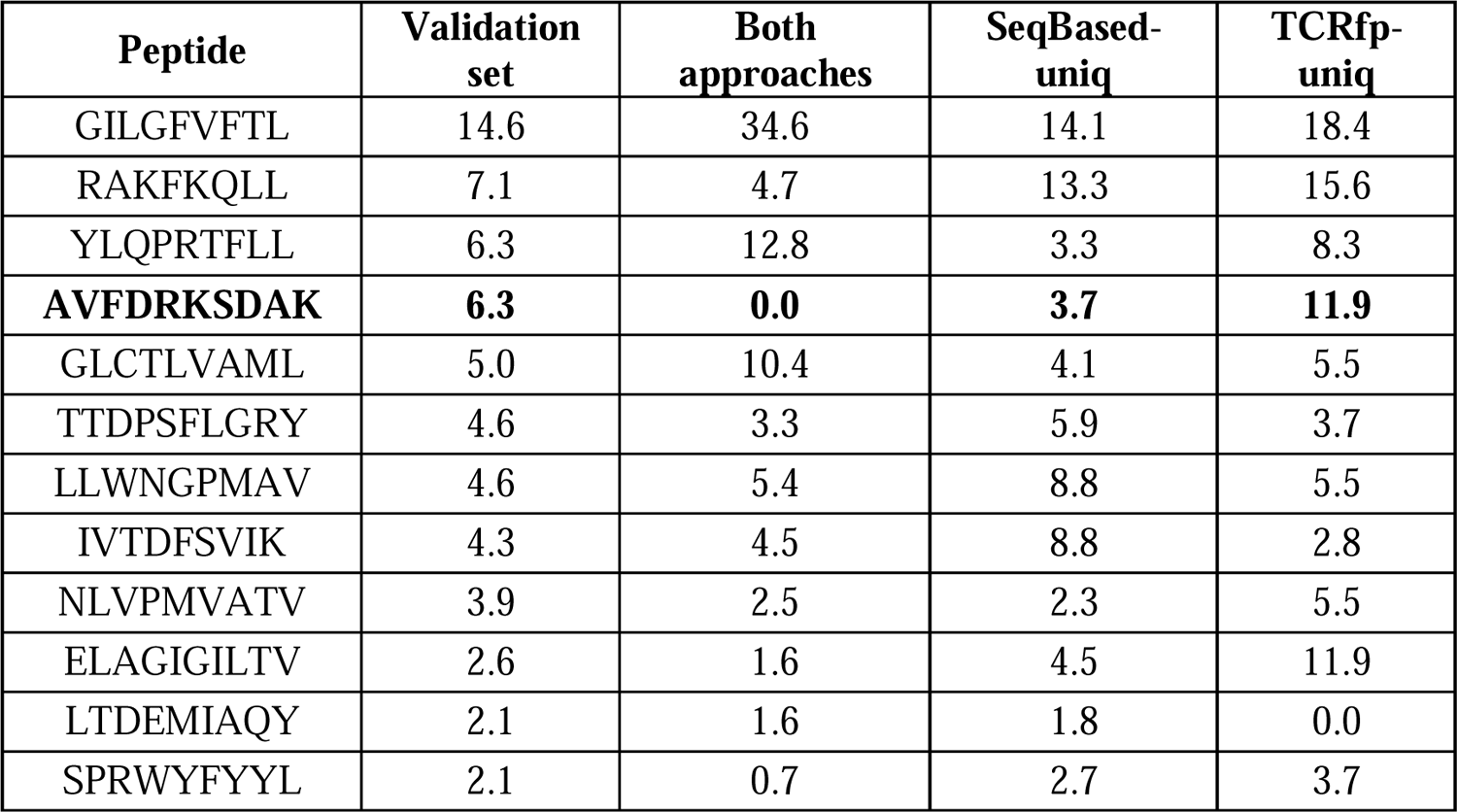
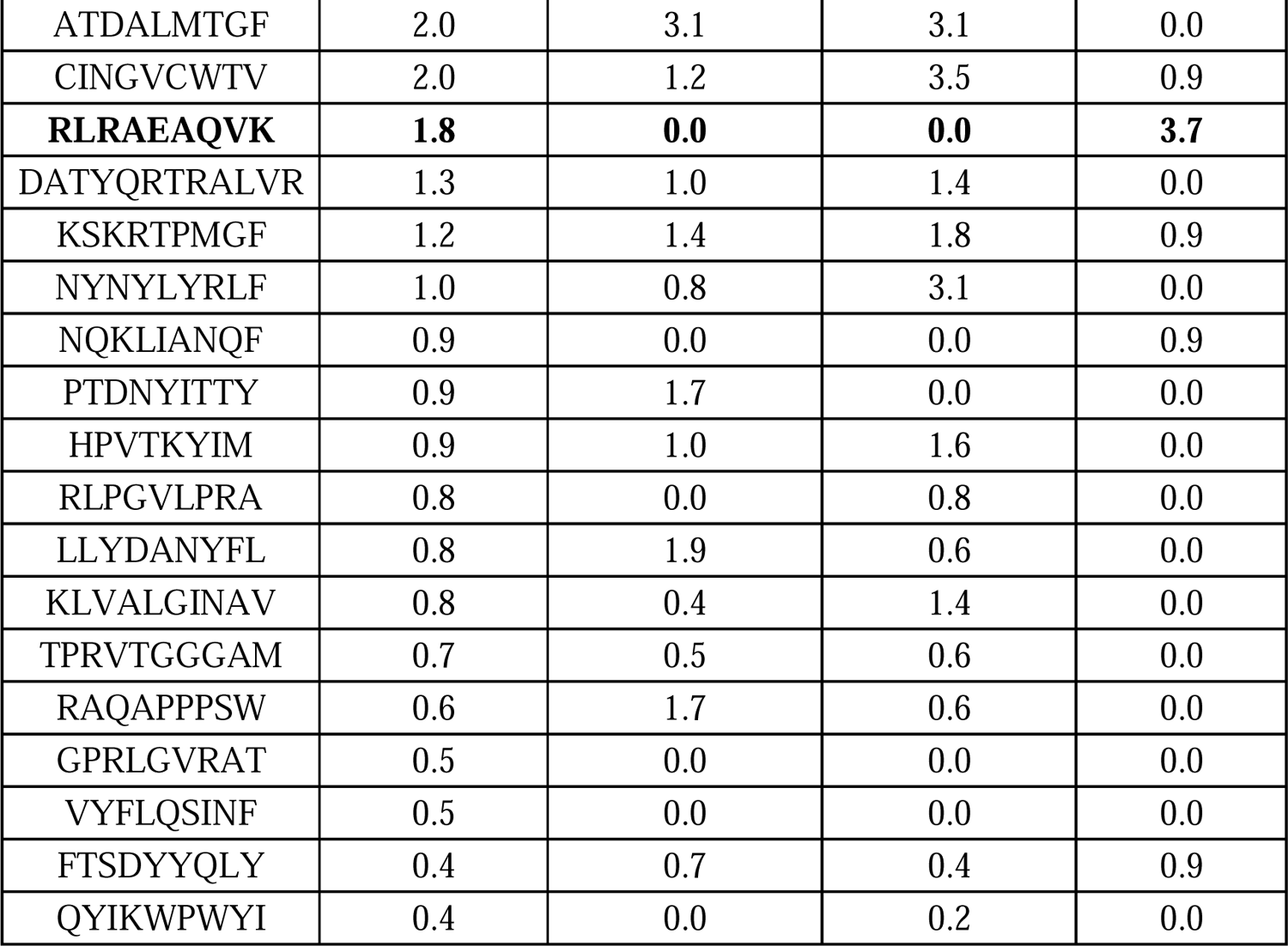
Representation of each peptide in the set of TCRs with specificity correctly predicted by: both approaches (column 3), uniquely by sequence-based (SeqBased-uniq, column 4) and uniquely by TCRfp (TCRfp-uniq, column 5). The representation is calculated as the ratio between the number of TCRs correctly paired binding a given peptide and the total number of TCRs correctly paired, at rank 1. The 30 most frequent peptides in the validation set are described and their respective frequency in the validation set is presented in column 2.

Altogether, both approaches, regardless of which one, are able to correctly predict 45% of the validation set. This value is quite interesting as it proves the importance of the complementarity of the sequence data with the structural and biophysiochemical information and supports the hypothesis that TCRfp can detect TCR similarity beyond a simple sequence similarity.

In summary, as mentioned above, TCRfp could correctly pair 88.7% of the TCRs at Rank 1 using a similarity threshold of 0.85. However, at this similarity threshold, only 19% of the TCRs of the validation set could be processed. Comparatively this BLOSUM62 sequence-based score was able to successfully pair 89.8% of the TCRs by setting up a threshold of 0.75, at which 22% of the TCRs could be processed. For the overall validation set, the sequence-based approach proved to perform generally better than our approach. Nevertheless, TCRfp performed better than the sequence-based score in particular cases, as for example, for pairing TCRs that recognize the RLRAEAQVK peptide. We have therefore combined the sequence-based score with TCRfp via a logistic regression (LR, see methods) and checked if a combination of both approaches would result in an increased predictive ability. We explored the performance of the three different methods, using three different threshold values (0.7, 0.8 and 0.9), for correctly pairing two TCRs binding to the same antigen (Table 6). We observed that, due to the different scoring functions used, the same similarity threshold has different impacts on the success rates of the approaches. For example, a similarity threshold of 0.7 seemed much more stringent for TCRfp, leading to a higher success rate than the other approaches, but at the cost of treating much less TCRs. For a fairer comparison, we selected a different threshold per each approach (Table 7). We then observed that using a combination of sequence based and TCRfp with such per-approach adjusted threshold was synergetic and resulted in an accuracy increase reaching 93%.

**Table 6.** Comparison of the TCRs correctly paired between TCRfp, sequence-based and a logistic regression (LR) approach for the probability thresholds 0.7, 0.8 and 0.9. The values noted in the Rank 1, Rank 2 and Rank 5 columns correspond to the peptide identity score % obtained with the validation set. Seq stands for sequence-based score and LR for logistic regression.

|  | Probability | Similarity calculated threshold | Rank1 | Paired TCRs | Rank2 | Paired TCRs | Rank5 | Paired TCRs | Total TCRs | TCRs clustered |
| --- | --- | --- | --- | --- | --- | --- | --- | --- | --- | --- |
| TCRfp | 0.7 | 0.77 | 60,07 | 725 | 64,13 | 774 | 68,85 | 831 | 1207 | 37,57% |
| Seq |  | 0.64 | 58,63 | 1179 | 63,45 | 1276 | 69,52 | 1298 | 2011 | 62,59% |
| LR |  | 0.66 | 60,00 | 1182 | 64,82 | 1277 | 70,61 | 1391 | 1970 | 61,31% |
| TCRfp | 0.8 | 0.8 | 75,48 | 631 | 78,47 | 656 | 81,70 | 683 | 836 | 26,02% |
| Seq |  | 0.66 | 65,48 | 1100 | 69,64 | 1170 | 75,30 | 1265 | 1680 | 52,29% |
| LR |  | 0.69 | 66,55 | 1110 | 71,04 | 1185 | 76,26 | 1272 | 1668 | 51,91% |
| TCRfp | 0.9 | 0.84 | 89,78 | 517 | 91,84 | 529 | 93,06 | 536 | 576 | 17,93% |
| Seq |  | 0.7 | 80,37 | 909 | 83,38 | 943 | 86,91 | 983 | 1131 | 35,20% |
| LR |  | 0.74 | 78,25 | 957 | 81,52 | 997 | 85,36 | 1044 | 1223 | 38,06% |

**Table 7.**
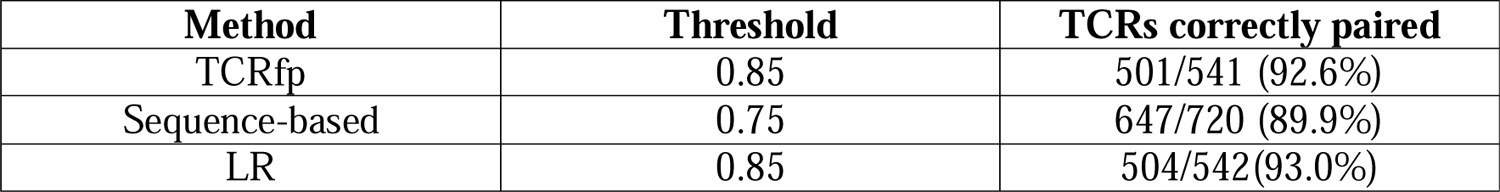
TCRs correctly paired using TCRfp, sequence-based and a logistic regression (LR) that combines TCRfp and the sequence-based score.

| Method | Threshold | TCRs correctly paired |
| --- | --- | --- |
| TCRfp | 0.85 | 501/541 (92.6%) |
| Sequence-based | 0.75 | 647/720 (89.9%) |
| LR | 0.85 | 504/542(93.0%) |

### 2.6. FPs are able to cluster experimental data

To assess the usefulness of TCRfp in processing clinical data, we applied it to a private set of 45 TCRs. For a further comparison, the sequence-based score and the logistic regression implemented (see the Methods section 3.8) were also applied to the private set. The results for the different methods can be seen in the Table 8.

**Table 8.** Comparison of the success rate in paring a TCR with another one showing the same specificity, using different approaches applied to a private set of TCRs. We compared the FP approach (FP) with the sequence-based approach and the logistic regression (LR), for the different ranks 1, 2, 5 and 10 without threshold and using a 0.7 threshold.

| Threshold | Rank | Method |  |  |
| --- | --- | --- | --- | --- |
|  |  | FP | Sequence-based | LR |
| no threshold | Rank 1 | 22.2 | 46.7 | 48.9 |
|  | Rank 2 | 35.6 | 60.0 | 62.2 |
|  | Rank 5 | 57.8 | 75.5 | 73.3 |
|  | Rank 10 | 73.3 | 77.8 | 77.8 |
| 0.7 | Rank 1 | 100 | 100 | 100 |
|  | Rank 2 | 100 | 100 | 100 |
|  | Rank 5 | 100 | 100 | 100 |
|  | Rank 10 | 100 | 100 | 100 |

As can be seen in Table 8, all methods were totally successful above the similarity threshold of 0.7. For the scores calculated without a threshold, the TCRfp provided the lowest success rate. However, the LR approach led to a substantially higher success rate than the sequence-based approach at Rank 1 and Rank 2.

## 4. Conclusion

This work introduced a new structural approach for TCR pairing and clustering based on ES5D FPs, called TCRfp, which involves the encoding of a 3D structure into a 1D vector. Although this unique approach was initially developed for small molecules, we hereby successfully applied the ES5D FPs on a substantial set of highly diverse TCRs. This novel approach has demonstrated the possibility of rapidly pairing TCRs in a way that correlates with their antigen specificity, underlining the importance of TCRs structural features for understanding their binding properties beyond purely sequence information. Importantly, while TCRfp proved competitive, the present study represents only a first exploration of the potential of structure-based TCR fingerprinting approaches in this context. These approaches offer a large number of possibilities of improvement and adaptation.

We also explored the possibility to provide a single generalized definition of the FP to improve the efficiency of the calculations using genetic algorithms. Genetic algorithms were applied over different training sets, using a similarity-based score (GA MATCH) or a distance-based score (GA MaxD). For both methods, the parameters were modified and tuned over different trials resulting in a better overall score and predictive ability for TCRfp optimized by the GA MATCH approach, although it was built using a smaller training set, which could have led to overfitting. Paradoxically, the predictive ability of TCRfp optimized by GA MaxD was lower, even though it was trained on a larger data set. The most probable reason is that the TCRfp MATCH-derived parameters were more accurate due to a more profound exploration of the genetic algorithms, and the different fitness used in the GA. The computational requirements of using bigger training slowed down the GA MaxD process and limited the exploration of TCRfp parameters. The fitness score used in GA MaxD, which takes into account an averaged value over a large dataset, probably does not offer sufficient variability in the fitness score as a function of the optimizable TCRfp parameters, making it less suitable for optimizing TCRfp performance. Of note, the approach using the original definition of the TCRfp based on centroids positioned on the tip of the loop, performed better than the GA-improved centroid definitions. Clearly, adapting the position of the centroid to each TCR proved more accurate to cluster TCRs in a way it that matches their specificity.

We also demonstrated that TCRfp is able to correctly pair TCRs which are not correctly matched with a sequence-based approach. The fact that our approach can predict TCR specificity when the sequence information is not sufficient, underlines the importance of the structure and biophyshicochemical properties of TCR loops for the TCR-pMHC interaction and TCR specificity determination.

Finally, we explored the ability of combining TCRfp with a sequence-based score using logistic regression. We found that the combined approach increased the accuracy of TCR specificity prediction. The enhanced predictive power of the new approach is in line with the importance of incorporating TCR structural parameters, as well as charges and lipophilicity information, in peptide specificity prediction. When applying the approach to an experimental dataset not used for the training, we observed an improvement in the performance of the logistic regression compared over the sequence-only approach. We thus demonstrated the ability of TCRfp to complement a sequence approach and provides additional meaningful information not encoded in sequence-based algorithms.

This work demonstrates the feasibility of rapid structure-based approach for TCR repertoire analysis, TCR clustering and potentially TCR specificity prediction, With possible clinical applications. TCRfp thus introduces a new class of approaches for TCR pairing and clustering that can shed some light on the complex structural mechanism underlying TCR-pMHC recognition.

## 3. Material and Methods

### 3.1. Data retrieval, cleaning, and curation

#### 3.1.1. TCR structures with known pMHC

To perform a preliminary analysis of our 3D-based approach, we used a set of 74 TCR structures with known pMHC (HLA-A2 restricted) taken from PDB as of September 2019. TCR-pMHC complexes were downloaded and then cleaned by removing pMHC, non-standard amino acids, water molecules and other TCR molecules in case several TCR dimers were present. In the end, we only retained the structure of one TCR pair (one α and one β chain). The PDB ID (PDBid) of the structures, the label of the retained chains, and the peptide (pMHC = pHLA-A2) can be seen in Supplementary Table 5. Peptides’ sequences were aligned based on their structural superposition using UCSF Chimera [36]. Since CDR loops of TCR are flexible and since we will not be able to use X-ray conformations in real case application, we decided to use the kinematic closure (KIC) method [37] of the Rosetta software version 3.11 [38] to create 100 different low energy conformations of the 6 CDR loops of each HLA-A*02 restricted TCR. The lowest energy conformation per TCR determined using Rosetta REF15 scoring function [39] was selected for further analysis.

#### 3.1.2. Development of a modelling pipeline based on Rosetta to create TCR models from sequences

To model TCR 3D structures models from their sequences a fast and automated TCR homology modelling pipeline was developed. The pipeline starts by reading the sequence information from the TRAV, TRAJ, TRBV, TRBJ genes as well as the CDR3 of each TCR chain. Then, the CDR3 sequences are aligned with the TRV and TRJ genes from the respective chains, and the full α and β chain sequences are reconstructed. Subsequently, the TCR sequences are converted into 3D structures by homology modelling using Rosetta version 3.11 [38] and the TCR modelling protocol described in Gowthaman et al [25]. Benchmarking of the Rosetta modelling protocol using a set of nonredundant TCR experimental structures showed that models are accurate and compare favourably to models from other available modelling methods [25]. Furthermore, in our own pipeline, 10 conformations per TCR were generated and ranked with the Rosetta REF15 scoring function [39] and the lowest-energy conformation was selected as the final TCR model. Finally, to detect and remove problematic models, all the final model conformations were subjected to distance-based quality controls. Indeed, by applying Rosetta protocol to model the TCR sequences from 10xGenomics we have observed that inaccurate TCRs models could be obtained (see results and discussion for further details). For this reason, we have developed filters based on the distances among specific conserved residues within the TCR, using the distances found among the conserved residues from experimental structures. To implement these filters, the TCRs were first renumbered according to IMGT [40] numbering using the ANARCI program [41] to keep the same residue number for constant residues across all the TCRs [41]. Based on [42], 4 conserved residues were selected for the filters (CYS23, LEU89, CYS104, TRP118). The distances between 9 combinations among them were calculated with UCSF Chimera [43] using a set of 92 experimental structures of TCR class I (Supplementary Table 6). For a model to be accepted as accurate, the distances must lie within the range defined as the average value in the experimental structures ± 3 times their standard deviation for each filter. The models that did not match the distance filter criteria were discarded.

### 3.2. Training and validation sets

#### 3.2.1. Training sets

CD8+ T cell sequences with known cognate pMHC were retrieved from the 10xGenomics database on the 25^th^ of November 2019 (CD8+ T cells from human Healthy Donors 1, 2, 3 and 4 (v1, Single Cell Immune Profiling Dataset by Cell Ranger 3.0.2, 10x Genomics, (2019, November 25)). The dataset was carefully cleaned by removing unreliable sequences. Additionally, all the retrieved T cell sequences lacking relevant information – i.e., a whole chain, a single gene, CDR3 data or the peptide specificity - were excluded. As the data used from 10xGenomics was obtained through Single-Cell Sequencing we had to construct the paired TCRs. During this process we encountered a high proportion of TCRs showing abnormal sequence count: TCRs with multiple α, β or both chains. Biologically, a fraction of T cells is known to express multiple α and β chains: two α and one β or vice versa [44]. To remove all possible ambiguities, we kept only TCR from T cells expressing a single α and a single β chain. After applying these stringent filters 14’479 TCR sequences with known pMHC were retained (Supplementary Table 1).

TCR sequences were then converted into 3D structures by applying the protocol previously described. Our TCR data set consists of 11’904 TCR models from 9 MHC and covers specificities for 33 different peptides (all the detailed data can be found in the Supplementary Table 1). Surprisingly, the majority of the TCRs found within our dataset are specific for the KLGGALQAK peptide (73.7%), followed by GILGFVFTL (7.7%), AVFDRKSDAK (5.8%) and RAKFKQLL (5.6%). The rest of the antigens account for 2% or less of representation from the total dataset.

### Creating TCR datasets to optimize TCRfp via heuristic search (MATCH)

To avoid over or under representation of a peptide in respect with the others we constructed 10 training sets. Each set was composed of groups of 9 TCRs, each group binding one of 13 different antigens, for a total of 117 TCRs per set (specified in the Supplementary Table 1).

### Optimizing TCRfp by maximizing the distance between TCRs (MaxD)

To mitigate the overfitting that could be generated small training sets, we examined optimizing the parameters of TCRfp to maximize the average distance between all TCRs considered. As this approach does not require an equal number of TCR binding the same peptide, we included all the TCR models data in a single dataset. Again, TCRs binding to several peptides were discarded beforehand to avoid biases in the scoring. However, it was possible to include peptides recognizing a single TCR, which were discarded in the previous MATCH training datasets, as the TCR specificity was not required in this new approach. To avoid a bias towards the peptide KLGGALQAK all the corresponding models were discarded from the training. The final training set consists of 2’931 TCRs that covered specificities for 32 different peptides.

#### 3.2.2. External test set

To assess the performance of our FP we used a different set that does not include any of the previous TCRs used for the training. This external test set was retrieved from the VDJdb curated database (data obtained in 2022). We modelled the TCR 3D structures using our TCR modeling pipeline, obtaining a total of 10’703 TCR structures to use as an external validation dataset for our FP scoring. To avoid a bias towards our results, all the redundant TCRs overlapping with the 10xGenomics database were excluded. Additionally, we discarded the TCRs recognizing the overrepresented peptide KLGGALQAK (53.6%) and the singleton TCRs that are impossible to pair (and so, useless for our test exercises). The final validation set consists of 3’213 TCRs, covering specificities for 200 different antigens. All the TCRs structures used in the final validation, including their genes composition and peptide specificity, are described in Supplementary Table 7.

### 3.3. Adaptation of the Electroshape algorithm for TCR models

We have applied Electroshape (ES5D) [28] to convert experimental and modelled TCR 3D structures into 1D vectors. The ES5D algorithm translates the 3D structure of the TCR in a 1D numerical vector, relying on 6 centroids. The latter are points of interest surrounding the molecule to describe (see latter for their definition of their positioning). In this approach, all centroids and atoms have 5D coordinates: the 3 Cartesian coordinates, the weighted atomic partial charges (4^th^ dimension) and the weighted lipophilicity (5^th^ dimension). The latter is defined as the atomic contribution to logP according to the WLOGP algorithm [45]. The weights of the atomic partial charges and lipophilicity are defined latter. To obtain the final 1D vector that defines the structural FP of a TCR, the average, standard deviation and third moment of the distances between all atoms (including hydrogen atoms) of the TCR’s CDR loops to each of the centroids were calculated, leading to a final vector of 18 values.

Once a FP has been caculated for each TCR, quantifying the similarity between two TCRs using ES5D boils down to calculating the distance between a pair of 1D vectors with a Manhattan distance-based score, which ranges from 0 for TCRs with totally dissimilar shapes to 1 for TCRs with perfectly identical shapes:

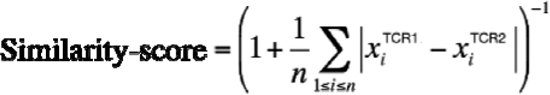

where *n* is the number of entries in the 1D vectors and *x* are the entry values of the vectors for each TCR.

### 3.4. TCR similarity scoring

To assess that our FP-based Similarity-score correlates with the likeliness of TCRs to share the same specificity, and thus that it can be used to apply the similarity principle, we have developed two other scoring systems based on the recognized peptide: the *sequence recapitulation* and the *peptide identity*.

The *sequence recapitulation* score estimates for each TCR within a set of TCRs with known specificity, if the closest TCRs among this set recognize peptide sequences with high degree of similarity. This is calculated by aligning the peptide sequence recognized by each TCR, taken as a reference, with the one of the closest TCR in the set according to our approach (excluding the TCR reference itself), and counting the number of identical residues in the alignment. The average value across all TCRs in the set represents the *sequence recapitulation* score. This score is particularly well adapted to test the approach on TCR-pMHC experimental structures, where several TCRs among the set are binding peptides baring a few point mutations.

The *peptide identity* score measures, for each TCR in a given set, how frequently the closest TCR according to TCRfp binds exactly the same peptide. This score is relevant when studying datasets with TCRs that recognize only a few unrelated peptides. Using a set of TCRs for which the cognate pMHC are known, the process involves identifying if they bind the same peptide (positive pair) or a different peptide (negative pair). The fraction of positive pairs among all pairs defines the peptide identity score expressed in %.

To translate TCR 3D structures into 1D fingerprint vectors, we initially placed each of the 6 centroids on the α carbons of the tip of the loops of one of the 6 CDRs (CDRs 1, 2, 3 from α chain and CDRs 1, 2, 3 from β chain; Figure 2). In the case of loops with an even sequence length, the tip of the loop was selected by taking the lowest from the two middle residues. As the tip of the loops are situated in flexible TCR regions and contact the pMHC, these represent strategic places to better describe the 5D shape of the TCRs. As mentioned earlier, the 4^th^ and 5^th^ dimensions of the atomic coordinates are defined as the weighted atomic partial charge and the weighted lipophilicity, respectively. To perform our first tests, after exploring a limited number of combinations (such as 25;4, 35;4 and 45;4), the charge and lipophilicity weighting parameters (C and P), were initially set to 25 and 4, respectively, since this combination was shown to perform the best. The C and P combination of values that were tested and finally used were inspired from previous studies with ES5D and small molecules [46] and a study performed in the lab (data not shown). These C and P values were later optimised by heuristic search, as described below.

### 3.5. Genetic algorithms with *peptide identity* scoring

We explored the possibility to define centroids as freely chosen points in space. Considering the infinite number of possible centroids positions in the 5D space and values for C and P, we searched for near-optimal solutions making use of genetic algorithms (GA). After superimposing all TCRs by centring them on the origin of the 3D space and aligning their main axes along the axes of the Cartesian space using the COOR ALIGN function of the CHARMM program, we apply our GA algorithms using the 10X genomics training sets.

Our GA performs as follows:

1. Initialization of the population: the optimizable genome, which characterises each individual of the population, was defined as the 5D coordinates of the six centroids, together with the C and P weighting parameters, for a total of 32 values to be optimized. The initial highly diverse population consisted in 400 individuals with a different genome chosen at random.
2. Fitness calculation: The fitness of each genome, was calculated as the average *peptide identity score* over the 10 sets of 117 TCRs, as defined above. Briefly, each genome was individually applied to each set, and a FP was generated for each TCR of the set. Thus, for each genome under consideration we obtained different TCR FPs. Genomes improving the overall TCRfp efficacy was characterised by increased values of the average *peptide identity score*.
3. Parent selection: The 50% best genomes (200), i.e., those showing the highest *peptide identity score,* were selected for the reproduction step to produce the next generation.
4. Crossover and/or mutation: The next generation offspring was built by applying mutations and/or crossovers to the genomes of the selected parents. For the crossover, a random number of centroids – between 1 and 6 - were exchanged between the selected parents (depending on the variant of algorithm used, the centroid can be entirely exchanged or just partly). The weighting parameters C and P could also be exchanged individually at random. From the two possible children generated this way, only one was randomly selected. A random number, between 2 and 4 (other values were explored but those were kept based on their higher performance), of genome entries were selected and randomly incremented by a random value ranging from −1 to 1. 200 children were generated this way before entering the population. Finally, the *peptide identity score* was estimated for each individual and the 200 ones with the worst scores were discarded to keep a population of 200 individuals. The GA was stopped when the fitness score of the best individual had not improved for at least 20 generations (Figure 12).

**Figure 12.**
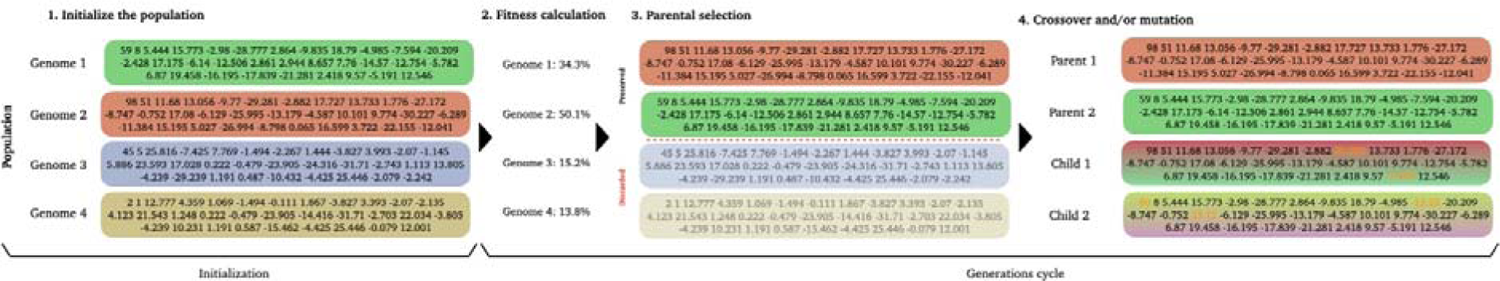
Overview of our genetic algorithm pipeline. 1. Initialize the population (for simplification, only 4 individuals are shown in this example). The genome of each individual is defined by randomly choosing the values of the 32 considered parameters (5 dimensions for each of the 6 centroids, plus 2 weighting parameters, C and P). 2. Fitness calculation. 3. Parent selection: the initial population is sorted based on the fitness scores. The upper half of the genomes (Genome 1 and Genome 3) are selected as parents for the following generation while the genomes 3 and 4 are discarded due to their lowest scores. 4. Crossover and/or mutation: each discarded individual is replaced by a child whose genome is constructed from the parent ones. For this step, the child genome is generated by combining those of two randomly selected parents (the proportion of each is also selected randomly). To increase the diversity of the population, we perform a mutation on the child genome by modifying random values in the vector. Once the offspring is generated, the GA continues at step 2 and reinitialize the process. The algorithm terminates when the fitness score does not improve over 40 generations.

Due to the large number of parameters to tune, finding the global optimum via this exploration would be considerably time consuming. We intended to perform a profound exploration of the best combinations of parameters to speed-up the search and to find near-optimal centroid values. This search was performed over 990 independent runs with different parameter combinations. In some runs, the centroids in the genomes were initially placed on the tip of the loop of each CDR (as specified for the original definition) and optimized starting from that position. To give more freedom to the algorithm to explore the search space, we also tested in some runs to initiate the centroid coordinates from random positions in the 3D space within a defined area of 30×30×30 Å^3^ (centred in the average middle of all the superimposed TCRs) and allowing them to move freely outside and inside that space. Analysing the outcome of two subsets of 350 runs each, we obtained an average of 36.98% ±1.19 *peptide identity score* when starting from the tip of the loop (corresponding to an improvement of 6.71% ±1.13 compared to the “strict” initial position), compared to a final 38.86% ±1.77 *peptide identity score when starting from random positions (corresponding to a* 9.02 ±1.67 improvement compared to the initial random genomes). We also increased the overall population size from 100 to 200 to test if it would improve the accuracy. Indeed, a population of 100 individual led to an average *peptide identity score* of 37.16% ±1.59 (improvement of 7.12% ±1.33 between the start and end of the runs) compared to 39.17% ±1.78 (improvement of 9.19% ±1.63 between the start and end of the runs) for a 200 population of 200 individuals, as measured over 350 runs each. Therefore, we decided to keep the initial random generation and the 200-individual population size for further improvements. As for the residues used to calculate the FP, the initial definition considered the 10 closest residues of the Cα of the tip of the loop. Eventually, we included the residues constituting the full length of each loop for the calculations, but no statistical difference was found for the algorithm and the original definition was kept.

Crossovers were performed by choosing a random position within a parent genome and replacing the values beyond that position with the corresponding ones taken from the other parent’s genome. We adapted the crossover to a centroid-based interchange to be in line with the biology behind the values. For this, we chose randomly a centroid and exchanged the corresponding 5 values between parent genomes. The two weighting parameters worked independently as two extra genes. As expected, the centroid-based crossover achieved a higher final *peptide identity score* (39.10% ±1.64) and improvement (9.1% ±1.76) compared to the initial crossover (37% ±1.38 and 7.2% ±1.59 respectively), averaged over 400 runs each. In addition, when exploring the centroid-based crossover performance, we tried to exchange the entire centroid or to break it at a random position to increase the variability. We generated 10 new runs for each approach and compared the average final *peptide identity score* as well as the improvement over the initial generations. In parallel, we explored the performance of exchanging several centroids at the same time. This test was performed using 10 more runs per each approach combining single and multiple centroid crossovers. We obtained a higher final *peptide identity score* and improvement (40.27%±2.20 and 8.21±2.41%, respectively) using multiple centroids than with the single-centroid approach (39.93% ±1.81; 7.79%±1.93). We only noticed a slight improvement when allowing to break centroids compared to consider them as a whole (40.82% ±2.06 peptide identity; 11.48% ±2.08 improvement) compared to considering them as a whole (39.72% ±2.43 peptide identity score; 10.65% ±1.4 improvement). There was no difference for the single-centroid approach. Therefore, we decided to keep the multiple crossovers for further improvement but to keep testing the entire centroid or the break.

Next, we assessed the effect of the amount of mutations performed on the offspring and their magnitude. We tested several ranges of values from which the algorithm could randomly choose the number of mutations applied to the offspring genome. The range performing better was from “1 to 2 mutations” (40.64%±1.95; 10.51%±1.92). The worst range was “5 to 15 mutations” (36.21%±1.22; 5.90%±1.32). For each of these ranges, we assessed different maximal incrementation values that the algorithm could choose to randomly subtract or add to the original gene value, i.e. 1, 2, 5, and 10. Of note, for both the weighting parameters C and P, the lowest range value led to better result (average 39.61%±1.43 *peptide identity score* with a 9.16%±1.7 improvement) compared with the highest value (38.30%±1.68 and 8.31%±1.88 improvement). Changing the mutation range for the centroids increased the peptide identity and the improvement from 36.35%±1.17 and 6.48%±1.04 for the lowest range, to 41.52%±1.54 and 11.35%±1.57 for a range of 10.

Additionally, some runs including the totality of the loop were tested for both TCRfp MATCH and TCRfp MaxD approaches instead of using only the mid residue and its adjacent ± 5 residues. without any noticeable increase in the scores. We kept trying different ranges on the further exploration as we were constantly calibrating new approaches but considering these values as the base of our algorithm. The details regarding the final algorithms can be found in the Supplementary Table 2.

Another parameter that we consider optimizing the GA was the fitness score itself. As previously described in section 3.4., the fitness *peptide identity score* described solely relies on the probability of a reference TCR and its closest one according to TCRfp to share a common pMHC target. Instead, we tested the score considering the 2 or 5 closest TCRs. The main goal was to test whether the GA would provide better results if a less restrictive fitness score was applied. The score, ranging from 0 to 1, was based on an average number of closest TCRs sharing an identical pMHC target than the reference TCR. Finally, like for the original *peptide identity score*, the average value over the whole set of TCRs was considered the fitness score of a given genome. To ease the score, a modification was incorporated to the 5 closest TCRs ranges. In that case, the fitness score was considered a binary score of 0 if none of the 5 closest TCRs was binding the same antigen as the reference TCR or 1 if at least one of the 5 closest TCRs was binding the same antigen. For comparison the three different new scoring protocols were tested using the parameters described in the Supplementary Table 2. We found that none of the GA using these new scoring schemes managed to surpass the best algorithms obtained with the original fitness score. The summary of the best centroids obtained from this test are also present in the Supplementary Table 3.

### 3.6. Genetic algorithms with *FP distance* (MaxD) scoring

The GA was modified by introducing another scoring system for the selection criteria. Instead of using the *peptide identity score*, the GA fitness was defined as the highest distance among all the FPs of the TCRs of the set that do not recognize the same pMHC. The GA objective was then to maximize this fitness. The principle of this fitness was to separate as much as possible the TCR that do not bind the same peptide, while others could remain close to each other. Unlike the *peptide identity score*, this new fitness did not require to match TCRs according to their specificity. As the results were not biased by the proportion of TCRs per antigen, a different training set was used. The latter was composed of 2831 TCR models that account for the specificity of 32 antigens. All the TCRs binding to the peptide KLGGALQAK were excluded from the pool of TCRs used by the GA to avoid an overrepresentation of this peptide in the results.

### 3.7. Comparison of TCRfp with a sequence-based approach

Most of the approaches analysing TCR repertoires and TCR specificities are based solely on their sequence. Contrarily to them, our approach encodes information regarding the biophysical and structural characteristics of the TCRs. To compare the performance of TCRfp with those of sequence-based approaches, we applied a sequence-based approach on the same validation set used for the validation of the TCRfp. BLOSUM matrices are usually used in sequence alignment algorithms to assess the similarity between sequence alignments. For this task, we calculated the BLOSUM62 score to assess the sequence similarity among the TCR sequences with an open gap penalty of −3 and an extension gap penalty of −1 for each TCR compared to the rest of TCRs of the validation set [35]. Of note, the scores given by this BLOSUM62 calculation can reach negative values. Hence, we normalized the BLOSUM score values so it would range from 0 to totally different TCR sequences to 1 for identical TCR sequences, like TCRfp. Using the *BLOSUM62 score*, it was possible to calculate *peptide identity scores* following the procedure defined in the section *2.4.2*, and compare with those obtained using TCRfp.

### 3.8. Logistic regression that combines the sequence-based score with TCRfp

We have implemented a logistic regression, based on the FP and on the *BLOSUM62 score*, to determine the probability of a TCR pair to bind the same peptide. The probability, *p*, of a TCR pair to recognize the same peptide was given by:

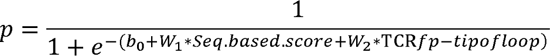

The bias term b0 and the weights Wn were determined using a set of 3213 TCRs (validation set, Supplementary Table 7), maximizing the likelihood that each TCR pair shares the same specificity. The accuracy of the logistic regression was determined by the area under the ROC curve (AUC) and by the % of correct pairs. The best model had a bias term b0 equals to −4.4545 while the weights W1 and W2 were 7.3739 and 0.5752, respectively. The AUC was only 0.66, emphasizing the difficulty to discriminate between TCRs that share the same specificity and TCRs that do not share the same specificity. The threshold of the classifier was set to 0.5, and we predicted TCRs sharing the same specificity if P>0.5. The regression trained on the 3213 TCRs was then applied to a private set of TCRs showing that the Logistic Regression is better than the BLOSUM score and or the FP alone.

## Supporting Information

Following supporting information available:

**Supplementary Table 1**: List of all the modeleld TCRs obtained from the 10xGenomics data base used in this project (10X id). The table contains information regarding all the genes and peptide specificity (pMHC = pHLA-A2) related to each TCR. The table also indicates if a 3D model was successfully obtained (tcr_model = Yes) and if that TCR was included within the TCRfp MATCH (match_set = Yes) or TCRfp MaxD (maxd_set = Yes) training sets.

**Supplementary Table 2**: Detailed GA parameters which led to the best genomes among all those tested for the TCRfp MATCH and TCRfp MaxD approaches. Main parameters: number of centroids and dimensions explored in each algorithm, initial size of the population and number of offspring genomes generated during each generation and the number of parents used from the main population to perform the next generation. Parameters of the initial generation: range of values for the C and P weighting parameters and X, Y, Z, c and p centroid coordinates, from within the initial genome parameters could be selected to generate a random initial population. Reproduction parameters: number of mutations that could be used when generating each child genome (the value is randomly chosen each time), the range value from which were selected the random value that would be subtracted or added to the parameter to be mutated depending if the parameter is the weighted C, weighted P, the X,Y,Z coordinates or the C and P values, the crossover type used for the algorithm (if a single centroid was swapped or multiple centroids) and the crossover point (if the entire centroid was swapped or a random portion of it).

**Supplementary Table 3:** Summary of the GA best algorithms. The table gives information regarding each algorithms’ approach used, if the initial generation was totally random or close to the tip of the loop (TOL), which training set was used, the maximum peptide identity score obtained corresponding to the best genome, the number of runs tested, and the mean final score obtained over all the runs. For each algorithm we provide the *peptide identity score* obtained for the top 5 centroids obtained by each algorithm along with the values for the C and P weighting parameters, and all the centroids’ coordinates.

**Supplementary Table 4:** TCR scores comparisons between the FP approach and the sequence-based approach. The table contains all the 10X ID (tcr_id) TCRs used in for the validation set, their peptide specificity (peptide), if the closest TCR obtained with the FP score (fp_MATCH = Yes) and the sequence-based score (BLOSUM_MATCH = Yes) are binding the same peptide, the respective antigens binding the closest TCRs (fp_peptide for the FP score peptide and BLOSUM_peptide for the sequence-based approach) and FP similarity (fp_score) and sequence similarity (BLOSUM_score) and the category according to the each approach MATCH (tag used for the colour visualization in the Figure 11).

**Supplementary Table 5:** Structural information of the PDB TCR class I structures used to calculate the preliminary results obtained from the Protein Data Bank (PDB). The columns contain the following information, from left to right: PDB ID, α chain, β chain and their cognate peptide (aligned among the rest of peptides).

**Supplementary Table 6:** List of all the IDs for the set of 92 experimental structures of TCR class I employed to calculate the distances for the distance-based filter.

**Supplementary Table 7:** List of all the TCRs modelled from the VDJ data base used as a validation set. The table contains the information regarding all the genes and peptide specificity (pMHC = pHLA-A2) related to each TCR.

**Supplementary** Figure 1: TCR ‘hierarchical’ trees built with ES5D FPs for the TCRfp MATCH approach applied to the 10 training sets of 117 TCRs. For each tree we can find the information regarding the peptide identity score obtained for the best algorithm, the pMHC-distance score, and the colour change value.

## Acknowledgements

Received: ((will be filled in by the editorial staff))

Revised: ((will be filled in by the editorial staff))

Published online: ((will be filled in by the editorial staff))

