## Supplementary Table 1 for "TCRfp: a new fingerprint-based approach for TCR repertoire analysis"

| tcr_id | TRAV | TRAJ | CDR3a | TRBV | TRBJ | CDR3b | Peptide | tcr_model | hs_set | maxd_set |
|---|---|---|---|---|---|---|---|---|---|---|
| 4 | TRAV30 | TRAJ20 | CGTEYPNDYKLSF | TRBV10-3 | TRBJ1-1 | CAIMIGTEAFF | GLCTLVAML | YES | YES | YES |
| 9 | TRAV1-2 | TRAJ20 | CAVREEDYKLSF | TRBV12-3 | TRBJ1-1 | CASSFRGPNLYTEAFF | KLQCVDLHV | YES | NO | YES |
| 14 | TRAV12-2 | TRAJ6 | CAASSGGSYIPTF | TRBV14 | TRBJ1-1 | CASSQDTSGSGLWTEAFF | ELAGIGILTV | NO | NO | NO |
| 15 | TRAV12-2 | TRAJ10 | CAVNPGGNKLTF | TRBV15 | TRBJ1-1 | CATSGGMGAEAFF | ELAGIGILTV | YES | NO | YES |
| 16 | TRAV1-2 | TRAJ6 | CAVRDKPPGPGGGSYIPTF | TRBV19 | TRBJ1-1 | CASSIHGAGTEAFF | GILGFVFTL | YES | NO | YES |
| 18 | TRAV12-2 | TRAJ42 | CAGDSGGGSQGNLIF | TRBV19 | TRBJ1-1 | CASSIRSNTEAFF | GILGFVFTL | YES | NO | YES |
| 25 | TRAV13-2 | TRAJ42 | CAENMGGGSQGNLIF | TRBV19 | TRBJ1-1 | CASSKRSGAEAFF | GILGFVFTL | YES | NO | YES |
| 30 | TRAV25 | TRAJ37 | CADPFSSNTGKLIF | TRBV19 | TRBJ1-1 | CASSTFSSGRTEAFF | GILGFVFTL | YES | NO | NO |
| 32 | TRAV27 | TRAJ42 | CAGGGSQGNLIF | TRBV19 | TRBJ1-1 | CASSIMALGRSEAFF | GILGFVFTL | YES | NO | YES |
| 33 | TRAV29DV5 | TRAJ39 | CAASGSSNAGNMLTF | TRBV19 | TRBJ1-1 | CASRAGREAFF | GILGFVFTL | YES | NO | YES |
| 34 | TRAV29DV5 | TRAJ53 | CAAISVGGSNYKLTF | TRBV19 | TRBJ1-1 | CASSSGLFSF | GILGFVFTL | YES | NO | YES |
| 35 | TRAV29DV5 | TRAJ53 | CAAQDSGGSNYKLTF | TRBV19 | TRBJ1-1 | CASSTYMTNTEAFF | LLDFVRFMGV | YES | NO | YES |
| 36 | TRAV35 | TRAJ31 | CAGHWDLGNNARLMF | TRBV19 | TRBJ1-1 | CASSVRHGTEAFF | GILGFVFTL | YES | YES | YES |
| 37 | TRAV35 | TRAJ42 | CAGAGDGGSQGNLIF | TRBV19 | TRBJ1-1 | CASSIRSTTEAFF | GILGFVFTL | YES | YES | YES |
| 39 | TRAV38-1 | TRAJ52 | CAFMKGAGGTSYGKLTF | TRBV19 | TRBJ1-1 | CASSPTSNTEAFF | GILGFVFTL | YES | YES | YES |
| 45 | TRAV8-1 | TRAJ52 | CAVMGAGGTSYGKLTF | TRBV19 | TRBJ1-1 | CASSIGSGEAFF | LLFGYPVYV | YES | YES | YES |
| 46 | TRAV8-4 | TRAJ20 | CAVSPNDYKLSF | TRBV19 | TRBJ1-1 | CASSIRSTTEAFF | GILGFVFTL | YES | YES | YES |
| 47 | TRAV8-4 | TRAJ37 | CAVSESNTGKLIF | TRBV19 | TRBJ1-1 | CASSIRAGYTEAFF | GILGFVFTL | YES | YES | YES |
| 51 | TRAV13-1 | TRAJ50 | CAAETSYDKVIF | TRBV24-1 | TRBJ1-1 | CATTYGDRRNTEAFF | GILGFVFTL | YES | NO | YES |
| 52 | TRAV12-2 | TRAJ13 | CAVFSGGYQKVTF | TRBV27 | TRBJ1-1 | CASSLGINTEAFF | ELAGIGILTV | YES | NO | YES |
| 53 | TRAV12-2 | TRAJ42 | CAVRIGSQGNLIF | TRBV27 | TRBJ1-1 | CASSLLQGTEAFF | IMDQVPFSV | YES | NO | YES |
| 54 | TRAV12-2 | TRAJ43 | CAVNNNNDMRF | TRBV27 | TRBJ1-1 | CASSFQGLGTEAFF | ELAGIGILTV | YES | YES | YES |
| 55 | TRAV12-2 | TRAJ52 | CAVNVGGTSYGKLTF | TRBV27 | TRBJ1-1 | CASSLSLGGTEAFF | ELAGIGILTV | YES | NO | YES |
| 56 | TRAV12-2 | TRAJ10 | CAVNKITGGGNKLTF | TRBV28 | TRBJ1-1 | CASKIQGSPEAFF | ELAGIGILTV | YES | NO | YES |
| 57 | TRAV12-2 | TRAJ18 | CAVSGRGSTLGRLYF | TRBV28 | TRBJ1-1 | CASSWGQMNTEAFF | ELAGIGILTV | YES | NO | YES |
| 58 | TRAV12-2 | TRAJ32 | CAVNGGGATNKLIF | TRBV28 | TRBJ1-1 | CAIPGPSNTEAFF | ELAGIGILTV | YES | NO | NO |
| 59 | TRAV21 | TRAJ39 | CAACNNAGNMLTF | TRBV29-1 | TRBJ1-1 | CSAGGNTEAFF | LLDFVRFMGV | YES | NO | YES |
| 60 | TRAV20 | TRAJ32 | CAVQAEGSYGGATNKLIF | TRBV30 | TRBJ1-1 | CAWSFLDVTAGTEAFF | GILGFVFTL | YES | NO | NO |
| 61 | TRAV12-2 | TRAJ35 | CAVGLGFGNVLHC | TRBV4-1 | TRBJ1-1 | CASSWEGNTEAFF | ELAGIGILTV | NO | NO | NO |
| 63 | TRAV12-2 | TRAJ45 | CAVGGGGADGLTF | TRBV4-2 | TRBJ1-1 | CASSQAQLNTEAFF | ELAGIGILTV | YES | YES | YES |
| 67 | TRAV8-6 | TRAJ28 | CAAISGAGSYQLTF | TRBV5-1 | TRBJ1-1 | CASSYGTGENTEAFF | RAKFKQLL | YES | NO | YES |
| 73 | TRAV13-2 | TRAJ45 | CADSGGGADGLTF | TRBV5-6 | TRBJ1-1 | CASSLRDGSEAFF | AVFDRKSDAK | YES | NO | YES |
| 75 | TRAV27 | TRAJ42 | CAGGGSQGNLIF | TRBV5-6 | TRBJ1-1 | CASSLRDGSEAFF | GILGFVFTL | YES | NO | YES |
| 78 | TRAV38-1 | TRAJ52 | CAFMKGAGGTSYGKLTF | TRBV5-6 | TRBJ1-1 | CASSLRDGSEAFF | RAKFKQLL | YES | NO | YES |
| 78 | TRAV38-1 | TRAJ52 | CAFMKGAGGTSYGKLTF | TRBV5-6 | TRBJ1-1 | CASSLRDGSEAFF | GILGFVFTL | YES | NO | YES |
| 84 | TRAV8-3 | TRAJ54 | CAAIKGAQKLVF | TRBV6-5 | TRBJ1-1 | CASSLNTEAFF | GILGFVFTL | YES | NO | YES |
| 84 | TRAV8-3 | TRAJ54 | CAAIKGAQKLVF | TRBV6-5 | TRBJ1-1 | CASSLNTEAFF | KLGGALQAK | YES | NO | YES |
| 85 | TRAV12-2 | TRAJ5 | CAVITGGRRALTF | TRBV6-6 | TRBJ1-1 | CASRLPGQGNTEAFF | SLFNTVATLY | NO | NO | NO |
| 90 | TRAV27 | TRAJ37 | CAGASNTGKLIF | TRBV7-9 | TRBJ1-1 | CASSLGDNTEAFF | GILGFVFTL | YES | NO | NO |
| 92 | TRAV38-1 | TRAJ52 | CAFMKGAGGTSYGKLTF | TRBV7-9 | TRBJ1-1 | CASSSLGGHGEAFF | GILGFVFTL | YES | NO | YES |
| 94 | TRAV12-2 | TRAJ20 | CAVKTHKLSF | TRBV9 | TRBJ1-1 | CASSDGGGGSEAFF | ELAGIGILTV | YES | NO | NO |
| 96 | TRAV26-1 | TRAJ17 | CILWVAAGNKLTF | TRBV9 | TRBJ1-1 | CASSVRSSMNTEAFF | AVFDRKSDAK | YES | NO | YES |
| 96 | TRAV26-1 | TRAJ17 | CILWVAAGNKLTF | TRBV9 | TRBJ1-1 | CASSVRSSMNTEAFF | IVTDFSVIK | YES | NO | YES |
| 98 | TRAV38-1 | TRAJ52 | CAFMKGAGGTSYGKLTF | TRBV9 | TRBJ1-1 | CASSVRSSMNTEAFF | GILGFVFTL | YES | NO | YES |
| 103 | TRAV27 | TRAJ42 | CAGGGSQGNLIF | TRBV11-2 | TRBJ1-2 | CASSWGGGSHYGYTF | GILGFVFTL | YES | NO | YES |
| 104 | TRAV38-2DV8 | TRAJ52 | CAWSSNAGGTSYGKLTF | TRBV11-2 | TRBJ1-2 | CASSWGGGSHYGYTF | GILGFVFTL | YES | NO | YES |
| 105 | TRAV38-2DV8 | TRAJ52 | CAYSPAAGGTSYGKLTF | TRBV11-2 | TRBJ1-2 | CASSWGGGSHYGYTF | GILGFVFTL | YES | NO | YES |
| 107 | TRAV27 | TRAJ42 | CAGGGSQGNLIF | TRBV12-3 | TRBJ1-2 | CASSLGDRAGGYTF | GILGFVFTL | YES | NO | YES |
| 112 | TRAV12-3 | TRAJ52 | CAMSVNAGGTSYGKLTF | TRBV19 | TRBJ1-2 | CASSIGLYGYTF | GILGFVFTL | YES | NO | YES |
| 118 | TRAV14DV4 | TRAJ18 | CAMSRGIDRGSTLGRLYF | TRBV19 | TRBJ1-2 | CASSIGVHGYTF | GILGFVFTL | YES | NO | YES |
| 119 | TRAV14DV4 | TRAJ49 | CAMRSTGNQFYF | TRBV19 | TRBJ1-2 | CASSIGVHGYTF | GILGFVFTL | YES | NO | YES |
| 121 | TRAV19 | TRAJ23 | CALSEAGFYNQGGKLIF | TRBV19 | TRBJ1-2 | CASSIYGGAGDGYTF | GILGFVFTL | YES | NO | YES |
| 123 | TRAV19 | TRAJ52 | CALMGNAGGTSYGKLTF | TRBV19 | TRBJ1-2 | CASSQGNYGYTF | GILGFVFTL | YES | NO | YES |
| 126 | TRAV21 | TRAJ52 | CAVRLEGPGGTSYGKLTF | TRBV19 | TRBJ1-2 | CASDTGGGYGYTF | GILGFVFTL | YES | NO | YES |
| 129 | TRAV27 | TRAJ42 | CAGGGSQGNLIF | TRBV19 | TRBJ1-2 | CASSIGSYGYTF | GILGFVFTL | YES | NO | YES |
| 131 | TRAV27 | TRAJ42 | CAGAGSQGNLIF | TRBV19 | TRBJ1-2 | CAGRGTDDYGYTF | GILGFVFTL | YES | NO | NO |
| 134 | TRAV29DV5 | TRAJ52 | CAASPNAGGTSYGKLTF | TRBV19 | TRBJ1-2 | CASSIGLYGYTF | GILGFVFTL | YES | NO | YES |
| 136 | TRAV36DV7 | TRAJ47 | CAVADMEYGNKLVF | TRBV19 | TRBJ1-2 | CASSTGTLYGYTF | GILGFVFTL | NO | NO | NO |
| 141 | TRAV38-1 | TRAJ52 | CAFMINAGGTSYGKLTF | TRBV19 | TRBJ1-2 | CASSIGYYGYTF | GILGFVFTL | YES | NO | YES |
| 142 | TRAV38-1 | TRAJ52 | CAFMTNAGGTSYGKLTF | TRBV19 | TRBJ1-2 | CASSQGAYGYTF | GILGFVFTL | YES | NO | YES |
| 143 | TRAV38-1 | TRAJ52 | CAFMTNAGGTSYGKLTF | TRBV19 | TRBJ1-2 | CASSIGVYGYTF | GILGFVFTL | YES | NO | YES |
| 144 | TRAV38-1 | TRAJ52 | CAFMIGAGGTSYGKLTF | TRBV19 | TRBJ1-2 | CASSIGSYGYTF | GILGFVFTL | YES | NO | YES |
| 145 | TRAV38-1 | TRAJ52 | CAFMINAGGTSYGKLTF | TRBV19 | TRBJ1-2 | CASSIGSYGYTF | GILGFVFTL | YES | NO | YES |
| 146 | TRAV38-1 | TRAJ52 | CAFMSNAGGTSYGKLTF | TRBV19 | TRBJ1-2 | CASSMGSYGYTF | GILGFVFTL | YES | NO | YES |
| 147 | TRAV38-1 | TRAJ52 | CAFMKDAGGTSYGKLTF | TRBV19 | TRBJ1-2 | CASSIGVYGYTF | GILGFVFTL | YES | NO | YES |
| 148 | TRAV38-1 | TRAJ52 | CAFMLDAGGTSYGKLTF | TRBV19 | TRBJ1-2 | CASSIGYYGYTF | GILGFVFTL | YES | NO | YES |
| 149 | TRAV38-1 | TRAJ52 | CAFMQNAGGTSYGKLTF | TRBV19 | TRBJ1-2 | CASSMGAYGYTF | GILGFVFTL | YES | NO | YES |
| 150 | TRAV38-1 | TRAJ52 | CAFMVGAGGTSYGKLTF | TRBV19 | TRBJ1-2 | CASSTGVYGYTF | GILGFVFTL | YES | NO | NO |
| 151 | TRAV38-1 | TRAJ52 | CAFMIGSGGTSYGKLTF | TRBV19 | TRBJ1-2 | CASSIGAYGYTF | GILGFVFTL | YES | NO | YES |
| 153 | TRAV38-1 | TRAJ52 | CAFMIGAGGTSYGKLTF | TRBV19 | TRBJ1-2 | CASSMGSYGYTF | GILGFVFTL | YES | NO | YES |
| 154 | TRAV38-1 | TRAJ52 | CAFMTNAGGTSYGKLTF | TRBV19 | TRBJ1-2 | CASSAGAYGYTF | GILGFVFTL | YES | NO | YES |
| 155 | TRAV38-1 | TRAJ52 | CAFMVNAGGTSYGKLTF | TRBV19 | TRBJ1-2 | CASSIGVYGYTF | GILGFVFTL | YES | NO | YES |
| 156 | TRAV38-1 | TRAJ52 | CAFMIGAGGTSYGKLTF | TRBV19 | TRBJ1-2 | CASSVGSYGYTF | GILGFVFTL | YES | NO | YES |
| 157 | TRAV38-1 | TRAJ52 | CAFMTNAGGTSYGKLTF | TRBV19 | TRBJ1-2 | CATSQGAYGYTF | GILGFVFTL | YES | NO | YES |
| 158 | TRAV38-1 | TRAJ52 | CAFMKGAGGTSYGKLTF | TRBV19 | TRBJ1-2 | CASSTGAYGYTF | GILGFVFTL | YES | NO | YES |
| 159 | TRAV38-1 | TRAJ52 | CAFMTNAGGTSYGKLTF | TRBV19 | TRBJ1-2 | CASSIGIYGYTF | GILGFVFTL | YES | NO | YES |
| 160 | TRAV38-1 | TRAJ52 | CAFMINAGGTSYGKLTF | TRBV19 | TRBJ1-2 | CASSIGAYGYTF | GILGFVFTL | YES | NO | NO |
| 161 | TRAV38-1 | TRAJ52 | CAFMVNAGGTSYGKLTF | TRBV19 | TRBJ1-2 | CASSIGSYGYTF | GILGFVFTL | YES | YES | YES |
| 162 | TRAV38-1 | TRAJ52 | CAFMTNAGGTSYGKLTF | TRBV19 | TRBJ1-2 | CASSQGSYGYTF | GILGFVFTL | YES | NO | YES |
| 169 | TRAV38-1 | TRAJ52 | CAFMIGAGGTSYGKLTF | TRBV19 | TRBJ1-2 | CASSIGYYGYTF | GILGFVFTL | YES | NO | YES |
| 170 | TRAV38-1 | TRAJ52 | CAFMIGAGGTSYGKLTF | TRBV19 | TRBJ1-2 | CASSIGVYGYTF | GILGFVFTL | YES | NO | NO |
| 171 | TRAV38-1 | TRAJ52 | CAFMVNAGGTSYGKLTF | TRBV19 | TRBJ1-2 | CASSIGAHGYTF | GILGFVFTL | YES | NO | YES |
| 172 | TRAV38-1 | TRAJ52 | CAFMKGAGGTSYGKLTF | TRBV19 | TRBJ1-2 | CASSVGSYGYTF | GILGFVFTL | YES | NO | YES |
| 173 | TRAV38-1 | TRAJ52 | CAFMKGAGGTSYGKLTF | TRBV19 | TRBJ1-2 | CASSTGSYGYTF | GILGFVFTL | YES | NO | YES |
| 174 | TRAV38-1 | TRAJ52 | CAFMRDAGGTSYGKLTF | TRBV19 | TRBJ1-2 | CASSIGVHGYTF | GILGFVFTL | YES | NO | YES |
| 175 | TRAV38-1 | TRAJ52 | CAFMVGAGGTSYGKLTF | TRBV19 | TRBJ1-2 | CASSTGIYGYTF | GILGFVFTL | YES | YES | YES |
| 176 | TRAV38-1 | TRAJ52 | CAFMNNAGGTSYGKLTF | TRBV19 | TRBJ1-2 | CASSIGVYGYTF | GILGFVFTL | YES | NO | YES |
| 177 | TRAV38-1 | TRAJ52 | CAFMTSPGGTSYGKLTF | TRBV19 | TRBJ1-2 | CASSIGVHGYTF | GILGFVFTL | YES | NO | YES |
| 178 | TRAV38-1 | TRAJ52 | CAFMVNAGGTSYGKLTF | TRBV19 | TRBJ1-2 | CASSIGVAGYTF | GILGFVFTL | YES | YES | YES |
| 179 | TRAV38-1 | TRAJ52 | CAFMINAGGTSYGKLTF | TRBV19 | TRBJ1-2 | CASSIGAWGYTF | GILGFVFTL | YES | NO | YES |
| 180 | TRAV38-1 | TRAJ52 | CAFMIGAGGTSYGKLTF | TRBV19 | TRBJ1-2 | CASSFGVHGYTF | GILGFVFTL | YES | NO | NO |
| 181 | TRAV38-1 | TRAJ52 | CAFMIGSGGTSYGKLTF | TRBV19 | TRBJ1-2 | CASSIGYYGYTF | GILGFVFTL | YES | YES | YES |
| 182 | TRAV38-1 | TRAJ52 | CAFMIGAGGTSYGKLTF | TRBV19 | TRBJ1-2 | CASSIGVWGYTF | GILGFVFTL | YES | NO | YES |
| 183 | TRAV38-1 | TRAJ52 | CAFMKGAGGTSYGKLTF | TRBV19 | TRBJ1-2 | CASSIGAYGYTF | GILGFVFTL | YES | NO | YES |
| 184 | TRAV38-1 | TRAJ52 | CAFMTGAGGTSYGKLTF | TRBV19 | TRBJ1-2 | CASSIGVYGYTF | GILGFVFTL | YES | NO | YES |
| 185 | TRAV38-1 | TRAJ52 | CAFMVDAGGTSYGKLTF | TRBV19 | TRBJ1-2 | CASSTGAYGYTF | GILGFVFTL | YES | NO | YES |
| 186 | TRAV38-2DV8 | TRAJ48 | CPSDYEKLTF | TRBV19 | TRBJ1-2 | CASSIGVHGYTF | GILGFVFTL | YES | NO | NO |
| 187 | TRAV38-2DV8 | TRAJ48 | CAYRTSILSNFGNEKLTF | TRBV19 | TRBJ1-2 | CASSQGSYGYTF | GILGFVFTL | YES | YES | YES |
| 189 | TRAV38-2DV8 | TRAJ52 | CAYSLSAGGTSYGKLTF | TRBV19 | TRBJ1-2 | CASSIGVYGYTF | GILGFVFTL | YES | NO | YES |
| 190 | TRAV38-2DV8 | TRAJ52 | CAYATNAGGTSYGKLTF | TRBV19 | TRBJ1-2 | CASSIGLYGYTF | GILGFVFTL | YES | NO | NO |
| 191 | TRAV38-2DV8 | TRAJ52 | CAYSHNAGGTSYGKLTF | TRBV19 | TRBJ1-2 | CASSIGVYGYTF | GILGFVFTL | YES | NO | YES |
| 192 | TRAV38-2DV8 | TRAJ52 | CAWSSNAGGTSYGKLTF | TRBV19 | TRBJ1-2 | CASSIGLYGYTF | GILGFVFTL | YES | NO | YES |
| 193 | TRAV38-2DV8 | TRAJ52 | CAYTPNAGGTSYGKLTF | TRBV19 | TRBJ1-2 | CASSIGIYGYTF | GILGFVFTL | YES | NO | YES |

| | | | | | | | | | | |
|---|---|---|---|---|---|---|---|---|---|---|
| 195 | TRAV38-2DV8 | TRAJ52 | CAYSVNAGGTSYGKLTF | TRBV19 | TRBJ1-2 | CASSIGLYGYTF | GILGFVFTL | YES | NO | YES |
| 196 | TRAV38-2DV8 | TRAJ52 | CAYMPNAGGTSYGKLTF | TRBV19 | TRBJ1-2 | CASSQGAYGYTF | GILGFVFTL | YES | NO | YES |
| 197 | TRAV38-2DV8 | TRAJ52 | CAYMTNAGGTSYGKLTF | TRBV19 | TRBJ1-2 | CASSEGSYGYTF | GILGFVFTL | YES | NO | YES |
| 198 | TRAV38-2DV8 | TRAJ52 | CAYMANAGGTSYGKLTF | TRBV19 | TRBJ1-2 | CASSIGVYGYTF | GILGFVFTL | YES | NO | YES |
| 199 | TRAV38-2DV8 | TRAJ52 | CAYSPAAGGTSYGKLTF | TRBV19 | TRBJ1-2 | CASSQGAYGYTF | GILGFVFTL | YES | NO | YES |
| 200 | TRAV38-2DV8 | TRAJ52 | CAYSYGAGGTSYGKLTF | TRBV19 | TRBJ1-2 | CASSIGLYGYTF | GILGFVFTL | YES | NO | NO |
| 201 | TRAV38-2DV8 | TRAJ52 | CAYMSNAGGTSYGKLTF | TRBV19 | TRBJ1-2 | CASSMGSYGYTF | GILGFVFTL | YES | NO | YES |
| 202 | TRAV38-2DV8 | TRAJ52 | CAYSANAGGTSYGKLTF | TRBV19 | TRBJ1-2 | CASSIGIYGYTF | GILGFVFTL | YES | NO | YES |
| 203 | TRAV38-2DV8 | TRAJ52 | CAYMVNAGGTSYGKLTF | TRBV19 | TRBJ1-2 | CASSLGSYGYTF | GILGFVFTL | YES | NO | YES |
| 204 | TRAV38-2DV8 | TRAJ58 | CAYRTKETSGSRLTF | TRBV19 | TRBJ1-2 | CASSIGSYGYTF | GILGFVFTL | YES | NO | YES |
| 207 | TRAV9-2 | TRAJ54 | CAPRALSGAQKLVF | TRBV19 | TRBJ1-2 | CASSTYSTGSGYTF | GILGFVFTL | YES | NO | YES |
| 213 | TRAV5 | TRAJ3 | CAEDSASKIIF | TRBV20-1 | TRBJ1-2 | CSARDETGNGYTF | GLCTLVAML | YES | YES | YES |
| 214 | TRAV5 | TRAJ31 | CAEDNNARLMF | TRBV20-1 | TRBJ1-2 | CSARDRTGNGYTF | GLCTLVAML | YES | YES | YES |
| 217 | TRAV12-2 | TRAJ22 | CAVNEGLGSARQLTF | TRBV24-1 | TRBJ1-2 | CATSDSSGGSGYTF | ELAGIGILTV | YES | NO | YES |
| 219 | TRAV12-2 | TRAJ43 | CAVNNDMRF | TRBV27 | TRBJ1-2 | CASSSPGTGAYGYTF | ELAGIGILTV | NO | NO | NO |
| 222 | TRAV27 | TRAJ47 | CAGGEYGNKLVF | TRBV27 | TRBJ1-2 | CASSPGRVYGYTF | GILGFVFTL | YES | NO | YES |
| 223 | TRAV12-3 | TRAJ39 | CAIGPGNMLTF | TRBV29-1 | TRBJ1-2 | CSVDLEANYGYTF | IVTDFSVIK | YES | NO | NO |
| 223 | TRAV12-3 | TRAJ39 | CAIGPGNMLTF | TRBV29-1 | TRBJ1-2 | CSVDLEANYGYTF | RTLNAWVKV | YES | NO | NO |
| 223 | TRAV12-3 | TRAJ39 | CAIGPGNMLTF | TRBV29-1 | TRBJ1-2 | CSVDLEANYGYTF | CLLGTYTQDV | YES | NO | NO |
| 223 | TRAV12-3 | TRAJ39 | CAIGPGNMLTF | TRBV29-1 | TRBJ1-2 | CSVDLEANYGYTF | FLASKIGRLV | YES | NO | NO |
| 224 | TRAV21 | TRAJ24 | CAVASTDSWGKLQF | TRBV29-1 | TRBJ1-2 | CSVSQSYGYTF | RTLNAWVKV | YES | YES | YES |
| 230 | TRAV17 | TRAJ24 | CATDSPTDSWGKLQF | TRBV4-2 | TRBJ1-2 | CASSQEQWSGETSNYGYTF | RTLNAWVKV | YES | YES | NO |
| 232 | TRAV12-2 | TRAJ11 | CAKGGSGYSTLTF | TRBV4-3 | TRBJ1-2 | CASSQEPGGAGGYTF | ELAGIGILTV | YES | NO | YES |
| 233 | TRAV12-2 | TRAJ13 | CAVGNSGGYQKVTF | TRBV4-3 | TRBJ1-2 | CASSQDHNRGQGYTF | ELAGIGILTV | YES | NO | YES |
| 234 | TRAV12-2 | TRAJ35 | CAVNRGFGNVLHC | TRBV5-1 | TRBJ1-2 | CASSLGTGDGYTF | ELAGIGILTV | NO | NO | NO |
| 237 | TRAV27 | TRAJ42 | CAGGGSQGNLIF | TRBV5-4 | TRBJ1-2 | CATTPGGSLSGYTF | GILGFVFTL | YES | NO | NO |
| 238 | TRAV27 | TRAJ42 | CAGEGGGSQGNLIF | TRBV5-4 | TRBJ1-2 | CASSLGTANYGYTF | GILGFVFTL | YES | NO | YES |
| 239 | TRAV38-2DV8 | TRAJ52 | CAFMINAGGTSYGKLTF | TRBV5-4 | TRBJ1-2 | CASSLGTANYGYTF | GILGFVFTL | YES | NO | NO |
| 241 | TRAV12-2 | TRAJ39 | CAVNAGNMLTF | TRBV6-1 | TRBJ1-2 | CASSDPIGGYGYTF | ELAGIGILTV | YES | NO | YES |
| 242 | TRAV12-2 | TRAJ41 | CAVNGGSNSGYALNF | TRBV6-3 | TRBJ1-2 | CASSYGTGNGYTF | ELAGIGILTV | YES | YES | YES |
| 243 | TRAV12-2 | TRAJ52 | CAGNAGGTSYGKLTF | TRBV6-3 | TRBJ1-2 | CASSWGQGSAYTF | ELAGIGILTV | YES | YES | YES |
| 244 | TRAV12-2 | TRAJ52 | CAVNLGLTAGGTSYGKLTF | TRBV6-4 | TRBJ1-2 | CASRAGTEISGYGYTF | ELAGIGILTV | YES | NO | YES |
| 247 | TRAV12-2 | TRAJ13 | CAPSNSGGYQKVTF | TRBV6-5 | TRBJ1-2 | CASSYGSGDGYTF | ELAGIGILTV | YES | NO | YES |
| 248 | TRAV13-2 | TRAJ11 | CAEYSGYSTLTF | TRBV6-5 | TRBJ1-2 | CASSSQGGFDGYTF | FLYALALLL | YES | YES | YES |
| 249 | TRAV17 | TRAJ11 | CATEGDSGYSTLTF | TRBV6-5 | TRBJ1-2 | CASSLQGGNYGYTF | FLYALALLL | YES | YES | YES |
| 250 | TRAV17 | TRAJ11 | CATEGNSGYSTLTF | TRBV6-5 | TRBJ1-2 | CASSPQGGGDGYTF | FLYALALLL | YES | YES | NO |
| 251 | TRAV17 | TRAJ11 | CATEGNSGYSTLTF | TRBV6-5 | TRBJ1-2 | CASTEQGGFHGYTF | FLYALALLL | YES | YES | YES |
| 252 | TRAV17 | TRAJ11 | CATEGNSGYSTLTF | TRBV6-5 | TRBJ1-2 | CASSRQGGNYGYTF | FLYALALLL | YES | YES | YES |
| 253 | TRAV17 | TRAJ11 | CATNGDSGYSTLTF | TRBV6-5 | TRBJ1-2 | CASSSQGGFDGYTF | FLYALALLL | YES | YES | YES |
| 254 | TRAV17 | TRAJ11 | CATEGDSGYSTLTF | TRBV6-5 | TRBJ1-2 | CASSYQGGNYGYTF | FLYALALLL | YES | YES | YES |
| 255 | TRAV17 | TRAJ11 | CATEGDSGYSTLTF | TRBV6-5 | TRBJ1-2 | CASSSQGGSYGYTF | FLYALALLL | YES | YES | YES |
| 256 | TRAV17 | TRAJ11 | CATEGDSGYSTLTF | TRBV6-5 | TRBJ1-2 | CASSSQGGGSGYTF | FLYALALLL | YES | YES | YES |
| 257 | TRAV17 | TRAJ11 | CATEGNSGYSTLTF | TRBV6-5 | TRBJ1-2 | CASSPLGGYYGYTF | FLYALALLL | YES | YES | YES |
| 262 | TRAV38-2DV8 | TRAJ52 | CAYSPAAGGTSYGKLTF | TRBV7-6 | TRBJ1-2 | CASSLAQGGALRGYTF | GILGFVFTL | YES | NO | YES |
| 263 | TRAV8-4 | TRAJ20 | CAVIQINDYKLSF | TRBV7-8 | TRBJ1-2 | CASSLGQDGYTF | IVTDFSVIK | YES | NO | NO |
| 263 | TRAV8-4 | TRAJ20 | CAVIQINDYKLSF | TRBV7-8 | TRBJ1-2 | CASSLGQDGYTF | SLFNTVATLY | YES | NO | NO |
| 263 | TRAV8-4 | TRAJ20 | CAVIQINDYKLSF | TRBV7-8 | TRBJ1-2 | CASSLGQDGYTF | FLASKIGRLV | YES | NO | NO |
| 265 | TRAV12-2 | TRAJ45 | CAAYSGGGADGLTF | TRBV9 | TRBJ1-2 | CASSPQTEYGYTF | ELAGIGILTV | YES | NO | YES |
| 266 | TRAV12-2 | TRAJ43 | CAVNSDMRF | TRBV12-3 | TRBJ1-3 | CASSSTGLGNTIYF | ELAGIGILTV | NO | NO | NO |
| 268 | TRAV26-1 | TRAJ33 | CIVRVEDSNYQLIW | TRBV16 | TRBJ1-3 | CASSQLFNLGQGFPGNTIYF | KTWGQYWQV | NO | NO | NO |
| 269 | TRAV12-3 | TRAJ42 | CAIGGDGGSQGNLIF | TRBV19 | TRBJ1-3 | CASSPRSGNTIYF | GILGFVFTL | YES | NO | YES |
| 270 | TRAV13-2 | TRAJ42 | CAENMGGGSQGNLIF | TRBV19 | TRBJ1-3 | CASSDRSGNTIYF | GILGFVFTL | YES | NO | NO |
| 271 | TRAV19 | TRAJ39 | CALSEAMNNAGNMLTF | TRBV19 | TRBJ1-3 | CASSIGHYSGNTIYF | GILGFVFTL | YES | NO | YES |
| 272 | TRAV20 | TRAJ30 | CVPVTLGVNRDDKIIF | TRBV19 | TRBJ1-3 | CASSQFFGNTIYF | GILGFVFTL | YES | NO | NO |
| 273 | TRAV38-2DV8 | TRAJ33 | CAYRSLPMDSNYQLIW | TRBV19 | TRBJ1-3 | CASSIYSGPGNTIYF | GILGFVFTL | YES | NO | YES |
| 274 | TRAV12-2 | TRAJ7 | CAVGGNNRLAF | TRBV2 | TRBJ1-3 | CASSEDFSGNTIYF | ELAGIGILTV | YES | NO | YES |
| 275 | TRAV12-3 | TRAJ27 | CAMKSTNAGKSTF | TRBV20-1 | TRBJ1-3 | CSARDRVGNTIYF | GLCTLVAML | YES | YES | YES |
| 279 | TRAV5 | TRAJ31 | CAEDNNARLMF | TRBV20-1 | TRBJ1-3 | CSARVGPTGNTIYF | GLCTLVAML | NO | NO | NO |
| 281 | TRAV5 | TRAJ31 | CAEDFNARLMF | TRBV20-1 | TRBJ1-3 | CSARIGVGNTIYF | GLCTLVAML | NO | NO | NO |
| 282 | TRAV5 | TRAJ31 | CAEDNNARLMF | TRBV20-1 | TRBJ1-3 | CSARSETGNTIYF | GLCTLVAML | YES | YES | YES |
| 285 | TRAV12-2 | TRAJ52 | CAVNLAGGTSYGKLTF | TRBV27 | TRBJ1-3 | CASSFAPGNTIYF | ELAGIGILTV | YES | NO | YES |
| 286 | TRAV12-2 | TRAJ28 | CAVNTYSGAGSYQLTF | TRBV28 | TRBJ1-3 | CASSSQGVGNTIYF | ELAGIGILTV | YES | NO | YES |
| 287 | TRAV12-2 | TRAJ54 | CAVGIIQGAQKLVF | TRBV28 | TRBJ1-3 | CASSLQGIGNTIYF | ELAGIGILTV | YES | YES | YES |
| 288 | TRAV12-2 | TRAJ8 | CAVNILGNTGFQKLVF | TRBV28 | TRBJ1-3 | CASSPMGVGGNTIYF | ELAGIGILTV | YES | YES | YES |
| 289 | TRAV12-2 | TRAJ43 | CAVNRGDMRF | TRBV3-1 | TRBJ1-3 | CASSDSGNTIYF | ELAGIGILTV | YES | NO | YES |
| 290 | TRAV12-2 | TRAJ35 | CAVGGFGNVLHC | TRBV4-1 | TRBJ1-3 | CASSQTRGAGNTIYF | ELAGIGILTV | NO | NO | NO |
| 293 | TRAV12-2 | TRAJ43 | CAVKGDMRF | TRBV6-3 | TRBJ1-3 | CASSYGEISGNTIYF | ELAGIGILTV | NO | NO | NO |
| 294 | TRAV12-2 | TRAJ52 | CAVPAGGTSYGKLTF | TRBV6-3 | TRBJ1-3 | CASSPGSGNTIYF | ELAGIGILTV | YES | NO | YES |
| 298 | TRAV38-2DV8 | TRAJ52 | CAYSHNAGGTSYGKLTF | TRBV6-5 | TRBJ1-3 | CASSYSTSGNTIYF | GILGFVFTL | YES | YES | YES |
| 299 | TRAV20 | TRAJ26 | CAVPDNYGQNFVF | TRBV6-6 | TRBJ1-3 | CASTTGVNTIYF | KLGGALQAK | YES | NO | NO |
| 299 | TRAV20 | TRAJ26 | CAVPDNYGQNFVF | TRBV6-6 | TRBJ1-3 | CASTTGVNTIYF | GILGFVFTL | YES | NO | NO |
| 308 | TRAV13-2 | TRAJ42 | CAENTGGGSQGNLIF | TRBV19 | TRBJ1-4 | CASSFRSSEKLFF | GILGFVFTL | YES | YES | YES |
| 309 | TRAV26-1 | TRAJ34 | CIVRVANTDKLIF | TRBV19 | TRBJ1-4 | CASSSFSANEKLFF | GILGFVFTL | YES | NO | YES |
| 311 | TRAV27 | TRAJ42 | CAGGGSQGNLIF | TRBV19 | TRBJ1-4 | CASSLQGANEKLFF | GILGFVFTL | YES | NO | YES |
| 313 | TRAV3 | TRAJ36 | CAVRDGTGANNLFF | TRBV19 | TRBJ1-4 | CASAGTGVEKLFF | GILGFVFTL | NO | NO | NO |
| 314 | TRAV35 | TRAJ42 | CAGGNYGGSQGNLIF | TRBV19 | TRBJ1-4 | CASSIRSDEKLFF | GILGFVFTL | YES | NO | YES |
| 315 | TRAV1-2 | TRAJ33 | CAVSDSNYQLIW | TRBV20-1 | TRBJ1-4 | CSAQSGPGEVEKLFF | GILGFVFTL | YES | NO | YES |
| 317 | TRAV9-2 | TRAJ13 | CALPSGGYQKVTF | TRBV27 | TRBJ1-4 | CASSFGQGAEKLFF | GILGFVFTL | YES | NO | YES |
| 320 | TRAV5 | TRAJ23 | CAESIGKLIF | TRBV29-1 | TRBJ1-4 | CSVGTGGTNEKLFF | GLCTLVAML | NO | NO | NO |
| 322 | TRAV5 | TRAJ37 | CAESTSGGKLIF | TRBV29-1 | TRBJ1-4 | CSVGSRGTNEKLFF | GLCTLVAML | NO | NO | NO |
| 323 | TRAV13-2 | TRAJ6 | CYQGGSYIPTF | TRBV4-3 | TRBJ1-4 | CASSQDRGGLGEKLFF | RTLNAWVKV | YES | NO | NO |
| 324 | TRAV39 | TRAJ58 | CAVVWGGSRLTF | TRBV6-6 | TRBJ1-4 | CASSYPEQGTSNEKLFF | GILGFVFTL | NO | NO | NO |
| 325 | TRAV3 | TRAJ45 | CAVAKDLQNSGGGADGLTF | TRBV7-9 | TRBJ1-4 | CASIDRPATNEKLFF | GILGFVFTL | NO | NO | NO |
| 328 | TRAV12-2 | TRAJ31 | CAVDNARLMF | TRBV12-3 | TRBJ1-5 | CASSFDQTNQPQHF | ELAGIGILTV | YES | NO | YES |
| 331 | TRAV1-2 | TRAJ6 | CAVRSSGGSYIPTF | TRBV15 | TRBJ1-5 | CATSTGDSNQPQHF | AVFDRKSDAK | YES | NO | NO |
| 335 | TRAV1-2 | TRAJ28 | CAVRDSDSGAGSYQLTF | TRBV19 | TRBJ1-5 | CASSARSGEPQHF | GILGFVFTL | YES | YES | YES |
| 338 | TRAV12-2 | TRAJ33 | CAVNTPPGDSNYQLIW | TRBV19 | TRBJ1-5 | CASAFYSNQPQHF | GILGFVFTL | YES | NO | YES |
| 340 | TRAV12-2 | TRAJ42 | CAVNQGGGSQGNLIF | TRBV19 | TRBJ1-5 | CASSIRSSEPQHF | GILGFVFTL | YES | NO | NO |
| 343 | TRAV12-3 | TRAJ42 | CAMSGDGGSQGNLIF | TRBV19 | TRBJ1-5 | CASSKRSNQPQHF | GILGFVFTL | YES | NO | YES |
| 344 | TRAV12-3 | TRAJ42 | CAMSGDGGSQGNLIF | TRBV19 | TRBJ1-5 | CASSWRSNQPQHF | GILGFVFTL | YES | NO | YES |
| 345 | TRAV12-3 | TRAJ52 | CAMRGQDAGGTSYGKLTF | TRBV19 | TRBJ1-5 | CASSVYSNQPQHF | GILGFVFTL | YES | NO | YES |
| 346 | TRAV12-3 | TRAJ52 | CAMRDTNAGGTSYGKLTF | TRBV19 | TRBJ1-5 | CASSILSNQPQHF | GILGFVFTL | YES | NO | YES |
| 347 | TRAV12-3 | TRAJ58 | CAMSGGGRETSGSRLTF | TRBV19 | TRBJ1-5 | CASSIRSNQPQHF | GILGFVFTL | YES | NO | YES |
| 349 | TRAV14DV4 | TRAJ33 | CAMREEGMDSNYQLIW | TRBV19 | TRBJ1-5 | CASSIFSGKPQHF | GILGFVFTL | YES | NO | YES |
| 350 | TRAV14DV4 | TRAJ52 | CAMREDPGGTSYGKLTF | TRBV19 | TRBJ1-5 | CASSWLSNQPQHF | GILGFVFTL | YES | NO | NO |
| 351 | TRAV14DV4 | TRAJ52 | CAMREDPGGTSYGKLTF | TRBV19 | TRBJ1-5 | CASSLYSNQPQHF | GILGFVFTL | YES | NO | YES |
| 352 | TRAV14DV4 | TRAJ52 | CAMRGDAGGTSYGKLTF | TRBV19 | TRBJ1-5 | CASSIYSNQPQHF | GILGFVFTL | YES | NO | YES |
| 353 | TRAV14DV4 | TRAJ52 | CAMRGDAGGTSYGKLTF | TRBV19 | TRBJ1-5 | CASSFYSNQPQHF | GILGFVFTL | YES | NO | YES |
| 354 | TRAV14DV4 | TRAJ52 | CAMREDPGGTSYGKLTF | TRBV19 | TRBJ1-5 | CASSTYSNQPQHF | GILGFVFTL | YES | NO | YES |
| 355 | TRAV17 | TRAJ53 | CATDGSSGGSNYKLTF | TRBV19 | TRBJ1-5 | CASSTSNEQPQHF | GILGFVFTL | YES | NO | YES |
| 357 | TRAV21 | TRAJ36 | CSPCKPEANNLFF | TRBV19 | TRBJ1-5 | CASSTHSGNQPQHF | GILGFVFTL | YES | NO | NO |
| 358 | TRAV22 | TRAJ33 | CAVVRGSMDSNYQLIW | TRBV19 | TRBJ1-5 | CASSIFTIEPQHF | GILGFVFTL | YES | NO | YES |
| 361 | TRAV23DV6 | TRAJ28 | CAASPRDQPLSGAGSYQLTF | TRBV19 | TRBJ1-5 | CASSIHSGNQPQHF | GILGFVFTL | NO | NO | NO |
| 365 | TRAV26-2 | TRAJ57 | CILSIQGGSEKLVF | TRBV19 | TRBJ1-5 | CASSPRSSQPQHF | GILGFVFTL | YES | NO | YES |

| | | | | | | | | | | |
|---|---|---|---|---|---|---|---|---|---|---|
| 366 | TRAV27 | TRAJ35 | CAGAVGGFGNVLHC | TRBV19 | TRBJ1-5 | CASSIYSNQPQHF | GILGFVFTL | NO | NO | NO |
| 367 | TRAV27 | TRAJ37 | CAGVSGNTGKLIF | TRBV19 | TRBJ1-5 | CASSTYSRQPQHF | GILGFVFTL | YES | YES | YES |
| 369 | TRAV27 | TRAJ42 | CAGGGSQGNLIF | TRBV19 | TRBJ1-5 | CASAFYSNQPQHF | GILGFVFTL | YES | NO | YES |
| 370 | TRAV27 | TRAJ42 | CAGGGQGGSQGNLIF | TRBV19 | TRBJ1-5 | CASSIYSNQPQHF | GILGFVFTL | YES | NO | NO |
| 371 | TRAV27 | TRAJ42 | CAGGGGGSQGNLIF | TRBV19 | TRBJ1-5 | CASSTSGANQPQHF | GILGFVFTL | YES | NO | YES |
| 372 | TRAV27 | TRAJ42 | CAGAGHGGSQGNLIF | TRBV19 | TRBJ1-5 | CASSYYSNQPQHF | GILGFVFTL | YES | NO | YES |
| 373 | TRAV27 | TRAJ42 | CAGAQGGGSQGNLIF | TRBV19 | TRBJ1-5 | CASSYYSNQPQHF | GILGFVFTL | YES | NO | YES |
| 374 | TRAV29DV5 | TRAJ27 | CAASGTNAGKSTF | TRBV19 | TRBJ1-5 | CASSVRSGWQPQHF | GILGFVFTL | YES | NO | YES |
| 377 | TRAV29DV5 | TRAJ50 | CAAKGTSYDKVIF | TRBV19 | TRBJ1-5 | CASSIHSNQPQHF | GILGFVFTL | YES | NO | YES |
| 379 | TRAV30 | TRAJ45 | CGTGVVSGGGADGLTF | TRBV19 | TRBJ1-5 | CASTVFQGGGQPQHF | GILGFVFTL | YES | NO | YES |
| 380 | TRAV30 | TRAJ6 | CGTGRGGSYIPTF | TRBV19 | TRBJ1-5 | CASREGHLEGQPQHF | GILGFVFTL | YES | NO | NO |
| 381 | TRAV35 | TRAJ42 | CAGGGDGGSQGNLIF | TRBV19 | TRBJ1-5 | CASSHRSGQPQHF | GILGFVFTL | YES | NO | YES |
| 382 | TRAV36DV7 | TRAJ47 | CAVADMEYGNKLVF | TRBV19 | TRBJ1-5 | CASSMDRFTNQPQHF | GILGFVFTL | NO | NO | NO |
| 385 | TRAV38-2DV8 | TRAJ26 | CAYRGPLDNYGQNFVF | TRBV19 | TRBJ1-5 | CASSIGGGNQPQHF | GILGFVFTL | YES | NO | YES |
| 387 | TRAV41 | TRAJ57 | CAAPITQGGSEKLVF | TRBV19 | TRBJ1-5 | CASSVYSNQPQHF | GILGFVFTL | YES | NO | YES |
| 390 | TRAV12-2 | TRAJ9 | CAVNTGGFKTIF | TRBV2 | TRBJ1-5 | CASTLGGGQPQHF | ELAGIGILTV | YES | NO | NO |
| 391 | TRAV20 | TRAJ42 | CAVHEGGSQGNLIF | TRBV20-1 | TRBJ1-5 | CSAREGNQPQHF | ELAGIGILTV | YES | NO | YES |
| 392 | TRAV27 | TRAJ42 | CAAGGSQGNLIF | TRBV20-1 | TRBJ1-5 | CSANRPGTEHSNQPQHF | GILGFVFTL | YES | NO | YES |
| 397 | TRAV12-2 | TRAJ31 | CAVTGARLMF | TRBV21-1 | TRBJ1-5 | CASSKARGNNQPQHF | ELAGIGILTV | YES | NO | YES |
| 403 | TRAV12-2 | TRAJ47 | CAVNDGNKLVF | TRBV27 | TRBJ1-5 | CASSLSWNQPQHF | ELAGIGILTV | YES | YES | YES |
| 405 | TRAV27 | TRAJ48 | CAGARHSFGNEKLTF | TRBV27 | TRBJ1-5 | CASSLYTRNQPQHF | IMDQVPFSV | YES | NO | YES |
| 407 | TRAV12-2 | TRAJ36 | CAVNIETGANNLFF | TRBV28 | TRBJ1-5 | CASRPTGLGQPQHF | ELAGIGILTV | YES | YES | YES |
| 408 | TRAV12-2 | TRAJ54 | CAEFQGAQKLVF | TRBV28 | TRBJ1-5 | CASSPTGLGQPQHF | ELAGIGILTV | YES | YES | YES |
| 409 | TRAV12-2 | TRAJ57 | CAVAQGGSEKLVF | TRBV28 | TRBJ1-5 | CASRGQGLGQPQHF | ELAGIGILTV | YES | NO | YES |
| 410 | TRAV12-3 | TRAJ41 | CAVTSNSNSGYALNF | TRBV28 | TRBJ1-5 | CASSPQGVGQPQHF | ELAGIGILTV | YES | NO | NO |
| 412 | TRAV12-2 | TRAJ46 | CAVNAPGSSGDKLTF | TRBV3-1 | TRBJ1-5 | CASSHFGLGQPQHF | ELAGIGILTV | YES | YES | YES |
| 413 | TRAV29DV5 | TRAJ49 | CAASAAGNQFYF | TRBV3-1 | TRBJ1-5 | CASSVGTGDHQPQHF | KLGGALQAK | YES | NO | NO |
| 413 | TRAV29DV5 | TRAJ49 | CAASAAGNQFYF | TRBV3-1 | TRBJ1-5 | CASSVGTGDHQPQHF | AVFDRKSDAK | YES | NO | NO |
| 413 | TRAV29DV5 | TRAJ49 | CAASAAGNQFYF | TRBV3-1 | TRBJ1-5 | CASSVGTGDHQPQHF | LLDFVRFMGV | YES | NO | NO |
| 414 | TRAV12-2 | TRAJ45 | CAVASGGADGLTF | TRBV30 | TRBJ1-5 | CAWGYTGGGQPQHF | ELAGIGILTV | NO | NO | NO |
| 415 | TRAV12-2 | TRAJ39 | CAVNAGNMLTF | TRBV4-1 | TRBJ1-5 | CASSQLGPGQPQHF | ELAGIGILTV | YES | YES | YES |
| 416 | TRAV12-3 | TRAJ33 | CAMSGDSNYQLIW | TRBV4-1 | TRBJ1-5 | CASSQDLVSQPQHF | RTLNAWVKV | YES | YES | YES |
| 418 | TRAV13-1 | TRAJ11 | CAAKEGYSTLTF | TRBV5-4 | TRBJ1-5 | CASSSTGGGEKDQPQHF | AVFDRKSDAK | YES | NO | NO |
| 418 | TRAV13-1 | TRAJ11 | CAAKEGYSTLTF | TRBV5-4 | TRBJ1-5 | CASSSTGGGEKDQPQHF | RLRAEAQVK | YES | NO | NO |
| 418 | TRAV13-1 | TRAJ11 | CAAKEGYSTLTF | TRBV5-4 | TRBJ1-5 | CASSSTGGGEKDQPQHF | RTLNAWVKV | YES | NO | NO |
| 418 | TRAV13-1 | TRAJ11 | CAAKEGYSTLTF | TRBV5-4 | TRBJ1-5 | CASSSTGGGEKDQPQHF | LLDFVRFMGV | YES | NO | NO |
| 422 | TRAV30 | TRAJ20 | CGTEYPNDYKLSF | TRBV5-4 | TRBJ1-5 | CASSSTGGGEKDQPQHF | GLCTLVAML | YES | YES | YES |
| 427 | TRAV26-2 | TRAJ45 | CILRGDSGGGADGLTF | TRBV6-1 | TRBJ1-5 | CASSDDPGFGQPQHF | KLGGALQAK | YES | NO | NO |
| 427 | TRAV26-2 | TRAJ45 | CILRGDSGGGADGLTF | TRBV6-1 | TRBJ1-5 | CASSDDPGFGQPQHF | GILGFVFTL | YES | NO | NO |
| 427 | TRAV26-2 | TRAJ45 | CILRGDSGGGADGLTF | TRBV6-1 | TRBJ1-5 | CASSDDPGFGQPQHF | AVFDRKSDAK | YES | NO | NO |
| 428 | TRAV12-2 | TRAJ15 | CAVKDQAGTALIF | TRBV6-3 | TRBJ1-5 | CASSYGTGQPQHF | ELAGIGILTV | YES | NO | YES |
| 429 | TRAV12-2 | TRAJ24 | CAVPTDSWGKLQF | TRBV6-3 | TRBJ1-5 | CASSPTGVGQPQHF | ELAGIGILTV | YES | NO | YES |
| 430 | TRAV12-2 | TRAJ27 | CAVPGGKSTF | TRBV6-3 | TRBJ1-5 | CASSYGGNQPQHF | ELAGIGILTV | YES | NO | NO |
| 434 | TRAV41 | TRAJ42 | CAVSSLGGGSQGNLIF | TRBV6-3 | TRBJ1-5 | CASSYWNQPQHF | ELAGIGILTV | YES | NO | YES |
| 435 | TRAV12-2 | TRAJ43 | CAAPSNNNDMRF | TRBV6-5 | TRBJ1-5 | CASSYGFNQPQHF | ELAGIGILTV | YES | NO | YES |
| 436 | TRAV12-2 | TRAJ45 | CAVAGGGADGLTF | TRBV6-5 | TRBJ1-5 | CASNPQGAGQPQHF | ELAGIGILTV | YES | YES | YES |
| 438 | TRAV41 | TRAJ42 | CAAPYGGSQGNLIF | TRBV6-6 | TRBJ1-5 | CASSYPGTGTSNQPQHF | GILGFVFTL | NO | NO | NO |
| 445 | TRAV38-2DV8 | TRAJ52 | CAYMVNAGGTSYGKLTF | TRBV13 | TRBJ1-6 | CASSLMRGGTYNSPLHF | GILGFVFTL | YES | NO | YES |
| 448 | TRAV12-2 | TRAJ48 | CAVNGFGNEKLTF | TRBV14 | TRBJ1-6 | CASSQEEGAYNSPLHF | ELAGIGILTV | NO | NO | NO |
| 449 | TRAV27 | TRAJ42 | CAGAYGGSQGNLIF | TRBV19 | TRBJ1-6 | CASSTRSDSPLHF | GILGFVFTL | YES | YES | YES |
| 451 | TRAV13-1 | TRAJ42 | CAYGGSQGNLIF | TRBV20-1 | TRBJ1-6 | CSARGGFLSPLHF | GILGFVFTL | YES | NO | YES |
| 454 | TRAV8-4 | TRAJ50 | CAVSVFPETSYDKVIF | TRBV20-1 | TRBJ1-6 | CSARESPGQGIFSPLHF | LLFGYPVYV | NO | NO | NO |
| 456 | TRAV17 | TRAJ11 | CATVRPYSGYSTLTF | TRBV27 | TRBJ1-6 | CASSLDRTSSPLHF | RTLNAWVKV | YES | YES | YES |
| 459 | TRAV12-2 | TRAJ36 | CAVSQTGANNLFF | TRBV28 | TRBJ1-6 | CASSPTGLGSPLHF | ELAGIGILTV | YES | NO | YES |
| 460 | TRAV12-2 | TRAJ30 | CAVNAGGDKIIF | TRBV30 | TRBJ1-6 | CAWSTGTGWPLHF | ELAGIGILTV | YES | YES | NO |
| 461 | TRAV19 | TRAJ17 | CALKLIKAAGNKLTF | TRBV30 | TRBJ1-6 | CAWTGTGKIGWDSPLHF | AVFDRKSDAK | YES | NO | NO |
| 461 | TRAV19 | TRAJ17 | CALKLIKAAGNKLTF | TRBV30 | TRBJ1-6 | CAWTGTGKIGWDSPLHF | GILGFVFTL | YES | NO | NO |
| 463 | TRAV9-2 | TRAJ20 | CALSDGNDYKLSF | TRBV4-3 | TRBJ1-6 | CASSQTGLAYNSPLHF | KLGGALQAK | YES | NO | NO |
| 463 | TRAV9-2 | TRAJ20 | CALSDGNDYKLSF | TRBV4-3 | TRBJ1-6 | CASSQTGLAYNSPLHF | SLFNTVATLY | YES | NO | NO |
| 464 | TRAV12-2 | TRAJ11 | CAVGANSGYSTLTF | TRBV5-1 | TRBJ1-6 | CASSLEGQGGSPLHF | ELAGIGILTV | YES | NO | YES |
| 466 | TRAV13-1 | TRAJ20 | CAPPSF | TRBV6-3 | TRBJ1-6 | CASSWGQGSPLHF | ELAGIGILTV | YES | NO | YES |
| 467 | TRAV12-2 | TRAJ13 | CAVGISGGYQKVTF | TRBV6-5 | TRBJ1-6 | CASSSGTGSPLHF | ELAGIGILTV | YES | NO | YES |
| 468 | TRAV12-2 | TRAJ30 | CAVKKGRDDKIIF | TRBV6-5 | TRBJ1-6 | CASSYLIATTFNSPLHF | ELAGIGILTV | YES | YES | YES |
| 469 | TRAV12-2 | TRAJ45 | CAVSTGGGADGLTF | TRBV6-5 | TRBJ1-6 | CASTPGQGGSPLHF | ELAGIGILTV | YES | YES | YES |
| 470 | TRAV12-2 | TRAJ29 | CASPSGNTPLVF | TRBV10-1 | TRBJ2-1 | CASSEVVLAGGQSEQFF | ELAGIGILTV | YES | NO | NO |
| 471 | TRAV12-1 | TRAJ45 | CVVNGGADGLTF | TRBV10-2 | TRBJ2-1 | CASRQSFNEQFF | GILGFVFTL | YES | NO | YES |
| 472 | TRAV12-2 | TRAJ23 | CAVTEGGKLIF | TRBV10-3 | TRBJ2-1 | CAISVVGGSYNEQFF | ELAGIGILTV | YES | YES | YES |
| 473 | TRAV12-2 | TRAJ5 | CARRDPTGRRALTF | TRBV10-3 | TRBJ2-1 | CAISSDSGDEQFF | YLNDHLEPWI | YES | NO | YES |
| 475 | TRAV8-6 | TRAJ20 | CAVSVGNDYKLSF | TRBV12-3 | TRBJ2-1 | CASSSPGGEQFF | GLCTLVAML | YES | YES | YES |
| 476 | TRAV14DV4 | TRAJ40 | CAMSGTSGTYKYIF | TRBV12-4 | TRBJ2-1 | CASSLLAPPPHNEQFF | KLGGALQAK | YES | NO | NO |
| 480 | TRAV12-3 | TRAJ34 | CAMSADTDKLIF | TRBV14 | TRBJ2-1 | CASSQVLYNEQFF | LLFGYPVYV | YES | YES | NO |
| 482 | TRAV12-2 | TRAJ35 | CAVNRGFGNVLHC | TRBV19 | TRBJ2-1 | CASSSGGGGQFF | ELAGIGILTV | NO | NO | NO |
| 483 | TRAV12-2 | TRAJ37 | CAVTPSNTGKLIF | TRBV19 | TRBJ2-1 | CASSTRAGGVGQFF | GILGFVFTL | YES | NO | NO |
| 484 | TRAV12-2 | TRAJ42 | CAVPGGGGSQGNLIF | TRBV19 | TRBJ2-1 | CASSMRSTNEQFF | GILGFVFTL | YES | NO | YES |
| 485 | TRAV12-2 | TRAJ42 | CAAGFMNYGGSQGNLIF | TRBV19 | TRBJ2-1 | CASSLTSVYNEQFF | KLGGALQAK | YES | NO | NO |
| 485 | TRAV12-2 | TRAJ42 | CAAGFMNYGGSQGNLIF | TRBV19 | TRBJ2-1 | CASSLTSVYNEQFF | RTLNAWVKV | YES | NO | NO |
| 485 | TRAV12-2 | TRAJ42 | CAAGFMNYGGSQGNLIF | TRBV19 | TRBJ2-1 | CASSLTSVYNEQFF | AVFDRKSDAK | YES | NO | NO |
| 485 | TRAV12-2 | TRAJ42 | CAAGFMNYGGSQGNLIF | TRBV19 | TRBJ2-1 | CASSLTSVYNEQFF | IVTDFSVIK | YES | NO | NO |
| 488 | TRAV13-2 | TRAJ42 | CAENIGGSQGNLIF | TRBV19 | TRBJ2-1 | CASSHYSVNEQFF | GILGFVFTL | YES | NO | YES |
| 491 | TRAV17 | TRAJ42 | CATDGGGGSQGNLIF | TRBV19 | TRBJ2-1 | CASSGLAGLNEQFF | GILGFVFTL | YES | NO | YES |
| 494 | TRAV19 | TRAJ13 | CALSELSGGYQKVTF | TRBV19 | TRBJ2-1 | CASSVLAGAYNEQFF | GILGFVFTL | YES | NO | YES |
| 495 | TRAV19 | TRAJ3 | CALSEPPGKSSASKIIF | TRBV19 | TRBJ2-1 | CASSTTSGGLNEQFF | GILGFVFTL | YES | NO | YES |
| 500 | TRAV24 | TRAJ37 | CASAGSGNTGKLIF | TRBV19 | TRBJ2-1 | CASSIRSAWEQFF | GILGFVFTL | YES | YES | NO |
| 502 | TRAV26-2 | TRAJ35 | CILREGFGNVLHC | TRBV19 | TRBJ2-1 | CASSMRSGSEQFF | GILGFVFTL | NO | NO | NO |
| 503 | TRAV27 | TRAJ37 | CAGAFGSSNTGKLIF | TRBV19 | TRBJ2-1 | CASSIRSSYEQFF | GILGFVFTL | YES | NO | YES |
| 506 | TRAV27 | TRAJ42 | CAGAGGGSQGNLIF | TRBV19 | TRBJ2-1 | CASSIRSSDEQFF | GILGFVFTL | YES | NO | YES |
| 507 | TRAV27 | TRAJ42 | CAGAGGGSQGNLIF | TRBV19 | TRBJ2-1 | CASSARSSNEQFF | GILGFVFTL | YES | NO | YES |
| 509 | TRAV35 | TRAJ42 | CAGQSSDFAYGGSQGNLIF | TRBV19 | TRBJ2-1 | CASSMVAALGEQFF | GILGFVFTL | YES | NO | YES |
| 513 | TRAV5 | TRAJ42 | CAEDYGGSQGNLIF | TRBV19 | TRBJ2-1 | CASSTRSSNEQFF | GILGFVFTL | YES | NO | YES |
| 516 | TRAV8-3 | TRAJ42 | CAVGAGGGSQGNLIF | TRBV19 | TRBJ2-1 | CASSMRASVEQFF | GILGFVFTL | YES | NO | YES |
| 517 | TRAV12-2 | TRAJ8 | CAVGFQKLVF | TRBV2 | TRBJ2-1 | CASSGIPGNEQFF | ELAGIGILTV | YES | NO | YES |
| 518 | TRAV12-1 | TRAJ30 | CVVNENDKIIF | TRBV20-1 | TRBJ2-1 | CSASKRLLNEQFF | LLFGYPVYV | NO | NO | NO |
| 519 | TRAV12-2 | TRAJ52 | CAPSAGKLTF | TRBV20-1 | TRBJ2-1 | CSASIAGVGNEQFF | ELAGIGILTV | NO | NO | NO |
| 521 | TRAV19 | TRAJ49 | CALSEATTGNQFYF | TRBV20-1 | TRBJ2-1 | CSAEWDRAYNEQFF | LLFGYPVYV | NO | NO | NO |
| 523 | TRAV3 | TRAJ26 | CAVKGNYGQNFVF | TRBV20-1 | TRBJ2-1 | CSAEWDRAYNEQFF | LLFGYPVYV | NO | NO | NO |
| 534 | TRAV12-2 | TRAJ10 | CAPGGGGNKLTF | TRBV24-1 | TRBJ2-1 | CATSDLFNNEQFF | ELAGIGILTV | YES | NO | YES |
| 535 | TRAV20 | TRAJ26 | CAVWPWGQNFVF | TRBV24-1 | TRBJ2-1 | CATSDFRQGDVLFWQFF | RTLNAWVKV | YES | NO | NO |
| 537 | TRAV12-2 | TRAJ6 | CAVNWEEQGGSYIPTF | TRBV27 | TRBJ2-1 | CASRVSGSLSYNEQFF | KLGGALQAK | YES | NO | NO |
| 537 | TRAV12-2 | TRAJ6 | CAVNWEEQGGSYIPTF | TRBV27 | TRBJ2-1 | CASRVSGSLSYNEQFF | AVFDRKSDAK | YES | NO | NO |
| 537 | TRAV12-2 | TRAJ6 | CAVNWEEQGGSYIPTF | TRBV27 | TRBJ2-1 | CASRVSGSLSYNEQFF | FLASKIGRLV | YES | NO | NO |
| 537 | TRAV12-2 | TRAJ6 | CAVNWEEQGGSYIPTF | TRBV27 | TRBJ2-1 | CASRVSGSLSYNEQFF | RTLNAWVKV | YES | NO | NO |
| 537 | TRAV12-2 | TRAJ6 | CAVNWEEQGGSYIPTF | TRBV27 | TRBJ2-1 | CASRVSGSLSYNEQFF | IVTDFSVIK | YES | NO | NO |
| 538 | TRAV2 | TRAJ22 | CAVEGSSGSARQLTF | TRBV27 | TRBJ2-1 | CASSSDGNEQFF | ELAGIGILTV | NO | NO | NO |
| 541 | TRAV12-2 | TRAJ9 | CAVNTGGFKTIF | TRBV28 | TRBJ2-1 | CASSLPMLYEQFF | ELAGIGILTV | YES | NO | NO |

| | | | | | | | | | | |
|---|---|---|---|---|---|---|---|---|---|---|
| 547 | TRAV29DV5 | TRAJ53 | CAASGRVSGGSNYKLTF | TRBV3-1 | TRBJ2-1 | CASSRLAGGWNEQFF | KLGGALQAK | YES | NO | NO |
| 547 | TRAV29DV5 | TRAJ53 | CAASGRVSGGSNYKLTF | TRBV3-1 | TRBJ2-1 | CASSRLAGGWNEQFF | AVFDRKSDAK | YES | NO | NO |
| 550 | TRAV17 | TRAJ37 | CATGLHGSGNTGKLIF | TRBV30 | TRBJ2-1 | CAWGSVRMISYNEQFF | RTLNAWVKV | YES | YES | NO |
| 552 | TRAV12-2 | TRAJ47 | CAVNGNKLVF | TRBV4-1 | TRBJ2-1 | CASSPGPLAGGDNEQFF | ELAGIGILTV | YES | NO | YES |
| 553 | TRAV38-2DV8 | TRAJ52 | CAYSPAAGGTSYGKLTF | TRBV4-1 | TRBJ2-1 | CASSLLWAEYNEQFF | GILGFVFTL | YES | YES | YES |
| 554 | TRAV38-2DV8 | TRAJ57 | CAYIGQGGSEKLVF | TRBV4-1 | TRBJ2-1 | CASSQDGFGLFNEQFF | GILGFVFTL | YES | NO | YES |
| 555 | TRAV5 | TRAJ10 | CAERVTGGGNKLTF | TRBV4-1 | TRBJ2-1 | CASSQDPGDEQFF | KLGGALQAK | YES | NO | NO |
| 558 | TRAV12-2 | TRAJ22 | CAVGAGSARQLTF | TRBV5-1 | TRBJ2-1 | CASSLSQGGNEQFF | ELAGIGILTV | YES | NO | YES |
| 563 | TRAV12-2 | TRAJ21 | CAVNKDFNKFYF | TRBV6-3 | TRBJ2-1 | CASSAGGSNEQFF | ELAGIGILTV | YES | NO | YES |
| 567 | TRAV12-2 | TRAJ31 | CAVSMDGLMF | TRBV6-4 | TRBJ2-1 | CASSGTQPGLGGEQFF | ELAGIGILTV | YES | NO | NO |
| 569 | TRAV23DV6 | TRAJ42 | CAADNYGGSQGNLIF | TRBV6-5 | TRBJ2-1 | CASSGLNEQFF | GILGFVFTL | NO | NO | NO |
| 570 | TRAV12-2 | TRAJ13 | CAVNIEGYQKVTF | TRBV6-6 | TRBJ2-1 | CASSDSSGGLYNEQFF | GILGFVFTL | NO | NO | NO |
| 576 | TRAV3 | TRAJ7 | CAVRDGNNRLAF | TRBV7-2 | TRBJ2-1 | CASSPRQVNYNEQFF | LLFGYPVYV | NO | NO | NO |
| 582 | TRAV30 | TRAJ26 | CGTHDNYGQNFVF | TRBV7-8 | TRBJ2-1 | CASSLANSYNEQFF | SLLMWITQV | YES | NO | YES |
| 584 | TRAV14DV4 | TRAJ5 | CAMRDPQGGRRALTF | TRBV7-9 | TRBJ2-1 | CASTGTSGGPTLRDEQFF | SLFNTVATLY | YES | NO | NO |
| 590 | TRAV35 | TRAJ49 | CAGHTGNQFYF | TRBV7-9 | TRBJ2-1 | CASSSGLAGGNEQFF | GILGFVFTL | YES | NO | NO |
| 591 | TRAV38-1 | TRAJ52 | CAFMKGAGGTSYGKLTF | TRBV7-9 | TRBJ2-1 | CASSPGHPSSYNEQFF | GILGFVFTL | YES | NO | YES |
| 592 | TRAV38-2DV8 | TRAJ52 | CAYSPAAGGTSYGKLTF | TRBV7-9 | TRBJ2-1 | CASSSGLAGVNEQFF | GILGFVFTL | YES | NO | YES |
| 593 | TRAV9-2 | TRAJ33 | CAREHMDSNYQLIW | TRBV7-9 | TRBJ2-1 | CASSQLGRGDNEQFF | AVFDRKSDAK | YES | NO | NO |
| 593 | TRAV9-2 | TRAJ33 | CAREHMDSNYQLIW | TRBV7-9 | TRBJ2-1 | CASSQLGRGDNEQFF | IVTDFSVIK | YES | NO | NO |
| 594 | TRAV1-2 | TRAJ36 | CAVRGTGANNLFF | TRBV9 | TRBJ2-1 | CASSPKDIRVYNEQFF | GILGFVFTL | YES | NO | YES |
| 596 | TRAV14DV4 | TRAJ52 | CAMREGDAGGTSYGKLTF | TRBV9 | TRBJ2-1 | CASSPSGSGNEQFF | YLLEMLWRL | YES | NO | YES |
| 601 | TRAV5 | TRAJ45 | CAEIFSGGGADGLTF | TRBV10-1 | TRBJ2-2 | CASNPASGGELFF | GILGFVFTL | YES | NO | YES |
| 602 | TRAV12-2 | TRAJ10 | CAVPLTGGGNKLTF | TRBV11-2 | TRBJ2-2 | CASSLGVAGTNTGELFF | GLCTLVAML | YES | YES | YES |
| 603 | TRAV12-3 | TRAJ27 | CAMKSTNAGKSTF | TRBV11-2 | TRBJ2-2 | CASSLGVAGTNTGELFF | IVTDFSVIK | YES | NO | NO |
| 603 | TRAV12-3 | TRAJ27 | CAMKSTNAGKSTF | TRBV11-2 | TRBJ2-2 | CASSLGVAGTNTGELFF | RLRAEAQVK | YES | NO | NO |
| 607 | TRAV12-2 | TRAJ42 | CAGGGDGGSQGNLIF | TRBV12-3 | TRBJ2-2 | CASSFSGNTGELFF | GILGFVFTL | YES | NO | YES |
| 611 | TRAV27 | TRAJ42 | CAGASGGSQGNLIF | TRBV12-3 | TRBJ2-2 | CASSFSGNTGELFF | GILGFVFTL | YES | NO | YES |
| 612 | TRAV27 | TRAJ42 | CAGGGSQGNLIF | TRBV12-3 | TRBJ2-2 | CASSFSGNTGELFF | GILGFVFTL | YES | NO | YES |
| 613 | TRAV38-1 | TRAJ52 | CAFMKGAGGTSYGKLTF | TRBV12-3 | TRBJ2-2 | CASSFSGNTGELFF | GILGFVFTL | YES | NO | YES |
| 614 | TRAV38-2DV8 | TRAJ52 | CAYSPAAGGTSYGKLTF | TRBV12-3 | TRBJ2-2 | CASSFSGGTGELFF | GILGFVFTL | YES | NO | YES |
| 615 | TRAV38-2DV8 | TRAJ52 | CAYSPAAGGTSYGKLTF | TRBV12-4 | TRBJ2-2 | CASSQGAGELFF | GILGFVFTL | YES | NO | YES |
| 616 | TRAV12-2 | TRAJ6 | CAVNLGSYIPTF | TRBV19 | TRBJ2-2 | CATSWGGSGELFF | ELAGIGILTV | YES | NO | YES |
| 618 | TRAV13-2 | TRAJ42 | CAENLGGGSQGNLIF | TRBV19 | TRBJ2-2 | CASSTRATGELFF | GILGFVFTL | YES | NO | YES |
| 619 | TRAV13-2 | TRAJ42 | CAENMGGGSQGNLIF | TRBV19 | TRBJ2-2 | CASSIRAAGELFF | GILGFVFTL | YES | YES | YES |
| 620 | TRAV13-2 | TRAJ42 | CAENKGGGSQGNLIF | TRBV19 | TRBJ2-2 | CASSIRSSGELFF | GILGFVFTL | YES | NO | NO |
| 623 | TRAV25 | TRAJ42 | CAANYGGSQGNLIF | TRBV19 | TRBJ2-2 | CASSGRSTGELFF | GILGFVFTL | YES | NO | YES |
| 624 | TRAV25 | TRAJ42 | CAGSGGGSQGNLIF | TRBV19 | TRBJ2-2 | CASSIRSTGELFF | GILGFVFTL | YES | NO | YES |
| 625 | TRAV25 | TRAJ42 | CAGNYGGSQGNLIF | TRBV19 | TRBJ2-2 | CASSVRSTGELFF | GILGFVFTL | YES | NO | YES |
| 627 | TRAV25 | TRAJ42 | CAGNYGGSQGNLIF | TRBV19 | TRBJ2-2 | CASSGRSTGELFF | GILGFVFTL | YES | NO | YES |
| 628 | TRAV27 | TRAJ37 | CAGAGNTGKLIF | TRBV19 | TRBJ2-2 | CASSGRSTGELFF | GILGFVFTL | YES | NO | YES |
| 629 | TRAV27 | TRAJ42 | CAGGDGGSQGNLIF | TRBV19 | TRBJ2-2 | CASSIRSTGELFF | GILGFVFTL | YES | YES | YES |
| 630 | TRAV27 | TRAJ42 | CAGASGGSQGNLIF | TRBV19 | TRBJ2-2 | CASSGRSTGELFF | GILGFVFTL | YES | NO | NO |
| 631 | TRAV27 | TRAJ42 | CAGAGGGSQGNLIF | TRBV19 | TRBJ2-2 | CASSTRSTGELFF | GILGFVFTL | YES | NO | YES |
| 632 | TRAV27 | TRAJ42 | CAGAGGGSQGNLIF | TRBV19 | TRBJ2-2 | CASSGRSTGELFF | GILGFVFTL | YES | NO | YES |
| 633 | TRAV27 | TRAJ42 | CAGAYGGSQGNLIF | TRBV19 | TRBJ2-2 | CASSGRSTGELFF | GILGFVFTL | YES | NO | YES |
| 634 | TRAV3 | TRAJ36 | CAVRDGTGANNLFF | TRBV19 | TRBJ2-2 | CASSISSTGELFF | GILGFVFTL | NO | NO | NO |
| 636 | TRAV5 | TRAJ26 | CAERQNFVF | TRBV19 | TRBJ2-2 | CASSISSTGELFF | GILGFVFTL | NO | NO | NO |
| 638 | TRAV8-1 | TRAJ27 | CAVNRTNAGKSTF | TRBV19 | TRBJ2-2 | CASSGRSTGELFF | GILGFVFTL | YES | NO | YES |
| 639 | TRAV12-1 | TRAJ12 | CVVNGGDSSYKLIF | TRBV2 | TRBJ2-2 | CASSEGQVSPGELFF | GLCTLVAML | YES | YES | YES |
| 641 | TRAV12-3 | TRAJ42 | CAMSGDGGSQGNLIF | TRBV2 | TRBJ2-2 | CASSLLLAGENTGELFF | GILGFVFTL | YES | NO | YES |
| 642 | TRAV13-1 | TRAJ50 | CAASETSYDKVIF | TRBV2 | TRBJ2-2 | CASSTPSQGTNTGELFF | RAKFKQLL | YES | YES | YES |
| 644 | TRAV12-2 | TRAJ48 | CAVNIGKLTF | TRBV20-1 | TRBJ2-2 | CSARDTGGGSGELFF | ELAGIGILTV | YES | NO | NO |
| 645 | TRAV12-2 | TRAJ22 | CAVGSSGSARQLTF | TRBV24-1 | TRBJ2-2 | CATSAGQGAFSGELFF | ELAGIGILTV | YES | YES | YES |
| 648 | TRAV8-3 | TRAJ31 | CAVGTARLMF | TRBV27 | TRBJ2-2 | CASSPHFQVDTGELFF | GILGFVFTL | YES | NO | NO |
| 653 | TRAV12-3 | TRAJ23 | CAMSADNQGGKLIF | TRBV4-2 | TRBJ2-2 | CASSQASGGSTNTGELFF | YLNDHLEPWI | YES | NO | YES |
| 656 | TRAV38-2DV8 | TRAJ58 | CAYRSAQETSGSRLTF | TRBV4-3 | TRBJ2-2 | CASSQGLLGGELFF | ELAGIGILTV | YES | YES | YES |
| 657 | TRAV12-3 | TRAJ48 | CAMAYSNFGNEKLTF | TRBV5-1 | TRBJ2-2 | CASSPGTPSLTGELFF | ELAGIGILTV | YES | NO | YES |
| 658 | TRAV1-2 | TRAJ35 | CAVGGPI | TRBV5-6 | TRBJ2-2 | CASTARALNTGELFF | LLDFVRFMGV | NO | NO | NO |
| 662 | TRAV12-2 | TRAJ45 | CAVDSGGGADGLTF | TRBV6-1 | TRBJ2-2 | CASSEGGVGAGELFF | ELAGIGILTV | YES | YES | YES |
| 664 | TRAV29DV5 | TRAJ31 | CAADDGRLMF | TRBV6-6 | TRBJ2-2 | CASSYGFPTSGGNTGELFF | GILGFVFTL | YES | NO | NO |
| 665 | TRAV38-2DV8 | TRAJ52 | CAYSPAAGGTSYGKLTF | TRBV7-2 | TRBJ2-2 | CASSFLIGELFF | GILGFVFTL | YES | NO | YES |
| 669 | TRAV12-2 | TRAJ42 | CAEGGGSQGNLIF | TRBV7-8 | TRBJ2-2 | CASSLAGLAGGELFF | ELAGIGILTV | YES | NO | YES |
| 670 | TRAV5 | TRAJ45 | CAEKEYADGLTF | TRBV7-8 | TRBJ2-2 | CASSSPEGQPLNTGELFF | GLCTLVAML | YES | YES | NO |
| 671 | TRAV34 | TRAJ3 | CGADSRYSSASKIIF | TRBV7-9 | TRBJ2-2 | CASSSRGADGELFF | RTLNAWVKV | YES | YES | YES |
| 675 | TRAV12-2 | TRAJ45 | CAVGGGGADGLTF | TRBV9 | TRBJ2-2 | CASSSSLAGGGELFF | ELAGIGILTV | YES | NO | YES |
| 676 | TRAV12-2 | TRAJ53 | CAVSGNYKLTF | TRBV9 | TRBJ2-2 | CASSEDTGELFF | YLLEMLWRL | YES | NO | YES |
| 678 | TRAV13-1 | TRAJ50 | CAASETSYDKVIF | TRBV9 | TRBJ2-2 | CASSSGLVSNTGELFF | RAKFKQLL | YES | YES | YES |
| 681 | TRAV30 | TRAJ20 | CGTEYPNDYKLSF | TRBV9 | TRBJ2-2 | CASSSGLVSNTGELFF | KLGGALQAK | YES | NO | NO |
| 683 | TRAV38-1 | TRAJ52 | CAFMINAGGTSYGKLTF | TRBV10-3 | TRBJ2-3 | CAISGAARDTQYF | GILGFVFTL | YES | NO | YES |
| 684 | TRAV12-2 | TRAJ23 | CAVKGGGKLIF | TRBV11-2 | TRBJ2-3 | CASSPEFEVWADTQYF | ELAGIGILTV | YES | YES | YES |
| 685 | TRAV8-1 | TRAJ20 | CAVTGNNDYKLSF | TRBV11-2 | TRBJ2-3 | CASSFTSGTTDTQYF | GLCTLVAML | YES | YES | YES |
| 686 | TRAV21 | TRAJ33 | CAVDSNYQLIW | TRBV12-5 | TRBJ2-3 | CASGLGTVASTDTQYF | KLGGALQAK | YES | NO | NO |
| 686 | TRAV21 | TRAJ33 | CAVDSNYQLIW | TRBV12-5 | TRBJ2-3 | CASGLGTVASTDTQYF | SLFNTVATLY | YES | NO | NO |
| 692 | TRAV12-1 | TRAJ42 | CVVNIGGGSQGNLIF | TRBV19 | TRBJ2-3 | CASSIRSTDTQYF | GILGFVFTL | YES | YES | YES |
| 693 | TRAV12-1 | TRAJ42 | CVVNIGGGSQGNLIF | TRBV19 | TRBJ2-3 | CASSIRSSDTQYF | GILGFVFTL | YES | NO | YES |
| 695 | TRAV12-2 | TRAJ42 | CAGGGDGGSQGNLIF | TRBV19 | TRBJ2-3 | CASSIRSTDTQYF | GILGFVFTL | YES | NO | YES |
| 696 | TRAV12-2 | TRAJ42 | CAVSLGGGSQGNLIF | TRBV19 | TRBJ2-3 | CASSIRSTDTQYF | GILGFVFTL | YES | NO | YES |
| 698 | TRAV12-2 | TRAJ5 | CAVPLAGTGRRALTF | TRBV19 | TRBJ2-3 | CASSILGVSTDTQYF | ELAGIGILTV | YES | NO | YES |
| 699 | TRAV12-3 | TRAJ42 | CAMSGDGGSQGNLIF | TRBV19 | TRBJ2-3 | CASSTRAGDTQYF | GILGFVFTL | YES | NO | YES |
| 703 | TRAV13-1 | TRAJ42 | CAASIGGGSQGNLIF | TRBV19 | TRBJ2-3 | CASSIRSTDTQYF | GILGFVFTL | YES | NO | YES |
| 704 | TRAV13-1 | TRAJ42 | CAASLGGGSQGNLIF | TRBV19 | TRBJ2-3 | CASSIRSTDTQYF | GILGFVFTL | YES | NO | YES |
| 705 | TRAV13-1 | TRAJ42 | CAASGGGSQGNLIF | TRBV19 | TRBJ2-3 | CASSMFGTDTQYF | GILGFVFTL | YES | NO | YES |
| 709 | TRAV13-2 | TRAJ42 | CAENKGGGSQGNLIF | TRBV19 | TRBJ2-3 | CASSIRSTDTQYF | GILGFVFTL | YES | NO | YES |
| 710 | TRAV13-2 | TRAJ42 | CAENTGGGSQGNLIF | TRBV19 | TRBJ2-3 | CASSWRSTDTQYF | GILGFVFTL | YES | NO | NO |
| 713 | TRAV16 | TRAJ43 | CALKNNDMRF | TRBV19 | TRBJ2-3 | CASSWRSTDTQYF | FLYALALLL | NO | NO | NO |
| 716 | TRAV17 | TRAJ42 | CATDGGGGSQGNLIF | TRBV19 | TRBJ2-3 | CATSQRSTDTQYF | GILGFVFTL | YES | NO | YES |
| 717 | TRAV17 | TRAJ42 | CATDGGGGSQGNLIF | TRBV19 | TRBJ2-3 | CASSPRSTDTQYF | GILGFVFTL | YES | NO | YES |
| 718 | TRAV19 | TRAJ47 | CALSYMEYGNKLVF | TRBV19 | TRBJ2-3 | CASSTRSTDTQYF | GILGFVFTL | YES | NO | YES |
| 720 | TRAV25 | TRAJ27 | CAGPTNAGKSTF | TRBV19 | TRBJ2-3 | CASSGISTDTQYF | GILGFVFTL | YES | NO | NO |
| 724 | TRAV25 | TRAJ42 | CAGNYGGSQGNLIF | TRBV19 | TRBJ2-3 | CASSGRSTDTQYF | GILGFVFTL | YES | NO | YES |
| 725 | TRAV25 | TRAJ42 | CAGNYGGSQGNLIF | TRBV19 | TRBJ2-3 | CASSTRSTDTQYF | GILGFVFTL | YES | NO | YES |
| 726 | TRAV25 | TRAJ42 | CAGNYGGSQGNLIF | TRBV19 | TRBJ2-3 | CASSQRSTDTQYF | GILGFVFTL | YES | NO | YES |
| 727 | TRAV25 | TRAJ42 | CAGNYGGSQGNLIF | TRBV19 | TRBJ2-3 | CASSPRSTDTQYF | GILGFVFTL | YES | NO | YES |
| 728 | TRAV25 | TRAJ42 | CAANYGGSQGNLIF | TRBV19 | TRBJ2-3 | CASSVRSTDTQYF | GILGFVFTL | YES | NO | YES |
| 729 | TRAV25 | TRAJ42 | CAATGGGSQGNLIF | TRBV19 | TRBJ2-3 | CASSMRSADTQYF | GILGFVFTL | YES | NO | YES |
| 730 | TRAV25 | TRAJ42 | CAGNYGGSQGNLIF | TRBV19 | TRBJ2-3 | CAASTRSTDTQYF | GILGFVFTL | YES | NO | NO |
| 731 | TRAV25 | TRAJ42 | CAGNYGGSQGNLIF | TRBV19 | TRBJ2-3 | CASSFRSTDTQYF | GILGFVFTL | YES | YES | YES |
| 732 | TRAV25 | TRAJ42 | CAGPEGGGSQGNLIF | TRBV19 | TRBJ2-3 | CASSGRSTDTQYF | GILGFVFTL | YES | NO | YES |
| 733 | TRAV25 | TRAJ42 | CAGPEEGGSQGNLIF | TRBV19 | TRBJ2-3 | CASSGRSTDTQYF | GILGFVFTL | YES | YES | YES |
| 734 | TRAV25 | TRAJ42 | CAGNYGGSQGNLIF | TRBV19 | TRBJ2-3 | CASSSRSTDTQYF | GILGFVFTL | YES | NO | YES |
| 735 | TRAV26-2 | TRAJ37 | CILPHAIYGSSNTGKLIF | TRBV19 | TRBJ2-3 | CASSVRSTDTQYF | GILGFVFTL | YES | YES | YES |
| 736 | TRAV27 | TRAJ37 | CAGANGSSNTGKLIF | TRBV19 | TRBJ2-3 | CASSIRSAYTQYF | GILGFVFTL | YES | YES | YES |
| 737 | TRAV27 | TRAJ37 | CAGAHGSSNTGKLIF | TRBV19 | TRBJ2-3 | CASSMDRGSADTQYF | GILGFVFTL | YES | NO | YES |

| | | | | | | | | | | |
|---|---|---|---|---|---|---|---|---|---|---|
| 738 | TRAV27 | TRAJ42 | CAGDGGGGSQGNLIF | TRBV19 | TRBJ2-3 | CASSHRSTDTQYF | GILGFVFTL | YES | NO | YES |
| 739 | TRAV27 | TRAJ42 | CAGDGGGGSQGNLIF | TRBV19 | TRBJ2-3 | CASSIRSLDTQYF | GILGFVFTL | YES | NO | YES |
| 740 | TRAV27 | TRAJ42 | CAGGGSQGNLIF | TRBV19 | TRBJ2-3 | CASSTRSTDTQYF | GILGFVFTL | YES | NO | NO |
| 744 | TRAV27 | TRAJ42 | CAGTGGGSQGNLIF | TRBV19 | TRBJ2-3 | CASSIRSTDTQYF | GILGFVFTL | YES | NO | YES |
| 745 | TRAV27 | TRAJ42 | CAGADGGSQGNLIF | TRBV19 | TRBJ2-3 | CASSQRSTDTQYF | GILGFVFTL | YES | YES | YES |
| 746 | TRAV27 | TRAJ42 | CAGAGGGSQGNLIF | TRBV19 | TRBJ2-3 | CASSTRSTDTQYF | GILGFVFTL | YES | NO | YES |
| 747 | TRAV27 | TRAJ42 | CAGAGGGSQGNLIF | TRBV19 | TRBJ2-3 | CASSIRSTDTQYF | GILGFVFTL | YES | NO | YES |
| 748 | TRAV27 | TRAJ42 | CAGAGGGSQGNLIF | TRBV19 | TRBJ2-3 | CASSQRSTDTQYF | GILGFVFTL | YES | NO | YES |
| 749 | TRAV27 | TRAJ42 | CAGTDGGSQGNLIF | TRBV19 | TRBJ2-3 | CASSPRSTDTQYF | GILGFVFTL | YES | NO | YES |
| 750 | TRAV27 | TRAJ42 | CAGTHGGSQGNLIF | TRBV19 | TRBJ2-3 | CASSVRSTDTQYF | GILGFVFTL | YES | NO | NO |
| 751 | TRAV27 | TRAJ42 | CAGAGGGSQGNLIF | TRBV19 | TRBJ2-3 | CASSFRSADTQYF | GILGFVFTL | YES | NO | YES |
| 752 | TRAV27 | TRAJ42 | CAGAGGGSQGNLIF | TRBV19 | TRBJ2-3 | CASSSRSTDTQYF | GILGFVFTL | YES | YES | YES |
| 753 | TRAV29DV5 | TRAJ43 | CAAPTGNNNDMRF | TRBV19 | TRBJ2-3 | CASSVRSTDTQYF | GILGFVFTL | YES | NO | YES |
| 754 | TRAV3 | TRAJ36 | CAVRDGTGANNLFF | TRBV19 | TRBJ2-3 | CASNPSSTDTQYF | GILGFVFTL | NO | NO | NO |
| 756 | TRAV30 | TRAJ27 | CGQTF | TRBV19 | TRBJ2-3 | CASSGRSTDTQYF | GILGFVFTL | YES | NO | NO |
| 758 | TRAV30 | TRAJ42 | CGTEMGGGSQGNLIF | TRBV19 | TRBJ2-3 | CASSIRSTDTQYF | GILGFVFTL | YES | NO | YES |
| 759 | TRAV30 | TRAJ42 | CGTVLEGGSQGNLIF | TRBV19 | TRBJ2-3 | CASSQRSTDTQYF | GILGFVFTL | YES | NO | YES |
| 760 | TRAV30 | TRAJ42 | CGTVFEGSQGNLIF | TRBV19 | TRBJ2-3 | CASSGRSTDTQYF | GILGFVFTL | YES | YES | NO |
| 761 | TRAV30 | TRAJ42 | CGTAMGGGSQGNLIF | TRBV19 | TRBJ2-3 | CASSMRSTDTQYF | GILGFVFTL | YES | YES | YES |
| 762 | TRAV30 | TRAJ42 | CGTDKGGGSQGNLIF | TRBV19 | TRBJ2-3 | CASSGRSTDTQYF | GILGFVFTL | YES | NO | YES |
| 763 | TRAV35 | TRAJ42 | CAGPAEVSGGSQGNLIF | TRBV19 | TRBJ2-3 | CASSIIALDTQYF | GILGFVFTL | YES | NO | YES |
| 767 | TRAV5 | TRAJ42 | CAEDYGGSQGNLIF | TRBV19 | TRBJ2-3 | CASSARSADTQYF | GILGFVFTL | YES | NO | YES |
| 768 | TRAV8-4 | TRAJ50 | CAPNKTSYDKVIF | TRBV19 | TRBJ2-3 | CASSRASTDTQYF | GILGFVFTL | YES | NO | YES |
| 770 | TRAV14DV4 | TRAJ39 | CAMREDNAGNMLTF | TRBV2 | TRBJ2-3 | CASSEKDRTVGTDTQYF | GILGFVFTL | YES | NO | NO |
| 771 | TRAV8-6 | TRAJ49 | CAVIHNTGNQFYF | TRBV2 | TRBJ2-3 | CASMGLTLRDTQYF | ELAGIGILTV | YES | YES | YES |
| 775 | TRAV27 | TRAJ42 | CAGGGSQGNLIF | TRBV20-1 | TRBJ2-3 | CSAPRGSGTIDTQYF | GILGFVFTL | YES | NO | YES |
| 777 | TRAV19 | TRAJ6 | CALSQVSGGSYIPTF | TRBV24-1 | TRBJ2-3 | CATSDYTSDTQYF | SLLMWITQV | YES | NO | YES |
| 778 | TRAV8-4 | TRAJ31 | CAVSENNNARLMF | TRBV24-1 | TRBJ2-3 | CATSEKLAVSTDTQYF | LLFGYPVYV | YES | YES | YES |
| 780 | TRAV12-2 | TRAJ31 | CAADNARLMF | TRBV27 | TRBJ2-3 | CASSIASLLAGGTDTQYF | ELAGIGILTV | YES | NO | NO |
| 783 | TRAV26-1 | TRAJ34 | CIVRVANTDKLIF | TRBV27 | TRBJ2-3 | CASSLGFTDTQYF | GILGFVFTL | YES | NO | YES |
| 784 | TRAV8-3 | TRAJ5 | CAVDPMDTGRRALTF | TRBV27 | TRBJ2-3 | CASSLGFTDTQYF | AVFDRKSDAK | YES | NO | NO |
| 784 | TRAV8-3 | TRAJ5 | CAVDPMDTGRRALTF | TRBV27 | TRBJ2-3 | CASSLGFTDTQYF | KLGGALQAK | YES | NO | NO |
| 784 | TRAV8-3 | TRAJ5 | CAVDPMDTGRRALTF | TRBV27 | TRBJ2-3 | CASSLGFTDTQYF | RLRAEAQVK | YES | NO | NO |
| 784 | TRAV8-3 | TRAJ5 | CAVDPMDTGRRALTF | TRBV27 | TRBJ2-3 | CASSLGFTDTQYF | IVTDFSVIK | YES | NO | NO |
| 787 | TRAV12-2 | TRAJ16 | CAASRGQKLLF | TRBV29-1 | TRBJ2-3 | CSVEDASGGTTDTQYF | ELAGIGILTV | NO | NO | NO |
| 791 | TRAV17 | TRAJ6 | CATEGASGGSYIPTF | TRBV30 | TRBJ2-3 | CAWVLGPAGDTQYF | KLGGALQAK | NO | NO | NO |
| 791 | TRAV17 | TRAJ6 | CATEGASGGSYIPTF | TRBV30 | TRBJ2-3 | CAWVLGPAGDTQYF | AVFDRKSDAK | NO | NO | NO |
| 791 | TRAV17 | TRAJ6 | CATEGASGGSYIPTF | TRBV30 | TRBJ2-3 | CAWVLGPAGDTQYF | GILGFVFTL | NO | NO | NO |
| 791 | TRAV17 | TRAJ6 | CATEGASGGSYIPTF | TRBV30 | TRBJ2-3 | CAWVLGPAGDTQYF | IVTDFSVIK | NO | NO | NO |
| 792 | TRAV29DV5 | TRAJ57 | CAAKGGSEKLVF | TRBV4-1 | TRBJ2-3 | CASSQRRTGPSSTDTQYF | RTLNAWVKV | YES | YES | YES |
| 795 | TRAV12-2 | TRAJ42 | CAVNYGGSQGNLIF | TRBV4-2 | TRBJ2-3 | CASSQDLLSGAQYF | ELAGIGILTV | YES | NO | NO |
| 796 | TRAV21 | TRAJ26 | CAVHGYGQNFVF | TRBV4-3 | TRBJ2-3 | CASSQEWLAVSTDTQYF | RLRAEAQVK | YES | NO | NO |
| 796 | TRAV21 | TRAJ26 | CAVHGYGQNFVF | TRBV4-3 | TRBJ2-3 | CASSQEWLAVSTDTQYF | SLFNTVATLY | YES | NO | NO |
| 797 | TRAV3 | TRAJ45 | CAVAKDLQNSGGGADGLTF | TRBV4-3 | TRBJ2-3 | CASSQEWLAVSTDTQYF | GILGFVFTL | YES | NO | YES |
| 798 | TRAV38-1 | TRAJ52 | CAFMINAGGTSYGKLTF | TRBV4-3 | TRBJ2-3 | CASSQEWLAVSTDTQYF | GILGFVFTL | YES | NO | YES |
| 799 | TRAV5 | TRAJ37 | CAEGGNTGKLIF | TRBV5-1 | TRBJ2-3 | CASSPGGLAGADTQYF | AVFDRKSDAK | YES | NO | NO |
| 801 | TRAV12-2 | TRAJ45 | CAVAVGVSGGGADGLTF | TRBV5-5 | TRBJ2-3 | CASSGIWDFTDTQYF | GILGFVFTL | YES | NO | YES |
| 802 | TRAV1-1 | TRAJ9 | CAVRPRGGFKTIF | TRBV6-1 | TRBJ2-3 | CASSITEGEDSVSTDTQYF | GILGFVFTL | YES | NO | NO |
| 803 | TRAV12-2 | TRAJ43 | CAVNTGDMRF | TRBV6-1 | TRBJ2-3 | CASSPGQGLDTQYF | ELAGIGILTV | YES | NO | YES |
| 804 | TRAV1-2 | TRAJ12 | CAVMDSSYKLIF | TRBV6-4 | TRBJ2-3 | CASSDAGGHTDTQYF | CLLWSFQTSA | YES | NO | YES |
| 806 | TRAV12-2 | TRAJ45 | CAVDSGGGADGLTF | TRBV6-5 | TRBJ2-3 | CASSSPANEGTDTQYF | ELAGIGILTV | NO | NO | NO |
| 808 | TRAV21 | TRAJ50 | CAVDLMKTSYDKVIF | TRBV6-6 | TRBJ2-3 | CASRTGLASTDTQYF | IVTDFSVIK | NO | NO | NO |
| 808 | TRAV21 | TRAJ50 | CAVDLMKTSYDKVIF | TRBV6-6 | TRBJ2-3 | CASRTGLASTDTQYF | KLGGALQAK | NO | NO | NO |
| 808 | TRAV21 | TRAJ50 | CAVDLMKTSYDKVIF | TRBV6-6 | TRBJ2-3 | CASRTGLASTDTQYF | AVFDRKSDAK | NO | NO | NO |
| 808 | TRAV21 | TRAJ50 | CAVDLMKTSYDKVIF | TRBV6-6 | TRBJ2-3 | CASRTGLASTDTQYF | FLASKIGRLV | NO | NO | NO |
| 808 | TRAV21 | TRAJ50 | CAVDLMKTSYDKVIF | TRBV6-6 | TRBJ2-3 | CASRTGLASTDTQYF | SLFNTVATLY | NO | NO | NO |
| 811 | TRAV38-2DV8 | TRAJ28 | CAYRSLSGAGSYQLTF | TRBV6-6 | TRBJ2-3 | CASSYPPSGVSTDTQYF | GILGFVFTL | NO | NO | NO |
| 816 | TRAV12-2 | TRAJ30 | CAVNGGDDKIIF | TRBV7-6 | TRBJ2-3 | CASSLAGGTDTQYF | KTWGQYWQV | YES | NO | YES |
| 822 | TRAV10 | TRAJ8 | CVVRSTGFQKLVF | TRBV9 | TRBJ2-3 | CASSVEGVGTSTDTQYF | AVFDRKSDAK | YES | NO | NO |
| 822 | TRAV10 | TRAJ8 | CVVRSTGFQKLVF | TRBV9 | TRBJ2-3 | CASSVEGVGTSTDTQYF | KLGGALQAK | YES | NO | NO |
| 823 | TRAV12-2 | TRAJ29 | CAPGGSGNTPLVF | TRBV9 | TRBJ2-3 | CASSEGGGALSTDTQYF | ELAGIGILTV | YES | NO | YES |
| 824 | TRAV12-2 | TRAJ45 | CAVLEGGGADGLTF | TRBV9 | TRBJ2-3 | CASSPGLAGGTQYF | ELAGIGILTV | YES | YES | YES |
| 825 | TRAV12-2 | TRAJ6 | CAGGGSYIPTF | TRBV9 | TRBJ2-3 | CASSSTDTQYF | ELAGIGILTV | YES | YES | YES |
| 828 | TRAV3 | TRAJ34 | CAVRDPHQGTDKLIF | TRBV9 | TRBJ2-3 | CASSVDGDTSTDTQYF | GILGFVFTL | NO | NO | NO |
| 829 | TRAV8-2 | TRAJ6 | CVVISSGGSYIPTF | TRBV9 | TRBJ2-3 | CASSPDQETSYTDTQYF | GILGFVFTL | YES | NO | YES |
| 830 | TRAV5 | TRAJ34 | CAESKPDKLIF | TRBV14 | TRBJ2-4 | CASSQSPGGIQYF | GLCTLVAML | NO | NO | NO |
| 831 | TRAV21 | TRAJ20 | CAVPLSNDYKLSF | TRBV19 | TRBJ2-4 | CASSRQGENIQYF | ELAGIGILTV | YES | NO | YES |
| 844 | TRAV12-2 | TRAJ5 | CAVDTGRRALTF | TRBV6-5 | TRBJ2-4 | CASTTSELAKNIQYF | ELAGIGILTV | YES | NO | YES |
| 846 | TRAV19 | TRAJ45 | CALLVAGGGADGLTF | TRBV10-2 | TRBJ2-5 | CASSERYEETQYF | LLDFVRFMGV | YES | NO | YES |
| 851 | TRAV38-2DV8 | TRAJ57 | CAYRSAESGSEKLVF | TRBV13 | TRBJ2-5 | CASSEGGVETQYF | KLGGALQAK | YES | NO | NO |
| 851 | TRAV38-2DV8 | TRAJ57 | CAYRSAESGSEKLVF | TRBV13 | TRBJ2-5 | CASSEGGVETQYF | GILGFVFTL | YES | NO | NO |
| 851 | TRAV38-2DV8 | TRAJ57 | CAYRSAESGSEKLVF | TRBV13 | TRBJ2-5 | CASSEGGVETQYF | AVFDRKSDAK | YES | NO | NO |
| 852 | TRAV27 | TRAJ42 | CAGAGSQGNLIF | TRBV14 | TRBJ2-5 | CASSQDRVRETQYF | GILGFVFTL | NO | NO | NO |
| 853 | TRAV5 | TRAJ36 | CAEAGANNLFF | TRBV14 | TRBJ2-5 | CASSQSPGGTQYF | GLCTLVAML | NO | NO | NO |
| 854 | TRAV5 | TRAJ4 | CAELKEKLIF | TRBV14 | TRBJ2-5 | CASSQSPGGTQYF | GLCTLVAML | NO | NO | NO |
| 857 | TRAV12-2 | TRAJ42 | CAVNIGGGSQGNLIF | TRBV19 | TRBJ2-5 | CASSIRSGETQYF | GILGFVFTL | YES | NO | YES |
| 858 | TRAV12-2 | TRAJ42 | CAGGLGGGSQGNLIF | TRBV19 | TRBJ2-5 | CASSIRASETQYF | GILGFVFTL | YES | NO | YES |
| 860 | TRAV12-3 | TRAJ42 | CAMSGDGGSQGNLIF | TRBV19 | TRBJ2-5 | CASSVRAGETQYF | GILGFVFTL | YES | NO | NO |
| 862 | TRAV13-1 | TRAJ23 | CAANNQGGKLIF | TRBV19 | TRBJ2-5 | CASSGRSGETQYF | GILGFVFTL | YES | NO | YES |
| 864 | TRAV13-1 | TRAJ26 | CAPEGGQNFVF | TRBV19 | TRBJ2-5 | CASSIRSGETQYF | GILGFVFTL | YES | NO | YES |
| 865 | TRAV13-1 | TRAJ37 | CAASEGSSNTGKLIF | TRBV19 | TRBJ2-5 | CASSIRSGETQYF | GILGFVFTL | YES | NO | YES |
| 866 | TRAV13-1 | TRAJ37 | CAAKSNTGKLIF | TRBV19 | TRBJ2-5 | CASSIRSSETQYF | GILGFVFTL | YES | NO | YES |
| 867 | TRAV13-1 | TRAJ37 | CAAPYSSNTGKLIF | TRBV19 | TRBJ2-5 | CASSPRAGWETQYF | GILGFVFTL | YES | NO | YES |
| 868 | TRAV13-1 | TRAJ37 | CAAISNTGKLIF | TRBV19 | TRBJ2-5 | CASSIRSGETQYF | GILGFVFTL | YES | NO | YES |
| 870 | TRAV13-1 | TRAJ42 | CAASGGGSQGNLIF | TRBV19 | TRBJ2-5 | CASSIRSQETQYF | GILGFVFTL | YES | NO | NO |
| 871 | TRAV13-1 | TRAJ42 | CAFGGSQGNLIF | TRBV19 | TRBJ2-5 | CASSIRSSETQYF | GILGFVFTL | YES | NO | YES |
| 872 | TRAV13-1 | TRAJ42 | CAASGGGSQGNLIF | TRBV19 | TRBJ2-5 | CASSGRSQETQYF | GILGFVFTL | YES | NO | YES |
| 873 | TRAV13-1 | TRAJ42 | CAEQGSQGNLIF | TRBV19 | TRBJ2-5 | CASSGRSGETQYF | GILGFVFTL | YES | YES | YES |
| 875 | TRAV13-1 | TRAJ42 | CAYGGSQRNLIF | TRBV19 | TRBJ2-5 | CASSFRSSETQYF | GILGFVFTL | YES | NO | YES |
| 876 | TRAV13-1 | TRAJ42 | CAASGSQGNLIF | TRBV19 | TRBJ2-5 | CASSIRSSETQYF | GILGFVFTL | YES | NO | YES |
| 877 | TRAV13-1 | TRAJ42 | CAGQGSQGNLIF | TRBV19 | TRBJ2-5 | CASSGRSGETQYF | GILGFVFTL | YES | NO | YES |
| 878 | TRAV13-1 | TRAJ42 | CALGGSQGNLIF | TRBV19 | TRBJ2-5 | CASSIRSSETQYF | GILGFVFTL | YES | NO | YES |
| 879 | TRAV13-2 | TRAJ42 | CAENIGGGSQGNLIF | TRBV19 | TRBJ2-5 | CASSFRSGETQYF | GILGFVFTL | YES | NO | YES |
| 880 | TRAV16 | TRAJ54 | CALNPLIQGAQKLVF | TRBV19 | TRBJ2-5 | CASSLRASGAQYF | GILGFVFTL | NO | NO | NO |
| 886 | TRAV29DV5 | TRAJ42 | CAARGGGGSQGNLIF | TRBV19 | TRBJ2-5 | CASSIRSGETQYF | GILGFVFTL | YES | NO | YES |
| 887 | TRAV6 | TRAJ42 | CALDMGGGSQGNLIF | TRBV19 | TRBJ2-5 | CASSSRAGFAQYF | GILGFVFTL | YES | NO | YES |
| 889 | TRAV8-3 | TRAJ42 | CAVGGGGGSQGNLIF | TRBV19 | TRBJ2-5 | CASSFRSSETQYF | GILGFVFTL | YES | NO | YES |
| 895 | TRAV8-3 | TRAJ20 | CAVGGLNDYKLSF | TRBV20-1 | TRBJ2-5 | CSARDDSGQGFGFLETQYF | GLCTLVAML | YES | YES | YES |
| 896 | TRAV12-2 | TRAJ43 | CADDMRF | TRBV27 | TRBJ2-5 | CASSLGYQGRETQYF | ELAGIGILTV | NO | NO | NO |
| 898 | TRAV12-2 | TRAJ8 | CAVSTGFQKLVF | TRBV27 | TRBJ2-5 | CASSLGETQYF | ELAGIGILTV | YES | YES | YES |
| 900 | TRAV12-2 | TRAJ45 | CAVNGGGADGLTF | TRBV28 | TRBJ2-5 | CASSLPGLAGEETQYF | ELAGIGILTV | YES | NO | NO |
| 904 | TRAV29DV5 | TRAJ52 | CAASSGGTSYGKLTF | TRBV3-1 | TRBJ2-5 | CASSPLWVAGGRETQYF | KLGGALQAK | YES | NO | NO |
| 904 | TRAV29DV5 | TRAJ52 | CAASSGGTSYGKLTF | TRBV3-1 | TRBJ2-5 | CASSPLWVAGGRETQYF | GILGFVFTL | YES | NO | NO |
| 907 | TRAV30 | TRAJ20 | CGTEYPNDYKLSF | TRBV4-3 | TRBJ2-5 | CASSLPGTVQETQYF | GLCTLVAML | YES | NO | YES |

| | | | | | | | | | | |
|---|---|---|---|---|---|---|---|---|---|---|
| 910 | TRAV12-1 | TRAJ45 | CVVNREVGGGGADGLTF | TRBV6-3 | TRBJ2-5 | CASGARPQETQYF | MLDLQPETT | YES | NO | NO |
| 916 | TRAV13-2 | TRAJ12 | CAEGGMDSSYKLIF | TRBV7-8 | TRBJ2-5 | CASSPLPRLAGEETQYF | SLFNTVATLY | YES | YES | YES |
| 918 | TRAV38-1 | TRAJ40 | CASMTSGTYKYIF | TRBV7-9 | TRBJ2-5 | CASSLEIAQETQYF | KTWGQYWQV | YES | NO | YES |
| 921 | TRAV19 | TRAJ29 | CALSESNSGNTPLVF | TRBV9 | TRBJ2-5 | CASSVDPAQETQYF | GILGFVFTL | YES | NO | YES |
| 922 | TRAV12-2 | TRAJ31 | CAVNNARLMF | TRBV10-3 | TRBJ2-6 | CAISEPGLLSGANVLTF | ELAGIGILTV | YES | YES | YES |
| 924 | TRAV3 | TRAJ45 | CAVAKDLQNSGGGADGLTF | TRBV15 | TRBJ2-6 | CATSYEDNSGANVLTF | GILGFVFTL | NO | NO | YES |
| 925 | TRAV13-1 | TRAJ37 | CATVPSNTGKLIF | TRBV19 | TRBJ2-6 | CASSMRSGANVLTF | GILGFVFTL | YES | NO | YES |
| 927 | TRAV10 | TRAJ50 | CVVSAGLGKTSYDKVIF | TRBV20-1 | TRBJ2-6 | CRYKGQGVSGANVLTF | GILGFVFTL | YES | NO | YES |
| 928 | TRAV20 | TRAJ32 | CAVHLIWEGGGATNKLIF | TRBV24-1 | TRBJ2-6 | CATSDFTDSGANVLTF | GILGFVFTL | YES | NO | YES |
| 935 | TRAV13-1 | TRAJ11 | CAAKEGYSTLTF | TRBV7-9 | TRBJ2-6 | CASSPVTGGGSGANVLTF | GILGFVFTL | YES | NO | YES |
| 943 | TRAV17 | TRAJ33 | CATDWDGYQLIW | TRBV11-2 | TRBJ2-7 | CASSSQVNYEQYF | FLASKIGRLV | YES | NO | YES |
| 944 | TRAV9-2 | TRAJ37 | CALDNTGKLIF | TRBV11-2 | TRBJ2-7 | CASSWNRASSYEQYF | GLCTLVAML | NO | NO | NO |
| 945 | TRAV12-2 | TRAJ4 | CAVRGFSGGYNKLIF | TRBV12-3 | TRBJ2-7 | CASSSPRGLEYF | MLDLQPETT | YES | NO | NO |
| 946 | TRAV13-1 | TRAJ50 | CAASETSYDKVIF | TRBV12-3 | TRBJ2-7 | CASSSLDVAGYEQYF | RAKFKQLL | YES | NO | YES |
| 947 | TRAV8-2 | TRAJ11 | CVVSDLRYSTLTF | TRBV12-4 | TRBJ2-7 | CASSPGTGLSYEQYF | FLASKIGRLV | YES | NO | YES |
| 949 | TRAV26-2 | TRAJ49 | CILRDFAGNQFYF | TRBV13 | TRBJ2-7 | CASSLGGSYEQYV | FLASKIGRLV | NO | NO | NO |
| 950 | TRAV5 | TRAJ29 | CAEMGGNTPLVF | TRBV13 | TRBJ2-7 | CASSLTGESYEQYF | LLDFVRFMGV | YES | NO | NO |
| 952 | TRAV12-2 | TRAJ54 | CAVVQGAQKLVF | TRBV14 | TRBJ2-7 | CASSYLTGTGAYEQYF | ELAGIGILTV | NO | NO | YES |
| 953 | TRAV5 | TRAJ45 | CAESTSGGGADGLTF | TRBV14 | TRBJ2-7 | CASSQSPGGMQYF | GLCTLVAML | NO | NO | NO |
| 954 | TRAV12-2 | TRAJ31 | CAVKNARLMF | TRBV15 | TRBJ2-7 | CATSRDPLNEGEQYF | ELAGIGILTV | YES | YES | YES |
| 957 | TRAV1-2 | TRAJ33 | CAVLDSNYQLIW | TRBV19 | TRBJ2-7 | CASSARSSTEQYF | GILGFVFTL | YES | NO | YES |
| 958 | TRAV1-2 | TRAJ33 | CAVADSNYQLIW | TRBV19 | TRBJ2-7 | CASSGRSGVEQYF | GILGFVFTL | YES | NO | YES |
| 961 | TRAV12-1 | TRAJ26 | CVVKAYGQNFVF | TRBV19 | TRBJ2-7 | CASSSRASYEQYF | GILGFVFTL | YES | NO | YES |
| 964 | TRAV12-1 | TRAJ40 | CVVNRWASGTYKYIF | TRBV19 | TRBJ2-7 | CASSARSSYEQYF | GILGFVFTL | YES | NO | YES |
| 965 | TRAV12-1 | TRAJ42 | CVVNIGGGSQGNLIF | TRBV19 | TRBJ2-7 | CASSIRSSYEQYF | GILGFVFTL | YES | NO | YES |
| 966 | TRAV12-1 | TRAJ42 | CVVNELDGGSQGNLIF | TRBV19 | TRBJ2-7 | CASSSRSSYEQYF | GILGFVFTL | YES | YES | YES |
| 967 | TRAV12-1 | TRAJ42 | CVVNIGGGSQGNLIF | TRBV19 | TRBJ2-7 | CASSLRSSYEQYF | GILGFVFTL | YES | NO | YES |
| 969 | TRAV12-1 | TRAJ50 | CVVTETSYDKVIF | TRBV19 | TRBJ2-7 | CASSARSSYEQYF | GILGFVFTL | YES | YES | YES |
| 971 | TRAV12-2 | TRAJ10 | CAPIGGGNKLTF | TRBV19 | TRBJ2-7 | CASSARSSYEQYF | GILGFVFTL | YES | YES | YES |
| 972 | TRAV12-2 | TRAJ10 | CAVPLTGGGNKLTF | TRBV19 | TRBJ2-7 | CASSIRSSYEQYF | GILGFVFTL | YES | NO | YES |
| 973 | TRAV12-2 | TRAJ12 | CASVGMDSSYKLIF | TRBV19 | TRBJ2-7 | CASSARSSYEQYF | GILGFVFTL | YES | NO | YES |
| 974 | TRAV12-2 | TRAJ21 | CAVNTPPFYF | TRBV19 | TRBJ2-7 | CASSIRSSYEQYF | ELAGIGILTV | YES | NO | NO |
| 975 | TRAV12-2 | TRAJ23 | CAAPSGNNQGGKLIF | TRBV19 | TRBJ2-7 | CASSVRSSYEQYF | GILGFVFTL | YES | NO | YES |
| 976 | TRAV12-2 | TRAJ3 | CAVDASKIIF | TRBV19 | TRBJ2-7 | CASRDLSSYEQYF | SLFNTVATLY | YES | YES | YES |
| 977 | TRAV12-2 | TRAJ37 | CAASQGSGNTGKLIF | TRBV19 | TRBJ2-7 | CASSQRSSYEQYF | GILGFVFTL | YES | NO | YES |
| 982 | TRAV12-2 | TRAJ42 | CAVNIGGGSQGNLIF | TRBV19 | TRBJ2-7 | CASSFRSSYEQYF | GILGFVFTL | YES | YES | YES |
| 983 | TRAV12-2 | TRAJ42 | CALNYGGSQGNLIF | TRBV19 | TRBJ2-7 | CASSVRSSYEQYF | GILGFVFTL | YES | NO | YES |
| 986 | TRAV12-2 | TRAJ42 | CAVNIGGGSQGNLIF | TRBV19 | TRBJ2-7 | CASSIRSGHEQYF | GILGFVFTL | YES | YES | YES |
| 987 | TRAV12-2 | TRAJ42 | CAVSGDGGSQGNLIF | TRBV19 | TRBJ2-7 | CASSTRSAYEQYF | GILGFVFTL | YES | NO | YES |
| 988 | TRAV12-2 | TRAJ42 | CALSPGGGSQGNLIF | TRBV19 | TRBJ2-7 | CASSIRSSYEQYF | GILGFVFTL | YES | NO | YES |
| 989 | TRAV12-2 | TRAJ42 | CAVNWGGGSQGNLIF | TRBV19 | TRBJ2-7 | CASSIRSSHEQYF | GILGFVFTL | YES | NO | YES |
| 990 | TRAV12-2 | TRAJ42 | CAVQGSQGNLIF | TRBV19 | TRBJ2-7 | CASSIRSAYEQYF | GILGFVFTL | YES | NO | NO |
| 991 | TRAV12-2 | TRAJ42 | CAVSSGGGSQGNLIF | TRBV19 | TRBJ2-7 | CASSIRSAYEQYF | GILGFVFTL | YES | NO | YES |
| 992 | TRAV12-2 | TRAJ43 | CAVGGDMRF | TRBV19 | TRBJ2-7 | CASSVRSSYEQYF | GILGFVFTL | NO | NO | NO |
| 994 | TRAV12-2 | TRAJ49 | CAATNQFYF | TRBV19 | TRBJ2-7 | CASSIRSSYEQYF | GILGFVFTL | NO | NO | YES |
| 1000 | TRAV12-3 | TRAJ42 | CAMSGDGGSQGNLIF | TRBV19 | TRBJ2-7 | CASSYRSSYEQYF | GILGFVFTL | YES | NO | NO |
| 1001 | TRAV12-3 | TRAJ42 | CAMSGDGGSQGNLIF | TRBV19 | TRBJ2-7 | CASSIRAAYEQYF | GILGFVFTL | YES | NO | YES |
| 1006 | TRAV13-1 | TRAJ42 | CAASMGGGSQGNLIF | TRBV19 | TRBJ2-7 | CASSIRSSYEQYF | GILGFVFTL | YES | NO | YES |
| 1007 | TRAV13-1 | TRAJ42 | CAGQGSQGNLIF | TRBV19 | TRBJ2-7 | CASSIRSSYEQYF | GILGFVFTL | YES | NO | YES |
| 1013 | TRAV13-2 | TRAJ30 | CADMNRDDKIIF | TRBV19 | TRBJ2-7 | CASSIRAAYEQYF | GILGFVFTL | YES | NO | YES |
| 1014 | TRAV13-2 | TRAJ42 | CAENMGGGSQGNLIF | TRBV19 | TRBJ2-7 | CASSIRSGNEQYF | GILGFVFTL | YES | YES | YES |
| 1016 | TRAV13-2 | TRAJ42 | CAENLAGGSQGNLIF | TRBV19 | TRBJ2-7 | CASSIRASYEQYF | GILGFVFTL | YES | NO | YES |
| 1017 | TRAV13-2 | TRAJ42 | CAENEGGGSQGNLIF | TRBV19 | TRBJ2-7 | CASSLYASYEQYF | GILGFVFTL | YES | YES | YES |
| 1018 | TRAV13-2 | TRAJ42 | CAEGGDGGSQGNLIF | TRBV19 | TRBJ2-7 | CASSTRSTYEQYF | GILGFVFTL | YES | YES | YES |
| 1022 | TRAV14DV4 | TRAJ41 | CAGGYALNF | TRBV19 | TRBJ2-7 | CASSTRSSYEQYF | GILGFVFTL | YES | NO | NO |
| 1023 | TRAV14DV4 | TRAJ42 | CASAGSQGNLIF | TRBV19 | TRBJ2-7 | CASSIRSSYEQYF | GILGFVFTL | YES | NO | NO |
| 1024 | TRAV14DV4 | TRAJ42 | CAMREGGGSQGNLIF | TRBV19 | TRBJ2-7 | CASSMRSGVEQYF | GILGFVFTL | YES | NO | YES |
| 1028 | TRAV16 | TRAJ38 | CALDRAGNNRKLIW | TRBV19 | TRBJ2-7 | CASSIRSSYEQYF | GILGFVFTL | NO | NO | NO |
| 1031 | TRAV17 | TRAJ28 | CATEGAYSGAGSYQLTF | TRBV19 | TRBJ2-7 | CASSIRSSYEQYF | GILGFVFTL | YES | NO | YES |
| 1035 | TRAV17 | TRAJ8 | CATANTGFQKLVF | TRBV19 | TRBJ2-7 | CASSIRSSYEQYF | GILGFVFTL | YES | NO | YES |
| 1036 | TRAV19 | TRAJ29 | CALSEPYSGNTPLVF | TRBV19 | TRBJ2-7 | CASSIRSSYEQYF | GILGFVFTL | YES | NO | YES |
| 1037 | TRAV19 | TRAJ33 | CALSESWTAMDSNYQLIW | TRBV19 | TRBJ2-7 | CASSTHPGNEQYF | GILGFVFTL | YES | NO | YES |
| 1043 | TRAV20 | TRAJ26 | CAVPDNYGQNFVF | TRBV19 | TRBJ2-7 | CASSARSSYEQYF | GILGFVFTL | YES | NO | YES |
| 1050 | TRAV21 | TRAJ34 | CAVDNTDKLIF | TRBV19 | TRBJ2-7 | CASSSRSSYEQYF | GILGFVFTL | YES | NO | NO |
| 1052 | TRAV21 | TRAJ39 | CAACNNAGNMLTF | TRBV19 | TRBJ2-7 | CASSIAPGSEQYF | LLDFVRFMGV | YES | NO | YES |
| 1053 | TRAV21 | TRAJ52 | CAVRIGGGTSYGKLTF | TRBV19 | TRBJ2-7 | CASSSRSGVEQYF | GILGFVFTL | YES | NO | YES |
| 1055 | TRAV22 | TRAJ37 | CAGSPSNTGKLIF | TRBV19 | TRBJ2-7 | CASSARSSYEQYF | GLCTLVAML | YES | NO | YES |
| 1072 | TRAV26-2 | TRAJ42 | CIPARGEYGGSQGNLIF | TRBV19 | TRBJ2-7 | CASSIRSSYEQYF | GILGFVFTL | YES | NO | NO |
| 1073 | TRAV26-2 | TRAJ43 | CILREFYNDMRF | TRBV19 | TRBJ2-7 | CASSARSSYEQYF | GILGFVFTL | YES | NO | YES |
| 1078 | TRAV27 | TRAJ20 | CAGAPSLFSNDYKLSF | TRBV19 | TRBJ2-7 | CASGLPYEQYF | GILGFVFTL | YES | NO | YES |
| 1079 | TRAV27 | TRAJ23 | CAGAHQGGKLIF | TRBV19 | TRBJ2-7 | CASSSRSAYEQYF | GILGFVFTL | YES | NO | YES |
| 1080 | TRAV27 | TRAJ23 | CAGDDQGGKLIF | TRBV19 | TRBJ2-7 | CASSIRSSYEQYF | GILGFVFTL | YES | YES | NO |
| 1081 | TRAV27 | TRAJ27 | CAGPPPNAGKSTF | TRBV19 | TRBJ2-7 | CASSMRSAYEQYF | GILGFVFTL | YES | NO | YES |
| 1082 | TRAV27 | TRAJ27 | CAGSNTNAGKSTF | TRBV19 | TRBJ2-7 | CASSIRSEYEQYF | GILGFVFTL | YES | NO | YES |
| 1083 | TRAV27 | TRAJ27 | CAGANTNAGKSTF | TRBV19 | TRBJ2-7 | CASSIRSEYEQYF | GILGFVFTL | YES | YES | YES |
| 1084 | TRAV27 | TRAJ30 | CAGAQITNRDDKIIF | TRBV19 | TRBJ2-7 | CASSIRSFYEQYF | GILGFVFTL | YES | NO | YES |
| 1085 | TRAV27 | TRAJ31 | CAGASSQGRLMF | TRBV19 | TRBJ2-7 | CASSIRSSYEQYF | GILGFVFTL | YES | NO | YES |
| 1093 | TRAV27 | TRAJ37 | CAGAGSSNTGKLIF | TRBV19 | TRBJ2-7 | CASSTRSGWEQYF | GILGFVFTL | YES | NO | YES |
| 1094 | TRAV27 | TRAJ37 | CAGAPGSSNTGKLIF | TRBV19 | TRBJ2-7 | CASSTRSAYEQYF | GILGFVFTL | YES | NO | YES |
| 1095 | TRAV27 | TRAJ37 | CAGGGSSNTGKLIF | TRBV19 | TRBJ2-7 | CASSIRSSYEQYF | GILGFVFTL | YES | NO | YES |
| 1096 | TRAV27 | TRAJ37 | CAGAVGSSNTGKLIF | TRBV19 | TRBJ2-7 | CASSIRSAYEQYF | GILGFVFTL | YES | NO | YES |
| 1097 | TRAV27 | TRAJ37 | CAGAVGSSNTGKLIF | TRBV19 | TRBJ2-7 | CASSIRSSYEQYF | GILGFVFTL | YES | NO | YES |
| 1098 | TRAV27 | TRAJ37 | CAGADGSSNTGKLIF | TRBV19 | TRBJ2-7 | CASSIRSAYEQYF | GILGFVFTL | YES | NO | YES |
| 1099 | TRAV27 | TRAJ37 | CAGASGSSNTGKLIF | TRBV19 | TRBJ2-7 | CASSTRSAYEQYF | GILGFVFTL | YES | NO | YES |
| 1100 | TRAV27 | TRAJ37 | CAGAEGSSNTGKLIF | TRBV19 | TRBJ2-7 | CASSIRSAYEQYF | GILGFVFTL | YES | NO | NO |
| 1101 | TRAV27 | TRAJ37 | CAGAGSSNTGKLIF | TRBV19 | TRBJ2-7 | CASSIRSAYEQYF | GILGFVFTL | YES | NO | YES |
| 1102 | TRAV27 | TRAJ37 | CAGANGSSNTGKLIF | TRBV19 | TRBJ2-7 | CASSIRSAYEQYF | GILGFVFTL | YES | NO | YES |
| 1103 | TRAV27 | TRAJ37 | CAGGGSGNTGKLIF | TRBV19 | TRBJ2-7 | CASSIRSAYEQYF | GILGFVFTL | YES | NO | YES |
| 1104 | TRAV27 | TRAJ37 | CAGAAGSSNTGKLIF | TRBV19 | TRBJ2-7 | CASSTRSAYEQYF | GILGFVFTL | YES | NO | YES |
| 1105 | TRAV27 | TRAJ37 | CAGAAGSSNTGKLIF | TRBV19 | TRBJ2-7 | CASSIRSSYEQYF | GILGFVFTL | YES | NO | YES |
| 1106 | TRAV27 | TRAJ37 | CAGGGSSNTGKLIF | TRBV19 | TRBJ2-7 | CASSVRSAYEQYF | GILGFVFTL | YES | NO | YES |
| 1108 | TRAV27 | TRAJ37 | CAGADGSGNTGKLIF | TRBV19 | TRBJ2-7 | CASSVRSAYEQYF | GILGFVFTL | YES | NO | YES |
| 1109 | TRAV27 | TRAJ37 | CAGAWNTGKLIF | TRBV19 | TRBJ2-7 | CASSIRSSYEQYF | GILGFVFTL | YES | NO | YES |
| 1110 | TRAV27 | TRAJ37 | CAGALGSSNTGKLIF | TRBV19 | TRBJ2-7 | CASSIRSAYEQYF | GILGFVFTL | YES | NO | NO |
| 1111 | TRAV27 | TRAJ37 | CAGALGSSNTGKLIF | TRBV19 | TRBJ2-7 | CASSTRSAYEQYF | GILGFVFTL | YES | NO | YES |
| 1112 | TRAV27 | TRAJ37 | CAGCHGSGNTGKLIF | TRBV19 | TRBJ2-7 | CASSSRSAYEQYF | GILGFVFTL | YES | NO | YES |
| 1113 | TRAV27 | TRAJ37 | CAGAHGSGNTGKLIF | TRBV19 | TRBJ2-7 | CASSIRSAYEQYF | GILGFVFTL | YES | NO | YES |
| 1114 | TRAV27 | TRAJ37 | CAGAFGSSNTGKLIF | TRBV19 | TRBJ2-7 | CASSIRSSYEQYF | GILGFVFTL | YES | NO | YES |
| 1115 | TRAV27 | TRAJ37 | CAGAGNIGKLIF | TRBV19 | TRBJ2-7 | CASSIRSSYEQYF | GILGFVFTL | YES | NO | YES |
| 1116 | TRAV27 | TRAJ37 | CAGARGSSNTGKLIF | TRBV19 | TRBJ2-7 | CASSIRSAYEQYF | GILGFVFTL | YES | NO | YES |
| 1117 | TRAV27 | TRAJ37 | CAGADGSSNTGKLIF | TRBV19 | TRBJ2-7 | CASSSRSAYEQYF | GILGFVFTL | YES | NO | YES |
| 1118 | TRAV27 | TRAJ37 | CAGGGSGNTGKLIF | TRBV19 | TRBJ2-7 | CASSIRSSYEQYF | GILGFVFTL | YES | NO | YES |
| 1123 | TRAV27 | TRAJ42 | CAAGGSQGNLIF | TRBV19 | TRBJ2-7 | CASSIRSAYEQYF | GILGFVFTL | YES | YES | YES |
| 1124 | TRAV27 | TRAJ42 | CAAGGSQGNLIF | TRBV19 | TRBJ2-7 | CASSIRSSYEQYF | GILGFVFTL | YES | NO | YES |

| | | | | | | | | | | |
|---|---|---|---|---|---|---|---|---|---|---|
| 1125 | TRAV27 | TRAJ42 | CAGPGSQGNLIF | TRBV19 | TRBJ2-7 | CASSIRSSYEQYF | GILGFVFTL | YES | NO | YES |
| 1126 | TRAV27 | TRAJ42 | CAGGGSQGNLIF | TRBV19 | TRBJ2-7 | CASSIRSAYEQYF | GILGFVFTL | YES | NO | YES |
| 1127 | TRAV27 | TRAJ42 | CAGGGSQGNLIF | TRBV19 | TRBJ2-7 | CASSTRSAYEQYF | GILGFVFTL | YES | NO | YES |
| 1128 | TRAV27 | TRAJ42 | CAGPGDGGSQGNLIF | TRBV19 | TRBJ2-7 | CASSIRSSYEQYF | GILGFVFTL | YES | NO | YES |
| 1129 | TRAV27 | TRAJ42 | CASGGSQGNLIF | TRBV19 | TRBJ2-7 | CASSVRSSYEQYF | GILGFVFTL | YES | NO | YES |
| 1130 | TRAV27 | TRAJ42 | CASGGSQGNLIF | TRBV19 | TRBJ2-7 | CASSIRSSYEQYF | GILGFVFTL | YES | NO | NO |
| 1131 | TRAV27 | TRAJ42 | CAGGGEGGSQGNLIF | TRBV19 | TRBJ2-7 | CASSTRSSYEQYF | GILGFVFTL | YES | NO | YES |
| 1132 | TRAV27 | TRAJ42 | CAGADGGGSQGNLIF | TRBV19 | TRBJ2-7 | CASSIRSGYEQYF | GILGFVFTL | YES | YES | YES |
| 1133 | TRAV27 | TRAJ42 | CAGAGSQGNLIF | TRBV19 | TRBJ2-7 | CASSTRSSYEQYF | GILGFVFTL | YES | NO | YES |
| 1134 | TRAV27 | TRAJ42 | CAGAMGGGSQGNLIF | TRBV19 | TRBJ2-7 | CASSIRASYEQYF | GILGFVFTL | YES | NO | YES |
| 1135 | TRAV27 | TRAJ42 | CAGGGSQGNLIF | TRBV19 | TRBJ2-7 | CASSDRSGTEQYF | GILGFVFTL | YES | YES | YES |
| 1136 | TRAV27 | TRAJ42 | CAGAKGGGSQGNLIF | TRBV19 | TRBJ2-7 | CASSIRSSYEQYF | GILGFVFTL | YES | NO | YES |
| 1147 | TRAV27 | TRAJ42 | CAGSGSQGNLIF | TRBV19 | TRBJ2-7 | CASSIRSSYEQYF | GILGFVFTL | YES | NO | YES |
| 1148 | TRAV27 | TRAJ42 | CAGEGGGSQGNLIF | TRBV19 | TRBJ2-7 | CASSIRSSYEQYF | GILGFVFTL | YES | NO | YES |
| 1149 | TRAV27 | TRAJ42 | CAGVEYGGSQGNLIF | TRBV19 | TRBJ2-7 | CASSVRSTYEQYF | GILGFVFTL | YES | NO | YES |
| 1150 | TRAV27 | TRAJ42 | CAAGGSQGNLIF | TRBV19 | TRBJ2-7 | CASSVRSSYEQYF | GILGFVFTL | YES | YES | NO |
| 1151 | TRAV27 | TRAJ42 | CAANYGGSQGNLIF | TRBV19 | TRBJ2-7 | CASSIRSSYEQYF | GILGFVFTL | YES | NO | YES |
| 1152 | TRAV27 | TRAJ42 | CAQAGSQGNLIF | TRBV19 | TRBJ2-7 | CASSIRSSYEQYF | GILGFVFTL | YES | NO | YES |
| 1153 | TRAV27 | TRAJ42 | CAGGGDGGSQGNLIF | TRBV19 | TRBJ2-7 | CASSIRSGVEQYF | GILGFVFTL | YES | NO | YES |
| 1154 | TRAV27 | TRAJ42 | CAGAGSQGNLIF | TRBV19 | TRBJ2-7 | CASSSRSAYEQYF | GILGFVFTL | YES | NO | YES |
| 1155 | TRAV27 | TRAJ42 | CASNYGGSQGNLIF | TRBV19 | TRBJ2-7 | CASSMRSSYEQYF | GILGFVFTL | YES | NO | YES |
| 1156 | TRAV27 | TRAJ42 | CAGNYGGSQGNLIF | TRBV19 | TRBJ2-7 | CASSTRSSYEQYF | GILGFVFTL | YES | NO | YES |
| 1157 | TRAV27 | TRAJ42 | CATNYGGSQGNLIF | TRBV19 | TRBJ2-7 | CASSIRSSYEQYF | GILGFVFTL | YES | NO | YES |
| 1158 | TRAV27 | TRAJ42 | CAGGGSQGNLIF | TRBV19 | TRBJ2-7 | CASSIRAAYEQYF | GILGFVFTL | YES | NO | YES |
| 1159 | TRAV27 | TRAJ42 | CAGGYGGSQGNLIF | TRBV19 | TRBJ2-7 | CASSIRSSHEQYF | GILGFVFTL | YES | NO | YES |
| 1160 | TRAV27 | TRAJ42 | CAGAGPGSQGNLIF | TRBV19 | TRBJ2-7 | CASSIRSSYEQYF | GILGFVFTL | YES | NO | NO |
| 1161 | TRAV27 | TRAJ42 | CAGSGSQGNLIF | TRBV19 | TRBJ2-7 | CASSIRSAYEQYF | GILGFVFTL | YES | NO | YES |
| 1162 | TRAV27 | TRAJ42 | CAVGGSQGNLIF | TRBV19 | TRBJ2-7 | CASSIRSSYEQYF | GILGFVFTL | YES | NO | YES |
| 1163 | TRAV27 | TRAJ42 | CAGALGGGSQGNLIF | TRBV19 | TRBJ2-7 | CASSSRAAYEQYF | GILGFVFTL | YES | NO | YES |
| 1164 | TRAV27 | TRAJ42 | CAAPGSQGNLIF | TRBV19 | TRBJ2-7 | CASSVRSSYEQYF | GILGFVFTL | YES | NO | YES |
| 1165 | TRAV27 | TRAJ42 | CSYGGSQGNLIF | TRBV19 | TRBJ2-7 | CASSVRSSYEQYF | GILGFVFTL | YES | NO | NO |
| 1166 | TRAV29DV5 | TRAJ21 | CAPLYNFNKFYF | TRBV19 | TRBJ2-7 | CASSTRSAYEQYF | GILGFVFTL | YES | NO | YES |
| 1167 | TRAV29DV5 | TRAJ24 | CAASTDSWGKLQF | TRBV19 | TRBJ2-7 | CASSIRGAYEQYF | GILGFVFTL | YES | NO | YES |
| 1170 | TRAV29DV5 | TRAJ42 | CAAREGGGSQGNLIF | TRBV19 | TRBJ2-7 | CASSIRSSNEQYF | GILGFVFTL | YES | NO | NO |
| 1174 | TRAV3 | TRAJ30 | CAVRDGENRDDKIIF | TRBV19 | TRBJ2-7 | CASSARSSYEQYF | GILGFVFTL | NO | NO | NO |
| 1175 | TRAV3 | TRAJ36 | CAVRDGTGANNLFF | TRBV19 | TRBJ2-7 | CASSGTSVHEQYF | GILGFVFTL | NO | NO | YES |
| 1176 | TRAV3 | TRAJ37 | CAVRDSSSGNTGKLIF | TRBV19 | TRBJ2-7 | CASSIRSSYEQYF | GILGFVFTL | NO | NO | NO |
| 1177 | TRAV3 | TRAJ4 | CAVRDPGGYNKLIF | TRBV19 | TRBJ2-7 | CASSARSSTEQYF | GILGFVFTL | NO | NO | NO |
| 1178 | TRAV3 | TRAJ42 | CAVSGSQGNLIF | TRBV19 | TRBJ2-7 | CASSIRSSYEQYF | GILGFVFTL | NO | NO | YES |
| 1179 | TRAV3 | TRAJ42 | CAPSAGGGSQGNLIF | TRBV19 | TRBJ2-7 | CASSVRSSYEQYF | GILGFVFTL | NO | NO | YES |
| 1182 | TRAV3 | TRAJ45 | CAVAKDLQNSGGGADGLTF | TRBV19 | TRBJ2-7 | CASSARSSSEQYF | GILGFVFTL | NO | NO | NO |
| 1187 | TRAV35 | TRAJ42 | CAGQLGGGSQGNLIF | TRBV19 | TRBJ2-7 | CASSMRSSYEQYF | GILGFVFTL | YES | NO | YES |
| 1188 | TRAV35 | TRAJ42 | CAGQLGGGSQGNLIF | TRBV19 | TRBJ2-7 | CASSTRSSYEQYF | GILGFVFTL | YES | NO | YES |
| 1189 | TRAV35 | TRAJ42 | CAGILYGGSQGNLIF | TRBV19 | TRBJ2-7 | CASSLRSSYEQYF | GILGFVFTL | YES | NO | YES |
| 1190 | TRAV38-1 | TRAJ28 | CAFMKQEVPGAGSYQLTF | TRBV19 | TRBJ2-7 | CASSISGTSAYYEQYF | GLCTLVAML | YES | YES | NO |
| 1191 | TRAV38-1 | TRAJ42 | CAFSRGHYGGSQGNLIF | TRBV19 | TRBJ2-7 | CASSIRSSYEQYF | GILGFVFTL | YES | NO | YES |
| 1194 | TRAV38-1 | TRAJ52 | CAFMKDAGGTSYGKLTF | TRBV19 | TRBJ2-7 | CASSTRSAYEQYF | GILGFVFTL | YES | NO | YES |
| 1195 | TRAV38-1 | TRAJ52 | CAFMKGAGGTSYGKLTF | TRBV19 | TRBJ2-7 | CASSIRSSYEQYF | GILGFVFTL | YES | NO | YES |
| 1196 | TRAV38-1 | TRAJ52 | CAFMINAGGTSYGKLTF | TRBV19 | TRBJ2-7 | CASSIRSSYEQYF | GILGFVFTL | YES | NO | YES |
| 1200 | TRAV38-1 | TRAJ52 | CAFMKGAGGTSYGKLTF | TRBV19 | TRBJ2-7 | CASSTRSSYEQYF | GILGFVFTL | YES | NO | NO |
| 1201 | TRAV38-1 | TRAJ52 | CAFMIGAGGTSYGKLTF | TRBV19 | TRBJ2-7 | CASSIRSSYEQYF | GILGFVFTL | YES | NO | YES |
| 1203 | TRAV38-2DV8 | TRAJ52 | CAYMSNAGGTSYGKLTF | TRBV19 | TRBJ2-7 | CASSIRSSYEQYF | GILGFVFTL | YES | NO | YES |
| 1204 | TRAV38-2DV8 | TRAJ52 | CAYSPAAGGTSYGKLTF | TRBV19 | TRBJ2-7 | CASSIRSSYEQYF | GILGFVFTL | YES | NO | YES |
| 1205 | TRAV38-2DV8 | TRAJ52 | CAFMKGAGGTSYGKLTF | TRBV19 | TRBJ2-7 | CASSIRSSYEQYF | GILGFVFTL | YES | NO | YES |
| 1206 | TRAV38-2DV8 | TRAJ57 | CAYRSQQGGSEKLVF | TRBV19 | TRBJ2-7 | CASSTRSSYEQYF | GILGFVFTL | YES | NO | YES |
| 1208 | TRAV39 | TRAJ42 | CAVDENGGSQGNLIF | TRBV19 | TRBJ2-7 | CASSIRSSYEQYF | GILGFVFTL | YES | NO | YES |
| 1215 | TRAV5 | TRAJ42 | CAESIGGGSQGNLIF | TRBV19 | TRBJ2-7 | CASSTRSGYEQYF | GILGFVFTL | YES | NO | YES |
| 1217 | TRAV5 | TRAJ6 | CAEMEGGSYIPTF | TRBV19 | TRBJ2-7 | CASSARSSYEQYF | GILGFVFTL | YES | NO | YES |
| 1218 | TRAV6 | TRAJ15 | CALKAGTALIF | TRBV19 | TRBJ2-7 | CASSIRASYEQYF | GILGFVFTL | YES | NO | YES |
| 1219 | TRAV8-1 | TRAJ11 | CAVNSGYSTLTF | TRBV19 | TRBJ2-7 | CASSIRSSYEQYF | GILGFVFTL | YES | NO | YES |
| 1220 | TRAV8-1 | TRAJ23 | CAVNEVEGGKLIF | TRBV19 | TRBJ2-7 | CASSTRSSVEQYF | GILGFVFTL | YES | NO | NO |
| 1221 | TRAV8-1 | TRAJ33 | CAVSMDSNYQLIW | TRBV19 | TRBJ2-7 | CASSTQVGYEQYF | RMFPNAPYL | YES | NO | YES |
| 1222 | TRAV8-1 | TRAJ42 | CAVNEGGGSQGNLIF | TRBV19 | TRBJ2-7 | CASSTRAAYEQYF | GILGFVFTL | YES | NO | YES |
| 1223 | TRAV8-1 | TRAJ42 | CAVNLYGGSQGNLIF | TRBV19 | TRBJ2-7 | CASSIRSSYEQYF | GILGFVFTL | YES | NO | YES |
| 1224 | TRAV8-1 | TRAJ42 | CAVNAGGGSQGNLIF | TRBV19 | TRBJ2-7 | CASSTRSSYEQYF | GILGFVFTL | YES | NO | YES |
| 1225 | TRAV8-1 | TRAJ42 | CAGDYGGSQGNLIF | TRBV19 | TRBJ2-7 | CASSIRSSYEQYF | GILGFVFTL | YES | NO | YES |
| 1226 | TRAV8-1 | TRAJ42 | CAVNAGGGSQGNLIF | TRBV19 | TRBJ2-7 | CASSVRSSYEQYF | GILGFVFTL | YES | NO | YES |
| 1228 | TRAV8-1 | TRAJ42 | CAVNGGGGSQGNLIF | TRBV19 | TRBJ2-7 | CASSMRAAYEQYF | GILGFVFTL | YES | NO | YES |
| 1229 | TRAV8-1 | TRAJ42 | CAVNLGGGSQGNLIF | TRBV19 | TRBJ2-7 | CASSSRASHEQYF | GILGFVFTL | YES | NO | YES |
| 1230 | TRAV8-1 | TRAJ42 | CAVNAGGGSQGNLIF | TRBV19 | TRBJ2-7 | CASSSRASYEQYF | GILGFVFTL | YES | NO | NO |
| 1231 | TRAV8-1 | TRAJ42 | CAVNAGGGSQGNLIF | TRBV19 | TRBJ2-7 | CASSTRASYEQYF | GILGFVFTL | YES | NO | YES |
| 1232 | TRAV8-1 | TRAJ42 | CAVNAGGGSQGNLIF | TRBV19 | TRBJ2-7 | CASSIRSSYEQYF | GILGFVFTL | YES | NO | YES |
| 1234 | TRAV8-3 | TRAJ11 | CAVGAGNSGYSTLTF | TRBV19 | TRBJ2-7 | CASSIRSAYEQYF | GILGFVFTL | YES | YES | YES |
| 1239 | TRAV8-3 | TRAJ37 | CAVVLGSGNTGKLIF | TRBV19 | TRBJ2-7 | CASSIRSAYEQYF | GILGFVFTL | YES | NO | YES |
| 1240 | TRAV8-3 | TRAJ42 | CAVGGGGGSQGNLIF | TRBV19 | TRBJ2-7 | CASSIRASYEQYF | GILGFVFTL | YES | NO | NO |
| 1241 | TRAV8-3 | TRAJ42 | CAVVEGGGSQGNLIF | TRBV19 | TRBJ2-7 | CASSIRSSYEQYF | GILGFVFTL | YES | NO | YES |
| 1243 | TRAV8-3 | TRAJ42 | CAVAGSQGNLIF | TRBV19 | TRBJ2-7 | CASSGRSGVEQYF | GILGFVFTL | YES | NO | YES |
| 1244 | TRAV8-3 | TRAJ42 | CAVPGSQGNLIF | TRBV19 | TRBJ2-7 | CASSLRASYEQYF | GILGFVFTL | YES | YES | YES |
| 1245 | TRAV8-3 | TRAJ42 | CAVGVGGSQGNLIF | TRBV19 | TRBJ2-7 | CASSIRSSYEQYF | GILGFVFTL | YES | YES | YES |
| 1246 | TRAV8-4 | TRAJ23 | CAVTSDPNQGGKLIF | TRBV19 | TRBJ2-7 | CASSIRSSYEQYF | GILGFVFTL | YES | NO | YES |
| 1247 | TRAV8-4 | TRAJ42 | CAEGGSQGNLIF | TRBV19 | TRBJ2-7 | CASSVRSSYEQYF | GILGFVFTL | YES | NO | YES |
| 1251 | TRAV8-6 | TRAJ42 | CAVTDGGSQGNLIF | TRBV19 | TRBJ2-7 | CASSIRSSYEQYF | GILGFVFTL | YES | NO | YES |
| 1252 | TRAV8-6 | TRAJ42 | CASGGSQGNLIF | TRBV19 | TRBJ2-7 | CASSIRSSYEQYF | GILGFVFTL | YES | YES | YES |
| 1253 | TRAV8-6 | TRAJ42 | CAVGGSQGNLIF | TRBV19 | TRBJ2-7 | CASSIRSSYEQYF | GILGFVFTL | YES | NO | YES |
| 1254 | TRAV8-6 | TRAJ42 | CAVGGSQGNLIF | TRBV19 | TRBJ2-7 | CASSIRSAYEQYF | GILGFVFTL | YES | YES | YES |
| 1255 | TRAV8-6 | TRAJ42 | CAVTGSQGNLIF | TRBV19 | TRBJ2-7 | CASSTRSSYEQYF | GILGFVFTL | YES | NO | YES |
| 1256 | TRAV8-6 | TRAJ42 | CAVGGSQGNLIF | TRBV19 | TRBJ2-7 | CASSVRSSYEQYF | GILGFVFTL | YES | NO | YES |
| 1260 | TRAV12-2 | TRAJ23 | CAVNIRGKLIF | TRBV20-1 | TRBJ2-7 | CSVPGGQGLNEQYF | ELAGIGILTV | NO | NO | NO |
| 1261 | TRAV13-1 | TRAJ12 | CAAKLSDSSYKLIF | TRBV20-1 | TRBJ2-7 | CSAMTSGSSYEQYF | KLGGALQAK | NO | NO | NO |
| 1261 | TRAV13-1 | TRAJ12 | CAAKLSDSSYKLIF | TRBV20-1 | TRBJ2-7 | CSAMTSGSSYEQYF | AVFDRKSDAK | NO | NO | NO |
| 1261 | TRAV13-1 | TRAJ12 | CAAKLSDSSYKLIF | TRBV20-1 | TRBJ2-7 | CSAMTSGSSYEQYF | FLASKIGRLV | NO | NO | NO |
| 1262 | TRAV27 | TRAJ42 | CAGGGSQGNLIF | TRBV20-1 | TRBJ2-7 | CSARDRLAGEYSEQYF | GILGFVFTL | NO | NO | NO |
| 1263 | TRAV27 | TRAJ42 | CAGGGSQGNLIF | TRBV20-1 | TRBJ2-7 | CSARDGRREYEQYF | GILGFVFTL | NO | NO | YES |
| 1267 | TRAV8-2 | TRAJ22 | CVGRKGSARQLTF | TRBV20-1 | TRBJ2-7 | CSARDLEVLSYEQYF | AVFDRKSDAK | YES | NO | YES |
| 1267 | TRAV8-2 | TRAJ22 | CVGRKGSARQLTF | TRBV20-1 | TRBJ2-7 | CSARDLEVLSYEQYF | SLFNTVATLY | YES | NO | YES |
| 1267 | TRAV8-2 | TRAJ22 | CVGRKGSARQLTF | TRBV20-1 | TRBJ2-7 | CSARDLEVLSYEQYF | RLRAEAQVK | YES | NO | YES |
| 1268 | TRAV9-2 | TRAJ42 | CALSRYGGSQGNLIF | TRBV20-1 | TRBJ2-7 | CSASNRESYEQYF | LLFGYPVYV | NO | NO | YES |
| 1269 | TRAV24 | TRAJ30 | CALRKVPDKIIF | TRBV21-1 | TRBJ2-7 | CASSKVWGTGRETYEQYF | LLDFVRFMGV | YES | NO | YES |
| 1272 | TRAV12-2 | TRAJ9 | CAVNTGGFKTIF | TRBV27 | TRBJ2-7 | CASSLMAGSYEQYF | GILGFVFTL | YES | NO | NO |
| 1273 | TRAV13-2 | TRAJ45 | CADSGGGADGLTF | TRBV27 | TRBJ2-7 | CASSERDRGYEQYF | LLFGYPVYV | YES | YES | YES |
| 1275 | TRAV27 | TRAJ42 | CAGGGSQGNLIF | TRBV27 | TRBJ2-7 | CASSLGRVSEQYF | GILGFVFTL | YES | NO | YES |
| 1276 | TRAV29DV5 | TRAJ28 | CAASSAGAGSYQLTF | TRBV27 | TRBJ2-7 | CASSLFGGTYEQYF | KLGGALQAK | YES | NO | NO |
| 1276 | TRAV29DV5 | TRAJ28 | CAASSAGAGSYQLTF | TRBV27 | TRBJ2-7 | CASSLFGGTYEQYF | AVFDRKSDAK | YES | NO | NO |
| 1277 | TRAV12-2 | TRAJ17 | CAVREAAGNKLTF | TRBV28 | TRBJ2-7 | CASRVQGLLMHYEQYF | ELAGIGILTV | YES | NO | YES |

| | | | | | | | | | | |
|---|---|---|---|---|---|---|---|---|---|---|
| 1278 | TRAV12-2 | TRAJ3 | CALGGYSSASKIIF | TRBV28 | TRBJ2-7 | CASTTTGTSSYEQYF | ELAGIGILTV | YES | NO | YES |
| 1280 | TRAV19 | TRAJ53 | CALSEAGGGGSNYKLTF | TRBV28 | TRBJ2-7 | CASSLTHTYEQYF | GILGFVFTL | YES | NO | NO |
| 1281 | TRAV12-2 | TRAJ31 | CAVLNNARLMF | TRBV29-1 | TRBJ2-7 | CSVDGSGGLYEQYF | ELAGIGILTV | YES | NO | YES |
| 1283 | TRAV12-3 | TRAJ53 | CAMTGSGGSNYKLTF | TRBV29-1 | TRBJ2-7 | CSVGTGDYEQYF | RAKFKQLL | YES | NO | NO |
| 1283 | TRAV12-3 | TRAJ53 | CAMTGSGGSNYKLTF | TRBV29-1 | TRBJ2-7 | CSVGTGDYEQYF | GILGFVFTL | YES | NO | NO |
| 1284 | TRAV17 | TRAJ53 | CATDASGGSNYKLTF | TRBV29-1 | TRBJ2-7 | CSVGTGDWGEQYF | GILGFVFTL | YES | NO | YES |
| 1285 | TRAV19 | TRAJ56 | CAPRNSKLTF | TRBV29-1 | TRBJ2-7 | CSVAGTTGYEQYF | KVLEYVIKV | NO | NO | YES |
| 1286 | TRAV27 | TRAJ42 | CAGGGSQGNLIF | TRBV29-1 | TRBJ2-7 | CSAGQGPYEQYF | RAKFKQLL | NO | NO | YES |
| 1287 | TRAV38-2DV8 | TRAJ52 | CALPSGGTSYGKLTF | TRBV29-1 | TRBJ2-7 | CSVEVAGHDEQYF | YLNDHLEPWI | YES | NO | YES |
| 1290 | TRAV1-1 | TRAJ15 | CAVRDSAGTALIF | TRBV3-1 | TRBJ2-7 | CASSLGTGGSYEQYF | AVFDRKSDAK | YES | NO | NO |
| 1293 | TRAV9-2 | TRAJ34 | CALYNTDKLIF | TRBV3-1 | TRBJ2-7 | CASSPVTGSSYEQYF | GLCTLVAML | YES | NO | YES |
| 1294 | TRAV12-2 | TRAJ23 | CAVNVRGKLIF | TRBV30 | TRBJ2-7 | CAWGGVQGINEQYF | ELAGIGILTV | YES | NO | YES |
| 1295 | TRAV12-2 | TRAJ54 | CAVNQGQGAQKLVF | TRBV30 | TRBJ2-7 | CAWSLPGLSYEQYF | ELAGIGILTV | YES | NO | YES |
| 1298 | TRAV12-2 | TRAJ45 | CAVSGGGADGLTF | TRBV4-2 | TRBJ2-7 | CASSHGRGNEQYF | ELAGIGILTV | YES | YES | YES |
| 1299 | TRAV19 | TRAJ53 | CALSEGHSGGSNYKLTF | TRBV4-2 | TRBJ2-7 | CASSRYLDTEQYF | LLFGYPVYV | YES | YES | YES |
| 1304 | TRAV3 | TRAJ31 | CAVRDENARLMF | TRBV4-3 | TRBJ2-7 | CASSQGLAGSEQYF | GILGFVFTL | YES | NO | YES |
| 1305 | TRAV12-2 | TRAJ35 | CAVSLGFGNVLHC | TRBV5-1 | TRBJ2-7 | CASSNPTGIYEQYF | ELAGIGILTV | NO | NO | YES |
| 1306 | TRAV4 | TRAJ42 | CLVGEGGGSQGNLIF | TRBV5-1 | TRBJ2-7 | CASSLGQGVAEQYF | RMFPNAPYL | NO | NO | NO |
| 1307 | TRAV12-2 | TRAJ33 | CAASNYQLIW | TRBV5-4 | TRBJ2-7 | CASSLTGGYEQYF | ELAGIGILTV | YES | NO | YES |
| 1308 | TRAV25 | TRAJ42 | CAGRYLNYGGSQGNLIF | TRBV5-4 | TRBJ2-7 | CASSLLPAGHEQYF | IMDQVPFSV | YES | NO | YES |
| 1310 | TRAV5 | TRAJ34 | CAESEGPTYTDKLIF | TRBV5-5 | TRBJ2-7 | CASSGPGGVDEQYF | IVTDFSVIK | YES | NO | NO |
| 1311 | TRAV12-1 | TRAJ43 | CVVNWPGDMRF | TRBV6-1 | TRBJ2-7 | CASKSERGTYEQYF | SLFNTVATLY | YES | NO | YES |
| 1311 | TRAV12-1 | TRAJ43 | CVVNWPGDMRF | TRBV6-1 | TRBJ2-7 | CASKSERGTYEQYF | KLGGALQAK | YES | NO | YES |
| 1312 | TRAV12-2 | TRAJ5 | CAVGAGRRALTF | TRBV6-1 | TRBJ2-7 | CATALGDSYEQYF | ELAGIGILTV | YES | NO | YES |
| 1313 | TRAV17 | TRAJ11 | CATEGDSGYSTLTF | TRBV6-1 | TRBJ2-7 | CASSEVVSTTYEQYF | FLYALALLL | YES | YES | YES |
| 1314 | TRAV4 | TRAJ26 | CLVGDIENYGQNFVF | TRBV6-1 | TRBJ2-7 | CASSEVQGSTYEQYV | KTWGQYWQV | NO | NO | YES |
| 1316 | TRAV12-2 | TRAJ27 | CAVGPNAGKSTF | TRBV6-3 | TRBJ2-7 | CASSPGQGYEQYF | ELAGIGILTV | YES | YES | YES |
| 1317 | TRAV12-2 | TRAJ9 | CAVNTGGFKTIF | TRBV6-3 | TRBJ2-7 | CASSYSTGYEQYF | ELAGIGILTV | YES | NO | YES |
| 1319 | TRAV12-2 | TRAJ32 | CAVNAPSATNKLIF | TRBV6-5 | TRBJ2-7 | CASSYGEGSYEQYF | ELAGIGILTV | YES | NO | YES |
| 1325 | TRAV12-2 | TRAJ33 | CAVINYQLIW | TRBV6-6 | TRBJ2-7 | CASSYYLAGTSIYEQYF | ELAGIGILTV | NO | NO | NO |
| 1328 | TRAV3 | TRAJ6 | CAVSAASGGSYIPTF | TRBV7-2 | TRBJ2-7 | CASSPRDRERGEQYF | AVFDRKSDAK | NO | NO | YES |
| 1329 | TRAV38-2DV8 | TRAJ28 | CAYRSARASGAGSYQLTF | TRBV7-2 | TRBJ2-7 | CASSFPPQLGAYEQYF | SLFNTVATLY | YES | YES | YES |
| 1330 | TRAV14DV4 | TRAJ39 | CAMREDNAGNMLTF | TRBV7-6 | TRBJ2-7 | CASSLQDRALSNDEQYF | GILGFVFTL | YES | NO | NO |
| 1331 | TRAV22 | TRAJ37 | CAGSPSNTGKLIF | TRBV7-9 | TRBJ2-7 | CASSTSEGGLFYEQYF | GILGFVFTL | YES | NO | YES |
| 1332 | TRAV27 | TRAJ42 | CAGGGQGGSQGNLIF | TRBV7-9 | TRBJ2-7 | CASSPDLGYEQYF | GILGFVFTL | YES | NO | YES |
| 1333 | TRAV3 | TRAJ45 | CAVAKDLQNSGGGADGLTF | TRBV7-9 | TRBJ2-7 | CASSQDPSRTYEQYF | GILGFVFTL | NO | NO | YES |
| 1334 | TRAV38-1 | TRAJ52 | CAFMISAGGTSYGKLTF | TRBV7-9 | TRBJ2-7 | CASSQDPSRTYEQYF | GILGFVFTL | NO | NO | NO |
| 1337 | TRAV12-2 | TRAJ30 | CAVNPDRDDKIIF | TRBV9 | TRBJ2-7 | CASSASALSYEQYF | ELAGIGILTV | YES | NO | YES |
| 1339 | TRAV21 | TRAJ20 | CAVSGDYKLSF | TRBV9 | TRBJ2-7 | CASSSRWGYEQYF | GILGFVFTL | YES | NO | YES |
| 1341 | TRAV1-2 | TRAJ20 | CAVRDRVGKLSF | TRBV10-1 | TRBJ1-1 | CASSETGKNTEAFF | KLGGALQAK | YES | NO | YES |
| 1342 | TRAV12-3 | TRAJ39 | CAIGPGNMLTF | TRBV10-1 | TRBJ1-1 | CASSESIGTGVNTEAFF | KLGGALQAK | YES | NO | NO |
| 1344 | TRAV13-1 | TRAJ18 | CAPRGERGSTLGRLYF | TRBV10-1 | TRBJ1-1 | CASRFSPGENTEAFF | KLGGALQAK | YES | NO | NO |
| 1344 | TRAV13-1 | TRAJ18 | CAPRGERGSTLGRLYF | TRBV10-1 | TRBJ1-1 | CASRFSPGENTEAFF | QYDPVAALF | YES | NO | NO |
| 1346 | TRAV27 | TRAJ26 | CAGAEYGQNFVF | TRBV10-1 | TRBJ1-1 | CASSAGQGSIGTEAFF | KLGGALQAK | YES | NO | NO |
| 1347 | TRAV29DV5 | TRAJ44 | CAARDTGTASKLTF | TRBV10-1 | TRBJ1-1 | CASSEGQGHTEAFF | KLGGALQAK | YES | NO | NO |
| 1348 | TRAV29DV5 | TRAJ45 | CAAISSGGGADGLTF | TRBV10-1 | TRBJ1-1 | CASSESGVTEAFF | KLGGALQAK | YES | NO | NO |
| 1349 | TRAV29DV5 | TRAJ45 | CAASATGGGADGLTF | TRBV10-1 | TRBJ1-1 | CASSESGVTEAFF | KLGGALQAK | YES | NO | NO |
| 1350 | TRAV3 | TRAJ43 | CAVNGQYNNDMRF | TRBV10-1 | TRBJ1-1 | CASSELRTLQVMNTEAFF | KLGGALQAK | NO | NO | NO |
| 1351 | TRAV41 | TRAJ42 | CAVSVYGGSQGNLIF | TRBV10-1 | TRBJ1-1 | CASSKSIIETGSNTEAFF | KLGGALQAK | YES | NO | NO |
| 1352 | TRAV8-3 | TRAJ6 | CAVGLSGGSYIPTF | TRBV10-1 | TRBJ1-1 | CASSDAGGLGEAFF | KLGGALQAK | YES | NO | NO |
| 1355 | TRAV21 | TRAJ5 | CAVKGRGDTGRRALTF | TRBV10-2 | TRBJ1-1 | CASSEQGSSEAFF | AVFDRKSDAK | YES | NO | NO |
| 1355 | TRAV21 | TRAJ5 | CAVKGRGDTGRRALTF | TRBV10-2 | TRBJ1-1 | CASSEQGSSEAFF | KLGGALQAK | YES | NO | NO |
| 1356 | TRAV8-2 | TRAJ29 | CVVSAGYSGNTPLVF | TRBV10-2 | TRBJ1-1 | CASSEDSNTEAFF | KLGGALQAK | YES | NO | NO |
| 1358 | TRAV8-6 | TRAJ22 | CAVTSLSSGSARQLTF | TRBV10-2 | TRBJ1-1 | CASSLTGGEGEAFF | KLGGALQAK | YES | NO | NO |
| 1359 | TRAV12-1 | TRAJ3 | CVVNGGASKIIF | TRBV10-3 | TRBJ1-1 | CATSPPGTGANTEAFF | KLGGALQAK | YES | NO | NO |
| 1360 | TRAV12-2 | TRAJ13 | CAVPSGGYQKVTF | TRBV10-3 | TRBJ1-1 | CAIRTLGGGDTEAFF | KLGGALQAK | YES | NO | NO |
| 1364 | TRAV17 | TRAJ45 | CATDAGGADGLTF | TRBV10-3 | TRBJ1-1 | CAISEGTEENTEAFF | KLGGALQAK | YES | NO | NO |
| 1365 | TRAV2 | TRAJ33 | CAVVRMDSNYQLIW | TRBV10-3 | TRBJ1-1 | CAISDQDKRALNTEAFF | KLGGALQAK | NO | NO | NO |
| 1366 | TRAV21 | TRAJ29 | CAVRNSGNTPLVF | TRBV10-3 | TRBJ1-1 | CAISESWGGAEAFF | KLGGALQAK | YES | NO | NO |
| 1367 | TRAV21 | TRAJ50 | CAEVKTSYDKVIF | TRBV10-3 | TRBJ1-1 | CAIRGFLGTGKNTEAFF | KLGGALQAK | YES | NO | NO |
| 1368 | TRAV24 | TRAJ40 | CAVHTSGTYKYIF | TRBV10-3 | TRBJ1-1 | CAISEGTEAFF | KLGGALQAK | YES | NO | NO |
| 1369 | TRAV27 | TRAJ24 | CAGYFSWGKLQF | TRBV10-3 | TRBJ1-1 | CAISGQNTEAFF | KLGGALQAK | YES | NO | NO |
| 1370 | TRAV29DV5 | TRAJ42 | CAAAGSQGNLIF | TRBV10-3 | TRBJ1-1 | CAISGQGEYTEAFF | KLGGALQAK | YES | NO | NO |
| 1371 | TRAV29DV5 | TRAJ45 | CAASAGGGADGLTF | TRBV10-3 | TRBJ1-1 | CAIGEAVRGQEAFF | KLGGALQAK | YES | NO | NO |
| 1372 | TRAV3 | TRAJ42 | CAVRDDLGSQGNLIF | TRBV10-3 | TRBJ1-1 | CAISDLTGGRTEAFF | KLGGALQAK | NO | NO | NO |
| 1373 | TRAV36DV7 | TRAJ53 | CAVEALNSGGSNYKLTF | TRBV10-3 | TRBJ1-1 | CAITRLIGTTDTEAFF | KLGGALQAK | NO | NO | NO |
| 1374 | TRAV38-1 | TRAJ47 | CAFMKLQPMEYGNKLVF | TRBV10-3 | TRBJ1-1 | CAIKGSATEAFF | KLGGALQAK | YES | NO | NO |
| 1375 | TRAV38-2DV8 | TRAJ23 | CAYSQGGKLIF | TRBV10-3 | TRBJ1-1 | CAISEPTGGHTEAFF | KLGGALQAK | YES | NO | NO |
| 1376 | TRAV8-3 | TRAJ42 | CAGEALHGGSQGNLIF | TRBV10-3 | TRBJ1-1 | CAISGQAGRNTEAFF | KLGGALQAK | YES | NO | NO |
| 1379 | TRAV24 | TRAJ11 | CAPSGYSTLTF | TRBV11-1 | TRBJ1-1 | CASSAEGAGNMNTEAFF | KLGGALQAK | YES | NO | NO |
| 1380 | TRAV38-2DV8 | TRAJ27 | CAYTNAGKSTF | TRBV11-1 | TRBJ1-1 | CASSLWAGTVNTEAFF | KLGGALQAK | YES | NO | NO |
| 1381 | TRAV1-2 | TRAJ11 | CAISGYSTLTF | TRBV11-2 | TRBJ1-1 | CASSLTEAFF | KLGGALQAK | YES | NO | NO |
| 1382 | TRAV10 | TRAJ40 | CVVSGPATSGTYKYIF | TRBV11-2 | TRBJ1-1 | CASSLGPSSTEAFF | KLGGALQAK | YES | NO | YES |
| 1386 | TRAV12-2 | TRAJ26 | CAVRGYGQNFVF | TRBV11-2 | TRBJ1-1 | CASSLDGGNTEAFF | KLGGALQAK | YES | NO | NO |
| 1387 | TRAV12-2 | TRAJ54 | CAAGAQKLVF | TRBV11-2 | TRBJ1-1 | CASSFGGDAEAFF | KLGGALQAK | YES | NO | NO |
| 1390 | TRAV13-1 | TRAJ5 | CAASQPGRRALTF | TRBV11-2 | TRBJ1-1 | CASSLGGSEAFF | KLGGALQAK | YES | NO | NO |
| 1391 | TRAV13-2 | TRAJ43 | CAENRIGNDMRF | TRBV11-2 | TRBJ1-1 | CASSLFQGSGTEAFF | KLGGALQAK | YES | NO | NO |
| 1393 | TRAV17 | TRAJ12 | CATVAKDSSYKLIF | TRBV11-2 | TRBJ1-1 | CASSLGWSTEAFF | KLGGALQAK | YES | NO | NO |
| 1394 | TRAV17 | TRAJ24 | CATDAGTTDSWGKLQF | TRBV11-2 | TRBJ1-1 | CASSLGYIRQGGFVTEAFF | KLGGALQAK | YES | NO | NO |
| 1395 | TRAV17 | TRAJ36 | CAQTGANNLFF | TRBV11-2 | TRBJ1-1 | CASSLFQGSGTEAFF | AVFDRKSDAK | YES | NO | NO |
| 1396 | TRAV17 | TRAJ48 | CATLDFGNEKLTF | TRBV11-2 | TRBJ1-1 | CASSFTGPNTEAFF | KLGGALQAK | YES | NO | NO |
| 1397 | TRAV19 | TRAJ42 | CALSEAGGSQGNLIF | TRBV11-2 | TRBJ1-1 | CASRIGGHTFLFGTEAFF | KLGGALQAK | YES | NO | NO |
| 1398 | TRAV19 | TRAJ48 | CALSEALVGNEKLTF | TRBV11-2 | TRBJ1-1 | CASSLFQGSGTEAFF | KLGGALQAK | YES | NO | NO |
| 1400 | TRAV29DV5 | TRAJ21 | CAASGNNFNKFYF | TRBV11-2 | TRBJ1-1 | CASSLKTVNTEAFF | KLGGALQAK | YES | NO | NO |
| 1401 | TRAV3 | TRAJ43 | CAVQYNNNDMRF | TRBV11-2 | TRBJ1-1 | CASSLGEGHMNTEAFF | RLRAEAQVK | YES | YES | YES |
| 1402 | TRAV38-2DV8 | TRAJ4 | CAYRSSFSGGYNKLIF | TRBV11-2 | TRBJ1-1 | CASSYKTGQGSMNTEAFF | KLGGALQAK | YES | NO | NO |
| 1407 | TRAV1-2 | TRAJ33 | CAVRDSNYQLIW | TRBV11-3 | TRBJ1-1 | CASSPRTGRAEAFF | KLGGALQAK | YES | NO | NO |
| 1408 | TRAV10 | TRAJ23 | CVVSDFYNQGGKLIF | TRBV11-3 | TRBJ1-1 | CASSLGENTEAFF | KLGGALQAK | YES | NO | NO |
| 1410 | TRAV12-1 | TRAJ4 | CVVHLPPYNKLIF | TRBV11-3 | TRBJ1-1 | CASSFESGGRREAFF | AVFDRKSDAK | YES | NO | NO |
| 1411 | TRAV12-3 | TRAJ35 | CAMKR | TRBV11-3 | TRBJ1-1 | CASSLELGLAEAFF | KLGGALQAK | NO | NO | NO |
| 1412 | TRAV13-1 | TRAJ11 | CAASIAGYSTLTF | TRBV11-3 | TRBJ1-1 | CASSLEQANTEAFF | KLGGALQAK | YES | NO | NO |
| 1415 | TRAV17 | TRAJ36 | CATDEGANNLFF | TRBV11-3 | TRBJ1-1 | CASRPVSLGGSNTEAFF | KLGGALQAK | YES | NO | NO |
| 1416 | TRAV21 | TRAJ22 | CAVSGGWAGSARQLTF | TRBV11-3 | TRBJ1-1 | CASSLGNTEAFF | KLGGALQAK | YES | NO | NO |
| 1417 | TRAV21 | TRAJ24 | CAVTDSWGKLQF | TRBV11-3 | TRBJ1-1 | CASSPPGRKNTEAFF | KLGGALQAK | YES | NO | YES |
| 1420 | TRAV29DV5 | TRAJ30 | CAASAGVDDKIIF | TRBV11-3 | TRBJ1-1 | CASSQWGTEAFF | KLGGALQAK | YES | NO | NO |
| 1421 | TRAV38-1 | TRAJ42 | CAFMKRFGGSQGNLIF | TRBV11-3 | TRBJ1-1 | CASSLSPATEAFF | KLGGALQAK | YES | NO | NO |
| 1422 | TRAV38-1 | TRAJ54 | CAFMSLIQGAQKLVF | TRBV11-3 | TRBJ1-1 | CASSLELGLAEAFF | KLGGALQAK | YES | NO | NO |
| 1422 | TRAV38-1 | TRAJ54 | CAFMSLIQGAQKLVF | TRBV11-3 | TRBJ1-1 | CASSLELGLAEAFF | AVFDRKSDAK | YES | NO | NO |
| 1423 | TRAV4 | TRAJ21 | CLVIYNFNKFYF | TRBV11-3 | TRBJ1-1 | CASSHNTGLVTEAFF | KLGGALQAK | YES | NO | NO |
| 1424 | TRAV8-1 | TRAJ8 | CAVNVKGFQKLVF | TRBV11-3 | TRBJ1-1 | CASSLDHGGGNTEAFF | KLGGALQAK | YES | YES | NO |
| 1425 | TRAV9-2 | TRAJ43 | CALINRNNDMRF | TRBV11-3 | TRBJ1-1 | CASHRGTGLRTEAFF | KLGGALQAK | YES | NO | YES |
| 1427 | TRAV13-1 | TRAJ13 | CAASRDQKVTF | TRBV12-3 | TRBJ1-1 | CASRGIGGASEAFF | KLGGALQAK | YES | NO | NO |
| 1430 | TRAV19 | TRAJ50 | CALSEAWDSPPDMKTSYDKVIF | TRBV12-3 | TRBJ1-1 | CASSAPKGWFTEAFF | KLGGALQAK | NO | NO | NO |
| 1431 | TRAV3 | TRAJ13 | CAVRDGGLGSGGYQKVTF | TRBV12-3 | TRBJ1-1 | CASSHRSGTEAFF | KLGGALQAK | NO | NO | NO |

| | | | | | | | | | | |
|---|---|---|---|---|---|---|---|---|---|---|
| 1432 | TRAV4 | TRAJ18 | CLVGDMEGGSTLGRLYF | TRBV12-3 | TRBJ1-1 | CASRYGTPGTEAFF | KLGGALQAK | YES | NO | NO |
| 1435 | TRAV1-1 | TRAJ29 | CAVSGNTPLVF | TRBV12-4 | TRBJ1-1 | CASRILPDRDGDTEAFF | KLGGALQAK | YES | NO | NO |
| 1436 | TRAV1-2 | TRAJ20 | CAVRHDYKLSF | TRBV12-4 | TRBJ1-1 | CASSTLRVGGRTEAFF | KLGGALQAK | YES | NO | NO |
| 1437 | TRAV12-2 | TRAJ49 | CAVILGNQFYF | TRBV12-4 | TRBJ1-1 | CASSSGNTEAFF | KLGGALQAK | YES | NO | NO |
| 1438 | TRAV13-2 | TRAJ18 | CAENIDRGSTLGRLYF | TRBV12-4 | TRBJ1-1 | CWGNTEAFF | KLGGALQAK | NO | NO | NO |
| 1442 | TRAV17 | TRAJ31 | CATATNNNARLMF | TRBV12-4 | TRBJ1-1 | CASRLGHRAKNTEAFF | KLGGALQAK | YES | NO | NO |
| 1443 | TRAV17 | TRAJ52 | CASRAGGTSYGKLTF | TRBV12-4 | TRBJ1-1 | CASSYPPNGRVEGAFF | KLGGALQAK | YES | NO | NO |
| 1444 | TRAV19 | TRAJ20 | CALSEAYLGDYKLSF | TRBV12-4 | TRBJ1-1 | CASRLGGYTEAFF | KLGGALQAK | YES | NO | NO |
| 1446 | TRAV19 | TRAJ56 | CALSEAPGANSKLTF | TRBV12-4 | TRBJ1-1 | CASSLAQGYAEAFF | KLGGALQAK | YES | NO | NO |
| 1447 | TRAV20 | TRAJ28 | CAVLYSGAGSYQLTF | TRBV12-4 | TRBJ1-1 | CASSFGSMNTEAFF | KLGGALQAK | YES | NO | NO |
| 1449 | TRAV24 | TRAJ49 | CAFFGTGNQFYF | TRBV12-4 | TRBJ1-1 | CASSLWVGSGEAFF | KLGGALQAK | YES | NO | YES |
| 1450 | TRAV26-1 | TRAJ52 | CIVAGGTSYGKLTF | TRBV12-4 | TRBJ1-1 | CASSLSTNTEAFF | KLGGALQAK | YES | NO | NO |
| 1451 | TRAV26-2 | TRAJ4 | CILRDIYNKLIF | TRBV12-4 | TRBJ1-1 | CASSLEGQGDTEAFF | KLGGALQAK | YES | NO | YES |
| 1453 | TRAV34 | TRAJ41 | CGAVSGYALNF | TRBV12-4 | TRBJ1-1 | CASTDRVETEAFF | KLGGALQAK | YES | NO | NO |
| 1454 | TRAV38-1 | TRAJ29 | CAFMKDYSGNTPLVF | TRBV12-4 | TRBJ1-1 | CASSLALILTETEAFF | KLGGALQAK | YES | NO | NO |
| 1457 | TRAV39 | TRAJ31 | CAVVNARLMF | TRBV12-4 | TRBJ1-1 | CASSSRGGSEAFF | KLGGALQAK | YES | NO | NO |
| 1458 | TRAV4 | TRAJ44 | CLVGDGTGTASKLTF | TRBV12-4 | TRBJ1-1 | CASSWGVGSTEAFF | KLGGALQAK | YES | NO | NO |
| 1460 | TRAV8-1 | TRAJ42 | CAVNAGGGSQGNLIF | TRBV12-4 | TRBJ1-1 | CASFDSMNTEAFF | KLGGALQAK | YES | NO | NO |
| 1462 | TRAV3 | TRAJ29 | CAVRDIGGNTPLVF | TRBV12-5 | TRBJ1-1 | CASGRQGENTEAFF | RLRAEAQVK | NO | NO | NO |
| 1463 | TRAV4 | TRAJ6 | CRAGGSYIPTF | TRBV12-5 | TRBJ1-1 | CASGALNTEAFF | KLGGALQAK | YES | NO | NO |
| 1465 | TRAV12-2 | TRAJ56 | CAVNPRLPGANSKLTF | TRBV13 | TRBJ1-1 | CASSLDGAFTEAFF | KLGGALQAK | YES | NO | NO |
| 1466 | TRAV14DV4 | TRAJ54 | CAMREGRGAQKLVF | TRBV13 | TRBJ1-1 | CASSLRLGGGTEAFF | KLGGALQAK | YES | NO | NO |
| 1467 | TRAV17 | TRAJ36 | CATVRQTGANNLFF | TRBV13 | TRBJ1-1 | CASSPTGDFGVGNTEAFF | KLGGALQAK | YES | NO | NO |
| 1469 | TRAV19 | TRAJ29 | CALSEAFGSGNTPLVF | TRBV13 | TRBJ1-1 | CASSHRTGGGSEAFF | KLGGALQAK | YES | NO | NO |
| 1470 | TRAV20 | TRAJ31 | CAVRTNNNARLMF | TRBV13 | TRBJ1-1 | CASRSQEGAEAFF | KLGGALQAK | YES | NO | NO |
| 1471 | TRAV26-2 | TRAJ58 | CGLETSGSRLTF | TRBV13 | TRBJ1-1 | CASSFGQANTEAFF | KLGGALQAK | YES | NO | NO |
| 1473 | TRAV36DV7 | TRAJ53 | CAVNSGGSNYKLTF | TRBV13 | TRBJ1-1 | CASSPGGQGRTEAFF | KLGGALQAK | NO | NO | NO |
| 1474 | TRAV8-3 | TRAJ36 | CAVGVGGANNLFF | TRBV13 | TRBJ1-1 | CASSSAEGLNTEAFF | KLGGALQAK | YES | NO | NO |
| 1477 | TRAV12-1 | TRAJ36 | CVVNLPTGANNLFF | TRBV14 | TRBJ1-1 | CASSQETVGTAIMNTEAFF | KLGGALQAK | NO | NO | NO |
| 1479 | TRAV2 | TRAJ36 | CAVEEAGANNLFF | TRBV14 | TRBJ1-1 | CASSQARGDTEAFF | KLGGALQAK | NO | NO | NO |
| 1480 | TRAV21 | TRAJ13 | CAPEGPSGGYQKVTF | TRBV14 | TRBJ1-1 | CASSQVSSYTEAFF | KLGGALQAK | NO | NO | NO |
| 1483 | TRAV8-1 | TRAJ26 | CAVPDNYGQNFVF | TRBV14 | TRBJ1-1 | CASSLVPGIAEAFF | KLGGALQAK | NO | NO | NO |
| 1485 | TRAV8-4 | TRAJ52 | CAVSDHIGAGGTSYGKLTF | TRBV14 | TRBJ1-1 | CASSQGFGVSRLNTEAFF | KLGGALQAK | NO | NO | NO |
| 1487 | TRAV12-1 | TRAJ48 | CVVNERGSNFGNEKLTF | TRBV15 | TRBJ1-1 | CATSRVRVAEAFF | KLGGALQAK | YES | NO | NO |
| 1487 | TRAV12-1 | TRAJ48 | CVVNERGSNFGNEKLTF | TRBV15 | TRBJ1-1 | CATSRVRVAEAFF | AVFDRKSDAK | YES | NO | NO |
| 1488 | TRAV13-1 | TRAJ17 | CAASEAAGNKLTF | TRBV15 | TRBJ1-1 | CATSTDSNTEAFF | KLGGALQAK | YES | NO | NO |
| 1489 | TRAV14DV4 | TRAJ5 | CAMREVDTGRRALTF | TRBV15 | TRBJ1-1 | CATSGGTGINTEAFF | KLGGALQAK | YES | NO | NO |
| 1490 | TRAV21 | TRAJ20 | CAVPLSNDYKLSF | TRBV15 | TRBJ1-1 | CATRPRGDTEAFF | KLGGALQAK | YES | NO | NO |
| 1491 | TRAV26-1 | TRAJ45 | CITGGGADGLTF | TRBV15 | TRBJ1-1 | CATSRDRLGRGKGTEAFF | KLGGALQAK | YES | NO | NO |
| 1493 | TRAV35 | TRAJ45 | CAGQRGMYSGGGADGLTF | TRBV15 | TRBJ1-1 | CATSIGTEIGYPEAFF | KLGGALQAK | YES | NO | NO |
| 1495 | TRAV5 | TRAJ20 | CAESFNDYKLSF | TRBV15 | TRBJ1-1 | CATSRDQGIRNTEAFF | KLGGALQAK | YES | NO | NO |
| 1496 | TRAV5 | TRAJ20 | CAESMNDYKLSF | TRBV15 | TRBJ1-1 | CATSRDQGTKNTEAFF | KLGGALQAK | YES | NO | NO |
| 1497 | TRAV8-6 | TRAJ30 | CAVSDLRDDKIIF | TRBV15 | TRBJ1-1 | CATSRDSGPNTEAFF | KLGGALQAK | YES | NO | NO |
| 1498 | TRAV2 | TRAJ41 | CAVEVPHSGYALNF | TRBV16 | TRBJ1-1 | CASSPGGTEAFF | KLGGALQAK | NO | NO | YES |
| 1499 | TRAV12-1 | TRAJ37 | CVVTGSGNTGKLIF | TRBV18 | TRBJ1-1 | CASSPRDRFMNTEAFF | KLGGALQAK | YES | NO | NO |
| 1500 | TRAV12-1 | TRAJ39 | CVGNNAGNMLTF | TRBV18 | TRBJ1-1 | CASSPPPGVSMNTEAFF | KLGGALQAK | YES | NO | NO |
| 1502 | TRAV13-1 | TRAJ49 | CAASTDTGNQFYF | TRBV18 | TRBJ1-1 | CASSPTGTSLSFF | KLGGALQAK | YES | NO | NO |
| 1503 | TRAV23DV6 | TRAJ58 | CAAREETSGSRLTF | TRBV18 | TRBJ1-1 | CASSPTGYMNTEAFF | RLRAEAQVK | YES | NO | YES |
| 1504 | TRAV27 | TRAJ26 | CAGIYGQNFVF | TRBV18 | TRBJ1-1 | CASSPPSRGRVNTEAFF | KLGGALQAK | YES | NO | NO |
| 1505 | TRAV1-1 | TRAJ24 | CAATDSWGKLQF | TRBV19 | TRBJ1-1 | CASSIDRGGNTEAFF | KLGGALQAK | YES | NO | NO |
| 1505 | TRAV1-1 | TRAJ24 | CAATDSWGKLQF | TRBV19 | TRBJ1-1 | CASSIDRGGNTEAFF | IVTDFSVIK | YES | NO | NO |
| 1505 | TRAV1-1 | TRAJ24 | CAATDSWGKLQF | TRBV19 | TRBJ1-1 | CASSIDRGGNTEAFF | AVFDRKSDAK | YES | NO | NO |
| 1512 | TRAV10 | TRAJ8 | CVVITGFQKLVF | TRBV19 | TRBJ1-1 | CASSITWTGNMNTEAFF | KLGGALQAK | YES | NO | NO |
| 1512 | TRAV10 | TRAJ8 | CVVITGFQKLVF | TRBV19 | TRBJ1-1 | CASSITWTGNMNTEAFF | AVFDRKSDAK | YES | NO | NO |
| 1513 | TRAV12-1 | TRAJ11 | CVVNGYSTLTF | TRBV19 | TRBJ1-1 | CASSIGRNTEAFF | KLGGALQAK | YES | NO | NO |
| 1514 | TRAV12-1 | TRAJ39 | CVVNGNNAGNMLTF | TRBV19 | TRBJ1-1 | CASSISRGYTEAFF | KLGGALQAK | YES | NO | NO |
| 1515 | TRAV12-1 | TRAJ5 | CVVNTGDTGRRALTF | TRBV19 | TRBJ1-1 | CASTGWEMNTEAFF | KLGGALQAK | YES | NO | NO |
| 1517 | TRAV12-2 | TRAJ30 | CAVMDRDDKIIF | TRBV19 | TRBJ1-1 | CATGQGAAEAFF | KLGGALQAK | YES | NO | NO |
| 1520 | TRAV12-2 | TRAJ53 | CAPRDEGSNYKLTF | TRBV19 | TRBJ1-1 | CASSLTALGTEAFF | KLGGALQAK | YES | NO | NO |
| 1522 | TRAV12-2 | TRAJ9 | CAVEARGGFKTIF | TRBV19 | TRBJ1-1 | CASSIVGGGEAFF | KLGGALQAK | YES | NO | NO |
| 1523 | TRAV12-2 | TRAJ9 | CAVNGPGGGGFKTIF | TRBV19 | TRBJ1-1 | CASSPTSNTEAFF | KLGGALQAK | YES | NO | NO |
| 1524 | TRAV12-3 | TRAJ31 | CAMSANNNARLMF | TRBV19 | TRBJ1-1 | CASSSPEVYRKF | KLGGALQAK | YES | NO | YES |
| 1527 | TRAV12-3 | TRAJ50 | CAVKTSYDKVIF | TRBV19 | TRBJ1-1 | CASSLAGNTEAFF | KLGGALQAK | YES | NO | NO |
| 1528 | TRAV13-1 | TRAJ13 | CAASLNSGGYQKVTF | TRBV19 | TRBJ1-1 | CATTTWDMNTEAFF | KLGGALQAK | YES | NO | YES |
| 1530 | TRAV13-1 | TRAJ43 | CAASSYNNNDMRF | TRBV19 | TRBJ1-1 | CASSILQQGEAFF | KLGGALQAK | YES | NO | NO |
| 1531 | TRAV13-1 | TRAJ45 | CAATPRGGADGLTF | TRBV19 | TRBJ1-1 | CATHRGGVNTEAFF | KLGGALQAK | YES | NO | YES |
| 1532 | TRAV13-1 | TRAJ50 | CALKTSYDKVIF | TRBV19 | TRBJ1-1 | CASSIAGWNTEAFF | KLGGALQAK | YES | NO | NO |
| 1534 | TRAV13-2 | TRAJ37 | CAETASNTGKLIF | TRBV19 | TRBJ1-1 | CASSMGPEGAFF | KLGGALQAK | YES | NO | NO |
| 1535 | TRAV13-2 | TRAJ8 | CAENGGFQKLVF | TRBV19 | TRBJ1-1 | CASSIEPGRTEAFF | KLGGALQAK | YES | NO | NO |
| 1536 | TRAV14DV4 | TRAJ12 | CAMRSMDSSYKLIF | TRBV19 | TRBJ1-1 | CASSLIPDQNTEAFF | KLGGALQAK | YES | NO | NO |
| 1537 | TRAV14DV4 | TRAJ12 | CAMRVGSSYKLIF | TRBV19 | TRBJ1-1 | CASSIGGIEAFF | KLGGALQAK | YES | NO | NO |
| 1538 | TRAV14DV4 | TRAJ23 | CAMREVIYNQGGKLIF | TRBV19 | TRBJ1-1 | CATSLGLNSEAFF | KLGGALQAK | YES | NO | YES |
| 1539 | TRAV14DV4 | TRAJ32 | CAMRERNYGGATNKLIF | TRBV19 | TRBJ1-1 | CASSITGDGLNTEAFF | KLGGALQAK | YES | NO | NO |
| 1540 | TRAV14DV4 | TRAJ37 | CAMREGSGNTGKLIF | TRBV19 | TRBJ1-1 | CASSMWGIGEAFF | KLGGALQAK | YES | NO | NO |
| 1542 | TRAV14DV4 | TRAJ42 | CAMREGDGGSQGNLIF | TRBV19 | TRBJ1-1 | CASSPLTAVNTEAFF | KLGGALQAK | YES | NO | NO |
| 1543 | TRAV14DV4 | TRAJ43 | CAMREGSNDMRF | TRBV19 | TRBJ1-1 | CASSIKGPFRTEAFF | KLGGALQAK | YES | NO | NO |
| 1544 | TRAV14DV4 | TRAJ49 | CAMREPRNTGNQFYF | TRBV19 | TRBJ1-1 | CAIHQGYTEAFF | KLGGALQAK | YES | NO | NO |
| 1545 | TRAV14DV4 | TRAJ6 | CAMREGLTGGSYIPTF | TRBV19 | TRBJ1-1 | CASSISLVGTEAFF | KLGGALQAK | YES | NO | NO |
| 1546 | TRAV16 | TRAJ29 | CALYNSGNTPLVF | TRBV19 | TRBJ1-1 | CASSIGGNTEAFF | KLGGALQAK | NO | NO | YES |
| 1548 | TRAV17 | TRAJ15 | CATDSRQAGTALIF | TRBV19 | TRBJ1-1 | CASSIESLGVTEAFF | KLGGALQAK | YES | NO | NO |
| 1549 | TRAV17 | TRAJ36 | CATDRTGANNLFF | TRBV19 | TRBJ1-1 | CASSTPYGETEAFF | KLGGALQAK | YES | NO | NO |
| 1550 | TRAV17 | TRAJ43 | CATDEVYNNDMRF | TRBV19 | TRBJ1-1 | CASRVGNTEAFF | KLGGALQAK | YES | NO | NO |
| 1552 | TRAV19 | TRAJ29 | CALSEAFSGNTPLVF | TRBV19 | TRBJ1-1 | CASSMSGPGTEAFF | AVFDRKSDAK | YES | NO | NO |
| 1553 | TRAV19 | TRAJ33 | CALSEVDTDSNYQLIW | TRBV19 | TRBJ1-1 | CASSIGRGSTEAFF | KLGGALQAK | YES | NO | NO |
| 1554 | TRAV19 | TRAJ34 | CALSERRNTDKLIF | TRBV19 | TRBJ1-1 | CASSEARGLNTEAFF | KLGGALQAK | YES | NO | NO |
| 1555 | TRAV19 | TRAJ36 | CALSESQTGANNLFF | TRBV19 | TRBJ1-1 | CAAGGGSEMNTEAFF | KLGGALQAK | YES | NO | NO |
| 1557 | TRAV19 | TRAJ5 | CALSEALGGRRALTF | TRBV19 | TRBJ1-1 | CASSILSTHTEAFF | KLGGALQAK | YES | NO | NO |
| 1558 | TRAV2 | TRAJ45 | CAVGGADGLTF | TRBV19 | TRBJ1-1 | CASSIGFRMNTEAFF | RLRAEAQVK | NO | NO | NO |
| 1559 | TRAV20 | TRAJ26 | CAVPQYGQNFVF | TRBV19 | TRBJ1-1 | CASRTLSQENTEAFF | KLGGALQAK | YES | NO | NO |
| 1560 | TRAV20 | TRAJ39 | CAVQAGHNAGNMLTF | TRBV19 | TRBJ1-1 | CASSYRNTEAFF | KLGGALQAK | YES | NO | NO |
| 1561 | TRAV21 | TRAJ37 | CAVVSGNTGKLIF | TRBV19 | TRBJ1-1 | CASSITGDTEAFF | KLGGALQAK | YES | NO | NO |
| 1563 | TRAV21 | TRAJ50 | CAVDLMKTSYDKVIF | TRBV19 | TRBJ1-1 | CASSITWTGNMNTEAFF | KLGGALQAK | YES | NO | NO |
| 1564 | TRAV22 | TRAJ28 | CAVEHPGAGSYQLTF | TRBV19 | TRBJ1-1 | CASSIDQGAEAFF | KLGGALQAK | YES | NO | NO |
| 1566 | TRAV24 | TRAJ20 | CAPINDYKLSF | TRBV19 | TRBJ1-1 | CASSPAVPGTEAFF | KLGGALQAK | YES | NO | NO |
| 1568 | TRAV25 | TRAJ20 | CAGTDYKLSF | TRBV19 | TRBJ1-1 | CASSLVWRGDVNTEAFF | KLGGALQAK | YES | NO | NO |
| 1569 | TRAV25 | TRAJ37 | CAGPNGSGNTGKLIF | TRBV19 | TRBJ1-1 | CASSIQGEWTEAFF | KLGGALQAK | YES | NO | NO |
| 1570 | TRAV25 | TRAJ50 | CAEKTSYDKVIF | TRBV19 | TRBJ1-1 | CASSIGGNTEAFF | KLGGALQAK | YES | NO | NO |
| 1571 | TRAV26-1 | TRAJ16 | CIVRDF | TRBV19 | TRBJ1-1 | CASSADRGGNTEAFF | KLGGALQAK | NO | NO | NO |
| 1572 | TRAV26-1 | TRAJ48 | CIVRVVLSNFGNEKLTF | TRBV19 | TRBJ1-1 | CASSMYTNTEAFF | KLGGALQAK | YES | NO | NO |
| 1576 | TRAV27 | TRAJ20 | CAGESLRDYKLSF | TRBV19 | TRBJ1-1 | CASSISTEAFF | KLGGALQAK | YES | NO | NO |
| 1577 | TRAV27 | TRAJ23 | CAGARYNQGGKLIF | TRBV19 | TRBJ1-1 | CASSSGTGALNTEAFF | KLGGALQAK | YES | NO | NO |
| 1580 | TRAV27 | TRAJ44 | CAGGVTGTASKLTF | TRBV19 | TRBJ1-1 | CASSTWDLNTEAFF | KLGGALQAK | YES | NO | NO |
| 1581 | TRAV27 | TRAJ48 | CAGSSNFGNEKLTF | TRBV19 | TRBJ1-1 | CASSIYLKGTEAFF | KLGGALQAK | YES | NO | NO |
| 1583 | TRAV29DV5 | TRAJ34 | CAASASNTDKLIF | TRBV19 | TRBJ1-1 | CASSWGGLRTEAFF | KLGGALQAK | YES | NO | NO |

| | | | | | | | | | | |
|---|---|---|---|---|---|---|---|---|---|---|
| 1584 | TRAV29DV5 | TRAJ41 | CAASENSGYALNF | TRBV19 | TRBJ1-1 | CASSILARDTEAFF | KLGGALQAK | YES | NO | NO |
| 1585 | TRAV29DV5 | TRAJ50 | CATKTSYDKVIF | TRBV19 | TRBJ1-1 | CAISRGGNTEAFF | KLGGALQAK | YES | NO | NO |
| 1587 | TRAV29DV5 | TRAJ6 | CAASARSGGSYIPTF | TRBV19 | TRBJ1-1 | CASSTYDGNTEAFF | KLGGALQAK | YES | NO | NO |
| 1588 | TRAV3 | TRAJ17 | CAVRDEKAAGNKLTF | TRBV19 | TRBJ1-1 | CASSIQGGYTEAFF | KLGGALQAK | NO | NO | NO |
| 1589 | TRAV3 | TRAJ18 | CAVRDGGGSTLGRLYF | TRBV19 | TRBJ1-1 | CATITGVTEAFF | KLGGALQAK | NO | NO | NO |
| 1590 | TRAV30 | TRAJ40 | CGTESTSGTYKYIF | TRBV19 | TRBJ1-1 | CASSISRSTEAFF | KLGGALQAK | YES | NO | NO |
| 1591 | TRAV35 | TRAJ40 | CAAPVGTYKYIF | TRBV19 | TRBJ1-1 | CASSMSGPGTEAFF | KLGGALQAK | YES | NO | NO |
| 1594 | TRAV38-2DV8 | TRAJ33 | CAYRSARGSNYQLIW | TRBV19 | TRBJ1-1 | CASTPAKGGVTEAFF | KLGGALQAK | YES | NO | NO |
| 1595 | TRAV38-2DV8 | TRAJ43 | CAYRSFVWSNNDMRF | TRBV19 | TRBJ1-1 | CASSIGRAEAFF | KLGGALQAK | YES | NO | NO |
| 1596 | TRAV38-2DV8 | TRAJ52 | CAYSSPGGTSYGKLTF | TRBV19 | TRBJ1-1 | CASSGGAGDGTEAFF | RLRAEAQVK | YES | YES | YES |
| 1597 | TRAV38-2DV8 | TRAJ58 | CASSGSRLTF | TRBV19 | TRBJ1-1 | CATRVSGAVTEAFF | KLGGALQAK | YES | NO | YES |
| 1598 | TRAV4 | TRAJ10 | CLVGEIFTGGGNKLTF | TRBV19 | TRBJ1-1 | CASQGRAPEAFF | KLGGALQAK | YES | NO | YES |
| 1599 | TRAV41 | TRAJ45 | CAVSLGGGADGLTF | TRBV19 | TRBJ1-1 | CASSIWSLRYTEAFF | KLGGALQAK | YES | NO | NO |
| 1600 | TRAV5 | TRAJ20 | CAENDYKLSF | TRBV19 | TRBJ1-1 | CASSNTRSNTEAFF | KLGGALQAK | YES | NO | NO |
| 1602 | TRAV6 | TRAJ21 | CAFSVVFNKFYF | TRBV19 | TRBJ1-1 | CASSIGKDMNTEAFF | KLGGALQAK | YES | NO | NO |
| 1602 | TRAV6 | TRAJ21 | CAFSVVFNKFYF | TRBV19 | TRBJ1-1 | CASSIGKDMNTEAFF | RLRAEAQVK | YES | NO | NO |
| 1604 | TRAV8-2 | TRAJ22 | CVGRKGSARQLTF | TRBV19 | TRBJ1-1 | CASSNQGTEAFF | KLGGALQAK | YES | NO | NO |
| 1608 | TRAV8-6 | TRAJ24 | CAATPLTTTDSWGKLQF | TRBV19 | TRBJ1-1 | CASSKGLTEAFF | KLGGALQAK | YES | NO | YES |
| 1611 | TRAV9-2 | TRAJ42 | CALSDLSGGSQGNLIF | TRBV19 | TRBJ1-1 | CASSIVGTGNTEAFF | KLGGALQAK | YES | NO | YES |
| 1612 | TRAV9-2 | TRAJ50 | CAEKTSYDKVIF | TRBV19 | TRBJ1-1 | CASSIAGYTEAFF | KLGGALQAK | YES | NO | NO |
| 1613 | TRAV9-2 | TRAJ56 | CALSGAGANSKLTF | TRBV19 | TRBJ1-1 | CASSRWEMNTEAFF | KLGGALQAK | YES | NO | NO |
| 1614 | TRAV12-1 | TRAJ21 | CVVNRNFNKFYF | TRBV2 | TRBJ1-1 | CASTKDPGREAFF | KLGGALQAK | YES | NO | NO |
| 1615 | TRAV12-1 | TRAJ33 | CVVNLLNYQLIW | TRBV2 | TRBJ1-1 | CASSDRGETEAFF | KLGGALQAK | YES | NO | NO |
| 1620 | TRAV19 | TRAJ32 | CALRPYGGATNKLIF | TRBV2 | TRBJ1-1 | CASFLGNTEAFF | KLGGALQAK | YES | NO | NO |
| 1621 | TRAV20 | TRAJ8 | CAVQAAGFQKLVF | TRBV2 | TRBJ1-1 | CATKGQLNTEAFF | KLGGALQAK | YES | NO | NO |
| 1622 | TRAV21 | TRAJ20 | CAVRYDYKLSF | TRBV2 | TRBJ1-1 | CASRQTKTNTEAFF | KLGGALQAK | YES | NO | NO |
| 1624 | TRAV22 | TRAJ3 | CAAGLYSSASKIIF | TRBV2 | TRBJ1-1 | CASSGFSHEEGTEAFF | KLGGALQAK | YES | NO | NO |
| 1625 | TRAV29DV5 | TRAJ28 | CAASPFSGAGSYQLTF | TRBV2 | TRBJ1-1 | CASSEMGVGAFF | KLGGALQAK | YES | NO | NO |
| 1626 | TRAV38-2DV8 | TRAJ45 | CAYRMYSGGGADGLTF | TRBV2 | TRBJ1-1 | CASSEAIVPTMNTEAFF | KLGGALQAK | YES | NO | NO |
| 1627 | TRAV38-2DV8 | TRAJ48 | CAYRAPRGNEKLTF | TRBV2 | TRBJ1-1 | CASSDRMNTEAFF | KLGGALQAK | YES | NO | NO |
| 1627 | TRAV38-2DV8 | TRAJ48 | CAYRAPRGNEKLTF | TRBV2 | TRBJ1-1 | CASSDRMNTEAFF | AVFDRKSDAK | YES | NO | NO |
| 1628 | TRAV39 | TRAJ45 | CAVLSGGGADGLTF | TRBV2 | TRBJ1-1 | CASIGHMNTEAFF | KLGGALQAK | YES | NO | NO |
| 1631 | TRAV9-2 | TRAJ21 | CALSVLDNFNKFYF | TRBV2 | TRBJ1-1 | CASPDSLGARDNTEAFF | KLGGALQAK | YES | NO | NO |
| 1633 | TRAV1-1 | TRAJ12 | CAVRPGDSSYKLIF | TRBV20-1 | TRBJ1-1 | CSAPDRASTEAFF | KLGGALQAK | YES | NO | NO |
| 1633 | TRAV1-1 | TRAJ12 | CAVRPGDSSYKLIF | TRBV20-1 | TRBJ1-1 | CSAPDRASTEAFF | AVFDRKSDAK | YES | NO | NO |
| 1637 | TRAV1-2 | TRAJ33 | CAVMDSNYQLIW | TRBV20-1 | TRBJ1-1 | CSARDRDTEAFF | KLGGALQAK | YES | NO | NO |
| 1638 | TRAV12-1 | TRAJ22 | CVVMVWQLTF | TRBV20-1 | TRBJ1-1 | CSARVPGELVTEAFF | KLGGALQAK | YES | NO | NO |
| 1640 | TRAV12-1 | TRAJ6 | CVVNFAPSSGGSYIPTF | TRBV20-1 | TRBJ1-1 | CSARDGRGENTEAFF | KLGGALQAK | YES | NO | NO |
| 1641 | TRAV12-2 | TRAJ21 | CAVQGNFNKFYF | TRBV20-1 | TRBJ1-1 | CSAEQLGVLNTEAFF | KLGGALQAK | YES | NO | NO |
| 1642 | TRAV12-2 | TRAJ48 | CAVFSGVNEKLTF | TRBV20-1 | TRBJ1-1 | CSALEGRAGMWNTEAFF | KLGGALQAK | YES | NO | NO |
| 1643 | TRAV12-3 | TRAJ32 | CAMISGATNKLIF | TRBV20-1 | TRBJ1-1 | CSARDPYEAFF | KLGGALQAK | YES | NO | NO |
| 1644 | TRAV13-1 | TRAJ12 | CAASRADSSYKLIF | TRBV20-1 | TRBJ1-1 | CSAPGIGLTEAFF | KLGGALQAK | YES | NO | NO |
| 1648 | TRAV14DV4 | TRAJ22 | CAMRRPISSGSARQLTF | TRBV20-1 | TRBJ1-1 | CSAWPWGGKRAEAFF | KLGGALQAK | YES | NO | NO |
| 1649 | TRAV14DV4 | TRAJ37 | CAMRGPFSSNTGKLIF | TRBV20-1 | TRBJ1-1 | CSAWDRLNTEAFF | KLGGALQAK | YES | NO | NO |
| 1649 | TRAV14DV4 | TRAJ37 | CAMRGPFSSNTGKLIF | TRBV20-1 | TRBJ1-1 | CSAWDRLNTEAFF | RLRAEAQVK | YES | NO | NO |
| 1649 | TRAV14DV4 | TRAJ37 | CAMRGPFSSNTGKLIF | TRBV20-1 | TRBJ1-1 | CSAWDRLNTEAFF | AVFDRKSDAK | YES | NO | NO |
| 1651 | TRAV14DV4 | TRAJ57 | CAMGTQGGSEKLVF | TRBV20-1 | TRBJ1-1 | CSAGDMNTEAFF | KLGGALQAK | YES | NO | NO |
| 1652 | TRAV17 | TRAJ30 | CATDVYRDDKIIF | TRBV20-1 | TRBJ1-1 | CSARGMNTEAFF | KLGGALQAK | YES | NO | NO |
| 1654 | TRAV19 | TRAJ4 | CALSISGGYNKLIF | TRBV20-1 | TRBJ1-1 | CSAPTATTSSVFTEAFF | KLGGALQAK | YES | NO | YES |
| 1655 | TRAV22 | TRAJ20 | CAVVPGTSNDYKLSF | TRBV20-1 | TRBJ1-1 | CSALSYRGRTEAFF | KLGGALQAK | YES | NO | NO |
| 1659 | TRAV4 | TRAJ40 | CLVGDPVGTYKYIF | TRBV20-1 | TRBJ1-1 | CSATVKSATEAFF | KLGGALQAK | NO | NO | YES |
| 1661 | TRAV5 | TRAJ12 | CAGMDSSYKLIF | TRBV20-1 | TRBJ1-1 | CSARGTSGTGIEAFF | KLGGALQAK | YES | NO | YES |
| 1663 | TRAV5 | TRAJ41 | CAESGNSGYALNF | TRBV20-1 | TRBJ1-1 | CSASSSTGEPEAFF | KLGGALQAK | NO | NO | NO |
| 1665 | TRAV6 | TRAJ30 | CALRDDKIIF | TRBV20-1 | TRBJ1-1 | CSARKWTEAFF | KLGGALQAK | YES | NO | NO |
| 1667 | TRAV8-3 | TRAJ30 | CAVLDDKIIF | TRBV20-1 | TRBJ1-1 | CSYRGLGTEAFF | KLGGALQAK | YES | NO | NO |
| 1668 | TRAV8-3 | TRAJ47 | CAVDTEYGNKLVF | TRBV20-1 | TRBJ1-1 | CSAPSGGLEAFF | KLGGALQAK | YES | NO | NO |
| 1669 | TRAV8-4 | TRAJ53 | CAVTLGGSNYKLTF | TRBV20-1 | TRBJ1-1 | CSARNPRGEDTEAFF | KLGGALQAK | YES | NO | YES |
| 1671 | TRAV17 | TRAJ47 | CATALLSHMEYGNKLVF | TRBV21-1 | TRBJ1-1 | CASSSSFATEAFF | KLGGALQAK | YES | NO | NO |
| 1673 | TRAV1-2 | TRAJ26 | CAVRDWGQNFVF | TRBV24-1 | TRBJ1-1 | CATRGGDRDFDTEAFF | KLGGALQAK | YES | NO | YES |
| 1674 | TRAV12-2 | TRAJ26 | CAPSRYGQNFVF | TRBV24-1 | TRBJ1-1 | CATSDFNGGLSEAFF | KLGGALQAK | YES | NO | NO |
| 1675 | TRAV12-2 | TRAJ28 | CAVSSGAGSYQLTF | TRBV24-1 | TRBJ1-1 | CATSDPNTEAFF | KLGGALQAK | YES | NO | NO |
| 1676 | TRAV12-2 | TRAJ5 | CAVNVDTGRRALTF | TRBV24-1 | TRBJ1-1 | CATRYSENTEAFF | KLGGALQAK | YES | NO | NO |
| 1677 | TRAV14DV4 | TRAJ4 | CAMRRPLGGYNKLIF | TRBV24-1 | TRBJ1-1 | CATSDGTGENTEAFF | KLGGALQAK | YES | NO | NO |
| 1678 | TRAV19 | TRAJ48 | CALSPVTNFGNEKLTF | TRBV24-1 | TRBJ1-1 | CATSGTGLNTEAFF | KLGGALQAK | YES | NO | NO |
| 1679 | TRAV20 | TRAJ41 | CAVQARNSGYALNF | TRBV24-1 | TRBJ1-1 | CATRISGFAAFF | KLGGALQAK | YES | NO | NO |
| 1680 | TRAV20 | TRAJ54 | CAVQVQIIQGAQKLVF | TRBV24-1 | TRBJ1-1 | CATSDLRNTEAFF | KLGGALQAK | YES | NO | NO |
| 1681 | TRAV21 | TRAJ30 | CAVRPMNRDDKIIF | TRBV24-1 | TRBJ1-1 | CATGQLNTEAFF | KLGGALQAK | YES | NO | NO |
| 1682 | TRAV21 | TRAJ43 | CAVWYMRF | TRBV24-1 | TRBJ1-1 | CATLLGNTEAFF | KLGGALQAK | NO | NO | NO |
| 1683 | TRAV5 | TRAJ43 | CAEFRGNNNDMRF | TRBV24-1 | TRBJ1-1 | CATMDSMNTEAFF | KLGGALQAK | YES | NO | NO |
| 1683 | TRAV5 | TRAJ43 | CAEFRGNNNDMRF | TRBV24-1 | TRBJ1-1 | CATMDSMNTEAFF | AVFDRKSDAK | YES | NO | NO |
| 1686 | TRAV9-2 | TRAJ26 | CALKDNYGQNFVF | TRBV24-1 | TRBJ1-1 | CATSDSKEQLNTEAFF | KLGGALQAK | YES | NO | NO |
| 1688 | TRAV12-2 | TRAJ24 | CAVNKSPSDSWGKFQF | TRBV25-1 | TRBJ1-1 | CASRLGQGEAFF | KLGGALQAK | YES | NO | NO |
| 1690 | TRAV13-2 | TRAJ30 | CAEILMNRDDKIIF | TRBV25-1 | TRBJ1-1 | CASSWGARRATEAFF | KLGGALQAK | YES | NO | NO |
| 1691 | TRAV13-2 | TRAJ52 | CAEVPAGGTSYGKLTF | TRBV25-1 | TRBJ1-1 | CASQTGLTEAFF | KLGGALQAK | YES | NO | NO |
| 1693 | TRAV29DV5 | TRAJ37 | CAASGTGNTGKLIF | TRBV25-1 | TRBJ1-1 | CASSETVIGTALEAFF | RLRAEAQVK | YES | YES | YES |
| 1695 | TRAV8-3 | TRAJ17 | CAIKAAGNKLTF | TRBV25-1 | TRBJ1-1 | CASRNGTVNTEAFF | KLGGALQAK | YES | NO | NO |
| 1696 | TRAV8-3 | TRAJ18 | CAVGAMGGSTLGRLYF | TRBV25-1 | TRBJ1-1 | CASSDPRTGWAFF | KLGGALQAK | YES | NO | NO |
| 1697 | TRAV8-3 | TRAJ20 | CAVGGNDYKLSF | TRBV25-1 | TRBJ1-1 | CASGGTLGMNTEAFF | KLGGALQAK | YES | NO | YES |
| 1699 | TRAV1-2 | TRAJ34 | CAVSSMITDKLIF | TRBV27 | TRBJ1-1 | CASRQQCWSGQGRCTEAFF | KLGGALQAK | YES | NO | NO |
| 1703 | TRAV12-1 | TRAJ23 | CVVNKIYNQGGKLIF | TRBV27 | TRBJ1-1 | CASSLTRAEMNTEAFF | KLGGALQAK | YES | NO | NO |
| 1704 | TRAV12-1 | TRAJ28 | CVVNRDHWGWELPTHF | TRBV27 | TRBJ1-1 | CASSSPSSEAFF | KLGGALQAK | NO | NO | NO |
| 1706 | TRAV12-2 | TRAJ33 | CAGTDSNYQLIW | TRBV27 | TRBJ1-1 | CASSLYSQDTEAFF | KLGGALQAK | YES | NO | NO |
| 1707 | TRAV12-2 | TRAJ49 | CAVTTGNQFYF | TRBV27 | TRBJ1-1 | CASSFGQGDFFTEAFF | KLGGALQAK | YES | NO | NO |
| 1709 | TRAV12-3 | TRAJ33 | CAMSGAPSVDSNYQLIW | TRBV27 | TRBJ1-1 | CASRPRDRGLSTEAFF | KLGGALQAK | YES | NO | NO |
| 1711 | TRAV12-3 | TRAJ39 | CAMRDNNAGNMLTF | TRBV27 | TRBJ1-1 | CASSGQMNTEAFF | KLGGALQAK | YES | NO | NO |
| 1712 | TRAV13-1 | TRAJ31 | CAASKGNNARLMF | TRBV27 | TRBJ1-1 | CASSLLGQKAFF | KLGGALQAK | YES | NO | NO |
| 1713 | TRAV13-1 | TRAJ52 | CAAAYGGTSYGKLTF | TRBV27 | TRBJ1-1 | CASSHTQGARTEAFF | KLGGALQAK | YES | YES | NO |
| 1714 | TRAV13-2 | TRAJ43 | CAERGRDDMRF | TRBV27 | TRBJ1-1 | CASSWTGGSLLNTEAFF | KLGGALQAK | YES | NO | YES |
| 1716 | TRAV14DV4 | TRAJ33 | CAMRESDSNYQLIW | TRBV27 | TRBJ1-1 | CASSLSAGMNTEAFF | KLGGALQAK | YES | NO | NO |
| 1717 | TRAV14DV4 | TRAJ48 | CASLPHFGNEKLTF | TRBV27 | TRBJ1-1 | CASSILTGTPTEAFF | KLGGALQAK | YES | NO | NO |
| 1718 | TRAV14DV4 | TRAJ49 | CAMRGPNTGNQFYF | TRBV27 | TRBJ1-1 | CASSFGAGVTEAFF | KLGGALQAK | YES | NO | NO |
| 1720 | TRAV17 | TRAJ20 | CATDGNDYKLSF | TRBV27 | TRBJ1-1 | CASSPEGQPNTEAFF | KLGGALQAK | YES | NO | NO |
| 1723 | TRAV17 | TRAJ40 | CATDGRGTYKYIF | TRBV27 | TRBJ1-1 | CASRWTGGDTEAFF | KLGGALQAK | YES | NO | YES |
| 1724 | TRAV17 | TRAJ49 | CATDTGNQFYF | TRBV27 | TRBJ1-1 | CASSKNRENTEAFF | KLGGALQAK | YES | NO | NO |
| 1725 | TRAV19 | TRAJ16 | CALSEF | TRBV27 | TRBJ1-1 | CASSPRGLVTEAFF | RLRAEAQVK | NO | NO | NO |
| 1726 | TRAV19 | TRAJ21 | CALSGAYNFNKFYF | TRBV27 | TRBJ1-1 | CASSLGPGGNTEAFF | KLGGALQAK | YES | NO | NO |
| 1727 | TRAV19 | TRAJ27 | CALSDLYNTNAGKSTF | TRBV27 | TRBJ1-1 | CASRSGGKNTEAFF | KLGGALQAK | YES | NO | NO |
| 1728 | TRAV19 | TRAJ30 | CALSTNRDDKIIF | TRBV27 | TRBJ1-1 | CASSRTLNTEAFF | KLGGALQAK | YES | NO | NO |
| 1731 | TRAV19 | TRAJ56 | CALTITGANSKLTF | TRBV27 | TRBJ1-1 | CASSTAGHNTEAFF | KLGGALQAK | YES | NO | NO |
| 1732 | TRAV20 | TRAJ11 | CAVQPRAMNSGYSTLTF | TRBV27 | TRBJ1-1 | CASSLIENTEAFF | KLGGALQAK | YES | NO | NO |
| 1733 | TRAV20 | TRAJ27 | CAVLSNAGKSTF | TRBV27 | TRBJ1-1 | CASSLLGGDTEAFF | RLRAEAQVK | YES | NO | YES |
| 1734 | TRAV20 | TRAJ34 | CAARIYNTDKLIF | TRBV27 | TRBJ1-1 | CASVGGGGTEAFF | KLGGALQAK | YES | NO | NO |
| 1735 | TRAV21 | TRAJ15 | CAVQGIQAGTALIF | TRBV27 | TRBJ1-1 | CASKGLRPEAFF | KLGGALQAK | YES | NO | NO |
| 1736 | TRAV21 | TRAJ29 | CAVPSLHSGNTPLVF | TRBV27 | TRBJ1-1 | CASISPSTEAFF | KLGGALQAK | YES | NO | NO |

| | | | | | | | | | | |
|---|---|---|---|---|---|---|---|---|---|---|
| 1737 | TRAV21 | TRAJ50 | CAVGGRTSYDKVIF | TRBV27 | TRBJ1-1 | CASSFSQATEAFF | KLGGALQAK | YES | NO | NO |
| 1738 | TRAV22 | TRAJ39 | CAVELEGNAGNMLTF | TRBV27 | TRBJ1-1 | CASCSLGGAEAFF | KLGGALQAK | YES | NO | NO |
| 1739 | TRAV22 | TRAJ42 | CAVEFVGYGGSQGNLIF | TRBV27 | TRBJ1-1 | CASSLIGFGGLNTEAFF | KLGGALQAK | YES | NO | NO |
| 1740 | TRAV26-2 | TRAJ53 | CILRDPNSGGSNYKLTF | TRBV27 | TRBJ1-1 | CATTAAGGLNTEAFF | KLGGALQAK | YES | NO | NO |
| 1744 | TRAV29DV5 | TRAJ41 | CAAPMDSGYALNF | TRBV27 | TRBJ1-1 | CASSFGRGNTEAFF | KLGGALQAK | YES | NO | NO |
| 1745 | TRAV29DV5 | TRAJ9 | CAASDSSGGFKTIF | TRBV27 | TRBJ1-1 | CASSSDLNTEAFF | KLGGALQAK | YES | NO | YES |
| 1747 | TRAV3 | TRAJ40 | CAVRDVATSGTYKYIF | TRBV27 | TRBJ1-1 | CASSLYPAMNTEAFF | KLGGALQAK | NO | NO | NO |
| 1750 | TRAV38-2DV8 | TRAJ43 | CAYRGDNDMRF | TRBV27 | TRBJ1-1 | CASSFVGGFMATEAFF | KLGGALQAK | YES | NO | NO |
| 1751 | TRAV39 | TRAJ27 | CAVDTNAGKSTF | TRBV27 | TRBJ1-1 | CASMTGSTEAFF | KLGGALQAK | YES | NO | NO |
| 1752 | TRAV4 | TRAJ18 | CLVGGGSTLGRLYF | TRBV27 | TRBJ1-1 | CASSWTGALNTEAFF | KLGGALQAK | YES | NO | NO |
| 1753 | TRAV41 | TRAJ48 | CAAPPSFGNEKLTF | TRBV27 | TRBJ1-1 | CASSLSGQGNTEAFF | KLGGALQAK | YES | NO | NO |
| 1754 | TRAV41 | TRAJ52 | CAVLPNAGGTSYGKLTF | TRBV27 | TRBJ1-1 | CASSPAGQGNTEAFF | KLGGALQAK | YES | NO | NO |
| 1757 | TRAV6 | TRAJ28 | CALGPGAGSYQLTF | TRBV27 | TRBJ1-1 | CASSLVGGDTEAFF | KLGGALQAK | YES | NO | NO |
| 1758 | TRAV6 | TRAJ28 | CALGPGAGSYQLTF | TRBV27 | TRBJ1-1 | CASSLAGGQTEAFF | KLGGALQAK | YES | NO | NO |
| 1761 | TRAV8-6 | TRAJ38 | CAVGAGNNRKLIW | TRBV27 | TRBJ1-1 | CASSLGQGNTEAFF | KLGGALQAK | YES | NO | NO |
| 1762 | TRAV12-1 | TRAJ49 | CVVNVENQFYF | TRBV28 | TRBJ1-1 | CASTEDRTGIVNTEAFF | KLGGALQAK | YES | NO | NO |
| 1763 | TRAV12-2 | TRAJ17 | CAVEAAGNKLTF | TRBV28 | TRBJ1-1 | CASSLGDPGTEAFF | KLGGALQAK | YES | NO | NO |
| 1764 | TRAV12-2 | TRAJ3 | CAVNRYSSASKIIF | TRBV28 | TRBJ1-1 | CASSLERGVSAAFF | KLGGALQAK | YES | NO | NO |
| 1765 | TRAV13-1 | TRAJ26 | CAAIDGQNFVF | TRBV28 | TRBJ1-1 | CASSLRQDMNTEAFF | KLGGALQAK | YES | NO | NO |
| 1766 | TRAV13-1 | TRAJ5 | CAASSAGRRALTF | TRBV28 | TRBJ1-1 | CASSFRGTEAFF | KLGGALQAK | YES | NO | NO |
| 1767 | TRAV13-2 | TRAJ11 | CAERDSGYSTLTF | TRBV28 | TRBJ1-1 | CASRAPGNTEAFF | KLGGALQAK | YES | NO | NO |
| 1771 | TRAV19 | TRAJ12 | CALSEAEMDSSYKLIF | TRBV28 | TRBJ1-1 | CASSLSGLSEAFF | KLGGALQAK | YES | NO | NO |
| 1772 | TRAV19 | TRAJ18 | CALSSDRGSTLGRLYF | TRBV28 | TRBJ1-1 | CASSLFGSKGGAFF | KLGGALQAK | YES | NO | NO |
| 1773 | TRAV20 | TRAJ52 | CAVRRAGGTSYGKLTF | TRBV28 | TRBJ1-1 | CASSPRWDDPVVNAEAFF | KLGGALQAK | YES | NO | NO |
| 1774 | TRAV3 | TRAJ26 | CAVKDYGQNFVF | TRBV28 | TRBJ1-1 | CASSFVGQGGTEAFF | KLGGALQAK | NO | NO | NO |
| 1775 | TRAV3 | TRAJ26 | CAVREPYGQNFVF | TRBV28 | TRBJ1-1 | CASRVEDRLNTEAFF | KLGGALQAK | NO | NO | NO |
| 1776 | TRAV38-2DV8 | TRAJ49 | CAYRSALTGNQFYF | TRBV28 | TRBJ1-1 | CASSPTGRVVPLPNTEAFF | KLGGALQAK | YES | NO | NO |
| 1780 | TRAV5 | TRAJ39 | CAEINAGNMLTF | TRBV28 | TRBJ1-1 | CASSLGGFDMNTEAFF | KLGGALQAK | YES | NO | NO |
| 1782 | TRAV8-3 | TRAJ49 | CAVGPNTGNQFYF | TRBV28 | TRBJ1-1 | CASSSNTEAFF | KLGGALQAK | YES | NO | NO |
| 1783 | TRAV8-6 | TRAJ4 | CAVSEVWGYNKLIF | TRBV28 | TRBJ1-1 | CASSLMTGMNTEAFF | KLGGALQAK | YES | NO | NO |
| 1784 | TRAV12-3 | TRAJ3 | CAMSASDASKIIF | TRBV29-1 | TRBJ1-1 | CSVHVPMNTEAFF | RLRAEAQVK | YES | YES | YES |
| 1785 | TRAV13-2 | TRAJ16 | CAENF | TRBV29-1 | TRBJ1-1 | CSVPREGNTEAFF | KLGGALQAK | NO | NO | NO |
| 1787 | TRAV14DV4 | TRAJ58 | CAMRDLGRGSRLTF | TRBV29-1 | TRBJ1-1 | CSVAGTFEAFF | KLGGALQAK | YES | NO | NO |
| 1789 | TRAV19 | TRAJ26 | CALSGANRNYGQNFVF | TRBV29-1 | TRBJ1-1 | CSVGFREMGATEAFF | KLGGALQAK | YES | NO | NO |
| 1793 | TRAV19 | TRAJ52 | CALSENAGGTSYGKLTF | TRBV29-1 | TRBJ1-1 | CSVPGGANTEAFF | KLGGALQAK | YES | NO | NO |
| 1794 | TRAV19 | TRAJ52 | CALSEASPGTSYGKLTF | TRBV29-1 | TRBJ1-1 | CSVGGASNTEAFF | KLGGALQAK | YES | NO | NO |
| 1795 | TRAV19 | TRAJ52 | CALSEGAGGTSYGKLTF | TRBV29-1 | TRBJ1-1 | CSVPPGGAGRAFF | KLGGALQAK | NO | NO | NO |
| 1796 | TRAV19 | TRAJ52 | CALSEATGGTSYGKLTF | TRBV29-1 | TRBJ1-1 | CSVPGTVNTEAFF | KLGGALQAK | NO | NO | NO |
| 1797 | TRAV19 | TRAJ54 | CALSASPVQGAQKLVF | TRBV29-1 | TRBJ1-1 | CSVEKNTEAFF | KLGGALQAK | YES | NO | NO |
| 1798 | TRAV21 | TRAJ21 | CAVRPGYNFNKFYF | TRBV29-1 | TRBJ1-1 | CSVRDSQNTEAFF | KLGGALQAK | YES | NO | NO |
| 1799 | TRAV21 | TRAJ22 | CAVSPVSSGSARQLTF | TRBV29-1 | TRBJ1-1 | CSVEWVGGPTEAFF | KLGGALQAK | YES | NO | NO |
| 1801 | TRAV21 | TRAJ24 | CAVVDSWGKLQF | TRBV29-1 | TRBJ1-1 | CSVEDFEGGSGTEAFF | KLGGALQAK | YES | NO | NO |
| 1802 | TRAV21 | TRAJ45 | CAVLGGGADGLTF | TRBV29-1 | TRBJ1-1 | CSFRIGGTEAFF | AVFDRKSDAK | NO | NO | NO |
| 1802 | TRAV21 | TRAJ45 | CAVLGGGADGLTF | TRBV29-1 | TRBJ1-1 | CSFRIGGTEAFF | IVTDFSVIK | NO | NO | NO |
| 1808 | TRAV27 | TRAJ20 | CAGAPSNDYKLSF | TRBV29-1 | TRBJ1-1 | CSVAATGNTEAFF | KLGGALQAK | YES | NO | NO |
| 1809 | TRAV27 | TRAJ20 | CAGISSNDYKLSF | TRBV29-1 | TRBJ1-1 | CSVAGTGNTEAFF | KLGGALQAK | YES | NO | NO |
| 1810 | TRAV27 | TRAJ20 | CAGVVTNDYKLSF | TRBV29-1 | TRBJ1-1 | CSVAGTGNTEAFF | KLGGALQAK | YES | NO | NO |
| 1811 | TRAV27 | TRAJ20 | CAGVESNDYKLSF | TRBV29-1 | TRBJ1-1 | CSVAAPGNTEAFF | KLGGALQAK | YES | NO | NO |
| 1811 | TRAV27 | TRAJ20 | CAGVESNDYKLSF | TRBV29-1 | TRBJ1-1 | CSVAAPGNTEAFF | RLRAEAQVK | YES | NO | NO |
| 1811 | TRAV27 | TRAJ20 | CAGVESNDYKLSF | TRBV29-1 | TRBJ1-1 | CSVAAPGNTEAFF | AVFDRKSDAK | YES | NO | NO |
| 1812 | TRAV27 | TRAJ20 | CAGPDVNDYKLSF | TRBV29-1 | TRBJ1-1 | CSVASVGNTEAFF | KLGGALQAK | YES | NO | YES |
| 1813 | TRAV27 | TRAJ20 | CAGVGSNDYKLSF | TRBV29-1 | TRBJ1-1 | CSVAGTGNTEAFF | KLGGALQAK | YES | NO | NO |
| 1814 | TRAV27 | TRAJ20 | CAGGPQNDYKLSF | TRBV29-1 | TRBJ1-1 | CSVAGVGNTEAFF | KLGGALQAK | YES | NO | NO |
| 1815 | TRAV27 | TRAJ20 | CAGVGSNDYKLSF | TRBV29-1 | TRBJ1-1 | CSVVGVGNTEAFF | KLGGALQAK | YES | NO | NO |
| 1816 | TRAV27 | TRAJ20 | CAGVASNDYKLSF | TRBV29-1 | TRBJ1-1 | CSVAGTGNTEAFF | KLGGALQAK | YES | NO | NO |
| 1817 | TRAV27 | TRAJ20 | CAGVPSNDYKLSF | TRBV29-1 | TRBJ1-1 | CSVVGVGNTEAFF | KLGGALQAK | YES | NO | NO |
| 1818 | TRAV27 | TRAJ20 | CAGVSSNDYKLSF | TRBV29-1 | TRBJ1-1 | CSVVGTGNTEAFF | KLGGALQAK | YES | NO | NO |
| 1819 | TRAV29DV5 | TRAJ28 | CAANSGAGSYQLTF | TRBV29-1 | TRBJ1-1 | CSVPGTGGLIAFF | KLGGALQAK | NO | NO | NO |
| 1821 | TRAV3 | TRAJ39 | CAVRRFNNAGNMLTF | TRBV29-1 | TRBJ1-1 | CSVEDRSGVTEAFF | KLGGALQAK | NO | NO | NO |
| 1822 | TRAV3 | TRAJ40 | CAVRDSASGTYKYIF | TRBV29-1 | TRBJ1-1 | CSVLQGYEAFF | KLGGALQAK | NO | NO | NO |
| 1824 | TRAV38-1 | TRAJ48 | CAFLFGNEKLTF | TRBV29-1 | TRBJ1-1 | CSVENLPEAFF | KLGGALQAK | YES | NO | NO |
| 1825 | TRAV38-2DV8 | TRAJ43 | CAYRSFGNNDMRF | TRBV29-1 | TRBJ1-1 | CSVNKRGAGGMNTEAFF | KLGGALQAK | YES | NO | YES |
| 1826 | TRAV4 | TRAJ10 | CLVGELGGGNKLTF | TRBV29-1 | TRBJ1-1 | CSVPSSNTEAFF | KLGGALQAK | YES | NO | NO |
| 1830 | TRAV8-2 | TRAJ30 | CVVSDLGKIIF | TRBV29-1 | TRBJ1-1 | CSVRARGYTEAFF | KLGGALQAK | YES | NO | NO |
| 1835 | TRAV9-2 | TRAJ53 | CALGGNYKLTF | TRBV29-1 | TRBJ1-1 | CSVDPGAEAFF | KLGGALQAK | YES | NO | NO |
| 1837 | TRAV1-2 | TRAJ8 | CAVRDADTGFQKLVF | TRBV3-1 | TRBJ1-1 | CASSQNGGAHRLMNTEAFF | KLGGALQAK | YES | NO | NO |
| 1838 | TRAV12-1 | TRAJ45 | CVARGVNSGGGADGLTF | TRBV3-1 | TRBJ1-1 | CASSQGGSFPAFF | KLGGALQAK | YES | NO | NO |
| 1839 | TRAV12-2 | TRAJ26 | CAVDEDNYGQNFVF | TRBV3-1 | TRBJ1-1 | CASSQRGNTEAFF | KLGGALQAK | YES | NO | NO |
| 1840 | TRAV12-2 | TRAJ26 | CAVDRDNYGQNFVF | TRBV3-1 | TRBJ1-1 | CASSQRGNTEAFF | KLGGALQAK | YES | NO | NO |
| 1841 | TRAV12-2 | TRAJ35 | CAVKRQGFGNVLHC | TRBV3-1 | TRBJ1-1 | CASSIQGSTEAFF | KLGGALQAK | NO | NO | NO |
| 1844 | TRAV17 | TRAJ52 | CAPDINAGGTSYGKLTF | TRBV3-1 | TRBJ1-1 | CASSQGGGGYLNTEAFF | KLGGALQAK | YES | NO | NO |
| 1850 | TRAV5 | TRAJ4 | CAEIPPPFSGGYNKLIF | TRBV3-1 | TRBJ1-1 | CASSQVTGGNTEAFF | KLGGALQAK | YES | NO | NO |
| 1853 | TRAV10 | TRAJ32 | CVVSARGGATNKLIF | TRBV30 | TRBJ1-1 | CAWSAGQGFTEAFF | KLGGALQAK | YES | NO | NO |
| 1854 | TRAV12-1 | TRAJ43 | CVVNTYNDMRF | TRBV30 | TRBJ1-1 | CAWSGGTTEAFF | KLGGALQAK | YES | NO | NO |
| 1855 | TRAV12-2 | TRAJ7 | CAVPSYGNNRLAF | TRBV30 | TRBJ1-1 | CAWSVGGGEAFF | KLGGALQAK | NO | NO | NO |
| 1856 | TRAV13-1 | TRAJ47 | CAASGNKLVF | TRBV30 | TRBJ1-1 | CAWNLGQLNTEAFF | KLGGALQAK | NO | NO | NO |
| 1857 | TRAV13-2 | TRAJ16 | CAEDTIDGQKLLF | TRBV30 | TRBJ1-1 | CAWSVTGNTEAFF | KLGGALQAK | NO | NO | NO |
| 1858 | TRAV13-2 | TRAJ40 | CAENKGGYKYIF | TRBV30 | TRBJ1-1 | CAWSVSAGRHTEAFF | KLGGALQAK | NO | NO | NO |
| 1860 | TRAV17 | TRAJ24 | CARDSWGKLQF | TRBV30 | TRBJ1-1 | CAWRGQDMNTEAFF | KLGGALQAK | YES | NO | NO |
| 1861 | TRAV19 | TRAJ52 | CALSEAYAGGTSYGKLTF | TRBV30 | TRBJ1-1 | CASRTGNLNTEAFF | AVFDRKSDAK | NO | NO | NO |
| 1862 | TRAV20 | TRAJ39 | CAVNPNNAGNMLTF | TRBV30 | TRBJ1-1 | CAWTRTGNTEAFF | KLGGALQAK | YES | NO | NO |
| 1863 | TRAV20 | TRAJ42 | CAVCPYGGSQGNLIF | TRBV30 | TRBJ1-1 | CAWSLSGGGEAFF | KLGGALQAK | NO | NO | NO |
| 1864 | TRAV21 | TRAJ43 | CAVRWRYNNNDMRF | TRBV30 | TRBJ1-1 | CAAEDPEWGAEAFF | KLGGALQAK | NO | NO | NO |
| 1865 | TRAV26-2 | TRAJ28 | CIPTGIYSGAGSYQLTF | TRBV30 | TRBJ1-1 | CAWKQGFLAEAFF | KLGGALQAK | NO | NO | NO |
| 1866 | TRAV38-1 | TRAJ35 | CAFMKHERDWGDFGNVLHC | TRBV30 | TRBJ1-1 | CASDRKNTEAFF | KLGGALQAK | NO | NO | NO |
| 1868 | TRAV5 | TRAJ31 | CAEYNARLMF | TRBV30 | TRBJ1-1 | CAWRISSEAFF | KLGGALQAK | NO | NO | NO |
| 1869 | TRAV5 | TRAJ39 | CAESIWNNAGNMLTF | TRBV30 | TRBJ1-1 | CAWSFGVDPGSEAFF | KLGGALQAK | YES | NO | NO |
| 1874 | TRAV1-2 | TRAJ10 | CAVEEGGGNKLTF | TRBV4-1 | TRBJ1-1 | CASSRQSNTEAFF | KLGGALQAK | YES | NO | NO |
| 1877 | TRAV12-1 | TRAJ49 | CVVTGNQFYF | TRBV4-1 | TRBJ1-1 | CASSQYHVEGSTEAFF | KLGGALQAK | YES | NO | NO |
| 1878 | TRAV12-2 | TRAJ31 | CAVNRVRDARLMF | TRBV4-1 | TRBJ1-1 | CASRVGNTEAFF | AVFDRKSDAK | YES | NO | NO |
| 1879 | TRAV12-3 | TRAJ34 | CAMSRLEGDKLIF | TRBV4-1 | TRBJ1-1 | CASRVGNTEAFF | KLGGALQAK | YES | NO | NO |
| 1880 | TRAV12-3 | TRAJ39 | CAIGPGNMLTF | TRBV4-1 | TRBJ1-1 | CASSQPYGQPNTEAFF | KLGGALQAK | YES | NO | NO |
| 1881 | TRAV13-1 | TRAJ4 | CAASSEAGGYNKLIF | TRBV4-1 | TRBJ1-1 | CASSQGQGATEAFF | KLGGALQAK | YES | NO | NO |
| 1882 | TRAV13-1 | TRAJ4 | CAASGAGGYNKLIF | TRBV4-1 | TRBJ1-1 | CASSQGTGGTEAFF | KLGGALQAK | YES | NO | YES |
| 1884 | TRAV13-2 | TRAJ45 | CAEAGGGADGLTF | TRBV4-1 | TRBJ1-1 | CASSSLGGVDTEAFF | KLGGALQAK | YES | NO | NO |
| 1885 | TRAV13-2 | TRAJ53 | CAENIDSGGSNYKLTF | TRBV4-1 | TRBJ1-1 | CASSQAPARNTEAFF | KLGGALQAK | YES | NO | NO |
| 1886 | TRAV13-2 | TRAJ7 | CAESVGYGNNRLAF | TRBV4-1 | TRBJ1-1 | CASRPSDRATEAFF | KLGGALQAK | YES | NO | NO |
| 1887 | TRAV14DV4 | TRAJ36 | CAITGANNLFF | TRBV4-1 | TRBJ1-1 | CASSQGHIGAVSMNTEAFF | KLGGALQAK | YES | NO | NO |
| 1888 | TRAV14DV4 | TRAJ5 | CAVYIMDTGRRALTF | TRBV4-1 | TRBJ1-1 | CASSEGRMNTEAFF | KLGGALQAK | YES | NO | NO |
| 1890 | TRAV17 | TRAJ20 | CAPDDYKLSF | TRBV4-1 | TRBJ1-1 | CASSRDRGDTEAFF | KLGGALQAK | YES | NO | NO |
| 1891 | TRAV19 | TRAJ26 | CALSEANYGQNFVF | TRBV4-1 | TRBJ1-1 | CASSHGTGGNTEAFF | KLGGALQAK | YES | NO | NO |
| 1892 | TRAV19 | TRAJ4 | CALSQAFSGGYNKLIF | TRBV4-1 | TRBJ1-1 | CASSQDSGGTLNTEAFF | KLGGALQAK | YES | NO | NO |
| 1896 | TRAV21 | TRAJ52 | CAVFNAGGTSYGKLTF | TRBV4-1 | TRBJ1-1 | CASSTYPTGNTEAFF | IVTDFSVIK | YES | NO | NO |
| 1896 | TRAV21 | TRAJ52 | CAVFNAGGTSYGKLTF | TRBV4-1 | TRBJ1-1 | CASSTYPTGNTEAFF | KLGGALQAK | YES | NO | NO |

| | | | | | | | | | | |
|---|---|---|---|---|---|---|---|---|---|---|
| 1898 | TRAV22 | TRAJ6 | CAVERRSGGSYIPTF | TRBV4-1 | TRBJ1-1 | CASSQGMTEAFF | KLGGALQAK | YES | NO | NO |
| 1899 | TRAV23DV6 | TRAJ45 | CAAVHSGGGADGLTF | TRBV4-1 | TRBJ1-1 | CASSPENRGAEAFF | KLGGALQAK | YES | NO | NO |
| 1901 | TRAV26-2 | TRAJ30 | CILIYMNRDDKIIF | TRBV4-1 | TRBJ1-1 | CASREGESNTEAFF | KLGGALQAK | YES | NO | YES |
| 1902 | TRAV27 | TRAJ40 | CAGISVSGTYKYIF | TRBV4-1 | TRBJ1-1 | CASSQGTGWDTEAFF | KLGGALQAK | YES | NO | NO |
| 1903 | TRAV29DV5 | TRAJ26 | CAASGDYGQNFVF | TRBV4-1 | TRBJ1-1 | CASSQAPLGNTEAFF | KLGGALQAK | YES | NO | NO |
| 1905 | TRAV29DV5 | TRAJ43 | CAASAWGNNDMRF | TRBV4-1 | TRBJ1-1 | CASSQGLPAGNTEAFF | KLGGALQAK | YES | NO | YES |
| 1907 | TRAV39 | TRAJ27 | CAVDMGAGKSTF | TRBV4-1 | TRBJ1-1 | CASSQQQGYGTEAFF | KLGGALQAK | YES | NO | NO |
| 1909 | TRAV5 | TRAJ20 | CAESMNDYKLSF | TRBV4-1 | TRBJ1-1 | CASSRPGQENTEAFF | KLGGALQAK | YES | NO | YES |
| 1914 | TRAV9-2 | TRAJ20 | CALDANDYKLSF | TRBV4-1 | TRBJ1-1 | CASSQTGLTNTEAFF | KLGGALQAK | YES | NO | YES |
| 1917 | TRAV1-2 | TRAJ17 | CAVEAAGNKLTF | TRBV4-2 | TRBJ1-1 | CASSQTTRQSNTEAFF | KLGGALQAK | YES | NO | YES |
| 1918 | TRAV1-2 | TRAJ23 | CAVHNQGGKLIF | TRBV4-2 | TRBJ1-1 | CASSQNRVGAEAFF | KLGGALQAK | YES | NO | YES |
| 1920 | TRAV17 | TRAJ54 | CATDPQGAQKLVF | TRBV4-2 | TRBJ1-1 | CASSQGGGWTEAFF | KLGGALQAK | YES | NO | NO |
| 1921 | TRAV19 | TRAJ12 | CALSEPMMDSSYKLIF | TRBV4-2 | TRBJ1-1 | CASSQAEGGNTEAFF | KLGGALQAK | YES | NO | NO |
| 1925 | TRAV36DV7 | TRAJ31 | CAVGNNNARLMF | TRBV4-2 | TRBJ1-1 | CASSPSQGLTEAFF | KLGGALQAK | NO | NO | YES |
| 1927 | TRAV1-2 | TRAJ34 | CAVRDSADKLIF | TRBV4-3 | TRBJ1-1 | CASSQDHQGGGTEAFF | KLGGALQAK | YES | NO | YES |
| 1928 | TRAV12-2 | TRAJ17 | CAVNEAAGNKLTF | TRBV4-3 | TRBJ1-1 | CASSQGYRLGTEAFF | KLGGALQAK | YES | NO | YES |
| 1930 | TRAV13-2 | TRAJ32 | CAENGGATNKLIF | TRBV4-3 | TRBJ1-1 | CASSQGQGNTEAFF | KLGGALQAK | YES | NO | NO |
| 1933 | TRAV21 | TRAJ21 | CAVTLYNFNKFYF | TRBV4-3 | TRBJ1-1 | CASSQGTGRNTEAFF | KLGGALQAK | YES | NO | NO |
| 1933 | TRAV21 | TRAJ21 | CAVTLYNFNKFYF | TRBV4-3 | TRBJ1-1 | CASSQGTGRNTEAFF | AVFDRKSDAK | YES | NO | NO |
| 1934 | TRAV24 | TRAJ52 | CAFIGDAGGTSYGKLTF | TRBV4-3 | TRBJ1-1 | CASSQDNGGMVAEAFF | KLGGALQAK | YES | NO | NO |
| 1937 | TRAV38-1 | TRAJ43 | CAFEWGNNNNDMRF | TRBV4-3 | TRBJ1-1 | CASSQGGQGATEAFF | KLGGALQAK | YES | NO | NO |
| 1938 | TRAV8-3 | TRAJ49 | CAVGAGTGNQFYF | TRBV4-3 | TRBJ1-1 | CASSQETGWGTEAFF | KLGGALQAK | YES | NO | NO |
| 1940 | TRAV9-2 | TRAJ17 | CALSDLKAAGNKLTF | TRBV4-3 | TRBJ1-1 | CASSQDLSRLNTEAFF | KLGGALQAK | YES | NO | NO |
| 1944 | TRAV1-2 | TRAJ39 | CAVRASNAGNMLTF | TRBV5-1 | TRBJ1-1 | CASSLAGQTPPWDTEAFF | KLGGALQAK | YES | NO | NO |
| 1946 | TRAV1-2 | TRAJ6 | CAVRSSGGSYIPTF | TRBV5-1 | TRBJ1-1 | CASSLTPNTEAFF | KLGGALQAK | YES | NO | NO |
| 1947 | TRAV12-1 | TRAJ43 | CVVGNNNDMRF | TRBV5-1 | TRBJ1-1 | CASSLVIRNEAFF | KLGGALQAK | YES | NO | NO |
| 1949 | TRAV13-1 | TRAJ8 | CAAITGFQKLVF | TRBV5-1 | TRBJ1-1 | CASSLRALNTEAFF | KLGGALQAK | YES | NO | NO |
| 1950 | TRAV13-2 | TRAJ15 | CAEIPKNQAGTALIF | TRBV5-1 | TRBJ1-1 | CASSDGTGNTEAFF | KLGGALQAK | YES | NO | NO |
| 1951 | TRAV13-2 | TRAJ4 | CAENIGTLSGGYNKLIF | TRBV5-1 | TRBJ1-1 | CASRTKQGATEAFF | KLGGALQAK | YES | NO | NO |
| 1952 | TRAV14DV4 | TRAJ22 | CAMRRPISSGSARQLTF | TRBV5-1 | TRBJ1-1 | CASSLGRGLNTEAFF | KLGGALQAK | YES | NO | NO |
| 1954 | TRAV14DV4 | TRAJ4 | CAMRRPSGGYNKLIF | TRBV5-1 | TRBJ1-1 | CASSLVWGAEPEAFF | KLGGALQAK | YES | NO | NO |
| 1955 | TRAV16 | TRAJ17 | CALSGRAAGNKLTF | TRBV5-1 | TRBJ1-1 | CASSLVDRAMNLNTEAFF | KLGGALQAK | NO | NO | NO |
| 1958 | TRAV17 | TRAJ36 | CAQTGANNLFF | TRBV5-1 | TRBJ1-1 | CASSLGGYRAEAFF | KLGGALQAK | YES | NO | NO |
| 1960 | TRAV20 | TRAJ20 | CAVPSSNDYKLSF | TRBV5-1 | TRBJ1-1 | CASSEGGGTEAFF | KLGGALQAK | YES | NO | NO |
| 1961 | TRAV20 | TRAJ20 | CAVRANDYKLSF | TRBV5-1 | TRBJ1-1 | CASSKGQASTEAFF | KLGGALQAK | YES | NO | NO |
| 1964 | TRAV30 | TRAJ17 | CGTLDAAGNKLTF | TRBV5-1 | TRBJ1-1 | CASSAGITNTEAFF | KLGGALQAK | YES | NO | NO |
| 1965 | TRAV38-1 | TRAJ41 | CAFRSGYALNF | TRBV5-1 | TRBJ1-1 | CASSLPGGALNTEAFF | KLGGALQAK | YES | NO | NO |
| 1966 | TRAV38-2DV8 | TRAJ32 | CAYRSEREVDYGGATNKLIF | TRBV5-1 | TRBJ1-1 | CASSPLRRLQPEAFF | KLGGALQAK | NO | NO | NO |
| 1969 | TRAV4 | TRAJ27 | CLVGDSAGKSTF | TRBV5-1 | TRBJ1-1 | CASSPHGQGYTEAFF | KLGGALQAK | NO | NO | NO |
| 1970 | TRAV5 | TRAJ45 | CAEIVRGGGADGLTF | TRBV5-1 | TRBJ1-1 | CASSADLGNTEAFF | KLGGALQAK | YES | NO | NO |
| 1971 | TRAV8-1 | TRAJ42 | CAEVYGGSQGNLIF | TRBV5-1 | TRBJ1-1 | CASSTTGDTEPWTEAFF | KLGGALQAK | YES | NO | NO |
| 1974 | TRAV1-2 | TRAJ15 | CAVRGLQAGTALIF | TRBV5-4 | TRBJ1-1 | CASSPTVNTEAFF | KLGGALQAK | YES | NO | NO |
| 1976 | TRAV14DV4 | TRAJ22 | CAMRGISSGSARQLTF | TRBV5-4 | TRBJ1-1 | CASSFWTGETEAFF | KLGGALQAK | YES | NO | NO |
| 1978 | TRAV17 | TRAJ42 | CATDSNYGGSQGNLIF | TRBV5-4 | TRBJ1-1 | CASSLAGGYTEAFF | RLRAEAQVK | YES | YES | YES |
| 1980 | TRAV20 | TRAJ40 | CAVQAWTYKYIF | TRBV5-4 | TRBJ1-1 | CASSLGTGAENTEAFF | KLGGALQAK | YES | NO | NO |
| 1981 | TRAV22 | TRAJ26 | CAVGDNYGQNFVF | TRBV5-4 | TRBJ1-1 | CASSGKAGADTEAFF | KLGGALQAK | YES | NO | NO |
| 1982 | TRAV25 | TRAJ49 | CAGGDTGNQFYF | TRBV5-4 | TRBJ1-1 | CASSWGVRDMNTEAFF | KLGGALQAK | YES | NO | NO |
| 1983 | TRAV26-2 | TRAJ52 | CILRDFIGAGGTSYGKLTF | TRBV5-4 | TRBJ1-1 | CASSLGGMEGTEAFF | KLGGALQAK | YES | NO | NO |
| 1986 | TRAV36DV7 | TRAJ56 | CASYTGANSKLTF | TRBV5-4 | TRBJ1-1 | CASSIRQRPGIRAFF | KLGGALQAK | NO | NO | NO |
| 1987 | TRAV38-2DV8 | TRAJ41 | CAYRSAADSGYALNF | TRBV5-4 | TRBJ1-1 | CASSLGLGKSVDTEAFF | KLGGALQAK | YES | NO | NO |
| 1988 | TRAV39 | TRAJ43 | CAVDMVNDMRF | TRBV5-4 | TRBJ1-1 | CASSLAWGLNTEAFF | KLGGALQAK | YES | NO | NO |
| 1991 | TRAV8-4 | TRAJ41 | CAVTPMISNSGYALNF | TRBV5-4 | TRBJ1-1 | CASSLGETEAFF | KLGGALQAK | YES | NO | NO |
| 1992 | TRAV8-6 | TRAJ21 | CAVSDRDNFNKFYF | TRBV5-4 | TRBJ1-1 | CASSRTENTEAFF | KLGGALQAK | YES | NO | NO |
| 1993 | TRAV1-1 | TRAJ12 | CAVHILDDSSYKLIF | TRBV5-5 | TRBJ1-1 | CASSSTGGTEAFF | KLGGALQAK | YES | NO | NO |
| 1995 | TRAV12-2 | TRAJ48 | CAVNYNFGNEKLTF | TRBV5-5 | TRBJ1-1 | CASSLAGGATEAFF | KLGGALQAK | YES | NO | NO |
| 1996 | TRAV13-2 | TRAJ16 | CAENIGDGQKLLF | TRBV5-5 | TRBJ1-1 | CASSSGTGGTEAFF | KLGGALQAK | NO | NO | NO |
| 1997 | TRAV26-2 | TRAJ48 | CILLDANFGNEKLTF | TRBV5-5 | TRBJ1-1 | CASSKGRQNTEAFF | KLGGALQAK | YES | NO | NO |
| 1998 | TRAV38-1 | TRAJ40 | CAFMSPSGTYKYIF | TRBV5-5 | TRBJ1-1 | CASSPTSGIPATEAFF | KLGGALQAK | YES | NO | NO |
| 1999 | TRAV13-1 | TRAJ16 | CAAIKVSDGQKLLF | TRBV6-5 | TRBJ2-5 | CASSYSYRGKETQYF | KLGGALQAK | NO | NO | NO |
| 2000 | TRAV5 | TRAJ37 | CAESRGNTGKLIF | TRBV5-5 | TRBJ1-1 | CASSPLDRGEAFF | KLGGALQAK | YES | NO | NO |
| 2001 | TRAV8-3 | TRAJ26 | CAVGDNYGQNFVF | TRBV5-5 | TRBJ1-1 | CASSLKGPIGTEAFF | KLGGALQAK | YES | NO | NO |
| 2002 | TRAV8-3 | TRAJ39 | CAVDLPENAGNMLTF | TRBV5-5 | TRBJ1-1 | CASSLAPWSGDTEAFF | KLGGALQAK | YES | NO | NO |
| 2003 | TRAV8-3 | TRAJ48 | CAVGGLVFGNEKLTF | TRBV5-5 | TRBJ1-1 | CASSLGSAEAFF | KLGGALQAK | YES | NO | NO |
| 2003 | TRAV8-3 | TRAJ48 | CAVGGLVFGNEKLTF | TRBV5-5 | TRBJ1-1 | CASSLGSAEAFF | AVFDRKSDAK | YES | NO | NO |
| 2004 | TRAV1-1 | TRAJ3 | CAVRDPPYSSASKIIF | TRBV5-6 | TRBJ1-1 | CASSLRDGSEAFF | KLGGALQAK | YES | NO | NO |
| 2006 | TRAV10 | TRAJ20 | CVVFNDYKLSF | TRBV5-6 | TRBJ1-1 | CASSLVGSEMNTEAFF | KLGGALQAK | YES | NO | NO |
| 2007 | TRAV10 | TRAJ7 | CVVSTFYGNNRLAF | TRBV5-6 | TRBJ1-1 | CASSLRDGSEAFF | KLGGALQAK | YES | NO | NO |
| 2008 | TRAV12-1 | TRAJ54 | CVAPLQGAQKLVF | TRBV5-6 | TRBJ1-1 | CASSLGVLTEAFF | RLRAEAQVK | YES | NO | YES |
| 2009 | TRAV13-1 | TRAJ11 | CAAKEGYSTLTF | TRBV5-6 | TRBJ1-1 | CASSLRDGSEAFF | KLGGALQAK | YES | NO | NO |
| 2010 | TRAV13-1 | TRAJ42 | CAARQASQGNLIF | TRBV5-6 | TRBJ1-1 | CASSLASGGGAAFF | KLGGALQAK | YES | NO | NO |
| 2011 | TRAV19 | TRAJ27 | CALSEANTNAGKSTF | TRBV5-6 | TRBJ1-1 | CASSLAGPNTEAFF | KLGGALQAK | YES | NO | NO |
| 2012 | TRAV20 | TRAJ52 | CAGWNAGGTSYGKLTF | TRBV5-6 | TRBJ1-1 | CASSFSFRENTEAFF | KLGGALQAK | YES | NO | NO |
| 2014 | TRAV29DV5 | TRAJ52 | CAASEPGGTSYGKLTF | TRBV5-6 | TRBJ1-1 | CASSGGVTEAFF | KLGGALQAK | YES | NO | NO |
| 2017 | TRAV38-2DV8 | TRAJ40 | CAIEDDSGTYKYIF | TRBV5-6 | TRBJ1-1 | CASSLRDGSEAFF | KLGGALQAK | YES | NO | NO |
| 2018 | TRAV38-2DV8 | TRAJ52 | CAYRSLGGTSYGKLTF | TRBV5-6 | TRBJ1-1 | CASRIGNTEAFF | KLGGALQAK | YES | NO | NO |
| 2019 | TRAV4 | TRAJ9 | CLVGDINTGGFKTIF | TRBV5-6 | TRBJ1-1 | CASSFSPTEAFF | KLGGALQAK | NO | NO | NO |
| 2021 | TRAV41 | TRAJ46 | CAPEIGSSGDKLTF | TRBV5-6 | TRBJ1-1 | CASGRLYRGTEAFF | KLGGALQAK | YES | NO | NO |
| 2024 | TRAV6 | TRAJ9 | CALDGNTGGFKTIF | TRBV5-6 | TRBJ1-1 | CASGQGNTEAFF | KLGGALQAK | YES | NO | NO |
| 2027 | TRAV9-2 | TRAJ40 | CALRRHGGTYKYIF | TRBV5-6 | TRBJ1-1 | CASSWNRDRYNTEAFF | KLGGALQAK | YES | NO | NO |
| 2028 | TRAV1-2 | TRAJ29 | CAVRDSGNTPLVF | TRBV5-8 | TRBJ1-1 | CASSLVDGNTEAFF | KLGGALQAK | YES | NO | NO |
| 2031 | TRAV17 | TRAJ6 | CATDDPRAAGGSYIPTF | TRBV5-8 | TRBJ1-1 | CASSLNGMNTEAFF | KLGGALQAK | YES | NO | NO |
| 2032 | TRAV19 | TRAJ13 | CALLSGGYQKVTF | TRBV5-8 | TRBJ1-1 | CASRYRDRMNTEAFF | KLGGALQAK | YES | NO | NO |
| 2034 | TRAV26-2 | TRAJ29 | CILRVTNSGNTPLVF | TRBV5-8 | TRBJ1-1 | CASLPDEVNTEAFF | KLGGALQAK | YES | NO | NO |
| 2035 | TRAV8-6 | TRAJ10 | CAVSEGLTGGGNKLTF | TRBV5-8 | TRBJ1-1 | CASSNRGLLAEAFF | KLGGALQAK | YES | NO | NO |
| 2036 | TRAV1-2 | TRAJ33 | CAVRDSNYQLIW | TRBV6-1 | TRBJ1-1 | CASSVGQGEGYTEAFF | KLGGALQAK | YES | NO | NO |
| 2037 | TRAV1-2 | TRAJ6 | CAGWTGGSYIPTF | TRBV6-1 | TRBJ1-1 | CASELTSREAFF | KLGGALQAK | YES | NO | NO |
| 2039 | TRAV12-1 | TRAJ6 | CVVNPSGGSYIPTF | TRBV6-1 | TRBJ1-1 | CASSEGQGWTEAFF | KLGGALQAK | YES | NO | NO |
| 2040 | TRAV12-2 | TRAJ48 | CAVDFGNEKLTF | TRBV6-1 | TRBJ1-1 | CASKEGTTGTEAFF | KLGGALQAK | NO | NO | NO |
| 2041 | TRAV12-2 | TRAJ53 | CAVMWRGGSNYKLTF | TRBV6-1 | TRBJ1-1 | CASSRNMNTEAFF | KLGGALQAK | YES | NO | NO |
| 2042 | TRAV12-3 | TRAJ20 | CAMSRKLSF | TRBV6-1 | TRBJ1-1 | CATIEGRMNTEAFF | KLGGALQAK | NO | NO | NO |
| 2045 | TRAV13-2 | TRAJ47 | CAENSLYGNKLVF | TRBV6-1 | TRBJ1-1 | CASSEGPDTEAFF | KLGGALQAK | YES | NO | NO |
| 2046 | TRAV14DV4 | TRAJ9 | CAMREARTGGFKTIF | TRBV6-1 | TRBJ1-1 | CASSVFLRGANTEAFF | KLGGALQAK | NO | NO | NO |
| 2049 | TRAV17 | TRAJ29 | CATDATLDTPLVF | TRBV6-1 | TRBJ1-1 | CATKGGSEAFF | KLGGALQAK | YES | NO | NO |
| 2051 | TRAV17 | TRAJ47 | CATVAIMEYGNKLVF | TRBV6-1 | TRBJ1-1 | CASREERKAFF | KLGGALQAK | YES | NO | NO |
| 2052 | TRAV17 | TRAJ56 | CATRFTGANSKLTF | TRBV6-1 | TRBJ1-1 | CASSEGVWNTEAFF | KLGGALQAK | YES | NO | NO |
| 2053 | TRAV19 | TRAJ52 | CALSENAGGTSYGKLTF | TRBV6-1 | TRBJ1-1 | CASSRDRRYMNTEAFF | KLGGALQAK | NO | NO | NO |
| 2054 | TRAV2 | TRAJ20 | CAVGPRDYKLSF | TRBV6-1 | TRBJ1-1 | CASSDIMGNTEAFF | KLGGALQAK | NO | NO | NO |
| 2056 | TRAV23DV6 | TRAJ33 | CAASRRSNYQLIW | TRBV6-1 | TRBJ1-1 | CASSDLPEAFF | KLGGALQAK | YES | NO | NO |
| 2058 | TRAV38-1 | TRAJ45 | CASHSGGGADGLTF | TRBV6-1 | TRBJ1-1 | CASSEVFGANTEAFF | RLRAEAQVK | YES | NO | NO |
| 2060 | TRAV4 | TRAJ5 | CPRQGRRALTF | TRBV6-1 | TRBJ1-1 | CASSDGDREIPEAFF | KLGGALQAK | YES | NO | NO |
| 2061 | TRAV5 | TRAJ39 | CAESSGNAGNMLTF | TRBV6-1 | TRBJ1-1 | CASSEAPPKTGGLGAFF | KLGGALQAK | YES | NO | NO |
| 2064 | TRAV8-4 | TRAJ29 | CAVGASGNTPLVF | TRBV6-1 | TRBJ1-1 | CASSKGHVNTEAFF | KLGGALQAK | YES | NO | NO |
| 2065 | TRAV1-2 | TRAJ4 | CAVFSGGYNKLIF | TRBV6-3 | TRBJ1-1 | CASSYRVSEAFF | KLGGALQAK | YES | NO | NO |
| 2067 | TRAV12-2 | TRAJ15 | CAVNDQAGTALIF | TRBV6-3 | TRBJ1-1 | CASSKQGATEAFF | KLGGALQAK | YES | NO | NO |

| | | | | | | | | | | |
|---|---|---|---|---|---|---|---|---|---|---|
| 2069 | TRAV13-1 | TRAJ20 | CAASGPNDYKLSF | TRBV6-3 | TRBJ1-1 | CASSYPQGGEAFF | KLGGALQAK | YES | NO | NO |
| 2070 | TRAV13-2 | TRAJ43 | CAENIENNDMRF | TRBV6-3 | TRBJ1-1 | CASSLGGQGWTEAFF | KLGGALQAK | YES | NO | NO |
| 2072 | TRAV17 | TRAJ24 | CATFLTDSWGKLQF | TRBV6-3 | TRBJ1-1 | CASSYSVTDNAEAFF | KLGGALQAK | YES | NO | NO |
| 2074 | TRAV24 | TRAJ52 | CAFVAGGTSYGKLTF | TRBV6-3 | TRBJ1-1 | CASSVKTGTGATEAFF | KLGGALQAK | YES | NO | NO |
| 2075 | TRAV25 | TRAJ53 | CAGTGSNYKLTF | TRBV6-3 | TRBJ1-1 | CASRGSMNTEAFF | KLGGALQAK | YES | NO | NO |
| 2076 | TRAV29DV5 | TRAJ40 | CAATGSGTYKYIF | TRBV6-3 | TRBJ1-1 | CASSQQGAEAFF | KLGGALQAK | YES | NO | NO |
| 2077 | TRAV4 | TRAJ27 | CLVGPVYNAGKSTF | TRBV6-3 | TRBJ1-1 | CASSSGTGANTEAFF | KLGGALQAK | NO | NO | NO |
| 2078 | TRAV4 | TRAJ45 | CLVGDSGGGADGLTF | TRBV6-3 | TRBJ1-1 | CASSWDRGPRTEAFF | KLGGALQAK | YES | NO | NO |
| 2080 | TRAV5 | TRAJ26 | CAPRGDNYGQNFVF | TRBV6-3 | TRBJ1-1 | CASSYSNAEAFF | KLGGALQAK | YES | NO | NO |
| 2082 | TRAV8-3 | TRAJ9 | CAVAPTGGFKTIF | TRBV6-3 | TRBJ1-1 | CASSYSSTGDSEAFF | KLGGALQAK | YES | NO | NO |
| 2083 | TRAV1-2 | TRAJ13 | CAVNSGGYQKVTF | TRBV6-4 | TRBJ1-1 | CASSDSGAGEAFF | KLGGALQAK | YES | NO | NO |
| 2084 | TRAV1-2 | TRAJ28 | CAVRAGGAGSYQLTF | TRBV6-4 | TRBJ1-1 | CASSDGRDGDTEAFF | KLGGALQAK | YES | NO | NO |
| 2085 | TRAV1-2 | TRAJ33 | CAVRDSNYQLIW | TRBV6-4 | TRBJ1-1 | CASSAGTGGTEAFF | KLGGALQAK | YES | NO | NO |
| 2086 | TRAV12-1 | TRAJ12 | CVVNGMDSSYKLIF | TRBV6-4 | TRBJ1-1 | CASSDSRTATEAFF | KLGGALQAK | YES | NO | NO |
| 2087 | TRAV13-2 | TRAJ30 | CAENIGDDKIIF | TRBV6-4 | TRBJ1-1 | CASSERWDTEAFF | KLGGALQAK | YES | NO | NO |
| 2089 | TRAV3 | TRAJ35 | CAVRDMAGFGNVLHC | TRBV6-4 | TRBJ1-1 | CASSVWEGTEAFF | KLGGALQAK | NO | NO | NO |
| 2091 | TRAV8-4 | TRAJ4 | CAVSATFSGGYNKLIF | TRBV6-4 | TRBJ1-1 | CASSDSYSPNTEAFF | KLGGALQAK | YES | NO | NO |
| 2093 | TRAV12-1 | TRAJ33 | CVVLDSNYQLIW | TRBV6-5 | TRBJ1-1 | CASSWDREMNTEAFF | KLGGALQAK | YES | NO | NO |
| 2094 | TRAV12-2 | TRAJ10 | CAVPRLTGGGNKLTF | TRBV6-5 | TRBJ1-1 | CASSYSKWGDWTEAFF | KLGGALQAK | YES | NO | NO |
| 2095 | TRAV12-2 | TRAJ16 | CAVIPF | TRBV6-5 | TRBJ1-1 | CASRLDSNTEAFF | KLGGALQAK | NO | NO | NO |
| 2098 | TRAV13-1 | TRAJ52 | CAAGGTSYGKLTF | TRBV6-5 | TRBJ1-1 | CASSYLDGNTEAFF | KLGGALQAK | YES | NO | NO |
| 2099 | TRAV13-1 | TRAJ6 | CAASGGSYIPTF | TRBV6-5 | TRBJ1-1 | CASSYAQGVSTEAFF | KLGGALQAK | YES | NO | NO |
| 2100 | TRAV13-2 | TRAJ30 | CAENSDDKIIF | TRBV6-5 | TRBJ1-1 | CASSYSTGGNTEAFF | KLGGALQAK | YES | NO | NO |
| 2101 | TRAV14DV4 | TRAJ3 | CAMREGGYSSASKIIF | TRBV6-5 | TRBJ1-1 | CASRPQGAVEAFF | KLGGALQAK | YES | NO | NO |
| 2102 | TRAV14DV4 | TRAJ40 | CAMREGQVTSGTYKYIF | TRBV6-5 | TRBJ1-1 | CASSLVELNTEAFF | KLGGALQAK | YES | NO | NO |
| 2104 | TRAV19 | TRAJ47 | CALGLMEYGNKLVF | TRBV6-5 | TRBJ1-1 | CASNPGQENTEAFF | KLGGALQAK | YES | NO | NO |
| 2105 | TRAV21 | TRAJ15 | CAVNFQAGTALIF | TRBV6-5 | TRBJ1-1 | CASTLRESEAFF | KLGGALQAK | YES | NO | NO |
| 2107 | TRAV21 | TRAJ35 | CAPI | TRBV6-5 | TRBJ1-1 | CASSLLGQTEAFF | KLGGALQAK | NO | NO | NO |
| 2108 | TRAV21 | TRAJ5 | CAVMDTGRRALTF | TRBV6-5 | TRBJ1-1 | CASSYSRTGGLNTEAFF | KLGGALQAK | YES | NO | NO |
| 2109 | TRAV21 | TRAJ52 | CAVMPDPNAGGTSYGKLTF | TRBV6-5 | TRBJ1-1 | CASSRAPNTEAFF | KLGGALQAK | YES | NO | NO |
| 2110 | TRAV21 | TRAJ54 | CAVSRMRGAQKLVF | TRBV6-5 | TRBJ1-1 | CASRPGQASEAFF | KLGGALQAK | YES | NO | NO |
| 2111 | TRAV24 | TRAJ21 | CAFHDNFNKFYF | TRBV6-5 | TRBJ1-1 | CASSYSTTSVLPSEAFF | KLGGALQAK | YES | NO | NO |
| 2113 | TRAV26-2 | TRAJ33 | CMYLMDSNYQLIW | TRBV6-5 | TRBJ1-1 | CASKTGVNTEAFF | KLGGALQAK | YES | NO | NO |
| 2115 | TRAV29DV5 | TRAJ23 | CAASFYNQGGKLIF | TRBV6-5 | TRBJ1-1 | CASSFDRAIGGTEAFF | KLGGALQAK | YES | NO | NO |
| 2116 | TRAV29DV5 | TRAJ42 | CAARPNYGGSQGNLIF | TRBV6-5 | TRBJ1-1 | CASSYPGLAEAFF | KLGGALQAK | YES | NO | NO |
| 2117 | TRAV3 | TRAJ26 | CAAPDNYGQNFVF | TRBV6-5 | TRBJ1-1 | CASRGGQGAVEAFF | KLGGALQAK | NO | NO | NO |
| 2119 | TRAV38-2DV8 | TRAJ32 | CAYNHGGATNKLIF | TRBV6-5 | TRBJ1-1 | CASSPRLMGTNTEAFF | KLGGALQAK | YES | NO | NO |
| 2120 | TRAV38-2DV8 | TRAJ39 | CGHNNAGNMLTF | TRBV6-5 | TRBJ1-1 | CASSFGTANTEAFF | KLGGALQAK | YES | NO | NO |
| 2121 | TRAV38-2DV8 | TRAJ43 | CAYRSAGDMRF | TRBV6-5 | TRBJ1-1 | CASSYLTGIGTEAFF | KLGGALQAK | YES | NO | NO |
| 2122 | TRAV38-2DV8 | TRAJ48 | CAYRSASFGNEKLTF | TRBV6-5 | TRBJ1-1 | CASSYGDTEAFF | KLGGALQAK | YES | NO | NO |
| 2124 | TRAV4 | TRAJ29 | CLVGDMAGNTPLVF | TRBV6-5 | TRBJ1-1 | CASSYIGVKTEAFF | KLGGALQAK | YES | NO | NO |
| 2125 | TRAV41 | TRAJ34 | CAVPTYNTDKLIF | TRBV6-5 | TRBJ1-1 | CASSYSGGENTEAFF | KLGGALQAK | YES | NO | NO |
| 2126 | TRAV41 | TRAJ45 | CAVRIQGGGADGLTF | TRBV6-5 | TRBJ1-1 | CASKTGLNTEAFF | KLGGALQAK | YES | NO | NO |
| 2128 | TRAV6 | TRAJ43 | CALGVTYNNNDMRF | TRBV6-5 | TRBJ1-1 | CASSYSEGANTEAFF | KLGGALQAK | YES | NO | NO |
| 2129 | TRAV8-3 | TRAJ49 | CAVARTGNQFYF | TRBV6-5 | TRBJ1-1 | CASSYIQGRNTEAFF | KLGGALQAK | YES | NO | NO |
| 2132 | TRAV9-2 | TRAJ10 | CALSEAGGGNKLTF | TRBV6-5 | TRBJ1-1 | CASSYSGGNTEAFF | KLGGALQAK | YES | NO | NO |
| 2133 | TRAV12-2 | TRAJ42 | CAVNDGYGGSQGNLIF | TRBV6-6 | TRBJ1-1 | CASSYPGVTMNTEAFF | KLGGALQAK | NO | NO | NO |
| 2135 | TRAV14DV4 | TRAJ3 | CAMRASIGSASKIIF | TRBV6-6 | TRBJ1-1 | CASSSLSRTGGLGTEAFF | KLGGALQAK | NO | NO | NO |
| 2137 | TRAV26-2 | TRAJ43 | CILKYNNNDMRF | TRBV6-6 | TRBJ1-1 | CASSRWDGMNTEAFF | KLGGALQAK | NO | NO | NO |
| 2138 | TRAV38-2DV8 | TRAJ50 | CAYRSVLTAASYDKVIF | TRBV6-6 | TRBJ1-1 | CASSYGELRMNTEAFF | KLGGALQAK | NO | NO | NO |
| 2139 | TRAV8-1 | TRAJ10 | CAVIFTGGGNKLTF | TRBV6-6 | TRBJ1-1 | CASEVNTEAFF | KLGGALQAK | NO | NO | NO |
| 2140 | TRAV8-2 | TRAJ45 | CVVSDGGSGGGADGLTF | TRBV6-6 | TRBJ1-1 | CASSDGRDTEAFF | KLGGALQAK | NO | NO | NO |
| 2144 | TRAV3 | TRAJ21 | CAVRDISYNFNKFYF | TRBV7-2 | TRBJ1-1 | CASIKDTGLGNTEAFF | KLGGALQAK | NO | NO | NO |
| 2145 | TRAV36DV7 | TRAJ45 | CAVRMYSGGGADGLTF | TRBV7-2 | TRBJ1-1 | CASRSRERGDEAFF | KLGGALQAK | NO | NO | NO |
| 2146 | TRAV8-2 | TRAJ40 | CVVSPGTYKYIF | TRBV7-2 | TRBJ1-1 | CASRGLNTEAFF | KLGGALQAK | NO | NO | NO |
| 2151 | TRAV14DV4 | TRAJ12 | CAMREEDSSYKLIF | TRBV7-3 | TRBJ1-1 | CASSPWNTEAFF | KLGGALQAK | NO | NO | NO |
| 2156 | TRAV26-2 | TRAJ58 | CILGPSETSGSRLTF | TRBV7-3 | TRBJ1-1 | CASSLNPVNTEAFF | AVFDRKSDAK | NO | NO | NO |
| 2156 | TRAV26-2 | TRAJ58 | CILGPSETSGSRLTF | TRBV7-3 | TRBJ1-1 | CASSLNPVNTEAFF | IVTDFSVIK | NO | NO | NO |
| 2159 | TRAV12-1 | TRAJ32 | CVVNGYGGATNKLIF | TRBV7-6 | TRBJ1-1 | CASSLSSGQGNTEAFF | KLGGALQAK | YES | NO | NO |
| 2160 | TRAV12-1 | TRAJ42 | CVVRLYYGGSQGNLIF | TRBV7-6 | TRBJ1-1 | CASSGTANMNTEAFF | KLGGALQAK | YES | NO | NO |
| 2162 | TRAV19 | TRAJ44 | CALFNPAGTASKLTF | TRBV7-6 | TRBJ1-1 | CASSLGQGRNTEAFF | KLGGALQAK | YES | NO | NO |
| 2163 | TRAV19 | TRAJ50 | CALRSLKTSYDKVIF | TRBV7-6 | TRBJ1-1 | CASSYGQGQNTEAFF | KLGGALQAK | YES | NO | NO |
| 2164 | TRAV19 | TRAJ52 | CALRAGGTSYGKLTF | TRBV7-6 | TRBJ1-1 | CASSSPQGVSNTEAFF | KLGGALQAK | YES | NO | NO |
| 2165 | TRAV20 | TRAJ58 | CAVQAREETSGSRLTF | TRBV7-6 | TRBJ1-1 | CASSLFPVAKAFF | KLGGALQAK | YES | NO | NO |
| 2166 | TRAV27 | TRAJ17 | CAGSMIKAAGNKLTF | TRBV7-6 | TRBJ1-1 | CASSPALGGVPEAFF | KLGGALQAK | YES | NO | NO |
| 2167 | TRAV3 | TRAJ8 | CAVRENTGFQKLVF | TRBV7-6 | TRBJ1-1 | CASSQDYGLNTEAFF | KLGGALQAK | NO | NO | NO |
| 2169 | TRAV8-2 | TRAJ15 | CVVSEGGGTALIF | TRBV7-6 | TRBJ1-1 | CASSFDRDLNTEAFF | KLGGALQAK | YES | NO | NO |
| 2171 | TRAV12-2 | TRAJ34 | CAVIRNTDKLIF | TRBV7-7 | TRBJ1-1 | CASSPLLQGNTEAFF | KLGGALQAK | YES | NO | NO |
| 2173 | TRAV8-4 | TRAJ45 | CAVSLSGGGADGLTF | TRBV7-7 | TRBJ1-1 | CASSTTMNTEAFF | RLRAEAQVK | YES | NO | YES |
| 2174 | TRAV12-1 | TRAJ27 | CVVNTNAGKSTF | TRBV7-8 | TRBJ1-1 | CASSPLPHMNTEAFF | KLGGALQAK | YES | NO | NO |
| 2176 | TRAV12-2 | TRAJ48 | CAGYISNFGNEKLTF | TRBV7-8 | TRBJ1-1 | CASSLEANTEAFF | KLGGALQAK | YES | NO | NO |
| 2178 | TRAV14DV4 | TRAJ49 | CAMRTNTGNQFYF | TRBV7-8 | TRBJ1-1 | CASSLGAGGAGNTEAFF | KLGGALQAK | YES | NO | NO |
| 2181 | TRAV25 | TRAJ40 | CAGPSSGTYKYIF | TRBV7-8 | TRBJ1-1 | CASSEYRGTEAFF | KLGGALQAK | YES | NO | NO |
| 2182 | TRAV27 | TRAJ30 | CAGVFRDDKIIF | TRBV7-8 | TRBJ1-1 | CASSFPDGNTEAFF | KLGGALQAK | YES | NO | NO |
| 2183 | TRAV29DV5 | TRAJ15 | CAAVNQAGTALIF | TRBV7-8 | TRBJ1-1 | CASSSRHEAFF | KLGGALQAK | YES | NO | NO |
| 2184 | TRAV29DV5 | TRAJ35 | CAASDGFGNVLHC | TRBV7-8 | TRBJ1-1 | CASSGVLGGMNTEAFF | KLGGALQAK | NO | NO | NO |
| 2186 | TRAV34 | TRAJ54 | CGADSPFPLQGAQKLVF | TRBV7-8 | TRBJ1-1 | CASSFFGNTEAFF | KLGGALQAK | YES | NO | NO |
| 2187 | TRAV38-2DV8 | TRAJ32 | CAYRPWSSGATNKLIF | TRBV7-8 | TRBJ1-1 | CASSHSTGSSTEAFF | KLGGALQAK | YES | NO | NO |
| 2188 | TRAV38-2DV8 | TRAJ33 | CAYRSTSSPWNSNYQLIW | TRBV7-8 | TRBJ1-1 | CASRPITGGGYTEAFF | KLGGALQAK | YES | NO | NO |
| 2189 | TRAV4 | TRAJ45 | CLVGVTPGGGADGLTF | TRBV7-8 | TRBJ1-1 | CASSLQEAFF | KLGGALQAK | YES | NO | NO |
| 2190 | TRAV8-1 | TRAJ17 | CAQYSIKAAGNKLTF | TRBV7-8 | TRBJ1-1 | CASSPGQGRGTEAFF | KLGGALQAK | YES | NO | NO |
| 2191 | TRAV1-1 | TRAJ26 | CAADLQNFVF | TRBV7-9 | TRBJ1-1 | CASSMTGPAEAFF | KLGGALQAK | YES | NO | NO |
| 2192 | TRAV1-1 | TRAJ5 | CAVRDTGRRALTF | TRBV7-9 | TRBJ1-1 | CASSLGDNTEAFF | KLGGALQAK | YES | NO | NO |
| 2193 | TRAV1-2 | TRAJ24 | CAVRGDSWGKFQF | TRBV7-9 | TRBJ1-1 | CASSLVGGSRAEAFF | KLGGALQAK | YES | NO | NO |
| 2194 | TRAV1-2 | TRAJ41 | CAVRTDSGYALNF | TRBV7-9 | TRBJ1-1 | CASSRQGAMNTEAFF | KLGGALQAK | YES | NO | NO |
| 2195 | TRAV12-1 | TRAJ15 | CVVTRGSGTALIF | TRBV7-9 | TRBJ1-1 | CASSAPDRGNTEAFF | KLGGALQAK | YES | NO | NO |
| 2196 | TRAV12-1 | TRAJ23 | CVCNQGGKLIF | TRBV7-9 | TRBJ1-1 | CASSLVSGPTEAFF | KLGGALQAK | YES | NO | NO |
| 2197 | TRAV12-2 | TRAJ24 | CAALDSWGKFQF | TRBV7-9 | TRBJ1-1 | CASSDRGGIGGSAFF | KLGGALQAK | YES | NO | NO |
| 2203 | TRAV14DV4 | TRAJ23 | CAMRLPGGKLIF | TRBV7-9 | TRBJ1-1 | CASSEGLGTEAFF | KLGGALQAK | YES | NO | NO |
| 2204 | TRAV14DV4 | TRAJ4 | CAMRRPSGGYNKLIF | TRBV7-9 | TRBJ1-1 | CASTFSFGQNTEAFF | KLGGALQAK | YES | NO | NO |
| 2205 | TRAV14DV4 | TRAJ4 | CAMRRPSGGYNKLIF | TRBV7-9 | TRBJ1-1 | CASSLASGRSTEAFF | AVFDRKSDAK | YES | NO | NO |
| 2205 | TRAV14DV4 | TRAJ4 | CAMRRPSGGYNKLIF | TRBV7-9 | TRBJ1-1 | CASSLASGRSTEAFF | KLGGALQAK | YES | NO | NO |
| 2205 | TRAV14DV4 | TRAJ4 | CAMRRPSGGYNKLIF | TRBV7-9 | TRBJ1-1 | CASSLASGRSTEAFF | IVTDFSVIK | YES | NO | NO |
| 2206 | TRAV14DV4 | TRAJ53 | CAMRSPGGSNYKLTF | TRBV7-9 | TRBJ1-1 | CASNSGLGTEAFF | KLGGALQAK | YES | NO | NO |
| 2206 | TRAV14DV4 | TRAJ53 | CAMRSPGGSNYKLTF | TRBV7-9 | TRBJ1-1 | CASNSGLGTEAFF | AVFDRKSDAK | YES | NO | NO |
| 2207 | TRAV17 | TRAJ11 | CATAMVGYSTLTF | TRBV7-9 | TRBJ1-1 | CASSPVAGLTLYLEAFF | KLGGALQAK | YES | NO | NO |
| 2210 | TRAV19 | TRAJ42 | CALSELNYGGSQGNLIF | TRBV7-9 | TRBJ1-1 | CASLGTPMNTEAFF | KLGGALQAK | YES | NO | NO |
| 2211 | TRAV19 | TRAJ52 | CALSAGGTSYGKLTF | TRBV7-9 | TRBJ1-1 | CASSFTLEAFF | KLGGALQAK | YES | NO | NO |
| 2212 | TRAV20 | TRAJ20 | CAVQAESNDYKLSF | TRBV7-9 | TRBJ1-1 | CASKTNTEAFF | KLGGALQAK | YES | NO | NO |
| 2213 | TRAV21 | TRAJ39 | CAALNNAGNMLTF | TRBV7-9 | TRBJ1-1 | CASSYLDGAGPEAFF | KLGGALQAK | YES | NO | NO |
| 2221 | TRAV29DV5 | TRAJ36 | CAATSQTGANNLFF | TRBV7-9 | TRBJ1-1 | CASSPGVNTEAFF | KLGGALQAK | YES | NO | NO |
| 2224 | TRAV3 | TRAJ41 | CAVRAYSGYALNF | TRBV7-9 | TRBJ1-1 | CASSLPGRGFFFNTEAFF | KLGGALQAK | YES | NO | NO |
| 2225 | TRAV38-1 | TRAJ7 | CALGNNRLAF | TRBV7-9 | TRBJ1-1 | CASSPTDAMNTEAFF | KLGGALQAK | YES | NO | NO |
| 2226 | TRAV4 | TRAJ42 | CPRGGGSQGNLIF | TRBV7-9 | TRBJ1-1 | CASSLALTGVGTEAFF | KLGGALQAK | NO | NO | NO |

| | | | | | | | | | | |
|---|---|---|---|---|---|---|---|---|---|---|
| 2227 | TRAV5 | TRAJ6 | CADMGGSYIPTF | TRBV7-9 | TRBJ1-1 | CASSLSEKLEAFF | KLGGALQAK | YES | NO | NO |
| 2229 | TRAV8-1 | TRAJ48 | CAVNAQETNFGNEKLTF | TRBV7-9 | TRBJ1-1 | CASSLASGRSTEAFF | KLGGALQAK | YES | NO | NO |
| 2233 | TRAV8-6 | TRAJ20 | CAVSGGDYKLSF | TRBV7-9 | TRBJ1-1 | CASSLAAGNTEAFF | KLGGALQAK | YES | NO | NO |
| 2234 | TRAV1-1 | TRAJ12 | CAVRALGDSSYKLIF | TRBV9 | TRBJ1-1 | CASSRGQGNEAFF | KLGGALQAK | YES | NO | NO |
| 2235 | TRAV1-1 | TRAJ6 | CVLSGGSYIPTF | TRBV9 | TRBJ1-1 | CASSGPGQLSTDTEAFF | KLGGALQAK | YES | NO | NO |
| 2236 | TRAV1-2 | TRAJ26 | CAVIGYGQNFVF | TRBV9 | TRBJ1-1 | CASSPIGGRDTEAFF | KLGGALQAK | YES | NO | NO |
| 2240 | TRAV1-2 | TRAJ40 | CAVRFTSGTYKYIF | TRBV9 | TRBJ1-1 | CASSAGTGGAASITEAFF | KLGGALQAK | YES | YES | NO |
| 2242 | TRAV10 | TRAJ33 | CVVSAVDSNYQLIW | TRBV9 | TRBJ1-1 | CASSDGGGGTEAFF | KLGGALQAK | YES | NO | NO |
| 2244 | TRAV10 | TRAJ4 | CVVRSGGYNKLIF | TRBV9 | TRBJ1-1 | CASSAGTGGAASITEAFF | KLGGALQAK | YES | NO | NO |
| 2245 | TRAV12-1 | TRAJ52 | CVVNMAYAGGTSYGKLTF | TRBV9 | TRBJ1-1 | CASSPGVGAEAFF | KLGGALQAK | YES | NO | NO |
| 2246 | TRAV12-2 | TRAJ38 | CAVTYAGNNRKLIW | TRBV9 | TRBJ1-1 | CASSPTGGNTEAFF | KLGGALQAK | YES | NO | NO |
| 2249 | TRAV13-2 | TRAJ20 | CAENGGSDYKLSF | TRBV9 | TRBJ1-1 | CASSVTNGGTEAFF | KLGGALQAK | YES | NO | NO |
| 2250 | TRAV13-2 | TRAJ52 | CAENIGAGGTSYGKLTF | TRBV9 | TRBJ1-1 | CASSVGGGTEAFF | KLGGALQAK | YES | NO | NO |
| 2254 | TRAV14DV4 | TRAJ28 | CAMRESGAGSYQLTF | TRBV9 | TRBJ1-1 | CASSGDRGLGTEAFF | KLGGALQAK | YES | NO | NO |
| 2255 | TRAV14DV4 | TRAJ39 | CAMREGNAGNMLTF | TRBV9 | TRBJ1-1 | CASSVDPSGGSGNTEAFF | KLGGALQAK | YES | NO | NO |
| 2256 | TRAV14DV4 | TRAJ49 | CAMREANTGNQFYF | TRBV9 | TRBJ1-1 | CASSVAGGEAEAFF | KLGGALQAK | YES | NO | NO |
| 2257 | TRAV17 | TRAJ40 | CATAPSGTYKYIF | TRBV9 | TRBJ1-1 | CASSVDTGFTEAFF | AVFDRKSDAK | YES | NO | NO |
| 2258 | TRAV17 | TRAJ40 | CATAPSGTYKYIF | TRBV9 | TRBJ1-1 | CASSVDTAYTEAFF | KLGGALQAK | YES | NO | NO |
| 2259 | TRAV17 | TRAJ57 | CATDQGGGSEKLVF | TRBV9 | TRBJ1-1 | CASSPWTGPLNTEAFF | KLGGALQAK | YES | NO | NO |
| 2260 | TRAV19 | TRAJ39 | CALSGENSNNAGNMLTF | TRBV9 | TRBJ1-1 | CASSPLGGEGELFMNTEAFF | KLGGALQAK | NO | NO | NO |
| 2261 | TRAV19 | TRAJ40 | CALSEASGTYKYIF | TRBV9 | TRBJ1-1 | CASSVSGLNTEAFF | KLGGALQAK | YES | NO | NO |
| 2266 | TRAV21 | TRAJ43 | CAVRLAIPSNNNDMRF | TRBV9 | TRBJ1-1 | CASSGGGYTEAFF | KLGGALQAK | YES | NO | NO |
| 2268 | TRAV25 | TRAJ27 | CAGFHTNAGKSTF | TRBV9 | TRBJ1-1 | CASSATGISTEAFF | KLGGALQAK | YES | NO | NO |
| 2269 | TRAV26-1 | TRAJ43 | CIVRANNNDMRF | TRBV9 | TRBJ1-1 | CASTRDITGGNTEAFF | KLGGALQAK | YES | NO | NO |
| 2270 | TRAV26-1 | TRAJ43 | CIVRDHNNDMRF | TRBV9 | TRBJ1-1 | CASSVVPMGTAGNTEAFF | KLGGALQAK | YES | NO | NO |
| 2272 | TRAV26-2 | TRAJ39 | CILRVPNAGNMLTF | TRBV9 | TRBJ1-1 | CASSYYEDRVSEAFF | KLGGALQAK | YES | NO | NO |
| 2273 | TRAV26-2 | TRAJ53 | CIRPSGGSNYKLTF | TRBV9 | TRBJ1-1 | CASSGGGNTEAFF | KLGGALQAK | YES | NO | NO |
| 2276 | TRAV27 | TRAJ53 | CAGGSNYKLTF | TRBV9 | TRBJ1-1 | CASSEGGQSREAFF | KLGGALQAK | YES | NO | NO |
| 2277 | TRAV29DV5 | TRAJ57 | CAASKTQGGSEKLVF | TRBV9 | TRBJ1-1 | CASSASVGGYTEAFF | KLGGALQAK | YES | NO | NO |
| 2280 | TRAV38-2DV8 | TRAJ39 | CAYRRGNAGNMLTF | TRBV9 | TRBJ1-1 | CASSVDGTMNTEAFF | KLGGALQAK | YES | NO | NO |
| 2281 | TRAV41 | TRAJ58 | CAVKEETSGSRLTF | TRBV9 | TRBJ1-1 | CASSGKQGCDTEAFF | KLGGALQAK | YES | NO | NO |
| 2283 | TRAV8-3 | TRAJ54 | CAVGQGAQKLVF | TRBV9 | TRBJ1-1 | CASSVGLGVTEAFF | KLGGALQAK | YES | NO | NO |
| 2284 | TRAV8-3 | TRAJ9 | CAVAPTGGFKTIF | TRBV9 | TRBJ1-1 | CASSDGGGGSEAFF | KLGGALQAK | YES | NO | NO |
| 2288 | TRAV12-1 | TRAJ42 | CVVNPFYGGSQGNLIF | TRBV10-2 | TRBJ1-2 | CASSEGSTYGYTF | KLGGALQAK | YES | NO | NO |
| 2289 | TRAV12-2 | TRAJ15 | CAVKEGQAGTALIF | TRBV10-2 | TRBJ1-2 | CASGDLGGYGYTF | KLGGALQAK | YES | NO | NO |
| 2293 | TRAV21 | TRAJ20 | CAVPSNDYKLSF | TRBV10-2 | TRBJ1-2 | CASTPRGSAGYTF | KLGGALQAK | YES | NO | NO |
| 2293 | TRAV21 | TRAJ20 | CAVPSNDYKLSF | TRBV10-2 | TRBJ1-2 | CASTPRGSAGYTF | AVFDRKSDAK | YES | NO | NO |
| 2294 | TRAV29DV5 | TRAJ35 | CAARRI | TRBV10-2 | TRBJ1-2 | CASSDSTGDGYTF | KLGGALQAK | NO | NO | NO |
| 2295 | TRAV29DV5 | TRAJ48 | CAVNFGNEKLTF | TRBV10-2 | TRBJ1-2 | CASSELLTGAFWGYTF | KLGGALQAK | YES | NO | NO |
| 2296 | TRAV10 | TRAJ6 | CVVSASRGSYIPTF | TRBV10-3 | TRBJ1-2 | CAISESGQEDNYGYTF | KLGGALQAK | YES | NO | NO |
| 2298 | TRAV14DV4 | TRAJ5 | CAMREYTGRRALTF | TRBV10-3 | TRBJ1-2 | CATSDRDNHLGYGYTF | KLGGALQAK | YES | NO | NO |
| 2299 | TRAV21 | TRAJ30 | CAVRGRDDKIIF | TRBV10-3 | TRBJ1-2 | CAISPGGVGYGYTF | KLGGALQAK | YES | NO | NO |
| 2300 | TRAV21 | TRAJ43 | CDTLDNNDMRF | TRBV10-3 | TRBJ1-2 | CAISESLTPPGLPLFNYGYTF | KLGGALQAK | NO | NO | NO |
| 2301 | TRAV21 | TRAJ49 | CAVSHQFYF | TRBV10-3 | TRBJ1-2 | CAISDQGTTYGYTF | KLGGALQAK | NO | NO | NO |
| 2302 | TRAV21 | TRAJ57 | CAARPHQGGSEKLVF | TRBV10-3 | TRBJ1-2 | CAISGPNYGYTF | KLGGALQAK | YES | NO | NO |
| 2303 | TRAV24 | TRAJ47 | CAHRGQEYGNKLVF | TRBV10-3 | TRBJ1-2 | CAISDGTVGGYTF | KLGGALQAK | YES | NO | NO |
| 2305 | TRAV5 | TRAJ34 | CAESRADKLIF | TRBV10-3 | TRBJ1-2 | CAISESDRTYGYTF | KLGGALQAK | YES | NO | NO |
| 2306 | TRAV1-1 | TRAJ29 | CAVRSQNSGNTPLVF | TRBV11-2 | TRBJ1-2 | CASSLVTGGRGYTF | KLGGALQAK | YES | NO | NO |
| 2307 | TRAV1-2 | TRAJ4 | CAVRGFSGGYNKLIF | TRBV11-2 | TRBJ1-2 | CASSGSFGTNYGYTF | KLGGALQAK | YES | NO | NO |
| 2308 | TRAV10 | TRAJ20 | CVVSPLNDYKLSF | TRBV11-2 | TRBJ1-2 | CASSLGDRIYTF | KLGGALQAK | YES | NO | NO |
| 2308 | TRAV10 | TRAJ20 | CVVSPLNDYKLSF | TRBV11-2 | TRBJ1-2 | CASSLGDRIYTF | AVFDRKSDAK | YES | NO | NO |
| 2312 | TRAV21 | TRAJ34 | CAVRKVSYNTDKLIF | TRBV11-2 | TRBJ1-2 | CASSLGGASNYGYTF | KLGGALQAK | YES | NO | NO |
| 2315 | TRAV27 | TRAJ57 | CAGAQGGSEKLVF | TRBV11-2 | TRBJ1-2 | CASSLALIPYGYTF | KLGGALQAK | YES | NO | NO |
| 2318 | TRAV29DV5 | TRAJ53 | CAASMGGGSNYKLTF | TRBV11-2 | TRBJ1-2 | CASSWTGGYTF | KLGGALQAK | YES | NO | NO |
| 2320 | TRAV3 | TRAJ30 | CAVRDSGDKIIF | TRBV11-2 | TRBJ1-2 | CASSWGGGSHYGYTF | KLGGALQAK | YES | NO | NO |
| 2321 | TRAV4 | TRAJ20 | CLVGDTDYKLSF | TRBV11-2 | TRBJ1-2 | CASSQETDLNYGYTF | KLGGALQAK | YES | NO | NO |
| 2322 | TRAV4 | TRAJ9 | CLVGDDGFKTIF | TRBV11-2 | TRBJ1-2 | CASSPLVEGNYGYTF | KLGGALQAK | YES | NO | NO |
| 2323 | TRAV41 | TRAJ45 | CAVLPGYSGGGADGLTF | TRBV11-2 | TRBJ1-2 | CASSLVWGGAYGYTF | KLGGALQAK | YES | NO | NO |
| 2324 | TRAV8-2 | TRAJ4 | CVVTFSGGYNKLIF | TRBV11-2 | TRBJ1-2 | CASSRRGEEGYTF | KLGGALQAK | YES | NO | NO |
| 2325 | TRAV8-2 | TRAJ9 | CVVSWTGGFKTIF | TRBV11-2 | TRBJ1-2 | CASSPGTAPGDYGYTF | KLGGALQAK | YES | NO | NO |
| 2326 | TRAV8-3 | TRAJ29 | CAVGEGGNTPLVF | TRBV11-2 | TRBJ1-2 | CASSLGTGDGYTF | KLGGALQAK | YES | NO | NO |
| 2328 | TRAV9-2 | TRAJ33 | CAREHMDSNYQLIW | TRBV11-2 | TRBJ1-2 | CASSHGSGSPYGYTF | KLGGALQAK | YES | NO | NO |
| 2329 | TRAV12-1 | TRAJ43 | CVVNEPPDMRF | TRBV11-3 | TRBJ1-2 | CASSSRGFYGYTF | KLGGALQAK | YES | NO | NO |
| 2332 | TRAV19 | TRAJ9 | CALSENTGGFKTIF | TRBV11-3 | TRBJ1-2 | CASSSPLREGYTF | KLGGALQAK | YES | NO | NO |
| 2333 | TRAV27 | TRAJ24 | CAGGSTDSWGKFQF | TRBV11-3 | TRBJ1-2 | CASSSQGGLYGYTF | KLGGALQAK | YES | NO | NO |
| 2335 | TRAV38-1 | TRAJ10 | CAAAGGGNKLTF | TRBV11-3 | TRBJ1-2 | CASSGRQGRGYTF | KLGGALQAK | YES | NO | NO |
| 2336 | TRAV8-3 | TRAJ45 | CAVGADSGGGADGLTF | TRBV11-3 | TRBJ1-2 | CASSPRQLLNYGYTF | KLGGALQAK | YES | NO | NO |
| 2337 | TRAV1-2 | TRAJ8 | CAVRDLYTGFQKLVF | TRBV12-3 | TRBJ1-2 | CASSFPSYGYTF | KLGGALQAK | YES | NO | NO |
| 2338 | TRAV12-3 | TRAJ39 | CAIGPGNMLTF | TRBV12-3 | TRBJ1-2 | CASSLGTDYGYTF | KLGGALQAK | YES | NO | NO |
| 2339 | TRAV13-1 | TRAJ10 | CAATGGGNKLTF | TRBV12-3 | TRBJ1-2 | CASSKDRLEGTF | KLGGALQAK | YES | NO | NO |
| 2340 | TRAV17 | TRAJ47 | CAPLSGNKLVF | TRBV12-3 | TRBJ1-2 | CASSYTNGYGYTF | KLGGALQAK | YES | NO | NO |
| 2341 | TRAV27 | TRAJ20 | CAGESADDYKLSF | TRBV12-3 | TRBJ1-2 | CASSLNNRQMITF | KLGGALQAK | YES | NO | NO |
| 2345 | TRAV36DV7 | TRAJ28 | CAVELAGAGSYQLTF | TRBV12-3 | TRBJ1-2 | CASSSWTASNYGYTF | KLGGALQAK | NO | NO | NO |
| 2346 | TRAV4 | TRAJ33 | CLVGDGNSNYQLIW | TRBV12-3 | TRBJ1-2 | CASSELGGYGYTF | KLGGALQAK | NO | NO | NO |
| 2347 | TRAV4 | TRAJ47 | CLVGDIGKLVF | TRBV12-3 | TRBJ1-2 | CASSPAGFYGYTF | KLGGALQAK | NO | NO | NO |
| 2348 | TRAV1-2 | TRAJ12 | CSQMDSSYKLIF | TRBV12-4 | TRBJ1-2 | CASSLSRTGRGYTF | KLGGALQAK | YES | NO | NO |
| 2349 | TRAV10 | TRAJ38 | CVVSVDAGNNRKLIW | TRBV12-4 | TRBJ1-2 | CASSGRYFYGYTF | KLGGALQAK | YES | NO | NO |
| 2351 | TRAV12-1 | TRAJ26 | CVVNPSGGQNFVF | TRBV12-4 | TRBJ1-2 | CASSLSPGTRRGYTF | KLGGALQAK | YES | NO | NO |
| 2352 | TRAV12-2 | TRAJ31 | CAVNNARLMF | TRBV12-4 | TRBJ1-2 | CASSSDPRYGYTF | KLGGALQAK | YES | NO | NO |
| 2354 | TRAV14DV4 | TRAJ47 | CAMREVEYGNKLVF | TRBV12-4 | TRBJ1-2 | CASSPTGYYGYTF | KLGGALQAK | YES | NO | NO |
| 2356 | TRAV19 | TRAJ45 | CALSDSGGGADGLTF | TRBV12-4 | TRBJ1-2 | CASSPGTINYGYTF | KLGGALQAK | YES | NO | NO |
| 2357 | TRAV24 | TRAJ52 | CAFIGDAGGTSYGKLTF | TRBV12-4 | TRBJ1-2 | CASSLVLQRIYGYTF | KLGGALQAK | YES | NO | NO |
| 2358 | TRAV27 | TRAJ24 | CAGVWADSWGKLQF | TRBV12-4 | TRBJ1-2 | CASSQLNTYGYTF | KLGGALQAK | YES | NO | NO |
| 2360 | TRAV29DV5 | TRAJ28 | CAASDGAGSYQLTF | TRBV12-4 | TRBJ1-2 | CASSLVPGPNGDYGYTF | RLRAEAQVK | YES | NO | NO |
| 2361 | TRAV29DV5 | TRAJ49 | CAASASGGQFYF | TRBV12-4 | TRBJ1-2 | CASSPGTNYGYTF | AVFDRKSDAK | YES | NO | NO |
| 2363 | TRAV35 | TRAJ40 | CAGRAGTYKYIF | TRBV12-4 | TRBJ1-2 | CASSSRTDTAYGYTF | KLGGALQAK | YES | NO | NO |
| 2364 | TRAV36DV7 | TRAJ54 | CAVIIQGAQKLVF | TRBV12-4 | TRBJ1-2 | CASSFWPGRNLLWGYTF | RLRAEAQVK | NO | NO | NO |
| 2365 | TRAV5 | TRAJ20 | CAETPNDYKLSF | TRBV12-4 | TRBJ1-2 | CASSLLGDYGYTF | KLGGALQAK | YES | NO | NO |
| 2366 | TRAV8-4 | TRAJ10 | CAVSDRGVTGGGNKLTF | TRBV12-4 | TRBJ1-2 | CASSLTSPSPDESLLSGLYGYTF | KLGGALQAK | NO | NO | NO |
| 2367 | TRAV8-6 | TRAJ49 | CAVSPTGNQFYF | TRBV12-4 | TRBJ1-2 | CASSLGEQGTF | KLGGALQAK | YES | NO | NO |
| 2370 | TRAV20 | TRAJ32 | CAVKDATNKLIF | TRBV12-5 | TRBJ1-2 | CASGLSQGFTF | KLGGALQAK | YES | NO | NO |
| 2373 | TRAV5 | TRAJ15 | CAESRGTALIF | TRBV12-5 | TRBJ1-2 | CASGSSGGAGYGYTF | KLGGALQAK | YES | NO | NO |
| 2374 | TRAV8-4 | TRAJ34 | CAVTSSYNTDKLIF | TRBV12-5 | TRBJ1-2 | CASGLVPRQGGNGYTF | KLGGALQAK | YES | NO | NO |
| 2376 | TRAV12-2 | TRAJ28 | CAVNRASGAGSYQLTF | TRBV13 | TRBJ1-2 | CASSSPGGEDTF | KLGGALQAK | YES | NO | NO |
| 2377 | TRAV19 | TRAJ30 | CALSEANRDDKIIF | TRBV13 | TRBJ1-2 | CASSLDRERYGYTF | KLGGALQAK | YES | NO | NO |
| 2379 | TRAV41 | TRAJ44 | CAVHTGTASKLTF | TRBV13 | TRBJ1-2 | CASSLFPNYGYTF | KLGGALQAK | YES | NO | NO |
| 2380 | TRAV8-1 | TRAJ37 | CAVGSGNTGKLIF | TRBV13 | TRBJ1-2 | CASSLVEATLNYGYTF | KLGGALQAK | YES | NO | NO |
| 2382 | TRAV8-4 | TRAJ3 | CAVSESLSSASKIIF | TRBV13 | TRBJ1-2 | CASSSGSYYGYTF | KLGGALQAK | YES | NO | NO |
| 2383 | TRAV12-2 | TRAJ15 | CAVNIPQAGTALIF | TRBV14 | TRBJ1-2 | CASSQVPVYGYTF | KLGGALQAK | NO | NO | NO |
| 2384 | TRAV12-3 | TRAJ39 | CAIGPGNMLTF | TRBV14 | TRBJ1-2 | CASSQVPGQGPNYGYTF | KLGGALQAK | NO | NO | NO |
| 2385 | TRAV13-2 | TRAJ53 | CAENWGSNYKLTF | TRBV14 | TRBJ1-2 | CASSQYLRPDYGYTF | KLGGALQAK | NO | NO | NO |
| 2386 | TRAV14DV4 | TRAJ5 | CAMRDPQGGRRALTF | TRBV14 | TRBJ1-2 | CASSQVPGQGPNYGYTF | KLGGALQAK | NO | NO | NO |
| 2387 | TRAV19 | TRAJ23 | CALSEAHQGGKLIF | TRBV14 | TRBJ1-2 | CASSQDHRGRISGYTF | KLGGALQAK | NO | NO | NO |
| 2388 | TRAV19 | TRAJ8 | CALSPRGFQKLVF | TRBV14 | TRBJ1-2 | CASSQVPGDIQGYTF | KLGGALQAK | NO | NO | NO |

| | | | | | | | | | |
|---|---|---|---|---|---|---|---|---|---|
| 2389 | TRAV20 | TRAJ15 | CAVQSNQAGTALIF | TRBV14 | TRBJ1-2 | CASSQSTGAYGYTF | KLGGALQAK | NO | NO | NO |
| 2390 | TRAV23DV6 | TRAJ20 | CAASGWYDYKLSF | TRBV14 | TRBJ1-2 | CASRDRGDYGYTF | KLGGALQAK | NO | NO | NO |
| 2393 | TRAV25 | TRAJ54 | CAGHVIQGAQKLVF | TRBV14 | TRBJ1-2 | CASSPFVSGSQTF | KLGGALQAK | NO | NO | NO |
| 2395 | TRAV29DV5 | TRAJ39 | CAASAINAGNMLTF | TRBV14 | TRBJ1-2 | CASSQVPGQGPNYGYTF | KLGGALQAK | NO | NO | NO |
| 2397 | TRAV4 | TRAJ20 | CLVGDDYKLSF | TRBV14 | TRBJ1-2 | CASSHTRLPNYGYTF | KLGGALQAK | NO | NO | NO |
| 2399 | TRAV8-1 | TRAJ24 | CAGMTTDSWGKLQF | TRBV14 | TRBJ1-2 | CASSQEGVHGYTF | KLGGALQAK | NO | NO | NO |
| 2404 | TRAV8-4 | TRAJ47 | CAVRNPMEYGNKLVF | TRBV14 | TRBJ1-2 | CASSQGQVGSGNYGYTF | KLGGALQAK | NO | NO | NO |
| 2405 | TRAV8-4 | TRAJ52 | CAVSALSPGGGTSYGKLTF | TRBV14 | TRBJ1-2 | CASSQVPGQGPNYGYTF | KLGGALQAK | NO | NO | NO |
| 2407 | TRAV12-2 | TRAJ6 | CAVKNPGGSYIPTF | TRBV15 | TRBJ1-2 | CATSRGGDRTKGYTF | KLGGALQAK | YES | NO | NO |
| 2409 | TRAV14DV4 | TRAJ20 | CAMRGSNDYKLSF | TRBV15 | TRBJ1-2 | CATSREEGDGYTF | KLGGALQAK | NO | NO | NO |
| 2410 | TRAV14DV4 | TRAJ44 | CAMREPNTGTASKLTF | TRBV15 | TRBJ1-2 | CATSRANRDGYTF | RLRAEAQVK | YES | NO | NO |
| 2412 | TRAV19 | TRAJ20 | CALSRNDYKLSF | TRBV15 | TRBJ1-2 | CATSREEGDGYTF | KLGGALQAK | YES | NO | NO |
| 2413 | TRAV21 | TRAJ52 | CAVIPLNAGGTSYGKLTF | TRBV15 | TRBJ1-2 | CATSSSSYGYTF | KLGGALQAK | YES | NO | NO |
| 2415 | TRAV27 | TRAJ42 | CAGGIYGGSQGNLIF | TRBV15 | TRBJ1-2 | CATSRDRVVGQGIDGYTF | KLGGALQAK | YES | NO | NO |
| 2416 | TRAV27 | TRAJ42 | CAGVEYGGSQGNLIF | TRBV15 | TRBJ1-2 | CATSRDRAGGANYGYTF | KLGGALQAK | YES | NO | NO |
| 2418 | TRAV29DV5 | TRAJ52 | CAASSTNAGGTSYGKLTF | TRBV15 | TRBJ1-2 | CATSSGYYGYTF | KLGGALQAK | YES | NO | NO |
| 2419 | TRAV5 | TRAJ8 | CAEEMNTGFQKLVF | TRBV15 | TRBJ1-2 | CATSRTGGRYGYTF | KLGGALQAK | YES | NO | NO |
| 2420 | TRAV8-1 | TRAJ3 | CAVNPEGASKIIF | TRBV15 | TRBJ1-2 | CATSTGRGQLYGYTF | KLGGALQAK | YES | NO | NO |
| 2423 | TRAV1-2 | TRAJ36 | CAVRSNAHQTGANNLFF | TRBV16 | TRBJ1-2 | CASSQSSGYGYTF | KLGGALQAK | YES | NO | NO |
| 2426 | TRAV12-1 | TRAJ6 | CVVIFLSSGGSYIPTF | TRBV18 | TRBJ1-2 | CASSERTGGSLLYGYTF | KLGGALQAK | YES | NO | NO |
| 2427 | TRAV14DV4 | TRAJ33 | CAMREHYSNYQLIW | TRBV18 | TRBJ1-2 | CASSPDSNYGYTF | RLRAEAQVK | YES | NO | YES |
| 2428 | TRAV17 | TRAJ57 | CATPVLRSEKLVF | TRBV18 | TRBJ1-2 | CASSPDRGLSYGYTF | KLGGALQAK | YES | NO | NO |
| 2429 | TRAV29DV5 | TRAJ52 | CAASAGGTSYGKLTF | TRBV18 | TRBJ1-2 | CASSPSTGGVYGYTF | KLGGALQAK | YES | NO | NO |
| 2430 | TRAV38-2DV8 | TRAJ22 | CAYRSSSGSARQLTF | TRBV18 | TRBJ1-2 | CASSPQWTLYGYTF | KLGGALQAK | YES | NO | NO |
| 2433 | TRAV1-1 | TRAJ29 | CAVSGNTPLVF | TRBV19 | TRBJ1-2 | CASSMGGYTYGYTF | KLGGALQAK | YES | NO | NO |
| 2435 | TRAV10 | TRAJ9 | CVVTPTGGFKTIF | TRBV19 | TRBJ1-2 | CASSMGLISTGDGYTF | KLGGALQAK | YES | NO | NO |
| 2436 | TRAV12-1 | TRAJ41 | CVVNSNSNSGYALNF | TRBV19 | TRBJ1-2 | CASTDRVGGYTF | KLGGALQAK | YES | NO | NO |
| 2437 | TRAV12-2 | TRAJ45 | CAVSGGGADGLTF | TRBV19 | TRBJ1-2 | CASSLGRAYGYTF | KLGGALQAK | YES | NO | NO |
| 2438 | TRAV12-2 | TRAJ49 | CAVNAGTGNQFYF | TRBV19 | TRBJ1-2 | CASTPRPGLLPPNYGYTF | KLGGALQAK | YES | NO | NO |
| 2439 | TRAV12-2 | TRAJ49 | CAVTGYSNTGNQFYF | TRBV19 | TRBJ1-2 | CASSMTGYSGYTF | KLGGALQAK | YES | NO | NO |
| 2442 | TRAV13-1 | TRAJ23 | CAALADNQGGKLIF | TRBV19 | TRBJ1-2 | CASALIQGVYTF | KLGGALQAK | YES | NO | NO |
| 2443 | TRAV13-1 | TRAJ40 | CAAPDGDTSGTYKYIF | TRBV19 | TRBJ1-2 | CASTPGHYGYTF | KLGGALQAK | YES | NO | NO |
| 2444 | TRAV13-1 | TRAJ52 | CAASNAGGTSYGKLTF | TRBV19 | TRBJ1-2 | CASSWGANYGYTF | KLGGALQAK | YES | NO | NO |
| 2445 | TRAV13-2 | TRAJ45 | CAERPSSGGGADGLTF | TRBV19 | TRBJ1-2 | CASSMGLISTGDGYTF | KLGGALQAK | YES | NO | NO |
| 2446 | TRAV14DV4 | TRAJ20 | CAMREGRGYKLSF | TRBV19 | TRBJ1-2 | CASSIEAGLYGYTF | KLGGALQAK | YES | NO | NO |
| 2448 | TRAV14DV4 | TRAJ23 | CAMRGGWGNQGGKLIF | TRBV19 | TRBJ1-2 | CASRDSLDYGYTF | KLGGALQAK | YES | NO | NO |
| 2449 | TRAV14DV4 | TRAJ8 | CAMREAAGNTGFQKLVF | TRBV19 | TRBJ1-2 | CASSITGTSNYGYTF | KLGGALQAK | YES | NO | NO |
| 2452 | TRAV17 | TRAJ22 | CATVPDDSARQLTF | TRBV19 | TRBJ1-2 | CASTYFPAYGYTF | KLGGALQAK | YES | NO | NO |
| 2453 | TRAV17 | TRAJ23 | CALFNQGGKLIF | TRBV19 | TRBJ1-2 | CASTPSGDGYTF | KLGGALQAK | YES | NO | NO |
| 2453 | TRAV17 | TRAJ23 | CALFNQGGKLIF | TRBV19 | TRBJ1-2 | CASTPSGDGYTF | AVFDRKSDAK | YES | NO | NO |
| 2455 | TRAV17 | TRAJ49 | CATDRSTGNQFYF | TRBV19 | TRBJ1-2 | CASSIDWPGIYGYTF | KLGGALQAK | YES | NO | NO |
| 2456 | TRAV17 | TRAJ53 | CATENSGGSNYKLTF | TRBV19 | TRBJ1-2 | CASSARTGVGYGYTF | AVFDRKSDAK | YES | NO | NO |
| 2456 | TRAV17 | TRAJ53 | CATENSGGSNYKLTF | TRBV19 | TRBJ1-2 | CASSARTGVGYGYTF | RLRAEAQVK | YES | NO | NO |
| 2458 | TRAV19 | TRAJ29 | CALSEPRSGNTPLVF | TRBV19 | TRBJ1-2 | CASSPGGGYGYTF | KLGGALQAK | YES | NO | NO |
| 2459 | TRAV19 | TRAJ43 | CALSEPRGVDMRF | TRBV19 | TRBJ1-2 | CASSMFTGQSYGYTF | KLGGALQAK | YES | NO | NO |
| 2461 | TRAV19 | TRAJ9 | CALSEAKGFKTIF | TRBV19 | TRBJ1-2 | CASSHAGTNYYGYTF | KLGGALQAK | YES | NO | NO |
| 2463 | TRAV20 | TRAJ20 | CAVRSEGSSDYKLSF | TRBV19 | TRBJ1-2 | CASSIGIGGNYGYTF | KLGGALQAK | YES | NO | NO |
| 2465 | TRAV20 | TRAJ9 | CAVLTNTGGFKTIF | TRBV19 | TRBJ1-2 | CASSITGFAPYGYTF | KLGGALQAK | YES | NO | NO |
| 2467 | TRAV21 | TRAJ17 | CAVTRVKAAGNKLTF | TRBV19 | TRBJ1-2 | CASSSAGGDGYTF | KLGGALQAK | YES | NO | NO |
| 2470 | TRAV21 | TRAJ52 | CAVISLLDAGGTSYGKLTF | TRBV19 | TRBJ1-2 | CASMTGARGDYGYTF | KLGGALQAK | YES | NO | NO |
| 2471 | TRAV21 | TRAJ56 | CAVESQANSKLTF | TRBV19 | TRBJ1-2 | CASRYGDGVLGYGYTF | KLGGALQAK | YES | NO | NO |
| 2472 | TRAV22 | TRAJ15 | CADNQAGTALIF | TRBV19 | TRBJ1-2 | CASSTGTGYGYTF | KLGGALQAK | YES | NO | NO |
| 2473 | TRAV24 | TRAJ28 | CAFSPSWSISGAGSYQLTF | TRBV19 | TRBJ1-2 | CASSISPGSYGYTF | KLGGALQAK | YES | NO | NO |
| 2476 | TRAV27 | TRAJ40 | CAGWGTYKYIF | TRBV19 | TRBJ1-2 | CASSIGGAGYTF | KLGGALQAK | YES | NO | NO |
| 2478 | TRAV29DV5 | TRAJ29 | CAASSGNTPLVF | TRBV19 | TRBJ1-2 | CASSITLADGYTF | KLGGALQAK | YES | NO | NO |
| 2479 | TRAV29DV5 | TRAJ31 | CAASAYARLMF | TRBV19 | TRBJ1-2 | CASSIETGGYYGYTF | KLGGALQAK | YES | NO | NO |
| 2480 | TRAV29DV5 | TRAJ42 | CAASAHYGGSQGNLIF | TRBV19 | TRBJ1-2 | CASSIKQGYGYTF | KLGGALQAK | YES | NO | NO |
| 2481 | TRAV3 | TRAJ23 | CAVRGTLINQGGKLIF | TRBV19 | TRBJ1-2 | CASRAKPSGTGFNYGYTF | KLGGALQAK | NO | NO | NO |
| 2482 | TRAV35 | TRAJ12 | CAGQTGLMDSSYKLIF | TRBV19 | TRBJ1-2 | CASMSIYGYTF | KLGGALQAK | YES | NO | NO |
| 2486 | TRAV5 | TRAJ49 | CAESIAGNQFYF | TRBV19 | TRBJ1-2 | CASRTTGGDGYTF | KLGGALQAK | YES | NO | NO |
| 2488 | TRAV8-1 | TRAJ27 | CAVNTNAGKSTF | TRBV19 | TRBJ1-2 | CASRLVLAGTGDGYTF | KLGGALQAK | YES | NO | NO |
| 2490 | TRAV8-3 | TRAJ21 | CAVGLFNFNKFYF | TRBV19 | TRBJ1-2 | CASSRRARGTPFGGYTF | KLGGALQAK | YES | NO | NO |
| 2491 | TRAV8-3 | TRAJ40 | CAVAPEGTYKYIF | TRBV19 | TRBJ1-2 | CASSVQGAYYGYTF | KLGGALQAK | YES | NO | NO |
| 2492 | TRAV8-4 | TRAJ14 | CAVSVGSTFIF | TRBV19 | TRBJ1-2 | CASSIDSLYGYTF | KLGGALQAK | YES | NO | NO |
| 2494 | TRAV9-2 | TRAJ49 | CALSDSNTGNQFYF | TRBV19 | TRBJ1-2 | CASSTQRDRELGYTF | KLGGALQAK | YES | NO | NO |
| 2496 | TRAV12-3 | TRAJ47 | CAMSAREYGNKLVF | TRBV2 | TRBJ1-2 | CAGIRTGTGPSPNYGYTF | KLGGALQAK | YES | NO | NO |
| 2497 | TRAV13-1 | TRAJ16 | CAASRASDGQKLLF | TRBV2 | TRBJ1-2 | CASSVIKNYGYTF | KLGGALQAK | NO | NO | NO |
| 2499 | TRAV17 | TRAJ13 | CATDAGGGYQKVTF | TRBV2 | TRBJ1-2 | CASSEARETATSNYGYTF | KLGGALQAK | YES | NO | NO |
| 2500 | TRAV17 | TRAJ48 | CATTFRSNFGNEKLTF | TRBV2 | TRBJ1-2 | CASSEVYENSNGYTF | KLGGALQAK | YES | NO | NO |
| 2503 | TRAV3 | TRAJ21 | CAVRDWNNFNKFYF | TRBV2 | TRBJ1-2 | CASSEGTGGILSYGYTF | KLGGALQAK | NO | NO | NO |
| 2504 | TRAV35 | TRAJ20 | CAGAADDYKLSF | TRBV2 | TRBJ1-2 | CASSPGQGWYGYTF | KLGGALQAK | YES | NO | NO |
| 2505 | TRAV35 | TRAJ49 | CAGPATGNQFYF | TRBV2 | TRBJ1-2 | CASSKDIGGNYGYTF | KLGGALQAK | YES | NO | NO |
| 2507 | TRAV38-2DV8 | TRAJ40 | CAYKVSGTYKYIF | TRBV2 | TRBJ1-2 | CASSEGWGYTF | KLGGALQAK | YES | NO | NO |
| 2508 | TRAV40 | TRAJ32 | CLLGGATNKLIF | TRBV2 | TRBJ1-2 | CASSGRMTFSGYTF | KLGGALQAK | NO | NO | NO |
| 2509 | TRAV40 | TRAJ35 | CSI | TRBV2 | TRBJ1-2 | CASSSASNGYTF | KLGGALQAK | NO | NO | NO |
| 2510 | TRAV41 | TRAJ33 | CAVRLDSNYQLIW | TRBV2 | TRBJ1-2 | CASSPGTAYGYTF | KLGGALQAK | YES | NO | NO |
| 2511 | TRAV8-3 | TRAJ6 | CAVVLYGGSYIPTF | TRBV2 | TRBJ1-2 | CASSEEGGGYTF | KLGGALQAK | YES | NO | NO |
| 2512 | TRAV1-1 | TRAJ30 | CAVRPRDDKIIF | TRBV20-1 | TRBJ1-2 | CSARDHRRGLYGYTF | KLGGALQAK | YES | NO | NO |
| 2513 | TRAV1-1 | TRAJ30 | CAVRPRDDKIIF | TRBV20-1 | TRBJ1-2 | CSARDSDRGIRDGYTF | KLGGALQAK | YES | NO | NO |
| 2515 | TRAV1-2 | TRAJ30 | CAVKGRDDKIIF | TRBV20-1 | TRBJ1-2 | CSARDPPAGRDGYTF | KLGGALQAK | YES | NO | NO |
| 2516 | TRAV1-2 | TRAJ33 | CAAMDSNYQLIW | TRBV20-1 | TRBJ1-2 | CSARTGQINGYTF | KLGGALQAK | YES | NO | NO |
| 2517 | TRAV1-2 | TRAJ34 | CAVREENTDKLIF | TRBV20-1 | TRBJ1-2 | CSARDLGRANGYTF | KLGGALQAK | YES | NO | NO |
| 2518 | TRAV10 | TRAJ49 | CVVASPTTGNQFYF | TRBV20-1 | TRBJ1-2 | CSARDHFAGGQANYGYTF | KLGGALQAK | YES | NO | NO |
| 2521 | TRAV12-1 | TRAJ42 | CVVRFGNYGGSQGNLIF | TRBV20-1 | TRBJ1-2 | CSARDVPGSIGGYTF | KLGGALQAK | YES | NO | NO |
| 2522 | TRAV12-2 | TRAJ49 | CAVNRGGNQFYF | TRBV20-1 | TRBJ1-2 | CSARTDRGRTF | AVFDRKSDAK | YES | NO | YES |
| 2523 | TRAV12-2 | TRAJ49 | CAVNRGGNQFYF | TRBV20-1 | TRBJ1-2 | CSARVPRGPGYTF | KLGGALQAK | YES | NO | NO |
| 2524 | TRAV12-2 | TRAJ49 | CAVNRGGNQFYF | TRBV20-1 | TRBJ1-2 | CSARVDRGYTF | KLGGALQAK | YES | NO | NO |
| 2525 | TRAV12-2 | TRAJ49 | CAVTSNTGNQFYF | TRBV20-1 | TRBJ1-2 | CSAGPEWGKDGYTF | KLGGALQAK | YES | NO | NO |
| 2527 | TRAV13-1 | TRAJ23 | CAARMIYNQGGKLIF | TRBV20-1 | TRBJ1-2 | CSAIIVGQYYGYTF | KLGGALQAK | YES | NO | NO |
| 2529 | TRAV14DV4 | TRAJ4 | CAMRPPFRGGYNKLIF | TRBV20-1 | TRBJ1-2 | CSARDLLAAGYGYTF | KLGGALQAK | YES | NO | NO |
| 2531 | TRAV16 | TRAJ54 | CALSPIQGAQKLVF | TRBV20-1 | TRBJ1-2 | CSARDRSLSGGYTF | KLGGALQAK | NO | NO | NO |
| 2533 | TRAV19 | TRAJ10 | CALSLLTGGGNKLTF | TRBV20-1 | TRBJ1-2 | CSAADRGIYGYTF | KLGGALQAK | YES | NO | NO |
| 2534 | TRAV19 | TRAJ4 | CALRISGGYNKLIF | TRBV20-1 | TRBJ1-2 | CSARDFDGYTF | KLGGALQAK | YES | NO | NO |
| 2535 | TRAV19 | TRAJ40 | CALSETSSGTYKYIF | TRBV20-1 | TRBJ1-2 | CSAPSPRHTIYGYTF | KLGGALQAK | YES | NO | NO |
| 2537 | TRAV20 | TRAJ53 | CAVQAPSQNSGGSNYKLTF | TRBV20-1 | TRBJ1-2 | CSAREGDPNYGYTF | KLGGALQAK | YES | NO | NO |
| 2538 | TRAV21 | TRAJ43 | CAVEGGNNDMRF | TRBV20-1 | TRBJ1-2 | CSASEQPYLQTGYTF | KLGGALQAK | YES | NO | NO |
| 2539 | TRAV21 | TRAJ48 | CAVLSLSNFGNEKLTF | TRBV20-1 | TRBJ1-2 | CSARTGRESTYYGYTF | KLGGALQAK | YES | NO | NO |
| 2542 | TRAV25 | TRAJ31 | CALPKGDARLMF | TRBV20-1 | TRBJ1-2 | CSARDRDRVYGYTF | KLGGALQAK | YES | NO | NO |
| 2544 | TRAV3 | TRAJ26 | CAVRDIERNYGQNFVF | TRBV20-1 | TRBJ1-2 | CSAKDPPGYTF | KLGGALQAK | NO | NO | NO |
| 2545 | TRAV3 | TRAJ6 | CAVRDEGGSYIPTF | TRBV20-1 | TRBJ1-2 | CSANTDRGYGYTF | KLGGALQAK | NO | NO | NO |
| 2546 | TRAV35 | TRAJ13 | CAGQLSGGYQKVTF | TRBV20-1 | TRBJ1-2 | CSARDQLREDNYGYTF | KLGGALQAK | YES | NO | NO |
| 2549 | TRAV5 | TRAJ22 | CAQVGSARQLTF | TRBV20-1 | TRBJ1-2 | CSLDRTPASNYGYTF | KLGGALQAK | YES | NO | NO |
| 2550 | TRAV5 | TRAJ4 | CAETSGGYNKLIF | TRBV20-1 | TRBJ1-2 | CSARDSILENYGYTF | KLGGALQAK | YES | NO | NO |
| 2551 | TRAV5 | TRAJ42 | CAENFGRGSQGNLIF | TRBV20-1 | TRBJ1-2 | CSARAGQGVDGYTF | KLGGALQAK | YES | NO | NO |

| | | | | | | | | | | |
|---|---|---|---|---|---|---|---|---|---|---|
| 2554 | TRAV8-3 | TRAJ4 | CAVGFSGGYNKLIF | TRBV20-1 | TRBJ1-2 | CSARVVWTAMGYGYTF | RLRAEAQVK | YES | NO | YES |
| 2557 | TRAV9-2 | TRAJ5 | CAAYTGRRALTF | TRBV20-1 | TRBJ1-2 | CSARGRGLDGYTF | KLGGALQAK | YES | NO | NO |
| 2558 | TRAV26-2 | TRAJ8 | CILRDAPRFQKLVF | TRBV21-1 | TRBJ1-2 | CASSKDGGTPTGYTF | KLGGALQAK | NO | NO | NO |
| 2560 | TRAV8-3 | TRAJ33 | CAVGDIVSNYQLIW | TRBV23-1 | TRBJ1-2 | CISAPALDQGVCYGYTF | KLGGALQAK | YES | NO | NO |
| 2561 | TRAV1-1 | TRAJ23 | CAAAAYNQGGKLIF | TRBV24-1 | TRBJ1-2 | CATSDPAGMTGGWHGYTF | KLGGALQAK | NO | NO | NO |
| 2566 | TRAV12-3 | TRAJ43 | CAMSWYNNNDMRF | TRBV24-1 | TRBJ1-2 | CATSDSGVNYGYTF | KLGGALQAK | YES | NO | NO |
| 2567 | TRAV13-1 | TRAJ54 | CAARGQGAQKLVF | TRBV24-1 | TRBJ1-2 | CATSNRQGVYGYTF | KLGGALQAK | YES | NO | NO |
| 2570 | TRAV21 | TRAJ40 | CAVRSSGTYKYIF | TRBV24-1 | TRBJ1-2 | CATSDGTGSNYGYTF | KLGGALQAK | YES | NO | NO |
| 2573 | TRAV27 | TRAJ39 | CAGVGNAGNMLTF | TRBV24-1 | TRBJ1-2 | CATRSSANYGYTF | KLGGALQAK | YES | NO | NO |
| 2574 | TRAV3 | TRAJ43 | CAVRDIDMRF | TRBV24-1 | TRBJ1-2 | CATRGSGTGGYGYTF | KLGGALQAK | NO | NO | NO |
| 2576 | TRAV4 | TRAJ34 | CLVGAMINTDKLIF | TRBV24-1 | TRBJ1-2 | CATSGETIDGYTF | KLGGALQAK | YES | NO | NO |
| 2578 | TRAV12-2 | TRAJ20 | CAAKNDYKLSF | TRBV25-1 | TRBJ1-2 | CASSEYRHWGNYGYTF | KLGGALQAK | YES | NO | NO |
| 2580 | TRAV24 | TRAJ31 | CASGGYNNARLMF | TRBV25-1 | TRBJ1-2 | CASSGRDRVWNNYGYTF | KLGGALQAK | YES | NO | NO |
| 2581 | TRAV3 | TRAJ21 | CAVRLYNFNKFYF | TRBV25-1 | TRBJ1-2 | CASIGEGTYNDYGYTF | KLGGALQAK | NO | NO | NO |
| 2582 | TRAV30 | TRAJ39 | CGTEGAGNMLTF | TRBV25-1 | TRBJ1-2 | CASSEYVKGYTF | KLGGALQAK | YES | NO | NO |
| 2583 | TRAV38-2DV8 | TRAJ29 | CAYRRPSGNTPLVF | TRBV25-1 | TRBJ1-2 | CASSDAGNYGYTF | KLGGALQAK | YES | NO | NO |
| 2586 | TRAV1-1 | TRAJ11 | CAVRSPGNSGYSTLTF | TRBV27 | TRBJ1-2 | CASSLSLPSGTVPSYGYTF | KLGGALQAK | YES | NO | NO |
| 2587 | TRAV1-1 | TRAJ34 | CAVRPGKYNTDKLIF | TRBV27 | TRBJ1-2 | CASSLAPSGNYGYTF | KLGGALQAK | YES | NO | NO |
| 2588 | TRAV1-2 | TRAJ20 | CAVRDRRDYKLSF | TRBV27 | TRBJ1-2 | CASRARTGGLGYTF | KLGGALQAK | YES | NO | NO |
| 2589 | TRAV1-2 | TRAJ28 | CAVPNSGAGSYQLTF | TRBV27 | TRBJ1-2 | CASKTGSYGYTF | KLGGALQAK | YES | NO | NO |
| 2592 | TRAV1-2 | TRAJ37 | CAVRDSHGSGNTGKLIF | TRBV27 | TRBJ1-2 | CASSPTGTATLSYTF | KLGGALQAK | YES | NO | NO |
| 2593 | TRAV1-2 | TRAJ42 | CAVRDYGGSQGNLIF | TRBV27 | TRBJ1-2 | CASSLGTANYGYTF | KLGGALQAK | YES | NO | NO |
| 2594 | TRAV10 | TRAJ17 | CVVSVAAGNKLTF | TRBV27 | TRBJ1-2 | CASTIRLGVYGYTF | KLGGALQAK | YES | NO | NO |
| 2595 | TRAV12-2 | TRAJ20 | CAVNKGYKLSF | TRBV27 | TRBJ1-2 | CASSLSWSGGDYGYTF | KLGGALQAK | YES | NO | NO |
| 2596 | TRAV12-2 | TRAJ34 | CAVNMSTDKLIF | TRBV27 | TRBJ1-2 | CASSLSPEGGGAGDGYTF | KLGGALQAK | YES | NO | NO |
| 2598 | TRAV12-2 | TRAJ49 | CASLNTGNQFYF | TRBV27 | TRBJ1-2 | CASSPQQGFTGYTF | KLGGALQAK | YES | NO | NO |
| 2601 | TRAV12-3 | TRAJ8 | CAMVPGFQKLVF | TRBV27 | TRBJ1-2 | CASSPTGTGDGYTF | KLGGALQAK | YES | NO | NO |
| 2602 | TRAV13-1 | TRAJ26 | CAASPGQNFVF | TRBV27 | TRBJ1-2 | CASSFLPLNYGYTF | KLGGALQAK | YES | NO | NO |
| 2604 | TRAV13-1 | TRAJ47 | CAPMGPKLVF | TRBV27 | TRBJ1-2 | CASSLTEDYGYTF | KLGGALQAK | YES | NO | NO |
| 2605 | TRAV13-1 | TRAJ49 | CAASPGNQFYF | TRBV27 | TRBJ1-2 | CASSLLPVNYGYTF | KLGGALQAK | YES | NO | NO |
| 2607 | TRAV14DV4 | TRAJ22 | CAMRRPISSGSARQLTF | TRBV27 | TRBJ1-2 | CASSLTGPDGYGYTF | KLGGALQAK | YES | NO | NO |
| 2608 | TRAV14DV4 | TRAJ24 | CAMREALKTDSWGKLQF | TRBV27 | TRBJ1-2 | CASTSGGFGGGYTF | KLGGALQAK | YES | NO | NO |
| 2610 | TRAV14DV4 | TRAJ36 | CAMREGWDQTGANNLFF | TRBV27 | TRBJ1-2 | CASSLTYRGLGYGYTF | KLGGALQAK | YES | NO | NO |
| 2613 | TRAV17 | TRAJ10 | CATGLTGGGNKLTF | TRBV27 | TRBJ1-2 | CASSLGGQNYGYTF | KLGGALQAK | YES | NO | NO |
| 2614 | TRAV17 | TRAJ10 | CAFGFTGGGNKLTF | TRBV27 | TRBJ1-2 | CASSLGGQNYGYTF | KLGGALQAK | YES | NO | NO |
| 2615 | TRAV17 | TRAJ43 | CATDLNNNDMRF | TRBV27 | TRBJ1-2 | CASSLMQGSYYGYTF | KLGGALQAK | YES | NO | NO |
| 2617 | TRAV19 | TRAJ13 | CALRPDSGGYQKVTF | TRBV27 | TRBJ1-2 | CASRPTGANYGYTF | KLGGALQAK | YES | NO | NO |
| 2618 | TRAV19 | TRAJ27 | CALSEGEGKSTF | TRBV27 | TRBJ1-2 | CASSLIGQRGYTF | KLGGALQAK | YES | YES | NO |
| 2619 | TRAV19 | TRAJ36 | CALSGTYQTGANNLFF | TRBV27 | TRBJ1-2 | CASSFLTGSYGYTF | KLGGALQAK | YES | NO | NO |
| 2620 | TRAV2 | TRAJ21 | CAVPGGHNFNKFYF | TRBV27 | TRBJ1-2 | CASSQRIILGEVYGYTF | KLGGALQAK | NO | NO | NO |
| 2621 | TRAV21 | TRAJ28 | CAVRASGAGSYQLTF | TRBV27 | TRBJ1-2 | CASSLGQGVYGYTF | KLGGALQAK | YES | NO | NO |
| 2622 | TRAV21 | TRAJ28 | CAAIGAGSYQLTF | TRBV27 | TRBJ1-2 | CASSFGGANYGYTF | KLGGALQAK | YES | NO | NO |
| 2624 | TRAV23DV6 | TRAJ37 | CASLESNTGKLIF | TRBV27 | TRBJ1-2 | CASMALMLQNYGYTF | KLGGALQAK | YES | NO | NO |
| 2626 | TRAV25 | TRAJ13 | CAGPVGGYQKVTF | TRBV27 | TRBJ1-2 | CASSLSNEGHTGYDYGYTF | KLGGALQAK | YES | NO | NO |
| 2629 | TRAV26-2 | TRAJ26 | CILRDRDNYGQNFVF | TRBV27 | TRBJ1-2 | CASSFRGVYGYTF | KLGGALQAK | YES | NO | NO |
| 2631 | TRAV27 | TRAJ22 | CAGASARQLTF | TRBV27 | TRBJ1-2 | CASSLYDSYGYTF | KLGGALQAK | YES | NO | NO |
| 2632 | TRAV27 | TRAJ36 | CSGANNLFF | TRBV27 | TRBJ1-2 | CASKGRGRINYGYTF | KLGGALQAK | NO | NO | NO |
| 2633 | TRAV27 | TRAJ40 | CAGPDSGTYKYIF | TRBV27 | TRBJ1-2 | CASRDGYYGYTF | KLGGALQAK | YES | NO | NO |
| 2634 | TRAV27 | TRAJ44 | CAGADGTASKLTF | TRBV27 | TRBJ1-2 | CASSLSGTWGEGYTF | RLRAEAQVK | YES | NO | NO |
| 2638 | TRAV34 | TRAJ52 | CGADPNAGGTSYGKLTF | TRBV27 | TRBJ1-2 | CASSFQSGQNFYGYTF | KLGGALQAK | YES | NO | NO |
| 2639 | TRAV35 | TRAJ45 | CAGRKYSGGGADGLTF | TRBV27 | TRBJ1-2 | CASSLYSRTLFYGYTF | KLGGALQAK | YES | NO | NO |
| 2640 | TRAV38-2DV8 | TRAJ57 | CAYRSAQGGSEKLVF | TRBV27 | TRBJ1-2 | CASSSQLALNYGYTF | KLGGALQAK | YES | NO | NO |
| 2642 | TRAV5 | TRAJ9 | CAESWGGGFKTIF | TRBV27 | TRBJ1-2 | CASSFSKGSGTDGYTF | KLGGALQAK | YES | NO | NO |
| 2643 | TRAV6 | TRAJ4 | CALSGGYNKLIF | TRBV27 | TRBJ1-2 | CASSSSPLQGTNYGYTF | KLGGALQAK | YES | NO | NO |
| 2648 | TRAV12-2 | TRAJ21 | CAGYNFNKFYF | TRBV28 | TRBJ1-2 | CASTTPPTGSGYTF | KLGGALQAK | YES | NO | NO |
| 2652 | TRAV29DV5 | TRAJ20 | CAASATNDYKLSF | TRBV28 | TRBJ1-2 | CASFRAVLYGYTF | KLGGALQAK | YES | NO | NO |
| 2653 | TRAV38-2DV8 | TRAJ32 | CAYRSFGGATNKLIF | TRBV28 | TRBJ1-2 | CASSSIREICSMCGYTF | RLRAEAQVK | YES | NO | YES |
| 2654 | TRAV9-2 | TRAJ48 | CALSEYGNEKLTF | TRBV28 | TRBJ1-2 | CASRLGTGGPSYGYTF | KLGGALQAK | YES | NO | NO |
| 2656 | TRAV1-2 | TRAJ18 | CAVRDDRGSTLGRLYF | TRBV29-1 | TRBJ1-2 | CSVDLEANYGYTF | KLGGALQAK | YES | NO | NO |
| 2658 | TRAV1-2 | TRAJ27 | CAVNAGKSTF | TRBV29-1 | TRBJ1-2 | CSVGVSGGYGYTF | KLGGALQAK | YES | NO | NO |
| 2664 | TRAV10 | TRAJ39 | CVVSNAGNMLTF | TRBV29-1 | TRBJ1-2 | CSVDLEANYGYTF | KLGGALQAK | YES | NO | NO |
| 2668 | TRAV12-1 | TRAJ53 | CVVGTWRGSNYKLTF | TRBV29-1 | TRBJ1-2 | CSVVLEGGVYGYTF | KLGGALQAK | YES | NO | NO |
| 2669 | TRAV12-2 | TRAJ45 | CAVYDSGGGADGLTF | TRBV29-1 | TRBJ1-2 | CSVPARAGYGYTF | KLGGALQAK | YES | NO | NO |
| 2670 | TRAV12-2 | TRAJ52 | CAVQPFAGGTSYGKLTF | TRBV29-1 | TRBJ1-2 | CSTRDRSYGYTF | KLGGALQAK | YES | NO | NO |
| 2672 | TRAV13-2 | TRAJ15 | CAENRDQAGTALIF | TRBV29-1 | TRBJ1-2 | CSVESKINYGYTF | KLGGALQAK | YES | NO | NO |
| 2677 | TRAV14DV4 | TRAJ23 | CASYYNQGGKLIF | TRBV29-1 | TRBJ1-2 | CSVDSHRTGYNYGYTF | KLGGALQAK | YES | NO | NO |
| 2680 | TRAV14DV4 | TRAJ9 | CAMREGPTGGFKTIF | TRBV29-1 | TRBJ1-2 | CSVDLEANYGYTF | KLGGALQAK | YES | NO | NO |
| 2683 | TRAV17 | TRAJ7 | CATEGAYGNNRLAF | TRBV29-1 | TRBJ1-2 | CSGWEGLYGYTF | KLGGALQAK | YES | NO | NO |
| 2686 | TRAV19 | TRAJ30 | CALSETDRDDKIIF | TRBV29-1 | TRBJ1-2 | CSVETTGLNYGYTF | KLGGALQAK | YES | NO | NO |
| 2687 | TRAV19 | TRAJ37 | CALSEFSSNTGKLIF | TRBV29-1 | TRBJ1-2 | CSVDLEANYGYTF | KLGGALQAK | YES | NO | NO |
| 2688 | TRAV19 | TRAJ39 | CALIYDNAGNMLTF | TRBV29-1 | TRBJ1-2 | CSVDLEANYGYTF | KLGGALQAK | YES | NO | NO |
| 2690 | TRAV20 | TRAJ49 | CAVQAKTGNQFYF | TRBV29-1 | TRBJ1-2 | CSVDLEANYGYTF | KLGGALQAK | YES | NO | NO |
| 2691 | TRAV21 | TRAJ37 | CAVLFNTGKLIF | TRBV29-1 | TRBJ1-2 | CSVDLEANYGYTF | KLGGALQAK | YES | NO | NO |
| 2694 | TRAV21 | TRAJ43 | CAVWYMRF | TRBV29-1 | TRBJ1-2 | CSVDLEANYGYTF | KLGGALQAK | NO | NO | NO |
| 2696 | TRAV23DV6 | TRAJ37 | CAASTYSNTGKLIF | TRBV29-1 | TRBJ1-2 | CSVEEWGRLGYGYTF | KLGGALQAK | YES | NO | NO |
| 2699 | TRAV26-2 | TRAJ22 | CILVLSGSARQLTF | TRBV29-1 | TRBJ1-2 | CSVGGDSVTNYGYTF | KLGGALQAK | NO | NO | NO |
| 2700 | TRAV26-2 | TRAJ40 | CIRGTYKYIF | TRBV29-1 | TRBJ1-2 | CSVSDKSYGYTF | KLGGALQAK | YES | NO | NO |
| 2701 | TRAV26-2 | TRAJ40 | CITGTYKYIF | TRBV29-1 | TRBJ1-2 | CSVGDRAYGYTF | KLGGALQAK | YES | NO | NO |
| 2702 | TRAV27 | TRAJ10 | CAGRGPGGGNKLTF | TRBV29-1 | TRBJ1-2 | CSVDLRGPGHGYTF | KLGGALQAK | YES | NO | NO |
| 2703 | TRAV27 | TRAJ22 | CAGQAPSGSARQLTF | TRBV29-1 | TRBJ1-2 | CSVDLRGPGHGYTF | KLGGALQAK | YES | NO | NO |
| 2703 | TRAV27 | TRAJ22 | CAGQAPSGSARQLTF | TRBV29-1 | TRBJ1-2 | CSVDLRGPGHGYTF | AVFDRKSDAK | YES | NO | NO |
| 2704 | TRAV27 | TRAJ29 | CAGEWQGNTPLVF | TRBV29-1 | TRBJ1-2 | CSVDLEANYGYTF | KLGGALQAK | YES | NO | NO |
| 2705 | TRAV29DV5 | TRAJ21 | CAAGRYNFNKFYF | TRBV29-1 | TRBJ1-2 | CSVDLEANYGYTF | KLGGALQAK | YES | NO | NO |
| 2707 | TRAV29DV5 | TRAJ34 | CAASPRGSDKLIF | TRBV29-1 | TRBJ1-2 | CSVDLEANYGYTF | KLGGALQAK | YES | NO | NO |
| 2708 | TRAV3 | TRAJ27 | CAVRDNAGKSTF | TRBV29-1 | TRBJ1-2 | CSVDLEANYGYTF | KLGGALQAK | NO | NO | NO |
| 2709 | TRAV30 | TRAJ37 | CGTASNTGKLIF | TRBV29-1 | TRBJ1-2 | CSVDLEANYGYTF | KLGGALQAK | YES | NO | NO |
| 2710 | TRAV34 | TRAJ3 | CGADSRYSSASKIIF | TRBV29-1 | TRBJ1-2 | CSVVGNYGYTF | KLGGALQAK | YES | NO | NO |
| 2710 | TRAV34 | TRAJ3 | CGADSRYSSASKIIF | TRBV29-1 | TRBJ1-2 | CSVVGNYGYTF | IVTDFSVIK | YES | NO | NO |
| 2710 | TRAV34 | TRAJ3 | CGADSRYSSASKIIF | TRBV29-1 | TRBJ1-2 | CSVVGNYGYTF | AVFDRKSDAK | YES | NO | NO |
| 2712 | TRAV39 | TRAJ54 | CAVEGIQGAQKLVF | TRBV29-1 | TRBJ1-2 | CSADHGNYGYTF | KLGGALQAK | YES | NO | NO |
| 2714 | TRAV5 | TRAJ37 | CAEISGNTGKLIF | TRBV29-1 | TRBJ1-2 | CSVEGLSALGKGYGYTF | KLGGALQAK | YES | NO | NO |
| 2715 | TRAV5 | TRAJ43 | CAETGNDMRF | TRBV29-1 | TRBJ1-2 | CSVDLEANYGYTF | KLGGALQAK | YES | NO | NO |
| 2716 | TRAV5 | TRAJ6 | CAGSGGSYIPTF | TRBV29-1 | TRBJ1-2 | CSVEVIGRGHGYTF | KLGGALQAK | YES | NO | NO |
| 2717 | TRAV8-2 | TRAJ22 | CVGRKGSARQLTF | TRBV29-1 | TRBJ1-2 | CSVDLEANYGYTF | KLGGALQAK | YES | NO | NO |
| 2718 | TRAV8-3 | TRAJ49 | CAVGAFRQFYF | TRBV29-1 | TRBJ1-2 | CSVDLEANYGYTF | KLGGALQAK | YES | NO | NO |
| 2719 | TRAV8-4 | TRAJ45 | CAVSYSGGGADGLTF | TRBV29-1 | TRBJ1-2 | CSVDLEANYGYTF | KLGGALQAK | YES | NO | NO |
| 2721 | TRAV12-1 | TRAJ10 | CVVSRGLTGGGNKLTF | TRBV3-1 | TRBJ1-2 | CASSPESLDGYTF | KLGGALQAK | YES | NO | NO |
| 2723 | TRAV12-2 | TRAJ10 | CAVNGGTGGGNKLTF | TRBV3-1 | TRBJ1-2 | CASSQLGNGYTF | KLGGALQAK | YES | NO | NO |
| 2727 | TRAV14DV4 | TRAJ34 | CAMRGYNTDKLIF | TRBV3-1 | TRBJ1-2 | CASSQGRDRHGYTF | KLGGALQAK | YES | NO | NO |
| 2728 | TRAV14DV4 | TRAJ4 | CAMRGPISGGYNKLIF | TRBV3-1 | TRBJ1-2 | CASSQGRDRHGYTF | AVFDRKSDAK | YES | NO | NO |
| 2728 | TRAV14DV4 | TRAJ4 | CAMRGPISGGYNKLIF | TRBV3-1 | TRBJ1-2 | CASSQGRDRHGYTF | IVTDFSVIK | YES | NO | NO |
| 2730 | TRAV2 | TRAJ31 | CAVETHNNARLMF | TRBV3-1 | TRBJ1-2 | CASSPPKGGENYGYTF | KLGGALQAK | NO | NO | NO |
| 2732 | TRAV21 | TRAJ33 | CAVRDSNYQLIW | TRBV3-1 | TRBJ1-2 | CASSPGQGSYGYTF | KLGGALQAK | YES | NO | NO |
| 2733 | TRAV21 | TRAJ54 | CAVEVQGAQKLVF | TRBV3-1 | TRBJ1-2 | CASSQGRDRHGYTF | KLGGALQAK | YES | NO | NO |

| | | | | | | | | | |
|---|---|---|---|---|---|---|---|---|---|
| 2734 | TRAV27 | TRAJ33 | CAGVMDSNYQLIW | TRBV3-1 | TRBJ1-2 | CASSQGRDRHGYTF | KLGGALQAK | YES | NO | NO |
| 2736 | TRAV29DV5 | TRAJ45 | CAASDMYSGGGADGLTF | TRBV3-1 | TRBJ1-2 | CASSQVGTVYGYTF | KLGGALQAK | YES | NO | NO |
| 2738 | TRAV38-2DV8 | TRAJ20 | CAYRSVYKLSF | TRBV3-1 | TRBJ1-2 | CASSQGRDRHGYTF | KLGGALQAK | YES | NO | NO |
| 2739 | TRAV41 | TRAJ57 | CAVRQGGSEKLVF | TRBV3-1 | TRBJ1-2 | CASSQDRTIWSYGYTF | KLGGALQAK | YES | NO | NO |
| 2740 | TRAV5 | TRAJ37 | CAEIRGGKLIF | TRBV3-1 | TRBJ1-2 | CASSQDHGGSSYGYTF | KLGGALQAK | YES | NO | NO |
| 2741 | TRAV8-1 | TRAJ13 | CAVSNSGGYQKVTF | TRBV3-1 | TRBJ1-2 | CASSSQGSNYGYTF | KLGGALQAK | YES | NO | NO |
| 2742 | TRAV8-3 | TRAJ36 | CAVGGETGANNLFF | TRBV3-1 | TRBJ1-2 | CASSQDSQHYGYTF | KLGGALQAK | YES | NO | NO |
| 2743 | TRAV8-3 | TRAJ42 | CAVDSNYGGSQGNLIF | TRBV3-1 | TRBJ1-2 | CASSSTGGGGYTF | KLGGALQAK | YES | NO | NO |
| 2744 | TRAV12-2 | TRAJ20 | CAVRSNDYKLSF | TRBV30 | TRBJ1-2 | CAWPGLSGYTF | KLGGALQAK | YES | YES | NO |
| 2745 | TRAV12-2 | TRAJ31 | CAVDLGLMF | TRBV30 | TRBJ1-2 | CAWSTTSRNYGYTF | KLGGALQAK | NO | NO | NO |
| 2746 | TRAV12-3 | TRAJ39 | CAIGPGNMLTF | TRBV30 | TRBJ1-2 | CAWGSGLGYGYTF | KLGGALQAK | YES | NO | NO |
| 2747 | TRAV14DV4 | TRAJ37 | CAMRGPYSSNTGKLIF | TRBV30 | TRBJ1-2 | CAWGSGLGYGYTF | AVFDRKSDAK | YES | NO | NO |
| 2747 | TRAV14DV4 | TRAJ37 | CAMRGPYSSNTGKLIF | TRBV30 | TRBJ1-2 | CAWGSGLGYGYTF | RLRAEAQVK | YES | NO | NO |
| 2748 | TRAV19 | TRAJ56 | CALSVGNTGANSKLTF | TRBV30 | TRBJ1-2 | CACPPEEGGHYGYTF | KLGGALQAK | NO | NO | NO |
| 2752 | TRAV38-2DV8 | TRAJ47 | CAYAWEYGNKLVF | TRBV30 | TRBJ1-2 | CAWSVLWSGRVYDYYGYTF | KLGGALQAK | YES | NO | NO |
| 2753 | TRAV8-1 | TRAJ34 | CAVNGLYNTDKLIF | TRBV30 | TRBJ1-2 | CAWSGGRMGGYTF | KLGGALQAK | NO | NO | NO |
| 2754 | TRAV8-2 | TRAJ4 | CVVSDLSGGYNKLIF | TRBV30 | TRBJ1-2 | CAWSVPPAASYGYTF | KLGGALQAK | NO | NO | NO |
| 2755 | TRAV1-2 | TRAJ28 | CAVRHSGAGSYQLTF | TRBV4-1 | TRBJ1-2 | CASSEGTDGYTF | KLGGALQAK | YES | YES | NO |
| 2759 | TRAV12-3 | TRAJ39 | CAIGPGNMLTF | TRBV4-1 | TRBJ1-2 | CASSLLTGPNYGYTF | KLGGALQAK | YES | NO | NO |
| 2761 | TRAV13-1 | TRAJ31 | CAANRVGSARLMF | TRBV4-1 | TRBJ1-2 | CASSQKTGQEDYGYTF | KLGGALQAK | YES | NO | NO |
| 2763 | TRAV13-2 | TRAJ16 | CAERGSDGQKLLF | TRBV4-1 | TRBJ1-2 | CASSQDFPGSDGYTF | KLGGALQAK | NO | NO | NO |
| 2764 | TRAV13-2 | TRAJ16 | CAEPWGGQKLLF | TRBV4-1 | TRBJ1-2 | CASSQVQGAQSSGYTF | KLGGALQAK | NO | NO | NO |
| 2768 | TRAV19 | TRAJ30 | CALIANRDDKIIF | TRBV4-1 | TRBJ1-2 | CASSQGVGQYGYTF | KLGGALQAK | YES | NO | NO |
| 2768 | TRAV19 | TRAJ30 | CALIANRDDKIIF | TRBV4-1 | TRBJ1-2 | CASSQGVGQYGYTF | RAKFKQLL | YES | NO | NO |
| 2769 | TRAV19 | TRAJ39 | CALSEDPGNAGNMLTF | TRBV4-1 | TRBJ1-2 | CASSPLTGANYGYTF | KLGGALQAK | YES | NO | NO |
| 2770 | TRAV20 | TRAJ30 | CAVNRDDKIIF | TRBV4-1 | TRBJ1-2 | CASSQGPGPGGYTF | KLGGALQAK | YES | NO | NO |
| 2771 | TRAV21 | TRAJ10 | CAVSGTTGGGNKLTF | TRBV4-1 | TRBJ1-2 | CASSQAGTGVSYGYTF | KLGGALQAK | YES | NO | NO |
| 2773 | TRAV26-1 | TRAJ4 | CIVVISGGYNKLIF | TRBV4-1 | TRBJ1-2 | CASSQGVWNYGYTF | KLGGALQAK | YES | NO | NO |
| 2774 | TRAV26-1 | TRAJ49 | CIAEASNTGNQFYF | TRBV4-1 | TRBJ1-2 | CASSQDLTGGYYGYTF | KLGGALQAK | YES | NO | NO |
| 2775 | TRAV27 | TRAJ26 | CAGDDNYGQNFVF | TRBV4-1 | TRBJ1-2 | CASSQLVQGADYGYTF | KLGGALQAK | YES | NO | NO |
| 2776 | TRAV27 | TRAJ54 | CAGVDQGAQKLVF | TRBV4-1 | TRBJ1-2 | CASSLLTGPNYGYTF | KLGGALQAK | YES | NO | NO |
| 2777 | TRAV29DV5 | TRAJ26 | CAASASYGQNFVF | TRBV4-1 | TRBJ1-2 | CASSQVGQVLHYGYTF | KLGGALQAK | YES | NO | NO |
| 2779 | TRAV29DV5 | TRAJ50 | CAASSYDKVIF | TRBV4-1 | TRBJ1-2 | CASSQGTLGSGYTF | KLGGALQAK | YES | NO | NO |
| 2780 | TRAV5 | TRAJ15 | CAEAGTALIF | TRBV4-1 | TRBJ1-2 | CASSQDRPRDYGYTF | KLGGALQAK | YES | NO | NO |
| 2781 | TRAV8-2 | TRAJ26 | CVYNYGQNFVF | TRBV4-1 | TRBJ1-2 | CASSAGLTNYGYTF | KLGGALQAK | YES | NO | NO |
| 2782 | TRAV8-2 | TRAJ50 | CVVSDQGASYDKVIF | TRBV4-1 | TRBJ1-2 | CASSHGPNYGYTF | KLGGALQAK | YES | NO | NO |
| 2784 | TRAV8-4 | TRAJ54 | CAVSVQGAQKLVF | TRBV4-1 | TRBJ1-2 | CASSQAWTGSYGYTF | AVFDRKSDAK | YES | NO | NO |
| 2784 | TRAV8-4 | TRAJ54 | CAVSVQGAQKLVF | TRBV4-1 | TRBJ1-2 | CASSQAWTGSYGYTF | KLGGALQAK | YES | NO | NO |
| 2786 | TRAV14DV4 | TRAJ5 | CAMRGYTGRRALTF | TRBV4-2 | TRBJ1-2 | CASSQEGATGSGYTF | KLGGALQAK | YES | NO | NO |
| 2787 | TRAV17 | TRAJ42 | CATFTGSQGNLIF | TRBV4-2 | TRBJ1-2 | CASSQAVAGYYGYTF | KLGGALQAK | YES | NO | NO |
| 2788 | TRAV19 | TRAJ52 | CALGGTSYGKLTF | TRBV4-2 | TRBJ1-2 | CASSQDEGTARGYTF | KLGGALQAK | YES | NO | NO |
| 2790 | TRAV24 | TRAJ6 | CASIASGGSYIPTF | TRBV4-2 | TRBJ1-2 | CASSLPLAVTF | KLGGALQAK | YES | NO | NO |
| 2791 | TRAV1-2 | TRAJ29 | CAVRSSGNTPLVF | TRBV4-3 | TRBJ1-2 | CASSQGTSVSGGYTF | KLGGALQAK | YES | NO | NO |
| 2792 | TRAV10 | TRAJ34 | CVGGGTDKLIF | TRBV4-3 | TRBJ1-2 | CASSQEGRGIYGYTF | KLGGALQAK | YES | NO | NO |
| 2793 | TRAV12-2 | TRAJ33 | CAVRSNYQLIW | TRBV4-3 | TRBJ1-2 | CASSQYVFQGSYGYTF | KLGGALQAK | YES | NO | NO |
| 2797 | TRAV14DV4 | TRAJ28 | CAMREPYSGAGSYQLTF | TRBV4-3 | TRBJ1-2 | CASSQDRQGISIGYTF | KLGGALQAK | YES | NO | NO |
| 2798 | TRAV14DV4 | TRAJ6 | CAMRGGEGGSYIPTF | TRBV4-3 | TRBJ1-2 | CASSFSGTGDLYGYTF | KLGGALQAK | YES | NO | NO |
| 2799 | TRAV17 | TRAJ11 | CATDGWSSGYSTLTF | TRBV4-3 | TRBJ1-2 | CASSHSGTGEYGYTF | KLGGALQAK | YES | NO | NO |
| 2803 | TRAV20 | TRAJ20 | CAVQALSNDYKLSF | TRBV4-3 | TRBJ1-2 | CASSPTVLEGYYGYTF | KLGGALQAK | YES | NO | NO |
| 2804 | TRAV21 | TRAJ21 | CAVKRSNFNKFYF | TRBV4-3 | TRBJ1-2 | CASSQDRGGYGYTF | KLGGALQAK | YES | NO | NO |
| 2806 | TRAV24 | TRAJ10 | CASVRRTGGGNKLTF | TRBV4-3 | TRBJ1-2 | CASSQGGKGEYLFNYGYTF | KLGGALQAK | YES | NO | NO |
| 2807 | TRAV26-1 | TRAJ23 | CIVRSWVFYNQGGKLIF | TRBV4-3 | TRBJ1-2 | CASSQEPDREGNYGYTF | KLGGALQAK | YES | NO | NO |
| 2808 | TRAV3 | TRAJ17 | CAVRDILKAAGNKLTF | TRBV4-3 | TRBJ1-2 | CASSQTLGGSYGYTF | AVFDRKSDAK | YES | NO | NO |
| 2809 | TRAV38-2DV8 | TRAJ54 | CAYRSTAIQGAQKLVF | TRBV4-3 | TRBJ1-2 | CASSQSQAWGRDGYTF | KLGGALQAK | YES | NO | NO |
| 2810 | TRAV39 | TRAJ58 | CAGTQETSGSRLTF | TRBV4-3 | TRBJ1-2 | CASSQVTYSYGYTF | KLGGALQAK | YES | NO | NO |
| 2812 | TRAV5 | TRAJ11 | CAEKRLNSGYSTLTF | TRBV4-3 | TRBJ1-2 | CASSQDPGSSYGYTF | KLGGALQAK | YES | NO | NO |
| 2814 | TRAV5 | TRAJ43 | CAETGNDMRF | TRBV4-3 | TRBJ1-2 | CASSQEGGVYGYTF | KLGGALQAK | YES | NO | NO |
| 2815 | TRAV1-1 | TRAJ20 | CASSSNDYKLSF | TRBV5-1 | TRBJ1-2 | CASSWGEQGLRGYTF | KLGGALQAK | YES | NO | NO |
| 2816 | TRAV1-2 | TRAJ34 | CAVGYTDKLIF | TRBV5-1 | TRBJ1-2 | CASMGTGFDGYTF | KLGGALQAK | YES | NO | NO |
| 2818 | TRAV12-1 | TRAJ4 | CVVKVAGGYNKLIF | TRBV5-1 | TRBJ1-2 | CASSLEGGSVYGYTF | KLGGALQAK | YES | NO | NO |
| 2822 | TRAV13-1 | TRAJ38 | CAATFNAGNNRKLIW | TRBV5-1 | TRBJ1-2 | CASSAENYGYTF | KLGGALQAK | YES | NO | NO |
| 2823 | TRAV13-1 | TRAJ6 | CAAGGSYIPTF | TRBV5-1 | TRBJ1-2 | CASSLGYEANPARGYGYTF | KLGGALQAK | YES | NO | NO |
| 2824 | TRAV14DV4 | TRAJ47 | CAMREREYGNKLVF | TRBV5-1 | TRBJ1-2 | CASSPGPGESGYGYTF | KLGGALQAK | YES | NO | NO |
| 2826 | TRAV21 | TRAJ4 | CAVIGFSGGYNKLIF | TRBV5-1 | TRBJ1-2 | CASGPERSERYGYTF | KLGGALQAK | YES | NO | NO |
| 2828 | TRAV21 | TRAJ54 | CAVRSSLIQGAQKLVF | TRBV5-1 | TRBJ1-2 | CASSFGQGENYGYTF | KLGGALQAK | YES | NO | NO |
| 2831 | TRAV27 | TRAJ30 | CAGELSDRDDKIIF | TRBV5-1 | TRBJ1-2 | CASSLHRGGFYGYTF | KLGGALQAK | YES | NO | NO |
| 2832 | TRAV29DV5 | TRAJ20 | CAASSDYKLSF | TRBV5-1 | TRBJ1-2 | CASSTGDRQVYYGYTF | KLGGALQAK | YES | NO | NO |
| 2833 | TRAV3 | TRAJ34 | CAVSGGRLDTDKLIF | TRBV5-1 | TRBJ1-2 | CASSLGERVGYGYTF | KLGGALQAK | YES | NO | NO |
| 2834 | TRAV39 | TRAJ54 | CAVVGGAQKLVF | TRBV5-1 | TRBJ1-2 | CASSLATSSNYGYTF | KLGGALQAK | YES | NO | NO |
| 2835 | TRAV4 | TRAJ31 | CLVGDFARLMF | TRBV5-1 | TRBJ1-2 | CASSLGQGVGYTF | KLGGALQAK | NO | NO | NO |
| 2837 | TRAV9-2 | TRAJ35 | CASDLI | TRBV5-1 | TRBJ1-2 | CASSPEGFSNYGYTF | KLGGALQAK | NO | NO | NO |
| 2839 | TRAV12-2 | TRAJ40 | CAADSGTYKYIF | TRBV5-4 | TRBJ1-2 | CASSLAVGGYTF | KLGGALQAK | YES | NO | NO |
| 2840 | TRAV12-3 | TRAJ32 | CAMNPAWGGATNKLIF | TRBV5-4 | TRBJ1-2 | CASTYLGGSLDGYTF | KLGGALQAK | YES | NO | NO |
| 2845 | TRAV19 | TRAJ26 | CALVRGGGQNFVF | TRBV5-4 | TRBJ1-2 | CASSLGTANYGYTF | KLGGALQAK | YES | NO | NO |
| 2846 | TRAV19 | TRAJ30 | CALSEGNRDDKIIF | TRBV5-4 | TRBJ1-2 | CASSWDRANGYTF | KLGGALQAK | YES | NO | NO |
| 2848 | TRAV19 | TRAJ5 | CALSEALGGRRALTF | TRBV5-4 | TRBJ1-2 | CASSLPSSNYGYTF | KLGGALQAK | YES | NO | NO |
| 2850 | TRAV21 | TRAJ24 | CAKTDSWGKLQF | TRBV5-4 | TRBJ1-2 | CASSLTAYGYTF | KLGGALQAK | YES | NO | NO |
| 2851 | TRAV21 | TRAJ42 | CAVSSGGSQGNLIF | TRBV5-4 | TRBJ1-2 | CASSLAPRAPLGYTF | KLGGALQAK | YES | NO | NO |
| 2852 | TRAV23DV6 | TRAJ45 | CAASQTGLTHSGGGADGLTF | TRBV5-4 | TRBJ1-2 | CASSLEALYGYTF | KLGGALQAK | NO | NO | NO |
| 2854 | TRAV26-2 | TRAJ23 | CILIPIYNQGGKLIF | TRBV5-4 | TRBJ1-2 | CASSLNGNYGYTF | KLGGALQAK | YES | NO | NO |
| 2855 | TRAV38-1 | TRAJ49 | CAFMKSFTGNQFYF | TRBV5-4 | TRBJ1-2 | CASSPEPGQGGYGYTF | KLGGALQAK | YES | NO | NO |
| 2857 | TRAV41 | TRAJ21 | CAVRWNFNKFYF | TRBV5-4 | TRBJ1-2 | CASSLLGGRYGYTF | KLGGALQAK | YES | NO | NO |
| 2858 | TRAV8-3 | TRAJ30 | CAVGAMNRDDKIIF | TRBV5-4 | TRBJ1-2 | CASSSGQTAQKHGYTF | KLGGALQAK | YES | NO | NO |
| 2860 | TRAV9-2 | TRAJ36 | CALNPGANNLFF | TRBV5-4 | TRBJ1-2 | CASSLGVTDLGYTF | KLGGALQAK | YES | NO | NO |
| 2862 | TRAV19 | TRAJ29 | CALSEANSGNTPLVF | TRBV5-5 | TRBJ1-2 | CASSLAWTGSFYGYTF | KLGGALQAK | YES | NO | NO |
| 2864 | TRAV23DV6 | TRAJ54 | CAATPPGTIQGAQKLVF | TRBV5-5 | TRBJ1-2 | CASSLLWGSGGFFYGYTF | KLGGALQAK | YES | NO | NO |
| 2869 | TRAV13-1 | TRAJ5 | CAANPPGRRALTF | TRBV5-6 | TRBJ1-2 | CASSLVRQGRKYGYTF | KLGGALQAK | YES | NO | NO |
| 2870 | TRAV14DV4 | TRAJ37 | CAMRGRWSSNTGKLIF | TRBV5-6 | TRBJ1-2 | CASSSSDQGRRGYTF | KLGGALQAK | YES | NO | NO |
| 2871 | TRAV17 | TRAJ30 | CATDGRGKIIF | TRBV5-6 | TRBJ1-2 | CASSSGPGAGGYGYTF | KLGGALQAK | YES | NO | NO |
| 2871 | TRAV17 | TRAJ30 | CATDGRGKIIF | TRBV5-6 | TRBJ1-2 | CASSSGPGAGGYGYTF | IVTDFSVIK | YES | NO | NO |
| 2872 | TRAV17 | TRAJ5 | CATDPDTGRRALTF | TRBV5-6 | TRBJ1-2 | CASSLLIREAYTF | KLGGALQAK | YES | NO | NO |
| 2874 | TRAV21 | TRAJ26 | CAVRDNYGQNFVF | TRBV5-6 | TRBJ1-2 | CASSLTTSYGYTF | KLGGALQAK | YES | NO | NO |
| 2875 | TRAV21 | TRAJ27 | CAVDTNAGKSTF | TRBV5-6 | TRBJ1-2 | CASGLFVVGTYGYTF | KLGGALQAK | YES | NO | NO |
| 2876 | TRAV21 | TRAJ31 | CAARRGARLMF | TRBV5-6 | TRBJ1-2 | CASSSTGVGYTF | KLGGALQAK | YES | NO | NO |
| 2876 | TRAV21 | TRAJ31 | CAARRGARLMF | TRBV5-6 | TRBJ1-2 | CASSSTGVGYTF | AVFDRKSDAK | YES | NO | NO |
| 2877 | TRAV21 | TRAJ39 | CAVRSGNAGNMLTF | TRBV5-6 | TRBJ1-2 | CASSLGRNYGYTF | KLGGALQAK | YES | NO | NO |
| 2878 | TRAV29DV5 | TRAJ6 | CAASALRASGGSYIPTF | TRBV5-6 | TRBJ1-2 | CASSFGPGRLYGYTF | KLGGALQAK | YES | NO | NO |
| 2881 | TRAV41 | TRAJ43 | CAARNDMRF | TRBV5-6 | TRBJ1-2 | CASSFRRDGGYGYTF | KLGGALQAK | NO | NO | NO |
| 2882 | TRAV41 | TRAJ43 | CAVRDHRYNNNDMRF | TRBV5-6 | TRBJ1-2 | CASSLTRRYTF | KLGGALQAK | YES | NO | NO |
| 2883 | TRAV6 | TRAJ34 | CALDGAGNTDKLIF | TRBV5-6 | TRBJ1-2 | CASSPGGGYTF | KLGGALQAK | YES | NO | NO |
| 2884 | TRAV8-2 | TRAJ46 | CVVSRGKSSGDKLTF | TRBV5-6 | TRBJ1-2 | CASSLGSSNYGYTF | KLGGALQAK | YES | NO | NO |
| 2886 | TRAV9-2 | TRAJ20 | CAPSDYKLSF | TRBV5-6 | TRBJ1-2 | CASSLPTQSYGYTF | KLGGALQAK | YES | NO | NO |
| 2888 | TRAV21 | TRAJ29 | CAVRQIIGNTPLVF | TRBV5-8 | TRBJ1-2 | CASSLGLGGYTF | KLGGALQAK | YES | NO | NO |
| 2889 | TRAV26-1 | TRAJ23 | CIVFYNQGGKLIF | TRBV5-8 | TRBJ1-2 | CASSYPGGYYGYTF | KLGGALQAK | YES | NO | NO |

| | | | | | | | | | | |
|---|---|---|---|---|---|---|---|---|---|---|
| 2890 | TRAV1-1 | TRAJ21 | CAVDNFNKFYF | TRBV6-1 | TRBJ1-2 | CASSEVEGHGYTF | KLGGALQAK | YES | NO | NO |
| 2891 | TRAV12-1 | TRAJ30 | CVVNKEDKIIF | TRBV6-1 | TRBJ1-2 | CASRTGGAGYTF | KLGGALQAK | YES | NO | NO |
| 2893 | TRAV12-2 | TRAJ41 | CAVIYSGYALNF | TRBV6-1 | TRBJ1-2 | CASTPGQEIYGYTF | KLGGALQAK | YES | NO | NO |
| 2896 | TRAV19 | TRAJ10 | CALSRPPITGGGNKLTF | TRBV6-1 | TRBJ1-2 | CASSDSIGYGYTF | KLGGALQAK | YES | NO | NO |
| 2898 | TRAV26-2 | TRAJ57 | CILPQGGSEKLVF | TRBV6-1 | TRBJ1-2 | CASSTNPYSANYGYTF | KLGGALQAK | YES | NO | NO |
| 2899 | TRAV3 | TRAJ11 | CAVRDIFSGYSTLTF | TRBV6-1 | TRBJ1-2 | CASSRPGQGVSYGYTF | KLGGALQAK | NO | NO | NO |
| 2902 | TRAV38-2DV8 | TRAJ33 | CAYRRMDSNYQLIW | TRBV6-1 | TRBJ1-2 | CASKFHGSHYGYTF | KLGGALQAK | YES | NO | NO |
| 2903 | TRAV39 | TRAJ43 | CAGEGNNNDMRF | TRBV6-1 | TRBJ1-2 | CAGEWDGSYGYTF | KLGGALQAK | YES | NO | NO |
| 2904 | TRAV41 | TRAJ31 | CAVFPNNARLMF | TRBV6-1 | TRBJ1-2 | CASIGRGGTDLGYTF | KLGGALQAK | YES | NO | NO |
| 2905 | TRAV8-1 | TRAJ4 | CAVNAPSFSGGYNKLIF | TRBV6-1 | TRBJ1-2 | CASSGGNILDGYTF | KLGGALQAK | YES | NO | NO |
| 2906 | TRAV1-1 | TRAJ30 | CAVRDGDRDDKIIF | TRBV6-3 | TRBJ1-2 | CASSYLGQGDAYGYTF | KLGGALQAK | YES | NO | NO |
| 2907 | TRAV1-2 | TRAJ13 | CAVMGYQKVTF | TRBV6-3 | TRBJ1-2 | CASSQQGAPYGYTF | KLGGALQAK | YES | NO | NO |
| 2908 | TRAV1-2 | TRAJ34 | CAVRDNTDKLIF | TRBV6-3 | TRBJ1-2 | CASSYWGTNYGYTF | KLGGALQAK | YES | NO | NO |
| 2909 | TRAV12-2 | TRAJ53 | CAVNGGSGGSNYKLTF | TRBV6-3 | TRBJ1-2 | CASSYSSRGNGYTF | KLGGALQAK | YES | NO | NO |
| 2910 | TRAV12-3 | TRAJ39 | CAIGPGNMLTF | TRBV6-3 | TRBJ1-2 | CASSQPLLRTSANYGYTF | KLGGALQAK | YES | NO | NO |
| 2911 | TRAV14DV4 | TRAJ45 | CAMSRDSGGGADGLTF | TRBV6-3 | TRBJ1-2 | CASSLSGTLLYGYTF | KLGGALQAK | YES | NO | NO |
| 2912 | TRAV17 | TRAJ9 | CATENTGGFKTIF | TRBV6-3 | TRBJ1-2 | CASSYSINAAVNYGYTF | KLGGALQAK | YES | NO | NO |
| 2913 | TRAV21 | TRAJ30 | CAVSMNRDDKIIF | TRBV6-3 | TRBJ1-2 | CASSGRTLSHYGYTF | KLGGALQAK | YES | NO | NO |
| 2915 | TRAV27 | TRAJ41 | CAGEGWNSGYALNF | TRBV6-3 | TRBJ1-2 | CASSYPFGYTF | KLGGALQAK | YES | NO | NO |
| 2916 | TRAV27 | TRAJ43 | CAGASHDMRF | TRBV6-3 | TRBJ1-2 | CASSQTGHYGYTF | KLGGALQAK | YES | NO | NO |
| 2918 | TRAV4 | TRAJ20 | CLVGGNNDYKLSF | TRBV6-3 | TRBJ1-2 | CASSSSTGTLYGYTF | KLGGALQAK | YES | NO | NO |
| 2919 | TRAV5 | TRAJ37 | CAESLASNTGKLIF | TRBV6-3 | TRBJ1-2 | CASSPTGGYGYTF | KLGGALQAK | YES | NO | NO |
| 2920 | TRAV8-3 | TRAJ15 | CAVGVLRGQAGTALIF | TRBV6-3 | TRBJ1-2 | CASRPGRADYGYTF | KLGGALQAK | YES | NO | NO |
| 2921 | TRAV1-1 | TRAJ32 | CAVRPRGGATNKLIF | TRBV6-4 | TRBJ1-2 | CASSDSGFGISDGYTF | KLGGALQAK | YES | NO | NO |
| 2922 | TRAV10 | TRAJ20 | CVVSPLADYKLSF | TRBV6-4 | TRBJ1-2 | CASSSTKNYGYTF | KLGGALQAK | YES | NO | NO |
| 2924 | TRAV19 | TRAJ39 | CALSENNAGNMLTF | TRBV6-4 | TRBJ1-2 | CASSDYPGAYGYTF | KLGGALQAK | YES | NO | NO |
| 2925 | TRAV19 | TRAJ43 | CALRYNNNDMRF | TRBV6-4 | TRBJ1-2 | CASSDTYGYTF | KLGGALQAK | YES | NO | NO |
| 2926 | TRAV19 | TRAJ48 | CALAEGGISNFGNEKLTF | TRBV6-4 | TRBJ1-2 | CASSDGENYGYTF | KLGGALQAK | YES | NO | NO |
| 2927 | TRAV6 | TRAJ40 | CALTNSGTYKYIF | TRBV6-4 | TRBJ1-2 | CASSETPQPFQNGYTF | RLRAEAQVK | YES | YES | YES |
| 2929 | TRAV1-1 | TRAJ12 | CAVFSDSSYKLIF | TRBV6-5 | TRBJ1-2 | CASSSGQDLNYGYTF | KLGGALQAK | YES | NO | NO |
| 2931 | TRAV12-1 | TRAJ43 | CVVNEGPSRNNDMRF | TRBV6-5 | TRBJ1-2 | CASSYDTANLANYGYTF | KLGGALQAK | YES | NO | NO |
| 2932 | TRAV12-2 | TRAJ35 | CAFRI | TRBV6-5 | TRBJ1-2 | CASRFTRDTENYGYTF | KLGGALQAK | NO | NO | NO |
| 2934 | TRAV13-2 | TRAJ38 | CAENIGAGNNRKLIW | TRBV6-5 | TRBJ1-2 | CASSPLEGADGYTF | KLGGALQAK | YES | NO | NO |
| 2935 | TRAV14DV4 | TRAJ11 | CAMRGSGYSTLTF | TRBV6-5 | TRBJ1-2 | CASSYLGGDAANYGYTF | KLGGALQAK | YES | NO | NO |
| 2936 | TRAV14DV4 | TRAJ42 | CAMRGSQGNLIF | TRBV6-5 | TRBJ1-2 | CASSNGGGRLYGYTF | KLGGALQAK | YES | NO | NO |
| 2937 | TRAV14DV4 | TRAJ58 | CAMREWGSGSRLTF | TRBV6-5 | TRBJ1-2 | CASSSQGSNYGYTF | KLGGALQAK | YES | NO | NO |
| 2938 | TRAV21 | TRAJ42 | CAVMNYGGSQGNLIF | TRBV6-5 | TRBJ1-2 | CASSTGLIYGYTF | KLGGALQAK | YES | NO | NO |
| 2941 | TRAV25 | TRAJ21 | CGPDNFNKFYF | TRBV6-5 | TRBJ1-2 | CASSYSSGVRGYTF | KLGGALQAK | YES | NO | NO |
| 2942 | TRAV26-2 | TRAJ36 | CILRDPQTGANNLFF | TRBV6-5 | TRBJ1-2 | CASSYGTAPTGYTF | KLGGALQAK | YES | NO | NO |
| 2943 | TRAV27 | TRAJ20 | CAGAGDDYKLSF | TRBV6-5 | TRBJ1-2 | CASSLSFGSPYHGYTF | KLGGALQAK | YES | NO | NO |
| 2944 | TRAV29DV5 | TRAJ42 | CAASVLHGGSQGNLIF | TRBV6-5 | TRBJ1-2 | CASSYGGLDGYTF | KLGGALQAK | YES | NO | NO |
| 2947 | TRAV35 | TRAJ40 | CAGSGTYKYIF | TRBV6-5 | TRBJ1-2 | CASSSGTGVYGYTF | KLGGALQAK | YES | NO | NO |
| 2948 | TRAV38-2DV8 | TRAJ31 | CAYRSGHARLMF | TRBV6-5 | TRBJ1-2 | CASRGQGASGYTF | KLGGALQAK | YES | NO | NO |
| 2949 | TRAV5 | TRAJ39 | CAESTNNAGNMLTF | TRBV6-5 | TRBJ1-2 | CASQGQMNYGYTF | KLGGALQAK | YES | NO | NO |
| 2951 | TRAV21 | TRAJ30 | CAVRPWRDDKIIF | TRBV6-6 | TRBJ1-2 | CASRESPGYTF | KLGGALQAK | NO | NO | NO |
| 2952 | TRAV21 | TRAJ4 | CAVSLSGGYNKLIF | TRBV6-6 | TRBJ1-2 | CASSYRGSRGYTF | KLGGALQAK | NO | NO | NO |
| 2954 | TRAV21 | TRAJ49 | CAVRPGTGNQFYF | TRBV6-6 | TRBJ1-2 | CASSYSPANYGYTF | KLGGALQAK | NO | NO | NO |
| 2956 | TRAV26-2 | TRAJ45 | CIRMYSGGGADGLTF | TRBV6-6 | TRBJ1-2 | CASSYSGGQGLRYTF | KLGGALQAK | NO | NO | NO |
| 2962 | TRAV19 | TRAJ20 | CALTAPNDYKLSF | TRBV7-2 | TRBJ1-2 | CASSLGIYYGYTF | KLGGALQAK | YES | NO | NO |
| 2963 | TRAV19 | TRAJ48 | CALSEALGNEKLTF | TRBV7-2 | TRBJ1-2 | CASSPHRSLSEINNYGYTF | KLGGALQAK | YES | NO | NO |
| 2964 | TRAV21 | TRAJ26 | CAVDNYGQNFVF | TRBV7-2 | TRBJ1-2 | CASSYRTGDRYGYTF | KLGGALQAK | NO | NO | NO |
| 2971 | TRAV12-3 | TRAJ33 | CAVGPDSNYQLIW | TRBV7-3 | TRBJ1-2 | CASSLTPGGNYGYTF | KLGGALQAK | NO | NO | NO |
| 2972 | TRAV12-3 | TRAJ54 | CAMSIQGAQKLVF | TRBV7-3 | TRBJ1-2 | CASSTRDITDRYTF | KLGGALQAK | NO | NO | NO |
| 2974 | TRAV21 | TRAJ33 | CAVTMDSNYQLIW | TRBV7-3 | TRBJ1-2 | CASSPYYSADGYTF | KLGGALQAK | NO | NO | NO |
| 2976 | TRAV22 | TRAJ8 | CADSGFQKLVF | TRBV7-3 | TRBJ1-2 | CASSFGTGNYGYTF | KLGGALQAK | YES | NO | NO |
| 2977 | TRAV26-2 | TRAJ47 | CILREYGNKLVF | TRBV7-3 | TRBJ1-2 | CASSSNPTGLYGYTF | KLGGALQAK | YES | NO | NO |
| 2978 | TRAV29DV5 | TRAJ53 | CAASDGGSNYKLTF | TRBV7-3 | TRBJ1-2 | CASSLTGQSQRVYGYTF | KLGGALQAK | NO | NO | NO |
| 2981 | TRAV35 | TRAJ28 | CAGHRDSGAGSYQLTF | TRBV7-3 | TRBJ1-2 | CASSLMGTSGYTF | KLGGALQAK | NO | NO | NO |
| 2983 | TRAV8-2 | TRAJ45 | CVVSDGGGADGLTF | TRBV7-3 | TRBJ1-2 | CASSPTGDGNYGYTF | KLGGALQAK | NO | NO | NO |
| 2984 | TRAV1-1 | TRAJ42 | CAVLGSQGNLIF | TRBV7-6 | TRBJ1-2 | CASSSRDREGGYTF | KLGGALQAK | YES | NO | NO |
| 2985 | TRAV12-1 | TRAJ13 | CVAMGSGGYQKVTF | TRBV7-6 | TRBJ1-2 | CASSLAPGTAGGYTF | KLGGALQAK | YES | NO | NO |
| 2989 | TRAV8-4 | TRAJ13 | CAVSFSGGYQKVTF | TRBV7-6 | TRBJ1-2 | CASSFGQGPLYVDGYTF | KLGGALQAK | YES | NO | NO |
| 2991 | TRAV8-6 | TRAJ57 | CAVSCPQGGSEKLVF | TRBV7-6 | TRBJ1-2 | CASSPGTGTTYGYTF | KLGGALQAK | YES | NO | NO |
| 2992 | TRAV8-6 | TRAJ58 | CAVYRPETSGSRLTF | TRBV7-6 | TRBJ1-2 | CASSDNAADYTF | KLGGALQAK | YES | NO | NO |
| 2994 | TRAV12-1 | TRAJ20 | CVVNLVNDYKLSF | TRBV7-8 | TRBJ1-2 | CASSLGTGNGYTF | KLGGALQAK | YES | NO | NO |
| 2995 | TRAV12-1 | TRAJ34 | CVVPGSYNTDKLIF | TRBV7-8 | TRBJ1-2 | CASSLATGEYPTNYGYTF | KLGGALQAK | YES | NO | NO |
| 2996 | TRAV12-1 | TRAJ49 | CVVNYPQFYF | TRBV7-8 | TRBJ1-2 | CASSLGDMTNYGYTF | KLGGALQAK | YES | NO | NO |
| 2996 | TRAV12-1 | TRAJ49 | CVVNYPQFYF | TRBV7-8 | TRBJ1-2 | CASSLGDMTNYGYTF | AVFDRKSDAK | YES | NO | NO |
| 3003 | TRAV19 | TRAJ44 | CALFNPAGTASKLTF | TRBV7-8 | TRBJ1-2 | CASSLGQDGYTF | KLGGALQAK | YES | NO | NO |
| 3004 | TRAV19 | TRAJ53 | CALSETRNSGGSNYKLTF | TRBV7-8 | TRBJ1-2 | CASSLGYGYTF | KLGGALQAK | YES | NO | NO |
| 3008 | TRAV25 | TRAJ32 | CAGNDYGGATNKLIF | TRBV7-8 | TRBJ1-2 | CASSLGQDGYTF | KLGGALQAK | YES | NO | NO |
| 3013 | TRAV29DV5 | TRAJ20 | CAAMSSNDYKLSF | TRBV7-8 | TRBJ1-2 | CASSLGSRPGYTF | KLGGALQAK | YES | NO | NO |
| 3016 | TRAV35 | TRAJ20 | CAGHDGNDYKLSF | TRBV7-8 | TRBJ1-2 | CASSLGYGYTF | AVFDRKSDAK | YES | NO | NO |
| 3019 | TRAV38-2DV8 | TRAJ49 | CAYRSAITGNQFYF | TRBV7-8 | TRBJ1-2 | CASSLVFQGAYGYTF | KLGGALQAK | YES | NO | NO |
| 3023 | TRAV8-3 | TRAJ43 | CAVGAEVRF | TRBV7-8 | TRBJ1-2 | CASSSGTMMNYGYTF | KLGGALQAK | NO | NO | NO |
| 3024 | TRAV8-4 | TRAJ20 | CAVIPINDYKLSF | TRBV7-8 | TRBJ1-2 | CASSLGQAYGYTF | AVFDRKSDAK | YES | NO | NO |
| 3026 | TRAV12-3 | TRAJ26 | CAMKDNYGQNFVF | TRBV7-9 | TRBJ1-2 | CASSLDAGVGGYTF | KLGGALQAK | YES | NO | NO |
| 3029 | TRAV13-2 | TRAJ28 | CAENGAGSYQLTF | TRBV7-9 | TRBJ1-2 | CASSLVLGQPNYGYTF | KLGGALQAK | YES | NO | NO |
| 3035 | TRAV14DV4 | TRAJ49 | CAMSPRSTGNQFYF | TRBV7-9 | TRBJ1-2 | CASTLGQGAPGYTF | KLGGALQAK | YES | NO | NO |
| 3036 | TRAV16 | TRAJ29 | CAHHSGNTPLVF | TRBV7-9 | TRBJ1-2 | CASSLDATHYGYTF | KLGGALQAK | NO | NO | NO |
| 3037 | TRAV16 | TRAJ6 | CALESGGSYIPTF | TRBV7-9 | TRBJ1-2 | CASSDWDRVVEGYTF | KLGGALQAK | NO | NO | NO |
| 3038 | TRAV17 | TRAJ31 | CATDAGARLMF | TRBV7-9 | TRBJ1-2 | CASSPTNGDGYTF | KLGGALQAK | YES | NO | NO |
| 3041 | TRAV19 | TRAJ9 | CALNGNTGGFKTIF | TRBV7-9 | TRBJ1-2 | CASSPSIGGQVIRGYTF | KLGGALQAK | YES | NO | NO |
| 3042 | TRAV21 | TRAJ20 | CAVNSNDYKLSF | TRBV7-9 | TRBJ1-2 | CASSDSYSYGYTF | KLGGALQAK | YES | NO | NO |
| 3043 | TRAV21 | TRAJ33 | CAVPDSNYQLIW | TRBV7-9 | TRBJ1-2 | CASSFAGGGGYTF | KLGGALQAK | YES | NO | NO |
| 3044 | TRAV21 | TRAJ43 | CAVVSNNNDMRF | TRBV7-9 | TRBJ1-2 | CASSDSYSYGYTF | KLGGALQAK | YES | NO | NO |
| 3044 | TRAV21 | TRAJ43 | CAVVSNNNDMRF | TRBV7-9 | TRBJ1-2 | CASSDSYSYGYTF | AVFDRKSDAK | YES | NO | NO |
| 3045 | TRAV21 | TRAJ43 | CAVRNNNNDMRF | TRBV7-9 | TRBJ1-2 | CASSDSYSNYGYTF | KLGGALQAK | YES | NO | NO |
| 3047 | TRAV26-2 | TRAJ20 | CILRDSNDYKLSF | TRBV7-9 | TRBJ1-2 | CASSSGGGGLDGYTF | KLGGALQAK | YES | NO | NO |
| 3053 | TRAV29DV5 | TRAJ43 | CAAKGSYNNNDMRF | TRBV7-9 | TRBJ1-2 | CASSLWEDGYTF | KLGGALQAK | YES | NO | NO |
| 3054 | TRAV35 | TRAJ27 | CAGQDTNAGKSTF | TRBV7-9 | TRBJ1-2 | CASSWQHGYTF | KLGGALQAK | YES | NO | NO |
| 3055 | TRAV38-2DV8 | TRAJ40 | CAYRSPTSGTYKYIF | TRBV7-9 | TRBJ1-2 | CASSTPGQGPNYGYTF | KLGGALQAK | YES | NO | NO |
| 3057 | TRAV4 | TRAJ49 | CLVGVTGNQFYF | TRBV7-9 | TRBJ1-2 | CASSSIGQGIRYGYTF | KLGGALQAK | YES | NO | NO |
| 3059 | TRAV5 | TRAJ23 | CAESIEGGKLIF | TRBV7-9 | TRBJ1-2 | CASSLVPGQANYGYTF | KLGGALQAK | YES | NO | NO |
| 3062 | TRAV8-2 | TRAJ49 | CVVSDRDGNQFYF | TRBV7-9 | TRBJ1-2 | CASSLAIGNYGYTF | KLGGALQAK | YES | NO | NO |
| 3066 | TRAV1-2 | TRAJ23 | CAVFFYNQGGKLIF | TRBV9 | TRBJ1-2 | CASSGDPGTGHLFNYGYTF | KLGGALQAK | YES | NO | NO |
| 3067 | TRAV1-2 | TRAJ41 | CAVRPGYALNF | TRBV9 | TRBJ1-2 | CASSVEAGLSYTF | KLGGALQAK | YES | NO | NO |
| 3068 | TRAV1-2 | TRAJ43 | CAVDVNDMRF | TRBV9 | TRBJ1-2 | CASSGGGGGINGYTF | KLGGALQAK | YES | NO | NO |
| 3069 | TRAV1-2 | TRAJ6 | CAVSSAGGSYIPTF | TRBV9 | TRBJ1-2 | CASNSGTYGYTF | KLGGALQAK | YES | NO | NO |
| 3071 | TRAV12-1 | TRAJ31 | CVLNDNARLMF | TRBV9 | TRBJ1-2 | CASSVVQGYGYTF | KLGGALQAK | YES | NO | NO |
| 3073 | TRAV12-1 | TRAJ43 | CVVTYNDMRF | TRBV9 | TRBJ1-2 | CASTGGGANGYTF | KLGGALQAK | YES | NO | NO |
| 3077 | TRAV12-3 | TRAJ53 | CAIGGSNYKLTF | TRBV9 | TRBJ1-2 | CASSLRTGAPHRYTF | KLGGALQAK | YES | NO | NO |
| 3078 | TRAV13-1 | TRAJ28 | CAASQGAGSYQLTF | TRBV9 | TRBJ1-2 | CASSVGTGLGDYGYTF | KLGGALQAK | YES | NO | NO |
| 3079 | TRAV14DV4 | TRAJ12 | CAMREGMDSSYKLIF | TRBV9 | TRBJ1-2 | CASADSYGYTF | KLGGALQAK | YES | NO | NO |

| | | | | | | | | | | |
|---|---|---|---|---|---|---|---|---|---|---|
| 3081 | TRAV14DV4 | TRAJ32 | CAMREGATNKLIF | TRBV9 | TRBJ1-2 | CASSPHAEGYTF | KLGGALQAK | YES | NO | NO |
| 3082 | TRAV14DV4 | TRAJ37 | CAMREGLSNTGKLIF | TRBV9 | TRBJ1-2 | CASSVGNRAGAYTF | KLGGALQAK | YES | NO | NO |
| 3084 | TRAV14DV4 | TRAJ52 | CAMRELAGGTSYGKLTF | TRBV9 | TRBJ1-2 | CASSRPERTNYGYTF | KLGGALQAK | YES | NO | NO |
| 3085 | TRAV14DV4 | TRAJ53 | CAMREGVGSNYKLTF | TRBV9 | TRBJ1-2 | CASSVTGMEYGYTF | KLGGALQAK | YES | YES | NO |
| 3086 | TRAV16 | TRAJ28 | CALNSAGSYQLTF | TRBV9 | TRBJ1-2 | CASRGFYGYTF | KLGGALQAK | NO | NO | NO |
| 3089 | TRAV19 | TRAJ13 | CALTSGGYQKVTF | TRBV9 | TRBJ1-2 | CASSVEKERASDYGYTF | KLGGALQAK | YES | NO | NO |
| 3093 | TRAV19 | TRAJ39 | CALKLGNAGNMLTF | TRBV9 | TRBJ1-2 | CASSSQSGSLYGYTF | KLGGALQAK | YES | NO | NO |
| 3095 | TRAV19 | TRAJ47 | CALSEAHEGNKLVF | TRBV9 | TRBJ1-2 | CASSVQAQGENYGYTF | KLGGALQAK | YES | NO | NO |
| 3096 | TRAV19 | TRAJ47 | CALSEAHGGNKLVF | TRBV9 | TRBJ1-2 | CASSVDPTGGNYGYTF | KLGGALQAK | YES | NO | NO |
| 3097 | TRAV19 | TRAJ52 | CALSEIGNAGGTSYGKLTF | TRBV9 | TRBJ1-2 | CASSVGQVFYSGTF | KLGGALQAK | NO | NO | NO |
| 3097 | TRAV19 | TRAJ52 | CALSEIGNAGGTSYGKLTF | TRBV9 | TRBJ1-2 | CASSVGQVFYSGTF | IVTDFSVIK | NO | NO | NO |
| 3098 | TRAV19 | TRAJ8 | CALSGAGFQKLVF | TRBV9 | TRBJ1-2 | CASSVVVETGSNYGYTF | KLGGALQAK | YES | NO | NO |
| 3099 | TRAV20 | TRAJ26 | CAVLGALSLNFVF | TRBV9 | TRBJ1-2 | CASSSGQGGYGYTF | KLGGALQAK | YES | NO | NO |
| 3100 | TRAV20 | TRAJ56 | CAVLTGANSKLTF | TRBV9 | TRBJ1-2 | CASSAAGGGDYGYTF | KLGGALQAK | YES | NO | NO |
| 3101 | TRAV21 | TRAJ27 | CAVRTNAGKSTF | TRBV9 | TRBJ1-2 | CASSNKAGGASGYTF | KLGGALQAK | YES | NO | NO |
| 3102 | TRAV22 | TRAJ28 | CGAGSYQLTF | TRBV9 | TRBJ1-2 | CASSVDGRSYTF | KLGGALQAK | YES | NO | NO |
| 3103 | TRAV29DV5 | TRAJ21 | CAATAFSRNFNKFYF | TRBV9 | TRBJ1-2 | CASSSQSGSLYGYTF | KLGGALQAK | YES | NO | NO |
| 3105 | TRAV29DV5 | TRAJ43 | CAASARNNDMRF | TRBV9 | TRBJ1-2 | CASSSQSGSLYGYTF | KLGGALQAK | YES | NO | NO |
| 3106 | TRAV29DV5 | TRAJ45 | CAASAGGGADGLTF | TRBV9 | TRBJ1-2 | CASNFVGGDYGYTF | KLGGALQAK | YES | NO | NO |
| 3107 | TRAV30 | TRAJ54 | CGTVQGAQKLVF | TRBV9 | TRBJ1-2 | CASSASGISNYGYTF | KLGGALQAK | YES | NO | NO |
| 3109 | TRAV38-1 | TRAJ39 | CAFMKNNNNAGNMLTF | TRBV9 | TRBJ1-2 | CASSEGQGNYGYTF | KLGGALQAK | YES | NO | NO |
| 3110 | TRAV38-2DV8 | TRAJ34 | CAADNTDKLIF | TRBV9 | TRBJ1-2 | CASSPPGIYGYTF | KLGGALQAK | YES | NO | NO |
| 3111 | TRAV38-2DV8 | TRAJ34 | CACGGGKLIF | TRBV9 | TRBJ1-2 | CASSDNQYRLNYGYTF | KLGGALQAK | YES | NO | NO |
| 3112 | TRAV38-2DV8 | TRAJ39 | CAHYGYFNNAGNMLTF | TRBV9 | TRBJ1-2 | CASSVASGTGGDTYGYTF | KLGGALQAK | YES | NO | NO |
| 3113 | TRAV39 | TRAJ33 | CAVDSNYQLIW | TRBV9 | TRBJ1-2 | CASRNEWGNYGYTF | KLGGALQAK | YES | NO | NO |
| 3116 | TRAV41 | TRAJ58 | CAGCEETSGSRLTF | TRBV9 | TRBJ1-2 | CASSVGGAAGSPKEGYTF | KLGGALQAK | YES | NO | NO |
| 3117 | TRAV8-3 | TRAJ11 | CAVGPAQSSGYSTLTF | TRBV9 | TRBJ1-2 | CASSVQAGGYGYTF | KLGGALQAK | YES | NO | NO |
| 3118 | TRAV8-3 | TRAJ27 | CAVFTNAGKSTF | TRBV9 | TRBJ1-2 | CASSAPAGIGNGYTF | KLGGALQAK | YES | NO | NO |
| 3123 | TRAV8-6 | TRAJ48 | CAVSESNFGNEKLTF | TRBV9 | TRBJ1-2 | CASSARDRGLNYGYTF | KLGGALQAK | YES | NO | NO |
| 3125 | TRAV13-1 | TRAJ45 | CAASNSGGADGLTF | TRBV10-2 | TRBJ1-3 | CASSTGLSGNTIYF | KLGGALQAK | YES | NO | NO |
| 3126 | TRAV29DV5 | TRAJ22 | CAASRGRGSARQLTF | TRBV10-2 | TRBJ1-3 | CASSEGPSRGNTIYF | KLGGALQAK | YES | NO | NO |
| 3127 | TRAV12-2 | TRAJ47 | CAVNIYGNKLVF | TRBV10-3 | TRBJ1-3 | CAISTGARPGNTIYF | KLGGALQAK | YES | NO | NO |
| 3128 | TRAV26-1 | TRAJ23 | CIVRLYQGGKLIF | TRBV10-3 | TRBJ1-3 | CAISHLGESGNTIYF | KLGGALQAK | YES | NO | NO |
| 3129 | TRAV29DV5 | TRAJ48 | CAAKDEKLTF | TRBV10-3 | TRBJ1-3 | CAISARQPWIGNTIYF | KLGGALQAK | YES | NO | NO |
| 3130 | TRAV10 | TRAJ33 | CVAWSDSNYQLIW | TRBV11-2 | TRBJ1-3 | CASSQTGSSGNTIYF | KLGGALQAK | YES | NO | NO |
| 3131 | TRAV14DV4 | TRAJ52 | CAMSANAGGTSYGKLTF | TRBV11-2 | TRBJ1-3 | CASSPYSSGNTIYF | KLGGALQAK | YES | NO | NO |
| 3132 | TRAV1-2 | TRAJ20 | CAVRGPYKLSF | TRBV12-3 | TRBJ1-3 | CASSPRTGGRDTIYF | KLGGALQAK | YES | NO | NO |
| 3135 | TRAV13-2 | TRAJ49 | CAENTGNQFYF | TRBV12-4 | TRBJ1-3 | CASSLTRAGNTIYF | KLGGALQAK | YES | NO | NO |
| 3141 | TRAV12-2 | TRAJ18 | CAVTFDRGSTLGRLYF | TRBV12-5 | TRBJ1-3 | CASAPSWPGNTIYF | KLGGALQAK | YES | NO | NO |
| 3143 | TRAV29DV5 | TRAJ23 | CAASGKIYNQGGKLIF | TRBV12-5 | TRBJ1-3 | CASGTPRSPSGNTIYF | KLGGALQAK | YES | NO | NO |
| 3144 | TRAV29DV5 | TRAJ31 | CAATASNARLMF | TRBV12-5 | TRBJ1-3 | CASGSGGSGNTIYF | KLGGALQAK | YES | NO | NO |
| 3145 | TRAV12-2 | TRAJ4 | CAVNTHGGYNKLIF | TRBV13 | TRBJ1-3 | CASSGEDIHSGNTIYF | KLGGALQAK | YES | NO | NO |
| 3145 | TRAV12-2 | TRAJ4 | CAVNTHGGYNKLIF | TRBV13 | TRBJ1-3 | CASSGEDIHSGNTIYF | AVFDRKSDAK | YES | NO | NO |
| 3146 | TRAV12-2 | TRAJ52 | CAVAGGTSYGKLTF | TRBV13 | TRBJ1-3 | CASSSGQPPRSGNTIYF | KLGGALQAK | YES | NO | NO |
| 3147 | TRAV17 | TRAJ12 | CATDVARDSSYKLIF | TRBV13 | TRBJ1-3 | CASSSTMGDTIYF | KLGGALQAK | YES | NO | NO |
| 3149 | TRAV29DV5 | TRAJ39 | CAAFGAGNMLTF | TRBV13 | TRBJ1-3 | CASSLEAGRPTSGNTIYF | KLGGALQAK | YES | NO | NO |
| 3150 | TRAV8-1 | TRAJ39 | CAVNAGNAGNMLTF | TRBV13 | TRBJ1-3 | CASSSSLQGTGNTIYF | KLGGALQAK | YES | NO | NO |
| 3152 | TRAV29DV5 | TRAJ48 | CAAHLSNFGNEKLTF | TRBV15 | TRBJ1-3 | CATSRDRGHGDTIYF | KLGGALQAK | YES | NO | NO |
| 3154 | TRAV12-1 | TRAJ33 | CVVNRDYQLIW | TRBV19 | TRBJ1-3 | CASSLTSPGNTIYF | KLGGALQAK | YES | NO | NO |
| 3155 | TRAV12-1 | TRAJ52 | CVVIPAAGGTSYGKLTF | TRBV19 | TRBJ1-3 | CASSIVAGGPNTIYF | KLGGALQAK | YES | NO | NO |
| 3156 | TRAV12-2 | TRAJ17 | CAVPGGFKAAGNKLTF | TRBV19 | TRBJ1-3 | CASSIGAGNTIYF | RLRAEAQVK | YES | NO | YES |
| 3157 | TRAV12-2 | TRAJ30 | CAVGDDKIIF | TRBV19 | TRBJ1-3 | CASVPFSGNTIYF | KLGGALQAK | YES | NO | NO |
| 3159 | TRAV13-2 | TRAJ52 | CAESRSNAGGTSYGKLTF | TRBV19 | TRBJ1-3 | CASSRRQEGNTIYF | KLGGALQAK | YES | NO | NO |
| 3162 | TRAV21 | TRAJ23 | CAIIFYNQGGKLIF | TRBV19 | TRBJ1-3 | CASESRDRASAIYF | KLGGALQAK | YES | NO | NO |
| 3163 | TRAV3 | TRAJ13 | CAVRDFNSGGYQKVTF | TRBV19 | TRBJ1-3 | CASSSPIRWFTGNTIYF | KLGGALQAK | NO | NO | NO |
| 3164 | TRAV3 | TRAJ20 | CAVRDLNDYKLSF | TRBV19 | TRBJ1-3 | CASSEDRDGNTIYF | KLGGALQAK | NO | NO | NO |
| 3165 | TRAV35 | TRAJ33 | CAGLDSNYQLIW | TRBV19 | TRBJ1-3 | CASSIAGFGNTIYF | KLGGALQAK | YES | NO | NO |
| 3166 | TRAV41 | TRAJ53 | CAVTLLGGSNYKLTF | TRBV19 | TRBJ1-3 | CASSFQGGLRGNTIYF | KLGGALQAK | YES | NO | NO |
| 3167 | TRAV8-3 | TRAJ35 | CAVGGFGNVLHC | TRBV19 | TRBJ1-3 | CASSILGTVGNTIYF | KLGGALQAK | NO | NO | NO |
| 3168 | TRAV17 | TRAJ5 | CATVDRFMDTGRRALTF | TRBV2 | TRBJ1-3 | CASGEKQGANTIYF | KLGGALQAK | YES | NO | NO |
| 3169 | TRAV8-3 | TRAJ50 | CAAQKTSYDKVIF | TRBV2 | TRBJ1-3 | CASSDWATGSSGNTIYF | KLGGALQAK | YES | NO | NO |
| 3174 | TRAV3 | TRAJ8 | CAVRGYEMNTGFQKLVF | TRBV20-1 | TRBJ1-3 | CSARALTGHESSGNTIYF | KLGGALQAK | NO | NO | NO |
| 3175 | TRAV39 | TRAJ40 | CAGTTSGTYKYIF | TRBV20-1 | TRBJ1-3 | CSAVALGRDSGNTIYF | KLGGALQAK | YES | NO | NO |
| 3179 | TRAV17 | TRAJ9 | CATDAGTGGFKTIF | TRBV24-1 | TRBJ1-3 | CATSDGNTIYF | KLGGALQAK | YES | NO | NO |
| 3183 | TRAV30 | TRAJ52 | CGTLGGTSYGKLTF | TRBV25-1 | TRBJ1-3 | CASSEGSSGNTIYF | KLGGALQAK | YES | NO | NO |
| 3184 | TRAV12-1 | TRAJ10 | CVGGGGGNKLTF | TRBV27 | TRBJ1-3 | CASSIVAGGGNTIYF | KLGGALQAK | YES | NO | NO |
| 3185 | TRAV12-1 | TRAJ15 | CVVPARRAGTALIF | TRBV27 | TRBJ1-3 | CASSLREPGLSGNTIYF | KLGGALQAK | YES | NO | NO |
| 3186 | TRAV13-2 | TRAJ13 | CAVTGGYQKVTF | TRBV27 | TRBJ1-3 | CASSFKSSGNTIYF | KLGGALQAK | YES | NO | NO |
| 3187 | TRAV4 | TRAJ9 | CLVGDINTGGFKTIF | TRBV27 | TRBJ1-3 | CASSSISGNTIYF | KLGGALQAK | NO | NO | NO |
| 3190 | TRAV12-2 | TRAJ20 | CAVKGLSF | TRBV28 | TRBJ1-3 | CASSSIIQVSGNTIYF | KLGGALQAK | NO | NO | NO |
| 3191 | TRAV38-2DV8 | TRAJ30 | CAYLDPRDDKIIF | TRBV28 | TRBJ1-3 | CASSFYRGGNTIYF | KLGGALQAK | YES | NO | NO |
| 3192 | TRAV38-2DV8 | TRAJ58 | CAYRSKETSGSRLTF | TRBV28 | TRBJ1-3 | CASSLYVSYPKNTIYF | KLGGALQAK | YES | NO | NO |
| 3194 | TRAV27 | TRAJ45 | CAGGGADGLTF | TRBV29-1 | TRBJ1-3 | CSVSDGYSGNTIYF | KLGGALQAK | YES | NO | NO |
| 3195 | TRAV38-2DV8 | TRAJ58 | CAYKGSRLTF | TRBV29-1 | TRBJ1-3 | CSVEDGQTSGNTIYF | KLGGALQAK | YES | NO | NO |
| 3196 | TRAV8-1 | TRAJ13 | CAVTRSGGYQKVTF | TRBV29-1 | TRBJ1-3 | CSGSEGNTIYF | KLGGALQAK | NO | NO | NO |
| 3197 | TRAV12-3 | TRAJ47 | CAMSAEYGNKLVF | TRBV3-1 | TRBJ1-3 | CASSQDLQGGNTIYF | KLGGALQAK | YES | NO | NO |
| 3198 | TRAV13-2 | TRAJ26 | CAEKPGQNFVF | TRBV3-1 | TRBJ1-3 | CASSPPPSGAKEGVRAGNTIYF | KLGGALQAK | NO | NO | NO |
| 3199 | TRAV14DV4 | TRAJ11 | CAMRGAHSGYSTLTF | TRBV3-1 | TRBJ1-3 | CASSPGQSSGNTIYF | KLGGALQAK | YES | NO | NO |
| 3200 | TRAV17 | TRAJ50 | CATDDWETSYDKVIF | TRBV3-1 | TRBJ1-3 | CASSQVESGNTIYF | KLGGALQAK | YES | NO | NO |
| 3202 | TRAV35 | TRAJ34 | CAGPDYNTDKLIF | TRBV3-1 | TRBJ1-3 | CASSRPDRVFGNTIYF | KLGGALQAK | YES | NO | NO |
| 3203 | TRAV41 | TRAJ42 | CAVLDYGGSQGNLIF | TRBV3-1 | TRBJ1-3 | CASSQVSTSPWAGPVSGNTIYF | RLRAEAQVK | NO | NO | NO |
| 3204 | TRAV8-3 | TRAJ13 | CAVEPSGGYQKVTF | TRBV3-1 | TRBJ1-3 | CASSHPMGGSGNTIYF | KLGGALQAK | YES | NO | NO |
| 3205 | TRAV12-1 | TRAJ28 | CVVNSRLSGAGSYQLTF | TRBV30 | TRBJ1-3 | CAWRTGPVGNTIYF | KLGGALQAK | YES | NO | NO |
| 3206 | TRAV12-2 | TRAJ43 | CAVTEYNNNDMRF | TRBV30 | TRBJ1-3 | CAWRPAGGNTIYF | KLGGALQAK | YES | NO | NO |
| 3207 | TRAV39 | TRAJ45 | CAVVLYSGGGADGLTF | TRBV30 | TRBJ1-3 | CALLSDSGNTIYF | KLGGALQAK | YES | NO | NO |
| 3208 | TRAV12-2 | TRAJ27 | CAVNGDTNAGKSTF | TRBV4-1 | TRBJ1-3 | CASRFDRGSGNTIYF | KLGGALQAK | YES | YES | NO |
| 3211 | TRAV2 | TRAJ31 | CASNNNARLMF | TRBV4-2 | TRBJ1-3 | CASSLNRELTGNTIYF | KLGGALQAK | NO | NO | NO |
| 3212 | TRAV5 | TRAJ36 | CAAPSGTGANNLFF | TRBV4-2 | TRBJ1-3 | CASSQDPGSGNTIYF | KLGGALQAK | YES | YES | NO |
| 3214 | TRAV1-2 | TRAJ23 | CAAHSWGQGGKLIF | TRBV5-1 | TRBJ1-3 | CASSAGTEAGNTIYF | KLGGALQAK | YES | NO | NO |
| 3215 | TRAV12-1 | TRAJ41 | CVVNRGDSGYALNF | TRBV5-1 | TRBJ1-3 | CASSLGQSSGNTIYF | KLGGALQAK | YES | NO | NO |
| 3216 | TRAV17 | TRAJ54 | CATVRYSSQKLVF | TRBV5-1 | TRBJ1-3 | CASSEDRATSGNTIYF | KLGGALQAK | YES | NO | NO |
| 3217 | TRAV26-2 | TRAJ49 | CIPLNTGNQFYF | TRBV5-1 | TRBJ1-3 | CASREGLNSGNTIYF | KLGGALQAK | YES | NO | NO |
| 3218 | TRAV27 | TRAJ33 | CAGIPDSNYQLIW | TRBV5-1 | TRBJ1-3 | CASSPDGLSGNTIYF | KLGGALQAK | YES | NO | NO |
| 3219 | TRAV29DV5 | TRAJ54 | CAASAVWIQGAQKLVF | TRBV5-1 | TRBJ1-3 | CASSPGERLTALSGNTIYF | KLGGALQAK | YES | NO | NO |
| 3220 | TRAV12-1 | TRAJ29 | CVVNLFSGNTPLVF | TRBV5-6 | TRBJ1-3 | CASTRTGFFDSGNTIYF | KLGGALQAK | YES | NO | NO |
| 3221 | TRAV13-2 | TRAJ37 | CAEIPHHHGSGNTGKLIF | TRBV5-6 | TRBJ1-3 | CASSLDSGNTIYF | KLGGALQAK | YES | NO | NO |
| 3223 | TRAV17 | TRAJ30 | CAPRRRDDKIIF | TRBV5-6 | TRBJ1-3 | CASSLVGGFSGNTIYF | KLGGALQAK | YES | NO | NO |
| 3224 | TRAV4 | TRAJ29 | CLVGDIYSGNTPLVF | TRBV5-6 | TRBJ1-3 | CASSPWPGNTIYF | KLGGALQAK | YES | NO | NO |
| 3225 | TRAV9-2 | TRAJ39 | CALAMNAGNMLTF | TRBV5-6 | TRBJ1-3 | CASSLGGNTIYF | KLGGALQAK | YES | NO | NO |
| 3226 | TRAV1-2 | TRAJ34 | CAVRDHTDKLIF | TRBV6-1 | TRBJ1-3 | CASSDKWGGSGNTIYF | KLGGALQAK | YES | NO | NO |
| 3227 | TRAV13-1 | TRAJ9 | CAASLDTGGFKTIF | TRBV6-1 | TRBJ1-3 | CASSDRAGNTIYF | KLGGALQAK | YES | NO | NO |
| 3228 | TRAV14DV4 | TRAJ57 | CAMRGPIQGGSEKLVF | TRBV6-1 | TRBJ1-3 | CASGLSAPGNTIYF | KLGGALQAK | YES | NO | NO |
| 3230 | TRAV14DV4 | TRAJ37 | CAMRGAFSSNTGKLIF | TRBV6-3 | TRBJ1-3 | CATCNVGNTIYF | AVFDRKSDAK | YES | NO | NO |
| 3230 | TRAV14DV4 | TRAJ37 | CAMRGAFSSNTGKLIF | TRBV6-3 | TRBJ1-3 | CATCNVGNTIYF | KLGGALQAK | YES | NO | NO |

| | | | | | | | | | | |
|---|---|---|---|---|---|---|---|---|---|---|
| 3233 | TRAV29DV5 | TRAJ33 | CAASPFQNSNYQLIW | TRBV6-5 | TRBJ1-3 | CASSFRETPSGNTIYF | KLGGALQAK | YES | NO | NO |
| 3234 | TRAV3 | TRAJ31 | CAVRDMVEGNARLMF | TRBV6-5 | TRBJ1-3 | CASRTQGVGNTIYF | KLGGALQAK | NO | NO | NO |
| 3235 | TRAV38-2DV8 | TRAJ54 | CAYRRYIQGAQKLVF | TRBV6-5 | TRBJ1-3 | CASSRVEDGNTIYF | KLGGALQAK | YES | NO | NO |
| 3236 | TRAV8-1 | TRAJ7 | CAVNGPHGNNRLAF | TRBV6-5 | TRBJ1-3 | CASSFLPGQGPPSGNTIYF | KLGGALQAK | YES | NO | NO |
| 3237 | TRAV8-3 | TRAJ43 | CAVGESDMRF | TRBV6-5 | TRBJ1-3 | CASGGGAGSGNTIYF | KLGGALQAK | YES | NO | NO |
| 3238 | TRAV8-4 | TRAJ43 | CAVLYNNNDMRF | TRBV6-5 | TRBJ1-3 | CASRRDFEGNTIYF | KLGGALQAK | YES | NO | NO |
| 3239 | TRAV12-2 | TRAJ45 | CAVNLMYSGGGADGLTF | TRBV6-6 | TRBJ1-3 | CASSIKEAGNTIYF | KLGGALQAK | YES | NO | NO |
| 3240 | TRAV19 | TRAJ28 | CALDSGAGSYQLTF | TRBV6-6 | TRBJ1-3 | CASEGRGSSGNTIYF | KLGGALQAK | YES | NO | NO |
| 3241 | TRAV21 | TRAJ12 | CAVLRMDSSYKLIF | TRBV6-6 | TRBJ1-3 | CASTPRRLGLFPSSGNTIYF | KLGGALQAK | NO | NO | NO |
| 3241 | TRAV21 | TRAJ12 | CAVLRMDSSYKLIF | TRBV6-6 | TRBJ1-3 | CASTPRRLGLFPSSGNTIYF | RLRAEAQVK | NO | NO | NO |
| 3242 | TRAV21 | TRAJ37 | CAVISNTGKLIF | TRBV7-2 | TRBJ1-3 | CASSYTQGLAGNTIYF | KLGGALQAK | YES | NO | NO |
| 3245 | TRAV12-2 | TRAJ4 | CAVTGSFSGGYNKLIF | TRBV7-6 | TRBJ1-3 | CASSLTQSSLSGNTIYF | KLGGALQAK | YES | NO | NO |
| 3246 | TRAV14DV4 | TRAJ17 | CAMREGLGAAGNKLTF | TRBV7-6 | TRBJ1-3 | CASSLATGGYGNTIYF | KLGGALQAK | YES | NO | NO |
| 3247 | TRAV5 | TRAJ44 | CAESTTGTASKLTF | TRBV7-6 | TRBJ1-3 | CASSLTGFSWKHHIF | KLGGALQAK | YES | NO | NO |
| 3249 | TRAV1-1 | TRAJ27 | CAVITNAGKSTF | TRBV7-8 | TRBJ1-3 | CASSLVGQSGNTIYF | KLGGALQAK | YES | NO | NO |
| 3251 | TRAV8-3 | TRAJ49 | CAVGARGNQFYF | TRBV7-8 | TRBJ1-3 | CASSQFLDSSGNTIYF | KLGGALQAK | YES | NO | NO |
| 3252 | TRAV1-2 | TRAJ28 | CAVSGYSGAGSYQLTF | TRBV7-9 | TRBJ1-3 | CASSLMVAGSSGNTIYF | KLGGALQAK | YES | NO | NO |
| 3253 | TRAV10 | TRAJ4 | CVVRAGGYNKLIF | TRBV7-9 | TRBJ1-3 | CASSFSLPAASDADRSGNTIYF | KLGGALQAK | NO | NO | NO |
| 3254 | TRAV12-2 | TRAJ42 | CAVIGSQGNLIF | TRBV7-9 | TRBJ1-3 | CASSGWVAGNTIYF | KLGGALQAK | YES | NO | NO |
| 3258 | TRAV14DV4 | TRAJ27 | CAMREGGQGAGKSTF | TRBV7-9 | TRBJ1-3 | CASSYSGVAGNTIYF | KLGGALQAK | YES | NO | NO |
| 3259 | TRAV17 | TRAJ58 | CATAGETSGSRLTF | TRBV7-9 | TRBJ1-3 | CASSLDWGNTIYF | KLGGALQAK | YES | NO | NO |
| 3260 | TRAV1-2 | TRAJ15 | CAVRRTALIF | TRBV9 | TRBJ1-3 | CASTGEGAPGNTIYF | KLGGALQAK | YES | YES | NO |
| 3262 | TRAV13-2 | TRAJ39 | CAENTDNAGNMLTF | TRBV9 | TRBJ1-3 | CASSFGTGDNTIYF | KLGGALQAK | YES | NO | NO |
| 3263 | TRAV14DV4 | TRAJ10 | CAMKGLTGGGNKLTF | TRBV9 | TRBJ1-3 | CASSVGRGGNTIYF | KLGGALQAK | YES | NO | NO |
| 3265 | TRAV17 | TRAJ30 | CATGMNRDDKIIF | TRBV9 | TRBJ1-3 | CASSVDGRVGNTIYF | KLGGALQAK | NO | NO | NO |
| 3266 | TRAV19 | TRAJ36 | CALSEAQTGANNLFF | TRBV9 | TRBJ1-3 | CASSHPQGVTSRSGNTIYF | KLGGALQAK | YES | NO | NO |
| 3267 | TRAV19 | TRAJ45 | CALSDWSGGGADGLTF | TRBV9 | TRBJ1-3 | CASSVGNTIYF | KLGGALQAK | YES | NO | NO |
| 3269 | TRAV30 | TRAJ23 | CGTEIYNQGGKLIF | TRBV9 | TRBJ1-3 | CASSVDRTSGNTIYF | KLGGALQAK | YES | NO | NO |
| 3270 | TRAV38-1 | TRAJ39 | CAFMKHENNAGNMLTF | TRBV9 | TRBJ1-3 | CASSARRAGGGNTIYF | KLGGALQAK | NO | NO | NO |
| 3274 | TRAV26-2 | TRAJ33 | CILIGDSNYQLIW | TRBV10-1 | TRBJ1-4 | CASSEGTVNEKLFF | KLGGALQAK | YES | NO | NO |
| 3276 | TRAV29DV5 | TRAJ27 | CAASATTNAGKSTF | TRBV10-2 | TRBJ1-4 | CASSEYGPEEKLFF | KLGGALQAK | YES | NO | NO |
| 3277 | TRAV1-2 | TRAJ6 | CAVGASGGSYIPTF | TRBV10-3 | TRBJ1-4 | CAAGGGGTEKLFF | KLGGALQAK | YES | NO | NO |
| 3279 | TRAV26-1 | TRAJ49 | CIVRFLFTGNQFYF | TRBV10-3 | TRBJ1-4 | CAIRGTANEKLFF | KLGGALQAK | YES | NO | NO |
| 3281 | TRAV17 | TRAJ16 | CATDAF | TRBV11-2 | TRBJ1-4 | CASSAPGTAREKLFF | KLGGALQAK | NO | NO | NO |
| 3282 | TRAV25 | TRAJ15 | CAARRQAGTALIF | TRBV11-2 | TRBJ1-4 | CASSLVVDGPAEKLFF | KLGGALQAK | YES | NO | NO |
| 3283 | TRAV26-1 | TRAJ45 | CIVSGGGADGLTF | TRBV11-2 | TRBJ1-4 | CASNPTGVGDEKLFF | KLGGALQAK | NO | NO | NO |
| 3283 | TRAV26-1 | TRAJ45 | CIVSGGGADGLTF | TRBV11-2 | TRBJ1-4 | CASNPTGVGDEKLFF | AVFDRKSDAK | NO | NO | NO |
| 3285 | TRAV1-2 | TRAJ3 | CAVRDRGSSASKIIF | TRBV11-3 | TRBJ1-4 | CASSGRKLFF | KLGGALQAK | YES | NO | NO |
| 3285 | TRAV1-2 | TRAJ3 | CAVRDRGSSASKIIF | TRBV11-3 | TRBJ1-4 | CASSGRKLFF | AVFDRKSDAK | YES | NO | NO |
| 3286 | TRAV5 | TRAJ33 | CAEWDSNYQLIW | TRBV11-3 | TRBJ1-4 | CASSGPGRTTNEKLFF | KLGGALQAK | YES | NO | NO |
| 3286 | TRAV5 | TRAJ33 | CAEWDSNYQLIW | TRBV11-3 | TRBJ1-4 | CASSGPGRTTNEKLFF | AVFDRKSDAK | YES | NO | NO |
| 3287 | TRAV12-1 | TRAJ31 | CVVNGYARLMF | TRBV12-3 | TRBJ1-4 | CASSPQEGGMNEKLFF | KLGGALQAK | YES | NO | NO |
| 3289 | TRAV13-2 | TRAJ24 | CAETDRTDSWGKFQF | TRBV12-3 | TRBJ1-4 | CASSFYVGQGATNEKLFF | KLGGALQAK | YES | NO | NO |
| 3291 | TRAV19 | TRAJ43 | CALSEDPPDMRF | TRBV12-4 | TRBJ1-4 | CASSAIGGEEKLFF | KLGGALQAK | YES | NO | NO |
| 3292 | TRAV20 | TRAJ42 | CAVRSYGGSQGNLIF | TRBV12-4 | TRBJ1-4 | CASSPMGTNEKLFF | KLGGALQAK | YES | NO | NO |
| 3294 | TRAV27 | TRAJ23 | CAGFYNQGGKLIF | TRBV12-4 | TRBJ1-4 | CASTPGGEKLFF | KLGGALQAK | YES | NO | NO |
| 3296 | TRAV8-1 | TRAJ3 | CAVSDYSSASKIIF | TRBV12-4 | TRBJ1-4 | CASSLAEKLFF | KLGGALQAK | YES | NO | NO |
| 3297 | TRAV12-2 | TRAJ11 | CAVKGLGYSTLTF | TRBV14 | TRBJ1-4 | CASSQSVGGLNEKLFF | KLGGALQAK | NO | NO | NO |
| 3299 | TRAV12-2 | TRAJ42 | CAVISFGSQGNLIF | TRBV14 | TRBJ1-4 | CASSQVPARKLFF | KLGGALQAK | NO | NO | NO |
| 3300 | TRAV12-2 | TRAJ52 | CAVGGTSYGKLTF | TRBV14 | TRBJ1-4 | CASSQGPGTVHEKLFF | KLGGALQAK | NO | NO | NO |
| 3301 | TRAV12-3 | TRAJ39 | CAIGPGNMLTF | TRBV14 | TRBJ1-4 | CASSQGPGTVHEKLFF | KLGGALQAK | NO | NO | NO |
| 3302 | TRAV14DV4 | TRAJ22 | CAMRRPISSGSARQLTF | TRBV14 | TRBJ1-4 | CASSHPGTRDEKLFF | KLGGALQAK | NO | NO | NO |
| 3309 | TRAV14DV4 | TRAJ30 | CAMREGVVTDDKIIF | TRBV15 | TRBJ1-4 | CATSLNVVRASNEKLFF | KLGGALQAK | NO | NO | NO |
| 3310 | TRAV29DV5 | TRAJ44 | CAARITGTASKLTF | TRBV15 | TRBJ1-4 | CATSNPDPLQGYRGEKLFF | KLGGALQAK | NO | NO | NO |
| 3312 | TRAV13-1 | TRAJ43 | CAAAYNNDMRF | TRBV16 | TRBJ1-4 | CASSQIEKLFF | KLGGALQAK | YES | NO | NO |
| 3313 | TRAV13-1 | TRAJ23 | CAASIGQGGKLIF | TRBV18 | TRBJ1-4 | CASSPYGYENEKLFF | KLGGALQAK | YES | NO | NO |
| 3315 | TRAV29DV5 | TRAJ39 | CAASADAGNMLTF | TRBV18 | TRBJ1-4 | CASSLGSTNEKLFF | KLGGALQAK | YES | NO | NO |
| 3316 | TRAV38-1 | TRAJ47 | CAFMTNMEYGNKLVF | TRBV18 | TRBJ1-4 | CASSPSSGTGGLATNEKLFF | KLGGALQAK | NO | NO | NO |
| 3317 | TRAV38-2DV8 | TRAJ40 | CAFQTSGTYKYIF | TRBV18 | TRBJ1-4 | CASSPKREVRVLEKLFF | KLGGALQAK | YES | NO | NO |
| 3318 | TRAV13-2 | TRAJ52 | CAVIGAGGTSYGKLTF | TRBV19 | TRBJ1-4 | CASSPAGGEKLFF | KLGGALQAK | YES | NO | NO |
| 3319 | TRAV17 | TRAJ57 | CATDAQRRAEKLVF | TRBV19 | TRBJ1-4 | CASSMGGFNEKLFF | KLGGALQAK | YES | NO | NO |
| 3320 | TRAV21 | TRAJ23 | CAVRRSIIGGKLIF | TRBV19 | TRBJ1-4 | CASRHRAGRDWMDEKLFF | KLGGALQAK | YES | NO | NO |
| 3321 | TRAV24 | TRAJ31 | CASWYARLMF | TRBV19 | TRBJ1-4 | CASSIQSEREKLFF | RLRAEAQVK | YES | YES | YES |
| 3323 | TRAV3 | TRAJ42 | CTIYGGSQGNLIF | TRBV19 | TRBJ1-4 | CASSGGQPNEKLFF | KLGGALQAK | NO | NO | NO |
| 3324 | TRAV3 | TRAJ48 | CAVRDPQAISNFGNEKLTF | TRBV19 | TRBJ1-4 | CASSASRAATNEKLFF | KLGGALQAK | NO | NO | NO |
| 3325 | TRAV38-1 | TRAJ57 | CAFMRFRGGSEKLVF | TRBV19 | TRBJ1-4 | CASSTLLQSREGSYEKLFF | KLGGALQAK | YES | NO | NO |
| 3326 | TRAV38-2DV8 | TRAJ21 | CAYPVYNFNKFYF | TRBV19 | TRBJ1-4 | CASSTIGTGGLEKLFF | KLGGALQAK | YES | NO | NO |
| 3326 | TRAV38-2DV8 | TRAJ21 | CAYPVYNFNKFYF | TRBV19 | TRBJ1-4 | CASSTIGTGGLEKLFF | AVFDRKSDAK | YES | NO | NO |
| 3327 | TRAV8-1 | TRAJ8 | CAVNAGFQKLVF | TRBV19 | TRBJ1-4 | CASRGTGIEKLFF | KLGGALQAK | YES | NO | NO |
| 3329 | TRAV9-2 | TRAJ5 | CALSGKDTGRRALTF | TRBV19 | TRBJ1-4 | CASNPGQSTNEKLFF | KLGGALQAK | YES | NO | NO |
| 3330 | TRAV12-1 | TRAJ11 | CVVKGYSTLTF | TRBV2 | TRBJ1-4 | CASSARVREKLFF | KLGGALQAK | YES | NO | NO |
| 3332 | TRAV14DV4 | TRAJ28 | CAMREGDLSGAGSYQLTF | TRBV2 | TRBJ1-4 | CASSYGEKLFF | KLGGALQAK | YES | NO | NO |
| 3333 | TRAV14DV4 | TRAJ56 | CAMRGDASTGANSKLTF | TRBV2 | TRBJ1-4 | CASSEMLGTNEKLFF | KLGGALQAK | YES | NO | NO |
| 3335 | TRAV3 | TRAJ6 | CAVRVPSGGSYIPTF | TRBV2 | TRBJ1-4 | CASRESLNEKLFF | KLGGALQAK | NO | NO | NO |
| 3336 | TRAV9-2 | TRAJ48 | CALSDVWGFGNEKLTF | TRBV2 | TRBJ1-4 | CASNAGTGHPNEKLFF | RLRAEAQVK | YES | NO | YES |
| 3337 | TRAV1-2 | TRAJ32 | CAVRGGYGGATNKLIF | TRBV20-1 | TRBJ1-4 | CSAREEGDTRRFF | KLGGALQAK | YES | NO | NO |
| 3338 | TRAV1-2 | TRAJ33 | CAVTDSNYQLIW | TRBV20-1 | TRBJ1-4 | CSARYREFEKLFF | KLGGALQAK | YES | NO | NO |
| 3339 | TRAV12-3 | TRAJ23 | CAMSVFYNQGGKLIF | TRBV20-1 | TRBJ1-4 | CSSPGGPNEKLFF | KLGGALQAK | YES | NO | NO |
| 3340 | TRAV12-3 | TRAJ57 | CAMSAVLSEKLVF | TRBV20-1 | TRBJ1-4 | CSARERWEGATNEKLFF | KLGGALQAK | YES | NO | NO |
| 3341 | TRAV13-1 | TRAJ39 | CAARNAGNMLTF | TRBV20-1 | TRBJ1-4 | CSSPGGPNEKLFF | KLGGALQAK | YES | NO | NO |
| 3343 | TRAV14DV4 | TRAJ22 | CAMRRPISSGSARQLTF | TRBV20-1 | TRBJ1-4 | CSSPGGPNEKLFF | KLGGALQAK | YES | NO | NO |
| 3343 | TRAV14DV4 | TRAJ22 | CAMRRPISSGSARQLTF | TRBV20-1 | TRBJ1-4 | CSSPGGPNEKLFF | AVFDRKSDAK | YES | NO | NO |
| 3344 | TRAV14DV4 | TRAJ22 | CAMRRPISSGSARQLTF | TRBV20-1 | TRBJ1-4 | CSAPRGTNEKLFF | KLGGALQAK | YES | NO | NO |
| 3347 | TRAV3 | TRAJ27 | CPLPTNAGKSTF | TRBV20-1 | TRBJ1-4 | CSARDYRTGDEKLFF | KLGGALQAK | NO | NO | NO |
| 3347 | TRAV3 | TRAJ27 | CPLPTNAGKSTF | TRBV20-1 | TRBJ1-4 | CSARDYRTGDEKLFF | AVFDRKSDAK | NO | NO | NO |
| 3349 | TRAV5 | TRAJ4 | CAETSPSFSGGYNKLIF | TRBV20-1 | TRBJ1-4 | CSSPGGPNEKLFF | KLGGALQAK | YES | NO | NO |
| 3351 | TRAV38-2DV8 | TRAJ8 | CNTGFQKLVF | TRBV21-1 | TRBJ1-4 | CASSYSQTFPGQGANEKLFF | KLGGALQAK | NO | NO | NO |
| 3354 | TRAV38-2DV8 | TRAJ53 | CAYRSLNSGGSNYKLTF | TRBV24-1 | TRBJ1-4 | CATRGPSEKLFF | KLGGALQAK | YES | NO | NO |
| 3355 | TRAV21 | TRAJ32 | CAVSYGGATNKLIF | TRBV25-1 | TRBJ1-4 | CASSNNRTGLFVYEKLFF | KLGGALQAK | YES | NO | NO |
| 3356 | TRAV27 | TRAJ49 | CAGAWNTGNQFYF | TRBV25-1 | TRBJ1-4 | CASSGGLGASEKLFF | KLGGALQAK | YES | NO | NO |
| 3357 | TRAV1-2 | TRAJ10 | CAVRVTSLTGGGNKLTF | TRBV27 | TRBJ1-4 | CASSFSGQGLTNEKLFF | KLGGALQAK | YES | NO | NO |
| 3358 | TRAV10 | TRAJ18 | CVVRGADRGSTLGRLYF | TRBV27 | TRBJ1-4 | CASSHLTEFNEKLFF | KLGGALQAK | YES | NO | NO |
| 3360 | TRAV12-2 | TRAJ52 | CAALPSNAGGTSYGKLTF | TRBV27 | TRBJ1-4 | CASSLYAGASEKLFF | KLGGALQAK | YES | NO | NO |
| 3361 | TRAV12-2 | TRAJ53 | CAVKADSGGSNYKLTF | TRBV27 | TRBJ1-4 | CASRLLGGRVEKLFF | KLGGALQAK | YES | NO | NO |
| 3363 | TRAV13-1 | TRAJ44 | CAAINTGTASKLTF | TRBV27 | TRBJ1-4 | CASSSPGPREKLFF | KLGGALQAK | YES | NO | NO |
| 3364 | TRAV14DV4 | TRAJ11 | CAMRLGYSTLTF | TRBV27 | TRBJ1-4 | CASSPGQGYEKLFF | KLGGALQAK | YES | NO | NO |
| 3365 | TRAV14DV4 | TRAJ23 | CAMRGWLNQGGKLIF | TRBV27 | TRBJ1-4 | CASSSGEKLFF | KLGGALQAK | YES | NO | NO |
| 3366 | TRAV17 | TRAJ53 | CAPLFSGGSNYKLTF | TRBV27 | TRBJ1-4 | CASGGATNEKLFF | KLGGALQAK | YES | NO | NO |
| 3368 | TRAV19 | TRAJ26 | CALSEDNYGQNFVF | TRBV27 | TRBJ1-4 | CASSPRDSPNEKLFF | KLGGALQAK | YES | NO | NO |
| 3369 | TRAV20 | TRAJ31 | CAVQEPNNNARLMF | TRBV27 | TRBJ1-4 | CASSLYAGASEKLFF | KLGGALQAK | YES | NO | NO |
| 3370 | TRAV25 | TRAJ32 | CAGNDYGGATNKLIF | TRBV27 | TRBJ1-4 | CASSLYAGASEKLFF | AVFDRKSDAK | YES | NO | NO |
| 3370 | TRAV25 | TRAJ32 | CAGNDYGGATNKLIF | TRBV27 | TRBJ1-4 | CASSLYAGASEKLFF | RLRAEAQVK | YES | NO | NO |
| 3370 | TRAV25 | TRAJ32 | CAGNDYGGATNKLIF | TRBV27 | TRBJ1-4 | CASSLYAGASEKLFF | IVTDFSVIK | YES | NO | NO |
| 3371 | TRAV27 | TRAJ23 | CAGVTQGGKLIF | TRBV27 | TRBJ1-4 | CASSLYAGASEKLFF | KLGGALQAK | YES | NO | NO |

| | | | | | | | | | | |
|---|---|---|---|---|---|---|---|---|---|---|
| 3372 | TRAV29DV5 | TRAJ57 | CVQGGSEKLVF | TRBV27 | TRBJ1-4 | CASSFFGGDEKLFF | KLGGALQAK | YES | NO | NO |
| 3374 | TRAV30 | TRAJ24 | CGALDSWGKLQF | TRBV27 | TRBJ1-4 | CASKTTGDSNEKLFF | KLGGALQAK | YES | NO | NO |
| 3375 | TRAV38-1 | TRAJ44 | CAFHVITGTASKLTF | TRBV27 | TRBJ1-4 | CASSWSQGVYEKLFF | KLGGALQAK | YES | NO | NO |
| 3377 | TRAV38-2DV8 | TRAJ22 | CAYRGSGSARQLTF | TRBV27 | TRBJ1-4 | CASSPGQGYEKLFF | KLGGALQAK | YES | NO | NO |
| 3378 | TRAV39 | TRAJ43 | CAVDNNNDMRF | TRBV27 | TRBJ1-4 | CASSLASYGGEKLFF | KLGGALQAK | YES | NO | NO |
| 3381 | TRAV8-2 | TRAJ47 | CAVTDGNKLVF | TRBV27 | TRBJ1-4 | CASSLGTHATNEKLFF | KLGGALQAK | YES | NO | NO |
| 3383 | TRAV8-6 | TRAJ54 | CAVSDYIIQGAQKLVF | TRBV27 | TRBJ1-4 | CASRPGGHEGEKLFF | KLGGALQAK | YES | NO | NO |
| 3384 | TRAV1-2 | TRAJ36 | CAVTPPETGANNLFF | TRBV28 | TRBJ1-4 | CASQPQGATSEKLFF | KLGGALQAK | YES | NO | NO |
| 3386 | TRAV12-3 | TRAJ34 | CAMSLTLSYNTDKLIF | TRBV28 | TRBJ1-4 | CASNAPGQGGEKLFF | KLGGALQAK | YES | NO | NO |
| 3388 | TRAV17 | TRAJ33 | CATDSDSNYQLIW | TRBV28 | TRBJ1-4 | CASSPRPTQLEKLFF | KLGGALQAK | YES | NO | NO |
| 3389 | TRAV29DV5 | TRAJ45 | CAASGGWGGLTF | TRBV28 | TRBJ1-4 | CASQDRGNEKLFF | KLGGALQAK | YES | NO | NO |
| 3391 | TRAV38-2DV8 | TRAJ49 | CAYRSPLTGNQFYF | TRBV28 | TRBJ1-4 | CASSLPRGGLFF | KLGGALQAK | YES | NO | NO |
| 3394 | TRAV12-3 | TRAJ28 | CAMSDSGAGSYQLTF | TRBV29-1 | TRBJ1-4 | CSVETNEKLFF | KLGGALQAK | YES | NO | NO |
| 3395 | TRAV17 | TRAJ43 | CATDSKGDMRF | TRBV29-1 | TRBJ1-4 | CSVVATNEKLFF | KLGGALQAK | YES | NO | NO |
| 3396 | TRAV27 | TRAJ37 | CAGAGNTGKLIF | TRBV29-1 | TRBJ1-4 | CSAAGQGPTNEKLFF | KLGGALQAK | YES | NO | NO |
| 3397 | TRAV5 | TRAJ9 | CAERTGGFKTIF | TRBV29-1 | TRBJ1-4 | CSVTEDEKLFF | KLGGALQAK | YES | NO | NO |
| 3399 | TRAV19 | TRAJ43 | CALSDLPNDMRF | TRBV3-1 | TRBJ1-4 | CASSTGTGVDEKLFF | KLGGALQAK | YES | NO | NO |
| 3400 | TRAV20 | TRAJ20 | CAVQADDYKLSF | TRBV3-1 | TRBJ1-4 | CASSPTGSNEKLFF | KLGGALQAK | YES | NO | NO |
| 3401 | TRAV26-1 | TRAJ33 | CIVRVTMDSNYQLIW | TRBV3-1 | TRBJ1-4 | CASSQDRGATDELFF | KLGGALQAK | YES | NO | NO |
| 3403 | TRAV12-2 | TRAJ21 | CAATPVYNFNKFYF | TRBV30 | TRBJ1-4 | CAWSLRIMDEKLFF | KLGGALQAK | NO | NO | NO |
| 3404 | TRAV13-2 | TRAJ15 | CAENIRSWQAGTALIF | TRBV30 | TRBJ1-4 | CAWAKPREIQVSGEKLFF | KLGGALQAK | NO | NO | NO |
| 3405 | TRAV23DV6 | TRAJ42 | CAASMNYGGSQGNLIF | TRBV30 | TRBJ1-4 | CAWSLGYGEDEKLFF | KLGGALQAK | YES | NO | NO |
| 3406 | TRAV12-1 | TRAJ36 | CVVIEGTGANNLFF | TRBV4-1 | TRBJ1-4 | CASSQGDWAFREKLFF | KLGGALQAK | YES | NO | NO |
| 3407 | TRAV12-2 | TRAJ31 | CAVNRVRDARLMF | TRBV4-1 | TRBJ1-4 | CASSQDHRMGGHEKLFF | KLGGALQAK | YES | NO | NO |
| 3408 | TRAV19 | TRAJ28 | CALGVGSGAGSYQLTF | TRBV4-1 | TRBJ1-4 | CASSQSKGQGNEKLFF | KLGGALQAK | YES | NO | NO |
| 3409 | TRAV20 | TRAJ8 | CAATGFQKLVF | TRBV4-1 | TRBJ1-4 | CASSQDAGVGEKLFF | KLGGALQAK | YES | NO | NO |
| 3410 | TRAV21 | TRAJ43 | CAVWYMRF | TRBV4-1 | TRBJ1-4 | CASSQDHRMGGHEKLFF | KLGGALQAK | NO | NO | NO |
| 3412 | TRAV38-2DV8 | TRAJ35 | CAYRSAI | TRBV4-1 | TRBJ1-4 | CASSQTGGPTNEKLFF | KLGGALQAK | NO | NO | NO |
| 3414 | TRAV8-1 | TRAJ38 | CAVKAGNNRKLIW | TRBV4-1 | TRBJ1-4 | CASSQTPGQGWLFF | KLGGALQAK | YES | NO | NO |
| 3416 | TRAV14DV4 | TRAJ54 | CAMREGKIQGAQKLVF | TRBV4-2 | TRBJ1-4 | CASSQGGQYNEKLFF | KLGGALQAK | YES | NO | NO |
| 3417 | TRAV26-2 | TRAJ5 | CILTPMDTGRRALTF | TRBV4-2 | TRBJ1-4 | CASSRGQGLSNEKLFF | KLGGALQAK | NO | NO | NO |
| 3419 | TRAV10 | TRAJ45 | CVVSSISGGGADGLTF | TRBV4-3 | TRBJ1-4 | CASSQDAAGEKLFF | KLGGALQAK | YES | NO | NO |
| 3424 | TRAV14DV4 | TRAJ37 | CAMRGPFSSNTGKLIF | TRBV4-3 | TRBJ1-4 | CASSQDHRMGGHEKLFF | RLRAEAQVK | YES | NO | YES |
| 3428 | TRAV19 | TRAJ50 | CALSVEKTSYDKVIF | TRBV4-3 | TRBJ1-4 | CASSQDAGGMTNEKLFF | KLGGALQAK | YES | NO | NO |
| 3430 | TRAV21 | TRAJ54 | CAVQGAQKLVF | TRBV4-3 | TRBJ1-4 | CASSQDHRMGGHEKLFF | KLGGALQAK | YES | NO | NO |
| 3431 | TRAV21 | TRAJ57 | CAVRQEGSEKLVF | TRBV4-3 | TRBJ1-4 | CASSQDMAGWGEKLFF | KLGGALQAK | YES | NO | NO |
| 3432 | TRAV24 | TRAJ52 | CAFIGDAGGTSYGKLTF | TRBV4-3 | TRBJ1-4 | CASSQDHRMGGHEKLFF | AVFDRKSDAK | YES | NO | NO |
| 3432 | TRAV24 | TRAJ52 | CAFIGDAGGTSYGKLTF | TRBV4-3 | TRBJ1-4 | CASSQDHRMGGHEKLFF | RLRAEAQVK | YES | NO | NO |
| 3432 | TRAV24 | TRAJ52 | CAFIGDAGGTSYGKLTF | TRBV4-3 | TRBJ1-4 | CASSQDHRMGGHEKLFF | IVTDFSVIK | YES | NO | NO |
| 3433 | TRAV24 | TRAJ52 | CAFIGDAGGTSYGKLTF | TRBV4-3 | TRBJ1-4 | CASSQDHRMGGQEKLFF | KLGGALQAK | YES | NO | NO |
| 3434 | TRAV29DV5 | TRAJ30 | CAASVQNRDDKIIF | TRBV4-3 | TRBJ1-4 | CASSQDHRMGGHEKLFF | KLGGALQAK | YES | NO | NO |
| 3435 | TRAV3 | TRAJ28 | CAVRDPSYSGAGSYQLTF | TRBV4-3 | TRBJ1-4 | CASSQKRTGVNEKLFF | KLGGALQAK | YES | NO | NO |
| 3437 | TRAV3 | TRAJ37 | CAVRDGGMGSNTGKLIF | TRBV4-3 | TRBJ1-4 | CASSQDHRMGGHEKLFF | KLGGALQAK | YES | NO | NO |
| 3439 | TRAV35 | TRAJ53 | CAGLEHSGGSNYKLTF | TRBV4-3 | TRBJ1-4 | CASSQARGKLFF | KLGGALQAK | YES | NO | NO |
| 3440 | TRAV38-1 | TRAJ34 | CAFVGGSTDKLIF | TRBV4-3 | TRBJ1-4 | CASSQYRGFEKLFF | KLGGALQAK | YES | NO | NO |
| 3442 | TRAV8-1 | TRAJ22 | CAVRSSGSARQLTF | TRBV4-3 | TRBJ1-4 | CASSQDHRMGGHEKLFF | KLGGALQAK | YES | NO | NO |
| 3443 | TRAV8-1 | TRAJ31 | CAVNEGNARLMF | TRBV4-3 | TRBJ1-4 | CASSQDHRMGGHEKLFF | KLGGALQAK | YES | NO | NO |
| 3445 | TRAV8-3 | TRAJ11 | CAVGGAWGYSTLTF | TRBV4-3 | TRBJ1-4 | CASSQDHRMGGHEKLFF | KLGGALQAK | YES | NO | NO |
| 3446 | TRAV12-1 | TRAJ40 | CVVNGDTSGTYKYIF | TRBV5-1 | TRBJ1-4 | CASSSGTHNEKLFF | KLGGALQAK | YES | NO | NO |
| 3447 | TRAV14DV4 | TRAJ49 | CAKREVSYTGNQFYF | TRBV5-1 | TRBJ1-4 | CASSLFSYNRDGNEKLFF | KLGGALQAK | YES | NO | NO |
| 3448 | TRAV21 | TRAJ21 | CAVPGLYNFNKFYF | TRBV5-1 | TRBJ1-4 | CASSSGQGAGEKLFF | KLGGALQAK | YES | NO | NO |
| 3449 | TRAV29DV5 | TRAJ28 | CAATNIATGAGSYQLTF | TRBV5-1 | TRBJ1-4 | CASSLALTGLPNEKLFF | KLGGALQAK | YES | NO | NO |
| 3450 | TRAV29DV5 | TRAJ42 | CAASAEGGSQGNLIF | TRBV5-1 | TRBJ1-4 | CASRKWSATNEKLFF | KLGGALQAK | YES | NO | NO |
| 3452 | TRAV30 | TRAJ36 | CGTESGANNLFF | TRBV5-1 | TRBJ1-4 | CASSLRGTGPDEKLFF | KLGGALQAK | YES | NO | NO |
| 3453 | TRAV30 | TRAJ36 | CGTEAGANNLFF | TRBV5-1 | TRBJ1-4 | CASSFDQGSSTNEKLFF | KLGGALQAK | YES | NO | NO |
| 3454 | TRAV34 | TRAJ4 | CGAAFFSGGYNKLIF | TRBV5-1 | TRBJ1-4 | CASSFTTNEKLFF | KLGGALQAK | YES | NO | NO |
| 3456 | TRAV39 | TRAJ43 | CAVDITADMRF | TRBV5-1 | TRBJ1-4 | CASGGSDEKLFF | KLGGALQAK | YES | NO | NO |
| 3457 | TRAV9-2 | TRAJ10 | CALTALTGGGNKLTF | TRBV5-1 | TRBJ1-4 | CASSPSTGGIGEKLFF | KLGGALQAK | YES | NO | NO |
| 3458 | TRAV1-2 | TRAJ31 | CAVQNNNARLMF | TRBV5-4 | TRBJ1-4 | CASSLGGGGVDEKLFF | KLGGALQAK | YES | NO | NO |
| 3460 | TRAV21 | TRAJ36 | CAVYPGANNLFF | TRBV5-4 | TRBJ1-4 | CASSLDQQGVGEKLFF | AVFDRKSDAK | YES | NO | NO |
| 3462 | TRAV34 | TRAJ6 | CGAFTSGGSYIPTF | TRBV5-4 | TRBJ1-4 | CATRVRGVEKLFF | KLGGALQAK | YES | NO | NO |
| 3463 | TRAV38-2DV8 | TRAJ44 | CALISDGTASKLTF | TRBV5-4 | TRBJ1-4 | CASSLMGFTNEKLFF | KLGGALQAK | YES | NO | NO |
| 3464 | TRAV25 | TRAJ28 | CAGFLITGAGSYQLTF | TRBV5-5 | TRBJ1-4 | CASSSTGTVSLEKLFF | KLGGALQAK | YES | NO | NO |
| 3465 | TRAV26-2 | TRAJ49 | CILKITGNQFYF | TRBV5-5 | TRBJ1-4 | CASSLELNSATNEKLFF | KLGGALQAK | YES | NO | NO |
| 3466 | TRAV13-1 | TRAJ26 | CAADGVWRYGQNFVF | TRBV5-6 | TRBJ1-4 | CASSLTGAPNEKLFF | KLGGALQAK | YES | NO | NO |
| 3468 | TRAV19 | TRAJ10 | CALSAPKRILTGGGNKLTF | TRBV5-6 | TRBJ1-4 | CASSPQPGGREKLFF | KLGGALQAK | YES | NO | NO |
| 3470 | TRAV19 | TRAJ20 | CALSRSNDYKLSF | TRBV5-6 | TRBJ1-4 | CASSLERFPDRVNEKLFF | KLGGALQAK | YES | NO | NO |
| 3475 | TRAV8-6 | TRAJ13 | CAVTSRSGGYQKVTF | TRBV5-6 | TRBJ1-4 | CASIAGPGTGGEKLFF | KLGGALQAK | YES | NO | NO |
| 3476 | TRAV12-2 | TRAJ18 | CAVNPDRGSTLGRLYF | TRBV6-1 | TRBJ1-4 | CASSEFIGSNEKLFF | KLGGALQAK | YES | NO | NO |
| 3477 | TRAV12-2 | TRAJ52 | CAALPSNAGGTSYGKLTF | TRBV6-1 | TRBJ1-4 | CASSEAWGTPLWGEKLFF | RLRAEAQVK | YES | NO | NO |
| 3477 | TRAV12-2 | TRAJ52 | CAALPSNAGGTSYGKLTF | TRBV6-1 | TRBJ1-4 | CASSEAWGTPLWGEKLFF | KLGGALQAK | YES | NO | NO |
| 3480 | TRAV19 | TRAJ20 | CALSDFNDYKLSF | TRBV6-1 | TRBJ1-4 | CASNQPGQGGFRLFF | KLGGALQAK | YES | NO | NO |
| 3481 | TRAV19 | TRAJ40 | CALGPTSGTYKYIF | TRBV6-1 | TRBJ1-4 | CASSERGTGIIWEKLFF | KLGGALQAK | YES | NO | NO |
| 3483 | TRAV21 | TRAJ20 | CAVSNDYKLSF | TRBV6-1 | TRBJ1-4 | CASSAYRDEKLFF | KLGGALQAK | YES | NO | NO |
| 3486 | TRAV13-2 | TRAJ21 | CADRWQFYF | TRBV6-3 | TRBJ1-4 | CASSRVTNEKLFF | KLGGALQAK | NO | NO | NO |
| 3490 | TRAV8-1 | TRAJ37 | CAVRIGSGNTGKLIF | TRBV6-3 | TRBJ1-4 | CASESGVELFF | KLGGALQAK | YES | NO | NO |
| 3491 | TRAV8-1 | TRAJ39 | CAVKVYNAGNMLTF | TRBV6-3 | TRBJ1-4 | CASESGVELFF | KLGGALQAK | YES | NO | NO |
| 3492 | TRAV1-2 | TRAJ33 | CAVQDSNYQLIW | TRBV6-4 | TRBJ1-4 | CASSEGRGSEKLFF | KLGGALQAK | YES | NO | NO |
| 3493 | TRAV13-1 | TRAJ20 | CAASNDYKLSF | TRBV6-5 | TRBJ1-4 | CATGQGGEKLFF | KLGGALQAK | NO | NO | NO |
| 3495 | TRAV14DV4 | TRAJ53 | CAMREDNSNSGGSNYKLTF | TRBV6-5 | TRBJ1-4 | CASSYSPFKNEKLFF | KLGGALQAK | NO | NO | NO |
| 3498 | TRAV25 | TRAJ23 | CAGDMIYNQGGKLIF | TRBV6-5 | TRBJ1-4 | CASSFGSGYEKLFF | KLGGALQAK | NO | NO | NO |
| 3499 | TRAV25 | TRAJ42 | CAGPLGGSQGNLIF | TRBV6-5 | TRBJ1-4 | CASSTTGGKLFF | KLGGALQAK | NO | NO | NO |
| 3500 | TRAV38-2DV8 | TRAJ45 | CALVWGGGADGLTF | TRBV6-5 | TRBJ1-4 | CASSHGTGAEKLFF | RLRAEAQVK | NO | NO | NO |
| 3501 | TRAV38-2DV8 | TRAJ56 | CAYRSGSAGANSKLTF | TRBV6-5 | TRBJ1-4 | CASSRLTATNEKLFF | KLGGALQAK | NO | NO | NO |
| 3502 | TRAV6 | TRAJ37 | CARYGGSSNTGKLIF | TRBV6-5 | TRBJ1-4 | CASSFDRGSEKLFF | KLGGALQAK | NO | NO | NO |
| 3504 | TRAV9-2 | TRAJ39 | CALRVNNAGNMLTF | TRBV6-5 | TRBJ1-4 | CASSYSWDTTNEKLFF | KLGGALQAK | NO | NO | NO |
| 3505 | TRAV36DV7 | TRAJ49 | CAVDTPNTGNQFYF | TRBV6-6 | TRBJ1-4 | CASSIYRGLNEKLFF | KLGGALQAK | NO | NO | NO |
| 3506 | TRAV1-2 | TRAJ33 | CAVLDSNYQLIW | TRBV7-2 | TRBJ1-4 | CASSSGQEVEKLFF | KLGGALQAK | YES | NO | NO |
| 3507 | TRAV1-2 | TRAJ4 | CAVDLVRYNKLIF | TRBV7-6 | TRBJ1-4 | CASSLRGSNEKLFF | KLGGALQAK | YES | NO | NO |
| 3512 | TRAV16 | TRAJ58 | CAPRGSRLTF | TRBV7-8 | TRBJ1-4 | CASSSAAGRINEKLFF | KLGGALQAK | NO | NO | NO |
| 3513 | TRAV8-6 | TRAJ21 | CAVSAYNFNKFYF | TRBV7-8 | TRBJ1-4 | CASTGGADEKLFF | KLGGALQAK | YES | NO | NO |
| 3514 | TRAV1-2 | TRAJ16 | CAASDGQKLLF | TRBV7-9 | TRBJ1-4 | CASGRLSGDRGWDNEKLFF | KLGGALQAK | NO | NO | NO |
| 3515 | TRAV1-2 | TRAJ34 | CAVRAPLYNTDKLIF | TRBV7-9 | TRBJ1-4 | CASSRVQGEKLFF | KLGGALQAK | YES | NO | NO |
| 3517 | TRAV12-1 | TRAJ28 | CVVNSWSGAGSYQLTF | TRBV7-9 | TRBJ1-4 | CASINRDTGGELFF | KLGGALQAK | YES | NO | NO |
| 3518 | TRAV12-1 | TRAJ6 | CVAEPSGGSYIPTF | TRBV7-9 | TRBJ1-4 | CASSLATTDGGGKLFF | KLGGALQAK | YES | NO | NO |
| 3519 | TRAV12-2 | TRAJ17 | CAATGKAAGNKLTF | TRBV7-9 | TRBJ1-4 | CASSRPLEKLFF | KLGGALQAK | YES | NO | NO |
| 3520 | TRAV13-1 | TRAJ50 | CAASETSYDKVIF | TRBV7-9 | TRBJ1-4 | CASSLVGGTDEKLFF | KLGGALQAK | YES | NO | NO |
| 3520 | TRAV13-1 | TRAJ50 | CAASETSYDKVIF | TRBV7-9 | TRBJ1-4 | CASSLVGGTDEKLFF | IVTDFSVIK | YES | NO | NO |
| 3521 | TRAV14DV4 | TRAJ57 | CAMREHLQGGSEKLVF | TRBV7-9 | TRBJ1-4 | CASSLVRESEKLFF | KLGGALQAK | YES | NO | NO |
| 3522 | TRAV14DV4 | TRAJ9 | CAMREGPTGGFKTIF | TRBV7-9 | TRBJ1-4 | CASSYQGGRGREKLFF | KLGGALQAK | YES | NO | NO |
| 3523 | TRAV2 | TRAJ12 | CAVNMDSSYKLIF | TRBV7-9 | TRBJ1-4 | CASSSPDRHNEKLFF | KLGGALQAK | NO | NO | NO |
| 3525 | TRAV20 | TRAJ5 | CAVHSLDTGRRALTF | TRBV7-9 | TRBJ1-4 | CASSLADASTNEKLFF | KLGGALQAK | YES | NO | NO |
| 3526 | TRAV21 | TRAJ43 | CAVWYMRF | TRBV7-9 | TRBJ1-4 | CASSRVQGWEKLFF | KLGGALQAK | NO | NO | NO |
| 3527 | TRAV24 | TRAJ36 | CAFLGGANNLFF | TRBV7-9 | TRBJ1-4 | CASSLVGGTDEKLFF | KLGGALQAK | YES | NO | NO |

| | | | | | | | | | | |
|---|---|---|---|---|---|---|---|---|---|---|
| 3528 | TRAV26-1 | TRAJ28 | CIVEISGAGSYQLTF | TRBV7-9 | TRBJ1-4 | CASSPPRGWVANEKLFF | KLGGALQAK | YES | NO | NO |
| 3529 | TRAV26-2 | TRAJ50 | CILRDVLKTSYDKVIF | TRBV7-9 | TRBJ1-4 | CASSLGDAEKLFF | KLGGALQAK | YES | NO | NO |
| 3530 | TRAV29DV5 | TRAJ26 | CAARPDNYGQNFVF | TRBV7-9 | TRBJ1-4 | CASCGGLNEKLFF | KLGGALQAK | YES | NO | NO |
| 3531 | TRAV3 | TRAJ17 | CAVRDGFKAAGNKLTF | TRBV7-9 | TRBJ1-4 | CASSRGTGSNEKLFF | KLGGALQAK | YES | NO | NO |
| 3532 | TRAV38-2DV8 | TRAJ24 | CAYRSGWGKLQF | TRBV7-9 | TRBJ1-4 | CASSLERVSEKLFF | KLGGALQAK | YES | NO | NO |
| 3533 | TRAV38-2DV8 | TRAJ29 | CAYWAGNTPLVF | TRBV7-9 | TRBJ1-4 | CASSPPGVSGEKLFF | KLGGALQAK | YES | NO | NO |
| 3534 | TRAV38-2DV8 | TRAJ43 | CAYRSADRDDMRF | TRBV7-9 | TRBJ1-4 | CASSLDRVPANEKLFF | KLGGALQAK | YES | NO | NO |
| 3535 | TRAV38-2DV8 | TRAJ48 | CAYRSSLGEKLTF | TRBV7-9 | TRBJ1-4 | CASSAGRNEKLFF | KLGGALQAK | YES | NO | NO |
| 3536 | TRAV8-2 | TRAJ43 | CVVSDHGDMRF | TRBV7-9 | TRBJ1-4 | CASSAPGTEGEKLFF | KLGGALQAK | YES | NO | NO |
| 3537 | TRAV12-1 | TRAJ33 | CVVMDSNYQLIW | TRBV9 | TRBJ1-4 | CASSVDPGNSEKLFF | KLGGALQAK | YES | NO | NO |
| 3539 | TRAV12-3 | TRAJ39 | CAIGPGNMLTF | TRBV9 | TRBJ1-4 | CASSGHGAGSSEKLFF | KLGGALQAK | YES | NO | NO |
| 3540 | TRAV14DV4 | TRAJ20 | CAMREGENDYKLSF | TRBV9 | TRBJ1-4 | CASSSGQGGKLFF | KLGGALQAK | YES | NO | NO |
| 3542 | TRAV19 | TRAJ26 | CALGPYGQNFVF | TRBV9 | TRBJ1-4 | CASSAGGGATNEKLFF | KLGGALQAK | YES | NO | NO |
| 3544 | TRAV19 | TRAJ39 | CALKLGNAGNMLTF | TRBV9 | TRBJ1-4 | CASSVNPNPIRDRDNEKLFF | KLGGALQAK | NO | NO | NO |
| 3545 | TRAV21 | TRAJ8 | CAVAPIEDTGFQKLVF | TRBV9 | TRBJ1-4 | CASSVQQGPNEKLFF | KLGGALQAK | YES | NO | NO |
| 3547 | TRAV38-2DV8 | TRAJ45 | CATGGGADGLTF | TRBV9 | TRBJ1-4 | CASSSWDRDEGKLFF | KLGGALQAK | YES | NO | NO |
| 3550 | TRAV13-2 | TRAJ5 | CAETDQRMDTGRRALTF | TRBV10-2 | TRBJ1-5 | CASSEGGHPFNQPQHF | KLGGALQAK | YES | NO | NO |
| 3551 | TRAV29DV5 | TRAJ34 | CAAPLDKLIF | TRBV10-2 | TRBJ1-5 | CASSPGTGGFQPQHF | KLGGALQAK | YES | NO | NO |
| 3553 | TRAV1-2 | TRAJ6 | CAVGSSGGSYIPTF | TRBV10-3 | TRBJ1-5 | CAISTGDSNQPQHF | KLGGALQAK | YES | NO | NO |
| 3554 | TRAV12-1 | TRAJ5 | CVVDPDTGRRALTF | TRBV10-3 | TRBJ1-5 | CAISEFSGTSNQPQHF | KLGGALQAK | YES | NO | NO |
| 3555 | TRAV12-2 | TRAJ52 | CAGGAGGTSYGKLTF | TRBV10-3 | TRBJ1-5 | CAISSGGNNQPQHF | KLGGALQAK | YES | NO | NO |
| 3558 | TRAV14DV4 | TRAJ22 | CAMRRPISSGSARQLTF | TRBV10-3 | TRBJ1-5 | CAISESEGAIQPQHF | KLGGALQAK | YES | NO | NO |
| 3559 | TRAV14DV4 | TRAJ52 | CAMTPRGGTSYGKLTF | TRBV10-3 | TRBJ1-5 | CAISISELQPQHF | KLGGALQAK | YES | NO | NO |
| 3561 | TRAV20 | TRAJ52 | CAVRRAGGTSYGKLTF | TRBV10-3 | TRBJ1-5 | CAISESEGAIQPQHF | KLGGALQAK | YES | NO | NO |
| 3564 | TRAV9-2 | TRAJ13 | CASVRNSGGYQKVTF | TRBV10-3 | TRBJ1-5 | CATLGGQGADQPQHF | KLGGALQAK | YES | NO | NO |
| 3566 | TRAV14DV4 | TRAJ22 | CAMRRPISSGSARQLTF | TRBV11-2 | TRBJ1-5 | CASSSRTGAVQPQHF | KLGGALQAK | YES | NO | NO |
| 3570 | TRAV1-2 | TRAJ34 | CAVRRTDKLIF | TRBV12-3 | TRBJ1-5 | CASSSKTGGDQPQHF | KLGGALQAK | YES | NO | NO |
| 3571 | TRAV10 | TRAJ6 | CVVSTRSPSGGSYIPTF | TRBV12-3 | TRBJ1-5 | CASSLGIWNRSGLIQPQHF | KLGGALQAK | YES | NO | NO |
| 3574 | TRAV29DV5 | TRAJ27 | CAAFRAGKSTF | TRBV12-3 | TRBJ1-5 | CASSTGGTPQHF | KLGGALQAK | YES | NO | NO |
| 3577 | TRAV1-1 | TRAJ26 | CAVRDSDYGQNFVF | TRBV12-4 | TRBJ1-5 | CASSQQGGQPQHF | KLGGALQAK | YES | NO | NO |
| 3578 | TRAV1-2 | TRAJ24 | CATGGTTDSWGKLQF | TRBV12-4 | TRBJ1-5 | CASSLRVGLDQPQHF | KLGGALQAK | YES | NO | NO |
| 3580 | TRAV10 | TRAJ15 | CVVGPETGTALIF | TRBV12-4 | TRBJ1-5 | CASSWTIAQPQHF | KLGGALQAK | YES | NO | NO |
| 3581 | TRAV12-1 | TRAJ43 | CVVISSEGDMRF | TRBV12-4 | TRBJ1-5 | CASSHGGGMGQPQHF | KLGGALQAK | YES | NO | NO |
| 3582 | TRAV12-3 | TRAJ9 | CAMSEGTGGFKTIF | TRBV12-4 | TRBJ1-5 | CASSFTSRVFGQPQHF | KLGGALQAK | YES | NO | NO |
| 3583 | TRAV13-1 | TRAJ6 | CAATRSGGSYIPTF | TRBV12-4 | TRBJ1-5 | CASSPALWVSNQPQHF | KLGGALQAK | YES | NO | NO |
| 3584 | TRAV14DV4 | TRAJ49 | CAMRVMDTGNQFYF | TRBV12-4 | TRBJ1-5 | CASSFYGGDQPQHF | KLGGALQAK | YES | NO | NO |
| 3585 | TRAV19 | TRAJ27 | CALSGLTNAGKSTF | TRBV12-4 | TRBJ1-5 | CASSLGQGTANQPQHF | KLGGALQAK | YES | NO | NO |
| 3588 | TRAV20 | TRAJ34 | CAVQASDNTDKLIF | TRBV12-4 | TRBJ1-5 | CASSYRFGSNQPQHF | KLGGALQAK | YES | NO | NO |
| 3589 | TRAV21 | TRAJ10 | CAVRFTGGGNKLTF | TRBV12-4 | TRBJ1-5 | CASSLGTVWYSNQPQHF | KLGGALQAK | YES | NO | NO |
| 3590 | TRAV26-1 | TRAJ6 | CIVRPTGGSYIPTF | TRBV12-4 | TRBJ1-5 | CASSPTGQVYNQPQHF | KLGGALQAK | YES | NO | NO |
| 3592 | TRAV35 | TRAJ29 | CAGLLNSGNTPLVF | TRBV12-4 | TRBJ1-5 | CASSGQGFNQPQHF | KLGGALQAK | YES | NO | NO |
| 3596 | TRAV8-6 | TRAJ37 | CAVSEWDNTGKLIF | TRBV12-4 | TRBJ1-5 | CASSGSDQAQHF | RLRAEAQVK | YES | YES | YES |
| 3597 | TRAV12-3 | TRAJ37 | CAMSVGSGNTGKLIF | TRBV12-5 | TRBJ1-5 | CASGRTDNSNQPQHF | KLGGALQAK | YES | NO | NO |
| 3598 | TRAV23DV6 | TRAJ41 | CAASLLSNSGYALNF | TRBV12-5 | TRBJ1-5 | CASGFQGINQPQHF | KLGGALQAK | YES | NO | NO |
| 3600 | TRAV26-2 | TRAJ29 | CILRLDSGNTPLVF | TRBV12-5 | TRBJ1-5 | CASGLYPGQPQHF | KLGGALQAK | YES | NO | NO |
| 3600 | TRAV26-2 | TRAJ29 | CILRLDSGNTPLVF | TRBV12-5 | TRBJ1-5 | CASGLYPGQPQHF | RLRAEAQVK | YES | NO | NO |
| 3604 | TRAV10 | TRAJ45 | CVVTYSGGGADGLTF | TRBV15 | TRBJ1-5 | CATSRDLGINQPQHF | KLGGALQAK | YES | NO | NO |
| 3605 | TRAV12-2 | TRAJ13 | CAVNKAGGYQKVTF | TRBV15 | TRBJ1-5 | CATSSRQGMKVSQPQHF | KLGGALQAK | YES | NO | NO |
| 3606 | TRAV13-2 | TRAJ11 | CAENIEGYSTLTF | TRBV15 | TRBJ1-5 | CATSPTGDQPQHF | KLGGALQAK | YES | NO | NO |
| 3609 | TRAV17 | TRAJ8 | CATDGSTGFQKLVF | TRBV15 | TRBJ1-5 | CATSTGDSNQPQHF | KLGGALQAK | YES | NO | NO |
| 3610 | TRAV19 | TRAJ27 | CALSEQETNAGKSTF | TRBV15 | TRBJ1-5 | CATSRDLGGGNQPQHF | KLGGALQAK | YES | NO | NO |
| 3611 | TRAV19 | TRAJ50 | CALSERLKTSYDKVIF | TRBV15 | TRBJ1-5 | CATSGLWDRGLGNQPQHF | KLGGALQAK | YES | NO | NO |
| 3612 | TRAV20 | TRAJ4 | CAVQGYGTGGYNKLIF | TRBV15 | TRBJ1-5 | CATSRDVTGSNQPQHF | KLGGALQAK | YES | NO | NO |
| 3613 | TRAV27 | TRAJ22 | CAGDFSSGSARQLTF | TRBV15 | TRBJ1-5 | CATSREVGVLQPQHF | KLGGALQAK | YES | NO | NO |
| 3614 | TRAV3 | TRAJ7 | CAVRDRGNNRLAF | TRBV15 | TRBJ1-5 | CATSRDRGTGNQPQHF | KLGGALQAK | NO | NO | NO |
| 3615 | TRAV38-2DV8 | TRAJ39 | CAYRSGNNAGNMLTF | TRBV15 | TRBJ1-5 | CATSPRQGQNQPQHF | KLGGALQAK | YES | NO | NO |
| 3616 | TRAV5 | TRAJ40 | CAGAKSTSGTYKYIF | TRBV15 | TRBJ1-5 | CATSSDRDQPQHF | KLGGALQAK | YES | NO | NO |
| 3619 | TRAV21 | TRAJ32 | CAVRRYGGATNKLIF | TRBV16 | TRBJ1-5 | CASSQSDLPQHF | KLGGALQAK | YES | NO | NO |
| 3620 | TRAV19 | TRAJ42 | CALSEPLWEGGSQGNLIF | TRBV18 | TRBJ1-5 | CASSPREGSIFQPQHF | KLGGALQAK | YES | NO | NO |
| 3621 | TRAV24 | TRAJ52 | CAFIGDAGGTSYGKLTF | TRBV18 | TRBJ1-5 | CASSQGDGVQPQHF | KLGGALQAK | YES | NO | NO |
| 3622 | TRAV26-2 | TRAJ40 | CILRDRTSGTYKYIF | TRBV18 | TRBJ1-5 | CASSPIRGNQPQHF | KLGGALQAK | YES | NO | NO |
| 3624 | TRAV38-2DV8 | TRAJ21 | CAYRSRVYNFNKFYF | TRBV18 | TRBJ1-5 | CASSPGPQDPFYSNQPQHF | KLGGALQAK | YES | NO | NO |
| 3626 | TRAV1-2 | TRAJ40 | CAPHRSGTYKYIF | TRBV19 | TRBJ1-5 | CASSPRDRGMEPQHF | KLGGALQAK | YES | NO | NO |
| 3627 | TRAV1-2 | TRAJ6 | CAVRASGGSYIPTF | TRBV19 | TRBJ1-5 | CATDTRDSNQPQHF | KLGGALQAK | YES | NO | NO |
| 3629 | TRAV12-1 | TRAJ8 | CVVNRVNTGFQKLVF | TRBV19 | TRBJ1-5 | CASSTWDSNQPQHF | KLGGALQAK | YES | NO | NO |
| 3630 | TRAV12-1 | TRAJ8 | CVVNTMNTGFQKLVF | TRBV19 | TRBJ1-5 | CASTTWDSNQPQHF | KLGGALQAK | YES | NO | NO |
| 3631 | TRAV12-2 | TRAJ27 | CAVTSNAGKSTF | TRBV19 | TRBJ1-5 | CASSRDGNQPQHF | KLGGALQAK | YES | NO | NO |
| 3634 | TRAV13-1 | TRAJ43 | CAASGGGDNDMRF | TRBV19 | TRBJ1-5 | CASSIGARGSNQPQHF | KLGGALQAK | YES | NO | NO |
| 3635 | TRAV14DV4 | TRAJ12 | CAMRALGGSSYKLIF | TRBV19 | TRBJ1-5 | CASSIDSGTSSPQHF | KLGGALQAK | YES | NO | NO |
| 3639 | TRAV14DV4 | TRAJ53 | CAMRDANSGGSNYKLTF | TRBV19 | TRBJ1-5 | CASSISGHTQPQHF | KLGGALQAK | YES | NO | NO |
| 3640 | TRAV17 | TRAJ23 | CATDHYNQGGKLIF | TRBV19 | TRBJ1-5 | CASDRASNQPQHF | KLGGALQAK | YES | NO | NO |
| 3641 | TRAV17 | TRAJ30 | CATDWNRDDKIIF | TRBV19 | TRBJ1-5 | CASSTGGFNQPQHF | KLGGALQAK | YES | NO | NO |
| 3642 | TRAV17 | TRAJ48 | CATERTNFGNEKLTF | TRBV19 | TRBJ1-5 | CASSIGSGVRSNQPQHF | KLGGALQAK | YES | NO | NO |
| 3643 | TRAV17 | TRAJ52 | CATDAGDAGGTSYGKLTF | TRBV19 | TRBJ1-5 | CASSIVGAGNQPQHF | KLGGALQAK | YES | NO | NO |
| 3644 | TRAV17 | TRAJ52 | CATDAGDAGGTSYGKLTF | TRBV19 | TRBJ1-5 | CASSLVGQGNQPQHF | KLGGALQAK | YES | NO | NO |
| 3645 | TRAV17 | TRAJ52 | CATDAGDAGGTSYGKLTF | TRBV19 | TRBJ1-5 | CASSIVGQGNQPQHF | KLGGALQAK | YES | NO | NO |
| 3647 | TRAV19 | TRAJ30 | CALTMNRDDKIIF | TRBV19 | TRBJ1-5 | CASSFNPPNQPQHF | KLGGALQAK | YES | NO | NO |
| 3651 | TRAV21 | TRAJ8 | CAVGVKMNTGFQKLVF | TRBV19 | TRBJ1-5 | CASSIPDRGEAQPQHF | KLGGALQAK | YES | NO | NO |
| 3653 | TRAV25 | TRAJ33 | CAAMDSNYQLIW | TRBV19 | TRBJ1-5 | CASSVGFTGGNQPQHF | KLGGALQAK | YES | NO | NO |
| 3654 | TRAV26-1 | TRAJ8 | CIVRVGRGFQKLVF | TRBV19 | TRBJ1-5 | CASSMRIFPQPQHF | KLGGALQAK | YES | NO | NO |
| 3655 | TRAV26-2 | TRAJ33 | CIRYNYQLIW | TRBV19 | TRBJ1-5 | CASSSEGTDNQPQHF | KLGGALQAK | YES | NO | NO |
| 3656 | TRAV27 | TRAJ40 | CAGVSSGTYKYIF | TRBV19 | TRBJ1-5 | CASSIASWAWGQPQHF | KLGGALQAK | YES | NO | NO |
| 3657 | TRAV29DV5 | TRAJ30 | CAAGGDDKIIF | TRBV19 | TRBJ1-5 | CASSITGGWRQPQHF | KLGGALQAK | YES | NO | NO |
| 3658 | TRAV29DV5 | TRAJ50 | CATKTSYDKVIF | TRBV19 | TRBJ1-5 | CASSTYSNQPQHF | KLGGALQAK | YES | NO | NO |
| 3659 | TRAV29DV5 | TRAJ52 | CAASEGGGTSYGKLTF | TRBV19 | TRBJ1-5 | CASSIGQGLNQPQHF | KLGGALQAK | YES | NO | NO |
| 3660 | TRAV3 | TRAJ17 | CAVRGMIKAAGNKLTF | TRBV19 | TRBJ1-5 | CASRGPSNQPQHF | KLGGALQAK | NO | NO | NO |
| 3663 | TRAV3 | TRAJ7 | CAVRGGVHGNNRLAF | TRBV19 | TRBJ1-5 | CASSISPNQPQHF | KLGGALQAK | NO | NO | NO |
| 3664 | TRAV35 | TRAJ52 | CAQRAGGTSYGKLTF | TRBV19 | TRBJ1-5 | CASSPAGQGDYQPQHF | KLGGALQAK | YES | NO | NO |
| 3665 | TRAV35 | TRAJ52 | CAGQLSDAGGTSYGKLTF | TRBV19 | TRBJ1-5 | CASSWDRVSNQPQHF | KLGGALQAK | YES | NO | NO |
| 3667 | TRAV4 | TRAJ28 | CLVGSGAGSYQLTF | TRBV19 | TRBJ1-5 | CASSGDRVGLELQPQHF | KLGGALQAK | NO | NO | NO |
| 3668 | TRAV41 | TRAJ30 | CAVFSGDDKIIF | TRBV19 | TRBJ1-5 | CASSSTGYGNQPQHF | KLGGALQAK | YES | NO | NO |
| 3670 | TRAV41 | TRAJ54 | CAVKSYQGAQKLVF | TRBV19 | TRBJ1-5 | CASSTDRYVRSNQPQHF | KLGGALQAK | YES | NO | NO |
| 3671 | TRAV6 | TRAJ43 | CALDNDMRF | TRBV19 | TRBJ1-5 | CASSLLPGNQPQHF | KLGGALQAK | NO | NO | NO |
| 3672 | TRAV8-2 | TRAJ23 | CVVSPERFYNQGGKLIF | TRBV19 | TRBJ1-5 | CASSIVGQPQHF | KLGGALQAK | YES | NO | NO |
| 3674 | TRAV8-3 | TRAJ24 | CAVGVISTDSWGKLQF | TRBV19 | TRBJ1-5 | CASSIGGGGNQPQHF | KLGGALQAK | YES | NO | NO |
| 3678 | TRAV12-1 | TRAJ23 | CVVNFGGKLIF | TRBV2 | TRBJ1-5 | CASSEAAENQPQHF | KLGGALQAK | YES | NO | NO |
| 3680 | TRAV16 | TRAJ21 | CALGDRYNFNKFYF | TRBV2 | TRBJ1-5 | CASSQGGGQPQHF | KLGGALQAK | NO | NO | NO |
| 3681 | TRAV21 | TRAJ33 | CAVEGWDSNYQLIW | TRBV2 | TRBJ1-5 | CASSEGQPQHF | KLGGALQAK | YES | NO | NO |
| 3682 | TRAV22 | TRAJ58 | CAVGVTSGSRLTF | TRBV2 | TRBJ1-5 | CASSPGQGYNQPQHF | KLGGALQAK | YES | NO | NO |
| 3683 | TRAV3 | TRAJ5 | CAVRDRYMDTGRRALTF | TRBV2 | TRBJ1-5 | CASSLGQGNQPQHF | KLGGALQAK | NO | NO | NO |
| 3685 | TRAV1-2 | TRAJ15 | CAVRVLNQAGTALIF | TRBV20-1 | TRBJ1-5 | CSARELTVQPQHF | KLGGALQAK | YES | NO | NO |
| 3687 | TRAV10 | TRAJ26 | CVVTPYGQNFVF | TRBV20-1 | TRBJ1-5 | CSAGTKRDRAQPQHF | KLGGALQAK | YES | NO | NO |
| 3688 | TRAV12-1 | TRAJ40 | CVVNPGRTYKYIF | TRBV20-1 | TRBJ1-5 | CSAKFPYSAAGQPQHF | KLGGALQAK | YES | NO | NO |
| 3689 | TRAV12-1 | TRAJ49 | CVANTGNQFYF | TRBV20-1 | TRBJ1-5 | CSARDLGQLNQPQHF | KLGGALQAK | YES | NO | NO |

| | | | | | | | | | | |
|---|---|---|---|---|---|---|---|---|---|---|
| 3691 | TRAV12-2 | TRAJ39 | CAVNAGNMLTF | TRBV20-1 | TRBJ1-5 | CSARDHRAEGQPQHF | KLGGALQAK | YES | NO | NO |
| 3694 | TRAV14DV4 | TRAJ17 | CAMREGSPIKAAGNKLTF | TRBV20-1 | TRBJ1-5 | CSARGGVAILNSNQPQHF | KLGGALQAK | YES | NO | NO |
| 3695 | TRAV14DV4 | TRAJ28 | CAMREGAYSGAGSYQLTF | TRBV20-1 | TRBJ1-5 | CSARGGDNQPQHF | KLGGALQAK | YES | NO | NO |
| 3697 | TRAV29DV5 | TRAJ29 | CAASAGSGNTPLVF | TRBV20-1 | TRBJ1-5 | CSARDDRGALSNQPQHF | KLGGALQAK | YES | NO | NO |
| 3699 | TRAV38-1 | TRAJ39 | CALIVDAGNMLTF | TRBV20-1 | TRBJ1-5 | CSARGPRGAGDYSNQPQHF | KLGGALQAK | YES | NO | NO |
| 3699 | TRAV38-1 | TRAJ39 | CALIVDAGNMLTF | TRBV20-1 | TRBJ1-5 | CSARGPRGAGDYSNQPQHF | AVFDRKSDAK | YES | NO | NO |
| 3700 | TRAV38-2DV8 | TRAJ50 | CAYRSADTSYDKVIF | TRBV20-1 | TRBJ1-5 | CSADTGTGGPNQPQHF | KLGGALQAK | YES | NO | NO |
| 3700 | TRAV38-2DV8 | TRAJ50 | CAYRSADTSYDKVIF | TRBV20-1 | TRBJ1-5 | CSADTGTGGPNQPQHF | AVFDRKSDAK | YES | NO | NO |
| 3701 | TRAV5 | TRAJ10 | CAEWGGGNKLTF | TRBV20-1 | TRBJ1-5 | CSAGDISNQPQHF | KLGGALQAK | YES | NO | NO |
| 3710 | TRAV1-2 | TRAJ32 | CAVRPLYGGATNKLIF | TRBV24-1 | TRBJ1-5 | CATSEGTVSNQPQHF | KLGGALQAK | YES | NO | NO |
| 3712 | TRAV14DV4 | TRAJ29 | CAMRGSASGNTPLVF | TRBV24-1 | TRBJ1-5 | CATSDNDRGHSQPQHF | KLGGALQAK | YES | NO | NO |
| 3713 | TRAV14DV4 | TRAJ49 | CAMREGPNTGNQFYF | TRBV24-1 | TRBJ1-5 | CATSDNDRGGGQPQHF | KLGGALQAK | YES | NO | NO |
| 3715 | TRAV29DV5 | TRAJ50 | CAAMKTSYDKVIF | TRBV24-1 | TRBJ1-5 | CATSDLPGPHFQPQHF | KLGGALQAK | YES | NO | NO |
| 3717 | TRAV5 | TRAJ4 | CAEFRFSGGYNKLIF | TRBV24-1 | TRBJ1-5 | CATSEGQGASQPQHF | KLGGALQAK | YES | NO | NO |
| 3719 | TRAV12-2 | TRAJ50 | CAAKTSYDKVIF | TRBV25-1 | TRBJ1-5 | CASSGHPGQGKPQHF | KLGGALQAK | YES | NO | NO |
| 3721 | TRAV3 | TRAJ26 | CAVINYGQNFVF | TRBV25-1 | TRBJ1-5 | CASSEPTGSQPQHF | KLGGALQAK | NO | NO | NO |
| 3722 | TRAV39 | TRAJ30 | CAVDMDPRDDKIIF | TRBV25-1 | TRBJ1-5 | CASSDSGEPQHF | KLGGALQAK | YES | NO | NO |
| 3723 | TRAV12-1 | TRAJ17 | CVVNDKAAGNKLTF | TRBV27 | TRBJ1-5 | CASSLGSSQPQHF | KLGGALQAK | NO | NO | NO |
| 3726 | TRAV12-3 | TRAJ3 | CASLPTGYSSASKIIF | TRBV27 | TRBJ1-5 | CASSLGVREAGSLNQPQHF | KLGGALQAK | YES | NO | NO |
| 3730 | TRAV13-1 | TRAJ44 | CAASISGTASKLTF | TRBV27 | TRBJ1-5 | CASSYQDGQPQHF | KLGGALQAK | YES | NO | NO |
| 3731 | TRAV13-1 | TRAJ8 | CAASGFQKLVF | TRBV27 | TRBJ1-5 | CASSLEGVEQPQHF | KLGGALQAK | YES | NO | NO |
| 3733 | TRAV13-2 | TRAJ6 | CAENGRGSYIPTF | TRBV27 | TRBJ1-5 | CASSHGQGDQPQHF | KLGGALQAK | YES | NO | NO |
| 3735 | TRAV14DV4 | TRAJ42 | CAMREPANMNYGGSQGNLIF | TRBV27 | TRBJ1-5 | CASRPQGFNQPQHF | KLGGALQAK | NO | NO | NO |
| 3736 | TRAV14DV4 | TRAJ5 | CAMREALPMDTGRRALTF | TRBV27 | TRBJ1-5 | CASSLIGESNQPQHF | KLGGALQAK | YES | NO | NO |
| 3737 | TRAV17 | TRAJ27 | CATDLNTNAGKSTF | TRBV27 | TRBJ1-5 | CASGTPNSNQPQHF | KLGGALQAK | YES | NO | NO |
| 3738 | TRAV19 | TRAJ13 | CALSEARFGYQKVTF | TRBV27 | TRBJ1-5 | CASSLSTGSNQPQHF | KLGGALQAK | NO | NO | NO |
| 3739 | TRAV19 | TRAJ24 | CALSEPEFRGKFQF | TRBV27 | TRBJ1-5 | CASSLDRVNQPQHF | KLGGALQAK | NO | NO | NO |
| 3740 | TRAV19 | TRAJ37 | CALTGSSNTGKLIF | TRBV27 | TRBJ1-5 | CASSLHGGGQPQHF | KLGGALQAK | YES | NO | NO |
| 3741 | TRAV19 | TRAJ54 | CALSEIQGAQKLVF | TRBV27 | TRBJ1-5 | CASGAANQPQHF | KLGGALQAK | YES | NO | NO |
| 3743 | TRAV21 | TRAJ28 | CAVSHSGAGSYQLTF | TRBV27 | TRBJ1-5 | CASSSLAGSNQPQHF | KLGGALQAK | YES | NO | NO |
| 3744 | TRAV21 | TRAJ42 | CAVPLYGGSQGNLIF | TRBV27 | TRBJ1-5 | CASSLAGLGNQPQHF | KLGGALQAK | YES | NO | NO |
| 3745 | TRAV21 | TRAJ48 | CAVLPENFGNEKLTF | TRBV27 | TRBJ1-5 | CASSPTTGPYSNQPQHF | KLGGALQAK | YES | NO | NO |
| 3746 | TRAV21 | TRAJ58 | CAVPNFRRTSGSRLTF | TRBV27 | TRBJ1-5 | CASSPGYIGGNQPQHF | KLGGALQAK | YES | YES | NO |
| 3747 | TRAV23DV6 | TRAJ13 | CAASRGGYQKVTF | TRBV27 | TRBJ1-5 | CASRYDSDSNQPQHF | KLGGALQAK | YES | NO | NO |
| 3748 | TRAV24 | TRAJ43 | CAFYNNDMRF | TRBV27 | TRBJ1-5 | CASSWTGGMWPQHF | KLGGALQAK | YES | NO | NO |
| 3752 | TRAV30 | TRAJ49 | CGTESNTGNQFYF | TRBV27 | TRBJ1-5 | CASSLLTGNQPQHF | KLGGALQAK | YES | NO | NO |
| 3753 | TRAV35 | TRAJ20 | CAGRRSNDYKLSF | TRBV27 | TRBJ1-5 | CASSLGDGSAQHF | KLGGALQAK | YES | NO | NO |
| 3754 | TRAV38-2DV8 | TRAJ4 | CAYRSAGAPFSGGYNKLIF | TRBV27 | TRBJ1-5 | CASSLLGSNQPQHF | KLGGALQAK | YES | NO | NO |
| 3756 | TRAV4 | TRAJ30 | CLVVSMNRDDKIIF | TRBV27 | TRBJ1-5 | CASSLSTANQPQHF | KLGGALQAK | YES | NO | NO |
| 3757 | TRAV41 | TRAJ54 | CAVQGAQKLVF | TRBV27 | TRBJ1-5 | CASSPEIGGNQPQHF | KLGGALQAK | YES | NO | NO |
| 3758 | TRAV5 | TRAJ43 | CAESMDNDMRF | TRBV27 | TRBJ1-5 | CASSWDRHKPQHF | KLGGALQAK | NO | NO | NO |
| 3759 | TRAV6 | TRAJ30 | CALDGGRDDKIIF | TRBV27 | TRBJ1-5 | CASSRGDNQPQHF | KLGGALQAK | YES | NO | NO |
| 3760 | TRAV8-1 | TRAJ40 | CAVPSSGTYKYIF | TRBV27 | TRBJ1-5 | CASSLGYGQEVEWPQHF | KLGGALQAK | YES | NO | NO |
| 3761 | TRAV8-2 | TRAJ12 | CVVSDFQWMDSSYKLIF | TRBV27 | TRBJ1-5 | CASRILGTGNQPQHF | KLGGALQAK | YES | NO | NO |
| 3762 | TRAV8-3 | TRAJ20 | CAVDDYKLSF | TRBV27 | TRBJ1-5 | CASSLSQNSNQPQHF | KLGGALQAK | YES | NO | NO |
| 3764 | TRAV8-4 | TRAJ45 | CAVSYSGGGADGLTF | TRBV27 | TRBJ1-5 | CASSWTGGMWPQHF | AVFDRKSDAK | YES | NO | NO |
| 3768 | TRAV19 | TRAJ21 | CALSEESYNFNKFYF | TRBV28 | TRBJ1-5 | CASSSLRGSNQPQHF | KLGGALQAK | YES | NO | NO |
| 3769 | TRAV20 | TRAJ32 | CAVQASGATNKLIF | TRBV28 | TRBJ1-5 | CASRETGGAQPQHF | KLGGALQAK | YES | NO | NO |
| 3770 | TRAV27 | TRAJ57 | CALTQGGSEKLVF | TRBV28 | TRBJ1-5 | CASSSPPRIYSNQPQHF | KLGGALQAK | YES | NO | NO |
| 3771 | TRAV8-4 | TRAJ8 | CAVSDRGVNTGFQKLVF | TRBV28 | TRBJ1-5 | CASSLENIRLQSNQPQHF | KLGGALQAK | YES | NO | NO |
| 3772 | TRAV12-2 | TRAJ31 | CAAGNNNARLMF | TRBV29-1 | TRBJ1-5 | CSVRQGSGNQPQHF | KLGGALQAK | YES | NO | NO |
| 3774 | TRAV27 | TRAJ53 | CAGDGGSNYKLTF | TRBV29-1 | TRBJ1-5 | CSVDRGNLKAQHF | KLGGALQAK | YES | NO | NO |
| 3775 | TRAV27 | TRAJ6 | CAGDWFGGSYIPTF | TRBV29-1 | TRBJ1-5 | CSVRQGSGNQPQHF | KLGGALQAK | YES | NO | NO |
| 3777 | TRAV38-2DV8 | TRAJ40 | CAYRSRTSGTYKYIF | TRBV29-1 | TRBJ1-5 | CSAMDSYQPQHF | KLGGALQAK | YES | NO | NO |
| 3779 | TRAV1-2 | TRAJ16 | CAVIDGQKLLF | TRBV3-1 | TRBJ1-5 | CASSQDMEPQHF | KLGGALQAK | NO | NO | NO |
| 3780 | TRAV12-2 | TRAJ32 | CAVNSGYGGATNKLIF | TRBV3-1 | TRBJ1-5 | CASSQERQVQPQHF | KLGGALQAK | YES | NO | NO |
| 3781 | TRAV12-2 | TRAJ33 | CAVLEDSNYQLIW | TRBV3-1 | TRBJ1-5 | CASSQGAEQWPQHF | KLGGALQAK | YES | NO | NO |
| 3784 | TRAV19 | TRAJ36 | CALSQLYSGANNLFF | TRBV3-1 | TRBJ1-5 | CASSQDLDRGPSGQPQHF | KLGGALQAK | YES | NO | NO |
| 3787 | TRAV29DV5 | TRAJ31 | CAASADARLMF | TRBV3-1 | TRBJ1-5 | CASSQDLDRGPSGQPQHF | KLGGALQAK | YES | NO | NO |
| 3789 | TRAV3 | TRAJ29 | CAVVGNTPLVF | TRBV3-1 | TRBJ1-5 | CASSQDLEDNQPQHF | KLGGALQAK | NO | NO | NO |
| 3791 | TRAV5 | TRAJ16 | CAESSVSDGQKLLF | TRBV3-1 | TRBJ1-5 | CASSLTDRVGQPQHF | AVFDRKSDAK | NO | NO | NO |
| 3793 | TRAV12-1 | TRAJ34 | CVVIPFTDKLIF | TRBV30 | TRBJ1-5 | CAWSSYGVSNQPQHF | KLGGALQAK | YES | NO | NO |
| 3794 | TRAV26-1 | TRAJ12 | CIVRLDSSYKLIF | TRBV30 | TRBJ1-5 | CAWTLGTGNQPQHF | KLGGALQAK | YES | NO | NO |
| 3796 | TRAV1-1 | TRAJ23 | CAWGGGKLIF | TRBV4-1 | TRBJ1-5 | CASSQGQGDQPQHF | KLGGALQAK | YES | NO | NO |
| 3797 | TRAV12-2 | TRAJ53 | CAVMGGGSNYKLTF | TRBV4-1 | TRBJ1-5 | CASSPPRYSNQPQHF | KLGGALQAK | YES | NO | NO |
| 3799 | TRAV12-3 | TRAJ39 | CAIGPGNMLTF | TRBV4-1 | TRBJ1-5 | CASSQDVARGNQPQHF | KLGGALQAK | YES | NO | NO |
| 3801 | TRAV13-1 | TRAJ43 | CAVSSVDNNDMRF | TRBV4-1 | TRBJ1-5 | CASSQVLGQANQPQHF | KLGGALQAK | YES | NO | NO |
| 3803 | TRAV19 | TRAJ43 | CALSEASNNDMRF | TRBV4-1 | TRBJ1-5 | CASSQAEGNFQPQHF | KLGGALQAK | YES | NO | NO |
| 3804 | TRAV19 | TRAJ57 | CALSEAKDEGSEKLVF | TRBV4-1 | TRBJ1-5 | CASSLSLHGNQPQHF | KLGGALQAK | YES | NO | NO |
| 3805 | TRAV23DV6 | TRAJ20 | CAARGYDYKLSF | TRBV4-1 | TRBJ1-5 | CASSPRDGFNQPQHF | KLGGALQAK | NO | NO | NO |
| 3806 | TRAV3 | TRAJ13 | CAVRASGGYQKVTF | TRBV4-1 | TRBJ1-5 | CASSQDGTAYSNQPQHF | KLGGALQAK | YES | NO | NO |
| 3808 | TRAV4 | TRAJ40 | CLRNSGTYKYIF | TRBV4-1 | TRBJ1-5 | CASSQDQQPQHF | KLGGALQAK | YES | NO | NO |
| 3810 | TRAV9-2 | TRAJ3 | CAPAYSSASKIIF | TRBV4-1 | TRBJ1-5 | CASSQDSTGSRGSNQPQHF | KLGGALQAK | YES | NO | NO |
| 3811 | TRAV12-1 | TRAJ31 | CVVPRNNNARLMF | TRBV4-2 | TRBJ1-5 | CASSQDRGLNQPQHF | KLGGALQAK | YES | NO | NO |
| 3812 | TRAV13-1 | TRAJ39 | CAARLGNMLTF | TRBV4-2 | TRBJ1-5 | CASSQGGPLSNQPQHF | KLGGALQAK | YES | NO | NO |
| 3813 | TRAV26-2 | TRAJ30 | CILRDRDDKIIF | TRBV4-2 | TRBJ1-5 | CASSQEGQLNQPQHF | KLGGALQAK | YES | NO | NO |
| 3815 | TRAV38-1 | TRAJ39 | CAFMSYNNAGNMLTF | TRBV4-2 | TRBJ1-5 | CASSHSVGSNQPQHF | KLGGALQAK | YES | NO | NO |
| 3816 | TRAV1-1 | TRAJ5 | CASTGRRALTF | TRBV4-3 | TRBJ1-5 | CASSQGTVLQPQHF | KLGGALQAK | YES | NO | NO |
| 3818 | TRAV1-2 | TRAJ5 | CASTGRRALTF | TRBV4-3 | TRBJ1-5 | CASSQGTVLQPQHF | KLGGALQAK | YES | NO | NO |
| 3820 | TRAV9-2 | TRAJ28 | CALSDPSGAGSYQLTF | TRBV4-3 | TRBJ1-5 | CASSQDRALNQPQHF | KLGGALQAK | YES | NO | NO |
| 3822 | TRAV13-2 | TRAJ15 | CAENPPSNQAGTALIF | TRBV5-1 | TRBJ1-5 | CASSAGTGIYQPQHF | KLGGALQAK | YES | NO | NO |
| 3823 | TRAV17 | TRAJ38 | CATDDDGNNRKLIW | TRBV5-1 | TRBJ1-5 | CASRRGWGDHQPQHF | KLGGALQAK | YES | NO | NO |
| 3825 | TRAV24 | TRAJ38 | CASDNAGNNRKLIW | TRBV5-1 | TRBJ1-5 | CASSLGQGVGQPQHF | KLGGALQAK | YES | NO | NO |
| 3826 | TRAV29DV5 | TRAJ53 | CAAKLKLTGEGSNYKLTF | TRBV5-1 | TRBJ1-5 | CASSLGSYSNQPQHF | KLGGALQAK | YES | NO | NO |
| 3827 | TRAV35 | TRAJ28 | CAGRKAGSYQLTF | TRBV5-1 | TRBJ1-5 | CASSLRGGQPQHF | KLGGALQAK | YES | NO | NO |
| 3828 | TRAV8-3 | TRAJ33 | CAVGADSNYQLIW | TRBV5-1 | TRBJ1-5 | CASSLGDSNQPQHF | KLGGALQAK | YES | NO | NO |
| 3830 | TRAV10 | TRAJ24 | CVFTDSWGKFQF | TRBV5-4 | TRBJ1-5 | CASSSGVGPQHF | KLGGALQAK | YES | NO | NO |
| 3833 | TRAV13-1 | TRAJ43 | CAANNNDMRF | TRBV5-4 | TRBJ1-5 | CASSSTGGGEKDQPQHF | KLGGALQAK | YES | NO | NO |
| 3838 | TRAV26-2 | TRAJ23 | CILIPIYNQGGKLIF | TRBV5-4 | TRBJ1-5 | CASSSTGGGEKDQPQHF | KLGGALQAK | YES | NO | NO |
| 3840 | TRAV38-2DV8 | TRAJ43 | CAYRSGNNNDMRF | TRBV5-4 | TRBJ1-5 | CASSLLNQPQHF | KLGGALQAK | YES | NO | NO |
| 3844 | TRAV1-2 | TRAJ15 | CAVRGVHQAGTALIF | TRBV5-5 | TRBJ1-5 | CASSPNSNQPQHF | KLGGALQAK | YES | NO | NO |
| 3846 | TRAV14DV4 | TRAJ47 | CAMREMEYGNKLVF | TRBV5-5 | TRBJ1-5 | CASSAQGALSFYSNQPQHF | KLGGALQAK | YES | NO | NO |
| 3847 | TRAV20 | TRAJ40 | CAVQENSGTYKYIF | TRBV5-5 | TRBJ1-5 | CASSLANQGYQPQHF | KLGGALQAK | YES | NO | NO |
| 3848 | TRAV29DV5 | TRAJ57 | CAASAVITQGGSEKLVF | TRBV5-5 | TRBJ1-5 | CASSLEGYSNQPQHF | KLGGALQAK | YES | NO | NO |
| 3852 | TRAV14DV4 | TRAJ4 | CAMRPPFRGGYNKLIF | TRBV5-6 | TRBJ1-5 | CASSSRDSQNQPQHF | AVFDRKSDAK | YES | NO | NO |
| 3853 | TRAV14DV4 | TRAJ4 | CAMRPPFRGGYNKLIF | TRBV5-6 | TRBJ1-5 | CVSSSRDSQNQPQHF | KLGGALQAK | YES | NO | NO |
| 3854 | TRAV14DV4 | TRAJ4 | CAMRPPFRGGYNKLIF | TRBV5-6 | TRBJ1-5 | CASSSRDSQNQPQHV | KLGGALQAK | NO | NO | NO |
| 3855 | TRAV17 | TRAJ45 | CAESGGGADGLTF | TRBV5-6 | TRBJ1-5 | CASSFWGPLRPQHF | KLGGALQAK | YES | NO | NO |
| 3859 | TRAV21 | TRAJ23 | CAVSPLYNQGGKLIF | TRBV5-6 | TRBJ1-5 | CASTKRGGTTSNQPQHF | KLGGALQAK | YES | NO | NO |
| 3861 | TRAV27 | TRAJ40 | CAGARGTYKYIF | TRBV5-6 | TRBJ1-5 | CASSFGRGYQPQHF | KLGGALQAK | YES | NO | NO |
| 3862 | TRAV29DV5 | TRAJ9 | CAASGRTGGFKTIF | TRBV5-6 | TRBJ1-5 | CASSFYRDGDHQPQHF | KLGGALQAK | YES | NO | NO |
| 3863 | TRAV3 | TRAJ32 | CAVRVLYGGATNKLIF | TRBV5-6 | TRBJ1-5 | CASSLEGGGLSNQPQHF | KLGGALQAK | YES | NO | NO |
| 3865 | TRAV41 | TRAJ32 | CAVHGGATNKLIF | TRBV5-6 | TRBJ1-5 | CASSLGLGGGRPQHF | KLGGALQAK | YES | NO | NO |
| 3866 | TRAV8-1 | TRAJ10 | CAVNAGITGGGNKLTF | TRBV5-6 | TRBJ1-5 | CASSRSLGNQPQHF | KLGGALQAK | YES | NO | NO |

| | | | | | | | | | |
|---|---|---|---|---|---|---|---|---|---|
| 3869 | TRAV1-2 | TRAJ20 | CAVRERDYKLSF | TRBV5-8 | TRBJ1-5 | CAYRGSNQPQHF | KLGGALQAK | YES | NO | NO |
| 3872 | TRAV21 | TRAJ12 | CAVRPRMDSSYKLIF | TRBV5-8 | TRBJ1-5 | CASSFSSGWDQPQHF | KLGGALQAK | YES | NO | NO |
| 3873 | TRAV25 | TRAJ24 | CAGRLVTDSWGKLQF | TRBV5-8 | TRBJ1-5 | CASSLAGAYSNQPQHF | KLGGALQAK | YES | NO | NO |
| 3874 | TRAV19 | TRAJ52 | CALGNAGGTSYGKLTF | TRBV6-1 | TRBJ1-5 | CASTIDGWGNQPQHF | KLGGALQAK | YES | NO | NO |
| 3875 | TRAV23DV6 | TRAJ43 | CAASGNHDMRF | TRBV6-1 | TRBJ1-5 | CASSEAPDGNQPQHF | KLGGALQAK | YES | NO | NO |
| 3877 | TRAV26-2 | TRAJ54 | CILRLYQGAQKLVF | TRBV6-1 | TRBJ1-5 | CASSWLGRVWGNQPQHF | KLGGALQAK | YES | NO | NO |
| 3878 | TRAV8-1 | TRAJ20 | CAVNAGDDYKLSF | TRBV6-1 | TRBJ1-5 | CASRGVQPQHF | KLGGALQAK | YES | NO | NO |
| 3879 | TRAV8-4 | TRAJ48 | CAVSGPFGNEKLTF | TRBV6-1 | TRBJ1-5 | CASRERDRGLNQPQHF | KLGGALQAK | YES | NO | NO |
| 3882 | TRAV12-2 | TRAJ24 | CAAPTDSWGKLQF | TRBV6-3 | TRBJ1-5 | CASSPGGHQPQHF | KLGGALQAK | YES | NO | NO |
| 3883 | TRAV12-2 | TRAJ24 | CAVNTDSWGKLQF | TRBV6-3 | TRBJ1-5 | CASSYGSNQPQHF | KLGGALQAK | YES | NO | NO |
| 3885 | TRAV13-1 | TRAJ23 | CAATSSYNQGGKLIF | TRBV6-3 | TRBJ1-5 | CASSPKALQLNSNQPQHF | KLGGALQAK | YES | NO | NO |
| 3886 | TRAV17 | TRAJ54 | CATDVQGAQKLVF | TRBV6-3 | TRBJ1-5 | CASSYISGGNQPQHF | KLGGALQAK | YES | NO | NO |
| 3888 | TRAV21 | TRAJ57 | CAVATQGGSEKLVF | TRBV6-3 | TRBJ1-5 | CASSYAVGNQPQHF | KLGGALQAK | YES | NO | NO |
| 3890 | TRAV27 | TRAJ26 | CAGAGDYGQNFVF | TRBV6-3 | TRBJ1-5 | CASSNRRGDQPQHF | KLGGALQAK | YES | NO | NO |
| 3892 | TRAV5 | TRAJ31 | CAESPNNARLMF | TRBV6-3 | TRBJ1-5 | CASSYSGTDTFYSNQPQHF | KLGGALQAK | YES | NO | NO |
| 3893 | TRAV5 | TRAJ45 | CAESITGVNLYSGGGADGLTF | TRBV6-3 | TRBJ1-5 | CASSYSTGADQPQHF | KLGGALQAK | NO | NO | NO |
| 3895 | TRAV8-4 | TRAJ20 | CAVIQINDYKLSF | TRBV6-3 | TRBJ1-5 | CASSYGTGQPQHF | KLGGALQAK | YES | NO | NO |
| 3897 | TRAV8-6 | TRAJ57 | CAVSYQGGSEKLVF | TRBV6-3 | TRBJ1-5 | CASSYSTGADQPQHF | KLGGALQAK | YES | NO | NO |
| 3900 | TRAV1-2 | TRAJ33 | CAVLDSNYQLIW | TRBV6-4 | TRBJ1-5 | CASSGTGNQPQHF | KLGGALQAK | YES | NO | NO |
| 3901 | TRAV2 | TRAJ30 | CAVPNRDDKIIF | TRBV6-4 | TRBJ1-5 | CASSDRTGTGRFNNQPQHF | KLGGALQAK | NO | NO | NO |
| 3902 | TRAV8-3 | TRAJ11 | CAVVMNSGYSTLTF | TRBV6-4 | TRBJ1-5 | CASSENSNQPQHF | KLGGALQAK | YES | NO | NO |
| 3904 | TRAV12-2 | TRAJ13 | CAVTQGYQKVTF | TRBV6-5 | TRBJ1-5 | CASSYLGEGQPQHF | KLGGALQAK | YES | NO | NO |
| 3905 | TRAV12-2 | TRAJ24 | CAALDSWGKFQF | TRBV6-5 | TRBJ1-5 | CASRTRGRDQPQHF | KLGGALQAK | YES | NO | NO |
| 3908 | TRAV12-3 | TRAJ48 | CAMSYTNFGNEKLTF | TRBV6-5 | TRBJ1-5 | CASSYSTGRPQHF | KLGGALQAK | YES | NO | NO |
| 3910 | TRAV14DV4 | TRAJ30 | CAMRERADDKIIF | TRBV6-5 | TRBJ1-5 | CASRSGPVNQPQHF | KLGGALQAK | YES | NO | NO |
| 3912 | TRAV14DV4 | TRAJ57 | CAMTVIPAKGGSEKLVF | TRBV6-5 | TRBJ1-5 | CASSIQPNSNQPQHF | KLGGALQAK | YES | NO | NO |
| 3913 | TRAV19 | TRAJ30 | CALSGKDDKIIF | TRBV6-5 | TRBJ1-5 | CASSYSPSFSNQPQHF | KLGGALQAK | YES | NO | NO |
| 3914 | TRAV19 | TRAJ5 | CALSEALGGRRALTF | TRBV6-5 | TRBJ1-5 | CASRSKTGSKQPQHF | AVFDRKSDAK | YES | NO | NO |
| 3914 | TRAV19 | TRAJ5 | CALSEALGGRRALTF | TRBV6-5 | TRBJ1-5 | CASRSKTGSKQPQHF | RLRAEAQVK | YES | NO | NO |
| 3916 | TRAV21 | TRAJ43 | CAVRYNNNDMRF | TRBV6-5 | TRBJ1-5 | CASSYGVGQPQHF | KLGGALQAK | YES | NO | NO |
| 3917 | TRAV21 | TRAJ52 | CAVMKDPNAGGTSYGKLTF | TRBV6-5 | TRBJ1-5 | CASSYAPNQPQHF | KLGGALQAK | YES | NO | NO |
| 3920 | TRAV38-1 | TRAJ30 | CAFDRDDKIIF | TRBV6-5 | TRBJ1-5 | CASSTQGGNQPQHF | KLGGALQAK | YES | NO | NO |
| 3921 | TRAV38-1 | TRAJ54 | CALIQGAQKLVF | TRBV6-5 | TRBJ1-5 | CASSPATGGHQPQHF | KLGGALQAK | YES | NO | NO |
| 3922 | TRAV38-2DV8 | TRAJ54 | CAYQGAQKLVF | TRBV6-5 | TRBJ1-5 | CASSYGGSSGPQHF | KLGGALQAK | YES | NO | NO |
| 3924 | TRAV8-2 | TRAJ26 | CVVSDLSNYGQNFVF | TRBV6-5 | TRBJ1-5 | CASSPSDRVVEPQHF | KLGGALQAK | YES | NO | NO |
| 3934 | TRAV17 | TRAJ54 | CATDGNQGAQKLVF | TRBV7-2 | TRBJ1-5 | CQQPRALPGQGFSSNQPQHF | KLGGALQAK | NO | NO | NO |
| 3935 | TRAV19 | TRAJ39 | CALSENNNAGNMLTF | TRBV7-2 | TRBJ1-5 | CASREGLGNNQPQHF | KLGGALQAK | YES | NO | NO |
| 3939 | TRAV13-1 | TRAJ18 | CAASRGDRGSTLGRLYF | TRBV7-6 | TRBJ1-5 | CASGEGQPQHF | KLGGALQAK | YES | NO | NO |
| 3942 | TRAV21 | TRAJ54 | CAVRSSLIQGAQKLVF | TRBV7-6 | TRBJ1-5 | CASSLAEGHQPQHF | KLGGALQAK | YES | NO | NO |
| 3943 | TRAV27 | TRAJ50 | CAGETSYDKVIF | TRBV7-6 | TRBJ1-5 | CASSLYAGGQPQHF | KLGGALQAK | YES | NO | NO |
| 3944 | TRAV5 | TRAJ12 | CAEPLMDSSYKLIF | TRBV7-6 | TRBJ1-5 | CASSLKTGAVNQPQHF | RLRAEAQVK | YES | YES | YES |
| 3948 | TRAV14DV4 | TRAJ53 | CAMRLYYSGGSNYKLTF | TRBV7-8 | TRBJ1-5 | CASSLDRTGGLGDRQPQHF | KLGGALQAK | YES | NO | NO |
| 3949 | TRAV19 | TRAJ26 | CALSEDNYGQNFVF | TRBV7-8 | TRBJ1-5 | CASSLSSRASNQPQHF | KLGGALQAK | YES | NO | NO |
| 3950 | TRAV19 | TRAJ29 | CALRNSGNTPLVF | TRBV7-8 | TRBJ1-5 | CASSLGPPNNQPQHF | KLGGALQAK | YES | NO | NO |
| 3952 | TRAV29DV5 | TRAJ52 | CAASALWGGGKLTF | TRBV7-8 | TRBJ1-5 | CASSFIPDRGNQPQHF | KLGGALQAK | NO | NO | NO |
| 3954 | TRAV10 | TRAJ4 | CVVRAGGYNKLIF | TRBV7-9 | TRBJ1-5 | CASSLTPGEYSNQPQHF | KLGGALQAK | YES | NO | NO |
| 3955 | TRAV12-3 | TRAJ37 | CAISSNTGKLIF | TRBV7-9 | TRBJ1-5 | CASSLSTGNSNQPQHF | KLGGALQAK | YES | NO | NO |
| 3956 | TRAV12-3 | TRAJ40 | CAMTGSNSGTYKYIF | TRBV7-9 | TRBJ1-5 | CASSLGTQPQHF | KLGGALQAK | YES | NO | NO |
| 3957 | TRAV13-1 | TRAJ30 | CAERDDKIIF | TRBV7-9 | TRBJ1-5 | CASSLMDSNQPQHF | KLGGALQAK | YES | NO | NO |
| 3959 | TRAV14DV4 | TRAJ22 | CAMRRPISSGSARQLTF | TRBV7-9 | TRBJ1-5 | CASSLGQGYQPQHF | KLGGALQAK | YES | NO | NO |
| 3960 | TRAV14DV4 | TRAJ24 | CAMRESDSWGKFQF | TRBV7-9 | TRBJ1-5 | CASSLVGAGENQPQHF | KLGGALQAK | YES | NO | NO |
| 3961 | TRAV14DV4 | TRAJ32 | CAMRGRYGGATNKLIF | TRBV7-9 | TRBJ1-5 | CASNSGVQPQHF | KLGGALQAK | YES | NO | NO |
| 3962 | TRAV14DV4 | TRAJ6 | CAMGPSGGSYIPTF | TRBV7-9 | TRBJ1-5 | CASSRGQINQPQHF | KLGGALQAK | YES | NO | NO |
| 3962 | TRAV14DV4 | TRAJ6 | CAMGPSGGSYIPTF | TRBV7-9 | TRBJ1-5 | CASSRGQINQPQHF | AVFDRKSDAK | YES | NO | NO |
| 3963 | TRAV17 | TRAJ13 | CATGEGYQKVTF | TRBV7-9 | TRBJ1-5 | CASSSNRDPIPQHF | KLGGALQAK | YES | NO | NO |
| 3964 | TRAV17 | TRAJ20 | CATTNDYKLSF | TRBV7-9 | TRBJ1-5 | CASSGALQDSNQPQHF | KLGGALQAK | YES | NO | NO |
| 3967 | TRAV20 | TRAJ30 | CAVQAKDDKIIF | TRBV7-9 | TRBJ1-5 | CASSFGVPHGWEDQPQHF | KLGGALQAK | YES | NO | NO |
| 3968 | TRAV26-2 | TRAJ43 | CILRDVEDMRF | TRBV7-9 | TRBJ1-5 | CASESDGTDEYSNQPQHF | KLGGALQAK | YES | NO | NO |
| 3969 | TRAV27 | TRAJ57 | CAGAWGAQGGSEKLVF | TRBV7-9 | TRBJ1-5 | CASSFGGYQPQHF | KLGGALQAK | YES | NO | NO |
| 3971 | TRAV30 | TRAJ57 | CGTEEVNKGGSEKLVF | TRBV7-9 | TRBJ1-5 | CASSSVRDRDRSQPQHF | KLGGALQAK | YES | NO | NO |
| 3973 | TRAV4 | TRAJ43 | CLVGEGYNNNDMRF | TRBV7-9 | TRBJ1-5 | CASSLVDTTNQPQHF | KLGGALQAK | YES | NO | NO |
| 3978 | TRAV8-6 | TRAJ45 | CAVIPQGDGLTF | TRBV7-9 | TRBJ1-5 | CASSFMDGGQPQHF | KLGGALQAK | YES | NO | NO |
| 3981 | TRAV13-1 | TRAJ48 | CAAKVYGNEKLTF | TRBV9 | TRBJ1-5 | CASSVGDSPPQPQHF | KLGGALQAK | YES | NO | NO |
| 3982 | TRAV14DV4 | TRAJ49 | CAMREDLTGNQFYF | TRBV9 | TRBJ1-5 | CASSVSGVGPQHF | KLGGALQAK | YES | NO | NO |
| 3985 | TRAV29DV5 | TRAJ45 | CAASDWAGGGADGLTF | TRBV9 | TRBJ1-5 | CASSATPLRQPQHF | KLGGALQAK | NO | NO | NO |
| 3986 | TRAV36DV7 | TRAJ37 | CAARHGSSNTGKLIF | TRBV9 | TRBJ1-5 | CASSVGSGRDQPQHF | KLGGALQAK | NO | NO | NO |
| 3988 | TRAV17 | TRAJ39 | CATGYNNAGNMLTF | TRBV10-1 | TRBJ1-6 | CASQGSNSPLHF | KLGGALQAK | YES | NO | NO |
| 3990 | TRAV41 | TRAJ48 | CAVRARGFGNEKLTF | TRBV10-3 | TRBJ1-6 | CAISKIWGLGSSPLHF | KLGGALQAK | YES | NO | NO |
| 3993 | TRAV12-2 | TRAJ45 | CAVNGAYSGGGADGLTF | TRBV11-1 | TRBJ1-6 | CASSPGLGSPLHF | KLGGALQAK | YES | NO | NO |
| 4000 | TRAV21 | TRAJ33 | CAVSNYQLIW | TRBV11-2 | TRBJ1-6 | CASSLAGWSYNSPLHF | KLGGALQAK | YES | NO | NO |
| 4003 | TRAV29DV5 | TRAJ49 | CAGTGNQFYF | TRBV11-2 | TRBJ1-6 | CASSFEQGNSPLHF | RLRAEAQVK | YES | NO | NO |
| 4003 | TRAV29DV5 | TRAJ49 | CAGTGNQFYF | TRBV11-2 | TRBJ1-6 | CASSFEQGNSPLHF | KLGGALQAK | YES | NO | NO |
| 4005 | TRAV17 | TRAJ26 | CATDSYGQNFVF | TRBV12-3 | TRBJ1-6 | CASSFGIDSPLHF | KLGGALQAK | YES | NO | NO |
| 4007 | TRAV24 | TRAJ52 | CARHPANAGGTSYGKLTF | TRBV12-4 | TRBJ1-6 | CASSLELGGQFPWDSPLHF | RLRAEAQVK | YES | YES | YES |
| 4008 | TRAV29DV5 | TRAJ45 | CAAKWGGGADGLTF | TRBV12-4 | TRBJ1-6 | CASSFNAALNSPLHF | KLGGALQAK | YES | NO | NO |
| 4011 | TRAV17 | TRAJ20 | CATDGDYKLSF | TRBV13 | TRBJ1-6 | CASRGSGNSPLHF | KLGGALQAK | YES | NO | NO |
| 4012 | TRAV19 | TRAJ49 | CALRAGNQFYF | TRBV13 | TRBJ1-6 | CASSLMRGGTYNSPLHF | AVFDRKSDAK | YES | NO | NO |
| 4013 | TRAV21 | TRAJ37 | CAVVLFGSSNTGKLIF | TRBV13 | TRBJ1-6 | CASSLAAGAVNSPLHF | KLGGALQAK | YES | NO | NO |
| 4014 | TRAV25 | TRAJ26 | CAGQEGQNFVF | TRBV13 | TRBJ1-6 | CASRIKGAWNSPLHF | KLGGALQAK | YES | NO | NO |
| 4016 | TRAV8-1 | TRAJ26 | CAVNAGNYGQNFVF | TRBV13 | TRBJ1-6 | CASSLGDRDDSPLHF | KLGGALQAK | YES | NO | NO |
| 4019 | TRAV12-1 | TRAJ28 | CVVWGYSGAGSYQLTF | TRBV14 | TRBJ1-6 | CASSQERNSPLHF | KLGGALQAK | NO | NO | NO |
| 4023 | TRAV21 | TRAJ43 | CAVWYMRF | TRBV15 | TRBJ1-6 | CATSTRTGLISPLHF | KLGGALQAK | NO | NO | NO |
| 4024 | TRAV24 | TRAJ39 | CAFMDNAGNMLTF | TRBV15 | TRBJ1-6 | CATSRDLGGSYNSPLHF | KLGGALQAK | YES | NO | NO |
| 4025 | TRAV38-1 | TRAJ43 | CAFIPLNDMRF | TRBV15 | TRBJ1-6 | CATSSSRDSNSPLHF | KLGGALQAK | YES | NO | NO |
| 4026 | TRAV21 | TRAJ54 | CAVRSSLIQGAQKLVF | TRBV18 | TRBJ1-6 | CASSPEGQTGSPLHF | KLGGALQAK | YES | NO | NO |
| 4027 | TRAV8-4 | TRAJ7 | CAVSDRYGNNRLAF | TRBV18 | TRBJ1-6 | CASSPPAGGQNSPLHF | KLGGALQAK | YES | NO | NO |
| 4028 | TRAV1-1 | TRAJ33 | CAVSHMDSNYQLIW | TRBV19 | TRBJ1-6 | CASGRDSQPNYNSPLHF | KLGGALQAK | YES | NO | NO |
| 4029 | TRAV12-1 | TRAJ37 | CVVNTGNTGKLIF | TRBV19 | TRBJ1-6 | CASRTGGNSPLHF | KLGGALQAK | YES | NO | NO |
| 4030 | TRAV12-1 | TRAJ8 | CVVNTGFQKLVF | TRBV19 | TRBJ1-6 | CASSVGPGESPLHF | KLGGALQAK | YES | NO | NO |
| 4032 | TRAV13-2 | TRAJ37 | CAVDSSNTGKLIF | TRBV19 | TRBJ1-6 | CASTNPGTTLHF | KLGGALQAK | YES | NO | NO |
| 4033 | TRAV17 | TRAJ52 | CATDALGAGGTSYGKLTF | TRBV19 | TRBJ1-6 | CASSIDVTGKGSPLHF | KLGGALQAK | YES | NO | NO |
| 4034 | TRAV19 | TRAJ23 | CALSEAGFYNQGGKLIF | TRBV19 | TRBJ1-6 | CASSIVSGDSYNSPLHF | KLGGALQAK | YES | NO | NO |
| 4035 | TRAV19 | TRAJ49 | CALSARTGNQFYF | TRBV19 | TRBJ1-6 | CASLDSSSRYNSPLHF | KLGGALQAK | YES | NO | NO |
| 4036 | TRAV20 | TRAJ20 | CAVLSNDYKLSF | TRBV19 | TRBJ1-6 | CASSSTGWGGSSPLHF | KLGGALQAK | YES | NO | NO |
| 4037 | TRAV21 | TRAJ38 | CAVSPVNAGNNRKLIW | TRBV19 | TRBJ1-6 | CASSMGRRSSYNSPLHF | KLGGALQAK | YES | NO | NO |
| 4038 | TRAV21 | TRAJ43 | CAVTPPGDMRF | TRBV19 | TRBJ1-6 | CASSIGQGQSPLHF | KLGGALQAK | YES | NO | NO |
| 4039 | TRAV26-1 | TRAJ40 | CIVRVASASGTYKYIF | TRBV19 | TRBJ1-6 | CASSHGTGYNSPLHF | KLGGALQAK | YES | NO | NO |
| 4040 | TRAV26-2 | TRAJ53 | CILIKPVWNSGGSNYKLTF | TRBV19 | TRBJ1-6 | CASSMGGSSYNSPLHF | KLGGALQAK | YES | NO | NO |
| 4041 | TRAV29DV5 | TRAJ34 | CAASVKTDKLIF | TRBV19 | TRBJ1-6 | CASTDRQGADSPLHF | KLGGALQAK | YES | NO | NO |
| 4042 | TRAV38-1 | TRAJ42 | CAFMNYGGSQGNLIF | TRBV19 | TRBJ1-6 | CASSIVPLGQGLGTLHF | KLGGALQAK | NO | NO | NO |
| 4045 | TRAV1-2 | TRAJ43 | CAVRDYNNNDMRF | TRBV2 | TRBJ1-6 | CASSKTEAGSPLHF | KLGGALQAK | YES | NO | NO |
| 4046 | TRAV12-1 | TRAJ12 | CVVNSMDSSYKLIF | TRBV2 | TRBJ1-6 | CASRGLGGTDSPLHF | KLGGALQAK | YES | NO | NO |
| 4047 | TRAV12-1 | TRAJ24 | CVVILTTDSWGKFQF | TRBV2 | TRBJ1-6 | CASSETGGVGSPLHF | KLGGALQAK | YES | NO | NO |
| 4049 | TRAV27 | TRAJ29 | CAGPSGNTPLVF | TRBV2 | TRBJ1-6 | CASTSQQAHDSPLHF | KLGGALQAK | YES | NO | NO |

| | | | | | | | | | | |
|---|---|---|---|---|---|---|---|---|---|---|
| 4050 | TRAV4 | TRAJ45 | CLVAEGSGGGADGLTF | TRBV2 | TRBJ1-6 | CASSEVTGAVNSPLHF | KLGGALQAK | YES | NO | NO |
| 4051 | TRAV8-4 | TRAJ50 | CAVSRNGPYDKVIF | TRBV2 | TRBJ1-6 | CASQQSGSSYNSPLHF | KLGGALQAK | YES | NO | NO |
| 4052 | TRAV1-2 | TRAJ33 | CAVMDSNYQLIW | TRBV20-1 | TRBJ1-6 | CSARRGTGGSPLHF | KLGGALQAK | YES | NO | NO |
| 4053 | TRAV1-2 | TRAJ33 | CASMDSNYQLIW | TRBV20-1 | TRBJ1-6 | CSLRTGEGSPLHF | KLGGALQAK | NO | NO | NO |
| 4056 | TRAV17 | TRAJ54 | CATDVLIQGAQKLVF | TRBV20-1 | TRBJ1-6 | CSARDGSVRVEYNSPLHF | KLGGALQAK | NO | NO | NO |
| 4058 | TRAV29DV5 | TRAJ27 | CAASEDTNAGKSTF | TRBV20-1 | TRBJ1-6 | CSAIHPRQSGSPLHF | KLGGALQAK | NO | NO | NO |
| 4061 | TRAV3 | TRAJ8 | CAVPEVDTGFQKLVF | TRBV20-1 | TRBJ1-6 | CSARDSQASSSYNSPLHF | KLGGALQAK | NO | NO | NO |
| 4062 | TRAV5 | TRAJ16 | CAESFKDGQKLLF | TRBV20-1 | TRBJ1-6 | CSATGTGYNSPLHF | KLGGALQAK | NO | NO | NO |
| 4068 | TRAV21 | TRAJ5 | CAVLGANAGRRALTF | TRBV25-1 | TRBJ1-6 | CASSDRDTDYNSPLHF | KLGGALQAK | YES | NO | NO |
| 4069 | TRAV9-2 | TRAJ39 | CALRDNNAGNMLTF | TRBV25-1 | TRBJ1-6 | CASSELEKGGNSPLHF | KLGGALQAK | YES | NO | NO |
| 4070 | TRAV1-2 | TRAJ31 | CAVRDNNARLMF | TRBV27 | TRBJ1-6 | CASSSGQGNSPLHF | KLGGALQAK | YES | NO | NO |
| 4071 | TRAV1-2 | TRAJ7 | CAVRVEGNNRLAF | TRBV27 | TRBJ1-6 | CASSLYGYNSPLHF | KLGGALQAK | YES | NO | NO |
| 4072 | TRAV12-1 | TRAJ27 | CVALSNAGKSTF | TRBV27 | TRBJ1-6 | CAIGNNSPLHF | KLGGALQAK | YES | NO | NO |
| 4073 | TRAV12-2 | TRAJ15 | CAVIQRSQAGTALIF | TRBV27 | TRBJ1-6 | CASRKTEFSYNSPLHF | KLGGALQAK | YES | NO | NO |
| 4074 | TRAV12-3 | TRAJ47 | CAMNREYGNKLVF | TRBV27 | TRBJ1-6 | CATSGLSYNSPLHF | KLGGALQAK | YES | NO | NO |
| 4075 | TRAV13-1 | TRAJ43 | CAPHNNDMRF | TRBV27 | TRBJ1-6 | CASKDRGDYSPLHF | KLGGALQAK | YES | NO | NO |
| 4076 | TRAV13-1 | TRAJ7 | CAASVYGNNRLAF | TRBV27 | TRBJ1-6 | CASSLQNSPLHF | KLGGALQAK | YES | NO | NO |
| 4077 | TRAV13-1 | TRAJ9 | CAALFTGGFKTIF | TRBV27 | TRBJ1-6 | CASTHQSGTGSPLHF | KLGGALQAK | YES | NO | NO |
| 4078 | TRAV13-2 | TRAJ15 | CAENIFQAGTALIF | TRBV27 | TRBJ1-6 | CASSLSKGLSSPLHF | KLGGALQAK | YES | NO | NO |
| 4079 | TRAV14DV4 | TRAJ53 | CAMRPLLGGSNYKLTF | TRBV27 | TRBJ1-6 | CASYPGTENNSPLHF | KLGGALQAK | YES | NO | NO |
| 4080 | TRAV14DV4 | TRAJ54 | CAMRQSKEGAQKLVF | TRBV27 | TRBJ1-6 | CASHRALYNSPLHF | AVFDRKSDAK | YES | NO | NO |
| 4080 | TRAV14DV4 | TRAJ54 | CAMRQSKEGAQKLVF | TRBV27 | TRBJ1-6 | CASHRALYNSPLHF | KLGGALQAK | YES | NO | NO |
| 4082 | TRAV17 | TRAJ45 | CATALMYSGGGADGLTF | TRBV27 | TRBJ1-6 | CASSPPPWGTNSPLHF | KLGGALQAK | YES | NO | NO |
| 4085 | TRAV19 | TRAJ43 | CALSEASNNDMRF | TRBV27 | TRBJ1-6 | CASSLSQFSPLHF | KLGGALQAK | YES | NO | NO |
| 4088 | TRAV21 | TRAJ17 | CAVDKAAGNKLTF | TRBV27 | TRBJ1-6 | CASSLLQGLDSPLHF | KLGGALQAK | YES | NO | NO |
| 4089 | TRAV21 | TRAJ34 | CAVYYGDKLIF | TRBV27 | TRBJ1-6 | CASSLRQGVGNSPLHF | KLGGALQAK | YES | NO | NO |
| 4090 | TRAV25 | TRAJ16 | CARPLVSDGQKLLF | TRBV27 | TRBJ1-6 | CASSLTESNSPLHF | KLGGALQAK | NO | NO | NO |
| 4091 | TRAV26-2 | TRAJ29 | CILFSGNTPLVF | TRBV27 | TRBJ1-6 | CASSLMMAGNSPLHF | KLGGALQAK | YES | NO | NO |
| 4093 | TRAV39 | TRAJ22 | CAATPYSSGSARQLTF | TRBV27 | TRBJ1-6 | CASSYQGGAYNSPLHF | KLGGALQAK | YES | NO | NO |
| 4095 | TRAV6 | TRAJ7 | CALAVYGNNRLAF | TRBV27 | TRBJ1-6 | CASSLPTGGQNSPLHF | KLGGALQAK | YES | NO | NO |
| 4096 | TRAV8-2 | TRAJ42 | CVVTVRTHGGSQGNLIF | TRBV27 | TRBJ1-6 | CASSPRGSPLHF | KLGGALQAK | YES | NO | NO |
| 4097 | TRAV8-3 | TRAJ20 | CAGLQNKWDYKLSF | TRBV27 | TRBJ1-6 | CASSLSPGVIFSYNSPLHF | KLGGALQAK | YES | NO | NO |
| 4098 | TRAV8-4 | TRAJ30 | CAVSERLWDDKIIF | TRBV27 | TRBJ1-6 | CASSLPGTGGSPLHF | KLGGALQAK | YES | NO | NO |
| 4100 | TRAV8-6 | TRAJ27 | CAVSDHTNAGKSTF | TRBV27 | TRBJ1-6 | CASKIVDEVGALHF | KLGGALQAK | YES | NO | NO |
| 4101 | TRAV13-1 | TRAJ3 | CAALSHASKIIF | TRBV28 | TRBJ1-6 | CASRPQQGHNSPLHF | KLGGALQAK | YES | NO | NO |
| 4101 | TRAV13-1 | TRAJ3 | CAALSHASKIIF | TRBV28 | TRBJ1-6 | CASRPQQGHNSPLHF | RLRAEAQVK | YES | NO | NO |
| 4104 | TRAV29DV5 | TRAJ32 | CAASADGATNKLIF | TRBV28 | TRBJ1-6 | CASSLRSLYNSPLHF | KLGGALQAK | YES | NO | NO |
| 4107 | TRAV8-4 | TRAJ45 | CAVTPDAGGGADGLTF | TRBV28 | TRBJ1-6 | CASSLGLKNSPLHF | KLGGALQAK | YES | NO | NO |
| 4108 | TRAV8-4 | TRAJ48 | CAVRDISNFGNEKLTF | TRBV28 | TRBJ1-6 | CASSRNRDNSPLHF | KLGGALQAK | YES | NO | NO |
| 4109 | TRAV9-2 | TRAJ18 | CALSVSDRGSTLGRLYF | TRBV28 | TRBJ1-6 | CASSLYGRKNSPLHF | KLGGALQAK | YES | NO | NO |
| 4110 | TRAV1-2 | TRAJ6 | CAATEGAAGGSYIPTF | TRBV3-1 | TRBJ1-6 | CASSEDWTGGYNSPLHF | KLGGALQAK | YES | NO | NO |
| 4111 | TRAV19 | TRAJ38 | CALSDPSHNAGNNRKLIW | TRBV3-1 | TRBJ1-6 | CASSQGQGTLPLHF | KLGGALQAK | YES | NO | NO |
| 4112 | TRAV41 | TRAJ42 | CGCMNYGGSQGNLIF | TRBV3-1 | TRBJ1-6 | CASSQVLLRVSSPLHF | KLGGALQAK | YES | NO | NO |
| 4113 | TRAV9-2 | TRAJ20 | CALDDYKLSF | TRBV3-1 | TRBJ1-6 | CASSQDRVKGLYNSPLHF | KLGGALQAK | YES | NO | NO |
| 4115 | TRAV12-3 | TRAJ47 | CAMSGYMEYGNKLVF | TRBV30 | TRBJ1-6 | CASKTDNYNSPLHF | KLGGALQAK | YES | NO | NO |
| 4116 | TRAV16 | TRAJ33 | CALFPHYQLIW | TRBV30 | TRBJ1-6 | CAWSANSPLHF | KLGGALQAK | NO | NO | NO |
| 4119 | TRAV25 | TRAJ52 | CAENAGGTSYGKLTF | TRBV30 | TRBJ1-6 | CAWSIGQPYSPLHF | KLGGALQAK | YES | NO | NO |
| 4121 | TRAV17 | TRAJ35 | CATDTI | TRBV4-1 | TRBJ1-6 | CASSQILPYNSPLHF | KLGGALQAK | NO | NO | NO |
| 4122 | TRAV26-2 | TRAJ13 | CIGAGMNSGGYQKVTF | TRBV4-1 | TRBJ1-6 | CASSQVGAGHNSPLHF | KLGGALQAK | YES | NO | NO |
| 4123 | TRAV27 | TRAJ39 | CAGAYINNAGNMLTF | TRBV4-1 | TRBJ1-6 | CASSQGLGGSPLHF | KLGGALQAK | YES | NO | NO |
| 4125 | TRAV38-2DV8 | TRAJ12 | CAYRSGDSSYKLIF | TRBV4-1 | TRBJ1-6 | CASSQGAGPNSPLHF | KLGGALQAK | YES | NO | NO |
| 4127 | TRAV8-2 | TRAJ9 | CVVSDNTGGFKTIF | TRBV4-1 | TRBJ1-6 | CASSQGTGVAFNSPLHF | KLGGALQAK | YES | NO | NO |
| 4128 | TRAV17 | TRAJ47 | CATGTMEYGNKLVF | TRBV4-2 | TRBJ1-6 | CASSQGGLGVSRLGSPLHF | KLGGALQAK | YES | NO | NO |
| 4129 | TRAV21 | TRAJ45 | CAVTQYSGGGADGLTF | TRBV4-2 | TRBJ1-6 | CASSQDNPLNSPLHF | KLGGALQAK | YES | NO | NO |
| 4131 | TRAV19 | TRAJ54 | CALSLIQGAQKLVF | TRBV4-3 | TRBJ1-6 | CASSQGHGQGAFSNSPLHF | KLGGALQAK | YES | NO | NO |
| 4132 | TRAV22 | TRAJ47 | CAVEPLYGNKLVF | TRBV4-3 | TRBJ1-6 | CASSQTSPGHGVSPLHF | KLGGALQAK | YES | NO | NO |
| 4133 | TRAV26-2 | TRAJ45 | CILSRQEGGADGLTF | TRBV4-3 | TRBJ1-6 | CASSQSGTYNSPLHF | KLGGALQAK | YES | NO | NO |
| 4134 | TRAV3 | TRAJ31 | CAVRDTIFGNNARLMF | TRBV4-3 | TRBJ1-6 | CASSPKTDSYNSPLHF | KLGGALQAK | YES | YES | NO |
| 4135 | TRAV4 | TRAJ33 | CLVGDMRGGSNYQLIW | TRBV4-3 | TRBJ1-6 | CASSQDTGTVNSPLHF | KLGGALQAK | YES | NO | NO |
| 4139 | TRAV38-2DV8 | TRAJ32 | CAYLWGGATNKLIF | TRBV5-1 | TRBJ1-6 | CASSPGTNQHSPLHF | KLGGALQAK | YES | NO | NO |
| 4140 | TRAV12-2 | TRAJ4 | CAAYSGGYNKLIF | TRBV5-4 | TRBJ1-6 | CASSLGSGKDNSPLHF | KLGGALQAK | YES | NO | NO |
| 4142 | TRAV25 | TRAJ44 | CAGIRSVENTGTASKLTF | TRBV5-4 | TRBJ1-6 | CASSLTGSGDSPLHF | KLGGALQAK | YES | NO | NO |
| 4144 | TRAV8-3 | TRAJ6 | CAVADLSGGSYIPTF | TRBV5-4 | TRBJ1-6 | CASSPGGSLHF | KLGGALQAK | YES | NO | NO |
| 4145 | TRAV12-2 | TRAJ26 | CAAHMDYGQNFVF | TRBV5-5 | TRBJ1-6 | CASSVDRWANSPLHF | KLGGALQAK | YES | NO | NO |
| 4146 | TRAV12-2 | TRAJ3 | CAVLRPSRSSASKIIF | TRBV5-5 | TRBJ1-6 | CASSFDGINSPLHF | KLGGALQAK | YES | NO | NO |
| 4149 | TRAV14DV4 | TRAJ33 | CAMRGSGNYQLIW | TRBV5-6 | TRBJ1-6 | CASSLESRLNSPLHF | KLGGALQAK | YES | NO | NO |
| 4152 | TRAV21 | TRAJ49 | CAWPNTGNQFYF | TRBV5-8 | TRBJ1-6 | CASSLVYGSGYSPLHF | KLGGALQAK | YES | NO | NO |
| 4153 | TRAV14DV4 | TRAJ43 | CAMREYDMRF | TRBV6-1 | TRBJ1-6 | CATRDANSPLHF | KLGGALQAK | YES | NO | NO |
| 4155 | TRAV8-3 | TRAJ39 | CAVGNNAGNMLTF | TRBV6-1 | TRBJ1-6 | CASSEAGDRYNSPLHF | KLGGALQAK | YES | NO | NO |
| 4156 | TRAV1-2 | TRAJ33 | CAVKDSNYQLIW | TRBV6-3 | TRBJ1-6 | CASSYSESGDSPLHF | KLGGALQAK | YES | NO | NO |
| 4157 | TRAV14DV4 | TRAJ21 | CAMRETYNFNKFYF | TRBV6-3 | TRBJ1-6 | CASSYSPDYNSPLHF | KLGGALQAK | YES | NO | NO |
| 4160 | TRAV13-1 | TRAJ9 | CAASGSGGFKTIF | TRBV6-5 | TRBJ1-6 | CASSYFLPGQRKTYSPLHF | KLGGALQAK | YES | NO | NO |
| 4161 | TRAV19 | TRAJ30 | CALSEATRDDKIIF | TRBV6-5 | TRBJ1-6 | CASSPRQGFSPLHF | KLGGALQAK | YES | NO | NO |
| 4163 | TRAV21 | TRAJ58 | CAAVGETSGSRLTF | TRBV6-5 | TRBJ1-6 | CASSYGVSPLHF | KLGGALQAK | YES | NO | NO |
| 4165 | TRAV39 | TRAJ49 | CAVDTFTGNQFYF | TRBV6-5 | TRBJ1-6 | CASSQTGRGDNSPLHF | KLGGALQAK | YES | NO | NO |
| 4167 | TRAV8-6 | TRAJ42 | CAVSDQSYYGGSQGNLIF | TRBV6-5 | TRBJ1-6 | CASRLQVNNSPLHF | KLGGALQAK | YES | NO | NO |
| 4169 | TRAV12-1 | TRAJ3 | CVVNTGASKIIF | TRBV6-6 | TRBJ1-6 | CASSEPGTNSPLHF | KLGGALQAK | NO | NO | NO |
| 4171 | TRAV21 | TRAJ17 | CAVRRMIKAAGNKLTF | TRBV6-6 | TRBJ1-6 | CASSPPGVSSPLHF | KLGGALQAK | NO | NO | NO |
| 4173 | TRAV20 | TRAJ34 | CAVQANTDKLIF | TRBV7-7 | TRBJ1-6 | CASSEDRGSPLHF | KLGGALQAK | YES | NO | NO |
| 4174 | TRAV8-2 | TRAJ12 | CVSRMDSSYKLIF | TRBV7-8 | TRBJ1-6 | CASSPPTGTDHRSPLHF | KLGGALQAK | YES | NO | NO |
| 4175 | TRAV1-2 | TRAJ15 | CAVRGQAGTALIF | TRBV7-9 | TRBJ1-6 | CASSAGQAGFSPLHF | KLGGALQAK | YES | NO | NO |
| 4178 | TRAV14DV4 | TRAJ23 | CAMRSQGGKLIF | TRBV7-9 | TRBJ1-6 | CASSQLEDRVGSPLHF | KLGGALQAK | YES | NO | NO |
| 4180 | TRAV21 | TRAJ16 | CAVMGSDGQKLLF | TRBV7-9 | TRBJ1-6 | CASRDGGFNSPLHF | KLGGALQAK | NO | NO | NO |
| 4184 | TRAV29DV5 | TRAJ40 | CAASASGTYKYIF | TRBV7-9 | TRBJ1-6 | CASSSIYRGYYSPLHF | KLGGALQAK | YES | NO | NO |
| 4188 | TRAV12-2 | TRAJ23 | CAVNSFYNQGGKLIF | TRBV9 | TRBJ1-6 | CASSEGQESFSPLHF | KLGGALQAK | NO | NO | NO |
| 4190 | TRAV13-1 | TRAJ6 | CAASGGSYIPTF | TRBV9 | TRBJ1-6 | CASSAGMSSYNSPLHF | KLGGALQAK | YES | NO | NO |
| 4191 | TRAV12-2 | TRAJ45 | CASDGGGADGLTF | TRBV10-1 | TRBJ2-1 | CASSFQGYNEQFF | KLGGALQAK | YES | NO | NO |
| 4193 | TRAV14DV4 | TRAJ22 | CAMRGSYSGSARQLTF | TRBV10-1 | TRBJ2-1 | CATRGGLENEQFF | KLGGALQAK | YES | NO | NO |
| 4196 | TRAV21 | TRAJ22 | CAVLPSSGSARQLTF | TRBV10-1 | TRBJ2-1 | CASSDGDSPYNEQFF | KLGGALQAK | YES | NO | NO |
| 4197 | TRAV4 | TRAJ45 | CLVETGGGADGLTF | TRBV10-1 | TRBJ2-1 | CASSQTSGGLNEQFF | KLGGALQAK | YES | NO | NO |
| 4198 | TRAV12-1 | TRAJ11 | CVVNIVGYSTLTF | TRBV10-2 | TRBJ2-1 | CASSESHGSTYEQFF | KLGGALQAK | NO | NO | NO |
| 4199 | TRAV12-1 | TRAJ26 | CVVSPNYGQNFVF | TRBV10-2 | TRBJ2-1 | CASSTGTTLYNEQFF | KLGGALQAK | YES | NO | NO |
| 4200 | TRAV12-2 | TRAJ43 | CAVTPGFNDMRF | TRBV10-2 | TRBJ2-1 | CASSEKPRGNNEQFF | KLGGALQAK | YES | NO | NO |
| 4201 | TRAV16 | TRAJ30 | CALSGVRDDKIIF | TRBV10-2 | TRBJ2-1 | CASSETSAYNEQFF | KLGGALQAK | NO | NO | NO |
| 4203 | TRAV21 | TRAJ5 | CAVRPRGDTGRRALTF | TRBV10-2 | TRBJ2-1 | CASGGQYNEQFF | KLGGALQAK | YES | NO | NO |
| 4204 | TRAV24 | TRAJ32 | CAFCGATNKLIF | TRBV10-2 | TRBJ2-1 | CASSGDRVTNEQFF | KLGGALQAK | YES | NO | NO |
| 4205 | TRAV27 | TRAJ6 | CARGSSGGSYIPTF | TRBV10-2 | TRBJ2-1 | CASSPSGGNEQFF | KLGGALQAK | YES | NO | NO |
| 4206 | TRAV29DV5 | TRAJ8 | CAASATGFQKLVF | TRBV10-2 | TRBJ2-1 | CASSESSPRGGSFF | KLGGALQAK | YES | NO | NO |
| 4208 | TRAV8-2 | TRAJ26 | CVVRDNYGQNFVF | TRBV10-2 | TRBJ2-1 | CASSYVQSSYNEQFF | KLGGALQAK | YES | NO | NO |
| 4211 | TRAV12-3 | TRAJ4 | CAIAPGGYNKLIF | TRBV10-3 | TRBJ2-1 | CAISEDRAPSNEQFF | KLGGALQAK | YES | NO | NO |
| 4212 | TRAV13-1 | TRAJ34 | CAAYNTDKLIF | TRBV10-3 | TRBJ2-1 | CAISDYAGYNEQFF | RLRAEAQVK | YES | NO | YES |
| 4213 | TRAV13-2 | TRAJ16 | CAENIPLSQKLLF | TRBV10-3 | TRBJ2-1 | CATSPPGIEQFF | KLGGALQAK | NO | NO | NO |
| 4219 | TRAV29DV5 | TRAJ49 | CAASSTTGNQFYF | TRBV10-3 | TRBJ2-1 | CAIRTGLAGHNEQFF | KLGGALQAK | YES | NO | NO |
| 4220 | TRAV39 | TRAJ28 | CAVDSEGAGSYQLTF | TRBV10-3 | TRBJ2-1 | CATRGSSGRAFFYEQFF | KLGGALQAK | YES | NO | NO |

| | | | | | | | | | | |
|---|---|---|---|---|---|---|---|---|---|---|
| 4222 | TRAV9-2 | TRAJ45 | CALLYSGGGADGLTF | TRBV10-3 | TRBJ2-1 | CAIGGDRYNEQFF | KLGGALQAK | YES | NO | NO |
| 4224 | TRAV3 | TRAJ24 | CAVNADSWGKLQF | TRBV11-1 | TRBJ2-1 | CVTGQGDINEQFF | KLGGALQAK | YES | NO | NO |
| 4226 | TRAV1-2 | TRAJ20 | CAVRDWGDYKLSF | TRBV11-2 | TRBJ2-1 | CASSPTGTSESYEQFF | KLGGALQAK | YES | NO | NO |
| 4227 | TRAV12-3 | TRAJ13 | CAMSFHSGGYQKVTF | TRBV11-2 | TRBJ2-1 | CASSPPPDNEQFF | KLGGALQAK | YES | NO | NO |
| 4231 | TRAV13-1 | TRAJ6 | CASRQSGGSYIPTF | TRBV11-2 | TRBJ2-1 | CASSLAGPAADRPYNEQFF | KLGGALQAK | YES | NO | NO |
| 4232 | TRAV14DV4 | TRAJ4 | CAMRGPISGGYNKLIF | TRBV11-2 | TRBJ2-1 | CASSLAALAGEHEQFF | KLGGALQAK | YES | NO | NO |
| 4233 | TRAV14DV4 | TRAJ8 | CAMNRNTGFQKLVF | TRBV11-2 | TRBJ2-1 | CASSLDIWTWDRSVYNEQFF | KLGGALQAK | NO | NO | NO |
| 4236 | TRAV17 | TRAJ18 | CATDALRGSTLGRLYF | TRBV11-2 | TRBJ2-1 | CASSETGTAFLNEQFF | KLGGALQAK | YES | NO | NO |
| 4237 | TRAV17 | TRAJ29 | CATGGNTPLVF | TRBV11-2 | TRBJ2-1 | CASSLAALAGEHEQFF | KLGGALQAK | YES | NO | NO |
| 4239 | TRAV19 | TRAJ32 | CALSGATNKLIF | TRBV11-2 | TRBJ2-1 | CASSLTQGSYNEQFF | KLGGALQAK | YES | NO | NO |
| 4241 | TRAV20 | TRAJ49 | CAVLVNTGNQFYF | TRBV11-2 | TRBJ2-1 | CASSLALRGSYNEQFF | KLGGALQAK | YES | NO | NO |
| 4244 | TRAV29DV5 | TRAJ3 | CAASSGYSSASKIIF | TRBV11-2 | TRBJ2-1 | CASSYSAGGRVNEQFF | RLRAEAQVK | YES | YES | YES |
| 4245 | TRAV29DV5 | TRAJ49 | CAASASGNQFYF | TRBV11-2 | TRBJ2-1 | CASSFWDNEQFF | KLGGALQAK | YES | NO | NO |
| 4246 | TRAV30 | TRAJ53 | CGTEAGSGGSNYKLTF | TRBV11-2 | TRBJ2-1 | CASSFGGPIYEQFF | KLGGALQAK | YES | NO | NO |
| 4248 | TRAV39 | TRAJ52 | CAVVSPNAGGTSYGKLTF | TRBV11-2 | TRBJ2-1 | CASSLYAPTNEQFF | KLGGALQAK | YES | NO | NO |
| 4249 | TRAV4 | TRAJ9 | CLVGDPGNTGGFKTIF | TRBV11-2 | TRBJ2-1 | CASSLGVTRYNEQFF | KLGGALQAK | YES | NO | NO |
| 4252 | TRAV1-2 | TRAJ33 | CAVRDSNYQLIW | TRBV11-3 | TRBJ2-1 | CASSFEDMGGEQFF | KLGGALQAK | YES | NO | NO |
| 4254 | TRAV12-2 | TRAJ35 | CAVIGFGNVLHC | TRBV11-3 | TRBJ2-1 | CASSSDGRDYHEQFF | KLGGALQAK | NO | NO | NO |
| 4256 | TRAV29DV5 | TRAJ28 | CAAIGAGSYQLTF | TRBV11-3 | TRBJ2-1 | CASRAFDPGTSGSSYNEQFF | KLGGALQAK | NO | NO | NO |
| 4259 | TRAV29DV5 | TRAJ42 | CAASAGGSQGNLIF | TRBV11-3 | TRBJ2-1 | CASSLEWGGNEQFF | KLGGALQAK | YES | NO | NO |
| 4260 | TRAV8-1 | TRAJ4 | CAVISGYNKLIF | TRBV11-3 | TRBJ2-1 | CASSLEAGSYNEQFF | KLGGALQAK | YES | NO | NO |
| 4261 | TRAV8-2 | TRAJ34 | CVVSDPYTDKLIF | TRBV11-3 | TRBJ2-1 | CASSPEGGGTYNEQFF | KLGGALQAK | YES | NO | NO |
| 4262 | TRAV8-4 | TRAJ42 | CAVSDLYGGSQGNLIF | TRBV11-3 | TRBJ2-1 | CASSPPGTSGSSSYNEQFF | KLGGALQAK | YES | NO | NO |
| 4266 | TRAV12-3 | TRAJ47 | CAMSEGYGNKLVF | TRBV12-3 | TRBJ2-1 | CASRYIAGGRNEQFF | KLGGALQAK | YES | YES | NO |
| 4267 | TRAV14DV4 | TRAJ22 | CAMRRPISSGSARQLTF | TRBV12-3 | TRBJ2-1 | CASSVGTGSYNEQFF | KLGGALQAK | YES | NO | NO |
| 4268 | TRAV19 | TRAJ44 | CALITGTASKLTF | TRBV12-3 | TRBJ2-1 | CASSFISSYNEQFF | KLGGALQAK | YES | NO | NO |
| 4270 | TRAV24 | TRAJ53 | CAFIMGGGSNYKLTF | TRBV12-3 | TRBJ2-1 | CASSLGPINEQFF | KLGGALQAK | YES | NO | NO |
| 4271 | TRAV25 | TRAJ24 | CAQTDSWGKLQF | TRBV12-3 | TRBJ2-1 | CASTQGTYYNEQFF | KLGGALQAK | YES | NO | NO |
| 4274 | TRAV29DV5 | TRAJ20 | CAASERNDYKLSF | TRBV12-3 | TRBJ2-1 | CASSFGTGSRNEQFF | AVFDRKSDAK | YES | NO | NO |
| 4274 | TRAV29DV5 | TRAJ20 | CAASERNDYKLSF | TRBV12-3 | TRBJ2-1 | CASSFGTGSRNEQFF | KLGGALQAK | YES | NO | NO |
| 4275 | TRAV29DV5 | TRAJ49 | CAAGESTGNQFYF | TRBV12-3 | TRBJ2-1 | CASSPARYNEQFF | KLGGALQAK | YES | NO | NO |
| 4276 | TRAV41 | TRAJ32 | CAVRPRADGGATNKLIF | TRBV12-3 | TRBJ2-1 | CASSFHADYNEQFF | KLGGALQAK | YES | NO | NO |
| 4277 | TRAV8-2 | TRAJ8 | CVVSVNTGFQKLVF | TRBV12-3 | TRBJ2-1 | CASSLNPTGDEQFF | KLGGALQAK | YES | NO | NO |
| 4278 | TRAV1-1 | TRAJ15 | CAVNQAGTALIF | TRBV12-4 | TRBJ2-1 | CASSLGGAGAWGYEQFF | KLGGALQAK | YES | NO | NO |
| 4279 | TRAV10 | TRAJ49 | CVVSVSGNQFYF | TRBV12-4 | TRBJ2-1 | CAIVLGTSAYNEQFF | KLGGALQAK | YES | NO | NO |
| 4283 | TRAV12-3 | TRAJ37 | CAPQRNTGKLIF | TRBV12-4 | TRBJ2-1 | CASTAVGIGSNEQFF | KLGGALQAK | YES | NO | NO |
| 4286 | TRAV13-1 | TRAJ43 | CADYMRF | TRBV12-4 | TRBJ2-1 | CASSSGTRVYEQFF | KLGGALQAK | NO | NO | NO |
| 4287 | TRAV13-1 | TRAJ6 | CAESGGSYIPTF | TRBV12-4 | TRBJ2-1 | CASSIGQSYNEQFF | KLGGALQAK | YES | NO | NO |
| 4290 | TRAV17 | TRAJ20 | CATDANDYKLSF | TRBV12-4 | TRBJ2-1 | CASSFGHNEQFF | KLGGALQAK | YES | NO | NO |
| 4291 | TRAV17 | TRAJ36 | CATAGANNLFF | TRBV12-4 | TRBJ2-1 | CASSFTGGPYNEQFF | KLGGALQAK | YES | NO | NO |
| 4292 | TRAV17 | TRAJ39 | CATDSNNAGNMLTF | TRBV12-4 | TRBJ2-1 | CASSLTDAEEQFF | KLGGALQAK | YES | NO | NO |
| 4293 | TRAV19 | TRAJ10 | CALSEGILTGGGNKLTF | TRBV12-4 | TRBJ2-1 | CASSLGGPYNEQFF | KLGGALQAK | YES | NO | NO |
| 4294 | TRAV19 | TRAJ10 | CALSEAFTGGGNKLTF | TRBV12-4 | TRBJ2-1 | CASSTLTIATLDNEQFF | KLGGALQAK | YES | NO | NO |
| 4295 | TRAV19 | TRAJ39 | CALSDYNNAGNMLTF | TRBV12-4 | TRBJ2-1 | CASSRYSGAANYNEQFF | KLGGALQAK | YES | NO | NO |
| 4296 | TRAV19 | TRAJ42 | CALSAYGGSQGNLIF | TRBV12-4 | TRBJ2-1 | CASSGGGASYNEQFF | KLGGALQAK | YES | NO | NO |
| 4298 | TRAV20 | TRAJ5 | CAVQAQGKMDTGRRALTF | TRBV12-4 | TRBJ2-1 | CASSLPKTGYNEQFF | KLGGALQAK | YES | NO | NO |
| 4299 | TRAV21 | TRAJ18 | CAVPDRGSTLGRLYF | TRBV12-4 | TRBJ2-1 | CASSLGYNEQFF | KLGGALQAK | YES | NO | NO |
| 4300 | TRAV21 | TRAJ30 | CAVAPNRDDKIIF | TRBV12-4 | TRBJ2-1 | CASSTQQPLYNEQFF | KLGGALQAK | YES | NO | NO |
| 4301 | TRAV21 | TRAJ43 | CAVNNDMRF | TRBV12-4 | TRBJ2-1 | CASRGRSYNEQFF | KLGGALQAK | NO | NO | NO |
| 4304 | TRAV26-2 | TRAJ41 | CILKSGYALNF | TRBV12-4 | TRBJ2-1 | CASSLSGGSYNEQFF | KLGGALQAK | YES | NO | NO |
| 4305 | TRAV29DV5 | TRAJ26 | CAASVGGQNFVF | TRBV12-4 | TRBJ2-1 | CASNPFPGVGNEQFF | KLGGALQAK | YES | NO | NO |
| 4307 | TRAV29DV5 | TRAJ54 | CARRPQGAQKLVF | TRBV12-4 | TRBJ2-1 | CASSWTGGYNEQFF | KLGGALQAK | YES | NO | NO |
| 4308 | TRAV38-1 | TRAJ42 | CALYPLLGNLIF | TRBV12-4 | TRBJ2-1 | CASSVILSPRGNEQFF | KLGGALQAK | YES | NO | NO |
| 4310 | TRAV41 | TRAJ34 | CAVSPYNTDKLIF | TRBV12-4 | TRBJ2-1 | CASSRGRGSSYNEQFF | KLGGALQAK | YES | NO | NO |
| 4311 | TRAV5 | TRAJ11 | CAEARSGYSTLTF | TRBV12-4 | TRBJ2-1 | CASSHLTDREQFF | KLGGALQAK | YES | NO | NO |
| 4313 | TRAV8-3 | TRAJ43 | CAVDNNNDMRF | TRBV12-4 | TRBJ2-1 | CASRTSGNEQFF | KLGGALQAK | YES | NO | NO |
| 4314 | TRAV8-6 | TRAJ36 | CAVSDQTGANNLFF | TRBV12-4 | TRBJ2-1 | CASSLARWQFF | KLGGALQAK | YES | NO | NO |
| 4315 | TRAV9-2 | TRAJ45 | CALSDGGGADGLTF | TRBV12-4 | TRBJ2-1 | CASTVTGNEQFF | KLGGALQAK | YES | YES | NO |
| 4317 | TRAV12-3 | TRAJ20 | CATTFGDYKLSF | TRBV12-5 | TRBJ2-1 | CASGSTLTGDYNEQFF | KLGGALQAK | YES | NO | NO |
| 4320 | TRAV29DV5 | TRAJ16 | CAASAF | TRBV12-5 | TRBJ2-1 | CASGWENNEQFF | KLGGALQAK | NO | NO | NO |
| 4321 | TRAV38-1 | TRAJ54 | CAFMSLIQGAQKLVF | TRBV12-5 | TRBJ2-1 | CASGSLQGFNEQFF | KLGGALQAK | YES | NO | NO |
| 4323 | TRAV4 | TRAJ43 | CLVGESVGNNNDMRF | TRBV12-5 | TRBJ2-1 | CASGAGTGTYNEQFF | KLGGALQAK | YES | NO | NO |
| 4325 | TRAV1-1 | TRAJ8 | CAVSLMNTGFQKLVF | TRBV13 | TRBJ2-1 | CASSLAGTGGNEQFF | KLGGALQAK | YES | NO | NO |
| 4326 | TRAV1-2 | TRAJ20 | CAVRRNDYKLSF | TRBV13 | TRBJ2-1 | CASSSSGGYNEQFF | RLRAEAQVK | YES | YES | YES |
| 4327 | TRAV12-2 | TRAJ18 | CAVMADRGSTLGRLYF | TRBV13 | TRBJ2-1 | CASSLALNNEQFF | KLGGALQAK | YES | NO | NO |
| 4328 | TRAV12-2 | TRAJ26 | CAARGKNYGQNFVF | TRBV13 | TRBJ2-1 | CASSLDPGVYNEQFF | KLGGALQAK | YES | NO | NO |
| 4329 | TRAV12-2 | TRAJ48 | CAVCLSNFGNEKLTF | TRBV13 | TRBJ2-1 | CASSAPGQGRQPTQFF | KLGGALQAK | YES | NO | NO |
| 4333 | TRAV17 | TRAJ45 | CATADSGGGADGLTF | TRBV13 | TRBJ2-1 | CASSLVEGGSQGAEPTYNEQFF | KLGGALQAK | NO | NO | NO |
| 4334 | TRAV19 | TRAJ15 | CALSEEPEAGTALIF | TRBV13 | TRBJ2-1 | CASSLDQEIIGNEQFF | KLGGALQAK | YES | NO | NO |
| 4335 | TRAV19 | TRAJ30 | CALSEAHGDDKIIF | TRBV13 | TRBJ2-1 | CASSSGGTGGPREQFF | KLGGALQAK | YES | NO | NO |
| 4340 | TRAV26-1 | TRAJ34 | CIVRLRCRTPTSSSL | TRBV13 | TRBJ2-1 | CASSLGGQGSNEQFF | KLGGALQAK | NO | NO | NO |
| 4341 | TRAV29DV5 | TRAJ45 | CAASGGSGGGADGLTF | TRBV13 | TRBJ2-1 | CASSLRLAGDIIVNEQFF | KLGGALQAK | YES | NO | NO |
| 4343 | TRAV36DV7 | TRAJ31 | CAVALWARLMF | TRBV13 | TRBJ2-1 | CASSLGGQGSNEQFF | KLGGALQAK | NO | NO | NO |
| 4345 | TRAV38-2DV8 | TRAJ53 | CAYQGGGSNYKLTF | TRBV13 | TRBJ2-1 | CASSLNGVNINEQFF | KLGGALQAK | YES | NO | NO |
| 4346 | TRAV38-2DV8 | TRAJ54 | CAYSPPQGAQKLVF | TRBV13 | TRBJ2-1 | CASSPGTSGSDEQFF | KLGGALQAK | YES | NO | NO |
| 4347 | TRAV8-1 | TRAJ37 | CAVTGGNTGKLIF | TRBV13 | TRBJ2-1 | CASSPGTSNEQFF | KLGGALQAK | YES | NO | NO |
| 4348 | TRAV8-1 | TRAJ42 | CAVRPRGSQGNLIF | TRBV13 | TRBJ2-1 | CASSSVGGVTNEQFF | KLGGALQAK | YES | NO | NO |
| 4349 | TRAV8-6 | TRAJ18 | CAVSGDRGSTLGRLYF | TRBV13 | TRBJ2-1 | CASSLKTGGNEQFF | KLGGALQAK | YES | NO | NO |
| 4352 | TRAV22 | TRAJ40 | CAVDLAFGTYKYIF | TRBV14 | TRBJ2-1 | CASSQDPHNEQFF | RLRAEAQVK | YES | YES | YES |
| 4354 | TRAV29DV5 | TRAJ34 | CAAESTYNTDKLIF | TRBV14 | TRBJ2-1 | CASSPTGGLYNEQFF | KLGGALQAK | YES | NO | NO |
| 4356 | TRAV38-2DV8 | TRAJ30 | CAYRSAGFMNRDDKIIF | TRBV14 | TRBJ2-1 | CASSQTQGGVSLYNEQFF | KLGGALQAK | YES | NO | NO |
| 4358 | TRAV8-2 | TRAJ34 | CVVSLVNTDKLIF | TRBV14 | TRBJ2-1 | CASSQEGGPPYNEQFF | KLGGALQAK | YES | NO | NO |
| 4359 | TRAV9-2 | TRAJ20 | CALSTDYKLSF | TRBV14 | TRBJ2-1 | CASSQVPSSYNEQFF | KLGGALQAK | YES | NO | NO |
| 4360 | TRAV1-2 | TRAJ26 | CAVRDYDNYGQNFVF | TRBV15 | TRBJ2-1 | CATSTSKGVAGDNEQFF | KLGGALQAK | YES | NO | NO |
| 4362 | TRAV10 | TRAJ38 | CVVSGYNAGNNRKLIW | TRBV15 | TRBJ2-1 | CATSIRLRPWQGGDEQFF | KLGGALQAK | YES | NO | NO |
| 4363 | TRAV12-1 | TRAJ34 | CVVNGDTDKLIF | TRBV15 | TRBJ2-1 | CATSSWTGGFDEQFF | KLGGALQAK | YES | NO | NO |
| 4364 | TRAV12-2 | TRAJ39 | CAVRNNNAGNMLTF | TRBV15 | TRBJ2-1 | CATSSNRDRGDNEQFF | KLGGALQAK | YES | NO | NO |
| 4365 | TRAV12-3 | TRAJ39 | CAIGPGNMLTF | TRBV15 | TRBJ2-1 | CATSRQTSIEQFF | KLGGALQAK | YES | NO | NO |
| 4368 | TRAV5 | TRAJ15 | CADFNQAGTALIF | TRBV15 | TRBJ2-1 | CATSRGQGVPNEQFF | KLGGALQAK | YES | NO | NO |
| 4369 | TRAV5 | TRAJ34 | CAESKADKLIF | TRBV15 | TRBJ2-1 | CATSSRLNNEQFF | KLGGALQAK | YES | NO | NO |
| 4371 | TRAV8-6 | TRAJ40 | CAVSPSGTYKYIF | TRBV15 | TRBJ2-1 | CATSSSGTSGGWSYNEQFF | KLGGALQAK | YES | NO | NO |
| 4373 | TRAV1-1 | TRAJ34 | CAVKGGKLIF | TRBV18 | TRBJ2-1 | CASNEDPGLAGDNEQFF | KLGGALQAK | YES | NO | NO |
| 4374 | TRAV1-2 | TRAJ20 | CAALYNDYKLSF | TRBV18 | TRBJ2-1 | CASSSPGQGLFEQFF | KLGGALQAK | YES | NO | NO |
| 4376 | TRAV12-2 | TRAJ15 | CAVNQAGTALIF | TRBV18 | TRBJ2-1 | CASSPPQGAVVEQFF | KLGGALQAK | YES | NO | NO |
| 4377 | TRAV12-3 | TRAJ52 | CAMSATAGGTSYGKLTF | TRBV18 | TRBJ2-1 | CASSPPQGGGTHNEQFF | KLGGALQAK | YES | NO | NO |
| 4378 | TRAV16 | TRAJ9 | CALSGTGGFKTIF | TRBV18 | TRBJ2-1 | CASSGTSGHNEQFF | KLGGALQAK | NO | NO | NO |
| 4379 | TRAV19 | TRAJ37 | CALNRGSSSNTGKLIF | TRBV18 | TRBJ2-1 | CASSPPTLTEQFF | KLGGALQAK | YES | NO | NO |
| 4380 | TRAV19 | TRAJ49 | CALSEPNTGNQFYF | TRBV18 | TRBJ2-1 | CASSPGGSYNEQFF | KLGGALQAK | YES | NO | NO |
| 4382 | TRAV36DV7 | TRAJ45 | CVPYSGGGADGLTF | TRBV18 | TRBJ2-1 | CASSPPTAYSYNEQFF | KLGGALQAK | NO | NO | NO |
| 4383 | TRAV38-1 | TRAJ36 | CAFDDQTGANNLFF | TRBV18 | TRBJ2-1 | CASSPDYNEQFF | KLGGALQAK | YES | NO | NO |
| 4384 | TRAV38-2DV8 | TRAJ37 | CAAHGSSNTGKLIF | TRBV18 | TRBJ2-1 | CASSPPSPGSYNEQFF | KLGGALQAK | YES | NO | NO |
| 4385 | TRAV38-2DV8 | TRAJ42 | CAYGGSQGNLIF | TRBV18 | TRBJ2-1 | CASSPGRPVETYNEQFF | KLGGALQAK | YES | NO | NO |
| 4387 | TRAV8-1 | TRAJ43 | CAVNGINNDMRF | TRBV18 | TRBJ2-1 | CASSSAGGTPYNEQFF | KLGGALQAK | YES | NO | NO |
| 4390 | TRAV12-1 | TRAJ23 | CVVNSGYNQGGKLIF | TRBV19 | TRBJ2-1 | CASSTAWTSGSYEQFF | KLGGALQAK | YES | NO | NO |

| | | | | | | | | | | |
|---|---|---|---|---|---|---|---|---|---|---|
| 4391 | TRAV12-1 | TRAJ30 | CVVNIAGNRDDKIIF | TRBV19 | TRBJ2-1 | CASSIGGHTYNEQFF | KLGGALQAK | YES | NO | NO |
| 4392 | TRAV12-1 | TRAJ4 | CVVDSGGYNKLIF | TRBV19 | TRBJ2-1 | CASSIRDREYNEQFF | KLGGALQAK | YES | NO | NO |
| 4394 | TRAV12-1 | TRAJ43 | CVVKDNNNDMRF | TRBV19 | TRBJ2-1 | CASVGWDNEQFF | KLGGALQAK | YES | NO | NO |
| 4396 | TRAV12-2 | TRAJ10 | CAVNIGLTGGGNKLTF | TRBV19 | TRBJ2-1 | CASSISLATYNEQFF | KLGGALQAK | YES | NO | NO |
| 4397 | TRAV12-2 | TRAJ15 | CAVNQAGTALIF | TRBV19 | TRBJ2-1 | CASSISSGVAYEQFF | KLGGALQAK | NO | NO | NO |
| 4399 | TRAV12-2 | TRAJ31 | CAGTGNNARLMF | TRBV19 | TRBJ2-1 | CASSISATGEQFF | KLGGALQAK | YES | NO | NO |
| 4400 | TRAV12-2 | TRAJ31 | CAVNRWNNNARLMF | TRBV19 | TRBJ2-1 | CATSRPREAYNEQFF | KLGGALQAK | YES | NO | NO |
| 4401 | TRAV12-2 | TRAJ33 | CAVKAWDSNYQLIW | TRBV19 | TRBJ2-1 | CASQTRGNNEQFF | KLGGALQAK | YES | NO | NO |
| 4402 | TRAV12-2 | TRAJ34 | CAVWRADKLIF | TRBV19 | TRBJ2-1 | CASSILGLAGGLNEQFF | KLGGALQAK | YES | NO | NO |
| 4405 | TRAV12-2 | TRAJ43 | CAVTLYNNNDMRF | TRBV19 | TRBJ2-1 | CASRDRGLNEQFF | KLGGALQAK | YES | NO | NO |
| 4406 | TRAV12-2 | TRAJ49 | CAVNMRLSNQFYF | TRBV19 | TRBJ2-1 | CASSWGSYNEQFF | KLGGALQAK | YES | NO | NO |
| 4407 | TRAV12-2 | TRAJ5 | CAVSDTGRRALTF | TRBV19 | TRBJ2-1 | CASSSGGSYNEQFF | KLGGALQAK | YES | NO | NO |
| 4408 | TRAV12-3 | TRAJ42 | CAMSAWGSQGNLIF | TRBV19 | TRBJ2-1 | CASSYGASDNEQFF | KLGGALQAK | YES | NO | NO |
| 4409 | TRAV13-1 | TRAJ32 | CAASQYGGATNKLIF | TRBV19 | TRBJ2-1 | CASSNKRDRGIDEQFF | KLGGALQAK | YES | NO | NO |
| 4411 | TRAV13-1 | TRAJ49 | CAASRISGNQFYF | TRBV19 | TRBJ2-1 | CASSPTGLVSYNEQFF | KLGGALQAK | YES | NO | NO |
| 4413 | TRAV14DV4 | TRAJ24 | CAMREDVDSWGKLQF | TRBV19 | TRBJ2-1 | CASSIGTSRNEQFF | KLGGALQAK | YES | NO | NO |
| 4414 | TRAV14DV4 | TRAJ30 | CAMRGGDDKIIF | TRBV19 | TRBJ2-1 | CASSIGLAGHSGDEQFF | KLGGALQAK | YES | NO | NO |
| 4418 | TRAV17 | TRAJ43 | CATELLLNNNDMRF | TRBV19 | TRBJ2-1 | CASSTSMIGGNEQFF | KLGGALQAK | YES | NO | NO |
| 4419 | TRAV17 | TRAJ47 | CATLLSYGNKLVF | TRBV19 | TRBJ2-1 | CASSPGTGGVNEQFF | KLGGALQAK | YES | NO | NO |
| 4420 | TRAV17 | TRAJ53 | CATENSGGSNYKLTF | TRBV19 | TRBJ2-1 | CASSIGGSSYNEQFF | KLGGALQAK | YES | NO | NO |
| 4421 | TRAV17 | TRAJ7 | CATDPLFGNNRLAF | TRBV19 | TRBJ2-1 | CASSTLNEQFF | KLGGALQAK | YES | NO | NO |
| 4423 | TRAV19 | TRAJ31 | CALSDRNNARLMF | TRBV19 | TRBJ2-1 | CAIAMGAYNEQFF | KLGGALQAK | YES | NO | NO |
| 4426 | TRAV19 | TRAJ47 | CALSEDYGNKLVF | TRBV19 | TRBJ2-1 | CASSSGGNNEQFF | KLGGALQAK | YES | NO | NO |
| 4429 | TRAV20 | TRAJ20 | CAVCSNDYKLSF | TRBV19 | TRBJ2-1 | CASTSGTLYNEQFF | KLGGALQAK | YES | NO | NO |
| 4431 | TRAV21 | TRAJ15 | CAAIPLNQAGTALIF | TRBV19 | TRBJ2-1 | CASSKTGNEQFF | KLGGALQAK | YES | NO | NO |
| 4433 | TRAV21 | TRAJ26 | CAVVDNYGQNFVF | TRBV19 | TRBJ2-1 | CASSSRGGNEQFF | KLGGALQAK | YES | NO | NO |
| 4434 | TRAV21 | TRAJ45 | CAVDYQGGADGLTF | TRBV19 | TRBJ2-1 | CASTKGTGGEQFF | KLGGALQAK | YES | NO | NO |
| 4435 | TRAV23DV6 | TRAJ40 | CAASGSGTYKYIF | TRBV19 | TRBJ2-1 | CASEIGQGLVNEQFF | KLGGALQAK | YES | NO | NO |
| 4438 | TRAV24 | TRAJ52 | CAFIGDAGGTSYGKLTF | TRBV19 | TRBJ2-1 | CASSIVGRTGFSYNEQFF | KLGGALQAK | YES | NO | NO |
| 4442 | TRAV26-2 | TRAJ34 | CILTLTDKLIF | TRBV19 | TRBJ2-1 | CASNRDSLAYNEQFF | KLGGALQAK | YES | YES | NO |
| 4443 | TRAV26-2 | TRAJ58 | CILRDEETSGSRLTF | TRBV19 | TRBJ2-1 | CASSLDGSSYNEQFF | KLGGALQAK | YES | NO | NO |
| 4444 | TRAV27 | TRAJ52 | CAGNHNAGGTSYGKLTF | TRBV19 | TRBJ2-1 | CASSIDSYYNEQFF | KLGGALQAK | YES | NO | NO |
| 4445 | TRAV27 | TRAJ9 | CAGPRTGGFKTIF | TRBV19 | TRBJ2-1 | CASGGWEGNEQFF | KLGGALQAK | YES | NO | NO |
| 4446 | TRAV29DV5 | TRAJ21 | CAARISYNFNKFYF | TRBV19 | TRBJ2-1 | CASSLGSSSYNEQFF | KLGGALQAK | YES | NO | NO |
| 4447 | TRAV29DV5 | TRAJ52 | CAASVRAGGTSYGKLTF | TRBV19 | TRBJ2-1 | CASSTLANEQFF | KLGGALQAK | YES | NO | NO |
| 4448 | TRAV3 | TRAJ13 | CAVRASGGYQKVTF | TRBV19 | TRBJ2-1 | CASSARGGSNIDEQFF | KLGGALQAK | NO | NO | NO |
| 4451 | TRAV3 | TRAJ43 | CAVRLYNNNDMRF | TRBV19 | TRBJ2-1 | CASSIGEIEQFF | KLGGALQAK | NO | NO | NO |
| 4452 | TRAV30 | TRAJ18 | CGTFDRGSTLGRLYF | TRBV19 | TRBJ2-1 | CASRTLAGGSSEQFF | KLGGALQAK | YES | NO | NO |
| 4453 | TRAV30 | TRAJ27 | CGTEENTNAGKSTF | TRBV19 | TRBJ2-1 | CASRVGQTNEQFF | KLGGALQAK | YES | NO | NO |
| 4454 | TRAV30 | TRAJ29 | CGVSGNTPLVF | TRBV19 | TRBJ2-1 | CASNAGVQNEQFF | KLGGALQAK | YES | NO | NO |
| 4457 | TRAV35 | TRAJ45 | CAGQLPSGGGADGLTF | TRBV19 | TRBJ2-1 | CASSFRNREGEQFF | KLGGALQAK | YES | NO | NO |
| 4458 | TRAV38-1 | TRAJ26 | CAFMKHADNYGQNFVF | TRBV19 | TRBJ2-1 | CASSIDRYNEQFF | KLGGALQAK | YES | NO | NO |
| 4460 | TRAV38-2DV8 | TRAJ42 | CAYRSAYGGSQGNLIF | TRBV19 | TRBJ2-1 | CASSRVAGNEQFF | KLGGALQAK | YES | NO | NO |
| 4461 | TRAV38-2DV8 | TRAJ50 | CAYRSASGMKTSYDKVIF | TRBV19 | TRBJ2-1 | CASSMWTSGNNEQFF | KLGGALQAK | YES | NO | NO |
| 4466 | TRAV5 | TRAJ42 | CAGGGSQGNLIF | TRBV19 | TRBJ2-1 | CASSISYEQFF | KLGGALQAK | YES | NO | NO |
| 4468 | TRAV8-1 | TRAJ32 | CAVIGYGGATNKLIF | TRBV19 | TRBJ2-1 | CASSDLLALREQFF | KLGGALQAK | YES | NO | NO |
| 4469 | TRAV8-1 | TRAJ38 | CAVHDAGNNRKLIW | TRBV19 | TRBJ2-1 | CASSYSGAGNMFGEQFF | KLGGALQAK | YES | NO | NO |
| 4470 | TRAV8-2 | TRAJ45 | CVVSDLVYSGGGADGLTF | TRBV19 | TRBJ2-1 | CASSIFDVGDNEQFF | KLGGALQAK | YES | NO | NO |
| 4471 | TRAV8-3 | TRAJ47 | CAVREYGNKLVF | TRBV19 | TRBJ2-1 | CASSIGQGGNEQFF | KLGGALQAK | YES | NO | NO |
| 4472 | TRAV8-3 | TRAJ49 | CAVGAFRQFYF | TRBV19 | TRBJ2-1 | CASGAGGLNEQFF | AVFDRKSDAK | YES | NO | NO |
| 4473 | TRAV8-4 | TRAJ13 | CAVSDLYSGGYQKVTF | TRBV19 | TRBJ2-1 | CASSMGQGGYNEQFF | KLGGALQAK | YES | NO | NO |
| 4476 | TRAV8-6 | TRAJ11 | CAVSPGYSTLTF | TRBV19 | TRBJ2-1 | CASSDSGERGEQFF | KLGGALQAK | YES | NO | NO |
| 4478 | TRAV9-2 | TRAJ3 | CALSVGHRYSSASKIIF | TRBV19 | TRBJ2-1 | CASSIGDTRYNEQFF | KLGGALQAK | YES | NO | NO |
| 4479 | TRAV9-2 | TRAJ54 | CALSGILTQGAQKLVF | TRBV19 | TRBJ2-1 | CASSIGRVADQFF | KLGGALQAK | YES | NO | NO |
| 4480 | TRAV1-2 | TRAJ20 | CAVSNDYKLSF | TRBV2 | TRBJ2-1 | CASSESSGGRNNEQFF | KLGGALQAK | YES | NO | NO |
| 4481 | TRAV10 | TRAJ10 | CVVSDGTGGGNKLTF | TRBV2 | TRBJ2-1 | CASSRDSHSYNEQFF | KLGGALQAK | YES | NO | NO |
| 4483 | TRAV10 | TRAJ45 | CVVSVLGGADGLTF | TRBV2 | TRBJ2-1 | CASSEAGAQLWSSYNEQFF | KLGGALQAK | YES | NO | NO |
| 4484 | TRAV12-1 | TRAJ12 | CVVSGDSSYKLIF | TRBV2 | TRBJ2-1 | CASSSPGLANNEQFF | KLGGALQAK | YES | NO | NO |
| 4485 | TRAV12-1 | TRAJ5 | CVVITGRRALTF | TRBV2 | TRBJ2-1 | CASTKGAVSSYNEQFF | KLGGALQAK | YES | NO | NO |
| 4486 | TRAV12-2 | TRAJ40 | CAVNPSGTYKYIF | TRBV2 | TRBJ2-1 | CASSPSGGDYNEQFF | KLGGALQAK | YES | NO | NO |
| 4487 | TRAV12-2 | TRAJ6 | CAVKNPAGGSYIPTF | TRBV2 | TRBJ2-1 | CASSSNQGLAGLHEQFF | KLGGALQAK | YES | NO | NO |
| 4488 | TRAV12-3 | TRAJ52 | CANSIAGGTSYGKLTF | TRBV2 | TRBJ2-1 | CASSEDTLTNEQFF | KLGGALQAK | YES | NO | NO |
| 4489 | TRAV13-1 | TRAJ9 | CAASRNTGGFKTIF | TRBV2 | TRBJ2-1 | CASSSTKYNEQFF | KLGGALQAK | YES | NO | NO |
| 4490 | TRAV13-2 | TRAJ39 | CAEHDNNAGNMLTF | TRBV2 | TRBJ2-1 | CASSETSGGYYNEQFF | KLGGALQAK | YES | NO | NO |
| 4491 | TRAV13-2 | TRAJ47 | CAEIPMEYGNKLVF | TRBV2 | TRBJ2-1 | CASSVGTTGFSYNEQFF | KLGGALQAK | YES | NO | NO |
| 4493 | TRAV17 | TRAJ24 | CATISTDSWGKFQF | TRBV2 | TRBJ2-1 | CASRQSGGSYNEQFF | KLGGALQAK | YES | NO | NO |
| 4494 | TRAV17 | TRAJ26 | CATGDVF | TRBV2 | TRBJ2-1 | CASSEWTRYNEQFF | AVFDRKSDAK | NO | NO | NO |
| 4494 | TRAV17 | TRAJ26 | CATGDVF | TRBV2 | TRBJ2-1 | CASSEWTRYNEQFF | KLGGALQAK | NO | NO | NO |
| 4495 | TRAV17 | TRAJ39 | CATDTNNAGNMLTF | TRBV2 | TRBJ2-1 | CASGTSGRNEQFF | KLGGALQAK | YES | NO | NO |
| 4496 | TRAV19 | TRAJ49 | CALSEDYTGNQFYF | TRBV2 | TRBJ2-1 | CASSDRGGGPNYNEQFF | KLGGALQAK | YES | NO | NO |
| 4498 | TRAV23DV6 | TRAJ56 | CAASPRTGANSKLTF | TRBV2 | TRBJ2-1 | CASSPDSYNEQFF | KLGGALQAK | YES | NO | NO |
| 4499 | TRAV27 | TRAJ39 | CAGGSNAGNMLTF | TRBV2 | TRBJ2-1 | CAQLAGGPYNEQFF | KLGGALQAK | YES | NO | NO |
| 4500 | TRAV29DV5 | TRAJ52 | CAAIEAGGTSYGKLTF | TRBV2 | TRBJ2-1 | CASVEGDEQFF | KLGGALQAK | YES | NO | NO |
| 4502 | TRAV30 | TRAJ32 | CGTETWGGATNKLIF | TRBV2 | TRBJ2-1 | CASSDLTFNEQFF | KLGGALQAK | YES | NO | NO |
| 4505 | TRAV39 | TRAJ20 | CAVNPSNDYKLSF | TRBV2 | TRBJ2-1 | CASSARLAGDFNEQFF | KLGGALQAK | YES | NO | NO |
| 4507 | TRAV5 | TRAJ3 | CAVASGYSSASKIIF | TRBV2 | TRBJ2-1 | CARVSGLGMYNEQFF | KLGGALQAK | YES | NO | NO |
| 4511 | TRAV1-1 | TRAJ43 | CAVRDNNNDMRF | TRBV20-1 | TRBJ2-1 | CSARDPSGSSYNEQFF | KLGGALQAK | NO | NO | NO |
| 4513 | TRAV1-2 | TRAJ33 | CAVMDSNYQLIW | TRBV20-1 | TRBJ2-1 | CSVPQTEDYNEQFF | KLGGALQAK | NO | NO | NO |
| 4516 | TRAV10 | TRAJ39 | CVVSAGYPGNMLTF | TRBV20-1 | TRBJ2-1 | CSARKGLADYNEQFF | KLGGALQAK | NO | NO | NO |
| 4517 | TRAV10 | TRAJ4 | CVVTDISGGYNKLIF | TRBV20-1 | TRBJ2-1 | CSASDPDNEQFF | KLGGALQAK | NO | NO | NO |
| 4520 | TRAV12-1 | TRAJ48 | CVVTTIFGNEKLTF | TRBV20-1 | TRBJ2-1 | CSARGLAGGYEQFF | KLGGALQAK | NO | NO | NO |
| 4521 | TRAV12-2 | TRAJ13 | CAVNHYSGGYQKVTF | TRBV20-1 | TRBJ2-1 | CSARGFGNHEQFF | KLGGALQAK | YES | NO | NO |
| 4525 | TRAV12-2 | TRAJ39 | CAIGPGNMLTF | TRBV20-1 | TRBJ2-1 | CSAPTGGGLYNEQFF | KLGGALQAK | NO | NO | NO |
| 4526 | TRAV12-2 | TRAJ44 | CAVNGDTGTASKLTF | TRBV20-1 | TRBJ2-1 | CSARDPLTSGHYNEQFF | KLGGALQAK | NO | NO | NO |
| 4528 | TRAV12-2 | TRAJ53 | CAVAGTGGSNYKLTF | TRBV20-1 | TRBJ2-1 | CSARGDRGGSSYNEQFF | KLGGALQAK | NO | NO | NO |
| 4529 | TRAV12-3 | TRAJ39 | CAIGPGNMLTF | TRBV20-1 | TRBJ2-1 | CSARDQRLAGGRVGNEQFF | KLGGALQAK | NO | NO | NO |
| 4532 | TRAV13-1 | TRAJ41 | CAASKGALNF | TRBV20-1 | TRBJ2-1 | CSARDKLAGTYNEQFF | KLGGALQAK | NO | NO | NO |
| 4534 | TRAV14DV4 | TRAJ18 | CAMREYDRGSTLGRLYF | TRBV20-1 | TRBJ2-1 | CSAPGQGVYNEQFF | KLGGALQAK | YES | NO | NO |
| 4535 | TRAV14DV4 | TRAJ22 | CAMRGPFSSARQLTF | TRBV20-1 | TRBJ2-1 | CSARDQRLAGGRVGNEQFF | KLGGALQAK | NO | NO | NO |
| 4536 | TRAV14DV4 | TRAJ24 | CAMRVWTDSWGKLQF | TRBV20-1 | TRBJ2-1 | CKYNEQFF | KLGGALQAK | NO | NO | NO |
| 4538 | TRAV16 | TRAJ12 | CALSWMDSSYKLIF | TRBV20-1 | TRBJ2-1 | CSARDLWGGPGNEQFF | KLGGALQAK | NO | NO | NO |
| 4545 | TRAV17 | TRAJ23 | CATAPYNQGGKLIF | TRBV20-1 | TRBJ2-1 | CSARGGLRDTYNEQFF | KLGGALQAK | NO | NO | NO |
| 4548 | TRAV19 | TRAJ18 | CALSVGRGSTLGRLYF | TRBV20-1 | TRBJ2-1 | CSAREQGATYNEQFF | KLGGALQAK | NO | NO | NO |
| 4552 | TRAV19 | TRAJ49 | CALSANGNQFYF | TRBV20-1 | TRBJ2-1 | CSAYPLAGTIDEQFF | KLGGALQAK | NO | NO | NO |
| 4553 | TRAV19 | TRAJ5 | CALSEALGGRRALTF | TRBV20-1 | TRBJ2-1 | CSASAGSGANNEQFF | KLGGALQAK | YES | NO | NO |
| 4554 | TRAV19 | TRAJ56 | CALRVTGANSKLTF | TRBV20-1 | TRBJ2-1 | CSARDIHGQGTYNEQFF | RLRAEAQVK | NO | NO | NO |
| 4555 | TRAV19 | TRAJ8 | CALSEARSTGFQKLVF | TRBV20-1 | TRBJ2-1 | CSARDIGVEQFF | KLGGALQAK | YES | NO | NO |
| 4556 | TRAV20 | TRAJ16 | CAVASF | TRBV20-1 | TRBJ2-1 | CSARGGGISYNEQFF | KLGGALQAK | NO | NO | NO |
| 4557 | TRAV20 | TRAJ30 | CAVQAGRDDKIIF | TRBV20-1 | TRBJ2-1 | CSARVPSYNEQFF | KLGGALQAK | NO | NO | NO |
| 4558 | TRAV20 | TRAJ42 | CAVQARGSQGNLIF | TRBV20-1 | TRBJ2-1 | CSARDALAGTYNEQFF | KLGGALQAK | NO | NO | NO |
| 4559 | TRAV21 | TRAJ23 | CAVRGGKLIF | TRBV20-1 | TRBJ2-1 | CSASFSPGQGMGTPYNEQFF | KLGGALQAK | NO | NO | NO |
| 4560 | TRAV21 | TRAJ24 | CETTDSWGKFQF | TRBV20-1 | TRBJ2-1 | CSARDRPGTGGFHEQFF | KLGGALQAK | NO | NO | NO |
| 4561 | TRAV21 | TRAJ28 | CAVSGIYSGAGSYQLTF | TRBV20-1 | TRBJ2-1 | CSAKLAGGHNEQFF | KLGGALQAK | NO | NO | NO |
| 4562 | TRAV21 | TRAJ35 | CAVSI | TRBV20-1 | TRBJ2-1 | CSASRLAGVYNEQFF | KLGGALQAK | NO | NO | NO |

| | | | | | | | | | | |
|---|---|---|---|---|---|---|---|---|---|---|
| 4564 | TRAV24 | TRAJ52 | CAFIGDAGGTSYGKLTF | TRBV20-1 | TRBJ2-1 | CSARDNNEQFF | KLGGALQAK | YES | NO | NO |
| 4565 | TRAV24 | TRAJ57 | CASDTQGGSEKLVF | TRBV20-1 | TRBJ2-1 | CSARDLSGGSYNEQFF | KLGGALQAK | NO | NO | NO |
| 4568 | TRAV26-2 | TRAJ42 | CILNYGGSQGNLIF | TRBV20-1 | TRBJ2-1 | CSASEGMGYNEQFF | KLGGALQAK | NO | NO | NO |
| 4569 | TRAV27 | TRAJ42 | CAGARSGSQGNLIF | TRBV20-1 | TRBJ2-1 | CSASGPGLYYNEQFF | KLGGALQAK | NO | NO | NO |
| 4569 | TRAV27 | TRAJ42 | CAGARSGSQGNLIF | TRBV20-1 | TRBJ2-1 | CSASGPGLYYNEQFF | AVFDRKSDAK | NO | NO | NO |
| 4570 | TRAV29DV5 | TRAJ37 | CAASGSSNTGKLIF | TRBV20-1 | TRBJ2-1 | CSAAKTSVYEQFF | KLGGALQAK | YES | NO | NO |
| 4572 | TRAV3 | TRAJ26 | CAVKGNYGQNFVF | TRBV20-1 | TRBJ2-1 | CSVWSGDSYNEQFF | KLGGALQAK | NO | NO | NO |
| 4573 | TRAV3 | TRAJ6 | CAVRDRGGGSYIPTF | TRBV20-1 | TRBJ2-1 | CSASLASGRDEQFF | KLGGALQAK | NO | NO | NO |
| 4574 | TRAV35 | TRAJ49 | CAGLGNQFYF | TRBV20-1 | TRBJ2-1 | CSARPTSSNEQFF | KLGGALQAK | NO | NO | NO |
| 4575 | TRAV36DV7 | TRAJ48 | CAVISNFGNEKLTF | TRBV20-1 | TRBJ2-1 | CSARETGYDGPYNEQFF | KLGGALQAK | NO | NO | NO |
| 4576 | TRAV38-1 | TRAJ54 | CAFMSLIQGAQKLVF | TRBV20-1 | TRBJ2-1 | CSARATSGTLNEQFF | KLGGALQAK | NO | NO | NO |
| 4577 | TRAV38-2DV8 | TRAJ28 | CAYPLGAGSYQLTF | TRBV20-1 | TRBJ2-1 | CSAWTSGGLSSYNEQFF | KLGGALQAK | NO | NO | NO |
| 4578 | TRAV38-2DV8 | TRAJ42 | CAYRSASYGGSQGNLIF | TRBV20-1 | TRBJ2-1 | CSATISGRPYNEQFF | KLGGALQAK | NO | NO | NO |
| 4580 | TRAV4 | TRAJ23 | CLVGGFYNQGGKLIF | TRBV20-1 | TRBJ2-1 | CSARTSGGDSNEQFF | KLGGALQAK | NO | NO | NO |
| 4581 | TRAV4 | TRAJ6 | CLVVAGGSYIPTF | TRBV20-1 | TRBJ2-1 | CSASWLAGFNEQFF | KLGGALQAK | YES | NO | NO |
| 4584 | TRAV5 | TRAJ42 | CAEYGGSQGNLIF | TRBV20-1 | TRBJ2-1 | CSALSTIQLNEQFF | KLGGALQAK | NO | NO | NO |
| 4585 | TRAV8-1 | TRAJ10 | CAVNAITGGGNKLTF | TRBV20-1 | TRBJ2-1 | CSAEWLGEQFF | AVFDRKSDAK | NO | NO | NO |
| 4585 | TRAV8-1 | TRAJ10 | CAVNAITGGGNKLTF | TRBV20-1 | TRBJ2-1 | CSAEWLGEQFF | KLGGALQAK | NO | NO | NO |
| 4586 | TRAV8-1 | TRAJ34 | CAVNGHNTDKLIF | TRBV20-1 | TRBJ2-1 | CSASDSYNEQFF | KLGGALQAK | YES | NO | NO |
| 4587 | TRAV8-2 | TRAJ13 | CVVSDLNLHSGGYQKVTF | TRBV20-1 | TRBJ2-1 | CSASNAGPRSEQFF | KLGGALQAK | YES | NO | NO |
| 4590 | TRAV8-3 | TRAJ37 | CAVGDSSNTGKLIF | TRBV20-1 | TRBJ2-1 | CSAREPGTIFYNEQFF | KLGGALQAK | NO | NO | NO |
| 4592 | TRAV8-3 | TRAJ53 | CAVGADRGSNYKLTF | TRBV20-1 | TRBJ2-1 | CSADKYNEQFF | KLGGALQAK | YES | NO | NO |
| 4596 | TRAV9-2 | TRAJ17 | CARAAGNKLTF | TRBV20-1 | TRBJ2-1 | CSARDGSYRTSGLNEQFF | KLGGALQAK | NO | NO | NO |
| 4604 | TRAV1-2 | TRAJ33 | CAPRMDSNYQLIW | TRBV24-1 | TRBJ2-1 | CATTGSNDEQFF | KLGGALQAK | YES | NO | NO |
| 4605 | TRAV1-2 | TRAJ6 | CAVRSSGGSYIPTF | TRBV24-1 | TRBJ2-1 | CATSDGRMVSYNEQFF | KLGGALQAK | YES | NO | NO |
| 4607 | TRAV12-1 | TRAJ12 | CVVNAVDSSYKLIF | TRBV24-1 | TRBJ2-1 | CATSERTGNEQFF | KLGGALQAK | YES | NO | NO |
| 4608 | TRAV12-1 | TRAJ20 | CVVNRGDYKLSF | TRBV24-1 | TRBJ2-1 | CATSDPWGGREWDNEQFF | KLGGALQAK | YES | NO | NO |
| 4612 | TRAV12-1 | TRAJ50 | CVVRVHDKVIF | TRBV24-1 | TRBJ2-1 | CATSELGGGLTDEQFF | KLGGALQAK | YES | NO | NO |
| 4613 | TRAV12-2 | TRAJ12 | CASVGMDSSYKLIF | TRBV24-1 | TRBJ2-1 | CATSDFRDSSNEQFF | KLGGALQAK | YES | NO | NO |
| 4614 | TRAV12-2 | TRAJ42 | CAVNIVYGGSQGNLIF | TRBV24-1 | TRBJ2-1 | CATSDTQKLDTYNEQFF | KLGGALQAK | YES | NO | NO |
| 4617 | TRAV12-3 | TRAJ9 | CAMSEDTGGFKTIF | TRBV24-1 | TRBJ2-1 | CATKRGGNNEQFF | KLGGALQAK | YES | NO | NO |
| 4620 | TRAV13-2 | TRAJ21 | CAENRNFNKFYF | TRBV24-1 | TRBJ2-1 | CATTSGSSYNEQFF | KLGGALQAK | YES | NO | NO |
| 4621 | TRAV13-2 | TRAJ43 | CAEISGDMRF | TRBV24-1 | TRBJ2-1 | CATRGGAGNEQFF | KLGGALQAK | YES | NO | NO |
| 4625 | TRAV14DV4 | TRAJ6 | CAMREPPASGGSYIPTF | TRBV24-1 | TRBJ2-1 | CATSDRDRGDEQFF | KLGGALQAK | YES | NO | NO |
| 4627 | TRAV17 | TRAJ23 | CATVLYNQGGKLIF | TRBV24-1 | TRBJ2-1 | CATSDGTGGYNEQFF | KLGGALQAK | YES | NO | NO |
| 4628 | TRAV19 | TRAJ26 | CALSENNYGQNFVF | TRBV24-1 | TRBJ2-1 | CATSDLRDREQFF | KLGGALQAK | YES | NO | NO |
| 4630 | TRAV21 | TRAJ26 | CAVNNYGQNFVF | TRBV24-1 | TRBJ2-1 | CATSEGTSGRYNEQFF | KLGGALQAK | YES | NO | NO |
| 4631 | TRAV21 | TRAJ5 | CAAKEATGRRALTF | TRBV24-1 | TRBJ2-1 | CATSDPDREYKDNEQFF | KLGGALQAK | YES | NO | NO |
| 4632 | TRAV22 | TRAJ23 | CAVGENQGGKLIF | TRBV24-1 | TRBJ2-1 | CATSDSGGSSYNEQFF | KLGGALQAK | YES | NO | NO |
| 4633 | TRAV26-2 | TRAJ20 | CILSNDYKLSF | TRBV24-1 | TRBJ2-1 | CATSRDHEQFF | KLGGALQAK | NO | NO | NO |
| 4634 | TRAV29DV5 | TRAJ33 | CAASGTDSNYQLIW | TRBV24-1 | TRBJ2-1 | CATSDLEGEQFF | KLGGALQAK | YES | NO | NO |
| 4637 | TRAV36DV7 | TRAJ48 | CAVRGANFGNEKLTF | TRBV24-1 | TRBJ2-1 | CATSLGNEQFF | KLGGALQAK | NO | NO | NO |
| 4638 | TRAV38-2DV8 | TRAJ49 | CAYEDPGNQFYF | TRBV24-1 | TRBJ2-1 | CATSGPGLDNEQFF | KLGGALQAK | YES | NO | NO |
| 4639 | TRAV38-2DV8 | TRAJ53 | CAYNGGSNYKLTF | TRBV24-1 | TRBJ2-1 | CATSDVGSWGRDEQFF | KLGGALQAK | YES | NO | NO |
| 4640 | TRAV4 | TRAJ43 | CLVTRDNNNDMRF | TRBV24-1 | TRBJ2-1 | CATSDFSGSYNEQFF | KLGGALQAK | YES | NO | NO |
| 4642 | TRAV8-2 | TRAJ11 | CVVSDSSGYSTLTF | TRBV24-1 | TRBJ2-1 | CATSDLSSGRGYNEQFF | KLGGALQAK | YES | NO | NO |
| 4643 | TRAV8-3 | TRAJ17 | CAVGELAAGNKLTF | TRBV24-1 | TRBJ2-1 | CATSDGRDDEQFF | KLGGALQAK | YES | NO | NO |
| 4645 | TRAV8-6 | TRAJ40 | CAAWGGTYKYIF | TRBV24-1 | TRBJ2-1 | CATSGGGPYNEQFF | KLGGALQAK | YES | NO | NO |
| 4646 | TRAV9-2 | TRAJ54 | CALSGLIQGAQKLVF | TRBV24-1 | TRBJ2-1 | CATSVDRGLYNEQFF | KLGGALQAK | YES | NO | NO |
| 4647 | TRAV10 | TRAJ41 | CVVSAGYALNF | TRBV25-1 | TRBJ2-1 | CASRSLQTFF | KLGGALQAK | YES | NO | NO |
| 4648 | TRAV12-2 | TRAJ26 | CAGIDNYGQNFVF | TRBV25-1 | TRBJ2-1 | CASSDRDRGHEQFF | KLGGALQAK | YES | NO | NO |
| 4649 | TRAV14DV4 | TRAJ20 | CAMREAFSNDYKLSF | TRBV25-1 | TRBJ2-1 | CASSEGLGLAGGEQFF | KLGGALQAK | YES | NO | NO |
| 4650 | TRAV16 | TRAJ8 | CALNFRTGFQKLVF | TRBV25-1 | TRBJ2-1 | CASSYTGQGLVNEQFF | KLGGALQAK | NO | NO | NO |
| 4652 | TRAV3 | TRAJ26 | CAVRDYGQNFVF | TRBV25-1 | TRBJ2-1 | CASIASGPYNEQFF | KLGGALQAK | NO | NO | NO |
| 4653 | TRAV3 | TRAJ49 | CAVRDSNTGNQFYF | TRBV25-1 | TRBJ2-1 | CASSVPGQNNEQFF | KLGGALQAK | NO | NO | NO |
| 4654 | TRAV1-1 | TRAJ23 | CAVSLHQGGKLIF | TRBV27 | TRBJ2-1 | CASSLRGLAMSSYNEQFF | KLGGALQAK | YES | NO | NO |
| 4655 | TRAV1-1 | TRAJ31 | CAVPNNARLMF | TRBV27 | TRBJ2-1 | CASSIEPYNEQFF | KLGGALQAK | YES | NO | NO |
| 4657 | TRAV1-2 | TRAJ27 | CAVIGNAGKSTF | TRBV27 | TRBJ2-1 | CASSIEPYNEQFF | KLGGALQAK | YES | NO | NO |
| 4657 | TRAV1-2 | TRAJ27 | CAVIGNAGKSTF | TRBV27 | TRBJ2-1 | CASSIEPYNEQFF | AVFDRKSDAK | YES | NO | NO |
| 4658 | TRAV1-2 | TRAJ27 | CAAGALNTNAGKSTF | TRBV27 | TRBJ2-1 | CASSFPGGQGYNEQFF | KLGGALQAK | YES | NO | NO |
| 4659 | TRAV1-2 | TRAJ33 | CAVKDSNYQLIW | TRBV27 | TRBJ2-1 | CASSNTRQGDEQFF | KLGGALQAK | YES | NO | NO |
| 4660 | TRAV10 | TRAJ6 | CVVSAWNPSGGSYIPTF | TRBV27 | TRBJ2-1 | CASSWPNGDEQFF | KLGGALQAK | YES | NO | NO |
| 4661 | TRAV12-1 | TRAJ32 | CVDANYGGATNKLIF | TRBV27 | TRBJ2-1 | CASSFWDTINEQFF | KLGGALQAK | YES | NO | NO |
| 4662 | TRAV12-1 | TRAJ42 | CVVNSRGYGGSQGNLIF | TRBV27 | TRBJ2-1 | CASSTSGPYNEQFF | KLGGALQAK | YES | NO | NO |
| 4663 | TRAV12-1 | TRAJ47 | CVVPHGNKLVF | TRBV27 | TRBJ2-1 | CASSLLGTQGGGDEQFF | KLGGALQAK | YES | NO | NO |
| 4664 | TRAV12-2 | TRAJ13 | CAGSGGYQKVTF | TRBV27 | TRBJ2-1 | CATGQTDNEQFF | KLGGALQAK | YES | NO | NO |
| 4665 | TRAV12-2 | TRAJ27 | CAVISAGKSTF | TRBV27 | TRBJ2-1 | CASNSGLDNNEQFF | KLGGALQAK | YES | NO | NO |
| 4666 | TRAV12-2 | TRAJ30 | CAARDDKIIF | TRBV27 | TRBJ2-1 | CASSLFGLAGDYNEQFF | KLGGALQAK | YES | NO | NO |
| 4668 | TRAV12-2 | TRAJ45 | CAVSGGGADGLTF | TRBV27 | TRBJ2-1 | CASANGLAGGEQFF | KLGGALQAK | YES | NO | NO |
| 4670 | TRAV12-3 | TRAJ53 | CAMSGGSNYKLTF | TRBV27 | TRBJ2-1 | CASSLEVQSRDEQFF | KLGGALQAK | YES | NO | NO |
| 4673 | TRAV13-2 | TRAJ15 | CAETNQAGTALIF | TRBV27 | TRBJ2-1 | CASSLGQVIDEQFF | KLGGALQAK | YES | NO | NO |
| 4675 | TRAV13-2 | TRAJ47 | CAESWEYGNKLVF | TRBV27 | TRBJ2-1 | CASGKSGTYGDEQFF | KLGGALQAK | YES | NO | NO |
| 4676 | TRAV13-2 | TRAJ58 | CAENQETSGSRLTF | TRBV27 | TRBJ2-1 | CASGLWNEQFF | KLGGALQAK | YES | NO | NO |
| 4679 | TRAV14DV4 | TRAJ29 | CAMRGLYSGNTPLVF | TRBV27 | TRBJ2-1 | CASSFPNLTPLIF | KLGGALQAK | YES | NO | NO |
| 4680 | TRAV14DV4 | TRAJ3 | CAMRGGYSSASKIIF | TRBV27 | TRBJ2-1 | CASSLSQTSGDTSRDEQFF | KLGGALQAK | YES | NO | NO |
| 4681 | TRAV14DV4 | TRAJ30 | CAMRNMNRDDKIIF | TRBV27 | TRBJ2-1 | CASSTPDRGRWGEQFF | KLGGALQAK | YES | NO | NO |
| 4682 | TRAV16 | TRAJ18 | CAGTDDGSTLGRLYF | TRBV27 | TRBJ2-1 | CASSLFPSYNEQFF | KLGGALQAK | NO | NO | NO |
| 4683 | TRAV17 | TRAJ20 | CATDADDYKLSF | TRBV27 | TRBJ2-1 | CASTPRDYNEQFF | KLGGALQAK | YES | NO | NO |
| 4684 | TRAV17 | TRAJ34 | CATDKRADKLIF | TRBV27 | TRBJ2-1 | CASRQGPNEQFF | KLGGALQAK | YES | NO | NO |
| 4686 | TRAV19 | TRAJ16 | CALSALDGQKLLF | TRBV27 | TRBJ2-1 | CASREKGHRRDNEQFF | KLGGALQAK | NO | NO | NO |
| 4687 | TRAV19 | TRAJ30 | CALSLNRDDKIIF | TRBV27 | TRBJ2-1 | CASSTGQIYNEQFF | KLGGALQAK | YES | NO | NO |
| 4688 | TRAV19 | TRAJ35 | CALSEGMAGFGNVLHC | TRBV27 | TRBJ2-1 | CASSLSGPSYNEQFF | KLGGALQAK | NO | NO | NO |
| 4691 | TRAV19 | TRAJ6 | CALSETASGGSYIPTF | TRBV27 | TRBJ2-1 | CASRRPTNEQFF | KLGGALQAK | YES | NO | NO |
| 4693 | TRAV20 | TRAJ58 | CAVQAWEETSGSRLTF | TRBV27 | TRBJ2-1 | CASRWAGGEQFF | KLGGALQAK | YES | NO | NO |
| 4694 | TRAV21 | TRAJ12 | CAVLMDSSYKLIF | TRBV27 | TRBJ2-1 | CASLKRGGSYNEQFF | AVFDRKSDAK | YES | NO | NO |
| 4695 | TRAV21 | TRAJ17 | CAVRPLIAAGNKLTF | TRBV27 | TRBJ2-1 | CASTQGAGVPYNEQFF | KLGGALQAK | YES | NO | NO |
| 4697 | TRAV21 | TRAJ43 | CAVSQKLYGRF | TRBV27 | TRBJ2-1 | CASRRTGGLNEQFF | KLGGALQAK | YES | NO | NO |
| 4698 | TRAV21 | TRAJ47 | CAVMEYGNKLVF | TRBV27 | TRBJ2-1 | CASTSPKTGVYNEQFF | KLGGALQAK | YES | NO | NO |
| 4699 | TRAV21 | TRAJ56 | CAVPSAYTGANSKLTF | TRBV27 | TRBJ2-1 | CASSTGTLGYNEQFF | KLGGALQAK | YES | NO | NO |
| 4700 | TRAV22 | TRAJ12 | CAVEDSSYKLIF | TRBV27 | TRBJ2-1 | CASSPTSGSLFTNEQFF | IVTDFSVIK | YES | NO | NO |
| 4701 | TRAV22 | TRAJ12 | CAVEDSSYKLIL | TRBV27 | TRBJ2-1 | CASSPTSGSLFTNEQFF | KLGGALQAK | NO | NO | NO |
| 4702 | TRAV22 | TRAJ21 | CAGRNNFNKFYF | TRBV27 | TRBJ2-1 | CASSPGQGTSYNEQFF | KLGGALQAK | YES | NO | NO |
| 4704 | TRAV24 | TRAJ42 | CAFYGGSQGNLIF | TRBV27 | TRBJ2-1 | CASSAGTGNNEQFF | KLGGALQAK | YES | NO | NO |
| 4705 | TRAV25 | TRAJ26 | CAGPDYGQNFVF | TRBV27 | TRBJ2-1 | CASSGQGAGYNEQFF | KLGGALQAK | YES | NO | NO |
| 4706 | TRAV26-1 | TRAJ34 | CIVRVYNTDKLIF | TRBV27 | TRBJ2-1 | CASSTRASGSDEQFF | KLGGALQAK | YES | NO | NO |
| 4707 | TRAV26-2 | TRAJ30 | CIRERMNRDDKIIF | TRBV27 | TRBJ2-1 | CASSPSTGGYEQFF | KLGGALQAK | YES | NO | NO |
| 4707 | TRAV26-2 | TRAJ30 | CIRERMNRDDKIIF | TRBV27 | TRBJ2-1 | CASSPSTGGYEQFF | AVFDRKSDAK | YES | NO | NO |
| 4708 | TRAV26-2 | TRAJ52 | CILRDEGAWAGGTSYGKLTF | TRBV27 | TRBJ2-1 | CASTTQGDSGQGLSAVGSYNEQFF | KLGGALQAK | NO | NO | NO |
| 4710 | TRAV27 | TRAJ20 | CAGVESNDYKLSF | TRBV27 | TRBJ2-1 | CASSPTSGSLFTNEQFF | KLGGALQAK | YES | NO | NO |
| 4711 | TRAV27 | TRAJ20 | CAGAAYKLSF | TRBV27 | TRBJ2-1 | CASSLYLQGRGEQFF | KLGGALQAK | YES | YES | NO |
| 4712 | TRAV27 | TRAJ20 | CAGEPSNDYKLSF | TRBV27 | TRBJ2-1 | CASSLSGSEGNEQFF | KLGGALQAK | YES | NO | NO |
| 4713 | TRAV27 | TRAJ48 | CAGAAAGFGNEKLTF | TRBV27 | TRBJ2-1 | CASSFGNEQFF | KLGGALQAK | YES | NO | NO |
| 4714 | TRAV29DV5 | TRAJ38 | CAAPKGNAGNNRKLIW | TRBV27 | TRBJ2-1 | CASSLWGASGDPFF | KLGGALQAK | YES | NO | NO |
| 4715 | TRAV3 | TRAJ26 | CAVRLHNYGQNFVF | TRBV27 | TRBJ2-1 | CASRVSGSLSYNEQFF | KLGGALQAK | NO | NO | NO |

| | | | | | | | | | | |
|---|---|---|---|---|---|---|---|---|---|---|
| 4716 | TRAV3 | TRAJ8 | CAVRSLTWKLVF | TRBV27 | TRBJ2-1 | CASSPTGGGYNEQFF | KLGGALQAK | NO | NO | NO |
| 4717 | TRAV30 | TRAJ58 | CGKGETSGSRLTF | TRBV27 | TRBJ2-1 | CAGSLAGTYNEQFF | KLGGALQAK | YES | NO | NO |
| 4719 | TRAV35 | TRAJ42 | CAGSYGGSQGNLIF | TRBV27 | TRBJ2-1 | CASRKGAPGINEQFF | KLGGALQAK | YES | NO | NO |
| 4721 | TRAV38-1 | TRAJ31 | CALNNARLMF | TRBV27 | TRBJ2-1 | CASSPLTEGDEQFF | KLGGALQAK | YES | NO | NO |
| 4722 | TRAV38-1 | TRAJ48 | CAFMKRLLNFGNEKLTF | TRBV27 | TRBJ2-1 | CASSLPGLAGGYNEQFF | KLGGALQAK | YES | NO | NO |
| 4724 | TRAV38-2DV8 | TRAJ29 | CAYGSGNTPLVF | TRBV27 | TRBJ2-1 | CASSFDRGYNEQFF | KLGGALQAK | YES | NO | NO |
| 4725 | TRAV38-2DV8 | TRAJ54 | CAYEGGAQKLVF | TRBV27 | TRBJ2-1 | CASSLFSGLADEQFF | KLGGALQAK | YES | NO | NO |
| 4726 | TRAV4 | TRAJ26 | CLVGDAWRDNYGQNFVF | TRBV27 | TRBJ2-1 | CASSLSSSGGPYNEQFF | KLGGALQAK | YES | NO | NO |
| 4727 | TRAV8-2 | TRAJ43 | CAVSDNNNDMRF | TRBV27 | TRBJ2-1 | CASKGLNYNEQFF | KLGGALQAK | YES | NO | NO |
| 4728 | TRAV8-2 | TRAJ43 | CVVSDSNDMRF | TRBV27 | TRBJ2-1 | CASSPSVHEQFF | KLGGALQAK | YES | NO | NO |
| 4729 | TRAV8-3 | TRAJ42 | CAVGRGSQGNLIF | TRBV27 | TRBJ2-1 | CASSPRLAFSSSYNEQFF | RLRAEAQVK | YES | NO | YES |
| 4730 | TRAV8-6 | TRAJ48 | CAVSANFGNEKLTF | TRBV27 | TRBJ2-1 | CASSVLRGNPYNEQFF | KLGGALQAK | YES | NO | NO |
| 4733 | TRAV9-2 | TRAJ11 | CALSDHSGYSTLTF | TRBV27 | TRBJ2-1 | CASSFTNNEQFF | KLGGALQAK | YES | NO | NO |
| 4734 | TRAV9-2 | TRAJ11 | CALSDHSGYSTLTF | TRBV27 | TRBJ2-1 | CASSPSDNEQFF | KLGGALQAK | YES | YES | NO |
| 4735 | TRAV1-2 | TRAJ13 | CAVRGTNSGGYQKVTF | TRBV28 | TRBJ2-1 | CASSLSSRLGVNEQFF | KLGGALQAK | YES | NO | NO |
| 4736 | TRAV10 | TRAJ40 | CVVEGSYKYIF | TRBV28 | TRBJ2-1 | CASGGSPISLVDNEQFF | KLGGALQAK | YES | NO | NO |
| 4737 | TRAV12-3 | TRAJ6 | CAVNWEEQGGSYIPTF | TRBV28 | TRBJ2-1 | CASSPSNEQFF | KLGGALQAK | YES | NO | NO |
| 4740 | TRAV14DV4 | TRAJ40 | CAMRINLKYIF | TRBV28 | TRBJ2-1 | CASRVEDRHSYNEQFF | KLGGALQAK | YES | NO | NO |
| 4741 | TRAV26-2 | TRAJ57 | CILVQGGSEKLVF | TRBV28 | TRBJ2-1 | CASSLGGFSSYNEQFF | KLGGALQAK | YES | NO | NO |
| 4744 | TRAV38-1 | TRAJ43 | CAFISNDMRF | TRBV28 | TRBJ2-1 | CASSFRGGGEQFF | KLGGALQAK | YES | NO | NO |
| 4745 | TRAV6 | TRAJ21 | CALDLYNFNKFYF | TRBV28 | TRBJ2-1 | CASSHSLGQYNEQFF | KLGGALQAK | YES | NO | NO |
| 4749 | TRAV12-1 | TRAJ39 | CVVNAFNNAGNMLTF | TRBV29-1 | TRBJ2-1 | CSVELTGVGDEQFF | KLGGALQAK | YES | NO | NO |
| 4750 | TRAV12-1 | TRAJ9 | CVVNTGGFKTIF | TRBV29-1 | TRBJ2-1 | CSGTGTSGRASNEQFF | KLGGALQAK | YES | NO | NO |
| 4751 | TRAV12-2 | TRAJ16 | CAVNRAPKLLF | TRBV29-1 | TRBJ2-1 | CSVELTGVGDEQFF | KLGGALQAK | NO | NO | NO |
| 4752 | TRAV12-2 | TRAJ17 | CAVPLTQGLTF | TRBV29-1 | TRBJ2-1 | CSVENPTGDNNEQFF | KLGGALQAK | YES | NO | NO |
| 4754 | TRAV12-2 | TRAJ47 | CAVEVEYGNKLVF | TRBV29-1 | TRBJ2-1 | CSVPGLAGIGDEQFF | AVFDRKSDAK | YES | NO | NO |
| 4756 | TRAV14DV4 | TRAJ32 | CAMRVGGATNKLIF | TRBV29-1 | TRBJ2-1 | CSVAGTPELYNEQFF | KLGGALQAK | YES | NO | NO |
| 4758 | TRAV17 | TRAJ52 | CATDALPNAGGTSYGKLTF | TRBV29-1 | TRBJ2-1 | CSVEDQNGGATYNEQFF | KLGGALQAK | YES | NO | NO |
| 4759 | TRAV19 | TRAJ18 | CALSEAFGRGSTLGRLYF | TRBV29-1 | TRBJ2-1 | CSVAGASGNEQFF | KLGGALQAK | YES | NO | NO |
| 4762 | TRAV19 | TRAJ43 | CALTVRNNNDMRF | TRBV29-1 | TRBJ2-1 | CSVDGLAGIYNEQFF | KLGGALQAK | YES | NO | NO |
| 4763 | TRAV19 | TRAJ52 | CALSEGGGGTSYGKLTF | TRBV29-1 | TRBJ2-1 | CSVPPGGANEQFF | KLGGALQAK | YES | YES | NO |
| 4766 | TRAV29DV5 | TRAJ31 | CAARLFDARLMF | TRBV29-1 | TRBJ2-1 | CSVESSEQFF | KLGGALQAK | YES | NO | NO |
| 4767 | TRAV29DV5 | TRAJ53 | CAASGGSNYKLTF | TRBV29-1 | TRBJ2-1 | CSVGSGVDQFF | KLGGALQAK | YES | NO | NO |
| 4768 | TRAV3 | TRAJ16 | CAVDDGQKLLF | TRBV29-1 | TRBJ2-1 | CSVLPLNNEQFF | KLGGALQAK | NO | NO | NO |
| 4770 | TRAV35 | TRAJ42 | CAGLPTMNYGGSQGNLIF | TRBV29-1 | TRBJ2-1 | CSVEDLRRSSYNEQFF | KLGGALQAK | YES | NO | NO |
| 4772 | TRAV8-1 | TRAJ45 | CAVNRGGGADGLTF | TRBV29-1 | TRBJ2-1 | CSVDHVPNEQFF | KLGGALQAK | YES | NO | NO |
| 4773 | TRAV1-2 | TRAJ15 | CAVSQAGTALIF | TRBV3-1 | TRBJ2-1 | CASSHLSGVRSYNEQFF | KLGGALQAK | YES | NO | NO |
| 4775 | TRAV10 | TRAJ31 | CVVSAVGANNARLMF | TRBV3-1 | TRBJ2-1 | CASSQDGGSSYNEQFF | KLGGALQAK | YES | NO | NO |
| 4776 | TRAV10 | TRAJ43 | CVVSAYHNNDMRF | TRBV3-1 | TRBJ2-1 | CASSSRLASSYNEQFF | KLGGALQAK | YES | NO | NO |
| 4777 | TRAV12-1 | TRAJ23 | CVVNRGYNQGGKLIF | TRBV3-1 | TRBJ2-1 | CASSQDASGYYNEQFF | KLGGALQAK | YES | NO | NO |
| 4779 | TRAV12-2 | TRAJ31 | CAVIQARLMF | TRBV3-1 | TRBJ2-1 | CASSQDTSGLYNEQFF | KLGGALQAK | YES | NO | NO |
| 4779 | TRAV12-2 | TRAJ31 | CAVIQARLMF | TRBV3-1 | TRBJ2-1 | CASSQDTSGLYNEQFF | AVFDRKSDAK | YES | NO | NO |
| 4780 | TRAV12-3 | TRAJ52 | CAMGAGGTSYGKLTF | TRBV3-1 | TRBJ2-1 | CASRTAGAPYNEQFF | KLGGALQAK | YES | NO | NO |
| 4781 | TRAV13-1 | TRAJ34 | CAALNTDKLIF | TRBV3-1 | TRBJ2-1 | CASSHTSASYNEQFF | KLGGALQAK | YES | NO | NO |
| 4786 | TRAV17 | TRAJ24 | CATPLDSWGKLQF | TRBV3-1 | TRBJ2-1 | CASSHRTTYNEQFF | KLGGALQAK | YES | NO | NO |
| 4788 | TRAV21 | TRAJ23 | CALIYNQGGKLIF | TRBV3-1 | TRBJ2-1 | CASSPRVWTYNEQFF | KLGGALQAK | YES | NO | NO |
| 4790 | TRAV21 | TRAJ7 | CAVPYGNNRLAF | TRBV3-1 | TRBJ2-1 | CASSQDRGGVPSYNEQFF | KLGGALQAK | YES | NO | NO |
| 4792 | TRAV26-2 | TRAJ12 | CILRDGMDSSYKLIF | TRBV3-1 | TRBJ2-1 | CASSQGLSGQGYNEQFF | KLGGALQAK | YES | NO | NO |
| 4793 | TRAV29DV5 | TRAJ52 | CAATDLTNAGGTSYGKLTF | TRBV3-1 | TRBJ2-1 | CASSRGGLAENEQFF | KLGGALQAK | YES | NO | NO |
| 4795 | TRAV38-2DV8 | TRAJ40 | CAYRSGGTYKYIF | TRBV3-1 | TRBJ2-1 | CASSPTGKSYNEQFF | KLGGALQAK | YES | NO | NO |
| 4796 | TRAV8-3 | TRAJ21 | CAVGDNFNKFYF | TRBV3-1 | TRBJ2-1 | CASSHLGGDRYMNEQFF | KLGGALQAK | YES | NO | NO |
| 4799 | TRAV9-2 | TRAJ30 | CALLNRDDKIIF | TRBV3-1 | TRBJ2-1 | CASSHLGGDRYMNEQFF | KLGGALQAK | YES | NO | NO |
| 4799 | TRAV9-2 | TRAJ30 | CALLNRDDKIIF | TRBV3-1 | TRBJ2-1 | CASSHLGGDRYMNEQFF | AVFDRKSDAK | YES | NO | NO |
| 4800 | TRAV1-1 | TRAJ27 | CAVNNAGKSTF | TRBV30 | TRBJ2-1 | CAWTPSGRANNEQFF | KLGGALQAK | YES | NO | NO |
| 4803 | TRAV12-2 | TRAJ30 | CAGGDKIIF | TRBV30 | TRBJ2-1 | CAWSLAYNEQFF | KLGGALQAK | NO | NO | NO |
| 4804 | TRAV12-2 | TRAJ54 | CAVIQGAQKLVF | TRBV30 | TRBJ2-1 | CAWSDSMWNEQFF | KLGGALQAK | NO | NO | NO |
| 4805 | TRAV12-3 | TRAJ49 | CAMYTGNQFYF | TRBV30 | TRBJ2-1 | CAWGGNNEQFF | KLGGALQAK | NO | NO | NO |
| 4808 | TRAV17 | TRAJ17 | CATDARAAGNKLTF | TRBV30 | TRBJ2-1 | CAWSRGQGDNEQFF | KLGGALQAK | YES | NO | NO |
| 4813 | TRAV29DV5 | TRAJ42 | CAASATPLFLYLEYLIF | TRBV30 | TRBJ2-1 | CAWETGSNEQFF | RLRAEAQVK | NO | NO | NO |
| 4816 | TRAV8-3 | TRAJ32 | CAVAGTGEGGATNKLIF | TRBV30 | TRBJ2-1 | CAWSRGGVDEQFF | KLGGALQAK | YES | NO | NO |
| 4820 | TRAV12-1 | TRAJ38 | CVVGDAGNNRKLIW | TRBV4-1 | TRBJ2-1 | CASSQALGSYNEQFF | KLGGALQAK | YES | NO | NO |
| 4821 | TRAV12-1 | TRAJ40 | CVVNSATSGTYKYIF | TRBV4-1 | TRBJ2-1 | CASSQGGTGLAGGPSYNEQFF | KLGGALQAK | NO | NO | NO |
| 4825 | TRAV13-1 | TRAJ24 | CAASSMADSWGKLQF | TRBV4-1 | TRBJ2-1 | CASSQEGASSYNEQFF | KLGGALQAK | YES | NO | NO |
| 4826 | TRAV13-1 | TRAJ4 | CGPGGYNKLIF | TRBV4-1 | TRBJ2-1 | CASSLAGTSGRDEQFF | KLGGALQAK | YES | NO | NO |
| 4827 | TRAV17 | TRAJ11 | CATAPGYSTLTF | TRBV4-1 | TRBJ2-1 | CASSQDGTGENEQFF | KLGGALQAK | YES | NO | NO |
| 4828 | TRAV17 | TRAJ17 | CATGKAAGNKLTF | TRBV4-1 | TRBJ2-1 | CASSQVPTSGNEQFF | KLGGALQAK | YES | NO | NO |
| 4830 | TRAV19 | TRAJ23 | CALSEPYNQGGKLIF | TRBV4-1 | TRBJ2-1 | CASSQEASGRGFNEQFF | KLGGALQAK | YES | NO | NO |
| 4831 | TRAV19 | TRAJ3 | CALSEVYSSASKIIF | TRBV4-1 | TRBJ2-1 | CASSQEDDGGAYNEQFF | KLGGALQAK | YES | NO | NO |
| 4833 | TRAV19 | TRAJ54 | CALSEAPGAQKLVF | TRBV4-1 | TRBJ2-1 | CASSQVSGTYNEQFF | KLGGALQAK | YES | NO | NO |
| 4835 | TRAV2 | TRAJ20 | CAVEEKADYKLSF | TRBV4-1 | TRBJ2-1 | CASSQDQGLGNEQFF | KLGGALQAK | NO | NO | NO |
| 4836 | TRAV24 | TRAJ13 | CAFCIIRGYQKVTF | TRBV4-1 | TRBJ2-1 | CASSQVVGADNEQFF | KLGGALQAK | YES | NO | NO |
| 4838 | TRAV27 | TRAJ54 | CAGARSQGAQKLVF | TRBV4-1 | TRBJ2-1 | CASSQTVPGSGRYNEQFF | KLGGALQAK | YES | NO | NO |
| 4839 | TRAV29DV5 | TRAJ18 | CAARGDRGSTLGRLYF | TRBV4-1 | TRBJ2-1 | CASSQGINEQFF | KLGGALQAK | YES | NO | NO |
| 4840 | TRAV29DV5 | TRAJ45 | CAARYSGGGADGLTF | TRBV4-1 | TRBJ2-1 | CASSQEPGLQYNEQFF | KLGGALQAK | YES | NO | NO |
| 4842 | TRAV4 | TRAJ37 | CLVGDGSGNTGKLIF | TRBV4-1 | TRBJ2-1 | CASSQVGNNEQFF | KLGGALQAK | YES | NO | NO |
| 4843 | TRAV4 | TRAJ43 | CLVGDGDNNDMRF | TRBV4-1 | TRBJ2-1 | CASSLSTGTFNNEQFF | KLGGALQAK | YES | NO | NO |
| 4845 | TRAV5 | TRAJ9 | CAERPLWAGGFKTIF | TRBV4-1 | TRBJ2-1 | CASSQEDSSYNEQFF | KLGGALQAK | YES | NO | NO |
| 4847 | TRAV8-1 | TRAJ10 | CAVIFTGGGNKLTF | TRBV4-1 | TRBJ2-1 | CASSLSGGNSYNEQFF | KLGGALQAK | YES | NO | NO |
| 4848 | TRAV8-1 | TRAJ9 | CAANTGGFKTIF | TRBV4-1 | TRBJ2-1 | CASSHSSGGGGEQFF | KLGGALQAK | YES | NO | NO |
| 4850 | TRAV1-2 | TRAJ24 | CAVGSDSWGKLQF | TRBV4-2 | TRBJ2-1 | CASSLTAGGPDEQFF | KLGGALQAK | YES | NO | NO |
| 4851 | TRAV1-2 | TRAJ33 | CAVLDSNYQLIW | TRBV4-2 | TRBJ2-1 | CASSQEAGGSYNEQFF | KLGGALQAK | YES | NO | NO |
| 4854 | TRAV30 | TRAJ53 | CGTEIDSGGSNYKLTF | TRBV4-2 | TRBJ2-1 | CASSQAYPDRGSYNEQFF | KLGGALQAK | YES | NO | NO |
| 4855 | TRAV8-4 | TRAJ20 | CAVSRSNDYKLSF | TRBV4-2 | TRBJ2-1 | CASSPLMQREGNEQFF | KLGGALQAK | YES | NO | NO |
| 4856 | TRAV9-2 | TRAJ26 | CALSDAESGQNFVF | TRBV4-2 | TRBJ2-1 | CASSQHTGSSYNEQFF | KLGGALQAK | YES | NO | NO |
| 4859 | TRAV12-1 | TRAJ40 | CVVNPPSGTYKYIF | TRBV4-3 | TRBJ2-1 | CASSQDIGGGYNNEQFF | KLGGALQAK | YES | NO | NO |
| 4862 | TRAV12-3 | TRAJ39 | CAIGPGNMLTF | TRBV4-3 | TRBJ2-1 | CASSQHGLAGDNEQFF | KLGGALQAK | YES | NO | NO |
| 4863 | TRAV12-3 | TRAJ39 | CAIGPGNMLTF | TRBV4-3 | TRBJ2-1 | CASSQDRPGGRPYNEQFF | KLGGALQAK | YES | NO | NO |
| 4864 | TRAV13-1 | TRAJ28 | CAARKSGAGSYQLTF | TRBV4-3 | TRBJ2-1 | CASSQGYFDGGTEYNEQFF | KLGGALQAK | YES | NO | NO |
| 4869 | TRAV2 | TRAJ5 | CAVEDDTGRRALTF | TRBV4-3 | TRBJ2-1 | CASSQESGIVYEQFF | RLRAEAQVK | NO | NO | NO |
| 4869 | TRAV2 | TRAJ5 | CAVEDDTGRRALTF | TRBV4-3 | TRBJ2-1 | CASSQESGIVYEQFF | KLGGALQAK | NO | NO | NO |
| 4871 | TRAV21 | TRAJ50 | CAVDLMKTSYDKVIF | TRBV4-3 | TRBJ2-1 | CASSHTSADEQFF | IVTDFSVIK | YES | YES | YES |
| 4873 | TRAV3 | TRAJ10 | CAVRDTFITGGGNKLTF | TRBV4-3 | TRBJ2-1 | CASSPSGLAGSYNEQFF | KLGGALQAK | YES | NO | NO |
| 4874 | TRAV3 | TRAJ11 | CAVTFIGYSTLTF | TRBV4-3 | TRBJ2-1 | CASSQDRPGGRPYNEQFF | KLGGALQAK | YES | NO | NO |
| 4876 | TRAV3 | TRAJ37 | CAVRDGGMGSNTGKLIF | TRBV4-3 | TRBJ2-1 | CASSPGQGHNEQFF | KLGGALQAK | YES | NO | NO |
| 4878 | TRAV36DV7 | TRAJ53 | CAPGVRGGSNYKLTF | TRBV4-3 | TRBJ2-1 | CASSQEGGEQFF | KLGGALQAK | NO | NO | NO |
| 4879 | TRAV4 | TRAJ11 | CLVAGGGSTLTF | TRBV4-3 | TRBJ2-1 | CASSQDPPGQLGNEQFF | KLGGALQAK | YES | NO | NO |
| 4881 | TRAV5 | TRAJ29 | CAESILEGNTPLVF | TRBV4-3 | TRBJ2-1 | CASSRWPTGGVEQFF | KLGGALQAK | YES | NO | NO |
| 4882 | TRAV8-1 | TRAJ34 | CASYYNTDKLIF | TRBV4-3 | TRBJ2-1 | CASSHTSADEQFF | KLGGALQAK | YES | NO | NO |
| 4882 | TRAV8-1 | TRAJ34 | CASYYNTDKLIF | TRBV4-3 | TRBJ2-1 | CASSHTSADEQFF | AVFDRKSDAK | YES | NO | NO |
| 4884 | TRAV12-1 | TRAJ15 | CVVGAGTALIF | TRBV5-1 | TRBJ2-1 | CASSLPPGYNEQFF | KLGGALQAK | YES | NO | NO |
| 4885 | TRAV12-1 | TRAJ22 | CVVVHSGSARQLTF | TRBV5-1 | TRBJ2-1 | CASSLELPHPDEQFF | KLGGALQAK | YES | NO | NO |
| 4887 | TRAV12-1 | TRAJ36 | CVVNLAGANNLFF | TRBV5-1 | TRBJ2-1 | CASSPDRAEEQFF | KLGGALQAK | YES | NO | NO |
| 4888 | TRAV12-3 | TRAJ39 | CAIGPGNMLTF | TRBV5-1 | TRBJ2-1 | CASSLLWEGPDEQFF | KLGGALQAK | YES | NO | NO |
| 4889 | TRAV13-1 | TRAJ20 | CAATADYKLSF | TRBV5-1 | TRBJ2-1 | CASSSGTGSYNEQFF | KLGGALQAK | YES | NO | NO |

| | | | | | | | | | | |
|---|---|---|---|---|---|---|---|---|---|---|
| 4890 | TRAV14DV4 | TRAJ11 | CAMRVSGYSTLTF | TRBV5-1 | TRBJ2-1 | CASSLAQFNEQFF | KLGGALQAK | YES | NO | NO |
| 4891 | TRAV14DV4 | TRAJ20 | CAMREGIGGNDYKLSF | TRBV5-1 | TRBJ2-1 | CASSLDYPYNEQFF | KLGGALQAK | YES | NO | NO |
| 4892 | TRAV14DV4 | TRAJ27 | CAMREGNTNAGKSTF | TRBV5-1 | TRBJ2-1 | CASSLGGGLGNEQFF | KLGGALQAK | YES | NO | NO |
| 4894 | TRAV16 | TRAJ21 | CALRDFNKFYF | TRBV5-1 | TRBJ2-1 | CATAGSLGYEQFF | KLGGALQAK | NO | NO | NO |
| 4895 | TRAV17 | TRAJ53 | CATVDNSGGSNYKLTF | TRBV5-1 | TRBJ2-1 | CASSQGNEQFF | KLGGALQAK | YES | NO | NO |
| 4896 | TRAV19 | TRAJ5 | CALSEALGGRRALTF | TRBV5-1 | TRBJ2-1 | CASSLEPWRGSSYNEQFF | KLGGALQAK | YES | NO | NO |
| 4898 | TRAV20 | TRAJ35 | CAVLGFGNVLHC | TRBV5-1 | TRBJ2-1 | CASSQRLAGSYNEQFF | KLGGALQAK | NO | NO | NO |
| 4899 | TRAV21 | TRAJ42 | CAVRLLNGGSQGNLIF | TRBV5-1 | TRBJ2-1 | CASSLLGGDYNEQFF | KLGGALQAK | YES | NO | NO |
| 4900 | TRAV22 | TRAJ22 | CAAGRISSGSARQLTF | TRBV5-1 | TRBJ2-1 | CASSLKRGEQFF | KLGGALQAK | YES | NO | NO |
| 4902 | TRAV22 | TRAJ30 | CAVESVDDKIIF | TRBV5-1 | TRBJ2-1 | CASRLSGGGYNEQFF | KLGGALQAK | YES | NO | NO |
| 4904 | TRAV26-2 | TRAJ32 | CILRDWAGGATNKLIF | TRBV5-1 | TRBJ2-1 | CASSSGLAGDYEQFF | KLGGALQAK | YES | NO | NO |
| 4907 | TRAV27 | TRAJ53 | CAGEGRGSNYKLTF | TRBV5-1 | TRBJ2-1 | CASSPVAPGGSSYNEQFF | KLGGALQAK | YES | NO | NO |
| 4913 | TRAV39 | TRAJ54 | CAVDIWQGAQKLVF | TRBV5-1 | TRBJ2-1 | CASSLVAGGYYNEQFF | KLGGALQAK | YES | NO | NO |
| 4915 | TRAV41 | TRAJ53 | CAVAGSGGSNYKLTF | TRBV5-1 | TRBJ2-1 | CASSLEASGPYNEQFF | RLRAEAQVK | YES | NO | YES |
| 4917 | TRAV8-3 | TRAJ33 | CAVGALTWDSNYQLIW | TRBV5-1 | TRBJ2-1 | CASSFAGNEQFF | KLGGALQAK | YES | NO | NO |
| 4918 | TRAV8-4 | TRAJ36 | CAVIQTGANNLFF | TRBV5-1 | TRBJ2-1 | CASSLVVVEDEQFF | KLGGALQAK | YES | NO | NO |
| 4920 | TRAV9-2 | TRAJ15 | CALSGSKAGTALIF | TRBV5-1 | TRBJ2-1 | CASVRTTDSYNEQFF | KLGGALQAK | YES | NO | NO |
| 4921 | TRAV1-2 | TRAJ27 | CAVRDSSGKSTF | TRBV5-4 | TRBJ2-1 | CATLQFGKHNEQFF | KLGGALQAK | YES | NO | NO |
| 4922 | TRAV10 | TRAJ4 | CVVSASGGYNKLIF | TRBV5-4 | TRBJ2-1 | CASGTNSYNEQFF | KLGGALQAK | YES | NO | NO |
| 4923 | TRAV12-1 | TRAJ41 | CVVNRYSGYALNF | TRBV5-4 | TRBJ2-1 | CASSLAGDPGEQFF | KLGGALQAK | YES | NO | NO |
| 4925 | TRAV13-1 | TRAJ11 | CAASRGYSTLTF | TRBV5-4 | TRBJ2-1 | CASSLGGGNNEQFF | AVFDRKSDAK | YES | NO | NO |
| 4926 | TRAV17 | TRAJ37 | CAPTSNTGKLIF | TRBV5-4 | TRBJ2-1 | CASSLGNEQFF | KLGGALQAK | YES | NO | NO |
| 4927 | TRAV17 | TRAJ45 | CATDGGADGLTF | TRBV5-4 | TRBJ2-1 | CASSGEGERGNEQFF | KLGGALQAK | YES | NO | NO |
| 4928 | TRAV17 | TRAJ49 | CATDAFLGNQFYF | TRBV5-4 | TRBJ2-1 | CASSPEGERNNEQFF | KLGGALQAK | YES | NO | NO |
| 4930 | TRAV20 | TRAJ29 | CAGPVSGNTPLVF | TRBV5-4 | TRBJ2-1 | CASSLGQGAYNEQFF | KLGGALQAK | YES | NO | NO |
| 4931 | TRAV20 | TRAJ34 | CAVQASNTDKLIF | TRBV5-4 | TRBJ2-1 | CASSLTSAAEFF | KLGGALQAK | YES | NO | NO |
| 4932 | TRAV20 | TRAJ34 | CAVQAGTDKLIF | TRBV5-4 | TRBJ2-1 | CASSARLAGSYNEQFF | KLGGALQAK | YES | NO | NO |
| 4933 | TRAV29DV5 | TRAJ45 | CAATTGHSGGGADGLTF | TRBV5-4 | TRBJ2-1 | CASSLGPRRGNEQFF | KLGGALQAK | YES | NO | NO |
| 4935 | TRAV8-2 | TRAJ15 | CVVSDVNQAGTALIF | TRBV5-4 | TRBJ2-1 | CASSSGQGRFDEQFF | KLGGALQAK | YES | NO | NO |
| 4936 | TRAV12-1 | TRAJ22 | CVVPTSGSARQLTF | TRBV5-5 | TRBJ2-1 | CASSQRGLAENEQFF | KLGGALQAK | YES | NO | NO |
| 4939 | TRAV26-2 | TRAJ40 | CILRDVSSGTYKYIF | TRBV5-5 | TRBJ2-1 | CASSLEAGGAYNEQFF | KLGGALQAK | YES | NO | NO |
| 4939 | TRAV26-2 | TRAJ40 | CILRDVSSGTYKYIF | TRBV5-5 | TRBJ2-1 | CASSLEAGGAYNEQFF | AVFDRKSDAK | YES | NO | NO |
| 4940 | TRAV8-3 | TRAJ45 | CAVGFSGGGADGLTF | TRBV5-5 | TRBJ2-1 | CASSLEAGGAYNEQFF | KLGGALQAK | YES | NO | NO |
| 4941 | TRAV1-2 | TRAJ31 | CAVRFNARLMF | TRBV5-6 | TRBJ2-1 | CASSLGRGGRNEQFF | KLGGALQAK | YES | NO | NO |
| 4942 | TRAV10 | TRAJ6 | CVVIRSGGSYIPTF | TRBV5-6 | TRBJ2-1 | CASSSIEIFNEQFF | KLGGALQAK | YES | NO | NO |
| 4944 | TRAV13-1 | TRAJ5 | CAATSMDTGRRALTF | TRBV5-6 | TRBJ2-1 | CASSTDIYNEQFF | KLGGALQAK | YES | NO | NO |
| 4947 | TRAV17 | TRAJ40 | CATAPSGTYKYIF | TRBV5-6 | TRBJ2-1 | CASSSTGLGSYNEQFF | KLGGALQAK | YES | NO | NO |
| 4948 | TRAV17 | TRAJ47 | CATDESKLVF | TRBV5-6 | TRBJ2-1 | CASRERGWVEQFF | KLGGALQAK | YES | NO | NO |
| 4950 | TRAV19 | TRAJ4 | CALSRWGGGYNKLIF | TRBV5-6 | TRBJ2-1 | CASSWRGAYNEQFF | KLGGALQAK | YES | NO | NO |
| 4951 | TRAV2 | TRAJ42 | CAVEDGYGGSQGNLIF | TRBV5-6 | TRBJ2-1 | CASSLGGTFYNEQFF | KLGGALQAK | NO | NO | NO |
| 4952 | TRAV21 | TRAJ30 | CAPDDKIIF | TRBV5-6 | TRBJ2-1 | CASTLAYLSEVGYNEQFF | KLGGALQAK | NO | NO | NO |
| 4953 | TRAV21 | TRAJ50 | CAVDLMKTSYDKVIF | TRBV5-6 | TRBJ2-1 | CASSLGINEQFF | KLGGALQAK | YES | NO | NO |
| 4958 | TRAV29DV5 | TRAJ45 | CAASSLGGGADGLTF | TRBV5-6 | TRBJ2-1 | CASTGGSSYNEQFF | KLGGALQAK | YES | NO | NO |
| 4959 | TRAV3 | TRAJ37 | CAVRDGGMGSNTGKLIF | TRBV5-6 | TRBJ2-1 | CASSLVGGEQFF | KLGGALQAK | NO | NO | NO |
| 4961 | TRAV40 | TRAJ28 | CLLRTGSYQLTF | TRBV5-6 | TRBJ2-1 | CASSHDTRNNEQFF | KLGGALQAK | NO | NO | NO |
| 4962 | TRAV8-1 | TRAJ29 | CAVISGNTPLVF | TRBV5-6 | TRBJ2-1 | CASNLGSYNEQFF | KLGGALQAK | YES | NO | NO |
| 4963 | TRAV8-2 | TRAJ39 | CVVSDIYNNAGNMLTF | TRBV5-6 | TRBJ2-1 | CASSRFRDSTNEQFF | KLGGALQAK | YES | NO | NO |
| 4964 | TRAV8-2 | TRAJ9 | CVVGNTGGFKTIF | TRBV5-6 | TRBJ2-1 | CASSLRAGSYNEQFF | KLGGALQAK | YES | NO | NO |
| 4966 | TRAV1-1 | TRAJ17 | CAVRDGAAGNKLTF | TRBV5-8 | TRBJ2-1 | CASSGGTSGSNEQFF | KLGGALQAK | YES | NO | NO |
| 4967 | TRAV1-2 | TRAJ42 | CAVRGLGGSQGNLIF | TRBV5-8 | TRBJ2-1 | CASSLVGEQFF | KLGGALQAK | YES | NO | NO |
| 4968 | TRAV12-2 | TRAJ32 | CAVNSGGATNKLIF | TRBV5-8 | TRBJ2-1 | CASSLGRTIPHEQFF | KLGGALQAK | YES | NO | NO |
| 4971 | TRAV27 | TRAJ45 | CAGVGGASGGGADGLTF | TRBV5-8 | TRBJ2-1 | CASSFGWYEQFF | KLGGALQAK | YES | NO | NO |
| 4973 | TRAV6 | TRAJ6 | CALDPPGGSYIPTF | TRBV5-8 | TRBJ2-1 | CASSYQTSGSIEQFF | KLGGALQAK | YES | NO | NO |
| 4976 | TRAV1-2 | TRAJ33 | CAVADSNYQLIW | TRBV6-1 | TRBJ2-1 | CASSEEAGEYNEQFF | KLGGALQAK | YES | NO | NO |
| 4977 | TRAV12-1 | TRAJ5 | CVVNSDTGRRALTF | TRBV6-1 | TRBJ2-1 | CASTTLRVSQFF | KLGGALQAK | YES | NO | NO |
| 4978 | TRAV12-2 | TRAJ9 | CAVTGGFKTIF | TRBV6-1 | TRBJ2-1 | CASSGGSGGLWDEQFF | KLGGALQAK | YES | NO | NO |
| 4979 | TRAV12-3 | TRAJ39 | CAIGPGNMLTF | TRBV6-1 | TRBJ2-1 | CASRDPGTTDYNEQFF | KLGGALQAK | YES | NO | NO |
| 4983 | TRAV27 | TRAJ44 | CAGADGTASKLTF | TRBV6-1 | TRBJ2-1 | CASSEVGGGYNEQFF | KLGGALQAK | YES | NO | NO |
| 4993 | TRAV9-2 | TRAJ23 | CALRPAVGNQGGKLIF | TRBV6-1 | TRBJ2-1 | CASTLAGGNNEQFF | KLGGALQAK | YES | NO | NO |
| 4995 | TRAV1-1 | TRAJ6 | CATLGLSGGSYIPTF | TRBV6-3 | TRBJ2-1 | CASSPPGSYNEQFF | KLGGALQAK | YES | NO | NO |
| 4997 | TRAV1-2 | TRAJ18 | CAVRDDDRGSTLGRLYF | TRBV6-3 | TRBJ2-1 | CASRGSNNEQFF | KLGGALQAK | YES | NO | NO |
| 4998 | TRAV1-2 | TRAJ33 | CAVMDSNYQLIW | TRBV6-3 | TRBJ2-1 | CASSYGTGGNEQFF | KLGGALQAK | YES | NO | NO |
| 4998 | TRAV1-2 | TRAJ33 | CAVMDSNYQLIW | TRBV6-3 | TRBJ2-1 | CASSYGTGGNEQFF | AVFDRKSDAK | YES | NO | NO |
| 4999 | TRAV10 | TRAJ17 | CVVSEGNKLTF | TRBV6-3 | TRBJ2-1 | CASSYSWTSGGVYNEQFF | KLGGALQAK | YES | NO | NO |
| 5000 | TRAV10 | TRAJ23 | CVVSAYNQGGKLIF | TRBV6-3 | TRBJ2-1 | CASSYSQINNEQFF | KLGGALQAK | YES | NO | NO |
| 5002 | TRAV10 | TRAJ42 | CVVRSYGGSQGNLIF | TRBV6-3 | TRBJ2-1 | CASIPWDRPYNEQFF | KLGGALQAK | YES | NO | NO |
| 5005 | TRAV12-1 | TRAJ40 | CVVNDDESGTYKYIF | TRBV6-3 | TRBJ2-1 | CASSYGRLAGGYVGNEQFF | KLGGALQAK | YES | NO | NO |
| 5007 | TRAV12-1 | TRAJ47 | CVVNRRGNKLVF | TRBV6-3 | TRBJ2-1 | CASTGLAGYNEQFF | KLGGALQAK | YES | NO | NO |
| 5008 | TRAV12-1 | TRAJ48 | CVVENPNFGNEKLTF | TRBV6-3 | TRBJ2-1 | CASSYGAGGYNEQFF | KLGGALQAK | YES | NO | NO |
| 5010 | TRAV12-2 | TRAJ13 | CAVNIGPGGYQKVTF | TRBV6-3 | TRBJ2-1 | CASSYSPGVAGGLVNEQFF | KLGGALQAK | YES | NO | NO |
| 5013 | TRAV12-3 | TRAJ52 | CAMSGTSYGKLTF | TRBV6-3 | TRBJ2-1 | CASSYGAGGYNEQFF | KLGGALQAK | YES | NO | NO |
| 5014 | TRAV12-3 | TRAJ54 | CAMSAGGAQKLVF | TRBV6-3 | TRBJ2-1 | CASSQNYNEQFF | KLGGALQAK | YES | NO | NO |
| 5016 | TRAV13-2 | TRAJ23 | CAGPNQGGKLIF | TRBV6-3 | TRBJ2-1 | CASSLSSGNEQFF | KLGGALQAK | YES | NO | NO |
| 5019 | TRAV14DV4 | TRAJ22 | CAMRGTFSGSARQLTF | TRBV6-3 | TRBJ2-1 | CASSYSLGLDFYNEQFF | KLGGALQAK | YES | NO | NO |
| 5021 | TRAV14DV4 | TRAJ29 | CAMRLGNTPLVF | TRBV6-3 | TRBJ2-1 | CASSYGAGGYNEQFF | KLGGALQAK | YES | NO | NO |
| 5025 | TRAV17 | TRAJ52 | CATDAFNAGGTSYGKLTF | TRBV6-3 | TRBJ2-1 | CASSYFRSGALYNEQFF | KLGGALQAK | YES | NO | NO |
| 5029 | TRAV2 | TRAJ3 | CAVDVDPYSSASKIIF | TRBV6-3 | TRBJ2-1 | CASLWLRGEYNEQFF | RLRAEAQVK | NO | NO | NO |
| 5030 | TRAV21 | TRAJ42 | CAVYWGGSQGNLIF | TRBV6-3 | TRBJ2-1 | CASTLGGLNEQFF | KLGGALQAK | YES | NO | NO |
| 5031 | TRAV22 | TRAJ39 | CAVDNAGNMLTF | TRBV6-3 | TRBJ2-1 | CASSLAWTYNEQFF | KLGGALQAK | YES | NO | NO |
| 5032 | TRAV25 | TRAJ18 | CAGAEVRGSTLGRLYF | TRBV6-3 | TRBJ2-1 | CASSYGAGGYNEQFF | KLGGALQAK | YES | NO | NO |
| 5034 | TRAV27 | TRAJ45 | CAGEGSGGGADGLTF | TRBV6-3 | TRBJ2-1 | CASWVANEQFF | KLGGALQAK | YES | NO | NO |
| 5035 | TRAV29DV5 | TRAJ26 | CAVRNYGQNFVF | TRBV6-3 | TRBJ2-1 | CASSYGTGSYEQFF | KLGGALQAK | YES | NO | NO |
| 5036 | TRAV29DV5 | TRAJ29 | CAATLNSGNTPLVF | TRBV6-3 | TRBJ2-1 | CASSSPELVGSYNEQFF | KLGGALQAK | YES | NO | NO |
| 5037 | TRAV29DV5 | TRAJ49 | CALPNTGNQFYF | TRBV6-3 | TRBJ2-1 | CASSYPTGLANNEQFF | KLGGALQAK | YES | NO | NO |
| 5038 | TRAV3 | TRAJ21 | CAVRDAYNFNKFYF | TRBV6-3 | TRBJ2-1 | CASNIEGSSYNEQFF | KLGGALQAK | NO | NO | NO |
| 5039 | TRAV3 | TRAJ30 | CAVRGYRDDKIIF | TRBV6-3 | TRBJ2-1 | CAIRGAGGSRRDEQFF | KLGGALQAK | NO | NO | NO |
| 5041 | TRAV35 | TRAJ42 | CAGQSGRVGGSQGNLIF | TRBV6-3 | TRBJ2-1 | CASSYGAGGYNEQFF | RLRAEAQVK | YES | NO | NO |
| 5042 | TRAV38-1 | TRAJ54 | CAFMSLIQGAQKLVF | TRBV6-3 | TRBJ2-1 | CASSYGAGGYNEQFF | KLGGALQAK | YES | NO | NO |
| 5045 | TRAV38-2DV8 | TRAJ48 | CAYRAPRGNEKLTF | TRBV6-3 | TRBJ2-1 | CASSYGAGGYNEQFF | KLGGALQAK | YES | NO | NO |
| 5047 | TRAV4 | TRAJ4 | CLVGDLPPFSGGYNKLIF | TRBV6-3 | TRBJ2-1 | CASSYKTGESYNEQFF | KLGGALQAK | YES | NO | NO |
| 5048 | TRAV41 | TRAJ40 | CAVSASGTYKYIF | TRBV6-3 | TRBJ2-1 | CASRASMSSYNEQFF | KLGGALQAK | YES | NO | NO |
| 5049 | TRAV41 | TRAJ43 | CAVRSRHDMRF | TRBV6-3 | TRBJ2-1 | CASSFPPSGGYNEQFF | KLGGALQAK | YES | NO | NO |
| 5051 | TRAV41 | TRAJ48 | CAVSPISNFGNEKLTF | TRBV6-3 | TRBJ2-1 | CASSSLAGPANEQFF | KLGGALQAK | NO | NO | NO |
| 5053 | TRAV8-3 | TRAJ6 | CASLGGSYIPTF | TRBV6-3 | TRBJ2-1 | CASSYGSGNEQFF | KLGGALQAK | YES | NO | NO |
| 5054 | TRAV8-4 | TRAJ20 | CAVIQINDYKLSF | TRBV6-3 | TRBJ2-1 | CASSYGAGGYNEQFF | KLGGALQAK | YES | NO | NO |
| 5055 | TRAV8-4 | TRAJ32 | CAVRSYGGATNKLIF | TRBV6-3 | TRBJ2-1 | CASSYPQGWEKRMGNEQFF | KLGGALQAK | YES | NO | NO |
| 5058 | TRAV1-2 | TRAJ12 | CAATDSSYKLIF | TRBV6-4 | TRBJ2-1 | CASSERGSYNEQFF | KLGGALQAK | YES | NO | NO |
| 5059 | TRAV1-2 | TRAJ12 | CAVMDSSYKLIF | TRBV6-4 | TRBJ2-1 | CASSDGTSGSNEQFF | KLGGALQAK | YES | NO | NO |
| 5060 | TRAV1-2 | TRAJ12 | CAVMDSSYKLIF | TRBV6-4 | TRBJ2-1 | CASSDGTGGYNEQFF | KLGGALQAK | YES | NO | NO |
| 5061 | TRAV1-2 | TRAJ33 | CAGQDSNYQLIW | TRBV6-4 | TRBJ2-1 | CATSPSGGTYNEQFF | KLGGALQAK | YES | NO | NO |
| 5062 | TRAV1-2 | TRAJ33 | CGPMDSNYQLIW | TRBV6-4 | TRBJ2-1 | CASSDGTGTYNEQFF | KLGGALQAK | YES | NO | NO |
| 5063 | TRAV1-2 | TRAJ33 | CAVRDSNYQLIW | TRBV6-4 | TRBJ2-1 | CASSDSTGGYNEQFF | KLGGALQAK | YES | NO | NO |
| 5064 | TRAV12-1 | TRAJ21 | CVGFNKFYF | TRBV6-4 | TRBJ2-1 | CASRLEGPHNEQFF | KLGGALQAK | NO | NO | NO |
| 5066 | TRAV12-3 | TRAJ10 | CAMSPGGVILTGGGNKLTF | TRBV6-4 | TRBJ2-1 | CASSEYSYNEQFF | KLGGALQAK | YES | NO | NO |

| | | | | | | | | | | |
|---|---|---|---|---|---|---|---|---|---|---|
| 5067 | TRAV13-1 | TRAJ26 | CAAKGYGQNFVF | TRBV6-4 | TRBJ2-1 | CASSDSLGSVEQFF | KLGGALQAK | YES | NO | NO |
| 5068 | TRAV13-1 | TRAJ28 | CAASQSGAGSYQLTF | TRBV6-4 | TRBJ2-1 | CASNNPYNEQFF | KLGGALQAK | YES | NO | NO |
| 5069 | TRAV13-1 | TRAJ43 | CAASHTVGDMRF | TRBV6-4 | TRBJ2-1 | CASSDSALRLEFF | KLGGALQAK | YES | NO | NO |
| 5071 | TRAV14DV4 | TRAJ9 | CAMREGATGGFKTIF | TRBV6-4 | TRBJ2-1 | CASSDPYNEQFF | KLGGALQAK | YES | NO | NO |
| 5072 | TRAV21 | TRAJ3 | CAVRKYSSASKIIF | TRBV6-4 | TRBJ2-1 | CASSDSIGGNEQFF | KLGGALQAK | YES | NO | NO |
| 5073 | TRAV22 | TRAJ38 | CAGNAGNNRKLIW | TRBV6-4 | TRBJ2-1 | CASSVGISYNEQFF | KLGGALQAK | YES | NO | NO |
| 5076 | TRAV41 | TRAJ54 | CAVPIQGAQKLVF | TRBV6-4 | TRBJ2-1 | CASSPREGSSYNEQFF | KLGGALQAK | YES | NO | NO |
| 5077 | TRAV8-6 | TRAJ37 | CAVSKDSSNTGKLIF | TRBV6-4 | TRBJ2-1 | CASSEGTRVGEQFF | KLGGALQAK | YES | NO | NO |
| 5078 | TRAV8-6 | TRAJ52 | CAVAGGTSYGKLTF | TRBV6-4 | TRBJ2-1 | CASRARHDSYNEQFF | KLGGALQAK | YES | NO | NO |
| 5079 | TRAV1-1 | TRAJ12 | CASYEDSSYKLIF | TRBV6-5 | TRBJ2-1 | CASSPLGGSSYNEQFF | KLGGALQAK | NO | NO | NO |
| 5080 | TRAV1-2 | TRAJ33 | CAVMDSNYQLIW | TRBV6-5 | TRBJ2-1 | CASSYGTSGPINEQFF | KLGGALQAK | NO | NO | NO |
| 5083 | TRAV12-2 | TRAJ15 | CAVGRGAGTALIF | TRBV6-5 | TRBJ2-1 | CASSTGLDEDEQFF | KLGGALQAK | NO | NO | NO |
| 5084 | TRAV12-2 | TRAJ6 | CAVNWEEQGGSYIPTF | TRBV6-5 | TRBJ2-1 | CASSTPGQGAGEQFF | KLGGALQAK | NO | NO | NO |
| 5085 | TRAV12-2 | TRAJ7 | CAVRYGNNRLAF | TRBV6-5 | TRBJ2-1 | CASSYSSSSGGSSYNEQFF | RAKFKQLL | NO | NO | NO |
| 5085 | TRAV12-2 | TRAJ7 | CAVRYGNNRLAF | TRBV6-5 | TRBJ2-1 | CASSYSSSSGGSSYNEQFF | KLGGALQAK | NO | NO | NO |
| 5086 | TRAV12-2 | TRAJ7 | CAVRYGNNRLAF | TRBV6-5 | TRBJ2-1 | CASSSLGAREQFF | KLGGALQAK | NO | NO | NO |
| 5087 | TRAV12-3 | TRAJ31 | CAMSESNARLMF | TRBV6-5 | TRBJ2-1 | CASSTTASSYNEQFF | KLGGALQAK | NO | NO | NO |
| 5088 | TRAV12-3 | TRAJ45 | CAMSRGGGADGLTF | TRBV6-5 | TRBJ2-1 | CASSTPGQGAGEQFF | KLGGALQAK | NO | NO | NO |
| 5088 | TRAV12-3 | TRAJ45 | CAMSRGGGADGLTF | TRBV6-5 | TRBJ2-1 | CASSTPGQGAGEQFF | AVFDRKSDAK | NO | NO | NO |
| 5088 | TRAV12-3 | TRAJ45 | CAMSRGGGADGLTF | TRBV6-5 | TRBJ2-1 | CASSTPGQGAGEQFF | IVTDFSVIK | NO | NO | NO |
| 5089 | TRAV13-1 | TRAJ43 | CAASGDNDMRF | TRBV6-5 | TRBJ2-1 | CASSTGLTHNEQFF | KLGGALQAK | NO | NO | NO |
| 5094 | TRAV20 | TRAJ7 | CAVQAGGNNRLAF | TRBV6-5 | TRBJ2-1 | CASSPDSNNEQFF | KLGGALQAK | NO | NO | NO |
| 5095 | TRAV21 | TRAJ43 | CAVWYMRF | TRBV6-5 | TRBJ2-1 | CASSPTSGTVNEQFF | KLGGALQAK | NO | NO | NO |
| 5096 | TRAV21 | TRAJ45 | CAVYSGGGADGLTF | TRBV6-5 | TRBJ2-1 | CASSFNGLAGENEQFF | KLGGALQAK | NO | NO | NO |
| 5097 | TRAV23DV6 | TRAJ8 | CAASNGNTGFQKLVF | TRBV6-5 | TRBJ2-1 | CASSYVNEQFF | KLGGALQAK | NO | NO | NO |
| 5099 | TRAV25 | TRAJ35 | CAGGSI | TRBV6-5 | TRBJ2-1 | CASSYYGTGAYNEQFF | KLGGALQAK | NO | NO | NO |
| 5103 | TRAV29DV5 | TRAJ49 | CAAAKEPTNTGNQFYF | TRBV6-5 | TRBJ2-1 | CASRYGGLSSYNEQFF | KLGGALQAK | NO | NO | NO |
| 5105 | TRAV35 | TRAJ53 | CAGQLESGGSNYKLTF | TRBV6-5 | TRBJ2-1 | CATGPGNEQFF | KLGGALQAK | NO | NO | NO |
| 5107 | TRAV38-2DV8 | TRAJ34 | CAAENTDKLIF | TRBV6-5 | TRBJ2-1 | CASSPQGARLDEQFF | KLGGALQAK | NO | NO | NO |
| 5109 | TRAV40 | TRAJ42 | CLLGDHGGSQGNLIF | TRBV6-5 | TRBJ2-1 | CASSYSGLAGYNEQFF | KLGGALQAK | NO | NO | NO |
| 5111 | TRAV6 | TRAJ11 | CASLYSTLTF | TRBV6-5 | TRBJ2-1 | CASSNGTGNEQFF | KLGGALQAK | NO | NO | NO |
| 5112 | TRAV8-2 | TRAJ47 | CVVSDLAKEYGNKLVF | TRBV6-5 | TRBJ2-1 | CASSYPGRGHNEQFF | KLGGALQAK | NO | NO | NO |
| 5113 | TRAV8-4 | TRAJ11 | CAVMGRGYSTLTF | TRBV6-5 | TRBJ2-1 | CASSSPKGSSYNEQFF | KLGGALQAK | NO | NO | NO |
| 5114 | TRAV9-2 | TRAJ28 | CALSSGAGSYQLTF | TRBV6-5 | TRBJ2-1 | CASSYSAGRDNEQFF | KLGGALQAK | NO | NO | NO |
| 5117 | TRAV12-2 | TRAJ42 | CAVRDYGGSQGNLIF | TRBV6-6 | TRBJ2-1 | CASAPGLAGNEQFF | KLGGALQAK | NO | NO | NO |
| 5117 | TRAV12-2 | TRAJ42 | CAVRDYGGSQGNLIF | TRBV6-6 | TRBJ2-1 | CASAPGLAGNEQFF | RAKFKQLL | NO | NO | NO |
| 5119 | TRAV19 | TRAJ42 | CALSGDYGGSQGNLIF | TRBV6-6 | TRBJ2-1 | CASSYSGTDYNEQFF | KLGGALQAK | NO | NO | NO |
| 5120 | TRAV2 | TRAJ9 | CAAPQTGGFKTIF | TRBV6-6 | TRBJ2-1 | CASSLPGTGNEQFF | KLGGALQAK | NO | NO | NO |
| 5121 | TRAV20 | TRAJ26 | CAVGGQNFVF | TRBV6-6 | TRBJ2-1 | CASSRPYNEQFF | KLGGALQAK | NO | NO | NO |
| 5123 | TRAV35 | TRAJ26 | CAGLDNYGQNFVF | TRBV6-6 | TRBJ2-1 | CASSPSGSYNEQFF | KLGGALQAK | NO | NO | NO |
| 5124 | TRAV8-1 | TRAJ15 | CAVNAWQAGTALIF | TRBV6-6 | TRBJ2-1 | CASSYQGQLPYNEQFF | KLGGALQAK | NO | NO | NO |
| 5125 | TRAV8-3 | TRAJ6 | CAVIRGSYIPTF | TRBV6-6 | TRBJ2-1 | CASSSEGGYNEQFF | KLGGALQAK | NO | NO | NO |
| 5131 | TRAV17 | TRAJ20 | CATDGNDYKLSF | TRBV7-2 | TRBJ2-1 | CASSLGPGNNEQFF | KLGGALQAK | YES | NO | NO |
| 5136 | TRAV29DV5 | TRAJ32 | CAASDDGGATNKLIF | TRBV7-2 | TRBJ2-1 | CASSLAGYNEQFF | KLGGALQAK | YES | NO | NO |
| 5140 | TRAV8-1 | TRAJ33 | CAVMFKDSNYQLIW | TRBV7-2 | TRBJ2-1 | CASSQHVRDWQFF | KLGGALQAK | YES | NO | NO |
| 5143 | TRAV9-2 | TRAJ7 | CAPSGGNNRLAF | TRBV7-2 | TRBJ2-1 | CASSLDLGGIQYNEQFF | KLGGALQAK | YES | NO | NO |
| 5145 | TRAV13-1 | TRAJ44 | CAAIDTGTASKLTF | TRBV7-3 | TRBJ2-1 | CASSLYGGMDYNEQFF | KLGGALQAK | YES | NO | NO |
| 5146 | TRAV13-1 | TRAJ45 | CAASTAGGGADGLTF | TRBV7-3 | TRBJ2-1 | CASSYPNNEQFF | KLGGALQAK | YES | NO | NO |
| 5147 | TRAV13-1 | TRAJ53 | CAAMDVGSNYKLTF | TRBV7-3 | TRBJ2-1 | CASSLVGGNEQFF | AVFDRKSDAK | YES | NO | NO |
| 5147 | TRAV13-1 | TRAJ53 | CAAMDVGSNYKLTF | TRBV7-3 | TRBJ2-1 | CASSLVGGNEQFF | KLGGALQAK | YES | NO | NO |
| 5150 | TRAV21 | TRAJ9 | CAVLDTGGFKTIF | TRBV7-3 | TRBJ2-1 | CASSFNLAGGDNEQFF | KLGGALQAK | YES | NO | NO |
| 5152 | TRAV38-2DV8 | TRAJ43 | CAYRSGRGNNDMRF | TRBV7-3 | TRBJ2-1 | CASSQSLAEGMNEQFF | KLGGALQAK | YES | NO | NO |
| 5153 | TRAV8-2 | TRAJ36 | CVLTGANNLFF | TRBV7-3 | TRBJ2-1 | CASSLGTAAYNEQFF | KLGGALQAK | YES | NO | NO |
| 5154 | TRAV8-3 | TRAJ31 | CAVRNNNARLMF | TRBV7-3 | TRBJ2-1 | CASSLMEYNEQFF | KLGGALQAK | YES | NO | NO |
| 5155 | TRAV38-2DV8 | TRAJ32 | CAYRRRGATNKLIF | TRBV7-4 | TRBJ2-1 | CASSFLLAGGPVDSSYNEQFF | KLGGALQAK | NO | NO | NO |
| 5156 | TRAV1-2 | TRAJ29 | CAAPSGNTPLVF | TRBV7-6 | TRBJ2-1 | CASSSSPGLAEGEQFF | KLGGALQAK | YES | NO | NO |
| 5157 | TRAV10 | TRAJ8 | CVVSENTGFQKLVF | TRBV7-6 | TRBJ2-1 | CASSVLYLNEQFF | KLGGALQAK | YES | NO | NO |
| 5158 | TRAV12-1 | TRAJ18 | CVVNNRDRGSTLGRLYF | TRBV7-6 | TRBJ2-1 | CASSLFRGGSYNEQFF | KLGGALQAK | YES | NO | NO |
| 5159 | TRAV12-2 | TRAJ26 | CAVVDNYGQNFVF | TRBV7-6 | TRBJ2-1 | CASSGSRAGGAGEQFF | KLGGALQAK | YES | NO | NO |
| 5160 | TRAV12-2 | TRAJ30 | CAVVRMNRDDKIIF | TRBV7-6 | TRBJ2-1 | CASSPVGGGYNEQFF | KLGGALQAK | YES | NO | NO |
| 5161 | TRAV12-2 | TRAJ45 | CAVISSGGGADGLTF | TRBV7-6 | TRBJ2-1 | CASSDTVDEQFF | KLGGALQAK | YES | NO | NO |
| 5162 | TRAV14DV4 | TRAJ30 | CAMRETFRDDKIIF | TRBV7-6 | TRBJ2-1 | CASSLAMREGAYNEQFF | KLGGALQAK | YES | NO | NO |
| 5164 | TRAV17 | TRAJ57 | CGGGSEKLVF | TRBV7-6 | TRBJ2-1 | CASSLAGAYEQFF | KLGGALQAK | YES | NO | NO |
| 5165 | TRAV19 | TRAJ24 | CALADSWGKFQF | TRBV7-6 | TRBJ2-1 | CASSLGVTPNEQFF | KLGGALQAK | YES | NO | NO |
| 5167 | TRAV19 | TRAJ37 | CALRLLSSNTGKLIF | TRBV7-6 | TRBJ2-1 | CASSLGLGTSNEQFF | KLGGALQAK | YES | NO | NO |
| 5170 | TRAV21 | TRAJ49 | CAVKGNQFYF | TRBV7-6 | TRBJ2-1 | CASSLLGDNEQFF | KLGGALQAK | YES | NO | NO |
| 5171 | TRAV29DV5 | TRAJ32 | CAASDDGGATNKLIF | TRBV7-6 | TRBJ2-1 | CASSSRGYNEQFF | RLRAEAQVK | YES | YES | YES |
| 5174 | TRAV39 | TRAJ45 | CAVPFRYSGGGADGLTF | TRBV7-6 | TRBJ2-1 | CASSFGTGIEQFF | KLGGALQAK | YES | NO | NO |
| 5175 | TRAV4 | TRAJ33 | CLVGDGKDSNYQLIW | TRBV7-6 | TRBJ2-1 | CASSLSLADNEQFF | KLGGALQAK | NO | NO | NO |
| 5176 | TRAV10 | TRAJ40 | CVVSATTSGTYKYIF | TRBV7-7 | TRBJ2-1 | CASSQGHYNEQFF | KLGGALQAK | YES | NO | NO |
| 5179 | TRAV8-4 | TRAJ45 | CAVTPGGGADGLTF | TRBV7-7 | TRBJ2-1 | CASSLFRGDNEQFF | KLGGALQAK | YES | NO | NO |
| 5181 | TRAV1-2 | TRAJ28 | CAVDVTGAGSYQLTF | TRBV7-8 | TRBJ2-1 | CASSLIEASGGFEQFF | KLGGALQAK | YES | NO | NO |
| 5182 | TRAV1-2 | TRAJ28 | CCGRYWGWELPTHF | TRBV7-8 | TRBJ2-1 | CASSLIEASGGFEQFF | KLGGALQAK | YES | NO | NO |
| 5184 | TRAV13-1 | TRAJ44 | CAAYTGTASKLTF | TRBV7-8 | TRBJ2-1 | CASSGTSGAYNEQFF | KLGGALQAK | YES | NO | NO |
| 5186 | TRAV14DV4 | TRAJ41 | CAMREGDSGYALNF | TRBV7-8 | TRBJ2-1 | CASSLGYEQFF | KLGGALQAK | YES | NO | NO |
| 5189 | TRAV17 | TRAJ9 | CATDHGDTGGFKTIF | TRBV7-8 | TRBJ2-1 | CASSFLAGGYNEQFF | KLGGALQAK | YES | NO | NO |
| 5190 | TRAV19 | TRAJ23 | CALSAPYNQGGKLIF | TRBV7-8 | TRBJ2-1 | CASSLLQGTYNEQFF | KLGGALQAK | YES | NO | NO |
| 5194 | TRAV26-2 | TRAJ48 | CILRDGGFGNEKLTF | TRBV7-8 | TRBJ2-1 | CASRQRGVSYNEQFF | KLGGALQAK | YES | NO | NO |
| 5195 | TRAV29DV5 | TRAJ35 | CAASDGFGNVLHC | TRBV7-8 | TRBJ2-1 | CASILGGSGGHMGNEQFF | KLGGALQAK | NO | NO | NO |
| 5206 | TRAV10 | TRAJ4 | CVVRSGGYNKLIF | TRBV7-9 | TRBJ2-1 | CASSLSLSGRGVEQFF | KLGGALQAK | YES | NO | NO |
| 5209 | TRAV12-1 | TRAJ30 | CVVITPKFRDDKIIF | TRBV7-9 | TRBJ2-1 | CASSSWTSGNNEQFF | KLGGALQAK | YES | NO | NO |
| 5212 | TRAV12-2 | TRAJ24 | CAVFISDSWGKLQF | TRBV7-9 | TRBJ2-1 | CASSLGGAGSSYNEQFF | KLGGALQAK | YES | NO | NO |
| 5213 | TRAV12-2 | TRAJ42 | CAVNRVGGSQGNLIF | TRBV7-9 | TRBJ2-1 | CASSLGPGEQFF | KLGGALQAK | YES | NO | NO |
| 5218 | TRAV12-3 | TRAJ39 | CAIGPGNMLTF | TRBV7-9 | TRBJ2-1 | CASSSGLAGENEQFF | KLGGALQAK | YES | NO | NO |
| 5219 | TRAV12-3 | TRAJ39 | CAIGPGNMLTF | TRBV7-9 | TRBJ2-1 | CASREGLSNEQFF | KLGGALQAK | YES | NO | NO |
| 5223 | TRAV13-1 | TRAJ45 | CAATPRGGADGLTF | TRBV7-9 | TRBJ2-1 | CASSNSLPSGVSYNEQFF | KLGGALQAK | YES | NO | NO |
| 5226 | TRAV13-2 | TRAJ49 | CAEIHTGNQFYF | TRBV7-9 | TRBJ2-1 | CASSRNEQFF | KLGGALQAK | YES | NO | NO |
| 5227 | TRAV13-2 | TRAJ49 | CAEIRGRDQFYF | TRBV7-9 | TRBJ2-1 | CASSSGLAGENEQFF | KLGGALQAK | YES | NO | NO |
| 5228 | TRAV14DV4 | TRAJ22 | CAMRRPISSGSARQLTF | TRBV7-9 | TRBJ2-1 | CASTGTSGGPTLRDEQFF | KLGGALQAK | YES | NO | NO |
| 5233 | TRAV17 | TRAJ10 | CATDPITGGGNKLTF | TRBV7-9 | TRBJ2-1 | CASSVLAGDHEQFF | KLGGALQAK | YES | NO | NO |
| 5235 | TRAV17 | TRAJ41 | CATDGNSGYALNF | TRBV7-9 | TRBJ2-1 | CASSSITSGRGGEQFF | KLGGALQAK | YES | NO | NO |
| 5236 | TRAV17 | TRAJ43 | CVGRGDMRF | TRBV7-9 | TRBJ2-1 | CASSADTVYNEQFF | KLGGALQAK | NO | NO | NO |
| 5237 | TRAV17 | TRAJ53 | CATDGRSGGSNYKLTF | TRBV7-9 | TRBJ2-1 | CASSQGPYNEQFF | KLGGALQAK | YES | NO | NO |
| 5238 | TRAV17 | TRAJ58 | CATDQTSGSRLTF | TRBV7-9 | TRBJ2-1 | CASVRDKYNEQFF | KLGGALQAK | YES | NO | NO |
| 5239 | TRAV19 | TRAJ12 | CALSEDMDSSYKLIF | TRBV7-9 | TRBJ2-1 | CASSHTSGVYNEQFF | KLGGALQAK | YES | NO | NO |
| 5240 | TRAV19 | TRAJ26 | CALSEAYYYGQNFVF | TRBV7-9 | TRBJ2-1 | CASSLEFTSGSHSYNEQFF | KLGGALQAK | YES | NO | NO |
| 5241 | TRAV19 | TRAJ37 | CALSSPGNTGKLIF | TRBV7-9 | TRBJ2-1 | CASSLDQGAQIRNEQFF | KLGGALQAK | YES | NO | NO |
| 5243 | TRAV19 | TRAJ43 | CALRPPGNNDMRF | TRBV7-9 | TRBJ2-1 | CASSFPGASYNEQFF | KLGGALQAK | YES | NO | NO |
| 5246 | TRAV19 | TRAJ5 | CALSEALGGRRALTF | TRBV7-9 | TRBJ2-1 | CASSLGPGEQFF | KLGGALQAK | YES | NO | NO |
| 5248 | TRAV2 | TRAJ10 | CAVGVTGGGNKLTF | TRBV7-9 | TRBJ2-1 | CASRKGGRADEQFF | KLGGALQAK | NO | NO | NO |
| 5251 | TRAV21 | TRAJ26 | CAVHGYGQNFVF | TRBV7-9 | TRBJ2-1 | CASSSGLAGVNEQFF | KLGGALQAK | YES | NO | NO |
| 5252 | TRAV21 | TRAJ39 | CAVTFNNAGNMLTF | TRBV7-9 | TRBJ2-1 | CASTGAWRGSSYNEQFF | KLGGALQAK | YES | NO | NO |
| 5259 | TRAV26-1 | TRAJ39 | CIVRVLSLNNNAGNMLTF | TRBV7-9 | TRBJ2-1 | CASSLTSGGAYDEQFF | KLGGALQAK | YES | NO | NO |
| 5264 | TRAV29DV5 | TRAJ4 | CAASFNSGGYNKLIF | TRBV7-9 | TRBJ2-1 | CASSLEREGDNEQFF | KLGGALQAK | YES | NO | NO |

| | | | | | | | | | | |
|---|---|---|---|---|---|---|---|---|---|---|
| 5266 | TRAV3 | TRAJ21 | CAVRERVRYNFNKFYF | TRBV7-9 | TRBJ2-1 | CASSLGATGEYNEQFF | KLGGALQAK | NO | NO | NO |
| 5268 | TRAV3 | TRAJ35 | CAVRDIKKAI | TRBV7-9 | TRBJ2-1 | CASKAGSGTEQFF | KLGGALQAK | NO | NO | NO |
| 5270 | TRAV3 | TRAJ38 | CAVRGFAGNNRKLIW | TRBV7-9 | TRBJ2-1 | CASSQLAGPFNEQFF | KLGGALQAK | NO | NO | NO |
| 5271 | TRAV3 | TRAJ39 | CAVRPLNNAGNMLTF | TRBV7-9 | TRBJ2-1 | CASTGTSGGPTLRDEQFF | KLGGALQAK | YES | NO | NO |
| 5273 | TRAV36DV7 | TRAJ42 | CALFGGSQGNLIF | TRBV7-9 | TRBJ2-1 | CASSTTGSSYNEQFF | KLGGALQAK | NO | NO | NO |
| 5274 | TRAV38-1 | TRAJ31 | CAFGFNNARLMF | TRBV7-9 | TRBJ2-1 | CASSSGLAGENEQFF | KLGGALQAK | YES | NO | NO |
| 5275 | TRAV38-2DV8 | TRAJ40 | CAYRDTSGTYKYIF | TRBV7-9 | TRBJ2-1 | CASSSPGESSYNEQFF | KLGGALQAK | YES | NO | NO |
| 5276 | TRAV38-2DV8 | TRAJ42 | CAYIYGGSQGNLIF | TRBV7-9 | TRBJ2-1 | CASSLTQGWYNEQFF | KLGGALQAK | YES | NO | NO |
| 5278 | TRAV38-2DV8 | TRAJ48 | CAYPSNEEKLTF | TRBV7-9 | TRBJ2-1 | CASSLGYGDSYNEQFF | KLGGALQAK | YES | NO | NO |
| 5279 | TRAV38-2DV8 | TRAJ57 | CAYSQGGSEKLVF | TRBV7-9 | TRBJ2-1 | CASSLAGGPADEQFF | KLGGALQAK | YES | NO | NO |
| 5280 | TRAV4 | TRAJ11 | CLVGDMRRGNSGYSTLTF | TRBV7-9 | TRBJ2-1 | CASSLAILTGHNEQFF | KLGGALQAK | YES | YES | NO |
| 5282 | TRAV5 | TRAJ24 | CADMPADSWGKLQF | TRBV7-9 | TRBJ2-1 | CASSQLGRGDNEQFF | KLGGALQAK | YES | NO | NO |
| 5285 | TRAV6 | TRAJ41 | CALGAGYALNF | TRBV7-9 | TRBJ2-1 | CASSPRPQDSYNEQFF | KLGGALQAK | YES | NO | NO |
| 5286 | TRAV8-1 | TRAJ21 | CAVNEPFNKFYF | TRBV7-9 | TRBJ2-1 | CASSSGLAGENEQFF | KLGGALQAK | YES | NO | NO |
| 5286 | TRAV8-1 | TRAJ21 | CAVNEPFNKFYF | TRBV7-9 | TRBJ2-1 | CASSSGLAGENEQFF | AVFDRKSDAK | YES | NO | NO |
| 5287 | TRAV8-1 | TRAJ34 | CAVTGSYNTDKLIF | TRBV7-9 | TRBJ2-1 | CASSSGLAGENEQFF | KLGGALQAK | YES | NO | NO |
| 5289 | TRAV8-1 | TRAJ6 | CAVLRTSGGSYIPTF | TRBV7-9 | TRBJ2-1 | CASSGRAGGPSEFF | KLGGALQAK | YES | NO | NO |
| 5292 | TRAV8-2 | TRAJ22 | CVGRKGSARQLTF | TRBV7-9 | TRBJ2-1 | CASSNSLPSGVSYNEQFF | KLGGALQAK | YES | NO | NO |
| 5293 | TRAV8-2 | TRAJ37 | CVVSDYSGSGNTGKLIF | TRBV7-9 | TRBJ2-1 | CASRGLPGTMENEQFF | KLGGALQAK | YES | NO | NO |
| 5295 | TRAV8-3 | TRAJ54 | CAVAFIQGAQKLVF | TRBV7-9 | TRBJ2-1 | CASSSLLAGAYNEQFF | KLGGALQAK | YES | NO | NO |
| 5296 | TRAV8-3 | TRAJ9 | CAVGLDTGGFKTIF | TRBV7-9 | TRBJ2-1 | CASTGTSGGPTLRDEQFF | KLGGALQAK | YES | NO | NO |
| 5297 | TRAV8-6 | TRAJ15 | CAVRRQAGTALIF | TRBV7-9 | TRBJ2-1 | CASSSGLAGENEQFF | KLGGALQAK | YES | NO | NO |
| 5298 | TRAV8-6 | TRAJ15 | CAVSHQAGTALIF | TRBV7-9 | TRBJ2-1 | CASRGQGRNNEQFF | KLGGALQAK | YES | NO | NO |
| 5300 | TRAV8-6 | TRAJ21 | CAVTLYNFNKFYF | TRBV7-9 | TRBJ2-1 | CASSLAIIGQGYGDEQFF | KLGGALQAK | YES | NO | NO |
| 5301 | TRAV9-2 | TRAJ16 | CALSEGGGQKLLF | TRBV7-9 | TRBJ2-1 | CASSLAYVSSYNEQFF | KLGGALQAK | NO | NO | NO |
| 5302 | TRAV1-1 | TRAJ23 | CAVTVRRNQGGKLIF | TRBV9 | TRBJ2-1 | CASSVGQGLSYNEQFF | KLGGALQAK | YES | NO | NO |
| 5303 | TRAV1-2 | TRAJ13 | CAVVGGYQKVTF | TRBV9 | TRBJ2-1 | CASSVEGGSKDNEQFF | KLGGALQAK | YES | NO | NO |
| 5304 | TRAV1-2 | TRAJ30 | CAVRRDDKIIF | TRBV9 | TRBJ2-1 | CASSAGTSGRVSYNEQFF | KLGGALQAK | YES | NO | NO |
| 5305 | TRAV1-2 | TRAJ5 | CAVRQDTGRRALTF | TRBV9 | TRBJ2-1 | CASSVASGRDEQFF | KLGGALQAK | YES | NO | NO |
| 5310 | TRAV12-3 | TRAJ20 | CAMSAHDYKLSF | TRBV9 | TRBJ2-1 | CASSVEGDANEQFF | KLGGALQAK | YES | NO | NO |
| 5311 | TRAV12-3 | TRAJ6 | CARGGSYIPTF | TRBV9 | TRBJ2-1 | CASSPQTSGGLNEQFF | KLGGALQAK | YES | NO | NO |
| 5312 | TRAV13-1 | TRAJ4 | CAASIRTGGYNKLIF | TRBV9 | TRBJ2-1 | CASSVEWTPSNEQFF | KLGGALQAK | YES | NO | NO |
| 5313 | TRAV14DV4 | TRAJ26 | CAIKNYGQNFVF | TRBV9 | TRBJ2-1 | CASSVEVGGLSNSYNEQFF | KLGGALQAK | YES | NO | NO |
| 5314 | TRAV14DV4 | TRAJ28 | CAMRPPYSGAGSYQLTF | TRBV9 | TRBJ2-1 | CASSPPGGSGGHYEQFF | KLGGALQAK | YES | NO | NO |
| 5315 | TRAV14DV4 | TRAJ53 | CAMKGGGSNYKLTF | TRBV9 | TRBJ2-1 | CASSAHRGGQGNEQFF | KLGGALQAK | YES | NO | NO |
| 5317 | TRAV17 | TRAJ54 | CATDSIQGAQKLVF | TRBV9 | TRBJ2-1 | CASSAGGAVYNEQFF | KLGGALQAK | YES | NO | NO |
| 5318 | TRAV17 | TRAJ57 | CATAQGGSEKLVF | TRBV9 | TRBJ2-1 | CASSVRLAGGVIDEQFF | KLGGALQAK | YES | NO | NO |
| 5319 | TRAV19 | TRAJ35 | CALSEGFGNVLHC | TRBV9 | TRBJ2-1 | CASSVETIPSPNEQFF | KLGGALQAK | NO | NO | NO |
| 5320 | TRAV19 | TRAJ37 | CALSGGENTGKLIF | TRBV9 | TRBJ2-1 | CASSVGGAGVSNEQFF | KLGGALQAK | YES | NO | NO |
| 5321 | TRAV19 | TRAJ43 | CALREPLYNNNDMRF | TRBV9 | TRBJ2-1 | CASSVDGYEQFF | KLGGALQAK | YES | NO | NO |
| 5322 | TRAV20 | TRAJ20 | CAVQVNDYKLSF | TRBV9 | TRBJ2-1 | CASSAPPGPYNEQFF | KLGGALQAK | YES | NO | NO |
| 5323 | TRAV20 | TRAJ42 | CAVQAQGSQGNLIF | TRBV9 | TRBJ2-1 | CASSHGGVDEQFF | KLGGALQAK | YES | NO | NO |
| 5324 | TRAV20 | TRAJ57 | CAVHSQGGSEKLVF | TRBV9 | TRBJ2-1 | CASSPRATVVDEQFF | KLGGALQAK | YES | NO | NO |
| 5326 | TRAV21 | TRAJ28 | CASVSYSGAGSYQLTF | TRBV9 | TRBJ2-1 | CASSETTGTGLYNEQFF | KLGGALQAK | YES | NO | NO |
| 5327 | TRAV21 | TRAJ43 | CAVGYNNNDMRF | TRBV9 | TRBJ2-1 | CASSVEDSSYNEQFF | KLGGALQAK | YES | NO | NO |
| 5328 | TRAV23DV6 | TRAJ29 | CAASRSTPLVF | TRBV9 | TRBJ2-1 | CASSVSLVNEQFF | KLGGALQAK | YES | NO | NO |
| 5329 | TRAV24 | TRAJ52 | CALNAGGTSYGKLTF | TRBV9 | TRBJ2-1 | CASSVDVFTNNEQFF | KLGGALQAK | YES | NO | NO |
| 5330 | TRAV26-2 | TRAJ36 | CILSQTGANNLFF | TRBV9 | TRBJ2-1 | CASSPEGLYYNEQFF | KLGGALQAK | YES | NO | NO |
| 5333 | TRAV29DV5 | TRAJ26 | CAASGFDGQNFVF | TRBV9 | TRBJ2-1 | CASSVTSVPYNEQFF | KLGGALQAK | YES | NO | NO |
| 5334 | TRAV29DV5 | TRAJ26 | CAASSDNYGQNFVF | TRBV9 | TRBJ2-1 | CASSVGAGEFNEQFF | KLGGALQAK | YES | NO | NO |
| 5335 | TRAV3 | TRAJ12 | CAVRDRKDSSYKLIF | TRBV9 | TRBJ2-1 | CASSVDFGGGLEQFF | KLGGALQAK | NO | NO | NO |
| 5336 | TRAV3 | TRAJ40 | CAVRDRLLTSGTYKYIF | TRBV9 | TRBJ2-1 | CASSVAHLINEQFF | RLRAEAQVK | NO | NO | NO |
| 5337 | TRAV30 | TRAJ23 | CGTEDNQGGKLIF | TRBV9 | TRBJ2-1 | CASSVEWTSGSEGYNEQFF | KLGGALQAK | YES | NO | NO |
| 5338 | TRAV35 | TRAJ43 | CAGLLPNNNDMRF | TRBV9 | TRBJ2-1 | CASSFPVGGLAGAGYNEQFF | KLGGALQAK | NO | NO | NO |
| 5339 | TRAV38-1 | TRAJ16 | CAFMKLWYDQKLLF | TRBV9 | TRBJ2-1 | CASSRDRHHNEQFF | KLGGALQAK | NO | NO | NO |
| 5341 | TRAV38-2DV8 | TRAJ30 | CAYRSAVDGGKIIF | TRBV9 | TRBJ2-1 | CASSWVGVPGNEQFF | KLGGALQAK | YES | NO | NO |
| 5342 | TRAV38-2DV8 | TRAJ47 | CAYRRGYGNKLVF | TRBV9 | TRBJ2-1 | CASSVSLVNEQFF | KLGGALQAK | YES | NO | NO |
| 5342 | TRAV38-2DV8 | TRAJ47 | CAYRRGYGNKLVF | TRBV9 | TRBJ2-1 | CASSVSLVNEQFF | AVFDRKSDAK | YES | NO | NO |
| 5342 | TRAV38-2DV8 | TRAJ47 | CAYRRGYGNKLVF | TRBV9 | TRBJ2-1 | CASSVSLVNEQFF | IVTDFSVIK | YES | NO | NO |
| 5344 | TRAV5 | TRAJ40 | CAETNSGTYKYIF | TRBV9 | TRBJ2-1 | CASSVGGTDFYNEQFF | KLGGALQAK | YES | NO | NO |
| 5346 | TRAV8-3 | TRAJ34 | CAVGYISYNTDKLIF | TRBV9 | TRBJ2-1 | CASSVVDNEQFF | KLGGALQAK | YES | NO | NO |
| 5348 | TRAV9-2 | TRAJ16 | CALEDGQKLLF | TRBV9 | TRBJ2-1 | CASSTGGAWNEQFF | KLGGALQAK | NO | NO | NO |
| 5349 | TRAV5 | TRAJ32 | CAINQGGGATNKLIF | TRBV10-1 | TRBJ2-2 | CASSPGRTGELFF | KLGGALQAK | YES | NO | NO |
| 5350 | TRAV13-1 | TRAJ32 | CAASDGGATNKLIF | TRBV10-2 | TRBJ2-2 | CASGVSGNTGELFF | KLGGALQAK | YES | NO | NO |
| 5356 | TRAV8-2 | TRAJ9 | CVVSDRGFKTIF | TRBV10-2 | TRBJ2-2 | CASIHTSGANTGELFF | KLGGALQAK | YES | NO | NO |
| 5357 | TRAV12-1 | TRAJ29 | CVVSPNSGNTPLVF | TRBV10-3 | TRBJ2-2 | CAIGFPSGGPWTGELFF | KLGGALQAK | YES | NO | NO |
| 5359 | TRAV12-2 | TRAJ45 | CAVNGYSGGGADGLTF | TRBV10-3 | TRBJ2-2 | CAISEGTSGYTGELFF | KLGGALQAK | YES | NO | NO |
| 5360 | TRAV12-2 | TRAJ47 | CAVNGNKLVF | TRBV10-3 | TRBJ2-2 | CAISDSPNTGELFF | KLGGALQAK | YES | NO | NO |
| 5364 | TRAV38-2DV8 | TRAJ21 | CAYRSAPFNKFYF | TRBV10-3 | TRBJ2-2 | CAISESGLAGGRSNTGELFF | KLGGALQAK | NO | NO | NO |
| 5365 | TRAV38-2DV8 | TRAJ49 | CAYRRAGNQFYF | TRBV10-3 | TRBJ2-2 | CAISESGILENTGELFF | KLGGALQAK | YES | NO | NO |
| 5366 | TRAV5 | TRAJ38 | CAELNAGNNRKLIW | TRBV10-3 | TRBJ2-2 | CAISEPTRQNVPNTGELFF | KLGGALQAK | YES | NO | NO |
| 5369 | TRAV1-1 | TRAJ13 | CAAGGGGYQKVTF | TRBV11-2 | TRBJ2-2 | CASSLSLSTGSTGELFF | KLGGALQAK | YES | NO | NO |
| 5371 | TRAV12-2 | TRAJ47 | CAVEVEYGNKLVF | TRBV11-2 | TRBJ2-2 | CASSLARGTGELFF | KLGGALQAK | YES | NO | NO |
| 5372 | TRAV13-1 | TRAJ40 | CAASILPGTYKYIF | TRBV11-2 | TRBJ2-2 | CATSAGTFHTGELFF | RLRAEAQVK | YES | YES | YES |
| 5373 | TRAV14DV4 | TRAJ13 | CAACISGGYQKVTF | TRBV11-2 | TRBJ2-2 | CASSLAGYTGELFF | KLGGALQAK | YES | NO | NO |
| 5374 | TRAV29DV5 | TRAJ26 | CAAAYGQNFVF | TRBV11-2 | TRBJ2-2 | CASSYPMDRSGELFF | KLGGALQAK | YES | NO | NO |
| 5377 | TRAV38-2DV8 | TRAJ52 | CASDAGGTSYGKLTF | TRBV11-2 | TRBJ2-2 | CASGTGELFF | KLGGALQAK | YES | NO | NO |
| 5378 | TRAV13-2 | TRAJ49 | CAENICRRHTGNQFYF | TRBV11-3 | TRBJ2-2 | CASSSQGTGYTGELFF | KLGGALQAK | YES | NO | NO |
| 5380 | TRAV14DV4 | TRAJ22 | CAMRRPISSGSARQLTF | TRBV11-3 | TRBJ2-2 | CASSPPGGTGDRVGELFF | KLGGALQAK | YES | NO | NO |
| 5381 | TRAV14DV4 | TRAJ22 | CAMRRPISSGSARQLTF | TRBV11-3 | TRBJ2-2 | CASSTRDSSTGELFF | KLGGALQAK | YES | NO | NO |
| 5382 | TRAV14DV4 | TRAJ22 | CAMRRPISSGSARQLTF | TRBV11-3 | TRBJ2-2 | CASSQSTGRAGELFF | KLGGALQAK | YES | NO | NO |
| 5383 | TRAV14DV4 | TRAJ31 | CAMREVGGGNARLMF | TRBV11-3 | TRBJ2-2 | CASRDRATSNTGELFF | KLGGALQAK | YES | NO | NO |
| 5387 | TRAV21 | TRAJ48 | CAAGYISNFGNEKLTF | TRBV11-3 | TRBJ2-2 | CASSAGGLAGPGELFF | KLGGALQAK | YES | NO | NO |
| 5391 | TRAV38-2DV8 | TRAJ29 | CAYRSVGTPLVF | TRBV11-3 | TRBJ2-2 | CASSLGVAGTNTGELFF | KLGGALQAK | YES | NO | NO |
| 5393 | TRAV12-1 | TRAJ37 | CVLATGKLIF | TRBV12-3 | TRBJ2-2 | CATHGGIDTGELFF | KLGGALQAK | YES | NO | NO |
| 5394 | TRAV14DV4 | TRAJ57 | CAMRESPGGSEKLVF | TRBV12-3 | TRBJ2-2 | CASSRTGLSGPGELFF | KLGGALQAK | YES | NO | NO |
| 5396 | TRAV16 | TRAJ32 | CALGCTGGATNKLIF | TRBV12-3 | TRBJ2-2 | CASNRGNTGELFF | KLGGALQAK | NO | NO | NO |
| 5397 | TRAV19 | TRAJ33 | CALSEAQNSNYQLIW | TRBV12-3 | TRBJ2-2 | CASSQYSGGAPGELFF | KLGGALQAK | YES | NO | NO |
| 5401 | TRAV29DV5 | TRAJ42 | CAEGFYGGSQGNLIF | TRBV12-3 | TRBJ2-2 | CASSFSGNTGELFF | KLGGALQAK | YES | NO | NO |
| 5402 | TRAV3 | TRAJ7 | CAVRDIIYGNNRLAF | TRBV12-3 | TRBJ2-2 | CASSFSGNTGELFF | KLGGALQAK | NO | NO | NO |
| 5403 | TRAV35 | TRAJ52 | CAGLANAGGTSYGKLTF | TRBV12-3 | TRBJ2-2 | CASSPGLAGAVSLGGELFF | KLGGALQAK | YES | NO | NO |
| 5410 | TRAV14DV4 | TRAJ22 | CAMRRPISSGSARQLTF | TRBV12-4 | TRBJ2-2 | CASSFGQGGYTGELFF | KLGGALQAK | YES | NO | NO |
| 5411 | TRAV2 | TRAJ13 | CAASGGYQKVTF | TRBV12-4 | TRBJ2-2 | CASRQGNTGELFF | KLGGALQAK | NO | NO | NO |
| 5413 | TRAV21 | TRAJ18 | CAFDRGSTLGRLYF | TRBV12-4 | TRBJ2-2 | CASSLSPNTGELFF | KLGGALQAK | YES | NO | NO |
| 5414 | TRAV21 | TRAJ40 | CAVRPSTSGTYKYIF | TRBV12-4 | TRBJ2-2 | CASSTRDRAPTGELFF | KLGGALQAK | YES | NO | NO |
| 5416 | TRAV27 | TRAJ36 | CAGLLKTGANNLFF | TRBV12-4 | TRBJ2-2 | CASSFLNTGELFF | KLGGALQAK | YES | NO | NO |
| 5417 | TRAV27 | TRAJ36 | CAGALVAGANNLFF | TRBV12-4 | TRBJ2-2 | CASSFVNTGELFF | KLGGALQAK | YES | NO | NO |
| 5418 | TRAV29DV5 | TRAJ49 | CAARNYDTGNQFYF | TRBV12-4 | TRBJ2-2 | CASSFPLALNTGELFF | KLGGALQAK | YES | NO | NO |
| 5419 | TRAV5 | TRAJ20 | CASINDYKLSF | TRBV12-4 | TRBJ2-2 | CASSLSGTGELFF | KLGGALQAK | YES | NO | NO |
| 5421 | TRAV10 | TRAJ48 | CVVREGSNFGNEKLTF | TRBV12-5 | TRBJ2-2 | CASGLVMLAGAGELFF | KLGGALQAK | YES | NO | NO |
| 5424 | TRAV27 | TRAJ23 | CAGALGNQGGKLIF | TRBV12-5 | TRBJ2-2 | CASRAGGNTGELFF | KLGGALQAK | YES | NO | NO |
| 5428 | TRAV26-1 | TRAJ10 | CIVRVAGGGGNKLTF | TRBV13 | TRBJ2-2 | CASSPAHTGELFF | KLGGALQAK | YES | NO | NO |
| 5428 | TRAV26-1 | TRAJ10 | CIVRVAGGGGNKLTF | TRBV13 | TRBJ2-2 | CASSPAHTGELFF | AVFDRKSDAK | YES | NO | NO |
| 5429 | TRAV26-1 | TRAJ10 | CIVRVPSGGGNKLTF | TRBV13 | TRBJ2-2 | CASSSPGENTGELFF | AVFDRKSDAK | NO | NO | NO |

| | | | | | | | | | | |
|---|---|---|---|---|---|---|---|---|---|---|
| 5429 | TRAV26-1 | TRAJ10 | CIVRVPSGGGNKLTF | TRBV13 | TRBJ2-2 | CASSSPGENTGELFF | KLGGALQAK | NO | NO | NO |
| 5430 | TRAV26-1 | TRAJ11 | CIVMGYSTLTF | TRBV13 | TRBJ2-2 | CASSPDKANTGELFF | KLGGALQAK | NO | NO | NO |
| 5434 | TRAV12-3 | TRAJ39 | CAIGPGNMLTF | TRBV14 | TRBJ2-2 | CASSLSAWAYSNTGELFF | KLGGALQAK | NO | NO | NO |
| 5435 | TRAV13-1 | TRAJ24 | CAARQTDSWGKFQF | TRBV14 | TRBJ2-2 | CASSLSAWAYSNTGELFF | KLGGALQAK | NO | NO | NO |
| 5436 | TRAV21 | TRAJ40 | CASHSGTYKYIF | TRBV14 | TRBJ2-2 | CASSQYLWGGELFF | KLGGALQAK | NO | NO | NO |
| 5437 | TRAV41 | TRAJ42 | CAVPMNYGGSQGNLIF | TRBV14 | TRBJ2-2 | CASSPPLIWDAGELFF | KLGGALQAK | NO | NO | NO |
| 5438 | TRAV1-2 | TRAJ30 | CAVRDYWDDKIIF | TRBV15 | TRBJ2-2 | CATSREVVGPGELFF | KLGGALQAK | YES | NO | NO |
| 5439 | TRAV1-2 | TRAJ33 | CAVRDSNYQLIW | TRBV15 | TRBJ2-2 | CATSTGQGETGELFF | KLGGALQAK | YES | NO | NO |
| 5440 | TRAV12-2 | TRAJ11 | CAVEASGGYSTLTF | TRBV15 | TRBJ2-2 | CATSESGGVGELFF | KLGGALQAK | YES | NO | NO |
| 5441 | TRAV13-1 | TRAJ44 | CAADNTGTASKLTF | TRBV15 | TRBJ2-2 | CATSSGANTGELFF | KLGGALQAK | YES | NO | NO |
| 5442 | TRAV16 | TRAJ47 | CALMEYGNKLVF | TRBV15 | TRBJ2-2 | CATSRDRTGEVVFF | KLGGALQAK | NO | NO | NO |
| 5444 | TRAV26-2 | TRAJ30 | CILNRDDKIIF | TRBV15 | TRBJ2-2 | CATSRDLGPGELFF | KLGGALQAK | YES | NO | NO |
| 5445 | TRAV27 | TRAJ41 | CAGENSGYALNF | TRBV15 | TRBJ2-2 | CATSRPRSLAVNTGELFF | KLGGALQAK | YES | NO | NO |
| 5446 | TRAV9-2 | TRAJ42 | CALNLPGVGSQGNLIF | TRBV16 | TRBJ2-2 | CASSPETGGNTGELFF | KLGGALQAK | YES | NO | NO |
| 5447 | TRAV1-2 | TRAJ42 | CAVRDEYGGSQGNLIF | TRBV18 | TRBJ2-2 | CASSPVGGSGELFF | KLGGALQAK | YES | NO | NO |
| 5449 | TRAV8-6 | TRAJ49 | CAVSGSGQFYF | TRBV18 | TRBJ2-2 | CASSPLREHTGELFF | KLGGALQAK | YES | NO | NO |
| 5450 | TRAV1-2 | TRAJ6 | CAVSTSGGSYIPTF | TRBV19 | TRBJ2-2 | CASSFSLEFTGELFF | KLGGALQAK | YES | NO | NO |
| 5451 | TRAV12-2 | TRAJ48 | CAVNINFGNEKLTF | TRBV19 | TRBJ2-2 | CASSWGPNTGELFF | KLGGALQAK | YES | NO | NO |
| 5455 | TRAV14DV4 | TRAJ22 | CAMRRPISSGSARQLTF | TRBV19 | TRBJ2-2 | CASSTQSGTASGELFF | KLGGALQAK | YES | NO | NO |
| 5456 | TRAV16 | TRAJ27 | CALGNAGKSTF | TRBV19 | TRBJ2-2 | CALRLPNTGELFF | KLGGALQAK | NO | NO | NO |
| 5457 | TRAV17 | TRAJ15 | CATDPYQAGTALIF | TRBV19 | TRBJ2-2 | CASSTHRDRGSGELFF | KLGGALQAK | YES | NO | NO |
| 5458 | TRAV17 | TRAJ52 | CATPTSYGKLTF | TRBV19 | TRBJ2-2 | CASTKSLGPNTGELFF | KLGGALQAK | YES | NO | NO |
| 5460 | TRAV19 | TRAJ24 | CALNDSWGKLQF | TRBV19 | TRBJ2-2 | CASSIGVRELFF | KLGGALQAK | YES | NO | NO |
| 5461 | TRAV19 | TRAJ29 | CALSGTRSGNTPLVF | TRBV19 | TRBJ2-2 | CASSGGQAAWGSVGSGELFF | KLGGALQAK | NO | NO | NO |
| 5462 | TRAV19 | TRAJ6 | CALSELGSGGSYIPTF | TRBV19 | TRBJ2-2 | CASMFTGELFF | KLGGALQAK | YES | NO | NO |
| 5463 | TRAV21 | TRAJ29 | CAVYSGNTPLVF | TRBV19 | TRBJ2-2 | CASSIVGNSELFF | KLGGALQAK | YES | NO | NO |
| 5465 | TRAV24 | TRAJ10 | CAFDSILTGGGNKLTF | TRBV19 | TRBJ2-2 | CASSILGQTIINTGELFF | KLGGALQAK | YES | NO | NO |
| 5466 | TRAV25 | TRAJ39 | CAGHQGNNAGNMLTF | TRBV19 | TRBJ2-2 | CASSMPGLVFPGELFF | KLGGALQAK | YES | NO | NO |
| 5468 | TRAV27 | TRAJ27 | CAGSWGTNAGKSTF | TRBV19 | TRBJ2-2 | CASRSRPGELFF | KLGGALQAK | YES | NO | NO |
| 5469 | TRAV29DV5 | TRAJ11 | CAASDGFGYSTLTF | TRBV19 | TRBJ2-2 | CASSIGGELFF | KLGGALQAK | YES | NO | NO |
| 5471 | TRAV36DV7 | TRAJ18 | CAAGFDRGSTLGRLYF | TRBV19 | TRBJ2-2 | CASSIFGNTGELFF | KLGGALQAK | NO | NO | NO |
| 5472 | TRAV38-2DV8 | TRAJ34 | CAYRTSASNTDKLIF | TRBV19 | TRBJ2-2 | CASSISYTGELFF | AVFDRKSDAK | YES | NO | NO |
| 5473 | TRAV39 | TRAJ33 | CAVEGDSNYQLIW | TRBV19 | TRBJ2-2 | CASSAGAAGELFF | KLGGALQAK | YES | NO | NO |
| 5474 | TRAV8-1 | TRAJ42 | CAVINYGGSQGNLIF | TRBV19 | TRBJ2-2 | CASSIPFGAGANTGELFF | KLGGALQAK | YES | NO | NO |
| 5476 | TRAV12-2 | TRAJ45 | CAVNVYSGGGADGLTF | TRBV2 | TRBJ2-2 | CASSEGTSGFTGELFF | KLGGALQAK | YES | NO | NO |
| 5477 | TRAV12-2 | TRAJ49 | CAVSPGNQFYF | TRBV2 | TRBJ2-2 | CASSETGAGDTGELFF | KLGGALQAK | YES | NO | NO |
| 5478 | TRAV12-3 | TRAJ21 | CAMKKYNFNKFYF | TRBV2 | TRBJ2-2 | CASRGAGELFF | KLGGALQAK | YES | NO | NO |
| 5478 | TRAV12-3 | TRAJ21 | CAMKKYNFNKFYF | TRBV2 | TRBJ2-2 | CASRGAGELFF | AVFDRKSDAK | YES | NO | NO |
| 5479 | TRAV13-1 | TRAJ50 | CAASETSYDKVIF | TRBV2 | TRBJ2-2 | CASRGAGELFF | KLGGALQAK | YES | NO | NO |
| 5480 | TRAV17 | TRAJ22 | CATEPYSGSARQLTF | TRBV2 | TRBJ2-2 | CASSDRAGELFF | KLGGALQAK | YES | NO | NO |
| 5483 | TRAV29DV5 | TRAJ40 | CAADATSGTYKYIF | TRBV2 | TRBJ2-2 | CASSGPSGNTGELFF | KLGGALQAK | YES | NO | NO |
| 5485 | TRAV41 | TRAJ49 | CAVEGTGNQFYF | TRBV2 | TRBJ2-2 | CASSRGGGVVDTGELFF | KLGGALQAK | NO | NO | NO |
| 5486 | TRAV5 | TRAJ20 | CAEKRNDYKLSF | TRBV2 | TRBJ2-2 | CASSPLNTGELFF | KLGGALQAK | YES | NO | NO |
| 5487 | TRAV8-3 | TRAJ40 | CAVGSLGASGTYKYIF | TRBV2 | TRBJ2-2 | CASSETPWTDGELFF | KLGGALQAK | YES | NO | NO |
| 5488 | TRAV8-3 | TRAJ9 | CAVYDGGFKTIF | TRBV2 | TRBJ2-2 | CASSEASGGATNTGELFF | KLGGALQAK | YES | NO | NO |
| 5489 | TRAV1-2 | TRAJ26 | CAVRRDYGQNFVF | TRBV20-1 | TRBJ2-2 | CSALDRATSNTGELFF | KLGGALQAK | YES | NO | NO |
| 5490 | TRAV12-2 | TRAJ11 | CAVNRGYSTLTF | TRBV20-1 | TRBJ2-2 | CSAGQGTGELFF | KLGGALQAK | YES | NO | NO |
| 5491 | TRAV12-2 | TRAJ15 | CAVNQAGTALIF | TRBV20-1 | TRBJ2-2 | CSAQAGGEMLPGELFF | KLGGALQAK | YES | NO | NO |
| 5494 | TRAV19 | TRAJ26 | CALSEAPDNYGQNFVF | TRBV20-1 | TRBJ2-2 | CSAREGGQGGIGELFF | KLGGALQAK | YES | NO | NO |
| 5496 | TRAV21 | TRAJ15 | CAVRRTQAGTALIF | TRBV20-1 | TRBJ2-2 | CSARAWDPNTGELFF | KLGGALQAK | YES | YES | NO |
| 5498 | TRAV39 | TRAJ43 | CAVARNNNDMRF | TRBV20-1 | TRBJ2-2 | CSARDMDISGELFF | KLGGALQAK | YES | NO | NO |
| 5499 | TRAV8-1 | TRAJ34 | CAVNADHNTDKLIF | TRBV20-1 | TRBJ2-2 | CSARAGTGNTGELFF | KLGGALQAK | YES | YES | NO |
| 5500 | TRAV9-2 | TRAJ29 | CALRDGTPLVF | TRBV20-1 | TRBJ2-2 | CSARDYSRTGELFF | KLGGALQAK | YES | NO | NO |
| 5501 | TRAV13-1 | TRAJ24 | CAASPFTTDSWGKLQF | TRBV21-1 | TRBJ2-2 | CASSKKIDRVLFWTGELFF | KLGGALQAK | YES | NO | NO |
| 5502 | TRAV21 | TRAJ43 | CAVWYMRF | TRBV21-1 | TRBJ2-2 | CASSKGKKGRGMDTGELFF | KLGGALQAK | NO | NO | NO |
| 5504 | TRAV1-1 | TRAJ20 | CAVRVFDKLSF | TRBV24-1 | TRBJ2-2 | CATSDLTDRAGGELFF | KLGGALQAK | YES | NO | NO |
| 5505 | TRAV1-2 | TRAJ33 | CAVLDSNYQLIW | TRBV24-1 | TRBJ2-2 | CATSDSQLTSGGETGELFF | KLGGALQAK | YES | NO | NO |
| 5506 | TRAV12-1 | TRAJ42 | CVVNGENYGGSQGNLIF | TRBV24-1 | TRBJ2-2 | CATSDWDTGELFF | KLGGALQAK | YES | NO | NO |
| 5510 | TRAV20 | TRAJ8 | CAVGMSTGFQKLVF | TRBV24-1 | TRBJ2-2 | CATSVRGVRTGELFF | KLGGALQAK | YES | NO | NO |
| 5511 | TRAV22 | TRAJ47 | CAVEPLYGNKLVF | TRBV24-1 | TRBJ2-2 | CATSDRLAGGELFF | KLGGALQAK | YES | NO | NO |
| 5511 | TRAV22 | TRAJ47 | CAVEPLYGNKLVF | TRBV24-1 | TRBJ2-2 | CATSDRLAGGELFF | AVFDRKSDAK | YES | NO | NO |
| 5511 | TRAV22 | TRAJ47 | CAVEPLYGNKLVF | TRBV24-1 | TRBJ2-2 | CATSDRLAGGELFF | RLRAEAQVK | YES | NO | NO |
| 5511 | TRAV22 | TRAJ47 | CAVEPLYGNKLVF | TRBV24-1 | TRBJ2-2 | CATSDRLAGGELFF | IVTDFSVIK | YES | NO | NO |
| 5514 | TRAV27 | TRAJ33 | CAGATDSNYQLIW | TRBV24-1 | TRBJ2-2 | CATSEGYTGELFF | KLGGALQAK | YES | NO | NO |
| 5515 | TRAV3 | TRAJ12 | CAVRDLDSSYKLIF | TRBV24-1 | TRBJ2-2 | CATSDRLAGGELFF | KLGGALQAK | NO | NO | NO |
| 5516 | TRAV4 | TRAJ33 | CLVGDMIVDSNYQLIW | TRBV24-1 | TRBJ2-2 | CATSDSIAGGPSTGELFF | KLGGALQAK | NO | NO | NO |
| 5517 | TRAV8-4 | TRAJ7 | CAVSEVGNNRLAF | TRBV24-1 | TRBJ2-2 | CATSGAEENTGELFF | KLGGALQAK | YES | NO | NO |
| 5519 | TRAV1-2 | TRAJ31 | CAVREWNNARLMF | TRBV25-1 | TRBJ2-2 | CASTPYTGELFF | KLGGALQAK | YES | NO | NO |
| 5520 | TRAV12-2 | TRAJ12 | CASVGMDSSYKLIF | TRBV25-1 | TRBJ2-2 | CASSELLAGMNTGELFF | KLGGALQAK | YES | NO | NO |
| 5522 | TRAV38-2DV8 | TRAJ30 | CAYRSNRDDKIIF | TRBV25-1 | TRBJ2-2 | CASSEWARGRGRYTGELFF | KLGGALQAK | YES | NO | NO |
| 5523 | TRAV12-1 | TRAJ20 | CVVSDYKLSF | TRBV27 | TRBJ2-2 | CASSQLTGTSPMELFF | KLGGALQAK | YES | NO | NO |
| 5524 | TRAV12-1 | TRAJ40 | CVVPSLTYKYIF | TRBV27 | TRBJ2-2 | CASSLFGAVGELFF | KLGGALQAK | YES | NO | NO |
| 5525 | TRAV12-2 | TRAJ9 | CAVKDGGFKTIF | TRBV27 | TRBJ2-2 | CASSFFAGGLGELFF | KLGGALQAK | YES | NO | NO |
| 5528 | TRAV13-1 | TRAJ15 | CAARVTAGTALIF | TRBV27 | TRBJ2-2 | CASSFGAGELFF | KLGGALQAK | YES | NO | NO |
| 5529 | TRAV13-1 | TRAJ16 | CAATPDGQKLLF | TRBV27 | TRBJ2-2 | CASSLLSRTGELFF | KLGGALQAK | NO | NO | NO |
| 5529 | TRAV13-1 | TRAJ16 | CAATPDGQKLLF | TRBV27 | TRBJ2-2 | CASSLLSRTGELFF | RLRAEAQVK | NO | NO | NO |
| 5530 | TRAV13-1 | TRAJ20 | CAASIPNDYKLSF | TRBV27 | TRBJ2-2 | CASTSGVNTGELFF | KLGGALQAK | NO | NO | NO |
| 5534 | TRAV14DV4 | TRAJ49 | CAMREVGNQFYF | TRBV27 | TRBJ2-2 | CASSLARGLGTGELFF | KLGGALQAK | YES | NO | NO |
| 5535 | TRAV14DV4 | TRAJ53 | CAMREGPGGSNYKLTF | TRBV27 | TRBJ2-2 | CASSLLGAGELFF | KLGGALQAK | YES | NO | NO |
| 5538 | TRAV19 | TRAJ15 | CALSENQAGTALIF | TRBV27 | TRBJ2-2 | CASSIRYTGELFF | KLGGALQAK | NO | NO | NO |
| 5540 | TRAV19 | TRAJ22 | CALSRLSGSARQLTF | TRBV27 | TRBJ2-2 | CASSALDSTGELFF | KLGGALQAK | YES | NO | NO |
| 5541 | TRAV19 | TRAJ24 | CALSEASDSWGKFQF | TRBV27 | TRBJ2-2 | CASTGGGAGVVGELFF | KLGGALQAK | YES | NO | NO |
| 5543 | TRAV19 | TRAJ6 | CALSEASSGGSYIPTF | TRBV27 | TRBJ2-2 | CASRRPTGELFF | KLGGALQAK | YES | NO | NO |
| 5544 | TRAV20 | TRAJ27 | CAVQAWDTNAGKSTF | TRBV27 | TRBJ2-2 | CASGSTGSGTGELFF | KLGGALQAK | YES | NO | NO |
| 5546 | TRAV25 | TRAJ48 | CAGAISNFGNEKLTF | TRBV27 | TRBJ2-2 | CASSSGTRLFSGELFF | KLGGALQAK | YES | NO | NO |
| 5547 | TRAV26-1 | TRAJ32 | CIVRGPHGGATNKLIF | TRBV27 | TRBJ2-2 | CASSFTEMIGELFF | KLGGALQAK | YES | NO | NO |
| 5548 | TRAV27 | TRAJ5 | CAGGHRHTGRRALTF | TRBV27 | TRBJ2-2 | CASSPLTGDTGELFF | KLGGALQAK | YES | NO | NO |
| 5551 | TRAV38-1 | TRAJ53 | CAFRPHSGGSNYKLTF | TRBV27 | TRBJ2-2 | CASGQGAYTGELFF | KLGGALQAK | YES | NO | NO |
| 5552 | TRAV38-2DV8 | TRAJ31 | CAYRVDNARLMF | TRBV27 | TRBJ2-2 | CASSLSMGGAGELFF | KLGGALQAK | YES | NO | NO |
| 5553 | TRAV4 | TRAJ20 | CLVGEIALSF | TRBV27 | TRBJ2-2 | CASSYAGGKNTGELFF | KLGGALQAK | YES | NO | NO |
| 5554 | TRAV41 | TRAJ47 | CAVTGNKLVF | TRBV27 | TRBJ2-2 | CASSLGQGALVPGELFF | KLGGALQAK | NO | NO | NO |
| 5555 | TRAV5 | TRAJ4 | CAENGGYNKLIF | TRBV27 | TRBJ2-2 | CASRRLAGNTGELFF | KLGGALQAK | YES | NO | NO |
| 5557 | TRAV8-3 | TRAJ20 | CAVGPFWDYKLSF | TRBV27 | TRBJ2-2 | CASSFGGGTGELFF | KLGGALQAK | YES | NO | NO |
| 5558 | TRAV8-3 | TRAJ37 | CAVVFSGNTGKLIF | TRBV27 | TRBJ2-2 | CASSLSGWSGTGELFF | KLGGALQAK | YES | NO | NO |
| 5560 | TRAV8-4 | TRAJ27 | CAVSYTNAGKSTF | TRBV27 | TRBJ2-2 | CASSLGTEWAAGELFF | KLGGALQAK | YES | NO | NO |
| 5561 | TRAV1-2 | TRAJ11 | CAVRERGYSTLTF | TRBV28 | TRBJ2-2 | CASEGAGGFRTGELFF | KLGGALQAK | YES | NO | NO |
| 5562 | TRAV13-1 | TRAJ11 | CAGYSTLTF | TRBV28 | TRBJ2-2 | CASSYTNTGELFF | KLGGALQAK | NO | NO | NO |
| 5564 | TRAV38-1 | TRAJ57 | CAFNQGGSEKLVF | TRBV28 | TRBJ2-2 | CASSPPGQGFGELFF | KLGGALQAK | YES | NO | NO |
| 5565 | TRAV8-1 | TRAJ39 | CAVNSNAGNMLTF | TRBV28 | TRBJ2-2 | CASTLITGELFF | KLGGALQAK | YES | NO | NO |
| 5566 | TRAV12-2 | TRAJ20 | CAVTGGNDYKLSF | TRBV29-1 | TRBJ2-2 | CSVQGANTGELFF | KLGGALQAK | YES | NO | NO |
| 5568 | TRAV13-1 | TRAJ3 | CAASRHSSASKIIF | TRBV29-1 | TRBJ2-2 | CSAVPGQGHTGELFF | KLGGALQAK | YES | NO | NO |
| 5569 | TRAV14DV4 | TRAJ22 | CAMRRPISSGSARQLTF | TRBV29-1 | TRBJ2-2 | CSVATDRGYTGELFF | KLGGALQAK | YES | NO | NO |
| 5570 | TRAV19 | TRAJ39 | CALRAYNAGNMLTF | TRBV29-1 | TRBJ2-2 | CSVEGTSGGDVTGELFF | KLGGALQAK | YES | NO | NO |
| 5571 | TRAV19 | TRAJ52 | CALNAGGTSYGKLTF | TRBV29-1 | TRBJ2-2 | CSARDTTTGELFF | KLGGALQAK | YES | NO | NO |

| | | | | | | | | | | |
|---|---|---|---|---|---|---|---|---|---|---|
| 5572 | TRAV2 | TRAJ24 | CAATTDSWGKFQF | TRBV29-1 | TRBJ2-2 | CSASIQGRAGELFF | KLGGALQAK | NO | NO | NO |
| 5574 | TRAV38-2DV8 | TRAJ23 | CAYKSAGKLIF | TRBV29-1 | TRBJ2-2 | CSVLTTGNTGELFF | KLGGALQAK | YES | NO | NO |
| 5575 | TRAV38-2DV8 | TRAJ32 | CASRNGGATNKLIF | TRBV29-1 | TRBJ2-2 | CSVVQPNTGELFF | KLGGALQAK | YES | NO | NO |
| 5577 | TRAV8-4 | TRAJ40 | CAVSAGTSGTYKYIF | TRBV29-1 | TRBJ2-2 | CSVAATTGELFF | KLGGALQAK | YES | NO | NO |
| 5578 | TRAV12-1 | TRAJ22 | CVVNSYSGSARQLTF | TRBV3-1 | TRBJ2-2 | CASSQVFRGPPGELFF | KLGGALQAK | YES | NO | NO |
| 5579 | TRAV12-2 | TRAJ15 | CAGGRGTALIF | TRBV3-1 | TRBJ2-2 | CASSQEEQGTGELFF | KLGGALQAK | YES | NO | NO |
| 5580 | TRAV12-3 | TRAJ35 | CAMKR | TRBV3-1 | TRBJ2-2 | CASSQVFRGPPGELFF | KLGGALQAK | NO | NO | NO |
| 5587 | TRAV5 | TRAJ16 | CAEEIF | TRBV3-1 | TRBJ2-2 | CASSQRGGELFF | KLGGALQAK | NO | NO | NO |
| 5588 | TRAV5 | TRAJ8 | CAEDQDTGFQKLVF | TRBV3-1 | TRBJ2-2 | CASSLPGQGAGELFF | KLGGALQAK | YES | NO | NO |
| 5589 | TRAV6 | TRAJ20 | CARPFNDYKLSF | TRBV3-1 | TRBJ2-2 | CASSHTGDTGELFF | KLGGALQAK | YES | NO | NO |
| 5591 | TRAV12-2 | TRAJ30 | CAVNRRDDKIIF | TRBV30 | TRBJ2-2 | CAWSVLVQGYTGELFF | KLGGALQAK | YES | NO | NO |
| 5592 | TRAV17 | TRAJ23 | CALFNQGGKLIF | TRBV30 | TRBJ2-2 | CAWSRGTGELFF | KLGGALQAK | YES | NO | NO |
| 5593 | TRAV19 | TRAJ41 | CALSEAQLGSGYALNF | TRBV30 | TRBJ2-2 | CAWSVGQGTDGELFF | KLGGALQAK | YES | NO | NO |
| 5594 | TRAV20 | TRAJ21 | CAVRDNFNKFYF | TRBV30 | TRBJ2-2 | CACLLLLNTGELFF | KLGGALQAK | YES | NO | NO |
| 5595 | TRAV29DV5 | TRAJ26 | CAASAFVDGQNFVF | TRBV30 | TRBJ2-2 | CAWRTQSWELFF | KLGGALQAK | YES | NO | NO |
| 5596 | TRAV38-1 | TRAJ57 | CAFSPQGGSEKLVF | TRBV30 | TRBJ2-2 | CAWSPNTGELFF | KLGGALQAK | YES | NO | NO |
| 5597 | TRAV8-1 | TRAJ33 | CAVNAGNYQLIW | TRBV30 | TRBJ2-2 | CAWSITGELFF | KLGGALQAK | YES | NO | NO |
| 5598 | TRAV1-2 | TRAJ27 | CAVRVGPRRTNAGKSTF | TRBV4-1 | TRBJ2-2 | CASSLHQTSGGNANTGELFF | KLGGALQAK | NO | NO | NO |
| 5599 | TRAV1-2 | TRAJ33 | CAPMDSNYQLIW | TRBV4-1 | TRBJ2-2 | CASSQEEIVSGELFF | KLGGALQAK | YES | NO | NO |
| 5600 | TRAV1-2 | TRAJ33 | CAAMDSNYQLIW | TRBV4-1 | TRBJ2-2 | CASSQDFGAAGELFF | KLGGALQAK | YES | NO | NO |
| 5601 | TRAV10 | TRAJ40 | CVVSGYKYIF | TRBV4-1 | TRBJ2-2 | CASSQTGTELAGANTGELFF | KLGGALQAK | NO | NO | NO |
| 5603 | TRAV13-1 | TRAJ23 | CAASGNNQGGKLIF | TRBV4-1 | TRBJ2-2 | CASSQVNTGELFF | KLGGALQAK | YES | NO | NO |
| 5605 | TRAV13-2 | TRAJ27 | CAENSSLAGKSTF | TRBV4-1 | TRBJ2-2 | CASSQDRGGPGELFF | KLGGALQAK | YES | NO | NO |
| 5606 | TRAV17 | TRAJ23 | CATELYNQGGKLIF | TRBV4-1 | TRBJ2-2 | CASSQHRGRTGELFF | KLGGALQAK | YES | NO | NO |
| 5609 | TRAV29DV5 | TRAJ49 | CAASDGNTGNQFYF | TRBV4-1 | TRBJ2-2 | CASSPYGGHGELFF | KLGGALQAK | YES | NO | NO |
| 5610 | TRAV4 | TRAJ43 | CLVGVYNNNDMRF | TRBV4-1 | TRBJ2-2 | CASSQEWLQTGYTGELFF | KLGGALQAK | YES | NO | NO |
| 5611 | TRAV8-2 | TRAJ45 | CVVSFYSGGGADGLTF | TRBV4-1 | TRBJ2-2 | CASSQAPGFDGELFF | KLGGALQAK | YES | NO | NO |
| 5616 | TRAV1-2 | TRAJ33 | CAVLDSNYQLIW | TRBV4-2 | TRBJ2-2 | CASSQEEGTRELFF | KLGGALQAK | YES | NO | NO |
| 5618 | TRAV13-1 | TRAJ35 | CAASRGFGNVLHC | TRBV4-2 | TRBJ2-2 | CASSQDQVGGAYLYTGELFF | KLGGALQAK | NO | NO | NO |
| 5619 | TRAV20 | TRAJ9 | CAVQAETGGFKTIF | TRBV4-2 | TRBJ2-2 | CASSPRLNRAGELFF | KLGGALQAK | YES | NO | NO |
| 5623 | TRAV12-2 | TRAJ29 | CAVSPGNTPLVF | TRBV4-3 | TRBJ2-2 | CASSQDTGGFAGELFF | KLGGALQAK | YES | NO | NO |
| 5624 | TRAV14DV4 | TRAJ15 | CAMREGSQAGTALIF | TRBV4-3 | TRBJ2-2 | CASSPGLAGGPDTGELFF | KLGGALQAK | YES | NO | NO |
| 5625 | TRAV19 | TRAJ48 | CALSEANFGNEKLTF | TRBV4-3 | TRBJ2-2 | CASSQDWVTGELFF | KLGGALQAK | YES | NO | NO |
| 5627 | TRAV21 | TRAJ31 | CAVNNNARLMF | TRBV4-3 | TRBJ2-2 | CASSQEAPGQGDTGELFF | KLGGALQAK | YES | NO | NO |
| 5628 | TRAV21 | TRAJ31 | CAVKLRDARLMF | TRBV4-3 | TRBJ2-2 | CASSQDYRLAGVAGELFF | KLGGALQAK | YES | NO | NO |
| 5629 | TRAV21 | TRAJ42 | CAVYYGGSQGNLIF | TRBV4-3 | TRBJ2-2 | CASSQDRGQGRSGELFF | KLGGALQAK | YES | NO | NO |
| 5633 | TRAV3 | TRAJ42 | CAVRADYGGSQGNLIF | TRBV4-3 | TRBJ2-2 | CASSQFQGAVTGELFF | KLGGALQAK | YES | NO | NO |
| 5635 | TRAV35 | TRAJ42 | CAGQSGRVGGSQGNLIF | TRBV4-3 | TRBJ2-2 | CASSFSGTGETGELFF | KLGGALQAK | YES | NO | NO |
| 5636 | TRAV5 | TRAJ26 | CAESPDNYGQNFVF | TRBV4-3 | TRBJ2-2 | CASSQAGSTGELFF | KLGGALQAK | YES | NO | NO |
| 5639 | TRAV12-2 | TRAJ15 | CAVNSGQAGTALIF | TRBV5-1 | TRBJ2-2 | CASSLGAGELFF | KLGGALQAK | YES | NO | NO |
| 5640 | TRAV12-2 | TRAJ42 | CAVGNLIF | TRBV5-1 | TRBJ2-2 | CASSLGDGGNTGELFF | KLGGALQAK | YES | NO | NO |
| 5642 | TRAV13-2 | TRAJ52 | CAENSLNAGGTSYGKLTF | TRBV5-1 | TRBJ2-2 | CASSAARSGTLNTGELFF | KLGGALQAK | YES | NO | NO |
| 5643 | TRAV16 | TRAJ43 | CALSIYNNNDMRF | TRBV5-1 | TRBJ2-2 | CASTGQGKANTGELFF | KLGGALQAK | NO | NO | NO |
| 5646 | TRAV25 | TRAJ41 | CAVNSGYALNF | TRBV5-1 | TRBJ2-2 | CASSSPRTSGSRGTGELFF | KLGGALQAK | YES | NO | NO |
| 5647 | TRAV8-1 | TRAJ43 | CAVNQNNNDMRF | TRBV5-1 | TRBJ2-2 | CASRETTGELFF | KLGGALQAK | YES | NO | NO |
| 5648 | TRAV8-6 | TRAJ52 | CAVSANAGGTSYGKLTF | TRBV5-1 | TRBJ2-2 | CASSPGLLSGELFF | KLGGALQAK | YES | NO | NO |
| 5651 | TRAV17 | TRAJ43 | CATDIINNNDMRF | TRBV5-4 | TRBJ2-2 | CASSLNTGELFF | KLGGALQAK | YES | NO | NO |
| 5653 | TRAV25 | TRAJ20 | CAGHLTNDYKLSF | TRBV5-4 | TRBJ2-2 | CASSLTRGMAGELFF | KLGGALQAK | YES | NO | NO |
| 5656 | TRAV10 | TRAJ15 | CVVSAVQAGTALIF | TRBV5-6 | TRBJ2-2 | CASSLAGGSNTGELFF | RLRAEAQVK | YES | YES | YES |
| 5657 | TRAV13-1 | TRAJ27 | CAASNEGTNAGKSTF | TRBV5-6 | TRBJ2-2 | CASTVYHGELFF | KLGGALQAK | YES | NO | NO |
| 5658 | TRAV14DV4 | TRAJ15 | CAMREGNPNQAGTALIF | TRBV5-6 | TRBJ2-2 | CASSQPGGGVNGELFF | KLGGALQAK | YES | NO | NO |
| 5659 | TRAV17 | TRAJ39 | CATGAYAGNMLTF | TRBV5-6 | TRBJ2-2 | CASRWDRSNTGELFF | KLGGALQAK | YES | NO | NO |
| 5660 | TRAV3 | TRAJ13 | CAVRVPTGGYQKVTF | TRBV5-6 | TRBJ2-2 | CASSLWQGTAGELFF | KLGGALQAK | NO | NO | NO |
| 5661 | TRAV38-1 | TRAJ49 | CAFMKQGNQFYF | TRBV5-6 | TRBJ2-2 | CASSSTGELFF | KLGGALQAK | YES | NO | NO |
| 5663 | TRAV4 | TRAJ3 | CLVGVSSASKIIF | TRBV5-6 | TRBJ2-2 | CASSPGQGPPKLFF | KLGGALQAK | NO | NO | NO |
| 5664 | TRAV6 | TRAJ34 | CALDGPGNTDKLIF | TRBV5-6 | TRBJ2-2 | CASSLIGELFF | KLGGALQAK | YES | NO | NO |
| 5667 | TRAV12-1 | TRAJ30 | CVVNRGDDKIIF | TRBV5-8 | TRBJ2-2 | CASSPLVDTGELFF | KLGGALQAK | YES | NO | NO |
| 5668 | TRAV14DV4 | TRAJ34 | CAMRELYNTDKLIF | TRBV5-8 | TRBJ2-2 | CASSFRQGDTGELFF | KLGGALQAK | YES | NO | NO |
| 5669 | TRAV17 | TRAJ6 | CATDDPRAAGGSYIPTF | TRBV5-8 | TRBJ2-2 | CASSFGGARTGELFF | KLGGALQAK | YES | NO | NO |
| 5671 | TRAV21 | TRAJ11 | CAVGSGYSTLTF | TRBV5-8 | TRBJ2-2 | CASSLYTSGGTGELFF | KLGGALQAK | YES | NO | NO |
| 5673 | TRAV41 | TRAJ47 | CAVNEYGNKLVF | TRBV5-8 | TRBJ2-2 | CASSFGGARTGELFF | KLGGALQAK | YES | NO | NO |
| 5675 | TRAV12-2 | TRAJ15 | CAVGAGTALIF | TRBV6-1 | TRBJ2-2 | CASSELAPNTGELFF | KLGGALQAK | YES | NO | NO |
| 5677 | TRAV24 | TRAJ40 | CAFPSGTYKYIF | TRBV6-1 | TRBJ2-2 | CASSEAWGATNTGELFF | KLGGALQAK | YES | NO | NO |
| 5677 | TRAV24 | TRAJ40 | CAFPSGTYKYIF | TRBV6-1 | TRBJ2-2 | CASSEAWGATNTGELFF | AVFDRKSDAK | YES | NO | NO |
| 5681 | TRAV8-3 | TRAJ20 | CAVGGPNDYKLSF | TRBV6-1 | TRBJ2-2 | CASGGTGGLTGELFF | KLGGALQAK | YES | NO | NO |
| 5682 | TRAV1-2 | TRAJ20 | CAVRLPNDYKLSF | TRBV6-3 | TRBJ2-2 | CASSNPRLDATGELFF | AVFDRKSDAK | YES | NO | NO |
| 5683 | TRAV12-1 | TRAJ5 | CVVFMDTGRRALTF | TRBV6-3 | TRBJ2-2 | CASRDGELFF | KLGGALQAK | YES | NO | NO |
| 5684 | TRAV13-1 | TRAJ23 | CAASLYNQGGKLIF | TRBV6-3 | TRBJ2-2 | CASSYINTGELFF | KLGGALQAK | YES | NO | NO |
| 5685 | TRAV13-1 | TRAJ36 | CAAETGANNLFF | TRBV6-3 | TRBJ2-2 | CASGTSGTAGELFF | KLGGALQAK | YES | NO | NO |
| 5687 | TRAV14DV4 | TRAJ54 | CAMREPLWGAQKLVF | TRBV6-3 | TRBJ2-2 | CASSLRDKPANTGELFF | KLGGALQAK | YES | NO | NO |
| 5689 | TRAV21 | TRAJ45 | CAVRPISGGGADGLTF | TRBV6-3 | TRBJ2-2 | CASSYFVDLTGELFF | KLGGALQAK | YES | NO | NO |
| 5690 | TRAV25 | TRAJ24 | CAGLLTTDSWGKLQF | TRBV6-3 | TRBJ2-2 | CASSPDTGELFF | KLGGALQAK | YES | NO | NO |
| 5691 | TRAV29DV5 | TRAJ53 | CAASANSGGSNYKLTF | TRBV6-3 | TRBJ2-2 | CASSYEISGEGKNTGELFF | KLGGALQAK | YES | NO | NO |
| 5692 | TRAV5 | TRAJ34 | CAERGNTDKLIF | TRBV6-3 | TRBJ2-2 | CASSYSGSGELFF | KLGGALQAK | YES | NO | NO |
| 5692 | TRAV5 | TRAJ34 | CAERGNTDKLIF | TRBV6-3 | TRBJ2-2 | CASSYSGSGELFF | RLRAEAQVK | YES | NO | NO |
| 5693 | TRAV1-1 | TRAJ30 | CAVPLNRDDKIIF | TRBV6-4 | TRBJ2-2 | CASTSANTGELFF | KLGGALQAK | YES | NO | NO |
| 5694 | TRAV1-2 | TRAJ20 | CAVRDRDYKLSF | TRBV6-4 | TRBJ2-2 | CASSRDTATGELFF | KLGGALQAK | YES | NO | NO |
| 5695 | TRAV1-2 | TRAJ33 | CAALDSNYQLIW | TRBV6-4 | TRBJ2-2 | CASSDGDNTGELFF | KLGGALQAK | YES | NO | NO |
| 5696 | TRAV1-2 | TRAJ33 | CAVMDSNYQLIW | TRBV6-4 | TRBJ2-2 | CASSDSTGELFF | KLGGALQAK | YES | NO | NO |
| 5698 | TRAV12-2 | TRAJ18 | CAVKRGSTLGRLYF | TRBV6-4 | TRBJ2-2 | CASSPENQGELFF | KLGGALQAK | YES | NO | NO |
| 5699 | TRAV8-1 | TRAJ10 | CAALTGGGNKLTF | TRBV6-4 | TRBJ2-2 | CATQQGLGTGELFF | AVFDRKSDAK | YES | NO | NO |
| 5699 | TRAV8-1 | TRAJ10 | CAALTGGGNKLTF | TRBV6-4 | TRBJ2-2 | CATQQGLGTGELFF | KLGGALQAK | YES | NO | NO |
| 5701 | TRAV10 | TRAJ15 | CVVSWDQAGTALIF | TRBV6-5 | TRBJ2-2 | CASTGAANTGELFF | KLGGALQAK | YES | NO | NO |
| 5702 | TRAV19 | TRAJ48 | CALSEANFGNEKLTF | TRBV6-5 | TRBJ2-2 | CASSYPGGGALNTGELFF | KLGGALQAK | YES | NO | NO |
| 5703 | TRAV19 | TRAJ49 | CALSDTGNQFYF | TRBV6-5 | TRBJ2-2 | CASSYDSGTGELFF | KLGGALQAK | YES | NO | NO |
| 5704 | TRAV21 | TRAJ11 | CAVTISGYSTLTF | TRBV6-5 | TRBJ2-2 | CASSYRLTVNTGELFF | KLGGALQAK | YES | NO | NO |
| 5705 | TRAV23DV6 | TRAJ20 | CAASRTFTNDYKLSF | TRBV6-5 | TRBJ2-2 | CASSYSAEQANTGELFF | KLGGALQAK | YES | NO | NO |
| 5706 | TRAV25 | TRAJ28 | CAHRDSGAGSYQLTF | TRBV6-5 | TRBJ2-2 | CASSPLGGYDGELFF | KLGGALQAK | YES | NO | NO |
| 5707 | TRAV26-1 | TRAJ52 | CFLDAGGTSYGKLTF | TRBV6-5 | TRBJ2-2 | CASSYPNTGELFF | AVFDRKSDAK | YES | NO | NO |
| 5707 | TRAV26-1 | TRAJ52 | CFLDAGGTSYGKLTF | TRBV6-5 | TRBJ2-2 | CASSYPNTGELFF | KLGGALQAK | YES | NO | NO |
| 5709 | TRAV29DV5 | TRAJ53 | CAASPIKHSGGSNYKLTF | TRBV6-5 | TRBJ2-2 | CASRSQGPGELFF | KLGGALQAK | YES | NO | NO |
| 5711 | TRAV3 | TRAJ43 | CAVRDMSMRF | TRBV6-5 | TRBJ2-2 | CASSVGTSGGELFF | KLGGALQAK | NO | NO | NO |
| 5712 | TRAV35 | TRAJ49 | CAGLKGNQFYF | TRBV6-5 | TRBJ2-2 | CASTGSGTGELFF | KLGGALQAK | YES | NO | NO |
| 5715 | TRAV13-2 | TRAJ18 | CAERGGGSTLGRLYF | TRBV7-2 | TRBJ2-2 | CASSLGVAGTNTGELFF | KLGGALQAK | YES | NO | NO |
| 5716 | TRAV21 | TRAJ40 | CAVGATSGTYKYIF | TRBV7-2 | TRBJ2-2 | CASSLGVAGTNTGELFF | KLGGALQAK | NO | NO | NO |
| 5718 | TRAV5 | TRAJ42 | CAESGGSQGNLIF | TRBV7-2 | TRBJ2-2 | CASSLRTQGTGELFF | KLGGALQAK | YES | NO | NO |
| 5719 | TRAV14DV4 | TRAJ47 | CAMREGGEYGNKLVF | TRBV7-3 | TRBJ2-2 | CASSRADLAGVPLGTGELFF | KLGGALQAK | NO | NO | NO |
| 5720 | TRAV14DV4 | TRAJ49 | CAMREAGNQFYF | TRBV7-3 | TRBJ2-2 | CASSAGTGGTGELFF | KLGGALQAK | YES | NO | NO |
| 5721 | TRAV20 | TRAJ31 | CAVVDTNARLMF | TRBV7-3 | TRBJ2-2 | CASSLYTGELFF | KLGGALQAK | NO | NO | NO |
| 5721 | TRAV20 | TRAJ31 | CAVVDTNARLMF | TRBV7-3 | TRBJ2-2 | CASSLYTGELFF | AVFDRKSDAK | NO | NO | NO |
| 5722 | TRAV29DV5 | TRAJ37 | CAASGNTGKLIF | TRBV7-3 | TRBJ2-2 | CASSLSGGNTGELFF | KLGGALQAK | NO | NO | NO |
| 5723 | TRAV29DV5 | TRAJ48 | CAASLFGNEKLTF | TRBV7-3 | TRBJ2-2 | CASSLRIFGAPGELFF | KLGGALQAK | NO | NO | NO |
| 5724 | TRAV6 | TRAJ49 | CALGGNQFYF | TRBV7-3 | TRBJ2-2 | CASSLGYGLAGTTGELFF | KLGGALQAK | NO | NO | NO |
| 5725 | TRAV1-2 | TRAJ22 | CAVSTPSGSARQLTF | TRBV7-6 | TRBJ2-2 | CASTQGLAGEWAGELFF | KLGGALQAK | YES | NO | NO |

| | | | | | | | | | | |
|---|---|---|---|---|---|---|---|---|---|---|
| 5726 | TRAV13-2 | TRAJ40 | CAENPGTSGTYKYIF | TRBV7-6 | TRBJ2-2 | CASNRGGTGELFF | KLGGALQAK | YES | NO | NO |
| 5728 | TRAV21 | TRAJ26 | CAGDNYGQNFVF | TRBV7-6 | TRBJ2-2 | CASSPGTVNTGELFF | KLGGALQAK | YES | NO | NO |
| 5729 | TRAV30 | TRAJ45 | CGTEPKRGGADGLTF | TRBV7-6 | TRBJ2-2 | CASTPSFGRNTGELFF | KLGGALQAK | YES | NO | NO |
| 5731 | TRAV38-2DV8 | TRAJ29 | CAYRSLFSGNTPLVF | TRBV7-6 | TRBJ2-2 | CASSLASAPGNRELFF | KLGGALQAK | YES | NO | NO |
| 5733 | TRAV10 | TRAJ15 | CVVSANQAGTALIF | TRBV7-7 | TRBJ2-2 | CASSLGHTGELFF | KLGGALQAK | YES | NO | NO |
| 5734 | TRAV25 | TRAJ20 | CAVLGSNDYKLSF | TRBV7-7 | TRBJ2-2 | CASSLAGDGTGELFF | KLGGALQAK | YES | NO | NO |
| 5738 | TRAV24 | TRAJ33 | CAFHPDSNYQLIW | TRBV7-8 | TRBJ2-2 | CASSVDGIAREGTGELFF | KLGGALQAK | YES | NO | NO |
| 5741 | TRAV27 | TRAJ54 | CAGVGAGAQKLVF | TRBV7-8 | TRBJ2-2 | CASSFPDNTGELFF | KLGGALQAK | YES | NO | NO |
| 5742 | TRAV29DV5 | TRAJ53 | CAAGYNSGGSNYKLTF | TRBV7-8 | TRBJ2-2 | CARKRGEDTGELFF | KLGGALQAK | YES | NO | NO |
| 5743 | TRAV35 | TRAJ26 | CAGLNYGQNFVF | TRBV7-8 | TRBJ2-2 | CASSLPANTGELFF | KLGGALQAK | YES | NO | NO |
| 5744 | TRAV4 | TRAJ9 | CLVALNTGGFKTIF | TRBV7-8 | TRBJ2-2 | CASSHYGDDIGELFF | KLGGALQAK | YES | NO | NO |
| 5748 | TRAV8-3 | TRAJ26 | CAVVLDNYGQNFVF | TRBV7-8 | TRBJ2-2 | CASSPRQGTGELFF | KLGGALQAK | YES | NO | NO |
| 5749 | TRAV8-3 | TRAJ4 | CAVGARTGGYNKLIF | TRBV7-8 | TRBJ2-2 | CASRSGTGSTGELFF | KLGGALQAK | YES | NO | NO |
| 5750 | TRAV1-1 | TRAJ31 | CASNNNARLMF | TRBV7-9 | TRBJ2-2 | CASTGTSSVSGELFF | AVFDRKSDAK | YES | NO | NO |
| 5751 | TRAV1-2 | TRAJ31 | CAVRDQRNARLMF | TRBV7-9 | TRBJ2-2 | CASSLGGGTGELFF | KLGGALQAK | YES | NO | NO |
| 5753 | TRAV10 | TRAJ43 | CVVSARYNNNDMRF | TRBV7-9 | TRBJ2-2 | CASSLSRPTGELFF | KLGGALQAK | YES | NO | NO |
| 5755 | TRAV12-1 | TRAJ28 | CVVNTFSGAGSYQLTF | TRBV7-9 | TRBJ2-2 | CASSLSGQGSGELFF | KLGGALQAK | YES | NO | NO |
| 5756 | TRAV12-1 | TRAJ28 | CVVNTFSGAGSYQLTF | TRBV7-9 | TRBJ2-2 | CASSRAGGGTGELFF | KLGGALQAK | YES | NO | NO |
| 5758 | TRAV12-2 | TRAJ43 | CAVESSDMRF | TRBV7-9 | TRBJ2-2 | CASSPGQGITGELFF | KLGGALQAK | YES | NO | NO |
| 5764 | TRAV14DV4 | TRAJ22 | CAMRRPISSGSARQLTF | TRBV7-9 | TRBJ2-2 | CASTGTSSVSGELFF | KLGGALQAK | YES | NO | NO |
| 5765 | TRAV16 | TRAJ13 | CALSSGGYQKVTF | TRBV7-9 | TRBJ2-2 | CASSPDYLAGGHFNTGELFF | KLGGALQAK | NO | NO | NO |
| 5765 | TRAV16 | TRAJ13 | CALSSGGYQKVTF | TRBV7-9 | TRBJ2-2 | CASSPDYLAGGHFNTGELFF | AVFDRKSDAK | NO | NO | NO |
| 5766 | TRAV16 | TRAJ7 | CALRYGNNRLAF | TRBV7-9 | TRBJ2-2 | CASGAGELFF | KLGGALQAK | NO | NO | NO |
| 5767 | TRAV19 | TRAJ37 | CALSEISSNTGKLIF | TRBV7-9 | TRBJ2-2 | CASSTTSGGPKSRELFF | KLGGALQAK | YES | NO | NO |
| 5768 | TRAV20 | TRAJ27 | CAVQARTNAGKSTF | TRBV7-9 | TRBJ2-2 | CASSLAKTANTGELFF | KLGGALQAK | YES | NO | NO |
| 5773 | TRAV25 | TRAJ45 | CAGLLPGGGADGLTF | TRBV7-9 | TRBJ2-2 | CASSLERLAGTVGELFF | KLGGALQAK | YES | NO | NO |
| 5774 | TRAV29DV5 | TRAJ33 | CAADSNYQLIW | TRBV7-9 | TRBJ2-2 | CASSLRDTDTGELFF | KLGGALQAK | YES | NO | NO |
| 5776 | TRAV38-1 | TRAJ20 | CASASSSNDYKLSF | TRBV7-9 | TRBJ2-2 | CASKREGVTGELFF | KLGGALQAK | YES | NO | NO |
| 5779 | TRAV5 | TRAJ24 | CAEKGSWGKFQF | TRBV7-9 | TRBJ2-2 | CASSFAGDGAGELFF | KLGGALQAK | YES | NO | NO |
| 5781 | TRAV8-2 | TRAJ48 | CVVRSNFGNEKLTF | TRBV7-9 | TRBJ2-2 | CASSLEGLYGGELFF | KLGGALQAK | YES | NO | NO |
| 5783 | TRAV8-4 | TRAJ39 | CAVSEGNMLTF | TRBV7-9 | TRBJ2-2 | CASSPSLGPNTGELFF | KLGGALQAK | YES | NO | NO |
| 5784 | TRAV8-4 | TRAJ42 | CAVSVEGSQGNLIF | TRBV7-9 | TRBJ2-2 | CASSLLAGAGAGELFF | KLGGALQAK | YES | NO | NO |
| 5785 | TRAV8-6 | TRAJ43 | CAVSDMRF | TRBV7-9 | TRBJ2-2 | CASRPHRGQSAGELFF | KLGGALQAK | NO | NO | NO |
| 5786 | TRAV8-6 | TRAJ54 | CAVSAWGAQKLVF | TRBV7-9 | TRBJ2-2 | CASSLDRTGGTGELFF | KLGGALQAK | YES | NO | NO |
| 5787 | TRAV1-1 | TRAJ3 | CAVSLYSSASKIIF | TRBV9 | TRBJ2-2 | CASSEDRHTGELFF | KLGGALQAK | YES | NO | NO |
| 5787 | TRAV1-1 | TRAJ3 | CAVSLYSSASKIIF | TRBV9 | TRBJ2-2 | CASSEDRHTGELFF | IVTDFSVIK | YES | NO | NO |
| 5787 | TRAV1-1 | TRAJ3 | CAVSLYSSASKIIF | TRBV9 | TRBJ2-2 | CASSEDRHTGELFF | AVFDRKSDAK | YES | NO | NO |
| 5788 | TRAV12-1 | TRAJ8 | CVVLAGFQKLVF | TRBV9 | TRBJ2-2 | CASTQRLTGELFF | KLGGALQAK | YES | NO | NO |
| 5790 | TRAV13-1 | TRAJ37 | CAASYSGNTGKLIF | TRBV9 | TRBJ2-2 | CASSVLGRGGANTGELFF | KLGGALQAK | YES | NO | NO |
| 5794 | TRAV19 | TRAJ50 | CALSGTRGTSYDKVIF | TRBV9 | TRBJ2-2 | CASSARMWERGSTGELFF | KLGGALQAK | YES | NO | NO |
| 5795 | TRAV24 | TRAJ42 | CAFSTGYGGSQGNLIF | TRBV9 | TRBJ2-2 | CASSVGDAGELFF | KLGGALQAK | YES | NO | NO |
| 5796 | TRAV25 | TRAJ21 | CAGQYNFNKFYF | TRBV9 | TRBJ2-2 | CASSNSGGAGTGELFF | KLGGALQAK | YES | NO | NO |
| 5800 | TRAV38-2DV8 | TRAJ45 | CALSGGGADGLTF | TRBV9 | TRBJ2-2 | CASSVPKEDSANTGELFF | KLGGALQAK | YES | NO | NO |
| 5801 | TRAV4 | TRAJ4 | CLVGPLGGYNKLIF | TRBV9 | TRBJ2-2 | CASSEGEYSGELFF | KLGGALQAK | YES | NO | NO |
| 5803 | TRAV8-4 | TRAJ20 | CAVIQINDYKLSF | TRBV9 | TRBJ2-2 | CASSVVAGELFF | KLGGALQAK | YES | NO | NO |
| 5804 | TRAV1-2 | TRAJ5 | CAVSRDTGRRALTF | TRBV10-1 | TRBJ2-3 | CASSRLAGAGDTQYF | KLGGALQAK | YES | NO | NO |
| 5806 | TRAV29DV5 | TRAJ30 | CAASARRDDKIIF | TRBV10-1 | TRBJ2-3 | CASSESEWGGTQYF | KLGGALQAK | YES | NO | NO |
| 5807 | TRAV3 | TRAJ34 | CAVRATDKLIF | TRBV10-1 | TRBJ2-3 | CASSELDSSTDTQYF | KLGGALQAK | NO | NO | NO |
| 5810 | TRAV8-4 | TRAJ49 | CAVSSNTGNQFYF | TRBV10-1 | TRBJ2-3 | CASKGAPGDTQYF | KLGGALQAK | YES | NO | NO |
| 5811 | TRAV12-1 | TRAJ27 | CVVFWTNAGKSTF | TRBV10-2 | TRBJ2-3 | CASSESFGGGVNTQYF | KLGGALQAK | YES | NO | NO |
| 5812 | TRAV2 | TRAJ30 | CAVEETMNRDDKIIF | TRBV10-2 | TRBJ2-3 | CASSQGEGLRTDTQYF | KLGGALQAK | NO | NO | NO |
| 5816 | TRAV1-2 | TRAJ24 | CAVSDSWGKLQF | TRBV10-3 | TRBJ2-3 | CAISESRGSLSDTQYF | KLGGALQAK | YES | NO | NO |
| 5817 | TRAV1-2 | TRAJ33 | CAVMDSNYQLIW | TRBV10-3 | TRBJ2-3 | CAISEAEPQDTQYF | KLGGALQAK | YES | NO | NO |
| 5819 | TRAV12-1 | TRAJ36 | CVVEAGANNLFF | TRBV10-3 | TRBJ2-3 | CAISEMAGGGTDTQYF | KLGGALQAK | YES | NO | NO |
| 5820 | TRAV19 | TRAJ33 | CALSENGVNSNYQLIW | TRBV10-3 | TRBJ2-3 | CAISDVSGEPTDTQYF | KLGGALQAK | YES | NO | NO |
| 5825 | TRAV21 | TRAJ49 | CAVSGHTGNQFYF | TRBV11-1 | TRBJ2-3 | CASTVSGSGADTQYF | KLGGALQAK | YES | NO | NO |
| 5826 | TRAV38-1 | TRAJ4 | CAFMFSGGYNKLIF | TRBV11-1 | TRBJ2-3 | CASSLGLAGPTDTQYF | KLGGALQAK | YES | NO | NO |
| 5828 | TRAV12-2 | TRAJ4 | CAVNRGYNKLIF | TRBV11-2 | TRBJ2-3 | CASSLVQVRVTDTQYF | RLRAEAQVK | YES | NO | YES |
| 5829 | TRAV13-1 | TRAJ12 | CAAKLSDSSYKLIF | TRBV11-2 | TRBJ2-3 | CASSFWGRNVDTQYF | KLGGALQAK | YES | NO | NO |
| 5833 | TRAV22 | TRAJ52 | CAVEGGGTSYGKLTF | TRBV11-2 | TRBJ2-3 | CASSLIGDTQYF | KLGGALQAK | YES | NO | NO |
| 5834 | TRAV23DV6 | TRAJ31 | CAASGNNARLMF | TRBV11-2 | TRBJ2-3 | CASSLGDFLRTDTQYF | KLGGALQAK | YES | NO | NO |
| 5837 | TRAV29DV5 | TRAJ57 | CAASLVTQGGSEKLVF | TRBV11-2 | TRBJ2-3 | CASSFWGRNVDTQYF | KLGGALQAK | YES | NO | NO |
| 5839 | TRAV38-1 | TRAJ27 | CAFNTNAGKSTF | TRBV11-2 | TRBJ2-3 | CASSLGWGPSEDTQYF | KLGGALQAK | YES | NO | NO |
| 5840 | TRAV5 | TRAJ30 | CADLNRDDKIIF | TRBV11-2 | TRBJ2-3 | CASSSGGAPDTQYF | KLGGALQAK | YES | NO | NO |
| 5842 | TRAV8-2 | TRAJ6 | CVVRSGGSYIPTF | TRBV11-2 | TRBJ2-3 | CASSQGTADTQYF | KLGGALQAK | YES | NO | NO |
| 5843 | TRAV9-2 | TRAJ41 | CALTGGYALNF | TRBV11-2 | TRBJ2-3 | CASSLDTQYF | KLGGALQAK | YES | NO | NO |
| 5844 | TRAV1-2 | TRAJ18 | CAVPADRGSTLGRLYF | TRBV11-3 | TRBJ2-3 | CASSLEVSGSLGTDTQYF | KLGGALQAK | YES | NO | NO |
| 5845 | TRAV1-2 | TRAJ29 | CAVRVLNSGNTPLVF | TRBV11-3 | TRBJ2-3 | CASSLELTGGLSDTQYF | KLGGALQAK | YES | NO | NO |
| 5846 | TRAV12-2 | TRAJ49 | CAVNGNQFYF | TRBV11-3 | TRBJ2-3 | CASSLDPVNTQYF | KLGGALQAK | YES | NO | NO |
| 5847 | TRAV17 | TRAJ54 | CATDGNQGAQKLVF | TRBV11-3 | TRBJ2-3 | CASSLAASDTQYF | KLGGALQAK | YES | NO | NO |
| 5848 | TRAV17 | TRAJ54 | CATDGNQGAQKLVF | TRBV11-3 | TRBJ2-3 | CASSLEATDTQYF | KLGGALQAK | YES | NO | NO |
| 5849 | TRAV17 | TRAJ54 | CATDGVAQKLVF | TRBV11-3 | TRBJ2-3 | CASSLGLAGPSTDTQYF | KLGGALQAK | YES | NO | NO |
| 5851 | TRAV29DV5 | TRAJ38 | CAASNAGNNRKLIW | TRBV11-3 | TRBJ2-3 | CASSFGTSNSDTQYF | KLGGALQAK | YES | NO | NO |
| 5852 | TRAV35 | TRAJ40 | CAAPVGTYKYIF | TRBV11-3 | TRBJ2-3 | CASRTPLDTQYF | KLGGALQAK | YES | NO | NO |
| 5853 | TRAV38-1 | TRAJ41 | CAFTQGDSGYALNF | TRBV11-3 | TRBJ2-3 | CASFGQHQDTQYF | KLGGALQAK | YES | NO | NO |
| 5855 | TRAV8-1 | TRAJ40 | CAVKYSGTYKYIF | TRBV11-3 | TRBJ2-3 | CASSLAAGQQYF | RLRAEAQVK | YES | NO | NO |
| 5856 | TRAV8-2 | TRAJ44 | CVVSPLTGTASKLTF | TRBV11-3 | TRBJ2-3 | CASSSRLDRTDTQYF | KLGGALQAK | YES | NO | NO |
| 5857 | TRAV8-2 | TRAJ48 | CVVSPFGNEKLTF | TRBV11-3 | TRBJ2-3 | CASSPRLAGGKGSTDTQYF | KLGGALQAK | YES | NO | NO |
| 5858 | TRAV8-4 | TRAJ20 | CAVIQINDYKLSF | TRBV11-3 | TRBJ2-3 | CASFGQHQDTQYF | KLGGALQAK | YES | NO | NO |
| 5861 | TRAV1-2 | TRAJ39 | CAVRDVAGNMLTF | TRBV12-3 | TRBJ2-3 | CASSFERGGPDTQYF | KLGGALQAK | YES | NO | NO |
| 5862 | TRAV10 | TRAJ43 | CVVTGYNNDMRF | TRBV12-3 | TRBJ2-3 | CASSSTDTQYF | KLGGALQAK | YES | NO | NO |
| 5863 | TRAV12-2 | TRAJ28 | CAVWGSGAGSYQLTF | TRBV12-3 | TRBJ2-3 | CASSWGTDTQYF | KLGGALQAK | YES | NO | NO |
| 5865 | TRAV13-2 | TRAJ11 | CAENGGYSTLTF | TRBV12-3 | TRBJ2-3 | CASSFGADTQYF | KLGGALQAK | YES | NO | NO |
| 5867 | TRAV19 | TRAJ11 | CALSNSGYSTLTF | TRBV12-3 | TRBJ2-3 | CASSGLATDTDTQYF | KLGGALQAK | YES | NO | NO |
| 5868 | TRAV20 | TRAJ53 | CAVQASGWDSGGSNYKLTF | TRBV12-3 | TRBJ2-3 | CASSFVRTVFYSTDTQYF | KLGGALQAK | YES | NO | NO |
| 5869 | TRAV21 | TRAJ40 | CAVRLTGTYKYIF | TRBV12-3 | TRBJ2-3 | CASSIEGGSTDTQYF | KLGGALQAK | YES | NO | NO |
| 5872 | TRAV29DV5 | TRAJ31 | CAASNNARLMF | TRBV12-3 | TRBJ2-3 | CASSQVGLSSTDTQYF | KLGGALQAK | YES | NO | NO |
| 5874 | TRAV38-2DV8 | TRAJ45 | CAYGGGADGLTF | TRBV12-3 | TRBJ2-3 | CASSLGQESSTDTQYF | KLGGALQAK | YES | NO | NO |
| 5877 | TRAV1-2 | TRAJ27 | CAVTTNAGKSTF | TRBV12-4 | TRBJ2-3 | CASSLLARGTDTQYF | KLGGALQAK | YES | NO | NO |
| 5880 | TRAV1-2 | TRAJ33 | CAVWVMDSNYQLIW | TRBV12-4 | TRBJ2-3 | CASSFFSQLTDTQYF | KLGGALQAK | YES | NO | NO |
| 5882 | TRAV10 | TRAJ6 | CVVRDVSGGSYIPTF | TRBV12-4 | TRBJ2-3 | CASSFWLADPSTDTQYF | KLGGALQAK | YES | NO | NO |
| 5883 | TRAV12-1 | TRAJ40 | CVVNKGTYKYIF | TRBV12-4 | TRBJ2-3 | CASSPMDTQYF | KLGGALQAK | YES | NO | NO |
| 5886 | TRAV12-2 | TRAJ30 | CAVYHDKIIF | TRBV12-4 | TRBJ2-3 | CASSPAGGGGTDTQYF | KLGGALQAK | YES | NO | NO |
| 5892 | TRAV13-1 | TRAJ47 | CAASYGNKLVF | TRBV12-4 | TRBJ2-3 | CASSPRAGGTDTQYF | KLGGALQAK | YES | NO | NO |
| 5898 | TRAV21 | TRAJ17 | CAVRPIKAAGNKLTF | TRBV12-4 | TRBJ2-3 | CASSFLVSNGADTQYF | KLGGALQAK | YES | NO | NO |
| 5900 | TRAV24 | TRAJ52 | CAFIGDAGGTSYGKLTF | TRBV12-4 | TRBJ2-3 | CASSPMDTQYF | KLGGALQAK | YES | NO | NO |
| 5902 | TRAV26-2 | TRAJ53 | CILRAPYVNSGGSNYKLTF | TRBV12-4 | TRBJ2-3 | CASSPMDTQYF | KLGGALQAK | YES | NO | NO |
| 5905 | TRAV29DV5 | TRAJ20 | CAASARNDYKLSF | TRBV12-4 | TRBJ2-3 | CASSSVLAGITDTQYF | KLGGALQAK | YES | NO | NO |
| 5907 | TRAV3 | TRAJ37 | CAVRDGGMGSNTGKLIF | TRBV12-4 | TRBJ2-3 | CASSPMDTQYF | RLRAEAQVK | NO | NO | NO |
| 5909 | TRAV38-1 | TRAJ54 | CAFMSLIQGAQKLVF | TRBV12-4 | TRBJ2-3 | CATKGEAGGTDTQYF | KLGGALQAK | YES | NO | NO |
| 5910 | TRAV38-1 | TRAJ54 | CAFMSLIQGAQKLVF | TRBV12-4 | TRBJ2-3 | CASSPMDTQYF | KLGGALQAK | YES | NO | NO |
| 5911 | TRAV41 | TRAJ48 | CAVSPDDFGNEKLTF | TRBV12-4 | TRBJ2-3 | CASTTQGPPVTPDTQYF | KLGGALQAK | YES | NO | NO |
| 5912 | TRAV5 | TRAJ24 | CAESGPDSWGKFQF | TRBV12-4 | TRBJ2-3 | CAVNGLAGPTDTQYF | KLGGALQAK | YES | NO | NO |
| 5918 | TRAV8-6 | TRAJ54 | CAVRPQEKLVF | TRBV12-4 | TRBJ2-3 | CASSPMDTQYF | KLGGALQAK | YES | NO | NO |

| | | | | | | | | | | |
|---|---|---|---|---|---|---|---|---|---|---|
| 5920 | TRAV9-2 | TRAJ45 | CALSEVYSGGGADGLTF | TRBV12-4 | TRBJ2-3 | CASSRGGVGDTQYF | KLGGALQAK | YES | NO | NO |
| 5922 | TRAV3 | TRAJ37 | CAVRDGGMGSNTGKLIF | TRBV12-5 | TRBJ2-3 | CASGLHVGTGGADTQYF | KLGGALQAK | NO | NO | NO |
| 5923 | TRAV1-2 | TRAJ28 | CAVRTAGSYQLTF | TRBV13 | TRBJ2-3 | CASSSPRGGGAGGTDTQYF | KLGGALQAK | YES | NO | NO |
| 5924 | TRAV13-1 | TRAJ45 | CAATPRGGADGLTF | TRBV13 | TRBJ2-3 | CASRGLAITDTQYF | KLGGALQAK | YES | NO | NO |
| 5927 | TRAV19 | TRAJ52 | CALSDLAGGTSYGKLTF | TRBV13 | TRBJ2-3 | CASSSAAGMSTDTQYF | KLGGALQAK | YES | NO | NO |
| 5929 | TRAV26-1 | TRAJ28 | CIVRVAYSGAGSYQLTF | TRBV13 | TRBJ2-3 | CASSFKPSGTDTQYF | KLGGALQAK | YES | NO | NO |
| 5932 | TRAV38-1 | TRAJ24 | CAFTDSWGKLQF | TRBV14 | TRBJ2-3 | CASSQESALASTDTQYF | KLGGALQAK | YES | NO | NO |
| 5933 | TRAV38-1 | TRAJ57 | CAFMKIQGGSEKLVF | TRBV14 | TRBJ2-3 | CASSLVERGWLTDTQYF | KLGGALQAK | YES | NO | NO |
| 5934 | TRAV38-2DV8 | TRAJ44 | CAYSPGTASKLTF | TRBV14 | TRBJ2-3 | CASSRHRGQDPTDTQYF | KLGGALQAK | YES | NO | NO |
| 5936 | TRAV8-2 | TRAJ43 | CVVSLRGNNDMRF | TRBV14 | TRBJ2-3 | CASSSMGGGTDTQYF | KLGGALQAK | YES | NO | NO |
| 5937 | TRAV8-4 | TRAJ32 | CAVRGYGGATNKLIF | TRBV14 | TRBJ2-3 | CASSQVNRGTDTQYF | KLGGALQAK | YES | NO | NO |
| 5938 | TRAV12-1 | TRAJ11 | CVVNGLGGYSTLTF | TRBV15 | TRBJ2-3 | CATSRVGADTQYF | KLGGALQAK | YES | NO | NO |
| 5939 | TRAV14DV4 | TRAJ22 | CAMSPGSARQLTF | TRBV15 | TRBJ2-3 | CATSRGLGAYTDTQYF | KLGGALQAK | YES | NO | NO |
| 5940 | TRAV14DV4 | TRAJ8 | CAMSAPLNTGFQKLVF | TRBV15 | TRBJ2-3 | CATSTGGAGGRTDTQYF | KLGGALQAK | YES | NO | NO |
| 5941 | TRAV19 | TRAJ52 | CALSEGEGGGTSYGKLTF | TRBV15 | TRBJ2-3 | CATSRGGGGDTQYF | KLGGALQAK | NO | NO | NO |
| 5942 | TRAV19 | TRAJ7 | CALSEEVDYGNNRLAF | TRBV15 | TRBJ2-3 | CATSLAESTDTQYF | KLGGALQAK | YES | NO | NO |
| 5943 | TRAV38-2DV8 | TRAJ49 | CAYSEWNTGNQFYF | TRBV15 | TRBJ2-3 | CATSRDAPPRVSTDTQYF | KLGGALQAK | YES | NO | NO |
| 5948 | TRAV1-1 | TRAJ33 | CAVSAMDSNYQLIW | TRBV18 | TRBJ2-3 | CASSPNRDINLDTQYF | KLGGALQAK | YES | NO | NO |
| 5949 | TRAV12-2 | TRAJ13 | CALSTGKVTF | TRBV18 | TRBJ2-3 | CASSPEDIGTDTQYF | KLGGALQAK | YES | NO | NO |
| 5950 | TRAV19 | TRAJ49 | CALSGDTGNQFYF | TRBV18 | TRBJ2-3 | CASSPSQGGNTDTQYF | KLGGALQAK | YES | NO | NO |
| 5951 | TRAV3 | TRAJ6 | CAVFLSGGSYIPTF | TRBV18 | TRBJ2-3 | CASSPTSTDTQYF | KLGGALQAK | NO | NO | NO |
| 5953 | TRAV1-1 | TRAJ30 | CAVRGGADDKIIF | TRBV19 | TRBJ2-3 | CASRGLAGGSTDTQYF | KLGGALQAK | YES | NO | NO |
| 5954 | TRAV1-2 | TRAJ33 | CASMDSNYQLIW | TRBV19 | TRBJ2-3 | CASSPGQGHTDTQYF | KLGGALQAK | YES | NO | NO |
| 5955 | TRAV12-1 | TRAJ12 | CVVNRDSSYKLIF | TRBV19 | TRBJ2-3 | CASSTAGVDTDTQYF | KLGGALQAK | YES | NO | NO |
| 5961 | TRAV14DV4 | TRAJ11 | CAMREDSGYSTLTF | TRBV19 | TRBJ2-3 | CASTTIWEGTLENTQYF | KLGGALQAK | YES | NO | NO |
| 5962 | TRAV14DV4 | TRAJ22 | CAMRRPISSGSARQLTF | TRBV19 | TRBJ2-3 | CASTRGGTQYF | KLGGALQAK | YES | NO | NO |
| 5963 | TRAV14DV4 | TRAJ30 | CAMREGWRDDKIIF | TRBV19 | TRBJ2-3 | CASSPGTGGTDTQYF | KLGGALQAK | YES | NO | NO |
| 5964 | TRAV14DV4 | TRAJ42 | CAMREAWGGGSQGNLIF | TRBV19 | TRBJ2-3 | CASSAGTGGTDTQYF | KLGGALQAK | YES | NO | NO |
| 5965 | TRAV14DV4 | TRAJ43 | CAMREGIHDMRF | TRBV19 | TRBJ2-3 | CASSIGLAGASTDTQYF | KLGGALQAK | YES | NO | NO |
| 5966 | TRAV14DV4 | TRAJ48 | CAMREGFGNEKLTF | TRBV19 | TRBJ2-3 | CASSWGAGGTDTQYF | KLGGALQAK | YES | YES | NO |
| 5967 | TRAV14DV4 | TRAJ52 | CAMRFAGGTSYGKLTF | TRBV19 | TRBJ2-3 | CASSSRSTDTQYF | KLGGALQAK | YES | NO | NO |
| 5968 | TRAV14DV4 | TRAJ9 | CAMREGQGGGFKTIF | TRBV19 | TRBJ2-3 | CARLPGQGVGDTQYF | KLGGALQAK | YES | NO | NO |
| 5972 | TRAV17 | TRAJ9 | CATVSSGGFKTIF | TRBV19 | TRBJ2-3 | CASSPWTGTDTQYF | KLGGALQAK | YES | NO | NO |
| 5973 | TRAV19 | TRAJ38 | CALSEAYAGNNRKLIW | TRBV19 | TRBJ2-3 | CASSIVWDRASLGTDTQYF | KLGGALQAK | YES | NO | NO |
| 5974 | TRAV19 | TRAJ39 | CALSEELENAGNMLTF | TRBV19 | TRBJ2-3 | CASSIDSPGLAGGPDTQYF | KLGGALQAK | YES | NO | NO |
| 5975 | TRAV19 | TRAJ47 | CALSDEYGNKLVF | TRBV19 | TRBJ2-3 | CASSASVAQYF | KLGGALQAK | YES | NO | NO |
| 5977 | TRAV2 | TRAJ21 | CASRGNFNKFYF | TRBV19 | TRBJ2-3 | CASSPGQGHTDTQYF | KLGGALQAK | NO | NO | NO |
| 5979 | TRAV2 | TRAJ31 | CAVDHPRARLMF | TRBV19 | TRBJ2-3 | CASSMTPLGEDTQYF | KLGGALQAK | NO | NO | NO |
| 5980 | TRAV20 | TRAJ35 | CAVQEGGFGNVLHC | TRBV19 | TRBJ2-3 | CASSIFHKGSTDTQYF | KLGGALQAK | NO | NO | NO |
| 5981 | TRAV20 | TRAJ4 | CSLFSGGYNKLIF | TRBV19 | TRBJ2-3 | CATEGDSADTQYF | KLGGALQAK | YES | NO | NO |
| 5982 | TRAV20 | TRAJ52 | CAVQYIAGGTSYGKLTF | TRBV19 | TRBJ2-3 | CASSLHPTSGTTDTQYF | KLGGALQAK | YES | NO | NO |
| 5983 | TRAV20 | TRAJ53 | CAVLGSGGSNYKLTF | TRBV19 | TRBJ2-3 | CASKSGRGLQDTQYF | KLGGALQAK | YES | NO | NO |
| 5984 | TRAV21 | TRAJ17 | CAVAVKAAGNKLTF | TRBV19 | TRBJ2-3 | CASSIGGTDTQYF | KLGGALQAK | YES | NO | NO |
| 5985 | TRAV21 | TRAJ23 | CAVIFYNQGGKLIF | TRBV19 | TRBJ2-3 | CASKVGASDTQYF | KLGGALQAK | YES | NO | NO |
| 5986 | TRAV21 | TRAJ57 | CAVPQGGSEKLVF | TRBV19 | TRBJ2-3 | CASRQGGTDTQYF | KLGGALQAK | YES | NO | NO |
| 5987 | TRAV26-1 | TRAJ52 | CIVRVGGTSYGKLTF | TRBV19 | TRBJ2-3 | CASSMSSWSADTQYF | KLGGALQAK | YES | NO | NO |
| 5988 | TRAV27 | TRAJ34 | CAGCPRYNTDKLIF | TRBV19 | TRBJ2-3 | CASSIVAILGTDTQYF | KLGGALQAK | YES | YES | NO |
| 5990 | TRAV29DV5 | TRAJ53 | CAADSGGSNYKLTF | TRBV19 | TRBJ2-3 | CASSIGDFYTDTQYF | KLGGALQAK | YES | NO | NO |
| 5991 | TRAV3 | TRAJ40 | CAVRDTEGTTSGTYKYIF | TRBV19 | TRBJ2-3 | CASRGVWGTDTQYF | KLGGALQAK | NO | NO | NO |
| 5992 | TRAV30 | TRAJ53 | CGTESSGGSNYKLTF | TRBV19 | TRBJ2-3 | CASSPLALQGPRDTQYF | KLGGALQAK | YES | NO | NO |
| 5993 | TRAV35 | TRAJ42 | CAGLPATYGGSQGNLIF | TRBV19 | TRBJ2-3 | CASSIMHPDTQYF | KLGGALQAK | YES | NO | NO |
| 5994 | TRAV38-2DV8 | TRAJ18 | CAYRSATDRGSTLGRLYF | TRBV19 | TRBJ2-3 | CASSWTGAFTDTQYF | RLRAEAQVK | YES | YES | YES |
| 5995 | TRAV38-2DV8 | TRAJ48 | CALRTLGNEKLTF | TRBV19 | TRBJ2-3 | CASSIWTDDTDTQYF | KLGGALQAK | YES | NO | NO |
| 5996 | TRAV4 | TRAJ43 | CLVGGLPLGNNDMRF | TRBV19 | TRBJ2-3 | CATGGPWADTQYF | KLGGALQAK | YES | NO | NO |
| 5997 | TRAV40 | TRAJ53 | CLLATGGGSNYKLTF | TRBV19 | TRBJ2-3 | CASGTYSGDTQYF | KLGGALQAK | NO | NO | NO |
| 5998 | TRAV8-2 | TRAJ12 | CVVSRMDSSYKLIF | TRBV19 | TRBJ2-3 | CASSMDRGSADTQYF | AVFDRKSDAK | YES | NO | NO |
| 5999 | TRAV8-2 | TRAJ8 | CVVTGFQKLVF | TRBV19 | TRBJ2-3 | CASSIDPSGPPSTDTQYF | KLGGALQAK | YES | NO | NO |
| 6002 | TRAV8-6 | TRAJ26 | CAVSDRDYGQNFVF | TRBV19 | TRBJ2-3 | CASSTTSGGLDTQYF | KLGGALQAK | YES | NO | NO |
| 6007 | TRAV12-2 | TRAJ29 | CAAPSGNTPLVF | TRBV2 | TRBJ2-3 | CASTAEGFHTDTQYF | KLGGALQAK | YES | NO | NO |
| 6008 | TRAV13-1 | TRAJ39 | CAANNAGNMLTF | TRBV2 | TRBJ2-3 | CASSVLYETDTQYF | KLGGALQAK | YES | NO | NO |
| 6009 | TRAV13-2 | TRAJ39 | CAENRGNNNAGNMLTF | TRBV2 | TRBJ2-3 | CASSDIRGGGTHLDTQYF | KLGGALQAK | YES | NO | NO |
| 6010 | TRAV17 | TRAJ34 | CAPEPYNTDKLIF | TRBV2 | TRBJ2-3 | CASSRGAGADTQYF | KLGGALQAK | YES | NO | NO |
| 6013 | TRAV19 | TRAJ57 | CALSGQEGGSEKLVF | TRBV2 | TRBJ2-3 | CASSRPGLAGDFGTDTQYF | KLGGALQAK | YES | NO | NO |
| 6016 | TRAV26-2 | TRAJ45 | CILRDRGGGADGLTF | TRBV2 | TRBJ2-3 | CASSVLIAGSGTDTQYF | KLGGALQAK | YES | NO | NO |
| 6020 | TRAV38-1 | TRAJ57 | CAEGQGGSEKLVF | TRBV2 | TRBJ2-3 | CASSDGHPVKGDTQYF | KLGGALQAK | YES | NO | NO |
| 6022 | TRAV4 | TRAJ44 | CLVGDTGTASKLTF | TRBV2 | TRBJ2-3 | CASSSSYLAGGRDTQYF | KLGGALQAK | YES | NO | NO |
| 6023 | TRAV5 | TRAJ15 | CAESVDNQAGTALIF | TRBV2 | TRBJ2-3 | CASSGLGYRGARADTQYF | KLGGALQAK | YES | NO | NO |
| 6024 | TRAV5 | TRAJ26 | CAESRGLDNYGQNFVF | TRBV2 | TRBJ2-3 | CASSVTSGGYSTDTQYF | KLGGALQAK | YES | NO | NO |
| 6025 | TRAV8-4 | TRAJ23 | CAVSGGNQGGKLIF | TRBV2 | TRBJ2-3 | CASSEQGLDTQYF | KLGGALQAK | YES | NO | NO |
| 6026 | TRAV9-2 | TRAJ40 | CALGGFGTYKYIF | TRBV2 | TRBJ2-3 | CASAPSGGATDTQYF | KLGGALQAK | YES | NO | NO |
| 6028 | TRAV1-1 | TRAJ13 | CAVRSYSGGYQKVTF | TRBV20-1 | TRBJ2-3 | CSAEGRGDRTDTQYF | KLGGALQAK | NO | NO | NO |
| 6029 | TRAV1-1 | TRAJ20 | CAVNIGNDYKLSF | TRBV20-1 | TRBJ2-3 | CSARDTSESTDTQYF | KLGGALQAK | YES | NO | NO |
| 6031 | TRAV1-2 | TRAJ16 | CAVRDSRDGQKLLF | TRBV20-1 | TRBJ2-3 | CSARDQDQRGSSTDTQYF | KLGGALQAK | NO | NO | NO |
| 6033 | TRAV1-2 | TRAJ39 | CAVRDYNNAGNMLTF | TRBV20-1 | TRBJ2-3 | CSASDVAGLYTDTQYF | KLGGALQAK | NO | NO | NO |
| 6035 | TRAV12-1 | TRAJ41 | CVVNPRNSGYALNF | TRBV20-1 | TRBJ2-3 | CSARVRGYRTDTQYF | KLGGALQAK | NO | NO | NO |
| 6036 | TRAV12-2 | TRAJ32 | CAVRGGATNKLIF | TRBV20-1 | TRBJ2-3 | CSARVRTGRGPQSSNTQYF | KLGGALQAK | NO | NO | NO |
| 6037 | TRAV12-2 | TRAJ4 | CAVKRGSGGYNKLIF | TRBV20-1 | TRBJ2-3 | CSADRDRGTDTQYF | KLGGALQAK | NO | NO | NO |
| 6040 | TRAV14DV4 | TRAJ53 | CAMREAPKLGGGSNYKLTF | TRBV20-1 | TRBJ2-3 | CSARDLASGSTDTQYF | KLGGALQAK | NO | NO | NO |
| 6042 | TRAV14DV4 | TRAJ9 | CAMRVNTGGFKTIF | TRBV20-1 | TRBJ2-3 | CSARGIFDNSDTQYF | AVFDRKSDAK | NO | NO | NO |
| 6042 | TRAV14DV4 | TRAJ9 | CAMRVNTGGFKTIF | TRBV20-1 | TRBJ2-3 | CSARGIFDNSDTQYF | KLGGALQAK | NO | NO | NO |
| 6043 | TRAV17 | TRAJ9 | CATGQTGGFKTIF | TRBV20-1 | TRBJ2-3 | CSARLSTLPADTQYF | KLGGALQAK | NO | NO | NO |
| 6045 | TRAV19 | TRAJ28 | CALTDYSGAGSYQLTF | TRBV20-1 | TRBJ2-3 | CSALPPLAGGHTDTQYF | AVFDRKSDAK | NO | NO | NO |
| 6046 | TRAV19 | TRAJ32 | CALSERDYGGATNKLIF | TRBV20-1 | TRBJ2-3 | CSARDLRGYTDTQYF | KLGGALQAK | NO | NO | NO |
| 6047 | TRAV19 | TRAJ40 | CALSEASGTYKYIF | TRBV20-1 | TRBJ2-3 | CSAPSGEGRDTQYF | KLGGALQAK | NO | NO | NO |
| 6048 | TRAV19 | TRAJ45 | CALSVFSGGGADGLTF | TRBV20-1 | TRBJ2-3 | CSARWGGGTDTQYF | KLGGALQAK | NO | NO | NO |
| 6053 | TRAV23DV6 | TRAJ34 | CAAKLRNTDKLIF | TRBV20-1 | TRBJ2-3 | CSASGYGTLKVTDTQYF | KLGGALQAK | NO | NO | NO |
| 6054 | TRAV25 | TRAJ20 | CASNDYKLSF | TRBV20-1 | TRBJ2-3 | CSARDLSSTDTQYF | KLGGALQAK | NO | NO | NO |
| 6055 | TRAV26-1 | TRAJ29 | CIVRVGLSGNTPLVF | TRBV20-1 | TRBJ2-3 | CSARETNTDTQYF | KLGGALQAK | NO | NO | NO |
| 6056 | TRAV26-1 | TRAJ53 | CIVRDPGGSNYKLTF | TRBV20-1 | TRBJ2-3 | CSASELPLGGGYTDTQYF | KLGGALQAK | NO | NO | NO |
| 6058 | TRAV27 | TRAJ11 | CAGALESGYSTLTF | TRBV20-1 | TRBJ2-3 | CSARELGADTQYF | KLGGALQAK | NO | NO | NO |
| 6059 | TRAV29DV5 | TRAJ36 | CAASSSTGANNLFF | TRBV20-1 | TRBJ2-3 | CSATSNGERTDTQYF | AVFDRKSDAK | NO | NO | NO |
| 6059 | TRAV29DV5 | TRAJ36 | CAASSSTGANNLFF | TRBV20-1 | TRBJ2-3 | CSATSNGERTDTQYF | RLRAEAQVK | NO | NO | NO |
| 6060 | TRAV3 | TRAJ39 | CAVLPGAGNMLTF | TRBV20-1 | TRBJ2-3 | CSASPSGVRFTDTQYF | KLGGALQAK | NO | NO | NO |
| 6062 | TRAV38-2DV8 | TRAJ23 | CASNQGGKLIF | TRBV20-1 | TRBJ2-3 | CSARDGASSDTQYF | KLGGALQAK | NO | NO | NO |
| 6063 | TRAV38-2DV8 | TRAJ42 | CAYVGGSQGNLIF | TRBV20-1 | TRBJ2-3 | CSARDLGLAGGLTVDTQYF | KLGGALQAK | NO | NO | NO |
| 6068 | TRAV41 | TRAJ43 | CAVRWRINDMRF | TRBV20-1 | TRBJ2-3 | CSASGTSSLVDTQYF | KLGGALQAK | NO | NO | NO |
| 6069 | TRAV5 | TRAJ10 | CAEIRDILTGGGNKLTF | TRBV20-1 | TRBJ2-3 | CSAMLGTSGSPPSTDTQYF | KLGGALQAK | NO | NO | NO |
| 6070 | TRAV5 | TRAJ42 | CAENWGGGSQGNLIF | TRBV20-1 | TRBJ2-3 | CSARDTWRRDPGTDTQYF | KLGGALQAK | NO | NO | NO |
| 6071 | TRAV5 | TRAJ43 | CAEYRDNDMRF | TRBV20-1 | TRBJ2-3 | CSAREDPQDTQYF | KLGGALQAK | NO | NO | NO |
| 6072 | TRAV6 | TRAJ5 | CALPRGRALTF | TRBV20-1 | TRBJ2-3 | CSARIADSTDTQYF | KLGGALQAK | NO | NO | NO |
| 6075 | TRAV8-2 | TRAJ9 | CALNTGGFKTIF | TRBV20-1 | TRBJ2-3 | CSAPGQGTDTQYF | KLGGALQAK | NO | NO | NO |
| 6076 | TRAV8-3 | TRAJ21 | CAVGGSYNFNKFYF | TRBV20-1 | TRBJ2-3 | CSASTTRTGGHWNTDTQYF | KLGGALQAK | NO | NO | NO |
| 6079 | TRAV9-2 | TRAJ26 | CALYYGQNFVF | TRBV20-1 | TRBJ2-3 | CSARNPGLASWSEDTQYF | KLGGALQAK | YES | NO | NO |
| 6082 | TRAV12-3 | TRAJ39 | CAIGPGNMLTF | TRBV21-1 | TRBJ2-3 | CQQQRNPGTNKRSTDTQYF | KLGGALQAK | YES | NO | NO |

| | | | | | | | | | | |
|---|---|---|---|---|---|---|---|---|---|---|
| 6086 | TRAV1-1 | TRAJ16 | CERSCCSDGQKLLF | TRBV24-1 | TRBJ2-3 | CATSGQGATDTQYF | KLGGALQAK | NO | NO | NO |
| 6090 | TRAV12-1 | TRAJ43 | CVVNIYNNNDMRF | TRBV24-1 | TRBJ2-3 | CATRTGSEDTQYF | KLGGALQAK | YES | NO | NO |
| 6091 | TRAV12-2 | TRAJ4 | CAVKPMFSGGYNKLIF | TRBV24-1 | TRBJ2-3 | CATSGTEGTDTQYF | KLGGALQAK | YES | NO | NO |
| 6091 | TRAV12-2 | TRAJ4 | CAVKPMFSGGYNKLIF | TRBV24-1 | TRBJ2-3 | CATSGTEGTDTQYF | RLRAEAQVK | YES | NO | NO |
| 6094 | TRAV14DV4 | TRAJ4 | CAMRGPISGGYNKLIF | TRBV24-1 | TRBJ2-3 | CATSRASTDTQYF | KLGGALQAK | YES | NO | NO |
| 6096 | TRAV19 | TRAJ58 | CALSEIPASGSRLTF | TRBV24-1 | TRBJ2-3 | CATSDRVLAGGADTQYF | KLGGALQAK | YES | NO | NO |
| 6097 | TRAV20 | TRAJ18 | CAVQADRGSTLGRLYF | TRBV24-1 | TRBJ2-3 | CATSDSWGGDTQYF | KLGGALQAK | YES | NO | NO |
| 6099 | TRAV25 | TRAJ35 | CAGSI | TRBV24-1 | TRBJ2-3 | CATRDSYTDTQYF | KLGGALQAK | NO | NO | NO |
| 6102 | TRAV3 | TRAJ39 | CAVRANDNAGNMLTF | TRBV24-1 | TRBJ2-3 | CATSDLSDTQYF | KLGGALQAK | NO | NO | NO |
| 6103 | TRAV38-2DV8 | TRAJ42 | CALRSQGNLIF | TRBV24-1 | TRBJ2-3 | CATKTGETDTQYF | KLGGALQAK | YES | NO | NO |
| 6104 | TRAV8-1 | TRAJ37 | CAVREVDRTGKLIF | TRBV24-1 | TRBJ2-3 | CATSRASTDTQYF | KLGGALQAK | YES | NO | NO |
| 6109 | TRAV19 | TRAJ28 | CALSGAGSYQLTF | TRBV25-1 | TRBJ2-3 | CASSEYSLTDTQYF | KLGGALQAK | YES | NO | NO |
| 6112 | TRAV1-1 | TRAJ24 | CAPKTDSWGKLQF | TRBV27 | TRBJ2-3 | CATMEPGQGPDTQYF | AVFDRKSDAK | YES | NO | NO |
| 6112 | TRAV1-1 | TRAJ24 | CAPKTDSWGKLQF | TRBV27 | TRBJ2-3 | CATMEPGQGPDTQYF | RLRAEAQVK | YES | NO | NO |
| 6114 | TRAV10 | TRAJ36 | CVVSDGFSGANNLFF | TRBV27 | TRBJ2-3 | CASSGLLSGTDTQYF | KLGGALQAK | YES | NO | NO |
| 6115 | TRAV12-1 | TRAJ26 | CVVTLGQNFVF | TRBV27 | TRBJ2-3 | CASSLGQGAIDTQYF | KLGGALQAK | YES | NO | NO |
| 6116 | TRAV12-1 | TRAJ49 | CVVNIEGTGNQFYF | TRBV27 | TRBJ2-3 | CASKPSNSRTDTQYF | KLGGALQAK | YES | NO | NO |
| 6118 | TRAV12-2 | TRAJ43 | CAVRVGDMRF | TRBV27 | TRBJ2-3 | CASSRAGSGEDTQYF | KLGGALQAK | YES | NO | NO |
| 6119 | TRAV12-2 | TRAJ49 | CAVNSNTGNQFYF | TRBV27 | TRBJ2-3 | CASSFTGSFTDTQYF | KLGGALQAK | YES | NO | NO |
| 6122 | TRAV13-1 | TRAJ45 | CAAYYSGGGADGLTF | TRBV27 | TRBJ2-3 | CASSFLAGGYGDTQYF | RLRAEAQVK | YES | NO | NO |
| 6122 | TRAV13-1 | TRAJ45 | CAAYYSGGGADGLTF | TRBV27 | TRBJ2-3 | CASSFLAGGYGDTQYF | KLGGALQAK | YES | NO | NO |
| 6122 | TRAV13-1 | TRAJ45 | CAAYYSGGGADGLTF | TRBV27 | TRBJ2-3 | CASSFLAGGYGDTQYF | AVFDRKSDAK | YES | NO | NO |
| 6129 | TRAV19 | TRAJ36 | CALSLGRGTGANNLFF | TRBV27 | TRBJ2-3 | CASARGTDTQYF | KLGGALQAK | YES | NO | NO |
| 6130 | TRAV19 | TRAJ53 | CALSGRYYGGSNYKLTF | TRBV27 | TRBJ2-3 | CATVRPSTDTQYF | KLGGALQAK | YES | NO | NO |
| 6131 | TRAV20 | TRAJ24 | CAVQPSWGKFQF | TRBV27 | TRBJ2-3 | CASTPGGGTDTQYF | KLGGALQAK | YES | NO | NO |
| 6132 | TRAV20 | TRAJ40 | CAVPTSGTYKYIF | TRBV27 | TRBJ2-3 | CATIPTGSTDTQYF | KLGGALQAK | YES | NO | NO |
| 6132 | TRAV20 | TRAJ40 | CAVPTSGTYKYIF | TRBV27 | TRBJ2-3 | CATIPTGSTDTQYF | AVFDRKSDAK | YES | NO | NO |
| 6135 | TRAV21 | TRAJ26 | CAVYNYGQNFVF | TRBV27 | TRBJ2-3 | CASSVSSTDTQYF | KLGGALQAK | YES | NO | NO |
| 6137 | TRAV21 | TRAJ4 | CAVPFPTGGYNKLIF | TRBV27 | TRBJ2-3 | CASSLMGITDTQYF | KLGGALQAK | YES | NO | NO |
| 6138 | TRAV21 | TRAJ56 | CAVKTGANSKLTF | TRBV27 | TRBJ2-3 | CASSSSAGLAYTDTQYF | KLGGALQAK | YES | NO | NO |
| 6139 | TRAV25 | TRAJ42 | CAGRGSQGNLIF | TRBV27 | TRBJ2-3 | CASSLYQGSSTDTQYF | KLGGALQAK | YES | NO | NO |
| 6140 | TRAV26-1 | TRAJ13 | CIVRPAGYQKVTF | TRBV27 | TRBJ2-3 | CASSSTSGSLLLTDTQYF | KLGGALQAK | YES | NO | NO |
| 6142 | TRAV26-2 | TRAJ12 | CIQDADSSYKLIF | TRBV27 | TRBJ2-3 | CASSHPTFIHTQYF | KLGGALQAK | YES | NO | NO |
| 6143 | TRAV26-2 | TRAJ30 | CILRDVIRDDKIIF | TRBV27 | TRBJ2-3 | CASSSWLGAPDTQYF | KLGGALQAK | YES | NO | NO |
| 6145 | TRAV27 | TRAJ52 | CAGVSWNAGGTSYGKLTF | TRBV27 | TRBJ2-3 | CASSLIAGGGDTQYF | KLGGALQAK | YES | NO | NO |
| 6146 | TRAV29DV5 | TRAJ53 | CAASGRGGSNYKLTF | TRBV27 | TRBJ2-3 | CASSLSLDIDTQYF | KLGGALQAK | YES | YES | NO |
| 6147 | TRAV3 | TRAJ20 | CAVRDTSNDYKLSF | TRBV27 | TRBJ2-3 | CASSFGEGLDTQYF | KLGGALQAK | NO | NO | NO |
| 6148 | TRAV38-2DV8 | TRAJ40 | CAYRTTSGTYKYIF | TRBV27 | TRBJ2-3 | CASRSWDPDTQYF | KLGGALQAK | YES | NO | NO |
| 6149 | TRAV38-2DV8 | TRAJ45 | CAYRSPNPYSGGGADGLTF | TRBV27 | TRBJ2-3 | CASSSGTRRTDTQYF | KLGGALQAK | YES | NO | NO |
| 6153 | TRAV8-3 | TRAJ33 | CAVASINYQLIW | TRBV27 | TRBJ2-3 | CASSPFWGGGHTDTQYF | KLGGALQAK | YES | NO | NO |
| 6155 | TRAV1-2 | TRAJ16 | CAVGHNGQKLLF | TRBV28 | TRBJ2-3 | CASSLRNLAELGDTQYF | KLGGALQAK | NO | NO | NO |
| 6156 | TRAV1-2 | TRAJ36 | CAVRPGANNLFF | TRBV28 | TRBJ2-3 | CASSPYSGSLTDTQYF | KLGGALQAK | YES | NO | NO |
| 6157 | TRAV12-2 | TRAJ53 | CAVLSASVSNYKLTF | TRBV28 | TRBJ2-3 | CASSLARGLSPDTQYF | KLGGALQAK | YES | NO | NO |
| 6159 | TRAV13-1 | TRAJ11 | CAARGGYSTLTF | TRBV28 | TRBJ2-3 | CASSLRTSGSFTDTQYF | RLRAEAQVK | YES | NO | NO |
| 6160 | TRAV14DV4 | TRAJ56 | CAMREEFGGTGANSKLTF | TRBV28 | TRBJ2-3 | CASSLGLGGDTQYF | KLGGALQAK | YES | NO | NO |
| 6161 | TRAV27 | TRAJ49 | CAGIDTGNQFYF | TRBV28 | TRBJ2-3 | CASKVVDTQYF | KLGGALQAK | YES | NO | NO |
| 6162 | TRAV38-2DV8 | TRAJ27 | CAYRSSPWGTNAGKSTF | TRBV28 | TRBJ2-3 | CASSFTYRDTQYF | KLGGALQAK | YES | NO | NO |
| 6163 | TRAV38-2DV8 | TRAJ49 | CAYRSSEGTGNQFYF | TRBV28 | TRBJ2-3 | CASSFGGWTDTQYF | KLGGALQAK | YES | NO | NO |
| 6168 | TRAV12-1 | TRAJ9 | CVVMGGGFKTIF | TRBV29-1 | TRBJ2-3 | CSVRWESTDTQYF | KLGGALQAK | YES | NO | NO |
| 6169 | TRAV14DV4 | TRAJ8 | CAMREGWNTGFQKLVF | TRBV29-1 | TRBJ2-3 | CSVETGGQGAWDTQYF | KLGGALQAK | YES | NO | NO |
| 6172 | TRAV17 | TRAJ9 | CATDGGTGGFKTIF | TRBV29-1 | TRBJ2-3 | CSVVGLAGGLIYTDTQYF | KLGGALQAK | YES | NO | NO |
| 6173 | TRAV19 | TRAJ34 | CALSELGYNTDKLIF | TRBV29-1 | TRBJ2-3 | CSVTFSSTDTQYF | KLGGALQAK | YES | NO | NO |
| 6174 | TRAV19 | TRAJ42 | CALSEARGSQGNLIF | TRBV29-1 | TRBJ2-3 | CSGNPGTSAATQYF | KLGGALQAK | YES | NO | NO |
| 6175 | TRAV19 | TRAJ58 | CALSEAVKTSGSRLTF | TRBV29-1 | TRBJ2-3 | CSVEYGLAGSTDTQYF | KLGGALQAK | YES | NO | NO |
| 6178 | TRAV21 | TRAJ22 | CAFSGSARQLTF | TRBV29-1 | TRBJ2-3 | CSVEDWTSGNTDTQYF | KLGGALQAK | YES | NO | NO |
| 6179 | TRAV26-1 | TRAJ36 | CIVPPQTGANNLFF | TRBV29-1 | TRBJ2-3 | CSVILTDSTDTQYF | KLGGALQAK | YES | NO | NO |
| 6180 | TRAV26-2 | TRAJ48 | CILRGEFGNEKLTF | TRBV29-1 | TRBJ2-3 | CSVEDSSTDTQYF | KLGGALQAK | YES | NO | NO |
| 6182 | TRAV41 | TRAJ48 | CAVRLHTRNFGNEKLTF | TRBV29-1 | TRBJ2-3 | CSVAGGGMDTQYF | KLGGALQAK | YES | NO | NO |
| 6184 | TRAV1-2 | TRAJ26 | CARSRYGQNFVF | TRBV3-1 | TRBJ2-3 | CASSPRDNTDTQYF | KLGGALQAK | YES | NO | NO |
| 6185 | TRAV1-2 | TRAJ31 | CAVRDARLMF | TRBV3-1 | TRBJ2-3 | CASSQEYRGSDTQYF | KLGGALQAK | YES | NO | NO |
| 6186 | TRAV12-3 | TRAJ45 | CAMSATRGGGADGLTF | TRBV3-1 | TRBJ2-3 | CASSLRRLTRFAGGSTDTQYF | KLGGALQAK | NO | NO | NO |
| 6187 | TRAV13-2 | TRAJ53 | CAENAGSNYKLTF | TRBV3-1 | TRBJ2-3 | CASSQWLRGLTDTQYF | KLGGALQAK | YES | NO | NO |
| 6188 | TRAV16 | TRAJ16 | CALPTPTSDGQKLLF | TRBV3-1 | TRBJ2-3 | CASSQDRVLGDTQYF | KLGGALQAK | NO | NO | NO |
| 6190 | TRAV23DV6 | TRAJ49 | CAARSAGNQFYF | TRBV3-1 | TRBJ2-3 | CASSQEYRGSDTQYF | KLGGALQAK | YES | NO | NO |
| 6192 | TRAV38-2DV8 | TRAJ17 | CAYRSYLMIKAAGNKLTF | TRBV3-1 | TRBJ2-3 | CASSLEQGARTDTQYF | KLGGALQAK | YES | NO | NO |
| 6195 | TRAV5 | TRAJ34 | CAERSLNNTDKLIF | TRBV3-1 | TRBJ2-3 | CASSPRPSGGHSTDTQYF | KLGGALQAK | YES | NO | NO |
| 6197 | TRAV8-4 | TRAJ42 | CAVSRDMNYGGSQGNLIF | TRBV3-1 | TRBJ2-3 | CASSQDLRGADTQYF | KLGGALQAK | YES | NO | NO |
| 6199 | TRAV14DV4 | TRAJ42 | CAMRNYGGSQGNLIF | TRBV30 | TRBJ2-3 | CAWSEGVAGFADTQYF | KLGGALQAK | YES | NO | NO |
| 6200 | TRAV17 | TRAJ30 | CATDRDDKIIF | TRBV30 | TRBJ2-3 | CAWSVQLSTDTQYF | KLGGALQAK | YES | NO | NO |
| 6201 | TRAV19 | TRAJ17 | CALSVIKAAGNKLTF | TRBV30 | TRBJ2-3 | CAWKNLFPRTDTQYF | KLGGALQAK | YES | NO | NO |
| 6201 | TRAV19 | TRAJ17 | CALSVIKAAGNKLTF | TRBV30 | TRBJ2-3 | CAWKNLFPRTDTQYF | AVFDRKSDAK | YES | NO | NO |
| 6202 | TRAV21 | TRAJ43 | CAVDNNNDMRF | TRBV30 | TRBJ2-3 | CAWKNLFPRTDTQYF | KLGGALQAK | NO | NO | NO |
| 6204 | TRAV38-2DV8 | TRAJ32 | CAYYGGATNKLIF | TRBV30 | TRBJ2-3 | CAWNWAGGSDTQYF | KLGGALQAK | YES | NO | NO |
| 6205 | TRAV8-1 | TRAJ36 | CAVNSGANNLFF | TRBV30 | TRBJ2-3 | CAWLSDTQYF | KLGGALQAK | YES | NO | NO |
| 6206 | TRAV1-2 | TRAJ9 | CAVISTGGFKTIF | TRBV4-1 | TRBJ2-3 | CASSQAAGGPPDTQYF | KLGGALQAK | YES | NO | NO |
| 6207 | TRAV12-1 | TRAJ3 | CVVNRYSSASKIIF | TRBV4-1 | TRBJ2-3 | CASSQQSGDTDTQYF | KLGGALQAK | YES | NO | NO |
| 6208 | TRAV12-1 | TRAJ37 | CVVNSFSSNTGKLIF | TRBV4-1 | TRBJ2-3 | CASSQYPGETDTQYF | KLGGALQAK | YES | NO | NO |
| 6208 | TRAV12-1 | TRAJ37 | CVVNSFSSNTGKLIF | TRBV4-1 | TRBJ2-3 | CASSQYPGETDTQYF | AVFDRKSDAK | YES | NO | NO |
| 6210 | TRAV13-2 | TRAJ44 | CAENTGTASKLTF | TRBV4-1 | TRBJ2-3 | CASSRLAESTDTQYF | KLGGALQAK | YES | NO | NO |
| 6213 | TRAV14DV4 | TRAJ37 | CAMRERRTGSGNTGKLIF | TRBV4-1 | TRBJ2-3 | CASSQDALAGTTDTQYF | KLGGALQAK | YES | NO | NO |
| 6214 | TRAV16 | TRAJ8 | CALGGLNTGFQKLVF | TRBV4-1 | TRBJ2-3 | CASSRVDTQYF | KLGGALQAK | NO | NO | NO |
| 6215 | TRAV17 | TRAJ49 | CATDDTGNQFYF | TRBV4-1 | TRBJ2-3 | CASTGQGATDTQYF | KLGGALQAK | NO | NO | NO |
| 6217 | TRAV17 | TRAJ9 | CATDEGGGFKTIF | TRBV4-1 | TRBJ2-3 | CASSQDMGSSKTQYF | KLGGALQAK | NO | NO | NO |
| 6218 | TRAV19 | TRAJ52 | CALRGGTSYGKLTF | TRBV4-1 | TRBJ2-3 | CASSSSGGSDTQYF | KLGGALQAK | YES | NO | NO |
| 6219 | TRAV21 | TRAJ33 | CAVDGDSNYQLIW | TRBV4-1 | TRBJ2-3 | CASSQYPGETDTQYF | KLGGALQAK | YES | NO | NO |
| 6220 | TRAV22 | TRAJ33 | CAVERDSNYQLIW | TRBV4-1 | TRBJ2-3 | CASSLYTSTDTQYF | KLGGALQAK | YES | NO | NO |
| 6221 | TRAV23DV6 | TRAJ4 | CAGGFGGGYNKLIF | TRBV4-1 | TRBJ2-3 | CASSQDDGLAGYSTDTQYF | KLGGALQAK | YES | NO | NO |
| 6222 | TRAV26-2 | TRAJ34 | CILRLSYNTDKLIF | TRBV4-1 | TRBJ2-3 | CASSQDIVASSGTDTQYF | KLGGALQAK | YES | NO | NO |
| 6223 | TRAV26-2 | TRAJ42 | CILINYGGSQGNLIF | TRBV4-1 | TRBJ2-3 | CASKGTVAAITDTQYF | KLGGALQAK | YES | NO | NO |
| 6226 | TRAV4 | TRAJ45 | CLVGDLTGGGADGLTF | TRBV4-1 | TRBJ2-3 | CASSQDDRGTDTQYF | KLGGALQAK | YES | NO | NO |
| 6228 | TRAV8-1 | TRAJ37 | CAVNAEGSGNTGKLIF | TRBV4-1 | TRBJ2-3 | CASSQVDRVTDTQYF | KLGGALQAK | YES | YES | NO |
| 6231 | TRAV8-4 | TRAJ48 | CAVSPGSFGNEKLTF | TRBV4-1 | TRBJ2-3 | CASSQGLASGGASTDTQYF | KLGGALQAK | YES | NO | NO |
| 6232 | TRAV8-6 | TRAJ52 | CAVTLTNAGGTSYGKLTF | TRBV4-1 | TRBJ2-3 | CASSQFTRLASTDTQYF | KLGGALQAK | YES | NO | NO |
| 6234 | TRAV12-2 | TRAJ4 | CAVNENTGGYNKLIF | TRBV4-2 | TRBJ2-3 | CASRALAGGVTDTQYF | KLGGALQAK | YES | NO | NO |
| 6236 | TRAV19 | TRAJ48 | CALSPTNFGNEKLTF | TRBV4-2 | TRBJ2-3 | CASSLPPDGTDTQYF | KLGGALQAK | YES | NO | NO |
| 6237 | TRAV29DV5 | TRAJ44 | CAAKTGTASKLTF | TRBV4-2 | TRBJ2-3 | CASSMTVSYTDTQYF | KLGGALQAK | YES | NO | NO |
| 6239 | TRAV38-2DV8 | TRAJ29 | CAYRSAWINTPLVF | TRBV4-2 | TRBJ2-3 | CASSRLALLTDTQYF | KLGGALQAK | YES | NO | NO |
| 6240 | TRAV4 | TRAJ20 | CLVGDDYKLSF | TRBV4-2 | TRBJ2-3 | CASSPDGTGPGDTQYF | KLGGALQAK | YES | NO | NO |
| 6241 | TRAV8-3 | TRAJ28 | CAVGASYSGAGSYQLTF | TRBV4-2 | TRBJ2-3 | CASSQDRGGGTDTQYF | KLGGALQAK | YES | NO | NO |
| 6242 | TRAV1-2 | TRAJ20 | CAVRDLSNDYKLSF | TRBV4-3 | TRBJ2-3 | CASSQGTGTDTQYF | KLGGALQAK | YES | NO | NO |
| 6243 | TRAV1-2 | TRAJ33 | CAVLDSNYQLIW | TRBV4-3 | TRBJ2-3 | CASSQEFRGQGADTQYF | KLGGALQAK | YES | NO | NO |
| 6248 | TRAV12-3 | TRAJ34 | CAMSRLEGDKLIF | TRBV4-3 | TRBJ2-3 | CASSQEGGFTDTQYF | RLRAEAQVK | YES | NO | NO |
| 6248 | TRAV12-3 | TRAJ34 | CAMSRLEGDKLIF | TRBV4-3 | TRBJ2-3 | CASSQEGGFTDTQYF | IVTDFSVIK | YES | NO | NO |
| 6251 | TRAV17 | TRAJ37 | CATDSNTGKLIF | TRBV4-3 | TRBJ2-3 | CASSQGQLSTDTQYF | KLGGALQAK | YES | NO | NO |

| | | | | | | | | | | |
|---|---|---|---|---|---|---|---|---|---|---|
| 6252 | TRAV19 | TRAJ16 | CALSARF | TRBV4-3 | TRBJ2-3 | CASSQDVQGTSTDTQYF | KLGGALQAK | NO | NO | NO |
| 6255 | TRAV21 | TRAJ24 | CAVSLSWGKFQF | TRBV4-3 | TRBJ2-3 | CASSQEGGFTDTQYF | KLGGALQAK | YES | NO | NO |
| 6257 | TRAV22 | TRAJ26 | CAVPDNYGQNFVF | TRBV4-3 | TRBJ2-3 | CASSQGDRAFTDTQYF | KLGGALQAK | YES | NO | NO |
| 6265 | TRAV9-2 | TRAJ33 | CALMDSNYQLIW | TRBV4-3 | TRBJ2-3 | CASSQVEGTDTQYF | KLGGALQAK | YES | NO | NO |
| 6267 | TRAV12-1 | TRAJ49 | CVVNIAGNQFYF | TRBV5-1 | TRBJ2-3 | CASSLLAGGSTDTQYF | KLGGALQAK | YES | NO | NO |
| 6269 | TRAV12-2 | TRAJ6 | CAVNWEEQGGSYIPTF | TRBV5-1 | TRBJ2-3 | CASRGQGVLDTQYF | KLGGALQAK | YES | NO | NO |
| 6270 | TRAV12-3 | TRAJ39 | CAIGPGNMLTF | TRBV5-1 | TRBJ2-3 | CASSMAWDRTDTQYF | KLGGALQAK | YES | NO | NO |
| 6271 | TRAV13-1 | TRAJ17 | CATPYRTAAGNKLTF | TRBV5-1 | TRBJ2-3 | CASSLASATDTQYF | KLGGALQAK | YES | NO | NO |
| 6273 | TRAV13-2 | TRAJ32 | CAESAPFPTNKLIF | TRBV5-1 | TRBJ2-3 | CASSPAGTSGKVWGTDTQYF | KLGGALQAK | NO | NO | NO |
| 6275 | TRAV13-2 | TRAJ8 | CAENTGFQKLVF | TRBV5-1 | TRBJ2-3 | CASKGASGATATDTQYF | KLGGALQAK | YES | NO | NO |
| 6277 | TRAV17 | TRAJ24 | CATASSTDSWGKLQF | TRBV5-1 | TRBJ2-3 | CASNLRTGGTDTQYF | KLGGALQAK | YES | NO | NO |
| 6278 | TRAV19 | TRAJ15 | CALSIDQAGTALIF | TRBV5-1 | TRBJ2-3 | CASSMWGGSTDTQYF | KLGGALQAK | YES | NO | NO |
| 6279 | TRAV19 | TRAJ20 | CALSEGSNDYKLSF | TRBV5-1 | TRBJ2-3 | CASSLKLADTQYF | KLGGALQAK | YES | NO | NO |
| 6280 | TRAV19 | TRAJ30 | CALSERTRDDKIIF | TRBV5-1 | TRBJ2-3 | CASSFSVRDRTTDTQYF | KLGGALQAK | YES | NO | NO |
| 6283 | TRAV21 | TRAJ24 | CAVSWTTDSWGKLQF | TRBV5-1 | TRBJ2-3 | CASSLAPSGVTQYF | KLGGALQAK | YES | NO | NO |
| 6285 | TRAV23DV6 | TRAJ26 | CAAFYGQNFVF | TRBV5-1 | TRBJ2-3 | CASSLWGGADTQYF | KLGGALQAK | YES | NO | NO |
| 6287 | TRAV29DV5 | TRAJ18 | CAASDRGSTLGRLYF | TRBV5-1 | TRBJ2-3 | CASSLYMQGQQITDTQYF | KLGGALQAK | YES | NO | NO |
| 6288 | TRAV29DV5 | TRAJ42 | CAAHDYGGSQGNLIF | TRBV5-1 | TRBJ2-3 | CASSLATTGTDTQYF | KLGGALQAK | YES | NO | NO |
| 6289 | TRAV29DV5 | TRAJ49 | CAASDTGNQFYF | TRBV5-1 | TRBJ2-3 | CASNLRTGGTDTQYF | KLGGALQAK | YES | NO | NO |
| 6290 | TRAV29DV5 | TRAJ53 | CAANIGGGSNYKLTF | TRBV5-1 | TRBJ2-3 | CASSFGTRGGMSDTQYF | KLGGALQAK | YES | NO | NO |
| 6291 | TRAV38-1 | TRAJ43 | CAFMYNNNDMRF | TRBV5-1 | TRBJ2-3 | CASSLGLEPSTDTQYF | KLGGALQAK | YES | NO | NO |
| 6292 | TRAV38-2DV8 | TRAJ44 | CAYRSILNTGGTASKLTF | TRBV5-1 | TRBJ2-3 | CASSWGTSGGRSTDTQYF | KLGGALQAK | YES | NO | NO |
| 6293 | TRAV4 | TRAJ22 | CLVGDRGFSSGSARQLTF | TRBV5-1 | TRBJ2-3 | CASRGQGVLDTQYF | KLGGALQAK | NO | NO | NO |
| 6294 | TRAV5 | TRAJ3 | CAETGSSASKIIF | TRBV5-1 | TRBJ2-3 | CASMDREGPDTQYF | KLGGALQAK | YES | NO | NO |
| 6298 | TRAV12-1 | TRAJ30 | CVVKNRDDKIIF | TRBV5-4 | TRBJ2-3 | CASSYGTEGTDTQYF | KLGGALQAK | YES | NO | NO |
| 6299 | TRAV12-1 | TRAJ6 | CVVNAAGGSYIPTF | TRBV5-4 | TRBJ2-3 | CASSLTGTPSTDTQYF | KLGGALQAK | YES | NO | NO |
| 6300 | TRAV12-2 | TRAJ5 | CALWGAPGRRALTF | TRBV5-4 | TRBJ2-3 | CASSRLAGESDTQYF | KLGGALQAK | YES | NO | NO |
| 6301 | TRAV12-2 | TRAJ53 | CAVNMVSGGSNYKLTF | TRBV5-4 | TRBJ2-3 | CASSSGTSKDTQYF | KLGGALQAK | YES | NO | NO |
| 6302 | TRAV12-2 | TRAJ7 | CAVIGGNNRLAF | TRBV5-4 | TRBJ2-3 | CASSLMTEGTQYF | KLGGALQAK | YES | NO | NO |
| 6303 | TRAV14DV4 | TRAJ9 | CAMREGLSTGGFKTIF | TRBV5-4 | TRBJ2-3 | CASSLGGYTDTQYF | KLGGALQAK | YES | NO | NO |
| 6304 | TRAV17 | TRAJ15 | CATDPDQAGTALIF | TRBV5-4 | TRBJ2-3 | CASSLVLTGADTQYF | KLGGALQAK | YES | NO | NO |
| 6305 | TRAV17 | TRAJ39 | CATDNNNAGNMLTF | TRBV5-4 | TRBJ2-3 | CASSLERSGAITDTQYF | KLGGALQAK | YES | NO | NO |
| 6306 | TRAV17 | TRAJ57 | CATDAGGSEKLVF | TRBV5-4 | TRBJ2-3 | CASSLPGRAPTDTQYF | KLGGALQAK | YES | NO | NO |
| 6307 | TRAV19 | TRAJ21 | CALSEPSYNFNKFYF | TRBV5-4 | TRBJ2-3 | CASSLLTSGGFTDTQYF | KLGGALQAK | YES | NO | NO |
| 6309 | TRAV24 | TRAJ30 | CALRKVPDKIIF | TRBV5-4 | TRBJ2-3 | CASSPRGTDTQYF | KLGGALQAK | YES | NO | NO |
| 6311 | TRAV3 | TRAJ37 | CAVRERFPGNTGKLIF | TRBV5-4 | TRBJ2-3 | CASSLSTDTQYF | KLGGALQAK | YES | NO | NO |
| 6312 | TRAV8-4 | TRAJ4 | CAVIPSSGGYNKLIF | TRBV5-4 | TRBJ2-3 | CASSAVLFGTEGDTQYF | IVTDFSVIK | YES | NO | NO |
| 6312 | TRAV8-4 | TRAJ4 | CAVIPSSGGYNKLIF | TRBV5-4 | TRBJ2-3 | CASSAVLFGTEGDTQYF | KLGGALQAK | YES | NO | NO |
| 6313 | TRAV8-4 | TRAJ50 | CAVSAEASYDKVIF | TRBV5-4 | TRBJ2-3 | CASSTPGLAGGPTDTQYF | KLGGALQAK | YES | NO | NO |
| 6315 | TRAV12-3 | TRAJ6 | CAMTLWGGSGGSYIPTF | TRBV5-5 | TRBJ2-3 | CASSLVEEAPTDTQYF | KLGGALQAK | YES | NO | NO |
| 6316 | TRAV13-2 | TRAJ8 | CAETSGFQKLVF | TRBV5-5 | TRBJ2-3 | CASPRLDRDTDTQYF | KLGGALQAK | YES | NO | NO |
| 6317 | TRAV19 | TRAJ17 | CALSGFKAAGNKLTF | TRBV5-5 | TRBJ2-3 | CASSTGQAFTDTQYF | KLGGALQAK | YES | NO | NO |
| 6321 | TRAV5 | TRAJ21 | CAEPYNFNKFYF | TRBV5-5 | TRBJ2-3 | CASSPGTATDTQYF | KLGGALQAK | YES | NO | NO |
| 6322 | TRAV12-2 | TRAJ12 | CAVNMYGDSSYKLIF | TRBV5-6 | TRBJ2-3 | CASSSTDTQYF | KLGGALQAK | YES | NO | NO |
| 6326 | TRAV13-1 | TRAJ53 | CAASMDSGGSNYKLTF | TRBV5-6 | TRBJ2-3 | CASSFWRGYTDTQYF | KLGGALQAK | YES | NO | NO |
| 6327 | TRAV14DV4 | TRAJ5 | CAMREMDTGRRALTF | TRBV5-6 | TRBJ2-3 | CASSLETGADTQYF | KLGGALQAK | YES | NO | NO |
| 6328 | TRAV14DV4 | TRAJ5 | CAMRPMDTGRRALTF | TRBV5-6 | TRBJ2-3 | CASSLETGPDTQYF | KLGGALQAK | YES | NO | NO |
| 6331 | TRAV29DV5 | TRAJ39 | CAASVDNNAGNMLTF | TRBV5-6 | TRBJ2-3 | CASSLAGRGSTDTQYF | KLGGALQAK | YES | YES | NO |
| 6333 | TRAV3 | TRAJ43 | CAVWKGRTDMRF | TRBV5-6 | TRBJ2-3 | CASRQILAANTDTQYF | RLRAEAQVK | NO | NO | NO |
| 6335 | TRAV38-2DV8 | TRAJ52 | CAYRRGRNAGGTSYGKLTF | TRBV5-6 | TRBJ2-3 | CASSRDRGEDTQYF | AVFDRKSDAK | YES | NO | NO |
| 6335 | TRAV38-2DV8 | TRAJ52 | CAYRRGRNAGGTSYGKLTF | TRBV5-6 | TRBJ2-3 | CASSRDRGEDTQYF | IVTDFSVIK | YES | NO | NO |
| 6336 | TRAV5 | TRAJ47 | CADELVF | TRBV5-6 | TRBJ2-3 | CASRQILAANTDTQYF | KLGGALQAK | YES | NO | NO |
| 6336 | TRAV5 | TRAJ47 | CADELVF | TRBV5-6 | TRBJ2-3 | CASRQILAANTDTQYF | RLRAEAQVK | YES | NO | NO |
| 6338 | TRAV12-1 | TRAJ26 | CVVNYGQNFVF | TRBV5-8 | TRBJ2-3 | CASSSLDSNGDTQYF | KLGGALQAK | YES | NO | NO |
| 6340 | TRAV38-2DV8 | TRAJ54 | CAYRSGQGAQKLVF | TRBV5-8 | TRBJ2-3 | CASSLNPTDTQYF | KLGGALQAK | YES | NO | NO |
| 6341 | TRAV1-2 | TRAJ12 | CAVMDSSYKLIF | TRBV6-1 | TRBJ2-3 | CASSELAGGPDTQYF | KLGGALQAK | YES | NO | NO |
| 6342 | TRAV1-2 | TRAJ20 | CAVLGNDYKLSF | TRBV6-1 | TRBJ2-3 | CASSGLAGGPDTQYF | KLGGALQAK | YES | NO | NO |
| 6344 | TRAV1-2 | TRAJ33 | CAVRDSNYQLIW | TRBV6-1 | TRBJ2-3 | CASSEVLDTTDTQYF | RLRAEAQVK | YES | NO | YES |
| 6345 | TRAV1-2 | TRAJ33 | CAVMDSNYQLIW | TRBV6-1 | TRBJ2-3 | CASSDGQATDTQYF | KLGGALQAK | YES | NO | NO |
| 6348 | TRAV10 | TRAJ4 | CVVSAQLTGGYNKLIF | TRBV6-1 | TRBJ2-3 | CASTVGIPLGTDTQYF | KLGGALQAK | YES | NO | NO |
| 6349 | TRAV12-2 | TRAJ23 | CAVFFYNQGGKLIF | TRBV6-1 | TRBJ2-3 | CASSEMMDTQYF | KLGGALQAK | YES | NO | NO |
| 6350 | TRAV12-2 | TRAJ6 | CAVNWEEQGGSYIPTF | TRBV6-1 | TRBJ2-3 | CASRTGLASTDTQYF | KLGGALQAK | YES | NO | NO |
| 6351 | TRAV13-1 | TRAJ15 | CAAIPNQAGTALIF | TRBV6-1 | TRBJ2-3 | CASSPTRDSALNTQYF | KLGGALQAK | YES | NO | NO |
| 6353 | TRAV14DV4 | TRAJ30 | CAMREGSRRRDDKIIF | TRBV6-1 | TRBJ2-3 | CASSNRGWGDTQYF | KLGGALQAK | YES | NO | NO |
| 6353 | TRAV14DV4 | TRAJ30 | CAMREGSRRRDDKIIF | TRBV6-1 | TRBJ2-3 | CASSNRGWGDTQYF | AVFDRKSDAK | YES | NO | NO |
| 6355 | TRAV19 | TRAJ22 | CALRILFSGSARQLTF | TRBV6-1 | TRBJ2-3 | CASSELADITDTQYF | KLGGALQAK | YES | NO | NO |
| 6356 | TRAV19 | TRAJ28 | CALSEAANSGAGSYQLTF | TRBV6-1 | TRBJ2-3 | CASSITEGEDSVSTDTQYF | AVFDRKSDAK | YES | NO | NO |
| 6356 | TRAV19 | TRAJ28 | CALSEAANSGAGSYQLTF | TRBV6-1 | TRBJ2-3 | CASSITEGEDSVSTDTQYF | KLGGALQAK | YES | NO | NO |
| 6357 | TRAV19 | TRAJ36 | CALTQWGRTGANNLFF | TRBV6-1 | TRBJ2-3 | CATPGQEADTQYF | KLGGALQAK | YES | NO | NO |
| 6359 | TRAV21 | TRAJ43 | CAVSGGNNDMRF | TRBV6-1 | TRBJ2-3 | CASRRGTTDTQYF | KLGGALQAK | YES | NO | NO |
| 6360 | TRAV24 | TRAJ45 | CAFKLYSGGGADGLTF | TRBV6-1 | TRBJ2-3 | CASSEWGQGRIDTQYF | KLGGALQAK | YES | NO | NO |
| 6360 | TRAV24 | TRAJ45 | CAFKLYSGGGADGLTF | TRBV6-1 | TRBJ2-3 | CASSEWGQGRIDTQYF | AVFDRKSDAK | YES | NO | NO |
| 6361 | TRAV38-2DV8 | TRAJ42 | CAFTNYGGSQGNLIF | TRBV6-1 | TRBJ2-3 | CASSATVITDTQYF | KLGGALQAK | YES | NO | NO |
| 6362 | TRAV8-1 | TRAJ26 | CAVNPYGQNFVF | TRBV6-1 | TRBJ2-3 | CASSEMDGRTDTQYF | KLGGALQAK | YES | NO | NO |
| 6363 | TRAV8-1 | TRAJ6 | CAVNDLSGGSYIPTF | TRBV6-1 | TRBJ2-3 | CASSDPGNTQYF | KLGGALQAK | YES | NO | NO |
| 6366 | TRAV1-2 | TRAJ33 | CAAMDSNYQLIW | TRBV6-3 | TRBJ2-3 | CASSYSAGPTDTQYF | AVFDRKSDAK | YES | NO | NO |
| 6366 | TRAV1-2 | TRAJ33 | CAAMDSNYQLIW | TRBV6-3 | TRBJ2-3 | CASSYSAGPTDTQYF | KLGGALQAK | YES | NO | NO |
| 6367 | TRAV12-1 | TRAJ11 | CVVNTLLSGYSTLTF | TRBV6-3 | TRBJ2-3 | CASSRVGADTQYF | KLGGALQAK | YES | NO | NO |
| 6368 | TRAV12-1 | TRAJ4 | CVVMGAGGYNKLIF | TRBV6-3 | TRBJ2-3 | CASSYSLPTSGEPSTDTQYF | KLGGALQAK | NO | NO | NO |
| 6369 | TRAV12-2 | TRAJ15 | CAVDQAGTALIF | TRBV6-3 | TRBJ2-3 | CASLRADTQYF | KLGGALQAK | YES | NO | NO |
| 6370 | TRAV12-2 | TRAJ45 | CAVWHLRGGGADGLTF | TRBV6-3 | TRBJ2-3 | CASSRRGDGTDTQYF | KLGGALQAK | YES | NO | NO |
| 6370 | TRAV12-2 | TRAJ45 | CAVWHLRGGGADGLTF | TRBV6-3 | TRBJ2-3 | CASSRRGDGTDTQYF | AVFDRKSDAK | YES | NO | NO |
| 6371 | TRAV12-2 | TRAJ6 | CAADEESGGSYIPTF | TRBV6-3 | TRBJ2-3 | CASSYGASTDTQYF | KLGGALQAK | YES | NO | NO |
| 6375 | TRAV13-1 | TRAJ11 | CAAISGYSTLTF | TRBV6-3 | TRBJ2-3 | CASCPSGPTDTQYF | KLGGALQAK | YES | NO | NO |
| 6376 | TRAV13-1 | TRAJ45 | CAATPRGGADGLTF | TRBV6-3 | TRBJ2-3 | CASRNIWATPAATDTQYF | KLGGALQAK | YES | NO | NO |
| 6381 | TRAV21 | TRAJ17 | CAVPGVKAAGNKLTF | TRBV6-3 | TRBJ2-3 | CASSFLGTRADTQYF | KLGGALQAK | YES | NO | NO |
| 6382 | TRAV21 | TRAJ43 | CAVWYMRF | TRBV6-3 | TRBJ2-3 | CASIPAGGPGDTQYF | IVTDFSVIK | NO | NO | NO |
| 6382 | TRAV21 | TRAJ43 | CAVWYMRF | TRBV6-3 | TRBJ2-3 | CASIPAGGPGDTQYF | AVFDRKSDAK | NO | NO | NO |
| 6385 | TRAV27 | TRAJ4 | CAGLISGGYNKLIF | TRBV6-3 | TRBJ2-3 | CASSYTNADTQYF | KLGGALQAK | YES | NO | NO |
| 6387 | TRAV29DV5 | TRAJ40 | CAARASPGTYKYIF | TRBV6-3 | TRBJ2-3 | CASSPPTSGSSTDTQYF | KLGGALQAK | YES | NO | NO |
| 6388 | TRAV29DV5 | TRAJ9 | CAASAGTGGFKTIF | TRBV6-3 | TRBJ2-3 | CATRWRLAGGPDTQYF | KLGGALQAK | YES | NO | NO |
| 6390 | TRAV3 | TRAJ8 | CAVREPGFQKLVF | TRBV6-3 | TRBJ2-3 | CASSYAPSGGVADTQYF | KLGGALQAK | NO | NO | NO |
| 6397 | TRAV8-4 | TRAJ39 | CAVTHAGNMLTF | TRBV6-3 | TRBJ2-3 | CASRSRLAGASTDTQYF | KLGGALQAK | YES | NO | NO |
| 6398 | TRAV1-1 | TRAJ33 | CATEGDSNYQLIW | TRBV6-4 | TRBJ2-3 | CASSDLTSGSHTQYF | KLGGALQAK | YES | NO | NO |
| 6399 | TRAV1-2 | TRAJ20 | CAVRDSDYKLSF | TRBV6-4 | TRBJ2-3 | CASSAGTGDTDTQYF | KLGGALQAK | YES | NO | NO |
| 6400 | TRAV1-2 | TRAJ33 | CAVRDSNYQLIW | TRBV6-4 | TRBJ2-3 | CASSASGGADTQYF | KLGGALQAK | YES | NO | NO |
| 6401 | TRAV1-2 | TRAJ33 | CAVRDSNYQLIW | TRBV6-4 | TRBJ2-3 | CASSVAGGTDTQYF | KLGGALQAK | YES | NO | NO |
| 6403 | TRAV1-2 | TRAJ33 | CAVMDSNYQLIW | TRBV6-4 | TRBJ2-3 | CASSLTGTDTQYF | KLGGALQAK | YES | NO | NO |
| 6405 | TRAV13-1 | TRAJ7 | CAAQEYGNNRLAF | TRBV6-4 | TRBJ2-3 | CASSDLAGSGTDTQYF | KLGGALQAK | YES | NO | NO |
| 6407 | TRAV13-2 | TRAJ33 | CAENGPNYQLIW | TRBV6-4 | TRBJ2-3 | CASSVYSGGTDTQYF | KLGGALQAK | YES | NO | NO |
| 6410 | TRAV2 | TRAJ4 | CAVGADGGYNKLIF | TRBV6-4 | TRBJ2-3 | CASSDSGTSGGGADTQYF | KLGGALQAK | NO | NO | NO |
| 6411 | TRAV3 | TRAJ40 | CAIFTSGTYKYIF | TRBV6-4 | TRBJ2-3 | CASSPGTGTIDTQYF | KLGGALQAK | NO | NO | NO |
| 6412 | TRAV3 | TRAJ41 | CAVRASGYALNF | TRBV6-4 | TRBJ2-3 | CASSDSKAGIMGTDTQYF | KLGGALQAK | NO | NO | NO |
| 6414 | TRAV1-2 | TRAJ31 | CAARGNNNARLMF | TRBV6-5 | TRBJ2-3 | CASSYVGGPGDTQYF | KLGGALQAK | NO | NO | NO |

| | | | | | | | | | | |
|---|---|---|---|---|---|---|---|---|---|---|
| 6416 | TRAV12-2 | TRAJ6 | CAVNQSGGSYIPTF | TRBV6-5 | TRBJ2-3 | CASSEGILDTDTQYF | KLGGALQAK | NO | NO | NO |
| 6419 | TRAV13-1 | TRAJ4 | CAAFSGGYNKLIF | TRBV6-5 | TRBJ2-3 | CASSYSGSDTQYF | KLGGALQAK | NO | NO | NO |
| 6422 | TRAV14DV4 | TRAJ42 | CAMREAWGGGSQGNLIF | TRBV6-5 | TRBJ2-3 | CASSYVGGPGDTQYF | KLGGALQAK | NO | NO | NO |
| 6423 | TRAV14DV4 | TRAJ42 | CAMRVYYGGSQGNLIF | TRBV6-5 | TRBJ2-3 | CASSYSLGTDTQYF | KLGGALQAK | NO | NO | NO |
| 6424 | TRAV14DV4 | TRAJ45 | CAMRTMYSGGGADGLTF | TRBV6-5 | TRBJ2-3 | CASSYSDSPTDTQYF | KLGGALQAK | NO | NO | NO |
| 6425 | TRAV14DV4 | TRAJ52 | CAITLNAGGTSYGKLTF | TRBV6-5 | TRBJ2-3 | CASSYLGGGDPFSTDTQYF | KLGGALQAK | NO | NO | NO |
| 6426 | TRAV14DV4 | TRAJ54 | CAMREGLRGAQKLVF | TRBV6-5 | TRBJ2-3 | CASSRLPGPIQYF | KLGGALQAK | NO | NO | NO |
| 6427 | TRAV16 | TRAJ31 | CALRRARLMF | TRBV6-5 | TRBJ2-3 | CASSPGTGGFTDTQYF | KLGGALQAK | NO | NO | NO |
| 6430 | TRAV21 | TRAJ22 | CAVLPSSGSARQLTF | TRBV6-5 | TRBJ2-3 | CASSTLDSPTDTQYF | KLGGALQAK | NO | NO | NO |
| 6430 | TRAV21 | TRAJ22 | CAVLPSSGSARQLTF | TRBV6-5 | TRBJ2-3 | CASSTLDSPTDTQYF | AVFDRKSDAK | NO | NO | NO |
| 6432 | TRAV21 | TRAJ38 | CAVNAGNNRKLIW | TRBV6-5 | TRBJ2-3 | CASSPDNWDSDTQYF | KLGGALQAK | NO | NO | NO |
| 6433 | TRAV21 | TRAJ54 | CAVRSSLIQGAQKLVF | TRBV6-5 | TRBJ2-3 | CASSYVGGPGDTQYF | KLGGALQAK | NO | NO | NO |
| 6433 | TRAV21 | TRAJ54 | CAVRSSLIQGAQKLVF | TRBV6-5 | TRBJ2-3 | CASSYVGGPGDTQYF | AVFDRKSDAK | NO | NO | NO |
| 6436 | TRAV25 | TRAJ32 | CAGNDYGGATNKLIF | TRBV6-5 | TRBJ2-3 | CASSLSGSTDTQYF | KLGGALQAK | NO | NO | NO |
| 6437 | TRAV25 | TRAJ37 | CAGNGNTGKLIF | TRBV6-5 | TRBJ2-3 | CASSYVGGPGDTQYF | RLRAEAQVK | NO | NO | NO |
| 6438 | TRAV27 | TRAJ20 | CAGASGDYKLSF | TRBV6-5 | TRBJ2-3 | CASSTSGTSTDTQYF | KLGGALQAK | NO | NO | NO |
| 6440 | TRAV38-2DV8 | TRAJ56 | CAYINTGANSKLTF | TRBV6-5 | TRBJ2-3 | CASSYWGLAGGTDTQYF | KLGGALQAK | NO | NO | NO |
| 6441 | TRAV4 | TRAJ12 | CLVAADSSYKLIF | TRBV6-5 | TRBJ2-3 | CASSYLTSGRFGADTQYF | KLGGALQAK | NO | NO | NO |
| 6442 | TRAV41 | TRAJ29 | CAVRAATGNTPLVF | TRBV6-5 | TRBJ2-3 | CASSLSGSTDTQYF | KLGGALQAK | NO | NO | NO |
| 6443 | TRAV41 | TRAJ54 | CAVQGAQKLVF | TRBV6-5 | TRBJ2-3 | CASSYVGGPGDTQYF | KLGGALQAK | NO | NO | NO |
| 6445 | TRAV5 | TRAJ10 | CAETRRVTGGGNKLTF | TRBV6-5 | TRBJ2-3 | CASSPGDTQYF | KLGGALQAK | NO | NO | NO |
| 6447 | TRAV8-1 | TRAJ45 | CAVNGYSGGGADGLTF | TRBV6-5 | TRBJ2-3 | CASSYATGDADTQYF | AVFDRKSDAK | NO | NO | NO |
| 6448 | TRAV8-1 | TRAJ45 | CAVNPYSGGGADGLTF | TRBV6-5 | TRBJ2-3 | CASSSGTGNLDTQYF | KLGGALQAK | NO | NO | NO |
| 6449 | TRAV8-1 | TRAJ45 | CAVNAYSGGGADGLTF | TRBV6-5 | TRBJ2-3 | CASSPGTGMEDTQYF | KLGGALQAK | NO | NO | NO |
| 6450 | TRAV8-4 | TRAJ31 | CAVMHGARLMF | TRBV6-5 | TRBJ2-3 | CASSYSSSTDTQYF | KLGGALQAK | NO | NO | NO |
| 6452 | TRAV13-1 | TRAJ39 | CAATQGNAGNMLTF | TRBV6-6 | TRBJ2-3 | CASSYLLAGETHTQYF | KLGGALQAK | NO | NO | NO |
| 6453 | TRAV14DV4 | TRAJ33 | CAMRSDSNYQLIW | TRBV6-6 | TRBJ2-3 | CASSQLAGEIDTQYF | KLGGALQAK | NO | NO | NO |
| 6454 | TRAV14DV4 | TRAJ34 | CAMRGYNTDKLIF | TRBV6-6 | TRBJ2-3 | CASFWTSGGSSTDTQYF | KLGGALQAK | NO | NO | NO |
| 6454 | TRAV14DV4 | TRAJ34 | CAMRGYNTDKLIF | TRBV6-6 | TRBJ2-3 | CASFWTSGGSSTDTQYF | AVFDRKSDAK | NO | NO | NO |
| 6456 | TRAV19 | TRAJ17 | CALSEALKAAGNKLTF | TRBV6-6 | TRBJ2-3 | CASSLRGTGTDTQYF | KLGGALQAK | NO | NO | NO |
| 6459 | TRAV23DV6 | TRAJ48 | CAAVTNFGNEKLTF | TRBV6-6 | TRBJ2-3 | CASSSRPGLAGYWVDTQYF | KLGGALQAK | NO | NO | NO |
| 6462 | TRAV8-2 | TRAJ27 | CVVSEFVKSTF | TRBV6-6 | TRBJ2-3 | CASSEGLAGVYTDTQYF | KLGGALQAK | NO | NO | NO |
| 6464 | TRAV39 | TRAJ52 | CAVVDAGGTSYGKLTF | TRBV6-9 | TRBJ2-3 | CASKNYSLYGTQYF | KLGGALQAK | YES | NO | NO |
| 6465 | TRAV1-2 | TRAJ13 | CAVRDGAGGYQKVTF | TRBV7-2 | TRBJ2-3 | CASANPVRIPDTDTQYF | KLGGALQAK | YES | NO | NO |
| 6467 | TRAV12-2 | TRAJ6 | CAFLGGSYIPTF | TRBV7-2 | TRBJ2-3 | CASSMDSTDTQYF | KLGGALQAK | YES | NO | NO |
| 6468 | TRAV12-3 | TRAJ50 | CAMRPSYDKVIF | TRBV7-2 | TRBJ2-3 | CASSERTGGNTDTQYF | KLGGALQAK | YES | NO | NO |
| 6472 | TRAV14DV4 | TRAJ37 | CAMREGGSSNTGKLIF | TRBV7-2 | TRBJ2-3 | CASSLLGYPMRDTQYF | KLGGALQAK | YES | NO | NO |
| 6476 | TRAV20 | TRAJ23 | CAVRSYNQGGKLIF | TRBV7-2 | TRBJ2-3 | CASSPRPITDTQYF | KLGGALQAK | YES | NO | NO |
| 6479 | TRAV3 | TRAJ37 | CAVWGSSNTGKLIF | TRBV7-2 | TRBJ2-3 | CASSLLAGAPDTQYF | KLGGALQAK | NO | NO | NO |
| 6480 | TRAV38-2DV8 | TRAJ45 | CAQMYSGGGADGLTF | TRBV7-2 | TRBJ2-3 | CASSFWGSGRDTQYF | KLGGALQAK | YES | NO | NO |
| 6480 | TRAV38-2DV8 | TRAJ45 | CAQMYSGGGADGLTF | TRBV7-2 | TRBJ2-3 | CASSFWGSGRDTQYF | AVFDRKSDAK | YES | NO | NO |
| 6484 | TRAV12-1 | TRAJ3 | CVVNDSSASKIIF | TRBV7-3 | TRBJ2-3 | CASSLDWTADTQYF | KLGGALQAK | YES | NO | NO |
| 6485 | TRAV12-2 | TRAJ13 | CAVKTGGYQKVTF | TRBV7-3 | TRBJ2-3 | CASSLDLSGGITDTQYF | AVFDRKSDAK | YES | NO | NO |
| 6485 | TRAV12-2 | TRAJ13 | CAVKTGGYQKVTF | TRBV7-3 | TRBJ2-3 | CASSLDLSGGITDTQYF | KLGGALQAK | YES | NO | NO |
| 6486 | TRAV12-2 | TRAJ31 | CAVNNNNARLMF | TRBV7-3 | TRBJ2-3 | CASSQGDFLITDTQYF | KLGGALQAK | YES | NO | NO |
| 6487 | TRAV13-1 | TRAJ26 | CAPRLNYGQNFVF | TRBV7-3 | TRBJ2-3 | CASSLRGRGADTQYF | KLGGALQAK | YES | NO | NO |
| 6488 | TRAV13-1 | TRAJ49 | CAAIRHGNQFYF | TRBV7-3 | TRBJ2-3 | CASSLVGGGTDTQYF | KLGGALQAK | YES | NO | NO |
| 6490 | TRAV19 | TRAJ49 | CALSEANTGNQFYF | TRBV7-3 | TRBJ2-3 | CASSAYRVDTQYF | KLGGALQAK | YES | NO | NO |
| 6491 | TRAV22 | TRAJ26 | CAVDPYGQNFVF | TRBV7-3 | TRBJ2-3 | CASSPLGSSTDTQYF | KLGGALQAK | YES | NO | NO |
| 6494 | TRAV3 | TRAJ35 | CAVRDPGFGNVLHC | TRBV7-3 | TRBJ2-3 | CASSALAGEDTQYF | KLGGALQAK | NO | NO | NO |
| 6495 | TRAV8-3 | TRAJ48 | CAVGGDFGNEKLTF | TRBV7-3 | TRBJ2-3 | CASSLGFGGPTDTQYF | KLGGALQAK | YES | NO | NO |
| 6496 | TRAV10 | TRAJ26 | CVVSASGQNFVF | TRBV7-6 | TRBJ2-3 | CASSPDPSGLTDTQYF | KLGGALQAK | NO | NO | NO |
| 6497 | TRAV13-2 | TRAJ36 | CAESLTGANNLFF | TRBV7-6 | TRBJ2-3 | CASSLVKGTDTQYF | KLGGALQAK | YES | NO | NO |
| 6498 | TRAV14DV4 | TRAJ48 | CAMREGRISNFGNEKLTF | TRBV7-6 | TRBJ2-3 | CASSLLASGSTDTQYF | KLGGALQAK | YES | NO | NO |
| 6499 | TRAV19 | TRAJ22 | CALSDGSARQLTF | TRBV7-6 | TRBJ2-3 | CASSSPEGTDTQYF | KLGGALQAK | YES | NO | NO |
| 6500 | TRAV19 | TRAJ26 | CALSEARNYGQNFVF | TRBV7-6 | TRBJ2-3 | CASSLGASRSTDTQYF | KLGGALQAK | YES | NO | NO |
| 6501 | TRAV19 | TRAJ31 | CALKHNNARLMF | TRBV7-6 | TRBJ2-3 | CASSPLAGVSDTQYF | KLGGALQAK | YES | NO | NO |
| 6502 | TRAV39 | TRAJ30 | CAVDRDDKIIF | TRBV7-6 | TRBJ2-3 | CASSIYPGTDTQYF | KLGGALQAK | YES | NO | NO |
| 6503 | TRAV5 | TRAJ6 | CAERSGGSYIPTF | TRBV7-6 | TRBJ2-3 | CASSRTGGTDTQYF | KLGGALQAK | YES | NO | NO |
| 6506 | TRAV13-2 | TRAJ53 | CAENSGGSNYKLTF | TRBV7-7 | TRBJ2-3 | CASSLDRGADTQYF | KLGGALQAK | YES | NO | NO |
| 6508 | TRAV41 | TRAJ28 | CAVPSSYSGAGSYQLTF | TRBV7-7 | TRBJ2-3 | CASSLSDRPSATDTQYF | KLGGALQAK | YES | NO | NO |
| 6509 | TRAV12-2 | TRAJ11 | CAVNKGYSTLTF | TRBV7-8 | TRBJ2-3 | CASSSSTSGYTDTQYF | KLGGALQAK | YES | NO | NO |
| 6511 | TRAV13-1 | TRAJ37 | CAASINYGSGNTGKLIF | TRBV7-8 | TRBJ2-3 | CASSLVGRGPSSDTQYF | KLGGALQAK | YES | NO | NO |
| 6513 | TRAV20 | TRAJ34 | CAVRGYNTDKLIF | TRBV7-8 | TRBJ2-3 | CASSDDRGGFLGTDTQYF | KLGGALQAK | YES | NO | NO |
| 6520 | TRAV8-1 | TRAJ32 | CAVRGGGATNKLIF | TRBV7-8 | TRBJ2-3 | CASSFPGQGSADTQYF | KLGGALQAK | YES | NO | NO |
| 6522 | TRAV8-2 | TRAJ57 | CVVRGFTQGGSEKLVF | TRBV7-8 | TRBJ2-3 | CASSPRLAGGPGSTDTQYF | KLGGALQAK | YES | NO | NO |
| 6523 | TRAV1-2 | TRAJ29 | CAVRDHSGNTPLVF | TRBV7-9 | TRBJ2-3 | CASSSRDGGTDTQYF | AVFDRKSDAK | YES | NO | YES |
| 6524 | TRAV1-2 | TRAJ39 | CAVRGVNAGNMLTF | TRBV7-9 | TRBJ2-3 | CASSFQGDGRDTQYF | KLGGALQAK | YES | NO | NO |
| 6527 | TRAV12-3 | TRAJ34 | CAMSADTDKLIF | TRBV7-9 | TRBJ2-3 | CASSLGTDTQYF | KLGGALQAK | YES | NO | NO |
| 6528 | TRAV12-3 | TRAJ39 | CAIGPGNMLTF | TRBV7-9 | TRBJ2-3 | CASSLTRTDTQYF | KLGGALQAK | YES | NO | NO |
| 6531 | TRAV13-1 | TRAJ53 | CAATWGGSNYKLTF | TRBV7-9 | TRBJ2-3 | CASSSRDGGTDTQYF | KLGGALQAK | YES | NO | NO |
| 6535 | TRAV14DV4 | TRAJ34 | CAMREGVDTDKLIF | TRBV7-9 | TRBJ2-3 | CASSSLDKGWGTDTQYF | KLGGALQAK | YES | NO | NO |
| 6537 | TRAV17 | TRAJ48 | CATDASGNEKLTF | TRBV7-9 | TRBJ2-3 | CASSRTGLVIGNTQYF | KLGGALQAK | YES | NO | NO |
| 6539 | TRAV19 | TRAJ26 | CALSDGQNFVF | TRBV7-9 | TRBJ2-3 | CASFTREGRGGYTQYF | KLGGALQAK | YES | NO | NO |
| 6541 | TRAV19 | TRAJ33 | CALSGVVVDSNYQLIW | TRBV7-9 | TRBJ2-3 | CASSGGQLTDTQYF | KLGGALQAK | YES | NO | NO |
| 6543 | TRAV21 | TRAJ31 | CAVNNARLMF | TRBV7-9 | TRBJ2-3 | CASSPVDFQNTDTQYF | KLGGALQAK | YES | NO | NO |
| 6544 | TRAV21 | TRAJ37 | CAVIRSGNTGKLIF | TRBV7-9 | TRBJ2-3 | CASSLGLAGLNTQYF | KLGGALQAK | YES | NO | NO |
| 6545 | TRAV21 | TRAJ56 | CAVNTGANSKLTF | TRBV7-9 | TRBJ2-3 | CASSLELAGPDTQYF | KLGGALQAK | YES | NO | NO |
| 6546 | TRAV22 | TRAJ8 | CALGFQKLVF | TRBV7-9 | TRBJ2-3 | CASSLGFGSPTSTDTQYF | KLGGALQAK | YES | NO | NO |
| 6552 | TRAV3 | TRAJ18 | CAVRDRGSTLGRLYF | TRBV7-9 | TRBJ2-3 | CASSLALPSGGGNTDTQYF | KLGGALQAK | NO | NO | NO |
| 6553 | TRAV3 | TRAJ27 | CAMGNTNAGKSTF | TRBV7-9 | TRBJ2-3 | CASSLPQEDTQYF | KLGGALQAK | NO | NO | NO |
| 6557 | TRAV38-2DV8 | TRAJ42 | CAYRIRGSQGNLIF | TRBV7-9 | TRBJ2-3 | CASSVGGADTQYF | KLGGALQAK | YES | NO | NO |
| 6558 | TRAV38-2DV8 | TRAJ6 | CAYRRGSYIPTF | TRBV7-9 | TRBJ2-3 | CASAVALAEPPDTQYF | KLGGALQAK | YES | NO | NO |
| 6559 | TRAV4 | TRAJ39 | CLVGDPPWAGNMLTF | TRBV7-9 | TRBJ2-3 | CASSFYEAPDTQYF | KLGGALQAK | YES | NO | NO |
| 6560 | TRAV5 | TRAJ30 | CAEPFMNRDDKIIF | TRBV7-9 | TRBJ2-3 | CASSPGLERSAATQYF | KLGGALQAK | YES | NO | NO |
| 6561 | TRAV8-2 | TRAJ45 | CVVSLGGGADGLTF | TRBV7-9 | TRBJ2-3 | CASSTRTSGMGDTQYF | KLGGALQAK | YES | NO | NO |
| 6564 | TRAV8-3 | TRAJ45 | CAVGGGADGLTF | TRBV7-9 | TRBJ2-3 | CASSLTRTDTQYF | KLGGALQAK | YES | NO | NO |
| 6565 | TRAV8-4 | TRAJ16 | CAVTAF | TRBV7-9 | TRBJ2-3 | CASSASLGGSTDTQYF | KLGGALQAK | NO | NO | NO |
| 6568 | TRAV1-1 | TRAJ23 | CAVLNQGGKLIF | TRBV9 | TRBJ2-3 | CASSVAGAQAGDTQYF | KLGGALQAK | YES | NO | NO |
| 6570 | TRAV1-2 | TRAJ28 | CAVPRSGAGSYQLTF | TRBV9 | TRBJ2-3 | CASSVDTSGGAGTDTQYF | KLGGALQAK | YES | NO | NO |
| 6572 | TRAV10 | TRAJ40 | CVVSAALTSGTYKYIF | TRBV9 | TRBJ2-3 | CASSAHWGSSTDTQYF | KLGGALQAK | YES | NO | NO |
| 6573 | TRAV12-1 | TRAJ17 | CVVNGGAAGNKLTF | TRBV9 | TRBJ2-3 | CASSVGVDSTDTQYF | KLGGALQAK | YES | NO | NO |
| 6574 | TRAV12-1 | TRAJ18 | CVVNLGGSTLGRLYF | TRBV9 | TRBJ2-3 | CASSASGGETDTQYF | KLGGALQAK | YES | NO | NO |
| 6576 | TRAV12-2 | TRAJ20 | CAVSGNDYKLSF | TRBV9 | TRBJ2-3 | CASTVSGEPSTDTQYF | KLGGALQAK | YES | NO | NO |
| 6579 | TRAV12-2 | TRAJ45 | CAVNRGGGADGLTF | TRBV9 | TRBJ2-3 | CASSVTWGLALDTQYF | KLGGALQAK | YES | NO | NO |
| 6580 | TRAV12-2 | TRAJ48 | CAVNSFGNEKLTF | TRBV9 | TRBJ2-3 | CASSVDRSTDTQYF | KLGGALQAK | YES | NO | NO |
| 6582 | TRAV13-1 | TRAJ11 | CAAKEGYSTLTF | TRBV9 | TRBJ2-3 | CASSPGTESTDTQYF | KLGGALQAK | YES | NO | NO |
| 6585 | TRAV13-2 | TRAJ40 | CAEWPPGTYKYIF | TRBV9 | TRBJ2-3 | CASSVVGGGTDTQYF | KLGGALQAK | YES | NO | NO |
| 6586 | TRAV13-2 | TRAJ53 | CADEGELSNYKLTF | TRBV9 | TRBJ2-3 | CASSVTGALDTQYF | KLGGALQAK | YES | NO | NO |
| 6587 | TRAV14DV4 | TRAJ11 | CAMREWLSGYSTLTF | TRBV9 | TRBJ2-3 | CASSVVGGADVDTQYF | KLGGALQAK | YES | NO | NO |
| 6589 | TRAV16 | TRAJ45 | CALWGGYSGGGADGLTF | TRBV9 | TRBJ2-3 | CASSAVAGGPTDTQYF | KLGGALQAK | NO | NO | NO |
| 6590 | TRAV17 | TRAJ33 | CATGGRDSNYQLIW | TRBV9 | TRBJ2-3 | CASSVAGGPTDTQYF | KLGGALQAK | YES | NO | NO |
| 6592 | TRAV17 | TRAJ57 | CATDRGGSEKLVF | TRBV9 | TRBJ2-3 | CASSLRLAGGPFDTQYF | KLGGALQAK | YES | NO | NO |
| 6593 | TRAV19 | TRAJ24 | CALSEETDSWGKLQF | TRBV9 | TRBJ2-3 | CASSAPGTDTQYF | KLGGALQAK | YES | NO | NO |

| | | | | | | | | | | |
|---|---|---|---|---|---|---|---|---|---|---|
| 6594 | TRAV19 | TRAJ33 | CALSEPNLDSNYQLIW | TRBV9 | TRBJ2-3 | CASSGRTVAADTQYF | KLGGALQAK | YES | NO | NO |
| 6595 | TRAV20 | TRAJ15 | CAVQAWSQAGTALIF | TRBV9 | TRBJ2-3 | CASSPSGAGTGTDTQYF | KLGGALQAK | YES | NO | NO |
| 6597 | TRAV21 | TRAJ33 | CAVRPVMDSNYQLIW | TRBV9 | TRBJ2-3 | CASSVTAGGADTQYF | KLGGALQAK | YES | NO | NO |
| 6601 | TRAV26-2 | TRAJ6 | CILRDWPGGSYIPTF | TRBV9 | TRBJ2-3 | CASSVAGTNPTDTQYF | KLGGALQAK | YES | NO | NO |
| 6603 | TRAV29DV5 | TRAJ39 | CAASDAGNMLTF | TRBV9 | TRBJ2-3 | CASSPTSGSGDTQYF | KLGGALQAK | YES | NO | NO |
| 6605 | TRAV30 | TRAJ26 | CGTAYGQNFVF | TRBV9 | TRBJ2-3 | CASSVNTATDTQYF | KLGGALQAK | YES | NO | NO |
| 6606 | TRAV30 | TRAJ31 | CGTEYNNARLMF | TRBV9 | TRBJ2-3 | CASSAEDGATDTQYF | KLGGALQAK | YES | NO | NO |
| 6607 | TRAV38-2DV8 | TRAJ28 | CAYIYSGAGSYQLTF | TRBV9 | TRBJ2-3 | CASSLEGGVADTQYF | KLGGALQAK | YES | NO | NO |
| 6608 | TRAV38-2DV8 | TRAJ29 | CALCGNTPLVF | TRBV9 | TRBJ2-3 | CASSDRLATTDTQYF | KLGGALQAK | YES | NO | NO |
| 6611 | TRAV38-2DV8 | TRAJ57 | CAYRSFGGGSEKLVF | TRBV9 | TRBJ2-3 | CASSVGGQGSTDTQYF | KLGGALQAK | YES | NO | NO |
| 6613 | TRAV4 | TRAJ39 | CLVGENNAGNMLTF | TRBV9 | TRBJ2-3 | CASSPLEYTDTQYF | KLGGALQAK | YES | NO | NO |
| 6614 | TRAV5 | TRAJ28 | CAESSSGAGSYQLTF | TRBV9 | TRBJ2-3 | CASSETGLAGVGTDTQYF | KLGGALQAK | YES | NO | NO |
| 6618 | TRAV8-3 | TRAJ9 | CAVGANTGGFKTIF | TRBV9 | TRBJ2-3 | CASSEGTSGPDTQYF | KLGGALQAK | YES | NO | NO |
| 6621 | TRAV9-2 | TRAJ48 | CALSVYFGNEKLTF | TRBV9 | TRBJ2-3 | CASSGLAGVYPQYF | KLGGALQAK | YES | NO | NO |
| 6623 | TRAV9-2 | TRAJ33 | CALRQDSNYQLIW | TRBV10-1 | TRBJ2-4 | CASSEGRGGKNIQYF | KLGGALQAK | YES | NO | NO |
| 6624 | TRAV12-1 | TRAJ9 | CVVNIATGGFKTIF | TRBV11-3 | TRBJ2-4 | CASSLVGGDIQYF | KLGGALQAK | YES | NO | NO |
| 6625 | TRAV39 | TRAJ43 | CAVDVNLYNNNDMRF | TRBV12-3 | TRBJ2-4 | CASRRTGGDLAKNIQYF | KLGGALQAK | YES | NO | NO |
| 6627 | TRAV12-2 | TRAJ43 | CAVNENNDMRF | TRBV12-4 | TRBJ2-4 | CASSAGTGDIQYF | KLGGALQAK | YES | NO | NO |
| 6628 | TRAV13-1 | TRAJ49 | CAASWGTGNQFYF | TRBV12-4 | TRBJ2-4 | CASSAGGDIQYF | RLRAEAQVK | YES | NO | NO |
| 6630 | TRAV26-2 | TRAJ49 | CILRDSLTGNQFYF | TRBV12-4 | TRBJ2-4 | CASSPTGLDIQYF | KLGGALQAK | YES | NO | NO |
| 6631 | TRAV38-2DV8 | TRAJ29 | CASSPSGNTPLVF | TRBV12-4 | TRBJ2-4 | CASSSGDRVENIQYF | KLGGALQAK | YES | NO | NO |
| 6632 | TRAV8-1 | TRAJ34 | CAVNTDKLIF | TRBV12-4 | TRBJ2-4 | CASSAALAKNIQYF | KLGGALQAK | YES | NO | NO |
| 6635 | TRAV27 | TRAJ29 | CAGNGNTPLVF | TRBV12-5 | TRBJ2-4 | CASGSSGGLAKNIQYF | KLGGALQAK | YES | YES | NO |
| 6636 | TRAV38-1 | TRAJ31 | CAFGFNNARLMF | TRBV12-5 | TRBJ2-4 | CASGLVEGDSKNIQYF | AVFDRKSDAK | YES | NO | NO |
| 6639 | TRAV9-2 | TRAJ7 | CALNGNNRLAF | TRBV13 | TRBJ2-4 | CASSLTGTPHQRLAKNIQYF | KLGGALQAK | NO | NO | NO |
| 6641 | TRAV25 | TRAJ50 | CAGKTSYDKVIF | TRBV14 | TRBJ2-4 | CASSQAYSGNKNIQYF | KLGGALQAK | NO | NO | NO |
| 6642 | TRAV8-6 | TRAJ3 | CAVSDYSSASKIIF | TRBV18 | TRBJ2-4 | CASSSDEELLLAKNIQYF | KLGGALQAK | YES | NO | NO |
| 6644 | TRAV12-1 | TRAJ20 | CVVTEDYKLSF | TRBV20-1 | TRBJ2-4 | CSAQLEAKNIQYF | KLGGALQAK | YES | NO | NO |
| 6645 | TRAV17 | TRAJ8 | CATDITGFQKLVF | TRBV20-1 | TRBJ2-4 | CSAPFGAKKNIQYF | KLGGALQAK | YES | NO | NO |
| 6647 | TRAV3 | TRAJ5 | CAVRDIITGRRALTF | TRBV20-1 | TRBJ2-4 | CSARPTRTFNIQYF | RLRAEAQVK | NO | NO | NO |
| 6649 | TRAV23DV6 | TRAJ21 | CAATNFNKFYF | TRBV27 | TRBJ2-4 | CASRTSGNKNIQYF | KLGGALQAK | YES | NO | NO |
| 6650 | TRAV29DV5 | TRAJ52 | CAASVYAGGTSYGKLTF | TRBV27 | TRBJ2-4 | CASSLPTLNGNIQYF | KLGGALQAK | YES | NO | NO |
| 6652 | TRAV8-6 | TRAJ31 | CAVSVPSNNARLMF | TRBV27 | TRBJ2-4 | CASRSGETGIIQYF | KLGGALQAK | YES | NO | NO |
| 6653 | TRAV38-2DV8 | TRAJ49 | CAYRSPLTGNQFYF | TRBV28 | TRBJ2-4 | CASSLPRGGSYF | KLGGALQAK | YES | NO | NO |
| 6654 | TRAV8-4 | TRAJ9 | CAVSHGGFKTIF | TRBV28 | TRBJ2-4 | CASSSPTGGQDKNIQYF | KLGGALQAK | YES | NO | NO |
| 6655 | TRAV23DV6 | TRAJ52 | CAATSFNAGGTSYGKLTF | TRBV29-1 | TRBJ2-4 | CSVAQGLAKNIQYF | KLGGALQAK | YES | NO | NO |
| 6656 | TRAV13-2 | TRAJ37 | CAENMSSGNTGKLIF | TRBV3-1 | TRBJ2-4 | CASSQESSLLAKNIQYF | KLGGALQAK | YES | NO | NO |
| 6661 | TRAV22 | TRAJ39 | CAVDHAGNMLTF | TRBV4-1 | TRBJ2-4 | CASSQPGVNIQYF | KLGGALQAK | YES | NO | NO |
| 6662 | TRAV25 | TRAJ40 | CAGPDSGTYKYIF | TRBV4-1 | TRBJ2-4 | CASSPGDIQYF | KLGGALQAK | YES | NO | NO |
| 6663 | TRAV38-2DV8 | TRAJ33 | CAYRSAGMDSNYQLIW | TRBV4-1 | TRBJ2-4 | CASSRGQNNAKNIQYF | KLGGALQAK | YES | NO | NO |
| 6665 | TRAV10 | TRAJ8 | CVVSRNTGFQKLVF | TRBV4-3 | TRBJ2-4 | CASSQEQGLAYIQYF | KLGGALQAK | NO | NO | NO |
| 6665 | TRAV10 | TRAJ8 | CVVSRNTGFQKLVF | TRBV4-3 | TRBJ2-4 | CASSQEQGLAYIQYF | AVFDRKSDAK | NO | NO | NO |
| 6667 | TRAV4 | TRAJ17 | CLVGDDVTAGNKLTF | TRBV4-3 | TRBJ2-4 | CASSQDFEPVKNIQYF | KLGGALQAK | YES | NO | NO |
| 6668 | TRAV41 | TRAJ30 | CAVRLGGMNRDDKIIF | TRBV5-1 | TRBJ2-4 | CASSYRTSGSNIQYF | KLGGALQAK | YES | NO | NO |
| 6669 | TRAV20 | TRAJ32 | CAVQDSGATNKLIF | TRBV5-4 | TRBJ2-4 | CASSTDSAENIQYF | KLGGALQAK | YES | NO | NO |
| 6671 | TRAV12-2 | TRAJ48 | CAVTDSNFGNEKLTF | TRBV5-6 | TRBJ2-4 | CASSLAQGLAKNIQYF | KLGGALQAK | YES | NO | NO |
| 6673 | TRAV1-2 | TRAJ33 | CAVRDSNYQLIW | TRBV6-1 | TRBJ2-4 | CASSEASGGANIQYF | KLGGALQAK | NO | NO | NO |
| 6675 | TRAV14DV4 | TRAJ4 | CAMRPPFSGGYNKLIF | TRBV6-1 | TRBJ2-4 | CASSDPDRIKNIQYF | AVFDRKSDAK | YES | NO | NO |
| 6675 | TRAV14DV4 | TRAJ4 | CAMRPPFSGGYNKLIF | TRBV6-1 | TRBJ2-4 | CASSDPDRIKNIQYF | KLGGALQAK | YES | NO | NO |
| 6676 | TRAV19 | TRAJ43 | CALSGENNDMRF | TRBV6-1 | TRBJ2-4 | CASRWGVLNIQYF | KLGGALQAK | YES | NO | NO |
| 6677 | TRAV8-4 | TRAJ20 | CAVIQINDYKLSF | TRBV6-1 | TRBJ2-4 | CASSDPDRIKNIQYF | KLGGALQAK | YES | NO | NO |
| 6682 | TRAV19 | TRAJ13 | CALSEASSGGYQKVTF | TRBV6-6 | TRBJ2-4 | CASSSIARVIQYF | KLGGALQAK | NO | NO | NO |
| 6684 | TRAV19 | TRAJ49 | CALSEGRRTGNQFYF | TRBV7-2 | TRBJ2-4 | CASSQTGTSGGVAKNIQYF | KLGGALQAK | YES | NO | NO |
| 6686 | TRAV5 | TRAJ11 | CAESPGYSTLTF | TRBV7-6 | TRBJ2-4 | CASSSGTGMDIQYF | KLGGALQAK | YES | NO | NO |
| 6688 | TRAV12-3 | TRAJ34 | CAMSAYTDKLIF | TRBV7-9 | TRBJ2-4 | CASSLSRWRQAKNIQYF | KLGGALQAK | YES | NO | NO |
| 6693 | TRAV21 | TRAJ58 | CAVPYWETSGSRLTF | TRBV7-9 | TRBJ2-4 | CASSQILQGKDIQYF | RLRAEAQVK | YES | NO | YES |
| 6694 | TRAV38-2DV8 | TRAJ42 | CASGVRGSQGNLIF | TRBV7-9 | TRBJ2-4 | CASSLAVGAKNIQYF | KLGGALQAK | YES | NO | NO |
| 6695 | TRAV38-2DV8 | TRAJ54 | CASEGAQKLVF | TRBV7-9 | TRBJ2-4 | CASSLGLDIQYF | KLGGALQAK | YES | NO | NO |
| 6696 | TRAV8-1 | TRAJ39 | CAVKVYNAGNMLTF | TRBV7-9 | TRBJ2-4 | CASSPSRDREFLYIQYF | KLGGALQAK | YES | NO | NO |
| 6699 | TRAV1-2 | TRAJ28 | CAVKYSGAGSYQLTF | TRBV9 | TRBJ2-4 | CASSGSSGTSKNIQYF | KLGGALQAK | YES | YES | NO |
| 6700 | TRAV19 | TRAJ43 | CALSEDDNNNDMRF | TRBV9 | TRBJ2-4 | CASSASGGAGNIQYF | KLGGALQAK | YES | NO | NO |
| 6702 | TRAV38-2DV8 | TRAJ42 | CAYRSRGYGGSQGNLIF | TRBV9 | TRBJ2-4 | CASSEGSAGLKNIQYF | KLGGALQAK | YES | NO | NO |
| 6705 | TRAV24 | TRAJ21 | CAFLGSGNFNKFYF | TRBV10-1 | TRBJ2-5 | CASSDLEPQETQYF | KLGGALQAK | YES | NO | NO |
| 6706 | TRAV8-1 | TRAJ40 | CAVKGGTYKYIF | TRBV10-1 | TRBJ2-5 | CASSELDEETQYF | KLGGALQAK | YES | NO | NO |
| 6707 | TRAV9-2 | TRAJ39 | CALSDGRGDNMLTF | TRBV10-1 | TRBJ2-5 | CASSRGGLAGEETQYF | KLGGALQAK | YES | NO | NO |
| 6708 | TRAV12-1 | TRAJ20 | CVVNSNDYKLSF | TRBV10-2 | TRBJ2-5 | CASSGGTGTDRDF | KLGGALQAK | YES | NO | NO |
| 6710 | TRAV13-1 | TRAJ33 | CAASNRDSNYQLIW | TRBV10-2 | TRBJ2-5 | CASTYSFETQYF | KLGGALQAK | YES | NO | NO |
| 6712 | TRAV19 | TRAJ29 | CALESGNTPLVF | TRBV10-2 | TRBJ2-5 | CASRVAAKANRETQYF | KLGGALQAK | YES | NO | NO |
| 6712 | TRAV19 | TRAJ29 | CALESGNTPLVF | TRBV10-2 | TRBJ2-5 | CASRVAAKANRETQYF | AVFDRKSDAK | YES | NO | NO |
| 6713 | TRAV26-1 | TRAJ38 | CIVRVARERAGNNRKLIW | TRBV10-2 | TRBJ2-5 | CASSDLLDGKETQYF | KLGGALQAK | YES | NO | NO |
| 6714 | TRAV27 | TRAJ52 | CIGGTSYGKLTF | TRBV10-2 | TRBJ2-5 | CASSDGFGETQYF | KLGGALQAK | YES | NO | NO |
| 6715 | TRAV41 | TRAJ43 | CAVWIYDMRF | TRBV10-2 | TRBJ2-5 | CASRVAAKANRETQYF | KLGGALQAK | YES | NO | NO |
| 6716 | TRAV5 | TRAJ10 | CAETLTGGGNKLTF | TRBV10-2 | TRBJ2-5 | CASKGRSGVGETQYF | KLGGALQAK | YES | NO | NO |
| 6717 | TRAV6 | TRAJ12 | CALDAGDSSYKLIF | TRBV10-3 | TRBJ2-5 | CAISGDSQETQYF | KLGGALQAK | YES | NO | NO |
| 6726 | TRAV13-1 | TRAJ26 | CAASRDGQNFVF | TRBV11-2 | TRBJ2-5 | CASSLDPGRGQETQYF | KLGGALQAK | YES | NO | NO |
| 6727 | TRAV13-1 | TRAJ36 | CAALRTGANNLFF | TRBV11-2 | TRBJ2-5 | CASSLAQETQYF | KLGGALQAK | YES | NO | NO |
| 6729 | TRAV17 | TRAJ22 | CATDRAGSARQLTF | TRBV11-2 | TRBJ2-5 | CASSLTGETQYF | KLGGALQAK | YES | NO | NO |
| 6730 | TRAV17 | TRAJ36 | CATDFVNLFF | TRBV11-2 | TRBJ2-5 | CASSTTRNQETQYF | KLGGALQAK | YES | NO | NO |
| 6731 | TRAV19 | TRAJ18 | CALPLGGSTLGRLYF | TRBV11-2 | TRBJ2-5 | CASSSPHRGLETQYF | KLGGALQAK | YES | NO | NO |
| 6735 | TRAV34 | TRAJ45 | CGAAPAEGGGADGLTF | TRBV11-2 | TRBJ2-5 | CASSLGFFGEVAQYF | KLGGALQAK | YES | NO | NO |
| 6737 | TRAV35 | TRAJ40 | CAGNSGTYKYIF | TRBV11-2 | TRBJ2-5 | CASSLGVGDGQETQYF | KLGGALQAK | YES | NO | NO |
| 6739 | TRAV38-2DV8 | TRAJ33 | CAYRSARGSNYQLIW | TRBV11-2 | TRBJ2-5 | CASSFWSGTQETQYF | KLGGALQAK | YES | NO | NO |
| 6739 | TRAV38-2DV8 | TRAJ33 | CAYRSARGSNYQLIW | TRBV11-2 | TRBJ2-5 | CASSFWSGTQETQYF | AVFDRKSDAK | YES | NO | NO |
| 6739 | TRAV38-2DV8 | TRAJ33 | CAYRSARGSNYQLIW | TRBV11-2 | TRBJ2-5 | CASSFWSGTQETQYF | IVTDFSVIK | YES | NO | NO |
| 6741 | TRAV10 | TRAJ20 | CVVRLVNDYKLSF | TRBV11-3 | TRBJ2-5 | CASSRTSGHQETQYF | KLGGALQAK | YES | NO | NO |
| 6743 | TRAV12-3 | TRAJ39 | CAIGPGNMLTF | TRBV11-3 | TRBJ2-5 | CASSQTGLVLGGQETQYF | AVFDRKSDAK | YES | NO | YES |
| 6746 | TRAV35 | TRAJ20 | CAGRDDYKLSF | TRBV11-3 | TRBJ2-5 | CASSLDLGQGKTQYF | KLGGALQAK | NO | NO | NO |
| 6748 | TRAV4 | TRAJ28 | CLVVFGAGSYQLTF | TRBV11-3 | TRBJ2-5 | CASSTQMETQYF | KLGGALQAK | NO | NO | NO |
| 6751 | TRAV8-4 | TRAJ44 | CAVTVTGTASKLTF | TRBV11-3 | TRBJ2-5 | CASSPPRVAGGVTQYF | KLGGALQAK | YES | NO | NO |
| 6753 | TRAV1-2 | TRAJ20 | CAVNHDYKLSF | TRBV12-3 | TRBJ2-5 | CASSSGLAGDVETQYF | KLGGALQAK | YES | NO | NO |
| 6754 | TRAV12-1 | TRAJ20 | CVVTDRSNDYKLSF | TRBV12-3 | TRBJ2-5 | CASSLLGGQETQYF | KLGGALQAK | YES | NO | NO |
| 6757 | TRAV24 | TRAJ24 | CASLTTDSWGKFQF | TRBV12-3 | TRBJ2-5 | CASSFGAGANQETQYF | KLGGALQAK | YES | NO | NO |
| 6758 | TRAV8-3 | TRAJ43 | CAVGAGNNDMRF | TRBV12-3 | TRBJ2-5 | CASATPEGETQYF | KLGGALQAK | YES | NO | NO |
| 6759 | TRAV8-3 | TRAJ43 | CAVGAGNNDMRF | TRBV12-3 | TRBJ2-5 | CASSPQEGETQYF | KLGGALQAK | YES | NO | NO |
| 6759 | TRAV8-3 | TRAJ43 | CAVGAGNNDMRF | TRBV12-3 | TRBJ2-5 | CASSPQEGETQYF | AVFDRKSDAK | YES | NO | NO |
| 6760 | TRAV8-3 | TRAJ43 | CAVGGGNNDMRF | TRBV12-3 | TRBJ2-5 | CASSPPEGETQYF | KLGGALQAK | YES | NO | NO |
| 6761 | TRAV8-3 | TRAJ43 | CAVGAGNNDMRF | TRBV12-3 | TRBJ2-5 | CASSYPEGETQYF | KLGGALQAK | YES | NO | NO |
| 6761 | TRAV8-3 | TRAJ43 | CAVGAGNNDMRF | TRBV12-3 | TRBJ2-5 | CASSYPEGETQYF | RLRAEAQVK | YES | NO | NO |
| 6762 | TRAV8-3 | TRAJ43 | CAVGAGNNDMRF | TRBV12-3 | TRBJ2-5 | CASSFPEGETQYF | KLGGALQAK | YES | NO | NO |
| 6763 | TRAV9-2 | TRAJ33 | CALSDVDSNYQLIW | TRBV12-3 | TRBJ2-5 | CASSPWAGQETQYF | KLGGALQAK | YES | NO | NO |
| 6764 | TRAV1-2 | TRAJ10 | CAVRGYILTGGGNKLTF | TRBV12-4 | TRBJ2-5 | CASSPARGGTETQYF | KLGGALQAK | YES | NO | NO |
| 6767 | TRAV13-1 | TRAJ45 | CLAGGGADGLTF | TRBV12-4 | TRBJ2-5 | CASSQGTGTQYF | KLGGALQAK | YES | NO | NO |
| 6769 | TRAV17 | TRAJ39 | CATWGAGNMLTF | TRBV12-4 | TRBJ2-5 | CASSKGGQETQYF | KLGGALQAK | YES | NO | NO |
| 6770 | TRAV17 | TRAJ43 | CAHAPLDDMRF | TRBV12-4 | TRBJ2-5 | CASRPSGGAETQYF | KLGGALQAK | YES | NO | NO |

| | | | | | | | | | |
|---|---|---|---|---|---|---|---|---|---|
| 6771 | TRAV17 | TRAJ43 | CATGWNNNDMRF | TRBV12-4 | TRBJ2-5 | CASGLAGFETQYF | KLGGALQAK | YES | NO | NO |
| 6772 | TRAV17 | TRAJ7 | CATVTYGNNRLAF | TRBV12-4 | TRBJ2-5 | CASSLDTEETQYF | KLGGALQAK | YES | NO | NO |
| 6774 | TRAV21 | TRAJ12 | CAVLDMDSSYKLIF | TRBV12-4 | TRBJ2-5 | CAISTSGETQYF | KLGGALQAK | YES | NO | NO |
| 6775 | TRAV21 | TRAJ17 | CAVRILAAGNKLTF | TRBV12-4 | TRBJ2-5 | CASSSRGSGAGETQYF | KLGGALQAK | YES | NO | NO |
| 6776 | TRAV24 | TRAJ40 | CASTSGTYKYIF | TRBV12-4 | TRBJ2-5 | CASSLTIQETQYF | KLGGALQAK | YES | NO | NO |
| 6777 | TRAV29DV5 | TRAJ39 | CAASESEGAGNMLTF | TRBV12-4 | TRBJ2-5 | CASRGTLAGWPPETQYF | KLGGALQAK | YES | NO | NO |
| 6778 | TRAV38-1 | TRAJ45 | CAFGGGADGLTF | TRBV12-4 | TRBJ2-5 | CASSRPGQGLQETQYF | KLGGALQAK | YES | NO | NO |
| 6779 | TRAV39 | TRAJ44 | CAVDSFFRTGTASKLTF | TRBV12-4 | TRBJ2-5 | CASSLGSGRYQETQYF | KLGGALQAK | YES | NO | NO |
| 6780 | TRAV8-4 | TRAJ38 | CAVSANAGNNRKLIW | TRBV12-4 | TRBJ2-5 | CASSYASGRGGTQYF | KLGGALQAK | YES | NO | NO |
| 6782 | TRAV22 | TRAJ48 | CALTNFGNEKLTF | TRBV12-5 | TRBJ2-5 | CASGLLGGWETQYF | KLGGALQAK | YES | NO | NO |
| 6785 | TRAV13-1 | TRAJ54 | CAASQGAQKLVF | TRBV13 | TRBJ2-5 | CASTEGGHETQYF | KLGGALQAK | YES | NO | NO |
| 6788 | TRAV19 | TRAJ17 | CALLKPAAGNKLTF | TRBV13 | TRBJ2-5 | CASSLIDQETQYF | KLGGALQAK | YES | NO | NO |
| 6789 | TRAV19 | TRAJ48 | CALSSNFGNEKLTF | TRBV13 | TRBJ2-5 | CASSLGLGGQEEETQYF | KLGGALQAK | YES | NO | NO |
| 6790 | TRAV21 | TRAJ58 | CAVRQETSGSRLTF | TRBV13 | TRBJ2-5 | CASSEDRFQETQYF | KLGGALQAK | YES | NO | NO |
| 6796 | TRAV14DV4 | TRAJ22 | CAMRRPISSGSARQLTF | TRBV14 | TRBJ2-5 | CASSQEWDPRETQYF | KLGGALQAK | NO | NO | NO |
| 6796 | TRAV14DV4 | TRAJ22 | CAMRRPISSGSARQLTF | TRBV14 | TRBJ2-5 | CASSQEWDPRETQYF | AVFDRKSDAK | NO | NO | NO |
| 6796 | TRAV14DV4 | TRAJ22 | CAMRRPISSGSARQLTF | TRBV14 | TRBJ2-5 | CASSQEWDPRETQYF | IVTDFSVIK | NO | NO | NO |
| 6798 | TRAV19 | TRAJ23 | CALSGIFQGGKLIF | TRBV14 | TRBJ2-5 | CASSLKHRTSGGDQETQYF | KLGGALQAK | NO | NO | NO |
| 6800 | TRAV24 | TRAJ3 | CAFNWYSSASKIIF | TRBV14 | TRBJ2-5 | CASSPIGGEALGETQYF | KLGGALQAK | NO | NO | NO |
| 6804 | TRAV1-2 | TRAJ18 | CAVRGDRGSTLGRLYF | TRBV15 | TRBJ2-5 | CATSRDLANQETQYF | KLGGALQAK | YES | NO | NO |
| 6806 | TRAV1-2 | TRAJ31 | CAVRDRGNNARLMF | TRBV15 | TRBJ2-5 | CATSSSGVQETQYF | KLGGALQAK | YES | NO | NO |
| 6808 | TRAV12-2 | TRAJ49 | CAVTLSGNQFYF | TRBV15 | TRBJ2-5 | CATSFRGQGGETQYF | KLGGALQAK | YES | NO | NO |
| 6810 | TRAV17 | TRAJ49 | CATDRLPGNQFYF | TRBV15 | TRBJ2-5 | CATRSAYQETQYF | KLGGALQAK | YES | NO | NO |
| 6811 | TRAV19 | TRAJ13 | CALSEGYQKVTF | TRBV15 | TRBJ2-5 | CATSRDSGSQETQYF | KLGGALQAK | YES | NO | NO |
| 6812 | TRAV19 | TRAJ24 | CALATDSWGKLQF | TRBV15 | TRBJ2-5 | CATSPDREYQETQYF | KLGGALQAK | YES | NO | NO |
| 6812 | TRAV19 | TRAJ24 | CALATDSWGKLQF | TRBV15 | TRBJ2-5 | CATSPDREYQETQYF | AVFDRKSDAK | YES | NO | NO |
| 6814 | TRAV8-2 | TRAJ38 | CVVSASDAGNNRKLIW | TRBV15 | TRBJ2-5 | CATSRVPRGLGETQYF | KLGGALQAK | YES | NO | NO |
| 6816 | TRAV14DV4 | TRAJ22 | CAMRRPISSGSARQLTF | TRBV18 | TRBJ2-5 | CASSPPGGQETQYF | RLRAEAQVK | YES | NO | NO |
| 6817 | TRAV14DV4 | TRAJ22 | CAMRRPISSGSARQLTF | TRBV18 | TRBJ2-5 | CASSQVFGDRGQETQYF | KLGGALQAK | YES | NO | NO |
| 6818 | TRAV14DV4 | TRAJ22 | CAMRRPISSGSARQLTF | TRBV18 | TRBJ2-5 | CASSPPARQETQYF | KLGGALQAK | YES | NO | NO |
| 6819 | TRAV14DV4 | TRAJ22 | CAMRRPISSGSARQLTF | TRBV18 | TRBJ2-5 | CASSLQGTQETQYF | KLGGALQAK | YES | NO | NO |
| 6821 | TRAV19 | TRAJ42 | CALSEVWGGSQGNLIF | TRBV18 | TRBJ2-5 | CASSPRTGDTETQYF | KLGGALQAK | YES | NO | NO |
| 6822 | TRAV22 | TRAJ10 | CAVERRWDGRLTGGGNKLTF | TRBV18 | TRBJ2-5 | CASSHGTLHF | KLGGALQAK | NO | NO | NO |
| 6825 | TRAV10 | TRAJ20 | CVVSAIDDYKLSF | TRBV19 | TRBJ2-5 | CASSIVTGEETQYF | KLGGALQAK | YES | NO | NO |
| 6826 | TRAV12-2 | TRAJ11 | CAVNNGGYSTLTF | TRBV19 | TRBJ2-5 | CASSKGGGEYGTQYF | KLGGALQAK | YES | NO | NO |
| 6827 | TRAV12-2 | TRAJ30 | CAVRGRDDKIIF | TRBV19 | TRBJ2-5 | CATGGDGYGETQYF | KLGGALQAK | YES | NO | NO |
| 6831 | TRAV14DV4 | TRAJ34 | CAMRGYNTDKLIF | TRBV19 | TRBJ2-5 | CASSNQDRGAQETQYF | KLGGALQAK | YES | NO | NO |
| 6832 | TRAV14DV4 | TRAJ43 | CAMREGFYNNDMRF | TRBV19 | TRBJ2-5 | CASSPGTGGTETQYF | KLGGALQAK | YES | NO | NO |
| 6836 | TRAV21 | TRAJ13 | CAVMNSGGYQKVTF | TRBV19 | TRBJ2-5 | CASSIDWRDRGEKIETQYF | AVFDRKSDAK | YES | NO | NO |
| 6836 | TRAV21 | TRAJ13 | CAVMNSGGYQKVTF | TRBV19 | TRBJ2-5 | CASSIDWRDRGEKIETQYF | KLGGALQAK | YES | NO | NO |
| 6838 | TRAV23DV6 | TRAJ45 | CAARYSGGGADGLTF | TRBV19 | TRBJ2-5 | CASSIGAGAQETQYF | KLGGALQAK | YES | NO | NO |
| 6842 | TRAV3 | TRAJ9 | CAEGGFKTIF | TRBV19 | TRBJ2-5 | CASSFGASGRKETQYF | KLGGALQAK | NO | NO | NO |
| 6843 | TRAV38-1 | TRAJ33 | CASLRPSNYQLIW | TRBV19 | TRBJ2-5 | CASREGETQYF | AVFDRKSDAK | YES | NO | NO |
| 6843 | TRAV38-1 | TRAJ33 | CASLRPSNYQLIW | TRBV19 | TRBJ2-5 | CASREGETQYF | KLGGALQAK | YES | NO | NO |
| 6844 | TRAV38-1 | TRAJ40 | CAFMKSPVTSGTYKYIF | TRBV19 | TRBJ2-5 | CASSIYHLAGLYETQYF | KLGGALQAK | YES | NO | NO |
| 6846 | TRAV5 | TRAJ3 | CAESGYSSASKIIF | TRBV19 | TRBJ2-5 | CASREGETQYF | KLGGALQAK | YES | NO | NO |
| 6847 | TRAV6 | TRAJ18 | CALDESNRGSTLGRLYF | TRBV19 | TRBJ2-5 | CASSIRLAGDQETQYF | KLGGALQAK | YES | NO | NO |
| 6849 | TRAV8-3 | TRAJ54 | CAVGAPGIQGAQKLVF | TRBV19 | TRBJ2-5 | CASREGETQYF | KLGGALQAK | YES | NO | NO |
| 6850 | TRAV9-2 | TRAJ52 | CALSRPGANAGGTSYGKLTF | TRBV19 | TRBJ2-5 | CASSIGWETQYF | KLGGALQAK | NO | NO | NO |
| 6851 | TRAV9-2 | TRAJ6 | CALSDAGGSYIPTF | TRBV19 | TRBJ2-5 | CASSIVVEKQETQYF | KLGGALQAK | YES | NO | NO |
| 6852 | TRAV1-2 | TRAJ15 | CALNQAGTALIF | TRBV2 | TRBJ2-5 | CASRARDYQETQYF | KLGGALQAK | YES | NO | NO |
| 6857 | TRAV17 | TRAJ3 | CATDYSSASKIIF | TRBV2 | TRBJ2-5 | CASVLLSSGRGQETQYF | KLGGALQAK | YES | NO | NO |
| 6858 | TRAV26-2 | TRAJ42 | CILSASYGGSQGNLIF | TRBV2 | TRBJ2-5 | CASRTPRGLAGLRETQYF | KLGGALQAK | YES | NO | NO |
| 6861 | TRAV1-2 | TRAJ33 | CAALDSNYQLIW | TRBV20-1 | TRBJ2-5 | CSAGSGTRGETQYF | KLGGALQAK | YES | NO | NO |
| 6861 | TRAV1-2 | TRAJ33 | CAALDSNYQLIW | TRBV20-1 | TRBJ2-5 | CSAGSGTRGETQYF | AVFDRKSDAK | YES | NO | NO |
| 6862 | TRAV1-2 | TRAJ33 | CAVMDSNYQLIW | TRBV20-1 | TRBJ2-5 | CSARLAGGPETQYF | KLGGALQAK | YES | NO | NO |
| 6863 | TRAV1-2 | TRAJ48 | CAVRGDFGNEKLTF | TRBV20-1 | TRBJ2-5 | CSAREGVRLVAGHQETQYF | KLGGALQAK | YES | NO | NO |
| 6865 | TRAV12-1 | TRAJ23 | CVVNRDYNQGGKLIF | TRBV20-1 | TRBJ2-5 | CSARDGMDLAPETQYF | KLGGALQAK | YES | NO | NO |
| 6866 | TRAV12-1 | TRAJ41 | CVVNKKNSGYALNF | TRBV20-1 | TRBJ2-5 | CSALAGGGETQYF | KLGGALQAK | NO | NO | NO |
| 6870 | TRAV13-1 | TRAJ57 | CAATQGGSEKLVF | TRBV20-1 | TRBJ2-5 | CSARLDRAGETQYF | KLGGALQAK | NO | NO | NO |
| 6871 | TRAV14DV4 | TRAJ18 | CAMRGPRGSTLGRLYF | TRBV20-1 | TRBJ2-5 | CSARDRAGKETQYF | AVFDRKSDAK | YES | NO | NO |
| 6875 | TRAV26-1 | TRAJ22 | CIVRASGSARQLTF | TRBV20-1 | TRBJ2-5 | CSARVINLGAGGLKETQYF | KLGGALQAK | YES | NO | NO |
| 6876 | TRAV26-1 | TRAJ57 | CIVRWTQGGSEKLVF | TRBV20-1 | TRBJ2-5 | CSARDVGEETQYF | KLGGALQAK | NO | NO | NO |
| 6878 | TRAV27 | TRAJ34 | CAGLTDKLIF | TRBV20-1 | TRBJ2-5 | CSANPSNRPFRTQYF | KLGGALQAK | YES | NO | NO |
| 6879 | TRAV29DV5 | TRAJ21 | CAASLTYNFNKFYF | TRBV20-1 | TRBJ2-5 | CSARVSSLAGGQETQYF | KLGGALQAK | NO | NO | NO |
| 6880 | TRAV5 | TRAJ23 | CALNTRGKLIF | TRBV20-1 | TRBJ2-5 | CSATWEGASGGETQYF | KLGGALQAK | YES | NO | NO |
| 6884 | TRAV12-1 | TRAJ40 | CVVNGEGTYKYIF | TRBV21-1 | TRBJ2-5 | CQQQTQPRDPVLRARHFRAPSETQYF | KLGGALQAK | NO | NO | NO |
| 6891 | TRAV12-2 | TRAJ3 | CAVYSSASKIIF | TRBV24-1 | TRBJ2-5 | CATSDGDKRARETQYF | KLGGALQAK | YES | NO | NO |
| 6892 | TRAV12-2 | TRAJ36 | CAVQTGANNLFF | TRBV24-1 | TRBJ2-5 | CATSDHGTGYDSETQYF | KLGGALQAK | YES | NO | NO |
| 6894 | TRAV17 | TRAJ53 | CATDAFPGSNYKLTF | TRBV24-1 | TRBJ2-5 | CATSDRTSGRAKETQYF | KLGGALQAK | YES | NO | NO |
| 6895 | TRAV19 | TRAJ40 | CALSGAPWSNKYIF | TRBV24-1 | TRBJ2-5 | CATSDKIAGGNQETQYF | KLGGALQAK | YES | NO | NO |
| 6896 | TRAV21 | TRAJ23 | CASDYNQGGKLIF | TRBV24-1 | TRBJ2-5 | CATSPGGQETQYF | KLGGALQAK | YES | NO | NO |
| 6898 | TRAV30 | TRAJ42 | CGTAQFYGGSQGNLIF | TRBV24-1 | TRBJ2-5 | CATSDGQTLETQYF | KLGGALQAK | YES | NO | NO |
| 6900 | TRAV39 | TRAJ30 | CAVDRNRKIIF | TRBV24-1 | TRBJ2-5 | CATSDPGTSGVETQYF | KLGGALQAK | YES | NO | NO |
| 6903 | TRAV1-2 | TRAJ42 | CAGLNYGGSQGNLIF | TRBV27 | TRBJ2-5 | CASSLDGTPGETQYF | KLGGALQAK | YES | NO | NO |
| 6904 | TRAV12-2 | TRAJ34 | CAAKLMYNTDKLIF | TRBV27 | TRBJ2-5 | CASSPYRGQQETQYF | KLGGALQAK | YES | NO | NO |
| 6906 | TRAV12-2 | TRAJ9 | CAVNPNTGGFKTIF | TRBV27 | TRBJ2-5 | CASSFSGGAFQETQYF | KLGGALQAK | YES | NO | NO |
| 6910 | TRAV13-2 | TRAJ8 | CAEIPMNTGFQKLVF | TRBV27 | TRBJ2-5 | CASSLFQAETQYF | KLGGALQAK | YES | NO | NO |
| 6912 | TRAV14DV4 | TRAJ53 | CAMRGVSGGSNYKLTF | TRBV27 | TRBJ2-5 | CASSPWTGGETQYF | KLGGALQAK | YES | NO | NO |
| 6913 | TRAV17 | TRAJ34 | CATVLNTDKLIF | TRBV27 | TRBJ2-5 | CASSRGHLQETQYF | KLGGALQAK | YES | NO | NO |
| 6915 | TRAV19 | TRAJ4 | CALSEAFSGGYNKLIF | TRBV27 | TRBJ2-5 | CASSLLGSGGQETQYF | KLGGALQAK | YES | NO | NO |
| 6916 | TRAV19 | TRAJ4 | CALSEAISGGYNKLIF | TRBV27 | TRBJ2-5 | CASSLLGSGVETQYF | KLGGALQAK | YES | NO | NO |
| 6919 | TRAV20 | TRAJ8 | CAVVGWENTGFQKLVF | TRBV27 | TRBJ2-5 | CASSLTGGGETQYF | KLGGALQAK | YES | NO | NO |
| 6920 | TRAV21 | TRAJ23 | CAVFTYNQGGKLIF | TRBV27 | TRBJ2-5 | CASTGRGAQETQYF | KLGGALQAK | YES | NO | NO |
| 6921 | TRAV21 | TRAJ49 | CAVRPGNQFYF | TRBV27 | TRBJ2-5 | CASSLSGTWGETQYF | KLGGALQAK | YES | NO | NO |
| 6922 | TRAV22 | TRAJ49 | CAVERDQFYF | TRBV27 | TRBJ2-5 | CASSQLGAWETQYF | KLGGALQAK | YES | NO | NO |
| 6926 | TRAV29DV5 | TRAJ20 | CAASPYDYKLSF | TRBV27 | TRBJ2-5 | CASSTTGQGGDKTQYF | KLGGALQAK | YES | NO | NO |
| 6927 | TRAV29DV5 | TRAJ23 | CAASDFQGGKLIF | TRBV27 | TRBJ2-5 | CASRPIGGRVVTQYF | KLGGALQAK | YES | NO | NO |
| 6928 | TRAV29DV5 | TRAJ57 | CAAIPQGGSEKLVF | TRBV27 | TRBJ2-5 | CASTGLAGGIGETQYF | KLGGALQAK | YES | NO | NO |
| 6929 | TRAV3 | TRAJ31 | CAVRDISGARLMF | TRBV27 | TRBJ2-5 | CASSVRTSGGQETQYF | KLGGALQAK | NO | NO | NO |
| 6931 | TRAV30 | TRAJ26 | CGTEPSDNYGQNFVF | TRBV27 | TRBJ2-5 | CASRPNPGGTQYF | KLGGALQAK | YES | NO | NO |
| 6932 | TRAV30 | TRAJ52 | CGTEIGGGTSYGKLTF | TRBV27 | TRBJ2-5 | CASGTETQYF | KLGGALQAK | YES | NO | NO |
| 6933 | TRAV34 | TRAJ23 | CGANGGNQGGKLIF | TRBV27 | TRBJ2-5 | CASSLSGTWGETQYF | KLGGALQAK | YES | NO | NO |
| 6934 | TRAV5 | TRAJ12 | CAESLMDSSYKLIF | TRBV27 | TRBJ2-5 | CASSLGGGTVRTQYF | KLGGALQAK | YES | NO | NO |
| 6935 | TRAV8-6 | TRAJ21 | CAVSDPNNFNKFYF | TRBV27 | TRBJ2-5 | CASRPRGYDETQYF | KLGGALQAK | YES | NO | NO |
| 6936 | TRAV8-6 | TRAJ4 | CAVISSGGYNKLIF | TRBV27 | TRBJ2-5 | CASSSLWEETQYF | KLGGALQAK | YES | NO | NO |
| 6938 | TRAV1-2 | TRAJ33 | CSSVDSNYQLIW | TRBV28 | TRBJ2-5 | CASESPGQGVRGTQYF | KLGGALQAK | YES | NO | NO |
| 6939 | TRAV12-1 | TRAJ52 | CVVKPNAGGTSYGKLTF | TRBV28 | TRBJ2-5 | CASSLYLGVGETQYF | KLGGALQAK | YES | NO | NO |
| 6941 | TRAV14DV4 | TRAJ37 | CAMREGDSSNTGKLIF | TRBV28 | TRBJ2-5 | CASTVGAGGLETQYF | KLGGALQAK | YES | NO | NO |
| 6942 | TRAV17 | TRAJ16 | CATDAWSDGQKLLF | TRBV28 | TRBJ2-5 | CASTVGDRAYQETQYF | KLGGALQAK | NO | NO | NO |
| 6943 | TRAV21 | TRAJ21 | CATYNFNKFYF | TRBV28 | TRBJ2-5 | CASSFETGGRTQYF | KLGGALQAK | YES | NO | NO |
| 6945 | TRAV29DV5 | TRAJ31 | CAASALGNNARLMF | TRBV28 | TRBJ2-5 | CASSLWGLAGGMGRTQYF | KLGGALQAK | YES | NO | NO |
| 6947 | TRAV12-2 | TRAJ56 | CAVIVGANSKLTF | TRBV29-1 | TRBJ2-5 | CSVEVAGGQETQYF | KLGGALQAK | YES | NO | NO |
| 6948 | TRAV27 | TRAJ23 | CAGEIYNQGGKLIF | TRBV29-1 | TRBJ2-5 | CSVEGEGGETQYF | KLGGALQAK | YES | NO | NO |

| | | | | | | | | | | |
|---|---|---|---|---|---|---|---|---|---|---|
| 6950 | TRAV3 | TRAJ40 | CAVRGFTSGTYKYIF | TRBV29-1 | TRBJ2-5 | CSVEHYQETQYF | KLGGALQAK | NO | NO | NO |
| 6951 | TRAV38-2DV8 | TRAJ48 | CAHFGNEKLTF | TRBV29-1 | TRBJ2-5 | CSVGSGIGGETQYF | KLGGALQAK | YES | NO | NO |
| 6953 | TRAV9-2 | TRAJ9 | CALSTGGFKTIF | TRBV29-1 | TRBJ2-5 | CSGLYQETQYF | KLGGALQAK | YES | NO | NO |
| 6955 | TRAV12-1 | TRAJ39 | CVVNAFNNAGNMLTF | TRBV3-1 | TRBJ2-5 | CASSQGPLGAGVQTQYF | KLGGALQAK | YES | NO | NO |
| 6956 | TRAV12-2 | TRAJ30 | CAVKRGDDKIIF | TRBV3-1 | TRBJ2-5 | CASSQGLAGIATQETQYF | KLGGALQAK | YES | NO | NO |
| 6957 | TRAV12-2 | TRAJ4 | CAVSAGGYNKLIF | TRBV3-1 | TRBJ2-5 | CASSQRTSAETQYF | KLGGALQAK | YES | NO | NO |
| 6959 | TRAV12-3 | TRAJ39 | CAIGPGNMLTF | TRBV3-1 | TRBJ2-5 | CASSHGKTQYF | KLGGALQAK | YES | NO | NO |
| 6960 | TRAV14DV4 | TRAJ26 | CAMCPVNYGQNFVF | TRBV3-1 | TRBJ2-5 | CASSQVGLAGGPETQYF | KLGGALQAK | YES | NO | NO |
| 6961 | TRAV17 | TRAJ56 | CATELYGANSKLTF | TRBV3-1 | TRBJ2-5 | CASTGLAGQETQYF | KLGGALQAK | YES | NO | NO |
| 6962 | TRAV2 | TRAJ15 | CAVTNQAGTALIF | TRBV3-1 | TRBJ2-5 | CASSPPTEETQYF | KLGGALQAK | NO | NO | NO |
| 6962 | TRAV2 | TRAJ15 | CAVTNQAGTALIF | TRBV3-1 | TRBJ2-5 | CASSPPTEETQYF | RLRAEAQVK | NO | NO | NO |
| 6963 | TRAV21 | TRAJ22 | CAVMGSRGSARQLTF | TRBV3-1 | TRBJ2-5 | CASSQDRDSPQETQYF | KLGGALQAK | YES | NO | NO |
| 6966 | TRAV27 | TRAJ40 | CADTTSGTYKYIF | TRBV3-1 | TRBJ2-5 | CASSHGKTQYF | KLGGALQAK | YES | NO | NO |
| 6968 | TRAV36DV7 | TRAJ45 | CAVGSHSGGGADGLTF | TRBV3-1 | TRBJ2-5 | CASSQETSGTNLGTQYF | KLGGALQAK | NO | NO | NO |
| 6970 | TRAV38-2DV8 | TRAJ47 | CAYRSAGEYGNKLVF | TRBV3-1 | TRBJ2-5 | CASNDRLFGTAYQETQYF | KLGGALQAK | YES | NO | NO |
| 6971 | TRAV41 | TRAJ54 | CAPTQGAQKLVF | TRBV3-1 | TRBJ2-5 | CAIAGGGAQYF | KLGGALQAK | YES | NO | NO |
| 6972 | TRAV5 | TRAJ39 | CAEMTFWAGNMLTF | TRBV3-1 | TRBJ2-5 | CASSPPSPKETQYF | KLGGALQAK | YES | NO | NO |
| 6977 | TRAV8-6 | TRAJ21 | CAVSAPDNFNKFYF | TRBV3-1 | TRBJ2-5 | CASSQGSGTYF | KLGGALQAK | YES | NO | NO |
| 6978 | TRAV14DV4 | TRAJ29 | CAMREGRKSGNTPLVF | TRBV30 | TRBJ2-5 | CAWSRGTGETQYF | KLGGALQAK | NO | NO | NO |
| 6979 | TRAV14DV4 | TRAJ42 | CAMREVGYGGSQGNLIF | TRBV30 | TRBJ2-5 | CAAMRVDQETQYF | KLGGALQAK | NO | NO | NO |
| 6981 | TRAV24 | TRAJ12 | CAFLMDSSYKLIF | TRBV30 | TRBJ2-5 | CAWSGRQGAGETQYF | KLGGALQAK | YES | NO | NO |
| 6982 | TRAV27 | TRAJ45 | CAITRSGGGADGLTF | TRBV30 | TRBJ2-5 | CAWDGGRGGETQYF | KLGGALQAK | YES | NO | NO |
| 6988 | TRAV17 | TRAJ49 | CATGRRNQFYF | TRBV4-1 | TRBJ2-5 | CASQSVAGDQETQYF | KLGGALQAK | NO | NO | NO |
| 6989 | TRAV22 | TRAJ9 | CAVEGFKTIF | TRBV4-1 | TRBJ2-5 | CASSQDGGAGQETQYF | KLGGALQAK | YES | NO | NO |
| 6990 | TRAV29DV5 | TRAJ20 | CAASGVGDYKLSF | TRBV4-1 | TRBJ2-5 | CASSHKPGQGSETQYF | KLGGALQAK | YES | NO | NO |
| 6991 | TRAV29DV5 | TRAJ29 | CAATTNSGNTPLVF | TRBV4-1 | TRBJ2-5 | CASSQLGWETQYF | KLGGALQAK | YES | NO | NO |
| 6992 | TRAV29DV5 | TRAJ45 | CAASAHAGGGADGLTF | TRBV4-1 | TRBJ2-5 | CASSHPIGGAETQYF | KLGGALQAK | YES | NO | NO |
| 6994 | TRAV3 | TRAJ28 | CAVRDPSYSGAGSYQLTF | TRBV4-1 | TRBJ2-5 | CASSQTSGGRETQYF | KLGGALQAK | YES | NO | NO |
| 6998 | TRAV8-4 | TRAJ23 | CAVSVGGGKLIF | TRBV4-1 | TRBJ2-5 | CASSQQETQYF | KLGGALQAK | YES | NO | NO |
| 7002 | TRAV21 | TRAJ18 | CAATRADRGSTLGRLYF | TRBV4-2 | TRBJ2-5 | CASSLDFDRGRLETQYF | KLGGALQAK | YES | NO | NO |
| 7003 | TRAV21 | TRAJ26 | CAVWDNYGQNFVF | TRBV4-2 | TRBJ2-5 | CASSQEWLAGGPGTQYF | KLGGALQAK | YES | NO | NO |
| 7006 | TRAV19 | TRAJ43 | CALSETANDMRF | TRBV4-3 | TRBJ2-5 | CASSQRRQITGGGETQYF | KLGGALQAK | YES | NO | NO |
| 7007 | TRAV24 | TRAJ53 | CAFPVGGGSNYKLTF | TRBV4-3 | TRBJ2-5 | CASSQGSRPQETQYF | KLGGALQAK | YES | NO | NO |
| 7008 | TRAV35 | TRAJ34 | CAGLSSDKLIF | TRBV4-3 | TRBJ2-5 | CASSQDRSGTSTETQYF | KLGGALQAK | YES | NO | NO |
| 7008 | TRAV35 | TRAJ34 | CAGLSSDKLIF | TRBV4-3 | TRBJ2-5 | CASSQDRSGTSTETQYF | AVFDRKSDAK | YES | NO | NO |
| 7009 | TRAV38-2DV8 | TRAJ57 | CAYRSGGGSEKLVF | TRBV4-3 | TRBJ2-5 | CASSQGTYQETQYF | KLGGALQAK | YES | NO | NO |
| 7011 | TRAV5 | TRAJ47 | CADELVF | TRBV4-3 | TRBJ2-5 | CASSQDRSGTSTETQYF | KLGGALQAK | YES | NO | NO |
| 7012 | TRAV1-2 | TRAJ33 | CAAMDSNYQLIW | TRBV5-1 | TRBJ2-5 | CASSLEWGGETQYF | KLGGALQAK | YES | NO | NO |
| 7013 | TRAV1-2 | TRAJ39 | CAGNAGNMLTF | TRBV5-1 | TRBJ2-5 | CASSLERTSARETQYF | KLGGALQAK | YES | NO | NO |
| 7014 | TRAV10 | TRAJ48 | CVVSFLDLSNFGNEKLTF | TRBV5-1 | TRBJ2-5 | CASSLYSGRIGGETQYF | KLGGALQAK | YES | NO | NO |
| 7015 | TRAV12-2 | TRAJ42 | CAVNFYGGSQGNLIF | TRBV5-1 | TRBJ2-5 | CASSLGQIEETQYF | KLGGALQAK | YES | NO | NO |
| 7017 | TRAV13-1 | TRAJ45 | CAASEGGADGLTF | TRBV5-1 | TRBJ2-5 | CASSLGQETQYF | KLGGALQAK | YES | NO | NO |
| 7018 | TRAV14DV4 | TRAJ36 | CAMGIPPPSTGANNLFF | TRBV5-1 | TRBJ2-5 | CASSLSQGWSETQYF | KLGGALQAK | YES | NO | NO |
| 7019 | TRAV17 | TRAJ22 | CATDERKGSARQLTF | TRBV5-1 | TRBJ2-5 | CASTLSLDPQETQYF | KLGGALQAK | YES | NO | NO |
| 7020 | TRAV17 | TRAJ8 | CATNTGFQKLVF | TRBV5-1 | TRBJ2-5 | CASRQGRQETQYF | KLGGALQAK | YES | NO | NO |
| 7023 | TRAV19 | TRAJ53 | CALSEAWDSGGSNYKLTF | TRBV5-1 | TRBJ2-5 | CASSLGYQGEGETQYF | KLGGALQAK | YES | NO | NO |
| 7024 | TRAV24 | TRAJ36 | CAFLGGANNLFF | TRBV5-1 | TRBJ2-5 | CASSLEWGGETQYF | AVFDRKSDAK | YES | NO | NO |
| 7025 | TRAV25 | TRAJ29 | CAGRNSGNTPLVF | TRBV5-1 | TRBJ2-5 | CASSLVGPGRELGETQYF | KLGGALQAK | YES | NO | NO |
| 7027 | TRAV3 | TRAJ6 | CAVRVASGGSYIPTF | TRBV5-1 | TRBJ2-5 | CASSPSGLLLAGGETQYF | KLGGALQAK | YES | NO | NO |
| 7030 | TRAV38-2DV8 | TRAJ17 | CAYLSAGNKLTF | TRBV5-1 | TRBJ2-5 | CASSPGTSGGETQYF | KLGGALQAK | YES | NO | NO |
| 7032 | TRAV38-2DV8 | TRAJ58 | CARTGGSRLTF | TRBV5-1 | TRBJ2-5 | CASSLGTGVKETQYF | KLGGALQAK | YES | NO | NO |
| 7033 | TRAV39 | TRAJ35 | CVVGFGNVLHC | TRBV5-1 | TRBJ2-5 | CASSLGYGGAGETQYF | KLGGALQAK | NO | NO | NO |
| 7035 | TRAV4 | TRAJ34 | CLVGDNTDKLIF | TRBV5-1 | TRBJ2-5 | CASSTYGTMEETQYF | RLRAEAQVK | YES | NO | YES |
| 7037 | TRAV6 | TRAJ13 | CALSSGGYQKVTF | TRBV5-1 | TRBJ2-5 | CASSSYGLTSGGAPEETQYF | KLGGALQAK | NO | NO | NO |
| 7038 | TRAV8-1 | TRAJ49 | CAVNPTGNQFYF | TRBV5-1 | TRBJ2-5 | CASSLPRDRVRETQYF | KLGGALQAK | YES | NO | NO |
| 7039 | TRAV8-3 | TRAJ35 | CALVGFGNVLHC | TRBV5-1 | TRBJ2-5 | CASSPGEYQETQYF | KLGGALQAK | NO | NO | NO |
| 7041 | TRAV8-3 | TRAJ43 | CAVGAFVTDMRF | TRBV5-1 | TRBJ2-5 | CASSSLSERGRETQYF | KLGGALQAK | YES | NO | NO |
| 7042 | TRAV19 | TRAJ34 | CALSEAYNTDKLIF | TRBV5-4 | TRBJ2-5 | CASSYNGRSLQETQYF | KLGGALQAK | YES | NO | NO |
| 7043 | TRAV38-2DV8 | TRAJ49 | CAYRSATGNQFYF | TRBV5-4 | TRBJ2-5 | CASSYGQGWLGQETQYF | KLGGALQAK | YES | NO | NO |
| 7046 | TRAV1-1 | TRAJ31 | CAVRDPLNNNARLMF | TRBV5-5 | TRBJ2-5 | CASSLGDRHEETQYF | KLGGALQAK | YES | NO | NO |
| 7047 | TRAV10 | TRAJ17 | CVVSDREAAGNKLTF | TRBV5-5 | TRBJ2-5 | CASSSGTSGGPQTQYF | KLGGALQAK | YES | NO | NO |
| 7048 | TRAV12-3 | TRAJ9 | CAMSYDTGGFKTIF | TRBV5-6 | TRBJ2-5 | CASSLGERQETQYF | KLGGALQAK | YES | NO | NO |
| 7049 | TRAV13-1 | TRAJ27 | CAAQDTNAGKSTF | TRBV5-6 | TRBJ2-5 | CASSLGSGETQYF | KLGGALQAK | YES | NO | NO |
| 7051 | TRAV14DV4 | TRAJ22 | CAMRRPISSGSARQLTF | TRBV5-6 | TRBJ2-5 | CASSFQGSPQETQYF | KLGGALQAK | YES | NO | NO |
| 7051 | TRAV14DV4 | TRAJ22 | CAMRRPISSGSARQLTF | TRBV5-6 | TRBJ2-5 | CASSFQGSPQETQYF | RLRAEAQVK | YES | NO | NO |
| 7052 | TRAV14DV4 | TRAJ22 | CAMRRPISSGSARQLTF | TRBV5-6 | TRBJ2-5 | CASSARQGQETQYF | KLGGALQAK | YES | NO | NO |
| 7053 | TRAV14DV4 | TRAJ22 | CAMRRPISSGSARQLTF | TRBV5-6 | TRBJ2-5 | CASSETSGQETQYF | KLGGALQAK | YES | NO | NO |
| 7055 | TRAV14DV4 | TRAJ5 | CAMRAMDTGRRALTF | TRBV5-6 | TRBJ2-5 | CASSPETGRETQYF | KLGGALQAK | YES | NO | NO |
| 7056 | TRAV19 | TRAJ24 | CALKSSWGKLQF | TRBV5-6 | TRBJ2-5 | CASMLAGSEETQYF | KLGGALQAK | YES | NO | NO |
| 7058 | TRAV2 | TRAJ21 | CAVEDVNNFNKFYF | TRBV5-6 | TRBJ2-5 | CASSSILAGAETQYF | KLGGALQAK | NO | NO | NO |
| 7060 | TRAV21 | TRAJ20 | CAVMGDGDYKLSF | TRBV5-6 | TRBJ2-5 | CASINVVGQGSETQYF | KLGGALQAK | YES | NO | NO |
| 7061 | TRAV3 | TRAJ37 | CAVRDGGMGSNTGKLIF | TRBV5-6 | TRBJ2-5 | CASSPETGGETQYF | KLGGALQAK | YES | NO | NO |
| 7062 | TRAV3 | TRAJ8 | CAVRDKSGFQKLVF | TRBV5-6 | TRBJ2-5 | CASSTGPTSGGETQYF | KLGGALQAK | YES | NO | NO |
| 7063 | TRAV8-4 | TRAJ30 | CAVSASGRDDKIIF | TRBV5-6 | TRBJ2-5 | CASSPRDPGETQYF | KLGGALQAK | YES | NO | NO |
| 7064 | TRAV12-2 | TRAJ42 | CAGWRTMNYGGSQGNLIF | TRBV5-8 | TRBJ2-5 | CASSLRDTTPAQYF | KLGGALQAK | YES | NO | NO |
| 7065 | TRAV12-3 | TRAJ27 | CAMSEPANAGKSTF | TRBV5-8 | TRBJ2-5 | CASSFRTGGETQYF | KLGGALQAK | YES | NO | NO |
| 7066 | TRAV13-1 | TRAJ42 | CAAGGSQGNLIF | TRBV5-8 | TRBJ2-5 | CASSLLAGVEETQYF | KLGGALQAK | YES | NO | NO |
| 7067 | TRAV1-1 | TRAJ27 | CAVRDTNAGKSTF | TRBV6-1 | TRBJ2-5 | CASSELAGGPETQYF | RLRAEAQVK | YES | NO | YES |
| 7069 | TRAV21 | TRAJ24 | CAVIGTTDSWGKLQF | TRBV6-1 | TRBJ2-5 | CASSGGYEVVTQYF | KLGGALQAK | YES | NO | NO |
| 7070 | TRAV21 | TRAJ34 | CAVTALYNTDKLIF | TRBV6-1 | TRBJ2-5 | CASTTRGGGTQETQYF | KLGGALQAK | YES | NO | NO |
| 7073 | TRAV1-2 | TRAJ33 | CAVRDRDYQLIW | TRBV6-3 | TRBJ2-5 | CASSTASAPETQYF | KLGGALQAK | YES | NO | NO |
| 7074 | TRAV12-2 | TRAJ49 | CAVSGNQFYF | TRBV6-3 | TRBJ2-5 | CASSYYHRGETQYF | KLGGALQAK | YES | NO | NO |
| 7075 | TRAV12-3 | TRAJ18 | CAMSEGGSTLGRLYF | TRBV6-3 | TRBJ2-5 | CASSYSLAGAEETQYF | KLGGALQAK | YES | NO | NO |
| 7076 | TRAV12-3 | TRAJ39 | CAMSAERSNMLTF | TRBV6-3 | TRBJ2-5 | CASSYGVNQETQYF | KLGGALQAK | YES | NO | NO |
| 7079 | TRAV14DV4 | TRAJ37 | CAMREGWSSNTGKLIF | TRBV6-3 | TRBJ2-5 | CASSGLAGGPYF | KLGGALQAK | YES | NO | NO |
| 7080 | TRAV17 | TRAJ21 | CATDSYNFNKFYF | TRBV6-3 | TRBJ2-5 | CASSGLERETQYF | KLGGALQAK | YES | NO | NO |
| 7081 | TRAV19 | TRAJ47 | CALGGMEYGNKLVF | TRBV6-3 | TRBJ2-5 | CASSYVPPSGNQETQYF | KLGGALQAK | YES | NO | NO |
| 7083 | TRAV29DV5 | TRAJ52 | CAARVSNAGGTSYGKLTF | TRBV6-3 | TRBJ2-5 | CASSLTTDGDAWETQYF | KLGGALQAK | YES | NO | NO |
| 7087 | TRAV5 | TRAJ43 | CAETPYNNNDMRF | TRBV6-3 | TRBJ2-5 | CASTRTSGGAWETQYF | KLGGALQAK | YES | NO | NO |
| 7088 | TRAV8-1 | TRAJ7 | CAVNAGGNNRLAF | TRBV6-3 | TRBJ2-5 | CASSSGQGLRAQYF | KLGGALQAK | YES | NO | NO |
| 7089 | TRAV8-3 | TRAJ48 | CAVGQGFGNEKLTF | TRBV6-3 | TRBJ2-5 | CASSYQTIGGSETQYF | KLGGALQAK | YES | NO | NO |
| 7090 | TRAV1-2 | TRAJ33 | CAFMDSNYQLIW | TRBV6-4 | TRBJ2-5 | CASSETSGGTETQYF | KLGGALQAK | YES | NO | NO |
| 7093 | TRAV29DV5 | TRAJ22 | CAASAPGSARQLTF | TRBV6-4 | TRBJ2-5 | CASSANRAETQYF | KLGGALQAK | YES | NO | NO |
| 7096 | TRAV12-2 | TRAJ7 | CAVRVGNNRLAF | TRBV6-5 | TRBJ2-5 | CASSYLGEETQYF | KLGGALQAK | NO | NO | NO |
| 7098 | TRAV13-1 | TRAJ53 | CAATSPSGGSNYKLTF | TRBV6-5 | TRBJ2-5 | CASSYRTRETQYF | KLGGALQAK | NO | NO | NO |
| 7099 | TRAV14DV4 | TRAJ23 | CAMREGGKLIF | TRBV6-5 | TRBJ2-5 | CASSSTLGTGGFETQYF | KLGGALQAK | NO | NO | NO |
| 7101 | TRAV17 | TRAJ11 | CATPSGYSTLTF | TRBV6-5 | TRBJ2-5 | CASSYSLLAAQETQYF | KLGGALQAK | NO | NO | NO |
| 7102 | TRAV17 | TRAJ30 | CATDLLNRDDKIIF | TRBV6-5 | TRBJ2-5 | CASTGEETQYF | KLGGALQAK | NO | NO | NO |
| 7105 | TRAV21 | TRAJ20 | CAVNSNDYKLSF | TRBV6-5 | TRBJ2-5 | CASGPSAQETQYF | KLGGALQAK | NO | NO | NO |
| 7105 | TRAV21 | TRAJ20 | CAVNSNDYKLSF | TRBV6-5 | TRBJ2-5 | CASGPSAQETQYF | IVTDFSVIK | NO | NO | NO |
| 7107 | TRAV26-1 | TRAJ52 | CIVRVLDSYGKLTF | TRBV6-5 | TRBJ2-5 | CASSYGRQETQYF | KLGGALQAK | NO | NO | NO |
| 7108 | TRAV3 | TRAJ17 | CAVRDILKAAGNKLTF | TRBV6-5 | TRBJ2-5 | CASSRTEETQYF | KLGGALQAK | YES | NO | NO |
| 7109 | TRAV3 | TRAJ34 | CAGYNTDKLIF | TRBV6-5 | TRBJ2-5 | CASSEGLAGGGGETQYF | KLGGALQAK | NO | NO | NO |
| 7110 | TRAV34 | TRAJ49 | CGAAMNTGNQFYF | TRBV6-5 | TRBJ2-5 | CASSYRRVQETQYF | KLGGALQAK | NO | NO | NO |

| | | | | | | | | | |
|---|---|---|---|---|---|---|---|---|---|
| 7111 | TRAV38-2DV8 | TRAJ22 | CSISGSARQLTF | TRBV6-5 | TRBJ2-5 | CASSIYPGEETQYF | KLGGALQAK | NO | NO | NO |
| 7113 | TRAV5 | TRAJ12 | CAVPFMDSSYKLIF | TRBV6-5 | TRBJ2-5 | CASSDRTGVGETQYF | KLGGALQAK | NO | NO | NO |
| 7114 | TRAV8-6 | TRAJ34 | CAVSDQYNTDKLIF | TRBV6-5 | TRBJ2-5 | CASIRSGTQYF | KLGGALQAK | NO | NO | NO |
| 7116 | TRAV21 | TRAJ34 | CAVRDNTDKLIF | TRBV6-6 | TRBJ2-5 | CATSYLTSGGYPQETQYF | KLGGALQAK | NO | NO | NO |
| 7117 | TRAV21 | TRAJ42 | CAAEGYGGSQGNLIF | TRBV6-6 | TRBJ2-5 | CASSYQAGPQETQYF | KLGGALQAK | NO | NO | NO |
| 7118 | TRAV8-3 | TRAJ26 | CAVAYGQNFVF | TRBV6-6 | TRBJ2-5 | CASRAGMYQETQYF | KLGGALQAK | NO | NO | NO |
| 7119 | TRAV8-6 | TRAJ39 | CAVSARNAGNMLTF | TRBV6-6 | TRBJ2-5 | CASQPESGGEETQYF | KLGGALQAK | NO | NO | NO |
| 7120 | TRAV12-2 | TRAJ6 | CAGVSGGSYIPTF | TRBV7-2 | TRBJ2-5 | CASSQYGGGDQYF | KLGGALQAK | YES | NO | NO |
| 7121 | TRAV13-2 | TRAJ42 | CAEQMNYGGSQGNLIF | TRBV7-2 | TRBJ2-5 | CASSLISGREETQYF | KLGGALQAK | YES | NO | NO |
| 7124 | TRAV29DV5 | TRAJ49 | CAASGNQFYF | TRBV7-3 | TRBJ2-5 | CASSLNSKETQYF | KLGGALQAK | NO | NO | NO |
| 7125 | TRAV4 | TRAJ29 | CLVGDSDSGNTPLVF | TRBV7-3 | TRBJ2-5 | CASSLNTGRKETQYF | KLGGALQAK | NO | NO | NO |
| 7126 | TRAV8-3 | TRAJ47 | CAVAPVRDGNKLVF | TRBV7-3 | TRBJ2-5 | CASSLGAGLRETQYF | KLGGALQAK | NO | NO | NO |
| 7127 | TRAV13-1 | TRAJ54 | CAASTGAQKLVF | TRBV7-6 | TRBJ2-5 | CASRQGRKETQYF | KLGGALQAK | YES | NO | NO |
| 7129 | TRAV23DV6 | TRAJ18 | CAASSRGSTLGRLYF | TRBV7-6 | TRBJ2-5 | CASSFGGGAPETQYF | KLGGALQAK | YES | NO | NO |
| 7131 | TRAV29DV5 | TRAJ20 | CAAKTLNDYKLSF | TRBV7-6 | TRBJ2-5 | CASSLTQGVGETQYF | KLGGALQAK | YES | NO | NO |
| 7137 | TRAV8-4 | TRAJ44 | CAVIVTTGTASKLTF | TRBV7-6 | TRBJ2-5 | CASSLAGPGLAGQETQYF | KLGGALQAK | YES | NO | NO |
| 7139 | TRAV1-1 | TRAJ28 | CAVAYSGAGSYQLTF | TRBV7-7 | TRBJ2-5 | CASSLGGNEETQYF | KLGGALQAK | YES | NO | NO |
| 7141 | TRAV13-2 | TRAJ12 | CAENMDSSYKLIF | TRBV7-7 | TRBJ2-5 | CASSLEAAKETQYF | KLGGALQAK | YES | NO | NO |
| 7143 | TRAV17 | TRAJ20 | CATDANDYKLSF | TRBV7-7 | TRBJ2-5 | CASSSTGIEETQYF | KLGGALQAK | YES | NO | NO |
| 7149 | TRAV13-1 | TRAJ23 | CAASSNQGGKLIF | TRBV7-8 | TRBJ2-5 | CASSVTDRAQYF | KLGGALQAK | YES | NO | NO |
| 7151 | TRAV19 | TRAJ17 | CALSVIKAAGNKLTF | TRBV7-8 | TRBJ2-5 | CASSPLPRLAGEETQYF | KLGGALQAK | YES | NO | NO |
| 7152 | TRAV19 | TRAJ21 | CALSEAGNFNKFYF | TRBV7-8 | TRBJ2-5 | CASSSEAGARETQYF | KLGGALQAK | YES | NO | NO |
| 7154 | TRAV26-1 | TRAJ40 | CIVREPGTYKYIF | TRBV7-8 | TRBJ2-5 | CASTPLGQGEETQYF | KLGGALQAK | YES | NO | NO |
| 7155 | TRAV38-2DV8 | TRAJ30 | CAYLDPRDDKIIF | TRBV7-8 | TRBJ2-5 | CASSPLPRLAGEETQYF | KLGGALQAK | YES | NO | NO |
| 7156 | TRAV38-2DV8 | TRAJ42 | CAYRSTFYGGSQGNLIF | TRBV7-8 | TRBJ2-5 | CASSLRLALLETQYF | KLGGALQAK | YES | NO | NO |
| 7159 | TRAV8-3 | TRAJ13 | CAVGAMGGYQKVTF | TRBV7-8 | TRBJ2-5 | CASSLDRLELQETQYF | KLGGALQAK | YES | NO | NO |
| 7160 | TRAV8-3 | TRAJ13 | CAVALRGGYQKVTF | TRBV7-8 | TRBJ2-5 | CASSLKETQYF | KLGGALQAK | YES | NO | NO |
| 7161 | TRAV8-6 | TRAJ29 | CAVSGNTPLVF | TRBV7-8 | TRBJ2-5 | CASSPLPRLAGEETQYF | KLGGALQAK | YES | NO | NO |
| 7164 | TRAV1-2 | TRAJ35 | CAVLGGFGNVLHC | TRBV7-9 | TRBJ2-5 | CASSLDDLQETQYF | KLGGALQAK | NO | NO | NO |
| 7165 | TRAV12-1 | TRAJ37 | CVVKEDSSNTGKLIF | TRBV7-9 | TRBJ2-5 | CASSPGLAGVGHF | KLGGALQAK | NO | NO | NO |
| 7166 | TRAV12-2 | TRAJ49 | CAVNLALYTGNQFYF | TRBV7-9 | TRBJ2-5 | CASSLVVGVQVIQYF | KLGGALQAK | YES | NO | NO |
| 7168 | TRAV13-1 | TRAJ53 | CAAPSGGSNYKLTF | TRBV7-9 | TRBJ2-5 | CASSFPTSGRGLAETQYF | KLGGALQAK | YES | NO | NO |
| 7169 | TRAV19 | TRAJ10 | CALSEKAGGGNKLTF | TRBV7-9 | TRBJ2-5 | CASRPDGSQETQYF | KLGGALQAK | YES | NO | NO |
| 7170 | TRAV19 | TRAJ23 | CALSEALIYNQGGKLIF | TRBV7-9 | TRBJ2-5 | CASRRLTQETQYF | KLGGALQAK | YES | NO | NO |
| 7171 | TRAV19 | TRAJ32 | CALSEAGGATNKLIF | TRBV7-9 | TRBJ2-5 | CASSPISRQETQYF | KLGGALQAK | NO | NO | NO |
| 7173 | TRAV19 | TRAJ43 | CALSDRVINNDMRF | TRBV7-9 | TRBJ2-5 | CASSGRGWETQYF | KLGGALQAK | YES | NO | NO |
| 7173 | TRAV19 | TRAJ43 | CALSDRVINNDMRF | TRBV7-9 | TRBJ2-5 | CASSGRGWETQYF | RLRAEAQVK | YES | NO | NO |
| 7174 | TRAV19 | TRAJ5 | CALGQDTGRRALTF | TRBV7-9 | TRBJ2-5 | CASSLEEGDAREAQYF | KLGGALQAK | YES | NO | NO |
| 7176 | TRAV21 | TRAJ4 | CAVRSGGYNKLIF | TRBV7-9 | TRBJ2-5 | CASSLVYGGNQETQYF | KLGGALQAK | YES | NO | NO |
| 7177 | TRAV24 | TRAJ36 | CAFTTGANNLFF | TRBV7-9 | TRBJ2-5 | CASSHRIGGETQYF | KLGGALQAK | YES | NO | NO |
| 7177 | TRAV24 | TRAJ36 | CAFTTGANNLFF | TRBV7-9 | TRBJ2-5 | CASSHRIGGETQYF | AVFDRKSDAK | YES | NO | NO |
| 7179 | TRAV26-2 | TRAJ39 | CILSGYNAGNMLTF | TRBV7-9 | TRBJ2-5 | CASSRTDRDRETQYF | KLGGALQAK | YES | NO | NO |
| 7181 | TRAV38-2DV8 | TRAJ29 | CAYRGNSGNTPLVF | TRBV7-9 | TRBJ2-5 | CASSLGLAGGPGETQYF | KLGGALQAK | YES | NO | NO |
| 7182 | TRAV38-2DV8 | TRAJ31 | CAYCFNARLMF | TRBV7-9 | TRBJ2-5 | CASSLAEYQETQYF | KLGGALQAK | YES | NO | NO |
| 7183 | TRAV38-2DV8 | TRAJ31 | CAYRSAFARLMF | TRBV7-9 | TRBJ2-5 | CASGGGPSETQYF | KLGGALQAK | YES | NO | NO |
| 7184 | TRAV8-3 | TRAJ26 | CAVGEDGQNFVF | TRBV7-9 | TRBJ2-5 | CASSWGGWGETQYF | KLGGALQAK | YES | NO | NO |
| 7185 | TRAV8-4 | TRAJ8 | CAVMRNTGFQKLVF | TRBV7-9 | TRBJ2-5 | CASSFVNQETQYF | KLGGALQAK | YES | NO | NO |
| 7187 | TRAV1-2 | TRAJ28 | CAVGTSGAGSYQLTF | TRBV9 | TRBJ2-5 | CASSGVTGDETQYF | KLGGALQAK | YES | NO | NO |
| 7188 | TRAV1-2 | TRAJ3 | CAVSGPGYSSASKIIF | TRBV9 | TRBJ2-5 | CASSSSSGGAQETQYF | KLGGALQAK | YES | NO | NO |
| 7189 | TRAV12-1 | TRAJ34 | CVVIVAADKLIF | TRBV9 | TRBJ2-5 | CASSAGTSEGTQYF | KLGGALQAK | YES | NO | NO |
| 7190 | TRAV12-2 | TRAJ27 | CAVNTNAGKSTF | TRBV9 | TRBJ2-5 | CASSQGQNQETQYF | KLGGALQAK | YES | NO | NO |
| 7192 | TRAV13-1 | TRAJ39 | CAASWGAGNMLTF | TRBV9 | TRBJ2-5 | CASSDRNAGTQYF | KLGGALQAK | YES | NO | NO |
| 7193 | TRAV13-2 | TRAJ56 | CAENTGANSKLTF | TRBV9 | TRBJ2-5 | CASSDSSETQYF | KLGGALQAK | YES | NO | NO |
| 7194 | TRAV14DV4 | TRAJ27 | CAMREGRTNAGKSTF | TRBV9 | TRBJ2-5 | CASSATSGSYVETQYF | KLGGALQAK | YES | NO | NO |
| 7195 | TRAV14DV4 | TRAJ8 | CAMRPMNTGFQKLVF | TRBV9 | TRBJ2-5 | CASGPGGAETQYF | KLGGALQAK | YES | NO | NO |
| 7198 | TRAV20 | TRAJ11 | CAVQMNSGYSTLTF | TRBV9 | TRBJ2-5 | CASSAGTSEGTQYF | KLGGALQAK | YES | NO | NO |
| 7199 | TRAV21 | TRAJ37 | CAVLSWGNTGKLIF | TRBV9 | TRBJ2-5 | CASSYTGTSGETQYF | KLGGALQAK | YES | NO | NO |
| 7201 | TRAV3 | TRAJ10 | CAVRDTWLTGGGNKLTF | TRBV9 | TRBJ2-5 | CASSAHQGLAGAGETQYF | KLGGALQAK | NO | NO | NO |
| 7202 | TRAV4 | TRAJ12 | CLVGDMASSSYKLIF | TRBV9 | TRBJ2-5 | CASSEPRLGLVETQYF | KLGGALQAK | YES | NO | NO |
| 7203 | TRAV4 | TRAJ31 | CLVGEDNARLMF | TRBV9 | TRBJ2-5 | CASSEAGGRETQYF | KLGGALQAK | NO | NO | NO |
| 7205 | TRAV8-1 | TRAJ12 | CAVNLDSSYKLIF | TRBV9 | TRBJ2-5 | CASNGAGGPETQYF | KLGGALQAK | YES | NO | NO |
| 7206 | TRAV23DV6 | TRAJ28 | CAAHRVDSGAGSYQLTF | TRBV10-1 | TRBJ2-6 | CASSRDLSGANVLTF | KLGGALQAK | YES | NO | NO |
| 7207 | TRAV27 | TRAJ38 | CAGYSINAGNNRKLIW | TRBV10-2 | TRBJ2-6 | CASSPGQVTGANVLTF | KLGGALQAK | YES | NO | NO |
| 7211 | TRAV8-2 | TRAJ23 | CVVSVSYNQGGKLIF | TRBV11-2 | TRBJ2-6 | CASSSPSGPLLSGANVLTF | KLGGALQAK | YES | NO | NO |
| 7212 | TRAV9-2 | TRAJ16 | CALSESWF | TRBV11-2 | TRBJ2-6 | CASSPKLTVSLGANVLTF | KLGGALQAK | NO | NO | NO |
| 7214 | TRAV21 | TRAJ58 | CAVGSVETSGSRLTF | TRBV11-3 | TRBJ2-6 | CASIGVPGANVLTF | KLGGALQAK | YES | NO | NO |
| 7216 | TRAV12-3 | TRAJ26 | CAMNNYGQNFVF | TRBV12-4 | TRBJ2-6 | CASSFANSGANVLTF | KLGGALQAK | YES | NO | NO |
| 7218 | TRAV38-2DV8 | TRAJ43 | CAYRRLNMRF | TRBV12-4 | TRBJ2-6 | CASSHRDRGRKGSGANVLTF | KLGGALQAK | NO | NO | NO |
| 7220 | TRAV26-2 | TRAJ42 | CILSYGGSQGNLIF | TRBV13 | TRBJ2-6 | CASSFQGGYSGANVLTF | KLGGALQAK | NO | NO | NO |
| 7223 | TRAV1-2 | TRAJ33 | CAVRDSNYQLIW | TRBV15 | TRBJ2-6 | CATSRESSGANVLTF | KLGGALQAK | YES | NO | NO |
| 7224 | TRAV21 | TRAJ17 | CAVRQTAAGNKLTF | TRBV15 | TRBJ2-6 | CATSYEDNSGANVLTF | KLGGALQAK | YES | NO | NO |
| 7225 | TRAV29DV5 | TRAJ24 | CAALRTDSWGKLQF | TRBV18 | TRBJ2-6 | CASSFGTSPGANVLTF | KLGGALQAK | YES | NO | NO |
| 7226 | TRAV38-2DV8 | TRAJ32 | CASYGGATNKLIF | TRBV18 | TRBJ2-6 | CASVSSELTSGANVLTF | KLGGALQAK | YES | NO | NO |
| 7229 | TRAV27 | TRAJ23 | CAGARYNQGGKLIF | TRBV19 | TRBJ2-6 | CASSITNSGANVLTF | KLGGALQAK | YES | NO | NO |
| 7230 | TRAV1-2 | TRAJ12 | CAVKMDSSYKLIF | TRBV2 | TRBJ2-6 | CASSGAGAMLGANVLTF | KLGGALQAK | YES | NO | NO |
| 7231 | TRAV25 | TRAJ40 | CAGPIPPFSGTYKYIF | TRBV2 | TRBJ2-6 | CASTPGGGSGANVLTF | KLGGALQAK | YES | NO | NO |
| 7232 | TRAV30 | TRAJ20 | CGTDSNDYKLSF | TRBV2 | TRBJ2-6 | CARQPLRGANVLTF | KLGGALQAK | YES | NO | NO |
| 7232 | TRAV30 | TRAJ20 | CGTDSNDYKLSF | TRBV2 | TRBJ2-6 | CARQPLRGANVLTF | RLRAEAQVK | YES | NO | NO |
| 7234 | TRAV12-1 | TRAJ23 | CVVGNYNQGGKLIF | TRBV20-1 | TRBJ2-6 | CSARRELDSGANVLTF | KLGGALQAK | YES | NO | NO |
| 7236 | TRAV12-3 | TRAJ39 | CAIGPGNMLTF | TRBV24-1 | TRBJ2-6 | CATSGAAYSGANVLTF | KLGGALQAK | YES | NO | NO |
| 7241 | TRAV26-2 | TRAJ11 | CILRDVGSGYSTLTF | TRBV24-1 | TRBJ2-6 | CATTTGDNSGANVLTF | KLGGALQAK | NO | NO | NO |
| 7243 | TRAV3 | TRAJ29 | CAVRDGGNTPLVF | TRBV24-1 | TRBJ2-6 | CATSEPTDATGYGANVLTF | KLGGALQAK | NO | NO | NO |
| 7244 | TRAV12-3 | TRAJ11 | CAMTRGRPTF | TRBV27 | TRBJ2-6 | CASSLHQNKYLSGANVLTF | KLGGALQAK | YES | NO | NO |
| 7245 | TRAV13-2 | TRAJ48 | CAESRNFGNEKLTF | TRBV27 | TRBJ2-6 | CASSSRTDRGKRFSGANVLTF | KLGGALQAK | NO | NO | NO |
| 7247 | TRAV8-6 | TRAJ5 | CAVNIRGWDTGRRALTF | TRBV27 | TRBJ2-6 | CASSTEYSGANVLTF | KLGGALQAK | NO | NO | NO |
| 7249 | TRAV17 | TRAJ34 | CATNLKDTDKLIF | TRBV28 | TRBJ2-6 | CASSLPDGTDLSGANVLTF | KLGGALQAK | YES | NO | NO |
| 7250 | TRAV8-3 | TRAJ20 | CAVGPTNDYKLSF | TRBV29-1 | TRBJ2-6 | CSVDDGNSGANVLTF | KLGGALQAK | YES | NO | NO |
| 7251 | TRAV1-2 | TRAJ33 | CAVKDSNYQLIW | TRBV3-1 | TRBJ2-6 | CASSQSVSGANVLTF | KLGGALQAK | YES | NO | NO |
| 7252 | TRAV38-1 | TRAJ28 | CAFSVSGAGSYQLTF | TRBV3-1 | TRBJ2-6 | CASSPKVTVFGANVLTF | KLGGALQAK | YES | NO | NO |
| 7254 | TRAV19 | TRAJ56 | CALSGGNTGANSKLTF | TRBV30 | TRBJ2-6 | CAGLGANVLTF | KLGGALQAK | NO | NO | NO |
| 7255 | TRAV22 | TRAJ28 | CAVARGAGSYQLTF | TRBV30 | TRBJ2-6 | CAWSGWNRDLAGANVLTF | KLGGALQAK | NO | NO | NO |
| 7256 | TRAV22 | TRAJ33 | CAVEQDSNYQLIW | TRBV30 | TRBJ2-6 | CAWALPPGANVLTF | KLGGALQAK | NO | NO | NO |
| 7257 | TRAV8-1 | TRAJ33 | CAVNSNYQLIW | TRBV30 | TRBJ2-6 | CAWTGDGANVLTF | KLGGALQAK | NO | NO | NO |
| 7260 | TRAV20 | TRAJ48 | CAVQHLPISNFGNEKLTF | TRBV4-1 | TRBJ2-6 | CASRFAQGSGANVLTF | KLGGALQAK | YES | NO | NO |
| 7270 | TRAV8-2 | TRAJ42 | CVVSDYGGSQGNLIF | TRBV5-1 | TRBJ2-6 | CAITVFSGANVLTF | KLGGALQAK | YES | NO | NO |
| 7271 | TRAV8-3 | TRAJ49 | CAVDLNTGNQFYF | TRBV5-4 | TRBJ2-6 | CASSGRAGPLLSGANVLTF | KLGGALQAK | YES | NO | NO |
| 7274 | TRAV38-2DV8 | TRAJ54 | CAYTGWPQGAQKLVF | TRBV5-6 | TRBJ2-6 | CASSLELAGESGANVLTF | KLGGALQAK | YES | NO | NO |
| 7275 | TRAV13-1 | TRAJ20 | CAASIFYDYKLSF | TRBV5-8 | TRBJ2-6 | CASSLPGQIISGANVLTF | KLGGALQAK | YES | NO | NO |
| 7277 | TRAV16 | TRAJ15 | CALVNQAGTALIF | TRBV6-3 | TRBJ2-6 | CASSYFLGSGAGPGANVLTF | KLGGALQAK | NO | NO | NO |
| 7279 | TRAV36DV7 | TRAJ43 | CAVGPYNNNDMRF | TRBV6-3 | TRBJ2-6 | CASRRSGANVLTF | KLGGALQAK | NO | NO | NO |
| 7280 | TRAV10 | TRAJ49 | CVVTLNQFYF | TRBV6-5 | TRBJ2-6 | CASSSPGANVLTF | KLGGALQAK | YES | NO | NO |
| 7282 | TRAV13-1 | TRAJ39 | CAASWGNAGNMLTF | TRBV6-5 | TRBJ2-6 | CATQGGRGGGANVLTF | KLGGALQAK | YES | NO | NO |
| 7283 | TRAV36DV7 | TRAJ40 | CAGRSSGTYKYIF | TRBV6-5 | TRBJ2-6 | CASSPGLETISGANVLTF | KLGGALQAK | NO | NO | NO |
| 7284 | TRAV5 | TRAJ10 | CAERPGYTGGGNKLTF | TRBV6-5 | TRBJ2-6 | CASSYSGANVLTF | KLGGALQAK | YES | NO | NO |

| | | | | | | | | | | |
|---|---|---|---|---|---|---|---|---|---|---|
| 7285 | TRAV8-3 | TRAJ6 | CAVVLHSYIPTF | TRBV6-5 | TRBJ2-6 | CASSYSLFGAGGSGANVLTF | RLRAEAQVK | NO | NO | NO |
| 7286 | TRAV8-6 | TRAJ53 | CAVSDGLGGGSNYKLTF | TRBV6-5 | TRBJ2-6 | CASSSGDGWGGANVLTF | KLGGALQAK | YES | NO | NO |
| 7287 | TRAV19 | TRAJ28 | CALSETAGAGSYQLTF | TRBV6-6 | TRBJ2-6 | CASSYSIPVDSGANVLTF | KLGGALQAK | YES | NO | NO |
| 7290 | TRAV3 | TRAJ26 | CAVRDNYGQNFVF | TRBV7-7 | TRBJ2-6 | CASSLGRRSGGANVLTF | KLGGALQAK | NO | NO | NO |
| 7291 | TRAV30 | TRAJ52 | CGTVNAGGTSYGKLTF | TRBV7-8 | TRBJ2-6 | CASSLYTISGANVLTF | KLGGALQAK | YES | NO | NO |
| 7295 | TRAV19 | TRAJ28 | CALGDSGAGSYQLTF | TRBV7-9 | TRBJ2-6 | CASREIQGSGANVLTF | KLGGALQAK | YES | NO | NO |
| 7295 | TRAV19 | TRAJ28 | CALGDSGAGSYQLTF | TRBV7-9 | TRBJ2-6 | CASREIQGSGANVLTF | AVFDRKSDAK | YES | NO | NO |
| 7295 | TRAV19 | TRAJ28 | CALGDSGAGSYQLTF | TRBV7-9 | TRBJ2-6 | CASREIQGSGANVLTF | RLRAEAQVK | YES | NO | NO |
| 7299 | TRAV1-2 | TRAJ9 | CAVLYTGGFKTIF | TRBV9 | TRBJ2-6 | CASSVEGAGGANVLTF | KLGGALQAK | YES | NO | NO |
| 7304 | TRAV21 | TRAJ42 | CAVSEGSQGNLIF | TRBV9 | TRBJ2-6 | CASSVLAGPYSGANVLTF | KLGGALQAK | YES | NO | NO |
| 7314 | TRAV12-2 | TRAJ45 | CAVSGGGGGADGLTF | TRBV10-2 | TRBJ2-7 | CASSGSGISYEQYF | KLGGALQAK | YES | NO | NO |
| 7317 | TRAV21 | TRAJ7 | CAVWSGGNNRLAF | TRBV10-2 | TRBJ2-7 | CASKDSGFSYEQYF | KLGGALQAK | YES | NO | NO |
| 7318 | TRAV22 | TRAJ15 | CAVPNQAGTALIF | TRBV10-2 | TRBJ2-7 | CASSGLADYEQYV | KLGGALQAK | NO | NO | NO |
| 7319 | TRAV3 | TRAJ44 | CAGTGTASKLTF | TRBV10-2 | TRBJ2-7 | CASSRQGEFYEQYF | KLGGALQAK | NO | NO | NO |
| 7320 | TRAV39 | TRAJ31 | CAVDYTRLMF | TRBV10-2 | TRBJ2-7 | CASSANSGGYEQYF | KLGGALQAK | YES | NO | NO |
| 7322 | TRAV5 | TRAJ24 | CAERTTTDSWGKLQF | TRBV10-2 | TRBJ2-7 | CASRGLQGANEQYF | KLGGALQAK | YES | NO | NO |
| 7323 | TRAV8-2 | TRAJ4 | CVVSDPEGTFSGGYNKLIF | TRBV10-2 | TRBJ2-7 | CASSESETGPYEQYV | KLGGALQAK | NO | NO | NO |
| 7327 | TRAV12-1 | TRAJ42 | CVVWGSQGNLIF | TRBV10-3 | TRBJ2-7 | CAIRGTSGVDEQYV | KLGGALQAK | NO | NO | NO |
| 7328 | TRAV13-2 | TRAJ4 | CAETPPGGYNKLIF | TRBV10-3 | TRBJ2-7 | CAIKTSGRAYEQYF | KLGGALQAK | YES | NO | NO |
| 7329 | TRAV14DV4 | TRAJ29 | CAMWAGNTPLVF | TRBV10-3 | TRBJ2-7 | CAISESVAIREQYV | KLGGALQAK | NO | NO | NO |
| 7331 | TRAV25 | TRAJ44 | CAGYTGTASKLTF | TRBV10-3 | TRBJ2-7 | CAISEGSGYEQYV | KLGGALQAK | NO | NO | NO |
| 7334 | TRAV13-2 | TRAJ53 | CAVQGSGGSNYKLTF | TRBV11-1 | TRBJ2-7 | CASSIQGAYEQYF | KLGGALQAK | YES | NO | NO |
| 7337 | TRAV1-1 | TRAJ29 | CAVFGNTPLVF | TRBV11-2 | TRBJ2-7 | CASSLDTSGSPYEQYF | KLGGALQAK | YES | NO | NO |
| 7338 | TRAV1-1 | TRAJ4 | CAVVFSGGYNKLIF | TRBV11-2 | TRBJ2-7 | CASSLDGTVTSYEQYV | KLGGALQAK | NO | NO | NO |
| 7340 | TRAV12-1 | TRAJ36 | CVSGANNLFF | TRBV11-2 | TRBJ2-7 | CASSLGAGIIYEQYF | KLGGALQAK | YES | NO | NO |
| 7341 | TRAV12-1 | TRAJ6 | CVVNPLTSGGSYIPTF | TRBV11-2 | TRBJ2-7 | CASSPGPTHEQYF | KLGGALQAK | YES | NO | NO |
| 7343 | TRAV12-3 | TRAJ54 | CAMSHLIIQGAQKLVF | TRBV11-2 | TRBJ2-7 | CASSLGQGLINYEQYF | KLGGALQAK | YES | NO | NO |
| 7344 | TRAV13-1 | TRAJ18 | CAASSGIADRGSTLGRLYF | TRBV11-2 | TRBJ2-7 | CASSRWTGVEEQYV | KLGGALQAK | NO | NO | NO |
| 7346 | TRAV13-2 | TRAJ13 | CAENSGGYQKVTF | TRBV11-2 | TRBJ2-7 | CASSHGQGYEQYF | KLGGALQAK | YES | NO | NO |
| 7347 | TRAV13-2 | TRAJ52 | CAEMPNAGGTSYGKLTF | TRBV11-2 | TRBJ2-7 | CASSWGHHYEQYF | KLGGALQAK | YES | NO | NO |
| 7350 | TRAV16 | TRAJ37 | CALSSMGSNTGKLIF | TRBV11-2 | TRBJ2-7 | CASSLVTSGETYEQYF | KLGGALQAK | NO | NO | NO |
| 7351 | TRAV17 | TRAJ11 | CATGGGYSTLTF | TRBV11-2 | TRBJ2-7 | CATKEEVTYEQYF | KLGGALQAK | YES | NO | NO |
| 7353 | TRAV17 | TRAJ20 | CATDDYKLSF | TRBV11-2 | TRBJ2-7 | CASSFSSGGDYEQYF | KLGGALQAK | YES | NO | NO |
| 7354 | TRAV17 | TRAJ36 | CATDARAGANNLFF | TRBV11-2 | TRBJ2-7 | CASSLGNSYEQYV | KLGGALQAK | NO | NO | NO |
| 7356 | TRAV21 | TRAJ4 | CAGGAFSGGYNKLIF | TRBV11-2 | TRBJ2-7 | CASSLVETGSYEQYF | KLGGALQAK | YES | NO | NO |
| 7358 | TRAV23DV6 | TRAJ45 | CAASDSGGGADGLTF | TRBV11-2 | TRBJ2-7 | CASSQQGQGSGYEQYV | KLGGALQAK | NO | NO | NO |
| 7359 | TRAV25 | TRAJ33 | CAGRDSNYQLIW | TRBV11-2 | TRBJ2-7 | CASSLDAGYEQYF | KLGGALQAK | YES | NO | NO |
| 7360 | TRAV26-2 | TRAJ23 | CILRDGGGKLIF | TRBV11-2 | TRBJ2-7 | CASSPGTAAYEQYF | KLGGALQAK | YES | NO | NO |
| 7363 | TRAV38-2DV8 | TRAJ31 | CAYRLGNNARLMF | TRBV11-2 | TRBJ2-7 | CASSARSGTNLYEQYF | KLGGALQAK | YES | NO | NO |
| 7364 | TRAV4 | TRAJ27 | CLVGDLNTNAGKSTF | TRBV11-2 | TRBJ2-7 | CASSLVIQEQYF | KLGGALQAK | YES | NO | NO |
| 7366 | TRAV40 | TRAJ56 | CLLRTGANSKLTF | TRBV11-2 | TRBJ2-7 | CASSLGTSGMGEQYF | KLGGALQAK | NO | NO | NO |
| 7367 | TRAV41 | TRAJ22 | CAVMPGSARQLTF | TRBV11-2 | TRBJ2-7 | CASSLFRSYEQYF | KLGGALQAK | NO | NO | NO |
| 7369 | TRAV1-2 | TRAJ21 | CAVNNFNKFYF | TRBV11-3 | TRBJ2-7 | CASSLDMSGSYEQYV | KLGGALQAK | NO | NO | NO |
| 7372 | TRAV12-3 | TRAJ30 | CAMSWGRDDKIIF | TRBV11-3 | TRBJ2-7 | CASSKGLTGNYEQYF | KLGGALQAK | YES | NO | NO |
| 7372 | TRAV12-3 | TRAJ30 | CAMSWGRDDKIIF | TRBV11-3 | TRBJ2-7 | CASSKGLTGNYEQYF | AVFDRKSDAK | YES | NO | NO |
| 7374 | TRAV13-1 | TRAJ53 | CAASIHSGGSNYKLTF | TRBV11-3 | TRBJ2-7 | CASSSRDKSYEQYV | KLGGALQAK | NO | NO | NO |
| 7375 | TRAV14DV4 | TRAJ21 | CAMSSLNKFYF | TRBV11-3 | TRBJ2-7 | CASSLLAGRGYEQYV | KLGGALQAK | NO | NO | NO |
| 7376 | TRAV14DV4 | TRAJ53 | CAMRSGGSNYKLTF | TRBV11-3 | TRBJ2-7 | CASSPMGLSYEQYV | KLGGALQAK | NO | NO | NO |
| 7380 | TRAV38-2DV8 | TRAJ20 | CAYRSVYKLSF | TRBV11-3 | TRBJ2-7 | CASSRTTYEQYV | KLGGALQAK | NO | NO | NO |
| 7381 | TRAV6 | TRAJ30 | CALDISPLDRDDKIIF | TRBV11-3 | TRBJ2-7 | CASSLVADYTYEQYV | KLGGALQAK | NO | NO | NO |
| 7382 | TRAV12-1 | TRAJ39 | CVVSLDNAGNMLTF | TRBV12-3 | TRBJ2-7 | CASRGQGVAQYV | KLGGALQAK | NO | NO | NO |
| 7385 | TRAV17 | TRAJ57 | CATQGGSEKLVF | TRBV12-3 | TRBJ2-7 | CASTRRTGDEQYF | KLGGALQAK | YES | NO | NO |
| 7387 | TRAV21 | TRAJ47 | CAVREYGNKLVF | TRBV12-3 | TRBJ2-7 | CASSWTSGSYEQYF | KLGGALQAK | YES | NO | NO |
| 7388 | TRAV29DV5 | TRAJ40 | CAASATSGTYKYIF | TRBV12-3 | TRBJ2-7 | CASSLLGNEQYF | KLGGALQAK | YES | YES | NO |
| 7393 | TRAV12-2 | TRAJ56 | CAVNTGANSKLTF | TRBV12-4 | TRBJ2-7 | CASSLYRGWYEQYF | KLGGALQAK | YES | NO | NO |
| 7394 | TRAV13-1 | TRAJ8 | CAAEGSFQKLVF | TRBV12-4 | TRBJ2-7 | CASSPLGPNYEQYF | KLGGALQAK | YES | NO | NO |
| 7397 | TRAV17 | TRAJ52 | CATDASTSYGKLTF | TRBV12-4 | TRBJ2-7 | CASSYIIIEQYF | KLGGALQAK | YES | NO | NO |
| 7400 | TRAV23DV6 | TRAJ31 | CAVPSSLRLMF | TRBV12-4 | TRBJ2-7 | CASSGAGTGPYEQYF | KLGGALQAK | YES | NO | NO |
| 7402 | TRAV24 | TRAJ20 | CAFQFSNDYKLSF | TRBV12-4 | TRBJ2-7 | CASSRSPGPLYEQYF | KLGGALQAK | YES | NO | NO |
| 7405 | TRAV26-2 | TRAJ42 | CIPNMNYGGSQGNLIF | TRBV12-4 | TRBJ2-7 | CASSFGQGGEQYF | KLGGALQAK | YES | NO | NO |
| 7406 | TRAV27 | TRAJ44 | CAGAIAGTASKLTF | TRBV12-4 | TRBJ2-7 | CASSSTGGYEQYF | KLGGALQAK | YES | NO | NO |
| 7408 | TRAV29DV5 | TRAJ30 | CAARHGFDDKIIF | TRBV12-4 | TRBJ2-7 | CASSLGTGSYEQYF | KLGGALQAK | YES | NO | NO |
| 7410 | TRAV4 | TRAJ30 | CLVGDPRDDKIIF | TRBV12-4 | TRBJ2-7 | CASRLGNHPPYEQYV | KLGGALQAK | NO | NO | NO |
| 7411 | TRAV8-1 | TRAJ17 | CAVRAAGNKLTF | TRBV12-4 | TRBJ2-7 | CASSLAGGSYEQYF | KLGGALQAK | YES | NO | NO |
| 7412 | TRAV8-1 | TRAJ9 | CAVIPTNTGGFKTIF | TRBV12-4 | TRBJ2-7 | CASGQGYEQYF | KLGGALQAK | YES | NO | NO |
| 7415 | TRAV8-4 | TRAJ54 | CAVQGAQKLVF | TRBV12-4 | TRBJ2-7 | CASSTRDRGSYEQYF | KLGGALQAK | YES | NO | NO |
| 7419 | TRAV25 | TRAJ37 | CAGLSSNTGKLIF | TRBV12-5 | TRBJ2-7 | CASGLWESYEQYF | KLGGALQAK | YES | NO | NO |
| 7421 | TRAV5 | TRAJ9 | CAESYTGGFKTIF | TRBV12-5 | TRBJ2-7 | CASGLTPPSRDRGTYEQYF | KLGGALQAK | YES | NO | NO |
| 7425 | TRAV13-1 | TRAJ8 | CAASRMNTGFQKLVF | TRBV13 | TRBJ2-7 | CASSLLGTGVVHEQYF | KLGGALQAK | YES | NO | NO |
| 7426 | TRAV14DV4 | TRAJ13 | CAMREGSGGYQKVTF | TRBV13 | TRBJ2-7 | CASSLRLAPYEQYV | KLGGALQAK | NO | NO | NO |
| 7427 | TRAV14DV4 | TRAJ17 | CAMRAIKAAGNKLTF | TRBV13 | TRBJ2-7 | CASSLTGESYEQYF | KLGGALQAK | YES | NO | NO |
| 7428 | TRAV14DV4 | TRAJ30 | CAMREGTGVNRDDKIIF | TRBV13 | TRBJ2-7 | CASSLSGNPYEQYV | KLGGALQAK | NO | NO | NO |
| 7429 | TRAV19 | TRAJ39 | CALSENNAGNMLTF | TRBV13 | TRBJ2-7 | CASSLGGVGSSYEQYF | KLGGALQAK | YES | NO | NO |
| 7430 | TRAV19 | TRAJ45 | CALSVRSGGGADGLTF | TRBV13 | TRBJ2-7 | CASSSRGQGGYEQYF | RLRAEAQVK | YES | NO | NO |
| 7432 | TRAV26-1 | TRAJ9 | CISRLPGGFKTIF | TRBV13 | TRBJ2-7 | CASGGALNYEQYF | KLGGALQAK | YES | NO | NO |
| 7435 | TRAV29DV5 | TRAJ57 | CAASPSWGGSEKLVF | TRBV13 | TRBJ2-7 | CASSSGQQAYEQYF | KLGGALQAK | YES | NO | NO |
| 7436 | TRAV30 | TRAJ57 | CGTGFTQGGSEKLVF | TRBV13 | TRBJ2-7 | CASSSPGTGISYEQYF | KLGGALQAK | YES | NO | NO |
| 7437 | TRAV38-2DV8 | TRAJ29 | CASCMISGNTPLVF | TRBV13 | TRBJ2-7 | CASTAVPNYEQYF | KLGGALQAK | YES | NO | NO |
| 7438 | TRAV38-2DV8 | TRAJ32 | CAYRSRRGATNKLIF | TRBV13 | TRBJ2-7 | CASSLAGTISYEQYV | KLGGALQAK | NO | NO | NO |
| 7439 | TRAV38-2DV8 | TRAJ45 | CAYRTYSGGGADGLTF | TRBV13 | TRBJ2-7 | CASRLQGAMGYEQYF | KLGGALQAK | YES | NO | NO |
| 7441 | TRAV6 | TRAJ13 | CALRNSGGYQKVTF | TRBV13 | TRBJ2-7 | CASSFGAGGPEQYF | KLGGALQAK | YES | NO | NO |
| 7442 | TRAV12-2 | TRAJ10 | CARRWILTGGGNKLTF | TRBV14 | TRBJ2-7 | CASSQDYEQYF | KLGGALQAK | NO | NO | NO |
| 7444 | TRAV13-1 | TRAJ41 | CAGDFSGYALNF | TRBV14 | TRBJ2-7 | CASSQQGPEQYF | KLGGALQAK | NO | NO | NO |
| 7445 | TRAV21 | TRAJ7 | CAVKSGNRLAF | TRBV14 | TRBJ2-7 | CASSFAGYEQYF | KLGGALQAK | NO | NO | NO |
| 7446 | TRAV22 | TRAJ34 | CASGSYNTDKLIF | TRBV14 | TRBJ2-7 | CASSQGYEQYF | KLGGALQAK | NO | NO | NO |
| 7447 | TRAV27 | TRAJ26 | CAAKFTYYGQNFVF | TRBV14 | TRBJ2-7 | CASSQGQGIHEQYF | KLGGALQAK | NO | NO | NO |
| 7448 | TRAV29DV5 | TRAJ49 | CAASAPGNQFYF | TRBV14 | TRBJ2-7 | CASSQAQGWYEQYF | KLGGALQAK | NO | NO | NO |
| 7449 | TRAV38-2DV8 | TRAJ36 | CAYWGSPGANNLFF | TRBV14 | TRBJ2-7 | CASSQELGTEQYF | KLGGALQAK | NO | NO | NO |
| 7451 | TRAV9-2 | TRAJ33 | CAREHMDSNYQLIW | TRBV14 | TRBJ2-7 | CASSSGREIWYEQYF | KLGGALQAK | NO | NO | NO |
| 7452 | TRAV9-2 | TRAJ34 | CALRSYNTDKLIF | TRBV14 | TRBJ2-7 | CASSHRTSGGAFPYEQYF | KLGGALQAK | NO | NO | NO |
| 7454 | TRAV1-2 | TRAJ33 | CAVMDSNYQLIW | TRBV15 | TRBJ2-7 | CATSRDSGHPDRQYF | KLGGALQAK | YES | NO | NO |
| 7455 | TRAV10 | TRAJ53 | CVVSADSGGSNYKLTF | TRBV15 | TRBJ2-7 | CATSRDRLGGGRYEQYF | KLGGALQAK | YES | NO | NO |
| 7456 | TRAV12-1 | TRAJ39 | CVVNGNNAGNMLTF | TRBV15 | TRBJ2-7 | CATSSRDRVGEQYF | KLGGALQAK | YES | NO | NO |
| 7457 | TRAV12-2 | TRAJ13 | CAVKTGGYQKVTF | TRBV15 | TRBJ2-7 | CATSRGRQGTLSYEQYF | KLGGALQAK | YES | NO | NO |
| 7458 | TRAV12-2 | TRAJ26 | CAVNSPYGQNFVF | TRBV15 | TRBJ2-7 | CATSRGAGFDEQYF | KLGGALQAK | YES | NO | NO |
| 7461 | TRAV19 | TRAJ33 | CASNLKGSNYQLIW | TRBV15 | TRBJ2-7 | CATSPGGGRDLYEQYF | KLGGALQAK | NO | NO | NO |
| 7461 | TRAV19 | TRAJ33 | CASNLKGSNYQLIW | TRBV15 | TRBJ2-7 | CATSPGGGRDLYEQYF | IVTDFSVIK | NO | NO | NO |
| 7463 | TRAV2 | TRAJ24 | CAPPGGTTDSWGKLQF | TRBV15 | TRBJ2-7 | CATSRDDIGTRDYEQYF | KLGGALQAK | NO | NO | NO |
| 7464 | TRAV22 | TRAJ49 | CAVRSTGNQFYF | TRBV15 | TRBJ2-7 | CATSRDVRSGGSYEQYF | KLGGALQAK | YES | NO | NO |
| 7467 | TRAV3 | TRAJ13 | CAVRDRGYQKVTF | TRBV15 | TRBJ2-7 | CATSRDRGTGSYEQYF | KLGGALQAK | NO | NO | NO |
| 7468 | TRAV3 | TRAJ22 | CAVRAAISSGSARQLTF | TRBV15 | TRBJ2-7 | CATSRTSASSYEQYF | KLGGALQAK | NO | NO | NO |
| 7469 | TRAV3 | TRAJ48 | CAVRDWGSNFGNEKLTF | TRBV15 | TRBJ2-7 | CATSRDRLAGSSYEQYF | KLGGALQAK | NO | NO | NO |
| 7470 | TRAV6 | TRAJ29 | CALAVAGNTPLVF | TRBV15 | TRBJ2-7 | CATSDSVKPYEQYF | KLGGALQAK | YES | NO | NO |
| 7471 | TRAV9-2 | TRAJ3 | CALKSSSASKIIF | TRBV15 | TRBJ2-7 | CATSTDLSGVTYEQYF | KLGGALQAK | YES | NO | NO |

| | | | | | | | | | | |
|---|---|---|---|---|---|---|---|---|---|---|
| 7473 | TRAV12-1 | TRAJ45 | CVVSSGGGADGLTF | TRBV18 | TRBJ2-7 | CASSPWTGRPYEQYV | KLGGALQAK | NO | NO | NO |
| 7476 | TRAV12-2 | TRAJ50 | CAVKGSYDKVIF | TRBV18 | TRBJ2-7 | CASSPPTGGKGEQYF | KLGGALQAK | YES | NO | NO |
| 7477 | TRAV12-2 | TRAJ58 | CAVAQETSGSRLTF | TRBV18 | TRBJ2-7 | CASSRQPSTYEQYF | KLGGALQAK | YES | NO | NO |
| 7478 | TRAV19 | TRAJ48 | CALSEANFGNEKLTF | TRBV18 | TRBJ2-7 | CASSPYTGTGPEQYF | KLGGALQAK | NO | NO | NO |
| 7479 | TRAV19 | TRAJ54 | CALSEAEGQGAQKLVF | TRBV18 | TRBJ2-7 | CASSLLAGLAYEQYV | KLGGALQAK | NO | NO | NO |
| 7480 | TRAV22 | TRAJ58 | CAVLETSGSRLTF | TRBV18 | TRBJ2-7 | CASSFRLARIYEQYF | KLGGALQAK | YES | NO | NO |
| 7481 | TRAV23DV6 | TRAJ23 | CAASVGDQGGKLIF | TRBV18 | TRBJ2-7 | CASSGGTGGPLEQYF | KLGGALQAK | NO | NO | NO |
| 7482 | TRAV24 | TRAJ30 | CAFRDDKIIF | TRBV18 | TRBJ2-7 | CASSPAGGTLYEQYF | KLGGALQAK | YES | NO | NO |
| 7483 | TRAV27 | TRAJ39 | CAGIAGNMLTF | TRBV18 | TRBJ2-7 | CASSPPIRGRARYEQYF | KLGGALQAK | YES | NO | NO |
| 7485 | TRAV1-1 | TRAJ21 | CAVSPGDNFNKFYF | TRBV19 | TRBJ2-7 | CASSIKGHYEQYF | KLGGALQAK | YES | NO | NO |
| 7486 | TRAV1-1 | TRAJ36 | CAALVGANNLFF | TRBV19 | TRBJ2-7 | CASSPPGRASYEQYF | KLGGALQAK | YES | NO | NO |
| 7488 | TRAV12-2 | TRAJ15 | CAVNQAGTALIF | TRBV19 | TRBJ2-7 | CASSIDGGVAYEQYF | KLGGALQAK | YES | NO | NO |
| 7489 | TRAV12-2 | TRAJ4 | CAVKPMFSGGYNKLIF | TRBV19 | TRBJ2-7 | CASSIRAAYEQYF | KLGGALQAK | YES | NO | NO |
| 7491 | TRAV12-2 | TRAJ42 | CAVNYNTGYGGSQGNLIF | TRBV19 | TRBJ2-7 | CASSISSGPAYEQYV | KLGGALQAK | NO | NO | NO |
| 7493 | TRAV12-3 | TRAJ21 | CAMSLYNFNKFYF | TRBV19 | TRBJ2-7 | CASGTSTNIYEQYF | KLGGALQAK | YES | NO | NO |
| 7495 | TRAV12-3 | TRAJ37 | CAMISSNTGKLIF | TRBV19 | TRBJ2-7 | CASSIRSSYEQYF | KLGGALQAK | YES | NO | NO |
| 7497 | TRAV12-3 | TRAJ52 | CAMSYINAGGTSYGKLTF | TRBV19 | TRBJ2-7 | CASSISSYEQYF | KLGGALQAK | YES | NO | NO |
| 7501 | TRAV14DV4 | TRAJ22 | CAMRVGSARQLTF | TRBV19 | TRBJ2-7 | CASSVSKARAFGEQYF | KLGGALQAK | YES | NO | NO |
| 7502 | TRAV14DV4 | TRAJ49 | CAMREGTWNQFYF | TRBV19 | TRBJ2-7 | CASSHGTGTYEQYF | KLGGALQAK | YES | NO | NO |
| 7503 | TRAV14DV4 | TRAJ9 | CAMRDGTGGFKTIF | TRBV19 | TRBJ2-7 | CASSIRRDVYYEQYF | KLGGALQAK | YES | NO | NO |
| 7504 | TRAV17 | TRAJ18 | CATANDRGSTLGRLYF | TRBV19 | TRBJ2-7 | CASSITGQHYEQYF | KLGGALQAK | YES | NO | NO |
| 7505 | TRAV17 | TRAJ29 | CATDAINSGNTPLVF | TRBV19 | TRBJ2-7 | CASTGTGGIYEQYF | KLGGALQAK | YES | NO | NO |
| 7508 | TRAV19 | TRAJ28 | CALSEAANSGAGSYQLTF | TRBV19 | TRBJ2-7 | CASSIVGTGSYEQYF | KLGGALQAK | YES | NO | NO |
| 7509 | TRAV19 | TRAJ39 | CALSGVNNNAGNMLTF | TRBV19 | TRBJ2-7 | CASSLSRSRGSYEQYF | KLGGALQAK | YES | NO | NO |
| 7510 | TRAV19 | TRAJ40 | CALQRGTYKYIF | TRBV19 | TRBJ2-7 | CASSARDTSYEQYF | KLGGALQAK | YES | NO | NO |
| 7511 | TRAV20 | TRAJ58 | CAVRGTSGSRLTF | TRBV19 | TRBJ2-7 | CASSMGWTGVVYEQYF | KLGGALQAK | YES | NO | NO |
| 7512 | TRAV20 | TRAJ8 | CAVTTGFQKLVF | TRBV19 | TRBJ2-7 | CASSPGMGPYEQYF | KLGGALQAK | YES | NO | NO |
| 7513 | TRAV21 | TRAJ17 | CAVYQGMIKAAGNKLTF | TRBV19 | TRBJ2-7 | CASSPGAGLSYEQYV | KLGGALQAK | NO | NO | NO |
| 7514 | TRAV21 | TRAJ33 | CAVFMDSNYQLIW | TRBV19 | TRBJ2-7 | CASSSGAYEQYF | KLGGALQAK | YES | NO | NO |
| 7516 | TRAV21 | TRAJ42 | CAVSGGSQGNLIF | TRBV19 | TRBJ2-7 | CASSIVGGVNEQYF | KLGGALQAK | YES | NO | NO |
| 7518 | TRAV24 | TRAJ39 | CASWGPYNAGNMLTF | TRBV19 | TRBJ2-7 | CASSIGDYEQYF | KLGGALQAK | YES | NO | NO |
| 7520 | TRAV25 | TRAJ31 | CAGGPLGNARLMF | TRBV19 | TRBJ2-7 | CASSLSSGTGLRV | KLGGALQAK | NO | NO | NO |
| 7521 | TRAV25 | TRAJ9 | CAGINTGGFKTIF | TRBV19 | TRBJ2-7 | CASSIGRGMDYEQYF | KLGGALQAK | YES | NO | NO |
| 7522 | TRAV26-2 | TRAJ49 | CILRDVSNTGNQFYF | TRBV19 | TRBJ2-7 | CASSILSGAYEQYF | KLGGALQAK | YES | NO | NO |
| 7524 | TRAV29DV5 | TRAJ29 | CAASAVLFEGNTPLVF | TRBV19 | TRBJ2-7 | CASSIDWHPWRAKGNYEQYF | RLRAEAQVK | NO | NO | NO |
| 7525 | TRAV29DV5 | TRAJ30 | CAAGLNRDDKIIF | TRBV19 | TRBJ2-7 | CASSIVGVPQTYEQYF | KLGGALQAK | YES | NO | NO |
| 7526 | TRAV29DV5 | TRAJ33 | CAASGRDSNYQLIW | TRBV19 | TRBJ2-7 | CASSIGTVGYEQYF | KLGGALQAK | YES | NO | NO |
| 7527 | TRAV29DV5 | TRAJ53 | CAASERSGGSNYKLTF | TRBV19 | TRBJ2-7 | CASSIVGSWEQYF | KLGGALQAK | YES | NO | NO |
| 7529 | TRAV30 | TRAJ36 | CGTAQTGANNLFF | TRBV19 | TRBJ2-7 | CASSPGLAYEQYF | KLGGALQAK | YES | NO | NO |
| 7530 | TRAV30 | TRAJ40 | CGTEMGASGTYKYIF | TRBV19 | TRBJ2-7 | CASSISAGGAYEQYF | KLGGALQAK | YES | NO | NO |
| 7531 | TRAV30 | TRAJ53 | CGTAHNSGGSNYKLTF | TRBV19 | TRBJ2-7 | CATSFTLSYEQYF | KLGGALQAK | YES | NO | NO |
| 7532 | TRAV38-2DV8 | TRAJ32 | CAISRGGATNKLIF | TRBV19 | TRBJ2-7 | CASSPLEGTVREQYF | KLGGALQAK | YES | NO | NO |
| 7533 | TRAV38-2DV8 | TRAJ56 | CAYRSGPTGANSKLTF | TRBV19 | TRBJ2-7 | CASSIDLLGYEQYF | KLGGALQAK | YES | NO | NO |
| 7538 | TRAV8-3 | TRAJ34 | CAVDTDKLIF | TRBV19 | TRBJ2-7 | CASSSRGPYEQYF | KLGGALQAK | YES | NO | NO |
| 7539 | TRAV8-3 | TRAJ53 | CAVGALSGGSNYKLTF | TRBV19 | TRBJ2-7 | CASSRDGAHEQYF | KLGGALQAK | YES | NO | NO |
| 7540 | TRAV8-3 | TRAJ57 | CAVGTQGGSEKLVF | TRBV19 | TRBJ2-7 | CASSTYSDLYEQYF | KLGGALQAK | YES | NO | NO |
| 7541 | TRAV9-2 | TRAJ45 | CALRFLGGGADGLTF | TRBV19 | TRBJ2-7 | CASSSRGGTYEQYF | RLRAEAQVK | YES | YES | YES |
| 7543 | TRAV1-1 | TRAJ13 | CAVRTDSGGYQKVTF | TRBV2 | TRBJ2-7 | CASRAGQSDYEQYF | KLGGALQAK | YES | NO | NO |
| 7545 | TRAV1-1 | TRAJ42 | CAVRDRVGSQGNLIF | TRBV2 | TRBJ2-7 | CASTPGTSGAGGEQYF | KLGGALQAK | YES | NO | NO |
| 7546 | TRAV1-2 | TRAJ39 | CAVRDNNAGNMLTF | TRBV2 | TRBJ2-7 | CASSDSGAGNEQYF | KLGGALQAK | YES | NO | NO |
| 7547 | TRAV12-1 | TRAJ20 | CVVNHDYKLSF | TRBV2 | TRBJ2-7 | CASRDTSYEQYV | KLGGALQAK | NO | NO | NO |
| 7548 | TRAV12-1 | TRAJ20 | CVVNKADYKLSF | TRBV2 | TRBJ2-7 | CASRRGGPYEQYF | KLGGALQAK | YES | NO | NO |
| 7549 | TRAV12-1 | TRAJ24 | CTDSWGKFQF | TRBV2 | TRBJ2-7 | CASSEVGGRGEQYF | KLGGALQAK | YES | NO | NO |
| 7551 | TRAV12-2 | TRAJ31 | CAVNMVARLMF | TRBV2 | TRBJ2-7 | CASSEASNAYEQYF | KLGGALQAK | YES | NO | NO |
| 7552 | TRAV12-2 | TRAJ31 | CAVNNARLMF | TRBV2 | TRBJ2-7 | CASTLSYEQYF | KLGGALQAK | YES | NO | NO |
| 7553 | TRAV12-2 | TRAJ43 | CAVNGGNNNDMRF | TRBV2 | TRBJ2-7 | CASRPQTGLTPRRQYF | KLGGALQAK | YES | NO | NO |
| 7554 | TRAV12-3 | TRAJ13 | CAMILSGGYQKVTF | TRBV2 | TRBJ2-7 | CASSTQTGGREQYF | KLGGALQAK | YES | NO | NO |
| 7555 | TRAV14DV4 | TRAJ37 | CAMRGAFSSNTGKLIF | TRBV2 | TRBJ2-7 | CASTEHSGGKYF | AVFDRKSDAK | YES | NO | YES |
| 7556 | TRAV17 | TRAJ40 | CATDSGTYKYIF | TRBV2 | TRBJ2-7 | CAGTPREGQGLREQYF | KLGGALQAK | YES | NO | NO |
| 7557 | TRAV20 | TRAJ34 | CAVQANTDKLIF | TRBV2 | TRBJ2-7 | CATTVPLSYEQYF | KLGGALQAK | YES | NO | NO |
| 7558 | TRAV21 | TRAJ10 | CAVRDTGGGNKLTF | TRBV2 | TRBJ2-7 | CASRAPPFHEQYV | KLGGALQAK | NO | NO | NO |
| 7559 | TRAV21 | TRAJ28 | CAVNPYSGAGSYQLTF | TRBV2 | TRBJ2-7 | CASSDGPAYEQYF | KLGGALQAK | YES | NO | NO |
| 7559 | TRAV21 | TRAJ28 | CAVNPYSGAGSYQLTF | TRBV2 | TRBJ2-7 | CASSDGPAYEQYF | AVFDRKSDAK | YES | NO | NO |
| 7564 | TRAV26-2 | TRAJ24 | CILRDPSTDSWGKLQF | TRBV2 | TRBJ2-7 | CASSGTEREGLTYEQYF | KLGGALQAK | YES | NO | NO |
| 7565 | TRAV27 | TRAJ26 | CAGIDNYGQNFVF | TRBV2 | TRBJ2-7 | CANQEGTGGSEQYF | KLGGALQAK | YES | NO | NO |
| 7566 | TRAV29DV5 | TRAJ29 | CAASAGAGNTPLVF | TRBV2 | TRBJ2-7 | CASAPPQGLILEQYF | KLGGALQAK | YES | NO | NO |
| 7567 | TRAV3 | TRAJ26 | CAVRDRVGQNFVF | TRBV2 | TRBJ2-7 | CASRDDRPAGSYEQYF | KLGGALQAK | NO | NO | NO |
| 7569 | TRAV41 | TRAJ32 | CAVRWADGGATNKLIF | TRBV2 | TRBJ2-7 | CASRAPNDKPYEQYV | KLGGALQAK | NO | NO | NO |
| 7570 | TRAV41 | TRAJ34 | CAAGRGGTDKLIF | TRBV2 | TRBJ2-7 | CASIGGIQYV | KLGGALQAK | NO | NO | NO |
| 7571 | TRAV5 | TRAJ6 | CAESSGGSYIPTF | TRBV2 | TRBJ2-7 | CASRGGQISYEQYF | KLGGALQAK | YES | NO | NO |
| 7573 | TRAV8-2 | TRAJ50 | CVVSETSYDKVIF | TRBV2 | TRBJ2-7 | CASSEFGTTTYEQYV | KLGGALQAK | NO | NO | NO |
| 7575 | TRAV1-1 | TRAJ23 | CAEDQGGKLIF | TRBV20-1 | TRBJ2-7 | CSARDGVGSGSTYEQYF | KLGGALQAK | NO | NO | NO |
| 7577 | TRAV1-2 | TRAJ20 | CAVRERDYKLSF | TRBV20-1 | TRBJ2-7 | CSALGLGTNEQYF | KLGGALQAK | YES | NO | NO |
| 7578 | TRAV1-2 | TRAJ22 | CAVRVRRQLTF | TRBV20-1 | TRBJ2-7 | CSARVSGTSGSSNEQYF | KLGGALQAK | NO | NO | NO |
| 7579 | TRAV1-2 | TRAJ26 | CAVKDNYGQNFVF | TRBV20-1 | TRBJ2-7 | CSASGLSSYEQYV | KLGGALQAK | NO | NO | NO |
| 7583 | TRAV1-2 | TRAJ33 | CAVQDSNYQLIW | TRBV20-1 | TRBJ2-7 | CSARTGEVEQYF | KLGGALQAK | YES | NO | NO |
| 7585 | TRAV1-2 | TRAJ33 | CAVRDSNYQLIW | TRBV20-1 | TRBJ2-7 | CSARDGRREYEQYF | AVFDRKSDAK | NO | NO | NO |
| 7588 | TRAV1-2 | TRAJ35 | CAVSGRGAGFGNVLHC | TRBV20-1 | TRBJ2-7 | CSARDPGQGYEQYF | KLGGALQAK | NO | NO | NO |
| 7590 | TRAV10 | TRAJ27 | CVGYTNAGKSTF | TRBV20-1 | TRBJ2-7 | CSARDGRREYEQYF | KLGGALQAK | YES | NO | NO |
| 7591 | TRAV12-1 | TRAJ10 | CVVTSFTGGGNKLTF | TRBV20-1 | TRBJ2-7 | CSAQKGADFSYEQYF | KLGGALQAK | NO | NO | NO |
| 7593 | TRAV12-1 | TRAJ23 | CVVTRTYNQGGKLIF | TRBV20-1 | TRBJ2-7 | CSARGGTEIPYEQYF | RLRAEAQVK | NO | NO | NO |
| 7594 | TRAV12-1 | TRAJ31 | CVVNVNARLMF | TRBV20-1 | TRBJ2-7 | CSARDRDRGYEQYF | KLGGALQAK | NO | NO | NO |
| 7595 | TRAV12-1 | TRAJ43 | CVVKDDMRF | TRBV20-1 | TRBJ2-7 | CSAREDERGVLYEQYF | KLGGALQAK | NO | NO | NO |
| 7596 | TRAV12-1 | TRAJ53 | CVVLWDSGGSNYKLTF | TRBV20-1 | TRBJ2-7 | CSASPPGQTIYEQYF | KLGGALQAK | NO | NO | NO |
| 7599 | TRAV12-2 | TRAJ17 | CAVVSAGNKLTF | TRBV20-1 | TRBJ2-7 | CSATLTGTSGGFSSYEQYF | KLGGALQAK | YES | NO | NO |
| 7600 | TRAV12-2 | TRAJ20 | CAGANNDYKLSF | TRBV20-1 | TRBJ2-7 | CSAPKAGSSYEQYF | KLGGALQAK | NO | NO | NO |
| 7601 | TRAV12-2 | TRAJ24 | CAVNTDSWGKLQF | TRBV20-1 | TRBJ2-7 | CSARDRLAGEYSEQYF | KLGGALQAK | NO | NO | NO |
| 7602 | TRAV12-2 | TRAJ33 | CAVGDSNYQLIW | TRBV20-1 | TRBJ2-7 | CSARAAGRQTSTYEQYF | KLGGALQAK | NO | NO | NO |
| 7603 | TRAV12-2 | TRAJ42 | CAAIRSQGNLIF | TRBV20-1 | TRBJ2-7 | CSARGVEGYEQYF | KLGGALQAK | YES | NO | NO |
| 7604 | TRAV12-2 | TRAJ46 | CAVKSSGDKLTF | TRBV20-1 | TRBJ2-7 | CSARDLEVLSYEQYF | KLGGALQAK | YES | NO | NO |
| 7605 | TRAV12-2 | TRAJ49 | CAVRTGNQFYF | TRBV20-1 | TRBJ2-7 | CSAIAGGYEQYF | KLGGALQAK | NO | NO | NO |
| 7606 | TRAV12-3 | TRAJ13 | CAMEGGYQKVTF | TRBV20-1 | TRBJ2-7 | CSASGASESYEQYF | KLGGALQAK | NO | NO | NO |
| 7608 | TRAV12-3 | TRAJ30 | CAMNGNNDKIIF | TRBV20-1 | TRBJ2-7 | CSARAVRVSVEQYF | KLGGALQAK | NO | NO | NO |
| 7609 | TRAV12-3 | TRAJ32 | CAMNPAWGGATNKLIF | TRBV20-1 | TRBJ2-7 | CSASPGDYEQYF | KLGGALQAK | NO | NO | NO |
| 7609 | TRAV12-3 | TRAJ32 | CAMNPAWGGATNKLIF | TRBV20-1 | TRBJ2-7 | CSASPGDYEQYF | AVFDRKSDAK | NO | NO | NO |
| 7609 | TRAV12-3 | TRAJ32 | CAMNPAWGGATNKLIF | TRBV20-1 | TRBJ2-7 | CSASPGDYEQYF | RLRAEAQVK | NO | NO | NO |
| 7610 | TRAV12-3 | TRAJ32 | CAMTPAWGGATNKLIF | TRBV20-1 | TRBJ2-7 | CSASPGDYEQYF | KLGGALQAK | NO | NO | NO |
| 7612 | TRAV12-3 | TRAJ39 | CAIGPGNMLTF | TRBV20-1 | TRBJ2-7 | CSARDLEVLSYEQYF | AVFDRKSDAK | YES | NO | NO |
| 7614 | TRAV12-3 | TRAJ39 | CAIGPGNMLTF | TRBV20-1 | TRBJ2-7 | CSARPNKLAGGYEQYF | KLGGALQAK | NO | NO | NO |
| 7615 | TRAV12-3 | TRAJ4 | CAMSPPMFSGGYNKLIF | TRBV20-1 | TRBJ2-7 | CSAMTSGSSYEQYF | KLGGALQAK | NO | NO | NO |
| 7616 | TRAV12-3 | TRAJ54 | CATEQGAQKLVF | TRBV20-1 | TRBJ2-7 | CSAGTSGESYEQYF | KLGGALQAK | NO | NO | NO |
| 7619 | TRAV13-2 | TRAJ45 | CADSGGGADGLTF | TRBV20-1 | TRBJ2-7 | CSARDGRREYEQYF | KLGGALQAK | NO | NO | NO |
| 7620 | TRAV14DV4 | TRAJ22 | CAMRRPISSGSARQLTF | TRBV20-1 | TRBJ2-7 | CSARDLEVLSYEQYF | KLGGALQAK | YES | NO | NO |
| 7622 | TRAV14DV4 | TRAJ48 | CAMRDPPMGGNEKLTF | TRBV20-1 | TRBJ2-7 | CSASGGAYEQYF | KLGGALQAK | NO | NO | NO |

| | | | | | | | | | | |
|---|---|---|---|---|---|---|---|---|---|---|
| 7626 | TRAV17 | TRAJ54 | CATDNQGAQKLVF | TRBV20-1 | TRBJ2-7 | CSARVRGKPYEQYV | KLGGALQAK | NO | NO | NO |
| 7631 | TRAV19 | TRAJ34 | CALSVYNTDKLIF | TRBV20-1 | TRBJ2-7 | CSARAPAGAYEQYF | KLGGALQAK | YES | NO | NO |
| 7632 | TRAV19 | TRAJ34 | CALSEEPYNTDKLIF | TRBV20-1 | TRBJ2-7 | CSASAGGTSVGYEQYF | KLGGALQAK | NO | NO | NO |
| 7633 | TRAV19 | TRAJ40 | CALSVTSGTYKYIF | TRBV20-1 | TRBJ2-7 | CSAIRSGGPYEQYF | KLGGALQAK | NO | NO | NO |
| 7634 | TRAV19 | TRAJ47 | CALEYGNKLVF | TRBV20-1 | TRBJ2-7 | CSALTGLIYHEQYF | KLGGALQAK | NO | NO | NO |
| 7635 | TRAV19 | TRAJ48 | CALSEPISNFGNEKLTF | TRBV20-1 | TRBJ2-7 | CSAPPPTRTALHEQYF | KLGGALQAK | NO | NO | NO |
| 7636 | TRAV19 | TRAJ53 | CALIGGSNYKLTF | TRBV20-1 | TRBJ2-7 | CSARVARDPAYEQYF | KLGGALQAK | NO | NO | NO |
| 7640 | TRAV20 | TRAJ45 | CAAHPPMYSGGGADGLTF | TRBV20-1 | TRBJ2-7 | CSADSGELSYEQYF | KLGGALQAK | YES | NO | NO |
| 7642 | TRAV21 | TRAJ20 | CAVMNDYKLSF | TRBV20-1 | TRBJ2-7 | CSAGRLAGISYEQYF | KLGGALQAK | YES | NO | NO |
| 7643 | TRAV21 | TRAJ36 | CAVYPGANNLFF | TRBV20-1 | TRBJ2-7 | CSAPRGGKYEQYF | KLGGALQAK | NO | NO | NO |
| 7645 | TRAV21 | TRAJ43 | CAVWYMRF | TRBV20-1 | TRBJ2-7 | CSARDLEVLSYEQYF | KLGGALQAK | NO | NO | NO |
| 7647 | TRAV21 | TRAJ48 | CAVRRFGNEKLTF | TRBV20-1 | TRBJ2-7 | CSADRDSSYEQYF | KLGGALQAK | NO | NO | NO |
| 7648 | TRAV23DV6 | TRAJ43 | CAASGNHDMRF | TRBV20-1 | TRBJ2-7 | CSASPGDYEQYF | RLRAEAQVK | NO | NO | NO |
| 7650 | TRAV24 | TRAJ45 | CAFRGGADGLTF | TRBV20-1 | TRBJ2-7 | CSTGTGYEQYF | KLGGALQAK | NO | NO | NO |
| 7654 | TRAV26-2 | TRAJ20 | CILRDSNDYKLSF | TRBV20-1 | TRBJ2-7 | CSARDLTGRIYEQYF | KLGGALQAK | NO | NO | NO |
| 7660 | TRAV27 | TRAJ33 | CAGVMDSNYQLIW | TRBV20-1 | TRBJ2-7 | CSARDLEVLSYEQYF | KLGGALQAK | YES | NO | NO |
| 7661 | TRAV27 | TRAJ44 | CAGADGTASKLTF | TRBV20-1 | TRBJ2-7 | CSASPTVKQSYEQYF | KLGGALQAK | YES | NO | NO |
| 7667 | TRAV29DV5 | TRAJ9 | CAASATGGFKTIF | TRBV20-1 | TRBJ2-7 | CSARNREGWGEQYF | KLGGALQAK | YES | NO | NO |
| 7668 | TRAV3 | TRAJ20 | CAVNDYKLSF | TRBV20-1 | TRBJ2-7 | CSADSGELSYEQYF | KLGGALQAK | NO | NO | NO |
| 7670 | TRAV3 | TRAJ26 | CAVRLHNYGQNFVF | TRBV20-1 | TRBJ2-7 | CSAPARSEPYEQYF | KLGGALQAK | NO | NO | NO |
| 7670 | TRAV3 | TRAJ26 | CAVRLHNYGQNFVF | TRBV20-1 | TRBJ2-7 | CSAPARSEPYEQYF | AVFDRKSDAK | NO | NO | NO |
| 7670 | TRAV3 | TRAJ26 | CAVRLHNYGQNFVF | TRBV20-1 | TRBJ2-7 | CSAPARSEPYEQYF | IVTDFSVIK | NO | NO | NO |
| 7671 | TRAV3 | TRAJ26 | CAVRLHNYGQNFVF | TRBV20-1 | TRBJ2-7 | CSARVSGTSGSSNEQYF | KLGGALQAK | NO | NO | NO |
| 7673 | TRAV3 | TRAJ37 | CAVRDGGMGSNTGKLIF | TRBV20-1 | TRBJ2-7 | CSARDLEVLSYEQYF | KLGGALQAK | NO | NO | NO |
| 7675 | TRAV30 | TRAJ24 | CGTERTDSWGKLQF | TRBV20-1 | TRBJ2-7 | CSAFPGPREAYEQYF | KLGGALQAK | NO | NO | NO |
| 7676 | TRAV34 | TRAJ44 | CGADFSSGAGTASKLTF | TRBV20-1 | TRBJ2-7 | CSATARGGTSGGKSYEQYF | KLGGALQAK | NO | NO | NO |
| 7677 | TRAV36DV7 | TRAJ54 | CAVSGIQGAQKLVF | TRBV20-1 | TRBJ2-7 | CSASDLGQGSNEQYF | KLGGALQAK | NO | NO | NO |
| 7678 | TRAV38-1 | TRAJ57 | CAFISPSQGGSEKLVF | TRBV20-1 | TRBJ2-7 | CSAREWDRRYEQYF | KLGGALQAK | NO | NO | NO |
| 7679 | TRAV38-2DV8 | TRAJ20 | CAYRSVYKLSF | TRBV20-1 | TRBJ2-7 | CSARDLEVLSYEQYF | KLGGALQAK | NO | NO | NO |
| 7680 | TRAV39 | TRAJ47 | CAVAMEYGNKLVF | TRBV20-1 | TRBJ2-7 | CSATYPRQGYYEQYF | KLGGALQAK | NO | NO | NO |
| 7681 | TRAV4 | TRAJ13 | CLVVSYSGGYQKVTF | TRBV20-1 | TRBJ2-7 | CSARQQGVLDEQYF | KLGGALQAK | NO | NO | NO |
| 7684 | TRAV41 | TRAJ45 | CAVPLNSGGGADGLTF | TRBV20-1 | TRBJ2-7 | CSAREALDEQYF | KLGGALQAK | NO | NO | NO |
| 7686 | TRAV5 | TRAJ43 | CAESIRNNDMRF | TRBV20-1 | TRBJ2-7 | CSARAYSSYEQYF | KLGGALQAK | YES | NO | NO |
| 7687 | TRAV8-1 | TRAJ8 | CAVNARTGFQKLVF | TRBV20-1 | TRBJ2-7 | CSARPNKLAGGYEQYF | KLGGALQAK | NO | NO | NO |
| 7688 | TRAV8-2 | TRAJ48 | CVVSYLISNFGNEKLTF | TRBV20-1 | TRBJ2-7 | CSAQWGDYEQYF | KLGGALQAK | YES | NO | NO |
| 7689 | TRAV8-3 | TRAJ45 | CAVGMYSGGGADGLTF | TRBV20-1 | TRBJ2-7 | CSATARGGTSGGKSYEQYF | KLGGALQAK | NO | NO | NO |
| 7690 | TRAV8-3 | TRAJ47 | CAVGEVRYGNKLVF | TRBV20-1 | TRBJ2-7 | CSARDSGTASYEQYF | KLGGALQAK | YES | NO | NO |
| 7691 | TRAV8-3 | TRAJ5 | CAVGPGGSYTGRRALTF | TRBV20-1 | TRBJ2-7 | CSASKPGPGTYEQYF | KLGGALQAK | YES | NO | NO |
| 7692 | TRAV8-4 | TRAJ16 | CAVSESHSDGQKLLF | TRBV20-1 | TRBJ2-7 | CSARAAGGQIYEQYF | KLGGALQAK | NO | NO | NO |
| 7693 | TRAV8-4 | TRAJ49 | CAVTLGGNQFYF | TRBV20-1 | TRBJ2-7 | CSAPGQGNYEQYF | KLGGALQAK | YES | NO | NO |
| 7702 | TRAV13-1 | TRAJ47 | CAPQIDGNKLVF | TRBV21-1 | TRBJ2-7 | CASSNRTGVHEQYV | KLGGALQAK | NO | NO | NO |
| 7711 | TRAV5 | TRAJ6 | CAEQGGGSYIPTF | TRBV24-1 | TRBJ2-7 | CATSGETESSYEQYF | KLGGALQAK | YES | NO | NO |
| 7713 | TRAV25 | TRAJ50 | CAGETSYDKVIF | TRBV25-1 | TRBJ2-7 | CASSEVGTYEQYF | KLGGALQAK | YES | NO | NO |
| 7714 | TRAV1-1 | TRAJ12 | CAGKGMDSSYKLIF | TRBV27 | TRBJ2-7 | CASSPTQGKYYEQYF | KLGGALQAK | YES | NO | NO |
| 7715 | TRAV1-1 | TRAJ33 | CAVFDSNYQLIW | TRBV27 | TRBJ2-7 | CASSTDSFSYEQYF | KLGGALQAK | YES | NO | NO |
| 7716 | TRAV1-1 | TRAJ33 | CAVRPPDSNYQLIW | TRBV27 | TRBJ2-7 | CASSTRDRFYEQYF | KLGGALQAK | YES | NO | NO |
| 7717 | TRAV1-2 | TRAJ18 | CAVRDDRGSTLGRLYF | TRBV27 | TRBJ2-7 | CASSPAGYEQYF | KLGGALQAK | YES | NO | NO |
| 7718 | TRAV10 | TRAJ47 | CVVSGMEYGNKLVF | TRBV27 | TRBJ2-7 | CASGSGSSSYEQYV | KLGGALQAK | NO | NO | NO |
| 7719 | TRAV10 | TRAJ49 | CVVSAFGQFYF | TRBV27 | TRBJ2-7 | CASSLRQTYEQYF | KLGGALQAK | YES | NO | NO |
| 7720 | TRAV12-1 | TRAJ18 | CVVNPTNRGSTLGRLYF | TRBV27 | TRBJ2-7 | CASSLTGLAGAEQYF | KLGGALQAK | YES | NO | NO |
| 7721 | TRAV12-1 | TRAJ39 | CVAENNAGNMLTF | TRBV27 | TRBJ2-7 | CASSQSTGGFYEQYV | KLGGALQAK | NO | NO | NO |
| 7722 | TRAV12-1 | TRAJ49 | CVVSTGNQFYF | TRBV27 | TRBJ2-7 | CASSVRGAFYEQYF | KLGGALQAK | YES | NO | NO |
| 7723 | TRAV12-2 | TRAJ15 | CAVANQAGTALIF | TRBV27 | TRBJ2-7 | CASSSSPGQWAYEQYF | KLGGALQAK | YES | NO | NO |
| 7724 | TRAV12-2 | TRAJ20 | CAVYPLINDYKLSF | TRBV27 | TRBJ2-7 | CASSLSNRGEQYF | KLGGALQAK | YES | NO | NO |
| 7726 | TRAV12-2 | TRAJ3 | CAVNGASKIIF | TRBV27 | TRBJ2-7 | CASGSGRLAEQYF | KLGGALQAK | YES | NO | NO |
| 7727 | TRAV12-2 | TRAJ43 | CAVNPGDMRF | TRBV27 | TRBJ2-7 | CASGTPGYEQYF | KLGGALQAK | YES | NO | NO |
| 7728 | TRAV12-2 | TRAJ45 | CAVKGTGASGGGADGLTF | TRBV27 | TRBJ2-7 | CASRGTGGLSYEQYF | KLGGALQAK | YES | NO | NO |
| 7729 | TRAV12-2 | TRAJ49 | CAAPNTGNQFYF | TRBV27 | TRBJ2-7 | CASSLGRNYEQYF | KLGGALQAK | YES | NO | NO |
| 7731 | TRAV12-2 | TRAJ9 | CAVRTGGFKTIF | TRBV27 | TRBJ2-7 | CASSIRGLAGPYEQYV | KLGGALQAK | NO | NO | NO |
| 7734 | TRAV12-3 | TRAJ47 | CAMTAMGEYGNKLVF | TRBV27 | TRBJ2-7 | CASSDYFSSYEQYF | KLGGALQAK | YES | NO | NO |
| 7737 | TRAV14DV4 | TRAJ22 | CAMRRPISSGSARQLTF | TRBV27 | TRBJ2-7 | CASSPRTGGQGYEQYV | KLGGALQAK | NO | NO | NO |
| 7739 | TRAV17 | TRAJ17 | CATDAFRTAGNKLTF | TRBV27 | TRBJ2-7 | CASAPGGPSYEQYF | KLGGALQAK | YES | NO | NO |
| 7740 | TRAV17 | TRAJ37 | CATAYINTGKLIF | TRBV27 | TRBJ2-7 | CATYRPGLEDEQYF | KLGGALQAK | YES | NO | NO |
| 7741 | TRAV17 | TRAJ4 | CATDASGGYNKLIF | TRBV27 | TRBJ2-7 | CASRWTGGYEQYF | KLGGALQAK | YES | NO | NO |
| 7742 | TRAV17 | TRAJ54 | CATSWGAQKLVF | TRBV27 | TRBJ2-7 | CASSETSGSAYEQYV | KLGGALQAK | NO | NO | NO |
| 7743 | TRAV17 | TRAJ56 | CATDSYLWDGANSKLTF | TRBV27 | TRBJ2-7 | CASSSPVGTHHSYEQYF | KLGGALQAK | YES | NO | NO |
| 7744 | TRAV19 | TRAJ21 | CALCGGGGRNFNKFYF | TRBV27 | TRBJ2-7 | CASSARDRAYYEQYF | KLGGALQAK | YES | NO | NO |
| 7745 | TRAV19 | TRAJ45 | CALHSGGGADGLTF | TRBV27 | TRBJ2-7 | CASSPRGGGYYEQYF | KLGGALQAK | YES | NO | NO |
| 7747 | TRAV19 | TRAJ56 | CALSGYTGANSKLTF | TRBV27 | TRBJ2-7 | CASSLSPGGSYEQYF | KLGGALQAK | YES | NO | NO |
| 7749 | TRAV20 | TRAJ22 | CAVLGGSARQLTF | TRBV27 | TRBJ2-7 | CASSHFPPPYEQYF | KLGGALQAK | YES | NO | NO |
| 7752 | TRAV21 | TRAJ26 | CAVSPIDNYGQNFVF | TRBV27 | TRBJ2-7 | CASSPGTNYLYEQYV | KLGGALQAK | NO | NO | NO |
| 7755 | TRAV21 | TRAJ40 | CAVGPGTYKYIF | TRBV27 | TRBJ2-7 | CASSPQGGSTPYEQYF | KLGGALQAK | YES | NO | NO |
| 7756 | TRAV21 | TRAJ54 | CAVWIQGAQKLVF | TRBV27 | TRBJ2-7 | CASSLAATYEQYF | KLGGALQAK | YES | NO | NO |
| 7758 | TRAV21 | TRAJ58 | CAVKETSGSRLTF | TRBV27 | TRBJ2-7 | CASSLLGGYEQYV | RLRAEAQVK | NO | NO | NO |
| 7759 | TRAV24 | TRAJ31 | CAFIKNNARLMF | TRBV27 | TRBJ2-7 | CASSRGTGFNEQYF | KLGGALQAK | YES | NO | NO |
| 7760 | TRAV24 | TRAJ48 | CACPYFGNEKLTF | TRBV27 | TRBJ2-7 | CASSSGLAGVAYEQYV | KLGGALQAK | NO | NO | NO |
| 7761 | TRAV25 | TRAJ28 | CAGSGAGSYQLTF | TRBV27 | TRBJ2-7 | CASSLERQSYEQYV | KLGGALQAK | NO | NO | NO |
| 7765 | TRAV26-1 | TRAJ54 | CIVRVFQIQGAQKLVF | TRBV27 | TRBJ2-7 | CASSLSGTGNYEQYV | KLGGALQAK | NO | NO | NO |
| 7766 | TRAV26-2 | TRAJ3 | CILRARYSSASKIIF | TRBV27 | TRBJ2-7 | CASSLTGSEQYF | KLGGALQAK | NO | NO | NO |
| 7767 | TRAV26-2 | TRAJ39 | CIRLAGNMLTF | TRBV27 | TRBJ2-7 | CASSSWDSSYEQYF | KLGGALQAK | YES | NO | NO |
| 7768 | TRAV27 | TRAJ45 | CAGALRGGADGLTF | TRBV27 | TRBJ2-7 | CAAWDRGSEQYF | KLGGALQAK | YES | NO | NO |
| 7769 | TRAV29DV5 | TRAJ18 | CAASDTARGSTLGRLYF | TRBV27 | TRBJ2-7 | CASSLTGTALNSYEQYF | KLGGALQAK | YES | NO | NO |
| 7770 | TRAV29DV5 | TRAJ52 | CAASAMRNYGKLTF | TRBV27 | TRBJ2-7 | CASSGLAGPLGEQYF | KLGGALQAK | YES | NO | NO |
| 7771 | TRAV3 | TRAJ23 | CVPKQNQGGKLIF | TRBV27 | TRBJ2-7 | CASSSGGDLYEQYF | KLGGALQAK | NO | NO | NO |
| 7773 | TRAV3 | TRAJ3 | CAVRDIGYSSASKIIF | TRBV27 | TRBJ2-7 | CASSLGIDYEQYV | KLGGALQAK | NO | NO | NO |
| 7776 | TRAV30 | TRAJ54 | CGTEGGAQKLVF | TRBV27 | TRBJ2-7 | CASSFQGGYEQYF | KLGGALQAK | YES | NO | NO |
| 7777 | TRAV34 | TRAJ9 | CGAEKDTGGFKTIF | TRBV27 | TRBJ2-7 | CASSFAGSYEQYF | KLGGALQAK | YES | NO | NO |
| 7778 | TRAV35 | TRAJ21 | CAGQSSGKFYF | TRBV27 | TRBJ2-7 | CASSRGTGEQYF | KLGGALQAK | YES | NO | NO |
| 7780 | TRAV38-1 | TRAJ28 | CAFFGSGAGSYQLTF | TRBV27 | TRBJ2-7 | CATWRGGREQYF | KLGGALQAK | YES | NO | NO |
| 7781 | TRAV41 | TRAJ28 | CAVLSGAGSYQLTF | TRBV27 | TRBJ2-7 | CASSLRGMSSYEQYF | KLGGALQAK | YES | NO | NO |
| 7782 | TRAV6 | TRAJ40 | CGGGTYKYIF | TRBV27 | TRBJ2-7 | CASSYDGSSYEQYF | KLGGALQAK | YES | NO | NO |
| 7783 | TRAV8-1 | TRAJ9 | CAVSTGGFKTIF | TRBV27 | TRBJ2-7 | CASSEQTGTNEQYF | KLGGALQAK | YES | NO | NO |
| 7786 | TRAV8-3 | TRAJ20 | CAVGDPNDYKLSF | TRBV27 | TRBJ2-7 | CASSSSDMGQKSYEQYF | KLGGALQAK | YES | NO | NO |
| 7787 | TRAV8-3 | TRAJ23 | CAVYQGGKLIF | TRBV27 | TRBJ2-7 | CASSFGGQLYEQYF | KLGGALQAK | YES | NO | NO |
| 7789 | TRAV8-6 | TRAJ8 | CAVSDKGFQKLVF | TRBV27 | TRBJ2-7 | CASSLSAGSSYEQYV | KLGGALQAK | NO | NO | NO |
| 7790 | TRAV9-2 | TRAJ12 | CALKMDSSYKLIF | TRBV27 | TRBJ2-7 | CASSFIGTGGEQYV | KLGGALQAK | NO | NO | NO |
| 7792 | TRAV9-2 | TRAJ48 | CATAGGFGNEKLTF | TRBV27 | TRBJ2-7 | CASSVRPPSGGEQYV | KLGGALQAK | NO | NO | NO |
| 7797 | TRAV12-2 | TRAJ13 | CAVNMRGGYQKVTF | TRBV28 | TRBJ2-7 | CASRPPGAVYEQYF | KLGGALQAK | YES | NO | NO |
| 7798 | TRAV12-2 | TRAJ20 | CAVNMRSDYKLSF | TRBV28 | TRBJ2-7 | CASSLSGTGSYEQYV | KLGGALQAK | NO | NO | NO |
| 7802 | TRAV13-1 | TRAJ43 | CAASITDNDMRF | TRBV28 | TRBJ2-7 | CASSFRGETYEQYF | KLGGALQAK | YES | NO | NO |
| 7803 | TRAV14DV4 | TRAJ33 | CAMREWSQDSNYQLIW | TRBV28 | TRBJ2-7 | CASSLEGTHEQYF | KLGGALQAK | YES | NO | NO |
| 7806 | TRAV17 | TRAJ33 | CATDGFYSNYQLIW | TRBV28 | TRBJ2-7 | CASRPTGTAALEQYF | KLGGALQAK | YES | NO | NO |
| 7807 | TRAV19 | TRAJ52 | CALSEAPGGTSYGKLTF | TRBV28 | TRBJ2-7 | CASSTRYEQYV | KLGGALQAK | NO | NO | NO |
| 7809 | TRAV22 | TRAJ30 | CTTLNRDDKIIF | TRBV28 | TRBJ2-7 | CASSSLGQILSYEQYF | KLGGALQAK | YES | NO | NO |

| | | | | | | | | | | |
|---|---|---|---|---|---|---|---|---|---|---|
| 7811 | TRAV25 | TRAJ10 | CAGITGGGNKLTF | TRBV28 | TRBJ2-7 | CASSLYFYSEQYV | KLGGALQAK | NO | NO | NO |
| 7812 | TRAV26-2 | TRAJ21 | CILSLTYNFNKFYF | TRBV28 | TRBJ2-7 | CASSPGRRSYEQYF | KLGGALQAK | YES | NO | NO |
| 7813 | TRAV26-2 | TRAJ23 | CILSPPYNQGGKLIF | TRBV28 | TRBJ2-7 | CASSLMAGTWFYEQYF | KLGGALQAK | YES | NO | NO |
| 7814 | TRAV27 | TRAJ13 | CAGATMNSGGYQKVTF | TRBV28 | TRBJ2-7 | CASRPAGLAYEQYF | KLGGALQAK | YES | NO | NO |
| 7815 | TRAV29DV5 | TRAJ40 | CAVSGTYKYIF | TRBV28 | TRBJ2-7 | CASSLYGGGGWEQYF | KLGGALQAK | YES | NO | NO |
| 7817 | TRAV3 | TRAJ16 | CAVRGRF | TRBV28 | TRBJ2-7 | CASSLDGSEGSEQYV | KLGGALQAK | NO | NO | NO |
| 7818 | TRAV3 | TRAJ33 | CAVRDLDSNYQLIW | TRBV28 | TRBJ2-7 | CASSFGPGLVYEQYV | KLGGALQAK | NO | NO | NO |
| 7819 | TRAV3 | TRAJ35 | CAVRDTI | TRBV28 | TRBJ2-7 | CASSLTDRVHEQYF | KLGGALQAK | NO | NO | NO |
| 7821 | TRAV5 | TRAJ31 | CAESKNNNARLMF | TRBV28 | TRBJ2-7 | CASSLSGYEQYF | KLGGALQAK | YES | NO | NO |
| 7821 | TRAV5 | TRAJ31 | CAESKNNNARLMF | TRBV28 | TRBJ2-7 | CASSLSGYEQYF | AVFDRKSDAK | YES | NO | NO |
| 7824 | TRAV9-2 | TRAJ30 | CALGNRDDKIIF | TRBV28 | TRBJ2-7 | CAIKSAGRSYEQYF | KLGGALQAK | YES | NO | NO |
| 7826 | TRAV12-1 | TRAJ49 | CVVAGNQFYF | TRBV29-1 | TRBJ2-7 | CSVSPTGTGTYEQYF | KLGGALQAK | YES | NO | NO |
| 7827 | TRAV13-1 | TRAJ20 | CAASGYDYKLSF | TRBV29-1 | TRBJ2-7 | CSVSASGGDEQYF | AVFDRKSDAK | YES | NO | NO |
| 7829 | TRAV14DV4 | TRAJ21 | CAMREYNFNKFYF | TRBV29-1 | TRBJ2-7 | CSVAGTSGGAYEQYF | KLGGALQAK | YES | NO | NO |
| 7829 | TRAV14DV4 | TRAJ21 | CAMREYNFNKFYF | TRBV29-1 | TRBJ2-7 | CSVAGTSGGAYEQYF | AVFDRKSDAK | YES | NO | NO |
| 7832 | TRAV21 | TRAJ49 | CAVPTGNQFYF | TRBV29-1 | TRBJ2-7 | CSVEDNSYEQYF | KLGGALQAK | YES | NO | NO |
| 7834 | TRAV22 | TRAJ42 | CAVMNYGGSQGNLIF | TRBV29-1 | TRBJ2-7 | CSVDGPSSYEQYF | KLGGALQAK | YES | NO | NO |
| 7836 | TRAV27 | TRAJ29 | CAGRLNSGNTPLVF | TRBV29-1 | TRBJ2-7 | CSVETGLGYEQYF | KLGGALQAK | YES | NO | NO |
| 7837 | TRAV29DV5 | TRAJ31 | CAASTNARLMF | TRBV29-1 | TRBJ2-7 | CSAQGGPFHEQYF | KLGGALQAK | YES | NO | NO |
| 7838 | TRAV29DV5 | TRAJ6 | CAAISGGSYIPTF | TRBV29-1 | TRBJ2-7 | CSVEDRGGYEQYF | KLGGALQAK | YES | NO | NO |
| 7840 | TRAV35 | TRAJ26 | CAGPTLINNYGQNFVF | TRBV29-1 | TRBJ2-7 | CSASAGTGGAGGTYEQYF | KLGGALQAK | YES | NO | NO |
| 7842 | TRAV38-2DV8 | TRAJ41 | CALGNSGYALNF | TRBV29-1 | TRBJ2-7 | CSVPGQSSYEQYF | KLGGALQAK | YES | NO | NO |
| 7843 | TRAV38-2DV8 | TRAJ47 | CAYRRRYMEYGNKLVF | TRBV29-1 | TRBJ2-7 | CSVVGGSFYEQYF | KLGGALQAK | YES | NO | NO |
| 7844 | TRAV38-2DV8 | TRAJ49 | CAYNTGNQFYF | TRBV29-1 | TRBJ2-7 | CSVGTGTYEQYF | KLGGALQAK | YES | NO | NO |
| 7846 | TRAV8-4 | TRAJ32 | CAVRNLDVDGGATNKLIF | TRBV29-1 | TRBJ2-7 | CSARQGMSYEQYF | KLGGALQAK | YES | NO | NO |
| 7847 | TRAV8-6 | TRAJ57 | CAVSDQGGSEKLVF | TRBV29-1 | TRBJ2-7 | CSVGGGNLYEQYF | KLGGALQAK | NO | NO | NO |
| 7850 | TRAV1-1 | TRAJ37 | CAVSNTGKLIF | TRBV3-1 | TRBJ2-7 | CASSQSGTREYEQYF | KLGGALQAK | YES | NO | NO |
| 7852 | TRAV12-1 | TRAJ34 | CVVNDNTDKLIF | TRBV3-1 | TRBJ2-7 | CASSQVLGTYEQYF | KLGGALQAK | YES | NO | NO |
| 7853 | TRAV12-1 | TRAJ47 | CGEYGNKLVF | TRBV3-1 | TRBJ2-7 | CASSPGAGTYEQYF | KLGGALQAK | YES | NO | NO |
| 7857 | TRAV13-2 | TRAJ43 | CAENNDMRF | TRBV3-1 | TRBJ2-7 | CASSPYIQGNEQYF | KLGGALQAK | NO | NO | NO |
| 7858 | TRAV14DV4 | TRAJ3 | CAMRGLLRSSASKIIF | TRBV3-1 | TRBJ2-7 | CASSPPRGALYHEQYF | KLGGALQAK | YES | NO | NO |
| 7860 | TRAV19 | TRAJ39 | CALSEALLAGNMLTF | TRBV3-1 | TRBJ2-7 | CASSQEPISYEQYF | KLGGALQAK | YES | NO | NO |
| 7861 | TRAV20 | TRAJ10 | CAVDVWGGGNKLTF | TRBV3-1 | TRBJ2-7 | CASSQPDNEQYF | KLGGALQAK | YES | NO | NO |
| 7862 | TRAV20 | TRAJ15 | CAVQANQAGTALIF | TRBV3-1 | TRBJ2-7 | CASSQFEGVYEQYF | KLGGALQAK | YES | NO | NO |
| 7864 | TRAV26-1 | TRAJ47 | CIVRLLMEYGNKLVF | TRBV3-1 | TRBJ2-7 | CASSRFAAERVPYEQYF | KLGGALQAK | YES | NO | NO |
| 7865 | TRAV29DV5 | TRAJ45 | CAASEGGGADGLTF | TRBV3-1 | TRBJ2-7 | CASGQGISYEQYF | KLGGALQAK | YES | NO | NO |
| 7866 | TRAV3 | TRAJ23 | CAVRVYNQGGKLIF | TRBV3-1 | TRBJ2-7 | CASSQEVRSSYEQYF | KLGGALQAK | NO | NO | NO |
| 7868 | TRAV38-2DV8 | TRAJ30 | CAYRRDDKIIF | TRBV3-1 | TRBJ2-7 | CASSQGLAASYEQYF | KLGGALQAK | NO | NO | NO |
| 7869 | TRAV5 | TRAJ42 | CIRGGNYGGSQGNLIF | TRBV3-1 | TRBJ2-7 | CASSLGTGGSYEQYF | KLGGALQAK | YES | NO | NO |
| 7870 | TRAV5 | TRAJ9 | CAELNTGGFKTIF | TRBV3-1 | TRBJ2-7 | CASSQVIEGRAYEQYF | KLGGALQAK | YES | NO | NO |
| 7871 | TRAV12-2 | TRAJ52 | CAAAGGTSYGKLTF | TRBV30 | TRBJ2-7 | CAWSVTSGRQYV | KLGGALQAK | NO | NO | NO |
| 7874 | TRAV17 | TRAJ53 | CATDPGSNYKLTF | TRBV30 | TRBJ2-7 | CAWGQNSYEQYF | KLGGALQAK | YES | NO | NO |
| 7875 | TRAV27 | TRAJ22 | CAGRVSGSARQLTF | TRBV30 | TRBJ2-7 | CAWSVGAGSNEQYV | KLGGALQAK | NO | NO | NO |
| 7876 | TRAV27 | TRAJ49 | CAGAYGNQFYF | TRBV30 | TRBJ2-7 | CAWSVNHEQYV | KLGGALQAK | NO | NO | NO |
| 7877 | TRAV4 | TRAJ13 | CLVEGGYQKVTF | TRBV30 | TRBJ2-7 | CAWMGTGTLYYEQYF | KLGGALQAK | NO | NO | NO |
| 7879 | TRAV1-1 | TRAJ17 | CAVRDTTKAAGNKLTF | TRBV4-1 | TRBJ2-7 | CASSLSDRASEQYF | KLGGALQAK | YES | NO | NO |
| 7879 | TRAV1-1 | TRAJ17 | CAVRDTTKAAGNKLTF | TRBV4-1 | TRBJ2-7 | CASSLSDRASEQYF | AVFDRKSDAK | YES | NO | NO |
| 7881 | TRAV12-2 | TRAJ39 | CAIGPGNMLTF | TRBV4-1 | TRBJ2-7 | CASSQGGRTGDYEQYF | KLGGALQAK | YES | NO | NO |
| 7882 | TRAV12-2 | TRAJ54 | CAVTQGAQKLVF | TRBV4-1 | TRBJ2-7 | CASSQSGGTSYEQYF | KLGGALQAK | YES | NO | NO |
| 7885 | TRAV13-1 | TRAJ42 | CAAHTGYGGSQGNLIF | TRBV4-1 | TRBJ2-7 | CASSQGTGSYEQYF | KLGGALQAK | YES | NO | NO |
| 7886 | TRAV13-1 | TRAJ54 | CAVQGAQKLVF | TRBV4-1 | TRBJ2-7 | CASSQGFSWEQYF | KLGGALQAK | YES | NO | NO |
| 7889 | TRAV13-2 | TRAJ49 | CAENGAGNQFYF | TRBV4-1 | TRBJ2-7 | CASSRTGTSYEQYF | KLGGALQAK | YES | NO | NO |
| 7891 | TRAV14DV4 | TRAJ42 | CAMRETFYGGSQGNLIF | TRBV4-1 | TRBJ2-7 | CASSQENSLIYEQYF | KLGGALQAK | YES | NO | NO |
| 7892 | TRAV14DV4 | TRAJ52 | CAMSNAGGTSYGKLTF | TRBV4-1 | TRBJ2-7 | CASSPQPGQGRNEQYV | KLGGALQAK | NO | NO | NO |
| 7893 | TRAV17 | TRAJ30 | CATDRRDDKIIF | TRBV4-1 | TRBJ2-7 | CASSQGTGSYEQYF | KLGGALQAK | YES | NO | NO |
| 7896 | TRAV21 | TRAJ10 | CAVYTGGGNKLTF | TRBV4-1 | TRBJ2-7 | CASSQNRDGPESEQYV | KLGGALQAK | NO | NO | NO |
| 7897 | TRAV21 | TRAJ40 | CAVRYTSGTYKYIF | TRBV4-1 | TRBJ2-7 | CASSQDRLAGGYEQYF | KLGGALQAK | YES | NO | NO |
| 7898 | TRAV21 | TRAJ52 | CAARAGGTSYGKLTF | TRBV4-1 | TRBJ2-7 | CASSQGGTGLGEQYF | KLGGALQAK | YES | NO | NO |
| 7901 | TRAV23DV6 | TRAJ37 | CAARFYDSSNTGKLIF | TRBV4-1 | TRBJ2-7 | CASSHGTRTYEQYF | KLGGALQAK | YES | NO | NO |
| 7903 | TRAV24 | TRAJ36 | CAFLGGANNLFF | TRBV4-1 | TRBJ2-7 | CASSQGDSRSYEQYF | KLGGALQAK | YES | NO | NO |
| 7904 | TRAV25 | TRAJ21 | CAGQYNFNKFYF | TRBV4-1 | TRBJ2-7 | CASSQGLLGYEQYF | KLGGALQAK | YES | NO | NO |
| 7905 | TRAV27 | TRAJ21 | CAGFVQVDFNKFYF | TRBV4-1 | TRBJ2-7 | CASSQGWDRRYEQYF | KLGGALQAK | YES | NO | NO |
| 7906 | TRAV27 | TRAJ57 | CAGQGGSEKLVF | TRBV4-1 | TRBJ2-7 | CASSLYTPGHFSDEQYF | KLGGALQAK | YES | NO | NO |
| 7908 | TRAV29DV5 | TRAJ30 | CAARILDDKIIF | TRBV4-1 | TRBJ2-7 | CASSQGRRGGSYEQYF | KLGGALQAK | YES | NO | NO |
| 7909 | TRAV29DV5 | TRAJ49 | CAARTGNQFYF | TRBV4-1 | TRBJ2-7 | CASSWQGTSYEQYF | KLGGALQAK | YES | NO | NO |
| 7910 | TRAV34 | TRAJ24 | CGADRTDSWGKLQF | TRBV4-1 | TRBJ2-7 | CASSQIQGQGAYEQYF | RLRAEAQVK | YES | NO | NO |
| 7910 | TRAV34 | TRAJ24 | CGADRTDSWGKLQF | TRBV4-1 | TRBJ2-7 | CASSQIQGQGAYEQYF | KLGGALQAK | YES | NO | NO |
| 7911 | TRAV34 | TRAJ40 | CGAGPSGTYKYIF | TRBV4-1 | TRBJ2-7 | CASSQNNQGRVEQYF | KLGGALQAK | NO | NO | NO |
| 7914 | TRAV38-1 | TRAJ54 | CAFMSLIQGAQKLVF | TRBV4-1 | TRBJ2-7 | CASSQGLSYEQYF | RLRAEAQVK | YES | YES | YES |
| 7915 | TRAV41 | TRAJ44 | CAVTGSGTASKLTF | TRBV4-1 | TRBJ2-7 | CASSQGPLSTEQYF | KLGGALQAK | YES | NO | NO |
| 7916 | TRAV5 | TRAJ45 | CAESKRRESGGGADGLTF | TRBV4-1 | TRBJ2-7 | CASSQGLLGYEQYF | KLGGALQAK | YES | NO | NO |
| 7918 | TRAV8-2 | TRAJ6 | CVVSDQGGSYIPTF | TRBV4-1 | TRBJ2-7 | CASSYGTSGTYEQYF | KLGGALQAK | YES | NO | NO |
| 7919 | TRAV8-3 | TRAJ11 | CAVGVMNSGYSTLTF | TRBV4-1 | TRBJ2-7 | CASSHPPQRGVSYEQYF | KLGGALQAK | YES | NO | NO |
| 7920 | TRAV8-3 | TRAJ7 | CAVGAQGNNRLAF | TRBV4-1 | TRBJ2-7 | CASSQTGTGSYEQYV | AVFDRKSDAK | NO | NO | NO |
| 7921 | TRAV8-6 | TRAJ17 | CAVSDKLVGAAGNKLTF | TRBV4-1 | TRBJ2-7 | CASSQHGGDYEQYF | KLGGALQAK | YES | NO | NO |
| 7924 | TRAV1-2 | TRAJ33 | CAVRDSNYQLIW | TRBV4-2 | TRBJ2-7 | CASSQEAESSYEQYF | KLGGALQAK | NO | NO | NO |
| 7925 | TRAV1-2 | TRAJ33 | CAVMDSNYQLIW | TRBV4-2 | TRBJ2-7 | CASSQESGGAGEQYF | KLGGALQAK | YES | NO | NO |
| 7926 | TRAV12-1 | TRAJ27 | CVVRTNAGKSTF | TRBV4-2 | TRBJ2-7 | CASSQKDRAPYEQYF | KLGGALQAK | NO | NO | NO |
| 7928 | TRAV21 | TRAJ12 | CAVIWMDSSYKLIF | TRBV4-2 | TRBJ2-7 | CASSPLGGGQNSYEQYF | KLGGALQAK | YES | NO | NO |
| 7929 | TRAV22 | TRAJ26 | CAVDNYGQNFVF | TRBV4-2 | TRBJ2-7 | CASSQERDYEQYF | KLGGALQAK | YES | NO | NO |
| 7930 | TRAV41 | TRAJ42 | CAVTYGGSQGNLIF | TRBV4-2 | TRBJ2-7 | CASSQQGGYEQYF | KLGGALQAK | YES | NO | NO |
| 7932 | TRAV1-1 | TRAJ5 | CASSNTGRRALTF | TRBV4-3 | TRBJ2-7 | CASSQDTGLGIYEQYF | KLGGALQAK | YES | NO | NO |
| 7933 | TRAV10 | TRAJ36 | CVVSGAGANNLFF | TRBV4-3 | TRBJ2-7 | CASSQDRRVSGGEQYF | KLGGALQAK | YES | NO | NO |
| 7934 | TRAV13-1 | TRAJ5 | CAASTGRRDTGRRALTF | TRBV4-3 | TRBJ2-7 | CASSQSSGSYEQYF | KLGGALQAK | YES | NO | NO |
| 7936 | TRAV14DV4 | TRAJ30 | CAMREGRDDKIIF | TRBV4-3 | TRBJ2-7 | CASSQDRGTGQYF | KLGGALQAK | YES | NO | NO |
| 7937 | TRAV14DV4 | TRAJ43 | CAMRGAIYNDMRF | TRBV4-3 | TRBJ2-7 | CASSIQTSGSYEQYF | KLGGALQAK | YES | NO | NO |
| 7942 | TRAV38-2DV8 | TRAJ26 | CAYRSPSGYGQNFVF | TRBV4-3 | TRBJ2-7 | CASSQETSTYEQYF | KLGGALQAK | YES | NO | NO |
| 7945 | TRAV1-2 | TRAJ21 | CAVRDNFNKFYF | TRBV5-1 | TRBJ2-7 | CASSLGSGTGGEQYV | KLGGALQAK | NO | NO | NO |
| 7946 | TRAV1-2 | TRAJ36 | CAVRAAAGANNLFF | TRBV5-1 | TRBJ2-7 | CASSFRGQGNEQYF | KLGGALQAK | YES | NO | NO |
| 7947 | TRAV12-1 | TRAJ8 | CVVNTGFQKLVF | TRBV5-1 | TRBJ2-7 | CASSLDGGDSYEQYF | KLGGALQAK | YES | NO | NO |
| 7948 | TRAV12-2 | TRAJ52 | CAVNHEGGTSYGKLTF | TRBV5-1 | TRBJ2-7 | CASSLQTGYSEDEQYF | KLGGALQAK | YES | NO | NO |
| 7950 | TRAV13-2 | TRAJ6 | CADASGGSYIPTF | TRBV5-1 | TRBJ2-7 | CASSLGAGMGEQYF | KLGGALQAK | YES | NO | NO |
| 7953 | TRAV17 | TRAJ45 | CATDGGGADGLTF | TRBV5-1 | TRBJ2-7 | CASSQDRLSYEQYF | KLGGALQAK | YES | NO | NO |
| 7954 | TRAV19 | TRAJ22 | CALSVISSGSARQLTF | TRBV5-1 | TRBJ2-7 | CASSLLGQPYEQYF | KLGGALQAK | YES | NO | NO |
| 7955 | TRAV19 | TRAJ5 | CALSEDGLMDTGRRALTF | TRBV5-1 | TRBJ2-7 | CASSLAPSGAYEQYF | KLGGALQAK | YES | NO | NO |
| 7956 | TRAV20 | TRAJ42 | CAGGTDGGSQGNLIF | TRBV5-1 | TRBJ2-7 | CASSPWGSGSSYEQYF | KLGGALQAK | YES | NO | NO |
| 7957 | TRAV21 | TRAJ10 | CAVITGGGNKLTF | TRBV5-1 | TRBJ2-7 | CASSSPKNEQYF | KLGGALQAK | YES | NO | NO |
| 7962 | TRAV26-1 | TRAJ37 | CIVRVPRFGSGNTGKLIF | TRBV5-1 | TRBJ2-7 | CASSGDINPYEQYF | KLGGALQAK | YES | NO | NO |
| 7963 | TRAV27 | TRAJ11 | CARHHYSGYSTLTF | TRBV5-1 | TRBJ2-7 | CASSFGEGVPYEQYF | KLGGALQAK | YES | NO | NO |
| 7965 | TRAV3 | TRAJ7 | CAVRDTFTAPGGNNRLAF | TRBV5-1 | TRBJ2-7 | CASSFSITYEQYV | KLGGALQAK | NO | NO | NO |
| 7966 | TRAV34 | TRAJ47 | CGADIWGIHGNKLVF | TRBV5-1 | TRBJ2-7 | CASSLPGSYEQYF | KLGGALQAK | YES | NO | NO |
| 7967 | TRAV35 | TRAJ20 | CAGRTNDYKLSF | TRBV5-1 | TRBJ2-7 | CASSLGTSGYYEQYF | KLGGALQAK | YES | NO | NO |
| 7968 | TRAV35 | TRAJ54 | CAGPFLIQGAQKLVF | TRBV5-1 | TRBJ2-7 | CASSLAADYEQYF | KLGGALQAK | YES | NO | NO |
| 7969 | TRAV35 | TRAJ54 | CAGPLLIQGAQKLVF | TRBV5-1 | TRBJ2-7 | CASSLAADYEQYF | KLGGALQAK | YES | NO | NO |

| | | | | | | | | | | |
|---|---|---|---|---|---|---|---|---|---|---|
| 7970 | TRAV38-2DV8 | TRAJ32 | CAFYRGRGATNKLIF | TRBV5-1 | TRBJ2-7 | CASSLASGVLPGQYF | KLGGALQAK | YES | NO | NO |
| 7971 | TRAV38-2DV8 | TRAJ33 | CAYGDSNYQLIW | TRBV5-1 | TRBJ2-7 | CASSLAADYEQYF | KLGGALQAK | YES | NO | NO |
| 7973 | TRAV39 | TRAJ24 | CAVDMRSWGKLQF | TRBV5-1 | TRBJ2-7 | CASGLDNVMTGYEQYF | RLRAEAQVK | YES | NO | YES |
| 7975 | TRAV41 | TRAJ50 | CASSHPGETSYDKVIF | TRBV5-1 | TRBJ2-7 | CASSLVQGLYEQYF | KLGGALQAK | YES | NO | NO |
| 7976 | TRAV8-1 | TRAJ44 | CAVRVNTGTASKLTF | TRBV5-1 | TRBJ2-7 | CASSLKVGGQGVSEQYF | KLGGALQAK | YES | NO | NO |
| 7976 | TRAV8-1 | TRAJ44 | CAVRVNTGTASKLTF | TRBV5-1 | TRBJ2-7 | CASSLKVGGQGVSEQYF | AVFDRKSDAK | YES | NO | NO |
| 7977 | TRAV8-2 | TRAJ34 | CVVSDVNTDKLIF | TRBV5-1 | TRBJ2-7 | CASSLFSGGDSSYEQYF | KLGGALQAK | YES | NO | NO |
| 7979 | TRAV8-3 | TRAJ16 | CAVGIDGQKLLF | TRBV5-1 | TRBJ2-7 | CASSLIGGRGVYEQYF | KLGGALQAK | NO | NO | NO |
| 7980 | TRAV8-4 | TRAJ11 | CAVSDIGYSTLTF | TRBV5-1 | TRBJ2-7 | CASSGTESYEQYF | KLGGALQAK | YES | NO | NO |
| 7981 | TRAV8-4 | TRAJ4 | CAVSDLEGFSGGYNKLIF | TRBV5-1 | TRBJ2-7 | CASSLAADYEQYF | KLGGALQAK | YES | YES | NO |
| 7982 | TRAV9-2 | TRAJ36 | CALTLTGGANNLFF | TRBV5-1 | TRBJ2-7 | CASSLGYEQYF | KLGGALQAK | YES | NO | NO |
| 7983 | TRAV10 | TRAJ43 | CVVSGDNDMRF | TRBV5-4 | TRBJ2-7 | CASSLELAGVHYEQYF | KLGGALQAK | YES | NO | NO |
| 7984 | TRAV12-1 | TRAJ28 | CVVRLSGAGSYQLTF | TRBV5-4 | TRBJ2-7 | CASSFGLGGEQYF | KLGGALQAK | YES | NO | NO |
| 7985 | TRAV12-2 | TRAJ38 | CAAHHKAGNNRKLIW | TRBV5-4 | TRBJ2-7 | CASSLEKVSGMPYEQYF | KLGGALQAK | YES | NO | NO |
| 7987 | TRAV12-3 | TRAJ43 | CAMSTYNNNDMRF | TRBV5-4 | TRBJ2-7 | CASSLSGGDEQYF | KLGGALQAK | YES | NO | NO |
| 7988 | TRAV12-3 | TRAJ6 | CAMSATGGSYIPTF | TRBV5-4 | TRBJ2-7 | CASSLGSAGAIPYEQYF | KLGGALQAK | YES | NO | NO |
| 7989 | TRAV13-1 | TRAJ11 | CAAKEGYSTLTF | TRBV5-4 | TRBJ2-7 | CASSPRDRERGEQYF | KLGGALQAK | YES | NO | NO |
| 7990 | TRAV13-1 | TRAJ11 | CAAKEGYSTLTF | TRBV5-4 | TRBJ2-7 | CASSYSGGSSYEQYF | KLGGALQAK | YES | NO | NO |
| 7991 | TRAV13-1 | TRAJ20 | CAAIRNDYKLSF | TRBV5-4 | TRBJ2-7 | CASSFDAAYEQYF | KLGGALQAK | YES | NO | NO |
| 7992 | TRAV13-2 | TRAJ30 | CAENSMNRDDKIIF | TRBV5-4 | TRBJ2-7 | CASSSRTGDSEQYF | KLGGALQAK | YES | NO | NO |
| 7993 | TRAV19 | TRAJ30 | CALSEGARDDKIIF | TRBV5-4 | TRBJ2-7 | CASSFSPSGAHEQYF | KLGGALQAK | YES | NO | NO |
| 7994 | TRAV19 | TRAJ4 | CALSDDVSGGYNKLIF | TRBV5-4 | TRBJ2-7 | CASRLSVDSGYEQYF | KLGGALQAK | YES | NO | NO |
| 7997 | TRAV36DV7 | TRAJ52 | CAVEAWFSEDGTSYGKLTF | TRBV5-4 | TRBJ2-7 | CASSFRQAVSEQYF | KLGGALQAK | NO | NO | NO |
| 7998 | TRAV38-1 | TRAJ15 | CAFTRPNQAGTALIF | TRBV5-4 | TRBJ2-7 | CASSQNPGQAYYEQYF | KLGGALQAK | YES | NO | NO |
| 7999 | TRAV41 | TRAJ21 | CAVSGYFNKFYF | TRBV5-4 | TRBJ2-7 | CASRNHVGQDYSYEQYF | KLGGALQAK | YES | NO | NO |
| 8000 | TRAV8-1 | TRAJ20 | CAVGRSNDYKLSF | TRBV5-4 | TRBJ2-7 | CASSKDSAYEQYF | KLGGALQAK | YES | NO | NO |
| 8002 | TRAV8-3 | TRAJ13 | CAVVGEGGYQKVTF | TRBV5-4 | TRBJ2-7 | CASSYNSYEQYF | KLGGALQAK | YES | NO | NO |
| 8003 | TRAV9-2 | TRAJ8 | CALDDFQKLVF | TRBV5-4 | TRBJ2-7 | CASSLGAGGPSYEQYV | KLGGALQAK | NO | NO | NO |
| 8006 | TRAV14DV4 | TRAJ4 | CAMREGMFSGGYNKLIF | TRBV5-5 | TRBJ2-7 | CASSLRSEGPTYEQYF | KLGGALQAK | YES | NO | NO |
| 8007 | TRAV16 | TRAJ54 | CALLQGAQKLVF | TRBV5-5 | TRBJ2-7 | CASSSDFYEQYF | KLGGALQAK | NO | NO | NO |
| 8009 | TRAV19 | TRAJ23 | CALSLRAYSGYNQGGKLIF | TRBV5-5 | TRBJ2-7 | CASRAGEHFYEQYF | KLGGALQAK | YES | NO | NO |
| 8010 | TRAV19 | TRAJ30 | CALVVRDDKIIF | TRBV5-5 | TRBJ2-7 | CASSFTDRGRDEQYF | KLGGALQAK | YES | NO | NO |
| 8011 | TRAV19 | TRAJ5 | CALSVLRDTGRRALTF | TRBV5-5 | TRBJ2-7 | CASSLEGNEQYF | KLGGALQAK | YES | NO | NO |
| 8012 | TRAV20 | TRAJ45 | CAAHPPMYSGGGADGLTF | TRBV5-5 | TRBJ2-7 | CASRAGEHFYEQYF | KLGGALQAK | YES | NO | NO |
| 8012 | TRAV20 | TRAJ45 | CAAHPPMYSGGGADGLTF | TRBV5-5 | TRBJ2-7 | CASRAGEHFYEQYF | AVFDRKSDAK | YES | NO | NO |
| 8014 | TRAV24 | TRAJ52 | CAFIGDAGGTSYGKLTF | TRBV5-5 | TRBJ2-7 | CASSNPYEQYV | KLGGALQAK | NO | NO | NO |
| 8015 | TRAV26-1 | TRAJ45 | CIVRVEGFHSGGGADGLTF | TRBV5-5 | TRBJ2-7 | CASSFFPTGGYHEQYF | KLGGALQAK | YES | NO | NO |
| 8016 | TRAV38-2DV8 | TRAJ39 | CAYRSSLNAGNMLTF | TRBV5-5 | TRBJ2-7 | CASSPPSYYEQYF | RLRAEAQVK | YES | YES | YES |
| 8017 | TRAV41 | TRAJ42 | CAVRFWGSQGNLIF | TRBV5-5 | TRBJ2-7 | CASSSTGLGEGIYEQYV | KLGGALQAK | NO | NO | NO |
| 8018 | TRAV41 | TRAJ49 | CSLNTGNQFYF | TRBV5-5 | TRBJ2-7 | CASSLGWVWRITGAYEQYF | KLGGALQAK | YES | NO | NO |
| 8019 | TRAV8-3 | TRAJ13 | CAVGGDSGGYQKVTF | TRBV5-5 | TRBJ2-7 | CASSDWGLSYEQYV | KLGGALQAK | NO | NO | NO |
| 8021 | TRAV12-2 | TRAJ23 | CAANLNQGGKLIF | TRBV5-6 | TRBJ2-7 | CASSLSGVPYEQYF | KLGGALQAK | YES | NO | NO |
| 8023 | TRAV12-2 | TRAJ48 | CAVNRRGFGNEKLTF | TRBV5-6 | TRBJ2-7 | CASSLEVRDGRAYEQYF | KLGGALQAK | YES | NO | NO |
| 8024 | TRAV13-2 | TRAJ28 | CAEIPSGAGSYQLTF | TRBV5-6 | TRBJ2-7 | CASSPGQGPYEQYF | KLGGALQAK | YES | NO | NO |
| 8025 | TRAV13-2 | TRAJ5 | CAENSVGRRALTF | TRBV5-6 | TRBJ2-7 | CASSLGEARGHEQYV | KLGGALQAK | NO | NO | NO |
| 8026 | TRAV14DV4 | TRAJ30 | CAMREWRDDKIIF | TRBV5-6 | TRBJ2-7 | CASSFGSGGRAGHEQYV | KLGGALQAK | NO | NO | NO |
| 8027 | TRAV14DV4 | TRAJ53 | CAMREDHSGGSNYKLTF | TRBV5-6 | TRBJ2-7 | CASSPGTAHEQYF | KLGGALQAK | YES | NO | NO |
| 8030 | TRAV21 | TRAJ26 | CAVPTPDNYGQNFVF | TRBV5-6 | TRBJ2-7 | CASSAVGTGDYEQYF | KLGGALQAK | YES | NO | NO |
| 8032 | TRAV24 | TRAJ37 | CALPRSGNTGKLIF | TRBV5-6 | TRBJ2-7 | CASSLIDTVWSLEQYF | KLGGALQAK | YES | NO | NO |
| 8033 | TRAV27 | TRAJ34 | CAGERTDKLIF | TRBV5-6 | TRBJ2-7 | CASSQGTGAYEQYV | KLGGALQAK | NO | NO | NO |
| 8034 | TRAV29DV5 | TRAJ45 | CAASGIGGADGLTF | TRBV5-6 | TRBJ2-7 | CASSLDPGEQYF | KLGGALQAK | YES | NO | NO |
| 8035 | TRAV29DV5 | TRAJ9 | CAAMNTGGFKTIF | TRBV5-6 | TRBJ2-7 | CASSPGTGPYEQYF | RLRAEAQVK | YES | NO | NO |
| 8037 | TRAV38-1 | TRAJ54 | CAFMSLIQGAQKLVF | TRBV5-6 | TRBJ2-7 | CASSFGGGGANEQYF | KLGGALQAK | YES | NO | NO |
| 8038 | TRAV38-1 | TRAJ54 | CAFMSLIQGAQKLVF | TRBV5-6 | TRBJ2-7 | CASSSDSTYEQYV | KLGGALQAK | NO | NO | NO |
| 8039 | TRAV41 | TRAJ21 | CAVRSGYNFNKFYF | TRBV5-6 | TRBJ2-7 | CASSLQVNIYEQYF | KLGGALQAK | YES | NO | NO |
| 8040 | TRAV41 | TRAJ34 | CAVRLYNTDKLIF | TRBV5-6 | TRBJ2-7 | CASSLFHEQYF | RLRAEAQVK | YES | NO | NO |
| 8041 | TRAV41 | TRAJ42 | CAVPRSQGNLIF | TRBV5-6 | TRBJ2-7 | CASSFVGSYEQYF | KLGGALQAK | YES | NO | NO |
| 8044 | TRAV12-1 | TRAJ49 | CVAYGNQFYF | TRBV5-8 | TRBJ2-7 | CASSLGGTYEQYF | KLGGALQAK | YES | NO | NO |
| 8045 | TRAV12-1 | TRAJ9 | CVVSGAGGFKTIF | TRBV5-8 | TRBJ2-7 | CASSLPRGQAYEQYF | KLGGALQAK | YES | NO | NO |
| 8046 | TRAV12-3 | TRAJ28 | CAMNSGAGSYQLTF | TRBV5-8 | TRBJ2-7 | CASSLPHYSDTGSYEQYF | KLGGALQAK | YES | NO | NO |
| 8048 | TRAV29DV5 | TRAJ26 | CAATYNYGQNFVF | TRBV5-8 | TRBJ2-7 | CASSPRETYEQYF | KLGGALQAK | YES | NO | NO |
| 8051 | TRAV5 | TRAJ18 | CAEVLSRGSTLGRLYF | TRBV5-8 | TRBJ2-7 | CASSVRSWEQYV | KLGGALQAK | NO | NO | NO |
| 8053 | TRAV9-2 | TRAJ32 | CALSDRAGLNYGGATNKLIF | TRBV5-8 | TRBJ2-7 | CASSEDSSGEQYF | RLRAEAQVK | NO | NO | NO |
| 8056 | TRAV1-2 | TRAJ33 | CAVMDSNYQLIW | TRBV6-1 | TRBJ2-7 | CASLLGTGGYEQYF | KLGGALQAK | YES | NO | NO |
| 8057 | TRAV12-1 | TRAJ49 | CVVNARNQFYF | TRBV6-1 | TRBJ2-7 | CASSEGQNTYEQYF | KLGGALQAK | YES | NO | NO |
| 8059 | TRAV13-1 | TRAJ49 | CAPSNTGNQFYF | TRBV6-1 | TRBJ2-7 | CASSRGALPVDEQYF | KLGGALQAK | YES | NO | NO |
| 8061 | TRAV14DV4 | TRAJ23 | CAMREGEERKLIF | TRBV6-1 | TRBJ2-7 | CAVWGSTTYEQYV | KLGGALQAK | NO | NO | NO |
| 8062 | TRAV19 | TRAJ49 | CALSDYTGNQFYF | TRBV6-1 | TRBJ2-7 | CASSEPGRYSSYEQYF | KLGGALQAK | YES | NO | NO |
| 8063 | TRAV20 | TRAJ57 | CAVTQGGSEKLVF | TRBV6-1 | TRBJ2-7 | CASSEYGAPYEQYF | KLGGALQAK | YES | NO | NO |
| 8065 | TRAV21 | TRAJ35 | CAVRGGFGNVLHC | TRBV6-1 | TRBJ2-7 | CASKGVVYEQYF | KLGGALQAK | NO | NO | NO |
| 8066 | TRAV24 | TRAJ15 | CALSEAGTALIF | TRBV6-1 | TRBJ2-7 | CASSDYEGYEQYV | KLGGALQAK | NO | NO | NO |
| 8067 | TRAV24 | TRAJ41 | CAPVSGYALNF | TRBV6-1 | TRBJ2-7 | CASSEGPLTGPYEQYF | RLRAEAQVK | YES | NO | YES |
| 8069 | TRAV26-2 | TRAJ52 | CILRDLNAGGTSYGKLTF | TRBV6-1 | TRBJ2-7 | CASSEVVSTTYEQYF | AVFDRKSDAK | YES | NO | NO |
| 8070 | TRAV29DV5 | TRAJ56 | CAASDTGANSKLTF | TRBV6-1 | TRBJ2-7 | CASSEDRGIRYEQYF | KLGGALQAK | YES | NO | NO |
| 8072 | TRAV3 | TRAJ31 | CAVRDTIFGNNARLMF | TRBV6-1 | TRBJ2-7 | CASSEKTGTGPHEQYF | KLGGALQAK | NO | NO | NO |
| 8072 | TRAV3 | TRAJ31 | CAVRDTIFGNNARLMF | TRBV6-1 | TRBJ2-7 | CASSEKTGTGPHEQYF | AVFDRKSDAK | NO | NO | NO |
| 8073 | TRAV38-2DV8 | TRAJ28 | CAYRSVYSGAGSYQLTF | TRBV6-1 | TRBJ2-7 | CASRHSYEQYF | KLGGALQAK | YES | NO | NO |
| 8075 | TRAV4 | TRAJ10 | CLVGAYTGGGNKLTF | TRBV6-1 | TRBJ2-7 | CASSGTDTYEQYF | KLGGALQAK | YES | NO | NO |
| 8076 | TRAV5 | TRAJ13 | CAEPSSGGYQKVTF | TRBV6-1 | TRBJ2-7 | CASSARDRVLYEQYF | KLGGALQAK | YES | NO | NO |
| 8077 | TRAV5 | TRAJ5 | CADLTGRRALTF | TRBV6-1 | TRBJ2-7 | CASSDSSGGYEQYF | KLGGALQAK | YES | NO | NO |
| 8079 | TRAV8-6 | TRAJ24 | CAVSDPSWGKLQF | TRBV6-1 | TRBJ2-7 | CASRDRSQYEQYF | KLGGALQAK | YES | NO | NO |
| 8081 | TRAV8-6 | TRAJ6 | CAVMGGSYIPTF | TRBV6-1 | TRBJ2-7 | CASHRGYPNEQYF | KLGGALQAK | YES | NO | NO |
| 8082 | TRAV1-1 | TRAJ42 | CALFYGGSQGNLIF | TRBV6-3 | TRBJ2-7 | CASSSDIGFYEQYV | KLGGALQAK | NO | NO | NO |
| 8083 | TRAV1-2 | TRAJ16 | CAVGSDGQKLLF | TRBV6-3 | TRBJ2-7 | CASSWAGENEQYV | KLGGALQAK | NO | NO | NO |
| 8085 | TRAV1-2 | TRAJ28 | CALPFSGAGSYQLTF | TRBV6-3 | TRBJ2-7 | CASRTGSGEQYF | KLGGALQAK | YES | NO | NO |
| 8086 | TRAV10 | TRAJ4 | CVVGDSGGYNKLIF | TRBV6-3 | TRBJ2-7 | CASSHGLAGVEQYF | KLGGALQAK | YES | NO | NO |
| 8087 | TRAV10 | TRAJ40 | CVVRSGTYKYIF | TRBV6-3 | TRBJ2-7 | CASSRGRGGEQYV | KLGGALQAK | NO | NO | NO |
| 8089 | TRAV12-2 | TRAJ54 | CAVGQVGQGAQKLVF | TRBV6-3 | TRBJ2-7 | CASTHLRSGGVYEQYF | KLGGALQAK | YES | NO | NO |
| 8093 | TRAV17 | TRAJ17 | CAAEAAGNKLTF | TRBV6-3 | TRBJ2-7 | CASSYDGLNEQYF | KLGGALQAK | YES | NO | NO |
| 8094 | TRAV19 | TRAJ47 | CALSETDYGNKLVF | TRBV6-3 | TRBJ2-7 | CASTSGAGHLTYEQYV | KLGGALQAK | NO | NO | NO |
| 8095 | TRAV2 | TRAJ17 | CPKAAGNKLTF | TRBV6-3 | TRBJ2-7 | CASSYSAGVPYEQYV | KLGGALQAK | NO | NO | NO |
| 8096 | TRAV20 | TRAJ7 | CAVQDYGNNRLAF | TRBV6-3 | TRBJ2-7 | CASSFGVGTESYEQYF | KLGGALQAK | YES | NO | NO |
| 8096 | TRAV20 | TRAJ7 | CAVQDYGNNRLAF | TRBV6-3 | TRBJ2-7 | CASSFGVGTESYEQYF | AVFDRKSDAK | YES | NO | NO |
| 8099 | TRAV26-2 | TRAJ33 | CILGTMDSNYQLIW | TRBV6-3 | TRBJ2-7 | CASSMGVYEQYV | KLGGALQAK | NO | NO | NO |
| 8102 | TRAV3 | TRAJ16 | CAVRDWF | TRBV6-3 | TRBJ2-7 | CASSYVGESYEQYF | KLGGALQAK | NO | NO | NO |
| 8103 | TRAV3 | TRAJ26 | CAVRDLWIGNYGQNFVF | TRBV6-3 | TRBJ2-7 | CASKGDRGAYEQYF | KLGGALQAK | NO | NO | NO |
| 8104 | TRAV3 | TRAJ31 | CAVRDPIGDNNARLMF | TRBV6-3 | TRBJ2-7 | CASSPGTSGNYEQYF | KLGGALQAK | NO | NO | NO |
| 8105 | TRAV4 | TRAJ26 | CLVGDESGGQNFVF | TRBV6-3 | TRBJ2-7 | CASRDRGYTYEQYV | KLGGALQAK | NO | NO | NO |
| 8106 | TRAV5 | TRAJ7 | CAELLYGNNRLAF | TRBV6-3 | TRBJ2-7 | CASRLAGGGEQYF | KLGGALQAK | YES | NO | NO |
| 8107 | TRAV6 | TRAJ40 | CALDIIGPSGTYKYIF | TRBV6-3 | TRBJ2-7 | CASKPGPTYEQYF | KLGGALQAK | YES | NO | NO |
| 8109 | TRAV8-3 | TRAJ47 | CAVGAQEYGNKLVF | TRBV6-3 | TRBJ2-7 | CASSYSTASGPYEQYV | KLGGALQAK | NO | NO | NO |
| 8110 | TRAV8-4 | TRAJ4 | CAVSDDSGGYNKLIF | TRBV6-3 | TRBJ2-7 | CASKPPGSSSYEQYF | KLGGALQAK | YES | NO | NO |
| 8113 | TRAV9-2 | TRAJ5 | CALTLITGRRALTF | TRBV6-3 | TRBJ2-7 | CASSYVLQTYEQYF | KLGGALQAK | YES | NO | NO |
| 8115 | TRAV4 | TRAJ45 | CLVGLIGFSGGGADGLTF | TRBV6-4 | TRBJ2-7 | CASNWGEQYF | KLGGALQAK | YES | NO | NO |

| | | | | | | | | | | |
|---|---|---|---|---|---|---|---|---|---|---|
| 8116 | TRAV1-1 | TRAJ4 | CAVSPWGGYNKLIF | TRBV6-5 | TRBJ2-7 | CASSYGRAYEQYF | KLGGALQAK | YES | NO | NO |
| 8116 | TRAV1-1 | TRAJ4 | CAVSPWGGYNKLIF | TRBV6-5 | TRBJ2-7 | CASSYGRAYEQYF | RLRAEAQVK | YES | NO | NO |
| 8116 | TRAV1-1 | TRAJ4 | CAVSPWGGYNKLIF | TRBV6-5 | TRBJ2-7 | CASSYGRAYEQYF | AVFDRKSDAK | YES | NO | NO |
| 8117 | TRAV1-2 | TRAJ34 | CAVRGEDTDKLIF | TRBV6-5 | TRBJ2-7 | CASSTRTDPPREQYF | KLGGALQAK | YES | NO | NO |
| 8118 | TRAV12-1 | TRAJ33 | CVVNTDSNYQLIW | TRBV6-5 | TRBJ2-7 | CASSTGGRHEQYF | KLGGALQAK | YES | NO | NO |
| 8120 | TRAV12-2 | TRAJ22 | CAVPPPKGSARQLTF | TRBV6-5 | TRBJ2-7 | CASSDTTGGFGEQYF | KLGGALQAK | YES | NO | NO |
| 8122 | TRAV13-1 | TRAJ10 | CAVLPGGGNKLTF | TRBV6-5 | TRBJ2-7 | CASSYSYEQYF | AVFDRKSDAK | YES | NO | NO |
| 8123 | TRAV13-1 | TRAJ10 | CAALPGGGNKLTF | TRBV6-5 | TRBJ2-7 | CASSYSYEQYF | KLGGALQAK | YES | NO | NO |
| 8124 | TRAV13-1 | TRAJ3 | CAATSYSSASKIIF | TRBV6-5 | TRBJ2-7 | CASSTLTSGSRSYEQYF | RLRAEAQVK | YES | YES | YES |
| 8125 | TRAV13-1 | TRAJ45 | CAASGAGGGADGLTF | TRBV6-5 | TRBJ2-7 | CASSYAHRGDMGEQYF | KLGGALQAK | YES | NO | NO |
| 8126 | TRAV13-1 | TRAJ5 | CAALDTGRRALTF | TRBV6-5 | TRBJ2-7 | CASSSGQGSYEQYV | KLGGALQAK | NO | NO | NO |
| 8127 | TRAV13-2 | TRAJ45 | CAERPSSGGGADGLTF | TRBV6-5 | TRBJ2-7 | CASTSGHEQYF | KLGGALQAK | YES | NO | NO |
| 8128 | TRAV14DV4 | TRAJ30 | CAMREGRMNRDDKIIF | TRBV6-5 | TRBJ2-7 | CASSFQTSTTLEQYF | KLGGALQAK | YES | NO | NO |
| 8129 | TRAV14DV4 | TRAJ36 | CAMREPDQTGANNLFF | TRBV6-5 | TRBJ2-7 | CASSYRRVQDEQYF | KLGGALQAK | YES | NO | NO |
| 8130 | TRAV14DV4 | TRAJ57 | CAMREGSQGGSEKLVF | TRBV6-5 | TRBJ2-7 | CASKTGSYEQYF | KLGGALQAK | YES | NO | NO |
| 8132 | TRAV19 | TRAJ32 | CALSEAQYGGATNKLIF | TRBV6-5 | TRBJ2-7 | CATSPNRAYEQYF | AVFDRKSDAK | YES | NO | NO |
| 8132 | TRAV19 | TRAJ32 | CALSEAQYGGATNKLIF | TRBV6-5 | TRBJ2-7 | CATSPNRAYEQYF | KLGGALQAK | YES | NO | NO |
| 8133 | TRAV21 | TRAJ15 | CAVRPVLKTGTALIF | TRBV6-5 | TRBJ2-7 | CASSYSPEVGEQYF | KLGGALQAK | YES | NO | NO |
| 8134 | TRAV21 | TRAJ49 | CAAYGGTGNQFYF | TRBV6-5 | TRBJ2-7 | CASSYGSGAYEQYV | KLGGALQAK | NO | NO | NO |
| 8135 | TRAV21 | TRAJ57 | CAVGRLTQGGSEKLVF | TRBV6-5 | TRBJ2-7 | CASTPWREYPLEQYF | KLGGALQAK | YES | NO | NO |
| 8137 | TRAV26-1 | TRAJ46 | CIAAGPESSGDKLTF | TRBV6-5 | TRBJ2-7 | CASSSSLFGRITSYEQYF | KLGGALQAK | YES | NO | NO |
| 8138 | TRAV27 | TRAJ26 | CAGPPYGQNFVF | TRBV6-5 | TRBJ2-7 | CASSLQGTGLYEQYF | KLGGALQAK | YES | NO | NO |
| 8141 | TRAV29DV5 | TRAJ29 | CAASRSGNTPLVF | TRBV6-5 | TRBJ2-7 | CASSYWGIGLAPTYEQYV | KLGGALQAK | NO | NO | NO |
| 8142 | TRAV29DV5 | TRAJ33 | CAASGRASNYQLIW | TRBV6-5 | TRBJ2-7 | CASSYSSGSYEQYF | KLGGALQAK | YES | NO | NO |
| 8143 | TRAV35 | TRAJ30 | CAGPYDKIIF | TRBV6-5 | TRBJ2-7 | CASSNRVDYEQYF | KLGGALQAK | YES | NO | NO |
| 8146 | TRAV4 | TRAJ33 | CLVGDGSNYQLIW | TRBV6-5 | TRBJ2-7 | CASSVVAGVGYEQYF | KLGGALQAK | NO | NO | NO |
| 8147 | TRAV41 | TRAJ57 | CALRNLDRGGSEKLVF | TRBV6-5 | TRBJ2-7 | CASSYGAGGYEQYV | KLGGALQAK | NO | NO | NO |
| 8149 | TRAV8-3 | TRAJ17 | CAVAAGNKLTF | TRBV6-5 | TRBJ2-7 | CASSPRTDEGQYF | KLGGALQAK | YES | NO | NO |
| 8150 | TRAV8-4 | TRAJ49 | CAVSFGNQFYF | TRBV6-5 | TRBJ2-7 | CASSRSGRERDEQYF | KLGGALQAK | YES | NO | NO |
| 8151 | TRAV12-2 | TRAJ29 | CAVNMKGNTPLVF | TRBV6-6 | TRBJ2-7 | CASSYSETYEQYF | KLGGALQAK | NO | NO | NO |
| 8153 | TRAV13-1 | TRAJ45 | CAASRSGGGADGLTF | TRBV6-6 | TRBJ2-7 | CASSSAGAGALPYEQYV | KLGGALQAK | NO | NO | NO |
| 8154 | TRAV13-2 | TRAJ50 | CAEKVTRYQTSYDKVIF | TRBV6-6 | TRBJ2-7 | CASSYYRDRVYEQYF | KLGGALQAK | NO | NO | NO |
| 8156 | TRAV14DV4 | TRAJ40 | CAMRETSGTYKYIF | TRBV6-6 | TRBJ2-7 | CASTNGGQSLV | KLGGALQAK | NO | NO | NO |
| 8157 | TRAV17 | TRAJ21 | CATEGGLYNFNKFYF | TRBV6-6 | TRBJ2-7 | CASSYGGQRASYEQYF | KLGGALQAK | NO | NO | NO |
| 8158 | TRAV21 | TRAJ49 | CAAHRDTGNQFYF | TRBV6-6 | TRBJ2-7 | CASTTPADGLSGTSGSSYEQYF | KLGGALQAK | NO | NO | NO |
| 8159 | TRAV23DV6 | TRAJ23 | CAASSNRYNQGGKLIF | TRBV6-6 | TRBJ2-7 | CASDAGTAYEQYF | KLGGALQAK | NO | NO | NO |
| 8160 | TRAV29DV5 | TRAJ31 | CAAPGSVGARLMF | TRBV6-6 | TRBJ2-7 | CASSEGRGEQYF | KLGGALQAK | NO | NO | NO |
| 8162 | TRAV36DV7 | TRAJ40 | CAVEPSGTYKYIF | TRBV6-6 | TRBJ2-7 | CASSYSGEQYF | KLGGALQAK | NO | NO | NO |
| 8163 | TRAV38-1 | TRAJ40 | CAFMGLSSGTYKYIF | TRBV6-6 | TRBJ2-7 | CASSYSPPDAVKYEQYF | KLGGALQAK | NO | NO | NO |
| 8165 | TRAV5 | TRAJ39 | CAEGTNNAGNMLTF | TRBV6-6 | TRBJ2-7 | CASSASDAYEQYF | KLGGALQAK | NO | NO | NO |
| 8168 | TRAV1-1 | TRAJ18 | CAVRRRGSTLGRLYF | TRBV7-2 | TRBJ2-7 | CASSPGTATYEQYF | KLGGALQAK | NO | NO | NO |
| 8171 | TRAV10 | TRAJ39 | CVVSAGAGNMLTF | TRBV7-2 | TRBJ2-7 | CASSETMSSYEQYV | KLGGALQAK | NO | NO | NO |
| 8172 | TRAV12-1 | TRAJ22 | CVVNLVGSARQLTF | TRBV7-2 | TRBJ2-7 | CASSLLGEGWHEQYF | KLGGALQAK | YES | NO | NO |
| 8173 | TRAV12-2 | TRAJ17 | CAVKGGAAGNKLTF | TRBV7-2 | TRBJ2-7 | CASSLDSAYEQYF | KLGGALQAK | NO | NO | NO |
| 8177 | TRAV13-2 | TRAJ52 | CAERDSNAGGTSYGKLTF | TRBV7-2 | TRBJ2-7 | CASSLPGYEQYF | KLGGALQAK | YES | NO | NO |
| 8180 | TRAV19 | TRAJ24 | CALSEDMTTDSWGKLQF | TRBV7-2 | TRBJ2-7 | CASSLAPSGGDEQYF | KLGGALQAK | NO | NO | NO |
| 8182 | TRAV19 | TRAJ52 | CALRADNAGGTSYGKLTF | TRBV7-2 | TRBJ2-7 | CASSSWGVGEQYV | KLGGALQAK | NO | NO | NO |
| 8183 | TRAV24 | TRAJ52 | CAFIGDAGGTSYGKLTF | TRBV7-2 | TRBJ2-7 | CASSLEPSYEQYF | KLGGALQAK | YES | NO | NO |
| 8184 | TRAV27 | TRAJ40 | CAGAPTSGTYKYIF | TRBV7-2 | TRBJ2-7 | CASSPGWLYEQYV | KLGGALQAK | NO | NO | NO |
| 8187 | TRAV38-1 | TRAJ57 | CAFMTGEQGGSEKLVF | TRBV7-2 | TRBJ2-7 | CASSLEWGLAGGAGYEQYV | KLGGALQAK | NO | NO | NO |
| 8192 | TRAV8-1 | TRAJ22 | CAVRSSGSARQLTF | TRBV7-2 | TRBJ2-7 | CASSLEPSYEQYF | IVTDFSVIK | NO | NO | NO |
| 8196 | TRAV29DV5 | TRAJ28 | CAASPEGAGSYQLTF | TRBV7-3 | TRBJ2-7 | CASSLYGGAGEQYF | KLGGALQAK | NO | NO | NO |
| 8199 | TRAV1-2 | TRAJ31 | CAVRSRDARLMF | TRBV7-6 | TRBJ2-7 | CASSLEAGGPSPLAYEQYF | KLGGALQAK | YES | NO | NO |
| 8200 | TRAV13-1 | TRAJ39 | CAATSRNAGNMLTF | TRBV7-6 | TRBJ2-7 | CASSLVRGVSSYEQYF | KLGGALQAK | YES | NO | NO |
| 8202 | TRAV13-1 | TRAJ54 | CAASTGGAQKLVF | TRBV7-6 | TRBJ2-7 | CASSLEQGTYEQYF | KLGGALQAK | YES | NO | NO |
| 8204 | TRAV21 | TRAJ57 | CAVIWQGGSEKLVF | TRBV7-6 | TRBJ2-7 | CASSPPRGYEQYF | KLGGALQAK | YES | NO | NO |
| 8206 | TRAV22 | TRAJ47 | CAVEPLYGNKLVF | TRBV7-6 | TRBJ2-7 | CASSPTQGAVNEQYF | KLGGALQAK | NO | NO | NO |
| 8207 | TRAV23DV6 | TRAJ58 | CAARKGTSGSRLTF | TRBV7-6 | TRBJ2-7 | CASSLIFGRQYF | KLGGALQAK | NO | NO | NO |
| 8209 | TRAV35 | TRAJ44 | CAGPGSDTGTASKLTF | TRBV7-6 | TRBJ2-7 | CASSDVKGSYEQYF | KLGGALQAK | YES | NO | NO |
| 8211 | TRAV38-1 | TRAJ29 | CAFMKQPLSGNTPLVF | TRBV7-6 | TRBJ2-7 | CASSYPGPYYEQYF | KLGGALQAK | YES | NO | NO |
| 8212 | TRAV41 | TRAJ40 | CAAEGTYKYIF | TRBV7-6 | TRBJ2-7 | CASSDSYEQYF | KLGGALQAK | YES | NO | NO |
| 8214 | TRAV8-6 | TRAJ57 | CAAPDQGGSEKLVF | TRBV7-6 | TRBJ2-7 | CASSWGQGNEQYV | KLGGALQAK | NO | NO | NO |
| 8215 | TRAV17 | TRAJ54 | CATGIQGAQKLVF | TRBV7-7 | TRBJ2-7 | CASSAWGLEYEQYF | KLGGALQAK | YES | NO | NO |
| 8217 | TRAV1-1 | TRAJ23 | CAVDKGGKLIF | TRBV7-8 | TRBJ2-7 | CASSSATGAYEQYF | KLGGALQAK | YES | NO | NO |
| 8218 | TRAV12-1 | TRAJ48 | CVVKVPLRNEKLTF | TRBV7-8 | TRBJ2-7 | CASSSATGAYEQYF | AVFDRKSDAK | YES | NO | NO |
| 8218 | TRAV12-1 | TRAJ48 | CVVKVPLRNEKLTF | TRBV7-8 | TRBJ2-7 | CASSSATGAYEQYF | KLGGALQAK | YES | NO | NO |
| 8227 | TRAV26-1 | TRAJ49 | CIVRVADTGNQFYF | TRBV7-8 | TRBJ2-7 | CASSLEEGRRDEQYF | KLGGALQAK | YES | NO | NO |
| 8228 | TRAV26-2 | TRAJ45 | CILRLGGGADGLTF | TRBV7-8 | TRBJ2-7 | CASSWGSGGETYEQYF | KLGGALQAK | YES | NO | NO |
| 8231 | TRAV29DV5 | TRAJ45 | CAAISGGGADGLTF | TRBV7-8 | TRBJ2-7 | CASSWTSYEQYF | KLGGALQAK | YES | NO | NO |
| 8232 | TRAV29DV5 | TRAJ6 | CAASSLSGSYIPTF | TRBV7-8 | TRBJ2-7 | CASRLSGSAYEQYF | KLGGALQAK | YES | NO | NO |
| 8232 | TRAV29DV5 | TRAJ6 | CAASSLSGSYIPTF | TRBV7-8 | TRBJ2-7 | CASRLSGSAYEQYF | RLRAEAQVK | YES | NO | NO |
| 8233 | TRAV38-2DV8 | TRAJ49 | CAYRRRGNQFYF | TRBV7-8 | TRBJ2-7 | CASSMREPEYGLEQYF | KLGGALQAK | YES | NO | NO |
| 8234 | TRAV39 | TRAJ31 | CAVDNGARLMF | TRBV7-8 | TRBJ2-7 | CASSYTRGGHEQYF | KLGGALQAK | YES | NO | NO |
| 8236 | TRAV8-1 | TRAJ20 | CAVNPVNDYKLSF | TRBV7-8 | TRBJ2-7 | CASSLGYEQYF | KLGGALQAK | YES | NO | NO |
| 8240 | TRAV1-2 | TRAJ20 | CAPESNDYKLSF | TRBV7-9 | TRBJ2-7 | CASSMGQASYEQYF | KLGGALQAK | YES | NO | NO |
| 8243 | TRAV12-2 | TRAJ10 | CAVNRNGGGNKLTF | TRBV7-9 | TRBJ2-7 | CASSSRDRGYEQYF | KLGGALQAK | YES | NO | NO |
| 8244 | TRAV12-2 | TRAJ17 | CAVNEAAGNKLTF | TRBV7-9 | TRBJ2-7 | CASSPSAGGAGNEQYF | KLGGALQAK | YES | NO | NO |
| 8245 | TRAV12-2 | TRAJ34 | CAVREYGKLIF | TRBV7-9 | TRBJ2-7 | CASSLVGGDEQYF | KLGGALQAK | YES | NO | NO |
| 8246 | TRAV12-2 | TRAJ45 | CAVSGGGADGLTF | TRBV7-9 | TRBJ2-7 | CAWSGTATPYEQYV | KLGGALQAK | NO | NO | NO |
| 8247 | TRAV12-2 | TRAJ48 | CAVNIGFGNEKLTF | TRBV7-9 | TRBJ2-7 | CASSSFGGTGTGSYEQYF | KLGGALQAK | YES | NO | NO |
| 8249 | TRAV12-2 | TRAJ54 | CAVQIQGAQKLVF | TRBV7-9 | TRBJ2-7 | CASWGLGLGEQYF | KLGGALQAK | YES | NO | NO |
| 8250 | TRAV12-3 | TRAJ31 | CAMSSNNNARLMF | TRBV7-9 | TRBJ2-7 | CASSLGLASYEQYF | KLGGALQAK | YES | NO | NO |
| 8255 | TRAV13-1 | TRAJ37 | CAASRGSSNTGKLIF | TRBV7-9 | TRBJ2-7 | CASSLGGQGSVEQYF | KLGGALQAK | YES | NO | NO |
| 8258 | TRAV14DV4 | TRAJ12 | CAMQDSSYKLIF | TRBV7-9 | TRBJ2-7 | CASSSWSGGASGEQYF | KLGGALQAK | YES | NO | NO |
| 8261 | TRAV14DV4 | TRAJ33 | CAMREGEDSNYQLIW | TRBV7-9 | TRBJ2-7 | CASSRTGEQYF | KLGGALQAK | YES | NO | NO |
| 8263 | TRAV14DV4 | TRAJ49 | CAMRSTGNQFYF | TRBV7-9 | TRBJ2-7 | CASSQDPSRTYEQYF | KLGGALQAK | NO | NO | NO |
| 8266 | TRAV14DV4 | TRAJ54 | CAMREVTQGAQKLVF | TRBV7-9 | TRBJ2-7 | CASSLRDGSYEQYF | KLGGALQAK | YES | NO | NO |
| 8268 | TRAV17 | TRAJ13 | CAQGYQKVTF | TRBV7-9 | TRBJ2-7 | CASSLGSRPSGGYEQYF | KLGGALQAK | YES | NO | NO |
| 8270 | TRAV19 | TRAJ22 | CALSEAGGSARQLTF | TRBV7-9 | TRBJ2-7 | CASSPQLAGGREQYF | KLGGALQAK | YES | NO | NO |
| 8271 | TRAV19 | TRAJ36 | CALSEALGRTGANNLFF | TRBV7-9 | TRBJ2-7 | CASSPRPQRGGYEQYF | KLGGALQAK | YES | NO | NO |
| 8277 | TRAV21 | TRAJ40 | CAVSPSTSGTYKYIF | TRBV7-9 | TRBJ2-7 | CASSLSGYYEQYF | KLGGALQAK | YES | NO | NO |
| 8278 | TRAV21 | TRAJ43 | CAVWYMRF | TRBV7-9 | TRBJ2-7 | CASKDGISYEQYF | KLGGALQAK | NO | NO | NO |
| 8279 | TRAV21 | TRAJ53 | CAPLGGGSNYKLTF | TRBV7-9 | TRBJ2-7 | CASSDPTVSGEQYF | KLGGALQAK | YES | NO | NO |
| 8280 | TRAV21 | TRAJ58 | CAVRGETSGSRLTF | TRBV7-9 | TRBJ2-7 | CASSPTGTSVYEQYF | KLGGALQAK | YES | NO | NO |
| 8282 | TRAV24 | TRAJ16 | CAFGRSGQKLLF | TRBV7-9 | TRBJ2-7 | CASSLVFWADNEQYF | KLGGALQAK | NO | NO | NO |
| 8283 | TRAV24 | TRAJ47 | CASMEYGNKLVF | TRBV7-9 | TRBJ2-7 | CASKDGISYEQYF | KLGGALQAK | YES | NO | NO |
| 8285 | TRAV26-1 | TRAJ54 | CIVRGADQGAQKLVF | TRBV7-9 | TRBJ2-7 | CASKDGISYEQYF | AVFDRKSDAK | YES | NO | NO |
| 8285 | TRAV26-1 | TRAJ54 | CIVRGADQGAQKLVF | TRBV7-9 | TRBJ2-7 | CASKDGISYEQYF | RLRAEAQVK | YES | NO | NO |
| 8288 | TRAV3 | TRAJ4 | CAVRDIGGYNKLIF | TRBV7-9 | TRBJ2-7 | CASSLLGTGVREQYF | KLGGALQAK | NO | NO | NO |
| 8289 | TRAV36DV7 | TRAJ41 | CAVEEPSGYALNF | TRBV7-9 | TRBJ2-7 | CASSNPGTAQREQYF | KLGGALQAK | NO | NO | NO |
| 8290 | TRAV38-1 | TRAJ52 | CAFMRGRGKLTF | TRBV7-9 | TRBJ2-7 | CASSFRDSYEQYF | KLGGALQAK | YES | NO | NO |
| 8291 | TRAV38-2DV8 | TRAJ23 | CAYRREGKLIF | TRBV7-9 | TRBJ2-7 | CASSSGGSYEQYF | KLGGALQAK | YES | NO | NO |
| 8292 | TRAV38-2DV8 | TRAJ42 | CAYYMNYGGSQGNLIF | TRBV7-9 | TRBJ2-7 | CASSLGGGLAGSGEQYF | KLGGALQAK | YES | NO | NO |
| 8293 | TRAV38-2DV8 | TRAJ44 | CAYRTTGTASKLTF | TRBV7-9 | TRBJ2-7 | CATARDRSEQYF | KLGGALQAK | YES | NO | NO |

| | | | | | | | | | | |
|---|---|---|---|---|---|---|---|---|---|---|
| 8296 | TRAV5 | TRAJ9 | CAEKDTGGFKTIF | TRBV7-9 | TRBJ2-7 | CASTGGDRTYYEQYF | KLGGALQAK | YES | NO | NO |
| 8297 | TRAV6 | TRAJ43 | CALGVTYNNNDMRF | TRBV7-9 | TRBJ2-7 | CASSGQVGYEQYF | KLGGALQAK | YES | NO | NO |
| 8298 | TRAV8-1 | TRAJ21 | CAVNEPFNKFYF | TRBV7-9 | TRBJ2-7 | CASSPSAGGAGNEQYF | KLGGALQAK | YES | NO | NO |
| 8302 | TRAV8-3 | TRAJ50 | CAVSKTSYDKVIF | TRBV7-9 | TRBJ2-7 | CASSSTMDIDYEQYV | KLGGALQAK | NO | NO | NO |
| 8303 | TRAV9-2 | TRAJ52 | CALSNAGGTSYGKLTF | TRBV7-9 | TRBJ2-7 | CASSGVAGGPYEQYV | KLGGALQAK | NO | NO | NO |
| 8305 | TRAV1-1 | TRAJ47 | CAVSELEYGNKLVF | TRBV9 | TRBJ2-7 | CASSQGGGEQYF | KLGGALQAK | YES | NO | NO |
| 8307 | TRAV1-2 | TRAJ28 | CAVSDSGAGSYQLTF | TRBV9 | TRBJ2-7 | CASSVGGSGGTYEQYF | KLGGALQAK | YES | NO | NO |
| 8310 | TRAV12-1 | TRAJ5 | CVVNPMDTGRRALTF | TRBV9 | TRBJ2-7 | CASSVGGGGAYEQYF | KLGGALQAK | YES | NO | NO |
| 8311 | TRAV12-2 | TRAJ16 | CAVNMDGQKLLF | TRBV9 | TRBJ2-7 | CASSVGESYEQYF | KLGGALQAK | NO | NO | NO |
| 8312 | TRAV12-2 | TRAJ47 | CAVDMEYGNKLVF | TRBV9 | TRBJ2-7 | CASSEGLAGGQGEQYF | KLGGALQAK | YES | NO | NO |
| 8315 | TRAV12-3 | TRAJ34 | CAPHPYNTDKLIF | TRBV9 | TRBJ2-7 | CASSPGGGKTYEQYF | KLGGALQAK | YES | NO | NO |
| 8327 | TRAV19 | TRAJ39 | CALSEAHEGNMLTF | TRBV9 | TRBJ2-7 | CASGIRKTGGEQYF | KLGGALQAK | YES | NO | NO |
| 8329 | TRAV19 | TRAJ43 | CALSEGHDMRF | TRBV9 | TRBJ2-7 | CASSGGQDPYEQYF | KLGGALQAK | YES | NO | NO |
| 8331 | TRAV19 | TRAJ52 | CALSENAGGTSYGKLTF | TRBV9 | TRBJ2-7 | CASSFTSGGYEQYF | AVFDRKSDAK | YES | NO | NO |
| 8331 | TRAV19 | TRAJ52 | CALSENAGGTSYGKLTF | TRBV9 | TRBJ2-7 | CASSFTSGGYEQYF | KLGGALQAK | YES | NO | NO |
| 8332 | TRAV19 | TRAJ58 | CALSVKETSGSRLTF | TRBV9 | TRBJ2-7 | CASSAGGAPLYEQYF | KLGGALQAK | YES | NO | NO |
| 8334 | TRAV21 | TRAJ16 | CAVIGSDGQKLLF | TRBV9 | TRBJ2-7 | CASSVGTSPYEQYF | KLGGALQAK | NO | NO | NO |
| 8339 | TRAV25 | TRAJ37 | CAGYSSGNTGKLIF | TRBV9 | TRBJ2-7 | CASSVEASGVYEQYF | KLGGALQAK | YES | NO | NO |
| 8340 | TRAV26-1 | TRAJ58 | CIVRVAEETSGSRLTF | TRBV9 | TRBJ2-7 | CASSVGIGAGYEQYF | KLGGALQAK | NO | NO | NO |
| 8342 | TRAV26-2 | TRAJ54 | CILAQKLVF | TRBV9 | TRBJ2-7 | CASSPQGTGLYEQYF | KLGGALQAK | NO | NO | NO |
| 8343 | TRAV29DV5 | TRAJ53 | CGGSNYKLTF | TRBV9 | TRBJ2-7 | CASSVDEQFSYEQYF | KLGGALQAK | YES | NO | NO |
| 8346 | TRAV3 | TRAJ45 | CAVRAHSGGGADGLTF | TRBV9 | TRBJ2-7 | CASSPGGGASYEQYF | RIAAWMATY | NO | NO | NO |
| 8348 | TRAV38-2DV8 | TRAJ49 | CAYRSAEVAGNQFYF | TRBV9 | TRBJ2-7 | CASSVRGQGPHEQYF | KLGGALQAK | YES | NO | NO |
| 8349 | TRAV39 | TRAJ49 | CAVDGNQFYF | TRBV9 | TRBJ2-7 | CASSVDEREQYF | KLGGALQAK | YES | NO | NO |
| 8350 | TRAV41 | TRAJ34 | CAVKATDKLIF | TRBV9 | TRBJ2-7 | CASSTQGVLTEQYF | KLGGALQAK | YES | NO | NO |
| 8352 | TRAV8-3 | TRAJ7 | CAVGGSLNYGNNRLAF | TRBV9 | TRBJ2-7 | CASSVAGTGGPGPSYEQYF | KLGGALQAK | YES | NO | NO |
| 8353 | TRAV9-2 | TRAJ6 | CALSISGGSYIPTF | TRBV9 | TRBJ2-7 | CASSPKGQGFEQYF | KLGGALQAK | YES | NO | NO |
| 8354 | TRAV16 | TRAJ11 | CALIDSSGYSTLTF | TRBV10-1 | TRBJ1-1 | CASSESWGTEAFF | AVFDRKSDAK | NO | NO | NO |
| 8356 | TRAV21 | TRAJ33 | CAVLMDSNYQLIW | TRBV10-2 | TRBJ1-1 | CASSDDGMNTEAFF | AVFDRKSDAK | YES | NO | YES |
| 8357 | TRAV26-1 | TRAJ27 | CIVRDLAGKSTF | TRBV10-2 | TRBJ1-1 | CASSESSQMNTEAFF | AVFDRKSDAK | YES | NO | YES |
| 8362 | TRAV3 | TRAJ4 | CAVSPFSGGYNKLIF | TRBV11-2 | TRBJ1-1 | CASSLEYGDIEAFF | AVFDRKSDAK | YES | NO | YES |
| 8363 | TRAV4 | TRAJ22 | CLVADLSSGSARQLTF | TRBV11-2 | TRBJ1-1 | CASSWEGFGTEAFF | AVFDRKSDAK | NO | NO | NO |
| 8366 | TRAV1-2 | TRAJ35 | CAVESFGNVLHC | TRBV12-3 | TRBJ1-1 | CASSGPLTGKGGEAFF | AVFDRKSDAK | NO | NO | NO |
| 8368 | TRAV27 | TRAJ20 | CAGANDYKLSF | TRBV12-3 | TRBJ1-1 | CASSFTGVNTEAFF | AVFDRKSDAK | YES | YES | YES |
| 8371 | TRAV27 | TRAJ24 | CAGAGSWGKLQF | TRBV12-4 | TRBJ1-1 | CASSLGVEGMNTEAFF | IVTDFSVIK | YES | NO | YES |
| 8372 | TRAV30 | TRAJ4 | CGTPISGGYNKLIF | TRBV12-4 | TRBJ1-1 | CASSFSERAFF | AVFDRKSDAK | YES | NO | YES |
| 8377 | TRAV12-1 | TRAJ21 | CVVYNFNKFYF | TRBV15 | TRBJ1-1 | CATSRAGGGTEAFF | AVFDRKSDAK | YES | NO | YES |
| 8380 | TRAV27 | TRAJ12 | CAGGMDSSYKLIF | TRBV18 | TRBJ1-1 | CASSPRGRRDTEAFF | AVFDRKSDAK | YES | NO | NO |
| 8382 | TRAV17 | TRAJ43 | CATDDPFNDMRF | TRBV19 | TRBJ1-1 | CASISTEAFF | AVFDRKSDAK | YES | NO | YES |
| 8383 | TRAV19 | TRAJ42 | CALSEALRNYGGSQGNLIF | TRBV19 | TRBJ1-1 | CASSLWDRVNTEAFF | AVFDRKSDAK | YES | NO | YES |
| 8384 | TRAV20 | TRAJ49 | CAVQGGNQFYF | TRBV19 | TRBJ1-1 | CASSMWGLLNTEAFF | AVFDRKSDAK | YES | NO | YES |
| 8387 | TRAV21 | TRAJ48 | CAVGANFGNEKLTF | TRBV19 | TRBJ1-1 | CASSWAGPHTEAFF | IVTDFSVIK | YES | YES | YES |
| 8388 | TRAV22 | TRAJ15 | CAVPPQANQAGTALIF | TRBV19 | TRBJ1-1 | CASSITGLGTEAFF | AVFDRKSDAK | YES | NO | YES |
| 8389 | TRAV3 | TRAJ5 | CAVSVDTGRRALTF | TRBV19 | TRBJ1-1 | CASRGWESNTEAFF | IVTDFSVIK | NO | NO | NO |
| 8392 | TRAV38-2DV8 | TRAJ31 | CAYYNARLMF | TRBV19 | TRBJ1-1 | CASRLSSRGMNTEAFF | AVFDRKSDAK | YES | YES | YES |
| 8394 | TRAV8-4 | TRAJ6 | CAVSDETGGSYIPTF | TRBV19 | TRBJ1-1 | CASSIGTGGTEAFF | AVFDRKSDAK | YES | NO | YES |
| 8395 | TRAV38-2DV8 | TRAJ45 | CAYRAYSGGGADGLTF | TRBV2 | TRBJ1-1 | CASSEALVPTWYTEAFF | AVFDRKSDAK | YES | NO | YES |
| 8400 | TRAV17 | TRAJ47 | CATDGVMEYGNKLVF | TRBV25-1 | TRBJ1-1 | CASRRQEYGTEAFF | AVFDRKSDAK | YES | NO | NO |
| 8401 | TRAV8-3 | TRAJ42 | CAVGDNYGGSQGNLIF | TRBV25-1 | TRBJ1-1 | CASRIGYTEAFF | AVFDRKSDAK | YES | NO | YES |
| 8402 | TRAV29DV5 | TRAJ49 | CAALFTGNQFYF | TRBV27 | TRBJ1-1 | CASNHGTGATEAFF | AVFDRKSDAK | YES | NO | YES |
| 8405 | TRAV12-1 | TRAJ20 | CVVNGNDYKLSF | TRBV28 | TRBJ1-1 | CASSLAAANTEAFF | IVTDFSVIK | YES | YES | YES |
| 8407 | TRAV29DV5 | TRAJ28 | CAAPSAYSGAGSYQLTF | TRBV28 | TRBJ1-1 | CASSLWRNTEAFF | AVFDRKSDAK | YES | NO | YES |
| 8410 | TRAV5 | TRAJ26 | CAESMGNYGQNFVF | TRBV29-1 | TRBJ1-1 | CSDLTGVDTEAFF | AVFDRKSDAK | YES | NO | NO |
| 8411 | TRAV12-1 | TRAJ4 | CVGESGGYNKLIF | TRBV3-1 | TRBJ1-1 | CASSQDVGSYTEAFF | AVFDRKSDAK | YES | NO | YES |
| 8412 | TRAV12-1 | TRAJ8 | CVVMNTGFQKLVF | TRBV30 | TRBJ1-1 | CAWSVGIQNTEAFF | IVTDFSVIK | NO | NO | NO |
| 8413 | TRAV17 | TRAJ20 | CATAHNDYKLSF | TRBV30 | TRBJ1-1 | CAWSDTGDTEAFF | IVTDFSVIK | YES | NO | YES |
| 8414 | TRAV38-2DV8 | TRAJ42 | CAYRSSPHGGSQGNLIF | TRBV30 | TRBJ1-1 | CAWSGGTTEAFF | AVFDRKSDAK | YES | NO | YES |
| 8415 | TRAV29DV5 | TRAJ48 | CAASLFGNEKLTF | TRBV4-1 | TRBJ1-1 | CASSQDRGGDTEAFF | IVTDFSVIK | YES | YES | YES |
| 8416 | TRAV8-1 | TRAJ40 | CAVNARGTYKYIF | TRBV4-1 | TRBJ1-1 | CASSQEEGTPNTEAFF | AVFDRKSDAK | YES | NO | YES |
| 8417 | TRAV8-4 | TRAJ27 | CAVSVSNAGKSTF | TRBV4-1 | TRBJ1-1 | CASSQVTGPYTEAFF | IVTDFSVIK | YES | NO | YES |
| 8420 | TRAV21 | TRAJ50 | CAVDLMKTSYDKVIF | TRBV4-3 | TRBJ1-1 | CASSQDRGSTEAFF | IVTDFSVIK | YES | YES | NO |
| 8421 | TRAV22 | TRAJ26 | CAVENYGQNFVF | TRBV4-3 | TRBJ1-1 | CASSYDSPTEAFF | IVTDFSVIK | YES | YES | YES |
| 8422 | TRAV1-2 | TRAJ40 | CAVRDTGTYKYIF | TRBV5-1 | TRBJ1-1 | CASSLTGTGDDTEAFF | AVFDRKSDAK | YES | NO | YES |
| 8423 | TRAV13-2 | TRAJ32 | CAETNYGGATNKLIF | TRBV5-1 | TRBJ1-1 | CASSWQPNTEAFF | AVFDRKSDAK | YES | NO | YES |
| 8427 | TRAV14DV4 | TRAJ20 | CAMSPSDDYKLSF | TRBV5-5 | TRBJ1-1 | CASSMWGRQLNTEAFF | AVFDRKSDAK | YES | YES | YES |
| 8429 | TRAV13-2 | TRAJ52 | CAENPGAGGTSYGKLTF | TRBV5-6 | TRBJ1-1 | CASSFGQGRTEAFF | AVFDRKSDAK | YES | NO | YES |
| 8430 | TRAV14DV4 | TRAJ45 | CAMSGYSGGGADGLTF | TRBV5-6 | TRBJ1-1 | CASSLPDRVDTEAFF | AVFDRKSDAK | YES | NO | NO |
| 8431 | TRAV21 | TRAJ57 | CAQGGSEKLVF | TRBV5-6 | TRBJ1-1 | CASSLAATHTEAFF | AVFDRKSDAK | YES | NO | YES |
| 8439 | TRAV12-1 | TRAJ40 | CVVDSGTYKYIF | TRBV6-1 | TRBJ1-1 | CASSNSPTEAFF | AVFDRKSDAK | YES | NO | YES |
| 8440 | TRAV21 | TRAJ39 | CASDSNNAGNMLTF | TRBV6-3 | TRBJ1-1 | CASSYGGNTEAFF | AVFDRKSDAK | YES | NO | NO |
| 8441 | TRAV21 | TRAJ50 | CAVDLMKTSYDKVIF | TRBV6-3 | TRBJ1-1 | CASSYGMGTEAFF | IVTDFSVIK | YES | NO | YES |
| 8444 | TRAV14DV4 | TRAJ4 | CAGDGGYNKLIF | TRBV7-2 | TRBJ1-1 | CASSLGNTEAFF | AVFDRKSDAK | YES | NO | NO |
| 8446 | TRAV12-2 | TRAJ29 | CAGSQGNTPLVF | TRBV7-3 | TRBJ1-1 | CASNTGQSAEAFF | AVFDRKSDAK | YES | NO | YES |
| 8451 | TRAV12-3 | TRAJ11 | CAMNSGYSTLTF | TRBV7-9 | TRBJ1-1 | CASSNTPLDEAFF | AVFDRKSDAK | YES | YES | YES |
| 8452 | TRAV17 | TRAJ39 | CATSAGNMLTF | TRBV7-9 | TRBJ1-1 | CASSLPDRGSRFGFF | AVFDRKSDAK | YES | NO | NO |
| 8453 | TRAV19 | TRAJ18 | CALDPSDRGSTLGRLYF | TRBV7-9 | TRBJ1-1 | CASSLEHTEAFF | AVFDRKSDAK | YES | NO | YES |
| 8456 | TRAV29DV5 | TRAJ48 | CATFGNEKLTF | TRBV7-9 | TRBJ1-1 | CASSLGFLDTGGLYGTEAFF | AVFDRKSDAK | NO | NO | NO |
| 8457 | TRAV6 | TRAJ40 | CALEGFPNYKYIF | TRBV7-9 | TRBJ1-1 | CASSPRGVTEAFF | AVFDRKSDAK | YES | YES | YES |
| 8460 | TRAV12-3 | TRAJ30 | CAMSWGRDDKIIF | TRBV9 | TRBJ1-1 | CASSVRSSMNTEAFF | AVFDRKSDAK | YES | NO | NO |
| 8464 | TRAV25 | TRAJ20 | CAGRSRSNDYKLSS | TRBV9 | TRBJ1-1 | CASSPSTGETAEAFF | AVFDRKSDAK | NO | NO | NO |
| 8466 | TRAV36DV7 | TRAJ54 | CAHLQGAQKLVF | TRBV9 | TRBJ1-1 | CASSEGDRVTEAFF | IVTDFSVIK | NO | NO | NO |
| 8472 | TRAV12-1 | TRAJ31 | CVVNMDNARLMF | TRBV11-2 | TRBJ1-2 | CASSWGGGSHYGYTF | IVTDFSVIK | YES | NO | YES |
| 8473 | TRAV12-2 | TRAJ12 | CAVNMYGDSSYKLIF | TRBV11-2 | TRBJ1-2 | CASSWGGGSHYGYTF | IVTDFSVIK | YES | NO | YES |
| 8477 | TRAV13-1 | TRAJ50 | CAASETSYDKVIF | TRBV11-2 | TRBJ1-2 | CASSWGGGSHYGYTF | IVTDFSVIK | YES | NO | YES |
| 8479 | TRAV14DV4 | TRAJ20 | CAMREGAGGNDYKLSF | TRBV11-2 | TRBJ1-2 | CASSLGTGARGYTF | IVTDFSVIK | YES | NO | YES |
| 8480 | TRAV19 | TRAJ49 | CALSEPNTGNQFYF | TRBV11-2 | TRBJ1-2 | CASSWGGGSHYGYTF | IVTDFSVIK | YES | YES | NO |
| 8483 | TRAV20 | TRAJ53 | CAVLLWGSGGSNYKLTF | TRBV11-2 | TRBJ1-2 | CASRSGQDYGYTF | IVTDFSVIK | YES | NO | YES |
| 8492 | TRAV3 | TRAJ37 | CAGGNTGKLIF | TRBV11-2 | TRBJ1-2 | CASSWGGGSHYGYTF | IVTDFSVIK | YES | NO | NO |
| 8495 | TRAV35 | TRAJ49 | CGCHTGNQFYF | TRBV11-2 | TRBJ1-2 | CASSHGSGSPYGYTF | IVTDFSVIK | YES | NO | NO |
| 8497 | TRAV38-1 | TRAJ52 | CAFMINAGGTSYGKLTF | TRBV11-2 | TRBJ1-2 | CASSWGGGSHYGYTF | IVTDFSVIK | YES | NO | YES |
| 8508 | TRAV20 | TRAJ40 | CAVQAYPGTYKYIF | TRBV12-5 | TRBJ1-2 | CASVATPYGYTF | AVFDRKSDAK | YES | NO | YES |
| 8509 | TRAV26-2 | TRAJ52 | CILRDLNAGGTSYGKLTF | TRBV12-5 | TRBJ1-2 | CASAPDGFFYGYTF | AVFDRKSDAK | YES | NO | NO |
| 8509 | TRAV26-2 | TRAJ52 | CILRDLNAGGTSYGKLTF | TRBV12-5 | TRBJ1-2 | CASAPDGFFYGYTF | RAKFKQLL | YES | NO | NO |
| 8513 | TRAV8-3 | TRAJ21 | CAVGDNFNKFYF | TRBV13 | TRBJ1-2 | CASTRDRPTLGYTF | AVFDRKSDAK | YES | YES | YES |
| 8514 | TRAV1-1 | TRAJ29 | CAVRSQNSGNTPLVF | TRBV14 | TRBJ1-2 | CASSQVPGQGPNYGYTF | AVFDRKSDAK | NO | NO | NO |
| 8516 | TRAV1-2 | TRAJ39 | CAVRARGANVAGNMLTF | TRBV19 | TRBJ1-2 | CASSMYGAANYGYTF | AVFDRKSDAK | YES | NO | YES |
| 8517 | TRAV12-2 | TRAJ24 | CAVIMTTDSWGKLQF | TRBV19 | TRBJ1-2 | CASSLTPGGGSYGYTF | AVFDRKSDAK | YES | YES | YES |
| 8526 | TRAV38-2DV8 | TRAJ30 | CAYRSSYRDDKIIF | TRBV19 | TRBJ1-2 | CASSIGGVGYTF | AVFDRKSDAK | YES | NO | YES |
| 8527 | TRAV1-2 | TRAJ4 | CAVRGSGGYNKLIF | TRBV2 | TRBJ1-2 | CASHATGAYGYTF | AVFDRKSDAK | YES | YES | YES |
| 8528 | TRAV1-1 | TRAJ33 | CAVADSNYQLIW | TRBV20-1 | TRBJ1-2 | CSARDFAGDYGYTF | AVFDRKSDAK | YES | NO | YES |
| 8529 | TRAV12-2 | TRAJ43 | CAVSGRF | TRBV20-1 | TRBJ1-2 | CSARWDGVPGGYTF | AVFDRKSDAK | NO | NO | NO |
| 8531 | TRAV21 | TRAJ50 | CAVDLMKTSYDKVIF | TRBV20-1 | TRBJ1-2 | CSARDRQDYGYTF | IVTDFSVIK | YES | NO | YES |
| 8533 | TRAV19 | TRAJ13 | CALSENPGGYQKVTF | TRBV24-1 | TRBJ1-2 | CATSDRDDSYGYTF | AVFDRKSDAK | YES | NO | YES |
| 8534 | TRAV21 | TRAJ50 | CAVDLMKTSYDKVIF | TRBV25-1 | TRBJ1-2 | CASSPPGDGYTF | IVTDFSVIK | YES | YES | YES |

| | | | | | | | | | | |
|---|---|---|---|---|---|---|---|---|---|---|
| 8535 | TRAV26-2 | TRAJ42 | CILRVDTLYGGSQGNLIF | TRBV25-1 | TRBJ1-2 | CASSEHMGGQGRVYGYTF | IVTDFSVIK | YES | YES | YES |
| 8536 | TRAV21 | TRAJ58 | CAAPPGETSGSRLTF | TRBV27 | TRBJ1-2 | CASRVVGVAGNYGYTF | AVFDRKSDAK | YES | NO | YES |
| 8538 | TRAV3 | TRAJ30 | CAVRDSMNRDDKIIF | TRBV27 | TRBJ1-2 | CASSFGGLNSLGGYGYTF | AVFDRKSDAK | NO | NO | NO |
| 8539 | TRAV35 | TRAJ34 | CAGLSLRSHSDKLIF | TRBV27 | TRBJ1-2 | CASSLTPGQGIYGYTF | IVTDFSVIK | YES | YES | YES |
| 8542 | TRAV2 | TRAJ6 | CAVSGGSYIPTF | TRBV28 | TRBJ1-2 | CASSSRTGYDGYTF | AVFDRKSDAK | NO | NO | NO |
| 8543 | TRAV21 | TRAJ50 | CAVDLMKTSYDKVIF | TRBV28 | TRBJ1-2 | CASSSRTGYDGYTF | IVTDFSVIK | YES | NO | YES |
| 8545 | TRAV1-2 | TRAJ16 | CAVRDGQKLLF | TRBV29-1 | TRBJ1-2 | CSVEDVVTGGPMNGYTF | AVFDRKSDAK | NO | NO | NO |
| 8546 | TRAV1-2 | TRAJ34 | CAVRDPNTDKLIF | TRBV29-1 | TRBJ1-2 | CSVDIELNYGYTF | AVFDRKSDAK | YES | NO | YES |
| 8547 | TRAV17 | TRAJ20 | CATDGDDYKLSF | TRBV29-1 | TRBJ1-2 | CSVDLRGPGHGYTF | AVFDRKSDAK | YES | NO | YES |
| 8548 | TRAV17 | TRAJ45 | CATWNSGGGADGLTF | TRBV29-1 | TRBJ1-2 | CSVDLEANYGYTF | AVFDRKSDAK | YES | NO | YES |
| 8549 | TRAV21 | TRAJ20 | CAVSPYKLSF | TRBV29-1 | TRBJ1-2 | CSVDLEANYGYTF | AVFDRKSDAK | YES | NO | YES |
| 8553 | TRAV35 | TRAJ38 | CAGQGNAGNNRKLIL | TRBV29-1 | TRBJ1-2 | CSVEDNGYGYTF | AVFDRKSDAK | NO | NO | NO |
| 8554 | TRAV13-2 | TRAJ52 | CAENAGGTSYGKLTF | TRBV3-1 | TRBJ1-2 | CASSQGRDRHGYTF | AVFDRKSDAK | YES | NO | YES |
| 8555 | TRAV3 | TRAJ35 | CAVRVGGNVLHC | TRBV30 | TRBJ1-2 | CAWSAATEYGYTF | AVFDRKSDAK | NO | NO | NO |
| 8559 | TRAV13-1 | TRAJ35 | CAARVRGFGNVLHC | TRBV4-1 | TRBJ1-2 | CASSPTPRDSYGYTF | AVFDRKSDAK | NO | NO | NO |
| 8562 | TRAV5 | TRAJ27 | CAEFDTNAGKSTF | TRBV4-2 | TRBJ1-2 | CASSQGGQGKGGYTF | AVFDRKSDAK | YES | NO | YES |
| 8564 | TRAV5 | TRAJ5 | CAENTGRRALTF | TRBV4-3 | TRBJ1-2 | CASSPTDGGYYGYTF | IVTDFSVIK | YES | NO | YES |
| 8565 | TRAV1-2 | TRAJ35 | CAVRGFGNVLHC | TRBV5-1 | TRBJ1-2 | CASSLDRGGGYTF | AVFDRKSDAK | NO | NO | NO |
| 8566 | TRAV12-1 | TRAJ20 | CVVRPNDYKLSF | TRBV5-1 | TRBJ1-2 | CASSIGTGGDINYGYTF | AVFDRKSDAK | YES | NO | YES |
| 8573 | TRAV3 | TRAJ27 | CAVRDDTNAGKSTF | TRBV5-5 | TRBJ1-2 | CASSADRAYGYTF | AVFDRKSDAK | YES | NO | YES |
| 8577 | TRAV19 | TRAJ24 | CALSEGLTTDSWGKLQF | TRBV5-8 | TRBJ1-2 | CASSWEDYGYTF | AVFDRKSDAK | YES | NO | YES |
| 8578 | TRAV21 | TRAJ38 | CAVHAGNNRKLIW | TRBV6-1 | TRBJ1-2 | CASIPTETYGYGYTF | AVFDRKSDAK | YES | NO | YES |
| 8579 | TRAV38-2DV8 | TRAJ49 | CAYLTGNQFYF | TRBV6-1 | TRBJ1-2 | CASSEQGDYGYTF | AVFDRKSDAK | YES | NO | YES |
| 8580 | TRAV1-2 | TRAJ33 | CAVKDSNYQLIW | TRBV6-3 | TRBJ1-2 | CASSYSTDSGYGYTF | AVFDRKSDAK | YES | NO | NO |
| 8581 | TRAV19 | TRAJ34 | CALLGDTDKLIF | TRBV6-3 | TRBJ1-2 | CASSYFLTGVSDGYTF | IVTDFSVIK | YES | YES | YES |
| 8582 | TRAV21 | TRAJ50 | CAVDLMKTSYDKVIF | TRBV6-5 | TRBJ1-2 | CASSYSGYGGYTF | IVTDFSVIK | YES | NO | YES |
| 8589 | TRAV29DV5 | TRAJ37 | CAATSSGNTGKLIF | TRBV7-8 | TRBJ1-2 | CASSPTGGGAYGYTF | AVFDRKSDAK | YES | NO | YES |
| 8591 | TRAV21 | TRAJ32 | CAVYGGATNKLIF | TRBV7-9 | TRBJ1-2 | CASSPDSGAFWPHGYTF | IVTDFSVIK | YES | NO | YES |
| 8594 | TRAV20 | TRAJ13 | CAVQAMNSGGYQKVTF | TRBV9 | TRBJ1-2 | CASSPASGRYKGYGYTF | AVFDRKSDAK | YES | NO | YES |
| 8596 | TRAV38-2DV8 | TRAJ29 | CAYSGNTPLVF | TRBV9 | TRBJ1-2 | CASSYFPLTSPNYGYTF | AVFDRKSDAK | YES | NO | YES |
| 8600 | TRAV3 | TRAJ13 | CAVTSRPSGGYQKVTF | TRBV14 | TRBJ1-3 | CASSQDLVVSSGNTIYF | AVFDRKSDAK | NO | NO | NO |
| 8601 | TRAV26-2 | TRAJ3 | CILRPYSSASKIIF | TRBV18 | TRBJ1-3 | CASSPVSQGANSGNTIYF | AVFDRKSDAK | YES | NO | YES |
| 8602 | TRAV12-2 | TRAJ30 | CAVKERDDKIIF | TRBV20-1 | TRBJ1-3 | CSARDVQLGNTIYF | AVFDRKSDAK | YES | NO | YES |
| 8603 | TRAV20 | TRAJ13 | CAVQARYQKVTF | TRBV20-1 | TRBJ1-3 | CSAPQGAGNTIYF | AVFDRKSDAK | YES | NO | YES |
| 8605 | TRAV8-3 | TRAJ40 | CAVGPLETYKYIF | TRBV27 | TRBJ1-3 | CASRPLTGSLSGNTIYF | AVFDRKSDAK | YES | NO | YES |
| 8617 | TRAV38-1 | TRAJ40 | CAFIGVTSGTYKYIF | TRBV19 | TRBJ1-4 | CASSIPRPTNEKLFF | AVFDRKSDAK | YES | NO | YES |
| 8620 | TRAV41 | TRAJ42 | CAVRGGSQGNLIF | TRBV2 | TRBJ1-4 | CASSEWVGINEKLFF | AVFDRKSDAK | YES | YES | NO |
| 8625 | TRAV1-2 | TRAJ27 | CAVRDRNTNAGKSTF | TRBV30 | TRBJ1-4 | CAWSPDRGRPNEKLFF | AVFDRKSDAK | YES | NO | YES |
| 8629 | TRAV27 | TRAJ42 | CAGAWNYGGSQGNLIF | TRBV4-3 | TRBJ1-4 | CASSQERLTTNEKLFF | AVFDRKSDAK | YES | YES | YES |
| 8632 | TRAV17 | TRAJ9 | CATDAGGFKTIF | TRBV5-6 | TRBJ1-4 | CASSLVEGGGEKLFF | IVTDFSVIK | YES | NO | NO |
| 8632 | TRAV17 | TRAJ9 | CATDAGGFKTIF | TRBV5-6 | TRBJ1-4 | CASSLVEGGGEKLFF | AVFDRKSDAK | YES | NO | NO |
| 8633 | TRAV1-2 | TRAJ33 | CAGMDSNYQLIW | TRBV6-1 | TRBJ1-4 | CASGGSTGTNEKLFF | AVFDRKSDAK | YES | NO | YES |
| 8634 | TRAV12-1 | TRAJ47 | CVVNDNKLVF | TRBV6-5 | TRBJ1-4 | CASSYQGPNEKLFF | AVFDRKSDAK | NO | NO | NO |
| 8635 | TRAV38-2DV8 | TRAJ30 | CAYRSAWLNRDDKIIF | TRBV6-5 | TRBJ1-4 | CASRRQETNEKLFF | AVFDRKSDAK | NO | NO | NO |
| 8636 | TRAV12-3 | TRAJ31 | CAMRNNNARLMF | TRBV6-6 | TRBJ1-4 | CASSSLRASTNEKLFF | AVFDRKSDAK | NO | NO | NO |
| 8637 | TRAV23DV6 | TRAJ22 | CAASKELGSARQLTF | TRBV6-6 | TRBJ1-4 | CASHFLRDDNEKLFF | AVFDRKSDAK | NO | NO | NO |
| 8638 | TRAV2 | TRAJ33 | CAVASNYQLIW | TRBV7-8 | TRBJ1-4 | CASSLVGQSNEKLFF | AVFDRKSDAK | NO | NO | NO |
| 8639 | TRAV38-1 | TRAJ26 | CAFMNNYGQNFVF | TRBV7-9 | TRBJ1-4 | CASSPDSYGEKLFF | AVFDRKSDAK | YES | NO | YES |
| 8643 | TRAV27 | TRAJ42 | CAGSILDGGSQGNLIF | TRBV10-2 | TRBJ1-5 | CASSESPSRGNQPQHF | AVFDRKSDAK | YES | NO | YES |
| 8645 | TRAV21 | TRAJ50 | CAVDLMKTSYDKVIF | TRBV13 | TRBJ1-5 | CASSFRGENQPQHF | IVTDFSVIK | YES | YES | YES |
| 8646 | TRAV5 | TRAJ42 | CAETLNYGGSQGNLIF | TRBV13 | TRBJ1-5 | CASSSGQGIHQPQHF | AVFDRKSDAK | YES | NO | YES |
| 8648 | TRAV1-2 | TRAJ23 | CAEDQGGKLIF | TRBV19 | TRBJ1-5 | CATGTGDSNQPQHF | AVFDRKSDAK | YES | NO | NO |
| 8649 | TRAV1-2 | TRAJ26 | CAVRDHYGQNFVF | TRBV19 | TRBJ1-5 | CASWTGDSNQPQHF | AVFDRKSDAK | YES | NO | YES |
| 8650 | TRAV1-2 | TRAJ33 | CAVMDSNYQLIW | TRBV19 | TRBJ1-5 | CASSSPEYQSRPQHF | AVFDRKSDAK | YES | NO | NO |
| 8653 | TRAV26-2 | TRAJ58 | CILRDVETSGSRLTF | TRBV19 | TRBJ1-5 | CASSIATYSNQPQHF | AVFDRKSDAK | YES | YES | YES |
| 8657 | TRAV26-1 | TRAJ26 | CIVRESGNYGQNFVF | TRBV24-1 | TRBJ1-5 | CATSDSLQGYSNQPQHF | AVFDRKSDAK | YES | NO | YES |
| 8658 | TRAV14DV4 | TRAJ20 | CAMREGVNDYKLSF | TRBV27 | TRBJ1-5 | CASSLSGTGIQPQHF | AVFDRKSDAK | YES | NO | YES |
| 8660 | TRAV21 | TRAJ50 | CAVDLMKTSYDKVIF | TRBV27 | TRBJ1-5 | CASGDRGWDQPQHF | IVTDFSVIK | YES | NO | NO |
| 8661 | TRAV26-2 | TRAJ42 | CILRDPGRNGGSQGNLIF | TRBV27 | TRBJ1-5 | CASSPFQSNQPQHF | AVFDRKSDAK | YES | NO | YES |
| 8662 | TRAV4 | TRAJ43 | CLVGGNDMRF | TRBV27 | TRBJ1-5 | CASSFRGSNQPQHF | AVFDRKSDAK | YES | NO | YES |
| 8663 | TRAV12-1 | TRAJ23 | CVVILLYNQGGKLIF | TRBV28 | TRBJ1-5 | CASSPTENQPQHF | AVFDRKSDAK | YES | NO | YES |
| 8668 | TRAV21 | TRAJ20 | CAVRFNDYKLSF | TRBV29-1 | TRBJ1-5 | CSVGTGGAGQPQHF | AVFDRKSDAK | YES | NO | YES |
| 8670 | TRAV13-1 | TRAJ49 | CAAFKPNTGNQFYF | TRBV4-1 | TRBJ1-5 | CASSPGGRGNQPQHF | AVFDRKSDAK | YES | NO | NO |
| 8677 | TRAV1-2 | TRAJ20 | CAVRERDYKLSF | TRBV6-3 | TRBJ1-5 | CATGHSNQPQHF | AVFDRKSDAK | YES | NO | NO |
| 8680 | TRAV12-3 | TRAJ45 | CAMSRGGGADGLTF | TRBV7-2 | TRBJ1-5 | CASSLAGVANQPQHF | AVFDRKSDAK | NO | NO | NO |
| 8684 | TRAV21 | TRAJ50 | CAVDLMKTSYDKVIF | TRBV7-9 | TRBJ1-5 | CASSLWGNQPQHF | IVTDFSVIK | YES | NO | YES |
| 8685 | TRAV8-1 | TRAJ9 | CAVLTYTGGFKTIF | TRBV7-9 | TRBJ1-5 | CASSPSGQPQHF | AVFDRKSDAK | YES | NO | NO |
| 8686 | TRAV8-6 | TRAJ6 | CAVSDGSGGSYIPTF | TRBV7-9 | TRBJ1-5 | CASSSVNSNQPQHF | IVTDFSVIK | YES | YES | YES |
| 8687 | TRAV12-1 | TRAJ31 | CVVNKNNNNARLMF | TRBV9 | TRBJ1-5 | CASSPEGKSNQPQHF | AVFDRKSDAK | YES | NO | YES |
| 8688 | TRAV39 | TRAJ40 | CAVDIFPSGTYKYIF | TRBV9 | TRBJ1-5 | CASSLSDNQPQHF | AVFDRKSDAK | NO | NO | NO |
| 8690 | TRAV1-2 | TRAJ6 | CAVRRGGSYIPTF | TRBV10-1 | TRBJ1-6 | CASSESGQGNSPLHF | AVFDRKSDAK | YES | NO | NO |
| 8691 | TRAV21 | TRAJ50 | CAVDLMKTSYDKVIF | TRBV10-2 | TRBJ1-6 | CISAPAVSLGVKGSNSPLHF | IVTDFSVIK | NO | NO | NO |
| 8693 | TRAV4 | TRAJ10 | CLVGEVLTGGGNKLTF | TRBV11-2 | TRBJ1-6 | CASSLREGSPLHF | AVFDRKSDAK | YES | YES | YES |
| 8694 | TRAV4 | TRAJ32 | CLVGVFYGGATNKLIF | TRBV12-4 | TRBJ1-6 | CASSMDRDYNSPLHF | AVFDRKSDAK | YES | NO | YES |
| 8695 | TRAV4 | TRAJ4 | CLVGASGGYNKLIF | TRBV14 | TRBJ1-6 | CASSQDFPASYNSPLHF | AVFDRKSDAK | NO | NO | NO |
| 8696 | TRAV12-2 | TRAJ49 | CAGFRTGNQFYF | TRBV19 | TRBJ1-6 | CASSKNRIQSSYNSPLHF | AVFDRKSDAK | YES | NO | YES |
| 8697 | TRAV21 | TRAJ50 | CAVDLMKTSYDKVIF | TRBV19 | TRBJ1-6 | CASSLRDTRNSLHF | IVTDFSVIK | YES | NO | NO |
| 8701 | TRAV13-2 | TRAJ40 | CAEIISGTYKYIF | TRBV27 | TRBJ1-6 | CASGWTGGNSPLHF | AVFDRKSDAK | YES | NO | YES |
| 8702 | TRAV14DV4 | TRAJ54 | CAMREVTRALFIQGAQKLVF | TRBV27 | TRBJ1-6 | CASSAGNSPLHF | AVFDRKSDAK | NO | NO | NO |
| 8712 | TRAV21 | TRAJ50 | CAVDLMKTSYDKVIF | TRBV4-3 | TRBJ1-6 | CASSQGTATYNSPLHF | IVTDFSVIK | YES | NO | YES |
| 8715 | TRAV29DV5 | TRAJ15 | CAASLNQAGTALIF | TRBV5-6 | TRBJ1-6 | CASSPETSGDSPLHF | AVFDRKSDAK | YES | NO | YES |
| 8716 | TRAV29DV5 | TRAJ54 | CAASGVGQGAQKLVF | TRBV5-6 | TRBJ1-6 | CASSQNSPLHF | IVTDFSVIK | YES | YES | YES |
| 8717 | TRAV12-3 | TRAJ39 | CAIGPGNMLTF | TRBV6-1 | TRBJ1-6 | CASSERTGMVAPLHF | AVFDRKSDAK | YES | NO | YES |
| 8718 | TRAV9-2 | TRAJ31 | CALSAYLGEYNARLMF | TRBV6-1 | TRBJ1-6 | CASTNEGPYSPLHF | AVFDRKSDAK | YES | NO | YES |
| 8720 | TRAV5 | TRAJ16 | CAESSVSDGQKLLF | TRBV6-5 | TRBJ1-6 | CASSYWGNSYNSPLHF | AVFDRKSDAK | NO | NO | NO |
| 8721 | TRAV23DV6 | TRAJ6 | CAASGGGSYIPTF | TRBV7-2 | TRBJ1-6 | CASSFMYQSNSPLHF | AVFDRKSDAK | NO | NO | NO |
| 8722 | TRAV19 | TRAJ24 | CALKDSWGKFQF | TRBV7-9 | TRBJ1-6 | CASSLAGDRGRNSPLHF | AVFDRKSDAK | YES | NO | YES |
| 8723 | TRAV24 | TRAJ16 | CAPF | TRBV7-9 | TRBJ1-6 | CASISVDGQINSPLHF | AVFDRKSDAK | NO | NO | NO |
| 8724 | TRAV1-1 | TRAJ23 | CAVRDYNQGGKLIF | TRBV9 | TRBJ1-6 | CASSAGTGFEGNSPLHF | AVFDRKSDAK | NO | NO | NO |
| 8738 | TRAV13-1 | TRAJ37 | CAADSSNTGKLIF | TRBV12-4 | TRBJ2-1 | CASSPGPSSYNEQFF | AVFDRKSDAK | YES | NO | YES |
| 8740 | TRAV38-1 | TRAJ12 | CAFMKSRELDSSYKLIF | TRBV12-4 | TRBJ2-1 | CASSFGPGWGNPSSYNEQFF | IVTDFSVIK | NO | NO | NO |
| 8743 | TRAV4 | TRAJ4 | CLVGETGGYNKLIF | TRBV14 | TRBJ2-1 | CASSPQYRDKNNEQFF | AVFDRKSDAK | YES | NO | YES |
| 8744 | TRAV21 | TRAJ50 | CAVDLMKTSYDKVIF | TRBV15 | TRBJ2-1 | CATSRGADSYNEQFF | IVTDFSVIK | YES | NO | YES |
| 8745 | TRAV25 | TRAJ23 | CAGVDNQGGKLIF | TRBV15 | TRBJ2-1 | CATSRDRVAGGYNEQFF | AVFDRKSDAK | YES | NO | YES |
| 8746 | TRAV20 | TRAJ30 | CAVQAENRDDKIIF | TRBV18 | TRBJ2-1 | CASSPGGGSSYNEQFF | AVFDRKSDAK | YES | NO | YES |
| 8748 | TRAV13-2 | TRAJ52 | CAENPINAGGTSYGKLTF | TRBV19 | TRBJ2-1 | CASRWGTDYNEQFF | AVFDRKSDAK | YES | YES | YES |
| 8750 | TRAV14DV4 | TRAJ42 | CAMREAFRGQGNLIF | TRBV19 | TRBJ2-1 | CASSQLQGPSFAEQFF | IVTDFSVIK | YES | NO | NO |
| 8752 | TRAV21 | TRAJ47 | CAVISVYGNKLVF | TRBV19 | TRBJ2-1 | CASSIGAGGYNEQFF | AVFDRKSDAK | YES | NO | YES |
| 8759 | TRAV4 | TRAJ8 | CLVGDRFVPMNTGFQKLVF | TRBV19 | TRBJ2-1 | CASSGGAVNEQFF | AVFDRKSDAK | YES | YES | YES |
| 8760 | TRAV8-1 | TRAJ33 | CAVTRIMDSNYQLIW | TRBV19 | TRBJ2-1 | CASSIDRAAGSYNEQFF | AVFDRKSDAK | YES | NO | NO |
| 8763 | TRAV17 | TRAJ24 | CATVPTDSWGKFQF | TRBV2 | TRBJ2-1 | CASRAGGSSYNEQFF | AVFDRKSDAK | YES | NO | YES |
| 8764 | TRAV17 | TRAJ57 | CATDQGGSEKLVF | TRBV2 | TRBJ2-1 | CARGGTGYNEQFF | AVFDRKSDAK | YES | NO | NO |
| 8765 | TRAV29DV5 | TRAJ42 | CAARYGGSQGNLIF | TRBV2 | TRBJ2-1 | CASTLGTSGPLGSYNEQFF | AVFDRKSDAK | YES | NO | YES |
| 8766 | TRAV30 | TRAJ33 | CGTERRLDSNYQLIW | TRBV2 | TRBJ2-1 | CASSDYRDTNNEQFF | AVFDRKSDAK | YES | NO | YES |

| | | | | | | | | | | |
|---|---|---|---|---|---|---|---|---|---|---|
| 8768 | TRAV1-2 | TRAJ33 | CAVLDSNYQLIW | TRBV20-1 | TRBJ2-1 | CSARDLSGGSYNEQFF | IVTDFSVIK | NO | NO | NO |
| 8769 | TRAV2 | TRAJ34 | CAVEDQGTDKLIF | TRBV20-1 | TRBJ2-1 | CSARDHNEGSYNEQFF | AVFDRKSDAK | NO | NO | NO |
| 8770 | TRAV20 | TRAJ42 | CAVKYFYGGSQGNLIF | TRBV20-1 | TRBJ2-1 | CSARDPRGDHNEQFF | AVFDRKSDAK | YES | NO | NO |
| 8777 | TRAV12-2 | TRAJ6 | CAVNIRGGSYIPTF | TRBV27 | TRBJ2-1 | CASSSTGTSGNRAEQFF | AVFDRKSDAK | YES | YES | YES |
| 8778 | TRAV21 | TRAJ50 | CAVDLMKTSYDKVIF | TRBV27 | TRBJ2-1 | CASRVSGSLSYNEQFF | IVTDFSVIK | YES | NO | YES |
| 8779 | TRAV21 | TRAJ50 | CAVDLMKTSYDKVIF | TRBV27 | TRBJ2-1 | CASSLSRREGHEQFF | IVTDFSVIK | YES | NO | YES |
| 8783 | TRAV26-1 | TRAJ42 | CIVRDPHYGGSQGNLIF | TRBV27 | TRBJ2-1 | CASSRLGGGVSSYNEQFF | AVFDRKSDAK | YES | NO | YES |
| 8785 | TRAV27 | TRAJ45 | CAGGSGGGADGLTF | TRBV27 | TRBJ2-1 | CASSSGTYYNEQFF | AVFDRKSDAK | YES | NO | YES |
| 8788 | TRAV20 | TRAJ10 | CAVQGILTGGGNKLTF | TRBV28 | TRBJ2-1 | CASSLYKSSYNEQFF | IVTDFSVIK | YES | NO | YES |
| 8790 | TRAV17 | TRAJ15 | CATEVNQAGTALIF | TRBV29-1 | TRBJ2-1 | CSAGLLGEQFF | AVFDRKSDAK | YES | NO | NO |
| 8793 | TRAV25 | TRAJ40 | CAGVSTSGTYKYIF | TRBV3-1 | TRBJ2-1 | CASSQDGLAGDNEQFF | IVTDFSVIK | YES | NO | YES |
| 8795 | TRAV8-6 | TRAJ58 | CAVSDSASGSRLTF | TRBV3-1 | TRBJ2-1 | CASSQVALGTSHNEQFF | AVFDRKSDAK | YES | NO | YES |
| 8796 | TRAV20 | TRAJ41 | CAVQANSGYALNF | TRBV30 | TRBJ2-1 | CAWNKRLAGSYNEQFF | AVFDRKSDAK | YES | NO | YES |
| 8797 | TRAV26-1 | TRAJ39 | CIVRVNAGNMLTF | TRBV30 | TRBJ2-1 | CAWSGRAGVSNEQFF | AVFDRKSDAK | NO | NO | NO |
| 8802 | TRAV8-3 | TRAJ21 | CAVGDNFNKFYF | TRBV4-1 | TRBJ2-1 | CASSLLWAEYNEQFF | AVFDRKSDAK | YES | NO | YES |
| 8804 | TRAV19 | TRAJ57 | CALLTTQGGSEKLVF | TRBV4-2 | TRBJ2-1 | CASSHPASGQFF | IVTDFSVIK | YES | NO | NO |
| 8808 | TRAV20 | TRAJ49 | CAVRAGNQFYF | TRBV4-3 | TRBJ2-1 | CASSQGVAGGQGVNEQFF | AVFDRKSDAK | YES | NO | YES |
| 8809 | TRAV21 | TRAJ50 | CAVDLMKTSYDKVIF | TRBV4-3 | TRBJ2-1 | CATRGVYEQFF | IVTDFSVIK | YES | NO | NO |
| 8811 | TRAV21 | TRAJ50 | CAVDLMKTSYDKVIF | TRBV5-1 | TRBJ2-1 | CASSLGPLAEPNNEQFF | IVTDFSVIK | YES | YES | YES |
| 8813 | TRAV12-2 | TRAJ13 | CAASGGYQKVTF | TRBV5-4 | TRBJ2-1 | CASSSFEGGDYNEQFF | AVFDRKSDAK | YES | NO | YES |
| 8814 | TRAV13-1 | TRAJ50 | CAAILRTSYDKVIF | TRBV5-4 | TRBJ2-1 | CASSLVEGRNEQFF | AVFDRKSDAK | YES | YES | YES |
| 8816 | TRAV23DV6 | TRAJ18 | CAAIYESRGSTLGRLYF | TRBV5-4 | TRBJ2-1 | CASSPPRVYSNGAGLAGVGWRNEQFF | AVFDRKSDAK | NO | NO | NO |
| 8818 | TRAV29DV5 | TRAJ39 | CAAEGNEAGNMLTF | TRBV5-5 | TRBJ2-1 | CASSPRTSHNEQFF | AVFDRKSDAK | YES | NO | YES |
| 8819 | TRAV21 | TRAJ20 | CAAERSNDYKLSF | TRBV5-6 | TRBJ2-1 | CASSLFAYNEQFF | AVFDRKSDAK | YES | NO | YES |
| 8820 | TRAV26-2 | TRAJ50 | CILRKAFFMKTSYDKVIF | TRBV5-6 | TRBJ2-1 | CASSLALAPRSYNEQFF | AVFDRKSDAK | YES | NO | NO |
| 8821 | TRAV12-2 | TRAJ49 | CAVMTGNQFYF | TRBV6-1 | TRBJ2-1 | CASRGGGSSYNEQFF | AVFDRKSDAK | YES | NO | YES |
| 8822 | TRAV21 | TRAJ39 | CAVNSNNAGNMLTF | TRBV6-1 | TRBJ2-1 | CASSEGWARNNEQFF | AVFDRKSDAK | YES | NO | YES |
| 8824 | TRAV14DV4 | TRAJ41 | CAMREGLNSGYALNF | TRBV6-3 | TRBJ2-1 | CASSPLLGGTYNEQFF | IVTDFSVIK | YES | NO | YES |
| 8827 | TRAV1-2 | TRAJ33 | CAVADSNYQLIW | TRBV6-4 | TRBJ2-1 | CASSDSIAGAYNEQFF | AVFDRKSDAK | YES | YES | YES |
| 8828 | TRAV1-2 | TRAJ33 | CAFMDSNYQLIW | TRBV6-4 | TRBJ2-1 | CASSDSPAGGNEQFF | AVFDRKSDAK | YES | NO | YES |
| 8830 | TRAV12-3 | TRAJ8 | CAMSSIMNTGFQKLVF | TRBV6-5 | TRBJ2-1 | CASSYGLTGSYNEQFF | AVFDRKSDAK | NO | NO | NO |
| 8834 | TRAV12-1 | TRAJ43 | CVGTANNNDMRF | TRBV6-6 | TRBJ2-1 | CAGSGQPYNEQFF | AVFDRKSDAK | NO | NO | NO |
| 8835 | TRAV17 | TRAJ45 | CATAGADSGGGADGLTF | TRBV6-6 | TRBJ2-1 | CASSYSGAAETSRNEQFF | AVFDRKSDAK | NO | NO | NO |
| 8836 | TRAV8-4 | TRAJ48 | CAVSDRPNFGNEKLTF | TRBV6-6 | TRBJ2-1 | CASSYLLAGLYNEQFF | AVFDRKSDAK | NO | NO | NO |
| 8839 | TRAV14DV4 | TRAJ11 | CAMREGRSGYSTLTF | TRBV7-6 | TRBJ2-1 | CASSLWERGASYNEQFF | IVTDFSVIK | YES | YES | YES |
| 8840 | TRAV14DV4 | TRAJ52 | CAMREGRGAGGTSYGKLTF | TRBV7-6 | TRBJ2-1 | CASSLGDTSYNEQFF | AVFDRKSDAK | YES | YES | NO |
| 8842 | TRAV12-2 | TRAJ7 | CAAIPYGNNRLAF | TRBV7-9 | TRBJ2-1 | CASSPPSGRAYEQFF | AVFDRKSDAK | NO | NO | NO |
| 8844 | TRAV19 | TRAJ32 | CALSGRGATNKLIF | TRBV7-9 | TRBJ2-1 | CASSLSESSYNEQFF | AVFDRKSDAK | YES | NO | YES |
| 8847 | TRAV21 | TRAJ50 | CAVDLMKTSYDKVIF | TRBV7-9 | TRBJ2-1 | CASSLGSGGYNEQFF | IVTDFSVIK | YES | NO | YES |
| 8850 | TRAV21 | TRAJ8 | CAVSNTGFQKLVF | TRBV7-9 | TRBJ2-1 | CASSLGVSSYNEQFF | AVFDRKSDAK | YES | NO | NO |
| 8854 | TRAV41 | TRAJ48 | CAVGLPVFGNEKLTF | TRBV7-9 | TRBJ2-1 | CASSLRVFHNEQFF | AVFDRKSDAK | YES | NO | YES |
| 8855 | TRAV5 | TRAJ7 | CAESRYYGNNRLAF | TRBV7-9 | TRBJ2-1 | CASSEEEALAGVDEQFF | AVFDRKSDAK | YES | NO | YES |
| 8858 | TRAV25 | TRAJ53 | CAGGGNSGGSNYKLTF | TRBV9 | TRBJ2-1 | CASSVGAGLNEQFF | AVFDRKSDAK | YES | NO | YES |
| 8865 | TRAV13-1 | TRAJ11 | CAASRGYSTLTF | TRBV12-3 | TRBJ2-2 | CASSFSGNTGELFF | AVFDRKSDAK | YES | YES | YES |
| 8869 | TRAV12-1 | TRAJ43 | CVVWRRF | TRBV12-4 | TRBJ2-2 | CATTGSSNTGELFF | AVFDRKSDAK | NO | NO | NO |
| 8871 | TRAV21 | TRAJ50 | CAVDLMKTSYDKVIF | TRBV12-5 | TRBJ2-2 | CASRAGGNTGELFF | IVTDFSVIK | YES | NO | YES |
| 8872 | TRAV21 | TRAJ50 | CAVDLMKTSYDKVIF | TRBV13 | TRBJ2-2 | CASSSPGENTGELFF | IVTDFSVIK | YES | NO | YES |
| 8873 | TRAV26-1 | TRAJ10 | CIVRVTEGGGNKLTF | TRBV13 | TRBJ2-2 | CASSFPGSTGELFF | AVFDRKSDAK | YES | NO | YES |
| 8876 | TRAV20 | TRAJ21 | CAVWYNFNKFYF | TRBV15 | TRBJ2-2 | CATSSRYLSGRTNTGELFF | IVTDFSVIK | YES | NO | YES |
| 8879 | TRAV3 | TRAJ34 | CAVRDTLYNTDKLIF | TRBV19 | TRBJ2-2 | CASSIELQGGTGELFF | IVTDFSVIK | NO | NO | NO |
| 8883 | TRAV12-1 | TRAJ21 | CVVLYNFNKFYF | TRBV20-1 | TRBJ2-2 | CSARARGLNTGELFF | AVFDRKSDAK | YES | YES | YES |
| 8884 | TRAV12-2 | TRAJ30 | CAVIDDKIIF | TRBV20-1 | TRBJ2-2 | CSARVENTGELFF | AVFDRKSDAK | YES | NO | YES |
| 8886 | TRAV21 | TRAJ50 | CAVDLMKTSYDKVIF | TRBV20-1 | TRBJ2-2 | CSAPNPNTGELFF | IVTDFSVIK | YES | NO | YES |
| 8889 | TRAV20 | TRAJ45 | CAAHPPMYSGGGADGLTF | TRBV24-1 | TRBJ2-2 | CATSDRLAGGELFF | IVTDFSVIK | YES | NO | YES |
| 8891 | TRAV27 | TRAJ33 | CAGTMDSNYQLIW | TRBV24-1 | TRBJ2-2 | CATSAGYTGELFF | AVFDRKSDAK | YES | NO | YES |
| 8893 | TRAV38-1 | TRAJ27 | CAFSSNAGKSTF | TRBV24-1 | TRBJ2-2 | CATSDRLAGGELFF | AVFDRKSDAK | YES | NO | YES |
| 8896 | TRAV1-2 | TRAJ20 | CAVKDDYKLSF | TRBV27 | TRBJ2-2 | CASGGGTGELFF | AVFDRKSDAK | YES | YES | YES |
| 8899 | TRAV13-1 | TRAJ11 | CAAKEGYSTLTF | TRBV28 | TRBJ2-2 | CASSLYSATGELFF | AVFDRKSDAK | YES | NO | YES |
| 8903 | TRAV14DV4 | TRAJ22 | CAMRRPISSGSARQLTF | TRBV29-1 | TRBJ2-2 | CSVAQEVAGPTGELFF | AVFDRKSDAK | YES | NO | YES |
| 8904 | TRAV19 | TRAJ26 | CALSDPIMAQNFVF | TRBV29-1 | TRBJ2-2 | CSVRAPLGELFF | AVFDRKSDAK | YES | NO | YES |
| 8905 | TRAV19 | TRAJ29 | CALEGNTPLVF | TRBV29-1 | TRBJ2-2 | CSSFDRVANTGELFF | AVFDRKSDAK | YES | NO | YES |
| 8906 | TRAV21 | TRAJ50 | CAVDLMKTSYDKVIF | TRBV29-1 | TRBJ2-2 | CSVPITGAGGVGELFF | IVTDFSVIK | YES | YES | YES |
| 8912 | TRAV21 | TRAJ50 | CAVDLMKTSYDKVIF | TRBV4-3 | TRBJ2-2 | CASSQGTGPVGTGELFF | IVTDFSVIK | YES | YES | YES |
| 8913 | TRAV21 | TRAJ50 | CAVDLMKTSYDKVIF | TRBV4-3 | TRBJ2-2 | CASRRSGLAPAGELFF | IVTDFSVIK | YES | YES | YES |
| 8916 | TRAV36DV7 | TRAJ54 | CAHLQGAQKLVF | TRBV5-6 | TRBJ2-2 | CASSLAGTGELFF | IVTDFSVIK | NO | NO | NO |
| 8918 | TRAV13-1 | TRAJ52 | CAADAGGTSYGKLTF | TRBV6-4 | TRBJ2-2 | CASSDGSTGELFF | IVTDFSVIK | YES | NO | YES |
| 8919 | TRAV12-2 | TRAJ45 | CAGSEGGGADGLTF | TRBV6-5 | TRBJ2-2 | CASTLRRGVDTGELFF | AVFDRKSDAK | YES | NO | YES |
| 8921 | TRAV21 | TRAJ50 | CAVDLMKTSYDKVIF | TRBV6-5 | TRBJ2-2 | CASSYPNTGELFF | IVTDFSVIK | YES | NO | YES |
| 8922 | TRAV26-2 | TRAJ40 | CIRRSGTYKYIF | TRBV6-5 | TRBJ2-2 | CASISPLGLASGTGELFF | IVTDFSVIK | YES | NO | YES |
| 8926 | TRAV20 | TRAJ11 | CAVQDSGYSTLTF | TRBV7-3 | TRBJ2-2 | CASSFMGGTGELFF | AVFDRKSDAK | NO | NO | NO |
| 8932 | TRAV21 | TRAJ50 | CAVDLMKTSYDKVIF | TRBV7-9 | TRBJ2-2 | CASSFRGGELFF | IVTDFSVIK | YES | NO | YES |
| 8935 | TRAV27 | TRAJ33 | CAGAPMDSNYQLIW | TRBV9 | TRBJ2-2 | CASSPRGTGELFF | AVFDRKSDAK | YES | YES | YES |
| 8936 | TRAV30 | TRAJ11 | CGTESGYSTLTF | TRBV9 | TRBJ2-2 | CASSVFDWASGRTGELFF | AVFDRKSDAK | YES | NO | YES |
| 8938 | TRAV21 | TRAJ50 | CAVDLMKTSYDKVIF | TRBV11-2 | TRBJ2-3 | CASSGTRADTQYF | IVTDFSVIK | YES | NO | YES |
| 8939 | TRAV17 | TRAJ32 | CATTYGGATNKLIF | TRBV12-3 | TRBJ2-3 | CASSSVGGGGDTQYF | IVTDFSVIK | YES | NO | YES |
| 8940 | TRAV12-2 | TRAJ11 | CAVKGLGYSTLTF | TRBV12-4 | TRBJ2-3 | CASSFPSPGDTQYF | AVFDRKSDAK | YES | NO | NO |
| 8945 | TRAV5 | TRAJ6 | CALSGGSYIPTF | TRBV15 | TRBJ2-3 | CATSRAAGGSTDTQYF | AVFDRKSDAK | YES | NO | YES |
| 8946 | TRAV13-1 | TRAJ43 | CAASGDNDMRF | TRBV19 | TRBJ2-3 | CASSILRSADTQYF | IVTDFSVIK | YES | YES | YES |
| 8947 | TRAV20 | TRAJ44 | CAVHLMDGTASKLTF | TRBV19 | TRBJ2-3 | CASSILNTDTQYF | AVFDRKSDAK | YES | NO | YES |
| 8949 | TRAV5 | TRAJ23 | CAEIGYNQGGKLIF | TRBV19 | TRBJ2-3 | CASSIRESTDTQYF | AVFDRKSDAK | YES | NO | YES |
| 8952 | TRAV1-2 | TRAJ6 | CAVSGGSYIPTF | TRBV20-1 | TRBJ2-3 | CSARDLDRDGTDTQYF | AVFDRKSDAK | NO | NO | NO |
| 8954 | TRAV13-2 | TRAJ36 | CAPKGGANNLFF | TRBV20-1 | TRBJ2-3 | CSASPAGSTDTQYF | AVFDRKSDAK | NO | NO | NO |
| 8957 | TRAV21 | TRAJ24 | CAVRSPHSWGKLQF | TRBV20-1 | TRBJ2-3 | CSASGLAGVLTDTQYF | AVFDRKSDAK | NO | NO | NO |
| 8958 | TRAV21 | TRAJ50 | CAVDLMKTSYDKVIF | TRBV20-1 | TRBJ2-3 | CSARDRWAGTSTDTQYF | IVTDFSVIK | NO | NO | NO |
| 8960 | TRAV25 | TRAJ52 | CAGAGGTSYGKLTF | TRBV20-1 | TRBJ2-3 | CSARDRLAGVADTQYF | AVFDRKSDAK | NO | NO | NO |
| 8962 | TRAV3 | TRAJ20 | CAVFNDYKLSF | TRBV20-1 | TRBJ2-3 | CSAPRGSGTIDTQYF | AVFDRKSDAK | NO | NO | NO |
| 8963 | TRAV3 | TRAJ27 | CPLPTNAGKSTF | TRBV20-1 | TRBJ2-3 | CSAPSGEGRDTQYF | AVFDRKSDAK | NO | NO | NO |
| 8964 | TRAV8-1 | TRAJ20 | CAVICNDYKLSF | TRBV20-1 | TRBJ2-3 | CSADGTHTPTDTQYF | AVFDRKSDAK | YES | NO | YES |
| 8970 | TRAV17 | TRAJ45 | CSGLSGGGADGLTF | TRBV27 | TRBJ2-3 | CASSFTGARTDTQYF | IVTDFSVIK | YES | NO | NO |
| 8971 | TRAV21 | TRAJ24 | CAVTDSWGKLQF | TRBV27 | TRBJ2-3 | CASKPTSTDTQYF | AVFDRKSDAK | YES | NO | YES |
| 8972 | TRAV21 | TRAJ50 | CAVDLMKTSYDKVIF | TRBV27 | TRBJ2-3 | CASSSSGPQTDTQYF | IVTDFSVIK | YES | NO | YES |
| 8973 | TRAV21 | TRAJ50 | CAVDLMKTSYDKVIF | TRBV27 | TRBJ2-3 | CASSPTSPTATDTQYF | IVTDFSVIK | YES | NO | YES |
| 8974 | TRAV38-1 | TRAJ43 | CAFMKLTYNNNDMRF | TRBV27 | TRBJ2-3 | CASSFGVSTDTQYF | AVFDRKSDAK | YES | NO | YES |
| 8976 | TRAV14DV4 | TRAJ45 | CAMREPYSGGGADGLTF | TRBV28 | TRBJ2-3 | CASSLLTSGGGSDTQYF | IVTDFSVIK | YES | NO | YES |
| 8977 | TRAV21 | TRAJ50 | CAVDLMKTSYDKVIF | TRBV28 | TRBJ2-3 | CASRPTSGGQADTQYF | IVTDFSVIK | YES | NO | YES |
| 8978 | TRAV38-2DV8 | TRAJ49 | CAYRSPLTGNQFYF | TRBV28 | TRBJ2-3 | CASTPPRGTQYF | AVFDRKSDAK | YES | NO | YES |
| 8981 | TRAV34 | TRAJ54 | CGADPIIQGAQKLVF | TRBV29-1 | TRBJ2-3 | CSVADGPGDTQYF | AVFDRKSDAK | YES | NO | YES |
| 8982 | TRAV35 | TRAJ49 | CAGHTGNQFYF | TRBV29-1 | TRBJ2-3 | CSVGSGEDSPQYF | IVTDFSVIK | YES | NO | NO |
| 8983 | TRAV30 | TRAJ15 | CGTPNQAGTALIF | TRBV3-1 | TRBJ2-3 | CASSQGDGTGTQYF | AVFDRKSDAK | YES | NO | YES |
| 8992 | TRAV19 | TRAJ5 | CALTTLGTGRRALTF | TRBV5-1 | TRBJ2-3 | CASSLASLAGNVDTQYF | IVTDFSVIK | YES | NO | YES |
| 8993 | TRAV27 | TRAJ18 | CAVDRGSTLGRLYF | TRBV5-1 | TRBJ2-3 | CASGRGTGGTDTQYF | AVFDRKSDAK | YES | NO | YES |
| 8995 | TRAV19 | TRAJ6 | CASSSGGSYIPTF | TRBV5-4 | TRBJ2-3 | CASSLARDIPRDF | IVTDFSVIK | YES | NO | NO |
| 8996 | TRAV2 | TRAJ21 | CAVGDFNKFYF | TRBV5-4 | TRBJ2-3 | CASSAWGTSSTDTQYF | IVTDFSVIK | NO | NO | NO |
| 8997 | TRAV41 | TRAJ45 | CAGSYSGGGADGLTF | TRBV5-4 | TRBJ2-3 | CASSLEAGYRDTDTQYF | AVFDRKSDAK | YES | NO | NO |

| | | | | | | | | | | |
|---|---|---|---|---|---|---|---|---|---|---|
| 8998 | TRAV12-2 | TRAJ21 | CAVSLYNFNKFYF | TRBV5-5 | TRBJ2-3 | CASVLWRASTDTQYF | AVFDRKSDAK | YES | NO | YES |
| 9002 | TRAV13-2 | TRAJ12 | CAENGGRDSSYKLIF | TRBV6-1 | TRBJ2-3 | CASSARTSTDTQYF | AVFDRKSDAK | YES | NO | YES |
| 9003 | TRAV29DV5 | TRAJ15 | CAASANPGTALIF | TRBV6-1 | TRBJ2-3 | CASSGLAGLFVTLGDTQYF | AVFDRKSDAK | YES | YES | YES |
| 9005 | TRAV38-2DV8 | TRAJ42 | CAYSGYGGSQGNLIF | TRBV6-1 | TRBJ2-3 | CASSATVIPDTQYF | IVTDFSVIK | YES | NO | YES |
| 9008 | TRAV36DV7 | TRAJ53 | CAVEVILNSGGSNYKLTF | TRBV6-3 | TRBJ2-3 | CASRNIWATPAATDTQYF | AVFDRKSDAK | NO | NO | NO |
| 9009 | TRAV41 | TRAJ48 | CAVSGGMGNEKLTF | TRBV6-3 | TRBJ2-3 | CASSFWTGGTDTQYF | IVTDFSVIK | YES | YES | YES |
| 9010 | TRAV1-2 | TRAJ12 | CAVLDSSYKLIF | TRBV6-4 | TRBJ2-3 | CASSPGGDRGADTQYF | AVFDRKSDAK | YES | NO | NO |
| 9013 | TRAV3 | TRAJ45 | CAGYPRETDGLTF | TRBV6-5 | TRBJ2-3 | CASSYVGGPGDTQYF | AVFDRKSDAK | NO | NO | NO |
| 9020 | TRAV14DV4 | TRAJ12 | CAMREGEDSSYKLIF | TRBV6-6 | TRBJ2-3 | CASRTGLASTDTQYF | IVTDFSVIK | NO | NO | NO |
| 9022 | TRAV17 | TRAJ24 | CATASSTDSWGKLQF | TRBV6-6 | TRBJ2-3 | CASRTGLASTDTQYF | IVTDFSVIK | NO | NO | NO |
| 9023 | TRAV19 | TRAJ17 | CALSVIKAAGNKLTF | TRBV6-6 | TRBJ2-3 | CASRTGLASTDTQYF | IVTDFSVIK | NO | NO | NO |
| 9024 | TRAV19 | TRAJ20 | CALKRSNDYKLSF | TRBV6-6 | TRBJ2-3 | CASRTGLASTDTQYF | IVTDFSVIK | NO | NO | NO |
| 9026 | TRAV21 | TRAJ20 | CAAFESNDYKLSF | TRBV6-6 | TRBJ2-3 | CASRTGLASTDTQYF | IVTDFSVIK | NO | NO | NO |
| 9028 | TRAV22 | TRAJ47 | CAVEPLYGNKLVF | TRBV6-6 | TRBJ2-3 | CASRTGLASTDTQYF | AVFDRKSDAK | NO | NO | NO |
| 9029 | TRAV27 | TRAJ35 | CAGTLLGFGNVLHC | TRBV6-6 | TRBJ2-3 | CASSLTLAGTGDTQYF | AVFDRKSDAK | NO | NO | NO |
| 9030 | TRAV3 | TRAJ12 | CAVRDLGDSSYKLIF | TRBV6-6 | TRBJ2-3 | CASRTGLASTDTQYF | IVTDFSVIK | NO | NO | NO |
| 9031 | TRAV3 | TRAJ12 | CAVRDIPASYKLIF | TRBV6-6 | TRBJ2-3 | CASRTGLASTDTQYF | IVTDFSVIK | NO | NO | NO |
| 9032 | TRAV3 | TRAJ37 | CAVRDGGMGSNTGKLIF | TRBV6-6 | TRBJ2-3 | CASRTGLASTDTQYF | IVTDFSVIK | NO | NO | NO |
| 9033 | TRAV38-1 | TRAJ37 | CAFTTGKLIF | TRBV6-6 | TRBJ2-3 | CASRTGLASTDTQYF | IVTDFSVIK | NO | NO | NO |
| 9035 | TRAV38-2DV8 | TRAJ30 | CVSYVSHDKIIF | TRBV6-6 | TRBJ2-3 | CASRTGLASTDTQYF | IVTDFSVIK | NO | NO | NO |
| 9038 | TRAV8-4 | TRAJ10 | CAVFTGGGNKLTF | TRBV6-6 | TRBJ2-3 | CASRTGLASTDTQYF | IVTDFSVIK | NO | NO | NO |
| 9041 | TRAV2 | TRAJ8 | CAVDVNTGFQKLVF | TRBV7-2 | TRBJ2-3 | CASSLGREGRMDTQYF | AVFDRKSDAK | NO | NO | NO |
| 9042 | TRAV8-1 | TRAJ28 | CAVIYSGAGSYQLTF | TRBV7-3 | TRBJ2-3 | CASSMGLAETDTQYF | AVFDRKSDAK | YES | NO | YES |
| 9045 | TRAV1-2 | TRAJ6 | CAVPVSGGSYIPTF | TRBV7-8 | TRBJ2-3 | CASSLFSASGSDTQYF | AVFDRKSDAK | YES | NO | YES |
| 9049 | TRAV8-2 | TRAJ21 | CVVSDRYNFNKFYF | TRBV7-9 | TRBJ2-3 | CASSSDSGVVADTQYF | AVFDRKSDAK | YES | NO | YES |
| 9051 | TRAV13-1 | TRAJ40 | CAASLSGTYKYIF | TRBV9 | TRBJ2-3 | CASSASGRVADTQYF | AVFDRKSDAK | YES | NO | YES |
| 9057 | TRAV12-3 | TRAJ39 | CAIGPGNMLTF | TRBV12-5 | TRBJ2-4 | CASGLVEGDSKNIQYF | AVFDRKSDAK | YES | NO | YES |
| 9067 | TRAV20 | TRAJ54 | CAVQARERGAQKLVF | TRBV4-3 | TRBJ2-4 | CASSQGTGSKNIQYF | AVFDRKSDAK | YES | NO | YES |
| 9069 | TRAV21 | TRAJ12 | CAVLRMDSSYKLIF | TRBV6-6 | TRBJ2-4 | CASSEASGGANIQYF | AVFDRKSDAK | NO | NO | NO |
| 9070 | TRAV23DV6 | TRAJ31 | CAASMIGARLMF | TRBV7-9 | TRBJ2-4 | CASRPWDLGLDLAKNIQYF | AVFDRKSDAK | YES | NO | NO |
| 9074 | TRAV13-1 | TRAJ52 | CAGTNAGGTSYGKLTF | TRBV11-2 | TRBJ2-5 | CASSLVGIEREGETQYF | IVTDFSVIK | YES | YES | YES |
| 9075 | TRAV21 | TRAJ50 | CAVDLMKTSYDKVIF | TRBV11-2 | TRBJ2-5 | CASSFWSGTQETQYF | IVTDFSVIK | YES | YES | YES |
| 9076 | TRAV4 | TRAJ39 | CLVGDYGNAGNMLTF | TRBV11-3 | TRBJ2-5 | CASSSRDYQETQYF | AVFDRKSDAK | YES | NO | YES |
| 9077 | TRAV9-2 | TRAJ50 | CALRETSYDKVIF | TRBV12-3 | TRBJ2-5 | CASSSGTSGGAGETQYF | RAKFKQLL | YES | NO | YES |
| 9080 | TRAV21 | TRAJ50 | CAVDLMKTSYDKVIF | TRBV12-4 | TRBJ2-5 | CASSLGQDKEAQYF | IVTDFSVIK | YES | YES | NO |
| 9081 | TRAV19 | TRAJ30 | CALRLRNRDDKIIF | TRBV13 | TRBJ2-5 | CASSQGLGRETQYF | AVFDRKSDAK | YES | NO | YES |
| 9083 | TRAV38-1 | TRAJ45 | CAFGVSTLYSGGGADGLTF | TRBV13 | TRBJ2-5 | CASSMRDPQETQYF | AVFDRKSDAK | YES | NO | YES |
| 9086 | TRAV4 | TRAJ13 | CLVGGYSGGYQKVTF | TRBV14 | TRBJ2-5 | CASSRDVQETQYF | AVFDRKSDAK | NO | NO | NO |
| 9088 | TRAV38-2DV8 | TRAJ48 | CASLTNFGNEKLTF | TRBV15 | TRBJ2-5 | CATSTAGTGLGETQYF | AVFDRKSDAK | YES | NO | NO |
| 9089 | TRAV12-1 | TRAJ44 | CVVNYQTGTASKLTF | TRBV19 | TRBJ2-5 | CASTTTGGETQYF | IVTDFSVIK | YES | NO | YES |
| 9092 | TRAV4 | TRAJ22 | CLVGALFSSGSARQLTF | TRBV2 | TRBJ2-5 | CASSEVGAVNQETQYF | AVFDRKSDAK | YES | NO | YES |
| 9093 | TRAV21 | TRAJ36 | CAVPTRSGANNLFF | TRBV20-1 | TRBJ2-5 | CSARDPPSDVGTQYF | IVTDFSVIK | YES | YES | YES |
| 9096 | TRAV21 | TRAJ50 | CAVDLMKTSYDKVIF | TRBV25-1 | TRBJ2-5 | CASSESDRAKETQYF | IVTDFSVIK | YES | NO | YES |
| 9104 | TRAV19 | TRAJ39 | CALSDLNNAGNMLTF | TRBV4-1 | TRBJ2-5 | CASSWVLAGTDETQYF | AVFDRKSDAK | YES | NO | YES |
| 9105 | TRAV21 | TRAJ50 | CAVDLMKTSYDKVIF | TRBV4-1 | TRBJ2-5 | CASSQQETQYF | IVTDFSVIK | YES | YES | YES |
| 9107 | TRAV8-4 | TRAJ13 | CAVSNSGGYQKVTF | TRBV4-1 | TRBJ2-5 | CASSQDRSGGGATQYF | IVTDFSVIK | YES | YES | YES |
| 9108 | TRAV1-2 | TRAJ12 | CAVLDSSYKLIF | TRBV4-2 | TRBJ2-5 | CASSQDRGGQETQYF | AVFDRKSDAK | NO | NO | NO |
| 9109 | TRAV20 | TRAJ58 | CAVQAGETSGSRLTF | TRBV4-2 | TRBJ2-5 | CAAGEMFGLGETQYF | AVFDRKSDAK | NO | NO | NO |
| 9113 | TRAV35 | TRAJ49 | CAGHTGNQFYF | TRBV5-1 | TRBJ2-5 | CASSLEWGGETQYF | IVTDFSVIK | YES | NO | YES |
| 9115 | TRAV3 | TRAJ37 | CAVRDGGMGSNTGKLIF | TRBV5-6 | TRBJ2-5 | CASSFQGSPQETQYF | AVFDRKSDAK | YES | NO | YES |
| 9116 | TRAV34 | TRAJ32 | CGARGGATNKLIF | TRBV6-1 | TRBJ2-5 | CASSETRIYMDSQETQYF | AVFDRKSDAK | YES | NO | YES |
| 9118 | TRAV17 | TRAJ6 | CATVASGGSYIPTF | TRBV6-3 | TRBJ2-5 | CASSYSQVAGELETQYF | AVFDRKSDAK | YES | NO | YES |
| 9119 | TRAV19 | TRAJ30 | CALSEAYDKIIF | TRBV6-3 | TRBJ2-5 | CASSYSQIVRGSGRQETQYF | AVFDRKSDAK | NO | NO | NO |
| 9120 | TRAV19 | TRAJ45 | CALSVFSGGGADGLTF | TRBV6-3 | TRBJ2-5 | CASSYSIGETQYF | AVFDRKSDAK | YES | NO | NO |
| 9121 | TRAV12-2 | TRAJ7 | CAVRYGNNRLAF | TRBV6-5 | TRBJ2-5 | CASSYRGAWETQYF | AVFDRKSDAK | NO | NO | NO |
| 9122 | TRAV4 | TRAJ33 | CLVGDGSNYQLIW | TRBV6-5 | TRBJ2-5 | CASSYEAGAGETQYF | AVFDRKSDAK | NO | NO | NO |
| 9128 | TRAV38-2DV8 | TRAJ44 | CAYRRGTGTASKLTF | TRBV7-8 | TRBJ2-5 | CASSPLPRLAGEETQYF | AVFDRKSDAK | YES | NO | YES |
| 9131 | TRAV13-1 | TRAJ50 | CAAAETSYDKVIF | TRBV7-9 | TRBJ2-5 | CASSFVDRAETQYF | IVTDFSVIK | YES | NO | NO |
| 9134 | TRAV38-1 | TRAJ48 | CAFTDFGNEKLTF | TRBV9 | TRBJ2-5 | CASSVGFSSGRGETQYF | IVTDFSVIK | YES | NO | YES |
| 9135 | TRAV8-3 | TRAJ21 | CAVGDNFNKFYF | TRBV10-1 | TRBJ2-6 | CASSRDLSGANVLTF | AVFDRKSDAK | YES | NO | YES |
| 9136 | TRAV13-2 | TRAJ37 | CAENLPSGGTGKLIF | TRBV18 | TRBJ2-6 | CASSPPFRAFLSGANVLTF | AVFDRKSDAK | YES | NO | YES |
| 9137 | TRAV21 | TRAJ50 | CAVDLMKTSYDKVIF | TRBV19 | TRBJ2-6 | CASSMSGANVLTF | IVTDFSVIK | YES | NO | YES |
| 9138 | TRAV13-2 | TRAJ49 | CAENRGGNQFYF | TRBV27 | TRBJ2-6 | CASSFSGANVLTF | AVFDRKSDAK | YES | NO | YES |
| 9142 | TRAV29DV5 | TRAJ38 | CAANGAGNNRKLIW | TRBV5-5 | TRBJ2-6 | CASSRQGDSGANVLTF | AVFDRKSDAK | YES | NO | YES |
| 9145 | TRAV21 | TRAJ50 | CAVDLMKTSYDKVIF | TRBV7-9 | TRBJ2-6 | CASSRSRDLANVLTF | IVTDFSVIK | YES | NO | YES |
| 9148 | TRAV8-3 | TRAJ21 | CAVGDNFNKFYF | TRBV9 | TRBJ2-6 | CASSVAVGTGSGANVLTF | AVFDRKSDAK | YES | NO | YES |
| 9149 | TRAV1-2 | TRAJ37 | CAVFSGNTGKLIF | TRBV10-1 | TRBJ2-7 | CASSESRAGEQYV | AVFDRKSDAK | NO | NO | NO |
| 9150 | TRAV26-2 | TRAJ39 | CILRDVNNAGNMLTF | TRBV10-2 | TRBJ2-7 | CASRGQGALMTYEQYF | AVFDRKSDAK | YES | YES | NO |
| 9153 | TRAV20 | TRAJ54 | CAVQAERQGAQKLVF | TRBV10-3 | TRBJ2-7 | CAISEPQGAVGEQYF | AVFDRKSDAK | YES | NO | YES |
| 9154 | TRAV21 | TRAJ32 | CAVKTEDGGATNKLIF | TRBV10-3 | TRBJ2-7 | CAISEGNSYEQYF | AVFDRKSDAK | YES | NO | YES |
| 9155 | TRAV22 | TRAJ20 | CAVGEYDYKLSF | TRBV10-3 | TRBJ2-7 | CASSESISYEQYF | AVFDRKSDAK | YES | YES | YES |
| 9162 | TRAV21 | TRAJ23 | CAVSGNQGGKLIF | TRBV11-2 | TRBJ2-7 | CASTLGSGSSYEQYF | AVFDRKSDAK | YES | NO | YES |
| 9163 | TRAV26-1 | TRAJ41 | CIVHSGYALNF | TRBV11-2 | TRBJ2-7 | CASSFGTASYEQYF | AVFDRKSDAK | YES | NO | YES |
| 9164 | TRAV26-2 | TRAJ36 | CILQTGANNLFF | TRBV11-2 | TRBJ2-7 | CASSFGGPYYEQYF | AVFDRKSDAK | YES | NO | YES |
| 9170 | TRAV38-1 | TRAJ54 | CAFMKARSQGAQKLVF | TRBV11-2 | TRBJ2-7 | CASSLDNSGPAYEQYF | AVFDRKSDAK | YES | NO | NO |
| 9172 | TRAV8-3 | TRAJ21 | CAVGDNFNKFYF | TRBV11-2 | TRBJ2-7 | CASSSGTAAYEQYF | AVFDRKSDAK | YES | YES | YES |
| 9173 | TRAV8-2 | TRAJ31 | CVVSDHHARLMF | TRBV11-3 | TRBJ2-7 | CASSEQDSAPYV | AVFDRKSDAK | NO | NO | NO |
| 9174 | TRAV12-2 | TRAJ23 | CAVKGGKLIF | TRBV12-3 | TRBJ2-7 | CASSSQGFEQYF | AVFDRKSDAK | YES | NO | YES |
| 9175 | TRAV29DV5 | TRAJ30 | CAAARDDKIIF | TRBV12-3 | TRBJ2-7 | CASSRGEGRGIQYF | AVFDRKSDAK | YES | NO | YES |
| 9176 | TRAV1-1 | TRAJ16 | CETLSSDGQKLLF | TRBV12-4 | TRBJ2-7 | CASSPRMSPYEQYF | AVFDRKSDAK | NO | NO | NO |
| 9177 | TRAV12-3 | TRAJ30 | CATRDDKIIF | TRBV12-4 | TRBJ2-7 | CASSPGTPYEQYF | AVFDRKSDAK | YES | YES | YES |
| 9178 | TRAV21 | TRAJ50 | CAVDLMKTSYDKVIF | TRBV12-4 | TRBJ2-7 | CASTNGSFGEQYF | IVTDFSVIK | YES | NO | YES |
| 9179 | TRAV25 | TRAJ42 | CAYYGGSQGNLIF | TRBV13 | TRBJ2-7 | CASSQLEDGHSYEQYF | AVFDRKSDAK | YES | NO | YES |
| 9185 | TRAV1-1 | TRAJ9 | CAVREVTGGFKTIF | TRBV19 | TRBJ2-7 | CASSPGQLYEQYF | AVFDRKSDAK | YES | NO | NO |
| 9187 | TRAV12-3 | TRAJ20 | CAIPGSNDYKLSF | TRBV19 | TRBJ2-7 | CASSISGGILYEQYF | AVFDRKSDAK | YES | NO | YES |
| 9189 | TRAV13-1 | TRAJ35 | CAARVRGFGNVLHC | TRBV19 | TRBJ2-7 | CASSIRSSYEQYF | AVFDRKSDAK | NO | NO | NO |
| 9190 | TRAV13-1 | TRAJ35 | CAARVRGFGNVLHC | TRBV19 | TRBJ2-7 | CASSARSSTEQYF | AVFDRKSDAK | NO | NO | NO |
| 9191 | TRAV20 | TRAJ48 | CAVQAYWVGNEKLTF | TRBV19 | TRBJ2-7 | CASSTRPGQGVEEQYF | AVFDRKSDAK | YES | NO | YES |
| 9202 | TRAV8-3 | TRAJ52 | CAVDPAGGTSYGKLTF | TRBV19 | TRBJ2-7 | CASSIGGNNEQYF | AVFDRKSDAK | YES | NO | YES |
| 9205 | TRAV21 | TRAJ50 | CAVDLMKTSYDKVIF | TRBV2 | TRBJ2-7 | CASSESLGATYEQYF | IVTDFSVIK | NO | NO | NO |
| 9206 | TRAV21 | TRAJ57 | CAVEGQGGSEKLVF | TRBV2 | TRBJ2-7 | CASSPSYEQYV | AVFDRKSDAK | NO | NO | NO |
| 9208 | TRAV3 | TRAJ26 | CAVRDGDYGQNFVF | TRBV2 | TRBJ2-7 | CASSGRDRYYEQYF | AVFDRKSDAK | NO | NO | NO |
| 9209 | TRAV5 | TRAJ8 | CADSNTGFQKLVF | TRBV2 | TRBJ2-7 | CASSARGGDSYEQYF | AVFDRKSDAK | YES | NO | YES |
| 9211 | TRAV12-1 | TRAJ57 | CVVTSQGGSEKLVF | TRBV20-1 | TRBJ2-7 | CSARLLDTNYEQYF | AVFDRKSDAK | YES | NO | YES |
| 9212 | TRAV12-2 | TRAJ43 | CAVNDNDMRF | TRBV20-1 | TRBJ2-7 | CSVPIMGYEQYF | AVFDRKSDAK | NO | NO | NO |
| 9214 | TRAV21 | TRAJ50 | CAVDLMKTSYDKVIF | TRBV20-1 | TRBJ2-7 | CSAPARSEPYEQYF | IVTDFSVIK | NO | NO | NO |
| 9215 | TRAV27 | TRAJ50 | CAGETSYDKVIF | TRBV20-1 | TRBJ2-7 | CSARDEAWASASYEQYF | AVFDRKSDAK | YES | NO | YES |
| 9219 | TRAV36DV7 | TRAJ42 | CAGGSQGNLIF | TRBV20-1 | TRBJ2-7 | CSARDNRGLGYEQYF | AVFDRKSDAK | NO | NO | NO |
| 9220 | TRAV5 | TRAJ20 | CAGSNDYKLSF | TRBV20-1 | TRBJ2-7 | CSAQAERSYEQYF | AVFDRKSDAK | NO | NO | NO |
| 9222 | TRAV23DV6 | TRAJ29 | CAAPIPGNTPLVF | TRBV24-1 | TRBJ2-7 | CATSEGTYEQYF | AVFDRKSDAK | YES | NO | YES |
| 9224 | TRAV12-1 | TRAJ8 | CVVRGTGFQKLVF | TRBV27 | TRBJ2-7 | CASSPPGSYEQYF | IVTDFSVIK | YES | NO | YES |
| 9226 | TRAV12-2 | TRAJ54 | CATRGAQKLVF | TRBV27 | TRBJ2-7 | CASSFSRNYEQYF | AVFDRKSDAK | YES | NO | YES |
| 9227 | TRAV13-1 | TRAJ33 | CAASSNYQLIW | TRBV27 | TRBJ2-7 | CASSYSGGAYEQYF | AVFDRKSDAK | YES | NO | YES |
| 9229 | TRAV14DV4 | TRAJ6 | CAFLTSGGSYIPTF | TRBV27 | TRBJ2-7 | CASSRSRGLEHYEQYF | AVFDRKSDAK | YES | NO | NO |

| | | | | | | | | | | |
|---|---|---|---|---|---|---|---|---|---|---|
| 9230 | TRAV17 | TRAJ26 | CATGDNYGQNFVF | TRBV27 | TRBJ2-7 | CASSDGQGLEQYF | AVFDRKSDAK | YES | NO | NO |
| 9232 | TRAV3 | TRAJ9 | CAVTSTGGFKTIF | TRBV27 | TRBJ2-7 | CASSDGTRRDEQYV | AVFDRKSDAK | NO | NO | NO |
| 9234 | TRAV36DV7 | TRAJ23 | CAATPFRYQGGKLIF | TRBV27 | TRBJ2-7 | CASSLIGTRDTSYEQYF | IVTDFSVIK | NO | NO | NO |
| 9236 | TRAV8-2 | TRAJ52 | CVVSEYGGTSYGKLTF | TRBV27 | TRBJ2-7 | CASRPGQLSEQYF | AVFDRKSDAK | YES | NO | YES |
| 9238 | TRAV13-1 | TRAJ9 | CAARTSGGFKTIF | TRBV28 | TRBJ2-7 | CASSYPRQGHPYEQYF | AVFDRKSDAK | YES | NO | NO |
| 9239 | TRAV17 | TRAJ20 | CATDEANDYKLSF | TRBV28 | TRBJ2-7 | CASSLATVSYEQYF | IVTDFSVIK | YES | NO | YES |
| 9240 | TRAV27 | TRAJ49 | CAGVSWSNQFYF | TRBV28 | TRBJ2-7 | CASSSGDRGLYEQYF | AVFDRKSDAK | YES | NO | NO |
| 9246 | TRAV12-1 | TRAJ39 | CVVNIRAGNMLTF | TRBV29-1 | TRBJ2-7 | CSVVMGYEQYF | AVFDRKSDAK | NO | NO | NO |
| 9247 | TRAV12-2 | TRAJ43 | CAVRGNNDMRF | TRBV29-1 | TRBJ2-7 | CSVEGGDRGYEQYF | IVTDFSVIK | YES | YES | YES |
| 9249 | TRAV20 | TRAJ8 | CAVQTLNTGFQKLVF | TRBV29-1 | TRBJ2-7 | CSVVLRGEDEQYF | IVTDFSVIK | YES | NO | YES |
| 9250 | TRAV27 | TRAJ34 | CAGGDNTDKLIF | TRBV29-1 | TRBJ2-7 | CSVQDGYEQYV | AVFDRKSDAK | NO | NO | NO |
| 9256 | TRAV21 | TRAJ50 | CAVDLMKTSYDKVIF | TRBV3-1 | TRBJ2-7 | CASSMQGKLYEQYF | IVTDFSVIK | YES | NO | YES |
| 9257 | TRAV21 | TRAJ50 | CAVDLMKTSYDKVIF | TRBV3-1 | TRBJ2-7 | CASSQVSLTGGYEQYF | IVTDFSVIK | YES | NO | YES |
| 9261 | TRAV21 | TRAJ41 | CAVRDSGYALNF | TRBV30 | TRBJ2-7 | CAWERVYEQYV | AVFDRKSDAK | NO | NO | NO |
| 9265 | TRAV12-2 | TRAJ49 | CAVGNTGNQFYF | TRBV4-1 | TRBJ2-7 | CASSQVIGRGYEQYF | AVFDRKSDAK | NO | NO | NO |
| 9268 | TRAV13-2 | TRAJ49 | CAENGNQFYF | TRBV4-1 | TRBJ2-7 | CASSRSGTGSYEQYF | AVFDRKSDAK | YES | NO | YES |
| 9270 | TRAV21 | TRAJ50 | CAVDLMKTSYDKVIF | TRBV4-1 | TRBJ2-7 | CASSQGGDGSYEQYF | IVTDFSVIK | YES | NO | NO |
| 9271 | TRAV23DV6 | TRAJ44 | CAARTTGTASKLTF | TRBV4-1 | TRBJ2-7 | CASSILAGSNEQYF | AVFDRKSDAK | YES | NO | YES |
| 9273 | TRAV4 | TRAJ20 | CLVGDDYKLSF | TRBV4-1 | TRBJ2-7 | CASSNGDEQYF | AVFDRKSDAK | YES | NO | YES |
| 9279 | TRAV8-4 | TRAJ45 | CAVSDQGGGADGLTF | TRBV5-1 | TRBJ2-7 | CASSGWTGGSYEQYF | AVFDRKSDAK | YES | NO | YES |
| 9280 | TRAV21 | TRAJ50 | CAVDLMKTSYDKVIF | TRBV5-4 | TRBJ2-7 | CASSLGRDRGDDEQYF | IVTDFSVIK | YES | NO | NO |
| 9284 | TRAV21 | TRAJ50 | CAVDLMKTSYDKVIF | TRBV5-5 | TRBJ2-7 | CASRYGQPHSYEQYF | IVTDFSVIK | YES | YES | YES |
| 9286 | TRAV1-1 | TRAJ20 | CAVRDDDYKLSF | TRBV5-6 | TRBJ2-7 | CASRFLAGGSHEQYF | AVFDRKSDAK | YES | NO | YES |
| 9287 | TRAV17 | TRAJ34 | CVTRPGTDKLIF | TRBV5-6 | TRBJ2-7 | CASSGGDITDHEQYV | IVTDFSVIK | NO | NO | NO |
| 9289 | TRAV19 | TRAJ20 | CALSTNDYKLSF | TRBV6-1 | TRBJ2-7 | CASSDGTGSFYEQYF | AVFDRKSDAK | YES | NO | YES |
| 9290 | TRAV19 | TRAJ44 | CALSEATAGTASKLTF | TRBV6-1 | TRBJ2-7 | CASRGSSAYEQYV | AVFDRKSDAK | NO | NO | NO |
| 9292 | TRAV21 | TRAJ50 | CAVDLMKTSYDKVIF | TRBV6-1 | TRBJ2-7 | CASRDSYEQYF | IVTDFSVIK | YES | YES | YES |
| 9295 | TRAV12-3 | TRAJ56 | CAMTWGANSKLTF | TRBV6-5 | TRBJ2-7 | CASSNRDRASYEQYF | IVTDFSVIK | YES | NO | YES |
| 9296 | TRAV5 | TRAJ17 | CAESMQKAAGNKLTF | TRBV6-5 | TRBJ2-7 | CASGEGGTTYEQYF | AVFDRKSDAK | YES | NO | YES |
| 9299 | TRAV17 | TRAJ35 | CATDRDVLHC | TRBV7-2 | TRBJ2-7 | CASSLGTGIYEQYV | AVFDRKSDAK | NO | NO | NO |
| 9300 | TRAV21 | TRAJ50 | CAVDLMKTSYDKVIF | TRBV7-2 | TRBJ2-7 | CASSLAPSGGDEQYF | IVTDFSVIK | YES | NO | NO |
| 9301 | TRAV21 | TRAJ50 | CAVDLMKTSYDKVIF | TRBV7-2 | TRBJ2-7 | CASSARGALAGGSYEQYF | IVTDFSVIK | NO | NO | NO |
| 9304 | TRAV20 | TRAJ37 | CAVLDSGNTGKLIF | TRBV7-3 | TRBJ2-7 | CASSPWSIWVAGSSYEQYF | AVFDRKSDAK | YES | NO | YES |
| 9305 | TRAV21 | TRAJ15 | CAVPNQAGTALIF | TRBV7-3 | TRBJ2-7 | CASSASGTTSEQYF | AVFDRKSDAK | NO | NO | NO |
| 9306 | TRAV8-3 | TRAJ21 | CAVGDNFNKFYF | TRBV7-3 | TRBJ2-7 | CASSLTRTRPLSYEQYF | AVFDRKSDAK | NO | NO | NO |
| 9307 | TRAV24 | TRAJ42 | CADYGGSQGNLIF | TRBV7-6 | TRBJ2-7 | CASSPTQGAVNEQYF | AVFDRKSDAK | YES | NO | YES |
| 9310 | TRAV21 | TRAJ50 | CAVDLMKTSYDKVIF | TRBV7-8 | TRBJ2-7 | CASSLQAGDGSYEQYF | IVTDFSVIK | YES | NO | NO |
| 9311 | TRAV4 | TRAJ4 | CLVAPGGYNKLIF | TRBV7-8 | TRBJ2-7 | CASSSQGALKREIPYEQYF | IVTDFSVIK | YES | NO | YES |
| 9312 | TRAV12-1 | TRAJ10 | CVVNGAGGGNKLTF | TRBV7-9 | TRBJ2-7 | CASSPVSSGLTYEQYF | AVFDRKSDAK | YES | YES | YES |
| 9314 | TRAV13-1 | TRAJ3 | CAAIRSASKIIF | TRBV7-9 | TRBJ2-7 | CASSLGQTGSYEQYF | AVFDRKSDAK | YES | NO | YES |
| 9324 | TRAV22 | TRAJ49 | CAWVENQFYF | TRBV9 | TRBJ2-7 | CASSVGSGPVEQYF | AVFDRKSDAK | YES | NO | YES |
| 9326 | TRAV27 | TRAJ45 | CAGVGGSGGGADGLTF | TRBV9 | TRBJ2-7 | CASSPGSDVGLYEQYF | AVFDRKSDAK | YES | YES | YES |
| 9330 | TRAV39 | TRAJ26 | CAVDFRDNYGQNFVF | TRBV9 | TRBJ2-7 | CASSLSSGTYEQYF | IVTDFSVIK | YES | YES | NO |
| 9332 | TRAV8-6 | TRAJ31 | CAVSDKVARLMF | TRBV9 | TRBJ2-7 | CASSVGGAGGTESYEQYF | AVFDRKSDAK | YES | NO | YES |
| 9333 | TRAV9-2 | TRAJ30 | CALLNRDDKIIF | TRBV9 | TRBJ2-7 | CASSLSSGTYEQYF | IVTDFSVIK | YES | NO | YES |
| 9334 | TRAV26-2 | TRAJ39 | CILPNNAGNMLTF | TRBV18 | TRBJ1-1 | CASSPLSGGTEAFF | AYAQKIFKI | NO | NO | NO |
| 9335 | TRAV12-1 | TRAJ42 | CVVARYGGSQGNLIF | TRBV4-3 | TRBJ1-1 | CASRPGTGGNTEAFF | CYTWNQMNL | YES | NO | YES |
| 9337 | TRAV25 | TRAJ43 | CAGPNNNDMRF | TRBV30 | TRBJ1-2 | CAWSGNDYGYTF | AYAQKIFKI | YES | YES | YES |
| 9340 | TRAV4 | TRAJ17 | CLVGDKAAGNKLTF | TRBV20-1 | TRBJ1-6 | CSAMGQGARGRRGYNSPLHF | CYTWNQMNL | NO | NO | NO |
| 9341 | TRAV29DV5 | TRAJ40 | CAASFSGTYKYIF | TRBV29-1 | TRBJ2-1 | CSVEPGRTGHLNEQFF | AYAQKIFKI | YES | YES | YES |
| 9343 | TRAV27 | TRAJ49 | CAGDLGNQFYF | TRBV30 | TRBJ2-1 | CAWSVSEQFF | AYAQKIFKI | YES | YES | YES |
| 9347 | TRAV4 | TRAJ4 | CLVGDPGGYNKLIF | TRBV20-1 | TRBJ2-5 | CSARVFGGDSQETQYF | AYAQKIFKI | YES | YES | YES |
| 9348 | TRAV12-1 | TRAJ30 | CVVDRDDKIIF | TRBV27 | TRBJ2-5 | CASSLGIAGGLGQTQYF | AYAQKIFKI | YES | YES | YES |
| 9349 | TRAV12-3 | TRAJ52 | CAIDLRSAGGTSYGKLTF | TRBV27 | TRBJ2-5 | CASSLSVIAGGRETQYF | CYTWNQMNL | YES | NO | YES |
| 9351 | TRAV38-2DV8 | TRAJ42 | CAYRSARGSQGNLIF | TRBV6-1 | TRBJ2-7 | CASSRRWGQGTWRGYEQYF | CYTWNQMNL | YES | NO | YES |
| 9352 | TRAV20 | TRAJ43 | CAVQADNDMRF | TRBV7-9 | TRBJ2-7 | CASSLRDRGHEQYF | AYAQKIFKI | YES | YES | YES |
| 9353 | TRAV12-2 | TRAJ21 | CAVRLDFNKFYF | TRBV20-1 | TRBJ2-1 | CSATTLVDSYNEQFF | QPRAPIRPI | NO | NO | NO |
| 9354 | TRAV12-2 | TRAJ48 | CAVCGSNFGNEKLTF | TRBV19 | TRBJ2-2 | CASSRQNTGELFF | RPHERNGFTVL | YES | NO | YES |
| 9355 | TRAV13-1 | TRAJ50 | CAASETSYDKVIF | TRBV10-3 | TRBJ1-1 | CAISGGTSRDTEAFF | RAKFKQLL | YES | NO | YES |
| 9356 | TRAV30 | TRAJ47 | CGTELYGNKLVF | TRBV11-2 | TRBJ1-1 | CASSLLRAGYTEAFF | RAKFKQLL | YES | YES | YES |
| 9364 | TRAV13-1 | TRAJ50 | CAPRETSYDKVIF | TRBV12-3 | TRBJ1-1 | CASSFLDRGDTEAFF | RAKFKQLL | YES | NO | YES |
| 9367 | TRAV8-6 | TRAJ39 | CAVSDRAGNMLTF | TRBV12-3 | TRBJ1-1 | CASSFLDRGPNTEAFF | RAKFKQLL | YES | NO | YES |
| 9385 | TRAV13-2 | TRAJ45 | CADSGGGADGLTF | TRBV27 | TRBJ1-1 | CASSVTGGDTEAFF | RAKFKQLL | YES | NO | YES |
| 9386 | TRAV13-1 | TRAJ12 | CAAARMDSSYKLIF | TRBV29-1 | TRBJ1-1 | CSAGGNTEAFF | RAKFKQLL | YES | NO | YES |
| 9390 | TRAV13-1 | TRAJ57 | CAAKEGGSEKLVF | TRBV29-1 | TRBJ1-1 | CSAGGNTEAFF | RAKFKQLL | NO | NO | NO |
| 9391 | TRAV38-2DV8 | TRAJ57 | CARAGGGSEKLVF | TRBV3-1 | TRBJ1-1 | CASSQIRGGTEAFF | RAKFKQLL | YES | NO | NO |
| 9402 | TRAV13-2 | TRAJ45 | CADEGEGGGADGLTF | TRBV4-1 | TRBJ1-1 | CASSLTGQMNTEAFF | RAKFKQLL | YES | NO | YES |
| 9403 | TRAV8-3 | TRAJ26 | CAVGESGGQNFVF | TRBV4-1 | TRBJ1-1 | CASSHWTGNTEAFF | RAKFKQLL | YES | NO | YES |
| 9405 | TRAV13-2 | TRAJ45 | CADS | TRBV5-1 | TRBJ1-1 | CASSYGTGENTEAFF | RAKFKQLL | YES | NO | YES |
| 9418 | TRAV12-1 | TRAJ34 | CVVDYTDKLIF | TRBV5-6 | TRBJ1-1 | CASSLRDGSEAFF | RAKFKQLL | YES | YES | YES |
| 9420 | TRAV12-1 | TRAJ50 | CVVTETSYDKVIF | TRBV5-6 | TRBJ1-1 | CASSLRDGSEAFF | RAKFKQLL | YES | YES | NO |
| 9428 | TRAV13-1 | TRAJ17 | CAATAAGNKLTF | TRBV5-6 | TRBJ1-1 | CASSLRDGSEAFF | RAKFKQLL | YES | NO | YES |
| 9429 | TRAV13-1 | TRAJ30 | CAALMNRDDKIIF | TRBV5-6 | TRBJ1-1 | CASSLRDGSEAFF | RAKFKQLL | YES | NO | YES |
| 9438 | TRAV13-2 | TRAJ45 | CADSGGGADGLTF | TRBV5-6 | TRBJ1-1 | CASSLREGSEAFF | RAKFKQLL | YES | YES | YES |
| 9439 | TRAV13-2 | TRAJ45 | CADS | TRBV5-6 | TRBJ1-1 | CASSLRDGSEAFF | RAKFKQLL | YES | NO | YES |
| 9442 | TRAV14DV4 | TRAJ30 | CAMREGSRRRDDKIIF | TRBV5-6 | TRBJ1-1 | CASSLRDGSEAFF | RAKFKQLL | YES | YES | YES |
| 9448 | TRAV17 | TRAJ22 | CATDRAGSARQLTF | TRBV5-6 | TRBJ1-1 | CASSLRDGSEAFF | RAKFKQLL | YES | NO | YES |
| 9455 | TRAV19 | TRAJ29 | CALRPFGGGNTPLVF | TRBV5-6 | TRBJ1-1 | CASSLRDGSEAFF | RAKFKQLL | YES | NO | YES |
| 9463 | TRAV26-1 | TRAJ17 | CILWVAAGNKLTF | TRBV5-6 | TRBJ1-1 | CASSLRDGSEAFF | RAKFKQLL | YES | NO | YES |
| 9467 | TRAV27 | TRAJ30 | CAGAQITNRDDKIIF | TRBV5-6 | TRBJ1-1 | CASSLRDGSEAFF | RAKFKQLL | YES | NO | YES |
| 9482 | TRAV5 | TRAJ34 | CAESEGPTYTDKLIF | TRBV5-6 | TRBJ1-1 | CASSLRDGSEAFF | RAKFKQLL | YES | NO | YES |
| 9488 | TRAV8-3 | TRAJ31 | CAVGTARLMF | TRBV5-6 | TRBJ1-1 | CASSLRDGSEAFF | RAKFKQLL | YES | NO | YES |
| 9501 | TRAV13-1 | TRAJ50 | CAAEETSYDKVIF | TRBV7-9 | TRBJ1-1 | CASSRLGQGAITEAFF | RAKFKQLL | YES | YES | YES |
| 9502 | TRAV13-1 | TRAJ50 | CAASETSYDKVIF | TRBV7-9 | TRBJ1-1 | CASSSLGGGNTEAFF | RAKFKQLL | YES | NO | YES |
| 9510 | TRAV10 | TRAJ27 | CVGYTNAGKSTF | TRBV11-2 | TRBJ1-2 | CASSWGGGSHYGYTF | RAKFKQLL | YES | NO | NO |
| 9512 | TRAV5 | TRAJ34 | CAESEGPTYTDKLIF | TRBV11-2 | TRBJ1-2 | CASSWGGGSHYGYTF | RAKFKQLL | YES | NO | YES |
| 9515 | TRAV13-1 | TRAJ50 | CAASETSYDKVIF | TRBV12-3 | TRBJ1-2 | CASSFTGGDLYGYTF | RAKFKQLL | YES | NO | YES |
| 9516 | TRAV13-1 | TRAJ50 | CAAMETSYDKVIF | TRBV12-3 | TRBJ1-2 | CASSLGDRAGGYTF | RAKFKQLL | YES | NO | YES |
| 9517 | TRAV13-1 | TRAJ50 | CAATETSYDKVIF | TRBV12-3 | TRBJ1-2 | CASSFTGTGSLGYTF | RAKFKQLL | YES | NO | YES |
| 9518 | TRAV13-1 | TRAJ50 | CAATETSYDKVIF | TRBV12-3 | TRBJ1-2 | CASSIGDTLNGYTF | RAKFKQLL | YES | YES | YES |
| 9519 | TRAV13-1 | TRAJ50 | CAATETSYDKVIF | TRBV12-3 | TRBJ1-2 | CASSFSGGASPPNYGYTF | RAKFKQLL | YES | NO | YES |
| 9520 | TRAV13-1 | TRAJ50 | CAGLETSYDKVIF | TRBV12-3 | TRBJ1-2 | CASSFSGTVRGGYTF | RAKFKQLL | YES | NO | NO |
| 9521 | TRAV13-1 | TRAJ50 | CAATETSYDKVIF | TRBV12-3 | TRBJ1-2 | CASSHTQPPPGRTGTYGYTF | RAKFKQLL | NO | NO | NO |
| 9522 | TRAV21 | TRAJ33 | CAVRMDSNYQLIW | TRBV12-3 | TRBJ1-2 | CASSLGDRAGGYTF | RAKFKQLL | YES | YES | YES |
| 9523 | TRAV26-2 | TRAJ50 | CILMETSYDKVIF | TRBV12-3 | TRBJ1-2 | CASSFSGANYGYTF | RAKFKQLL | YES | NO | YES |
| 9527 | TRAV13-2 | TRAJ45 | CADSGGGADGLTF | TRBV12-5 | TRBJ1-2 | CASAPDGFFYGYTF | RAKFKQLL | YES | YES | YES |
| 9529 | TRAV29DV5 | TRAJ20 | CAAEVMGDYKLSF | TRBV14 | TRBJ1-2 | CASSPHRGLEGYTF | RAKFKQLL | NO | NO | NO |
| 9531 | TRAV13-1 | TRAJ50 | CAASETSYDKVIF | TRBV18 | TRBJ1-2 | CASSSGEYGYTF | RAKFKQLL | YES | NO | YES |
| 9540 | TRAV2 | TRAJ26 | CAVEDNYGQNFVF | TRBV29-1 | TRBJ1-2 | CSTWGTGLNGYTF | RAKFKQLL | NO | NO | NO |
| 9541 | TRAV38-2DV8 | TRAJ29 | CAYRGSDSGNTPLVF | TRBV30 | TRBJ1-2 | CAWNKYGSLTYGYTF | RAKFKQLL | NO | NO | NO |
| 9543 | TRAV12-2 | TRAJ10 | CAPIGGGNKLTF | TRBV4-1 | TRBJ1-2 | CASSTSTGTGYGYTF | RAKFKQLL | YES | NO | YES |
| 9544 | TRAV13-1 | TRAJ20 | CAASESSNDYKLSF | TRBV4-1 | TRBJ1-2 | CASSPGTDYGYTF | RAKFKQLL | YES | NO | YES |
| 9545 | TRAV13-1 | TRAJ50 | CAAGETSYDKVIF | TRBV4-1 | TRBJ1-2 | CASSQDRSGEGYTF | RAKFKQLL | YES | YES | YES |
| 9548 | TRAV19 | TRAJ28 | CALGDSGAGSYQLTF | TRBV4-1 | TRBJ1-2 | CASSTSTGTGYGYTF | RAKFKQLL | YES | NO | YES |

| | | | | | | | | | | |
|---|---|---|---|---|---|---|---|---|---|---|
| 9552 | TRAV27 | TRAJ26 | CAGGDNYGQNFVF | TRBV4-1 | TRBJ1-2 | CASSTSTGTGYGYTF | RAKFKQLL | YES | NO | YES |
| 9554 | TRAV5 | TRAJ6 | CAEMEGGSYIPTF | TRBV4-1 | TRBJ1-2 | CASSRDRDYGYTF | RAKFKQLL | YES | NO | YES |
| 9562 | TRAV13-1 | TRAJ50 | CAASETSYDKVIF | TRBV5-4 | TRBJ1-2 | CASRLRDSYTF | RAKFKQLL | YES | NO | YES |
| 9577 | TRAV36DV7 | TRAJ36 | CAVEAEQTGANNLFF | TRBV9 | TRBJ1-2 | CASSVGAGLYGYTF | RAKFKQLL | NO | NO | NO |
| 9578 | TRAV5 | TRAJ30 | CAESLWDDKIIF | TRBV9 | TRBJ1-2 | CASSKWDSRNGYTF | RAKFKQLL | YES | NO | YES |
| 9584 | TRAV3 | TRAJ44 | CAVRDYGASKLTF | TRBV24-1 | TRBJ1-3 | CATSDWPKGDTIYF | RAKFKQLL | NO | NO | NO |
| 9587 | TRAV19 | TRAJ40 | CALSEASGTYKYIF | TRBV4-1 | TRBJ1-3 | CASSQTRGAGNTIYF | RAKFKQLL | YES | NO | YES |
| 9590 | TRAV5 | TRAJ30 | CAESLWDDKIIF | TRBV4-1 | TRBJ1-3 | CASSQTRGAGNTIYF | RAKFKQLL | YES | NO | NO |
| 9592 | TRAV13-1 | TRAJ50 | CAEAETSYDKVIF | TRBV7-9 | TRBJ1-3 | CASSPLGGPKDTIYF | RAKFKQLL | YES | YES | YES |
| 9594 | TRAV13-1 | TRAJ50 | CAAMETSYDKVIF | TRBV11-2 | TRBJ1-4 | CASSFHGGNEKLFF | RAKFKQLL | YES | NO | YES |
| 9595 | TRAV13-1 | TRAJ50 | CAAMETSYDKVIF | TRBV12-3 | TRBJ1-4 | CASSLGSATNEKLFF | RAKFKQLL | YES | NO | YES |
| 9596 | TRAV13-1 | TRAJ50 | CAARETSYDKVIF | TRBV12-3 | TRBJ1-4 | CASSLQGGEKLFF | RAKFKQLL | YES | YES | YES |
| 9597 | TRAV13-1 | TRAJ50 | CAATETSYDKVIF | TRBV12-3 | TRBJ1-4 | CASGPPGGEKLFF | RAKFKQLL | YES | NO | YES |
| 9600 | TRAV10 | TRAJ27 | CVGYTNAGKSTF | TRBV15 | TRBJ1-4 | CATSRSGQGEKLFF | RAKFKQLL | YES | NO | NO |
| 9601 | TRAV13-1 | TRAJ50 | CAASETSYDKVIF | TRBV15 | TRBJ1-4 | CATSRSGQGEKLFF | RAKFKQLL | YES | NO | YES |
| 9606 | TRAV13-1 | TRAJ50 | CAAMETSYDKVIF | TRBV19 | TRBJ1-4 | CASSLQGANEKLFF | RAKFKQLL | YES | NO | YES |
| 9607 | TRAV13-1 | TRAJ50 | CAAMETSYDKVIF | TRBV19 | TRBJ1-4 | CASSSQGANEKLFF | RAKFKQLL | YES | NO | YES |
| 9608 | TRAV2 | TRAJ36 | CAVEEAGANNLFF | TRBV19 | TRBJ1-4 | CASSSQGANEKLFF | RAKFKQLL | NO | NO | NO |
| 9610 | TRAV26-2 | TRAJ53 | CILREKSGGSNYKLTF | TRBV19 | TRBJ1-4 | CASSIVLGLGEKLFF | RAKFKQLL | YES | NO | NO |
| 9611 | TRAV5 | TRAJ30 | CAESLWDDKIIF | TRBV19 | TRBJ1-4 | CASSTIGTGGLEKLFF | RAKFKQLL | YES | NO | YES |
| 9614 | TRAV13-1 | TRAJ50 | CAAMETSYDKVIF | TRBV25-1 | TRBJ1-4 | CASSEQGQGEKLFF | RAKFKQLL | YES | YES | YES |
| 9615 | TRAV38-2DV8 | TRAJ56 | CAYSGDNSKLTF | TRBV27 | TRBJ1-4 | CASSSRLGRYNEKLFF | RAKFKQLL | YES | NO | YES |
| 9617 | TRAV13-1 | TRAJ50 | CAALETSYDKVIF | TRBV28 | TRBJ1-4 | CASTRQGANEKLFF | RAKFKQLL | YES | NO | YES |
| 9619 | TRAV21 | TRAJ12 | CALREDSSYKLIF | TRBV4-3 | TRBJ1-4 | CASSQGGEVGEKLFF | RAKFKQLL | YES | NO | YES |
| 9621 | TRAV12-3 | TRAJ9 | CAMISNTGGFKTIF | TRBV5-4 | TRBJ1-4 | CASSFSVGENEKLFF | RAKFKQLL | YES | NO | NO |
| 9623 | TRAV8-3 | TRAJ5 | CAVDPMDTGRRALTF | TRBV5-4 | TRBJ1-4 | CASSFSVGENEKLFF | RAKFKQLL | YES | NO | YES |
| 9627 | TRAV5 | TRAJ50 | CAERETSYDKVIF | TRBV7-9 | TRBJ1-4 | CASSLGQGKEATNEKLFF | RAKFKQLL | YES | NO | YES |
| 9628 | TRAV13-1 | TRAJ50 | CAAGETSYDKVIF | TRBV12-3 | TRBJ1-5 | CASRSGDSNQPQHF | RAKFKQLL | YES | NO | YES |
| 9629 | TRAV13-1 | TRAJ34 | CAASEGAPDKLIF | TRBV12-4 | TRBJ1-5 | CASSLVAGASNQPQHF | RAKFKQLL | YES | YES | YES |
| 9630 | TRAV13-1 | TRAJ50 | CAAVETSYDKVIF | TRBV12-5 | TRBJ1-5 | CASGSSESGDQPQHF | RAKFKQLL | YES | NO | NO |
| 9631 | TRAV13-1 | TRAJ50 | CAAMETSYDKVIF | TRBV15 | TRBJ1-5 | CATSTGDSNQPQHF | RAKFKQLL | YES | NO | YES |
| 9636 | TRAV13-1 | TRAJ50 | CAAMETSYDKVIF | TRBV19 | TRBJ1-5 | CASSWLSNQPQHF | RAKFKQLL | YES | YES | YES |
| 9644 | TRAV38-2DV8 | TRAJ47 | CAYRDEGNKLVF | TRBV27 | TRBJ1-5 | CASSSRTESPHQPQHF | RAKFKQLL | YES | NO | YES |
| 9646 | TRAV34 | TRAJ40 | CGADGGGTYKYIF | TRBV29-1 | TRBJ1-5 | CSYSAGEDQPQHF | RAKFKQLL | YES | NO | YES |
| 9647 | TRAV13-1 | TRAJ57 | CAAKEGGSEKLVF | TRBV3-1 | TRBJ1-5 | CASSLGLSNQPQHF | RAKFKQLL | YES | NO | YES |
| 9649 | TRAV38-2DV8 | TRAJ49 | CAYRSARETGNQFYF | TRBV4-1 | TRBJ1-5 | CASSRSLHSNQPQHF | RAKFKQLL | YES | YES | YES |
| 9652 | TRAV13-1 | TRAJ57 | CAAKEGGSEKLVF | TRBV5-4 | TRBJ1-5 | CASSSTGGGEKDQPQHF | RAKFKQLL | YES | NO | YES |
| 9654 | TRAV26-2 | TRAJ50 | CILRETSYDKVIF | TRBV5-5 | TRBJ1-5 | CASRPGVGNQPQHF | RAKFKQLL | YES | NO | YES |
| 9657 | TRAV38-1 | TRAJ48 | CAFMTNFGNEKLTF | TRBV6-3 | TRBJ1-5 | CASDRQDFYSNQPQHF | RAKFKQLL | YES | YES | YES |
| 9659 | TRAV13-1 | TRAJ50 | CAASETSYDKVIF | TRBV7-9 | TRBJ1-5 | CASSLGTQPQHF | RAKFKQLL | YES | NO | YES |
| 9662 | TRAV13-1 | TRAJ50 | CAASETSYDKVIF | TRBV12-3 | TRBJ1-6 | CASGQLSYNSPLHF | RAKFKQLL | YES | NO | YES |
| 9663 | TRAV26-2 | TRAJ50 | CILRETSYDKVIF | TRBV12-5 | TRBJ1-6 | CASALLSSYNSPLHF | RAKFKQLL | YES | NO | YES |
| 9676 | TRAV13-1 | TRAJ50 | CAALETSYDKVIF | TRBV3-1 | TRBJ1-6 | CASSQADSLNSPLHF | RAKFKQLL | YES | NO | YES |
| 9680 | TRAV13-2 | TRAJ45 | CADSGGGADGLTF | TRBV5-6 | TRBJ1-6 | CASSLMRGGTYNSPLHF | RAKFKQLL | YES | NO | NO |
| 9682 | TRAV13-1 | TRAJ50 | CAASETSYDKVIF | TRBV7-9 | TRBJ1-6 | CASSSLGTAKSPLHF | RAKFKQLL | YES | NO | YES |
| 9683 | TRAV13-1 | TRAJ50 | CAASDTSYDKVIF | TRBV7-9 | TRBJ1-6 | CASSFLGQKNSPLHF | RAKFKQLL | YES | NO | YES |
| 9684 | TRAV13-1 | TRAJ50 | CAASETSYDKVIF | TRBV7-9 | TRBJ1-6 | CASSLLGDPNSPLHF | RAKFKQLL | YES | NO | YES |
| 9693 | TRAV13-1 | TRAJ50 | CAASETSYDKVIF | TRBV12-3 | TRBJ2-1 | CASSFSGATHNEQFF | RAKFKQLL | YES | YES | YES |
| 9694 | TRAV13-1 | TRAJ50 | CAATETSYDKVIF | TRBV12-3 | TRBJ2-1 | CASLLVGAGGAEQFF | RAKFKQLL | YES | NO | YES |
| 9695 | TRAV13-1 | TRAJ50 | CAASETSYDKVIF | TRBV12-3 | TRBJ2-1 | CASLLVGAGGAEQFF | RAKFKQLL | YES | NO | YES |
| 9699 | TRAV10 | TRAJ27 | CVGYTNAGKSTF | TRBV12-4 | TRBJ2-1 | CASGRLSYNEQFF | RAKFKQLL | YES | NO | YES |
| 9700 | TRAV12-3 | TRAJ10 | CAIITGGGNKLTF | TRBV12-4 | TRBJ2-1 | CASSAGTSGFGYNEQFF | RAKFKQLL | YES | NO | NO |
| 9701 | TRAV2 | TRAJ21 | CAVEDYNKFYF | TRBV12-4 | TRBJ2-1 | CASGRLSYNEQFF | RAKFKQLL | NO | NO | NO |
| 9702 | TRAV29DV5 | TRAJ13 | CAASEDLPGGYQKVTF | TRBV12-5 | TRBJ2-1 | CASGSCAGTGSYCEQFF | RAKFKQLL | YES | NO | YES |
| 9712 | TRAV1-2 | TRAJ33 | CAVSDSNYQLIW | TRBV20-1 | TRBJ2-1 | CSARGGTGELNEQFF | RAKFKQLL | YES | NO | YES |
| 9723 | TRAV8-2 | TRAJ42 | CVVSDEYGGSQGNLIF | TRBV27 | TRBJ2-1 | CASSLRSVYNEQFF | RAKFKQLL | YES | NO | YES |
| 9737 | TRAV8-6 | TRAJ33 | CAVSERSNYQLIW | TRBV30 | TRBJ2-1 | CAWSGRAGVSNEQFF | RAKFKQLL | NO | NO | NO |
| 9742 | TRAV13-1 | TRAJ50 | CAASETSYDKVIF | TRBV4-1 | TRBJ2-1 | CASSPLAGIYNEQFF | RAKFKQLL | YES | NO | YES |
| 9744 | TRAV17 | TRAJ22 | CATDRAGSARQLTF | TRBV4-1 | TRBJ2-1 | CASSLGGTNEQFF | RAKFKQLL | YES | NO | YES |
| 9750 | TRAV24 | TRAJ12 | CAWMDSSYKLIF | TRBV4-1 | TRBJ2-1 | CASSLLAGEYNEQFF | RAKFKQLL | YES | YES | NO |
| 9754 | TRAV5 | TRAJ33 | CAETQDSNYQLIW | TRBV4-1 | TRBJ2-1 | CASSLLWAEYNEQFF | RAKFKQLL | YES | NO | YES |
| 9761 | TRAV13-1 | TRAJ50 | CAASETSYDKVIF | TRBV5-1 | TRBJ2-1 | CASSSPAGGQYNEQFF | RAKFKQLL | YES | NO | YES |
| 9765 | TRAV35 | TRAJ24 | CAGTDSWGKFQF | TRBV5-4 | TRBJ2-1 | CASSSSSGVHEQFF | RAKFKQLL | YES | NO | YES |
| 9768 | TRAV13-2 | TRAJ45 | CADSGGGADGLTF | TRBV5-6 | TRBJ2-1 | CASSSTLYGSYNEQFF | RAKFKQLL | YES | NO | YES |
| 9776 | TRAV13-2 | TRAJ42 | CAENEVGSQGNLIF | TRBV7-3 | TRBJ2-1 | CASSFYAGSTNEQFF | RAKFKQLL | YES | NO | YES |
| 9785 | TRAV26-2 | TRAJ50 | CILRETSYDKVIF | TRBV7-9 | TRBJ2-1 | CASSSGLAGVNEQFF | RAKFKQLL | YES | NO | YES |
| 9790 | TRAV20 | TRAJ35 | CAVQGVGFGNVLHC | TRBV9 | TRBJ2-1 | CASSVDLAGGREQFF | RAKFKQLL | NO | NO | NO |
| 9795 | TRAV26-2 | TRAJ50 | CILRETSYDKVIF | TRBV11-2 | TRBJ2-2 | CASSLGVAGTNTGELFF | RAKFKQLL | YES | YES | YES |
| 9796 | TRAV1-1 | TRAJ15 | CAVRDSAGTALIF | TRBV12-3 | TRBJ2-2 | CASSFSGNTGELFF | RAKFKQLL | YES | NO | YES |
| 9798 | TRAV10 | TRAJ8 | CVVRNMNTGFQKLVF | TRBV12-3 | TRBJ2-2 | CASSFSGNTGELFF | RAKFKQLL | YES | NO | YES |
| 9800 | TRAV12-1 | TRAJ50 | CVVTETSYDKVIF | TRBV12-3 | TRBJ2-2 | CASSFSGNTGELFF | RAKFKQLL | YES | NO | NO |
| 9806 | TRAV13-1 | TRAJ50 | CAARETSYDKVIF | TRBV12-3 | TRBJ2-2 | CASRKQGSDTGELFF | RAKFKQLL | YES | YES | YES |
| 9807 | TRAV13-1 | TRAJ50 | CVGRETSYDKVIF | TRBV12-3 | TRBJ2-2 | CASSSSGAGELFF | RAKFKQLL | YES | NO | NO |
| 9808 | TRAV13-1 | TRAJ50 | CAASKTSYDKVIF | TRBV12-3 | TRBJ2-2 | CASSFSGNTGELFF | RAKFKQLL | YES | YES | YES |
| 9809 | TRAV13-1 | TRAJ50 | CAASETSYDKVIF | TRBV12-3 | TRBJ2-2 | CASSRPGQGARTGELFF | RAKFKQLL | YES | NO | YES |
| 9811 | TRAV13-2 | TRAJ45 | CADSGGGADGLTF | TRBV12-3 | TRBJ2-2 | CASSFSGNTGELFF | RAKFKQLL | YES | NO | YES |
| 9817 | TRAV26-1 | TRAJ45 | CIVRVIVSGGGADGLTF | TRBV12-3 | TRBJ2-2 | CASSFSGNTGELFF | RAKFKQLL | YES | NO | YES |
| 9821 | TRAV29DV5 | TRAJ44 | CAASVSIWTGTASKLTF | TRBV12-3 | TRBJ2-2 | CASSFSGNTGELFF | RAKFKQLL | YES | NO | YES |
| 9824 | TRAV38-2DV8 | TRAJ40 | CAYRSLTSGTYKYIF | TRBV12-3 | TRBJ2-2 | CASSFSGNTGELFF | RAKFKQLL | YES | NO | YES |
| 9825 | TRAV38-2DV8 | TRAJ52 | CAYSPAAGGTSYGKLTF | TRBV12-3 | TRBJ2-2 | CASSFSGNTGELFF | RAKFKQLL | YES | NO | YES |
| 9828 | TRAV41 | TRAJ34 | CAVTAYNTDKLIF | TRBV12-3 | TRBJ2-2 | CASSFSGNTGELFF | RAKFKQLL | YES | YES | YES |
| 9829 | TRAV41 | TRAJ42 | CAVSIGAYGGSQGNLIF | TRBV12-3 | TRBJ2-2 | CASSFSGNTGELFF | RAKFKQLL | YES | YES | YES |
| 9830 | TRAV41 | TRAJ57 | CAVLGTQGGSEKLVF | TRBV12-3 | TRBJ2-2 | CASSFSGGTGELFF | RAKFKQLL | YES | NO | NO |
| 9835 | TRAV9-2 | TRAJ20 | CALSDYKLSF | TRBV12-3 | TRBJ2-2 | CASSFSGNTGELFF | RAKFKQLL | YES | NO | YES |
| 9837 | TRAV13-1 | TRAJ50 | CAASETSYDKVIF | TRBV12-4 | TRBJ2-2 | CASSRSGGAGELFF | RAKFKQLL | YES | NO | YES |
| 9838 | TRAV13-2 | TRAJ50 | CAERETSYDKVIF | TRBV12-4 | TRBJ2-2 | CASSFTGAEGTGELFF | RAKFKQLL | YES | NO | YES |
| 9841 | TRAV26-1 | TRAJ8 | CIVMDDFQKLVF | TRBV14 | TRBJ2-2 | CASSQATNTGELFF | RAKFKQLL | NO | NO | NO |
| 9843 | TRAV10 | TRAJ27 | CVGYTNAGKSTF | TRBV18 | TRBJ2-2 | CASSPPGGGNTGELFF | RAKFKQLL | YES | NO | YES |
| 9847 | TRAV13-1 | TRAJ50 | CAAMETSYDKVIF | TRBV18 | TRBJ2-2 | CASSPPGGGNTGELFF | RAKFKQLL | YES | NO | YES |
| 9851 | TRAV26-2 | TRAJ44 | CKAGTASKLTF | TRBV18 | TRBJ2-2 | CASSPPGGGNTGELFF | RAKFKQLL | YES | NO | NO |
| 9856 | TRAV14DV4 | TRAJ17 | CAMRAIKAAGNKLTF | TRBV19 | TRBJ2-2 | CASSQRSTGELFF | RAKFKQLL | YES | YES | YES |
| 9857 | TRAV19 | TRAJ5 | CALSEVMDTGRRALTF | TRBV19 | TRBJ2-2 | CASSPRTDLSAPNTGELFF | RAKFKQLL | YES | NO | YES |
| 9867 | TRAV1-2 | TRAJ20 | CAVRDPGRDYKLSF | TRBV20-1 | TRBJ2-2 | CSARGHETTGELFF | RAKFKQLL | YES | NO | YES |
| 9868 | TRAV10 | TRAJ48 | CVPRNFGNEKLTF | TRBV27 | TRBJ2-2 | CASSPHFQVDTGELFF | RAKFKQLL | YES | NO | YES |
| 9871 | TRAV13-1 | TRAJ50 | CAASETSYDKVIF | TRBV28 | TRBJ2-2 | CASSLYSATGELFF | RAKFKQLL | YES | NO | YES |
| 9872 | TRAV9-2 | TRAJ50 | CALRETSYDKVIF | TRBV28 | TRBJ2-2 | CASSLYSATGELFF | RAKFKQLL | YES | NO | YES |
| 9873 | TRAV13-2 | TRAJ45 | CADSGGGADGLTF | TRBV29-1 | TRBJ2-2 | CSVGTRTTGELFF | RAKFKQLL | YES | NO | YES |
| 9874 | TRAV12-2 | TRAJ33 | CAVNIRDSNYQLIW | TRBV3-1 | TRBJ2-2 | CASSYSGGAGELFF | RAKFKQLL | YES | NO | YES |
| 9880 | TRAV13-2 | TRAJ45 | CADSGGGADGLTF | TRBV4-1 | TRBJ2-2 | CASSQDRGGPGELFF | RAKFKQLL | YES | NO | NO |
| 9886 | TRAV13-2 | TRAJ3 | CAENPSSASKIIF | TRBV4-2 | TRBJ2-2 | CASSQRPSEVGELFF | RAKFKQLL | YES | NO | YES |
| 9889 | TRAV22 | TRAJ23 | CAVDNQGGKLIF | TRBV4-2 | TRBJ2-2 | CASSQRPSEVGELFF | RAKFKQLL | YES | NO | YES |
| 9906 | TRAV3 | TRAJ20 | CAVRALNDYKLSF | TRBV6-1 | TRBJ2-2 | CASIALTGELFF | RAKFKQLL | NO | NO | NO |
| 9908 | TRAV13-2 | TRAJ45 | CADSGGGADGLTF | TRBV6-3 | TRBJ2-2 | CASSNPRLDATGELFF | RAKFKQLL | YES | NO | YES |
| 9913 | TRAV13-2 | TRAJ45 | CADSGGGADGLTF | TRBV7-2 | TRBJ2-2 | CASSLGVAGTNTGELFF | RAKFKQLL | YES | NO | YES |

| | | | | | | | | | | |
|---|---|---|---|---|---|---|---|---|---|---|
| 9916 | TRAV12-2 | TRAJ43 | CAVSQRF | TRBV7-3 | TRBJ2-2 | CASSLGTSGGTGELFF | RAKFKQLL | NO | NO | NO |
| 9917 | TRAV13-1 | TRAJ21 | CAPWDFNKFYF | TRBV7-3 | TRBJ2-2 | CASSLQGGTGELFF | RAKFKQLL | NO | NO | NO |
| 9922 | TRAV26-2 | TRAJ45 | CILRGDSGGGADGLTF | TRBV7-3 | TRBJ2-2 | CASSLGTSGGTGELFF | RAKFKQLL | NO | NO | NO |
| 9924 | TRAV12-2 | TRAJ42 | CADESGGSQGNLIF | TRBV7-9 | TRBJ2-2 | CASSPRTGNTGELFF | RAKFKQLL | YES | NO | YES |
| 9925 | TRAV13-2 | TRAJ50 | CAEGETSYDKVIF | TRBV7-9 | TRBJ2-2 | CASSFTGGTGELFF | RAKFKQLL | YES | NO | YES |
| 9927 | TRAV13-1 | TRAJ50 | CAAMETSYDKVIF | TRBV9 | TRBJ2-2 | CASSSGLVSNTGELFF | RAKFKQLL | YES | NO | YES |
| 9928 | TRAV13-1 | TRAJ50 | CAAMETSYDKVIF | TRBV9 | TRBJ2-2 | CASSVARTGTNTGELFF | RAKFKQLL | YES | NO | YES |
| 9929 | TRAV17 | TRAJ58 | CATDGETSGSRLTF | TRBV9 | TRBJ2-2 | CASSVAYYLGENTGELFF | RAKFKQLL | YES | NO | YES |
| 9932 | TRAV13-2 | TRAJ45 | CADSGGGADGLTF | TRBV10-3 | TRBJ2-3 | CAINPLGSSSKKGTQYF | RAKFKQLL | YES | YES | YES |
| 9937 | TRAV9-2 | TRAJ50 | CALMETSYDKVIF | TRBV12-3 | TRBJ2-3 | CASSFPTEGLGATDTQYF | RAKFKQLL | YES | NO | YES |
| 9940 | TRAV12-2 | TRAJ11 | CAVSEYSTLTF | TRBV12-5 | TRBJ2-3 | CASGLNPQERSTDTQYF | RAKFKQLL | YES | NO | NO |
| 9950 | TRAV14DV4 | TRAJ18 | CAMREDDRGSTLGRLYF | TRBV20-1 | TRBJ2-3 | CSAGQGLAGGPGTDTQYF | RAKFKQLL | NO | NO | NO |
| 9958 | TRAV29DV5 | TRAJ37 | CAASNTGKLIF | TRBV27 | TRBJ2-3 | CASSLGFTDTQYF | RAKFKQLL | YES | YES | YES |
| 9960 | TRAV13-1 | TRAJ50 | CAASETSYDKVIF | TRBV28 | TRBJ2-3 | CASSLTDTQYF | RAKFKQLL | YES | NO | NO |
| 9973 | TRAV29DV5 | TRAJ54 | CAAGTQGAQKLVF | TRBV4-1 | TRBJ2-3 | CASSQRIGRGGPRNTDTQYF | RAKFKQLL | NO | NO | NO |
| 9980 | TRAV8-3 | TRAJ29 | CAVGEGTPLVF | TRBV4-2 | TRBJ2-3 | CASSPRLSTDTQYF | RAKFKQLL | YES | NO | NO |
| 9983 | TRAV1-2 | TRAJ33 | CAVGDSNYQLIW | TRBV6-1 | TRBJ2-3 | CASSDGARGGTDTQYF | RAKFKQLL | YES | NO | YES |
| 9987 | TRAV12-2 | TRAJ29 | CAVRPGNTPLVF | TRBV6-4 | TRBJ2-3 | CASSYTLWETDTQYF | RAKFKQLL | YES | NO | YES |
| 9992 | TRAV13-2 | TRAJ45 | CADSGGGADGLTF | TRBV7-2 | TRBJ2-3 | CASAHPRASGSTDTQYF | RAKFKQLL | YES | NO | YES |
| 10001 | TRAV13-2 | TRAJ45 | CADSGGGADGLTF | TRBV7-9 | TRBJ2-3 | CASSSRDGGTDTQYF | RAKFKQLL | YES | NO | YES |
| 10002 | TRAV21 | TRAJ8 | CAVMMNTGFQKLVF | TRBV7-9 | TRBJ2-3 | CASSPSGGAADTQYF | RAKFKQLL | YES | NO | YES |
| 10003 | TRAV9-2 | TRAJ34 | CALSEAGPDKLIF | TRBV7-9 | TRBJ2-3 | CASSSRDGGTDTQYF | RAKFKQLL | YES | NO | YES |
| 10013 | TRAV9-2 | TRAJ50 | CALRETSYDKVIF | TRBV12-3 | TRBJ2-5 | CASSGGVSGGAGETQYF | RAKFKQLL | YES | YES | YES |
| 10014 | TRAV9-2 | TRAJ50 | CALRETSYDKVIF | TRBV12-3 | TRBJ2-5 | CASSFPSGDGTQYF | RAKFKQLL | YES | NO | YES |
| 10026 | TRAV38-2DV8 | TRAJ43 | CARSWGYDMRF | TRBV3-1 | TRBJ2-5 | CASSHRPGLNQETQYF | RAKFKQLL | YES | NO | NO |
| 10027 | TRAV26-2 | TRAJ48 | CILREFTFGNEKLTF | TRBV4-1 | TRBJ2-5 | CASSQLSPLAGGLETQYF | RAKFKQLL | YES | NO | YES |
| 10041 | TRAV13-2 | TRAJ45 | CADSGGGADGLTF | TRBV7-3 | TRBJ2-5 | CASSSRGGQETQYF | RAKFKQLL | YES | NO | YES |
| 10042 | TRAV29DV5 | TRAJ36 | CAAREDQTGANNLFF | TRBV7-3 | TRBJ2-5 | CASSSRGGQETQYF | RAKFKQLL | NO | NO | NO |
| 10045 | TRAV13-1 | TRAJ50 | CAARETSYDKVIF | TRBV7-9 | TRBJ2-5 | CASSLVGTKRETQYF | RAKFKQLL | YES | NO | YES |
| 10064 | TRAV13-1 | TRAJ50 | CAASETSYDKVIF | TRBV11-2 | TRBJ2-7 | CASRGSVYEQYF | RAKFKQLL | YES | NO | YES |
| 10069 | TRAV13-1 | TRAJ50 | CAASETSYDKVIF | TRBV12-3 | TRBJ2-7 | CASRPGTSDSYEQYF | RAKFKQLL | YES | NO | YES |
| 10071 | TRAV21 | TRAJ57 | CAVAQGGSEKLVF | TRBV12-3 | TRBJ2-7 | CASSLWGDYEQYF | RAKFKQLL | YES | NO | YES |
| 10072 | TRAV26-1 | TRAJ30 | CIVREGDRDDKIIF | TRBV12-3 | TRBJ2-7 | CASSLWGDYEQYF | RAKFKQLL | YES | NO | YES |
| 10085 | TRAV13-1 | TRAJ50 | CAAMETSYDKVIF | TRBV19 | TRBJ2-7 | CASSIRSSYEQYF | RAKFKQLL | YES | NO | YES |
| 10087 | TRAV13-2 | TRAJ45 | CADSGGGADGLTF | TRBV19 | TRBJ2-7 | CASSPRSSYEQYF | RAKFKQLL | YES | NO | YES |
| 10089 | TRAV5 | TRAJ34 | CAESEGPTYTDKLIF | TRBV19 | TRBJ2-7 | CASSIRSSYEQYF | RAKFKQLL | YES | NO | YES |
| 10092 | TRAV13-2 | TRAJ45 | CADSGGGADGLTF | TRBV2 | TRBJ2-7 | CASSEGPGQGYEQYF | RAKFKQLL | YES | NO | YES |
| 10110 | TRAV30 | TRAJ18 | CGTESLRGSTLGRLYF | TRBV27 | TRBJ2-7 | CASSLSGRPYEQYF | RAKFKQLL | YES | NO | NO |
| 10115 | TRAV17 | TRAJ43 | CATDAYNNDMRF | TRBV28 | TRBJ2-7 | CASSPEFSRGGLGEQYF | RAKFKQLL | YES | NO | YES |
| 10123 | TRAV12-1 | TRAJ31 | CVVNIDNARLMF | TRBV29-1 | TRBJ2-7 | CSTGQGPYEQYF | RAKFKQLL | NO | NO | NO |
| 10124 | TRAV12-1 | TRAJ31 | CVVNIDNARLMF | TRBV29-1 | TRBJ2-7 | CSVGQGPYEQYF | RAKFKQLL | NO | NO | NO |
| 10126 | TRAV12-2 | TRAJ53 | CAMTFSGGSNYKLTF | TRBV29-1 | TRBJ2-7 | CSVGTGDYEQYF | RAKFKQLL | YES | NO | YES |
| 10129 | TRAV12-3 | TRAJ53 | CAMSGSGGSNYKLTF | TRBV29-1 | TRBJ2-7 | CSVGSGDYEQYF | RAKFKQLL | YES | NO | YES |
| 10130 | TRAV12-3 | TRAJ53 | CAMTFSGGSNYKLTF | TRBV29-1 | TRBJ2-7 | CSVGTGDYEQYF | RAKFKQLL | YES | NO | NO |
| 10134 | TRAV13-2 | TRAJ45 | CADSGGGADGLTF | TRBV29-1 | TRBJ2-7 | CSVGSGEDNEQYF | RAKFKQLL | YES | NO | YES |
| 10139 | TRAV22 | TRAJ23 | CAVDPSIYNQGGKLIF | TRBV29-1 | TRBJ2-7 | CSVGTGDWGEQYF | RAKFKQLL | YES | NO | YES |
| 10141 | TRAV27 | TRAJ54 | CAGAKGAQKLVF | TRBV29-1 | TRBJ2-7 | CSVGSGDYEQYF | RAKFKQLL | YES | NO | YES |
| 10152 | TRAV5 | TRAJ6 | CAEMEGGSYIPTF | TRBV3-1 | TRBJ2-7 | CASSLGTGGSYEQYF | RAKFKQLL | YES | NO | YES |
| 10154 | TRAV9-2 | TRAJ43 | CALSESGGDDMRF | TRBV3-1 | TRBJ2-7 | CASSPLTSGSYEQYF | RAKFKQLL | YES | NO | YES |
| 10155 | TRAV8-6 | TRAJ9 | CAVSDQNGGFKTIF | TRBV4-1 | TRBJ2-7 | CASSLTGEDEQYF | RAKFKQLL | YES | NO | NO |
| 10162 | TRAV13-2 | TRAJ36 | CAETGGANNLFF | TRBV5-1 | TRBJ2-7 | CASSLNGGNSYEQYF | RAKFKQLL | YES | NO | YES |
| 10164 | TRAV13-2 | TRAJ45 | CADSGGGADGLTF | TRBV5-1 | TRBJ2-7 | CASSRDRSSYEQYF | RAKFKQLL | YES | NO | YES |
| 10166 | TRAV13-1 | TRAJ50 | CAATETSYDKVIF | TRBV5-4 | TRBJ2-7 | CASKRQGSLSYEQYF | RAKFKQLL | YES | NO | YES |
| 10168 | TRAV27 | TRAJ43 | CAGGFLNNNDMRF | TRBV5-4 | TRBJ2-7 | CASSLISGPPYEQYF | RAKFKQLL | YES | NO | YES |
| 10183 | TRAV2 | TRAJ23 | CAVPYNQGGKLIF | TRBV5-5 | TRBJ2-7 | CASSGPGGVDEQYF | RAKFKQLL | NO | NO | NO |
| 10190 | TRAV5 | TRAJ6 | CAEMEGGSYIPTF | TRBV5-6 | TRBJ2-7 | CASSLVGAPRREQYF | RAKFKQLL | YES | NO | NO |
| 10191 | TRAV8-6 | TRAJ28 | CAAISGAGSYQLTF | TRBV5-6 | TRBJ2-7 | CASSPSYEQYF | RAKFKQLL | YES | NO | YES |
| 10192 | TRAV13-1 | TRAJ50 | CAASETSYDKVIF | TRBV6-1 | TRBJ2-7 | CASSEVVSTTYEQYF | RAKFKQLL | YES | NO | YES |
| 10203 | TRAV13-1 | TRAJ50 | CAASETSYDKVIF | TRBV7-9 | TRBJ2-7 | CASSFVDRVYEQYF | RAKFKQLL | YES | NO | YES |
| 10204 | TRAV13-2 | TRAJ50 | CAEMETSYDKVIF | TRBV7-9 | TRBJ2-7 | CASTSVQRQGGDASYEQYF | RAKFKQLL | YES | NO | YES |
| 10207 | TRAV27 | TRAJ50 | CAGAKETSYDKVIF | TRBV7-9 | TRBJ2-7 | CASSSAAAYEQYF | RAKFKQLL | YES | NO | YES |
| 10211 | TRAV21 | TRAJ40 | CAVVGSGTYKYIF | TRBV29-1 | TRBJ2-7 | CSVVTFYYEQYF | IPSINVHHY | YES | NO | YES |
| 10212 | TRAV12-2 | TRAJ31 | CAVNNARLMF | TRBV4-3 | TRBJ2-3 | CASSQGGGGTDTQYF | SLFNTVATL | YES | NO | YES |
| 10213 | TRAV12-2 | TRAJ43 | CAVRNNDMRF | TRBV7-3 | TRBJ2-3 | CASSPLGQDTQYF | SLFNTVATL | YES | NO | YES |
| 10215 | TRAV21 | TRAJ31 | CAARRGARLMF | TRBV5-6 | TRBJ1-5 | CASSSTGGGEKDQPQHF | KLGGALQAK | YES | NO | NO |
| 10216 | TRAV19 | TRAJ40 | CALRTYKYIF | TRBV7-2 | TRBJ2-5 | CASGYWKLAGGPQETQYF | KLGGALQAK | YES | NO | NO |
| 10217 | TRAV39 | TRAJ58 | CAGSTSGSRLTF | TRBV20-1 | TRBJ2-7 | CSATYEQYF | KLGGALQAK | NO | NO | NO |
| 10218 | TRAV12-2 | TRAJ52 | CAVEAGGTSYGKLTF | TRBV20-1 | TRBJ1-5 | CSASDVGLNQPQHF | KLGGALQAK | YES | NO | NO |
| 10219 | TRAV19 | TRAJ24 | CALSEADDSWGKLQF | TRBV27 | TRBJ2-2 | CASSLYGNLGTGELFF | KLGGALQAK | YES | NO | NO |
| 10220 | TRAV1-1 | TRAJ28 | CERSGYSGAGSYQLTF | TRBV7-9 | TRBJ2-7 | CASSLELASYEQYF | KLGGALQAK | YES | NO | NO |
| 10221 | TRAV25 | TRAJ42 | CAGNYGGSQGNLIF | TRBV19 | TRBJ2-2 | CASSSRSTGELFF | GILGFVFTL | YES | NO | YES |
| 10222 | TRAV12-3 | TRAJ18 | CAMSSADRGSTLGRLYF | TRBV9 | TRBJ2-3 | CASSTKAGEYADTQYF | KLGGALQAK | YES | NO | NO |
| 10223 | TRAV9-2 | TRAJ52 | CALRNGGTSYGKLTF | TRBV7-7 | TRBJ2-6 | CASSRGGIASGANVLTF | KLGGALQAK | YES | NO | NO |
| 10224 | TRAV27 | TRAJ26 | CAGVRYGQNFVF | TRBV4-3 | TRBJ2-2 | CASSQDLAGNTGELFF | RAKFKQLL | YES | NO | YES |
| 10226 | TRAV13-1 | TRAJ57 | CAASFQGGSEKLVF | TRBV10-3 | TRBJ2-4 | CAISPSRSKNIQYF | AVFDRKSDAK | YES | NO | YES |
| 10227 | TRAV24 | TRAJ23 | CAFPYNQGGKLIF | TRBV15 | TRBJ1-5 | CATSREFQGAQPQHF | KLGGALQAK | YES | NO | NO |
| 10228 | TRAV8-3 | TRAJ42 | CAVGTDYGGSQGNLIF | TRBV2 | TRBJ2-3 | CASSEIGQGGTDTQYF | KLGGALQAK | YES | NO | NO |
| 10229 | TRAV35 | TRAJ37 | CAGQTGNTGKLIF | TRBV19 | TRBJ2-3 | CASSVRSADTQYF | GILGFVFTL | YES | NO | YES |
| 10230 | TRAV2 | TRAJ13 | CAVEDLNSGGYQKVTF | TRBV28 | TRBJ2-7 | CASKRTATYEQYF | KLGGALQAK | NO | NO | NO |
| 10231 | TRAV24 | TRAJ53 | CALNSGGSNYKLTF | TRBV4-3 | TRBJ2-1 | CASTPSGASYNEQFF | KLGGALQAK | YES | NO | NO |
| 10232 | TRAV13-1 | TRAJ50 | CAASETSYDKVIF | TRBV12-5 | TRBJ2-7 | CASGLAQFGEQYF | RAKFKQLL | YES | NO | YES |
| 10233 | TRAV12-2 | TRAJ47 | CAVNYGNKLVF | TRBV20-1 | TRBJ1-1 | CSASAGLAEAFF | ELAGIGILTV | YES | YES | YES |
| 10234 | TRAV4 | TRAJ6 | CLVGDASGGSYIPTF | TRBV20-1 | TRBJ2-1 | CSASGQSGVYNEQFF | AVFDRKSDAK | NO | NO | NO |
| 10235 | TRAV17 | TRAJ30 | CATDRDDKIIF | TRBV9 | TRBJ1-1 | CASSVRSSMNTEAFF | KLGGALQAK | YES | NO | NO |
| 10236 | TRAV14DV4 | TRAJ9 | CAMREGPVKTIF | TRBV3-1 | TRBJ2-7 | CASSQDRVLAGAYEQYF | KLGGALQAK | YES | NO | NO |
| 10237 | TRAV12-3 | TRAJ52 | CARSTSYGKLTF | TRBV4-1 | TRBJ2-7 | CASSQDGNRGRDEQYF | KLGGALQAK | YES | NO | NO |
| 10238 | TRAV34 | TRAJ30 | CGADIDDKIIF | TRBV19 | TRBJ2-1 | CASSASVLGVNEQFF | KLGGALQAK | YES | NO | NO |
| 10239 | TRAV12-2 | TRAJ43 | CAVRNDMRF | TRBV28 | TRBJ1-2 | CASSPQGQGNGYTF | ELAGIGILTV | NO | NO | NO |
| 10240 | TRAV12-2 | TRAJ16 | CAVTDRF | TRBV5-4 | TRBJ2-4 | CASSWDGAGNIQYF | ELAGIGILTV | NO | NO | NO |
| 10241 | TRAV13-2 | TRAJ45 | CADSGGGADGLTF | TRBV19 | TRBJ2-1 | CASSIRSGYEQFF | GILGFVFTL | YES | NO | YES |
| 10242 | TRAV1-2 | TRAJ20 | CAVRERDYKLSF | TRBV19 | TRBJ1-5 | CASKEGNQPQHF | KLGGALQAK | YES | NO | NO |
| 10243 | TRAV12-2 | TRAJ34 | CAVIFTDKLIF | TRBV12-4 | TRBJ2-1 | CASGSGGGNEQFF | AVFDRKSDAK | YES | NO | YES |
| 10244 | TRAV8-3 | TRAJ24 | CAVGALGDSWGKLQF | TRBV9 | TRBJ2-6 | CASSVVSGANVLTF | KLGGALQAK | YES | NO | NO |
| 10245 | TRAV14DV4 | TRAJ56 | CAMREGLFTGANSKLTF | TRBV14 | TRBJ2-2 | CASSPNLDTGELFF | KLGGALQAK | NO | NO | NO |
| 10246 | TRAV19 | TRAJ17 | CALSEAYGLKAAGNKLTF | TRBV10-3 | TRBJ2-3 | CAIRDVSSTDTQYF | KLGGALQAK | YES | NO | NO |
| 10247 | TRAV14DV4 | TRAJ54 | CAMRGLIQGAQKLVF | TRBV7-9 | TRBJ2-5 | CASSSAGTGGQETQYF | KLGGALQAK | YES | NO | NO |
| 10248 | TRAV5 | TRAJ13 | CAESVSGGYQKVTF | TRBV5-1 | TRBJ2-3 | CASSFPGTPTREDTQYF | KLGGALQAK | YES | NO | NO |
| 10249 | TRAV12-2 | TRAJ53 | CAVNGAESSGGSNYKLTF | TRBV9 | TRBJ1-6 | CASSVGGVSPLHF | YLLEMLWRL | YES | NO | YES |
| 10250 | TRAV5 | TRAJ45 | CAESYSGGGADGLTF | TRBV14 | TRBJ1-1 | CASSPLGAYTGFF | KLGGALQAK | NO | NO | NO |
| 10251 | TRAV36DV7 | TRAJ35 | CAGAI | TRBV24-1 | TRBJ1-2 | CATSTGTRNYGYTF | KLGGALQAK | NO | NO | NO |
| 10252 | TRAV13-1 | TRAJ42 | CAASGGGSQGNLIF | TRBV19 | TRBJ2-3 | CASSIRSQDTQYF | GILGFVFTL | YES | NO | YES |
| 10253 | TRAV17 | TRAJ11 | CATVGNSGYSTLTF | TRBV6-5 | TRBJ2-1 | CASSLQGGNEQFF | FLYALALLL | NO | NO | NO |
| 10254 | TRAV8-3 | TRAJ42 | CAVGGGGGSQGNLIF | TRBV19 | TRBJ2-1 | CASSSRSGTEQFF | GILGFVFTL | YES | NO | YES |
| 10255 | TRAV17 | TRAJ53 | CATLNSGGSNYKLTF | TRBV11-2 | TRBJ2-7 | CASSFGTGGYEQYF | KLGGALQAK | YES | NO | NO |

| | | | | | | | | | | |
|---|---|---|---|---|---|---|---|---|---|---|
| 10256 | TRAV14DV4 | TRAJ17 | CAMRASEQAAGNKLTF | TRBV19 | TRBJ2-7 | CASSIRSSYEQYF | GILGFVFTL | YES | NO | YES |
| 10257 | TRAV1-2 | TRAJ10 | CAVRGTGGGNKLTF | TRBV2 | TRBJ1-2 | CASSDAPGAINRIF | KLGGALQAK | NO | NO | NO |
| 10258 | TRAV12-1 | TRAJ34 | CVVLNTDKLIF | TRBV7-2 | TRBJ2-1 | CASSFGEQFF | GILGFVFTL | YES | NO | YES |
| 10259 | TRAV21 | TRAJ27 | CAVRPNTNAGKSTF | TRBV7-2 | TRBJ2-3 | CASSSDRGIGTDTQYF | KLGGALQAK | YES | NO | NO |
| 10260 | TRAV8-1 | TRAJ24 | CAVTWRTTDSWGKLQF | TRBV20-1 | TRBJ2-7 | CSARDLAKYEQYF | KLGGALQAK | NO | NO | NO |
| 10261 | TRAV36DV7 | TRAJ52 | CAPNAGGTSYGKLTF | TRBV7-9 | TRBJ2-7 | CASSGGLAPPYEQYF | KLGGALQAK | NO | NO | NO |
| 10262 | TRAV29DV5 | TRAJ45 | CASYSGGGADGLTF | TRBV28 | TRBJ2-2 | CASRRLAGGAGELFF | AVFDRKSDAK | YES | NO | NO |
| 10263 | TRAV22 | TRAJ37 | CADGSSNTGKLIF | TRBV4-2 | TRBJ1-1 | CATSSGGTEAFF | RLRAEAQVK | YES | NO | YES |
| 10264 | TRAV13-1 | TRAJ45 | CAASESGGGADGLTF | TRBV7-2 | TRBJ2-7 | CASSPSSYEQYF | KLGGALQAK | YES | NO | NO |
| 10265 | TRAV1-1 | TRAJ30 | CAVRGGNRDDKIIF | TRBV10-3 | TRBJ2-1 | CAISESGDYEQFF | KLGGALQAK | YES | NO | NO |
| 10266 | TRAV14DV4 | TRAJ48 | CAMRDPNFGNEKLTF | TRBV20-1 | TRBJ2-1 | CSVFGGRSYNEQFF | KLGGALQAK | NO | NO | NO |
| 10267 | TRAV3 | TRAJ34 | CAVRLYNTDKLIF | TRBV12-4 | TRBJ2-5 | CASLLEGAEGETQYF | KLGGALQAK | NO | NO | NO |
| 10268 | TRAV29DV5 | TRAJ37 | CAASSDSSNTGKLIF | TRBV6-1 | TRBJ1-1 | CASSVSMNTEAFF | KLGGALQAK | YES | NO | NO |
| 10269 | TRAV13-2 | TRAJ42 | CAENMGGGSQGNLIF | TRBV19 | TRBJ2-5 | CASSSRSGETQYF | GILGFVFTL | YES | NO | YES |
| 10270 | TRAV21 | TRAJ45 | CAVSGGGADGLTF | TRBV28 | TRBJ1-1 | CASSPNKLMNTEAFF | KLGGALQAK | YES | NO | NO |
| 10271 | TRAV8-3 | TRAJ20 | CAVGATNDYKLSF | TRBV12-3 | TRBJ1-1 | CASSLGPNTEAFF | KLGGALQAK | YES | NO | NO |
| 10272 | TRAV19 | TRAJ48 | CALSEALHFGNEKLTF | TRBV6-3 | TRBJ2-1 | CASSYLAGSSVYNEQFF | KLGGALQAK | YES | NO | NO |
| 10273 | TRAV27 | TRAJ24 | CAGPDSWGKLQF | TRBV10-3 | TRBJ2-1 | CAISESLAGRTYNEQFF | KLGGALQAK | YES | NO | NO |
| 10275 | TRAV14DV4 | TRAJ57 | CAMREGQGGGSEKLVF | TRBV19 | TRBJ1-5 | CASTSGVYQPQHF | KLGGALQAK | YES | NO | NO |
| 10276 | TRAV26-2 | TRAJ42 | CILMNYGGSQGNLIF | TRBV4-3 | TRBJ2-3 | CASSYGLAGGPWDTDTQYF | KLGGALQAK | YES | NO | NO |
| 10277 | TRAV19 | TRAJ34 | CALKVLGKLIF | TRBV30 | TRBJ1-1 | CAWSGLTITEAFF | IVTDFSVIK | YES | NO | NO |
| 10278 | TRAV19 | TRAJ47 | CALSDVEYGNKLVF | TRBV5-1 | TRBJ1-5 | CASSLDGSNQPQHF | KLGGALQAK | YES | NO | NO |
| 10279 | TRAV24 | TRAJ3 | CAFGSSSASKIIF | TRBV10-3 | TRBJ1-6 | CAISAGDSPLHF | KLGGALQAK | YES | NO | NO |
| 10280 | TRAV12-2 | TRAJ4 | CADSSYMFSGGYNKLIF | TRBV28 | TRBJ1-6 | CASTERGQGDSPLHF | KLGGALQAK | YES | NO | NO |
| 10281 | TRAV8-4 | TRAJ43 | CAVSAWSADMRF | TRBV2 | TRBJ1-2 | CARRQGSYGYTF | KLGGALQAK | YES | NO | NO |
| 10282 | TRAV12-2 | TRAJ13 | CAASGGYQKVTF | TRBV15 | TRBJ2-3 | CATSSATPPGLAIPDTQYF | KLGGALQAK | YES | NO | NO |
| 10284 | TRAV35 | TRAJ52 | CAGAGGTSYGKLTF | TRBV27 | TRBJ1-6 | CASSYTGYSSPLHF | KLGGALQAK | YES | NO | NO |
| 10286 | TRAV8-3 | TRAJ23 | CAVGNQGGKLIF | TRBV20-1 | TRBJ2-3 | CSAIVGRPYFTDTQYF | AVFDRKSDAK | NO | NO | NO |
| 10287 | TRAV19 | TRAJ56 | CALSEFRGANSKLTF | TRBV5-1 | TRBJ1-2 | CASSYLARRVGYTF | KLGGALQAK | YES | NO | NO |
| 10289 | TRAV13-1 | TRAJ41 | CAASPFDSGYALNF | TRBV4-3 | TRBJ1-3 | CASSQGFWTFGNTIYF | KLGGALQAK | YES | NO | NO |
| 10290 | TRAV1-2 | TRAJ23 | CAGAYNQGGKLIF | TRBV12-4 | TRBJ1-2 | CASSLEGGLLYGYTF | KLGGALQAK | YES | NO | NO |
| 10291 | TRAV13-2 | TRAJ39 | CAENPNAGNMLTF | TRBV6-6 | TRBJ2-3 | CASSPMGDTQYF | KLGGALQAK | NO | NO | NO |
| 10292 | TRAV12-2 | TRAJ17 | CAVSIKAAGNKLTF | TRBV6-5 | TRBJ1-2 | CASSPGMGSGYTF | ELAGIGILTV | YES | NO | YES |
| 10293 | TRAV12-1 | TRAJ3 | CVVSSYSSASKIIF | TRBV19 | TRBJ2-7 | CASSPRGLSSYEQYF | KLGGALQAK | YES | NO | NO |
| 10294 | TRAV9-2 | TRAJ26 | CALSDPQRNYGQNFVF | TRBV7-8 | TRBJ2-7 | CASSLTSEGYEQYF | KLGGALQAK | YES | NO | NO |
| 10295 | TRAV29DV5 | TRAJ34 | CAAPRGGTDKLIF | TRBV29-1 | TRBJ2-7 | CSVFGGRLYEQYF | KLGGALQAK | YES | NO | NO |
| 10296 | TRAV8-3 | TRAJ42 | CAVGGYGGSQGNLIF | TRBV19 | TRBJ2-7 | CASSGRSGYEQYF | GILGFVFTL | YES | NO | YES |
| 10297 | TRAV4 | TRAJ44 | CLVGVGGTASKLTF | TRBV7-3 | TRBJ1-2 | CASSLTANYGYTF | AYAQKIFKI | NO | NO | NO |
| 10298 | TRAV1-1 | TRAJ11 | CAVRDGGSGYSTLTF | TRBV11-2 | TRBJ1-2 | CASSWGGGSHYGYTF | IVTDFSVIK | YES | NO | YES |
| 10299 | TRAV25 | TRAJ42 | CAGGSGGSQGNLIF | TRBV19 | TRBJ2-2 | CASSMRSTGELFF | GILGFVFTL | YES | NO | YES |
| 10300 | TRAV20 | TRAJ54 | CAVCRVQGAQKLVF | TRBV27 | TRBJ1-6 | CASSWRGRNSPLHF | KLGGALQAK | YES | NO | NO |
| 10301 | TRAV27 | TRAJ37 | CAGAEGSGNTGKLIF | TRBV19 | TRBJ2-7 | CASSIRSAYEQYF | GILGFVFTL | YES | NO | YES |
| 10302 | TRAV13-1 | TRAJ53 | CAASFVERSNYKLTF | TRBV6-3 | TRBJ2-5 | CASEEQGGLETQYF | AVFDRKSDAK | YES | NO | YES |
| 10303 | TRAV12-3 | TRAJ49 | CAMKGNQFYF | TRBV4-3 | TRBJ2-7 | CASSRLPGLAGVGNHEQYF | KLGGALQAK | YES | NO | NO |
| 10304 | TRAV12-2 | TRAJ32 | CAMNPAWGGATNKLIF | TRBV20-1 | TRBJ2-7 | CSASPGDYEQYF | KLGGALQAK | NO | NO | NO |
| 10305 | TRAV17 | TRAJ47 | CATGSMEYGNKLVF | TRBV9 | TRBJ2-2 | CASRMGDINTGELFF | IVTDFSVIK | YES | NO | YES |
| 10306 | TRAV13-1 | TRAJ48 | CAANEKLTF | TRBV6-5 | TRBJ2-7 | CASSYGTGGTYEQYF | KLGGALQAK | YES | NO | NO |
| 10309 | TRAV8-6 | TRAJ29 | CAVSVPLGNTPLVF | TRBV11-3 | TRBJ2-1 | CASSPRTSGIHNEQFF | KLGGALQAK | YES | NO | NO |
| 10310 | TRAV13-1 | TRAJ21 | CAASSWVNFNKFYF | TRBV7-2 | TRBJ2-1 | CASSFQGEAGLNEQFF | KLGGALQAK | YES | NO | NO |
| 10311 | TRAV3 | TRAJ31 | CAVRGSDSARLMF | TRBV6-5 | TRBJ1-1 | CASTQGANTEAFF | KLGGALQAK | NO | NO | NO |
| 10312 | TRAV8-3 | TRAJ45 | CAVGAGSGGGADGLTF | TRBV29-1 | TRBJ1-2 | CSVESRNYGYTF | AVFDRKSDAK | YES | YES | YES |
| 10313 | TRAV16 | TRAJ26 | CALLDNYGQNFVF | TRBV7-8 | TRBJ1-1 | CASSLERGNTEAFF | KLGGALQAK | NO | NO | NO |
| 10314 | TRAV12-1 | TRAJ41 | CVVNSGYALNF | TRBV27 | TRBJ2-1 | CASSFTGAGNEQFF | KLGGALQAK | YES | NO | NO |
| 10315 | TRAV12-2 | TRAJ35 | CAVAI | TRBV20-1 | TRBJ1-1 | CSARQGAEAFF | ELAGIGILTV | NO | NO | NO |
| 10316 | TRAV12-2 | TRAJ52 | CAAASGGTSYGKLTF | TRBV11-2 | TRBJ1-2 | CASSWGGGSHYGYTF | IVTDFSVIK | YES | NO | YES |
| 10317 | TRAV10 | TRAJ20 | CVVSGWSDYKLSF | TRBV11-3 | TRBJ2-5 | CASSSPNQETQYF | AVFDRKSDAK | YES | YES | YES |
| 10318 | TRAV26-2 | TRAJ33 | CIRIAGDSNYQLIW | TRBV27 | TRBJ2-7 | CASSLGILYEQYF | KLGGALQAK | YES | NO | NO |
| 10319 | TRAV14DV4 | TRAJ13 | CAMEAGGYQKVTF | TRBV28 | TRBJ2-7 | CASSPRLAESSYEQYF | KLGGALQAK | YES | YES | NO |
| 10320 | TRAV19 | TRAJ48 | CALSERGNFGNEKLTF | TRBV24-1 | TRBJ2-1 | CATSGPTSREGDNEQFF | KLGGALQAK | YES | NO | NO |
| 10321 | TRAV14DV4 | TRAJ43 | CAMRDSYNNNDMRF | TRBV6-5 | TRBJ2-1 | CASSPSLGTGGGLQFF | AVFDRKSDAK | NO | NO | NO |
| 10322 | TRAV12-2 | TRAJ54 | CAAQGAQKLVF | TRBV4-3 | TRBJ1-6 | CASSQGLVSGSPLHF | ELAGIGILTV | YES | YES | YES |
| 10323 | TRAV8-4 | TRAJ7 | CAVSPTYGNNRLAF | TRBV6-3 | TRBJ2-1 | CASSYLTSGGASEQFF | KLGGALQAK | YES | NO | NO |
| 10324 | TRAV26-2 | TRAJ50 | CILAETSYDKVIF | TRBV7-9 | TRBJ2-6 | CASSLTFSGANVLTF | RAKFKQLL | YES | NO | YES |
| 10325 | TRAV12-2 | TRAJ45 | CASNSGGGADGLTF | TRBV5-1 | TRBJ2-3 | CASSSEAGLAATDTQYF | ELAGIGILTV | YES | NO | YES |
| 10326 | TRAV5 | TRAJ6 | CAEMEGGSYIPTF | TRBV4-1 | TRBJ1-1 | CASSQGTVNTEAFF | KLGGALQAK | YES | NO | NO |
| 10327 | TRAV10 | TRAJ7 | CVVSRQEQQTRF | TRBV20-1 | TRBJ2-7 | CSARDINRGSYEQYF | KLGGALQAK | NO | NO | NO |
| 10328 | TRAV12-1 | TRAJ48 | CVGPRTF | TRBV28 | TRBJ1-5 | CASSLTGLGQPQHF | ELAGIGILTV | YES | NO | NO |
| 10329 | TRAV12-3 | TRAJ48 | CAMSPSNFGNEKLTF | TRBV14 | TRBJ1-3 | CASSQDFVLYSGNTIYF | ELAGIGILTV | NO | NO | NO |
| 10330 | TRAV17 | TRAJ5 | CATDAGDTGRRALTF | TRBV7-9 | TRBJ2-7 | CASTPPLASGRYYEQYF | KLGGALQAK | YES | NO | NO |
| 10331 | TRAV6 | TRAJ37 | CAPSSNTGKLIF | TRBV7-9 | TRBJ2-3 | CASSFAVSTDTQYF | KLGGALQAK | YES | NO | NO |
| 10332 | TRAV29DV5 | TRAJ47 | CAASAGEYGNKLVF | TRBV4-1 | TRBJ1-1 | CASSQEYSGTWTEAFF | KLGGALQAK | YES | NO | NO |
| 10333 | TRAV21 | TRAJ42 | CAVRAWNGGSQGNLIF | TRBV13 | TRBJ2-1 | CASLGGGPYNEQFF | KLGGALQAK | YES | NO | NO |
| 10334 | TRAV40 | TRAJ34 | CLRGVDNTDKLIF | TRBV19 | TRBJ2-7 | CASSIRSSYEQYF | KLGGALQAK | NO | NO | NO |
| 10335 | TRAV24 | TRAJ40 | CASIAGTYKYIF | TRBV9 | TRBJ2-6 | CASSVVSGANVLTF | KLGGALQAK | YES | NO | NO |
| 10336 | TRAV12-3 | TRAJ42 | CAMSGDGGSQGNLIF | TRBV21-1 | TRBJ2-7 | CQQQKSYEQYF | GILGFVFTL | YES | NO | NO |
| 10337 | TRAV38-1 | TRAJ49 | CAFMIGTGNQFYF | TRBV19 | TRBJ2-1 | CASSISGSSYNEQFF | KLGGALQAK | YES | NO | NO |
| 10338 | TRAV29DV5 | TRAJ49 | CAATLTGNQFYF | TRBV10-3 | TRBJ1-1 | CAISESILWVVRMNTEAFF | RLRAEAQVK | YES | NO | YES |
| 10340 | TRAV14DV4 | TRAJ34 | CAMSASYNTDKLIF | TRBV4-3 | TRBJ1-1 | CASTYAVTGNTEAFF | KLGGALQAK | YES | NO | NO |
| 10341 | TRAV23DV6 | TRAJ12 | CAARRRGMDSSYKLIF | TRBV19 | TRBJ1-5 | CASSINSNQPQHF | GILGFVFTL | YES | NO | YES |
| 10342 | TRAV3 | TRAJ23 | CAVRDIGNQGGKLIF | TRBV7-9 | TRBJ2-3 | CASSHSSGGVADTQYF | KLGGALQAK | NO | NO | NO |
| 10344 | TRAV38-2DV8 | TRAJ47 | CAYRSGYGYGNKLVF | TRBV4-3 | TRBJ1-2 | CASSQDGEPYGYTF | KLGGALQAK | YES | NO | NO |
| 10345 | TRAV29DV5 | TRAJ33 | CAALDSNYQLIW | TRBV11-2 | TRBJ1-2 | CASSWGGGSHYGYTF | IVTDFSVIK | YES | NO | YES |
| 10347 | TRAV35 | TRAJ33 | CAGLSVMDSNYQLIW | TRBV4-1 | TRBJ2-3 | CASSHLRFTQYF | KLGGALQAK | YES | YES | NO |
| 10348 | TRAV19 | TRAJ35 | CALSEAEVGFGNVLHC | TRBV9 | TRBJ1-2 | CASSARDTTNYGYTF | KLGGALQAK | NO | NO | NO |
| 10349 | TRAV26-2 | TRAJ15 | CILRAPGAGTALIF | TRBV19 | TRBJ1-2 | CASSIGVYGYTF | GILGFVFTL | YES | NO | YES |
| 10350 | TRAV12-1 | TRAJ41 | CVVSDSNSGYALNF | TRBV5-1 | TRBJ1-5 | CASSPGTGPNQPQHF | GILGFVFTL | YES | NO | NO |
| 10351 | TRAV26-2 | TRAJ45 | CILSGGGGADGLTF | TRBV7-9 | TRBJ1-4 | CASSDWGNEKLFF | KLGGALQAK | YES | NO | NO |
| 10353 | TRAV22 | TRAJ48 | CAVDGNEKLTF | TRBV27 | TRBJ2-7 | CASSFYVGGPYEQYF | KLGGALQAK | YES | NO | NO |
| 10354 | TRAV5 | TRAJ28 | CAEVYSGAGSYQLTF | TRBV18 | TRBJ2-1 | CASSPPGSDEQFF | KLGGALQAK | YES | NO | NO |
| 10355 | TRAV13-2 | TRAJ45 | CADSGGGADGLTF | TRBV27 | TRBJ2-3 | CASKGTDTQYF | KLGGALQAK | YES | NO | NO |
| 10356 | TRAV12-2 | TRAJ45 | CAVVEGGGADGLTF | TRBV27 | TRBJ1-2 | CASSLSPFYSYGYTF | ELAGIGILTV | YES | NO | YES |
| 10357 | TRAV13-1 | TRAJ49 | CAASMTGNQFYF | TRBV12-5 | TRBJ2-1 | CASGPLGGYNEQFF | KLGGALQAK | YES | NO | NO |
| 10358 | TRAV1-2 | TRAJ33 | CAKRSNYQLIW | TRBV6-3 | TRBJ2-1 | CASSPTSFGEQFF | KLGGALQAK | YES | NO | NO |
| 10359 | TRAV8-1 | TRAJ39 | CAVNADNNAGNMLTF | TRBV7-2 | TRBJ2-7 | CASSEQGYEQYF | AVFDRKSDAK | NO | NO | NO |
| 10360 | TRAV8-6 | TRAJ17 | CAVTMAAGNKLTF | TRBV7-9 | TRBJ1-1 | CASTVGSTEAFF | KLGGALQAK | YES | NO | NO |
| 10361 | TRAV24 | TRAJ42 | CAFRGGGSQGNLIF | TRBV19 | TRBJ2-7 | CASGTSGAYEQYF | KLGGALQAK | YES | NO | NO |
| 10362 | TRAV8-3 | TRAJ44 | CAVGNTGTASKLTF | TRBV14 | TRBJ2-2 | CASSQFTGESSGELFF | KLGGALQAK | NO | NO | NO |
| 10363 | TRAV13-2 | TRAJ33 | CAVPKVDSNYQLIW | TRBV4-1 | TRBJ2-7 | CASSQAQASSYEQYF | KLGGALQAK | YES | NO | NO |
| 10364 | TRAV25 | TRAJ42 | CAGNYGGSQGNLIF | TRBV19 | TRBJ2-2 | CASSKRSTGELFF | GILGFVFTL | YES | NO | YES |
| 10365 | TRAV1-2 | TRAJ12 | CAPMDSSYKLIF | TRBV6-6 | TRBJ2-3 | CASSGTSDPADTQYF | KLGGALQAK | NO | NO | NO |
| 10366 | TRAV1-2 | TRAJ33 | CAVRDSNYQLIW | TRBV27 | TRBJ2-2 | CASSFHGGELFF | KLGGALQAK | YES | NO | NO |
| 10367 | TRAV29DV5 | TRAJ22 | CAASGGLSARQLTF | TRBV6-3 | TRBJ1-1 | CASSYGGGTEAFF | AVFDRKSDAK | YES | NO | YES |
| 10368 | TRAV14DV4 | TRAJ57 | CAMREPLTQGGSEKLVF | TRBV11-2 | TRBJ1-2 | CASSWGGGSHYGYTF | IVTDFSVIK | YES | YES | YES |
| 10369 | TRAV29DV5 | TRAJ40 | CAAGTYKYIF | TRBV9 | TRBJ2-3 | CASSPEGRPGDTQYF | KLGGALQAK | YES | NO | NO |

| | | | | | | | | | | |
|---|---|---|---|---|---|---|---|---|---|---|
| 10370 | TRAV3 | TRAJ40 | CAVRDIASGTYKYIF | TRBV5-5 | TRBJ2-3 | CASSLGGRGVTDTQYF | KLGGALQAK | NO | NO | NO |
| 10371 | TRAV14DV4 | TRAJ27 | CAMRESNAGKSTF | TRBV7-2 | TRBJ2-7 | CASSLTSETSYEQYF | RAKFKQLL | YES | NO | YES |
| 10372 | TRAV24 | TRAJ28 | CAFPWGAGSYQLTF | TRBV24-1 | TRBJ2-3 | CATMVTYF | RAKFKQLL | NO | NO | NO |
| 10373 | TRAV6 | TRAJ5 | CVPMDTGRRALTF | TRBV4-3 | TRBJ1-2 | CASSQDSPTNYGYTF | KLGGALQAK | YES | NO | NO |
| 10374 | TRAV3 | TRAJ5 | CAVRDIDTGRRALTF | TRBV5-5 | TRBJ1-2 | CASSPRDTYYGYTF | KLGGALQAK | YES | NO | NO |
| 10375 | TRAV12-2 | TRAJ42 | CAVNLGGGSQGNLIF | TRBV19 | TRBJ2-7 | CASSIRSTYEQYF | GILGFVFTL | YES | NO | YES |
| 10376 | TRAV19 | TRAJ26 | CALSEAGNYGQNFVF | TRBV12-4 | TRBJ2-1 | CASSLGGPGLYNEQFF | KLGGALQAK | YES | NO | NO |
| 10377 | TRAV12-2 | TRAJ12 | CAVAGEDSSYKLIF | TRBV7-9 | TRBJ2-3 | CASSKKATDTQYF | KLGGALQAK | YES | NO | NO |
| 10378 | TRAV17 | TRAJ41 | CATDASSGYALNF | TRBV7-2 | TRBJ1-6 | CASSLAWDLNSPLHF | KLGGALQAK | NO | NO | NO |
| 10379 | TRAV12-2 | TRAJ22 | CAVRDSSGSARQLTF | TRBV4-1 | TRBJ1-1 | CASSQGTGNTEAFF | KLGGALQAK | YES | NO | NO |
| 10380 | TRAV3 | TRAJ46 | CAVRAIKSSGDKLTF | TRBV20-1 | TRBJ2-1 | CSAPGRYYNEQFF | KLGGALQAK | NO | NO | NO |
| 10381 | TRAV2 | TRAJ16 | CAVEDF | TRBV19 | TRBJ2-2 | CASTRTEPNTGELFF | KLGGALQAK | NO | NO | NO |
| 10382 | TRAV16 | TRAJ20 | CALRPSNDYKLSF | TRBV18 | TRBJ1-2 | CASSPGQGIHGYTF | KLGGALQAK | NO | NO | NO |
| 10383 | TRAV26-1 | TRAJ26 | CIVRESQNFVF | TRBV4-3 | TRBJ2-3 | CASSQDADTDTQYF | AVFDRKSDAK | YES | NO | YES |
| 10384 | TRAV12-2 | TRAJ45 | CAVSQGGADGLTF | TRBV4-2 | TRBJ1-1 | CASSLTGGAEAFF | ELAGIGILTV | YES | NO | YES |
| 10385 | TRAV21 | TRAJ26 | CAPRNYGQNFVF | TRBV27 | TRBJ2-6 | CASRNLAPLHSGANVLTF | KLGGALQAK | YES | NO | NO |
| 10386 | TRAV5 | TRAJ7 | CAEIIYGNNRLAF | TRBV5-5 | TRBJ2-3 | CASSSRQSEADTQYF | KLGGALQAK | YES | NO | NO |
| 10387 | TRAV35 | TRAJ48 | CAGPISNFGNEKLTF | TRBV29-1 | TRBJ1-1 | CSAASGTEAFF | KLGGALQAK | YES | NO | NO |
| 10389 | TRAV8-2 | TRAJ20 | CVVSDLGYKLSF | TRBV6-3 | TRBJ2-1 | CASRTSGGAYNEQFF | KLGGALQAK | YES | NO | NO |
| 10390 | TRAV41 | TRAJ47 | CAVIGGYGNKLVF | TRBV2 | TRBJ2-3 | CASSDLADTDTQYF | KLGGALQAK | YES | NO | NO |
| 10391 | TRAV12-3 | TRAJ30 | CAMQSPRDDKIIF | TRBV24-1 | TRBJ2-6 | CATSDSYSGANVLTF | AVFDRKSDAK | YES | NO | YES |
| 10392 | TRAV3 | TRAJ7 | CAVRDIIYGNNRLAF | TRBV7-6 | TRBJ2-1 | CASSLVEGGLAGSYNEQFF | KLGGALQAK | NO | NO | NO |
| 10393 | TRAV12-2 | TRAJ32 | CAMNPAWGGATNKLIF | TRBV9 | TRBJ2-7 | CASSVGDAQYEQYF | KLGGALQAK | YES | NO | NO |
| 10394 | TRAV3 | TRAJ37 | CAGGNTGKLIF | TRBV7-2 | TRBJ1-5 | CASSLGQGASQPQHF | GLCTLVAML | NO | NO | NO |
| 10395 | TRAV13-2 | TRAJ42 | CAERYNYGGSQGNLIF | TRBV6-4 | TRBJ1-2 | CASSDTGQGLYGYTF | AVFDRKSDAK | YES | NO | YES |
| 10396 | TRAV8-4 | TRAJ20 | CAVNSNDYKLSF | TRBV7-9 | TRBJ1-1 | CASSLGGTVEAFF | KLGGALQAK | YES | NO | NO |
| 10397 | TRAV19 | TRAJ32 | CALSGNYGGATNKLIF | TRBV7-2 | TRBJ2-3 | CASSLGVGADTDTQYF | KLGGALQAK | YES | NO | NO |
| 10398 | TRAV5 | TRAJ42 | CAESTPNGGSQGNLIF | TRBV4-3 | TRBJ2-3 | CASSPTQGAGDTQYF | AVFDRKSDAK | YES | NO | YES |
| 10399 | TRAV1-2 | TRAJ20 | CAVNSDYKLSF | TRBV20-1 | TRBJ2-2 | CSAGQGPNTGELFF | KLGGALQAK | YES | NO | NO |
| 10400 | TRAV5 | TRAJ41 | CAVSFSGYALNF | TRBV18 | TRBJ2-1 | CASSLAGSSGNEQFF | KLGGALQAK | YES | NO | NO |
| 10401 | TRAV8-3 | TRAJ28 | CAVGAPGAGSYQLTF | TRBV3-1 | TRBJ2-7 | CASSQSAGPRSYEQYF | KLGGALQAK | YES | NO | NO |
| 10402 | TRAV29DV5 | TRAJ43 | CAASALRNNDMRF | TRBV27 | TRBJ2-3 | CASSLDRVAADTQYF | KLGGALQAK | YES | NO | NO |
| 10403 | TRAV27 | TRAJ26 | CAGVNNYGQNFVF | TRBV28 | TRBJ1-4 | CASSLCPLPTVCYEKLFF | KLGGALQAK | YES | NO | NO |
| 10404 | TRAV14DV4 | TRAJ22 | CAMREPASGSARQLTF | TRBV6-3 | TRBJ1-5 | CASSYSKGGPYQPQHF | KLGGALQAK | YES | NO | NO |
| 10405 | TRAV26-2 | TRAJ52 | CILRDYRRGAGGTSYGKLTF | TRBV28 | TRBJ1-4 | CASSWRGASGKLFF | KLGGALQAK | NO | NO | NO |
| 10406 | TRAV8-3 | TRAJ34 | CAVGAGNTDKLIF | TRBV9 | TRBJ2-3 | CASSVAPDTQYF | KLGGALQAK | YES | NO | NO |
| 10407 | TRAV3 | TRAJ13 | CAVSASGGYQKVTF | TRBV20-1 | TRBJ2-3 | CSAIPWTGSTDTQYF | KLGGALQAK | NO | NO | NO |
| 10408 | TRAV9-2 | TRAJ49 | CALIIPPADTGNQFYF | TRBV6-5 | TRBJ2-5 | CASSKRGVRWAQYF | KLGGALQAK | NO | NO | NO |
| 10409 | TRAV38-1 | TRAJ52 | CAFMVNAGGTSYGKLTF | TRBV19 | TRBJ1-2 | CASSTGSYGYTF | GILGFVFTL | YES | NO | YES |
| 10410 | TRAV8-1 | TRAJ3 | CAVNPEGASKIIF | TRBV5-1 | TRBJ2-7 | CASSLDVGRAASYEQYF | KLGGALQAK | YES | NO | NO |
| 10412 | TRAV26-2 | TRAJ42 | CILSVYYGGSQGNLIF | TRBV4-1 | TRBJ1-2 | CASSQELAGGNANYGYTF | KLGGALQAK | YES | NO | NO |
| 10413 | TRAV3 | TRAJ6 | CAVRAPSGGSYIPTF | TRBV13 | TRBJ2-7 | CASSLGTSGSSYEQYF | IVTDFSVIK | NO | NO | NO |
| 10414 | TRAV19 | TRAJ24 | CALSETTTDSWGKLQF | TRBV6-3 | TRBJ1-1 | CASRDQMNTEAFF | KLGGALQAK | YES | NO | NO |
| 10415 | TRAV12-2 | TRAJ37 | CAVGGTGKLIF | TRBV19 | TRBJ2-7 | CASGLSGDEQYF | ELAGIGILTV | YES | YES | YES |
| 10416 | TRAV12-2 | TRAJ40 | CAVTSGTYKYIF | TRBV7-8 | TRBJ2-1 | CASSLGMRGGWDEQFF | KLGGALQAK | YES | NO | NO |
| 10418 | TRAV12-1 | TRAJ12 | CVVNLQMDSSYKLIF | TRBV7-3 | TRBJ2-5 | CASSVFGGGETQYF | KLGGALQAK | NO | NO | NO |
| 10419 | TRAV25 | TRAJ44 | CAGPLTGTASKLTF | TRBV9 | TRBJ2-7 | CASSVGGTGVYEQYF | KLGGALQAK | YES | NO | NO |
| 10420 | TRAV12-3 | TRAJ8 | CAMQGFQKLVF | TRBV20-1 | TRBJ2-1 | CSARVSGGGYNEQFF | KLGGALQAK | NO | NO | NO |
| 10421 | TRAV22 | TRAJ40 | CAVERASGTYKYIF | TRBV11-1 | TRBJ2-1 | CASSSLAGGRDEQFF | KLGGALQAK | YES | NO | NO |
| 10422 | TRAV19 | TRAJ56 | CALSVRWGANSKLTF | TRBV5-4 | TRBJ1-2 | CASSLGGRGSYGYTF | KLGGALQAK | YES | NO | NO |
| 10423 | TRAV21 | TRAJ39 | CAAALYNNAGNMLTF | TRBV19 | TRBJ1-5 | CASSIYSNQPQHF | GILGFVFTL | YES | NO | YES |
| 10424 | TRAV22 | TRAJ49 | CAARTGNQFYF | TRBV4-1 | TRBJ1-1 | CASSQEELVLAFF | KLGGALQAK | YES | NO | NO |
| 10425 | TRAV5 | TRAJ42 | CAPGGSQGNLIF | TRBV6-6 | TRBJ1-1 | CASSPTGLEAFF | KLGGALQAK | NO | NO | NO |
| 10426 | TRAV4 | TRAJ8 | CLSYSFQKLVF | TRBV5-4 | TRBJ1-6 | CASSLFAGERSPLHF | KLGGALQAK | YES | NO | NO |
| 10427 | TRAV41 | TRAJ52 | CAVSNAGGTSYGKLTF | TRBV6-5 | TRBJ2-7 | CASSSTTLLAGGYEQYF | KLGGALQAK | YES | NO | NO |
| 10428 | TRAV10 | TRAJ33 | CVVLGVMDSNYQLIW | TRBV18 | TRBJ1-1 | CASSLDPGQGWTNTEAFF | GILGFVFTL | YES | NO | YES |
| 10429 | TRAV3 | TRAJ33 | CALDSNYQLIW | TRBV2 | TRBJ1-4 | CASRRQGNEKLFF | KLGGALQAK | NO | NO | NO |
| 10430 | TRAV14DV4 | TRAJ11 | CAMRELAEYSTLTF | TRBV29-1 | TRBJ2-7 | CSASGVRDSLYEQYF | AVFDRKSDAK | YES | NO | NO |
| 10431 | TRAV8-4 | TRAJ13 | CAVSVPNSGGYQKVTF | TRBV5-1 | TRBJ1-3 | CASSLDGLVTGNTIYF | KLGGALQAK | YES | NO | NO |
| 10432 | TRAV8-6 | TRAJ17 | CAVSQAAGNKLTF | TRBV7-9 | TRBJ2-7 | CASSRRLAGYYEQYF | KLGGALQAK | YES | NO | NO |
| 10433 | TRAV13-1 | TRAJ11 | CAASIGSGYSTLTF | TRBV14 | TRBJ1-4 | CASSQAAGDNEKLFF | KLGGALQAK | NO | NO | NO |
| 10434 | TRAV12-2 | TRAJ31 | CAGNNARLMF | TRBV28 | TRBJ1-5 | CASRGTGLGNQPQHF | ELAGIGILTV | YES | NO | YES |
| 10435 | TRAV22 | TRAJ4 | CAVRGVAAGGYNKLIF | TRBV28 | TRBJ2-3 | CASSLIGVGEFTDTQYF | KLGGALQAK | YES | NO | NO |
| 10436 | TRAV6 | TRAJ32 | CALDGGATNKLIF | TRBV5-6 | TRBJ2-5 | CASSLAGGGLPEETQYF | KLGGALQAK | YES | NO | NO |
| 10437 | TRAV12-2 | TRAJ23 | CAVILFYNQGGKLIF | TRBV6-3 | TRBJ1-2 | CASSTPRGGYTF | MLDLQPETT | YES | NO | YES |
| 10438 | TRAV29DV5 | TRAJ53 | CALYGSGGSNYKLTF | TRBV9 | TRBJ1-3 | CASSVEAGASSGNTIYF | AVFDRKSDAK | NO | NO | NO |
| 10439 | TRAV12-2 | TRAJ52 | CAASHSGTSYGKLTF | TRBV27 | TRBJ2-2 | CASSLSLAGELFF | ELAGIGILTV | NO | NO | NO |
| 10440 | TRAV29DV5 | TRAJ20 | CAASESNYKLSF | TRBV10-3 | TRBJ2-3 | CAITSVSGGLDTDTQYF | AVFDRKSDAK | YES | NO | NO |
| 10441 | TRAV8-1 | TRAJ45 | CAVISDSGGGADGLTF | TRBV7-9 | TRBJ2-1 | CASSSLGEFYNEQFF | KLGGALQAK | YES | NO | NO |
| 10442 | TRAV27 | TRAJ40 | CAGYPSGTYKYIF | TRBV20-1 | TRBJ2-1 | CSAPPSGGAHEQFF | KLGGALQAK | NO | NO | NO |
| 10443 | TRAV5 | TRAJ30 | CAESLWDDKIIF | TRBV11-2 | TRBJ1-2 | CASSWGGGSHYGYTF | IVTDFSVIK | YES | YES | YES |
| 10444 | TRAV21 | TRAJ48 | CAVKTLGNEKLTF | TRBV5-1 | TRBJ2-5 | CASSWTTSGGQETQYF | RAKFKQLL | YES | NO | YES |
| 10445 | TRAV19 | TRAJ6 | CALSEAGTGGSYIPTF | TRBV19 | TRBJ1-1 | CASSSFLGAEAFF | GILGFVFTL | YES | NO | YES |
| 10446 | TRAV19 | TRAJ56 | CALSELNYTGANSKLTF | TRBV19 | TRBJ2-3 | CASSSHSGRANTDTQYF | RAKFKQLL | YES | NO | YES |
| 10447 | TRAV16 | TRAJ49 | CALSDTNTGNQFYF | TRBV4-1 | TRBJ1-6 | CASSQEKGTAVPNSPLHF | KLGGALQAK | NO | NO | NO |
| 10448 | TRAV12-1 | TRAJ13 | CVVNGNSGGYQKVTF | TRBV4-2 | TRBJ2-3 | CASSDLAGDPTDTQYF | GILGFVFTL | YES | NO | YES |
| 10449 | TRAV9-2 | TRAJ21 | CALSAILYNFNKFYF | TRBV5-6 | TRBJ1-6 | CASSHRAGALHF | KLGGALQAK | YES | NO | NO |
| 10450 | TRAV14DV4 | TRAJ12 | CAIRMDSSYKLIF | TRBV7-9 | TRBJ2-2 | CASSSQTSGSTGELFF | KLGGALQAK | YES | NO | NO |
| 10451 | TRAV1-2 | TRAJ12 | CAVMDSSYKLIF | TRBV6-4 | TRBJ2-3 | CASSEGDTDTQYF | KLGGALQAK | YES | NO | NO |
| 10452 | TRAV12-1 | TRAJ52 | CVVNIGTLNAGGTSYGKLTF | TRBV10-3 | TRBJ2-7 | CAISEGVWTYEQYF | YLLEMLWRL | NO | NO | NO |
| 10453 | TRAV9-2 | TRAJ21 | CAPNWNFNKFYF | TRBV19 | TRBJ2-5 | CASSISQGAQETQYF | KLGGALQAK | YES | NO | NO |
| 10455 | TRAV22 | TRAJ21 | CAVVKYNFNKFYF | TRBV6-3 | TRBJ1-5 | CASSRGTGRIGNQPQHF | KLGGALQAK | YES | NO | NO |
| 10456 | TRAV8-3 | TRAJ45 | CAVGGYSGGGADGLTF | TRBV18 | TRBJ1-2 | CASSPETGAYGYTF | AVFDRKSDAK | YES | NO | YES |
| 10457 | TRAV9-2 | TRAJ53 | CALESSNYKLTF | TRBV7-3 | TRBJ2-5 | CASSSGALPRGETQYF | RLRAEAQVK | NO | NO | NO |
| 10458 | TRAV27 | TRAJ42 | CAAGGSQGNLIF | TRBV19 | TRBJ2-7 | CASSTRSAYEQYF | GILGFVFTL | YES | NO | YES |
| 10459 | TRAV8-1 | TRAJ45 | CAVNRAGGGADGLTF | TRBV2 | TRBJ2-1 | CASSEASGGFYNEQFF | IVTDFSVIK | YES | YES | YES |
| 10460 | TRAV30 | TRAJ20 | CGTEYPNDYKLSF | TRBV11-2 | TRBJ1-2 | CASSWGGGSHYGYTF | IVTDFSVIK | YES | NO | NO |
| 10461 | TRAV29DV5 | TRAJ39 | CAASASNAGNMLTF | TRBV29-1 | TRBJ2-1 | CSVVGVGYNEQFF | KLGGALQAK | YES | NO | NO |
| 10462 | TRAV26-2 | TRAJ30 | CILRDGDDKIIF | TRBV6-6 | TRBJ1-6 | CASSYGGENNSPLHF | KLGGALQAK | NO | NO | NO |
| 10463 | TRAV17 | TRAJ53 | CATAGPNSGGSNYKLTF | TRBV5-6 | TRBJ1-6 | CASSFRAPFDSPLHF | FLASKIGRLV | YES | NO | YES |
| 10464 | TRAV1-2 | TRAJ29 | CAPRGGNTPLVF | TRBV7-9 | TRBJ2-5 | CASSLGTSGRETQYF | KLGGALQAK | YES | NO | NO |
| 10465 | TRAV24 | TRAJ11 | CARSGYSTLTF | TRBV6-1 | TRBJ2-7 | CASGQSPSGMEYEQYF | KLGGALQAK | YES | NO | NO |
| 10466 | TRAV8-4 | TRAJ31 | CAVSDPGNNARLMF | TRBV5-6 | TRBJ2-7 | CASSFGVTYEQYF | KLGGALQAK | YES | NO | NO |
| 10467 | TRAV29DV5 | TRAJ45 | CAASGDSGGGADGLTF | TRBV11-2 | TRBJ2-5 | CASSLEFVVGETQYF | KLGGALQAK | YES | NO | NO |
| 10468 | TRAV1-2 | TRAJ20 | CAVRAPGLNDYKLSF | TRBV5-6 | TRBJ1-6 | CASNEQFSHSPLHF | KLGGALQAK | YES | NO | NO |
| 10469 | TRAV23DV6 | TRAJ40 | CAASSFSGTYKYIF | TRBV6-5 | TRBJ2-1 | CASSLTPPAVAGNEQFF | ELAGIGILTV | NO | NO | NO |
| 10470 | TRAV13-1 | TRAJ9 | CAATGGFKTIF | TRBV12-5 | TRBJ2-5 | CASGLKQGLETQYF | KLGGALQAK | YES | NO | NO |
| 10471 | TRAV14DV4 | TRAJ26 | CAMTNNYGQNFVF | TRBV2 | TRBJ1-1 | CASRGLDRVNTEAFF | KLGGALQAK | YES | NO | NO |
| 10472 | TRAV12-2 | TRAJ35 | CAVLGFGNVLHC | TRBV12-3 | TRBJ2-1 | CASSPTGFF | AVFDRKSDAK | NO | NO | NO |
| 10473 | TRAV38-1 | TRAJ52 | CASSGGTSYGKLTF | TRBV7-2 | TRBJ2-3 | CASILEMTSGADTQYF | AVFDRKSDAK | YES | NO | NO |
| 10474 | TRAV1-2 | TRAJ33 | CAVRDSNYQLIW | TRBV6-6 | TRBJ2-7 | CASSYETGAYEQYF | AVFDRKSDAK | NO | NO | NO |
| 10475 | TRAV27 | TRAJ41 | CAGALDSGYALNF | TRBV5-6 | TRBJ1-1 | CASSLETANTEAFF | SLFNTVATLY | YES | YES | YES |
| 10476 | TRAV9-2 | TRAJ7 | CALLNYGNNRLAF | TRBV7-8 | TRBJ2-7 | CASSLVRDRAYEQYF | AVFDRKSDAK | YES | NO | YES |
| 10477 | TRAV14DV4 | TRAJ28 | CAMREATSGAGSYQLTF | TRBV7-9 | TRBJ2-1 | CASSLGGSSYNEQFF | KLGGALQAK | YES | NO | NO |

| | | | | | | | | | | |
|---|---|---|---|---|---|---|---|---|---|---|
| 10478 | TRAV12-3 | TRAJ56 | CAMSALRGGANSKLTF | TRBV7-9 | TRBJ2-7 | CASSWDLTSYEQYF | KLGGALQAK | YES | NO | NO |
| 10479 | TRAV19 | TRAJ48 | CALSEYWTNEKLTF | TRBV20-1 | TRBJ2-1 | CSARLPRLAGNEQFF | RLRAEAQVK | YES | YES | YES |
| 10480 | TRAV12-1 | TRAJ22 | CVVLSGSARQLTF | TRBV7-9 | TRBJ2-1 | CASSGGPRLAGNEQFF | KLGGALQAK | YES | NO | NO |
| 10481 | TRAV3 | TRAJ33 | CAVRDWDSNYQLIW | TRBV7-9 | TRBJ2-2 | CASSYTAGELFF | KLGGALQAK | YES | NO | NO |
| 10482 | TRAV19 | TRAJ52 | CALTLNAGGTSYGKLTF | TRBV6-5 | TRBJ2-2 | CASSYSVRGTGELFF | KLGGALQAK | YES | NO | NO |
| 10483 | TRAV19 | TRAJ32 | CALSDRYGGATNKLIF | TRBV6-3 | TRBJ1-1 | CASTPIVGGHHTEAFF | KLGGALQAK | YES | NO | NO |
| 10484 | TRAV12-2 | TRAJ41 | CAVKDSGYALNF | TRBV28 | TRBJ1-5 | CASSLFWVFSGTASNQPQHF | KLGGALQAK | NO | NO | NO |
| 10485 | TRAV1-2 | TRAJ34 | CAVPWSTDKLIF | TRBV4-3 | TRBJ1-2 | CASSQGTQDGYTF | KLGGALQAK | YES | NO | NO |
| 10486 | TRAV19 | TRAJ21 | CALSLKYNFNKFYF | TRBV12-3 | TRBJ2-7 | CASSLARTEDNYEQYF | KLGGALQAK | YES | NO | NO |
| 10487 | TRAV12-2 | TRAJ24 | CAVKTDSWGKLQF | TRBV13 | TRBJ2-7 | CASSQWGSGVYEQYF | LLFGYPVYV | YES | YES | YES |
| 10488 | TRAV27 | TRAJ39 | CADNAGNMLTF | TRBV6-3 | TRBJ2-1 | CASRETYNEQFF | KLGGALQAK | YES | NO | NO |
| 10489 | TRAV8-1 | TRAJ17 | CAVNARAAGNKLTF | TRBV12-5 | TRBJ2-5 | CASGSAVRGGLETQYF | AVFDRKSDAK | YES | YES | YES |
| 10490 | TRAV5 | TRAJ11 | CAESMGYSTLTF | TRBV12-4 | TRBJ2-7 | CASSLVGGEQYF | KLGGALQAK | YES | NO | NO |
| 10491 | TRAV17 | TRAJ23 | CATDGLYNQGGKLIF | TRBV2 | TRBJ1-6 | CASSESADNSPLHF | KLGGALQAK | YES | NO | NO |
| 10492 | TRAV5 | TRAJ41 | CAENLPNSGYALNF | TRBV7-8 | TRBJ2-1 | CASSPPGGNEQFF | IVTDFSVIK | YES | NO | YES |
| 10493 | TRAV8-3 | TRAJ21 | CAVGDNFNKFYF | TRBV12-3 | TRBJ2-7 | CASSSLDVAGYEQYF | AVFDRKSDAK | YES | NO | YES |
| 10494 | TRAV20 | TRAJ36 | CAVQAILGRTGANNLFF | TRBV27 | TRBJ2-7 | CASSLGTAYEQYF | KLGGALQAK | YES | NO | NO |
| 10495 | TRAV17 | TRAJ30 | CATGPNRDDKIIF | TRBV12-4 | TRBJ2-2 | CASLGRRGGLFF | AVFDRKSDAK | YES | NO | YES |
| 10496 | TRAV12-2 | TRAJ28 | CAVRNSGAGSYQLTF | TRBV3-1 | TRBJ2-7 | CASSQDFTYEQYF | KLGGALQAK | YES | NO | NO |
| 10497 | TRAV5 | TRAJ28 | CAEAYSGAGSYQLTF | TRBV24-1 | TRBJ1-1 | CATSEMTGTEAFF | KLGGALQAK | YES | NO | NO |
| 10498 | TRAV12-2 | TRAJ22 | CAGPREARQLTF | TRBV20-1 | TRBJ1-5 | CSATPTGVGQPQHF | ELAGIGILTV | YES | NO | YES |
| 10499 | TRAV12-3 | TRAJ54 | CAMMGAQKLVF | TRBV20-1 | TRBJ2-1 | CSAHNEQFF | AVFDRKSDAK | NO | NO | NO |
| 10500 | TRAV17 | TRAJ39 | CATFPGDAGNMLTF | TRBV11-1 | TRBJ2-3 | CASSLTAAEWTQYF | KLGGALQAK | YES | NO | NO |
| 10501 | TRAV38-1 | TRAJ52 | CAFMTNAGGTSYGKLTF | TRBV19 | TRBJ1-2 | CASSAGSYGYTF | GILGFVFTL | YES | YES | YES |
| 10502 | TRAV38-1 | TRAJ52 | CAFMVNAGGTSYGKLTF | TRBV19 | TRBJ1-2 | CASSQGSYGYTF | GILGFVFTL | YES | NO | YES |
| 10503 | TRAV3 | TRAJ39 | CAVRDVITDNAGNMLTF | TRBV6-6 | TRBJ2-7 | CASSPTGSSYEQYF | IVTDFSVIK | NO | NO | NO |
| 10504 | TRAV38-2DV8 | TRAJ31 | CAYRVDNARLMF | TRBV7-3 | TRBJ1-1 | CASSPVPGGTEAFF | KLGGALQAK | NO | NO | NO |
| 10505 | TRAV8-1 | TRAJ42 | CAVNEGGGSQGNLIF | TRBV19 | TRBJ2-7 | CASSMRSSYEQYF | GILGFVFTL | YES | NO | YES |
| 10506 | TRAV24 | TRAJ15 | CASLGAGTALIF | TRBV15 | TRBJ2-5 | CATSRVKGIETQYF | KLGGALQAK | YES | NO | NO |
| 10507 | TRAV13-1 | TRAJ11 | CAARQDSGYSTLTF | TRBV19 | TRBJ1-3 | CASRLRQGYGNTIYF | KLGGALQAK | YES | NO | NO |
| 10508 | TRAV1-2 | TRAJ13 | CASLGGYQKVTF | TRBV7-9 | TRBJ2-1 | CASSPEGDEQFF | KLGGALQAK | YES | NO | NO |
| 10509 | TRAV27 | TRAJ42 | CAGANGGSQGNLIF | TRBV19 | TRBJ2-2 | CASSVRSAGELFF | GILGFVFTL | YES | NO | YES |
| 10510 | TRAV12-2 | TRAJ42 | CAPPRRGSQGNLIF | TRBV7-2 | TRBJ2-7 | CASSLILHEQYF | KLGGALQAK | NO | NO | NO |
| 10511 | TRAV8-3 | TRAJ43 | CAVGALSNDMRF | TRBV27 | TRBJ2-7 | CASSFGGYEQYF | KLGGALQAK | YES | NO | NO |
| 10512 | TRAV19 | TRAJ57 | CALSDTGGSEKLVF | TRBV5-5 | TRBJ1-6 | CASSYASNSPLHF | KLGGALQAK | YES | NO | NO |
| 10513 | TRAV1-2 | TRAJ20 | CAVNSRDYKLSF | TRBV12-5 | TRBJ1-5 | CASGIQGIEPQHF | KLGGALQAK | YES | NO | NO |
| 10514 | TRAV12-2 | TRAJ37 | CAVMGGNTGKLIF | TRBV5-1 | TRBJ1-1 | CASSLKGAMNTEAFF | KLGGALQAK | YES | NO | NO |
| 10515 | TRAV12-2 | TRAJ5 | CAVIAAYTGRRALTF | TRBV7-2 | TRBJ2-1 | CASSLGPDNEQFF | KLGGALQAK | YES | NO | NO |
| 10516 | TRAV1-2 | TRAJ20 | CAVRENYKLSF | TRBV9 | TRBJ2-7 | CASSAPDGSYEQYF | KLGGALQAK | YES | NO | NO |
| 10517 | TRAV12-2 | TRAJ42 | CAFSMGGGSQGNLIF | TRBV19 | TRBJ2-3 | CASSIRSTDTQYF | GILGFVFTL | YES | NO | YES |
| 10518 | TRAV29DV5 | TRAJ34 | CAASASTDKLIF | TRBV20-1 | TRBJ2-7 | CSATDSYEQYF | KLGGALQAK | NO | NO | NO |
| 10519 | TRAV25 | TRAJ37 | CAVPGNTGKLIF | TRBV7-3 | TRBJ1-1 | CASSLTSGNTEAFF | AVFDRKSDAK | NO | NO | NO |
| 10520 | TRAV19 | TRAJ6 | CALIPAGGSYIPTF | TRBV27 | TRBJ2-2 | CASSTQTGSNTGELFF | KLGGALQAK | YES | NO | NO |
| 10521 | TRAV20 | TRAJ54 | CAVQAQIQGAQKLVF | TRBV18 | TRBJ2-7 | CASSPGQEVYEQYF | KLGGALQAK | YES | NO | NO |
| 10522 | TRAV38-1 | TRAJ26 | CAFYGQNFVF | TRBV4-3 | TRBJ1-2 | CASSQDGGYYGYTF | KLGGALQAK | YES | NO | NO |
| 10523 | TRAV41 | TRAJ54 | CAGDIQGAQKLVF | TRBV19 | TRBJ1-5 | CASSIAPWHRNNQPQHF | RLRAEAQVK | YES | NO | NO |
| 10524 | TRAV12-1 | TRAJ38 | CVVNWGNAGNNRKLIW | TRBV7-2 | TRBJ2-3 | CASSPLAGVTDTQYF | AVFDRKSDAK | YES | NO | YES |
| 10525 | TRAV14DV4 | TRAJ48 | CAMREDNFGNEKLTF | TRBV7-3 | TRBJ2-7 | CASSHGQGREQYF | KLGGALQAK | NO | NO | NO |
| 10526 | TRAV6 | TRAJ5 | CALPAGGRRALTF | TRBV5-8 | TRBJ2-1 | CASTVQGGNEQFF | KLGGALQAK | YES | NO | NO |
| 10527 | TRAV41 | TRAJ10 | CAASITGGGNKLTF | TRBV16 | TRBJ1-1 | CASSRSTEAFF | KLGGALQAK | NO | NO | NO |
| 10528 | TRAV5 | TRAJ6 | CAETLAPGGSYIPTF | TRBV20-1 | TRBJ2-3 | CSALGGTQYF | GILGFVFTL | NO | NO | NO |
| 10529 | TRAV1-2 | TRAJ20 | CAVRNDYKLSF | TRBV7-2 | TRBJ2-4 | CASSSGQGMDIQYF | KLGGALQAK | YES | NO | NO |
| 10530 | TRAV27 | TRAJ53 | CAGEAGGSNYKLTF | TRBV20-1 | TRBJ2-1 | CSASGLAAYNEQFF | KLGGALQAK | YES | NO | NO |
| 10531 | TRAV29DV5 | TRAJ42 | CAASAQGGSQGNLIF | TRBV5-5 | TRBJ2-5 | CASSLGGGQETQYF | KLGGALQAK | YES | NO | NO |
| 10532 | TRAV5 | TRAJ49 | CAEYTTGNQFYF | TRBV7-9 | TRBJ2-1 | CASSLGQINEQFF | KLGGALQAK | YES | NO | NO |
| 10533 | TRAV22 | TRAJ53 | CAVERLGGSNYKLTF | TRBV9 | TRBJ1-1 | CASSPGPMNTEAFF | IVTDFSVIK | YES | NO | YES |
| 10534 | TRAV14DV4 | TRAJ32 | CAMREGGSGATNKLIF | TRBV7-3 | TRBJ2-1 | CASSFLAGGFNEQFF | KLGGALQAK | YES | NO | NO |
| 10535 | TRAV12-2 | TRAJ32 | CAVPYGGATNKLIF | TRBV6-3 | TRBJ2-3 | CASRQGWLLMADTQYF | ELAGIGILTV | YES | YES | YES |
| 10536 | TRAV19 | TRAJ39 | CALSEAQHNAGNMLTF | TRBV7-9 | TRBJ2-5 | CASSPGTPIGTQYF | KLGGALQAK | YES | NO | NO |
| 10537 | TRAV12-2 | TRAJ52 | CAVNRSSYGKLTF | TRBV6-5 | TRBJ2-7 | CASNPGLAGGPHEQYF | LLFGYPVYV | YES | YES | YES |
| 10538 | TRAV5 | TRAJ6 | CAEMEGGSYIPTF | TRBV11-2 | TRBJ1-2 | CASSWGGGSHYGYTF | IVTDFSVIK | YES | YES | YES |
| 10539 | TRAV12-1 | TRAJ6 | CVVWGGSYIPTF | TRBV28 | TRBJ2-2 | CASSFPPDLNTGELFF | AVFDRKSDAK | YES | NO | YES |
| 10540 | TRAV12-2 | TRAJ29 | CAVTGNTPLVF | TRBV29-1 | TRBJ2-5 | CSVTPSGEQTQYF | KLGGALQAK | YES | NO | NO |
| 10541 | TRAV19 | TRAJ49 | CALSEADTGNQFYF | TRBV7-9 | TRBJ2-2 | CASSPRGEYTGELFF | RAKFKQLL | YES | NO | YES |
| 10542 | TRAV12-2 | TRAJ45 | CGVSGGGADGLTF | TRBV6-3 | TRBJ2-7 | CASTDSPGLAGGYEQYF | ELAGIGILTV | YES | YES | YES |
| 10543 | TRAV10 | TRAJ39 | CVVSANNAGNMLTF | TRBV11-3 | TRBJ2-2 | CASSVHRGSGGELFF | KLGGALQAK | YES | NO | NO |
| 10544 | TRAV8-3 | TRAJ9 | CAVALRAGGFKTIF | TRBV29-1 | TRBJ2-7 | CSVEVPWTGTYEQYF | KLGGALQAK | YES | NO | NO |
| 10545 | TRAV17 | TRAJ11 | CATAGDSGYSTLTF | TRBV6-5 | TRBJ1-2 | CASSPQGGNYGYTF | FLYALALLL | YES | YES | YES |
| 10546 | TRAV26-2 | TRAJ58 | CILQETSGSRLTF | TRBV11-2 | TRBJ1-1 | CASSSGTGEPNTEAFF | KLGGALQAK | YES | NO | NO |
| 10547 | TRAV8-6 | TRAJ4 | CAVRSFSGGYNKLIF | TRBV19 | TRBJ1-4 | CASSPTTGGRNEKLFF | KLGGALQAK | YES | NO | NO |
| 10548 | TRAV8-6 | TRAJ49 | CAVSDRGNQFYF | TRBV5-1 | TRBJ2-1 | CASSLDLAVDNEQFF | KLGGALQAK | YES | NO | NO |
| 10549 | TRAV21 | TRAJ37 | CALGSSNTGKLIF | TRBV18 | TRBJ2-3 | CASSPKILAYTDTQYF | KLGGALQAK | YES | NO | NO |
| 10550 | TRAV13-1 | TRAJ42 | CAASMGGGSQGNLIF | TRBV5-1 | TRBJ1-2 | CASSLGQGVFYTF | IVTDFSVIK | YES | NO | NO |
| 10551 | TRAV16 | TRAJ57 | CALQGGSEKLVF | TRBV5-1 | TRBJ1-3 | CASSLVGVGGSGNTIYF | KLGGALQAK | NO | NO | NO |
| 10552 | TRAV13-2 | TRAJ34 | CAERPYNTDKLIF | TRBV7-2 | TRBJ1-4 | CASSSPPGTGEVLFF | KLGGALQAK | YES | NO | NO |
| 10553 | TRAV1-2 | TRAJ6 | CAVGRGSYIPTF | TRBV7-9 | TRBJ2-3 | CASSFSQDPQYF | IVTDFSVIK | YES | YES | YES |
| 10554 | TRAV12-1 | TRAJ39 | CGAAGNMLTF | TRBV11-2 | TRBJ1-2 | CASSWGGGSHYGYTF | IVTDFSVIK | YES | NO | NO |
| 10555 | TRAV3 | TRAJ21 | CAVRDMLGFNKFYF | TRBV6-4 | TRBJ2-2 | CASSPTGSDTGELFF | GILGFVFTL | NO | NO | NO |
| 10556 | TRAV24 | TRAJ4 | CAPEGYNKLIF | TRBV11-2 | TRBJ2-7 | CASSLGQGPYEQYF | KLGGALQAK | YES | NO | NO |
| 10557 | TRAV13-2 | TRAJ52 | CAEKRYAGGTSYGKLTF | TRBV4-1 | TRBJ1-1 | CASSQGQGNTEAFF | AVFDRKSDAK | YES | NO | YES |
| 10558 | TRAV21 | TRAJ58 | CAVRETSGSRLTF | TRBV9 | TRBJ2-3 | CASSVEEGGPTDTQYF | KLGGALQAK | YES | NO | NO |
| 10559 | TRAV8-3 | TRAJ56 | CASGWGANSKLTF | TRBV20-1 | TRBJ2-3 | CSGASGRHTQYF | KLGGALQAK | NO | NO | NO |
| 10560 | TRAV9-2 | TRAJ33 | CALSHMDSNYQLIW | TRBV7-9 | TRBJ1-5 | CASSLFGTGGDQPQHF | KLGGALQAK | YES | NO | NO |
| 10561 | TRAV10 | TRAJ21 | CVVSARDNFNKFYF | TRBV27 | TRBJ1-3 | CASSFGGRGLGFSGNTIYF | GILGFVFTL | YES | NO | YES |
| 10562 | TRAV8-3 | TRAJ28 | CAVGTYSGAGSYQLTF | TRBV13 | TRBJ1-1 | CASSLGVGGGVGTEAFF | AVFDRKSDAK | YES | YES | YES |
| 10563 | TRAV19 | TRAJ3 | CALSEDSSASKIIF | TRBV28 | TRBJ2-7 | CASEPGQGPGGWSYEQYF | SLFNTVATLY | YES | YES | YES |
| 10564 | TRAV1-2 | TRAJ33 | CAVVDSNYQLIW | TRBV20-1 | TRBJ2-1 | CSARVLAGGIHEQFF | KLGGALQAK | YES | NO | NO |
| 10565 | TRAV8-2 | TRAJ53 | CVVSDLGSNYKLTF | TRBV11-2 | TRBJ1-2 | CASSWGGGSHYGYTF | IVTDFSVIK | YES | NO | YES |
| 10566 | TRAV13-1 | TRAJ50 | CAASETSYDKVIF | TRBV11-2 | TRBJ2-3 | CASSLTALGRGTDTQYF | AVFDRKSDAK | YES | NO | YES |
| 10567 | TRAV12-1 | TRAJ6 | CVVNIGYIPTF | TRBV12-3 | TRBJ2-7 | CASSPNANEQYF | KLGGALQAK | YES | NO | NO |
| 10568 | TRAV13-1 | TRAJ48 | CETFGNEKLTF | TRBV7-9 | TRBJ2-7 | CASSLEGPYEQYF | KLGGALQAK | YES | NO | NO |
| 10569 | TRAV13-1 | TRAJ20 | CAASRGRLSF | TRBV4-3 | TRBJ2-3 | CASSYLAGSTDTQYF | KLGGALQAK | YES | NO | NO |
| 10570 | TRAV4 | TRAJ36 | CLVGDETGANNLFF | TRBV27 | TRBJ2-3 | CASSFMSTDTQYF | IVTDFSVIK | YES | NO | NO |
| 10571 | TRAV17 | TRAJ11 | CATVGNSGYSTLTF | TRBV6-5 | TRBJ1-2 | CASSLQGGIDGYTF | FLYALALLL | YES | YES | YES |
| 10572 | TRAV41 | TRAJ45 | CAVPGGGADGLTF | TRBV5-5 | TRBJ2-3 | CASSLATSGWWDTQYF | KLGGALQAK | YES | NO | NO |
| 10573 | TRAV39 | TRAJ20 | CAVDKIDYKLSF | TRBV5-5 | TRBJ1-5 | CASSWGAARGNQPQHF | KLGGALQAK | YES | NO | NO |
| 10574 | TRAV27 | TRAJ42 | CAGAGVGYGGSQGNLIF | TRBV19 | TRBJ2-7 | CASSIRSAYEQYF | GILGFVFTL | YES | NO | YES |
| 10575 | TRAV8-1 | TRAJ10 | CAVNEVRGNKLTF | TRBV7-8 | TRBJ2-1 | CASTPTGTYNEQFF | KLGGALQAK | YES | NO | NO |
| 10577 | TRAV12-2 | TRAJ40 | CAVNKGTYKYIF | TRBV6-3 | TRBJ1-5 | CASSYWGFNQPQHF | RAKFKQLL | YES | NO | YES |
| 10578 | TRAV22 | TRAJ42 | CAVDRDYGGSQGNLIF | TRBV4-3 | TRBJ1-3 | CASSQDELTGAGNTIYF | KLGGALQAK | YES | NO | NO |
| 10579 | TRAV25 | TRAJ42 | CKGGSQGNLIF | TRBV28 | TRBJ2-7 | CASSFQGLGYEQYF | ELAGIGILTV | YES | NO | YES |
| 10580 | TRAV1-2 | TRAJ20 | CAVSNDYKLSF | TRBV9 | TRBJ2-1 | CASSVGLAGSNEQFF | AVFDRKSDAK | YES | NO | NO |
| 10581 | TRAV12-1 | TRAJ20 | CVVSVLNDYKLSF | TRBV28 | TRBJ2-7 | CASTPWAGGTYEQYF | KLGGALQAK | NO | NO | NO |
| 10582 | TRAV12-2 | TRAJ54 | CAVNPFQGGAQKLVF | TRBV6-6 | TRBJ2-3 | CASSYNLAGGPPDTQYF | AVFDRKSDAK | NO | NO | NO |

| | | | | | | | | | | |
|---|---|---|---|---|---|---|---|---|---|---|
| 10583 | TRAV12-2 | TRAJ24 | CAVSTDSWGKLQF | TRBV20-1 | TRBJ2-7 | CSARARLAGEFSEQYF | KLGGALQAK | YES | NO | NO |
| 10584 | TRAV8-3 | TRAJ13 | CAVVQMNSGGYQKVTF | TRBV28 | TRBJ2-1 | CATRPAGYNEQFF | KLGGALQAK | YES | NO | NO |
| 10585 | TRAV8-4 | TRAJ43 | CAVSDGNNDMRF | TRBV6-6 | TRBJ2-6 | CASSSLGPQGSGANVLTF | RAKFKQLL | YES | NO | YES |
| 10586 | TRAV35 | TRAJ49 | CAGHTGNQSYF | TRBV11-2 | TRBJ1-2 | CASSWGGGSHYGYTF | IVTDFSVIK | YES | NO | YES |
| 10587 | TRAV9-2 | TRAJ36 | CALIDQTGANNLFF | TRBV21-1 | TRBJ2-7 | CASPDDGELHEQYF | IVTDFSVIK | YES | YES | YES |
| 10589 | TRAV8-2 | TRAJ27 | CVVRPAGKSTF | TRBV27 | TRBJ2-5 | CASSLFLAAQETQYF | AVFDRKSDAK | YES | NO | YES |
| 10590 | TRAV25 | TRAJ32 | CAGPKGGATNKLIF | TRBV28 | TRBJ2-1 | CASSLRGRLRHYNEQFF | KLGGALQAK | YES | NO | NO |
| 10591 | TRAV1-2 | TRAJ33 | CAPMDSNYQLIW | TRBV20-1 | TRBJ1-2 | CSASEGAGEVYGYTF | KLGGALQAK | YES | NO | NO |
| 10592 | TRAV12-2 | TRAJ42 | CAVNGGSQGNLIF | TRBV2 | TRBJ2-7 | CASTIFWTGTYYEQYF | KLGGALQAK | YES | NO | NO |
| 10593 | TRAV5 | TRAJ13 | CAEKEWGYQKVTF | TRBV6-5 | TRBJ1-1 | CASTVQQGATEAFF | KLGGALQAK | YES | NO | NO |
| 10594 | TRAV35 | TRAJ49 | CAGHTGNQFYF | TRBV6-5 | TRBJ2-7 | CASSLGDSSYEQYF | RAKFKQLL | YES | NO | YES |
| 10595 | TRAV22 | TRAJ42 | CAENYGGSQGNLIF | TRBV4-1 | TRBJ2-7 | CASSQGQGGEQYF | KLGGALQAK | YES | NO | NO |
| 10596 | TRAV23DV6 | TRAJ17 | CAATSGAAGNKLTF | TRBV5-1 | TRBJ2-6 | CASSPGTGGWFIFSGANVLTF | RLRAEAQVK | NO | NO | NO |
| 10597 | TRAV26-1 | TRAJ35 | CIVRPAGFGNVLHC | TRBV7-9 | TRBJ2-7 | CASSLTGTPHEQYF | KLGGALQAK | NO | NO | NO |
| 10598 | TRAV12-2 | TRAJ5 | CAPGAGRRALTF | TRBV13 | TRBJ2-1 | CASSLLNEQFF | ELAGIGILTV | YES | YES | YES |
| 10599 | TRAV21 | TRAJ40 | CAVDSGTYKYIF | TRBV7-8 | TRBJ2-1 | CASSPRTSEQFF | KLGGALQAK | YES | NO | NO |
| 10600 | TRAV38-1 | TRAJ26 | CAFMKTYDYGQNFVF | TRBV28 | TRBJ2-1 | CASSRLDNYNEQFF | KLGGALQAK | YES | NO | NO |
| 10602 | TRAV19 | TRAJ24 | CALSEAGTDSWGKLQF | TRBV6-5 | TRBJ1-2 | CASSSTLQDRANYGYTF | KLGGALQAK | YES | NO | NO |
| 10603 | TRAV19 | TRAJ20 | CALSDYDYKLSF | TRBV4-1 | TRBJ2-5 | CASSQAPGETQYF | AVFDRKSDAK | YES | NO | YES |
| 10604 | TRAV8-4 | TRAJ43 | CAVSDGVNNNDMRF | TRBV5-4 | TRBJ1-1 | CASSPGGAEAFF | KLGGALQAK | YES | NO | NO |
| 10605 | TRAV25 | TRAJ11 | CAGVYSGYSTLTF | TRBV29-1 | TRBJ2-2 | CSSRMENTGELFF | KLGGALQAK | YES | NO | NO |
| 10606 | TRAV13-1 | TRAJ44 | CAADNTGTASKLTF | TRBV6-5 | TRBJ2-3 | CASSYSMSGGTDTQYF | KLGGALQAK | NO | NO | NO |
| 10607 | TRAV14DV4 | TRAJ22 | CAMREGQAGSARQLTF | TRBV11-2 | TRBJ1-2 | CASSWGGGSHYGYTF | IVTDFSVIK | YES | YES | YES |
| 10608 | TRAV19 | TRAJ20 | CALSEATWLRDYKLSF | TRBV7-8 | TRBJ2-7 | CASSLNVEQYF | RLRAEAQVK | YES | YES | YES |
| 10609 | TRAV38-1 | TRAJ54 | CAFMKHAIIIQGAQKLVF | TRBV28 | TRBJ2-7 | CASSLWRALEQYF | RAKFKQLL | YES | NO | YES |
| 10610 | TRAV12-2 | TRAJ10 | CAVNRPLTGGGNKLTF | TRBV3-1 | TRBJ2-7 | CASSQATSGIFVYEQYF | ELAGIGILTV | YES | YES | NO |
| 10611 | TRAV3 | TRAJ20 | CAVREMSDYKLSF | TRBV28 | TRBJ2-3 | CASRGVFGAGLTDTQYF | KLGGALQAK | NO | NO | NO |
| 10612 | TRAV12-2 | TRAJ10 | CAVPITGGGNKLTF | TRBV6-3 | TRBJ2-3 | CASSYSSFGTDTQYF | ELAGIGILTV | YES | NO | YES |
| 10613 | TRAV8-3 | TRAJ42 | CAVGMNYGGSQGNLIF | TRBV13 | TRBJ1-2 | CASSGVLGSDGYTF | AVFDRKSDAK | YES | NO | YES |
| 10614 | TRAV27 | TRAJ27 | CAGSNTNAGKSTF | TRBV19 | TRBJ2-7 | CASSIRSSYEQYF | GILGFVFTL | YES | NO | YES |
| 10615 | TRAV8-6 | TRAJ53 | CAVSDRGSNYKLTF | TRBV27 | TRBJ1-1 | CASSLSRGNTEAFF | AVFDRKSDAK | YES | NO | YES |
| 10616 | TRAV36DV7 | TRAJ40 | CAVRPSGTYKYIF | TRBV7-3 | TRBJ1-2 | CASSFYPGQDYGYTF | KLGGALQAK | NO | NO | NO |
| 10617 | TRAV1-1 | TRAJ21 | CAPYNFNKFYF | TRBV12-3 | TRBJ2-1 | CASSPGREGYNEQFF | AYAQKIFKI | YES | YES | YES |
| 10618 | TRAV2 | TRAJ9 | CAVESRGGFKTIF | TRBV5-1 | TRBJ1-2 | CASSLVWFGQGVGTAGYTF | KLGGALQAK | NO | NO | NO |
| 10619 | TRAV38-2DV8 | TRAJ57 | CAYRTPQGGSEKLVF | TRBV13 | TRBJ1-1 | CASSTETGNTEAFF | KLGGALQAK | YES | NO | NO |
| 10620 | TRAV17 | TRAJ7 | CATGHYGNNRLAF | TRBV25-1 | TRBJ1-2 | CASSESKGYTF | KLGGALQAK | YES | NO | NO |
| 10621 | TRAV35 | TRAJ49 | CAGHTGNQFYF | TRBV9 | TRBJ2-2 | CASSSGLVSNTGELFF | IVTDFSVIK | YES | NO | YES |
| 10622 | TRAV39 | TRAJ42 | CAVDEGGGSQGNLIF | TRBV19 | TRBJ2-3 | CASSIRSTDTQYF | GILGFVFTL | YES | NO | YES |
| 10623 | TRAV12-1 | TRAJ30 | CVVNHFRDDKIIF | TRBV4-3 | TRBJ1-4 | CASSQTGFPTTNEKLFF | KLGGALQAK | YES | NO | NO |
| 10624 | TRAV2 | TRAJ31 | CAVGPNNNARLMF | TRBV6-3 | TRBJ2-7 | CASSYPGTGEYEQYF | IVTDFSVIK | NO | NO | NO |
| 10625 | TRAV12-1 | TRAJ23 | CVVNFMIYNQGGKLIF | TRBV7-9 | TRBJ1-1 | CASSLVGAGGNTEAFF | KLGGALQAK | YES | NO | NO |
| 10626 | TRAV14DV4 | TRAJ24 | CAMREPPTDSWGKLQF | TRBV13 | TRBJ1-3 | CASSLGQQPGPGNTIYF | KLGGALQAK | YES | NO | NO |
| 10627 | TRAV12-2 | TRAJ21 | CAVNGGEFYF | TRBV9 | TRBJ2-1 | CASSAPGGTSYNEQFF | KLGGALQAK | YES | NO | NO |
| 10628 | TRAV8-3 | TRAJ7 | CAVAYYGNNRLAF | TRBV6-3 | TRBJ2-2 | CASSYSGGTNTGELFF | KLGGALQAK | YES | NO | NO |
| 10629 | TRAV12-2 | TRAJ36 | CAVDGTGANNLFF | TRBV5-1 | TRBJ2-1 | CASSLAPSGEQFF | ELAGIGILTV | YES | NO | YES |
| 10630 | TRAV3 | TRAJ30 | CAVRDGNRDDKIIF | TRBV29-1 | TRBJ2-3 | CSVELTTGTDTQYF | KLGGALQAK | NO | NO | NO |
| 10631 | TRAV38-1 | TRAJ32 | CAFMRDYGGATNKLIF | TRBV11-2 | TRBJ2-3 | CASSPFAGGPADTQYF | KLGGALQAK | YES | NO | NO |
| 10632 | TRAV38-1 | TRAJ48 | CAFKTYEKLTF | TRBV9 | TRBJ2-6 | CASSVRDSILPSGANVLTF | KLGGALQAK | YES | NO | NO |
| 10633 | TRAV21 | TRAJ43 | CAVQRYNNNDMRF | TRBV7-6 | TRBJ2-2 | CASSLVAGELFF | KLGGALQAK | YES | NO | NO |
| 10634 | TRAV12-1 | TRAJ16 | CVVKGLTPFRDGQKLLF | TRBV9 | TRBJ2-1 | CASSSPSSSYNEQFF | KLGGALQAK | NO | NO | NO |
| 10635 | TRAV21 | TRAJ27 | CAGGNTNAGKSTF | TRBV7-9 | TRBJ1-2 | CASSIFGQGNYGYTF | AVFDRKSDAK | YES | NO | YES |
| 10636 | TRAV13-1 | TRAJ45 | CAASRGKGGGADGLTF | TRBV6-5 | TRBJ2-7 | CASSYSTGTVDYEQYF | GILGFVFTL | YES | NO | YES |
| 10637 | TRAV13-1 | TRAJ49 | CAATYTGNQFYF | TRBV11-2 | TRBJ1-5 | CASSLGRTNQPQHF | GILGFVFTL | YES | NO | YES |
| 10638 | TRAV13-1 | TRAJ43 | CAASSGDMRF | TRBV11-2 | TRBJ1-2 | CASSWGGGSHYGYTF | IVTDFSVIK | YES | NO | YES |
| 10639 | TRAV12-2 | TRAJ4 | CAVPSGGYNKLIF | TRBV12-5 | TRBJ1-3 | CASGLDSSSGNTIYF | AVFDRKSDAK | YES | NO | YES |
| 10640 | TRAV3 | TRAJ42 | CALTYGGSQGNLIF | TRBV19 | TRBJ2-7 | CASSLRAGIEQYF | GILGFVFTL | NO | NO | NO |
| 10641 | TRAV6 | TRAJ4 | CALHYSGGYNKLIF | TRBV5-5 | TRBJ2-1 | CASSPSAGGAYNEQFF | KLGGALQAK | YES | NO | NO |
| 10642 | TRAV9-2 | TRAJ28 | CAPTSTWGWELPTHF | TRBV19 | TRBJ2-1 | CASSISLAAEQFF | LLDFVRFMGV | NO | NO | NO |
| 10643 | TRAV5 | TRAJ18 | CAEGFDRGSTLGRLYF | TRBV7-2 | TRBJ2-3 | CASRPIPGAYAQYF | KLGGALQAK | YES | NO | NO |
| 10644 | TRAV24 | TRAJ6 | CAWASGGSYIPTF | TRBV20-1 | TRBJ2-3 | CSARDTGVSDTQYF | RAKFKQLL | NO | NO | NO |
| 10645 | TRAV3 | TRAJ39 | CAVRDIGNAGNMLTF | TRBV29-1 | TRBJ2-5 | CSPMDRQETQYF | KLGGALQAK | NO | NO | NO |
| 10646 | TRAV41 | TRAJ44 | CAVRGTGTASKLTF | TRBV6-5 | TRBJ2-7 | CASRQLRGLMAYEQYF | ELAGIGILTV | YES | NO | NO |
| 10647 | TRAV22 | TRAJ50 | CAVRTSYDKVIF | TRBV9 | TRBJ2-7 | CASSSAGADYEQYF | KLGGALQAK | YES | NO | NO |
| 10648 | TRAV13-1 | TRAJ45 | CAASIRRSSGGGADGLTF | TRBV9 | TRBJ1-2 | CASSHIGQGYTF | KLGGALQAK | YES | NO | NO |
| 10649 | TRAV19 | TRAJ48 | CALSEAPNFGNEKLTF | TRBV3-1 | TRBJ1-5 | CASSQDSPGIRDNQPQHF | KLGGALQAK | YES | NO | NO |
| 10650 | TRAV38-2DV8 | TRAJ29 | CAYRTYSGNTPLVF | TRBV28 | TRBJ1-6 | CASSFGGPRGPLHF | AVFDRKSDAK | YES | NO | NO |
| 10651 | TRAV10 | TRAJ12 | CVVRGWMDSSYKLIF | TRBV6-5 | TRBJ2-6 | CASSLNRGTGANVLTF | KLGGALQAK | YES | NO | NO |
| 10652 | TRAV8-4 | TRAJ43 | CAVSENDMRF | TRBV6-1 | TRBJ2-5 | CASSEGTDPQETQYF | KLGGALQAK | YES | NO | NO |
| 10653 | TRAV12-2 | TRAJ45 | CAVKDSGGGADGLTF | TRBV27 | TRBJ1-5 | CASSPQGLGQPQHF | ELAGIGILTV | YES | YES | YES |
| 10654 | TRAV27 | TRAJ27 | CAGANTNAGKSTF | TRBV19 | TRBJ2-7 | CASSTRSSYEQYF | GILGFVFTL | YES | NO | YES |
| 10655 | TRAV41 | TRAJ34 | CAVSSYNTDKLIF | TRBV5-1 | TRBJ2-5 | CASSLDGSVETQYF | AVFDRKSDAK | YES | NO | NO |
| 10656 | TRAV1-2 | TRAJ33 | CAVTDSNYQLIW | TRBV20-1 | TRBJ2-7 | CSAYLQGSSYEQYF | KLGGALQAK | NO | NO | NO |
| 10657 | TRAV35 | TRAJ42 | CAGPTHNMNYGGSQGNLIF | TRBV19 | TRBJ1-5 | CASSYYSNQPQHF | GILGFVFTL | YES | NO | YES |
| 10658 | TRAV25 | TRAJ20 | CAGLRDYKLSF | TRBV5-4 | TRBJ2-1 | CASRADSYNEQFF | KLGGALQAK | YES | NO | NO |
| 10659 | TRAV12-2 | TRAJ34 | CAVSPDKLIF | TRBV20-1 | TRBJ1-5 | CSASAGNQPQHF | ELAGIGILTV | YES | YES | YES |
| 10660 | TRAV12-2 | TRAJ43 | CAVKGNNDMRF | TRBV6-5 | TRBJ2-7 | CASSYSALAYEQYF | ELAGIGILTV | YES | YES | NO |
| 10661 | TRAV38-1 | TRAJ42 | CAFSRGHYGGSQGNLIF | TRBV25-1 | TRBJ2-7 | CASSDSRRGTGTYEQYF | KLGGALQAK | YES | NO | NO |
| 10662 | TRAV27 | TRAJ57 | CAGAQGGSEKLVF | TRBV6-5 | TRBJ2-7 | CASSDGFPDSTYEQYF | KLGGALQAK | YES | NO | NO |
| 10663 | TRAV12-2 | TRAJ10 | CAATGGGGNKLTF | TRBV9 | TRBJ2-2 | CASSVGGRTGELFF | IMDQVPFSV | YES | NO | YES |
| 10664 | TRAV35 | TRAJ45 | CAGQWDSGGGADGLTF | TRBV5-6 | TRBJ2-4 | CASSSEGQVAKNIQYF | KLGGALQAK | YES | NO | NO |
| 10665 | TRAV12-3 | TRAJ48 | CAMSAMVGNEKLTF | TRBV14 | TRBJ2-3 | CASSHVGTGWDLRTDTQYF | KLGGALQAK | YES | NO | NO |
| 10667 | TRAV29DV5 | TRAJ45 | CAASEPLYSGGGADGLTF | TRBV7-9 | TRBJ2-1 | CASSDPTSGGPYNEQFF | RAKFKQLL | YES | NO | YES |
| 10668 | TRAV8-3 | TRAJ27 | CAVASPNNTNAGKSTF | TRBV27 | TRBJ2-3 | CASSLGFTDTQYF | KLGGALQAK | YES | NO | NO |
| 10669 | TRAV12-3 | TRAJ48 | CAMSASNFGNEKLTF | TRBV7-2 | TRBJ1-3 | CASSSIRGPLSGNTIYF | GILGFVFTL | YES | NO | YES |
| 10670 | TRAV12-3 | TRAJ48 | CAMSGSNFGNEKLTF | TRBV29-1 | TRBJ2-2 | CSVEKGTGELFF | ELAGIGILTV | YES | NO | NO |
| 10671 | TRAV17 | TRAJ8 | CATDGGFQKLVF | TRBV9 | TRBJ2-2 | CASSYPGGYPNTGELFF | RLRAEAQVK | YES | NO | YES |
| 10672 | TRAV17 | TRAJ37 | CATDGSSSNTGKLIF | TRBV19 | TRBJ2-1 | CASSIYSGGYNEQFF | GILGFVFTL | YES | NO | YES |
| 10673 | TRAV12-2 | TRAJ32 | CAAPGGATNKLIF | TRBV6-5 | TRBJ2-3 | CASSYSTPSWTDTQYF | ELAGIGILTV | NO | NO | NO |
| 10674 | TRAV20 | TRAJ4 | CAVQPFSGGYNKLIF | TRBV18 | TRBJ2-1 | CASSLQGAPTYNEQFF | KLGGALQAK | YES | NO | NO |
| 10675 | TRAV8-4 | TRAJ13 | CAVSTVPPSGGYQKVTF | TRBV11-2 | TRBJ1-2 | CASSWGGGSHYGYTF | IVTDFSVIK | YES | NO | YES |
| 10676 | TRAV35 | TRAJ49 | CAGHTGNQFYF | TRBV3-1 | TRBJ2-1 | CASSHLGGDRYMNEQFF | IVTDFSVIK | YES | YES | YES |
| 10677 | TRAV8-3 | TRAJ20 | CAVGALGDYKLSF | TRBV6-5 | TRBJ1-1 | CASSQDRGLVDTEAFF | KLGGALQAK | YES | NO | NO |
| 10678 | TRAV12-2 | TRAJ9 | CAVNPTGGFKTIF | TRBV6-3 | TRBJ2-7 | CASCAGMGNEQYF | ELAGIGILTV | YES | NO | NO |
| 10679 | TRAV14DV4 | TRAJ39 | CAMREGIGNAGNMLTF | TRBV11-2 | TRBJ1-2 | CASSWGGGSHYGYTF | IVTDFSVIK | YES | NO | YES |
| 10680 | TRAV29DV5 | TRAJ43 | CAASETNNDMRF | TRBV7-9 | TRBJ2-2 | CASSWGMTGELFF | KLGGALQAK | YES | NO | NO |
| 10681 | TRAV10 | TRAJ34 | CVVSAPHTDKLIF | TRBV28 | TRBJ2-2 | CASQVDTFTDGGELFF | KLGGALQAK | YES | NO | NO |
| 10682 | TRAV13-2 | TRAJ28 | CAGIGAGSYQLTF | TRBV9 | TRBJ1-1 | CASSAPRGTQAFF | KLGGALQAK | YES | NO | NO |
| 10683 | TRAV12-2 | TRAJ29 | CAVTGNTPLVF | TRBV20-1 | TRBJ1-2 | CSARVGIPNYGYTF | KLGGALQAK | YES | NO | NO |
| 10684 | TRAV27 | TRAJ42 | CAGASGGSQGNLIF | TRBV29-1 | TRBJ1-5 | CSVGTGGAGQPQHF | GILGFVFTL | YES | NO | YES |
| 10686 | TRAV29DV5 | TRAJ18 | CAASDRGSTLGRLYF | TRBV7-9 | TRBJ1-2 | CASSPISSEYGYTF | KLGGALQAK | YES | NO | NO |
| 10687 | TRAV8-3 | TRAJ54 | CAVGPVQGAQKLVF | TRBV6-3 | TRBJ2-1 | CASSYSVGAAPYNEQFF | KLGGALQAK | YES | NO | NO |
| 10688 | TRAV38-2DV8 | TRAJ33 | CAYRSEYEWDSNYQLIW | TRBV5-1 | TRBJ2-7 | CASRAGGSYEQYF | KLGGALQAK | YES | NO | NO |
| 10689 | TRAV1-1 | TRAJ12 | CAVSRMDSSYKLIF | TRBV2 | TRBJ1-3 | CASSGGANSGNTIYF | KLGGALQAK | YES | NO | NO |
| 10690 | TRAV14DV4 | TRAJ23 | CAMREVYNQGGKLIF | TRBV7-9 | TRBJ1-4 | CASGTPGGEKLFF | KLGGALQAK | YES | NO | NO |

| | | | | | | | | | |
|---|---|---|---|---|---|---|---|---|---|
| 10691 | TRAV8-1 | TRAJ42 | CAVNAGGGSQGNLIF | TRBV12-5 | TRBJ1-2 | CASAPDGFFYGYTF | GILGFVFTL | YES | NO | YES |
| 10692 | TRAV38-2DV8 | TRAJ28 | CAYRDSYSGAGSYQLTF | TRBV19 | TRBJ2-5 | CASSIPGRFQETQYF | KLGGALQAK | YES | NO | NO |
| 10693 | TRAV25 | TRAJ42 | CAGNYGGSQGNLIF | TRBV19 | TRBJ2-2 | CASSARSTGELFF | GILGFVFTL | YES | NO | YES |
| 10694 | TRAV12-3 | TRAJ47 | CAIYMEYGNKLVF | TRBV20-1 | TRBJ2-7 | CSARDINLDTYEQYF | KLGGALQAK | NO | NO | NO |
| 10695 | TRAV4 | TRAJ5 | CLVGDDRDTGRRALTF | TRBV7-9 | TRBJ2-4 | CASSLGTLAKNIQYF | LLFGYPVYV | YES | YES | YES |
| 10696 | TRAV29DV5 | TRAJ52 | CAASGGTSYGKLTF | TRBV25-1 | TRBJ2-7 | CASRGEKAYEQYF | FLYALALLL | YES | YES | YES |
| 10697 | TRAV41 | TRAJ49 | CAKPGGNQFYF | TRBV12-5 | TRBJ1-3 | CASGLIPGGPGNTIYF | AVFDRKSDAK | YES | YES | YES |
| 10698 | TRAV17 | TRAJ23 | CATDAYNQGGKLIF | TRBV6-3 | TRBJ1-5 | CASGLGVQDLNQPQHF | KLGGALQAK | YES | NO | NO |
| 10699 | TRAV8-3 | TRAJ20 | CAVGPLNDYKLSF | TRBV11-2 | TRBJ2-5 | CATSSTGLKTQYF | RLRAEAQVK | YES | YES | YES |
| 10700 | TRAV8-4 | TRAJ57 | CAVSDLTQGGSEKLVF | TRBV19 | TRBJ1-2 | CATQDGRGYTF | KLGGALQAK | YES | NO | NO |
| 10701 | TRAV38-1 | TRAJ52 | CAFMMGAGGTSYGKLTF | TRBV21-1 | TRBJ1-5 | CASKRGMSNQPQHF | GILGFVFTL | YES | NO | YES |
| 10702 | TRAV13-1 | TRAJ11 | CAAENSGYSTLTF | TRBV3-1 | TRBJ2-7 | CASSQDWTEQYF | KLGGALQAK | YES | NO | NO |
| 10703 | TRAV5 | TRAJ43 | CAESNGMRF | TRBV7-2 | TRBJ2-7 | CASSRSSYEQYF | AVFDRKSDAK | NO | NO | NO |
| 10704 | TRAV10 | TRAJ37 | CVVTGSSNTGKLIF | TRBV12-4 | TRBJ2-5 | CASSFVVQETQYF | KLGGALQAK | YES | NO | NO |
| 10705 | TRAV1-2 | TRAJ7 | CAVREVSRLAF | TRBV2 | TRBJ1-1 | CASSEPPQGRNTEAFF | RLRAEAQVK | YES | NO | NO |
| 10706 | TRAV16 | TRAJ15 | CALNQAGTALIF | TRBV5-4 | TRBJ2-7 | CASSPRGQSYEQYF | GILGFVFTL | NO | NO | NO |
| 10707 | TRAV12-2 | TRAJ43 | CAVSNNNDMRF | TRBV4-1 | TRBJ1-5 | CASSQDRWLSGQPQHF | ELAGIGILTV | YES | NO | YES |
| 10708 | TRAV36DV7 | TRAJ43 | CAVYNNNDMRF | TRBV28 | TRBJ2-1 | CASSLLRGAKGGRPSYNEQFF | KLGGALQAK | NO | NO | NO |
| 10709 | TRAV41 | TRAJ42 | CAVANGSQGNLIF | TRBV4-3 | TRBJ1-2 | CASSGYWGLGYGYTF | KLGGALQAK | YES | NO | NO |
| 10710 | TRAV12-3 | TRAJ3 | CADSGYSSASKIIF | TRBV7-2 | TRBJ2-1 | CASSLGLAGPIGNEQFF | AVFDRKSDAK | YES | NO | NO |
| 10712 | TRAV29DV5 | TRAJ29 | CAARGKGSLGGNTPLVF | TRBV7-6 | TRBJ1-1 | CASSSGQMNTEAFF | KLGGALQAK | YES | NO | NO |
| 10713 | TRAV35 | TRAJ34 | CAGQLVFDTDKLIF | TRBV12-3 | TRBJ1-2 | CASSYLSTNGYTF | RAKFKQLL | YES | YES | YES |
| 10714 | TRAV8-6 | TRAJ7 | CAVSDGGNNRLAF | TRBV7-9 | TRBJ1-3 | CASRSGTGVSGNTIYF | KLGGALQAK | YES | NO | NO |
| 10715 | TRAV39 | TRAJ48 | CAVTNLFGNEKLTF | TRBV7-8 | TRBJ2-4 | CASSEWRERTKNIQYF | KLGGALQAK | NO | NO | NO |
| 10716 | TRAV12-1 | TRAJ43 | CVVISYNDMRF | TRBV3-1 | TRBJ2-3 | CASSQPGAFFTDTQYF | AVFDRKSDAK | YES | NO | YES |
| 10717 | TRAV17 | TRAJ29 | CATGNSGNTPLVF | TRBV12-5 | TRBJ2-2 | CASGWDRPNTGELFF | KLGGALQAK | YES | NO | NO |
| 10718 | TRAV41 | TRAJ26 | CAVEEDNYGQNFVF | TRBV5-1 | TRBJ2-1 | CASSLEGSHSYNEQFF | KLGGALQAK | YES | NO | NO |
| 10719 | TRAV8-4 | TRAJ43 | CAVRSPGNNDMRF | TRBV7-9 | TRBJ2-7 | CASSLGLSGLGYEQYF | KLGGALQAK | YES | NO | NO |
| 10720 | TRAV38-2DV8 | TRAJ57 | CAYRTITQGGSEKLVF | TRBV6-1 | TRBJ1-2 | CASSEFGRNYGYTF | KLGGALQAK | YES | NO | NO |
| 10721 | TRAV27 | TRAJ42 | CAGAGSQGNLIF | TRBV28 | TRBJ2-2 | CASSLYSATGELFF | AVFDRKSDAK | YES | NO | YES |
| 10722 | TRAV1-2 | TRAJ6 | CAVRGWGGSYIPTF | TRBV20-1 | TRBJ2-2 | CSARQGTTGELFF | RAKFKQLL | YES | YES | YES |
| 10723 | TRAV8-1 | TRAJ21 | CAVSLFEYNFNKFYF | TRBV5-1 | TRBJ1-1 | CASSQGGGLGTEAFF | RLRAEAQVK | YES | NO | YES |
| 10724 | TRAV23DV6 | TRAJ34 | CAASRGTDKLIF | TRBV29-1 | TRBJ2-7 | CSVEDGPPGEGGYEQYF | KLGGALQAK | YES | NO | NO |
| 10725 | TRAV2 | TRAJ22 | CAVEDRGSARQLTF | TRBV25-1 | TRBJ2-2 | CASSPTRGRKNTGELFF | KLGGALQAK | NO | NO | NO |
| 10726 | TRAV14DV4 | TRAJ4 | CAIGFGGYNKLIF | TRBV7-8 | TRBJ2-1 | CASSSGSGSPEMGDEQFF | KLGGALQAK | YES | NO | NO |
| 10727 | TRAV17 | TRAJ30 | CATDPIMNRDDKIIF | TRBV29-1 | TRBJ1-1 | CSVVLAQGHDTEAFF | KLGGALQAK | YES | NO | NO |
| 10728 | TRAV14DV4 | TRAJ5 | CAMREGLGTGRRALTF | TRBV4-3 | TRBJ2-1 | CASSQEVSSGGSNEQFF | AVFDRKSDAK | YES | NO | YES |
| 10729 | TRAV17 | TRAJ10 | CATDTGGGNKLTF | TRBV13 | TRBJ1-1 | CASSPPRREVEAFF | KLGGALQAK | YES | NO | NO |
| 10730 | TRAV12-1 | TRAJ39 | CVVNPNNAGNMLTF | TRBV28 | TRBJ2-7 | CASSLSGYEQYF | KLGGALQAK | YES | NO | NO |
| 10731 | TRAV38-2DV8 | TRAJ43 | CAYRMRF | TRBV7-6 | TRBJ2-7 | CASSPGVGLSYEQYF | KLGGALQAK | NO | NO | NO |
| 10732 | TRAV14DV4 | TRAJ20 | CAMRELLDYKLSF | TRBV9 | TRBJ2-3 | CASSQTSGSGDTQYF | AVFDRKSDAK | YES | NO | YES |
| 10733 | TRAV21 | TRAJ26 | CAVIEDYGQNFVF | TRBV12-4 | TRBJ1-1 | CASSLSFGDTEAFF | KLGGALQAK | YES | NO | NO |
| 10734 | TRAV19 | TRAJ54 | CALRNIIQGAQKLVF | TRBV7-3 | TRBJ2-5 | CASSPYGWAKETQYF | KLGGALQAK | NO | NO | NO |
| 10735 | TRAV8-3 | TRAJ29 | CAVATWNSGNTPLVF | TRBV6-4 | TRBJ1-5 | CASTLQASISNQPQHF | KLGGALQAK | YES | NO | NO |
| 10736 | TRAV38-1 | TRAJ54 | CAFQGRIQGAQKLVF | TRBV10-3 | TRBJ1-1 | CAISESPGPEKNTEAFF | KLGGALQAK | YES | NO | NO |
| 10737 | TRAV12-2 | TRAJ4 | CAVNGGYNKLIF | TRBV7-9 | TRBJ1-5 | CASSLVGRGGNQPQHF | KLGGALQAK | YES | NO | NO |
| 10738 | TRAV38-1 | TRAJ34 | CAFMDYYNTDKLIF | TRBV19 | TRBJ2-1 | CASTLTSGSYFSYNEQFF | KLGGALQAK | YES | NO | NO |
| 10739 | TRAV8-3 | TRAJ35 | CAVLI | TRBV7-2 | TRBJ2-7 | CASSEEGPPWEQYF | KLGGALQAK | NO | NO | NO |
| 10740 | TRAV38-1 | TRAJ49 | CAFNGNTGNQFYF | TRBV4-3 | TRBJ2-5 | CASSQEWTGLQETQYF | KLGGALQAK | YES | NO | NO |
| 10741 | TRAV22 | TRAJ45 | CAVYSGGGADGLTF | TRBV30 | TRBJ1-1 | CAWSVVRKGTEAFF | KLGGALQAK | YES | NO | NO |
| 10742 | TRAV19 | TRAJ56 | CALSEAWANSKLTF | TRBV18 | TRBJ1-2 | CASSPQWRQHTNYGYTF | RLRAEAQVK | YES | NO | YES |
| 10743 | TRAV23DV6 | TRAJ20 | CAASRSNDYKLSF | TRBV5-6 | TRBJ1-1 | CASSLRDGSEAFF | RAKFKQLL | YES | NO | YES |
| 10744 | TRAV17 | TRAJ7 | CATDAYGNNRLAF | TRBV7-3 | TRBJ1-1 | CASSLAGAGTEAFF | SLFNTVATLY | NO | NO | NO |
| 10745 | TRAV21 | TRAJ30 | CAVNFRDDKIIF | TRBV20-1 | TRBJ2-2 | CSASQGLNTGELFF | KTWGQYWQV | YES | NO | YES |
| 10746 | TRAV14DV4 | TRAJ39 | CAMRTPNNAGNMLTF | TRBV24-1 | TRBJ2-2 | CATSSVTGELFF | KLGGALQAK | YES | NO | NO |
| 10747 | TRAV22 | TRAJ37 | CAVVLDGSSNTGKLIF | TRBV6-5 | TRBJ2-2 | CASSYGQGTGELFF | KLGGALQAK | YES | NO | NO |
| 10748 | TRAV8-1 | TRAJ34 | CAVNDYNTDKLIF | TRBV5-4 | TRBJ2-4 | CASSPLGAKNIQYF | KLGGALQAK | YES | NO | NO |
| 10749 | TRAV17 | TRAJ4 | CATRLLFSGGYNKLIF | TRBV28 | TRBJ2-3 | CASSLPGAGSSTDTQYF | KLGGALQAK | YES | NO | NO |
| 10750 | TRAV12-1 | TRAJ33 | CVVNRDSNYQLIW | TRBV10-3 | TRBJ1-4 | CAISGGLQGQEKLFF | KLGGALQAK | YES | NO | NO |
| 10751 | TRAV1-2 | TRAJ28 | CAALWGWELPTHF | TRBV28 | TRBJ2-3 | CASSLYRGADTQYF | KLGGALQAK | NO | NO | NO |
| 10753 | TRAV3 | TRAJ26 | CAVRDYDNYGQNFVF | TRBV6-3 | TRBJ2-7 | CASSYEVPTLYEQYF | KLGGALQAK | NO | NO | NO |
| 10754 | TRAV13-1 | TRAJ28 | CAAIESGAGSYQLTF | TRBV27 | TRBJ1-1 | CASSTGTEAFF | KLGGALQAK | YES | NO | NO |
| 10755 | TRAV24 | TRAJ6 | CASQASGGSYIPTF | TRBV10-3 | TRBJ1-4 | CAISGGGNEKLFF | KLGGALQAK | YES | NO | NO |
| 10756 | TRAV12-3 | TRAJ54 | CAALEVQGAQKLVF | TRBV28 | TRBJ1-2 | CASSLWARKDRGLEGYTF | RLRAEAQVK | YES | YES | YES |
| 10757 | TRAV13-2 | TRAJ52 | CAEMAGGTSYGKLTF | TRBV12-5 | TRBJ2-7 | CASGFSGTEQYF | KLGGALQAK | YES | NO | NO |
| 10758 | TRAV38-2DV8 | TRAJ48 | CAPLSNFGNEKLTF | TRBV7-9 | TRBJ2-3 | CAGARVDTQYF | KLGGALQAK | YES | NO | NO |
| 10759 | TRAV1-2 | TRAJ6 | CAVRGRAGGSYIPTF | TRBV4-3 | TRBJ2-5 | CASSQGTGQGETQYF | CYTWNQMNL | YES | NO | YES |
| 10760 | TRAV36DV7 | TRAJ41 | CAVGGYALNF | TRBV7-2 | TRBJ1-1 | CASSLVHSLWGLSTEAFF | KLGGALQAK | NO | NO | NO |
| 10761 | TRAV1-1 | TRAJ34 | CAVSPYNTDKLIF | TRBV4-3 | TRBJ2-3 | CASSQGPAGLTDTQYF | KLGGALQAK | YES | NO | NO |
| 10762 | TRAV26-1 | TRAJ27 | CIVFNTNAGKSTF | TRBV6-3 | TRBJ2-1 | CASSYFFQGPLYEQFF | KLGGALQAK | YES | NO | NO |
| 10763 | TRAV12-3 | TRAJ34 | CAIGPGNTDKLIF | TRBV20-1 | TRBJ2-7 | CSARPERELYEQYF | KLGGALQAK | NO | NO | NO |
| 10764 | TRAV36DV7 | TRAJ38 | CAVEVNAGNNRKLIW | TRBV19 | TRBJ2-7 | CASSIRSSYEQYF | GILGFVFTL | NO | NO | NO |
| 10765 | TRAV12-3 | TRAJ54 | CAMSLIQGAQKLVF | TRBV5-1 | TRBJ2-6 | CASSVYRGLGANVLTF | KLGGALQAK | YES | NO | NO |
| 10766 | TRAV12-3 | TRAJ11 | CAMDLVNSGYSTLTF | TRBV27 | TRBJ2-7 | CASSSGTSGAYEQYF | KLGGALQAK | YES | NO | NO |
| 10767 | TRAV20 | TRAJ52 | CAGTAGGTSYGKLTF | TRBV10-2 | TRBJ1-1 | CASTPGYMNTEAFF | KLGGALQAK | YES | NO | NO |
| 10768 | TRAV30 | TRAJ22 | CGTASSGSARQLTF | TRBV14 | TRBJ2-7 | CASSQTGGKTYEQYF | AVFDRKSDAK | NO | NO | NO |
| 10769 | TRAV23DV6 | TRAJ5 | CAAWEGTGRRALTF | TRBV11-2 | TRBJ1-2 | CASSWGGGSHYGYTF | IVTDFSVIK | YES | NO | YES |
| 10771 | TRAV12-1 | TRAJ36 | CVVTPGANNLFF | TRBV28 | TRBJ2-5 | CASSSPETQYF | KLGGALQAK | YES | NO | NO |
| 10772 | TRAV29DV5 | TRAJ48 | CAAKGGNEKLTF | TRBV7-2 | TRBJ2-7 | CASSLTGGMREQYF | RAKFKQLL | YES | NO | YES |
| 10773 | TRAV36DV7 | TRAJ42 | CAVESQGNLIF | TRBV2 | TRBJ1-1 | CASAGNTEAFF | KLGGALQAK | NO | NO | NO |
| 10774 | TRAV12-2 | TRAJ23 | CAGQGGKLIF | TRBV20-1 | TRBJ1-4 | CSAALGLGEKLFF | ELAGIGILTV | YES | NO | YES |
| 10775 | TRAV13-1 | TRAJ42 | CAASNPPGSQGNLIF | TRBV7-3 | TRBJ1-1 | CASSLTRQGGTEAFF | AVFDRKSDAK | YES | NO | YES |
| 10776 | TRAV12-3 | TRAJ57 | CATQGGSEKLVF | TRBV13 | TRBJ2-1 | CASSPHEVGLGDFRYNEQFF | KLGGALQAK | NO | NO | NO |
| 10777 | TRAV29DV5 | TRAJ40 | CAASAGTYKYIF | TRBV4-3 | TRBJ2-5 | CASSQELGPRETQYF | KLGGALQAK | NO | NO | NO |
| 10778 | TRAV12-2 | TRAJ8 | CAVTGGFQKLVF | TRBV14 | TRBJ1-5 | CASSHTGLGQPQHF | ELAGIGILTV | NO | NO | NO |
| 10780 | TRAV19 | TRAJ42 | CALSDSYYGGSQGNLIF | TRBV9 | TRBJ1-6 | CASSPLRDRGNSPLHF | KLGGALQAK | NO | NO | NO |
| 10781 | TRAV13-2 | TRAJ45 | CAENGGGGADGLTF | TRBV6-6 | TRBJ1-1 | CASSTTAGDTEAFF | KLGGALQAK | NO | NO | NO |
| 10782 | TRAV25 | TRAJ52 | CAGPPGVAGGTSYGKLTF | TRBV4-1 | TRBJ2-3 | CASSRASRQGSTDTQYF | AVFDRKSDAK | YES | NO | YES |
| 10783 | TRAV1-1 | TRAJ26 | CAAYDNYGQNFVF | TRBV6-5 | TRBJ2-7 | CASSSSRRGGRAYEQYF | KLGGALQAK | YES | NO | NO |
| 10784 | TRAV8-6 | TRAJ37 | CAVSSDNTGKLIF | TRBV19 | TRBJ2-7 | CASSIRSNYEQYF | GILGFVFTL | YES | NO | YES |
| 10785 | TRAV12-3 | TRAJ8 | CAMFGFQKLVF | TRBV24-1 | TRBJ2-2 | CATSDLGSNGELFF | KLGGALQAK | YES | NO | NO |
| 10786 | TRAV24 | TRAJ54 | CALIQGAQKLVF | TRBV5-6 | TRBJ1-4 | CASSYRGQPRREKLFF | KLGGALQAK | YES | NO | NO |
| 10787 | TRAV22 | TRAJ37 | CAGPPGSSNTGKLIF | TRBV19 | TRBJ2-7 | CASSIRSSYEQYF | GILGFVFTL | YES | NO | NO |
| 10788 | TRAV17 | TRAJ30 | CATEGRDDKIIF | TRBV11-1 | TRBJ2-7 | CASSLWTGNQYF | KLGGALQAK | YES | NO | NO |
| 10789 | TRAV16 | TRAJ49 | CAPGNQFYF | TRBV11-2 | TRBJ1-2 | CASSWGGGSHYGYTF | IVTDFSVIK | NO | NO | NO |
| 10790 | TRAV39 | TRAJ38 | CAGPNAGNNRKLIW | TRBV4-1 | TRBJ2-5 | CASSHPPRLAGGYQETQYF | KLGGALQAK | YES | NO | NO |
| 10791 | TRAV22 | TRAJ26 | CAVDNYGQNFVF | TRBV4-2 | TRBJ2-7 | CASSQELSYEQYF | KLGGALQAK | YES | NO | NO |
| 10792 | TRAV12-2 | TRAJ45 | CAVEGGADGLTF | TRBV24-1 | TRBJ2-5 | CATSDPSGTGVETQYF | ELAGIGILTV | YES | NO | YES |
| 10793 | TRAV19 | TRAJ9 | CALSGTYTGGFKTIF | TRBV7-9 | TRBJ2-5 | CASSLAPGQGPIKETQYF | KLGGALQAK | YES | NO | NO |
| 10794 | TRAV1-1 | TRAJ29 | CAVDSTPLVF | TRBV25-1 | TRBJ1-2 | CASSEAGAFGYTF | KLGGALQAK | YES | NO | NO |
| 10795 | TRAV1-1 | TRAJ12 | CAVSRMDSSYKLIF | TRBV19 | TRBJ2-7 | CASSHLLSTVDYEQYF | KLGGALQAK | YES | NO | NO |
| 10796 | TRAV8-4 | TRAJ37 | CAVSAPSGNTGKLIF | TRBV11-2 | TRBJ2-5 | CASSLGDFFSQETQYF | AVFDRKSDAK | YES | NO | YES |
| 10797 | TRAV29DV5 | TRAJ44 | CAASTTGTASKLTF | TRBV4-1 | TRBJ2-3 | CASSPSRSGADTQYF | RAKFKQLL | YES | NO | YES |
| 10798 | TRAV29DV5 | TRAJ58 | CAASEIGGSRLTF | TRBV4-3 | TRBJ2-2 | CASSRVQATTGELFF | KLGGALQAK | YES | NO | NO |

| | | | | | | | | | | |
|---|---|---|---|---|---|---|---|---|---|---|
| 10799 | TRAV14DV4 | TRAJ58 | CAMRTGETSGSRLTF | TRBV27 | TRBJ2-7 | CASSLSGWTYEQYF | KLGGALQAK | YES | NO | NO |
| 10800 | TRAV38-2DV8 | TRAJ33 | CAFSGSNYQLIW | TRBV7-9 | TRBJ2-3 | CASSRLTSTDTQYF | KLGGALQAK | YES | NO | NO |
| 10801 | TRAV1-2 | TRAJ6 | CASPRGSYIPTF | TRBV6-1 | TRBJ2-2 | CASYTPENTGELFF | IVTDFSVIK | YES | NO | YES |
| 10804 | TRAV1-2 | TRAJ20 | CAVRERDYKLSF | TRBV9 | TRBJ2-1 | CASSVEASGERWLFF | KLGGALQAK | YES | NO | NO |
| 10805 | TRAV8-6 | TRAJ30 | CAVSEKNRDDKIIF | TRBV12-3 | TRBJ1-4 | CASSPRGEKLFF | KLGGALQAK | YES | NO | NO |
| 10806 | TRAV17 | TRAJ29 | CATDLFSGNTPLVF | TRBV11-2 | TRBJ2-1 | CASSLGKNEQFF | KLGGALQAK | YES | NO | NO |
| 10807 | TRAV12-2 | TRAJ28 | CAVNSVPWGWDPPQLTF | TRBV28 | TRBJ2-1 | CASSRKTGANEQFF | LLFGYPVYV | YES | NO | NO |
| 10808 | TRAV5 | TRAJ13 | CADDWGGYQKVTF | TRBV12-4 | TRBJ1-1 | CASSSSSQENTEAFF | KLGGALQAK | YES | NO | NO |
| 10809 | TRAV27 | TRAJ42 | CAGAGGGSQGNLIF | TRBV19 | TRBJ2-3 | CASSGRSTDTQYF | GILGFVFTL | YES | NO | YES |
| 10810 | TRAV19 | TRAJ40 | CALSEAYYPGTYKYIF | TRBV27 | TRBJ1-5 | CASSLLVGSQPQHF | KLGGALQAK | YES | NO | NO |
| 10811 | TRAV12-2 | TRAJ16 | CAVSDGQKLLF | TRBV25-1 | TRBJ2-3 | CASSELNGMDTQYF | KLGGALQAK | NO | NO | NO |
| 10812 | TRAV38-1 | TRAJ56 | CALHTGANSKLTF | TRBV11-2 | TRBJ1-2 | CASSWGGGSHYGYTF | IVTDFSVIK | YES | NO | NO |
| 10813 | TRAV8-6 | TRAJ42 | CAVSGSQGNLIF | TRBV19 | TRBJ2-7 | CASSLRSSYEQYF | GILGFVFTL | YES | NO | YES |
| 10814 | TRAV38-2DV8 | TRAJ57 | CAYNRPQGGSEKLVF | TRBV5-1 | TRBJ1-1 | CASSFGQGANTEAFF | KLGGALQAK | YES | NO | NO |
| 10815 | TRAV12-2 | TRAJ26 | CAVDNYGQNFVF | TRBV6-5 | TRBJ2-3 | CASSYGVLSTDTQYF | ELAGIGILTV | NO | NO | NO |
| 10816 | TRAV35 | TRAJ30 | CAGQYRDDKIIF | TRBV19 | TRBJ2-7 | CASSIGLGEQYF | GILGFVFTL | YES | NO | YES |
| 10817 | TRAV8-3 | TRAJ27 | CAVGSNTNAGKSTF | TRBV12-3 | TRBJ1-2 | CASSSGANYGYTF | KLGGALQAK | YES | NO | NO |
| 10818 | TRAV25 | TRAJ20 | CAVSIPTDYKLSF | TRBV3-1 | TRBJ2-1 | CASSPDLVHEQFF | KLGGALQAK | YES | NO | NO |
| 10819 | TRAV22 | TRAJ58 | CAVVPRGTSGSRLTF | TRBV18 | TRBJ1-5 | CASSFLIGGNQPQHF | KLGGALQAK | YES | NO | NO |
| 10820 | TRAV12-3 | TRAJ45 | CAIMRGADGLTF | TRBV4-2 | TRBJ2-6 | CASSAGLIGANVLTF | ELAGIGILTV | YES | NO | NO |
| 10821 | TRAV12-2 | TRAJ45 | CAPSGGGADGLTF | TRBV28 | TRBJ1-1 | CASSPIGYTEAFF | ELAGIGILTV | YES | NO | YES |
| 10822 | TRAV12-2 | TRAJ9 | CAVNTGGFKTIF | TRBV14 | TRBJ1-2 | CASSQEGGNSPYGYTF | KLGGALQAK | NO | NO | NO |
| 10823 | TRAV5 | TRAJ41 | CAESPNSGYALNF | TRBV11-1 | TRBJ2-1 | CASSLGQGRNEQFF | KLGGALQAK | YES | NO | NO |
| 10824 | TRAV26-2 | TRAJ13 | CILFSGGYQKVTF | TRBV6-3 | TRBJ2-1 | CASRTDRGDEQFF | AVFDRKSDAK | YES | NO | YES |
| 10825 | TRAV13-1 | TRAJ20 | CAASMDDYKLSF | TRBV27 | TRBJ1-1 | CASTSRWGTEAFF | KLGGALQAK | YES | NO | NO |
| 10826 | TRAV12-2 | TRAJ31 | CAVNNARLMF | TRBV4-2 | TRBJ2-7 | CASSPSAQGNEQYF | ELAGIGILTV | YES | YES | YES |
| 10827 | TRAV21 | TRAJ28 | CAVTSDSGAGSYQLTF | TRBV19 | TRBJ2-1 | CASSISLAAEQFF | LLDFVRFMGV | YES | NO | YES |
| 10828 | TRAV27 | TRAJ42 | CAGAGSQGNLIF | TRBV9 | TRBJ2-7 | CASSALSGRAIDEQYF | GILGFVFTL | YES | NO | YES |
| 10829 | TRAV5 | TRAJ34 | CAESGGTDKLIF | TRBV5-1 | TRBJ2-5 | CASSDDLETQYF | KLGGALQAK | YES | NO | NO |
| 10830 | TRAV12-2 | TRAJ24 | CAALDSWGKFQF | TRBV19 | TRBJ1-5 | CASSARSGEPQHF | GILGFVFTL | YES | NO | NO |
| 10831 | TRAV12-3 | TRAJ53 | CAMRNGGSNYKLTF | TRBV28 | TRBJ1-3 | CASSLSGQRVSGNTIYF | KLGGALQAK | YES | NO | NO |
| 10832 | TRAV16 | TRAJ26 | CALIDNYGQNFVF | TRBV7-8 | TRBJ1-3 | CASSLERGAAGNTIYF | KLGGALQAK | NO | NO | NO |
| 10833 | TRAV1-2 | TRAJ13 | CAVRSGGYQKVTF | TRBV5-6 | TRBJ2-1 | CASSLKRDTYNEQFF | KLGGALQAK | YES | NO | NO |
| 10834 | TRAV1-2 | TRAJ33 | CAVCDSNYQLIW | TRBV6-3 | TRBJ2-7 | CASSYSDNYEQYF | KLGGALQAK | YES | NO | NO |
| 10835 | TRAV19 | TRAJ17 | CALKLIKAAGNKLTF | TRBV6-5 | TRBJ1-4 | CASSPGGLNEKLFF | KLGGALQAK | NO | NO | NO |
| 10836 | TRAV13-2 | TRAJ6 | CAENRGSYIPTF | TRBV7-9 | TRBJ2-3 | CASSTHRLGTSGGPPDTQYF | KLGGALQAK | NO | NO | NO |
| 10838 | TRAV12-3 | TRAJ42 | CARNYGGSQGNLIF | TRBV14 | TRBJ2-1 | CASSLQGSGYNEQFF | KLGGALQAK | YES | NO | NO |
| 10839 | TRAV3 | TRAJ4 | CAVPFSGGYNKLIF | TRBV7-6 | TRBJ2-1 | CASSLISGRPEQFF | KLGGALQAK | NO | NO | NO |
| 10840 | TRAV17 | TRAJ54 | CATRAQGAQKLVF | TRBV9 | TRBJ2-7 | CASSPGLAAMYEQYF | KLGGALQAK | YES | NO | NO |
| 10841 | TRAV21 | TRAJ30 | CAVIMNRDDKIIF | TRBV7-2 | TRBJ2-1 | CASSGWVYNEQFF | KLGGALQAK | YES | NO | NO |
| 10842 | TRAV21 | TRAJ44 | CAVRRGGRNTGTASKLTF | TRBV9 | TRBJ1-3 | CASSLSGAGNTIYF | KLGGALQAK | YES | NO | NO |
| 10843 | TRAV1-2 | TRAJ24 | CALPTDSWGKLQF | TRBV28 | TRBJ2-7 | CASSPLRGTGDYEQYF | KLGGALQAK | YES | NO | NO |
| 10844 | TRAV14DV4 | TRAJ57 | CAMRDPQGGSEKLVF | TRBV7-9 | TRBJ2-4 | CASSSTSGRFQNIQYF | KLGGALQAK | YES | NO | NO |
| 10845 | TRAV41 | TRAJ58 | CAAAQETSGSRLTF | TRBV19 | TRBJ1-1 | CASSIEGGTTEAFF | KLGGALQAK | YES | NO | NO |
| 10846 | TRAV17 | TRAJ58 | CATDQETSGSRLTF | TRBV7-3 | TRBJ2-7 | CASSPGGLNEQYF | KLGGALQAK | NO | NO | NO |
| 10847 | TRAV12-2 | TRAJ33 | CAFSDSNYQLIW | TRBV10-3 | TRBJ2-1 | CATGTGTGDNEQFF | AVFDRKSDAK | YES | NO | NO |
| 10848 | TRAV8-4 | TRAJ31 | CAVRPNARLMF | TRBV7-8 | TRBJ2-7 | CASSPGQGGYEQYF | AVFDRKSDAK | YES | NO | YES |
| 10849 | TRAV8-3 | TRAJ4 | CAVGAGGYNKLIF | TRBV5-6 | TRBJ1-2 | CASSGTGANYGYTF | KLGGALQAK | YES | NO | NO |
| 10850 | TRAV13-1 | TRAJ47 | CAAYQEYGNKLVF | TRBV4-1 | TRBJ2-7 | CASATGTGGNEQYF | AVFDRKSDAK | YES | YES | NO |
| 10851 | TRAV13-1 | TRAJ21 | CAAFDFNKFYF | TRBV29-1 | TRBJ1-2 | CSVEGRAQGYTF | KLGGALQAK | YES | NO | NO |
| 10853 | TRAV14DV4 | TRAJ30 | CAMRDWDDKIIF | TRBV6-3 | TRBJ2-1 | CASTLTGLADTYNEQFF | KLGGALQAK | YES | NO | NO |
| 10854 | TRAV19 | TRAJ20 | CARATNDYKLSF | TRBV19 | TRBJ1-2 | CASSSAGPYGYTF | GILGFVFTL | YES | NO | NO |
| 10855 | TRAV1-2 | TRAJ33 | CAVMDSNYQLIW | TRBV20-1 | TRBJ2-5 | CSARDPGGRETQYF | KLGGALQAK | YES | NO | NO |
| 10856 | TRAV21 | TRAJ54 | CAVYGIQGAQKLVF | TRBV13 | TRBJ1-5 | CASSLGLEGTNQPQHF | KTWGQYWQV | YES | NO | YES |
| 10857 | TRAV13-1 | TRAJ20 | CAASPGDYKLSF | TRBV19 | TRBJ2-3 | CASTPGGIDTQYF | KLGGALQAK | YES | NO | NO |
| 10858 | TRAV12-2 | TRAJ17 | CAVNKDAGNKLTF | TRBV6-5 | TRBJ2-1 | CASSYLTGTSGEQFF | AVFDRKSDAK | NO | NO | NO |
| 10859 | TRAV1-2 | TRAJ18 | CAVREGDRGSTLGRLYF | TRBV7-9 | TRBJ2-5 | CASSLVGGQETQYF | KLGGALQAK | YES | NO | NO |
| 10860 | TRAV1-2 | TRAJ6 | CAVKSSGGSYIPTF | TRBV10-3 | TRBJ1-5 | CAIGTGDTNQPQHF | KLGGALQAK | YES | NO | NO |
| 10861 | TRAV21 | TRAJ42 | CAVWSYGGSQGNLIF | TRBV4-3 | TRBJ1-1 | CASSHGQGRGNTEAFF | KLGGALQAK | YES | NO | NO |
| 10862 | TRAV1-2 | TRAJ33 | CAAVNSNYQLIW | TRBV4-3 | TRBJ2-1 | CASSQDGESSYNEQFF | KLGGALQAK | YES | NO | NO |
| 10863 | TRAV5 | TRAJ49 | CAESIWNTGNQFYF | TRBV20-1 | TRBJ2-1 | CSALIRDYNEQFF | KLGGALQAK | NO | NO | NO |
| 10864 | TRAV38-2DV8 | TRAJ54 | CAYRRGAQKLVF | TRBV11-2 | TRBJ1-1 | CASSFPRTKNTEAFF | KLGGALQAK | YES | NO | NO |
| 10865 | TRAV13-2 | TRAJ34 | CAENMRNTDKLIF | TRBV19 | TRBJ2-2 | CASSSFGGRELFF | GILGFVFTL | YES | NO | YES |
| 10866 | TRAV34 | TRAJ21 | CGADYNFNKFYF | TRBV27 | TRBJ2-7 | CASSLLRTGGNEQYF | KLGGALQAK | YES | NO | NO |
| 10867 | TRAV29DV5 | TRAJ52 | CAASAPGGTSYGKLTF | TRBV9 | TRBJ2-7 | CASSVAASGSYEQYF | KLGGALQAK | YES | NO | NO |
| 10868 | TRAV12-1 | TRAJ34 | CVAYNTDKLIF | TRBV7-2 | TRBJ2-1 | CASSLGEQFF | KLGGALQAK | YES | NO | NO |
| 10869 | TRAV35 | TRAJ49 | CAGHTGNQFYF | TRBV11-2 | TRBJ1-6 | CASSTTGPQTPLHF | IVTDFSVIK | YES | NO | YES |
| 10870 | TRAV6 | TRAJ8 | CALIGTGFQKLVF | TRBV12-5 | TRBJ2-1 | CASGLPLAGGPYPYNEQFF | KLGGALQAK | YES | NO | NO |
| 10871 | TRAV19 | TRAJ31 | CALSEKEARLMF | TRBV7-9 | TRBJ2-7 | CASSLERFYEQYF | KLGGALQAK | YES | NO | NO |
| 10872 | TRAV21 | TRAJ37 | CAVSLSGNTGKLIF | TRBV4-1 | TRBJ2-1 | CASSQGPYNEQFF | KLGGALQAK | YES | NO | NO |
| 10873 | TRAV12-2 | TRAJ48 | CAGFFGNEKLTF | TRBV6-6 | TRBJ2-7 | CASRIAHEQYF | KLGGALQAK | NO | NO | NO |
| 10874 | TRAV27 | TRAJ40 | CAGLTSGTYKYIF | TRBV11-2 | TRBJ2-3 | CASSLDGEGYGAFTDTQYF | KLGGALQAK | YES | NO | NO |
| 10875 | TRAV38-1 | TRAJ38 | CAFMKPPSAGNNRKLIW | TRBV4-3 | TRBJ1-1 | CASSQVPGADTEAFF | KLGGALQAK | YES | NO | NO |
| 10876 | TRAV8-3 | TRAJ43 | CAVGGNNNDMRF | TRBV20-1 | TRBJ2-7 | CSARDYLNSYEQYF | GILGFVFTL | YES | NO | YES |
| 10877 | TRAV8-4 | TRAJ8 | CAVGTGFQKLVF | TRBV7-9 | TRBJ1-1 | CASSLPVEVNTEAFF | KLGGALQAK | YES | NO | NO |
| 10878 | TRAV12-1 | TRAJ11 | CVVSGYSTLTF | TRBV19 | TRBJ2-2 | CASSTMTGSTGELFF | KLGGALQAK | YES | NO | NO |
| 10879 | TRAV12-2 | TRAJ34 | CAVNSGTDKLIF | TRBV28 | TRBJ1-5 | CASRFISLGQPQHF | ELAGIGILTV | YES | NO | YES |
| 10880 | TRAV13-1 | TRAJ34 | CAASPFSYNTDKLIF | TRBV27 | TRBJ2-7 | CASSFGQLHIEYF | KLGGALQAK | YES | NO | NO |
| 10881 | TRAV12-2 | TRAJ11 | CAVENSGYSTLTF | TRBV6-1 | TRBJ1-1 | CASRGVLGGQGETEAFF | KLGGALQAK | YES | NO | NO |
| 10882 | TRAV8-3 | TRAJ26 | CAVGDNYGQNFVF | TRBV6-5 | TRBJ2-5 | CASGGETQYF | KLGGALQAK | NO | NO | NO |
| 10883 | TRAV12-2 | TRAJ45 | CAVNIGGGADGLTF | TRBV6-5 | TRBJ2-7 | CASSSGQGNEQYF | ELAGIGILTV | YES | NO | YES |
| 10884 | TRAV29DV5 | TRAJ53 | CAASASGGSNYKLTF | TRBV7-9 | TRBJ2-3 | CAIKRLSTDTQYF | KLGGALQAK | YES | NO | NO |
| 10885 | TRAV22 | TRAJ40 | CAVPTTSGTYKYIF | TRBV5-4 | TRBJ2-2 | CASSFGTPTTGELFF | KLGGALQAK | YES | NO | NO |
| 10886 | TRAV27 | TRAJ37 | CAGAHGSSNTGKLIF | TRBV19 | TRBJ2-7 | CASSIRSSYEQYF | GILGFVFTL | YES | YES | YES |
| 10887 | TRAV35 | TRAJ44 | CAGSHPDTGTASKLTF | TRBV27 | TRBJ1-6 | CASSLLGSPLHF | KLGGALQAK | YES | NO | NO |
| 10888 | TRAV21 | TRAJ39 | CAVRNNAGNMLTF | TRBV30 | TRBJ2-1 | CAWSVSASEQFF | LLDFVRFMGV | NO | NO | NO |
| 10889 | TRAV8-3 | TRAJ39 | CAVYYNAGNMLTF | TRBV12-4 | TRBJ1-6 | CASSFLGLDGSPLHF | AVFDRKSDAK | YES | NO | YES |
| 10890 | TRAV13-2 | TRAJ45 | CAENSGGGADGLTF | TRBV9 | TRBJ1-1 | CASSVVSGGTEAFF | KLGGALQAK | YES | NO | NO |
| 10891 | TRAV26-2 | TRAJ54 | CILIQGAQKLVF | TRBV10-3 | TRBJ2-5 | CAISELFLAGPMRGTQYF | KLGGALQAK | YES | NO | NO |
| 10892 | TRAV38-2DV8 | TRAJ28 | CAYQAPGAGSYQLTF | TRBV13 | TRBJ2-3 | CASSVLAGYTDTQYF | KLGGALQAK | YES | NO | NO |
| 10893 | TRAV19 | TRAJ48 | CALSEAQANFGNEKLTF | TRBV4-2 | TRBJ2-1 | CASSQEQASNEQFF | IVTDFSVIK | YES | NO | YES |
| 10894 | TRAV13-1 | TRAJ53 | CAAKGSNYKLTF | TRBV5-1 | TRBJ2-3 | CASSFSGGAEGDTQYF | KLGGALQAK | YES | NO | NO |
| 10895 | TRAV5 | TRAJ24 | CADMPADSWGKLQF | TRBV4-1 | TRBJ1-5 | CASSQEGGAQPQHF | KLGGALQAK | YES | NO | NO |
| 10896 | TRAV26-1 | TRAJ58 | CIVIETSGSRLTF | TRBV20-1 | TRBJ2-6 | CSGEGAGANVLTF | KLGGALQAK | NO | NO | NO |
| 10897 | TRAV12-2 | TRAJ20 | CAVRRMRDYKLSF | TRBV7-9 | TRBJ2-7 | CASSNGTGGVYEQYF | KLGGALQAK | YES | NO | NO |
| 10898 | TRAV29DV5 | TRAJ37 | CAAKGSSNTGKLIF | TRBV19 | TRBJ2-7 | CASSIRSSYEQYF | GILGFVFTL | YES | NO | YES |
| 10899 | TRAV1-2 | TRAJ31 | CAVSDNNARLMF | TRBV27 | TRBJ2-3 | CASSSLAGGLTDTQYF | KLGGALQAK | YES | NO | NO |
| 10900 | TRAV8-6 | TRAJ39 | CALNAGNMLTF | TRBV19 | TRBJ2-7 | CASSPQGAYEQYF | KLGGALQAK | YES | NO | NO |
| 10901 | TRAV12-2 | TRAJ45 | CAVPNSGGGADGLTF | TRBV9 | TRBJ2-1 | CASSPGLAGGEQFF | ELAGIGILTV | YES | YES | YES |
| 10902 | TRAV13-1 | TRAJ30 | CAASPRRDDKIIF | TRBV5-1 | TRBJ2-7 | CASSFIPAELRYEQYF | KLGGALQAK | YES | NO | NO |
| 10903 | TRAV13-1 | TRAJ50 | CAASETSYDKVIF | TRBV19 | TRBJ1-2 | CASSIGVYGYTF | GILGFVFTL | YES | NO | YES |
| 10904 | TRAV12-2 | TRAJ45 | CAAYSGGGADGLTF | TRBV9 | TRBJ2-7 | CASSASGVGYEQYF | KLGGALQAK | YES | NO | NO |
| 10905 | TRAV8-1 | TRAJ37 | CAVNAPGNTGKLIF | TRBV6-3 | TRBJ2-1 | CASSYGPGLAGRSSYNEQFF | KLGGALQAK | NO | NO | NO |
| 10906 | TRAV34 | TRAJ43 | CGAGRVDDMRF | TRBV11-2 | TRBJ1-2 | CASSWGGGSHYGYTF | IVTDFSVIK | YES | NO | YES |

| | | | | | | | | | | |
|---|---|---|---|---|---|---|---|---|---|---|
| 10907 | TRAV19 | TRAJ9 | CALSEAGGFKTIF | TRBV11-2 | TRBJ1-2 | CASSWGGGSHYGYTF | IVTDFSVIK | YES | NO | NO |
| 10909 | TRAV12-2 | TRAJ29 | CAVDSGNTPLVF | TRBV11-2 | TRBJ2-1 | CASSLDTLAGGQFF | ELAGIGILTV | YES | NO | NO |
| 10910 | TRAV4 | TRAJ3 | CLVGDPFYSSASKIIF | TRBV7-2 | TRBJ2-7 | CASSSPFSASSYEQYF | AVFDRKSDAK | NO | NO | NO |
| 10911 | TRAV12-2 | TRAJ41 | CAVKAGYALNF | TRBV11-2 | TRBJ1-2 | CASSWGGGSHYGYTF | IVTDFSVIK | YES | NO | YES |
| 10912 | TRAV19 | TRAJ49 | CALSELYTGNQFYF | TRBV5-1 | TRBJ2-3 | CASRRGPVGDTQYF | KLGGALQAK | YES | NO | NO |
| 10913 | TRAV12-2 | TRAJ33 | CAVSNYQLIW | TRBV11-1 | TRBJ2-1 | CASTSGDNEQFF | ELAGIGILTV | YES | YES | YES |
| 10914 | TRAV13-1 | TRAJ42 | CALTGGGSQGNLIF | TRBV19 | TRBJ2-3 | CASSIRSGDTQYF | GILGFVFTL | YES | NO | YES |
| 10915 | TRAV35 | TRAJ49 | CAGHTGNQFYF | TRBV5-1 | TRBJ2-3 | CASSPGGLAGADTQYF | IVTDFSVIK | YES | NO | YES |
| 10916 | TRAV30 | TRAJ20 | CGTEYPNDYKLSF | TRBV9 | TRBJ2-7 | CASSLSSGTYEQYF | IVTDFSVIK | YES | YES | YES |
| 10917 | TRAV19 | TRAJ37 | CAHSSNTGKLIF | TRBV6-5 | TRBJ2-1 | CASSYARNYNEQFF | KLGGALQAK | NO | NO | NO |
| 10918 | TRAV8-3 | TRAJ28 | CAVGARRNSGAGSYQLTF | TRBV4-1 | TRBJ2-3 | CASSQEPLTDTQYF | KLGGALQAK | YES | NO | NO |
| 10919 | TRAV25 | TRAJ15 | CAVVGQAGTALIF | TRBV3-1 | TRBJ2-1 | CASSQVRIYNEQFF | KLGGALQAK | YES | NO | NO |
| 10920 | TRAV29DV5 | TRAJ45 | CAATPSGGGADGLTF | TRBV7-8 | TRBJ1-5 | CASSFTLAAYSNQPQHF | KLGGALQAK | YES | NO | NO |
| 10921 | TRAV12-2 | TRAJ24 | CAVSDSWGKLQF | TRBV28 | TRBJ2-4 | CASTNTRESAKNIQYF | KLGGALQAK | YES | YES | NO |
| 10922 | TRAV30 | TRAJ20 | CGTEINDYKLSF | TRBV2 | TRBJ2-1 | CARQGTSGSNEQFF | KLGGALQAK | YES | NO | NO |
| 10923 | TRAV20 | TRAJ26 | CAVQAHYGQNFVF | TRBV19 | TRBJ2-1 | CASHSVTNEQFF | KLGGALQAK | YES | NO | NO |
| 10924 | TRAV1-2 | TRAJ20 | CAVRERDYKLSF | TRBV19 | TRBJ1-5 | CASRQSNQPQHF | RLRAEAQVK | YES | NO | YES |
| 10925 | TRAV14DV4 | TRAJ28 | CAMSDLYSGAGSYQLTF | TRBV5-1 | TRBJ2-7 | CASSADSPSYEQYF | KLGGALQAK | YES | NO | NO |
| 10926 | TRAV21 | TRAJ54 | CAVGGAQKLVF | TRBV11-1 | TRBJ1-1 | CASSLWDRGWTEAFF | KLGGALQAK | YES | NO | NO |
| 10927 | TRAV12-3 | TRAJ23 | CAMSGHNQGGKLIF | TRBV9 | TRBJ2-7 | CASSVDGKTEQYF | KLGGALQAK | YES | NO | NO |
| 10928 | TRAV13-2 | TRAJ26 | CAENENYGQNFVF | TRBV3-1 | TRBJ2-7 | CASSLGQGGYEQYF | KLGGALQAK | YES | NO | NO |
| 10929 | TRAV38-1 | TRAJ53 | CAFMKLYSGGSNYKLTF | TRBV5-5 | TRBJ1-5 | CASSLVRGGNQPQHF | AVFDRKSDAK | YES | YES | YES |
| 10930 | TRAV13-2 | TRAJ42 | CAENIGGGSQGNLIF | TRBV19 | TRBJ2-7 | CASSIRSSYEQYF | GILGFVFTL | YES | NO | NO |
| 10931 | TRAV13-2 | TRAJ42 | CAENMGGGSQGNLIF | TRBV19 | TRBJ2-5 | CASSVRAGETQYF | GILGFVFTL | YES | NO | YES |
| 10932 | TRAV20 | TRAJ44 | CAVYRSGTASKLTF | TRBV15 | TRBJ2-1 | CATRKTGNEQFF | KLGGALQAK | YES | NO | NO |
| 10933 | TRAV20 | TRAJ26 | CAVQWTIWSHGQNFVF | TRBV20-1 | TRBJ2-1 | CSARDDYLGNWNEQFF | KLGGALQAK | NO | NO | NO |
| 10934 | TRAV1-2 | TRAJ31 | CAVRDGARLMF | TRBV7-2 | TRBJ1-2 | CASSAGTGVYGYTF | KLGGALQAK | NO | NO | NO |
| 10935 | TRAV13-2 | TRAJ42 | CAENNGGGSQGNLIF | TRBV19 | TRBJ2-1 | CASSIRSVHEQFF | GILGFVFTL | YES | YES | YES |
| 10936 | TRAV8-3 | TRAJ43 | CAVLNDMRF | TRBV20-1 | TRBJ1-1 | CSANGGDTEAFF | YLLEMLWRL | NO | NO | NO |
| 10937 | TRAV14DV4 | TRAJ20 | CAMEGDYKLSF | TRBV7-6 | TRBJ1-6 | CASSSKEGVLLHF | KLGGALQAK | YES | NO | NO |
| 10938 | TRAV14DV4 | TRAJ40 | CAMRELPTTSGTYKYIF | TRBV5-6 | TRBJ1-1 | CASSFARRVNTEAFF | KLGGALQAK | YES | NO | NO |
| 10939 | TRAV27 | TRAJ17 | CAIPQKAAGNKLTF | TRBV28 | TRBJ1-1 | CASSFPQFVTGLPEAFF | ELAGIGILTV | YES | NO | YES |
| 10940 | TRAV26-2 | TRAJ21 | CILRVRGFNKFYF | TRBV20-1 | TRBJ1-4 | CSAREGGILFF | KLGGALQAK | NO | NO | NO |
| 10941 | TRAV1-2 | TRAJ20 | CAVNSDYKLSF | TRBV20-1 | TRBJ2-2 | CSAREEAGDDTGELFF | AVFDRKSDAK | NO | NO | NO |
| 10942 | TRAV4 | TRAJ9 | CLVGVHTGGFKTIF | TRBV20-1 | TRBJ2-7 | CSAIPQVRAHEQYF | KLGGALQAK | YES | NO | NO |
| 10943 | TRAV12-2 | TRAJ31 | CAVDARLMF | TRBV6-6 | TRBJ2-2 | CASSPLSRQGGTGELFF | ELAGIGILTV | NO | NO | NO |
| 10944 | TRAV1-2 | TRAJ22 | CAVSGYISSGSARQLTF | TRBV10-2 | TRBJ2-7 | CASRETGEGYEQYF | RLRAEAQVK | YES | YES | YES |
| 10945 | TRAV27 | TRAJ9 | CAGWTGGFKTIF | TRBV28 | TRBJ2-7 | CASRPGTGDYEQYF | KLGGALQAK | YES | NO | NO |
| 10946 | TRAV13-2 | TRAJ11 | CAENSGYSTLTF | TRBV30 | TRBJ2-1 | CASQATTTYNEQFF | KLGGALQAK | YES | NO | NO |
| 10947 | TRAV1-2 | TRAJ4 | CAVNGYNKLIF | TRBV27 | TRBJ1-1 | CASSLSGGVTEAFF | KLGGALQAK | YES | NO | NO |
| 10948 | TRAV38-2DV8 | TRAJ29 | CAYRSLSGNTPLVF | TRBV7-2 | TRBJ2-2 | CASSLDQDTFLGVVTGELFF | YLLEMLWRL | NO | NO | NO |
| 10949 | TRAV12-2 | TRAJ49 | CAVNFITGNQFYF | TRBV28 | TRBJ2-3 | CASSVGGPPPDTQYF | KLGGALQAK | YES | NO | NO |
| 10950 | TRAV14DV4 | TRAJ52 | CAMRESNAGGTSYGKLTF | TRBV5-1 | TRBJ1-2 | CASSLYGTVESNYGYTF | KLGGALQAK | YES | NO | NO |
| 10951 | TRAV24 | TRAJ57 | CAFIQGGSEKLVF | TRBV15 | TRBJ2-7 | CATSRDRLAGSNYEQYF | KLGGALQAK | YES | NO | NO |
| 10952 | TRAV36DV7 | TRAJ48 | CAVGNFGNEKLTF | TRBV15 | TRBJ1-5 | CATSRDDGQGYQPQHF | KLGGALQAK | NO | NO | NO |
| 10953 | TRAV1-2 | TRAJ9 | CALNTGGFKTIF | TRBV28 | TRBJ2-1 | CASSWGQSISGVFNNEQFF | RLRAEAQVK | YES | NO | NO |
| 10954 | TRAV14DV4 | TRAJ52 | CAMREYTSNAGGTSYGKLTF | TRBV7-6 | TRBJ2-5 | CASSYVGGAGETQYF | KLGGALQAK | NO | NO | NO |
| 10955 | TRAV12-2 | TRAJ42 | CAVPLGGGSQGNLIF | TRBV19 | TRBJ1-1 | CASSIRAATEAFF | GILGFVFTL | YES | NO | YES |
| 10956 | TRAV1-1 | TRAJ29 | CAVRTRNSGNTPLVF | TRBV5-6 | TRBJ1-5 | CASSLRGLGQPQHF | ELAGIGILTV | YES | NO | YES |
| 10957 | TRAV13-1 | TRAJ9 | CAASITGGFKTIF | TRBV6-5 | TRBJ1-1 | CASSKQGALTEAFF | KLGGALQAK | YES | NO | NO |
| 10958 | TRAV13-2 | TRAJ21 | CAETWNNFNKFYF | TRBV5-6 | TRBJ2-1 | CASSLGPITGLPYNEQFF | GILGFVFTL | YES | NO | YES |
| 10959 | TRAV8-3 | TRAJ30 | CAVVRDDKIIF | TRBV7-2 | TRBJ1-2 | CASSSDWTGGAYYGYTF | KLGGALQAK | NO | NO | NO |
| 10960 | TRAV35 | TRAJ42 | CAGKNYGGSQGNLIF | TRBV2 | TRBJ2-3 | CASSPFPSTDTQYF | FLASKIGRLV | YES | NO | NO |
| 10961 | TRAV1-2 | TRAJ29 | CAVRSVSSGNTPLVF | TRBV7-9 | TRBJ1-1 | CASSWGGPNTEAFF | KLGGALQAK | YES | NO | NO |
| 10962 | TRAV9-2 | TRAJ29 | CALEPGNTPLVF | TRBV20-1 | TRBJ2-3 | CSAWGIQDTQYF | KLGGALQAK | NO | NO | NO |
| 10963 | TRAV14DV4 | TRAJ4 | CAMRELLISGGYNKLIF | TRBV27 | TRBJ2-5 | CASSSPSGLAGQETQYF | RTLNAWVKV | YES | YES | YES |
| 10964 | TRAV8-3 | TRAJ21 | CAVGDNFNKFYF | TRBV29-1 | TRBJ2-7 | CSVSASGGDEQYF | KLGGALQAK | YES | NO | NO |
| 10965 | TRAV38-1 | TRAJ33 | CAFMILWDSNYQLIW | TRBV19 | TRBJ2-1 | CASSTGGDRAGYNEQFF | KLGGALQAK | YES | NO | NO |
| 10966 | TRAV19 | TRAJ50 | CALSLTSYDKVIF | TRBV7-9 | TRBJ2-5 | CASSKTTEFTQEETQYF | KLGGALQAK | YES | NO | NO |
| 10967 | TRAV19 | TRAJ54 | CALGGAQKLVF | TRBV12-4 | TRBJ2-1 | CASSFGGSGRLYNEQFF | KLGGALQAK | YES | NO | NO |
| 10968 | TRAV29DV5 | TRAJ49 | CAASVTNTGNQFYF | TRBV19 | TRBJ2-7 | CASSIRSTYEQYF | GILGFVFTL | YES | NO | YES |
| 10969 | TRAV38-1 | TRAJ28 | CAFMNSGAGSYQLTF | TRBV11-2 | TRBJ1-2 | CASSWGGGSHYGYTF | IVTDFSVIK | YES | YES | YES |
| 10970 | TRAV29DV5 | TRAJ45 | CAASGGGADGLTF | TRBV3-1 | TRBJ2-7 | CASSQVKLAGEQYF | RAKFKQLL | YES | NO | NO |
| 10971 | TRAV19 | TRAJ33 | CALSEASLVDSNYQLIW | TRBV19 | TRBJ2-1 | CASSPTSGGYNEQFF | GILGFVFTL | YES | NO | YES |
| 10972 | TRAV19 | TRAJ49 | CALSEAEYQFYF | TRBV7-6 | TRBJ2-2 | CASSSDRGTGELFF | AVFDRKSDAK | YES | NO | YES |
| 10973 | TRAV39 | TRAJ40 | CAVESGTYKYIF | TRBV9 | TRBJ2-7 | CASSADRGGDEQYF | RAKFKQLL | YES | NO | YES |
| 10974 | TRAV12-2 | TRAJ22 | CAVGGSARQLTF | TRBV2 | TRBJ1-3 | CASSEGQTGNTIYF | ELAGIGILTV | YES | YES | YES |
| 10975 | TRAV29DV5 | TRAJ5 | CAASETGRRALTF | TRBV11-2 | TRBJ2-7 | CASSLGDGNYEQYF | KLGGALQAK | YES | NO | NO |
| 10976 | TRAV12-2 | TRAJ50 | CAVQNAPMKTSYDKVIF | TRBV7-2 | TRBJ1-4 | CASSLGTGEKLFF | KLGGALQAK | YES | NO | NO |
| 10977 | TRAV5 | TRAJ3 | CAEYSSASKIIF | TRBV14 | TRBJ2-3 | CASSQSPGGTQYF | GLCTLVAML | YES | NO | YES |
| 10978 | TRAV1-1 | TRAJ28 | CAVRDQDTL | TRBV7-9 | TRBJ2-7 | CASSLELASYEQYF | KLGGALQAK | YES | NO | NO |
| 10979 | TRAV19 | TRAJ3 | CALSVPGYSSASKIIF | TRBV4-3 | TRBJ2-1 | CASSQDGYNEQFF | KLGGALQAK | YES | NO | NO |
| 10980 | TRAV12-2 | TRAJ45 | CAPYSGGGADGLTF | TRBV28 | TRBJ2-1 | CASSFPGTSGIDEQFF | KLGGALQAK | YES | NO | NO |
| 10981 | TRAV13-2 | TRAJ45 | CADNSGGGADGLTF | TRBV11-1 | TRBJ2-1 | CASSSLANEQFF | KLGGALQAK | YES | NO | NO |
| 10982 | TRAV8-3 | TRAJ47 | CAVGGYMEYGNKLVF | TRBV20-1 | TRBJ2-3 | CSASPRGAWYTDTQYF | RLRAEAQVK | YES | NO | YES |
| 10984 | TRAV17 | TRAJ33 | CATDGDSNYQLIW | TRBV27 | TRBJ2-7 | CASSLTGGSYEQYF | KLGGALQAK | YES | NO | NO |
| 10985 | TRAV19 | TRAJ30 | CALSEAADDKIIF | TRBV3-1 | TRBJ2-3 | CASSLTSLGSGSTDTQYF | AVFDRKSDAK | YES | NO | YES |
| 10986 | TRAV21 | TRAJ31 | CAVSLYARLMF | TRBV7-8 | TRBJ1-1 | CASSLYGGGRENLRTEAFF | KLGGALQAK | YES | NO | NO |
| 10987 | TRAV4 | TRAJ20 | CLVVIRSNDYKLSF | TRBV7-9 | TRBJ1-1 | CASSLTGTEAFF | KLGGALQAK | YES | NO | NO |
| 10988 | TRAV21 | TRAJ33 | CAVRGSNYQLIW | TRBV5-4 | TRBJ2-7 | CASSFFRIIGRGGYEQYF | KLGGALQAK | YES | NO | NO |
| 10989 | TRAV1-2 | TRAJ33 | CAVRDSNYQLIW | TRBV20-1 | TRBJ1-5 | CSAQWTPGGNQPQHF | KLGGALQAK | YES | NO | NO |
| 10990 | TRAV22 | TRAJ45 | CAPGGGADGLTF | TRBV4-1 | TRBJ1-4 | CASSPVSGDEKLFF | KLGGALQAK | YES | NO | NO |
| 10992 | TRAV39 | TRAJ49 | CAVDTGTGNQFYF | TRBV27 | TRBJ2-7 | CASRRGWSDHEQYF | KLGGALQAK | YES | NO | NO |
| 10993 | TRAV19 | TRAJ17 | CALRGEAAGNKLTF | TRBV24-1 | TRBJ2-2 | CATSDEDRGFTGELFF | KLGGALQAK | YES | NO | NO |
| 10994 | TRAV12-3 | TRAJ11 | CAMNRYSTLTF | TRBV2 | TRBJ2-7 | CASSEGSSYEQYF | RAKFKQLL | YES | NO | YES |
| 10995 | TRAV3 | TRAJ29 | CAVRDINRGNTPLVF | TRBV13 | TRBJ2-7 | CASSLGSYEQYF | KLGGALQAK | NO | NO | NO |
| 10997 | TRAV19 | TRAJ54 | CALITIQGAQKLVF | TRBV7-9 | TRBJ1-1 | CASSFLFHTEAFF | RLRAEAQVK | YES | YES | YES |
| 10998 | TRAV35 | TRAJ49 | CAGQDTGNQFYF | TRBV27 | TRBJ1-6 | CASSLWGISPLHF | RLRAEAQVK | YES | NO | YES |
| 10999 | TRAV27 | TRAJ49 | CAVYSLGQFYF | TRBV29-1 | TRBJ1-1 | CSAATGNTEAFF | KLGGALQAK | NO | NO | NO |
| 11000 | TRAV24 | TRAJ53 | CAFGGGSNYKLTF | TRBV5-1 | TRBJ2-2 | CASSRTVWSITGELFF | AVFDRKSDAK | YES | YES | NO |
| 11001 | TRAV8-2 | TRAJ52 | CVVRGGRAGGTSYGKLTF | TRBV6-5 | TRBJ2-5 | CASSRQGSWLETQYF | KLGGALQAK | NO | NO | NO |
| 11002 | TRAV1-2 | TRAJ40 | CAATTSGTYKYIF | TRBV10-3 | TRBJ1-2 | CAISESGTGSVFDGYTF | AVFDRKSDAK | YES | YES | YES |
| 11003 | TRAV8-1 | TRAJ32 | CAVISYGGATNKLIF | TRBV5-5 | TRBJ2-2 | CASSLGGGTGELFF | KTWGQYWQV | YES | NO | YES |
| 11004 | TRAV35 | TRAJ49 | CAGHTGNQFYF | TRBV11-2 | TRBJ1-5 | CASSSNQGDNQPQHF | IVTDFSVIK | YES | NO | YES |
| 11005 | TRAV12-2 | TRAJ48 | CAVTYRLSGLTF | TRBV4-1 | TRBJ2-2 | CASSPTQGATGELFF | IVTDFSVIK | YES | NO | NO |
| 11006 | TRAV19 | TRAJ30 | CALSEARDDKIIF | TRBV7-9 | TRBJ2-1 | CASSSEGGGEQFF | KLGGALQAK | YES | NO | NO |
| 11007 | TRAV19 | TRAJ54 | CALRKTQGAQKLVF | TRBV11-2 | TRBJ1-2 | CASSWGGGSHYGYTF | IVTDFSVIK | YES | NO | YES |
| 11008 | TRAV14DV4 | TRAJ57 | CATQGGSEKLVF | TRBV13 | TRBJ2-3 | CASSPAVGFLTDTQYF | KLGGALQAK | YES | NO | NO |
| 11009 | TRAV13-2 | TRAJ29 | CAEISGNTPLVF | TRBV11-2 | TRBJ2-6 | CASSAGAAGANVLTF | KLGGALQAK | YES | NO | NO |
| 11010 | TRAV23DV6 | TRAJ27 | CAAYRMGKSTF | TRBV14 | TRBJ2-1 | CASSTRDRGWTSGYNEQFF | KLGGALQAK | YES | NO | NO |
| 11011 | TRAV12-2 | TRAJ39 | CAVNNTGNMLTF | TRBV6-5 | TRBJ2-1 | CASKNSQGTGNEQFF | ELAGIGILTV | NO | NO | NO |
| 11012 | TRAV23DV6 | TRAJ42 | CAADNYGGSQGNLIF | TRBV19 | TRBJ2-3 | CASSIRSTDTQYF | GILGFVFTL | YES | NO | YES |
| 11013 | TRAV25 | TRAJ42 | CAGNYGGSQGNLIF | TRBV19 | TRBJ1-4 | CASSIRSDEKLFF | GILGFVFTL | YES | NO | YES |
| 11014 | TRAV27 | TRAJ43 | CAGVWNDMRF | TRBV7-2 | TRBJ1-1 | CASSLVLIGQGYTEAFF | KLGGALQAK | NO | NO | NO |

| | | | | | | | | | | |
|---|---|---|---|---|---|---|---|---|---|---|
| 11015 | TRAV21 | TRAJ57 | CAVFSFQGGSEKLVF | TRBV20-1 | TRBJ1-1 | CSASPRVNVHTEAFF | AVFDRKSDAK | YES | NO | YES |
| 11017 | TRAV17 | TRAJ57 | CATDAQGGSEKLVF | TRBV12-4 | TRBJ2-7 | CASSLALGTGRYSSYEQYF | KLGGALQAK | YES | NO | NO |
| 11018 | TRAV1-2 | TRAJ3 | CAGVYSSASKIIF | TRBV6-4 | TRBJ2-1 | CASSDYGLAGEQFF | KLGGALQAK | YES | NO | NO |
| 11019 | TRAV12-1 | TRAJ43 | CVANNDMRF | TRBV7-2 | TRBJ2-3 | CASSLAGDKSLTDTQYF | KLGGALQAK | NO | NO | NO |
| 11020 | TRAV12-2 | TRAJ35 | CAVSI | TRBV27 | TRBJ2-1 | CASSGRAGTGVQFF | ELAGIGILTV | NO | NO | NO |
| 11021 | TRAV9-2 | TRAJ22 | CALKGSGSARQLTF | TRBV18 | TRBJ2-5 | CASSPLTAGGVF | RLRAEAQVK | YES | NO | NO |
| 11022 | TRAV14DV4 | TRAJ53 | CAMILASNYKLTF | TRBV12-4 | TRBJ2-5 | CASSRSGGLQETQYF | KLGGALQAK | YES | NO | NO |
| 11023 | TRAV8-3 | TRAJ8 | CAVGAGTGFQKLVF | TRBV6-3 | TRBJ2-7 | CASSNHLWGRNEQYF | KLGGALQAK | YES | NO | NO |
| 11025 | TRAV13-1 | TRAJ50 | CAASETSYDKVIF | TRBV7-2 | TRBJ2-3 | CASSLLAGVSADTQYF | RAKFKQLL | YES | NO | YES |
| 11026 | TRAV19 | TRAJ32 | CALSEGGGATNKLIF | TRBV25-1 | TRBJ2-1 | CASSPGVGTYNEQFF | KLGGALQAK | YES | NO | NO |
| 11027 | TRAV12-3 | TRAJ56 | CAMTLTGANSKLTF | TRBV14 | TRBJ1-3 | CASSQGSFSGNTIYF | IVTDFSVIK | NO | NO | NO |
| 11028 | TRAV24 | TRAJ37 | CASAGSGNTGKLIF | TRBV19 | TRBJ1-1 | CASSTRSGTEAFF | GILGFVFTL | YES | NO | YES |
| 11029 | TRAV8-4 | TRAJ29 | CAISGNTPLVF | TRBV6-3 | TRBJ1-6 | CASSSLGQRIPNSPLHF | KLGGALQAK | YES | NO | NO |
| 11030 | TRAV29DV5 | TRAJ28 | CAASSAGAGSYQLTF | TRBV27 | TRBJ2-1 | CASSLDSYNEQFF | KLGGALQAK | YES | NO | NO |
| 11031 | TRAV12-2 | TRAJ45 | CAVNGGGADGLTF | TRBV6-3 | TRBJ2-1 | CASSPGVLASNEQFF | ELAGIGILTV | YES | NO | YES |
| 11032 | TRAV27 | TRAJ48 | CAGGPLNFGNEKLTF | TRBV15 | TRBJ2-7 | CATSSLRQGTYEQYF | KLGGALQAK | YES | NO | NO |
| 11033 | TRAV12-2 | TRAJ39 | CAAGGAGNMLTF | TRBV6-5 | TRBJ1-3 | CASSLGEGNTIYF | ELAGIGILTV | YES | YES | YES |
| 11034 | TRAV29DV5 | TRAJ50 | CAATGTSYDKVIF | TRBV19 | TRBJ1-5 | CASSYHSNQPQHF | GILGFVFTL | YES | YES | YES |
| 11035 | TRAV4 | TRAJ47 | CLGSEYGNKLVF | TRBV6-6 | TRBJ1-2 | CASSPDSGIGGNYGYTF | KLGGALQAK | NO | NO | NO |
| 11036 | TRAV19 | TRAJ28 | CALGDSGAGSYQLTF | TRBV11-2 | TRBJ1-2 | CASSWGGGSHYGYTF | IVTDFSVIK | YES | YES | YES |
| 11037 | TRAV20 | TRAJ52 | CAVPPGEGGTSYGKLTF | TRBV5-1 | TRBJ2-7 | CASSLFASGLAGYEQYF | RLRAEAQVK | YES | NO | YES |
| 11038 | TRAV12-3 | TRAJ5 | CAMKTGRRALTF | TRBV27 | TRBJ2-3 | CASSLSIDSTDTQYF | KLGGALQAK | YES | NO | NO |
| 11039 | TRAV38-2DV8 | TRAJ34 | CAYNTDKLIF | TRBV4-1 | TRBJ2-1 | CASSISRVYNEQFF | RAKFKQLL | YES | NO | YES |
| 11040 | TRAV8-1 | TRAJ52 | CAVNAGGTSYGKLTF | TRBV20-1 | TRBJ2-7 | CSADGGPTTPNEQYF | KLGGALQAK | NO | NO | NO |
| 11041 | TRAV29DV5 | TRAJ39 | CAASGYNAGNMLTF | TRBV21-1 | TRBJ1-5 | CASSKGLGGDQPQHF | AVFDRKSDAK | YES | NO | YES |
| 11043 | TRAV3 | TRAJ3 | CAVRDSSSASKIIF | TRBV27 | TRBJ2-3 | CASTETRSDADTQYF | KLGGALQAK | NO | NO | NO |
| 11044 | TRAV12-2 | TRAJ31 | CAVLNARLMF | TRBV6-5 | TRBJ2-5 | CASSSGGARETQYF | ELAGIGILTV | NO | NO | NO |
| 11045 | TRAV12-2 | TRAJ33 | CAVNRDYQLIW | TRBV20-1 | TRBJ2-1 | CSARPSLAGGYNEQFF | AVFDRKSDAK | YES | NO | YES |
| 11046 | TRAV12-3 | TRAJ11 | CAMGSGYSTLTF | TRBV6-1 | TRBJ1-1 | CASSPNGLNTEAFF | KLGGALQAK | YES | NO | NO |
| 11047 | TRAV38-2DV8 | TRAJ52 | CAYGTNAGGTSYGKLTF | TRBV19 | TRBJ1-2 | CASSIGVYGYTF | GILGFVFTL | YES | NO | YES |
| 11048 | TRAV21 | TRAJ16 | CAVF | TRBV13 | TRBJ1-2 | CASSLAGFPDGYTF | AVFDRKSDAK | NO | NO | NO |
| 11049 | TRAV25 | TRAJ24 | CAGATDSWGKLQF | TRBV11-2 | TRBJ1-2 | CASSWGGGSHYGYTF | IVTDFSVIK | YES | YES | YES |
| 11050 | TRAV13-1 | TRAJ27 | CAATNTNAGKSTF | TRBV7-6 | TRBJ2-1 | CASSPTTGEDNEQFF | RAKFKQLL | YES | YES | NO |
| 11051 | TRAV23DV6 | TRAJ27 | CAAKGSGAGKSTF | TRBV7-2 | TRBJ2-1 | CASSLGILAGGLSDSYNEQFF | KLGGALQAK | NO | NO | NO |
| 11052 | TRAV19 | TRAJ36 | CALSEAPGANNLFF | TRBV4-3 | TRBJ1-2 | CASSQEGESNYGYTF | KLGGALQAK | YES | NO | NO |
| 11053 | TRAV19 | TRAJ27 | CALTSNTNAGKSTF | TRBV5-4 | TRBJ2-6 | CASSLRDRLITGANVLTF | KLGGALQAK | YES | NO | NO |
| 11054 | TRAV38-2DV8 | TRAJ43 | CGFYNNNDMRF | TRBV6-3 | TRBJ2-3 | CATGTDTQYF | KLGGALQAK | YES | NO | NO |
| 11055 | TRAV38-2DV8 | TRAJ52 | CAYWGGTSYGKLTF | TRBV20-1 | TRBJ2-1 | CSASGYIVNEQFF | KLGGALQAK | YES | NO | NO |
| 11056 | TRAV1-2 | TRAJ4 | CAVRSFSGGYNKLIF | TRBV7-2 | TRBJ2-7 | CASSLAGDSYEQYF | GILGFVFTL | YES | YES | YES |
| 11057 | TRAV12-2 | TRAJ24 | CAVTTDSWGKLQF | TRBV20-1 | TRBJ2-2 | CSARPRTSGSGAGELFF | ELAGIGILTV | YES | YES | YES |
| 11058 | TRAV12-2 | TRAJ13 | CAVKTGGYQKVTF | TRBV4-2 | TRBJ2-2 | CASSQDASGGAGELFF | KLGGALQAK | YES | NO | NO |
| 11060 | TRAV12-2 | TRAJ49 | CAVNGNQFYF | TRBV4-1 | TRBJ2-7 | CASSQEEESLGYEQYF | ELAGIGILTV | YES | YES | NO |
| 11061 | TRAV14DV4 | TRAJ45 | CAMREGRKSGGGADGLTF | TRBV11-2 | TRBJ2-7 | CASSLQVIYEQYF | KLGGALQAK | YES | NO | NO |
| 11062 | TRAV29DV5 | TRAJ43 | CAASASNNNDMRF | TRBV19 | TRBJ2-1 | CASSMVSQGEQFF | GILGFVFTL | YES | NO | YES |
| 11063 | TRAV16 | TRAJ54 | CALRYQGAQKLVF | TRBV27 | TRBJ1-1 | CASSLSGLGTEAFF | KLGGALQAK | NO | NO | NO |
| 11064 | TRAV10 | TRAJ4 | CVVDESGGYNKLIF | TRBV4-3 | TRBJ2-7 | CASSQDSAGGREQYF | AVFDRKSDAK | YES | NO | YES |
| 11065 | TRAV38-2DV8 | TRAJ48 | CAYVSNFGNEKLTF | TRBV12-5 | TRBJ2-1 | CASGWNLDEQFF | KLGGALQAK | YES | NO | NO |
| 11066 | TRAV16 | TRAJ58 | CALSEETSGSRLTF | TRBV2 | TRBJ2-2 | CASTTLWGGIGAGELFF | KLGGALQAK | NO | NO | NO |
| 11067 | TRAV17 | TRAJ20 | CATVSDYKLSF | TRBV5-1 | TRBJ2-7 | CASGDSPNEQYF | KLGGALQAK | YES | NO | NO |
| 11068 | TRAV5 | TRAJ13 | CAESIGSYQKVTF | TRBV3-1 | TRBJ1-2 | CASSQDGRTYGYTF | IVTDFSVIK | YES | YES | YES |
| 11069 | TRAV12-2 | TRAJ47 | CAAYSMEYGNKLVF | TRBV12-4 | TRBJ1-3 | CASSLGTTGNTIYF | KLGGALQAK | YES | NO | NO |
| 11070 | TRAV14DV4 | TRAJ5 | CAMREVGRRALTF | TRBV11-2 | TRBJ2-2 | CASSLGVAGTNTGELFF | KLGGALQAK | YES | NO | NO |
| 11071 | TRAV27 | TRAJ37 | CAGASGSSNTGKLIF | TRBV19 | TRBJ2-7 | CASSIRAAYEQYF | GILGFVFTL | YES | YES | YES |
| 11072 | TRAV8-4 | TRAJ17 | CAVRRGAAGNKLTF | TRBV7-2 | TRBJ1-5 | CASSVLGLGNQPQHF | KLGGALQAK | YES | NO | NO |
| 11073 | TRAV1-2 | TRAJ17 | CAVEAAGNKLTF | TRBV4-2 | TRBJ2-6 | CASSDGGSNSGANVLTF | AVFDRKSDAK | YES | YES | YES |
| 11074 | TRAV2 | TRAJ42 | CAFTGGSQGNLIF | TRBV29-1 | TRBJ1-3 | CSVMDPMSGYTIYF | ELAGIGILTV | NO | NO | NO |
| 11076 | TRAV12-2 | TRAJ33 | CAVINYQLIW | TRBV7-9 | TRBJ2-1 | CASSLDRDASYNEQFF | ELAGIGILTV | YES | NO | YES |
| 11077 | TRAV13-2 | TRAJ42 | CAERGGGGSQGNLIF | TRBV19 | TRBJ2-5 | CASSIRASETQYF | GILGFVFTL | YES | NO | YES |
| 11078 | TRAV22 | TRAJ53 | CAVATMGSGGSNYKLTF | TRBV7-2 | TRBJ2-2 | CASSFVGGGTGAGELFF | KLGGALQAK | YES | NO | NO |
| 11079 | TRAV1-2 | TRAJ11 | CAVGLVGYSTLTF | TRBV28 | TRBJ2-6 | CASRPGAGGNVLTF | KLGGALQAK | YES | NO | NO |
| 11080 | TRAV17 | TRAJ58 | CATDVGGETSGSRLTF | TRBV2 | TRBJ1-1 | CARYRHLGEAFF | KLGGALQAK | YES | NO | NO |
| 11081 | TRAV38-1 | TRAJ52 | CAFWAGGTSYGKLTF | TRBV11-2 | TRBJ2-5 | CASSFPGLAMETQYF | AVFDRKSDAK | YES | NO | YES |
| 11082 | TRAV22 | TRAJ30 | CAVGNRDDKIIF | TRBV9 | TRBJ1-6 | CASSVMLDSPLHF | KLGGALQAK | NO | NO | NO |
| 11083 | TRAV12-2 | TRAJ21 | CAVPYYNFNKFYF | TRBV27 | TRBJ2-2 | CASSLSRKGELFF | AVFDRKSDAK | YES | NO | YES |
| 11084 | TRAV38-1 | TRAJ45 | CAFRAMGGADGLTF | TRBV12-5 | TRBJ2-1 | CASGLVLSWGSRTSGRLDEQFF | KLGGALQAK | NO | NO | NO |
| 11085 | TRAV38-2DV8 | TRAJ53 | CSNYKLTF | TRBV7-9 | TRBJ2-7 | CASSWDRVLYEQYF | SLLMWITQV | NO | NO | NO |
| 11086 | TRAV8-4 | TRAJ48 | CAVSDISNFGNEKLTF | TRBV11-2 | TRBJ1-1 | CASSLGDRNTEAFF | LLFGYPVYV | YES | YES | YES |
| 11087 | TRAV19 | TRAJ17 | CALSEAQAAGNKLTF | TRBV12-3 | TRBJ1-4 | CASSLAGWKLFF | KLGGALQAK | YES | NO | NO |
| 11088 | TRAV10 | TRAJ8 | CVVSENTGFQKLVF | TRBV7-6 | TRBJ2-7 | CASSLGGGDYEQYF | RLRAEAQVK | YES | NO | YES |
| 11089 | TRAV14DV4 | TRAJ33 | CAMREGQDSNYQLIW | TRBV3-1 | TRBJ1-4 | CASSQERGLPNEKLFF | IVTDFSVIK | YES | YES | YES |
| 11090 | TRAV25 | TRAJ41 | CAGHLSGYALNF | TRBV6-3 | TRBJ2-1 | CATSTGTTSVNEQFF | KLGGALQAK | YES | NO | NO |
| 11091 | TRAV35 | TRAJ58 | CAGQLKRETSGSRLTF | TRBV7-2 | TRBJ1-5 | CASSAYALYSNQPQHF | KLGGALQAK | YES | NO | NO |
| 11092 | TRAV4 | TRAJ11 | CLLNSGYSTLTF | TRBV4-2 | TRBJ1-1 | CASSQDQGQDTEAFF | KLGGALQAK | YES | NO | NO |
| 11093 | TRAV30 | TRAJ37 | CGTEDMKSSGNTGKLIF | TRBV7-2 | TRBJ2-1 | CASSLSAGGYNEQFF | IVTDFSVIK | YES | NO | YES |
| 11094 | TRAV8-6 | TRAJ42 | CAVSGSQGNLIF | TRBV19 | TRBJ2-7 | CASSIRSSYEQYF | GILGFVFTL | YES | NO | YES |
| 11095 | TRAV12-2 | TRAJ52 | CAAGARAGGTSYGKLTF | TRBV12-3 | TRBJ2-7 | CASSGGLAGAPYEQYF | KLGGALQAK | YES | NO | NO |
| 11096 | TRAV36DV7 | TRAJ31 | CAVALWARLMF | TRBV11-2 | TRBJ1-2 | CASSWGGGSHYGYTF | IVTDFSVIK | NO | NO | NO |
| 11097 | TRAV21 | TRAJ23 | CAVLYNQGGKLIF | TRBV12-5 | TRBJ2-3 | CASGFGANTWDTQYF | KLGGALQAK | YES | NO | NO |
| 11098 | TRAV12-2 | TRAJ10 | CAVATGGGNKLTF | TRBV13 | TRBJ2-1 | CASSLLSGTSAYNEQFF | KLGGALQAK | YES | NO | NO |
| 11099 | TRAV13-1 | TRAJ47 | CAASIRPMEYGNKLVF | TRBV19 | TRBJ1-3 | CASSTTGTPGNTIYF | GILGFVFTL | YES | NO | YES |
| 11100 | TRAV25 | TRAJ31 | CAGPISARLMF | TRBV7-9 | TRBJ2-1 | CASSTTGFSYNEQFF | KLGGALQAK | YES | NO | NO |
| 11101 | TRAV27 | TRAJ26 | CAGAEYGQNFVF | TRBV7-9 | TRBJ1-1 | CASREGGGGTEAFF | KLGGALQAK | YES | NO | NO |
| 11102 | TRAV3 | TRAJ43 | CAVIRWGDMRF | TRBV11-2 | TRBJ1-2 | CASSWGGGSHYGYTF | IVTDFSVIK | YES | YES | YES |
| 11103 | TRAV8-1 | TRAJ43 | CAVNGFDMRF | TRBV14 | TRBJ2-3 | CASSQDMGWGTDTQYF | FLASKIGRLV | YES | NO | YES |
| 11104 | TRAV1-1 | TRAJ9 | CAVKGTGGFKTIF | TRBV28 | TRBJ1-5 | CASRSQGALYSNQPQHF | KLGGALQAK | YES | NO | NO |
| 11105 | TRAV1-1 | TRAJ29 | CAVSALENTPLVF | TRBV7-2 | TRBJ2-6 | CASSPGLKLSGANVLTF | KLGGALQAK | YES | NO | NO |
| 11106 | TRAV25 | TRAJ42 | CAGNGGGSQGNLIF | TRBV19 | TRBJ2-2 | CASSGRSTGELFF | GILGFVFTL | YES | NO | YES |
| 11107 | TRAV5 | TRAJ42 | CADLGSQGNLIF | TRBV6-1 | TRBJ2-7 | CASSEGWHSYEQYF | KLGGALQAK | YES | NO | NO |
| 11108 | TRAV14DV4 | TRAJ28 | CAMRTSGAGSYQLTF | TRBV7-9 | TRBJ1-6 | CASSPGQGNSPLHF | KLGGALQAK | YES | NO | NO |
| 11109 | TRAV12-2 | TRAJ5 | CATGGTGRRALTF | TRBV4-2 | TRBJ1-2 | CASSQEGGDGYTF | ELAGIGILTV | YES | NO | YES |
| 11110 | TRAV21 | TRAJ54 | CAVYQGAQKLVF | TRBV6-3 | TRBJ1-5 | CASSYTADQPQHF | KLGGALQAK | YES | NO | NO |
| 11111 | TRAV12-2 | TRAJ31 | CAVNGNARLMF | TRBV20-1 | TRBJ1-1 | CSLIQGLGTEAFF | ELAGIGILTV | YES | NO | YES |
| 11112 | TRAV1-2 | TRAJ33 | CAVLDSNYQLIW | TRBV20-1 | TRBJ2-1 | CSAREIRGEGIHEQFF | KLGGALQAK | NO | NO | NO |
| 11113 | TRAV12-2 | TRAJ43 | CAVGIMWDDMRF | TRBV11-2 | TRBJ2-7 | CASSAPGTGHYEQYF | ELAGIGILTV | YES | YES | YES |
| 11114 | TRAV19 | TRAJ43 | CALSNNDMRF | TRBV27 | TRBJ2-7 | CASSPARGRVEQYF | KLGGALQAK | YES | NO | NO |
| 11115 | TRAV12-3 | TRAJ49 | CAMSTNTGNQFYF | TRBV6-5 | TRBJ2-2 | CASSYFSPLAVNTGELFF | KLGGALQAK | YES | NO | NO |
| 11116 | TRAV12-3 | TRAJ49 | CAMSYTGNQFYF | TRBV7-6 | TRBJ2-1 | CASSPGGDEQFF | KLGGALQAK | YES | NO | NO |
| 11117 | TRAV10 | TRAJ33 | CVVSSDSNYQLIW | TRBV10-3 | TRBJ1-6 | CAISGAGLRYNSPLHF | MLDLQPETT | YES | NO | YES |
| 11118 | TRAV1-2 | TRAJ34 | CAVEYTDKLIF | TRBV7-2 | TRBJ2-7 | CASSSEGRMTYEQYF | KLGGALQAK | YES | NO | NO |
| 11119 | TRAV12-2 | TRAJ35 | CAASI | TRBV27 | TRBJ2-3 | CASSLGTDTQYF | ELAGIGILTV | NO | NO | NO |
| 11120 | TRAV27 | TRAJ37 | CAGADDTGKLIF | TRBV19 | TRBJ2-7 | CASSIRSSYEQYF | GILGFVFTL | YES | NO | NO |
| 11121 | TRAV8-4 | TRAJ47 | CAVSTEYGNKLVF | TRBV6-1 | TRBJ2-7 | CASSPWTGARWEQYF | AVFDRKSDAK | YES | NO | YES |
| 11122 | TRAV4 | TRAJ11 | CLVGDMNSGYSTLTF | TRBV11-2 | TRBJ2-1 | CASSLDLIPAGGSTPYNEQFF | KLGGALQAK | NO | NO | NO |
| 11123 | TRAV13-2 | TRAJ52 | CAENIVGAGGTSYGKLTF | TRBV27 | TRBJ2-3 | CASSPRGWNGLSTDTQYF | RLRAEAQVK | YES | YES | YES |

| | | | | | | | | | | |
|---|---|---|---|---|---|---|---|---|---|---|
| 11124 | TRAV1-2 | TRAJ33 | CAVSDSNYQLIW | TRBV4-2 | TRBJ1-2 | CASSQDTPDSNYGYTF | KLGGALQAK | YES | NO | NO |
| 11125 | TRAV8-3 | TRAJ28 | CAVVSGAGSYQLTF | TRBV30 | TRBJ2-2 | CAWSITDTGELFF | KLGGALQAK | YES | NO | NO |
| 11126 | TRAV41 | TRAJ58 | CAVKETSGSRLTF | TRBV12-4 | TRBJ2-6 | CASSLGGSGANVLTF | KLGGALQAK | YES | NO | NO |
| 11127 | TRAV12-2 | TRAJ26 | CAVPFGNYGQNFVF | TRBV28 | TRBJ1-5 | CASSPTGVGQPQHF | ELAGIGILTV | YES | NO | YES |
| 11128 | TRAV1-2 | TRAJ17 | CAVGMIKAAGNKLTF | TRBV20-1 | TRBJ2-3 | CSALGGTQYF | GILGFVFTL | NO | NO | NO |
| 11129 | TRAV14DV4 | TRAJ23 | CAMREGHQGGKLIF | TRBV27 | TRBJ1-5 | CASSLGTGENQPQHF | KLGGALQAK | NO | NO | NO |
| 11130 | TRAV12-1 | TRAJ12 | CVVNGRWMDSSYKLIF | TRBV12-4 | TRBJ2-2 | CASSLSTDGPISNTGELFF | KLGGALQAK | YES | NO | NO |
| 11131 | TRAV1-2 | TRAJ26 | CAVRDNYGQNFVF | TRBV5-4 | TRBJ1-1 | CASSLGGTEAFF | KLGGALQAK | YES | NO | NO |
| 11132 | TRAV26-2 | TRAJ58 | CILRDSEETSGSRLTF | TRBV4-1 | TRBJ2-2 | CASSRTGGINTGELFF | KLGGALQAK | YES | NO | NO |
| 11133 | TRAV3 | TRAJ23 | CAVASYNQGGKLIF | TRBV18 | TRBJ2-3 | CASSPSLYLTDTQYF | KLGGALQAK | NO | NO | NO |
| 11134 | TRAV3 | TRAJ8 | CAVRDGVNTGFQKLVF | TRBV2 | TRBJ1-1 | CASSEGVGLQGWNTEAFF | KLGGALQAK | NO | NO | NO |
| 11135 | TRAV14DV4 | TRAJ45 | CAMTKRGGGGADGLTF | TRBV27 | TRBJ2-7 | CASSLSPSGSYEQYF | AVFDRKSDAK | YES | NO | YES |
| 11136 | TRAV14DV4 | TRAJ3 | CAMRGIYSSASKIIF | TRBV2 | TRBJ2-1 | CASSEASGGFYNEQFF | IVTDFSVIK | YES | YES | YES |
| 11137 | TRAV13-1 | TRAJ6 | CAAFGGSYIPTF | TRBV7-3 | TRBJ2-3 | CASSLTPGTDTQYF | AVFDRKSDAK | YES | NO | YES |
| 11138 | TRAV24 | TRAJ44 | CAHDTGTASKLTF | TRBV6-3 | TRBJ2-7 | CASSYSDETYEQYF | KLGGALQAK | YES | NO | NO |
| 11139 | TRAV5 | TRAJ31 | CAERGEWARLMF | TRBV19 | TRBJ1-5 | CASSIDDGTLINQPQHF | RLRAEAQVK | YES | NO | YES |
| 11140 | TRAV13-2 | TRAJ45 | CADSGGGADGLTF | TRBV19 | TRBJ2-7 | CASSAQQGLSSYEQYF | RAKFKQLL | YES | NO | NO |
| 11141 | TRAV12-2 | TRAJ17 | CAVKAAGNKLTF | TRBV11-2 | TRBJ1-2 | CASSWGGGSHYGYTF | IVTDFSVIK | YES | NO | YES |
| 11142 | TRAV13-2 | TRAJ42 | CAEIALQNYGGSQGNLIF | TRBV4-1 | TRBJ1-2 | CASSQEGGLDYGYTF | GILGFVFTL | YES | NO | YES |
| 11143 | TRAV12-2 | TRAJ30 | CAVNIPSRDDKIIF | TRBV12-3 | TRBJ2-7 | CASNMGPYEQYF | AVFDRKSDAK | YES | NO | YES |
| 11144 | TRAV8-1 | TRAJ42 | CAVNAGGGSQGNLIF | TRBV19 | TRBJ2-7 | CASSIRAAYEQYF | GILGFVFTL | YES | NO | YES |
| 11145 | TRAV12-2 | TRAJ41 | CAVNPSGYALNF | TRBV12-4 | TRBJ1-5 | CASSLGGLGQPQHF | ELAGIGILTV | YES | YES | YES |
| 11146 | TRAV17 | TRAJ11 | CATQRGYSTLTF | TRBV11-2 | TRBJ2-3 | CASSPPWAAPRTQYF | KLGGALQAK | YES | NO | NO |
| 11147 | TRAV17 | TRAJ43 | CATDAKNNNDMRF | TRBV19 | TRBJ1-1 | CATRHLNTEAFF | KLGGALQAK | YES | NO | NO |
| 11149 | TRAV12-2 | TRAJ34 | CAVNLNTDKLIF | TRBV28 | TRBJ1-3 | CASSSQGLGNTIYF | ELAGIGILTV | YES | NO | YES |
| 11150 | TRAV23DV6 | TRAJ15 | CAATNQAGTALIF | TRBV4-1 | TRBJ2-3 | CASSPDLDTQYF | KLGGALQAK | YES | NO | NO |
| 11151 | TRAV9-2 | TRAJ58 | CALTRETSGSRLTF | TRBV4-1 | TRBJ2-3 | CASSQGRVLPSRTDTQYF | KLGGALQAK | YES | NO | NO |
| 11152 | TRAV12-2 | TRAJ23 | CAGPYNQGGKLIF | TRBV19 | TRBJ2-5 | CASSIRSTETQYF | GILGFVFTL | YES | NO | YES |
| 11153 | TRAV35 | TRAJ49 | CGCHTGNQFYF | TRBV12-3 | TRBJ2-2 | CASSFSGNTGELFF | IVTDFSVIK | YES | NO | NO |
| 11154 | TRAV12-2 | TRAJ23 | CAVGGGKLIF | TRBV6-3 | TRBJ1-5 | CASNKGMNQPQHF | ELAGIGILTV | YES | NO | YES |
| 11155 | TRAV27 | TRAJ42 | CAGDYGGSQGNLIF | TRBV20-1 | TRBJ2-3 | CSALPPLAGGHTDTQYF | KLGGALQAK | NO | NO | NO |
| 11156 | TRAV27 | TRAJ42 | CAGGGSQGNLIF | TRBV20-1 | TRBJ2-7 | CSARDLTAGSYEQYF | GILGFVFTL | YES | NO | YES |
| 11157 | TRAV39 | TRAJ40 | CAVDIMGTYKYIF | TRBV9 | TRBJ2-2 | CASRGLAGEAGELFF | KLGGALQAK | YES | NO | NO |
| 11158 | TRAV27 | TRAJ58 | CAGEKETSGSRLTF | TRBV19 | TRBJ2-7 | CASSIRSSYEQYF | GILGFVFTL | YES | NO | YES |
| 11159 | TRAV12-2 | TRAJ23 | CAVNQGGKLIF | TRBV12-4 | TRBJ1-3 | CASGDSGNTIYF | ELAGIGILTV | YES | NO | YES |
| 11161 | TRAV29DV5 | TRAJ45 | CAASAGGGADGLTF | TRBV4-1 | TRBJ2-1 | CASSQRRGRHNEQFF | KLGGALQAK | YES | NO | NO |
| 11162 | TRAV29DV5 | TRAJ42 | CAATGGSQGNLIF | TRBV18 | TRBJ1-2 | CASSPRDRTGGYTF | MLDLQPETT | YES | NO | YES |
| 11163 | TRAV8-2 | TRAJ29 | CVVSAHWWNTPLVF | TRBV10-3 | TRBJ1-1 | CAIKVTENTEAFF | LLDFVRFMGV | YES | NO | YES |
| 11164 | TRAV12-2 | TRAJ13 | CAVNRGYQKVTF | TRBV10-3 | TRBJ1-1 | CAISESVNTEAFF | QPRAPIRPI | YES | NO | YES |
| 11165 | TRAV22 | TRAJ52 | CVMCNAGGTSYGKLTF | TRBV10-3 | TRBJ1-6 | CAISVRSRSPLHF | KLGGALQAK | YES | NO | NO |
| 11166 | TRAV13-2 | TRAJ30 | CAEMLDDKIIF | TRBV19 | TRBJ1-1 | CATQGLGTEAFF | KLGGALQAK | YES | NO | NO |
| 11167 | TRAV29DV5 | TRAJ20 | CAASDVDYKLSF | TRBV19 | TRBJ2-7 | CASSIRSSYEQYF | GILGFVFTL | YES | NO | YES |
| 11168 | TRAV1-2 | TRAJ28 | CAVRDEYSGAGSYQLTF | TRBV6-1 | TRBJ2-7 | CASSDLGQGHEQYF | KLGGALQAK | YES | NO | NO |
| 11169 | TRAV27 | TRAJ37 | CAGAAGSSNTGKLIF | TRBV11-2 | TRBJ1-2 | CASSWGGGSHYGYTF | IVTDFSVIK | YES | NO | YES |
| 11170 | TRAV17 | TRAJ56 | CATDGGTGANSKLTF | TRBV7-2 | TRBJ2-1 | CASSSPCCSHSYNEQFF | KLGGALQAK | YES | NO | NO |
| 11171 | TRAV21 | TRAJ33 | CAVRMDSNYQLIW | TRBV4-3 | TRBJ1-1 | CASSQGAETGSTEAFF | KLGGALQAK | YES | NO | NO |
| 11172 | TRAV14DV4 | TRAJ11 | CAWFPMNSGYSTLTF | TRBV7-9 | TRBJ2-1 | CASSLGARLAGYNEQFF | KLGGALQAK | YES | NO | NO |
| 11173 | TRAV9-2 | TRAJ33 | CALSFMDSNYQLIW | TRBV7-9 | TRBJ1-1 | CASSRHGQGNTEAFF | AVFDRKSDAK | YES | NO | YES |
| 11174 | TRAV17 | TRAJ20 | CATDANDYKLSF | TRBV5-1 | TRBJ2-5 | CASSIGQGETQYF | FLASKIGRLV | YES | NO | YES |
| 11175 | TRAV8-3 | TRAJ20 | CAVVANDYKLSF | TRBV12-3 | TRBJ2-1 | CASSVGPGNEQFF | RAKFKQLL | YES | NO | YES |
| 11176 | TRAV24 | TRAJ49 | CAFAGNQFYF | TRBV15 | TRBJ1-2 | CATSRDKGADGYTF | IVTDFSVIK | YES | NO | YES |
| 11177 | TRAV13-2 | TRAJ11 | CAETNQLYSTLTF | TRBV30 | TRBJ2-1 | CAWSDEASYNEQFF | KLGGALQAK | YES | NO | NO |
| 11178 | TRAV2 | TRAJ24 | CAVPMTTDSWGKLQF | TRBV7-9 | TRBJ2-1 | CASSFHRRPFDGYNEQFF | KLGGALQAK | NO | NO | NO |
| 11179 | TRAV14DV4 | TRAJ6 | CAMGPSGGSYIPTF | TRBV11-2 | TRBJ1-2 | CASSWGGGSHYGYTF | IVTDFSVIK | YES | NO | YES |
| 11181 | TRAV4 | TRAJ10 | CLVGDVVTGGGNKLTF | TRBV4-3 | TRBJ1-2 | CASSQETGYGYTF | KLGGALQAK | YES | NO | NO |
| 11182 | TRAV9-2 | TRAJ11 | CALRPNSGYSTLTF | TRBV7-2 | TRBJ1-5 | CASSSGTGWLGSNQPQHF | KLGGALQAK | NO | NO | NO |
| 11183 | TRAV41 | TRAJ22 | CAVRLLLLSGSARQLTF | TRBV20-1 | TRBJ2-7 | CSAHRQGRTYEQYF | RLRAEAQVK | NO | NO | NO |
| 11184 | TRAV14DV4 | TRAJ8 | CAMRVNTGFQKLVF | TRBV28 | TRBJ2-3 | CASTGPRSKTDTQYF | KLGGALQAK | YES | NO | NO |
| 11185 | TRAV21 | TRAJ45 | CAGHSGGGADGLTF | TRBV19 | TRBJ1-1 | CASRLVSMNTEAFF | KLGGALQAK | YES | NO | NO |
| 11186 | TRAV8-3 | TRAJ21 | CAVGDNFNKFYF | TRBV10-1 | TRBJ2-7 | CASSRDASSYEQYF | AVFDRKSDAK | YES | NO | YES |
| 11187 | TRAV19 | TRAJ53 | CALSEARSGGSNYKLTF | TRBV19 | TRBJ2-1 | CASSMGQGSYNEQFF | KLGGALQAK | YES | NO | NO |
| 11188 | TRAV12-1 | TRAJ13 | CVVGMNSGGYQKVTF | TRBV7-9 | TRBJ1-1 | CASSPRATEAFF | KLGGALQAK | YES | NO | NO |
| 11189 | TRAV12-2 | TRAJ52 | CAVYYGWAGGTSYGKLTF | TRBV6-3 | TRBJ2-1 | CASSISEYNEQFF | AYAQKIFKI | YES | YES | YES |
| 11190 | TRAV19 | TRAJ20 | CALRYGSNDYKLSF | TRBV18 | TRBJ1-4 | CASSPYTDRDTATNEKLFF | KLGGALQAK | YES | NO | NO |
| 11191 | TRAV17 | TRAJ43 | CATDWNNDMRF | TRBV5-8 | TRBJ2-1 | CASSFPGLYNEQFF | KLGGALQAK | YES | NO | NO |
| 11192 | TRAV12-2 | TRAJ42 | CAVNQGGGSQGNLIF | TRBV19 | TRBJ2-3 | CASSIRSADTQYF | GILGFVFTL | YES | NO | YES |
| 11193 | TRAV17 | TRAJ58 | CATLAETSGSRLTF | TRBV6-6 | TRBJ2-1 | CASSLGDRLPYNEQFF | KLGGALQAK | NO | NO | NO |
| 11194 | TRAV12-2 | TRAJ3 | CAARALYSSASKIIF | TRBV28 | TRBJ2-7 | CASKNPGTEIYEQYF | KLGGALQAK | YES | NO | NO |
| 11195 | TRAV22 | TRAJ50 | CAVKTSYDKVIF | TRBV20-1 | TRBJ1-2 | CSAYTGGGYTF | KLGGALQAK | YES | NO | NO |
| 11196 | TRAV27 | TRAJ49 | CAGKNTGNQFYF | TRBV7-2 | TRBJ1-2 | CASSLTSRGGNGYTF | KLGGALQAK | NO | NO | NO |
| 11197 | TRAV13-1 | TRAJ49 | CAASIWNGNQFYF | TRBV7-2 | TRBJ2-1 | CASSRWTSEDEQFF | KLGGALQAK | YES | NO | NO |
| 11198 | TRAV12-2 | TRAJ7 | CAVSEGNNRLAF | TRBV5-1 | TRBJ2-3 | CASSSGMGTDTQYF | KLGGALQAK | YES | NO | NO |
| 11199 | TRAV19 | TRAJ9 | CALSEANTGGFKTIF | TRBV11-2 | TRBJ1-2 | CASSWGGGSHYGYTF | IVTDFSVIK | YES | NO | NO |
| 11201 | TRAV24 | TRAJ53 | CAEGGSNYKLTF | TRBV5-1 | TRBJ1-1 | CASSTTGGSTEAFF | KLGGALQAK | YES | NO | NO |
| 11203 | TRAV2 | TRAJ21 | CAVEDPIYNFNKFYF | TRBV27 | TRBJ2-2 | CASSPSRPWGPHSNTGELFF | KLGGALQAK | NO | NO | NO |
| 11204 | TRAV13-2 | TRAJ42 | CAENLGGGSQGNLIF | TRBV11-2 | TRBJ2-2 | CASSLGVAGTNTGELFF | GILGFVFTL | YES | NO | YES |
| 11205 | TRAV24 | TRAJ49 | CAFTRFGGNQFYF | TRBV5-1 | TRBJ2-1 | CASSLTDRLNEQFF | KLGGALQAK | YES | NO | NO |
| 11206 | TRAV1-1 | TRAJ47 | CAVSLMEYGNKLVF | TRBV9 | TRBJ1-2 | CASSAGGNGYTF | KLGGALQAK | YES | NO | NO |
| 11207 | TRAV9-2 | TRAJ52 | CALRGGGTSYGKLTF | TRBV4-2 | TRBJ2-7 | CASSRPVTSGRHEQYF | KLGGALQAK | YES | NO | NO |
| 11208 | TRAV1-2 | TRAJ33 | CAGMDSNYQLIW | TRBV15 | TRBJ2-2 | CATSRDDPGTAPGELFF | KLGGALQAK | YES | NO | NO |
| 11209 | TRAV12-3 | TRAJ26 | CAMSGGTWGYGQNFVF | TRBV7-7 | TRBJ2-3 | CASSPGTSGGADTQYF | KLGGALQAK | YES | NO | NO |
| 11210 | TRAV38-1 | TRAJ49 | CASNTGNQFYF | TRBV9 | TRBJ1-6 | CASSGAGQENNSPLHF | SLLMWITQV | YES | NO | NO |
| 11211 | TRAV38-1 | TRAJ30 | CAFMFNRDDKIIF | TRBV3-1 | TRBJ2-5 | CASSQRRRGRETQYF | KLGGALQAK | YES | NO | NO |
| 11212 | TRAV12-2 | TRAJ28 | CAVKGPGAGSYQLTF | TRBV9 | TRBJ1-1 | CASSGGTGGTEAFF | KLGGALQAK | YES | NO | NO |
| 11213 | TRAV1-2 | TRAJ49 | CAVRGRAGNQFYF | TRBV4-1 | TRBJ2-5 | CASSQGEGRETQYF | AVFDRKSDAK | YES | NO | YES |
| 11214 | TRAV29DV5 | TRAJ47 | CAASDGNKLVF | TRBV25-1 | TRBJ2-3 | CASSSRGRSTDTQYF | KLGGALQAK | YES | NO | NO |
| 11215 | TRAV38-2DV8 | TRAJ31 | CAYWASRPNNARLMF | TRBV7-9 | TRBJ2-1 | CASSLGGPPVNEQFF | KLGGALQAK | YES | NO | NO |
| 11216 | TRAV1-1 | TRAJ20 | CAVRSDYKLSF | TRBV11-1 | TRBJ2-3 | CASSLPTDTQYF | RLRAEAQVK | YES | YES | YES |
| 11217 | TRAV21 | TRAJ45 | CAVHSGGGADGLTF | TRBV6-5 | TRBJ2-5 | CASSWLQGSEQYF | KLGGALQAK | NO | NO | NO |
| 11218 | TRAV12-1 | TRAJ11 | CVVNREGYSTLTF | TRBV2 | TRBJ1-1 | CASSESVLGLNTEAFF | GLCTLVAML | YES | YES | YES |
| 11219 | TRAV14DV4 | TRAJ21 | CAMRDVVLNFNKFYF | TRBV28 | TRBJ2-2 | CASSGLASTGELFF | KLGGALQAK | YES | NO | NO |
| 11220 | TRAV8-1 | TRAJ42 | CAVFRGGSQGNLIF | TRBV13 | TRBJ2-7 | CASSFLAGGNYEQYF | KLGGALQAK | YES | NO | NO |
| 11221 | TRAV38-2DV8 | TRAJ42 | CAYRADYGGSQGNLIF | TRBV19 | TRBJ2-5 | CASSIDRTNQETQYF | KLGGALQAK | YES | NO | NO |
| 11222 | TRAV26-1 | TRAJ32 | CIVRVSYGGATNKLIF | TRBV4-1 | TRBJ2-2 | CASSQEEAGTGELFF | KLGGALQAK | YES | NO | NO |
| 11223 | TRAV29DV5 | TRAJ57 | CAASAQGGSEKLVF | TRBV6-5 | TRBJ2-7 | CASSYSGRAHEQYF | KLGGALQAK | YES | NO | NO |
| 11224 | TRAV26-2 | TRAJ43 | CILGDKYNNNDMRF | TRBV27 | TRBJ1-2 | CASTRVGGDYGYTF | KLGGALQAK | YES | NO | NO |
| 11225 | TRAV16 | TRAJ52 | CALSSNAGGTSYGKLTF | TRBV7-2 | TRBJ2-1 | CASSFGEQFF | KLGGALQAK | NO | NO | NO |
| 11226 | TRAV25 | TRAJ42 | CAGTYGGSQGNLIF | TRBV19 | TRBJ1-5 | CASSGRSSQPQHF | GILGFVFTL | YES | NO | YES |
| 11227 | TRAV12-1 | TRAJ12 | CVVNGGDSSYKLIF | TRBV11-2 | TRBJ1-2 | CASSWGGGSHYGYTF | IVTDFSVIK | YES | YES | YES |
| 11229 | TRAV12-2 | TRAJ45 | CAVSGGADGLTF | TRBV12-4 | TRBJ2-1 | CASSLGGNEQFF | ELAGIGILTV | YES | NO | YES |
| 11230 | TRAV19 | TRAJ49 | CALSEAAGNQFYF | TRBV5-5 | TRBJ2-7 | CASSLDPDRVFREQYF | KLGGALQAK | YES | NO | NO |
| 11231 | TRAV8-1 | TRAJ45 | CAVKSGGADGLTF | TRBV30 | TRBJ1-1 | CALVGLNTEAFF | RLRAEAQVK | YES | NO | YES |
| 11232 | TRAV38-1 | TRAJ45 | CAFRKWGGADGLTF | TRBV13 | TRBJ1-5 | CASSQYGNQPQHF | KVLEYVIKV | NO | NO | NO |
| 11233 | TRAV12-3 | TRAJ33 | CAMKDSNYQLIW | TRBV4-2 | TRBJ1-2 | CASSQEEGPVGYTF | KLGGALQAK | YES | NO | NO |

| | | | | | | | | | | |
|---|---|---|---|---|---|---|---|---|---|---|
| 11234 | TRAV20 | TRAJ8 | CALNTGFQKLVF | TRBV11-2 | TRBJ2-1 | CASSLVTSGIIEQFF | KLGGALQAK | YES | NO | NO |
| 11235 | TRAV1-2 | TRAJ33 | CAVLDSNYQLIW | TRBV20-1 | TRBJ2-3 | CSASRGRESDTQYF | KLGGALQAK | NO | NO | NO |
| 11236 | TRAV29DV5 | TRAJ37 | CAASPLDDGSSNTGKLIF | TRBV4-3 | TRBJ2-3 | CASSQESPREAADTQYF | KLGGALQAK | YES | NO | NO |
| 11237 | TRAV8-1 | TRAJ54 | CAVNARGAQKLVF | TRBV18 | TRBJ2-1 | CASSPDLRGPNEQFF | KLGGALQAK | YES | NO | NO |
| 11238 | TRAV17 | TRAJ54 | CATDRIQGAQKLVF | TRBV5-4 | TRBJ2-4 | CASSYDRVKNIQYF | KLGGALQAK | YES | NO | NO |
| 11239 | TRAV3 | TRAJ33 | CAHPDSNYQLIW | TRBV27 | TRBJ2-2 | CASSLRLGNTGELFF | KLGGALQAK | NO | NO | NO |
| 11240 | TRAV2 | TRAJ36 | CAVETGANNLFF | TRBV20-1 | TRBJ2-1 | CSAPPSSGRSYNEQFF | KLGGALQAK | NO | NO | NO |
| 11241 | TRAV29DV5 | TRAJ57 | CAARETQGGSEKLVF | TRBV13 | TRBJ2-5 | CASSLGQAETQYF | KLGGALQAK | YES | NO | NO |
| 11242 | TRAV29DV5 | TRAJ31 | CAADDGRLMF | TRBV30 | TRBJ1-6 | CAWTGTGKIGWDSPLHF | RAKFKQLL | NO | NO | NO |
| 11243 | TRAV35 | TRAJ42 | CAGPSYYGGSQGNLIF | TRBV27 | TRBJ2-3 | CASGQGTDTQYF | KLGGALQAK | YES | NO | NO |
| 11244 | TRAV9-2 | TRAJ42 | CALSDRVFNYGGSQGNLIF | TRBV10-3 | TRBJ1-2 | CAISEGIGQNYGYTF | KLGGALQAK | YES | NO | NO |
| 11245 | TRAV8-6 | TRAJ54 | CAVSQWAQKLVF | TRBV5-4 | TRBJ2-1 | CASSLGLAGLNEQFF | KLGGALQAK | YES | NO | NO |
| 11247 | TRAV13-1 | TRAJ24 | CAAKSISTDSWGKLQF | TRBV3-1 | TRBJ2-5 | CASSQGVAGGWGTQYF | KLGGALQAK | YES | NO | NO |
| 11248 | TRAV38-1 | TRAJ27 | CAFISVNAGKSTF | TRBV7-2 | TRBJ1-1 | CASSYCCGTEAFF | KLGGALQAK | YES | NO | NO |
| 11249 | TRAV9-2 | TRAJ28 | CALSGYSGAGSYQLTF | TRBV6-1 | TRBJ2-7 | CASSEGGGLPSYEQYF | QYDPVAALF | YES | NO | YES |
| 11250 | TRAV12-2 | TRAJ12 | CAVNREDSSYKLIF | TRBV20-1 | TRBJ2-5 | CSAPTEEGSLQETQYF | KLGGALQAK | YES | NO | NO |
| 11251 | TRAV8-1 | TRAJ13 | CAVDSNSGGYQKVTF | TRBV7-9 | TRBJ2-5 | CASSYLLTGTSRQETQYF | CYTWNQMNL | YES | NO | YES |
| 11252 | TRAV38-2DV8 | TRAJ53 | CAYIESGGSNYKLTF | TRBV4-1 | TRBJ2-7 | CASSQDRQGGDEQYF | AVFDRKSDAK | YES | YES | YES |
| 11253 | TRAV22 | TRAJ56 | CAGTGANSKLTF | TRBV6-6 | TRBJ1-5 | CASRSRQGYQPQHF | KLGGALQAK | NO | NO | NO |
| 11254 | TRAV12-1 | TRAJ6 | CVVEGGSYIPTF | TRBV7-2 | TRBJ1-6 | CASSLKAREAYNSPLHF | KLGGALQAK | YES | NO | NO |
| 11255 | TRAV41 | TRAJ48 | CAVSLNFGNEKLTF | TRBV7-2 | TRBJ2-1 | CASSWGDYNEQFF | YLLEMLWRL | YES | NO | YES |
| 11256 | TRAV12-2 | TRAJ22 | CAVGGQLTF | TRBV14 | TRBJ2-7 | CASSQANYEQYF | KLGGALQAK | NO | NO | NO |
| 11257 | TRAV12-1 | TRAJ37 | CVVKRSSNTGKLIF | TRBV9 | TRBJ1-6 | CASSAGDWISDNSPLHF | IVTDFSVIK | YES | YES | YES |
| 11258 | TRAV22 | TRAJ24 | CAVHDSWGKLQF | TRBV3-1 | TRBJ2-1 | CASSQGTSFFEQFF | KLGGALQAK | YES | NO | NO |
| 11259 | TRAV9-2 | TRAJ33 | CALSDLDSNYQLIW | TRBV7-2 | TRBJ2-3 | CASSLYTDTQYF | RMFPNAPYL | YES | NO | YES |
| 11260 | TRAV27 | TRAJ41 | CADSNSGYALNF | TRBV4-3 | TRBJ2-7 | CASSQGGGQGYEQYF | AVFDRKSDAK | YES | NO | NO |
| 11261 | TRAV12-2 | TRAJ5 | CAVGGTGRRALTF | TRBV4-3 | TRBJ1-1 | CASSQAEGAGTEAFF | ELAGIGILTV | YES | YES | YES |
| 11263 | TRAV8-4 | TRAJ43 | CAVSDGNNDMRF | TRBV10-2 | TRBJ1-2 | CASRGGTGDNYGYTF | KLGGALQAK | YES | NO | NO |
| 11264 | TRAV12-2 | TRAJ41 | CAAGSGYALNF | TRBV7-9 | TRBJ1-2 | CASSFRSGDGYTF | KLGGALQAK | YES | NO | NO |
| 11265 | TRAV41 | TRAJ58 | CAVRLGETSGSRLTF | TRBV11-2 | TRBJ2-5 | CASSLGDFFTQETQYF | KLGGALQAK | YES | NO | NO |
| 11266 | TRAV26-1 | TRAJ17 | CILVVAAGNKLTF | TRBV11-2 | TRBJ1-2 | CASSWGGGSHYGYTF | IVTDFSVIK | YES | NO | YES |
| 11267 | TRAV19 | TRAJ57 | CALSEPQGGSEKLVF | TRBV9 | TRBJ2-5 | CASSPPGRGVVQETQYF | KLGGALQAK | YES | NO | NO |
| 11268 | TRAV1-2 | TRAJ10 | CAVRAGGGNKLTF | TRBV20-1 | TRBJ2-2 | CSARDLGGPNTGELFF | AVFDRKSDAK | YES | NO | YES |
| 11269 | TRAV13-1 | TRAJ28 | CYSTWRLSCLLLCRDSAGAGSYQLTF | TRBV9 | TRBJ2-2 | CASSANRDTEDTGELFF | KLGGALQAK | NO | NO | NO |
| 11270 | TRAV21 | TRAJ21 | CAAEYNFNKFYF | TRBV12-3 | TRBJ2-1 | CASSLGWGVNEQFF | KLGGALQAK | YES | NO | NO |
| 11271 | TRAV8-4 | TRAJ31 | CAVSDPRNNARLMF | TRBV19 | TRBJ1-2 | CASSARTGDGYTF | KLGGALQAK | YES | NO | NO |
| 11272 | TRAV13-1 | TRAJ22 | CAASGAARQLTF | TRBV7-2 | TRBJ1-5 | CATGYRFNQPQHF | AVFDRKSDAK | YES | NO | NO |
| 11273 | TRAV26-1 | TRAJ45 | CIVRAPWYSGGGADGLTF | TRBV4-2 | TRBJ1-1 | CASSPPGGGNTEAFF | KLGGALQAK | YES | NO | NO |
| 11275 | TRAV12-3 | TRAJ36 | CAMGPKGANNLFF | TRBV3-1 | TRBJ1-1 | CASSHLDRGRTEAFF | KLGGALQAK | YES | NO | NO |
| 11276 | TRAV24 | TRAJ8 | CAQTGFQKLVF | TRBV6-5 | TRBJ2-7 | CASSLTSGKPYEQYF | KLGGALQAK | YES | NO | NO |
| 11277 | TRAV13-2 | TRAJ4 | CAPGGYNKLIF | TRBV7-2 | TRBJ2-2 | CASSLGGLAGVGTGELFF | KLGGALQAK | NO | NO | NO |
| 11278 | TRAV8-3 | TRAJ33 | CAVVAGDSNYQLIW | TRBV27 | TRBJ2-3 | CASSIGLLTDTQYF | KLGGALQAK | YES | NO | NO |
| 11279 | TRAV22 | TRAJ58 | CAVERRETSGSRLTF | TRBV7-2 | TRBJ2-7 | CASSLGRAYEQYF | KLGGALQAK | NO | NO | NO |
| 11280 | TRAV12-2 | TRAJ23 | CAVSPGKLIF | TRBV18 | TRBJ2-3 | CASSPQDSPTDTQYF | KLGGALQAK | YES | NO | NO |
| 11281 | TRAV12-2 | TRAJ45 | CAVPYSGGGADGLTF | TRBV25-1 | TRBJ2-7 | CASSPHPGTGGYEQYF | ELAGIGILTV | YES | YES | YES |
| 11282 | TRAV13-2 | TRAJ45 | CADSGGGADGLTF | TRBV11-2 | TRBJ1-2 | CASSWGGGSHYGYTF | IVTDFSVIK | YES | NO | YES |
| 11283 | TRAV8-2 | TRAJ42 | CVVSDRGSQGNLIF | TRBV5-6 | TRBJ2-1 | CASSGGTSGYNEQFF | IVTDFSVIK | YES | NO | YES |
| 11284 | TRAV12-2 | TRAJ43 | CAASYNNDMRF | TRBV24-1 | TRBJ1-5 | CATSVPTGHNQPQHF | KLGGALQAK | YES | NO | NO |
| 11285 | TRAV19 | TRAJ52 | CALSAAGGTSYGKLTF | TRBV28 | TRBJ2-2 | CASSLYSATGELFF | AVFDRKSDAK | YES | YES | YES |
| 11286 | TRAV38-1 | TRAJ44 | CAFLNTGTASKLTF | TRBV4-3 | TRBJ2-1 | CASSQAHPWRRSYNEQFF | KLGGALQAK | YES | NO | NO |
| 11287 | TRAV12-2 | TRAJ17 | CAVKNAGNKLTF | TRBV7-2 | TRBJ2-7 | CASSLNDFYEQYF | ELAGIGILTV | YES | NO | YES |
| 11288 | TRAV12-2 | TRAJ40 | CAVKYPSGTYKYIF | TRBV6-5 | TRBJ1-2 | CASNDRGNGYTF | GILGFVFTL | YES | NO | YES |
| 11289 | TRAV19 | TRAJ54 | CALSEASGIQGAQKLVF | TRBV15 | TRBJ1-6 | CATSRDVVFGTGGGPYNSPLHF | KLGGALQAK | NO | NO | NO |
| 11290 | TRAV13-1 | TRAJ35 | CAARARGFGNVLHC | TRBV28 | TRBJ2-2 | CASSLYSATGELFF | AVFDRKSDAK | NO | NO | NO |
| 11291 | TRAV1-1 | TRAJ37 | CAATSNTGKLIF | TRBV12-4 | TRBJ2-3 | CASRTREGRTDTQYF | RAKFKQLL | YES | YES | YES |
| 11292 | TRAV14DV4 | TRAJ9 | CAMREEGGGFKTIF | TRBV7-9 | TRBJ2-1 | CASSSIAEQFF | KLGGALQAK | YES | NO | NO |
| 11294 | TRAV6 | TRAJ37 | CALWPFSGNTGKLIF | TRBV12-4 | TRBJ1-5 | CASSPRYSNQPQHF | KLGGALQAK | YES | NO | NO |
| 11295 | TRAV24 | TRAJ52 | CAFNAGGTSYGKLTF | TRBV11-2 | TRBJ2-1 | CASSPDWTSFNEQFF | KLGGALQAK | YES | NO | NO |
| 11296 | TRAV38-2DV8 | TRAJ39 | CAYRSLIRNAGNMLTF | TRBV11-2 | TRBJ2-1 | CASSQYGTSSVNEQFF | KLGGALQAK | YES | NO | NO |
| 11297 | TRAV10 | TRAJ20 | CVVSDINDYKLSF | TRBV9 | TRBJ2-3 | CASSLTSGGATDTQYF | KLGGALQAK | YES | NO | NO |
| 11298 | TRAV8-3 | TRAJ54 | CAVAVIQGAQKLVF | TRBV7-2 | TRBJ2-7 | CASSPRDRERGEQYV | KLGGALQAK | NO | NO | NO |
| 11299 | TRAV12-2 | TRAJ15 | CAANQAGTALIF | TRBV4-3 | TRBJ1-5 | CASSQGVLLGQPQHF | ELAGIGILTV | YES | YES | YES |
| 11300 | TRAV20 | TRAJ42 | CAVRMGGGGSQGNLIF | TRBV28 | TRBJ2-2 | CASSLLAHTGELFF | MLDLQPETT | YES | NO | NO |
| 11301 | TRAV12-2 | TRAJ24 | CAATDSWGKLQF | TRBV28 | TRBJ2-3 | CASTELTGTGHTDTQYF | KLGGALQAK | YES | NO | NO |
| 11302 | TRAV25 | TRAJ28 | CAGIYSGAGSYQLTF | TRBV27 | TRBJ2-1 | CASTPGTSAYNEQFF | KLGGALQAK | YES | NO | NO |
| 11303 | TRAV29DV5 | TRAJ54 | CAAKARWGAQKLVF | TRBV5-1 | TRBJ1-2 | CASSLPGQDYGYTF | RLRAEAQVK | YES | NO | YES |
| 11304 | TRAV3 | TRAJ5 | CAVRAPDTGRRALTF | TRBV24-1 | TRBJ2-3 | CATSEGDRVGEIRTDTQYF | KLGGALQAK | NO | NO | NO |
| 11305 | TRAV29DV5 | TRAJ36 | CAASAMIKTGANNLFF | TRBV7-9 | TRBJ2-7 | CASSDILAGVGEQYF | KLGGALQAK | YES | NO | NO |
| 11306 | TRAV5 | TRAJ4 | CAETPLFSGGYNKLIF | TRBV12-5 | TRBJ1-1 | CASGLDRMNTEAFF | KLGGALQAK | YES | NO | NO |
| 11307 | TRAV6 | TRAJ6 | CALDSPGGSYIPTF | TRBV25-1 | TRBJ2-7 | CATGQGAGYEQYF | KLGGALQAK | YES | NO | NO |
| 11308 | TRAV4 | TRAJ22 | CLVLSSGSARQLTF | TRBV4-3 | TRBJ2-2 | CASSFGTGAGGELFF | CYTWNQMNL | NO | NO | NO |
| 11309 | TRAV12-1 | TRAJ24 | CVVNSWGKLQF | TRBV28 | TRBJ2-3 | CASSNKVADTQYF | MLDLQPETT | YES | NO | YES |
| 11310 | TRAV12-2 | TRAJ29 | CAPYSGNTPLVF | TRBV24-1 | TRBJ2-1 | CATSEQGQSLTTYNEQFF | ELAGIGILTV | YES | YES | NO |
| 11311 | TRAV12-2 | TRAJ10 | CAVKGVTGGGNKLTF | TRBV9 | TRBJ2-3 | CASSVAGTTDTQYF | GLCTLVAML | YES | YES | YES |
| 11312 | TRAV12-2 | TRAJ57 | CAAQGGSEKLVF | TRBV9 | TRBJ2-1 | CASSVEGNEQFF | KLGGALQAK | YES | NO | NO |
| 11313 | TRAV8-6 | TRAJ52 | CAVSDYAGGTSYGKLTF | TRBV19 | TRBJ2-7 | CATQRDETYEQYF | KLGGALQAK | YES | NO | NO |
| 11314 | TRAV21 | TRAJ54 | CAVKIQGAQKLVF | TRBV3-1 | TRBJ1-1 | CASSQDGGDETPNTEAFF | CYTWNQMNL | YES | NO | YES |
| 11315 | TRAV21 | TRAJ57 | CAVAQGGSEKLVF | TRBV5-1 | TRBJ1-1 | CASSLRGRLLLTRTEAFF | KLGGALQAK | YES | NO | NO |
| 11316 | TRAV8-3 | TRAJ15 | CAVGAGTALIF | TRBV11-2 | TRBJ1-2 | CASSWGGGSHYGYTF | IVTDFSVIK | YES | NO | YES |
| 11317 | TRAV8-1 | TRAJ7 | CAAYGNNRLAF | TRBV13 | TRBJ1-2 | CASRRTTESTHGYTF | KLGGALQAK | YES | NO | NO |
| 11318 | TRAV12-3 | TRAJ53 | CATGDSGGSNYKLTF | TRBV13 | TRBJ1-4 | CASSLVGVNEKLFF | KLGGALQAK | YES | NO | NO |
| 11319 | TRAV8-3 | TRAJ33 | CAVASDSNYQLIW | TRBV6-5 | TRBJ1-3 | CASSYSSGNTIYF | KLGGALQAK | YES | NO | NO |
| 11320 | TRAV25 | TRAJ13 | CAVDSGGYQKVTF | TRBV4-2 | TRBJ1-1 | CASSQEARKGRVTEAFF | KLGGALQAK | YES | NO | NO |
| 11321 | TRAV1-2 | TRAJ26 | CAVRDNYGQNFVF | TRBV5-4 | TRBJ2-3 | CASSSSDTQYF | KLGGALQAK | YES | NO | NO |
| 11322 | TRAV12-2 | TRAJ52 | CAVNPHAPGGTSYGKLTF | TRBV28 | TRBJ1-5 | CASSLVGGLGQPQHF | KLGGALQAK | YES | NO | NO |
| 11323 | TRAV29DV5 | TRAJ16 | CAARADGQKLLF | TRBV20-1 | TRBJ1-6 | CSARDTRQLSSYNSPLHF | KLGGALQAK | NO | NO | NO |
| 11324 | TRAV10 | TRAJ32 | CVVSRWGWGGATNKLIF | TRBV11-1 | TRBJ2-5 | CASSPRLAGAETQYF | AVFDRKSDAK | YES | NO | YES |
| 11325 | TRAV14DV4 | TRAJ20 | CAMREGDRNDYKLSF | TRBV9 | TRBJ2-2 | CASSVGREWTGELFF | KLGGALQAK | YES | NO | NO |
| 11326 | TRAV10 | TRAJ26 | CVVSDNYGQNFVF | TRBV27 | TRBJ1-3 | CASSLSWGAGNTIYF | KLGGALQAK | YES | NO | NO |
| 11327 | TRAV5 | TRAJ22 | CAEIVGQLTF | TRBV7-9 | TRBJ2-7 | CASSPGGLENEQYF | KLGGALQAK | YES | NO | NO |
| 11328 | TRAV8-2 | TRAJ21 | CVVIYNFNKFYF | TRBV4-1 | TRBJ2-1 | CASSKLAGGLKNEQFF | KLGGALQAK | YES | NO | NO |
| 11329 | TRAV30 | TRAJ42 | CGTADGGSQGNLIF | TRBV19 | TRBJ2-4 | CASSIRSDDIQYF | GILGFVFTL | YES | NO | YES |
| 11330 | TRAV1-2 | TRAJ33 | CAAMDSNYQLIW | TRBV20-1 | TRBJ2-1 | CSAADSNLSYNEQFF | KLGGALQAK | NO | NO | NO |
| 11331 | TRAV12-3 | TRAJ45 | CAMSSPGWGGGADGLTF | TRBV27 | TRBJ1-2 | CASSPTGTAYGYTF | KLGGALQAK | YES | NO | NO |
| 11332 | TRAV8-3 | TRAJ45 | CAVGPDSGGGADGLTF | TRBV13 | TRBJ2-7 | CASSSDSYEQYF | KLGGALQAK | YES | NO | NO |
| 11333 | TRAV8-4 | TRAJ33 | CAVSQDSNYQLIW | TRBV11-2 | TRBJ1-2 | CASSWGGGSHYGYTF | IVTDFSVIK | YES | YES | YES |
| 11334 | TRAV20 | TRAJ37 | CAVQDPGSSNTGKLIF | TRBV12-4 | TRBJ2-3 | CASSLYGVGDTQYF | KLGGALQAK | YES | NO | NO |
| 11335 | TRAV1-2 | TRAJ41 | CAVRGSANSGYALNF | TRBV6-5 | TRBJ1-1 | CATSDRGGTEAFF | AVFDRKSDAK | YES | NO | YES |
| 11336 | TRAV12-2 | TRAJ28 | CAAGAGSYQLTF | TRBV6-3 | TRBJ1-5 | CASSQGQGQPQHF | ELAGIGILTV | YES | NO | YES |
| 11337 | TRAV1-2 | TRAJ20 | CAVNEGDYKLSF | TRBV20-1 | TRBJ2-1 | CSARHLSGIEQFF | SLFNTVATLY | NO | NO | NO |
| 11338 | TRAV36DV7 | TRAJ40 | CAVTTSGTYKYIF | TRBV4-1 | TRBJ2-3 | CASSRRGPWVTKGTDTQYF | RLRAEAQVK | NO | NO | NO |
| 11339 | TRAV24 | TRAJ52 | CAFNAGGTSYGKLTF | TRBV7-2 | TRBJ1-5 | CASSLDYYSNQPQHF | AVFDRKSDAK | YES | NO | YES |
| 11340 | TRAV36DV7 | TRAJ40 | CAVGGWAITSGTYKYIF | TRBV12-4 | TRBJ2-7 | CASSRDHSYEQYF | KLGGALQAK | NO | NO | NO |
| 11341 | TRAV12-1 | TRAJ40 | CVVNMDPSGTYKYIF | TRBV28 | TRBJ1-2 | CASRPTGGASLGYTF | RAKFKQLL | YES | YES | YES |

| | | | | | | | | | | |
|---|---|---|---|---|---|---|---|---|---|---|
| 11342 | TRAV14DV4 | TRAJ33 | CAMRPLGMDSNYQLIW | TRBV19 | TRBJ2-7 | CASSTHGADEQYF | GILGFVFTL | YES | NO | YES |
| 11343 | TRAV8-6 | TRAJ6 | CAVSLLSGGSYIPTF | TRBV14 | TRBJ1-4 | CASSQPGAGFGEKLFF | KLGGALQAK | YES | NO | NO |
| 11344 | TRAV19 | TRAJ42 | CALRDGGSQGNLIF | TRBV2 | TRBJ1-1 | CASTLTGVNTEAFF | KLGGALQAK | YES | NO | NO |
| 11345 | TRAV8-3 | TRAJ42 | CAVGGDGGSQGNLIF | TRBV19 | TRBJ2-7 | CASSIRSSYEQYF | GILGFVFTL | YES | NO | YES |
| 11346 | TRAV8-6 | TRAJ17 | CAVKIKAAGNKLTF | TRBV13 | TRBJ2-5 | CASSSTGGKETQYF | AVFDRKSDAK | YES | NO | YES |
| 11348 | TRAV19 | TRAJ41 | CALSEAYEYALNF | TRBV20-1 | TRBJ2-1 | CSPKEGNNEQFF | AVFDRKSDAK | NO | NO | NO |
| 11349 | TRAV39 | TRAJ34 | CAVDDSDKLIF | TRBV12-4 | TRBJ2-3 | CASSHSGGTDTQYF | KLGGALQAK | YES | NO | NO |
| 11351 | TRAV24 | TRAJ16 | CLLTSDGQKLLF | TRBV4-2 | TRBJ2-1 | CASSQDIRTGNNEQFF | KLGGALQAK | NO | NO | NO |
| 11352 | TRAV1-1 | TRAJ12 | CAGIQDSSYKLIF | TRBV20-1 | TRBJ2-1 | CSARAKRGLFNEQFF | KLGGALQAK | NO | NO | NO |
| 11353 | TRAV29DV5 | TRAJ28 | CAASAPKPGSYQLTF | TRBV10-2 | TRBJ2-7 | CASRETGVYEQYF | KLGGALQAK | YES | NO | NO |
| 11354 | TRAV10 | TRAJ27 | CVGYTNAGKSTF | TRBV19 | TRBJ2-7 | CASSIRASYEQYF | GILGFVFTL | YES | NO | YES |
| 11355 | TRAV1-2 | TRAJ8 | CAVRGLNTGFQKLVF | TRBV14 | TRBJ1-2 | CASSQQGSYGYTF | RAKFKQLL | NO | NO | NO |
| 11356 | TRAV19 | TRAJ41 | CALSVFSNSGYALNF | TRBV20-1 | TRBJ2-2 | CSARSGTGELFF | KLGGALQAK | YES | NO | NO |
| 11357 | TRAV24 | TRAJ12 | CAWMDSSYKLIF | TRBV19 | TRBJ2-5 | CASSSGSLETQYF | KLGGALQAK | YES | NO | NO |
| 11358 | TRAV12-1 | TRAJ48 | CVVTPSNFGNEKLTF | TRBV5-6 | TRBJ2-5 | CASSLAGLAGETQYF | ELAGIGILTV | YES | YES | YES |
| 11359 | TRAV8-2 | TRAJ30 | CVVSDYRDDKIIF | TRBV27 | TRBJ1-1 | CASTQNRENTEAFF | KLGGALQAK | YES | NO | NO |
| 11360 | TRAV20 | TRAJ39 | CAVRVAGNMLTF | TRBV30 | TRBJ2-4 | CAWSVQTPTQHF | KLGGALQAK | YES | NO | NO |
| 11361 | TRAV3 | TRAJ4 | CAVRDPYQTGGYNKLIF | TRBV13 | TRBJ2-3 | CASSSPERGRGTDTQYF | KLGGALQAK | NO | NO | NO |
| 11362 | TRAV12-2 | TRAJ34 | CAVRPDKLIF | TRBV2 | TRBJ2-1 | CASREREGLAVYNEQFF | ELAGIGILTV | YES | NO | YES |
| 11363 | TRAV12-2 | TRAJ28 | CAVIAPGAGSYQLTF | TRBV6-3 | TRBJ2-2 | CASSYSILDTGELFF | KLGGALQAK | YES | NO | NO |
| 11364 | TRAV17 | TRAJ48 | CATPDFYSNFGNEKLTF | TRBV5-5 | TRBJ2-5 | CASSGGITGEETQYF | IVTDFSVIK | YES | NO | YES |
| 11365 | TRAV12-2 | TRAJ48 | CAVTGSNFGNEKLTF | TRBV20-1 | TRBJ1-2 | CSARPGEQGGDGYTF | ELAGIGILTV | YES | NO | YES |
| 11366 | TRAV8-3 | TRAJ20 | CAVGTDYKLSF | TRBV13 | TRBJ1-2 | CASSTDTFGYGYTF | KLGGALQAK | YES | NO | NO |
| 11367 | TRAV13-1 | TRAJ35 | CAARVRGFGNVLHC | TRBV28 | TRBJ1-6 | CASRPQQGHNSPLHF | AVFDRKSDAK | NO | NO | NO |
| 11368 | TRAV29DV5 | TRAJ52 | CAASAKGGTSYGKLTF | TRBV7-8 | TRBJ1-1 | CASSLRVTNPQGKGTEAFF | KLGGALQAK | YES | NO | NO |
| 11369 | TRAV29DV5 | TRAJ36 | CAAETGANNLFF | TRBV19 | TRBJ1-2 | CASSATGTGVGYTF | KLGGALQAK | YES | NO | NO |
| 11370 | TRAV19 | TRAJ17 | CALSEIAAGNKLTF | TRBV4-3 | TRBJ1-2 | CASSQDWQAKGAGYTF | KLGGALQAK | YES | NO | NO |
| 11371 | TRAV6 | TRAJ53 | CALDPSFSGGSNYKLTF | TRBV18 | TRBJ2-1 | CASSPRTPYNEQFF | KLGGALQAK | YES | NO | NO |
| 11372 | TRAV8-4 | TRAJ47 | CAVEYGNKLVF | TRBV6-5 | TRBJ2-7 | CASSTYASGAISYEQYF | IVTDFSVIK | YES | NO | YES |
| 11373 | TRAV3 | TRAJ21 | CAVRDPYNFNKFYF | TRBV20-1 | TRBJ2-1 | CSVRGGGISYNEQFF | AVFDRKSDAK | NO | NO | NO |
| 11374 | TRAV21 | TRAJ39 | CAVRGNNAGNMLTF | TRBV7-3 | TRBJ2-1 | CASSLIPGPYNEQFF | RAKFKQLL | YES | NO | YES |
| 11375 | TRAV19 | TRAJ28 | CALTDYSGAGSYQLTF | TRBV19 | TRBJ1-1 | CASSIRSGTEAFF | GILGFVFTL | YES | NO | YES |
| 11376 | TRAV13-2 | TRAJ42 | CAENLGGGSQGNLIF | TRBV19 | TRBJ2-1 | CASSIRAASYNEQFF | GILGFVFTL | YES | NO | YES |
| 11377 | TRAV8-3 | TRAJ34 | CAVGAAYNTDKLIF | TRBV7-9 | TRBJ1-3 | CASSLVGVKAGNTIYF | KLGGALQAK | YES | NO | NO |
| 11378 | TRAV12-2 | TRAJ45 | CAVNLGSGGGADGLTF | TRBV7-9 | TRBJ1-1 | CASSPSGNTEAFF | KLGGALQAK | YES | NO | NO |
| 11379 | TRAV8-4 | TRAJ36 | CAVSDQGANNLFF | TRBV6-3 | TRBJ2-2 | CASSYYGWGTGELFF | KLGGALQAK | YES | NO | NO |
| 11380 | TRAV8-6 | TRAJ8 | CAVSGGIGFQKLVF | TRBV5-1 | TRBJ2-4 | CASSFKGPGAKNIQYF | KLGGALQAK | YES | NO | NO |
| 11381 | TRAV12-2 | TRAJ24 | CAVTPTDSWGKLQF | TRBV6-3 | TRBJ1-5 | CASSYGMGQPQHF | ELAGIGILTV | YES | YES | YES |
| 11382 | TRAV21 | TRAJ57 | CAVKGGSEKLVF | TRBV5-1 | TRBJ2-3 | CASSLVAHTQYF | KLGGALQAK | YES | NO | NO |
| 11383 | TRAV17 | TRAJ41 | CATDSNSGYALNF | TRBV6-3 | TRBJ2-5 | CASSYSIGETQYF | AVFDRKSDAK | YES | NO | YES |
| 11384 | TRAV8-3 | TRAJ28 | CAVGGYRGAGSYQLTF | TRBV6-5 | TRBJ1-5 | CASSLEPTANQPQHF | KVLEYVIKV | YES | NO | YES |
| 11385 | TRAV39 | TRAJ58 | CAVDMYETSGSRLTF | TRBV6-3 | TRBJ2-3 | CASGYWSGDTQYF | RLRAEAQVK | YES | YES | YES |
| 11386 | TRAV29DV5 | TRAJ53 | CAAIGSNYKLTF | TRBV11-2 | TRBJ1-2 | CASSWGGGSHYGYTF | IVTDFSVIK | YES | NO | YES |
| 11387 | TRAV20 | TRAJ53 | CAVPLTSGGGSNYKLTF | TRBV5-8 | TRBJ2-1 | CASSPTVTYNEQFF | KLGGALQAK | YES | NO | NO |
| 11388 | TRAV1-2 | TRAJ33 | CAVQDSNYQLIW | TRBV11-2 | TRBJ1-2 | CASSWGGGSHYGYTF | IVTDFSVIK | YES | YES | YES |
| 11389 | TRAV2 | TRAJ16 | CAVRPCF | TRBV27 | TRBJ1-5 | CASSLAGKGAQHF | KLGGALQAK | NO | NO | NO |
| 11390 | TRAV35 | TRAJ37 | CAGQGVPAGGNTGKLIF | TRBV19 | TRBJ2-3 | CASSIVSTDTQYF | GILGFVFTL | YES | NO | NO |
| 11391 | TRAV34 | TRAJ47 | CGAAYGNKLVF | TRBV4-2 | TRBJ2-2 | CASSQDPPPTANTGELFF | AVFDRKSDAK | YES | YES | YES |
| 11392 | TRAV39 | TRAJ13 | CAVVPSGGYQKVTF | TRBV9 | TRBJ1-5 | CASSVGAGSGQPQHF | KLGGALQAK | YES | NO | NO |
| 11393 | TRAV35 | TRAJ42 | CASRLVGGSQGNLIF | TRBV9 | TRBJ2-1 | CASSTSGYSSYNEQFF | KLGGALQAK | YES | NO | NO |
| 11394 | TRAV22 | TRAJ10 | CAGPVTGGGNKLTF | TRBV16 | TRBJ1-1 | CASSQSTEAFF | KLGGALQAK | YES | NO | NO |
| 11395 | TRAV25 | TRAJ42 | CAGNYGGSQGNLIF | TRBV19 | TRBJ2-2 | CASSTRSTGELFF | GILGFVFTL | YES | NO | YES |
| 11396 | TRAV12-2 | TRAJ54 | CAVRGGAQKLVF | TRBV9 | TRBJ2-2 | CASSVGSSGIYGELFF | IVTDFSVIK | YES | NO | YES |
| 11397 | TRAV14DV4 | TRAJ54 | CAMRGLQGAQKLVF | TRBV18 | TRBJ2-3 | CASSLEDRESTDTQYF | KLGGALQAK | YES | NO | NO |
| 11398 | TRAV1-2 | TRAJ20 | CAVRERDYKLSF | TRBV19 | TRBJ1-5 | CAIRLGNQPQHF | KLGGALQAK | YES | NO | NO |
| 11399 | TRAV41 | TRAJ42 | CAVSIGAYGGSQGNLIF | TRBV7-6 | TRBJ2-2 | CASSLLAENTGELFF | KLGGALQAK | YES | NO | NO |
| 11400 | TRAV8-3 | TRAJ22 | CAVGAVGSARQLTF | TRBV19 | TRBJ1-1 | CASSIPRQGRHTEAFF | KLGGALQAK | YES | NO | NO |
| 11401 | TRAV12-1 | TRAJ11 | CVVSPGYSTLTF | TRBV14 | TRBJ1-2 | CASSQSGTGYGYTF | AVFDRKSDAK | NO | NO | NO |
| 11402 | TRAV13-1 | TRAJ10 | CAALPGGGNKLTF | TRBV6-3 | TRBJ2-7 | CASSFSYEQYF | AVFDRKSDAK | YES | NO | YES |
| 11403 | TRAV14DV4 | TRAJ13 | CAMRDAPGGYQKVTF | TRBV5-1 | TRBJ2-5 | CASSFSRARRETQYF | KLGGALQAK | YES | NO | NO |
| 11404 | TRAV38-2DV8 | TRAJ40 | CAYRARVSGTYKYIF | TRBV7-2 | TRBJ1-2 | CASSLVGQGRDYGYTF | KLGGALQAK | YES | NO | NO |
| 11405 | TRAV14DV4 | TRAJ43 | CAMREGAYNNNDMRF | TRBV4-3 | TRBJ2-2 | CASSQEFWVLGELFF | LLFGYPVYV | YES | YES | YES |
| 11406 | TRAV9-2 | TRAJ6 | CALTLSGGSYIPTF | TRBV4-2 | TRBJ2-3 | CASSQEVVLSTDTQYF | KLGGALQAK | YES | NO | NO |
| 11407 | TRAV12-1 | TRAJ49 | CVLSDTGNQFYF | TRBV7-2 | TRBJ2-7 | CASSASYEQYF | KLGGALQAK | YES | NO | NO |
| 11408 | TRAV12-2 | TRAJ49 | CAVPGSTGNQFYF | TRBV6-3 | TRBJ1-5 | CASSFGFGQPQHF | ELAGIGILTV | YES | NO | YES |
| 11409 | TRAV4 | TRAJ20 | CLVGDGDYKLSF | TRBV24-1 | TRBJ2-1 | CATSDFVGPALGSYNEQFF | KLGGALQAK | YES | NO | NO |
| 11410 | TRAV8-4 | TRAJ39 | CAVRREAGNMLTF | TRBV7-9 | TRBJ1-1 | CASSLGRVEGGTEAFF | KLGGALQAK | YES | NO | NO |
| 11411 | TRAV12-2 | TRAJ8 | CAADPGFQKLVF | TRBV2 | TRBJ2-1 | CASGMSGGAENEQFF | ELAGIGILTV | YES | YES | YES |
| 11412 | TRAV3 | TRAJ36 | CAVRDGTGANNLFF | TRBV19 | TRBJ1-2 | CASTTVQGGYTF | GILGFVFTL | NO | NO | NO |
| 11413 | TRAV1-2 | TRAJ28 | CAVPPGAGSYQLTF | TRBV24-1 | TRBJ2-7 | CATSDGTPDSDPYEQYF | KLGGALQAK | YES | NO | NO |
| 11414 | TRAV4 | TRAJ47 | CLVGDGWEYGNKLVF | TRBV20-1 | TRBJ2-2 | CSASGQGLNTGELFF | KLGGALQAK | NO | NO | NO |
| 11415 | TRAV29DV5 | TRAJ57 | CAASPSQGGSEKLVF | TRBV7-6 | TRBJ2-4 | CASSHLNGYIQYF | KLGGALQAK | YES | NO | NO |
| 11416 | TRAV35 | TRAJ48 | CAGLDFGNEKLTF | TRBV7-9 | TRBJ1-1 | CASSLATSPLMNTEAFF | ELAGIGILTV | YES | NO | YES |
| 11417 | TRAV14DV4 | TRAJ54 | CAMREDIIQGAQKLVF | TRBV27 | TRBJ2-7 | CASSLAGTGYEQYF | KLGGALQAK | YES | NO | NO |
| 11418 | TRAV1-2 | TRAJ26 | CAVIRYGQNFVF | TRBV27 | TRBJ1-1 | CASSLKQSPHTEAFF | RAKFKQLL | YES | NO | YES |
| 11419 | TRAV12-2 | TRAJ31 | CAVNSNARLMF | TRBV4-3 | TRBJ1-2 | CASSPTGGRYGYTF | MLDLQPETT | YES | NO | YES |
| 11420 | TRAV41 | TRAJ33 | CAVRVDSNYQLIW | TRBV4-3 | TRBJ2-7 | CASSQAGGLFEQYF | AVFDRKSDAK | YES | NO | NO |
| 11421 | TRAV23DV6 | TRAJ28 | CAAITTPYSGAGSYQLTF | TRBV27 | TRBJ2-3 | CASSLRRTDTQYF | AVFDRKSDAK | YES | NO | YES |
| 11422 | TRAV19 | TRAJ48 | CALSEAWSNFGNEKLTF | TRBV7-2 | TRBJ2-7 | CASSPTSGGVYEQYF | KLGGALQAK | YES | NO | NO |
| 11423 | TRAV1-2 | TRAJ32 | CAGDYGGATNKLIF | TRBV11-2 | TRBJ1-2 | CASSWGGGSHYGYTF | IVTDFSVIK | YES | NO | YES |
| 11424 | TRAV5 | TRAJ24 | CAELADSWGKLQF | TRBV20-1 | TRBJ1-2 | CSAPGQGYNYGYTF | KLGGALQAK | YES | NO | NO |
| 11425 | TRAV1-2 | TRAJ23 | CAVPYNQGGKLIF | TRBV6-6 | TRBJ2-5 | CASSTLAGGQETQYF | GILGFVFTL | NO | NO | NO |
| 11426 | TRAV26-1 | TRAJ23 | CIVFYNQGGKLIF | TRBV19 | TRBJ2-7 | CASSTRSSYEQYF | GILGFVFTL | YES | YES | YES |
| 11427 | TRAV12-1 | TRAJ20 | CVVNDYKLSF | TRBV12-3 | TRBJ1-2 | CASSLAPGIHYGYTF | KLGGALQAK | YES | NO | NO |
| 11428 | TRAV36DV7 | TRAJ48 | CAVTNNFGNEKLTF | TRBV6-3 | TRBJ1-2 | CASSYWPDYGYTF | KLGGALQAK | NO | NO | NO |
| 11429 | TRAV23DV6 | TRAJ33 | CAASPTGMDSNYQLIW | TRBV19 | TRBJ1-5 | CASSIHTANQPQHF | GILGFVFTL | YES | NO | YES |
| 11430 | TRAV12-2 | TRAJ52 | CAVNLAGGTSYGKLTF | TRBV27 | TRBJ1-3 | CASSPGLGNTIYF | ELAGIGILTV | YES | YES | NO |
| 11431 | TRAV5 | TRAJ31 | CAEDNNARLMF | TRBV20-1 | TRBJ1-2 | CSARDSTGNGYTF | GLCTLVAML | YES | YES | YES |
| 11432 | TRAV12-2 | TRAJ4 | CAVSFSGGYNKLIF | TRBV2 | TRBJ1-5 | CASSDGGLGQPQHF | ELAGIGILTV | YES | YES | YES |
| 11433 | TRAV35 | TRAJ52 | CARTSYGKLTF | TRBV7-3 | TRBJ2-2 | CASSRRGQPNTGELFF | KLGGALQAK | NO | NO | NO |
| 11434 | TRAV12-1 | TRAJ18 | CVVNFALGRLYF | TRBV7-9 | TRBJ2-2 | CASSLVGTSATGELFF | AVFDRKSDAK | YES | NO | YES |
| 11435 | TRAV17 | TRAJ46 | CATAVLLYGSGDKLTF | TRBV27 | TRBJ2-7 | CASSSGTSGSYEQYF | KLGGALQAK | YES | NO | NO |
| 11436 | TRAV3 | TRAJ16 | CAVNRYDGQKLLF | TRBV7-9 | TRBJ1-1 | CASSLQSTEAFF | KLGGALQAK | NO | NO | NO |
| 11438 | TRAV13-1 | TRAJ33 | CAADSNYQLIW | TRBV7-9 | TRBJ2-7 | CASSLARGMPYEQYF | KLGGALQAK | YES | NO | NO |
| 11439 | TRAV12-2 | TRAJ29 | CAVGHSGNTPLVF | TRBV28 | TRBJ2-7 | CASSLGQGSSYEQYF | KLGGALQAK | YES | NO | NO |
| 11440 | TRAV13-2 | TRAJ33 | CAERPSMDSNYQLIW | TRBV4-1 | TRBJ2-1 | CASSQGSAGGNEQFF | KLGGALQAK | YES | NO | NO |
| 11441 | TRAV41 | TRAJ33 | CAVISYMDSNYQLIW | TRBV28 | TRBJ2-7 | CATDPGQGSYEQYF | KLGGALQAK | YES | NO | NO |
| 11442 | TRAV5 | TRAJ8 | CAENEGFQKLVF | TRBV12-4 | TRBJ2-3 | CASSFTDTQYF | KLGGALQAK | YES | NO | NO |
| 11443 | TRAV12-2 | TRAJ38 | CAVRNAGNNRKLIW | TRBV28 | TRBJ1-2 | CASSFSLSGGYTF | ELAGIGILTV | YES | YES | YES |
| 11444 | TRAV13-2 | TRAJ42 | CAEIPNYGGSQGNLIF | TRBV5-1 | TRBJ1-1 | CASSPKHHTEAFF | KLGGALQAK | YES | NO | NO |
| 11445 | TRAV12-2 | TRAJ49 | CAWKSNQFYF | TRBV4-2 | TRBJ2-2 | CASSQELLSSGELFF | ELAGIGILTV | YES | NO | YES |
| 11447 | TRAV29DV5 | TRAJ31 | CAASDLARLMF | TRBV4-1 | TRBJ2-5 | CASSQAGRVVTQYF | KLGGALQAK | YES | NO | NO |
| 11448 | TRAV20 | TRAJ45 | CAVQALMYSGGGADGLTF | TRBV7-8 | TRBJ2-1 | CASSFLAGGYNEQFF | IVTDFSVIK | YES | NO | YES |
| 11449 | TRAV38-2DV8 | TRAJ43 | CAYRISLNNNDMRF | TRBV19 | TRBJ1-1 | CASSVHGGANTEAFF | GILGFVFTL | YES | NO | YES |

| | | | | | | | | | | |
|---|---|---|---|---|---|---|---|---|---|---|
| 11450 | TRAV13-1 | TRAJ43 | CAASINNDMRF | TRBV10-3 | TRBJ1-4 | CAISELRGRKEKLFF | KLGGALQAK | YES | NO | NO |
| 11451 | TRAV8-1 | TRAJ15 | CAVNVNQAGTALIF | TRBV20-1 | TRBJ2-2 | CSAREGTGGTTGELFF | KLGGALQAK | YES | NO | NO |
| 11452 | TRAV35 | TRAJ49 | CAGHTGNQFYV | TRBV11-2 | TRBJ1-2 | CASSWGGGSHYGYTF | IVTDFSVIK | NO | NO | NO |
| 11453 | TRAV27 | TRAJ26 | CAGAPYGQNFVF | TRBV19 | TRBJ1-1 | CASTPDWDTEAFF | AVFDRKSDAK | YES | NO | YES |
| 11454 | TRAV3 | TRAJ8 | CAVRGQGTGFQKLVF | TRBV6-3 | TRBJ1-5 | CASSYTPERALPNQPQHF | KLGGALQAK | NO | NO | NO |
| 11455 | TRAV14DV4 | TRAJ10 | CAMREGITGGGNKLTF | TRBV7-9 | TRBJ2-7 | CASSPPGTAGSEQYF | KLGGALQAK | YES | YES | NO |
| 11456 | TRAV3 | TRAJ13 | CAVSSNSGGYQKVTF | TRBV7-8 | TRBJ2-3 | CASNMGGLLSTDTQYF | ELAGIGILTV | NO | NO | NO |
| 11457 | TRAV3 | TRAJ33 | CAVRPPMDSNYQLIW | TRBV29-1 | TRBJ2-3 | CSVVNTGVWDTQYF | RAKFKQLL | NO | NO | NO |
| 11458 | TRAV8-4 | TRAJ56 | CAVSLTGANSKLTF | TRBV5-1 | TRBJ1-4 | CASSGTGRYEKLFF | AVFDRKSDAK | YES | NO | YES |
| 11459 | TRAV12-2 | TRAJ31 | CAVKNARLMF | TRBV20-1 | TRBJ2-2 | CSARDLGSGTGELFF | ELAGIGILTV | YES | YES | YES |
| 11461 | TRAV38-1 | TRAJ48 | CAFMKLLTNFGNEKLTF | TRBV20-1 | TRBJ2-7 | CSAYPWDRVAYEQYF | KLGGALQAK | YES | NO | NO |
| 11462 | TRAV38-1 | TRAJ40 | CAFTSGTYKYIF | TRBV19 | TRBJ1-5 | CASSIVQGSNQPQHF | SLFNTVATLY | YES | YES | YES |
| 11463 | TRAV29DV5 | TRAJ29 | CAASGGNTPLVF | TRBV11-2 | TRBJ2-1 | CASSPGQGYNEQFF | KLGGALQAK | YES | NO | NO |
| 11464 | TRAV12-3 | TRAJ9 | CAMSATGGFKTIF | TRBV27 | TRBJ2-7 | CASSLYGLGYEQYF | KLGGALQAK | YES | NO | NO |
| 11465 | TRAV39 | TRAJ26 | CAVDFRDNYGQNFVF | TRBV11-2 | TRBJ1-2 | CASSWGGGSHYGYTF | IVTDFSVIK | YES | NO | YES |
| 11466 | TRAV1-2 | TRAJ33 | CASMDSNYQLIW | TRBV20-1 | TRBJ2-2 | CSARGGGGPENTGELFF | KLGGALQAK | YES | NO | NO |
| 11467 | TRAV35 | TRAJ49 | CAGHTGNHFYF | TRBV11-2 | TRBJ1-2 | CASSWGGGSHYGYTF | IVTDFSVIK | YES | YES | YES |
| 11469 | TRAV13-1 | TRAJ4 | CAATSGTGGYNKLIF | TRBV4-1 | TRBJ2-7 | CASSQGLSNEQYF | KLGGALQAK | YES | NO | NO |
| 11470 | TRAV13-1 | TRAJ13 | CAASIAGGYQKVTF | TRBV7-9 | TRBJ2-1 | CASSLGQGVFYEQFF | AVFDRKSDAK | YES | NO | NO |
| 11471 | TRAV12-1 | TRAJ33 | CVVGGMDSNYQLIW | TRBV4-1 | TRBJ1-6 | CASSLTTGIINSPLHF | KLGGALQAK | YES | NO | NO |
| 11472 | TRAV19 | TRAJ20 | CALSENRGDYKLSF | TRBV10-3 | TRBJ2-3 | CAIRELPTRTDTQYF | KLGGALQAK | YES | NO | NO |
| 11473 | TRAV13-1 | TRAJ45 | CAAQGKSGGGADGLTF | TRBV25-1 | TRBJ2-7 | CASSDSYEQYF | KLGGALQAK | YES | NO | NO |
| 11474 | TRAV35 | TRAJ49 | CAGHTGNQFYF | TRBV11-2 | TRBJ2-7 | CASSLEGQLFYEQYF | IVTDFSVIK | YES | YES | YES |
| 11475 | TRAV38-2DV8 | TRAJ49 | CAYFPGNQFYF | TRBV7-9 | TRBJ2-1 | CASSPKTSVTYNEQFF | KLGGALQAK | YES | NO | NO |
| 11476 | TRAV5 | TRAJ43 | CAEGDNDMRF | TRBV12-3 | TRBJ1-4 | CASSLGGGVLFF | KTWGQYWQV | YES | NO | NO |
| 11477 | TRAV12-1 | TRAJ42 | CVVTHYGGSQGNLIF | TRBV2 | TRBJ2-4 | CASSDELDRGRALAKNIQYF | KLGGALQAK | NO | NO | NO |
| 11479 | TRAV4 | TRAJ24 | CLVGDSWGKLQF | TRBV20-1 | TRBJ2-7 | CSAPGLADSYEQYF | KLGGALQAK | YES | NO | NO |
| 11480 | TRAV19 | TRAJ49 | CALRAGNQFYF | TRBV19 | TRBJ1-4 | CASSSDSLKLFF | KLGGALQAK | YES | NO | NO |
| 11481 | TRAV38-1 | TRAJ52 | CAFMLGAGGTSYGKLTF | TRBV7-6 | TRBJ1-1 | CASSFLTGNTEAFF | GILGFVFTL | YES | NO | YES |
| 11482 | TRAV29DV5 | TRAJ54 | CAAEGAQKLVF | TRBV7-8 | TRBJ2-7 | CASSSGLADQDLYEQYF | KLGGALQAK | YES | NO | NO |
| 11483 | TRAV38-2DV8 | TRAJ30 | CAYWTNRDDKIIF | TRBV7-8 | TRBJ2-7 | CASSLHRSSSSYEQYF | IVTDFSVIK | YES | NO | YES |
| 11484 | TRAV1-2 | TRAJ17 | CAVLGAAGNKLTF | TRBV5-1 | TRBJ2-1 | CASGDNEQFF | KLGGALQAK | YES | NO | NO |
| 11485 | TRAV12-2 | TRAJ54 | CAVGVQGAQKLVF | TRBV9 | TRBJ1-1 | CASSVDSGAGTEAFF | ELAGIGILTV | YES | NO | YES |
| 11486 | TRAV10 | TRAJ28 | CVVTPYSGAGSYQLTF | TRBV5-1 | TRBJ1-1 | CASSLVHRGHANTEAFF | KLGGALQAK | YES | NO | NO |
| 11488 | TRAV12-1 | TRAJ6 | CVVPSGSYIPTF | TRBV7-8 | TRBJ2-2 | CASSPQGHTGELFF | AVFDRKSDAK | YES | NO | YES |
| 11489 | TRAV3 | TRAJ22 | CAVRDVVSGSARQLTF | TRBV7-8 | TRBJ2-7 | CASRLDGTSSHEQYF | KLGGALQAK | NO | NO | NO |
| 11490 | TRAV14DV4 | TRAJ48 | CAMREGRVSNFGNEKLTF | TRBV7-2 | TRBJ1-5 | CASSRDYYSNQPQHF | KLGGALQAK | YES | NO | NO |
| 11491 | TRAV12-2 | TRAJ45 | CAVSGGADGLTF | TRBV4-2 | TRBJ1-6 | CASSQGGSPLHF | ELAGIGILTV | YES | NO | YES |
| 11492 | TRAV2 | TRAJ4 | CAVPFSGGYNKLIF | TRBV6-3 | TRBJ2-1 | CASSKTGTSSYNEQFF | AVFDRKSDAK | NO | NO | NO |
| 11493 | TRAV23DV6 | TRAJ33 | CAASMDSNYQLIW | TRBV11-2 | TRBJ1-2 | CASSWGGGSHYGYTF | IVTDFSVIK | YES | YES | YES |
| 11494 | TRAV5 | TRAJ12 | CAERPVDSSYKLIF | TRBV4-3 | TRBJ2-1 | CASSHGGTGRIDNEQFF | KLGGALQAK | YES | NO | NO |
| 11495 | TRAV21 | TRAJ34 | CAVRPVDKLIF | TRBV12-3 | TRBJ1-2 | CASSTGTAYGYTF | KLGGALQAK | YES | NO | NO |
| 11496 | TRAV21 | TRAJ6 | CAVRTAKGGSYIPTF | TRBV9 | TRBJ1-1 | CASSVGHTEAFF | AVFDRKSDAK | YES | NO | YES |
| 11497 | TRAV8-4 | TRAJ13 | CAVSLANSGGYQKVTF | TRBV20-1 | TRBJ2-1 | CSARGDYSFHEQFF | KLGGALQAK | NO | NO | NO |
| 11498 | TRAV5 | TRAJ39 | CAADNNAGNMLTF | TRBV7-3 | TRBJ2-7 | CASSLSGLDEQYF | RAKFKQLL | NO | NO | NO |
| 11500 | TRAV17 | TRAJ39 | CATDANNAGNMLTF | TRBV4-3 | TRBJ2-1 | CASSQVASDEQFF | KLGGALQAK | YES | NO | NO |
| 11502 | TRAV21 | TRAJ53 | CAGNSGGSNYKLTF | TRBV12-3 | TRBJ2-1 | CASRLDWHDNEQFF | KLGGALQAK | YES | NO | NO |
| 11503 | TRAV13-1 | TRAJ11 | CAAKEGYSTLTF | TRBV9 | TRBJ1-1 | CASSEGDRVTEAFF | IVTDFSVIK | YES | NO | YES |
| 11504 | TRAV1-2 | TRAJ33 | CAVVDSNYQLIW | TRBV6-1 | TRBJ2-2 | CASSEEIGDFPGELFF | AVFDRKSDAK | YES | YES | YES |
| 11505 | TRAV13-1 | TRAJ16 | CAAPF | TRBV5-1 | TRBJ2-1 | CASSLERSLISYNEQFF | AVFDRKSDAK | NO | NO | NO |
| 11506 | TRAV38-2DV8 | TRAJ53 | CAYKGSGGSNYKLTF | TRBV20-1 | TRBJ2-3 | CSATGLAGGEFTDTQYF | KLGGALQAK | YES | NO | NO |
| 11507 | TRAV19 | TRAJ33 | CALSDFLDSNYQLIW | TRBV4-1 | TRBJ2-5 | CASSQLVAGALGETQYF | KLGGALQAK | NO | NO | NO |
| 11508 | TRAV20 | TRAJ48 | CAVQAVANFGNEKLTF | TRBV4-2 | TRBJ2-4 | CASSQDRVTWGGIAKNIQYF | KLGGALQAK | NO | NO | NO |
| 11509 | TRAV8-4 | TRAJ13 | CAVSSLWGYQKVTF | TRBV11-2 | TRBJ2-3 | CASSLPLSRTTDTQYF | AVFDRKSDAK | YES | NO | YES |
| 11510 | TRAV1-1 | TRAJ21 | CAVLNNFNKFYF | TRBV20-1 | TRBJ2-2 | CSARNTGELFF | KLGGALQAK | YES | NO | NO |
| 11511 | TRAV25 | TRAJ8 | CAGKMNTGFQKLVF | TRBV20-1 | TRBJ2-1 | CSARQMDYNEQFF | KLGGALQAK | NO | NO | NO |
| 11512 | TRAV35 | TRAJ53 | CAGREGSNYKLTF | TRBV29-1 | TRBJ1-1 | CSVERDMGPDTEAFF | KLGGALQAK | YES | NO | NO |
| 11513 | TRAV12-3 | TRAJ23 | CAMYHGGKLIF | TRBV7-9 | TRBJ2-1 | CASSSPGAGPNNEQFF | KLGGALQAK | YES | NO | NO |
| 11514 | TRAV21 | TRAJ33 | CAVIKDSNYQLIW | TRBV2 | TRBJ2-3 | CASSEGWQGANTQYF | KLGGALQAK | YES | NO | NO |
| 11515 | TRAV12-2 | TRAJ29 | CAFDSGNTPLVF | TRBV27 | TRBJ1-1 | CASSPGQGIRPGEAFF | KLGGALQAK | YES | NO | NO |
| 11516 | TRAV19 | TRAJ16 | CALSEMF | TRBV6-3 | TRBJ2-1 | CASSYGGTSGGPFF | CYTWNQMNL | NO | NO | NO |
| 11517 | TRAV29DV5 | TRAJ53 | CAGCGGSNYKLTF | TRBV4-1 | TRBJ1-6 | CASSQRGGVYNSPLHF | RLRAEAQVK | YES | YES | YES |
| 11518 | TRAV1-2 | TRAJ20 | CAVRGRDYKLSF | TRBV3-1 | TRBJ2-5 | CASSQPVAARGAQYF | AVFDRKSDAK | YES | NO | YES |
| 11519 | TRAV35 | TRAJ42 | CAGSGGSQGNLIF | TRBV3-1 | TRBJ1-2 | CASSQETYYGYTF | KLGGALQAK | YES | NO | NO |
| 11521 | TRAV12-1 | TRAJ39 | CVVNKNNAGNMLTF | TRBV6-3 | TRBJ1-6 | CASSYFGQRNSPLHF | KLGGALQAK | YES | NO | NO |
| 11522 | TRAV5 | TRAJ26 | CAETPWNYGQNFVF | TRBV15 | TRBJ2-4 | CATSRDRIAGWDLLAKNIQYF | KLGGALQAK | NO | NO | NO |
| 11523 | TRAV1-2 | TRAJ16 | CAVGSDGQKLLF | TRBV2 | TRBJ2-5 | CASSEKGGRETQYF | FLASKIGRLV | NO | NO | NO |
| 11524 | TRAV21 | TRAJ28 | CAADYSGAGSYQLTF | TRBV7-9 | TRBJ2-7 | CASSLGFGGREAYEQYF | RMFPNAPYL | YES | NO | YES |
| 11525 | TRAV38-1 | TRAJ57 | CAFILTQGGSEKLVF | TRBV5-4 | TRBJ2-3 | CASSTGLLTDTQYF | KLGGALQAK | YES | NO | NO |
| 11526 | TRAV8-1 | TRAJ53 | CAVNHPNYKLTF | TRBV11-1 | TRBJ2-1 | CASSLIGILYNEQFF | KLGGALQAK | YES | NO | NO |
| 11527 | TRAV29DV5 | TRAJ32 | CAAPYTPGGATNKLIF | TRBV7-9 | TRBJ2-2 | CASSLCCSPNTGELFF | KLGGALQAK | YES | NO | NO |
| 11528 | TRAV8-4 | TRAJ47 | CAVGSMEYGNKLVF | TRBV4-3 | TRBJ2-7 | CASSQDMGGIDEQYF | KLGGALQAK | YES | NO | NO |
| 11529 | TRAV14DV4 | TRAJ52 | CAMSSNAGGTSYGKLTF | TRBV11-2 | TRBJ1-2 | CASSWGGGSHYGYTF | IVTDFSVIK | YES | NO | YES |
| 11530 | TRAV17 | TRAJ49 | CATYTGNQFYF | TRBV7-9 | TRBJ1-2 | CASSLAAGMGGYTF | KLGGALQAK | YES | NO | NO |
| 11531 | TRAV14DV4 | TRAJ43 | CAMRERNDMRF | TRBV27 | TRBJ2-7 | CASSPYLDIAGYEQYF | KLGGALQAK | YES | NO | NO |
| 11532 | TRAV13-1 | TRAJ34 | CAATPYNTDKLIF | TRBV7-3 | TRBJ1-1 | CASSRFGDTEAFF | KLGGALQAK | YES | NO | NO |
| 11533 | TRAV9-2 | TRAJ54 | CALSDRSQGAQKLVF | TRBV11-2 | TRBJ2-5 | CASSLVGGFGFETQYF | KLGGALQAK | YES | NO | NO |
| 11534 | TRAV27 | TRAJ18 | CAGAGRGSTLGRLYF | TRBV20-1 | TRBJ2-7 | CSASPGDYEQYF | KLGGALQAK | NO | NO | NO |
| 11535 | TRAV38-2DV8 | TRAJ40 | CAYRSSAGGSGTYKYIF | TRBV27 | TRBJ1-2 | CASSLYQGSNYGYTF | RLRAEAQVK | YES | YES | YES |
| 11536 | TRAV24 | TRAJ52 | CAFMGYAGGTSYGKLTF | TRBV12-3 | TRBJ2-2 | CASSVRVSGELFF | KLGGALQAK | YES | NO | NO |
| 11537 | TRAV13-1 | TRAJ9 | CAASDTGGFKTIF | TRBV5-1 | TRBJ2-7 | CASSRFWVGPYEQYF | KLGGALQAK | YES | NO | NO |
| 11538 | TRAV13-1 | TRAJ31 | CAAYDNNARLMF | TRBV28 | TRBJ1-5 | CASSFGQPQHF | KLGGALQAK | YES | NO | NO |
| 11539 | TRAV12-2 | TRAJ50 | CAVNGVKTSYDKVIF | TRBV7-9 | TRBJ2-1 | CASSFSYQGPYMFF | KLGGALQAK | YES | NO | NO |
| 11540 | TRAV35 | TRAJ42 | CAGQVNGGSQGNLIF | TRBV19 | TRBJ2-3 | CASSTYSIDTQYF | GILGFVFTL | YES | YES | NO |
| 11541 | TRAV12-1 | TRAJ31 | CVVNMDNARLMF | TRBV11-2 | TRBJ2-3 | CASSLGDFLRTDTQYF | RAKFKQLL | YES | NO | YES |
| 11544 | TRAV25 | TRAJ28 | CIRSGAGSYQLTF | TRBV12-4 | TRBJ1-5 | CASSLQGGFSNQPQHF | KLGGALQAK | YES | NO | NO |
| 11545 | TRAV12-2 | TRAJ35 | CAVRGFGNVLHC | TRBV7-2 | TRBJ2-5 | CASTPPRLVRWETQYF | KLGGALQAK | NO | NO | NO |
| 11546 | TRAV5 | TRAJ13 | CADSSGGYQKVTF | TRBV20-1 | TRBJ2-2 | CSANRRASGRNTGELFF | KLGGALQAK | YES | NO | NO |
| 11547 | TRAV26-1 | TRAJ50 | CIVLGGMKTSYDKVIF | TRBV9 | TRBJ2-7 | CASSLSSGTYEQYF | IVTDFSVIK | YES | NO | YES |
| 11548 | TRAV25 | TRAJ11 | CAAFYSTLTF | TRBV12-4 | TRBJ2-7 | CASSSPWGQGPHEQYF | GILGFVFTL | YES | NO | YES |
| 11549 | TRAV12-2 | TRAJ48 | CAVKRSNFGNEKLTF | TRBV3-1 | TRBJ2-2 | CASSQGFYLSGWDTGELFF | ELAGIGILTV | YES | YES | YES |
| 11550 | TRAV8-3 | TRAJ37 | CAVVVGSSNTGKLIF | TRBV19 | TRBJ2-7 | CASSIRSSYEQYF | GILGFVFTL | YES | NO | NO |
| 11551 | TRAV26-2 | TRAJ13 | CTLSGGYQKVTF | TRBV12-4 | TRBJ2-5 | CASSSPTSGGQETQYF | AVFDRKSDAK | YES | NO | NO |
| 11552 | TRAV2 | TRAJ26 | CAVEDDNYGQNFVF | TRBV19 | TRBJ2-2 | CASSTGQATGELFF | AVFDRKSDAK | NO | NO | NO |
| 11553 | TRAV14DV4 | TRAJ11 | CAMNSGYSTLTF | TRBV7-3 | TRBJ2-1 | CASSLSGQGRGEQFF | RAKFKQLL | YES | NO | YES |
| 11554 | TRAV19 | TRAJ30 | CALCKGDDKIIF | TRBV19 | TRBJ2-2 | CASSMRSTGELFF | GILGFVFTL | YES | NO | YES |
| 11555 | TRAV20 | TRAJ44 | CAVRTTGTASKLTF | TRBV11-2 | TRBJ1-2 | CASSWGGGSHYGYTF | IVTDFSVIK | YES | NO | YES |
| 11556 | TRAV8-6 | TRAJ8 | CAVMDTGFQKLVF | TRBV7-9 | TRBJ2-7 | CASSFLGDRDEQYF | KLGGALQAK | YES | NO | NO |
| 11557 | TRAV10 | TRAJ6 | CVVSEGGGSYIPTF | TRBV18 | TRBJ1-3 | CASSPGKDRFSGNTIYF | KLGGALQAK | YES | NO | NO |
| 11558 | TRAV14DV4 | TRAJ49 | CAMRVGTGNQFYF | TRBV7-2 | TRBJ1-1 | CASSFVGVGTEAFF | KLGGALQAK | YES | NO | NO |
| 11559 | TRAV16 | TRAJ40 | CALSPLSGTYKYIF | TRBV7-9 | TRBJ2-1 | CASSLGGLGNEQFF | KLGGALQAK | NO | NO | NO |
| 11560 | TRAV13-1 | TRAJ30 | CAARLNRDDKIIF | TRBV6-6 | TRBJ1-5 | CASSAVTEYSNQPQHF | KLGGALQAK | NO | NO | NO |
| 11562 | TRAV22 | TRAJ52 | CAVVDGTSYGKLTF | TRBV12-5 | TRBJ2-3 | CASGFRTDTQYF | KLGGALQAK | YES | NO | NO |
| 11564 | TRAV12-2 | TRAJ29 | CAVPSGNTPLVF | TRBV2 | TRBJ2-2 | CASSEYGGGVAGELFF | ELAGIGILTV | YES | NO | YES |

| | | | | | | | | | | |
|---|---|---|---|---|---|---|---|---|---|---|
| 11565 | TRAV13-1 | TRAJ4 | CAATVSGGYNKLIF | TRBV27 | TRBJ2-1 | CASSLWGPDSNNEQFF | RAKFKQLL | YES | NO | YES |
| 11566 | TRAV17 | TRAJ29 | CATDKFSGNTPLVF | TRBV11-2 | TRBJ2-1 | CASSLGKNEQFF | KLGGALQAK | YES | NO | NO |
| 11567 | TRAV12-2 | TRAJ41 | CAASNSGYALNF | TRBV27 | TRBJ2-2 | CASSSPSGSGTGELFF | ELAGIGILTV | YES | NO | YES |
| 11568 | TRAV4 | TRAJ45 | CLARSGGGADGLTF | TRBV6-5 | TRBJ2-1 | CASSQGQGSYFEQFF | KLGGALQAK | NO | NO | NO |
| 11569 | TRAV38-2DV8 | TRAJ57 | CALGGGSEKLVF | TRBV11-2 | TRBJ1-2 | CASSWGGGSHYGYTF | IVTDFSVIK | YES | NO | NO |
| 11570 | TRAV4 | TRAJ8 | CLVGESFQKLVF | TRBV11-2 | TRBJ2-5 | CASGRHRTAQETQYF | KLGGALQAK | YES | NO | NO |
| 11571 | TRAV3 | TRAJ33 | CAVSSGDSNYQLIW | TRBV4-1 | TRBJ1-1 | CASSQGMNTEAFF | KLGGALQAK | YES | NO | NO |
| 11573 | TRAV10 | TRAJ30 | CVVRMRDDKIIF | TRBV28 | TRBJ1-2 | CASSPRTKSFLSYGYTF | KLGGALQAK | YES | NO | NO |
| 11574 | TRAV8-3 | TRAJ42 | CANTGSQGNLIF | TRBV19 | TRBJ2-7 | CASSDRSGTEQYF | GILGFVFTL | YES | YES | YES |
| 11575 | TRAV39 | TRAJ35 | CAVGEGFGNVLHC | TRBV5-1 | TRBJ1-2 | CASSLRGASAGYTF | KLGGALQAK | NO | NO | NO |
| 11576 | TRAV14DV4 | TRAJ37 | CAMSSYSGNTGKLIF | TRBV12-4 | TRBJ2-1 | CASSYGTSDEQFF | RAKFKQLL | YES | NO | YES |
| 11577 | TRAV1-2 | TRAJ4 | CAVHNPGGYNKLIF | TRBV9 | TRBJ2-2 | CASSVAAGNTGELFF | KLGGALQAK | YES | NO | NO |
| 11578 | TRAV12-1 | TRAJ38 | CVVNRAGNNRKLIW | TRBV24-1 | TRBJ2-1 | CATSDTHQGYQQFF | KLGGALQAK | YES | NO | NO |
| 11579 | TRAV29DV5 | TRAJ17 | CAARDYEAAGNKLTF | TRBV5-4 | TRBJ2-6 | CASSSQGAGANVLTF | AVFDRKSDAK | YES | NO | YES |
| 11580 | TRAV8-6 | TRAJ6 | CAVSDTSGGSYIPTF | TRBV19 | TRBJ1-5 | CASSMGQGTLRNQPQHF | AVFDRKSDAK | YES | NO | NO |
| 11581 | TRAV38-2DV8 | TRAJ40 | CAYSSSGTYKYIF | TRBV7-9 | TRBJ2-1 | CASSLVVTEQFF | KLGGALQAK | YES | NO | NO |
| 11582 | TRAV12-2 | TRAJ48 | CAVSSNFGNEKLTF | TRBV2 | TRBJ2-7 | CASSTDYEQYF | KLGGALQAK | YES | NO | NO |
| 11583 | TRAV27 | TRAJ42 | CAGGGSQGNLIF | TRBV19 | TRBJ1-2 | CASSTGTLYGYTF | GILGFVFTL | YES | NO | YES |
| 11584 | TRAV8-4 | TRAJ37 | CAVPGGNTGKLIF | TRBV19 | TRBJ2-7 | CASSIRSSYEQYF | GILGFVFTL | YES | NO | YES |
| 11585 | TRAV12-2 | TRAJ33 | CAVSAPFMDSNYQLIW | TRBV28 | TRBJ2-2 | CASSPRLAGTGELFF | KLGGALQAK | YES | NO | NO |
| 11586 | TRAV3 | TRAJ18 | CAVRDRGSTLGRLYF | TRBV4-2 | TRBJ2-5 | CASSQDGKAQYF | KLGGALQAK | YES | NO | NO |
| 11587 | TRAV24 | TRAJ42 | CAFVDGSQGNLIF | TRBV19 | TRBJ2-3 | CASSIGTDTQYF | GILGFVFTL | YES | NO | YES |
| 11588 | TRAV39 | TRAJ43 | CAVDINNDMRF | TRBV9 | TRBJ2-2 | CASSVDGGDPPWELFF | KLGGALQAK | YES | NO | NO |
| 11589 | TRAV13-2 | TRAJ45 | CADSGGGADGLTF | TRBV6-6 | TRBJ2-5 | CASSYISTGAPTQYF | RAKFKQLL | NO | NO | NO |
| 11590 | TRAV13-1 | TRAJ28 | CAAGGYSGAGSYQLTF | TRBV12-3 | TRBJ2-7 | CASSLVLGQVYEQYF | GILGFVFTL | NO | NO | NO |
| 11591 | TRAV8-4 | TRAJ4 | CAVSDRGYNKLIF | TRBV7-2 | TRBJ1-1 | CASSFEEETEAFF | AVFDRKSDAK | YES | NO | YES |
| 11592 | TRAV12-3 | TRAJ8 | CAMKGNTGFQKLVF | TRBV14 | TRBJ2-3 | CASSQEMGGPDTQYF | AVFDRKSDAK | YES | NO | YES |
| 11593 | TRAV24 | TRAJ27 | CAFKRNTNAGKSTF | TRBV11-2 | TRBJ1-5 | CASSFWPGPNQPQHF | KLGGALQAK | YES | NO | NO |
| 11594 | TRAV27 | TRAJ40 | CAGERASGTYKYIF | TRBV7-9 | TRBJ2-1 | CASSVSERELRNEQFF | KLGGALQAK | YES | NO | NO |
| 11595 | TRAV3 | TRAJ35 | CAAPEGGFGNVLHC | TRBV5-1 | TRBJ2-2 | CASSLAGLAVNTGELFF | KLGGALQAK | NO | NO | NO |
| 11596 | TRAV3 | TRAJ35 | CAVSVGFGNVLHC | TRBV5-6 | TRBJ2-3 | CASREPGTYTDTQYF | KLGGALQAK | NO | NO | NO |
| 11597 | TRAV12-2 | TRAJ24 | CAVNPGGKLQF | TRBV4-3 | TRBJ2-2 | CASSQAQIQTGELFF | ELAGIGILTV | YES | NO | NO |
| 11598 | TRAV25 | TRAJ37 | CAGVGSSNTGKLIF | TRBV20-1 | TRBJ2-7 | CSATALGPLDEQYF | KLGGALQAK | NO | NO | NO |
| 11599 | TRAV27 | TRAJ24 | CAGVRGWGKLQF | TRBV7-3 | TRBJ2-1 | CASSLRGLAGNEQFF | KLGGALQAK | YES | NO | NO |
| 11600 | TRAV1-2 | TRAJ20 | CAVRERDYKLSF | TRBV6-3 | TRBJ1-5 | CASAVANQPQHF | KLGGALQAK | YES | NO | NO |
| 11602 | TRAV35 | TRAJ28 | CAGRTWAGSYQLTF | TRBV6-5 | TRBJ1-2 | CASSYRGSRGYTF | KLGGALQAK | YES | NO | NO |
| 11603 | TRAV8-6 | TRAJ34 | CAVRYNTDKLIF | TRBV29-1 | TRBJ1-2 | CSVPGTVYGYTF | KLGGALQAK | YES | NO | NO |
| 11604 | TRAV19 | TRAJ54 | CALTPLIIQGAQKLVF | TRBV28 | TRBJ2-3 | CASSLTGGDTQYF | LLFGYPVYV | YES | YES | YES |
| 11605 | TRAV38-1 | TRAJ34 | CAFFGGSTDKLIF | TRBV6-5 | TRBJ2-7 | CASSNTGRREQYF | LLDFVRFMGV | YES | NO | YES |
| 11606 | TRAV36DV7 | TRAJ54 | CAHLQGAQKLVF | TRBV28 | TRBJ2-2 | CASSLYSATGELFF | AVFDRKSDAK | NO | NO | NO |
| 11607 | TRAV19 | TRAJ45 | CALRDLYSGGGADGLTF | TRBV11-2 | TRBJ1-1 | CASSFGQNTEAFF | KLGGALQAK | YES | NO | NO |
| 11608 | TRAV34 | TRAJ28 | CGADSKTGSYQLTF | TRBV28 | TRBJ2-1 | CASSFLPRAYNEQFF | AVFDRKSDAK | YES | NO | YES |
| 11609 | TRAV1-2 | TRAJ12 | CAVFLMDSSYKLIF | TRBV20-1 | TRBJ2-7 | CSARDLGLAGDEQYF | AYAQKIFKI | NO | NO | NO |
| 11610 | TRAV8-4 | TRAJ15 | CAVTPWEQAGTALIF | TRBV5-1 | TRBJ2-1 | CASSLDAYNEQFF | KLGGALQAK | YES | NO | NO |
| 11611 | TRAV14DV4 | TRAJ23 | CAMREGPGKLIF | TRBV27 | TRBJ2-7 | CASRRDGSYEQYF | KLGGALQAK | YES | NO | NO |
| 11612 | TRAV21 | TRAJ53 | CAVNSGGSNYKLTF | TRBV6-1 | TRBJ1-2 | CASSVTGPYGYTF | KLGGALQAK | YES | NO | NO |
| 11613 | TRAV13-1 | TRAJ20 | CAASGYDYKLSF | TRBV11-2 | TRBJ1-2 | CASSWGGGSHYGYTF | IVTDFSVIK | YES | NO | YES |
| 11614 | TRAV12-2 | TRAJ49 | CAVNGGNQFYF | TRBV4-2 | TRBJ1-1 | CASSLEMGAGTEAFF | ELAGIGILTV | YES | NO | YES |
| 11615 | TRAV39 | TRAJ58 | CAVDRWETSGSRLTF | TRBV6-1 | TRBJ2-2 | CASSEAPGRRTGELFF | KLGGALQAK | YES | NO | NO |
| 11616 | TRAV35 | TRAJ26 | CAGHGGQNFVF | TRBV11-2 | TRBJ1-2 | CASSPGGGSSFGYTF | IVTDFSVIK | YES | YES | YES |
| 11617 | TRAV12-2 | TRAJ53 | CAVNPGWGGSNYKLTF | TRBV9 | TRBJ2-5 | CASSVTGQLQETQYF | ELAGIGILTV | YES | YES | YES |
| 11618 | TRAV19 | TRAJ34 | CALSSYNTDKLIF | TRBV13 | TRBJ2-7 | CASSSLLAGGLEQYF | KLGGALQAK | YES | NO | NO |
| 11619 | TRAV29DV5 | TRAJ30 | CAASAGVDDKIIF | TRBV27 | TRBJ2-1 | CASSFGLGGAYNEQFF | AVFDRKSDAK | YES | YES | YES |
| 11620 | TRAV13-2 | TRAJ54 | CAEDLIQGAQKLVF | TRBV4-3 | TRBJ2-1 | CASSQGREAGGPAGYNEQFF | KLGGALQAK | NO | NO | NO |
| 11621 | TRAV12-3 | TRAJ23 | CAIFYNQGGKLIF | TRBV20-1 | TRBJ1-1 | CSASSYMNTEAFF | KLGGALQAK | YES | NO | NO |
| 11622 | TRAV5 | TRAJ20 | CAESKELSF | TRBV15 | TRBJ1-1 | CATSSKTGLTEAFF | KLGGALQAK | NO | NO | NO |
| 11623 | TRAV19 | TRAJ9 | CALRGNTGGFKTIF | TRBV27 | TRBJ2-1 | CASSLSGATYNEQFF | RTLNAWVKV | YES | NO | NO |
| 11624 | TRAV19 | TRAJ32 | CALSEAGDYGGATNKLIF | TRBV7-2 | TRBJ2-3 | CASRLAGRMDTQYF | KLGGALQAK | YES | NO | NO |
| 11626 | TRAV29DV5 | TRAJ11 | CAASFGLRGYSTLTF | TRBV9 | TRBJ1-1 | CASSIIDRSGTEAFF | IMDQVPFSV | YES | NO | YES |
| 11627 | TRAV41 | TRAJ10 | CAASITGGGNKLTF | TRBV9 | TRBJ2-7 | CASSVVDAGFEQYF | AVFDRKSDAK | NO | NO | NO |
| 11628 | TRAV41 | TRAJ33 | CAVRMDSNYQLIW | TRBV3-1 | TRBJ2-1 | CASSQDPSGRWYNEQFF | KLGGALQAK | YES | NO | NO |
| 11629 | TRAV1-2 | TRAJ20 | CAVRERDYKLSF | TRBV9 | TRBJ1-2 | CASSAMDRGADGYTF | KLGGALQAK | YES | NO | NO |
| 11630 | TRAV38-2DV8 | TRAJ41 | CAYRSLYALNF | TRBV19 | TRBJ1-6 | CASSSGTGGYNSPLHF | IVTDFSVIK | YES | YES | NO |
| 11631 | TRAV14DV4 | TRAJ10 | CAILTGGGNKLTF | TRBV10-3 | TRBJ1-2 | CAISESNKGTPGYTF | KLGGALQAK | YES | NO | NO |
| 11632 | TRAV13-2 | TRAJ45 | CADSGGGADGLTF | TRBV4-1 | TRBJ2-1 | CASSQGSAGGNEQFF | AVFDRKSDAK | YES | YES | YES |
| 11633 | TRAV8-4 | TRAJ26 | CAVSGDYGQNFVF | TRBV29-1 | TRBJ2-7 | CSVESGRTDEQYF | IPSINVHHY | YES | NO | YES |
| 11634 | TRAV14DV4 | TRAJ41 | CAMRGHSGYALNF | TRBV7-2 | TRBJ1-5 | CASSLAGRTANQPQHF | KLGGALQAK | YES | NO | NO |
| 11635 | TRAV38-1 | TRAJ43 | CAFMKPGNNDMRF | TRBV9 | TRBJ2-3 | CASSVAGQDTDTQYF | KLGGALQAK | YES | NO | NO |
| 11636 | TRAV30 | TRAJ42 | CGTGLGGGSQGNLIF | TRBV19 | TRBJ2-1 | CASSIRSSNEQFF | GILGFVFTL | YES | NO | YES |
| 11637 | TRAV27 | TRAJ42 | CSTGGSQGNLIF | TRBV19 | TRBJ2-7 | CASSIRSSYEQYF | GILGFVFTL | YES | NO | NO |
| 11638 | TRAV38-1 | TRAJ23 | CAPLPQGGKLIF | TRBV27 | TRBJ1-1 | CASSWPGGTEAFF | KLGGALQAK | YES | NO | NO |
| 11639 | TRAV13-2 | TRAJ26 | CAENDNYGQNFVF | TRBV9 | TRBJ2-1 | CASSATGTENNEQFF | KLGGALQAK | YES | NO | NO |
| 11640 | TRAV30 | TRAJ48 | CGTDPNFGNEKLTF | TRBV10-3 | TRBJ2-1 | CAISESGSGGPGSRFF | KLGGALQAK | NO | NO | NO |
| 11641 | TRAV12-2 | TRAJ48 | CAVKRSNFGNEKLTF | TRBV6-5 | TRBJ1-1 | CASSYTPVSMNTEAFF | ELAGIGILTV | YES | YES | YES |
| 11642 | TRAV35 | TRAJ53 | CAGPLGGGSNYKLTF | TRBV12-3 | TRBJ2-3 | CASSYPFGVGPDTQYF | KLGGALQAK | YES | NO | NO |
| 11643 | TRAV8-6 | TRAJ49 | CAVTAANTGNQFYF | TRBV5-1 | TRBJ2-2 | CASSFTAGSANTGELFF | KLGGALQAK | YES | NO | NO |
| 11644 | TRAV12-2 | TRAJ48 | CAVKGSTF | TRBV15 | TRBJ1-1 | CATSGSSGTEAFF | ELAGIGILTV | YES | NO | YES |
| 11645 | TRAV8-1 | TRAJ42 | CAVNTNGGSQGNLIF | TRBV19 | TRBJ2-7 | CASSIRSSYEQYF | GILGFVFTL | YES | NO | YES |
| 11646 | TRAV24 | TRAJ52 | CAFNAGGTSYGKLTF | TRBV7-8 | TRBJ1-5 | CASSVDSYSNQPQHF | KLGGALQAK | YES | NO | NO |
| 11647 | TRAV19 | TRAJ41 | CALSVEAGYALNF | TRBV5-6 | TRBJ1-1 | CATGTVNTEAFF | KLGGALQAK | YES | NO | NO |
| 11648 | TRAV1-2 | TRAJ34 | CAVRDLGNTDKLIF | TRBV25-1 | TRBJ1-5 | CASSESPVNQPQHF | KLGGALQAK | YES | NO | NO |
| 11649 | TRAV14DV4 | TRAJ52 | CAMRLINAGGTSYGKLTF | TRBV20-1 | TRBJ1-2 | CSARTGTNYGYTF | KLGGALQAK | YES | NO | NO |
| 11650 | TRAV20 | TRAJ56 | CAVQAWGANSKLTF | TRBV28 | TRBJ2-7 | CASSWGSYEQYF | KLGGALQAK | YES | NO | NO |
| 11651 | TRAV38-2DV8 | TRAJ40 | CAIEDDSGTYKYIF | TRBV27 | TRBJ2-3 | CASSLKTSGRGDTQYF | KLGGALQAK | YES | NO | NO |
| 11652 | TRAV12-2 | TRAJ43 | CAVNAGNDMRF | TRBV6-3 | TRBJ1-2 | CASSYGQGWGYTF | ELAGIGILTV | YES | NO | YES |
| 11653 | TRAV8-1 | TRAJ22 | CAVNAEGSARQLTF | TRBV4-3 | TRBJ2-1 | CASSQGLSGSSYNEQFF | KLGGALQAK | YES | NO | NO |
| 11654 | TRAV1-2 | TRAJ32 | CAVDRGATNKLIF | TRBV6-3 | TRBJ2-2 | CASARTTGELFF | KLGGALQAK | YES | NO | NO |
| 11655 | TRAV12-2 | TRAJ35 | CAVSI | TRBV4-2 | TRBJ1-1 | CASSPPGEEAFF | ELAGIGILTV | NO | NO | NO |
| 11656 | TRAV1-2 | TRAJ11 | CAVNRYSTLTF | TRBV20-1 | TRBJ2-7 | CSASLGRGYEQYF | AVFDRKSDAK | NO | NO | NO |
| 11657 | TRAV8-3 | TRAJ27 | CAVGAPAGKSTF | TRBV7-9 | TRBJ2-3 | CASSYLQDFTDTQYF | KLGGALQAK | YES | NO | NO |
| 11658 | TRAV21 | TRAJ58 | CAVDGTSGSRLTF | TRBV7-2 | TRBJ2-3 | CATQLGSFTDTQYF | KLGGALQAK | YES | NO | NO |
| 11659 | TRAV12-2 | TRAJ48 | CAVSGNEKLTF | TRBV9 | TRBJ1-3 | CASSVYLSGNTIYF | ELAGIGILTV | NO | NO | NO |
| 11660 | TRAV23DV6 | TRAJ48 | CAAYNFGNEKLTF | TRBV12-5 | TRBJ1-2 | CASGLVRGQGNGYTF | KLGGALQAK | YES | NO | NO |
| 11661 | TRAV19 | TRAJ36 | CALSEGQTGANNLFF | TRBV19 | TRBJ1-5 | CASSWGGLSNQPQHF | ELAGIGILTV | YES | NO | YES |
| 11662 | TRAV12-2 | TRAJ42 | CAVKRGSQGNLIF | TRBV28 | TRBJ1-5 | CASSLAWGQPQHF | ELAGIGILTV | YES | YES | YES |
| 11663 | TRAV26-2 | TRAJ17 | CILRDVIKAAGNKLTF | TRBV27 | TRBJ1-2 | CASSSVRGPNYGYTF | RAKFKQLL | YES | NO | YES |
| 11664 | TRAV8-6 | TRAJ42 | CAVQGSQGNLIF | TRBV19 | TRBJ2-7 | CASSSRSSYEQYF | GILGFVFTL | YES | NO | YES |
| 11665 | TRAV12-1 | TRAJ27 | CVVNEDTNAGKSTF | TRBV6-3 | TRBJ1-1 | CASREGVTEAFF | KLGGALQAK | YES | NO | NO |
| 11666 | TRAV38-1 | TRAJ44 | CAFMKLTGTASKLTF | TRBV5-1 | TRBJ2-1 | CASSFSGANPYNEQFF | AVFDRKSDAK | YES | NO | YES |
| 11667 | TRAV12-2 | TRAJ42 | CAARNYGGSQGNLIF | TRBV18 | TRBJ1-1 | CASSHNSEYTEAFF | KLGGALQAK | YES | NO | NO |
| 11668 | TRAV5 | TRAJ5 | CAEAVMDTGRRALTF | TRBV7-9 | TRBJ2-7 | CASSLADGGFSYEQYF | KLGGALQAK | YES | NO | NO |
| 11669 | TRAV26-2 | TRAJ43 | CILRESWNNDMRF | TRBV20-1 | TRBJ1-1 | CSARDGTTEAFF | KLGGALQAK | YES | NO | NO |
| 11670 | TRAV23DV6 | TRAJ28 | CAAHRVDSGAGSYQLTF | TRBV6-5 | TRBJ1-2 | CASSALGGSYGYTF | FLYALALLL | YES | YES | NO |
| 11671 | TRAV29DV5 | TRAJ57 | CAASRFFPIWEKLVF | TRBV5-6 | TRBJ2-3 | CASSLARAGGFNTQYF | KLGGALQAK | YES | NO | NO |

| | | | | | | | | | | |
|---|---|---|---|---|---|---|---|---|---|---|
| 11672 | TRAV2 | TRAJ8 | CAVNTGFQKLVF | TRBV10-3 | TRBJ2-3 | CAIGLAGDTQYF | KLGGALQAK | NO | NO | NO |
| 11673 | TRAV19 | TRAJ3 | CALSDPGYSSASKIIF | TRBV12-5 | TRBJ2-1 | CASGLNIDGDEQFF | KLGGALQAK | YES | NO | NO |
| 11674 | TRAV5 | TRAJ3 | CAEYSSASKIIF | TRBV14 | TRBJ2-5 | CASSQSPGGTQYF | GLCTLVAML | NO | NO | NO |
| 11675 | TRAV8-3 | TRAJ24 | CAVGGPRSDSWGKLQF | TRBV6-6 | TRBJ2-1 | CASSNPLEGIDEQFF | KLGGALQAK | NO | NO | NO |
| 11676 | TRAV5 | TRAJ32 | CADHGGATNKLIF | TRBV4-1 | TRBJ2-7 | CASSTGAGVNEQYF | AVFDRKSDAK | YES | NO | YES |
| 11678 | TRAV12-1 | TRAJ6 | CVVESGGSYIPTF | TRBV19 | TRBJ2-7 | CASSARSSYEQYF | GILGFVFTL | YES | NO | YES |
| 11679 | TRAV14DV4 | TRAJ21 | CAMREADFNKFYF | TRBV5-6 | TRBJ2-7 | CASSWSSAGGPSYEQYF | IVTDFSVIK | YES | NO | YES |
| 11680 | TRAV8-6 | TRAJ42 | CAGAGSQGNLIF | TRBV19 | TRBJ2-7 | CASSIRSSYEQYF | GILGFVFTL | YES | NO | NO |
| 11681 | TRAV5 | TRAJ6 | CAEMEGGSYIPTF | TRBV29-1 | TRBJ1-4 | CSVGSRGTNEKLFF | GLCTLVAML | YES | YES | YES |
| 11682 | TRAV20 | TRAJ41 | CAVQGLNSGYALNF | TRBV20-1 | TRBJ1-1 | CSAPRRTTEAFF | KLGGALQAK | YES | NO | NO |
| 11683 | TRAV29DV5 | TRAJ21 | CAATAFSRNFNKFYF | TRBV9 | TRBJ2-6 | CASSVAVGTGSGANVLTF | AVFDRKSDAK | YES | NO | YES |
| 11684 | TRAV1-2 | TRAJ20 | CAGLTGDYKLSF | TRBV6-3 | TRBJ2-3 | CASRGGWEADTQYF | KLGGALQAK | YES | NO | NO |
| 11685 | TRAV39 | TRAJ42 | CAVDMAGGSQGNLIF | TRBV19 | TRBJ2-7 | CASSFRSGIEQYF | GILGFVFTL | YES | NO | YES |
| 11686 | TRAV39 | TRAJ26 | CAVDFRDNYGQNFVF | TRBV5-6 | TRBJ1-1 | CASSLRDGSEAFF | RAKFKQLL | YES | YES | YES |
| 11687 | TRAV12-2 | TRAJ36 | CAVIGANNLFF | TRBV30 | TRBJ1-3 | CASRIGNGNTIYF | KLGGALQAK | YES | NO | NO |
| 11688 | TRAV38-1 | TRAJ56 | CAFMKLAGANSKLTF | TRBV3-1 | TRBJ1-6 | CASKSEKGAGGSPLHF | KLGGALQAK | YES | NO | NO |
| 11689 | TRAV1-2 | TRAJ32 | CAVEDYGGATNKLIF | TRBV10-1 | TRBJ2-7 | CASNRDLSSYEQYF | KLGGALQAK | YES | NO | NO |
| 11690 | TRAV13-1 | TRAJ43 | CAPRAPTRNNDMRF | TRBV15 | TRBJ2-2 | CATSSPGGYTGELFF | KLGGALQAK | YES | NO | NO |
| 11691 | TRAV12-3 | TRAJ42 | CAMNYGGSQGNLIF | TRBV3-1 | TRBJ1-2 | CASSQDWGTVYGYTF | KLGGALQAK | YES | NO | NO |
| 11692 | TRAV13-1 | TRAJ4 | CAAILPSGGYNKLIF | TRBV20-1 | TRBJ1-2 | CSARDRTGNGYTF | GLCTLVAML | YES | YES | YES |
| 11693 | TRAV27 | TRAJ42 | CAGNYGGSQGNLIF | TRBV19 | TRBJ2-7 | CASSLRSSYEQYF | GILGFVFTL | YES | NO | YES |
| 11694 | TRAV29DV5 | TRAJ49 | CAASTGNQFYF | TRBV27 | TRBJ2-3 | CASSGGASPYTQYF | KLGGALQAK | YES | NO | NO |
| 11695 | TRAV27 | TRAJ42 | CAGAPGGGSQGNLIF | TRBV19 | TRBJ2-7 | CASSIRSSYEQYF | GILGFVFTL | YES | NO | YES |
| 11696 | TRAV26-2 | TRAJ44 | CILSPAGTASKLTF | TRBV11-2 | TRBJ2-1 | CASSLGQQGVNRDWYNEQFF | AVFDRKSDAK | NO | NO | NO |
| 11697 | TRAV39 | TRAJ10 | CAARGVLTGGGNKLTF | TRBV9 | TRBJ2-7 | CASSRTGVEQYF | AVFDRKSDAK | YES | NO | YES |
| 11698 | TRAV8-3 | TRAJ26 | CAVGARDYGQNFVF | TRBV7-3 | TRBJ2-2 | CASSLGTSGGTGELFF | RAKFKQLL | YES | YES | YES |
| 11699 | TRAV35 | TRAJ31 | CAGPSYARLMF | TRBV19 | TRBJ1-2 | CASSIDHRNGYTF | KLGGALQAK | YES | NO | NO |
| 11700 | TRAV30 | TRAJ20 | CGTEYPNDYKLSF | TRBV7-9 | TRBJ2-5 | CASSFVDRAETQYF | RAKFKQLL | YES | NO | NO |
| 11701 | TRAV8-6 | TRAJ27 | CAVSSGPGKSTF | TRBV6-3 | TRBJ1-5 | CASSYNPRTEVSNQPQHF | GILGFVFTL | YES | NO | YES |
| 11702 | TRAV13-2 | TRAJ50 | CAERETSYDKVIF | TRBV7-9 | TRBJ1-1 | CASSSLGGHGEAFF | RAKFKQLL | YES | NO | YES |
| 11703 | TRAV8-6 | TRAJ42 | CAVTDGGSQGNLIF | TRBV19 | TRBJ2-7 | CASSARSSYEQYF | GILGFVFTL | YES | NO | YES |
| 11704 | TRAV12-3 | TRAJ40 | CAMIGRTSGTYKYIF | TRBV7-9 | TRBJ2-1 | CASSLARQGDEQFF | KLGGALQAK | YES | NO | NO |
| 11705 | TRAV17 | TRAJ39 | CATGYNNAGNMLTF | TRBV27 | TRBJ1-2 | CASSLGGQNYGYTF | KLGGALQAK | YES | NO | NO |
| 11706 | TRAV13-2 | TRAJ45 | CADSGGGADGLTF | TRBV5-4 | TRBJ1-5 | CASSSTGGGEKDQPQHF | RAKFKQLL | YES | NO | YES |
| 11707 | TRAV8-3 | TRAJ21 | CAVGDNFNKFYF | TRBV3-1 | TRBJ2-7 | CASSPVTGSSYEQYF | GLCTLVAML | YES | YES | YES |
| 11708 | TRAV4 | TRAJ3 | CLVGDPFYSSASKIIF | TRBV12-3 | TRBJ2-2 | CASSFSGNTGELFF | RAKFKQLL | YES | NO | YES |
| 11709 | TRAV17 | TRAJ36 | CATDSLTGANNLFF | TRBV2 | TRBJ2-1 | CASVWQGAYNEQFF | KLGGALQAK | YES | NO | NO |
| 11710 | TRAV22 | TRAJ20 | CAVEPPNDYKLSF | TRBV27 | TRBJ2-1 | CASSYFGNPVQFF | KLGGALQAK | YES | NO | NO |
| 11711 | TRAV25 | TRAJ42 | CAGNYGGSQGNLIF | TRBV19 | TRBJ2-3 | CASSVRSSDTQYF | GILGFVFTL | YES | NO | YES |
| 11712 | TRAV12-2 | TRAJ37 | CAGSGGNTGKLIF | TRBV12-4 | TRBJ2-1 | CASGRLSYNEQFF | RAKFKQLL | YES | NO | YES |
| 11713 | TRAV21 | TRAJ41 | CAVISNSGYALNF | TRBV18 | TRBJ2-3 | CASTTTGNTDTQYF | GLCTLVAML | YES | NO | YES |
| 11714 | TRAV10 | TRAJ9 | CVVTLNTGGFKTIF | TRBV9 | TRBJ2-5 | CASSGDPTGLAPQMAQYF | KLGGALQAK | YES | NO | NO |
| 11715 | TRAV23DV6 | TRAJ48 | CAAPPVSNFGNEKLTF | TRBV7-9 | TRBJ1-2 | CASSLAGRNYGYTF | KLGGALQAK | NO | NO | NO |
| 11716 | TRAV14DV4 | TRAJ50 | CAMREGSKTSYDKVIF | TRBV6-5 | TRBJ1-5 | CASSYSMGPNQPQHF | KLGGALQAK | YES | NO | NO |
| 11717 | TRAV22 | TRAJ45 | CAVLYSGGGADGLTF | TRBV7-8 | TRBJ2-7 | CASSLLGGDYEQYF | KLGGALQAK | YES | NO | NO |
| 11718 | TRAV9-2 | TRAJ20 | CALIEGDDYKLSF | TRBV12-3 | TRBJ1-2 | CASSLGGGYGYTF | KLGGALQAK | YES | NO | NO |
| 11719 | TRAV27 | TRAJ42 | CAGAGSQGNLIF | TRBV13 | TRBJ2-1 | CASSLTTGGRNEQFF | GILGFVFTL | YES | NO | YES |
| 11721 | TRAV8-3 | TRAJ5 | CAVDPMDTGRRALTF | TRBV19 | TRBJ1-5 | CASSIYSNQPQHF | GILGFVFTL | YES | NO | YES |
| 11722 | TRAV35 | TRAJ42 | CAGQLGGGSQGNLIF | TRBV19 | TRBJ2-7 | CASSQRSSYEQYF | GILGFVFTL | YES | NO | YES |
| 11723 | TRAV13-1 | TRAJ11 | CAAKEGYSTLTF | TRBV29-1 | TRBJ1-1 | CSAGGNTEAFF | RAKFKQLL | YES | NO | YES |
| 11724 | TRAV21 | TRAJ28 | CAVPYSGAGSYQLTF | TRBV9 | TRBJ2-2 | CASSGSTSQGGDTGELFF | KLGGALQAK | YES | NO | NO |
| 11725 | TRAV8-1 | TRAJ17 | CAVTGAAGNKLTF | TRBV9 | TRBJ2-1 | CASSHRVGNEQFF | KLGGALQAK | YES | NO | NO |
| 11726 | TRAV27 | TRAJ17 | CAPHKAAGNKLTF | TRBV19 | TRBJ1-5 | CASSKRSNQPQHF | GILGFVFTL | YES | NO | YES |
| 11727 | TRAV39 | TRAJ43 | CAVGRNNNDMRF | TRBV20-1 | TRBJ2-2 | CSARDFTNTGELFF | KLGGALQAK | YES | NO | NO |
| 11728 | TRAV17 | TRAJ40 | CATASTSGTYKYIF | TRBV4-1 | TRBJ2-3 | CASSQEWLAVSTDTQYF | RAKFKQLL | YES | NO | YES |
| 11729 | TRAV4 | TRAJ13 | CLVGDAYSGGYQKVTF | TRBV12-3 | TRBJ2-2 | CASSFSGNTGELFF | RAKFKQLL | NO | NO | NO |
| 11730 | TRAV8-4 | TRAJ40 | CAVSVTTSGTYKYIF | TRBV20-1 | TRBJ2-7 | CSASGWVRQGAFYEQYF | RLRAEAQVK | NO | NO | NO |
| 11731 | TRAV12-2 | TRAJ45 | CAVGGGADGLTF | TRBV27 | TRBJ2-7 | CASSLGGLGGSEQYF | ELAGIGILTV | YES | YES | YES |
| 11732 | TRAV35 | TRAJ42 | CAGQLGGGSQGNLIF | TRBV19 | TRBJ2-5 | CASSIRSSETQYF | GILGFVFTL | YES | NO | YES |
| 11733 | TRAV1-2 | TRAJ33 | CAVEDSNYQLIW | TRBV6-1 | TRBJ1-1 | CASSERQEGNTEAFF | KLGGALQAK | YES | NO | NO |
| 11734 | TRAV16 | TRAJ16 | CALGFAADGQKLLF | TRBV19 | TRBJ1-1 | CASRTGNTEAFF | GILGFVFTL | NO | NO | NO |
| 11735 | TRAV38-1 | TRAJ26 | CAFNERDNYGQNFVF | TRBV10-1 | TRBJ2-1 | CASSRDSSANEQFF | KLGGALQAK | YES | NO | NO |
| 11736 | TRAV21 | TRAJ29 | CAVRPDHTPLVF | TRBV27 | TRBJ2-5 | CASSAIPGTSYQETQYF | KLGGALQAK | YES | NO | NO |
| 11737 | TRAV27 | TRAJ42 | CAGAGSQGNLIF | TRBV12-4 | TRBJ2-1 | CASGRLSYNEQFF | RAKFKQLL | YES | NO | YES |
| 11738 | TRAV27 | TRAJ42 | CAGAPGGGSQGNLIF | TRBV19 | TRBJ1-2 | CASSLTVLYGYTF | GILGFVFTL | YES | NO | YES |
| 11739 | TRAV1-2 | TRAJ20 | CAAYSSNDYKLSF | TRBV15 | TRBJ1-5 | CATSRATGGGPQHF | KLGGALQAK | YES | NO | NO |
| 11740 | TRAV29DV5 | TRAJ42 | CAASGGSQGNLIF | TRBV12-3 | TRBJ2-3 | CASSLMGGRTDTQYF | KLGGALQAK | YES | NO | NO |
| 11741 | TRAV1-2 | TRAJ20 | CAVRDGDYKLSF | TRBV12-3 | TRBJ2-2 | CASSFSGNTGELFF | KLGGALQAK | YES | NO | NO |
| 11742 | TRAV17 | TRAJ36 | CATGTQTGANNLFF | TRBV11-2 | TRBJ2-3 | CASLPSGTDTQYF | KLGGALQAK | YES | NO | NO |
| 11743 | TRAV35 | TRAJ42 | CAPAGDGGSQGNLIF | TRBV19 | TRBJ1-5 | CASSPRSSQPQHF | GILGFVFTL | YES | YES | YES |
| 11744 | TRAV17 | TRAJ47 | CATAQYGNKLVF | TRBV5-6 | TRBJ1-1 | CASSLRDGSEAFF | RAKFKQLL | YES | NO | YES |
| 11745 | TRAV10 | TRAJ27 | CVGYTNAGKSTF | TRBV5-6 | TRBJ1-1 | CASSLRDGSEAFF | RAKFKQLL | YES | NO | YES |
| 11746 | TRAV24 | TRAJ15 | CAFNQAGTALIF | TRBV7-9 | TRBJ1-2 | CASSSLSGGYTF | RLRAEAQVK | YES | YES | YES |
| 11747 | TRAV13-1 | TRAJ42 | CAASGGGSQGNLIF | TRBV19 | TRBJ2-3 | CASSGRSTDTQYF | GILGFVFTL | YES | NO | YES |
| 11748 | TRAV3 | TRAJ43 | CAVRAPLYNNNDMRF | TRBV6-4 | TRBJ2-1 | CASSDHPVSPPGEQFF | KLGGALQAK | NO | NO | NO |
| 11749 | TRAV12-2 | TRAJ42 | CAVNTGGGSQGNLIF | TRBV19 | TRBJ1-4 | CASSIRSDEKLFF | GILGFVFTL | YES | NO | YES |
| 11750 | TRAV35 | TRAJ42 | CAGPSGNMNYGGSQGNLIF | TRBV19 | TRBJ2-1 | CASSIVALDEQFF | GILGFVFTL | YES | NO | NO |
| 11751 | TRAV5 | TRAJ20 | CAERKDDYKLSF | TRBV7-9 | TRBJ1-1 | CGSDRADTEAFF | KLGGALQAK | YES | NO | NO |
| 11752 | TRAV39 | TRAJ30 | CAGADDKIIF | TRBV12-4 | TRBJ2-3 | CASRASATQYF | SLFNTVATLY | YES | YES | YES |
| 11753 | TRAV30 | TRAJ42 | CGTDPLGYGGSQGNLIF | TRBV19 | TRBJ2-3 | CASSARSSDTQYF | GILGFVFTL | YES | YES | YES |
| 11754 | TRAV26-1 | TRAJ39 | CIVRAANNAGNMLTF | TRBV13 | TRBJ2-4 | CASRWKGSGAKNIQYF | IVTDFSVIK | YES | YES | YES |
| 11755 | TRAV38-1 | TRAJ33 | CAFTSDSNYQLIW | TRBV4-1 | TRBJ2-1 | CASSLLWAEYNEQFF | RAKFKQLL | YES | NO | YES |
| 11756 | TRAV1-2 | TRAJ29 | CAVRVPLENTPLVF | TRBV7-7 | TRBJ2-3 | CASSLSLTGGTDTQYF | KLGGALQAK | YES | NO | NO |
| 11757 | TRAV13-1 | TRAJ42 | CAYGGSQGNLIF | TRBV11-2 | TRBJ1-2 | CASSWGGGSHYGYTF | IVTDFSVIK | YES | NO | YES |
| 11758 | TRAV29DV5 | TRAJ34 | CAASGQNTDKLIF | TRBV9 | TRBJ2-5 | CASSVEAVPETQYF | KLGGALQAK | YES | NO | NO |
| 11759 | TRAV12-1 | TRAJ28 | CAPSGAGSYQLTF | TRBV27 | TRBJ2-3 | CASSLLGGYTDTQYF | KLGGALQAK | YES | NO | NO |
| 11760 | TRAV8-6 | TRAJ53 | CAVSGSGGSNYKLTF | TRBV29-1 | TRBJ2-3 | CTTSSGEADTQYF | RAKFKQLL | YES | NO | NO |
| 11761 | TRAV27 | TRAJ42 | CAGGTGGGSQGNLIF | TRBV19 | TRBJ2-7 | CASSIRASYEQYF | GILGFVFTL | YES | NO | YES |
| 11762 | TRAV20 | TRAJ29 | CAVFGSGNTPLVF | TRBV6-1 | TRBJ2-7 | CASSLTGPPYEQYF | KLGGALQAK | YES | NO | NO |
| 11763 | TRAV12-1 | TRAJ48 | CVVNSRVNFGNEKLTF | TRBV6-3 | TRBJ2-3 | CASSQLARIDTQYF | KLGGALQAK | YES | NO | NO |
| 11764 | TRAV24 | TRAJ49 | CAFNTGNQFYF | TRBV3-1 | TRBJ2-2 | CASSPRGSGGVTGELFF | KLGGALQAK | YES | NO | NO |
| 11765 | TRAV5 | TRAJ29 | CAESIDGGNTPLVF | TRBV14 | TRBJ1-1 | CASSQYPRGGNTEAFF | RAKFKQLL | NO | NO | NO |
| 11766 | TRAV39 | TRAJ48 | CAGPLDFGNEKLTF | TRBV2 | TRBJ2-7 | CASRELTGFSYEQYF | KLGGALQAK | YES | NO | NO |
| 11767 | TRAV26-1 | TRAJ21 | CIAPVPLNFNKFYF | TRBV29-1 | TRBJ1-2 | CSVEGSEDYGYTF | IPSINVHHY | YES | NO | YES |
| 11768 | TRAV17 | TRAJ6 | CATEGASGGSYIPTF | TRBV6-5 | TRBJ1-2 | CASSYQGGNYGYTF | FLYALALLL | YES | YES | YES |
| 11769 | TRAV8-4 | TRAJ49 | CAVSGNTGNQFYF | TRBV15 | TRBJ2-3 | CATSREPGLATDTQYF | AVFDRKSDAK | YES | NO | YES |
| 11770 | TRAV35 | TRAJ54 | CAGPIQGAQKLVF | TRBV19 | TRBJ1-5 | CASSLIEGLSWAQHF | KLGGALQAK | YES | NO | NO |
| 11771 | TRAV8-1 | TRAJ37 | CASFTGKLIF | TRBV11-2 | TRBJ2-7 | CASSTGTAAYEQYF | KLGGALQAK | YES | NO | NO |
| 11772 | TRAV13-2 | TRAJ53 | CAETHSGGSNYKLTF | TRBV29-1 | TRBJ1-5 | CSYSAGEDQPQHF | RAKFKQLL | YES | NO | YES |
| 11773 | TRAV20 | TRAJ32 | CAVRGATNKLIF | TRBV14 | TRBJ2-3 | CASSQEDSWGDTQYF | KLGGALQAK | YES | NO | NO |
| 11774 | TRAV30 | TRAJ42 | CGTDTGGGSQGNLIF | TRBV19 | TRBJ2-3 | CASSGRSTDTQYF | GILGFVFTL | YES | NO | YES |
| 11775 | TRAV12-2 | TRAJ27 | CAVYTNAGKSTF | TRBV11-2 | TRBJ2-2 | CASSSDPGAGELFF | KLGGALQAK | YES | NO | NO |
| 11776 | TRAV35 | TRAJ28 | CAGRKAGSLPTHF | TRBV5-1 | TRBJ1-5 | CASSLRGGQPQHF | KLGGALQAK | YES | NO | NO |
| 11777 | TRAV19 | TRAJ6 | CALSEAGVTGGSYIPTF | TRBV19 | TRBJ1-1 | CASSIFLGAEAFF | GILGFVFTL | YES | NO | YES |

| | | | | | | | | | | |
|---|---|---|---|---|---|---|---|---|---|---|
| 11778 | TRAV8-3 | TRAJ48 | CAVVPEYISNFGNEKLTF | TRBV4-1 | TRBJ2-7 | CASSPSTVSAEQYF | RAKFKQLL | YES | NO | YES |
| 11779 | TRAV29DV5 | TRAJ29 | CAARIGNTPLVF | TRBV4-1 | TRBJ2-3 | CASTPIESSTDTQYF | AVFDRKSDAK | YES | NO | YES |
| 11780 | TRAV13-2 | TRAJ39 | CAESSLDNMLTF | TRBV4-1 | TRBJ1-1 | CASSHGTVETEAFF | GILGFVFTL | YES | NO | NO |
| 11781 | TRAV12-2 | TRAJ15 | CAVNMRQAGTALIF | TRBV2 | TRBJ1-1 | CASSDPTRGTEAFF | KLGGALQAK | YES | NO | NO |
| 11782 | TRAV13-2 | TRAJ28 | CAENRYGSGAGSYQLTF | TRBV4-1 | TRBJ1-5 | CASSQDGWAGDQPQHF | KLGGALQAK | YES | NO | NO |
| 11783 | TRAV19 | TRAJ53 | CALSEEGGSNYKLTF | TRBV20-1 | TRBJ1-6 | CSAFPGRYNSPLHF | KLGGALQAK | NO | NO | NO |
| 11784 | TRAV36DV7 | TRAJ47 | CAVDGNKLVF | TRBV20-1 | TRBJ1-2 | CSAKEGPTGGYGYTF | KLGGALQAK | NO | NO | NO |
| 11785 | TRAV8-6 | TRAJ52 | CAVSDGDAGGTSYGKLTF | TRBV19 | TRBJ1-1 | CASSTSTGSEAFF | AVFDRKSDAK | YES | NO | YES |
| 11786 | TRAV13-2 | TRAJ45 | CADSGGGADGLTF | TRBV6-1 | TRBJ2-7 | CASSPSRLGTGFYEQYF | RAKFKQLL | YES | NO | YES |
| 11787 | TRAV13-2 | TRAJ50 | CAEMETSYDKVIF | TRBV7-9 | TRBJ2-6 | CASSEGGSGANVLTF | RAKFKQLL | YES | YES | YES |
| 11788 | TRAV36DV7 | TRAJ8 | CAVDNDGGGFQKLVF | TRBV5-4 | TRBJ2-1 | CASRTSGSSYNEQFF | KLGGALQAK | NO | NO | NO |
| 11789 | TRAV19 | TRAJ9 | CALSEAIRNTGGFKTIF | TRBV7-9 | TRBJ1-2 | CASSPRGDDYGYTF | KLGGALQAK | YES | NO | NO |
| 11790 | TRAV12-3 | TRAJ30 | CAMSWGRDDKIIF | TRBV12-3 | TRBJ2-2 | CASSFSGNTGELFF | RAKFKQLL | YES | YES | NO |
| 11791 | TRAV1-2 | TRAJ33 | CAVEDSNYQLIW | TRBV6-4 | TRBJ2-5 | CASSDSAGGQETQYF | KLGGALQAK | YES | NO | NO |
| 11792 | TRAV5 | TRAJ27 | CAEWDAGKSTF | TRBV7-8 | TRBJ1-1 | CASSLEAGNTEAFF | KLGGALQAK | YES | NO | NO |
| 11793 | TRAV17 | TRAJ22 | CATDRAGSARQLTF | TRBV11-2 | TRBJ1-2 | CASSWGGGSHYGYTF | KLGGALQAK | YES | NO | NO |
| 11794 | TRAV39 | TRAJ27 | CAGTNAGKSTF | TRBV7-3 | TRBJ2-1 | CASSFLAGVGYNEQFF | KLGGALQAK | YES | NO | NO |
| 11795 | TRAV29DV5 | TRAJ42 | CAASAGGGSQGNLIF | TRBV19 | TRBJ2-7 | CASSIRASYEQYF | GILGFVFTL | YES | NO | YES |
| 11796 | TRAV5 | TRAJ6 | CAEMEGGSYIPTF | TRBV4-2 | TRBJ1-5 | CASSQGSNQPQHF | RAKFKQLL | YES | NO | YES |
| 11797 | TRAV39 | TRAJ15 | CAVDGFRSRQAGTALIF | TRBV10-3 | TRBJ1-2 | CAISPPRVGANYGYTF | RLRAEAQVK | YES | YES | YES |
| 11798 | TRAV13-1 | TRAJ50 | CAASETSYDKVIF | TRBV25-1 | TRBJ1-4 | CASSEQGQGEKLFF | RAKFKQLL | YES | NO | YES |
| 11799 | TRAV26-1 | TRAJ27 | CIVRVVERTVSHTNAGKSTF | TRBV7-9 | TRBJ2-7 | CASSLAYFGGSYEQYF | AVFDRKSDAK | NO | NO | NO |
| 11800 | TRAV9-2 | TRAJ20 | CALVDYKLSF | TRBV6-6 | TRBJ1-1 | CASVTRDRGPNTEAFF | KLGGALQAK | NO | NO | NO |
| 11801 | TRAV17 | TRAJ24 | CATVSTDSWGKFQF | TRBV2 | TRBJ2-1 | CASRLAGGQLNEQFF | KLGGALQAK | YES | NO | NO |
| 11802 | TRAV5 | TRAJ6 | CAEMEGGSYIPTF | TRBV19 | TRBJ2-7 | CASSIRSSYEQYF | RAKFKQLL | YES | NO | YES |
| 11803 | TRAV8-4 | TRAJ45 | CAVSGHARGADGLTF | TRBV14 | TRBJ1-2 | CASSLGTILSYTF | KLGGALQAK | NO | NO | NO |
| 11804 | TRAV17 | TRAJ39 | CATGFNNAGNMLTF | TRBV20-1 | TRBJ2-5 | CSAPTSGAPETQYF | KLGGALQAK | YES | NO | NO |
| 11805 | TRAV13-2 | TRAJ45 | CADSGGGADGLTF | TRBV2 | TRBJ2-7 | CASSGRTGSSYEQYF | GILGFVFTL | YES | NO | YES |
| 11806 | TRAV5 | TRAJ6 | CAEMEGGSYIPTF | TRBV7-8 | TRBJ2-2 | CASSSPEGQPLNTGELFF | GLCTLVAML | YES | YES | YES |
| 11807 | TRAV13-1 | TRAJ23 | CAASNQGGKLIF | TRBV5-5 | TRBJ2-7 | CASSPWTGDYEQYF | KLGGALQAK | YES | NO | NO |
| 11808 | TRAV17 | TRAJ45 | CATDPSGGGADGLTF | TRBV27 | TRBJ1-2 | CASSPSDWNGYTF | KLGGALQAK | YES | NO | NO |
| 11809 | TRAV8-1 | TRAJ9 | CAVDTGGFKTIF | TRBV14 | TRBJ2-7 | CASSPGSPGYEQYF | KLGGALQAK | NO | NO | NO |
| 11810 | TRAV9-2 | TRAJ30 | CALLNRDDKIIF | TRBV5-1 | TRBJ1-1 | CASSYGTGENTEAFF | RAKFKQLL | YES | NO | NO |
| 11811 | TRAV1-2 | TRAJ34 | CAVRDGVSYNTDKLIF | TRBV20-1 | TRBJ2-6 | CSAMTSEGSGANVLTF | KLGGALQAK | YES | NO | NO |
| 11812 | TRAV35 | TRAJ42 | CAGQSEEANYGGSQGNLIF | TRBV19 | TRBJ2-7 | CASSSRSQYEQYF | GILGFVFTL | YES | NO | YES |
| 11813 | TRAV13-1 | TRAJ42 | CAASMGGGSQGNLIF | TRBV20-1 | TRBJ2-7 | CSASPGDYEQYF | GILGFVFTL | NO | NO | NO |
| 11814 | TRAV2 | TRAJ7 | CPRYGNNRLAF | TRBV7-8 | TRBJ2-2 | CASSPPIAGGYTGELFF | AVFDRKSDAK | NO | NO | NO |
| 11815 | TRAV19 | TRAJ27 | CALSEDTNAGKSTF | TRBV12-3 | TRBJ2-2 | CASSFSGNTGELFF | RAKFKQLL | YES | NO | YES |
| 11816 | TRAV20 | TRAJ23 | CAVLPSYNQGGKLIF | TRBV6-1 | TRBJ2-7 | CASSEFPPPYEQYF | KLGGALQAK | YES | NO | NO |
| 11817 | TRAV8-3 | TRAJ5 | CAVDPMDTGRRALTF | TRBV12-3 | TRBJ2-2 | CASSFSGNTGELFF | RAKFKQLL | YES | NO | YES |
| 11818 | TRAV1-2 | TRAJ33 | CAVRAHSNYQLIW | TRBV7-9 | TRBJ2-1 | CASSSGLAGLNEQFF | KLGGALQAK | YES | NO | NO |
| 11819 | TRAV10 | TRAJ45 | CVVSADGGGADGLTF | TRBV5-4 | TRBJ2-1 | CASSLGASGGRYNEQFF | KLGGALQAK | YES | YES | NO |
| 11820 | TRAV38-2DV8 | TRAJ35 | CALGFGNVLHC | TRBV7-3 | TRBJ2-1 | CASSLTLGGSRNEQFF | KLGGALQAK | NO | NO | NO |
| 11821 | TRAV38-1 | TRAJ52 | CAFMKGAGGTSYGKLTF | TRBV19 | TRBJ1-5 | CASSPRSNQPQHF | GILGFVFTL | YES | NO | YES |
| 11822 | TRAV8-1 | TRAJ42 | CAVNAGGGSQGNLIF | TRBV19 | TRBJ2-3 | CASSPRSTDTQYF | GILGFVFTL | YES | NO | YES |
| 11823 | TRAV19 | TRAJ49 | CALSEATGNQFYF | TRBV7-8 | TRBJ2-1 | CASSLGMRGGWDEQFF | KLGGALQAK | YES | NO | NO |
| 11824 | TRAV17 | TRAJ42 | CATDAGGGSQGNLIF | TRBV19 | TRBJ2-1 | CASMELAGLNEQFF | GILGFVFTL | YES | NO | YES |
| 11825 | TRAV12-2 | TRAJ31 | CASFSDARLMF | TRBV12-4 | TRBJ1-1 | CASGGGAAFF | IVTDFSVIK | YES | NO | YES |
| 11826 | TRAV19 | TRAJ57 | CALRGQGGSEKLVF | TRBV7-9 | TRBJ2-1 | CASSLRGVDEQFF | KLGGALQAK | YES | NO | NO |
| 11827 | TRAV26-2 | TRAJ52 | CILRDLNAGGTSYGKLTF | TRBV4-1 | TRBJ2-3 | CASSQVWRGTDTQYF | KLGGALQAK | YES | NO | NO |
| 11828 | TRAV4 | TRAJ32 | CLVGDLSGATNKLIF | TRBV4-2 | TRBJ2-2 | CASSQRPSEVGELFF | RAKFKQLL | YES | NO | YES |
| 11829 | TRAV38-2DV8 | TRAJ40 | CAYPIGTGTYKYIF | TRBV19 | TRBJ2-2 | CASSLVGGELFF | RAKFKQLL | YES | YES | YES |
| 11830 | TRAV21 | TRAJ28 | CAVRVSL | TRBV11-3 | TRBJ2-1 | CASSSHGAGAYNEQFF | RAKFKQLL | YES | NO | NO |
| 11831 | TRAV19 | TRAJ32 | CALSEANYGGATNKLIF | TRBV11-1 | TRBJ2-7 | CASSLSRDSYYEQYF | KLGGALQAK | YES | NO | NO |
| 11832 | TRAV27 | TRAJ42 | CAGAGGGSQGNLIF | TRBV19 | TRBJ2-3 | CASSMRSADTQYF | GILGFVFTL | YES | NO | YES |
| 11833 | TRAV26-2 | TRAJ50 | CILTETSYDKVIF | TRBV7-9 | TRBJ2-7 | CASASLGGGLYEQYF | RAKFKQLL | YES | NO | YES |
| 11834 | TRAV12-2 | TRAJ45 | CAGGNSGGGADGLTF | TRBV29-1 | TRBJ2-2 | CSASRGSVELFF | KLGGALQAK | NO | NO | NO |
| 11835 | TRAV12-2 | TRAJ42 | CAVNQGGGSQGNLIF | TRBV19 | TRBJ1-2 | CASSFRAPYGYTF | GILGFVFTL | YES | NO | YES |
| 11836 | TRAV12-3 | TRAJ54 | CAMTSQGAQKLVF | TRBV9 | TRBJ2-3 | CASSVYKDILGADTQYF | IVTDFSVIK | YES | NO | YES |
| 11837 | TRAV29DV5 | TRAJ44 | CAASDPGTASKLTF | TRBV7-9 | TRBJ1-2 | CASSLALTAEIHGYTF | QYDPVAALF | YES | NO | YES |
| 11838 | TRAV25 | TRAJ42 | CAGSYGGSQGNLIF | TRBV19 | TRBJ2-3 | CASSTRSTDTQYF | GILGFVFTL | YES | NO | YES |
| 11839 | TRAV17 | TRAJ39 | CATDDNAGNMLTF | TRBV7-9 | TRBJ2-7 | CASSLRGIDEQYF | KLGGALQAK | YES | NO | NO |
| 11840 | TRAV38-1 | TRAJ52 | CAFMKGAGGTSYGKLTF | TRBV19 | TRBJ2-7 | CASSARSSYEQYF | GILGFVFTL | YES | NO | NO |
| 11841 | TRAV25 | TRAJ42 | CAGSNGGSQGNLIF | TRBV19 | TRBJ2-3 | CASSTRSADTQYF | GILGFVFTL | YES | NO | YES |
| 11842 | TRAV21 | TRAJ26 | CAVHGYGQNFVF | TRBV12-3 | TRBJ2-2 | CASSFSGNTGELFF | RAKFKQLL | YES | NO | YES |
| 11843 | TRAV19 | TRAJ4 | CALSERVSGGYNKLIF | TRBV13 | TRBJ2-3 | CASSLKGQGSDTQYF | IVTDFSVIK | YES | YES | YES |
| 11844 | TRAV25 | TRAJ40 | CGSGTYKYIF | TRBV27 | TRBJ2-7 | CASSLRPGRAYEQYF | KLGGALQAK | YES | NO | NO |
| 11845 | TRAV27 | TRAJ30 | CAGMNRDDKIIF | TRBV4-2 | TRBJ2-7 | CASSHVIGQGGYEQYF | RAKFKQLL | YES | NO | YES |
| 11846 | TRAV1-2 | TRAJ33 | CAAMDSNYQLIW | TRBV6-5 | TRBJ2-5 | CASSYSSGGGETQYF | KLGGALQAK | NO | NO | NO |
| 11847 | TRAV22 | TRAJ41 | CAVVDSGYALNF | TRBV3-1 | TRBJ2-6 | CASSPLIGANVLTF | KLGGALQAK | YES | NO | NO |
| 11848 | TRAV1-1 | TRAJ13 | CAVSPNSGGYQKVTF | TRBV7-9 | TRBJ2-3 | CASSLPDTQYF | KLGGALQAK | YES | NO | NO |
| 11849 | TRAV19 | TRAJ32 | CALAGGATNKLIF | TRBV5-4 | TRBJ1-2 | CASSFNADYGYTF | KLGGALQAK | YES | NO | NO |
| 11850 | TRAV20 | TRAJ53 | CAVSGGSNYKLTF | TRBV12-3 | TRBJ2-2 | CASSFSGNTGELFF | RAKFKQLL | YES | NO | NO |
| 11851 | TRAV17 | TRAJ11 | CATEGDSGYSTLTF | TRBV9 | TRBJ1-4 | CASSVGGHLNEKLFF | FLYALALLL | YES | YES | YES |
| 11852 | TRAV14DV4 | TRAJ53 | CALLFLSGGSNYKLTF | TRBV6-5 | TRBJ1-2 | CASSNGGGRLYGYTF | KLGGALQAK | YES | NO | NO |
| 11853 | TRAV24 | TRAJ33 | CAFKDSNYQLIW | TRBV5-1 | TRBJ2-7 | CASNAFDYEQYF | KLGGALQAK | YES | NO | NO |
| 11854 | TRAV26-2 | TRAJ5 | CILRMDTGRRALTF | TRBV7-2 | TRBJ1-1 | CASSLATGPGGVRNTEAFF | KLGGALQAK | NO | NO | NO |
| 11855 | TRAV26-1 | TRAJ47 | CIVRDMEYGNKLVF | TRBV7-3 | TRBJ2-3 | CASSLVVAGGPGTDTQYF | RAKFKQLL | YES | NO | YES |
| 11856 | TRAV12-2 | TRAJ9 | CALNTGGFKTIF | TRBV4-2 | TRBJ2-1 | CASSQETSGGASQFF | ELAGIGILTV | YES | NO | NO |
| 11858 | TRAV27 | TRAJ20 | CAGPGSNDYKLSF | TRBV6-1 | TRBJ2-1 | CASSVDVGLYNEQFF | KLGGALQAK | YES | NO | NO |
| 11859 | TRAV17 | TRAJ53 | CATDASGGSNYKLTF | TRBV12-3 | TRBJ2-2 | CASSFSGNTGELFF | RAKFKQLL | YES | NO | YES |
| 11860 | TRAV12-1 | TRAJ43 | CVVNRGSNDMRF | TRBV10-3 | TRBJ1-5 | CAISEPGTNQPQHF | RLRAEAQVK | YES | YES | NO |
| 11861 | TRAV38-1 | TRAJ52 | CAFMKGAGGTSYGKLTF | TRBV30 | TRBJ2-2 | CAWSPKTSDTGELFF | GILGFVFTL | YES | NO | YES |
| 11862 | TRAV19 | TRAJ28 | CALSEGAGAGSYQLTF | TRBV4-1 | TRBJ1-1 | CASSQGDTEDPEAFF | KLGGALQAK | YES | NO | NO |
| 11863 | TRAV12-2 | TRAJ39 | CAVDNNAGNMLTF | TRBV7-9 | TRBJ1-1 | CASSWGVNPEAFF | KLGGALQAK | YES | NO | NO |
| 11864 | TRAV5 | TRAJ34 | CAESEGPTYTDKLIF | TRBV7-3 | TRBJ2-2 | CASSLGTSGGTGELFF | RAKFKQLL | YES | NO | YES |
| 11865 | TRAV5 | TRAJ6 | CAEMEGGNYIPTF | TRBV4-2 | TRBJ2-2 | CASSQRPSEVGELFF | RAKFKQLL | YES | YES | YES |
| 11866 | TRAV13-2 | TRAJ45 | CADSGGGADGLTF | TRBV19 | TRBJ2-7 | CASSIRSSYEQYF | GILGFVFTL | YES | NO | YES |
| 11867 | TRAV6 | TRAJ17 | CALLEMVIKAAGNKLTF | TRBV28 | TRBJ1-5 | CASSTDRAAQPQHF | RTLNAWVKV | YES | YES | YES |
| 11868 | TRAV13-1 | TRAJ50 | CAASETSYDKVIF | TRBV12-3 | TRBJ2-3 | CASSLIDTQYF | RAKFKQLL | YES | NO | YES |
| 11869 | TRAV5 | TRAJ26 | CAESMDNYGQNFVF | TRBV4-2 | TRBJ2-2 | CASSQRPSEVGELFF | RAKFKQLL | YES | NO | YES |
| 11870 | TRAV13-1 | TRAJ42 | CAASGGGSQGNLIF | TRBV19 | TRBJ2-7 | CASSARSSYEQYF | GILGFVFTL | YES | NO | NO |
| 11871 | TRAV12-3 | TRAJ28 | CAMNRAGSYQLTF | TRBV19 | TRBJ2-1 | CASSIDHNQGYNEQFF | AVFDRKSDAK | YES | NO | YES |
| 11872 | TRAV1-2 | TRAJ31 | CAVKRARLMF | TRBV27 | TRBJ1-1 | CASTFGPEAFF | KLGGALQAK | YES | NO | NO |
| 11873 | TRAV27 | TRAJ41 | CAGHNSGYALNF | TRBV19 | TRBJ2-7 | CASRLGGEQYF | KLGGALQAK | YES | NO | NO |
| 11874 | TRAV12-2 | TRAJ15 | CAVPQAGTALIF | TRBV19 | TRBJ1-1 | CASSPGALNTEAFF | ELAGIGILTV | YES | NO | YES |
| 11875 | TRAV12-2 | TRAJ39 | CAVNAGNMLTF | TRBV15 | TRBJ2-7 | CATSRGQGNEQYF | ELAGIGILTV | YES | YES | YES |
| 11876 | TRAV12-1 | TRAJ4 | CVVNMVSGGYNKLIF | TRBV7-8 | TRBJ2-2 | CASSLRLGAGANTGELFF | KLGGALQAK | YES | NO | NO |
| 11877 | TRAV1-2 | TRAJ12 | CAVMDSSYKLIF | TRBV6-3 | TRBJ2-5 | CASSGQEQETQYF | AVFDRKSDAK | YES | NO | YES |
| 11878 | TRAV41 | TRAJ34 | CAVGPFYNTDKLIF | TRBV6-5 | TRBJ2-1 | CASRGQGSYNEQFF | KLGGALQAK | NO | NO | NO |
| 11879 | TRAV12-2 | TRAJ7 | CAVREGNNRLAF | TRBV6-5 | TRBJ2-7 | CASSSLGADEQYF | KLGGALQAK | YES | NO | NO |
| 11880 | TRAV38-2DV8 | TRAJ13 | CAYTGGYQKVTF | TRBV9 | TRBJ2-2 | CASSEKGLADTGELFF | KLGGALQAK | YES | NO | NO |
| 11881 | TRAV13-1 | TRAJ41 | CAAGDGNALNF | TRBV12-4 | TRBJ2-2 | CASSRGTGELFF | AVFDRKSDAK | YES | NO | NO |
| 11882 | TRAV13-2 | TRAJ53 | CAETHSGGSNYKLTF | TRBV12-4 | TRBJ2-7 | CASSPSSSYEQYF | RAKFKQLL | YES | NO | YES |

| | | | | | | | | | | |
|---|---|---|---|---|---|---|---|---|---|---|
| 11883 | TRAV27 | TRAJ42 | CAGPGDGGSQGNLIF | TRBV12-3 | TRBJ2-7 | CASSLWGDYEQYF | GILGFVFTL | YES | NO | YES |
| 11884 | TRAV12-2 | TRAJ7 | CAVRVGTTDSLL | TRBV19 | TRBJ2-3 | CASSPLSTDTQYF | GILGFVFTL | YES | NO | NO |
| 11885 | TRAV21 | TRAJ17 | CAVRQTAAGNKLTF | TRBV15 | TRBJ1-2 | CATRRNRGNTYGYTF | KLGGALQAK | YES | NO | NO |
| 11886 | TRAV14DV4 | TRAJ49 | CAMREWNTGNQFYF | TRBV4-2 | TRBJ2-7 | CASSPGTSSIYEQYF | GILGFVFTL | YES | NO | YES |
| 11887 | TRAV1-2 | TRAJ16 | CAVRPSRF | TRBV7-9 | TRBJ2-1 | CASSPGLAGGDEQFF | KLGGALQAK | NO | NO | NO |
| 11888 | TRAV19 | TRAJ42 | CALSEAGRSQGNLIF | TRBV10-1 | TRBJ2-7 | CASSRDASSYEQYF | KLGGALQAK | YES | NO | NO |
| 11889 | TRAV13-1 | TRAJ50 | CAAGETSYDKVIF | TRBV18 | TRBJ1-2 | CASSPWPDGGSNYGYTF | RAKFKQLL | YES | NO | YES |
| 11890 | TRAV38-2DV8 | TRAJ57 | CAYRVGGGSWGKTQGGSEKLVF | TRBV19 | TRBJ2-2 | CASSSSGGAGELFF | GILGFVFTL | NO | NO | NO |
| 11891 | TRAV16 | TRAJ43 | CALKNNDMRF | TRBV19 | TRBJ2-7 | CASSARSSYEQYF | GILGFVFTL | NO | NO | NO |
| 11892 | TRAV23DV6 | TRAJ43 | CAAQYNDMRF | TRBV5-1 | TRBJ1-2 | CASSQGERGYTF | AVFDRKSDAK | YES | NO | YES |
| 11893 | TRAV20 | TRAJ33 | CAVQARFDSNYQLIW | TRBV29-1 | TRBJ2-7 | CSVEGWGGFYEQYF | KLGGALQAK | YES | NO | NO |
| 11894 | TRAV8-1 | TRAJ3 | CAVNLYSSASKIIF | TRBV5-1 | TRBJ1-1 | CASSFGDHEAFF | KLGGALQAK | YES | NO | NO |
| 11895 | TRAV29DV5 | TRAJ49 | CAASASGGQFYF | TRBV12-3 | TRBJ2-2 | CASSFSGNTGELFF | RAKFKQLL | YES | NO | YES |
| 11896 | TRAV16 | TRAJ30 | CALEGDDKIIF | TRBV20-1 | TRBJ2-3 | CSARWGGGTDTQYF | KLGGALQAK | NO | NO | NO |
| 11897 | TRAV27 | TRAJ42 | CAGAGSQGNLIF | TRBV29-1 | TRBJ2-1 | CSLQGGRYNEQFF | GILGFVFTL | YES | NO | YES |
| 11898 | TRAV13-1 | TRAJ26 | CAAVDNYGQNFVF | TRBV29-1 | TRBJ2-1 | CSGGQGPDNEQFF | RAKFKQLL | YES | NO | YES |
| 11899 | TRAV16 | TRAJ10 | CALPTGGGNKLTF | TRBV5-6 | TRBJ2-1 | CASSLRQAYNEQFF | KLGGALQAK | NO | NO | NO |
| 11900 | TRAV9-2 | TRAJ34 | CTSVLCITPTSSSL | TRBV3-1 | TRBJ2-7 | CASSPVTGSSYEQYF | GLCTLVAML | NO | NO | NO |
| 11901 | TRAV12-2 | TRAJ39 | CAVDSNAGNMLTF | TRBV28 | TRBJ2-5 | CASSFSSETQYF | KLGGALQAK | YES | NO | NO |
| 11903 | TRAV8-3 | TRAJ7 | CAVGLYYGNNRLAF | TRBV3-1 | TRBJ2-7 | CASSPVTGSSYEQYF | GLCTLVAML | YES | YES | YES |
| 11904 | TRAV21 | TRAJ33 | CAVTWSNYQLIW | TRBV30 | TRBJ1-1 | CAWSLKPNTEAFF | KLGGALQAK | NO | NO | NO |
| 11905 | TRAV12-3 | TRAJ42 | CAMSGEGGSQGNLIF | TRBV19 | TRBJ1-5 | CASSIRSNQPQHF | GILGFVFTL | YES | NO | YES |
| 11906 | TRAV8-3 | TRAJ23 | CAVGSIYNQGGKLIF | TRBV5-6 | TRBJ1-1 | CASSLRDGSEAFF | RAKFKQLL | YES | YES | YES |
| 11907 | TRAV12-2 | TRAJ13 | CAAKVTF | TRBV19 | TRBJ2-7 | CASSARSSYEQYF | GILGFVFTL | YES | NO | NO |
| 11908 | TRAV13-1 | TRAJ21 | CAAKKYNFNKFYF | TRBV2 | TRBJ2-5 | CASRGQQETQYF | KLGGALQAK | YES | NO | NO |
| 11909 | TRAV26-1 | TRAJ12 | CIVSYKLIF | TRBV20-1 | TRBJ2-1 | CSARLAGLTYNEQFF | GILGFVFTL | NO | NO | NO |
| 11910 | TRAV12-1 | TRAJ11 | CVVNGNSGYSTLTF | TRBV15 | TRBJ2-4 | CATSRVGADIQYF | KLGGALQAK | YES | NO | NO |
| 11911 | TRAV19 | TRAJ36 | CALTPGLGANNLFF | TRBV9 | TRBJ2-3 | CASSVNRAGGVTDTQYF | KLGGALQAK | YES | NO | NO |
| 11912 | TRAV19 | TRAJ5 | CALTSGGGRRALTF | TRBV29-1 | TRBJ2-1 | CSVEDPGSGGEQWQFF | KLGGALQAK | YES | NO | NO |
| 11913 | TRAV27 | TRAJ42 | CAGGGSQGNLIF | TRBV5-1 | TRBJ2-5 | CASSLEWGGETQYF | KLGGALQAK | YES | NO | NO |
| 11914 | TRAV36DV7 | TRAJ57 | CAVEAREGGSEKLVF | TRBV5-1 | TRBJ1-2 | CASKLSFNYGYTF | KLGGALQAK | NO | NO | NO |
| 11915 | TRAV26-2 | TRAJ43 | CILREFYNDMRF | TRBV7-3 | TRBJ2-5 | CASSSRGGQETQYF | RAKFKQLL | NO | NO | NO |
| 11916 | TRAV14DV4 | TRAJ27 | CAMREGQGTNAGKSTF | TRBV18 | TRBJ2-1 | CASSLAQGHNEQFF | KLGGALQAK | YES | YES | NO |
| 11917 | TRAV16 | TRAJ13 | CALLIYGGYQKVTF | TRBV12-3 | TRBJ2-2 | CASSFSGNTGELFF | RAKFKQLL | NO | NO | NO |
| 11918 | TRAV25 | TRAJ42 | CAGPQDGGSQGNLIF | TRBV19 | TRBJ1-5 | CASSYYSNQPQHF | GILGFVFTL | YES | NO | YES |
| 11919 | TRAV13-1 | TRAJ50 | CAASETSYDKVIF | TRBV7-9 | TRBJ1-1 | CASSLLGQPNTEAFF | RAKFKQLL | YES | NO | YES |
| 11920 | TRAV13-1 | TRAJ34 | CAASRDTDKLIF | TRBV4-1 | TRBJ2-2 | CASSQQQGADGELFF | RAKFKQLL | YES | NO | NO |
| 11921 | TRAV1-2 | TRAJ33 | CAVLDSNYQLIW | TRBV6-5 | TRBJ2-7 | CASKKLAGDSYEQYF | KLGGALQAK | YES | NO | NO |
| 11922 | TRAV27 | TRAJ27 | CAGDSPAGKSTF | TRBV19 | TRBJ2-7 | CASSIRSSYEQYF | GILGFVFTL | YES | NO | YES |
| 11923 | TRAV29DV5 | TRAJ40 | CAASASGTYKYIF | TRBV19 | TRBJ2-7 | CASSARSSYEQYF | GILGFVFTL | YES | NO | YES |
| 11924 | TRAV29DV5 | TRAJ24 | CAAFLLQLGEIPVWSRDP | TRBV6-3 | TRBJ2-7 | CASGMTGLTSEQYF | KLGGALQAK | NO | NO | NO |
| 11925 | TRAV12-2 | TRAJ27 | CAVRTGKSTF | TRBV6-5 | TRBJ2-6 | CASKSPGQGRSGANVLTF | KLGGALQAK | YES | NO | NO |
| 11926 | TRAV13-1 | TRAJ50 | CARGETSYDKVIF | TRBV12-4 | TRBJ2-7 | CASSSSGVGFYEQYF | RAKFKQLL | YES | YES | YES |
| 11927 | TRAV12-2 | TRAJ53 | CAVVEFGGSNYKLTF | TRBV4-1 | TRBJ1-1 | CASSQGLGENTEAFF | RAKFKQLL | YES | NO | YES |
| 11928 | TRAV21 | TRAJ8 | CAVRGIISFQKLVF | TRBV5-1 | TRBJ1-4 | CASSLGQGAVNEKLFF | KLGGALQAK | YES | NO | NO |
| 11929 | TRAV5 | TRAJ42 | CAEDYGGSQGNLIF | TRBV19 | TRBJ2-1 | CASSSRSANEQFF | GILGFVFTL | YES | NO | YES |
| 11930 | TRAV13-1 | TRAJ53 | CAAIDLGSNYKLTF | TRBV7-2 | TRBJ2-1 | CASSLVGGNEQFF | KLGGALQAK | YES | NO | NO |
| 11931 | TRAV12-1 | TRAJ47 | CVVPNPEYGNKLVF | TRBV4-1 | TRBJ1-5 | CASSQDPGYSNQPQHF | KLGGALQAK | YES | NO | NO |
| 11932 | TRAV1-2 | TRAJ10 | CAVKGLTGGGNKLTF | TRBV7-9 | TRBJ2-7 | CASSTSEGGLFYEQYF | KLGGALQAK | YES | NO | NO |
| 11933 | TRAV13-1 | TRAJ11 | CAAKEGYSTLTF | TRBV12-3 | TRBJ2-2 | CASSFSGNTGELFF | RAKFKQLL | YES | NO | YES |
| 11934 | TRAV1-2 | TRAJ12 | CAVMDSSYKLIF | TRBV6-4 | TRBJ2-3 | CASSVGVTDTQYF | KLGGALQAK | YES | NO | NO |
| 11935 | TRAV21 | TRAJ58 | CAVETSGSRLTF | TRBV2 | TRBJ1-1 | CASRAGATEAFF | KLGGALQAK | YES | NO | NO |
| 11936 | TRAV38-1 | TRAJ53 | CAFMKHVSSGGSNYKLTF | TRBV27 | TRBJ2-2 | CASSPHFQVDTGELFF | RAKFKQLL | YES | NO | YES |
| 11937 | TRAV9-2 | TRAJ50 | CALRETSYDKVIF | TRBV12-3 | TRBJ1-1 | CASSFLDRGPNTEAFF | RAKFKQLL | YES | YES | YES |
| 11938 | TRAV9-2 | TRAJ41 | CALRVNSGYALNF | TRBV11-2 | TRBJ2-3 | CASSSDGGLGTDTQYF | KLGGALQAK | YES | NO | NO |
| 11939 | TRAV10 | TRAJ41 | CVVNQDSGYALNF | TRBV6-5 | TRBJ2-2 | CASSYSGSGELFF | KLGGALQAK | YES | NO | NO |
| 11940 | TRAV6 | TRAJ20 | CALGVDYKLSF | TRBV7-9 | TRBJ2-5 | CASTPGVSNQETQYF | KLGGALQAK | YES | NO | NO |
| 11941 | TRAV39 | TRAJ48 | CAPSVGGNEKLTF | TRBV7-7 | TRBJ2-7 | CASSLVPWTAKFF | KLGGALQAK | YES | NO | NO |
| 11942 | TRAV17 | TRAJ11 | CATEGGSGYSTLTF | TRBV6-5 | TRBJ1-2 | CASSYQGGGKGGYTF | FLYALALLL | YES | YES | YES |
| 11943 | TRAV13-1 | TRAJ50 | CAASEDIGNDKVIF | TRBV3-1 | TRBJ1-2 | CASSYPLGARWRGYTF | RAKFKQLL | YES | NO | YES |
| 11944 | TRAV13-1 | TRAJ50 | CAASETSYDKVIF | TRBV12-3 | TRBJ1-2 | CASSPGTNYGYTF | RAKFKQLL | YES | NO | YES |
| 11945 | TRAV17 | TRAJ42 | CATDDGGSQGNLIF | TRBV27 | TRBJ2-1 | CASRKGGPNNEQFF | KLGGALQAK | YES | NO | NO |
| 11946 | TRAV35 | TRAJ49 | CAGHTGNQFYF | TRBV7-9 | TRBJ2-3 | CASSLDTQYF | IVTDFSVIK | YES | NO | YES |
| 11947 | TRAV1-2 | TRAJ26 | CAVRGNYGQNFVF | TRBV7-9 | TRBJ2-7 | CASSSALAGPNEQYF | IVTDFSVIK | YES | NO | YES |
| 11948 | TRAV12-3 | TRAJ13 | CAFHSGGYQKVTF | TRBV11-2 | TRBJ2-2 | CASSLGVAGTNTGELFF | KLGGALQAK | YES | NO | NO |
| 11949 | TRAV20 | TRAJ27 | CAVQAIRANAGKSTF | TRBV27 | TRBJ2-7 | CASSLAGTGNEQYF | KLGGALQAK | YES | NO | NO |
| 11950 | TRAV12-3 | TRAJ42 | CAMSGDGGSQGNLIF | TRBV19 | TRBJ1-5 | CASSPRSNQPQHF | GILGFVFTL | YES | NO | NO |
| 11951 | TRAV10 | TRAJ42 | CVVSAEGSQGNLIF | TRBV5-4 | TRBJ1-5 | CASSLHPREAQHF | KLGGALQAK | YES | NO | NO |
| 11952 | TRAV29DV5 | TRAJ49 | CAASSGTGNQFYF | TRBV5-6 | TRBJ1-1 | CASSLRDGSEAFF | RAKFKQLL | YES | YES | YES |
| 11953 | TRAV13-1 | TRAJ42 | CAARTMEGSQGNLIF | TRBV9 | TRBJ2-7 | CASSVAPGEYEQYF | KLGGALQAK | YES | NO | NO |
| 11954 | TRAV12-1 | TRAJ40 | CVVNSPSGTYKYIF | TRBV3-1 | TRBJ2-1 | CASSPYSGGLYNEQFF | RLRAEAQVK | YES | YES | YES |
| 11955 | TRAV23DV6 | TRAJ42 | CAASMNYGGSQGNLIF | TRBV12-4 | TRBJ1-1 | CASSRPEPLNTEAFF | KLGGALQAK | YES | NO | NO |
| 11956 | TRAV14DV4 | TRAJ15 | CAMRETNQAGTALIF | TRBV12-3 | TRBJ2-2 | CASSFSGNTGELFF | RAKFKQLL | YES | NO | YES |
| 11957 | TRAV8-3 | TRAJ52 | CAVASAGGTSYGKLTF | TRBV16 | TRBJ2-7 | CASSPPKGRGSYEQYF | AYAQKIFKI | YES | YES | YES |
| 11958 | TRAV1-2 | TRAJ33 | CAGTDSNYQLIW | TRBV6-1 | TRBJ1-1 | CASSEFRGGASEAFF | KLGGALQAK | YES | NO | NO |
| 11959 | TRAV1-2 | TRAJ8 | CAAPLGGFQKLVF | TRBV10-3 | TRBJ1-5 | CAISHPGQGGQPQHF | KLGGALQAK | YES | NO | NO |
| 11960 | TRAV8-3 | TRAJ21 | CAVGDNFNKFYF | TRBV11-2 | TRBJ2-1 | CASSLGQQGVNRDWYNEQFF | KLGGALQAK | NO | NO | NO |
| 11961 | TRAV39 | TRAJ52 | CAGGLENAGGTSYGKLTF | TRBV19 | TRBJ1-5 | CASSRLSNQPQHF | GILGFVFTL | YES | NO | YES |
| 11962 | TRAV1-2 | TRAJ26 | CAVRDHYGQNFVF | TRBV14 | TRBJ2-1 | CASSVRQTYNEQFF | KLGGALQAK | YES | NO | NO |
| 11963 | TRAV22 | TRAJ5 | CAVEGMDTGRRALTF | TRBV4-1 | TRBJ2-5 | CASSLAMARKAQYF | KLGGALQAK | YES | NO | NO |
| 11964 | TRAV1-2 | TRAJ33 | CAVRDSNYQLIW | TRBV6-1 | TRBJ2-3 | CASSEKSGGPDTQYF | KLGGALQAK | YES | NO | NO |
| 11965 | TRAV13-1 | TRAJ42 | CAATPYYGGSQGNLIF | TRBV12-3 | TRBJ2-2 | CASSFSGNTGELFF | RAKFKQLL | YES | NO | YES |
| 11966 | TRAV4 | TRAJ30 | CLVGDKGDDKIIF | TRBV9 | TRBJ1-2 | CASSFFRSGGSLNYGYTF | KLGGALQAK | YES | NO | NO |
| 11967 | TRAV8-1 | TRAJ39 | CAVNRNAGNMLTF | TRBV19 | TRBJ2-7 | CASSIRSSYEQYF | GILGFVFTL | YES | NO | YES |
| 11968 | TRAV27 | TRAJ37 | CAGAHGSSNTGKLIF | TRBV19 | TRBJ1-1 | CASSIRSGTEAFF | GILGFVFTL | YES | NO | YES |
| 11969 | TRAV5 | TRAJ24 | CAESGPDSWGKFQF | TRBV19 | TRBJ1-2 | CASSIGIYGYTF | GILGFVFTL | YES | NO | YES |
| 11970 | TRAV5 | TRAJ36 | CAERRSDGANNLFF | TRBV4-1 | TRBJ1-5 | CASSQDRGSNQPQHF | KLGGALQAK | YES | NO | NO |
| 11971 | TRAV5 | TRAJ42 | CAESSRF | TRBV6-3 | TRBJ2-7 | CASSEGAFSYEQYF | KLGGALQAK | YES | NO | NO |
| 11972 | TRAV30 | TRAJ20 | CGTEYPNDYKLSF | TRBV7-8 | TRBJ2-1 | CASSLGGNNEQFF | GLCTLVAML | YES | YES | YES |
| 11973 | TRAV6 | TRAJ31 | CSWGNARLMF | TRBV27 | TRBJ1-5 | CASMRGADNQPQHF | KLGGALQAK | YES | NO | NO |
| 11974 | TRAV13-1 | TRAJ36 | CAASEALEANNLFF | TRBV12-4 | TRBJ2-7 | CASRVSGGEEQYF | KLGGALQAK | YES | NO | NO |
| 11975 | TRAV29DV5 | TRAJ44 | CAASARRIDTGTASKLTF | TRBV13 | TRBJ2-7 | CASRHRGAPSYEQYF | RLRAEAQVK | YES | YES | YES |
| 11976 | TRAV38-2DV8 | TRAJ57 | CAYRSAESGSGKLVF | TRBV13 | TRBJ2-5 | CASSEGGVETQYF | AVFDRKSDAK | YES | NO | YES |
| 11977 | TRAV38-1 | TRAJ52 | CAFMTNAGGTSYGKLTF | TRBV19 | TRBJ1-2 | CATSQGSYGYTF | GILGFVFTL | YES | NO | YES |
| 11978 | TRAV13-1 | TRAJ50 | CAASETSYDKVIF | TRBV7-9 | TRBJ2-5 | CASSFVDRAETQYF | RAKFKQLL | YES | NO | YES |
| 11979 | TRAV29DV5 | TRAJ28 | CAAITGAGSYQLTF | TRBV7-9 | TRBJ1-2 | CASSDSGGHYGYTF | KLGGALQAK | YES | NO | NO |
| 11980 | TRAV8-1 | TRAJ39 | CAVNLTNNAGNMLTF | TRBV19 | TRBJ2-3 | CASSGRSTDTQYF | GILGFVFTL | YES | NO | NO |
| 11981 | TRAV17 | TRAJ44 | CATGFHTGTASKLTF | TRBV3-1 | TRBJ2-3 | CASSQVPGQGWARTDTQYF | KLGGALQAK | YES | NO | NO |
| 11982 | TRAV1-1 | TRAJ39 | CAVRDAHNNAGNMLTF | TRBV9 | TRBJ2-3 | CASSVTSSRGGTDTQYF | KLGGALQAK | YES | NO | NO |
| 11983 | TRAV1-2 | TRAJ33 | CAVMDSNYQLIW | TRBV20-1 | TRBJ1-1 | CSARTGLETEAFF | KLGGALQAK | YES | NO | NO |
| 11984 | TRAV12-1 | TRAJ27 | CVVETNAGKSTF | TRBV15 | TRBJ2-7 | CATSRVRVDEQYF | KLGGALQAK | YES | NO | NO |
| 11985 | TRAV29DV5 | TRAJ31 | CAASTNARLMF | TRBV29-1 | TRBJ2-1 | CSLQGGRYNEQFF | KLGGALQAK | YES | NO | NO |
| 11987 | TRAV26-2 | TRAJ50 | CILRETSYDKVIF | TRBV7-9 | TRBJ1-4 | CASSPRQGWTNEKLFF | GILGFVFTL | YES | NO | YES |
| 11988 | TRAV5 | TRAJ34 | CAESEGPTYTDKLIF | TRBV27 | TRBJ2-7 | CASSLFGGTYEQYF | RAKFKQLL | YES | NO | YES |

| | | | | | | | | | | |
|---|---|---|---|---|---|---|---|---|---|---|
| 11989 | TRAV13-1 | TRAJ49 | CAAKKGNQFYF | TRBV12-3 | TRBJ1-4 | CASGPPGGEKLFF | RAKFKQLL | YES | NO | YES |
| 11990 | TRAV1-2 | TRAJ33 | CAAMDSNYQLIW | TRBV20-1 | TRBJ2-2 | CSARQQGDTGELFF | KLGGALQAK | YES | NO | NO |
| 11991 | TRAV13-1 | TRAJ50 | CAASETSYDKVIF | TRBV7-9 | TRBJ1-4 | CASSFQGAPDEKLFF | RAKFKQLL | YES | NO | YES |
| 11992 | TRAV1-1 | TRAJ49 | CAVSNTGNQFYF | TRBV9 | TRBJ2-7 | CASSVGGLAGEEQYF | AVFDRKSDAK | YES | NO | YES |
| 11993 | TRAV13-1 | TRAJ50 | CAASETSYDKVIF | TRBV7-9 | TRBJ2-7 | CASSLTDGGEQYF | RAKFKQLL | YES | YES | YES |
| 11994 | TRAV36DV7 | TRAJ34 | CAVLLFNTDKLIF | TRBV20-1 | TRBJ2-7 | CSARGPGRGEQYF | KLGGALQAK | NO | NO | NO |
| 11995 | TRAV13-1 | TRAJ50 | CAASETSYDKVIF | TRBV7-9 | TRBJ2-7 | CASSLAGSYEQYF | RAKFKQLL | YES | NO | YES |
| 11996 | TRAV24 | TRAJ40 | CASIAGTYKYIF | TRBV12-4 | TRBJ2-1 | CASGRLSYNEQFF | RAKFKQLL | YES | YES | YES |
| 11997 | TRAV25 | TRAJ42 | CAGNYGGSQGNLIF | TRBV19 | TRBJ1-5 | CASSTRSHEPQHF | GILGFVFTL | YES | YES | YES |
| 11998 | TRAV8-6 | TRAJ11 | CAVSSGYSTLTF | TRBV10-2 | TRBJ2-7 | CASSLPRDSFSYEQYF | KLGGALQAK | YES | NO | NO |
| 11999 | TRAV35 | TRAJ42 | CAGHLYGGSQGNLIF | TRBV7-9 | TRBJ1-1 | CASSLWTGGVRDTEAFF | KLGGALQAK | YES | NO | NO |
| 12000 | TRAV1-2 | TRAJ5 | CAVRDPGTGRRALTF | TRBV12-3 | TRBJ2-2 | CASSFSGNTGELFF | RAKFKQLL | YES | NO | NO |
| 12001 | TRAV14DV4 | TRAJ56 | CAMREGWANSKLTF | TRBV7-3 | TRBJ2-3 | CASSLTGDGPDTQYF | KLGGALQAK | YES | NO | NO |
| 12002 | TRAV13-1 | TRAJ50 | CAASETSYDKVIF | TRBV29-1 | TRBJ1-1 | CSAGGNTEAFF | RAKFKQLL | YES | NO | YES |
| 12003 | TRAV8-1 | TRAJ41 | CAVNADSGYALNF | TRBV6-6 | TRBJ1-1 | CATYVGGNTEAFF | KLGGALQAK | NO | NO | NO |
| 12004 | TRAV8-4 | TRAJ16 | CAVSGGGQKLLF | TRBV20-1 | TRBJ2-7 | CSAPGQGSSEQYF | KLGGALQAK | NO | NO | NO |
| 12005 | TRAV5 | TRAJ10 | CAEMTGGGNKLTF | TRBV15 | TRBJ2-7 | CATGTGISYEQYF | IVTDFSVIK | YES | YES | YES |
| 12006 | TRAV14DV4 | TRAJ50 | CAMRETETSYDKVIF | TRBV2 | TRBJ2-7 | CASSFLAPGDLLLYEQYF | KLGGALQAK | YES | NO | NO |
| 12007 | TRAV12-2 | TRAJ37 | CAVNPLEGNTGKLIF | TRBV19 | TRBJ2-7 | CASSHLLSTVDYEQYF | KLGGALQAK | YES | NO | NO |
| 12008 | TRAV41 | TRAJ49 | CAVSNTGNQFYF | TRBV12-3 | TRBJ1-1 | CASSFLDRGPNTEAFF | RAKFKQLL | YES | NO | YES |
| 12009 | TRAV35 | TRAJ42 | CAGTIAGGSQGNLIF | TRBV19 | TRBJ1-5 | CASSIVSNQPQHF | GILGFVFTL | YES | NO | YES |
| 12010 | TRAV3 | TRAJ34 | CAVRTSILYNTDKLIF | TRBV20-1 | TRBJ1-5 | CSARDLQGGNQPQHF | KLGGALQAK | NO | NO | NO |
| 12011 | TRAV13-1 | TRAJ11 | CAAKEGYSTLTF | TRBV7-3 | TRBJ2-5 | CASSSRGGQETQYF | KLGGALQAK | NO | NO | NO |
| 12012 | TRAV12-2 | TRAJ39 | CAVRRAGNMLTF | TRBV12-4 | TRBJ2-7 | CASTDRTRLNEQYF | KLGGALQAK | YES | NO | NO |
| 12013 | TRAV19 | TRAJ45 | CALSVFSGGGADGLTF | TRBV20-1 | TRBJ1-6 | CSAVGPPLHF | KLGGALQAK | NO | NO | NO |
| 12014 | TRAV1-2 | TRAJ33 | CAVRDSNYQLIW | TRBV4-2 | TRBJ2-3 | CASSQERGGGTDTQYF | KLGGALQAK | YES | NO | NO |
| 12015 | TRAV35 | TRAJ23 | CAGRIYNQGGKLIF | TRBV4-1 | TRBJ1-6 | CASSLGRDSPLHF | KLGGALQAK | YES | NO | NO |
| 12016 | TRAV13-1 | TRAJ43 | CAALAWRAYNNNDMRF | TRBV2 | TRBJ1-2 | CASSEGGQYDDYGYTF | RLRAEAQVK | YES | NO | YES |
| 12017 | TRAV1-2 | TRAJ33 | CAVEDSNYQLIW | TRBV6-4 | TRBJ2-1 | CASSDRASGGADEQFF | KLGGALQAK | YES | NO | NO |
| 12018 | TRAV8-4 | TRAJ3 | CAVGSSASKIIF | TRBV28 | TRBJ1-1 | CASSPPGTGEAQAFF | KLGGALQAK | YES | NO | NO |
| 12019 | TRAV13-2 | TRAJ13 | CAEKPYSGGYQKVTF | TRBV4-1 | TRBJ2-7 | CASSLGQGTYEQYF | KLGGALQAK | YES | NO | NO |
| 12020 | TRAV8-4 | TRAJ11 | CAVSGYSTLTF | TRBV5-6 | TRBJ1-1 | CASSLRDGSEAFF | RAKFKQLL | YES | NO | NO |
| 12021 | TRAV13-1 | TRAJ39 | CAASFNNAGNMLTF | TRBV6-1 | TRBJ2-4 | CASNRLATNLAKNIQYF | KLGGALQAK | YES | NO | NO |
| 12022 | TRAV38-2DV8 | TRAJ40 | CAYRSDSGTYKYIF | TRBV4-1 | TRBJ1-2 | CASSTSTGTGYGYTF | RAKFKQLL | YES | NO | YES |
| 12023 | TRAV13-1 | TRAJ50 | CAASETSYDKVIF | TRBV7-9 | TRBJ1-1 | CASSSLGDRNTEAFF | RAKFKQLL | YES | NO | YES |
| 12024 | TRAV26-2 | TRAJ47 | CILRDWERKYGNKLVF | TRBV5-1 | TRBJ2-5 | CASSAGTGRVETQYF | KLGGALQAK | YES | NO | NO |
| 12025 | TRAV27 | TRAJ42 | CAENYGGSQGNLIF | TRBV19 | TRBJ2-7 | CASSIRSSYEQYF | GILGFVFTL | YES | NO | YES |
| 12026 | TRAV9-2 | TRAJ7 | CALSVNGNNRLAF | TRBV7-6 | TRBJ2-1 | CASSLGTAGFEQFF | KLGGALQAK | YES | NO | NO |
| 12027 | TRAV13-1 | TRAJ50 | CAASETSYDKVIF | TRBV7-9 | TRBJ2-7 | CASSLVGAEPYEQYF | RAKFKQLL | YES | NO | YES |
| 12028 | TRAV1-2 | TRAJ33 | CAVMDSNYQLIW | TRBV5-4 | TRBJ1-5 | CASSSTGGGEKDQPQHF | KLGGALQAK | YES | NO | NO |
| 12029 | TRAV17 | TRAJ28 | CATFNSGAGSYQLTF | TRBV12-3 | TRBJ1-2 | CASSFESGLGYGYTF | KLGGALQAK | YES | NO | NO |
| 12030 | TRAV14DV4 | TRAJ57 | CAMREAPQGGSEKLVF | TRBV20-1 | TRBJ1-3 | CSAQGGNTIYF | KLGGALQAK | YES | NO | NO |
| 12031 | TRAV13-2 | TRAJ45 | CADSGGGADGLTF | TRBV5-6 | TRBJ2-1 | CASSLTLAAYNEQFF | RAKFKQLL | YES | NO | YES |
| 12032 | TRAV25 | TRAJ42 | CAGNYGGSQGNLIF | TRBV19 | TRBJ2-3 | CASSIRSSDTQYF | GILGFVFTL | YES | NO | YES |
| 12033 | TRAV13-2 | TRAJ23 | CALEGYNQGGKLIF | TRBV9 | TRBJ2-7 | CASSARQGGSEQYF | KLGGALQAK | YES | NO | NO |
| 12034 | TRAV17 | TRAJ11 | CATEGDSGYSTLTF | TRBV6-5 | TRBJ1-2 | CASSPQGGGYGYTF | FLYALALLL | YES | YES | YES |
| 12036 | TRAV19 | TRAJ34 | CALSGGLATDKLIF | TRBV29-1 | TRBJ1-2 | CSVRRGDEGYTF | KLGGALQAK | YES | NO | NO |
| 12037 | TRAV13-1 | TRAJ42 | CAASNRGSQGNLIF | TRBV3-1 | TRBJ2-3 | CASSQVPGAAGEPTDTQYF | KLGGALQAK | YES | NO | NO |
| 12038 | TRAV12-2 | TRAJ28 | CAVNIGTSGAGSYQLTF | TRBV29-1 | TRBJ2-7 | CSAGQGPYEQYF | RAKFKQLL | NO | NO | NO |
| 12039 | TRAV17 | TRAJ33 | CATARSSMDSNYQLIW | TRBV19 | TRBJ1-5 | CATAPHSGNQPQHF | GILGFVFTL | YES | NO | NO |
| 12040 | TRAV14DV4 | TRAJ49 | CAMREGPNTGNQFYF | TRBV10-1 | TRBJ2-7 | CASSRDASSYEQYF | KLGGALQAK | YES | NO | NO |
| 12041 | TRAV13-2 | TRAJ45 | CADSGGGADGLTF | TRBV5-1 | TRBJ2-5 | CASSQQGQPQETQYF | RAKFKQLL | YES | NO | YES |
| 12042 | TRAV12-1 | TRAJ45 | CVGTGGADGLTF | TRBV11-2 | TRBJ2-7 | CASSFAGSGYEQYF | KLGGALQAK | YES | NO | NO |
| 12043 | TRAV14DV4 | TRAJ49 | CAMREGSGYTGNQFYF | TRBV29-1 | TRBJ2-7 | CSAGQGPYEQYF | RAKFKQLL | NO | NO | NO |
| 12044 | TRAV19 | TRAJ17 | CALKLIKAAGNKLTF | TRBV30 | TRBJ1-6 | CAWSHPSPLHF | KLGGALQAK | YES | NO | NO |
| 12045 | TRAV19 | TRAJ21 | CALSEAGLYNFNKFYF | TRBV19 | TRBJ1-6 | CASSIIQGHSYNSPLHF | KLGGALQAK | YES | NO | NO |
| 12046 | TRAV13-1 | TRAJ6 | CAASITGGSYIPTF | TRBV5-4 | TRBJ2-7 | CASSETHRGEQYF | KLGGALQAK | YES | NO | NO |
| 12047 | TRAV25 | TRAJ13 | CAGSRGGYQKVTF | TRBV7-6 | TRBJ2-5 | CASSLGGVLGETQYF | KLGGALQAK | YES | NO | NO |
| 12048 | TRAV23DV6 | TRAJ40 | CAASPGTYKYIF | TRBV27 | TRBJ2-1 | CASSWGDSYNEQFF | KLGGALQAK | YES | NO | NO |
| 12049 | TRAV19 | TRAJ44 | CALSINTGTASKLTF | TRBV7-9 | TRBJ2-5 | CASSLTGYQTQYF | KLGGALQAK | YES | NO | NO |
| 12050 | TRAV4 | TRAJ30 | CLVVEGNRDDKIIF | TRBV12-3 | TRBJ2-1 | CASREWGGSVNEQFF | RAKFKQLL | YES | NO | NO |
| 12051 | TRAV13-2 | TRAJ12 | CAENGGDSSYKLIF | TRBV12-4 | TRBJ2-5 | CASRPSGGAETQYF | KLGGALQAK | YES | NO | NO |
| 12052 | TRAV5 | TRAJ43 | CAEDNNNDMRF | TRBV9 | TRBJ2-7 | CASSVTGTFGYEQYF | GLCTLVAML | YES | NO | YES |
| 12053 | TRAV12-2 | TRAJ40 | CAVNRGASGTYKYIF | TRBV4-1 | TRBJ2-7 | CASRVASGFGEQYF | AVFDRKSDAK | YES | YES | YES |
| 12054 | TRAV12-2 | TRAJ28 | CAVNGGPYSGAGSYQLTF | TRBV11-2 | TRBJ1-1 | CASSLGDTEAFF | KLGGALQAK | YES | NO | NO |
| 12055 | TRAV38-2DV8 | TRAJ43 | CAYRSAYDNDMRF | TRBV11-2 | TRBJ2-2 | CASSLGVAGTNTGELFF | KLGGALQAK | YES | NO | NO |
| 12056 | TRAV17 | TRAJ23 | CATDQGGKLIF | TRBV2 | TRBJ1-2 | CASSPGGAWNGYTF | KLGGALQAK | YES | NO | NO |
| 12057 | TRAV19 | TRAJ30 | CALSEAGADKIIF | TRBV9 | TRBJ1-2 | CASSGQGTYGYTF | KLGGALQAK | YES | NO | NO |
| 12058 | TRAV1-2 | TRAJ33 | CAVVDSNYQLIW | TRBV28 | TRBJ2-3 | CASSRGDTHTDTQYF | KLGGALQAK | YES | NO | NO |
| 12059 | TRAV35 | TRAJ42 | CAGQLGGGSQGNLIF | TRBV19 | TRBJ2-7 | CASSSRSSYEQYF | GILGFVFTL | YES | NO | YES |
| 12060 | TRAV19 | TRAJ44 | CALSPPRGTASKLTF | TRBV6-6 | TRBJ2-1 | CASSYSTSGSGRLNEQFF | KLGGALQAK | NO | NO | NO |
| 12061 | TRAV30 | TRAJ20 | CGTEYPNDYKLSF | TRBV4-1 | TRBJ2-7 | CASSQGQGISYEQYF | GLCTLVAML | NO | NO | NO |
| 12062 | TRAV35 | TRAJ42 | CAGSGDGGSQGNLIF | TRBV19 | TRBJ2-5 | CASSIRSGATQYF | GILGFVFTL | YES | NO | YES |
| 12063 | TRAV8-4 | TRAJ56 | CAVSERRIGANSKLTF | TRBV11-1 | TRBJ2-7 | CASSFGTAPSYEQYF | RAKFKQLL | YES | NO | YES |
| 12064 | TRAV1-2 | TRAJ33 | CAVRDKDYQLIW | TRBV24-1 | TRBJ2-2 | CATSEATDTGELFF | KLGGALQAK | YES | NO | NO |
| 12065 | TRAV10 | TRAJ30 | CVVSRDDKIIF | TRBV4-1 | TRBJ1-2 | CASSQGQGQDYGYTF | RAKFKQLL | YES | NO | YES |
| 12066 | TRAV14DV4 | TRAJ18 | CAMREDDRDSTLGRLYF | TRBV20-1 | TRBJ2-3 | CSAGQGLAGGPGTDTQYF | RAKFKQLL | NO | NO | NO |
| 12067 | TRAV1-2 | TRAJ33 | CASMDSNYQLIW | TRBV15 | TRBJ1-4 | CATSRVTEEKLFF | KLGGALQAK | YES | NO | NO |
| 12068 | TRAV29DV5 | TRAJ40 | CASYLSGTYKYIF | TRBV4-1 | TRBJ2-3 | CASSQVGGAKDTQYF | KLGGALQAK | YES | NO | NO |
| 12069 | TRAV1-2 | TRAJ33 | CACMDSNYQLIW | TRBV20-1 | TRBJ1-1 | CSARTGDTEAFF | KLGGALQAK | YES | NO | NO |
| 12070 | TRAV13-1 | TRAJ50 | CAASETSYDKVIF | TRBV29-1 | TRBJ2-3 | CSVGSGEDSPQYF | RAKFKQLL | YES | NO | NO |
| 12071 | TRAV24 | TRAJ23 | CAPFQGKLIF | TRBV11-2 | TRBJ2-7 | CASSSGTAAYEQYF | RLRAEAQVK | YES | NO | NO |
| 12072 | TRAV13-2 | TRAJ42 | CAENMGGGSQGNLIF | TRBV19 | TRBJ2-3 | CASSTRSGDTQYF | GILGFVFTL | YES | NO | YES |
| 12073 | TRAV13-1 | TRAJ42 | CAASGSQGNLIF | TRBV29-1 | TRBJ2-3 | CSVGSGEDSPQYF | RAKFKQLL | YES | NO | NO |
| 12074 | TRAV2 | TRAJ15 | CAVDQAGTALIF | TRBV7-9 | TRBJ1-1 | CASSLDRGVNTEAFF | KLGGALQAK | NO | NO | NO |
| 12075 | TRAV5 | TRAJ31 | CAEDNNARLMF | TRBV20-1 | TRBJ1-3 | CSARPQPVGNTIYF | GLCTLVAML | YES | YES | YES |
| 12076 | TRAV22 | TRAJ47 | CAVDRAIEYGNKLVF | TRBV25-1 | TRBJ2-2 | CASSVGTGELFF | RTLNAWVKV | YES | YES | YES |
| 12077 | TRAV27 | TRAJ37 | CAGGGSSNTGKLIF | TRBV7-9 | TRBJ2-7 | CASSSAAAYEQYF | RAKFKQLL | YES | NO | YES |
| 12078 | TRAV13-2 | TRAJ42 | CAENNGGGSQGNLIF | TRBV19 | TRBJ2-1 | CASSIRSGNEQFF | GILGFVFTL | YES | NO | YES |
| 12079 | TRAV17 | TRAJ20 | CATDESNDYKLSF | TRBV15 | TRBJ2-1 | CATSIEAGGGNEQFF | KLGGALQAK | YES | NO | NO |
| 12080 | TRAV19 | TRAJ26 | CALSEAENYGQNFVF | TRBV27 | TRBJ1-2 | CASGYPGQSNYGYTF | KLGGALQAK | YES | NO | NO |
| 12081 | TRAV8-3 | TRAJ5 | CAVDPMDTGRRALTF | TRBV11-2 | TRBJ2-2 | CASSLGVAGTNTGELFF | KLGGALQAK | YES | NO | NO |
| 12082 | TRAV13-1 | TRAJ31 | CAASITARLMF | TRBV7-9 | TRBJ2-3 | CASSLILYTDTQYF | KLGGALQAK | YES | NO | NO |
| 12083 | TRAV12-1 | TRAJ28 | CVVNSFSGAGSYQLTF | TRBV7-9 | TRBJ2-2 | CASSPLQGANTGELFF | KLGGALQAK | YES | NO | NO |
| 12084 | TRAV1-2 | TRAJ33 | CAVRDSNYQLIW | TRBV9 | TRBJ2-7 | CASSLSSGTYEQYF | KLGGALQAK | YES | NO | NO |
| 12085 | TRAV12-2 | TRAJ8 | CAVNTGFQKLVF | TRBV27 | TRBJ2-1 | CASSLSGNEQFF | ELAGIGILTV | YES | NO | YES |
| 12086 | TRAV12-2 | TRAJ15 | CAVNIRQAGTALIF | TRBV2 | TRBJ1-1 | CASSGGDRGTEAFF | KLGGALQAK | YES | NO | NO |
| 12087 | TRAV17 | TRAJ54 | CATDGGGAQKLVF | TRBV7-9 | TRBJ1-5 | CASSSSLAGDQPQHF | RAKFKQLL | YES | NO | YES |
| 12088 | TRAV29DV5 | TRAJ26 | CAPLQIYNYGQNFVF | TRBV19 | TRBJ1-2 | CASSLGLYGYTF | GILGFVFTL | YES | NO | YES |
| 12089 | TRAV1-2 | TRAJ33 | CAVMDSNYQLIW | TRBV6-6 | TRBJ1-3 | CASTAAISGNTIYF | KLGGALQAK | YES | NO | NO |
| 12090 | TRAV13-2 | TRAJ30 | CAVMNRDDKIIF | TRBV28 | TRBJ2-1 | CASSLMGGTYNEQFF | KLGGALQAK | YES | NO | NO |
| 12091 | TRAV12-3 | TRAJ33 | CANYQLIW | TRBV19 | TRBJ2-7 | CASSIRSSYEQYF | GILGFVFTL | NO | NO | NO |
| 12092 | TRAV12-1 | TRAJ40 | CVVIAPSGTYKYIF | TRBV19 | TRBJ2-7 | CASSARSSYEQYF | GILGFVFTL | YES | NO | YES |
| 12093 | TRAV13-1 | TRAJ50 | CAATETSYDKVIF | TRBV7-9 | TRBJ1-3 | CASSFLGTGGASNLISGNTIYF | RAKFKQLL | NO | NO | NO |

| | | | | | | | | | | |
|---|---|---|---|---|---|---|---|---|---|---|
| 12094 | TRAV8-3 | TRAJ17 | CAVKAAGNKLTF | TRBV12-4 | TRBJ2-1 | CASGRLSYNEQFF | RAKFKQLL | YES | NO | YES |
| 12095 | TRAV12-3 | TRAJ42 | CALSPGGGSQGNLIF | TRBV19 | TRBJ2-7 | CASSIRSSYEQYF | GILGFVFTL | YES | NO | YES |
| 12096 | TRAV12-2 | TRAJ42 | CAVTGDGGSQGNLIF | TRBV19 | TRBJ2-7 | CASSIRSSYEQYF | GILGFVFTL | YES | NO | YES |
| 12097 | TRAV10 | TRAJ41 | CVVSGPLFSNSNSGYALNF | TRBV20-1 | TRBJ2-1 | CSARDLPSGGDNEQFF | KLGGALQAK | NO | NO | NO |
| 12098 | TRAV24 | TRAJ11 | CALGYSTLTF | TRBV11-1 | TRBJ2-5 | CASSGRGGGPAETQYF | KLGGALQAK | YES | NO | NO |
| 12099 | TRAV13-1 | TRAJ50 | CAAMETSYDKVIF | TRBV9 | TRBJ2-1 | CASSIPNSGAVNEQFF | KLGGALQAK | YES | NO | NO |
| 12100 | TRAV13-1 | TRAJ23 | CAASGGKLIF | TRBV30 | TRBJ1-1 | CAWNLGQVNTEAFF | AVFDRKSDAK | YES | NO | NO |
| 12101 | TRAV13-1 | TRAJ50 | CAAMETSYDKVIF | TRBV5-4 | TRBJ2-1 | CASSLADEQFF | RAKFKQLL | YES | NO | YES |
| 12102 | TRAV26-1 | TRAJ39 | CIVRVNAGNMLTF | TRBV9 | TRBJ2-2 | CASSVRVASMTGELFF | KLGGALQAK | YES | NO | NO |
| 12103 | TRAV12-1 | TRAJ28 | CVVNMFSGAGSYQLTF | TRBV7-9 | TRBJ2-2 | CASSPRTGNTGELFF | IVTDFSVIK | YES | YES | YES |
| 12104 | TRAV27 | TRAJ42 | CALNYGGSQGNLIF | TRBV19 | TRBJ2-7 | CASSIRSSYEQYF | GILGFVFTL | YES | NO | YES |
| 12105 | TRAV12-3 | TRAJ42 | CAMSGDGGSQGNLIF | TRBV19 | TRBJ2-5 | CASSMRAGETQYF | GILGFVFTL | YES | NO | YES |
| 12106 | TRAV19 | TRAJ26 | CALIRAGGQNFVF | TRBV5-4 | TRBJ1-2 | CASSLGTANYGYTF | KLGGALQAK | YES | NO | NO |
| 12107 | TRAV12-2 | TRAJ31 | CAVGWNNNARLMF | TRBV20-1 | TRBJ1-4 | CSARGISRPYGDEKLFF | RAKFKQLL | YES | NO | YES |
| 12108 | TRAV5 | TRAJ6 | CAEMEGGSYIPTF | TRBV7-9 | TRBJ2-5 | CASSFVDRAETQYF | RAKFKQLL | YES | NO | YES |
| 12109 | TRAV19 | TRAJ29 | CALSAFSGNTPLVF | TRBV20-1 | TRBJ2-7 | CSASEQTGGYEQYF | KLGGALQAK | NO | NO | NO |
| 12110 | TRAV8-1 | TRAJ4 | CAGRFSGGYNKLIF | TRBV15 | TRBJ1-2 | CATSTGRGQLYGYTF | AVFDRKSDAK | YES | NO | NO |
| 12111 | TRAV19 | TRAJ7 | CALRSPGNNRLAF | TRBV20-1 | TRBJ2-5 | CSARGFSGITGQETQYF | KLGGALQAK | NO | NO | NO |
| 12112 | TRAV5 | TRAJ30 | CAENRDDKIIF | TRBV29-1 | TRBJ1-2 | CSADWSTGGANYGYTF | KLGGALQAK | YES | NO | NO |
| 12113 | TRAV29DV5 | TRAJ35 | CAASDGFGNVLHC | TRBV7-9 | TRBJ1-4 | CASSLVGGTDEKLFF | RAKFKQLL | NO | NO | NO |
| 12114 | TRAV19 | TRAJ45 | CALSVFSGGGADGLTF | TRBV11-2 | TRBJ2-2 | CASSLGVAGTNTGELFF | AVFDRKSDAK | YES | YES | YES |
| 12115 | TRAV27 | TRAJ42 | CAGGGSQGNLIF | TRBV20-1 | TRBJ1-2 | CSARDRTGNGYTF | GILGFVFTL | YES | YES | YES |
| 12116 | TRAV19 | TRAJ40 | CAPRTSGTYKYIF | TRBV19 | TRBJ1-1 | CASSIFGQVDAFF | GILGFVFTL | YES | NO | NO |
| 12117 | TRAV25 | TRAJ42 | CAGQYGGSQGNLIF | TRBV19 | TRBJ2-3 | CASSMRSSDTQYF | GILGFVFTL | YES | YES | YES |
| 12118 | TRAV29DV5 | TRAJ58 | CAASTSGSRLTF | TRBV7-9 | TRBJ2-1 | CASSLNPGNPNEQFF | RAKFKQLL | YES | NO | YES |
| 12119 | TRAV13-2 | TRAJ42 | CAENSGGGSQGNLIF | TRBV19 | TRBJ2-5 | CASSIRAAETQYF | GILGFVFTL | YES | NO | YES |
| 12120 | TRAV5 | TRAJ6 | CAEMEGGSYIPTF | TRBV9 | TRBJ2-7 | CASSVTGTFGYEQYF | GLCTLVAML | YES | YES | NO |
| 12121 | TRAV12-2 | TRAJ7 | CAVSGNNRLAF | TRBV14 | TRBJ1-1 | CASSQVGAFMTEAFF | AVFDRKSDAK | NO | NO | NO |
| 12122 | TRAV13-1 | TRAJ11 | CAASRGYSTLTF | TRBV5-4 | TRBJ2-7 | CASSLGGGINEQYF | KLGGALQAK | YES | NO | NO |
| 12123 | TRAV12-2 | TRAJ4 | CAVEPPGGYNKLIF | TRBV6-1 | TRBJ1-1 | CASSESSEAFF | IVTDFSVIK | YES | NO | YES |
| 12124 | TRAV1-2 | TRAJ33 | CAVLDSNYQLIW | TRBV3-1 | TRBJ2-3 | CASNLGQGATDTQYF | RLRAEAQVK | YES | NO | YES |
| 12125 | TRAV35 | TRAJ49 | CAGHTGNQFYF | TRBV12-3 | TRBJ1-1 | CASSFLDRGPNTEAFF | RAKFKQLL | YES | NO | YES |
| 12126 | TRAV19 | TRAJ39 | CALSEANNAGNMLTF | TRBV12-3 | TRBJ2-2 | CASSFSGNTGELFF | RAKFKQLL | YES | NO | YES |
| 12127 | TRAV30 | TRAJ26 | CGTAGDNYGQNFVF | TRBV5-4 | TRBJ2-7 | CASSLNGVYMEQYF | AVFDRKSDAK | YES | NO | YES |
| 12128 | TRAV36DV7 | TRAJ7 | CAVVGNRLAF | TRBV20-1 | TRBJ2-1 | CSARGDRGGSSYNEQFF | KLGGALQAK | NO | NO | NO |
| 12129 | TRAV1-2 | TRAJ12 | CAVMDSSYKLIF | TRBV19 | TRBJ2-7 | CASSIRSSYEQYF | GILGFVFTL | YES | NO | YES |
| 12130 | TRAV5 | TRAJ6 | CAEMEGGSYIPTF | TRBV12-3 | TRBJ1-1 | CASSFLDRGPNTEAFF | RAKFKQLL | YES | NO | NO |
| 12131 | TRAV24 | TRAJ47 | CAHGGNKLVF | TRBV11-2 | TRBJ2-7 | CASSPGTALSYEQYF | KLGGALQAK | YES | NO | NO |
| 12132 | TRAV26-1 | TRAJ15 | CIVRVWGQAGTALIF | TRBV4-1 | TRBJ2-1 | CASSQERGPYNEQFF | IVTDFSVIK | YES | YES | YES |
| 12133 | TRAV29DV5 | TRAJ15 | CAASAQAGTALIF | TRBV14 | TRBJ2-7 | CASSQVLFAYEQYF | KLGGALQAK | NO | NO | NO |
| 12134 | TRAV13-1 | TRAJ47 | CAASEEYGNKLVF | TRBV4-2 | TRBJ2-2 | CASSQRPSEVGELFF | RAKFKQLL | YES | NO | YES |
| 12135 | TRAV4 | TRAJ42 | CLVEFEGSQGNLIF | TRBV2 | TRBJ2-7 | CASRGGASYEQYF | KLGGALQAK | NO | NO | NO |
| 12136 | TRAV12-3 | TRAJ54 | CATAQGAQKLVF | TRBV11-2 | TRBJ1-2 | CASSFGGPNYGYTF | KLGGALQAK | YES | NO | NO |
| 12137 | TRAV1-2 | TRAJ33 | CAVLDSNYQLIW | TRBV6-1 | TRBJ2-7 | CASKGVVYEQYF | KLGGALQAK | YES | NO | NO |
| 12138 | TRAV12-2 | TRAJ47 | CAVRGAEYGNKLVF | TRBV18 | TRBJ2-7 | CASSPTGSVAEQYF | GILGFVFTL | YES | NO | YES |
| 12139 | TRAV13-2 | TRAJ49 | CAGTEISNQFYF | TRBV20-1 | TRBJ1-2 | CSARMVGGHNYGYTF | KLGGALQAK | YES | NO | NO |
| 12140 | TRAV35 | TRAJ42 | CAGAGDGGSQGNLIF | TRBV4-3 | TRBJ2-3 | CASSQEWLAVSTDTQYF | GILGFVFTL | YES | NO | NO |
| 12141 | TRAV8-6 | TRAJ4 | CAVSSFSGGYNKLIF | TRBV14 | TRBJ1-2 | CASSPGTVYGYTF | KLGGALQAK | NO | NO | NO |
| 12142 | TRAV26-1 | TRAJ5 | CIGCWEGRRALTF | TRBV9 | TRBJ2-1 | CASSSPGTRVNEQFF | RAKFKQLL | YES | NO | NO |
| 12143 | TRAV14DV4 | TRAJ56 | CAMREGGGANSKLTF | TRBV7-9 | TRBJ2-7 | CASSLVGVRYEQYF | KLGGALQAK | YES | NO | NO |
| 12144 | TRAV13-1 | TRAJ49 | CAAGSANTGNQFYF | TRBV9 | TRBJ1-2 | CASSVNPGQGANYGYTF | KLGGALQAK | YES | NO | NO |
| 12145 | TRAV19 | TRAJ39 | CALSEAGSNAGNMLTF | TRBV7-9 | TRBJ2-7 | CASSFREKYEQYF | KLGGALQAK | YES | NO | NO |
| 12146 | TRAV8-1 | TRAJ33 | CAVSPIDMDSNYQLIW | TRBV19 | TRBJ1-5 | CASSTHSGNQPQHF | GILGFVFTL | YES | NO | YES |
| 12147 | TRAV9-2 | TRAJ49 | CALSDPNTGNQFYF | TRBV20-1 | TRBJ2-2 | CSARDIERTGELFF | KLGGALQAK | YES | NO | NO |
| 12148 | TRAV1-2 | TRAJ33 | CAVTDSNYQLIW | TRBV20-1 | TRBJ2-5 | CSARDPSGGDQGTQYF | KLGGALQAK | YES | NO | NO |
| 12149 | TRAV8-3 | TRAJ28 | CAQEAGSYQLTF | TRBV11-2 | TRBJ2-2 | CASSLGGANTGELFF | KLGGALQAK | YES | NO | NO |
| 12150 | TRAV38-1 | TRAJ45 | CAFGGGADGLTF | TRBV13 | TRBJ2-1 | CASSPEFRAFEQFF | KLGGALQAK | YES | NO | NO |
| 12151 | TRAV30 | TRAJ28 | CGTAVADSGAGSYQLTF | TRBV5-1 | TRBJ2-5 | CASSPEGQGASAQYF | KLGGALQAK | YES | NO | NO |
| 12152 | TRAV8-3 | TRAJ11 | CAVGPTDPGYSTLTF | TRBV12-3 | TRBJ2-2 | CASSFSGNTGELFF | RAKFKQLL | YES | YES | YES |
| 12153 | TRAV12-2 | TRAJ30 | CAFRMNRDDKIIF | TRBV4-1 | TRBJ1-2 | CASSQDRSGEGYTF | RAKFKQLL | YES | NO | YES |
| 12154 | TRAV13-1 | TRAJ21 | CAAFDFNKFYF | TRBV12-3 | TRBJ2-2 | CASSFSGNTGELFF | RAKFKQLL | YES | NO | YES |
| 12155 | TRAV8-4 | TRAJ15 | CAVRERANQAGTALIF | TRBV11-2 | TRBJ1-2 | CASSWGGGSHYGYTF | IVTDFSVIK | YES | NO | YES |
| 12156 | TRAV20 | TRAJ39 | CAVLDNNAGNMLTF | TRBV6-5 | TRBJ1-1 | CASSYVGGASRAFF | KLGGALQAK | YES | NO | NO |
| 12157 | TRAV29DV5 | TRAJ26 | CAAPGNYGQNFVF | TRBV6-5 | TRBJ1-1 | CASLPGNGAEAFF | RAKFKQLL | YES | YES | YES |
| 12158 | TRAV5 | TRAJ40 | CGEGTYKYIF | TRBV19 | TRBJ1-5 | CASSIEDKDQPQHF | AVFDRKSDAK | YES | NO | YES |
| 12159 | TRAV8-1 | TRAJ28 | CAVNVDSGAGSYQLTF | TRBV7-9 | TRBJ2-1 | CASSGPPRGPYNEQFF | KLGGALQAK | YES | NO | NO |
| 12160 | TRAV13-1 | TRAJ50 | CAGGETSYDKVIF | TRBV12-3 | TRBJ2-2 | CASSFTGGRTGELFF | RAKFKQLL | YES | NO | NO |
| 12161 | TRAV1-2 | TRAJ33 | CAVSDSNYQLIW | TRBV3-1 | TRBJ2-1 | CASSQELEVYNEQFF | RAKFKQLL | YES | YES | YES |
| 12162 | TRAV12-1 | TRAJ37 | CVVPRQSSNTGKLIF | TRBV20-1 | TRBJ1-1 | CSAREGFSTEAFF | KLGGALQAK | YES | NO | NO |
| 12163 | TRAV1-2 | TRAJ12 | CAPRMDSSYKLIF | TRBV19 | TRBJ2-7 | CASSIRSSYEQYF | GILGFVFTL | YES | NO | YES |
| 12164 | TRAV6 | TRAJ8 | CARGFQKLVF | TRBV27 | TRBJ2-3 | CASSPDDTTDTQYF | KLGGALQAK | YES | NO | NO |
| 12165 | TRAV12-3 | TRAJ40 | CAMSGKTSGTYKYIF | TRBV20-1 | TRBJ2-7 | CSASSLTSGGAPEQYF | KLGGALQAK | NO | NO | NO |
| 12166 | TRAV12-3 | TRAJ53 | CAMSESGGSNYKLTF | TRBV29-1 | TRBJ2-7 | CSVGSGDYEQYF | RAKFKQLL | YES | NO | YES |
| 12167 | TRAV24 | TRAJ36 | CAFLGGANNLFF | TRBV9 | TRBJ1-2 | CASSADPSQGDGNYGYTF | KLGGALQAK | YES | NO | NO |
| 12168 | TRAV4 | TRAJ36 | CLVGDIRTGANNLFF | TRBV11-2 | TRBJ2-1 | CASSLVFTGGAGNEQFF | KLGGALQAK | YES | NO | NO |
| 12169 | TRAV26-2 | TRAJ49 | CILRLFRNTGNQFYF | TRBV9 | TRBJ2-7 | CASSVESGSAYEQYF | KLGGALQAK | NO | NO | NO |
| 12170 | TRAV19 | TRAJ58 | CALTLPLTSGSRLTF | TRBV19 | TRBJ2-3 | CASSSRTGPDTQYF | KLGGALQAK | YES | NO | NO |
| 12171 | TRAV13-1 | TRAJ50 | CAASETSYDKVIF | TRBV12-3 | TRBJ1-2 | CASSYTGGLRYGYTF | RAKFKQLL | YES | NO | YES |
| 12172 | TRAV27 | TRAJ42 | CAGPLAGGSQGNLIF | TRBV19 | TRBJ1-5 | CASSIYSNQPQHF | GILGFVFTL | YES | NO | YES |
| 12173 | TRAV23DV6 | TRAJ31 | CAATDNNARLMF | TRBV9 | TRBJ2-6 | CASSVAVGTGSGANVLTF | KLGGALQAK | YES | NO | NO |
| 12174 | TRAV39 | TRAJ54 | CAVNQGAQKLVF | TRBV19 | TRBJ2-5 | CASSGRSGETQYF | GILGFVFTL | YES | NO | YES |
| 12175 | TRAV19 | TRAJ31 | CALSEAYNNARLMF | TRBV5-4 | TRBJ1-1 | CASSFNGQTNTEAFF | KLGGALQAK | YES | NO | NO |
| 12176 | TRAV25 | TRAJ42 | CAMNYGGSQGNLIF | TRBV19 | TRBJ2-3 | CASSTRSTDTQYF | GILGFVFTL | YES | NO | YES |
| 12177 | TRAV26-1 | TRAJ17 | CILWVAAGNKLTF | TRBV11-3 | TRBJ2-1 | CASSLGLAGGGGIEQFF | ELAGIGILTV | YES | YES | YES |
| 12178 | TRAV13-2 | TRAJ15 | CAETEAGTALIF | TRBV12-3 | TRBJ2-7 | CASSWGYEQYF | KLGGALQAK | YES | NO | NO |
| 12179 | TRAV29DV5 | TRAJ28 | CAASSAGAGSYQLTF | TRBV5-6 | TRBJ1-1 | CASSLRDGSEAFF | RAKFKQLL | YES | YES | YES |
| 12180 | TRAV19 | TRAJ17 | CALKLIKAAGNKLTF | TRBV4-1 | TRBJ1-2 | CASSTSTGTGYGYTF | RAKFKQLL | YES | NO | NO |
| 12181 | TRAV10 | TRAJ45 | CVVMGGADGLTF | TRBV30 | TRBJ2-1 | CAWSDDRGYNEQFF | KLGGALQAK | NO | NO | NO |
| 12182 | TRAV8-3 | TRAJ32 | CAGGGATNKLIF | TRBV15 | TRBJ2-3 | CATSRTSGLDTQYF | KLGGALQAK | YES | NO | NO |
| 12183 | TRAV1-2 | TRAJ23 | CAVQEGKLIF | TRBV29-1 | TRBJ1-2 | CSALRQGLLEYTF | KLGGALQAK | YES | NO | NO |
| 12184 | TRAV12-3 | TRAJ23 | CASYNQGGKLIF | TRBV19 | TRBJ2-7 | CASSIRSSYEQYF | GILGFVFTL | YES | NO | YES |
| 12185 | TRAV13-1 | TRAJ50 | CAASETSYDKVIF | TRBV12-5 | TRBJ1-1 | CASGLVESAGTEAFF | RAKFKQLL | YES | NO | YES |
| 12186 | TRAV39 | TRAJ43 | CAVDNNNDMRF | TRBV6-5 | TRBJ2-1 | CASSYSSYGGHNEQFF | RTLNAWVKV | NO | NO | NO |
| 12187 | TRAV29DV5 | TRAJ44 | CAASETGTASKLTF | TRBV10-3 | TRBJ2-5 | CAISGWDRQETQYF | KLGGALQAK | YES | NO | NO |
| 12188 | TRAV8-3 | TRAJ42 | CAVLNYGGSQGNLIF | TRBV7-9 | TRBJ2-7 | CASSLFGTDEQYF | KLGGALQAK | YES | NO | NO |
| 12189 | TRAV1-1 | TRAJ5 | CAVRDTGRRALTF | TRBV12-3 | TRBJ2-3 | CASTELAGGQWLDTQYF | KLGGALQAK | YES | NO | NO |
| 12190 | TRAV12-3 | TRAJ45 | CAMSDSGGGADGLTF | TRBV20-1 | TRBJ2-1 | CSARDRVNYNEQFF | KLGGALQAK | NO | NO | NO |
| 12191 | TRAV35 | TRAJ50 | CAGIYDKVIF | TRBV11-2 | TRBJ1-1 | CASSFGGRAEAFF | KLGGALQAK | YES | NO | NO |
| 12192 | TRAV38-2DV8 | TRAJ52 | CAYSPAAGGTSYGKLTF | TRBV20-1 | TRBJ1-1 | CSARTGDTEAFF | GILGFVFTL | YES | NO | YES |
| 12193 | TRAV12-2 | TRAJ36 | CASQVGPNNLFF | TRBV4-1 | TRBJ2-3 | CASSQGTDTQYF | KLGGALQAK | YES | NO | NO |
| 12194 | TRAV26-1 | TRAJ41 | CIVRGFSNSGYALNF | TRBV19 | TRBJ2-1 | CASSIYSRGDEQFF | GILGFVFTL | YES | YES | YES |
| 12195 | TRAV1-1 | TRAJ21 | CAVFNFNKFYF | TRBV15 | TRBJ1-2 | CATSRDGPGHEVTANYGYTF | KLGGALQAK | NO | NO | NO |
| 12196 | TRAV22 | TRAJ26 | CAPRRNYGQNFVF | TRBV18 | TRBJ2-7 | CASSPGQGGYEQYF | IMDQVPFSV | YES | NO | YES |
| 12197 | TRAV12-2 | TRAJ49 | CAVNAGNQFYF | TRBV11-2 | TRBJ2-2 | CASSLGFLTGELFF | ELAGIGILTV | YES | YES | YES |

| | | | | | | | | | | |
|---|---|---|---|---|---|---|---|---|---|---|
| 12198 | TRAV21 | TRAJ27 | CAVPFNAGKSTF | TRBV5-6 | TRBJ2-1 | CASSLWTGGHNEQFF | KLGGALQAK | YES | NO | NO |
| 12199 | TRAV27 | TRAJ37 | CAGAHGSSNTGKLIF | TRBV19 | TRBJ1-1 | CASYDFTEAFF | GILGFVFTL | YES | NO | YES |
| 12200 | TRAV3 | TRAJ45 | CAVAKDLQNSGGGADGLTF | TRBV19 | TRBJ1-1 | CASSVRHGTEAFF | GILGFVFTL | NO | NO | NO |
| 12201 | TRAV19 | TRAJ20 | CALSDPYFKLSF | TRBV9 | TRBJ1-1 | CASSVKGTMNTEAFF | KLGGALQAK | YES | NO | NO |
| 12202 | TRAV24 | TRAJ42 | CAFGGGGSQGNLIF | TRBV19 | TRBJ2-3 | CASSIRSVDTQYF | GILGFVFTL | YES | NO | YES |
| 12203 | TRAV27 | TRAJ28 | CAGARGGSGAGSYQLTF | TRBV10-3 | TRBJ1-1 | CAISAYSRNTEAFF | KLGGALQAK | YES | NO | NO |
| 12204 | TRAV29DV5 | TRAJ49 | CAASASGGQFYF | TRBV19 | TRBJ1-1 | CASRAGREAFF | GILGFVFTL | YES | NO | YES |
| 12206 | TRAV30 | TRAJ28 | CGTEMNSGAGSYQLTF | TRBV7-9 | TRBJ2-7 | CASSFWTGPTYEQYF | KLGGALQAK | NO | NO | NO |
| 12207 | TRAV29DV5 | TRAJ49 | CAATNTGNQFYF | TRBV7-9 | TRBJ2-6 | CASSPVTGGGSGANVLTF | KLGGALQAK | YES | NO | NO |
| 12208 | TRAV38-2DV8 | TRAJ40 | CAYTGTSGTYKYIF | TRBV5-1 | TRBJ1-2 | CASSLSGPGYGYTF | KLGGALQAK | YES | NO | NO |
| 12209 | TRAV12-2 | TRAJ36 | CAVNDPGANNLFF | TRBV6-4 | TRBJ1-1 | CASSDSWGTEAFF | KLGGALQAK | YES | NO | NO |
| 12210 | TRAV41 | TRAJ49 | CAHFSGNQFYF | TRBV27 | TRBJ1-5 | CASRGAGQPQHF | KLGGALQAK | NO | NO | NO |
| 12211 | TRAV12-2 | TRAJ31 | CAVRNNNARLMF | TRBV19 | TRBJ1-1 | CASSPGGGSEAFF | AVFDRKSDAK | YES | NO | YES |
| 12212 | TRAV29DV5 | TRAJ34 | CAAYPEGLYNTDKLIF | TRBV5-4 | TRBJ2-3 | CASSFNGQGATDTQYF | RAKFKQLL | YES | YES | YES |
| 12213 | TRAV12-1 | TRAJ20 | CVVNYDYKLSF | TRBV11-2 | TRBJ2-2 | CASSLERGTGELFF | KLGGALQAK | YES | NO | NO |
| 12214 | TRAV8-2 | TRAJ34 | CVVSVGNTDKLIF | TRBV12-4 | TRBJ2-7 | CASSLGTGYYEQYF | KLGGALQAK | YES | NO | NO |
| 12215 | TRAV38-2DV8 | TRAJ43 | CAYRSFGNNDMRF | TRBV12-3 | TRBJ2-7 | CASSSLDVAGYEQYF | RAKFKQLL | YES | NO | YES |
| 12216 | TRAV29DV5 | TRAJ52 | CAAIYSGTSYGKLTF | TRBV2 | TRBJ2-5 | CASSLSGNRQETQYF | KLGGALQAK | YES | NO | NO |
| 12217 | TRAV9-2 | TRAJ33 | CAREHMDSNYQLIW | TRBV7-9 | TRBJ2-6 | CASSPVTGGGSGANVLTF | KLGGALQAK | YES | NO | NO |
| 12218 | TRAV19 | TRAJ49 | CALSEAHTGNQFYF | TRBV9 | TRBJ1-2 | CASSVRTTTGVGDYGYTF | AVFDRKSDAK | YES | NO | YES |
| 12219 | TRAV35 | TRAJ38 | CAGQGSAGNNRKLIW | TRBV29-1 | TRBJ1-2 | CSVEDLAYGYTF | KLGGALQAK | YES | NO | NO |
| 12220 | TRAV19 | TRAJ57 | CALSEALQGGSEKLVF | TRBV20-1 | TRBJ2-3 | CSATHLELTTDTQYF | KLGGALQAK | NO | NO | NO |
| 12221 | TRAV1-2 | TRAJ33 | CAAVDSNYQLIW | TRBV6-1 | TRBJ2-3 | CASSEEGGRGADTQYF | KLGGALQAK | YES | NO | NO |
| 12222 | TRAV21 | TRAJ20 | CAVVVNDYKLSF | TRBV6-4 | TRBJ2-2 | CATQQGLGTGELFF | AVFDRKSDAK | YES | NO | NO |
| 12223 | TRAV12-2 | TRAJ24 | CAVNTDSWGKLQF | TRBV5-6 | TRBJ1-1 | CASSLRDGSEAFF | RAKFKQLL | YES | NO | YES |
| 12224 | TRAV29DV5 | TRAJ33 | CAAREHSNYQLIW | TRBV7-2 | TRBJ1-5 | CASSPWDREPQHF | KLGGALQAK | YES | NO | NO |
| 12225 | TRAV1-2 | TRAJ33 | CAVIESNYQLIW | TRBV11-2 | TRBJ2-3 | CASSRLATTDTQYF | KLGGALQAK | YES | NO | NO |
| 12226 | TRAV38-2DV8 | TRAJ28 | CAYRSPYSGAGSYQLTF | TRBV12-3 | TRBJ2-1 | CASSIKLRDSNEQFF | KLGGALQAK | YES | NO | NO |
| 12227 | TRAV13-1 | TRAJ9 | CAASEDGGFKTIF | TRBV4-1 | TRBJ2-2 | CASSQVQGAVGELFF | RAKFKQLL | YES | NO | NO |
| 12228 | TRAV19 | TRAJ18 | CALIWGSTLGRLYF | TRBV20-1 | TRBJ2-1 | CSARVAKGYNEQFF | KLGGALQAK | NO | NO | NO |
| 12229 | TRAV12-3 | TRAJ23 | CAMSAYDQGGKLIF | TRBV7-9 | TRBJ2-5 | CASSPRLGTETQYF | AVFDRKSDAK | YES | NO | YES |
| 12230 | TRAV19 | TRAJ34 | CAPYNTDKLIF | TRBV27 | TRBJ1-2 | CASSLAGSSYGYTF | KLGGALQAK | YES | NO | NO |
| 12231 | TRAV29DV5 | TRAJ42 | CAVIGGSQGNLIF | TRBV4-2 | TRBJ2-2 | CASSPNPGTNTGELFF | KLGGALQAK | YES | NO | NO |
| 12232 | TRAV25 | TRAJ44 | CAGHGTASKLTF | TRBV19 | TRBJ2-3 | CASSGRSTDTQYF | GILGFVFTL | YES | NO | YES |
| 12233 | TRAV5 | TRAJ33 | CAETQDSNYQLIW | TRBV21-1 | TRBJ2-4 | CASSMGDRSMGNIQYF | RAKFKQLL | YES | NO | YES |
| 12234 | TRAV27 | TRAJ42 | CAGAGSQGNLIF | TRBV19 | TRBJ2-7 | CASSIRSAYEQYF | GILGFVFTL | YES | NO | YES |
| 12235 | TRAV39 | TRAJ31 | CAVGQNNARLMF | TRBV20-1 | TRBJ2-2 | CSARDIERTGELFF | KLGGALQAK | YES | NO | NO |
| 12236 | TRAV13-1 | TRAJ50 | CAASETSYDKVIF | TRBV12-3 | TRBJ2-7 | CASQPHGTGDHEQYF | RAKFKQLL | YES | NO | YES |
| 12237 | TRAV13-1 | TRAJ42 | CAASGGGSQGNLIF | TRBV12-3 | TRBJ2-2 | CASSFSGNTGELFF | RAKFKQLL | YES | NO | YES |
| 12238 | TRAV3 | TRAJ6 | CAVRRTSGGSYIPTF | TRBV27 | TRBJ2-7 | CASSPGTPFYYEQYF | KLGGALQAK | NO | NO | NO |
| 12239 | TRAV12-1 | TRAJ26 | CVVGLDNYGQNFVF | TRBV15 | TRBJ2-2 | CATSRVESGTGELFF | KLGGALQAK | YES | NO | NO |
| 12240 | TRAV1-2 | TRAJ24 | CAVTRTDSWGKFQF | TRBV20-1 | TRBJ1-3 | CSAPGEGYGNTIYF | KLGGALQAK | YES | NO | NO |
| 12241 | TRAV12-2 | TRAJ31 | CAVIVARLMF | TRBV3-1 | TRBJ2-6 | CASSRTTSGGSGANVLTF | KLGGALQAK | YES | NO | NO |
| 12242 | TRAV25 | TRAJ20 | CAGLNDYKLSF | TRBV5-4 | TRBJ2-1 | CASRPDSYNEQFF | KLGGALQAK | YES | NO | NO |
| 12243 | TRAV19 | TRAJ6 | CALGGSYIPTF | TRBV11-2 | TRBJ2-7 | CASSSDQLDRGSYEQYF | RTLNAWVKV | YES | YES | YES |
| 12244 | TRAV13-2 | TRAJ29 | CAENKREAGNTPLVF | TRBV7-3 | TRBJ2-2 | CASSLGTSGGTGELFF | RAKFKQLL | YES | NO | YES |
| 12245 | TRAV13-1 | TRAJ47 | CAAGEGNKLVF | TRBV12-4 | TRBJ2-2 | CASSQGAGELFF | KLGGALQAK | YES | NO | NO |
| 12246 | TRAV13-1 | TRAJ39 | CAALRNTAGNMLTF | TRBV20-1 | TRBJ2-5 | CSARDYSGPTSGTQYF | KLGGALQAK | YES | NO | NO |
| 12247 | TRAV17 | TRAJ6 | CATEGASGGSYIPTF | TRBV19 | TRBJ2-7 | CASSIRSSYEQYF | GILGFVFTL | YES | NO | YES |
| 12248 | TRAV13-2 | TRAJ53 | CAETHSGGSNYKLTF | TRBV9 | TRBJ2-2 | CASSSGLVSNTGELFF | GLCTLVAML | YES | YES | YES |
| 12249 | TRAV20 | TRAJ24 | CAVWIREATDSWGKLQF | TRBV2 | TRBJ2-1 | CASLLLLNNEQFF | LLFGYPVYV | YES | YES | YES |
| 12251 | TRAV1-2 | TRAJ20 | CAVVQGDYKLSF | TRBV20-1 | TRBJ2-1 | CSARDLPSGGDNEQFF | KLGGALQAK | NO | NO | NO |
| 12252 | TRAV29DV5 | TRAJ13 | CAASPHSGGYQKVTF | TRBV19 | TRBJ2-3 | CASSVRSTDTQYF | GILGFVFTL | YES | NO | YES |
| 12253 | TRAV21 | TRAJ5 | CAVMDTGRRALTF | TRBV6-5 | TRBJ1-1 | CASSYSWDRGINTEAFF | KTWGQYWQV | YES | NO | YES |
| 12254 | TRAV13-1 | TRAJ35 | CAARVRGFGNVLHC | TRBV4-1 | TRBJ2-3 | CASTPIESSTDTQYF | KLGGALQAK | NO | NO | NO |
| 12255 | TRAV21 | TRAJ17 | CAVSRIKAAGNKLTF | TRBV19 | TRBJ1-5 | CASSIYSNQPQHF | GILGFVFTL | YES | YES | YES |
| 12256 | TRAV36DV7 | TRAJ54 | CAHLQGAQKLVF | TRBV4-2 | TRBJ2-2 | CASSQRPSEVGELFF | RAKFKQLL | NO | NO | NO |
| 12257 | TRAV12-2 | TRAJ7 | CAVLYGNNRLAF | TRBV6-5 | TRBJ2-4 | CASSYSASSGTKNIQYF | AVFDRKSDAK | YES | YES | YES |
| 12258 | TRAV8-3 | TRAJ42 | CAVGGSGGSQGNLIF | TRBV19 | TRBJ2-7 | CASSVRSSYEQYF | GILGFVFTL | YES | NO | YES |
| 12259 | TRAV2 | TRAJ29 | CAVEDHNTPLVF | TRBV7-7 | TRBJ2-1 | CASSLDSSGGAYNEQFF | KLGGALQAK | NO | NO | NO |
| 12260 | TRAV17 | TRAJ53 | CATDASGGNNYKLTF | TRBV29-1 | TRBJ2-7 | CSVGTGDWGEQYF | RAKFKQLL | YES | NO | NO |
| 12261 | TRAV14DV4 | TRAJ21 | CAMREASSRYNFNKFYF | TRBV5-6 | TRBJ2-3 | CASSLDLRQGRDTQYF | KLGGALQAK | YES | NO | NO |
| 12262 | TRAV35 | TRAJ42 | CASNYGGSQGNLIF | TRBV19 | TRBJ2-3 | CASSMRSSDTQYF | GILGFVFTL | YES | NO | YES |
| 12263 | TRAV6 | TRAJ27 | CALDAPTANAGKSTF | TRBV11-2 | TRBJ2-7 | CASSPGTALYEQYF | IVTDFSVIK | YES | YES | YES |
| 12264 | TRAV12-2 | TRAJ30 | CAVKDKIIF | TRBV27 | TRBJ1-5 | CASSPIFRNSNQPQHF | AYAQKIFKI | NO | NO | NO |
| 12265 | TRAV13-1 | TRAJ31 | CAAREEGDARLMF | TRBV19 | TRBJ2-5 | CASSIRSGETQYF | GILGFVFTL | YES | NO | YES |
| 12266 | TRAV39 | TRAJ48 | CAVDNFGNEKLTF | TRBV6-1 | TRBJ1-1 | CASSEGTGGTEAFF | KLGGALQAK | YES | NO | NO |
| 12267 | TRAV13-1 | TRAJ50 | CAASETSYDKVIF | TRBV12-3 | TRBJ1-2 | CASSFWPGRNLLWGYTF | RAKFKQLL | YES | NO | YES |
| 12268 | TRAV38-2DV8 | TRAJ52 | CAWSSNAGGTSYGKLTF | TRBV6-1 | TRBJ2-1 | CASSEGGFTFYNEQFF | GLCTLVAML | YES | YES | YES |
| 12269 | TRAV35 | TRAJ42 | CAGPSSSFNYGGSQGNLIF | TRBV19 | TRBJ1-5 | CASSIYSNQPQHF | GILGFVFTL | YES | NO | YES |
| 12270 | TRAV27 | TRAJ33 | CAGAGNYQLIW | TRBV12-3 | TRBJ1-1 | CASSFLDRGPNTEAFF | RAKFKQLL | YES | NO | NO |
| 12271 | TRAV38-2DV8 | TRAJ57 | CAYRSQQGGSEKLVF | TRBV9 | TRBJ2-2 | CASSPRDSGVTGELFF | GILGFVFTL | YES | NO | YES |
| 12272 | TRAV26-2 | TRAJ22 | CILRDGGPGQLTF | TRBV6-6 | TRBJ2-7 | CASSYTGGPNEQYF | KLGGALQAK | NO | NO | NO |
| 12273 | TRAV19 | TRAJ6 | CALSEAGTGGSYIPTF | TRBV19 | TRBJ1-5 | CASSIFGGQPQHF | GILGFVFTL | YES | NO | YES |
| 12274 | TRAV5 | TRAJ34 | CAESEGPTYTDKLIF | TRBV7-9 | TRBJ2-5 | CASRPPAAAADKTQYF | KLGGALQAK | YES | NO | NO |
| 12275 | TRAV12-2 | TRAJ6 | CAVNMRASGGSYIPTF | TRBV11-2 | TRBJ2-3 | CASSLVRTSGGAISDTQYF | KLGGALQAK | YES | NO | NO |
| 12276 | TRAV5 | TRAJ33 | CAETQDSNYQLIW | TRBV12-3 | TRBJ2-2 | CASSFSGNTGELFF | RAKFKQLL | YES | NO | YES |
| 12277 | TRAV27 | TRAJ17 | CAGLDEKAAGNKLTF | TRBV5-1 | TRBJ2-2 | CASSLGLTTGELFF | KLGGALQAK | YES | NO | NO |
| 12278 | TRAV38-1 | TRAJ52 | CAFMKGAGGTSYGKLTF | TRBV28 | TRBJ2-1 | CASSLMGGTYNEQFF | KLGGALQAK | YES | NO | NO |
| 12279 | TRAV8-6 | TRAJ42 | CAVGGSQGNLIF | TRBV19 | TRBJ2-7 | CASSIRSSLEQYF | GILGFVFTL | YES | NO | YES |
| 12280 | TRAV13-1 | TRAJ35 | CAARVRGFGNVLHC | TRBV12-4 | TRBJ2-1 | CASGRLSYNEQFF | RAKFKQLL | NO | NO | NO |
| 12281 | TRAV27 | TRAJ42 | CAGAGGGSQGNLIF | TRBV19 | TRBJ2-3 | CASSKRSTDTQYF | GILGFVFTL | YES | NO | YES |
| 12282 | TRAV5 | TRAJ5 | CAEIEDTGRRALTF | TRBV9 | TRBJ2-7 | CASSVDGEPYEQYF | KLGGALQAK | YES | NO | NO |
| 12283 | TRAV12-2 | TRAJ30 | CAGGDDKIIF | TRBV20-1 | TRBJ1-5 | CSASRTGLGQPQHF | ELAGIGILTV | YES | YES | YES |
| 12284 | TRAV12-2 | TRAJ28 | CAVTGSYQLTF | TRBV24-1 | TRBJ2-2 | CATSEGGTGELFF | RAKFKQLL | YES | NO | YES |
| 12285 | TRAV13-1 | TRAJ50 | CAASETSYDKVIF | TRBV12-3 | TRBJ1-2 | CASSSDPRYGYTF | RAKFKQLL | YES | NO | YES |
| 12286 | TRAV9-2 | TRAJ3 | CARAPLSGSSASKIIF | TRBV7-2 | TRBJ2-5 | CASSLGTSGGETQYF | GILGFVFTL | YES | NO | YES |
| 12287 | TRAV14DV4 | TRAJ9 | CAMREGQGGGFKTIF | TRBV12-3 | TRBJ2-4 | CASSTPQGNIQYF | KLGGALQAK | YES | NO | NO |
| 12288 | TRAV20 | TRAJ49 | CAAHGNQFYF | TRBV6-5 | TRBJ1-2 | CASSAQGGNYGYTF | FLYALALLL | YES | YES | YES |
| 12289 | TRAV1-2 | TRAJ33 | CAVLDSNYQLIW | TRBV6-1 | TRBJ2-6 | CASSPKIGTGANVLTF | KLGGALQAK | YES | NO | NO |
| 12290 | TRAV1-2 | TRAJ4 | CAVRSPFSGGYNKLIF | TRBV14 | TRBJ1-1 | CASSQERGATEAFF | KLGGALQAK | NO | NO | NO |
| 12291 | TRAV1-2 | TRAJ33 | CAVKDSNYQLIW | TRBV4-1 | TRBJ2-2 | CASSQEGAVSGELFF | KLGGALQAK | YES | NO | NO |
| 12292 | TRAV8-3 | TRAJ42 | CAGPLVNYGGSQGNLIF | TRBV19 | TRBJ2-7 | CASSIRSAYEQYF | GILGFVFTL | YES | NO | YES |
| 12293 | TRAV12-3 | TRAJ15 | CAVSRGQAGTALIF | TRBV30 | TRBJ2-1 | CAWSLAGGEQFF | KLGGALQAK | YES | NO | NO |
| 12294 | TRAV21 | TRAJ43 | CAVWADMRF | TRBV7-8 | TRBJ2-3 | CASSPGDTRTDTQYF | AVFDRKSDAK | NO | NO | NO |
| 12295 | TRAV1-2 | TRAJ16 | CARWSDGQKLLF | TRBV6-1 | TRBJ1-5 | CASSSARGISNQPQHF | KLGGALQAK | NO | NO | NO |
| 12296 | TRAV8-1 | TRAJ18 | CAVNAQGGGSTLGRLYF | TRBV19 | TRBJ2-5 | CASSIHGLETQYF | GILGFVFTL | YES | NO | YES |
| 12297 | TRAV12-2 | TRAJ9 | CAVNAGGFKTIF | TRBV4-2 | TRBJ2-2 | CASSQEGNTGELFF | ELAGIGILTV | YES | NO | NO |
| 12298 | TRAV6 | TRAJ34 | CALDGPGNTDKLIF | TRBV5-6 | TRBJ2-2 | CASSSTGELFF | KLGGALQAK | YES | NO | NO |
| 12299 | TRAV12-1 | TRAJ20 | CVAFFDYKLSF | TRBV9 | TRBJ1-2 | CASSVGNDYTF | KLGGALQAK | YES | NO | NO |
| 12300 | TRAV38-2DV8 | TRAJ29 | CAHHGGNTPLVF | TRBV11-2 | TRBJ2-7 | CASSGWDRPSVIYEQYF | KLGGALQAK | YES | NO | NO |
| 12301 | TRAV25 | TRAJ26 | CAGQEGQNFVF | TRBV19 | TRBJ2-3 | CASSPRSTDTQYF | GILGFVFTL | YES | NO | YES |
| 12302 | TRAV10 | TRAJ26 | CVVSPDNYGQNFVF | TRBV11-2 | TRBJ2-7 | CASSFARGHEQYF | KLGGALQAK | YES | NO | NO |
| 12303 | TRAV29DV5 | TRAJ31 | CAASSNARLMF | TRBV29-1 | TRBJ2-1 | CSVQGGSYNEQFF | KLGGALQAK | YES | NO | NO |

| | | | | | | | | | | |
|---|---|---|---|---|---|---|---|---|---|---|
| 12304 | TRAV12-2 | TRAJ42 | CAVNMGGGSQGNLIF | TRBV19 | TRBJ2-7 | CASSKRSSYEQYF | GILGFVFTL | YES | NO | YES |
| 12305 | TRAV12-2 | TRAJ26 | CAVNNYGQNFVF | TRBV11-1 | TRBJ1-5 | CASSSPAEGGQPQHF | AVFDRKSDAK | YES | NO | YES |
| 12306 | TRAV22 | TRAJ41 | CAVERNSGYALNF | TRBV4-1 | TRBJ2-1 | CASSQGLYNEQFF | KLGGALQAK | YES | NO | NO |
| 12307 | TRAV36DV7 | TRAJ52 | CASGSNAGGTSYGKLTF | TRBV6-3 | TRBJ2-3 | CASSPLGKAQYF | KLGGALQAK | NO | NO | NO |
| 12308 | TRAV12-3 | TRAJ53 | CAMSRSGGSNYKLTF | TRBV29-1 | TRBJ2-7 | CSVGSGDYEQYF | RAKFKQLL | YES | NO | YES |
| 12309 | TRAV14DV4 | TRAJ30 | CAMREGGLDRDDKIIF | TRBV13 | TRBJ1-2 | CASPPTGALGGYTF | KLGGALQAK | YES | NO | NO |
| 12310 | TRAV23DV6 | TRAJ43 | CAASGNHDMRF | TRBV12-3 | TRBJ2-2 | CASSFSGNTGELFF | RAKFKQLL | YES | NO | NO |
| 12311 | TRAV21 | TRAJ36 | CAVRQKQTGANNLFF | TRBV6-1 | TRBJ1-4 | CASSPLIRGLNEKLFF | AVFDRKSDAK | YES | NO | YES |
| 12312 | TRAV35 | TRAJ38 | CAGQGSAGNNRKLIW | TRBV29-1 | TRBJ1-2 | CSVELLDYGYTF | KLGGALQAK | YES | NO | NO |
| 12313 | TRAV13-1 | TRAJ4 | CAATPQGYNKLIF | TRBV4-2 | TRBJ2-1 | CASSPQSGKYNEQFF | RLRAEAQVK | YES | NO | YES |
| 12314 | TRAV8-3 | TRAJ12 | CAVGRMDSSYKLIF | TRBV28 | TRBJ2-7 | CASGLAGGLNEQYF | KLGGALQAK | YES | NO | NO |
| 12315 | TRAV29DV5 | TRAJ54 | CPQGAQKLVF | TRBV12-3 | TRBJ2-2 | CASSFSGNTGELFF | RAKFKQLL | YES | NO | NO |
| 12316 | TRAV1-2 | TRAJ33 | CAVSDSNYQLIW | TRBV20-1 | TRBJ2-1 | CSAGNSGGYNEQFF | KLGGALQAK | NO | NO | NO |
| 12317 | TRAV5 | TRAJ37 | CAEWGSGNTGKLIF | TRBV9 | TRBJ2-7 | CASSVAGGTEQYF | KLGGALQAK | YES | NO | NO |
| 12318 | TRAV9-2 | TRAJ31 | CALKEARLMF | TRBV12-4 | TRBJ2-7 | CASRVSGGEEQYF | AVFDRKSDAK | YES | YES | YES |
| 12319 | TRAV20 | TRAJ29 | CAVLSNSGNTPLVF | TRBV6-5 | TRBJ2-7 | CASSVGQGPYEQYF | KLGGALQAK | YES | NO | NO |
| 12320 | TRAV17 | TRAJ40 | CATASTSGTYKYIF | TRBV13 | TRBJ2-1 | CASSLGGQGSNEQFF | RAKFKQLL | YES | NO | NO |
| 12321 | TRAV29DV5 | TRAJ53 | CAASRLGGSNYKLTF | TRBV16 | TRBJ2-7 | CASSRDRGVEQYF | KLGGALQAK | YES | NO | NO |
| 12322 | TRAV38-1 | TRAJ52 | CAFMKDAGGTSYGKLTF | TRBV4-1 | TRBJ1-2 | CASSTSTGTGYGYTF | GILGFVFTL | YES | NO | YES |
| 12323 | TRAV38-2DV8 | TRAJ54 | CASQPFQGAQKLVF | TRBV4-1 | TRBJ2-2 | CASSNLDGTGELFF | RAKFKQLL | YES | NO | NO |
| 12324 | TRAV26-1 | TRAJ39 | CIVRVNAGNMLTF | TRBV29-1 | TRBJ2-1 | CSVGSGDHGEQFF | RAKFKQLL | YES | NO | YES |
| 12325 | TRAV17 | TRAJ52 | CATAGGTSYGKLTF | TRBV12-3 | TRBJ2-2 | CASSFSGNTGELFF | RAKFKQLL | YES | NO | YES |
| 12326 | TRAV13-1 | TRAJ39 | CAPGEGNMLTF | TRBV12-4 | TRBJ2-2 | CASSQGTGELFF | KLGGALQAK | YES | NO | NO |
| 12327 | TRAV25 | TRAJ32 | CAGGATNKLIF | TRBV5-4 | TRBJ2-1 | CASRPGFPSYNEQFF | KLGGALQAK | YES | NO | NO |
| 12328 | TRAV12-1 | TRAJ50 | CVVTETSYDKVIF | TRBV13 | TRBJ2-3 | CASSSPGLAVDTQYF | RAKFKQLL | YES | NO | YES |
| 12329 | TRAV21 | TRAJ11 | CAVTISGYSTLTF | TRBV12-3 | TRBJ2-2 | CASSFSGNTGELFF | RAKFKQLL | YES | NO | YES |
| 12330 | TRAV20 | TRAJ29 | CAVQAELSGNTPLVF | TRBV6-6 | TRBJ2-2 | CASSIPVGDHTGELFF | KLGGALQAK | YES | NO | NO |
| 12331 | TRAV12-2 | TRAJ24 | CAVIDSWGKLQF | TRBV7-9 | TRBJ2-1 | CASIRTSGNDNEQFF | KLGGALQAK | YES | NO | NO |
| 12332 | TRAV27 | TRAJ42 | CAGGGSQGNLIF | TRBV5-5 | TRBJ2-7 | CASSGPGGVDEQYF | RAKFKQLL | YES | NO | YES |
| 12333 | TRAV3 | TRAJ45 | CAVAKDLQNSGGGADGLTF | TRBV11-1 | TRBJ1-1 | CASSPEDRVGDNTEAFF | GILGFVFTL | YES | NO | YES |
| 12336 | TRAV29DV5 | TRAJ40 | CAASSTSGTYKYIF | TRBV11-2 | TRBJ1-2 | CASSLDGQGPLYGYTF | KLGGALQAK | YES | NO | NO |
| 12337 | TRAV8-2 | TRAJ3 | CVVSEGASKIIF | TRBV6-6 | TRBJ2-1 | CASSLGDRLPYNEQFF | KLGGALQAK | NO | NO | NO |
| 12338 | TRAV38-1 | TRAJ52 | CAFMINAGGTSYGKLTF | TRBV19 | TRBJ1-2 | CASSIGVYGYTF | GILGFVFTL | YES | NO | YES |
| 12339 | TRAV12-1 | TRAJ9 | CVAENTGGFKTIF | TRBV15 | TRBJ2-3 | CATSRVRADTQYF | KLGGALQAK | YES | NO | NO |
| 12340 | TRAV9-2 | TRAJ31 | CAPFRGRLMF | TRBV11-2 | TRBJ2-7 | CASSSGTAAYEQYF | KLGGALQAK | YES | NO | NO |
| 12341 | TRAV8-6 | TRAJ37 | CAVTSNTGKLIF | TRBV19 | TRBJ2-7 | CASSFRSSYEQYF | GILGFVFTL | YES | NO | YES |
| 12342 | TRAV13-1 | TRAJ20 | CAASGYDYKLSF | TRBV12-3 | TRBJ2-2 | CASSFSGNTGELFF | RAKFKQLL | YES | NO | YES |
| 12343 | TRAV13-2 | TRAJ52 | CAEMYDGTSYGKLTF | TRBV9 | TRBJ2-5 | CASSAFSGAVETQYF | KLGGALQAK | YES | NO | NO |
| 12344 | TRAV1-2 | TRAJ37 | CAVQGNTGKLIF | TRBV20-1 | TRBJ2-3 | CSAPGQGLSEGPTDTQYF | AVFDRKSDAK | YES | NO | YES |
| 12345 | TRAV41 | TRAJ30 | CAAPLDDKIIF | TRBV12-3 | TRBJ2-2 | CASSFSGNTGELFF | RAKFKQLL | YES | NO | NO |
| 12346 | TRAV12-2 | TRAJ12 | CAVNMYGDSSYKLIF | TRBV12-3 | TRBJ1-1 | CASSFLDRGPNTEAFF | RAKFKQLL | YES | NO | YES |
| 12347 | TRAV26-2 | TRAJ50 | CITQETSYDKVIF | TRBV12-3 | TRBJ2-1 | CASRRTDDHEQFF | RAKFKQLL | YES | NO | NO |
| 12348 | TRAV19 | TRAJ44 | CALSGMTNTGTASKLTF | TRBV7-8 | TRBJ1-3 | CASSLAKGAGNTIYF | KLGGALQAK | YES | NO | NO |
| 12349 | TRAV13-1 | TRAJ11 | CAAKEGYSTLTL | TRBV5-4 | TRBJ1-5 | CASSSTGGGEKDQPQHF | KLGGALQAK | NO | NO | NO |
| 12350 | TRAV12-3 | TRAJ53 | CAMSYSGGSNYKLTF | TRBV29-1 | TRBJ2-7 | CSVGSGDYEQYF | RAKFKQLL | YES | NO | NO |
| 12351 | TRAV25 | TRAJ40 | CAAGTYKYIF | TRBV27 | TRBJ2-7 | CASSLRAGRAYEQYF | KLGGALQAK | YES | NO | NO |
| 12352 | TRAV38-1 | TRAJ34 | CAFMPPYNTDKLIF | TRBV12-3 | TRBJ2-2 | CASSFSGNTGELFF | RAKFKQLL | YES | NO | YES |
| 12353 | TRAV12-3 | TRAJ4 | CAIRSGGYNKLIF | TRBV2 | TRBJ2-7 | CASSEGQAYEQYF | AVFDRKSDAK | YES | NO | YES |
| 12354 | TRAV1-2 | TRAJ31 | CAVWGPWGRLMF | TRBV20-1 | TRBJ2-7 | CSARDKGGGAREQYF | KLGGALQAK | NO | NO | NO |
| 12355 | TRAV21 | TRAJ40 | CAVRPTSGTYKYIF | TRBV12-4 | TRBJ2-5 | CASSPGQPETQYF | KLGGALQAK | YES | NO | NO |
| 12356 | TRAV19 | TRAJ15 | CALRSSQAGTALIF | TRBV6-1 | TRBJ2-1 | CASSGQTRNSRDEQFF | KLGGALQAK | YES | NO | NO |
| 12357 | TRAV25 | TRAJ34 | CAGFGNTDKLIF | TRBV10-1 | TRBJ2-2 | CASSRDSASTGELFF | KLGGALQAK | YES | NO | NO |
| 12358 | TRAV22 | TRAJ18 | CAVEFLDRGSTLGRLYF | TRBV19 | TRBJ2-3 | CASSIISVDTQYF | GILGFVFTL | YES | NO | YES |
| 12359 | TRAV12-2 | TRAJ6 | CAVRDGGSYIPTF | TRBV15 | TRBJ2-7 | CATSRDGGHEQYF | AVFDRKSDAK | YES | NO | YES |
| 12360 | TRAV12-2 | TRAJ28 | CAVTRPMGAGSYQLTF | TRBV20-1 | TRBJ2-1 | CSGHPNEQFF | KLGGALQAK | NO | NO | NO |
| 12361 | TRAV4 | TRAJ28 | CLVGEGSGAGSYQLTF | TRBV7-9 | TRBJ2-1 | CASSLSDSLYNEQFF | KLGGALQAK | YES | NO | NO |
| 12362 | TRAV16 | TRAJ43 | CALSKNNNDMRF | TRBV20-1 | TRBJ2-2 | CSARDFDLTGELFF | KLGGALQAK | NO | NO | NO |
| 12363 | TRAV24 | TRAJ42 | CASPGSQGNLIF | TRBV9 | TRBJ2-7 | CASSADRFPYEQYF | AVFDRKSDAK | YES | NO | YES |
| 12364 | TRAV19 | TRAJ9 | CALSEAKGTGGFKTIF | TRBV30 | TRBJ1-6 | CAWRGGPYNSPLHF | KLGGALQAK | NO | NO | NO |
| 12365 | TRAV9-2 | TRAJ54 | CALMIQGAQKLVF | TRBV7-6 | TRBJ2-7 | CASSFGPAAGRSSYEQYF | KLGGALQAK | YES | NO | NO |
| 12366 | TRAV19 | TRAJ49 | CALRAGNQFYF | TRBV12-3 | TRBJ2-2 | CASSFSGNTGELFF | RAKFKQLL | YES | NO | YES |
| 12367 | TRAV29DV5 | TRAJ47 | CAAGEYGNKLVF | TRBV19 | TRBJ2-7 | CASSVRSSYEQYF | GILGFVFTL | YES | YES | YES |
| 12368 | TRAV19 | TRAJ23 | CALSEGVGKLIF | TRBV19 | TRBJ2-7 | CASSIRSSYEQYF | GILGFVFTL | YES | NO | NO |
| 12369 | TRAV12-2 | TRAJ26 | CAVNNYGQNFVF | TRBV6-3 | TRBJ1-2 | CASSLPANIYAYGYTF | KLGGALQAK | YES | NO | NO |
| 12370 | TRAV6 | TRAJ37 | CALGGYGSSNTGKLIF | TRBV6-5 | TRBJ2-2 | CASRLAGAPSTGELFF | AVFDRKSDAK | YES | YES | NO |
| 12371 | TRAV12-2 | TRAJ9 | CAVGRDTGGFKTIF | TRBV19 | TRBJ2-2 | CASSIFGAGELFF | GILGFVFTL | YES | NO | NO |
| 12372 | TRAV12-2 | TRAJ27 | CAAYNAGKSTF | TRBV11-2 | TRBJ2-5 | CASSLGSGAQETQYF | KLGGALQAK | YES | NO | NO |
| 12373 | TRAV27 | TRAJ42 | CAGAQGGGSQGNLIF | TRBV12-3 | TRBJ2-2 | CASSFSGNTGELFF | GILGFVFTL | YES | NO | YES |
| 12374 | TRAV19 | TRAJ45 | CALSEGGGADGLTF | TRBV4-2 | TRBJ2-2 | CASSQGETTGELFF | KLGGALQAK | YES | NO | NO |
| 12375 | TRAV27 | TRAJ50 | CAGARETSYDKVIF | TRBV7-9 | TRBJ2-7 | CASSSTGPSSYEQYF | RAKFKQLL | YES | YES | YES |
| 12376 | TRAV1-2 | TRAJ33 | CAVKDSNYQLIW | TRBV20-1 | TRBJ1-2 | CSARTGEAGYTF | AVFDRKSDAK | YES | NO | YES |
| 12377 | TRAV40 | TRAJ34 | CLLGTDKLIF | TRBV12-3 | TRBJ2-2 | CASSFSGNTGELFF | RAKFKQLL | NO | NO | NO |
| 12378 | TRAV27 | TRAJ26 | CAGDRLGGQNFVF | TRBV5-5 | TRBJ2-7 | CASSGPGGVDEQYF | RAKFKQLL | YES | NO | YES |
| 12379 | TRAV8-1 | TRAJ4 | CAVIVLSGGYNKLIF | TRBV27 | TRBJ2-1 | CASSLREDEQFF | KLGGALQAK | YES | NO | NO |
| 12380 | TRAV8-4 | TRAJ34 | CAVSESYNTDKLIF | TRBV5-4 | TRBJ1-1 | CASSLNGGGGTTDEAFF | KLGGALQAK | YES | NO | NO |
| 12381 | TRAV12-1 | TRAJ15 | CVVNQAGTALIF | TRBV19 | TRBJ2-7 | CASSTRSSYEQYF | GILGFVFTL | YES | NO | YES |
| 12382 | TRAV13-1 | TRAJ42 | CAYGGSQGNLIF | TRBV12-3 | TRBJ2-2 | CASSFSGNTGELFF | RAKFKQLL | YES | YES | YES |
| 12383 | TRAV20 | TRAJ32 | CAVRGATNKLIF | TRBV11-2 | TRBJ2-2 | CASSLGVAGTNTGELFF | KLGGALQAK | YES | NO | NO |
| 12384 | TRAV5 | TRAJ43 | CAESDNNNDMRF | TRBV5-5 | TRBJ2-3 | CASSLEGQGATTDTQYF | AVFDRKSDAK | YES | NO | YES |
| 12385 | TRAV13-1 | TRAJ50 | CAAMETSYDKVIF | TRBV7-9 | TRBJ2-1 | CASSSGLAGLNEQFF | RAKFKQLL | YES | YES | YES |
| 12386 | TRAV26-1 | TRAJ29 | CIVRVAGNSGNTPLVF | TRBV4-1 | TRBJ1-1 | CASSQDSGVVTFF | KLGGALQAK | YES | NO | NO |
| 12387 | TRAV17 | TRAJ11 | CATEGLSGYSTLTF | TRBV6-5 | TRBJ1-2 | CASSYQGGNYGYTF | FLYALALLL | YES | YES | YES |
| 12388 | TRAV17 | TRAJ36 | CATGDQTGANNLFF | TRBV11-2 | TRBJ2-1 | CASLRSGGDEQFF | KLGGALQAK | YES | NO | NO |
| 12389 | TRAV12-3 | TRAJ17 | CAMSEAAGNKLTF | TRBV2 | TRBJ1-5 | CASRIPGTGRDQPQHF | KLGGALQAK | YES | NO | NO |
| 12390 | TRAV5 | TRAJ31 | CAGRGNNARLMF | TRBV19 | TRBJ1-2 | CASSIGSYGYTF | GILGFVFTL | YES | NO | NO |
| 12391 | TRAV17 | TRAJ11 | CATEGNSGYSTLTF | TRBV6-5 | TRBJ1-2 | CASSWQGGNGYTF | FLYALALLL | YES | YES | YES |
| 12392 | TRAV19 | TRAJ22 | CALPSSGSARQLTF | TRBV11-2 | TRBJ2-3 | CASSLLGDTQYF | KLGGALQAK | YES | NO | NO |
| 12393 | TRAV38-1 | TRAJ52 | CAFAGGTSYGKLTF | TRBV12-3 | TRBJ2-2 | CASSFSGNTGELFF | RAKFKQLL | YES | NO | YES |
| 12394 | TRAV10 | TRAJ45 | CVVQGGADGLTF | TRBV30 | TRBJ2-1 | CAWSWDRGYNEQFF | KLGGALQAK | YES | NO | NO |
| 12395 | TRAV13-2 | TRAJ50 | CAETETSYDKVIF | TRBV7-9 | TRBJ2-3 | CASSLTELGHRADTQYF | RAKFKQLL | YES | NO | YES |
| 12396 | TRAV8-2 | TRAJ45 | CVVTERGGADGLTF | TRBV7-9 | TRBJ2-3 | CASSPGTPNTQYF | KLGGALQAK | YES | NO | NO |
| 12397 | TRAV25 | TRAJ45 | CAGPRGGGADGLTF | TRBV11-2 | TRBJ1-2 | CASSFGGPTYGYTF | KLGGALQAK | YES | NO | NO |
| 12398 | TRAV8-3 | TRAJ43 | CAVGAKNNDMRF | TRBV9 | TRBJ2-7 | CASSPGGTGTYEQYF | KLGGALQAK | YES | NO | NO |
| 12399 | TRAV38-2DV8 | TRAJ40 | CAYMDSGTYKYIF | TRBV28 | TRBJ2-3 | CASSPGPGWRTDTQYF | GLCTLVAML | YES | YES | YES |
| 12400 | TRAV14DV4 | TRAJ27 | CASPRRNTNAGKSTF | TRBV4-1 | TRBJ2-3 | CASSQDPRPDTQYF | KLGGALQAK | YES | NO | NO |
| 12401 | TRAV8-3 | TRAJ47 | CAVGPPMEYGNKLVF | TRBV10-3 | TRBJ1-3 | CAINPGTGSGNTIYF | KLGGALQAK | YES | NO | NO |
| 12402 | TRAV17 | TRAJ29 | CAQVMNSGNTPLVF | TRBV3-1 | TRBJ2-1 | CASSQGAYAIDEQFF | KLGGALQAK | YES | NO | NO |
| 12403 | TRAV19 | TRAJ37 | CALSESGSSNTGKLIF | TRBV12-5 | TRBJ1-1 | CASGLAVNTEAFF | KLGGALQAK | YES | NO | NO |
| 12404 | TRAV13-1 | TRAJ47 | CAASEYGNKLVF | TRBV4-1 | TRBJ1-2 | CASSQTPTGSNYGYTF | RAKFKQLL | YES | NO | YES |
| 12405 | TRAV27 | TRAJ42 | CAGAQGGGSQGNLIF | TRBV6-3 | TRBJ2-7 | CASGMTGLTSEQYF | GILGFVFTL | YES | NO | YES |
| 12406 | TRAV5 | TRAJ31 | CAEDNNARLMF | TRBV20-1 | TRBJ1-2 | CSARTGTGNSYTF | GLCTLVAML | YES | YES | YES |
| 12407 | TRAV20 | TRAJ8 | CAVQADTGFQKLVF | TRBV5-1 | TRBJ2-7 | CASSLYGDYEQYF | KLGGALQAK | YES | NO | NO |
| 12408 | TRAV29DV5 | TRAJ34 | CAAIGGGTDKLIF | TRBV5-1 | TRBJ2-1 | CASSLAGYNEQFF | KLGGALQAK | YES | NO | NO |
| 12409 | TRAV14DV4 | TRAJ48 | CAMREFGNEKLTF | TRBV29-1 | TRBJ2-7 | CSVVYRAYEQYF | KLGGALQAK | YES | NO | NO |

| | | | | | | | | | | |
|---|---|---|---|---|---|---|---|---|---|---|
| 12410 | TRAV13-1 | TRAJ20 | CAASLNDYKLSF | TRBV5-4 | TRBJ2-2 | CASSFDLADTGELFF | KLGGALQAK | YES | NO | NO |
| 12411 | TRAV1-2 | TRAJ33 | CAATDSNYQLIW | TRBV20-1 | TRBJ2-1 | CSARDLAGNEQFF | KLGGALQAK | NO | NO | NO |
| 12412 | TRAV17 | TRAJ53 | CATDSGGSNYKLTF | TRBV29-1 | TRBJ2-7 | CSVGAGEDDEQYF | RAKFKQLL | NO | NO | NO |
| 12413 | TRAV17 | TRAJ31 | CATDRNNNARLMF | TRBV11-1 | TRBJ1-2 | CASSLSTGSWGYTF | KLGGALQAK | YES | NO | NO |
| 12414 | TRAV1-2 | TRAJ33 | CAVMDSNYQLIW | TRBV29-1 | TRBJ2-2 | CSVGGGGDTGELFF | KLGGALQAK | YES | NO | NO |
| 12415 | TRAV17 | TRAJ11 | CATEGNSGYSTLTF | TRBV6-5 | TRBJ1-2 | CATAGQGGNGYTF | FLYALALLL | YES | NO | NO |
| 12416 | TRAV14DV4 | TRAJ15 | CAMREATGTALIF | TRBV19 | TRBJ2-7 | CASSARSSYEQYF | GILGFVFTL | YES | NO | YES |
| 12417 | TRAV14DV4 | TRAJ41 | CAMSPGGSGYALNF | TRBV10-2 | TRBJ1-2 | CASSESGIGGYTF | KLGGALQAK | YES | NO | NO |
| 12418 | TRAV27 | TRAJ37 | CAGAAGSSNTGKLIF | TRBV19 | TRBJ2-7 | CASSIRSAYEQYF | GILGFVFTL | YES | NO | YES |
| 12419 | TRAV12-3 | TRAJ43 | CAMSEGNNNDMRF | TRBV5-1 | TRBJ1-2 | CASSLPLGDYGYTF | KLGGALQAK | YES | NO | NO |
| 12420 | TRAV3 | TRAJ45 | CAVAKDLQNSGGGADGLTF | TRBV19 | TRBJ2-7 | CASSIRASYEQYF | GILGFVFTL | NO | NO | NO |
| 12421 | TRAV12-1 | TRAJ23 | CVVNQGGKLIF | TRBV7-2 | TRBJ1-2 | CASSSDHRGGYTF | KLGGALQAK | YES | NO | NO |
| 12422 | TRAV8-3 | TRAJ28 | CALRSINPSGAGSYQLTF | TRBV10-3 | TRBJ2-3 | CAISEPQLSYTDTQYF | RLRAEAQVK | YES | NO | YES |
| 12423 | TRAV27 | TRAJ42 | CAGGGSQGNLIF | TRBV19 | TRBJ2-5 | CASSFRSSETQYF | GILGFVFTL | YES | YES | YES |
| 12424 | TRAV29DV5 | TRAJ28 | CAASSAGAGSYQLTF | TRBV29-1 | TRBJ1-1 | CSAGGNTEAFF | RAKFKQLL | NO | NO | NO |
| 12425 | TRAV12-1 | TRAJ33 | CVVNRDYQLIW | TRBV12-5 | TRBJ2-7 | CASGLVQYEQYF | KLGGALQAK | YES | NO | NO |
| 12426 | TRAV12-2 | TRAJ23 | CAVIDQGGKLIF | TRBV7-9 | TRBJ2-7 | CASIGTSPRYEQYF | KLGGALQAK | YES | NO | NO |
| 12427 | TRAV21 | TRAJ48 | CAVRYLSNFGNEKLTF | TRBV12-3 | TRBJ2-2 | CASSFSGNTGELFF | RAKFKQLL | YES | NO | YES |
| 12428 | TRAV12-2 | TRAJ57 | CAVWEGGSEKLVF | TRBV9 | TRBJ2-2 | CASSSGLVSNTGELFF | KLGGALQAK | YES | YES | NO |
| 12430 | TRAV39 | TRAJ28 | CAVDILSHPSSGAGSYQLTF | TRBV10-3 | TRBJ2-1 | CAISKAVFFSYNEQFF | KLGGALQAK | NO | NO | NO |
| 12431 | TRAV13-1 | TRAJ52 | CAAHPRNTNAGGTSYGKLTF | TRBV11-2 | TRBJ2-1 | CASSLELGRNEQFF | KLGGALQAK | NO | NO | NO |
| 12432 | TRAV26-1 | TRAJ50 | CIVRETSYDKVIF | TRBV12-4 | TRBJ2-2 | CASSSTQGPGELFF | RAKFKQLL | YES | NO | YES |
| 12434 | TRAV23DV6 | TRAJ31 | CAATDNNARLMF | TRBV12-3 | TRBJ2-2 | CASSFSGNTGELFF | RAKFKQLL | YES | NO | YES |
| 12435 | TRAV27 | TRAJ37 | CAGASGSSNTGKLIF | TRBV5-6 | TRBJ1-1 | CASSLRDGSEAFF | RAKFKQLL | YES | NO | YES |
| 12436 | TRAV6 | TRAJ6 | CARTGGSYIPTF | TRBV6-5 | TRBJ2-5 | CASSYAETQYF | GILGFVFTL | NO | NO | NO |
| 12437 | TRAV13-1 | TRAJ50 | CAASETSYDKVIF | TRBV7-9 | TRBJ1-3 | CASSLLGAENTIYF | RAKFKQLL | YES | NO | YES |
| 12438 | TRAV1-1 | TRAJ12 | CAVSRMDSSYKLIF | TRBV7-9 | TRBJ1-3 | CASSPLGGPKDTIYF | RAKFKQLL | YES | NO | YES |
| 12439 | TRAV13-2 | TRAJ45 | CADSGGGADGLTF | TRBV5-6 | TRBJ2-1 | CASSFTGGRSSYNEQFF | RAKFKQLL | YES | NO | YES |
| 12440 | TRAV12-1 | TRAJ43 | CVVNTGSNDMRF | TRBV10-3 | TRBJ1-5 | CAISEAGADRPQHF | KLGGALQAK | YES | NO | NO |
| 12441 | TRAV17 | TRAJ31 | CATDGGNNARLMF | TRBV30 | TRBJ2-7 | CAWSWGSGGEQYF | KLGGALQAK | YES | NO | NO |
| 12442 | TRAV8-4 | TRAJ40 | CAVSQDSSGTYKYIF | TRBV5-1 | TRBJ2-1 | CASSHGLNTYNEQFF | KLGGALQAK | YES | NO | NO |
| 12443 | TRAV13-1 | TRAJ11 | CAAKEGYSTLTF | TRBV4-1 | TRBJ2-1 | CASSLLWAEYNEQFF | RAKFKQLL | YES | NO | YES |
| 12444 | TRAV12-2 | TRAJ9 | CAVWSGGFKTIF | TRBV12-3 | TRBJ2-7 | CASSSLDVAGYEQYF | RAKFKQLL | YES | NO | NO |
| 12445 | TRAV8-3 | TRAJ54 | CAGSIQGAQKLVF | TRBV3-1 | TRBJ2-5 | CASSQGGGSETQYF | KLGGALQAK | YES | NO | NO |
| 12446 | TRAV35 | TRAJ40 | CAGFHSGTYKYIF | TRBV7-2 | TRBJ1-1 | CASSLPRTPGRNTEAFF | KLGGALQAK | YES | NO | NO |
| 12447 | TRAV38-2DV8 | TRAJ45 | CAYPPRGGADGLTF | TRBV5-4 | TRBJ1-1 | CASSLGVNTEAFF | KLGGALQAK | YES | NO | NO |
| 12448 | TRAV8-3 | TRAJ24 | CALDSWGKLQF | TRBV20-1 | TRBJ2-1 | CSARDPDVPGGNEQFF | KLGGALQAK | NO | NO | NO |
| 12449 | TRAV35 | TRAJ42 | CAGSGDGGSQGNLIF | TRBV7-9 | TRBJ2-1 | CASSFGSDYNEQFF | GILGFVFTL | YES | NO | YES |
| 12451 | TRAV27 | TRAJ42 | CAGAGGGSQGNLIF | TRBV19 | TRBJ2-3 | CASSMRSTDTQYF | GILGFVFTL | YES | NO | YES |
| 12452 | TRAV5 | TRAJ42 | CAEDSGNLIF | TRBV9 | TRBJ2-6 | CASSVAVGTGSGANVLTF | GLCTLVAML | YES | NO | NO |
| 12453 | TRAV19 | TRAJ33 | CALSEASRMDSNYQLIW | TRBV19 | TRBJ1-1 | CASSTHGGGNTEAFF | GILGFVFTL | YES | NO | YES |
| 12454 | TRAV13-1 | TRAJ50 | CAATETSYDKVIF | TRBV29-1 | TRBJ1-2 | CSVEDLAYGYTF | RAKFKQLL | YES | NO | YES |
| 12455 | TRAV1-2 | TRAJ33 | CAVKDSNYQLIW | TRBV20-1 | TRBJ1-1 | CSVLSGDREINTEAFF | KLGGALQAK | YES | NO | NO |
| 12456 | TRAV27 | TRAJ42 | CAGAGQRGSQGNLIF | TRBV19 | TRBJ1-5 | CASSTYSNQPQHF | GILGFVFTL | YES | NO | YES |
| 12457 | TRAV27 | TRAJ30 | CAGMNRDDKIIF | TRBV20-1 | TRBJ1-3 | CSARDRVGNTIYF | GLCTLVAML | YES | YES | YES |
| 12458 | TRAV17 | TRAJ39 | CATGYNNAGNMLTF | TRBV27 | TRBJ1-2 | CASSSSGQTYGYTF | KLGGALQAK | YES | NO | NO |
| 12459 | TRAV3 | TRAJ29 | CAVRDTHSGNTPLVF | TRBV18 | TRBJ2-5 | CASSPAGFEETQYF | KLGGALQAK | NO | NO | NO |
| 12460 | TRAV1-2 | TRAJ33 | CAVMDSNYQLIW | TRBV6-1 | TRBJ2-1 | CASTSGLAELYNEQFF | KLGGALQAK | YES | NO | NO |
| 12461 | TRAV19 | TRAJ49 | CALRAGNQFYF | TRBV19 | TRBJ2-7 | CASSIRSSYEQYF | GILGFVFTL | YES | NO | YES |
| 12462 | TRAV8-3 | TRAJ45 | CAVGRGISGGGADGLTF | TRBV9 | TRBJ1-2 | CASSLGQGKVNTF | KLGGALQAK | YES | NO | NO |
| 12463 | TRAV1-2 | TRAJ28 | CAVRADSGAGSYQLTF | TRBV20-1 | TRBJ1-3 | CSARDRVGNTIYF | GLCTLVAML | YES | YES | YES |
| 12464 | TRAV27 | TRAJ42 | CAYSYGGSQGNLIF | TRBV19 | TRBJ2-3 | CASSMRSSDTQYF | GILGFVFTL | YES | NO | YES |
| 12465 | TRAV29DV5 | TRAJ23 | CAASFYNQGGKLIF | TRBV19 | TRBJ1-2 | CASSIGVHGYTF | GILGFVFTL | YES | NO | YES |
| 12466 | TRAV35 | TRAJ29 | CAGDRNSGNTPLVF | TRBV5-4 | TRBJ2-2 | CASSLATGWTGELFF | KLGGALQAK | YES | NO | NO |
| 12467 | TRAV21 | TRAJ30 | CAVEIDDKIIF | TRBV3-1 | TRBJ2-7 | CASSPISGSSYEQYF | KLGGALQAK | YES | NO | NO |
| 12468 | TRAV23DV6 | TRAJ34 | CAASIAYNTDKLIF | TRBV5-4 | TRBJ1-5 | CASSFNGQSNQPQHF | KLGGALQAK | YES | NO | NO |
| 12469 | TRAV9-2 | TRAJ15 | CALSGASGTALIF | TRBV7-8 | TRBJ2-3 | CASSLGGGGLTDTQYF | KLGGALQAK | YES | NO | NO |
| 12470 | TRAV12-2 | TRAJ15 | CAVSANQAGTALIF | TRBV7-9 | TRBJ2-7 | CASSWAGEQYF | KLGGALQAK | YES | NO | NO |
| 12471 | TRAV12-2 | TRAJ45 | CAPYSGGGADGLTF | TRBV19 | TRBJ2-7 | CASSIDGDRVYEQYF | IVTDFSVIK | YES | YES | YES |
| 12472 | TRAV13-1 | TRAJ12 | CAAARIDSSYKLIF | TRBV29-1 | TRBJ2-3 | CSVGSGEDSPQYF | RAKFKQLL | YES | NO | NO |
| 12473 | TRAV30 | TRAJ49 | CGTPHTGNQFYF | TRBV11-2 | TRBJ1-2 | CASSPGGPGYGYTF | KLGGALQAK | YES | NO | NO |
| 12474 | TRAV13-2 | TRAJ39 | CADPGNMLTF | TRBV7-9 | TRBJ1-1 | CASSFSQNTEAFF | RAKFKQLL | YES | NO | YES |
| 12475 | TRAV14DV4 | TRAJ4 | CAMRASPSGGYNKLIF | TRBV4-2 | TRBJ2-2 | CASSQERGDAGELFF | AVFDRKSDAK | YES | NO | YES |
| 12476 | TRAV3 | TRAJ42 | CAVTYGGSQGNLIF | TRBV19 | TRBJ2-7 | CASSTRAAYEQYF | GILGFVFTL | NO | NO | NO |
| 12477 | TRAV13-1 | TRAJ50 | CAASETSYDKVIF | TRBV7-9 | TRBJ2-4 | CASTRLGGLDIQYF | RAKFKQLL | YES | NO | YES |
| 12478 | TRAV1-2 | TRAJ12 | CAGMDSSYKLIF | TRBV6-1 | TRBJ2-5 | CASSELAGGQETQYF | KLGGALQAK | YES | NO | NO |
| 12479 | TRAV26-2 | TRAJ50 | CILRETSYDKVIF | TRBV18 | TRBJ2-2 | CASSPPGGGNTGELFF | RAKFKQLL | YES | NO | YES |
| 12480 | TRAV8-3 | TRAJ40 | CAVATTSGTYKYIF | TRBV9 | TRBJ1-2 | CASSVGGQYYGYTF | KLGGALQAK | YES | NO | NO |
| 12481 | TRAV21 | TRAJ4 | CAVSLSGGYNKLIF | TRBV6-3 | TRBJ2-7 | CASGMTGLTSEQYF | KLGGALQAK | YES | NO | NO |
| 12482 | TRAV1-2 | TRAJ33 | CAVMDSNYQLIW | TRBV6-5 | TRBJ2-1 | CASSFTSGGANEQFF | KLGGALQAK | NO | NO | NO |
| 12484 | TRAV17 | TRAJ10 | CATALTGGGNKLTF | TRBV27 | TRBJ1-2 | CASSMGGQTYGYTF | KLGGALQAK | YES | NO | NO |
| 12485 | TRAV38-2DV8 | TRAJ52 | CAYSPAAGGTSYGKLTF | TRBV12-4 | TRBJ2-5 | CASSVTSGGRETQYF | GILGFVFTL | YES | NO | YES |
| 12486 | TRAV14DV4 | TRAJ9 | CAMRERMLGGGFKTIF | TRBV14 | TRBJ2-3 | CASSQSPGGTQYF | GLCTLVAML | YES | NO | NO |
| 12487 | TRAV17 | TRAJ11 | CATAGNSGYSTLTF | TRBV6-5 | TRBJ2-1 | CASEGQGGNEQFF | FLYALALLL | NO | NO | NO |
| 12488 | TRAV19 | TRAJ33 | CALSRPVDSNYQLIW | TRBV4-1 | TRBJ1-5 | CASSQDRGYQPQHF | KLGGALQAK | YES | NO | NO |
| 12489 | TRAV17 | TRAJ10 | CAKGLTGGGNKLTF | TRBV27 | TRBJ1-2 | CASSPGGQNYGYTF | KLGGALQAK | YES | NO | NO |
| 12490 | TRAV36DV7 | TRAJ54 | CAYLQGAQKLVF | TRBV7-6 | TRBJ1-1 | CASSLVSTEAFF | RAKFKQLL | NO | NO | NO |
| 12491 | TRAV16 | TRAJ23 | CALLSQGPGPGRKAYLRT | TRBV7-9 | TRBJ2-3 | CASSPLGLAGGPGEDTQYF | RAKFKQLL | NO | NO | NO |
| 12492 | TRAV8-3 | TRAJ21 | CAVGLFNFNKFYF | TRBV20-1 | TRBJ1-3 | CSARDRVGNTIYF | GLCTLVAML | YES | YES | YES |
| 12493 | TRAV12-1 | TRAJ45 | CVVTRRGGADGLTF | TRBV2 | TRBJ1-5 | CASSNRDRGRQPQHF | AVFDRKSDAK | YES | NO | YES |
| 12494 | TRAV13-1 | TRAJ35 | CAARVRGFGNVLHC | TRBV6-4 | TRBJ1-6 | CASSDSGTGGSPLHF | KLGGALQAK | NO | NO | NO |
| 12495 | TRAV13-2 | TRAJ45 | CADSGGGADGLTF | TRBV13 | TRBJ1-6 | CASSLMRGGTYNSPLHF | RAKFKQLL | YES | NO | YES |
| 12496 | TRAV26-2 | TRAJ28 | CTDSGAGSYQLTF | TRBV5-6 | TRBJ1-1 | CASSLRDGSEAFF | RAKFKQLL | YES | NO | NO |
| 12497 | TRAV29DV5 | TRAJ20 | CAATETNDYKLSF | TRBV2 | TRBJ2-2 | CASSTPRQGSNTGELFF | RAKFKQLL | YES | YES | YES |
| 12498 | TRAV22 | TRAJ27 | CAVVTNAGKSTF | TRBV7-6 | TRBJ2-1 | CASSVLTSGGEQFF | AVFDRKSDAK | YES | YES | YES |
| 12499 | TRAV41 | TRAJ58 | CAVGVEETSGSRLTF | TRBV19 | TRBJ1-1 | CASSIFYRAEAFF | GILGFVFTL | YES | NO | YES |
| 12500 | TRAV13-1 | TRAJ50 | CAASETSYDKVIF | TRBV7-9 | TRBJ2-5 | CASLRLGIDQETQYF | RAKFKQLL | YES | NO | NO |
| 12501 | TRAV12-2 | TRAJ40 | CAAPTPTSGTYKYIF | TRBV7-9 | TRBJ1-5 | CASSLGTQPQHF | KLGGALQAK | YES | NO | NO |
| 12502 | TRAV29DV5 | TRAJ40 | CAAFYSGTYKYIF | TRBV7-9 | TRBJ2-3 | CASSSRDGGTDTQYF | KLGGALQAK | YES | NO | NO |
| 12503 | TRAV19 | TRAJ12 | CALSKMDSSYKLIF | TRBV18 | TRBJ2-1 | CASSLPPRSGAAYNEQFF | KLGGALQAK | YES | NO | NO |
| 12504 | TRAV1-2 | TRAJ16 | CAVREVSQKLLF | TRBV7-9 | TRBJ2-1 | CASSSGLAGRNEQFF | KLGGALQAK | NO | NO | NO |
| 12505 | TRAV39 | TRAJ43 | CAGEGNNNDMRF | TRBV7-9 | TRBJ1-6 | CASSLLGGGPNSPLHF | RAKFKQLL | YES | NO | YES |
| 12506 | TRAV25 | TRAJ42 | CAGKGGGSQGNLIF | TRBV19 | TRBJ1-1 | CASSIRSTEGAFF | GILGFVFTL | YES | NO | YES |
| 12507 | TRAV1-2 | TRAJ12 | CAVRDDSSYKLIF | TRBV9 | TRBJ2-1 | CASSVASGLVAYNEQFF | KLGGALQAK | YES | NO | NO |
| 12508 | TRAV13-1 | TRAJ50 | CAASETSYDKVIF | TRBV5-4 | TRBJ2-5 | CASRGGTATYQETQYF | RAKFKQLL | YES | YES | YES |
| 12509 | TRAV27 | TRAJ42 | CAGAGSQGNLIF | TRBV19 | TRBJ1-2 | CASSIGLYGYTF | GILGFVFTL | YES | NO | YES |
| 12510 | TRAV1-2 | TRAJ20 | CAVSLLNDYKLSF | TRBV9 | TRBJ2-5 | CASSTGQVTQETQYF | GLCTLVAML | YES | YES | NO |
| 12511 | TRAV12-3 | TRAJ9 | CARGGGFKTIF | TRBV12-3 | TRBJ2-2 | CASSFSGNTGELFF | RAKFKQLL | YES | NO | NO |
| 12512 | TRAV13-1 | TRAJ50 | CAAGETSYDKVIF | TRBV7-9 | TRBJ1-6 | CASSLLGGGPNSPLHF | RAKFKQLL | YES | NO | YES |
| 12513 | TRAV30 | TRAJ30 | CGTTNRDDKIIF | TRBV5-1 | TRBJ2-1 | CASSPSSGAVDEQFF | RTLNAWVKV | YES | YES | YES |
| 12514 | TRAV35 | TRAJ49 | CAGHTGNQFYF | TRBV20-1 | TRBJ2-3 | CSARLSTLPADTQYF | KLGGALQAK | NO | NO | NO |
| 12515 | TRAV5 | TRAJ6 | CAEMEGGSYIPTF | TRBV9 | TRBJ1-1 | CASSVRSSMNTEAFF | RAKFKQLL | YES | NO | YES |
| 12516 | TRAV19 | TRAJ49 | CALRAGNQFYF | TRBV12-4 | TRBJ2-1 | CASGRLSYNEQFF | RAKFKQLL | YES | NO | YES |
| 12517 | TRAV12-2 | TRAJ31 | CAVKARLMF | TRBV20-1 | TRBJ2-5 | CSARDSRGTRQETQYF | KLGGALQAK | NO | NO | NO |

| | | | | | | | | | | |
|---|---|---|---|---|---|---|---|---|---|---|
| 12518 | TRAV13-1 | TRAJ20 | CAAKPDYKLSF | TRBV2 | TRBJ2-3 | CASSTMGLAGGLTDTQYF | KLGGALQAK | YES | NO | NO |
| 12519 | TRAV21 | TRAJ13 | CAVMNSGGYQKVTF | TRBV5-5 | TRBJ2-7 | CASSGPGGVDEQYF | RAKFKQLL | YES | NO | YES |
| 12520 | TRAV12-1 | TRAJ42 | CVVNIGGGSQGNLIF | TRBV12-3 | TRBJ1-1 | CASSFLDRGPNTEAFF | RAKFKQLL | YES | NO | NO |
| 12521 | TRAV1-2 | TRAJ33 | CAVRDRNYQLIW | TRBV6-1 | TRBJ2-1 | CASSEEAGGYNEQFF | KLGGALQAK | YES | NO | NO |
| 12522 | TRAV9-2 | TRAJ20 | CALRTANDYKLSF | TRBV6-6 | TRBJ2-5 | CASSYQATGPQETQYF | KLGGALQAK | NO | NO | NO |
| 12523 | TRAV5 | TRAJ31 | CAETFNARLMF | TRBV6-6 | TRBJ2-2 | CASSYGFPTSGGNTGELFF | RAKFKQLL | YES | NO | YES |
| 12524 | TRAV13-1 | TRAJ50 | CAPRETSYDKVIF | TRBV5-1 | TRBJ2-1 | CASSLPDYNEQFF | RAKFKQLL | YES | NO | YES |
| 12525 | TRAV13-1 | TRAJ20 | CAASSKLSF | TRBV11-2 | TRBJ1-2 | CASSRDANYGYTF | KLGGALQAK | NO | NO | NO |
| 12526 | TRAV12-1 | TRAJ29 | CVVNIVDRSGNTPLVF | TRBV7-9 | TRBJ2-2 | CASSGRASGTYTGELFF | KLGGALQAK | YES | NO | NO |
| 12527 | TRAV12-2 | TRAJ26 | CAVNAVDNYGQNFVF | TRBV10-1 | TRBJ2-3 | CASSKDSSTDTQYF | KLGGALQAK | YES | NO | NO |
| 12528 | TRAV1-2 | TRAJ24 | CAVGSWGKLQF | TRBV19 | TRBJ2-3 | CASSARAGITQYF | GILGFVFTL | YES | YES | YES |
| 12529 | TRAV27 | TRAJ37 | CAGGGSSNTGKLIF | TRBV19 | TRBJ2-1 | CASSIRSSYEQFF | GILGFVFTL | YES | NO | YES |
| 12530 | TRAV5 | TRAJ6 | CAEMEGGSYIPTF | TRBV4-1 | TRBJ1-2 | CASSTSTGTGYGYTF | RAKFKQLL | YES | NO | NO |
| 12531 | TRAV13-2 | TRAJ8 | CAETYSNTGFQKLVF | TRBV4-2 | TRBJ2-3 | CASSQDRAPTDTQYF | AVFDRKSDAK | YES | YES | YES |
| 12532 | TRAV13-1 | TRAJ37 | CAVVPGNTGKLIF | TRBV4-2 | TRBJ2-2 | CASSQRPSEVGELFF | RAKFKQLL | YES | NO | YES |
| 12533 | TRAV29DV5 | TRAJ40 | CAATPGTYKYIF | TRBV5-6 | TRBJ1-1 | CASSLRDGSEAFF | RAKFKQLL | YES | NO | YES |
| 12534 | TRAV2 | TRAJ6 | CAVEDQGSGGSYIPTF | TRBV19 | TRBJ2-1 | CASSLPLGSTYNEQFF | KLGGALQAK | NO | NO | NO |
| 12535 | TRAV29DV5 | TRAJ34 | CAAGFGTDKLIF | TRBV4-1 | TRBJ2-1 | CASSLLWAEYNEQFF | RAKFKQLL | YES | NO | YES |
| 12536 | TRAV13-1 | TRAJ50 | CAAAETSYDKVIF | TRBV7-9 | TRBJ2-7 | CASSLRGIDEQYF | RAKFKQLL | YES | NO | YES |
| 12537 | TRAV6 | TRAJ24 | CAPAPMTTDSWGKLQF | TRBV5-6 | TRBJ2-2 | CASSLIGELFF | KLGGALQAK | YES | YES | NO |
| 12538 | TRAV1-2 | TRAJ28 | CAVRDSRGTL | TRBV6-1 | TRBJ2-5 | CASSASREAGETQYF | KLGGALQAK | YES | NO | NO |
| 12539 | TRAV1-2 | TRAJ33 | CAVSDSNYQLIW | TRBV4-1 | TRBJ2-6 | CASSQDSGTGANVLTF | KLGGALQAK | YES | NO | NO |
| 12540 | TRAV25 | TRAJ28 | CAGPPYSGAGSYQLTF | TRBV6-5 | TRBJ1-2 | CASSYSHSNYGYTF | KLGGALQAK | YES | NO | NO |
| 12541 | TRAV12-3 | TRAJ16 | CAMESQKLLF | TRBV20-1 | TRBJ2-3 | CSARGSGGDPTDTQYF | RAKFKQLL | NO | NO | NO |
| 12542 | TRAV9-2 | TRAJ21 | CALQYNFNKFYF | TRBV5-6 | TRBJ1-1 | CASSLRDGSEAFF | RAKFKQLL | YES | NO | YES |
| 12543 | TRAV2 | TRAJ4 | CAVEDLFSGGYNKLIF | TRBV29-1 | TRBJ2-1 | CSVEDRRPDEQFF | KLGGALQAK | NO | NO | NO |
| 12544 | TRAV12-1 | TRAJ10 | CVVSGLTGGGNKLTF | TRBV5-1 | TRBJ2-1 | CASSLERGSYNEQFF | KLGGALQAK | YES | NO | NO |
| 12545 | TRAV25 | TRAJ49 | CAGGDTGNQFYF | TRBV12-3 | TRBJ2-2 | CASSFSGNTGELFF | RAKFKQLL | YES | NO | YES |
| 12546 | TRAV1-1 | TRAJ24 | CAVRTTDSWGKLQF | TRBV5-6 | TRBJ2-7 | CASSSYSYEQYF | AVFDRKSDAK | YES | NO | YES |
| 12547 | TRAV13-1 | TRAJ7 | CAASIWEQQTRF | TRBV29-1 | TRBJ1-1 | CSAGGNTEAFF | RAKFKQLL | YES | NO | NO |
| 12548 | TRAV29DV5 | TRAJ53 | CAASALSSGGSNYKLTF | TRBV7-9 | TRBJ2-3 | CASSSRDGGTDTQYF | KLGGALQAK | YES | NO | NO |
| 12549 | TRAV17 | TRAJ22 | CATDRAGSARQLIF | TRBV11-2 | TRBJ2-5 | CASSLTGETQYF | KLGGALQAK | YES | NO | NO |
| 12550 | TRAV12-2 | TRAJ42 | CAVSEGGGSQGNLIF | TRBV19 | TRBJ2-7 | CASSIRSSYEQYF | GILGFVFTL | YES | NO | NO |
| 12551 | TRAV16 | TRAJ52 | CALSAGGTSYGKLTF | TRBV7-6 | TRBJ1-1 | CASSFQPGMGTEAFF | KLGGALQAK | NO | NO | NO |
| 12552 | TRAV30 | TRAJ34 | CGTPKRPTSSSL | TRBV27 | TRBJ2-2 | CASSGTGELFF | KLGGALQAK | YES | NO | NO |
| 12553 | TRAV5 | TRAJ34 | CAESEGPTYTDKLIF | TRBV7-2 | TRBJ2-7 | CASSSPFSASSYEQYF | RAKFKQLL | NO | NO | NO |
| 12554 | TRAV12-2 | TRAJ43 | CAVNIGDMRF | TRBV27 | TRBJ1-1 | CAVRRVPGTEAFF | KLGGALQAK | YES | NO | NO |
| 12555 | TRAV29DV5 | TRAJ37 | CAATVSGNTGKLIF | TRBV5-1 | TRBJ1-1 | CASSYNTGLTRLGMNTEAFF | KLGGALQAK | NO | NO | NO |
| 12556 | TRAV9-2 | TRAJ43 | CALSDATNNDMRF | TRBV27 | TRBJ2-5 | CASSLMGSATQYF | KLGGALQAK | YES | NO | NO |
| 12557 | TRAV1-2 | TRAJ20 | CAVPNDYKLSF | TRBV4-1 | TRBJ2-5 | CASSQTPPAGETQYF | KLGGALQAK | YES | NO | NO |
| 12558 | TRAV8-3 | TRAJ45 | CAVYSGGGADGLTF | TRBV12-4 | TRBJ2-7 | CASSLGTSYEQYF | KLGGALQAK | YES | NO | NO |
| 12559 | TRAV13-2 | TRAJ45 | CADSGGGADGLTF | TRBV5-5 | TRBJ2-7 | CASSGPGGVDEQYF | RAKFKQLL | YES | NO | YES |
| 12560 | TRAV8-4 | TRAJ11 | CAVSDSDSGYSTLTF | TRBV6-5 | TRBJ1-1 | CASSFPYQGDTEAFF | LLFGYPVYV | YES | YES | NO |
| 12561 | TRAV22 | TRAJ32 | CAVEQDYGGATNKLIF | TRBV7-6 | TRBJ2-1 | CASSLGTAGFEQFF | AVFDRKSDAK | YES | NO | YES |
| 12562 | TRAV12-2 | TRAJ9 | CAVTHTGGFKTIF | TRBV18 | TRBJ2-2 | CASSPPGGGNTGELFF | RAKFKQLL | YES | NO | NO |
| 12563 | TRAV21 | TRAJ40 | CAVEAPGSGTYKYIF | TRBV27 | TRBJ2-3 | CASRVDRGGDTQYF | KLGGALQAK | YES | NO | NO |
| 12564 | TRAV21 | TRAJ9 | CVFKTIF | TRBV7-9 | TRBJ2-7 | CASSFLDSGDYEQYF | KLGGALQAK | YES | NO | NO |
| 12565 | TRAV19 | TRAJ42 | CALSENGRSQGNLIF | TRBV5-6 | TRBJ1-1 | CASSLRDGSEAFF | RAKFKQLL | YES | YES | YES |
| 12566 | TRAV35 | TRAJ42 | CAGPHHMNYGGSQGNLIF | TRBV19 | TRBJ1-5 | CASSVYSNQPQHF | GILGFVFTL | YES | NO | YES |
| 12567 | TRAV8-2 | TRAJ8 | CVVSENTGFQKLVF | TRBV5-6 | TRBJ1-1 | CASSLRDGSEAFF | RAKFKQLL | YES | NO | YES |
| 12568 | TRAV4 | TRAJ33 | CLVENVDSNYQLIW | TRBV9 | TRBJ1-1 | CASSVRPRTPENTEAFF | KLGGALQAK | NO | NO | NO |
| 12569 | TRAV19 | TRAJ28 | CALKGSGAGSYQLTF | TRBV11-1 | TRBJ2-7 | CASSIRDRAFGEQYF | KLGGALQAK | YES | NO | NO |
| 12570 | TRAV38-1 | TRAJ52 | CAFMVGAGGTSYGKLTF | TRBV19 | TRBJ1-2 | CASSIGIYGYTF | GILGFVFTL | YES | NO | NO |
| 12571 | TRAV8-6 | TRAJ42 | CAVTDGGSQGNLIF | TRBV12-3 | TRBJ2-2 | CASSFSGNTGELFF | RAKFKQLL | YES | NO | YES |
| 12572 | TRAV26-2 | TRAJ34 | CILYNTDKLIF | TRBV2 | TRBJ2-1 | CASSDRDYLIHEQFF | KLGGALQAK | YES | NO | NO |
| 12573 | TRAV13-1 | TRAJ50 | CAASETSYDKVIF | TRBV12-3 | TRBJ1-2 | CASSPGDRGSGYTF | RAKFKQLL | YES | NO | YES |
| 12574 | TRAV5 | TRAJ26 | CAESIDGQNFVF | TRBV6-1 | TRBJ2-1 | CASSEAPREQFF | KLGGALQAK | YES | NO | NO |
| 12575 | TRAV3 | TRAJ35 | CAVRDIGFGNVLHC | TRBV5-6 | TRBJ1-1 | CASSPTTGHTEAFF | KLGGALQAK | NO | NO | NO |
| 12576 | TRAV27 | TRAJ42 | CGYGGSQGNLIF | TRBV19 | TRBJ2-7 | CASSIRSSYEQYF | GILGFVFTL | YES | NO | NO |
| 12577 | TRAV1-2 | TRAJ7 | CAVRERGNNRLAF | TRBV10-3 | TRBJ2-7 | CAISESLTGNYEQYF | KLGGALQAK | YES | NO | NO |
| 12578 | TRAV25 | TRAJ42 | CAGSYGGSQGNLIF | TRBV19 | TRBJ2-3 | CASSVRSTDTQYF | GILGFVFTL | YES | NO | YES |
| 12579 | TRAV41 | TRAJ42 | CAAPYGGSQGNLIF | TRBV7-9 | TRBJ2-3 | CASSSRDGGTDTQYF | GILGFVFTL | YES | NO | YES |
| 12580 | TRAV1-1 | TRAJ40 | CAVRDWASGTYKYIF | TRBV6-5 | TRBJ1-3 | CASSYSPTQGENTIYF | AVFDRKSDAK | YES | NO | NO |
| 12581 | TRAV8-6 | TRAJ6 | CAAWAPASGGSYIPTF | TRBV9 | TRBJ1-2 | CASSQTGVGYTF | KLGGALQAK | YES | NO | NO |
| 12582 | TRAV14DV4 | TRAJ3 | CAMREGESSASKIIF | TRBV29-1 | TRBJ1-2 | CSAGLQYGYTF | KLGGALQAK | NO | NO | NO |
| 12583 | TRAV27 | TRAJ20 | CAGAVNDYKLSF | TRBV29-1 | TRBJ2-7 | CSAGQGPYEQYF | RAKFKQLL | NO | NO | NO |
| 12584 | TRAV22 | TRAJ20 | CAVGEYDYKLSF | TRBV5-6 | TRBJ1-1 | CASSLRDGSEAFF | RAKFKQLL | YES | NO | YES |
| 12585 | TRAV13-1 | TRAJ3 | CAASLTGYSSASKIIF | TRBV7-9 | TRBJ2-2 | CASSPLGGKNTGELFF | KLGGALQAK | YES | NO | NO |
| 12586 | TRAV12-2 | TRAJ24 | CAALDSWGKFQF | TRBV19 | TRBJ2-7 | CASSIRSSYEQYF | GILGFVFTL | YES | NO | YES |
| 12587 | TRAV8-3 | TRAJ6 | CAVGATGGSYIPTF | TRBV12-4 | TRBJ1-5 | CASSSHGNQPQHF | KLGGALQAK | YES | NO | NO |
| 12588 | TRAV13-1 | TRAJ50 | CAASETSYDKVIF | TRBV19 | TRBJ2-7 | CASSTRSSVEQYF | GILGFVFTL | YES | NO | YES |
| 12589 | TRAV5 | TRAJ37 | CAEGGNTGKLIF | TRBV12-3 | TRBJ2-5 | CASSFPSGDGTQYF | RAKFKQLL | YES | YES | YES |
| 12590 | TRAV39 | TRAJ54 | CAVDRGAQKLVF | TRBV10-3 | TRBJ2-5 | CAIRTSGVETQYF | KLGGALQAK | YES | NO | NO |
| 12591 | TRAV27 | TRAJ49 | CAGGPDTGNQFYF | TRBV6-5 | TRBJ1-1 | CASRSQGSTEAFF | KLGGALQAK | YES | NO | NO |
| 12592 | TRAV8-1 | TRAJ24 | CAVNAMGTTDSWGKLQF | TRBV28 | TRBJ2-3 | CASSLPTGTGDTDTQYF | KLGGALQAK | YES | NO | NO |
| 12593 | TRAV17 | TRAJ45 | CATARGADGLTF | TRBV5-1 | TRBJ1-2 | CASSLAGGRTGDYGYTF | KLGGALQAK | YES | NO | NO |
| 12594 | TRAV19 | TRAJ38 | CALSEFEGAGNNRKLIW | TRBV4-2 | TRBJ2-3 | CASSRLGDTQYF | KLGGALQAK | YES | NO | NO |
| 12595 | TRAV1-2 | TRAJ33 | CAVMDSNYQLIW | TRBV20-1 | TRBJ2-1 | CSARNREAEQFF | KLGGALQAK | NO | NO | NO |
| 12596 | TRAV8-3 | TRAJ49 | CAVVGTGNQFYF | TRBV12-4 | TRBJ2-7 | CASSNSWGSYEQYF | GLCTLVAML | YES | NO | YES |
| 12597 | TRAV13-1 | TRAJ50 | CAASETSYDKVIF | TRBV7-9 | TRBJ2-3 | CASSLLGGPDTQYF | RAKFKQLL | YES | NO | YES |
| 12598 | TRAV8-3 | TRAJ21 | CAVGDNFNKFYF | TRBV9 | TRBJ2-2 | CASSSGLVSNTGELFF | GLCTLVAML | YES | YES | YES |
| 12599 | TRAV19 | TRAJ49 | CALRAGNQFYF | TRBV4-1 | TRBJ1-3 | CASSQTRGAGNTIYF | RAKFKQLL | YES | NO | YES |
| 12600 | TRAV8-4 | TRAJ31 | CAVSDHSNNARLMF | TRBV4-1 | TRBJ2-1 | CASSLGGTNEQFF | RAKFKQLL | YES | NO | NO |
| 12601 | TRAV19 | TRAJ48 | CALSGYFGNEKLTF | TRBV6-3 | TRBJ2-7 | CASTPLRGDRPYEQYF | KLGGALQAK | YES | NO | NO |
| 12602 | TRAV19 | TRAJ13 | CATSGGYQKVTF | TRBV9 | TRBJ1-1 | CASSALAGNTEAFF | RAKFKQLL | YES | YES | YES |
| 12603 | TRAV30 | TRAJ20 | CGTEYPNDYKLSF | TRBV27 | TRBJ2-7 | CASSLSGRPYEQYF | RAKFKQLL | YES | YES | YES |
| 12604 | TRAV8-3 | TRAJ31 | CAVGTARLMF | TRBV6-5 | TRBJ2-7 | CASSLRFPDIGEQYF | RAKFKQLL | YES | NO | YES |
| 12605 | TRAV26-1 | TRAJ34 | CIVRVANPTDKLIF | TRBV9 | TRBJ1-1 | CASSVRSSMNTEAFF | KLGGALQAK | YES | NO | NO |
| 12606 | TRAV12-2 | TRAJ29 | CATPSGNTPLVF | TRBV29-1 | TRBJ1-2 | CSVEEGHYGYTF | KLGGALQAK | YES | NO | NO |
| 12607 | TRAV22 | TRAJ32 | CAVYGGATNKLIF | TRBV2 | TRBJ1-1 | CASSVGEAFF | KLGGALQAK | YES | NO | NO |
| 12608 | TRAV5 | TRAJ28 | CAESIGYSGAGSYQLTF | TRBV5-1 | TRBJ1-6 | CASSLEVGESLHF | KLGGALQAK | YES | NO | NO |
| 12609 | TRAV14DV4 | TRAJ29 | CAMREDTSGNTPLVF | TRBV7-8 | TRBJ2-7 | CASSHPPTGAYEQYF | KLGGALQAK | YES | NO | NO |
| 12610 | TRAV6 | TRAJ28 | CAFNCWELPTHF | TRBV7-6 | TRBJ2-5 | CASSLGGVLGETQYF | RLRAEAQVK | YES | NO | NO |
| 12611 | TRAV17 | TRAJ36 | CATDSLTGANNLFF | TRBV12-4 | TRBJ2-7 | CASSPSSSYEQYF | KLGGALQAK | YES | NO | NO |
| 12612 | TRAV27 | TRAJ48 | CAASNPNGKLTF | TRBV19 | TRBJ2-7 | CASSVRSSYEQYF | GILGFVFTL | YES | NO | YES |
| 12613 | TRAV19 | TRAJ18 | CALSEANRGSTLGRLYF | TRBV20-1 | TRBJ2-3 | CSAIGGGREDTDTQYF | KLGGALQAK | NO | NO | NO |
| 12614 | TRAV12-2 | TRAJ33 | CAVNMKDSNYQLIW | TRBV27 | TRBJ2-7 | CASSLDKATEQYF | KLGGALQAK | YES | NO | NO |
| 12615 | TRAV14DV4 | TRAJ49 | CAMRSTGNQFYF | TRBV19 | TRBJ2-7 | CASSIRSSYEQYF | GILGFVFTL | YES | NO | YES |
| 12616 | TRAV1-2 | TRAJ38 | CAVRDRNAGNNRKLIW | TRBV28 | TRBJ2-3 | CASGRGGGPDTQYF | KLGGALQAK | YES | NO | NO |
| 12617 | TRAV21 | TRAJ39 | CAALNNAGNMLTF | TRBV19 | TRBJ2-1 | CASSIALASEQFF | LLDFVRFMGV | YES | NO | YES |
| 12618 | TRAV23DV6 | TRAJ33 | CAASMDSNYQLIW | TRBV5-1 | TRBJ2-5 | CASSLGYGGAGETQYF | KLGGALQAK | YES | NO | NO |
| 12619 | TRAV12-3 | TRAJ9 | CARGGGFKTIF | TRBV7-9 | TRBJ1-5 | CASSLADVGGPQHF | RAKFKQLL | YES | NO | NO |
| 12620 | TRAV17 | TRAJ20 | CATDRYNDYKLSF | TRBV7-9 | TRBJ1-6 | CASSPGWGTDHSPLHF | KLGGALQAK | YES | NO | NO |
| 12621 | TRAV19 | TRAJ29 | CALSESNSGNTPLVF | TRBV20-1 | TRBJ1-1 | CSAWTGTEAFF | GLCTLVAML | YES | NO | YES |

| | | | | | | | | | | |
|---|---|---|---|---|---|---|---|---|---|---|
| 12622 | TRAV8-4 | TRAJ10 | CAVTGGGNKLTF | TRBV20-1 | TRBJ2-7 | CSAPGQGVAEQYF | KLGGALQAK | NO | NO | NO |
| 12623 | TRAV12-3 | TRAJ37 | CAMNSNTGKLIF | TRBV4-2 | TRBJ2-2 | CASSQRPSEVGELFF | RAKFKQLL | YES | YES | YES |
| 12624 | TRAV14DV4 | TRAJ11 | CAMREGRLSGYSTLTF | TRBV9 | TRBJ2-2 | CASSVVQENTGELFF | KLGGALQAK | YES | NO | NO |
| 12625 | TRAV38-1 | TRAJ53 | CALNSGGSNYKLTF | TRBV6-6 | TRBJ2-1 | CASSYLGPRDEQFF | KLGGALQAK | NO | NO | NO |
| 12626 | TRAV12-3 | TRAJ42 | CAMSGDGGSQGNLIF | TRBV19 | TRBJ2-7 | CASSTRSAYEQYF | GILGFVFTL | YES | YES | YES |
| 12627 | TRAV17 | TRAJ45 | CATAAQGGADGLTF | TRBV6-3 | TRBJ2-3 | CASSLASSADTQYF | RAKFKQLL | YES | NO | YES |
| 12628 | TRAV25 | TRAJ40 | CGSGTYKYIF | TRBV12-3 | TRBJ2-5 | CASSGGVSGGAGETQYF | RAKFKQLL | YES | NO | NO |
| 12629 | TRAV12-1 | TRAJ37 | CVVTHITRGNTGKLIF | TRBV7-9 | TRBJ2-7 | CASSPGAYEQYF | KLGGALQAK | YES | NO | NO |
| 12630 | TRAV8-1 | TRAJ23 | CAINQGGKLIF | TRBV6-5 | TRBJ1-2 | CASSSQGGFDGYTF | FLYALALLL | YES | YES | NO |
| 12631 | TRAV1-2 | TRAJ4 | CASLGGYNKLIF | TRBV7-9 | TRBJ2-2 | CASSQQGADGELFF | KLGGALQAK | YES | NO | NO |
| 12632 | TRAV35 | TRAJ26 | CAGPLNYGQNFVF | TRBV5-6 | TRBJ2-5 | CASSLPGQGLETQYF | AVFDRKSDAK | YES | NO | YES |
| 12633 | TRAV1-1 | TRAJ39 | CAVQAGNMLTF | TRBV5-4 | TRBJ2-7 | CASTRGDSYEQYF | KLGGALQAK | YES | NO | NO |
| 12634 | TRAV5 | TRAJ43 | CAVGNNDMRF | TRBV7-9 | TRBJ2-7 | CASSPRPGGYEQYF | KLGGALQAK | YES | NO | NO |
| 12636 | TRAV29DV5 | TRAJ45 | CAGGGADGLTF | TRBV5-1 | TRBJ1-5 | CASSLGVLQPQHF | KLGGALQAK | YES | NO | NO |
| 12637 | TRAV34 | TRAJ49 | CGAAMNTGNQFYF | TRBV15 | TRBJ2-7 | CATSRGLAGAYEQYF | KLGGALQAK | YES | NO | NO |
| 12638 | TRAV1-2 | TRAJ33 | CAVKDSNYQLIW | TRBV4-2 | TRBJ2-1 | CASSQEEGSGWVLFF | RLRAEAQVK | NO | NO | NO |
| 12639 | TRAV8-2 | TRAJ3 | CVLYSSASKIIF | TRBV20-1 | TRBJ2-7 | CSAPGQGTYEQYF | KLGGALQAK | NO | NO | NO |
| 12640 | TRAV10 | TRAJ7 | CVVTRGYGNNRLAF | TRBV6-4 | TRBJ2-3 | CASSLTSSADTQYF | KLGGALQAK | YES | NO | NO |
| 12641 | TRAV29DV5 | TRAJ15 | CAADQAGTALIF | TRBV5-4 | TRBJ1-2 | CASRLRDSYTF | RAKFKQLL | YES | YES | YES |
| 12642 | TRAV13-1 | TRAJ11 | CAAWSPEYSTLTF | TRBV19 | TRBJ2-5 | CASSMIGETQYF | GILGFVFTL | YES | NO | YES |
| 12643 | TRAV27 | TRAJ42 | CAGGGSQGNLIF | TRBV19 | TRBJ1-1 | CASKSGLGGTTEAFF | GILGFVFTL | YES | NO | YES |
| 12644 | TRAV1-2 | TRAJ9 | CAVRSNTGGFKTIF | TRBV7-9 | TRBJ2-7 | CASSSQTSGGSYEQYF | KLGGALQAK | YES | NO | NO |
| 12645 | TRAV21 | TRAJ32 | CAAVGAYGGATNKLIF | TRBV28 | TRBJ2-5 | CASSSPYRIQETQYF | KLGGALQAK | YES | YES | NO |
| 12646 | TRAV1-2 | TRAJ33 | CAAMDSNYQLIW | TRBV7-9 | TRBJ2-7 | CASSSAAAYEQYF | RAKFKQLL | YES | NO | YES |
| 12647 | TRAV27 | TRAJ20 | CAGARSNDYKLSF | TRBV27 | TRBJ2-7 | CASSLRDPGSNPYEQYF | KLGGALQAK | YES | NO | NO |
| 12648 | TRAV12-1 | TRAJ39 | CVVNGAGNMLTF | TRBV29-1 | TRBJ2-7 | CSVWTPRSQTYEQYF | KLGGALQAK | NO | NO | NO |
| 12649 | TRAV3 | TRAJ12 | CAVRDSEGDSSYKLIF | TRBV4-1 | TRBJ2-7 | CASSQAPAGAGHEQYF | RAKFKQLL | YES | YES | YES |
| 12650 | TRAV13-1 | TRAJ7 | CAASPLPNNRLAF | TRBV5-4 | TRBJ2-6 | CASSLAGGRGRSGANVLTF | KLGGALQAK | YES | NO | NO |
| 12651 | TRAV12-3 | TRAJ42 | CAMSGDGGSQGNLIF | TRBV19 | TRBJ2-7 | CASSARSSYEQYF | GILGFVFTL | YES | NO | YES |
| 12652 | TRAV1-2 | TRAJ33 | CAVMGSNYQLIW | TRBV6-4 | TRBJ1-6 | CASTDGVENSPLHF | RAKFKQLL | YES | YES | YES |
| 12653 | TRAV12-3 | TRAJ42 | CAMSGGGGSQGNLIF | TRBV19 | TRBJ2-3 | CASSGRSTDTQYF | GILGFVFTL | YES | NO | YES |
| 12654 | TRAV22 | TRAJ37 | CAGSPSNTGKLIF | TRBV19 | TRBJ1-5 | CASSIHSGNQPQHF | GILGFVFTL | YES | NO | NO |
| 12655 | TRAV13-1 | TRAJ11 | CAAKEGYSTLTF | TRBV11-2 | TRBJ2-2 | CASSLGVAGTNTGELFF | KLGGALQAK | YES | NO | NO |
| 12656 | TRAV29DV5 | TRAJ43 | CAASENNDMRF | TRBV5-5 | TRBJ2-7 | CASTPDWGGTYEQYF | KLGGALQAK | YES | NO | NO |
| 12657 | TRAV35 | TRAJ45 | CAGLHLSGGGADGLTF | TRBV6-1 | TRBJ2-7 | CASSDSWGSSYEQYF | KLGGALQAK | YES | NO | NO |
| 12658 | TRAV38-1 | TRAJ34 | CAFMKFYNTDKLIF | TRBV14 | TRBJ2-1 | CASSQDRLAGGYNEQFF | KLGGALQAK | YES | NO | NO |
| 12661 | TRAV13-1 | TRAJ50 | CAASETSYDKVIF | TRBV12-3 | TRBJ1-1 | CASSFTGQANTEAFF | RAKFKQLL | YES | NO | YES |
| 12662 | TRAV5 | TRAJ13 | CAEPSGGYQKVTF | TRBV18 | TRBJ2-7 | CASSRMDGPAYEQYF | KLGGALQAK | YES | NO | NO |
| 12663 | TRAV19 | TRAJ26 | CALTLNYGQNFVF | TRBV7-8 | TRBJ2-7 | CASRYGGPSDEQYF | KLGGALQAK | YES | YES | NO |
| 12664 | TRAV12-1 | TRAJ29 | CVVNPPGNTPLVF | TRBV12-3 | TRBJ2-2 | CASSFSGNTGELFF | RAKFKQLL | YES | YES | YES |
| 12665 | TRAV5 | TRAJ4 | CAELKEKLIF | TRBV20-1 | TRBJ2-3 | CSARWGGGTDTQYF | GLCTLVAML | NO | NO | NO |
| 12666 | TRAV1-2 | TRAJ33 | CAVTDSNYQLIW | TRBV15 | TRBJ2-3 | CATRREAGGPDTQYF | KLGGALQAK | YES | NO | NO |
| 12668 | TRAV26-1 | TRAJ43 | CIVREINNDMRF | TRBV4-1 | TRBJ1-2 | CASSPTPTANYGYTF | RAKFKQLL | YES | NO | YES |
| 12669 | TRAV21 | TRAJ26 | CAVRPNYGQNFVF | TRBV9 | TRBJ1-4 | CASSSTTATNEKLFF | KLGGALQAK | YES | YES | NO |
| 12670 | TRAV19 | TRAJ13 | CALSEGNSGGYQKVTF | TRBV10-2 | TRBJ2-7 | CASSESWGPYEQYF | AVFDRKSDAK | YES | NO | NO |
| 12671 | TRAV12-2 | TRAJ53 | CAVMWDSGGSNYKLTF | TRBV12-4 | TRBJ2-1 | CASSTVGALYNEQFF | KLGGALQAK | YES | NO | NO |
| 12672 | TRAV13-1 | TRAJ11 | CAAKEGYSTLTF | TRBV20-1 | TRBJ2-6 | CRYKGQGVSGANVLTF | GILGFVFTL | YES | NO | NO |
| 12673 | TRAV6 | TRAJ43 | CALVRINDMRF | TRBV29-1 | TRBJ2-7 | CSAGQGPYEQYF | RAKFKQLL | NO | NO | NO |
| 12674 | TRAV21 | TRAJ32 | CAVRNYGGATNKLIF | TRBV27 | TRBJ2-7 | CASSLSGRPYEQYF | RAKFKQLL | YES | NO | YES |
| 12675 | TRAV13-1 | TRAJ20 | CAASGYDYKLSF | TRBV7-9 | TRBJ2-7 | CASSSALAGPNEQYF | RLRAEAQVK | YES | NO | YES |
| 12676 | TRAV13-2 | TRAJ45 | CADSGGGADGLTF | TRBV5-6 | TRBJ1-5 | CASSLTEGGRNQPQHF | RAKFKQLL | YES | NO | YES |
| 12677 | TRAV1-2 | TRAJ33 | CAVRGSNYQLIW | TRBV7-9 | TRBJ2-1 | CASSTGLAGTNEQFF | KLGGALQAK | YES | NO | NO |
| 12678 | TRAV2 | TRAJ26 | CAVEDRDNYGQNFVF | TRBV19 | TRBJ2-7 | CASSIRSSYEQYF | GILGFVFTL | NO | NO | NO |
| 12679 | TRAV21 | TRAJ4 | CAVSRFSGGYNKLIF | TRBV5-1 | TRBJ2-7 | CASRHSREQYF | KLGGALQAK | YES | NO | NO |
| 12680 | TRAV21 | TRAJ27 | CAVKRAGKSTF | TRBV27 | TRBJ1-5 | CASSSGQGWEQPQHF | AVFDRKSDAK | YES | NO | NO |
| 12681 | TRAV17 | TRAJ40 | CATAPTSGTYKYIF | TRBV2 | TRBJ2-3 | CASIRTRDPKAQYF | AVFDRKSDAK | YES | NO | NO |
| 12682 | TRAV12-1 | TRAJ31 | CVVNMDNARLMF | TRBV19 | TRBJ2-3 | CASSGRSTDTQYF | GILGFVFTL | YES | NO | YES |
| 12683 | TRAV12-2 | TRAJ37 | CAVHGSSNTGKLIF | TRBV5-1 | TRBJ2-7 | CASSFTYGPGGHEQYF | KLGGALQAK | YES | NO | NO |
| 12684 | TRAV13-1 | TRAJ12 | CAASMDSSYKLIF | TRBV9 | TRBJ1-1 | CASSVRSSMNTEAFF | KLGGALQAK | YES | NO | NO |
| 12685 | TRAV35 | TRAJ49 | CAGHTGNQFYF | TRBV12-3 | TRBJ2-7 | CASSSLDVAGYEQYF | RAKFKQLL | YES | YES | YES |
| 12686 | TRAV21 | TRAJ57 | CAVRNQGGSEKLVF | TRBV19 | TRBJ1-2 | CASSLGAYGYTF | GILGFVFTL | YES | NO | YES |
| 12687 | TRAV1-1 | TRAJ37 | CAVIEDTGKLIF | TRBV7-9 | TRBJ1-1 | CASSLVLGRETEAFF | RAKFKQLL | YES | NO | YES |
| 12688 | TRAV13-1 | TRAJ40 | CAATSPSSGTYKYIF | TRBV5-4 | TRBJ2-7 | CASREGVGEQYF | KLGGALQAK | YES | NO | NO |
| 12689 | TRAV29DV5 | TRAJ20 | CAASAMIIDYKLSF | TRBV6-6 | TRBJ2-2 | CASSYGFPTSGGNTGELFF | RAKFKQLL | YES | NO | YES |
| 12690 | TRAV22 | TRAJ20 | CAVRFNDYKLSF | TRBV7-2 | TRBJ2-2 | CASSPPGLVGGELFF | GLCTLVAML | YES | YES | NO |
| 12691 | TRAV4 | TRAJ13 | CLVGGRSGGYQKVTF | TRBV12-4 | TRBJ2-3 | CAVNGLAGPTDTQYF | KLGGALQAK | NO | NO | NO |
| 12692 | TRAV1-2 | TRAJ33 | CAVLDSNYQLIW | TRBV9 | TRBJ1-1 | CASSSGRLTNTEAFF | GLCTLVAML | YES | YES | YES |
| 12693 | TRAV13-2 | TRAJ34 | CAENEDTDKLIF | TRBV11-1 | TRBJ2-2 | CASSPGQGIAGELFF | KLGGALQAK | YES | NO | NO |
| 12694 | TRAV9-2 | TRAJ50 | CALSETSYDKVIF | TRBV12-3 | TRBJ2-2 | CASRVGAGGPDGELFF | RAKFKQLL | YES | NO | YES |
| 12695 | TRAV17 | TRAJ45 | CATAAGGGADGLTF | TRBV7-9 | TRBJ2-1 | CASSSAGPYTRRTRQGGQFF | KLGGALQAK | NO | NO | NO |
| 12696 | TRAV13-2 | TRAJ45 | CADSGGGADGLTF | TRBV20-1 | TRBJ2-7 | CSALWTDSYEQYF | RAKFKQLL | NO | NO | NO |
| 12697 | TRAV12-3 | TRAJ48 | CAMARSNFGNEKLTF | TRBV19 | TRBJ2-3 | CASSIFYSSTDTQYF | GILGFVFTL | YES | NO | YES |
| 12698 | TRAV13-2 | TRAJ39 | CAENNAGNMLTF | TRBV15 | TRBJ2-1 | CATSRLASSYNEQFF | KLGGALQAK | YES | NO | NO |
| 12699 | TRAV13-2 | TRAJ45 | CAENGGGGADGLTF | TRBV6-5 | TRBJ2-5 | CASSYTPTAQETQYF | KLGGALQAK | NO | NO | NO |
| 12700 | TRAV29DV5 | TRAJ49 | CAAPTGNQFYF | TRBV4-1 | TRBJ1-3 | CASSQTRGAGNTIYF | RAKFKQLL | YES | NO | NO |
| 12701 | TRAV1-2 | TRAJ12 | CAALDSSYKLIF | TRBV6-1 | TRBJ1-5 | CASSEAPDGNQPQHF | KLGGALQAK | YES | NO | NO |
| 12702 | TRAV14DV4 | TRAJ52 | CAMRGDAGGTSYGKLTF | TRBV12-3 | TRBJ2-2 | CASSFSGNTGELFF | RAKFKQLL | YES | NO | YES |
| 12703 | TRAV17 | TRAJ37 | CATDFSGNTGKLIF | TRBV11-2 | TRBJ2-3 | CASSLYGRTTDTQYF | KLGGALQAK | YES | NO | NO |
| 12704 | TRAV13-1 | TRAJ50 | CAAMETSYDKVIF | TRBV11-2 | TRBJ1-2 | CASSWGGGSHYGYTF | RAKFKQLL | YES | NO | YES |
| 12705 | TRAV13-1 | TRAJ49 | CAVHTGNQFYF | TRBV12-3 | TRBJ2-2 | CASSFSGNTGELFF | RAKFKQLL | YES | NO | YES |
| 12706 | TRAV8-1 | TRAJ13 | CAVNARSGGYQKVTF | TRBV11-2 | TRBJ2-3 | CASLISGTDTQYF | KLGGALQAK | YES | NO | NO |
| 12707 | TRAV5 | TRAJ39 | CAEDAGNMLTF | TRBV19 | TRBJ2-3 | CASSIRSTDTQYF | GILGFVFTL | YES | NO | YES |
| 12708 | TRAV8-2 | TRAJ15 | CVVSGEGWAAGTALIF | TRBV4-1 | TRBJ2-1 | CASSLLWAEYNEQFF | RAKFKQLL | YES | NO | YES |
| 12709 | TRAV25 | TRAJ42 | CAATGGGSQGNLIF | TRBV19 | TRBJ1-2 | CASSIGVHGYTF | GILGFVFTL | YES | NO | YES |
| 12710 | TRAV38-1 | TRAJ52 | CAFMKGAGGTSYGKLTF | TRBV4-3 | TRBJ1-6 | CASSQEIGFSPLHF | GILGFVFTL | YES | NO | NO |
| 12711 | TRAV19 | TRAJ43 | CALSGANNNDMRF | TRBV5-6 | TRBJ2-5 | CASSFPGLADRQYF | KLGGALQAK | YES | NO | NO |
| 12712 | TRAV29DV5 | TRAJ40 | CAASTTSGTYKYIF | TRBV11-2 | TRBJ1-2 | CASSLDGQGPLYGYTF | KLGGALQAK | YES | NO | NO |
| 12713 | TRAV13-1 | TRAJ20 | CAASGYDYKLSF | TRBV29-1 | TRBJ2-7 | CSVGTGDWGEQYF | RAKFKQLL | YES | NO | YES |
| 12714 | TRAV13-2 | TRAJ53 | CAETHSGGSNYNLTF | TRBV29-1 | TRBJ2-1 | CSVGSGDHGEQFF | RAKFKQLL | YES | YES | YES |
| 12715 | TRAV12-1 | TRAJ54 | CVVCSLQGAQKLVF | TRBV12-3 | TRBJ2-2 | CASSFSGNTGELFF | RAKFKQLL | YES | YES | YES |
| 12716 | TRAV27 | TRAJ37 | CAGNSGNTGKLIF | TRBV19 | TRBJ2-7 | CASSSRSGYEQYF | GILGFVFTL | YES | YES | YES |
| 12717 | TRAV27 | TRAJ40 | CAGGTGTYKYIF | TRBV5-4 | TRBJ2-1 | CASSRRLAVYNEQFF | KLGGALQAK | YES | NO | NO |
| 12718 | TRAV13-2 | TRAJ45 | CADSGGSADGLTF | TRBV5-6 | TRBJ1-1 | CASSLRDGSEAFF | RAKFKQLL | YES | NO | YES |
| 12719 | TRAV19 | TRAJ23 | CAFLYNQGGKLIF | TRBV12-4 | TRBJ2-4 | CASSPGSRGNIQYF | KLGGALQAK | YES | NO | NO |
| 12720 | TRAV12-1 | TRAJ6 | CVVTPTSGGSYIPTF | TRBV7-2 | TRBJ2-5 | CASILPQGTQYF | KLGGALQAK | YES | NO | NO |
| 12721 | TRAV8-3 | TRAJ26 | CAVAYNYGQNFVF | TRBV5-5 | TRBJ1-2 | CASSLASNLGYGYTF | RLRAEAQVK | YES | YES | YES |
| 12722 | TRAV13-2 | TRAJ45 | CADSGGGADGLTF | TRBV20-1 | TRBJ1-2 | CSARDRTGNGYTF | RAKFKQLL | YES | NO | YES |
| 12723 | TRAV1-2 | TRAJ33 | CAVINSNYQLIW | TRBV20-1 | TRBJ2-3 | CSARDLLAGDTDTQYF | KLGGALQAK | NO | NO | NO |
| 12724 | TRAV38-1 | TRAJ49 | CAFMKREVGGNQFYF | TRBV10-3 | TRBJ2-1 | CAISIASYNEQFF | AYAQKIFKI | YES | YES | YES |
| 12725 | TRAV5 | TRAJ8 | CAEIRDTGFQKLVF | TRBV3-1 | TRBJ1-2 | CASSHYRRAEGGAGYTF | RAKFKQLL | YES | NO | YES |
| 12726 | TRAV13-1 | TRAJ50 | CAASETSYDKVIF | TRBV18 | TRBJ1-5 | CASSPRQGDQPQHF | RAKFKQLL | YES | YES | YES |
| 12727 | TRAV41 | TRAJ52 | CAAAGGTSYGKLTF | TRBV2 | TRBJ1-5 | CASSERIPNQPQHF | KLGGALQAK | YES | NO | NO |
| 12728 | TRAV12-1 | TRAJ23 | CVVYYGGKLIF | TRBV12-3 | TRBJ2-2 | CASSFSGNTGELFF | RAKFKQLL | YES | NO | YES |
| 12729 | TRAV29DV5 | TRAJ34 | CAASATRGTDKLIF | TRBV6-1 | TRBJ1-6 | CASSEALDPTGSSPLHF | KLGGALQAK | YES | NO | NO |

| | | | | | | | | | | |
|---|---|---|---|---|---|---|---|---|---|---|
| 12730 | TRAV17 | TRAJ11 | CATEGDSGYSTLTF | TRBV5-1 | TRBJ2-5 | CASSLEWGGETQYF | FLYALALLL | YES | YES | NO |
| 12731 | TRAV17 | TRAJ20 | CATDGNDYKLSF | TRBV2 | TRBJ2-3 | CAKGSSGGPHTQYF | KLGGALQAK | YES | NO | NO |
| 12732 | TRAV36DV7 | TRAJ52 | CAVAGGTSYGKLTF | TRBV19 | TRBJ2-7 | CASSLDSSYEQYF | KLGGALQAK | NO | NO | NO |
| 12733 | TRAV12-2 | TRAJ4 | CAVALFSGGYNKLIF | TRBV7-6 | TRBJ2-1 | CASSLYAPGTGVYNEQFF | KLGGALQAK | YES | NO | NO |
| 12734 | TRAV20 | TRAJ47 | CAVQAPGNKLVF | TRBV15 | TRBJ1-2 | CATSRVGEHGYTF | AVFDRKSDAK | YES | YES | YES |
| 12735 | TRAV12-3 | TRAJ31 | CAMKSANNARLMF | TRBV7-9 | TRBJ2-7 | CASSFAGDEQYF | RTLNAWVKV | YES | YES | YES |
| 12736 | TRAV35 | TRAJ45 | CAGPLRTRSGGGADGLTF | TRBV20-1 | TRBJ2-7 | CSARDLAGGREQYF | KLGGALQAK | NO | NO | NO |
| 12737 | TRAV1-2 | TRAJ8 | CAVRDEGNTGFQKLVF | TRBV20-1 | TRBJ2-7 | CSARVWGANYEQYF | KLGGALQAK | NO | NO | NO |
| 12738 | TRAV13-1 | TRAJ4 | CAASGAGGYNKLIF | TRBV20-1 | TRBJ2-7 | CSARDGRREYEQYF | KLGGALQAK | YES | NO | NO |
| 12739 | TRAV22 | TRAJ37 | CAGSPSNTGKLIF | TRBV12-4 | TRBJ2-1 | CASGRLSYNEQFF | RAKFKQLL | YES | NO | NO |
| 12740 | TRAV1-2 | TRAJ36 | CAVRDRTGANNLFF | TRBV6-1 | TRBJ2-7 | CASSPSRLGTGFYEQYF | KLGGALQAK | YES | NO | NO |
| 12741 | TRAV12-1 | TRAJ15 | CVVNGGQAGTALIF | TRBV7-2 | TRBJ2-7 | CASSFDPSRGEQYF | RLRAEAQVK | NO | NO | NO |
| 12742 | TRAV20 | TRAJ37 | CAVQDRGGSGNTGKLIF | TRBV20-1 | TRBJ1-2 | CSARSEGRYTF | KLGGALQAK | YES | NO | NO |
| 12743 | TRAV36DV7 | TRAJ7 | CAVDGNRLAF | TRBV20-1 | TRBJ2-1 | CSARGDRGGSSYNEQFF | KLGGALQAK | NO | NO | NO |
| 12744 | TRAV12-1 | TRAJ29 | CVVKVGAPLVF | TRBV24-1 | TRBJ2-5 | CATSDPKGDQETQYF | KLGGALQAK | YES | NO | NO |
| 12745 | TRAV5 | TRAJ31 | CAEDNNARLMF | TRBV20-1 | TRBJ1-5 | CSASRPYSNQPQHF | GLCTLVAML | YES | YES | YES |
| 12746 | TRAV1-2 | TRAJ33 | CAVMDSNYQLIW | TRBV6-1 | TRBJ2-1 | CASSELAGANEQFF | KLGGALQAK | YES | NO | NO |
| 12747 | TRAV8-3 | TRAJ7 | CAVFLYGNNRLAF | TRBV4-2 | TRBJ2-2 | CASSQRPSEVGELFF | RAKFKQLL | YES | NO | YES |
| 12748 | TRAV13-1 | TRAJ49 | CAAQPPTGNQFYF | TRBV11-2 | TRBJ2-5 | CASSFLTGQGETQYF | RAKFKQLL | YES | NO | YES |
| 12749 | TRAV12-2 | TRAJ49 | CAVKPLTTNTGNQFYF | TRBV10-3 | TRBJ1-1 | CAISEPTFMDTEAFF | KLGGALQAK | YES | NO | NO |
| 12750 | TRAV27 | TRAJ42 | CAGNYGGSQGNLIF | TRBV19 | TRBJ2-7 | CASSMRSSHEQYF | GILGFVFTL | YES | NO | NO |
| 12751 | TRAV13-1 | TRAJ45 | CAASTYSGGGADGLTF | TRBV6-6 | TRBJ2-1 | CASSPFSDEQFF | KLGGALQAK | NO | NO | NO |
| 12752 | TRAV13-2 | TRAJ53 | CAETHSGGSNYKLTF | TRBV19 | TRBJ2-7 | CASSARSSYEQYF | GILGFVFTL | YES | NO | YES |
| 12753 | TRAV12-1 | TRAJ20 | CVVNRDYKLSF | TRBV11-2 | TRBJ2-2 | CASSLARGTGELFF | KLGGALQAK | YES | NO | NO |
| 12754 | TRAV12-2 | TRAJ15 | CAVWGQAGTALIF | TRBV29-1 | TRBJ1-6 | CSVEEGGGSPLHF | KLGGALQAK | NO | NO | NO |
| 12755 | TRAV3 | TRAJ36 | CAVRDGTGANNLFF | TRBV19 | TRBJ2-3 | CASSTGGTDTQYF | GILGFVFTL | NO | NO | NO |
| 12756 | TRAV25 | TRAJ26 | CAWNPDNYGQNFVF | TRBV5-4 | TRBJ2-5 | CASSFNGVRETQYF | KLGGALQAK | YES | NO | NO |
| 12757 | TRAV17 | TRAJ45 | CATVPYFSPSGGGADGLTF | TRBV7-9 | TRBJ1-1 | CASSWDRDTEAFF | KLGGALQAK | YES | NO | NO |
| 12758 | TRAV5 | TRAJ6 | CAVMEGGSYIPTF | TRBV4-2 | TRBJ2-2 | CASSQRPSEVGELFF | RAKFKQLL | YES | NO | YES |
| 12759 | TRAV12-2 | TRAJ11 | CAVKVTGYSTLTF | TRBV5-6 | TRBJ2-2 | CASSLLGHNTGELFF | KLGGALQAK | YES | NO | NO |
| 12760 | TRAV26-1 | TRAJ38 | CIVRTAGNNRKLIW | TRBV19 | TRBJ1-5 | CASSMGGLQPQHF | KLGGALQAK | YES | NO | NO |
| 12761 | TRAV8-3 | TRAJ36 | CAPGETGANNLFF | TRBV19 | TRBJ1-5 | CASSFYSNQPQHF | GILGFVFTL | YES | NO | YES |
| 12762 | TRAV12-2 | TRAJ41 | CAAYQGRRDSGYALNF | TRBV20-1 | TRBJ2-7 | CSVGGYSGGYEQYF | KLGGALQAK | YES | NO | NO |
| 12763 | TRAV1-2 | TRAJ17 | CAVRASKAAGNKLTF | TRBV5-6 | TRBJ1-1 | CASSLRDGSEAFF | RAKFKQLL | YES | NO | YES |
| 12764 | TRAV35 | TRAJ38 | CAGQGNAGNNRKLIW | TRBV29-1 | TRBJ1-2 | CSVEDNGYGYTF | KLGGALQAK | YES | NO | NO |
| 12765 | TRAV21 | TRAJ26 | CAVHGYGQNFVF | TRBV4-2 | TRBJ2-1 | CASSHPALGNYDQLSYNEQFF | KLGGALQAK | NO | NO | NO |
| 12766 | TRAV29DV5 | TRAJ41 | CAEKPGSGYALNF | TRBV7-8 | TRBJ2-5 | CASSLTPGGETQYF | KLGGALQAK | YES | NO | NO |
| 12767 | TRAV1-2 | TRAJ33 | CAVSDSNYQLIW | TRBV30 | TRBJ2-2 | CAWRTGFGDTGELFF | KLGGALQAK | YES | NO | NO |
| 12768 | TRAV29DV5 | TRAJ32 | CAASSSGGATNKLIF | TRBV4-2 | TRBJ2-7 | CASSGGGYEQYF | KLGGALQAK | YES | NO | NO |
| 12769 | TRAV12-2 | TRAJ40 | CAPRGSGTYKYIF | TRBV24-1 | TRBJ2-1 | CATSETGGVDEQFF | AYAQKIFKI | YES | YES | YES |
| 12770 | TRAV12-2 | TRAJ45 | CARRGGADGLTF | TRBV5-6 | TRBJ2-7 | CASSLALAGPYEQYF | ELAGIGILTV | YES | YES | NO |
| 12771 | TRAV17 | TRAJ44 | CATDRGTASKLTF | TRBV10-3 | TRBJ2-3 | CAINPLGSSSKKGTQYF | KLGGALQAK | YES | NO | NO |
| 12772 | TRAV38-1 | TRAJ52 | CAFMINAGGTSYGKLTF | TRBV19 | TRBJ1-2 | CASSIGSHGYTF | GILGFVFTL | YES | NO | YES |
| 12773 | TRAV12-1 | TRAJ34 | CVPHNTDKLIF | TRBV6-3 | TRBJ2-7 | CASSYSFAGGIDEQYF | KLGGALQAK | YES | NO | NO |
| 12774 | TRAV8-3 | TRAJ21 | CAVGDNFNKFYF | TRBV7-9 | TRBJ1-1 | CASSSLGGHGEAFF | RAKFKQLL | YES | NO | YES |
| 12775 | TRAV14DV4 | TRAJ26 | CAMRDDDNYGQNFVF | TRBV12-3 | TRBJ2-2 | CASSFSGNTGELFF | RAKFKQLL | YES | NO | YES |
| 12776 | TRAV12-3 | TRAJ26 | CAMKDYGQNFVF | TRBV6-1 | TRBJ1-1 | CASSVSSEWNTEAFF | KLGGALQAK | YES | NO | NO |
| 12777 | TRAV27 | TRAJ42 | CAGGGSQGNLIF | TRBV4-2 | TRBJ2-2 | CASSQRPSEVGELFF | RAKFKQLL | YES | NO | YES |
| 12778 | TRAV13-2 | TRAJ45 | CADSGGGADGLTF | TRBV5-6 | TRBJ1-1 | CASSLGASMNTEAFF | RAKFKQLL | YES | NO | YES |
| 12779 | TRAV13-1 | TRAJ50 | CAAMETSYDKVIF | TRBV9 | TRBJ2-2 | CASSTRTGTETNTGELFF | RAKFKQLL | YES | NO | YES |
| 12780 | TRAV5 | TRAJ42 | CAEVKGGGSQGNLIF | TRBV19 | TRBJ2-7 | CASSIRAAYEQYF | GILGFVFTL | YES | NO | NO |
| 12781 | TRAV26-1 | TRAJ43 | CIVREGNLDDMRF | TRBV7-9 | TRBJ2-3 | CASSPLAGAAIHTQYF | RAKFKQLL | YES | YES | YES |
| 12782 | TRAV17 | TRAJ4 | CATDGFSGGYNKLIF | TRBV27 | TRBJ2-2 | CASSLAQGAGGELFF | AYAQKIFKI | YES | YES | YES |
| 12783 | TRAV29DV5 | TRAJ15 | CAASQQAGTALIF | TRBV14 | TRBJ2-7 | CASSQVLWAYEQYF | KLGGALQAK | NO | NO | NO |
| 12784 | TRAV17 | TRAJ50 | CAEGGMKTSYDKVIF | TRBV29-1 | TRBJ2-1 | CSVQGGRYNEQFF | KLGGALQAK | YES | NO | NO |
| 12785 | TRAV3 | TRAJ31 | CAVRGYNNARLMF | TRBV28 | TRBJ1-2 | CASSKTGIYGYTF | KLGGALQAK | YES | NO | NO |
| 12786 | TRAV1-2 | TRAJ33 | CAIMDSNYQLIW | TRBV20-1 | TRBJ2-7 | CSARPLKGEIEQYF | RLRAEAQVK | NO | NO | NO |
| 12787 | TRAV17 | TRAJ23 | CATDPGRGKLIF | TRBV9 | TRBJ2-3 | CASSAGGGPDTQYF | KLGGALQAK | YES | NO | NO |
| 12788 | TRAV12-1 | TRAJ39 | CGAAGNMLTF | TRBV7-9 | TRBJ2-6 | CASSPVTGGGSGANVLTF | AVFDRKSDAK | YES | NO | NO |
| 12789 | TRAV12-3 | TRAJ13 | CAFHSGGYQKVTF | TRBV12-4 | TRBJ2-7 | CASSLWGPRTYEQYF | KLGGALQAK | YES | NO | NO |
| 12790 | TRAV16 | TRAJ26 | CALNNYGQNFVF | TRBV19 | TRBJ2-1 | CASSMVGSSTYNEQFF | KLGGALQAK | NO | NO | NO |
| 12791 | TRAV13-2 | TRAJ45 | CADSGGGADGLTF | TRBV7-3 | TRBJ2-2 | CASSLGTSGGTGELFF | RAKFKQLL | NO | NO | NO |
| 12792 | TRAV8-3 | TRAJ28 | CAVGALYSGAGSYQLTF | TRBV7-6 | TRBJ2-7 | CASSSLTGLHEQYF | KLGGALQAK | YES | NO | NO |
| 12793 | TRAV26-1 | TRAJ52 | CIAHFGGTSYGKLTF | TRBV5-6 | TRBJ2-3 | CASSLVAGGMADTQYF | KLGGALQAK | YES | NO | NO |
| 12794 | TRAV9-2 | TRAJ50 | CALSETSYDKVIF | TRBV12-3 | TRBJ2-1 | CASGPGQGEYNEQFF | RAKFKQLL | YES | NO | YES |
| 12795 | TRAV30 | TRAJ20 | CGTEYPNDYKLSF | TRBV12-3 | TRBJ2-2 | CASSFSGGTGELFF | RAKFKQLL | YES | NO | YES |
| 12796 | TRAV13-1 | TRAJ23 | CAASANQGGKLIF | TRBV7-6 | TRBJ2-7 | CASSSGPNYEQYF | KLGGALQAK | YES | NO | NO |
| 12797 | TRAV8-3 | TRAJ3 | CAVGPWYSSASKIIF | TRBV5-1 | TRBJ1-1 | CASSSEGGSTEAFF | KLGGALQAK | YES | NO | NO |
| 12798 | TRAV19 | TRAJ48 | CALSEALSNFGNEKLTF | TRBV19 | TRBJ1-5 | CASSIHSGNQPQHF | GILGFVFTL | YES | NO | YES |
| 12799 | TRAV12-3 | TRAJ21 | CAMRKYNFNKFYF | TRBV2 | TRBJ2-1 | CASRVRDEQFF | AVFDRKSDAK | YES | NO | YES |
| 12800 | TRAV12-2 | TRAJ45 | CAVNMEVSRGGGADGLTF | TRBV28 | TRBJ1-5 | CASTLGGLGQPQHF | ELAGIGILTV | YES | YES | NO |
| 12801 | TRAV26-2 | TRAJ40 | CIRGTYKYIF | TRBV12-3 | TRBJ2-2 | CASSFSGNTGELFF | RAKFKQLL | YES | NO | NO |
| 12802 | TRAV2 | TRAJ42 | CGRQMLLFTTVLWAYYGGSQGNLIF | TRBV5-6 | TRBJ1-1 | CASSLRDGSEAFF | RAKFKQLL | NO | NO | NO |
| 12803 | TRAV3 | TRAJ23 | CAVRPGGKLIF | TRBV11-2 | TRBJ2-7 | CASSVGTALYEQYF | KLGGALQAK | YES | NO | NO |
| 12804 | TRAV12-3 | TRAJ54 | CAMIQGAQKLVF | TRBV6-6 | TRBJ2-1 | CASTGGGSGVYNEQFF | KLGGALQAK | NO | NO | NO |
| 12805 | TRAV27 | TRAJ42 | CAGAHGGSQGNLIF | TRBV19 | TRBJ2-3 | CASSGRSADTQYF | GILGFVFTL | YES | NO | YES |
| 12806 | TRAV5 | TRAJ37 | CAERSNTGKLIF | TRBV4-2 | TRBJ2-2 | CASSQGAGGLGGELFF | AVFDRKSDAK | YES | YES | YES |
| 12807 | TRAV12-2 | TRAJ11 | CAVNTNSGYSTLTF | TRBV10-3 | TRBJ2-3 | CAISPPDSSTQYF | AVFDRKSDAK | YES | NO | YES |
| 12808 | TRAV13-2 | TRAJ45 | CADSGGGADGLTF | TRBV7-9 | TRBJ2-6 | CASREIQGSGANVLTF | RAKFKQLL | YES | NO | YES |
| 12809 | TRAV1-2 | TRAJ33 | CAVVNSNYQLIW | TRBV20-1 | TRBJ2-1 | CSGGERLEGNEQFF | KLGGALQAK | NO | NO | NO |
| 12810 | TRAV1-2 | TRAJ33 | CAVTDSNYQLIW | TRBV6-4 | TRBJ2-1 | CASRGGDSHNEQFF | KLGGALQAK | YES | NO | NO |
| 12811 | TRAV14DV4 | TRAJ53 | CAMSGGSNYKLTF | TRBV30 | TRBJ1-1 | CAWTWGTLNTEAFF | KLGGALQAK | YES | NO | NO |
| 12812 | TRAV12-1 | TRAJ28 | CVVNSFSGAGSYQLTF | TRBV7-9 | TRBJ2-2 | CASSNSAGELFF | KLGGALQAK | YES | NO | NO |
| 12813 | TRAV26-1 | TRAJ8 | CIVMDDFQKLVF | TRBV12-3 | TRBJ2-2 | CASSFSGNTGELFF | RAKFKQLL | YES | NO | YES |
| 12814 | TRAV41 | TRAJ34 | CAARWDKLIF | TRBV20-1 | TRBJ2-5 | CSARARKGGGLETQYF | KLGGALQAK | YES | NO | NO |
| 12815 | TRAV12-2 | TRAJ7 | CAGASWGNNRLAF | TRBV6-5 | TRBJ2-3 | CATRPPSTDTQYF | KLGGALQAK | NO | NO | NO |
| 12816 | TRAV13-2 | TRAJ43 | CAENSKGHDNDMRF | TRBV27 | TRBJ2-7 | CASSLSGRPYEQYF | RAKFKQLL | YES | NO | YES |
| 12817 | TRAV41 | TRAJ57 | CAVLGTQGGSEKLVF | TRBV5-1 | TRBJ2-1 | CASFRGAAYNEQFF | RAKFKQLL | YES | NO | YES |
| 12818 | TRAV12-2 | TRAJ7 | CAVRYGNNRLAF | TRBV6-5 | TRBJ2-4 | CASSYSASGGIKNIQYF | KLGGALQAK | YES | YES | NO |
| 12819 | TRAV1-2 | TRAJ33 | CAGLDSNYQLIW | TRBV6-5 | TRBJ2-5 | CASSYMMGQGGETQYF | KLGGALQAK | NO | NO | NO |
| 12820 | TRAV12-2 | TRAJ42 | CAVNEGGGSQGNLIF | TRBV19 | TRBJ2-7 | CASSPRSSYEQYF | GILGFVFTL | YES | NO | NO |
| 12821 | TRAV22 | TRAJ37 | CAGSPSNTGKLIF | TRBV19 | TRBJ1-5 | CASSLYSNQPQHF | GILGFVFTL | YES | NO | NO |
| 12822 | TRAV13-1 | TRAJ50 | CAASETSYDKVIF | TRBV7-9 | TRBJ1-1 | CASSRLGQGAITEAFF | RAKFKQLL | YES | NO | YES |
| 12823 | TRAV8-3 | TRAJ13 | CAVSHSGGYQKVTF | TRBV19 | TRBJ2-3 | CASSMRSADTQYF | GILGFVFTL | YES | NO | YES |
| 12824 | TRAV13-1 | TRAJ47 | CAASKGHLAYGNKLVF | TRBV18 | TRBJ2-2 | CASSPPGGGNTGELFF | RAKFKQLL | YES | NO | YES |
| 12825 | TRAV38-2DV8 | TRAJ48 | CAYRSVTSNFGNEKLTF | TRBV6-5 | TRBJ2-1 | CASSDPPGSSYNEQFF | GILGFVFTL | NO | NO | NO |
| 12827 | TRAV17 | TRAJ28 | CATDLNPGAGSYQLTF | TRBV7-9 | TRBJ2-7 | CASSLTGDEQYF | KLGGALQAK | YES | NO | NO |
| 12828 | TRAV8-2 | TRAJ9 | CVVSGGGFKTIF | TRBV6-1 | TRBJ2-3 | CASSAQTGEADTQYF | KLGGALQAK | YES | NO | NO |
| 12829 | TRAV1-2 | TRAJ33 | CAVGDSNYQLIW | TRBV20-1 | TRBJ2-7 | CSARGQTDYEQYF | KLGGALQAK | NO | NO | NO |
| 12830 | TRAV12-1 | TRAJ48 | CVVNGAKFGNEKLTF | TRBV15 | TRBJ1-6 | CATSRILGSPLHF | KLGGALQAK | YES | NO | NO |
| 12831 | TRAV38-1 | TRAJ52 | CAFMVGAGGTSYGKLTF | TRBV19 | TRBJ2-7 | CASSIRSSYEQYF | GILGFVFTL | YES | NO | YES |
| 12832 | TRAV8-3 | TRAJ40 | CAVHTSGTYKYIF | TRBV11-2 | TRBJ2-3 | CASSFSAGTTDTQYF | KLGGALQAK | YES | NO | NO |
| 12833 | TRAV4 | TRAJ4 | CLVPDGAGGYNKLIF | TRBV11-2 | TRBJ2-7 | CASSLETGYEQYF | KLGGALQAK | YES | NO | NO |
| 12835 | TRAV13-2 | TRAJ42 | CAENGGGGSQGNLIF | TRBV19 | TRBJ2-3 | CASSYRATDTQYF | GILGFVFTL | YES | NO | YES |

| | | | | | | | | | | |
|---|---|---|---|---|---|---|---|---|---|---|
| 12836 | TRAV13-1 | TRAJ50 | CAAMETSYDKVIF | TRBV5-4 | TRBJ2-1 | CASSHRQGVVLDEQFF | RAKFKQLL | YES | NO | YES |
| 12837 | TRAV8-2 | TRAJ24 | CVVSDVSWGKLQF | TRBV19 | TRBJ2-2 | CASSSYTGELFF | SLFNTVATL | YES | NO | YES |
| 12838 | TRAV19 | TRAJ8 | CALRTGFQKLVF | TRBV7-9 | TRBJ2-7 | CASSLDRAYHYEQYF | RLRAEAQVK | YES | NO | YES |
| 12839 | TRAV13-1 | TRAJ42 | CAASGGGSQGNLIF | TRBV19 | TRBJ1-5 | CASSKRSQEPQHF | GILGFVFTL | YES | NO | YES |
| 12840 | TRAV27 | TRAJ37 | CAGAVGSSNTGKLIF | TRBV25-1 | TRBJ2-5 | CASSDPGIQETQYF | GILGFVFTL | YES | NO | NO |
| 12841 | TRAV2 | TRAJ27 | CGRQMLLFTTSAANAGKSTF | TRBV27 | TRBJ2-1 | CASRTGGVNNEQFF | KLGGALQAK | NO | NO | NO |
| 12842 | TRAV35 | TRAJ43 | CAGQQTDMRF | TRBV7-3 | TRBJ1-1 | CASSLMGNTEAFF | KLGGALQAK | NO | NO | NO |
| 12843 | TRAV19 | TRAJ52 | CALFNAGGTSYGKLTF | TRBV19 | TRBJ2-7 | CASSLDSSYEQYF | KLGGALQAK | YES | NO | NO |
| 12844 | TRAV13-1 | TRAJ35 | CAARVRGFGNVLHC | TRBV12-3 | TRBJ2-2 | CASSFSGNTGELFF | RAKFKQLL | NO | NO | NO |
| 12845 | TRAV12-1 | TRAJ20 | CVVNRDYRLSF | TRBV11-2 | TRBJ2-2 | CASSLARGTGELFF | KLGGALQAK | YES | NO | NO |
| 12846 | TRAV5 | TRAJ30 | CAESLWDDKIIF | TRBV7-9 | TRBJ2-3 | CASSFQGDGRDTQYF | RAKFKQLL | YES | NO | YES |
| 12847 | TRAV8-2 | TRAJ44 | CVVDTGTASKLTF | TRBV4-1 | TRBJ2-3 | CASKVVTESTDTQYF | KLGGALQAK | YES | NO | NO |
| 12848 | TRAV12-3 | TRAJ48 | CAMSDGNEKLTF | TRBV7-9 | TRBJ1-5 | CASSSGWGGVQPQHF | KLGGALQAK | YES | NO | NO |
| 12849 | TRAV1-2 | TRAJ33 | CAVRDSNYQLIW | TRBV4-2 | TRBJ2-3 | CASSQERGGPDTQYF | CLLGTYTQDV | YES | NO | YES |
| 12850 | TRAV13-1 | TRAJ50 | CAASETSYDKVIF | TRBV20-1 | TRBJ1-3 | CSARALPVGNTIYF | GLCTLVAML | YES | YES | NO |
| 12851 | TRAV8-3 | TRAJ11 | CAVASGYSTLTF | TRBV13 | TRBJ1-1 | CASKVAEHTEAFF | KLGGALQAK | YES | NO | NO |
| 12853 | TRAV17 | TRAJ28 | CATDAPPELPTHF | TRBV10-3 | TRBJ2-1 | CAISESRASGDNNEQFF | KLGGALQAK | NO | NO | NO |
| 12854 | TRAV12-2 | TRAJ42 | CAVEGSQGNLIF | TRBV7-9 | TRBJ1-5 | CASSYGQLNQPQHF | RAKFKQLL | YES | NO | YES |
| 12855 | TRAV5 | TRAJ6 | CAEMEGGSYIPTF | TRBV4-2 | TRBJ1-5 | CASSQEGQLNQPQHF | RAKFKQLL | YES | NO | YES |
| 12856 | TRAV27 | TRAJ42 | CAGGGSQGNLIF | TRBV6-1 | TRBJ2-7 | CASSVSRNYEQYF | KLGGALQAK | YES | NO | NO |
| 12857 | TRAV26-1 | TRAJ34 | CIVREVGDKLIF | TRBV29-1 | TRBJ2-5 | CSVEGPLPAGTLGTQYF | RAKFKQLL | YES | NO | YES |
| 12858 | TRAV13-1 | TRAJ42 | CAYGGSQGNLIF | TRBV6-3 | TRBJ1-2 | CASSYSINAAVNYGYTF | GILGFVFTL | YES | NO | YES |
| 12859 | TRAV12-2 | TRAJ15 | CAVNIRGTALIF | TRBV20-1 | TRBJ2-6 | CSAFTGVVGSGANVLTF | AVFDRKSDAK | YES | YES | YES |
| 12860 | TRAV1-2 | TRAJ33 | CAVMDSNYQLIW | TRBV20-1 | TRBJ2-5 | CSARFGRGETQYF | GLCTLVAML | YES | YES | NO |
| 12861 | TRAV5 | TRAJ31 | CAATDNNARLMF | TRBV4-2 | TRBJ2-2 | CASSQRPSEVGELFF | RAKFKQLL | YES | NO | YES |
| 12862 | TRAV19 | TRAJ57 | CALSEAKDEGSEKLVF | TRBV19 | TRBJ2-7 | CASSARSSYEQYF | GILGFVFTL | YES | NO | YES |
| 12863 | TRAV13-1 | TRAJ50 | CAAMETSYDKVIF | TRBV12-3 | TRBJ2-2 | CASSFSGNTGELFF | RAKFKQLL | YES | NO | YES |
| 12864 | TRAV38-1 | TRAJ43 | CAFLYKVNNNDMRF | TRBV24-1 | TRBJ2-7 | CATSSGQYPYEQYF | KLGGALQAK | YES | NO | NO |
| 12865 | TRAV13-2 | TRAJ45 | CADSGGGADGLTF | TRBV5-6 | TRBJ2-1 | CASSLGLAAAYNEQFF | RAKFKQLL | YES | YES | YES |
| 12866 | TRAV35 | TRAJ48 | CAGQWASNFGNEKLTF | TRBV2 | TRBJ1-4 | CASESGGGAEKLFF | KLGGALQAK | YES | NO | NO |
| 12868 | TRAV1-2 | TRAJ33 | CAVMDSNYQLIW | TRBV20-1 | TRBJ2-2 | CSAQGGLESGELFF | KLGGALQAK | YES | NO | NO |
| 12869 | TRAV27 | TRAJ42 | CAMNYGGSQGNLIF | TRBV19 | TRBJ2-7 | CASSIRSSHEQYF | GILGFVFTL | YES | NO | YES |
| 12870 | TRAV13-1 | TRAJ49 | CAAGSANTGNQFYF | TRBV12-3 | TRBJ2-2 | CASSFSGNTGELFF | RAKFKQLL | YES | YES | NO |
| 12871 | TRAV26-1 | TRAJ34 | CIVRVNTDKLIF | TRBV7-9 | TRBJ2-1 | CASSSAGPYTRRTRQGGQFF | KLGGALQAK | NO | NO | NO |
| 12872 | TRAV16 | TRAJ28 | CARSGAGSYQLTF | TRBV18 | TRBJ1-1 | CASSPTGPEAFF | KLGGALQAK | NO | NO | NO |
| 12873 | TRAV13-1 | TRAJ50 | CAAAETSYDKVIF | TRBV7-6 | TRBJ2-7 | CASSLKGGRHEQYF | RLRAEAQVK | YES | YES | YES |
| 12874 | TRAV13-1 | TRAJ11 | CAAKEGYSTLTF | TRBV4-1 | TRBJ2-5 | CASSDLAGYQETQYF | RAKFKQLL | YES | NO | YES |
| 12875 | TRAV13-1 | TRAJ33 | CAAIQDFNYQLIW | TRBV7-9 | TRBJ2-7 | CASSLSSYEQYF | KLGGALQAK | YES | NO | NO |
| 12876 | TRAV8-1 | TRAJ37 | CAVVHGSSNTGKLIF | TRBV19 | TRBJ2-7 | CASSIRSSYEQYF | GILGFVFTL | YES | NO | YES |
| 12877 | TRAV2 | TRAJ21 | CAVEDLDNFNKFYF | TRBV4-2 | TRBJ2-6 | CASSQTSSSGANVLTF | AVFDRKSDAK | NO | NO | NO |
| 12878 | TRAV21 | TRAJ28 | CAVDPPPYSGAGSYQLTF | TRBV7-9 | TRBJ1-1 | CASSLPEGALNTEAFF | KLGGALQAK | YES | NO | NO |
| 12879 | TRAV13-1 | TRAJ50 | CAASETSYDKVIF | TRBV3-1 | TRBJ2-2 | CASSYSGGAGELFF | RAKFKQLL | YES | NO | YES |
| 12880 | TRAV38-2DV8 | TRAJ48 | CAYSVTINFGNEKLTF | TRBV9 | TRBJ1-5 | CASSEGKGRDSNQPQHF | KLGGALQAK | YES | NO | NO |
| 12881 | TRAV21 | TRAJ45 | CAVRAYSGGGADGLTF | TRBV18 | TRBJ2-5 | CASSPFQREETQYF | KLGGALQAK | YES | NO | NO |
| 12882 | TRAV14DV4 | TRAJ21 | CAMNNFNKFYF | TRBV20-1 | TRBJ1-5 | CSARAYGDTSKHSNQPQHF | KLGGALQAK | YES | NO | NO |
| 12883 | TRAV9-2 | TRAJ33 | CAREHMDSNYQLIW | TRBV4-2 | TRBJ2-2 | CASSQRPSEVGELFF | RAKFKQLL | YES | NO | YES |
| 12884 | TRAV24 | TRAJ10 | CAPAGGGGNKLTF | TRBV7-9 | TRBJ2-3 | CASSLAQASTDTQYF | KLGGALQAK | YES | NO | NO |
| 12885 | TRAV12-1 | TRAJ44 | CVVHTGTASKLTF | TRBV27 | TRBJ1-4 | CASSLYSPLAYSWNEKLFF | KLGGALQAK | YES | NO | NO |
| 12886 | TRAV27 | TRAJ42 | CAGGGGGSQGNLIF | TRBV19 | TRBJ1-5 | CASSISGANQPQHF | GILGFVFTL | YES | NO | YES |
| 12887 | TRAV14DV4 | TRAJ9 | CAMREGENGGFKTIF | TRBV7-9 | TRBJ1-4 | CASSLVGGTDEKLFF | RAKFKQLL | YES | NO | NO |
| 12888 | TRAV20 | TRAJ28 | CAVSGAGSYQLTF | TRBV5-4 | TRBJ2-7 | CASSLGAGDTLEQYF | KLGGALQAK | YES | NO | NO |
| 12889 | TRAV17 | TRAJ24 | CATADSWGKFQF | TRBV5-5 | TRBJ2-2 | CASSKLRTGYTGELFF | KLGGALQAK | YES | NO | NO |
| 12890 | TRAV13-1 | TRAJ50 | CAASETSYDKVIF | TRBV12-3 | TRBJ1-5 | CASSLAHNQPQHF | RAKFKQLL | YES | NO | NO |
| 12891 | TRAV13-1 | TRAJ50 | CAARETSYDKVIF | TRBV12-3 | TRBJ2-2 | CASSFSLDKPTNTGELFF | RAKFKQLL | YES | NO | YES |
| 12892 | TRAV30 | TRAJ30 | CGTDRDDKIIF | TRBV10-2 | TRBJ1-1 | CASIFARGEAFF | KLGGALQAK | YES | NO | NO |
| 12893 | TRAV12-1 | TRAJ12 | CVVNPVDSSYKLIF | TRBV12-4 | TRBJ2-1 | CASSLSGGSYNEQFF | KLGGALQAK | YES | NO | NO |
| 12894 | TRAV12-2 | TRAJ47 | CAVKEYGNKLVF | TRBV4-1 | TRBJ2-3 | CASSPSELADTQYF | ELAGIGILTV | YES | YES | YES |
| 12895 | TRAV5 | TRAJ42 | CAGGGSGGSQGNLIF | TRBV19 | TRBJ1-1 | CASSWRSDTEAFF | GILGFVFTL | YES | YES | YES |
| 12896 | TRAV13-1 | TRAJ50 | CAASETSYDKVIF | TRBV7-9 | TRBJ2-1 | CASSVTGPGDNEQFF | RAKFKQLL | YES | NO | YES |
| 12897 | TRAV35 | TRAJ49 | CAGPPNTGNQFYF | TRBV19 | TRBJ2-2 | CASSIYGFTGELFF | KLGGALQAK | YES | NO | NO |
| 12898 | TRAV19 | TRAJ6 | CALSEVSGGSYIPTF | TRBV5-1 | TRBJ2-7 | CASRGLLSNEQYF | KLGGALQAK | YES | NO | NO |
| 12899 | TRAV13-1 | TRAJ50 | CAASETSYDKVIF | TRBV12-4 | TRBJ2-2 | CASSFSGNTGELFF | RAKFKQLL | YES | NO | YES |
| 12900 | TRAV5 | TRAJ48 | CAERIPNFGNEKLTF | TRBV19 | TRBJ2-7 | CASSIRSSYEQYF | GILGFVFTL | YES | NO | NO |
| 12901 | TRAV8-6 | TRAJ23 | CAVSGLNNQGGKLIF | TRBV19 | TRBJ2-7 | CASSIRSSYEQYF | GILGFVFTL | YES | NO | YES |
| 12902 | TRAV12-1 | TRAJ17 | CVVMIKAAGNKLTF | TRBV28 | TRBJ1-2 | CASSLDSSYGYTF | AVFDRKSDAK | YES | NO | YES |
| 12903 | TRAV17 | TRAJ8 | CATDGSTGFQKLVF | TRBV15 | TRBJ1-3 | CATRSQGSYFSGNTIYF | KLGGALQAK | YES | NO | NO |
| 12904 | TRAV12-3 | TRAJ42 | CAMSGDGGSQGNLIF | TRBV19 | TRBJ1-5 | CASSHRSNQPQHF | GILGFVFTL | YES | NO | YES |
| 12905 | TRAV24 | TRAJ36 | CAFLGGANNLFF | TRBV19 | TRBJ2-3 | CASSTRSTDTQYF | GILGFVFTL | YES | NO | YES |
| 12906 | TRAV9-2 | TRAJ42 | CALYYGGSQGNLIF | TRBV27 | TRBJ2-1 | CASRQGAPLNEQFF | KLGGALQAK | YES | NO | NO |
| 12907 | TRAV17 | TRAJ53 | CATDAGSNYKLTF | TRBV6-3 | TRBJ1-2 | CASSDTGPANGYTF | KLGGALQAK | YES | NO | NO |
| 12908 | TRAV12-2 | TRAJ49 | CAGTAGNQFYF | TRBV6-6 | TRBJ2-2 | CASLRDGELFF | KLGGALQAK | YES | NO | NO |
| 12909 | TRAV9-2 | TRAJ6 | CALSDWESGGSYIPTF | TRBV7-6 | TRBJ1-4 | CASSQRPKGSEKLFF | AVFDRKSDAK | YES | NO | YES |
| 12910 | TRAV40 | TRAJ56 | CLLGASGANSKLTF | TRBV20-1 | TRBJ2-1 | CSATSAGGSYNEQFF | KLGGALQAK | NO | NO | NO |
| 12911 | TRAV8-3 | TRAJ16 | CAVVAF | TRBV6-4 | TRBJ2-1 | CASSDSGSEQFF | KLGGALQAK | NO | NO | NO |
| 12912 | TRAV25 | TRAJ45 | CAGRGGGADGLTF | TRBV6-6 | TRBJ2-1 | CAVGLILEQFF | KLGGALQAK | NO | NO | NO |
| 12913 | TRAV3 | TRAJ49 | CAVRDTRYTGNQFYF | TRBV30 | TRBJ1-1 | CAWSDPGGLPWAFF | KLGGALQAK | NO | NO | NO |
| 12914 | TRAV13-1 | TRAJ35 | CAARVRGFGNVLHC | TRBV5-4 | TRBJ1-5 | CASSSTGGGEKDQPQHF | KLGGALQAK | NO | NO | NO |
| 12915 | TRAV27 | TRAJ26 | CAGLTDGQNFVF | TRBV12-4 | TRBJ1-2 | CASSPDSNGYTF | KLGGALQAK | YES | NO | NO |
| 12916 | TRAV13-2 | TRAJ45 | CADPGGGADGLTF | TRBV5-6 | TRBJ1-1 | CASSLRDGSEAFF | RAKFKQLL | YES | NO | YES |
| 12917 | TRAV41 | TRAJ33 | CAVEHMDSNYQLIW | TRBV4-2 | TRBJ2-7 | CASSQVLGGGLNEQYF | RAKFKQLL | YES | NO | YES |
| 12918 | TRAV9-2 | TRAJ20 | CALSPNDYKLSF | TRBV28 | TRBJ2-5 | CASSLQGTQETQYF | KLGGALQAK | YES | NO | NO |
| 12919 | TRAV5 | TRAJ6 | CAEMEGGSYIPTF | TRBV6-1 | TRBJ2-3 | CASSITEGEDSVSTDTQYF | RAKFKQLL | YES | NO | YES |
| 12920 | TRAV25 | TRAJ49 | CAGGDTGNQFYF | TRBV5-4 | TRBJ1-5 | CASSSTGGGEKDQPQHF | KLGGALQAK | YES | NO | NO |
| 12921 | TRAV19 | TRAJ30 | CALSVNRDDKIIF | TRBV27 | TRBJ2-1 | CASSPGQIYNEQFF | KLGGALQAK | YES | NO | NO |
| 12922 | TRAV13-1 | TRAJ11 | CAASRGYSTLTF | TRBV5-4 | TRBJ1-5 | CASSSTGGGEKDQPQHF | KLGGALQAK | YES | NO | NO |
| 12923 | TRAV30 | TRAJ20 | CGTEINDYKLSF | TRBV20-1 | TRBJ1-1 | CSAWTGTEAFF | GLCTLVAML | YES | YES | YES |
| 12924 | TRAV13-1 | TRAJ50 | CAASETSYDKVIF | TRBV7-9 | TRBJ2-1 | CASSSGLAGVNEQFF | KLGGALQAK | YES | NO | NO |
| 12925 | TRAV24 | TRAJ18 | CAFMVRGSTLGRLYF | TRBV28 | TRBJ2-3 | CASSFLAGTDTQYF | KLGGALQAK | YES | NO | NO |
| 12926 | TRAV13-1 | TRAJ50 | CAATETSYDKVIF | TRBV12-3 | TRBJ1-6 | CASSYTGSSNSPLHF | RAKFKQLL | YES | NO | YES |
| 12927 | TRAV5 | TRAJ50 | CAESETSYDKVIF | TRBV5-1 | TRBJ1-1 | CASRAVEGKRGAEAFF | RAKFKQLL | YES | NO | YES |
| 12928 | TRAV29DV5 | TRAJ40 | CAARWIGSGTYKYIF | TRBV5-6 | TRBJ2-5 | CASSLARTDKETQYF | RAKFKQLL | YES | NO | YES |
| 12929 | TRAV1-2 | TRAJ26 | CAVRDHYGQNFVF | TRBV7-9 | TRBJ2-1 | CASSSGLAGGNEQFF | KLGGALQAK | YES | NO | NO |
| 12930 | TRAV13-2 | TRAJ45 | CADSGGGADGLTF | TRBV4-1 | TRBJ1-5 | CAAGSSGNQPQHF | RAKFKQLL | YES | NO | NO |
| 12931 | TRAV1-2 | TRAJ33 | CARMDSNYQLIW | TRBV6-3 | TRBJ2-7 | CASSASPDRWGEQYF | KLGGALQAK | YES | NO | NO |
| 12932 | TRAV13-1 | TRAJ20 | CAASGYDYKLSF | TRBV5-6 | TRBJ1-1 | CASSLRDGSEAFF | RAKFKQLL | YES | NO | YES |
| 12933 | TRAV8-2 | TRAJ4 | CVVSEAGGYNKLIF | TRBV5-1 | TRBJ1-1 | CASSLGSGWEAFF | KLGGALQAK | YES | NO | NO |
| 12934 | TRAV13-2 | TRAJ52 | CAERVGAGGTSYGKLTF | TRBV6-3 | TRBJ2-2 | CASSYGFGGHNTGELFF | KLGGALQAK | YES | NO | NO |
| 12935 | TRAV25 | TRAJ42 | CAGNYGGSQGNLIF | TRBV5-6 | TRBJ1-1 | CASSLRDGSEAFF | RAKFKQLL | YES | NO | YES |
| 12936 | TRAV8-3 | TRAJ50 | CAVGEDKVIF | TRBV4-1 | TRBJ1-1 | CASSHWQTNTEAFF | RAKFKQLL | YES | NO | YES |
| 12937 | TRAV1-2 | TRAJ33 | CAALDSNYQLIW | TRBV7-9 | TRBJ1-5 | CASSLGGDPGVGPQHF | IVTDFSVIK | YES | NO | YES |
| 12938 | TRAV1-2 | TRAJ33 | CAVRDSNYQLIW | TRBV20-1 | TRBJ2-6 | CSASPGAPSGANVLTF | IVTDFSVIK | YES | YES | YES |
| 12939 | TRAV12-2 | TRAJ15 | CAVNQAGTALIF | TRBV20-1 | TRBJ2-1 | CSAFARDRVYEQFF | KLGGALQAK | NO | NO | NO |
| 12940 | TRAV26-2 | TRAJ32 | CILRGNYGGATNKLIF | TRBV6-1 | TRBJ1-5 | CASRITGESDQPQHF | RAKFKQLL | YES | NO | NO |
| 12941 | TRAV23DV6 | TRAJ34 | CAATESYNTDKLIF | TRBV5-4 | TRBJ1-1 | CASSWNGQGKDSEAFF | AVFDRKSDAK | YES | NO | YES |

| | | | | | | | | | | |
|---|---|---|---|---|---|---|---|---|---|---|
| 12942 | TRAV8-1 | TRAJ42 | CAVDADYGGSQGNLIF | TRBV19 | TRBJ2-7 | CASSIRASVEQYF | GILGFVFTL | YES | YES | YES |
| 12943 | TRAV41 | TRAJ33 | CAASMDSNYQLIW | TRBV10-3 | TRBJ1-2 | CAISELDRGHGYTF | IVTDFSVIK | YES | NO | YES |
| 12944 | TRAV13-1 | TRAJ42 | CAASMGGGSQGNLIF | TRBV19 | TRBJ1-2 | CASSQGAYGYTF | GILGFVFTL | YES | NO | YES |
| 12945 | TRAV13-1 | TRAJ50 | CAAMETSYDKVIF | TRBV29-1 | TRBJ2-1 | CSVGSGDHGEQFF | RAKFKQLL | YES | YES | YES |
| 12946 | TRAV5 | TRAJ27 | CAEWDAGKSTF | TRBV11-2 | TRBJ2-2 | CASSLGVAGTNTGELFF | KLGGALQAK | YES | NO | NO |
| 12947 | TRAV27 | TRAJ33 | CAGVMDSNYQLIW | TRBV6-6 | TRBJ1-1 | CASTPGRAGSGVAFF | KLGGALQAK | NO | NO | NO |
| 12948 | TRAV8-6 | TRAJ47 | CAVSDMEYGNKLVF | TRBV7-3 | TRBJ2-1 | CASSLIVAGGYNEQFF | RAKFKQLL | YES | NO | YES |
| 12949 | TRAV13-1 | TRAJ50 | CAASETSYDKVIF | TRBV12-3 | TRBJ1-2 | CASRPTGDYGYTF | RAKFKQLL | YES | NO | YES |
| 12950 | TRAV13-1 | TRAJ50 | CAAVETSYDKVIF | TRBV7-9 | TRBJ1-6 | CASSPIGEVSPLHF | RAKFKQLL | YES | YES | NO |
| 12951 | TRAV1-2 | TRAJ33 | CAAMDSNYQLIW | TRBV6-1 | TRBJ1-3 | CASSEATGSGNTIYF | KLGGALQAK | YES | NO | NO |
| 12952 | TRAV13-2 | TRAJ29 | CAENKREAGNTPLVF | TRBV19 | TRBJ2-7 | CASSIRSSYEQYF | GILGFVFTL | YES | NO | YES |
| 12953 | TRAV21 | TRAJ12 | CAVRMDSSYKLIF | TRBV29-1 | TRBJ2-1 | CSVGAGEGEQFF | RAKFKQLL | YES | YES | YES |
| 12954 | TRAV23DV6 | TRAJ28 | CAAHRVDSGAGSYQLTF | TRBV7-9 | TRBJ2-3 | CASSSRDGGTDTQYF | KLGGALQAK | YES | NO | NO |
| 12955 | TRAV39 | TRAJ42 | CAVDEGGGSQGNLIF | TRBV6-1 | TRBJ1-1 | CASSEGTVTEAFF | GILGFVFTL | NO | NO | NO |
| 12956 | TRAV23DV6 | TRAJ43 | CAASGNHDMRL | TRBV6-6 | TRBJ1-1 | CASSTTAGDTEAFF | RAKFKQLL | NO | NO | NO |
| 12957 | TRAV19 | TRAJ30 | CALSEPTRDDKIIF | TRBV6-1 | TRBJ2-5 | CASSEEPGGETQYF | KLGGALQAK | YES | YES | NO |
| 12958 | TRAV8-1 | TRAJ36 | CAVQPGANNLFF | TRBV29-1 | TRBJ1-2 | CSVELIGYTF | KLGGALQAK | YES | NO | NO |
| 12959 | TRAV1-2 | TRAJ23 | CAVKEVNNQGGKLIF | TRBV2 | TRBJ2-3 | CASRIRTSVADTQYF | KLGGALQAK | YES | NO | NO |
| 12960 | TRAV27 | TRAJ42 | CAGGGSQGNLIF | TRBV4-1 | TRBJ2-1 | CASSLLWAEYNEQFF | RAKFKQLL | YES | YES | NO |
| 12961 | TRAV35 | TRAJ54 | CAGQLGAQKLVF | TRBV18 | TRBJ1-3 | CASLQYPGNTIYF | KLGGALQAK | NO | NO | NO |
| 12962 | TRAV19 | TRAJ9 | CALSEANTGGFKTIF | TRBV29-1 | TRBJ2-7 | CSVGTGDWGEQYF | RAKFKQLL | YES | NO | NO |
| 12963 | TRAV38-2DV8 | TRAJ33 | CALVDSNYQLIW | TRBV4-1 | TRBJ1-2 | CASSQDGRGGYGYTF | KLGGALQAK | YES | NO | NO |
| 12964 | TRAV19 | TRAJ16 | CALSESDGQKLLF | TRBV4-2 | TRBJ2-2 | CASSQRPSEVGELFF | RAKFKQLL | NO | NO | NO |
| 12965 | TRAV8-1 | TRAJ40 | CAAYFSGTYKYIF | TRBV2 | TRBJ2-7 | CASSFNLADSYEQYF | KLGGALQAK | YES | NO | NO |
| 12966 | TRAV14DV4 | TRAJ27 | CAMREGWPNTNAGKSTF | TRBV19 | TRBJ2-5 | CASSIRAAETQYF | GILGFVFTL | YES | NO | YES |
| 12967 | TRAV9-2 | TRAJ28 | CALSEGGGILWGWELPTHF | TRBV29-1 | TRBJ1-1 | CSVEGRSGLPNTEAFF | AVFDRKSDAK | NO | NO | NO |
| 12968 | TRAV26-2 | TRAJ30 | CILSELNRDDKIIF | TRBV27 | TRBJ2-6 | CASTTHLVVGGPGANVLTF | KLGGALQAK | NO | NO | NO |
| 12969 | TRAV12-1 | TRAJ37 | CVVISNTGKLIF | TRBV7-9 | TRBJ2-2 | CAIPGRATGELFF | KLGGALQAK | YES | NO | NO |
| 12970 | TRAV29DV5 | TRAJ15 | CAADQAGTALIF | TRBV7-9 | TRBJ1-2 | CASSQTGGPLGGYTF | KLGGALQAK | YES | NO | NO |
| 12971 | TRAV19 | TRAJ42 | CALSALMNYGGSQGNLIF | TRBV4-1 | TRBJ2-3 | CASSQDPAGALTDTQYF | KLGGALQAK | YES | NO | NO |
| 12972 | TRAV36DV7 | TRAJ43 | CAVEWGGRF | TRBV7-9 | TRBJ1-3 | CATELSPGTVGNTIYF | KLGGALQAK | NO | NO | NO |
| 12973 | TRAV12-2 | TRAJ10 | CALVTGGGNKLTF | TRBV25-1 | TRBJ2-1 | CASRSLQTFF | KLGGALQAK | YES | NO | NO |
| 12974 | TRAV1-2 | TRAJ29 | CAVRVPLENTPLVF | TRBV12-3 | TRBJ2-3 | CASSFGVSGGTDTQYF | KLGGALQAK | YES | NO | NO |
| 12975 | TRAV8-3 | TRAJ20 | CAVGASNDYKLSF | TRBV7-9 | TRBJ2-6 | CASSPVTGGGSGANVLTF | KLGGALQAK | YES | NO | NO |
| 12976 | TRAV12-2 | TRAJ29 | CAPSSGGNTPLVF | TRBV29-1 | TRBJ1-2 | CSVEGAGQGQLLYGYTF | KLGGALQAK | YES | NO | NO |
| 12977 | TRAV12-2 | TRAJ12 | CASVGMDSSYKLIF | TRBV19 | TRBJ1-1 | CASSTRSGTEAFF | GILGFVFTL | YES | NO | YES |
| 12978 | TRAV21 | TRAJ32 | CAINGGATNKLIF | TRBV5-6 | TRBJ2-7 | CASSPDGSYEQYF | AVFDRKSDAK | YES | NO | YES |
| 12979 | TRAV12-3 | TRAJ42 | CAMSGEGGSQGNLIF | TRBV19 | TRBJ1-5 | CASSTRSNQPQHF | GILGFVFTL | YES | NO | YES |
| 12980 | TRAV13-2 | TRAJ45 | CADSGGGADGLTF | TRBV27 | TRBJ2-7 | CASSLSGRPYEQYF | RAKFKQLL | YES | YES | NO |
| 12981 | TRAV12-2 | TRAJ15 | CAVRVNQAGTALIF | TRBV21-1 | TRBJ2-1 | CASSRGGLSYNEQFF | GILGFVFTL | YES | YES | YES |
| 12982 | TRAV21 | TRAJ47 | CAASPMEYGNKLVF | TRBV5-1 | TRBJ2-1 | CASSLPLAGAYNEQFF | AYAQKIFKI | YES | YES | YES |
| 12983 | TRAV12-3 | TRAJ57 | CAMSQGGSEKLVF | TRBV3-1 | TRBJ2-1 | CASSHLGGDRYMNEQFF | KLGGALQAK | YES | NO | NO |
| 12984 | TRAV3 | TRAJ42 | CAGPGSQGNLIF | TRBV19 | TRBJ2-7 | CASSVRSSYEQYF | GILGFVFTL | NO | NO | NO |
| 12985 | TRAV1-2 | TRAJ12 | CAVIRGMDSSYKLIF | TRBV19 | TRBJ2-7 | CASSIVGTGTYEQYF | GILGFVFTL | YES | YES | YES |
| 12986 | TRAV1-1 | TRAJ13 | CAVRGVRESGGYQKVTF | TRBV27 | TRBJ2-7 | CASTGTGGEHEQYF | KLGGALQAK | YES | NO | NO |
| 12987 | TRAV8-6 | TRAJ54 | CAVSDQEGAQKLVF | TRBV7-3 | TRBJ1-1 | CASSLIALGNTEAFF | RAKFKQLL | YES | NO | YES |
| 12988 | TRAV19 | TRAJ31 | CALSGVNNARLMF | TRBV12-3 | TRBJ2-3 | CASSSQGDTQYF | KLGGALQAK | YES | NO | NO |
| 12989 | TRAV1-2 | TRAJ9 | CAVRDRTGGFKTIF | TRBV29-1 | TRBJ2-1 | CSVEDRRPDEQFF | KLGGALQAK | YES | NO | NO |
| 12990 | TRAV5 | TRAJ33 | CAETQDSNYQLIW | TRBV4-1 | TRBJ1-5 | CASSQDGWAGDQPQHF | RAKFKQLL | YES | NO | NO |
| 12991 | TRAV8-3 | TRAJ56 | CAVLGGANSKLTF | TRBV19 | TRBJ2-1 | CASSFLATDEQFF | GILGFVFTL | YES | NO | YES |
| 12992 | TRAV13-1 | TRAJ47 | CAAGDGNKLVF | TRBV12-4 | TRBJ2-2 | CASSQGTGELFF | KLGGALQAK | YES | NO | NO |
| 12993 | TRAV41 | TRAJ49 | CAVLTGNQFYF | TRBV30 | TRBJ2-5 | CAWSLTTPETQYF | KLGGALQAK | NO | NO | NO |
| 12994 | TRAV25 | TRAJ42 | CAGTHGGSQGNLIF | TRBV19 | TRBJ1-6 | CASSPRSDSPLHF | GILGFVFTL | YES | NO | YES |
| 12995 | TRAV19 | TRAJ56 | CALSEVNTGANSKLTF | TRBV6-5 | TRBJ2-1 | CASSFSTSGLIDEQFF | KLGGALQAK | NO | NO | NO |
| 12996 | TRAV27 | TRAJ42 | CAGAGSQGNLIF | TRBV4-1 | TRBJ2-1 | CASSLLWAEYNEQFF | RAKFKQLL | YES | YES | YES |
| 12997 | TRAV13-2 | TRAJ42 | CAEKNYGGSQGNLIF | TRBV29-1 | TRBJ1-2 | CSADGGGGYTF | KLGGALQAK | YES | NO | NO |
| 12998 | TRAV38-2DV8 | TRAJ41 | CAYKRANSGYALNF | TRBV4-1 | TRBJ2-7 | CASSLEVPSYEQYF | KLGGALQAK | YES | NO | NO |
| 12999 | TRAV27 | TRAJ42 | CAGSGSQGNLIF | TRBV19 | TRBJ1-2 | CASSIGVHGYTF | GILGFVFTL | YES | NO | YES |
| 13000 | TRAV5 | TRAJ34 | CAESEGPTYTDKLIF | TRBV7-2 | TRBJ2-7 | CASSPRDRERGEQYF | RAKFKQLL | NO | NO | NO |
| 13001 | TRAV19 | TRAJ26 | CALHNYGQNFVF | TRBV11-2 | TRBJ1-1 | CASSSGPTEAFF | IVTDFSVIK | YES | YES | YES |
| 13002 | TRAV8-6 | TRAJ42 | CAVTDGGSQGNLIF | TRBV19 | TRBJ1-2 | CASSQGAYGYTF | GILGFVFTL | YES | NO | YES |
| 13003 | TRAV8-3 | TRAJ7 | CAVFLYGNNRLAF | TRBV4-1 | TRBJ2-1 | CASSLLWAEYNEQFF | RAKFKQLL | YES | NO | YES |
| 13004 | TRAV19 | TRAJ40 | CALSEGTYKYIF | TRBV28 | TRBJ2-2 | CASSSRGRRDPGELFF | KLGGALQAK | YES | NO | NO |
| 13005 | TRAV5 | TRAJ20 | CAEIRANDYKLSF | TRBV5-1 | TRBJ1-2 | CASSLGQRYTF | KLGGALQAK | YES | NO | NO |
| 13006 | TRAV12-3 | TRAJ9 | CARGGGFKTIF | TRBV7-9 | TRBJ2-3 | CASSSRDGGTDTQYF | KLGGALQAK | YES | NO | NO |
| 13007 | TRAV23DV6 | TRAJ34 | CAASEAYNTDKLIF | TRBV5-4 | TRBJ2-3 | CASSFNGGAIDTQYF | RLRAEAQVK | YES | YES | YES |
| 13008 | TRAV1-2 | TRAJ4 | CAVTSGGYNKLIF | TRBV5-1 | TRBJ2-7 | CASSFFTRGGEDEQYF | RLRAEAQVK | YES | YES | YES |
| 13009 | TRAV1-2 | TRAJ10 | CAVRAFTGGGNKLTF | TRBV7-8 | TRBJ2-4 | CASSLEGLAGAGIQYF | KLGGALQAK | YES | NO | NO |
| 13010 | TRAV12-1 | TRAJ31 | CVVNRDNARLMF | TRBV29-1 | TRBJ2-7 | CSAGQGPYEQYF | RAKFKQLL | NO | NO | NO |
| 13011 | TRAV8-4 | TRAJ43 | CAVSVNDMRF | TRBV7-8 | TRBJ1-2 | CASSSELTGTVYGYTF | KLGGALQAK | YES | NO | NO |
| 13012 | TRAV12-3 | TRAJ11 | CAMNIMNSGYSTLTF | TRBV5-1 | TRBJ2-5 | CASSYFSGGETQYF | KLGGALQAK | YES | NO | NO |
| 13013 | TRAV13-1 | TRAJ20 | CAASEGLNDYKLSF | TRBV3-1 | TRBJ1-1 | CASSQAMGTEAFF | RAKFKQLL | YES | NO | YES |
| 13014 | TRAV17 | TRAJ6 | CATEGASGGSYIPTF | TRBV27 | TRBJ2-7 | CASSLSGRPYEQYF | RAKFKQLL | YES | NO | YES |
| 13015 | TRAV35 | TRAJ28 | CAGQLRELVLSATGAGSYQLTF | TRBV2 | TRBJ2-1 | CASLGQAYNEQFF | IVTDFSVIK | NO | NO | NO |
| 13016 | TRAV27 | TRAJ42 | CAGGGSQGNLIF | TRBV14 | TRBJ2-3 | CASSQSPGGTQYF | GILGFVFTL | YES | NO | YES |
| 13017 | TRAV8-3 | TRAJ31 | CAVGTARLMF | TRBV27 | TRBJ1-6 | CASSLPGTGGSPLHF | RAKFKQLL | YES | NO | YES |
| 13018 | TRAV38-1 | TRAJ52 | CAFMINAGGTSYGKLTF | TRBV4-1 | TRBJ1-2 | CASSTSTGTGYGYTF | RAKFKQLL | YES | NO | YES |
| 13019 | TRAV19 | TRAJ39 | CALSAYNAGNMLTF | TRBV11-2 | TRBJ2-1 | CASSLAGGPSYNEQFF | KLGGALQAK | YES | NO | NO |
| 13020 | TRAV13-2 | TRAJ45 | CADSGGGADGLTF | TRBV29-1 | TRBJ2-3 | CSVGSGEDSPQYF | RAKFKQLL | YES | NO | NO |
| 13021 | TRAV12-2 | TRAJ57 | CAVNDRGSEKLVF | TRBV19 | TRBJ2-7 | CASSARSSYEQYF | GILGFVFTL | YES | NO | YES |
| 13022 | TRAV8-4 | TRAJ45 | CAVRGRSGGGADGLTF | TRBV7-9 | TRBJ2-3 | CASSLVRPPTDTQYF | KLGGALQAK | YES | NO | NO |
| 13023 | TRAV12-1 | TRAJ17 | CVSAAGNKLTF | TRBV6-4 | TRBJ2-1 | CASKDKEGTGYNEQFF | AVFDRKSDAK | YES | NO | YES |
| 13024 | TRAV26-2 | TRAJ50 | CILRETSYDKVIF | TRBV12-3 | TRBJ2-2 | CASTDVRRVDGELFF | RAKFKQLL | YES | NO | YES |
| 13025 | TRAV13-1 | TRAJ50 | CAASETSYDKVIF | TRBV27 | TRBJ1-1 | CASSPIPVVTEAFF | RAKFKQLL | YES | NO | YES |
| 13026 | TRAV26-1 | TRAJ20 | CIVRASGDYKLSF | TRBV19 | TRBJ2-1 | CASSPRSGVEQFF | GILGFVFTL | YES | NO | YES |
| 13027 | TRAV3 | TRAJ36 | CAVRDGTGANNLFF | TRBV19 | TRBJ2-4 | CASAGTGVNIQYF | GILGFVFTL | NO | NO | NO |
| 13028 | TRAV27 | TRAJ37 | CAGAHGSGNTGKLIF | TRBV4-1 | TRBJ1-3 | CASSQTRGAGNTIYF | RAKFKQLL | YES | NO | YES |
| 13029 | TRAV8-3 | TRAJ27 | CAYNTNAGKSTF | TRBV19 | TRBJ1-6 | CATDMGGIRGVSPLHF | AVFDRKSDAK | YES | NO | NO |
| 13030 | TRAV8-1 | TRAJ10 | CAALTGGGNKLTF | TRBV6-1 | TRBJ2-2 | CATQDGPGLGELFF | KLGGALQAK | YES | NO | NO |
| 13031 | TRAV9-2 | TRAJ50 | CALRETSYDKVIF | TRBV18 | TRBJ1-4 | CASSPRDRGGSNEKLFF | RAKFKQLL | YES | NO | YES |
| 13032 | TRAV1-2 | TRAJ20 | CAVRDQDYKLSF | TRBV6-1 | TRBJ2-3 | CASSESQGGTDTQYF | KLGGALQAK | YES | NO | NO |
| 13033 | TRAV35 | TRAJ41 | CAGQTNSGYALNF | TRBV20-1 | TRBJ2-5 | CSARDDSGQGFGFLETQYF | GLCTLVAML | NO | NO | NO |
| 13034 | TRAV27 | TRAJ37 | CAGAGASNTGKLIF | TRBV19 | TRBJ2-7 | CASSIRSSYEQYF | GILGFVFTL | YES | NO | YES |
| 13035 | TRAV25 | TRAJ42 | CAGTYGGSQGNLIF | TRBV19 | TRBJ2-3 | CAASSRSTDTQYF | GILGFVFTL | YES | NO | YES |
| 13036 | TRAV13-1 | TRAJ53 | CAARHRSGSNYKLTF | TRBV20-1 | TRBJ1-1 | CSARALLRDTEAFF | AVFDRKSDAK | YES | YES | YES |
| 13037 | TRAV22 | TRAJ45 | CAPDGRMYSGGGADGLTF | TRBV27 | TRBJ2-3 | CASSTTRMDTQYF | AVFDRKSDAK | YES | NO | YES |
| 13038 | TRAV21 | TRAJ26 | CAVHGYGQNFVF | TRBV29-1 | TRBJ2-1 | CSVGSGDHGEQFF | RAKFKQLL | YES | NO | YES |
| 13039 | TRAV35 | TRAJ42 | CAGGGYGGSQGNLIF | TRBV19 | TRBJ1-1 | CASSIRSDTEAFF | GILGFVFTL | YES | NO | YES |
| 13040 | TRAV12-3 | TRAJ42 | CAMGGGGGSQGNLIF | TRBV19 | TRBJ2-5 | CASSFRSSETQYF | GILGFVFTL | YES | YES | NO |
| 13041 | TRAV29DV5 | TRAJ53 | CAASVVSGGSNYKLTF | TRBV27 | TRBJ2-7 | CASSSLPTSFYEQYF | KLGGALQAK | YES | NO | NO |
| 13042 | TRAV14DV4 | TRAJ52 | CAMRRPNAGGTSYGKLTF | TRBV4-1 | TRBJ1-2 | CASSTSTGTGYGYTF | RAKFKQLL | YES | NO | YES |
| 13043 | TRAV1-2 | TRAJ13 | CASDSGGYQKVTF | TRBV20-1 | TRBJ2-3 | CSAQPGLADTDTQYF | KLGGALQAK | NO | NO | NO |
| 13044 | TRAV17 | TRAJ52 | CATGARANAGGTSYGKLTF | TRBV7-7 | TRBJ2-2 | CASSPTSGGFRELFF | AVFDRKSDAK | YES | NO | YES |
| 13045 | TRAV10 | TRAJ50 | CVVSAGLGKTSYDKVIF | TRBV12-3 | TRBJ2-7 | CASSSLDVAGYEQYF | RAKFKQLL | YES | NO | YES |

| | | | | | | | | | | |
|---|---|---|---|---|---|---|---|---|---|---|
| 13046 | TRAV30 | TRAJ37 | CGTEPGNTGKLIF | TRBV19 | TRBJ2-3 | CASSIRSTDTQYF | GILGFVFTL | YES | YES | YES |
| 13047 | TRAV13-2 | TRAJ53 | CAETHSGGSNYKLTF | TRBV5-6 | TRBJ1-1 | CASSLRDGSEAFF | RAKFKQLL | YES | NO | YES |
| 13049 | TRAV3 | TRAJ4 | CAVRDFSGGYNKLIF | TRBV2 | TRBJ2-1 | CASRDVYEQFF | KLGGALQAK | NO | NO | NO |
| 13050 | TRAV13-1 | TRAJ50 | CAAVETSYDKVIF | TRBV7-9 | TRBJ1-4 | CASSLEGTNEKLFF | RAKFKQLL | YES | NO | NO |
| 13051 | TRAV13-1 | TRAJ10 | CAAPTGGGNKLTF | TRBV7-8 | TRBJ1-2 | CASSFTGSSNYGYTF | KLGGALQAK | YES | NO | NO |
| 13052 | TRAV5 | TRAJ30 | CAESLWDDKIIF | TRBV4-1 | TRBJ2-3 | CASSQGAGIGRTDTQYF | RAKFKQLL | YES | NO | YES |
| 13053 | TRAV3 | TRAJ12 | CAVLWMDSSYKLIF | TRBV29-1 | TRBJ2-7 | CSVVWAEQYF | IVTDFSVIK | NO | NO | NO |
| 13054 | TRAV21 | TRAJ37 | CAVGPTGKLIF | TRBV7-9 | TRBJ2-5 | CASSEDRGYQETQYF | KLGGALQAK | NO | NO | NO |
| 13056 | TRAV26-2 | TRAJ49 | CIPADNTGNQFYF | TRBV19 | TRBJ2-3 | CASRIASRGGDTQYF | AVFDRKSDAK | YES | YES | YES |
| 13057 | TRAV26-2 | TRAJ50 | CILMETSYDKVIF | TRBV12-3 | TRBJ2-2 | CASSSSSTGELFF | RAKFKQLL | YES | NO | YES |
| 13058 | TRAV8-1 | TRAJ28 | CAAAGAGSYQLTF | TRBV6-1 | TRBJ2-7 | CASKSERGTYEQYF | KLGGALQAK | YES | NO | NO |
| 13059 | TRAV8-3 | TRAJ50 | CAVGAETSYDKVIF | TRBV4-2 | TRBJ2-2 | CASSQRPSEVGELFF | RAKFKQLL | YES | NO | YES |
| 13060 | TRAV8-4 | TRAJ26 | CAVSDDYGQNFVF | TRBV5-1 | TRBJ2-4 | CASRTSTGQGNIQYF | KLGGALQAK | YES | NO | NO |
| 13061 | TRAV12-2 | TRAJ43 | CAVKNNDMRF | TRBV4-1 | TRBJ2-2 | CASSQDGLLGELFF | ELAGIGILTV | YES | NO | YES |
| 13062 | TRAV12-2 | TRAJ30 | CAVNKRDDKIIF | TRBV20-1 | TRBJ2-7 | CSARDLGSTEIYEQYF | KLGGALQAK | NO | NO | NO |
| 13063 | TRAV8-3 | TRAJ30 | CAVTRDDKIIF | TRBV5-6 | TRBJ1-1 | CASSLRDGSEAFF | RAKFKQLL | YES | NO | YES |
| 13064 | TRAV27 | TRAJ42 | CAGAGGGSQGNLIF | TRBV19 | TRBJ2-3 | CASSIRSADTQYF | GILGFVFTL | YES | YES | YES |
| 13065 | TRAV38-1 | TRAJ52 | CAFMTNAGGTSYGKLTF | TRBV19 | TRBJ1-2 | CATSTGSYGYTF | GILGFVFTL | YES | NO | YES |
| 13066 | TRAV12-2 | TRAJ4 | CAVGAGGYNKLIF | TRBV11-3 | TRBJ2-2 | CASSLGASAGELFF | ELAGIGILTV | YES | YES | YES |
| 13067 | TRAV35 | TRAJ53 | CAGPPGSNYKLTF | TRBV6-6 | TRBJ2-7 | CASSGAGAGYEQYF | KLGGALQAK | NO | NO | NO |
| 13068 | TRAV8-4 | TRAJ17 | CAVSVVKAAGNKLTF | TRBV12-4 | TRBJ1-2 | CASSPGTNYGYTF | KLGGALQAK | YES | NO | NO |
| 13069 | TRAV1-2 | TRAJ33 | CAVMDSNYQLIW | TRBV20-1 | TRBJ2-7 | CSARAGTGLDEQYF | KLGGALQAK | NO | NO | NO |
| 13070 | TRAV29DV5 | TRAJ33 | CAASADSNYQLIW | TRBV21-1 | TRBJ2-4 | CAPLVAGGGIGTKNIQYF | KLGGALQAK | YES | NO | NO |
| 13071 | TRAV13-1 | TRAJ29 | CAASSPGNTPLVF | TRBV7-2 | TRBJ2-7 | CASSLVVPYEQYF | KLGGALQAK | NO | NO | NO |
| 13072 | TRAV24 | TRAJ28 | CALYAYSGAGSYQLTF | TRBV7-9 | TRBJ2-3 | CASSLVWGLSTDTQYF | KLGGALQAK | YES | NO | NO |
| 13073 | TRAV25 | TRAJ10 | CASFTGGGNKLTF | TRBV19 | TRBJ1-1 | CASSSTGGGVTEAFF | GILGFVFTL | YES | NO | YES |
| 13074 | TRAV17 | TRAJ7 | CALRRGYGNNRLAF | TRBV28 | TRBJ2-2 | CASSSWDIQTNTGELFF | ELAGIGILTV | YES | YES | YES |
| 13075 | TRAV29DV5 | TRAJ54 | CAAGTQGAQKLVF | TRBV4-1 | TRBJ2-5 | CASSHSGGDQETQYF | RAKFKQLL | YES | NO | YES |
| 13076 | TRAV13-1 | TRAJ50 | CAASETSYDKVIF | TRBV12-4 | TRBJ2-2 | CASSFTGAEGTGELFF | RAKFKQLL | YES | NO | YES |
| 13077 | TRAV13-2 | TRAJ45 | CAENGGGGADGLTF | TRBV29-1 | TRBJ2-7 | CSVGSGDYEQYF | RAKFKQLL | YES | NO | YES |
| 13078 | TRAV16 | TRAJ42 | CALMPPAGYGGSQGNLIF | TRBV5-1 | TRBJ2-7 | CASSLRDRVHEQYF | KLGGALQAK | NO | NO | NO |
| 13079 | TRAV13-1 | TRAJ11 | CAAKEGYSTLTF | TRBV7-9 | TRBJ2-5 | CASSLVGTKRETQYF | RAKFKQLL | YES | NO | YES |
| 13080 | TRAV19 | TRAJ26 | CALSDYGQNFVF | TRBV5-4 | TRBJ1-1 | CASGLNTEAFF | RAKFKQLL | YES | NO | NO |
| 13081 | TRAV35 | TRAJ26 | CAGPPLIDNYGQNFVF | TRBV12-3 | TRBJ2-7 | CASSLTLSSYEQYF | KLGGALQAK | YES | NO | NO |
| 13082 | TRAV16 | TRAJ26 | CALGGYGQNFVF | TRBV7-8 | TRBJ1-5 | CASSLAGNRQGARSQPQHF | KLGGALQAK | NO | NO | NO |
| 13083 | TRAV13-1 | TRAJ50 | CAAMETSYDKVIF | TRBV5-6 | TRBJ1-1 | CASSLRDGSEAFF | RAKFKQLL | YES | NO | YES |
| 13084 | TRAV8-3 | TRAJ48 | CAVGSYFGNEKLTF | TRBV11-2 | TRBJ2-5 | CATQTAQGTEQTQYF | KLGGALQAK | YES | NO | NO |
| 13085 | TRAV24 | TRAJ47 | CAPGARKLVF | TRBV11-2 | TRBJ2-7 | CASSPGTALSYEQYF | KLGGALQAK | YES | NO | NO |
| 13086 | TRAV19 | TRAJ34 | CALSVLLYNTDKLIF | TRBV20-1 | TRBJ2-7 | CSARSYGGGSSYEQYF | KLGGALQAK | YES | NO | NO |
| 13087 | TRAV26-2 | TRAJ23 | CILYMIYNQGGKLIF | TRBV5-4 | TRBJ2-3 | CASSFNGGGTDTQYF | KLGGALQAK | YES | NO | NO |
| 13088 | TRAV25 | TRAJ49 | CAGQNRGIGANQFYF | TRBV5-4 | TRBJ1-1 | CASSLSPGQGFRNTEAFF | KLGGALQAK | YES | NO | NO |
| 13089 | TRAV12-3 | TRAJ53 | CAMRDRNSGGSNYKLTF | TRBV13 | TRBJ2-2 | CASSSDRDNTGELFF | GILGFVFTL | NO | NO | NO |
| 13090 | TRAV1-2 | TRAJ27 | CAVRTNAGKSTF | TRBV4-2 | TRBJ2-7 | CASSQSWGDSYEQYF | KLGGALQAK | NO | NO | NO |
| 13091 | TRAV12-2 | TRAJ3 | CAALYSSASKIIF | TRBV19 | TRBJ2-7 | CASSIRSSYEQYF | GILGFVFTL | YES | NO | YES |
| 13092 | TRAV21 | TRAJ57 | CAVNYQGGSEKLVF | TRBV19 | TRBJ1-1 | CASSIRSTTEAFF | GILGFVFTL | YES | NO | YES |
| 13093 | TRAV9-2 | TRAJ50 | CALSETSYDKVIF | TRBV12-3 | TRBJ1-6 | CASSELQGAQDISPLHF | RAKFKQLL | YES | NO | YES |
| 13094 | TRAV25 | TRAJ20 | CAGPVLNDYKLSF | TRBV20-1 | TRBJ1-1 | CKPGMNTEAFF | KLGGALQAK | YES | NO | NO |
| 13095 | TRAV36DV7 | TRAJ57 | CASGGGSEKLVF | TRBV19 | TRBJ2-3 | CASSIRSTDTQYF | GILGFVFTL | NO | NO | NO |
| 13096 | TRAV12-2 | TRAJ4 | CAVNEAGGYNKLIF | TRBV6-5 | TRBJ2-7 | CASSYMGLGAYEQYF | ELAGIGILTV | YES | YES | YES |
| 13097 | TRAV12-2 | TRAJ26 | CAVNTGLFGQNFVF | TRBV6-5 | TRBJ2-1 | CASTSGYNEQFF | KLGGALQAK | NO | NO | NO |
| 13098 | TRAV13-1 | TRAJ50 | CAASETSYDKVIF | TRBV12-3 | TRBJ1-2 | CASHPRTPIYGYTF | RAKFKQLL | YES | NO | YES |
| 13099 | TRAV8-3 | TRAJ21 | CAVGDNFNKSYF | TRBV12-3 | TRBJ2-2 | CASSFSGNTGELFF | RAKFKQLL | YES | NO | YES |
| 13101 | TRAV13-1 | TRAJ11 | CAAKEGYSTLTF | TRBV19 | TRBJ1-2 | CASSGGGVYGYTF | KLGGALQAK | YES | NO | NO |
| 13102 | TRAV27 | TRAJ24 | CAGSTTDSWGKLQF | TRBV27 | TRBJ1-1 | CASYDRGTEAFF | KLGGALQAK | YES | NO | NO |
| 13103 | TRAV9-2 | TRAJ34 | CALPYNTDKLIF | TRBV11-2 | TRBJ2-1 | CASSLGAFENEQFF | KLGGALQAK | YES | NO | NO |
| 13104 | TRAV8-3 | TRAJ37 | CAADHGSGNTGKLIF | TRBV27 | TRBJ2-1 | CASSLDRDLSSYNEQFF | KLGGALQAK | YES | NO | NO |
| 13105 | TRAV17 | TRAJ9 | CATDPPEGGFKTIF | TRBV11-2 | TRBJ2-5 | CASSPRVGETQYF | KLGGALQAK | YES | NO | NO |
| 13106 | TRAV17 | TRAJ24 | CATDAATTDSWGKFQF | TRBV7-9 | TRBJ1-1 | CASSIGMGDWAFF | KLGGALQAK | YES | NO | NO |
| 13107 | TRAV1-2 | TRAJ6 | CAVRSSGGSYIPTF | TRBV12-3 | TRBJ2-2 | CASSFSGNTGELFF | RAKFKQLL | YES | NO | YES |
| 13108 | TRAV26-2 | TRAJ56 | CILSQGAVIELTF | TRBV7-9 | TRBJ1-2 | CASSLAGQGYGYTF | KLGGALQAK | YES | NO | NO |
| 13109 | TRAV13-2 | TRAJ45 | CADSGGGADGLTF | TRBV5-1 | TRBJ1-2 | CASSLMSQEFRYGYTF | RAKFKQLL | YES | NO | YES |
| 13110 | TRAV12-3 | TRAJ42 | CAMSGDGGSQGNLIF | TRBV19 | TRBJ1-5 | CASSARSNQPQHF | GILGFVFTL | YES | YES | NO |
| 13111 | TRAV12-2 | TRAJ48 | CAVGSNFGNEKLTF | TRBV20-1 | TRBJ2-5 | CSASVDSQYF | KLGGALQAK | NO | NO | NO |
| 13112 | TRAV22 | TRAJ11 | CAVAGYSTLTF | TRBV12-3 | TRBJ2-1 | CASSPTSNEQFF | KLGGALQAK | YES | NO | NO |
| 13113 | TRAV5 | TRAJ11 | CAEISSTLTF | TRBV19 | TRBJ1-2 | CASTLEGSYGYTF | AVFDRKSDAK | YES | NO | YES |
| 13114 | TRAV13-1 | TRAJ50 | CAAMETSYDKVIF | TRBV18 | TRBJ2-2 | CASSPRKDSTGELFF | RAKFKQLL | YES | YES | YES |
| 13115 | TRAV5 | TRAJ31 | CAEDSNARLMF | TRBV20-1 | TRBJ1-3 | CSARSRVGNTIYF | GLCTLVAML | YES | YES | YES |
| 13116 | TRAV20 | TRAJ48 | CAALLDFGNEKLTF | TRBV4-1 | TRBJ2-1 | CASSLLWAEYNEQFF | RAKFKQLL | YES | NO | YES |
| 13117 | TRAV38-1 | TRAJ34 | CAFMPPYNTDKLIF | TRBV7-3 | TRBJ1-1 | CASSLGQGAINTEAFF | KLGGALQAK | NO | NO | NO |
| 13118 | TRAV13-1 | TRAJ50 | CAASETSYDKVIF | TRBV12-3 | TRBJ2-2 | CASIVRNDRDPTGELFF | RAKFKQLL | YES | NO | YES |
| 13119 | TRAV12-2 | TRAJ45 | CAVNKGGGGGADGLTF | TRBV12-4 | TRBJ2-1 | CASRPLLAGGDEQFF | KLGGALQAK | YES | NO | NO |
| 13120 | TRAV13-2 | TRAJ45 | CADSGGGADGLTF | TRBV5-6 | TRBJ2-4 | CASSSEGQVAKNIQYF | RAKFKQLL | YES | NO | NO |
| 13121 | TRAV14DV4 | TRAJ9 | CAMREGQGGGFKTIF | TRBV12-3 | TRBJ2-3 | CAATRAGSPDTQYF | KLGGALQAK | YES | NO | NO |
| 13122 | TRAV3 | TRAJ34 | CAVRDSRRYNTDKLIF | TRBV19 | TRBJ2-1 | CASSIDGSGGNEQFF | KLGGALQAK | NO | NO | NO |
| 13123 | TRAV3 | TRAJ9 | CAVRDDTGGFKTIF | TRBV9 | TRBJ2-2 | CASSGEGGTTNTGELFF | GILGFVFTL | NO | NO | NO |
| 13124 | TRAV4 | TRAJ15 | CLVDEAGTALIF | TRBV20-1 | TRBJ1-1 | CSASQGLMNTEAFF | ELAGIGILTV | NO | NO | NO |
| 13125 | TRAV29DV5 | TRAJ49 | CAASDGNQFYF | TRBV7-9 | TRBJ2-1 | CASPYLTGHEQFF | KLGGALQAK | YES | NO | NO |
| 13126 | TRAV8-3 | TRAJ21 | CAVGDNFNKFYF | TRBV7-9 | TRBJ1-3 | CASSPLGGPKDTIYF | RAKFKQLL | YES | NO | YES |
| 13127 | TRAV13-2 | TRAJ45 | CADSGGGADGLTF | TRBV7-8 | TRBJ2-2 | CASSSPEGQPLNTGELFF | GLCTLVAML | YES | YES | YES |
| 13128 | TRAV17 | TRAJ54 | CATHPIQGAQKLVF | TRBV2 | TRBJ1-2 | CASSGGITVYGYTF | GILGFVFTL | YES | NO | YES |
| 13129 | TRAV6 | TRAJ43 | CALGNNDMRF | TRBV27 | TRBJ1-5 | CASSFGADNQPQHF | KLGGALQAK | YES | NO | NO |
| 13130 | TRAV8-3 | TRAJ37 | CAVGPVHGSSNTGKLIF | TRBV14 | TRBJ2-7 | CASSLGLNTYEQYF | KLGGALQAK | NO | NO | NO |
| 13131 | TRAV10 | TRAJ27 | CVGYTNAGKSTF | TRBV7-9 | TRBJ2-1 | CASSFDKTEQFF | RAKFKQLL | YES | NO | YES |
| 13132 | TRAV17 | TRAJ10 | CATGLTGGGNKLTF | TRBV19 | TRBJ2-7 | CASSIRSSYEQYF | GILGFVFTL | YES | NO | YES |
| 13133 | TRAV8-2 | TRAJ3 | CVVSDPESSASKIIF | TRBV7-2 | TRBJ1-2 | CASSLNSGLDGYTF | KLGGALQAK | YES | NO | NO |
| 13134 | TRAV12-3 | TRAJ30 | CAMSWGRDDKIIF | TRBV7-9 | TRBJ2-7 | CASSKGLTGNYEQYF | KLGGALQAK | YES | NO | NO |
| 13135 | TRAV1-2 | TRAJ36 | CAPDQTGANNLFF | TRBV7-9 | TRBJ2-7 | CASSSPPTPYEQYF | KLGGALQAK | YES | NO | NO |
| 13136 | TRAV13-2 | TRAJ53 | CAETHSGGSNYKLTF | TRBV13 | TRBJ2-7 | CASSLPSGMLYEQYF | RAKFKQLL | YES | NO | YES |
| 13137 | TRAV9-2 | TRAJ30 | CALLNRDDKIIF | TRBV7-9 | TRBJ2-7 | CASSFVDRVYEQYF | RAKFKQLL | YES | NO | YES |
| 13138 | TRAV29DV5 | TRAJ28 | CAARKGAGSYQLTF | TRBV7-8 | TRBJ2-1 | CASSLRRVEQFF | KLGGALQAK | YES | NO | NO |
| 13139 | TRAV35 | TRAJ49 | CAGHTGNQFYF | TRBV19 | TRBJ2-1 | CASSIRSGYEQFF | GILGFVFTL | YES | NO | YES |
| 13140 | TRAV35 | TRAJ42 | CAGAGDGGSQGNLIF | TRBV6-3 | TRBJ2-3 | CASSYSSGGADTQYF | GILGFVFTL | YES | NO | NO |
| 13141 | TRAV12-3 | TRAJ32 | CAMNPAWGGATNKLIF | TRBV9 | TRBJ1-1 | CASSALAGNTEAFF | AVFDRKSDAK | YES | NO | YES |
| 13142 | TRAV35 | TRAJ6 | CAGPTPSGGSYIPTF | TRBV15 | TRBJ2-6 | CATSYEDNSGANVLTF | KLGGALQAK | YES | NO | NO |
| 13143 | TRAV35 | TRAJ36 | CAVRLQTGANNLFF | TRBV12-4 | TRBJ2-1 | CASSFSAEQFF | KLGGALQAK | YES | NO | NO |
| 13144 | TRAV19 | TRAJ49 | CALRAGNQFYF | TRBV9 | TRBJ2-7 | CASSVDSGSDYEQYF | KLGGALQAK | YES | NO | NO |
| 13145 | TRAV29DV5 | TRAJ31 | CAASTNARLMF | TRBV19 | TRBJ2-7 | CASSIRSSYEQYF | GILGFVFTL | YES | NO | YES |
| 13146 | TRAV41 | TRAJ42 | CAVEVGGSQGNLIF | TRBV19 | TRBJ2-2 | CASTKAEYKTGELFF | KLGGALQAK | YES | NO | NO |
| 13147 | TRAV13-2 | TRAJ45 | CADSGGGADGLTF | TRBV5-6 | TRBJ2-7 | CASSLIQTGQALEQYF | RAKFKQLL | YES | NO | YES |
| 13148 | TRAV8-4 | TRAJ34 | CAVTHTDKLIF | TRBV5-4 | TRBJ2-1 | CASSSRAEQFF | KLGGALQAK | YES | NO | NO |
| 13149 | TRAV12-3 | TRAJ20 | CAMRENDYKLSF | TRBV7-2 | TRBJ2-7 | CASSPSYSKHYEQYF | KLGGALQAK | YES | NO | NO |
| 13150 | TRAV17 | TRAJ18 | CATDDYDRGSTLGRLYF | TRBV19 | TRBJ2-1 | CASKGTSGGVNEQFF | GILGFVFTL | YES | YES | NO |
| 13151 | TRAV36DV7 | TRAJ17 | CAARMIKAAGNKLTF | TRBV19 | TRBJ2-7 | CASSIRASYEQYF | GILGFVFTL | NO | NO | NO |
| 13152 | TRAV5 | TRAJ6 | CAEMEGGSYIPTF | TRBV4-2 | TRBJ1-5 | CASSQEGQINQPQHF | RAKFKQLL | NO | NO | NO |

| | | | | | | | | | | |
|---|---|---|---|---|---|---|---|---|---|---|
| 13153 | TRAV13-1 | TRAJ35 | CAARVRGFGNVLHC | TRBV2 | TRBJ2-7 | CASAPPQGLILEQYF | KLGGALQAK | NO | NO | NO |
| 13154 | TRAV1-2 | TRAJ31 | CAVNRDNARLMF | TRBV12-3 | TRBJ2-2 | CASRKQGSDTGELFF | RAKFKQLL | YES | NO | YES |
| 13155 | TRAV13-2 | TRAJ50 | CAERETSYDKVIF | TRBV7-9 | TRBJ2-4 | CASSHSEGAENIQYF | RAKFKQLL | NO | NO | NO |
| 13156 | TRAV38-1 | TRAJ52 | CAFMINAGGTSYGKLTF | TRBV11-2 | TRBJ2-3 | CASLISGTDTQYF | KLGGALQAK | YES | NO | NO |
| 13157 | TRAV1-2 | TRAJ8 | CAEGFQKLVF | TRBV12-4 | TRBJ2-7 | CASSFSSGLGYEQYF | AVFDRKSDAK | YES | NO | YES |
| 13158 | TRAV19 | TRAJ30 | CALIANRDDKIIF | TRBV12-3 | TRBJ1-1 | CASSFLDRGPNTEAFF | RAKFKQLL | YES | NO | YES |
| 13159 | TRAV3 | TRAJ5 | CAVRPVDTGRRALTF | TRBV5-4 | TRBJ1-2 | CASDRTPKSQDGYTF | KLGGALQAK | YES | NO | NO |
| 13160 | TRAV12-2 | TRAJ6 | CAVNTAGGSYIPTF | TRBV14 | TRBJ1-6 | CASSQHSGSPLHF | KLGGALQAK | NO | NO | NO |
| 13161 | TRAV25 | TRAJ42 | CAGGDGGSQGNLIF | TRBV19 | TRBJ2-3 | CASSVRSTDTQYF | GILGFVFTL | YES | NO | YES |
| 13162 | TRAV35 | TRAJ49 | CAGHTGNQFYF | TRBV2 | TRBJ2-2 | CASSTPRQGSNTGELFF | RAKFKQLL | YES | NO | YES |
| 13163 | TRAV5 | TRAJ31 | CAEDNNARLMF | TRBV20-1 | TRBJ1-3 | CSARDREFGNTIYF | GLCTLVAML | YES | YES | YES |
| 13164 | TRAV13-1 | TRAJ42 | CAATGGGSQGNLIF | TRBV19 | TRBJ2-5 | CASSYRSQETQYF | GILGFVFTL | YES | NO | YES |
| 13165 | TRAV26-2 | TRAJ13 | CILRAPSGGYQKVTF | TRBV5-1 | TRBJ2-3 | CASTPVGGRGTDTQYF | AVFDRKSDAK | YES | NO | YES |
| 13166 | TRAV13-1 | TRAJ50 | CAAMETSYDKVIF | TRBV6-3 | TRBJ2-7 | CASGMTGLTSEQYF | RAKFKQLL | YES | NO | YES |
| 13167 | TRAV12-2 | TRAJ45 | CAGGNSGGGADGLTF | TRBV29-1 | TRBJ1-1 | CSAGGNTEAFF | KLGGALQAK | NO | NO | NO |
| 13168 | TRAV6 | TRAJ40 | CARTSGTYKYIF | TRBV7-2 | TRBJ2-2 | CASSLAGLAAWDGELFF | KLGGALQAK | YES | NO | NO |
| 13169 | TRAV8-4 | TRAJ27 | CAVTSTNAGKSTF | TRBV4-1 | TRBJ2-7 | CASSQVTGVLYEQYF | KLGGALQAK | NO | NO | NO |
| 13170 | TRAV38-2DV8 | TRAJ52 | CAYSSNAGGTSYGKLTF | TRBV19 | TRBJ1-2 | CASSIGLYGYTF | GILGFVFTL | YES | YES | NO |
| 13171 | TRAV8-4 | TRAJ44 | CAVSVGTASKLTF | TRBV4-1 | TRBJ1-2 | CASSQDVRLDGYTF | KLGGALQAK | YES | NO | NO |
| 13172 | TRAV21 | TRAJ37 | CAVSGSSNTGKLIF | TRBV5-1 | TRBJ2-7 | CASRGLLSNEQYF | KLGGALQAK | YES | NO | NO |
| 13173 | TRAV12-1 | TRAJ53 | CVARGSNYKLTF | TRBV2 | TRBJ2-6 | CASSELSSGANVLTF | KLGGALQAK | YES | NO | NO |
| 13174 | TRAV12-1 | TRAJ44 | CVVRFTGTASKLTF | TRBV6-1 | TRBJ2-3 | CASRAGQGRTDTQYF | KLGGALQAK | YES | NO | NO |
| 13175 | TRAV17 | TRAJ17 | CATDAWAAGNKLTF | TRBV7-3 | TRBJ2-5 | CASSLKPRSGGASETQYF | KLGGALQAK | NO | NO | NO |
| 13176 | TRAV14DV4 | TRAJ52 | CAMRDLDGGGTSYGKLTF | TRBV5-1 | TRBJ2-1 | CASSPTHRYNEQFF | KLGGALQAK | YES | NO | NO |
| 13177 | TRAV22 | TRAJ37 | CAGSPSNTGKLIF | TRBV7-2 | TRBJ2-7 | CASSPRDRERGEQYF | GILGFVFTL | NO | NO | NO |
| 13178 | TRAV8-4 | TRAJ30 | CAVSGRKIIF | TRBV11-2 | TRBJ1-2 | CASSFAGDRGLGSYGYTF | KLGGALQAK | YES | NO | NO |
| 13179 | TRAV17 | TRAJ22 | CATDRAGSARQLTF | TRBV20-1 | TRBJ1-2 | CSARDRTGNGYTF | GLCTLVAML | YES | YES | YES |
| 13180 | TRAV20 | TRAJ16 | CAVQAVGSDGQKLLF | TRBV19 | TRBJ2-4 | CASSIAHAGGLAKNIQYF | KLGGALQAK | NO | NO | NO |
| 13181 | TRAV13-1 | TRAJ50 | CAASETSYDKVIF | TRBV7-9 | TRBJ2-5 | CASSLLGTDQETQYF | RAKFKQLL | YES | NO | YES |
| 13182 | TRAV13-1 | TRAJ15 | CAASIPNQAGTALIF | TRBV12-3 | TRBJ2-2 | CASSFSGNTGELFF | RAKFKQLL | YES | NO | YES |
| 13183 | TRAV29DV5 | TRAJ52 | CAASSGGTSYGKLTF | TRBV27 | TRBJ2-2 | CASSPHFQVDTGELFF | RAKFKQLL | YES | NO | YES |
| 13184 | TRAV8-2 | TRAJ6 | CVTSLSGGSYIPTF | TRBV5-6 | TRBJ1-1 | CASSLRDGSEAFF | RAKFKQLL | YES | NO | YES |
| 13185 | TRAV12-1 | TRAJ7 | CVVNLEVFAYYGNNRLAF | TRBV7-9 | TRBJ2-7 | CASSLGTVSYEQYF | GILGFVFTL | YES | NO | YES |
| 13186 | TRAV26-1 | TRAJ27 | CIADNTNAGKSTF | TRBV29-1 | TRBJ2-6 | CSVQGAPGANVLTF | KLGGALQAK | YES | NO | NO |
| 13187 | TRAV8-2 | TRAJ34 | CVVSDVNTDKLIF | TRBV20-1 | TRBJ1-3 | CSARSRVGNTIYF | GLCTLVAML | YES | YES | YES |
| 13188 | TRAV12-1 | TRAJ30 | CVVNRDRDDKIIF | TRBV4-1 | TRBJ2-7 | CASSQARDRRYEQYF | KLGGALQAK | YES | NO | NO |
| 13189 | TRAV8-1 | TRAJ15 | CAVNAVSNQAGTALIF | TRBV5-6 | TRBJ1-1 | CASSLRDGSEAFF | RAKFKQLL | YES | NO | YES |
| 13190 | TRAV4 | TRAJ10 | CLVGYTGGGNKLTF | TRBV5-6 | TRBJ1-1 | CASSLRDGSEAFF | RAKFKQLL | YES | NO | NO |
| 13191 | TRAV13-2 | TRAJ45 | CADSGGGADGLTF | TRBV6-4 | TRBJ2-1 | CATSPSGGTYNEQFF | RAKFKQLL | YES | NO | YES |
| 13192 | TRAV39 | TRAJ42 | CAVDEGGGSQGNLIF | TRBV13 | TRBJ2-5 | CASSEGGVETQYF | GILGFVFTL | YES | NO | YES |
| 13193 | TRAV38-2DV8 | TRAJ31 | CAYNNARLMF | TRBV12-4 | TRBJ2-1 | CASGRLSYNEQFF | RAKFKQLL | YES | NO | YES |
| 13194 | TRAV38-2DV8 | TRAJ58 | CAYRSKETSGSRLTF | TRBV18 | TRBJ2-2 | CASSPPGGGNTGELFF | RAKFKQLL | YES | YES | YES |
| 13195 | TRAV3 | TRAJ20 | CAVFNDYKLSF | TRBV4-2 | TRBJ2-2 | CASSQRPSEVGELFF | RAKFKQLL | YES | NO | YES |
| 13196 | TRAV13-2 | TRAJ45 | CADSGGGADGLTF | TRBV5-6 | TRBJ2-1 | CASSLGGGGTYNEQFF | RAKFKQLL | YES | NO | YES |
| 13197 | TRAV34 | TRAJ28 | CGAHGGSGAGSYQLTF | TRBV19 | TRBJ2-7 | CASSTLAGGYYEQYF | AVFDRKSDAK | YES | NO | YES |
| 13198 | TRAV3 | TRAJ21 | CAVRDISNFNKFYF | TRBV19 | TRBJ1-1 | CASSTSDRGNTEAFF | KLGGALQAK | NO | NO | NO |
| 13199 | TRAV39 | TRAJ43 | CAVDRNDMRF | TRBV20-1 | TRBJ2-3 | CSARPTSVSPASTDTQYF | KLGGALQAK | NO | NO | NO |
| 13200 | TRAV19 | TRAJ45 | CALTGGGADGLTF | TRBV5-5 | TRBJ2-7 | CASSGPGGVDEQYF | RAKFKQLL | NO | NO | NO |
| 13201 | TRAV12-2 | TRAJ10 | CAVPLTGGGNKLTF | TRBV19 | TRBJ2-7 | CASSTRSSYEQYF | GLCTLVAML | YES | NO | YES |
| 13202 | TRAV19 | TRAJ26 | CALSPAVTTLTFVF | TRBV20-1 | TRBJ1-6 | CSAPSYRDRDNSPLHF | AVFDRKSDAK | NO | NO | NO |
| 13203 | TRAV19 | TRAJ45 | CALSVFSGGGADGLTF | TRBV30 | TRBJ2-2 | CAWSVGQGTDGELFF | KLGGALQAK | YES | NO | NO |
| 13204 | TRAV14DV4 | TRAJ30 | CAMREGSRRRDDKIIF | TRBV4-1 | TRBJ1-2 | CASSTSTGTGYGYTF | RAKFKQLL | YES | NO | YES |
| 13205 | TRAV1-2 | TRAJ20 | CAVRDGDYKLSF | TRBV20-1 | TRBJ2-7 | CSARELADYEQYF | KLGGALQAK | NO | NO | NO |
| 13206 | TRAV27 | TRAJ42 | CAGATGGGSQGNLIF | TRBV19 | TRBJ1-1 | CASSIRSSFEAFF | GILGFVFTL | YES | NO | YES |
| 13207 | TRAV12-2 | TRAJ43 | CAVRVGDMRF | TRBV5-4 | TRBJ1-5 | CASSSTGGGEKDQPQHF | KLGGALQAK | YES | NO | NO |
| 13208 | TRAV8-2 | TRAJ12 | CVVSRMDSSYKLIF | TRBV7-2 | TRBJ2-7 | CASSLQGVSYEQYF | KLGGALQAK | YES | NO | NO |
| 13209 | TRAV35 | TRAJ54 | CAGPRYAQGAQKLVF | TRBV27 | TRBJ1-5 | CASSFQGLGQPQHF | ELAGIGILTV | NO | NO | NO |
| 13211 | TRAV12-2 | TRAJ8 | CAVNAGFQKLVF | TRBV7-6 | TRBJ1-1 | CASSLGVEAFF | ELAGIGILTV | YES | NO | YES |
| 13212 | TRAV3 | TRAJ43 | CAVIRWGDMRF | TRBV5-6 | TRBJ1-1 | CASSLRDGSEAFF | RAKFKQLL | YES | NO | YES |
| 13213 | TRAV13-1 | TRAJ8 | CAALEPGFQKLVF | TRBV19 | TRBJ2-3 | CASSPLSTDTQYF | GILGFVFTL | YES | YES | YES |
| 13214 | TRAV10 | TRAJ27 | CVGYTNAGKSTF | TRBV19 | TRBJ1-1 | CASRAGREAFF | GILGFVFTL | YES | YES | YES |
| 13215 | TRAV26-2 | TRAJ26 | CILRDVRGQNFVF | TRBV20-1 | TRBJ1-3 | CSAFTGEGNTIYF | RAKFKQLL | NO | NO | NO |
| 13216 | TRAV12-3 | TRAJ52 | CAMRDTNAGGTSYGKLTF | TRBV19 | TRBJ2-1 | CASSMRSGSEQFF | GILGFVFTL | YES | NO | YES |
| 13217 | TRAV1-2 | TRAJ15 | CAVRQAGTALIF | TRBV7-9 | TRBJ2-3 | CASSLGLAGTDTQYF | KLGGALQAK | YES | NO | NO |
| 13218 | TRAV1-2 | TRAJ45 | CAVRDGGGADGLTF | TRBV7-9 | TRBJ2-3 | CASSSGLAGAGTQYF | KLGGALQAK | YES | NO | NO |
| 13219 | TRAV1-2 | TRAJ33 | CAAMDSNYQLIW | TRBV19 | TRBJ2-2 | CASKGLGADTGELFF | KLGGALQAK | YES | NO | NO |
| 13220 | TRAV12-2 | TRAJ45 | CAVTRGADGLTF | TRBV11-2 | TRBJ2-5 | CASSRTTGQAETQYF | KLGGALQAK | YES | NO | NO |
| 13221 | TRAV19 | TRAJ17 | CALKLIKAAGNKLTF | TRBV19 | TRBJ1-1 | CASSIRSTTEAFF | GILGFVFTL | YES | NO | YES |
| 13222 | TRAV4 | TRAJ36 | CLVPTGANNLFF | TRBV20-1 | TRBJ2-7 | CSARDLTAGSYEQYF | GILGFVFTL | NO | NO | NO |
| 13223 | TRAV36DV7 | TRAJ27 | CAVGPNAGKSTF | TRBV11-2 | TRBJ2-3 | CASSRAGRSTDTQYF | KLGGALQAK | NO | NO | NO |
| 13224 | TRAV38-1 | TRAJ57 | CAFMKHGGGSEKLVF | TRBV14 | TRBJ1-1 | CASSHYFGTEAFF | KLGGALQAK | NO | NO | NO |
| 13225 | TRAV1-2 | TRAJ8 | CAVKTGFQKLVF | TRBV10-3 | TRBJ2-7 | CAISGGRAGEYAYEQYF | KLGGALQAK | YES | NO | NO |
| 13226 | TRAV1-2 | TRAJ33 | CAVRDSNYQLIW | TRBV20-1 | TRBJ1-1 | CSVEQTNTEAFF | KLGGALQAK | YES | NO | NO |
| 13227 | TRAV19 | TRAJ45 | CALSEAGGMGGGADGLTF | TRBV12-3 | TRBJ2-2 | CASSFSGNTGELFF | RAKFKQLL | YES | NO | YES |
| 13228 | TRAV12-1 | TRAJ37 | CVVNIPFNPGKLIF | TRBV4-1 | TRBJ2-1 | CASSLLWAEYNEQFF | RAKFKQLL | YES | YES | YES |
| 13229 | TRAV9-2 | TRAJ40 | CALRLEGGSGTYKYIF | TRBV7-8 | TRBJ1-2 | CASSLEVGLGGYTF | GILGFVFTL | YES | NO | YES |
| 13230 | TRAV13-1 | TRAJ50 | CAATETSYDKVIF | TRBV29-1 | TRBJ2-1 | CSVGSGDHGEQFF | RAKFKQLL | YES | NO | NO |
| 13231 | TRAV27 | TRAJ42 | CAGGGSQGNLSF | TRBV19 | TRBJ2-7 | CASSIRSSYEQYF | GILGFVFTL | YES | NO | YES |
| 13232 | TRAV17 | TRAJ40 | CATDAGILSGTYKYIF | TRBV9 | TRBJ1-6 | CASSVRVGSPLHF | KLGGALQAK | YES | NO | NO |
| 13233 | TRAV38-1 | TRAJ39 | CAFMKRNAGNMLTF | TRBV5-6 | TRBJ1-1 | CASSLRDGSEAFF | RAKFKQLL | YES | NO | YES |
| 13234 | TRAV27 | TRAJ34 | CAGELTPTSSSL | TRBV5-6 | TRBJ1-1 | CASSLRDGSEAFF | RAKFKQLL | YES | NO | YES |
| 13235 | TRAV21 | TRAJ26 | CAVHGYGQNFVF | TRBV7-3 | TRBJ2-2 | CASSLGTSGGTGELFF | RAKFKQLL | YES | YES | YES |
| 13236 | TRAV3 | TRAJ36 | CAVRDGTGANNLFF | TRBV19 | TRBJ2-2 | CASSITSTGELFF | GILGFVFTL | NO | NO | NO |
| 13237 | TRAV3 | TRAJ6 | CAVSAASGGSYIPTF | TRBV12-5 | TRBJ1-1 | CASGLVESAGTEAFF | RAKFKQLL | NO | NO | NO |
| 13238 | TRAV3 | TRAJ34 | CAVRGFIGTDKLIF | TRBV9 | TRBJ2-1 | CASSEEEGGWEQFF | KLGGALQAK | NO | NO | NO |
| 13239 | TRAV9-2 | TRAJ43 | CALSGKDMRF | TRBV6-1 | TRBJ2-5 | CASSELIGIRPGPAATKF | KLGGALQAK | NO | NO | NO |
| 13240 | TRAV12-1 | TRAJ31 | CVVNIDNARLMF | TRBV29-1 | TRBJ2-3 | CSVGSGEDSPQYF | RAKFKQLL | YES | NO | NO |
| 13241 | TRAV35 | TRAJ28 | CAGRKAGSYQLTF | TRBV19 | TRBJ2-1 | CASSIDHNQGYNEQFF | KLGGALQAK | YES | NO | NO |
| 13242 | TRAV35 | TRAJ42 | CAGQEGGGSQGNLIF | TRBV19 | TRBJ2-7 | CASSLRSSYEQYF | GILGFVFTL | YES | NO | YES |
| 13243 | TRAV12-3 | TRAJ36 | CAFGGANNLFF | TRBV7-9 | TRBJ2-7 | CASSPTGAYEQYF | KLGGALQAK | YES | NO | NO |
| 13244 | TRAV8-1 | TRAJ26 | CAAHGQNFVF | TRBV12-5 | TRBJ2-3 | CASGQELAGTDTQYF | KLGGALQAK | YES | NO | NO |
| 13245 | TRAV12-2 | TRAJ24 | CAVKTDSWGKLQF | TRBV4-1 | TRBJ1-1 | CASSPGELVPNTEAFF | ELAGIGILTV | YES | NO | YES |
| 13246 | TRAV27 | TRAJ3 | CAGGYSSASKIIF | TRBV6-5 | TRBJ2-7 | CASSPSLSYEQYF | RAKFKQLL | YES | NO | YES |
| 13247 | TRAV29DV5 | TRAJ31 | CAASTNARLMF | TRBV29-1 | TRBJ2-1 | CSVQGGAYNEQFF | KLGGALQAK | YES | NO | NO |
| 13248 | TRAV27 | TRAJ43 | CAGYNNNDMRF | TRBV18 | TRBJ2-1 | CASSPVGVGEQFF | GILGFVFTL | YES | NO | YES |
| 13249 | TRAV9-2 | TRAJ28 | CALSPYSGAGSYQLTF | TRBV7-8 | TRBJ2-3 | CASSSPEIGRSTDTQYF | KLGGALQAK | YES | NO | NO |
| 13250 | TRAV26-1 | TRAJ52 | CIVNDAGTSYGKLTF | TRBV16 | TRBJ1-1 | CASSQELSRRNTEAFF | KLGGALQAK | YES | NO | NO |
| 13251 | TRAV1-2 | TRAJ33 | CAVMDSNYQLIW | TRBV20-1 | TRBJ1-1 | CSASGTTDTEAFF | KLGGALQAK | YES | NO | NO |
| 13252 | TRAV38-2DV8 | TRAJ52 | CAYMTNAGGTSYGKLTF | TRBV7-9 | TRBJ2-3 | CASSSRDGGTDTQYF | GILGFVFTL | YES | NO | YES |
| 13253 | TRAV1-2 | TRAJ33 | CAVSNSNYQLIW | TRBV6-1 | TRBJ2-3 | CASSEAAGGGTDTQYF | KLGGALQAK | YES | NO | NO |
| 13254 | TRAV25 | TRAJ42 | CAAVYGGSQGNLIF | TRBV27 | TRBJ2-7 | CASRRGWPGNEQYF | KLGGALQAK | YES | NO | NO |
| 13255 | TRAV13-2 | TRAJ42 | CAERDGGGSQGNLIF | TRBV19 | TRBJ2-5 | CASSIRAAETQYF | GILGFVFTL | YES | NO | YES |
| 13256 | TRAV20 | TRAJ33 | CALSGDSNYQLIW | TRBV7-9 | TRBJ2-3 | CASSSPPLDTQYF | KLGGALQAK | YES | NO | NO |
| 13257 | TRAV12-2 | TRAJ42 | CAVNLGGGSQGNLIF | TRBV2 | TRBJ2-5 | CASSQQWEETQYF | GILGFVFTL | YES | NO | YES |

| | | | | | | | | | | |
|---|---|---|---|---|---|---|---|---|---|---|
| 13258 | TRAV17 | TRAJ11 | CATEGNSGYSTLTF | TRBV6-5 | TRBJ1-2 | CASSYSGGYYGYTF | FLYALALLL | YES | YES | YES |
| 13259 | TRAV35 | TRAJ57 | CAGPTLTQGGSEKLVF | TRBV6-5 | TRBJ1-3 | CASSYGVQGNTIYF | KLGGALQAK | YES | NO | NO |
| 13260 | TRAV17 | TRAJ39 | CATDPRANAGNMLTF | TRBV4-1 | TRBJ1-4 | CASGYEKLFF | RLRAEAQVK | YES | YES | NO |
| 13261 | TRAV5 | TRAJ31 | CAEDSNARLMF | TRBV11-2 | TRBJ2-2 | CASSLGVAGTNTGELFF | GLCTLVAML | YES | YES | YES |
| 13262 | TRAV17 | TRAJ52 | CATAKYNAGGTSYGKLTF | TRBV19 | TRBJ1-2 | CASSIGSHGYTF | GILGFVFTL | YES | NO | YES |
| 13263 | TRAV27 | TRAJ42 | CAGGGSQGNLIF | TRBV7-9 | TRBJ2-3 | CASSSRDGGTDTQYF | GILGFVFTL | YES | NO | YES |
| 13264 | TRAV24 | TRAJ20 | CAFPFDDYKLSF | TRBV5-5 | TRBJ2-7 | CASSLARTAYEQYF | AVFDRKSDAK | YES | NO | YES |
| 13265 | TRAV13-2 | TRAJ42 | CAENTGGGSQGNLIF | TRBV19 | TRBJ1-2 | CASSIGLYGYTF | GILGFVFTL | YES | YES | YES |
| 13266 | TRAV21 | TRAJ12 | CAVLPMDSSYKLIF | TRBV5-4 | TRBJ1-6 | CASSVQGLGNSPLHF | KLGGALQAK | YES | NO | NO |
| 13267 | TRAV38-1 | TRAJ52 | CAFMTSPGGTSYGKLTF | TRBV11-2 | TRBJ2-3 | CASSLGDFLRTDTQYF | GILGFVFTL | YES | NO | YES |
| 13268 | TRAV2 | TRAJ38 | CGRQMLLFTTVLYTKAGNNRKLIW | TRBV6-4 | TRBJ2-1 | CASKTGDNEQFF | KLGGALQAK | NO | NO | NO |
| 13269 | TRAV14DV4 | TRAJ22 | CAMKGSARQLTF | TRBV11-2 | TRBJ2-3 | CASLISGTDTQYF | KLGGALQAK | YES | NO | NO |
| 13270 | TRAV10 | TRAJ38 | CVVTDFNAGNNRKLIW | TRBV5-6 | TRBJ1-2 | CASSPGGGYTF | KLGGALQAK | YES | NO | NO |
| 13271 | TRAV38-2DV8 | TRAJ58 | CAYRSATRETSGSRLTF | TRBV5-1 | TRBJ2-7 | CASSFVTASYEQYF | KLGGALQAK | YES | NO | NO |
| 13272 | TRAV12-3 | TRAJ34 | CAMSYVNTDKLIF | TRBV10-3 | TRBJ2-3 | CAISNVDTGVGADTQYF | KLGGALQAK | YES | NO | NO |
| 13273 | TRAV38-2DV8 | TRAJ42 | CAYVGGSQGNLIF | TRBV12-3 | TRBJ2-2 | CASSFSGNTGELFF | RAKFKQLL | YES | NO | YES |
| 13274 | TRAV29DV5 | TRAJ34 | CAASEGNTDKLIF | TRBV20-1 | TRBJ1-1 | CSAEAEAFF | KLGGALQAK | NO | NO | NO |
| 13275 | TRAV13-1 | TRAJ50 | CAASETSYDKVIF | TRBV4-1 | TRBJ2-1 | CASSLLAGEYNEQFF | RAKFKQLL | YES | NO | YES |
| 13276 | TRAV14DV4 | TRAJ28 | CAMSAYSGAGSYQLTF | TRBV5-5 | TRBJ2-7 | CASSLGTLPYEQYF | KLGGALQAK | YES | NO | NO |
| 13277 | TRAV27 | TRAJ42 | CAGGGSQGNLIF | TRBV21-1 | TRBJ2-4 | CISVPAAWGTGRWETFSTSAP | GILGFVFTL | NO | NO | NO |
| 13278 | TRAV13-1 | TRAJ42 | CAASGGGSQGNLIF | TRBV7-3 | TRBJ2-1 | CASSFYAGSTNEQFF | RAKFKQLL | YES | NO | YES |
| 13279 | TRAV27 | TRAJ42 | CAGSGSQGNLIF | TRBV19 | TRBJ1-2 | CASSQGAYGYTF | GILGFVFTL | YES | NO | YES |
| 13280 | TRAV5 | TRAJ50 | CAETETSYDKVIF | TRBV7-9 | TRBJ2-3 | CASRPRGTTDTQYF | RAKFKQLL | YES | NO | NO |
| 13281 | TRAV21 | TRAJ33 | CAVLMDSNYQLIW | TRBV10-2 | TRBJ1-1 | CASSEDGMNTEAFF | KLGGALQAK | YES | NO | NO |
| 13282 | TRAV8-1 | TRAJ50 | CAVRETSYDKVIF | TRBV18 | TRBJ2-3 | CASSPWADVVTDTQYF | RAKFKQLL | YES | NO | YES |
| 13283 | TRAV12-3 | TRAJ42 | CAMSGDGGSQGNLIF | TRBV19 | TRBJ1-5 | CASSIRSSEPQHF | GILGFVFTL | YES | NO | YES |
| 13284 | TRAV14DV4 | TRAJ9 | CAMREGLNTGGFKTIF | TRBV5-4 | TRBJ2-3 | CASSFGGYTDTQYF | KLGGALQAK | YES | NO | NO |
| 13285 | TRAV25 | TRAJ42 | CAITYGGSQGNLIF | TRBV19 | TRBJ2-3 | CASSTRSTDTQYF | GILGFVFTL | YES | NO | YES |
| 13286 | TRAV20 | TRAJ40 | CAVRSSGTYKYIF | TRBV9 | TRBJ1-1 | CASSVEGGNTEAFF | KLGGALQAK | YES | NO | NO |
| 13287 | TRAV14DV4 | TRAJ7 | CAMIRPSGNNRLAF | TRBV6-5 | TRBJ1-2 | CASSYQGGNYGYTF | FLYALALLL | YES | YES | YES |
| 13288 | TRAV12-2 | TRAJ57 | CAVWEGGSEKLVF | TRBV9 | TRBJ1-1 | CASSVEGRMAAEAFF | KLGGALQAK | YES | NO | NO |
| 13289 | TRAV1-2 | TRAJ33 | CAVMDSNYQLIW | TRBV6-1 | TRBJ2-6 | CASSGGASGANVLTF | KLGGALQAK | YES | NO | NO |
| 13290 | TRAV38-2DV8 | TRAJ45 | CAYSSGGGADGLTF | TRBV5-1 | TRBJ2-5 | CASSFGTDALYQETQYF | KLGGALQAK | YES | NO | NO |
| 13291 | TRAV16 | TRAJ10 | CALVTFTGGGNKLTF | TRBV5-4 | TRBJ1-5 | CASSSTGGGEKDQPQHF | GILGFVFTL | NO | NO | NO |
| 13292 | TRAV1-2 | TRAJ33 | CAAIDSNYQLIW | TRBV20-1 | TRBJ2-5 | CSVPGGETETQYF | KLGGALQAK | YES | NO | NO |
| 13293 | TRAV13-2 | TRAJ42 | CAENIGGGSQGNLIF | TRBV19 | TRBJ2-3 | CASSSRATDTQYF | GILGFVFTL | YES | NO | YES |
| 13294 | TRAV38-1 | TRAJ52 | CAFMKGAGGTSYGKLTF | TRBV7-9 | TRBJ2-3 | CASSSRDGGTDTQYF | GILGFVFTL | YES | NO | YES |
| 13295 | TRAV21 | TRAJ26 | CAVPNYGQNFVF | TRBV7-3 | TRBJ2-5 | CASSSRGGQETQYF | RAKFKQLL | NO | NO | NO |
| 13296 | TRAV8-3 | TRAJ9 | CAVGAAGGFKTIF | TRBV5-6 | TRBJ1-1 | CASSLRDGSEAFF | RAKFKQLL | YES | NO | NO |
| 13297 | TRAV17 | TRAJ53 | CATDRGSNYKLTF | TRBV13 | TRBJ2-1 | CASSPGGGNEQFF | KLGGALQAK | YES | NO | NO |
| 13298 | TRAV21 | TRAJ17 | CALGAAGNKLTF | TRBV12-3 | TRBJ2-1 | CASSVSGGGYNEQFF | AVFDRKSDAK | YES | NO | YES |
| 13299 | TRAV20 | TRAJ29 | CAVQDEFSGNTPLVF | TRBV6-6 | TRBJ1-1 | CASSYPQGTGTEAFF | KLGGALQAK | NO | NO | NO |
| 13300 | TRAV21 | TRAJ24 | CAVEDSWGKLQF | TRBV7-9 | TRBJ1-2 | CASSLGTGDGYGYTF | IMDQVPFSV | YES | NO | NO |
| 13301 | TRAV2 | TRAJ43 | CAVEDYNDMRF | TRBV12-3 | TRBJ2-1 | CASSREGEQFF | KLGGALQAK | NO | NO | NO |
| 13302 | TRAV8-3 | TRAJ12 | CAVGRMDSSYKLIF | TRBV5-6 | TRBJ1-1 | CASSLRDGSEAFF | RAKFKQLL | YES | NO | YES |
| 13303 | TRAV41 | TRAJ42 | CAIYYGGSQGNLIF | TRBV27 | TRBJ2-3 | CASRQGAPLNTQYF | KLGGALQAK | YES | NO | NO |
| 13304 | TRAV8-4 | TRAJ21 | CAVSGSYNFNKFYF | TRBV27 | TRBJ1-2 | CASSPRTGVYGYTF | KLGGALQAK | YES | NO | NO |
| 13305 | TRAV10 | TRAJ27 | CVGYTNAGKSTF | TRBV6-3 | TRBJ2-7 | CASGMTGLTSEQYF | KLGGALQAK | YES | NO | NO |
| 13306 | TRAV9-2 | TRAJ54 | CALTYQGAQKLVF | TRBV27 | TRBJ2-3 | CASRPRGATDTQYF | KLGGALQAK | YES | NO | NO |
| 13307 | TRAV19 | TRAJ20 | CALSEDNDYKLSF | TRBV10-3 | TRBJ2-5 | CAISESVAGEQETQYF | RAKFKQLL | YES | NO | YES |
| 13308 | TRAV5 | TRAJ31 | CAEDNNARLMF | TRBV4-2 | TRBJ2-2 | CASSQRPSEVGELFF | RAKFKQLL | YES | NO | YES |
| 13309 | TRAV1-2 | TRAJ33 | CAVKNSNYQLIW | TRBV6-4 | TRBJ2-6 | CASSDSGEGANVLTF | KLGGALQAK | YES | NO | NO |
| 13310 | TRAV1-2 | TRAJ33 | CAVLDSNYQLIW | TRBV27 | TRBJ2-7 | CASSFYEQYF | AVFDRKSDAK | YES | NO | NO |
| 13311 | TRAV29DV5 | TRAJ28 | CAANYGAGSYQLTF | TRBV4-2 | TRBJ1-3 | CASSQVVGNTIYF | GILGFVFTL | YES | NO | YES |
| 13312 | TRAV1-2 | TRAJ33 | CAQIDSNYQLFW | TRBV6-4 | TRBJ2-2 | CASSATSGVAGELFF | KLGGALQAK | YES | NO | NO |
| 13313 | TRAV13-1 | TRAJ23 | CAASINQGGKLIF | TRBV7-6 | TRBJ2-7 | CASSRGPNYEQYF | KLGGALQAK | YES | NO | NO |
| 13314 | TRAV5 | TRAJ36 | CAESLQTGANNLFF | TRBV20-1 | TRBJ2-1 | CSARDFQGLAYNEQFF | KLGGALQAK | NO | NO | NO |
| 13315 | TRAV16 | TRAJ53 | CALSYSGGSNYKLTF | TRBV10-3 | TRBJ2-1 | CAISESDAGLYNEQFF | AVFDRKSDAK | NO | NO | NO |
| 13316 | TRAV12-2 | TRAJ4 | CAVPSGGYNKLIF | TRBV19 | TRBJ2-7 | CASSGTSGGYEQYF | GILGFVFTL | YES | NO | YES |
| 13317 | TRAV13-2 | TRAJ12 | CAENMGDSSYKLIF | TRBV12-4 | TRBJ2-7 | CASRPLGGTEEQYF | KLGGALQAK | YES | NO | NO |
| 13318 | TRAV8-6 | TRAJ36 | CAVSDPQTGANNLFF | TRBV5-4 | TRBJ2-7 | CASSFRGPGYEQYF | AVFDRKSDAK | YES | NO | YES |
| 13319 | TRAV13-1 | TRAJ50 | CAASETSYDKVIF | TRBV12-3 | TRBJ1-2 | CASTQQGSYGYTF | RAKFKQLL | YES | YES | YES |
| 13320 | TRAV21 | TRAJ50 | CAVKTSYDKVIF | TRBV7-9 | TRBJ2-1 | CASSSRLAGAEDEQFF | KLGGALQAK | YES | NO | NO |
| 13321 | TRAV1-2 | TRAJ12 | CAVLDSSYKLIF | TRBV6-4 | TRBJ2-3 | CASSDSSTDTQYF | AVFDRKSDAK | YES | NO | YES |
| 13322 | TRAV1-2 | TRAJ33 | CAVRDSNYQLIW | TRBV6-1 | TRBJ2-6 | CASDESPGANVLTF | KLGGALQAK | YES | NO | NO |
| 13323 | TRAV13-1 | TRAJ40 | CAASPPGYIF | TRBV6-6 | TRBJ2-5 | CASSYGRGSGETQYF | KLGGALQAK | NO | NO | NO |
| 13324 | TRAV13-2 | TRAJ45 | CAENGGGGADGLTF | TRBV12-3 | TRBJ1-4 | CASSLGSATNEKLFF | RAKFKQLL | YES | NO | YES |
| 13325 | TRAV12-3 | TRAJ9 | CAMKGGFKTIF | TRBV15 | TRBJ2-1 | CAILQGTANEQFF | AVFDRKSDAK | YES | NO | NO |
| 13326 | TRAV14DV4 | TRAJ27 | CAMREGRANAGKSTF | TRBV6-1 | TRBJ2-3 | CASTAGTASTDTQYF | KLGGALQAK | YES | NO | NO |
| 13327 | TRAV29DV5 | TRAJ49 | CAASQRGDQFYF | TRBV5-1 | TRBJ2-7 | CASSLAQGFHEQYF | AVFDRKSDAK | YES | NO | YES |
| 13328 | TRAV14DV4 | TRAJ45 | CAMRGDSGGGADGLTF | TRBV12-4 | TRBJ2-1 | CASGRLSYNEQFF | RAKFKQLL | YES | NO | YES |
| 13329 | TRAV13-1 | TRAJ22 | CAASGGQLTF | TRBV30 | TRBJ1-1 | CAWNLGQVYTEAFF | KLGGALQAK | YES | NO | NO |
| 13330 | TRAV12-3 | TRAJ32 | CAMNPAWGGATNKLIF | TRBV19 | TRBJ2-7 | CASSARSSYEQYF | GILGFVFTL | YES | NO | NO |
| 13331 | TRAV5 | TRAJ6 | CAEMEGGSYIPTF | TRBV4-2 | TRBJ1-5 | CASSSTGGGEKDQPQHF | RAKFKQLL | YES | NO | YES |
| 13332 | TRAV13-1 | TRAJ50 | CAAAETSYDKVIF | TRBV19 | TRBJ2-2 | CASSWIAANTGELFF | RAKFKQLL | YES | NO | YES |
| 13333 | TRAV13-1 | TRAJ21 | CAPWGFNKFYF | TRBV7-3 | TRBJ2-3 | CASSLMATDTQYF | RAKFKQLL | YES | NO | YES |
| 13334 | TRAV3 | TRAJ20 | CAVRALNDYKLSF | TRBV19 | TRBJ2-7 | CASSARSSYEQYF | GILGFVFTL | NO | NO | NO |
| 13335 | TRAV29DV5 | TRAJ33 | CAARGGSNYQLIW | TRBV4-2 | TRBJ1-5 | CASSQDHLGPNQPQHF | KLGGALQAK | YES | NO | NO |
| 13336 | TRAV24 | TRAJ29 | CASPYSGNTPLVF | TRBV11-2 | TRBJ2-7 | CASSVGGPSYEQYF | KLGGALQAK | YES | NO | NO |
| 13337 | TRAV12-3 | TRAJ56 | CAMPGANSKLTF | TRBV6-5 | TRBJ2-1 | CASSLNHLSSYNEQFF | KLGGALQAK | NO | NO | NO |
| 13339 | TRAV3 | TRAJ36 | CAVRDGTGANNLFF | TRBV19 | TRBJ2-1 | CASNSRQGRQFF | GILGFVFTL | NO | NO | NO |
| 13340 | TRAV1-2 | TRAJ15 | CAVRGVHQAGTALIF | TRBV5-5 | TRBJ1-1 | CASSLRDGSEAFF | KLGGALQAK | YES | NO | NO |
| 13341 | TRAV1-1 | TRAJ35 | CAVRGGFGNVLHC | TRBV27 | TRBJ2-7 | CASSLSGRPYEQYF | RAKFKQLL | NO | NO | NO |
| 13342 | TRAV21 | TRAJ20 | CAVRYDYKLSF | TRBV7-9 | TRBJ2-1 | CASSYPHEQFF | KLGGALQAK | YES | NO | NO |
| 13343 | TRAV17 | TRAJ24 | CATDDLGLTTDSWGKFQF | TRBV19 | TRBJ1-5 | CASSIFGSSNQPQHF | GILGFVFTL | YES | NO | YES |
| 13344 | TRAV1-2 | TRAJ31 | CAVENNNARLMF | TRBV4-2 | TRBJ2-1 | CASSQDPGQGAHEQFF | KLGGALQAK | YES | NO | NO |
| 13345 | TRAV17 | TRAJ53 | CATDAKGGSNYKLTF | TRBV29-1 | TRBJ2-1 | CSVGSGDGGEQFF | RAKFKQLL | YES | NO | YES |
| 13346 | TRAV34 | TRAJ49 | CGADPNTGNQFYF | TRBV30 | TRBJ1-1 | CAWRIDRPYEAFF | KLGGALQAK | NO | NO | NO |
| 13347 | TRAV8-3 | TRAJ21 | CAVGDNFNKFYF | TRBV5-1 | TRBJ1-1 | CASSYGTGENTEAFF | RAKFKQLL | YES | NO | YES |
| 13348 | TRAV13-2 | TRAJ45 | CADSGGGADGLTF | TRBV5-1 | TRBJ2-7 | CASSYKGLAITTYEQYF | RAKFKQLL | YES | NO | YES |
| 13349 | TRAV27 | TRAJ42 | CAGPGDGGSQGNLIF | TRBV6-3 | TRBJ2-7 | CASSYSLQGGLDEQYF | GILGFVFTL | YES | NO | YES |
| 13350 | TRAV1-2 | TRAJ33 | CAVLDSNYQLIW | TRBV6-3 | TRBJ1-5 | CASSYTDNQPQHF | KLGGALQAK | YES | NO | NO |
| 13351 | TRAV9-2 | TRAJ9 | CALTLNTGGFKTIF | TRBV10-3 | TRBJ2-1 | CAISSRDGRTDNEQFF | KLGGALQAK | YES | NO | NO |
| 13352 | TRAV1-2 | TRAJ31 | CAVRDRSNARLMF | TRBV29-1 | TRBJ1-1 | CSDWEGGSTEAFF | AVFDRKSDAK | YES | NO | YES |
| 13354 | TRAV12-2 | TRAJ52 | CAVNGRDRGTSYGKLTF | TRBV3-1 | TRBJ2-5 | CASSMTSGSFQTQYF | KLGGALQAK | YES | YES | NO |
| 13355 | TRAV20 | TRAJ34 | CAVPNPYNTDKLIF | TRBV3-1 | TRBJ1-1 | CASSQETGAEAFF | AVFDRKSDAK | YES | NO | YES |
| 13356 | TRAV1-2 | TRAJ31 | CAVRDPRARLMF | TRBV9 | TRBJ2-2 | CASSVELGGQGATTGELFF | KLGGALQAK | YES | NO | NO |
| 13357 | TRAV38-2DV8 | TRAJ31 | CAYRSAWENNARLMF | TRBV9 | TRBJ1-2 | CASSGGAGNNYGYTF | KLGGALQAK | YES | NO | NO |
| 13358 | TRAV12-1 | TRAJ17 | CVVNFPQAAGNKLTF | TRBV4-1 | TRBJ2-5 | CASSQVLLAGGLVPEETQYF | KLGGALQAK | NO | NO | NO |
| 13359 | TRAV3 | TRAJ12 | CAVRSGGMDSSYKLIF | TRBV6-3 | TRBJ1-2 | CATLLGVDYTF | KLGGALQAK | NO | NO | NO |
| 13360 | TRAV8-3 | TRAJ40 | CAVGARSGTYKYIF | TRBV12-3 | TRBJ2-3 | CASSLILTDGTDTQYF | KLGGALQAK | YES | NO | NO |
| 13362 | TRAV24 | TRAJ39 | CAFDRNNAGNMLTF | TRBV4-3 | TRBJ2-3 | CASSLGSGGTDTQYF | KLGGALQAK | YES | NO | NO |
| 13363 | TRAV21 | TRAJ22 | CAVLDVGSARQLTF | TRBV7-6 | TRBJ2-3 | CASSFVEATSGSSAQYF | KLGGALQAK | YES | NO | NO |
| 13364 | TRAV4 | TRAJ45 | CLVGDKGGGADGLTF | TRBV7-9 | TRBJ2-6 | CASSLSGAGANVLTF | KLGGALQAK | YES | NO | NO |

| | | | | | | | | | | |
|---|---|---|---|---|---|---|---|---|---|---|
| 13365 | TRAV1-2 | TRAJ12 | CAFKMDSSYKLIF | TRBV9 | TRBJ2-1 | CASSPGLAVHNEQFF | KLGGALQAK | YES | NO | NO |
| 13366 | TRAV1-2 | TRAJ31 | CAVRDNNNNARLMF | TRBV6-3 | TRBJ2-7 | CASSSPRDEQYV | KLGGALQAK | NO | NO | NO |
| 13367 | TRAV23DV6 | TRAJ58 | CAAMEETSGSRLTF | TRBV5-6 | TRBJ1-2 | CASSLIGYGYTF | KLGGALQAK | NO | NO | NO |
| 13368 | TRAV39 | TRAJ34 | CAVVDNTDKLIF | TRBV6-5 | TRBJ2-1 | CASSYSGGNEQFF | KLGGALQAK | NO | NO | NO |
| 13369 | TRAV8-1 | TRAJ35 | CAVKGVGFGNVLHC | TRBV19 | TRBJ1-2 | CASRNSGQGVDGYTF | KLGGALQAK | NO | NO | NO |
| 13370 | TRAV25 | TRAJ43 | CAGFSNNDMRF | TRBV19 | TRBJ1-1 | CATGYNNPAMTEAFF | KLGGALQAK | YES | NO | NO |
| 13371 | TRAV13-2 | TRAJ33 | CAENEHSNYQLIW | TRBV12-4 | TRBJ2-7 | CASTATGFYEQYF | KLGGALQAK | YES | NO | NO |
| 13372 | TRAV1-2 | TRAJ20 | CAVKTNDYKLSF | TRBV12-5 | TRBJ1-1 | CASGLVPENTEAFF | KLGGALQAK | YES | NO | NO |
| 13373 | TRAV16 | TRAJ28 | CALAAYSGAGSYQLTF | TRBV7-2 | TRBJ1-2 | CASSSSLGPYGYTF | AVFDRKSDAK | NO | NO | NO |
| 13374 | TRAV10 | TRAJ34 | CVVSLFDTDKLIF | TRBV11-2 | TRBJ2-5 | CASSHGLAGGETQYF | KLGGALQAK | YES | NO | NO |
| 13375 | TRAV4 | TRAJ23 | CLVGNQGGKLIF | TRBV11-2 | TRBJ2-3 | CASSPSNPSTDTQYF | KLGGALQAK | NO | NO | NO |
| 13376 | TRAV19 | TRAJ40 | CALSGAPSGTYKYIF | TRBV29-1 | TRBJ1-2 | CSVDLEANYGYTF | KLGGALQAK | YES | NO | NO |
| 13377 | TRAV25 | TRAJ20 | CAGRPINDYKLSF | TRBV4-3 | TRBJ1-5 | CASSQVYEGYQPQHF | AVFDRKSDAK | YES | NO | YES |
| 13378 | TRAV17 | TRAJ52 | CATDGADAGGTSYGKLTF | TRBV6-3 | TRBJ2-1 | CASSYGAGGYNEQFF | KLGGALQAK | YES | NO | NO |
| 13379 | TRAV3 | TRAJ6 | CAVRDIRAGGSYIPTF | TRBV27 | TRBJ1-2 | CASSQAPGPAYGYTF | KLGGALQAK | NO | NO | NO |
| 13380 | TRAV21 | TRAJ57 | CAVTPAQGGSEKLVF | TRBV6-5 | TRBJ1-5 | CASSYDGGNQPQHF | KLGGALQAK | YES | NO | NO |
| 13381 | TRAV35 | TRAJ54 | CAGGIIQGAQKLVF | TRBV7-9 | TRBJ1-2 | CASSPGQGSYGYTF | KLGGALQAK | YES | NO | NO |
| 13382 | TRAV12-2 | TRAJ52 | CAVNFRSSYGKLTF | TRBV27 | TRBJ2-7 | CASSSLIRGAVGEQYF | KLGGALQAK | YES | NO | NO |
| 13383 | TRAV25 | TRAJ9 | CAGLNTGGFKTIF | TRBV10-2 | TRBJ1-2 | CASSESVDGRIINRYTF | KLGGALQAK | YES | NO | NO |
| 13384 | TRAV29DV5 | TRAJ45 | CAAVASGGGADGLTF | TRBV10-1 | TRBJ1-1 | CASSDSGITEAFF | KLGGALQAK | YES | NO | NO |
| 13385 | TRAV21 | TRAJ49 | CAVSTGNQFYF | TRBV19 | TRBJ2-7 | CATRQGYEQYF | KLGGALQAK | YES | NO | NO |
| 13386 | TRAV12-2 | TRAJ54 | CAAGAQKLVF | TRBV4-3 | TRBJ2-3 | CASSQEGGFTDTQYF | KLGGALQAK | YES | NO | NO |
| 13387 | TRAV12-1 | TRAJ40 | CVVNPPSGTYKYIF | TRBV4-3 | TRBJ2-1 | CASSQDLGGGGGNEQFF | KLGGALQAK | YES | NO | NO |
| 13388 | TRAV14DV4 | TRAJ9 | CAMRAYTGGFKTIF | TRBV7-8 | TRBJ1-4 | CASSLSQVTNEKLFF | KLGGALQAK | YES | NO | NO |
| 13389 | TRAV27 | TRAJ53 | CAGESGGSNYKLTF | TRBV7-3 | TRBJ1-1 | CASSYGTGWGTEAFF | KLGGALQAK | NO | NO | NO |
| 13390 | TRAV8-4 | TRAJ49 | CAVRAGNQFYF | TRBV7-6 | TRBJ2-2 | CASSPGRNTGELFF | KLGGALQAK | YES | NO | NO |
| 13391 | TRAV19 | TRAJ57 | CALSDESQGGSEKLVF | TRBV19 | TRBJ1-1 | CASSMGWGPAFF | AVFDRKSDAK | YES | NO | YES |
| 13392 | TRAV12-1 | TRAJ49 | CVVYTGNQFYF | TRBV7-9 | TRBJ2-2 | CASSPLAPTGELFF | KLGGALQAK | YES | NO | NO |
| 13393 | TRAV3 | TRAJ24 | CAVRDNKSWGKFQF | TRBV27 | TRBJ2-1 | CASSLYGTSGSSYNEQFF | KLGGALQAK | NO | NO | NO |
| 13394 | TRAV17 | TRAJ22 | CATADISGSARQLTF | TRBV10-3 | TRBJ2-2 | CAISALGGVIGELFF | KLGGALQAK | YES | NO | NO |
| 13395 | TRAV41 | TRAJ57 | CAHPQGGSEKLVF | TRBV5-1 | TRBJ2-7 | CASSLNGGTYEQYV | AVFDRKSDAK | NO | NO | NO |
| 13396 | TRAV19 | TRAJ57 | CALSLPQGGSEKLVF | TRBV19 | TRBJ1-5 | CASSIALGGWQPQHF | KLGGALQAK | YES | NO | NO |
| 13397 | TRAV3 | TRAJ34 | CAVRDGGGTDKLIF | TRBV3-1 | TRBJ2-2 | CASSLRAGVGTGELFF | KLGGALQAK | NO | NO | NO |
| 13398 | TRAV13-1 | TRAJ40 | CAADSSTSGTYKYIF | TRBV4-1 | TRBJ1-1 | CASSQQGATEAFF | KLGGALQAK | YES | NO | NO |
| 13399 | TRAV27 | TRAJ15 | CAGDSGQAGTALIF | TRBV19 | TRBJ2-1 | CASSINLGLAGVWSYNEQFF | KLGGALQAK | NO | NO | NO |
| 13400 | TRAV14DV4 | TRAJ50 | CARWRKTSYDKVIF | TRBV29-1 | TRBJ1-1 | CSVVVTEAFF | KLGGALQAK | YES | NO | NO |
| 13401 | TRAV1-2 | TRAJ23 | CAVSWRNQGGKLIF | TRBV6-5 | TRBJ2-1 | CASSYSIAGVYNEQFF | KLGGALQAK | NO | NO | NO |
| 13402 | TRAV14DV4 | TRAJ5 | CAMQDTGRRALTF | TRBV14 | TRBJ1-3 | CASSQASHQTRNTIYF | KLGGALQAK | NO | NO | NO |
| 13403 | TRAV13-1 | TRAJ45 | CAASRPGGGADGLTF | TRBV20-1 | TRBJ1-1 | CSAWDRLNTEAFF | KLGGALQAK | YES | NO | NO |
| 13404 | TRAV24 | TRAJ11 | CAPQDSTLTF | TRBV19 | TRBJ1-1 | CASTVGGTEAFF | KLGGALQAK | YES | NO | NO |
| 13405 | TRAV14DV4 | TRAJ48 | CAMREGYNFGNEKLTF | TRBV13 | TRBJ2-2 | CASSRIEQANTGELFF | KLGGALQAK | NO | NO | NO |
| 13406 | TRAV1-2 | TRAJ26 | CAVRDNPYGQNFVF | TRBV6-6 | TRBJ2-5 | CASRPGLAGAGGIVSEETQYF | RLRAEAQVK | NO | NO | NO |
| 13407 | TRAV12-1 | TRAJ23 | CVVNGNQGGKLIF | TRBV4-1 | TRBJ2-3 | CASSPFAGSTDTQYF | AVFDRKSDAK | YES | NO | YES |
| 13408 | TRAV21 | TRAJ52 | CAVLNAGGTSYGKLTF | TRBV28 | TRBJ1-5 | CASSLSSNQPQHF | KLGGALQAK | YES | NO | NO |
| 13409 | TRAV8-6 | TRAJ39 | CAVSEQNNAGNMLTF | TRBV5-6 | TRBJ1-5 | CASSLGGSNQPQHF | KLGGALQAK | YES | NO | NO |
| 13410 | TRAV29DV5 | TRAJ56 | CAARPTGANSKLTF | TRBV7-9 | TRBJ2-2 | CASIGLSGAGELFF | IVTDFSVIK | YES | NO | NO |
| 13411 | TRAV5 | TRAJ23 | CAESIDNQGGKLIF | TRBV29-1 | TRBJ1-2 | CSVDLEANYGYTF | KLGGALQAK | YES | NO | NO |
| 13412 | TRAV14DV4 | TRAJ29 | CAMRASGNTPLVF | TRBV10-2 | TRBJ1-2 | CASSETVSALNGYTF | KLGGALQAK | YES | NO | NO |
| 13413 | TRAV17 | TRAJ15 | CATDARQAGTALIF | TRBV4-1 | TRBJ2-5 | CASSQASRGWETQYF | KLGGALQAK | YES | NO | NO |
| 13414 | TRAV22 | TRAJ15 | CAVSSPLNQAGTALIF | TRBV5-5 | TRBJ1-1 | CASSLEGGKAFF | KLGGALQAK | YES | NO | NO |
| 13415 | TRAV12-1 | TRAJ44 | CVVNTLTGTASKLTF | TRBV7-9 | TRBJ2-1 | CASSSPGLAGGSSYNEQFF | KLGGALQAK | YES | NO | NO |
| 13416 | TRAV2 | TRAJ31 | CAVEGIYNARLMF | TRBV6-6 | TRBJ1-2 | CASSYSFFEGAEGEHGYTF | KLGGALQAK | NO | NO | NO |
| 13417 | TRAV19 | TRAJ33 | CALSEDSYGDSNYQLIW | TRBV20-1 | TRBJ1-6 | CSARDPTRTVGAYNSPLHF | AVFDRKSDAK | NO | NO | NO |
| 13418 | TRAV3 | TRAJ43 | CAVLLYNNNDMRF | TRBV20-1 | TRBJ2-7 | CSAPQKRGGSYEQYF | AVFDRKSDAK | NO | NO | NO |
| 13419 | TRAV35 | TRAJ28 | CAGSYSGAGSYQLTF | TRBV6-1 | TRBJ2-2 | CASSTFRVKHTGELFF | KLGGALQAK | YES | NO | NO |
| 13420 | TRAV1-2 | TRAJ22 | CAVRDMGSARQLTF | TRBV7-2 | TRBJ1-4 | CASSPPGAGEKLFF | KLGGALQAK | YES | NO | NO |
| 13421 | TRAV1-2 | TRAJ34 | CASYNTDKLIF | TRBV27 | TRBJ1-1 | CASRGQGTTEAFF | KLGGALQAK | YES | NO | NO |
| 13422 | TRAV9-2 | TRAJ28 | CALSEGLTF | TRBV11-2 | TRBJ2-3 | CASSLSGGGQSTDTQYF | KLGGALQAK | YES | NO | NO |
| 13423 | TRAV3 | TRAJ13 | CAVRDFRHSGGYQKVTF | TRBV7-7 | TRBJ1-1 | CASSVPTGDIEAFF | KLGGALQAK | NO | NO | NO |
| 13424 | TRAV17 | TRAJ10 | CATDSTGGGNKLTF | TRBV12-4 | TRBJ2-2 | CASSLVHLWTGELFF | KLGGALQAK | YES | NO | NO |
| 13425 | TRAV27 | TRAJ37 | CAGSSNTGKLIF | TRBV7-9 | TRBJ1-1 | CASSLASGRSTEAFF | KLGGALQAK | YES | NO | NO |
| 13426 | TRAV25 | TRAJ33 | CAGEYMDSNYQLIW | TRBV3-1 | TRBJ2-3 | CASSQDFGGTSTDTQYF | KLGGALQAK | YES | NO | NO |
| 13427 | TRAV3 | TRAJ23 | CAVRERNNQGGKLIF | TRBV11-3 | TRBJ2-3 | CASRTPLDTQYF | KLGGALQAK | YES | NO | NO |
| 13428 | TRAV5 | TRAJ33 | CAESIADSNYQLIW | TRBV14 | TRBJ1-1 | CASSQEGHIEAFF | KLGGALQAK | NO | NO | NO |
| 13429 | TRAV4 | TRAJ8 | CLVGEQGFQKLVF | TRBV5-6 | TRBJ1-1 | CASSPDLGTEAFF | KLGGALQAK | YES | NO | NO |
| 13430 | TRAV12-1 | TRAJ11 | CVVAPWQGYSTLTF | TRBV12-3 | TRBJ1-1 | CASRDGTEAFF | KLGGALQAK | YES | NO | NO |
| 13431 | TRAV8-3 | TRAJ42 | CAVGAGYGGSQGNLIF | TRBV19 | TRBJ1-5 | CASREQGTHSNQPQHF | KLGGALQAK | YES | NO | NO |
| 13432 | TRAV25 | TRAJ28 | CAGRSYSGAGSYQLTF | TRBV7-9 | TRBJ2-6 | CASSSRTEIFSGANVLTF | KLGGALQAK | YES | NO | NO |
| 13433 | TRAV41 | TRAJ6 | CAVSLGGSYIPTF | TRBV3-1 | TRBJ1-3 | CASSPNPTQGTSGNTIYF | KLGGALQAK | YES | NO | NO |
| 13434 | TRAV36DV7 | TRAJ48 | CAVVPNFGNEKLTF | TRBV19 | TRBJ2-2 | CASRRLASDTGELFF | KLGGALQAK | NO | NO | NO |
| 13435 | TRAV14DV4 | TRAJ5 | CAMREMDTGRRALTF | TRBV5-6 | TRBJ2-5 | CASSPETGGETQYF | KLGGALQAK | YES | NO | NO |
| 13436 | TRAV14DV4 | TRAJ4 | CAMRADPGYNKLIF | TRBV25-1 | TRBJ2-1 | CASSVTGIYNEQFF | KLGGALQAK | YES | NO | NO |
| 13437 | TRAV19 | TRAJ37 | CALSEAQFNGSSNTGKLIF | TRBV5-5 | TRBJ1-1 | CASSLSSLGLPEAFF | KLGGALQAK | YES | NO | NO |
| 13438 | TRAV12-3 | TRAJ58 | CAMSAASGSRLTF | TRBV24-1 | TRBJ1-2 | CATSDSFIRTDRKWDGYTF | KLGGALQAK | YES | NO | NO |
| 13439 | TRAV8-4 | TRAJ12 | CAVSALLMDSSYKLIF | TRBV5-8 | TRBJ2-7 | CASSLPGSYEQYF | KLGGALQAK | YES | NO | NO |
| 13440 | TRAV17 | TRAJ30 | CATENRDDKIIF | TRBV5-6 | TRBJ2-1 | CASSSGLARNEQFF | KLGGALQAK | YES | NO | NO |
| 13441 | TRAV17 | TRAJ10 | CATEAYAGGGNKLTF | TRBV7-9 | TRBJ1-3 | CASSLAADRVFGNTIYF | IVTDFSVIK | YES | NO | YES |
| 13442 | TRAV21 | TRAJ30 | CAVNRDDKIIF | TRBV7-9 | TRBJ2-3 | CASSLAVAAWSTDTQYF | KLGGALQAK | YES | NO | NO |
| 13443 | TRAV39 | TRAJ45 | CAAAGGVYSGGGADGLTF | TRBV7-6 | TRBJ2-7 | CASSLAERGGGEQYF | KLGGALQAK | YES | NO | NO |
| 13444 | TRAV9-2 | TRAJ21 | CALSERDVYNFNKFYF | TRBV6-6 | TRBJ1-1 | CASSSDSEAFF | KLGGALQAK | NO | NO | NO |
| 13445 | TRAV20 | TRAJ28 | CAVRAGAGSYQLTF | TRBV11-2 | TRBJ2-5 | CASSLGTLATSETQYF | KLGGALQAK | YES | NO | NO |
| 13446 | TRAV1-2 | TRAJ34 | CVTLPAADKLIF | TRBV4-3 | TRBJ1-4 | CASSQELAGTGPDEKLFF | KLGGALQAK | YES | NO | NO |
| 13447 | TRAV39 | TRAJ40 | CAGWSGTYKYIF | TRBV7-2 | TRBJ1-1 | CASSLVLGTTEAFF | KLGGALQAK | NO | NO | NO |
| 13448 | TRAV24 | TRAJ42 | CAFKRDYGGSQGNLIF | TRBV7-8 | TRBJ1-1 | CASSKRTGQNTEAFF | KLGGALQAK | YES | NO | NO |
| 13449 | TRAV5 | TRAJ11 | CAESPSGYSTLTF | TRBV11-2 | TRBJ2-2 | CASSLKTGELFF | KLGGALQAK | YES | NO | NO |
| 13450 | TRAV8-1 | TRAJ22 | CAVRSSGSARQLTF | TRBV7-2 | TRBJ2-1 | CASSHQARYDLFNEQFF | KLGGALQAK | YES | NO | NO |
| 13451 | TRAV14DV4 | TRAJ20 | CAMRELNDYKLSF | TRBV7-2 | TRBJ1-2 | CASSRMGARDGYTF | KLGGALQAK | YES | NO | NO |
| 13452 | TRAV8-2 | TRAJ39 | CVVSDHNNAGNMLTF | TRBV2 | TRBJ2-7 | CASSEEWGEIYEQYF | KLGGALQAK | YES | NO | NO |
| 13453 | TRAV25 | TRAJ36 | CAGILYQTGANNLFF | TRBV27 | TRBJ2-2 | CASSLLWGELFF | KLGGALQAK | YES | NO | NO |
| 13454 | TRAV1-2 | TRAJ32 | CAVRVLIGGATNKLIF | TRBV20-1 | TRBJ1-4 | CSARYRGIKEKLFF | KLGGALQAK | YES | NO | NO |
| 13455 | TRAV19 | TRAJ53 | CAFGGSNYKLTF | TRBV6-1 | TRBJ1-1 | CASNAAGTAFNTEAFF | KLGGALQAK | YES | NO | NO |
| 13456 | TRAV2 | TRAJ15 | CAVEDDQAGTALIF | TRBV4-1 | TRBJ1-1 | CASSQELRDRVSEAFF | KLGGALQAK | NO | NO | NO |
| 13457 | TRAV38-2DV8 | TRAJ33 | CAYTSNSNYQLIW | TRBV11-2 | TRBJ2-5 | CASSEGTSWVETQYF | KLGGALQAK | YES | NO | NO |
| 13458 | TRAV8-2 | TRAJ22 | CVGRKGSARQLTF | TRBV3-1 | TRBJ1-5 | CASSHYQPQHF | KLGGALQAK | YES | NO | NO |
| 13459 | TRAV24 | TRAJ49 | CASPNTGNQFYF | TRBV12-5 | TRBJ1-5 | CASGLAGQTSNQPQHF | KLGGALQAK | YES | NO | NO |
| 13460 | TRAV3 | TRAJ5 | CAVRAYTGRRALTF | TRBV4-1 | TRBJ1-4 | CASSQTTGEGEKLFF | KLGGALQAK | YES | NO | NO |
| 13461 | TRAV19 | TRAJ26 | CALSEDNYGQNFVF | TRBV7-6 | TRBJ2-5 | CASSLVGTSGRTQETQYF | KLGGALQAK | YES | NO | NO |
| 13462 | TRAV4 | TRAJ40 | CLVAYSGTYKYIF | TRBV7-9 | TRBJ2-1 | CASSYPGLQSNEQFF | AVFDRKSDAK | YES | NO | YES |
| 13463 | TRAV27 | TRAJ44 | CAGLYTGTASKLTF | TRBV14 | TRBJ1-3 | CASRRRTGNTIYF | KLGGALQAK | NO | NO | NO |
| 13464 | TRAV5 | TRAJ37 | CAETPGSSNTGKLIF | TRBV19 | TRBJ2-1 | CASSLPAQSSGRFFYEQFF | KLGGALQAK | YES | NO | NO |
| 13465 | TRAV19 | TRAJ52 | CALSEADAGGTSYGKLTF | TRBV9 | TRBJ2-1 | CASSSGRASDEQFF | KLGGALQAK | YES | NO | NO |
| 13466 | TRAV8-1 | TRAJ9 | CAVNANTGGFKTIF | TRBV29-1 | TRBJ2-7 | CSVAGGKNYEQYF | KLGGALQAK | NO | NO | NO |
| 13467 | TRAV12-2 | TRAJ26 | CAVNDNYGQNFVF | TRBV5-1 | TRBJ2-7 | CASSSSGSADEQYV | KLGGALQAK | NO | NO | NO |
| 13468 | TRAV8-3 | TRAJ50 | CAVEGSYDKVIF | TRBV7-9 | TRBJ2-7 | CASSRVSGQYF | KLGGALQAK | YES | NO | NO |

| | | | | | | | | | | |
|---|---|---|---|---|---|---|---|---|---|---|
| 13469 | TRAV41 | TRAJ42 | CAVRYLRSQGNLIF | TRBV20-1 | TRBJ1-6 | CSAHGQGAYSPLHF | KLGGALQAK | YES | NO | NO |
| 13470 | TRAV21 | TRAJ42 | CAVDGGSQGNLIF | TRBV19 | TRBJ1-2 | CASREGQAYGYTF | KLGGALQAK | YES | NO | NO |
| 13471 | TRAV27 | TRAJ52 | CAAPNAGGTSYGKLTF | TRBV11-2 | TRBJ2-3 | CASSLDYFGVFSTDTQYF | KLGGALQAK | YES | NO | NO |
| 13472 | TRAV10 | TRAJ4 | CVVRSGGYNKLIF | TRBV6-5 | TRBJ2-7 | CASSYSAGEDYEQYF | KLGGALQAK | YES | NO | NO |
| 13473 | TRAV35 | TRAJ26 | CAGQGNYGQNFVF | TRBV6-5 | TRBJ2-1 | CASSRYNEQFF | KLGGALQAK | NO | NO | NO |
| 13474 | TRAV39 | TRAJ54 | CAVDSAGAQKLVF | TRBV10-3 | TRBJ1-1 | CAIRETAMNTEAFF | KLGGALQAK | YES | NO | NO |
| 13475 | TRAV1-2 | TRAJ24 | CAVSATDSWGKFQF | TRBV4-1 | TRBJ1-5 | CASSHPDLNQPQHF | KLGGALQAK | YES | NO | NO |
| 13476 | TRAV23DV6 | TRAJ52 | CAATGEGGGTSYGKLTF | TRBV10-2 | TRBJ2-1 | CASSDGGGGYYNEQFF | KLGGALQAK | YES | NO | NO |
| 13477 | TRAV19 | TRAJ48 | CALWRNFGNEKLTF | TRBV4-1 | TRBJ2-3 | CASSQEGLGQGGPDTQYF | KLGGALQAK | YES | NO | NO |
| 13478 | TRAV1-2 | TRAJ15 | CAVRDGPAGQAGTALIF | TRBV18 | TRBJ2-5 | CASSPASGAQETQYF | KLGGALQAK | YES | NO | NO |
| 13479 | TRAV8-3 | TRAJ39 | CAVGAVGNAGNMLTF | TRBV4-1 | TRBJ1-5 | CASSQEFGQSNQPQHF | KLGGALQAK | YES | NO | NO |
| 13480 | TRAV3 | TRAJ44 | CAVINTGTASKLTF | TRBV5-1 | TRBJ2-7 | CASSWDRAYEQYF | KLGGALQAK | YES | NO | NO |
| 13481 | TRAV19 | TRAJ17 | CALSRGAAGNKLTF | TRBV29-1 | TRBJ1-2 | CSVDLEANYGYTF | KLGGALQAK | YES | NO | NO |
| 13482 | TRAV8-3 | TRAJ23 | CAVGEGGKLIF | TRBV5-4 | TRBJ2-3 | CASSWGDTQYF | AVFDRKSDAK | YES | NO | YES |
| 13483 | TRAV14DV4 | TRAJ49 | CAMREDFTGNQFYF | TRBV25-1 | TRBJ2-1 | CASIERGSYNEQFF | KLGGALQAK | YES | NO | NO |
| 13484 | TRAV35 | TRAJ57 | CAGQPPWVAQGGSEKLVF | TRBV7-2 | TRBJ2-7 | CASSLMILYEQYF | KLGGALQAK | YES | NO | NO |
| 13485 | TRAV19 | TRAJ24 | CALSAATDSWGKFQF | TRBV20-1 | TRBJ2-3 | CSARDFSGSTDTQYF | RLRAEAQVK | YES | NO | YES |
| 13486 | TRAV8-6 | TRAJ20 | CAVRANDYKLSF | TRBV4-3 | TRBJ1-1 | CASSQGSEAFF | GILGFVFTL | YES | NO | YES |
| 13487 | TRAV12-3 | TRAJ52 | CAILNAGGTSYGKLTF | TRBV27 | TRBJ1-2 | CASSNAGTGDGWDYGYTF | KLGGALQAK | YES | NO | NO |
| 13488 | TRAV1-2 | TRAJ28 | CAVMTGAGSYQLTF | TRBV6-3 | TRBJ2-3 | CASRDPRDTQYF | KLGGALQAK | YES | NO | NO |
| 13489 | TRAV9-2 | TRAJ48 | CALTSPSNFGNEKLTF | TRBV2 | TRBJ2-7 | CASSDRIYEQYV | AVFDRKSDAK | NO | NO | NO |
| 13491 | TRAV23DV6 | TRAJ24 | CAASPTDSWGKLQF | TRBV9 | TRBJ1-2 | CASSADRTTDYGYTF | KLGGALQAK | YES | NO | NO |
| 13492 | TRAV17 | TRAJ15 | CATDLGGTALIF | TRBV11-2 | TRBJ2-1 | CASSLIRGQGGLRDRGEQFF | KLGGALQAK | NO | NO | NO |
| 13493 | TRAV23DV6 | TRAJ13 | CAAIYSGGYQKVTF | TRBV7-6 | TRBJ2-1 | CASSSRTADNEQFF | KLGGALQAK | YES | NO | NO |
| 13494 | TRAV29DV5 | TRAJ29 | CAAPGNTPLVF | TRBV10-2 | TRBJ1-5 | CASSPGTGGYQPQHF | KLGGALQAK | YES | NO | NO |
| 13495 | TRAV25 | TRAJ40 | CAGQTGTYKYIF | TRBV27 | TRBJ1-5 | CASSLGQGVQPQHF | KLGGALQAK | YES | NO | NO |
| 13496 | TRAV41 | TRAJ48 | CAVRSVSNFGNEKLTF | TRBV7-8 | TRBJ1-5 | CASSLAGPQRYSNQPQHF | KLGGALQAK | YES | NO | NO |
| 13497 | TRAV8-3 | TRAJ36 | CAVPHVGTGANNLFF | TRBV6-5 | TRBJ2-1 | CASSYSLAGGYNEQFF | KLGGALQAK | NO | NO | NO |
| 13498 | TRAV26-1 | TRAJ36 | CIVRVAMGANNLFF | TRBV6-3 | TRBJ1-5 | CASSYGSPIQPQHF | KLGGALQAK | YES | NO | NO |
| 13499 | TRAV29DV5 | TRAJ53 | CAASSFGGSNYKLTF | TRBV7-9 | TRBJ2-1 | CASRRGGGQGFHNEQFF | AVFDRKSDAK | YES | NO | YES |
| 13500 | TRAV14DV4 | TRAJ4 | CAMRPPFRGGYNKLIF | TRBV5-4 | TRBJ2-1 | CASSGLAIEQFF | KLGGALQAK | YES | NO | NO |
| 13501 | TRAV14DV4 | TRAJ4 | CAMRRPSGGYNKLIF | TRBV7-9 | TRBJ2-1 | CASTGTSGGPTLRDEQFF | KLGGALQAK | YES | NO | NO |
| 13502 | TRAV19 | TRAJ58 | CALSEVEETSGSRLTF | TRBV7-8 | TRBJ2-2 | CASSTGLAGGPDTGELFF | KLGGALQAK | YES | YES | NO |
| 13503 | TRAV12-2 | TRAJ34 | CAVRSDTDKLIF | TRBV4-1 | TRBJ1-6 | CASSQLGEGSSPLHF | KLGGALQAK | YES | NO | NO |
| 13504 | TRAV8-2 | TRAJ9 | CVVSRGFKTIF | TRBV9 | TRBJ1-3 | CASSVGEGTGGNTIYF | KLGGALQAK | YES | NO | NO |
| 13505 | TRAV8-3 | TRAJ35 | CAVSI | TRBV11-2 | TRBJ1-1 | CASSSEGMMNTEAFF | KLGGALQAK | NO | NO | NO |
| 13506 | TRAV19 | TRAJ21 | CALSEGRDFNKFYF | TRBV4-3 | TRBJ2-6 | CASSQDLGSGANVLTF | KLGGALQAK | YES | NO | NO |
| 13507 | TRAV19 | TRAJ52 | CALSEVAGGTSYGKLTF | TRBV20-1 | TRBJ2-1 | CSALRYNEQFF | KLGGALQAK | NO | NO | NO |
| 13508 | TRAV38-1 | TRAJ48 | CAFMKHGNFGNEKLTF | TRBV11-3 | TRBJ2-1 | CASSLTFAPYNEQFF | KLGGALQAK | YES | NO | NO |
| 13509 | TRAV12-2 | TRAJ39 | CAGNAGNMLTF | TRBV7-8 | TRBJ1-1 | CASSLVSPNTEAFF | KLGGALQAK | YES | NO | NO |
| 13510 | TRAV17 | TRAJ52 | CATPAGGTSYGKLTF | TRBV7-9 | TRBJ2-3 | CASSSTRQGTDTQYF | KLGGALQAK | YES | NO | NO |
| 13511 | TRAV19 | TRAJ24 | CALAPTTDSWGKFQF | TRBV19 | TRBJ1-1 | CASGWFAEAFF | KLGGALQAK | YES | NO | NO |
| 13512 | TRAV26-2 | TRAJ58 | CILRETSGSRLTF | TRBV20-1 | TRBJ1-5 | CSASGGTHQPQHF | KLGGALQAK | YES | NO | NO |
| 13513 | TRAV16 | TRAJ42 | CALAHDGSQGNLIF | TRBV20-1 | TRBJ2-1 | CSARMVGALQFF | KLGGALQAK | NO | NO | NO |
| 13514 | TRAV5 | TRAJ42 | CAELGGSQGNLIF | TRBV6-5 | TRBJ2-7 | CASSHEVSYEQYF | KLGGALQAK | YES | NO | NO |
| 13515 | TRAV26-1 | TRAJ52 | CIVNVPGGTSYGKLTF | TRBV12-4 | TRBJ1-1 | CASSLTGNTEAFF | KLGGALQAK | YES | NO | NO |
| 13516 | TRAV20 | TRAJ4 | CAVQVSGGYNKLIF | TRBV6-3 | TRBJ1-6 | CASSYTGTDSPLHF | AVFDRKSDAK | YES | NO | YES |
| 13517 | TRAV17 | TRAJ31 | CATDAFANARLMF | TRBV3-1 | TRBJ1-2 | CASRGRTTNYGYTF | KLGGALQAK | YES | NO | NO |
| 13519 | TRAV4 | TRAJ34 | CLVGEGNTDKLIF | TRBV2 | TRBJ2-7 | CASSEEISGSYEQYF | KLGGALQAK | YES | NO | NO |
| 13520 | TRAV5 | TRAJ31 | CAERAGDARLMF | TRBV28 | TRBJ2-5 | CASTAQETQYF | KLGGALQAK | YES | NO | NO |
| 13521 | TRAV12-3 | TRAJ49 | CAGTGNQFYF | TRBV10-3 | TRBJ2-1 | CAVNPAGAKTYNEQFF | KLGGALQAK | YES | NO | NO |
| 13522 | TRAV25 | TRAJ41 | CAGGDSGYALNF | TRBV6-1 | TRBJ2-1 | CASSTLMPPVQFF | KLGGALQAK | YES | NO | NO |
| 13523 | TRAV1-2 | TRAJ28 | CAVRYGAGSYQLTF | TRBV9 | TRBJ2-1 | CASSVVPGTDNEQFF | KLGGALQAK | YES | NO | NO |
| 13524 | TRAV22 | TRAJ13 | CAVGRGGYQKVTF | TRBV11-2 | TRBJ2-7 | CASSLDFFDRTYEQYV | AVFDRKSDAK | NO | NO | NO |
| 13525 | TRAV38-2DV8 | TRAJ31 | CAYRGLNNNARLMF | TRBV5-4 | TRBJ1-1 | CASSQPENTEAFF | KLGGALQAK | YES | NO | NO |
| 13526 | TRAV12-2 | TRAJ15 | CAVFNQAGTALIF | TRBV7-9 | TRBJ1-1 | CASSSVGTEAFF | KLGGALQAK | YES | NO | NO |
| 13527 | TRAV17 | TRAJ44 | CATDRGGKLTF | TRBV10-3 | TRBJ2-2 | CAISAAGTSGGYTGELFF | KLGGALQAK | YES | NO | NO |
| 13528 | TRAV1-2 | TRAJ9 | CAVRDRAGGFKTIF | TRBV7-2 | TRBJ2-7 | CASSLQTSGGEQYF | KLGGALQAK | YES | NO | NO |
| 13530 | TRAV8-6 | TRAJ54 | CAVSAIQGAQKLVF | TRBV5-1 | TRBJ1-1 | CASSLGRGLNTEAFF | KLGGALQAK | YES | NO | NO |
| 13531 | TRAV38-2DV8 | TRAJ23 | CAYRSALEGKLIF | TRBV19 | TRBJ1-1 | CASSMGPNTEAFF | KLGGALQAK | YES | NO | NO |
| 13532 | TRAV1-2 | TRAJ21 | CAVRDQGNFNKFYF | TRBV20-1 | TRBJ1-3 | CSARDLHRPGNTIYF | KLGGALQAK | YES | NO | NO |
| 13533 | TRAV29DV5 | TRAJ34 | CAASRYNTDKLIF | TRBV7-9 | TRBJ2-1 | CASSLVGGQGVANEQFF | AVFDRKSDAK | YES | NO | YES |
| 13534 | TRAV21 | TRAJ54 | CAVEVQGAQKLVF | TRBV29-1 | TRBJ1-2 | CSVDLEANYGYTF | KLGGALQAK | YES | NO | NO |
| 13535 | TRAV21 | TRAJ36 | CAVPDQTGANNLFF | TRBV2 | TRBJ2-5 | CASSEPLQDYQETQYF | KLGGALQAK | YES | NO | NO |
| 13536 | TRAV20 | TRAJ53 | CAVQVAGGSNYKLTF | TRBV4-1 | TRBJ1-3 | CASSPRTGGNTIYF | KLGGALQAK | YES | NO | NO |
| 13538 | TRAV5 | TRAJ18 | CAESGRGSTLGRLYF | TRBV4-1 | TRBJ2-7 | CASSPKTGKGEQYF | KLGGALQAK | YES | NO | NO |
| 13539 | TRAV3 | TRAJ35 | CAVYGFGNVLHC | TRBV6-6 | TRBJ2-1 | CASSILAGSNEQFF | KLGGALQAK | NO | NO | NO |
| 13540 | TRAV14DV4 | TRAJ4 | CAMRPPFSGGYNKLIF | TRBV4-3 | TRBJ1-4 | CASSQDHRMGGHEKLFF | KLGGALQAK | YES | NO | NO |
| 13541 | TRAV12-2 | TRAJ20 | CAVMGNDYKLSF | TRBV27 | TRBJ2-2 | CASSFSPGSTGELFF | KLGGALQAK | YES | NO | NO |
| 13542 | TRAV38-2DV8 | TRAJ30 | CAYSDLDDKIIF | TRBV19 | TRBJ1-2 | CASSILGTANYGYTF | KLGGALQAK | YES | NO | NO |
| 13543 | TRAV29DV5 | TRAJ17 | CAASAQAAGNKLTF | TRBV7-6 | TRBJ1-1 | CASSPPGYTEAFF | KLGGALQAK | YES | NO | NO |
| 13544 | TRAV8-4 | TRAJ30 | CAVSDHRDDKIIF | TRBV4-1 | TRBJ1-6 | CASTLALGDSGKAPLHF | KLGGALQAK | YES | NO | NO |
| 13545 | TRAV12-1 | TRAJ49 | CVVERLLATTGNQFYF | TRBV7-9 | TRBJ2-3 | CASSHRVADTQYF | KLGGALQAK | YES | NO | NO |
| 13547 | TRAV12-1 | TRAJ52 | CVAFGTSYGKLTF | TRBV5-1 | TRBJ1-2 | CASSFRTGETNYGYTF | KLGGALQAK | YES | NO | NO |
| 13548 | TRAV12-2 | TRAJ18 | CAGTDRGSTLGRLYF | TRBV13 | TRBJ2-5 | CASSYAGQHQETQYF | AVFDRKSDAK | YES | NO | YES |
| 13549 | TRAV35 | TRAJ49 | CAGQSGNQFYF | TRBV4-1 | TRBJ1-2 | CASSSVQGTGYGYTF | KLGGALQAK | YES | NO | NO |
| 13550 | TRAV1-2 | TRAJ45 | CAVTILYSGGGADGLTF | TRBV20-1 | TRBJ1-1 | CSARVGLGTEAFF | KLGGALQAK | YES | NO | NO |
| 13551 | TRAV13-1 | TRAJ52 | CAASKDAGGTSYGKLTF | TRBV7-6 | TRBJ2-7 | CASSLQQGVYEQYF | KLGGALQAK | YES | NO | NO |
| 13552 | TRAV8-6 | TRAJ40 | CAVSDNWGTYKYIF | TRBV27 | TRBJ2-4 | CASSPDLGDIQYF | AVFDRKSDAK | YES | NO | YES |
| 13553 | TRAV39 | TRAJ43 | CAVVNNDMRF | TRBV4-1 | TRBJ2-7 | CASSQGRLVRGGYEQYF | KLGGALQAK | YES | NO | NO |
| 13554 | TRAV17 | TRAJ45 | CATDAYSGGGADGLTF | TRBV13 | TRBJ2-5 | CASSTYGETQYF | KLGGALQAK | YES | NO | NO |
| 13555 | TRAV21 | TRAJ20 | CARPGDYKLSF | TRBV5-1 | TRBJ2-1 | CASSLWLAGGYNEQFF | KLGGALQAK | YES | NO | NO |
| 13556 | TRAV12-2 | TRAJ56 | CAPYSGANSKLTF | TRBV11-2 | TRBJ2-7 | CASSSGTRPTEQYV | KLGGALQAK | NO | NO | NO |
| 13557 | TRAV9-2 | TRAJ28 | CALSDAGAGSYQLTF | TRBV12-3 | TRBJ2-2 | CASRDDRSRNTGELFF | AVFDRKSDAK | YES | NO | YES |
| 13558 | TRAV14DV4 | TRAJ20 | CAMSPRSNDYKLSF | TRBV12-4 | TRBJ2-3 | CASSRGPDTQYF | KLGGALQAK | YES | NO | NO |
| 13559 | TRAV19 | TRAJ24 | CALSEPDSWGKFQF | TRBV5-6 | TRBJ2-1 | CASSLAVKAGGPKDEQFF | KLGGALQAK | YES | NO | NO |
| 13560 | TRAV38-2DV8 | TRAJ53 | CAYRSGSGGSNYKLTF | TRBV5-5 | TRBJ2-4 | CASSSEQGLRNIQYF | KLGGALQAK | YES | NO | NO |
| 13561 | TRAV22 | TRAJ33 | CAVERWDSNYQLIW | TRBV7-9 | TRBJ1-6 | CASSLSPGGNSPLHF | KLGGALQAK | YES | NO | NO |
| 13562 | TRAV21 | TRAJ45 | CAVWAPGGGADGLTF | TRBV19 | TRBJ1-2 | CASSIGNSGYTF | KLGGALQAK | YES | NO | NO |
| 13563 | TRAV29DV5 | TRAJ31 | CAASYNARLMF | TRBV27 | TRBJ1-2 | CASSPNPDIRNTF | KLGGALQAK | YES | NO | NO |
| 13564 | TRAV26-2 | TRAJ49 | CILRVGYQFYF | TRBV9 | TRBJ2-2 | CASSVGGLAGGVNTGELFF | KLGGALQAK | YES | NO | NO |
| 13565 | TRAV8-1 | TRAJ13 | CAVLTGGYQKVTF | TRBV12-5 | TRBJ2-2 | CAHTDPNPTGELFF | KLGGALQAK | YES | NO | NO |
| 13566 | TRAV8-3 | TRAJ6 | CAVISGGSYIPTF | TRBV20-1 | TRBJ1-2 | CSARDGDRGLYGYTF | KLGGALQAK | YES | NO | NO |
| 13567 | TRAV38-1 | TRAJ48 | CAFVAGFGNEKLTF | TRBV7-2 | TRBJ1-3 | CASSFGGNTIYF | KLGGALQAK | YES | NO | NO |
| 13568 | TRAV5 | TRAJ13 | CAGMGGYQKVTF | TRBV29-1 | TRBJ2-5 | CSVEFARYQETQYF | AVFDRKSDAK | YES | NO | YES |
| 13569 | TRAV13-1 | TRAJ38 | CAASLNAGNNRKLIW | TRBV27 | TRBJ1-1 | CASSLSVRGSTEAFF | MLDLQPETT | YES | NO | YES |
| 13570 | TRAV19 | TRAJ56 | CALSAGANSKLTF | TRBV9 | TRBJ1-1 | CASSPWTGPLNTEAFF | KLGGALQAK | YES | NO | NO |
| 13571 | TRAV14DV4 | TRAJ24 | CAMREGLTTDSWGKLQF | TRBV10-3 | TRBJ2-3 | CAISEDGQQLGDTQYF | KLGGALQAK | YES | NO | NO |
| 13572 | TRAV8-4 | TRAJ54 | CAVSGPKLVF | TRBV4-1 | TRBJ1-2 | CASSRAAGGVDGYTF | KLGGALQAK | YES | NO | NO |
| 13573 | TRAV1-2 | TRAJ17 | CAVRDGAAGNKLTF | TRBV7-9 | TRBJ1-1 | CASSLARTVNTEAFF | KLGGALQAK | YES | NO | NO |
| 13574 | TRAV25 | TRAJ45 | CAGLLPGGGADGLTF | TRBV5-1 | TRBJ2-5 | CASSIREDFQETQYF | KLGGALQAK | YES | NO | NO |
| 13575 | TRAV14DV4 | TRAJ13 | CAMREVNSGGYQKVTF | TRBV15 | TRBJ2-3 | CATSRATSGRAPDTQYF | KLGGALQAK | YES | NO | NO |
| 13576 | TRAV29DV5 | TRAJ48 | CAASVGDFGNEKLTF | TRBV4-3 | TRBJ1-4 | CASSQDHRMGGHEKLFF | KLGGALQAK | YES | NO | NO |
| 13577 | TRAV19 | TRAJ52 | CALSEAGAGGTSYGKLTF | TRBV12-4 | TRBJ2-2 | CASSKIEAAGELFF | KLGGALQAK | YES | NO | NO |

| | | | | | | | | | | |
|---|---|---|---|---|---|---|---|---|---|---|
| 13578 | TRAV13-2 | TRAJ45 | CAESKGGADGLTF | TRBV4-1 | TRBJ2-1 | CASSQGVHEQFF | KLGGALQAK | YES | NO | NO |
| 13579 | TRAV13-1 | TRAJ10 | CAFLPGGGNKLTF | TRBV6-3 | TRBJ2-7 | CASSYSYEQYF | KLGGALQAK | YES | NO | NO |
| 13580 | TRAV38-1 | TRAJ58 | CAFRETSGSRLTF | TRBV5-6 | TRBJ1-2 | CASSLLGTNYGYTF | KLGGALQAK | YES | NO | NO |
| 13581 | TRAV12-1 | TRAJ49 | CVVNTGNQFYF | TRBV7-2 | TRBJ2-5 | CASSLAFRRGGETQYF | KLGGALQAK | YES | NO | NO |
| 13582 | TRAV13-2 | TRAJ28 | CAENLGGSGAGSYQLTF | TRBV7-9 | TRBJ1-2 | CASSFTDYGYTF | KLGGALQAK | YES | NO | NO |
| 13583 | TRAV5 | TRAJ23 | CAETLTYNQGGKLIF | TRBV4-1 | TRBJ2-2 | CASSHSQGATGELFF | KLGGALQAK | YES | NO | NO |
| 13584 | TRAV8-4 | TRAJ45 | CAVSAFSGGGADGLTF | TRBV7-9 | TRBJ1-2 | CASSHRYFYGYTF | KLGGALQAK | YES | NO | NO |
| 13585 | TRAV16 | TRAJ34 | CALSGLDTDKLIF | TRBV20-1 | TRBJ1-2 | CSAPPAGGRQGYTF | KLGGALQAK | NO | NO | NO |
| 13586 | TRAV12-2 | TRAJ18 | CAVKDRGSTLGRLYF | TRBV7-2 | TRBJ2-1 | CASSSPPGLAGSHEQFF | KLGGALQAK | YES | NO | NO |
| 13587 | TRAV34 | TRAJ45 | CGAGLYSGGGADGLTF | TRBV4-1 | TRBJ1-1 | CASSSRMNTEAFF | KLGGALQAK | YES | NO | NO |
| 13588 | TRAV26-2 | TRAJ31 | CILHLNNARLMF | TRBV4-1 | TRBJ1-1 | CASSPHPMNTEAFF | KLGGALQAK | YES | NO | NO |
| 13589 | TRAV41 | TRAJ28 | CAVRYSGAGSYQLTF | TRBV7-3 | TRBJ2-7 | CASSLGLGSSYEQYF | KLGGALQAK | NO | NO | NO |
| 13590 | TRAV3 | TRAJ28 | CAVRDGSGAGSYQLTF | TRBV10-2 | TRBJ1-1 | CASRSGWNTEAFF | KLGGALQAK | NO | NO | NO |
| 13591 | TRAV13-2 | TRAJ47 | CADPGQYGNKLVF | TRBV7-8 | TRBJ2-1 | CASSLSGGTNEQFF | KLGGALQAK | YES | NO | NO |
| 13592 | TRAV26-2 | TRAJ13 | CILRDTGGYQKVTF | TRBV15 | TRBJ1-6 | CATSRDPQDNSPLHF | RLRAEAQVK | YES | NO | YES |
| 13593 | TRAV17 | TRAJ26 | CATDDNYGQNFVF | TRBV3-1 | TRBJ1-3 | CASSRQPSGNTIYF | KLGGALQAK | YES | NO | NO |
| 13594 | TRAV12-2 | TRAJ40 | CAVKATSGTYKYIF | TRBV29-1 | TRBJ2-1 | CSVEFSSGGVI | KLGGALQAK | NO | NO | NO |
| 13595 | TRAV41 | TRAJ33 | CAVWTGMDSNYQLIW | TRBV2 | TRBJ2-7 | CASSDAGSGYEQYV | KLGGALQAK | NO | NO | NO |
| 13596 | TRAV8-1 | TRAJ35 | CAVNAI | TRBV14 | TRBJ2-7 | CASSLYYEQYF | KLGGALQAK | NO | NO | NO |
| 13597 | TRAV17 | TRAJ9 | CATDATGGFKTIF | TRBV6-5 | TRBJ2-2 | CASSLMTGLGDTGELFF | KLGGALQAK | YES | NO | NO |
| 13598 | TRAV5 | TRAJ31 | CAETANNNARLMF | TRBV19 | TRBJ1-1 | CASSIGLNTEAFF | KLGGALQAK | YES | NO | NO |
| 13599 | TRAV8-6 | TRAJ53 | CAVSDQNSGGSNYKLTF | TRBV27 | TRBJ2-1 | CASTPRRTLEQFF | KLGGALQAK | YES | NO | NO |
| 13600 | TRAV35 | TRAJ45 | CAGALRLNLVSGGGADGLTF | TRBV6-5 | TRBJ2-2 | CASSSTGGSGELFF | KLGGALQAK | NO | NO | NO |
| 13601 | TRAV26-2 | TRAJ57 | CILRQGGSEKLVF | TRBV7-6 | TRBJ2-3 | CASSLAPGTDTQYF | KLGGALQAK | YES | NO | NO |
| 13602 | TRAV35 | TRAJ57 | CAGPPAGTQGGSEKLVF | TRBV19 | TRBJ1-6 | CASNDRGHSPLHF | KLGGALQAK | YES | NO | NO |
| 13603 | TRAV17 | TRAJ20 | CATDRDDYKLSF | TRBV6-3 | TRBJ2-5 | CASRRESQETQYF | KLGGALQAK | YES | NO | NO |
| 13604 | TRAV8-3 | TRAJ45 | CAVGAVGSGGGADGLTF | TRBV6-6 | TRBJ2-7 | CASGTAPYEQYF | KLGGALQAK | NO | NO | NO |
| 13605 | TRAV10 | TRAJ3 | CVVTSSSASKIIF | TRBV12-4 | TRBJ2-1 | CASSGTSGYNEQFF | KLGGALQAK | YES | NO | NO |
| 13606 | TRAV25 | TRAJ34 | CAGRNTDKLIF | TRBV5-1 | TRBJ1-1 | CASSLGRGLNTEAFF | KLGGALQAK | YES | NO | NO |
| 13607 | TRAV8-1 | TRAJ6 | CAASGGSYIPTF | TRBV11-2 | TRBJ2-1 | CASSPLDGEQFF | RLRAEAQVK | YES | YES | YES |
| 13608 | TRAV12-2 | TRAJ30 | CAVIGRDDKIIF | TRBV15 | TRBJ2-1 | CAIKIAEAYNEQFF | KLGGALQAK | YES | NO | NO |
| 13609 | TRAV12-2 | TRAJ48 | CAEGTPNFGNEKLTF | TRBV29-1 | TRBJ1-2 | CSVEGGAFGGSNYGYTF | KLGGALQAK | YES | NO | NO |
| 13610 | TRAV13-1 | TRAJ33 | CAASPDSNYQLIW | TRBV7-8 | TRBJ1-2 | CASSRDDAGGGYGYTF | KLGGALQAK | YES | NO | NO |
| 13611 | TRAV12-2 | TRAJ54 | CAGGGGAQKLVF | TRBV12-4 | TRBJ1-1 | CASSFSATEAFF | KLGGALQAK | YES | NO | NO |
| 13612 | TRAV14DV4 | TRAJ42 | CAMREAWGGGSQGNLIF | TRBV7-9 | TRBJ2-3 | CASSLTRTDTQYF | KLGGALQAK | YES | NO | NO |
| 13613 | TRAV3 | TRAJ23 | CAVSGMIYNQGGKLIF | TRBV12-5 | TRBJ1-1 | CASGSGGRTEAFF | KLGGALQAK | NO | NO | NO |
| 13614 | TRAV38-1 | TRAJ52 | CAFMHLNTGGTSYGKLTF | TRBV9 | TRBJ1-1 | CASRGRDRTSTEAFF | KLGGALQAK | YES | NO | NO |
| 13615 | TRAV26-2 | TRAJ49 | CILRDYGNQFYF | TRBV9 | TRBJ2-1 | CASSVGGTGYNEQFF | KLGGALQAK | YES | NO | NO |
| 13616 | TRAV17 | TRAJ11 | CATDSGGYSTLTF | TRBV7-9 | TRBJ1-1 | CASSLQPSGGANTEAFF | KLGGALQAK | YES | NO | NO |
| 13617 | TRAV8-1 | TRAJ20 | CAVKDYKLSF | TRBV10-2 | TRBJ2-7 | CASSLGQGYEQYF | KLGGALQAK | YES | NO | NO |
| 13618 | TRAV19 | TRAJ42 | CALSEALGYGGSQGNLIF | TRBV19 | TRBJ2-4 | CASTPVGQGAKNIQYF | KLGGALQAK | YES | NO | NO |
| 13619 | TRAV12-1 | TRAJ4 | CVVNDPGGYNKLIF | TRBV12-4 | TRBJ1-3 | CASSLQRRGENTIYF | KLGGALQAK | YES | NO | NO |
| 13620 | TRAV12-3 | TRAJ49 | CAMRGLGNQFYF | TRBV28 | TRBJ2-7 | CASSSPGTAHSYEQYV | RTLNAWVKV | NO | NO | NO |
| 13621 | TRAV9-2 | TRAJ39 | CALRDNNAGNMLTF | TRBV12-4 | TRBJ1-1 | CASSPMGGLAEAFF | KLGGALQAK | YES | NO | NO |
| 13622 | TRAV8-3 | TRAJ34 | CAVGFPDNTDKLIF | TRBV24-1 | TRBJ1-1 | CATLGQENTEAFF | KLGGALQAK | YES | NO | NO |
| 13623 | TRAV24 | TRAJ52 | CAFSSAGGTSYGKLTF | TRBV12-3 | TRBJ2-1 | CASPSGGYNEQFF | KLGGALQAK | YES | NO | NO |
| 13624 | TRAV13-2 | TRAJ47 | CAENMEYGNKLVF | TRBV7-8 | TRBJ1-1 | CASSVTGTEAFF | KLGGALQAK | YES | NO | NO |
| 13625 | TRAV21 | TRAJ43 | CAVSNNNDMRF | TRBV6-5 | TRBJ1-6 | CASSYSGTGASPLHF | KLGGALQAK | YES | NO | NO |
| 13626 | TRAV12-3 | TRAJ47 | CAMSYFPEYGNKLVF | TRBV4-1 | TRBJ2-3 | CASSQEPTSGGPTDTQYF | AVFDRKSDAK | YES | NO | YES |
| 13627 | TRAV2 | TRAJ49 | CAVEWGGTGNQFYF | TRBV5-4 | TRBJ2-1 | CASSFTGLGNEQFF | KLGGALQAK | NO | NO | NO |
| 13628 | TRAV14DV4 | TRAJ20 | CAMRERTNDYKLSF | TRBV13 | TRBJ2-5 | CASKGGGETQYF | KLGGALQAK | YES | NO | NO |
| 13629 | TRAV29DV5 | TRAJ39 | CAASDAGNMLTF | TRBV11-2 | TRBJ2-5 | CASRGPVETQYF | KLGGALQAK | YES | NO | NO |
| 13630 | TRAV17 | TRAJ26 | CATDNNYGQNFVF | TRBV5-1 | TRBJ2-1 | CASSLVAGGRDEQFF | KLGGALQAK | YES | NO | NO |
| 13631 | TRAV12-3 | TRAJ33 | CAMRHSNYQLIW | TRBV7-6 | TRBJ2-1 | CASSSDYNEQFF | KLGGALQAK | YES | NO | NO |
| 13632 | TRAV21 | TRAJ21 | CAVTFEDFNKFYF | TRBV12-4 | TRBJ2-6 | CASSAGTGYGANVLTF | KLGGALQAK | YES | NO | NO |
| 13633 | TRAV12-3 | TRAJ48 | CAMSLSNFGNEKLTF | TRBV20-1 | TRBJ2-2 | CSASETSKGTGELFF | KLGGALQAK | YES | NO | NO |
| 13634 | TRAV29DV5 | TRAJ34 | CAAPLDKLIF | TRBV7-2 | TRBJ1-2 | CASSLTGPNYGYTF | KLGGALQAK | YES | NO | NO |
| 13635 | TRAV14DV4 | TRAJ22 | CAMRRPISSGSARQLTF | TRBV5-6 | TRBJ1-5 | CASSSRDSQNQPQHF | KLGGALQAK | YES | NO | NO |
| 13636 | TRAV35 | TRAJ48 | CAGQLGVGNEKLTF | TRBV6-3 | TRBJ2-2 | CASSYSRDLENTGELFF | KLGGALQAK | YES | NO | NO |
| 13637 | TRAV38-1 | TRAJ52 | CAFLFTNAGGTSYGKLTF | TRBV6-6 | TRBJ1-1 | CASSHYAVEAFF | KLGGALQAK | NO | NO | NO |
| 13639 | TRAV10 | TRAJ10 | CVVSHTTGGGNKLTF | TRBV19 | TRBJ1-5 | CASSHWQISNQPQHF | KLGGALQAK | YES | NO | NO |
| 13640 | TRAV21 | TRAJ31 | CAVNNNARLMF | TRBV27 | TRBJ2-1 | CASNTGTGIYNEQFF | KLGGALQAK | YES | NO | NO |
| 13641 | TRAV12-2 | TRAJ31 | CAVNDGARLMF | TRBV12-5 | TRBJ2-4 | CASGLVEGDSKNIQYF | KLGGALQAK | YES | NO | NO |
| 13642 | TRAV19 | TRAJ52 | CALSDPNAGGTSYGKLTF | TRBV4-3 | TRBJ2-3 | CASSQARSGRHTQYF | KLGGALQAK | YES | NO | NO |
| 13643 | TRAV13-1 | TRAJ45 | CAASSGGGADGLTF | TRBV12-3 | TRBJ2-5 | CASSMGGGIIFRTQYF | KLGGALQAK | YES | NO | NO |
| 13644 | TRAV29DV5 | TRAJ57 | CAASGIQGGSEKLVF | TRBV7-2 | TRBJ2-7 | CASSMGLRGYEQYF | KLGGALQAK | NO | NO | NO |
| 13645 | TRAV14DV4 | TRAJ52 | CAMRGSGGTSYGKLTF | TRBV12-5 | TRBJ2-5 | CASGSGVAGAYQETQYF | KLGGALQAK | YES | NO | NO |
| 13646 | TRAV1-2 | TRAJ4 | CAVISGGYNKLIF | TRBV10-1 | TRBJ2-7 | CASSAGTGGGGQYF | KLGGALQAK | YES | NO | NO |
| 13648 | TRAV14DV4 | TRAJ32 | CAMREDGGATNKLIF | TRBV12-4 | TRBJ1-2 | CASSFNIKGLNYGYTF | KLGGALQAK | YES | NO | NO |
| 13649 | TRAV14DV4 | TRAJ29 | CAMREGRNTPLVF | TRBV29-1 | TRBJ1-2 | CSVPLQGEYGYTF | KLGGALQAK | YES | NO | NO |
| 13650 | TRAV12-3 | TRAJ3 | CAMSSEYSSASKIIF | TRBV20-1 | TRBJ2-3 | CSAGSDLAGVVTDTQYF | KLGGALQAK | NO | NO | NO |
| 13651 | TRAV1-2 | TRAJ22 | CAVPGLISSGSARQLTF | TRBV18 | TRBJ1-2 | CASSPTDRANGYTF | KLGGALQAK | YES | NO | NO |
| 13652 | TRAV27 | TRAJ35 | CAGEAFGNVLHC | TRBV19 | TRBJ1-1 | CASRPGIEGSEAFF | KLGGALQAK | NO | NO | NO |
| 13653 | TRAV14DV4 | TRAJ23 | CAMREPGYNQGGKLIF | TRBV7-6 | TRBJ2-3 | CASSLVSGTALGTQYF | KLGGALQAK | YES | NO | NO |
| 13654 | TRAV13-2 | TRAJ15 | CAVSDQAGTALIF | TRBV5-1 | TRBJ1-1 | CASSLAGGTEAFF | KLGGALQAK | YES | NO | NO |
| 13655 | TRAV26-2 | TRAJ11 | CILISGYSTLTF | TRBV9 | TRBJ2-3 | CASSVATLGSTDTQYF | KLGGALQAK | YES | NO | NO |
| 13656 | TRAV3 | TRAJ31 | CAVRAGAGDARLMF | TRBV20-1 | TRBJ1-2 | CSARDPGTNYGYTF | KLGGALQAK | NO | NO | NO |
| 13657 | TRAV1-2 | TRAJ33 | CAVRDSNYQLIW | TRBV4-1 | TRBJ2-3 | CASSQFPLGEGTDTQYF | KLGGALQAK | YES | NO | NO |
| 13658 | TRAV35 | TRAJ39 | CAGHPRTGNMLTF | TRBV7-9 | TRBJ2-1 | CASSSAGISYNEQFF | KLGGALQAK | YES | NO | NO |
| 13659 | TRAV1-2 | TRAJ29 | CATLSGNTPLVF | TRBV12-4 | TRBJ2-5 | CASSVLPGQGDETQYF | KLGGALQAK | YES | NO | NO |
| 13660 | TRAV1-1 | TRAJ30 | CAVVRDDKIIF | TRBV27 | TRBJ2-7 | CASSLGGGRIPYEQYV | KLGGALQAK | NO | NO | NO |
| 13661 | TRAV6 | TRAJ9 | CALDANVGGFKTIF | TRBV5-6 | TRBJ1-1 | CASSMGNTEAFF | KLGGALQAK | YES | NO | NO |
| 13662 | TRAV14DV4 | TRAJ45 | CAMREEFPYSGGGADGLTF | TRBV11-1 | TRBJ1-1 | CASSPSLRTEAFF | KLGGALQAK | YES | NO | NO |
| 13663 | TRAV24 | TRAJ6 | CAFRSGGSYIPTF | TRBV6-4 | TRBJ1-2 | CASSEGQGANYGYTF | AVFDRKSDAK | YES | NO | YES |
| 13664 | TRAV8-6 | TRAJ4 | CAVSDAGGYNKLIF | TRBV6-3 | TRBJ2-5 | CASSWGETQYF | KLGGALQAK | YES | NO | NO |
| 13665 | TRAV10 | TRAJ52 | CVVTNAGGTSYGKLTF | TRBV5-6 | TRBJ1-6 | CASSLKGTYNSPLHF | KLGGALQAK | YES | NO | NO |
| 13666 | TRAV29DV5 | TRAJ26 | CAASGKNNYGQNFVF | TRBV12-3 | TRBJ1-1 | CASSFSVWDRPNTEAFF | KLGGALQAK | YES | NO | NO |
| 13667 | TRAV12-2 | TRAJ49 | CAGRNTGNQFYF | TRBV4-1 | TRBJ2-7 | CASSQVLAGGPYEQYF | AVFDRKSDAK | YES | NO | YES |
| 13668 | TRAV17 | TRAJ26 | CATDAHNYGQNFVF | TRBV13 | TRBJ1-2 | CASSGGQGVMDGYTF | KLGGALQAK | YES | NO | NO |
| 13669 | TRAV12-1 | TRAJ11 | CVVNRQAGYSTLTF | TRBV2 | TRBJ2-3 | CASSERSGGDTDTQYF | KLGGALQAK | YES | NO | NO |
| 13670 | TRAV14DV4 | TRAJ8 | CAMREGGDTGFQKLVF | TRBV4-1 | TRBJ2-5 | CASSQDRTWETQYF | KLGGALQAK | YES | NO | NO |
| 13671 | TRAV14DV4 | TRAJ48 | CAMREDESNFGNEKLTF | TRBV5-1 | TRBJ2-4 | CASSSLAGLQRIQYF | KLGGALQAK | YES | NO | NO |
| 13672 | TRAV25 | TRAJ9 | CAGEHTGGFKTIF | TRBV5-5 | TRBJ1-1 | CASSWGTGEPEAFF | KLGGALQAK | YES | NO | NO |
| 13673 | TRAV41 | TRAJ42 | CAVGGSQGNLIF | TRBV7-6 | TRBJ2-2 | CASSLLNTGELFF | KLGGALQAK | YES | NO | NO |
| 13674 | TRAV38-2DV8 | TRAJ56 | CAYRSPRGNSKLTF | TRBV4-2 | TRBJ2-7 | CASSHIMDRDHEQYF | KLGGALQAK | NO | NO | NO |
| 13675 | TRAV25 | TRAJ52 | CAGRAGGTSYGKLTF | TRBV6-3 | TRBJ2-7 | CASSWTSDYEQYV | KLGGALQAK | NO | NO | NO |
| 13676 | TRAV8-6 | TRAJ33 | CAVGSNYQLIW | TRBV12-4 | TRBJ2-2 | CASSPGLANTGELFF | KLGGALQAK | YES | NO | NO |
| 13677 | TRAV8-3 | TRAJ49 | CAVGAFRQFYF | TRBV4-3 | TRBJ1-4 | CASSQDHRMGGHEKLFF | KLGGALQAK | YES | NO | NO |
| 13678 | TRAV8-3 | TRAJ29 | CAVGADSGNTPLVF | TRBV18 | TRBJ1-4 | CASSPGGVSEKLFF | KLGGALQAK | YES | NO | NO |
| 13679 | TRAV20 | TRAJ21 | CAVQADFNKFYF | TRBV19 | TRBJ2-7 | CASSPRTGNYEQYF | KLGGALQAK | YES | NO | NO |
| 13680 | TRAV25 | TRAJ15 | CAGSDQAGTALIF | TRBV10-3 | TRBJ1-1 | CAISETGGRTEAFF | KLGGALQAK | YES | NO | NO |
| 13681 | TRAV21 | TRAJ37 | CAVSGNTGKLIF | TRBV4-1 | TRBJ1-2 | CASSVIPGQGSYGYTF | KLGGALQAK | YES | NO | NO |
| 13682 | TRAV1-2 | TRAJ32 | CAVWRYGGATNKLIF | TRBV7-3 | TRBJ1-1 | CASKGGLGTEAFF | KLGGALQAK | NO | NO | NO |
| 13683 | TRAV39 | TRAJ57 | CAVDKSGSEKLVF | TRBV7-2 | TRBJ2-7 | CASSLVSFGDEQYV | KLGGALQAK | NO | NO | NO |

| | | | | | | | | | | |
|---|---|---|---|---|---|---|---|---|---|---|
| 13684 | TRAV26-2 | TRAJ36 | CILPQTGANNLFF | TRBV15 | TRBJ1-5 | CATRTGGNQPQHF | KLGGALQAK | YES | NO | NO |
| 13685 | TRAV27 | TRAJ35 | CAGAI | TRBV18 | TRBJ2-1 | CASSRDGYNEQFF | KLGGALQAK | NO | NO | NO |
| 13686 | TRAV29DV5 | TRAJ48 | CAASDTINEKLTF | TRBV5-5 | TRBJ1-2 | CASSLGGLGYGYTF | KLGGALQAK | YES | NO | NO |
| 13687 | TRAV41 | TRAJ34 | CAVKGSSGYNTDKLIF | TRBV27 | TRBJ2-7 | CASSLGTVGYEQYF | KLGGALQAK | YES | NO | NO |
| 13688 | TRAV2 | TRAJ27 | CAVEDNTNAGKSTF | TRBV19 | TRBJ2-2 | CASSIILGNTGELFF | KLGGALQAK | NO | NO | NO |
| 13689 | TRAV8-2 | TRAJ34 | CVLNTDKLIF | TRBV20-1 | TRBJ2-7 | CSASDRATYEQYF | KLGGALQAK | NO | NO | NO |
| 13690 | TRAV21 | TRAJ9 | CAVSVHTGGFKTIF | TRBV7-9 | TRBJ2-1 | CASSLGLGLDNEQFF | KLGGALQAK | YES | NO | NO |
| 13691 | TRAV2 | TRAJ42 | CAVPPYGGSQGNLIF | TRBV7-9 | TRBJ2-3 | CASSSTTPSGRNTDTQYF | KLGGALQAK | NO | NO | NO |
| 13692 | TRAV19 | TRAJ28 | CALSEEYSGAGSYQLTF | TRBV9 | TRBJ2-1 | CASSLPRPGGSYNEQFF | KLGGALQAK | YES | NO | NO |
| 13693 | TRAV13-2 | TRAJ54 | CAENRRDIQGAQKLVF | TRBV5-1 | TRBJ1-1 | CASSQHRATEAFF | AVFDRKSDAK | YES | NO | YES |
| 13694 | TRAV29DV5 | TRAJ54 | CAAGHIIQGAQKLVF | TRBV7-9 | TRBJ2-7 | CASSPGPQGQYF | KLGGALQAK | YES | NO | NO |
| 13696 | TRAV26-1 | TRAJ40 | CIVGSGTYKYIF | TRBV4-1 | TRBJ1-5 | CASSQDSGVHQPQHF | AVFDRKSDAK | YES | NO | YES |
| 13697 | TRAV26-1 | TRAJ40 | CIVRVVYPGTYKYIF | TRBV14 | TRBJ2-5 | CASSPGLEKTQYF | KLGGALQAK | NO | NO | NO |
| 13698 | TRAV19 | TRAJ58 | CALRRETSGSRLTF | TRBV2 | TRBJ2-1 | CASGRLAGGYNEQFF | KLGGALQAK | YES | NO | NO |
| 13699 | TRAV24 | TRAJ52 | CAFFNAGGTSYGKLTF | TRBV10-2 | TRBJ1-1 | CASGQVREAFF | KLGGALQAK | YES | NO | NO |
| 13700 | TRAV13-2 | TRAJ28 | CAENSDGSYQLTF | TRBV4-3 | TRBJ1-3 | CASSADRASDTIYF | AVFDRKSDAK | YES | NO | NO |
| 13701 | TRAV14DV4 | TRAJ37 | CAMRGPFSSNTGKLIF | TRBV2 | TRBJ1-1 | CASWASLGGGGEAFF | KLGGALQAK | YES | NO | NO |
| 13702 | TRAV27 | TRAJ49 | CAGLAGGNQFYF | TRBV27 | TRBJ2-3 | CASTGLPGTQYF | KLGGALQAK | YES | NO | NO |
| 13703 | TRAV8-6 | TRAJ42 | CAVSDSASYGGSQGNLIF | TRBV9 | TRBJ1-4 | CASSEGGDPYEKLFF | KLGGALQAK | YES | NO | NO |
| 13704 | TRAV13-2 | TRAJ52 | CAENPSNAGGTSYGKLTF | TRBV5-4 | TRBJ2-1 | CASPGSVSNEQFF | KLGGALQAK | YES | NO | NO |
| 13705 | TRAV35 | TRAJ33 | CAGRDSNYQLIW | TRBV4-1 | TRBJ2-1 | CASSPGTFNEQFF | KLGGALQAK | YES | NO | NO |
| 13706 | TRAV8-3 | TRAJ15 | CAGANQAGTALIF | TRBV12-4 | TRBJ1-2 | CASSSDRQGAVTGYTF | KLGGALQAK | YES | NO | NO |
| 13707 | TRAV16 | TRAJ32 | CALIKRHGGATNKLIF | TRBV20-1 | TRBJ2-2 | CSARDQGGYTGELFF | KLGGALQAK | NO | NO | NO |
| 13709 | TRAV19 | TRAJ12 | CALSEGGDSSYKLIF | TRBV7-9 | TRBJ1-4 | CASSLSPGDSEKLFF | KLGGALQAK | YES | NO | NO |
| 13710 | TRAV21 | TRAJ48 | CAVTPNFGNEKLTF | TRBV10-3 | TRBJ2-3 | CAINNGPLAPPDTQYF | KLGGALQAK | YES | YES | NO |
| 13711 | TRAV20 | TRAJ58 | CAVQGVGSRLTF | TRBV4-1 | TRBJ1-1 | CASSPDPDRNTEAFF | KLGGALQAK | YES | NO | NO |
| 13712 | TRAV24 | TRAJ52 | CALNAGGTSYGKLTF | TRBV6-5 | TRBJ1-5 | CASSQDESHYQPQHF | KLGGALQAK | YES | NO | NO |
| 13713 | TRAV14DV4 | TRAJ23 | CAMREAPRGKLIF | TRBV4-3 | TRBJ1-1 | CASSQDLGGGTEAFF | KLGGALQAK | YES | NO | NO |
| 13714 | TRAV1-1 | TRAJ31 | CASNNNARLMF | TRBV11-2 | TRBJ2-3 | CASRDPTDTQYF | KLGGALQAK | YES | NO | NO |
| 13715 | TRAV19 | TRAJ40 | CALTVTTSGTYKYIF | TRBV7-9 | TRBJ2-7 | CASSRTPESSYEQYV | KLGGALQAK | NO | NO | NO |
| 13716 | TRAV8-1 | TRAJ30 | CAVTPNRDDKIIF | TRBV6-5 | TRBJ1-2 | CASSYSSGGYGYTF | KLGGALQAK | YES | NO | NO |
| 13717 | TRAV17 | TRAJ40 | CATDARGTYKYIF | TRBV27 | TRBJ2-7 | CASSLTPPGQGFYEQYF | KLGGALQAK | YES | NO | NO |
| 13718 | TRAV27 | TRAJ33 | CADMDSNYQLIW | TRBV4-1 | TRBJ2-1 | CASSQWAGDYNEQFF | KLGGALQAK | YES | NO | NO |
| 13719 | TRAV6 | TRAJ21 | CAFSVVFNKFYF | TRBV19 | TRBJ2-1 | CASSIGGSSYNEQFF | KLGGALQAK | YES | NO | NO |
| 13720 | TRAV4 | TRAJ17 | CLVGTTHRIKAAGNKLTF | TRBV6-5 | TRBJ2-7 | CASNNRGRSYEQYF | KLGGALQAK | YES | NO | NO |
| 13721 | TRAV8-1 | TRAJ44 | CAVGTGTASKLTF | TRBV4-3 | TRBJ1-1 | CASSPSTPGNTEAFF | KLGGALQAK | YES | NO | NO |
| 13722 | TRAV6 | TRAJ35 | CALDMWK | TRBV19 | TRBJ1-1 | CASSTNRGLFSGDTEAFF | KLGGALQAK | NO | NO | NO |
| 13723 | TRAV5 | TRAJ42 | CAENGGSQGNLIF | TRBV9 | TRBJ1-1 | CASSRPSGMNTEAFF | KLGGALQAK | YES | NO | NO |
| 13724 | TRAV39 | TRAJ18 | CAVVPDRGSTLGRLYF | TRBV5-1 | TRBJ2-7 | CASVRDNSYEQYF | KLGGALQAK | YES | NO | NO |
| 13725 | TRAV27 | TRAJ47 | CAGGSWEYGNKLVF | TRBV10-3 | TRBJ2-2 | CAISETKNTGELFF | KLGGALQAK | YES | NO | NO |
| 13726 | TRAV39 | TRAJ31 | CASPERKNARLMF | TRBV11-2 | TRBJ1-2 | CASSLGRNTLRNYGYTF | KLGGALQAK | YES | NO | NO |
| 13727 | TRAV14DV4 | TRAJ4 | CAMRGPFSGGYNKLIF | TRBV6-3 | TRBJ2-1 | CASILDSLSNEQFF | KLGGALQAK | YES | NO | NO |
| 13728 | TRAV19 | TRAJ5 | CALSEALGGRRALTF | TRBV7-8 | TRBJ2-5 | CASYPVPGLKTQYF | KLGGALQAK | YES | NO | NO |
| 13729 | TRAV14DV4 | TRAJ34 | CAMREHNTDKLIF | TRBV19 | TRBJ2-7 | CASKAGTGEYEQYF | KLGGALQAK | YES | NO | NO |
| 13730 | TRAV16 | TRAJ53 | CALVGRSNYKLTF | TRBV11-2 | TRBJ2-1 | CASSFGTAEEQFF | KLGGALQAK | NO | NO | NO |
| 13731 | TRAV8-1 | TRAJ42 | CAVNAGSQGNLIF | TRBV6-6 | TRBJ2-5 | CATEGLAEWGQQETQYF | KLGGALQAK | NO | NO | NO |
| 13732 | TRAV2 | TRAJ20 | CAVVDDYKLSF | TRBV3-1 | TRBJ2-7 | CASSQDLYEQYF | KLGGALQAK | NO | NO | NO |
| 13733 | TRAV13-2 | TRAJ15 | CAENIRNNQAGTALIF | TRBV13 | TRBJ1-4 | CASSPGTLDEKLFF | KLGGALQAK | YES | NO | NO |
| 13734 | TRAV14DV4 | TRAJ9 | CAMREGPGGGGFKTIF | TRBV24-1 | TRBJ2-1 | CATSATVYNEQFF | KLGGALQAK | YES | NO | NO |
| 13735 | TRAV38-2DV8 | TRAJ44 | CAYDTGTASKLTF | TRBV20-1 | TRBJ1-1 | CSARDFDRGLATEAFF | IVTDFSVIK | YES | NO | YES |
| 13736 | TRAV8-3 | TRAJ43 | CAVAFADMRF | TRBV27 | TRBJ2-6 | CASSSRDLDGSGANVLTF | IVTDFSVIK | NO | NO | NO |
| 13737 | TRAV41 | TRAJ45 | CAVSSPSGGGADGLTF | TRBV12-4 | TRBJ1-1 | CASSFFGGYTEAFF | KLGGALQAK | YES | NO | NO |
| 13738 | TRAV17 | TRAJ23 | CALFNQGGKLIF | TRBV6-5 | TRBJ1-5 | CASRSKTGSKQPQHF | KLGGALQAK | YES | NO | NO |
| 13739 | TRAV22 | TRAJ32 | CAVYKEDYGGATNKLIF | TRBV19 | TRBJ2-7 | CASSSWTGAYEQYF | KLGGALQAK | YES | NO | NO |
| 13740 | TRAV17 | TRAJ28 | CATATRGAGSYQLTF | TRBV7-9 | TRBJ1-4 | CASSPDNGEKLFF | KLGGALQAK | YES | NO | NO |
| 13741 | TRAV19 | TRAJ43 | CALSEAGNNNDMRF | TRBV4-2 | TRBJ1-5 | CASSQGGRINQPQHF | KLGGALQAK | YES | NO | NO |
| 13742 | TRAV8-6 | TRAJ44 | CAVSVRAGTASKLTF | TRBV11-3 | TRBJ2-2 | CASSLDSGFGELFF | KLGGALQAK | YES | NO | NO |
| 13743 | TRAV12-2 | TRAJ35 | CAGRYGNVLHC | TRBV7-6 | TRBJ2-2 | CASSLQGTGELFF | KLGGALQAK | NO | NO | NO |
| 13744 | TRAV1-2 | TRAJ15 | CAVPNQAGTALIF | TRBV27 | TRBJ2-1 | CASSPESSYNEQFF | KLGGALQAK | YES | NO | NO |
| 13745 | TRAV12-1 | TRAJ15 | CVAHQAGTALIF | TRBV5-4 | TRBJ2-2 | CASSGTGSETGELFF | KLGGALQAK | YES | NO | NO |
| 13746 | TRAV41 | TRAJ33 | CAARISQMDSNYQLIW | TRBV9 | TRBJ2-2 | CASSVEPNTGELFF | KLGGALQAK | YES | NO | NO |
| 13747 | TRAV14DV4 | TRAJ45 | CAMRTRSGGADGLTF | TRBV28 | TRBJ2-2 | CASSRDRASFTGELFF | KLGGALQAK | YES | NO | NO |
| 13749 | TRAV38-2DV8 | TRAJ57 | CAYRSFGGGSEKLVF | TRBV24-1 | TRBJ2-2 | CATSDEGQITGELFF | KLGGALQAK | YES | NO | NO |
| 13750 | TRAV26-2 | TRAJ37 | CILRFGSSNTGKLIF | TRBV7-9 | TRBJ2-7 | CAATLGGGYEQYF | KLGGALQAK | YES | NO | NO |
| 13751 | TRAV25 | TRAJ16 | CAGRSDGQKLLF | TRBV11-2 | TRBJ2-1 | CASSPTGQGYNEQFF | KLGGALQAK | NO | NO | NO |
| 13752 | TRAV35 | TRAJ54 | CAGGRIQGAQKLVF | TRBV7-9 | TRBJ1-2 | CASSPGTGNYGYTF | KLGGALQAK | YES | NO | NO |
| 13753 | TRAV8-1 | TRAJ40 | CAVILPITTSGTYKYIF | TRBV27 | TRBJ2-2 | CASSLLGGTGELFF | KLGGALQAK | YES | NO | NO |
| 13754 | TRAV21 | TRAJ58 | CAVRGETSGSRLTF | TRBV12-4 | TRBJ1-4 | CASSLTDLSTNEKLFF | KLGGALQAK | YES | NO | NO |
| 13755 | TRAV5 | TRAJ26 | CAEKVSYGQNFVF | TRBV11-2 | TRBJ1-1 | CASSAPGQMNTEAFF | KLGGALQAK | YES | NO | NO |
| 13756 | TRAV1-2 | TRAJ20 | CAGTFNDYKLSF | TRBV7-2 | TRBJ2-1 | CASSPSVSEQFF | KLGGALQAK | YES | NO | NO |
| 13757 | TRAV12-2 | TRAJ28 | CAVNTYSGAGSYQLTF | TRBV9 | TRBJ1-4 | CASSVALGQEGEKLFF | KLGGALQAK | YES | NO | NO |
| 13758 | TRAV2 | TRAJ32 | CAVGWGGATNKLIF | TRBV30 | TRBJ1-2 | CAWSVGVGQLDGYTF | KLGGALQAK | NO | NO | NO |
| 13759 | TRAV12-2 | TRAJ33 | CAVTNSNYQLIW | TRBV25-1 | TRBJ2-7 | CASSKMGQGSEQYF | KLGGALQAK | YES | NO | NO |
| 13760 | TRAV29DV5 | TRAJ48 | CAAGPRNFGNEKLTF | TRBV4-3 | TRBJ2-1 | CASSQEASGTYNEQFF | KLGGALQAK | YES | NO | NO |
| 13761 | TRAV8-1 | TRAJ6 | CALSGGSYIPTF | TRBV11-2 | TRBJ2-4 | CASSLGTKTIAKNIQYF | KLGGALQAK | YES | NO | NO |
| 13762 | TRAV13-1 | TRAJ6 | CAASMGGGSYIPTF | TRBV6-5 | TRBJ2-2 | CASSPTSGSFTGELFF | KLGGALQAK | YES | NO | NO |
| 13763 | TRAV12-3 | TRAJ54 | CASGQGAQKLVF | TRBV3-1 | TRBJ2-1 | CASSPRTSGVANEQFF | KLGGALQAK | YES | NO | NO |
| 13764 | TRAV17 | TRAJ15 | CATDNQAGTALIF | TRBV11-2 | TRBJ1-1 | CASSLDGTGTEAFF | KLGGALQAK | YES | NO | NO |
| 13765 | TRAV12-1 | TRAJ6 | CVVNAGGSYIPTF | TRBV14 | TRBJ1-1 | CASPPEEGLLAEAFF | KLGGALQAK | NO | NO | NO |
| 13766 | TRAV8-2 | TRAJ3 | CVVTDYSSASKIIF | TRBV10-2 | TRBJ1-2 | CASSELRAGQDGYTF | KLGGALQAK | YES | NO | NO |
| 13767 | TRAV23DV6 | TRAJ23 | CAASSSYNQGGKLIF | TRBV6-3 | TRBJ2-1 | CASSYGTAIGVTF | KLGGALQAK | NO | NO | NO |
| 13768 | TRAV17 | TRAJ42 | CATDRGGGSQGNLIF | TRBV27 | TRBJ2-7 | CASSLSGGINEQYF | KLGGALQAK | YES | NO | NO |
| 13769 | TRAV29DV5 | TRAJ17 | CAARGIAAGNKLTF | TRBV7-2 | TRBJ2-1 | CASSPEPGLAGDNEQFF | KLGGALQAK | YES | NO | NO |
| 13770 | TRAV19 | TRAJ16 | CALSEAVSDGQKLLF | TRBV11-2 | TRBJ1-4 | CASSLDFWTGLPNEKLFF | KLGGALQAK | NO | NO | NO |
| 13771 | TRAV12-2 | TRAJ5 | CAVNGGTGRRALTF | TRBV19 | TRBJ1-5 | CASTTWDFNQPQHF | KLGGALQAK | YES | NO | NO |
| 13772 | TRAV5 | TRAJ49 | CAEAWAGNQFYF | TRBV4-3 | TRBJ1-6 | CASSHGNSPLHF | KLGGALQAK | YES | NO | NO |
| 13773 | TRAV27 | TRAJ39 | CAGARLDNAGNMLTF | TRBV10-3 | TRBJ2-3 | CAISALAGGTDTQYF | KLGGALQAK | YES | NO | NO |
| 13774 | TRAV27 | TRAJ42 | CAGVIRGSQGNLIF | TRBV4-3 | TRBJ1-1 | CASSQDLNRGALEAFF | KLGGALQAK | YES | NO | NO |
| 13775 | TRAV13-2 | TRAJ23 | CAENVGQGGKLIF | TRBV12-4 | TRBJ2-7 | CASSDGTSYEQYV | KLGGALQAK | NO | NO | NO |
| 13777 | TRAV38-2DV8 | TRAJ45 | CACSGGGADGLTF | TRBV9 | TRBJ1-2 | CASSVAGTEINYGYTF | KLGGALQAK | YES | NO | NO |
| 13778 | TRAV19 | TRAJ34 | CALSEALPRDKLIF | TRBV12-5 | TRBJ1-1 | CASSIWAGTEAFF | KLGGALQAK | YES | NO | NO |
| 13779 | TRAV25 | TRAJ52 | CAGLDLMNAGGTSYGKLTF | TRBV5-1 | TRBJ1-2 | CASSLYGTGVDGYTF | KLGGALQAK | YES | NO | NO |
| 13780 | TRAV38-2DV8 | TRAJ40 | CAYTSGTYKYIF | TRBV6-5 | TRBJ1-4 | CASSYTGVPEKLFF | KLGGALQAK | NO | NO | NO |
| 13781 | TRAV9-2 | TRAJ47 | CASMMEYGNKLVF | TRBV7-8 | TRBJ1-4 | CASSLLRAATNEKLFF | KLGGALQAK | YES | NO | NO |
| 13782 | TRAV13-1 | TRAJ20 | CAASPNDYKLSF | TRBV7-6 | TRBJ1-1 | CASSDPGYEAFF | KLGGALQAK | YES | NO | NO |
| 13783 | TRAV12-3 | TRAJ49 | CAMSGWTGNQFYF | TRBV5-4 | TRBJ2-1 | CASSLAVIRGRRNEQFF | KLGGALQAK | YES | NO | NO |
| 13784 | TRAV20 | TRAJ26 | CAVGSGYGQNFVF | TRBV7-9 | TRBJ1-2 | CASSLIGSNYGYTF | KLGGALQAK | YES | NO | NO |
| 13785 | TRAV8-1 | TRAJ20 | CAVNTNDYKLSF | TRBV5-1 | TRBJ2-4 | CASSSGLAGGAKNIQYF | KLGGALQAK | YES | NO | NO |
| 13786 | TRAV12-2 | TRAJ50 | CAVNRLASYDKVIF | TRBV29-1 | TRBJ2-1 | CSVEDVRDEQFF | KLGGALQAK | YES | NO | NO |
| 13787 | TRAV19 | TRAJ9 | CALLRDGGFKTIF | TRBV20-1 | TRBJ2-3 | CSAQQTGTTDTQYF | KLGGALQAK | NO | NO | NO |
| 13788 | TRAV13-2 | TRAJ43 | CAENDMRF | TRBV4-1 | TRBJ2-3 | CASSPTASGSYTQYF | KLGGALQAK | NO | NO | NO |
| 13789 | TRAV12-3 | TRAJ11 | CAMSGSGYSTLTF | TRBV11-2 | TRBJ2-1 | CASSLAGGLEEQFF | KLGGALQAK | YES | NO | NO |
| 13790 | TRAV35 | TRAJ45 | CAGRAGGGADGLTF | TRBV7-8 | TRBJ1-1 | CASSIPGSPGNTEAFF | KLGGALQAK | YES | NO | NO |
| 13791 | TRAV4 | TRAJ30 | CLVLDDKIIF | TRBV9 | TRBJ2-3 | CASSVLAGGSTDTQYF | AVFDRKSDAK | YES | YES | YES |

| | | | | | | | | | | |
|---|---|---|---|---|---|---|---|---|---|---|
| 13792 | TRAV14DV4 | TRAJ22 | CAMRRPISSGSARQLTF | TRBV6-3 | TRBJ2-1 | CASSPGLAGDNEQFF | KLGGALQAK | YES | NO | NO |
| 13793 | TRAV39 | TRAJ42 | CAVVLSLSTLMNYGGSQGNLIF | TRBV6-4 | TRBJ1-6 | CASSDSSPGGRNSPLHF | KLGGALQAK | NO | NO | NO |
| 13794 | TRAV29DV5 | TRAJ31 | CAASAADNARLMF | TRBV24-1 | TRBJ2-3 | CATSDLSDADTQYF | KLGGALQAK | YES | NO | NO |
| 13795 | TRAV21 | TRAJ4 | CAVKPGGYNKLIF | TRBV20-1 | TRBJ2-2 | CSAQDRRSTGELFF | KLGGALQAK | YES | NO | NO |
| 13796 | TRAV14DV4 | TRAJ33 | CAMRGLDSNYQLIW | TRBV30 | TRBJ1-1 | CAWSVFGQGMNTEAFF | KLGGALQAK | YES | NO | NO |
| 13797 | TRAV5 | TRAJ9 | CAESPDTGGFKTIF | TRBV7-9 | TRBJ2-2 | CASSHTSGEGTGELFF | KLGGALQAK | YES | NO | NO |
| 13798 | TRAV13-2 | TRAJ39 | CAENNAGNMLTF | TRBV7-9 | TRBJ2-1 | CASRGGGDEQFF | KLGGALQAK | YES | NO | NO |
| 13799 | TRAV19 | TRAJ56 | CATTGANSKLTF | TRBV19 | TRBJ1-5 | CASSSGIGTGGSQPQHF | KLGGALQAK | YES | NO | NO |
| 13800 | TRAV39 | TRAJ48 | CAVDPTNFGNEKLTF | TRBV6-1 | TRBJ2-1 | CASSEALFELVYNEQFF | KLGGALQAK | YES | NO | NO |
| 13801 | TRAV30 | TRAJ38 | CGTEYFTGNNRKLIW | TRBV2 | TRBJ1-5 | CASSEGRTLNQPQHF | KLGGALQAK | YES | NO | NO |
| 13802 | TRAV8-4 | TRAJ8 | CAVSDWDTGFQKLVF | TRBV13 | TRBJ1-6 | CASRLSDYNSPLHF | KLGGALQAK | YES | NO | NO |
| 13803 | TRAV41 | TRAJ30 | CAVLLGGAMNRDDKIIF | TRBV29-1 | TRBJ2-1 | CSVALRAAGYNEQFF | AVFDRKSDAK | YES | NO | NO |
| 13804 | TRAV29DV5 | TRAJ42 | CAAGGSQGNLIF | TRBV10-3 | TRBJ2-1 | CAISAVGGAGPGNEQFF | KLGGALQAK | YES | NO | NO |
| 13806 | TRAV22 | TRAJ49 | CAGPNTGNQFYF | TRBV11-2 | TRBJ2-1 | CASSLDTFGTSGRGEQFF | KLGGALQAK | YES | NO | NO |
| 13807 | TRAV20 | TRAJ37 | CAVQAGSSNTGKLIF | TRBV27 | TRBJ2-7 | CASSYGAAYEQYF | KLGGALQAK | YES | NO | NO |
| 13808 | TRAV12-3 | TRAJ28 | CAMSDIGAGSYQLTF | TRBV4-1 | TRBJ2-1 | CASSQGPEQFF | KLGGALQAK | YES | NO | NO |
| 13809 | TRAV8-6 | TRAJ37 | CAVGHLLGSSNTGKLIF | TRBV7-9 | TRBJ1-1 | CASSLMGQAIMNTEAFF | KLGGALQAK | YES | NO | NO |
| 13810 | TRAV14DV4 | TRAJ22 | CAMRRPISSGSARQLTF | TRBV10-2 | TRBJ2-2 | CASSESAPTGGTGELFF | KLGGALQAK | YES | NO | NO |
| 13811 | TRAV3 | TRAJ13 | CAVRDLGPGSGGYQKVTF | TRBV29-1 | TRBJ1-1 | CSVPWGFTEAFF | KLGGALQAK | NO | NO | NO |
| 13812 | TRAV1-2 | TRAJ27 | CAVVNTNAGKSTF | TRBV9 | TRBJ2-5 | CASSVAWRGEDTQYF | KLGGALQAK | YES | NO | NO |
| 13813 | TRAV3 | TRAJ3 | CAVRARSASKIIF | TRBV19 | TRBJ2-5 | CASSMETQYF | AVFDRKSDAK | NO | NO | NO |
| 13814 | TRAV17 | TRAJ57 | CATAQGGSEKLVF | TRBV5-1 | TRBJ1-1 | CASSLAPGGNTEAFF | KLGGALQAK | YES | NO | NO |
| 13815 | TRAV8-3 | TRAJ29 | CALFRNSGNTPLVF | TRBV3-1 | TRBJ2-7 | CASSQGTYEQYF | KLGGALQAK | YES | NO | NO |
| 13816 | TRAV12-1 | TRAJ4 | CVVNIFSGGYNKLIF | TRBV7-9 | TRBJ2-7 | CASSLGDYEQYV | KLGGALQAK | NO | NO | NO |
| 13817 | TRAV36DV7 | TRAJ48 | CALLISNFGNEKLTF | TRBV5-1 | TRBJ2-1 | CASSILTAYNEQFF | KLGGALQAK | NO | NO | NO |
| 13818 | TRAV13-2 | TRAJ26 | CAENKDNYGQNFVF | TRBV20-1 | TRBJ2-1 | CSVTQGSSYNEQFF | KLGGALQAK | YES | NO | NO |
| 13819 | TRAV1-2 | TRAJ5 | CAVRDHTGRRALTF | TRBV7-3 | TRBJ2-2 | CASSTDSYTGELFF | AVFDRKSDAK | NO | NO | NO |
| 13820 | TRAV35 | TRAJ42 | CAGLGTYGGSQGNLIF | TRBV12-4 | TRBJ1-1 | CASRQNGWEGSTEAFF | AVFDRKSDAK | YES | NO | NO |
| 13821 | TRAV19 | TRAJ20 | CALRSDYKLSF | TRBV7-2 | TRBJ2-1 | CASSVGGRGNEQFF | KLGGALQAK | YES | NO | NO |
| 13822 | TRAV13-1 | TRAJ16 | CAASDRF | TRBV6-4 | TRBJ2-1 | CASSDERFYNEQFF | KLGGALQAK | NO | NO | NO |
| 13823 | TRAV19 | TRAJ30 | CALSEDPTNRDDKIIF | TRBV9 | TRBJ1-1 | CASSVGSGGVMNTEAFF | KLGGALQAK | YES | NO | NO |
| 13824 | TRAV12-2 | TRAJ26 | CAADLDNYGQNFVF | TRBV3-1 | TRBJ1-1 | CASSQRGNTEAFF | KLGGALQAK | YES | NO | NO |
| 13825 | TRAV16 | TRAJ17 | CAPPGRAAGNKLTF | TRBV19 | TRBJ1-4 | CASSNRDTLLATNEKLFF | KLGGALQAK | NO | NO | NO |
| 13826 | TRAV3 | TRAJ33 | CAVSPRSTGNYQLIW | TRBV3-1 | TRBJ2-5 | CASSPFRLAGGQETQYF | KLGGALQAK | NO | NO | NO |
| 13827 | TRAV14DV4 | TRAJ9 | CAMREIVWTGGFKTIF | TRBV11-2 | TRBJ1-4 | CASSFATANEKLFF | KLGGALQAK | YES | NO | NO |
| 13828 | TRAV14DV4 | TRAJ41 | CAMRVRDSGYALNF | TRBV5-6 | TRBJ2-3 | CASSGWGSGSTDTQYF | KLGGALQAK | YES | NO | NO |
| 13829 | TRAV29DV5 | TRAJ28 | CAASARAGSYQLTF | TRBV5-6 | TRBJ1-5 | CASSLKTGFSNQPQHF | KLGGALQAK | YES | NO | NO |
| 13830 | TRAV19 | TRAJ4 | CALSVAFSGGYNKLIF | TRBV7-9 | TRBJ2-5 | CASRGNHGDNQETQYF | KLGGALQAK | YES | NO | NO |
| 13831 | TRAV21 | TRAJ29 | CAVFSGNTPLVF | TRBV19 | TRBJ2-1 | CASSIASGGRGKFF | KLGGALQAK | YES | NO | NO |
| 13832 | TRAV20 | TRAJ15 | CAVQAWQAGTALIF | TRBV5-1 | TRBJ1-5 | CASSLSGTGADQPQHF | KLGGALQAK | YES | NO | NO |
| 13833 | TRAV34 | TRAJ43 | CGAAYNNNDMRF | TRBV4-3 | TRBJ2-1 | CASSPGQGWNEQFF | KLGGALQAK | YES | NO | NO |
| 13834 | TRAV8-3 | TRAJ47 | CALPVEYGNKLVF | TRBV10-2 | TRBJ2-7 | CASSDLGVYEQYF | KLGGALQAK | YES | NO | NO |
| 13835 | TRAV21 | TRAJ34 | CAVGLLDTDKLIF | TRBV20-1 | TRBJ2-7 | CSARGQGVYEQYV | KLGGALQAK | NO | NO | NO |
| 13836 | TRAV19 | TRAJ58 | CALSGSRLTF | TRBV20-1 | TRBJ1-2 | CSARDSRQGIGYGYTF | KLGGALQAK | YES | NO | NO |
| 13837 | TRAV8-3 | TRAJ26 | CAVDNYGQNFVF | TRBV19 | TRBJ2-2 | CASSTGTSGGELFF | KLGGALQAK | YES | NO | NO |
| 13838 | TRAV8-6 | TRAJ11 | CAVSDRYSTLTF | TRBV12-4 | TRBJ2-1 | CASSLASGTSNYNEQFF | KLGGALQAK | YES | NO | NO |
| 13839 | TRAV19 | TRAJ42 | CALDALGQGNLIF | TRBV6-1 | TRBJ2-1 | CASTPSQGAFNEQFF | KLGGALQAK | YES | NO | NO |
| 13841 | TRAV8-1 | TRAJ6 | CAVNSPGGSYIPTF | TRBV10-3 | TRBJ2-3 | CAISGTSGEISTDTQYF | KLGGALQAK | YES | NO | NO |
| 13842 | TRAV19 | TRAJ28 | CALSEAGGSGAGSYQLTF | TRBV12-3 | TRBJ2-5 | CASSMTGNQETQYF | KLGGALQAK | YES | NO | NO |
| 13843 | TRAV10 | TRAJ49 | CVVSGANTGNQFYF | TRBV13 | TRBJ2-7 | CASSLGRSTYEQYF | KLGGALQAK | YES | NO | NO |
| 13844 | TRAV38-2DV8 | TRAJ42 | CAYRSAYGYGGSQGNLIF | TRBV7-9 | TRBJ2-6 | CASSSRTEIFSGANVLTF | KLGGALQAK | YES | NO | NO |
| 13845 | TRAV12-3 | TRAJ39 | CAIGPGNMLTF | TRBV29-1 | TRBJ2-7 | CSVDGQGAGEQYF | KLGGALQAK | YES | NO | NO |
| 13846 | TRAV1-2 | TRAJ31 | CAVGISTSNPSGARLMF | TRBV12-5 | TRBJ2-7 | CASGAAGTAYEQYF | KLGGALQAK | YES | NO | NO |
| 13847 | TRAV12-1 | TRAJ41 | CVVTPISGYALNF | TRBV9 | TRBJ1-2 | CASSEGAQGAGYYGYTF | KLGGALQAK | YES | NO | NO |
| 13848 | TRAV41 | TRAJ47 | CAVRFEYGNKLVF | TRBV19 | TRBJ2-5 | CASSIVGRGPARETQYF | KLGGALQAK | YES | NO | NO |
| 13849 | TRAV12-2 | TRAJ12 | CAVAMDSSYKLIF | TRBV5-4 | TRBJ2-3 | CASSGLASEDTQYF | KLGGALQAK | YES | NO | NO |
| 13850 | TRAV13-2 | TRAJ49 | CAENFDTGNQFYF | TRBV19 | TRBJ2-7 | CASSIGQHYEQYF | KLGGALQAK | YES | NO | NO |
| 13851 | TRAV29DV5 | TRAJ34 | CAGGYNTDKLIF | TRBV5-5 | TRBJ2-2 | CASSISGDLGGELFF | AVFDRKSDAK | YES | NO | YES |
| 13852 | TRAV38-2DV8 | TRAJ50 | CAYRSNVGRTSYDKVIF | TRBV27 | TRBJ2-3 | CASSLSNPDTQYF | KLGGALQAK | YES | NO | NO |
| 13853 | TRAV26-1 | TRAJ39 | CIVRANNAGNMLTF | TRBV7-8 | TRBJ1-2 | CASSLGYGYTF | KLGGALQAK | YES | NO | NO |
| 13854 | TRAV13-2 | TRAJ48 | CAETRGNEKLTF | TRBV9 | TRBJ1-4 | CASSGAPTDNNEKLFF | KLGGALQAK | YES | NO | NO |
| 13855 | TRAV19 | TRAJ9 | CALSGSLSGFKTIF | TRBV4-1 | TRBJ2-3 | CASSLLRDTQYF | KLGGALQAK | YES | NO | NO |
| 13856 | TRAV1-2 | TRAJ33 | CAVRAMDSNYQLIW | TRBV6-3 | TRBJ2-7 | CASTHGSGLSYEQYF | KLGGALQAK | YES | YES | NO |
| 13857 | TRAV25 | TRAJ40 | CAVSGTYKYIF | TRBV20-1 | TRBJ2-1 | CSAGRGTSSFYNEQFF | KLGGALQAK | NO | NO | NO |
| 13858 | TRAV12-2 | TRAJ9 | CAVNPGGFKTIF | TRBV19 | TRBJ2-7 | CASSMGPGFEQYV | ELAGIGILTV | NO | NO | NO |
| 13859 | TRAV24 | TRAJ7 | CASFDYGNNRLAF | TRBV27 | TRBJ2-7 | CASSFAPQGNYEQYF | KLGGALQAK | YES | NO | NO |
| 13860 | TRAV12-3 | TRAJ39 | CAVDNNAGNMLTF | TRBV7-2 | TRBJ1-1 | CASSLVTLTNTEAFF | KLGGALQAK | YES | NO | NO |
| 13861 | TRAV1-2 | TRAJ33 | CALWDYQLIW | TRBV12-5 | TRBJ2-1 | CASGLVRGSYNEQFF | KLGGALQAK | YES | NO | NO |
| 13862 | TRAV14DV4 | TRAJ4 | CAMRGPISGGYNKLIF | TRBV3-1 | TRBJ1-5 | CASSLTDRVGQPQHF | KLGGALQAK | YES | NO | NO |
| 13863 | TRAV17 | TRAJ45 | CATDARSGGGADGLTF | TRBV4-1 | TRBJ2-7 | CASSQDIRTGGYEQYF | KLGGALQAK | YES | NO | NO |
| 13864 | TRAV25 | TRAJ21 | CAGLGNFNKFYF | TRBV6-5 | TRBJ1-2 | CASSPTYYGYTF | RLRAEAQVK | YES | NO | YES |
| 13865 | TRAV2 | TRAJ47 | CGRQMLLFTTVLWRSREEYGNKLVF | TRBV12-4 | TRBJ2-4 | CASSFGASAKNIQYF | KLGGALQAK | NO | NO | NO |
| 13866 | TRAV12-2 | TRAJ52 | CAVHNAGGTSYGKLTF | TRBV3-1 | TRBJ1-5 | CASSQGVGQGNQPQHF | KLGGALQAK | YES | NO | NO |
| 13867 | TRAV21 | TRAJ52 | CAVLLGTSYGKLTF | TRBV5-1 | TRBJ2-1 | CASSLVAGYEQFF | KLGGALQAK | YES | NO | NO |
| 13868 | TRAV24 | TRAJ6 | CAFITSGGSYIPTF | TRBV7-6 | TRBJ1-5 | CASSLDRGSYSNQPQHF | KLGGALQAK | YES | NO | NO |
| 13869 | TRAV4 | TRAJ18 | CLVGDNRGSTLGRLYF | TRBV5-1 | TRBJ2-3 | CASSVAGGPAVTDTQYF | KLGGALQAK | YES | NO | NO |
| 13870 | TRAV12-1 | TRAJ8 | CGLRSFQKLVF | TRBV6-9 | TRBJ1-1 | CASRFQGLTEAFF | KLGGALQAK | YES | NO | NO |
| 13871 | TRAV19 | TRAJ24 | CALIRADSWGKFQF | TRBV10-3 | TRBJ1-6 | CAISEGEGVSYNSPLHF | KLGGALQAK | YES | NO | NO |
| 13872 | TRAV30 | TRAJ57 | CGTEWAGGSEKLVF | TRBV5-1 | TRBJ2-2 | CASSFFYNTGELFF | KLGGALQAK | YES | NO | NO |
| 13873 | TRAV29DV5 | TRAJ28 | CAASSYSGAGSYQLTF | TRBV12-4 | TRBJ2-7 | CASSGQLGSYEQYF | KLGGALQAK | YES | NO | NO |
| 13874 | TRAV8-3 | TRAJ4 | CAVGADISGGYNKLIF | TRBV5-1 | TRBJ2-6 | CASSLALGGPGANVLTF | KLGGALQAK | YES | NO | NO |
| 13875 | TRAV27 | TRAJ42 | CAGTPRGSQGNLIF | TRBV27 | TRBJ2-7 | CASSFLAGGLSYEQYF | KLGGALQAK | YES | NO | NO |
| 13876 | TRAV17 | TRAJ52 | CATVLNAGGTSYGKLTF | TRBV6-6 | TRBJ2-7 | CASSPRTSGRVDYEQYV | KLGGALQAK | NO | NO | NO |
| 13877 | TRAV19 | TRAJ40 | CALSDATSGTYKYIF | TRBV9 | TRBJ2-2 | CASSVLRGGLNTGELFF | KLGGALQAK | YES | NO | NO |
| 13878 | TRAV19 | TRAJ54 | CALSASPIQGAQKLVF | TRBV29-1 | TRBJ1-1 | CSVERNTEAFF | KLGGALQAK | YES | NO | NO |
| 13879 | TRAV6 | TRAJ30 | CALGPGKDDKIIF | TRBV12-3 | TRBJ1-2 | CASSLGTDYGYTF | KLGGALQAK | YES | NO | NO |
| 13880 | TRAV14DV4 | TRAJ22 | CAMRGTFSGSARQLTF | TRBV29-1 | TRBJ2-5 | CSVRTSGGLETQYF | KLGGALQAK | YES | NO | NO |
| 13881 | TRAV12-2 | TRAJ15 | CAVTNNQAGTALIF | TRBV4-3 | TRBJ1-6 | CASSQDSGAGSPLHF | AVFDRKSDAK | YES | NO | YES |
| 13882 | TRAV13-2 | TRAJ39 | CAENNAGNMLTF | TRBV11-1 | TRBJ2-7 | CASSLETGTAYEQYF | KLGGALQAK | YES | NO | NO |
| 13883 | TRAV17 | TRAJ57 | CATGYAQGGSEKLVF | TRBV30 | TRBJ2-1 | CASPGGASSYNEQFF | KLGGALQAK | NO | NO | NO |
| 13884 | TRAV25 | TRAJ34 | CAGPGTDKLIF | TRBV29-1 | TRBJ2-1 | CSVLSGVTYNEQFF | KLGGALQAK | YES | NO | NO |
| 13885 | TRAV38-1 | TRAJ31 | CAFGFNNARLMF | TRBV6-3 | TRBJ2-1 | CASSYGAGGYNEQFF | KLGGALQAK | YES | NO | NO |
| 13886 | TRAV1-2 | TRAJ31 | CAVRDSNNARLMF | TRBV19 | TRBJ1-6 | CASSPGGMAPLHF | KLGGALQAK | YES | NO | NO |
| 13887 | TRAV35 | TRAJ16 | CAGKQLSDGQKLLF | TRBV25-1 | TRBJ2-1 | CASSDGDSSYNEQFF | KLGGALQAK | NO | NO | NO |
| 13888 | TRAV38-2DV8 | TRAJ58 | CAYRSAETSGSRLTF | TRBV7-8 | TRBJ2-1 | CASSLHRGYLNEQFF | KLGGALQAK | YES | NO | NO |
| 13889 | TRAV12-1 | TRAJ27 | CVVFWTNAGKSTF | TRBV5-6 | TRBJ1-5 | CASSSRDSQNQPQHF | KLGGALQAK | YES | NO | NO |
| 13890 | TRAV1-2 | TRAJ29 | CAVRSSGNTPLVF | TRBV20-1 | TRBJ1-4 | CSARTGTDFNEKLFF | KLGGALQAK | YES | NO | NO |
| 13891 | TRAV13-1 | TRAJ49 | CAASHTNTGNQFYF | TRBV7-8 | TRBJ2-1 | CASSSPGKSSYNEQFF | KLGGALQAK | YES | NO | NO |
| 13892 | TRAV12-1 | TRAJ42 | CVVLNYGGSQGNLIF | TRBV7-9 | TRBJ1-2 | CASSLGTGDGYTF | KLGGALQAK | YES | NO | NO |
| 13893 | TRAV12-2 | TRAJ43 | CAAEGTDDMRF | TRBV10-2 | TRBJ2-7 | CASREGLRGWEQYF | KLGGALQAK | YES | NO | NO |
| 13894 | TRAV8-6 | TRAJ7 | CAVGGEQQTRF | TRBV12-4 | TRBJ2-3 | CASSPMDTQYF | KLGGALQAK | YES | NO | NO |
| 13895 | TRAV12-2 | TRAJ7 | CAVSLTIYGNNRLAF | TRBV29-1 | TRBJ2-1 | CSSTEYNEQFF | KLGGALQAK | YES | NO | NO |
| 13896 | TRAV22 | TRAJ27 | CAVSDAGKSTF | TRBV5-4 | TRBJ2-7 | CASSFGQGAGEQYV | KLGGALQAK | NO | NO | NO |
| 13897 | TRAV8-1 | TRAJ22 | CAVRSSGSARQLTF | TRBV7-6 | TRBJ2-7 | CASSLGTESYEQYF | KLGGALQAK | YES | NO | NO |

| | | | | | | | | | | |
|---|---|---|---|---|---|---|---|---|---|---|
| 13898 | TRAV29DV5 | TRAJ49 | CAASAFGNQFYF | TRBV27 | TRBJ1-4 | CASSQTEFATNEKLFF | KLGGALQAK | YES | NO | NO |
| 13899 | TRAV39 | TRAJ34 | CAVEAFYTDKLIF | TRBV9 | TRBJ2-1 | CASSPSPAGASTNEQFF | KLGGALQAK | YES | NO | NO |
| 13900 | TRAV13-1 | TRAJ23 | CAALYNQGGKLIF | TRBV27 | TRBJ2-1 | CASSLMGIGSYNEQFF | KLGGALQAK | YES | NO | NO |
| 13901 | TRAV26-1 | TRAJ52 | CIVRVNAGGTSYGKLTF | TRBV19 | TRBJ1-6 | CASSIGFSPLHF | KLGGALQAK | YES | NO | NO |
| 13902 | TRAV13-2 | TRAJ50 | CAENPQTSYDKVIF | TRBV20-1 | TRBJ2-5 | CSANSWTSGEETQYF | KLGGALQAK | YES | NO | NO |
| 13903 | TRAV6 | TRAJ10 | CALDSLGGGNKLTF | TRBV5-6 | TRBJ1-2 | CASSIGLLDGYTF | KLGGALQAK | YES | NO | NO |
| 13904 | TRAV29DV5 | TRAJ29 | CAASAPTPLVF | TRBV12-4 | TRBJ2-1 | CASSLLAGGGYNEQFF | KLGGALQAK | YES | NO | NO |
| 13905 | TRAV12-1 | TRAJ49 | CVVNRRYPGNQFYF | TRBV5-1 | TRBJ2-5 | CASSLGAQETQYF | KLGGALQAK | YES | NO | NO |
| 13906 | TRAV14DV4 | TRAJ5 | CAMRYDTGRRALTF | TRBV7-3 | TRBJ2-2 | CASSPHDGITGELFF | KLGGALQAK | NO | NO | NO |
| 13907 | TRAV14DV4 | TRAJ50 | CAMREPARLMKTSYDKVIF | TRBV20-1 | TRBJ1-5 | CSASAGFSDQPQHF | KLGGALQAK | YES | NO | NO |
| 13908 | TRAV3 | TRAJ31 | CAVRDMYNARLMF | TRBV11-2 | TRBJ2-7 | CASSPPGQLSYEQYF | AVFDRKSDAK | YES | NO | YES |
| 13909 | TRAV25 | TRAJ24 | CAGMTTDSWGKFQF | TRBV10-1 | TRBJ2-3 | CASSRGQGSTDTQYF | KLGGALQAK | YES | NO | NO |
| 13910 | TRAV14DV4 | TRAJ4 | CAMREGPFSGGYNKLIF | TRBV9 | TRBJ2-5 | CASSPSGLGVLARETQYF | KLGGALQAK | YES | NO | NO |
| 13912 | TRAV22 | TRAJ24 | CAVERISTDSWGKLQF | TRBV27 | TRBJ2-7 | CASSPSSSYEQYV | KLGGALQAK | NO | NO | NO |
| 13913 | TRAV12-2 | TRAJ15 | CAVNGAGTALIF | TRBV19 | TRBJ2-7 | CASSISLGTAYEQYF | KLGGALQAK | YES | NO | NO |
| 13914 | TRAV36DV7 | TRAJ39 | CAVGANAGNMLTF | TRBV5-1 | TRBJ1-1 | CASSAIQGARLLNTEAFF | KLGGALQAK | NO | NO | NO |
| 13915 | TRAV12-1 | TRAJ27 | CVVFWTNAGKSTF | TRBV5-1 | TRBJ1-2 | CASSLMSQGNYGYTF | KLGGALQAK | YES | NO | NO |
| 13916 | TRAV1-1 | TRAJ27 | CEGEGNAGKSTF | TRBV3-1 | TRBJ1-2 | CASSQVTGGPNYGYTF | KLGGALQAK | YES | NO | NO |
| 13917 | TRAV29DV5 | TRAJ53 | CAASRGSNYKLTF | TRBV29-1 | TRBJ1-1 | CSVEQGVTEAFF | KLGGALQAK | YES | NO | NO |
| 13918 | TRAV38-1 | TRAJ42 | CAFSGSQGNLIF | TRBV7-9 | TRBJ2-7 | CASSFYRGLSGSYEQYV | KLGGALQAK | NO | NO | NO |
| 13919 | TRAV21 | TRAJ20 | CAVMGSNDYKLSF | TRBV19 | TRBJ2-1 | CASSAMGQHNEQFF | KLGGALQAK | YES | NO | NO |
| 13920 | TRAV27 | TRAJ44 | CAGADGTASKLTF | TRBV13 | TRBJ2-7 | CASSQLEDGHSYEQYF | KLGGALQAK | YES | NO | NO |
| 13921 | TRAV22 | TRAJ27 | CAVPYHAGKSTF | TRBV5-1 | TRBJ2-7 | CASSLYGADEQYF | KLGGALQAK | YES | NO | NO |
| 13922 | TRAV19 | TRAJ50 | CALKTSYDKVIF | TRBV7-9 | TRBJ2-2 | CASSLAREVPKTGELFF | KLGGALQAK | YES | YES | NO |
| 13923 | TRAV12-1 | TRAJ52 | CVVSTFNAGGTSYGKLTF | TRBV29-1 | TRBJ1-1 | CSNIEMNTEAFF | KLGGALQAK | YES | NO | NO |
| 13924 | TRAV19 | TRAJ58 | CALWGETSGSRLTF | TRBV5-4 | TRBJ2-5 | CASSLSGGPETQYF | KLGGALQAK | YES | NO | NO |
| 13925 | TRAV19 | TRAJ39 | CALSGPLKAGNMLTF | TRBV5-1 | TRBJ1-4 | CASSSSMGDMDEKLFF | KLGGALQAK | YES | NO | NO |
| 13926 | TRAV38-2DV8 | TRAJ28 | CAFSGAGSYQLTF | TRBV2 | TRBJ2-1 | CASSEGGRRYNEQFF | KLGGALQAK | YES | NO | NO |
| 13927 | TRAV1-2 | TRAJ6 | CAVRDTGGSYIPTF | TRBV15 | TRBJ2-1 | CATSGTGTSGYNEQFF | KLGGALQAK | YES | NO | NO |
| 13928 | TRAV8-4 | TRAJ48 | CAVSGPADFGNEKLTF | TRBV19 | TRBJ1-2 | CASSFVPNYGYTF | KLGGALQAK | YES | NO | NO |
| 13929 | TRAV35 | TRAJ45 | CAGLGTRNLIGGGADGLTF | TRBV7-9 | TRBJ2-1 | CASTPREAGDWYEQFF | KLGGALQAK | YES | NO | NO |
| 13930 | TRAV29DV5 | TRAJ36 | CAASSSSGANNLFF | TRBV20-1 | TRBJ2-3 | CSATDNRGRTDTQYF | KLGGALQAK | NO | NO | NO |
| 13931 | TRAV12-1 | TRAJ41 | CVERGSNSGYALNF | TRBV7-9 | TRBJ2-5 | CASTAYQETQYF | KLGGALQAK | YES | NO | NO |
| 13932 | TRAV20 | TRAJ16 | CAVQASDGQKLLF | TRBV19 | TRBJ2-7 | CASSAGTGNEQYF | KLGGALQAK | NO | NO | NO |
| 13933 | TRAV5 | TRAJ40 | CAIGAGTYKYIF | TRBV27 | TRBJ2-2 | CASRISTGELFF | KLGGALQAK | YES | NO | NO |
| 13934 | TRAV19 | TRAJ30 | CALSEAGRDDKIIF | TRBV5-1 | TRBJ1-3 | CASSLPDSGNTIYF | KLGGALQAK | YES | NO | NO |
| 13935 | TRAV19 | TRAJ42 | CALSELNYGGSQGNLIF | TRBV7-9 | TRBJ1-5 | CASSSGQGPNQPQHF | KLGGALQAK | YES | NO | NO |
| 13936 | TRAV19 | TRAJ57 | CAPTQGGSEKLVF | TRBV7-9 | TRBJ2-7 | CASSSGNSYEQYF | KLGGALQAK | YES | NO | NO |
| 13937 | TRAV20 | TRAJ57 | CAVTQGGSEKLVF | TRBV5-6 | TRBJ2-7 | CASSPWTGLLSEQYF | KLGGALQAK | YES | NO | NO |
| 13938 | TRAV14DV4 | TRAJ40 | CAMRGGSGTYKYIF | TRBV7-9 | TRBJ2-1 | CASSFGLAGLGDEQFF | KLGGALQAK | YES | NO | NO |
| 13939 | TRAV8-2 | TRAJ23 | CVPREPGRKAYLRT | TRBV29-1 | TRBJ1-1 | CSVEDGAIGTEAFF | KLGGALQAK | NO | NO | NO |
| 13940 | TRAV35 | TRAJ47 | CAGQSYGNKLVF | TRBV6-3 | TRBJ1-5 | CASSPGQGYQPQHF | KLGGALQAK | YES | NO | NO |
| 13941 | TRAV8-6 | TRAJ50 | CAVSDPSGTSYDKVIF | TRBV7-8 | TRBJ1-1 | CASRRTGGERSEAFF | KLGGALQAK | YES | NO | NO |
| 13942 | TRAV1-2 | TRAJ27 | CAVRSTTNAGKSTF | TRBV24-1 | TRBJ1-1 | CATSDKGGLKAFF | KLGGALQAK | YES | NO | NO |
| 13943 | TRAV14DV4 | TRAJ39 | CAMREGFNNAGNMLTF | TRBV6-3 | TRBJ1-1 | CASSYEEITMNTEAFF | KLGGALQAK | YES | NO | NO |
| 13944 | TRAV10 | TRAJ42 | CVVSARNYGGSQGNLIF | TRBV19 | TRBJ1-5 | CASSTTGGYSNQPQHF | KLGGALQAK | YES | NO | NO |
| 13945 | TRAV8-3 | TRAJ41 | CAVGTERPNSGYALNF | TRBV19 | TRBJ1-1 | CASSPNRGLVTEAFF | KLGGALQAK | YES | NO | NO |
| 13946 | TRAV12-2 | TRAJ22 | CAAEVVSSGSARQLTF | TRBV6-3 | TRBJ1-6 | CASSYSGTAGSPLHF | KLGGALQAK | YES | NO | NO |
| 13947 | TRAV27 | TRAJ45 | CAGAGSGGGADGLTF | TRBV6-6 | TRBJ2-2 | CASSYERGGPPNTGELFF | KLGGALQAK | YES | NO | NO |
| 13948 | TRAV17 | TRAJ54 | CATDARLQGAQKLVF | TRBV7-8 | TRBJ2-2 | CASSLESGVVTGELFF | KLGGALQAK | YES | NO | NO |
| 13949 | TRAV26-1 | TRAJ41 | CIVRGISNSGYALNF | TRBV4-1 | TRBJ1-4 | CASSQDLPDGEKLFF | KLGGALQAK | NO | NO | NO |
| 13950 | TRAV12-1 | TRAJ31 | CVVNMGNNARLMF | TRBV27 | TRBJ1-4 | CASSVLPGSEKLFF | KLGGALQAK | YES | NO | NO |
| 13951 | TRAV19 | TRAJ57 | CALSGLTQGGSEKLVF | TRBV6-3 | TRBJ2-5 | CASSGLAGGPYF | KLGGALQAK | YES | NO | NO |
| 13953 | TRAV13-2 | TRAJ45 | CAERARSGGGADGLTF | TRBV4-3 | TRBJ1-2 | CASSRTHGTGKGYTF | KLGGALQAK | YES | NO | NO |
| 13954 | TRAV16 | TRAJ8 | CALSSTETGFQKLVF | TRBV20-1 | TRBJ2-7 | CSASGGTAVPYEQYF | KLGGALQAK | NO | NO | NO |
| 13955 | TRAV19 | TRAJ43 | CALSGYSRGDMRF | TRBV11-1 | TRBJ2-7 | CASSPWDSRETYEQYF | KLGGALQAK | YES | NO | NO |
| 13956 | TRAV13-1 | TRAJ49 | CAALFYF | TRBV5-4 | TRBJ1-1 | CASSWDTNTEAFF | KLGGALQAK | NO | NO | NO |
| 13957 | TRAV26-2 | TRAJ8 | CILRDRNTGFQKLVF | TRBV9 | TRBJ2-3 | CASSDTASGDTDTQYF | KLGGALQAK | YES | NO | NO |
| 13958 | TRAV5 | TRAJ43 | CAANDMRF | TRBV4-1 | TRBJ1-1 | CASSQEGGGWGTEAFF | KLGGALQAK | NO | NO | NO |
| 13959 | TRAV1-2 | TRAJ20 | CAVRDPPPGDYKLSF | TRBV20-1 | TRBJ2-7 | CSARLQGNEQYF | KLGGALQAK | NO | NO | NO |
| 13960 | TRAV8-4 | TRAJ43 | CAVSLRGDMRF | TRBV20-1 | TRBJ2-4 | CSARARVLGWWGGRSQNIQYF | KLGGALQAK | NO | NO | NO |
| 13961 | TRAV14DV4 | TRAJ37 | CAMRGPYSSNTGKLIF | TRBV30 | TRBJ2-1 | CAWQGLSEQFF | KLGGALQAK | NO | NO | NO |
| 13962 | TRAV38-1 | TRAJ44 | CAFAGTASKLTF | TRBV3-1 | TRBJ1-3 | CASSPPPSGAKEGVRAGNTIYF | AVFDRKSDAK | NO | NO | NO |
| 13963 | TRAV29DV5 | TRAJ42 | CAARDLGGSQGNLIF | TRBV4-1 | TRBJ2-1 | CASSQDIGTGGVNEQFF | KLGGALQAK | YES | NO | NO |
| 13964 | TRAV26-1 | TRAJ24 | CIAITDSWGKLQF | TRBV2 | TRBJ2-3 | CASEEPGGPDTQYF | KLGGALQAK | YES | NO | NO |
| 13965 | TRAV8-2 | TRAJ48 | CVVSESVGRGNEKLTF | TRBV27 | TRBJ1-2 | CASSWYGGTNYGYTF | FLASKIGRLV | YES | NO | YES |
| 13966 | TRAV27 | TRAJ32 | CAGAMGKYGGATNKLIF | TRBV4-3 | TRBJ2-2 | CASSPDSVNTGELFF | KLGGALQAK | YES | NO | NO |
| 13967 | TRAV13-1 | TRAJ49 | CAGPNTGNQFYF | TRBV7-8 | TRBJ2-3 | CASSLMWLAGAADTQYF | KLGGALQAK | YES | NO | NO |
| 13968 | TRAV17 | TRAJ54 | CARQGAQKLVF | TRBV6-6 | TRBJ1-2 | CATRSATGDNYGYTF | KLGGALQAK | NO | NO | NO |
| 13969 | TRAV17 | TRAJ43 | CATDRIRNDMRF | TRBV7-9 | TRBJ1-5 | CASSFHRDSFNQPQHF | KLGGALQAK | YES | NO | NO |
| 13970 | TRAV17 | TRAJ36 | CAYDQTGANNLFF | TRBV19 | TRBJ1-1 | CASSLGTGNTEAFF | KLGGALQAK | YES | NO | NO |
| 13971 | TRAV26-1 | TRAJ24 | CIVRVTTDSWGKLQF | TRBV7-9 | TRBJ1-6 | CASSLDSYNSPLHF | KLGGALQAK | YES | NO | NO |
| 13972 | TRAV25 | TRAJ37 | CAGWGNTGKLIF | TRBV14 | TRBJ1-1 | CASSQDWGLGHTEAFF | KLGGALQAK | NO | NO | NO |
| 13974 | TRAV21 | TRAJ48 | CAVEDFGNEKLTF | TRBV27 | TRBJ1-1 | CASSRTGLNTEAFF | AVFDRKSDAK | YES | NO | YES |
| 13975 | TRAV12-2 | TRAJ37 | CAVEHGSGNTGKLIF | TRBV27 | TRBJ2-7 | CASSLFPRLYEQYF | KLGGALQAK | YES | NO | NO |
| 13976 | TRAV19 | TRAJ45 | CALSQERGGGADGLTF | TRBV13 | TRBJ1-3 | CASGWDPRSGNTIYF | KLGGALQAK | YES | NO | NO |
| 13977 | TRAV17 | TRAJ54 | CATDGIQGAQKLVF | TRBV11-3 | TRBJ2-1 | CASSLESSTEQFF | KLGGALQAK | YES | NO | NO |
| 13978 | TRAV29DV5 | TRAJ36 | CAASEAGANNLFF | TRBV14 | TRBJ1-1 | CASSQGGVTEAFF | KLGGALQAK | NO | NO | NO |
| 13979 | TRAV17 | TRAJ47 | CATDVNEYGNKLVF | TRBV19 | TRBJ1-1 | CASSISGQVEAFF | KLGGALQAK | YES | NO | NO |
| 13980 | TRAV21 | TRAJ45 | CAVYSGGGADGLTF | TRBV2 | TRBJ2-1 | CASRERELGWFEQFF | KLGGALQAK | YES | NO | NO |
| 13981 | TRAV27 | TRAJ3 | CAGVLYSSASKIIF | TRBV7-9 | TRBJ2-3 | CASSPPEDHVATDTQYF | KLGGALQAK | YES | NO | NO |
| 13982 | TRAV29DV5 | TRAJ34 | CAASAVNDKLIF | TRBV29-1 | TRBJ1-2 | CSVDLANGYTF | KLGGALQAK | YES | NO | NO |
| 13983 | TRAV10 | TRAJ39 | CVVSRVSNAGNMLTF | TRBV5-5 | TRBJ2-2 | CASSPSTSGSPDGELFF | KLGGALQAK | YES | NO | NO |
| 13984 | TRAV19 | TRAJ28 | CALSEAEYSGAGSYQLTF | TRBV4-1 | TRBJ2-1 | CASSLSPPPDEQFF | AVFDRKSDAK | YES | NO | YES |
| 13985 | TRAV10 | TRAJ31 | CVVSYARLMF | TRBV7-2 | TRBJ2-5 | CASSVTSGGLETQYF | RLRAEAQVK | YES | YES | YES |
| 13986 | TRAV8-6 | TRAJ39 | CAVILGNAGNMLTF | TRBV7-3 | TRBJ2-2 | CASSPLSNTGELFF | KLGGALQAK | NO | NO | NO |
| 13987 | TRAV14DV4 | TRAJ16 | CAMRDLGQKLLF | TRBV12-4 | TRBJ2-3 | CASRAGGAADTQYF | KLGGALQAK | NO | NO | NO |
| 13988 | TRAV1-1 | TRAJ28 | CAVRADPGAGSYQLTF | TRBV13 | TRBJ1-2 | CASSLTGQGPGYGYTF | KLGGALQAK | YES | NO | NO |
| 13989 | TRAV17 | TRAJ31 | CATDAEDARLMF | TRBV25-1 | TRBJ2-2 | CASSETPGLAGELFF | KLGGALQAK | YES | NO | NO |
| 13990 | TRAV8-3 | TRAJ17 | CAVGAPPAAGNKLTF | TRBV27 | TRBJ2-3 | CASRTRQGVLDTQYF | KLGGALQAK | YES | NO | NO |
| 13991 | TRAV12-3 | TRAJ21 | CAMRLRYNFNKFYF | TRBV7-8 | TRBJ2-7 | CASSPRLAGGYEQYV | KLGGALQAK | NO | NO | NO |
| 13992 | TRAV14DV4 | TRAJ37 | CAMREGGSGNTGKLIF | TRBV5-6 | TRBJ2-2 | CASSLLGKTGELFF | KLGGALQAK | YES | NO | NO |
| 13993 | TRAV19 | TRAJ31 | CALSEAWNNARLMF | TRBV5-4 | TRBJ2-2 | CASSRTRANTGELFF | KLGGALQAK | YES | NO | NO |
| 13994 | TRAV35 | TRAJ49 | CAGSNTGNQFYF | TRBV27 | TRBJ2-7 | CASSPSPGTGTYEQYF | KLGGALQAK | YES | NO | NO |
| 13995 | TRAV12-3 | TRAJ52 | CAMSPNAGGTSYGKLTF | TRBV20-1 | TRBJ1-1 | CSARESRGYTEAFF | KLGGALQAK | YES | NO | NO |
| 13996 | TRAV14DV4 | TRAJ15 | CAMREGQAGTALIF | TRBV19 | TRBJ1-2 | CASSIEAAGGLGGYTF | KLGGALQAK | YES | NO | NO |
| 13997 | TRAV8-1 | TRAJ35 | CAVLPGAGFGNVLHC | TRBV18 | TRBJ2-1 | CASSLRSGTGRPNNEQFF | AVFDRKSDAK | NO | NO | NO |
| 13998 | TRAV8-4 | TRAJ20 | CAVSVSDKLSF | TRBV4-1 | TRBJ1-2 | CASSQEDRPLYGYTF | KLGGALQAK | YES | NO | NO |
| 13999 | TRAV8-2 | TRAJ42 | CVVSDFFQGQGNLIF | TRBV7-2 | TRBJ2-7 | CASSHLGGRDEQYF | KLGGALQAK | NO | NO | NO |
| 14000 | TRAV8-3 | TRAJ42 | CAVFVGDGGSQGNLIF | TRBV2 | TRBJ2-1 | CASSERREGNEQFF | KLGGALQAK | YES | NO | NO |
| 14001 | TRAV19 | TRAJ58 | CALKKTSGSRLTF | TRBV5-4 | TRBJ1-1 | CASSPGNLPEAFF | KLGGALQAK | YES | NO | NO |
| 14002 | TRAV13-1 | TRAJ45 | CAASIWPGGGADGLTF | TRBV7-2 | TRBJ1-1 | CASSFSPGATEAFF | KLGGALQAK | YES | NO | NO |
| 14003 | TRAV21 | TRAJ21 | CAVVDRNFNKFYF | TRBV11-2 | TRBJ2-3 | CASSFSGGAGDTQYF | KLGGALQAK | YES | NO | NO |
| 14004 | TRAV12-2 | TRAJ39 | CAVNAGNMLTF | TRBV7-9 | TRBJ1-1 | CASSSVATEAFF | KLGGALQAK | YES | NO | NO |

| | | | | | | | | | | |
|---|---|---|---|---|---|---|---|---|---|---|
| 14005 | TRAV29DV5 | TRAJ40 | CAASLFSGTYKYIF | TRBV12-3 | TRBJ2-7 | CATGSGNEQYF | KLGGALQAK | YES | NO | NO |
| 14006 | TRAV26-2 | TRAJ33 | CILRLAVGNYQLIW | TRBV4-1 | TRBJ2-7 | CASSQDRGGTQYF | KLGGALQAK | YES | NO | NO |
| 14007 | TRAV29DV5 | TRAJ20 | CAAMSSNDYKLSF | TRBV11-3 | TRBJ2-5 | CASSQTGLVLGGQETQYF | KLGGALQAK | YES | NO | NO |
| 14008 | TRAV36DV7 | TRAJ52 | CAVRINAGGTSYGKLTF | TRBV6-1 | TRBJ1-5 | CASRRDSQGYSNQPQHF | AVFDRKSDAK | NO | NO | NO |
| 14009 | TRAV13-2 | TRAJ21 | CAETRSGVNKFYF | TRBV27 | TRBJ1-5 | CASSDRGQPQHF | KLGGALQAK | YES | NO | NO |
| 14010 | TRAV12-2 | TRAJ21 | CAVNQGNFNKFYF | TRBV13 | TRBJ1-1 | CASSLLSMNTEAFF | KLGGALQAK | YES | NO | NO |
| 14011 | TRAV1-2 | TRAJ34 | CAVFVDTDKLIF | TRBV24-1 | TRBJ2-3 | CATSDFGTGGGLAELTDTQYF | KLGGALQAK | NO | NO | NO |
| 14012 | TRAV8-3 | TRAJ27 | CAVGAGTNAGKSTF | TRBV19 | TRBJ2-1 | CASSITLGGITGNEQFF | KLGGALQAK | YES | NO | NO |
| 14013 | TRAV3 | TRAJ20 | CAVNDYKLSF | TRBV4-1 | TRBJ2-5 | CASSPGQGSAPETQYF | KLGGALQAK | YES | NO | NO |
| 14014 | TRAV35 | TRAJ54 | CAGQGAQKLVF | TRBV10-2 | TRBJ2-7 | CASSLAGVLEQYF | IVTDFSVIK | YES | NO | YES |
| 14015 | TRAV3 | TRAJ27 | CAVRDGNAGKSTF | TRBV3-1 | TRBJ2-1 | CASRFAGDHSYNEQFF | KLGGALQAK | NO | NO | NO |
| 14016 | TRAV29DV5 | TRAJ9 | CAARDNTGGFKTIF | TRBV27 | TRBJ1-1 | CASSLWVGGKVEAFF | KLGGALQAK | YES | NO | NO |
| 14017 | TRAV41 | TRAJ32 | CAVADFFSGATNKLIF | TRBV4-3 | TRBJ1-6 | CASSQDFREGKNSPLHF | KLGGALQAK | YES | NO | NO |
| 14018 | TRAV5 | TRAJ13 | CAVGSGGYQKVTF | TRBV15 | TRBJ2-2 | CASDREGSGELFF | KLGGALQAK | YES | NO | NO |
| 14019 | TRAV17 | TRAJ45 | CATDYSGGGADGLTF | TRBV20-1 | TRBJ1-2 | CSARTVTYGYTF | KLGGALQAK | YES | NO | NO |
| 14020 | TRAV9-2 | TRAJ47 | CALSGEYGNKLVF | TRBV9 | TRBJ2-1 | CASSDGGGSYNEQFF | KLGGALQAK | YES | NO | NO |
| 14021 | TRAV1-2 | TRAJ24 | CAVRGPTDSWGKLQF | TRBV27 | TRBJ1-6 | CASSFSMDSPAYSPLHF | KLGGALQAK | YES | NO | NO |
| 14022 | TRAV19 | TRAJ57 | CALSEAVGFEDRGGSEKLVF | TRBV19 | TRBJ1-1 | CASSWSSNTEAFF | KLGGALQAK | NO | NO | NO |
| 14023 | TRAV20 | TRAJ47 | CAVQGTYGNKLVF | TRBV4-2 | TRBJ1-1 | CASSQGGENTEAFF | KLGGALQAK | YES | NO | NO |
| 14024 | TRAV29DV5 | TRAJ27 | CAASVGAGKSTF | TRBV6-5 | TRBJ1-2 | CASSYWESSYGYTF | KLGGALQAK | YES | NO | NO |
| 14025 | TRAV14DV4 | TRAJ4 | CAMRGPISGGYNKLIF | TRBV15 | TRBJ2-7 | CATSASWDRVHEQYF | KLGGALQAK | YES | NO | NO |
| 14026 | TRAV5 | TRAJ6 | CAESFTTSGGSYIPTF | TRBV5-1 | TRBJ2-1 | CASRGGQGYNEQFF | AVFDRKSDAK | YES | NO | YES |
| 14027 | TRAV8-1 | TRAJ39 | CAFLRHNAGNMLTF | TRBV29-1 | TRBJ2-6 | CSVWTAISSGANVLTF | KLGGALQAK | YES | NO | NO |
| 14028 | TRAV17 | TRAJ15 | CATDYNQAGTALIF | TRBV9 | TRBJ1-1 | CASSVGGTEAFF | KLGGALQAK | YES | NO | NO |
| 14029 | TRAV26-1 | TRAJ42 | CIPWGSQGNLIF | TRBV7-8 | TRBJ1-5 | CASSLVRGSNQPQHF | KLGGALQAK | YES | NO | NO |
| 14030 | TRAV13-2 | TRAJ45 | CAENSPGGGADGLTF | TRBV4-1 | TRBJ1-5 | CASSQVGWNQPQHF | KLGGALQAK | YES | NO | NO |
| 14031 | TRAV8-1 | TRAJ28 | CAVNPLSYQLTF | TRBV20-1 | TRBJ2-1 | CSAEVTGGVDEQFF | KLGGALQAK | NO | NO | NO |
| 14032 | TRAV35 | TRAJ34 | CAGQNYNTDKLIF | TRBV7-6 | TRBJ2-7 | CASSLVSSGPRTYEQYF | KLGGALQAK | YES | NO | NO |
| 14033 | TRAV14DV4 | TRAJ12 | CAMREGRSDSSYKLIF | TRBV6-1 | TRBJ2-5 | CASSVYRDSHETQYF | KLGGALQAK | YES | NO | NO |
| 14034 | TRAV27 | TRAJ31 | CAGAVLNNARLMF | TRBV5-1 | TRBJ1-2 | CASSLKKQTGGSNYGYTF | KLGGALQAK | YES | NO | NO |
| 14035 | TRAV25 | TRAJ52 | CAGPDAGGTSYGKLTF | TRBV24-1 | TRBJ2-5 | CATSDPRTSGSVRRWETQYF | KLGGALQAK | NO | NO | NO |
| 14036 | TRAV3 | TRAJ37 | CAVRDGGMGSNTGKLIF | TRBV7-2 | TRBJ2-7 | CASSLEPSYEQYF | AVFDRKSDAK | NO | NO | NO |
| 14037 | TRAV14DV4 | TRAJ53 | CAMRDVVFGSNYKLTF | TRBV25-1 | TRBJ1-6 | CASSEFGDRGLSYNSPLHF | KLGGALQAK | YES | NO | NO |
| 14038 | TRAV24 | TRAJ31 | CASPNNNARLMF | TRBV10-3 | TRBJ2-7 | CAISERGGVNEQYF | KLGGALQAK | YES | NO | NO |
| 14039 | TRAV35 | TRAJ33 | CAGQPLMDSNYQLIW | TRBV18 | TRBJ1-1 | CASSPGRTAQAEAFF | KLGGALQAK | YES | NO | NO |
| 14040 | TRAV8-3 | TRAJ52 | CAALISNAGGTSYGKLTF | TRBV3-1 | TRBJ2-3 | CASSQDSSLTSGSSTDTQYF | KLGGALQAK | NO | NO | NO |
| 14041 | TRAV13-2 | TRAJ41 | CAEIAANSNSGYALNF | TRBV7-6 | TRBJ1-1 | CASSPALGGVPEAFF | KLGGALQAK | YES | NO | NO |
| 14042 | TRAV20 | TRAJ21 | CAVSNFNKFYF | TRBV7-8 | TRBJ2-2 | CASSLNLGQANTGELFF | KLGGALQAK | YES | NO | NO |
| 14043 | TRAV30 | TRAJ48 | CGTERSNFGNEKLTF | TRBV29-1 | TRBJ1-1 | CSVTGIQPEAFF | KLGGALQAK | YES | NO | NO |
| 14044 | TRAV3 | TRAJ28 | CAVRVSGAGSYQLTF | TRBV2 | TRBJ2-2 | CARARDRELFF | KLGGALQAK | NO | NO | NO |
| 14045 | TRAV41 | TRAJ45 | CAVTPRAGGGADGLTF | TRBV29-1 | TRBJ1-2 | CSVDLEANYGYTF | KLGGALQAK | YES | NO | NO |
| 14046 | TRAV4 | TRAJ8 | CLVGDRNTGFQKLVF | TRBV30 | TRBJ1-2 | CAWSATYASTIYGYTF | KLGGALQAK | YES | NO | NO |
| 14047 | TRAV20 | TRAJ48 | CAVQAWDFGNEKLTF | TRBV14 | TRBJ2-6 | CASSQDGGDLSSGANVLTF | KLGGALQAK | NO | NO | NO |
| 14048 | TRAV41 | TRAJ54 | CAAFRTIIQGAQKLVF | TRBV6-1 | TRBJ2-3 | CASSERKFETSTDTQYF | KLGGALQAK | YES | NO | NO |
| 14049 | TRAV4 | TRAJ27 | CLAGSNAGKSTF | TRBV6-6 | TRBJ1-2 | CASSYSEGLADGYTF | AYAQKIFKI | NO | NO | NO |
| 14050 | TRAV29DV5 | TRAJ22 | CAALSSGSARQLTF | TRBV19 | TRBJ2-3 | CASSTTFTLADTQYF | KLGGALQAK | YES | NO | NO |
| 14051 | TRAV8-2 | TRAJ36 | CVVSEGGANNPFF | TRBV10-3 | TRBJ1-1 | CAIPHARPMNTEAFF | KLGGALQAK | YES | NO | NO |
| 14052 | TRAV29DV5 | TRAJ32 | CAASDDGGATNKLIF | TRBV7-2 | TRBJ2-1 | CASSLPGYNEQFF | KLGGALQAK | YES | NO | NO |
| 14053 | TRAV38-2DV8 | TRAJ45 | CAFVYSGGGADGLTF | TRBV12-4 | TRBJ1-2 | CASSLVWAPRDYGYTF | KLGGALQAK | YES | NO | NO |
| 14054 | TRAV3 | TRAJ20 | CAVRDYKLSF | TRBV9 | TRBJ1-3 | CASSVDTPPGNTIYF | KLGGALQAK | NO | NO | NO |
| 14055 | TRAV14DV4 | TRAJ54 | CAMRRGGAQKLVF | TRBV7-9 | TRBJ2-2 | CASQSLARNTGELFF | KLGGALQAK | YES | NO | NO |
| 14056 | TRAV12-2 | TRAJ11 | CAVTVGYSTLTF | TRBV4-3 | TRBJ2-3 | CASSQEGGFTDTQYF | KLGGALQAK | YES | NO | NO |
| 14057 | TRAV8-6 | TRAJ52 | CAVSDQGAGGTSYGKLTF | TRBV20-1 | TRBJ1-6 | CSARGNRQGLGSPLHF | KLGGALQAK | NO | NO | NO |
| 14058 | TRAV36DV7 | TRAJ47 | CAVGEEYGNKLVF | TRBV4-3 | TRBJ2-6 | CASSEFTGANVLTF | KLGGALQAK | NO | NO | NO |
| 14059 | TRAV12-3 | TRAJ33 | CAMSDSNYQLIW | TRBV18 | TRBJ2-2 | CASSPPSAGELFF | KLGGALQAK | YES | NO | NO |
| 14060 | TRAV8-1 | TRAJ28 | CAVRYSGAGSYQLTF | TRBV7-2 | TRBJ1-1 | CASSWEGRLGTEAFF | KLGGALQAK | YES | NO | NO |
| 14061 | TRAV12-2 | TRAJ33 | CAVDSNYQLIW | TRBV27 | TRBJ1-1 | CASSFRAFF | KLGGALQAK | NO | NO | NO |
| 14062 | TRAV25 | TRAJ16 | CAGLFMWF | TRBV20-1 | TRBJ2-1 | CSARDLRGPYNEQFF | KLGGALQAK | NO | NO | NO |
| 14063 | TRAV21 | TRAJ24 | CAVTDSWGKLQF | TRBV10-3 | TRBJ2-3 | CAIRSGGGRTDTQYF | KLGGALQAK | YES | NO | NO |
| 14064 | TRAV3 | TRAJ35 | CAVKGFGNVLHC | TRBV5-6 | TRBJ1-5 | CASSVTGGRSNQPQHF | KLGGALQAK | NO | NO | NO |
| 14065 | TRAV16 | TRAJ36 | CALSGGANNLFF | TRBV12-4 | TRBJ2-1 | CASSFPGLISYNEQFF | KLGGALQAK | NO | NO | NO |
| 14066 | TRAV14DV4 | TRAJ52 | CAMREGFVPGGTSYGKLTF | TRBV9 | TRBJ1-1 | CASSVGASPCDFF | KLGGALQAK | YES | NO | NO |
| 14067 | TRAV24 | TRAJ53 | CAFPNSGGSNYKLTF | TRBV15 | TRBJ2-1 | CATGGGLAGVHPYNEQFF | KLGGALQAK | YES | NO | NO |
| 14068 | TRAV17 | TRAJ11 | CATDAGSTLTF | TRBV7-9 | TRBJ1-2 | CASSPLNRDGYTF | KLGGALQAK | YES | NO | NO |
| 14069 | TRAV8-1 | TRAJ26 | CAVNAGQNFVF | TRBV9 | TRBJ2-5 | CASSVGISAGVGQETQYF | KLGGALQAK | YES | NO | NO |
| 14070 | TRAV3 | TRAJ6 | CALESPSGGSYIPTF | TRBV13 | TRBJ2-2 | CASSPNTGELFF | KLGGALQAK | NO | NO | NO |
| 14071 | TRAV21 | TRAJ52 | CAVTNAGGTSYGKLTF | TRBV19 | TRBJ2-2 | CASSLMGGSNTGELFF | KLGGALQAK | YES | NO | NO |
| 14072 | TRAV5 | TRAJ15 | CAESSGTALIF | TRBV25-1 | TRBJ1-1 | CASSVLNTEAFF | KLGGALQAK | YES | NO | NO |
| 14073 | TRAV12-1 | TRAJ16 | CVVNINSGGWF | TRBV5-1 | TRBJ2-1 | CASSEGYNEQFF | KLGGALQAK | NO | NO | NO |
| 14074 | TRAV1-2 | TRAJ24 | CAVSDSWGKLQF | TRBV19 | TRBJ1-1 | CASSMTGTGGTEAFF | GILGFVFTL | YES | NO | YES |
| 14075 | TRAV41 | TRAJ17 | CAVKAAGNKLTF | TRBV4-1 | TRBJ2-2 | CASSQTAQNTGELFF | KLGGALQAK | YES | NO | NO |
| 14076 | TRAV13-2 | TRAJ10 | CAEMGGTGGGNKLTF | TRBV7-3 | TRBJ2-3 | CASSLLSTDTQYF | KLGGALQAK | YES | NO | NO |
| 14077 | TRAV27 | TRAJ54 | CAGRFQGAQKLVF | TRBV9 | TRBJ1-1 | CASSVGGGTEAFF | KLGGALQAK | YES | NO | NO |
| 14078 | TRAV1-2 | TRAJ40 | CAVSVLSGTYKYIF | TRBV5-1 | TRBJ2-7 | CASSLRYDSYEQYF | KLGGALQAK | YES | NO | NO |
| 14079 | TRAV1-2 | TRAJ20 | CAVTLNDYKLSF | TRBV5-1 | TRBJ2-7 | CASSLQDRAYEQYF | KLGGALQAK | YES | NO | NO |
| 14080 | TRAV17 | TRAJ3 | CATDPYSSASKIIF | TRBV4-1 | TRBJ2-2 | CASSQDDLGSTPIYTGELFF | KLGGALQAK | NO | NO | NO |
| 14081 | TRAV39 | TRAJ57 | CAVDPAQGGSEKLVF | TRBV10-3 | TRBJ2-3 | CAISESWRFSTDTQYF | KLGGALQAK | YES | NO | NO |
| 14082 | TRAV1-1 | TRAJ7 | CAVYYGNNRLAF | TRBV27 | TRBJ1-1 | CASSLRAMNTEAFF | KLGGALQAK | YES | NO | NO |
| 14083 | TRAV13-2 | TRAJ20 | CAENGSNDYKLSF | TRBV27 | TRBJ2-7 | CASSFTGGSYEQYF | KLGGALQAK | YES | NO | NO |
| 14084 | TRAV1-2 | TRAJ17 | CAVNNQRAAGNKLTF | TRBV6-3 | TRBJ1-5 | CASSYRGLGNQPQHF | KLGGALQAK | YES | NO | NO |
| 14085 | TRAV13-1 | TRAJ49 | CAAQLGDTGNQFYF | TRBV7-9 | TRBJ1-1 | CASSLTPGQANTEAFF | KLGGALQAK | YES | NO | NO |
| 14086 | TRAV1-2 | TRAJ5 | CAVRLDTGRRALTF | TRBV5-1 | TRBJ1-1 | CASSLAPRTVGMNTEAFF | KLGGALQAK | YES | NO | NO |
| 14087 | TRAV8-4 | TRAJ13 | CAVSDRSGGYQKVTF | TRBV7-8 | TRBJ2-2 | CASSPGTGKAGELFF | KLGGALQAK | YES | NO | NO |
| 14088 | TRAV19 | TRAJ37 | CALSEPSNTGKLIF | TRBV20-1 | TRBJ2-7 | CSARDQGMPYEQYF | KLGGALQAK | YES | NO | NO |
| 14089 | TRAV27 | TRAJ44 | CAANTGTASKLTF | TRBV11-1 | TRBJ1-1 | CASSLETGYTEAFF | KLGGALQAK | YES | NO | NO |
| 14090 | TRAV23DV6 | TRAJ29 | CAASRSTPLVF | TRBV27 | TRBJ2-7 | CASSLSQGTYEQYF | KLGGALQAK | YES | NO | NO |
| 14091 | TRAV6 | TRAJ32 | CALDGGATNKLIF | TRBV13 | TRBJ1-1 | CASSTRGGTLHTEAFF | KLGGALQAK | YES | NO | NO |
| 14092 | TRAV19 | TRAJ39 | CALSVHNAGNMLTF | TRBV4-1 | TRBJ2-1 | CASSQLGLAGAEQFF | KLGGALQAK | YES | NO | NO |
| 14093 | TRAV13-2 | TRAJ42 | CAENSDGGSQGNLIF | TRBV4-1 | TRBJ2-7 | CASSQVIGREQYF | KLGGALQAK | YES | NO | NO |
| 14094 | TRAV9-2 | TRAJ17 | CALIKIKAAGNKLTF | TRBV20-1 | TRBJ2-5 | CSSARTGGVQETQYF | KLGGALQAK | YES | NO | NO |
| 14095 | TRAV8-4 | TRAJ42 | CAVGNYGGSQGNLIF | TRBV10-1 | TRBJ2-1 | CASSLELTGSYNEQFF | KLGGALQAK | YES | NO | NO |
| 14096 | TRAV41 | TRAJ42 | CAVEGSQGNLIF | TRBV25-1 | TRBJ2-2 | CASSEWNTGELFF | AVFDRKSDAK | YES | NO | YES |
| 14097 | TRAV22 | TRAJ20 | CAVGGNDYKLSF | TRBV4-1 | TRBJ2-1 | CASSRLTGVVFNEQFF | KLGGALQAK | YES | NO | NO |
| 14098 | TRAV4 | TRAJ27 | CLVGDIANAGKSTF | TRBV4-1 | TRBJ2-5 | CASSQVLTSSQETQYF | KLGGALQAK | YES | NO | NO |
| 14099 | TRAV12-2 | TRAJ8 | CAVNGGFQKLVF | TRBV4-2 | TRBJ2-1 | CASSQDVPNEQFF | KLGGALQAK | YES | NO | NO |
| 14100 | TRAV1-1 | TRAJ16 | CAVRDAF | TRBV6-1 | TRBJ1-1 | CAGAQNTEAFF | KLGGALQAK | NO | NO | NO |
| 14101 | TRAV12-1 | TRAJ32 | CVVNVIYGGATNKLIF | TRBV11-2 | TRBJ2-3 | CASSLDTPWGFNTDTQYF | KLGGALQAK | YES | NO | NO |
| 14102 | TRAV17 | TRAJ45 | CATWNSGGGADGLTF | TRBV7-8 | TRBJ2-5 | CASYPVPGLKTQYF | KLGGALQAK | YES | NO | NO |
| 14103 | TRAV17 | TRAJ47 | CATGTMEYGNKLVF | TRBV3-1 | TRBJ2-5 | CASSQYPRLAGEKNQETQYF | AVFDRKSDAK | NO | NO | NO |
| 14104 | TRAV19 | TRAJ50 | CALSELGTSYDKVIF | TRBV4-1 | TRBJ2-1 | CASSQDRGPANEQFF | RLRAEAQVK | NO | NO | NO |
| 14105 | TRAV12-3 | TRAJ39 | CAIGPGNMLTF | TRBV4-1 | TRBJ2-3 | CASSQGVGLAGAPSTDTQYF | KLGGALQAK | NO | NO | NO |
| 14106 | TRAV29DV5 | TRAJ26 | CAASGNGQNFVF | TRBV15 | TRBJ2-5 | CATSRDRANWETQYF | KLGGALQAK | YES | NO | NO |
| 14107 | TRAV9-2 | TRAJ16 | CALKDGQKLLF | TRBV20-1 | TRBJ2-3 | CSARRLQGELDTQYF | KLGGALQAK | NO | NO | NO |
| 14108 | TRAV21 | TRAJ34 | CAVSKSYNTDKLIF | TRBV20-1 | TRBJ1-1 | CSATNPSGSLDTEAFF | KLGGALQAK | YES | NO | NO |

| | | | | | | | | | |
|---|---|---|---|---|---|---|---|---|---|
| 14109 | TRAV29DV5 | TRAJ10 | CAAPIPGGGNKLTF | TRBV7-9 | TRBJ1-1 | CASTKDRSEAFF | IVTDFSVIK | YES | NO | YES |
| 14110 | TRAV13-1 | TRAJ10 | CAVLPGGGNKLTF | TRBV4-1 | TRBJ1-1 | CASSHRQGAGTEAFF | KLGGALQAK | YES | NO | NO |
| 14111 | TRAV34 | TRAJ32 | CGADKYGGATNKLIF | TRBV10-3 | TRBJ2-4 | CAISGGTVESKNIQYF | KLGGALQAK | YES | NO | NO |
| 14112 | TRAV12-3 | TRAJ43 | CAMSAGNNNDMRF | TRBV6-1 | TRBJ2-7 | CASSDRDRVEQYF | KLGGALQAK | YES | NO | NO |
| 14113 | TRAV35 | TRAJ43 | CAGGRIDNDMRF | TRBV6-1 | TRBJ2-1 | CASSHGGISYNEQFF | KLGGALQAK | YES | NO | NO |
| 14114 | TRAV8-4 | TRAJ20 | CAVIQINDYKLSF | TRBV7-8 | TRBJ1-2 | CASSLGYGYTF | KLGGALQAK | YES | NO | NO |
| 14115 | TRAV22 | TRAJ48 | CAVVSNFGNEKLTF | TRBV5-5 | TRBJ2-7 | CASSGDWNLDEQYV | KLGGALQAK | NO | NO | NO |
| 14116 | TRAV12-1 | TRAJ8 | CVVNFGTGFQKLVF | TRBV7-2 | TRBJ1-1 | CASSLAQVQYTEAFF | KLGGALQAK | YES | NO | NO |
| 14117 | TRAV13-2 | TRAJ28 | CAENKGSGAGSYQLTF | TRBV19 | TRBJ2-7 | CASSPGTGGYEQYV | KLGGALQAK | NO | NO | NO |
| 14118 | TRAV3 | TRAJ32 | CAPHYGGATNKLIF | TRBV30 | TRBJ1-2 | CAWSPGNYGYTF | KLGGALQAK | YES | NO | NO |
| 14119 | TRAV4 | TRAJ41 | CLVGAKASNSGYALNF | TRBV7-2 | TRBJ2-1 | CASSLSISAYNEQFF | AVFDRKSDAK | YES | NO | YES |
| 14120 | TRAV4 | TRAJ32 | CLVGGSGATNKLIF | TRBV2 | TRBJ2-1 | CASSVGYGTGPPGGQFF | KLGGALQAK | YES | NO | NO |
| 14121 | TRAV38-1 | TRAJ32 | CAFMSFGGATNKLIF | TRBV13 | TRBJ2-1 | CASSSSQEDNEQFF | KLGGALQAK | YES | NO | NO |
| 14122 | TRAV12-1 | TRAJ45 | CVVNNRGADGLTF | TRBV29-1 | TRBJ2-1 | CSVVWTSRRYEQFF | KLGGALQAK | YES | YES | NO |
| 14123 | TRAV39 | TRAJ43 | CAVDPYNNNDMRF | TRBV10-3 | TRBJ2-2 | CAISGRGTNTGELFF | KLGGALQAK | YES | YES | NO |
| 14124 | TRAV3 | TRAJ6 | CAVRDTSSGGSYIPTF | TRBV7-2 | TRBJ2-7 | CASSAGTSGSSYEQYV | KLGGALQAK | NO | NO | NO |
| 14125 | TRAV3 | TRAJ40 | CAVRSFSGTYKYIF | TRBV3-1 | TRBJ1-2 | CASSQRGQETLYGYTF | KLGGALQAK | NO | NO | NO |
| 14126 | TRAV35 | TRAJ44 | CAGQERDTGTASKLTF | TRBV27 | TRBJ2-5 | CASRRRDGTQYF | KLGGALQAK | YES | NO | NO |
| 14127 | TRAV1-2 | TRAJ13 | CAVRDQGGYQKVTF | TRBV28 | TRBJ1-5 | CASRPVEGNQPQHF | KLGGALQAK | YES | NO | NO |
| 14128 | TRAV36DV7 | TRAJ48 | CAVGLISNFGNEKLTF | TRBV12-4 | TRBJ1-2 | CASSLDDRRNYGYTF | KLGGALQAK | NO | NO | NO |
| 14129 | TRAV1-2 | TRAJ20 | CAVREGDYKLSF | TRBV6-4 | TRBJ2-3 | CASSGTSGATDTQYF | KLGGALQAK | YES | YES | NO |
| 14130 | TRAV14DV4 | TRAJ12 | CAMRDRDSSYKLIF | TRBV2 | TRBJ2-2 | CASTNPGLAGPSGTGELFF | KLGGALQAK | YES | NO | NO |
| 14131 | TRAV8-1 | TRAJ11 | CAVSGYSTLTF | TRBV5-1 | TRBJ2-7 | CASSLGQGVYEQYF | KLGGALQAK | YES | NO | NO |
| 14132 | TRAV1-2 | TRAJ28 | CAGRYSGAGSYQLTF | TRBV2 | TRBJ1-1 | CASSESLGVNTEAFF | KLGGALQAK | YES | NO | NO |
| 14133 | TRAV29DV5 | TRAJ40 | CAAYGTSGTYKYIF | TRBV19 | TRBJ1-2 | CASSIVGSNYGYTF | KLGGALQAK | YES | NO | NO |
| 14134 | TRAV16 | TRAJ4 | CALRFSGGYNKLIF | TRBV7-3 | TRBJ2-3 | CASSLGPPGTSTDTQYF | KLGGALQAK | NO | NO | NO |
| 14135 | TRAV24 | TRAJ40 | CAFPSGTYKYIF | TRBV5-4 | TRBJ1-4 | CASSLDQQGVGEKLFF | KLGGALQAK | YES | NO | NO |
| 14136 | TRAV9-2 | TRAJ42 | CALSDPRDGGSQGNLIF | TRBV6-6 | TRBJ2-3 | CASNPRIRARTDTQYF | KLGGALQAK | NO | NO | NO |
| 14137 | TRAV19 | TRAJ57 | CALSEAFSQGGSEKLVF | TRBV9 | TRBJ2-3 | CASSDLSHSGRADTQYF | KLGGALQAK | YES | NO | NO |
| 14138 | TRAV17 | TRAJ48 | CATDPANFGNEKLTF | TRBV7-6 | TRBJ2-1 | CASSLVTGTGASYNEQFF | KLGGALQAK | YES | NO | NO |
| 14139 | TRAV12-1 | TRAJ42 | CVVNIVGGSQGNLIF | TRBV6-5 | TRBJ1-2 | CASSYSGGNGYTF | KLGGALQAK | YES | NO | NO |
| 14140 | TRAV26-1 | TRAJ11 | CIVRVDNSGYSTLTF | TRBV7-2 | TRBJ2-4 | CASSLMIGAGEAKNIQYF | KLGGALQAK | YES | NO | NO |
| 14141 | TRAV21 | TRAJ12 | CAVLMDSSYKLIF | TRBV7-9 | TRBJ2-1 | CASSNSLPSGVSYNEQFF | KLGGALQAK | YES | NO | NO |
| 14142 | TRAV1-2 | TRAJ28 | CAVRRSGAGSYQLTF | TRBV9 | TRBJ2-1 | CASSVGISGGPGQFEQFF | KLGGALQAK | YES | NO | NO |
| 14143 | TRAV21 | TRAJ33 | CAVTLDSNYQLIW | TRBV7-2 | TRBJ2-7 | CASSPNGPHEQYV | KLGGALQAK | NO | NO | NO |
| 14144 | TRAV17 | TRAJ52 | CATDASTSYGKLTF | TRBV19 | TRBJ1-1 | CASNPWDMNTEAFF | KLGGALQAK | YES | NO | NO |
| 14145 | TRAV17 | TRAJ17 | CAALDGIKAAGNKLTF | TRBV19 | TRBJ1-1 | CASKTGGALNTEAFF | KLGGALQAK | YES | NO | NO |
| 14146 | TRAV1-2 | TRAJ11 | CAVRDLGGYSTLTF | TRBV27 | TRBJ2-5 | CASSPLAGDGKTQYF | KLGGALQAK | YES | NO | NO |
| 14147 | TRAV24 | TRAJ33 | CAHQGDSNYQLIW | TRBV4-2 | TRBJ2-1 | CASSQELGSSYNEQFF | KLGGALQAK | YES | NO | NO |
| 14148 | TRAV17 | TRAJ6 | CVAGGSYIPTF | TRBV12-4 | TRBJ1-4 | CASSLASIGEKLFF | KLGGALQAK | YES | NO | NO |
| 14149 | TRAV38-2DV8 | TRAJ11 | CAYRSEISSTLTF | TRBV7-9 | TRBJ2-7 | CASSLASFDEQYF | KLGGALQAK | YES | NO | NO |
| 14150 | TRAV19 | TRAJ34 | CALSELDTDKLIF | TRBV6-5 | TRBJ2-7 | CASSYGAGERYEQYF | KLGGALQAK | YES | NO | NO |
| 14151 | TRAV8-6 | TRAJ49 | CAVSDRVVNTGNQFYF | TRBV30 | TRBJ1-2 | CAWSVLSDFNYGYTF | KLGGALQAK | YES | NO | NO |
| 14152 | TRAV17 | TRAJ54 | CATDDPQGAQKLVF | TRBV6-5 | TRBJ2-7 | CASSDGTGLNEQYF | KLGGALQAK | YES | NO | NO |
| 14153 | TRAV1-1 | TRAJ26 | CAVLYGQNFVF | TRBV27 | TRBJ2-1 | CASSLQALAGGGEQFF | KLGGALQAK | YES | NO | NO |
| 14154 | TRAV35 | TRAJ26 | CAGFSYGQNFVF | TRBV27 | TRBJ1-1 | CASSITFKANEAFF | KLGGALQAK | YES | NO | NO |
| 14155 | TRAV13-1 | TRAJ41 | CAASIWSNSGYALNF | TRBV19 | TRBJ1-2 | CASSIAGGGYGYTF | KLGGALQAK | YES | NO | NO |
| 14156 | TRAV1-1 | TRAJ42 | CAVLGSQGNLIF | TRBV20-1 | TRBJ1-1 | CSAMTGLNTEAFF | KLGGALQAK | YES | NO | NO |
| 14157 | TRAV12-2 | TRAJ16 | CAVRDGQKLLF | TRBV29-1 | TRBJ1-3 | CSVEQGESGNTIYF | KLGGALQAK | NO | NO | NO |
| 14158 | TRAV29DV5 | TRAJ49 | CAASGDQFYF | TRBV27 | TRBJ2-5 | CASSSDRIGETQYF | KLGGALQAK | YES | NO | NO |
| 14159 | TRAV8-6 | TRAJ43 | CAVSSYNNNDMRF | TRBV7-6 | TRBJ1-6 | CASSMGQDSPLHF | KLGGALQAK | YES | NO | NO |
| 14160 | TRAV1-2 | TRAJ15 | CAVTLNQAGTALIF | TRBV9 | TRBJ1-2 | CASSVDPWDTSGYTF | KLGGALQAK | YES | NO | NO |
| 14161 | TRAV24 | TRAJ16 | CASITDGQKLLF | TRBV6-6 | TRBJ1-1 | CASSPLLWGMNTEAFF | KLGGALQAK | NO | NO | NO |
| 14162 | TRAV26-1 | TRAJ34 | CIVIRGGKLIF | TRBV19 | TRBJ2-2 | CASIERGLAGGSTGELFF | KLGGALQAK | YES | NO | NO |
| 14163 | TRAV1-2 | TRAJ20 | CAVLNDYKLSF | TRBV27 | TRBJ2-5 | CASSRTGNQETQYF | KLGGALQAK | YES | NO | NO |
| 14164 | TRAV21 | TRAJ48 | CAAPPGFGNEKLTF | TRBV19 | TRBJ2-1 | CASSTGGGRYNEQFF | KLGGALQAK | YES | NO | NO |
| 14165 | TRAV1-2 | TRAJ6 | CAVRDSGAGGSYIPTF | TRBV7-6 | TRBJ1-5 | CASSESTGTGGGQPQHF | KLGGALQAK | YES | NO | NO |
| 14166 | TRAV1-2 | TRAJ42 | CAVLNYGGSQGNLIF | TRBV5-4 | TRBJ1-2 | CASAAGTGGDYGYTF | KLGGALQAK | YES | NO | NO |
| 14167 | TRAV35 | TRAJ17 | CAGQVLGAAGNKLTF | TRBV10-2 | TRBJ2-7 | CASSDISPTYEQYV | KLGGALQAK | NO | NO | NO |
| 14168 | TRAV8-3 | TRAJ20 | CAVGPGGDYKLSF | TRBV7-8 | TRBJ2-7 | CASSTSQTGDFAYEQYV | KLGGALQAK | NO | NO | NO |
| 14169 | TRAV12-2 | TRAJ40 | CAVKGSGTYKYIF | TRBV7-3 | TRBJ1-1 | CASSPHPGSTEAFF | KLGGALQAK | YES | NO | NO |
| 14170 | TRAV1-2 | TRAJ32 | CAVLGGATNKLIF | TRBV7-2 | TRBJ1-6 | CASSSRTGIYNSPLHF | AVFDRKSDAK | YES | NO | NO |
| 14171 | TRAV8-6 | TRAJ34 | CAVRGRVSTDKLIF | TRBV4-1 | TRBJ1-1 | CASSQDQGITEAFF | KLGGALQAK | YES | NO | NO |
| 14172 | TRAV2 | TRAJ11 | CAVGGYSTLTF | TRBV4-1 | TRBJ1-2 | CASSQENRDYGYTF | KLGGALQAK | NO | NO | NO |
| 14173 | TRAV39 | TRAJ45 | CAVDYYSGGGADGLTF | TRBV4-1 | TRBJ2-2 | CASSQRLGPEQGGTGELFF | LLFGYPVYV | YES | YES | YES |
| 14174 | TRAV1-2 | TRAJ33 | CAVVDSNYQLIW | TRBV20-1 | TRBJ1-1 | CSARDRGDTEAFF | KLGGALQAK | YES | NO | NO |
| 14175 | TRAV13-2 | TRAJ52 | CAETTNAGGTSYGKLTF | TRBV19 | TRBJ2-5 | CASSPGGPETQYF | KLGGALQAK | YES | NO | NO |
| 14176 | TRAV20 | TRAJ23 | CAVQSVYNQGGKLIF | TRBV6-3 | TRBJ2-1 | CASIEALAGSYNEQFF | KLGGALQAK | YES | NO | NO |
| 14177 | TRAV21 | TRAJ20 | CAVLANDYKLSF | TRBV12-3 | TRBJ2-1 | CASRLAGDEQFF | KLGGALQAK | YES | NO | NO |
| 14178 | TRAV14DV4 | TRAJ9 | CAMRAGGFKTIF | TRBV30 | TRBJ2-3 | CAWSVLGSTDTQYF | KLGGALQAK | YES | NO | NO |
| 14179 | TRAV8-6 | TRAJ53 | CAVSDVGGSNYKLTF | TRBV19 | TRBJ2-1 | CASSIYTADEQFF | KLGGALQAK | YES | NO | NO |
| 14180 | TRAV8-1 | TRAJ15 | CAVNIQAGTALIF | TRBV20-1 | TRBJ1-2 | CSARDSDYGYTF | KLGGALQAK | YES | NO | NO |
| 14181 | TRAV20 | TRAJ28 | CAASGAGSYQLTF | TRBV5-6 | TRBJ2-2 | CASSSSNTGELFF | AVFDRKSDAK | YES | YES | YES |
| 14182 | TRAV21 | TRAJ12 | CAVGGDSSYKLIF | TRBV20-1 | TRBJ2-1 | CSAPRTSSYNEQFF | KLGGALQAK | NO | NO | NO |
| 14183 | TRAV21 | TRAJ52 | CAADPLHLNAGGTSYGKLTF | TRBV5-6 | TRBJ1-1 | CASNPETGINTEAFF | KLGGALQAK | NO | NO | NO |
| 14184 | TRAV25 | TRAJ42 | CAGTYYGGSQGNLIF | TRBV4-1 | TRBJ1-5 | CASSQDRTQGQPQHF | KLGGALQAK | YES | NO | NO |
| 14185 | TRAV1-2 | TRAJ33 | CAIMDSNYQLIW | TRBV3-1 | TRBJ2-2 | CASSPRTGDTGELFF | KLGGALQAK | YES | NO | NO |
| 14186 | TRAV1-1 | TRAJ30 | CAVRDYKNRDDKIIF | TRBV20-1 | TRBJ2-2 | CSATQGPGELFF | KLGGALQAK | YES | NO | NO |
| 14187 | TRAV8-4 | TRAJ20 | CAVIQINDYKLSF | TRBV12-4 | TRBJ2-3 | CASSPMDTQYF | KLGGALQAK | YES | NO | NO |
| 14188 | TRAV16 | TRAJ31 | CALNNNARLMF | TRBV27 | TRBJ2-5 | CASSLYPYSETQYF | KLGGALQAK | NO | NO | NO |
| 14189 | TRAV36DV7 | TRAJ20 | CAVYNDYKLSF | TRBV12-5 | TRBJ2-2 | CASGLDMAGGELFF | AVFDRKSDAK | NO | NO | NO |
| 14190 | TRAV10 | TRAJ37 | CVVSALGYGNTGKLIF | TRBV15 | TRBJ2-4 | CATSRGPTSGTGSAKNIQYF | KLGGALQAK | NO | NO | NO |
| 14191 | TRAV29DV5 | TRAJ21 | CAARATYNFNKFYF | TRBV10-3 | TRBJ2-7 | CASKPRARQGSSYEQYF | KLGGALQAK | YES | NO | NO |
| 14192 | TRAV20 | TRAJ28 | CAVLDSGAGSYQLTF | TRBV27 | TRBJ1-2 | CASSPGIAYGYTF | KLGGALQAK | YES | NO | NO |
| 14193 | TRAV2 | TRAJ32 | CAVEDGGATNKLIF | TRBV2 | TRBJ2-5 | CASSALGESQETQYF | AVFDRKSDAK | NO | NO | NO |
| 14194 | TRAV8-2 | TRAJ34 | CVVSDRRTDKLIF | TRBV6-5 | TRBJ2-3 | CASIPSGVPSTDTQYF | KLGGALQAK | NO | NO | NO |
| 14196 | TRAV12-3 | TRAJ42 | CASMNYGGSQGNLIF | TRBV12-4 | TRBJ1-5 | CASSLVGTLNQPQHF | KLGGALQAK | YES | NO | NO |
| 14197 | TRAV8-1 | TRAJ36 | CAVTEANNLFF | TRBV19 | TRBJ2-5 | CASSMTMLRGRIWETQYF | KLGGALQAK | YES | NO | NO |
| 14198 | TRAV29DV5 | TRAJ34 | CAASAFLLTDKLIF | TRBV7-9 | TRBJ2-2 | CASSRPGQEPGELFF | KLGGALQAK | YES | YES | NO |
| 14199 | TRAV38-2DV8 | TRAJ33 | CAFLFGNYQLIW | TRBV9 | TRBJ2-7 | CASSVDNHYEQYF | KLGGALQAK | YES | NO | NO |
| 14200 | TRAV40 | TRAJ54 | CLLGEMIGQGAQKLVF | TRBV5-6 | TRBJ1-5 | CASSSLQQPQHF | KLGGALQAK | NO | NO | NO |
| 14201 | TRAV38-2DV8 | TRAJ48 | CAYRSNFGNEKLTF | TRBV7-2 | TRBJ1-6 | CASRPSMWGNSPLHF | KLGGALQAK | NO | NO | NO |
| 14202 | TRAV35 | TRAJ58 | CAGPDRQTSGSRLTF | TRBV19 | TRBJ1-5 | CASTGQGLGNQPQHF | KLGGALQAK | YES | NO | NO |
| 14203 | TRAV5 | TRAJ42 | CALIGSQGNLIF | TRBV5-8 | TRBJ2-7 | CASSLADTYEQYV | KLGGALQAK | NO | NO | NO |
| 14204 | TRAV30 | TRAJ30 | CGTAPHRDDKIIF | TRBV19 | TRBJ2-1 | CASSLDTSVYNEQFF | KLGGALQAK | YES | NO | NO |
| 14205 | TRAV40 | TRAJ43 | CLLGDESNNNDMRF | TRBV7-9 | TRBJ1-2 | CASSYGYGYTF | KLGGALQAK | NO | NO | NO |
| 14206 | TRAV19 | TRAJ48 | CALSDPNFGNEKLTF | TRBV20-1 | TRBJ2-7 | CSARVDSSYEQYF | KLGGALQAK | YES | NO | NO |
| 14207 | TRAV24 | TRAJ52 | CAFIGDAGGTSYGKLTF | TRBV4-3 | TRBJ1-2 | CASSQTLGGSYGYTF | KLGGALQAK | YES | NO | NO |
| 14208 | TRAV1-2 | TRAJ33 | CAVMDSNYQLIW | TRBV10-3 | TRBJ2-1 | CAISETSGSYNEQFF | KLGGALQAK | YES | NO | NO |
| 14209 | TRAV29DV5 | TRAJ5 | CATRDTGRRALTF | TRBV12-4 | TRBJ1-4 | CASSGFEFQEKLFF | KLGGALQAK | YES | NO | NO |
| 14211 | TRAV1-1 | TRAJ42 | CAVRDGGSQGNLIF | TRBV12-3 | TRBJ2-3 | CASRSGPQDTQYF | KLGGALQAK | YES | NO | NO |
| 14212 | TRAV12-2 | TRAJ44 | CAVDTGTASKLTF | TRBV5-1 | TRBJ2-7 | CASSLYGTGNLYEQYF | KLGGALQAK | YES | NO | NO |
| 14213 | TRAV38-1 | TRAJ54 | CAFMSLIQGAQKLVF | TRBV7-9 | TRBJ1-1 | CASSPIGGNTEAFF | KLGGALQAK | YES | NO | NO |
| 14214 | TRAV1-1 | TRAJ34 | CAVNPTDKLIF | TRBV20-1 | TRBJ2-3 | CSGAGVPSTDTQYF | KLGGALQAK | NO | NO | NO |

| | | | | | | | | | | |
|---|---|---|---|---|---|---|---|---|---|---|
| 14215 | TRAV13-2 | TRAJ53 | CAENSGGSNYKLTF | TRBV7-9 | TRBJ2-7 | CASKDGISYEQYF | KLGGALQAK | YES | NO | NO |
| 14216 | TRAV29DV5 | TRAJ41 | CAASANSGYALNF | TRBV11-2 | TRBJ1-1 | CASSFHRNAEAFF | AVFDRKSDAK | YES | NO | YES |
| 14217 | TRAV20 | TRAJ22 | CAVQALWGSARQLTF | TRBV29-1 | TRBJ2-3 | CSVYSGMDTQYF | KLGGALQAK | YES | NO | NO |
| 14218 | TRAV3 | TRAJ50 | CAALKTSYDKVIF | TRBV5-1 | TRBJ2-2 | CASSYGLADYTTGELFF | KLGGALQAK | YES | NO | NO |
| 14219 | TRAV14DV4 | TRAJ40 | CAMRVFTKYIF | TRBV13 | TRBJ1-6 | CASSPDGDNSPLHF | KLGGALQAK | YES | NO | NO |
| 14220 | TRAV17 | TRAJ52 | CAKGKAGGTSYGKLTF | TRBV19 | TRBJ2-7 | CASTAGQGGTYEQYF | KLGGALQAK | YES | NO | NO |
| 14221 | TRAV8-3 | TRAJ27 | CAVVSPNAGKSTF | TRBV6-5 | TRBJ2-3 | CASSYSGGRGPTDTQYF | KLGGALQAK | NO | NO | NO |
| 14222 | TRAV25 | TRAJ48 | CAGPGLTNFGNEKLTF | TRBV10-2 | TRBJ1-5 | CASSEWGTGDQPQHF | KLGGALQAK | YES | NO | NO |
| 14223 | TRAV8-2 | TRAJ4 | CVVSISGGYNKLIF | TRBV11-2 | TRBJ2-2 | CASYRPGAGELFF | KLGGALQAK | YES | NO | NO |
| 14224 | TRAV19 | TRAJ39 | CALSGRAGNMLTF | TRBV7-9 | TRBJ1-2 | CASSLVLLGDYGYTF | KLGGALQAK | YES | NO | NO |
| 14225 | TRAV1-1 | TRAJ12 | CAVRDGWDSSYKLIF | TRBV9 | TRBJ2-2 | CASSVETNTGELFF | KLGGALQAK | YES | NO | NO |
| 14226 | TRAV3 | TRAJ40 | CAVTTSGTYKYIF | TRBV3-1 | TRBJ1-4 | CASSLKISWTNEKLFF | KLGGALQAK | NO | NO | NO |
| 14227 | TRAV38-1 | TRAJ22 | CACLSGSARQLTF | TRBV12-4 | TRBJ2-3 | CASSLNIVGGGSTDTQYF | KLGGALQAK | YES | NO | NO |
| 14228 | TRAV4 | TRAJ26 | CLVGDVDNYGQNFVF | TRBV7-9 | TRBJ2-3 | CASSIDTDTQYF | KLGGALQAK | YES | NO | NO |
| 14229 | TRAV29DV5 | TRAJ40 | CAASEVMASGTYKYIF | TRBV7-2 | TRBJ2-5 | CASSLMRLQETQYF | KLGGALQAK | YES | NO | NO |
| 14230 | TRAV14DV4 | TRAJ34 | CAMRERLYNTDKLIF | TRBV4-3 | TRBJ2-1 | CASSQPSSVEQFF | KLGGALQAK | YES | NO | NO |
| 14231 | TRAV8-3 | TRAJ6 | CASLGGSYIPTF | TRBV9 | TRBJ2-2 | CASRVGLADTGELFF | KLGGALQAK | YES | NO | NO |
| 14232 | TRAV1-2 | TRAJ26 | CAVRGHNYGQNFVF | TRBV5-1 | TRBJ1-6 | CASSLDDNSPLHF | KLGGALQAK | YES | NO | NO |
| 14233 | TRAV12-3 | TRAJ54 | CAMSIQGAQKLVF | TRBV19 | TRBJ1-1 | CASSIGKDMNTEAFF | KLGGALQAK | YES | NO | NO |
| 14234 | TRAV35 | TRAJ49 | CAGQLTGNQFYF | TRBV27 | TRBJ1-2 | CASRPNPGSTYGYTF | KLGGALQAK | YES | NO | NO |
| 14235 | TRAV8-6 | TRAJ17 | CAVSEAAGNKLTF | TRBV7-8 | TRBJ1-1 | CASSLGGAKNTEAFF | KLGGALQAK | YES | NO | NO |
| 14236 | TRAV8-3 | TRAJ20 | CAVGANDYKLSF | TRBV10-1 | TRBJ2-1 | CATGTSGNEQFF | KLGGALQAK | YES | NO | NO |
| 14237 | TRAV8-4 | TRAJ20 | CAVIQINDYKLSF | TRBV20-1 | TRBJ2-7 | CSARDLEVLSYEQYF | KLGGALQAK | YES | NO | NO |
| 14238 | TRAV12-1 | TRAJ43 | CVVTYQYNNNDMRF | TRBV19 | TRBJ1-1 | CASSLGQGGTEAFF | AVFDRKSDAK | YES | NO | YES |
| 14239 | TRAV26-1 | TRAJ38 | CIVRVPYAGNNRKLIW | TRBV15 | TRBJ1-3 | CATSRDLGTGNTIYF | KLGGALQAK | YES | NO | NO |
| 14240 | TRAV20 | TRAJ54 | CAVQPQIQGAQKLVF | TRBV7-9 | TRBJ2-1 | CASSLSYNEQFF | KLGGALQAK | YES | NO | NO |
| 14241 | TRAV14DV4 | TRAJ40 | CAMRASGTYKYIF | TRBV9 | TRBJ1-1 | CASSVEGRGTEAFF | KLGGALQAK | YES | NO | NO |
| 14242 | TRAV10 | TRAJ45 | CVVSDLAGGGADGLTF | TRBV14 | TRBJ1-2 | CASSQDWDSTYGYTF | KLGGALQAK | NO | NO | NO |
| 14243 | TRAV14DV4 | TRAJ49 | CAMRDIIGNQFYF | TRBV20-1 | TRBJ2-7 | CSARDWGGGGDLYEQYF | KLGGALQAK | NO | NO | NO |
| 14244 | TRAV14DV4 | TRAJ9 | CAMSYRGGGFKTIF | TRBV27 | TRBJ2-1 | CASSIRQGPNEQFF | KLGGALQAK | YES | NO | NO |
| 14245 | TRAV8-3 | TRAJ53 | CAVWNSGGSNYKLTF | TRBV6-5 | TRBJ1-2 | CASSYSHRQGNYGYTF | KLGGALQAK | YES | NO | NO |
| 14246 | TRAV1-2 | TRAJ12 | CAVMDSSYKLIF | TRBV6-1 | TRBJ2-3 | CASSEGTGHTDTQYF | KLGGALQAK | YES | NO | NO |
| 14248 | TRAV13-1 | TRAJ23 | CAAYNQGGKLIF | TRBV6-5 | TRBJ2-7 | CASSIQGPDEQYF | AVFDRKSDAK | YES | NO | YES |
| 14249 | TRAV29DV5 | TRAJ49 | CAASANTGNQFYF | TRBV7-9 | TRBJ2-6 | CASSLDRQAMTF | KLGGALQAK | YES | NO | NO |
| 14250 | TRAV29DV5 | TRAJ40 | CAASATISGTYKYIF | TRBV7-9 | TRBJ2-3 | CASSTVPAAPTDTQYF | RLRAEAQVK | YES | YES | NO |
| 14251 | TRAV29DV5 | TRAJ29 | CAASVGSGNTPLVF | TRBV7-2 | TRBJ1-1 | CASRLGVFGGTEAFF | KLGGALQAK | YES | NO | NO |
| 14252 | TRAV14DV4 | TRAJ49 | CAMREGPTGNQFYF | TRBV2 | TRBJ2-5 | CASSDQGRQETQYF | KLGGALQAK | YES | NO | NO |
| 14253 | TRAV16 | TRAJ39 | CALSGPAGNMLTF | TRBV29-1 | TRBJ2-1 | CSSPDPNRPYNEQFF | KLGGALQAK | NO | NO | NO |
| 14254 | TRAV27 | TRAJ28 | CAGYSGAGSYQLTF | TRBV11-2 | TRBJ1-4 | CASSFGTGATNEKLFF | KLGGALQAK | YES | NO | NO |
| 14255 | TRAV14DV4 | TRAJ53 | CAMRSPGGSNYKLTF | TRBV7-2 | TRBJ2-3 | CASSPSDGRTDTQYF | KLGGALQAK | YES | NO | NO |
| 14256 | TRAV29DV5 | TRAJ45 | CAASVRGGADGLTF | TRBV4-1 | TRBJ2-5 | CASSQPSGGSQYF | KLGGALQAK | YES | NO | NO |
| 14257 | TRAV26-2 | TRAJ52 | CILRDVGGGTSYGKLTF | TRBV19 | TRBJ1-1 | CASSIGGLLNTEAFF | KLGGALQAK | YES | NO | NO |
| 14258 | TRAV26-1 | TRAJ24 | CIVMATTDSWGKLQF | TRBV9 | TRBJ2-1 | CASTTGLAGVEQFF | KLGGALQAK | YES | NO | NO |
| 14259 | TRAV19 | TRAJ53 | CALSTEGGSNYKLTF | TRBV4-3 | TRBJ1-1 | CASSQDMSPGTEAFF | KLGGALQAK | YES | NO | NO |
| 14260 | TRAV30 | TRAJ42 | CGTERVGGSQGNLIF | TRBV27 | TRBJ2-7 | CASSLSGGSYEQYF | KLGGALQAK | YES | YES | NO |
| 14261 | TRAV26-1 | TRAJ54 | CIVRGADQGAQKLVF | TRBV29-1 | TRBJ1-2 | CSVDLEANYGYTF | KLGGALQAK | YES | NO | NO |
| 14262 | TRAV3 | TRAJ26 | CAVRDILGNYGQNFVF | TRBV10-1 | TRBJ2-3 | CASSEVGLAEDTQYF | KLGGALQAK | NO | NO | NO |
| 14263 | TRAV29DV5 | TRAJ27 | CAANTNAGKSTF | TRBV10-1 | TRBJ2-3 | CASSSTDTQYF | KLGGALQAK | YES | NO | NO |
| 14264 | TRAV8-2 | TRAJ33 | CVVSGGDSNYQLIW | TRBV12-4 | TRBJ1-1 | CASSLGGWGAFF | KLGGALQAK | YES | NO | NO |
| 14265 | TRAV3 | TRAJ36 | CAVRESGGANNLFF | TRBV30 | TRBJ2-5 | CAWDSGQGFLETQYF | KLGGALQAK | NO | NO | NO |
| 14266 | TRAV14DV4 | TRAJ48 | CAMRGLSNFGNEKLTF | TRBV19 | TRBJ1-5 | CASSIDRGGGLGNQPQHF | KLGGALQAK | YES | NO | NO |
| 14267 | TRAV14DV4 | TRAJ4 | CAMRGPISGGYNKLIF | TRBV11-2 | TRBJ2-5 | CASSYMRGAYQETQYF | KLGGALQAK | YES | NO | NO |
| 14268 | TRAV8-1 | TRAJ7 | CAVNEGNNRLAF | TRBV5-1 | TRBJ1-1 | CASSRENEQGEAFF | KLGGALQAK | YES | NO | NO |
| 14269 | TRAV1-2 | TRAJ39 | CAGGGAGNMLTF | TRBV10-3 | TRBJ2-3 | CAIRTTPPGHSTDTQYF | KLGGALQAK | YES | NO | NO |
| 14270 | TRAV10 | TRAJ39 | CVVRNNAGNMLTF | TRBV5-8 | TRBJ2-3 | CASSLGSSGRSPFGTDTQYF | KLGGALQAK | NO | NO | NO |
| 14271 | TRAV13-1 | TRAJ48 | CAASGGGNFGNEKLTF | TRBV19 | TRBJ2-5 | CASSIGGGLETQYF | KLGGALQAK | YES | NO | NO |
| 14273 | TRAV14DV4 | TRAJ9 | CAMREGSGGFKTIF | TRBV9 | TRBJ1-2 | CASSSTGTGDGYTF | AVFDRKSDAK | YES | NO | NO |
| 14274 | TRAV17 | TRAJ34 | CATGLSYNTDKLIF | TRBV7-6 | TRBJ2-1 | CASSLATGNEQFF | KLGGALQAK | YES | NO | NO |
| 14275 | TRAV8-1 | TRAJ45 | CAVRGGADGLTF | TRBV16 | TRBJ1-5 | CASRSLGAGLYSNQPQHF | KLGGALQAK | YES | NO | NO |
| 14276 | TRAV12-1 | TRAJ24 | CVVPPVWGKFQF | TRBV12-3 | TRBJ2-1 | CASSSTNEQFF | KLGGALQAK | YES | NO | NO |
| 14277 | TRAV19 | TRAJ9 | CALSGATGGFKTIF | TRBV7-9 | TRBJ1-2 | CASSDGGGADALVRDGYTF | KLGGALQAK | YES | NO | NO |
| 14279 | TRAV14DV4 | TRAJ33 | CAMREMDSNYQLIW | TRBV9 | TRBJ2-1 | CASSPTGPEQFF | KLGGALQAK | YES | NO | NO |
| 14280 | TRAV13-1 | TRAJ4 | CAAGDSGGYNKLIF | TRBV19 | TRBJ1-1 | CASNPGGGFTEAFF | KLGGALQAK | YES | NO | NO |
| 14281 | TRAV8-3 | TRAJ10 | CAVADTGGGNKLTF | TRBV19 | TRBJ1-5 | CASSILSNQPQHF | KLGGALQAK | YES | NO | NO |
| 14282 | TRAV12-1 | TRAJ27 | CVVNVYNTNAGKSTF | TRBV12-3 | TRBJ1-2 | CASTGGGDGYTF | KLGGALQAK | YES | NO | NO |
| 14283 | TRAV12-3 | TRAJ39 | CAIGPGNMLTF | TRBV9 | TRBJ1-1 | CASSLSQGALNTEAFF | KLGGALQAK | YES | NO | NO |
| 14284 | TRAV12-3 | TRAJ13 | CAMRLINSGGYQKVTF | TRBV19 | TRBJ2-5 | CASSIVPAETQYF | AVFDRKSDAK | YES | NO | YES |
| 14285 | TRAV26-2 | TRAJ26 | CIRLDNYGQNFVF | TRBV6-5 | TRBJ1-2 | CASSYIPGQAYGYTF | KLGGALQAK | YES | NO | NO |
| 14286 | TRAV17 | TRAJ37 | CATDAWYSNTGKLIF | TRBV20-1 | TRBJ2-3 | CSARDPHTDTQYF | KLGGALQAK | NO | NO | NO |
| 14287 | TRAV30 | TRAJ17 | CGTAIKAAGNKLTF | TRBV19 | TRBJ2-7 | CASSIGTSVHEQYF | KLGGALQAK | YES | NO | NO |
| 14288 | TRAV13-2 | TRAJ8 | CAETTPGNTGFQKLVF | TRBV2 | TRBJ2-7 | CASSEVASGTYEQYF | KLGGALQAK | YES | NO | NO |
| 14289 | TRAV26-1 | TRAJ49 | CIAGFNNQFYF | TRBV7-9 | TRBJ2-4 | CASSLVRGANIQYF | KLGGALQAK | YES | NO | NO |
| 14290 | TRAV8-3 | TRAJ6 | CAASSGGSYIPTF | TRBV10-3 | TRBJ1-3 | CAIRVPTTESGGNTIYF | KLGGALQAK | YES | NO | NO |
| 14291 | TRAV12-1 | TRAJ52 | CAPRSGTSYGKLTF | TRBV5-1 | TRBJ1-1 | CASSLGGYRAEAFF | KLGGALQAK | YES | NO | NO |
| 14292 | TRAV26-2 | TRAJ30 | CIPEGNRDDKIIF | TRBV19 | TRBJ1-2 | CASSPTGLGFYGYTF | KLGGALQAK | YES | NO | NO |
| 14293 | TRAV14DV4 | TRAJ54 | CAMREGLFIQGAQKLVF | TRBV19 | TRBJ2-4 | CASSKGQGLNIQYF | KLGGALQAK | YES | NO | NO |
| 14294 | TRAV6 | TRAJ24 | CAIYDSWGKLQF | TRBV11-2 | TRBJ2-1 | CASSLALAIEQFF | KLGGALQAK | YES | NO | NO |
| 14295 | TRAV2 | TRAJ13 | CAVEASGGYQKVTF | TRBV12-4 | TRBJ2-5 | CASSFSQGAFSETQYF | KLGGALQAK | NO | NO | NO |
| 14296 | TRAV17 | TRAJ53 | CATDAGSNYKLTF | TRBV13 | TRBJ2-3 | CASSSRPADTQYF | KLGGALQAK | YES | NO | NO |
| 14297 | TRAV38-2DV8 | TRAJ15 | CAYQAGTALIF | TRBV4-3 | TRBJ1-6 | CASSQSSVNSPLHF | KLGGALQAK | YES | NO | NO |
| 14298 | TRAV1-1 | TRAJ32 | CAVNYGGATNKLIF | TRBV7-6 | TRBJ2-7 | CASSPQRMGDEQYF | KLGGALQAK | YES | NO | NO |
| 14299 | TRAV13-1 | TRAJ26 | CAADGVWRYGQNFVF | TRBV20-1 | TRBJ1-6 | CSVAPDRFNSPLHF | KLGGALQAK | YES | NO | NO |
| 14300 | TRAV27 | TRAJ15 | CAGALWQAGTALIF | TRBV11-2 | TRBJ2-1 | CASSPGLEGSNEQFF | KLGGALQAK | YES | NO | NO |
| 14301 | TRAV27 | TRAJ26 | CAGDLDYGQNFVF | TRBV7-9 | TRBJ2-1 | CASSSRSREVEQFF | KLGGALQAK | YES | NO | NO |
| 14302 | TRAV12-1 | TRAJ49 | CVVNYNQFYF | TRBV20-1 | TRBJ2-3 | CSALEYLGQSTDTQYF | KLGGALQAK | NO | NO | NO |
| 14303 | TRAV13-2 | TRAJ26 | CAENDNYGQNFVF | TRBV19 | TRBJ2-5 | CASSIEGAHEETQYF | KLGGALQAK | YES | NO | NO |
| 14304 | TRAV4 | TRAJ49 | CLVGVRTGNQFYF | TRBV20-1 | TRBJ2-1 | CSARDRDGYNEQFF | KLGGALQAK | NO | NO | NO |
| 14305 | TRAV12-2 | TRAJ15 | CAVNQAGTALIF | TRBV18 | TRBJ2-1 | CASSPKLRRDNEQFF | KLGGALQAK | YES | NO | NO |
| 14306 | TRAV8-3 | TRAJ7 | CAVGAEGNNRLAF | TRBV18 | TRBJ2-1 | CASSSGQGAGYNEQFF | KLGGALQAK | YES | NO | NO |
| 14307 | TRAV23DV6 | TRAJ12 | CAASEVDSSYKLIF | TRBV13 | TRBJ1-5 | CASSLQGGLNQPQHF | KLGGALQAK | NO | NO | NO |
| 14308 | TRAV8-1 | TRAJ13 | CAASSGGYQKVTF | TRBV4-3 | TRBJ1-1 | CASREIQGGTEAFF | KLGGALQAK | YES | NO | NO |
| 14309 | TRAV2 | TRAJ11 | CAVKGYSTLTF | TRBV27 | TRBJ1-1 | CASTASMNTEAFF | KLGGALQAK | NO | NO | NO |
| 14310 | TRAV8-3 | TRAJ4 | CAVGSGGYNKLIF | TRBV15 | TRBJ2-1 | CATRDDEDIRSYNEQFF | KLGGALQAK | YES | NO | NO |
| 14311 | TRAV26-2 | TRAJ53 | CILRGSNYKLTF | TRBV4-1 | TRBJ1-1 | CASSAGAWTEAFF | KLGGALQAK | YES | NO | NO |
| 14312 | TRAV12-1 | TRAJ42 | CVVNLDPVDGGSQGNLIF | TRBV2 | TRBJ2-7 | CASSVFREPYEQYF | KLGGALQAK | YES | NO | NO |
| 14313 | TRAV5 | TRAJ50 | CAESMETSYDKVIF | TRBV20-1 | TRBJ1-5 | CSARLSGSYQPQHF | KLGGALQAK | YES | NO | NO |
| 14314 | TRAV17 | TRAJ36 | CAQTGANNLFF | TRBV4-3 | TRBJ1-4 | CASSQDHRMGGHEKLFF | KLGGALQAK | YES | NO | NO |
| 14315 | TRAV19 | TRAJ9 | CALSEAKGFKTIF | TRBV9 | TRBJ2-3 | CASSIYTDTQYF | KLGGALQAK | YES | NO | NO |
| 14316 | TRAV10 | TRAJ34 | CVVRAYNTDKLIF | TRBV5-6 | TRBJ2-2 | CASSLFGGRDTGELFF | KLGGALQAK | YES | NO | NO |
| 14317 | TRAV13-1 | TRAJ43 | CAASYDDMRF | TRBV12-4 | TRBJ1-1 | CASSEGYGMNTEAFF | KLGGALQAK | YES | NO | NO |
| 14318 | TRAV14DV4 | TRAJ4 | CAMRPPFRGGYNKLIF | TRBV7-9 | TRBJ1-2 | CASSSTGTGPNYGYTF | KLGGALQAK | YES | NO | NO |
| 14319 | TRAV21 | TRAJ12 | CAPRRMDSSYKLIF | TRBV13 | TRBJ1-2 | CASSLTAYGYTF | KLGGALQAK | YES | NO | NO |
| 14320 | TRAV1-2 | TRAJ37 | CAVRAGNTGKLIF | TRBV7-9 | TRBJ2-7 | CASTPPRTDPYEQYF | KLGGALQAK | YES | NO | NO |
| 14321 | TRAV14DV4 | TRAJ22 | CAMRRPISSGSARQLTF | TRBV11-3 | TRBJ2-2 | CASSSNKLAGGPSGELFF | KLGGALQAK | YES | NO | NO |

| | | | | | | | | | | |
|---|---|---|---|---|---|---|---|---|---|---|
| 14322 | TRAV19 | TRAJ48 | CALSEAIMNFGNEKLTF | TRBV6-1 | TRBJ1-1 | CASYWLRDRDPEAFF | KLGGALQAK | YES | NO | NO |
| 14323 | TRAV22 | TRAJ11 | CAVDEYSTLTF | TRBV29-1 | TRBJ2-2 | CSVFQENTGELFF | AVFDRKSDAK | YES | YES | YES |
| 14324 | TRAV17 | TRAJ26 | CATAWGSYGQNFVF | TRBV15 | TRBJ2-5 | CATSIPLAGDQETQYF | RLRAEAQVK | YES | NO | YES |
| 14325 | TRAV13-1 | TRAJ50 | CAATPKTSYDKVIF | TRBV19 | TRBJ1-2 | CASSTTGGPYGYTF | KLGGALQAK | YES | NO | NO |
| 14326 | TRAV14DV4 | TRAJ22 | CAMRRPISSGSARQLTF | TRBV29-1 | TRBJ2-2 | CSVGRDRGPTGELFF | KLGGALQAK | YES | NO | NO |
| 14327 | TRAV24 | TRAJ12 | CAFYSSYKLIF | TRBV11-2 | TRBJ1-1 | CASSLVTGLNTEAFF | KLGGALQAK | YES | NO | NO |
| 14328 | TRAV8-6 | TRAJ20 | CAVSLGDYKLSF | TRBV29-1 | TRBJ2-2 | CSVEGFGRAGTGELFF | KLGGALQAK | YES | NO | NO |
| 14329 | TRAV8-1 | TRAJ36 | CAVAGANNLFF | TRBV6-6 | TRBJ2-5 | CASSYVPGRPETQYF | RLRAEAQVK | NO | NO | NO |
| 14330 | TRAV13-1 | TRAJ49 | CAARRGLAGNQFYF | TRBV9 | TRBJ2-6 | CASSAREQLVPSISGANVLTF | KLGGALQAK | NO | NO | NO |
| 14331 | TRAV1-2 | TRAJ20 | CAVEDYKLSF | TRBV14 | TRBJ1-3 | CASSRPSSGNTIYF | RLRAEAQVK | NO | NO | NO |
| 14332 | TRAV25 | TRAJ40 | CAVTSGTYKYIF | TRBV12-4 | TRBJ2-2 | CASTPWGGAGGELFF | KLGGALQAK | YES | NO | NO |
| 14333 | TRAV1-2 | TRAJ20 | CAVSAVNDYKLSF | TRBV20-1 | TRBJ2-1 | CGGLGGTSGRNEQFF | AVFDRKSDAK | NO | NO | NO |
| 14334 | TRAV14DV4 | TRAJ48 | CAMREHYFGNEKLTF | TRBV5-1 | TRBJ2-1 | CASSWDSPGEFF | KLGGALQAK | YES | NO | NO |
| 14335 | TRAV24 | TRAJ28 | CAFDSGAGSYQLTF | TRBV28 | TRBJ2-3 | CASISADTQYF | KLGGALQAK | YES | NO | NO |
| 14336 | TRAV3 | TRAJ12 | CAVRDDRDSSYKLIF | TRBV7-9 | TRBJ2-7 | CASSTQGGEQYF | KLGGALQAK | YES | NO | NO |
| 14338 | TRAV8-3 | TRAJ17 | CALKAAGNKLTF | TRBV25-1 | TRBJ1-1 | CASRDGTANTEAFF | KLGGALQAK | YES | YES | NO |
| 14339 | TRAV12-2 | TRAJ50 | CAVKTSYDKVIF | TRBV19 | TRBJ2-7 | CASKPDGSKNEQYF | KLGGALQAK | YES | NO | NO |
| 14340 | TRAV1-1 | TRAJ23 | CAVMIYNQGGKLIF | TRBV7-9 | TRBJ2-2 | CASSLAGTSRTGELFF | KLGGALQAK | YES | NO | NO |
| 14341 | TRAV8-3 | TRAJ47 | CAVDTEYGNKLVF | TRBV6-5 | TRBJ2-1 | CASSPYMTSGSVSEQFF | KLGGALQAK | NO | NO | NO |
| 14342 | TRAV25 | TRAJ13 | CVGGYQKVTF | TRBV19 | TRBJ2-1 | CASSEQYNEQFF | KLGGALQAK | YES | NO | NO |
| 14343 | TRAV8-4 | TRAJ6 | CAVSESGGSYIPTF | TRBV27 | TRBJ1-5 | CASSLGGSNQPQHF | AVFDRKSDAK | NO | NO | NO |
| 14344 | TRAV13-2 | TRAJ54 | CAENIFSGAQKLVF | TRBV9 | TRBJ2-7 | CASSVGTGNYEQYF | KLGGALQAK | YES | NO | NO |
| 14345 | TRAV19 | TRAJ30 | CALSENEDDKIIF | TRBV9 | TRBJ2-3 | CASRLGGASTDTQYF | KLGGALQAK | YES | NO | NO |
| 14346 | TRAV3 | TRAJ29 | CAVRDRHSGNTPLVF | TRBV13 | TRBJ2-7 | CASSSGGAAEQYF | KLGGALQAK | NO | NO | NO |
| 14347 | TRAV22 | TRAJ6 | CAGGGSGGSYIPTF | TRBV6-3 | TRBJ1-1 | CASSPFNTEAFF | KLGGALQAK | YES | NO | NO |
| 14348 | TRAV8-4 | TRAJ40 | CAVRTSGTYKYIF | TRBV7-3 | TRBJ2-3 | CASSLGTTSTDTQYF | KLGGALQAK | YES | NO | NO |
| 14349 | TRAV12-2 | TRAJ50 | CAVNIHAGRTSYDKVIF | TRBV7-7 | TRBJ2-1 | CASSSYGQGGYNEQFF | AVFDRKSDAK | YES | YES | YES |
| 14350 | TRAV4 | TRAJ34 | CLVGPYNTDKLIF | TRBV20-1 | TRBJ1-1 | CSARDLGTPNTEAFF | AVFDRKSDAK | NO | NO | NO |
| 14351 | TRAV22 | TRAJ41 | CAVPVYALNF | TRBV20-1 | TRBJ1-2 | CSAKTGGYGYTF | KLGGALQAK | YES | NO | NO |
| 14352 | TRAV26-1 | TRAJ30 | CIVRYFTNRDDKIIF | TRBV4-1 | TRBJ2-3 | CASSPWGSGGAGDTQYF | KLGGALQAK | YES | NO | NO |
| 14353 | TRAV12-1 | TRAJ44 | CVVEYTGTASKLTF | TRBV5-5 | TRBJ2-2 | CASSSPPMGGELFF | KLGGALQAK | YES | NO | NO |
| 14354 | TRAV30 | TRAJ43 | CGTEWANNNDMRF | TRBV19 | TRBJ1-1 | CASSMGDPWAFF | KLGGALQAK | YES | NO | NO |
| 14355 | TRAV14DV4 | TRAJ48 | CAMREDPNFGNEKLTF | TRBV7-3 | TRBJ2-2 | CASSLGQVGLAGAPNTGELFF | KLGGALQAK | NO | NO | NO |
| 14356 | TRAV29DV5 | TRAJ32 | CAASTWLDGGATNKLIF | TRBV12-4 | TRBJ1-1 | CASSLGLGYTEAFF | KLGGALQAK | YES | NO | NO |
| 14357 | TRAV5 | TRAJ12 | CAEEGMDSSYKLIF | TRBV5-1 | TRBJ2-7 | CASSSADGSYEQYF | KLGGALQAK | YES | NO | NO |
| 14358 | TRAV29DV5 | TRAJ52 | CASGGTSYGKLTF | TRBV7-9 | TRBJ2-2 | CASSDDMRQDPGELFF | KLGGALQAK | YES | NO | NO |
| 14359 | TRAV8-1 | TRAJ6 | CAGAGGSYIPTF | TRBV24-1 | TRBJ2-1 | CATSDGTGVYNEQFF | KLGGALQAK | YES | NO | NO |
| 14360 | TRAV17 | TRAJ31 | CAMGNNARLMF | TRBV9 | TRBJ2-1 | CASSVAPGPNEQFF | KLGGALQAK | YES | NO | NO |
| 14361 | TRAV20 | TRAJ37 | CAVLGNTGKLIF | TRBV27 | TRBJ2-7 | CASSSGQGIGEQYF | KLGGALQAK | YES | NO | NO |
| 14362 | TRAV20 | TRAJ57 | CAVRGGSEKLVF | TRBV7-2 | TRBJ2-1 | CASSLVGQSYNEQFF | KLGGALQAK | YES | NO | NO |
| 14363 | TRAV8-6 | TRAJ30 | CAFMNRDDKIIF | TRBV20-1 | TRBJ1-2 | CSAGPISDDNYGYTF | KLGGALQAK | YES | NO | NO |
| 14364 | TRAV26-2 | TRAJ41 | CILRANSGYALNF | TRBV6-1 | TRBJ2-7 | CASSEALHGGYRSYEQYF | KLGGALQAK | YES | NO | NO |
| 14365 | TRAV30 | TRAJ26 | CGTTEDNYGQNFVF | TRBV5-4 | TRBJ2-5 | CASSYGLAQQETQYF | KLGGALQAK | YES | NO | NO |
| 14366 | TRAV38-1 | TRAJ52 | CAFMKAGGTSYGKLTF | TRBV19 | TRBJ1-5 | CASNPGGLHQPQHF | AVFDRKSDAK | YES | NO | YES |
| 14367 | TRAV21 | TRAJ47 | CAVVSMEYGNKLVF | TRBV11-2 | TRBJ1-1 | CASSLFQGSGTEAFF | KLGGALQAK | YES | NO | NO |
| 14368 | TRAV38-2DV8 | TRAJ45 | CAFGGGADGLTF | TRBV2 | TRBJ2-2 | CASSDEGGLTITGELFF | KLGGALQAK | YES | NO | NO |
| 14369 | TRAV21 | TRAJ16 | CAVKHPLDGQKLLF | TRBV4-3 | TRBJ2-1 | CASSQGFLAGAPEQFF | KLGGALQAK | NO | NO | NO |
| 14370 | TRAV4 | TRAJ4 | CLVGDTSGYNKLIF | TRBV11-2 | TRBJ2-3 | CASSLSTGVGPDTQYF | KLGGALQAK | YES | NO | NO |
| 14371 | TRAV19 | TRAJ20 | CALSEGYSNDYKLSF | TRBV20-1 | TRBJ2-7 | CSARRTDDYEQYF | KLGGALQAK | YES | NO | NO |
| 14372 | TRAV12-3 | TRAJ45 | CAMSSYSGGGADGLTF | TRBV19 | TRBJ2-2 | CASSTITGELFF | KLGGALQAK | YES | NO | NO |
| 14373 | TRAV24 | TRAJ44 | CARYTGTASKLTF | TRBV6-6 | TRBJ2-5 | CASSQGTGAGETQYF | KLGGALQAK | NO | NO | NO |
| 14374 | TRAV14DV4 | TRAJ48 | CAMRAAFGNEKLTF | TRBV12-4 | TRBJ2-6 | CASSLGTGVSGANVLTF | KLGGALQAK | YES | NO | NO |
| 14375 | TRAV29DV5 | TRAJ23 | CAAPDFYNQGGKLIF | TRBV4-1 | TRBJ1-3 | CASSPTKGGSGNTIYF | KLGGALQAK | YES | NO | NO |
| 14376 | TRAV38-2DV8 | TRAJ48 | CAYRAPRGNEKLTF | TRBV11-2 | TRBJ2-7 | CASSYGTGTYEQYF | KLGGALQAK | YES | NO | NO |
| 14377 | TRAV20 | TRAJ45 | CAVPRIQRGGADGLTF | TRBV4-1 | TRBJ2-7 | CASSPQGASYEQYF | KLGGALQAK | YES | NO | NO |
| 14378 | TRAV19 | TRAJ10 | CALSEFWWAGGGNKLTF | TRBV7-8 | TRBJ1-6 | CASSLVAAQLHTEENSPLHF | KLGGALQAK | NO | NO | NO |
| 14379 | TRAV5 | TRAJ42 | CADTDYGGSQGNLIF | TRBV3-1 | TRBJ1-2 | CASSHTTGSPFYGYTF | KLGGALQAK | YES | NO | NO |
| 14380 | TRAV13-2 | TRAJ43 | CAEFNNNDMRF | TRBV7-8 | TRBJ2-1 | CASSTRDRAYNEQFF | KLGGALQAK | YES | NO | NO |
| 14381 | TRAV29DV5 | TRAJ43 | CAASATYNDMRF | TRBV7-9 | TRBJ2-4 | CASSTDRGGTKNIQYF | KLGGALQAK | YES | NO | NO |
| 14382 | TRAV3 | TRAJ33 | CAVRDSNYQLIW | TRBV5-6 | TRBJ2-1 | CASRDLGPEQFF | KLGGALQAK | YES | NO | NO |
| 14383 | TRAV20 | TRAJ15 | CAVRTDNQAGTALIF | TRBV18 | TRBJ2-1 | CASSPSSNEQFF | KLGGALQAK | YES | NO | NO |
| 14384 | TRAV27 | TRAJ26 | CAGLDNYGQNFVF | TRBV20-1 | TRBJ2-3 | CSVANTPFPADTQYF | KLGGALQAK | NO | NO | NO |
| 14385 | TRAV12-1 | TRAJ23 | CVVWIYNQGGKLIF | TRBV4-1 | TRBJ1-1 | CASSLSGTETEAFF | KLGGALQAK | YES | NO | NO |
| 14386 | TRAV5 | TRAJ31 | CADRDNNARLMF | TRBV3-1 | TRBJ2-1 | CASSQVDSSYNEQFF | IVTDFSVIK | YES | NO | YES |
| 14387 | TRAV35 | TRAJ16 | CAGQYTNSDGQKLLF | TRBV7-9 | TRBJ2-1 | CASREGLSNEQFF | KLGGALQAK | NO | NO | NO |
| 14388 | TRAV12-1 | TRAJ6 | CVVNTGGSYIPTF | TRBV12-3 | TRBJ1-1 | CASTDLNTEAFF | KLGGALQAK | YES | NO | NO |
| 14389 | TRAV19 | TRAJ48 | CALRFISNFGNEKLTF | TRBV7-9 | TRBJ2-3 | CASSLKADSRTDTQYF | KLGGALQAK | YES | NO | NO |
| 14390 | TRAV3 | TRAJ29 | CAVRDGAGNTPLVF | TRBV4-1 | TRBJ2-5 | CASSQTLAGGPWETQYF | KLGGALQAK | YES | NO | NO |
| 14391 | TRAV26-1 | TRAJ57 | CIVRVAYDEGGSEKLVF | TRBV2 | TRBJ1-1 | CASSVSVNTEAFF | KLGGALQAK | YES | NO | NO |
| 14392 | TRAV5 | TRAJ41 | CAESTPNSGYALNF | TRBV24-1 | TRBJ1-1 | CATMDTMNTEAFF | KLGGALQAK | YES | NO | NO |
| 14393 | TRAV25 | TRAJ42 | CAGLNGDYGGSQGNLIF | TRBV18 | TRBJ2-7 | CASSPHPPPMGEQYF | KLGGALQAK | YES | NO | NO |
| 14394 | TRAV17 | TRAJ20 | CAINPNDYKLSF | TRBV4-1 | TRBJ2-7 | CASSQDGSSYYEQYF | KLGGALQAK | YES | NO | NO |
| 14395 | TRAV21 | TRAJ54 | CAVRIQGAQKLVF | TRBV27 | TRBJ1-1 | CASSRGGGGLEAFF | KLGGALQAK | YES | NO | NO |
| 14396 | TRAV13-1 | TRAJ31 | CAASSSNNARLMF | TRBV7-2 | TRBJ2-1 | CASSLVGSYNEQFF | KLGGALQAK | YES | NO | NO |
| 14397 | TRAV26-2 | TRAJ54 | CILRGVLGAQKLVF | TRBV19 | TRBJ2-7 | CASSTVDRNSYEQYV | KLGGALQAK | NO | NO | NO |
| 14398 | TRAV19 | TRAJ28 | CALSEETSGAGSYQLTF | TRBV29-1 | TRBJ2-6 | CSVVSGDSGANVLTF | KLGGALQAK | YES | NO | NO |
| 14399 | TRAV25 | TRAJ37 | CAAPSSNTGKLIF | TRBV27 | TRBJ2-1 | CASSLDRYNEQFF | KLGGALQAK | YES | NO | NO |
| 14400 | TRAV39 | TRAJ34 | CAVANTDKLIF | TRBV4-1 | TRBJ2-7 | CASSQDRVSYEQYF | KLGGALQAK | YES | NO | NO |
| 14401 | TRAV3 | TRAJ35 | CAVRDIEGFGNVLHC | TRBV6-1 | TRBJ1-1 | CASSDRVGGMNTEAFF | KLGGALQAK | NO | NO | NO |
| 14402 | TRAV17 | TRAJ27 | CATDAGYTNAGKSTF | TRBV11-2 | TRBJ1-1 | CASSYTGGNTEAFF | KLGGALQAK | YES | NO | NO |
| 14403 | TRAV22 | TRAJ30 | CAVGGRDDKIIF | TRBV12-4 | TRBJ1-1 | CASRLWAGTEAFF | KLGGALQAK | YES | NO | NO |
| 14404 | TRAV14DV4 | TRAJ53 | CAIPEIESGGSNYKLTF | TRBV19 | TRBJ1-1 | CASSMGTGFFMNTEAFF | AVFDRKSDAK | YES | NO | YES |
| 14405 | TRAV14DV4 | TRAJ45 | CAMRERSGGGADGLTF | TRBV6-5 | TRBJ1-1 | CASSYSRKAHTEAFF | KLGGALQAK | YES | NO | NO |
| 14406 | TRAV22 | TRAJ13 | CAEGSGGYQKVTF | TRBV5-1 | TRBJ1-2 | CASSTLSGANYGYTF | KLGGALQAK | YES | NO | NO |
| 14407 | TRAV12-1 | TRAJ40 | CVSHSGTYKYIF | TRBV4-3 | TRBJ2-1 | CASSQGSTNNEQFF | KLGGALQAK | YES | NO | NO |
| 14408 | TRAV38-2DV8 | TRAJ33 | CAYRSSMDSNYQLIW | TRBV19 | TRBJ2-3 | CASSWGLQGNTQYF | AVFDRKSDAK | YES | NO | YES |
| 14409 | TRAV12-1 | TRAJ23 | CVVNIRGKLIF | TRBV7-9 | TRBJ2-6 | CASSLGESGANVLTF | KLGGALQAK | YES | NO | NO |
| 14410 | TRAV36DV7 | TRAJ33 | CALHSNYQLIW | TRBV6-5 | TRBJ2-3 | CASSRQSADTQYF | AVFDRKSDAK | NO | NO | NO |
| 14411 | TRAV3 | TRAJ6 | CAVRGGSYIPTF | TRBV10-2 | TRBJ1-5 | CASRLQGLDQPQHF | KLGGALQAK | NO | NO | NO |
| 14412 | TRAV21 | TRAJ26 | CAVKHYGQNFVF | TRBV12-4 | TRBJ2-2 | CASSSEGAGELFF | KLGGALQAK | YES | NO | NO |
| 14413 | TRAV17 | TRAJ45 | CATDLTNSGGGADGLTF | TRBV19 | TRBJ2-7 | CASSMAGYEQYF | KLGGALQAK | YES | NO | NO |
| 14414 | TRAV5 | TRAJ49 | CADGNQFYF | TRBV20-1 | TRBJ1-1 | CSALGGVGVRENTEAFF | KLGGALQAK | NO | NO | NO |
| 14415 | TRAV8-2 | TRAJ45 | CVVSDWSGGGADGLTF | TRBV7-2 | TRBJ2-5 | CASSFIVGGQETQYF | KLGGALQAK | YES | NO | NO |
| 14416 | TRAV14DV4 | TRAJ5 | CAMRDPQGGRRALTF | TRBV7-9 | TRBJ2-1 | CASSSRLAVYNEQFF | KLGGALQAK | YES | NO | NO |
| 14417 | TRAV17 | TRAJ23 | CATGYYNQGGKLIF | TRBV20-1 | TRBJ2-1 | CSARDRPQETYNEQFF | KLGGALQAK | NO | NO | NO |
| 14418 | TRAV29DV5 | TRAJ33 | CAASDSNYQLIW | TRBV7-9 | TRBJ1-2 | CASSPAGGYSYGYTF | KLGGALQAK | YES | NO | NO |
| 14419 | TRAV8-3 | TRAJ40 | CAVGGSGSGTYKYIF | TRBV5-4 | TRBJ2-1 | CASSLLGATEQFF | KLGGALQAK | YES | NO | NO |
| 14420 | TRAV21 | TRAJ48 | CAFISNFGNEKLTF | TRBV6-3 | TRBJ1-1 | CASSYGQGYTEAFF | KLGGALQAK | YES | NO | NO |
| 14421 | TRAV26-2 | TRAJ39 | CILSNNAGNMLTF | TRBV5-1 | TRBJ2-1 | CASSLKGGTVLYNEQFF | KLGGALQAK | YES | NO | NO |
| 14422 | TRAV12-1 | TRAJ43 | CVVAYNNNDMRF | TRBV6-1 | TRBJ1-2 | CASSEEPGYGYTF | KLGGALQAK | YES | NO | NO |
| 14423 | TRAV21 | TRAJ41 | CAVNSNSGYALNF | TRBV7-2 | TRBJ1-3 | CASSLVGTGPPGNTIYF | KLGGALQAK | YES | NO | NO |
| 14424 | TRAV17 | TRAJ23 | CATDENQGGKLIF | TRBV9 | TRBJ2-5 | CASSVAGEETQYF | KLGGALQAK | YES | NO | NO |
| 14425 | TRAV36DV7 | TRAJ45 | CAVEAADSGGGADGLTF | TRBV13 | TRBJ1-1 | CASSLEGTSSEAFF | KLGGALQAK | NO | NO | NO |
| 14426 | TRAV8-3 | TRAJ47 | CAAMEYGNKLVF | TRBV6-5 | TRBJ2-3 | CASSVGPSTDTQYF | KLGGALQAK | NO | NO | NO |

| | | | | | | | | | | |
|---|---|---|---|---|---|---|---|---|---|---|
| 14427 | TRAV19 | TRAJ52 | CALSETNAGGTSYGKLTF | TRBV4-1 | TRBJ2-7 | CASSHPQGTSYEQYF | KLGGALQAK | YES | NO | NO |
| 14428 | TRAV29DV5 | TRAJ34 | CAASFHTDKLIF | TRBV7-8 | TRBJ1-2 | CASSLSLGTTNYGYTF | KLGGALQAK | YES | NO | NO |
| 14429 | TRAV25 | TRAJ28 | CAGSGAGSYQLTF | TRBV7-9 | TRBJ1-2 | CASSSNQGRSYGYTF | KLGGALQAK | YES | NO | NO |
| 14430 | TRAV41 | TRAJ34 | CAVSLGWNTDKLIF | TRBV27 | TRBJ1-5 | CASSTLSYNQPQHF | KLGGALQAK | YES | NO | NO |
| 14431 | TRAV34 | TRAJ45 | CGAANSGGGADGLTF | TRBV19 | TRBJ2-1 | CASSKYGGNEQFF | KLGGALQAK | YES | NO | NO |
| 14432 | TRAV17 | TRAJ20 | CATDNNDYKLSF | TRBV7-9 | TRBJ1-1 | CASSLDVGDTEAFF | KLGGALQAK | YES | NO | NO |
| 14433 | TRAV14DV4 | TRAJ50 | CAMSLKTSYDKVIF | TRBV7-9 | TRBJ2-7 | CASKDGISYEQYF | KLGGALQAK | YES | NO | NO |
| 14434 | TRAV38-2DV8 | TRAJ48 | CAYRSADFGNEKLTF | TRBV4-1 | TRBJ2-7 | CASSPGLLTYEQYF | KLGGALQAK | YES | NO | NO |
| 14435 | TRAV1-2 | TRAJ15 | CAVRLNQAGTALIF | TRBV3-1 | TRBJ2-3 | CASSQDLATWVQYF | KLGGALQAK | YES | NO | NO |
| 14436 | TRAV4 | TRAJ23 | CLVGETYNQGGKLIF | TRBV16 | TRBJ2-3 | CASGQGVTDTQYF | KLGGALQAK | NO | NO | NO |
| 14437 | TRAV13-1 | TRAJ42 | CAASIDGGSQGNLIF | TRBV20-1 | TRBJ2-1 | CSARSLYDYPNEQFF | KLGGALQAK | YES | NO | NO |
| 14438 | TRAV19 | TRAJ26 | CALRAYYGQNFVF | TRBV3-1 | TRBJ2-7 | CASSQEGAGGIHEQYF | KLGGALQAK | YES | NO | NO |
| 14439 | TRAV26-1 | TRAJ17 | CIVRDEIKAAGNKLTF | TRBV19 | TRBJ2-7 | CASSIVDAGLYEQYF | KLGGALQAK | YES | NO | NO |
| 14440 | TRAV34 | TRAJ20 | CGADLNDYKLSF | TRBV27 | TRBJ2-5 | CASSSFTGTETQYF | KLGGALQAK | YES | NO | NO |
| 14441 | TRAV4 | TRAJ29 | CLVGDNGSGNTPLVF | TRBV7-9 | TRBJ2-1 | CASSLEAGLNNEQFF | KLGGALQAK | YES | YES | NO |
| 14442 | TRAV29DV5 | TRAJ42 | CAARFGGSQGNLIF | TRBV7-6 | TRBJ2-1 | CASSSPLAGRFNEQFF | KLGGALQAK | YES | NO | NO |
| 14443 | TRAV13-2 | TRAJ39 | CAENLWNAGNMLTF | TRBV12-3 | TRBJ2-3 | CASSLLGGGITDTQYF | KLGGALQAK | YES | NO | NO |
| 14444 | TRAV38-2DV8 | TRAJ49 | CAYRSAYTGNQFYF | TRBV7-8 | TRBJ1-2 | CASSLGQDGYTF | KLGGALQAK | YES | NO | NO |
| 14445 | TRAV10 | TRAJ48 | CVVSANFGNEKLTF | TRBV2 | TRBJ2-3 | CASSERAGVMDTQYF | KLGGALQAK | YES | NO | NO |
| 14447 | TRAV12-2 | TRAJ45 | CAVYGSGGGADGLTF | TRBV6-5 | TRBJ1-1 | CASSPGNTEAFF | AVFDRKSDAK | YES | YES | YES |
| 14448 | TRAV13-1 | TRAJ20 | CAASLSNDYKLSF | TRBV27 | TRBJ2-1 | CASQGASNEQFF | KLGGALQAK | YES | NO | NO |
| 14449 | TRAV19 | TRAJ6 | CALRDASGGSYIPTF | TRBV4-1 | TRBJ2-3 | CASSQRARAPDTQYF | KLGGALQAK | YES | NO | NO |
| 14450 | TRAV29DV5 | TRAJ47 | CAASVMEYGNKLVF | TRBV5-6 | TRBJ2-5 | CASSLPLARASQYF | AVFDRKSDAK | YES | NO | NO |
| 14451 | TRAV13-1 | TRAJ43 | CAATTYNNNDMRF | TRBV19 | TRBJ1-1 | CASSIGGGNTEAFF | KLGGALQAK | YES | NO | NO |
| 14452 | TRAV19 | TRAJ34 | CALSGGSYNTDKLIF | TRBV5-1 | TRBJ2-1 | CASSLRGDEQFF | KLGGALQAK | YES | NO | NO |
| 14453 | TRAV27 | TRAJ24 | CAGVTTDSWGKFQF | TRBV9 | TRBJ1-1 | CASSEDGELSNTEAFF | KLGGALQAK | YES | NO | NO |
| 14454 | TRAV3 | TRAJ8 | CASFTGFQKLVF | TRBV6-5 | TRBJ1-4 | CASSYAGPSEKLFF | KLGGALQAK | NO | NO | NO |
| 14455 | TRAV12-2 | TRAJ13 | CAVEEGYQKVTF | TRBV4-1 | TRBJ1-2 | CASSQGQGDGYTF | KLGGALQAK | YES | NO | NO |
| 14456 | TRAV12-1 | TRAJ43 | CVVNRGWDMRF | TRBV3-1 | TRBJ2-6 | CASSQEKSRVGANVLTF | KLGGALQAK | YES | NO | NO |
| 14457 | TRAV19 | TRAJ37 | CALSETYSSNTGKLIF | TRBV7-9 | TRBJ1-1 | CASSRPTLNTEAFF | KLGGALQAK | YES | NO | NO |
| 14458 | TRAV24 | TRAJ15 | CASFNQAGTALIF | TRBV27 | TRBJ1-5 | CASSLSRNQPQHF | KLGGALQAK | YES | NO | NO |
| 14459 | TRAV8-1 | TRAJ33 | CAVVDSNYQLIW | TRBV11-2 | TRBJ1-2 | CASSSTGLGYTF | KLGGALQAK | YES | NO | NO |
| 14460 | TRAV21 | TRAJ26 | CAVTYYGQNFVF | TRBV12-4 | TRBJ2-1 | CASSPSASGSNDEQFF | KLGGALQAK | YES | NO | NO |
| 14461 | TRAV12-2 | TRAJ35 | CAEIGFGNVLHC | TRBV9 | TRBJ2-5 | CASSVGDGGSGGKETQYF | KLGGALQAK | NO | NO | NO |
| 14462 | TRAV27 | TRAJ26 | CAGDENYGQNFVF | TRBV4-1 | TRBJ1-2 | CASSQVTQGAGYGYTF | KLGGALQAK | YES | NO | NO |
| 14463 | TRAV20 | TRAJ23 | CAVEQGGKLIF | TRBV12-3 | TRBJ2-1 | CASSLASGTEQFF | KLGGALQAK | YES | NO | NO |
| 14464 | TRAV5 | TRAJ23 | CAERVYNQGGKLIF | TRBV19 | TRBJ2-3 | CASTPGQTITDTQYF | KLGGALQAK | YES | NO | NO |
| 14465 | TRAV12-2 | TRAJ11 | CAVGGYSTLTF | TRBV5-6 | TRBJ1-5 | CASSFSGEYQPQHF | KLGGALQAK | YES | NO | NO |
| 14466 | TRAV12-2 | TRAJ49 | CAVNIGGNQFYF | TRBV18 | TRBJ2-3 | CASSQGTGEDTQYF | KLGGALQAK | YES | NO | NO |
| 14467 | TRAV24 | TRAJ20 | CASANDYKLSF | TRBV20-1 | TRBJ1-2 | CSARELGQGAFYGYTF | KLGGALQAK | YES | NO | NO |
| 14469 | TRAV39 | TRAJ33 | CAVEKDSNYQLIW | TRBV7-9 | TRBJ2-2 | CAMGPNTGELFF | KLGGALQAK | YES | NO | NO |
| 14471 | TRAV14DV4 | TRAJ45 | CAMREMYSGGGADGLTF | TRBV20-1 | TRBJ2-7 | CSARDRQGFYEQYF | KLGGALQAK | NO | NO | NO |
| 14472 | TRAV13-2 | TRAJ33 | CAERGIMDSNYQLIW | TRBV5-1 | TRBJ2-1 | CASSLGDNEQFF | AVFDRKSDAK | YES | YES | YES |
| 14473 | TRAV19 | TRAJ37 | CALSEMIGSGNTGKLIF | TRBV13 | TRBJ2-1 | CASGTSVYNEQFF | KLGGALQAK | YES | NO | NO |
| 14474 | TRAV8-4 | TRAJ20 | CAVIQINDYKLSF | TRBV5-6 | TRBJ1-5 | CASSSRDSQNQPQHF | KLGGALQAK | YES | NO | NO |
| 14475 | TRAV12-1 | TRAJ46 | CVVNGRGSSGDKLTF | TRBV27 | TRBJ1-5 | CASSWAGSQPQHF | KLGGALQAK | YES | NO | NO |
| 14476 | TRAV4 | TRAJ15 | CLVGDGQAGTALIF | TRBV4-1 | TRBJ1-2 | CASSQPGQDGINYGYTF | KLGGALQAK | NO | NO | NO |
| 14477 | TRAV27 | TRAJ28 | CAEMGGAGSYQLTF | TRBV6-5 | TRBJ1-1 | CASKDIMNTEAFF | KLGGALQAK | YES | NO | NO |
| 14478 | TRAV12-3 | TRAJ42 | CAMMTGGSQGNLIF | TRBV20-1 | TRBJ1-1 | CSARNRGWTEAFF | KLGGALQAK | YES | NO | NO |
| 14479 | TRAV21 | TRAJ27 | CAVKDTNAGKSTF | TRBV20-1 | TRBJ2-1 | CSAREAGVWFNEQFF | RLRAEAQVK | NO | NO | NO |
| 14480 | TRAV5 | TRAJ32 | CAESAGATNKLIF | TRBV12-4 | TRBJ2-3 | CASSLDPGQGWDTQYF | KLGGALQAK | YES | NO | NO |
| 14481 | TRAV34 | TRAJ26 | CGADIGNYGQNFVF | TRBV6-5 | TRBJ2-1 | CASSYGTYSYNEQFF | KLGGALQAK | NO | NO | NO |
| 14482 | TRAV21 | TRAJ29 | CAVGSGNTPLVF | TRBV7-6 | TRBJ2-1 | CASSQAFSPGLAGASYNEQFF | KLGGALQAK | NO | NO | NO |
| 14483 | TRAV8-3 | TRAJ53 | CAVLRGSFNSGGSNYKLTF | TRBV5-1 | TRBJ2-7 | CASRPGRADYEQYF | KLGGALQAK | YES | NO | NO |
| 14484 | TRAV8-6 | TRAJ30 | CAVSLRNRDDKIIF | TRBV20-1 | TRBJ2-3 | CSARDHFSGGYTDTQYF | AVFDRKSDAK | NO | NO | NO |
| 14485 | TRAV3 | TRAJ36 | CAVRDGTGANNLFF | TRBV19 | TRBJ2-1 | CASSIGVNEQFF | KLGGALQAK | NO | NO | NO |
| 14486 | TRAV12-1 | TRAJ39 | CVVKTDNAGNMLTF | TRBV5-1 | TRBJ1-6 | CASSPGTENYNSPLHF | KLGGALQAK | YES | NO | NO |
| 14487 | TRAV1-2 | TRAJ47 | CAVPQEYGNKLVF | TRBV24-1 | TRBJ2-2 | CATSDRGFSTGELFF | KLGGALQAK | YES | NO | NO |
| 14488 | TRAV17 | TRAJ43 | CATEDDMRF | TRBV7-9 | TRBJ1-1 | CASSLASANTEAFF | KLGGALQAK | NO | NO | NO |
| 14489 | TRAV3 | TRAJ41 | CAVRDMGPGYALNF | TRBV29-1 | TRBJ1-2 | CSVDDRGGYTF | KLGGALQAK | NO | NO | NO |
| 14490 | TRAV4 | TRAJ41 | CLVGEAMNSNSGYALNF | TRBV7-9 | TRBJ1-1 | CASSHSGGRVNTEAFF | AVFDRKSDAK | YES | NO | NO |
| 14491 | TRAV35 | TRAJ18 | CAGQDRGSTLGRLYF | TRBV19 | TRBJ1-1 | CASTQGGSNTEAFF | KLGGALQAK | YES | NO | NO |
| 14492 | TRAV8-4 | TRAJ57 | CAVSADQGGSEKLVF | TRBV2 | TRBJ1-1 | CASSDRMNTEAFF | KLGGALQAK | YES | NO | NO |
| 14493 | TRAV29DV5 | TRAJ47 | CAASATLGEYGNKLVF | TRBV4-3 | TRBJ1-1 | CASSQDRGLSTEAFF | KLGGALQAK | YES | NO | NO |
| 14494 | TRAV24 | TRAJ22 | CAFTISSGSARQLTF | TRBV15 | TRBJ2-7 | CATSRAPGTSGKQSIYEQYF | KLGGALQAK | NO | NO | NO |
| 14495 | TRAV35 | TRAJ42 | CAGQWGSQGNLIF | TRBV6-1 | TRBJ1-5 | CASSEHQPQHF | KLGGALQAK | YES | NO | NO |
| 14496 | TRAV3 | TRAJ34 | CAVRDMRNTDKLIF | TRBV11-3 | TRBJ2-4 | CASSLELDIQYF | KLGGALQAK | YES | NO | NO |
| 14497 | TRAV27 | TRAJ23 | CAGGDNQGGKLIF | TRBV6-5 | TRBJ2-1 | CASRPLAGAYNEQFF | KLGGALQAK | NO | NO | NO |
| 14498 | TRAV29DV5 | TRAJ45 | CAAMGGADGLTF | TRBV12-4 | TRBJ2-1 | CASMDRPYNEQFF | KLGGALQAK | YES | NO | NO |
| 14499 | TRAV41 | TRAJ52 | CAVRSHAGGTSYGKLTF | TRBV9 | TRBJ2-1 | CASSVFGEDNEQFF | KLGGALQAK | YES | NO | NO |
| 14500 | TRAV13-2 | TRAJ17 | CAENDKAAGNKLTF | TRBV13 | TRBJ1-3 | CASSFRGSGNTIYF | KLGGALQAK | YES | NO | NO |
| 14502 | TRAV19 | TRAJ54 | CALSEDVLQGAQKLVF | TRBV7-3 | TRBJ2-3 | CASSPHQDRVTDTQYF | KLGGALQAK | YES | NO | NO |
| 14503 | TRAV19 | TRAJ34 | CALSEAQNTDKLIF | TRBV19 | TRBJ1-5 | CASSDLNSRQPQHF | KLGGALQAK | YES | NO | NO |
| 14504 | TRAV26-2 | TRAJ28 | CILRDLGGAGSYQLTF | TRBV19 | TRBJ1-1 | CASSITGNTEAFF | KLGGALQAK | YES | NO | NO |
| 14505 | TRAV27 | TRAJ41 | CAGDGDSGYALNF | TRBV7-9 | TRBJ2-1 | CASSKLAGGPDEQFF | AVFDRKSDAK | YES | NO | YES |
| 14506 | TRAV5 | TRAJ23 | CAVSWGQGGKLIF | TRBV7-9 | TRBJ2-3 | CASSIPLAESTDTQYF | KLGGALQAK | YES | NO | NO |
| 14507 | TRAV25 | TRAJ23 | CAGQGNQGGKLIF | TRBV9 | TRBJ1-3 | CASSARGQGGNTIYF | KLGGALQAK | YES | NO | NO |
| 14508 | TRAV3 | TRAJ36 | CAVSPGANNLFF | TRBV11-2 | TRBJ1-1 | CASSLPSTGPSEIEAFF | KLGGALQAK | YES | NO | NO |
| 14509 | TRAV3 | TRAJ17 | CAVRGSIKAAGNKLTF | TRBV2 | TRBJ2-1 | CASRSSGIPNEQFF | KLGGALQAK | NO | NO | NO |
| 14510 | TRAV12-2 | TRAJ26 | CAVMDYGQNFVF | TRBV11-2 | TRBJ1-2 | CASSLGEGGGYGYTF | YLNDHLEPWI | YES | NO | NO |
| 14511 | TRAV14DV4 | TRAJ30 | CAMREGLDDKIIF | TRBV19 | TRBJ2-2 | CASSITGQGSTGELFF | KLGGALQAK | YES | NO | NO |
| 14512 | TRAV5 | TRAJ23 | CAESMGNQGGKLIF | TRBV7-9 | TRBJ1-1 | CASSPERGRAEAFF | KLGGALQAK | YES | NO | NO |
| 14513 | TRAV38-1 | TRAJ11 | CAFMKPLNSGYSTLTF | TRBV4-1 | TRBJ2-4 | CASSHGRTNIQYF | KLGGALQAK | YES | NO | NO |
| 14514 | TRAV29DV5 | TRAJ27 | CAAQANAGKSTF | TRBV2 | TRBJ2-7 | CASSSGSSYEQYF | KLGGALQAK | YES | NO | NO |
| 14515 | TRAV12-1 | TRAJ29 | CVPHSGNTPLVF | TRBV5-4 | TRBJ1-2 | CASTPPGGAGYGYTF | KLGGALQAK | YES | NO | NO |
| 14516 | TRAV38-2DV8 | TRAJ41 | CAYRSALNSGYALNF | TRBV18 | TRBJ2-1 | CASSPPIPRRWDEQFF | KLGGALQAK | YES | NO | NO |
| 14517 | TRAV1-2 | TRAJ21 | CAVRFPTRFNKFYF | TRBV9 | TRBJ2-2 | CASSAPDRNTGELFF | KLGGALQAK | YES | YES | NO |
| 14518 | TRAV8-1 | TRAJ33 | CAVIVDSNYQLIW | TRBV7-9 | TRBJ2-7 | CASSSGLSYEQYF | KLGGALQAK | YES | NO | NO |
| 14519 | TRAV17 | TRAJ20 | CATDRDDYKLSF | TRBV19 | TRBJ2-1 | CASSDSGERGEQFF | KLGGALQAK | YES | NO | NO |
| 14520 | TRAV12-1 | TRAJ4 | CVVILLEFSGGYNKLIF | TRBV5-1 | TRBJ2-1 | CASSFSLYNEQFF | KLGGALQAK | YES | NO | NO |
| 14521 | TRAV36DV7 | TRAJ47 | CAVELEYGNKLVF | TRBV4-1 | TRBJ1-2 | CASSADPLGYGYTF | AVFDRKSDAK | NO | NO | NO |
| 14522 | TRAV13-1 | TRAJ23 | CAARSHQGGKLIF | TRBV4-2 | TRBJ2-1 | CASSQGLADNNEQFF | KLGGALQAK | YES | NO | NO |
| 14523 | TRAV8-1 | TRAJ36 | CAVQTGANNLFF | TRBV11-2 | TRBJ1-2 | CASSLGGYRGYTF | KLGGALQAK | YES | NO | NO |
| 14524 | TRAV4 | TRAJ34 | CLVVGEADKLIF | TRBV6-1 | TRBJ2-4 | CASSETPGALNIQYF | AVFDRKSDAK | NO | NO | NO |
| 14525 | TRAV19 | TRAJ4 | CALSEGLSGGYNKLIF | TRBV27 | TRBJ2-1 | CASSLIGGSGEQFF | KLGGALQAK | YES | NO | NO |
| 14526 | TRAV14DV4 | TRAJ47 | CAMREGQGNKLVF | TRBV13 | TRBJ1-1 | CASSLLQPFGTEAFF | KLGGALQAK | YES | NO | NO |
| 14527 | TRAV29DV5 | TRAJ53 | CAALQGSNYKLTF | TRBV9 | TRBJ1-2 | CASSVGSRGVYGYTF | KLGGALQAK | YES | NO | NO |
| 14528 | TRAV17 | TRAJ31 | CATGNNNARLMF | TRBV7-9 | TRBJ2-5 | CASRQLALGETQYF | AVFDRKSDAK | YES | NO | YES |
| 14529 | TRAV19 | TRAJ52 | CALSGGAGGTSYGKLTF | TRBV7-2 | TRBJ1-5 | CASSLVHGSNQPQHF | KLGGALQAK | YES | NO | NO |
| 14530 | TRAV35 | TRAJ49 | CAGYTGNQFYF | TRBV7-9 | TRBJ1-4 | CASSFGGPPLFF | KLGGALQAK | YES | NO | NO |
| 14531 | TRAV8-3 | TRAJ20 | CAVGSNDYKLSF | TRBV7-2 | TRBJ1-2 | CASSELPGGYTF | KLGGALQAK | YES | NO | NO |
| 14532 | TRAV19 | TRAJ33 | CALSEVRSNYQLIW | TRBV6-5 | TRBJ2-1 | CATYPTGAPVDEQFF | KLGGALQAK | NO | NO | NO |
| 14533 | TRAV26-2 | TRAJ54 | CILFQGAQKLVF | TRBV27 | TRBJ1-1 | CASSSTTGGNTEAFF | KLGGALQAK | YES | NO | NO |
| 14534 | TRAV14DV4 | TRAJ33 | CAMREDSNYQLIW | TRBV7-8 | TRBJ2-1 | CASSLGVGRSSYNEQFF | KLGGALQAK | YES | NO | NO |

| | | | | | | | | | | |
|---|---|---|---|---|---|---|---|---|---|---|
| 14535 | TRAV25 | TRAJ47 | CAGPKEYGNKLVF | TRBV6-4 | TRBJ2-1 | CASSERGEVYNEQFF | KLGGALQAK | YES | NO | NO |
| 14536 | TRAV6 | TRAJ23 | CALNQGGKLIF | TRBV20-1 | TRBJ2-1 | CSARSRETVYNEQFF | KLGGALQAK | NO | NO | NO |
| 14537 | TRAV14DV4 | TRAJ4 | CAMRRPSGGYNKLIF | TRBV9 | TRBJ2-3 | CASSVDGRGGADTQYF | KLGGALQAK | YES | NO | NO |
| 14538 | TRAV21 | TRAJ49 | CAEDTGNQFYF | TRBV27 | TRBJ1-6 | CASSRETGNNSPLHF | KLGGALQAK | YES | NO | NO |
| 14539 | TRAV8-1 | TRAJ4 | CAVKWFSGGYNKLIF | TRBV28 | TRBJ1-3 | CASGGDLSGNTIYF | KLGGALQAK | YES | NO | NO |
| 14540 | TRAV1-1 | TRAJ21 | CAVRDGYNFNKFYF | TRBV20-1 | TRBJ1-1 | CSARDQSNTEAFF | KLGGALQAK | YES | NO | NO |
| 14541 | TRAV8-4 | TRAJ44 | CAVSEAGTASKLTF | TRBV11-2 | TRBJ2-1 | CASSTWTSGIYNEQFF | KLGGALQAK | YES | NO | NO |
| 14542 | TRAV1-2 | TRAJ10 | CAVPGGGNKLTF | TRBV4-1 | TRBJ2-5 | CASSQDGTETYQETQYF | KLGGALQAK | YES | NO | NO |
| 14543 | TRAV12-1 | TRAJ9 | CVVNAGGFKTIF | TRBV10-2 | TRBJ2-1 | CASSEEEQFF | KLGGALQAK | YES | NO | NO |
| 14544 | TRAV13-1 | TRAJ9 | CAASTGGFKTIF | TRBV29-1 | TRBJ2-5 | CSVEGRLAEETQYF | KLGGALQAK | YES | NO | NO |
| 14546 | TRAV17 | TRAJ40 | CATAPTSGTYKYIF | TRBV5-1 | TRBJ2-3 | CASSLGQGADTQYF | KLGGALQAK | YES | NO | NO |
| 14547 | TRAV12-1 | TRAJ15 | CVVNDQAGTALIF | TRBV3-1 | TRBJ2-1 | CASNTEYNEQFF | KLGGALQAK | YES | NO | NO |
| 14548 | TRAV26-1 | TRAJ41 | CIALPPGYALNF | TRBV27 | TRBJ1-5 | CASSLGVNQPQHF | KLGGALQAK | NO | NO | NO |
| 14549 | TRAV16 | TRAJ39 | CALGAGNMLTF | TRBV7-3 | TRBJ2-1 | CASSGAGTSGRNEQFF | KLGGALQAK | NO | NO | NO |
| 14550 | TRAV38-1 | TRAJ50 | CAFKTSYDKVIF | TRBV27 | TRBJ1-1 | CASSFAGTYEAFF | KLGGALQAK | YES | NO | NO |
| 14551 | TRAV9-2 | TRAJ40 | CALSEEGTYKYIF | TRBV11-3 | TRBJ2-5 | CASSSISGKQETQYF | KLGGALQAK | YES | NO | NO |
| 14552 | TRAV12-2 | TRAJ45 | CAVNWPWESGGGADGLTF | TRBV6-6 | TRBJ2-7 | CASSYMQGKGYEQYF | KLGGALQAK | NO | NO | NO |
| 14553 | TRAV24 | TRAJ15 | CAFQTVAGTALIF | TRBV7-9 | TRBJ2-1 | CASSWKQGIEQFF | KLGGALQAK | YES | NO | NO |
| 14554 | TRAV27 | TRAJ54 | CAYFQGAQKLVF | TRBV7-2 | TRBJ2-7 | CASSSGLASYEQYF | AVFDRKSDAK | YES | NO | YES |
| 14555 | TRAV1-2 | TRAJ39 | CAGYNAGNMLTF | TRBV7-9 | TRBJ1-1 | CASSPIQDTEAFF | KLGGALQAK | YES | NO | NO |
| 14556 | TRAV5 | TRAJ31 | CAEADNNARLMF | TRBV19 | TRBJ1-3 | CASSIDLQRNTIYF | KLGGALQAK | YES | NO | NO |
| 14557 | TRAV8-2 | TRAJ32 | CVVTLYYGGATNKLIF | TRBV10-3 | TRBJ2-3 | CAISEYGAKDTDTQYF | KLGGALQAK | YES | NO | NO |
| 14558 | TRAV34 | TRAJ40 | CGADTSGTYKYIF | TRBV29-1 | TRBJ1-2 | CSVDRDRGQFDYGYTF | KLGGALQAK | YES | NO | NO |
| 14559 | TRAV8-4 | TRAJ13 | CAVSYSGGYQKVTF | TRBV20-1 | TRBJ1-1 | CSARDEIRHTEAFF | KLGGALQAK | YES | NO | NO |
| 14560 | TRAV39 | TRAJ54 | CAGAPGAQKLVF | TRBV5-1 | TRBJ2-1 | CASSLDSYEQFF | KLGGALQAK | YES | NO | NO |
| 14561 | TRAV19 | TRAJ13 | CALSEAYGGYQKVTF | TRBV6-3 | TRBJ2-7 | CASSYGTSGRAYYYEQYV | KLGGALQAK | NO | NO | NO |
| 14562 | TRAV19 | TRAJ48 | CALSEAEGNEKLTF | TRBV12-4 | TRBJ2-1 | CASSFPLAGTLMNEQFF | KLGGALQAK | YES | NO | NO |
| 14563 | TRAV38-2DV8 | TRAJ32 | CAYRRAYGGATNKLIF | TRBV2 | TRBJ2-1 | CASTEGTYNEQFF | KLGGALQAK | YES | NO | NO |
| 14564 | TRAV12-2 | TRAJ52 | CAPAAGGTSYGKLTF | TRBV20-1 | TRBJ2-5 | CSARGLEETQYF | KLGGALQAK | YES | NO | NO |
| 14565 | TRAV12-3 | TRAJ39 | CAIGPGNMLTF | TRBV5-4 | TRBJ2-3 | CASSPRGTDTQYF | KLGGALQAK | YES | NO | NO |
| 14566 | TRAV12-1 | TRAJ20 | CVVNGNDYKLSF | TRBV12-3 | TRBJ1-1 | CASGPYDRADTEAFF | KLGGALQAK | YES | NO | NO |
| 14567 | TRAV41 | TRAJ42 | CALTTYGGSQGNLIF | TRBV2 | TRBJ2-7 | CASSGGTGPYEQYF | KLGGALQAK | YES | NO | NO |
| 14568 | TRAV19 | TRAJ22 | CALDFSSGSARQLTF | TRBV19 | TRBJ1-2 | CASRDSLDYGYTF | KLGGALQAK | YES | NO | NO |
| 14569 | TRAV9-2 | TRAJ33 | CALMPPMDSNYQLIW | TRBV11-2 | TRBJ1-1 | CASSLVVRGTEAFF | KLGGALQAK | YES | NO | NO |
| 14570 | TRAV12-2 | TRAJ50 | CAGKTSYDKVIF | TRBV11-3 | TRBJ1-1 | CASSPVNGRDTEAFF | KLGGALQAK | YES | NO | NO |
| 14571 | TRAV17 | TRAJ18 | CATANDRGSTLGRLYF | TRBV7-9 | TRBJ1-3 | CASSDSRPAGNTIYF | KLGGALQAK | YES | NO | NO |
| 14572 | TRAV27 | TRAJ13 | CAGFNSGGYQKVTF | TRBV27 | TRBJ1-4 | CASSLYAGASEKLFF | KLGGALQAK | YES | NO | NO |
| 14573 | TRAV25 | TRAJ50 | CAEKTSYDKVIF | TRBV19 | TRBJ2-1 | CASSIAGYYEQFF | KLGGALQAK | YES | NO | NO |
| 14574 | TRAV12-1 | TRAJ47 | CVVNVEYGNKLVF | TRBV7-8 | TRBJ1-2 | CASSLGQDGYTF | KLGGALQAK | YES | NO | NO |
| 14575 | TRAV3 | TRAJ37 | CALSWHSGNTGKLIF | TRBV7-9 | TRBJ2-7 | CASSWSGGSSYEQYV | KLGGALQAK | NO | NO | NO |
| 14577 | TRAV13-1 | TRAJ34 | CAASRGNTDKLIF | TRBV4-1 | TRBJ2-1 | CASSLLSGRVNEQFF | KLGGALQAK | YES | NO | NO |
| 14578 | TRAV8-3 | TRAJ33 | CALQSSMDSNYQLIW | TRBV7-9 | TRBJ2-1 | CASSYSAPEQFF | KLGGALQAK | YES | NO | NO |
| 14579 | TRAV25 | TRAJ40 | CAGPSQTFSGTYKYIF | TRBV2 | TRBJ2-2 | CASKGDTGELFF | KLGGALQAK | YES | NO | NO |
| 14580 | TRAV3 | TRAJ36 | CAVETGANNLFF | TRBV30 | TRBJ2-1 | CAQGENEQFF | KLGGALQAK | NO | NO | NO |
| 14581 | TRAV19 | TRAJ4 | CALHGGYNKLIF | TRBV5-6 | TRBJ1-1 | CASSGTGSMNTEAFF | KLGGALQAK | YES | NO | NO |
| 14582 | TRAV17 | TRAJ53 | CATVENSGGSNYKLTF | TRBV5-1 | TRBJ2-1 | CASSFGNEQFF | KLGGALQAK | YES | NO | NO |
| 14583 | TRAV36DV7 | TRAJ50 | CAGGTSYDKVIF | TRBV11-3 | TRBJ2-1 | CASSGWGQRNNEQFF | KLGGALQAK | NO | NO | NO |
| 14584 | TRAV8-6 | TRAJ31 | CAVSDKVARLMF | TRBV20-1 | TRBJ2-5 | CSADPVDSTLEEETQYF | KLGGALQAK | NO | NO | NO |
| 14585 | TRAV1-2 | TRAJ4 | CAAFSGGYNKLIF | TRBV7-2 | TRBJ1-2 | CASSPLSGSGYTF | KLGGALQAK | YES | NO | NO |
| 14586 | TRAV22 | TRAJ10 | CAVEIWLTGGGNKLTF | TRBV5-4 | TRBJ1-1 | CASSSDRVYTEAFF | IVTDFSVIK | YES | YES | YES |
| 14587 | TRAV29DV5 | TRAJ42 | CAASADYGGSQGNLIF | TRBV12-3 | TRBJ2-2 | CASSPLQGRGEAGELFF | KLGGALQAK | YES | NO | NO |
| 14588 | TRAV20 | TRAJ48 | CAVQSFGNEKLTF | TRBV7-2 | TRBJ2-1 | CASSLPGYNEQFF | KLGGALQAK | YES | NO | NO |
| 14589 | TRAV8-6 | TRAJ52 | CAVSERGAGGTSYGKLTF | TRBV19 | TRBJ2-1 | CATTSAEVRNEQFF | KLGGALQAK | YES | NO | NO |
| 14590 | TRAV29DV5 | TRAJ56 | CAASEAGANSKLTF | TRBV9 | TRBJ2-2 | CASSEFSRELFF | KLGGALQAK | YES | NO | NO |
| 14591 | TRAV39 | TRAJ57 | CAVPQTQGGSEKLVF | TRBV9 | TRBJ1-1 | CASSVATMDSLMNTEAFF | KLGGALQAK | YES | NO | NO |
| 14592 | TRAV19 | TRAJ56 | CALSGSTGANSKLTF | TRBV12-4 | TRBJ2-7 | CATGRGGYEQYF | KLGGALQAK | YES | NO | NO |
| 14593 | TRAV12-1 | TRAJ34 | CVVNIVRLIF | TRBV7-3 | TRBJ2-3 | CASSWGTNTDTQYF | KLGGALQAK | YES | NO | NO |
| 14594 | TRAV29DV5 | TRAJ23 | CAASAIQGGKLIF | TRBV2 | TRBJ2-7 | CASSDWGQGGYEQYF | KLGGALQAK | YES | NO | NO |
| 14595 | TRAV35 | TRAJ27 | CAGHPESGKSTF | TRBV29-1 | TRBJ1-1 | CSVTTGVNTEAFF | KLGGALQAK | YES | NO | NO |
| 14596 | TRAV19 | TRAJ20 | CALRGSNDYKLSF | TRBV27 | TRBJ1-1 | CASSLSGSLTNTEAFF | KLGGALQAK | YES | NO | NO |
| 14597 | TRAV24 | TRAJ40 | CAFPSGTYKYIF | TRBV7-9 | TRBJ2-7 | CASKDGISYEQYF | KLGGALQAK | YES | NO | NO |
| 14598 | TRAV9-2 | TRAJ41 | CALSDLRSNSGYALNF | TRBV7-9 | TRBJ2-7 | CASSLAYLPLEQYV | KLGGALQAK | NO | NO | NO |
| 14599 | TRAV29DV5 | TRAJ4 | CAASDSGGYNKLIF | TRBV2 | TRBJ1-5 | CASSEALGQTKQGNQPQHF | KLGGALQAK | YES | NO | NO |
| 14600 | TRAV8-6 | TRAJ11 | CAVSEYGYSTLTF | TRBV20-1 | TRBJ2-1 | CSARTSGGGTRTILSYNEQFF | KLGGALQAK | NO | NO | NO |
| 14601 | TRAV41 | TRAJ49 | CATPTGNQFYF | TRBV10-3 | TRBJ1-4 | CAIGDRLYEKLFF | KLGGALQAK | YES | NO | NO |
| 14602 | TRAV35 | TRAJ29 | CAGQRPGNTPLVF | TRBV7-3 | TRBJ2-7 | CASSLSGGGSYEQYF | KLGGALQAK | NO | NO | NO |
| 14603 | TRAV38-1 | TRAJ28 | CAFMKDGYSGAGSYQLTF | TRBV12-4 | TRBJ1-1 | CASTKKWVQGPNTEAFF | KLGGALQAK | YES | NO | NO |
| 14604 | TRAV25 | TRAJ37 | CAAGGNTGKLIF | TRBV19 | TRBJ1-2 | CASSILGNGYTF | KLGGALQAK | NO | NO | NO |
| 14605 | TRAV12-2 | TRAJ42 | CAVLGGNLIF | TRBV9 | TRBJ1-1 | CASSAGNTEAFF | KLGGALQAK | YES | NO | NO |
| 14606 | TRAV8-6 | TRAJ27 | CAVTPYNTNAGKSTF | TRBV19 | TRBJ1-3 | CASSGFSSSGNTIYF | KLGGALQAK | YES | NO | NO |
| 14607 | TRAV25 | TRAJ49 | CAGLPNTGNQFYF | TRBV2 | TRBJ2-5 | CASSEVRWSASSETQYF | KLGGALQAK | YES | NO | NO |
| 14608 | TRAV13-1 | TRAJ45 | CAASESGGGADGLTF | TRBV7-2 | TRBJ2-1 | CASSLGYEQFF | KLGGALQAK | YES | NO | NO |
| 14609 | TRAV8-1 | TRAJ17 | CAVSAAGNKLTF | TRBV5-4 | TRBJ1-2 | CASSLDRGQGANGYTF | KLGGALQAK | YES | NO | NO |
| 14610 | TRAV24 | TRAJ26 | CALYRDNYGQNFVF | TRBV5-1 | TRBJ2-1 | CASSLAPGLPNEQFF | KLGGALQAK | YES | NO | NO |
| 14611 | TRAV8-6 | TRAJ43 | CAVIVNNNDMRF | TRBV20-1 | TRBJ2-7 | CSARSQGIYEQYF | KLGGALQAK | NO | NO | NO |
| 14612 | TRAV26-1 | TRAJ23 | CIVRDAGGKLIF | TRBV7-3 | TRBJ1-2 | CASSFGELDSRGYTF | KLGGALQAK | YES | NO | NO |
| 14613 | TRAV29DV5 | TRAJ45 | CAAPRTPGGGADGLTF | TRBV9 | TRBJ2-2 | CASSVSGGITGELFF | AVFDRKSDAK | YES | NO | YES |
| 14614 | TRAV13-1 | TRAJ31 | CAASADARLMF | TRBV6-5 | TRBJ2-5 | CASSYVAGGQETQYF | KLGGALQAK | NO | NO | NO |
| 14615 | TRAV29DV5 | TRAJ58 | CAATGSGSRLTF | TRBV7-9 | TRBJ1-1 | CASSLGFREGTEAFF | KLGGALQAK | YES | NO | NO |
| 14616 | TRAV20 | TRAJ49 | CAVRSNTGNQFYF | TRBV12-4 | TRBJ1-2 | CASSWDAGHGYTF | KLGGALQAK | YES | NO | NO |
| 14617 | TRAV24 | TRAJ33 | CYSNYQLIW | TRBV30 | TRBJ1-2 | CAWSATGDYGYTF | KLGGALQAK | NO | NO | NO |
| 14618 | TRAV29DV5 | TRAJ26 | CAALGQNFVF | TRBV19 | TRBJ2-1 | CASSISGTGGFYEQFF | KLGGALQAK | YES | NO | NO |
| 14619 | TRAV8-3 | TRAJ34 | CAVGPYNTDKLIF | TRBV7-7 | TRBJ1-5 | CASSLSLGQPQHF | KLGGALQAK | YES | NO | NO |
| 14620 | TRAV8-3 | TRAJ43 | CAVGGFDDMRF | TRBV12-4 | TRBJ2-2 | CASSLDSGNTGELFF | KLGGALQAK | YES | NO | NO |
| 14621 | TRAV3 | TRAJ28 | CAGDSGAGSYQLTF | TRBV7-3 | TRBJ2-3 | CASSLLAGGRDTQYF | KLGGALQAK | NO | NO | NO |
| 14622 | TRAV19 | TRAJ42 | CALSELNYGGSQGNLIF | TRBV7-2 | TRBJ2-2 | CASSLVPLTSGSTGELFF | KLGGALQAK | NO | NO | NO |
| 14623 | TRAV20 | TRAJ23 | CAVPPYNQGGKLIF | TRBV7-9 | TRBJ1-2 | CASSALEAGYGYTF | KLGGALQAK | YES | NO | NO |
| 14624 | TRAV21 | TRAJ47 | CAVREYGNKLVF | TRBV3-1 | TRBJ2-7 | CASSSPTSGTYEQYF | KLGGALQAK | YES | NO | NO |
| 14625 | TRAV29DV5 | TRAJ28 | CGLSGAGSYQLTF | TRBV5-1 | TRBJ2-1 | CASSSGTGPYNEQFF | KLGGALQAK | YES | NO | NO |
| 14626 | TRAV19 | TRAJ28 | CALTWAGSYQLTF | TRBV20-1 | TRBJ2-1 | CSASAGEQFF | KLGGALQAK | NO | NO | NO |
| 14627 | TRAV12-3 | TRAJ52 | CAMISANAGGTSYGKLTF | TRBV9 | TRBJ1-6 | CASSATGGPNSPLHF | KLGGALQAK | YES | NO | NO |
| 14628 | TRAV3 | TRAJ18 | CAVRADRGSTLGRLYF | TRBV19 | TRBJ1-1 | CASSIGTVNTEAFF | AVFDRKSDAK | NO | NO | NO |
| 14629 | TRAV19 | TRAJ4 | CALSEGVSGGYNKLIF | TRBV27 | TRBJ2-7 | CASSLLGQGTGEQYF | AVFDRKSDAK | YES | NO | YES |
| 14630 | TRAV14DV4 | TRAJ20 | CAMREAPGKLSF | TRBV6-1 | TRBJ2-7 | CASSEVDRVAEKQYF | KLGGALQAK | YES | NO | NO |
| 14631 | TRAV1-2 | TRAJ9 | CAVRDEDGGFKTIF | TRBV4-1 | TRBJ2-3 | CASSQGVGLAGAPSTDTQYF | KLGGALQAK | NO | NO | NO |
| 14632 | TRAV29DV5 | TRAJ22 | CAASFAASGSARQLTF | TRBV6-3 | TRBJ2-1 | CASSYGAGGYNEQFF | KLGGALQAK | YES | NO | NO |
| 14633 | TRAV12-1 | TRAJ41 | CVVNPPNSGYALNF | TRBV6-5 | TRBJ1-3 | CASLGAGNTIYF | KLGGALQAK | YES | NO | NO |
| 14634 | TRAV19 | TRAJ42 | CAYYGGSQGNLIF | TRBV29-1 | TRBJ2-1 | CSTLAGGYNEQFF | KLGGALQAK | YES | NO | NO |
| 14635 | TRAV2 | TRAJ22 | CAVDPLSSGSARQLTF | TRBV12-4 | TRBJ1-5 | CASSFGGSGNQPQHF | KLGGALQAK | NO | NO | NO |
| 14636 | TRAV5 | TRAJ10 | CAEITGGGNKLTF | TRBV7-9 | TRBJ1-2 | CASSKAAGDYGYTF | AVFDRKSDAK | YES | NO | YES |
| 14637 | TRAV8-6 | TRAJ56 | CAVSPGANSKLTF | TRBV11-2 | TRBJ2-7 | CASSFRGGGMDEQYF | KLGGALQAK | YES | NO | NO |
| 14638 | TRAV8-2 | TRAJ49 | CVVSDVDGNQFYF | TRBV7-9 | TRBJ1-2 | CASSLAVGNYGYTF | KLGGALQAK | YES | NO | NO |
| 14639 | TRAV17 | TRAJ58 | CATDALGSRLTF | TRBV5-8 | TRBJ2-3 | CASSGTEVTDTQYF | KLGGALQAK | YES | NO | NO |
| 14640 | TRAV39 | TRAJ44 | CAVDLTASKLTF | TRBV2 | TRBJ2-3 | CASSVWGRSTDTQYF | KLGGALQAK | YES | NO | NO |

| | | | | | | | | | | |
|---|---|---|---|---|---|---|---|---|---|---|
| 14641 | TRAV12-2 | TRAJ37 | CAVNPSNTGKLIF | TRBV29-1 | TRBJ2-3 | CSVRLGFTDTQYF | KLGGALQAK | YES | NO | NO |
| 14642 | TRAV1-2 | TRAJ33 | CAVWDSNYQLIW | TRBV4-2 | TRBJ1-5 | CASSQEGEGFQPQHF | KLGGALQAK | YES | NO | NO |
| 14643 | TRAV35 | TRAJ44 | CAGLNTGTASKLTF | TRBV5-1 | TRBJ2-1 | CASSLSWTVPEQFF | KLGGALQAK | YES | NO | NO |
| 14644 | TRAV17 | TRAJ45 | CRDSGGGADGLTF | TRBV7-8 | TRBJ1-5 | CASSLNPQGIQPQHF | AVFDRKSDAK | YES | NO | NO |
| 14645 | TRAV1-2 | TRAJ34 | CAVRPATSSSL | TRBV11-2 | TRBJ2-7 | CASSLDPGTYEQYF | KLGGALQAK | YES | NO | NO |
| 14646 | TRAV8-1 | TRAJ39 | CAVKASLEFDAGNMLTF | TRBV5-5 | TRBJ2-5 | CASLQRLEETQYF | KLGGALQAK | YES | NO | NO |
| 14647 | TRAV26-1 | TRAJ16 | CIVSTGQKLLF | TRBV7-9 | TRBJ2-1 | CASSQEGDEQFF | KLGGALQAK | NO | NO | NO |
| 14648 | TRAV21 | TRAJ40 | CAVSSTSGTYKYIF | TRBV7-3 | TRBJ2-2 | CASSSETAVSGELFF | KLGGALQAK | NO | NO | NO |
| 14649 | TRAV8-3 | TRAJ34 | CAVGPSYNTDKLIF | TRBV10-2 | TRBJ1-2 | CASSNQGHDGYTF | KLGGALQAK | YES | NO | NO |
| 14650 | TRAV12-2 | TRAJ28 | CAVRRWEGGAGSYQLTF | TRBV11-2 | TRBJ2-2 | CASSWSSGELFF | KLGGALQAK | YES | NO | NO |
| 14652 | TRAV12-1 | TRAJ33 | CVVMDSNYQLIW | TRBV7-2 | TRBJ2-3 | CASSLVAGDLGDTQYF | KLGGALQAK | YES | NO | NO |
| 14653 | TRAV8-4 | TRAJ52 | CAVSQAGGTSYGKLTF | TRBV9 | TRBJ2-3 | CASSGVSDTQYF | KLGGALQAK | YES | NO | NO |
| 14654 | TRAV19 | TRAJ6 | CALSEAGPWEGSYIPTF | TRBV6-5 | TRBJ2-3 | CASSASEGDTQYF | KLGGALQAK | NO | NO | NO |
| 14655 | TRAV20 | TRAJ35 | CAVQAGGFGNVLHC | TRBV7-2 | TRBJ2-5 | CASSLGSLLAGVF | KLGGALQAK | NO | NO | NO |
| 14656 | TRAV8-3 | TRAJ47 | CATSYGNKLVF | TRBV20-1 | TRBJ1-2 | CSARSQDGYTF | KLGGALQAK | YES | NO | NO |
| 14657 | TRAV12-1 | TRAJ53 | CVVEGGSNYKLTF | TRBV19 | TRBJ2-7 | CASSAGQGGYEQYF | KLGGALQAK | YES | NO | NO |
| 14658 | TRAV8-3 | TRAJ47 | CAVKYGNKLVF | TRBV6-3 | TRBJ2-1 | CASTVGLAENAYYNEQFF | KLGGALQAK | YES | NO | NO |
| 14659 | TRAV4 | TRAJ32 | CLVGDDYGGATNKLIF | TRBV5-1 | TRBJ1-1 | CASSPTTSGQPEAFF | KLGGALQAK | YES | NO | NO |
| 14660 | TRAV38-1 | TRAJ28 | CAFMIAYSGAGSYQLTF | TRBV20-1 | TRBJ2-1 | CSAKGLAGVEVNEQFF | KLGGALQAK | NO | NO | NO |
| 14661 | TRAV8-4 | TRAJ44 | CAVMRDTGTASKLTF | TRBV20-1 | TRBJ2-7 | CSATPTPGQFYEQYF | KLGGALQAK | NO | NO | NO |
| 14663 | TRAV8-3 | TRAJ43 | CAVGTNDMRF | TRBV7-9 | TRBJ2-3 | CASKNGGQLYTDTQYF | KLGGALQAK | YES | NO | NO |
| 14664 | TRAV12-1 | TRAJ49 | CVVNPNTGNQFYF | TRBV7-9 | TRBJ2-2 | CASSLVLNTGELFF | KLGGALQAK | YES | NO | NO |
| 14665 | TRAV8-4 | TRAJ20 | CAVSGSRDYKLSF | TRBV6-1 | TRBJ2-7 | CASSFQTSGGTILPYEQYF | KLGGALQAK | YES | NO | NO |
| 14666 | TRAV14DV4 | TRAJ22 | CAMRRPISSGSARQLTF | TRBV11-2 | TRBJ2-3 | CASSFSLAGGPSTDTQYF | KLGGALQAK | YES | NO | NO |
| 14667 | TRAV30 | TRAJ21 | CGTESYNFNKFYF | TRBV15 | TRBJ2-7 | CATSRDRIGGDEQYF | AVFDRKSDAK | YES | NO | YES |
| 14668 | TRAV19 | TRAJ52 | CALRTAAGGTSYGKLTF | TRBV9 | TRBJ1-1 | CASSVGGENTEAFF | KLGGALQAK | YES | YES | NO |
| 14669 | TRAV20 | TRAJ12 | CAVPMDSSYKLIF | TRBV20-1 | TRBJ2-3 | CSARDPDNPDTQYF | KLGGALQAK | NO | NO | NO |
| 14671 | TRAV5 | TRAJ37 | CAEASGNTGKLIF | TRBV4-1 | TRBJ1-1 | CASSQGMGQGSGNTEAFF | KTWGQYWQV | NO | NO | NO |
| 14672 | TRAV19 | TRAJ15 | CAPPAGTALIF | TRBV10-3 | TRBJ1-4 | CAISEPGAPNEKLFF | KLGGALQAK | YES | NO | NO |
| 14673 | TRAV12-2 | TRAJ43 | CAVNVLGMRF | TRBV7-2 | TRBJ2-7 | CASSLEPSYEQYF | AVFDRKSDAK | NO | NO | NO |
| 14674 | TRAV25 | TRAJ57 | CAVPLITQGGSEKLVF | TRBV6-3 | TRBJ2-1 | CASSGQGVIGEQFF | RLRAEAQVK | YES | YES | YES |
| 14675 | TRAV39 | TRAJ40 | CAVDSGTYKYIF | TRBV18 | TRBJ1-2 | CASSPPGANYGYTF | KLGGALQAK | YES | NO | NO |
| 14676 | TRAV22 | TRAJ48 | CAVALTREISNFGNEKLTF | TRBV27 | TRBJ2-2 | CASSLSGQGVSVSTGELFF | KLGGALQAK | YES | NO | NO |
| 14677 | TRAV21 | TRAJ21 | CAVVGLYNFNKFYF | TRBV19 | TRBJ2-3 | CASRGTSAKDTQYF | AVFDRKSDAK | YES | NO | YES |
| 14678 | TRAV8-6 | TRAJ17 | CAVFRAAGNKLTF | TRBV27 | TRBJ1-1 | CASSSFTGGTEAFF | KLGGALQAK | YES | NO | NO |
| 14679 | TRAV10 | TRAJ6 | CVVTRRASGGSYIPTF | TRBV4-3 | TRBJ2-1 | CASSQGSLAGGPNEQFF | KLGGALQAK | YES | NO | NO |
| 14680 | TRAV6 | TRAJ39 | CALSAGNMLTF | TRBV7-9 | TRBJ2-1 | CASRPARGYEQFF | KLGGALQAK | YES | NO | NO |
| 14681 | TRAV1-1 | TRAJ12 | CAFSGGSSYKLIF | TRBV10-3 | TRBJ2-5 | CAISESPMSGGEKETQYF | KLGGALQAK | YES | NO | NO |
| 14682 | TRAV8-3 | TRAJ39 | CAASEAGNMLTF | TRBV4-1 | TRBJ2-7 | CASSQTGTGSHEQYV | KLGGALQAK | NO | NO | NO |
| 14683 | TRAV19 | TRAJ48 | CALSEVGFGNEKLTF | TRBV12-5 | TRBJ2-7 | CASGPQGAYEQYF | KLGGALQAK | YES | NO | NO |
| 14684 | TRAV21 | TRAJ6 | CAVRASGGSYIPTF | TRBV27 | TRBJ2-7 | CASSPQTMTSGRVYEQYV | KLGGALQAK | NO | NO | NO |
| 14685 | TRAV27 | TRAJ57 | CAGATQGGSEKLVF | TRBV7-9 | TRBJ1-2 | CASSLVFAEYGYTF | AVFDRKSDAK | YES | NO | YES |
| 14686 | TRAV13-2 | TRAJ31 | CAENNNARLMF | TRBV19 | TRBJ2-1 | CASSLSGYNEQFF | KLGGALQAK | YES | NO | NO |
| 14687 | TRAV12-1 | TRAJ23 | CVVNGRGKLIF | TRBV7-9 | TRBJ2-6 | CASSLTESGANVLTF | AVFDRKSDAK | YES | NO | YES |
| 14688 | TRAV25 | TRAJ27 | CAHTNAGKSTF | TRBV7-9 | TRBJ2-7 | CASSHIRDYEQYF | KLGGALQAK | YES | NO | NO |
| 14689 | TRAV3 | TRAJ30 | CAVRPQMNRDDKIIF | TRBV7-3 | TRBJ2-7 | CASSLTSDGYYEQYF | KLGGALQAK | NO | NO | NO |
| 14690 | TRAV12-2 | TRAJ45 | CAVNRGGGADGLTF | TRBV19 | TRBJ1-2 | CASWGVPYGYTF | KLGGALQAK | YES | NO | NO |
| 14691 | TRAV12-1 | TRAJ40 | CVVNPGTYKYIF | TRBV7-2 | TRBJ2-1 | CASSLVYSSSYNEQFF | KLGGALQAK | YES | NO | NO |
| 14692 | TRAV12-2 | TRAJ29 | CAGNTPLVF | TRBV27 | TRBJ1-5 | CASSLGLGQPQHF | ELAGIGILTV | NO | NO | NO |
| 14693 | TRAV12-1 | TRAJ45 | CVVNVDSGGGADGLTF | TRBV6-6 | TRBJ1-5 | CASSLTWELRF | KLGGALQAK | NO | NO | NO |
| 14694 | TRAV38-2DV8 | TRAJ48 | CAYRAPRGNEKLTF | TRBV19 | TRBJ1-4 | CASSQGSEKLFF | KLGGALQAK | YES | NO | NO |
| 14695 | TRAV9-2 | TRAJ13 | CALSDPKVTF | TRBV3-1 | TRBJ2-5 | CASSQGLAGANQETQYF | KLGGALQAK | YES | NO | NO |
| 14696 | TRAV1-1 | TRAJ22 | CAVSPGGTARQLTF | TRBV27 | TRBJ1-5 | CASSSYTGGLNQPQHF | KLGGALQAK | YES | NO | NO |
| 14697 | TRAV27 | TRAJ40 | CAGGRGTYKYIF | TRBV20-1 | TRBJ2-7 | CSARDVHRGIEQYF | KLGGALQAK | NO | NO | NO |
| 14699 | TRAV19 | TRAJ13 | CALRPSGGYQKVTF | TRBV29-1 | TRBJ2-1 | CSASTDEQFF | KLGGALQAK | YES | NO | NO |
| 14700 | TRAV1-2 | TRAJ6 | CAVSDASGGSYIPTF | TRBV19 | TRBJ1-1 | CASSISTGGNTEAFF | KLGGALQAK | YES | NO | NO |
| 14701 | TRAV19 | TRAJ22 | CALYPLSGSARQLTF | TRBV4-1 | TRBJ1-1 | CASSRPGQENTEAFF | KLGGALQAK | YES | NO | NO |
| 14702 | TRAV16 | TRAJ17 | CAPRGKAAGNKLTF | TRBV4-1 | TRBJ1-1 | CASSQVPGRGGNTEAFF | KLGGALQAK | NO | NO | NO |
| 14703 | TRAV17 | TRAJ43 | CATVFYNNNDMRF | TRBV19 | TRBJ2-1 | CASQTQSSYNEQFF | KLGGALQAK | YES | NO | NO |
| 14704 | TRAV12-1 | TRAJ12 | CVVLDSSYKLIF | TRBV6-5 | TRBJ1-5 | CASRSKTGSKQPQHF | KLGGALQAK | YES | NO | NO |
| 14705 | TRAV12-2 | TRAJ37 | CAGGDSGNTGKLIF | TRBV7-9 | TRBJ2-7 | CASSPKTGYYEQYF | KLGGALQAK | YES | NO | NO |
| 14706 | TRAV17 | TRAJ42 | CATDADYGGSQGNLIF | TRBV5-6 | TRBJ1-6 | CASSLDGQVWSPLHF | KLGGALQAK | YES | NO | NO |
| 14707 | TRAV27 | TRAJ31 | CAGNNARLMF | TRBV5-1 | TRBJ1-6 | CASSLDPRGSSYNSPLHF | AVFDRKSDAK | YES | NO | YES |
| 14708 | TRAV16 | TRAJ27 | CALLTNAGKSTF | TRBV11-2 | TRBJ2-2 | CASSLGTGTGELFF | KLGGALQAK | NO | NO | NO |
| 14709 | TRAV3 | TRAJ22 | CAVRAYSSGSARQLTF | TRBV25-1 | TRBJ2-2 | CASSEDLWGIMNTGELFF | KLGGALQAK | NO | NO | NO |
| 14710 | TRAV21 | TRAJ57 | CAVRRKQGGSEKLVF | TRBV19 | TRBJ1-5 | CASSMYVNQPQHF | KLGGALQAK | YES | NO | NO |
| 14711 | TRAV9-2 | TRAJ30 | CALPMNRDDKIIF | TRBV2 | TRBJ2-1 | CASSDFIVSEQFF | KLGGALQAK | YES | NO | NO |
| 14712 | TRAV38-1 | TRAJ58 | CAFMRETSGSRLTF | TRBV5-1 | TRBJ2-1 | CASSLAAAGANEQFF | KLGGALQAK | YES | NO | NO |
| 14713 | TRAV12-3 | TRAJ39 | CAIGPGNMLTF | TRBV6-1 | TRBJ2-7 | CASSERDRATYEQYF | KLGGALQAK | YES | NO | NO |
| 14714 | TRAV27 | TRAJ13 | CAGARSGGYQKVTF | TRBV4-1 | TRBJ2-1 | CASSPPNTYEQFF | KLGGALQAK | YES | NO | NO |
| 14715 | TRAV8-2 | TRAJ45 | CVVSAEHSGGGADGLTF | TRBV7-9 | TRBJ1-4 | CASSLTGGEKLFF | KLGGALQAK | YES | NO | NO |
| 14716 | TRAV4 | TRAJ32 | CLVGPLNLNYGGATNKLIF | TRBV5-1 | TRBJ2-2 | CASTGSGRELIGELFF | KLGGALQAK | YES | NO | NO |
| 14717 | TRAV36DV7 | TRAJ30 | CAVERDDKIIF | TRBV5-1 | TRBJ1-2 | CASRTGLDGYTF | KLGGALQAK | NO | NO | NO |
| 14718 | TRAV29DV5 | TRAJ45 | CAATDSGGGADGLTF | TRBV6-3 | TRBJ2-1 | CASSRYNEQFF | KLGGALQAK | YES | NO | NO |
| 14719 | TRAV8-1 | TRAJ5 | CAAVLDTGRRALTF | TRBV5-6 | TRBJ2-7 | CASSPGQAPYEQYF | KLGGALQAK | YES | NO | NO |
| 14721 | TRAV13-1 | TRAJ49 | CAASAGSNQFYF | TRBV30 | TRBJ2-1 | CAWSVGGVNVFF | KLGGALQAK | NO | NO | NO |
| 14722 | TRAV13-2 | TRAJ37 | CAENKAPSNTGKLIF | TRBV4-1 | TRBJ1-5 | CASSHWHGTVLNQPQHF | CYTWNQMNL | YES | NO | YES |
| 14723 | TRAV19 | TRAJ43 | CALMENNNDMRF | TRBV7-9 | TRBJ2-4 | CASSPTGGPKNIQYF | KLGGALQAK | YES | NO | NO |
| 14724 | TRAV3 | TRAJ39 | CAVRDPNNAGNMLTF | TRBV27 | TRBJ2-1 | CASGTTGFYNEQFF | KLGGALQAK | NO | NO | NO |
| 14725 | TRAV1-1 | TRAJ12 | CAVRPPMDSSYKLIF | TRBV7-9 | TRBJ1-1 | CASSLGGGMNTEAFF | KLGGALQAK | YES | NO | NO |
| 14726 | TRAV17 | TRAJ24 | CATDPTDSWGKFQF | TRBV6-3 | TRBJ2-1 | CASTPGGRSNEQFF | KLGGALQAK | YES | NO | NO |
| 14727 | TRAV22 | TRAJ34 | CAVELLYNTDKLIF | TRBV18 | TRBJ2-1 | CASSPEGSYNEQFF | KLGGALQAK | YES | NO | NO |
| 14728 | TRAV17 | TRAJ45 | CATGRGGGADGLTF | TRBV19 | TRBJ2-7 | CASSPGVSYEQYF | KLGGALQAK | YES | NO | NO |
| 14729 | TRAV1-2 | TRAJ27 | CAVRSNAGKSTF | TRBV4-1 | TRBJ1-4 | CASSPGASNEKLFF | KLGGALQAK | YES | NO | NO |
| 14730 | TRAV2 | TRAJ29 | CAVEDRRNTPLVF | TRBV7-2 | TRBJ2-3 | CASSPPVWEMTDTQYF | KLGGALQAK | NO | NO | NO |
| 14731 | TRAV34 | TRAJ44 | CGADIAMNAGTASKLTF | TRBV12-4 | TRBJ1-1 | CASIRGRGYTEAFF | KLGGALQAK | YES | NO | NO |
| 14732 | TRAV19 | TRAJ9 | CALSEAGAGGFKTIF | TRBV7-9 | TRBJ2-3 | CASSPSAWADTQYF | KLGGALQAK | YES | NO | NO |
| 14733 | TRAV21 | TRAJ54 | CAVRSSLIQGAQKLVF | TRBV30 | TRBJ2-1 | CAWSVSGATDEQFF | AVFDRKSDAK | YES | NO | YES |
| 14734 | TRAV20 | TRAJ10 | CAVRPLTGGGNKLTF | TRBV30 | TRBJ2-1 | CAWNPGPSYGEQFF | KLGGALQAK | YES | NO | NO |
| 14735 | TRAV6 | TRAJ39 | CARGNAGNMLTF | TRBV5-8 | TRBJ2-2 | CASSPGVTGELFF | KLGGALQAK | YES | NO | NO |
| 14736 | TRAV39 | TRAJ33 | CAVDTLRHDSNYQLIW | TRBV24-1 | TRBJ1-2 | CATSDFRGQHYGYTF | KLGGALQAK | YES | NO | NO |
| 14737 | TRAV14DV4 | TRAJ52 | CAMREPAGGTSYGKLTF | TRBV4-3 | TRBJ2-7 | CASSQDVAGGLHEQYV | KLGGALQAK | NO | NO | NO |
| 14738 | TRAV29DV5 | TRAJ50 | CAAKTSYDKVIF | TRBV19 | TRBJ1-1 | CASSIAGNTEAFF | KLGGALQAK | YES | NO | NO |
| 14739 | TRAV19 | TRAJ52 | CALSANAGGTSYGKLTF | TRBV6-3 | TRBJ2-1 | CASRPINSYNEQFF | KLGGALQAK | YES | NO | NO |
| 14740 | TRAV29DV5 | TRAJ31 | CAASATNNARLMF | TRBV24-1 | TRBJ1-5 | CATAQGMGQPQHF | KLGGALQAK | YES | NO | NO |
| 14741 | TRAV13-2 | TRAJ11 | CAENRGRSGYSTLTF | TRBV19 | TRBJ2-1 | CASSITLAGGHLYNEQFF | KLGGALQAK | YES | NO | NO |
| 14742 | TRAV1-2 | TRAJ23 | CAVRFRGKLIF | TRBV4-1 | TRBJ2-5 | CASSQDSGEETQYF | AVFDRKSDAK | YES | NO | NO |
| 14743 | TRAV17 | TRAJ23 | CATVPVYNQGGKLIF | TRBV19 | TRBJ2-2 | CASSMTGLNTGELFF | KLGGALQAK | YES | NO | NO |
| 14744 | TRAV8-2 | TRAJ47 | CVASYGNKLVF | TRBV4-1 | TRBJ1-1 | CASSPSGGLGTEAFF | KLGGALQAK | YES | NO | NO |
| 14745 | TRAV24 | TRAJ43 | CASRESNDMRF | TRBV7-6 | TRBJ2-5 | CASEASGLAGLGTQYF | KLGGALQAK | YES | NO | NO |
| 14746 | TRAV19 | TRAJ49 | CALSELSNTGNQFYF | TRBV10-3 | TRBJ2-5 | CAISEGTSGLETQYF | KLGGALQAK | YES | NO | NO |
| 14747 | TRAV24 | TRAJ47 | CAAPMEYGNKLVF | TRBV9 | TRBJ1-2 | CASSSGGTYGYTF | KLGGALQAK | YES | NO | NO |
| 14748 | TRAV39 | TRAJ33 | CAVDSNYQLIW | TRBV20-1 | TRBJ2-7 | CSARDLEVLSYEQYF | KLGGALQAK | YES | NO | NO |
| 14749 | TRAV20 | TRAJ58 | CARYETSGSRLTF | TRBV7-9 | TRBJ1-5 | CASSLGATHQPQHF | KLGGALQAK | YES | NO | NO |

| | | | | | | | | | | |
|---|---|---|---|---|---|---|---|---|---|---|
| 14750 | TRAV20 | TRAJ48 | CAVQASSNFGNEKLTF | TRBV4-3 | TRBJ2-1 | CASSQARSNEQFF | KLGGALQAK | YES | NO | NO |
| 14751 | TRAV26-1 | TRAJ29 | CIVRIVFSGNTPLVF | TRBV29-1 | TRBJ1-2 | CSVEFWGRYGYTF | KLGGALQAK | YES | NO | NO |
| 14752 | TRAV8-3 | TRAJ37 | CAAQDGSSNTGKLIF | TRBV5-1 | TRBJ2-3 | CASSLASVSSTDTQYF | KLGGALQAK | YES | NO | NO |
| 14753 | TRAV26-2 | TRAJ29 | CILNSGNTPLVF | TRBV6-3 | TRBJ1-2 | CASSYSNGYVNGYTF | KLGGALQAK | YES | NO | NO |
| 14754 | TRAV1-1 | TRAJ11 | CAVSKNSGYSTLTF | TRBV5-6 | TRBJ2-3 | CASSPLGGTQYF | KLGGALQAK | YES | NO | NO |
| 14755 | TRAV8-2 | TRAJ42 | CVVREPGGSQGNLIF | TRBV7-6 | TRBJ2-6 | CASSYPPLESGANVLTF | KLGGALQAK | YES | NO | NO |
| 14756 | TRAV29DV5 | TRAJ30 | CAASLRDDKIIF | TRBV11-1 | TRBJ2-1 | CASSAPRGPGTNEQFF | KLGGALQAK | YES | NO | NO |
| 14757 | TRAV12-2 | TRAJ12 | CAVISIYKLIF | TRBV11-3 | TRBJ2-3 | CASSEQDSTDTQYF | KLGGALQAK | YES | NO | NO |
| 14758 | TRAV26-2 | TRAJ48 | CILRGFGNEKLTF | TRBV12-4 | TRBJ2-1 | CASSFGTVYNEQFF | KLGGALQAK | YES | NO | NO |
| 14759 | TRAV26-1 | TRAJ40 | CIVRVASASGTYKYIF | TRBV27 | TRBJ2-2 | CASSLQNTGELFF | KLGGALQAK | NO | NO | NO |
| 14760 | TRAV13-1 | TRAJ20 | CAAQDYKLSF | TRBV12-3 | TRBJ2-3 | CASSGTSGGPEPPSTDTQYF | KLGGALQAK | NO | NO | NO |
| 14761 | TRAV35 | TRAJ32 | CAGQLEVMNYGGATNKLIF | TRBV4-3 | TRBJ2-1 | CASSQDDGGYNEQFF | KLGGALQAK | YES | NO | NO |
| 14762 | TRAV8-3 | TRAJ40 | CAAFTSGTYKYIF | TRBV6-5 | TRBJ2-3 | CASSYVGGPGDTQYF | KLGGALQAK | NO | NO | NO |
| 14763 | TRAV5 | TRAJ12 | CAVSDSSYKLIF | TRBV2 | TRBJ1-1 | CASSEPQDTEAFF | KLGGALQAK | YES | NO | NO |
| 14764 | TRAV1-2 | TRAJ40 | CAVRGFTSGTYKYIF | TRBV9 | TRBJ1-1 | CASSPDDTEAFF | QPRAPIRPI | YES | NO | YES |
| 14765 | TRAV1-2 | TRAJ6 | CAVEPGGSYIPTF | TRBV7-9 | TRBJ2-1 | CASSSPSGLADYNEQFF | KLGGALQAK | YES | NO | NO |
| 14766 | TRAV29DV5 | TRAJ43 | CAASQNNDMRF | TRBV30 | TRBJ1-1 | CAWSYGTEAFF | KLGGALQAK | YES | NO | NO |
| 14767 | TRAV29DV5 | TRAJ42 | CAASSYGGSQGNLIF | TRBV7-9 | TRBJ2-1 | CASSPLYGNEQFF | KLGGALQAK | YES | NO | NO |
| 14768 | TRAV41 | TRAJ48 | CAVSPPNFGNEKLTF | TRBV19 | TRBJ1-2 | CASSEGGVYGYTF | KLGGALQAK | YES | NO | NO |
| 14769 | TRAV19 | TRAJ43 | CALSEPPINNNDMRF | TRBV20-1 | TRBJ1-5 | CSARDSGLYQPQHF | KLGGALQAK | NO | NO | NO |
| 14770 | TRAV4 | TRAJ42 | CLVGVYGGSQGNLIF | TRBV6-4 | TRBJ2-3 | CASRPTSGHTDTQYF | KLGGALQAK | NO | NO | NO |
| 14771 | TRAV29DV5 | TRAJ57 | CAASSDQGGSEKLVF | TRBV3-1 | TRBJ1-1 | CASSQDEGLNTEAFF | KLGGALQAK | YES | NO | NO |
| 14772 | TRAV8-1 | TRAJ43 | CAVSYNNNDMRF | TRBV9 | TRBJ2-1 | CASSASFYGSGMEEQFF | KLGGALQAK | YES | NO | NO |
| 14773 | TRAV41 | TRAJ48 | CAVRSQGFGNEKLTF | TRBV14 | TRBJ1-2 | CASSQDRGFNYGYTF | KLGGALQAK | NO | NO | NO |
| 14774 | TRAV8-2 | TRAJ21 | CVVSDLVDFNKFYF | TRBV19 | TRBJ1-4 | CVAHGTANEKLFF | KLGGALQAK | YES | NO | NO |
| 14775 | TRAV26-2 | TRAJ45 | CILRDSSGGADGLTF | TRBV28 | TRBJ1-1 | CASSLPYPGVGAFF | KLGGALQAK | YES | NO | NO |
| 14776 | TRAV30 | TRAJ30 | CGTEIRRDDKIIF | TRBV19 | TRBJ1-6 | CASSIGTNSPLHF | KLGGALQAK | YES | NO | NO |
| 14777 | TRAV38-2DV8 | TRAJ41 | CAYKGLNSGYALNF | TRBV11-2 | TRBJ2-3 | CASSIHSTDTQYF | KLGGALQAK | YES | NO | NO |
| 14778 | TRAV1-2 | TRAJ13 | CAVLNSGGYQKVTF | TRBV2 | TRBJ2-5 | CASSEGTSGQETQYF | KLGGALQAK | YES | NO | NO |
| 14779 | TRAV8-4 | TRAJ54 | CAVSIQGAQKLVF | TRBV24-1 | TRBJ2-5 | CATSDPASGALQETQYF | KLGGALQAK | YES | NO | NO |
| 14780 | TRAV35 | TRAJ15 | CAGPHYNQAGTALIF | TRBV5-5 | TRBJ1-1 | CASSLDGLGAFF | SLFNTVATLY | YES | YES | NO |
| 14781 | TRAV12-1 | TRAJ47 | CVVNRGTEYGNKLVF | TRBV6-3 | TRBJ2-3 | CASSYSPQGKGSTDTQYF | KLGGALQAK | YES | NO | NO |
| 14782 | TRAV4 | TRAJ29 | CLVGDPPGNTPLVF | TRBV20-1 | TRBJ2-7 | CSARDWDSRTETSYEQYF | KLGGALQAK | NO | NO | NO |
| 14783 | TRAV27 | TRAJ37 | CAGPPASGNTGKLIF | TRBV5-1 | TRBJ1-4 | CASQTWGAQGEKLFF | AVFDRKSDAK | YES | NO | YES |
| 14784 | TRAV8-3 | TRAJ48 | CAVAPSNFGNEKLTF | TRBV12-4 | TRBJ1-4 | CASRPSAGGKGDEKLFF | KLGGALQAK | YES | NO | NO |
| 14785 | TRAV21 | TRAJ8 | CAVTGFQKLVF | TRBV4-1 | TRBJ2-1 | CASSQEGVAGGYNEQFF | KLGGALQAK | YES | NO | NO |
| 14786 | TRAV14DV4 | TRAJ32 | CATWGGGYGGATNKLIF | TRBV6-5 | TRBJ1-3 | CASSRGTGAFSGNTIYF | KLGGALQAK | YES | NO | NO |
| 14787 | TRAV29DV5 | TRAJ40 | CAASSGTYKYIF | TRBV7-2 | TRBJ2-2 | CASGPLGSGELFF | KLGGALQAK | YES | NO | NO |
| 14788 | TRAV1-2 | TRAJ40 | CAVRERGSGTYKYIF | TRBV10-3 | TRBJ1-2 | CAISEEWGSYGYTF | KLGGALQAK | YES | NO | NO |
| 14789 | TRAV19 | TRAJ34 | CALRPSWGNTDKLIF | TRBV11-2 | TRBJ2-1 | CASSLVLAGGNEQFF | KLGGALQAK | YES | NO | NO |
| 14790 | TRAV8-3 | TRAJ24 | CAVGEYLDSWGKLQF | TRBV11-2 | TRBJ2-1 | CASSPGLEGSNEQFF | KLGGALQAK | YES | NO | NO |
| 14791 | TRAV8-2 | TRAJ22 | CVVRIALLGSARQLTF | TRBV11-2 | TRBJ2-1 | CASSLAASGPTGDEQFF | AVFDRKSDAK | YES | NO | YES |
| 14792 | TRAV8-2 | TRAJ47 | CVVRGSMEYGNKLVF | TRBV30 | TRBJ1-1 | CAWSVDNMNTEAFF | KLGGALQAK | YES | NO | NO |
| 14793 | TRAV8-3 | TRAJ52 | CAVGSNAGGTSYGKLTF | TRBV19 | TRBJ2-1 | CASSWTSGRSNEQFF | KLGGALQAK | YES | NO | NO |
| 14794 | TRAV27 | TRAJ20 | CAGPDVNDYKLSF | TRBV6-3 | TRBJ2-1 | CASSYGAGGYNEQFF | KLGGALQAK | YES | NO | NO |
| 14795 | TRAV8-2 | TRAJ8 | CVVSDLGTGFQKLVF | TRBV2 | TRBJ2-5 | CASSEGLAGKYQETQYF | KLGGALQAK | YES | NO | NO |
| 14796 | TRAV13-1 | TRAJ34 | CAASLDTDKLIF | TRBV18 | TRBJ2-7 | CASSPADREFGWPYEQYV | KLGGALQAK | NO | NO | NO |
| 14797 | TRAV26-1 | TRAJ44 | CIVRVGGTASKLTF | TRBV4-1 | TRBJ2-5 | CASSQDSGEETQYF | KLGGALQAK | YES | NO | NO |
| 14798 | TRAV2 | TRAJ44 | CVRYTGTASKLTF | TRBV2 | TRBJ1-1 | CASTGLIEAFF | KLGGALQAK | NO | NO | NO |
| 14799 | TRAV29DV5 | TRAJ48 | CAAISNFGNEKLTF | TRBV9 | TRBJ1-5 | CASSPADGLNQPQHF | KLGGALQAK | NO | NO | NO |
| 14800 | TRAV8-4 | TRAJ48 | CAVSDRFGNEKLTF | TRBV9 | TRBJ1-1 | CASSAFRTDNTIENTEAFF | AVFDRKSDAK | YES | NO | NO |
| 14801 | TRAV12-2 | TRAJ42 | CAVTGGSQGNLIF | TRBV7-9 | TRBJ1-6 | CASSFGPPLHF | KLGGALQAK | YES | NO | NO |
| 14802 | TRAV1-1 | TRAJ15 | CAVRDWAGTALIF | TRBV19 | TRBJ1-5 | CASSIGGVGNQPQHF | KLGGALQAK | YES | NO | NO |
| 14803 | TRAV27 | TRAJ54 | CAVAGIQGAQKLVF | TRBV5-6 | TRBJ1-1 | CASSLGVFEAFF | AVFDRKSDAK | YES | NO | YES |
| 14804 | TRAV27 | TRAJ33 | CAGVMDSNYQLIW | TRBV24-1 | TRBJ2-2 | CATSRGLTGELFF | KLGGALQAK | YES | NO | NO |
| 14805 | TRAV19 | TRAJ16 | CALSEGISDGQKLLF | TRBV4-1 | TRBJ2-7 | CASSQDFGVYEQYF | KLGGALQAK | NO | NO | NO |
| 14806 | TRAV29DV5 | TRAJ54 | CAASGWGAQKLVF | TRBV14 | TRBJ2-1 | CASSQRPFPAGGDSYNEQFF | KLGGALQAK | NO | NO | NO |
| 14807 | TRAV17 | TRAJ23 | CATDNQGGKLIF | TRBV12-4 | TRBJ2-1 | CASSSTALSYNEQFF | KLGGALQAK | YES | NO | NO |
| 14808 | TRAV19 | TRAJ24 | CALSFDSWGKLQF | TRBV20-1 | TRBJ2-7 | CSARDRRQGVGYEQYF | KLGGALQAK | NO | NO | NO |
| 14809 | TRAV38-1 | TRAJ47 | CAFMKLQPMEYGNKLVF | TRBV11-2 | TRBJ2-7 | CASSFGSTGGIYEQYV | AVFDRKSDAK | NO | NO | NO |
| 14810 | TRAV8-1 | TRAJ9 | CAVRAGSTGGFKTIF | TRBV7-9 | TRBJ1-1 | CASSTITGENTEAFF | KLGGALQAK | YES | NO | NO |
| 14811 | TRAV12-2 | TRAJ41 | CAVNLGNSGYALNF | TRBV4-1 | TRBJ1-1 | CASSQGRLRTEAFF | KLGGALQAK | YES | NO | NO |
| 14812 | TRAV8-1 | TRAJ39 | CAVKGAGNMLTF | TRBV12-3 | TRBJ1-2 | CASGEWTKANYGYTF | KLGGALQAK | YES | NO | NO |
| 14813 | TRAV25 | TRAJ30 | CAGFLRNRDDKIIF | TRBV19 | TRBJ2-1 | CASSPGTTYNEQFF | KLGGALQAK | YES | NO | NO |
| 14814 | TRAV19 | TRAJ57 | CAPSFITQGGSEKLVF | TRBV4-1 | TRBJ2-1 | CASSLIAGGPNEQFF | KLGGALQAK | YES | NO | NO |
| 14815 | TRAV20 | TRAJ24 | CAVRPTDSWGKLQF | TRBV9 | TRBJ2-1 | CASSVDRELAYNEQFF | KLGGALQAK | YES | NO | NO |
| 14816 | TRAV13-1 | TRAJ4 | CAASEPGSGGYNKLIF | TRBV13 | TRBJ2-6 | CASSTGQGAPGANVLTF | KLGGALQAK | YES | NO | NO |
| 14817 | TRAV14DV4 | TRAJ48 | CAMREASGRVGNEKLTF | TRBV9 | TRBJ2-5 | CASSPPREEETQYF | KLGGALQAK | YES | NO | NO |
| 14818 | TRAV8-1 | TRAJ10 | CAVNDILTGGGNKLTF | TRBV28 | TRBJ2-1 | CASSSAEQAGVYNEQFF | KLGGALQAK | YES | NO | NO |
| 14819 | TRAV17 | TRAJ22 | CATGTGSARQLTF | TRBV7-2 | TRBJ2-1 | CASSLRSGGGGAYNEQFF | KLGGALQAK | YES | NO | NO |
| 14820 | TRAV13-1 | TRAJ11 | CAAKRNSGYSTLTF | TRBV11-3 | TRBJ1-2 | CASSLRTGQLNNYGYTF | KLGGALQAK | YES | NO | NO |
| 14821 | TRAV4 | TRAJ30 | CLVGDTNRDDKIIF | TRBV11-2 | TRBJ2-1 | CASSQTFGGAYNEQFF | AVFDRKSDAK | YES | NO | YES |
| 14822 | TRAV5 | TRAJ15 | CAVRNQAGTALIF | TRBV3-1 | TRBJ1-5 | CASSQESTSQPQHF | KLGGALQAK | YES | NO | NO |
| 14823 | TRAV1-2 | TRAJ33 | CAVLDSNYQLIW | TRBV20-1 | TRBJ2-1 | CSQTSGDSYNEQFF | KLGGALQAK | NO | NO | NO |
| 14824 | TRAV13-1 | TRAJ45 | CAASLGGGADGLTF | TRBV27 | TRBJ1-1 | CASSPSGDEAFF | KLGGALQAK | YES | NO | NO |
| 14825 | TRAV8-1 | TRAJ49 | CAVYTGNQFYF | TRBV30 | TRBJ2-3 | CAGTAGAGTDTQYF | RTLNAWVKV | NO | NO | NO |
| 14826 | TRAV25 | TRAJ28 | CARVSYSGAGSYQLTF | TRBV9 | TRBJ2-5 | CASSGHRGVGETQYF | KLGGALQAK | YES | NO | NO |
| 14827 | TRAV35 | TRAJ43 | CAGHWNNDMRF | TRBV11-2 | TRBJ1-6 | CASSLGGVSPLHF | KLGGALQAK | YES | NO | NO |
| 14828 | TRAV14DV4 | TRAJ50 | CAMREGLETSYDKVIF | TRBV20-1 | TRBJ2-3 | CSARSLTPILAGGASTDTQYF | KLGGALQAK | NO | NO | NO |
| 14829 | TRAV3 | TRAJ28 | CAVRDRYSGAGSYQLTF | TRBV7-9 | TRBJ1-1 | CASSLEAGGDQAFF | KLGGALQAK | NO | NO | NO |
| 14830 | TRAV19 | TRAJ53 | CALKGDSGGSNYKLTF | TRBV5-6 | TRBJ1-1 | CASSLEPPGTEAFF | KLGGALQAK | YES | NO | NO |
| 14831 | TRAV12-2 | TRAJ6 | CAVNIPRSGGSYIPTF | TRBV7-6 | TRBJ2-1 | CASRTGVEQFF | KLGGALQAK | YES | NO | NO |
| 14832 | TRAV13-2 | TRAJ26 | CAENYGQNFVF | TRBV12-3 | TRBJ2-7 | CASREGLGRTHEQYV | KLGGALQAK | NO | NO | NO |
| 14833 | TRAV1-2 | TRAJ29 | CAVDSGNTPLVF | TRBV14 | TRBJ2-5 | CASSQEYPGQETQYF | KLGGALQAK | NO | NO | NO |
| 14834 | TRAV17 | TRAJ53 | CATVGSGGSNYKLTF | TRBV6-1 | TRBJ1-3 | CASQDRRSGNTIYF | KLGGALQAK | YES | NO | NO |
| 14835 | TRAV1-2 | TRAJ33 | CAVRDRAKSNYQLIW | TRBV7-9 | TRBJ2-1 | CASSLGESGRAHEQFF | RLRAEAQVK | YES | YES | YES |
| 14836 | TRAV29DV5 | TRAJ58 | CAARETSGSRLTF | TRBV7-8 | TRBJ2-3 | CASSLTSGSTDTQYF | KLGGALQAK | YES | NO | NO |
| 14837 | TRAV9-2 | TRAJ8 | CALAMNTGFQKLVF | TRBV7-9 | TRBJ2-7 | CASRDGGVASYEQYV | KLGGALQAK | NO | NO | NO |
| 14838 | TRAV12-1 | TRAJ47 | CVVNELDKLVF | TRBV6-6 | TRBJ1-2 | CASSYLAGDGYTF | KLGGALQAK | NO | NO | NO |
| 14839 | TRAV38-1 | TRAJ48 | CAFNFGNEKLTF | TRBV7-9 | TRBJ2-7 | CASSLALAGQEQYF | KLGGALQAK | YES | NO | NO |
| 14840 | TRAV24 | TRAJ52 | CAFIGDAGGTSYGKLTF | TRBV6-3 | TRBJ2-1 | CASSYGAGGYNEQFF | KLGGALQAK | YES | NO | NO |
| 14841 | TRAV12-1 | TRAJ49 | CVVTNTGNQFYF | TRBV7-8 | TRBJ2-7 | CASSPVRYEQYF | KLGGALQAK | YES | NO | NO |
| 14842 | TRAV13-1 | TRAJ48 | CAASMGRGRFGNEKLTF | TRBV20-1 | TRBJ2-3 | CSARDISDTQYF | KLGGALQAK | YES | NO | NO |
| 14843 | TRAV13-1 | TRAJ45 | CAAKRSGGGADGLTF | TRBV7-6 | TRBJ2-4 | CASSLGARNLAKNIQYF | KLGGALQAK | YES | NO | NO |
| 14844 | TRAV1-2 | TRAJ33 | CAVRAMDSNYQLIW | TRBV6-3 | TRBJ2-7 | CATSASSGISYEQYV | KLGGALQAK | NO | NO | NO |
| 14845 | TRAV30 | TRAJ36 | CGTESGANNLFF | TRBV5-1 | TRBJ1-2 | CASSFSVEETANYGYTF | KLGGALQAK | YES | NO | NO |
| 14846 | TRAV25 | TRAJ10 | CAGLTGGGNKLTF | TRBV11-2 | TRBJ1-1 | CASSLDLWGRNTEAFF | KLGGALQAK | YES | NO | NO |
| 14847 | TRAV27 | TRAJ54 | CAGVQGAQKLVF | TRBV11-2 | TRBJ1-2 | CASSISWGGYGYTF | KLGGALQAK | YES | NO | NO |
| 14848 | TRAV12-2 | TRAJ33 | CAVTDSNYQLIW | TRBV7-9 | TRBJ2-2 | CASSPGTGDSLGELFF | KLGGALQAK | YES | NO | NO |
| 14849 | TRAV8-6 | TRAJ54 | CAVSEIQGAQKLVF | TRBV20-1 | TRBJ1-1 | CSAAQGKGVYTEAFF | KLGGALQAK | YES | NO | NO |
| 14850 | TRAV12-1 | TRAJ9 | CVVTHTGGFKTIF | TRBV20-1 | TRBJ1-4 | CSARGLEDRATTNEKLFF | KLGGALQAK | NO | NO | NO |
| 14851 | TRAV22 | TRAJ33 | CAVKDSNYQLIW | TRBV10-2 | TRBJ2-3 | CASRQGSDTQYF | KLGGALQAK | YES | NO | NO |
| 14852 | TRAV4 | TRAJ37 | CLVGDSGNTGKLIF | TRBV15 | TRBJ2-2 | CAVHPGMNTGELFF | KLGGALQAK | NO | NO | NO |
| 14853 | TRAV12-2 | TRAJ49 | CAVTTGNQFYF | TRBV12-4 | TRBJ2-5 | CASFSGGEETQYF | RLRAEAQVK | YES | NO | YES |

| | | | | | | | | | | |
|---|---|---|---|---|---|---|---|---|---|---|
| 14854 | TRAV8-3 | TRAJ48 | CAVGATLGNEKLTF | TRBV20-1 | TRBJ2-7 | CSAREQDTQLYEQYF | KLGGALQAK | YES | NO | NO |
| 14855 | TRAV38-2DV8 | TRAJ28 | CAYLISYSGAGSYQLTF | TRBV4-1 | TRBJ1-5 | CASSQETGPAPQHF | KLGGALQAK | YES | NO | NO |
| 14856 | TRAV4 | TRAJ5 | CLVGEDTGRRALTF | TRBV11-2 | TRBJ2-3 | CASSYSGGALDTQYF | KLGGALQAK | YES | NO | NO |
| 14857 | TRAV38-2DV8 | TRAJ43 | CASYNNNDMRF | TRBV27 | TRBJ2-1 | CASSLPGRNLAGEQFF | KLGGALQAK | YES | NO | NO |
| 14858 | TRAV9-2 | TRAJ16 | CALSDQNGQKLLF | TRBV4-1 | TRBJ2-1 | CASSQDLSLAGYNEQFF | KLGGALQAK | NO | NO | NO |
| 14859 | TRAV38-1 | TRAJ54 | CAFMSLIQGAQKLVF | TRBV3-1 | TRBJ2-5 | CASSHGKTQYF | KLGGALQAK | YES | NO | NO |
| 14860 | TRAV1-2 | TRAJ31 | CAVKNNNARLMF | TRBV7-8 | TRBJ1-2 | CASSPSPRQGGKGGYTF | KLGGALQAK | YES | NO | NO |
| 14861 | TRAV35 | TRAJ30 | CAGLDRDDKIIF | TRBV19 | TRBJ2-7 | CASSSTPYEQYF | KLGGALQAK | YES | NO | NO |
| 14862 | TRAV24 | TRAJ8 | CAFPNMNTGFQKLVF | TRBV10-3 | TRBJ1-1 | CAISQSVNTEAFF | KLGGALQAK | YES | NO | NO |
| 14863 | TRAV5 | TRAJ40 | CAEVRTSGTYKYIF | TRBV9 | TRBJ2-5 | CASSAGGTGSETQYF | KLGGALQAK | YES | NO | NO |
| 14865 | TRAV12-3 | TRAJ49 | CAMSEGQFYF | TRBV12-4 | TRBJ1-2 | CASRAISYYGYTF | KLGGALQAK | YES | NO | NO |
| 14866 | TRAV1-2 | TRAJ20 | CAVRMDYKLSF | TRBV7-9 | TRBJ2-1 | CASSPEDGPYNEQFF | KLGGALQAK | YES | NO | NO |
| 14867 | TRAV5 | TRAJ34 | CAGDGGGDKLIF | TRBV4-1 | TRBJ2-3 | CASRISRGSTDTQYF | IVTDFSVIK | YES | NO | YES |
| 14868 | TRAV12-3 | TRAJ13 | CAMMAGGYQKVTF | TRBV29-1 | TRBJ1-1 | CSSESGTSEAFF | KLGGALQAK | YES | NO | NO |
| 14869 | TRAV12-2 | TRAJ41 | CAVDRRGSNSGYALNF | TRBV2 | TRBJ2-3 | CASSELTDTQYF | KLGGALQAK | YES | NO | NO |
| 14870 | TRAV13-2 | TRAJ22 | CAESPISGSARQLTF | TRBV11-1 | TRBJ2-7 | CASSFVGGSYEQYF | KLGGALQAK | YES | NO | NO |
| 14871 | TRAV29DV5 | TRAJ34 | CAASARDWNTDKLIF | TRBV5-1 | TRBJ2-7 | CASTTGVSYEQYF | KLGGALQAK | YES | NO | NO |
| 14872 | TRAV12-3 | TRAJ39 | CAIGPGNMLTF | TRBV15 | TRBJ1-5 | CATSGGMGQPQHF | ELAGIGILTV | YES | NO | YES |
| 14873 | TRAV13-1 | TRAJ40 | CAARSGTYKYIF | TRBV11-1 | TRBJ2-5 | CASSPPGAQETQYF | KLGGALQAK | YES | NO | NO |
| 14874 | TRAV14DV4 | TRAJ54 | CGTQGAQKLVF | TRBV4-1 | TRBJ2-6 | CASSQVQLSGANVLTF | KLGGALQAK | YES | NO | NO |
| 14875 | TRAV1-1 | TRAJ23 | CALVLNQGGKLIF | TRBV18 | TRBJ2-1 | CASSPPFEREDNEQFF | KLGGALQAK | YES | NO | NO |
| 14876 | TRAV27 | TRAJ27 | CAGALPNAGKSTF | TRBV4-1 | TRBJ1-1 | CASSQFQNTEAFF | KLGGALQAK | YES | NO | NO |
| 14877 | TRAV27 | TRAJ40 | CAGEPSGTYKYIF | TRBV5-6 | TRBJ1-1 | CASSDTGDGNTEAFF | KLGGALQAK | YES | NO | NO |
| 14878 | TRAV12-2 | TRAJ23 | CAVNRLHNQGGKLIF | TRBV4-3 | TRBJ1-5 | CASSLSLGGTQPQHF | KLGGALQAK | YES | NO | NO |
| 14879 | TRAV24 | TRAJ52 | CAFIGDAGGTSYGKLTF | TRBV20-1 | TRBJ2-7 | CSARDLEVLSYEQYF | KLGGALQAK | YES | NO | NO |
| 14880 | TRAV22 | TRAJ12 | CAVEDSSYKLIF | TRBV2 | TRBJ2-7 | CASSENWSGEQYV | KLGGALQAK | NO | NO | NO |
| 14881 | TRAV29DV5 | TRAJ54 | CAASALQGAQKLVF | TRBV7-9 | TRBJ1-2 | CASSLGRGNLYGYTF | KLGGALQAK | YES | NO | NO |
| 14882 | TRAV22 | TRAJ23 | CAVGYNQGGKLIF | TRBV15 | TRBJ1-1 | CATSRANTEAFF | KLGGALQAK | YES | NO | NO |
| 14883 | TRAV4 | TRAJ26 | CLVADNYGQNFVF | TRBV12-4 | TRBJ1-2 | CASSLVDRANYGYTF | KLGGALQAK | YES | NO | NO |
| 14884 | TRAV22 | TRAJ36 | CAVDPETGANNLFF | TRBV5-4 | TRBJ2-2 | CASSFNVGDTGELFF | KLGGALQAK | YES | NO | NO |
| 14885 | TRAV17 | TRAJ36 | CAQTGANNLFF | TRBV3-1 | TRBJ1-1 | CASSPLPRRGPNTEAFF | KLGGALQAK | YES | NO | NO |
| 14886 | TRAV8-3 | TRAJ43 | CAVNNNDMRF | TRBV7-8 | TRBJ2-1 | CASSDPGRNNEQFF | KLGGALQAK | YES | NO | NO |
| 14887 | TRAV10 | TRAJ24 | CVVSPLMIADSWGKFQF | TRBV7-2 | TRBJ1-1 | CASSLAMGNTEAFF | KLGGALQAK | YES | NO | NO |
| 14888 | TRAV12-3 | TRAJ22 | CAMRGRISSGSARQLTF | TRBV13 | TRBJ1-4 | CASSLVRDEGVPTNEKLFF | KLGGALQAK | YES | NO | NO |
| 14889 | TRAV8-4 | TRAJ53 | CAVSDWGSNYKLTF | TRBV7-2 | TRBJ2-3 | CASSELGTSGNTDTQYF | KLGGALQAK | YES | NO | NO |
| 14890 | TRAV4 | TRAJ18 | CLVGEDRGSTLGRLYF | TRBV6-1 | TRBJ1-3 | CATLGDRQASGNTIYF | KLGGALQAK | YES | NO | NO |
| 14891 | TRAV17 | TRAJ16 | CATALNRF | TRBV20-1 | TRBJ2-4 | CSAGTPDQNIQYF | KLGGALQAK | NO | NO | NO |
| 14892 | TRAV12-1 | TRAJ39 | CVVLNVNAGNMLTF | TRBV15 | TRBJ1-6 | CATSRERGGGNSPLHF | KLGGALQAK | YES | NO | NO |
| 14893 | TRAV14DV4 | TRAJ30 | CAMIRGGHDKIIF | TRBV6-1 | TRBJ2-7 | CASSVQGARGYEQYF | KLGGALQAK | YES | NO | NO |
| 14894 | TRAV4 | TRAJ40 | CLVGGTYKYIF | TRBV11-3 | TRBJ2-1 | CASSLVAVADNEQFF | KLGGALQAK | YES | NO | NO |
| 14895 | TRAV26-2 | TRAJ15 | CILRDSNQAGTALIF | TRBV29-1 | TRBJ2-1 | CSAKPPGPQSYNEQFF | KLGGALQAK | YES | NO | NO |
| 14896 | TRAV35 | TRAJ41 | CAGLFSGYALNF | TRBV7-3 | TRBJ1-2 | CASSLEPPGRGFAGYTF | KLGGALQAK | YES | NO | NO |
| 14897 | TRAV26-1 | TRAJ26 | CIAHGQNFVF | TRBV27 | TRBJ2-2 | CASKLGGSTGELFF | KLGGALQAK | NO | NO | NO |
| 14898 | TRAV1-2 | TRAJ34 | CAVRSTDKLIF | TRBV7-9 | TRBJ2-6 | CASSLVQEVGANVLTF | KLGGALQAK | YES | NO | NO |
| 14900 | TRAV5 | TRAJ40 | CAASGTYKYIF | TRBV6-6 | TRBJ2-2 | CASLDPGQTSGELFF | KLGGALQAK | YES | NO | NO |
| 14901 | TRAV8-1 | TRAJ49 | CAAQTGNQFYF | TRBV9 | TRBJ1-1 | CASSEGQGNTEAFF | KLGGALQAK | YES | NO | NO |
| 14902 | TRAV38-1 | TRAJ42 | CAFRGSQGNLIF | TRBV6-5 | TRBJ1-5 | CASSYSIGNQPQHF | KLGGALQAK | YES | NO | NO |
| 14903 | TRAV22 | TRAJ43 | CAVDLIVYDMRF | TRBV19 | TRBJ2-7 | CASSPSGIYEQYF | KLGGALQAK | YES | NO | NO |
| 14904 | TRAV29DV5 | TRAJ12 | CAASGADSSYKLIF | TRBV20-1 | TRBJ1-2 | CSATPRWGGYTF | KLGGALQAK | YES | NO | NO |
| 14905 | TRAV8-1 | TRAJ20 | CAVNPVNDYKLSF | TRBV7-9 | TRBJ1-2 | CASSPGTGNYGYTF | AVFDRKSDAK | YES | YES | YES |
| 14906 | TRAV12-3 | TRAJ39 | CAIGPGNMLTF | TRBV7-3 | TRBJ2-7 | CASSPSGGGVDEQYF | KLGGALQAK | NO | NO | NO |
| 14907 | TRAV27 | TRAJ35 | CAGDLGFGNVLHC | TRBV19 | TRBJ1-2 | CASSMGSLGYTF | KLGGALQAK | NO | NO | NO |
| 14908 | TRAV14DV4 | TRAJ18 | CAMDRGSTLGRLYF | TRBV7-9 | TRBJ2-1 | CASSFHAGGRGHEQFF | KLGGALQAK | YES | YES | NO |
| 14909 | TRAV19 | TRAJ54 | CALRPIIQGAQKLVF | TRBV4-1 | TRBJ1-4 | CASSGPRQGNEKLFF | KLGGALQAK | YES | NO | NO |
| 14910 | TRAV30 | TRAJ37 | CGTETGSSNTGKLIF | TRBV7-9 | TRBJ2-7 | CASLAPGISEQYV | KLGGALQAK | NO | NO | NO |
| 14911 | TRAV13-1 | TRAJ15 | CAVFNQAGTALIF | TRBV7-3 | TRBJ2-5 | CASSLVGGHQETQYF | KLGGALQAK | NO | NO | NO |
| 14912 | TRAV20 | TRAJ47 | CAVLTMEYGNKLVF | TRBV12-3 | TRBJ2-1 | CASSLGAASGLYYNEQFF | KLGGALQAK | YES | NO | NO |
| 14913 | TRAV17 | TRAJ30 | CATERMNRDDKIIF | TRBV19 | TRBJ2-7 | CASSAGTGGPYEQYF | KLGGALQAK | YES | NO | NO |
| 14914 | TRAV19 | TRAJ39 | CALSAYAGNMLTF | TRBV28 | TRBJ2-7 | CASSFRQAAPYEQYV | KLGGALQAK | NO | NO | NO |
| 14915 | TRAV22 | TRAJ35 | CAVAGFGNVLHC | TRBV5-5 | TRBJ2-7 | CASSNPYEQYV | KLGGALQAK | NO | NO | NO |
| 14916 | TRAV17 | TRAJ9 | CATDDTGGFKTIF | TRBV6-1 | TRBJ2-3 | CATIDWGGEDTQYF | AVFDRKSDAK | YES | NO | NO |
| 14917 | TRAV29DV5 | TRAJ36 | CAASVTGANNLFF | TRBV7-9 | TRBJ2-7 | CASSPPLWQIYEQYF | KLGGALQAK | YES | NO | NO |
| 14918 | TRAV14DV4 | TRAJ54 | CAMRDVIQGAQKLVF | TRBV7-9 | TRBJ2-7 | CASSPQGSYEQYF | KLGGALQAK | YES | NO | NO |
| 14919 | TRAV26-2 | TRAJ43 | CILSYADNDMRF | TRBV6-1 | TRBJ1-1 | CASNGFGVEAFF | KLGGALQAK | YES | NO | NO |
| 14920 | TRAV12-2 | TRAJ33 | CAVRDSNYQLIW | TRBV9 | TRBJ2-7 | CASSVATSGEQYF | KLGGALQAK | YES | NO | NO |
| 14921 | TRAV8-1 | TRAJ42 | CAVTIVLEGGSQGNLIF | TRBV7-6 | TRBJ2-5 | CASSLYQETQYF | KVLEYVIKV | YES | NO | YES |
| 14922 | TRAV12-3 | TRAJ21 | CATQRNNFNKFYF | TRBV12-4 | TRBJ1-1 | CASSFSGVGTEAFF | KLGGALQAK | YES | NO | NO |
| 14923 | TRAV14DV4 | TRAJ30 | CAMREGNRDDKIIF | TRBV5-1 | TRBJ2-5 | CASSQGTTNQETQYF | KLGGALQAK | YES | NO | NO |
| 14924 | TRAV29DV5 | TRAJ7 | CAASADGNNRLAF | TRBV27 | TRBJ1-5 | CASSLSGGYQPQHF | KLGGALQAK | NO | NO | NO |
| 14925 | TRAV14DV4 | TRAJ34 | CAMREGWGTDKLIF | TRBV19 | TRBJ2-5 | CASSPGTGGTETQYF | KLGGALQAK | YES | NO | NO |
| 14926 | TRAV41 | TRAJ49 | CAVSPFTGNQFYF | TRBV6-1 | TRBJ1-2 | CASSDWGQGYGYTF | KLGGALQAK | YES | NO | NO |
| 14927 | TRAV24 | TRAJ32 | CAFPASYGGATNKLIF | TRBV13 | TRBJ2-1 | CASSPAVGPMNEQFF | KLGGALQAK | YES | NO | NO |
| 14928 | TRAV8-3 | TRAJ24 | CAVGTDSWGKLQF | TRBV24-1 | TRBJ2-1 | CATSESLAGGGYNEQFF | AVFDRKSDAK | YES | NO | YES |
| 14929 | TRAV12-2 | TRAJ15 | CAVNINQAGTALIF | TRBV7-2 | TRBJ2-5 | CASSLVRQGASTIGTQYF | KLGGALQAK | YES | NO | NO |
| 14930 | TRAV3 | TRAJ48 | CAVTDLLGNEKLTF | TRBV7-8 | TRBJ1-6 | CASSLEGNSYNSPLHF | KLGGALQAK | NO | NO | NO |
| 14931 | TRAV13-2 | TRAJ20 | CAESSNDYKLSF | TRBV6-5 | TRBJ1-1 | CASSYGGSTEAFF | KLGGALQAK | YES | NO | NO |
| 14932 | TRAV8-3 | TRAJ28 | CAVGAYSGAGSYQLTF | TRBV7-9 | TRBJ2-2 | CASSSGDSFFTGELFF | KLGGALQAK | YES | NO | NO |
| 14933 | TRAV8-3 | TRAJ13 | CAVYPMNSGGYQKVTF | TRBV20-1 | TRBJ1-2 | CSASPRTGGIPNYGYTF | KLGGALQAK | YES | NO | NO |
| 14934 | TRAV26-2 | TRAJ33 | CILRLLDSNYQLIW | TRBV15 | TRBJ2-1 | CATSRVRGPEAYNEQFF | AVFDRKSDAK | YES | NO | YES |
| 14935 | TRAV8-2 | TRAJ20 | CVVIPSNDYKLSF | TRBV5-6 | TRBJ2-5 | CASSLGPTSGRETQYF | KLGGALQAK | YES | NO | NO |
| 14936 | TRAV38-2DV8 | TRAJ50 | CAYRKTSYDKVIF | TRBV6-3 | TRBJ2-5 | CASSYSILAGGQETQYF | KLGGALQAK | YES | NO | NO |
| 14937 | TRAV14DV4 | TRAJ18 | CAMRGPRGSTLGRLYF | TRBV4-3 | TRBJ1-4 | CASSQDHRMGGHEKLFF | KLGGALQAK | YES | NO | NO |
| 14938 | TRAV12-2 | TRAJ41 | CAVNSIYSGYALNF | TRBV19 | TRBJ1-1 | CASSMSQTGNTEAFF | KLGGALQAK | YES | NO | NO |
| 14939 | TRAV12-3 | TRAJ39 | CAIGPGNMLTF | TRBV20-1 | TRBJ2-3 | CSARKGEPPSTDTQYF | KLGGALQAK | YES | NO | NO |
| 14940 | TRAV9-2 | TRAJ12 | CALRDSSYKLIF | TRBV7-9 | TRBJ1-1 | CASSLERLEAFF | KLGGALQAK | YES | NO | NO |
| 14941 | TRAV29DV5 | TRAJ31 | CAASAHQDRLMF | TRBV7-8 | TRBJ2-1 | CASSLGLAGGNSYNEQFF | KLGGALQAK | YES | NO | NO |
| 14942 | TRAV4 | TRAJ29 | CLVGGPHSGNTPLVF | TRBV5-5 | TRBJ1-1 | CASSLGGPGGGNTEAFF | KLGGALQAK | YES | NO | NO |
| 14943 | TRAV12-2 | TRAJ49 | CAVKLVGNQFYF | TRBV20-1 | TRBJ2-1 | CSARDLLVASYNEQFF | KLGGALQAK | NO | NO | NO |
| 14944 | TRAV13-2 | TRAJ23 | CAETHYNQGGKLIF | TRBV7-2 | TRBJ2-1 | CASSPSPYNEQFF | KLGGALQAK | YES | NO | NO |
| 14945 | TRAV9-2 | TRAJ52 | CALSGSPDAGGTSYGKLTF | TRBV19 | TRBJ2-2 | CASKGLAGAPGELFF | KLGGALQAK | YES | NO | NO |
| 14946 | TRAV3 | TRAJ42 | CAVRPPGGGSQGNLIF | TRBV19 | TRBJ1-2 | CASSFDTPPVYGYTF | AVFDRKSDAK | NO | NO | NO |
| 14947 | TRAV1-1 | TRAJ5 | CAVRDRDTGRRALTF | TRBV12-4 | TRBJ2-5 | CASSHIAGQETQYF | FLASKIGRLV | YES | NO | YES |
| 14948 | TRAV8-4 | TRAJ34 | CAVSFNTDKLIF | TRBV6-3 | TRBJ2-1 | CASSYSGGLDEQFF | KLGGALQAK | YES | NO | NO |
| 14949 | TRAV1-2 | TRAJ41 | CAVRVLTAYRYALNF | TRBV19 | TRBJ2-3 | CASSPGTSGRAEDTQYF | KLGGALQAK | YES | NO | NO |
| 14950 | TRAV3 | TRAJ38 | CAVRGLNAGNNRKLIW | TRBV3-1 | TRBJ2-2 | CASINGLANTGELFF | KLGGALQAK | NO | NO | NO |
| 14951 | TRAV1-2 | TRAJ27 | CAVKGHAGKSTF | TRBV5-4 | TRBJ2-7 | CASRGDPGEHEQYV | KLGGALQAK | NO | NO | NO |
| 14952 | TRAV5 | TRAJ29 | CAFSGNTPLVF | TRBV10-3 | TRBJ2-2 | CAITGLAMNTGELFF | AVFDRKSDAK | YES | YES | YES |
| 14953 | TRAV39 | TRAJ47 | CAVDMEYGNKLVF | TRBV15 | TRBJ2-7 | CATSRDISIGSYEQYF | KLGGALQAK | YES | NO | NO |
| 14954 | TRAV9-2 | TRAJ23 | CALRGRKAYLRT | TRBV29-1 | TRBJ2-1 | CSVLAGSSYNEQFF | KLGGALQAK | NO | NO | NO |
| 14955 | TRAV1-2 | TRAJ20 | CAGSNDYKLSF | TRBV5-4 | TRBJ1-1 | CASGDGTANTEAFF | KLGGALQAK | YES | NO | NO |
| 14956 | TRAV35 | TRAJ16 | CAGPDGQKLLF | TRBV27 | TRBJ1-1 | CASSSITRFTEAFF | KLGGALQAK | NO | NO | NO |
| 14957 | TRAV10 | TRAJ42 | CVVSARDGGSQGNLIF | TRBV4-3 | TRBJ2-1 | CASSQQTSAYNEQFF | KLGGALQAK | YES | NO | NO |
| 14958 | TRAV8-4 | TRAJ58 | CAVSDQETSGSRLTF | TRBV13 | TRBJ2-5 | CASSSTGSRQETQYF | FLASKIGRLV | YES | NO | YES |
| 14959 | TRAV25 | TRAJ53 | CAGPNSGGSNYKLTF | TRBV7-9 | TRBJ2-2 | CASSLVGWANQPPGELFF | KLGGALQAK | YES | NO | NO |

| | | | | | | | | | | |
|---|---|---|---|---|---|---|---|---|---|---|
| 14960 | TRAV29DV5 | TRAJ57 | CAASAAGGSEKLVF | TRBV11-2 | TRBJ2-6 | CASSSRLTGDYGANVLTF | KLGGALQAK | YES | NO | NO |
| 14961 | TRAV35 | TRAJ40 | CAGHFTSGTYKYIF | TRBV19 | TRBJ1-2 | CASGQGVFGYTF | KLGGALQAK | YES | NO | NO |
| 14962 | TRAV13-2 | TRAJ45 | CAENSRGGGADGLTF | TRBV7-7 | TRBJ2-2 | CASSLSSGITGELFF | KLGGALQAK | YES | NO | NO |
| 14963 | TRAV27 | TRAJ20 | CAVTRSNDYKLSF | TRBV20-1 | TRBJ2-1 | CSAGGGATYNEQFF | KLGGALQAK | NO | NO | NO |
| 14964 | TRAV13-2 | TRAJ50 | CAEKQDDKVIF | TRBV6-5 | TRBJ2-2 | CASRYTGATGELFF | KLGGALQAK | YES | NO | NO |
| 14965 | TRAV29DV5 | TRAJ47 | CAASGYGNKLVF | TRBV20-1 | TRBJ2-1 | CSATSGGAPHNEQFF | KLGGALQAK | NO | NO | NO |
| 14966 | TRAV12-1 | TRAJ44 | CVVSITGTASKLTF | TRBV19 | TRBJ2-1 | CASTGTGVYEQFF | AVFDRKSDAK | YES | NO | YES |
| 14967 | TRAV17 | TRAJ24 | CATTPTDSWGKLQF | TRBV6-5 | TRBJ2-2 | CASSYSSAGGLLAGELFF | KLGGALQAK | YES | NO | NO |
| 14968 | TRAV8-1 | TRAJ41 | CAVTSNSGYALNF | TRBV2 | TRBJ2-1 | CASSLVAGGYNEQFF | KLGGALQAK | YES | NO | NO |
| 14969 | TRAV29DV5 | TRAJ58 | CAATSGSRLTF | TRBV7-9 | TRBJ1-2 | CASSFPGTGEAYGYTF | KLGGALQAK | YES | NO | NO |
| 14970 | TRAV29DV5 | TRAJ40 | CAASFTSGTYKYIF | TRBV7-2 | TRBJ2-2 | CASSLQNTGELFF | KLGGALQAK | YES | NO | NO |
| 14971 | TRAV1-2 | TRAJ29 | CAVPEVNTPLVF | TRBV7-9 | TRBJ1-6 | CASSLGQGGAPLHF | KLGGALQAK | YES | NO | NO |
| 14972 | TRAV8-1 | TRAJ5 | CAVNLGRRALTF | TRBV6-3 | TRBJ1-1 | CASSIMNTEAFF | KLGGALQAK | YES | NO | NO |
| 14973 | TRAV12-3 | TRAJ39 | CAIGPGNMLTF | TRBV20-1 | TRBJ1-1 | CSAWDRLNTEAFF | KLGGALQAK | YES | NO | NO |
| 14974 | TRAV1-1 | TRAJ37 | CALGSGNTGKLIF | TRBV19 | TRBJ2-7 | CASSSGVYEQYF | KLGGALQAK | YES | NO | NO |
| 14975 | TRAV27 | TRAJ58 | CAGLDSSGSRLTF | TRBV27 | TRBJ1-4 | CASSLSGTWGEKLFF | KLGGALQAK | YES | NO | NO |
| 14976 | TRAV1-2 | TRAJ29 | CAVRDKRNTPLVF | TRBV15 | TRBJ2-5 | CATSRGSGQYQETQYF | KLGGALQAK | YES | NO | NO |
| 14977 | TRAV8-1 | TRAJ22 | CAVRSSGSARQLTF | TRBV5-5 | TRBJ2-7 | CASNPSGTGTNYEQYF | KLGGALQAK | YES | NO | NO |
| 14978 | TRAV14DV4 | TRAJ41 | CAMREGAGYALNF | TRBV19 | TRBJ2-1 | CASSILAGYYNEQFF | KLGGALQAK | YES | NO | NO |
| 14980 | TRAV26-1 | TRAJ39 | CIVRANNAGNMLTF | TRBV19 | TRBJ1-2 | CASSARTGVGYGYTF | AVFDRKSDAK | YES | YES | NO |
| 14981 | TRAV24 | TRAJ52 | CAFIGDAGGTSYGKLTF | TRBV19 | TRBJ2-5 | CASIEGETQYF | KLGGALQAK | YES | NO | NO |
| 14982 | TRAV8-4 | TRAJ49 | CAVRGNTGNQFYF | TRBV5-6 | TRBJ1-5 | CASSLFDQPQHF | KLGGALQAK | YES | NO | NO |
| 14983 | TRAV17 | TRAJ3 | CATDGSSASKIIF | TRBV9 | TRBJ2-2 | CASMAENTGELFF | KLGGALQAK | YES | NO | NO |
| 14984 | TRAV13-1 | TRAJ45 | CAASRGGADGLTF | TRBV6-3 | TRBJ2-1 | CASSSVSLEQFF | KLGGALQAK | YES | NO | NO |
| 14985 | TRAV14DV4 | TRAJ39 | CARDNAGNMLTF | TRBV7-8 | TRBJ2-3 | CASSSNDRGPADTQYF | KLGGALQAK | YES | NO | NO |
| 14986 | TRAV38-2DV8 | TRAJ58 | CAYRSAPETSGSRLTF | TRBV30 | TRBJ2-3 | CAWRTGIGTDTQYF | KLGGALQAK | YES | NO | NO |
| 14987 | TRAV10 | TRAJ53 | CVVSGWGGSNYKLTF | TRBV20-1 | TRBJ1-1 | CSASLGGGPDMNTEAFF | KLGGALQAK | YES | NO | NO |
| 14988 | TRAV29DV5 | TRAJ37 | CAASALGNTGKLIF | TRBV7-9 | TRBJ2-1 | CASSSGLAGENEQFF | KLGGALQAK | YES | NO | NO |
| 14989 | TRAV12-2 | TRAJ16 | CAVIADGQKLLF | TRBV4-1 | TRBJ1-1 | CASSQTKYTEAFF | KLGGALQAK | NO | NO | NO |
| 14990 | TRAV25 | TRAJ56 | CAGLTGANSKLTF | TRBV27 | TRBJ2-7 | CASSWGSSYEQYF | KLGGALQAK | YES | NO | NO |
| 14991 | TRAV41 | TRAJ58 | CAVSQIRTSGSRLTF | TRBV4-1 | TRBJ1-5 | CASSQLGTGYQPQHF | KLGGALQAK | YES | NO | NO |
| 14992 | TRAV17 | TRAJ53 | CATPGGSNYKLTF | TRBV7-4 | TRBJ1-2 | CASSPRQVGYGYTF | KLGGALQAK | YES | NO | NO |
| 14993 | TRAV20 | TRAJ39 | CAVQENNAGNMLTF | TRBV29-1 | TRBJ1-2 | CSVEEVLVGGYTF | KLGGALQAK | YES | NO | NO |
| 14994 | TRAV17 | TRAJ57 | CATDGWGGSEKLVF | TRBV7-2 | TRBJ2-3 | CASSLNIVVGPSDTQYF | KLGGALQAK | YES | NO | NO |
| 14995 | TRAV30 | TRAJ24 | CGTAFRTDSWGKLQF | TRBV19 | TRBJ2-7 | CASSIGGANQPYEQYF | KLGGALQAK | YES | NO | NO |
| 14996 | TRAV1-2 | TRAJ33 | CAVSDSNYQLIW | TRBV6-3 | TRBJ1-5 | CASSSAGGHQPPQHF | RLRAEAQVK | YES | NO | YES |
| 14997 | TRAV35 | TRAJ49 | CAGANTGNQFYF | TRBV7-2 | TRBJ1-2 | CASSWTQGYSGYTF | KLGGALQAK | NO | NO | NO |
| 14998 | TRAV14DV4 | TRAJ43 | CAMREVTHNDMRF | TRBV5-1 | TRBJ2-1 | CASSSNRDRVYNEQFF | KLGGALQAK | YES | NO | NO |
| 14999 | TRAV10 | TRAJ20 | CVVIAPNDYKLSF | TRBV5-1 | TRBJ1-1 | CASSLDSGVHTEAFF | KLGGALQAK | YES | NO | NO |
| 15000 | TRAV21 | TRAJ52 | CAGGLINAGGTSYGKLTF | TRBV20-1 | TRBJ2-7 | CSARDWSSYEQYV | KLGGALQAK | NO | NO | NO |
| 15002 | TRAV17 | TRAJ20 | CALGSNDYKLSF | TRBV6-4 | TRBJ1-5 | CASSEYSGGGQPQHF | KLGGALQAK | YES | NO | NO |
| 15003 | TRAV38-1 | TRAJ11 | CAFIPNSGYSTLTF | TRBV6-3 | TRBJ1-5 | CASSTPWGNQPQHF | KLGGALQAK | YES | NO | NO |
| 15004 | TRAV12-2 | TRAJ41 | CAVSNSGYALNF | TRBV12-4 | TRBJ2-7 | CASSLADRVYEQYF | KLGGALQAK | YES | NO | NO |
| 15005 | TRAV13-2 | TRAJ16 | CAENKAWTDGQKLLF | TRBV4-1 | TRBJ2-7 | CASSLKGEGIYEQYF | KLGGALQAK | NO | NO | NO |
| 15006 | TRAV24 | TRAJ20 | CAINDYKLSF | TRBV19 | TRBJ2-7 | CASSTSSYEQYF | KLGGALQAK | YES | NO | NO |
| 15007 | TRAV17 | TRAJ20 | CATDRSNDYKLSF | TRBV27 | TRBJ2-1 | CASSLAANYEQFF | KLGGALQAK | YES | NO | NO |
| 15008 | TRAV5 | TRAJ42 | CAEVNGGSQGNLIF | TRBV20-1 | TRBJ2-7 | CSANPTPSYEQYF | KLGGALQAK | NO | NO | NO |
| 15009 | TRAV1-2 | TRAJ39 | CAVRDIVPNAGNMLTF | TRBV4-3 | TRBJ1-1 | CASARASMNTEAFF | KLGGALQAK | YES | NO | NO |
| 15010 | TRAV1-2 | TRAJ15 | CAVRDAQAGTALIF | TRBV4-3 | TRBJ2-1 | CASSQDVGLAGTNEQFF | KLGGALQAK | YES | NO | NO |
| 15011 | TRAV8-3 | TRAJ39 | CAVGDGNAGNMLTF | TRBV27 | TRBJ1-6 | CASRRTGQSYNSPLHF | KLGGALQAK | YES | NO | NO |
| 15012 | TRAV5 | TRAJ5 | CAEREDTGRRALTF | TRBV6-1 | TRBJ1-1 | CASSDLGTGHTEAFF | KLGGALQAK | YES | NO | NO |
| 15013 | TRAV21 | TRAJ12 | CAPGPDSSYKLIF | TRBV20-1 | TRBJ2-3 | CSARVPAGASTDTQYF | KLGGALQAK | NO | NO | NO |
| 15014 | TRAV25 | TRAJ11 | CAVDSGYSTLTF | TRBV4-1 | TRBJ2-7 | CASSPPDSYEQYF | KLGGALQAK | YES | NO | NO |
| 15015 | TRAV20 | TRAJ23 | CAVQAYNQGGKLIF | TRBV5-5 | TRBJ2-1 | CASSLVVTGLEQFF | KLGGALQAK | YES | NO | NO |
| 15016 | TRAV13-1 | TRAJ52 | CSAGGTSYGKLTF | TRBV27 | TRBJ2-3 | CASSPLDGRADTQYF | RLRAEAQVK | YES | NO | NO |
| 15017 | TRAV27 | TRAJ36 | CAGARTGANNLFF | TRBV21-1 | TRBJ2-2 | CQQRLGGLGTGELFF | KLGGALQAK | YES | NO | NO |
| 15018 | TRAV27 | TRAJ44 | CAGADGTASKLTF | TRBV12-4 | TRBJ2-7 | CASSYIIIEQYF | KLGGALQAK | YES | NO | NO |
| 15019 | TRAV13-1 | TRAJ3 | CAAPMDYSSASKIIF | TRBV6-5 | TRBJ1-2 | CASSFGPNYGYTF | KLGGALQAK | YES | NO | NO |
| 15020 | TRAV29DV5 | TRAJ28 | CAVTYSGAGSYQLTF | TRBV9 | TRBJ2-7 | CASSVGGGDYEQYF | KLGGALQAK | YES | NO | NO |
| 15021 | TRAV8-3 | TRAJ45 | CAVGAPQGGADGLTF | TRBV4-3 | TRBJ1-4 | CASSQDHRMGGHEKLFF | KLGGALQAK | YES | NO | NO |
| 15022 | TRAV29DV5 | TRAJ26 | CAASASNYGQNFVF | TRBV18 | TRBJ2-7 | CASSFGRGYEQYV | KLGGALQAK | NO | NO | NO |
| 15023 | TRAV5 | TRAJ3 | CAETRSSASKIIF | TRBV20-1 | TRBJ2-7 | CSGTGTDNSYEQYF | KLGGALQAK | NO | NO | NO |
| 15024 | TRAV23DV6 | TRAJ42 | CAASPDYGGSQGNLIF | TRBV19 | TRBJ1-3 | CASSMATGLSGNTIYF | KLGGALQAK | YES | NO | NO |
| 15025 | TRAV14DV4 | TRAJ52 | CAMREGYAGGTSYGKLTF | TRBV7-9 | TRBJ1-1 | CASSQNRGLNTEAFF | KLGGALQAK | YES | NO | NO |
| 15026 | TRAV20 | TRAJ8 | CAVQLNTGFQKLVF | TRBV7-9 | TRBJ1-1 | CASSLFDIGEAFF | KLGGALQAK | YES | NO | NO |
| 15027 | TRAV12-3 | TRAJ57 | CAIRNRGGSEKLVF | TRBV28 | TRBJ1-1 | CASSLGGQGRNTEAFF | KLGGALQAK | YES | NO | NO |
| 15028 | TRAV8-4 | TRAJ39 | CAVSDPNNAGNMLTF | TRBV19 | TRBJ2-7 | CASSWNDYEQYF | KLGGALQAK | YES | NO | NO |
| 15029 | TRAV8-6 | TRAJ13 | CATDSGGYQKVTF | TRBV5-1 | TRBJ2-7 | CASRQGATYEQYF | KLGGALQAK | YES | NO | NO |
| 15030 | TRAV29DV5 | TRAJ40 | CAASALTGTYKYIF | TRBV24-1 | TRBJ2-3 | CATRIGGGAADTQYF | KLGGALQAK | YES | NO | NO |
| 15031 | TRAV25 | TRAJ29 | CAGTNSGNTPLVF | TRBV6-3 | TRBJ2-1 | CASSYGAGGYNEQFF | KLGGALQAK | YES | NO | NO |
| 15032 | TRAV17 | TRAJ53 | CATEYSGGSNYKLTF | TRBV7-6 | TRBJ2-7 | CASSLALSYEQYF | IVTDFSVIK | YES | YES | YES |
| 15033 | TRAV8-1 | TRAJ24 | CAVNAQVSWGKLQF | TRBV4-3 | TRBJ2-4 | CASSRETGGAKNIQYF | KLGGALQAK | YES | NO | NO |
| 15034 | TRAV1-2 | TRAJ30 | CAVRGRDDKIIF | TRBV4-1 | TRBJ2-7 | CASSQAIAGGSYEQYF | KLGGALQAK | YES | NO | NO |
| 15035 | TRAV36DV7 | TRAJ11 | CAVAGYSTLTF | TRBV12-3 | TRBJ1-5 | CASSLGGSGPQHF | KLGGALQAK | NO | NO | NO |
| 15036 | TRAV1-1 | TRAJ21 | CAVRDHGTLYNFNKFYF | TRBV12-3 | TRBJ2-1 | CASSSPGGGPSYNEQFF | KLGGALQAK | YES | NO | NO |
| 15037 | TRAV8-2 | TRAJ40 | CVVSPTSGTYKYIF | TRBV6-3 | TRBJ2-5 | CASSQVDRALETQYF | KLGGALQAK | YES | NO | NO |
| 15038 | TRAV21 | TRAJ6 | CAVRLQFSGGSYIPTF | TRBV11-3 | TRBJ2-7 | CASSETSGPSYEQYF | KLGGALQAK | YES | NO | NO |
| 15039 | TRAV27 | TRAJ53 | CAGAGGGSNYKLTF | TRBV5-1 | TRBJ2-3 | CASSRYSGGFGDTQYF | KLGGALQAK | YES | NO | NO |
| 15040 | TRAV17 | TRAJ23 | CATAPYNQGGKLIF | TRBV20-1 | TRBJ2-1 | CSARDRDRATYNEQFF | KLGGALQAK | NO | NO | NO |
| 15041 | TRAV21 | TRAJ40 | CAVLPTSGTYKYIF | TRBV6-1 | TRBJ2-7 | CASSDLHEQYF | KLGGALQAK | YES | NO | NO |
| 15042 | TRAV8-3 | TRAJ44 | CAVGAGTASKLTF | TRBV6-3 | TRBJ2-1 | CASSLLNPATPNNEQFF | KLGGALQAK | YES | NO | NO |
| 15043 | TRAV12-3 | TRAJ49 | CAMRASSNTGNQFYF | TRBV6-5 | TRBJ2-1 | CASGLLWGQEQFF | KLGGALQAK | NO | NO | NO |
| 15044 | TRAV8-6 | TRAJ29 | CAVSDGYGNTPLVF | TRBV11-2 | TRBJ1-1 | CASSLDFPEAFF | KLGGALQAK | YES | NO | NO |
| 15045 | TRAV12-2 | TRAJ42 | CAVMDDGGSQGNLIF | TRBV12-4 | TRBJ2-5 | CASSSPYGTADYQETQYF | KLGGALQAK | YES | NO | NO |
| 15046 | TRAV14DV4 | TRAJ16 | CAMREHQDGQKLLF | TRBV5-1 | TRBJ1-2 | CASSLEGEGYTF | AVFDRKSDAK | NO | NO | NO |
| 15047 | TRAV38-2DV8 | TRAJ42 | CAYRSPSRWGGSQGNLIF | TRBV5-1 | TRBJ2-1 | CASSLGGADEQFF | AVFDRKSDAK | YES | YES | YES |
| 15048 | TRAV19 | TRAJ21 | CALSDDLYNFNKFYF | TRBV20-1 | TRBJ2-1 | CSASGLAVNEQFF | AVFDRKSDAK | NO | NO | NO |
| 15049 | TRAV8-3 | TRAJ42 | CAVRGSQGNLIF | TRBV27 | TRBJ1-2 | CASSLSQGADYGYTF | KLGGALQAK | YES | NO | NO |
| 15050 | TRAV34 | TRAJ48 | CGAGAIWYEKLTF | TRBV6-6 | TRBJ1-1 | CASRAGVNTEAFF | KLGGALQAK | NO | NO | NO |
| 15051 | TRAV13-1 | TRAJ29 | CAAKPEAGNTPLVF | TRBV7-9 | TRBJ1-2 | CASSKKGGGASGYTF | KLGGALQAK | YES | NO | NO |
| 15053 | TRAV19 | TRAJ20 | CALSEAIHNDYKLSF | TRBV19 | TRBJ1-4 | CASSILGARAGEKLFF | KLGGALQAK | YES | NO | NO |
| 15054 | TRAV13-2 | TRAJ53 | CAETLGSNYKLTF | TRBV9 | TRBJ2-2 | CASSAPGGEGVDTGELFF | KLGGALQAK | YES | NO | NO |
| 15055 | TRAV12-1 | TRAJ13 | CVVNIQDSGGYQKVTF | TRBV20-1 | TRBJ2-7 | CSARGRLAYEQYF | KLGGALQAK | NO | NO | NO |
| 15056 | TRAV13-1 | TRAJ43 | CAASRVANNNDMRF | TRBV6-3 | TRBJ2-1 | CASSYGAGGYNEQFF | KLGGALQAK | YES | NO | NO |
| 15057 | TRAV1-2 | TRAJ37 | CAVSNTGKLIF | TRBV7-9 | TRBJ1-2 | CASSYQQGAYDGYTF | KLGGALQAK | YES | NO | NO |
| 15058 | TRAV27 | TRAJ42 | CAGVNYGGSQGNLIF | TRBV2 | TRBJ2-2 | CASRWTLNTGELFF | KLGGALQAK | YES | NO | NO |
| 15059 | TRAV3 | TRAJ11 | CAVSGYSTLTF | TRBV19 | TRBJ2-2 | CASSIDQGGELFF | AVFDRKSDAK | NO | NO | NO |
| 15060 | TRAV19 | TRAJ30 | CALIRGDDKIIF | TRBV7-9 | TRBJ1-3 | CASSPSRDRGSGNTIYF | KLGGALQAK | YES | NO | NO |
| 15061 | TRAV38-2DV8 | TRAJ30 | CAYRSIRDDKIIF | TRBV19 | TRBJ1-5 | CASIEWQGSNQPQHF | KLGGALQAK | YES | NO | NO |
| 15062 | TRAV21 | TRAJ48 | CAVRPGFTSNFGNEKLTF | TRBV7-8 | TRBJ1-1 | CASSLMGQLNTEAFF | KLGGALQAK | YES | NO | NO |
| 15063 | TRAV23DV6 | TRAJ52 | CAAGGAGGTSYGKLTF | TRBV6-3 | TRBJ2-1 | CASRWDSSYNEQFF | KLGGALQAK | YES | NO | NO |
| 15064 | TRAV9-2 | TRAJ16 | CALSSLWF | TRBV7-9 | TRBJ1-1 | CASSLATGTEAFF | KLGGALQAK | NO | NO | NO |
| 15065 | TRAV29DV5 | TRAJ42 | CAASDENGGSQGNLIF | TRBV15 | TRBJ1-2 | CATSRVDDRVDGYTF | KLGGALQAK | YES | NO | NO |
| 15066 | TRAV21 | TRAJ41 | CAVGEPNSGYALNF | TRBV29-1 | TRBJ1-5 | CSVESGQGGQPQHF | KLGGALQAK | YES | NO | NO |

| | | | | | | | | | | |
|---|---|---|---|---|---|---|---|---|---|---|
| 15067 | TRAV29DV5 | TRAJ45 | CAASTGADGLTF | TRBV7-9 | TRBJ2-1 | CASTQWSGGSYNEQFF | KLGGALQAK | YES | NO | NO |
| 15068 | TRAV1-1 | TRAJ24 | CAPKTDSWGKLQF | TRBV7-2 | TRBJ2-7 | CASSLEPSYEQYF | KLGGALQAK | YES | NO | NO |
| 15069 | TRAV24 | TRAJ24 | CAFSGSDSWGKLQF | TRBV20-1 | TRBJ2-5 | CSARDRGAGAQYF | KLGGALQAK | YES | NO | NO |
| 15070 | TRAV14DV4 | TRAJ28 | CAMREVAYSGAGSYQLTF | TRBV12-4 | TRBJ2-7 | CASSLDSREQYV | KLGGALQAK | NO | NO | NO |
| 15071 | TRAV29DV5 | TRAJ16 | CAASAF | TRBV12-5 | TRBJ2-1 | CASGLENNEQFF | KLGGALQAK | NO | NO | NO |
| 15072 | TRAV19 | TRAJ24 | CALSEALSMTTDSWGKFQF | TRBV19 | TRBJ1-1 | CASSIPGTKGSLNTEAFF | KLGGALQAK | YES | NO | NO |
| 15073 | TRAV1-2 | TRAJ39 | CAVKTAAGNMLTF | TRBV10-3 | TRBJ2-5 | CAISESGLAGGETQYF | KLGGALQAK | YES | NO | NO |
| 15074 | TRAV20 | TRAJ26 | CAVLRNYGQNFVF | TRBV6-3 | TRBJ2-2 | CASSSTKTAYTTGELFF | KLGGALQAK | YES | NO | NO |
| 15075 | TRAV19 | TRAJ33 | CALSEASADSNYQLIW | TRBV9 | TRBJ2-4 | CASSRLGFGSKNIQYF | KLGGALQAK | NO | NO | NO |
| 15076 | TRAV26-1 | TRAJ36 | CIVRVAYQTGANNLFF | TRBV6-3 | TRBJ2-7 | CASTKLAQSSYEQYF | KLGGALQAK | YES | NO | NO |
| 15077 | TRAV19 | TRAJ30 | CALTLARDDKIIF | TRBV29-1 | TRBJ2-1 | CSVQLHEQFF | KLGGALQAK | YES | NO | NO |
| 15078 | TRAV12-1 | TRAJ49 | CVVSYNTGNQFYF | TRBV12-4 | TRBJ2-5 | CASSLGDRQETQYF | KLGGALQAK | YES | NO | NO |
| 15079 | TRAV19 | TRAJ47 | CALSGNKLVF | TRBV6-5 | TRBJ1-3 | CASSTVGSSGNTIYF | RTLNAWVKV | YES | YES | YES |
| 15080 | TRAV19 | TRAJ16 | CALNSWADGQKLLF | TRBV7-9 | TRBJ2-6 | CASSLKAGWGSGANVLTF | KLGGALQAK | NO | NO | NO |
| 15081 | TRAV1-2 | TRAJ17 | CAVKAAGNKLTF | TRBV5-1 | TRBJ1-3 | CASSLGTNSGNTIYF | KLGGALQAK | YES | NO | NO |
| 15082 | TRAV1-2 | TRAJ11 | CATNSGYSTLTF | TRBV2 | TRBJ2-3 | CASSALDSGDLTDTQYF | KLGGALQAK | YES | NO | NO |
| 15083 | TRAV12-1 | TRAJ39 | CVVNGGNAGNMLTF | TRBV4-1 | TRBJ2-7 | CASSQLTGVYEQYF | KLGGALQAK | YES | NO | NO |
| 15084 | TRAV26-2 | TRAJ52 | CILRDENAGGTSYGKLTF | TRBV27 | TRBJ1-5 | CASSLYWGASNQPQHF | KLGGALQAK | NO | NO | NO |
| 15085 | TRAV27 | TRAJ40 | CAGAGYIF | TRBV19 | TRBJ2-1 | CASSFSAGGPYEQFF | KLGGALQAK | YES | NO | NO |
| 15086 | TRAV8-2 | TRAJ44 | CVVSVAGTASKLTF | TRBV4-1 | TRBJ2-2 | CASSQVWPLATTGELFF | KLGGALQAK | YES | NO | NO |
| 15087 | TRAV29DV5 | TRAJ41 | CAASGEGSGYALNF | TRBV13 | TRBJ1-4 | CASSLGQGRKLFF | KLGGALQAK | YES | NO | NO |
| 15088 | TRAV9-2 | TRAJ24 | CALSDPADSWGKFQF | TRBV19 | TRBJ2-3 | CASSIGLAGEVDTQYF | KLGGALQAK | YES | NO | NO |
| 15089 | TRAV1-2 | TRAJ8 | CAVTGFQKLVF | TRBV6-5 | TRBJ1-2 | CASSAKDGNYGYTF | KLGGALQAK | YES | NO | NO |
| 15090 | TRAV12-2 | TRAJ30 | CAVNRDDKIIF | TRBV6-5 | TRBJ1-1 | CASSLTGAEAFF | KLGGALQAK | YES | NO | NO |
| 15091 | TRAV38-2DV8 | TRAJ42 | CAYRDGGSQGNLIF | TRBV4-3 | TRBJ2-3 | CASSQDKVRGLASGKTDTQYF | KLGGALQAK | NO | NO | NO |
| 15092 | TRAV13-1 | TRAJ7 | CAASRGNNRLAF | TRBV6-5 | TRBJ2-1 | CASGSTAYNEQFF | KLGGALQAK | NO | NO | NO |
| 15093 | TRAV19 | TRAJ30 | CALRGYRDDKIIF | TRBV11-1 | TRBJ2-1 | CASSLRGAYNEQFF | KLGGALQAK | YES | NO | NO |
| 15094 | TRAV8-6 | TRAJ22 | CAVSDPESSGSARQLTF | TRBV14 | TRBJ1-2 | CASSYRQNYGYTF | KLGGALQAK | NO | NO | NO |
| 15095 | TRAV12-3 | TRAJ11 | CAMTREMNSGYSTLTF | TRBV7-9 | TRBJ2-1 | CASSQDRGWYNEQFF | KLGGALQAK | YES | NO | NO |
| 15096 | TRAV17 | TRAJ43 | CATENFYNNNDMRF | TRBV10-3 | TRBJ1-6 | CAIKPGQNSPLHF | KLGGALQAK | YES | NO | NO |
| 15097 | TRAV12-1 | TRAJ49 | CVVTITGNQFYF | TRBV6-1 | TRBJ1-2 | CASSAPRGYGYTF | KLGGALQAK | YES | NO | NO |
| 15098 | TRAV19 | TRAJ39 | CALSEADAGNMLTF | TRBV9 | TRBJ2-7 | CASSVGEGTPHSYEQYF | KLGGALQAK | YES | NO | NO |
| 15099 | TRAV17 | TRAJ32 | CATPYGGATNKLIF | TRBV7-3 | TRBJ2-1 | CASSLSGEQFF | KLGGALQAK | YES | NO | NO |
| 15100 | TRAV21 | TRAJ57 | CAVNSQGGSEKLVF | TRBV5-1 | TRBJ2-7 | CASSPDTLLTYEQYF | KLGGALQAK | YES | NO | NO |
| 15101 | TRAV8-4 | TRAJ42 | CAVSDDGGSQGNLIF | TRBV6-3 | TRBJ2-2 | CASSYTFLSLGGQDDTGELFF | KLGGALQAK | NO | NO | NO |
| 15102 | TRAV8-4 | TRAJ47 | CAVSGPEMEYGNKLVF | TRBV7-6 | TRBJ1-1 | CASSLPFSEAFF | KLGGALQAK | YES | NO | NO |
| 15103 | TRAV12-3 | TRAJ49 | CAMRDTGNQFYF | TRBV4-1 | TRBJ1-2 | CASSQAGAVYGYTF | KLGGALQAK | YES | NO | NO |
| 15104 | TRAV14DV4 | TRAJ26 | CAMRENNYGQNFVF | TRBV7-9 | TRBJ1-1 | CASSLAPRNEAFF | KLGGALQAK | YES | NO | NO |
| 15105 | TRAV24 | TRAJ52 | CAFIGDAGGTSYGKLTF | TRBV19 | TRBJ2-7 | CASSFTGGRYEQYF | KLGGALQAK | YES | NO | NO |
| 15106 | TRAV14DV4 | TRAJ37 | CAMRGPYSSNTGKLIF | TRBV6-1 | TRBJ2-3 | CASSEGDTQYF | KLGGALQAK | YES | NO | NO |
| 15107 | TRAV3 | TRAJ4 | CAVRDAFSGGYNKLIF | TRBV12-5 | TRBJ1-4 | CASGFAGDEKLFF | KLGGALQAK | NO | NO | NO |
| 15108 | TRAV27 | TRAJ48 | CAGVFGNEKLTF | TRBV7-8 | TRBJ1-6 | CASSYGGTINSPLHF | AVFDRKSDAK | YES | NO | YES |
| 15109 | TRAV4 | TRAJ27 | CLVGAYNTNAGKSTF | TRBV6-3 | TRBJ2-7 | CASSYGQGAYEQYF | KLGGALQAK | NO | NO | NO |
| 15110 | TRAV1-2 | TRAJ36 | CAVRDAGANNLFF | TRBV7-9 | TRBJ2-7 | CASSSGTEVPEQYF | KLGGALQAK | YES | NO | NO |
| 15111 | TRAV35 | TRAJ49 | CAGQIFTGNQFYF | TRBV5-1 | TRBJ1-1 | CASSATGGDSEAFF | KLGGALQAK | YES | NO | NO |
| 15112 | TRAV20 | TRAJ39 | CAVQARNNAGNMLTF | TRBV9 | TRBJ1-2 | CASSTPGQGARGYTF | KLGGALQAK | YES | YES | NO |
| 15113 | TRAV1-2 | TRAJ30 | CAVRFTRMNRDDKIIF | TRBV4-1 | TRBJ2-5 | CASSPTSGQETQYF | KLGGALQAK | YES | NO | NO |
| 15114 | TRAV8-3 | TRAJ48 | CAVGARGFGNEKLTF | TRBV25-1 | TRBJ2-2 | CASSDEGVGDDGELFF | KLGGALQAK | YES | NO | NO |
| 15115 | TRAV17 | TRAJ26 | CATDYGQNFVF | TRBV9 | TRBJ1-4 | CASSPNGVSVEKLFF | KLGGALQAK | YES | NO | NO |
| 15116 | TRAV14DV4 | TRAJ43 | CAMREGNNNDMRF | TRBV7-6 | TRBJ2-2 | CASSLPVTYTGELFF | KLGGALQAK | YES | NO | NO |
| 15117 | TRAV8-6 | TRAJ27 | CAVHIHDTNAGKSTF | TRBV19 | TRBJ2-1 | CASSIVNGNEQFF | KLGGALQAK | YES | NO | NO |
| 15118 | TRAV13-1 | TRAJ8 | CAVHMNTGFQKLVF | TRBV4-1 | TRBJ2-5 | CASSQGGGQETQYF | KLGGALQAK | YES | NO | NO |
| 15119 | TRAV19 | TRAJ52 | CALSPSNAGGTSYGKLTF | TRBV4-2 | TRBJ2-3 | CASSQDFFRGLPSSDTQYF | KLGGALQAK | YES | NO | NO |
| 15120 | TRAV8-1 | TRAJ23 | CAVNINNQGGKLIF | TRBV7-3 | TRBJ2-3 | CASSLNLDPSTDTQYF | KLGGALQAK | YES | NO | NO |
| 15121 | TRAV38-2DV8 | TRAJ43 | CAYRENDMRF | TRBV19 | TRBJ2-1 | CASSMVAGVYNEQFF | KLGGALQAK | YES | NO | NO |
| 15122 | TRAV17 | TRAJ45 | CATDAWGGGADGLTF | TRBV5-4 | TRBJ1-4 | CASSLGAGALNEKLFF | KLGGALQAK | YES | NO | NO |
| 15123 | TRAV3 | TRAJ21 | CAVRGLYNFNKFYF | TRBV7-8 | TRBJ2-1 | CASSSGSYNEQFF | AVFDRKSDAK | NO | NO | NO |
| 15124 | TRAV27 | TRAJ34 | CAGEDNTDKLIF | TRBV7-8 | TRBJ2-7 | CASSLRRGLWSYEQYF | KLGGALQAK | YES | NO | NO |
| 15125 | TRAV19 | TRAJ42 | CALSEATYGGSQGNLIF | TRBV9 | TRBJ2-7 | CASSVAPQESYEQYF | KLGGALQAK | YES | NO | NO |
| 15126 | TRAV9-2 | TRAJ29 | CALSDFSENTPLVF | TRBV24-1 | TRBJ2-1 | CATSDLQGGGPFNEQFF | KLGGALQAK | YES | NO | NO |
| 15127 | TRAV24 | TRAJ23 | CARREGGKLIF | TRBV5-6 | TRBJ2-5 | CASSLSFLGRETQYF | KLGGALQAK | YES | NO | NO |
| 15128 | TRAV8-1 | TRAJ23 | CAGMIYNQGGKLIF | TRBV4-1 | TRBJ2-7 | CASSQDTGEQYF | KLGGALQAK | YES | NO | NO |
| 15129 | TRAV41 | TRAJ52 | CAVKFLAGGTSYGKLTF | TRBV7-6 | TRBJ2-2 | CASSDYRGDFAGELFF | KLGGALQAK | YES | NO | NO |
| 15130 | TRAV26-2 | TRAJ48 | CIQHNFGNEKLTF | TRBV11-3 | TRBJ2-2 | CASSPTGELFF | KLGGALQAK | YES | NO | NO |
| 15131 | TRAV8-4 | TRAJ21 | CAVGLYNFNKFYF | TRBV14 | TRBJ1-2 | CASSQGQGAYGYTF | KLGGALQAK | NO | NO | NO |
| 15132 | TRAV36DV7 | TRAJ47 | CAVEMREYGNKLVF | TRBV19 | TRBJ2-1 | CASRYGLYNEQFF | KLGGALQAK | NO | NO | NO |
| 15133 | TRAV35 | TRAJ28 | CAGPSGAGSYQLTF | TRBV19 | TRBJ1-1 | CASSILGMSTEAFF | KLGGALQAK | YES | NO | NO |
| 15134 | TRAV26-2 | TRAJ27 | CILAWGNTNAGKSTF | TRBV7-9 | TRBJ2-5 | CASSLAVAETQYF | KLGGALQAK | NO | NO | NO |
| 15135 | TRAV8-4 | TRAJ58 | CAVSDQLASGSRLTF | TRBV7-9 | TRBJ2-2 | CASSDHRWGNTGELFF | KLGGALQAK | YES | NO | NO |
| 15136 | TRAV39 | TRAJ58 | CAAGASGSRLTF | TRBV7-6 | TRBJ2-7 | CASSLASRGAEQYF | KLGGALQAK | YES | NO | NO |
| 15137 | TRAV4 | TRAJ30 | CLVVTNRDDKIIF | TRBV6-4 | TRBJ2-1 | CASSGTPPFGEQFF | KLGGALQAK | YES | NO | NO |
| 15138 | TRAV27 | TRAJ28 | CAAVEPGAGSYQLTF | TRBV27 | TRBJ1-1 | CASSWGGPDTEAFF | KLGGALQAK | YES | NO | NO |
| 15139 | TRAV1-2 | TRAJ33 | CAVLDSNYQLIW | TRBV20-1 | TRBJ2-1 | CSARDPGSNYNEQFF | KLGGALQAK | NO | NO | NO |
| 15140 | TRAV29DV5 | TRAJ37 | CAARHGSSNTGKLIF | TRBV5-5 | TRBJ2-3 | CASSSQPLAGKDTQYF | KLGGALQAK | YES | NO | NO |
| 15141 | TRAV26-2 | TRAJ58 | CIPEETSGSRLTF | TRBV16 | TRBJ1-6 | CASRPTDYNSPLHF | KLGGALQAK | YES | NO | NO |
| 15142 | TRAV2 | TRAJ37 | CAVEDKKPSNTGKLIF | TRBV4-1 | TRBJ1-1 | CASAGPGTGSTEAFF | KLGGALQAK | NO | NO | NO |
| 15143 | TRAV27 | TRAJ44 | CAGADGTASKLTF | TRBV6-5 | TRBJ2-3 | CASSYATGDADTQYF | KLGGALQAK | NO | NO | NO |
| 15144 | TRAV8-3 | TRAJ40 | CAVGWKSGSGTYKYIF | TRBV12-4 | TRBJ2-3 | CASSLAHGSEDTQYF | KLGGALQAK | YES | YES | NO |
| 15145 | TRAV12-2 | TRAJ9 | CAVKGGGFKTIF | TRBV3-1 | TRBJ2-2 | CASSQLPLDTELFF | AVFDRKSDAK | YES | NO | NO |
| 15146 | TRAV5 | TRAJ10 | CAEFTAGGGNKLTF | TRBV5-1 | TRBJ1-6 | CASSSGSVKSSYNSPLHF | KLGGALQAK | YES | NO | NO |
| 15147 | TRAV26-2 | TRAJ39 | CILRDSDAGNMLTF | TRBV11-2 | TRBJ2-7 | CASSTGTGGLIEQYF | KLGGALQAK | YES | NO | NO |
| 15148 | TRAV5 | TRAJ33 | CAEGGSNYQLIW | TRBV7-9 | TRBJ1-1 | CASSIDLEAFF | KLGGALQAK | YES | NO | NO |
| 15149 | TRAV12-1 | TRAJ20 | CVVRISNDYKLSF | TRBV6-3 | TRBJ2-7 | CASSLKGVSSYEQYF | RLRAEAQVK | YES | YES | YES |
| 15150 | TRAV16 | TRAJ50 | CALKTSYDKVIF | TRBV19 | TRBJ1-1 | CASSIAGYVEAFF | KLGGALQAK | NO | NO | NO |
| 15151 | TRAV8-4 | TRAJ52 | CAVPAGGTSYGKLTF | TRBV6-5 | TRBJ2-1 | CASSPLVGTYNEQFF | KLGGALQAK | NO | NO | NO |
| 15152 | TRAV13-2 | TRAJ48 | CAENPLSNFGNEKLTF | TRBV4-1 | TRBJ2-1 | CASSQGSEQFF | AVFDRKSDAK | YES | NO | YES |
| 15153 | TRAV19 | TRAJ40 | CALSEATGTYKYIF | TRBV20-1 | TRBJ2-7 | CSARDHPPSGYYEQYF | KLGGALQAK | YES | NO | NO |
| 15154 | TRAV24 | TRAJ52 | CAFIGDAGGTSYGKLTF | TRBV7-9 | TRBJ2-7 | CASKDGISYEQYF | AVFDRKSDAK | YES | NO | YES |
| 15155 | TRAV17 | TRAJ53 | CATVQNSGGSNYKLTF | TRBV5-1 | TRBJ2-1 | CASSLGNEQFF | KLGGALQAK | YES | NO | NO |
| 15156 | TRAV21 | TRAJ37 | CAVTHSGNTGKLIF | TRBV19 | TRBJ2-7 | CASSIGEGLSYEQYF | KLGGALQAK | YES | NO | NO |
| 15157 | TRAV9-2 | TRAJ56 | CALTSLSPLNTGANSKLTF | TRBV13 | TRBJ2-1 | CASSSGLDKYNEQFF | KLGGALQAK | YES | NO | NO |
| 15158 | TRAV17 | TRAJ49 | CATYRNQFYF | TRBV29-1 | TRBJ2-3 | CSVMGRSITDTQYF | AVFDRKSDAK | YES | NO | YES |
| 15159 | TRAV17 | TRAJ36 | CATDFKANNLFF | TRBV12-4 | TRBJ2-1 | CASTSGRSNEQFF | KLGGALQAK | YES | NO | NO |
| 15160 | TRAV1-2 | TRAJ4 | CAAVSGGYNKLIF | TRBV5-4 | TRBJ1-4 | CASSARDQDEKLFF | KLGGALQAK | YES | NO | NO |
| 15161 | TRAV39 | TRAJ40 | CAVGDGSGTYKYIF | TRBV18 | TRBJ2-1 | CASSPPGADPYNEQFF | KLGGALQAK | YES | NO | NO |
| 15162 | TRAV14DV4 | TRAJ53 | CADSGGSNYKLTF | TRBV10-3 | TRBJ2-3 | CAISDGQGVWTDTQYF | AVFDRKSDAK | YES | NO | NO |
| 15163 | TRAV12-1 | TRAJ21 | CVVNFNKFYF | TRBV27 | TRBJ1-5 | CASSYGRGGQPQHF | AVFDRKSDAK | YES | NO | YES |
| 15164 | TRAV27 | TRAJ54 | CAVTQGAQKLVF | TRBV6-6 | TRBJ2-3 | CASASGSTQYF | KLGGALQAK | NO | NO | NO |
| 15165 | TRAV17 | TRAJ15 | CATDQAGTALIF | TRBV4-1 | TRBJ2-1 | CASSPAPGVDEQFF | KLGGALQAK | YES | NO | NO |
| 15166 | TRAV10 | TRAJ18 | CVVSGPDRGSTLGRLYF | TRBV27 | TRBJ2-7 | CASSLSMIDFYEQYF | KLGGALQAK | YES | NO | NO |
| 15167 | TRAV13-1 | TRAJ49 | CAARAHGDNQFYF | TRBV7-8 | TRBJ2-1 | CASSLGLAGGNEQFF | KLGGALQAK | YES | NO | NO |
| 15168 | TRAV21 | TRAJ11 | CAVDSGYSTLTF | TRBV5-6 | TRBJ2-3 | CASSLVSVADTQYF | KLGGALQAK | YES | NO | NO |
| 15169 | TRAV12-2 | TRAJ30 | CAVNPLRDDKIIF | TRBV27 | TRBJ2-1 | CASSFGPNNEQFF | ELAGIGILTV | YES | YES | YES |
| 15170 | TRAV12-2 | TRAJ4 | CAVNTVGGYNKLIF | TRBV13 | TRBJ1-1 | CASGQDVEAFF | KLGGALQAK | YES | NO | NO |

| | | | | | | | | | | |
|---|---|---|---|---|---|---|---|---|---|---|
| 15171 | TRAV13-1 | TRAJ50 | CAASNKKTSYDKVIF | TRBV6-5 | TRBJ1-2 | CASSYGLPYGYTF | KLGGALQAK | YES | NO | NO |
| 15172 | TRAV29DV5 | TRAJ37 | CAASSSNTGKLIF | TRBV12-4 | TRBJ2-1 | CASRDRGPYNEQFF | KLGGALQAK | YES | NO | NO |
| 15173 | TRAV20 | TRAJ13 | CAVQAGGYQKVTF | TRBV7-9 | TRBJ1-5 | CASSSNRGFQPQHF | KLGGALQAK | YES | NO | NO |
| 15174 | TRAV26-2 | TRAJ43 | CILRGGIYNNNDMRF | TRBV7-9 | TRBJ1-1 | CASSPVPEENTEAFF | KLGGALQAK | YES | NO | NO |
| 15175 | TRAV1-2 | TRAJ24 | CAVRGTDSWGKLQF | TRBV7-2 | TRBJ2-1 | CASRPSSQGLAGGYNEQFF | KLGGALQAK | YES | NO | NO |
| 15176 | TRAV34 | TRAJ32 | CGADNYNKLIF | TRBV4-1 | TRBJ1-4 | CASSQTGPPGEKLFF | KLGGALQAK | YES | NO | NO |
| 15177 | TRAV29DV5 | TRAJ35 | CAARQI | TRBV5-1 | TRBJ1-2 | CASSLGDGYTF | KLGGALQAK | NO | NO | NO |
| 15178 | TRAV38-1 | TRAJ54 | CASQIQGAQKLVF | TRBV7-9 | TRBJ2-3 | CASSVAKGRTGAAQSTDTQYF | KLGGALQAK | NO | NO | NO |
| 15179 | TRAV17 | TRAJ31 | CATGLYNNARLMF | TRBV4-1 | TRBJ1-5 | CASSQGDWIRDGTYSNQPQHF | KLGGALQAK | NO | NO | NO |
| 15180 | TRAV12-1 | TRAJ41 | CVVKINSGYALNF | TRBV5-8 | TRBJ2-4 | CASSLVTSGLNIQYF | KLGGALQAK | YES | NO | NO |
| 15181 | TRAV12-3 | TRAJ33 | CASHSMDSNYQLIW | TRBV5-6 | TRBJ2-3 | CASSSRRVSSTDTQYF | KLGGALQAK | YES | NO | NO |
| 15182 | TRAV25 | TRAJ52 | CAGPGVGGTSYGKLTF | TRBV7-6 | TRBJ2-7 | CASSLEAVTTYEQYF | KLGGALQAK | YES | NO | NO |
| 15183 | TRAV5 | TRAJ45 | CAESRYSGGGADGLTF | TRBV20-1 | TRBJ2-1 | CSARSEGAGQFF | KLGGALQAK | YES | NO | NO |
| 15184 | TRAV27 | TRAJ12 | CAGEMDSSYKLIF | TRBV12-3 | TRBJ2-1 | CASTLFGEQFF | KLGGALQAK | YES | NO | NO |
| 15185 | TRAV3 | TRAJ22 | CAVRALSAGSARQLTF | TRBV27 | TRBJ1-5 | CASSPEYSYSNQPQHF | KLGGALQAK | NO | NO | NO |
| 15186 | TRAV4 | TRAJ39 | CLVGPNNAGNMLTF | TRBV2 | TRBJ2-2 | CASMGIDTGELFF | KLGGALQAK | YES | NO | NO |
| 15187 | TRAV14DV4 | TRAJ43 | CAMREGSNDMRF | TRBV19 | TRBJ2-3 | CASSTGLAGDNTDTQYF | KLGGALQAK | YES | NO | NO |
| 15188 | TRAV12-2 | TRAJ28 | CAVTYSGAGSYQLTF | TRBV20-1 | TRBJ2-1 | CSAPLAGGPDEQFF | KLGGALQAK | NO | NO | NO |
| 15189 | TRAV17 | TRAJ57 | CATDFQGGSEKLVF | TRBV25-1 | TRBJ2-5 | CASSATYQETQYF | KLGGALQAK | YES | NO | NO |
| 15190 | TRAV14DV4 | TRAJ52 | CAMREDAGGTSYGKLTF | TRBV13 | TRBJ1-1 | CASSSHLGSWGTEAFF | YLLEMLWRL | YES | NO | NO |
| 15191 | TRAV38-2DV8 | TRAJ48 | CAYREKLTF | TRBV7-9 | TRBJ2-7 | CASSLADRGLSEQYF | KLGGALQAK | YES | NO | NO |
| 15192 | TRAV8-4 | TRAJ45 | CAVRRYSGGGADGLTF | TRBV24-1 | TRBJ1-2 | CATSGLSASGGYTF | KLGGALQAK | YES | NO | NO |
| 15193 | TRAV20 | TRAJ39 | CAVRGNAGNMLTF | TRBV12-4 | TRBJ1-1 | CASSLTGQYTEAFF | KLGGALQAK | YES | NO | NO |
| 15194 | TRAV1-2 | TRAJ29 | CAALNSGNTPLVF | TRBV25-1 | TRBJ1-2 | CASSEGTGGLGYGYTF | KLGGALQAK | YES | NO | NO |
| 15195 | TRAV8-3 | TRAJ47 | CAVGYGNKLVF | TRBV5-4 | TRBJ1-2 | CASSQNPGQFYGYTF | KLGGALQAK | YES | NO | NO |
| 15196 | TRAV12-2 | TRAJ17 | CAVNVIKAAGNKLTF | TRBV25-1 | TRBJ2-1 | CASQFEEGGVDEQFF | KLGGALQAK | YES | NO | NO |
| 15197 | TRAV17 | TRAJ15 | CATDANQAGTALIF | TRBV3-1 | TRBJ2-2 | CASSGTPGLAYTGELFF | KLGGALQAK | YES | NO | NO |
| 15198 | TRAV1-1 | TRAJ12 | CAGVWMDSSYKLIF | TRBV7-6 | TRBJ2-5 | CASSGVSQRETQYF | KLGGALQAK | YES | NO | NO |
| 15199 | TRAV12-1 | TRAJ23 | CVPGDRQGGKLIF | TRBV2 | TRBJ2-1 | CASRLAGYNEQFF | KLGGALQAK | YES | NO | NO |
| 15200 | TRAV17 | TRAJ11 | CATDRDSGYSTLTF | TRBV6-5 | TRBJ1-2 | CASSYLGGEGYTF | KLGGALQAK | YES | NO | NO |
| 15201 | TRAV8-6 | TRAJ47 | CAVSDGNKLVF | TRBV11-2 | TRBJ2-7 | CASSPNAGTGETYEQYF | KLGGALQAK | YES | NO | NO |
| 15202 | TRAV13-1 | TRAJ30 | CAASRLVGDRDDKIIF | TRBV5-1 | TRBJ2-1 | CASSSTGGTHNEQFF | KLGGALQAK | YES | NO | NO |
| 15203 | TRAV29DV5 | TRAJ23 | CAAVSYNQGGKLIF | TRBV7-8 | TRBJ1-4 | CASSLSLSNEKLFF | KLGGALQAK | YES | NO | NO |
| 15204 | TRAV3 | TRAJ28 | CAVRDPILGSYQLTF | TRBV3-1 | TRBJ2-4 | CASSHTGGPTLAKNIQYF | KLGGALQAK | NO | NO | NO |
| 15206 | TRAV38-2DV8 | TRAJ48 | CAYREANFGNEKLTF | TRBV27 | TRBJ1-1 | CASSLSGGTEAFF | KLGGALQAK | YES | NO | NO |
| 15207 | TRAV3 | TRAJ37 | CAVRVSSNTGKLIF | TRBV11-2 | TRBJ2-1 | CASSLVDEQFF | KLGGALQAK | YES | NO | NO |
| 15208 | TRAV14DV4 | TRAJ54 | CAMRGDITQGAQKLVF | TRBV19 | TRBJ2-5 | CASSHRTGEETQYF | KLGGALQAK | YES | NO | NO |
| 15209 | TRAV29DV5 | TRAJ41 | CAASNSNSGYALNF | TRBV27 | TRBJ2-1 | CASSFWGATYNEQFF | KLGGALQAK | YES | NO | NO |
| 15210 | TRAV8-3 | TRAJ31 | CAVGRNNNARLMF | TRBV11-2 | TRBJ2-1 | CASSLIGGANEQFF | KLGGALQAK | YES | NO | NO |
| 15211 | TRAV8-4 | TRAJ34 | CAVSRAYNTDKLIF | TRBV2 | TRBJ2-2 | CASSEAHWDRGTGELFF | KLGGALQAK | YES | NO | NO |
| 15212 | TRAV24 | TRAJ41 | CAFNSNSGYALNF | TRBV25-1 | TRBJ1-1 | CASSGSPGLTEAFF | KLGGALQAK | YES | NO | NO |
| 15213 | TRAV23DV6 | TRAJ58 | CAASPSGSRLTF | TRBV10-3 | TRBJ2-1 | CAIRRRQDSYNEQFF | KLGGALQAK | NO | NO | NO |
| 15214 | TRAV8-4 | TRAJ20 | CAVIQINDYKLSF | TRBV12-4 | TRBJ1-3 | CASSLTRAGNTIYF | KLGGALQAK | YES | NO | NO |
| 15215 | TRAV38-2DV8 | TRAJ43 | CAYRTYDMRF | TRBV12-3 | TRBJ2-1 | CASSFGGREIGSYNEQFF | KLGGALQAK | YES | NO | NO |
| 15216 | TRAV12-2 | TRAJ54 | CAVIIQGAQKLVF | TRBV29-1 | TRBJ1-1 | CSVEQMNTEAFF | KLGGALQAK | YES | YES | NO |
| 15217 | TRAV29DV5 | TRAJ24 | CAAREDSWGKLQF | TRBV28 | TRBJ2-2 | CASSFLGAGELFF | KLGGALQAK | YES | NO | NO |
| 15218 | TRAV25 | TRAJ26 | CAGYLYGQNFVF | TRBV6-5 | TRBJ1-2 | CASSYGKIYGYTF | KLGGALQAK | YES | NO | NO |
| 15219 | TRAV19 | TRAJ35 | CALII | TRBV24-1 | TRBJ1-5 | CATSEGQSQPQHF | KLGGALQAK | NO | NO | NO |
| 15220 | TRAV14DV4 | TRAJ33 | CAMSDSNYQLIW | TRBV6-5 | TRBJ2-1 | CASREPVGRNEQFF | KLGGALQAK | NO | NO | NO |
| 15221 | TRAV12-1 | TRAJ49 | CVVNVYTGNQFYF | TRBV7-9 | TRBJ2-2 | CASSSTGSPGELFF | AVFDRKSDAK | YES | YES | YES |
| 15222 | TRAV21 | TRAJ48 | CAVRPYFGNEKLTF | TRBV4-1 | TRBJ1-1 | CASSPLKATQNTEAFF | KLGGALQAK | YES | NO | NO |
| 15223 | TRAV13-2 | TRAJ17 | CAETSRKAAGNKLTF | TRBV9 | TRBJ2-1 | CASRGTSGRSSYNEQFF | KLGGALQAK | YES | NO | NO |
| 15224 | TRAV29DV5 | TRAJ21 | CAARISYNFNKFYF | TRBV19 | TRBJ1-1 | CASSIWDFSTEAFF | KLGGALQAK | YES | NO | NO |
| 15225 | TRAV26-2 | TRAJ48 | CILGVPNFGNEKLTF | TRBV29-1 | TRBJ2-1 | CSGIAGNYNEQFF | KLGGALQAK | YES | NO | NO |
| 15226 | TRAV3 | TRAJ34 | CAVRDFPNTDKLIF | TRBV5-6 | TRBJ2-3 | CASSLFSDTQYF | AVFDRKSDAK | NO | NO | NO |
| 15227 | TRAV13-1 | TRAJ40 | CAASGTYKYIF | TRBV5-1 | TRBJ2-7 | CASSLGLAVDEQYF | KLGGALQAK | YES | NO | NO |
| 15228 | TRAV4 | TRAJ10 | CLVGDPPPGGGNKLTF | TRBV27 | TRBJ2-1 | CASSYRDSTLYYNEQFF | KLGGALQAK | YES | NO | NO |
| 15229 | TRAV8-1 | TRAJ26 | CAVNPYGQNFVF | TRBV7-2 | TRBJ2-3 | CASSPSDGRTDTQYF | KLGGALQAK | YES | NO | NO |
| 15230 | TRAV17 | TRAJ53 | CATENSGGSNYKLTF | TRBV4-1 | TRBJ2-7 | CASSFGTGAYEQYF | KLGGALQAK | YES | NO | NO |
| 15231 | TRAV12-1 | TRAJ39 | CVVNNNAGNMLTF | TRBV11-2 | TRBJ2-3 | CASSLDPNTDTQYF | KLGGALQAK | YES | NO | NO |
| 15232 | TRAV30 | TRAJ45 | CGTEDSGGGADGLTF | TRBV19 | TRBJ1-5 | CASSISQGRKYSNQPQHF | KLGGALQAK | YES | NO | NO |
| 15233 | TRAV8-6 | TRAJ34 | CAVTGWDTDKLIF | TRBV19 | TRBJ1-1 | CASSIQTGGTEAFF | AVFDRKSDAK | YES | NO | YES |
| 15234 | TRAV13-2 | TRAJ47 | CAESGHGNKLVF | TRBV2 | TRBJ2-5 | CASIWTSGSQETQYF | KLGGALQAK | YES | NO | NO |
| 15235 | TRAV25 | TRAJ37 | CAGPNSSNTGKLIF | TRBV7-6 | TRBJ2-3 | CASSLVWTRPASTDTQYF | KLGGALQAK | YES | NO | NO |
| 15236 | TRAV25 | TRAJ43 | CAGVNNNDMRF | TRBV3-1 | TRBJ2-1 | CASSQVVYNEQFF | KLGGALQAK | YES | NO | NO |
| 15237 | TRAV29DV5 | TRAJ28 | CAASAGSGAGSYQLTF | TRBV7-9 | TRBJ1-2 | CASSLMGGYTF | KLGGALQAK | YES | NO | NO |
| 15238 | TRAV21 | TRAJ42 | CAANYGGSQGNLIF | TRBV2 | TRBJ2-1 | CATPPGRNYNEQFF | KLGGALQAK | YES | NO | NO |
| 15239 | TRAV12-1 | TRAJ40 | CVEGASGTYKYIF | TRBV12-4 | TRBJ1-1 | CASSFPLQGQGEAFF | KLGGALQAK | YES | NO | NO |
| 15240 | TRAV17 | TRAJ7 | CATEIYGNNRLAF | TRBV5-6 | TRBJ2-2 | CASSTDRGGTGELFF | AVFDRKSDAK | YES | NO | NO |
| 15241 | TRAV17 | TRAJ48 | CAVSMLGNEKLTF | TRBV29-1 | TRBJ1-2 | CSVAQNDYGYTF | KLGGALQAK | YES | NO | NO |
| 15242 | TRAV12-3 | TRAJ24 | CAMMTTDSWGKFQF | TRBV12-4 | TRBJ2-3 | CASSPMDTQYF | KLGGALQAK | YES | NO | NO |
| 15243 | TRAV40 | TRAJ48 | CLAGSNFGNEKLTF | TRBV7-9 | TRBJ1-6 | CASSLPNSPLHF | KLGGALQAK | NO | NO | NO |
| 15244 | TRAV19 | TRAJ22 | CALSEAQGSARQLTF | TRBV19 | TRBJ1-3 | CASSIALVGSGNTIYF | KLGGALQAK | YES | NO | NO |
| 15245 | TRAV1-1 | TRAJ35 | CAPEGFGNVLHC | TRBV27 | TRBJ1-1 | CASVSTEAFF | KLGGALQAK | NO | NO | NO |
| 15246 | TRAV3 | TRAJ37 | CAVRDGGMGSNTGKLIF | TRBV11-2 | TRBJ1-1 | CASSLFQGSGTEAFF | KLGGALQAK | YES | NO | NO |
| 15247 | TRAV16 | TRAJ28 | CALHSGAGSYQLTF | TRBV5-1 | TRBJ1-1 | CASHGGGTEAFF | RLRAEAQVK | NO | NO | NO |
| 15248 | TRAV17 | TRAJ30 | CARGGGDDKIIF | TRBV6-1 | TRBJ1-5 | CASSEQGASNQPQHF | KLGGALQAK | YES | NO | NO |
| 15249 | TRAV12-2 | TRAJ3 | CAVGYSGYSSASKIIF | TRBV25-1 | TRBJ2-5 | CASSEGISNQETQYF | KLGGALQAK | YES | NO | NO |
| 15250 | TRAV1-2 | TRAJ24 | CAVSKPPLRLWTTDSWGKLQF | TRBV19 | TRBJ2-2 | CASSIDMGLTGELFF | KLGGALQAK | NO | NO | NO |
| 15251 | TRAV8-3 | TRAJ47 | CAVGAGGKLVF | TRBV7-2 | TRBJ2-7 | CASSSAGYEQYF | AVFDRKSDAK | YES | NO | YES |
| 15252 | TRAV29DV5 | TRAJ42 | CAASARYGGSQGNLIF | TRBV9 | TRBJ2-3 | CASSVDGGTQYF | KLGGALQAK | YES | NO | NO |
| 15253 | TRAV30 | TRAJ57 | CGTGQGGSEKLVF | TRBV7-9 | TRBJ1-3 | CASSLNPRVGNTIYF | KLGGALQAK | YES | NO | NO |
| 15254 | TRAV4 | TRAJ26 | CLVGAGNYGQNFVF | TRBV7-2 | TRBJ2-6 | CASSLPAGRFSGANVLTF | KLGGALQAK | NO | NO | NO |
| 15255 | TRAV14DV4 | TRAJ5 | CAMRDPQGGRRALTF | TRBV3-1 | TRBJ2-5 | CASSHGKTQYF | RTLNAWVKV | YES | YES | YES |
| 15256 | TRAV8-1 | TRAJ31 | CAVIAGARLMF | TRBV12-5 | TRBJ2-1 | CASGFDWNNEQFF | KLGGALQAK | YES | NO | NO |
| 15257 | TRAV8-1 | TRAJ9 | CAVNASPGGFKTIF | TRBV4-1 | TRBJ2-5 | CASSHPAGGEETQYF | AVFDRKSDAK | YES | NO | NO |
| 15258 | TRAV12-2 | TRAJ30 | CAVLNRDDKIIF | TRBV15 | TRBJ2-3 | CATSRITSSTQYF | KLGGALQAK | YES | NO | NO |
| 15259 | TRAV13-2 | TRAJ9 | CAENTGGFKTIF | TRBV27 | TRBJ2-3 | CASSEGDTQYF | KLGGALQAK | YES | NO | NO |
| 15260 | TRAV35 | TRAJ20 | CAGHDGNDYKLSF | TRBV5-1 | TRBJ1-2 | CASSLETENYGYTF | IVTDFSVIK | YES | YES | NO |
| 15261 | TRAV19 | TRAJ20 | CALSADLDDYKLSF | TRBV12-3 | TRBJ2-1 | CASRGLAGEPSYNEQFF | KLGGALQAK | YES | NO | NO |
| 15262 | TRAV1-1 | TRAJ4 | CAVRGGYDYNKLIF | TRBV10-2 | TRBJ1-1 | CATHERGNTEAFF | KLGGALQAK | YES | NO | NO |
| 15263 | TRAV8-1 | TRAJ52 | CAVNGVGAGGTSYGKLTF | TRBV27 | TRBJ2-7 | CASSWGVSTYF | KLGGALQAK | YES | NO | NO |
| 15264 | TRAV23DV6 | TRAJ24 | CAAMSDSWGKFQF | TRBV10-2 | TRBJ2-7 | CASSEDGDYEQYV | KLGGALQAK | NO | NO | NO |
| 15265 | TRAV5 | TRAJ6 | CAESKEGPGGSYIPTF | TRBV24-1 | TRBJ2-3 | CATSPTETDTQYF | KLGGALQAK | YES | NO | NO |
| 15266 | TRAV12-2 | TRAJ39 | CAVSRAGNMLTF | TRBV4-3 | TRBJ2-1 | CASSQDSGVAYEQFF | KLGGALQAK | NO | NO | NO |
| 15267 | TRAV4 | TRAJ3 | CLVGAPGYSSASKIIF | TRBV6-1 | TRBJ2-5 | CASSVVGANQETQYF | KLGGALQAK | YES | NO | NO |
| 15268 | TRAV5 | TRAJ26 | CAPRGDNYGQNFVF | TRBV5-6 | TRBJ1-4 | CASIAGPGTGGEKLFF | KLGGALQAK | YES | NO | NO |
| 15269 | TRAV25 | TRAJ40 | CAGTTSGTYKYIF | TRBV5-1 | TRBJ2-1 | CASSLGQGDWGEQFF | KLGGALQAK | YES | NO | NO |
| 15270 | TRAV22 | TRAJ52 | CAVRPSNAGGTSYGKLTF | TRBV18 | TRBJ2-3 | CASSPRASGTSTDTQYF | KLGGALQAK | YES | NO | NO |
| 15271 | TRAV19 | TRAJ16 | CALSGVF | TRBV4-1 | TRBJ1-2 | CASSQAADRGNYGYTF | KLGGALQAK | NO | NO | NO |
| 15272 | TRAV29DV5 | TRAJ28 | CAASAAGAGSYQLTF | TRBV11-2 | TRBJ2-2 | CASSLAAGQGNGGDTGELFF | KLGGALQAK | NO | NO | NO |
| 15273 | TRAV1-2 | TRAJ30 | CAVRGDDKIIF | TRBV19 | TRBJ1-1 | CASSIPQKAFF | KLGGALQAK | YES | NO | NO |
| 15274 | TRAV1-2 | TRAJ31 | CAVPNNNARLMF | TRBV29-1 | TRBJ1-4 | CSVEEAGGNEKLFF | AVFDRKSDAK | YES | YES | YES |
| 15275 | TRAV29DV5 | TRAJ52 | CAASDFHAGGTSYGKLTF | TRBV9 | TRBJ2-5 | CASSVGAFQETQYF | KLGGALQAK | YES | NO | NO |

| | | | | | | | | | |
|---|---|---|---|---|---|---|---|---|---|
| 15276 | TRAV29DV5 | TRAJ50 | CAANSMKTSYDKVIF | TRBV19 | TRBJ1-1 | CASSMGVNTEAFF | KLGGALQAK | YES | NO | NO |
| 15277 | TRAV20 | TRAJ17 | CAVKRAAGNKLTF | TRBV5-1 | TRBJ1-1 | CQSSNTEAFF | KLGGALQAK | YES | NO | NO |
| 15278 | TRAV25 | TRAJ20 | CAANNDYKLSF | TRBV7-9 | TRBJ2-2 | CASRARGQGPTGELFF | KLGGALQAK | YES | NO | NO |
| 15279 | TRAV17 | TRAJ36 | CATDLQTGANNLFF | TRBV19 | TRBJ1-4 | CASRVQMENEKLFF | KLGGALQAK | YES | NO | NO |
| 15280 | TRAV25 | TRAJ26 | CAAADNYGQNFVF | TRBV30 | TRBJ2-3 | CAWSSGEGTDTQYF | AVFDRKSDAK | YES | NO | NO |
| 15281 | TRAV29DV5 | TRAJ23 | CAAPATYNQGGKLIF | TRBV18 | TRBJ2-7 | CASSPIQMDQSGSYEQYF | KLGGALQAK | YES | NO | NO |
| 15282 | TRAV19 | TRAJ56 | CALSEARSGANSKLTF | TRBV9 | TRBJ2-3 | CASSARLDTQYF | KLGGALQAK | YES | NO | NO |
| 15283 | TRAV35 | TRAJ48 | CAGQPHHISNFGNEKLTF | TRBV19 | TRBJ2-1 | CASSIFSAGGKDEQFF | GILGFVFTL | YES | NO | YES |
| 15284 | TRAV13-2 | TRAJ39 | CAENMRAGNMLTF | TRBV6-5 | TRBJ1-5 | CASSSFMGNQPQHF | KLGGALQAK | YES | NO | NO |
| 15285 | TRAV26-1 | TRAJ54 | CIVRVADPIQGAQKLVF | TRBV9 | TRBJ2-5 | CASSVNWAGPPQETQYF | AVFDRKSDAK | YES | NO | YES |
| 15286 | TRAV1-2 | TRAJ33 | CAVMDSNYQLIW | TRBV27 | TRBJ1-1 | CASSLAGGQTEAFF | KLGGALQAK | YES | NO | NO |
| 15287 | TRAV8-3 | TRAJ43 | CAVVSDNNNDMRF | TRBV7-9 | TRBJ1-2 | CASRKETGLLYGYTF | KLGGALQAK | YES | NO | NO |
| 15288 | TRAV19 | TRAJ33 | CALSETMDSNYQLIW | TRBV11-2 | TRBJ2-5 | CASSLIFPGDLPHQETQYF | KLGGALQAK | YES | NO | NO |
| 15289 | TRAV3 | TRAJ40 | CAVRATSGTYKYIF | TRBV6-3 | TRBJ2-1 | CASIGRQLFF | KLGGALQAK | NO | NO | NO |
| 15290 | TRAV41 | TRAJ30 | CAALDDKIIF | TRBV27 | TRBJ2-1 | CASSFGLLAGGNEQFF | KLGGALQAK | YES | NO | NO |
| 15291 | TRAV9-2 | TRAJ43 | CALRIAPDMRF | TRBV12-4 | TRBJ1-2 | CASRSYRNYGYTF | KLGGALQAK | YES | NO | NO |
| 15292 | TRAV8-3 | TRAJ30 | CAHLNRDDKIIF | TRBV20-1 | TRBJ1-4 | CSARDFRSPTNEKLFF | KLGGALQAK | YES | NO | NO |
| 15293 | TRAV8-1 | TRAJ20 | CAVREGDYKLSF | TRBV7-7 | TRBJ2-1 | CASSPTRLGWHNEQFF | KLGGALQAK | YES | NO | NO |
| 15294 | TRAV12-2 | TRAJ48 | CAVNVNFGNEKLTF | TRBV5-1 | TRBJ2-7 | CASSGWGGLAGDLFHEQYF | KLGGALQAK | YES | YES | NO |
| 15295 | TRAV8-1 | TRAJ33 | CAVKGVSNYQLIW | TRBV2 | TRBJ2-1 | CASSVIAGGPYNEQFF | KLGGALQAK | YES | NO | NO |
| 15296 | TRAV24 | TRAJ26 | CAYNNYGQNFVF | TRBV9 | TRBJ2-5 | CASSPGSRREETQYF | KLGGALQAK | YES | NO | NO |
| 15297 | TRAV12-3 | TRAJ40 | CAMCAPGTYKYIF | TRBV19 | TRBJ2-7 | CASSTGTGLPYEQYF | RLRAEAQVK | YES | YES | YES |
| 15298 | TRAV8-4 | TRAJ29 | CAVSEAGNTPLVF | TRBV6-3 | TRBJ2-7 | CASRGYPSYEQYV | KLGGALQAK | NO | NO | NO |
| 15299 | TRAV14DV4 | TRAJ22 | CAMRRPISSGSARQLTF | TRBV12-5 | TRBJ2-2 | CASGLAGVGELFF | AVFDRKSDAK | YES | NO | YES |
| 15300 | TRAV19 | TRAJ32 | CALSEESGEVIF | TRBV10-1 | TRBJ2-1 | CASSDFGISGTYNEQFF | KLGGALQAK | YES | NO | NO |
| 15301 | TRAV21 | TRAJ28 | CAVDGRYSGAGSYQLTF | TRBV5-1 | TRBJ1-1 | CASSLDRGTEAFF | KLGGALQAK | YES | NO | NO |
| 15302 | TRAV38-2DV8 | TRAJ34 | CAYRSARFTDKLIF | TRBV4-2 | TRBJ2-2 | CASSPSLDAGTGELFF | KLGGALQAK | YES | NO | NO |
| 15303 | TRAV3 | TRAJ45 | CAVREGSGGGADGLTF | TRBV4-1 | TRBJ2-7 | CASSQDRGVRQYF | KLGGALQAK | YES | NO | NO |
| 15304 | TRAV40 | TRAJ42 | CLLVDYGGSQGNLIF | TRBV7-2 | TRBJ2-2 | CASSFVMFHTGELFF | KLGGALQAK | NO | NO | NO |
| 15305 | TRAV12-1 | TRAJ17 | CVAGGAAGNKLTF | TRBV7-3 | TRBJ2-7 | CASSPRDSPFEQYF | KLGGALQAK | NO | NO | NO |
| 15306 | TRAV12-3 | TRAJ9 | CAMKPATGGFKTIF | TRBV27 | TRBJ2-3 | CASSPDWDTQYF | KLGGALQAK | YES | NO | NO |
| 15307 | TRAV38-2DV8 | TRAJ30 | CAYRSSIEIIF | TRBV6-3 | TRBJ2-1 | CASSYGAGGYNEQFF | KLGGALQAK | YES | NO | NO |
| 15308 | TRAV29DV5 | TRAJ54 | CAAQAGYQGAQKLVF | TRBV12-5 | TRBJ2-4 | CASGSSFGDNIQYF | KLGGALQAK | YES | NO | NO |
| 15309 | TRAV1-2 | TRAJ9 | CAVRGTGGFKTIF | TRBV5-5 | TRBJ1-2 | CASSSEQGTQRGYTF | KLGGALQAK | YES | NO | NO |
| 15310 | TRAV27 | TRAJ34 | CAGYHNTDKLIF | TRBV10-3 | TRBJ1-1 | CAILGNSGTEAFF | AVFDRKSDAK | YES | NO | NO |
| 15311 | TRAV8-6 | TRAJ21 | CAVSANNFNKFYF | TRBV20-1 | TRBJ1-2 | CSARRTGALYGYTF | KLGGALQAK | YES | NO | NO |
| 15312 | TRAV21 | TRAJ41 | CAVINSNSGYALNF | TRBV14 | TRBJ1-1 | CASSPPLFRDTEAFF | KLGGALQAK | NO | NO | NO |
| 15313 | TRAV4 | TRAJ34 | CLVVGTDKLIF | TRBV15 | TRBJ2-7 | CATRGQGVWEQYF | KLGGALQAK | YES | NO | NO |
| 15314 | TRAV8-3 | TRAJ20 | CAVGGGDYKLSF | TRBV19 | TRBJ1-5 | CASSTGTSGVQPQHF | RLRAEAQVK | YES | YES | YES |
| 15315 | TRAV21 | TRAJ9 | CAVRRDTGGFKTIF | TRBV13 | TRBJ1-6 | CASSLGRTENSPLHF | KLGGALQAK | YES | NO | NO |
| 15316 | TRAV3 | TRAJ20 | CAVRGSNDYKLSF | TRBV29-1 | TRBJ2-1 | CSVETGYSYNEQFF | KLGGALQAK | NO | NO | NO |
| 15317 | TRAV8-2 | TRAJ16 | CVVTRF | TRBV14 | TRBJ1-2 | CASSQEGEPNYGYTF | KLGGALQAK | NO | NO | NO |
| 15318 | TRAV29DV5 | TRAJ40 | CAADTSGTYKYIF | TRBV7-9 | TRBJ1-1 | CASSATQGATEAFF | IVTDFSVIK | YES | NO | YES |
| 15319 | TRAV12-2 | TRAJ48 | CAVRSNFGNEKLTF | TRBV29-1 | TRBJ2-3 | CSVELAGSTDTQYF | AVFDRKSDAK | YES | NO | YES |
| 15320 | TRAV23DV6 | TRAJ40 | CAASSGTYKYIF | TRBV7-8 | TRBJ2-1 | CASSPEGPIYNEQFF | KLGGALQAK | YES | NO | NO |
| 15321 | TRAV3 | TRAJ16 | CAVRPDGQKLLF | TRBV24-1 | TRBJ1-1 | CATSDSVSGEAFF | KLGGALQAK | NO | NO | NO |
| 15322 | TRAV19 | TRAJ41 | CALSEGSGYALNF | TRBV4-1 | TRBJ1-5 | CASSQEFRADQPQHF | KLGGALQAK | YES | NO | NO |
| 15323 | TRAV12-1 | TRAJ11 | CVVNEGSTLTF | TRBV3-1 | TRBJ2-3 | CASSQGLAGVLADTQYF | KLGGALQAK | YES | NO | NO |
| 15324 | TRAV21 | TRAJ24 | CAVKPDSWGKLQF | TRBV4-1 | TRBJ1-2 | CASSQIPDRDYGYTF | KLGGALQAK | YES | NO | NO |
| 15325 | TRAV29DV5 | TRAJ22 | CAAGISGSARQLTF | TRBV7-2 | TRBJ2-7 | CASSLAWGTLPPYEQYV | KLGGALQAK | NO | NO | NO |
| 15326 | TRAV21 | TRAJ34 | CAVIGWTDKLIF | TRBV20-1 | TRBJ2-1 | CSAEFPGAVYNEQFF | KLGGALQAK | NO | NO | NO |
| 15327 | TRAV27 | TRAJ57 | CAGSQGGSEKLVF | TRBV19 | TRBJ1-2 | CASSSLHYGYTF | KLGGALQAK | YES | NO | NO |
| 15328 | TRAV12-2 | TRAJ9 | CAVNFNTGGFKTIF | TRBV3-1 | TRBJ2-1 | CASSPGLAGGYEQFF | KLGGALQAK | YES | NO | NO |
| 15329 | TRAV17 | TRAJ12 | CATDLMDSSYKLIF | TRBV24-1 | TRBJ2-6 | CATSDRPGGANVLTF | KLGGALQAK | YES | NO | NO |
| 15330 | TRAV1-2 | TRAJ24 | CAVRDDRDSWGKLQF | TRBV20-1 | TRBJ1-3 | CSARPDRAVGNTIYF | KLGGALQAK | YES | NO | NO |
| 15331 | TRAV24 | TRAJ49 | CAFPNTGNQFYF | TRBV7-8 | TRBJ1-4 | CASSSTLFNVEEKLFF | KLGGALQAK | YES | NO | NO |
| 15332 | TRAV13-1 | TRAJ36 | CAATPPRGANNLFF | TRBV12-3 | TRBJ1-4 | CASGPVQDLTGINEKLFF | KLGGALQAK | YES | NO | NO |
| 15333 | TRAV41 | TRAJ30 | CAVWAYRDDKIIF | TRBV20-1 | TRBJ2-7 | CSARDKYSYEQYV | KLGGALQAK | NO | NO | NO |
| 15334 | TRAV1-2 | TRAJ16 | CAVGSDGQKLLF | TRBV5-1 | TRBJ2-5 | CASGLEGRWQETQYF | KLGGALQAK | NO | NO | NO |
| 15335 | TRAV12-3 | TRAJ34 | CAMSSTDKLIF | TRBV12-4 | TRBJ2-7 | CASSFPGTAPYEQYF | KLGGALQAK | YES | NO | NO |
| 15336 | TRAV29DV5 | TRAJ54 | CAGWGGGAQKLVF | TRBV30 | TRBJ2-1 | CAWTSGLGNEQFF | KLGGALQAK | YES | NO | NO |
| 15337 | TRAV12-2 | TRAJ53 | CAVNSGGSNYKLTF | TRBV4-3 | TRBJ2-1 | CASSQDNWSDYNEQFF | KLGGALQAK | YES | NO | NO |
| 15338 | TRAV22 | TRAJ48 | CAVELLAGNEKLTF | TRBV7-2 | TRBJ2-7 | CASSFRSATHTYEQYF | KLGGALQAK | YES | NO | NO |
| 15339 | TRAV12-1 | TRAJ31 | CVVSPYARLMF | TRBV14 | TRBJ2-2 | CASTLTGSTGELFF | KLGGALQAK | NO | NO | NO |
| 15340 | TRAV26-1 | TRAJ45 | CIVRVPMYSGGGADGLTF | TRBV27 | TRBJ2-1 | CASSLTSGNNEQFF | KLGGALQAK | NO | NO | NO |
| 15341 | TRAV5 | TRAJ4 | CADLSGGYNKLIF | TRBV18 | TRBJ1-1 | CASSPPHRGTEAFF | KLGGALQAK | YES | NO | NO |
| 15342 | TRAV14DV4 | TRAJ43 | CAMREGANNDMRF | TRBV7-9 | TRBJ2-3 | CASSALGTSLADTQYF | KLGGALQAK | YES | NO | NO |
| 15343 | TRAV4 | TRAJ37 | CLVGVYPSNTGKLIF | TRBV21-1 | TRBJ2-6 | CASSKAETFPSSGANVLTF | KLGGALQAK | NO | NO | NO |
| 15344 | TRAV12-2 | TRAJ20 | CAASNDYKLSF | TRBV15 | TRBJ1-1 | CATGWGNTEAFF | KLGGALQAK | YES | NO | NO |
| 15345 | TRAV39 | TRAJ30 | CAVDMPMNRDDKIIF | TRBV6-5 | TRBJ1-5 | CASSYTWLGDQPQHF | KLGGALQAK | YES | NO | NO |
| 15346 | TRAV1-2 | TRAJ6 | CAVSTSGGSYIPTF | TRBV4-1 | TRBJ2-6 | CASSERDYLDSGANVLTF | KLGGALQAK | YES | NO | NO |
| 15347 | TRAV19 | TRAJ17 | CALSEAKAAGNKLTF | TRBV5-1 | TRBJ2-1 | CASSLWEGRSYNEQFF | KLGGALQAK | YES | NO | NO |
| 15349 | TRAV39 | TRAJ42 | CAVDPNYGGSQGNLIF | TRBV6-6 | TRBJ2-7 | CASFFPYEQYF | KLGGALQAK | NO | NO | NO |
| 15350 | TRAV29DV5 | TRAJ24 | CAASSTDSWGKFQF | TRBV4-1 | TRBJ2-1 | CASRTGQGAFEQFF | KLGGALQAK | YES | NO | NO |
| 15351 | TRAV12-3 | TRAJ50 | CAMKKTSYDKVIF | TRBV7-9 | TRBJ1-1 | CASSSGQGAKTEAFF | KLGGALQAK | YES | NO | NO |
| 15352 | TRAV26-1 | TRAJ38 | CIVRPRYNAGNNRKLIW | TRBV11-2 | TRBJ2-5 | CASSLSSGGAWETQYF | KLGGALQAK | YES | NO | NO |
| 15353 | TRAV27 | TRAJ43 | CAGARNNNDMRF | TRBV6-3 | TRBJ2-7 | CASSYHMRNYEQYF | KLGGALQAK | YES | NO | NO |
| 15354 | TRAV9-2 | TRAJ26 | CALSEKNYGQNFVF | TRBV4-3 | TRBJ2-5 | CASSQEFGGRGLEETQYF | AVFDRKSDAK | YES | NO | YES |
| 15355 | TRAV20 | TRAJ15 | CAVPGNQAGTALIF | TRBV9 | TRBJ2-3 | CASSAFSGTTDTQYF | AVFDRKSDAK | YES | NO | YES |
| 15356 | TRAV9-2 | TRAJ43 | CALSDYDNNDMRF | TRBV15 | TRBJ1-6 | CATRSTGTLITDSPLHF | KLGGALQAK | YES | NO | NO |
| 15357 | TRAV8-3 | TRAJ15 | CAVVGTALIF | TRBV9 | TRBJ1-2 | CASSADKLYGYTF | KLGGALQAK | YES | NO | NO |
| 15358 | TRAV17 | TRAJ52 | CATDAGDAGGTSYGKLTF | TRBV19 | TRBJ1-5 | CASSIIGSGNQPQHF | KLGGALQAK | YES | NO | NO |
| 15359 | TRAV1-2 | TRAJ31 | CAVSNNARLMF | TRBV12-3 | TRBJ2-5 | CASSLKMAIQETQYF | KLGGALQAK | YES | NO | NO |
| 15360 | TRAV12-3 | TRAJ31 | CAMSEFLRVYARLMF | TRBV9 | TRBJ1-1 | CASSGQGANTEAFF | KLGGALQAK | YES | NO | NO |
| 15361 | TRAV17 | TRAJ29 | CATDVLGNTPLVF | TRBV5-4 | TRBJ2-5 | CASSLVGRPSVYQETQYF | KLGGALQAK | YES | NO | NO |
| 15362 | TRAV29DV5 | TRAJ40 | CAASARNSGTYKYIF | TRBV7-9 | TRBJ2-3 | CASSFEVGGSTDTQYF | KLGGALQAK | YES | NO | NO |
| 15363 | TRAV8-3 | TRAJ37 | CVEDRPGNTGKLIF | TRBV6-1 | TRBJ2-1 | CASSEAGTNYYNEQFF | KLGGALQAK | YES | NO | NO |
| 15364 | TRAV29DV5 | TRAJ47 | CAASSMEYGNKLVF | TRBV12-3 | TRBJ2-3 | CASSYLSASTDTQYF | KLGGALQAK | YES | NO | NO |
| 15365 | TRAV19 | TRAJ20 | CALSGGERSNDYKLSF | TRBV9 | TRBJ2-2 | CASSARGLAGELFF | KLGGALQAK | YES | NO | NO |
| 15366 | TRAV24 | TRAJ40 | CALTGTYKYIF | TRBV5-4 | TRBJ2-3 | CASSLPTSGGDTQYF | KLGGALQAK | YES | NO | NO |
| 15367 | TRAV1-2 | TRAJ26 | CAVFLYGQNFVF | TRBV12-4 | TRBJ2-3 | CASHPQSGSWSDTQYF | KLGGALQAK | YES | NO | NO |
| 15368 | TRAV38-1 | TRAJ56 | CAFMKPYTGANSKLTF | TRBV9 | TRBJ2-7 | CASSVGQDYEQYF | KLGGALQAK | YES | NO | NO |
| 15369 | TRAV23DV6 | TRAJ7 | CAASPYGNNRLAF | TRBV20-1 | TRBJ1-4 | CSSPGGPNEKLFF | RLRAEAQVK | YES | YES | YES |
| 15370 | TRAV20 | TRAJ17 | CAVQAGIKAAGNKLTF | TRBV7-9 | TRBJ1-1 | CASSSPQSGNTEAFF | KLGGALQAK | YES | NO | NO |
| 15371 | TRAV17 | TRAJ45 | CATDSGGGADGLTF | TRBV5-8 | TRBJ2-2 | CASTEDRADTGELFF | KLGGALQAK | YES | NO | NO |
| 15372 | TRAV17 | TRAJ30 | CATVTRNRDDKIIF | TRBV19 | TRBJ2-7 | CASSSGTGDVYEQYF | KLGGALQAK | YES | NO | NO |
| 15373 | TRAV8-3 | TRAJ28 | CAVGGRAGSYQLTF | TRBV9 | TRBJ2-3 | CASSWIDTQYF | KLGGALQAK | YES | NO | NO |
| 15374 | TRAV12-2 | TRAJ16 | CAVNVPSDGQKLLF | TRBV12-4 | TRBJ2-2 | CASAPHGGPINTGELFF | KLGGALQAK | NO | NO | NO |
| 15375 | TRAV38-2DV8 | TRAJ52 | CAYRRGRNAGGTSYGKLTF | TRBV30 | TRBJ1-2 | CAWGSGLGYGYTF | KLGGALQAK | YES | NO | NO |
| 15376 | TRAV30 | TRAJ8 | CGTEIVQKLVF | TRBV7-9 | TRBJ1-6 | CASNPTGAYNSPLHF | KLGGALQAK | YES | NO | NO |
| 15377 | TRAV19 | TRAJ34 | CALGRDTDKLIF | TRBV27 | TRBJ2-1 | CASRLRTGNYNEQFF | KLGGALQAK | YES | NO | NO |
| 15378 | TRAV9-2 | TRAJ20 | CALSEPPVENDYKLSF | TRBV19 | TRBJ1-3 | CASSRWTIYF | KLGGALQAK | YES | NO | NO |
| 15379 | TRAV6 | TRAJ22 | CALLIISSGSARQLTF | TRBV7-8 | TRBJ2-3 | CASSLNGGLLVRTDTQYF | KLGGALQAK | YES | NO | NO |
| 15380 | TRAV1-2 | TRAJ12 | CAVMDSSYKLIF | TRBV4-3 | TRBJ2-6 | CASSQVAGSGANVLTF | AVFDRKSDAK | YES | NO | NO |

| | | | | | | | | | | |
|---|---|---|---|---|---|---|---|---|---|---|
| 15381 | TRAV21 | TRAJ37 | CAVLSGNTGKLIF | TRBV12-4 | TRBJ2-2 | CASSLGTWELFF | KLGGALQAK | YES | NO | NO |
| 15382 | TRAV13-2 | TRAJ11 | CAESGYSTLTF | TRBV6-6 | TRBJ1-2 | CASALEGVSYGYTF | KLGGALQAK | NO | NO | NO |
| 15383 | TRAV3 | TRAJ41 | CAVSAANSNSGYALNF | TRBV7-9 | TRBJ2-7 | CASSGQTAFPSLRSEQYF | KLGGALQAK | NO | NO | NO |
| 15384 | TRAV38-2DV8 | TRAJ21 | CAYRSGVYHNFNKFYF | TRBV27 | TRBJ2-2 | CASSLGGTGELFF | KLGGALQAK | YES | NO | NO |
| 15385 | TRAV41 | TRAJ48 | CAAPYFGNEKLTF | TRBV7-9 | TRBJ2-7 | CASSSPVGYEQYF | KLGGALQAK | YES | NO | NO |
| 15386 | TRAV13-2 | TRAJ40 | CADTASGTYKYIF | TRBV12-4 | TRBJ2-2 | CASSFPGELFF | KLGGALQAK | YES | NO | NO |
| 15387 | TRAV14DV4 | TRAJ30 | CAMRRLNRDDKIIF | TRBV9 | TRBJ2-1 | CASSVVESGNYNEQFF | KLGGALQAK | YES | NO | NO |
| 15388 | TRAV8-3 | TRAJ29 | CAVGLSSGNTPLVF | TRBV2 | TRBJ2-1 | CASSPSAGVPYNEQFF | KLGGALQAK | YES | NO | NO |
| 15389 | TRAV27 | TRAJ24 | CAGGDPTDSWGKLQF | TRBV19 | TRBJ1-5 | CASSSARTQHF | KLGGALQAK | YES | NO | NO |
| 15390 | TRAV9-2 | TRAJ57 | CALSELGGSEKLVF | TRBV13 | TRBJ2-1 | CASSLGTSGIGNEQFF | KLGGALQAK | YES | NO | NO |
| 15391 | TRAV12-2 | TRAJ4 | CAVNRHGGYNKLIF | TRBV13 | TRBJ1-1 | CASSGDRGVDAFF | KLGGALQAK | YES | NO | NO |
| 15392 | TRAV17 | TRAJ43 | CATDRDNNDMRF | TRBV4-1 | TRBJ2-1 | CASSQDWGGSYNEQFF | KLGGALQAK | YES | NO | NO |
| 15393 | TRAV17 | TRAJ47 | CATPGTMEYGNKLVF | TRBV12-4 | TRBJ1-6 | CASSLAPGGSNSPLHF | KLGGALQAK | YES | NO | NO |
| 15394 | TRAV9-2 | TRAJ30 | CALSDLGDRDDKIIF | TRBV14 | TRBJ2-2 | CASSQVFGSSNTGELFF | KLGGALQAK | NO | NO | NO |
| 15395 | TRAV12-1 | TRAJ17 | CVVKAAGNKLTF | TRBV27 | TRBJ1-5 | CASSYSDSNQPQHF | KLGGALQAK | YES | NO | NO |
| 15396 | TRAV38-2DV8 | TRAJ29 | CAYRGLNSGNTPLVF | TRBV4-1 | TRBJ2-7 | CASSQKEGGGSYEQYV | KLGGALQAK | NO | NO | NO |
| 15397 | TRAV14DV4 | TRAJ34 | CAMREGWRNTDKLIF | TRBV7-2 | TRBJ1-5 | CASRIGNQPQHF | KLGGALQAK | YES | NO | NO |
| 15398 | TRAV25 | TRAJ47 | CAGYYMEYGNKLVF | TRBV5-4 | TRBJ2-1 | CASSNLGTGNFF | KLGGALQAK | YES | NO | NO |
| 15399 | TRAV24 | TRAJ10 | CAFNAGGGNKLTF | TRBV3-1 | TRBJ2-1 | CASSQGRTGYNEQFF | KLGGALQAK | YES | NO | NO |
| 15400 | TRAV35 | TRAJ16 | CAGQLPRF | TRBV27 | TRBJ2-2 | CASRLAGSTGELFF | KLGGALQAK | NO | NO | NO |
| 15401 | TRAV12-2 | TRAJ54 | CAVNTGAQKLVF | TRBV29-1 | TRBJ2-7 | CLTGAYEQYF | AVFDRKSDAK | NO | NO | NO |
| 15402 | TRAV27 | TRAJ23 | CAGGHTYNQGGKLIF | TRBV2 | TRBJ2-1 | CATRDGLSDEQFF | KLGGALQAK | YES | NO | NO |
| 15403 | TRAV14DV4 | TRAJ4 | CAMRPPFRGGYNKLIF | TRBV15 | TRBJ2-4 | CATSRGPTSGTGSAKNIQYF | KLGGALQAK | NO | NO | NO |
| 15404 | TRAV12-1 | TRAJ20 | CVVTPDYKLSF | TRBV20-1 | TRBJ1-1 | CSARMGDTEAFF | KLGGALQAK | YES | NO | NO |
| 15405 | TRAV14DV4 | TRAJ52 | CATLAGGTSYGKLTF | TRBV4-1 | TRBJ2-7 | CASSQYPTSGTSEHEQYF | KLGGALQAK | YES | NO | NO |
| 15406 | TRAV8-3 | TRAJ37 | CAFGSSNTGKLIF | TRBV6-6 | TRBJ2-3 | CASSPRDTQYF | KLGGALQAK | NO | NO | NO |
| 15407 | TRAV21 | TRAJ42 | CAVGGSQGNLIF | TRBV13 | TRBJ1-5 | CASSLALNQPQHF | KLGGALQAK | NO | NO | NO |
| 15408 | TRAV26-2 | TRAJ31 | CILRDRYDARLMF | TRBV5-6 | TRBJ1-1 | CASSLVPSSGTEAFF | KLGGALQAK | NO | NO | NO |
| 15409 | TRAV14DV4 | TRAJ57 | CAMRETQGGSEKLVF | TRBV5-5 | TRBJ1-2 | CASSLASVYGYTF | KLGGALQAK | YES | NO | NO |
| 15410 | TRAV14DV4 | TRAJ49 | CAMREGWTGNQFYF | TRBV19 | TRBJ2-3 | CASSWGTGGTDTQYF | KLGGALQAK | YES | NO | NO |
| 15411 | TRAV17 | TRAJ49 | CATPVGNQFYF | TRBV6-3 | TRBJ1-1 | CASSYGTEAFF | KLGGALQAK | YES | NO | NO |
| 15412 | TRAV38-2DV8 | TRAJ42 | CAFTYGGSQGNLIF | TRBV7-9 | TRBJ1-4 | CASSRGGQGINEKLFF | KLGGALQAK | YES | NO | NO |
| 15414 | TRAV19 | TRAJ43 | CALRPKNMRF | TRBV11-2 | TRBJ2-1 | CASSLEDRSWNNEQFF | KLGGALQAK | YES | NO | NO |
| 15415 | TRAV8-2 | TRAJ28 | CVVSSYSGAGSYQLTF | TRBV5-8 | TRBJ2-7 | CASSFGDRSLSYEQYF | KLGGALQAK | YES | NO | NO |
| 15416 | TRAV38-1 | TRAJ53 | CAFNGGSNYKLTF | TRBV6-6 | TRBJ2-2 | CASSLGGYMSGELFF | KLGGALQAK | YES | NO | NO |
| 15417 | TRAV14DV4 | TRAJ28 | CAMREIGAGSYQLTF | TRBV7-9 | TRBJ1-5 | CASSLGPSNQPQHF | KLGGALQAK | YES | NO | NO |
| 15418 | TRAV13-2 | TRAJ28 | CAENNIGAGSYQLTF | TRBV13 | TRBJ2-1 | CASSYPTSGSYNEQFF | KLGGALQAK | YES | NO | NO |
| 15419 | TRAV8-2 | TRAJ26 | CVVADNYGQNFVF | TRBV27 | TRBJ2-7 | CASSSTWTSGRYEQYV | KLGGALQAK | NO | NO | NO |
| 15420 | TRAV3 | TRAJ26 | CAVVSYGQNFVF | TRBV27 | TRBJ2-1 | CASSPGPGVNEQFF | KLGGALQAK | NO | NO | NO |
| 15421 | TRAV22 | TRAJ36 | CAVYRTGANNLFF | TRBV7-9 | TRBJ2-2 | CASSPFSGAGELFF | KLGGALQAK | YES | NO | NO |
| 15422 | TRAV5 | TRAJ4 | CADESIGGYNKLIF | TRBV4-3 | TRBJ2-1 | CASSQDDRRGSYNEQFF | AVFDRKSDAK | YES | NO | YES |
| 15423 | TRAV19 | TRAJ34 | CALSGALTPDKLIF | TRBV7-8 | TRBJ2-1 | CASSSGTSGSIEQFF | KLGGALQAK | YES | NO | NO |
| 15424 | TRAV1-2 | TRAJ10 | CAVRGGGNKLTF | TRBV5-5 | TRBJ1-2 | CASIGQDGPDYGYTF | KLGGALQAK | YES | NO | NO |
| 15425 | TRAV12-1 | TRAJ21 | CVVKVGYNFNKFYF | TRBV3-1 | TRBJ2-3 | CASSLMEGLSTSTDTQYF | KLGGALQAK | YES | NO | NO |
| 15426 | TRAV19 | TRAJ9 | CALSDFNTGGFKTIF | TRBV9 | TRBJ2-3 | CASSDSGGSTDTQYF | KLGGALQAK | YES | NO | NO |
| 15427 | TRAV20 | TRAJ53 | CAVQVGGSNYKLTF | TRBV19 | TRBJ2-1 | CASSILLAGLGEQFF | KLGGALQAK | YES | NO | NO |
| 15428 | TRAV35 | TRAJ41 | CAGQLGPNSGYALNF | TRBV7-9 | TRBJ1-1 | CASSPPGSTEAFF | KLGGALQAK | YES | NO | NO |
| 15429 | TRAV38-1 | TRAJ53 | CASLNSGGSNYKLTF | TRBV4-1 | TRBJ2-7 | CASSQDRVYEQYF | KLGGALQAK | YES | NO | NO |
| 15430 | TRAV16 | TRAJ7 | CLLVTGNRLAF | TRBV19 | TRBJ2-7 | CASSTGGGYEQYV | AVFDRKSDAK | NO | NO | NO |
| 15431 | TRAV17 | TRAJ43 | CATDLDNNDMRF | TRBV4-1 | TRBJ2-1 | CASSQDWGGSYNEQFF | KLGGALQAK | YES | NO | NO |
| 15432 | TRAV5 | TRAJ24 | CATRTDSWGKLQF | TRBV12-4 | TRBJ2-1 | CASSDPGGSYNEQFF | KLGGALQAK | YES | NO | NO |
| 15433 | TRAV12-2 | TRAJ15 | CAVNQAGTALIF | TRBV4-2 | TRBJ2-7 | CASSQDTGVAYEQYF | KLGGALQAK | NO | NO | NO |
| 15434 | TRAV17 | TRAJ43 | CATGWNNNDMRF | TRBV11-2 | TRBJ2-7 | CASSSSGFYEQYF | KLGGALQAK | YES | NO | NO |
| 15435 | TRAV12-2 | TRAJ20 | CGSGDYKLSF | TRBV7-2 | TRBJ2-6 | CASMAVGANVLTF | ELAGIGILTV | NO | NO | NO |
| 15436 | TRAV12-2 | TRAJ45 | CAANAGGADGLTF | TRBV7-3 | TRBJ1-1 | CASSSLTNTEAFF | KLGGALQAK | NO | NO | NO |
| 15437 | TRAV1-1 | TRAJ42 | CAVRVDGGSQGNLIF | TRBV5-4 | TRBJ1-1 | CASSLVANTEAFF | KLGGALQAK | YES | NO | NO |
| 15438 | TRAV13-1 | TRAJ40 | CAATGTYKYIF | TRBV7-8 | TRBJ2-4 | CASSLGPWGVKNIQYF | KLGGALQAK | YES | NO | NO |
| 15439 | TRAV1-2 | TRAJ43 | CAVRDEGNDMRF | TRBV20-1 | TRBJ1-1 | CSARDQLAGSTEAFF | KLGGALQAK | YES | NO | NO |
| 15440 | TRAV4 | TRAJ11 | CLVGDAGYSTLTF | TRBV12-3 | TRBJ2-5 | CASSFTTSDETQYF | KLGGALQAK | YES | NO | NO |
| 15441 | TRAV26-1 | TRAJ36 | CIVRVAGQTGANNLFF | TRBV25-1 | TRBJ2-2 | CASSESPRSATGELFF | KLGGALQAK | YES | NO | NO |
| 15442 | TRAV4 | TRAJ37 | CLVGDEYGSSNTGKLIF | TRBV11-2 | TRBJ2-1 | CASSLTRTPSSYNEQFF | KLGGALQAK | NO | NO | NO |
| 15443 | TRAV27 | TRAJ58 | CAGVYLETSGSRLTF | TRBV6-6 | TRBJ2-1 | CASGRPINNEQFF | KLGGALQAK | NO | NO | NO |
| 15444 | TRAV19 | TRAJ21 | CALSELDNFNKFYF | TRBV5-1 | TRBJ2-7 | CASSLEVPSGGSNEQYF | KLGGALQAK | YES | NO | NO |
| 15445 | TRAV8-6 | TRAJ12 | CAVRRDSSYKLIF | TRBV19 | TRBJ1-1 | CASSIRGAPEAFF | KLGGALQAK | YES | NO | NO |
| 15446 | TRAV26-2 | TRAJ48 | CIRGGSNFGNEKLTF | TRBV9 | TRBJ2-3 | CASSVETSGTSTDTQYF | KLGGALQAK | YES | NO | NO |
| 15447 | TRAV12-1 | TRAJ26 | CVVNGDYGQNFVF | TRBV4-1 | TRBJ2-1 | CASSRQGENNEQFF | KLGGALQAK | YES | NO | NO |
| 15448 | TRAV19 | TRAJ45 | CALSEARSGGGADGLTF | TRBV19 | TRBJ1-5 | CASSYGLYSNQPQHF | KLGGALQAK | YES | NO | NO |
| 15449 | TRAV12-2 | TRAJ50 | CAVKTSYDKVIF | TRBV12-3 | TRBJ2-5 | CASRLLGGRTLETQYF | KLGGALQAK | YES | NO | NO |
| 15450 | TRAV5 | TRAJ21 | CAEITAFNKFYF | TRBV6-1 | TRBJ1-1 | CASRLGTEMNTEAFF | KLGGALQAK | YES | NO | NO |
| 15451 | TRAV4 | TRAJ36 | CLVGFQTGANNLFF | TRBV9 | TRBJ2-1 | CASSVEGGSSYNEQFF | KLGGALQAK | YES | NO | NO |
| 15452 | TRAV29DV5 | TRAJ41 | CAASDPNSGYALNF | TRBV19 | TRBJ2-3 | CASSDAPGTGASTDTQYF | KLGGALQAK | YES | NO | NO |
| 15453 | TRAV35 | TRAJ27 | CAGHDAGKSTF | TRBV6-5 | TRBJ1-1 | CASSYSGGDTEAFF | KLGGALQAK | YES | NO | NO |
| 15454 | TRAV3 | TRAJ43 | CAVRAPGASYNNNDMRF | TRBV3-1 | TRBJ2-5 | CASSQPRTGETQYF | KLGGALQAK | NO | NO | NO |
| 15455 | TRAV9-2 | TRAJ8 | CALSEDTGFQKLVF | TRBV11-1 | TRBJ2-7 | CASSSLQSGSYEQYF | KLGGALQAK | YES | NO | NO |
| 15456 | TRAV1-2 | TRAJ35 | CAVLGFGNVLHC | TRBV19 | TRBJ1-1 | CASSAGGTEAFF | KLGGALQAK | NO | NO | NO |
| 15457 | TRAV3 | TRAJ29 | CAVRDIAGNTPLVF | TRBV6-3 | TRBJ2-2 | CASSQTGGELFF | KLGGALQAK | NO | NO | NO |
| 15458 | TRAV12-3 | TRAJ39 | CAIGPGNMLTF | TRBV27 | TRBJ2-1 | CASSLPWTGAKDNEQFF | KLGGALQAK | YES | NO | NO |
| 15459 | TRAV22 | TRAJ45 | CAAYSGGGADGLTF | TRBV4-3 | TRBJ2-1 | CASSPRLAGEYTSSYNEQFF | KLGGALQAK | NO | NO | NO |
| 15460 | TRAV20 | TRAJ57 | CAVRTQGGSEKLVF | TRBV6-5 | TRBJ1-2 | CASRQKNRDGYTF | AVFDRKSDAK | YES | NO | NO |
| 15461 | TRAV13-1 | TRAJ44 | CAASITGTASKLTF | TRBV19 | TRBJ2-7 | CASSHGTGTYEQYF | KLGGALQAK | YES | NO | NO |
| 15462 | TRAV22 | TRAJ44 | CAVTGTASKLTF | TRBV7-9 | TRBJ2-7 | CASSLATSEQYV | KLGGALQAK | NO | NO | NO |
| 15463 | TRAV13-2 | TRAJ40 | CAETPGGTYKYIF | TRBV7-6 | TRBJ2-3 | CASSLAPPDTQYF | KLGGALQAK | YES | NO | NO |
| 15464 | TRAV24 | TRAJ9 | CASSNTGGFKTIF | TRBV6-5 | TRBJ2-3 | CASSYNDGLTDTQYF | KLGGALQAK | NO | NO | NO |
| 15465 | TRAV29DV5 | TRAJ45 | CAASLGGGADGLTF | TRBV18 | TRBJ2-7 | CASSPTTFLYEQYF | KLGGALQAK | YES | NO | NO |
| 15466 | TRAV25 | TRAJ37 | CAGGGSGNTGKLIF | TRBV7-2 | TRBJ2-1 | CASSLDLGGSPYNEQFF | KLGGALQAK | YES | NO | NO |
| 15467 | TRAV5 | TRAJ10 | CAGEWTGGGNKLTF | TRBV4-1 | TRBJ2-7 | CASSQDSSGIYEQYF | KLGGALQAK | YES | NO | NO |
| 15468 | TRAV1-2 | TRAJ33 | CAPMDSNYQLIW | TRBV6-4 | TRBJ2-1 | CASSEAGGEAYNEQFF | KLGGALQAK | YES | NO | NO |
| 15469 | TRAV12-3 | TRAJ39 | CAIGPGNMLTF | TRBV7-9 | TRBJ2-1 | CASSRAGATYNEQFF | KLGGALQAK | YES | NO | NO |
| 15470 | TRAV14DV4 | TRAJ4 | CAMRPPFRGGYNKLIF | TRBV20-1 | TRBJ2-7 | CSARTGTAVYEQYF | AVFDRKSDAK | NO | NO | NO |
| 15471 | TRAV8-3 | TRAJ48 | CAVGTNFGNEKLTF | TRBV7-7 | TRBJ2-1 | CASSSTSGDYNEQFF | AVFDRKSDAK | YES | NO | YES |
| 15472 | TRAV19 | TRAJ4 | CALSEPMVGYNKLIF | TRBV2 | TRBJ2-1 | CASSEIRKTKPSFF | KLGGALQAK | YES | NO | NO |
| 15473 | TRAV17 | TRAJ17 | CATGGGAAGNKLTF | TRBV4-2 | TRBJ2-2 | CASSRGGSGNTGELFF | KLGGALQAK | YES | NO | NO |
| 15474 | TRAV24 | TRAJ45 | CASSSGGGADGLTF | TRBV27 | TRBJ2-1 | CASSRLNNEQFF | KLGGALQAK | YES | NO | NO |
| 15475 | TRAV27 | TRAJ31 | CAGANARLMF | TRBV12-3 | TRBJ1-1 | CASSYRTGGTEAFF | KLGGALQAK | YES | NO | NO |
| 15476 | TRAV26-2 | TRAJ43 | CIGWRGNDMRF | TRBV7-3 | TRBJ2-3 | CASSLLVGDTQYF | KLGGALQAK | YES | NO | NO |
| 15477 | TRAV13-2 | TRAJ23 | CAENNNQGGKLIF | TRBV7-2 | TRBJ2-7 | CASSPETGGSYEQYF | KLGGALQAK | YES | NO | NO |
| 15478 | TRAV17 | TRAJ23 | CATDGNQGGKLIF | TRBV10-1 | TRBJ2-5 | CASSDSGTGGETQYF | KLGGALQAK | YES | NO | NO |
| 15479 | TRAV12-2 | TRAJ13 | CAVTSGGYQKVTF | TRBV19 | TRBJ2-1 | CASSIGAGDEQFF | AVFDRKSDAK | YES | NO | YES |
| 15480 | TRAV26-1 | TRAJ26 | CIVIVGGQNFVF | TRBV7-9 | TRBJ2-7 | CASKDGISYEQYF | KLGGALQAK | YES | NO | NO |
| 15481 | TRAV1-2 | TRAJ41 | CAGMTNSGYALNF | TRBV7-9 | TRBJ1-2 | CASSDGQGANYGYTF | KLGGALQAK | YES | NO | NO |
| 15482 | TRAV8-2 | TRAJ37 | CVVSVISSNTGKLIF | TRBV27 | TRBJ1-2 | CASKTYSTGGNGYTF | KLGGALQAK | YES | NO | NO |
| 15483 | TRAV12-1 | TRAJ44 | CVVTRTGTASKLTF | TRBV6-1 | TRBJ1-1 | CASSDFAGLPGYTEAFF | KLGGALQAK | YES | NO | NO |
| 15484 | TRAV19 | TRAJ13 | CALSEYRLRSGGYQKVTF | TRBV6-5 | TRBJ1-5 | CASSSTGGGNQPQHF | KLGGALQAK | YES | NO | NO |
| 15485 | TRAV13-1 | TRAJ39 | CAASIYAGNMLTF | TRBV12-5 | TRBJ1-5 | CASGAKQPQHF | KLGGALQAK | YES | NO | NO |

| | | | | | | | | | | |
|---|---|---|---|---|---|---|---|---|---|---|
| 15486 | TRAV14DV4 | TRAJ4 | CAMRGPISGGYNKLIF | TRBV12-5 | TRBJ2-7 | CASGSPVREGSYEQYV | KLGGALQAK | NO | NO | NO |
| 15487 | TRAV12-2 | TRAJ7 | CAVTPYGNNRLAF | TRBV5-4 | TRBJ2-2 | CASSLWTLNTGELFF | KLGGALQAK | YES | YES | NO |
| 15488 | TRAV26-2 | TRAJ52 | CILRDPHNAGGTSYGKLTF | TRBV2 | TRBJ1-1 | CASSEGRGIGTEAFF | KLGGALQAK | NO | NO | NO |
| 15489 | TRAV17 | TRAJ37 | CATDSPGNTGKLIF | TRBV5-1 | TRBJ1-2 | CASRPPGRDYGYTF | KLGGALQAK | YES | YES | NO |
| 15491 | TRAV35 | TRAJ52 | CAGPLSGGTSYGKLTF | TRBV6-1 | TRBJ2-1 | CASIQPTEQFF | KLGGALQAK | YES | NO | NO |
| 15492 | TRAV8-4 | TRAJ30 | CAVRTNRDDKIIF | TRBV7-8 | TRBJ2-7 | CASSLDRASSYEQYV | KLGGALQAK | NO | NO | NO |
| 15493 | TRAV12-2 | TRAJ9 | CAVNPYTGGFKTIF | TRBV3-1 | TRBJ1-1 | CASSQEEGGSPEAFF | KLGGALQAK | YES | NO | NO |
| 15494 | TRAV12-2 | TRAJ45 | CAVNRIRTGGGADGLTF | TRBV9 | TRBJ1-1 | CASSVVTGGLGTEAFF | KLGGALQAK | YES | NO | NO |
| 15495 | TRAV26-1 | TRAJ27 | CMRVNNTNAGKSTF | TRBV7-2 | TRBJ2-5 | CASRGGLQETQYF | KLGGALQAK | YES | NO | NO |
| 15496 | TRAV29DV5 | TRAJ42 | CAASARIYGGSQGNLIF | TRBV4-1 | TRBJ1-5 | CASSQGFSNQPQHF | KLGGALQAK | YES | NO | NO |
| 15497 | TRAV12-3 | TRAJ34 | CAMSPYNTDKLIF | TRBV5-4 | TRBJ2-2 | CASSSTGANTGELFF | KLGGALQAK | YES | NO | NO |
| 15498 | TRAV14DV4 | TRAJ40 | CAMRETSGTYKYIF | TRBV20-1 | TRBJ2-3 | CSARGISTDTQYF | KLGGALQAK | NO | NO | NO |
| 15499 | TRAV8-6 | TRAJ54 | CAVRPIIQGAQKLVF | TRBV4-3 | TRBJ1-5 | CASSQDEGSNQPQHF | KLGGALQAK | YES | NO | NO |
| 15500 | TRAV1-2 | TRAJ34 | CAVWGYNTDKLIF | TRBV9 | TRBJ1-1 | CASSGRQGDEAFF | AVFDRKSDAK | YES | NO | NO |
| 15501 | TRAV35 | TRAJ17 | CAGQYLKAAGNKLTF | TRBV19 | TRBJ1-1 | CASSIGSNTEAFF | KLGGALQAK | YES | NO | NO |
| 15502 | TRAV12-2 | TRAJ31 | CAVNNARLMF | TRBV4-3 | TRBJ1-3 | CASSQAHGTMGNTIYF | KLGGALQAK | YES | NO | NO |
| 15503 | TRAV13-2 | TRAJ44 | CAENPHFSGTASKLTF | TRBV4-1 | TRBJ2-3 | CASSARLADTQYF | KLGGALQAK | YES | NO | NO |
| 15504 | TRAV9-2 | TRAJ54 | CALSDQGAQKLVF | TRBV19 | TRBJ2-1 | CASSIISNNEQFF | KLGGALQAK | YES | NO | NO |
| 15505 | TRAV12-2 | TRAJ39 | CAIGPGNMLTF | TRBV5-1 | TRBJ2-1 | CATRPDSYNEQFF | KLGGALQAK | YES | NO | NO |
| 15506 | TRAV41 | TRAJ33 | CAVADSNYQLIW | TRBV7-9 | TRBJ1-2 | CASSSGTLGDYGYTF | KLGGALQAK | YES | NO | NO |
| 15507 | TRAV8-2 | TRAJ50 | CVVSQKKTSYDKVIF | TRBV4-1 | TRBJ1-5 | CASSQQQGARSNQPQHF | AVFDRKSDAK | YES | NO | YES |
| 15509 | TRAV16 | TRAJ40 | CALSGFTSGTYKYIF | TRBV7-9 | TRBJ2-7 | CASSFGGGETYEQYF | KLGGALQAK | NO | NO | NO |
| 15510 | TRAV12-3 | TRAJ31 | CAMSAFSNARLMF | TRBV29-1 | TRBJ1-2 | CSVDLEANYGYTF | KLGGALQAK | YES | NO | NO |
| 15511 | TRAV12-1 | TRAJ41 | CVVNIANSGYALNF | TRBV12-4 | TRBJ2-3 | CASSFRGGEAIHTDTQYF | KLGGALQAK | YES | NO | NO |
| 15512 | TRAV39 | TRAJ48 | CAATNFGNEKLTF | TRBV4-3 | TRBJ2-1 | CASSQDVGTYNEQFF | KLGGALQAK | YES | NO | NO |
| 15513 | TRAV4 | TRAJ9 | CLVGDGTGGFKTIF | TRBV10-1 | TRBJ2-1 | CATRGGLENEQFF | KLGGALQAK | YES | NO | NO |
| 15514 | TRAV21 | TRAJ57 | CAVRGQGGSEKLVF | TRBV7-9 | TRBJ2-7 | CASSDSLGDRGSYEQYF | KLGGALQAK | YES | NO | NO |
| 15515 | TRAV8-6 | TRAJ24 | CAVSDPKTDSWGKFQF | TRBV7-9 | TRBJ1-6 | CASSATGTGRGSPLHF | KLGGALQAK | YES | NO | NO |
| 15516 | TRAV1-1 | TRAJ20 | CAVRDHNDYKLSF | TRBV18 | TRBJ2-1 | CASSPPSGFSYNEQFF | KLGGALQAK | YES | NO | NO |
| 15517 | TRAV41 | TRAJ32 | CAVTLSPHGGATNKLIF | TRBV20-1 | TRBJ1-6 | CSATRRDRILTGYNSPLHF | KLGGALQAK | YES | NO | NO |
| 15518 | TRAV19 | TRAJ17 | CALSEAIKAAGNKLTF | TRBV4-3 | TRBJ2-3 | CASSQDSGSADTQYF | KLGGALQAK | NO | NO | NO |
| 15519 | TRAV25 | TRAJ28 | CAGLGAGSYQLTF | TRBV6-3 | TRBJ2-5 | CASSSGLEGETQYF | KLGGALQAK | YES | NO | NO |
| 15520 | TRAV22 | TRAJ36 | CAAQRPGANNLFF | TRBV4-1 | TRBJ2-7 | CASSPQPGILSYEQYF | KLGGALQAK | YES | NO | NO |
| 15521 | TRAV25 | TRAJ31 | CAGPNARLMF | TRBV19 | TRBJ2-1 | CASSIGGVVGSDYNEQFF | KLGGALQAK | YES | NO | NO |
| 15522 | TRAV9-2 | TRAJ49 | CALTYTGNQFYF | TRBV9 | TRBJ2-2 | CASRQGNTGELFF | KLGGALQAK | YES | NO | NO |
| 15523 | TRAV8-6 | TRAJ57 | CAVSEDGGGSEKLVF | TRBV20-1 | TRBJ2-1 | CSATDRFF | KLGGALQAK | NO | NO | NO |
| 15524 | TRAV29DV5 | TRAJ42 | CAASASRGGSQGNLIF | TRBV6-6 | TRBJ2-3 | CASNERFFLSSTDTQYF | KLGGALQAK | NO | NO | NO |
| 15525 | TRAV13-1 | TRAJ23 | CAASSQDNQGGKLIF | TRBV2 | TRBJ2-2 | CASRTPGLAPGELFF | KLGGALQAK | YES | NO | NO |
| 15526 | TRAV4 | TRAJ41 | CLVGAGALNF | TRBV29-1 | TRBJ1-1 | CSVPDSQVPEAFF | KLGGALQAK | YES | NO | NO |
| 15527 | TRAV19 | TRAJ30 | CALSEVNRDDKIIF | TRBV12-4 | TRBJ2-1 | CASSSEALGRGSYNEQFF | KLGGALQAK | YES | NO | NO |
| 15528 | TRAV13-2 | TRAJ54 | CAESAVQGAQKLVF | TRBV7-8 | TRBJ1-1 | CASSVQATEAFF | KLGGALQAK | YES | NO | NO |
| 15529 | TRAV14DV4 | TRAJ39 | CAMRETNAGNMLTF | TRBV20-1 | TRBJ2-1 | CSARAHYGDEQFF | KLGGALQAK | NO | NO | NO |
| 15530 | TRAV20 | TRAJ32 | CAVTTYGGATNKLIF | TRBV19 | TRBJ2-3 | CASSAGARNTQYF | KLGGALQAK | YES | NO | NO |
| 15531 | TRAV21 | TRAJ7 | CAVRPGEQQTRF | TRBV7-8 | TRBJ2-7 | CASRLSGSAYEQYF | KLGGALQAK | YES | NO | NO |
| 15532 | TRAV20 | TRAJ47 | CAVQTFEYGNKLVF | TRBV10-3 | TRBJ2-1 | CAISGESHEQFF | KLGGALQAK | YES | NO | NO |
| 15533 | TRAV4 | TRAJ4 | CLVGDIGSGGYNKLIF | TRBV7-9 | TRBJ2-2 | CASSRSDGDTGELFF | KLGGALQAK | YES | NO | NO |
| 15534 | TRAV12-2 | TRAJ45 | CAPVSGGGADGLTF | TRBV6-6 | TRBJ2-1 | CASSYRVGEQFF | KLGGALQAK | NO | NO | NO |
| 15535 | TRAV9-2 | TRAJ53 | CALIGGSNYKLTF | TRBV19 | TRBJ1-5 | CASGYSNQPQHF | KLGGALQAK | YES | NO | NO |
| 15536 | TRAV9-2 | TRAJ28 | CALSAYSGAGSYQLTF | TRBV4-1 | TRBJ1-5 | CASSQDPQGRQPQHF | KLGGALQAK | YES | NO | NO |
| 15537 | TRAV8-3 | TRAJ48 | CAANFGNEKLTF | TRBV13 | TRBJ2-1 | CASSSLAGGRLNNEQFF | AVFDRKSDAK | YES | NO | YES |
| 15538 | TRAV13-1 | TRAJ34 | CAASINTDKLIF | TRBV20-1 | TRBJ2-7 | CSAREWGQRGEQYF | KLGGALQAK | YES | NO | NO |
| 15539 | TRAV27 | TRAJ44 | CAGADGTASKLTF | TRBV4-3 | TRBJ1-1 | CASSQGQGNTEAFF | KLGGALQAK | YES | NO | NO |
| 15540 | TRAV1-2 | TRAJ20 | CAVREGDYKLSF | TRBV6-4 | TRBJ2-1 | CASSDSAGGYNEQFF | KLGGALQAK | YES | NO | NO |
| 15541 | TRAV17 | TRAJ9 | CATTGGFKTIF | TRBV20-1 | TRBJ2-3 | CSATSNGERTDTQYF | KLGGALQAK | NO | NO | NO |
| 15542 | TRAV9-2 | TRAJ17 | CALSDSEAAGNKLTF | TRBV19 | TRBJ2-6 | CASSIDARDSGANVLTF | KLGGALQAK | YES | NO | NO |
| 15543 | TRAV21 | TRAJ33 | CAVLRDSNYQLIW | TRBV14 | TRBJ2-1 | CASSQEAPPIYNEQFF | KLGGALQAK | YES | NO | NO |
| 15544 | TRAV1-2 | TRAJ49 | CAAKNTGNQFYF | TRBV20-1 | TRBJ1-5 | CSAIGTGGGQPQHF | KLGGALQAK | YES | NO | NO |
| 15545 | TRAV12-1 | TRAJ41 | CVVNSNSGYALNF | TRBV7-9 | TRBJ2-3 | CASSQTSGGTDTQYF | KLGGALQAK | YES | NO | NO |
| 15546 | TRAV14DV4 | TRAJ48 | CAMRGGFGNEKLTF | TRBV27 | TRBJ2-3 | CASSLLTGQGSGTQYF | RLRAEAQVK | YES | YES | YES |
| 15547 | TRAV29DV5 | TRAJ45 | CAAYAGGGADGLTF | TRBV9 | TRBJ1-5 | CASGDSFSNQPQHF | KLGGALQAK | NO | NO | NO |
| 15548 | TRAV29DV5 | TRAJ22 | CAASSTGSARQLTF | TRBV27 | TRBJ1-2 | CASSPTGGSGYTF | KLGGALQAK | YES | NO | NO |
| 15549 | TRAV22 | TRAJ35 | CAVNEFTGFGNVLHC | TRBV6-3 | TRBJ1-5 | CASSYEREQPQHF | KLGGALQAK | NO | NO | NO |
| 15550 | TRAV14DV4 | TRAJ9 | CAMREAESGGFKTIF | TRBV9 | TRBJ2-3 | CASSPLAGGSVDTQYF | KLGGALQAK | YES | NO | NO |
| 15551 | TRAV29DV5 | TRAJ36 | CAADLQTGANNLFF | TRBV27 | TRBJ2-7 | CASMKQGSYEQYF | KLGGALQAK | YES | NO | NO |
| 15552 | TRAV13-2 | TRAJ42 | CAENTPITGSQGNLIF | TRBV4-1 | TRBJ1-1 | CASSQEHRANTEAFF | KLGGALQAK | YES | NO | NO |
| 15553 | TRAV1-1 | TRAJ38 | CAVWYAGNNRKLIW | TRBV4-1 | TRBJ1-1 | CASSQAAGGNTEAFF | KLGGALQAK | YES | YES | NO |
| 15554 | TRAV12-3 | TRAJ47 | CAMEYGNKLVF | TRBV29-1 | TRBJ1-2 | CSSGGTYGYTF | KLGGALQAK | YES | NO | NO |
| 15555 | TRAV25 | TRAJ52 | CAGGRIAGGTSYGKLTF | TRBV19 | TRBJ2-1 | CASSGGGWNEQFF | KLGGALQAK | YES | NO | NO |
| 15556 | TRAV3 | TRAJ41 | CAVRDSRARNF | TRBV4-3 | TRBJ2-1 | CASSQDHAASYNEQFF | KLGGALQAK | YES | NO | NO |
| 15557 | TRAV22 | TRAJ35 | CAVDGGFGNVLHC | TRBV18 | TRBJ1-1 | CASSPVDTEAFF | KLGGALQAK | NO | NO | NO |
| 15558 | TRAV14DV4 | TRAJ22 | CAMREGPLYSARQLTF | TRBV7-8 | TRBJ1-4 | CASSLVEDEKLFF | KLGGALQAK | YES | NO | NO |
| 15559 | TRAV12-3 | TRAJ39 | CAIGPGNMLTF | TRBV6-3 | TRBJ2-3 | CASALAGGTDTQYF | KLGGALQAK | YES | NO | NO |
| 15560 | TRAV25 | TRAJ26 | CAGRHYGQNFVF | TRBV12-4 | TRBJ2-5 | CASSLGAGGAGETQYF | RLRAEAQVK | YES | YES | NO |
| 15561 | TRAV19 | TRAJ30 | CALSEGNRDDKIIF | TRBV10-2 | TRBJ2-7 | CASSEMASGSREQYV | KLGGALQAK | NO | NO | NO |
| 15562 | TRAV12-3 | TRAJ13 | CAIPSGGYQKVTF | TRBV7-9 | TRBJ2-7 | CASGGTETYEQYF | KLGGALQAK | YES | NO | NO |
| 15563 | TRAV12-1 | TRAJ26 | CVVCDYGQNFVF | TRBV19 | TRBJ1-1 | CASIRDGQGGTEAFF | AVFDRKSDAK | YES | YES | YES |
| 15564 | TRAV35 | TRAJ9 | CAGSNTGGFKTIF | TRBV11-2 | TRBJ1-4 | CASKGDIGEKLFF | KLGGALQAK | YES | NO | NO |
| 15565 | TRAV5 | TRAJ39 | CALTGNAGNMLTF | TRBV6-3 | TRBJ1-1 | CASSYPRQGFEAFF | KLGGALQAK | YES | NO | NO |
| 15566 | TRAV19 | TRAJ48 | CALSDDSNFGNEKLTF | TRBV6-1 | TRBJ2-7 | CASSEVFGTVLEQYF | KLGGALQAK | YES | NO | NO |
| 15567 | TRAV29DV5 | TRAJ32 | CAASDDGGATNKLIF | TRBV7-2 | TRBJ2-1 | CASSTSGYNEQFF | KLGGALQAK | YES | NO | NO |
| 15568 | TRAV17 | TRAJ8 | CAPSNTGFQKLVF | TRBV27 | TRBJ2-5 | CASSLGETQYF | KLGGALQAK | YES | NO | NO |
| 15569 | TRAV1-2 | TRAJ40 | CAVNGYKYIF | TRBV7-6 | TRBJ2-5 | CASSLGVGDVQETQYF | AVFDRKSDAK | YES | NO | YES |
| 15570 | TRAV27 | TRAJ24 | CAGAFPTDSWGKLQF | TRBV12-5 | TRBJ1-5 | CASSPPNQPQHF | KLGGALQAK | YES | NO | NO |
| 15571 | TRAV14DV4 | TRAJ9 | CAMRDSYTGGFKTIF | TRBV20-1 | TRBJ2-7 | CSATGTASASYEQYF | KLGGALQAK | NO | NO | NO |
| 15572 | TRAV27 | TRAJ37 | CAGGSSNTGKLIF | TRBV4-1 | TRBJ1-5 | CASSQDWFVRGPNQPQHF | AVFDRKSDAK | YES | NO | YES |
| 15573 | TRAV35 | TRAJ49 | CAGLGNQFYF | TRBV20-1 | TRBJ2-1 | CSARGDSRNEQFF | KLGGALQAK | YES | NO | NO |
| 15574 | TRAV9-2 | TRAJ12 | CASGMDSSYKLIF | TRBV12-4 | TRBJ2-7 | CASSSPDRGLDEQYF | KLGGALQAK | YES | NO | NO |
| 15575 | TRAV1-2 | TRAJ31 | CAVRLNNARLMF | TRBV12-4 | TRBJ2-3 | CASGVGGSSTDTQYF | KLGGALQAK | YES | NO | NO |
| 15576 | TRAV9-2 | TRAJ37 | CALFASGNTGKLIF | TRBV19 | TRBJ1-2 | CAIAGPSRGYTF | KLGGALQAK | YES | NO | NO |
| 15577 | TRAV22 | TRAJ12 | CAVEDSSYKLIF | TRBV4-3 | TRBJ1-4 | CASSQDHRMGGHEKLFF | KLGGALQAK | YES | NO | NO |
| 15578 | TRAV9-2 | TRAJ20 | CALIEDYKLSF | TRBV7-9 | TRBJ2-1 | CASRPQGVYNEQFF | KLGGALQAK | YES | NO | NO |
| 15579 | TRAV17 | TRAJ50 | CARVGKTSYDKVIF | TRBV27 | TRBJ2-7 | CASRGQGYV | KLGGALQAK | NO | NO | NO |
| 15580 | TRAV12-1 | TRAJ26 | CVVNRIRDNYGQNFVF | TRBV19 | TRBJ1-2 | CASTFGTVYGYTF | KLGGALQAK | YES | NO | NO |
| 15581 | TRAV25 | TRAJ47 | CAGPGGNKLVF | TRBV30 | TRBJ1-1 | CAWSVRRQGAFF | KLGGALQAK | YES | NO | NO |
| 15582 | TRAV20 | TRAJ22 | CAVQAWVLGTSGSARQLTF | TRBV5-8 | TRBJ2-1 | CASSPDWSSHNEQFF | KLGGALQAK | YES | NO | NO |
| 15583 | TRAV8-1 | TRAJ32 | CAVNSWYGGATNKLIF | TRBV19 | TRBJ2-1 | CASSDPTSGGYNEQFF | KLGGALQAK | YES | NO | NO |
| 15584 | TRAV8-1 | TRAJ13 | CAVFSGGYQKVTF | TRBV7-9 | TRBJ2-2 | CASSWGAPNTGELFF | KLGGALQAK | YES | NO | NO |
| 15585 | TRAV12-1 | TRAJ43 | CVVNDNDMRF | TRBV5-4 | TRBJ1-2 | CATSSGNYGYTF | KLGGALQAK | YES | NO | NO |
| 15586 | TRAV13-1 | TRAJ40 | CAAGRGTYKYIF | TRBV12-4 | TRBJ2-1 | CASSAPGLAVNEQFF | AVFDRKSDAK | YES | NO | YES |
| 15587 | TRAV3 | TRAJ36 | CAVRDWQTGANNLFF | TRBV2 | TRBJ2-7 | CASSEEDNSYEQYV | KLGGALQAK | NO | NO | NO |
| 15588 | TRAV14DV4 | TRAJ8 | CAMSRQGFQKLVF | TRBV11-2 | TRBJ2-1 | CASSLVAGVMDEQFF | KLGGALQAK | YES | NO | NO |
| 15589 | TRAV5 | TRAJ13 | CAESRGYQKVTF | TRBV5-8 | TRBJ1-2 | CASIEKGYGYTF | KLGGALQAK | YES | NO | NO |
| 15590 | TRAV1-2 | TRAJ20 | CAVRVNDYKLSF | TRBV2 | TRBJ2-7 | CASSEAPQHYEQYV | KLGGALQAK | NO | NO | NO |
| 15591 | TRAV1-2 | TRAJ33 | CAVRDGGNYQLIW | TRBV27 | TRBJ1-3 | CASSPGDFFRDTIYF | KLGGALQAK | YES | NO | NO |

| | | | | | | | | | | |
|---|---|---|---|---|---|---|---|---|---|---|
| 15592 | TRAV8-6 | TRAJ54 | CAVTPPIQGAQKLVF | TRBV4-3 | TRBJ2-1 | CASSPGLAGDEQFF | KLGGALQAK | YES | NO | NO |
| 15593 | TRAV3 | TRAJ49 | CAVREDGNTLLGGYTGNQFYF | TRBV19 | TRBJ1-1 | CASSQEGRWNTEAFF | KLGGALQAK | NO | NO | NO |
| 15594 | TRAV17 | TRAJ54 | CATDPFQGAQKLVF | TRBV19 | TRBJ1-6 | CASSKGRLPNSPLHF | KLGGALQAK | YES | NO | NO |
| 15595 | TRAV12-2 | TRAJ45 | CAVNERGGADGLTF | TRBV12-3 | TRBJ2-1 | CAGATGTTYNEQFF | KLGGALQAK | YES | NO | NO |
| 15596 | TRAV27 | TRAJ41 | CAGTPASRGYALNF | TRBV13 | TRBJ1-1 | CASIGEMNTEAFF | KLGGALQAK | YES | NO | NO |
| 15597 | TRAV13-2 | TRAJ34 | CAENLPNTDKLIF | TRBV5-1 | TRBJ2-3 | CASSGWTSGTSTDTQYF | KLGGALQAK | YES | NO | NO |
| 15598 | TRAV41 | TRAJ40 | CAVFSGTYKYIF | TRBV25-1 | TRBJ1-2 | CASSCTGTGYGYTF | KLGGALQAK | YES | NO | NO |
| 15599 | TRAV38-2DV8 | TRAJ12 | CAYRPMDSSYKLIF | TRBV20-1 | TRBJ2-3 | CSARDLQAGGPADTQYF | KLGGALQAK | NO | NO | NO |
| 15600 | TRAV4 | TRAJ33 | CLVGDSSNYQLIW | TRBV18 | TRBJ1-2 | CASSPGTDPFNYGYTF | KLGGALQAK | NO | NO | NO |
| 15601 | TRAV13-2 | TRAJ31 | CAEINARLMF | TRBV4-1 | TRBJ2-2 | CASSQDRDRELFF | KLGGALQAK | YES | NO | NO |
| 15602 | TRAV14DV4 | TRAJ4 | CAMRRPSGGYNKLIF | TRBV12-4 | TRBJ2-1 | CASSRTESGNEQFF | KLGGALQAK | YES | NO | NO |
| 15603 | TRAV8-6 | TRAJ23 | CAVSDPYYNQGGKLIF | TRBV7-3 | TRBJ2-1 | CASSFTDGDNEQFF | KLGGALQAK | YES | NO | NO |
| 15604 | TRAV25 | TRAJ53 | CAGLGGGSNYKLTF | TRBV10-3 | TRBJ1-1 | CAISDPTDLNTEAFF | KLGGALQAK | YES | NO | NO |
| 15605 | TRAV8-3 | TRAJ57 | CASPQGGSEKLVF | TRBV7-9 | TRBJ1-4 | CASRFAGGNEKLFF | KLGGALQAK | YES | NO | NO |
| 15606 | TRAV30 | TRAJ52 | CGTEANAGGTSYGKLTF | TRBV19 | TRBJ1-5 | CASSMIGALNQPQHF | KLGGALQAK | YES | NO | NO |
| 15607 | TRAV29DV5 | TRAJ26 | CAAFNNYGQNFVF | TRBV4-3 | TRBJ1-2 | CASSQVGTGHGYTF | KLGGALQAK | YES | NO | NO |
| 15608 | TRAV38-2DV8 | TRAJ52 | CAYMWGGTSYGKLTF | TRBV12-4 | TRBJ1-5 | CASRWDSYNQPQHF | KLGGALQAK | YES | NO | NO |
| 15609 | TRAV10 | TRAJ40 | CVVSDTSGTYKYIF | TRBV27 | TRBJ2-1 | CASSFLGGRNPNEQFF | KLGGALQAK | YES | NO | NO |
| 15610 | TRAV25 | TRAJ21 | CAGVYNFNKFYF | TRBV9 | TRBJ2-2 | CASSHSGGAGTGELFF | KLGGALQAK | YES | NO | NO |
| 15611 | TRAV8-6 | TRAJ49 | CAVTPRNQFYF | TRBV4-1 | TRBJ2-6 | CASSQDRVGAGANVLTF | KLGGALQAK | YES | NO | NO |
| 15612 | TRAV19 | TRAJ54 | CALSEAQGAQKLVF | TRBV3-1 | TRBJ2-5 | CASSLPVEGTETQYF | KLGGALQAK | YES | NO | NO |
| 15613 | TRAV29DV5 | TRAJ40 | CAASGSGTYKYIF | TRBV19 | TRBJ2-3 | CASSPPLGADTQYF | KLGGALQAK | YES | NO | NO |
| 15614 | TRAV12-1 | TRAJ54 | CVVNRIIQGAQKLVF | TRBV7-8 | TRBJ2-1 | CASSPSRMSSYNEQFF | KLGGALQAK | YES | NO | NO |
| 15615 | TRAV1-2 | TRAJ34 | CAVRWGTTDKLIF | TRBV5-5 | TRBJ1-1 | CASSLPPPGGGAFF | KLGGALQAK | YES | NO | NO |
| 15616 | TRAV1-2 | TRAJ31 | CAVRDNARLMF | TRBV7-9 | TRBJ2-7 | CASSSGTGAYEQYF | KLGGALQAK | YES | NO | NO |
| 15617 | TRAV29DV5 | TRAJ43 | CAARLNDMRF | TRBV4-3 | TRBJ1-5 | CASSQDLRSNQPQHF | KLGGALQAK | YES | NO | NO |
| 15618 | TRAV5 | TRAJ20 | CAELNDYKLSF | TRBV5-1 | TRBJ2-1 | CASSHAGTGAGEQFF | KLGGALQAK | YES | NO | NO |
| 15619 | TRAV21 | TRAJ45 | CAVPGGGADGLTF | TRBV2 | TRBJ1-6 | CASSVSTPGQGLFYNSPLHF | KLGGALQAK | NO | NO | NO |
| 15620 | TRAV19 | TRAJ5 | CALSEESWRALTF | TRBV7-2 | TRBJ2-1 | CASSLGVIAGGLNEQFF | KLGGALQAK | YES | NO | NO |
| 15621 | TRAV3 | TRAJ28 | CAVRNSGAGSYQLTF | TRBV20-1 | TRBJ2-5 | CSARVRDRATHKETQYF | KLGGALQAK | NO | NO | NO |
| 15622 | TRAV24 | TRAJ21 | CAFCGNFNKFYF | TRBV7-2 | TRBJ2-7 | CASSFGQQLEQYV | KLGGALQAK | NO | NO | NO |
| 15623 | TRAV12-2 | TRAJ35 | CAVIGFGNVLHC | TRBV29-1 | TRBJ2-1 | CSGSKKGVAEQFF | KLGGALQAK | NO | NO | NO |
| 15624 | TRAV12-1 | TRAJ10 | CVVRAGGGNKLTF | TRBV9 | TRBJ2-1 | CASSPDSLNEQFF | KLGGALQAK | YES | NO | NO |
| 15625 | TRAV38-2DV8 | TRAJ31 | CAYRSAFDARLMF | TRBV7-2 | TRBJ2-7 | CASSLAPTSGRASEQYV | KLGGALQAK | NO | NO | NO |
| 15626 | TRAV12-1 | TRAJ31 | CVVNALNNNARLMF | TRBV7-9 | TRBJ1-2 | CASGFSGTGPLFYGYTF | KLGGALQAK | YES | NO | NO |
| 15627 | TRAV12-2 | TRAJ44 | CAARFTGTASKLTF | TRBV19 | TRBJ1-3 | CASSPVSGNTIYF | KLGGALQAK | YES | NO | NO |
| 15628 | TRAV20 | TRAJ40 | CAVPTLRGGTYKYIF | TRBV29-1 | TRBJ2-1 | CSVLRTGGLDEQFF | KLGGALQAK | YES | NO | NO |
| 15629 | TRAV12-2 | TRAJ54 | CAVRVQGAQKLVF | TRBV5-1 | TRBJ1-2 | CASSLDRGTGGYTF | KLGGALQAK | YES | NO | NO |
| 15630 | TRAV12-2 | TRAJ42 | CAFQGGGSQGNLIF | TRBV6-5 | TRBJ1-1 | CASSYSPRKGYEAFF | KLGGALQAK | YES | NO | NO |
| 15631 | TRAV8-3 | TRAJ7 | CAVESPYGNNRLAF | TRBV28 | TRBJ2-7 | CASSLNISYEQYF | KLGGALQAK | YES | NO | NO |
| 15632 | TRAV14DV4 | TRAJ39 | CAMREPIKNAGNMLTF | TRBV7-9 | TRBJ1-5 | CASSLVGPGSNQPQHF | KLGGALQAK | YES | NO | NO |
| 15633 | TRAV12-3 | TRAJ43 | CAMSWQAFPENDMRF | TRBV12-5 | TRBJ2-2 | CASGLPQGDSSGELFF | KLGGALQAK | YES | NO | NO |
| 15634 | TRAV25 | TRAJ26 | CAGRYGQNFVF | TRBV27 | TRBJ2-5 | CASSYQGYQETQYF | KLGGALQAK | YES | NO | NO |
| 15635 | TRAV10 | TRAJ37 | CVVSFLNGSGNTGKLIF | TRBV6-6 | TRBJ2-2 | CASRTGDYRRAGELFF | KLGGALQAK | YES | NO | NO |
| 15636 | TRAV1-2 | TRAJ20 | CAVRDGDYKLSF | TRBV6-1 | TRBJ2-7 | CASSEESSGGYEQYF | KLGGALQAK | YES | NO | NO |
| 15637 | TRAV38-2DV8 | TRAJ26 | CAYRSEDNYGQNFVF | TRBV9 | TRBJ2-2 | CASSAGTGFFSGELFF | KLGGALQAK | YES | NO | NO |
| 15638 | TRAV29DV5 | TRAJ49 | CAASLLGNQFYF | TRBV4-1 | TRBJ2-3 | CASSHGAGDTDTQYF | KLGGALQAK | YES | NO | NO |
| 15639 | TRAV24 | TRAJ52 | CAFIGDAGGTSYGKLTF | TRBV14 | TRBJ1-4 | CASSQGPGTVHEKLFF | KLGGALQAK | NO | NO | NO |
| 15640 | TRAV21 | TRAJ40 | CAGSTSGTYKYIF | TRBV5-6 | TRBJ2-7 | CASSLAAGWQYV | KLGGALQAK | NO | NO | NO |
| 15641 | TRAV26-1 | TRAJ26 | CIAGYGQNFVF | TRBV11-2 | TRBJ2-5 | CASSLGKRQGRGQETQYF | KLGGALQAK | YES | NO | NO |
| 15642 | TRAV5 | TRAJ40 | CAEKASGTYKYIF | TRBV12-4 | TRBJ2-2 | CASSLGENTGELFF | AVFDRKSDAK | YES | NO | YES |
| 15643 | TRAV19 | TRAJ20 | CALRLYNDYKLSF | TRBV7-9 | TRBJ1-1 | CASSPKGQGIVTEAFF | KLGGALQAK | YES | NO | NO |
| 15644 | TRAV1-1 | TRAJ30 | CAVRHRDDKIIF | TRBV2 | TRBJ1-2 | CASGKTGGLTNYGYTF | KLGGALQAK | YES | NO | NO |
| 15645 | TRAV17 | TRAJ58 | CAPSGGTSGSRLTF | TRBV19 | TRBJ2-1 | CASSSMTGLAGGSYNEQFF | KLGGALQAK | YES | NO | NO |
| 15646 | TRAV19 | TRAJ30 | CALSRDDKIIF | TRBV11-3 | TRBJ2-1 | CASKPRLSSYNEQFF | KLGGALQAK | YES | NO | NO |
| 15647 | TRAV1-2 | TRAJ34 | CAVHNTDKLIF | TRBV6-5 | TRBJ1-3 | CASSLGQGLGNTIYF | KLGGALQAK | YES | NO | NO |
| 15648 | TRAV25 | TRAJ31 | CAGVNARLMF | TRBV12-3 | TRBJ2-3 | CASSFSGRATDTQYF | AVFDRKSDAK | YES | YES | YES |
| 15649 | TRAV1-1 | TRAJ31 | CAVEDNARLMF | TRBV3-1 | TRBJ2-5 | CASSQVSEQETQYF | KLGGALQAK | YES | NO | NO |
| 15650 | TRAV38-1 | TRAJ28 | CAFMKPHSGAGSYQLTF | TRBV20-1 | TRBJ2-5 | CSVADRGETQYF | KLGGALQAK | YES | NO | NO |
| 15651 | TRAV17 | TRAJ42 | CATAPMNYGGSQGNLIF | TRBV11-2 | TRBJ2-7 | CASSFSFPAYEQYV | KLGGALQAK | NO | NO | NO |
| 15652 | TRAV19 | TRAJ10 | CALSEALTGGGNKLTF | TRBV29-1 | TRBJ2-7 | CSVEDGGTAPYEQYF | KLGGALQAK | YES | NO | NO |
| 15653 | TRAV19 | TRAJ39 | CALSEARNAGNMLTF | TRBV15 | TRBJ2-7 | CATSRDQAPGEQYF | KLGGALQAK | YES | NO | NO |
| 15654 | TRAV20 | TRAJ9 | CAVYTGGFKTIF | TRBV9 | TRBJ1-2 | CASSPPGGTGGADGYTF | KLGGALQAK | YES | NO | NO |
| 15655 | TRAV21 | TRAJ36 | CAVNQTGANNLFF | TRBV7-9 | TRBJ2-4 | CASSRLTSANIQYF | KLGGALQAK | YES | NO | NO |
| 15656 | TRAV39 | TRAJ31 | CAVDGGNARLMF | TRBV7-6 | TRBJ2-5 | CASSLASLGETQYF | KLGGALQAK | YES | NO | NO |
| 15657 | TRAV12-2 | TRAJ50 | CAVARKTSYDKVIF | TRBV6-3 | TRBJ2-5 | CASNGRGIQETQYF | KLGGALQAK | YES | YES | NO |
| 15658 | TRAV8-2 | TRAJ18 | CVVSDLNRGSTLGRLYF | TRBV20-1 | TRBJ2-7 | CSTNSASGDYEQYF | KLGGALQAK | NO | NO | NO |
| 15659 | TRAV12-1 | TRAJ42 | CVVNRGSQGNLIF | TRBV12-4 | TRBJ1-4 | CASSFIQGRDEKLFF | KLGGALQAK | YES | NO | NO |
| 15660 | TRAV22 | TRAJ12 | CAVKGWVDSSYKLIF | TRBV24-1 | TRBJ1-2 | CATSDEDTGYGYTF | KLGGALQAK | YES | NO | NO |
| 15661 | TRAV19 | TRAJ54 | CALSGPRIQGAQKLVF | TRBV4-1 | TRBJ2-1 | CASSQESGLAHNEQFF | FLASKIGRLV | YES | NO | YES |
| 15662 | TRAV17 | TRAJ44 | CATDVTGTASKLTF | TRBV5-1 | TRBJ2-1 | CASKDPSGDPFYNEQFF | KLGGALQAK | YES | NO | NO |
| 15663 | TRAV12-2 | TRAJ49 | CAVNSPGNQFYF | TRBV7-9 | TRBJ1-1 | CASSVLGQGRSEAFF | KLGGALQAK | YES | NO | NO |
| 15664 | TRAV35 | TRAJ49 | CAGPGTGNQFYF | TRBV7-9 | TRBJ1-2 | CASSFSYTGHYGYTF | KLGGALQAK | YES | NO | NO |
| 15665 | TRAV9-2 | TRAJ47 | CALSGRGNKLVF | TRBV10-3 | TRBJ1-2 | CAISGGQIVPNGYTF | KLGGALQAK | YES | NO | NO |
| 15666 | TRAV8-4 | TRAJ31 | CAVRNNARLMF | TRBV4-3 | TRBJ2-1 | CASSQVGLAELYNEQFF | KLGGALQAK | YES | NO | NO |
| 15667 | TRAV38-1 | TRAJ26 | CAFSKYTNYGQNFVF | TRBV6-3 | TRBJ2-5 | CASSYSQETQYF | KLGGALQAK | YES | YES | NO |
| 15668 | TRAV12-2 | TRAJ12 | CAVLDSSYKLIF | TRBV7-3 | TRBJ2-1 | CASSVTFGVARYEQFF | KLGGALQAK | YES | NO | NO |
| 15669 | TRAV1-2 | TRAJ23 | CAVRAGGKLIF | TRBV29-1 | TRBJ2-1 | CSVEEGGLGLFF | KLGGALQAK | NO | NO | NO |
| 15670 | TRAV41 | TRAJ43 | CAVDASNDMRF | TRBV19 | TRBJ2-1 | CASSINPPGQGSYNEQFF | KLGGALQAK | YES | NO | NO |
| 15671 | TRAV8-3 | TRAJ45 | CAVGSIGSGGGADGLTF | TRBV6-5 | TRBJ2-7 | CASGRSPYEQYF | KLGGALQAK | YES | NO | NO |
| 15672 | TRAV4 | TRAJ32 | CLVAGGGATNKLIF | TRBV7-6 | TRBJ1-1 | CASSSGTGFPHNTEAFF | KLGGALQAK | YES | NO | NO |
| 15673 | TRAV17 | TRAJ26 | CATDADYGQNFVF | TRBV19 | TRBJ1-5 | CASSVLRDQPQHF | KLGGALQAK | YES | NO | NO |
| 15674 | TRAV38-2DV8 | TRAJ42 | CAYRSPTVVRNLIF | TRBV7-9 | TRBJ2-7 | CASSSSGTKDYEQYF | KLGGALQAK | YES | NO | NO |
| 15675 | TRAV17 | TRAJ47 | CATDGRMEYGNKLVF | TRBV5-1 | TRBJ1-3 | CASSLEGPGSSGNTIYF | KLGGALQAK | YES | NO | NO |
| 15676 | TRAV29DV5 | TRAJ30 | CAASYDKIIF | TRBV12-3 | TRBJ2-1 | CASSLQGLAVEQFF | KLGGALQAK | YES | NO | NO |
| 15677 | TRAV12-3 | TRAJ21 | CAMSAEYNFNKFYF | TRBV6-1 | TRBJ2-1 | CATLRQGAYNEQFF | KLGGALQAK | YES | NO | NO |
| 15678 | TRAV2 | TRAJ40 | CAPHPSGTYKYIF | TRBV19 | TRBJ2-1 | CASTRDGYNEQFF | KLGGALQAK | NO | NO | NO |
| 15679 | TRAV17 | TRAJ53 | CATKNSGGSNYKLTF | TRBV19 | TRBJ1-2 | CASSARTGVGYGYTF | KLGGALQAK | YES | NO | NO |
| 15680 | TRAV8-1 | TRAJ28 | CAVNPLYSGAGSYQLTF | TRBV7-8 | TRBJ2-2 | CASSLGSGSLPGELFF | KLGGALQAK | YES | NO | NO |
| 15681 | TRAV12-2 | TRAJ34 | CAVRRNTDKLIF | TRBV7-9 | TRBJ2-7 | CASSPKTDSYEQYF | KLGGALQAK | YES | NO | NO |
| 15682 | TRAV8-1 | TRAJ30 | CAVVGGRDDKIIF | TRBV20-1 | TRBJ2-5 | CSARDPEYGETQYF | KLGGALQAK | YES | NO | NO |
| 15683 | TRAV19 | TRAJ37 | CALGHGSSNTGKLIF | TRBV7-2 | TRBJ2-3 | CASSLLISQNTDTQYF | AVFDRKSDAK | YES | NO | YES |
| 15684 | TRAV12-3 | TRAJ40 | CAMSFTSGTYKYIF | TRBV6-1 | TRBJ2-5 | CASSEGDPQETQYF | KLGGALQAK | YES | NO | NO |
| 15685 | TRAV4 | TRAJ11 | CLVGELGYSTLTF | TRBV12-4 | TRBJ2-3 | CASSLATSTDTQYF | AVFDRKSDAK | YES | NO | YES |
| 15686 | TRAV21 | TRAJ47 | CAVVSMEYGNKLVF | TRBV12-4 | TRBJ1-5 | CASSLSASNQPQHF | KLGGALQAK | YES | NO | NO |
| 15687 | TRAV12-1 | TRAJ18 | CVVNRDRGSTLGRLYF | TRBV20-1 | TRBJ1-5 | CSARSGTGGFDQPQHF | KLGGALQAK | YES | NO | NO |
| 15688 | TRAV9-2 | TRAJ32 | CALSEGYGGATNKLIF | TRBV7-9 | TRBJ1-1 | CASTLPRDTEAFF | KLGGALQAK | YES | NO | NO |
| 15689 | TRAV3 | TRAJ7 | CAVRDYGNNRLAF | TRBV4-1 | TRBJ1-5 | CASSNDRDSNQPQHF | KLGGALQAK | YES | NO | NO |
| 15690 | TRAV12-1 | TRAJ33 | CVVMDSNYQLIW | TRBV18 | TRBJ2-5 | CASSPEGLQETQYF | KLGGALQAK | YES | NO | NO |
| 15691 | TRAV13-1 | TRAJ49 | CAASINRNTGNQFYF | TRBV7-6 | TRBJ2-5 | CASSLGWTGQETQYF | KLGGALQAK | YES | NO | NO |
| 15692 | TRAV25 | TRAJ29 | CAGTNSGNTPLVF | TRBV19 | TRBJ1-1 | CASQQGAVTEAFF | KLGGALQAK | YES | NO | NO |
| 15693 | TRAV14DV4 | TRAJ53 | CAMREGGGSNYKLTF | TRBV27 | TRBJ2-2 | CASSPYLTLGELFF | KLGGALQAK | YES | NO | NO |
| 15694 | TRAV3 | TRAJ26 | CAVRDILGQNFVF | TRBV19 | TRBJ1-1 | CASRSGTGGTNTEAFF | KLGGALQAK | NO | NO | NO |
| 15695 | TRAV8-4 | TRAJ8 | CAVRADGFQKLVF | TRBV7-9 | TRBJ1-5 | CASSLVRGGGDQPQHF | KLGGALQAK | YES | NO | NO |

| | | | | | | | | | | |
|---|---|---|---|---|---|---|---|---|---|---|
| 15696 | TRAV14DV4 | TRAJ16 | CAMREGWF | TRBV12-4 | TRBJ2-5 | CASSPGGREQETQYF | KLGGALQAK | NO | NO | NO |
| 15697 | TRAV30 | TRAJ22 | CGNLHFGSARQLTF | TRBV12-4 | TRBJ2-2 | CASSLWEADTGELFF | KLGGALQAK | YES | NO | NO |
| 15698 | TRAV13-1 | TRAJ8 | CAARRGFQKLVF | TRBV9 | TRBJ2-1 | CASSEDGSSYNEQFF | KLGGALQAK | YES | NO | NO |
| 15699 | TRAV14DV4 | TRAJ9 | CAMREADTGGFKTIF | TRBV12-4 | TRBJ2-5 | CASSLRARLETQYF | KLGGALQAK | YES | NO | NO |
| 15700 | TRAV8-3 | TRAJ45 | CAVGGGADGLTF | TRBV6-4 | TRBJ2-2 | CASSDSNTGELFF | KLGGALQAK | YES | NO | NO |
| 15701 | TRAV38-2DV8 | TRAJ23 | CAYRSLYQGGKLIF | TRBV7-9 | TRBJ1-1 | CASSLDLGTEAFF | KLGGALQAK | YES | YES | NO |
| 15702 | TRAV27 | TRAJ45 | CAGVGGSGGGADGLTF | TRBV29-1 | TRBJ2-2 | CSVATDRGYTGELFF | KLGGALQAK | YES | NO | NO |
| 15703 | TRAV12-3 | TRAJ43 | CAMMYNNNDMRF | TRBV3-1 | TRBJ1-1 | CASFGDTEAFF | KLGGALQAK | YES | NO | NO |
| 15704 | TRAV14DV4 | TRAJ33 | CAMRMDSNYQLIW | TRBV25-1 | TRBJ1-5 | CASSEYTGSNQPQHF | KLGGALQAK | YES | NO | NO |
| 15705 | TRAV10 | TRAJ20 | CVVSIGSNDYKLSF | TRBV27 | TRBJ2-7 | CASRGELYEQYF | IVTDFSVIK | YES | NO | YES |
| 15706 | TRAV29DV5 | TRAJ52 | CAASLANAGGTSYGKLTF | TRBV4-3 | TRBJ2-1 | CASSQDRRVASGGNYNEQFF | KLGGALQAK | NO | NO | NO |
| 15707 | TRAV8-1 | TRAJ40 | CAVGLGTYKYIF | TRBV30 | TRBJ1-4 | CAWSVGQAPEKLFF | KLGGALQAK | YES | NO | NO |
| 15708 | TRAV14DV4 | TRAJ28 | CAMREHSGAGSYQLTF | TRBV6-5 | TRBJ2-3 | CASSTPGNTQYF | KLGGALQAK | NO | NO | NO |
| 15709 | TRAV21 | TRAJ23 | CAVIKGGKLIF | TRBV6-3 | TRBJ2-7 | CASSYSILAGLYEQYV | AVFDRKSDAK | NO | NO | NO |
| 15710 | TRAV1-1 | TRAJ30 | CAVRDDKIIF | TRBV27 | TRBJ2-3 | CASSQPGQGWQDTQYF | KLGGALQAK | YES | NO | NO |
| 15711 | TRAV26-2 | TRAJ52 | CILTANAGGTSYGKLTF | TRBV16 | TRBJ2-7 | CASSHGQFSLYEQYF | AVFDRKSDAK | YES | NO | YES |
| 15712 | TRAV1-2 | TRAJ33 | CAVEASDSNYQLIW | TRBV4-1 | TRBJ1-1 | CASSHRAGTAGTEAFF | KLGGALQAK | YES | NO | NO |
| 15713 | TRAV1-2 | TRAJ28 | CAVSPASDVAGSYQLTF | TRBV19 | TRBJ1-2 | CASTGGPNYGYTF | KLGGALQAK | YES | NO | NO |
| 15714 | TRAV17 | TRAJ30 | CATDSSRDDKIIF | TRBV19 | TRBJ1-1 | CASSSGTGVTEAFF | KLGGALQAK | YES | NO | NO |
| 15715 | TRAV26-1 | TRAJ49 | CIVRTNTGNQFYF | TRBV10-2 | TRBJ1-2 | CASSESFLSPTYGYTF | KLGGALQAK | YES | NO | NO |
| 15717 | TRAV1-1 | TRAJ28 | CAAGGYSGAGSYQLTF | TRBV7-9 | TRBJ2-7 | CASSSGTRTYEQYF | KLGGALQAK | YES | NO | NO |
| 15718 | TRAV8-3 | TRAJ40 | CAVEIPHTSGTYKYIF | TRBV10-3 | TRBJ2-3 | CAIKRESTDTQYF | KLGGALQAK | YES | NO | NO |
| 15719 | TRAV22 | TRAJ53 | CAVDRSGGSNYKLTF | TRBV20-1 | TRBJ1-1 | CSAFRRGMNTEAFF | KLGGALQAK | YES | NO | NO |
| 15720 | TRAV19 | TRAJ37 | CALSEEKGSSNTGKLIF | TRBV4-1 | TRBJ2-3 | CASSQDMGQGTDTQYF | KLGGALQAK | YES | NO | NO |
| 15721 | TRAV19 | TRAJ16 | CALSDPF | TRBV5-6 | TRBJ1-3 | CASSSFTGMNSGNTIYF | KLGGALQAK | NO | NO | NO |
| 15722 | TRAV26-1 | TRAJ52 | CIVTSSYLAGGTSYGKLTF | TRBV6-4 | TRBJ1-6 | CASSDDRSYNSPLHF | KLGGALQAK | YES | NO | NO |
| 15723 | TRAV19 | TRAJ6 | CALSEPSGGSYIPTF | TRBV2 | TRBJ2-4 | CASSDRSAKNIQYF | KLGGALQAK | YES | NO | NO |
| 15724 | TRAV12-1 | TRAJ8 | CVVKGAGFQKLVF | TRBV4-3 | TRBJ2-1 | CASSQIGSYNEQFF | KLGGALQAK | YES | NO | NO |
| 15725 | TRAV8-1 | TRAJ13 | CAVNAPGGYQKVTF | TRBV19 | TRBJ2-3 | CASSPGRGVADTQYF | KLGGALQAK | YES | NO | NO |
| 15726 | TRAV12-3 | TRAJ5 | CASDTGRRALTF | TRBV10-2 | TRBJ1-1 | CAIEDGMNTEAFF | KLGGALQAK | YES | NO | NO |
| 15727 | TRAV1-2 | TRAJ11 | CALMDSGYSTLTF | TRBV6-3 | TRBJ1-3 | CASVTGPGNTIYF | KLGGALQAK | YES | NO | NO |
| 15728 | TRAV1-2 | TRAJ33 | CAVMDSNYQLIW | TRBV20-1 | TRBJ1-1 | CSAGQGDTEAFF | KLGGALQAK | YES | NO | NO |
| 15729 | TRAV14DV4 | TRAJ41 | CAMREGGWDSGYALNF | TRBV7-3 | TRBJ2-1 | CASSPAGAELYNEQFF | KLGGALQAK | YES | NO | NO |
| 15730 | TRAV19 | TRAJ33 | CALSEVSNYQLIW | TRBV9 | TRBJ2-1 | CASSVEPSGYNEQFF | KLGGALQAK | YES | NO | NO |
| 15731 | TRAV29DV5 | TRAJ45 | CAASGTGGGADGLTF | TRBV4-2 | TRBJ1-2 | CASSHGQGNYGYTF | KLGGALQAK | YES | NO | NO |
| 15732 | TRAV1-2 | TRAJ48 | CAVRSISNFGNEKLTF | TRBV6-1 | TRBJ2-1 | CASSEARQAVLNEQFF | KLGGALQAK | YES | NO | NO |
| 15733 | TRAV38-1 | TRAJ30 | CAFHVNRDDKIIF | TRBV19 | TRBJ1-1 | CASSILSPSTEAFF | KLGGALQAK | YES | NO | NO |
| 15734 | TRAV1-2 | TRAJ4 | CAPGPFLSGGYNKLIF | TRBV14 | TRBJ1-1 | CASSPLGGREAFF | KLGGALQAK | NO | NO | NO |
| 15735 | TRAV27 | TRAJ41 | CAGPNSNSGYALNF | TRBV4-2 | TRBJ1-2 | CASSQETGYGYTF | AVFDRKSDAK | YES | NO | YES |
| 15736 | TRAV22 | TRAJ54 | CAVEGTIQGAQKLVF | TRBV27 | TRBJ2-7 | CASSYKQGYEQYF | KLGGALQAK | YES | NO | NO |
| 15737 | TRAV12-1 | TRAJ39 | CVVNGGSAAGNMLTF | TRBV15 | TRBJ2-2 | CATSRERFTGELFF | KLGGALQAK | YES | NO | NO |
| 15738 | TRAV1-1 | TRAJ8 | CAVSTGFQKLVF | TRBV27 | TRBJ2-1 | CASSSDPGQGASYNEQFF | KLGGALQAK | YES | NO | NO |
| 15739 | TRAV19 | TRAJ56 | CALSGSGTGANSKLTF | TRBV4-1 | TRBJ2-5 | CASSQGGGRGQETQYF | KLGGALQAK | YES | NO | NO |
| 15740 | TRAV5 | TRAJ5 | CAEISVDTGRRALTF | TRBV15 | TRBJ1-5 | CATSTYPTGEILVNQPQHF | KLGGALQAK | YES | NO | NO |
| 15741 | TRAV8-3 | TRAJ36 | CAVGAGDQTGANNLFF | TRBV10-3 | TRBJ2-6 | CAISEPHNVLTF | KLGGALQAK | YES | NO | NO |
| 15742 | TRAV19 | TRAJ23 | CALMYNQGGKLIF | TRBV4-1 | TRBJ1-2 | CASSQGDRNDVYGYTF | KLGGALQAK | YES | NO | NO |
| 15743 | TRAV26-2 | TRAJ52 | CILRDSGGTSYGKLTF | TRBV20-1 | TRBJ2-1 | CSAREVNTNEQFF | KLGGALQAK | NO | NO | NO |
| 15744 | TRAV14DV4 | TRAJ35 | CAMREARGFGNVLHC | TRBV6-5 | TRBJ2-2 | CASSYSNSLTGELFF | KLGGALQAK | NO | NO | NO |
| 15745 | TRAV21 | TRAJ23 | CAVPFQGGKLIF | TRBV4-2 | TRBJ1-4 | CASSLLDEKLFF | KLGGALQAK | YES | NO | NO |
| 15746 | TRAV12-1 | TRAJ3 | CVVNNFYSSASKIIF | TRBV4-3 | TRBJ1-2 | CASSQAHPTGKDGYTF | KLGGALQAK | YES | NO | NO |
| 15747 | TRAV29DV5 | TRAJ49 | CAATGNQFYF | TRBV7-9 | TRBJ1-2 | CASSLGNYGYTF | KLGGALQAK | YES | NO | NO |
| 15748 | TRAV3 | TRAJ43 | CAVRGKNNNDMRF | TRBV20-1 | TRBJ2-3 | CSARSRTSGGPPVGDTQYF | KLGGALQAK | NO | NO | NO |
| 15749 | TRAV19 | TRAJ8 | CALSEARFQKLVF | TRBV4-1 | TRBJ1-1 | CASSQDRQALSTEAFF | KLGGALQAK | YES | NO | NO |
| 15750 | TRAV22 | TRAJ49 | CAGLNTGNQFYF | TRBV19 | TRBJ1-2 | CASSTLTGANGYTF | KLGGALQAK | YES | NO | NO |
| 15751 | TRAV8-2 | TRAJ52 | CVVSEIHAGGTSYGKLTF | TRBV7-6 | TRBJ2-5 | CASSLLFGGEGQETQYF | KLGGALQAK | YES | NO | NO |
| 15752 | TRAV12-2 | TRAJ30 | CAVNVLNRDDKIIF | TRBV11-2 | TRBJ1-4 | CASSLTWDEKLFF | KLGGALQAK | YES | NO | NO |
| 15753 | TRAV13-1 | TRAJ47 | CAAREYGNKLVF | TRBV3-1 | TRBJ1-2 | CASSQDRGGPEGLNYGYTF | KLGGALQAK | YES | NO | NO |
| 15754 | TRAV12-1 | TRAJ49 | CVVNTGNQFYF | TRBV12-4 | TRBJ2-7 | CAATGTVSYEQYF | KLGGALQAK | YES | NO | NO |
| 15755 | TRAV21 | TRAJ45 | CAVDSGGGADGLTF | TRBV6-5 | TRBJ1-5 | CASSDFRYQPQHF | KLGGALQAK | YES | NO | NO |
| 15756 | TRAV13-2 | TRAJ28 | CAENPYSGAGSYQLTF | TRBV19 | TRBJ2-1 | CASRRGTVYNEQFF | FLASKIGRLV | YES | NO | YES |
| 15757 | TRAV3 | TRAJ26 | CAVRDYYGQNFVF | TRBV27 | TRBJ1-1 | CASSSSGQFEAFF | KLGGALQAK | NO | NO | NO |
| 15758 | TRAV3 | TRAJ20 | CAVRVFNDYKLSF | TRBV29-1 | TRBJ1-1 | CSAGTGGPNTEAFF | KLGGALQAK | NO | NO | NO |
| 15759 | TRAV19 | TRAJ10 | CALSGLTGGGNKLTF | TRBV19 | TRBJ2-1 | CASSMRTGAHNEQFF | KLGGALQAK | YES | NO | NO |
| 15760 | TRAV19 | TRAJ49 | CALSETGNQFYF | TRBV6-4 | TRBJ2-1 | CASSPLAVPYNEQFF | KLGGALQAK | YES | NO | NO |
| 15761 | TRAV1-2 | TRAJ11 | CAVRDSNSGYSTLTF | TRBV4-1 | TRBJ1-6 | CASSQERAGGVSPLHF | KLGGALQAK | YES | NO | NO |
| 15762 | TRAV27 | TRAJ52 | CAGVVQAGGTSYGKLTF | TRBV9 | TRBJ2-1 | CASSGVAYNEQFF | KLGGALQAK | YES | NO | NO |
| 15763 | TRAV14DV4 | TRAJ50 | CAMREAQTSYDKVIF | TRBV6-3 | TRBJ2-3 | CASSSRQGPRTDTQYF | KLGGALQAK | YES | NO | NO |
| 15764 | TRAV12-3 | TRAJ39 | CAIGPGNMLTF | TRBV4-1 | TRBJ2-2 | CAIVGDTGELFF | KLGGALQAK | YES | NO | NO |
| 15765 | TRAV13-1 | TRAJ42 | CAAVGVYGGSQGNLIF | TRBV9 | TRBJ2-5 | CASSVGETQYF | KLGGALQAK | YES | NO | NO |
| 15766 | TRAV21 | TRAJ57 | CAVSPPKLTQGGSEKLVF | TRBV7-6 | TRBJ1-1 | CASSSPLDRAGSAFF | KLGGALQAK | YES | NO | NO |
| 15767 | TRAV8-1 | TRAJ31 | CAVKAGNNARLMF | TRBV29-1 | TRBJ1-2 | CSVVRADGYTF | KLGGALQAK | YES | NO | NO |
| 15768 | TRAV29DV5 | TRAJ40 | CAAKPSGTYKYIF | TRBV4-1 | TRBJ1-1 | CASSQGRGAEAFF | KLGGALQAK | YES | NO | NO |
| 15769 | TRAV3 | TRAJ23 | CAVRDINYNQGGKLIF | TRBV3-1 | TRBJ2-1 | CASSEKSSGAGYNEQFF | KLGGALQAK | NO | NO | NO |
| 15770 | TRAV1-1 | TRAJ26 | CAVRWDYGQNFVF | TRBV6-5 | TRBJ1-2 | CATLISGNNYGYTF | KLGGALQAK | YES | NO | NO |
| 15771 | TRAV20 | TRAJ30 | CAVQAGGDKIIF | TRBV19 | TRBJ1-5 | CASSTSGYNQPQHF | KLGGALQAK | YES | NO | NO |
| 15772 | TRAV19 | TRAJ21 | CALSEAGNFNKFYF | TRBV19 | TRBJ2-5 | CASSSTGTSDGETQYF | KLGGALQAK | YES | NO | NO |
| 15773 | TRAV29DV5 | TRAJ29 | CAARYSGNTPLVF | TRBV7-9 | TRBJ2-3 | CASSPETPGGESGGTDTQYF | KLGGALQAK | NO | NO | NO |
| 15774 | TRAV29DV5 | TRAJ45 | CAASVGGGADGLTF | TRBV5-6 | TRBJ2-1 | CASIGGTGYNEQFF | KLGGALQAK | YES | NO | NO |
| 15775 | TRAV5 | TRAJ41 | CAENSNSGYALNF | TRBV18 | TRBJ2-7 | CASSPRDRVLEQYF | KLGGALQAK | YES | NO | NO |
| 15776 | TRAV27 | TRAJ23 | CAGPTYNQGGKLIF | TRBV4-1 | TRBJ2-2 | CASSQEGPGLFTGELFF | KLGGALQAK | YES | NO | NO |
| 15777 | TRAV4 | TRAJ49 | CLVGDIGNQFYF | TRBV5-1 | TRBJ1-2 | CASSLTTGGFNYGYTF | KLGGALQAK | YES | NO | NO |
| 15778 | TRAV35 | TRAJ54 | CAGGRIQGAQKLVF | TRBV12-4 | TRBJ2-5 | CASSLWDGQETQYF | AVFDRKSDAK | YES | NO | YES |
| 15779 | TRAV8-6 | TRAJ49 | CAVSDRSGNQFYF | TRBV12-4 | TRBJ1-1 | CASSFYNTEAFF | KLGGALQAK | YES | NO | NO |
| 15780 | TRAV8-1 | TRAJ17 | CAVKGRAAGNKLTF | TRBV7-9 | TRBJ2-1 | CASSLSGLGDYNEQFF | KLGGALQAK | YES | NO | NO |
| 15781 | TRAV3 | TRAJ34 | CAVRDIGGTDKLIF | TRBV2 | TRBJ2-7 | CASREGGTSYEQYF | KLGGALQAK | NO | NO | NO |
| 15782 | TRAV25 | TRAJ31 | CAGPPWNNNARLMF | TRBV6-6 | TRBJ2-2 | CASLRSGELFF | KLGGALQAK | YES | NO | NO |
| 15783 | TRAV8-6 | TRAJ5 | CAVTNADTGRRALTF | TRBV9 | TRBJ2-7 | CASSEGGEFGDEQYV | KLGGALQAK | NO | NO | NO |
| 15784 | TRAV12-3 | TRAJ42 | CAMSLYGGSQGNLIF | TRBV20-1 | TRBJ2-1 | CSARPTSSPLLNEQFF | KLGGALQAK | NO | NO | NO |
| 15785 | TRAV12-3 | TRAJ39 | CAIGPGNMLTF | TRBV14 | TRBJ1-5 | CASSQTGLGSQPQHF | KLGGALQAK | NO | NO | NO |
| 15786 | TRAV17 | TRAJ15 | CATDASQAGTALIF | TRBV4-1 | TRBJ2-7 | CASSQGLASSYEQYV | KLGGALQAK | NO | NO | NO |
| 15787 | TRAV29DV5 | TRAJ27 | CAVVNAGKSTF | TRBV5-4 | TRBJ2-1 | CASSISGTILNEQFF | AVFDRKSDAK | YES | NO | YES |
| 15788 | TRAV5 | TRAJ37 | CAESITGKLIF | TRBV7-2 | TRBJ2-7 | CASSLEPSYEQYF | KLGGALQAK | NO | NO | NO |
| 15789 | TRAV8-3 | TRAJ53 | CAVGGSGGSNYKLTF | TRBV19 | TRBJ2-6 | CASSMRAGRPGANVLTF | KLGGALQAK | YES | NO | NO |
| 15790 | TRAV12-2 | TRAJ45 | CALDSGGGADGLTF | TRBV6-3 | TRBJ2-7 | CASSYGVQFEQYV | KLGGALQAK | NO | NO | NO |
| 15791 | TRAV20 | TRAJ45 | CAVRGGGADGLTF | TRBV7-8 | TRBJ1-1 | CASSLVGQGYTEAFF | KLGGALQAK | YES | NO | NO |
| 15792 | TRAV19 | TRAJ34 | CALSEELNTDKLIF | TRBV4-2 | TRBJ1-4 | CASSQVRSLGQENEKLFF | KLGGALQAK | YES | NO | NO |
| 15793 | TRAV24 | TRAJ26 | CASPDNYGQNFVF | TRBV19 | TRBJ1-1 | CASSLGTEGGTEAFF | KLGGALQAK | YES | NO | NO |
| 15794 | TRAV8-1 | TRAJ37 | CAVSSGNTGKLIF | TRBV13 | TRBJ2-7 | CASSYGLAESYEQYF | KLGGALQAK | YES | NO | NO |
| 15795 | TRAV8-6 | TRAJ53 | CAVGHEDGGSNYKLTF | TRBV7-9 | TRBJ2-7 | CASSPVFRTEQYF | KLGGALQAK | YES | NO | NO |
| 15796 | TRAV17 | TRAJ49 | CATDFGQFYF | TRBV7-2 | TRBJ2-4 | CASSLGIDIQYF | KLGGALQAK | YES | NO | NO |
| 15797 | TRAV29DV5 | TRAJ36 | CAAQLPGTGANNLFF | TRBV20-1 | TRBJ1-4 | CSVNQILGQPNEKLFF | KLGGALQAK | YES | NO | NO |
| 15798 | TRAV19 | TRAJ36 | CALSEAGGRGANNLFF | TRBV11-2 | TRBJ2-3 | CAIGQRGGLDTQYF | KLGGALQAK | YES | NO | NO |
| 15799 | TRAV4 | TRAJ23 | CLVGPFYNQGGKLIF | TRBV7-8 | TRBJ2-1 | CASSPGTGMDEQFF | KLGGALQAK | NO | NO | NO |
| 15800 | TRAV38-2DV8 | TRAJ30 | CAYRAGGDDKIIF | TRBV9 | TRBJ1-1 | CASSAPEGLETEAFF | KLGGALQAK | YES | NO | NO |

| | | | | | | | | | | |
|---|---|---|---|---|---|---|---|---|---|---|
| 15801 | TRAV12-3 | TRAJ39 | CAIGPGNMLTF | TRBV14 | TRBJ1-1 | CASSLVPGIAEAFF | KLGGALQAK | NO | NO | NO |
| 15802 | TRAV19 | TRAJ23 | CALSPFYNQGGKLIF | TRBV7-9 | TRBJ1-1 | CASSSSQGPEAFF | KLGGALQAK | YES | NO | NO |
| 15803 | TRAV9-2 | TRAJ26 | CALIDQNFVF | TRBV7-9 | TRBJ2-7 | CASLTRDRGFDEQYF | KLGGALQAK | YES | NO | NO |
| 15804 | TRAV3 | TRAJ34 | CAVRDGVFWDNTDKLIF | TRBV27 | TRBJ1-2 | CASSLLGGGLSYGYTF | KLGGALQAK | NO | NO | NO |
| 15805 | TRAV25 | TRAJ41 | CAGQNSGYALNF | TRBV19 | TRBJ1-1 | CASSVDRGLNTEAFF | KLGGALQAK | YES | NO | NO |
| 15806 | TRAV36DV7 | TRAJ27 | CAAGNTNAGKSTF | TRBV19 | TRBJ2-1 | CASTATGQGEQFF | KLGGALQAK | NO | NO | NO |
| 15807 | TRAV21 | TRAJ21 | CAVWDNFNKFYF | TRBV4-1 | TRBJ1-1 | CASSRPGTEAFF | KLGGALQAK | YES | NO | NO |
| 15808 | TRAV29DV5 | TRAJ20 | CAATVDYKLSF | TRBV5-4 | TRBJ2-2 | CASSLTGTRGRAGELFF | KLGGALQAK | YES | NO | NO |
| 15809 | TRAV26-2 | TRAJ40 | CTRTSGTYKYIF | TRBV4-1 | TRBJ2-4 | CASSQRDEENIQYF | KLGGALQAK | YES | NO | NO |
| 15810 | TRAV38-2DV8 | TRAJ52 | CAYRRGRNAGGTSYGKLTF | TRBV9 | TRBJ2-6 | CASSARPGTGTGANVLTF | KLGGALQAK | YES | NO | NO |
| 15811 | TRAV29DV5 | TRAJ40 | CAADTSGTYKYIF | TRBV7-9 | TRBJ1-2 | CASSLDLRTVNYGYTF | KLGGALQAK | YES | NO | NO |
| 15812 | TRAV27 | TRAJ40 | CAGAGSGTYKYIF | TRBV4-2 | TRBJ2-7 | CASSQGQVAYEQYF | KLGGALQAK | YES | NO | NO |
| 15813 | TRAV12-2 | TRAJ48 | CAGGGLFGNEKLTF | TRBV28 | TRBJ1-2 | CASSADRGLYGYTF | KLGGALQAK | YES | NO | NO |
| 15814 | TRAV17 | TRAJ20 | CATDGGDYKLSF | TRBV12-4 | TRBJ1-5 | CASTPDFGLSNQPQHF | AVFDRKSDAK | YES | NO | YES |
| 15815 | TRAV19 | TRAJ5 | CALSEGTGRRALTF | TRBV4-3 | TRBJ2-3 | CASSQDNRGATDTQYF | KLGGALQAK | YES | NO | NO |
| 15816 | TRAV24 | TRAJ45 | CANSLAGGADGLTF | TRBV20-1 | TRBJ2-1 | CSAPPIRAGSYNEQFF | KLGGALQAK | NO | NO | NO |
| 15817 | TRAV21 | TRAJ49 | CAVQNTGNQFYF | TRBV19 | TRBJ2-1 | CASSIGLAGYNEQFF | KLGGALQAK | YES | NO | NO |
| 15818 | TRAV3 | TRAJ7 | CAPDGGEQQTRF | TRBV20-1 | TRBJ2-3 | CSAFLDRISSGREKYPPDTQYF | KLGGALQAK | NO | NO | NO |
| 15819 | TRAV13-1 | TRAJ41 | CAASINSGYALNF | TRBV7-9 | TRBJ1-4 | CASSSDAGFSAEKLFF | KLGGALQAK | YES | NO | NO |
| 15820 | TRAV30 | TRAJ48 | CGGPYLNFGNEKLTF | TRBV19 | TRBJ1-1 | CASSSGTGGTEAFF | KLGGALQAK | YES | NO | NO |
| 15821 | TRAV27 | TRAJ16 | CAGWLAWGQKLLF | TRBV12-5 | TRBJ1-3 | CASGDQGLPGNTIYF | RLRAEAQVK | NO | NO | NO |
| 15822 | TRAV19 | TRAJ17 | CALSEALKAAGNKLTF | TRBV4-3 | TRBJ1-4 | CASSQDHRMGGHEKLFF | KLGGALQAK | YES | NO | NO |
| 15823 | TRAV29DV5 | TRAJ47 | CAATGGVYGNKLVF | TRBV13 | TRBJ2-6 | CASSGQGALVSGANVLTF | KLGGALQAK | YES | NO | NO |
| 15824 | TRAV29DV5 | TRAJ48 | CAASVGGNEKLTF | TRBV20-1 | TRBJ1-5 | CSAYGPFGSQHF | KLGGALQAK | YES | NO | NO |
| 15825 | TRAV13-1 | TRAJ39 | CAASNNAGNMLTF | TRBV12-3 | TRBJ2-5 | CASSKLADYQETQYF | KLGGALQAK | YES | NO | NO |
| 15826 | TRAV29DV5 | TRAJ52 | CAASARGNAGGTSYGKLTF | TRBV6-5 | TRBJ2-1 | CASSYGSYNEQFF | AVFDRKSDAK | NO | NO | NO |
| 15827 | TRAV8-4 | TRAJ41 | CAVTGYSNSGYALNF | TRBV10-3 | TRBJ2-2 | CAISPGTSGRNTGELFF | KLGGALQAK | YES | NO | NO |
| 15828 | TRAV38-1 | TRAJ47 | CALMEYGNKLVF | TRBV7-9 | TRBJ2-1 | CASSFFGGDSYNEQFF | KLGGALQAK | YES | NO | NO |
| 15829 | TRAV1-2 | TRAJ41 | CGGGDSGYALNF | TRBV5-5 | TRBJ2-3 | CASSTGQAFTDTQYF | KLGGALQAK | YES | NO | NO |
| 15830 | TRAV4 | TRAJ13 | CLVDSGGYQKVTF | TRBV2 | TRBJ2-1 | CASSDIQYNEQFF | KLGGALQAK | NO | NO | NO |
| 15831 | TRAV8-2 | TRAJ49 | CVVSDRNTGNQFYF | TRBV28 | TRBJ2-3 | CASSTTSGGSSTDTQYF | KLGGALQAK | YES | NO | NO |
| 15832 | TRAV35 | TRAJ20 | CAGSSSNDYKLSF | TRBV6-3 | TRBJ2-2 | CASSLILTSRELFF | KLGGALQAK | YES | NO | NO |
| 15833 | TRAV1-2 | TRAJ23 | CAPTWNQGGKLIF | TRBV6-6 | TRBJ2-1 | CANGVGGEGGQFF | KLGGALQAK | NO | NO | NO |
| 15834 | TRAV1-1 | TRAJ39 | CAVADNNAGNMLTF | TRBV7-9 | TRBJ1-2 | CASSPEMAANYGYTF | KLGGALQAK | YES | NO | NO |
| 15835 | TRAV20 | TRAJ34 | CAVQAENTDKLIF | TRBV5-1 | TRBJ2-2 | CASGEYPGNTGELFF | KLGGALQAK | YES | NO | NO |
| 15836 | TRAV1-2 | TRAJ13 | CAAKNSGGYQKVTF | TRBV4-1 | TRBJ1-1 | CASSQDDRGSFTEAFF | KLGGALQAK | YES | NO | NO |
| 15837 | TRAV29DV5 | TRAJ37 | CAASVGSSNTGKLIF | TRBV3-1 | TRBJ1-1 | CASSPTLGQMNTEAFF | KLGGALQAK | YES | NO | NO |
| 15838 | TRAV12-1 | TRAJ42 | CVAIGGSQGNLIF | TRBV2 | TRBJ1-2 | CASSDRTYGYTF | KLGGALQAK | YES | NO | NO |
| 15839 | TRAV35 | TRAJ49 | CAGREAGNQFYF | TRBV10-3 | TRBJ1-6 | CAITGGNSPLHF | KLGGALQAK | YES | NO | NO |
| 15840 | TRAV12-3 | TRAJ11 | CAMTREMNSGYSTLTF | TRBV7-2 | TRBJ2-7 | CASSLEPSYEQYF | KLGGALQAK | NO | NO | NO |
| 15841 | TRAV12-1 | TRAJ5 | CVVNMGTGRRALTF | TRBV4-3 | TRBJ2-1 | CASSQDGSGLAGSNEQFF | KLGGALQAK | YES | NO | NO |
| 15842 | TRAV1-2 | TRAJ45 | CAVYSGGGADGLTF | TRBV25-1 | TRBJ1-2 | CASSGEQGSYGYTF | KLGGALQAK | YES | NO | NO |
| 15843 | TRAV8-1 | TRAJ11 | CAVKVAGYSTLTF | TRBV5-4 | TRBJ2-2 | CASSLGNTGELFF | KLGGALQAK | YES | NO | NO |
| 15844 | TRAV12-1 | TRAJ49 | CVVSNTGNQFYF | TRBV7-9 | TRBJ2-5 | CASRRAGDWETQYF | KLGGALQAK | YES | NO | NO |
| 15845 | TRAV29DV5 | TRAJ49 | CAARTRTGNQFYF | TRBV20-1 | TRBJ1-5 | CSASDRDRGFYSNQPQHF | KLGGALQAK | YES | NO | NO |
| 15846 | TRAV12-2 | TRAJ50 | CATTETSYDKVIF | TRBV10-2 | TRBJ1-5 | CASSETASGNQPQHF | KLGGALQAK | YES | NO | NO |
| 15847 | TRAV8-1 | TRAJ26 | CAVNPYGQNFVF | TRBV6-1 | TRBJ2-3 | CASSEQDRRTDTQYF | KLGGALQAK | YES | NO | NO |
| 15848 | TRAV13-2 | TRAJ39 | CAENMYNAGNMLTF | TRBV9 | TRBJ1-2 | CASRGTDANYGYTF | KLGGALQAK | YES | NO | NO |
| 15849 | TRAV35 | TRAJ22 | CAGFSGSARQLTF | TRBV27 | TRBJ2-2 | CASTPNNRNTGELFF | KLGGALQAK | YES | NO | NO |
| 15850 | TRAV17 | TRAJ52 | CATDGGGGTSYGKLTF | TRBV4-1 | TRBJ2-7 | CASSQVEGGLNEQYF | KLGGALQAK | YES | NO | NO |
| 15851 | TRAV19 | TRAJ47 | CALSSLSKLVF | TRBV18 | TRBJ1-2 | CASSPPDRLYGYTF | KLGGALQAK | YES | NO | NO |
| 15852 | TRAV8-2 | TRAJ48 | CVVRGFGNEKLTF | TRBV6-5 | TRBJ1-2 | CASSDPPTDWSRGLYGYTF | KLGGALQAK | YES | NO | NO |
| 15853 | TRAV3 | TRAJ8 | CAVSREGFQKLVF | TRBV3-1 | TRBJ1-2 | CASSQTGTGDYGYTF | KLGGALQAK | NO | NO | NO |
| 15854 | TRAV12-1 | TRAJ8 | CVVTEMNTGFQKLVF | TRBV6-1 | TRBJ1-1 | CASRTGAAEAFF | KLGGALQAK | NO | NO | NO |
| 15855 | TRAV12-3 | TRAJ39 | CAIGPGNMLTF | TRBV7-9 | TRBJ1-5 | CASSLAGQGRQPQHF | KLGGALQAK | YES | NO | NO |
| 15856 | TRAV24 | TRAJ40 | CALLTTSGTYKYIF | TRBV9 | TRBJ2-1 | CASSVEVLAGLILTQSNEQFF | KLGGALQAK | NO | NO | NO |
| 15857 | TRAV12-3 | TRAJ30 | CAMADDKIIF | TRBV7-9 | TRBJ1-1 | CASSSSGTGGTEAFF | KLGGALQAK | YES | NO | NO |
| 15858 | TRAV12-2 | TRAJ28 | CAATGAYSGAGSYQLTF | TRBV7-9 | TRBJ2-2 | CASTFGGELFF | KLGGALQAK | YES | NO | NO |
| 15859 | TRAV29DV5 | TRAJ48 | CAASAGFGNEKLTF | TRBV29-1 | TRBJ2-1 | CSVTDSSYNEQFF | KLGGALQAK | YES | NO | NO |
| 15860 | TRAV12-1 | TRAJ40 | CVVNSAGTYKYIF | TRBV9 | TRBJ2-7 | CASSPPTSYEQYF | KLGGALQAK | YES | NO | NO |
| 15861 | TRAV8-3 | TRAJ13 | CAVEGGYQKVTF | TRBV5-8 | TRBJ2-1 | CASSAQGARGQFF | KLGGALQAK | YES | NO | NO |
| 15862 | TRAV10 | TRAJ49 | CVVSAIGNQFYF | TRBV12-4 | TRBJ2-3 | CASTRTSGKDTQYF | KLGGALQAK | YES | NO | NO |
| 15863 | TRAV13-2 | TRAJ4 | CAPRKLIF | TRBV12-4 | TRBJ1-5 | CASSFGGSGNQPQHF | KLGGALQAK | YES | NO | NO |
| 15864 | TRAV8-1 | TRAJ3 | CAVNAYSSASKIIF | TRBV2 | TRBJ1-2 | CASSELLTKGTIANYGYTF | KLGGALQAK | YES | NO | NO |
| 15865 | TRAV20 | TRAJ42 | CAVQVSRRYGGSQGNLIF | TRBV27 | TRBJ2-1 | CASSPSGGGYNEQFF | KLGGALQAK | YES | NO | NO |
| 15866 | TRAV12-3 | TRAJ39 | CAIGPGNMLTF | TRBV27 | TRBJ1-1 | CASSLSEGTLNTEAFF | KLGGALQAK | YES | YES | NO |
| 15867 | TRAV2 | TRAJ31 | CAVNNARLMF | TRBV27 | TRBJ2-1 | CASRGGISGNNEQFF | KLGGALQAK | NO | NO | NO |
| 15868 | TRAV25 | TRAJ52 | CAGDNAGGTSYGKLTF | TRBV6-3 | TRBJ1-1 | CASSYSNAEAFF | AVFDRKSDAK | YES | YES | YES |
| 15869 | TRAV3 | TRAJ43 | CAVRLAYNNNDMRF | TRBV4-1 | TRBJ1-2 | CASSQNTRTDGYTF | KLGGALQAK | YES | NO | NO |
| 15870 | TRAV38-1 | TRAJ41 | CAFMNSGYALNF | TRBV9 | TRBJ2-1 | CASSAGTSGDNEQFF | KLGGALQAK | YES | NO | NO |
| 15871 | TRAV14DV4 | TRAJ6 | CAMREDTGGSYIPTF | TRBV7-9 | TRBJ2-5 | CASSPDRRQETQYF | KLGGALQAK | YES | NO | NO |
| 15872 | TRAV17 | TRAJ47 | CSMEYGNKLVF | TRBV12-4 | TRBJ2-6 | CASSRPRIGANVLTF | KLGGALQAK | YES | NO | NO |
| 15873 | TRAV41 | TRAJ43 | CAVRRPNNNDMRF | TRBV9 | TRBJ2-7 | CASSVEGGPGTSGGRSSYEQYF | AVFDRKSDAK | NO | NO | NO |
| 15874 | TRAV19 | TRAJ54 | CALSDSPVQGAQKLVF | TRBV29-1 | TRBJ1-1 | CSVEGNTEAFF | KLGGALQAK | NO | NO | NO |
| 15875 | TRAV12-1 | TRAJ54 | CVVNIGQGAQKLVF | TRBV4-1 | TRBJ2-7 | CASSQGAGTGFTYEQYF | KLGGALQAK | YES | NO | NO |
| 15876 | TRAV26-2 | TRAJ57 | CILSQQIQGGSEKLVF | TRBV6-5 | TRBJ1-4 | CASSYWTPATNEKLFF | KLGGALQAK | NO | NO | NO |
| 15877 | TRAV8-6 | TRAJ42 | CAPGGSQGNLIF | TRBV7-3 | TRBJ2-1 | CASSLTSGGAVGNEQFF | KLGGALQAK | YES | NO | NO |
| 15878 | TRAV8-3 | TRAJ43 | CAVGPNNNDMRF | TRBV11-1 | TRBJ1-1 | CASSLTTGRNTEAFF | KLGGALQAK | YES | NO | NO |
| 15879 | TRAV3 | TRAJ32 | CAVRDLNYGGATNKLIF | TRBV12-3 | TRBJ1-5 | CASSSEDSSNQPQHF | KLGGALQAK | NO | NO | NO |
| 15880 | TRAV13-2 | TRAJ45 | CAENNLGGGADGLTF | TRBV5-4 | TRBJ2-1 | CASSYSGSNEQFF | KLGGALQAK | YES | NO | NO |
| 15881 | TRAV26-1 | TRAJ58 | CIVRVAMKTSGSRLTF | TRBV7-3 | TRBJ1-1 | CASSLMGGANTEAFF | AVFDRKSDAK | NO | NO | NO |
| 15882 | TRAV17 | TRAJ45 | CASVPYSGGGADGLTF | TRBV6-3 | TRBJ1-2 | CASGTPWDKQTF | KLGGALQAK | YES | NO | NO |
| 15883 | TRAV38-2DV8 | TRAJ28 | CAYTFYGAGSYQLTF | TRBV6-5 | TRBJ2-7 | CASSNYPDRGPSPYYEQYF | KLGGALQAK | YES | NO | NO |
| 15884 | TRAV12-2 | TRAJ26 | CAVNYGQNFVF | TRBV4-1 | TRBJ2-1 | CASSQDPHGGDYNEQFF | ELAGIGILTV | YES | NO | YES |
| 15885 | TRAV34 | TRAJ49 | CGADSRNQFYF | TRBV12-5 | TRBJ2-1 | CASGLQHLREPRNEQFF | KLGGALQAK | YES | NO | NO |
| 15886 | TRAV5 | TRAJ35 | CAEIGFGNVLHC | TRBV19 | TRBJ1-2 | CASQYTQGPGSGYTF | KLGGALQAK | NO | NO | NO |
| 15888 | TRAV21 | TRAJ36 | CAVYGGANNLFF | TRBV19 | TRBJ1-5 | CASMGGSNQPQHF | KLGGALQAK | YES | NO | NO |
| 15889 | TRAV17 | TRAJ36 | CAQTGANNLFF | TRBV12-4 | TRBJ2-3 | CASSPMDTQYF | KLGGALQAK | YES | NO | NO |
| 15890 | TRAV12-1 | TRAJ45 | CVVNRYSGGGADGLTF | TRBV10-2 | TRBJ2-7 | CASSPNPQWEQYF | KLGGALQAK | YES | NO | NO |
| 15891 | TRAV5 | TRAJ8 | CAETTGFQKLVF | TRBV9 | TRBJ2-1 | CASSATGTRNSNEQFF | KLGGALQAK | YES | NO | NO |
| 15892 | TRAV29DV5 | TRAJ44 | CAASEVTGTASKLTF | TRBV3-1 | TRBJ1-2 | CASSLRAGFYGYTF | KLGGALQAK | YES | NO | NO |
| 15893 | TRAV13-2 | TRAJ17 | CAESSLKAAGNKLTF | TRBV29-1 | TRBJ1-2 | CSVDTVWGHGYTF | KLGGALQAK | YES | NO | NO |
| 15894 | TRAV17 | TRAJ29 | CATDANTPLVF | TRBV4-1 | TRBJ1-5 | CASSQVGGQPQHF | AVFDRKSDAK | NO | NO | NO |
| 15895 | TRAV38-1 | TRAJ44 | CAFAGTASKLTF | TRBV5-4 | TRBJ1-2 | CASSLALGSAGYTF | KLGGALQAK | YES | NO | NO |
| 15896 | TRAV17 | TRAJ40 | CATAPSGTYKYIF | TRBV7-9 | TRBJ1-5 | CASSLAGYSNQPQHF | KLGGALQAK | YES | NO | NO |
| 15897 | TRAV27 | TRAJ6 | CAGASGGSYIPTF | TRBV15 | TRBJ2-3 | CATTGGSNTDTQYF | KLGGALQAK | YES | NO | NO |
| 15898 | TRAV13-2 | TRAJ52 | CAEMLGAGGTSYGKLTF | TRBV10-3 | TRBJ2-2 | CAIDSNTGELFF | KLGGALQAK | YES | NO | NO |
| 15899 | TRAV29DV5 | TRAJ49 | CAYWGGNQFYF | TRBV6-5 | TRBJ2-3 | CASSYVGGPGDTQYF | KLGGALQAK | NO | NO | NO |
| 15900 | TRAV29DV5 | TRAJ28 | CAALSGAGSYQLTF | TRBV19 | TRBJ2-3 | CASSIYGGGNTDTQYF | KLGGALQAK | YES | NO | NO |
| 15901 | TRAV14DV4 | TRAJ34 | CAMRGPSYNTDKLIF | TRBV5-6 | TRBJ2-1 | CASSLVREYNEQFF | KLGGALQAK | YES | NO | NO |
| 15902 | TRAV12-3 | TRAJ53 | CARALGNYKLTF | TRBV12-4 | TRBJ1-6 | CASSLSGGGFSPLHF | KLGGALQAK | YES | NO | NO |
| 15903 | TRAV39 | TRAJ45 | CAVQPRGGGADGLTF | TRBV10-2 | TRBJ2-1 | CASSGASGMGGSYNEQFF | KLGGALQAK | YES | NO | NO |
| 15904 | TRAV21 | TRAJ43 | CAVWYMRF | TRBV18 | TRBJ1-1 | CASSPRSVPTEAFF | KLGGALQAK | NO | NO | NO |
| 15905 | TRAV22 | TRAJ45 | CAVDIPSGGGADGLTF | TRBV3-1 | TRBJ2-7 | CASSQDLGYEQYV | KLGGALQAK | NO | NO | NO |

| | | | | | | | | | | |
|---|---|---|---|---|---|---|---|---|---|---|
| 15907 | TRAV16 | TRAJ21 | CAKYNFNKFYF | TRBV7-2 | TRBJ2-7 | CASSTGTTEAYEQYF | KLGGALQAK | NO | NO | NO |
| 15908 | TRAV39 | TRAJ30 | CAVDRDDKIIF | TRBV10-3 | TRBJ1-1 | CAISESISARINRLEAFF | KLGGALQAK | YES | NO | NO |
| 15909 | TRAV1-2 | TRAJ12 | CAAMDSSYKLIF | TRBV6-1 | TRBJ2-7 | CASSERTGESYEQYF | KLGGALQAK | YES | NO | NO |
| 15910 | TRAV20 | TRAJ37 | CAVRSGSSNTGKLIF | TRBV4-1 | TRBJ1-6 | CASSQDRAGSPLHF | KLGGALQAK | YES | NO | NO |
| 15911 | TRAV9-2 | TRAJ58 | CALRKETSGSRLTF | TRBV27 | TRBJ2-7 | CASTPMGGRYEQYF | KLGGALQAK | YES | NO | NO |
| 15912 | TRAV14DV4 | TRAJ5 | CAMREGVDTGRRALTF | TRBV6-1 | TRBJ1-1 | CASSEFMGMNTEAFF | KLGGALQAK | NO | NO | NO |
| 15913 | TRAV12-1 | TRAJ45 | CVVPTRTVGGADGLTF | TRBV4-1 | TRBJ2-7 | CASSQAGTGLKF | KLGGALQAK | NO | NO | NO |
| 15914 | TRAV41 | TRAJ30 | CAVRSVFGNRDDKIIF | TRBV5-1 | TRBJ2-7 | CASSFSVEQYF | KLGGALQAK | YES | NO | NO |
| 15915 | TRAV20 | TRAJ49 | CAVHGNQFYF | TRBV7-9 | TRBJ2-1 | CASSIHEQFF | KLGGALQAK | YES | NO | NO |
| 15916 | TRAV17 | TRAJ47 | CATLFPMEYGNKLVF | TRBV18 | TRBJ1-2 | CASSLVGNYGYTF | KLGGALQAK | YES | NO | NO |
| 15917 | TRAV1-2 | TRAJ41 | CAVKDSGYALNF | TRBV12-4 | TRBJ2-7 | CASSLGGQATYEQYV | KLGGALQAK | NO | NO | NO |
| 15918 | TRAV19 | TRAJ11 | CALREVSGYSTLTF | TRBV20-1 | TRBJ2-5 | CSARGYSGLQETQYF | KLGGALQAK | YES | NO | NO |
| 15919 | TRAV4 | TRAJ9 | CLVGTTGGFKTIF | TRBV7-9 | TRBJ2-4 | CASSPVLRAKNIQYF | KLGGALQAK | YES | NO | NO |
| 15920 | TRAV17 | TRAJ56 | CATDDGANSKLTF | TRBV7-9 | TRBJ2-2 | CASSFAPGGGYTGELFF | KLGGALQAK | YES | NO | NO |
| 15921 | TRAV1-1 | TRAJ30 | CAVMNRDDKIIF | TRBV9 | TRBJ2-7 | CASSGLGQTYEQYF | KLGGALQAK | YES | NO | NO |
| 15922 | TRAV1-2 | TRAJ20 | CAVAQFYKLSF | TRBV27 | TRBJ2-2 | CASSPSSWDRPGTGELFF | RLRAEAQVK | YES | NO | YES |
| 15923 | TRAV39 | TRAJ41 | CAVDEWGSGYALNF | TRBV4-1 | TRBJ2-4 | CASSQRLSGRRKNIQYF | RLRAEAQVK | YES | NO | YES |
| 15924 | TRAV1-2 | TRAJ33 | CAVKVNYQLIW | TRBV6-6 | TRBJ1-2 | CASSWTDDYGYTF | KLGGALQAK | NO | NO | NO |
| 15925 | TRAV22 | TRAJ28 | CAVDFGAGSYQLTF | TRBV7-2 | TRBJ2-3 | CASRNRGGGAHNTQYF | KLGGALQAK | YES | NO | NO |
| 15926 | TRAV12-1 | TRAJ33 | CVVTTDSNYQLIW | TRBV7-9 | TRBJ2-3 | CASLEEGTDTQYF | KLGGALQAK | YES | NO | NO |
| 15927 | TRAV19 | TRAJ23 | CALSEVIYNQGGKLIF | TRBV24-1 | TRBJ1-1 | CATQTDWNTEAFF | KLGGALQAK | YES | NO | NO |
| 15928 | TRAV26-1 | TRAJ21 | CIAVYNFNKFYF | TRBV4-1 | TRBJ1-4 | CASSQVPLSTRGCEKLFF | KLGGALQAK | YES | NO | NO |
| 15929 | TRAV22 | TRAJ52 | CAVPGGTSYGKLTF | TRBV27 | TRBJ2-1 | CASFSGGSYNEQFF | KLGGALQAK | YES | NO | NO |
| 15930 | TRAV9-2 | TRAJ47 | CALMEYGNKLVF | TRBV27 | TRBJ2-3 | CASSHGWLAGGEDTQYF | KLGGALQAK | YES | NO | NO |
| 15931 | TRAV29DV5 | TRAJ8 | CAASMDTGFQKLVF | TRBV6-1 | TRBJ1-5 | CASSEWTGWGSNQPQHF | KLGGALQAK | YES | NO | NO |
| 15932 | TRAV8-1 | TRAJ4 | CAVAMFSGGYNKLIF | TRBV30 | TRBJ1-2 | CAWRGVTFSNYGYTF | KLGGALQAK | NO | NO | NO |
| 15933 | TRAV21 | TRAJ13 | CAVRPGGYQKVTF | TRBV19 | TRBJ1-5 | CASKGSRDSFNQPQHF | KLGGALQAK | YES | NO | NO |
| 15934 | TRAV12-2 | TRAJ33 | CAVYMDSNYQLIW | TRBV6-1 | TRBJ2-7 | CASSEEHISYEQYV | AVFDRKSDAK | NO | NO | NO |
| 15935 | TRAV35 | TRAJ58 | CAGQLTSGSRLTF | TRBV6-6 | TRBJ2-2 | CASSYYPDSGELFF | KLGGALQAK | YES | NO | NO |
| 15936 | TRAV8-1 | TRAJ42 | CAVNGGSQGNLIF | TRBV20-1 | TRBJ2-7 | CSARLGGQQYF | KLGGALQAK | NO | NO | NO |
| 15937 | TRAV19 | TRAJ26 | CALSEPQGYGQNFVF | TRBV6-6 | TRBJ2-5 | CASSYSMGGAGETQYF | KLGGALQAK | NO | NO | NO |
| 15938 | TRAV35 | TRAJ52 | CAGRPAGGTSYGKLTF | TRBV2 | TRBJ2-7 | CASSATGLFSPYEQYV | KLGGALQAK | NO | NO | NO |
| 15939 | TRAV13-1 | TRAJ3 | CAASGSASKIIF | TRBV9 | TRBJ2-3 | CASSVGGPSTDTQYF | KLGGALQAK | YES | NO | NO |
| 15940 | TRAV12-1 | TRAJ47 | CVVNVEYGNKLVF | TRBV20-1 | TRBJ2-7 | CSARGLAGFVEQYF | KLGGALQAK | NO | NO | NO |
| 15941 | TRAV8-6 | TRAJ52 | CAVSDSDAGGTSYGKLTF | TRBV19 | TRBJ2-3 | CASSARGGASDTQYF | KLGGALQAK | YES | NO | NO |
| 15942 | TRAV13-1 | TRAJ33 | CAASPFMDSNYQLIW | TRBV3-1 | TRBJ2-1 | CASSPDENEQFF | KLGGALQAK | YES | NO | NO |
| 15943 | TRAV14DV4 | TRAJ22 | CAMRRPISSGSARQLTF | TRBV6-3 | TRBJ2-1 | CASSYGAGGYNEQFF | AVFDRKSDAK | YES | NO | YES |
| 15944 | TRAV8-3 | TRAJ45 | CAVGDSGGGADGLTF | TRBV5-1 | TRBJ2-3 | CASSLGTSGGGDTQYF | KLGGALQAK | YES | NO | NO |
| 15945 | TRAV17 | TRAJ39 | CATDAYNNAGNMLTF | TRBV9 | TRBJ2-3 | CASTSDGTSYTDTQYF | KLGGALQAK | YES | NO | NO |
| 15946 | TRAV17 | TRAJ58 | CATDAWETSGSRLTF | TRBV12-4 | TRBJ2-1 | CASSWTSGYNEQFF | KLGGALQAK | YES | NO | NO |
| 15947 | TRAV21 | TRAJ48 | CAVSGNEKLTF | TRBV7-3 | TRBJ2-3 | CASSLVGAGTDTQYF | KLGGALQAK | YES | NO | NO |
| 15948 | TRAV14DV4 | TRAJ48 | CAMRVPPEFGNEKLTF | TRBV9 | TRBJ1-2 | CASSVAARYTF | KLGGALQAK | YES | NO | NO |
| 15949 | TRAV41 | TRAJ49 | CAPSTGNQFYF | TRBV10-3 | TRBJ2-4 | CAISEDSDIQYF | KLGGALQAK | NO | NO | NO |
| 15950 | TRAV14DV4 | TRAJ36 | CAMSLGAGANNLFF | TRBV12-4 | TRBJ1-5 | CASSPDREDQPQHF | KLGGALQAK | YES | NO | NO |
| 15951 | TRAV9-2 | TRAJ44 | CALSDNTGTASKLTF | TRBV12-3 | TRBJ2-3 | CASSLNGDTQYF | KLGGALQAK | YES | NO | NO |
| 15952 | TRAV21 | TRAJ43 | CAVVSNNNDMRF | TRBV5-1 | TRBJ2-7 | CASSETSGPPYEQYF | KLGGALQAK | NO | NO | NO |
| 15953 | TRAV29DV5 | TRAJ15 | CAASDSQAGTALIF | TRBV4-2 | TRBJ1-5 | CASSQSRGLSNQPQHF | KLGGALQAK | YES | NO | NO |
| 15954 | TRAV12-3 | TRAJ11 | CAMTRSGYSTLTF | TRBV9 | TRBJ1-3 | CASSPDAGNTIYF | KLGGALQAK | YES | NO | NO |
| 15955 | TRAV41 | TRAJ58 | CAVRPWMTSGSRLTF | TRBV3-1 | TRBJ2-1 | CASSPTSGANLNEQFF | KLGGALQAK | YES | NO | NO |
| 15956 | TRAV5 | TRAJ42 | CAVGGSQGNLIF | TRBV7-6 | TRBJ2-2 | CASSLGTANTGELFF | KLGGALQAK | YES | NO | NO |
| 15957 | TRAV8-4 | TRAJ33 | CAVSDPSNYQLIW | TRBV19 | TRBJ1-2 | CASSPSGGTNYGYTF | KLGGALQAK | YES | NO | NO |
| 15958 | TRAV24 | TRAJ52 | CARLGAGGTSYGKLTF | TRBV19 | TRBJ1-1 | CASSQGMNTEAFF | KLGGALQAK | YES | NO | NO |
| 15959 | TRAV8-3 | TRAJ45 | CAVGTWSGGGADGLTF | TRBV7-9 | TRBJ2-7 | CASSSDSYEQYF | KLGGALQAK | YES | NO | NO |
| 15960 | TRAV25 | TRAJ52 | CAGEQGVGGTSYGKLTF | TRBV19 | TRBJ1-1 | CASSIVGAWNTEAFF | KLGGALQAK | YES | NO | NO |
| 15962 | TRAV17 | TRAJ23 | CATVLYNQGGKLIF | TRBV6-5 | TRBJ1-6 | CASSYSAPTSKSPLHF | KLGGALQAK | YES | NO | NO |
| 15963 | TRAV14DV4 | TRAJ22 | CAMRRPISSGSARQLTF | TRBV6-5 | TRBJ2-7 | CASSPVVPYEQYF | KLGGALQAK | YES | NO | NO |
| 15964 | TRAV26-2 | TRAJ45 | CILRDPVSGGGADGLTF | TRBV15 | TRBJ1-5 | CATSRGTSNQPQHF | KLGGALQAK | YES | NO | NO |
| 15965 | TRAV41 | TRAJ54 | CAVEEGAQKLVF | TRBV5-1 | TRBJ1-1 | CASSPRVEGAWYTEAFF | KLGGALQAK | YES | NO | NO |
| 15966 | TRAV1-2 | TRAJ20 | CAVRRSNDYKLSF | TRBV2 | TRBJ2-3 | CASSETSGSLGDTQYF | AVFDRKSDAK | YES | NO | YES |
| 15967 | TRAV17 | TRAJ49 | CATDPDTGNQFYF | TRBV7-8 | TRBJ2-7 | CASSLVGLSYEQYV | KLGGALQAK | NO | NO | NO |
| 15968 | TRAV24 | TRAJ52 | CAFIGDAGGTSYGKLTF | TRBV14 | TRBJ2-1 | CASSQDPPLWGQTNEQFF | KLGGALQAK | YES | NO | NO |
| 15969 | TRAV12-1 | TRAJ39 | CVVSLVAGNMLTF | TRBV9 | TRBJ2-3 | CASSPSLLSREGHPDTQYF | KLGGALQAK | YES | NO | NO |
| 15970 | TRAV12-2 | TRAJ10 | CARRTGGGNKLTF | TRBV6-5 | TRBJ1-5 | CASSEEGTPSNQPQHF | AVFDRKSDAK | YES | NO | NO |
| 15971 | TRAV5 | TRAJ43 | CAEKDNDMRF | TRBV6-5 | TRBJ1-5 | CASSAYSGTGPQPQHF | KLGGALQAK | YES | NO | NO |
| 15972 | TRAV24 | TRAJ6 | CAQHSSGGSYIPTF | TRBV4-1 | TRBJ1-3 | CASSQEIRGSGNTIYF | KLGGALQAK | YES | NO | NO |
| 15973 | TRAV19 | TRAJ17 | CALSEATKAAGNKLTF | TRBV27 | TRBJ2-2 | CASSWGLAGGGTGELFF | KLGGALQAK | YES | NO | NO |
| 15974 | TRAV36DV7 | TRAJ44 | CAVGPLSTGTASKLTF | TRBV5-1 | TRBJ1-1 | CASGTGGTRTMNTEAFF | KLGGALQAK | NO | NO | NO |
| 15975 | TRAV29DV5 | TRAJ34 | CAASDTDKLIF | TRBV7-8 | TRBJ2-2 | CASSFNFGTGGRTGELFF | KLGGALQAK | YES | NO | NO |
| 15976 | TRAV13-1 | TRAJ49 | CAASAGNQFYF | TRBV4-1 | TRBJ2-3 | CASSQALGGGYTQYF | KLGGALQAK | YES | NO | NO |
| 15977 | TRAV5 | TRAJ16 | CAESPLGQKLLF | TRBV5-1 | TRBJ2-2 | CASSFRPGEAGELFF | KLGGALQAK | NO | NO | NO |
| 15978 | TRAV8-4 | TRAJ52 | CAVRANAGGTSYGKLTF | TRBV24-1 | TRBJ1-1 | CATSAGLNTEAFF | KLGGALQAK | YES | NO | NO |
| 15980 | TRAV19 | TRAJ22 | CALDFSSGSARQLTF | TRBV4-3 | TRBJ1-4 | CASSQDHRMGGHEKLFF | AVFDRKSDAK | YES | NO | NO |
| 15981 | TRAV3 | TRAJ8 | CAVREEGFQKLVF | TRBV5-4 | TRBJ2-7 | CASSLGVGPITYEQYV | KLGGALQAK | NO | NO | NO |
| 15983 | TRAV8-3 | TRAJ22 | CAVGSGGQLTF | TRBV19 | TRBJ1-4 | CASAVGAGDKLFF | KLGGALQAK | YES | NO | NO |
| 15984 | TRAV8-6 | TRAJ4 | CAVSLAGGYNKLIF | TRBV29-1 | TRBJ1-4 | CSVAAGADEKLFF | KLGGALQAK | YES | NO | NO |
| 15985 | TRAV12-3 | TRAJ26 | CAMSPLPNGQNFVF | TRBV13 | TRBJ1-1 | CASSLGTRGGAPEAFF | RLRAEAQVK | YES | YES | YES |
| 15988 | TRAV41 | TRAJ17 | CAVRFDLAAGNKLTF | TRBV27 | TRBJ2-1 | CASSFWGAGEQFF | KLGGALQAK | YES | NO | NO |
| 15989 | TRAV8-3 | TRAJ22 | CAGGSARQLTF | TRBV20-1 | TRBJ1-2 | CSARGLHAPNYGYTF | KLGGALQAK | YES | NO | NO |
| 15990 | TRAV5 | TRAJ8 | CAGRMNTGFQKLVF | TRBV9 | TRBJ1-5 | CASSADRDRGQPQHF | KLGGALQAK | YES | NO | NO |
| 15991 | TRAV14DV4 | TRAJ43 | CAMREVVNNDMRF | TRBV7-6 | TRBJ1-6 | CASSLRTGSSSYNSPLHF | KLGGALQAK | YES | NO | NO |
| 15992 | TRAV13-1 | TRAJ40 | CAATGTYKYIF | TRBV28 | TRBJ2-2 | CASSLSRATGELFF | KLGGALQAK | YES | NO | NO |
| 15993 | TRAV12-1 | TRAJ39 | CVVNNNAGNMLTF | TRBV11-1 | TRBJ2-4 | CASSLYDRVVKNIQYF | KLGGALQAK | YES | NO | NO |
| 15994 | TRAV8-1 | TRAJ20 | CAVIPNDYKLSF | TRBV6-3 | TRBJ2-2 | CASDKSQGSPGELFF | KLGGALQAK | YES | NO | NO |
| 15995 | TRAV17 | TRAJ17 | CATDAEAAGNKLTF | TRBV5-1 | TRBJ1-2 | CASSLAEGFYGYTF | KLGGALQAK | YES | YES | NO |
| 15996 | TRAV1-2 | TRAJ27 | CAVRDTNAGKSTF | TRBV5-1 | TRBJ1-1 | CASSLEQGAFTEAFF | KLGGALQAK | YES | NO | NO |
| 15997 | TRAV12-2 | TRAJ34 | CAVWRADKLIF | TRBV20-1 | TRBJ1-1 | CSARDKLGIPEAFF | KLGGALQAK | YES | NO | NO |
| 15998 | TRAV1-1 | TRAJ24 | CAPKTDSWGKLQF | TRBV13 | TRBJ2-3 | CASSMGQGGNTQYF | KLGGALQAK | YES | NO | NO |
| 15999 | TRAV20 | TRAJ39 | CAVLNAGNMLTF | TRBV27 | TRBJ2-4 | CASSFGPKNIQYF | KLGGALQAK | YES | NO | NO |
| 16000 | TRAV20 | TRAJ26 | CAVQVYSRNYGQNFVF | TRBV19 | TRBJ2-3 | CASSILGRTDTQYF | KLGGALQAK | YES | NO | NO |
| 16001 | TRAV10 | TRAJ20 | CVVMFRSNDYKLSF | TRBV27 | TRBJ2-3 | CASTPLGGGLNTQYF | KLGGALQAK | YES | NO | NO |
| 16002 | TRAV29DV5 | TRAJ50 | CAAKDSYDKVIF | TRBV19 | TRBJ1-5 | CASSGYSNQPQHF | KLGGALQAK | YES | NO | NO |
| 16003 | TRAV29DV5 | TRAJ40 | CAASSTSGTYKYIF | TRBV19 | TRBJ1-2 | CASSIDMWQFGF | KLGGALQAK | YES | NO | NO |
| 16004 | TRAV26-2 | TRAJ43 | CILTKLNDMRF | TRBV5-1 | TRBJ1-3 | CASRTDGFSGNTIYF | KLGGALQAK | YES | NO | NO |
| 16005 | TRAV13-1 | TRAJ23 | CAATPHPFQGGKLIF | TRBV25-1 | TRBJ2-2 | CASSESPRSATGELFF | KLGGALQAK | YES | NO | NO |
| 16006 | TRAV26-2 | TRAJ23 | CILRDDGYNQGGKLIF | TRBV4-1 | TRBJ2-5 | CASSQTSGKIQETQYF | KLGGALQAK | NO | NO | NO |
| 16007 | TRAV9-2 | TRAJ28 | CALDSGAGSYQLTF | TRBV7-8 | TRBJ2-7 | CASSHGLAGGDLGYEQYV | KLGGALQAK | NO | NO | NO |
| 16008 | TRAV3 | TRAJ33 | CAVRDWSNYQLIW | TRBV27 | TRBJ2-7 | CASSSTDSYEQYF | KLGGALQAK | NO | NO | NO |
| 16009 | TRAV24 | TRAJ17 | CAFKAAGNKLTF | TRBV4-1 | TRBJ2-2 | CASSQVLGGDTGELFF | KLGGALQAK | YES | NO | NO |
| 16010 | TRAV1-2 | TRAJ10 | CAVRSIGGGGNKLTF | TRBV5-4 | TRBJ2-1 | CASSFQPSSYNEQFF | KLGGALQAK | YES | NO | NO |
| 16011 | TRAV5 | TRAJ45 | CAAGGGADGLTF | TRBV11-3 | TRBJ2-7 | CASSFSGGSYEQYF | KLGGALQAK | YES | NO | NO |
| 16012 | TRAV3 | TRAJ35 | CAVNGFGNVLHC | TRBV27 | TRBJ2-2 | CASSSQGNTGELFF | KLGGALQAK | NO | NO | NO |
| 16013 | TRAV13-2 | TRAJ4 | CAENGGGYNKLIF | TRBV4-1 | TRBJ1-2 | CASSQGGDGYTF | KLGGALQAK | YES | NO | NO |
| 16014 | TRAV14DV4 | TRAJ53 | CAMREREASGGSNYKLTF | TRBV27 | TRBJ1-2 | CASSFFRNYGYTF | KLGGALQAK | YES | NO | NO |
| 16015 | TRAV25 | TRAJ33 | CAGPGMDSNYQLIW | TRBV30 | TRBJ1-1 | CAWSVLGKATEAFF | KLGGALQAK | YES | YES | NO |

| | | | | | | | | | | |
|---|---|---|---|---|---|---|---|---|---|---|
| 16016 | TRAV12-2 | TRAJ49 | CAPSTGNQFYF | TRBV5-6 | TRBJ2-1 | CASSLSHSYNEQFF | KLGGALQAK | YES | NO | NO |
| 16017 | TRAV30 | TRAJ32 | CGTEFHYGGATNKLIF | TRBV4-3 | TRBJ2-1 | CASSQDGGGYNEQFF | KLGGALQAK | YES | NO | NO |
| 16018 | TRAV8-4 | TRAJ13 | CAVLSGGYQKVTF | TRBV6-3 | TRBJ1-5 | CASSYSNQFGHQPQHF | KLGGALQAK | YES | NO | NO |
| 16019 | TRAV25 | TRAJ40 | CAPTSGTYKYIF | TRBV7-8 | TRBJ2-7 | CASSLAVAGQVSYEQYV | KLGGALQAK | NO | NO | NO |
| 16021 | TRAV8-6 | TRAJ52 | CAVSVSDAGGTSYGKLTF | TRBV19 | TRBJ1-5 | CASSIGGYSNQPQHF | KLGGALQAK | YES | NO | NO |
| 16022 | TRAV1-2 | TRAJ6 | CAVEGGSYIPTF | TRBV7-9 | TRBJ2-2 | CASSGITGGDTGELFF | KLGGALQAK | YES | NO | NO |
| 16023 | TRAV29DV5 | TRAJ28 | CGAGSYQLTF | TRBV7-9 | TRBJ1-5 | CASSLTDNQPQHF | KLGGALQAK | YES | NO | NO |
| 16024 | TRAV8-6 | TRAJ39 | CAVSVRRGAGNMLTF | TRBV12-4 | TRBJ1-1 | CASSFATEAFF | KLGGALQAK | YES | NO | NO |
| 16025 | TRAV38-2DV8 | TRAJ43 | CAYSNNNDMRF | TRBV4-1 | TRBJ1-4 | CASSQDRLNEKLFF | KLGGALQAK | YES | NO | NO |
| 16026 | TRAV8-4 | TRAJ48 | CAVSGINEKLTF | TRBV13 | TRBJ2-1 | CASSLAGVNEQFF | LLDFVRFMGV | YES | NO | YES |
| 16027 | TRAV5 | TRAJ8 | CAMNTGFQKLVF | TRBV25-1 | TRBJ1-1 | CASSEMNTEAFF | KLGGALQAK | YES | NO | NO |
| 16028 | TRAV13-2 | TRAJ54 | CAETVQGAQKLVF | TRBV4-1 | TRBJ2-7 | CASSQGRGPYEQYF | KLGGALQAK | YES | NO | NO |
| 16029 | TRAV38-2DV8 | TRAJ49 | CAYRSTGNQFYF | TRBV7-9 | TRBJ2-7 | CASSLQGRPRDEQYF | KLGGALQAK | YES | NO | NO |
| 16030 | TRAV8-2 | TRAJ44 | CVVRGHPYTGTASKLTF | TRBV7-9 | TRBJ2-1 | CASSTGATEQFF | KLGGALQAK | YES | NO | NO |
| 16031 | TRAV21 | TRAJ18 | CAVSLDRGSTLGRLYF | TRBV7-6 | TRBJ2-2 | CASSLGTANTGELFF | KLGGALQAK | YES | NO | NO |
| 16032 | TRAV17 | TRAJ53 | CATDPSLSGGSNYKLTF | TRBV20-1 | TRBJ2-5 | CSARVAGPKGETQYF | KLGGALQAK | NO | NO | NO |
| 16033 | TRAV25 | TRAJ38 | CAGGAGNNRKLIW | TRBV9 | TRBJ2-2 | CASSVTATATGELFF | KLGGALQAK | YES | NO | NO |
| 16034 | TRAV8-4 | TRAJ17 | CAVTTYKAAGNKLTF | TRBV4-1 | TRBJ2-3 | CASSQAGGDTQYF | KLGGALQAK | YES | NO | NO |
| 16035 | TRAV5 | TRAJ26 | CAENYGQNFVF | TRBV5-4 | TRBJ2-1 | CASSPTYNEQFF | KLGGALQAK | YES | NO | NO |
| 16036 | TRAV8-3 | TRAJ20 | CAVAAYKLSF | TRBV27 | TRBJ2-5 | CASRPAGEETQYF | KLGGALQAK | YES | NO | NO |
| 16037 | TRAV4 | TRAJ22 | CLVGAVPFSGSARQLTF | TRBV24-1 | TRBJ2-1 | CATTSGTYDNEQFF | KLGGALQAK | NO | NO | NO |
| 16038 | TRAV9-2 | TRAJ17 | CALSDLWAAGNKLTF | TRBV29-1 | TRBJ2-1 | CSGVAPDSEQFF | KLGGALQAK | YES | NO | NO |
| 16039 | TRAV3 | TRAJ6 | CAVRDTGGGSYIPTF | TRBV7-9 | TRBJ2-2 | CASSLLRDIPPGELFF | KLGGALQAK | YES | NO | NO |
| 16040 | TRAV19 | TRAJ21 | CALREGFNKFYF | TRBV15 | TRBJ1-5 | CATSRDRGGTANQPQHF | KLGGALQAK | YES | NO | NO |
| 16041 | TRAV3 | TRAJ37 | CAVRDGGMGSNTGKLIF | TRBV11-2 | TRBJ2-5 | CASRGPVETQYF | KLGGALQAK | YES | NO | NO |
| 16042 | TRAV5 | TRAJ13 | CAETEGGYQKVTF | TRBV6-1 | TRBJ2-3 | CASTPAGGPTDF | KLGGALQAK | NO | NO | NO |
| 16043 | TRAV24 | TRAJ40 | CAHTKRTTSGTYKYIF | TRBV2 | TRBJ2-1 | CASEGLAGLNEQFF | KLGGALQAK | YES | NO | NO |
| 16044 | TRAV25 | TRAJ52 | CAGPYAGGTSYGKLTF | TRBV19 | TRBJ2-1 | CASTPPGANEQFF | KLGGALQAK | YES | NO | NO |
| 16045 | TRAV14DV4 | TRAJ16 | CAMRVNRF | TRBV19 | TRBJ1-5 | CASSAGTGYQPQHF | KLGGALQAK | NO | NO | NO |
| 16046 | TRAV26-2 | TRAJ20 | CILMDDYKLSF | TRBV13 | TRBJ2-1 | CASSLQGVAYNEQFF | KLGGALQAK | YES | NO | NO |
| 16047 | TRAV3 | TRAJ39 | CAVRDGNNAGNMLTF | TRBV13 | TRBJ2-1 | CASSSVGEQFF | KLGGALQAK | NO | NO | NO |
| 16048 | TRAV22 | TRAJ48 | CAVDESNFGNEKLTF | TRBV3-1 | TRBJ2-7 | CASSQDPGYEQYF | KLGGALQAK | YES | NO | NO |
| 16049 | TRAV8-1 | TRAJ13 | CAASYQKVTF | TRBV12-4 | TRBJ2-3 | CASNAQGISDTQYF | KLGGALQAK | YES | NO | NO |
| 16050 | TRAV20 | TRAJ42 | CAVQDFYGGSQGNLIF | TRBV7-8 | TRBJ1-1 | CASSSGTGLNTEAFF | AVFDRKSDAK | YES | NO | NO |
| 16051 | TRAV12-3 | TRAJ39 | CAIGPGNMLTF | TRBV25-1 | TRBJ1-2 | CASSGRDRVWNNYGYTF | KLGGALQAK | YES | NO | NO |
| 16052 | TRAV35 | TRAJ58 | CAGRGGETSGSRLTF | TRBV7-7 | TRBJ2-1 | CASSLLGDSYNEQFF | KLGGALQAK | YES | NO | NO |
| 16053 | TRAV26-1 | TRAJ50 | CIAPETSYDKVIF | TRBV7-9 | TRBJ2-1 | CASSLGLAGPYNEQFF | KLGGALQAK | YES | NO | NO |
| 16054 | TRAV14DV4 | TRAJ9 | CAMRENNGGFKTIF | TRBV10-2 | TRBJ2-7 | CASSETAVGNEQYF | KLGGALQAK | YES | NO | NO |
| 16055 | TRAV26-1 | TRAJ26 | CIASFYGQNFVF | TRBV7-9 | TRBJ2-2 | CASSKAPQANTGELFF | KLGGALQAK | YES | NO | NO |
| 16056 | TRAV12-1 | TRAJ9 | CVVDLGGFKTIF | TRBV4-1 | TRBJ1-2 | CASSQPYRSDYGYTF | KLGGALQAK | YES | NO | NO |
| 16057 | TRAV6 | TRAJ24 | CALDEDKWDSWGKLQF | TRBV4-1 | TRBJ1-1 | CASSPPGQANTEAFF | KLGGALQAK | YES | NO | NO |
| 16058 | TRAV12-1 | TRAJ49 | CVVNIPPGNQFYF | TRBV9 | TRBJ1-4 | CASSTTVTTTNEKLFF | IVTDFSVIK | YES | NO | YES |
| 16059 | TRAV21 | TRAJ12 | CAVIMDSSYKLIF | TRBV7-2 | TRBJ2-2 | CASSLNRDTATNTGELFF | KLGGALQAK | YES | NO | NO |
| 16060 | TRAV29DV5 | TRAJ32 | CAATGGATNKLIF | TRBV24-1 | TRBJ2-1 | CATSDPNTDRTSGTDYNEQFF | KLGGALQAK | NO | NO | NO |
| 16061 | TRAV19 | TRAJ36 | CALSEGETGANNLFF | TRBV6-5 | TRBJ2-7 | CASSGRTGWNEQYF | KLGGALQAK | YES | NO | NO |
| 16062 | TRAV26-2 | TRAJ48 | CILQPSHFGNEKLTF | TRBV19 | TRBJ2-3 | CASSLRTSGTDTQYF | KLGGALQAK | YES | NO | NO |
| 16063 | TRAV24 | TRAJ28 | CAFVSGAGSYQLTF | TRBV4-3 | TRBJ2-3 | CASSQEAGTASPITDTQYF | KLGGALQAK | YES | NO | NO |
| 16064 | TRAV12-1 | TRAJ33 | CVVIPPQDSNYQLIW | TRBV20-1 | TRBJ1-2 | CSARERGNNYGYTF | KLGGALQAK | YES | NO | NO |
| 16065 | TRAV29DV5 | TRAJ31 | CAATMEDARLMF | TRBV24-1 | TRBJ2-1 | CATSDFSSGDYNEQFF | KLGGALQAK | YES | NO | NO |
| 16066 | TRAV12-1 | TRAJ40 | CVVRVGYKYIF | TRBV7-6 | TRBJ2-3 | CASSRTTSGEDTDTQYF | KLGGALQAK | YES | NO | NO |
| 16067 | TRAV27 | TRAJ20 | CAGGPQNDYKLSF | TRBV11-2 | TRBJ2-5 | CASRGPVETQYF | KLGGALQAK | YES | NO | NO |
| 16069 | TRAV19 | TRAJ57 | CALTGVQGGSEKLVF | TRBV19 | TRBJ1-1 | CASSIFSGPGIEAFF | GILGFVFTL | YES | NO | YES |
| 16070 | TRAV4 | TRAJ30 | CLVLDDKIIF | TRBV24-1 | TRBJ1-1 | CATSEGEGTEAFF | KLGGALQAK | YES | NO | NO |
| 16071 | TRAV1-2 | TRAJ28 | CAVRYSGAGSYQLTF | TRBV3-1 | TRBJ2-5 | CASSQSQGDGETQYF | KLGGALQAK | NO | NO | NO |
| 16072 | TRAV13-1 | TRAJ47 | CAASWGEYGNKLVF | TRBV13 | TRBJ1-2 | CASSTTGGVGGYTF | KLGGALQAK | YES | NO | NO |
| 16073 | TRAV16 | TRAJ53 | CALSALGGSNYKLTF | TRBV20-1 | TRBJ2-7 | CSARFPSSYEQYF | KLGGALQAK | NO | NO | NO |
| 16074 | TRAV24 | TRAJ52 | CAFIGDAGGTSYGKLTF | TRBV27 | TRBJ2-2 | CASSQLTGTSPMELFF | KLGGALQAK | YES | NO | NO |
| 16075 | TRAV12-3 | TRAJ39 | CAIGPGNMLTF | TRBV7-9 | TRBJ2-7 | CASSLLGTGVREQYF | KLGGALQAK | YES | NO | NO |
| 16076 | TRAV19 | TRAJ29 | CALSELNSGNTPLVF | TRBV7-9 | TRBJ1-1 | CASSFPESLVTGGDPTEAFF | KLGGALQAK | NO | NO | NO |
| 16077 | TRAV1-2 | TRAJ33 | CAALGDSNYQLIW | TRBV9 | TRBJ2-1 | CASSVDGASYNEQFF | KLGGALQAK | YES | NO | NO |
| 16078 | TRAV20 | TRAJ31 | CAVQAGGRLMF | TRBV5-1 | TRBJ2-2 | CASSPPNSPFELFF | KLGGALQAK | YES | NO | NO |
| 16079 | TRAV14DV4 | TRAJ4 | CAMRGPISGGYNKLIF | TRBV10-3 | TRBJ1-2 | CAISDQASYGYTF | KLGGALQAK | YES | NO | NO |
| 16080 | TRAV8-4 | TRAJ23 | CAVSGQGGKLIF | TRBV13 | TRBJ2-5 | CASSLGPVIQYF | IMDQVPFSV | YES | NO | NO |
| 16081 | TRAV1-1 | TRAJ30 | CAGRDDKIIF | TRBV6-3 | TRBJ2-7 | CASKPGQSSYEQYV | KLGGALQAK | NO | NO | NO |
| 16082 | TRAV1-1 | TRAJ23 | CAGNQGGKLIF | TRBV20-1 | TRBJ2-1 | CSARSRENPDNEQFF | KLGGALQAK | NO | NO | NO |
| 16083 | TRAV6 | TRAJ26 | CALDPRNYGQNFVF | TRBV4-1 | TRBJ1-3 | CASSQGSSGNTIYF | KLGGALQAK | YES | NO | NO |
| 16084 | TRAV24 | TRAJ49 | CASSNTGNQFYF | TRBV12-3 | TRBJ2-5 | CASSFLAGMETQYF | KLGGALQAK | YES | NO | NO |
| 16085 | TRAV8-4 | TRAJ20 | CAVIQINDYKLSF | TRBV6-5 | TRBJ2-5 | CASSTTGTGTQYF | KLGGALQAK | NO | NO | NO |
| 16086 | TRAV34 | TRAJ34 | CGARDWSDKLIF | TRBV29-1 | TRBJ1-1 | CSVEGRGADTEAFF | KLGGALQAK | YES | NO | NO |
| 16087 | TRAV8-1 | TRAJ28 | CAVNAYSGAGSYQLTF | TRBV6-1 | TRBJ1-1 | CATGRANTEAFF | KLGGALQAK | YES | NO | NO |
| 16088 | TRAV24 | TRAJ17 | CAFGPVKAAGNKLTF | TRBV19 | TRBJ2-7 | CASSIAGSHEQYF | KLGGALQAK | YES | NO | NO |
| 16089 | TRAV21 | TRAJ17 | CAVLGAAGNKLTF | TRBV7-9 | TRBJ1-5 | CASSFSSSNQPQHF | KLGGALQAK | YES | NO | NO |
| 16090 | TRAV8-6 | TRAJ11 | CAVSESGYSTLTF | TRBV11-2 | TRBJ2-1 | CASPGQGYNEQFF | KLGGALQAK | YES | NO | NO |
| 16091 | TRAV8-1 | TRAJ20 | CAVRLTNDYKLSF | TRBV6-5 | TRBJ2-3 | CASGSPGTQYF | KLGGALQAK | NO | NO | NO |
| 16092 | TRAV27 | TRAJ20 | CAGYNDYKLSF | TRBV7-3 | TRBJ1-1 | CASSLGENTEAFF | KLGGALQAK | NO | NO | NO |
| 16093 | TRAV13-2 | TRAJ16 | CAENMGDGQKLLF | TRBV6-5 | TRBJ1-2 | CASSYGGDGYTF | KLGGALQAK | NO | NO | NO |
| 16094 | TRAV29DV5 | TRAJ23 | CAASLRGTKLIF | TRBV27 | TRBJ2-2 | CASSLAALSGGELFF | KLGGALQAK | YES | NO | NO |
| 16095 | TRAV17 | TRAJ15 | CATDQAGTALIF | TRBV15 | TRBJ2-5 | CATSSTDRVGQETQYF | KLGGALQAK | YES | NO | NO |
| 16096 | TRAV29DV5 | TRAJ6 | CAASAGESGGSYIPTF | TRBV7-9 | TRBJ2-1 | CASSSPLPGTGVNEQFF | KLGGALQAK | YES | NO | NO |
| 16097 | TRAV17 | TRAJ24 | CASTDSWGKLQF | TRBV20-1 | TRBJ2-3 | CSARDRVGADYTDTQYF | KLGGALQAK | NO | NO | NO |
| 16098 | TRAV14DV4 | TRAJ20 | CAMREGGRYDYKLSF | TRBV3-1 | TRBJ1-3 | CASSQVPQGLSGNTIYF | AVFDRKSDAK | YES | NO | YES |
| 16099 | TRAV29DV5 | TRAJ45 | CAASVRGGADGLTF | TRBV9 | TRBJ1-1 | CASSVDTGFTEAFF | KLGGALQAK | YES | NO | NO |
| 16100 | TRAV17 | TRAJ20 | CATGKYLDYKLSF | TRBV11-2 | TRBJ2-7 | CASSLDLPGSYEQYF | KLGGALQAK | YES | NO | NO |
| 16101 | TRAV12-2 | TRAJ52 | CAVKGGGGTSYGKLTF | TRBV19 | TRBJ2-1 | CASSLPGRAYNEQFF | KLGGALQAK | YES | NO | NO |
| 16103 | TRAV19 | TRAJ17 | CALSEVKAAGNKLTF | TRBV19 | TRBJ2-1 | CASSGSTDPPYNEQFF | KLGGALQAK | YES | NO | NO |
| 16104 | TRAV12-3 | TRAJ58 | CAMGLLETSGSRLTF | TRBV5-1 | TRBJ2-7 | CASSPVAGGDYEQYF | KLGGALQAK | YES | NO | NO |
| 16105 | TRAV19 | TRAJ49 | CALSEWTGNQFYF | TRBV7-9 | TRBJ1-5 | CASSFEQGGEAQHF | KLGGALQAK | YES | NO | NO |
| 16106 | TRAV38-2DV8 | TRAJ48 | CAYREPNFGNEKLTF | TRBV27 | TRBJ2-7 | CASSLYGAAYEQYV | KLGGALQAK | NO | NO | NO |
| 16107 | TRAV1-2 | TRAJ4 | CAVRHSGGYNKLIF | TRBV25-1 | TRBJ1-5 | CASSERQGDQPQHF | KLGGALQAK | YES | NO | NO |
| 16108 | TRAV20 | TRAJ5 | CAVRDTGRRALTF | TRBV24-1 | TRBJ2-2 | CATSDRSQGVNTGELFF | KLGGALQAK | YES | NO | NO |
| 16109 | TRAV1-2 | TRAJ16 | CAVINEAYGQKLLF | TRBV10-2 | TRBJ2-3 | CASSESTSGGVTDTQYF | KLGGALQAK | NO | NO | NO |
| 16110 | TRAV9-2 | TRAJ58 | CALSRETSGSRLTF | TRBV20-1 | TRBJ1-5 | CSASIPGQGVFQPQHF | RLRAEAQVK | YES | YES | NO |
| 16111 | TRAV10 | TRAJ20 | CVVSEDKLSF | TRBV3-1 | TRBJ2-5 | CASSPLTSGGAEETQYF | KLGGALQAK | YES | NO | NO |
| 16112 | TRAV8-6 | TRAJ24 | CAVADSWGKFQF | TRBV20-1 | TRBJ2-1 | CSARRRQGVEQFF | KLGGALQAK | NO | NO | NO |
| 16113 | TRAV26-1 | TRAJ24 | CIVNSGWGKLQF | TRBV12-4 | TRBJ2-7 | CASSLRDSRDGEQYF | KLGGALQAK | YES | NO | NO |
| 16114 | TRAV3 | TRAJ29 | CAVRDEGDSGNTPLVF | TRBV10-1 | TRBJ2-2 | CASSERPGLAPGELFF | KLGGALQAK | NO | NO | NO |
| 16115 | TRAV21 | TRAJ26 | CAVDNYGQNFVF | TRBV4-3 | TRBJ1-4 | CASSQDHRMGGHEKLFF | KLGGALQAK | YES | NO | NO |
| 16116 | TRAV3 | TRAJ37 | CAVRDGGMGSNTGKLIF | TRBV21-1 | TRBJ2-3 | CQQQRNPGTNKRSTDTQYF | KLGGALQAK | NO | NO | NO |
| 16117 | TRAV1-2 | TRAJ36 | CAVRETGANNLFF | TRBV4-1 | TRBJ2-7 | CASSQGSSGPDEQYF | KLGGALQAK | YES | NO | NO |
| 16118 | TRAV3 | TRAJ34 | CAVRDGGTDKLIF | TRBV20-1 | TRBJ1-5 | CSAETGNLPGNQPQHF | KLGGALQAK | NO | NO | NO |
| 16119 | TRAV19 | TRAJ52 | CALRGGTSYGKLTF | TRBV15 | TRBJ2-5 | CATREDINQETQYF | KLGGALQAK | YES | NO | NO |
| 16120 | TRAV38-1 | TRAJ40 | CAFMNPAGTYKYIF | TRBV4-1 | TRBJ1-1 | CASSLQLGENTEAFF | KLGGALQAK | YES | NO | NO |
| 16121 | TRAV35 | TRAJ28 | CAGRVISYQLTF | TRBV2 | TRBJ2-3 | CASSVLPADTQYF | KLGGALQAK | YES | NO | NO |
| 16123 | TRAV19 | TRAJ34 | CALRVAYNTDKLIF | TRBV5-5 | TRBJ2-3 | CASSGLLVDTQYF | KLGGALQAK | YES | NO | NO |

| | | | | | | | | | | |
|---|---|---|---|---|---|---|---|---|---|---|
| 16124 | TRAV9-2 | TRAJ21 | CALSESLNFNKFYF | TRBV6-6 | TRBJ2-7 | CASSRPPVASGSYEQYV | KLGGALQAK | NO | NO | NO |
| 16125 | TRAV1-1 | TRAJ28 | CAVRDLGYQLTF | TRBV7-2 | TRBJ2-5 | CASSLAESETQYF | KLGGALQAK | YES | NO | NO |
| 16126 | TRAV1-2 | TRAJ42 | CAVNYGGSQGNLIF | TRBV7-9 | TRBJ1-4 | CASSLGEGIEELFF | KLGGALQAK | YES | NO | NO |
| 16127 | TRAV12-1 | TRAJ3 | CVVNPLDPYSSASKIIF | TRBV9 | TRBJ1-1 | CASSGDTMNTEAFF | KLGGALQAK | YES | NO | NO |
| 16128 | TRAV8-6 | TRAJ32 | CAVSPYGGATNKLIF | TRBV19 | TRBJ1-4 | CASSPYGGSANEKLFF | KLGGALQAK | YES | NO | NO |
| 16129 | TRAV19 | TRAJ53 | CALTLHLTSGGSNYKLTF | TRBV13 | TRBJ1-2 | CASSIGLAYGYTF | KLGGALQAK | YES | NO | NO |
| 16130 | TRAV41 | TRAJ54 | CAVRRAQKLVF | TRBV5-1 | TRBJ2-3 | CASSWARVTSTDTQYF | KLGGALQAK | YES | NO | NO |
| 16131 | TRAV14DV4 | TRAJ47 | CAMREVQYGNKLVF | TRBV12-4 | TRBJ1-5 | CASRAGTNQPQHF | KLGGALQAK | YES | NO | NO |
| 16132 | TRAV25 | TRAJ39 | CAGRNNAGNMLTF | TRBV24-1 | TRBJ1-5 | CATSETSRGSQHF | RLRAEAQVK | YES | YES | YES |
| 16133 | TRAV8-3 | TRAJ10 | CAGYILTGGGNKLTF | TRBV27 | TRBJ1-1 | CASRSSWGGATEAFF | KLGGALQAK | YES | NO | NO |
| 16134 | TRAV1-2 | TRAJ4 | CAVRVPSGGYNKLIF | TRBV2 | TRBJ2-5 | CASIGESGELQETQYF | KLGGALQAK | YES | NO | NO |
| 16135 | TRAV17 | TRAJ21 | CATVLFDNFNKFYF | TRBV30 | TRBJ2-2 | CAWTHRGKDGELFF | KLGGALQAK | YES | NO | NO |
| 16136 | TRAV35 | TRAJ57 | CAGQGGSEKLVF | TRBV10-3 | TRBJ2-7 | CAPNRDSPYEQYF | KLGGALQAK | YES | NO | NO |
| 16137 | TRAV3 | TRAJ8 | CAVRDITETGFQKLVF | TRBV9 | TRBJ2-1 | CASSVAFEGMGSYNEQFF | KLGGALQAK | NO | NO | NO |
| 16138 | TRAV3 | TRAJ20 | CAVRDMGNDYKLSF | TRBV7-9 | TRBJ2-5 | CASSTWDSQETQYF | KLGGALQAK | NO | NO | NO |
| 16140 | TRAV4 | TRAJ45 | CLVGDGTGGGADGLTF | TRBV11-2 | TRBJ2-1 | CASSLGLVTFF | KLGGALQAK | YES | NO | NO |
| 16141 | TRAV3 | TRAJ34 | CAVRTYNTDKLIF | TRBV11-2 | TRBJ2-7 | CASSFSGTVAEQYF | KLGGALQAK | YES | NO | NO |
| 16142 | TRAV8-1 | TRAJ32 | CAVRGGATNKLIF | TRBV5-6 | TRBJ2-2 | CASSGANTGELFF | KLGGALQAK | YES | NO | NO |
| 16143 | TRAV4 | TRAJ23 | CLVGEGNQGGKLIF | TRBV20-1 | TRBJ2-7 | CSARERYEMGEQYF | AVFDRKSDAK | NO | NO | NO |
| 16144 | TRAV19 | TRAJ35 | CALMEGFGNVLHC | TRBV12-4 | TRBJ1-1 | CASTVALMEIAFF | KLGGALQAK | NO | NO | NO |
| 16145 | TRAV3 | TRAJ34 | CAVRGMDTDKLIF | TRBV15 | TRBJ2-2 | CATSYLTGANTGELFF | KLGGALQAK | NO | NO | NO |
| 16146 | TRAV5 | TRAJ35 | CAELIGNVLHC | TRBV7-2 | TRBJ1-2 | CASSSGVGYTF | KLGGALQAK | NO | NO | NO |
| 16147 | TRAV17 | TRAJ53 | CATENSGGSNYKLTF | TRBV5-1 | TRBJ2-5 | CASSLSSGIEETQYF | KLGGALQAK | YES | NO | NO |
| 16148 | TRAV14DV4 | TRAJ37 | CAMRGPFSSNTGKLIF | TRBV12-4 | TRBJ1-1 | CASSLTAYIVNTEAFF | KLGGALQAK | YES | NO | NO |
| 16149 | TRAV8-2 | TRAJ24 | CVVSDQPDSWGKFQF | TRBV19 | TRBJ2-2 | CASSIESRNGELFF | KLGGALQAK | YES | NO | NO |
| 16151 | TRAV14DV4 | TRAJ54 | CAMRDGQGAQKLVF | TRBV20-1 | TRBJ2-3 | CSASRTSGLTDTQYF | KLGGALQAK | NO | NO | NO |
| 16152 | TRAV17 | TRAJ6 | CATDGGSYIPTF | TRBV7-9 | TRBJ1-4 | CASSLRQGVPNEKLFF | KLGGALQAK | YES | NO | NO |
| 16153 | TRAV14DV4 | TRAJ21 | CAMREGDNKFYF | TRBV19 | TRBJ2-5 | CASSIGGLAGATQETQYF | KLGGALQAK | YES | NO | NO |
| 16154 | TRAV21 | TRAJ50 | CAVKTSYDKVIF | TRBV24-1 | TRBJ2-2 | CATSDSGTGGTGELFF | AVFDRKSDAK | YES | NO | YES |
| 16155 | TRAV1-2 | TRAJ32 | CAVRDEYGGATNKLIF | TRBV6-3 | TRBJ2-1 | CASSSKREALGEQFF | KLGGALQAK | YES | NO | NO |
| 16156 | TRAV26-2 | TRAJ48 | CILRVNFGNEKLTF | TRBV6-5 | TRBJ2-1 | CASSIIGLQGNEQFF | KLGGALQAK | NO | NO | NO |
| 16157 | TRAV39 | TRAJ47 | CAVEYGNKLVF | TRBV20-1 | TRBJ2-7 | CSAFPEERQGPFMEEQYF | KLGGALQAK | NO | NO | NO |
| 16158 | TRAV19 | TRAJ48 | CALSESNFGNEKLTF | TRBV5-6 | TRBJ1-2 | CASSPWALTYNYGYTF | KLGGALQAK | YES | NO | NO |
| 16159 | TRAV36DV7 | TRAJ45 | CAVSGGGADGLTF | TRBV6-5 | TRBJ2-3 | CASGAEGQGSDTQYF | KLGGALQAK | NO | NO | NO |
| 16160 | TRAV35 | TRAJ47 | CGPMEYGNKLVF | TRBV3-1 | TRBJ1-2 | CASSQTGENLYGYTF | KLGGALQAK | YES | NO | NO |
| 16161 | TRAV3 | TRAJ24 | CAVRGIMTTDSWGKFQF | TRBV10-1 | TRBJ2-2 | CASRESSNTGELFF | KLGGALQAK | NO | NO | NO |
| 16162 | TRAV13-2 | TRAJ5 | CAEISMDTGRRALTF | TRBV19 | TRBJ2-7 | CASSIFESEQYF | KLGGALQAK | YES | NO | NO |
| 16163 | TRAV1-1 | TRAJ26 | CAVREPSYGQNFVF | TRBV7-9 | TRBJ1-5 | CASSLNYNQPQHF | KLGGALQAK | YES | NO | NO |
| 16164 | TRAV17 | TRAJ54 | CATDQGAQKLVF | TRBV24-1 | TRBJ2-1 | CATRDSGNNEQFF | KLGGALQAK | YES | NO | NO |
| 16165 | TRAV12-3 | TRAJ31 | CAVNRVRDARLMF | TRBV4-1 | TRBJ1-1 | CASRVGNTEAFF | KLGGALQAK | YES | NO | NO |
| 16166 | TRAV38-2DV8 | TRAJ49 | CAYTNTGNQFYF | TRBV5-5 | TRBJ2-1 | CASSLYTSGYNEQFF | KLGGALQAK | YES | NO | NO |
| 16167 | TRAV5 | TRAJ31 | CAETYMNNARLMF | TRBV5-1 | TRBJ2-2 | CASSGTDFNTGELFF | KLGGALQAK | YES | NO | NO |
| 16168 | TRAV35 | TRAJ29 | CAGQIISGNTPLVF | TRBV27 | TRBJ2-1 | CASSLSTNEQFF | KLGGALQAK | YES | NO | NO |
| 16169 | TRAV13-1 | TRAJ53 | CAASLGGSNYKLTF | TRBV4-3 | TRBJ1-2 | CASSQARAYGYTF | KLGGALQAK | YES | NO | NO |
| 16170 | TRAV17 | TRAJ37 | CATASVSSNTGKLIF | TRBV19 | TRBJ1-1 | CATEAGTGNTEAFF | KLGGALQAK | YES | NO | NO |
| 16171 | TRAV8-3 | TRAJ4 | CAVGDFSFGGYNKLIF | TRBV27 | TRBJ1-5 | CASSQASNQPQHF | KLGGALQAK | NO | NO | NO |
| 16172 | TRAV14DV4 | TRAJ33 | CAMRVESNYQLIW | TRBV5-4 | TRBJ1-1 | CASSLGEMNTEAFF | KLGGALQAK | YES | NO | NO |
| 16173 | TRAV35 | TRAJ58 | CAGHGETSGSRLTF | TRBV18 | TRBJ2-1 | CASSVAGGNEQFF | RLRAEAQVK | YES | YES | YES |
| 16174 | TRAV13-1 | TRAJ22 | CAASIRAGSSGSARQLTF | TRBV4-1 | TRBJ1-6 | CASSQGGNSPLHF | KLGGALQAK | YES | NO | NO |
| 16175 | TRAV17 | TRAJ49 | CATEAGNLFYF | TRBV4-1 | TRBJ1-1 | CASSQGQGYTEAFF | KLGGALQAK | NO | NO | NO |
| 16176 | TRAV17 | TRAJ9 | CATDPNTGGFKTIF | TRBV27 | TRBJ2-1 | CASISPGYNEQFF | KLGGALQAK | YES | NO | NO |
| 16177 | TRAV17 | TRAJ28 | CATDRSPQYSGAGSYQLTF | TRBV12-3 | TRBJ1-1 | CASSLSVHLGGNTEAFF | KLGGALQAK | YES | NO | NO |
| 16178 | TRAV21 | TRAJ43 | CAVWYMRF | TRBV5-5 | TRBJ1-1 | CASSPLDRGEAFF | KLGGALQAK | NO | NO | NO |
| 16179 | TRAV12-1 | TRAJ34 | CVVNPQTYNTDKLIF | TRBV19 | TRBJ1-1 | CASSLGTGWTEAFF | KLGGALQAK | YES | NO | NO |
| 16180 | TRAV36DV7 | TRAJ39 | CAVDFNNAGNMLTF | TRBV16 | TRBJ1-2 | CASSQQTGYGYTF | KLGGALQAK | NO | NO | NO |
| 16181 | TRAV12-2 | TRAJ42 | CAVGRGGSQGNLIF | TRBV12-4 | TRBJ1-2 | CASSLATAYYGYTF | KLGGALQAK | YES | NO | NO |
| 16182 | TRAV25 | TRAJ35 | CAGPTGFGNVLHC | TRBV6-3 | TRBJ2-7 | CASSYPSPGQGVNEQYF | KLGGALQAK | NO | NO | NO |
| 16183 | TRAV3 | TRAJ10 | CAVRDPAGGGNKLTF | TRBV11-3 | TRBJ1-1 | CASSSTTAIRDTEAFF | KLGGALQAK | YES | NO | NO |
| 16184 | TRAV12-1 | TRAJ36 | CVVSQTGANNLFF | TRBV5-1 | TRBJ2-7 | CASSTGPRDRGYEQYV | KLGGALQAK | NO | NO | NO |
| 16185 | TRAV14DV4 | TRAJ58 | CAMREEETSGSRLTF | TRBV4-3 | TRBJ2-4 | CASSQVRGINIQYF | KLGGALQAK | YES | NO | NO |
| 16186 | TRAV21 | TRAJ57 | CAVLTQGGSEKLVF | TRBV4-2 | TRBJ2-5 | CASSQMGTQYF | KLGGALQAK | YES | NO | NO |
| 16187 | TRAV12-1 | TRAJ6 | CVVRTTAGGSYIPTF | TRBV7-9 | TRBJ2-6 | CASSSGESGANVLTF | KLGGALQAK | YES | NO | NO |
| 16188 | TRAV1-1 | TRAJ23 | CAVEIQGGKLIF | TRBV7-9 | TRBJ2-2 | CASSSSGTEFHTGELFF | KLGGALQAK | YES | NO | NO |
| 16189 | TRAV12-2 | TRAJ49 | CAGLTGNQFYF | TRBV20-1 | TRBJ1-1 | CSVQTGRAFF | KLGGALQAK | NO | NO | NO |
| 16190 | TRAV13-2 | TRAJ24 | CADLPLTDSWGKFQF | TRBV4-1 | TRBJ2-3 | CASSQDMGGRQYF | KLGGALQAK | YES | NO | NO |
| 16191 | TRAV3 | TRAJ37 | CAVRDGGMGSNTGKLIF | TRBV13 | TRBJ2-3 | CASSALSGDSTSTDTQYF | KLGGALQAK | NO | NO | NO |
| 16192 | TRAV27 | TRAJ33 | CAGVWSNYQLIW | TRBV7-9 | TRBJ1-4 | CATRGLSNEKLFF | KLGGALQAK | YES | NO | NO |
| 16193 | TRAV13-1 | TRAJ11 | CASPRRGYSTLTF | TRBV6-3 | TRBJ2-2 | CASSPSGRFGELFF | KLGGALQAK | YES | NO | NO |
| 16194 | TRAV26-2 | TRAJ57 | CILRDFPGGSEKLVF | TRBV19 | TRBJ2-1 | CASSISSYNEQFF | KLGGALQAK | YES | NO | NO |
| 16195 | TRAV26-2 | TRAJ54 | CILLQGAQKLVF | TRBV27 | TRBJ1-2 | CASSPQPDKYYGYTF | KLGGALQAK | YES | NO | NO |
| 16196 | TRAV8-6 | TRAJ36 | CAVSDTQTGANNLFF | TRBV18 | TRBJ1-6 | CASSPGQGSDNSPLHF | KLGGALQAK | YES | NO | NO |
| 16197 | TRAV8-4 | TRAJ17 | CAVNHKAAGNKLTF | TRBV5-6 | TRBJ2-6 | CASSLAGVSGANVLTF | KLGGALQAK | YES | NO | NO |
| 16198 | TRAV3 | TRAJ42 | CAVRDERGSQGNLIF | TRBV7-2 | TRBJ2-7 | CASSDDRGPFYEQYV | KLGGALQAK | NO | NO | NO |
| 16199 | TRAV27 | TRAJ42 | CAGVGGGSQGNLIF | TRBV19 | TRBJ2-7 | CASSIRATYEQYF | GILGFVFTL | YES | NO | YES |
| 16200 | TRAV8-3 | TRAJ15 | CAVVPNTNQAGTALIF | TRBV20-1 | TRBJ1-1 | CSARSWGVEAFF | KLGGALQAK | YES | NO | NO |
| 16202 | TRAV24 | TRAJ52 | CAFIGDAGGTSYGKLTF | TRBV27 | TRBJ2-1 | CASSFKGGAGYNEQFF | KLGGALQAK | YES | NO | NO |
| 16203 | TRAV21 | TRAJ36 | CAVRTGANNLFF | TRBV11-2 | TRBJ2-7 | CASSTGLYEQYF | KLGGALQAK | YES | NO | NO |
| 16204 | TRAV23DV6 | TRAJ17 | CAASPAAGNKLTF | TRBV29-1 | TRBJ2-7 | CSVAPVVSYEQYF | KLGGALQAK | NO | NO | NO |
| 16205 | TRAV25 | TRAJ33 | CAGLDSNYQLIW | TRBV9 | TRBJ2-7 | CASSPRGWDEQYF | KLGGALQAK | YES | NO | NO |
| 16206 | TRAV8-3 | TRAJ45 | CAVGPQPGGGADGLTF | TRBV13 | TRBJ2-1 | CASTLGASGGNEQFF | KLGGALQAK | YES | NO | NO |
| 16207 | TRAV14DV4 | TRAJ54 | CAMRAEIQGAQKLVF | TRBV5-1 | TRBJ2-7 | CASSLESAGGEQYF | AVFDRKSDAK | YES | NO | YES |
| 16208 | TRAV1-2 | TRAJ17 | CAVRDEAAGNKLTF | TRBV12-4 | TRBJ2-1 | CASGGNEQFF | KLGGALQAK | YES | NO | NO |
| 16209 | TRAV25 | TRAJ34 | CAGDLWNTDKLIF | TRBV25-1 | TRBJ2-3 | CASSSTVYQLTDTQYF | KLGGALQAK | YES | NO | NO |
| 16210 | TRAV29DV5 | TRAJ26 | CAAKDNYGQNFVF | TRBV9 | TRBJ2-1 | CASSADLYNEQFF | KLGGALQAK | YES | NO | NO |
| 16211 | TRAV12-3 | TRAJ26 | CAMSAWDNYGQNFVF | TRBV2 | TRBJ2-2 | CASRGGELFF | KLGGALQAK | YES | NO | NO |
| 16212 | TRAV13-2 | TRAJ44 | CAENTGTASKLTF | TRBV9 | TRBJ2-5 | CASSGQGSETQYF | KLGGALQAK | YES | NO | NO |
| 16213 | TRAV12-3 | TRAJ13 | CAMSESGGYQKVTF | TRBV4-1 | TRBJ2-3 | CASSPLVDSDSTDTQYF | KLGGALQAK | YES | NO | NO |
| 16214 | TRAV12-2 | TRAJ39 | CAVKNNAGNMLTF | TRBV7-9 | TRBJ1-5 | CASSEQGAGSNQPQHF | KLGGALQAK | YES | NO | NO |
| 16215 | TRAV21 | TRAJ41 | CAPGGSGYALNF | TRBV12-4 | TRBJ2-7 | CASSPSSYEQYF | KLGGALQAK | YES | NO | NO |
| 16216 | TRAV12-1 | TRAJ4 | CVVNYPGGYNKLIF | TRBV29-1 | TRBJ1-1 | CSVVDTEAFF | KLGGALQAK | NO | NO | NO |
| 16217 | TRAV17 | TRAJ30 | CATSTGDDKIIF | TRBV27 | TRBJ2-1 | CASSLSAYNEQFF | KLGGALQAK | YES | NO | NO |
| 16218 | TRAV17 | TRAJ21 | CVPLGDNFNKFYF | TRBV7-9 | TRBJ2-5 | CASSFPGSETQYF | AVFDRKSDAK | YES | NO | NO |
| 16219 | TRAV10 | TRAJ15 | CVVSLNQAGTALIF | TRBV24-1 | TRBJ2-3 | CATSGPHTDTQYF | KLGGALQAK | YES | NO | NO |
| 16220 | TRAV36DV7 | TRAJ42 | CAVELDGGSQGNLIF | TRBV5-1 | TRBJ2-7 | CASSSIGAYEQYF | KLGGALQAK | NO | NO | NO |
| 16221 | TRAV38-1 | TRAJ43 | CAFMPGNYNNNDMRF | TRBV7-6 | TRBJ2-3 | CASSLRSGAGVAQIPQYF | KLGGALQAK | YES | NO | NO |
| 16222 | TRAV12-2 | TRAJ49 | CAVREAGNQFYF | TRBV11-3 | TRBJ2-7 | CASSDHPYEQYV | KLGGALQAK | NO | NO | NO |
| 16223 | TRAV8-6 | TRAJ54 | CAVSDPQGAQKLVF | TRBV7-6 | TRBJ1-3 | CASGTQNTIYF | AVFDRKSDAK | YES | NO | YES |
| 16224 | TRAV38-2DV8 | TRAJ44 | CAFTHLTGTASKLTF | TRBV13 | TRBJ2-1 | CASSPPGTSGFYEQFF | KLGGALQAK | YES | NO | NO |
| 16225 | TRAV13-2 | TRAJ39 | CAETDNAGNMLTF | TRBV6-5 | TRBJ2-7 | CASSFTVLGDEQYF | KLGGALQAK | YES | NO | NO |
| 16226 | TRAV19 | TRAJ57 | CALSSITQGGSEKLVF | TRBV9 | TRBJ2-1 | CASSVLENNEQFF | KLGGALQAK | YES | NO | NO |
| 16227 | TRAV21 | TRAJ53 | CAVIGDSGGSNYKLTF | TRBV19 | TRBJ1-5 | CASSIVGQPQHF | AVFDRKSDAK | YES | NO | YES |
| 16228 | TRAV22 | TRAJ23 | CAVVEGGKLIF | TRBV9 | TRBJ2-1 | CASSVEDRVNEQFF | KLGGALQAK | YES | NO | NO |
| 16229 | TRAV29DV5 | TRAJ48 | CAARVGEKLTF | TRBV28 | TRBJ2-3 | CASSLWLKDFAGGGDTQYF | KLGGALQAK | YES | NO | NO |
| 16230 | TRAV12-1 | TRAJ4 | CVVTLLSGGYNKLIF | TRBV6-5 | TRBJ2-1 | CASSYHSSGEGPNEQFF | KLGGALQAK | NO | NO | NO |

| | | | | | | | | | | |
|---|---|---|---|---|---|---|---|---|---|---|
| 16231 | TRAV1-2 | TRAJ33 | CAVADSNYQLIW | TRBV6-6 | TRBJ2-2 | CASSYRGEAGELFF | KLGGALQAK | YES | NO | NO |
| 16232 | TRAV38-2DV8 | TRAJ30 | CAYRSSIEIIF | TRBV4-1 | TRBJ1-1 | CASSQNLGQGVAEAFF | KLGGALQAK | YES | NO | NO |
| 16233 | TRAV27 | TRAJ13 | CAGAAGGYQKVTF | TRBV4-1 | TRBJ1-5 | CASSHVNQPQHF | KLGGALQAK | YES | NO | NO |
| 16234 | TRAV20 | TRAJ31 | CAVQAHARLMF | TRBV27 | TRBJ2-3 | CASRPLTGRVHDTQYF | KLGGALQAK | YES | NO | NO |
| 16235 | TRAV35 | TRAJ31 | CAGPAGARLMF | TRBV7-9 | TRBJ1-2 | CASSLYVNNGYTF | AVFDRKSDAK | YES | NO | YES |
| 16236 | TRAV35 | TRAJ37 | CAGQGPLRNTGKLIF | TRBV29-1 | TRBJ1-2 | CSPDGYTF | KLGGALQAK | NO | NO | NO |
| 16237 | TRAV25 | TRAJ39 | CAANMNNNAGNMLTF | TRBV12-4 | TRBJ1-5 | CASSLSNSNQPQHF | KLGGALQAK | YES | NO | NO |
| 16238 | TRAV1-2 | TRAJ33 | CAPLNSNYQLIW | TRBV20-1 | TRBJ2-2 | CSARDLGHDTGELFF | KLGGALQAK | YES | NO | NO |
| 16239 | TRAV12-2 | TRAJ37 | CAVTGKLIF | TRBV12-4 | TRBJ1-1 | CASSFGVGTEAFF | KLGGALQAK | YES | NO | NO |
| 16240 | TRAV35 | TRAJ26 | CAGPRDNYGQNFVF | TRBV27 | TRBJ2-7 | CASSSGTNSYEQYF | KLGGALQAK | YES | NO | NO |
| 16241 | TRAV4 | TRAJ42 | CLVGENYGGSQGNLIF | TRBV6-3 | TRBJ1-5 | CASSYGVGNQPQHF | KLGGALQAK | NO | NO | NO |
| 16242 | TRAV16 | TRAJ57 | CALSGFSQGGSEKLVF | TRBV11-1 | TRBJ2-1 | CASSWAYNEQFF | KLGGALQAK | NO | NO | NO |
| 16243 | TRAV19 | TRAJ53 | CVLRSGGSNYKLTF | TRBV13 | TRBJ2-6 | CASSLGLSGANVLTF | KLGGALQAK | NO | NO | NO |
| 16244 | TRAV13-1 | TRAJ33 | CAASNIRGRDSNYQLIW | TRBV6-3 | TRBJ2-7 | CASSWTENYEQYV | KLGGALQAK | NO | NO | NO |
| 16245 | TRAV24 | TRAJ52 | CAFIGDAGGTSYGKLTF | TRBV4-1 | TRBJ2-3 | CASSQYPGETDTQYF | KLGGALQAK | YES | NO | NO |
| 16246 | TRAV1-2 | TRAJ34 | CAVRAYNTDKLIF | TRBV9 | TRBJ2-3 | CASSVVGGATDTQYF | KLGGALQAK | YES | NO | NO |
| 16247 | TRAV1-1 | TRAJ31 | CAVRWEARLMF | TRBV5-6 | TRBJ2-2 | CASSFGGTGELFF | KLGGALQAK | YES | NO | NO |
| 16248 | TRAV13-1 | TRAJ36 | CAAIEGAGANNLFF | TRBV12-4 | TRBJ2-3 | CASSLAGGNTDTQYF | KLGGALQAK | YES | NO | NO |
| 16249 | TRAV38-1 | TRAJ33 | CAFMKNPDSNYQLIW | TRBV11-2 | TRBJ1-1 | CASSLDSFGARGPNTEAFF | KLGGALQAK | YES | NO | NO |
| 16250 | TRAV12-2 | TRAJ54 | CAHLQGAQKLVF | TRBV29-1 | TRBJ1-5 | CSVLGGVSNQPQHF | KLGGALQAK | YES | NO | NO |
| 16251 | TRAV13-1 | TRAJ23 | CAASRYNQGGKLIF | TRBV5-6 | TRBJ2-7 | CASSLISPYEQYF | KLGGALQAK | YES | NO | NO |
| 16252 | TRAV19 | TRAJ29 | CALSEASSGNTPLVF | TRBV7-8 | TRBJ2-1 | CASSLWPSGTNEQFF | KLGGALQAK | YES | NO | NO |
| 16253 | TRAV12-1 | TRAJ45 | CVVITGADGLTF | TRBV7-9 | TRBJ2-5 | CASSLIPPETQYF | KLGGALQAK | YES | NO | NO |
| 16254 | TRAV24 | TRAJ36 | CASPGTGANNLFF | TRBV27 | TRBJ2-1 | CASSMTSGRYNEQFF | KLGGALQAK | YES | NO | NO |
| 16255 | TRAV13-2 | TRAJ37 | CAESYSNTGKLIF | TRBV27 | TRBJ1-1 | CASSLAGGEAFF | KLGGALQAK | YES | NO | NO |
| 16256 | TRAV19 | TRAJ4 | CALSEGVSGGYNKLIF | TRBV27 | TRBJ1-5 | CASSLLGQGLDQPQHF | KLGGALQAK | YES | NO | NO |
| 16257 | TRAV13-1 | TRAJ45 | CAASTGGGADGLTF | TRBV11-2 | TRBJ2-1 | CASSSPERFGYNEQFF | KLGGALQAK | YES | NO | NO |
| 16258 | TRAV8-4 | TRAJ50 | CAVSDTSYDKVIF | TRBV20-1 | TRBJ2-7 | CSARGLSSYEQYF | KLGGALQAK | YES | NO | NO |
| 16259 | TRAV14DV4 | TRAJ54 | CAMRERFLQGAQKLVF | TRBV20-1 | TRBJ2-1 | CSAEGLAGVRANNEQFF | KLGGALQAK | NO | NO | NO |
| 16260 | TRAV12-2 | TRAJ41 | CAVTSVQNSNSGYALNF | TRBV19 | TRBJ1-1 | CASSMDEGSTEAFF | KLGGALQAK | YES | NO | NO |
| 16261 | TRAV29DV5 | TRAJ27 | CAASRKNTNAGKSTF | TRBV7-9 | TRBJ1-1 | CASSLGLWGLDTEAFF | KLGGALQAK | YES | NO | NO |
| 16262 | TRAV4 | TRAJ28 | CLVGVPGAGSYQLTF | TRBV5-1 | TRBJ2-1 | CASSSPGTYNEQFF | KLGGALQAK | NO | NO | NO |
| 16263 | TRAV6 | TRAJ28 | CALPAYSGAGSYQLTF | TRBV6-3 | TRBJ2-7 | CASSYSGGGDSGTYEQYV | KLGGALQAK | NO | NO | NO |
| 16264 | TRAV1-2 | TRAJ33 | CAVKDSNYQLIW | TRBV15 | TRBJ2-2 | CATSKGSDTGELFF | KLGGALQAK | YES | NO | NO |
| 16265 | TRAV27 | TRAJ52 | CAGDLGYGKLTF | TRBV7-9 | TRBJ2-1 | CASSLVIVDLGANEQFF | KLGGALQAK | YES | NO | NO |
| 16266 | TRAV8-1 | TRAJ20 | CAVRNDYKLSF | TRBV19 | TRBJ2-2 | CATSNNRGPGELFF | KLGGALQAK | YES | NO | NO |
| 16267 | TRAV14DV4 | TRAJ22 | CAMRRPISSGSARQLTF | TRBV14 | TRBJ1-4 | CASSQPGTDPNEKLFF | KLGGALQAK | NO | NO | NO |
| 16268 | TRAV19 | TRAJ53 | CALRNNSGGSNYKLTF | TRBV7-9 | TRBJ1-5 | CASSSSGTTKQPQHF | KLGGALQAK | YES | NO | NO |
| 16269 | TRAV17 | TRAJ49 | CATPTGNQFYF | TRBV9 | TRBJ2-1 | CASSVGQGLVYEQFF | KLGGALQAK | YES | NO | NO |
| 16270 | TRAV12-3 | TRAJ39 | CAIGPGNMLTF | TRBV7-2 | TRBJ2-3 | CASSPRLGLLSTDTQYF | KLGGALQAK | YES | NO | NO |
| 16271 | TRAV38-2DV8 | TRAJ49 | CAYRSACNQFYF | TRBV11-2 | TRBJ2-3 | CASSLVCTDTQYF | AVFDRKSDAK | YES | NO | YES |
| 16272 | TRAV1-2 | TRAJ31 | CAVRKDNNARLMF | TRBV4-1 | TRBJ2-3 | CASSQDRGQASQYF | KLGGALQAK | YES | NO | NO |
| 16273 | TRAV14DV4 | TRAJ49 | CAMREGDIGNQFYF | TRBV20-1 | TRBJ2-1 | CSARDYLRPLDNEQFF | KLGGALQAK | NO | NO | NO |
| 16274 | TRAV21 | TRAJ37 | CAVRSYQGKLIF | TRBV7-2 | TRBJ2-7 | CASSSGYHEQYF | KLGGALQAK | YES | NO | NO |
| 16275 | TRAV25 | TRAJ50 | CAEKTSYDKVIF | TRBV19 | TRBJ1-2 | CASSARTGVGYGYTF | KLGGALQAK | YES | NO | NO |
| 16276 | TRAV8-2 | TRAJ22 | CVGRKGSARQLTF | TRBV30 | TRBJ1-2 | CAWGSGLGYGYTF | KLGGALQAK | NO | NO | NO |
| 16277 | TRAV36DV7 | TRAJ52 | CAGANAGGTSYGKLTF | TRBV19 | TRBJ2-2 | CASSGGTNPGELFF | AVFDRKSDAK | NO | NO | NO |
| 16278 | TRAV12-3 | TRAJ54 | CAMSEQGAQKLVF | TRBV6-5 | TRBJ2-7 | CASSPGGPAEQYF | KLGGALQAK | YES | NO | NO |
| 16279 | TRAV3 | TRAJ30 | CAVREIGRDDKIIF | TRBV5-6 | TRBJ2-3 | CASSLDQPGGADTQYF | KLGGALQAK | NO | NO | NO |
| 16280 | TRAV27 | TRAJ27 | CAGVQNTNAGKSTF | TRBV7-9 | TRBJ1-1 | CASTWTDTEAFF | KLGGALQAK | YES | NO | NO |
| 16281 | TRAV17 | TRAJ47 | CATDAGASNGNKLVF | TRBV9 | TRBJ1-4 | CASSATGVQKLFF | KLGGALQAK | YES | NO | NO |
| 16282 | TRAV26-1 | TRAJ26 | CIAPSYGQNFVF | TRBV2 | TRBJ2-3 | CASRPPGREEDTQYF | KLGGALQAK | YES | NO | NO |
| 16283 | TRAV27 | TRAJ50 | CAGPDSKTSYDKVIF | TRBV19 | TRBJ2-7 | CATLGTGGYYEQYF | KLGGALQAK | YES | NO | NO |
| 16284 | TRAV19 | TRAJ36 | CALSLGSGANNLFF | TRBV14 | TRBJ2-1 | CASSLVIAGPYQFF | KLGGALQAK | YES | NO | NO |
| 16285 | TRAV13-2 | TRAJ12 | CAENSMDSSYKLIF | TRBV19 | TRBJ1-1 | CASRDGMADTEAFF | KLGGALQAK | YES | NO | NO |
| 16286 | TRAV8-4 | TRAJ10 | CAVRGARGGNKLTF | TRBV12-3 | TRBJ2-1 | CASRVNNEQFF | KLGGALQAK | YES | NO | NO |
| 16287 | TRAV12-2 | TRAJ13 | CAVNMRGGYQKVTF | TRBV7-9 | TRBJ2-3 | CASSFTSGSTDTQYF | KLGGALQAK | YES | NO | NO |
| 16288 | TRAV12-1 | TRAJ53 | CVVNPPAGSNYKLTF | TRBV12-4 | TRBJ2-4 | CASSASLGVDLAKNIQYF | KLGGALQAK | YES | NO | NO |
| 16289 | TRAV1-1 | TRAJ11 | CAVQGNSGYSTLTF | TRBV6-3 | TRBJ2-3 | CASSGLADSTDTQYF | KLGGALQAK | YES | NO | NO |
| 16290 | TRAV8-6 | TRAJ56 | CAVSDRTGANSKLTF | TRBV19 | TRBJ1-1 | CASSMTGFTEAFF | KLGGALQAK | YES | NO | NO |
| 16291 | TRAV35 | TRAJ48 | CAGQRESNFGNEKLTF | TRBV9 | TRBJ2-7 | CASSVEIGAYEQYF | KLGGALQAK | YES | NO | NO |
| 16292 | TRAV8-3 | TRAJ7 | CAVGTTYGNNRLAF | TRBV3-1 | TRBJ2-5 | CASSQNDGRQTQETQYF | KLGGALQAK | YES | NO | NO |
| 16294 | TRAV12-3 | TRAJ39 | CAIGPGNMLTF | TRBV7-8 | TRBJ2-5 | CASYPVPGLKTQYF | KLGGALQAK | YES | NO | NO |
| 16295 | TRAV8-4 | TRAJ32 | CAVSGGFGGATNKLIF | TRBV12-4 | TRBJ2-3 | CASSLFRRETIHTDTQYF | KLGGALQAK | YES | NO | NO |
| 16296 | TRAV16 | TRAJ11 | CALGNSGYSTLTF | TRBV7-2 | TRBJ2-2 | CASRGHTGELFF | KLGGALQAK | NO | NO | NO |
| 16297 | TRAV1-2 | TRAJ24 | CAVRDESWGKFQF | TRBV30 | TRBJ1-2 | CAWSVPQGAHYGYTF | KLGGALQAK | NO | NO | NO |
| 16298 | TRAV24 | TRAJ26 | CAFNNYGQNFVF | TRBV9 | TRBJ2-3 | CASSPGSKREGTQYF | KLGGALQAK | YES | NO | NO |
| 16299 | TRAV14DV4 | TRAJ23 | CAMREGLRGNQGGKLIF | TRBV20-1 | TRBJ2-1 | CSALGLSRDLSSYNEQFF | KLGGALQAK | NO | NO | NO |
| 16300 | TRAV29DV5 | TRAJ42 | CAASALRGGSQGNLIF | TRBV11-3 | TRBJ2-1 | CASSLSLGGNEQFF | KLGGALQAK | YES | NO | NO |
| 16301 | TRAV14DV4 | TRAJ32 | CAMREGRDYGGATNKLIF | TRBV29-1 | TRBJ2-1 | CSGVLAPSYNEQFF | KLGGALQAK | YES | NO | NO |
| 16302 | TRAV13-1 | TRAJ34 | CAAGYGTDKLIF | TRBV7-9 | TRBJ1-1 | CASSFGSEAFF | KLGGALQAK | YES | NO | NO |
| 16303 | TRAV6 | TRAJ49 | CALDYSPTGNQFYF | TRBV3-1 | TRBJ2-1 | CASSQDLGGVSYNEQFF | KLGGALQAK | YES | NO | NO |
| 16304 | TRAV8-3 | TRAJ34 | CAVDNTDKLIF | TRBV11-2 | TRBJ1-1 | CASSSNRDRGTKDTEAFF | KLGGALQAK | YES | NO | NO |
| 16305 | TRAV14DV4 | TRAJ50 | CAMRERKTSYDKVIF | TRBV9 | TRBJ2-5 | CASSVGQPGETQYF | KLGGALQAK | YES | NO | NO |
| 16306 | TRAV25 | TRAJ26 | CAGADNYGQNFVF | TRBV29-1 | TRBJ1-6 | CSVEVGDNSPLHF | KLGGALQAK | NO | NO | NO |
| 16307 | TRAV38-1 | TRAJ54 | CAFMSLIQGAQKLVF | TRBV7-9 | TRBJ2-1 | CASSNSLPSGVSYNEQFF | KLGGALQAK | YES | NO | NO |
| 16308 | TRAV13-2 | TRAJ13 | CAESPGYQKVTF | TRBV7-9 | TRBJ2-1 | CASRIGTSLDNEQFF | KLGGALQAK | YES | YES | NO |
| 16309 | TRAV10 | TRAJ39 | CVVSARGRNAGNMLTF | TRBV13 | TRBJ2-5 | CASSLSPSTGYYSQETQYF | KLGGALQAK | YES | NO | NO |
| 16310 | TRAV4 | TRAJ33 | CLVGADSNYQLIW | TRBV6-3 | TRBJ2-1 | CASSNRLAGITYNEQFF | KLGGALQAK | NO | NO | NO |
| 16311 | TRAV21 | TRAJ34 | CAVSLLNTDKLIF | TRBV19 | TRBJ1-5 | CASSLGSNQPQHF | KLGGALQAK | YES | NO | NO |
| 16312 | TRAV17 | TRAJ26 | CATDDNYGQNFVF | TRBV6-6 | TRBJ2-1 | CASSSPGTGNEQFF | KLGGALQAK | NO | NO | NO |
| 16313 | TRAV8-1 | TRAJ30 | CAVNSNRDDKIIF | TRBV27 | TRBJ2-3 | CASSPRQGEYYTDTQYF | KLGGALQAK | YES | NO | NO |
| 16314 | TRAV8-4 | TRAJ43 | CAVSDGGNNDMRF | TRBV20-1 | TRBJ2-2 | CSARDRLAFDTGELFF | KLGGALQAK | YES | NO | NO |
| 16315 | TRAV14DV4 | TRAJ29 | CAMREGGRSGNTPLVF | TRBV4-1 | TRBJ2-7 | CASSQGKLLSYEQYF | KLGGALQAK | YES | NO | NO |
| 16316 | TRAV8-3 | TRAJ42 | CAVAPGGGSQGNLIF | TRBV24-1 | TRBJ2-7 | CATSGLTDYEQYV | KLGGALQAK | NO | NO | NO |
| 16317 | TRAV10 | TRAJ31 | CVVSLNNARLMF | TRBV19 | TRBJ1-5 | CASTGGAYNQPQHF | KLGGALQAK | YES | NO | NO |
| 16318 | TRAV29DV5 | TRAJ52 | CAASTTNAGGTSYGKLTF | TRBV20-1 | TRBJ1-2 | CSVRRGYYGYTF | KLGGALQAK | YES | NO | NO |
| 16319 | TRAV20 | TRAJ42 | CAVQVGSGGSQGNLIF | TRBV7-6 | TRBJ2-1 | CASSPGASYNEQFF | KLGGALQAK | YES | NO | NO |
| 16320 | TRAV12-3 | TRAJ53 | CAMSGSGGSNYKLTF | TRBV19 | TRBJ2-3 | CASSILLVNTDTQYF | KLGGALQAK | YES | NO | NO |
| 16321 | TRAV12-1 | TRAJ4 | CVVTGGYNKLIF | TRBV12-3 | TRBJ2-3 | CASSPTDRATDTQYF | AVFDRKSDAK | YES | NO | YES |
| 16322 | TRAV25 | TRAJ16 | CAVDDGQKLLF | TRBV19 | TRBJ1-2 | CATTNGGRGYTF | AVFDRKSDAK | NO | NO | NO |
| 16323 | TRAV13-1 | TRAJ9 | CAATNTGGFKTIF | TRBV28 | TRBJ1-6 | CASSLGPTGGSPLHF | KLGGALQAK | YES | NO | NO |
| 16324 | TRAV12-3 | TRAJ40 | CAMTSGTYKYIF | TRBV20-1 | TRBJ2-2 | CSARPLVSAGELFF | KLGGALQAK | YES | NO | NO |
| 16325 | TRAV10 | TRAJ40 | CVVISGTYKYIF | TRBV6-6 | TRBJ1-4 | CASSSLGQGSVNEKLFF | KLGGALQAK | NO | NO | NO |
| 16326 | TRAV12-3 | TRAJ12 | CAMGSMDSSYKLIF | TRBV14 | TRBJ1-5 | CASSQDAGQPQHF | KLGGALQAK | NO | NO | NO |
| 16327 | TRAV22 | TRAJ42 | CAVYGGSQGNLIF | TRBV7-9 | TRBJ2-1 | CASSLLAGGLGEQFF | KLGGALQAK | YES | NO | NO |
| 16328 | TRAV30 | TRAJ42 | CGWGGSQGNLIF | TRBV9 | TRBJ2-7 | CASSVGSSLHEQYF | KLGGALQAK | YES | NO | NO |
| 16329 | TRAV12-2 | TRAJ43 | CAVPRGF | TRBV19 | TRBJ1-1 | CASSIGRKNTEAFF | KLGGALQAK | NO | NO | NO |
| 16330 | TRAV3 | TRAJ35 | CAVRDPSGFGNVLHC | TRBV7-9 | TRBJ2-1 | CASSSSGPYNEQFF | KLGGALQAK | NO | NO | NO |
| 16331 | TRAV29DV5 | TRAJ49 | CAAKNTGNQFYF | TRBV4-1 | TRBJ1-1 | CASSQGGGLTEAFF | KLGGALQAK | YES | NO | NO |
| 16332 | TRAV29DV5 | TRAJ49 | CAASPTGNQFYF | TRBV29-1 | TRBJ2-6 | CSVADRGSSSGANVLTF | KLGGALQAK | YES | NO | NO |
| 16333 | TRAV25 | TRAJ9 | CAGLITGGFKTIF | TRBV15 | TRBJ2-5 | CATSRDLFETQYF | KLGGALQAK | YES | NO | NO |
| 16334 | TRAV17 | TRAJ58 | CATEAVWETSGSRLTF | TRBV19 | TRBJ2-7 | CASSMGTGLDEQYF | KLGGALQAK | YES | NO | NO |
| 16335 | TRAV13-1 | TRAJ47 | CAVVEYGNKLVF | TRBV12-3 | TRBJ2-3 | CASSPGYTQYF | KLGGALQAK | YES | NO | NO |

| | | | | | | | | | | |
|---|---|---|---|---|---|---|---|---|---|---|
| 16336 | TRAV41 | TRAJ30 | CAVRDDKIIF | TRBV4-3 | TRBJ1-3 | CASSQDRTSGNTIYF | AVFDRKSDAK | YES | NO | NO |
| 16337 | TRAV17 | TRAJ18 | CATDSDRGSTLGRLYF | TRBV27 | TRBJ1-1 | CASSPRQVPNTEAFF | KLGGALQAK | YES | NO | NO |
| 16338 | TRAV34 | TRAJ52 | CGADDESFSAGGTSYGKLTF | TRBV19 | TRBJ2-1 | CASRRTGGSYNEQFF | KLGGALQAK | NO | NO | NO |
| 16339 | TRAV19 | TRAJ26 | CALSEGIDNYGQNFVF | TRBV2 | TRBJ1-6 | CAGGTGYNSPLHF | KLGGALQAK | YES | NO | NO |
| 16340 | TRAV5 | TRAJ9 | CAESYTGGFKTIF | TRBV13 | TRBJ2-3 | CASSSGTVTDTQYF | KLGGALQAK | YES | NO | NO |
| 16341 | TRAV5 | TRAJ39 | CAETNAGNMLTF | TRBV9 | TRBJ2-7 | CASSADGEQYF | KLGGALQAK | YES | NO | NO |
| 16342 | TRAV4 | TRAJ42 | CLVGDLLYGGSQGNLIF | TRBV6-1 | TRBJ1-1 | CASLGKIGPMNTEAFF | KLGGALQAK | NO | NO | NO |
| 16343 | TRAV23DV6 | TRAJ5 | CAASPMDTGRRALTF | TRBV7-9 | TRBJ1-4 | CASGEGAANEKLFF | KLGGALQAK | YES | NO | NO |
| 16344 | TRAV3 | TRAJ43 | CAVRDYNNNDMRF | TRBV6-5 | TRBJ2-1 | CASSSDTGLAVDEQFF | KLGGALQAK | YES | NO | NO |
| 16345 | TRAV12-3 | TRAJ20 | CAAVNDYKLSF | TRBV13 | TRBJ2-5 | CASSLGDRTQYF | KLGGALQAK | YES | NO | NO |
| 16346 | TRAV12-2 | TRAJ39 | CAVHNAGNMLTF | TRBV5-1 | TRBJ2-3 | CASSFSGEADTQYF | KLGGALQAK | YES | NO | NO |
| 16347 | TRAV21 | TRAJ13 | CAVNSGGYQKVTF | TRBV3-1 | TRBJ2-1 | CASRGISGNNNEQFF | KLGGALQAK | YES | NO | NO |
| 16348 | TRAV25 | TRAJ20 | CAGFFDYKLSF | TRBV5-1 | TRBJ2-7 | CASSLSPRRTQDGGASYEQYF | KLGGALQAK | NO | NO | NO |
| 16349 | TRAV27 | TRAJ29 | CAGNGNTPLVF | TRBV6-3 | TRBJ2-1 | CASSYGAGGYNEQFF | KLGGALQAK | YES | NO | NO |
| 16350 | TRAV21 | TRAJ22 | CAGISSGSARQLTF | TRBV16 | TRBJ2-1 | CASGRLDIEQFF | KLGGALQAK | YES | NO | NO |
| 16351 | TRAV25 | TRAJ18 | CAGQYRGSTLGRLYF | TRBV27 | TRBJ2-1 | CASSLLPGSPDNEQFF | KLGGALQAK | YES | NO | NO |
| 16352 | TRAV8-4 | TRAJ49 | CAVIQRGNQFYF | TRBV7-9 | TRBJ1-1 | CASSSPRGGRITEAFF | KLGGALQAK | YES | NO | NO |
| 16353 | TRAV12-3 | TRAJ52 | CAMPRGTSYGKLTF | TRBV20-1 | TRBJ2-1 | CSARDFNSGGSNYNEQFF | KLGGALQAK | NO | NO | NO |
| 16354 | TRAV34 | TRAJ24 | CGADRGTGWGKFQF | TRBV14 | TRBJ2-1 | CASSQVLAGMWNEQFF | AVFDRKSDAK | YES | NO | YES |
| 16355 | TRAV13-1 | TRAJ23 | CAARLIYNQGGKLIF | TRBV12-4 | TRBJ2-1 | CASSLTGQREGTHNEQFF | KLGGALQAK | YES | NO | NO |
| 16356 | TRAV3 | TRAJ33 | CAVRDIKNSNYQLIW | TRBV30 | TRBJ2-3 | CAWSVLGSTDTQYF | KLGGALQAK | YES | NO | NO |
| 16357 | TRAV19 | TRAJ49 | CALSEFGTGNQFYF | TRBV7-2 | TRBJ2-3 | CASSSLAGYAQYF | KLGGALQAK | YES | NO | NO |
| 16358 | TRAV25 | TRAJ11 | CAGPNSGYSTLTF | TRBV19 | TRBJ2-1 | CASSTKPVWSSYNEQFF | KLGGALQAK | YES | NO | NO |
| 16359 | TRAV9-2 | TRAJ45 | CALINSGGGADGLTF | TRBV7-6 | TRBJ2-3 | CASSLAGGVDTQYF | KTWGQYWQV | YES | NO | YES |
| 16360 | TRAV13-2 | TRAJ9 | CAENTGGFKTIF | TRBV6-3 | TRBJ2-3 | CASSYRAGVADTQYF | KLGGALQAK | YES | NO | NO |
| 16361 | TRAV35 | TRAJ26 | CAGQNYGQNFVF | TRBV5-8 | TRBJ1-5 | CASTPLGGQPQHF | KLGGALQAK | YES | NO | NO |
| 16362 | TRAV20 | TRAJ4 | CAVQDENGGYNKLIF | TRBV5-1 | TRBJ1-1 | CASSGTVMNTEAFF | KLGGALQAK | YES | NO | NO |
| 16363 | TRAV22 | TRAJ34 | CAVGRNTDKLIF | TRBV19 | TRBJ1-1 | CASSNVGGIWTEAFF | KLGGALQAK | YES | NO | NO |
| 16364 | TRAV30 | TRAJ28 | CGTKTSGAGSYQLTF | TRBV7-8 | TRBJ1-1 | CASSLLVRANPNTEAFF | KLGGALQAK | YES | NO | NO |
| 16365 | TRAV2 | TRAJ45 | CAVRGGADGLTF | TRBV4-1 | TRBJ1-6 | CASSQDKGRDSPLHF | KLGGALQAK | NO | NO | NO |
| 16366 | TRAV1-2 | TRAJ33 | CAVRDSNYQLIW | TRBV6-3 | TRBJ2-5 | CASSDNREWETQYF | KLGGALQAK | YES | NO | NO |
| 16367 | TRAV16 | TRAJ22 | CAFLSGSARQLTF | TRBV5-1 | TRBJ1-1 | CASSFGANTEAFF | KLGGALQAK | NO | NO | NO |
| 16368 | TRAV27 | TRAJ48 | CAGTFGNEKLTF | TRBV7-6 | TRBJ2-6 | CASSSSGPFNVLTF | KLGGALQAK | YES | NO | NO |
| 16369 | TRAV29DV5 | TRAJ53 | CAASAGSGGSNYKLTF | TRBV13 | TRBJ1-5 | CASSPRHEDQPQHF | KLGGALQAK | NO | NO | NO |
| 16370 | TRAV20 | TRAJ32 | CAVQANYGGATNKLIF | TRBV7-9 | TRBJ2-2 | CASSLSPGGELFF | KLGGALQAK | YES | NO | NO |
| 16371 | TRAV12-2 | TRAJ20 | CAVIPDYKLSF | TRBV29-1 | TRBJ2-1 | CSVEVAGDNEQFF | KLGGALQAK | YES | NO | NO |
| 16372 | TRAV1-2 | TRAJ7 | CAGDYGNNRLAF | TRBV5-1 | TRBJ1-1 | CASRQTGIGYTEAFF | KLGGALQAK | YES | NO | NO |
| 16373 | TRAV35 | TRAJ45 | CAGTGGGADGLTF | TRBV4-1 | TRBJ2-7 | CASSPRAGGFYEQYF | KLGGALQAK | YES | NO | NO |
| 16374 | TRAV13-2 | TRAJ40 | CAENKGGSGTYKYIF | TRBV3-1 | TRBJ2-7 | CASSQDSGTGGYYEQYF | KLGGALQAK | YES | NO | NO |
| 16375 | TRAV21 | TRAJ21 | CAVRPGYNFNKFYF | TRBV5-1 | TRBJ2-3 | CASNEGTGGTDTQYF | KLGGALQAK | YES | NO | NO |
| 16376 | TRAV8-6 | TRAJ32 | CAVIPYGGATNKLIF | TRBV5-6 | TRBJ1-1 | CASSPGTGSTEAFF | RLRAEAQVK | YES | YES | YES |
| 16377 | TRAV17 | TRAJ21 | CATVGYNFNKFYF | TRBV4-1 | TRBJ1-5 | CASSPVTGFNQPQHF | KLGGALQAK | YES | NO | NO |
| 16378 | TRAV17 | TRAJ56 | CATDGWAGANSKLTF | TRBV27 | TRBJ2-3 | CASSSTGGSLITDTQYF | KLGGALQAK | YES | NO | NO |
| 16379 | TRAV1-2 | TRAJ45 | CAVRKGGGADGLTF | TRBV10-3 | TRBJ2-2 | CAINLGGTNTGELFF | KLGGALQAK | YES | NO | NO |
| 16380 | TRAV29DV5 | TRAJ39 | CAAYNNAGNMLTF | TRBV7-9 | TRBJ1-6 | CASSSNTSPLHF | KLGGALQAK | YES | NO | NO |
| 16381 | TRAV12-3 | TRAJ42 | CAMSRGGGSQGNLIF | TRBV5-1 | TRBJ2-7 | CASSGGDSSSYEQYV | KLGGALQAK | NO | NO | NO |
| 16382 | TRAV39 | TRAJ33 | CAGEMDSNYQLIW | TRBV12-4 | TRBJ2-5 | CASSLVSGGEETQYF | KLGGALQAK | YES | NO | NO |
| 16383 | TRAV35 | TRAJ45 | CAGRAGGGADGLTF | TRBV7-8 | TRBJ1-1 | CASSLPGGEDTEAFF | KLGGALQAK | YES | NO | NO |
| 16384 | TRAV8-4 | TRAJ39 | CAVSDSVNAGNMLTF | TRBV7-9 | TRBJ2-7 | CASKDGISYEQYF | KLGGALQAK | YES | NO | NO |
| 16385 | TRAV26-2 | TRAJ52 | CILDGGTSYGKLTF | TRBV5-1 | TRBJ2-5 | CASSWEGEEETQYF | KLGGALQAK | YES | NO | NO |
| 16386 | TRAV10 | TRAJ31 | CVVSARNNARLMF | TRBV6-6 | TRBJ2-5 | CASSYLLTGSLEETQYF | KLGGALQAK | NO | NO | NO |
| 16387 | TRAV4 | TRAJ20 | CLVGAVNDYKLSF | TRBV3-1 | TRBJ2-5 | CASSQDPGQGFNETQYF | KLGGALQAK | YES | NO | NO |
| 16388 | TRAV12-3 | TRAJ20 | CAMSVADYKLSF | TRBV19 | TRBJ2-1 | CASSDLIQGVFFEQFF | KLGGALQAK | YES | NO | NO |
| 16389 | TRAV19 | TRAJ43 | CALSEENNNDMRF | TRBV10-3 | TRBJ2-3 | CAISDSGGRTDTQYF | KLGGALQAK | YES | NO | NO |
| 16390 | TRAV38-1 | TRAJ50 | CAFMKIDKVIF | TRBV6-5 | TRBJ1-1 | CASSYETGITEAFF | KLGGALQAK | YES | NO | NO |
| 16391 | TRAV29DV5 | TRAJ45 | CAASAGWGGGADGLTF | TRBV15 | TRBJ1-2 | CATSRVGQTPYGYTF | KLGGALQAK | YES | NO | NO |
| 16392 | TRAV38-2DV8 | TRAJ57 | CAANQGGSEKLVF | TRBV18 | TRBJ2-3 | CASSPRQGLTDTQYF | KLGGALQAK | YES | NO | NO |
| 16393 | TRAV1-2 | TRAJ24 | CAVMTTDSWGKFQF | TRBV2 | TRBJ2-2 | CATISGVSGELFF | KLGGALQAK | YES | NO | NO |
| 16394 | TRAV29DV5 | TRAJ8 | CAAVGGTGFQKLVF | TRBV6-5 | TRBJ2-1 | CASSPSGSYNEQFF | KLGGALQAK | NO | NO | NO |
| 16395 | TRAV41 | TRAJ57 | CAVRLITQGGSEKLVF | TRBV11-2 | TRBJ2-7 | CASSGYHEQYV | KLGGALQAK | NO | NO | NO |
| 16396 | TRAV12-2 | TRAJ8 | CAVNTGFQKLVF | TRBV12-4 | TRBJ2-2 | CASSLGQGTSGGGINTGELFF | KLGGALQAK | NO | NO | NO |
| 16397 | TRAV17 | TRAJ23 | CATDLLYNQGGKLIF | TRBV13 | TRBJ2-1 | CASSSNRRGGEQFF | KLGGALQAK | YES | NO | NO |
| 16398 | TRAV38-1 | TRAJ58 | CYEETSGSRLTF | TRBV7-9 | TRBJ2-1 | CASSLFIGGYNEQFF | KLGGALQAK | YES | NO | NO |
| 16400 | TRAV24 | TRAJ44 | CAYHTGTASKLTF | TRBV4-1 | TRBJ1-4 | CASSEGQSTNEKLFF | KLGGALQAK | YES | NO | NO |
| 16401 | TRAV38-1 | TRAJ48 | CAFMRGFGNEKLTF | TRBV7-8 | TRBJ1-5 | CASSFAPRRLPQPQHF | KLGGALQAK | YES | NO | NO |
| 16402 | TRAV25 | TRAJ37 | CAGNGNTGKLIF | TRBV7-4 | TRBJ2-2 | CASSYTPSGGENTGELFF | KLGGALQAK | YES | NO | NO |
| 16403 | TRAV12-2 | TRAJ39 | CAVNGAGNMLTF | TRBV19 | TRBJ2-7 | CASSISAGVGYEQYV | KLGGALQAK | NO | NO | NO |
| 16404 | TRAV39 | TRAJ27 | CAVVSAGKSTF | TRBV12-3 | TRBJ1-2 | CASRAGTGYGYTF | KLGGALQAK | YES | NO | NO |
| 16405 | TRAV12-3 | TRAJ7 | CATNYGNNRLAF | TRBV7-2 | TRBJ2-3 | CASSLAAGATDTQYF | KLGGALQAK | YES | NO | NO |
| 16406 | TRAV17 | TRAJ54 | CAQQGVQGAQKLVF | TRBV11-3 | TRBJ2-1 | CASSLLGGGNEQFF | KLGGALQAK | YES | NO | NO |
| 16407 | TRAV8-1 | TRAJ12 | CAVNALRDSSYKLIF | TRBV6-4 | TRBJ2-1 | CASSERIAGVDEQFF | KLGGALQAK | YES | NO | NO |
| 16408 | TRAV38-2DV8 | TRAJ54 | CAYRSLIQGAQKLVF | TRBV5-1 | TRBJ1-1 | CASSFMLNDRGAEAFF | KLGGALQAK | YES | NO | NO |
| 16409 | TRAV21 | TRAJ18 | CAVSPQSLYRGSTLGRLYF | TRBV19 | TRBJ1-3 | CASSIGVSGNTIYF | KLGGALQAK | YES | NO | NO |
| 16410 | TRAV12-2 | TRAJ41 | CAVQGLSNSGYALNF | TRBV20-1 | TRBJ2-3 | CSAPKDGDTQYF | KLGGALQAK | NO | NO | NO |
| 16411 | TRAV1-1 | TRAJ42 | CAAGWVSQGNLIF | TRBV27 | TRBJ2-1 | CASSPSYNEQFF | RLRAEAQVK | YES | NO | YES |
| 16412 | TRAV4 | TRAJ28 | CLGSGAGSYQLTF | TRBV3-1 | TRBJ2-1 | CASSQASGVINEQFF | KLGGALQAK | YES | NO | NO |
| 16413 | TRAV27 | TRAJ48 | CAGAQDFGNEKLTF | TRBV7-9 | TRBJ2-5 | CASSPQTGPNQETQYF | KLGGALQAK | YES | NO | NO |
| 16414 | TRAV8-6 | TRAJ32 | CAVRIMNYGGATNKLIF | TRBV20-1 | TRBJ1-2 | CSARDFQGYTF | KLGGALQAK | YES | NO | NO |
| 16415 | TRAV12-2 | TRAJ23 | CAVNMLNQGGKLIF | TRBV2 | TRBJ1-1 | CASNRPGHELVLNTEAFF | KLGGALQAK | YES | NO | NO |
| 16416 | TRAV20 | TRAJ58 | CAVQATETSGSRLTF | TRBV9 | TRBJ2-1 | CASSGYNEQFF | KLGGALQAK | YES | NO | NO |
| 16417 | TRAV17 | TRAJ9 | CATNTGGFKTIF | TRBV15 | TRBJ1-2 | CATSVGQGGWDLGYTF | KLGGALQAK | YES | NO | NO |
| 16418 | TRAV19 | TRAJ34 | CALSEATYNTDKLIF | TRBV12-4 | TRBJ2-3 | CASSLGGERGTDTQYF | KLGGALQAK | YES | NO | NO |
| 16419 | TRAV8-1 | TRAJ39 | CAANAGNMLTF | TRBV30 | TRBJ2-1 | CAWKPLLANEQFF | KLGGALQAK | YES | NO | NO |
| 16420 | TRAV19 | TRAJ48 | CALSEENFGNEKLTF | TRBV7-6 | TRBJ2-2 | CASSLASSAGELFF | KLGGALQAK | YES | NO | NO |
| 16421 | TRAV19 | TRAJ24 | CALSESTDSWGKFQF | TRBV12-5 | TRBJ1-1 | CASGLSDRENTEAFF | KLGGALQAK | YES | NO | NO |
| 16422 | TRAV17 | TRAJ22 | CATDIGGSARQLTF | TRBV6-5 | TRBJ2-3 | CASSAIGGTDTQYF | KLGGALQAK | NO | NO | NO |
| 16423 | TRAV29DV5 | TRAJ33 | CAARDDSNYQLIW | TRBV12-4 | TRBJ2-5 | CASSLVTDSQETQYF | KLGGALQAK | YES | NO | NO |
| 16424 | TRAV12-1 | TRAJ15 | CVVNQAGTALIF | TRBV5-1 | TRBJ1-1 | CASSKGTEAFF | KLGGALQAK | YES | NO | NO |
| 16425 | TRAV29DV5 | TRAJ28 | CAARAAGAGSYQLTF | TRBV7-9 | TRBJ2-5 | CASSSPGASAVQETQYF | KLGGALQAK | YES | NO | NO |
| 16426 | TRAV27 | TRAJ36 | CAVLTGANNLFF | TRBV30 | TRBJ2-1 | CAGTLAGAYNEQFF | KLGGALQAK | YES | NO | NO |
| 16427 | TRAV26-2 | TRAJ54 | CILRRQKLVF | TRBV5-1 | TRBJ2-7 | CASSLDRTGPYEQYF | KLGGALQAK | YES | NO | NO |
| 16428 | TRAV1-2 | TRAJ16 | CAVRDRSGQKLLF | TRBV12-4 | TRBJ2-1 | CASSLGGEQFF | KLGGALQAK | NO | NO | NO |
| 16429 | TRAV1-2 | TRAJ6 | CAVMGGGSYIPTF | TRBV3-1 | TRBJ2-1 | CASSQPRLAGDGNEQFF | KLGGALQAK | YES | NO | NO |
| 16430 | TRAV16 | TRAJ53 | CALSGGSNYKLTF | TRBV20-1 | TRBJ2-1 | CSANSGTSPNEQFF | KLGGALQAK | NO | NO | NO |
| 16431 | TRAV2 | TRAJ39 | CAGTDNNAGNMLTF | TRBV28 | TRBJ2-1 | CASTVAAYNEQFF | KLGGALQAK | NO | NO | NO |
| 16432 | TRAV12-2 | TRAJ30 | CAVYNRDDKIIF | TRBV20-1 | TRBJ1-2 | CSARGGENYGYTF | KLGGALQAK | YES | NO | NO |
| 16433 | TRAV13-2 | TRAJ39 | CAENTLTF | TRBV4-1 | TRBJ2-7 | CASSKTGSGSYEQYF | KLGGALQAK | YES | NO | NO |
| 16434 | TRAV1-2 | TRAJ6 | CAVRDWGGSYIPTF | TRBV6-1 | TRBJ1-6 | CASSEVGTGGLIGSPLHF | KLGGALQAK | YES | NO | NO |
| 16435 | TRAV39 | TRAJ47 | CAVEYGNKLVF | TRBV12-4 | TRBJ1-5 | CASSSLTGPHQPQHF | KLGGALQAK | YES | NO | NO |
| 16436 | TRAV12-2 | TRAJ50 | CAVDDVKTSYDKVIF | TRBV20-1 | TRBJ2-5 | CSAPGGLETQYF | KLGGALQAK | YES | NO | NO |
| 16437 | TRAV24 | TRAJ45 | CAFPGGGADGLTF | TRBV3-1 | TRBJ2-7 | CASTSGLAAIYEQYF | KLGGALQAK | YES | NO | NO |
| 16438 | TRAV17 | TRAJ17 | CATDAWAAGNKLTF | TRBV7-3 | TRBJ2-5 | CASGNGGEETQYF | KLGGALQAK | NO | NO | NO |
| 16439 | TRAV34 | TRAJ49 | CGAESNTGNQFYF | TRBV6-5 | TRBJ2-1 | CAARSGYNEQFF | KLGGALQAK | NO | NO | NO |
| 16440 | TRAV19 | TRAJ53 | CALSEAGSGGSNYKLTF | TRBV10-2 | TRBJ1-5 | CASSGKYSNQPQHF | KLGGALQAK | YES | NO | NO |

| | | | | | | | | | | |
|---|---|---|---|---|---|---|---|---|---|---|
| 16441 | TRAV20 | TRAJ26 | CAVQAWGQNFVF | TRBV19 | TRBJ2-5 | CASSNPGEGETQYF | KLGGALQAK | YES | NO | NO |
| 16442 | TRAV4 | TRAJ8 | CLVGSSMNTGFQKLVF | TRBV11-3 | TRBJ2-1 | CASSRSGFGGRGYNEQFF | KLGGALQAK | YES | NO | NO |
| 16443 | TRAV12-1 | TRAJ23 | CVVGLGGKLIF | TRBV4-1 | TRBJ1-5 | CASSQAWSDSVMGQPQHF | KLGGALQAK | YES | NO | NO |
| 16444 | TRAV12-1 | TRAJ29 | CVVDSGNTPLVF | TRBV2 | TRBJ1-1 | CARGGFNRNTEAFF | AVFDRKSDAK | YES | NO | NO |
| 16445 | TRAV16 | TRAJ45 | CALSTYSGGGADGLTF | TRBV7-2 | TRBJ1-1 | CASSLPSGPGSEAFF | KLGGALQAK | NO | NO | NO |
| 16446 | TRAV17 | TRAJ49 | CATDAPNTGNQFYF | TRBV2 | TRBJ1-1 | CASSESVGMNTEAFF | KLGGALQAK | YES | NO | NO |
| 16447 | TRAV30 | TRAJ29 | CGTEFSGNTPLVF | TRBV27 | TRBJ2-1 | CASSVGTSGRGNEQFF | KLGGALQAK | YES | NO | NO |
| 16448 | TRAV14DV4 | TRAJ12 | CAMRDAWMDSSYKLIF | TRBV6-5 | TRBJ2-5 | CASHDPRAQETQYF | KLGGALQAK | NO | NO | NO |
| 16449 | TRAV22 | TRAJ44 | CAVVTGTASKLTF | TRBV7-2 | TRBJ1-1 | CASSLVPPGQGAFFTEAFF | KLGGALQAK | YES | NO | NO |
| 16450 | TRAV35 | TRAJ45 | CAGSAFSGGGADGLTF | TRBV12-4 | TRBJ2-7 | CASSPIRAMGSYEQYF | KLGGALQAK | YES | NO | NO |
| 16451 | TRAV8-2 | TRAJ20 | CVVIESNDYKLSF | TRBV19 | TRBJ1-1 | CASSIRGDLNTEAFF | AVFDRKSDAK | YES | NO | YES |
| 16452 | TRAV1-2 | TRAJ50 | CAVRSTSYDKVIF | TRBV6-3 | TRBJ2-7 | CASSYVQSSYEQYV | KLGGALQAK | NO | NO | NO |
| 16453 | TRAV21 | TRAJ52 | CAVILGGAGGTSYGKLTF | TRBV12-4 | TRBJ2-5 | CASSLSQWETQYF | KLGGALQAK | YES | NO | NO |
| 16454 | TRAV22 | TRAJ33 | CAVYGSNYQLIW | TRBV28 | TRBJ2-2 | CASSFGVLELFF | KLGGALQAK | YES | NO | NO |
| 16455 | TRAV19 | TRAJ34 | CALSEHDNTDKLIF | TRBV12-4 | TRBJ2-7 | CASSDSKGIYEQYV | KLGGALQAK | NO | NO | NO |
| 16456 | TRAV14DV4 | TRAJ7 | CAMRHHYGNNRLAF | TRBV11-2 | TRBJ1-1 | CASSLTENTEAFF | KLGGALQAK | YES | NO | NO |
| 16457 | TRAV12-1 | TRAJ34 | CVVNQDNTDKLIF | TRBV2 | TRBJ1-6 | CASSPEGQTYNSPLHF | KLGGALQAK | YES | NO | NO |
| 16458 | TRAV13-2 | TRAJ49 | CAENITPTGNQFYF | TRBV4-3 | TRBJ1-6 | CASSQDSMYNSPLHF | KLGGALQAK | YES | NO | NO |
| 16459 | TRAV14DV4 | TRAJ54 | CAMREGPEREKLVF | TRBV4-1 | TRBJ2-3 | CASREVGGLDTDTQYF | KLGGALQAK | YES | NO | NO |
| 16460 | TRAV8-1 | TRAJ15 | CAVNDQAGTALIF | TRBV20-1 | TRBJ2-7 | CSAGGDRSYEQYF | AVFDRKSDAK | NO | NO | NO |
| 16461 | TRAV38-1 | TRAJ52 | CAFMSLAGGTSYGKLTF | TRBV7-2 | TRBJ2-7 | CASIPGLAGSYEQYV | KLGGALQAK | NO | NO | NO |
| 16462 | TRAV21 | TRAJ58 | CAVIQETSGSRLTF | TRBV13 | TRBJ2-5 | CASSLRTSGLQETQYF | KLGGALQAK | YES | NO | NO |
| 16463 | TRAV12-2 | TRAJ26 | CAVDADNYGQNFVF | TRBV3-1 | TRBJ1-1 | CASSQRGNTEAFF | KLGGALQAK | YES | NO | NO |
| 16464 | TRAV17 | TRAJ57 | CATVSLTQGGSEKLVF | TRBV30 | TRBJ2-7 | CAWSPTGSEQYV | AVFDRKSDAK | NO | NO | NO |
| 16465 | TRAV8-1 | TRAJ22 | CAVNGPVSSGSARQLTF | TRBV9 | TRBJ2-1 | CASSPTDHNEQFF | KLGGALQAK | YES | NO | NO |
| 16466 | TRAV12-2 | TRAJ39 | CAIGPGNMLTF | TRBV4-1 | TRBJ1-1 | CASRVGNTEAFF | IVTDFSVIK | YES | YES | YES |
| 16467 | TRAV25 | TRAJ28 | CAGLPYSGAGSYQLTF | TRBV14 | TRBJ1-2 | CASSPIYNYGYTF | KLGGALQAK | NO | NO | NO |
| 16468 | TRAV14DV4 | TRAJ36 | CAMREGHGANNLFF | TRBV10-3 | TRBJ2-7 | CAISHTDRGPTYEQYF | KLGGALQAK | YES | NO | NO |
| 16469 | TRAV19 | TRAJ11 | CALSVIMNSGYSTLTF | TRBV12-4 | TRBJ2-2 | CASSQWTSGGHTGELFF | KLGGALQAK | YES | NO | NO |
| 16470 | TRAV26-1 | TRAJ43 | CIVRVANDMRF | TRBV6-5 | TRBJ1-1 | CSAGQGARRAFF | KLGGALQAK | YES | NO | NO |
| 16471 | TRAV12-1 | TRAJ37 | CVVNTGSSNTGKLIF | TRBV4-1 | TRBJ2-1 | CASSQEAGTGYNEQFF | KLGGALQAK | YES | NO | NO |
| 16472 | TRAV1-2 | TRAJ16 | CAVRLF | TRBV4-1 | TRBJ1-2 | CASSQDSARVDYGYTF | KLGGALQAK | NO | NO | NO |
| 16473 | TRAV25 | TRAJ26 | CAGADNYGQNFVF | TRBV27 | TRBJ1-2 | CASSPYRTYGYTF | KLGGALQAK | YES | NO | NO |
| 16474 | TRAV38-1 | TRAJ26 | CAFMKHLNYGQNFVF | TRBV9 | TRBJ2-7 | CASSVGTSGGYEQYF | KLGGALQAK | YES | NO | NO |
| 16475 | TRAV1-2 | TRAJ39 | CAVREDAGNMLTF | TRBV12-5 | TRBJ2-2 | CASGPGVAHLTGELFF | KLGGALQAK | YES | NO | NO |
| 16476 | TRAV1-2 | TRAJ41 | CAVRDLTGYALNF | TRBV19 | TRBJ2-2 | CASLWGDTGELFF | KLGGALQAK | YES | NO | NO |
| 16477 | TRAV29DV5 | TRAJ43 | CAASAGMNDMRF | TRBV12-4 | TRBJ2-2 | CASSLVHLWTGELFF | KLGGALQAK | YES | NO | NO |
| 16478 | TRAV19 | TRAJ30 | CALSGFRDDKIIF | TRBV19 | TRBJ2-5 | CASSISVGETQYF | KLGGALQAK | YES | NO | NO |
| 16479 | TRAV14DV4 | TRAJ56 | CAMREVPNTGANSKLTF | TRBV12-3 | TRBJ1-1 | CASGPGGSTEAFF | KLGGALQAK | YES | NO | NO |
| 16480 | TRAV16 | TRAJ49 | CAPRKLTGNQFYF | TRBV19 | TRBJ2-7 | CASSILLAGRTYEQYF | KLGGALQAK | NO | NO | NO |
| 16481 | TRAV1-2 | TRAJ10 | CACLTGGGNKLTF | TRBV3-1 | TRBJ2-5 | CASSQRTRGGETQYF | KLGGALQAK | YES | NO | NO |
| 16482 | TRAV17 | TRAJ20 | CATDPNDYKLSF | TRBV5-4 | TRBJ2-1 | CASTTSGRANEQFF | KLGGALQAK | YES | NO | NO |
| 16483 | TRAV29DV5 | TRAJ22 | CAAKSSGSARQLTF | TRBV4-1 | TRBJ1-4 | CASSQGSNEKLFF | KLGGALQAK | YES | NO | NO |
| 16484 | TRAV30 | TRAJ45 | CGTERGGGADGLTF | TRBV4-1 | TRBJ1-1 | CASSQWTGGEAEAFF | KLGGALQAK | YES | NO | NO |
| 16485 | TRAV9-2 | TRAJ48 | CALSDPIFGNEKLTF | TRBV2 | TRBJ1-1 | CASRAFF | KLGGALQAK | NO | NO | NO |
| 16486 | TRAV26-2 | TRAJ26 | CILRDGINYGQNFVF | TRBV27 | TRBJ2-5 | CASSLDSTPRETQYF | KLGGALQAK | YES | NO | NO |
| 16487 | TRAV5 | TRAJ3 | CAESGYSSASKIIF | TRBV3-1 | TRBJ1-2 | CASSLTGMSEREDYTF | KLGGALQAK | YES | NO | NO |
| 16488 | TRAV13-2 | TRAJ29 | CAENKNSGNTPLVF | TRBV27 | TRBJ2-1 | CASSILDREYEQFF | KLGGALQAK | YES | NO | NO |
| 16489 | TRAV20 | TRAJ5 | CAVQRDTGRRALTF | TRBV3-1 | TRBJ2-7 | CASSQDGGSYEQYF | AVFDRKSDAK | YES | NO | YES |
| 16490 | TRAV30 | TRAJ53 | CGTANSGGSNYKLTF | TRBV19 | TRBJ2-2 | CASSIWTGNTGELFF | KLGGALQAK | YES | NO | NO |
| 16491 | TRAV40 | TRAJ52 | CLLIFGRRGTSYGKLTF | TRBV4-1 | TRBJ1-2 | CASSQKQTPSNYGYTF | KLGGALQAK | NO | NO | NO |
| 16493 | TRAV8-3 | TRAJ16 | CAVGEVGQKLLF | TRBV11-2 | TRBJ2-5 | CASSFWQGQETQYF | KLGGALQAK | NO | NO | NO |
| 16494 | TRAV12-3 | TRAJ40 | CAMRAPSGTYKYIF | TRBV6-3 | TRBJ2-7 | CASTEEGRDGSYEQYV | RLRAEAQVK | NO | NO | NO |
| 16495 | TRAV16 | TRAJ36 | CALRLNGANNLFF | TRBV2 | TRBJ2-5 | CASKEVAGAQETQYF | RTLNAWVKV | NO | NO | NO |
| 16496 | TRAV20 | TRAJ52 | CAVQARGAGGTSYGKLTF | TRBV4-1 | TRBJ2-1 | CASSQDPPGTSGGAEQFF | KLGGALQAK | YES | NO | NO |
| 16497 | TRAV1-1 | TRAJ15 | CAVRDPQGQAGTALIF | TRBV10-2 | TRBJ2-7 | CATLAVDSYEQYF | KLGGALQAK | YES | NO | NO |
| 16498 | TRAV17 | TRAJ15 | CATGEAGTALIF | TRBV4-1 | TRBJ1-2 | CASSQKQTPSNYGYTF | AVFDRKSDAK | YES | NO | YES |
| 16499 | TRAV4 | TRAJ37 | CLVGDGGSSNTGKLIF | TRBV24-1 | TRBJ2-1 | CATSERYGRVYNEQFF | KLGGALQAK | YES | NO | NO |
| 16500 | TRAV13-1 | TRAJ10 | CAVLEGGGNKLTF | TRBV6-3 | TRBJ1-2 | CASSYSYGYTF | KLGGALQAK | YES | NO | NO |
| 16501 | TRAV1-2 | TRAJ33 | CAVGDSNYQLIW | TRBV30 | TRBJ1-4 | CAWSVGQVNEKLFF | KLGGALQAK | YES | NO | NO |
| 16502 | TRAV21 | TRAJ17 | CAVKAAGNKLTF | TRBV27 | TRBJ1-1 | CASSPIDRNEAFF | KLGGALQAK | YES | NO | NO |
| 16503 | TRAV12-2 | TRAJ30 | CAVNEANRDDKIIF | TRBV27 | TRBJ2-7 | CASSFARGYEQYF | KLGGALQAK | YES | NO | NO |
| 16504 | TRAV19 | TRAJ29 | CALSESDSGNTPLVF | TRBV7-9 | TRBJ1-2 | CASSSNGDHQMGIYGYTF | KLGGALQAK | YES | NO | NO |
| 16505 | TRAV1-2 | TRAJ39 | CAVRNNAGNMLTF | TRBV20-1 | TRBJ1-2 | CSARVPGSVYGYTF | KLGGALQAK | YES | NO | NO |
| 16506 | TRAV19 | TRAJ40 | CALTTSGTYKYIF | TRBV7-2 | TRBJ2-1 | CASSLGFGTRVEQFF | KLGGALQAK | NO | NO | NO |
| 16507 | TRAV3 | TRAJ30 | CAVSVTMNRDDKIIF | TRBV10-2 | TRBJ2-3 | CASSTGLVPSTDTQYF | KLGGALQAK | NO | NO | NO |
| 16508 | TRAV26-1 | TRAJ50 | CIVRVVETSYDKVIF | TRBV5-6 | TRBJ2-1 | CASSQLLAGYNEQFF | KLGGALQAK | YES | NO | NO |
| 16509 | TRAV19 | TRAJ27 | CALSEARTNAGKSTF | TRBV7-8 | TRBJ1-3 | CASSPSLNSGNTIYF | KLGGALQAK | YES | NO | NO |
| 16510 | TRAV3 | TRAJ37 | CAVRDGGMGSNTGKLIF | TRBV12-5 | TRBJ1-1 | CASGLNTRYVGGTEAFF | KLGGALQAK | NO | NO | NO |
| 16511 | TRAV17 | TRAJ22 | CATDESGSARQLTF | TRBV5-1 | TRBJ2-7 | CASSWTTGTAYEQYF | KLGGALQAK | NO | NO | NO |
| 16512 | TRAV13-1 | TRAJ49 | CAASIGNTGNQFYF | TRBV4-1 | TRBJ1-3 | CASSQVSGGFGNTIYF | KLGGALQAK | YES | NO | NO |
| 16513 | TRAV29DV5 | TRAJ34 | CAACYNTDKLIF | TRBV12-4 | TRBJ2-1 | CASSLNPGGQNNEQFF | KLGGALQAK | YES | NO | NO |
| 16514 | TRAV1-2 | TRAJ12 | CAVRGEDSSYKLIF | TRBV30 | TRBJ1-2 | CAWSVPSYYGYTF | KLGGALQAK | NO | NO | NO |
| 16515 | TRAV14DV4 | TRAJ42 | CAMREGRGSQGNLIF | TRBV7-3 | TRBJ1-4 | CASSSLDPTPVSEKLFF | KLGGALQAK | NO | NO | NO |
| 16516 | TRAV19 | TRAJ44 | CALSDLNTGTASKLTF | TRBV9 | TRBJ2-1 | CASSVVSSAYNEQFF | KLGGALQAK | YES | NO | NO |
| 16517 | TRAV19 | TRAJ30 | CALPNRDDKIIF | TRBV13 | TRBJ1-1 | CASSLRGTEAFF | KLGGALQAK | YES | NO | NO |
| 16518 | TRAV27 | TRAJ42 | CAGDGGSQGNLIF | TRBV6-1 | TRBJ1-3 | CASSKDRATGNTIYF | KLGGALQAK | YES | NO | NO |
| 16519 | TRAV1-1 | TRAJ38 | CAVISAGNNRKLIW | TRBV12-4 | TRBJ2-2 | CASSLRPPDTGELFF | KLGGALQAK | YES | NO | NO |
| 16520 | TRAV27 | TRAJ28 | CAARYSGAGSYQLTF | TRBV7-9 | TRBJ2-1 | CASSLSGSSYNEQFF | KLGGALQAK | YES | NO | NO |
| 16521 | TRAV12-2 | TRAJ4 | CAVGSGGYNKLIF | TRBV20-1 | TRBJ1-1 | CSARPRLSTEAFF | KLGGALQAK | YES | NO | NO |
| 16522 | TRAV26-2 | TRAJ40 | CILKLAGTYKYIF | TRBV29-1 | TRBJ1-1 | CSTPGTNTEAFF | KLGGALQAK | NO | NO | NO |
| 16523 | TRAV29DV5 | TRAJ44 | CAARITGTASKLTF | TRBV7-9 | TRBJ1-1 | CASSLMGDEAFF | KLGGALQAK | YES | NO | NO |
| 16524 | TRAV8-1 | TRAJ28 | CAVKWPSGAGSYQLTF | TRBV12-3 | TRBJ2-1 | CASRDYNEQFF | KLGGALQAK | YES | NO | NO |
| 16525 | TRAV34 | TRAJ49 | CGAVRTPQFYF | TRBV7-8 | TRBJ2-1 | CASRPLLAGEYYNEQFF | KLGGALQAK | YES | NO | NO |
| 16526 | TRAV10 | TRAJ42 | CVVSPNYGGSQGNLIF | TRBV2 | TRBJ2-5 | CASRPHQETQYF | KLGGALQAK | YES | NO | NO |
| 16527 | TRAV17 | TRAJ4 | CATGPGGYNKLIF | TRBV19 | TRBJ2-1 | CASSIPLAGGLAYNEQFF | KLGGALQAK | YES | NO | NO |
| 16528 | TRAV17 | TRAJ42 | CATPYGGSQGNLIF | TRBV7-9 | TRBJ2-5 | CASSHIPGRETQYF | KLGGALQAK | YES | NO | NO |
| 16529 | TRAV22 | TRAJ26 | CAARGYGQNFVF | TRBV7-6 | TRBJ1-1 | CASSLGQAAEAFF | KLGGALQAK | YES | NO | NO |
| 16530 | TRAV29DV5 | TRAJ32 | CAASDDGGATNKLIF | TRBV7-2 | TRBJ2-1 | CASSFSGYNEQFF | KLGGALQAK | YES | NO | NO |
| 16531 | TRAV27 | TRAJ35 | CAGAGAGGFGNVLHC | TRBV6-5 | TRBJ1-1 | CASSTWAGDTEAFF | KLGGALQAK | NO | NO | NO |
| 16532 | TRAV3 | TRAJ33 | CAVRGDSNYQLIW | TRBV7-9 | TRBJ1-5 | CASSLNYNQPQHF | AVFDRKSDAK | YES | NO | YES |
| 16533 | TRAV12-3 | TRAJ52 | CAMISANAGGTSYGKLTF | TRBV4-1 | TRBJ1-2 | CASSQGLAYGYTF | KLGGALQAK | YES | NO | NO |
| 16534 | TRAV19 | TRAJ47 | CALSGEYGNKLVF | TRBV12-3 | TRBJ2-1 | CASSPIAGGPYNEQFF | KLGGALQAK | YES | NO | NO |
| 16535 | TRAV20 | TRAJ40 | CAVPAAGSGTYKYIF | TRBV4-1 | TRBJ1-1 | CASSQGPGLGDTEAFF | KLGGALQAK | YES | NO | NO |
| 16536 | TRAV16 | TRAJ50 | CALSGRKTSYDKVIF | TRBV6-6 | TRBJ2-3 | CASSYAGGTDTQYF | KLGGALQAK | NO | NO | NO |
| 16537 | TRAV12-3 | TRAJ48 | CAMSPSNFGNEKLTF | TRBV2 | TRBJ1-3 | CAIPRRAVSGNTIYF | ELAGIGILTV | YES | NO | NO |
| 16538 | TRAV8-4 | TRAJ42 | CAVSELFGGSQGNLIF | TRBV5-6 | TRBJ2-7 | CASSLAYTSGSYYEQYV | KLGGALQAK | NO | NO | NO |
| 16539 | TRAV6 | TRAJ41 | CALVTNSNSGYALNF | TRBV4-1 | TRBJ1-4 | CASSQTVGLGEKLFF | KLGGALQAK | YES | NO | NO |
| 16540 | TRAV13-1 | TRAJ3 | CAASWSSASKIIF | TRBV12-5 | TRBJ1-4 | CASGYGTGSNEKLFF | AVFDRKSDAK | YES | NO | NO |
| 16541 | TRAV25 | TRAJ21 | CAGQVIDSFNKFYF | TRBV24-1 | TRBJ1-1 | CATSGVAEAFF | KLGGALQAK | YES | NO | NO |
| 16542 | TRAV3 | TRAJ7 | CAVRDVDGNNRLAF | TRBV7-8 | TRBJ2-1 | CASSFARVPGDEQFF | KLGGALQAK | NO | NO | NO |
| 16543 | TRAV12-2 | TRAJ21 | CAVNDGFNKFYF | TRBV7-8 | TRBJ1-1 | CASSLGRVSTEAFF | KLGGALQAK | YES | NO | NO |
| 16544 | TRAV24 | TRAJ8 | CAFSTGFQKLVF | TRBV12-4 | TRBJ2-7 | CASSSGTGAYEQYV | KLGGALQAK | NO | NO | NO |
| 16545 | TRAV17 | TRAJ52 | CATEGNAGGTSYGKLTF | TRBV7-6 | TRBJ2-7 | CASSWTGGDTYEQYF | KLGGALQAK | YES | NO | NO |

| | | | | | | | | | | |
|---|---|---|---|---|---|---|---|---|---|---|
| 16546 | TRAV3 | TRAJ12 | CAVRDTSSYKLIF | TRBV30 | TRBJ2-7 | CAWMKGLGYEQYV | KLGGALQAK | NO | NO | NO |
| 16547 | TRAV14DV4 | TRAJ43 | CAMREGFNNNDMRF | TRBV19 | TRBJ2-5 | CASSPGTGGTETQYF | KLGGALQAK | YES | NO | NO |
| 16548 | TRAV12-1 | TRAJ13 | CVCPSGGYQKVTF | TRBV19 | TRBJ1-1 | CASSRGGGRVNTEAFF | KLGGALQAK | YES | NO | NO |
| 16549 | TRAV27 | TRAJ26 | CAGRHDNYGQNFVF | TRBV9 | TRBJ2-7 | CASSVGGREQYF | KLGGALQAK | YES | NO | NO |
| 16550 | TRAV12-2 | TRAJ57 | CAVTQGGSEKLVF | TRBV4-1 | TRBJ2-7 | CASSQKIGGYEQYV | KLGGALQAK | NO | NO | NO |
| 16551 | TRAV38-1 | TRAJ28 | CAFMKPTSWAGSYQLTF | TRBV19 | TRBJ1-2 | CASSMGDGYTF | KLGGALQAK | YES | NO | NO |
| 16552 | TRAV14DV4 | TRAJ48 | CAMRDRNFGNEKLTF | TRBV19 | TRBJ2-2 | CASSITGTGALTNTGELFF | KLGGALQAK | YES | NO | NO |
| 16553 | TRAV12-3 | TRAJ52 | CALNKGGKLTF | TRBV12-4 | TRBJ2-6 | CASSLVHVAGANVLTF | KLGGALQAK | YES | NO | NO |
| 16554 | TRAV35 | TRAJ37 | CAGRSNTGKLIF | TRBV11-2 | TRBJ1-2 | CASSLLVADNYGYTF | KLGGALQAK | YES | NO | NO |
| 16555 | TRAV17 | TRAJ39 | CATDARNNAGNMLTF | TRBV29-1 | TRBJ2-1 | CSVDVRVADEQFF | KLGGALQAK | YES | NO | NO |
| 16556 | TRAV3 | TRAJ48 | CAVRVISNFGNEKLTF | TRBV14 | TRBJ2-5 | CASSQGGQGALETQYF | KLGGALQAK | NO | NO | NO |
| 16557 | TRAV9-2 | TRAJ35 | CALSARIRGSI | TRBV6-5 | TRBJ1-2 | CASSGFMGPNYGYTF | KLGGALQAK | NO | NO | NO |
| 16558 | TRAV20 | TRAJ45 | CAVQAVFLYSGGGADGLTF | TRBV7-9 | TRBJ1-1 | CASSLGQGDTEAFF | KLGGALQAK | YES | NO | NO |
| 16559 | TRAV35 | TRAJ33 | CAGPLPRDSNYQLIW | TRBV4-1 | TRBJ1-2 | CASSLGQGTTNGYTF | KLGGALQAK | YES | NO | NO |
| 16560 | TRAV3 | TRAJ47 | CAVREGYGNKLVF | TRBV14 | TRBJ2-3 | CASSQDRGSTDTQYF | AVFDRKSDAK | NO | NO | NO |
| 16561 | TRAV21 | TRAJ24 | CAVTTDSWGKLQF | TRBV5-6 | TRBJ1-5 | CASSFGTGAQPQHF | IVTDFSVIK | YES | NO | YES |
| 16562 | TRAV12-3 | TRAJ4 | CAMRMFSGGYNKLIF | TRBV27 | TRBJ1-5 | CASTDGTGGLNQPQHF | KLGGALQAK | NO | NO | NO |
| 16564 | TRAV8-6 | TRAJ9 | CAVSDRGSSGGFKTIF | TRBV4-1 | TRBJ1-1 | CASSHVRENTEAFF | KLGGALQAK | YES | NO | NO |
| 16566 | TRAV3 | TRAJ33 | CAVRDYSNYQLIW | TRBV6-5 | TRBJ1-5 | CASSYGTGNQPQHF | KLGGALQAK | NO | NO | NO |
| 16567 | TRAV38-1 | TRAJ54 | CVFMSLIQGAQKLVF | TRBV7-9 | TRBJ2-7 | CASKDGISYEQYF | KLGGALQAK | YES | NO | NO |
| 16568 | TRAV5 | TRAJ42 | CAESIDGGSQGNLIF | TRBV7-8 | TRBJ2-1 | CASSFFAGEKNEQFF | KLGGALQAK | YES | NO | NO |
| 16569 | TRAV41 | TRAJ54 | CAVNSGAQKLVF | TRBV3-1 | TRBJ2-5 | CASSQDPEPIRQTQYF | KLGGALQAK | YES | NO | NO |
| 16570 | TRAV20 | TRAJ29 | CAVGVHSGNTPLVF | TRBV10-1 | TRBJ2-1 | CASSATGLGNNEQFF | KLGGALQAK | YES | NO | NO |
| 16571 | TRAV1-2 | TRAJ23 | CAVRDLGNQGGKLIF | TRBV12-4 | TRBJ1-1 | CASSLAGRGGRNTEAFF | KLGGALQAK | YES | NO | NO |
| 16572 | TRAV24 | TRAJ11 | CALNSGYSTLTF | TRBV7-9 | TRBJ2-3 | CASSFTSGTTDTQYF | KLGGALQAK | YES | NO | NO |
| 16573 | TRAV1-2 | TRAJ20 | CAVKNDYKLSF | TRBV9 | TRBJ1-2 | CASSVEGLTGDYGYTF | KLGGALQAK | YES | NO | NO |
| 16574 | TRAV17 | TRAJ30 | CATELGPRKIIF | TRBV13 | TRBJ1-2 | CASSFVRGYTF | KLGGALQAK | YES | NO | NO |
| 16575 | TRAV12-3 | TRAJ43 | CAMSDNDMRF | TRBV27 | TRBJ1-1 | CASSSNTEAFF | KLGGALQAK | YES | NO | NO |
| 16576 | TRAV26-2 | TRAJ54 | CILRDFLQGAQKLVF | TRBV5-1 | TRBJ2-1 | CASSPEAGALFNEQFF | KLGGALQAK | YES | NO | NO |
| 16577 | TRAV30 | TRAJ49 | CGTEPTGNQFYF | TRBV5-6 | TRBJ1-2 | CASSLLGLKARGYTF | KLGGALQAK | YES | NO | NO |
| 16578 | TRAV29DV5 | TRAJ49 | CAATVTGNQFYF | TRBV30 | TRBJ2-2 | CAWKLLVSTGELFF | KLGGALQAK | YES | NO | NO |
| 16579 | TRAV38-2DV8 | TRAJ23 | CAYRSALAFPQGGKLIF | TRBV18 | TRBJ1-6 | CASSRGAGGGSPLHF | KLGGALQAK | YES | NO | NO |
| 16580 | TRAV1-2 | TRAJ33 | CAVIHHSNYQLIW | TRBV19 | TRBJ1-2 | CASSARTGVGYGYTF | KLGGALQAK | YES | NO | NO |
| 16581 | TRAV3 | TRAJ44 | CAVRTGTASKLTF | TRBV27 | TRBJ2-1 | CASSHSDRIINEQFF | AVFDRKSDAK | NO | NO | NO |
| 16582 | TRAV1-2 | TRAJ24 | CAVSALTDSWGKFQF | TRBV9 | TRBJ2-7 | CASSVPDRNQNYEQYF | AVFDRKSDAK | YES | NO | YES |
| 16583 | TRAV14DV4 | TRAJ52 | CAMREYYAGGTSYGKLTF | TRBV15 | TRBJ1-4 | CATSRDRVAEKLFF | ELAGIGILTV | YES | NO | YES |
| 16584 | TRAV20 | TRAJ54 | CAVQARDGAQKLVF | TRBV11-1 | TRBJ1-1 | CASSFLNTEAFF | KLGGALQAK | YES | NO | NO |
| 16585 | TRAV36DV7 | TRAJ11 | CAVWGASGYSTLTF | TRBV9 | TRBJ2-3 | CASSTQGLAESKYF | KLGGALQAK | NO | NO | NO |
| 16586 | TRAV8-3 | TRAJ44 | CAGEPGTASKLTF | TRBV27 | TRBJ1-2 | CASSGPTGTGGYGYTF | KLGGALQAK | YES | NO | NO |
| 16587 | TRAV12-1 | TRAJ47 | CVVRAGYGNKLVF | TRBV24-1 | TRBJ2-2 | CATNVANTGELFF | KLGGALQAK | YES | NO | NO |
| 16588 | TRAV14DV4 | TRAJ49 | CAMREGTWNQFYF | TRBV6-1 | TRBJ1-4 | CASSVGVGGPYEKLFF | KLGGALQAK | YES | NO | NO |
| 16589 | TRAV27 | TRAJ23 | CAGADYNQGGKLIF | TRBV6-3 | TRBJ2-7 | CASSSTYEQYV | KLGGALQAK | NO | NO | NO |
| 16590 | TRAV38-2DV8 | TRAJ49 | CAYRSAFKPGNQFYF | TRBV6-1 | TRBJ2-1 | CASSEEGVAGVFNEQFF | AVFDRKSDAK | YES | NO | NO |
| 16591 | TRAV21 | TRAJ58 | CAVKETSGSRLTF | TRBV19 | TRBJ1-6 | CASSIGNSPLHF | KLGGALQAK | YES | NO | NO |
| 16592 | TRAV14DV4 | TRAJ42 | CAMREGMYGGSQGNLIF | TRBV5-1 | TRBJ2-3 | CASSFLGDKHTDTQYF | KLGGALQAK | YES | NO | NO |
| 16593 | TRAV5 | TRAJ8 | CAEKGFQKLVF | TRBV5-1 | TRBJ2-1 | CASSFTSGDSYNEQFF | KLGGALQAK | YES | NO | NO |
| 16594 | TRAV38-2DV8 | TRAJ48 | CAYNFGNEKLTF | TRBV7-9 | TRBJ2-3 | CASSLCCWSHRGLHTDTQYF | KLGGALQAK | NO | NO | NO |
| 16595 | TRAV41 | TRAJ34 | CAVSSPYNTDKLIF | TRBV4-1 | TRBJ2-2 | CASSYWTSGGPGAAGELFF | KLGGALQAK | YES | NO | NO |
| 16596 | TRAV12-2 | TRAJ30 | CAVGDDKIIF | TRBV12-5 | TRBJ2-7 | CASGLGSGVYEQYF | KLGGALQAK | YES | NO | NO |
| 16597 | TRAV12-2 | TRAJ31 | CAVNSWNNNARLMF | TRBV20-1 | TRBJ1-1 | CSASPRGYTEAFF | KLGGALQAK | YES | NO | NO |
| 16598 | TRAV22 | TRAJ58 | CAGETSGSRLTF | TRBV6-5 | TRBJ2-7 | CASSPAWGLEQYV | KLGGALQAK | NO | NO | NO |
| 16599 | TRAV34 | TRAJ37 | CGADIRSSNTGKLIF | TRBV19 | TRBJ1-1 | CASRDGRIEAFF | KLGGALQAK | NO | NO | NO |
| 16600 | TRAV5 | TRAJ34 | CAEKPDTDKLIF | TRBV7-9 | TRBJ2-3 | CASSLDRDTQYF | KLGGALQAK | YES | NO | NO |
| 16601 | TRAV8-6 | TRAJ20 | CAVSDGGKLSF | TRBV29-1 | TRBJ2-5 | CSEGSPQETQYF | KLGGALQAK | YES | NO | NO |
| 16602 | TRAV13-1 | TRAJ7 | CAVGYGNNRLAF | TRBV12-4 | TRBJ2-5 | CASSLNGLEETQYF | KLGGALQAK | YES | NO | NO |
| 16604 | TRAV12-1 | TRAJ45 | CVVDSGGGADGLTF | TRBV4-2 | TRBJ2-7 | CASSPDLGTYEQYF | KLGGALQAK | YES | NO | NO |
| 16606 | TRAV29DV5 | TRAJ41 | CAADSGYALNF | TRBV19 | TRBJ1-5 | CASNWSPGIDQPQHF | KLGGALQAK | YES | NO | NO |
| 16607 | TRAV20 | TRAJ42 | CAVLGSQGNLIF | TRBV4-3 | TRBJ2-2 | CASSQLNLGLAGVVSGELFF | KLGGALQAK | NO | NO | NO |
| 16608 | TRAV24 | TRAJ49 | CALLNTGNQFYF | TRBV4-3 | TRBJ1-5 | CASSQDRALNQPQHF | KLGGALQAK | YES | NO | NO |
| 16609 | TRAV12-2 | TRAJ45 | CAVSGGGADGLTF | TRBV29-1 | TRBJ2-1 | CSDRPSGITNEQFF | KLGGALQAK | YES | NO | NO |
| 16610 | TRAV19 | TRAJ6 | CALSEVQASGGSYIPTF | TRBV7-9 | TRBJ1-5 | CASSLWAQVAEDSNQPQHF | AVFDRKSDAK | YES | NO | NO |
| 16611 | TRAV8-6 | TRAJ20 | CAVSPLAGDYKLSF | TRBV5-1 | TRBJ2-3 | CASSFRGDTQYF | KLGGALQAK | YES | NO | NO |
| 16612 | TRAV19 | TRAJ33 | CALSDGMDSNYQLIW | TRBV19 | TRBJ1-1 | CASSIDQGGRVYEAFF | KLGGALQAK | YES | NO | NO |
| 16613 | TRAV21 | TRAJ54 | CAVQGAQKLVF | TRBV10-3 | TRBJ2-7 | CAISTHELAGYEQYV | KLGGALQAK | NO | NO | NO |
| 16614 | TRAV13-1 | TRAJ21 | CAASGYNFNKFYF | TRBV24-1 | TRBJ2-2 | CATSDHETWTDANTGELFF | KLGGALQAK | YES | NO | NO |
| 16615 | TRAV13-1 | TRAJ45 | CAASIRRGADGLTF | TRBV12-4 | TRBJ2-2 | CASSLEVLEEREGDGELFF | KLGGALQAK | YES | NO | NO |
| 16616 | TRAV14DV4 | TRAJ54 | CAMREVTQGAQKLVF | TRBV7-9 | TRBJ2-7 | CASALRDGSYEQYF | KLGGALQAK | YES | NO | NO |
| 16617 | TRAV12-1 | TRAJ29 | CVVNMKGNTPLVF | TRBV20-1 | TRBJ1-1 | CSARDLARHTEAFF | KLGGALQAK | YES | NO | NO |
| 16618 | TRAV6 | TRAJ43 | CALDVPRF | TRBV2 | TRBJ1-1 | CASSEAGGLGLTEAFF | KLGGALQAK | NO | NO | NO |
| 16619 | TRAV17 | TRAJ13 | CATGPMNSGGYQKVTF | TRBV19 | TRBJ2-5 | CASSTRHQETQYF | KLGGALQAK | YES | NO | NO |
| 16620 | TRAV2 | TRAJ33 | CAVDKDSNYQLIW | TRBV9 | TRBJ2-2 | CASSELGTGELFF | KLGGALQAK | NO | NO | NO |
| 16621 | TRAV12-2 | TRAJ37 | CAVNPGNTGKLIF | TRBV19 | TRBJ1-2 | CASSVILGGGGYTF | RLRAEAQVK | YES | YES | YES |
| 16622 | TRAV17 | TRAJ28 | CAAPHSGAGSYQLTF | TRBV10-3 | TRBJ1-2 | CAISESNTGTDYGYTF | KLGGALQAK | YES | NO | NO |
| 16623 | TRAV9-2 | TRAJ34 | CALDTDKLIF | TRBV29-1 | TRBJ1-1 | CSAPPQGRTEAFF | KLGGALQAK | YES | NO | NO |
| 16624 | TRAV5 | TRAJ45 | CAESMYSGGGADGLTF | TRBV3-1 | TRBJ2-3 | CASSPIPGGSDTQYF | KLGGALQAK | YES | NO | NO |
| 16625 | TRAV17 | TRAJ7 | CATDDYGNNRLAF | TRBV12-4 | TRBJ2-5 | CASSSSRGQGTQETQYF | AVFDRKSDAK | YES | NO | YES |
| 16626 | TRAV4 | TRAJ6 | CLVALSGGSYIPTF | TRBV29-1 | TRBJ2-3 | CSVPVVDTQYF | KLGGALQAK | YES | NO | NO |
| 16627 | TRAV1-2 | TRAJ20 | CAVSEYDYKLSF | TRBV20-1 | TRBJ1-2 | CSAPGQSRGYTF | KLGGALQAK | YES | NO | NO |
| 16628 | TRAV4 | TRAJ31 | CLVGEPGVLNNARLMF | TRBV4-3 | TRBJ1-1 | CASSQAVLNTEAFF | KLGGALQAK | NO | NO | NO |
| 16629 | TRAV12-2 | TRAJ12 | CAVNNRRDSSYKLIF | TRBV27 | TRBJ1-5 | CASSFSSGGNQPQHF | KLGGALQAK | YES | NO | NO |
| 16630 | TRAV8-3 | TRAJ33 | CAVGAGESNYQLIW | TRBV29-1 | TRBJ1-2 | CSVLDQGANYGYTF | KLGGALQAK | YES | NO | NO |
| 16631 | TRAV19 | TRAJ26 | CALSEDNYGQNFVF | TRBV7-9 | TRBJ2-1 | CASSSLLGGHEQFF | KLGGALQAK | YES | NO | NO |
| 16632 | TRAV12-2 | TRAJ28 | CAVPFFSGAGSYQLTF | TRBV24-1 | TRBJ1-2 | CATSGGHGYTF | KLGGALQAK | YES | NO | NO |
| 16633 | TRAV24 | TRAJ32 | CALLGGATNKLIF | TRBV19 | TRBJ2-2 | CASSIVVTGELFF | AVFDRKSDAK | YES | NO | YES |
| 16634 | TRAV12-2 | TRAJ33 | CAVTDSNYQLIW | TRBV7-9 | TRBJ1-2 | CASSPDRAITNYGYTF | KLGGALQAK | YES | NO | NO |
| 16635 | TRAV12-2 | TRAJ43 | CAVTRNNNDMRF | TRBV11-2 | TRBJ1-6 | CASSLGQGNSPLHF | KLGGALQAK | YES | NO | NO |
| 16636 | TRAV2 | TRAJ13 | CAVEDQSGGYQKVTF | TRBV4-3 | TRBJ1-5 | CASSQDRMSNQPQHF | KLGGALQAK | NO | NO | NO |
| 16637 | TRAV5 | TRAJ27 | CADTNAGKSTF | TRBV19 | TRBJ1-4 | CASSIGVAGEKLFF | KLGGALQAK | YES | NO | NO |
| 16638 | TRAV8-4 | TRAJ52 | CAVTHRNAGGTSYGKLTF | TRBV3-1 | TRBJ1-2 | CASSHPETVRGYTF | KLGGALQAK | YES | NO | NO |
| 16639 | TRAV12-1 | TRAJ4 | CVGYSGGYNKLIF | TRBV4-1 | TRBJ1-3 | CASRAGTEIAGNTIYF | KLGGALQAK | YES | NO | NO |
| 16640 | TRAV13-2 | TRAJ42 | CAEMKGQGNLIF | TRBV7-9 | TRBJ2-3 | CASSSAGSGGPTDTQYF | KLGGALQAK | YES | NO | NO |
| 16641 | TRAV36DV7 | TRAJ48 | CAVANFGNEKLTF | TRBV19 | TRBJ1-1 | CASRPTLWSANTEAFF | KLGGALQAK | NO | NO | NO |
| 16642 | TRAV13-2 | TRAJ39 | CAEKGNNAGNMLTF | TRBV5-1 | TRBJ1-1 | CASSLASGTEAFF | KLGGALQAK | YES | NO | NO |
| 16643 | TRAV35 | TRAJ52 | CAGGTSYGKLTF | TRBV7-9 | TRBJ1-1 | CASTDWGSTEAFF | KLGGALQAK | YES | NO | NO |
| 16644 | TRAV9-2 | TRAJ31 | CALFSDNARLMF | TRBV11-3 | TRBJ1-1 | CASSKRQSMNTEAFF | KLGGALQAK | YES | NO | NO |
| 16645 | TRAV1-2 | TRAJ28 | CAVSDSGAGSYQLTF | TRBV5-6 | TRBJ2-2 | CASSEGLAGEGLAGELFF | KLGGALQAK | YES | NO | NO |
| 16646 | TRAV22 | TRAJ43 | CAVEGGMRF | TRBV12-3 | TRBJ1-5 | CASSLGAGQPQHF | KLGGALQAK | NO | NO | NO |
| 16647 | TRAV13-2 | TRAJ29 | CAETLTGNTPLVF | TRBV4-1 | TRBJ2-7 | CASSQVRGSYEQYF | AVFDRKSDAK | YES | NO | YES |
| 16648 | TRAV12-1 | TRAJ32 | CVVAAYATNKLIF | TRBV7-9 | TRBJ1-5 | CASSRTEPTQPQHF | AVFDRKSDAK | YES | NO | YES |
| 16649 | TRAV38-2DV8 | TRAJ38 | CAYRSPQGAGNNRKLIW | TRBV7-2 | TRBJ2-1 | CASSLVGGLGNNEQFF | KLGGALQAK | YES | NO | NO |
| 16650 | TRAV29DV5 | TRAJ28 | CAAPSGAGSYQLTF | TRBV4-1 | TRBJ2-5 | CASSHGLAGEETQYF | KLGGALQAK | YES | NO | NO |
| 16651 | TRAV4 | TRAJ4 | CPLFSGGYNKLIF | TRBV5-6 | TRBJ1-4 | CASSFLSGDTGNEKLFF | KLGGALQAK | YES | NO | NO |
| 16652 | TRAV19 | TRAJ20 | CALSEEFNDYKLSF | TRBV12-3 | TRBJ2-1 | CASRGLAGVGVLSGYNEQFF | AVFDRKSDAK | NO | NO | NO |
| 16653 | TRAV29DV5 | TRAJ52 | CAASGYWGKLTF | TRBV7-2 | TRBJ2-1 | CASSLDRAVSSYNEQFF | KLGGALQAK | YES | NO | NO |

| | | | | | | | | | | |
|---|---|---|---|---|---|---|---|---|---|---|
| 16654 | TRAV10 | TRAJ47 | CVATEYGNKLVF | TRBV12-4 | TRBJ1-5 | CASSLGVEGNSNQPQHF | KLGGALQAK | YES | NO | NO |
| 16655 | TRAV9-2 | TRAJ37 | CALSVGSSNTGKLIF | TRBV9 | TRBJ2-2 | CASSESGTNTGELFF | KLGGALQAK | YES | NO | NO |
| 16656 | TRAV14DV4 | TRAJ4 | CAMREWFSGGYNKLIF | TRBV15 | TRBJ1-2 | CATSRDDSDRDPGGYTF | KLGGALQAK | YES | NO | NO |
| 16657 | TRAV4 | TRAJ28 | CLVGDAYSGAGSYQLTF | TRBV5-4 | TRBJ2-7 | CASSGETYEQYF | KLGGALQAK | NO | NO | NO |
| 16658 | TRAV17 | TRAJ23 | CATARYNQGGKLIF | TRBV24-1 | TRBJ2-1 | CATSDGLAGGYNEQFF | KLGGALQAK | YES | NO | NO |
| 16659 | TRAV36DV7 | TRAJ46 | CAGKTSGDKLTF | TRBV9 | TRBJ2-1 | CASSPTGGNNEQFF | KLGGALQAK | NO | NO | NO |
| 16660 | TRAV29DV5 | TRAJ31 | CAASLNNARLMF | TRBV4-3 | TRBJ2-5 | CASSQDLGLAGGETQYF | KLGGALQAK | YES | NO | NO |
| 16661 | TRAV29DV5 | TRAJ20 | CAASDRDYKLSF | TRBV6-1 | TRBJ1-5 | CASSGTGNQPQHF | KLGGALQAK | YES | NO | NO |
| 16662 | TRAV1-1 | TRAJ36 | CAVIAGANNLFF | TRBV30 | TRBJ1-3 | CAWSGPAGNTIYF | KLGGALQAK | YES | NO | NO |
| 16663 | TRAV26-1 | TRAJ28 | CIVRVVEGAGSYQLTF | TRBV5-5 | TRBJ2-7 | CASSLFYSSYEQYF | KLGGALQAK | YES | NO | NO |
| 16664 | TRAV8-6 | TRAJ30 | CAVSPGGNDKIIF | TRBV29-1 | TRBJ2-2 | CSVLLHTGELFF | KLGGALQAK | NO | NO | NO |
| 16665 | TRAV39 | TRAJ57 | CAVDTSGSEKLVF | TRBV2 | TRBJ1-5 | CASSDIRGLGSNQPQHF | AVFDRKSDAK | YES | NO | YES |
| 16666 | TRAV4 | TRAJ35 | CLVGMI | TRBV7-9 | TRBJ1-1 | CASSSTRDRAGYEAFF | KLGGALQAK | NO | NO | NO |
| 16667 | TRAV12-1 | TRAJ40 | CVVISTTSGTYKYIF | TRBV11-2 | TRBJ2-3 | CASSSSQEALTDTQYF | KLGGALQAK | YES | NO | NO |
| 16668 | TRAV12-1 | TRAJ13 | CVVPSGGYQKVTF | TRBV12-4 | TRBJ1-1 | CASSSPGQGDTEAFF | KLGGALQAK | YES | NO | NO |
| 16669 | TRAV8-3 | TRAJ49 | CAVDPTGNQFYF | TRBV7-9 | TRBJ1-2 | CASSLAGNYGYTF | KLGGALQAK | YES | NO | NO |
| 16670 | TRAV29DV5 | TRAJ20 | CAASATNDYKLSF | TRBV12-3 | TRBJ2-3 | CASILRASDTQYF | KLGGALQAK | YES | NO | NO |
| 16671 | TRAV1-2 | TRAJ15 | CAVRDHQAGTALIF | TRBV3-1 | TRBJ2-2 | CASSQGAGRDPTGELFF | KLGGALQAK | YES | NO | NO |
| 16672 | TRAV26-1 | TRAJ20 | CIVRVGHKLSF | TRBV19 | TRBJ2-1 | CASSTIPGRGNEQFF | KLGGALQAK | YES | NO | NO |
| 16673 | TRAV27 | TRAJ53 | CAGAGSGGSNYKLTF | TRBV7-8 | TRBJ2-5 | CASSPGTSGSTQYF | KLGGALQAK | YES | NO | NO |
| 16674 | TRAV19 | TRAJ31 | CALSEYYNARLMF | TRBV2 | TRBJ2-3 | CASSEVSTDTQYF | KLGGALQAK | YES | NO | NO |
| 16675 | TRAV12-2 | TRAJ33 | CAAKDSNYQLIW | TRBV5-1 | TRBJ2-6 | CASSSGTSGIRGTGANVLTF | KLGGALQAK | NO | NO | NO |
| 16676 | TRAV29DV5 | TRAJ33 | CAASALSNYQLIW | TRBV29-1 | TRBJ2-7 | CSVEEQGNYEQYF | KLGGALQAK | YES | NO | NO |
| 16677 | TRAV2 | TRAJ40 | CAVSANSGTYKYIF | TRBV3-1 | TRBJ1-6 | CASSHPGGQAVSSYNSPLHF | KLGGALQAK | NO | NO | NO |
| 16678 | TRAV13-2 | TRAJ17 | CAEKRWAAGNKLTF | TRBV20-1 | TRBJ2-7 | CSASSRVAYEQYF | KLGGALQAK | NO | NO | NO |
| 16679 | TRAV12-2 | TRAJ43 | CAVVYNNNDMRF | TRBV4-3 | TRBJ2-1 | CASSQDSGGVNEQFF | KLGGALQAK | YES | NO | NO |
| 16680 | TRAV36DV7 | TRAJ48 | CAVGFGNEKLTF | TRBV4-3 | TRBJ1-1 | CASSQDPENTEAFF | AVFDRKSDAK | NO | NO | NO |
| 16681 | TRAV10 | TRAJ42 | CVVSGGSQGNLIF | TRBV4-2 | TRBJ1-2 | CASSQEGTGNYGYTF | KLGGALQAK | YES | NO | NO |
| 16683 | TRAV25 | TRAJ44 | CAGNTGTASKLTF | TRBV5-5 | TRBJ2-1 | CASSLEAGGAYNEQFF | KLGGALQAK | YES | NO | NO |
| 16684 | TRAV41 | TRAJ54 | CAVLIQGAQKLVF | TRBV19 | TRBJ1-6 | CASSKNRQGSSYNSPLHF | KLGGALQAK | YES | NO | NO |
| 16685 | TRAV1-2 | TRAJ28 | CAVTYSGAGSYQLTF | TRBV19 | TRBJ2-1 | CASSIVAGGLDEQFF | KLGGALQAK | YES | NO | NO |
| 16686 | TRAV24 | TRAJ40 | CAFDPTSGTYKYIF | TRBV27 | TRBJ1-2 | CASSPLTGTGGGYTF | KLGGALQAK | YES | NO | NO |
| 16687 | TRAV29DV5 | TRAJ40 | CAASGTTSGTYKYIF | TRBV7-9 | TRBJ2-1 | CASNGGQGYEQFF | KLGGALQAK | YES | NO | NO |
| 16688 | TRAV17 | TRAJ4 | CATDPFSGGYNKLIF | TRBV5-1 | TRBJ2-1 | CASSLGLADYYNEQFF | KLGGALQAK | YES | NO | NO |
| 16689 | TRAV21 | TRAJ35 | CAVGRI | TRBV11-2 | TRBJ2-1 | CASSPLGLGGGEQFF | KLGGALQAK | NO | NO | NO |
| 16690 | TRAV19 | TRAJ49 | CALSGPNTGNQFYF | TRBV13 | TRBJ1-5 | CASSLGILGGSLEPQHF | KVLEYVIKV | YES | NO | NO |
| 16691 | TRAV14DV4 | TRAJ5 | CAMRDPQGGRRALTF | TRBV6-5 | TRBJ2-3 | CASSYVGGPGDTQYF | KLGGALQAK | NO | NO | NO |
| 16692 | TRAV1-2 | TRAJ28 | CAVRDLPGSYQLTF | TRBV14 | TRBJ1-3 | CASSQDEGQDGPGNTIYF | KLGGALQAK | NO | NO | NO |
| 16693 | TRAV19 | TRAJ28 | CALRTYSGAGSYQLTF | TRBV4-2 | TRBJ1-1 | CASSQTRGLISEAFF | KLGGALQAK | YES | NO | NO |
| 16694 | TRAV29DV5 | TRAJ34 | CAASARDTDKLIF | TRBV5-6 | TRBJ2-1 | CASSRTDSYNEQFF | KLGGALQAK | YES | NO | NO |
| 16695 | TRAV5 | TRAJ45 | CAETHSGGGADGLTF | TRBV29-1 | TRBJ1-2 | CSVETDYGYTF | KLGGALQAK | NO | NO | NO |
| 16696 | TRAV13-1 | TRAJ5 | CAASMSRGRALTF | TRBV7-9 | TRBJ2-4 | CASSLGESIAKNIQYF | KLGGALQAK | YES | NO | NO |
| 16697 | TRAV19 | TRAJ23 | CALSEALGLGKLQGGKLIF | TRBV3-1 | TRBJ2-1 | CASSQGKRDEQFF | KLGGALQAK | YES | NO | NO |
| 16698 | TRAV19 | TRAJ38 | CALSGAINAGNNRKLIW | TRBV7-9 | TRBJ2-1 | CASSRTGTGGGNEQFF | KLGGALQAK | YES | NO | NO |
| 16699 | TRAV19 | TRAJ58 | CALSEARETSGSRLTF | TRBV9 | TRBJ2-1 | CASSVRPGGAGNEQFF | KLGGALQAK | YES | NO | NO |
| 16700 | TRAV26-2 | TRAJ45 | CILRALASKMYSGGGADGLTF | TRBV7-2 | TRBJ2-3 | CASSPRVFSPGTDTQYF | KLGGALQAK | NO | NO | NO |
| 16701 | TRAV17 | TRAJ43 | CATGPSPNDMRF | TRBV12-4 | TRBJ1-5 | CASSLALEANQPQHF | KLGGALQAK | YES | NO | NO |
| 16702 | TRAV5 | TRAJ40 | CAELFTSGTYKYIF | TRBV4-3 | TRBJ2-2 | CASSQVTGLSDGELFF | AVFDRKSDAK | YES | NO | YES |
| 16703 | TRAV1-2 | TRAJ33 | CAVRDSNYQLIW | TRBV20-1 | TRBJ1-1 | CSARSLGGTEAFF | KLGGALQAK | YES | NO | NO |
| 16704 | TRAV17 | TRAJ26 | CATDADYGQNFVF | TRBV7-6 | TRBJ1-5 | CASSSGPGEVQPQHF | KLGGALQAK | YES | NO | NO |
| 16705 | TRAV29DV5 | TRAJ20 | CAASPSSNDYKLSF | TRBV7-8 | TRBJ2-2 | CASSLGLPGLEGNTGELFF | AVFDRKSDAK | YES | NO | YES |
| 16706 | TRAV20 | TRAJ9 | CAVRIYTGGFKTIF | TRBV4-3 | TRBJ2-2 | CASSQDVGPDSLNTGELFF | KLGGALQAK | YES | NO | NO |
| 16707 | TRAV1-2 | TRAJ41 | CAVRASNSGYALNF | TRBV20-1 | TRBJ2-7 | CSAAGTGTPYEQYV | KLGGALQAK | NO | NO | NO |
| 16708 | TRAV8-1 | TRAJ40 | CAVGGPGTYKYIF | TRBV5-6 | TRBJ2-3 | CASSSGQNTDTQYF | KLGGALQAK | YES | NO | NO |
| 16709 | TRAV27 | TRAJ39 | CAGGGAGNMLTF | TRBV7-7 | TRBJ1-3 | CASSLAGSGNTIYF | KLGGALQAK | YES | NO | NO |
| 16710 | TRAV26-2 | TRAJ43 | CILRDGFSPSNNNDMRF | TRBV20-1 | TRBJ2-3 | CSAGGLAGRDTQYF | KLGGALQAK | YES | NO | NO |
| 16711 | TRAV29DV5 | TRAJ13 | CAASDSGGYQKVTF | TRBV12-4 | TRBJ1-2 | CASSSRVSEYYGYTF | KLGGALQAK | YES | NO | NO |
| 16712 | TRAV12-2 | TRAJ9 | CAVSRAGGFKTIF | TRBV19 | TRBJ1-5 | CASSIDEGFWISNQPQHF | KLGGALQAK | YES | NO | NO |
| 16713 | TRAV3 | TRAJ15 | CAVRDPITANQAGTALIF | TRBV19 | TRBJ1-1 | CASINYGAGLNTEAFF | KLGGALQAK | NO | NO | NO |
| 16714 | TRAV13-2 | TRAJ11 | CAGSTNSGYSTLTF | TRBV19 | TRBJ1-2 | CASSDGPNYGYTF | KLGGALQAK | YES | NO | NO |
| 16715 | TRAV19 | TRAJ37 | CAKSNTGKLIF | TRBV12-4 | TRBJ1-2 | CASRRRDRGFNYGYTF | KLGGALQAK | YES | NO | NO |
| 16716 | TRAV38-2DV8 | TRAJ18 | CAYRDSTLGRLYF | TRBV7-9 | TRBJ2-5 | CASSRTEGVETQYF | KLGGALQAK | YES | NO | NO |
| 16717 | TRAV19 | TRAJ12 | CALSARMDSSYKLIF | TRBV19 | TRBJ2-2 | CASSITAGELFF | KLGGALQAK | YES | NO | NO |
| 16718 | TRAV23DV6 | TRAJ37 | CAASIGGYGSGNTGKLIF | TRBV7-3 | TRBJ1-1 | CASSLLENTEAFF | KLGGALQAK | NO | NO | NO |
| 16719 | TRAV24 | TRAJ52 | CAFSRDSGTSYGKLTF | TRBV19 | TRBJ1-5 | CASSIGRLQPQHF | KLGGALQAK | YES | NO | NO |
| 16720 | TRAV1-1 | TRAJ37 | CAVRVGSGNTGKLIF | TRBV9 | TRBJ1-1 | CASSVSSGGPEAFF | AVFDRKSDAK | YES | NO | NO |
| 16721 | TRAV35 | TRAJ42 | CAGQSGRVGGSQGNLIF | TRBV20-1 | TRBJ2-1 | CSASSSGGAEDEQFF | KLGGALQAK | YES | NO | NO |
| 16722 | TRAV19 | TRAJ40 | CALSELSLSGTYKYIF | TRBV4-1 | TRBJ1-6 | CASSRGTESSYNSPLHF | KLGGALQAK | YES | NO | NO |
| 16723 | TRAV24 | TRAJ17 | CALFPLPIKAAGNKLTF | TRBV4-2 | TRBJ1-5 | CASSQFRAWQPQHF | KLGGALQAK | YES | NO | NO |
| 16724 | TRAV29DV5 | TRAJ54 | CAAIPLQGAQKLVF | TRBV13 | TRBJ1-4 | CASSEDRGADYEKLFF | KLGGALQAK | YES | NO | NO |
| 16725 | TRAV1-2 | TRAJ33 | CAVRDSNYQLIW | TRBV20-1 | TRBJ1-1 | CSAFLDNTEAFF | KLGGALQAK | YES | NO | NO |
| 16726 | TRAV19 | TRAJ49 | CALSGTGNQFYF | TRBV19 | TRBJ2-3 | CASSMFLDTDTQYF | KLGGALQAK | YES | NO | NO |
| 16727 | TRAV39 | TRAJ33 | CAVVGMDSNYQLIW | TRBV19 | TRBJ2-1 | CASSIYKSSYNEQFF | KLGGALQAK | YES | NO | NO |
| 16728 | TRAV13-2 | TRAJ30 | CAENINRDDKIIF | TRBV5-1 | TRBJ2-1 | CASSWTRYNEQFF | KLGGALQAK | YES | NO | NO |
| 16729 | TRAV19 | TRAJ58 | CALNSLTPRLTF | TRBV20-1 | TRBJ1-2 | CSAFPKGTHTF | KLGGALQAK | YES | NO | NO |
| 16730 | TRAV25 | TRAJ37 | CAGTPGNTGKLIF | TRBV6-3 | TRBJ1-1 | CASSYLWNTEAFF | AVFDRKSDAK | YES | YES | NO |
| 16731 | TRAV8-1 | TRAJ27 | CAVSTNAGKSTF | TRBV30 | TRBJ2-2 | CAWSVTNTGELFF | KLGGALQAK | YES | NO | NO |
| 16732 | TRAV17 | TRAJ40 | CATDPQYSGTYKYIF | TRBV19 | TRBJ1-1 | CASSSGTANTEAFF | KLGGALQAK | YES | NO | NO |
| 16733 | TRAV34 | TRAJ48 | CGAATPNFGNEKLTF | TRBV19 | TRBJ2-3 | CASRSGMSTDTQYF | KLGGALQAK | YES | NO | NO |
| 16734 | TRAV1-2 | TRAJ13 | CAAGPSGGYQKVTF | TRBV7-2 | TRBJ1-2 | CASSLVLPGQGIGYGYTF | KLGGALQAK | YES | NO | NO |
| 16735 | TRAV14DV4 | TRAJ30 | CAMREGMNRDDKIIF | TRBV4-3 | TRBJ2-4 | CASSQDLTAGAKNIQYF | KLGGALQAK | YES | NO | NO |
| 16736 | TRAV1-2 | TRAJ22 | CAVRGRGSARQLTF | TRBV19 | TRBJ1-2 | CASWGIWGKQGAPGYTF | KLGGALQAK | YES | NO | NO |
| 16737 | TRAV38-2DV8 | TRAJ42 | CAYNTPHGGGSQGNLIF | TRBV7-8 | TRBJ2-7 | CASSLDSGFEQYV | KLGGALQAK | NO | NO | NO |
| 16738 | TRAV3 | TRAJ36 | CAVRDTGTGANNLFF | TRBV6-5 | TRBJ1-3 | CASSPTGGSGNTIYF | KLGGALQAK | NO | NO | NO |
| 16739 | TRAV1-1 | TRAJ21 | CAVYNFNKFYF | TRBV27 | TRBJ1-1 | CASSLRAMNTEAFF | KLGGALQAK | YES | NO | NO |
| 16740 | TRAV12-3 | TRAJ50 | CAMSAGTSYDKVIF | TRBV20-1 | TRBJ1-2 | CSARVHRYNYGYTF | KLGGALQAK | YES | NO | NO |
| 16741 | TRAV1-1 | TRAJ23 | CAVRDNQGGKLIF | TRBV19 | TRBJ2-5 | CASSLTAGAAETQYF | KLGGALQAK | YES | NO | NO |
| 16742 | TRAV21 | TRAJ24 | CAGTDSWGKLQF | TRBV4-2 | TRBJ2-3 | CASSQDYPRLAGTDTQYF | KLGGALQAK | YES | NO | NO |
| 16743 | TRAV25 | TRAJ30 | CAGRGGRDDKIIF | TRBV19 | TRBJ1-2 | CASSIGGRGYTF | KLGGALQAK | YES | NO | NO |
| 16744 | TRAV29DV5 | TRAJ47 | CAAREYGNKLVF | TRBV6-5 | TRBJ2-2 | CASSPGTGENTGELFF | KLGGALQAK | YES | NO | NO |
| 16745 | TRAV8-2 | TRAJ22 | CVGRKGSARQLTF | TRBV10-3 | TRBJ2-3 | CAISESRGSLSDTQYF | KLGGALQAK | YES | NO | NO |
| 16746 | TRAV25 | TRAJ9 | CAGPIAGGFKTIF | TRBV19 | TRBJ2-1 | CASSMWTGDNEQFF | IVTDFSVIK | YES | NO | NO |
| 16747 | TRAV5 | TRAJ40 | CAEEYSGTYKYIF | TRBV30 | TRBJ1-1 | CAWSVERTEAFF | KLGGALQAK | YES | NO | NO |
| 16748 | TRAV14DV4 | TRAJ27 | CAMREFYAGKSTF | TRBV6-3 | TRBJ2-7 | CASSTPDGGRTYEQYV | KLGGALQAK | NO | NO | NO |
| 16749 | TRAV4 | TRAJ29 | CLVGDNGNTPLVF | TRBV4-2 | TRBJ1-1 | CASSQYTVDTEAFF | KLGGALQAK | YES | NO | NO |
| 16750 | TRAV8-2 | TRAJ43 | CVVSDLVDDMRF | TRBV6-3 | TRBJ2-5 | CASSGIETQYF | KLGGALQAK | YES | NO | NO |
| 16751 | TRAV13-1 | TRAJ30 | CAVLNRDDKIIF | TRBV11-2 | TRBJ1-4 | CASSTTGGNEKLFF | KLGGALQAK | YES | NO | NO |
| 16752 | TRAV12-2 | TRAJ23 | CAVTPFGGGGKLIF | TRBV5-1 | TRBJ1-6 | CASSLEATGDNSPLHF | KLGGALQAK | YES | NO | NO |
| 16753 | TRAV29DV5 | TRAJ49 | CAASESGNQFYF | TRBV12-4 | TRBJ2-3 | CASSLAVSTDTQYF | KLGGALQAK | YES | NO | NO |
| 16754 | TRAV29DV5 | TRAJ6 | CAANFGGSYIPTF | TRBV5-1 | TRBJ2-3 | CASSLGLAGGLMDTQYF | KLGGALQAK | YES | NO | NO |
| 16755 | TRAV8-2 | TRAJ49 | CVVSGYTGNQFYF | TRBV30 | TRBJ2-3 | CAWSVLGSTDTQYF | KLGGALQAK | YES | NO | NO |
| 16756 | TRAV21 | TRAJ20 | CAVDRSNDYKLSF | TRBV19 | TRBJ1-1 | CASKLTTMTEAFF | KLGGALQAK | YES | NO | NO |
| 16757 | TRAV16 | TRAJ17 | CALSDQFKAAGNKLTF | TRBV6-4 | TRBJ1-6 | CASSDSQGSYNSPLHF | KLGGALQAK | NO | NO | NO |
| 16758 | TRAV8-3 | TRAJ54 | CAVGPGAQKLVF | TRBV11-2 | TRBJ1-1 | CASSPTGTGTEAFF | KLGGALQAK | YES | NO | NO |

| | | | | | | | | | | |
|---|---|---|---|---|---|---|---|---|---|---|
| 16759 | TRAV25 | TRAJ50 | CAVNMKTSYDKVIF | TRBV19 | TRBJ2-1 | CASSRDSYNEQFF | KLGGALQAK | YES | NO | NO |
| 16760 | TRAV13-2 | TRAJ20 | CAERGDYKLSF | TRBV6-5 | TRBJ1-1 | CASSLQDMNTEAFF | KLGGALQAK | YES | NO | NO |
| 16761 | TRAV12-2 | TRAJ33 | CAGGGGDSNYQLIW | TRBV20-1 | TRBJ1-5 | CSARDGVSPQHF | KLGGALQAK | YES | NO | NO |
| 16762 | TRAV8-4 | TRAJ43 | CAVSYNNNDMRF | TRBV30 | TRBJ2-1 | CAWTEGRGVGEQFF | KLGGALQAK | NO | NO | NO |
| 16763 | TRAV13-2 | TRAJ54 | CAENIGQGAQKLVF | TRBV2 | TRBJ2-2 | CASYGAETGELFF | KLGGALQAK | YES | NO | NO |
| 16764 | TRAV13-2 | TRAJ52 | CAERDAGGTSYGKLTF | TRBV5-1 | TRBJ2-5 | CASSLARTEETQYF | KLGGALQAK | YES | NO | NO |
| 16765 | TRAV8-3 | TRAJ53 | CAVGAGSGGSNYKLTF | TRBV7-7 | TRBJ1-2 | CASSPLGYGYTF | KLGGALQAK | YES | NO | NO |
| 16766 | TRAV3 | TRAJ5 | CAVRGVMDTGRRALTF | TRBV27 | TRBJ2-2 | CASSVVQTNTGELFF | KLGGALQAK | NO | NO | NO |
| 16767 | TRAV10 | TRAJ40 | CVVRFSGTYKYIF | TRBV5-1 | TRBJ2-1 | CASSSQELVVHNEQFF | KLGGALQAK | YES | NO | NO |
| 16768 | TRAV13-1 | TRAJ48 | CAALSNFGNEKLTF | TRBV6-3 | TRBJ2-4 | CASSYPRVGSKNIQYF | KLGGALQAK | YES | NO | NO |
| 16769 | TRAV29DV5 | TRAJ22 | CAAISSGSARQLTF | TRBV10-3 | TRBJ2-2 | CAISDAGLATGELFF | KLGGALQAK | YES | NO | NO |
| 16770 | TRAV29DV5 | TRAJ44 | CAASSTGTASKLTF | TRBV6-1 | TRBJ2-7 | CASRETYKYEQYV | KLGGALQAK | NO | NO | NO |
| 16772 | TRAV4 | TRAJ3 | CLVGDIGYSSASKIIF | TRBV12-4 | TRBJ2-3 | CASTLEDSTDTQYF | KLGGALQAK | YES | NO | NO |
| 16773 | TRAV17 | TRAJ34 | CATVVSFNTDKLIF | TRBV6-3 | TRBJ1-6 | CASTLTVVGNSPLHF | AVFDRKSDAK | YES | NO | YES |
| 16774 | TRAV5 | TRAJ32 | CAEWSGGATNKLIF | TRBV7-9 | TRBJ2-1 | CASSLGSNEQFF | KLGGALQAK | YES | NO | NO |
| 16775 | TRAV8-3 | TRAJ49 | CAVGGNQFYF | TRBV11-3 | TRBJ2-4 | CASSGDIQYF | KLGGALQAK | YES | NO | NO |
| 16776 | TRAV29DV5 | TRAJ34 | CAASAEYNTDKLIF | TRBV7-9 | TRBJ1-1 | CASSYAPGEAFF | KLGGALQAK | YES | NO | NO |
| 16777 | TRAV12-1 | TRAJ18 | CVVINPVDRGSTLGRLYF | TRBV7-7 | TRBJ2-2 | CASSTQGLAGVNGELFF | KLGGALQAK | YES | NO | NO |
| 16778 | TRAV26-2 | TRAJ30 | CILNRDDKIIF | TRBV20-1 | TRBJ2-3 | CSALGTGGFVTDTQYF | KLGGALQAK | NO | NO | NO |
| 16779 | TRAV9-2 | TRAJ52 | CALRGTSYGKLTF | TRBV28 | TRBJ1-2 | CASSPPTGVGYGYTF | KLGGALQAK | YES | NO | NO |
| 16780 | TRAV21 | TRAJ36 | CAVYPGANNLFF | TRBV6-5 | TRBJ2-7 | CASSLQGTGLYEQYF | KLGGALQAK | YES | NO | NO |
| 16781 | TRAV12-2 | TRAJ45 | CAVNRGIYSGGGADGLTF | TRBV6-3 | TRBJ2-1 | CASSPGLAGDNEQFF | AVFDRKSDAK | YES | NO | YES |
| 16782 | TRAV8-2 | TRAJ22 | CVGRKGSARQLTF | TRBV7-2 | TRBJ2-7 | CASSLEPSYEQYF | KLGGALQAK | YES | NO | NO |
| 16783 | TRAV39 | TRAJ53 | CAVADSGGSNYKLTF | TRBV12-4 | TRBJ2-3 | CASSSRTSSTDTQYF | KLGGALQAK | YES | NO | NO |
| 16784 | TRAV8-6 | TRAJ15 | CAVNGQAGTALIF | TRBV5-1 | TRBJ1-1 | CASVSGNEAFF | AVFDRKSDAK | YES | NO | YES |
| 16785 | TRAV3 | TRAJ23 | CAVRDRVYNQGGKLIF | TRBV7-2 | TRBJ2-3 | CASSSRGEVRTDTQYF | KLGGALQAK | NO | NO | NO |
| 16786 | TRAV8-1 | TRAJ43 | CAVEMGNNNDMRF | TRBV30 | TRBJ2-5 | CAWSSGRGETQYF | KLGGALQAK | YES | NO | NO |
| 16787 | TRAV8-2 | TRAJ22 | CVGRKGSARQLTF | TRBV19 | TRBJ1-5 | CASSIGGGSYQPQHF | KLGGALQAK | YES | NO | NO |
| 16788 | TRAV1-1 | TRAJ29 | CAVEGNTPLVF | TRBV7-6 | TRBJ2-1 | CASSPPSPGLALNEQFF | KLGGALQAK | YES | NO | NO |
| 16789 | TRAV20 | TRAJ54 | CAVQAGGAQKLVF | TRBV27 | TRBJ2-1 | CASSFQTGNEQFF | KLGGALQAK | YES | NO | NO |
| 16791 | TRAV12-3 | TRAJ26 | CAILNYGQNFVF | TRBV7-2 | TRBJ1-3 | CASSLEITSGNTIYF | KLGGALQAK | YES | NO | NO |
| 16792 | TRAV23DV6 | TRAJ56 | CAASTTGANSKLTF | TRBV12-4 | TRBJ1-2 | CASRVTGGNGYTF | KLGGALQAK | YES | NO | NO |
| 16793 | TRAV12-1 | TRAJ33 | CVVNVGYQLIW | TRBV3-1 | TRBJ2-1 | CASSPPPLSGGYTEQFF | KLGGALQAK | YES | NO | NO |
| 16794 | TRAV8-1 | TRAJ13 | CAVNAGGYQKVTF | TRBV27 | TRBJ2-1 | CASSLAGGNNEQFF | IVTDFSVIK | YES | NO | YES |
| 16795 | TRAV38-2DV8 | TRAJ33 | CAYMDSNYQLIW | TRBV7-9 | TRBJ1-2 | CASETVHYGYTF | KLGGALQAK | YES | NO | NO |
| 16796 | TRAV1-1 | TRAJ21 | CAVSPGDNFNKFYF | TRBV6-1 | TRBJ1-1 | CASREQGNTEAFF | KLGGALQAK | YES | NO | NO |
| 16797 | TRAV12-2 | TRAJ49 | CAVNSNTGNQFYF | TRBV7-6 | TRBJ2-4 | CASSSWTGVAKNIQYF | KLGGALQAK | YES | NO | NO |
| 16798 | TRAV35 | TRAJ37 | CAGRARNTGKLIF | TRBV19 | TRBJ1-5 | CASSPGTGVDPQHF | KLGGALQAK | YES | NO | NO |
| 16799 | TRAV24 | TRAJ52 | CAFIGDAGGTSYGKLTF | TRBV15 | TRBJ2-1 | CATSRGGYEQFF | KLGGALQAK | YES | NO | NO |
| 16800 | TRAV8-4 | TRAJ43 | CAVSGLTNDMRF | TRBV6-5 | TRBJ1-1 | CASKIGGKNTEAFF | KLGGALQAK | YES | NO | NO |
| 16801 | TRAV8-4 | TRAJ40 | CAVWSSGTYKYIF | TRBV7-2 | TRBJ1-2 | CASSWGDGNYGYTF | KLGGALQAK | YES | NO | NO |
| 16802 | TRAV12-1 | TRAJ9 | CVVNGYTGGFKTIF | TRBV9 | TRBJ1-5 | CASSVSGGGQPQHF | KLGGALQAK | NO | NO | NO |
| 16803 | TRAV24 | TRAJ24 | CASDDSWGKFQF | TRBV15 | TRBJ2-2 | CATSGTGITTGELFF | KLGGALQAK | YES | NO | NO |
| 16804 | TRAV13-2 | TRAJ13 | CAENMNSGGYQKVTF | TRBV6-3 | TRBJ2-7 | CASSYDSSYEQYV | KLGGALQAK | NO | NO | NO |
| 16805 | TRAV20 | TRAJ33 | CAVQIWDSNYQLIW | TRBV14 | TRBJ2-1 | CASSEPLVNEQFF | KLGGALQAK | YES | NO | NO |
| 16806 | TRAV21 | TRAJ42 | CAVLNYGGSQGNLIF | TRBV5-1 | TRBJ1-4 | CASTGTGNEKLFF | KLGGALQAK | YES | NO | NO |
| 16807 | TRAV27 | TRAJ31 | CAGVNNARLMF | TRBV12-3 | TRBJ2-2 | CASSLAPGQGGFLVTGELFF | KLGGALQAK | NO | NO | NO |
| 16808 | TRAV19 | TRAJ26 | CALIDNYGQNFVF | TRBV12-4 | TRBJ1-2 | CASSLDNYGYTF | KLGGALQAK | YES | NO | NO |
| 16809 | TRAV19 | TRAJ39 | CALSPDNNAGNMLTF | TRBV7-7 | TRBJ2-1 | CASSLRLADFFEQFF | KLGGALQAK | YES | NO | NO |
| 16810 | TRAV8-3 | TRAJ40 | CAVGEAGTYKYIF | TRBV11-1 | TRBJ2-3 | CASSLEGQVADTQYF | KLGGALQAK | YES | NO | NO |
| 16811 | TRAV19 | TRAJ48 | CALTVRTNFGNEKLTF | TRBV19 | TRBJ1-5 | CASSIAGQGYSNQPQHF | KLGGALQAK | YES | NO | NO |
| 16812 | TRAV3 | TRAJ37 | CAVRDGGMGSNTGKLIF | TRBV7-9 | TRBJ2-1 | CASSNSLPSGVSYNEQFF | KLGGALQAK | NO | NO | NO |
| 16813 | TRAV14DV4 | TRAJ17 | CAMRDKTPAAGNKLTF | TRBV19 | TRBJ2-3 | CASNPRPRPSYTDTQYF | KLGGALQAK | YES | NO | NO |
| 16814 | TRAV17 | TRAJ40 | CATDAEMSGTYKYIF | TRBV20-1 | TRBJ2-1 | CSARLLGSNEQFF | KLGGALQAK | NO | NO | NO |
| 16815 | TRAV8-1 | TRAJ29 | CAVKNSGNTPLVF | TRBV6-5 | TRBJ1-2 | CASSGQGAAGYTF | KLGGALQAK | YES | NO | NO |
| 16816 | TRAV12-1 | TRAJ45 | CVVTDSGGGADGLTF | TRBV30 | TRBJ1-3 | CAWNGGSGNTIYF | KLGGALQAK | YES | NO | NO |
| 16817 | TRAV8-4 | TRAJ48 | CAVRGISNFGNEKLTF | TRBV27 | TRBJ2-7 | CASSTHLEGGYEQYV | KLGGALQAK | NO | NO | NO |
| 16818 | TRAV3 | TRAJ8 | CAVRAFMNTGFQKLVF | TRBV7-9 | TRBJ2-5 | CASSLDLVRSQETQYF | KLGGALQAK | YES | NO | NO |
| 16819 | TRAV19 | TRAJ52 | CALSDSGGTSYGKLTF | TRBV19 | TRBJ2-2 | CASSGTGELFF | KLGGALQAK | YES | NO | NO |
| 16820 | TRAV6 | TRAJ31 | CALDHNARLMF | TRBV20-1 | TRBJ2-1 | CSARDLAGGTYNEQFF | KLGGALQAK | NO | NO | NO |
| 16821 | TRAV14DV4 | TRAJ42 | CAMREGWGYGGSQGNLIF | TRBV3-1 | TRBJ1-4 | CASSPGTGGTEKLFF | KLGGALQAK | YES | NO | NO |
| 16822 | TRAV17 | TRAJ47 | CATDGIMEYGNKLVF | TRBV6-1 | TRBJ1-1 | CASRENREAFF | KLGGALQAK | YES | NO | NO |
| 16823 | TRAV8-4 | TRAJ34 | CAVSGYNTDKLIF | TRBV6-3 | TRBJ2-1 | CAAGGASYNEQFF | KLGGALQAK | YES | NO | NO |
| 16824 | TRAV39 | TRAJ6 | CALASALPSGGSYIPTF | TRBV4-1 | TRBJ2-4 | CASSRPSAKNIQYF | RLRAEAQVK | YES | YES | YES |
| 16825 | TRAV12-2 | TRAJ36 | CAVRNNLFF | TRBV12-3 | TRBJ1-3 | CASNFDSSGNTIYF | KLGGALQAK | NO | NO | NO |
| 16826 | TRAV12-1 | TRAJ12 | CVVNHRDSSYKLIF | TRBV4-3 | TRBJ2-1 | CASSQDEVREQFF | KLGGALQAK | YES | NO | NO |
| 16827 | TRAV1-2 | TRAJ27 | CAVRGPRAGKSTF | TRBV27 | TRBJ2-7 | CASNPAGEMYEQYF | KLGGALQAK | YES | NO | NO |
| 16828 | TRAV17 | TRAJ10 | CATGILTGGGNKLTF | TRBV6-5 | TRBJ2-1 | CASSYGLAGPSYNEQFF | KLGGALQAK | NO | NO | NO |
| 16829 | TRAV27 | TRAJ21 | CAGRWFNFNKFYF | TRBV7-9 | TRBJ2-3 | CASSPGTVQSDTQYF | KLGGALQAK | YES | NO | NO |
| 16830 | TRAV14DV4 | TRAJ9 | CAMRGGGFKTIF | TRBV25-1 | TRBJ2-5 | CASSWGQAYPETQYF | KLGGALQAK | YES | NO | NO |
| 16831 | TRAV19 | TRAJ42 | CALSEAPYGGSQGNLIF | TRBV7-9 | TRBJ1-5 | CASSPGQVQPQHF | KLGGALQAK | YES | NO | NO |
| 16832 | TRAV13-2 | TRAJ4 | CAGISGGYNKLIF | TRBV27 | TRBJ2-1 | CASSLRRNEQFF | AVFDRKSDAK | YES | NO | YES |
| 16833 | TRAV3 | TRAJ34 | CAVRAYTDKLIF | TRBV20-1 | TRBJ2-1 | CSARGATSGNEQFF | KLGGALQAK | NO | NO | NO |
| 16834 | TRAV12-2 | TRAJ22 | CAVTSVIGSARQLTF | TRBV7-6 | TRBJ1-3 | CASSYRRAGAISSGNTIYF | KLGGALQAK | YES | NO | NO |
| 16835 | TRAV1-2 | TRAJ42 | CAVNHGGSQGNLIF | TRBV3-1 | TRBJ2-1 | CASSQDMDEQFF | AVFDRKSDAK | YES | NO | YES |
| 16836 | TRAV1-2 | TRAJ4 | CAVPSGGYNKLIF | TRBV20-1 | TRBJ2-2 | CSARENTGELFF | RLRAEAQVK | YES | YES | YES |
| 16837 | TRAV1-2 | TRAJ33 | CAVFGSNYQLIW | TRBV7-8 | TRBJ2-3 | CASSFPGQGSADTQYF | KLGGALQAK | YES | NO | NO |
| 16838 | TRAV4 | TRAJ8 | CLVGDLFMNTGFQKLVF | TRBV27 | TRBJ2-3 | CASSDKGADTQYF | KLGGALQAK | YES | NO | NO |
| 16839 | TRAV26-2 | TRAJ40 | CILRDLPGTYKYIF | TRBV18 | TRBJ2-1 | CASSPPDRADEQFF | KLGGALQAK | YES | NO | NO |
| 16840 | TRAV12-2 | TRAJ45 | CAPSGGGADGLTF | TRBV12-3 | TRBJ2-4 | CASSSDEHENIQYF | ELAGIGILTV | YES | NO | NO |
| 16841 | TRAV2 | TRAJ13 | CAVDSSGGYQKVTF | TRBV7-9 | TRBJ2-2 | CASSLPAFGGAGELFF | KLGGALQAK | NO | NO | NO |
| 16842 | TRAV1-1 | TRAJ26 | CAVKSYGQNFVF | TRBV12-4 | TRBJ2-3 | CASSRGSQVPDTQYF | KLGGALQAK | YES | NO | NO |
| 16843 | TRAV26-1 | TRAJ39 | CIVQAILGAGNMLTF | TRBV7-6 | TRBJ1-5 | CASSDSTGQPQHF | KLGGALQAK | YES | NO | NO |
| 16844 | TRAV13-2 | TRAJ16 | CAEGRF | TRBV27 | TRBJ2-1 | CATGLAGVNNEQFF | KLGGALQAK | NO | NO | NO |
| 16845 | TRAV29DV5 | TRAJ23 | CAASVIKGGKLIF | TRBV7-3 | TRBJ2-3 | CASSSTWEGGGDTQYF | KLGGALQAK | YES | NO | NO |
| 16847 | TRAV12-2 | TRAJ43 | CAVNDMRF | TRBV7-9 | TRBJ2-1 | CASSPGPGNEQFF | ELAGIGILTV | NO | NO | NO |
| 16848 | TRAV27 | TRAJ49 | CAGYYTGNQFYF | TRBV20-1 | TRBJ2-3 | CSAREKTSARTDTQYF | KLGGALQAK | YES | NO | NO |
| 16849 | TRAV10 | TRAJ45 | CVVTASGGGADGLTF | TRBV6-3 | TRBJ2-3 | CASSPNFRTQYF | KLGGALQAK | YES | NO | NO |
| 16850 | TRAV8-2 | TRAJ34 | CVVRPEYTDKLIF | TRBV7-9 | TRBJ2-1 | CASVVRGAYNEQFF | KLGGALQAK | YES | NO | NO |
| 16851 | TRAV35 | TRAJ36 | CAGPTAGANNLFF | TRBV7-8 | TRBJ1-2 | CASSPYVFGEMTYGYTF | KLGGALQAK | YES | NO | NO |
| 16852 | TRAV4 | TRAJ29 | CLVGDPGNTPLVF | TRBV14 | TRBJ2-7 | CASSQGGGYEQYF | KLGGALQAK | NO | NO | NO |
| 16853 | TRAV12-1 | TRAJ43 | CVVNRFNNDMRF | TRBV6-5 | TRBJ2-5 | CASSYSPYQETQYF | KLGGALQAK | NO | NO | NO |
| 16854 | TRAV29DV5 | TRAJ42 | CAASVGYGGSQGNLIF | TRBV9 | TRBJ2-1 | CASSDSAYDYNEQFF | KLGGALQAK | YES | NO | NO |
| 16855 | TRAV19 | TRAJ45 | CALSESEMYSGGGADGLTF | TRBV18 | TRBJ1-1 | CASSPQEFENTEAFF | KLGGALQAK | YES | NO | NO |
| 16856 | TRAV12-2 | TRAJ40 | CAAHSGTYKYIF | TRBV12-3 | TRBJ2-5 | CASSFGSQETQYF | KLGGALQAK | YES | NO | NO |
| 16857 | TRAV19 | TRAJ22 | CALVRGSARQLTF | TRBV27 | TRBJ2-1 | CASSLGDYYNEQFF | AVFDRKSDAK | YES | NO | YES |
| 16858 | TRAV26-1 | TRAJ30 | CIVRAGSNRDDKIIF | TRBV11-2 | TRBJ2-1 | CASSLHVLNEQFF | KLGGALQAK | YES | NO | NO |
| 16859 | TRAV25 | TRAJ41 | CAGDSNSGYALNF | TRBV10-2 | TRBJ2-3 | CASSSSGGITDTQYF | KLGGALQAK | YES | NO | NO |
| 16860 | TRAV25 | TRAJ17 | CMIKAAGNKLTF | TRBV10-3 | TRBJ1-5 | CAISESLSQPQHF | KLGGALQAK | YES | NO | NO |
| 16861 | TRAV17 | TRAJ22 | CATEVFPGSARQLTF | TRBV4-1 | TRBJ2-4 | CASSHLTGLAKNIQYF | KLGGALQAK | YES | NO | NO |
| 16862 | TRAV1-2 | TRAJ29 | CAVRDIGEGNTPLVF | TRBV10-3 | TRBJ1-5 | CAISESGKGATANQPQHF | KLGGALQAK | YES | NO | NO |
| 16863 | TRAV6 | TRAJ41 | CALNSGYALNF | TRBV7-2 | TRBJ2-1 | CASSPNYNEQFF | KLGGALQAK | YES | NO | NO |
| 16864 | TRAV21 | TRAJ16 | CAVRF | TRBV15 | TRBJ2-2 | CATSRDFETEWNTGELFF | KLGGALQAK | NO | NO | NO |
| 16865 | TRAV39 | TRAJ40 | CAGESGTYKYIF | TRBV3-1 | TRBJ2-7 | CASSRPNEQYF | KLGGALQAK | YES | NO | NO |

| | | | | | | | | | | |
|---|---|---|---|---|---|---|---|---|---|---|
| 16866 | TRAV8-3 | TRAJ44 | CAVGAPHTGTASKLTF | TRBV5-4 | TRBJ2-7 | CASSLAQSYEQYF | KLGGALQAK | YES | NO | NO |
| 16867 | TRAV14DV4 | TRAJ39 | CAMREINNAGNMLTF | TRBV12-4 | TRBJ2-2 | CASSASPGLAISLDTGELFF | KLGGALQAK | NO | NO | NO |
| 16869 | TRAV19 | TRAJ29 | CALSEVDSGNTPLVF | TRBV7-9 | TRBJ2-7 | CASSPPGEGEQYF | AVFDRKSDAK | YES | NO | YES |
| 16870 | TRAV14DV4 | TRAJ41 | CAMREANSGYALNF | TRBV20-1 | TRBJ1-4 | CSASGDREVSGEKLFF | KLGGALQAK | YES | NO | NO |
| 16871 | TRAV3 | TRAJ31 | CAVRDLNNARLMF | TRBV5-6 | TRBJ2-1 | CASSLGGQQFF | KLGGALQAK | NO | NO | NO |
| 16872 | TRAV22 | TRAJ56 | CAVEAGANSKLTF | TRBV7-9 | TRBJ2-6 | CASSLDWTGANVLTF | KLGGALQAK | YES | NO | NO |
| 16873 | TRAV22 | TRAJ44 | CAGAPLTGTASKLTF | TRBV7-9 | TRBJ2-3 | CASSASSGGAYYTQYF | KLGGALQAK | YES | NO | NO |
| 16874 | TRAV3 | TRAJ40 | CAVRDTAGTYKYIF | TRBV3-1 | TRBJ2-7 | CASSQDRSGGGYEQYF | KLGGALQAK | NO | NO | NO |
| 16875 | TRAV12-1 | TRAJ50 | CVVNTPNTLKTSYDKVIF | TRBV7-6 | TRBJ2-7 | CASSLGQGLAYEQYF | KLGGALQAK | NO | NO | NO |
| 16876 | TRAV12-3 | TRAJ37 | CAMAGSGNTGKLIF | TRBV7-9 | TRBJ2-7 | CASSDSLGDRGSYEQYF | KLGGALQAK | YES | NO | NO |
| 16877 | TRAV12-1 | TRAJ45 | CVGRGGGADGLTF | TRBV6-5 | TRBJ1-2 | CASSYGTRSGYTF | KLGGALQAK | YES | NO | NO |
| 16878 | TRAV14DV4 | TRAJ49 | CAMRELRTGNQFYF | TRBV7-9 | TRBJ1-1 | CASSATGLTNTEAFF | KLGGALQAK | YES | NO | NO |
| 16879 | TRAV8-3 | TRAJ52 | CAALSAGGTSYGKLTF | TRBV27 | TRBJ2-1 | CASIPLAGVRDEQFF | AVFDRKSDAK | YES | NO | YES |
| 16880 | TRAV6 | TRAJ26 | CALEGNYGQNFVF | TRBV7-2 | TRBJ2-3 | CASSQGTILTDTQYF | KLGGALQAK | YES | NO | NO |
| 16881 | TRAV3 | TRAJ22 | CAVRDPGSARQLTF | TRBV5-1 | TRBJ2-2 | CASSEPGQGYTGELFF | KLGGALQAK | YES | NO | NO |
| 16883 | TRAV13-2 | TRAJ39 | CAETHNAGNMLTF | TRBV7-9 | TRBJ2-3 | CASSWHSSTDTQYF | AVFDRKSDAK | YES | NO | YES |
| 16884 | TRAV8-3 | TRAJ15 | CAVPNQAGTALIF | TRBV2 | TRBJ2-7 | CASSDWDSSYEQYF | KLGGALQAK | YES | NO | NO |
| 16885 | TRAV1-1 | TRAJ42 | CALNGGSQGNLIF | TRBV28 | TRBJ2-4 | CATGGSGNIQYF | KLGGALQAK | YES | NO | NO |
| 16886 | TRAV25 | TRAJ36 | CAGLGETGANNLFF | TRBV27 | TRBJ1-2 | CASSGLVDNGYTF | KLGGALQAK | YES | NO | NO |
| 16887 | TRAV8-2 | TRAJ48 | CVVSDLDFGNEKLTF | TRBV6-5 | TRBJ2-5 | CASSYSAGLETQYF | KLGGALQAK | NO | NO | NO |
| 16888 | TRAV5 | TRAJ12 | CAEMDSSYKLIF | TRBV2 | TRBJ1-1 | CASIPDMNTEAFF | KLGGALQAK | YES | NO | NO |
| 16889 | TRAV8-1 | TRAJ20 | CAVNPINDYKLSF | TRBV7-8 | TRBJ2-1 | CASSLGYEQFF | KLGGALQAK | YES | NO | NO |
| 16890 | TRAV29DV5 | TRAJ42 | CAASGLGGSQGNLIF | TRBV27 | TRBJ2-7 | CASSFEGPSYEQYF | KLGGALQAK | YES | NO | NO |
| 16891 | TRAV19 | TRAJ36 | CALSGVQTGANNLFF | TRBV4-3 | TRBJ2-1 | CASSQDPGLGSNEQFF | KLGGALQAK | YES | NO | NO |
| 16892 | TRAV13-1 | TRAJ42 | CAASRPYGGSQGNLIF | TRBV4-1 | TRBJ2-2 | CASSQATYTNTGELFF | AVFDRKSDAK | YES | YES | YES |
| 16893 | TRAV8-3 | TRAJ15 | CAVVILSRQAGTALIF | TRBV19 | TRBJ1-2 | CATTDGTGELEGYTF | KLGGALQAK | YES | NO | NO |
| 16894 | TRAV12-3 | TRAJ39 | CAIGPGNMLTF | TRBV7-9 | TRBJ2-2 | CASSPPRDRGTGELFF | KLGGALQAK | YES | NO | NO |
| 16895 | TRAV13-2 | TRAJ40 | CAWSGTYKYIF | TRBV7-3 | TRBJ2-3 | CASSLTAGGPADTQYF | KLGGALQAK | YES | NO | NO |
| 16896 | TRAV27 | TRAJ34 | CAGDNTDKLIF | TRBV7-3 | TRBJ2-3 | CASSPSYGTQYF | KLGGALQAK | YES | NO | NO |
| 16897 | TRAV12-2 | TRAJ10 | CAVTGGGNKLTF | TRBV4-3 | TRBJ1-4 | CASSQDHRMGGHEKLFF | KLGGALQAK | YES | NO | NO |
| 16898 | TRAV4 | TRAJ43 | CLVGESHDMRF | TRBV7-2 | TRBJ2-7 | CASSLGANRFYEQYF | KLGGALQAK | NO | NO | NO |
| 16899 | TRAV29DV5 | TRAJ37 | CAASGSSNTGKLIF | TRBV7-9 | TRBJ2-7 | CASSSTDQQYV | KLGGALQAK | NO | NO | NO |
| 16900 | TRAV13-1 | TRAJ47 | CAAPQEYGNKLVF | TRBV12-4 | TRBJ2-7 | CASSPGLATYEQYF | KLGGALQAK | YES | NO | NO |
| 16901 | TRAV21 | TRAJ15 | CAVSSQAGTALIF | TRBV19 | TRBJ1-5 | CASSIDEGGNQPQHF | KLGGALQAK | YES | NO | NO |
| 16902 | TRAV27 | TRAJ42 | CADNYGGSQGNLIF | TRBV19 | TRBJ2-3 | CASSISGQGTDTQYF | KLGGALQAK | YES | NO | NO |
| 16903 | TRAV1-2 | TRAJ4 | CAVFSGGYNKLIF | TRBV27 | TRBJ1-3 | CASSFGGAGNTIYF | KLGGALQAK | YES | NO | NO |
| 16904 | TRAV5 | TRAJ21 | CAESKTFNKFYF | TRBV4-1 | TRBJ2-7 | CASSQGENEQYF | KLGGALQAK | YES | NO | NO |
| 16905 | TRAV13-1 | TRAJ49 | CAAQTGNQFYF | TRBV15 | TRBJ2-1 | CATSLGTGGWDEQFF | KLGGALQAK | YES | NO | NO |
| 16906 | TRAV9-2 | TRAJ52 | CALSRPGANAGGTSYGKLTF | TRBV7-2 | TRBJ2-7 | CASSLEPSYEQYF | AVFDRKSDAK | NO | NO | NO |
| 16907 | TRAV8-3 | TRAJ13 | CAVEGGYQKVTF | TRBV7-8 | TRBJ1-2 | CASSLGYGYTF | KLGGALQAK | YES | NO | NO |
| 16908 | TRAV5 | TRAJ42 | CAVLNYGGSQGNLIF | TRBV5-1 | TRBJ2-3 | CASSATSGSTDTQYF | KLGGALQAK | YES | NO | NO |
| 16909 | TRAV20 | TRAJ40 | CAVRPGTYKYIF | TRBV30 | TRBJ2-2 | CAWSDGSVGELFF | KLGGALQAK | YES | NO | NO |
| 16910 | TRAV24 | TRAJ16 | CACAADGQKLLF | TRBV19 | TRBJ1-1 | CASNPGSLNTEAFF | KLGGALQAK | NO | NO | NO |
| 16911 | TRAV39 | TRAJ52 | CAVPNAGGTSYGKLTF | TRBV19 | TRBJ1-5 | CASSMVGHQPQHF | KLGGALQAK | YES | NO | NO |
| 16912 | TRAV6 | TRAJ20 | CAPLDFDDYKLSF | TRBV13 | TRBJ2-6 | CASSLRLAEISGANVLTF | KLGGALQAK | NO | NO | NO |
| 16913 | TRAV22 | TRAJ48 | CAVQNFGNEKLTF | TRBV19 | TRBJ2-2 | CASSITGVTGELFF | KLGGALQAK | YES | NO | NO |
| 16914 | TRAV4 | TRAJ26 | CLVGGGQNFVF | TRBV7-8 | TRBJ2-7 | CASSLSEGPTGPYEQYV | KLGGALQAK | NO | NO | NO |
| 16915 | TRAV8-3 | TRAJ45 | CAVGPLYSGGGADGLTF | TRBV18 | TRBJ2-1 | CASSPGLAGGPLEQFF | KLGGALQAK | YES | NO | NO |
| 16916 | TRAV13-1 | TRAJ45 | CAASQFLVYSGGGADGLTF | TRBV19 | TRBJ2-1 | CASSQGGNEQFF | KLGGALQAK | YES | NO | NO |
| 16917 | TRAV19 | TRAJ5 | CALSEALGGRRALTF | TRBV7-9 | TRBJ2-7 | CASKDGISYEQYF | KLGGALQAK | YES | NO | NO |
| 16918 | TRAV22 | TRAJ45 | CAGGDSGGGADGLTF | TRBV25-1 | TRBJ2-1 | CASSERTPPPYEQFF | KLGGALQAK | NO | NO | NO |
| 16919 | TRAV4 | TRAJ45 | CLVGEYSGGGADGLTF | TRBV4-1 | TRBJ1-1 | CASSQGTGGTEAFF | KLGGALQAK | YES | NO | NO |
| 16920 | TRAV22 | TRAJ20 | CAGTNDYKLSF | TRBV9 | TRBJ1-3 | CASSVSFFSGNTIYF | KLGGALQAK | NO | NO | NO |
| 16921 | TRAV29DV5 | TRAJ48 | CAASATNFGNEKLTF | TRBV6-3 | TRBJ2-7 | CASSPGGADFTTYEQYV | AVFDRKSDAK | NO | NO | NO |
| 16922 | TRAV13-1 | TRAJ26 | CAASNYGQNFVF | TRBV27 | TRBJ1-2 | CASSLSIGGAYGYTF | KLGGALQAK | YES | NO | NO |
| 16923 | TRAV12-1 | TRAJ9 | CVADTGGFKTIF | TRBV5-4 | TRBJ1-2 | CASSDLTDYGYTF | KLGGALQAK | YES | NO | NO |
| 16924 | TRAV17 | TRAJ54 | CATDPFQGAQKLVF | TRBV5-1 | TRBJ2-5 | CASSPGTSGKETQYF | KLGGALQAK | YES | NO | NO |
| 16925 | TRAV13-2 | TRAJ48 | CADPVGNEKLTF | TRBV12-4 | TRBJ1-2 | CASSLSLVGGSDGYTF | KLGGALQAK | YES | NO | NO |
| 16926 | TRAV29DV5 | TRAJ45 | CAAREGLYSGGGADGLTF | TRBV10-3 | TRBJ1-1 | CAISETGVAPNTEAFF | KLGGALQAK | YES | NO | NO |
| 16927 | TRAV12-2 | TRAJ32 | CAVKRYGGATNKLIF | TRBV7-3 | TRBJ2-2 | CASSRTAGLGANTGELFF | KLGGALQAK | NO | NO | NO |
| 16928 | TRAV12-3 | TRAJ23 | CAMSGGKLIF | TRBV18 | TRBJ1-2 | CASSPATADGYTF | KLGGALQAK | YES | NO | NO |
| 16929 | TRAV39 | TRAJ20 | CAVNPYKLSF | TRBV11-2 | TRBJ2-3 | CASSLTPGLAGGSTDTQYF | KLGGALQAK | YES | NO | NO |
| 16930 | TRAV12-1 | TRAJ53 | CVVNMDSGGSNYKLTF | TRBV4-3 | TRBJ2-2 | CASSQWGTGELFF | KLGGALQAK | YES | NO | NO |
| 16931 | TRAV12-2 | TRAJ5 | CAVTGGRRALTF | TRBV27 | TRBJ2-3 | CASSLSGPGQGFADTQYF | KLGGALQAK | YES | NO | NO |
| 16932 | TRAV12-2 | TRAJ26 | CVRTGQNFVF | TRBV10-2 | TRBJ2-2 | CASSELGLTGELFF | AVFDRKSDAK | YES | NO | NO |
| 16933 | TRAV14DV4 | TRAJ22 | CAMRRPISSGSARQLTF | TRBV6-5 | TRBJ2-5 | CASSEGLAGGGGETQYF | KLGGALQAK | NO | NO | NO |
| 16934 | TRAV19 | TRAJ30 | CALKHNRDDKIIF | TRBV20-1 | TRBJ2-3 | CSARRAGTGVFTDTQYF | KLGGALQAK | YES | NO | NO |
| 16935 | TRAV13-2 | TRAJ42 | CAETDGGSQGNLIF | TRBV14 | TRBJ1-3 | CASRRRTGNTIYF | KLGGALQAK | NO | NO | NO |
| 16936 | TRAV35 | TRAJ42 | CAGQSGRVGGSQGNLIF | TRBV7-9 | TRBJ2-7 | CASKDGISYEQYF | KLGGALQAK | YES | NO | NO |
| 16937 | TRAV12-1 | TRAJ3 | CVASRGASKIIF | TRBV5-4 | TRBJ1-2 | CASTSDRDYGYTF | KLGGALQAK | YES | NO | NO |
| 16938 | TRAV12-1 | TRAJ50 | CVVSLKTSYDKVIF | TRBV6-5 | TRBJ2-7 | CASSWNDRLQSYEQYF | KLGGALQAK | YES | NO | NO |
| 16939 | TRAV26-2 | TRAJ42 | CIPDGGSQGNLIF | TRBV27 | TRBJ2-2 | CASSDTGTSNTGELFF | KLGGALQAK | YES | NO | NO |
| 16940 | TRAV9-2 | TRAJ13 | CALSDLGGYQKVTF | TRBV14 | TRBJ2-1 | CASSPDNPLALPYNEQFF | KLGGALQAK | YES | NO | NO |
| 16941 | TRAV12-1 | TRAJ48 | CVVPYFGNEKLTF | TRBV4-1 | TRBJ1-1 | CASSQYRVDYTEAFF | KLGGALQAK | YES | NO | NO |
| 16942 | TRAV3 | TRAJ21 | CAALLYNFNKFYF | TRBV6-3 | TRBJ2-1 | CASSPVLVNEQFF | KLGGALQAK | NO | NO | NO |
| 16943 | TRAV30 | TRAJ21 | CGTAGGNFNKFYF | TRBV19 | TRBJ1-2 | CASTPGGPGYGYTF | RLRAEAQVK | YES | NO | YES |
| 16944 | TRAV38-2DV8 | TRAJ54 | CAYSFQGAQKLVF | TRBV6-3 | TRBJ2-1 | CASSYGAGGYNEQFF | AVFDRKSDAK | YES | NO | YES |
| 16945 | TRAV34 | TRAJ48 | CGAANFGNEKLTF | TRBV29-1 | TRBJ2-7 | CSVELGDSYEQYF | KLGGALQAK | YES | NO | NO |
| 16946 | TRAV5 | TRAJ23 | CAESGYNQGGKLIF | TRBV6-1 | TRBJ2-2 | CASSEAYMNTGELFF | KLGGALQAK | YES | NO | NO |
| 16947 | TRAV3 | TRAJ34 | CAVRPDNTDKLIF | TRBV2 | TRBJ2-1 | CASSPSWYNEQFF | KLGGALQAK | NO | NO | NO |
| 16948 | TRAV8-3 | TRAJ33 | CAVGADSNYQLIW | TRBV9 | TRBJ2-7 | CASSAYWAGKYEQYF | KLGGALQAK | YES | NO | NO |
| 16949 | TRAV6 | TRAJ24 | CALEATDSWGKLQF | TRBV6-1 | TRBJ2-1 | CASSVDSNSYNEQFF | KLGGALQAK | YES | NO | NO |
| 16950 | TRAV29DV5 | TRAJ27 | CAASAAGKSTF | TRBV20-1 | TRBJ2-1 | CSAREGVYEQFF | KLGGALQAK | YES | NO | NO |
| 16951 | TRAV5 | TRAJ31 | CAEINNNARLMF | TRBV6-5 | TRBJ1-2 | CASSYSIRDGYTF | KLGGALQAK | YES | NO | NO |
| 16952 | TRAV13-2 | TRAJ28 | CAENRSGAGSYQLTF | TRBV7-9 | TRBJ2-5 | CASSSTEETQYF | KLGGALQAK | YES | NO | NO |
| 16953 | TRAV17 | TRAJ54 | CATDLQGAQKLVF | TRBV11-1 | TRBJ1-1 | CASSSGTDEAFF | KLGGALQAK | YES | NO | NO |
| 16954 | TRAV17 | TRAJ7 | CATVGWRGNNRLAF | TRBV7-2 | TRBJ2-5 | CASAPSGETQYF | KLGGALQAK | YES | NO | NO |
| 16955 | TRAV8-1 | TRAJ48 | CAVQNEKLTF | TRBV20-1 | TRBJ2-3 | CSARAHTDGADTQYF | KLGGALQAK | NO | NO | NO |
| 16956 | TRAV19 | TRAJ40 | CALSEAGTYKYIF | TRBV19 | TRBJ1-1 | CASSIGNTEAFF | KLGGALQAK | YES | NO | NO |
| 16957 | TRAV14DV4 | TRAJ52 | CAMREGGYAGGTSYGKLTF | TRBV27 | TRBJ2-1 | CASSLLGGGHNEQFF | KLGGALQAK | YES | NO | NO |
| 16958 | TRAV5 | TRAJ47 | CAESLPMEYGNKLVF | TRBV4-3 | TRBJ2-1 | CASSQVLVGRDSYNEQFF | KLGGALQAK | YES | YES | NO |
| 16959 | TRAV1-2 | TRAJ40 | CAVKDSGTYKYIF | TRBV2 | TRBJ2-7 | CASSGLGGSSYEQYF | KLGGALQAK | YES | NO | NO |
| 16960 | TRAV29DV5 | TRAJ56 | CAASGTGANSKLTF | TRBV5-4 | TRBJ2-1 | CASSGLAIEQFF | KLGGALQAK | YES | NO | NO |
| 16961 | TRAV8-1 | TRAJ42 | CAVNWDGGSQGNLIF | TRBV20-1 | TRBJ2-7 | CSARVGYEQYF | KLGGALQAK | NO | NO | NO |
| 16962 | TRAV29DV5 | TRAJ34 | CAASGDTDKLIF | TRBV7-9 | TRBJ1-2 | CASSAVEDYGYTF | KLGGALQAK | YES | NO | NO |
| 16963 | TRAV17 | TRAJ23 | CATGFYNQGGKLIF | TRBV27 | TRBJ1-1 | CASSSGQGGTEAFF | KLGGALQAK | YES | NO | NO |
| 16964 | TRAV29DV5 | TRAJ39 | CAASGNAGNMLTF | TRBV5-4 | TRBJ2-7 | CASSLGTGSYEQYF | KLGGALQAK | YES | NO | NO |
| 16965 | TRAV12-3 | TRAJ39 | CAIGPGNMLTF | TRBV14 | TRBJ1-1 | CASSQGGVTEAFF | KLGGALQAK | NO | NO | NO |
| 16966 | TRAV9-2 | TRAJ32 | CALRDGGATNKLIF | TRBV20-1 | TRBJ2-1 | CSARDSYSGTYNEQFF | KLGGALQAK | NO | NO | NO |
| 16967 | TRAV14DV4 | TRAJ47 | CAMREGKYGNKLVF | TRBV4-3 | TRBJ1-1 | CASSQDEGQGAEAFF | AVFDRKSDAK | YES | NO | YES |
| 16968 | TRAV25 | TRAJ24 | CGRMTTDSWGKFQF | TRBV19 | TRBJ1-1 | CASSTSMGEAFF | KLGGALQAK | YES | NO | NO |
| 16969 | TRAV12-2 | TRAJ58 | CAVTKTSGSRLTF | TRBV9 | TRBJ1-1 | CASSDNRDRGGGTEAFF | KLGGALQAK | YES | NO | NO |
| 16970 | TRAV12-2 | TRAJ28 | CAVESGAGSYQLTF | TRBV12-5 | TRBJ1-1 | CASGYNTEAFF | ELAGIGILTV | YES | NO | NO |
| 16971 | TRAV12-2 | TRAJ48 | CAVTPISNFGNEKLTF | TRBV13 | TRBJ1-1 | CASSLFLPAGNTEAFF | KLGGALQAK | YES | NO | NO |

| | | | | | | | | | | |
|---|---|---|---|---|---|---|---|---|---|---|
| 16972 | TRAV12-2 | TRAJ43 | CAVDRYNNNDMRF | TRBV12-5 | TRBJ2-3 | CASDTRQADTQYF | KLGGALQAK | YES | NO | NO |
| 16973 | TRAV38-2DV8 | TRAJ31 | CAYRSASNNARLMF | TRBV2 | TRBJ1-5 | CASSPVGGYQPQHF | KLGGALQAK | YES | NO | NO |
| 16974 | TRAV24 | TRAJ33 | CAASGNYQLIW | TRBV10-1 | TRBJ2-1 | CASSVDQLGYNEQFF | AVFDRKSDAK | YES | NO | YES |
| 16975 | TRAV14DV4 | TRAJ4 | CAMRGPISGGYNKLIF | TRBV11-3 | TRBJ2-1 | CASSSERDAFDNEQFF | KLGGALQAK | YES | NO | NO |
| 16976 | TRAV36DV7 | TRAJ44 | CVLNTGTASKLTF | TRBV19 | TRBJ2-1 | CASSHGGRYNEQFF | KLGGALQAK | NO | NO | NO |
| 16977 | TRAV17 | TRAJ33 | CATDGEDSNYQLIW | TRBV3-1 | TRBJ1-2 | CASSQEGTGNYGYTF | KLGGALQAK | YES | NO | NO |
| 16978 | TRAV35 | TRAJ24 | CAGYSLTDSWGKLQF | TRBV7-2 | TRBJ2-3 | CASSFQDTQYF | KLGGALQAK | YES | NO | NO |
| 16979 | TRAV17 | TRAJ53 | CATENSGGSNYKLTF | TRBV11-2 | TRBJ2-5 | CASRGPVETQYF | KLGGALQAK | YES | NO | NO |
| 16980 | TRAV5 | TRAJ15 | CAESNSNQAGTALIF | TRBV6-3 | TRBJ1-2 | CASRGGQSLGYTF | KLGGALQAK | YES | NO | NO |
| 16981 | TRAV19 | TRAJ10 | CALSEDILTGGGNKLTF | TRBV7-9 | TRBJ2-3 | CASSSRLAGGLSDTQYF | KLGGALQAK | YES | NO | NO |
| 16982 | TRAV14DV4 | TRAJ22 | CAMKDPGSARQLTF | TRBV10-3 | TRBJ2-7 | CAIGDYEQYV | AVFDRKSDAK | NO | NO | NO |
| 16983 | TRAV12-2 | TRAJ50 | CAVNPTSYDKVIF | TRBV27 | TRBJ1-2 | CASSFGGASGYTF | ELAGIGILTV | YES | YES | YES |
| 16984 | TRAV20 | TRAJ39 | CAVGNNAGNMLTF | TRBV4-1 | TRBJ2-1 | CASSQGTSNNEQFF | KLGGALQAK | YES | NO | NO |
| 16985 | TRAV13-2 | TRAJ9 | CAETVNTGGFKTIF | TRBV5-1 | TRBJ2-7 | CASSLGTDYEQYF | KLGGALQAK | YES | NO | NO |
| 16986 | TRAV9-2 | TRAJ52 | CALAGGTSYGKLTF | TRBV2 | TRBJ2-5 | CASSESGFDSETQYF | KLGGALQAK | YES | NO | NO |
| 16987 | TRAV10 | TRAJ34 | CVVSGYNTDKLIF | TRBV5-6 | TRBJ2-1 | CASSLFGGRYNEQFF | KLGGALQAK | YES | NO | NO |
| 16988 | TRAV17 | TRAJ43 | CATPGGYNNNDMRF | TRBV6-3 | TRBJ2-2 | CASSYGHNTGELFF | KLGGALQAK | YES | NO | NO |
| 16989 | TRAV12-2 | TRAJ11 | CAVNSGYSTLTF | TRBV7-6 | TRBJ2-3 | CASSWQLAGPHTDTQYF | KLGGALQAK | YES | NO | NO |
| 16990 | TRAV14DV4 | TRAJ22 | CAMRRPISSGSARQLTF | TRBV12-4 | TRBJ1-4 | CASSLGSGRLMEKLFF | KLGGALQAK | YES | NO | NO |
| 16991 | TRAV8-4 | TRAJ21 | CAVKANFNKFYF | TRBV20-1 | TRBJ1-6 | CSAPKDGPLHF | KLGGALQAK | YES | NO | NO |
| 16992 | TRAV12-1 | TRAJ30 | CVVNTGDKIIF | TRBV7-9 | TRBJ1-5 | CASSLYWDQPQHF | KLGGALQAK | YES | NO | NO |
| 16993 | TRAV34 | TRAJ57 | CGADGGGSEKLVF | TRBV5-1 | TRBJ2-7 | CASSYPPSGGVAYEQYF | KLGGALQAK | YES | NO | NO |
| 16994 | TRAV14DV4 | TRAJ28 | CAMREPAYSGAGSYQLTF | TRBV4-2 | TRBJ1-6 | CASSQGGYQGRNSPLHF | KLGGALQAK | YES | NO | NO |
| 16995 | TRAV8-2 | TRAJ34 | CVVSDDGTDKLIF | TRBV19 | TRBJ2-1 | CASSIAPGGLAGIEQFF | KLGGALQAK | YES | NO | NO |
| 16996 | TRAV3 | TRAJ18 | CAVRDKWLRGSTLGRLYF | TRBV27 | TRBJ2-5 | CASSHYKYNTGGQETQYF | KLGGALQAK | NO | NO | NO |
| 16997 | TRAV17 | TRAJ42 | CATDEGGSQGNLIF | TRBV12-3 | TRBJ2-5 | CASTAKQDFQETQYF | KLGGALQAK | YES | NO | NO |
| 16998 | TRAV8-3 | TRAJ8 | CAVRMNTGFQKLVF | TRBV27 | TRBJ2-3 | CASSLQGHSTDTQYF | KLGGALQAK | YES | NO | NO |
| 16999 | TRAV14DV4 | TRAJ48 | CAMREGISNFGNEKLTF | TRBV24-1 | TRBJ2-5 | CATSDRGEGIETQYF | KLGGALQAK | YES | NO | NO |
| 17000 | TRAV8-4 | TRAJ20 | CAVSDVGDYKLSF | TRBV20-1 | TRBJ2-1 | CSARFSGGGRSEQFF | KLGGALQAK | NO | NO | NO |
| 17001 | TRAV8-1 | TRAJ42 | CAVNVGRSQGNLIF | TRBV24-1 | TRBJ2-1 | CATSSTDDYKHYNEQFF | KLGGALQAK | YES | NO | NO |
| 17002 | TRAV17 | TRAJ43 | CATPLNNDMRF | TRBV7-9 | TRBJ1-4 | CASSLGGEKLFF | KLGGALQAK | YES | NO | NO |
| 17003 | TRAV9-2 | TRAJ41 | CALSNSNSGYALNF | TRBV7-9 | TRBJ2-1 | CASSDLMGLALDEQFF | KLGGALQAK | YES | NO | NO |
| 17004 | TRAV12-2 | TRAJ13 | CAVASFYQKVTF | TRBV3-1 | TRBJ2-7 | CASSQDPGTGTDEQYF | AVFDRKSDAK | YES | NO | YES |
| 17005 | TRAV30 | TRAJ40 | CGTESVLPFSGTYKYIF | TRBV12-4 | TRBJ2-5 | CASSWVMETQYF | KLGGALQAK | YES | NO | NO |
| 17006 | TRAV21 | TRAJ52 | CAVRPAGGTSYGKLTF | TRBV13 | TRBJ2-1 | CASSLGGYEQFF | KLGGALQAK | YES | NO | NO |
| 17007 | TRAV8-2 | TRAJ43 | CVVSDRINDMRF | TRBV5-1 | TRBJ1-5 | CASSLGLNQPQHF | KLGGALQAK | YES | NO | NO |
| 17008 | TRAV8-3 | TRAJ42 | CARGGSQGNLIF | TRBV10-2 | TRBJ2-7 | CASAPGEQPYEQYF | KLGGALQAK | YES | NO | NO |
| 17009 | TRAV8-4 | TRAJ37 | CAVSDSGNTGKLIF | TRBV25-1 | TRBJ1-1 | CASSEGDRGGWNTEAFF | KLGGALQAK | YES | NO | NO |
| 17010 | TRAV38-2DV8 | TRAJ48 | CAYRAPRGNEKLTF | TRBV7-2 | TRBJ2-7 | CASSLAFGVEQYV | KLGGALQAK | NO | NO | NO |
| 17011 | TRAV12-1 | TRAJ15 | CVAGNQAGTALIF | TRBV7-2 | TRBJ2-5 | CASSLGGPGETQYF | KLGGALQAK | YES | NO | NO |
| 17012 | TRAV8-1 | TRAJ45 | CAVIGGSGGGADGLTF | TRBV7-8 | TRBJ1-1 | CASSALQGWMTEAFF | KLGGALQAK | YES | NO | NO |
| 17013 | TRAV19 | TRAJ29 | CALSEAFSGNTPLVF | TRBV11-2 | TRBJ2-5 | CASRGPVETQYF | KLGGALQAK | YES | NO | NO |
| 17014 | TRAV27 | TRAJ37 | CAGVGSSNTGKLIF | TRBV29-1 | TRBJ1-1 | CSVRPPGIF | KLGGALQAK | NO | NO | NO |
| 17015 | TRAV26-2 | TRAJ37 | CILSSGNTGKLIF | TRBV15 | TRBJ1-1 | CATSRQGATEAFF | KLGGALQAK | YES | NO | NO |
| 17016 | TRAV30 | TRAJ42 | CGTDHLLNYGGSQGNLIF | TRBV12-4 | TRBJ1-4 | CASSLEVDNEKLFF | KLGGALQAK | YES | NO | NO |
| 17017 | TRAV13-1 | TRAJ48 | CAASVFFGNEKLTF | TRBV5-6 | TRBJ2-7 | CASSDSYEQYF | KLGGALQAK | YES | NO | NO |
| 17019 | TRAV8-3 | TRAJ4 | CAVGPSGGYNKLIF | TRBV27 | TRBJ2-3 | CASSPPGPVVTDTQYF | KLGGALQAK | YES | NO | NO |
| 17020 | TRAV12-2 | TRAJ43 | CAATVTGDMRF | TRBV15 | TRBJ1-2 | CATSREVVDDGYTF | KLGGALQAK | YES | NO | NO |
| 17021 | TRAV29DV5 | TRAJ43 | CAASARDNDMRF | TRBV7-2 | TRBJ1-1 | CASSQLGQGHTEAFF | KLGGALQAK | NO | NO | NO |
| 17022 | TRAV1-2 | TRAJ6 | CAVRDTSTGGSYIPTF | TRBV4-2 | TRBJ2-3 | CASSSRGGLAGDSTDTQYF | KLGGALQAK | YES | NO | NO |
| 17023 | TRAV5 | TRAJ24 | CAETDSWGKLQF | TRBV12-3 | TRBJ1-4 | CASSFVESNEKLFF | KLGGALQAK | YES | NO | NO |
| 17024 | TRAV12-1 | TRAJ5 | CVIWDTGRRALTF | TRBV4-1 | TRBJ1-1 | CASSSGPEAFF | AVFDRKSDAK | YES | NO | YES |
| 17025 | TRAV25 | TRAJ26 | CAARDNYGQNFVF | TRBV9 | TRBJ2-1 | CASSVLAGDNEQFF | KLGGALQAK | YES | NO | NO |
| 17026 | TRAV10 | TRAJ39 | CVVSALSAGNMLTF | TRBV6-5 | TRBJ1-5 | CASSLTRDSSPQHF | KLGGALQAK | YES | NO | NO |
| 17027 | TRAV3 | TRAJ30 | CAVRPMNRDDKIIF | TRBV3-1 | TRBJ2-7 | CASSQFRTSLSYEQYF | KLGGALQAK | NO | NO | NO |
| 17028 | TRAV39 | TRAJ48 | CAVDMGGLSNFGNEKLTF | TRBV7-7 | TRBJ2-2 | CASSAFVLAANTGELFF | KLGGALQAK | YES | NO | NO |
| 17030 | TRAV8-6 | TRAJ9 | CAGGFKTIF | TRBV6-1 | TRBJ2-2 | CASSEPQTSGNTGELFF | KLGGALQAK | YES | NO | NO |
| 17031 | TRAV3 | TRAJ43 | CAVRAYNNNDMRF | TRBV24-1 | TRBJ2-3 | CATSVSGGADTQYF | KLGGALQAK | NO | NO | NO |
| 17032 | TRAV35 | TRAJ43 | CAGSLSNNDMRF | TRBV6-5 | TRBJ1-2 | CASSYGRSYGYTF | KLGGALQAK | YES | NO | NO |
| 17033 | TRAV29DV5 | TRAJ40 | CAAISSSGTYKYIF | TRBV20-1 | TRBJ1-1 | CSAPDRASTEAFF | KLGGALQAK | YES | NO | NO |
| 17034 | TRAV1-2 | TRAJ29 | CAVRSSGNTPLVF | TRBV14 | TRBJ1-2 | CASSQVPGQGPNYGYTF | KLGGALQAK | NO | NO | NO |
| 17035 | TRAV16 | TRAJ28 | CAHAYSGAGSYQLTF | TRBV29-1 | TRBJ1-1 | CSVVTGGTLNTEAFF | KLGGALQAK | NO | NO | NO |
| 17036 | TRAV8-1 | TRAJ22 | CAVRSSGSARQLTF | TRBV3-1 | TRBJ1-2 | CASSQGRDRHGYTF | KLGGALQAK | YES | NO | NO |
| 17037 | TRAV13-2 | TRAJ26 | CAEPDNYGQNFVF | TRBV11-2 | TRBJ2-3 | CASSISGGASDTQYF | KLGGALQAK | YES | NO | NO |
| 17038 | TRAV14DV4 | TRAJ40 | CAMRESTSGTYKYIF | TRBV9 | TRBJ2-1 | CASSVGAVTYNEQFF | KLGGALQAK | YES | NO | NO |
| 17039 | TRAV29DV5 | TRAJ12 | CAASHSSYKLIF | TRBV27 | TRBJ1-1 | CASSFELNTEAFF | KLGGALQAK | YES | NO | NO |
| 17040 | TRAV12-3 | TRAJ41 | CAMKDSGYALNF | TRBV11-2 | TRBJ1-4 | CASSSMLTEKLFF | KLGGALQAK | YES | NO | NO |
| 17041 | TRAV38-1 | TRAJ29 | CAFMNTPLVF | TRBV11-2 | TRBJ2-3 | CASSLGHDLADTQYF | KLGGALQAK | YES | NO | NO |
| 17042 | TRAV21 | TRAJ57 | CAGPFQGGSEKLVF | TRBV19 | TRBJ2-2 | CASSATGGPTGELFF | KLGGALQAK | YES | NO | NO |
| 17043 | TRAV8-3 | TRAJ23 | CAVGAYQGGKLIF | TRBV7-9 | TRBJ2-2 | CASSPGQGMSGELFF | KLGGALQAK | YES | NO | NO |
| 17044 | TRAV19 | TRAJ12 | CALGRGWMDSSYKLIF | TRBV14 | TRBJ1-2 | CASSQDPQVVGYTF | KLGGALQAK | NO | NO | NO |
| 17045 | TRAV21 | TRAJ57 | CAAELVF | TRBV7-3 | TRBJ2-1 | CASSGAGTSGRNEQFF | KLGGALQAK | YES | NO | NO |
| 17046 | TRAV29DV5 | TRAJ45 | CAASGYSGGGADGLTF | TRBV7-9 | TRBJ1-1 | CASSLELDEAFF | KLGGALQAK | YES | NO | NO |
| 17047 | TRAV14DV4 | TRAJ4 | CAMRPPFRGGYNKLIF | TRBV6-5 | TRBJ2-1 | CASSTGLTHNEQFF | KLGGALQAK | NO | NO | NO |
| 17048 | TRAV4 | TRAJ39 | CLVGDSNAGNMLTF | TRBV5-1 | TRBJ1-4 | CASSLGIMTASTNEKLFF | KLGGALQAK | YES | NO | NO |
| 17049 | TRAV14DV4 | TRAJ21 | CAMREGPNNFNKFYF | TRBV5-6 | TRBJ2-1 | CASSASDNLDNEQFF | KLGGALQAK | YES | NO | NO |
| 17050 | TRAV41 | TRAJ43 | CAVRSYNNNDMRF | TRBV6-6 | TRBJ1-3 | CASSRREASGNTIYF | KLGGALQAK | YES | NO | NO |
| 17051 | TRAV23DV6 | TRAJ27 | CAATPNTNAGKSTF | TRBV11-2 | TRBJ1-4 | CASSLDQWRGQREKLFF | KLGGALQAK | YES | NO | NO |
| 17052 | TRAV21 | TRAJ43 | CAVWYMRF | TRBV5-6 | TRBJ2-5 | CASSFQGSPQETQYF | KLGGALQAK | NO | NO | NO |
| 17053 | TRAV5 | TRAJ15 | CAESTSPNQAGTALIF | TRBV7-9 | TRBJ2-1 | CASSFPRIGDNEQFF | KLGGALQAK | YES | NO | NO |
| 17054 | TRAV12-1 | TRAJ7 | CVRGSYGNNRLAF | TRBV7-2 | TRBJ1-1 | CASRSTGSPEAFF | KLGGALQAK | NO | NO | NO |
| 17055 | TRAV21 | TRAJ37 | CAQHGSGNTGKLIF | TRBV7-9 | TRBJ2-7 | CASSSTLIHYEQYF | KLGGALQAK | YES | NO | NO |
| 17056 | TRAV20 | TRAJ33 | CAVRADSNYQLIW | TRBV3-1 | TRBJ2-3 | CASSPSEWGGGDTDTQYF | IVTDFSVIK | YES | YES | YES |
| 17057 | TRAV16 | TRAJ33 | CALGMDSNYQLIW | TRBV30 | TRBJ2-3 | CAWSAFSGTDTQYF | KLGGALQAK | NO | NO | NO |
| 17058 | TRAV26-2 | TRAJ58 | CILTETSGSRLTF | TRBV2 | TRBJ2-7 | CASTGTGVSYEQYF | KLGGALQAK | YES | NO | NO |
| 17059 | TRAV39 | TRAJ44 | CAAVTGTASKLTF | TRBV20-1 | TRBJ1-1 | CSARVFYTFFRGGFKSAEAFF | KLGGALQAK | NO | NO | NO |
| 17060 | TRAV1-2 | TRAJ26 | CAVKPSGGQNFVF | TRBV12-3 | TRBJ1-1 | CASSPQAFF | KLGGALQAK | NO | NO | NO |
| 17061 | TRAV12-2 | TRAJ50 | CAVNGGKTSYDKVIF | TRBV24-1 | TRBJ2-2 | CATDSGLNTGELFF | KLGGALQAK | YES | NO | NO |
| 17062 | TRAV5 | TRAJ13 | CAESIAGYQKVTF | TRBV7-9 | TRBJ1-1 | CASSSGRDTEAFF | KLGGALQAK | YES | NO | NO |
| 17063 | TRAV17 | TRAJ24 | CATAAALTTDSWGKLQF | TRBV5-4 | TRBJ2-1 | CASSWGQGDEQFF | KLGGALQAK | YES | NO | NO |
| 17064 | TRAV29DV5 | TRAJ26 | CAASEDYGQNFVF | TRBV18 | TRBJ2-5 | CASSPSGTEETQYF | KLGGALQAK | YES | NO | NO |
| 17065 | TRAV9-2 | TRAJ16 | CALSDPPYSLSDGQKLLF | TRBV5-1 | TRBJ2-7 | CASSLGRSQNEQYF | KLGGALQAK | NO | NO | NO |
| 17066 | TRAV12-2 | TRAJ45 | CAVNTYSGGGADGLTF | TRBV7-9 | TRBJ1-1 | CASRLFGTEAFF | KLGGALQAK | YES | NO | NO |
| 17067 | TRAV12-1 | TRAJ49 | CVVPYGGNQFYF | TRBV24-1 | TRBJ2-2 | CATTGTGTGELFF | KLGGALQAK | YES | NO | NO |
| 17068 | TRAV21 | TRAJ26 | CAVIPDDNYGQNFVF | TRBV29-1 | TRBJ1-2 | CSVDLEANYGYTF | KLGGALQAK | YES | NO | NO |
| 17069 | TRAV39 | TRAJ49 | CAGLKSNTGNQFYF | TRBV19 | TRBJ1-5 | CASSPWTGGSNQPQHF | KLGGALQAK | YES | NO | NO |
| 17070 | TRAV35 | TRAJ56 | CAGPSSGGANSKLTF | TRBV27 | TRBJ2-5 | CASSLFPRVETQYF | KLGGALQAK | YES | NO | NO |
| 17071 | TRAV23DV6 | TRAJ52 | CAARSNAGGTSYGKLTF | TRBV19 | TRBJ2-3 | CASSLGSFTDTQYF | KLGGALQAK | YES | NO | NO |
| 17072 | TRAV27 | TRAJ33 | CAGPSDSNYQLIW | TRBV24-1 | TRBJ1-6 | CATSAGLSSPLHF | KLGGALQAK | YES | NO | NO |
| 17073 | TRAV21 | TRAJ24 | CAVKDSWGKFQF | TRBV11-3 | TRBJ1-1 | CASSPPGQMNTEAFF | KLGGALQAK | YES | NO | NO |
| 17074 | TRAV8-6 | TRAJ29 | CAVSENSGNTPLVF | TRBV9 | TRBJ2-7 | CASSSQWSSYEQYF | KLGGALQAK | YES | NO | NO |
| 17075 | TRAV8-1 | TRAJ34 | CAVSTGEDKLIF | TRBV10-2 | TRBJ2-7 | CASSEISGGSEQYF | AVFDRKSDAK | YES | NO | YES |
| 17076 | TRAV14DV4 | TRAJ26 | CAMRDPGQNFVF | TRBV9 | TRBJ1-5 | CASSVGSDGSNQPQHF | KLGGALQAK | YES | NO | NO |
| 17077 | TRAV14DV4 | TRAJ48 | CAMRTRSNFGNEKLTF | TRBV7-3 | TRBJ1-1 | CASSTVVGGTEAFF | KLGGALQAK | NO | NO | NO |

| | | | | | | | | | | |
|---|---|---|---|---|---|---|---|---|---|---|
| 17078 | TRAV13-2 | TRAJ52 | CAETSGGTSYGKLTF | TRBV7-2 | TRBJ2-7 | CASSLAVNYEQYV | KLGGALQAK | NO | NO | NO |
| 17079 | TRAV12-3 | TRAJ45 | CAMSGGGADGLTF | TRBV13 | TRBJ1-1 | CASSSGGQRATEAFF | KLGGALQAK | YES | NO | NO |
| 17080 | TRAV19 | TRAJ40 | CALSESSPGTYKYIF | TRBV24-1 | TRBJ1-2 | CATKGTESNYGYTF | KLGGALQAK | YES | NO | NO |
| 17081 | TRAV13-2 | TRAJ47 | CAEMGEYGNKLVF | TRBV5-4 | TRBJ2-7 | CASSGGQGDDEQYF | KLGGALQAK | YES | NO | NO |
| 17082 | TRAV14DV4 | TRAJ26 | CAMREGNYGQNFVF | TRBV6-5 | TRBJ1-1 | CASSGQFMNTEAFF | KLGGALQAK | YES | NO | NO |
| 17083 | TRAV38-2DV8 | TRAJ54 | CAYYQGAQKLVF | TRBV20-1 | TRBJ2-1 | CSAHTPAFGRTSSYNEQFF | KLGGALQAK | NO | NO | NO |
| 17084 | TRAV19 | TRAJ54 | CALSGEIIQGAQKLVF | TRBV5-1 | TRBJ1-1 | CASSLTGDTEAFF | KLGGALQAK | YES | NO | NO |
| 17085 | TRAV17 | TRAJ9 | CATPRRTGGFKTIF | TRBV5-6 | TRBJ1-4 | CASSLGPTGGDEKLFF | KLGGALQAK | YES | NO | NO |
| 17086 | TRAV8-3 | TRAJ52 | CAVEDNAGGTSYGKLTF | TRBV5-1 | TRBJ2-5 | CASRYLIVDQETQYF | KLGGALQAK | YES | NO | NO |
| 17087 | TRAV8-6 | TRAJ44 | CAVTLGDTGTASKLTF | TRBV7-6 | TRBJ2-7 | CASSQGLGTDSYEQYF | KLGGALQAK | YES | NO | NO |
| 17088 | TRAV1-2 | TRAJ33 | CAVRDSNYQLIW | TRBV7-8 | TRBJ2-1 | CASSLIEASGGFEQFF | KLGGALQAK | YES | NO | NO |
| 17089 | TRAV38-1 | TRAJ40 | CAFMNPTSGTYKYIF | TRBV10-1 | TRBJ1-2 | CASSDFETPDYYGYTF | KLGGALQAK | YES | NO | NO |
| 17090 | TRAV14DV4 | TRAJ22 | CAMRRPISSGSARQLTF | TRBV12-4 | TRBJ2-2 | CASSGREDNTGELFF | KLGGALQAK | YES | NO | NO |
| 17091 | TRAV19 | TRAJ3 | CALSEAYSSASKIIF | TRBV5-6 | TRBJ2-2 | CASSLLPGTSETGELFF | KLGGALQAK | YES | NO | NO |
| 17092 | TRAV14DV4 | TRAJ11 | CAMTMGTLTF | TRBV11-2 | TRBJ1-3 | CASSPFNVGAGNTIYF | KLGGALQAK | YES | NO | NO |
| 17093 | TRAV17 | TRAJ53 | CATPTYSGGSNYKLTF | TRBV5-6 | TRBJ2-7 | CASSPGQGVSEQYF | KLGGALQAK | YES | NO | NO |
| 17094 | TRAV4 | TRAJ45 | CLVGDTSLSRGGADGLTF | TRBV4-1 | TRBJ1-2 | CASSQDLGQGIFYGYTF | KLGGALQAK | YES | NO | NO |
| 17095 | TRAV21 | TRAJ26 | CAVMDNYGQNFVF | TRBV5-5 | TRBJ2-5 | CASSLRDGFPQTQYF | KLGGALQAK | YES | NO | NO |
| 17096 | TRAV20 | TRAJ20 | CALIRGDYKLSF | TRBV7-9 | TRBJ1-1 | CASSLARGVTEAFF | KLGGALQAK | YES | NO | NO |
| 17097 | TRAV25 | TRAJ20 | CAAYSNDYKLSF | TRBV5-1 | TRBJ2-1 | CASSLGTGAIYNEQFF | KLGGALQAK | YES | NO | NO |
| 17098 | TRAV8-6 | TRAJ30 | CAVSDSEVYRDDKIIF | TRBV12-5 | TRBJ1-1 | CASGLGNPGSGVTEAFF | KLGGALQAK | YES | NO | NO |
| 17099 | TRAV13-2 | TRAJ53 | CAENTGGGGSNYKLTF | TRBV4-1 | TRBJ2-1 | CASSQDWVLAGTQEQFF | KLGGALQAK | YES | NO | NO |
| 17100 | TRAV12-2 | TRAJ15 | CAVNLWNQAGTALIF | TRBV9 | TRBJ2-5 | CASSEDGAEETQYF | KLGGALQAK | YES | NO | NO |
| 17101 | TRAV24 | TRAJ34 | CASPFYTDKLIF | TRBV7-8 | TRBJ2-1 | CASSLDTSSYNEQFF | KLGGALQAK | YES | NO | NO |
| 17102 | TRAV24 | TRAJ16 | CASWF | TRBV7-9 | TRBJ1-1 | CASSHTGGYTEAFF | KLGGALQAK | NO | NO | NO |
| 17103 | TRAV41 | TRAJ42 | CAVFDGGSQGNLIF | TRBV19 | TRBJ2-1 | CASTRGGSSYNEQFF | KLGGALQAK | YES | NO | NO |
| 17104 | TRAV1-2 | TRAJ31 | CAVRDDNNARLMF | TRBV12-5 | TRBJ1-2 | CASGLGPAGYTF | KLGGALQAK | YES | NO | NO |
| 17105 | TRAV22 | TRAJ40 | CAVDESGTYKYIF | TRBV2 | TRBJ2-7 | CAISSGYEQYF | KLGGALQAK | YES | NO | NO |
| 17106 | TRAV20 | TRAJ40 | CAVQEDTTTSGTYKYIF | TRBV9 | TRBJ1-2 | CASSEGTVNYGYTF | AVFDRKSDAK | YES | NO | YES |
| 17107 | TRAV1-2 | TRAJ28 | CAVRRAPL | TRBV12-5 | TRBJ1-5 | CASAPAGGTDPIYSNQPQHF | KLGGALQAK | NO | NO | NO |
| 17108 | TRAV8-6 | TRAJ52 | CAVSEMSGTSYGKLTF | TRBV2 | TRBJ2-1 | CASSLGQLSYNEQFF | KLGGALQAK | YES | NO | NO |
| 17109 | TRAV26-1 | TRAJ8 | CIVRVTGFQKLVF | TRBV27 | TRBJ2-5 | CASSLGTGSMGETQYF | RLRAEAQVK | YES | YES | YES |
| 17110 | TRAV1-2 | TRAJ8 | CAVRPGGMNTGFQKLVF | TRBV12-5 | TRBJ2-3 | CASGLPQGGSFVTDTQYF | KLGGALQAK | YES | NO | NO |
| 17112 | TRAV14DV4 | TRAJ45 | CAMRERGGADGLTF | TRBV19 | TRBJ2-7 | CASSITGNSFEQYV | KLGGALQAK | NO | NO | NO |
| 17113 | TRAV14DV4 | TRAJ7 | CAMRGVKGNNRLAF | TRBV2 | TRBJ1-1 | CASSEEVSTEAFF | KLGGALQAK | YES | NO | NO |
| 17114 | TRAV17 | TRAJ31 | CATDDARLMF | TRBV5-4 | TRBJ1-1 | CASSPGQREAFF | KLGGALQAK | YES | NO | NO |
| 17115 | TRAV14DV4 | TRAJ20 | CAMRVNDYKLSF | TRBV11-3 | TRBJ2-1 | CASSLDGAGGPWDEQFF | KLGGALQAK | YES | NO | NO |
| 17116 | TRAV3 | TRAJ29 | CAVRDNSGNTPLVF | TRBV4-1 | TRBJ1-1 | CASSQRQGLGNTEAFF | KLGGALQAK | YES | NO | NO |
| 17117 | TRAV12-3 | TRAJ31 | CAMSANNNARLMF | TRBV19 | TRBJ1-2 | CASSARTGVGYGYTF | KLGGALQAK | YES | NO | NO |
| 17118 | TRAV9-2 | TRAJ3 | CALSDPPRGSSASKIIF | TRBV19 | TRBJ2-3 | CASSIKTSGDPDTQYF | KLGGALQAK | YES | NO | NO |
| 17119 | TRAV21 | TRAJ6 | CAVPTGGSYIPTF | TRBV11-2 | TRBJ2-7 | CASSLGSNEQYV | KLGGALQAK | NO | NO | NO |
| 17120 | TRAV12-2 | TRAJ16 | CAVGPLQRF | TRBV4-1 | TRBJ2-2 | CASSQAMGTGELFF | KLGGALQAK | NO | NO | NO |
| 17121 | TRAV23DV6 | TRAJ23 | CAAYYNQGGKLIF | TRBV27 | TRBJ1-1 | CASFRGFTEAFF | KLGGALQAK | YES | NO | NO |
| 17122 | TRAV5 | TRAJ11 | CAESGYSTLTF | TRBV7-9 | TRBJ1-2 | CASSLAGQGAHGYTF | KLGGALQAK | YES | NO | NO |
| 17123 | TRAV12-2 | TRAJ37 | CAVDSGNTGKLIF | TRBV12-4 | TRBJ1-2 | CASSLVLQRIYGYTF | AVFDRKSDAK | YES | NO | YES |
| 17124 | TRAV17 | TRAJ26 | CATDDNYGQNFVF | TRBV5-4 | TRBJ2-5 | CASSLAETQYF | KLGGALQAK | YES | NO | NO |
| 17125 | TRAV29DV5 | TRAJ52 | CAASDTSYGKLTF | TRBV19 | TRBJ1-1 | CASSVDRGYNTEAFF | KLGGALQAK | YES | NO | NO |
| 17126 | TRAV12-2 | TRAJ43 | CAVNINDMRF | TRBV24-1 | TRBJ2-2 | CATGAGSNTGELFF | KLGGALQAK | YES | NO | NO |
| 17127 | TRAV12-2 | TRAJ24 | CAVTHGGRKTTDSWGKFQF | TRBV7-9 | TRBJ2-1 | CASSLVKGQGGDEQFF | KLGGALQAK | YES | NO | NO |
| 17128 | TRAV1-1 | TRAJ32 | CAVRDDGGATNKLIF | TRBV4-1 | TRBJ2-1 | CASSRAVDNEQFF | KLGGALQAK | YES | NO | NO |
| 17129 | TRAV13-1 | TRAJ32 | CAASDGGATNKLIF | TRBV10-2 | TRBJ2-2 | CAIGTSGNTGELFF | KLGGALQAK | YES | NO | NO |
| 17130 | TRAV12-2 | TRAJ28 | CAVIVLAYSGAGSYQLTF | TRBV11-2 | TRBJ2-7 | CASSLVSGGDYEQYV | KLGGALQAK | NO | NO | NO |
| 17131 | TRAV19 | TRAJ56 | CALSEAAGANSKLTF | TRBV4-1 | TRBJ1-2 | CASSHRMGPDYGYTF | KLGGALQAK | YES | NO | NO |
| 17132 | TRAV30 | TRAJ26 | CGTEDNYGQNFVF | TRBV5-4 | TRBJ1-1 | CASTLREGLNTEAFF | KLGGALQAK | YES | NO | NO |
| 17133 | TRAV29DV5 | TRAJ48 | CAASAGFGNEKLTF | TRBV29-1 | TRBJ2-1 | CSGTGSSYNEQFF | KLGGALQAK | YES | NO | NO |
| 17134 | TRAV25 | TRAJ29 | CAGNSGNTPLVF | TRBV19 | TRBJ2-1 | CASSIGNYNEQFF | KLGGALQAK | YES | NO | NO |
| 17135 | TRAV12-1 | TRAJ15 | CVVNQAGTALIF | TRBV10-3 | TRBJ2-1 | CATSGQGAPEQFF | KLGGALQAK | YES | NO | NO |
| 17136 | TRAV12-2 | TRAJ6 | CAVNRSGGSYIPTF | TRBV9 | TRBJ1-5 | CASSEGDSNQPQHF | KLGGALQAK | YES | NO | NO |
| 17137 | TRAV25 | TRAJ32 | CAGPGYGGATNKLIF | TRBV7-9 | TRBJ2-1 | CASSLEGGLAAYNEQFF | AVFDRKSDAK | YES | YES | YES |
| 17138 | TRAV35 | TRAJ30 | CAGLENRDDKIIF | TRBV5-1 | TRBJ2-7 | CASSFAPGRGYEQYV | KLGGALQAK | NO | NO | NO |
| 17139 | TRAV8-4 | TRAJ36 | CAVSDQVAGANNLFF | TRBV11-2 | TRBJ2-5 | CASSLASGGFETQYF | KLGGALQAK | YES | NO | NO |
| 17140 | TRAV13-1 | TRAJ44 | CAASKSAGTASKLTF | TRBV18 | TRBJ1-5 | CASSHSGDQPQHF | AVFDRKSDAK | YES | NO | NO |
| 17141 | TRAV27 | TRAJ21 | CAGADFNKFYF | TRBV3-1 | TRBJ2-3 | CASSQDRANTDTQYF | KLGGALQAK | YES | NO | NO |
| 17142 | TRAV5 | TRAJ34 | CAERLPYNTDKLIF | TRBV7-6 | TRBJ2-1 | CASSLDRLGDNEQFF | AVFDRKSDAK | YES | NO | YES |
| 17143 | TRAV22 | TRAJ45 | CAAGGGGADGLTF | TRBV10-3 | TRBJ1-1 | CAISGMNTEAFF | KLGGALQAK | YES | NO | NO |
| 17144 | TRAV27 | TRAJ37 | CAGGSGNTGKLIF | TRBV2 | TRBJ2-1 | CASSEAMSAYEQFF | KLGGALQAK | YES | NO | NO |
| 17145 | TRAV26-2 | TRAJ58 | CIFLRGETSGSRLTF | TRBV19 | TRBJ2-2 | CASSIWENTGELFF | KLGGALQAK | YES | NO | NO |
| 17146 | TRAV26-2 | TRAJ54 | CILFQGAQKLVF | TRBV27 | TRBJ2-7 | CASSPTLAGTYEQYF | KLGGALQAK | YES | NO | NO |
| 17147 | TRAV14DV4 | TRAJ11 | CAMREVNSGYSTLTF | TRBV6-3 | TRBJ2-1 | CASSYGAGGYNEQFF | KLGGALQAK | YES | NO | NO |
| 17148 | TRAV38-2DV8 | TRAJ53 | CASGSNYKLTF | TRBV7-9 | TRBJ1-1 | CASSSNDRYGTEAFF | KLGGALQAK | YES | NO | NO |
| 17149 | TRAV17 | TRAJ23 | CATEGNQGGKLIF | TRBV27 | TRBJ2-2 | CASSWGPGELFF | SLFNTVATLY | YES | YES | YES |
| 17150 | TRAV8-6 | TRAJ49 | CAVSLTGNQFYF | TRBV7-9 | TRBJ1-2 | CASSLPLSQRDYGYTF | AVFDRKSDAK | YES | NO | NO |
| 17151 | TRAV21 | TRAJ43 | CAVTGNNDMRF | TRBV12-5 | TRBJ1-3 | CASRSSGNTIYF | KLGGALQAK | YES | NO | NO |
| 17152 | TRAV8-4 | TRAJ54 | CAVSVQGAQKLVF | TRBV6-3 | TRBJ2-1 | CASSYGAGGYNEQFF | KLGGALQAK | YES | NO | NO |
| 17153 | TRAV12-1 | TRAJ39 | CVVNDGGNAGNMLTF | TRBV19 | TRBJ1-5 | CASSMSVGFSNQPQHF | KLGGALQAK | YES | NO | NO |
| 17154 | TRAV8-1 | TRAJ39 | CAVNAGNMLTF | TRBV25-1 | TRBJ1-2 | CASSEQGGGYTF | AVFDRKSDAK | YES | NO | YES |
| 17155 | TRAV13-2 | TRAJ20 | CAENGGSNDYKLSF | TRBV6-1 | TRBJ1-1 | CASREGMTEAFF | KLGGALQAK | YES | NO | NO |
| 17156 | TRAV13-1 | TRAJ44 | CAASIAGTASKLTF | TRBV27 | TRBJ1-1 | CASSLHQVDTEAFF | KLGGALQAK | YES | NO | NO |
| 17157 | TRAV39 | TRAJ30 | CAVVPMNRDDKIIF | TRBV12-4 | TRBJ2-4 | CASSSAAFVNIQYF | KLGGALQAK | YES | YES | NO |
| 17158 | TRAV23DV6 | TRAJ58 | CAASSQLETSGSRLTF | TRBV27 | TRBJ1-2 | CASKGVRNYGYTF | KLGGALQAK | YES | NO | NO |
| 17159 | TRAV26-2 | TRAJ23 | CILRLYNQGGKLIF | TRBV6-5 | TRBJ1-5 | CASSYNGQLNQPQHF | KLGGALQAK | YES | NO | NO |
| 17160 | TRAV12-2 | TRAJ4 | CAVGSGGYNKLIF | TRBV4-1 | TRBJ2-4 | CASSPDSDRGNIQYF | KLGGALQAK | YES | NO | NO |
| 17161 | TRAV19 | TRAJ36 | CALRAHESILYQTGANNLFF | TRBV19 | TRBJ2-1 | CASTEAGGPGNEQFF | KLGGALQAK | NO | NO | NO |
| 17162 | TRAV12-3 | TRAJ34 | CAMGGNTDKLIF | TRBV11-2 | TRBJ2-5 | CASSLGTSGTETQYF | KLGGALQAK | YES | NO | NO |
| 17163 | TRAV29DV5 | TRAJ45 | CAALPAGGGADGLTF | TRBV29-1 | TRBJ1-3 | CSVDGLLTGEGGNTIYF | KLGGALQAK | YES | NO | NO |
| 17164 | TRAV25 | TRAJ42 | CAGTGSQGNLIF | TRBV7-8 | TRBJ1-1 | CASSRFGQGLVGTEAFF | KLGGALQAK | YES | NO | NO |
| 17165 | TRAV8-6 | TRAJ13 | CAAFSGGYQKVTF | TRBV20-1 | TRBJ2-1 | CSPGQGPYNEQFF | KLGGALQAK | YES | NO | NO |
| 17166 | TRAV1-2 | TRAJ39 | CAVRDDNNAGNMLTF | TRBV11-1 | TRBJ1-2 | CASSYGAGGDGYTF | KLGGALQAK | YES | NO | NO |
| 17167 | TRAV14DV4 | TRAJ49 | CAMREVDTGNQFYF | TRBV9 | TRBJ2-5 | CASNRGGPIETQYF | KLGGALQAK | YES | NO | NO |
| 17168 | TRAV5 | TRAJ18 | CAESGGRGSTLGRLYF | TRBV5-1 | TRBJ2-3 | CASSFTSGGDTDTQYF | KLGGALQAK | YES | NO | NO |
| 17169 | TRAV14DV4 | TRAJ4 | CAMRPPFRGGYNKLIF | TRBV20-1 | TRBJ2-7 | CSARDLEVLSYEQYF | KLGGALQAK | YES | NO | NO |
| 17170 | TRAV29DV5 | TRAJ28 | CAASAVNSGAGSYQLTF | TRBV7-9 | TRBJ2-5 | CASSFREPSSQYF | KLGGALQAK | YES | NO | NO |
| 17171 | TRAV3 | TRAJ20 | CAVRPLSNDYKLSF | TRBV7-9 | TRBJ2-7 | CASSLMDGDLYEQYV | KLGGALQAK | NO | NO | NO |
| 17173 | TRAV8-3 | TRAJ10 | CAVGVTGGGNKLTF | TRBV2 | TRBJ2-1 | CASRPDPTGFFGNEQFF | KLGGALQAK | YES | NO | NO |
| 17174 | TRAV5 | TRAJ17 | CAEAAAGNKLTF | TRBV7-2 | TRBJ2-2 | CASSPLTSGITGELFF | KLGGALQAK | YES | NO | NO |
| 17175 | TRAV38-1 | TRAJ40 | CAFILTTTSGTYKYIF | TRBV2 | TRBJ2-5 | CASRGWETQYF | KLGGALQAK | YES | NO | NO |
| 17176 | TRAV22 | TRAJ34 | CAVPPYNTDKLIF | TRBV4-3 | TRBJ2-3 | CASSQGVGGNTDTQYF | KLGGALQAK | YES | NO | NO |
| 17177 | TRAV4 | TRAJ13 | CLAPSGGYQKVTF | TRBV4-1 | TRBJ2-1 | CASSPQGGTPNEQFF | KLGGALQAK | NO | NO | NO |
| 17178 | TRAV8-1 | TRAJ41 | CAVPLPGYALNF | TRBV6-3 | TRBJ2-1 | CASSYLRGSYNEQFF | KLGGALQAK | YES | NO | NO |
| 17179 | TRAV13-1 | TRAJ49 | CAELTGNQFYF | TRBV11-3 | TRBJ2-1 | CASSLGGGPYNEQFF | KLGGALQAK | YES | NO | NO |
| 17180 | TRAV27 | TRAJ28 | CAGQRGAGSYQLTF | TRBV30 | TRBJ1-5 | CAWSWTTGGNQPQHF | KLGGALQAK | YES | NO | NO |
| 17181 | TRAV3 | TRAJ43 | CAVRAYNNNDMRF | TRBV20-1 | TRBJ2-5 | CSARESPPPIEETQYF | KLGGALQAK | YES | NO | NO |
| 17182 | TRAV1-1 | TRAJ29 | CAVRSQNSGNTPLVF | TRBV27 | TRBJ2-1 | CASGGADEQFF | KLGGALQAK | YES | NO | NO |
| 17183 | TRAV1-1 | TRAJ34 | CAADNTDKLIF | TRBV9 | TRBJ2-5 | CASSEDFRKQETQYF | KLGGALQAK | YES | NO | NO |

| | | | | | | | | | | |
|---|---|---|---|---|---|---|---|---|---|---|
| 17184 | TRAV9-2 | TRAJ45 | CARRTPRYSGGGADGLTF | TRBV5-1 | TRBJ2-7 | CASSGPTGAYEQYF | KLGGALQAK | YES | NO | NO |
| 17185 | TRAV17 | TRAJ33 | CATPRRDSNYQLIW | TRBV6-5 | TRBJ2-1 | CASSYAISYNEQFF | KLGGALQAK | NO | NO | NO |
| 17186 | TRAV14DV4 | TRAJ4 | CAMRPAFSGGYNKLIF | TRBV5-8 | TRBJ1-2 | CASSTGLNYGYTF | KLGGALQAK | YES | NO | NO |
| 17187 | TRAV19 | TRAJ34 | CAPHITPTSSSL | TRBV4-3 | TRBJ2-2 | CASTAGVETGELFF | KLGGALQAK | YES | NO | NO |
| 17188 | TRAV1-2 | TRAJ33 | CAVREDSNYQLIW | TRBV7-9 | TRBJ1-5 | CASSLGGGEQPQHF | KLGGALQAK | YES | NO | NO |
| 17189 | TRAV12-2 | TRAJ47 | CAVMEYGNKLVF | TRBV19 | TRBJ2-7 | CASSPAGLAYEQYF | KLGGALQAK | YES | NO | NO |
| 17190 | TRAV12-1 | TRAJ49 | CVVNHNTGNQFYF | TRBV6-1 | TRBJ1-2 | CASSEVAESSYGYTF | KLGGALQAK | YES | NO | NO |
| 17191 | TRAV19 | TRAJ8 | CALRVGTGFQKLVF | TRBV7-2 | TRBJ2-7 | CASSLVTDSYEQYV | KLGGALQAK | NO | NO | NO |
| 17192 | TRAV6 | TRAJ6 | CALASGGSYIPTF | TRBV7-9 | TRBJ1-2 | CASSLSEDNYGYTF | KLGGALQAK | YES | NO | NO |
| 17193 | TRAV23DV6 | TRAJ10 | CAASNRLTGGGNKLTF | TRBV5-4 | TRBJ1-2 | CASLLQGGGYTF | KLGGALQAK | YES | NO | NO |
| 17194 | TRAV12-2 | TRAJ57 | CAVNAEKLVF | TRBV29-1 | TRBJ1-1 | CSVVGPNTEAFF | KLGGALQAK | YES | NO | NO |
| 17195 | TRAV19 | TRAJ32 | CALSELGGATNKLIF | TRBV7-8 | TRBJ1-6 | CASSLVGLPSSYNSPLHF | KLGGALQAK | YES | NO | NO |
| 17196 | TRAV8-6 | TRAJ26 | CAVIRTYGQNFVF | TRBV27 | TRBJ1-1 | CASSFTEPGTEAFF | KLGGALQAK | YES | NO | NO |
| 17197 | TRAV8-4 | TRAJ48 | CAVTLNFGNEKLTF | TRBV4-1 | TRBJ2-1 | CASSQKRVNEQFF | KLGGALQAK | YES | NO | NO |
| 17198 | TRAV12-2 | TRAJ34 | CAVKRNTDKLIF | TRBV2 | TRBJ2-4 | CASKGPENIQYF | KLGGALQAK | YES | NO | NO |
| 17199 | TRAV29DV5 | TRAJ30 | CAASAVGDKIIF | TRBV4-1 | TRBJ1-1 | CASSQPLGGSTEAFF | KLGGALQAK | YES | NO | NO |
| 17200 | TRAV19 | TRAJ49 | CALKENTGNQFYF | TRBV20-1 | TRBJ2-2 | CSASDTVDLAGELFF | AVFDRKSDAK | YES | NO | NO |
| 17201 | TRAV17 | TRAJ23 | CATDAFMIYNQGGKLIF | TRBV15 | TRBJ2-5 | CATSGEGQQQETQYF | KLGGALQAK | YES | NO | NO |
| 17202 | TRAV8-3 | TRAJ52 | CAVGADAGGTSYGKLTF | TRBV12-5 | TRBJ1-3 | CASGFFPLGLGNTIYF | KLGGALQAK | YES | NO | NO |
| 17203 | TRAV26-1 | TRAJ49 | CIVSPRTGNQFYF | TRBV5-5 | TRBJ1-5 | CASSSQGARNQPQHF | KLGGALQAK | YES | NO | NO |
| 17204 | TRAV14DV4 | TRAJ45 | CAMRELYSGGGADGLTF | TRBV7-6 | TRBJ1-1 | CASSSPGGGLNTEAFF | KLGGALQAK | YES | NO | NO |
| 17205 | TRAV13-1 | TRAJ20 | CAASWSDYKLSF | TRBV2 | TRBJ2-4 | CASNRLAGGPAKNIQYF | KLGGALQAK | YES | NO | NO |
| 17206 | TRAV12-2 | TRAJ9 | CAVPLNTGGFKTIF | TRBV5-4 | TRBJ2-5 | CASSLELAGGEGETQYF | KLGGALQAK | YES | NO | NO |
| 17207 | TRAV38-2DV8 | TRAJ45 | CAYRSLESGGGADGLTF | TRBV9 | TRBJ1-2 | CASSVAVSSFYGYTF | KLGGALQAK | YES | NO | NO |
| 17208 | TRAV21 | TRAJ53 | CAVNSGGSNYKLTF | TRBV2 | TRBJ1-1 | CASSERLGMNTEAFF | KLGGALQAK | YES | NO | NO |
| 17209 | TRAV19 | TRAJ27 | CALSEALAGKSTF | TRBV4-3 | TRBJ2-1 | CASSQSANLPGTLSNEQFF | KLGGALQAK | YES | NO | NO |
| 17210 | TRAV38-2DV8 | TRAJ48 | CAYRAPRGNEKLTF | TRBV9 | TRBJ2-7 | CASSPVSGSYEQYF | KLGGALQAK | YES | NO | NO |
| 17211 | TRAV12-3 | TRAJ53 | CAMSYRSGGSNYKLTF | TRBV7-2 | TRBJ1-2 | CASSPRGVGDGYTF | KLGGALQAK | YES | NO | NO |
| 17212 | TRAV30 | TRAJ40 | CGATLTSGTYKYIF | TRBV6-5 | TRBJ2-1 | CASSFLSEDRVNEQFF | KLGGALQAK | NO | NO | NO |
| 17213 | TRAV8-3 | TRAJ27 | CAVGEGDTNAGKSTF | TRBV3-1 | TRBJ2-1 | CASSPNRANEQFF | AVFDRKSDAK | YES | NO | YES |
| 17214 | TRAV8-3 | TRAJ42 | CAVGAGGGSQGNLIF | TRBV9 | TRBJ2-1 | CASSEAGGPYNEQFF | KLGGALQAK | YES | NO | NO |
| 17215 | TRAV8-2 | TRAJ43 | CVVSDPGDMRF | TRBV30 | TRBJ1-1 | CAWSEILNTEAFF | KLGGALQAK | YES | NO | NO |
| 17216 | TRAV8-3 | TRAJ4 | CAVGASPLSGGYNKLIF | TRBV11-2 | TRBJ1-2 | CASRAGKANYGYTF | KLGGALQAK | YES | NO | NO |
| 17217 | TRAV12-2 | TRAJ52 | CADAGGTSYGKLTF | TRBV4-1 | TRBJ2-1 | CASSQVKGGDEQFF | KLGGALQAK | YES | NO | NO |
| 17218 | TRAV4 | TRAJ24 | CLVGPPQTDSWGKFQF | TRBV10-2 | TRBJ2-2 | CASGTSGDTGELFF | KLGGALQAK | YES | NO | NO |
| 17219 | TRAV9-2 | TRAJ33 | CAPGNYQLIW | TRBV4-1 | TRBJ1-2 | CASSQELQSYGYTF | KLGGALQAK | YES | NO | NO |
| 17220 | TRAV35 | TRAJ7 | CAGTTDGNNRLAF | TRBV4-1 | TRBJ1-2 | CASSPGGHGYTF | KLGGALQAK | YES | NO | NO |
| 17221 | TRAV12-2 | TRAJ5 | CAVTGRRALTF | TRBV28 | TRBJ1-5 | CASSFFRDTIGNQPQHF | KLGGALQAK | YES | NO | NO |
| 17222 | TRAV19 | TRAJ40 | CALMGSPGTYKYIF | TRBV29-1 | TRBJ1-1 | CSVPYTPGTEAFF | KLGGALQAK | YES | NO | NO |
| 17223 | TRAV12-3 | TRAJ34 | CAMRAWNNTDKLIF | TRBV5-1 | TRBJ2-3 | CASRSVLGGDTQYF | KLGGALQAK | YES | NO | NO |
| 17224 | TRAV1-2 | TRAJ34 | CTTMTYTDKLIF | TRBV11-2 | TRBJ2-5 | CASSLATSGGYQETQYF | KLGGALQAK | YES | NO | NO |
| 17225 | TRAV10 | TRAJ45 | CVVSALPSGGGADGLTF | TRBV7-8 | TRBJ1-5 | CASSSSRAASNQPQHF | KLGGALQAK | YES | NO | NO |
| 17226 | TRAV21 | TRAJ41 | CAVRPPNSGYALNF | TRBV29-1 | TRBJ2-2 | CSVVAWSGELFF | KLGGALQAK | NO | NO | NO |
| 17227 | TRAV22 | TRAJ31 | CAVVANNARLMF | TRBV5-5 | TRBJ1-1 | CASAGPEAFF | KLGGALQAK | YES | NO | NO |
| 17228 | TRAV14DV4 | TRAJ52 | CAMREGAGGTSYGKLTF | TRBV19 | TRBJ1-2 | CASSTGHYGYTF | KLGGALQAK | YES | NO | NO |
| 17229 | TRAV4 | TRAJ3 | CLVGSWYSSASKIIF | TRBV30 | TRBJ2-2 | CAWAPTGVAGELFF | KLGGALQAK | NO | NO | NO |
| 17230 | TRAV13-1 | TRAJ27 | CAASGNAGKSTF | TRBV27 | TRBJ1-1 | CASSPGNTEAFF | RLRAEAQVK | YES | NO | NO |
| 17231 | TRAV38-2DV8 | TRAJ29 | CAENSGNTPLVF | TRBV5-8 | TRBJ1-6 | CASSWTGGANSPLHF | KLGGALQAK | YES | NO | NO |
| 17232 | TRAV41 | TRAJ52 | CAVRPPNAGGTSYGKLTF | TRBV6-5 | TRBJ1-6 | CASSYGDWDRHNSPLHF | KLGGALQAK | YES | NO | NO |
| 17233 | TRAV1-2 | TRAJ40 | CAGSGTYKYIF | TRBV5-1 | TRBJ2-5 | CASSRTGPGETQYF | KLGGALQAK | YES | NO | NO |
| 17234 | TRAV38-2DV8 | TRAJ34 | CAYRSLNTDKLIF | TRBV7-9 | TRBJ1-6 | CASRQGGTSPLHF | IVTDFSVIK | YES | NO | YES |
| 17235 | TRAV14DV4 | TRAJ40 | CAMREGPASGTYKYIF | TRBV4-3 | TRBJ2-3 | CASSQVGGRATDTQYF | KLGGALQAK | YES | NO | NO |
| 17236 | TRAV1-2 | TRAJ9 | CAVREAGGFKTIF | TRBV19 | TRBJ2-1 | CASSLLAGSDEQFF | AVFDRKSDAK | YES | NO | NO |
| 17237 | TRAV26-2 | TRAJ34 | CILRDASYNTDKLIF | TRBV3-1 | TRBJ2-4 | CASSDGLAGEAKNIQYF | KLGGALQAK | YES | NO | NO |
| 17238 | TRAV38-2DV8 | TRAJ31 | CAYRRFNNARLMF | TRBV4-1 | TRBJ2-7 | CASSHMYLYYEQYF | KLGGALQAK | YES | NO | NO |
| 17239 | TRAV19 | TRAJ42 | CALSDSMNYGGSQGNLIF | TRBV12-4 | TRBJ2-5 | CASSLDTVGPTGGEETQYF | KLGGALQAK | YES | NO | NO |
| 17240 | TRAV12-1 | TRAJ42 | CVVNADYGGSQGNLIF | TRBV6-1 | TRBJ2-7 | CASSDPQGAGNEQYF | KLGGALQAK | YES | NO | NO |
| 17241 | TRAV1-2 | TRAJ17 | CAPRAAGNKLTF | TRBV9 | TRBJ2-1 | CASSFTEGNEQFF | KLGGALQAK | YES | NO | NO |
| 17242 | TRAV5 | TRAJ40 | CAEEGASSGTYKYIF | TRBV7-9 | TRBJ1-1 | CASSVNRGEAFF | KLGGALQAK | YES | NO | NO |
| 17243 | TRAV1-2 | TRAJ37 | CAVRDTGKLIF | TRBV7-9 | TRBJ2-7 | CASSTRDGPYEQYF | KLGGALQAK | YES | NO | NO |
| 17244 | TRAV1-2 | TRAJ20 | CAVRGGNDYKLSF | TRBV19 | TRBJ2-3 | CASGAWADTQYF | KLGGALQAK | YES | NO | NO |
| 17245 | TRAV17 | TRAJ16 | CATDPSDGQKLLF | TRBV4-1 | TRBJ1-5 | CASSQWDSNQPQHF | KLGGALQAK | NO | NO | NO |
| 17246 | TRAV13-1 | TRAJ45 | CAATPRGGADGLTF | TRBV2 | TRBJ1-1 | CASSDRMNTEAFF | KLGGALQAK | YES | NO | NO |
| 17247 | TRAV12-2 | TRAJ3 | CAVPNSSASKIIF | TRBV6-5 | TRBJ2-1 | CAIGLAGANEQFF | KLGGALQAK | NO | NO | NO |
| 17248 | TRAV8-3 | TRAJ27 | CAADGNAGKSTF | TRBV11-2 | TRBJ2-1 | CASSPSLGTRYNEQFF | KLGGALQAK | YES | NO | NO |
| 17249 | TRAV10 | TRAJ40 | CVVSAYPGTYKYIF | TRBV7-2 | TRBJ2-7 | CASSPGTNFHEQYV | KLGGALQAK | NO | NO | NO |
| 17250 | TRAV8-3 | TRAJ13 | CAVAEGSGGYQKVTF | TRBV7-7 | TRBJ2-5 | CASSRGQGWETQYF | KLGGALQAK | YES | NO | NO |
| 17251 | TRAV8-6 | TRAJ13 | CAVSKVGGYQKVTF | TRBV18 | TRBJ2-5 | CASSLQGTQETQYF | KLGGALQAK | YES | NO | NO |
| 17252 | TRAV29DV5 | TRAJ43 | CAATRYNNNDMRF | TRBV12-3 | TRBJ1-5 | CASTREGAGWSAQHF | KLGGALQAK | YES | NO | NO |
| 17253 | TRAV13-1 | TRAJ50 | CAASSGTSYDKVIF | TRBV7-9 | TRBJ2-3 | CASSSVDGDTQYF | KLGGALQAK | YES | NO | NO |
| 17254 | TRAV9-2 | TRAJ43 | CALSDLNDMRF | TRBV7-9 | TRBJ1-2 | CASSHPPRWDGYTF | KLGGALQAK | YES | NO | NO |
| 17255 | TRAV21 | TRAJ8 | CAVPLTGFQKLVF | TRBV7-3 | TRBJ2-1 | CASSLISGYNEQFF | KLGGALQAK | YES | NO | NO |
| 17256 | TRAV22 | TRAJ45 | CAASGGGADGLTF | TRBV4-1 | TRBJ2-7 | CASSQATEQYF | KLGGALQAK | YES | NO | NO |
| 17257 | TRAV8-1 | TRAJ9 | CAVNTGGFKTIF | TRBV19 | TRBJ1-5 | CASTPGEVGNQPQHF | KLGGALQAK | YES | NO | NO |
| 17258 | TRAV6 | TRAJ49 | CALPRPNTGNQFYF | TRBV24-1 | TRBJ1-5 | CATSPGIHQPQHF | KLGGALQAK | YES | NO | NO |
| 17259 | TRAV20 | TRAJ34 | CAVQARTDKLIF | TRBV29-1 | TRBJ2-1 | CSGATGLSYNEQFF | KLGGALQAK | YES | NO | NO |
| 17260 | TRAV1-2 | TRAJ40 | CAVRPPTSGTYKYIF | TRBV6-6 | TRBJ1-4 | CASIQGTNEKLFF | KLGGALQAK | NO | NO | NO |
| 17261 | TRAV8-1 | TRAJ49 | CAVNYQFYF | TRBV7-8 | TRBJ1-2 | CASSLFEVNYGYTF | KLGGALQAK | NO | NO | NO |
| 17262 | TRAV13-1 | TRAJ42 | CAAGNDGGSQGNLIF | TRBV24-1 | TRBJ1-5 | CATSDLVANQPQHF | KLGGALQAK | YES | NO | NO |
| 17263 | TRAV3 | TRAJ16 | CAVRDGISDGQKLLF | TRBV9 | TRBJ1-1 | CASSADRFPNTEAFF | KLGGALQAK | NO | NO | NO |
| 17264 | TRAV20 | TRAJ34 | CAVQVISRNTDKLIF | TRBV9 | TRBJ1-5 | CASSVGSITPQHF | KLGGALQAK | YES | NO | NO |
| 17265 | TRAV14DV4 | TRAJ38 | CAMRGILGAGNNRKLIW | TRBV7-9 | TRBJ1-1 | CASSFNTEAFF | IVTDFSVIK | YES | NO | YES |
| 17266 | TRAV8-3 | TRAJ41 | CAVEQDSGYALNF | TRBV7-9 | TRBJ2-3 | CASSYPSGANKPTDTQYF | KLGGALQAK | YES | NO | NO |
| 17267 | TRAV20 | TRAJ24 | CAVQAGDSWGKFQF | TRBV18 | TRBJ2-1 | CASSPEQQGHEQFF | KLGGALQAK | YES | NO | NO |
| 17268 | TRAV6 | TRAJ37 | CALPSNTGKLIF | TRBV13 | TRBJ2-1 | CASSLNHEQFF | KLGGALQAK | YES | NO | NO |
| 17269 | TRAV1-1 | TRAJ33 | CAVRAHMDSNYQLIW | TRBV5-4 | TRBJ2-5 | CASSPGLAGAQETQYF | KLGGALQAK | YES | NO | NO |
| 17270 | TRAV21 | TRAJ57 | CAPSQGGSEKLVF | TRBV20-1 | TRBJ2-1 | CSARGDGAIINEQFF | KLGGALQAK | NO | NO | NO |
| 17271 | TRAV2 | TRAJ32 | CAVEDGGGATNKLIF | TRBV19 | TRBJ2-1 | CASRMDSYRNNEQFF | KLGGALQAK | NO | NO | NO |
| 17272 | TRAV14DV4 | TRAJ22 | CAMRGISSGSARQLTF | TRBV4-3 | TRBJ1-4 | CASSQDHRMGGHEKLFF | KLGGALQAK | YES | YES | NO |
| 17273 | TRAV10 | TRAJ7 | CVVSDQLWEQQTRF | TRBV20-1 | TRBJ1-5 | CSAGDRYSNQPQHF | KLGGALQAK | YES | NO | NO |
| 17274 | TRAV1-2 | TRAJ34 | CAVRGTYNTDKLIF | TRBV5-1 | TRBJ2-2 | CASSSTGELFF | FLASKIGRLV | YES | NO | YES |
| 17275 | TRAV12-1 | TRAJ9 | CVVNILNTGGFKTIF | TRBV25-1 | TRBJ2-7 | CASSEVGLSYEQYF | KLGGALQAK | YES | NO | NO |
| 17276 | TRAV13-2 | TRAJ16 | CAETGGQKLLF | TRBV19 | TRBJ1-1 | CASSTQGSGTEAFF | KLGGALQAK | NO | NO | NO |
| 17277 | TRAV34 | TRAJ49 | CGADGGNQFYF | TRBV18 | TRBJ2-5 | CASSPPGVQETQYF | KLGGALQAK | YES | NO | NO |
| 17279 | TRAV8-3 | TRAJ40 | CAVVSGTYKYIF | TRBV18 | TRBJ1-1 | CASSPEAGNMNTEAFF | AVFDRKSDAK | YES | NO | YES |
| 17280 | TRAV20 | TRAJ32 | CAGGGGATNKLIF | TRBV24-1 | TRBJ2-2 | CATSETGVENTGELFF | KLGGALQAK | YES | NO | NO |
| 17281 | TRAV27 | TRAJ20 | CAGAESNDYKLSF | TRBV29-1 | TRBJ1-1 | CSVAGTGNTEAFF | KLGGALQAK | YES | NO | NO |
| 17282 | TRAV30 | TRAJ3 | CGTFPPYSSASKIIF | TRBV6-3 | TRBJ1-1 | CASSYFGGTEAFF | KLGGALQAK | YES | NO | NO |
| 17283 | TRAV8-3 | TRAJ45 | CAVGRGGADGLTF | TRBV11-2 | TRBJ2-1 | CASSQARVSYNEQFF | KLGGALQAK | YES | NO | NO |
| 17284 | TRAV21 | TRAJ40 | CAVEFATSGTYKYIF | TRBV7-9 | TRBJ2-2 | CASSLDNAGELFF | KLGGALQAK | YES | NO | NO |
| 17285 | TRAV1-2 | TRAJ6 | CAVRGGSYIPTF | TRBV11-2 | TRBJ2-7 | CASSLGGGRVLYEQYV | KLGGALQAK | NO | NO | NO |
| 17286 | TRAV4 | TRAJ12 | CLVGGPMDSSYKLIF | TRBV29-1 | TRBJ2-1 | CSETSGTGFYEQFF | KLGGALQAK | YES | NO | NO |
| 17287 | TRAV17 | TRAJ5 | CAHSLDTGRRALTF | TRBV5-6 | TRBJ2-6 | CASSGGQVTGANVLTF | KLGGALQAK | YES | NO | NO |
| 17288 | TRAV38-2DV8 | TRAJ57 | CAYRSAGGGGSEKLVF | TRBV2 | TRBJ1-5 | CASSAIGTGVQPQHF | KLGGALQAK | YES | NO | NO |

| | | | | | | | | | | |
|---|---|---|---|---|---|---|---|---|---|---|
| 17289 | TRAV34 | TRAJ42 | CGADGYVYGGSQGNLIF | TRBV18 | TRBJ2-1 | CASSPRSGPSYNEQFF | KLGGALQAK | YES | NO | NO |
| 17290 | TRAV20 | TRAJ8 | CAVHPDRDTGFQKLVF | TRBV19 | TRBJ1-5 | CASSMMGQPQHF | KLGGALQAK | YES | NO | NO |
| 17291 | TRAV12-2 | TRAJ49 | CAVNTGNQFYF | TRBV28 | TRBJ2-2 | CASQGVGSTGELFF | KLGGALQAK | YES | NO | NO |
| 17292 | TRAV19 | TRAJ43 | CALRGTYNNNDMRF | TRBV11-2 | TRBJ2-1 | CASSPRASGGRWHEQFF | KLGGALQAK | YES | NO | NO |
| 17293 | TRAV4 | TRAJ42 | CLVGEDERGSQGNLIF | TRBV10-3 | TRBJ2-1 | CAISDLVGQGNEQFF | KLGGALQAK | NO | NO | NO |
| 17294 | TRAV29DV5 | TRAJ33 | CAATDMDSNYQLIW | TRBV4-1 | TRBJ2-5 | CASSYKGLGGQETQYF | KLGGALQAK | YES | NO | NO |
| 17295 | TRAV19 | TRAJ49 | CALSEASNTGNQFYF | TRBV7-7 | TRBJ2-1 | CASSLGLVDEQFF | KLGGALQAK | YES | NO | NO |
| 17296 | TRAV8-3 | TRAJ28 | CAVVPLTGAGSYQLTF | TRBV20-1 | TRBJ2-7 | CSASPRPDPYEQYF | KLGGALQAK | NO | NO | NO |
| 17297 | TRAV3 | TRAJ33 | CAVRDGDSNYQLIW | TRBV30 | TRBJ1-1 | CAWSVHRNTEAFF | KLGGALQAK | YES | NO | NO |
| 17298 | TRAV1-1 | TRAJ26 | CAVKSNYGQNFVF | TRBV20-1 | TRBJ1-1 | CSARGPRGGNTEAFF | KLGGALQAK | YES | NO | NO |
| 17299 | TRAV17 | TRAJ58 | CATDARETSGSRLTF | TRBV6-5 | TRBJ1-2 | CASSYHHRGDGYTF | KLGGALQAK | YES | NO | NO |
| 17300 | TRAV12-1 | TRAJ20 | CVVNKDYKLSF | TRBV12-4 | TRBJ2-2 | CASSFALAGGPGTGELFF | KLGGALQAK | YES | NO | NO |
| 17301 | TRAV13-2 | TRAJ47 | CAEKPVMEYGNKLVF | TRBV6-1 | TRBJ2-1 | CASRPTLSSYNEQFF | KLGGALQAK | YES | NO | NO |
| 17302 | TRAV6 | TRAJ39 | CALDGDAGNMLTF | TRBV5-6 | TRBJ2-1 | CASSLYSVYNEQFF | AVFDRKSDAK | YES | NO | YES |
| 17303 | TRAV1-2 | TRAJ31 | CAVSGDNARLMF | TRBV27 | TRBJ2-7 | CASSLSWGGRGEQYV | KLGGALQAK | NO | NO | NO |
| 17304 | TRAV19 | TRAJ37 | CALSEASASNTGKLIF | TRBV15 | TRBJ2-4 | CASMGWGSKNIQYF | KLGGALQAK | YES | NO | NO |
| 17305 | TRAV2 | TRAJ36 | CAVEAGANNLFF | TRBV5-1 | TRBJ2-5 | CASSAGTSGGETQYF | KLGGALQAK | NO | NO | NO |
| 17306 | TRAV8-3 | TRAJ45 | CAAYHRYSGGGADGLTF | TRBV6-3 | TRBJ1-2 | CASSQQQGVLYGYTF | AVFDRKSDAK | YES | NO | YES |
| 17307 | TRAV13-2 | TRAJ53 | CAENSGGSNYKLTF | TRBV28 | TRBJ2-2 | CASSLQAGETGELFF | KLGGALQAK | YES | NO | NO |
| 17308 | TRAV8-2 | TRAJ8 | CVVDDTGFQKLVF | TRBV7-2 | TRBJ2-1 | CASSLLLGLADKGQFF | KLGGALQAK | YES | NO | NO |
| 17309 | TRAV1-2 | TRAJ33 | CAVRDSNYQLIW | TRBV20-1 | TRBJ1-1 | CSASPGQVEAFF | KLGGALQAK | YES | NO | NO |
| 17310 | TRAV8-4 | TRAJ12 | CAVSDPMDSSYKLIF | TRBV28 | TRBJ2-2 | CASSLGDSYTGELFF | KLGGALQAK | YES | NO | NO |
| 17311 | TRAV38-2DV8 | TRAJ57 | CAYSSRGGSEKLVF | TRBV3-1 | TRBJ1-2 | CASSQVALGQGNYGYTF | KLGGALQAK | YES | NO | NO |
| 17312 | TRAV29DV5 | TRAJ34 | CAASYNTDKLIF | TRBV7-9 | TRBJ2-7 | CASSLATGYEQYV | KLGGALQAK | NO | NO | NO |
| 17313 | TRAV13-1 | TRAJ43 | CAASIMGNNNDMRF | TRBV10-2 | TRBJ2-1 | CATGTSGTDNEQFF | KLGGALQAK | YES | NO | NO |
| 17314 | TRAV1-2 | TRAJ34 | CAVKNTDKLIF | TRBV6-3 | TRBJ2-7 | CASSYTDRSYEQYV | AVFDRKSDAK | NO | NO | NO |
| 17315 | TRAV2 | TRAJ26 | CAVDNYGQNFVF | TRBV14 | TRBJ2-3 | CASSQDSGGSQDTQYF | KLGGALQAK | NO | NO | NO |
| 17316 | TRAV39 | TRAJ45 | CAVRGGGADGLTF | TRBV7-9 | TRBJ2-7 | CASSSSGRLHEQYV | KLGGALQAK | NO | NO | NO |
| 17317 | TRAV3 | TRAJ11 | CAVRDEYSTLTF | TRBV7-9 | TRBJ2-5 | CASSLSERWETQYF | KLGGALQAK | NO | NO | NO |
| 17318 | TRAV12-2 | TRAJ13 | CAVNGGYQKVTF | TRBV19 | TRBJ2-2 | CASSIPNGGELFF | KLGGALQAK | YES | NO | NO |
| 17319 | TRAV19 | TRAJ57 | CALRRTQGGSEKLVF | TRBV7-2 | TRBJ2-1 | CASSSKSGTSLGYNEQFF | KLGGALQAK | NO | NO | NO |
| 17320 | TRAV12-1 | TRAJ20 | CVVNSPSDYKLSF | TRBV20-1 | TRBJ2-3 | CSALSSGGRTDTQYF | KLGGALQAK | NO | NO | NO |
| 17321 | TRAV24 | TRAJ50 | CASSPRTSYDKVIF | TRBV6-1 | TRBJ1-5 | CASSDPGLVPQHF | KLGGALQAK | YES | NO | NO |
| 17322 | TRAV41 | TRAJ35 | CALI | TRBV20-1 | TRBJ2-2 | CSATRITSGRAGGELFF | KLGGALQAK | NO | NO | NO |
| 17323 | TRAV39 | TRAJ58 | CAVDFETSGSRLTF | TRBV30 | TRBJ1-2 | CAWRGSTGQGWGYGYTF | KLGGALQAK | YES | NO | NO |
| 17324 | TRAV12-2 | TRAJ28 | CAASGAGSYQLTF | TRBV7-9 | TRBJ2-4 | CASSSARGGRDIQYF | AVFDRKSDAK | YES | YES | YES |
| 17325 | TRAV12-2 | TRAJ4 | CAGGGGYNKLIF | TRBV24-1 | TRBJ1-6 | CATSDSGSNSPLHF | KLGGALQAK | YES | NO | NO |
| 17326 | TRAV24 | TRAJ32 | CAFGTWGATNKLIF | TRBV19 | TRBJ1-6 | CATGTANSPLHF | KLGGALQAK | YES | NO | NO |
| 17327 | TRAV24 | TRAJ52 | CAFIGDAGGTSYGKLTF | TRBV10-2 | TRBJ2-2 | CASSDLVIGELFF | KLGGALQAK | YES | NO | NO |
| 17328 | TRAV29DV5 | TRAJ54 | CAALGAQKLVF | TRBV13 | TRBJ2-2 | CASSLGTGFRGELFF | KLGGALQAK | NO | NO | NO |
| 17329 | TRAV38-1 | TRAJ54 | CAFMSLIQGAQKLVF | TRBV4-1 | TRBJ1-2 | CASSREWGTIYGYTF | KLGGALQAK | YES | NO | NO |
| 17330 | TRAV1-2 | TRAJ20 | CAVRGGDYKLSF | TRBV19 | TRBJ2-6 | CASSINEGSGANVLTF | KLGGALQAK | YES | NO | NO |
| 17331 | TRAV41 | TRAJ32 | CAVRSRDYGGATNKLIF | TRBV6-5 | TRBJ2-1 | CASSYGGAASYNEQFF | KLGGALQAK | NO | NO | NO |
| 17332 | TRAV14DV4 | TRAJ17 | CAMREQVFKAAGNKLTF | TRBV7-6 | TRBJ1-2 | CASSPTDRANYGYTF | KLGGALQAK | YES | NO | NO |
| 17333 | TRAV38-2DV8 | TRAJ57 | CAYRSGSEKLVF | TRBV11-2 | TRBJ2-7 | CASSLAAGGFYEQYV | KLGGALQAK | NO | NO | NO |
| 17334 | TRAV12-2 | TRAJ15 | CAVPANQAGTALIF | TRBV7-2 | TRBJ2-1 | CASSLVGGGAYNEQFF | KLGGALQAK | YES | NO | NO |
| 17335 | TRAV36DV7 | TRAJ57 | CAPQGGSEKLVF | TRBV20-1 | TRBJ2-1 | CSASERGFEGPYNEQFF | KLGGALQAK | NO | NO | NO |
| 17336 | TRAV20 | TRAJ53 | CAVHSGYSGGSNYKLTF | TRBV9 | TRBJ2-5 | CASSAGTAARTQYF | KLGGALQAK | YES | NO | NO |
| 17337 | TRAV29DV5 | TRAJ52 | CAASRANAGGTSYGKLTF | TRBV4-1 | TRBJ1-1 | CASSQDKVSWTGVSGGMNTEAFF | KLGGALQAK | NO | NO | NO |
| 17338 | TRAV19 | TRAJ27 | CALSAANAGKSTF | TRBV18 | TRBJ1-1 | CASSIYRTGDLNTEAFF | KLGGALQAK | YES | NO | NO |
| 17339 | TRAV12-1 | TRAJ20 | CVVNDYKLSF | TRBV7-2 | TRBJ1-4 | CASSLTGGFSTNEKLFF | KLGGALQAK | NO | NO | NO |
| 17340 | TRAV13-2 | TRAJ30 | CAENRDRDDKIIF | TRBV5-4 | TRBJ2-3 | CASSLRGSDTQYF | KLGGALQAK | YES | NO | NO |
| 17341 | TRAV17 | TRAJ20 | CATDYNDYKLSF | TRBV5-6 | TRBJ2-7 | CASSLEVQGGPEQYF | KLGGALQAK | YES | NO | NO |
| 17342 | TRAV19 | TRAJ36 | CALRGANGAGANNLFF | TRBV24-1 | TRBJ2-3 | CATSGGRTDTQYF | KLGGALQAK | YES | NO | NO |
| 17343 | TRAV1-2 | TRAJ17 | CAVSKAAGNKLTF | TRBV6-1 | TRBJ1-3 | CASSEARTDGNTIYF | KLGGALQAK | YES | NO | NO |
| 17344 | TRAV25 | TRAJ26 | CAGRDYGQNFVF | TRBV24-1 | TRBJ2-1 | CATSDPLAGGGYNEQFF | KLGGALQAK | YES | NO | NO |
| 17345 | TRAV14DV4 | TRAJ49 | CAMREGKSVTTGNQFYF | TRBV19 | TRBJ1-1 | CATEVGGLAEAFF | AVFDRKSDAK | YES | NO | NO |
| 17346 | TRAV22 | TRAJ48 | CAVGFGNEKLTF | TRBV9 | TRBJ2-1 | CASSGVSTYNEQFF | KLGGALQAK | YES | NO | NO |
| 17347 | TRAV4 | TRAJ5 | CLVGDIGDTGRRALTF | TRBV11-2 | TRBJ2-1 | CASTNLAGYDEQFF | KLGGALQAK | YES | NO | NO |
| 17348 | TRAV8-1 | TRAJ39 | CAVNFDAGNMLTF | TRBV7-8 | TRBJ2-1 | CASSPGIPEGVF | KLGGALQAK | YES | NO | NO |
| 17349 | TRAV19 | TRAJ44 | CALSDPGTASKLTF | TRBV5-1 | TRBJ2-2 | CASSLTGELFF | KLGGALQAK | YES | NO | NO |
| 17350 | TRAV29DV5 | TRAJ29 | CAASDPSGNTPLVF | TRBV11-3 | TRBJ1-1 | CASSLQSGTGELVNTEAFF | KLGGALQAK | YES | NO | NO |
| 17351 | TRAV20 | TRAJ45 | CAVHWGGADGLTF | TRBV2 | TRBJ1-1 | CASRDRGRAEAFF | KLGGALQAK | YES | NO | NO |
| 17352 | TRAV20 | TRAJ21 | CAVDDFNKFYF | TRBV6-5 | TRBJ2-2 | CASTVGGPGTGELFF | KLGGALQAK | YES | NO | NO |
| 17353 | TRAV17 | TRAJ41 | CATDRGNSGYALNF | TRBV7-8 | TRBJ2-5 | CASSLVEVETQYF | KLGGALQAK | YES | NO | NO |
| 17354 | TRAV1-1 | TRAJ17 | CAVRDPQAAGNKLTF | TRBV5-1 | TRBJ1-2 | CASSSTDRNYGYTF | AVFDRKSDAK | YES | NO | YES |
| 17355 | TRAV8-3 | TRAJ33 | CAVEGDSNYQLIW | TRBV7-7 | TRBJ2-1 | CASSEVGPGEQFF | KLGGALQAK | YES | NO | NO |
| 17356 | TRAV1-2 | TRAJ32 | CAVANYGGATNKLIF | TRBV10-2 | TRBJ1-3 | CASSDREPGNTIYF | KLGGALQAK | YES | NO | NO |
| 17357 | TRAV19 | TRAJ54 | CALSEALIQGAQKLVF | TRBV5-5 | TRBJ2-6 | CASSLEWPGANVLTF | KLGGALQAK | YES | NO | NO |
| 17358 | TRAV4 | TRAJ52 | CLVGVKAGGTSYGKLTF | TRBV19 | TRBJ2-1 | CASSIGQGAYEQFF | AVFDRKSDAK | YES | YES | YES |
| 17359 | TRAV27 | TRAJ20 | CAGPSSNDYKLSF | TRBV29-1 | TRBJ1-1 | CSVAGTGNTEAFF | KLGGALQAK | YES | NO | NO |
| 17360 | TRAV10 | TRAJ24 | CVVMQDSWGKLQF | TRBV9 | TRBJ2-3 | CASSGVAGLTDTQYF | KLGGALQAK | YES | NO | NO |
| 17361 | TRAV14DV4 | TRAJ53 | CAMREGVLGSGGSNYKLTF | TRBV4-3 | TRBJ2-3 | CASSQDGGAGNFADTQYF | KLGGALQAK | YES | NO | NO |
| 17362 | TRAV1-2 | TRAJ4 | CAVRDQGLSGGYNKLIF | TRBV4-1 | TRBJ1-2 | CASSQAGSGYTF | AVFDRKSDAK | YES | NO | YES |
| 17363 | TRAV8-6 | TRAJ49 | CAVTLNTGNQFYF | TRBV6-6 | TRBJ2-7 | CASTQAGLAGFEQYV | KLGGALQAK | NO | NO | NO |
| 17364 | TRAV20 | TRAJ45 | CAVRSHSGGGADGLTF | TRBV6-1 | TRBJ2-7 | CASSEYLSGSSYEQYF | KLGGALQAK | YES | NO | NO |
| 17365 | TRAV29DV5 | TRAJ26 | CAASANNYGQNFVF | TRBV19 | TRBJ1-1 | CATNRGFTEAFF | AVFDRKSDAK | YES | NO | NO |
| 17366 | TRAV19 | TRAJ11 | CALNSGYSTLTF | TRBV27 | TRBJ2-6 | CASSLRTGEAGANVLTF | KLGGALQAK | YES | NO | NO |
| 17368 | TRAV12-2 | TRAJ4 | CAVLSGGYNKLIF | TRBV11-3 | TRBJ2-2 | CASSLPAGRVSAGELFF | RLRAEAQVK | YES | NO | YES |
| 17369 | TRAV1-1 | TRAJ24 | CAVTDSWGKLQF | TRBV7-8 | TRBJ1-2 | CASSLGYGYTF | KLGGALQAK | YES | NO | NO |
| 17370 | TRAV1-2 | TRAJ32 | CAVKSYGGATNKLIF | TRBV25-1 | TRBJ1-2 | CASRDSANYGYTF | KLGGALQAK | YES | NO | NO |
| 17371 | TRAV19 | TRAJ42 | CALSEAWRSQGNLIF | TRBV30 | TRBJ2-1 | CAWATGTGGSGYEQFF | KLGGALQAK | NO | NO | NO |
| 17372 | TRAV13-2 | TRAJ23 | CAENYQGGKLIF | TRBV7-6 | TRBJ1-3 | CASSRTGNTIYF | KLGGALQAK | YES | NO | NO |
| 17373 | TRAV17 | TRAJ39 | CATEDAGNMLTF | TRBV11-2 | TRBJ2-7 | CASSPGPMDEQYF | KLGGALQAK | YES | NO | NO |
| 17374 | TRAV12-3 | TRAJ30 | CAMSAQVGWDDKIIF | TRBV13 | TRBJ2-5 | CASSLTLEYETQYF | KLGGALQAK | YES | NO | NO |
| 17375 | TRAV1-2 | TRAJ33 | CAVMDSNYQLIW | TRBV6-4 | TRBJ2-1 | CASSDSSGGNEQFF | KLGGALQAK | YES | NO | NO |
| 17376 | TRAV5 | TRAJ29 | CAEPFSGNTPLVF | TRBV19 | TRBJ1-1 | CTSSAGDTEAFF | KLGGALQAK | YES | NO | NO |
| 17377 | TRAV29DV5 | TRAJ42 | CAASGGSQGNLIF | TRBV27 | TRBJ2-7 | CASSLWAGSYEQYF | KLGGALQAK | YES | NO | NO |
| 17378 | TRAV12-2 | TRAJ52 | CAVNIASSNAGGTSYGKLTF | TRBV19 | TRBJ1-1 | CASSPWGYTEAFF | KLGGALQAK | NO | NO | NO |
| 17379 | TRAV14DV4 | TRAJ32 | CAMRIYGGATNKLIF | TRBV12-4 | TRBJ2-7 | CASSLSPWEQYF | KLGGALQAK | YES | NO | NO |
| 17380 | TRAV19 | TRAJ54 | CALSEGALIQGAQKLVF | TRBV27 | TRBJ2-5 | CASRRQIDQETQYF | KLGGALQAK | YES | NO | NO |
| 17381 | TRAV8-6 | TRAJ57 | CAVSLTQGGSEKLVF | TRBV6-5 | TRBJ2-7 | CASSYQGHYEQYF | KLGGALQAK | YES | NO | NO |
| 17382 | TRAV13-1 | TRAJ33 | CASYTPDSNYQLIW | TRBV4-1 | TRBJ2-7 | CASSQYRGSYEQYF | KLGGALQAK | YES | NO | NO |
| 17383 | TRAV20 | TRAJ43 | CAASNDMRF | TRBV3-1 | TRBJ2-1 | CASSQDVGGNSYNEQFF | KLGGALQAK | NO | NO | NO |
| 17385 | TRAV1-1 | TRAJ9 | CAATGGFKTIF | TRBV12-4 | TRBJ2-1 | CASSLPASGGATDNEQFF | KLGGALQAK | YES | NO | NO |
| 17386 | TRAV21 | TRAJ43 | CAVVYMRF | TRBV12-4 | TRBJ1-2 | CASSLWPDYGYTF | KLGGALQAK | NO | NO | NO |
| 17387 | TRAV8-1 | TRAJ28 | CAVKTISGAGSYQLTF | TRBV19 | TRBJ2-1 | CASRNRGYEQFF | KLGGALQAK | YES | YES | NO |
| 17389 | TRAV14DV4 | TRAJ20 | CALLLNDYKLSF | TRBV20-1 | TRBJ1-1 | CSARDFGTGGTEAFF | KLGGALQAK | YES | NO | NO |
| 17390 | TRAV12-1 | TRAJ36 | CVVNRRGGANNLFF | TRBV5-5 | TRBJ1-3 | CASSVVRGEAGNTIYF | KLGGALQAK | YES | NO | NO |
| 17391 | TRAV26-2 | TRAJ48 | CILRDGGILGNEKLTF | TRBV6-5 | TRBJ2-3 | CASSPGQGFDTQYF | KLGGALQAK | NO | NO | NO |
| 17392 | TRAV12-3 | TRAJ4 | CAMAVLESGGYNKLIF | TRBV20-1 | TRBJ2-1 | CSARDLPGGYEQFF | KLGGALQAK | NO | NO | NO |
| 17393 | TRAV30 | TRAJ48 | CGTIGNEKLTF | TRBV19 | TRBJ2-7 | CASSAGQGSYEQYF | KLGGALQAK | YES | NO | NO |
| 17394 | TRAV29DV5 | TRAJ22 | CAATFSGSARQLTF | TRBV4-1 | TRBJ2-3 | CASSQPGATDTQYF | KLGGALQAK | YES | NO | NO |
| 17395 | TRAV29DV5 | TRAJ20 | CAASPNDYKLSF | TRBV12-3 | TRBJ2-7 | CASSLLGSYEQYF | KLGGALQAK | YES | NO | NO |

| | | | | | | | | | | |
|---|---|---|---|---|---|---|---|---|---|---|
| 17397 | TRAV26-2 | TRAJ33 | CILRPMDSNYQLIW | TRBV4-3 | TRBJ1-1 | CASSQDGGNTEAFF | KLGGALQAK | YES | NO | NO |
| 17398 | TRAV12-1 | TRAJ15 | CVVLNQAGTALIF | TRBV20-1 | TRBJ2-3 | CSARDRDRQRDTQYF | KLGGALQAK | NO | NO | NO |
| 17399 | TRAV5 | TRAJ36 | CAGRTGANNLFF | TRBV7-9 | TRBJ1-2 | CASSSSSETVIYGYTF | KLGGALQAK | YES | NO | NO |
| 17400 | TRAV8-1 | TRAJ39 | CAAINNAGNMLTF | TRBV4-1 | TRBJ2-1 | CASSHQTSSYNEQFF | KLGGALQAK | YES | NO | NO |
| 17401 | TRAV20 | TRAJ33 | CAVQAADSNYQLIW | TRBV5-4 | TRBJ2-1 | CASSSGTSSYNEQFF | KLGGALQAK | YES | NO | NO |
| 17402 | TRAV24 | TRAJ48 | CASSAFGNEKLTF | TRBV4-1 | TRBJ2-3 | CASSQEFGFRVDTQYF | KLGGALQAK | YES | NO | NO |
| 17403 | TRAV35 | TRAJ41 | CAGSLTLSNSGYALNF | TRBV29-1 | TRBJ2-1 | CSVRLAGGKNEQFF | KLGGALQAK | YES | NO | NO |
| 17404 | TRAV4 | TRAJ36 | CLVVGTGANNLFF | TRBV4-3 | TRBJ1-1 | CASSQGTVLNTEAFF | KLGGALQAK | YES | NO | NO |
| 17405 | TRAV29DV5 | TRAJ54 | CAASSSGAQKLVF | TRBV6-1 | TRBJ2-4 | CASSARDGNIQYF | KLGGALQAK | YES | NO | NO |
| 17406 | TRAV35 | TRAJ43 | CAGQGYNNNDMRF | TRBV7-9 | TRBJ1-2 | CASSLVQGVVGYTF | KLGGALQAK | YES | NO | NO |
| 17407 | TRAV14DV4 | TRAJ4 | CAMRRPSGGYNKLIF | TRBV11-2 | TRBJ1-2 | CASSLEPTDRDSYGYTF | KLGGALQAK | YES | YES | NO |
| 17408 | TRAV1-2 | TRAJ33 | CAGLDSNYQLIW | TRBV20-1 | TRBJ2-7 | CSGLAGEPYEQYF | KLGGALQAK | NO | NO | NO |
| 17409 | TRAV8-3 | TRAJ53 | CAVGTPNSGGSNYKLTF | TRBV9 | TRBJ2-1 | CASSSGLAGASDEQFF | KLGGALQAK | YES | NO | NO |
| 17410 | TRAV25 | TRAJ20 | CAGPSSNDYKLSF | TRBV18 | TRBJ1-1 | CAVMEGSNEAFF | KLGGALQAK | YES | NO | NO |
| 17411 | TRAV29DV5 | TRAJ29 | CAASGTGNTPLVF | TRBV30 | TRBJ1-2 | CAWSTGGGYGYTF | KLGGALQAK | YES | NO | NO |
| 17412 | TRAV29DV5 | TRAJ29 | CAASAALGNTPLVF | TRBV12-3 | TRBJ1-1 | CASSFYEAFF | KLGGALQAK | YES | NO | NO |
| 17413 | TRAV14DV4 | TRAJ24 | CAMREMMTTDSWGKLQF | TRBV9 | TRBJ2-1 | CASSVDDGEYYNEQFF | KLGGALQAK | YES | NO | NO |
| 17414 | TRAV12-2 | TRAJ52 | CAVMGAGGTSYGKLTF | TRBV12-5 | TRBJ1-1 | CASGPPRDRARNTEAFF | KLGGALQAK | YES | NO | NO |
| 17415 | TRAV19 | TRAJ17 | CALSEAVEAAGNKLTF | TRBV19 | TRBJ1-1 | CASSMGTGLKAFF | KLGGALQAK | YES | NO | NO |
| 17416 | TRAV8-3 | TRAJ35 | CVRI | TRBV6-5 | TRBJ2-3 | CASSRIGPDTQYF | KLGGALQAK | NO | NO | NO |
| 17417 | TRAV17 | TRAJ27 | CATDLFYTGKSTF | TRBV27 | TRBJ2-1 | CASKGLADNEQFF | KLGGALQAK | YES | NO | NO |
| 17418 | TRAV35 | TRAJ53 | CAGPWGGSNYKLTF | TRBV6-1 | TRBJ2-7 | CASSSHPRTRVYEQYV | KLGGALQAK | NO | NO | NO |
| 17419 | TRAV12-2 | TRAJ5 | CAVNGALTF | TRBV20-1 | TRBJ2-2 | CSALYRDGWNTGELFF | KLGGALQAK | NO | NO | NO |
| 17420 | TRAV9-2 | TRAJ42 | CALRRYGGSQGNLIF | TRBV7-9 | TRBJ1-2 | CASSLAIGATPDSNYGYTF | KLGGALQAK | YES | NO | NO |
| 17421 | TRAV13-1 | TRAJ33 | CAASKDSNYQLIW | TRBV11-2 | TRBJ2-5 | CASSFYGTQETQYF | KLGGALQAK | YES | NO | NO |
| 17422 | TRAV1-2 | TRAJ34 | CAVNTDKLIF | TRBV6-6 | TRBJ2-1 | CASRTTGFGFNEQFF | KLGGALQAK | NO | NO | NO |
| 17423 | TRAV3 | TRAJ28 | CAVRLAPYSGAGSYQLTF | TRBV7-9 | TRBJ1-5 | CASSTGQGKNQPQHF | KLGGALQAK | NO | NO | NO |
| 17424 | TRAV8-4 | TRAJ3 | CAVSAGYSSASKIIF | TRBV12-4 | TRBJ2-1 | CASSFHEREMNNEQFF | LLFGYPVYV | YES | YES | YES |
| 17425 | TRAV26-2 | TRAJ47 | CILSPMEYGNKLVF | TRBV6-1 | TRBJ2-7 | CASSELGYEQYV | AVFDRKSDAK | NO | NO | NO |
| 17426 | TRAV14DV4 | TRAJ9 | CAMREGGGGFKTIF | TRBV5-4 | TRBJ1-1 | CASSPRGEGHVEAFF | KLGGALQAK | YES | NO | NO |
| 17427 | TRAV3 | TRAJ20 | CAVRDIEGGDYKLSF | TRBV14 | TRBJ2-1 | CASSPLAGGQDEQFF | KLGGALQAK | NO | NO | NO |
| 17428 | TRAV14DV4 | TRAJ22 | CAMREGSARQLTF | TRBV19 | TRBJ2-4 | CASSTGFNIQYF | KLGGALQAK | YES | NO | NO |
| 17429 | TRAV1-2 | TRAJ4 | CAFSAGGYNKLIF | TRBV7-9 | TRBJ2-7 | CASSPSGSYEQYF | KLGGALQAK | YES | NO | NO |
| 17430 | TRAV3 | TRAJ40 | CAVTSTSGTYKYIF | TRBV7-9 | TRBJ2-7 | CASSPPGQGYWEQYV | KLGGALQAK | NO | NO | NO |
| 17431 | TRAV19 | TRAJ44 | CALSINAGTASKLTF | TRBV27 | TRBJ2-2 | CASSDRGPGTGELFF | KLGGALQAK | YES | NO | NO |
| 17432 | TRAV1-2 | TRAJ17 | CAPEAAGNKLTF | TRBV27 | TRBJ1-4 | CASSFDYNEKLFF | KLGGALQAK | YES | NO | NO |
| 17433 | TRAV13-1 | TRAJ50 | CAPTVYSYDKVIF | TRBV12-4 | TRBJ1-6 | CASTLGSGNSPLHF | KLGGALQAK | YES | NO | NO |
| 17434 | TRAV12-2 | TRAJ50 | CAVKTSYDKVIF | TRBV20-1 | TRBJ1-1 | CSARDSRANTEAFF | KLGGALQAK | YES | NO | NO |
| 17435 | TRAV14DV4 | TRAJ7 | CAMWAHKGNNRLAF | TRBV12-4 | TRBJ1-1 | CASSLIGGTEAFF | KLGGALQAK | YES | NO | NO |
| 17436 | TRAV1-2 | TRAJ38 | CAVHFNAGNNRKLIW | TRBV4-1 | TRBJ1-1 | CASSQDWAAEAFF | KLGGALQAK | YES | NO | NO |
| 17437 | TRAV20 | TRAJ52 | CAVQANAGGTSYGKLTF | TRBV19 | TRBJ2-2 | CASSIGRLAGVTGELFF | KLGGALQAK | YES | NO | NO |
| 17438 | TRAV1-2 | TRAJ18 | CAVRGRGSTLGRLYF | TRBV12-4 | TRBJ1-2 | CASSLSQPGAGTANYGYTF | KLGGALQAK | YES | NO | NO |
| 17439 | TRAV23DV6 | TRAJ28 | CAASKLITGAGSYQLTF | TRBV7-9 | TRBJ1-1 | CASSPGEKNTEAFF | KLGGALQAK | YES | NO | NO |
| 17441 | TRAV1-2 | TRAJ13 | CAALNSGGYQKVTF | TRBV9 | TRBJ2-2 | CASSVASGELFF | KLGGALQAK | YES | NO | NO |
| 17442 | TRAV13-2 | TRAJ40 | CAETSSTSGTYKYIF | TRBV6-3 | TRBJ2-7 | CASSYGGSYEQYF | KLGGALQAK | YES | NO | NO |
| 17443 | TRAV38-2DV8 | TRAJ49 | CAFNTGNQFYF | TRBV30 | TRBJ2-3 | CAWSVLSYTDTQYF | KLGGALQAK | NO | NO | NO |
| 17444 | TRAV12-3 | TRAJ52 | CAMSADAGGTSYGKLTF | TRBV27 | TRBJ2-1 | CASSLLGQQFF | KLGGALQAK | YES | NO | NO |
| 17445 | TRAV17 | TRAJ45 | CAPQYSGGGADGLTF | TRBV12-4 | TRBJ1-4 | CASSGDEKLFF | KLGGALQAK | YES | NO | NO |
| 17446 | TRAV36DV7 | TRAJ48 | CAVVVPNFGNEKLTF | TRBV7-8 | TRBJ1-2 | CASSLAWQPDGYTF | KLGGALQAK | NO | NO | NO |
| 17447 | TRAV9-2 | TRAJ26 | CALSDPGDNYGQNFVF | TRBV4-3 | TRBJ2-2 | CASSQYDSSGELFF | AVFDRKSDAK | YES | NO | YES |
| 17448 | TRAV17 | TRAJ48 | CATKDFGNEKLTF | TRBV5-6 | TRBJ2-5 | CASSLIGTGGETQYF | KLGGALQAK | YES | NO | NO |
| 17449 | TRAV24 | TRAJ53 | CASGGSNYKLTF | TRBV4-1 | TRBJ1-1 | CASSQVQGAPTEAFF | KLGGALQAK | YES | YES | NO |
| 17450 | TRAV26-1 | TRAJ27 | CIVSGDNTNAGKSTF | TRBV4-1 | TRBJ1-6 | CASSRLRDGGGYNSPLHF | RLRAEAQVK | YES | NO | NO |
| 17451 | TRAV2 | TRAJ42 | CAVEVTWGGSQGNLIF | TRBV9 | TRBJ2-2 | CASSPPGANTGELFF | KLGGALQAK | NO | NO | NO |
| 17452 | TRAV1-2 | TRAJ21 | CAVRIFNKFYF | TRBV3-1 | TRBJ2-1 | CASSQASGAGEKRSSYNEQFF | KLGGALQAK | NO | NO | NO |
| 17453 | TRAV13-1 | TRAJ52 | CAASNRSNAGGTSYGKLTF | TRBV5-6 | TRBJ2-1 | CASSEITSGVYNEQFF | KLGGALQAK | YES | NO | NO |
| 17454 | TRAV29DV5 | TRAJ40 | CAARLMGTYKYIF | TRBV12-4 | TRBJ2-3 | CANREGGTDTQYF | KLGGALQAK | YES | NO | NO |
| 17455 | TRAV26-2 | TRAJ43 | CILRDEVNNNDMRF | TRBV3-1 | TRBJ2-1 | CASSVSSYGLAAYNEQFF | KLGGALQAK | YES | NO | NO |
| 17456 | TRAV19 | TRAJ57 | CALSEAYTQGGSEKLVF | TRBV6-6 | TRBJ2-5 | CASSYSERGAAHETQYF | KLGGALQAK | NO | NO | NO |
| 17457 | TRAV8-3 | TRAJ23 | CAVGAGIYNQGGKLIF | TRBV20-1 | TRBJ1-1 | CSASNRGLAEAFF | KLGGALQAK | YES | NO | NO |
| 17458 | TRAV4 | TRAJ40 | CLQTTSGTYKYIF | TRBV5-1 | TRBJ1-4 | CASSLGQGPNEKLFF | KLGGALQAK | YES | NO | NO |
| 17459 | TRAV12-1 | TRAJ33 | CVALDSNYQLIW | TRBV5-1 | TRBJ1-2 | CASSLGQGPDGYTF | KLGGALQAK | YES | NO | NO |
| 17461 | TRAV21 | TRAJ26 | CAVRYYYGQNFVF | TRBV7-8 | TRBJ2-5 | CASSSTGTFDETQYF | KLGGALQAK | YES | NO | NO |
| 17462 | TRAV13-2 | TRAJ52 | CAELSNAGGTSYGKLTF | TRBV6-1 | TRBJ1-2 | CASSTQGANYGYTF | KLGGALQAK | YES | NO | NO |
| 17463 | TRAV8-1 | TRAJ6 | CAVNAPGGSYIPTF | TRBV5-1 | TRBJ1-4 | CASSSGQGGEKLFF | KLGGALQAK | YES | NO | NO |
| 17464 | TRAV12-2 | TRAJ49 | CAVPYHTGNQFYF | TRBV7-9 | TRBJ2-5 | CASSLGQAIETQYF | KLGGALQAK | YES | NO | NO |
| 17465 | TRAV12-2 | TRAJ48 | CAVNPEPGNFGNEKLTF | TRBV5-4 | TRBJ2-2 | CASSFWDFRTGELFF | KLGGALQAK | YES | NO | NO |
| 17466 | TRAV39 | TRAJ26 | CAVDSDNYGQNFVF | TRBV5-4 | TRBJ2-2 | CASSWGTGELFF | KLGGALQAK | YES | NO | NO |
| 17467 | TRAV1-2 | TRAJ10 | CARSGLTGGGNKLTF | TRBV7-2 | TRBJ2-7 | CASSLVGGGDEQYF | AVFDRKSDAK | NO | NO | NO |
| 17469 | TRAV14DV4 | TRAJ43 | CAMREGWGFNNNDMRF | TRBV9 | TRBJ2-1 | CASSARPGLAGVGRDEQFF | KLGGALQAK | YES | NO | NO |
| 17470 | TRAV12-1 | TRAJ23 | CVVNNNQGGKLIF | TRBV2 | TRBJ2-5 | CASSEMWRSVEETQYF | KLGGALQAK | YES | NO | NO |
| 17471 | TRAV38-1 | TRAJ41 | CAQLPPISNSGYALNF | TRBV19 | TRBJ2-7 | CASSYGYEQYF | AVFDRKSDAK | YES | NO | NO |
| 17472 | TRAV17 | TRAJ15 | CTTGTALIF | TRBV9 | TRBJ2-1 | CASSVAGVGHNEQFF | KLGGALQAK | NO | NO | NO |
| 17473 | TRAV17 | TRAJ53 | CATDESGGSNYKLTF | TRBV27 | TRBJ1-2 | CASSFARNYGYTF | KLGGALQAK | YES | NO | NO |
| 17474 | TRAV4 | TRAJ37 | CLVGDGSSNTGKLIF | TRBV4-1 | TRBJ1-4 | CASSQDRGIIEKLFF | KLGGALQAK | YES | NO | NO |
| 17475 | TRAV19 | TRAJ12 | CALSGGDSSYKLIF | TRBV7-3 | TRBJ1-1 | CASSLGFLGGTEAFF | KLGGALQAK | NO | NO | NO |
| 17476 | TRAV19 | TRAJ40 | CALSGQGTYKYIF | TRBV7-9 | TRBJ2-5 | CASSPTAQETQYF | KLGGALQAK | YES | NO | NO |
| 17477 | TRAV27 | TRAJ40 | CAATTSGTYKYIF | TRBV20-1 | TRBJ2-1 | CSASPLSGSTYNEQFF | KLGGALQAK | NO | NO | NO |
| 17478 | TRAV30 | TRAJ12 | CGTEMDSSYKLIF | TRBV7-8 | TRBJ1-1 | CASSPHQGLVTEAFF | KLGGALQAK | YES | NO | NO |
| 17479 | TRAV19 | TRAJ26 | CALSEVNYGQNFVF | TRBV11-2 | TRBJ2-5 | CASSGRQGAGRGAQYF | KLGGALQAK | YES | NO | NO |
| 17480 | TRAV13-1 | TRAJ45 | CAASAYSGGGADGLTF | TRBV12-4 | TRBJ2-3 | CASSPVVDTQYF | KLGGALQAK | YES | NO | NO |
| 17481 | TRAV8-6 | TRAJ50 | CAPLGPVIF | TRBV12-3 | TRBJ2-1 | CASRISWAGEVESEQFF | KLGGALQAK | NO | NO | NO |
| 17482 | TRAV12-1 | TRAJ27 | CVVTEDAGKSTF | TRBV7-9 | TRBJ2-7 | CASSGSYEQYF | KLGGALQAK | YES | NO | NO |
| 17483 | TRAV17 | TRAJ33 | CATDSNYQLIW | TRBV19 | TRBJ2-7 | CATGTAPSYEQYF | KLGGALQAK | YES | NO | NO |
| 17484 | TRAV4 | TRAJ26 | CLVATNYGQNFVF | TRBV19 | TRBJ2-1 | CASSMSVWGTANNEQFF | KLGGALQAK | YES | NO | NO |
| 17485 | TRAV9-2 | TRAJ30 | CALSDPMNRDDKIIF | TRBV7-9 | TRBJ1-1 | CASSLAYGTPEAFF | KLGGALQAK | YES | NO | NO |
| 17486 | TRAV19 | TRAJ6 | CALSPLITSGGSYIPTF | TRBV7-3 | TRBJ1-2 | CASSLSSGGMDYGYTF | KLGGALQAK | NO | NO | NO |
| 17487 | TRAV12-2 | TRAJ6 | CAVKGVGGSYIPTF | TRBV6-1 | TRBJ2-2 | CASGGGGTGELFF | KLGGALQAK | YES | NO | NO |
| 17488 | TRAV12-1 | TRAJ39 | CVVNDAGNMLTF | TRBV18 | TRBJ2-3 | CASSPPGTDTQYF | KLGGALQAK | YES | NO | NO |
| 17489 | TRAV35 | TRAJ29 | CAGQWGNTPLVF | TRBV29-1 | TRBJ2-1 | CSVESILAGGGPNEQFF | AVFDRKSDAK | YES | NO | YES |
| 17490 | TRAV13-2 | TRAJ11 | CAEGDGTLTF | TRBV4-3 | TRBJ1-4 | CASSQDPVRSNEKLFF | KLGGALQAK | YES | NO | NO |
| 17491 | TRAV4 | TRAJ40 | CLVGDASGTYKYIF | TRBV12-4 | TRBJ2-1 | CASSYLTGGWNEQFF | KLGGALQAK | YES | NO | NO |
| 17492 | TRAV12-1 | TRAJ49 | CVVMTGNQFYF | TRBV12-4 | TRBJ2-7 | CASSAYLHEQYV | KLGGALQAK | NO | NO | NO |
| 17493 | TRAV14DV4 | TRAJ30 | CAMRKTNRDDKIIF | TRBV5-8 | TRBJ2-5 | CASSSGLGPRETQYF | KLGGALQAK | YES | NO | NO |
| 17494 | TRAV19 | TRAJ54 | CALRVYQGAQKLVF | TRBV6-3 | TRBJ1-2 | CASSTLGDGVRGYTF | KLGGALQAK | YES | NO | NO |
| 17495 | TRAV29DV5 | TRAJ45 | CAASASGGGADGLTF | TRBV7-9 | TRBJ2-5 | CASSPPSEAQYF | KLGGALQAK | YES | NO | NO |
| 17496 | TRAV29DV5 | TRAJ26 | CAASFYTSDNYGQNFVF | TRBV14 | TRBJ2-3 | CASSQVVLYEDTQYF | KLGGALQAK | YES | NO | NO |
| 17497 | TRAV17 | TRAJ47 | CATEYGNKLVF | TRBV7-6 | TRBJ1-1 | CASSPTGGYTEAFF | KLGGALQAK | YES | NO | NO |
| 17498 | TRAV2 | TRAJ42 | CAVERAYGGSQGNLIF | TRBV7-6 | TRBJ2-1 | CASSIAARGSSYNEQFF | KLGGALQAK | NO | NO | NO |
| 17499 | TRAV13-1 | TRAJ6 | CAATHRSGGSYIPTF | TRBV4-1 | TRBJ1-5 | CASSHSVQGNQPQHF | AVFDRKSDAK | YES | NO | YES |
| 17500 | TRAV1-2 | TRAJ4 | CAVRGDLSGGYNKLIF | TRBV4-3 | TRBJ2-4 | CASSQSHGAGDIQYF | AVFDRKSDAK | NO | NO | NO |
| 17501 | TRAV19 | TRAJ52 | CALSEAAGGTSYGKLTF | TRBV29-1 | TRBJ1-1 | CSVPGSANTEAFF | KLGGALQAK | YES | NO | NO |
| 17502 | TRAV17 | TRAJ43 | CATDDDMRF | TRBV20-1 | TRBJ1-1 | CSAIWEEKGENTEAFF | KLGGALQAK | NO | NO | NO |
| 17503 | TRAV17 | TRAJ48 | CATPLNFGNEKLTF | TRBV4-1 | TRBJ1-4 | CASRDKAAQDEKLFF | KLGGALQAK | YES | NO | NO |

| | | | | | | | | | | |
|---|---|---|---|---|---|---|---|---|---|---|
| 17504 | TRAV19 | TRAJ54 | CALSEDGQGAQKLVF | TRBV5-1 | TRBJ2-4 | CASSLGWGAKNIQYF | KLGGALQAK | YES | NO | NO |
| 17505 | TRAV14DV4 | TRAJ7 | CAMREEHGRLAF | TRBV12-4 | TRBJ2-7 | CASSFGREQYF | KLGGALQAK | YES | NO | NO |
| 17506 | TRAV19 | TRAJ31 | CALSAGWPDNARLMF | TRBV28 | TRBJ2-6 | CASSTPPGQGSLGANVLTF | AVFDRKSDAK | YES | NO | YES |
| 17507 | TRAV38-2DV8 | TRAJ43 | CAYGPKGHDMRF | TRBV9 | TRBJ1-3 | CASSAAGASSGNTIYF | KLGGALQAK | NO | NO | NO |
| 17508 | TRAV24 | TRAJ40 | CASKSGTYKYIF | TRBV10-3 | TRBJ2-3 | CAISDQGRTDTQYF | KLGGALQAK | YES | NO | NO |
| 17509 | TRAV8-6 | TRAJ53 | CAVTGGSGGSNYKLTF | TRBV27 | TRBJ1-1 | CASSLSGGMNTEAFF | AVFDRKSDAK | YES | NO | YES |
| 17510 | TRAV12-1 | TRAJ40 | CVVSLTSGTYKYIF | TRBV7-9 | TRBJ1-1 | CASSNYRLTENTEAFF | KLGGALQAK | YES | NO | NO |
| 17511 | TRAV20 | TRAJ26 | CAVQSYGQNFVF | TRBV2 | TRBJ2-7 | CASREGTGNEQYF | KLGGALQAK | YES | NO | NO |
| 17512 | TRAV26-2 | TRAJ45 | CILRDGGGADGLTF | TRBV4-1 | TRBJ1-2 | CASSPTVLYGYTF | KLGGALQAK | YES | NO | NO |
| 17513 | TRAV35 | TRAJ49 | CAGQNTGNQFYF | TRBV27 | TRBJ2-3 | CASSPSGGTGTDTQYF | KLGGALQAK | YES | NO | NO |
| 17514 | TRAV1-2 | TRAJ37 | CAVGSGNTGKLIF | TRBV7-9 | TRBJ2-3 | CASSKWGGGSTDTQYF | KLGGALQAK | YES | NO | NO |
| 17515 | TRAV19 | TRAJ57 | CALSENEGGGSEKLVF | TRBV12-4 | TRBJ1-1 | CASSFGGVRNTEAFF | KLGGALQAK | YES | NO | NO |
| 17516 | TRAV13-1 | TRAJ6 | CAASTSGGSYIPTF | TRBV12-4 | TRBJ2-1 | CASSQLRTSLDEQFF | KLGGALQAK | YES | NO | NO |
| 17517 | TRAV5 | TRAJ23 | CAEKSHNQGGKLIF | TRBV24-1 | TRBJ2-2 | CATSDGLNTGELFF | KLGGALQAK | YES | NO | NO |
| 17518 | TRAV12-2 | TRAJ49 | CAVNMDTGNQFYF | TRBV27 | TRBJ2-7 | CASSPHSYEQYF | ELAGIGILTV | YES | YES | YES |
| 17519 | TRAV3 | TRAJ27 | CAVRDSNLNAGKSTF | TRBV7-9 | TRBJ2-1 | CASLSGTKGFGEQFF | KLGGALQAK | NO | NO | NO |
| 17520 | TRAV21 | TRAJ34 | CAVLGDKLIF | TRBV29-1 | TRBJ2-1 | CSVDTYNEQFF | KLGGALQAK | YES | NO | NO |
| 17521 | TRAV17 | TRAJ15 | CATDQAGTALIF | TRBV7-9 | TRBJ1-6 | CASSLVPTNSPLHF | KLGGALQAK | YES | NO | NO |
| 17522 | TRAV19 | TRAJ30 | CALSEPDRDDKIIF | TRBV12-4 | TRBJ2-7 | CASSLSGQSSYEQYF | KLGGALQAK | YES | NO | NO |
| 17523 | TRAV8-6 | TRAJ6 | CAVRSGGSYIPTF | TRBV2 | TRBJ2-7 | CATAPRERSYEQYF | KLGGALQAK | YES | NO | NO |
| 17524 | TRAV8-6 | TRAJ26 | CAVSDNYGQNFVF | TRBV4-1 | TRBJ1-2 | CASSQNRDRGIYGYTF | KLGGALQAK | YES | NO | NO |
| 17525 | TRAV12-1 | TRAJ39 | CVVNKGGNAGNMLTF | TRBV20-1 | TRBJ2-1 | CSAYRDSYNEQFF | KLGGALQAK | NO | NO | NO |
| 17526 | TRAV12-1 | TRAJ43 | CVVINNDMRF | TRBV6-3 | TRBJ1-1 | CASSRRQLTNTEAFF | KLGGALQAK | YES | NO | NO |
| 17527 | TRAV19 | TRAJ8 | CALSEARRTGFQKLVF | TRBV20-1 | TRBJ2-7 | CSARVAGYYEQYF | KLGGALQAK | YES | NO | NO |
| 17528 | TRAV14DV4 | TRAJ15 | CAMSHFNQAGTALIF | TRBV9 | TRBJ2-3 | CASSVEEYTDTQYF | KLGGALQAK | YES | NO | NO |
| 17529 | TRAV4 | TRAJ36 | CLVGDIDGRTGANNLFF | TRBV19 | TRBJ2-1 | CASNPEPGNEQFF | KLGGALQAK | YES | NO | NO |
| 17530 | TRAV12-1 | TRAJ49 | CVVLLTGNQFYF | TRBV5-5 | TRBJ2-2 | CASSGLGVGNTGELFF | KLGGALQAK | YES | NO | NO |
| 17531 | TRAV35 | TRAJ56 | CAGHPLTGANSKLTF | TRBV5-1 | TRBJ2-7 | CASWDRGGRYEQYF | KLGGALQAK | YES | NO | NO |
| 17532 | TRAV25 | TRAJ17 | CAGPPWAAGNKLTF | TRBV24-1 | TRBJ2-1 | CATSDRTSGSFNEQFF | KLGGALQAK | YES | NO | NO |
| 17533 | TRAV2 | TRAJ3 | CAGPYSSASKIIF | TRBV20-1 | TRBJ1-2 | CSARTGGDYGYTF | KLGGALQAK | NO | NO | NO |
| 17534 | TRAV16 | TRAJ10 | CALSLTGGGNKLTF | TRBV27 | TRBJ2-7 | CASSLSSGGQGEQYF | KLGGALQAK | NO | NO | NO |
| 17535 | TRAV24 | TRAJ36 | CAPGVTGANNLFF | TRBV19 | TRBJ1-1 | CASSIGVATEAFF | KLGGALQAK | YES | NO | NO |
| 17536 | TRAV12-2 | TRAJ12 | CAVSNKLIF | TRBV7-9 | TRBJ2-4 | CASSLPRDNQCF | KLGGALQAK | NO | NO | NO |
| 17537 | TRAV8-1 | TRAJ39 | CAVNLPGNNAGNMLTF | TRBV7-9 | TRBJ2-7 | CASSLGPYEQYV | KLGGALQAK | NO | NO | NO |
| 17538 | TRAV19 | TRAJ5 | CALSEYLDTGRRALTF | TRBV5-6 | TRBJ2-3 | CASSLELGPDTQYF | AVFDRKSDAK | YES | NO | YES |
| 17539 | TRAV12-2 | TRAJ57 | CAPGETQGGSEKLVF | TRBV19 | TRBJ1-1 | CASSPNGPGAEAFF | KLGGALQAK | YES | NO | NO |
| 17540 | TRAV13-1 | TRAJ45 | CAATPRGGADGLTF | TRBV13 | TRBJ2-1 | CASSLYRVRISGAYNEQFF | KLGGALQAK | YES | NO | NO |
| 17541 | TRAV12-3 | TRAJ49 | CAMAGDTGNQFYF | TRBV12-4 | TRBJ2-1 | CASIPPGRSRDEQFF | AVFDRKSDAK | YES | YES | YES |
| 17542 | TRAV3 | TRAJ20 | CAVREQTNDYKLSF | TRBV19 | TRBJ1-1 | CASSTEPGLNTEAFF | IVTDFSVIK | NO | NO | NO |
| 17543 | TRAV27 | TRAJ8 | CAANTGFQKLVF | TRBV4-3 | TRBJ1-5 | CASSQDRSPSQPQHF | AVFDRKSDAK | YES | NO | YES |
| 17544 | TRAV12-1 | TRAJ49 | CVVNTGNQFYF | TRBV12-4 | TRBJ1-2 | CASTGTGEYYGYTF | KLGGALQAK | YES | NO | NO |
| 17545 | TRAV12-1 | TRAJ5 | CVVKSTGRRALTF | TRBV27 | TRBJ1-2 | CASSFGPGTGVGYTF | KLGGALQAK | YES | NO | NO |
| 17546 | TRAV9-2 | TRAJ54 | CALLWGAQKLVF | TRBV5-1 | TRBJ2-7 | CASSPMRGALLEQYF | RLRAEAQVK | YES | YES | YES |
| 17547 | TRAV27 | TRAJ20 | CAAVTNDYKLSF | TRBV19 | TRBJ2-1 | CASSIDLNNEQFF | KLGGALQAK | YES | NO | NO |
| 17548 | TRAV14DV4 | TRAJ6 | CAMRPPSGGSYIPTF | TRBV7-9 | TRBJ2-3 | CASTSDDTQYF | KLGGALQAK | YES | NO | NO |
| 17549 | TRAV6 | TRAJ31 | CALDGNARLMF | TRBV7-9 | TRBJ1-1 | CASSLSFGVRNTEAFF | KLGGALQAK | YES | NO | NO |
| 17550 | TRAV12-1 | TRAJ4 | CVVPSGGYNKLIF | TRBV19 | TRBJ2-3 | CASSIQGGIPDTQYF | KLGGALQAK | YES | NO | NO |
| 17551 | TRAV12-2 | TRAJ31 | CAVNRVRDARLMF | TRBV12-4 | TRBJ2-1 | CASSLNPGLATYNEQFF | KLGGALQAK | YES | NO | NO |
| 17552 | TRAV14DV4 | TRAJ49 | CAMREITGNQFYF | TRBV2 | TRBJ2-1 | CASRTGLAGPQDEQFF | KLGGALQAK | YES | NO | NO |
| 17553 | TRAV22 | TRAJ20 | CAVERSDYKLSF | TRBV14 | TRBJ2-3 | CASRTVSTDTQYF | AVFDRKSDAK | YES | NO | YES |
| 17554 | TRAV12-1 | TRAJ41 | CVETNSGYALNF | TRBV5-5 | TRBJ2-1 | CASSIEDGGYNEQFF | KLGGALQAK | YES | NO | NO |
| 17555 | TRAV12-1 | TRAJ49 | CVAAYGTGNQFYF | TRBV5-1 | TRBJ2-7 | CASSPGRSYEQYF | KLGGALQAK | YES | NO | NO |
| 17556 | TRAV8-3 | TRAJ23 | CAVGPYNQGGKLIF | TRBV2 | TRBJ2-5 | CASSDQREPAETQYF | AVFDRKSDAK | YES | NO | YES |
| 17557 | TRAV1-2 | TRAJ33 | CAVGFMDSNYQLIW | TRBV20-1 | TRBJ1-1 | CSAYPASNTEAFF | KLGGALQAK | NO | NO | NO |
| 17558 | TRAV17 | TRAJ39 | CATDNAGNMLTF | TRBV10-1 | TRBJ1-1 | CASSEYRRADTEAFF | KLGGALQAK | YES | NO | NO |
| 17559 | TRAV10 | TRAJ4 | CVVRSGGYNKLIF | TRBV19 | TRBJ1-4 | CASRALGQRNEKLFF | KLGGALQAK | YES | NO | NO |
| 17560 | TRAV12-1 | TRAJ33 | CVVNKGGSNYQLIW | TRBV5-5 | TRBJ2-1 | CASSTYGEGLGLTYNEQFF | KLGGALQAK | YES | NO | NO |
| 17561 | TRAV25 | TRAJ39 | CAGINAGNMLTF | TRBV4-1 | TRBJ1-2 | CASSLFQGRSIANYGYTF | KLGGALQAK | YES | NO | NO |
| 17562 | TRAV8-3 | TRAJ45 | CAVGGAYSGGGADGLTF | TRBV6-6 | TRBJ1-1 | CASSYSGTGNTEAFF | KLGGALQAK | NO | NO | NO |
| 17563 | TRAV3 | TRAJ20 | CAVRDMKDDYKLSF | TRBV11-3 | TRBJ2-5 | CASSYSGTGARETQYF | KLGGALQAK | YES | NO | NO |
| 17564 | TRAV17 | TRAJ43 | CATDGNNDMRF | TRBV6-5 | TRBJ2-7 | CASSWTSGGAGYEQYF | KLGGALQAK | YES | NO | NO |
| 17565 | TRAV10 | TRAJ38 | CVVSFNAGNNRKLIW | TRBV7-2 | TRBJ2-3 | CASSLGASGGDTQYF | KLGGALQAK | YES | NO | NO |
| 17566 | TRAV35 | TRAJ11 | CAGPGYSTLTF | TRBV19 | TRBJ1-1 | CASSYQGAGTEAFF | KLGGALQAK | YES | NO | NO |
| 17567 | TRAV29DV5 | TRAJ17 | CAASALGPLKAAGNKLTF | TRBV10-2 | TRBJ2-1 | CASSHTLSSYNEQFF | KLGGALQAK | YES | NO | NO |
| 17568 | TRAV14DV4 | TRAJ9 | CAMREEDTGGFKTIF | TRBV12-3 | TRBJ2-1 | CASSFTGSSYNEQFF | KLGGALQAK | YES | NO | NO |
| 17569 | TRAV13-2 | TRAJ42 | CAENINGYGGSQGNLIF | TRBV7-9 | TRBJ2-2 | CASSLGDGELFF | KLGGALQAK | YES | NO | NO |
| 17570 | TRAV17 | TRAJ37 | CATVPFFSSNTGKLIF | TRBV20-1 | TRBJ2-3 | CSARGGVRTTPHEETSTDTQYF | KLGGALQAK | NO | NO | NO |
| 17571 | TRAV12-1 | TRAJ40 | CVVIKFETSGTYKYIF | TRBV20-1 | TRBJ2-2 | CSAAGQANTGELFF | KLGGALQAK | YES | NO | NO |
| 17572 | TRAV8-6 | TRAJ20 | CAVSRSNDYKLSF | TRBV29-1 | TRBJ2-5 | CSVGTAQETQYF | KLGGALQAK | YES | NO | NO |
| 17573 | TRAV17 | TRAJ41 | CATDSGYALNF | TRBV12-4 | TRBJ2-2 | CASSLGDRGQGGELFF | KLGGALQAK | YES | NO | NO |
| 17574 | TRAV24 | TRAJ13 | CACSGGYQKVTF | TRBV2 | TRBJ2-5 | CASSRGPGQGEETQYF | KLGGALQAK | YES | NO | NO |
| 17575 | TRAV14DV4 | TRAJ54 | CAMREVLGGAQKLVF | TRBV7-2 | TRBJ2-1 | CASSSTGGVANEQFF | AVFDRKSDAK | YES | NO | YES |
| 17576 | TRAV38-1 | TRAJ47 | CASMEYGNKLVF | TRBV7-9 | TRBJ1-1 | CASSFLTAENTEAFF | KLGGALQAK | YES | NO | NO |
| 17577 | TRAV1-2 | TRAJ20 | CAVRPFNDYKLSF | TRBV2 | TRBJ2-3 | CASEGGGASTTDTQYF | KLGGALQAK | YES | NO | NO |
| 17578 | TRAV38-1 | TRAJ53 | CAFRGSGGSNYKLTF | TRBV6-4 | TRBJ1-5 | CASSDREGSNQPQHF | KLGGALQAK | YES | NO | NO |
| 17579 | TRAV3 | TRAJ11 | CAVSFDSGYSTLTF | TRBV7-8 | TRBJ2-6 | CASSLGGLWTEYDSGANVLTF | KLGGALQAK | NO | NO | NO |
| 17580 | TRAV12-1 | TRAJ26 | CVVNTVYGQNFVF | TRBV6-1 | TRBJ2-4 | CASSEPSDIAKNIQYF | KLGGALQAK | YES | NO | NO |
| 17581 | TRAV4 | TRAJ33 | CLCGDSNYQLIW | TRBV12-5 | TRBJ2-1 | CASGFGSGSGDEQFF | KLGGALQAK | NO | NO | NO |
| 17582 | TRAV30 | TRAJ48 | CGTEIGGFGNEKLTF | TRBV5-1 | TRBJ2-1 | CASSPRVSYNEQFF | KLGGALQAK | YES | NO | NO |
| 17583 | TRAV21 | TRAJ40 | CAFTSGTYKYIF | TRBV3-1 | TRBJ2-2 | CASSQEARGRSNTGELFF | AVFDRKSDAK | YES | YES | YES |
| 17584 | TRAV29DV5 | TRAJ40 | CAASAAGTYKYIF | TRBV11-3 | TRBJ2-1 | CASSIAGGYEQFF | KLGGALQAK | YES | NO | NO |
| 17585 | TRAV14DV4 | TRAJ20 | CAMRGPSNDYKLSF | TRBV27 | TRBJ1-1 | CASSRPGFATEAFF | KLGGALQAK | YES | NO | NO |
| 17586 | TRAV12-3 | TRAJ31 | CAMSADARLMF | TRBV4-2 | TRBJ2-7 | CASSQETGGEQYV | KLGGALQAK | NO | NO | NO |
| 17587 | TRAV38-2DV8 | TRAJ23 | CASLIGNQGGKLIF | TRBV7-9 | TRBJ1-5 | CASSPGTISNQPQHF | KLGGALQAK | YES | NO | NO |
| 17588 | TRAV5 | TRAJ42 | CAERGGSQGNLIF | TRBV7-8 | TRBJ2-2 | CASSSRGTENTGELFF | KLGGALQAK | YES | NO | NO |
| 17589 | TRAV4 | TRAJ34 | CLVGRYNTDKLIF | TRBV12-4 | TRBJ1-1 | CASSNSLGDEAFF | KLGGALQAK | YES | NO | NO |
| 17590 | TRAV35 | TRAJ45 | CAGPNSGGGADGLTF | TRBV4-1 | TRBJ2-1 | CASSQHTEFYHEQFF | KLGGALQAK | YES | NO | NO |
| 17591 | TRAV41 | TRAJ43 | CAVRRYNNNDMRF | TRBV4-3 | TRBJ2-2 | CASSQERFRPGELFF | KLGGALQAK | YES | NO | NO |
| 17592 | TRAV9-2 | TRAJ41 | CALSDRVWIDSGYALNF | TRBV4-3 | TRBJ2-1 | CASSPTGGGSYNEQFF | KLGGALQAK | YES | NO | NO |
| 17593 | TRAV20 | TRAJ26 | CAVRESNYGQNFVF | TRBV27 | TRBJ1-1 | CASSTNTEAFF | KLGGALQAK | YES | NO | NO |
| 17594 | TRAV13-1 | TRAJ41 | CAAPPDSNSGYALNF | TRBV7-6 | TRBJ2-1 | CASSPRGAWFDEQFF | KLGGALQAK | YES | NO | NO |
| 17595 | TRAV21 | TRAJ26 | CAVKANYGQNFVF | TRBV5-4 | TRBJ2-1 | CASSLVQGGPLVEQFF | KLGGALQAK | YES | NO | NO |
| 17596 | TRAV5 | TRAJ34 | CAENLVLNTDKLIF | TRBV6-6 | TRBJ1-1 | CASSSTGLNTEAFF | KLGGALQAK | NO | NO | NO |
| 17597 | TRAV38-2DV8 | TRAJ34 | CAYKNDTDKLIF | TRBV9 | TRBJ2-5 | CASSVGTDLYQETQYF | KLGGALQAK | YES | NO | NO |
| 17598 | TRAV12-3 | TRAJ16 | CAISSSDGQKLLF | TRBV6-1 | TRBJ2-2 | CASSVIKENTGELFF | KLGGALQAK | NO | NO | NO |
| 17599 | TRAV12-2 | TRAJ3 | CAVGGGASKIIF | TRBV24-1 | TRBJ2-3 | CATSDPDRITDTQYF | KLGGALQAK | YES | NO | NO |
| 17600 | TRAV20 | TRAJ32 | CAVQASIYGGATNKLIF | TRBV19 | TRBJ1-5 | CASSPWLGDRQPQHF | KLGGALQAK | YES | NO | NO |
| 17601 | TRAV1-2 | TRAJ16 | CAVLGGQKLLF | TRBV12-4 | TRBJ2-3 | CASRIRGVPDTQYF | KLGGALQAK | NO | NO | NO |
| 17602 | TRAV12-1 | TRAJ30 | CVVNPYRDDKIIF | TRBV5-1 | TRBJ2-7 | CASSLGLAGTYEQYF | KLGGALQAK | YES | NO | NO |
| 17603 | TRAV1-2 | TRAJ20 | CAPVSNDYKLSF | TRBV27 | TRBJ2-2 | CASSLGPMNTGELFF | KLGGALQAK | YES | NO | NO |
| 17604 | TRAV38-2DV8 | TRAJ49 | CAYRSFSGNQFYF | TRBV20-1 | TRBJ1-2 | CSASRRDIISGYTF | KLGGALQAK | YES | NO | NO |
| 17605 | TRAV5 | TRAJ9 | CAERGGTGGFKTIF | TRBV3-1 | TRBJ1-5 | CASSQGEGYSNQPQHF | KLGGALQAK | YES | NO | NO |
| 17606 | TRAV8-3 | TRAJ21 | CAVGADNFNKFYF | TRBV5-4 | TRBJ2-3 | CASSVGMAGGTDTQYF | KLGGALQAK | YES | NO | NO |
| 17607 | TRAV12-1 | TRAJ49 | CVVTGNQFYF | TRBV10-3 | TRBJ2-7 | CAISESITGFYEQYF | AVFDRKSDAK | NO | NO | NO |

| | | | | | | | | | | |
|---|---|---|---|---|---|---|---|---|---|---|
| 17608 | TRAV21 | TRAJ26 | CAVLYYGQNFVF | TRBV6-3 | TRBJ1-2 | CASSSNRAGYTF | KLGGALQAK | YES | NO | NO |
| 17609 | TRAV8-4 | TRAJ49 | CAVSGTGNQFYF | TRBV11-3 | TRBJ1-3 | CASSLLISSGNTIYF | KLGGALQAK | YES | NO | NO |
| 17610 | TRAV8-6 | TRAJ16 | CAVSDTLSDGQKLLF | TRBV6-1 | TRBJ2-1 | CASSELRYNEQFF | KLGGALQAK | NO | NO | NO |
| 17611 | TRAV19 | TRAJ52 | CALIGGTSYGKLTF | TRBV6-1 | TRBJ1-4 | CASYRGITNEKLFF | KLGGALQAK | YES | NO | NO |
| 17612 | TRAV21 | TRAJ34 | CAVRYGTDKLIF | TRBV7-9 | TRBJ2-7 | CASSSTSKGQGSYEQYF | KLGGALQAK | YES | NO | NO |
| 17613 | TRAV12-3 | TRAJ39 | CAIGPGNMLTF | TRBV4-1 | TRBJ2-5 | CASSHFLTEETQYF | KLGGALQAK | YES | NO | NO |
| 17614 | TRAV8-2 | TRAJ20 | CVVSDLYDYKLSF | TRBV24-1 | TRBJ2-1 | CATSGNQLGTGAYNEQFF | KLGGALQAK | YES | NO | NO |
| 17615 | TRAV5 | TRAJ49 | CAEKKLTGNQFYF | TRBV20-1 | TRBJ2-1 | CSARGSGLERSSYNEQFF | KLGGALQAK | YES | NO | NO |
| 17616 | TRAV19 | TRAJ48 | CALSESISNFGNEKLTF | TRBV7-9 | TRBJ1-1 | CASSPREEAFF | KLGGALQAK | YES | NO | NO |
| 17617 | TRAV38-2DV8 | TRAJ48 | CAPNFGNEKLTF | TRBV3-1 | TRBJ2-1 | CASSQDGGIAYNEQFF | KLGGALQAK | YES | NO | NO |
| 17618 | TRAV29DV5 | TRAJ48 | CAASAGFGNEKLTF | TRBV29-1 | TRBJ2-1 | CSTAESSYNEQFF | KLGGALQAK | YES | NO | NO |
| 17619 | TRAV19 | TRAJ52 | CALVNAGGTSYGKLTF | TRBV12-4 | TRBJ1-2 | CKRTETGKTNYGYTF | KLGGALQAK | YES | NO | NO |
| 17620 | TRAV17 | TRAJ47 | CATDGYGNKLVF | TRBV18 | TRBJ1-2 | CASSPPASFPYGYTF | KLGGALQAK | YES | NO | NO |
| 17621 | TRAV1-2 | TRAJ33 | CAVMDSNYQLIW | TRBV20-1 | TRBJ1-1 | CSARAGDTEAFF | KLGGALQAK | YES | NO | NO |
| 17622 | TRAV8-4 | TRAJ52 | CAVLQGAGGTSYGKLTF | TRBV27 | TRBJ2-3 | CASSPGLVLSDTQYF | KLGGALQAK | YES | NO | NO |
| 17623 | TRAV20 | TRAJ6 | CAVQGTGGSYIPTF | TRBV5-1 | TRBJ2-1 | CASSWGQGNNEQFF | KLGGALQAK | YES | NO | NO |
| 17624 | TRAV12-2 | TRAJ9 | CAVPYTGGFKTIF | TRBV7-9 | TRBJ2-5 | CASSKGDQGETQYF | KLGGALQAK | YES | NO | NO |
| 17625 | TRAV17 | TRAJ39 | CATNNNAGNMLTF | TRBV4-1 | TRBJ2-5 | CASSQVLEGETQYF | IVTDFSVIK | YES | NO | YES |
| 17626 | TRAV5 | TRAJ20 | CAEMRSNDYKLSF | TRBV19 | TRBJ1-1 | CASSIRMNTEAFF | KLGGALQAK | YES | NO | NO |
| 17627 | TRAV38-1 | TRAJ28 | CAFITSGAGSYQLTF | TRBV7-9 | TRBJ2-1 | CASSGRGTSGRNEQFF | KLGGALQAK | YES | NO | NO |
| 17628 | TRAV36DV7 | TRAJ53 | CGGSNYKLTF | TRBV7-9 | TRBJ2-1 | CASSKEPGLEFF | KLGGALQAK | NO | NO | NO |
| 17629 | TRAV22 | TRAJ43 | CAVEPSRNNDMRF | TRBV5-4 | TRBJ2-7 | CASSLYGYEQYV | KLGGALQAK | NO | NO | NO |
| 17630 | TRAV38-2DV8 | TRAJ50 | CAYGTSYDKVIF | TRBV12-5 | TRBJ2-1 | CASGLEGQGAGEQFF | KLGGALQAK | YES | NO | NO |
| 17631 | TRAV20 | TRAJ45 | CAAYTGGGADGLTF | TRBV5-4 | TRBJ1-2 | CASSGLDKANYGYTF | KLGGALQAK | YES | NO | NO |
| 17632 | TRAV29DV5 | TRAJ23 | CAAYIYNQGGKLIF | TRBV7-9 | TRBJ2-3 | CASSGGLAGGTDTQYF | KLGGALQAK | YES | NO | NO |
| 17633 | TRAV10 | TRAJ43 | CVVSPKFNNNDMRF | TRBV30 | TRBJ1-6 | CAWRAPDNSPLHF | KLGGALQAK | NO | NO | NO |
| 17634 | TRAV19 | TRAJ3 | CALSDIRDSSASKIIF | TRBV5-4 | TRBJ2-1 | CASSLDGTEYYNEQFF | KLGGALQAK | YES | NO | NO |
| 17635 | TRAV6 | TRAJ30 | CALDIGDDKIIF | TRBV20-1 | TRBJ2-5 | CSARDGDQGRGDNQETQYF | KLGGALQAK | NO | NO | NO |
| 17636 | TRAV8-1 | TRAJ39 | CAVKRAGNMLTF | TRBV6-5 | TRBJ2-3 | CASSPAGFSGGATDTQYF | KLGGALQAK | NO | NO | NO |
| 17637 | TRAV13-1 | TRAJ34 | CAAKTDKLIF | TRBV27 | TRBJ1-2 | CASSYGTDYGYTF | KLGGALQAK | YES | NO | NO |
| 17638 | TRAV19 | TRAJ29 | CALSEAFSGNTPLVF | TRBV19 | TRBJ1-2 | CASSARTGVGYGYTF | KLGGALQAK | YES | NO | NO |
| 17640 | TRAV26-2 | TRAJ57 | CILKWGGSEKLVF | TRBV4-1 | TRBJ1-2 | CASSQEKFLPGGYTF | KLGGALQAK | YES | NO | NO |
| 17641 | TRAV16 | TRAJ40 | CALRFLGTYKYIF | TRBV25-1 | TRBJ2-1 | CASIPDGYNEQFF | KLGGALQAK | NO | NO | NO |
| 17642 | TRAV12-1 | TRAJ28 | CVVNHPLWGWELPTHF | TRBV6-5 | TRBJ2-3 | CASSSGTGNLDTQYF | KLGGALQAK | NO | NO | NO |
| 17643 | TRAV12-2 | TRAJ13 | CAVNRSGGYQKVTF | TRBV4-1 | TRBJ2-1 | CASSQAGDNEQFF | KLGGALQAK | YES | NO | NO |
| 17644 | TRAV5 | TRAJ23 | CAESKHNQGGKLIF | TRBV11-2 | TRBJ2-3 | CASSFIGSTDTQYF | KLGGALQAK | YES | NO | NO |
| 17645 | TRAV30 | TRAJ49 | CGTERNTGNQFYF | TRBV7-9 | TRBJ2-7 | CASKDGISYEQYF | KLGGALQAK | YES | NO | NO |
| 17646 | TRAV8-3 | TRAJ10 | CAVDPRILTGGGNKLTF | TRBV5-6 | TRBJ2-3 | CASSPGGPYFTDTQYF | KLGGALQAK | YES | NO | NO |
| 17647 | TRAV14DV4 | TRAJ33 | CAMREGRMDSNYQLIW | TRBV3-1 | TRBJ2-3 | CASSQDSGYPTDTQYF | KLGGALQAK | YES | NO | NO |
| 17648 | TRAV8-3 | TRAJ34 | CAVGEGDKLIF | TRBV10-3 | TRBJ2-5 | CAITISGLQETQYF | KLGGALQAK | YES | NO | NO |
| 17649 | TRAV1-1 | TRAJ21 | CAVRDFNKFYF | TRBV7-2 | TRBJ2-3 | CASSLVGGRASDTQYF | KLGGALQAK | YES | NO | NO |
| 17650 | TRAV14DV4 | TRAJ53 | CAMREALLWGGSNYKLTF | TRBV7-2 | TRBJ2-1 | CASSSKREENEQFF | KLGGALQAK | YES | NO | NO |
| 17651 | TRAV19 | TRAJ35 | CALSEGGSI | TRBV7-7 | TRBJ2-1 | CASSLLAGSMGEQFF | KLGGALQAK | NO | NO | NO |
| 17652 | TRAV17 | TRAJ11 | CATDSGYSTLTF | TRBV7-8 | TRBJ2-1 | CASSAPSGSHYNEQFF | KLGGALQAK | YES | NO | NO |
| 17653 | TRAV13-1 | TRAJ29 | CAANWIGNTPLVF | TRBV5-1 | TRBJ1-1 | CASSLVQGMNTEAFF | KLGGALQAK | YES | NO | NO |
| 17654 | TRAV12-2 | TRAJ7 | CALDGNNRLAF | TRBV24-1 | TRBJ2-1 | CATSGLGGADEQFF | KLGGALQAK | YES | NO | NO |
| 17655 | TRAV12-3 | TRAJ39 | CAIGPGNMLTF | TRBV19 | TRBJ1-1 | CASSTGWTEAFF | KLGGALQAK | YES | NO | NO |
| 17656 | TRAV12-1 | TRAJ21 | CVVNTGFNKFYF | TRBV3-1 | TRBJ1-1 | CASSLTTDWAEAFF | KLGGALQAK | YES | NO | NO |
| 17657 | TRAV12-2 | TRAJ20 | CAVNRDYKLSF | TRBV29-1 | TRBJ2-3 | CSVAGTSGRGPDTQYF | ELAGIGILTV | YES | YES | YES |
| 17658 | TRAV12-1 | TRAJ41 | CVVNKDSGYALNF | TRBV5-1 | TRBJ2-1 | CASSIGLAGANEQFF | KLGGALQAK | YES | NO | NO |
| 17659 | TRAV3 | TRAJ34 | CAVRDVRSYNTDKLIF | TRBV7-9 | TRBJ1-2 | CASSLLSGDPERYTF | KLGGALQAK | NO | NO | NO |
| 17660 | TRAV12-3 | TRAJ32 | CAMVGDGGATNKLIF | TRBV6-5 | TRBJ2-1 | CASSYSGSNEQFF | KLGGALQAK | NO | NO | NO |
| 17661 | TRAV29DV5 | TRAJ33 | CAASAGDSNYQLIW | TRBV24-1 | TRBJ2-7 | CATSDQGTEQYF | KLGGALQAK | YES | NO | NO |
| 17662 | TRAV4 | TRAJ36 | CLVGDPRTGANNLFF | TRBV5-4 | TRBJ1-1 | CASSLAVGMNTEAFF | KLGGALQAK | YES | NO | NO |
| 17663 | TRAV5 | TRAJ27 | CAEFGYNTNAGKSTF | TRBV5-6 | TRBJ2-1 | CASSSARDNEQFF | KLGGALQAK | YES | NO | NO |
| 17664 | TRAV3 | TRAJ5 | CAVRDRVMDTGRRALTF | TRBV2 | TRBJ1-5 | CASRPGQTNQPQHF | KLGGALQAK | NO | NO | NO |
| 17666 | TRAV1-2 | TRAJ5 | CAVSPDTGRRALTF | TRBV29-1 | TRBJ1-2 | CSVDIRQGFNNYGYTF | AVFDRKSDAK | YES | NO | YES |
| 17667 | TRAV5 | TRAJ43 | CAESLRNNDMRF | TRBV12-5 | TRBJ2-2 | CASGQGQGVTGELFF | KLGGALQAK | YES | NO | NO |
| 17668 | TRAV29DV5 | TRAJ33 | CAASADSNYQLIW | TRBV5-1 | TRBJ2-7 | CASSLEPSYEQYF | KLGGALQAK | YES | NO | NO |
| 17669 | TRAV20 | TRAJ26 | CAVRQNYGQNFVF | TRBV4-1 | TRBJ2-1 | CASSQALSGYNEQFF | KLGGALQAK | YES | NO | NO |
| 17670 | TRAV8-2 | TRAJ10 | CVSSWGGGGNKLTF | TRBV20-1 | TRBJ2-1 | CSAVEGTSGGAYNEQFF | KLGGALQAK | NO | NO | NO |
| 17671 | TRAV35 | TRAJ42 | CAGQSGRVGGSQGNLIF | TRBV7-9 | TRBJ1-1 | CASGGAIDTEAFF | KLGGALQAK | YES | NO | NO |
| 17672 | TRAV9-2 | TRAJ24 | CALSDGDSWGKFQF | TRBV12-4 | TRBJ1-3 | CASSSLKLSGNTIYF | KLGGALQAK | YES | NO | NO |
| 17673 | TRAV5 | TRAJ43 | CAEILLNDMRF | TRBV5-1 | TRBJ1-1 | CASSTGQGNTEAFF | KLGGALQAK | YES | NO | NO |
| 17674 | TRAV1-2 | TRAJ38 | CAVWYAGNNRKLIW | TRBV6-5 | TRBJ1-1 | CASSYGENTEAFF | KLGGALQAK | YES | NO | NO |
| 17675 | TRAV17 | TRAJ21 | CATDWGGKFYF | TRBV12-4 | TRBJ1-1 | CASSFLSATEAFF | KLGGALQAK | YES | NO | NO |
| 17676 | TRAV12-1 | TRAJ42 | CVGPNYGGSQGNLIF | TRBV9 | TRBJ2-1 | CASSVEAGWDEQFF | KLGGALQAK | YES | NO | NO |
| 17677 | TRAV19 | TRAJ5 | CALSEALGGRRALTF | TRBV19 | TRBJ1-2 | CASSISPGFYGYTF | KLGGALQAK | YES | NO | NO |
| 17678 | TRAV8-4 | TRAJ23 | CAVSDYNQGGKLIF | TRBV5-6 | TRBJ2-1 | CASSLREQGRYNEQFF | KLGGALQAK | YES | NO | NO |
| 17679 | TRAV41 | TRAJ53 | CAVRGLRGSNYKLTF | TRBV19 | TRBJ1-5 | CASNMQGGHQPQHF | KLGGALQAK | YES | NO | NO |
| 17680 | TRAV20 | TRAJ4 | CAVQDLISGGYNKLIF | TRBV19 | TRBJ2-1 | CASSIDSGGSYNEQFF | KLGGALQAK | YES | NO | NO |
| 17681 | TRAV1-2 | TRAJ34 | CAVYTDKLIF | TRBV4-1 | TRBJ2-7 | CASSRGGGLYEQYF | RLRAEAQVK | YES | YES | YES |
| 17682 | TRAV20 | TRAJ39 | CAVRDNAGNMLTF | TRBV19 | TRBJ1-1 | CASSITPLNTEAFF | KLGGALQAK | YES | NO | NO |
| 17685 | TRAV1-2 | TRAJ36 | CACQTGANNLFF | TRBV6-1 | TRBJ1-1 | CASTPGTGPTEAFF | KLGGALQAK | YES | NO | NO |
| 17686 | TRAV35 | TRAJ48 | CAGPSAKLTF | TRBV9 | TRBJ2-2 | CASSPTGTGELFF | KLGGALQAK | YES | NO | NO |
| 17687 | TRAV24 | TRAJ49 | CAFIITGNQFYF | TRBV9 | TRBJ2-7 | CASSVMGSPYEQYF | KLGGALQAK | YES | NO | NO |
| 17688 | TRAV8-1 | TRAJ7 | CAVRDGNNRLAF | TRBV6-4 | TRBJ2-1 | CASSDGLNEQFF | KLGGALQAK | YES | NO | NO |
| 17689 | TRAV12-1 | TRAJ42 | CVVNRWGGGSQGNLIF | TRBV25-1 | TRBJ1-1 | CASSDRQGVTEAFF | KLGGALQAK | YES | NO | NO |
| 17690 | TRAV21 | TRAJ37 | CAARLAGNTGKLIF | TRBV19 | TRBJ1-1 | CASSVQGATEAFF | KLGGALQAK | YES | NO | NO |
| 17691 | TRAV8-3 | TRAJ47 | CAVVYMEYGNKLVF | TRBV7-9 | TRBJ2-7 | CASSLGYEQYF | KLGGALQAK | YES | NO | NO |
| 17692 | TRAV25 | TRAJ52 | CAGANAGGTSYGKLTF | TRBV29-1 | TRBJ2-5 | CSVEDRGGGETQYF | KLGGALQAK | YES | NO | NO |
| 17693 | TRAV17 | TRAJ12 | CATEKDSSYKLIF | TRBV5-1 | TRBJ2-3 | CASSPGDQSTDTQYF | KLGGALQAK | YES | NO | NO |
| 17694 | TRAV40 | TRAJ53 | CLLISGGSNYKLTF | TRBV12-4 | TRBJ2-2 | CASSTTLYTGELFF | KLGGALQAK | NO | NO | NO |
| 17695 | TRAV26-1 | TRAJ30 | CIEDDKIIF | TRBV2 | TRBJ1-5 | CASTTYFGGNQPQHF | KLGGALQAK | NO | NO | NO |
| 17696 | TRAV8-4 | TRAJ49 | CAVSATGNQFYF | TRBV15 | TRBJ1-2 | CATSRVGDSHNYGYTF | KLGGALQAK | YES | NO | NO |
| 17697 | TRAV12-2 | TRAJ10 | CAVSPTGGGNKLTF | TRBV4-2 | TRBJ1-3 | CASSQGVSPRGGNTIYF | KLGGALQAK | YES | NO | NO |
| 17698 | TRAV12-1 | TRAJ26 | CVVIHNYGQNFVF | TRBV4-1 | TRBJ1-5 | CASSQDPGSNQPQHF | KLGGALQAK | YES | NO | NO |
| 17699 | TRAV1-1 | TRAJ33 | CAVRRSNYQLIW | TRBV29-1 | TRBJ2-5 | CSVGTGTGVETQYF | KLGGALQAK | YES | NO | NO |
| 17700 | TRAV19 | TRAJ52 | CALSEAGGGTSYGKLTF | TRBV29-1 | TRBJ2-3 | CSVPPASGGAGDTQYF | KLGGALQAK | YES | NO | NO |
| 17701 | TRAV13-2 | TRAJ45 | CAETGEYSGGGADGLTF | TRBV9 | TRBJ2-2 | CASSVAGGGTGELFF | KLGGALQAK | YES | NO | NO |
| 17702 | TRAV12-3 | TRAJ39 | CAIGPGNMLTF | TRBV5-6 | TRBJ2-7 | CASSGLAARPTGYYEQYV | KLGGALQAK | NO | NO | NO |
| 17703 | TRAV12-2 | TRAJ43 | CAVKRANNDMRF | TRBV7-3 | TRBJ1-1 | CASSLLPGRITEAFF | AVFDRKSDAK | NO | NO | NO |
| 17705 | TRAV38-1 | TRAJ52 | CAFMSHPAGGGTSYGKLTF | TRBV27 | TRBJ1-1 | CASSFSRGLGTEAFF | KLGGALQAK | YES | NO | NO |
| 17706 | TRAV29DV5 | TRAJ40 | CAARPLGSGTYKYIF | TRBV4-1 | TRBJ1-1 | CASSQDPQGAVNTEAFF | KLGGALQAK | YES | NO | NO |
| 17707 | TRAV12-2 | TRAJ20 | CAVSGYKLSF | TRBV20-1 | TRBJ2-1 | CSAMEGSYNEQFF | AVFDRKSDAK | YES | NO | YES |
| 17708 | TRAV4 | TRAJ32 | CLLDYGGATNKLIF | TRBV4-1 | TRBJ1-2 | CASNPGRGVSYTF | KLGGALQAK | YES | NO | NO |
| 17709 | TRAV8-3 | TRAJ30 | CAVYVRDRDDKIIF | TRBV5-1 | TRBJ1-1 | CASSGGQGSEAFF | KLGGALQAK | YES | NO | NO |
| 17710 | TRAV38-2DV8 | TRAJ54 | CALCLIQGAQKLVF | TRBV7-9 | TRBJ2-3 | CASSLDGMGRDTQYF | KLGGALQAK | YES | NO | NO |
| 17711 | TRAV19 | TRAJ52 | CALSDSNAGGTSYGKLTF | TRBV13 | TRBJ1-1 | CASSTEKGAEAFF | KLGGALQAK | YES | NO | NO |
| 17712 | TRAV3 | TRAJ6 | CAVITGGGSYIPTF | TRBV30 | TRBJ2-7 | CAWSVQASGEQYV | KLGGALQAK | NO | NO | NO |
| 17713 | TRAV12-2 | TRAJ52 | CAVHLNAGGTSYGKLTF | TRBV19 | TRBJ1-5 | CASSIWGISGQPQHF | ELAGIGILTV | YES | NO | YES |
| 17714 | TRAV19 | TRAJ45 | CALSENSGGGADGLTF | TRBV5-6 | TRBJ2-4 | CASGPADRGFDIQYF | KLGGALQAK | YES | NO | NO |
| 17715 | TRAV1-2 | TRAJ15 | CAVITQAGTALIF | TRBV5-1 | TRBJ2-3 | CASSLLTGRGEAAQYF | KLGGALQAK | YES | NO | NO |
| 17716 | TRAV12-3 | TRAJ39 | CAIGPGNMLTF | TRBV27 | TRBJ2-2 | CASSLQNTGELFF | KLGGALQAK | NO | NO | NO |

| | | | | | | | | | |
|---|---|---|---|---|---|---|---|---|---|
| 17717 | TRAV12-2 | TRAJ39 | CAVKNDGDNNAGNMLTF | TRBV5-1 | TRBJ2-7 | CASSLDPGSPKYV | KLGGALQAK | NO | NO | NO |
| 17718 | TRAV1-2 | TRAJ5 | CAPDTGRRALTF | TRBV4-3 | TRBJ1-4 | CASSPEGDIANEKLFF | KLGGALQAK | YES | NO | NO |
| 17719 | TRAV17 | TRAJ37 | CATASNTGKLIF | TRBV11-2 | TRBJ2-7 | CASRSIYEQYV | KLGGALQAK | NO | NO | NO |
| 17720 | TRAV12-2 | TRAJ6 | CVRRAGGSYIPTF | TRBV19 | TRBJ2-1 | CASSFGGGEQFF | ELAGIGILTV | YES | NO | NO |
| 17721 | TRAV36DV7 | TRAJ20 | CAVNSNDYKLSF | TRBV5-1 | TRBJ2-4 | CASSLSGNIQYF | KLGGALQAK | NO | NO | NO |
| 17722 | TRAV21 | TRAJ35 | CAVRRVGFGNVLHC | TRBV24-1 | TRBJ2-7 | CATMSAGGEQYF | KLGGALQAK | NO | NO | NO |
| 17723 | TRAV13-2 | TRAJ4 | CAESKPGGYNKLIF | TRBV20-1 | TRBJ2-7 | CSARTPDRDLVYEQYF | KLGGALQAK | NO | NO | NO |
| 17724 | TRAV8-2 | TRAJ22 | CVGRKGSARQLTF | TRBV7-9 | TRBJ2-7 | CASKDGISYEQYF | KLGGALQAK | YES | NO | NO |
| 17725 | TRAV19 | TRAJ15 | CALSEATQAGTALIF | TRBV6-6 | TRBJ2-2 | CAITGVGTGELFF | KLGGALQAK | YES | NO | NO |
| 17726 | TRAV20 | TRAJ20 | CAVQGSVASDYKLSF | TRBV4-1 | TRBJ2-7 | CASSLLAEYEQYF | AVFDRKSDAK | YES | NO | YES |
| 17727 | TRAV8-6 | TRAJ49 | CAVSDRNQFYF | TRBV11-2 | TRBJ2-7 | CASSLDSAYEQYF | KLGGALQAK | YES | NO | NO |
| 17728 | TRAV19 | TRAJ49 | CALSEALGNQFYF | TRBV15 | TRBJ2-2 | CATSRDVGAGNTGELFF | AVFDRKSDAK | YES | NO | YES |
| 17729 | TRAV4 | TRAJ17 | CLVGDIIKAAGNKLTF | TRBV9 | TRBJ2-1 | CASSLREGYEQFF | KLGGALQAK | YES | NO | NO |
| 17730 | TRAV25 | TRAJ36 | CAGELLGANNLFF | TRBV5-5 | TRBJ2-7 | CASSLASDLYEQYV | AVFDRKSDAK | NO | NO | NO |
| 17731 | TRAV8-2 | TRAJ23 | CVVSGPGGKLIF | TRBV27 | TRBJ1-4 | CASSPSGTWGEKLFF | KLGGALQAK | YES | NO | NO |
| 17732 | TRAV14DV4 | TRAJ21 | CAMRGAIYNFNKFYF | TRBV20-1 | TRBJ2-7 | CSASLKEGREIYEQYF | KLGGALQAK | NO | NO | NO |
| 17733 | TRAV24 | TRAJ52 | CAFIGDAGGTSYGKLTF | TRBV4-3 | TRBJ2-3 | CASSQDHASGGALDTQYF | KLGGALQAK | YES | NO | NO |
| 17734 | TRAV26-2 | TRAJ49 | CILRPSNTGNQFYF | TRBV4-1 | TRBJ1-2 | CASSQGGTGIYGYTF | KLGGALQAK | YES | NO | NO |
| 17735 | TRAV3 | TRAJ26 | CAVRSYGQNFVF | TRBV20-1 | TRBJ2-1 | CSASTSGGSSYNEQFF | KLGGALQAK | NO | NO | NO |
| 17736 | TRAV21 | TRAJ26 | CAALDNYGQNFVF | TRBV6-3 | TRBJ2-2 | CASSYKREYTGELFF | KLGGALQAK | YES | NO | NO |
| 17737 | TRAV20 | TRAJ54 | CAVKIQGAQKLVF | TRBV7-2 | TRBJ2-2 | CASSLVGPGELFF | KLGGALQAK | YES | NO | NO |
| 17738 | TRAV1-2 | TRAJ31 | CAVKGARLMF | TRBV7-9 | TRBJ1-2 | CASSSSFTSVTANYGYTF | IVTDFSVIK | YES | YES | YES |
| 17739 | TRAV9-2 | TRAJ58 | CALSETSGSRLTF | TRBV10-3 | TRBJ2-1 | CATQRRGGYYNEQFF | KLGGALQAK | YES | NO | NO |
| 17740 | TRAV4 | TRAJ9 | CLVGGGTGGFKTIF | TRBV29-1 | TRBJ2-1 | CSVEWLNNEQFF | IPSINVHHY | YES | NO | NO |
| 17741 | TRAV12-1 | TRAJ41 | CVVNDGLAPNSNSGYALNF | TRBV6-1 | TRBJ2-7 | CASMWSDAYEQYF | KLGGALQAK | YES | NO | NO |
| 17742 | TRAV26-1 | TRAJ53 | CTRNGGSNYKLTF | TRBV19 | TRBJ1-1 | CASSAGLILGDTEAFF | KLGGALQAK | YES | NO | NO |
| 17743 | TRAV1-1 | TRAJ31 | CAVTGNNARLMF | TRBV4-3 | TRBJ1-2 | CASSPGTGDAYGYTF | KLGGALQAK | YES | NO | NO |
| 17744 | TRAV29DV5 | TRAJ49 | CAGSNTGNQFYF | TRBV7-9 | TRBJ2-1 | CASSHPWAGSYNEQFF | KLGGALQAK | YES | NO | NO |
| 17745 | TRAV8-2 | TRAJ49 | CVVSPGGNQFYF | TRBV20-1 | TRBJ2-7 | CSATPSGLAGAYEQYV | KLGGALQAK | NO | NO | NO |
| 17746 | TRAV19 | TRAJ39 | CALSEFNNAGNMLTF | TRBV27 | TRBJ2-1 | CASSLGDVSYNEQFF | KLGGALQAK | YES | NO | NO |
| 17747 | TRAV8-2 | TRAJ13 | CVVSDLNSGGYQKVTF | TRBV7-9 | TRBJ1-6 | CASSLKNGGPGGIGLHF | KLGGALQAK | YES | NO | NO |
| 17748 | TRAV25 | TRAJ45 | CAAYSGGGADGLTF | TRBV7-9 | TRBJ1-1 | CASSLGLVTEAFF | KLGGALQAK | YES | NO | NO |
| 17749 | TRAV14DV4 | TRAJ42 | CAMREGVEYYGGSQGNLIF | TRBV7-9 | TRBJ2-1 | CASSFEREQFF | KLGGALQAK | YES | NO | NO |
| 17750 | TRAV12-2 | TRAJ20 | CAVRRSNDYKLSF | TRBV6-3 | TRBJ2-2 | CASTPLRLMNTGELFF | KLGGALQAK | YES | NO | NO |
| 17751 | TRAV4 | TRAJ11 | CLVGDWENSGYSTLTF | TRBV7-6 | TRBJ2-3 | CASSLESTSGSPDTQYF | KLGGALQAK | NO | NO | NO |
| 17752 | TRAV3 | TRAJ52 | CAVRDRNAGGTSYGKLTF | TRBV5-6 | TRBJ2-5 | CASSLGPSTGSETQYF | KLGGALQAK | YES | NO | NO |
| 17753 | TRAV12-2 | TRAJ47 | CAVKLMEYGNKLVF | TRBV19 | TRBJ1-1 | CASIRGAGTEAFF | KLGGALQAK | YES | NO | NO |
| 17754 | TRAV1-2 | TRAJ28 | CALPAYSGAGSYQLTF | TRBV3-1 | TRBJ2-5 | CASSRGLAGSHQETQYF | KLGGALQAK | YES | NO | NO |
| 17755 | TRAV8-1 | TRAJ30 | CAVNVVVDDKIIF | TRBV20-1 | TRBJ2-2 | CSASGSGVGTGELFF | KLGGALQAK | YES | NO | NO |
| 17756 | TRAV9-2 | TRAJ53 | CALIPDSNYKLTF | TRBV10-1 | TRBJ2-1 | CASNDRGMFNEQFF | CLLWSFQTSA | YES | NO | YES |
| 17757 | TRAV20 | TRAJ42 | CAVQANDGGSQGNLIF | TRBV19 | TRBJ2-1 | CASSDVLGGQFF | KLGGALQAK | YES | NO | NO |
| 17758 | TRAV23DV6 | TRAJ47 | CAASMGEYGNKLVF | TRBV19 | TRBJ1-1 | CASSSSMNTEAFF | KLGGALQAK | YES | NO | NO |
| 17759 | TRAV27 | TRAJ13 | CAGFNSGGYQKVTF | TRBV12-3 | TRBJ2-7 | CASSLEGAGEQYV | KLGGALQAK | NO | NO | NO |
| 17760 | TRAV8-3 | TRAJ48 | CAVGARSNFGNEKLTF | TRBV20-1 | TRBJ1-5 | CSARDGIYSNQPQHF | KLGGALQAK | YES | NO | NO |
| 17761 | TRAV14DV4 | TRAJ23 | CAMREGIYNQGGKLIF | TRBV5-1 | TRBJ2-7 | CASSLLGLGEQYV | KLGGALQAK | NO | NO | NO |
| 17762 | TRAV16 | TRAJ29 | CALIRGNTPLVF | TRBV28 | TRBJ1-2 | CASSPPAVKFYGYTF | KLGGALQAK | NO | NO | NO |
| 17763 | TRAV17 | TRAJ58 | CAVARETSGSRLTF | TRBV19 | TRBJ2-1 | CASTITSGSSYNEQFF | KLGGALQAK | YES | NO | NO |
| 17764 | TRAV8-2 | TRAJ22 | CVGRKGSARQLTF | TRBV11-2 | TRBJ2-5 | CASRGPVETQYF | KLGGALQAK | YES | NO | NO |
| 17765 | TRAV26-2 | TRAJ48 | CILRGGSNFGNEKLTF | TRBV5-1 | TRBJ1-2 | CASSFITWLDSRNYGYTF | KLGGALQAK | YES | NO | NO |
| 17766 | TRAV12-1 | TRAJ18 | CVVNTDRGSTLGRLYF | TRBV24-1 | TRBJ2-2 | CATSDLPGQSNTGELFF | KLGGALQAK | YES | NO | NO |
| 17767 | TRAV38-1 | TRAJ31 | CAFMNNNARLMF | TRBV7-8 | TRBJ1-1 | CASREVNTEAFF | KLGGALQAK | YES | NO | NO |
| 17768 | TRAV17 | TRAJ10 | CATDTGGGNKLTF | TRBV19 | TRBJ1-5 | CASNPGTNNQPQHF | KLGGALQAK | YES | NO | NO |
| 17769 | TRAV13-2 | TRAJ49 | CAETSGNQFYF | TRBV19 | TRBJ2-7 | CASSPGSSYEQYF | KLGGALQAK | YES | NO | NO |
| 17770 | TRAV3 | TRAJ29 | CAVRDEGNTPLVF | TRBV10-3 | TRBJ2-1 | CAISETSGGGSNEQFF | KLGGALQAK | NO | NO | NO |
| 17771 | TRAV38-1 | TRAJ54 | CAFMSLIQGAQKLVF | TRBV3-1 | TRBJ2-2 | CASSQVFRGPPGELFF | KLGGALQAK | YES | NO | NO |
| 17772 | TRAV30 | TRAJ20 | CGTEVNDYKLSF | TRBV6-6 | TRBJ1-5 | CASRPAAPYQPQHF | KLGGALQAK | NO | NO | NO |
| 17773 | TRAV12-1 | TRAJ5 | CVVNGDTGRRALTF | TRBV19 | TRBJ1-1 | CASSSWTVNTEAFF | KLGGALQAK | YES | NO | NO |
| 17774 | TRAV8-1 | TRAJ44 | CAVKTGTASKLTF | TRBV4-1 | TRBJ1-1 | CASSHATGGTNTEAFF | AVFDRKSDAK | YES | NO | YES |
| 17775 | TRAV12-3 | TRAJ39 | CAIGPGNMLTF | TRBV24-1 | TRBJ2-3 | CATSQIPGTSGSSDTQYF | RLRAEAQVK | YES | NO | YES |
| 17776 | TRAV35 | TRAJ28 | CAGQRTYSGAGSYQLTF | TRBV7-2 | TRBJ1-6 | CASRNLINSPLHF | KLGGALQAK | YES | NO | NO |
| 17777 | TRAV12-2 | TRAJ45 | CAVDSGGGADGLTF | TRBV18 | TRBJ2-7 | CASSPWGSPSYEQYF | LLFGYPVYV | YES | YES | YES |
| 17778 | TRAV39 | TRAJ32 | CAPGGATNKLIF | TRBV7-9 | TRBJ2-1 | CASSERRGDEQFF | KLGGALQAK | YES | NO | NO |
| 17779 | TRAV39 | TRAJ11 | CAVDILGYSTLTF | TRBV5-6 | TRBJ1-5 | CASSLIDSNQPQHF | KLGGALQAK | YES | NO | NO |
| 17780 | TRAV13-2 | TRAJ5 | CAEESMDTGRRALTF | TRBV19 | TRBJ1-1 | CASSVYSTEAFF | KLGGALQAK | YES | NO | NO |
| 17781 | TRAV27 | TRAJ20 | CAGVSSNDYKLSF | TRBV29-1 | TRBJ1-1 | CSVSGTGNTEAFF | KLGGALQAK | YES | NO | NO |
| 17782 | TRAV25 | TRAJ30 | CAGPRWRDDKIIF | TRBV27 | TRBJ2-7 | CASSPGGIHEQYF | KLGGALQAK | YES | NO | NO |
| 17783 | TRAV13-1 | TRAJ8 | CAASNTGFQKLVF | TRBV12-4 | TRBJ1-1 | CASKTPSRGPHTEAFF | AVFDRKSDAK | YES | NO | YES |
| 17784 | TRAV1-2 | TRAJ12 | CAVWMDSSYKLIF | TRBV14 | TRBJ1-4 | CASSQEAADNEKLFF | NLVPMVATV | NO | NO | NO |
| 17785 | TRAV12-1 | TRAJ30 | CVVNRDDKIIF | TRBV9 | TRBJ1-2 | CASSGGNRRDTEKGYTF | KLGGALQAK | YES | NO | NO |
| 17786 | TRAV20 | TRAJ26 | CAVQAYRDNYGQNFVF | TRBV10-3 | TRBJ2-2 | CAISESNSNTGELFF | KLGGALQAK | YES | NO | NO |
| 17787 | TRAV14DV4 | TRAJ40 | CAMRRAGTYKYIF | TRBV7-9 | TRBJ1-4 | CASSWGDRGRSEKLFF | KLGGALQAK | YES | NO | NO |
| 17788 | TRAV19 | TRAJ48 | CALSERRNEKLTF | TRBV19 | TRBJ1-1 | CASIDRVSVAEAFF | KLGGALQAK | YES | NO | NO |
| 17789 | TRAV3 | TRAJ20 | CAVRDGAPTYDYKLSF | TRBV6-5 | TRBJ2-3 | CASSYSIPGQGKDTQYF | KLGGALQAK | YES | NO | NO |
| 17790 | TRAV17 | TRAJ21 | CASDFNKFYF | TRBV15 | TRBJ2-5 | CATSSDRRPGEETQYF | KLGGALQAK | YES | NO | NO |
| 17791 | TRAV16 | TRAJ39 | CALSSNAGNMLTF | TRBV16 | TRBJ2-7 | CASSQSTSYEQYV | KLGGALQAK | NO | NO | NO |
| 17792 | TRAV14DV4 | TRAJ52 | CAMREVPRRAGGTSYGKLTF | TRBV7-9 | TRBJ2-7 | CASSLGSYEQYV | KLGGALQAK | NO | NO | NO |
| 17793 | TRAV12-1 | TRAJ4 | CVGYSGGYNKLIF | TRBV4-1 | TRBJ2-3 | CASSHVVSGGSTDTQYF | AVFDRKSDAK | YES | NO | YES |
| 17794 | TRAV17 | TRAJ31 | CATDPNNNARLMF | TRBV4-1 | TRBJ1-2 | CASSQDSKDRGLDGYTF | KLGGALQAK | YES | NO | NO |
| 17795 | TRAV5 | TRAJ32 | CASGGATNKLIF | TRBV6-5 | TRBJ1-1 | CASSYSENTEAFF | KLGGALQAK | YES | NO | NO |
| 17796 | TRAV36DV7 | TRAJ48 | CAVVPLPDNFGNEKLTF | TRBV12-4 | TRBJ2-1 | CASSLLSSGRVNEQFF | KLGGALQAK | NO | NO | NO |
| 17797 | TRAV12-2 | TRAJ9 | CAVKTGFKTIF | TRBV5-1 | TRBJ1-1 | CASSAGQGGNTEAFF | KLGGALQAK | YES | NO | NO |
| 17798 | TRAV24 | TRAJ47 | CASPYGNKLVF | TRBV4-1 | TRBJ2-3 | CASSPNLSTDTQYF | KLGGALQAK | YES | NO | NO |
| 17799 | TRAV41 | TRAJ42 | CAVRPGGSQGNLIF | TRBV9 | TRBJ1-3 | CASSTNWDSGNTIYF | KLGGALQAK | YES | NO | NO |
| 17800 | TRAV3 | TRAJ43 | CAVRDPLFNNDMRF | TRBV29-1 | TRBJ2-1 | CSALLAGDFDEQFF | AVFDRKSDAK | NO | NO | NO |
| 17801 | TRAV3 | TRAJ42 | CAVRDSPYGGSQGNLIF | TRBV6-1 | TRBJ2-5 | CASSEEQSGGPQETQYF | KLGGALQAK | NO | NO | NO |
| 17802 | TRAV10 | TRAJ20 | CVVRPLNDYKLSF | TRBV12-4 | TRBJ2-2 | CASSPPQGLITGELFF | KLGGALQAK | YES | NO | NO |
| 17803 | TRAV1-1 | TRAJ28 | CAVRGASGAGSYQLTF | TRBV9 | TRBJ1-2 | CASSVVPDSSQAYGYTF | KLGGALQAK | YES | NO | NO |
| 17804 | TRAV1-2 | TRAJ12 | CAVSGEGSSSYKLIF | TRBV9 | TRBJ2-2 | CASSVATSGTGELFF | KLGGALQAK | YES | NO | NO |
| 17805 | TRAV13-2 | TRAJ11 | CAENMNSGYSTLTF | TRBV5-8 | TRBJ2-2 | CASSLGGGGELFF | KLGGALQAK | YES | NO | NO |
| 17806 | TRAV8-1 | TRAJ24 | CAVTADSWGKLQF | TRBV20-1 | TRBJ2-1 | CSALAGNEQFF | KLGGALQAK | NO | NO | NO |
| 17807 | TRAV20 | TRAJ37 | CAVPIPLVPGKLIF | TRBV5-1 | TRBJ1-4 | CASSPQGGEEKLFF | KLGGALQAK | YES | NO | NO |
| 17808 | TRAV12-2 | TRAJ20 | CAVHNDYKLSF | TRBV5-1 | TRBJ1-6 | CASSLSGTRNSPLHF | KLGGALQAK | YES | NO | NO |
| 17809 | TRAV8-3 | TRAJ27 | CAVGAADAGKSTF | TRBV30 | TRBJ2-3 | CAWKGGPVSTDTQYF | KLGGALQAK | YES | NO | NO |
| 17810 | TRAV22 | TRAJ49 | CAVVNPGNQFYF | TRBV7-2 | TRBJ2-1 | CASSSMAVEQFF | KLGGALQAK | YES | NO | NO |
| 17811 | TRAV30 | TRAJ24 | CGTVTTDSWGKLQF | TRBV7-6 | TRBJ1-2 | CASSLGLPHGYTF | KLGGALQAK | YES | NO | NO |
| 17812 | TRAV23DV6 | TRAJ42 | CAASKKNYGGSQGNLIF | TRBV5-1 | TRBJ1-2 | CASSWGPATLSHYGYTF | AVFDRKSDAK | YES | YES | YES |
| 17813 | TRAV25 | TRAJ34 | CADQAINTDKLIF | TRBV27 | TRBJ1-5 | CASSLAGGGQPQHF | KLGGALQAK | YES | NO | NO |
| 17814 | TRAV1-2 | TRAJ31 | CAVSRLNNARLMF | TRBV20-1 | TRBJ2-2 | CSVEAGVHTGELFF | KLGGALQAK | YES | NO | NO |
| 17816 | TRAV25 | TRAJ52 | CAGRAGGTSYGKLTF | TRBV9 | TRBJ2-1 | CASSASGTFLGNEQFF | KLGGALQAK | YES | NO | NO |
| 17817 | TRAV29DV5 | TRAJ53 | CAAGPSLGGSNYKLTF | TRBV19 | TRBJ2-2 | CASSISYTGELFF | KLGGALQAK | YES | NO | NO |
| 17818 | TRAV12-3 | TRAJ39 | CAIGPGNMLTF | TRBV3-1 | TRBJ1-3 | CASSQGLLTERGYTIYF | KLGGALQAK | YES | NO | NO |
| 17820 | TRAV13-2 | TRAJ45 | CAESRGGGADGLTF | TRBV11-2 | TRBJ2-5 | CASSSDYIMGETQYF | KLGGALQAK | YES | NO | NO |
| 17821 | TRAV17 | TRAJ31 | CGENNNARLMF | TRBV7-2 | TRBJ2-1 | CASNEDRATYNEQFF | KLGGALQAK | YES | NO | NO |
| 17822 | TRAV3 | TRAJ8 | CAVRDLGMYMNTGFQKLVF | TRBV6-3 | TRBJ2-1 | CASSYSGTNNEQFF | KLGGALQAK | NO | NO | NO |

| | | | | | | | | | | |
|---|---|---|---|---|---|---|---|---|---|---|
| 17823 | TRAV8-1 | TRAJ24 | CAVRPPTDSWGKFQF | TRBV5-1 | TRBJ2-1 | CASSLLSGDTYSYNEQFF | KLGGALQAK | YES | NO | NO |
| 17824 | TRAV13-2 | TRAJ43 | CAENNDMRF | TRBV6-5 | TRBJ2-6 | CASSYSGGTGANVLTF | KLGGALQAK | NO | NO | NO |
| 17825 | TRAV21 | TRAJ41 | CAGRSGYALNF | TRBV6-6 | TRBJ1-2 | CASSFEGALGYTF | KLGGALQAK | NO | NO | NO |
| 17827 | TRAV41 | TRAJ35 | CASSFGNVLHC | TRBV7-2 | TRBJ2-2 | CASSQIPGAGELFF | KLGGALQAK | NO | NO | NO |
| 17829 | TRAV1-2 | TRAJ33 | CAVSDSNYQLIW | TRBV20-1 | TRBJ2-3 | CSARDFSGDVDTQYF | AVFDRKSDAK | NO | NO | NO |
| 17830 | TRAV8-4 | TRAJ49 | CAVSETGNQFYF | TRBV18 | TRBJ1-1 | CASSPPGRNTEAFF | KLGGALQAK | YES | NO | NO |
| 17831 | TRAV39 | TRAJ45 | CAVDTSSGGGADGLTF | TRBV7-9 | TRBJ1-1 | CASSEGTAPEAFF | KLGGALQAK | YES | NO | NO |
| 17832 | TRAV8-4 | TRAJ21 | CAGAYNFNKFYF | TRBV11-2 | TRBJ2-3 | CASSLTGTLTKVGSDTQYF | KLGGALQAK | YES | NO | NO |
| 17833 | TRAV41 | TRAJ58 | CAVRSGETSGSRLTF | TRBV5-1 | TRBJ1-2 | CASSFRVDNSYGYTF | KLGGALQAK | YES | NO | NO |
| 17834 | TRAV38-2DV8 | TRAJ32 | CAYDGGATNKLIF | TRBV20-1 | TRBJ1-1 | CSAAGVLGMNTEAFF | KLGGALQAK | YES | NO | NO |
| 17835 | TRAV12-1 | TRAJ30 | CVVKNRDDKIIF | TRBV7-6 | TRBJ2-4 | CASSLAGDIQYF | KLGGALQAK | YES | NO | NO |
| 17836 | TRAV23DV6 | TRAJ44 | CAASDSGTASKLTF | TRBV7-2 | TRBJ1-2 | CASSFPPDRGRVYYGYTF | KLGGALQAK | NO | NO | NO |
| 17837 | TRAV5 | TRAJ30 | CAEITNRDDKIIF | TRBV9 | TRBJ1-6 | CASSPGGGPGALHF | KLGGALQAK | NO | NO | NO |
| 17838 | TRAV2 | TRAJ10 | CAVAAGGGNKLTF | TRBV7-9 | TRBJ2-2 | CASRNPGTSGTTGELFF | KLGGALQAK | NO | NO | NO |
| 17839 | TRAV1-2 | TRAJ20 | CAVISNDYKLSF | TRBV20-1 | TRBJ2-3 | CSARDPASRTDTQYF | KLGGALQAK | YES | NO | NO |
| 17840 | TRAV19 | TRAJ27 | CALSAGTNAGKSTF | TRBV2 | TRBJ2-7 | CASSEAGGEQYV | KLGGALQAK | NO | NO | NO |
| 17841 | TRAV13-2 | TRAJ45 | CAENSGGGADGLTF | TRBV4-3 | TRBJ1-4 | CASSQDHRMGGHEKLFF | KLGGALQAK | YES | NO | NO |
| 17842 | TRAV8-3 | TRAJ37 | CAVGHSGNTGKLIF | TRBV29-1 | TRBJ2-2 | CSVLRDTGELFF | AVFDRKSDAK | YES | NO | YES |
| 17843 | TRAV38-2DV8 | TRAJ32 | CACGATNKLIF | TRBV19 | TRBJ1-4 | CASSWVGRVLFF | KLGGALQAK | YES | NO | NO |
| 17844 | TRAV8-1 | TRAJ40 | CAVNARTSGTYKYIF | TRBV12-4 | TRBJ1-5 | CASSSTGWGQPQHF | KLGGALQAK | YES | NO | NO |
| 17845 | TRAV24 | TRAJ52 | CAFIGDAGGTSYGKLTF | TRBV2 | TRBJ2-6 | CASTPGGGSGANVLTF | KLGGALQAK | YES | NO | NO |
| 17846 | TRAV38-2DV8 | TRAJ38 | CAYSNFAGNNRKLIW | TRBV9 | TRBJ2-1 | CASSVIGSSNEQFF | KLGGALQAK | YES | NO | NO |
| 17847 | TRAV8-6 | TRAJ29 | CAVSISLGNTPLVF | TRBV5-6 | TRBJ1-1 | CASSARDRGSEAFF | KLGGALQAK | YES | NO | NO |
| 17848 | TRAV19 | TRAJ23 | CALSEVYNQGGKLIF | TRBV3-1 | TRBJ2-3 | CASSQEGDPSTDTQYF | KLGGALQAK | YES | NO | NO |
| 17849 | TRAV10 | TRAJ40 | CVVPGTYKYIF | TRBV6-1 | TRBJ2-7 | CASSEELAEYYEQYV | KLGGALQAK | NO | NO | NO |
| 17850 | TRAV8-2 | TRAJ20 | CVVTLNDYKLSF | TRBV6-1 | TRBJ2-2 | CASSEGYRALLRTGELFF | KLGGALQAK | YES | NO | NO |
| 17851 | TRAV38-1 | TRAJ49 | CAFMKHVAGNQFYF | TRBV27 | TRBJ2-5 | CASSFLSQETQYF | KLGGALQAK | YES | NO | NO |
| 17852 | TRAV19 | TRAJ20 | CALSERKDDYKLSF | TRBV28 | TRBJ2-1 | CASSLRTGEIDLGKFF | KLGGALQAK | YES | NO | NO |
| 17853 | TRAV13-2 | TRAJ23 | CATIWGGKLIF | TRBV12-4 | TRBJ1-2 | CASRMIDRDYGYTF | KLGGALQAK | YES | NO | NO |
| 17854 | TRAV8-3 | TRAJ42 | CAVGEGYGGSQGNLIF | TRBV7-6 | TRBJ1-1 | CASSSSDREMNTEAFF | KLGGALQAK | YES | NO | NO |
| 17855 | TRAV10 | TRAJ45 | CVVSEGYSGGGADGLTF | TRBV10-3 | TRBJ2-7 | CATGQGPSYEQYF | KLGGALQAK | YES | NO | NO |
| 17856 | TRAV1-1 | TRAJ13 | CAVRDRSGGYQKVTF | TRBV10-1 | TRBJ2-1 | CASSESTRRGRVGEQFF | KLGGALQAK | YES | NO | NO |
| 17857 | TRAV29DV5 | TRAJ44 | CAATGTASKLTF | TRBV4-1 | TRBJ2-5 | CASSHPAGGEETQYF | KLGGALQAK | YES | NO | NO |
| 17858 | TRAV17 | TRAJ11 | CATAMVGYSTLTF | TRBV7-8 | TRBJ1-2 | CASSLGYGYTF | KLGGALQAK | YES | NO | NO |
| 17859 | TRAV1-2 | TRAJ33 | CAVLDSNYQLIW | TRBV20-1 | TRBJ2-1 | CSARSGVEHNEQFF | KLGGALQAK | NO | NO | NO |
| 17860 | TRAV5 | TRAJ20 | CAESRNDYKLSF | TRBV7-2 | TRBJ1-2 | CATPNRGFDGYTF | KLGGALQAK | YES | NO | NO |
| 17861 | TRAV38-2DV8 | TRAJ18 | CAYMADRGSTLGRLYF | TRBV27 | TRBJ2-5 | CASSFRAQYF | KLGGALQAK | YES | NO | NO |
| 17862 | TRAV27 | TRAJ32 | CAGGGNYGGATNKLIF | TRBV27 | TRBJ2-1 | CASSLSSLYEQFF | KLGGALQAK | YES | NO | NO |
| 17863 | TRAV27 | TRAJ37 | CAGRMYGSSNTGKLIF | TRBV9 | TRBJ2-1 | CASSDIKRDYNEQFF | KLGGALQAK | YES | NO | NO |
| 17864 | TRAV29DV5 | TRAJ45 | CAASAFGGGADGLTF | TRBV7-9 | TRBJ2-1 | CASSNSLPSGVSYNEQFF | KLGGALQAK | YES | NO | NO |
| 17865 | TRAV22 | TRAJ43 | CAVESGYNNDMRF | TRBV15 | TRBJ2-1 | CATSRGRDLDNEQFF | KLGGALQAK | YES | NO | NO |
| 17866 | TRAV29DV5 | TRAJ28 | CAARYSGAGSYQLTF | TRBV27 | TRBJ1-1 | CASSLGRNTEAFF | KLGGALQAK | YES | NO | NO |
| 17867 | TRAV23DV6 | TRAJ26 | CAASGRDNYGQNFVF | TRBV9 | TRBJ2-1 | CASSVSDGGYNEQFF | KLGGALQAK | YES | NO | NO |
| 17868 | TRAV20 | TRAJ34 | CAPRSYNTDKLIF | TRBV4-1 | TRBJ2-2 | CASSPGRGTFAGELFF | KLGGALQAK | YES | NO | NO |
| 17869 | TRAV17 | TRAJ54 | CATRGGLQGAQKLVF | TRBV13 | TRBJ1-1 | CASSDFGRTEAFF | KLGGALQAK | YES | NO | NO |
| 17870 | TRAV8-2 | TRAJ30 | CVVRSRDDKIIF | TRBV2 | TRBJ1-6 | CASSEGGFSYNSPLHF | KLGGALQAK | YES | NO | NO |
| 17871 | TRAV13-2 | TRAJ9 | CAENWDTGGFKTIF | TRBV12-4 | TRBJ1-1 | CASSLDPQGGTEAFF | KLGGALQAK | YES | NO | NO |
| 17872 | TRAV17 | TRAJ13 | CATDSYFSGGYQKVTF | TRBV7-6 | TRBJ1-5 | CASSPDSHSNQPQHF | KLGGALQAK | YES | NO | NO |
| 17873 | TRAV20 | TRAJ37 | CAVHSSGNTGKLIF | TRBV10-3 | TRBJ2-7 | CAIREEVSYEQYV | KLGGALQAK | NO | NO | NO |
| 17874 | TRAV3 | TRAJ20 | CAVRDPPNDYKLSF | TRBV19 | TRBJ1-5 | CASDPGTDNQPQHF | KLGGALQAK | NO | NO | NO |
| 17875 | TRAV1-2 | TRAJ26 | CAVKRDYGQNFVF | TRBV4-3 | TRBJ1-2 | CASSHPGQGRGYTF | KLGGALQAK | YES | NO | NO |
| 17876 | TRAV10 | TRAJ18 | CVVSERGSTLGRLYF | TRBV4-1 | TRBJ2-7 | CASSQGVSSGGGYEQYF | KLGGALQAK | YES | NO | NO |
| 17877 | TRAV22 | TRAJ18 | CAVEEGPRGSTLGRLYF | TRBV7-6 | TRBJ2-5 | CASSFRTGQETQYF | KLGGALQAK | YES | NO | NO |
| 17878 | TRAV38-1 | TRAJ43 | CASRGGDNNDMRF | TRBV6-5 | TRBJ1-6 | CASSYWGNSYNSPLHF | KLGGALQAK | YES | NO | NO |
| 17879 | TRAV8-2 | TRAJ22 | CVGRKGSARQLTF | TRBV12-4 | TRBJ2-3 | CASSPMDTQYF | KLGGALQAK | YES | NO | NO |
| 17880 | TRAV12-1 | TRAJ48 | CVVNRARFLGNEKLTF | TRBV20-1 | TRBJ1-3 | CSARGREAGNTIYF | KLGGALQAK | YES | NO | NO |
| 17881 | TRAV12-3 | TRAJ52 | CAMTGGTSYGKLTF | TRBV6-3 | TRBJ2-2 | CASSYSLAGVRDRHNTGELFF | KLGGALQAK | NO | NO | NO |
| 17882 | TRAV19 | TRAJ24 | CALSEATDSWGKLQF | TRBV19 | TRBJ2-4 | CASSNIQGAPNIQYF | KLGGALQAK | YES | NO | NO |
| 17883 | TRAV29DV5 | TRAJ22 | CASFSSGSARQLTF | TRBV7-2 | TRBJ1-1 | CASSLGPNTEAFF | KLGGALQAK | YES | NO | NO |
| 17884 | TRAV3 | TRAJ36 | CAVRVRVQTGANNLFF | TRBV10-3 | TRBJ2-3 | CAISESVRVVSDTQYF | KLGGALQAK | NO | NO | NO |
| 17885 | TRAV10 | TRAJ6 | CVVSATGGSYIPTF | TRBV5-6 | TRBJ2-5 | CASSNGGAQETQYF | AVFDRKSDAK | YES | YES | YES |
| 17886 | TRAV26-2 | TRAJ3 | CILSGYSSASKIIF | TRBV4-2 | TRBJ1-2 | CASSQGTVYGYTF | KLGGALQAK | YES | NO | NO |
| 17887 | TRAV12-1 | TRAJ12 | CVVIPGMDSSYKLIF | TRBV5-6 | TRBJ1-2 | CASSLETHNYGYTF | KLGGALQAK | YES | NO | NO |
| 17888 | TRAV27 | TRAJ52 | CAGVYAGGTSYGKLTF | TRBV7-8 | TRBJ1-3 | CASSQGTVSGNTIYF | KLGGALQAK | YES | NO | NO |
| 17889 | TRAV12-2 | TRAJ52 | CAVKEAGGTSYGKLTF | TRBV5-1 | TRBJ2-1 | CASSIWAGDPYNEQFF | KLGGALQAK | YES | NO | NO |
| 17890 | TRAV8-1 | TRAJ45 | CAARSGGGADGLTF | TRBV13 | TRBJ1-2 | CASQIRDRGNYGYTF | KLGGALQAK | YES | NO | NO |
| 17891 | TRAV19 | TRAJ45 | CALSEGYSGGGADGLTF | TRBV19 | TRBJ2-7 | CASSIGTGGIYEQYF | KLGGALQAK | YES | NO | NO |
| 17892 | TRAV29DV5 | TRAJ20 | CAASGASNDYKLSF | TRBV7-8 | TRBJ1-5 | CATLGQGEVEPQHF | KLGGALQAK | NO | NO | NO |
| 17893 | TRAV6 | TRAJ29 | CALSGNTPLVF | TRBV12-4 | TRBJ2-2 | CASSLRSGRVGTGELFF | KLGGALQAK | YES | NO | NO |
| 17894 | TRAV12-1 | TRAJ40 | CVVDDSGTYKYIF | TRBV6-5 | TRBJ2-5 | CASSSPRTSWETQYF | KLGGALQAK | NO | NO | NO |
| 17895 | TRAV24 | TRAJ21 | CALYNFNKFYF | TRBV20-1 | TRBJ2-2 | CSARDGTTSGIDTGELFF | KLGGALQAK | YES | NO | NO |
| 17896 | TRAV29DV5 | TRAJ53 | CAASVGGGSNYKLTF | TRBV19 | TRBJ2-3 | CASSNGQGSDTQYF | KLGGALQAK | YES | NO | NO |
| 17897 | TRAV14DV4 | TRAJ40 | CAMRDVGTYKYIF | TRBV4-1 | TRBJ2-1 | CASSQVFERVYNEQFF | KLGGALQAK | YES | NO | NO |
| 17898 | TRAV13-2 | TRAJ28 | CAENIGSGAGSYQLTF | TRBV9 | TRBJ1-5 | CASSATGQWSGQPQHF | KLGGALQAK | YES | NO | NO |
| 17899 | TRAV2 | TRAJ41 | CAVEDSSGYALNF | TRBV6-3 | TRBJ1-2 | CASSWIQRPDYGYTF | KLGGALQAK | NO | NO | NO |
| 17900 | TRAV8-6 | TRAJ57 | CAVSDLMAGQGGSEKLVF | TRBV15 | TRBJ2-1 | CATSRGGYEQFF | KLGGALQAK | YES | NO | NO |
| 17901 | TRAV29DV5 | TRAJ31 | CAASVNNARLMF | TRBV4-1 | TRBJ2-2 | CASSQVGLGTGELFF | KLGGALQAK | YES | NO | NO |
| 17902 | TRAV14DV4 | TRAJ49 | CAMRETGNQFYF | TRBV4-3 | TRBJ1-4 | CASSQDHRMGGHEKLFF | KLGGALQAK | YES | YES | NO |
| 17903 | TRAV3 | TRAJ8 | CAVRWNTGFQKLVF | TRBV19 | TRBJ1-1 | CASSGTEAHSYTEAFF | KLGGALQAK | NO | NO | NO |
| 17905 | TRAV14DV4 | TRAJ4 | CAMREGMFSGGYNKLIF | TRBV3-1 | TRBJ1-2 | CASSQGRDRHGYTF | KLGGALQAK | YES | NO | NO |
| 17906 | TRAV38-1 | TRAJ43 | CAFMRPYNNNDMRF | TRBV2 | TRBJ1-2 | CASSEGIEGWGYTF | KLGGALQAK | YES | NO | NO |
| 17907 | TRAV19 | TRAJ17 | CALSEAAAAGNKLTF | TRBV10-3 | TRBJ2-1 | CAISESVRGGRDEQFF | KLGGALQAK | YES | NO | NO |
| 17908 | TRAV25 | TRAJ11 | CAGPSPQSGYSTLTF | TRBV5-1 | TRBJ2-7 | CASSHGTSETYEQYV | KLGGALQAK | NO | NO | NO |
| 17909 | TRAV24 | TRAJ15 | CAFANQAGTALIF | TRBV19 | TRBJ1-3 | CASSIWTGLSGNTIYF | KLGGALQAK | YES | NO | NO |
| 17910 | TRAV12-1 | TRAJ3 | CVVNGYSSASKIIF | TRBV12-4 | TRBJ2-7 | CASSLQGDEQYV | KLGGALQAK | NO | NO | NO |
| 17911 | TRAV13-2 | TRAJ45 | CAENMYSGGGADGLTF | TRBV19 | TRBJ1-3 | CASSWVSGNTIYF | KLGGALQAK | YES | NO | NO |
| 17912 | TRAV1-2 | TRAJ39 | CAAPVDGGNMLTF | TRBV9 | TRBJ2-3 | CASSVVGSTDTQYF | KLGGALQAK | YES | NO | NO |
| 17913 | TRAV12-1 | TRAJ31 | CVVNTRARLMF | TRBV7-9 | TRBJ2-6 | CASSITESGANVLTF | KLGGALQAK | YES | NO | NO |
| 17914 | TRAV24 | TRAJ41 | CALSNSGYALNF | TRBV20-1 | TRBJ2-1 | CSAEPGGISDSSYNEQFF | KLGGALQAK | NO | NO | NO |
| 17915 | TRAV25 | TRAJ26 | CAYLEDNYGQNFVF | TRBV14 | TRBJ1-2 | CASSRKWRTDSYGYTF | KLGGALQAK | NO | NO | NO |
| 17916 | TRAV29DV5 | TRAJ6 | CAASSLSGSYIPTF | TRBV20-1 | TRBJ1-2 | CSAPEWDRGLENYGYTF | AVFDRKSDAK | YES | NO | YES |
| 17917 | TRAV14DV4 | TRAJ21 | CAMRGLYNFNKFYF | TRBV9 | TRBJ2-5 | CASSESSGNLKETQYF | KLGGALQAK | YES | NO | NO |
| 17918 | TRAV13-1 | TRAJ40 | CAATGTYKYIF | TRBV7-9 | TRBJ2-1 | CASSLRDRVNEQFF | KLGGALQAK | YES | NO | NO |
| 17919 | TRAV5 | TRAJ29 | CAESTSGNTPLVF | TRBV12-4 | TRBJ1-2 | CASSLYDYGYTF | KLGGALQAK | YES | NO | NO |
| 17921 | TRAV21 | TRAJ41 | CAVRDSNSGYALNF | TRBV12-4 | TRBJ2-3 | CASSHFEGLDTQYF | KLGGALQAK | YES | NO | NO |
| 17922 | TRAV29DV5 | TRAJ22 | CAASLSRQLTF | TRBV7-9 | TRBJ2-6 | CASSFLAGANVLTF | KLGGALQAK | YES | NO | NO |
| 17923 | TRAV12-2 | TRAJ53 | CAVSGGSNYKLTF | TRBV29-1 | TRBJ2-1 | CSGRQGGRNEQFF | KLGGALQAK | YES | NO | NO |
| 17924 | TRAV12-1 | TRAJ26 | CVVNRGNYGQNFVF | TRBV20-1 | TRBJ2-7 | CSASSGGGYEQYF | KLGGALQAK | NO | NO | NO |
| 17925 | TRAV19 | TRAJ39 | CALSEGAAGNMLTF | TRBV15 | TRBJ2-7 | CATSRDHNAWYEQYV | KLGGALQAK | NO | NO | NO |
| 17927 | TRAV19 | TRAJ7 | CALSGFGGNNRLAF | TRBV18 | TRBJ1-2 | CASSPDLNNDPNYGYTF | KLGGALQAK | YES | NO | NO |
| 17928 | TRAV24 | TRAJ57 | CALTDQGGSEKLVF | TRBV7-9 | TRBJ2-2 | CASSPRDRRAGELFF | KLGGALQAK | YES | NO | NO |
| 17929 | TRAV26-2 | TRAJ45 | CILRDVPGGGADGLTF | TRBV13 | TRBJ2-7 | CASTEGATESSYEQYF | KLGGALQAK | YES | NO | NO |
| 17930 | TRAV13-1 | TRAJ27 | CAASIPYAGKSTF | TRBV11-2 | TRBJ1-1 | CASSPGGGKGDTEAFF | KLGGALQAK | YES | NO | NO |
| 17931 | TRAV35 | TRAJ40 | CAGPTSGTYKYIF | TRBV6-5 | TRBJ1-1 | CASSYTPETTEAFF | KLGGALQAK | YES | NO | NO |

| | | | | | | | | | | |
|---|---|---|---|---|---|---|---|---|---|---|
| 17932 | TRAV12-3 | TRAJ50 | CAMSAVVKTSYDKVIF | TRBV13 | TRBJ2-6 | CASSLGGVGANVLTF | KLGGALQAK | YES | NO | NO |
| 17933 | TRAV1-2 | TRAJ20 | CAVNDYKLSF | TRBV20-1 | TRBJ2-1 | CSAIGLAGGSQQFF | KLGGALQAK | NO | NO | NO |
| 17934 | TRAV13-2 | TRAJ6 | CAEASGGSYIPTF | TRBV29-1 | TRBJ2-5 | CSVEEGGAGGLETQYF | KLGGALQAK | YES | NO | NO |
| 17935 | TRAV10 | TRAJ53 | CVVSEGSGGSNYKLTF | TRBV19 | TRBJ2-1 | CASSIVGGGYAVYNEQFF | KLGGALQAK | YES | NO | NO |
| 17936 | TRAV26-2 | TRAJ53 | CILRDLGGSNYKLTF | TRBV5-6 | TRBJ1-1 | CASSSMTGSNTEAFF | KLGGALQAK | NO | NO | NO |
| 17937 | TRAV23DV6 | TRAJ17 | CAASIRAAGNKLTF | TRBV6-3 | TRBJ2-1 | CASSYHEMTGTVEQFF | KLGGALQAK | YES | NO | NO |
| 17938 | TRAV16 | TRAJ50 | CALSGRKTSYDKVIF | TRBV10-2 | TRBJ1-1 | CASSESPITDGELNTEAFF | KLGGALQAK | NO | NO | NO |
| 17939 | TRAV29DV5 | TRAJ50 | CAASAIKTSYDKVIF | TRBV27 | TRBJ1-2 | CASSFGSSYGYTF | AVFDRKSDAK | YES | YES | YES |
| 17940 | TRAV13-2 | TRAJ22 | CAENIGSARQLTF | TRBV7-9 | TRBJ2-3 | CASSWATGADTQYF | KLGGALQAK | YES | NO | NO |
| 17941 | TRAV14DV4 | TRAJ49 | CAMKSWTGNQFYF | TRBV9 | TRBJ2-2 | CASSSPGTGWTGELFF | KLGGALQAK | YES | NO | NO |
| 17942 | TRAV4 | TRAJ23 | CLVSQGGKLIF | TRBV12-4 | TRBJ2-5 | CASSVPRDRDYLETQYF | KLGGALQAK | NO | NO | NO |
| 17943 | TRAV19 | TRAJ33 | CALSDMDSNYQLIW | TRBV7-9 | TRBJ1-5 | CASSSGHYSNQPQHF | KLGGALQAK | YES | NO | NO |
| 17944 | TRAV14DV4 | TRAJ42 | CAMNGGSQGNLIF | TRBV4-1 | TRBJ2-3 | CASSQVWPGPSTDTQYF | KLGGALQAK | YES | NO | NO |
| 17945 | TRAV9-2 | TRAJ20 | CALSDGDYKLSF | TRBV12-4 | TRBJ2-5 | CASSGTPQETQYF | KLGGALQAK | YES | NO | NO |
| 17946 | TRAV8-2 | TRAJ11 | CVVSESAGYSTLTF | TRBV4-1 | TRBJ1-5 | CASSQGRGSNQPQHF | KLGGALQAK | YES | NO | NO |
| 17947 | TRAV14DV4 | TRAJ37 | CAMRGPFSSNTGKLIF | TRBV11-2 | TRBJ2-7 | CASSLEFAEPSYEQYF | KLGGALQAK | YES | NO | NO |
| 17948 | TRAV19 | TRAJ26 | CALSVNYGQNFVF | TRBV19 | TRBJ1-1 | CASSTIGISLNLTEAFF | KLGGALQAK | YES | NO | NO |
| 17949 | TRAV35 | TRAJ8 | CAGQEGNTGFQKLVF | TRBV7-9 | TRBJ1-4 | CASSLALGGFDEKLFF | KLGGALQAK | YES | NO | NO |
| 17950 | TRAV12-1 | TRAJ26 | CVVNMGSASRDNYGQNFVF | TRBV19 | TRBJ1-1 | CASTSGNTEAFF | KLGGALQAK | YES | YES | NO |
| 17951 | TRAV5 | TRAJ37 | CAEISGNTGKLIF | TRBV20-1 | TRBJ1-2 | CSATGTGILDGYTF | AVFDRKSDAK | YES | NO | YES |
| 17952 | TRAV25 | TRAJ42 | CAGDPNYGGSQGNLIF | TRBV20-1 | TRBJ1-2 | CSATGTGYNYGYTF | KLGGALQAK | YES | NO | NO |
| 17953 | TRAV21 | TRAJ57 | CAVWGQGGSEKLVF | TRBV27 | TRBJ2-5 | CASSLQAGLTIETQYF | AVFDRKSDAK | YES | NO | YES |
| 17954 | TRAV1-2 | TRAJ8 | CAVDTGFQKLVF | TRBV9 | TRBJ2-2 | CASSGWIGTENTGELFF | AVFDRKSDAK | YES | NO | YES |
| 17955 | TRAV4 | TRAJ34 | CLVGGGFDTDKLIF | TRBV6-1 | TRBJ1-1 | CASTRQGRTEAFF | KLGGALQAK | NO | NO | NO |
| 17956 | TRAV1-1 | TRAJ37 | CAGGGNTGKLIF | TRBV19 | TRBJ2-2 | CASSSLSGGRAFGELFF | KLGGALQAK | YES | NO | NO |
| 17957 | TRAV21 | TRAJ11 | CAVMKGYSTLTF | TRBV6-5 | TRBJ2-1 | CASRIGTGGEEQFF | KLGGALQAK | NO | NO | NO |
| 17958 | TRAV8-4 | TRAJ36 | CAVTETGANNLFF | TRBV7-8 | TRBJ2-7 | CASSTERGEAYEQYF | LLFGYPVYV | NO | NO | NO |
| 17959 | TRAV5 | TRAJ15 | CADPNQAGTALIF | TRBV13 | TRBJ1-3 | CASSPRASGNTIYF | KLGGALQAK | YES | NO | NO |
| 17960 | TRAV8-1 | TRAJ17 | CAVNAHQAAGNKLTF | TRBV19 | TRBJ2-1 | CASSIGGAGEQFF | KLGGALQAK | YES | NO | NO |
| 17961 | TRAV8-3 | TRAJ47 | CAVGWMEYGNKLVF | TRBV24-1 | TRBJ2-2 | CATSDFRGRDTGELFF | KLGGALQAK | YES | NO | NO |
| 17962 | TRAV5 | TRAJ13 | CAESEGGYQKVTF | TRBV20-1 | TRBJ2-2 | CSVGWTESGELFF | KLGGALQAK | YES | NO | NO |
| 17963 | TRAV21 | TRAJ37 | CAAIEISGNTGKLIF | TRBV2 | TRBJ1-6 | CARGLDRGLYNSPLHF | KLGGALQAK | YES | NO | NO |
| 17964 | TRAV12-1 | TRAJ44 | CVVIITGTASKLTF | TRBV6-1 | TRBJ2-5 | CASSEAPWNSGGAVGAQETQYF | KLGGALQAK | NO | NO | NO |
| 17966 | TRAV6 | TRAJ35 | CALESVGGGFGNVLHC | TRBV12-4 | TRBJ2-5 | CASSLNTETQYF | KLGGALQAK | NO | NO | NO |
| 17967 | TRAV13-2 | TRAJ52 | CAERLKTGGTSYGKLTF | TRBV7-2 | TRBJ2-7 | CASSSQTLLGQGAYEQYF | KLGGALQAK | YES | NO | NO |
| 17968 | TRAV24 | TRAJ52 | CAFIPGGTSYGKLTF | TRBV4-1 | TRBJ2-1 | CASSQSGTYNEQFF | KLGGALQAK | YES | NO | NO |
| 17969 | TRAV8-3 | TRAJ16 | CAVGADGQKLLF | TRBV7-6 | TRBJ1-2 | CASSLQGGDYGYTF | KLGGALQAK | NO | NO | NO |
| 17970 | TRAV8-1 | TRAJ18 | CAVGRGSTLGRLYF | TRBV6-6 | TRBJ2-2 | CASTTDRGSNTGELFF | KLGGALQAK | YES | NO | NO |
| 17971 | TRAV1-1 | TRAJ33 | CAVSEGNYQLIW | TRBV5-6 | TRBJ2-1 | CASSLDLAGGSNEQFF | KLGGALQAK | YES | NO | NO |
| 17972 | TRAV9-2 | TRAJ23 | CALIYNQGGKLIF | TRBV12-5 | TRBJ2-1 | CASGLAGQGPYNEQFF | KLGGALQAK | YES | NO | NO |
| 17973 | TRAV6 | TRAJ31 | CALDARLMF | TRBV19 | TRBJ2-2 | CASSIANTGELFF | KLGGALQAK | NO | NO | NO |
| 17974 | TRAV21 | TRAJ12 | CAVRCRWSSYKLIF | TRBV7-2 | TRBJ2-1 | CASSLKGLPQEGDEQFF | KLGGALQAK | YES | NO | NO |
| 17975 | TRAV9-2 | TRAJ54 | CALREGAQKLVF | TRBV5-4 | TRBJ2-4 | CASSWEGLAGAGKNIQYF | KLGGALQAK | YES | NO | NO |
| 17976 | TRAV17 | TRAJ23 | CATAPYNQGGKLIF | TRBV20-1 | TRBJ2-1 | CSARDSYSGTYNEQFF | KLGGALQAK | NO | NO | NO |
| 17977 | TRAV8-6 | TRAJ26 | CAVSDNYGQNFVF | TRBV20-1 | TRBJ2-1 | CSASRARYNEQFF | KLGGALQAK | YES | NO | NO |
| 17978 | TRAV35 | TRAJ20 | CAGHDGNDYKLSF | TRBV7-8 | TRBJ1-2 | CASSLGQAYGYTF | KLGGALQAK | YES | NO | NO |
| 17979 | TRAV8-3 | TRAJ48 | CAVVMNFGNEKLTF | TRBV19 | TRBJ2-1 | CASLDRNYNEQFF | KLGGALQAK | YES | NO | NO |
| 17980 | TRAV25 | TRAJ43 | CAGGDMRF | TRBV19 | TRBJ2-1 | CASNPTSGGSWNEQFF | KLGGALQAK | NO | NO | NO |
| 17981 | TRAV21 | TRAJ30 | CAVNRDDKIIF | TRBV29-1 | TRBJ1-2 | CSARVLGGYTF | KLGGALQAK | YES | NO | NO |
| 17982 | TRAV8-4 | TRAJ48 | CAVSGYFGNEKLTF | TRBV6-6 | TRBJ2-3 | CASFWTSGGSSTDTQYF | KLGGALQAK | NO | NO | NO |
| 17983 | TRAV12-1 | TRAJ13 | CVVNDPGGYQKVTF | TRBV20-1 | TRBJ2-5 | CSARETSLQETQYF | KLGGALQAK | NO | NO | NO |
| 17984 | TRAV26-1 | TRAJ54 | CIVRGADQGAQKLVF | TRBV7-9 | TRBJ2-1 | CASTGTSGGPTLRDEQFF | KLGGALQAK | YES | NO | NO |
| 17985 | TRAV29DV5 | TRAJ52 | CAASSPANAGGTSYGKLTF | TRBV4-1 | TRBJ1-1 | CASSQDFRNTEAFF | KLGGALQAK | YES | NO | NO |
| 17986 | TRAV29DV5 | TRAJ50 | CAVKTSYDKVIF | TRBV19 | TRBJ1-1 | CASSMAANTEAFF | KLGGALQAK | YES | NO | NO |
| 17987 | TRAV27 | TRAJ26 | CAGAPIDYGQNFVF | TRBV7-9 | TRBJ2-1 | CASDTVAGYNEQFF | KLGGALQAK | YES | NO | NO |
| 17988 | TRAV16 | TRAJ4 | CALLLSGGYNKLIF | TRBV10-3 | TRBJ1-5 | CAISESGRGFYFDQPQHF | KLGGALQAK | NO | NO | NO |
| 17989 | TRAV29DV5 | TRAJ29 | CAASARGSGNTPLVF | TRBV13 | TRBJ2-2 | CASSLGAGELFF | KLGGALQAK | YES | NO | NO |
| 17990 | TRAV39 | TRAJ48 | CAAGNNFGNEKLTF | TRBV7-2 | TRBJ1-5 | CASSLLLPGTSFFHQPQHF | KLGGALQAK | YES | NO | NO |
| 17991 | TRAV29DV5 | TRAJ54 | CAASFQGAQKLVF | TRBV5-5 | TRBJ1-5 | CASSLGRGNQPQHF | KLGGALQAK | YES | NO | NO |
| 17992 | TRAV8-4 | TRAJ20 | CAVIQINDYKLSF | TRBV4-1 | TRBJ2-1 | CASSRVPGLAGGPPYNEQFF | KLGGALQAK | NO | NO | NO |
| 17993 | TRAV29DV5 | TRAJ39 | CAASEDNAGNMLTF | TRBV6-3 | TRBJ1-6 | CASVLGTGVSPLHF | KLGGALQAK | YES | NO | NO |
| 17994 | TRAV38-2DV8 | TRAJ48 | CAYRAPRGNEKLTF | TRBV3-1 | TRBJ1-2 | CASSQGRDRHGYTF | KLGGALQAK | YES | NO | NO |
| 17995 | TRAV14DV4 | TRAJ12 | CAMSEMDSSYKLIF | TRBV6-1 | TRBJ2-7 | CASRYLAGGPGEQYV | KLGGALQAK | NO | NO | NO |
| 17996 | TRAV13-2 | TRAJ21 | CAQDNFNKFYF | TRBV4-1 | TRBJ1-1 | CASSQGQGLLEAFF | KLGGALQAK | YES | NO | NO |
| 17997 | TRAV25 | TRAJ56 | CAGFLLGGSELTF | TRBV24-1 | TRBJ2-1 | CATTGEGNEQFF | AVFDRKSDAK | YES | NO | NO |
| 17998 | TRAV12-3 | TRAJ13 | CAMSFSGGYQKVTF | TRBV7-6 | TRBJ1-3 | CASSPPTTDFSGNTIYF | KLGGALQAK | YES | NO | NO |
| 17999 | TRAV17 | TRAJ49 | CATDPHTGNQFYF | TRBV25-1 | TRBJ2-5 | CASSRGGEETQYF | AVFDRKSDAK | YES | NO | YES |
| 18000 | TRAV21 | TRAJ37 | CAVSAGGNTGKLIF | TRBV7-9 | TRBJ1-2 | CASSFTAMENTF | KLGGALQAK | YES | NO | NO |
| 18001 | TRAV8-6 | TRAJ11 | CAVAEGYSTLTF | TRBV15 | TRBJ1-2 | CATSTGQLNYGYTF | KLGGALQAK | YES | NO | NO |
| 18002 | TRAV1-2 | TRAJ15 | CAVRDGSQAGTALIF | TRBV7-9 | TRBJ1-1 | CASSLGSGTEAFF | KLGGALQAK | YES | NO | NO |
| 18003 | TRAV2 | TRAJ28 | CAVEDLRDSGAGSYQLTF | TRBV4-1 | TRBJ2-5 | CASSQNRETQYF | AVFDRKSDAK | NO | NO | NO |
| 18004 | TRAV36DV7 | TRAJ48 | CAGRGNFGNEKLTF | TRBV6-5 | TRBJ1-1 | CASSYKYRENTEAFF | KLGGALQAK | NO | NO | NO |
| 18005 | TRAV35 | TRAJ49 | CAGPTGNQFYF | TRBV9 | TRBJ1-1 | CASSTLDTEAFF | KLGGALQAK | YES | NO | NO |
| 18006 | TRAV12-2 | TRAJ56 | CAVTSLGANSKLTF | TRBV19 | TRBJ2-7 | CASSIDLGVAYEQYF | KLGGALQAK | YES | NO | NO |
| 18007 | TRAV5 | TRAJ36 | CAGNAIQTGANNLFF | TRBV19 | TRBJ1-1 | CASRTTGAGDTEAFF | AVFDRKSDAK | YES | NO | YES |
| 18008 | TRAV19 | TRAJ54 | CALSDAIIQGAQKLVF | TRBV5-6 | TRBJ1-2 | CASSLSHPHGYTF | KLGGALQAK | NO | NO | NO |
| 18009 | TRAV5 | TRAJ48 | CAELSNFGNEKLTF | TRBV29-1 | TRBJ2-1 | CSVETHNEQFF | KLGGALQAK | YES | NO | NO |
| 18010 | TRAV1-2 | TRAJ26 | CAVRPNYGQNFVF | TRBV9 | TRBJ2-2 | CASNGNTANTGELFF | KLGGALQAK | YES | NO | NO |
| 18011 | TRAV1-2 | TRAJ33 | CAVMDSNYQLIW | TRBV4-2 | TRBJ2-2 | CASSQESLHTGELFF | KLGGALQAK | YES | NO | NO |
| 18012 | TRAV12-2 | TRAJ48 | CAVSSNFGNEKLTF | TRBV6-1 | TRBJ2-5 | CASRGLPNQETQYF | KLGGALQAK | YES | NO | NO |
| 18013 | TRAV1-2 | TRAJ10 | CAVREGLTGGGNKLTF | TRBV4-1 | TRBJ1-5 | CASSQYPGSEGGNQPQHF | KLGGALQAK | YES | NO | NO |
| 18014 | TRAV35 | TRAJ13 | CAGQNSGGYQKVTF | TRBV7-8 | TRBJ2-7 | CASSLVSGTGIYEQYF | KLGGALQAK | YES | NO | NO |
| 18015 | TRAV19 | TRAJ40 | CALSDLSGTYKYIF | TRBV7-9 | TRBJ2-6 | CASSLSQGEAGANVLTF | AVFDRKSDAK | YES | NO | YES |
| 18016 | TRAV17 | TRAJ49 | CATGSITGNQFYF | TRBV20-1 | TRBJ2-2 | CSALGQGNRIHANTGELFF | KLGGALQAK | YES | NO | NO |
| 18017 | TRAV1-2 | TRAJ39 | CAALFAGNMLTF | TRBV24-1 | TRBJ2-2 | CATSEPDRATNSGELFF | KLGGALQAK | YES | NO | NO |
| 18018 | TRAV12-2 | TRAJ29 | CAVNGGGNTPLVF | TRBV11-1 | TRBJ2-2 | CASSLATGELFF | KLGGALQAK | YES | NO | NO |
| 18019 | TRAV16 | TRAJ28 | CAPTSGAGSYQLTF | TRBV9 | TRBJ2-2 | CASSVRPGTLLGGELFF | KLGGALQAK | NO | NO | NO |
| 18020 | TRAV35 | TRAJ45 | CAGPTGADGLTF | TRBV12-4 | TRBJ1-6 | CASSFGSYNSPLHF | KLGGALQAK | YES | NO | NO |
| 18021 | TRAV16 | TRAJ11 | CALSGLNSGYSTLTF | TRBV6-3 | TRBJ1-1 | CASRKTLNTEAFF | KLGGALQAK | NO | NO | NO |
| 18022 | TRAV8-6 | TRAJ44 | CAVSDPSGAGTASKLTF | TRBV5-1 | TRBJ2-7 | CASSWGTYEQYF | KLGGALQAK | YES | NO | NO |
| 18023 | TRAV23DV6 | TRAJ48 | CAARFGNEKLTF | TRBV6-3 | TRBJ2-2 | CAEGPWGGELFF | KLGGALQAK | YES | NO | NO |
| 18024 | TRAV12-2 | TRAJ39 | CAANAGNMLTF | TRBV11-2 | TRBJ1-1 | CASSLGGGTEAFF | ELAGIGILTV | YES | YES | YES |
| 18026 | TRAV34 | TRAJ3 | CGADSIASKIIF | TRBV6-5 | TRBJ2-2 | CASSTGVGELFF | KLGGALQAK | YES | NO | NO |
| 18027 | TRAV27 | TRAJ21 | CAGMIYNFNKFYF | TRBV19 | TRBJ2-5 | CASSILGGGETQYF | KLGGALQAK | YES | NO | NO |
| 18028 | TRAV35 | TRAJ42 | CAGHVYGGSQGNLIF | TRBV27 | TRBJ1-5 | CASSLLTGGVSSVQPQHF | KLGGALQAK | YES | NO | NO |
| 18029 | TRAV21 | TRAJ45 | CALRADGLTF | TRBV6-5 | TRBJ2-6 | CASSYRVFGSGANVLTF | KLGGALQAK | YES | NO | NO |
| 18030 | TRAV38-2DV8 | TRAJ57 | CAYRSAKTQGGSEKLVF | TRBV7-8 | TRBJ2-1 | CASSSFLAGEVYNEQFF | KLGGALQAK | YES | NO | NO |
| 18031 | TRAV8-3 | TRAJ17 | CAVLKAAGNKLTF | TRBV20-1 | TRBJ1-1 | CSAWDRLNTEAFF | KLGGALQAK | YES | NO | NO |
| 18032 | TRAV13-1 | TRAJ23 | CAASIGQGGKLIF | TRBV3-1 | TRBJ1-2 | CASSQGRDRHGYTF | RLRAEAQVK | YES | NO | YES |
| 18033 | TRAV1-2 | TRAJ6 | CAVSPGGSYIPTF | TRBV5-5 | TRBJ1-1 | CASSLDLGRGTEAFF | KLGGALQAK | YES | NO | NO |
| 18034 | TRAV19 | TRAJ50 | CASKTSYDKVIF | TRBV4-1 | TRBJ2-2 | CASSQDRISSRSNTGELFF | KLGGALQAK | YES | NO | NO |
| 18035 | TRAV13-2 | TRAJ43 | CAENGNNNDMRF | TRBV7-6 | TRBJ2-5 | CASSRNYGQETQYF | KLGGALQAK | YES | NO | NO |
| 18036 | TRAV12-3 | TRAJ27 | CAMSDTNAGKSTF | TRBV6-1 | TRBJ2-7 | CASSARVDYEQYF | KLGGALQAK | YES | NO | NO |
| 18037 | TRAV14DV4 | TRAJ22 | CAMRRPISSGSARQLTF | TRBV4-1 | TRBJ2-1 | CASSQDPVTSGGRSYEQFF | KTWGQYWQV | YES | NO | YES |

| | | | | | | | | | | |
|---|---|---|---|---|---|---|---|---|---|---|
| 18038 | TRAV13-1 | TRAJ30 | CAASEGGDDKIIF | TRBV10-3 | TRBJ2-5 | CATSAGTSGGPQETQYF | KLGGALQAK | YES | NO | NO |
| 18039 | TRAV5 | TRAJ8 | CAEHKILNTGFQKLVF | TRBV4-1 | TRBJ2-5 | CASSQTSGGRETQYF | KLGGALQAK | YES | NO | NO |
| 18040 | TRAV38-2DV8 | TRAJ34 | CAYRGIYNTDKLIF | TRBV20-1 | TRBJ2-1 | CSVFDRHTYNEQFF | KLGGALQAK | NO | NO | NO |
| 18041 | TRAV26-2 | TRAJ52 | CILRSGGTSYGKLTF | TRBV7-9 | TRBJ2-1 | CASSLWPYYNEQFF | KLGGALQAK | YES | NO | NO |
| 18042 | TRAV27 | TRAJ45 | CAGAESGGGADGLTF | TRBV4-3 | TRBJ1-1 | CASSQVAGFTEAFF | KLGGALQAK | YES | NO | NO |
| 18043 | TRAV21 | TRAJ39 | CAVAPLTIINAGNMLTF | TRBV19 | TRBJ2-2 | CASSIASNTGELFF | KLGGALQAK | YES | NO | NO |
| 18044 | TRAV16 | TRAJ58 | CALGETSGSRLTF | TRBV7-7 | TRBJ2-1 | CASSLAGRNEQFF | KLGGALQAK | NO | NO | NO |
| 18045 | TRAV17 | TRAJ20 | CATDGNDYKLSF | TRBV19 | TRBJ2-2 | CASSIGYFANTGELFF | KLGGALQAK | YES | NO | NO |
| 18046 | TRAV22 | TRAJ28 | CAVEPGYSGAGSYQLTF | TRBV7-2 | TRBJ2-7 | CASSQPSGYEQYV | KLGGALQAK | NO | NO | NO |
| 18047 | TRAV12-2 | TRAJ18 | CAVSRGSTLGRLYF | TRBV11-3 | TRBJ1-2 | CASSFPDVFDGYTF | KLGGALQAK | YES | NO | NO |
| 18048 | TRAV9-2 | TRAJ6 | CALTEGAGGSYIPTF | TRBV10-1 | TRBJ2-3 | CASSEYSGIGTDTQYF | KLGGALQAK | YES | NO | NO |
| 18049 | TRAV9-2 | TRAJ56 | CALRDTGANSKLTF | TRBV7-9 | TRBJ2-3 | CASWTGLADTQYF | KLGGALQAK | YES | NO | NO |
| 18050 | TRAV29DV5 | TRAJ29 | CAASGPGNTPLVF | TRBV12-4 | TRBJ2-5 | CASSLLDGETQYF | KLGGALQAK | YES | NO | NO |
| 18051 | TRAV8-1 | TRAJ40 | CAVTSGTYKYIF | TRBV7-9 | TRBJ2-7 | CASSSSDKYEQYF | KLGGALQAK | YES | NO | NO |
| 18052 | TRAV17 | TRAJ30 | CATENRDDKIIF | TRBV9 | TRBJ1-2 | CASRGSRGSYGYTF | KLGGALQAK | YES | NO | NO |
| 18053 | TRAV1-2 | TRAJ39 | CAVRGVNAGNMLTF | TRBV6-6 | TRBJ2-7 | CASSPEGQGSYEQYV | KLGGALQAK | NO | NO | NO |
| 18054 | TRAV13-2 | TRAJ28 | CAENLFSGAGSYQLTF | TRBV19 | TRBJ2-7 | CASSASGGSYEQYF | IVTDFSVIK | YES | NO | YES |
| 18055 | TRAV35 | TRAJ37 | CAGPHGSSNTGKLIF | TRBV13 | TRBJ1-5 | CASSLGGVQPQHF | KLGGALQAK | YES | NO | NO |
| 18056 | TRAV13-2 | TRAJ43 | CAENMYDMRF | TRBV5-1 | TRBJ1-1 | CASSLEGQRYTEAFF | AVFDRKSDAK | YES | NO | YES |
| 18057 | TRAV41 | TRAJ34 | CAVRADNTDKLIF | TRBV4-1 | TRBJ2-1 | CASSQEDWYNEQFF | KTWGQYWQV | YES | NO | NO |
| 18058 | TRAV27 | TRAJ57 | CAGEEGGSEKLVF | TRBV9 | TRBJ2-7 | CASSANSGGLYEQYF | KLGGALQAK | YES | NO | NO |
| 18059 | TRAV17 | TRAJ7 | CATDGGRLAF | TRBV3-1 | TRBJ2-2 | CASSYRVGDTGELFF | KLGGALQAK | YES | NO | NO |
| 18060 | TRAV13-1 | TRAJ28 | CAAIDNSGAGSYQLTF | TRBV6-3 | TRBJ2-2 | CASSYSIPGELFF | KLGGALQAK | YES | NO | NO |
| 18061 | TRAV1-2 | TRAJ33 | CAVKGMDSNYQLIW | TRBV2 | TRBJ1-5 | CASSEALLRLTGATNQPQHF | KLGGALQAK | NO | NO | NO |
| 18062 | TRAV34 | TRAJ40 | CGADHSGTYKYIF | TRBV24-1 | TRBJ2-7 | CATSDPKTDEQYF | RPPIFIRRL | YES | NO | YES |
| 18063 | TRAV26-2 | TRAJ30 | CILNRDDKIIF | TRBV11-3 | TRBJ1-5 | CASSPGYSNQPQHF | KLGGALQAK | YES | NO | NO |
| 18064 | TRAV17 | TRAJ23 | CATAPYNQGGKLIF | TRBV20-1 | TRBJ2-1 | CSARVRTQGTYNEQFF | KLGGALQAK | NO | NO | NO |
| 18065 | TRAV20 | TRAJ24 | CATQGGTTDSWGKLQF | TRBV7-6 | TRBJ1-1 | CASSLGQGRNTEAFF | KLGGALQAK | YES | NO | NO |
| 18066 | TRAV17 | TRAJ52 | CATDAPRGSALGYGKLTF | TRBV10-3 | TRBJ1-1 | CAISAGTGAFF | KLGGALQAK | YES | YES | NO |
| 18067 | TRAV3 | TRAJ39 | CAVRDSNNAGNMLTF | TRBV9 | TRBJ2-7 | CASSKGLAGAEQYV | KLGGALQAK | NO | NO | NO |
| 18068 | TRAV4 | TRAJ34 | CLVGALDNTDKLIF | TRBV9 | TRBJ2-4 | CASSLTGGRDIQYF | KLGGALQAK | YES | NO | NO |
| 18069 | TRAV12-2 | TRAJ52 | CAGGYGAGGTSYGKLTF | TRBV5-6 | TRBJ2-7 | CASSWTQGSEQYF | KLGGALQAK | YES | NO | NO |
| 18070 | TRAV8-6 | TRAJ41 | CAVRSNSGYALNF | TRBV6-1 | TRBJ2-1 | CASSEGNYNEQFF | KLGGALQAK | YES | NO | NO |
| 18071 | TRAV12-2 | TRAJ57 | CAVINQGGSEKLVF | TRBV13 | TRBJ2-5 | CASSSEGGGETQYF | KLGGALQAK | YES | NO | NO |
| 18072 | TRAV17 | TRAJ54 | CATERGAQKLVF | TRBV5-1 | TRBJ1-1 | CASSFQAAGGTEAFF | KLGGALQAK | YES | NO | NO |
| 18073 | TRAV8-3 | TRAJ18 | CAVEKEGRGSTLGRLYF | TRBV19 | TRBJ2-1 | CASTPGLALNNEQFF | KLGGALQAK | YES | NO | NO |
| 18074 | TRAV9-2 | TRAJ10 | CALNSFGGGNKLTF | TRBV3-1 | TRBJ2-1 | CASSQDPAGASYNEQFF | KLGGALQAK | YES | NO | NO |
| 18075 | TRAV1-2 | TRAJ40 | CAVRDSGTYKYIF | TRBV7-2 | TRBJ2-1 | CASNSGGVSEQFF | KLGGALQAK | YES | NO | NO |
| 18076 | TRAV38-2DV8 | TRAJ23 | CAYRSGSNQGGKLIF | TRBV7-6 | TRBJ2-4 | CASSLEKASGGLAKNIQYF | AVFDRKSDAK | YES | NO | YES |
| 18077 | TRAV16 | TRAJ31 | CALSNNNARLMF | TRBV4-1 | TRBJ2-1 | CASSPLAGGPGYNEQFF | KLGGALQAK | NO | NO | NO |
| 18078 | TRAV19 | TRAJ34 | CALGSINTDKLIF | TRBV7-8 | TRBJ2-7 | CASSPTSGSHEQYV | KLGGALQAK | NO | NO | NO |
| 18079 | TRAV29DV5 | TRAJ23 | CAAKGGPDNQGGKLIF | TRBV4-1 | TRBJ2-7 | CASSQYGISEQYF | KLGGALQAK | YES | NO | NO |
| 18080 | TRAV3 | TRAJ37 | CAVRDIDGSGNTGKLIF | TRBV6-1 | TRBJ2-1 | CASSEGLAGGYEQFF | KLGGALQAK | NO | NO | NO |
| 18081 | TRAV41 | TRAJ53 | CAASFLGGSNYKLTF | TRBV12-4 | TRBJ2-3 | CASRPLLAETGASTDTQYF | KLGGALQAK | YES | NO | NO |
| 18082 | TRAV8-3 | TRAJ43 | CAVGAPDNNNDMRF | TRBV10-2 | TRBJ2-7 | CASSGQGQSYEQYV | KLGGALQAK | NO | NO | NO |
| 18083 | TRAV1-2 | TRAJ26 | CAFDNYGQNFVF | TRBV7-2 | TRBJ1-5 | CASSLDDNNQPQHF | KLGGALQAK | YES | NO | NO |
| 18084 | TRAV8-4 | TRAJ57 | CAVSDQGGSEKLVF | TRBV5-1 | TRBJ2-7 | CASSGGQGLFYEQYF | KLGGALQAK | YES | NO | NO |
| 18085 | TRAV1-2 | TRAJ20 | CAVSFSNDYKLSF | TRBV7-9 | TRBJ2-2 | CASSSTASGNTGELFF | KLGGALQAK | YES | NO | NO |
| 18086 | TRAV19 | TRAJ17 | CALSEAWAAGNKLTF | TRBV19 | TRBJ1-1 | CASSTLGTEAFF | KLGGALQAK | YES | NO | NO |
| 18087 | TRAV14DV4 | TRAJ53 | CAMRDESGGSNYKLTF | TRBV12-4 | TRBJ1-3 | CASSERFSGNTIYF | KLGGALQAK | YES | NO | NO |
| 18088 | TRAV24 | TRAJ40 | CASPAPTSGTYKYIF | TRBV7-9 | TRBJ1-1 | CASSSGWTEAFF | KLGGALQAK | YES | NO | NO |
| 18089 | TRAV4 | TRAJ43 | CLVGDNDMRF | TRBV7-3 | TRBJ2-2 | CASSLRTANTGELFF | KLGGALQAK | NO | NO | NO |
| 18090 | TRAV17 | TRAJ28 | CATDGSGAGSYQLTF | TRBV7-6 | TRBJ1-1 | CASRSSLTEAFF | KLGGALQAK | YES | NO | NO |
| 18091 | TRAV26-2 | TRAJ12 | CILMDSSYKLIF | TRBV2 | TRBJ2-2 | CASSVGVAGELFF | KLGGALQAK | YES | NO | NO |
| 18092 | TRAV19 | TRAJ53 | CALSETNSGGSNYKLTF | TRBV27 | TRBJ2-1 | CASSRRGSSYNEQFF | KLGGALQAK | YES | NO | NO |
| 18093 | TRAV25 | TRAJ13 | CAGSIHSGGYQKVTF | TRBV4-1 | TRBJ2-7 | CASSQDSRTGAVYEQYF | KLGGALQAK | YES | NO | NO |
| 18094 | TRAV17 | TRAJ23 | CATVLYNQGGKLIF | TRBV24-1 | TRBJ2-1 | CATSEGTGGYNEQFF | KLGGALQAK | YES | NO | NO |
| 18095 | TRAV3 | TRAJ17 | CAVRGLKAAGNKLTF | TRBV24-1 | TRBJ2-2 | CATSDHDRAGELFF | KLGGALQAK | NO | NO | NO |
| 18096 | TRAV8-2 | TRAJ22 | CVGRKGSARQLTF | TRBV12-3 | TRBJ2-1 | CASSAGAQSSYNEQFF | KLGGALQAK | YES | NO | NO |
| 18097 | TRAV39 | TRAJ50 | CAVDMAETSYDKVIF | TRBV27 | TRBJ2-1 | CASSLSSNEQFF | KLGGALQAK | YES | NO | NO |
| 18098 | TRAV12-2 | TRAJ23 | CAVNRINNQGGKLIF | TRBV7-2 | TRBJ1-2 | CASSLDRVGSYGYTF | KLGGALQAK | NO | NO | NO |
| 18099 | TRAV17 | TRAJ15 | CATDGEAGTALIF | TRBV4-1 | TRBJ2-7 | CASSQGRGAELYEQYF | AVFDRKSDAK | YES | NO | YES |
| 18100 | TRAV12-2 | TRAJ40 | CAVDPTSGTYKYIF | TRBV6-3 | TRBJ1-1 | CASSYARGRGTEAFF | KLGGALQAK | YES | NO | NO |
| 18101 | TRAV35 | TRAJ34 | CAGQRGGLSTDKLIF | TRBV6-3 | TRBJ1-6 | CAGSQGMRSPLHF | KLGGALQAK | YES | NO | NO |
| 18102 | TRAV3 | TRAJ37 | CAVIPTHGSSNTGKLIF | TRBV24-1 | TRBJ2-5 | CATSEFDRDGETQYF | KLGGALQAK | NO | NO | NO |
| 18103 | TRAV35 | TRAJ54 | CAGQLGQGAQKLVF | TRBV11-3 | TRBJ1-2 | CASSLRGDYGYTF | KLGGALQAK | YES | NO | NO |
| 18104 | TRAV17 | TRAJ30 | CATARLRDDKIIF | TRBV12-4 | TRBJ1-1 | CASSPTGPNTEAFF | KLGGALQAK | YES | NO | NO |
| 18105 | TRAV8-4 | TRAJ57 | CAVRSQGGSEKLVF | TRBV29-1 | TRBJ2-1 | CSASDRGGWGQFF | KLGGALQAK | YES | NO | NO |
| 18106 | TRAV12-1 | TRAJ34 | CVVNARGVRSADKLIF | TRBV4-1 | TRBJ1-5 | CASSQTPNQPQHF | KLGGALQAK | YES | NO | NO |
| 18107 | TRAV12-1 | TRAJ23 | CVVAPFYNQGGKLIF | TRBV13 | TRBJ2-2 | CASSNGLAGTGELFF | KLGGALQAK | NO | NO | NO |
| 18108 | TRAV1-1 | TRAJ16 | CAVRDGQKLLF | TRBV20-1 | TRBJ2-7 | CSARVASSYEQYF | KLGGALQAK | NO | NO | NO |
| 18109 | TRAV12-2 | TRAJ53 | CASGGSNYKLTF | TRBV19 | TRBJ1-1 | CASSIQIGNTEAFF | KLGGALQAK | YES | NO | NO |
| 18110 | TRAV12-3 | TRAJ15 | CAMSNQAGTALIF | TRBV30 | TRBJ1-1 | CAWSGVTEAFF | KLGGALQAK | NO | NO | NO |
| 18111 | TRAV39 | TRAJ45 | CAVSGGGGADGLTF | TRBV5-1 | TRBJ2-1 | CASIAGGVLSNEQFF | KLGGALQAK | YES | NO | NO |
| 18112 | TRAV12-2 | TRAJ11 | CAVTGYSTLTF | TRBV3-1 | TRBJ2-7 | CASSQDGRYEQYF | KLGGALQAK | YES | NO | NO |
| 18113 | TRAV8-4 | TRAJ23 | CAVSDVYNQGGKLIF | TRBV5-5 | TRBJ2-7 | CASSFCTQSMEFCEQYV | KLGGALQAK | NO | NO | NO |
| 18114 | TRAV12-1 | TRAJ22 | CVVTESGSARQLTF | TRBV7-9 | TRBJ2-2 | CASSLERKAGELFF | KLGGALQAK | YES | NO | NO |
| 18115 | TRAV12-3 | TRAJ9 | CAMSVGYTGGFKTIF | TRBV27 | TRBJ2-7 | CASSEVWDRGYEQYF | KLGGALQAK | YES | NO | NO |
| 18116 | TRAV3 | TRAJ39 | CAVRGLNNAGNMLTF | TRBV20-1 | TRBJ2-2 | CSARDLLAGGYTGELFF | KLGGALQAK | NO | NO | NO |
| 18117 | TRAV39 | TRAJ32 | CAVWTPSYGGATNKLIF | TRBV4-1 | TRBJ1-6 | CASSQERRYNSPLHF | KLGGALQAK | YES | NO | NO |
| 18118 | TRAV9-2 | TRAJ20 | CAREGVYDYKLSF | TRBV19 | TRBJ2-2 | CASSMGLAENTGELFF | RLRAEAQVK | YES | NO | YES |
| 18119 | TRAV12-2 | TRAJ17 | CAVNALEIKAAGNKLTF | TRBV12-4 | TRBJ2-5 | CASSFTETQYF | AVFDRKSDAK | YES | NO | YES |
| 18120 | TRAV34 | TRAJ11 | CGADPDSGYSTLTF | TRBV7-8 | TRBJ2-5 | CASSSTGYPQYF | AVFDRKSDAK | YES | NO | NO |
| 18121 | TRAV12-2 | TRAJ34 | CAVLLYNTDKLIF | TRBV6-5 | TRBJ1-1 | CASSYGEGTEAFF | KLGGALQAK | YES | NO | NO |
| 18122 | TRAV4 | TRAJ44 | CLVGDMDGTASKLTF | TRBV27 | TRBJ2-1 | CASSFSLAGYNEQFF | KLGGALQAK | YES | NO | NO |
| 18123 | TRAV3 | TRAJ35 | CAVRDEGGFGNVLHC | TRBV19 | TRBJ2-1 | CASSIELRLAGVPNEQFF | KLGGALQAK | NO | NO | NO |
| 18124 | TRAV19 | TRAJ47 | CALSEPRLEYGNKLVF | TRBV27 | TRBJ1-1 | CASSLRWDRGSNTEAFF | KLGGALQAK | YES | NO | NO |
| 18125 | TRAV26-1 | TRAJ31 | CIVRVADNARLMF | TRBV6-5 | TRBJ1-2 | CASLSTWDGYTF | KLGGALQAK | YES | NO | NO |
| 18126 | TRAV8-6 | TRAJ12 | CAVKSRMDSSYKLIF | TRBV13 | TRBJ1-2 | CASSGGQEDYGYTF | KLGGALQAK | YES | NO | NO |
| 18127 | TRAV24 | TRAJ44 | CAYFTGTASKLTF | TRBV9 | TRBJ1-2 | CASSARDRPIYGYTF | KLGGALQAK | YES | NO | NO |
| 18128 | TRAV17 | TRAJ45 | CATGGRGGGADGLTF | TRBV13 | TRBJ2-5 | CASSGGGDRSGRTQYF | KLGGALQAK | YES | NO | NO |
| 18129 | TRAV6 | TRAJ40 | CALVSTTSGTYKYIF | TRBV7-8 | TRBJ1-1 | CASSADMGTEAFF | KLGGALQAK | YES | NO | NO |
| 18130 | TRAV29DV5 | TRAJ42 | CAASDYGGSQGNLIF | TRBV6-1 | TRBJ2-2 | CASSEYYSTGELFF | KLGGALQAK | YES | NO | NO |
| 18131 | TRAV19 | TRAJ40 | CALSENAGTYKYIF | TRBV19 | TRBJ2-2 | CASTPGQGAGELFF | KLGGALQAK | YES | NO | NO |
| 18132 | TRAV12-3 | TRAJ36 | CAGTGANNLFF | TRBV7-9 | TRBJ2-2 | CASGLASTGELFF | KLGGALQAK | YES | NO | NO |
| 18133 | TRAV21 | TRAJ54 | CAVYGGAQKLVF | TRBV5-4 | TRBJ2-1 | CASSLDASLNEQFF | KLGGALQAK | YES | NO | NO |
| 18134 | TRAV35 | TRAJ54 | CAGGSIRQGAQKLVF | TRBV7-9 | TRBJ1-2 | CASSSGQGNYGYTF | KLGGALQAK | YES | NO | NO |
| 18135 | TRAV1-2 | TRAJ12 | CAVRDQADSSYKLIF | TRBV7-8 | TRBJ1-2 | CASSIIRESNYGYTF | AVFDRKSDAK | YES | NO | YES |
| 18136 | TRAV20 | TRAJ42 | CAVPNYGGSQGNLIF | TRBV4-3 | TRBJ2-1 | CASSQDSPLATIYNEQFF | KLGGALQAK | YES | NO | NO |
| 18137 | TRAV14DV4 | TRAJ11 | CAMREVNSGYSTLTF | TRBV14 | TRBJ2-3 | CASSPQGAYLDTQYF | KLGGALQAK | YES | NO | NO |
| 18138 | TRAV6 | TRAJ47 | CALNMEYGNKLVF | TRBV6-5 | TRBJ2-5 | CASREGTGIRETQYF | KLGGALQAK | NO | NO | NO |
| 18139 | TRAV1-2 | TRAJ23 | CAVRARGGKLIF | TRBV7-9 | TRBJ1-1 | CASSLSESGSTEAFF | KLGGALQAK | YES | NO | NO |
| 18140 | TRAV3 | TRAJ32 | CAVRDIRYGGATNKLIF | TRBV12-3 | TRBJ1-5 | CASSSEDSGDQPQHF | KLGGALQAK | NO | NO | NO |
| 18141 | TRAV19 | TRAJ9 | CALSEFLTGGFKTIF | TRBV27 | TRBJ2-1 | CASWTGTSAEDYNEQFF | KLGGALQAK | YES | NO | NO |

| | | | | | | | | | | |
|---|---|---|---|---|---|---|---|---|---|---|
| 18142 | TRAV22 | TRAJ40 | CAVTSTSGTYKYIF | TRBV19 | TRBJ1-1 | CASSIGQGPCCFF | KLGGALQAK | YES | NO | NO |
| 18143 | TRAV29DV5 | TRAJ28 | CAASEREGYYSGAGSYQLTF | TRBV6-6 | TRBJ2-3 | CASSYSRGLGVMDPSTDTQYF | KLGGALQAK | NO | NO | NO |
| 18144 | TRAV26-1 | TRAJ26 | CIVRGDNYGQNFVF | TRBV19 | TRBJ2-7 | CASSMTGGSYEQYF | KLGGALQAK | YES | NO | NO |
| 18145 | TRAV27 | TRAJ34 | CAGYNTDKLIF | TRBV24-1 | TRBJ1-1 | CATSVVITEAFF | KLGGALQAK | YES | NO | NO |
| 18146 | TRAV1-1 | TRAJ27 | CANGDTNAGKSTF | TRBV6-4 | TRBJ1-2 | CASRRDNNYGYTF | KLGGALQAK | YES | NO | NO |
| 18147 | TRAV12-2 | TRAJ48 | CAVDFGNEKLTF | TRBV29-1 | TRBJ1-2 | CSVGGTEYGYTF | KLGGALQAK | YES | NO | NO |
| 18148 | TRAV8-6 | TRAJ15 | CAVSDSEAGTALIF | TRBV6-6 | TRBJ2-1 | CASSYPLAGGGSYNEQFF | KLGGALQAK | NO | NO | NO |
| 18149 | TRAV6 | TRAJ20 | CARPSDYKLSF | TRBV6-5 | TRBJ1-1 | CASSSDSGTEAFF | KLGGALQAK | YES | NO | NO |
| 18150 | TRAV13-1 | TRAJ3 | CAAPYSSASKIIF | TRBV7-8 | TRBJ2-3 | CASSIGTSLARDTQYF | KLGGALQAK | YES | NO | NO |
| 18151 | TRAV27 | TRAJ43 | CAGAAGNDMRF | TRBV11-2 | TRBJ2-1 | CASSLGRGPGNEQFF | KLGGALQAK | YES | NO | NO |
| 18152 | TRAV14DV4 | TRAJ53 | CAMRPSYSGGSNYKLTF | TRBV7-8 | TRBJ1-6 | CASSLATGGYSPLHF | KLGGALQAK | YES | NO | NO |
| 18153 | TRAV38-2DV8 | TRAJ35 | CAYRSAI | TRBV4-1 | TRBJ2-1 | CASSQGPGLLNEQFF | KLGGALQAK | NO | NO | NO |
| 18154 | TRAV12-1 | TRAJ9 | CVVNPGGFKTIF | TRBV20-1 | TRBJ1-1 | CSASAMNTEAFF | KLGGALQAK | YES | NO | NO |
| 18155 | TRAV4 | TRAJ3 | CLVGDIGYSSASKIIF | TRBV7-6 | TRBJ2-3 | CASSLSEGVDTQYF | KLGGALQAK | NO | NO | NO |
| 18156 | TRAV3 | TRAJ13 | CAVRVAGGYQKVTF | TRBV4-1 | TRBJ2-2 | CASSQEGLRNTGELFF | AVFDRKSDAK | YES | NO | YES |
| 18157 | TRAV8-3 | TRAJ17 | CAVGAYNAPRGNKLTF | TRBV4-3 | TRBJ2-2 | CASSQDPLASGELFF | KLGGALQAK | YES | NO | NO |
| 18158 | TRAV25 | TRAJ56 | CAGPRYTGANSKLTF | TRBV7-3 | TRBJ2-7 | CASSLIQESEQYF | KLGGALQAK | YES | NO | NO |
| 18159 | TRAV29DV5 | TRAJ17 | CAATGIAAGNKLTF | TRBV20-1 | TRBJ2-6 | CSARNPAGAGANVLTF | KLGGALQAK | YES | NO | NO |
| 18160 | TRAV38-2DV8 | TRAJ42 | CAYRSETGGSQGNLIF | TRBV7-2 | TRBJ2-7 | CASSLGLAGVYEQYV | RLRAEAQVK | NO | NO | NO |
| 18161 | TRAV19 | TRAJ32 | CALRGSGGATNKLIF | TRBV29-1 | TRBJ1-1 | CSAHDRGLNTEAFF | KLGGALQAK | NO | NO | NO |
| 18162 | TRAV17 | TRAJ18 | CATVTDRGSTLGRLYF | TRBV5-1 | TRBJ2-1 | CASSLAESSYNEQFF | KLGGALQAK | YES | NO | NO |
| 18163 | TRAV12-2 | TRAJ52 | CAVNLNAGGTSYGKLTF | TRBV19 | TRBJ2-1 | CASSRGLYNEQFF | KLGGALQAK | YES | NO | NO |
| 18164 | TRAV39 | TRAJ58 | CGTSGSRLTF | TRBV4-1 | TRBJ2-3 | CASSQVLAGVADTQYF | KLGGALQAK | YES | NO | NO |
| 18165 | TRAV20 | TRAJ36 | CAVQPEGANNLFF | TRBV2 | TRBJ1-5 | CASSAVYSNQPQHF | KLGGALQAK | YES | NO | NO |
| 18166 | TRAV19 | TRAJ58 | CALSAFSGSRLTF | TRBV7-2 | TRBJ2-1 | CASSLVTGSEQFF | KLGGALQAK | YES | NO | NO |
| 18167 | TRAV14DV4 | TRAJ11 | CAMREGNSGYSTLTF | TRBV7-3 | TRBJ2-5 | CASSLAGGPLETQYF | KLGGALQAK | NO | NO | NO |
| 18168 | TRAV35 | TRAJ28 | CAGQFDSGAGSYQLTF | TRBV19 | TRBJ2-2 | CASSLLVTGELFF | KLGGALQAK | YES | NO | NO |
| 18169 | TRAV41 | TRAJ43 | CAGLYNNNDMRF | TRBV2 | TRBJ2-2 | CASEAGELFF | KLGGALQAK | YES | NO | NO |
| 18170 | TRAV5 | TRAJ21 | CAVLGYNFNKFYF | TRBV5-6 | TRBJ1-4 | CASSFPGDREKLFF | KLGGALQAK | YES | NO | NO |
| 18171 | TRAV35 | TRAJ34 | CAGQLGRYNTDKLIF | TRBV24-1 | TRBJ2-1 | CATSDLPGAYNEQFF | KLGGALQAK | YES | NO | NO |
| 18172 | TRAV1-1 | TRAJ28 | CAVRPGAGSYQLTF | TRBV19 | TRBJ1-1 | CASTQRGHTEAFF | KLGGALQAK | YES | NO | NO |
| 18173 | TRAV38-2DV8 | TRAJ32 | CAYRGPNYGGATNKLIF | TRBV4-1 | TRBJ2-5 | CASSQWRSGGGAGKETQYF | KLGGALQAK | YES | NO | NO |
| 18174 | TRAV23DV6 | TRAJ30 | CAASWGVDDKIIF | TRBV7-2 | TRBJ1-6 | CASSWDGTGHVGSPLHF | KLGGALQAK | NO | NO | NO |
| 18175 | TRAV29DV5 | TRAJ43 | CATNDMRF | TRBV10-2 | TRBJ1-5 | CASSPGQGGNQPQHF | KLGGALQAK | NO | NO | NO |
| 18176 | TRAV16 | TRAJ28 | CALTYSGAGSYQLTF | TRBV7-9 | TRBJ2-1 | CASRQNSGSSYNEQFF | KLGGALQAK | NO | NO | NO |
| 18177 | TRAV30 | TRAJ45 | CGTDSSSGGGADGLTF | TRBV4-1 | TRBJ2-3 | CASSQGTWKTSTDTQYF | KLGGALQAK | YES | NO | NO |
| 18178 | TRAV23DV6 | TRAJ37 | CAADSHGSSNTGKLIF | TRBV7-6 | TRBJ2-1 | CASSLTGNEQFF | KLGGALQAK | YES | NO | NO |
| 18179 | TRAV21 | TRAJ50 | CAVRPQLETSYDKVIF | TRBV4-1 | TRBJ2-3 | CASSPRVGDLTQYF | KLGGALQAK | YES | NO | NO |
| 18180 | TRAV21 | TRAJ37 | CAVRGLGGNTGKLIF | TRBV6-5 | TRBJ2-3 | CASSYADTQYF | RLRAEAQVK | NO | NO | NO |
| 18181 | TRAV23DV6 | TRAJ42 | CAAGMNYGGSQGNLIF | TRBV7-2 | TRBJ2-6 | CASSLIFIRLAGGVFGANVLTF | KLGGALQAK | NO | NO | NO |
| 18182 | TRAV4 | TRAJ18 | CLVGDRGSTLGRLYF | TRBV4-1 | TRBJ2-7 | CASSPRLEAYEQYF | KLGGALQAK | YES | NO | NO |
| 18183 | TRAV17 | TRAJ20 | CATHLNDYKLSF | TRBV29-1 | TRBJ2-1 | CSVDASDRGEQFF | KLGGALQAK | YES | NO | NO |
| 18184 | TRAV17 | TRAJ40 | CATPQVTSGTYKYIF | TRBV6-1 | TRBJ2-4 | CASTPRTAKNIQYF | KLGGALQAK | YES | NO | NO |
| 18185 | TRAV36DV7 | TRAJ52 | CAVGANAGGTSYGKLTF | TRBV25-1 | TRBJ1-5 | CASSWSNQPQHF | KLGGALQAK | NO | NO | NO |
| 18186 | TRAV14DV4 | TRAJ28 | CASAYSGAGSYQLTF | TRBV27 | TRBJ2-1 | CASSLRWTNEQFF | KLGGALQAK | YES | NO | NO |
| 18187 | TRAV14DV4 | TRAJ50 | CAMREGETSYDKVIF | TRBV6-5 | TRBJ2-7 | CASSSPRRGEYEQYF | KLGGALQAK | YES | NO | NO |
| 18188 | TRAV27 | TRAJ52 | CAGATAGGTSYGKLTF | TRBV18 | TRBJ2-7 | CASSPLGPPYEQYF | KLGGALQAK | YES | NO | NO |
| 18189 | TRAV23DV6 | TRAJ28 | CAASMLISGAGSYQLTF | TRBV27 | TRBJ2-7 | CASSAGVAQGEQYF | KLGGALQAK | YES | NO | NO |
| 18190 | TRAV29DV5 | TRAJ54 | CAASARLIQGAQKLVF | TRBV5-1 | TRBJ2-4 | CASSLELAGLGLAKNIQYF | KLGGALQAK | YES | NO | NO |
| 18191 | TRAV29DV5 | TRAJ53 | CAAMMSGGSNYKLTF | TRBV7-2 | TRBJ1-5 | CASSLALGGHQPQHF | KLGGALQAK | NO | NO | NO |
| 18192 | TRAV10 | TRAJ33 | CVVSAIEGMDSNYQLIW | TRBV2 | TRBJ2-1 | CASSEFASVSYNEQFF | RLRAEAQVK | YES | YES | YES |
| 18193 | TRAV12-2 | TRAJ45 | CAVTISGGGADGLTF | TRBV2 | TRBJ2-5 | CASSELQETQYF | KLGGALQAK | YES | NO | NO |
| 18194 | TRAV40 | TRAJ42 | CLLGDLGGGSQGNLIF | TRBV7-9 | TRBJ2-5 | CASSSTTSGRGWETQYF | KLGGALQAK | NO | NO | NO |
| 18195 | TRAV8-2 | TRAJ42 | CVVSDLGSQGNLIF | TRBV7-9 | TRBJ2-7 | CASSFPSVSYEQYF | KLGGALQAK | YES | NO | NO |
| 18196 | TRAV27 | TRAJ52 | CADIDAGGTSYGKLTF | TRBV4-1 | TRBJ2-1 | CASSLAAGFEQFF | KLGGALQAK | YES | NO | NO |
| 18197 | TRAV10 | TRAJ35 | CVVRSFGNVLHC | TRBV30 | TRBJ2-6 | CAWSRRSGANVLTF | KLGGALQAK | NO | NO | NO |
| 18198 | TRAV29DV5 | TRAJ32 | CAASDDGGATNKLIF | TRBV7-2 | TRBJ2-1 | CASTTGYNEQFF | KLGGALQAK | YES | NO | NO |
| 18199 | TRAV16 | TRAJ31 | CALNNARLMF | TRBV5-4 | TRBJ2-3 | CASSFHSGTDTQYF | KLGGALQAK | NO | NO | NO |
| 18200 | TRAV3 | TRAJ39 | CAVRGLSGLDNNAGNMLTF | TRBV20-1 | TRBJ1-5 | CSARLSGGGNQPQHF | KLGGALQAK | NO | NO | NO |
| 18201 | TRAV12-2 | TRAJ52 | CAVNGDAGGTSYGKLTF | TRBV7-8 | TRBJ2-7 | CASSPGASYEQYF | KLGGALQAK | YES | YES | NO |
| 18202 | TRAV13-2 | TRAJ30 | CAENNMNRDDKIIF | TRBV20-1 | TRBJ2-7 | CSALSDRDYEQYF | KLGGALQAK | YES | NO | NO |
| 18203 | TRAV19 | TRAJ34 | CALSWDTDKLIF | TRBV5-1 | TRBJ2-3 | CASSPYGTSYTDTQYF | KLGGALQAK | YES | NO | NO |
| 18204 | TRAV1-2 | TRAJ10 | CAVELAFTGGGNKLTF | TRBV14 | TRBJ1-2 | CASSQVDSYGYTF | AYAQKIFKI | YES | YES | YES |
| 18205 | TRAV19 | TRAJ49 | CALSARDTGNQFYF | TRBV13 | TRBJ1-1 | CASSWGTEAFF | KLGGALQAK | YES | NO | NO |
| 18206 | TRAV20 | TRAJ18 | CAVQAERGSTLGRLYF | TRBV6-6 | TRBJ1-5 | CASTSPPGQPQHF | KLGGALQAK | NO | NO | NO |
| 18207 | TRAV12-1 | TRAJ43 | CVVNGWVHNDMRF | TRBV11-3 | TRBJ2-2 | CASSLAPGLAGTGELFF | KLGGALQAK | YES | NO | NO |
| 18208 | TRAV27 | TRAJ17 | CAGAAGGDNKLTF | TRBV12-4 | TRBJ1-4 | CASSLGSGQAEKLFF | KLGGALQAK | YES | NO | NO |
| 18209 | TRAV21 | TRAJ23 | CAVRPGGKLIF | TRBV29-1 | TRBJ1-2 | CSVEGDWRQGASGYTF | KLGGALQAK | YES | NO | NO |
| 18210 | TRAV12-2 | TRAJ39 | CAIGPGNMLTF | TRBV3-1 | TRBJ2-7 | CASSQGGVEQYF | KLGGALQAK | YES | NO | NO |
| 18211 | TRAV25 | TRAJ26 | CAGRGNYGQNFVF | TRBV7-6 | TRBJ1-2 | CASSSATLGGRIYGYTF | KLGGALQAK | YES | NO | NO |
| 18212 | TRAV8-1 | TRAJ39 | CAVNAANAGNMLTF | TRBV11-2 | TRBJ2-3 | CASRTPPSLSSTDTQYF | AYAQKIFKI | YES | YES | YES |
| 18213 | TRAV6 | TRAJ49 | CAIGNQFYF | TRBV29-1 | TRBJ2-2 | CSVESAASSGELFF | KLGGALQAK | NO | NO | NO |
| 18214 | TRAV29DV5 | TRAJ42 | CAASVGGGSQGNLIF | TRBV12-4 | TRBJ2-5 | CASSSFLAGAKTQYF | KLGGALQAK | YES | NO | NO |
| 18215 | TRAV13-1 | TRAJ9 | CAASINGGFKTIF | TRBV7-6 | TRBJ1-5 | CASSPGGGRDQPQHF | KLGGALQAK | YES | NO | NO |
| 18216 | TRAV29DV5 | TRAJ40 | CAASWDPTSGTYKYIF | TRBV11-2 | TRBJ2-1 | CASSLVAGGQNEQFF | KLGGALQAK | YES | NO | NO |
| 18217 | TRAV27 | TRAJ26 | CAGAGNYGQNFVF | TRBV29-1 | TRBJ2-7 | CSVEEGNRGGYEQYF | KLGGALQAK | YES | NO | NO |
| 18218 | TRAV1-2 | TRAJ35 | CAVREGGFGNVLHC | TRBV7-9 | TRBJ2-5 | CASSDTPGLAGGEETQYF | KLGGALQAK | NO | NO | NO |
| 18219 | TRAV13-1 | TRAJ34 | CAASNGDTDKLIF | TRBV12-4 | TRBJ2-7 | CASSVGGTHYEQYF | KLGGALQAK | YES | NO | NO |
| 18220 | TRAV9-2 | TRAJ34 | CALSGTDKLIF | TRBV9 | TRBJ1-1 | CASSPRGTVNAEAFF | KLGGALQAK | YES | NO | NO |
| 18221 | TRAV27 | TRAJ20 | CAGVGSNDYKLSF | TRBV29-1 | TRBJ1-1 | CSVSGTGNTEAFF | KLGGALQAK | YES | NO | NO |
| 18222 | TRAV19 | TRAJ53 | CALTASGGSNYKLTF | TRBV6-1 | TRBJ2-1 | CASRASGRNEQFF | KLGGALQAK | YES | NO | NO |
| 18223 | TRAV17 | TRAJ22 | CATDAGTSGSARQLTF | TRBV19 | TRBJ1-5 | CASSIGIPRWPQHF | KLGGALQAK | YES | NO | NO |
| 18224 | TRAV12-3 | TRAJ35 | CAMTPTPQDGFGNVLHC | TRBV20-1 | TRBJ2-5 | CSASFFRDQETQYF | AVFDRKSDAK | NO | NO | NO |
| 18225 | TRAV29DV5 | TRAJ13 | CAASAPRGYQKVTF | TRBV7-3 | TRBJ2-2 | CASSSTDNTGELFF | KLGGALQAK | YES | NO | NO |
| 18226 | TRAV6 | TRAJ28 | CALGSGAGSYQLTF | TRBV27 | TRBJ2-3 | CASSGYTDTQYF | KLGGALQAK | YES | NO | NO |
| 18227 | TRAV8-6 | TRAJ54 | CAVSDPQGAQKLVF | TRBV19 | TRBJ2-7 | CASSITGYSEQYF | KLGGALQAK | YES | NO | NO |
| 18228 | TRAV12-2 | TRAJ40 | CAVNPGTYKYIF | TRBV4-1 | TRBJ1-4 | CASSTGGEKLFF | KLGGALQAK | YES | NO | NO |
| 18229 | TRAV1-2 | TRAJ28 | CAARGGSYQLTF | TRBV20-1 | TRBJ2-1 | CSVPPRDRYNEQFF | KLGGALQAK | NO | NO | NO |
| 18230 | TRAV12-2 | TRAJ18 | CAVGRGSTLGRLYF | TRBV7-9 | TRBJ1-4 | CASSSRPQGTNEKLFF | KLGGALQAK | YES | NO | NO |
| 18231 | TRAV12-1 | TRAJ45 | CVAERGGADGLTF | TRBV3-1 | TRBJ2-7 | CASSPGLGAYEQYV | KLGGALQAK | NO | NO | NO |
| 18232 | TRAV9-2 | TRAJ6 | CALRDAGGSYIPTF | TRBV6-3 | TRBJ2-7 | CASSTFEEILIGYEQYF | KLGGALQAK | YES | NO | NO |
| 18233 | TRAV29DV5 | TRAJ39 | CAASGNNAGNMLTF | TRBV6-6 | TRBJ1-1 | CASSPGGHTAAFF | KLGGALQAK | NO | NO | NO |
| 18234 | TRAV1-2 | TRAJ35 | CAVPI | TRBV7-9 | TRBJ2-2 | CASTGTSSVSGELFF | KLGGALQAK | NO | NO | NO |
| 18235 | TRAV17 | TRAJ29 | CATDAYSGNTPLVF | TRBV4-1 | TRBJ2-1 | CASSQDSGGGDEQFF | KLGGALQAK | YES | NO | NO |
| 18236 | TRAV14DV4 | TRAJ9 | CAMREGSTGGFKTIF | TRBV4-1 | TRBJ1-1 | CASSLKRDRETEAFF | KLGGALQAK | YES | NO | NO |
| 18237 | TRAV29DV5 | TRAJ46 | CAAGGDKLTF | TRBV7-6 | TRBJ1-3 | CASSADSIRNTIYF | KLGGALQAK | YES | NO | NO |
| 18238 | TRAV10 | TRAJ45 | CVVRETAGGGADGLTF | TRBV7-9 | TRBJ2-3 | CASSLAITSGSDTQYF | KLGGALQAK | YES | NO | NO |
| 18239 | TRAV23DV6 | TRAJ10 | CAASRKLTGGGNKLTF | TRBV5-1 | TRBJ1-6 | CASSEGGYSYNSPLHF | KLGGALQAK | YES | NO | NO |
| 18240 | TRAV13-1 | TRAJ50 | CAARTMKTSYDKVIF | TRBV6-5 | TRBJ2-1 | CASSIVGGGYNEQFF | KLGGALQAK | NO | NO | NO |
| 18241 | TRAV22 | TRAJ26 | CAVGNYGQNFVF | TRBV4-1 | TRBJ1-4 | CASSQEGEKLFF | SLFNTVATL | YES | NO | YES |
| 18242 | TRAV13-1 | TRAJ9 | CAASILARTGGFKTIF | TRBV19 | TRBJ1-2 | CASSNQGGRTF | KLGGALQAK | YES | NO | NO |
| 18243 | TRAV27 | TRAJ27 | CAGAFDTNAGKSTF | TRBV10-3 | TRBJ2-7 | CAISDYYEQYF | KLGGALQAK | YES | NO | NO |
| 18244 | TRAV39 | TRAJ45 | CAVDIGGGADGLTF | TRBV5-1 | TRBJ2-1 | CASSLVGYEQFF | KLGGALQAK | YES | NO | NO |
| 18245 | TRAV12-2 | TRAJ31 | CAVTKARLMF | TRBV29-1 | TRBJ2-2 | CSVVHGELFF | KLGGALQAK | NO | NO | NO |

| | | | | | | | | | | |
|---|---|---|---|---|---|---|---|---|---|---|
| 18246 | TRAV20 | TRAJ44 | CAGGTGTASKLTF | TRBV4-1 | TRBJ2-2 | CASSRTGELFF | KLGGALQAK | YES | NO | NO |
| 18247 | TRAV12-1 | TRAJ41 | CVVKAGSSGYALNF | TRBV12-4 | TRBJ2-2 | CASTGGRGLDTGELFF | KLGGALQAK | YES | NO | NO |
| 18248 | TRAV29DV5 | TRAJ53 | CAIGSNYKLTF | TRBV27 | TRBJ1-1 | CASSLQGNTEAFF | KLGGALQAK | YES | NO | NO |
| 18249 | TRAV1-1 | TRAJ29 | CAVVNTPLVF | TRBV15 | TRBJ2-6 | CATSNTGYSGANVLTF | KLGGALQAK | YES | NO | NO |
| 18250 | TRAV20 | TRAJ17 | CAVGRAAGNKLTF | TRBV9 | TRBJ2-1 | CASSVLVGGSYNEQFF | AVFDRKSDAK | YES | NO | NO |
| 18251 | TRAV14DV4 | TRAJ11 | CAMRETSGYSTLTF | TRBV6-4 | TRBJ1-2 | CASSEGRGYTF | KLGGALQAK | YES | NO | NO |
| 18252 | TRAV14DV4 | TRAJ49 | CAMSFWGGNQFYF | TRBV9 | TRBJ2-5 | CASSVELAGGPQTQYF | KLGGALQAK | YES | NO | NO |
| 18253 | TRAV12-2 | TRAJ53 | CAGGGSNYKLTF | TRBV6-1 | TRBJ2-1 | CASRGGAGGAGGNEQFF | AVFDRKSDAK | YES | NO | YES |
| 18254 | TRAV20 | TRAJ6 | CAVPGAPGGSYIPTF | TRBV19 | TRBJ1-1 | CASITTGANTEAFF | KLGGALQAK | YES | NO | NO |
| 18255 | TRAV13-2 | TRAJ26 | CAENSVDNYGQNFVF | TRBV6-6 | TRBJ2-1 | CASQTSGSDEQFF | KLGGALQAK | NO | NO | NO |
| 18256 | TRAV8-3 | TRAJ34 | CAVATYNTDKLIF | TRBV14 | TRBJ2-7 | CASSPIAAEDLYEQYV | KLGGALQAK | NO | NO | NO |
| 18257 | TRAV4 | TRAJ33 | CLVGDGYQLIW | TRBV11-2 | TRBJ2-7 | CASSQDFGYEQYV | KLGGALQAK | NO | NO | NO |
| 18258 | TRAV13-1 | TRAJ43 | CAASGDNDMRF | TRBV11-2 | TRBJ2-3 | CASSLSRGHNTDTQYF | KLGGALQAK | YES | NO | NO |
| 18259 | TRAV34 | TRAJ53 | CGADIGSGGSNYKLTF | TRBV4-1 | TRBJ2-7 | CASSRTTTPSYEQYF | KLGGALQAK | YES | NO | NO |
| 18260 | TRAV12-3 | TRAJ26 | CAMSDGQNFVF | TRBV4-1 | TRBJ1-3 | CASSQVLNSAIYF | KLGGALQAK | YES | NO | NO |
| 18261 | TRAV12-3 | TRAJ34 | CAMSYNTDKLIF | TRBV27 | TRBJ2-7 | CASKSFPGDPDLGEQYF | KLGGALQAK | YES | NO | NO |
| 18262 | TRAV19 | TRAJ10 | CALSALTGGGNKLTF | TRBV7-2 | TRBJ2-1 | CASSLGTSGSTLFDEQFF | KLGGALQAK | YES | NO | NO |
| 18263 | TRAV3 | TRAJ24 | CAGPGVDSWGKLQF | TRBV19 | TRBJ1-4 | CASSTYRGPTNEKLFF | AVFDRKSDAK | NO | NO | NO |
| 18264 | TRAV13-2 | TRAJ40 | CAEIHTSGTYKYIF | TRBV27 | TRBJ2-7 | CASSLTGGYEQYF | KLGGALQAK | YES | NO | NO |
| 18265 | TRAV4 | TRAJ44 | CLVGEINTGTASKLTF | TRBV7-9 | TRBJ2-1 | CASSSGLAGENEQFF | KLGGALQAK | YES | NO | NO |
| 18266 | TRAV4 | TRAJ30 | CLVGEGDDKIIF | TRBV5-6 | TRBJ2-7 | CASSLGGGPYEQYV | KLGGALQAK | NO | NO | NO |
| 18267 | TRAV8-4 | TRAJ28 | CAVSDSGAGSYQLTF | TRBV13 | TRBJ2-7 | CASSSRQTDYEQYF | KLGGALQAK | YES | NO | NO |
| 18268 | TRAV2 | TRAJ45 | CAVEDLIGGADGLTF | TRBV6-1 | TRBJ2-2 | CASSVYRGNTGELFF | KLGGALQAK | NO | NO | NO |
| 18269 | TRAV16 | TRAJ58 | CALMEEGETSGSRLTF | TRBV7-9 | TRBJ1-1 | CASSPPAGGNTEAFF | KLGGALQAK | NO | NO | NO |
| 18270 | TRAV9-2 | TRAJ11 | CALSEGYSTLTF | TRBV7-8 | TRBJ1-1 | CASISPGQNTEAFF | KLGGALQAK | YES | NO | NO |
| 18271 | TRAV14DV4 | TRAJ33 | CAMRKDSNYQLIW | TRBV27 | TRBJ1-1 | CASSFIGSDWNTEAFF | KLGGALQAK | YES | NO | NO |
| 18272 | TRAV21 | TRAJ18 | CAVKGDDRGSTLGRLYF | TRBV20-1 | TRBJ2-7 | CSATARGGTSGGKSYEQYF | KLGGALQAK | NO | NO | NO |
| 18273 | TRAV30 | TRAJ6 | CGTEMASGGSYIPTF | TRBV29-1 | TRBJ1-2 | CSVDLEANYGYTF | KLGGALQAK | YES | NO | NO |
| 18274 | TRAV13-2 | TRAJ45 | CAEKAPKGGGADGLTF | TRBV5-5 | TRBJ1-1 | CASSLQGAPEAFF | KLGGALQAK | YES | NO | NO |
| 18275 | TRAV29DV5 | TRAJ48 | CAASVGLGGNEKLTF | TRBV11-2 | TRBJ1-1 | CASSLDLETGDTEAFF | KLGGALQAK | YES | NO | NO |
| 18276 | TRAV13-1 | TRAJ39 | CAASSWNNAGNMLTF | TRBV7-3 | TRBJ1-1 | CASSATGGAGHTEAFF | KLGGALQAK | NO | NO | NO |
| 18277 | TRAV12-2 | TRAJ15 | CAVQAGTALIF | TRBV5-4 | TRBJ2-4 | CASSSRGAKNIQYF | KLGGALQAK | YES | NO | NO |
| 18278 | TRAV13-1 | TRAJ21 | CAASIYNFNKFYF | TRBV5-6 | TRBJ2-2 | CASSLLGQNTGELFF | IVTDFSVIK | YES | YES | YES |
| 18279 | TRAV8-3 | TRAJ43 | CAVTDMRF | TRBV6-5 | TRBJ2-5 | CASSYTGVVETQYF | KLGGALQAK | NO | NO | NO |
| 18280 | TRAV23DV6 | TRAJ49 | CAASIDTGNQFYF | TRBV7-9 | TRBJ1-4 | CASSSWTANEKLFF | KLGGALQAK | YES | NO | NO |
| 18281 | TRAV25 | TRAJ8 | CAGSGTGFQKLVF | TRBV27 | TRBJ1-2 | CASSPGNGANYGYTF | KLGGALQAK | YES | NO | NO |
| 18282 | TRAV13-2 | TRAJ43 | CAENGAVYNNNDMRF | TRBV7-9 | TRBJ1-1 | CASSPGQNTEAFF | KLGGALQAK | YES | NO | NO |
| 18283 | TRAV25 | TRAJ39 | CAGLSFVDAGNMLTF | TRBV11-3 | TRBJ2-1 | CASSLWGRHNEQFF | KLGGALQAK | YES | NO | NO |
| 18284 | TRAV14DV4 | TRAJ37 | CAMREGWSSNTGKLIF | TRBV6-5 | TRBJ2-3 | CASSYVGGPGDTQYF | KLGGALQAK | NO | NO | NO |
| 18285 | TRAV25 | TRAJ52 | CAGLRAGGTSYGKLTF | TRBV27 | TRBJ2-6 | CASSFSGANVLTF | KLGGALQAK | YES | NO | NO |
| 18286 | TRAV23DV6 | TRAJ42 | CAASNIGGSQGNLIF | TRBV19 | TRBJ1-5 | CASSMGGAGNQPQHF | KLGGALQAK | YES | NO | NO |
| 18287 | TRAV4 | TRAJ31 | CLVGDVNNNARLMF | TRBV5-1 | TRBJ2-3 | CASSPPDSFTQYF | KLGGALQAK | NO | NO | NO |
| 18288 | TRAV13-1 | TRAJ48 | CAASKFRGFGNEKLTF | TRBV7-9 | TRBJ2-1 | CASSLAGLSDEQFF | KLGGALQAK | YES | NO | NO |
| 18289 | TRAV13-1 | TRAJ28 | CAATSRPLILWGWELPTHF | TRBV5-5 | TRBJ1-1 | CASSFSVNTEAFF | KLGGALQAK | NO | NO | NO |
| 18290 | TRAV13-1 | TRAJ44 | CAASKTGTASKLTF | TRBV7-9 | TRBJ2-3 | CASSKRGGQTDTQYF | KLGGALQAK | YES | NO | NO |
| 18291 | TRAV19 | TRAJ16 | CALSEVFWSDGQKLLF | TRBV3-1 | TRBJ2-6 | CASTLDGRTSGANVLTF | KLGGALQAK | NO | NO | NO |
| 18292 | TRAV21 | TRAJ5 | CAVRRTGRRALTF | TRBV19 | TRBJ1-4 | CASSSPVTDPQNEKLFF | KLGGALQAK | YES | NO | NO |
| 18293 | TRAV8-3 | TRAJ49 | CAVGARGNQFYF | TRBV7-9 | TRBJ2-4 | CASSPSRDREFLYIQYF | KLGGALQAK | YES | NO | NO |
| 18294 | TRAV35 | TRAJ27 | CAGLGDTNAGKSTF | TRBV7-6 | TRBJ2-1 | CASSLSWNEQFF | KLGGALQAK | YES | NO | NO |
| 18295 | TRAV21 | TRAJ20 | CAVSSNDYKLSF | TRBV6-6 | TRBJ2-3 | CASSYYSGGSTDTQYF | KLGGALQAK | NO | NO | NO |
| 18296 | TRAV8-6 | TRAJ23 | CAVSANMIYNQGGKLIF | TRBV5-1 | TRBJ2-5 | CASSFRLAGEETQYF | AVFDRKSDAK | YES | NO | YES |
| 18297 | TRAV19 | TRAJ34 | CALSEGRDTDKLIF | TRBV4-1 | TRBJ2-1 | CASSQGPSNNEQFF | KLGGALQAK | YES | NO | NO |
| 18298 | TRAV20 | TRAJ8 | CAAGGDGFQKLVF | TRBV12-4 | TRBJ2-3 | CASSQSDNDDYTQYF | KLGGALQAK | YES | NO | NO |
| 18299 | TRAV1-2 | TRAJ30 | CAVSNRDDKIIF | TRBV4-1 | TRBJ1-1 | CASQDAVGTEAFF | AVFDRKSDAK | YES | NO | YES |
| 18300 | TRAV12-2 | TRAJ29 | CAPEGGNTPLVF | TRBV30 | TRBJ2-2 | CAWSETSGSLNTGELFF | KLGGALQAK | YES | NO | NO |
| 18301 | TRAV9-2 | TRAJ53 | CALSDLGSGGSNYKLTF | TRBV7-9 | TRBJ2-7 | CASRPGLNYEQYF | KLGGALQAK | YES | NO | NO |
| 18302 | TRAV12-2 | TRAJ49 | CAVNRNSNQFYF | TRBV4-1 | TRBJ2-7 | CASSSGGYEQYF | KLGGALQAK | YES | NO | NO |
| 18304 | TRAV20 | TRAJ30 | CAGPRYDKIIF | TRBV29-1 | TRBJ1-2 | CSVFSRLYGYTF | KLGGALQAK | YES | NO | NO |
| 18305 | TRAV24 | TRAJ52 | CAFIGDAGGSSYGKLTF | TRBV4-3 | TRBJ1-4 | CASSQDHRMGGHEKLFF | KLGGALQAK | YES | NO | NO |
| 18306 | TRAV29DV5 | TRAJ24 | CAARGTTDSWGKLQF | TRBV6-1 | TRBJ2-1 | CAINNEQFF | KLGGALQAK | NO | NO | NO |
| 18307 | TRAV8-3 | TRAJ34 | CAVGAPNTDKLIF | TRBV10-1 | TRBJ2-7 | CASSMETDPYEQYF | KLGGALQAK | YES | NO | NO |
| 18308 | TRAV20 | TRAJ23 | CAVQGHYNQGGKLIF | TRBV5-5 | TRBJ1-5 | CASSDQTPGNQPQHF | KLGGALQAK | YES | NO | NO |
| 18309 | TRAV39 | TRAJ29 | CAVDRSGTPLVF | TRBV4-1 | TRBJ2-3 | CASSQGGTGEKRDTQYF | KLGGALQAK | YES | NO | NO |
| 18310 | TRAV19 | TRAJ54 | CALSEGAQKLVF | TRBV13 | TRBJ2-2 | CASVRRLATGELFF | KLGGALQAK | YES | NO | NO |
| 18311 | TRAV27 | TRAJ30 | CAGVWRDDKIIF | TRBV10-3 | TRBJ2-1 | CATLQESSYNEQFF | KLGGALQAK | YES | NO | NO |
| 18312 | TRAV1-2 | TRAJ6 | CAVSVSGGSYIPTF | TRBV20-1 | TRBJ2-1 | CSAAEGLSDEQFF | KLGGALQAK | NO | NO | NO |
| 18313 | TRAV38-2DV8 | TRAJ30 | CAQFTMNRDDKIIF | TRBV20-1 | TRBJ2-3 | CSARAYGTSTTSTDTQYF | KTWGQYWQV | YES | NO | NO |
| 18314 | TRAV22 | TRAJ23 | CAVDPDTQGGKLIF | TRBV5-1 | TRBJ2-7 | CASSPSTGSYEQYV | KLGGALQAK | NO | NO | NO |
| 18315 | TRAV35 | TRAJ44 | CAGNTGTASKLTF | TRBV5-1 | TRBJ1-1 | CASSFYPGTGRPNTEAFF | KLGGALQAK | YES | NO | NO |
| 18316 | TRAV35 | TRAJ43 | CAGHSTPYNNNDMRF | TRBV7-8 | TRBJ2-4 | CASSLGSGSPSANIQYF | KLGGALQAK | YES | NO | NO |
| 18317 | TRAV13-1 | TRAJ41 | CAASIQDSGYALNF | TRBV2 | TRBJ1-5 | CASSEFGRTFINQPQHF | KLGGALQAK | YES | NO | NO |
| 18318 | TRAV13-2 | TRAJ13 | CAENSGGYQKVTF | TRBV11-2 | TRBJ1-1 | CASSLSTEAFF | KLGGALQAK | YES | NO | NO |
| 18319 | TRAV19 | TRAJ37 | CALSEFSSNTGKLIF | TRBV10-1 | TRBJ1-4 | CASSDSNGINEKLFF | KLGGALQAK | YES | NO | NO |
| 18320 | TRAV12-2 | TRAJ47 | CAVKGKLVF | TRBV7-9 | TRBJ1-1 | CASSLAAENEAFF | KLGGALQAK | NO | NO | NO |
| 18321 | TRAV27 | TRAJ27 | CAGRHNTNAGKSTF | TRBV29-1 | TRBJ2-2 | CSVVGTGELFF | KLGGALQAK | YES | NO | NO |
| 18322 | TRAV25 | TRAJ42 | CAGGDDGGSQGNLIF | TRBV15 | TRBJ2-1 | CATSRDQVLAPSYNEQFF | KLGGALQAK | YES | NO | NO |
| 18323 | TRAV39 | TRAJ30 | CAVDNPLWDDKIIF | TRBV24-1 | TRBJ1-5 | CATSDRRGGGNQPQHF | KLGGALQAK | YES | NO | NO |
| 18324 | TRAV35 | TRAJ53 | CAGQGGGSNYKLTF | TRBV7-8 | TRBJ1-2 | CASSPGQGVHGYTF | AVFDRKSDAK | YES | NO | YES |
| 18325 | TRAV19 | TRAJ53 | CALSEASGGSNYKLTF | TRBV4-1 | TRBJ2-3 | CASSQDRVTDTQYF | KLGGALQAK | YES | NO | NO |
| 18326 | TRAV12-2 | TRAJ50 | CAFETSYDKVIF | TRBV18 | TRBJ1-5 | CASSPLSPGENSNQPQHF | KLGGALQAK | YES | NO | NO |
| 18327 | TRAV9-2 | TRAJ12 | CALSDRMDSSYKLIF | TRBV13 | TRBJ1-1 | CASSLVGGLNTEAFF | KLGGALQAK | YES | NO | NO |
| 18328 | TRAV1-2 | TRAJ45 | CAPMYSGGGADGLTF | TRBV2 | TRBJ2-1 | CASPGTGSYNEQFF | KLGGALQAK | YES | NO | NO |
| 18329 | TRAV19 | TRAJ30 | CALSEANRDDKIIF | TRBV7-9 | TRBJ2-3 | CASSLGDYGPDTQYF | KLGGALQAK | YES | NO | NO |
| 18330 | TRAV26-2 | TRAJ39 | CILRDFNNAGNMLTF | TRBV4-1 | TRBJ2-1 | CASSQISASYNEQFF | KLGGALQAK | YES | NO | NO |
| 18331 | TRAV1-2 | TRAJ9 | CAVREDTGGFKTIF | TRBV10-3 | TRBJ1-6 | CAISEMTGSVSPLHF | AVFDRKSDAK | YES | NO | NO |
| 18332 | TRAV24 | TRAJ16 | CAFNSDGQKLLF | TRBV12-4 | TRBJ2-3 | CASSTPLEDTQYF | AVFDRKSDAK | NO | NO | NO |
| 18333 | TRAV8-3 | TRAJ39 | CAVSRGKNAGNMLTF | TRBV27 | TRBJ1-5 | CASRGSDRGLQGPQHF | KLGGALQAK | YES | NO | NO |
| 18334 | TRAV13-1 | TRAJ43 | CAASRARDDMRF | TRBV11-3 | TRBJ2-1 | CASSTPGDNEQFF | AVFDRKSDAK | YES | NO | YES |
| 18335 | TRAV23DV6 | TRAJ8 | CAGPTGFQKLVF | TRBV4-1 | TRBJ2-7 | CASSQVLDRGFYEQYF | KLGGALQAK | YES | NO | NO |
| 18336 | TRAV8-3 | TRAJ47 | CAVVFMEYGNKLVF | TRBV7-9 | TRBJ2-7 | CASSGQVGYEQYF | KLGGALQAK | YES | NO | NO |
| 18337 | TRAV14DV4 | TRAJ53 | CAIRKGSNYKLTF | TRBV11-3 | TRBJ2-5 | CASSFNSDSIQETQYF | RLRAEAQVK | YES | NO | YES |
| 18338 | TRAV19 | TRAJ31 | CALSGYNARLMF | TRBV7-9 | TRBJ1-1 | CASSPLTLLGNTEAFF | KLGGALQAK | YES | NO | NO |
| 18339 | TRAV13-2 | TRAJ9 | CAVNTGGFKTIF | TRBV3-1 | TRBJ2-5 | CASSQDWGGETQYF | KLGGALQAK | YES | NO | NO |
| 18340 | TRAV1-2 | TRAJ34 | CAPTNTDKLIF | TRBV5-1 | TRBJ2-1 | CASSEASNEQFF | KLGGALQAK | YES | NO | NO |
| 18341 | TRAV35 | TRAJ40 | CAGQPSGTYKYIF | TRBV5-1 | TRBJ1-1 | CASSWGQGADEAFF | KLGGALQAK | YES | NO | NO |
| 18342 | TRAV10 | TRAJ20 | CVVKSSNDYKLSF | TRBV27 | TRBJ2-1 | CASSDSAGLALNEQFF | AVFDRKSDAK | YES | NO | YES |
| 18343 | TRAV12-1 | TRAJ41 | CVVTQAHSGYALNF | TRBV5-1 | TRBJ1-5 | CASSLGFEGNQPQHF | KLGGALQAK | YES | NO | NO |
| 18344 | TRAV39 | TRAJ24 | CAVVSRFDSWGKLQF | TRBV6-5 | TRBJ2-5 | CASSDTGQDEETQYF | KLGGALQAK | NO | NO | NO |
| 18345 | TRAV19 | TRAJ39 | CALGRYNNAGNMLTF | TRBV4-1 | TRBJ1-1 | CASSQDNQQGGTEAFF | KLGGALQAK | YES | NO | NO |
| 18346 | TRAV25 | TRAJ32 | CAGNDYGGATNKLIF | TRBV5-6 | TRBJ1-5 | CASSSRDSQNQPQHF | KLGGALQAK | YES | NO | NO |
| 18347 | TRAV17 | TRAJ22 | CASKQLTF | TRBV7-9 | TRBJ2-1 | CASSLLPVQEPNNEQFF | KLGGALQAK | YES | NO | NO |
| 18348 | TRAV17 | TRAJ28 | CATDSGAGSYQLTF | TRBV7-2 | TRBJ2-5 | CASSSTGTGLFETQYF | KLGGALQAK | YES | NO | NO |
| 18349 | TRAV12-2 | TRAJ10 | CAVNKITGGGNKLTF | TRBV29-1 | TRBJ2-5 | CSVEETGQQETQYF | KLGGALQAK | YES | NO | NO |
| 18350 | TRAV1-2 | TRAJ29 | CAVRDNSGNTPLVF | TRBV24-1 | TRBJ2-5 | CATSDNQGSLSTQYF | KLGGALQAK | YES | NO | NO |

| | | | | | | | | | | |
|---|---|---|---|---|---|---|---|---|---|---|
| 18351 | TRAV4 | TRAJ32 | CLVGGGGATNKLIF | TRBV4-3 | TRBJ1-1 | CASSQEDRGLTEAFF | KLGGALQAK | YES | NO | NO |
| 18352 | TRAV12-2 | TRAJ31 | CAVKTGARLMF | TRBV20-1 | TRBJ2-5 | CSASWGLAGALGGQETQYF | KLGGALQAK | YES | NO | NO |
| 18353 | TRAV38-2DV8 | TRAJ49 | CAYISGNQFYF | TRBV6-5 | TRBJ1-2 | CASRGTDYGYTF | KLGGALQAK | YES | NO | NO |
| 18354 | TRAV12-3 | TRAJ47 | CAMNTNGNKLVF | TRBV19 | TRBJ1-2 | CATSGTVYGYTF | RLRAEAQVK | YES | YES | YES |
| 18355 | TRAV4 | TRAJ31 | CLVGDIDNNARLMF | TRBV7-2 | TRBJ1-5 | CASSRGRGGYQPQHF | AVFDRKSDAK | NO | NO | NO |
| 18356 | TRAV21 | TRAJ48 | CAVPNFGNEKLTF | TRBV2 | TRBJ2-2 | CASSFTGSDTGELFF | KLGGALQAK | YES | NO | NO |
| 18357 | TRAV14DV4 | TRAJ22 | CAMREGLSGSARQLTF | TRBV13 | TRBJ2-1 | CASSFSGGTLYNEQFF | AVFDRKSDAK | YES | NO | YES |
| 18358 | TRAV1-2 | TRAJ10 | CAVRDDTGGGNKLTF | TRBV10-1 | TRBJ2-7 | CASSQARTSEQYF | KLGGALQAK | YES | NO | NO |
| 18359 | TRAV13-2 | TRAJ42 | CAETPPYGGSQGNLIF | TRBV4-1 | TRBJ2-2 | CASSQWRLLGQGSTGELFF | KLGGALQAK | YES | NO | NO |
| 18360 | TRAV25 | TRAJ36 | CGTSGANNLFF | TRBV10-2 | TRBJ2-2 | CASSLGTPNTGELFF | KLGGALQAK | YES | NO | NO |
| 18361 | TRAV38-2DV8 | TRAJ43 | CAYRSAGDMRF | TRBV11-3 | TRBJ2-5 | CASKLRGEETQYF | KLGGALQAK | YES | NO | NO |
| 18362 | TRAV35 | TRAJ49 | CAGQLWNTGNQFYF | TRBV24-1 | TRBJ2-2 | CATTDKNTGELFF | KLGGALQAK | YES | NO | NO |
| 18363 | TRAV17 | TRAJ40 | CATDRTSGTYKYIF | TRBV27 | TRBJ2-7 | CASSFGDSHEQYF | KLGGALQAK | YES | NO | NO |
| 18364 | TRAV14DV4 | TRAJ45 | CAMREGSGGGADGLTF | TRBV7-8 | TRBJ2-5 | CASSLRAEYLETQYF | KLGGALQAK | YES | NO | NO |
| 18365 | TRAV1-2 | TRAJ9 | CAVNDGDGGFKTIF | TRBV5-1 | TRBJ1-1 | CASSLLGEGRSEAFF | KLGGALQAK | YES | NO | NO |
| 18366 | TRAV35 | TRAJ41 | CAGQDGNSGYALNF | TRBV29-1 | TRBJ1-2 | CSVQAPRGGYTF | KLGGALQAK | YES | NO | NO |
| 18367 | TRAV8-4 | TRAJ33 | CAVSVQGSNYQLIW | TRBV12-4 | TRBJ2-7 | CASSLRFRQSYEQYF | KLGGALQAK | YES | NO | NO |
| 18368 | TRAV13-1 | TRAJ45 | CAASETGGGADGLTF | TRBV7-8 | TRBJ1-1 | CASSSPPGTGGGNTEAFF | KLGGALQAK | YES | NO | NO |
| 18369 | TRAV1-1 | TRAJ26 | CAVRAPDNYGQNFVF | TRBV5-1 | TRBJ2-5 | CASSPLPQETQYF | KLGGALQAK | YES | NO | NO |
| 18370 | TRAV8-1 | TRAJ33 | CAVNAHSNYQLIW | TRBV27 | TRBJ2-5 | CASSPSQGAKETQYF | KLGGALQAK | YES | NO | NO |
| 18371 | TRAV30 | TRAJ52 | CGTVPSGGTSYGKLTF | TRBV27 | TRBJ2-3 | CASSLRGGTQYF | AVFDRKSDAK | YES | YES | YES |
| 18372 | TRAV27 | TRAJ47 | CAGAQYGNKLVF | TRBV6-1 | TRBJ2-7 | CASTSGVNYEQYF | KLGGALQAK | YES | NO | NO |
| 18373 | TRAV14DV4 | TRAJ37 | CAMRGPFSSNTGKLIF | TRBV7-2 | TRBJ2-7 | CASSLEPSYEQYF | KLGGALQAK | NO | NO | NO |
| 18374 | TRAV30 | TRAJ54 | CGTEEGPGAQKLVF | TRBV4-1 | TRBJ1-5 | CASSQEIGGNQPQHF | KLGGALQAK | YES | YES | NO |
| 18375 | TRAV8-1 | TRAJ33 | CAVNARSNYQLIW | TRBV14 | TRBJ1-2 | CASSQWDDYGYTF | KLGGALQAK | NO | NO | NO |
| 18376 | TRAV12-3 | TRAJ37 | CAMSSSNTGKLIF | TRBV7-6 | TRBJ1-1 | CASSLSSGRTEAFF | AVFDRKSDAK | YES | NO | YES |
| 18377 | TRAV5 | TRAJ37 | CAESNWFGSSNTGKLIF | TRBV12-4 | TRBJ2-3 | CASSLAGDSPTDTQYF | KLGGALQAK | YES | NO | NO |
| 18378 | TRAV8-3 | TRAJ7 | CAVGAGNNRLAF | TRBV5-6 | TRBJ1-1 | CASSFGVGGVGSAFF | KLGGALQAK | YES | NO | NO |
| 18379 | TRAV21 | TRAJ36 | CAVYPGANNLFF | TRBV15 | TRBJ1-6 | CATSSSRDSNSPLHF | KLGGALQAK | YES | NO | NO |
| 18380 | TRAV12-2 | TRAJ54 | CAVKDRGAQKLVF | TRBV6-1 | TRBJ2-1 | CASSEWTGDYNEQFF | KLGGALQAK | YES | NO | NO |
| 18381 | TRAV20 | TRAJ52 | CAVRRAGGTSYGKLTF | TRBV27 | TRBJ1-5 | CASSLSTANQPQHF | KLGGALQAK | YES | NO | NO |
| 18382 | TRAV21 | TRAJ37 | CAAFSNTGKLIF | TRBV7-3 | TRBJ2-1 | CASSFSQNEQFF | KLGGALQAK | YES | YES | NO |
| 18383 | TRAV8-6 | TRAJ58 | CAVTGETSGSRLTF | TRBV5-4 | TRBJ2-3 | CASSLDTDQDTQYF | KLGGALQAK | YES | NO | NO |
| 18384 | TRAV29DV5 | TRAJ17 | CAAAGVGAAGNKLTF | TRBV12-4 | TRBJ2-3 | CASSLVGTPDTQYF | KLGGALQAK | YES | NO | NO |
| 18385 | TRAV29DV5 | TRAJ39 | CAASGNAGNMLTF | TRBV5-5 | TRBJ2-1 | CASSGLGLAVFNEQFF | KLGGALQAK | YES | NO | NO |
| 18386 | TRAV27 | TRAJ27 | CAVSAGKSTF | TRBV7-9 | TRBJ1-2 | CASSLALVGVAGYTF | KLGGALQAK | YES | NO | NO |
| 18387 | TRAV17 | TRAJ48 | CATANFGNEKLTF | TRBV27 | TRBJ2-7 | CASSLGNNYEQYV | KLGGALQAK | NO | NO | NO |
| 18388 | TRAV5 | TRAJ28 | CADPSGAGSYQLTF | TRBV6-3 | TRBJ1-1 | CASRDRVEPEAFF | KLGGALQAK | YES | NO | NO |
| 18389 | TRAV9-2 | TRAJ34 | CALGVYNTDKLIF | TRBV2 | TRBJ1-1 | CASSPTGDPGAFF | KLGGALQAK | YES | NO | NO |
| 18390 | TRAV14DV4 | TRAJ11 | CAMREVNSGYSTLTF | TRBV7-9 | TRBJ2-1 | CASSNSLPSGVSYNEQFF | KLGGALQAK | YES | NO | NO |
| 18391 | TRAV12-3 | TRAJ48 | CAMSLSNFGNEKLTF | TRBV27 | TRBJ1-2 | CASSPGTGILYGYTF | ELAGIGILTV | YES | NO | YES |
| 18392 | TRAV10 | TRAJ20 | CVVSALDDYKLSF | TRBV29-1 | TRBJ2-3 | CSVDLEGATTDTQYF | KLGGALQAK | YES | NO | NO |
| 18393 | TRAV1-2 | TRAJ27 | CAVQTNAGKSTF | TRBV3-1 | TRBJ1-2 | CASSSRDYFNYGYTF | RLRAEAQVK | YES | YES | YES |
| 18395 | TRAV35 | TRAJ23 | CAGHNQGGKLIF | TRBV12-4 | TRBJ2-7 | CASSLWTGEHEQYF | KLGGALQAK | YES | NO | NO |
| 18396 | TRAV21 | TRAJ43 | CAVEDNNDMRF | TRBV19 | TRBJ2-1 | CASSRGADEQFF | KLGGALQAK | YES | NO | NO |
| 18398 | TRAV30 | TRAJ49 | CGTENSGNQFYF | TRBV7-9 | TRBJ1-5 | CASSLVVMTGKSNQPQHF | KLGGALQAK | YES | NO | NO |
| 18399 | TRAV25 | TRAJ52 | CAAKDENAGGTSYGKLTF | TRBV12-3 | TRBJ1-1 | CASSLVPVDTEAFF | KLGGALQAK | YES | NO | NO |
| 18400 | TRAV29DV5 | TRAJ36 | CAASPDQTGANNLFF | TRBV12-3 | TRBJ2-5 | CASSLGGEETQYF | KLGGALQAK | YES | NO | NO |
| 18401 | TRAV21 | TRAJ43 | CAVNNNNNDMRF | TRBV7-6 | TRBJ1-1 | CASSRSGSTEAFF | KLGGALQAK | YES | NO | NO |
| 18403 | TRAV8-1 | TRAJ35 | CAARAPI | TRBV5-6 | TRBJ1-1 | CASSLWATEAFF | KLGGALQAK | NO | NO | NO |
| 18404 | TRAV26-2 | TRAJ47 | CILRYMEYGNKLVF | TRBV4-1 | TRBJ1-4 | CASSKDGRRRSEKLFF | KLGGALQAK | NO | NO | NO |
| 18405 | TRAV17 | TRAJ7 | CATVPPTYGNNRLAF | TRBV19 | TRBJ1-5 | CASSIGQGDQPQHF | KLGGALQAK | YES | NO | NO |
| 18406 | TRAV21 | TRAJ33 | CAVSSMDSNYQLIW | TRBV7-9 | TRBJ1-1 | CASSPQVNTEAFF | KLGGALQAK | YES | NO | NO |
| 18407 | TRAV14DV4 | TRAJ44 | CAMRSVPRTASKLTF | TRBV27 | TRBJ1-5 | CASSLGGAQPQHF | ELAGIGILTV | YES | YES | YES |
| 18408 | TRAV34 | TRAJ26 | CGADGDYGQNFVF | TRBV7-2 | TRBJ2-7 | CASSLVVQRAGGEQYF | KLGGALQAK | NO | NO | NO |
| 18409 | TRAV39 | TRAJ52 | CAVGSNAGGTSYGKLTF | TRBV5-1 | TRBJ1-1 | CASSPRGTRNTEAFF | KLGGALQAK | YES | NO | NO |
| 18410 | TRAV26-1 | TRAJ45 | CIEGADGLTF | TRBV4-1 | TRBJ1-1 | CASSHPLGGTDTEAFF | KLGGALQAK | NO | NO | NO |
| 18411 | TRAV35 | TRAJ33 | CAGPNSRYQLIW | TRBV5-1 | TRBJ2-2 | CASNGGETNTGELFF | KLGGALQAK | YES | NO | NO |
| 18412 | TRAV3 | TRAJ45 | CAVRDRTVARGGADGLTF | TRBV7-6 | TRBJ1-2 | CASSPGASGYTF | KLGGALQAK | NO | NO | NO |
| 18413 | TRAV5 | TRAJ34 | CAESKGNTDKLIF | TRBV12-4 | TRBJ2-7 | CASSPVAGLPGEQYF | KLGGALQAK | YES | NO | NO |
| 18414 | TRAV19 | TRAJ29 | CALSDTSGNTPLVF | TRBV7-9 | TRBJ2-1 | CASSLGTGGYNEQFF | KLGGALQAK | YES | NO | NO |
| 18415 | TRAV17 | TRAJ56 | CATDPNTGANSKLTF | TRBV4-1 | TRBJ2-6 | CASAQESGANVLTF | KLGGALQAK | YES | NO | NO |
| 18416 | TRAV19 | TRAJ18 | CALSEASSGGSTLGRLYF | TRBV30 | TRBJ2-1 | CAWIPAGVYNEQFF | KLGGALQAK | YES | NO | NO |
| 18417 | TRAV1-2 | TRAJ29 | CAYSGNTPLVF | TRBV7-2 | TRBJ2-3 | CASSSRLAGVYSDTQYF | KLGGALQAK | YES | NO | NO |
| 18418 | TRAV14DV4 | TRAJ49 | CAMREGDGNQFYF | TRBV9 | TRBJ1-6 | CASSARSWGSYNSPLHF | KLGGALQAK | YES | NO | NO |
| 18419 | TRAV24 | TRAJ53 | CAPSGGSNYKLTF | TRBV19 | TRBJ2-1 | CASSTGQGQFF | KLGGALQAK | YES | NO | NO |
| 18420 | TRAV17 | TRAJ48 | CATDANFGNEKLTF | TRBV29-1 | TRBJ2-2 | CSVPGETAGELFF | KLGGALQAK | YES | NO | NO |
| 18421 | TRAV12-3 | TRAJ39 | CAIGPGNMLTF | TRBV12-4 | TRBJ1-6 | CASSLGTARGPLHF | KLGGALQAK | YES | NO | NO |
| 18422 | TRAV12-2 | TRAJ48 | CAVNENFGNEKLTF | TRBV7-3 | TRBJ2-1 | CASSLKTSAAEQFF | AVFDRKSDAK | YES | NO | YES |
| 18423 | TRAV14DV4 | TRAJ37 | CAMRGPYSSNTGKLIF | TRBV5-4 | TRBJ1-2 | CASSLPSSNYGYTF | AVFDRKSDAK | YES | NO | YES |
| 18424 | TRAV14DV4 | TRAJ42 | CAMDWLGSQGNLIF | TRBV5-6 | TRBJ2-5 | CASSLAGRGGSQYF | KLGGALQAK | YES | NO | NO |
| 18426 | TRAV8-4 | TRAJ48 | CAVSGNFGNEKLTF | TRBV29-1 | TRBJ1-2 | CSVEDTDYGYTF | KLGGALQAK | YES | NO | NO |
| 18427 | TRAV29DV5 | TRAJ32 | CAASDDGGATNKLIF | TRBV7-2 | TRBJ2-1 | CASTRGYNEQFF | KLGGALQAK | YES | NO | NO |
| 18428 | TRAV14DV4 | TRAJ49 | CAMREDLFDTGNQFYF | TRBV19 | TRBJ2-4 | CASSAPYWQEVNIQYF | KLGGALQAK | YES | NO | NO |
| 18429 | TRAV20 | TRAJ26 | CAVPPYGQNFVF | TRBV6-5 | TRBJ1-1 | CASSSPQDLEAFF | AVFDRKSDAK | YES | NO | YES |
| 18430 | TRAV27 | TRAJ39 | CAGDWSNAGNMLTF | TRBV12-4 | TRBJ2-1 | CASSPSPGLAGDEQFF | KLGGALQAK | YES | NO | NO |
| 18431 | TRAV10 | TRAJ27 | CVVDNTNAGKSTF | TRBV18 | TRBJ1-2 | CASSTGTDYGYTF | KLGGALQAK | YES | NO | NO |
| 18432 | TRAV19 | TRAJ57 | CALSGGGKITQGGSEKLVF | TRBV27 | TRBJ2-2 | CASSLGQNTGELFF | KLGGALQAK | YES | NO | NO |
| 18433 | TRAV38-1 | TRAJ43 | CAFTGYNNNDMRF | TRBV7-9 | TRBJ2-1 | CASSLGTFSYNEQFF | KLGGALQAK | YES | NO | NO |
| 18434 | TRAV29DV5 | TRAJ24 | CAASGTTDSWGKLQF | TRBV7-9 | TRBJ1-1 | CASSSSLGWEAFF | KLGGALQAK | YES | NO | NO |
| 18435 | TRAV29DV5 | TRAJ31 | CAARSSNARLMF | TRBV14 | TRBJ2-5 | CASSQGRETQYF | KLGGALQAK | NO | NO | NO |
| 18436 | TRAV1-2 | TRAJ20 | CAVRDYKLSF | TRBV7-6 | TRBJ2-3 | CASSTGLAESTDTQYF | KLGGALQAK | YES | NO | NO |
| 18437 | TRAV3 | TRAJ35 | CAVRDPVGFGNVLHC | TRBV29-1 | TRBJ2-7 | CSVVGGPYEQYF | KLGGALQAK | NO | NO | NO |
| 18439 | TRAV3 | TRAJ5 | CAVRDRYMDTGRRALTF | TRBV2 | TRBJ1-2 | CASSVGQGAPGYTF | KLGGALQAK | NO | NO | NO |
| 18440 | TRAV8-4 | TRAJ50 | CAVSGETSYDKVIF | TRBV12-4 | TRBJ2-7 | CASSFGASYEQYV | AVFDRKSDAK | NO | NO | NO |
| 18441 | TRAV8-6 | TRAJ30 | CAVSDRGNRDDKIIF | TRBV20-1 | TRBJ2-5 | CSTPQGVFQYF | KLGGALQAK | NO | NO | NO |
| 18442 | TRAV8-4 | TRAJ15 | CAVSGSQAGTALIF | TRBV24-1 | TRBJ2-3 | CATSDRTSGPADTQYF | KLGGALQAK | YES | NO | NO |
| 18443 | TRAV29DV5 | TRAJ39 | CAASSGNMLTF | TRBV6-5 | TRBJ2-1 | CASSYKVYNEQFF | KLGGALQAK | NO | NO | NO |
| 18444 | TRAV13-1 | TRAJ53 | CAASNPPLVRSGGSNYKLTF | TRBV4-1 | TRBJ2-7 | CASSQPPRGEQYF | KLGGALQAK | NO | NO | NO |
| 18445 | TRAV12-3 | TRAJ7 | CAMKEWGNNRLAF | TRBV12-4 | TRBJ2-5 | CASSTGLAGGQETQYF | KLGGALQAK | YES | NO | NO |
| 18446 | TRAV12-2 | TRAJ43 | CAGHNNNDMRF | TRBV5-1 | TRBJ1-1 | CASEVGQGILTEAFF | KLGGALQAK | YES | NO | NO |
| 18447 | TRAV3 | TRAJ43 | CAVRDPHPLYNNNDMRF | TRBV27 | TRBJ2-5 | CASSLRDSQETQYF | AVFDRKSDAK | NO | NO | NO |
| 18448 | TRAV35 | TRAJ37 | CAGRSTGKLIF | TRBV27 | TRBJ2-3 | CASSLMLAGTTDTQYF | AVFDRKSDAK | YES | NO | YES |
| 18449 | TRAV20 | TRAJ45 | CAVQAWSGGGADGLTF | TRBV5-6 | TRBJ2-7 | CASSLGQAYEQYV | KLGGALQAK | NO | NO | NO |
| 18450 | TRAV4 | TRAJ10 | CLVVILTGGGNKLTF | TRBV7-9 | TRBJ1-1 | CASSSMNTEAFF | KLGGALQAK | YES | NO | NO |
| 18451 | TRAV21 | TRAJ43 | CAVRIFERNNDMRF | TRBV7-8 | TRBJ1-2 | CASSAGQPEGDYGYTF | KLGGALQAK | YES | NO | NO |
| 18452 | TRAV8-3 | TRAJ43 | CAVGADDMRF | TRBV12-4 | TRBJ2-1 | CASSSSLEQFF | KLGGALQAK | YES | NO | NO |
| 18453 | TRAV19 | TRAJ23 | CALSEAGYNQGGKLIF | TRBV28 | TRBJ1-5 | CASSLEGDSNQPQHF | KLGGALQAK | YES | NO | NO |
| 18454 | TRAV12-2 | TRAJ20 | CAVTGDYKLSF | TRBV10-2 | TRBJ2-4 | CASSDLGGVGKNIQYF | LLFGYPVYV | YES | YES | YES |
| 18455 | TRAV8-2 | TRAJ47 | CVVSDRDGNKLVF | TRBV7-9 | TRBJ1-2 | CASSLAVGNYGYTF | KLGGALQAK | YES | NO | NO |
| 18456 | TRAV5 | TRAJ6 | CAELASGGSYIPTF | TRBV20-1 | TRBJ2-1 | CSARDMAGVASDEQFF | AVFDRKSDAK | NO | NO | NO |
| 18457 | TRAV20 | TRAJ23 | CAALYNQGGKLIF | TRBV10-2 | TRBJ1-5 | CARGDGHQPQHF | KLGGALQAK | YES | NO | NO |
| 18458 | TRAV22 | TRAJ43 | CVNDMRF | TRBV12-4 | TRBJ1-2 | CASGGMGDGYTF | KLGGALQAK | NO | NO | NO |
| 18459 | TRAV25 | TRAJ16 | CAGQLQSGQKLLF | TRBV10-2 | TRBJ2-7 | CASRDSYYEQYF | KLGGALQAK | NO | NO | NO |

| | | | | | | | | | | |
|---|---|---|---|---|---|---|---|---|---|---|
| 18460 | TRAV14DV4 | TRAJ12 | CAMRRPSSYKLIF | TRBV11-3 | TRBJ1-3 | CASSLTGGYGNTIYF | KLGGALQAK | YES | NO | NO |
| 18461 | TRAV38-2DV8 | TRAJ28 | CAYRDSGAGSYQLTF | TRBV7-9 | TRBJ1-1 | CASTQGSTGAAFF | KLGGALQAK | YES | NO | NO |
| 18462 | TRAV12-2 | TRAJ26 | CAVKGYGQNFVF | TRBV9 | TRBJ2-7 | CASSVSGGDPYEQYF | KLGGALQAK | YES | NO | NO |
| 18463 | TRAV8-6 | TRAJ49 | CAVRHPFNQFYF | TRBV5-5 | TRBJ1-1 | CASSLDRVNTEAFF | KLGGALQAK | YES | NO | NO |
| 18464 | TRAV12-2 | TRAJ45 | CAVNIPGGGADGLTF | TRBV7-2 | TRBJ2-5 | CASSFTSGRPTETQYF | KLGGALQAK | YES | NO | NO |
| 18465 | TRAV13-1 | TRAJ6 | CAAIKSGGSYIPTF | TRBV12-4 | TRBJ1-5 | CASSFTLWVSNQPQHF | KLGGALQAK | YES | NO | NO |
| 18466 | TRAV38-2DV8 | TRAJ53 | CAYRSNYKLTF | TRBV7-9 | TRBJ2-1 | CASRSLLGEEQFF | KLGGALQAK | YES | NO | NO |
| 18467 | TRAV19 | TRAJ40 | CALCNIRTYKYIF | TRBV20-1 | TRBJ1-1 | CSARDLETEAFF | KLGGALQAK | YES | NO | NO |
| 18468 | TRAV12-3 | TRAJ20 | CAMSDYKLSF | TRBV12-4 | TRBJ1-4 | CASSLTRSTATNEKLFF | KLGGALQAK | YES | NO | NO |
| 18469 | TRAV20 | TRAJ52 | CAVRRAGGTSYGKLTF | TRBV3-1 | TRBJ2-7 | CASSPRGTYEQYF | KLGGALQAK | YES | NO | NO |
| 18470 | TRAV19 | TRAJ53 | CALMGGGSNYKLTF | TRBV11-2 | TRBJ1-6 | CASTFVYGVDSPLHF | AVFDRKSDAK | YES | NO | NO |
| 18471 | TRAV14DV4 | TRAJ54 | CAMRGPYRIQGAQKLVF | TRBV20-1 | TRBJ2-1 | CSAIGDSPFHEQFF | KLGGALQAK | NO | NO | NO |
| 18472 | TRAV8-4 | TRAJ20 | CAVIQINDYKLSF | TRBV24-1 | TRBJ1-2 | CATSEKGAGDGNGYTF | KLGGALQAK | YES | NO | NO |
| 18473 | TRAV38-1 | TRAJ28 | CGLHPF | TRBV19 | TRBJ1-5 | CASSMGSYQPQHF | KLGGALQAK | NO | NO | NO |
| 18474 | TRAV29DV5 | TRAJ28 | CAASEDSGAGSYQLTF | TRBV6-5 | TRBJ1-3 | CASSYSFMIKQNTIYF | KLGGALQAK | YES | NO | NO |
| 18475 | TRAV1-2 | TRAJ18 | CAVRQDRGSTLGRLYF | TRBV10-1 | TRBJ2-2 | CASSLNRAGELFF | KLGGALQAK | YES | NO | NO |
| 18476 | TRAV16 | TRAJ29 | CALKTGNTPLVF | TRBV12-4 | TRBJ1-2 | CASSLWTTNYGYTF | KLGGALQAK | NO | NO | NO |
| 18477 | TRAV20 | TRAJ17 | CAVQAVVAAGNKLTF | TRBV4-1 | TRBJ2-1 | CASSRGYNEQFF | KLGGALQAK | YES | NO | NO |
| 18478 | TRAV8-3 | TRAJ7 | CAVGALYGNNRLAF | TRBV3-1 | TRBJ2-5 | CASSHPSTSGGAGETQYF | KLGGALQAK | YES | NO | NO |
| 18479 | TRAV8-3 | TRAJ42 | CAVEGSQGNLIF | TRBV5-1 | TRBJ2-2 | CASSLVSGRKWSTGELFF | KLGGALQAK | YES | NO | NO |
| 18480 | TRAV8-2 | TRAJ26 | CVVSDRGGQNFVF | TRBV4-1 | TRBJ2-7 | CASSQDVAGSAYEQYV | KLGGALQAK | NO | NO | NO |
| 18481 | TRAV20 | TRAJ41 | CAVQALSGSGYALNF | TRBV30 | TRBJ2-7 | CAWSVPQAPGEQYV | KLGGALQAK | NO | NO | NO |
| 18482 | TRAV29DV5 | TRAJ20 | CAASANDYKLSF | TRBV3-1 | TRBJ2-2 | CASSQLSGGGTGELFF | AVFDRKSDAK | YES | NO | YES |
| 18483 | TRAV22 | TRAJ45 | CAGNPSGGGADGLTF | TRBV5-6 | TRBJ2-7 | CASSFGQGTYEQYV | KLGGALQAK | NO | NO | NO |
| 18484 | TRAV12-2 | TRAJ35 | CAATLRGFGNVLHC | TRBV7-9 | TRBJ1-2 | CASSLDPGTGVYGYTF | KLGGALQAK | NO | NO | NO |
| 18486 | TRAV8-1 | TRAJ40 | CAVILDSGTYKYIF | TRBV7-9 | TRBJ1-2 | CASSLLADGNYGYTF | KLGGALQAK | YES | NO | NO |
| 18487 | TRAV13-1 | TRAJ18 | CAASMNRGSTLGRLYF | TRBV7-8 | TRBJ2-1 | CASSLVAWLAENNEQFF | KLGGALQAK | YES | NO | NO |
| 18488 | TRAV3 | TRAJ13 | CAESSGGYQKVTF | TRBV27 | TRBJ2-3 | CASTDSGDTQYF | KLGGALQAK | NO | NO | NO |
| 18489 | TRAV24 | TRAJ48 | CAFVSNFGNEKLTF | TRBV19 | TRBJ2-1 | CASSIGLVNEQFF | KLGGALQAK | YES | NO | NO |
| 18490 | TRAV19 | TRAJ34 | CALSEGSYNTDKLIF | TRBV11-2 | TRBJ1-4 | CASSSGDRTNEKLFF | KLGGALQAK | YES | NO | NO |
| 18491 | TRAV2 | TRAJ6 | CAVEETGGSYIPTF | TRBV2 | TRBJ2-6 | CASGSLESGANVLTF | KLGGALQAK | NO | NO | NO |
| 18492 | TRAV21 | TRAJ34 | CAVRLGNTDKLIF | TRBV10-1 | TRBJ2-3 | CASSTGFGTDTQYF | KLGGALQAK | YES | NO | NO |
| 18493 | TRAV13-2 | TRAJ49 | CAEIRGRDQFYF | TRBV27 | TRBJ2-1 | CASSPTSGSLFTNEQFF | KLGGALQAK | YES | NO | NO |
| 18494 | TRAV26-2 | TRAJ31 | CILRDNARLMF | TRBV11-3 | TRBJ2-3 | CASSFRTSGATDTQYF | KLGGALQAK | YES | NO | NO |
| 18495 | TRAV2 | TRAJ26 | CASRDNYGQNFVF | TRBV15 | TRBJ1-1 | CATSSSQEATEAFF | KLGGALQAK | NO | NO | NO |
| 18496 | TRAV8-6 | TRAJ17 | CAVSDPDQAAGNKLTF | TRBV7-6 | TRBJ2-7 | CASSLLPGADEQYV | KLGGALQAK | NO | NO | NO |
| 18497 | TRAV1-2 | TRAJ34 | CAAYTNTDKLIF | TRBV4-2 | TRBJ2-1 | CASSQVGQRGGLLNEQFF | AVFDRKSDAK | YES | YES | YES |
| 18498 | TRAV26-1 | TRAJ50 | CIVRVGVTSYDKVIF | TRBV20-1 | TRBJ2-2 | CSASMRGTNTGELFF | RLRAEAQVK | YES | YES | YES |
| 18499 | TRAV17 | TRAJ7 | CATDARENYGNNRLAF | TRBV27 | TRBJ1-5 | CASSLGTGWGNQPQHF | KLGGALQAK | YES | NO | NO |
| 18500 | TRAV5 | TRAJ39 | CAEISNAGNMLTF | TRBV29-1 | TRBJ2-7 | CSVETSGTGERVVPYEQYF | KLGGALQAK | YES | NO | NO |
| 18501 | TRAV29DV5 | TRAJ45 | CAASVYSGGGADGLTF | TRBV20-1 | TRBJ2-5 | CSARDRAGKETQYF | KLGGALQAK | YES | NO | NO |
| 18502 | TRAV12-2 | TRAJ53 | CAGRGSNYKLTF | TRBV11-3 | TRBJ1-1 | CASRYTGNTEAFF | KLGGALQAK | YES | NO | NO |
| 18503 | TRAV12-1 | TRAJ40 | CVVNIIQGTYKYIF | TRBV7-2 | TRBJ2-3 | CASTLTVTDTQYF | KLGGALQAK | YES | NO | NO |
| 18504 | TRAV1-2 | TRAJ12 | CAVRDPWSSYKLIF | TRBV4-1 | TRBJ1-1 | CASSQGDTEAFF | KLGGALQAK | YES | NO | NO |
| 18505 | TRAV1-2 | TRAJ32 | CAVRDLNYGGATNKLIF | TRBV10-3 | TRBJ2-3 | CAISGGSGGAQDTQYF | KLGGALQAK | YES | NO | NO |
| 18506 | TRAV13-1 | TRAJ17 | CAASIVAAGNKLTF | TRBV2 | TRBJ1-2 | CASFGKSNYGYTF | KLGGALQAK | YES | NO | NO |
| 18507 | TRAV9-2 | TRAJ36 | CALGTGANNLFF | TRBV11-2 | TRBJ2-7 | CASSLSNPYEQYV | KLGGALQAK | NO | NO | NO |
| 18508 | TRAV29DV5 | TRAJ22 | CAVLSSGSARQLTF | TRBV4-3 | TRBJ2-2 | CASSQYIENTGELFF | KLGGALQAK | YES | NO | NO |
| 18509 | TRAV14DV4 | TRAJ22 | CAMRRPISSGSARQLTF | TRBV14 | TRBJ2-5 | CASSSRDNQETQYF | KLGGALQAK | NO | NO | NO |
| 18510 | TRAV30 | TRAJ28 | CGTEDPYSGAGSYQLTF | TRBV18 | TRBJ2-3 | CASSPLDPTDTQYF | KLGGALQAK | YES | NO | NO |
| 18511 | TRAV14DV4 | TRAJ9 | CAMREAYTGGFKTIF | TRBV27 | TRBJ1-2 | CASSLDGTGGSGYTF | KLGGALQAK | YES | NO | NO |
| 18512 | TRAV4 | TRAJ30 | CLVVSMNRDDKIIF | TRBV3-1 | TRBJ1-2 | CASSQGRDRHGYTF | KLGGALQAK | YES | NO | NO |
| 18513 | TRAV12-1 | TRAJ42 | CVVTAYGGSQGNLIF | TRBV12-3 | TRBJ1-1 | CASSTTTGNTEAFF | KLGGALQAK | YES | NO | NO |
| 18514 | TRAV21 | TRAJ41 | CAVGNPNSGYALNF | TRBV4-3 | TRBJ1-2 | CASSFEQSYGYTF | KLGGALQAK | YES | NO | NO |
| 18515 | TRAV19 | TRAJ53 | CALSDAGGSNYKLTF | TRBV11-1 | TRBJ2-7 | CASSLLAGTHEQYF | KLGGALQAK | YES | NO | NO |
| 18516 | TRAV3 | TRAJ31 | CAVRDRNNARLMF | TRBV15 | TRBJ2-7 | CATTKDRGTGSYEQYF | AVFDRKSDAK | NO | NO | NO |
| 18517 | TRAV22 | TRAJ56 | CAVGDRGANSKLTF | TRBV7-9 | TRBJ2-3 | CASSPIADHTQYF | KLGGALQAK | YES | NO | NO |
| 18518 | TRAV13-1 | TRAJ39 | CAANNNAGNMLTF | TRBV11-2 | TRBJ1-1 | CASSLLNTEAFF | KLGGALQAK | YES | NO | NO |
| 18519 | TRAV27 | TRAJ33 | CAGVDSNYQLIW | TRBV4-1 | TRBJ1-2 | CASSQVGGIDLGYTF | KLGGALQAK | YES | NO | NO |
| 18520 | TRAV22 | TRAJ26 | CAVERHGQNFVF | TRBV24-1 | TRBJ2-2 | CATSDLTGELFF | KLGGALQAK | YES | NO | NO |
| 18521 | TRAV17 | TRAJ20 | CATDANDYKLSF | TRBV5-1 | TRBJ2-5 | CASSSGGTGGETQYF | KLGGALQAK | YES | NO | NO |
| 18522 | TRAV24 | TRAJ45 | CACSDSGGGADGLTF | TRBV5-1 | TRBJ2-3 | CASSLETSTSTDTQYF | KLGGALQAK | YES | NO | NO |
| 18523 | TRAV16 | TRAJ39 | CALDNAGNMLTF | TRBV11-2 | TRBJ2-1 | CASSQPWTGGSSYNEQFF | KLGGALQAK | NO | NO | NO |
| 18524 | TRAV1-2 | TRAJ27 | CAPTNTNAGKSTF | TRBV11-2 | TRBJ2-1 | CASSPGQGTHTPPNEQFF | KLGGALQAK | YES | NO | NO |
| 18525 | TRAV19 | TRAJ52 | CALSDLDTGANAGGTSYGKLTF | TRBV19 | TRBJ2-6 | CASSMQPVGANVLTF | KLGGALQAK | NO | NO | NO |
| 18526 | TRAV4 | TRAJ45 | CLVVAHSGGGADGLTF | TRBV11-2 | TRBJ2-5 | CASLGQGYQETQYF | KLGGALQAK | YES | NO | NO |
| 18527 | TRAV8-3 | TRAJ36 | CAVKTGANNLFF | TRBV19 | TRBJ2-7 | CASSAGLTYEQYV | KLGGALQAK | NO | NO | NO |
| 18528 | TRAV9-2 | TRAJ45 | CALSFLSGGGADGLTF | TRBV6-1 | TRBJ2-1 | CASSTYRVRVSESNEQFF | KLGGALQAK | YES | NO | NO |
| 18529 | TRAV9-2 | TRAJ47 | CALSVAEYGNKLVF | TRBV27 | TRBJ2-1 | CASSFRDRYNNEQFF | KLGGALQAK | YES | NO | NO |
| 18530 | TRAV27 | TRAJ43 | CAGGPRF | TRBV3-1 | TRBJ1-3 | CASSQDGSSGNTIYF | KLGGALQAK | NO | NO | NO |
| 18531 | TRAV29DV5 | TRAJ28 | CAASAPGSYQLTF | TRBV27 | TRBJ2-7 | CASSLAGVKYEQYF | KLGGALQAK | YES | NO | NO |
| 18532 | TRAV13-1 | TRAJ48 | CAASSGGNEKLTF | TRBV6-5 | TRBJ2-7 | CASSFGTGGWYEQYF | KLGGALQAK | YES | NO | NO |
| 18533 | TRAV23DV6 | TRAJ54 | CAARIQGAQKLVF | TRBV10-3 | TRBJ1-1 | CAISEGNRGVTEAFF | KLGGALQAK | YES | NO | NO |
| 18534 | TRAV6 | TRAJ39 | CALDLRQNNAGNMLTF | TRBV5-1 | TRBJ2-3 | CASSLVSVGLAWDTQYF | KLGGALQAK | YES | NO | NO |
| 18535 | TRAV25 | TRAJ24 | CAGGDSWGKLQF | TRBV7-6 | TRBJ1-6 | CASSLGGDSPLHF | KLGGALQAK | YES | NO | NO |
| 18536 | TRAV35 | TRAJ31 | CAGPNARLMF | TRBV9 | TRBJ1-2 | CASSVGGAGYTF | KLGGALQAK | YES | NO | NO |
| 18537 | TRAV20 | TRAJ28 | CAPSHQLTF | TRBV4-3 | TRBJ1-4 | CASSQDHRMGGHEKLFF | KLGGALQAK | YES | NO | NO |
| 18538 | TRAV26-2 | TRAJ40 | CILSPRTYKYIF | TRBV7-9 | TRBJ1-5 | CASSSGTVNNQPQHF | KLGGALQAK | YES | NO | NO |
| 18539 | TRAV8-6 | TRAJ20 | CAVRDYKLSF | TRBV29-1 | TRBJ1-1 | CSVMSGPEAFF | KLGGALQAK | YES | NO | NO |
| 18540 | TRAV8-2 | TRAJ29 | CVVPISGNTPLVF | TRBV7-9 | TRBJ1-5 | CASSPFRTGEQPQHF | KLGGALQAK | YES | NO | NO |
| 18541 | TRAV13-2 | TRAJ34 | CAVASYNTDKLIF | TRBV19 | TRBJ1-2 | CASSVGQGAFGYTF | KLGGALQAK | YES | NO | NO |
| 18542 | TRAV24 | TRAJ52 | CAFIGDAGGTSYGKLTF | TRBV7-9 | TRBJ2-4 | CASSPSRDREFLYIQYF | KLGGALQAK | YES | NO | NO |
| 18543 | TRAV1-2 | TRAJ42 | CAVYHGGSQGNLIF | TRBV7-2 | TRBJ1-1 | CASSLHPYTEAFF | KLGGALQAK | YES | NO | NO |
| 18544 | TRAV27 | TRAJ52 | CAGINAGGTSYGKLTF | TRBV27 | TRBJ1-4 | CASQGFANEKLFF | KLGGALQAK | YES | NO | NO |
| 18545 | TRAV1-2 | TRAJ33 | CALQGNYQLIW | TRBV7-8 | TRBJ1-1 | CASSLGLQNTEAFF | KLGGALQAK | YES | NO | NO |
| 18546 | TRAV8-6 | TRAJ57 | CAVSYQGGSEKLVF | TRBV6-3 | TRBJ2-1 | CASSYGAGGYNEQFF | KLGGALQAK | YES | NO | NO |
| 18547 | TRAV14DV4 | TRAJ42 | CAMRVDGGSQGNLIF | TRBV9 | TRBJ2-1 | CASRSGSSYNEQFF | KLGGALQAK | YES | NO | NO |
| 18548 | TRAV12-1 | TRAJ20 | CVVNDYKLSF | TRBV11-2 | TRBJ2-3 | CASRKTSGGVSTDTQYF | KLGGALQAK | YES | NO | NO |
| 18549 | TRAV26-1 | TRAJ26 | CIVRDHNYGQNFVF | TRBV4-1 | TRBJ1-1 | CASSQDFSGAIMNTEAFF | KLGGALQAK | YES | NO | NO |
| 18550 | TRAV12-1 | TRAJ13 | CVVRDSGGYQKVTF | TRBV4-1 | TRBJ1-1 | CASSQVQGPNTEAFF | KLGGALQAK | YES | NO | NO |
| 18551 | TRAV1-2 | TRAJ6 | CAVRDRGGSYIPTF | TRBV20-1 | TRBJ2-3 | CSARDKWTQDTQYF | KLGGALQAK | NO | NO | NO |
| 18552 | TRAV12-2 | TRAJ36 | CAVTGANNLFF | TRBV10-1 | TRBJ2-1 | CASSPGANNEQFF | KLGGALQAK | YES | NO | NO |
| 18553 | TRAV1-2 | TRAJ41 | CAPVGRNSGYALNF | TRBV20-1 | TRBJ2-7 | CSARGSGTYEQYF | KLGGALQAK | YES | NO | NO |
| 18554 | TRAV29DV5 | TRAJ15 | CAASDNQAGTALIF | TRBV7-9 | TRBJ2-7 | CASSSDARGYEQYF | KLGGALQAK | YES | NO | NO |
| 18555 | TRAV12-2 | TRAJ47 | CAVYKYGNKLVF | TRBV4-3 | TRBJ2-2 | CASSQDHFQGTGGELFF | KLGGALQAK | YES | NO | NO |
| 18556 | TRAV29DV5 | TRAJ40 | CAVTSGTYKYIF | TRBV10-3 | TRBJ2-3 | CAISESLAGGPGTQYF | KLGGALQAK | YES | NO | NO |
| 18557 | TRAV5 | TRAJ42 | CAESDDGGSQGNLIF | TRBV20-1 | TRBJ1-1 | CSASSQGKAVNTEAFF | KLGGALQAK | YES | NO | NO |
| 18558 | TRAV8-3 | TRAJ23 | CAVDNQGGKLIF | TRBV19 | TRBJ2-3 | CASSPPPSPRRVWGTDTQYF | KLGGALQAK | NO | NO | NO |
| 18559 | TRAV8-1 | TRAJ42 | CAVRGGSQGNLIF | TRBV7-8 | TRBJ2-7 | CASSSTDADEQYF | KLGGALQAK | YES | NO | NO |
| 18560 | TRAV13-2 | TRAJ5 | CAENNTGRRALTF | TRBV7-9 | TRBJ2-1 | CASSLFDNEQFF | KLGGALQAK | YES | NO | NO |
| 18561 | TRAV13-2 | TRAJ30 | CAENSHRDDKIIF | TRBV19 | TRBJ1-1 | CASRLSWGTEAFF | KLGGALQAK | YES | NO | NO |
| 18562 | TRAV8-6 | TRAJ53 | CAVSDSGGSNYKLTF | TRBV7-9 | TRBJ1-3 | CASSLQSSGNTIYF | KLGGALQAK | YES | NO | NO |
| 18563 | TRAV17 | TRAJ47 | CATVHIMEYGNKLVF | TRBV6-1 | TRBJ1-6 | CASRGEGNSYNSPLHF | KLGGALQAK | YES | NO | NO |
| 18564 | TRAV14DV4 | TRAJ57 | CAMREGYLFQGGSEKLVF | TRBV27 | TRBJ2-3 | CASSPPGQGWSDTQYF | KLGGALQAK | YES | NO | NO |

| | | | | | | | | | | |
|---|---|---|---|---|---|---|---|---|---|---|
| 18565 | TRAV29DV5 | TRAJ43 | CAASAYVNDMRF | TRBV7-9 | TRBJ2-7 | CASKDGISYEQYF | KLGGALQAK | YES | NO | NO |
| 18566 | TRAV19 | TRAJ12 | CALSEARDSSYKLIF | TRBV5-1 | TRBJ2-1 | CASSLDSSYNEQFF | KLGGALQAK | YES | NO | NO |
| 18567 | TRAV5 | TRAJ23 | CAEYPIYNQGGKLIF | TRBV4-2 | TRBJ2-3 | CASSQFPSGGTDTQYF | KLGGALQAK | YES | NO | NO |
| 18568 | TRAV27 | TRAJ43 | CAGLYNNNDMRF | TRBV24-1 | TRBJ2-1 | CATSDLPQDEQFF | KLGGALQAK | YES | NO | NO |
| 18569 | TRAV29DV5 | TRAJ47 | CAATGPYGNKLVF | TRBV7-9 | TRBJ2-2 | CASSSPLNWGAGGELFF | KLGGALQAK | YES | NO | NO |
| 18570 | TRAV14DV4 | TRAJ10 | CAMRGLILTGGGNKLTF | TRBV9 | TRBJ2-1 | CASSPGLGGTYNEQFF | KLGGALQAK | YES | NO | NO |
| 18571 | TRAV9-2 | TRAJ23 | CALSDHPLEGGKLIF | TRBV20-1 | TRBJ2-3 | CRFSGRAAQSDTQYF | KLGGALQAK | NO | NO | NO |
| 18572 | TRAV39 | TRAJ23 | CAVLQGGKLIF | TRBV6-5 | TRBJ1-2 | CASRGGIHYGYTF | KLGGALQAK | YES | NO | NO |
| 18573 | TRAV12-1 | TRAJ47 | CVVSMEYGNKLVF | TRBV12-5 | TRBJ2-1 | CASGLVQLSNEQFF | KLGGALQAK | YES | NO | NO |
| 18574 | TRAV8-4 | TRAJ20 | CAVIQINDYKLSF | TRBV7-8 | TRBJ1-2 | CASSLGQAYGYTF | KLGGALQAK | YES | NO | NO |
| 18575 | TRAV14DV4 | TRAJ22 | CAMRRPISSGSARQLTF | TRBV27 | TRBJ2-1 | CASRTASSYNEQFF | KLGGALQAK | YES | NO | NO |
| 18576 | TRAV27 | TRAJ28 | CAGVPFSGAGSYQLTF | TRBV7-8 | TRBJ2-3 | CASSLEGGGSTDTQYF | KLGGALQAK | YES | NO | NO |
| 18577 | TRAV12-2 | TRAJ57 | CAVNWGGSEKLVF | TRBV2 | TRBJ2-3 | CASSDGGGATDTQYF | KLGGALQAK | YES | NO | NO |
| 18578 | TRAV13-2 | TRAJ13 | CAEMDSGGYQKVTF | TRBV6-5 | TRBJ1-1 | CASSYEGNTEAFF | KLGGALQAK | YES | NO | NO |
| 18579 | TRAV9-2 | TRAJ47 | CALLDMEYGNKLVF | TRBV7-2 | TRBJ2-1 | CASSGNSGVPGNEQFF | KLGGALQAK | YES | NO | NO |
| 18580 | TRAV12-1 | TRAJ49 | CVVNMGPTGNQFYF | TRBV6-3 | TRBJ1-6 | CASSRGSYNSPLHF | KLGGALQAK | YES | NO | NO |
| 18581 | TRAV13-1 | TRAJ48 | CAASREKLTF | TRBV19 | TRBJ2-1 | CASSILGGGYNEQFF | KLGGALQAK | YES | NO | NO |
| 18584 | TRAV12-1 | TRAJ27 | CVVNVGANAGKSTF | TRBV11-2 | TRBJ2-1 | CASSFLGNEQFF | KLGGALQAK | YES | NO | NO |
| 18585 | TRAV1-2 | TRAJ28 | CAVRDRSQPGAGSYQLTF | TRBV4-2 | TRBJ2-5 | CASSRTKGGKETQYF | IVTDFSVIK | YES | NO | YES |
| 18586 | TRAV29DV5 | TRAJ42 | CAASDYGGSQGNLIF | TRBV6-4 | TRBJ1-1 | CASSEFMGNTEAFF | KLGGALQAK | YES | NO | NO |
| 18588 | TRAV13-1 | TRAJ54 | CAASISRAIPQGAQKLVF | TRBV7-9 | TRBJ2-2 | CASSFWTDTGELFF | AVFDRKSDAK | YES | NO | YES |
| 18589 | TRAV14DV4 | TRAJ57 | CAMREDITQGGSEKLVF | TRBV5-1 | TRBJ2-5 | CASSLWPSRQETQYF | KLGGALQAK | YES | NO | NO |
| 18590 | TRAV20 | TRAJ54 | CAVGAQKLVF | TRBV4-1 | TRBJ2-4 | CASSPWGVIAKNIQYF | KLGGALQAK | YES | NO | NO |
| 18591 | TRAV20 | TRAJ48 | CAVQGSNFGNEKLTF | TRBV7-8 | TRBJ2-7 | CASSSSGNYEQYV | KLGGALQAK | NO | NO | NO |
| 18592 | TRAV21 | TRAJ20 | CAPLRGDYKLSF | TRBV6-3 | TRBJ2-1 | CASNPGGNEQFF | KLGGALQAK | YES | NO | NO |
| 18593 | TRAV9-2 | TRAJ7 | CALSGYGNNRLAF | TRBV6-5 | TRBJ2-3 | CASSYQGGTGTDTQYF | KLGGALQAK | NO | NO | NO |
| 18594 | TRAV35 | TRAJ34 | CAGYLRELIF | TRBV29-1 | TRBJ2-7 | CSVTEGFGEQYF | KLGGALQAK | YES | NO | NO |
| 18595 | TRAV12-2 | TRAJ47 | CAVEYGNKLVF | TRBV12-3 | TRBJ1-2 | CASSFRQGHTF | KLGGALQAK | YES | NO | NO |
| 18596 | TRAV38-2DV8 | TRAJ32 | CAYRSAQGYGGATNKLIF | TRBV4-3 | TRBJ2-5 | CASSQRSRGRVETQYF | KLGGALQAK | YES | NO | NO |
| 18597 | TRAV27 | TRAJ40 | CAGSGATTSGTYKYIF | TRBV7-9 | TRBJ1-3 | CASSLSKHWEISGNTIYF | KLGGALQAK | YES | NO | NO |
| 18598 | TRAV8-1 | TRAJ32 | CAVIQGWYGGATNKLIF | TRBV28 | TRBJ1-6 | CASSPRVLSYNSPLHF | KLGGALQAK | YES | NO | NO |
| 18599 | TRAV29DV5 | TRAJ29 | CAASFSGNTPLVF | TRBV27 | TRBJ1-2 | CASSFGGVDGYTF | AVFDRKSDAK | YES | YES | YES |
| 18600 | TRAV17 | TRAJ43 | CATDDMRF | TRBV18 | TRBJ1-2 | CASSPPGAPNYGYTF | KLGGALQAK | NO | NO | NO |
| 18601 | TRAV39 | TRAJ53 | CAVDKSRSSGGSNYKLTF | TRBV19 | TRBJ2-1 | CASSIGTSGNNEQFF | KLGGALQAK | YES | NO | NO |
| 18602 | TRAV12-2 | TRAJ17 | CAATGKAAGNKLTF | TRBV19 | TRBJ1-2 | CASREGQAYGYTF | KLGGALQAK | YES | NO | NO |
| 18603 | TRAV17 | TRAJ48 | CATLSNFGNEKLTF | TRBV20-1 | TRBJ2-1 | CSARDWGFGDYNEQFF | KLGGALQAK | NO | NO | NO |
| 18604 | TRAV17 | TRAJ40 | CATDEGTYKYIF | TRBV5-1 | TRBJ2-7 | CASSEGQGAGYEQYF | KLGGALQAK | NO | NO | NO |
| 18605 | TRAV12-3 | TRAJ39 | CAIGPGNMLTF | TRBV7-9 | TRBJ1-4 | CASSPGLEEKLFF | KLGGALQAK | YES | NO | NO |
| 18606 | TRAV35 | TRAJ45 | CAGRKGGGADGLTF | TRBV7-8 | TRBJ1-1 | CASSLPGTPANTEAFF | KLGGALQAK | YES | NO | NO |
| 18607 | TRAV21 | TRAJ30 | CAVRPTNDKIIF | TRBV6-1 | TRBJ2-1 | CASSGSSYNEQFF | KLGGALQAK | YES | NO | NO |
| 18608 | TRAV21 | TRAJ20 | CAVRPNDYKLSF | TRBV6-6 | TRBJ2-3 | CASSYQGDTQYF | KLGGALQAK | NO | NO | NO |
| 18609 | TRAV5 | TRAJ40 | CAGSYSGTYKYIF | TRBV13 | TRBJ1-5 | CASSLAGNNQPQHF | KLGGALQAK | YES | NO | NO |
| 18610 | TRAV20 | TRAJ50 | CAVQAWETSYDKVIF | TRBV6-3 | TRBJ2-1 | CASSYNHRVHEQFF | KLGGALQAK | YES | NO | NO |
| 18611 | TRAV22 | TRAJ45 | CAVLGGGADGLTF | TRBV19 | TRBJ1-5 | CASSIWTSDQPQHF | IVTDFSVIK | YES | YES | YES |
| 18612 | TRAV29DV5 | TRAJ52 | CAVNAGGTSYGKLTF | TRBV5-1 | TRBJ2-3 | CASSGGTGSTDTQYF | KLGGALQAK | YES | NO | NO |
| 18613 | TRAV13-2 | TRAJ49 | CAEIRGRDQFYF | TRBV4-3 | TRBJ2-3 | CASSQEGGFTDTQYF | KLGGALQAK | YES | NO | NO |
| 18614 | TRAV1-2 | TRAJ33 | CAGLLDSNYQLIW | TRBV7-9 | TRBJ1-1 | CASMGQLSEAFF | KLGGALQAK | YES | NO | NO |
| 18615 | TRAV19 | TRAJ52 | CALSEVLNAGGTSYGKLTF | TRBV19 | TRBJ2-3 | CASSQSAGGIFNTQYF | KLGGALQAK | YES | YES | NO |
| 18616 | TRAV41 | TRAJ54 | CAVRIQGAQKLVF | TRBV7-2 | TRBJ2-7 | CASSLRQPGLELYEQYF | KLGGALQAK | YES | NO | NO |
| 18617 | TRAV9-2 | TRAJ30 | CALSDRMNRDDKIIF | TRBV7-9 | TRBJ1-6 | CASSLAFGDSPLHF | KLGGALQAK | YES | NO | NO |
| 18618 | TRAV19 | TRAJ57 | CAPEPNFQGGSEKLVF | TRBV24-1 | TRBJ2-1 | CATSDWLAGDEQFF | KLGGALQAK | YES | NO | NO |
| 18619 | TRAV27 | TRAJ54 | CAGAFWQGAQKLVF | TRBV3-1 | TRBJ2-1 | CASSQAGQRLAGALNPTQSYNEQFF | AVFDRKSDAK | NO | NO | NO |
| 18620 | TRAV29DV5 | TRAJ28 | CAATPRGAGSYQLTF | TRBV11-2 | TRBJ1-1 | CASSLFTEAFF | KLGGALQAK | YES | NO | NO |
| 18621 | TRAV30 | TRAJ49 | CGTRRGNQFYF | TRBV11-2 | TRBJ2-7 | CASSLAGTELYEQYV | KLGGALQAK | NO | NO | NO |
| 18622 | TRAV8-6 | TRAJ39 | CAVSDGGAGNMLTF | TRBV19 | TRBJ2-1 | CASSRQGPYNEQFF | KLGGALQAK | YES | NO | NO |
| 18623 | TRAV26-2 | TRAJ40 | CIARSGTYKYIF | TRBV19 | TRBJ1-2 | CASSMGSGQSYGYTF | KLGGALQAK | YES | NO | NO |
| 18624 | TRAV12-2 | TRAJ30 | CAVKDRDDKIIF | TRBV20-1 | TRBJ2-7 | CSARPKRGVEQYF | KLGGALQAK | NO | NO | NO |
| 18625 | TRAV8-6 | TRAJ45 | CAVSRGADGLTF | TRBV20-1 | TRBJ2-7 | CSARDSRFTYEQYF | KLGGALQAK | NO | NO | NO |
| 18626 | TRAV12-3 | TRAJ47 | CAMGREYGNKLVF | TRBV6-3 | TRBJ2-3 | CASIPAGGPGDTQYF | KLGGALQAK | YES | NO | NO |
| 18627 | TRAV1-2 | TRAJ31 | CAVYNNARLMF | TRBV11-2 | TRBJ1-5 | CASSPGDSGNQPQHF | KLGGALQAK | YES | NO | NO |
| 18628 | TRAV38-1 | TRAJ54 | CAFMSLIQGAQKLVF | TRBV9 | TRBJ2-7 | CASSVGGSGGTYEQYF | KLGGALQAK | YES | NO | NO |
| 18629 | TRAV14DV4 | TRAJ50 | CAMRGTKTSYDKVIF | TRBV19 | TRBJ1-1 | CASSIGRSTEAFF | KLGGALQAK | YES | NO | NO |
| 18630 | TRAV21 | TRAJ6 | CAASPGASGGSYIPTF | TRBV7-2 | TRBJ1-6 | CASSLADSISPLHF | KLGGALQAK | NO | NO | NO |
| 18631 | TRAV24 | TRAJ33 | CAFFDSNYQLIW | TRBV7-9 | TRBJ2-7 | CASSWTGGYEQYF | AVFDRKSDAK | YES | NO | YES |
| 18632 | TRAV38-2DV8 | TRAJ23 | CAYGDDNQGGKLIF | TRBV6-5 | TRBJ2-6 | CASTPWADYNSGANVLTF | KLGGALQAK | YES | NO | NO |
| 18633 | TRAV29DV5 | TRAJ56 | CAASDTGANSKLTF | TRBV4-1 | TRBJ1-1 | CASSPGLPAHTEAFF | KLGGALQAK | YES | NO | NO |
| 18634 | TRAV1-1 | TRAJ23 | CAVRDRDYNQGGKLIF | TRBV7-9 | TRBJ2-2 | CASSARVWPGELFF | KLGGALQAK | YES | NO | NO |
| 18635 | TRAV21 | TRAJ28 | CAVRPPGPGLGVTNSLS | TRBV14 | TRBJ2-7 | CASSPLNDLDEQYF | KLGGALQAK | NO | NO | NO |
| 18636 | TRAV10 | TRAJ48 | CVVSLPRFGNEKLTF | TRBV7-3 | TRBJ2-3 | CASSVLAGPYTDTQYF | KLGGALQAK | YES | NO | NO |
| 18637 | TRAV3 | TRAJ28 | CAVRDPGAGSYQLTF | TRBV5-6 | TRBJ1-4 | CASRRLTAKNEKLFF | KLGGALQAK | NO | NO | NO |
| 18638 | TRAV21 | TRAJ52 | CAVAVNAGGTSYGKLTF | TRBV4-1 | TRBJ2-1 | CASSQSPLGDYNEQFF | KLGGALQAK | YES | NO | NO |
| 18639 | TRAV21 | TRAJ33 | CAVIGSNYQLIW | TRBV27 | TRBJ1-5 | CASSESNPTGSWPQHF | AVFDRKSDAK | YES | NO | YES |
| 18640 | TRAV12-2 | TRAJ8 | CAPWGGSTGFQKLVF | TRBV6-3 | TRBJ2-3 | CASSYGTSSTDTQYF | KLGGALQAK | YES | NO | NO |
| 18641 | TRAV35 | TRAJ53 | CAGQLGSNYKLTF | TRBV12-4 | TRBJ1-2 | CASVGMLARTEANYGYTF | KLGGALQAK | YES | NO | NO |
| 18642 | TRAV13-1 | TRAJ13 | CAAVFGGYQKVTF | TRBV5-6 | TRBJ2-2 | CASSLMWTPPGGELFF | KLGGALQAK | YES | NO | NO |
| 18643 | TRAV29DV5 | TRAJ49 | CAASAGNQFYF | TRBV19 | TRBJ1-2 | CASSIAVGMPIGYTF | KLGGALQAK | YES | NO | NO |
| 18644 | TRAV12-1 | TRAJ3 | CVVNMETAYSSASKIIF | TRBV11-2 | TRBJ2-1 | CASSLARTSGTLTYNEQFF | KLGGALQAK | YES | NO | NO |
| 18645 | TRAV1-2 | TRAJ15 | CALRPNQAGTALIF | TRBV4-1 | TRBJ1-1 | CASSSGTGNTEAFF | KLGGALQAK | YES | NO | NO |
| 18646 | TRAV14DV4 | TRAJ37 | CAMRGPYSSNTGKLIF | TRBV5-4 | TRBJ2-3 | CASSLSTDTQYF | RLRAEAQVK | YES | YES | YES |
| 18647 | TRAV16 | TRAJ33 | CALIMDSNYQLIW | TRBV12-4 | TRBJ1-1 | CASSPQGKAFF | KLGGALQAK | NO | NO | NO |
| 18648 | TRAV8-4 | TRAJ20 | CAVIQINDYKLSF | TRBV19 | TRBJ1-1 | CASFGDRADGEAFF | KLGGALQAK | YES | NO | NO |
| 18649 | TRAV24 | TRAJ30 | CASSGNRDDKIIF | TRBV19 | TRBJ1-2 | CASSIGYTLNYGYTF | KLGGALQAK | YES | NO | NO |
| 18650 | TRAV27 | TRAJ34 | CAGAQSRNTDKLIF | TRBV19 | TRBJ2-7 | CASSITGGFLSYEQYF | KLGGALQAK | YES | NO | NO |
| 18651 | TRAV8-6 | TRAJ30 | CAVSDPTNRDDKIIF | TRBV29-1 | TRBJ1-1 | CSVTKKRGFMNTEAFF | KLGGALQAK | YES | NO | NO |
| 18652 | TRAV36DV7 | TRAJ45 | CAVDIYSGGGADGLTF | TRBV7-9 | TRBJ2-2 | CASSLEGLGKTNTGELFF | KLGGALQAK | NO | NO | NO |
| 18654 | TRAV12-1 | TRAJ49 | CVVNRAGNQFYF | TRBV20-1 | TRBJ1-2 | CSAREQEANYGYTF | KLGGALQAK | YES | NO | NO |
| 18655 | TRAV12-2 | TRAJ17 | CAVTLKAAGNKLTF | TRBV6-5 | TRBJ1-2 | CASSYGTAPTGYTF | KLGGALQAK | YES | NO | NO |
| 18656 | TRAV35 | TRAJ39 | CAGRAENAGNMLTF | TRBV5-1 | TRBJ1-1 | CASSLAGTAEAFF | KLGGALQAK | YES | NO | NO |
| 18657 | TRAV3 | TRAJ44 | CAVRVVTGTASKLTF | TRBV5-5 | TRBJ1-5 | CASSLEGEYAQPQHF | KLGGALQAK | NO | NO | NO |
| 18658 | TRAV38-2DV8 | TRAJ57 | CAYRSAYEGGSEKLVF | TRBV5-4 | TRBJ1-1 | CASSLYGRGGNTEAFF | KLGGALQAK | YES | NO | NO |
| 18659 | TRAV29DV5 | TRAJ32 | CAASVGGATNKLIF | TRBV12-4 | TRBJ2-1 | CAIMGGRWDEQFF | KLGGALQAK | YES | NO | NO |
| 18660 | TRAV29DV5 | TRAJ52 | CAASARAGGTSYGKLTF | TRBV7-8 | TRBJ1-5 | CASSSRRGYSNQPQHF | KLGGALQAK | YES | NO | NO |
| 18661 | TRAV35 | TRAJ11 | CAGQRGYSTLTF | TRBV7-9 | TRBJ1-6 | CASSQGQTGADGSPLHF | KLGGALQAK | YES | NO | NO |
| 18662 | TRAV12-2 | TRAJ42 | CAVISFGSQGNLIF | TRBV6-3 | TRBJ2-1 | CASSYGAGGYNEQFF | KLGGALQAK | YES | NO | NO |
| 18663 | TRAV13-2 | TRAJ13 | CADNSGGYQKVTF | TRBV7-9 | TRBJ2-1 | CASSLGGSVLLAGGNEQFF | KLGGALQAK | YES | NO | NO |
| 18664 | TRAV41 | TRAJ52 | CAADTITSSLHGGTSYGKLTF | TRBV7-8 | TRBJ1-1 | CASSLQQGRKAFF | KLGGALQAK | NO | NO | NO |
| 18666 | TRAV14DV4 | TRAJ34 | CAMREGTDKLIF | TRBV19 | TRBJ2-3 | CASSRSGLAGSNTDTQYF | KLGGALQAK | YES | NO | NO |
| 18667 | TRAV41 | TRAJ40 | CAAQSTSGTYKYIF | TRBV7-2 | TRBJ2-7 | CASSFRDRRNYEQYF | KLGGALQAK | YES | NO | NO |
| 18668 | TRAV8-1 | TRAJ42 | CAVRGSQGNLIF | TRBV9 | TRBJ2-7 | CASSVGTGEGTYEQYF | KLGGALQAK | YES | NO | NO |
| 18669 | TRAV4 | TRAJ37 | CLVGDIIFGSGNTGKLIF | TRBV5-1 | TRBJ1-2 | CASSLAETGDGYTF | KLGGALQAK | YES | NO | NO |
| 18670 | TRAV5 | TRAJ3 | CAESRDYSSASKIIF | TRBV5-6 | TRBJ2-3 | CASSLSSPGHPTDTQYF | KLGGALQAK | YES | NO | NO |
| 18671 | TRAV29DV5 | TRAJ16 | CAAISLSDGQKLLF | TRBV7-9 | TRBJ1-2 | CASSFQGPASFYGYTF | KLGGALQAK | NO | NO | NO |
| 18672 | TRAV3 | TRAJ5 | CAVRVPNTGRRALTF | TRBV20-1 | TRBJ2-3 | CSAKTGLAGPDTDTQYF | KLGGALQAK | NO | NO | NO |
| 18673 | TRAV13-2 | TRAJ8 | CAETTGFQKLVF | TRBV5-1 | TRBJ2-3 | CASRGSGDSTSTDTQYF | KLGGALQAK | YES | NO | NO |

| | | | | | | | | | | |
|---|---|---|---|---|---|---|---|---|---|---|
| 18674 | TRAV8-6 | TRAJ34 | CAVSGAYNTDKLIF | TRBV19 | TRBJ2-1 | CASSLTSGGTNEQFF | KLGGALQAK | YES | NO | NO |
| 18675 | TRAV19 | TRAJ4 | CALSAFSGGYNKLIF | TRBV4-3 | TRBJ2-5 | CASSQESSNWETQYF | KLGGALQAK | YES | NO | NO |
| 18676 | TRAV24 | TRAJ44 | CALDTGTASKLTF | TRBV6-3 | TRBJ2-1 | CASSYPGAQDEQFF | KLGGALQAK | YES | NO | NO |
| 18677 | TRAV41 | TRAJ42 | CAVVGHGGSQGNLIF | TRBV28 | TRBJ2-1 | CASSPHRLAQFF | KLGGALQAK | YES | NO | NO |
| 18678 | TRAV1-2 | TRAJ36 | CAVLTNNLFF | TRBV7-8 | TRBJ2-1 | CASSLVGRSDNEQFF | KLGGALQAK | YES | NO | NO |
| 18679 | TRAV25 | TRAJ48 | CEGLNFGNEKLTF | TRBV6-5 | TRBJ2-2 | CASRTAGGPTGELFF | KLGGALQAK | YES | NO | NO |
| 18680 | TRAV6 | TRAJ13 | CARPNSGGYQKVTF | TRBV20-1 | TRBJ2-7 | CSANWDTNIYEQYF | KLGGALQAK | YES | NO | NO |
| 18681 | TRAV3 | TRAJ37 | CAVRDGGMGSNTGKLIF | TRBV19 | TRBJ2-1 | CAGGWGDEQFF | KLGGALQAK | NO | NO | NO |
| 18682 | TRAV5 | TRAJ40 | CAESIPSGTYKYIF | TRBV6-3 | TRBJ2-2 | CAILTNTGELFF | KLGGALQAK | YES | NO | NO |
| 18683 | TRAV1-2 | TRAJ29 | CAGGPGNTPLVF | TRBV12-3 | TRBJ2-3 | CASTRSGRTDTQYF | KLGGALQAK | YES | NO | NO |
| 18684 | TRAV23DV6 | TRAJ22 | CAAVPSGSARQLTF | TRBV11-2 | TRBJ2-5 | CASSPFYMETQYF | KLGGALQAK | NO | NO | NO |
| 18685 | TRAV22 | TRAJ48 | CAGGYNFGNEKLTF | TRBV13 | TRBJ2-1 | CASSTSGGVFREQFF | KLGGALQAK | YES | NO | NO |
| 18686 | TRAV2 | TRAJ37 | CAIFDSGNTGKLIF | TRBV5-1 | TRBJ2-1 | CASSLPNEQFF | KLGGALQAK | NO | NO | NO |
| 18687 | TRAV17 | TRAJ11 | CATDGWSSGYSTLTF | TRBV18 | TRBJ2-5 | CASSPPGGQETQYF | AVFDRKSDAK | YES | NO | YES |
| 18688 | TRAV12-3 | TRAJ39 | CAIGPGNMLTF | TRBV19 | TRBJ2-7 | CASKGTSYEQYF | KLGGALQAK | YES | NO | NO |
| 18689 | TRAV17 | TRAJ36 | CATREDQTGANNLFF | TRBV4-1 | TRBJ1-4 | CASSQGPGGMKEKLFF | AVFDRKSDAK | YES | NO | YES |
| 18690 | TRAV19 | TRAJ23 | CALNQGGKLIF | TRBV10-1 | TRBJ2-1 | CASRGAGVDNEQFF | KLGGALQAK | YES | NO | NO |
| 18691 | TRAV8-4 | TRAJ21 | CAVSEGNFNKFYF | TRBV4-2 | TRBJ2-3 | CASSQIAGVGSVPGTDTQYF | KLGGALQAK | NO | NO | NO |
| 18692 | TRAV10 | TRAJ45 | CVVSARGGADGLTF | TRBV5-1 | TRBJ2-3 | CASSLPLAGGSTDTQYF | KLGGALQAK | YES | NO | NO |
| 18693 | TRAV8-6 | TRAJ34 | CAVSETTDKLIF | TRBV12-4 | TRBJ1-1 | CASSNAGTYTEAFF | KLGGALQAK | YES | NO | NO |
| 18695 | TRAV10 | TRAJ39 | CVVSAGAGNMLTF | TRBV7-2 | TRBJ2-7 | CASSATMSSYEQYV | KLGGALQAK | NO | NO | NO |
| 18696 | TRAV12-1 | TRAJ26 | CVVLLGQNFVF | TRBV7-9 | TRBJ2-1 | CASSSSSSHEQFF | KLGGALQAK | YES | NO | NO |
| 18697 | TRAV8-6 | TRAJ34 | CAVSQTTHTDKLIF | TRBV7-7 | TRBJ1-3 | CASSPLGDSDSGNTIYF | KLGGALQAK | YES | NO | NO |
| 18698 | TRAV12-2 | TRAJ48 | CAEGDFGNEKLTF | TRBV13 | TRBJ1-1 | CASSISTGGFTEAFF | KLGGALQAK | YES | NO | NO |
| 18700 | TRAV10 | TRAJ20 | CVVSAPVDYKLSF | TRBV4-3 | TRBJ2-3 | CASSQEVLAGPRVDTQYF | AVFDRKSDAK | YES | NO | NO |
| 18701 | TRAV8-3 | TRAJ24 | CAVGANPTDSWGKLQF | TRBV5-4 | TRBJ1-2 | CASTVQADYGYTF | KLGGALQAK | YES | NO | NO |
| 18702 | TRAV5 | TRAJ42 | CAESMNYGGSQGNLIF | TRBV20-1 | TRBJ2-1 | CSATRTNYNEQFF | KLGGALQAK | YES | NO | NO |
| 18703 | TRAV17 | TRAJ20 | CATDDDYKLSF | TRBV7-2 | TRBJ2-2 | CASSAWAGANTGELFF | AVFDRKSDAK | NO | NO | NO |
| 18704 | TRAV5 | TRAJ15 | CAEKGQAGTALIF | TRBV5-1 | TRBJ2-3 | CASFYRDTQYF | KLGGALQAK | YES | NO | NO |
| 18705 | TRAV12-2 | TRAJ10 | CAVNTIGAGGGNKLTF | TRBV19 | TRBJ1-5 | CASSTGGRGNQPQHF | KLGGALQAK | YES | YES | NO |
| 18706 | TRAV4 | TRAJ26 | CLVVAHYGQNFVF | TRBV6-5 | TRBJ2-3 | CASTFRDRGTDTQYF | KLGGALQAK | NO | NO | NO |
| 18707 | TRAV12-1 | TRAJ34 | CVVNRFDIYNTDKLIF | TRBV20-1 | TRBJ2-3 | CSARGAPGSNTDTQYF | AVFDRKSDAK | NO | NO | NO |
| 18708 | TRAV25 | TRAJ49 | CAGPGDTGNQFYF | TRBV30 | TRBJ1-1 | CAWSVQRGSEAFF | KLGGALQAK | YES | NO | NO |
| 18709 | TRAV1-2 | TRAJ37 | CAVPPDSGNTGKLIF | TRBV7-8 | TRBJ1-1 | CASSPRAGTNTEAFF | KLGGALQAK | YES | NO | NO |
| 18710 | TRAV12-3 | TRAJ26 | CAMSGWNYGQNFVF | TRBV19 | TRBJ1-1 | CASSIGTAAEAFF | AVFDRKSDAK | YES | NO | NO |
| 18711 | TRAV14DV4 | TRAJ4 | CAMRPPFSGGYNKLIF | TRBV7-2 | TRBJ2-7 | CASSLEPSYEQYF | KLGGALQAK | YES | NO | NO |
| 18712 | TRAV3 | TRAJ23 | CAVRDVKTWNQGGKLIF | TRBV19 | TRBJ1-1 | CASSLGHRTEAFF | KLGGALQAK | NO | NO | NO |
| 18713 | TRAV8-3 | TRAJ45 | CAVGASGGGADGLTF | TRBV6-3 | TRBJ1-1 | CASSYRAGGLTFF | KLGGALQAK | YES | NO | NO |
| 18714 | TRAV22 | TRAJ58 | CAVGSLETSGSRLTF | TRBV5-4 | TRBJ2-1 | CASSSQAGLTYNEQFF | KLGGALQAK | YES | NO | NO |
| 18715 | TRAV1-1 | TRAJ13 | CAVYGSGGYQKVTF | TRBV19 | TRBJ1-3 | CAATNRLGNTIYF | KLGGALQAK | YES | NO | NO |
| 18716 | TRAV1-2 | TRAJ9 | CAVREDTGGFKTIF | TRBV20-1 | TRBJ1-6 | CSARDPREVNSPLHF | KLGGALQAK | YES | NO | NO |
| 18717 | TRAV1-2 | TRAJ15 | CAVSNQAGTALIF | TRBV9 | TRBJ2-5 | CASSGQGVVETQYF | QPRAPIRPI | YES | NO | YES |
| 18718 | TRAV25 | TRAJ10 | CAVPGGGNKLTF | TRBV12-4 | TRBJ2-3 | CASSEMTDTQYF | KLGGALQAK | YES | NO | NO |
| 18719 | TRAV12-1 | TRAJ22 | CVVYEGGSARQLTF | TRBV19 | TRBJ1-2 | CASRNPAGVVYGYTF | KLGGALQAK | YES | NO | NO |
| 18720 | TRAV5 | TRAJ40 | CADISSGTYKYIF | TRBV30 | TRBJ2-2 | CAWSTETGPGELFF | KLGGALQAK | YES | NO | NO |
| 18721 | TRAV29DV5 | TRAJ22 | CAASGDASGSARQLTF | TRBV4-1 | TRBJ2-5 | CASSQDTKTQYF | KLGGALQAK | YES | NO | NO |
| 18722 | TRAV12-1 | TRAJ37 | CVVNSNGNTGKLIF | TRBV3-1 | TRBJ2-1 | CASSPRTSGAYNEQFF | KLGGALQAK | YES | NO | NO |
| 18723 | TRAV26-2 | TRAJ45 | CILRELSGGGADGLTF | TRBV18 | TRBJ2-1 | CASSATEQFF | KLGGALQAK | YES | NO | NO |
| 18724 | TRAV16 | TRAJ9 | CALYTGGFKTIF | TRBV11-2 | TRBJ2-1 | CASSSQAGGRENEQFF | KLGGALQAK | NO | NO | NO |
| 18725 | TRAV12-1 | TRAJ4 | CVVNIRGYNKLIF | TRBV7-9 | TRBJ1-4 | CASSLGGGDNEKLFF | KLGGALQAK | YES | NO | NO |
| 18726 | TRAV26-2 | TRAJ52 | CILRNAGGTSYGKLTF | TRBV27 | TRBJ2-7 | CASSLMGGGEQYF | KLGGALQAK | YES | NO | NO |
| 18727 | TRAV8-3 | TRAJ56 | CAVVRDTGANSKLTF | TRBV5-1 | TRBJ2-3 | CASSYRTSGINTQYF | KLGGALQAK | YES | NO | NO |
| 18729 | TRAV17 | TRAJ13 | CATDRSGGGGGYQKVTF | TRBV27 | TRBJ1-5 | CASSLGTYSNQPQHF | KLGGALQAK | YES | NO | NO |
| 18730 | TRAV3 | TRAJ49 | CAVRDNGNQFYF | TRBV7-2 | TRBJ2-7 | CASSLDSAHEQYV | KLGGALQAK | NO | NO | NO |
| 18731 | TRAV35 | TRAJ42 | CAGQKGSQGNLIF | TRBV12-3 | TRBJ2-2 | CASRDSNTGELFF | KLGGALQAK | YES | NO | NO |
| 18732 | TRAV17 | TRAJ40 | CATDSGTYKYIF | TRBV18 | TRBJ2-5 | CASSPSADQETQYF | KLGGALQAK | YES | NO | NO |
| 18733 | TRAV38-1 | TRAJ28 | CAFTHAYSGAGSYQLTF | TRBV7-6 | TRBJ2-7 | CASSLGPNYEQYF | KLGGALQAK | YES | NO | NO |
| 18734 | TRAV12-2 | TRAJ42 | CAVNTYGGSQGNLIF | TRBV12-4 | TRBJ1-4 | CASTQHRDSGEKLFF | KLGGALQAK | YES | NO | NO |
| 18735 | TRAV29DV5 | TRAJ20 | CAASEGSNDYKLSF | TRBV12-4 | TRBJ2-5 | CASSFSGEGETQYF | KLGGALQAK | YES | NO | NO |
| 18736 | TRAV12-3 | TRAJ33 | CAMSAASNYQLIW | TRBV24-1 | TRBJ2-3 | CATSDPGTGFTDTQYF | KLGGALQAK | YES | NO | NO |
| 18737 | TRAV12-1 | TRAJ7 | CVVIAYGNNRLAF | TRBV24-1 | TRBJ2-4 | CATSAVDRGKNIQYF | KLGGALQAK | YES | NO | NO |
| 18738 | TRAV19 | TRAJ40 | CALINTSGTYKYIF | TRBV27 | TRBJ1-2 | CASSTADRWYGYTF | KLGGALQAK | YES | NO | NO |
| 18739 | TRAV14DV4 | TRAJ23 | CAMRGFYNQGGKLIF | TRBV7-9 | TRBJ2-3 | CASTPSSGGLTDTQYF | KLGGALQAK | YES | NO | NO |
| 18741 | TRAV12-1 | TRAJ5 | CVVRGDTGRRALTF | TRBV19 | TRBJ1-1 | CASSPHTMNTEAFF | KLGGALQAK | YES | NO | NO |
| 18742 | TRAV9-2 | TRAJ5 | CALSVGGPDTGRRALTF | TRBV7-9 | TRBJ1-4 | CASSSSYSHEKLFF | KLGGALQAK | YES | NO | NO |
| 18743 | TRAV25 | TRAJ26 | CAGTLLRYGQNFVF | TRBV5-4 | TRBJ2-1 | CASSLGLAGTYNEQFF | KLGGALQAK | YES | NO | NO |
| 18744 | TRAV8-1 | TRAJ43 | CAVNAGDMRF | TRBV27 | TRBJ2-7 | CASSLRGGRNPYEQYF | KLGGALQAK | YES | NO | NO |
| 18745 | TRAV9-2 | TRAJ30 | CAREGGDDKIIF | TRBV11-2 | TRBJ2-3 | CASSLDLRTSGGIYTDTQYF | KLGGALQAK | NO | NO | NO |
| 18746 | TRAV24 | TRAJ40 | CAYSSGTYKYIF | TRBV3-1 | TRBJ1-1 | CASSRGTGGHTEAFF | KLGGALQAK | YES | NO | NO |
| 18747 | TRAV2 | TRAJ40 | CAVERGSGTYKYIF | TRBV27 | TRBJ2-1 | CASSSPFSYNEQFF | KLGGALQAK | NO | NO | NO |
| 18748 | TRAV38-1 | TRAJ58 | CAFGASGSRLTF | TRBV13 | TRBJ1-1 | CASSQTQGVTEAFF | KLGGALQAK | YES | NO | NO |
| 18749 | TRAV1-2 | TRAJ9 | CAVRDQDGGFKTIF | TRBV10-2 | TRBJ2-2 | CASSAPSQNTGELFF | KLGGALQAK | YES | NO | NO |
| 18750 | TRAV27 | TRAJ45 | CAGEGSGGGADGLTF | TRBV19 | TRBJ1-6 | CASNAGTGGYSPLHF | KLGGALQAK | YES | NO | NO |
| 18751 | TRAV14DV4 | TRAJ16 | CAMREISDGQKLLF | TRBV5-5 | TRBJ2-5 | CASSFIGSETQYF | KLGGALQAK | NO | NO | NO |
| 18752 | TRAV20 | TRAJ20 | CAVETRTNDYKLSF | TRBV12-4 | TRBJ1-4 | CASSLVAEYNEKLFF | KLGGALQAK | YES | NO | NO |
| 18754 | TRAV13-1 | TRAJ28 | CAAFRDSGAGSYQLTF | TRBV7-9 | TRBJ2-2 | CASSLRGGYTGELFF | KLGGALQAK | YES | NO | NO |
| 18755 | TRAV29DV5 | TRAJ20 | CAASANDYKLSF | TRBV12-4 | TRBJ1-4 | CASSSIEKLFF | KLGGALQAK | YES | NO | NO |
| 18756 | TRAV14DV4 | TRAJ43 | CAMREGLRNNDMRF | TRBV19 | TRBJ2-7 | CASSPGTGYEQYF | KLGGALQAK | YES | NO | NO |
| 18757 | TRAV22 | TRAJ30 | CAVERDDKIIF | TRBV5-4 | TRBJ2-5 | CASSGPTGIQETQYF | KLGGALQAK | YES | NO | NO |
| 18758 | TRAV29DV5 | TRAJ27 | CAASTNTNAGKSTF | TRBV7-9 | TRBJ2-1 | CASSLGLNEQFF | KLGGALQAK | YES | NO | NO |
| 18759 | TRAV19 | TRAJ20 | CALSEQSNDYKLSF | TRBV27 | TRBJ2-3 | CASTTRRFGKGATDTQYF | KLGGALQAK | YES | NO | NO |
| 18760 | TRAV39 | TRAJ44 | CAAVTGTASKLTF | TRBV27 | TRBJ1-2 | CASSPFTGVGYGYTF | KLGGALQAK | YES | NO | NO |
| 18761 | TRAV17 | TRAJ3 | CATDSGYSSASKIIF | TRBV27 | TRBJ1-5 | CASSPGQTSNQPQHF | KLGGALQAK | YES | NO | NO |
| 18762 | TRAV6 | TRAJ31 | CALWFFEGARLMF | TRBV7-8 | TRBJ1-5 | CASSLEYRTGVLNQPQHF | KLGGALQAK | YES | NO | NO |
| 18763 | TRAV8-3 | TRAJ9 | CAVGADTGGFKTIF | TRBV12-3 | TRBJ1-2 | CASSHLYYDSSYGYTF | AVFDRKSDAK | YES | NO | NO |
| 18764 | TRAV12-2 | TRAJ33 | CAAFNYQLIW | TRBV4-1 | TRBJ1-1 | CASSPGQGGTEAFF | ELAGIGILTV | YES | NO | YES |
| 18765 | TRAV12-1 | TRAJ23 | CVVNHRGKLIF | TRBV7-9 | TRBJ2-6 | CASSMTESGANVLTF | KLGGALQAK | YES | NO | NO |
| 18766 | TRAV12-1 | TRAJ39 | CVVNSAGNMLTF | TRBV20-1 | TRBJ1-2 | CSARDQDRGFGYTF | KLGGALQAK | YES | NO | NO |
| 18767 | TRAV1-2 | TRAJ33 | CAVRDSNYQLIW | TRBV4-1 | TRBJ2-2 | CASSSQRNTGELFF | KLGGALQAK | YES | NO | NO |
| 18769 | TRAV9-2 | TRAJ6 | CALSDSGGSYIPTF | TRBV7-9 | TRBJ2-7 | CASSPQTGKRLYEQYF | AVFDRKSDAK | YES | NO | YES |
| 18770 | TRAV1-2 | TRAJ11 | CAVAMNSGYSTLTF | TRBV7-9 | TRBJ2-2 | CASSLTNRGRGTGELFF | KLGGALQAK | YES | NO | NO |
| 18771 | TRAV13-1 | TRAJ27 | CAASNAGKSTF | TRBV6-6 | TRBJ2-3 | CASRQTTSTDTQYF | KLGGALQAK | NO | NO | NO |
| 18772 | TRAV4 | TRAJ45 | CLVGDNIGGGADGLTF | TRBV7-9 | TRBJ1-5 | CASSLAGGNQPQHF | KLGGALQAK | YES | NO | NO |
| 18773 | TRAV13-1 | TRAJ20 | CAAGDYKLSF | TRBV11-3 | TRBJ2-5 | CASSSEPGTGVQETQYF | KLGGALQAK | YES | NO | NO |
| 18774 | TRAV8-3 | TRAJ12 | CAVVPDSSYKLIF | TRBV7-2 | TRBJ2-1 | CASSAGQGAVDEQFF | KLGGALQAK | YES | NO | NO |
| 18775 | TRAV3 | TRAJ44 | CAVRSDTGTASKLTF | TRBV20-1 | TRBJ1-2 | CSARVVLSRGYGYTF | KLGGALQAK | NO | NO | NO |
| 18776 | TRAV1-2 | TRAJ32 | CAFYGGATNKLIF | TRBV3-1 | TRBJ1-3 | CASSQDDDSGNTIYF | KLGGALQAK | YES | NO | NO |
| 18777 | TRAV22 | TRAJ50 | CAVEPVTSYDKVIF | TRBV4-3 | TRBJ1-2 | CASSQDGPGVGANYGYTF | KLGGALQAK | YES | NO | NO |
| 18778 | TRAV1-1 | TRAJ23 | CAVRAQGGKLIF | TRBV7-9 | TRBJ1-2 | CASSLGESGTGPHGYTF | KLGGALQAK | YES | NO | NO |
| 18779 | TRAV10 | TRAJ34 | CVVSYNTDKLIF | TRBV6-5 | TRBJ1-1 | CASSPLGGRTEAFF | KLGGALQAK | YES | NO | NO |
| 18780 | TRAV39 | TRAJ37 | CAVMQYPGNTGKLIF | TRBV7-9 | TRBJ2-1 | CASSLVDSGSSYNEQFF | KLGGALQAK | YES | NO | NO |
| 18781 | TRAV19 | TRAJ49 | CALSEAFDTGNQFYF | TRBV4-3 | TRBJ2-6 | CASSLRRTGASSGANVLTF | KLGGALQAK | YES | NO | NO |
| 18782 | TRAV27 | TRAJ39 | CAGYVRENAGNMLTF | TRBV3-1 | TRBJ2-2 | CASSQAEDTGELFF | KLGGALQAK | YES | NO | NO |
| 18783 | TRAV1-2 | TRAJ33 | CAGTDSNYQLIW | TRBV15 | TRBJ2-6 | CATSRDNSGANVLTF | KLGGALQAK | YES | NO | NO |

| | | | | | | | | | | |
|---|---|---|---|---|---|---|---|---|---|---|
| 18784 | TRAV21 | TRAJ37 | CAVLFNTGKLIF | TRBV7-3 | TRBJ2-1 | CASSLGREQFF | KLGGALQAK | YES | NO | NO |
| 18785 | TRAV24 | TRAJ36 | CAFIPVVSQTGANNLFF | TRBV12-4 | TRBJ1-2 | CASNPGTGRYGYTF | KLGGALQAK | YES | NO | NO |
| 18786 | TRAV8-3 | TRAJ49 | CAVGTNTGNQFYF | TRBV4-3 | TRBJ2-1 | CASSQVQGQGFWEQFF | KLGGALQAK | YES | NO | NO |
| 18787 | TRAV13-1 | TRAJ31 | CAASILNARLMF | TRBV7-9 | TRBJ2-1 | CASGLAGNNEQFF | KLGGALQAK | YES | NO | NO |
| 18788 | TRAV41 | TRAJ57 | CAAQGGSEKLVF | TRBV27 | TRBJ1-2 | CASSSQGTDLGYTF | KLGGALQAK | YES | NO | NO |
| 18789 | TRAV8-3 | TRAJ42 | CAVGATDYGGSQGNLIF | TRBV12-3 | TRBJ2-7 | CASSLGQGARNYEQYF | KLGGALQAK | YES | YES | NO |
| 18790 | TRAV12-2 | TRAJ40 | CAVNSNKYIF | TRBV5-4 | TRBJ2-1 | CASSGLAPEQFF | KLGGALQAK | YES | NO | NO |
| 18791 | TRAV38-1 | TRAJ36 | CAFPSPGANNLFF | TRBV7-9 | TRBJ2-1 | CASMAGTYNEQFF | KLGGALQAK | YES | NO | NO |
| 18792 | TRAV1-2 | TRAJ30 | CAVSLVGRDKIIF | TRBV9 | TRBJ1-4 | CASKGVLDEKLFF | KLGGALQAK | YES | NO | NO |
| 18793 | TRAV14DV4 | TRAJ37 | CAMRGPFSSNTGKLIF | TRBV2 | TRBJ1-6 | CASSGTSPLHF | KLGGALQAK | YES | NO | NO |
| 18794 | TRAV8-1 | TRAJ30 | CAVNAGRDDKIIF | TRBV13 | TRBJ2-1 | CASSSLAVSYNEQFF | KLGGALQAK | YES | NO | NO |
| 18795 | TRAV14DV4 | TRAJ58 | CAMRELETSGSRLTF | TRBV9 | TRBJ2-2 | CASSPLGLGRTGELFF | KLGGALQAK | YES | YES | NO |
| 18796 | TRAV1-1 | TRAJ30 | CAGVGDKIIF | TRBV4-2 | TRBJ2-3 | CASRGGRGLDTQYF | KLGGALQAK | YES | NO | NO |
| 18797 | TRAV3 | TRAJ36 | CAVRDIGGTGANNLFF | TRBV7-9 | TRBJ2-5 | CASSSDRRQETQYF | AVFDRKSDAK | NO | NO | NO |
| 18798 | TRAV41 | TRAJ49 | CAVTPTNTGNQFYF | TRBV5-1 | TRBJ2-7 | CASSPGTGSYEQYF | KLGGALQAK | YES | NO | NO |
| 18799 | TRAV17 | TRAJ8 | CATDVGGFQKLVF | TRBV28 | TRBJ2-1 | CASSFVGDSSYNEQFF | KLGGALQAK | YES | NO | NO |
| 18800 | TRAV14DV4 | TRAJ45 | CAMREGTGGGADGLTF | TRBV18 | TRBJ1-4 | CASSPFATNEKLFF | KLGGALQAK | YES | NO | NO |
| 18801 | TRAV29DV5 | TRAJ26 | CAASVSNYGQNFVF | TRBV4-1 | TRBJ1-4 | CASSQVAGEREKLFF | KLGGALQAK | YES | NO | NO |
| 18802 | TRAV3 | TRAJ28 | CAVRDSGAGSYQLTF | TRBV3-1 | TRBJ1-1 | CASSRALGWGLDTEAFF | AVFDRKSDAK | NO | NO | NO |
| 18803 | TRAV22 | TRAJ29 | CAVEPGISGNTPLVF | TRBV5-6 | TRBJ2-5 | CASSLPPGSGGARETQYF | KLGGALQAK | YES | NO | NO |
| 18804 | TRAV27 | TRAJ9 | CAGAPLHTGGFKTIF | TRBV27 | TRBJ1-1 | CASSFSQGAVLKEAFF | KLGGALQAK | YES | NO | NO |
| 18805 | TRAV38-2DV8 | TRAJ56 | CAYLGGANSKLTF | TRBV15 | TRBJ2-1 | CATSRDLSGTNYNEQFF | KLGGALQAK | YES | NO | NO |
| 18806 | TRAV12-3 | TRAJ17 | CAMRRAAGNKLTF | TRBV20-1 | TRBJ2-3 | CSARDRIGGRLTDTQYF | KLGGALQAK | YES | NO | NO |
| 18807 | TRAV26-2 | TRAJ27 | CILRSNTNAGKSTF | TRBV7-9 | TRBJ2-2 | CAERRQITNTGELFF | KLGGALQAK | YES | NO | NO |
| 18808 | TRAV38-2DV8 | TRAJ40 | CAYGASGTYKYIF | TRBV27 | TRBJ2-5 | CASSPYPGETQYF | KLGGALQAK | YES | NO | NO |
| 18809 | TRAV9-2 | TRAJ42 | CALSPYYGGSQGNLIF | TRBV4-1 | TRBJ2-2 | CASSHPSGGQNTGELFF | KLGGALQAK | YES | NO | NO |
| 18810 | TRAV8-6 | TRAJ41 | CAVSDPTTNSGYALNF | TRBV5-8 | TRBJ2-7 | CASSPSPDTYEQYF | KLGGALQAK | YES | NO | NO |
| 18811 | TRAV12-2 | TRAJ42 | CAVSNYGGSQGNLIF | TRBV6-3 | TRBJ1-1 | CASSRDTEAFF | KLGGALQAK | YES | NO | NO |
| 18812 | TRAV8-3 | TRAJ7 | CAVGPLGVPIYGNNRLAF | TRBV6-1 | TRBJ1-6 | CASSERRVPFGDSPLHF | KLGGALQAK | YES | NO | NO |
| 18813 | TRAV14DV4 | TRAJ4 | CAMREGSGGYNKLIF | TRBV7-9 | TRBJ2-3 | CASSGGDRPTTDTQYF | KLGGALQAK | YES | NO | NO |
| 18814 | TRAV4 | TRAJ13 | CLVGDMEARRYQKVTF | TRBV12-4 | TRBJ1-1 | CASTVSGTGAWNTEAFF | KLGGALQAK | YES | NO | NO |
| 18815 | TRAV12-1 | TRAJ34 | CVVMLLSYNTDKLIF | TRBV19 | TRBJ1-1 | CASSIGVNTEAFF | KLGGALQAK | YES | NO | NO |
| 18816 | TRAV8-6 | TRAJ38 | CAVRETNAGNNRKLIW | TRBV7-9 | TRBJ2-5 | CASSLVVVEQETQYF | KLGGALQAK | YES | NO | NO |
| 18817 | TRAV8-1 | TRAJ7 | CAVNVYGNNRLAF | TRBV12-4 | TRBJ1-1 | CASSFISPPQTEAFF | KLGGALQAK | YES | NO | NO |
| 18818 | TRAV12-2 | TRAJ33 | CAVNGWDSNYQLIW | TRBV19 | TRBJ2-7 | CASMAGGTYEQYF | KLGGALQAK | YES | NO | NO |
| 18819 | TRAV24 | TRAJ23 | CAFPYYNQGGKLIF | TRBV5-1 | TRBJ2-2 | CASSSVTGGTGELFF | KLGGALQAK | YES | NO | NO |
| 18820 | TRAV13-1 | TRAJ24 | CAASRGGSWGKLQF | TRBV29-1 | TRBJ2-6 | CSVEGATGANVLTF | AVFDRKSDAK | YES | NO | NO |
| 18821 | TRAV38-2DV8 | TRAJ45 | CARISGGGADGLTF | TRBV6-6 | TRBJ1-2 | CASSYSQGWRNYGYTF | KLGGALQAK | NO | NO | NO |
| 18822 | TRAV17 | TRAJ7 | CATDLYYGNNRLAF | TRBV2 | TRBJ1-5 | CASSEAESQPQHF | AVFDRKSDAK | YES | NO | YES |
| 18823 | TRAV10 | TRAJ31 | CVVSRGNNNARLMF | TRBV7-9 | TRBJ2-3 | CASSGSGGSTDTQYF | KLGGALQAK | YES | NO | NO |
| 18824 | TRAV17 | TRAJ28 | CATVGAGSYQLTF | TRBV7-7 | TRBJ2-7 | CASSLAAGAYEQYF | KLGGALQAK | YES | NO | NO |
| 18825 | TRAV5 | TRAJ32 | CAENDGGATNKLIF | TRBV15 | TRBJ2-2 | CATSTPGLAGHTGELFF | KLGGALQAK | YES | NO | NO |
| 18826 | TRAV29DV5 | TRAJ32 | CAASDDGGATNKLIF | TRBV7-2 | TRBJ2-1 | CASSLGGYNEQFF | KLGGALQAK | YES | NO | NO |
| 18827 | TRAV8-2 | TRAJ47 | CVVSESGEYGNKLVF | TRBV4-3 | TRBJ2-1 | CASSQADSSYNEQFF | KLGGALQAK | YES | NO | NO |
| 18828 | TRAV1-2 | TRAJ16 | CAVRDF | TRBV9 | TRBJ2-7 | CASSVVGESYEQYF | KLGGALQAK | NO | NO | NO |
| 18829 | TRAV8-3 | TRAJ24 | CAVGDSGTTDSWGKFQF | TRBV7-8 | TRBJ1-2 | CASSLGQDGYTF | KLGGALQAK | YES | NO | NO |
| 18830 | TRAV8-6 | TRAJ32 | CAVSDPGGATNKLIF | TRBV19 | TRBJ1-4 | CASRPSDLPREKLFF | KLGGALQAK | YES | NO | NO |
| 18831 | TRAV14DV4 | TRAJ45 | CAMREGPGGADGLTF | TRBV9 | TRBJ2-5 | CASSESTTSGSGETQYF | KLGGALQAK | YES | NO | NO |
| 18832 | TRAV27 | TRAJ47 | CAGDEYGNKLVF | TRBV7-8 | TRBJ1-2 | CASSLEAGGGYTF | KLGGALQAK | YES | NO | NO |
| 18833 | TRAV5 | TRAJ50 | CAESPMKTSYDKVIF | TRBV5-5 | TRBJ2-2 | CASSLHVAGDTGELFF | KLGGALQAK | YES | NO | NO |
| 18834 | TRAV38-1 | TRAJ31 | CAFGFNNARLMF | TRBV29-1 | TRBJ2-5 | CSVEGSGGFQETQYF | KLGGALQAK | YES | NO | NO |
| 18835 | TRAV26-1 | TRAJ40 | CIVRAEARTYKYIF | TRBV27 | TRBJ1-5 | CASSDGPNQPQHF | KLGGALQAK | YES | NO | NO |
| 18836 | TRAV36DV7 | TRAJ52 | CAGANAGGTSYGKLTF | TRBV19 | TRBJ2-1 | CASKMLNEQFF | KLGGALQAK | NO | NO | NO |
| 18837 | TRAV39 | TRAJ34 | CAVDSEVNTDKLIF | TRBV5-1 | TRBJ2-1 | CASSWTGSYNEQFF | KLGGALQAK | YES | NO | NO |
| 18838 | TRAV3 | TRAJ21 | CAVRSNFNKFYF | TRBV11-3 | TRBJ2-1 | CASIGGDNEQFF | KLGGALQAK | YES | NO | NO |
| 18839 | TRAV19 | TRAJ37 | CALSEFSSNTGKLIF | TRBV6-3 | TRBJ2-1 | CASSYGAGGYNEQFF | KLGGALQAK | YES | NO | NO |
| 18840 | TRAV12-2 | TRAJ42 | CAVGGSQGNLIF | TRBV19 | TRBJ2-7 | CASSMAGGDGEQYF | KLGGALQAK | YES | NO | NO |
| 18841 | TRAV14DV4 | TRAJ34 | CAMREGPSYNTDKLIF | TRBV7-9 | TRBJ1-6 | CASSLVIPHGTRWDSPLHF | KLGGALQAK | YES | NO | NO |
| 18842 | TRAV13-1 | TRAJ3 | CAAGTKIIF | TRBV7-9 | TRBJ1-1 | CASSLANTEAFF | KLGGALQAK | YES | NO | NO |
| 18843 | TRAV12-2 | TRAJ31 | CAFDNARLMF | TRBV11-2 | TRBJ1-1 | CASSSWTGMNTEAFF | ELAGIGILTV | YES | NO | YES |
| 18844 | TRAV13-1 | TRAJ52 | CAAPSNAGGTSYGKLTF | TRBV2 | TRBJ1-1 | CASSDPGTGRTEAFF | KLGGALQAK | YES | YES | NO |
| 18845 | TRAV38-1 | TRAJ45 | CAFMTPGRGADGLTF | TRBV11-2 | TRBJ2-7 | CASRNWDGTYEQYV | KLGGALQAK | NO | NO | NO |
| 18846 | TRAV12-2 | TRAJ20 | CAVTGDDYKLSF | TRBV7-9 | TRBJ1-5 | CASSSLQSEPQHF | IVTDFSVIK | YES | YES | YES |
| 18847 | TRAV34 | TRAJ54 | CGAVFRNPIQGAQKLVF | TRBV7-6 | TRBJ2-1 | CASSLIGGFTNNEQFF | AVFDRKSDAK | YES | NO | YES |
| 18848 | TRAV13-2 | TRAJ11 | CAENIGYSTLTF | TRBV19 | TRBJ1-1 | CASMGPIEAFF | AYAQKIFKI | YES | YES | YES |
| 18849 | TRAV12-2 | TRAJ9 | CAVYTGGFKTIF | TRBV6-1 | TRBJ2-1 | CASSVLHGLGNEQFF | KLGGALQAK | YES | NO | NO |
| 18850 | TRAV1-2 | TRAJ20 | CAVRDFGDYKLSF | TRBV9 | TRBJ1-1 | CASSVVKPQTGGYTEAFF | KLGGALQAK | YES | NO | NO |
| 18851 | TRAV8-2 | TRAJ13 | CVVSDRAGGYQKVTF | TRBV7-8 | TRBJ2-2 | CASSPGTGRNGELFF | KLGGALQAK | YES | NO | NO |
| 18852 | TRAV25 | TRAJ50 | CAGHTPILKTSYDKVIF | TRBV10-3 | TRBJ1-6 | CAINQGGTDNSPLHF | KLGGALQAK | YES | NO | NO |
| 18853 | TRAV14DV4 | TRAJ40 | CAMREGQVTSGTYKYIF | TRBV5-1 | TRBJ2-7 | CASSGLGEQYF | KLGGALQAK | YES | NO | NO |
| 18854 | TRAV14DV4 | TRAJ34 | CAMREGSTDKLIF | TRBV9 | TRBJ2-1 | CASSVAGGRGNNEQFF | KLGGALQAK | YES | NO | NO |
| 18855 | TRAV35 | TRAJ29 | CAGRPLSGNTPLVF | TRBV9 | TRBJ2-1 | CASSVLASGSHEQFF | KLGGALQAK | YES | NO | NO |
| 18856 | TRAV22 | TRAJ33 | CAVELAGSNYQLIW | TRBV12-4 | TRBJ2-7 | CASHSGTGEEQYF | KLGGALQAK | YES | NO | NO |
| 18857 | TRAV1-2 | TRAJ39 | CAVRDWNAGNMLTF | TRBV6-1 | TRBJ2-1 | CASSEASYGATYNEQFF | KLGGALQAK | YES | NO | NO |
| 18858 | TRAV35 | TRAJ28 | CAGQSYSGAGSYQLTF | TRBV25-1 | TRBJ2-6 | CASSSTGIGANVLTF | KLGGALQAK | YES | NO | NO |
| 18859 | TRAV1-2 | TRAJ8 | CAVRRNTGFQKLVF | TRBV12-5 | TRBJ2-6 | CASGGAGTGGANVLTF | KLGGALQAK | YES | NO | NO |
| 18860 | TRAV39 | TRAJ41 | CAVVLGYALNF | TRBV11-2 | TRBJ2-7 | CASSLTRTEQYV | KLGGALQAK | NO | NO | NO |
| 18861 | TRAV29DV5 | TRAJ43 | CAASASNDMRF | TRBV18 | TRBJ1-1 | CASSPNLVGLNTEAFF | KLGGALQAK | YES | NO | NO |
| 18862 | TRAV26-2 | TRAJ27 | CILRGINTNAGKSTF | TRBV13 | TRBJ2-1 | CASSLELGQGRYNEQFF | KLGGALQAK | YES | NO | NO |
| 18863 | TRAV17 | TRAJ12 | CATVPMDSSYKLIF | TRBV7-9 | TRBJ1-4 | CASSTNQQGMGEKLFF | KLGGALQAK | YES | NO | NO |
| 18864 | TRAV9-2 | TRAJ52 | CALSADAGGTSYGKLTF | TRBV4-1 | TRBJ1-2 | CASGHWGASGYTF | KLGGALQAK | YES | NO | NO |
| 18865 | TRAV12-2 | TRAJ31 | CAVDNARLMF | TRBV11-2 | TRBJ2-5 | CASTPGVAGGPGETQYF | ELAGIGILTV | YES | NO | YES |
| 18866 | TRAV3 | TRAJ6 | CAVRDLSGGSYIPTF | TRBV6-5 | TRBJ2-2 | CASSPDTGELFF | KLGGALQAK | NO | NO | NO |
| 18867 | TRAV1-2 | TRAJ39 | CAQNAGNMLTF | TRBV9 | TRBJ2-7 | CASSVGFRGHEQYF | KLGGALQAK | YES | NO | NO |
| 18868 | TRAV19 | TRAJ23 | CALSEPLVYNQGGKLIF | TRBV20-1 | TRBJ1-2 | CSAPLGDHYGYTF | KLGGALQAK | YES | NO | NO |
| 18869 | TRAV1-1 | TRAJ20 | CAVSLSNDYKLSF | TRBV12-4 | TRBJ2-3 | CASSIAGSTDTQYF | KLGGALQAK | YES | NO | NO |
| 18870 | TRAV8-4 | TRAJ6 | CAVSEPGGSYIPTF | TRBV7-2 | TRBJ2-1 | CASSSRVDEQFF | KLGGALQAK | YES | NO | NO |
| 18871 | TRAV13-2 | TRAJ24 | CAENLMTTDSWGKLQF | TRBV19 | TRBJ1-1 | CASRGGGLTEAFF | KLGGALQAK | YES | NO | NO |
| 18872 | TRAV5 | TRAJ41 | CAESTLSGYALNF | TRBV20-1 | TRBJ2-5 | CSAPTSGQETQYF | KLGGALQAK | YES | NO | NO |
| 18873 | TRAV22 | TRAJ40 | CAVEEASGTYKYIF | TRBV20-1 | TRBJ2-1 | CSARQGYQFF | KLGGALQAK | NO | NO | NO |
| 18874 | TRAV12-1 | TRAJ34 | CVVNDNTDKLIF | TRBV2 | TRBJ2-5 | CASSRGPGQGEETQYF | AVFDRKSDAK | YES | NO | YES |
| 18875 | TRAV14DV4 | TRAJ49 | CAMETSTGNQFYF | TRBV6-5 | TRBJ1-4 | CASSYSGRSNEKLFF | KLGGALQAK | NO | NO | NO |
| 18876 | TRAV3 | TRAJ29 | CAVRALLGNTPLVF | TRBV19 | TRBJ2-2 | CASTTYTGELFF | KLGGALQAK | NO | NO | NO |
| 18877 | TRAV21 | TRAJ58 | CAVPRETSGSRLTF | TRBV7-9 | TRBJ2-5 | CASSLAVTPISGTQYF | KLGGALQAK | YES | NO | NO |
| 18878 | TRAV12-1 | TRAJ53 | CVVLRGSNYKLTF | TRBV14 | TRBJ2-2 | CASMRQGSAGELFF | KLGGALQAK | NO | NO | NO |
| 18879 | TRAV9-2 | TRAJ17 | CALSDWAGNKLTF | TRBV6-6 | TRBJ2-1 | CASSYRDIDNEQFF | KLGGALQAK | NO | NO | NO |
| 18880 | TRAV38-1 | TRAJ29 | CAFMKHWGSGNTPLVF | TRBV9 | TRBJ2-1 | CASSLPNRDPHNEQFF | KLGGALQAK | YES | NO | NO |
| 18881 | TRAV20 | TRAJ22 | CAVPKVSGSARQLTF | TRBV20-1 | TRBJ2-3 | CSADGLSTDTQYF | KLGGALQAK | NO | NO | NO |
| 18882 | TRAV12-2 | TRAJ53 | CAVRSGGSNYKLTF | TRBV7-9 | TRBJ1-1 | CASSLVLNTEAFF | KLGGALQAK | YES | NO | NO |
| 18883 | TRAV6 | TRAJ35 | CAPMPLI | TRBV5-1 | TRBJ2-3 | CASSLTSGAGDTQYF | KLGGALQAK | NO | NO | NO |
| 18884 | TRAV3 | TRAJ28 | CAVSQPGAGSYQLTF | TRBV7-9 | TRBJ1-3 | CASSHPQENTIYF | KLGGALQAK | NO | NO | NO |
| 18885 | TRAV27 | TRAJ20 | CAGSDYKLSF | TRBV19 | TRBJ2-1 | CASSALAGGYNEQFF | KLGGALQAK | YES | NO | NO |
| 18886 | TRAV29DV5 | TRAJ36 | CAGDGANNLFF | TRBV6-4 | TRBJ1-2 | CASSENPGQNYGYTF | KLGGALQAK | YES | NO | NO |
| 18887 | TRAV17 | TRAJ9 | CATALRGGFKTIF | TRBV19 | TRBJ1-5 | CASNSWTGDQPQHF | KLGGALQAK | YES | NO | NO |

| | | | | | | | | | | |
|---|---|---|---|---|---|---|---|---|---|---|
| 18888 | TRAV14DV4 | TRAJ4 | CAMSHISGGYNKLIF | TRBV6-5 | TRBJ2-5 | CASSYWSGPPGETQYF | KLGGALQAK | NO | NO | NO |
| 18889 | TRAV3 | TRAJ20 | CAVREQSSNDYKLSF | TRBV5-1 | TRBJ1-1 | CASSATFTNTEAFF | KLGGALQAK | YES | NO | NO |
| 18890 | TRAV24 | TRAJ52 | CAFIGDAGGTSYGKLTF | TRBV19 | TRBJ2-7 | CASSIGLAGGSLRPKSYEQYF | KLGGALQAK | NO | NO | NO |
| 18891 | TRAV27 | TRAJ20 | CAGVIYDYKLSF | TRBV3-1 | TRBJ1-2 | CASSQDFGTADGYTF | KLGGALQAK | YES | NO | NO |
| 18892 | TRAV21 | TRAJ57 | CAGETQGGSEKLVF | TRBV13 | TRBJ1-1 | CASPLPGQGTAFF | AVFDRKSDAK | NO | NO | NO |
| 18893 | TRAV38-1 | TRAJ54 | CAFMSLIQGAQKLVF | TRBV6-1 | TRBJ1-4 | CASSEAWGTPLWGEKLFF | KLGGALQAK | YES | NO | NO |
| 18894 | TRAV12-3 | TRAJ8 | CAMIDTGFQKLVF | TRBV2 | TRBJ2-5 | CASKVFGDGQETQYF | KLGGALQAK | YES | NO | NO |
| 18895 | TRAV35 | TRAJ49 | CAGSNTGNQFYF | TRBV27 | TRBJ2-1 | CASSPLSGRYNEQFF | KLGGALQAK | YES | NO | NO |
| 18896 | TRAV29DV5 | TRAJ48 | CAARGPNFGNEKLTF | TRBV15 | TRBJ2-7 | CATSRSGLAGVYSYEQYV | AVFDRKSDAK | NO | NO | NO |
| 18897 | TRAV19 | TRAJ58 | CALSEAGETSGSRLTF | TRBV19 | TRBJ2-3 | CASSPFAGGMSTDTQYF | KLGGALQAK | YES | NO | NO |
| 18898 | TRAV5 | TRAJ32 | CAEIPYGGATNKLIF | TRBV20-1 | TRBJ2-1 | CSARELLREQFF | KLGGALQAK | YES | NO | NO |
| 18899 | TRAV5 | TRAJ40 | CAESNSGTYKYIF | TRBV20-1 | TRBJ2-6 | CSGVSLQESSGANVLTF | AVFDRKSDAK | YES | NO | YES |
| 18900 | TRAV3 | TRAJ15 | CAVRVDQAGTALIF | TRBV11-3 | TRBJ2-3 | CASSFRLAGAGDTQYF | KLGGALQAK | NO | NO | NO |
| 18901 | TRAV9-2 | TRAJ18 | CALIVGRGSTLGRLYF | TRBV24-1 | TRBJ1-2 | CATSDLGRRDGYTF | KLGGALQAK | YES | NO | NO |
| 18902 | TRAV19 | TRAJ4 | CALSELSFSGGYNKLIF | TRBV5-6 | TRBJ1-5 | CASSLWPMGQDHQPQHF | KLGGALQAK | YES | NO | NO |
| 18903 | TRAV8-1 | TRAJ49 | CAAYTGNQFYF | TRBV7-2 | TRBJ1-3 | CASSPWMGRGNTIYF | KLGGALQAK | YES | NO | NO |
| 18904 | TRAV1-1 | TRAJ34 | CAVDNTDKLIF | TRBV29-1 | TRBJ2-1 | CSVEGSGGHEQFF | KLGGALQAK | YES | NO | NO |
| 18905 | TRAV9-2 | TRAJ31 | CALGDARLMF | TRBV7-9 | TRBJ1-1 | CASSLEPGGRNTEAFF | KLGGALQAK | YES | NO | NO |
| 18906 | TRAV13-2 | TRAJ41 | CAEGANSGYALNF | TRBV5-6 | TRBJ1-1 | CASSSGGTEAFF | KLGGALQAK | YES | NO | NO |
| 18907 | TRAV5 | TRAJ7 | CAELLYGNNRLAF | TRBV5-1 | TRBJ2-5 | CASRHEAGEGPETQYF | KLGGALQAK | YES | NO | NO |
| 18908 | TRAV23DV6 | TRAJ9 | CAASNQTNTGGFKTIF | TRBV7-9 | TRBJ2-7 | CASSGASYEQYF | KLGGALQAK | YES | NO | NO |
| 18909 | TRAV8-3 | TRAJ52 | CADGAGGTSYGKLTF | TRBV12-4 | TRBJ2-1 | CASSLGRESGEQFF | KLGGALQAK | YES | NO | NO |
| 18910 | TRAV38-2DV8 | TRAJ20 | CASLSSNDYKLSF | TRBV12-4 | TRBJ2-7 | CASSSDGYEQYF | KLGGALQAK | YES | NO | NO |
| 18911 | TRAV12-1 | TRAJ12 | CVVNAVDSSYKLIF | TRBV6-3 | TRBJ2-1 | CASSYGAGGYNEQFF | KLGGALQAK | YES | NO | NO |
| 18912 | TRAV20 | TRAJ15 | CAVQAGQAGTALIF | TRBV20-1 | TRBJ2-1 | CSARSRGSYNEQFF | KLGGALQAK | NO | NO | NO |
| 18913 | TRAV19 | TRAJ49 | CALSETPNTGNQFYF | TRBV3-1 | TRBJ2-7 | CASSQGGGGGYEQYF | KLGGALQAK | YES | NO | NO |
| 18914 | TRAV9-2 | TRAJ58 | CALSKETSGSRLTF | TRBV20-1 | TRBJ2-2 | CSAIRGLAGTNTGELFF | KLGGALQAK | YES | NO | NO |
| 18915 | TRAV12-2 | TRAJ18 | CAVSRGSTLGRLYF | TRBV6-1 | TRBJ1-3 | CASRGISGNTIYF | AVFDRKSDAK | YES | NO | YES |
| 18916 | TRAV12-3 | TRAJ11 | CAMSGSGYSTLTF | TRBV10-3 | TRBJ2-3 | CAISESRGSLSDTQYF | KLGGALQAK | YES | NO | NO |
| 18917 | TRAV12-3 | TRAJ49 | CAMLSGNQFYF | TRBV27 | TRBJ2-7 | CASSPTTVHYEQYV | KLGGALQAK | NO | NO | NO |
| 18918 | TRAV38-1 | TRAJ40 | CAFMKPYTTSGTYKYIF | TRBV12-4 | TRBJ2-3 | CASSSPLAGHGDTQYF | KLGGALQAK | YES | NO | NO |
| 18919 | TRAV27 | TRAJ16 | CAARF | TRBV6-6 | TRBJ2-3 | CASSYPNPTDTQYF | KLGGALQAK | NO | NO | NO |
| 18921 | TRAV21 | TRAJ39 | CAVDNNAGNMLTF | TRBV4-1 | TRBJ1-2 | CASSQEDRLPYGYTF | KLGGALQAK | YES | NO | NO |
| 18923 | TRAV10 | TRAJ43 | CVVSAIEDNDMRF | TRBV4-3 | TRBJ1-2 | CASSQDTFKGHAYGYTF | KLGGALQAK | YES | NO | NO |
| 18924 | TRAV12-1 | TRAJ21 | CVAFKYNFNKFYF | TRBV12-4 | TRBJ1-6 | CASRSDSSYNSPLHF | KLGGALQAK | YES | NO | NO |
| 18925 | TRAV13-1 | TRAJ41 | CAVPNSGYALNF | TRBV24-1 | TRBJ2-1 | CATSDWSASGLDDNEQFF | KLGGALQAK | YES | NO | NO |
| 18926 | TRAV19 | TRAJ31 | CALSNNARLMF | TRBV7-6 | TRBJ1-1 | CASSLGYLELNTEAFF | KLGGALQAK | YES | NO | NO |
| 18927 | TRAV19 | TRAJ54 | CALRGQGAQKLVF | TRBV7-7 | TRBJ2-7 | CASSLERSREQYF | KLGGALQAK | YES | NO | NO |
| 18928 | TRAV29DV5 | TRAJ34 | CAARDEYTDKLIF | TRBV5-5 | TRBJ2-7 | CASRGLAGSYEQYF | KLGGALQAK | YES | NO | NO |
| 18929 | TRAV17 | TRAJ28 | CATDASNSGAGSYQLTF | TRBV5-1 | TRBJ2-2 | CASSLGTGITGELFF | KLGGALQAK | YES | NO | NO |
| 18930 | TRAV22 | TRAJ52 | CAVANAGGTSYGKLTF | TRBV5-4 | TRBJ2-7 | CASSLGQGGYEQYV | KLGGALQAK | NO | NO | NO |
| 18931 | TRAV5 | TRAJ40 | CAESRTYKYIF | TRBV11-2 | TRBJ2-3 | CASSPLAGEGTDTQYF | KLGGALQAK | YES | NO | NO |
| 18932 | TRAV1-1 | TRAJ21 | CAVRDGNNFNKFYF | TRBV20-1 | TRBJ1-1 | CSARDYTNTEAFF | KLGGALQAK | YES | NO | NO |
| 18933 | TRAV8-3 | TRAJ47 | CAVGLMEYGNKLVF | TRBV7-6 | TRBJ1-5 | CASSPRFRNQPQHF | KLGGALQAK | YES | NO | NO |
| 18934 | TRAV38-2DV8 | TRAJ37 | CAYLPGNTGKLIF | TRBV6-5 | TRBJ2-2 | CASRPQGARGELFF | KLGGALQAK | YES | NO | NO |
| 18935 | TRAV12-1 | TRAJ22 | CVVIASGSARQLTF | TRBV7-9 | TRBJ1-2 | CASSAPGEGGYTF | KLGGALQAK | YES | NO | NO |
| 18936 | TRAV13-1 | TRAJ16 | CAASSDGQKLLF | TRBV7-3 | TRBJ2-1 | CASSLIAGEKVNEQFF | KLGGALQAK | NO | NO | NO |
| 18937 | TRAV13-2 | TRAJ9 | CAENTGGFKTIF | TRBV7-9 | TRBJ2-7 | CASSWTSGEQYV | KLGGALQAK | NO | NO | NO |
| 18938 | TRAV30 | TRAJ28 | CGTEEGTVPGAGSYQLTF | TRBV4-2 | TRBJ2-5 | CASSHGLGWETQYF | AVFDRKSDAK | YES | NO | YES |
| 18939 | TRAV10 | TRAJ31 | CVVSAGARLMF | TRBV4-2 | TRBJ1-4 | CASSQILQGYEKLFF | KLGGALQAK | YES | NO | NO |
| 18940 | TRAV19 | TRAJ47 | CALSEGMEYGNKLVF | TRBV9 | TRBJ2-5 | CASSGSDGQAKWETQYF | KLGGALQAK | YES | NO | NO |
| 18941 | TRAV12-3 | TRAJ44 | CAMRITGTASKLTF | TRBV24-1 | TRBJ2-1 | CATMDDEYNEQFF | KLGGALQAK | YES | NO | NO |
| 18942 | TRAV41 | TRAJ43 | CAAGNDMRF | TRBV6-5 | TRBJ2-3 | CASRPRPGQASTDTQYF | KLGGALQAK | NO | NO | NO |
| 18943 | TRAV8-3 | TRAJ52 | CAVGRQGAGGTSYGKLTF | TRBV27 | TRBJ1-5 | CASRHSGVNQPQHF | AVFDRKSDAK | YES | NO | YES |
| 18944 | TRAV27 | TRAJ23 | CAGYPPGLWGKLIF | TRBV19 | TRBJ1-1 | CASSIVSGDTEAFF | KLGGALQAK | YES | NO | NO |
| 18945 | TRAV20 | TRAJ41 | CAVQDLYSGYALNF | TRBV11-3 | TRBJ2-2 | CASSLGEPATGELFF | KLGGALQAK | YES | NO | NO |
| 18946 | TRAV14DV4 | TRAJ6 | CAMRERKGGSYIPTF | TRBV5-1 | TRBJ1-3 | CASSLQGDTIYF | KLGGALQAK | YES | NO | NO |
| 18947 | TRAV19 | TRAJ12 | CALSEGMDSSYKLIF | TRBV4-1 | TRBJ1-1 | CASSSGTGQVMNTEAFF | KLGGALQAK | YES | NO | NO |
| 18948 | TRAV19 | TRAJ56 | CALSETTGANSKLTF | TRBV19 | TRBJ1-1 | CASSMDMWTEAFF | KLGGALQAK | YES | NO | NO |
| 18949 | TRAV1-2 | TRAJ24 | CAVRDNWGKLQF | TRBV6-5 | TRBJ2-1 | CASTQRAGGRIEQFF | AVFDRKSDAK | NO | NO | NO |
| 18950 | TRAV1-2 | TRAJ24 | CAVRDHPDSWGKLQF | TRBV3-1 | TRBJ2-1 | CASSQDPGQGSNEQFF | KLGGALQAK | YES | NO | NO |
| 18951 | TRAV14DV4 | TRAJ57 | CAMREAWGGSEKLVF | TRBV9 | TRBJ2-1 | CASNPAGGVYNEQFF | KLGGALQAK | YES | NO | NO |
| 18952 | TRAV24 | TRAJ30 | CAFIRDDKIIF | TRBV19 | TRBJ1-1 | CAISSGTGATEAFF | KLGGALQAK | YES | NO | NO |
| 18953 | TRAV8-3 | TRAJ41 | CAVGALDSGYALNF | TRBV20-1 | TRBJ1-2 | CSAPGQGVGYTF | AVFDRKSDAK | YES | NO | YES |
| 18954 | TRAV19 | TRAJ34 | CALTLDNTDKLIF | TRBV5-6 | TRBJ1-1 | CASSLVGSTEAFF | KLGGALQAK | YES | NO | NO |
| 18955 | TRAV19 | TRAJ17 | CALSEAIKAAGNKLTF | TRBV7-6 | TRBJ1-5 | CASSLETGAYQPQHF | KLGGALQAK | YES | NO | NO |
| 18956 | TRAV19 | TRAJ48 | CALSNFGNEKLTF | TRBV28 | TRBJ2-7 | CASSRDNSYEQYV | RLRAEAQVK | NO | NO | NO |
| 18957 | TRAV8-1 | TRAJ10 | CAVNEILTGGGNKLTF | TRBV19 | TRBJ2-1 | CASSIPNENEQFF | KLGGALQAK | YES | NO | NO |
| 18958 | TRAV8-3 | TRAJ34 | CAVGASYNTDKLIF | TRBV15 | TRBJ2-3 | CATSKLAGGGTDTQYF | KLGGALQAK | YES | NO | NO |
| 18959 | TRAV12-2 | TRAJ37 | CAVRGSSNTGKLIF | TRBV4-3 | TRBJ1-3 | CASSQEETAVSGNTIYF | ELAGIGILTV | YES | NO | YES |
| 18960 | TRAV4 | TRAJ23 | CLAGEGGKLIF | TRBV7-9 | TRBJ2-7 | CASTHSGTLTYEQYF | KLGGALQAK | NO | NO | NO |
| 18961 | TRAV19 | TRAJ41 | CALSEDASGYALNF | TRBV5-4 | TRBJ1-3 | CASSWGLSGNTIYF | KLGGALQAK | YES | NO | NO |
| 18963 | TRAV13-2 | TRAJ43 | CAENMAHYNNDMRF | TRBV9 | TRBJ2-1 | CASSPSGGAGNEQFF | KLGGALQAK | YES | NO | NO |
| 18964 | TRAV34 | TRAJ57 | CGAPSTQGGSEKLVF | TRBV10-3 | TRBJ2-1 | CAISLSGRGNEQFF | KLGGALQAK | YES | NO | NO |
| 18965 | TRAV38-2DV8 | TRAJ40 | CAYRTTSGTYKYIF | TRBV4-2 | TRBJ1-6 | CASSPGEGPHSPLHF | KLGGALQAK | YES | NO | NO |
| 18966 | TRAV13-2 | TRAJ16 | CAEISTDGQKLLF | TRBV9 | TRBJ2-7 | CASSVGQGSYEQYF | KLGGALQAK | NO | NO | NO |
| 18967 | TRAV16 | TRAJ21 | CALGGYNFNKFYF | TRBV19 | TRBJ2-3 | CASRDRVSDSDTQYF | KLGGALQAK | NO | NO | NO |
| 18968 | TRAV12-3 | TRAJ23 | CAMRNNQGGKLIF | TRBV19 | TRBJ1-5 | CASSIALSSNQPQHF | KLGGALQAK | YES | NO | NO |
| 18969 | TRAV12-1 | TRAJ23 | CVVNSAGQGGKLIF | TRBV4-1 | TRBJ2-7 | CASSQTPGEQYF | RLRAEAQVK | YES | NO | YES |
| 18970 | TRAV12-1 | TRAJ28 | CVVNRGYSGAGSYQLTF | TRBV5-1 | TRBJ2-7 | CASSLAGGTYEQYF | KLGGALQAK | YES | NO | NO |
| 18972 | TRAV4 | TRAJ27 | CLVGDIAAGKSTF | TRBV12-4 | TRBJ2-3 | CASSLRVFTGVGTDTQYF | KLGGALQAK | YES | YES | NO |
| 18973 | TRAV12-2 | TRAJ50 | CAVTPKTSYDKVIF | TRBV20-1 | TRBJ2-3 | CSARAGRGLDTDTQYF | KLGGALQAK | NO | NO | NO |
| 18974 | TRAV36DV7 | TRAJ40 | CAVVRTSGTYKYIF | TRBV12-4 | TRBJ1-2 | CASSPDLRAPDTF | KLGGALQAK | NO | NO | NO |
| 18975 | TRAV8-6 | TRAJ53 | CAVIFLSGGSNYKLTF | TRBV5-4 | TRBJ1-3 | CASSSRGSSGNTIYF | KLGGALQAK | YES | NO | NO |
| 18976 | TRAV25 | TRAJ33 | CAGSRSNYQLIW | TRBV6-3 | TRBJ2-7 | CASGTGWSYEQYV | AVFDRKSDAK | NO | NO | NO |
| 18977 | TRAV2 | TRAJ32 | CAVEDGGATNKLIF | TRBV12-4 | TRBJ2-2 | CASTSAGPNTGELFF | KLGGALQAK | NO | NO | NO |
| 18978 | TRAV10 | TRAJ8 | CVVTLVGFQKLVF | TRBV7-3 | TRBJ2-3 | CASSSGLARTDTQYF | KLGGALQAK | YES | NO | NO |
| 18979 | TRAV21 | TRAJ10 | CAVEGLTGGGNKLTF | TRBV20-1 | TRBJ2-7 | CSARDHTERGGIYEQYF | KLGGALQAK | NO | NO | NO |
| 18980 | TRAV5 | TRAJ17 | CAESMEEAAGNKLTF | TRBV19 | TRBJ2-7 | CASMGQSYEQYF | KLGGALQAK | YES | NO | NO |
| 18981 | TRAV12-1 | TRAJ9 | CVVRNTGGFKTIF | TRBV5-1 | TRBJ1-1 | CASSLLGQAVMNTEAFF | KLGGALQAK | YES | NO | NO |
| 18982 | TRAV8-2 | TRAJ22 | CVGRKGSARQLTF | TRBV6-3 | TRBJ2-3 | CASIPAGGPGDTQYF | KLGGALQAK | YES | NO | NO |
| 18983 | TRAV25 | TRAJ13 | CAGVTLNSGGYQKVTF | TRBV6-5 | TRBJ1-1 | CASSYSMMRNTEAFF | KLGGALQAK | YES | NO | NO |
| 18984 | TRAV23DV6 | TRAJ45 | CAASTVYSGGGADGLTF | TRBV27 | TRBJ1-6 | CASSFYPAGGNSPLHF | KLGGALQAK | YES | NO | NO |
| 18985 | TRAV35 | TRAJ28 | CAGQPEYSGAGSYQLTF | TRBV5-1 | TRBJ2-3 | CASSRGQGADTQYF | KLGGALQAK | YES | NO | NO |
| 18986 | TRAV3 | TRAJ37 | CAVRDGGMGSNTGKLIF | TRBV29-1 | TRBJ1-2 | CSVDLEANYGYTF | KLGGALQAK | NO | NO | NO |
| 18987 | TRAV1-2 | TRAJ30 | CAVDRDDKIIF | TRBV12-4 | TRBJ2-3 | CASSHREDDTDTQYF | KLGGALQAK | YES | NO | NO |
| 18988 | TRAV16 | TRAJ18 | CALSGAFGRGSTLGRLYF | TRBV27 | TRBJ1-5 | CASQDRGNGNQPQHF | KLGGALQAK | NO | NO | NO |
| 18989 | TRAV1-2 | TRAJ36 | CAPSGANNLFF | TRBV27 | TRBJ1-6 | CASSWDRPENSPLHF | KLGGALQAK | YES | NO | NO |
| 18990 | TRAV19 | TRAJ23 | CALSQPIYNQGGKLIF | TRBV12-4 | TRBJ2-1 | CASSLAGGNNEQFF | KLGGALQAK | YES | NO | NO |
| 18991 | TRAV8-4 | TRAJ52 | CAAFWGKLTF | TRBV14 | TRBJ2-1 | CASSRVPTGEQFF | KLGGALQAK | YES | NO | NO |
| 18992 | TRAV13-1 | TRAJ43 | CAAQNNDMRF | TRBV27 | TRBJ1-2 | CASSFGTGAGYTF | KLGGALQAK | YES | NO | NO |
| 18993 | TRAV12-2 | TRAJ29 | CAVASGNTPLVF | TRBV2 | TRBJ1-1 | CASSEVEGGTEAFF | KLGGALQAK | YES | NO | NO |
| 18994 | TRAV12-1 | TRAJ6 | CVVNPHGSYIPTF | TRBV19 | TRBJ1-1 | CASNRDPLDTEAFF | KLGGALQAK | YES | NO | NO |
| 18995 | TRAV26-2 | TRAJ12 | CIPWMDSSYKLIF | TRBV27 | TRBJ2-7 | CASGGGGPYEQYF | AVFDRKSDAK | YES | NO | NO |

| | | | | | | | | | | |
|---|---|---|---|---|---|---|---|---|---|---|
| 18996 | TRAV29DV5 | TRAJ20 | CAASPSNDYKLSF | TRBV10-3 | TRBJ1-2 | CAISESTSPYGYTF | KLGGALQAK | YES | NO | NO |
| 18997 | TRAV26-2 | TRAJ57 | CILETQGGSEKLVF | TRBV7-9 | TRBJ2-5 | CASSPAPGQGEDSAQYF | KLGGALQAK | YES | NO | NO |
| 18998 | TRAV5 | TRAJ23 | CAEIGDQGGKLIF | TRBV6-5 | TRBJ1-1 | CASGITEAFF | KLGGALQAK | YES | NO | NO |
| 18999 | TRAV27 | TRAJ27 | CAGALWDTNAGKSTF | TRBV10-2 | TRBJ2-7 | CASSDGQGPNEQYF | KLGGALQAK | YES | NO | NO |
| 19000 | TRAV5 | TRAJ33 | CAEMADSNYQLIW | TRBV6-5 | TRBJ2-5 | CASSYGEAQETQYF | KLGGALQAK | NO | NO | NO |
| 19001 | TRAV21 | TRAJ12 | CAVKEEDSSYKLIF | TRBV9 | TRBJ2-1 | CASSPGPHEQFF | KLGGALQAK | YES | NO | NO |
| 19002 | TRAV3 | TRAJ4 | CAVRDLSGGYNKLIF | TRBV24-1 | TRBJ2-3 | CATRDGATDTQYF | KLGGALQAK | NO | NO | NO |
| 19003 | TRAV8-6 | TRAJ54 | CAVTSRGHQGAQKLVF | TRBV24-1 | TRBJ2-1 | CATSKKGWANEQFF | KLGGALQAK | YES | NO | NO |
| 19004 | TRAV12-1 | TRAJ37 | CVAHGSSNTGKLIF | TRBV29-1 | TRBJ1-2 | CSVDLEANYGYTF | KLGGALQAK | YES | NO | NO |
| 19005 | TRAV13-2 | TRAJ5 | CAEGAMDTGRRALTF | TRBV19 | TRBJ1-1 | CASSIFETEAFF | KLGGALQAK | YES | NO | NO |
| 19006 | TRAV1-2 | TRAJ31 | CAVRGDNARLMF | TRBV12-4 | TRBJ1-1 | CASSLQEAFF | KLGGALQAK | YES | NO | NO |
| 19007 | TRAV13-2 | TRAJ45 | CAENDSGGGADGLTF | TRBV11-2 | TRBJ2-5 | CASSLGVSGGYQETQYF | KLGGALQAK | YES | NO | NO |
| 19008 | TRAV1-1 | TRAJ22 | CAVKSSGSARQLTF | TRBV7-9 | TRBJ2-7 | CASSLPTQGGSYEQYF | KLGGALQAK | YES | NO | NO |
| 19009 | TRAV12-1 | TRAJ3 | CVAYSSASKIIF | TRBV20-1 | TRBJ2-2 | CSASWPGANTGELFF | KLGGALQAK | YES | NO | NO |
| 19010 | TRAV8-3 | TRAJ53 | CAVGGLSGGSNYKLTF | TRBV7-8 | TRBJ2-1 | CASSYNRDVGRNEQFF | AVFDRKSDAK | YES | NO | NO |
| 19011 | TRAV14DV4 | TRAJ28 | CAMREADPAIVGSYQLTF | TRBV5-1 | TRBJ2-3 | CASTQEAALAGAGTDTQYF | AVFDRKSDAK | YES | NO | YES |
| 19012 | TRAV14DV4 | TRAJ21 | CAMRDLYNFNKFYF | TRBV12-4 | TRBJ1-2 | CASRATGPDVYGYTF | KLGGALQAK | YES | NO | NO |
| 19013 | TRAV14DV4 | TRAJ11 | CAMREGSGYSTLTF | TRBV15 | TRBJ1-1 | CATSQPGVNTEAFF | KLGGALQAK | YES | NO | NO |
| 19014 | TRAV39 | TRAJ32 | CALGGATNKLIF | TRBV7-9 | TRBJ2-5 | CASRAMETQYF | KLGGALQAK | YES | NO | NO |
| 19015 | TRAV1-2 | TRAJ32 | CAVRDAGRFYGGATNKLIF | TRBV6-6 | TRBJ2-4 | CASSIGTELAKNIQYF | KLGGALQAK | NO | NO | NO |
| 19016 | TRAV1-2 | TRAJ18 | CAVAQRRGSTLGRLYF | TRBV20-1 | TRBJ2-7 | CSARPEPNTYEQYF | KLGGALQAK | NO | NO | NO |
| 19017 | TRAV12-3 | TRAJ31 | CAMSADARLMF | TRBV7-9 | TRBJ2-2 | CASRTPSGYTGELFF | KLGGALQAK | YES | NO | NO |
| 19018 | TRAV12-1 | TRAJ41 | CVVNKYVAGYALNF | TRBV7-2 | TRBJ2-3 | CASSLDLSFTDTQYF | KLGGALQAK | YES | NO | NO |
| 19019 | TRAV29DV5 | TRAJ26 | CAARGIGGQNFVF | TRBV7-3 | TRBJ1-1 | CASSVGTGLSNTEAFF | KLGGALQAK | NO | NO | NO |
| 19020 | TRAV12-1 | TRAJ50 | CVVNLGTSYDKVIF | TRBV27 | TRBJ2-1 | CASSLSWTGGSYNEQFF | KLGGALQAK | YES | NO | NO |
| 19021 | TRAV36DV7 | TRAJ48 | CAVSNFGNEKLTF | TRBV29-1 | TRBJ2-1 | CSVEGPRLEFSYNEQFF | KLGGALQAK | NO | NO | NO |
| 19022 | TRAV25 | TRAJ41 | CAPGGYALNF | TRBV24-1 | TRBJ2-1 | CATSGGRSGTSGSYNEQFF | KLGGALQAK | YES | NO | NO |
| 19023 | TRAV41 | TRAJ34 | CAVRGATWNTDKLIF | TRBV29-1 | TRBJ2-5 | CSVEDRGDYETQYF | KLGGALQAK | YES | NO | NO |
| 19024 | TRAV24 | TRAJ45 | CAFIYPGGGADGLTF | TRBV9 | TRBJ1-2 | CASRGQGGGYTF | KLGGALQAK | YES | NO | NO |
| 19025 | TRAV12-2 | TRAJ52 | CAVNGLNAGGTSYGKLTF | TRBV20-1 | TRBJ1-1 | CSARDQGLPWGEAFF | KLGGALQAK | YES | NO | NO |
| 19026 | TRAV19 | TRAJ27 | CALSEGNAGKSTF | TRBV7-9 | TRBJ2-3 | CASSLGRASNTDTQYF | KLGGALQAK | YES | NO | NO |
| 19027 | TRAV36DV7 | TRAJ50 | CAVPKTSYDKVIF | TRBV6-5 | TRBJ2-5 | CASSYSWSGTSLETQYF | KLGGALQAK | NO | NO | NO |
| 19028 | TRAV35 | TRAJ45 | CAGGLVSGGGADGLTF | TRBV30 | TRBJ1-2 | CAWSPGQGPDGYTF | KLGGALQAK | YES | NO | NO |
| 19029 | TRAV40 | TRAJ34 | CLLGEGNTDKLIF | TRBV19 | TRBJ1-1 | CASSIQGVNTEAFF | KLGGALQAK | NO | NO | NO |
| 19030 | TRAV25 | TRAJ58 | CAGIWETSGSRLTF | TRBV19 | TRBJ1-2 | CASSILGALSYGYTF | KLGGALQAK | YES | NO | NO |
| 19031 | TRAV29DV5 | TRAJ17 | CAASDGEAAGNKLTF | TRBV5-1 | TRBJ1-2 | CASSLGLTGLAYGYTF | KLGGALQAK | YES | NO | NO |
| 19032 | TRAV21 | TRAJ50 | CAVDLMKTSYDKVIF | TRBV6-5 | TRBJ2-3 | CASRTGLASTDTQYF | IVTDFSVIK | NO | NO | NO |
| 19033 | TRAV5 | TRAJ42 | CAENEGGGSQGNLIF | TRBV19 | TRBJ2-7 | CASSSRAGGEQYF | GILGFVFTL | YES | NO | YES |
| 19034 | TRAV8-3 | TRAJ21 | CAVGDNFNKFYF | TRBV28 | TRBJ2-2 | CASSLYSATGELFF | AVFDRKSDAK | YES | NO | YES |
| 19035 | TRAV27 | TRAJ42 | CAGGGSQGNLIF | TRBV19 | TRBJ2-7 | CASSIRSSYEQYF | GILGFVFTL | YES | NO | YES |
| 19036 | TRAV13-1 | TRAJ35 | CAARVRGFGNVLHC | TRBV28 | TRBJ2-2 | CASSLYSATGELFF | AVFDRKSDAK | NO | NO | NO |
| 19037 | TRAV38-1 | TRAJ52 | CAFMKGAGGTSYGKLTF | TRBV19 | TRBJ1-2 | CASSWGAYGYTF | GILGFVFTL | YES | NO | YES |
| 19038 | TRAV27 | TRAJ42 | CAGGGSQGNLIF | TRBV19 | TRBJ2-7 | CASSVRSSYEQYF | GILGFVFTL | YES | NO | YES |
| 19039 | TRAV23DV6 | TRAJ42 | CAADNYGGSQGNLIF | TRBV19 | TRBJ1-2 | CASSIYGGAGDGYTF | GILGFVFTL | YES | NO | YES |
| 19040 | TRAV21 | TRAJ39 | CAAINNAGNMLTF | TRBV19 | TRBJ2-1 | CASSISLAAEQFF | LLDFVRFMGV | YES | NO | NO |
| 19041 | TRAV35 | TRAJ49 | CAGHTGNQFYF | TRBV11-2 | TRBJ1-2 | CASSWGGGSHYGYTF | IVTDFSVIK | YES | NO | YES |
| 19042 | TRAV21 | TRAJ26 | CAVHGYGQNFVF | TRBV4-3 | TRBJ2-3 | CASSQEWLAVSTDTQYF | AVFDRKSDAK | YES | NO | NO |
| 19043 | TRAV8-2 | TRAJ12 | CVVSRMDSSYKLIF | TRBV19 | TRBJ2-3 | CASSMDRGSADTQYF | KLGGALQAK | YES | NO | NO |
| 19045 | TRAV1-2 | TRAJ6 | CAVRSSGGSYIPTF | TRBV15 | TRBJ1-5 | CATSTGDSNQPQHF | KLGGALQAK | YES | NO | NO |
| 19047 | TRAV19 | TRAJ17 | CALKLIKAAGNKLTF | TRBV30 | TRBJ1-6 | CAWTGTGKIGWDSPLHF | KLGGALQAK | YES | NO | NO |
| 19048 | TRAV21 | TRAJ26 | CAVHGYGQNFVF | TRBV4-3 | TRBJ2-3 | CASSQEWLAVSTDTQYF | KLGGALQAK | YES | NO | NO |
| 19049 | TRAV29DV5 | TRAJ42 | CAARGGGGSQGNLIF | TRBV19 | TRBJ2-7 | CASSIRASYEQYF | GILGFVFTL | YES | NO | YES |
| 19050 | TRAV27 | TRAJ37 | CAGAHGSSNTGKLIF | TRBV19 | TRBJ2-7 | CASSIRSAYEQYF | GILGFVFTL | YES | NO | NO |
| 19051 | TRAV38-1 | TRAJ52 | CAFMKDAGGTSYGKLTF | TRBV19 | TRBJ1-2 | CASSIGSYGYTF | GILGFVFTL | YES | NO | YES |
| 19052 | TRAV5 | TRAJ48 | CAALSPNFGNEKLTF | TRBV13 | TRBJ2-7 | CASSRTGTSYEQYF | LLDFVRFMGV | YES | NO | YES |
| 19053 | TRAV14DV4 | TRAJ49 | CAMTRSNTGNQFYF | TRBV14 | TRBJ2-3 | CASSGRHVDTKDTQYF | IVTDFSVIK | YES | NO | YES |
| 19055 | TRAV16 | TRAJ38 | CALDRAGNNRKLIW | TRBV21-1 | TRBJ1-5 | CASSYRVDSNQPQHF | GILGFVFTL | NO | NO | NO |
| 19057 | TRAV12-2 | TRAJ47 | CAVEVEYGNKLVF | TRBV29-1 | TRBJ2-1 | CSVPGLAGIGDEQFF | KLGGALQAK | YES | NO | NO |
| 19058 | TRAV13-1 | TRAJ20 | CAASGYDYKLSF | TRBV29-1 | TRBJ2-7 | CSVSASGGDEQYF | KLGGALQAK | YES | NO | NO |
| 19059 | TRAV4 | TRAJ28 | CLVGDMRSGAGSYQLTF | TRBV20-1 | TRBJ2-7 | CSARDINRGSYEQYF | KLGGALQAK | NO | NO | NO |
| 19060 | TRAV1-2 | TRAJ20 | CAVRLPNDYKLSF | TRBV6-3 | TRBJ2-2 | CASSNPRLDATGELFF | KLGGALQAK | YES | NO | NO |
| 19062 | TRAV27 | TRAJ42 | CAGAGSQGNLIF | TRBV19 | TRBJ2-7 | CASSIRSSYEQYF | GILGFVFTL | YES | NO | YES |
| 19063 | TRAV9-2 | TRAJ33 | CAREHMDSNYQLIW | TRBV7-9 | TRBJ2-1 | CASSQLGRGDNEQFF | KLGGALQAK | YES | NO | NO |
| 19065 | TRAV13-1 | TRAJ37 | CAVVPGNTGKLIF | TRBV19 | TRBJ1-1 | CASSIRSGESRAFF | GILGFVFTL | YES | NO | YES |
| 19066 | TRAV38-1 | TRAJ52 | CAFMTNAGGTSYGKLTF | TRBV19 | TRBJ2-7 | CASSIRSSYEQYF | GILGFVFTL | YES | NO | YES |
| 19067 | TRAV14DV4 | TRAJ41 | CAMREGLNSGYALNF | TRBV19 | TRBJ2-1 | CASRQRDRPNNEQFF | IVTDFSVIK | YES | NO | YES |
| 19069 | TRAV27 | TRAJ17 | CAPHKAAGNKLTF | TRBV10-3 | TRBJ1-4 | CAAGGGTEKLFF | KLGGALQAK | YES | NO | NO |
| 19071 | TRAV27 | TRAJ37 | CAGASGSSNTGKLIF | TRBV19 | TRBJ2-7 | CASSIRSSYEQYF | GILGFVFTL | YES | NO | YES |
| 19072 | TRAV25 | TRAJ42 | CAGNYGGSQGNLIF | TRBV19 | TRBJ2-2 | CASSIRSTGELFF | GILGFVFTL | YES | NO | YES |
| 19073 | TRAV19 | TRAJ17 | CALKLIKAAGNKLTF | TRBV30 | TRBJ1-6 | CAWTGTGKIGWDSPLHF | IVTDFSVIK | YES | NO | NO |
| 19074 | TRAV27 | TRAJ42 | CAGASGGGSQGNLIF | TRBV19 | TRBJ2-1 | CASSIRSGYEQFF | GILGFVFTL | YES | NO | YES |
| 19075 | TRAV20 | TRAJ32 | CAVFFEGGATNKLIF | TRBV19 | TRBJ1-2 | CASSQGSWGYTF | GILGFVFTL | YES | NO | YES |
| 19076 | TRAV35 | TRAJ43 | CAGLYNNNDMRF | TRBV5-1 | TRBJ1-2 | CASSLGQGVFYTF | IVTDFSVIK | YES | NO | YES |
| 19077 | TRAV5 | TRAJ37 | CAEGGNTGKLIF | TRBV5-1 | TRBJ2-3 | CASSPGGLAGADTQYF | KLGGALQAK | YES | NO | NO |
| 19078 | TRAV38-2DV8 | TRAJ52 | CAYMSNAGGTSYGKLTF | TRBV19 | TRBJ1-2 | CASSFGSYGYTF | GILGFVFTL | YES | YES | YES |
| 19079 | TRAV3 | TRAJ28 | CAVRDMSGARGAGSYQLTF | TRBV30 | TRBJ2-3 | CAWVLGPAGDTQYF | KLGGALQAK | NO | NO | NO |
| 19081 | TRAV8-4 | TRAJ48 | CAVSDQISGNEKLTF | TRBV19 | TRBJ1-5 | CASSIHKNQPQHF | GILGFVFTL | YES | NO | YES |
| 19082 | TRAV8-3 | TRAJ21 | CAVGDNFNKFYF | TRBV28 | TRBJ1-1 | CASSLRSAAGWPFF | AVFDRKSDAK | YES | NO | NO |
| 19083 | TRAV38-2DV8 | TRAJ29 | CAYRSVGTPLVF | TRBV11-3 | TRBJ2-3 | CASSLGAGKVTDTQYF | KLGGALQAK | YES | NO | NO |
| 19084 | TRAV27 | TRAJ37 | CAGAIGSSNTGKLIF | TRBV19 | TRBJ2-7 | CASSIRSSYEQYF | GILGFVFTL | YES | NO | YES |
| 19085 | TRAV12-2 | TRAJ12 | CASVGMDSSYKLIF | TRBV24-1 | TRBJ2-1 | CATSDFRDSSNEQFF | AVFDRKSDAK | YES | NO | NO |
| 19086 | TRAV13-2 | TRAJ42 | CAENIGGGSQGNLIF | TRBV19 | TRBJ1-1 | CASSIRSGTEAFF | GILGFVFTL | YES | NO | YES |
| 19087 | TRAV21 | TRAJ7 | CAARPSF | TRBV19 | TRBJ2-7 | CASSIRSSYEQYF | GILGFVFTL | YES | NO | NO |
| 19088 | TRAV23DV6 | TRAJ43 | CAASGNHDMRF | TRBV6-6 | TRBJ1-1 | CASSTTAGDTEAFF | KLGGALQAK | NO | NO | NO |
| 19089 | TRAV12-3 | TRAJ42 | CAMSGDGGSQGNLIF | TRBV19 | TRBJ2-5 | CASSIRAGETQYF | GILGFVFTL | YES | NO | YES |
| 19090 | TRAV12-2 | TRAJ27 | CAVGDTNAGKSTF | TRBV27 | TRBJ2-3 | CASTRLAGAGTDTQYF | AVFDRKSDAK | YES | NO | NO |
| 19092 | TRAV12-2 | TRAJ42 | CAVQGSQGNLIF | TRBV19 | TRBJ2-7 | CASSIRSSYEQYF | GILGFVFTL | YES | NO | YES |
| 19093 | TRAV4 | TRAJ5 | CLSPMDTGRRALTF | TRBV10-3 | TRBJ1-5 | CATSTGDSNQPQHF | KLGGALQAK | YES | NO | NO |
| 19094 | TRAV23DV6 | TRAJ43 | CAASGNHDMRF | TRBV6-6 | TRBJ1-1 | CASSTTAGDTEAFF | AVFDRKSDAK | NO | NO | NO |
| 19095 | TRAV27 | TRAJ40 | CAGARLRGTYKYIF | TRBV12-4 | TRBJ1-4 | CASTLTGATNEKLFF | KLGGALQAK | YES | NO | NO |
| 19096 | TRAV13-2 | TRAJ45 | CADSGGGADGLTF | TRBV5-6 | TRBJ1-1 | CASSLRDGSEAFF | RAKFKQLL | YES | NO | NO |
| 19097 | TRAV10 | TRAJ16 | CVVSANSDGQKLLF | TRBV9 | TRBJ2-6 | CASSVVSGANVLTF | KLGGALQAK | NO | NO | NO |
| 19098 | TRAV8-3 | TRAJ7 | CAVFLYGNNRLAF | TRBV29-1 | TRBJ2-7 | CSVSASGGDEQYF | KLGGALQAK | YES | NO | NO |
| 19102 | TRAV16 | TRAJ26 | CALTGNYGQNFVF | TRBV19 | TRBJ2-7 | CASSIRSSYEQYF | GILGFVFTL | NO | NO | NO |
| 19103 | TRAV5 | TRAJ37 | CAESTGKLIF | TRBV29-1 | TRBJ1-4 | CSVGTGGTNEKLFF | GLCTLVAML | YES | YES | YES |
| 19105 | TRAV12-2 | TRAJ45 | CAVAVGVSGGGADGLTF | TRBV19 | TRBJ1-2 | CASSQGAYGYTF | GILGFVFTL | YES | NO | YES |
| 19106 | TRAV23DV6 | TRAJ33 | CAASMDSNYQLIW | TRBV10-3 | TRBJ2-7 | CASSESISYEQYF | KLGGALQAK | YES | NO | NO |
| 19108 | TRAV27 | TRAJ42 | CAAGGSQGNLIF | TRBV19 | TRBJ2-1 | CASSIRSSYEQFF | GILGFVFTL | YES | NO | YES |
| 19110 | TRAV38-1 | TRAJ52 | CAFMIGAGGTSYGKLTF | TRBV19 | TRBJ1-2 | CASSTGSYGYTF | GILGFVFTL | YES | NO | NO |
| 19111 | TRAV19 | TRAJ11 | CALSDNEDSTLTF | TRBV19 | TRBJ2-1 | CASSPLAVTYNEQFF | GILGFVFTL | YES | NO | YES |
| 19113 | TRAV21 | TRAJ26 | CAVHGYGQNFVF | TRBV4-3 | TRBJ2-3 | CASSQEWLAVSTDTQYF | IVTDFSVIK | YES | NO | NO |
| 19115 | TRAV8-3 | TRAJ21 | CAVGDNFNKFYF | TRBV11-2 | TRBJ1-2 | CASSWGGGSHYGYTF | IVTDFSVIK | YES | NO | YES |
| 19118 | TRAV12-2 | TRAJ10 | CAPIGGGNKLTF | TRBV28 | TRBJ1-2 | CASSLPTANYGYTF | KLGGALQAK | YES | NO | NO |
| 19119 | TRAV17 | TRAJ53 | CATDGSSGGSNYKLTF | TRBV19 | TRBJ1-2 | CASSIGLYGYTF | GILGFVFTL | YES | YES | YES |

| | | | | | | | | | | |
|---|---|---|---|---|---|---|---|---|---|---|
| 19122 | TRAV19 | TRAJ32 | CALSEANYGGATNKLIF | TRBV7-9 | TRBJ2-3 | CASSPGPAKDTQYF | KLGGALQAK | YES | NO | NO |
| 19123 | TRAV8-3 | TRAJ21 | CAVGDNFNKFYF | TRBV12-3 | TRBJ2-2 | CASSFSGNTGELFF | AVFDRKSDAK | YES | NO | YES |
| 19125 | TRAV27 | TRAJ37 | CAGAGNTGKLIF | TRBV19 | TRBJ2-7 | CASSIRSSYEQYF | GILGFVFTL | YES | NO | YES |
| 19126 | TRAV3 | TRAJ40 | CAIFTSGTYKYIF | TRBV6-4 | TRBJ2-3 | CASSPGTGTIDTQYF | AVFDRKSDAK | NO | NO | NO |
| 19127 | TRAV13-1 | TRAJ50 | CAASETSYDKVIF | TRBV12-3 | TRBJ2-2 | CASSFSGNTGELFF | RAKFKQLL | YES | NO | YES |
| 19128 | TRAV25 | TRAJ42 | CAGSSGGSQGNLIF | TRBV19 | TRBJ2-3 | CASSPRSTDTQYF | GILGFVFTL | YES | NO | YES |
| 19129 | TRAV3 | TRAJ6 | CAVSAASGGSYIPTF | TRBV7-2 | TRBJ2-7 | CASSPRDRERGEQYF | KLGGALQAK | NO | NO | NO |
| 19132 | TRAV39 | TRAJ42 | CAVDEGGGSQGNLIF | TRBV19 | TRBJ2-3 | CASSARAGITQYF | GILGFVFTL | YES | YES | YES |
| 19133 | TRAV26-2 | TRAJ50 | CILRETSYDKVIF | TRBV12-4 | TRBJ2-1 | CASGRLSYNEQFF | RAKFKQLL | YES | NO | YES |
| 19134 | TRAV30 | TRAJ20 | CGTEYPNDYKLSF | TRBV9 | TRBJ2-2 | CASSSGLVSNTGELFF | GLCTLVAML | YES | NO | NO |
| 19135 | TRAV22 | TRAJ37 | CAGSPSNTGKLIF | TRBV19 | TRBJ2-7 | CASSARSSYEQYF | GILGFVFTL | YES | NO | NO |
| 19136 | TRAV13-1 | TRAJ42 | CAASMGGGSQGNLIF | TRBV19 | TRBJ2-3 | CASSIRSTDTQYF | GILGFVFTL | YES | NO | YES |
| 19137 | TRAV27 | TRAJ42 | CAGGGSQGNLIF | TRBV21-1 | TRBJ2-4 | CASSMGDRSMGNIQYF | GILGFVFTL | YES | NO | YES |
| 19138 | TRAV13-1 | TRAJ50 | CAATETSYDKVIF | TRBV29-1 | TRBJ1-1 | CSAGGNTEAFF | RAKFKQLL | YES | NO | YES |
| 19139 | TRAV9-2 | TRAJ15 | CALNQAGTALIF | TRBV9 | TRBJ2-3 | CASSANWRGSTDTQYF | KLGGALQAK | YES | NO | NO |
| 19140 | TRAV12-1 | TRAJ50 | CVVTETSYDKVIF | TRBV12-3 | TRBJ1-1 | CASSFLDRGPNTEAFF | RAKFKQLL | YES | YES | NO |
| 19141 | TRAV3 | TRAJ45 | CAVAKDLQNSGGGADGLTF | TRBV19 | TRBJ2-7 | CASSARSSTEQYF | GILGFVFTL | NO | NO | NO |
| 19142 | TRAV13-2 | TRAJ53 | CAETHSGGSNYKLTF | TRBV29-1 | TRBJ2-1 | CSVGSGDHGEQFF | RAKFKQLL | YES | NO | NO |
| 19143 | TRAV5 | TRAJ6 | CAEMEGGSYIPTF | TRBV4-2 | TRBJ2-2 | CASSQRPSEVGELFF | RAKFKQLL | YES | NO | YES |
| 19144 | TRAV29DV5 | TRAJ37 | CAASNTGKLIF | TRBV27 | TRBJ2-2 | CASSPHFQVDTGELFF | RAKFKQLL | YES | NO | YES |
| 19145 | TRAV13-1 | TRAJ34 | CAASEGAPDKLIF | TRBV12-4 | TRBJ2-5 | CASSLLGVSAQETQYF | RAKFKQLL | YES | NO | YES |
| 19146 | TRAV13-1 | TRAJ42 | CAYGGSQGNLIF | TRBV19 | TRBJ2-5 | CASSFRSSETQYF | GILGFVFTL | YES | NO | YES |
| 19147 | TRAV27 | TRAJ42 | CAGAGSQGNLIF | TRBV19 | TRBJ2-7 | CASSTRSAYEQYF | GILGFVFTL | YES | NO | YES |
| 19148 | TRAV13-2 | TRAJ45 | CADSGGGADGLTF | TRBV9 | TRBJ1-1 | CASSVRSSMNTEAFF | RAKFKQLL | YES | NO | YES |
| 19149 | TRAV5 | TRAJ34 | CAESEGPTYTDKLIF | TRBV5-5 | TRBJ2-7 | CASSGPGGVDEQYF | RAKFKQLL | YES | NO | NO |
| 19150 | TRAV38-1 | TRAJ52 | CAFMKGAGGTSYGKLTF | TRBV19 | TRBJ1-2 | CASSIGVHGYTF | GILGFVFTL | YES | NO | NO |
| 19151 | TRAV20 | TRAJ42 | CAVQAEGGGSQGNLIF | TRBV19 | TRBJ2-1 | CASSIALASEQFF | LLDFVRFMGV | YES | NO | YES |
| 19152 | TRAV12-1 | TRAJ31 | CVVNMDNARLMF | TRBV29-1 | TRBJ2-7 | CSAGQGPYEQYF | RAKFKQLL | NO | NO | NO |
| 19153 | TRAV13-1 | TRAJ50 | CAASETSYDKVIF | TRBV7-9 | TRBJ1-4 | CASSLVGGTDEKLFF | RAKFKQLL | YES | NO | NO |
| 19154 | TRAV13-1 | TRAJ50 | CAAAETSYDKVIF | TRBV3-1 | TRBJ2-2 | CASSYSGGAGELFF | RAKFKQLL | YES | YES | YES |
| 19155 | TRAV12-3 | TRAJ9 | CARGGGFKTIF | TRBV7-9 | TRBJ2-6 | CASSPVTGGGSGANVLTF | KLGGALQAK | YES | NO | NO |
| 19156 | TRAV13-1 | TRAJ50 | CAAMETSYDKVIF | TRBV15 | TRBJ1-4 | CATSRSGQGEKLFF | RAKFKQLL | YES | NO | YES |
| 19157 | TRAV9-2 | TRAJ34 | CALSEAGPDKLIF | TRBV12-4 | TRBJ2-7 | CASSPSSSYEQYF | RAKFKQLL | YES | NO | YES |
| 19158 | TRAV26-1 | TRAJ17 | CILWVAAGNKLTF | TRBV9 | TRBJ1-1 | CASSVRSSMNTEAFF | KLGGALQAK | YES | NO | NO |
| 19159 | TRAV13-1 | TRAJ50 | CAASETSYDKVIF | TRBV12-3 | TRBJ1-3 | CASSRTDGNTIYF | RAKFKQLL | YES | NO | YES |
| 19160 | TRAV13-1 | TRAJ57 | CAAKEGGSEKLVF | TRBV27 | TRBJ2-7 | CASSLSGRPYEQYF | RAKFKQLL | YES | NO | NO |
| 19161 | TRAV13-1 | TRAJ50 | CAASETSYDKVIF | TRBV7-9 | TRBJ2-3 | CASSSRDGGTDTQYF | RAKFKQLL | YES | NO | YES |
| 19162 | TRAV13-1 | TRAJ50 | CAASETSYDKVIF | TRBV18 | TRBJ2-2 | CASSPPGGGNTGELFF | RAKFKQLL | YES | NO | YES |
| 19163 | TRAV13-1 | TRAJ11 | CAAKEGYSTLTF | TRBV5-4 | TRBJ1-5 | CASSSTGGGEKDQPQHF | KLGGALQAK | YES | NO | NO |
| 19164 | TRAV26-2 | TRAJ43 | CILREFYNDMRF | TRBV7-3 | TRBJ2-2 | CASSLGTSGGTGELFF | RAKFKQLL | NO | NO | NO |
| 19165 | TRAV13-1 | TRAJ12 | CAAARMDSSYKLIF | TRBV29-1 | TRBJ2-3 | CSVGSGEDSPQYF | RAKFKQLL | YES | NO | NO |
| 19166 | TRAV27 | TRAJ42 | CAGGGSQGNLIF | TRBV19 | TRBJ2-7 | CASSTRSSYEQYF | GILGFVFTL | YES | NO | YES |
| 19167 | TRAV38-2DV8 | TRAJ40 | CAYRSLTSGTYKYIF | TRBV13 | TRBJ2-5 | CASSPRGGVEKTQYF | KLGGALQAK | YES | NO | NO |
| 19168 | TRAV29DV5 | TRAJ31 | CAADDGRLMF | TRBV6-6 | TRBJ2-2 | CASSYGFPTSGGNTGELFF | RAKFKQLL | YES | NO | NO |
| 19169 | TRAV24 | TRAJ36 | CAFLGGANNLFF | TRBV5-1 | TRBJ2-5 | CASSLEWGGETQYF | KLGGALQAK | YES | NO | NO |
| 19170 | TRAV38-2DV8 | TRAJ52 | CAYSPAAGGTSYGKLTF | TRBV19 | TRBJ1-2 | CASSIGLYGYTF | GILGFVFTL | YES | NO | NO |
| 19171 | TRAV5 | TRAJ33 | CAETQDSNYQLIW | TRBV4-1 | TRBJ1-2 | CASSTSTGTGYGYTF | RAKFKQLL | YES | YES | YES |
| 19172 | TRAV17 | TRAJ53 | CATDASGGSNYKLTF | TRBV29-1 | TRBJ2-7 | CSVGTGDWGEQYF | RAKFKQLL | YES | NO | NO |
| 19173 | TRAV26-2 | TRAJ50 | CILMETSYDKVIF | TRBV12-3 | TRBJ2-2 | CASSFSGGTGELFF | RAKFKQLL | YES | NO | YES |
| 19174 | TRAV27 | TRAJ42 | CAGAQGGGSQGNLIF | TRBV19 | TRBJ2-7 | CASSIRAAYEQYF | GILGFVFTL | YES | NO | YES |
| 19175 | TRAV2 | TRAJ40 | CGRQMLLFTTVLWRTGTSGTYKYIF | TRBV29-1 | TRBJ2-3 | CSVGSGEDSPQYF | RAKFKQLL | NO | NO | NO |
| 19177 | TRAV1-2 | TRAJ29 | CAVRDYGGNTPLVF | TRBV9 | TRBJ2-7 | CASSVDSGSDYEQYF | KLGGALQAK | YES | NO | NO |
| 19178 | TRAV8-3 | TRAJ31 | CAVGTARLMF | TRBV27 | TRBJ2-2 | CASSPHFQVDTGELFF | RAKFKQLL | NO | NO | NO |
| 19179 | TRAV10 | TRAJ27 | CVGYTNAGKSTF | TRBV5-5 | TRBJ2-7 | CASSGPGGVDEQYF | RAKFKQLL | YES | NO | YES |
| 19180 | TRAV12-2 | TRAJ42 | CADESGGSQGNLIF | TRBV19 | TRBJ1-4 | CASSSQGANEKLFF | RAKFKQLL | YES | NO | NO |
| 19181 | TRAV26-2 | TRAJ52 | CILRDLNAGGTSYGKLTF | TRBV6-1 | TRBJ2-7 | CASSEVVSTTYEQYF | KLGGALQAK | YES | NO | NO |
| 19182 | TRAV22 | TRAJ53 | CAVATMGSGGSNYKLTF | TRBV19 | TRBJ2-7 | CASSSFLLSEQYF | GILGFVFTL | YES | NO | YES |
| 19183 | TRAV41 | TRAJ57 | CAVLGTQGGSEKLVF | TRBV4-1 | TRBJ2-1 | CASSLLWAEYNEQFF | RAKFKQLL | YES | NO | YES |
| 19184 | TRAV12-1 | TRAJ43 | CVVNDMRF | TRBV19 | TRBJ1-4 | CASSIFQNEKLFF | GILGFVFTL | NO | NO | NO |
| 19185 | TRAV8-3 | TRAJ42 | CALPGSQGNLIF | TRBV19 | TRBJ2-7 | CASSIRASYEQYF | GILGFVFTL | YES | NO | YES |
| 19186 | TRAV13-1 | TRAJ50 | CAAGETSYDKVIF | TRBV12-3 | TRBJ2-2 | CASIYSGNGELFF | RAKFKQLL | YES | NO | YES |
| 19187 | TRAV13-1 | TRAJ9 | CAASEGGFKTIF | TRBV11-2 | TRBJ2-7 | CASSLGTGENEQYF | RAKFKQLL | YES | NO | NO |
| 19188 | TRAV12-3 | TRAJ53 | CAMSGSGGSNYKLTF | TRBV29-1 | TRBJ2-7 | CSVGAGDYEQYF | RAKFKQLL | YES | NO | YES |
| 19189 | TRAV12-1 | TRAJ11 | CVVNREGYSTLTF | TRBV6-1 | TRBJ2-1 | CASSEGGFTFYNEQFF | GLCTLVAML | YES | YES | YES |
| 19190 | TRAV13-1 | TRAJ50 | CAAMETSYDKVIF | TRBV12-3 | TRBJ1-2 | CASSYTGGLRYGYTF | RAKFKQLL | YES | NO | NO |
| 19191 | TRAV27 | TRAJ42 | CAGAGGGSQGNLIF | TRBV19 | TRBJ2-3 | CASSMRSSDTQYF | GILGFVFTL | YES | NO | YES |
| 19192 | TRAV12-2 | TRAJ10 | CAVPLTGGGNKLTF | TRBV11-2 | TRBJ1-1 | CASSLNRGDPEAFF | GLCTLVAML | YES | YES | YES |
| 19193 | TRAV13-1 | TRAJ50 | CAASETSYDKVIF | TRBV11-2 | TRBJ2-3 | CASLISGTDTQYF | RAKFKQLL | YES | NO | YES |
| 19194 | TRAV1-1 | TRAJ15 | CAVRDSAGTALIF | TRBV3-1 | TRBJ2-7 | CASSLGTGGSYEQYF | KLGGALQAK | YES | NO | NO |
| 19195 | TRAV5 | TRAJ31 | CAEDSNARLMF | TRBV20-1 | TRBJ1-3 | CSARDRVGNTIYF | GLCTLVAML | YES | YES | YES |
| 19196 | TRAV13-2 | TRAJ29 | CAENKREAGNTPLVF | TRBV5-5 | TRBJ1-5 | CASRPGVGNQPQHF | KLGGALQAK | YES | NO | NO |
| 19197 | TRAV8-3 | TRAJ40 | CASRSGTYKYIF | TRBV9 | TRBJ1-2 | CASSVNPGQGANYGYTF | KLGGALQAK | YES | NO | NO |
| 19199 | TRAV27 | TRAJ37 | CAGGGSSNTGKLIF | TRBV19 | TRBJ2-7 | CASSIRSAYEQYF | GILGFVFTL | YES | NO | YES |
| 19200 | TRAV12-2 | TRAJ40 | CAASGGTYKYIF | TRBV9 | TRBJ1-1 | CASSSGRLTNTEAFF | GLCTLVAML | YES | NO | NO |
| 19201 | TRAV17 | TRAJ40 | CATASTSGTYKYIF | TRBV4-1 | TRBJ2-1 | CASSLLAGEYNEQFF | RAKFKQLL | YES | NO | YES |
| 19203 | TRAV26-2 | TRAJ30 | CIFVRNRDDKIIF | TRBV19 | TRBJ2-7 | CASSIRSSYEQYF | GILGFVFTL | YES | NO | NO |
| 19204 | TRAV12-3 | TRAJ27 | CAMKSTNAGKSTF | TRBV11-2 | TRBJ2-2 | CASSLGVAGTNTGELFF | KLGGALQAK | YES | NO | NO |
| 19205 | TRAV8-2 | TRAJ24 | CVVRPGGDSWGKLQF | TRBV5-6 | TRBJ1-1 | CASSFDSKAFF | KLGGALQAK | YES | NO | NO |
| 19206 | TRAV5 | TRAJ42 | CAENQGGGSQGNLIF | TRBV19 | TRBJ1-1 | CASSTRSGTEAFF | GILGFVFTL | YES | NO | YES |
| 19207 | TRAV12-3 | TRAJ9 | CARGGGFKTIF | TRBV7-9 | TRBJ2-6 | CASSPVTGGGSGANVLTF | AVFDRKSDAK | YES | NO | NO |
| 19209 | TRAV29DV5 | TRAJ43 | CAASVYRGYNNNDMRF | TRBV19 | TRBJ1-5 | CASSPRSNQPQHF | GILGFVFTL | YES | NO | YES |
| 19211 | TRAV13-1 | TRAJ50 | CAASETSYDKVIF | TRBV13 | TRBJ1-6 | CASSLMRGGTYNSPLHF | RAKFKQLL | YES | NO | YES |
| 19212 | TRAV19 | TRAJ49 | CALRAGNQFYF | TRBV13 | TRBJ1-6 | CASSLMRGGTYNSPLHF | KLGGALQAK | YES | NO | NO |
| 19215 | TRAV38-1 | TRAJ52 | CAFMKGAGGTSYGKLTF | TRBV19 | TRBJ1-2 | CASSIGSHGYTF | GILGFVFTL | YES | NO | YES |
| 19216 | TRAV5 | TRAJ8 | CAEDLQKLVF | TRBV20-1 | TRBJ1-3 | CSARALPVGNTIYF | GLCTLVAML | YES | YES | YES |
| 19217 | TRAV2 | TRAJ24 | CAVEGPTDSWGKFQF | TRBV6-6 | TRBJ2-3 | CASSPAGGTDTQYF | KLGGALQAK | NO | NO | NO |
| 19218 | TRAV26-1 | TRAJ8 | CIVMDDFQKLVF | TRBV11-3 | TRBJ2-1 | CASSSHGAGAYNEQFF | RAKFKQLL | YES | NO | YES |
| 19219 | TRAV1-2 | TRAJ39 | CAVRGYNAGNMLTF | TRBV7-9 | TRBJ2-1 | CASSSGLAGVNEQFF | KLGGALQAK | YES | NO | NO |
| 19220 | TRAV21 | TRAJ47 | CAVRRDGNKLVF | TRBV19 | TRBJ2-5 | CASSMRAGETQYF | GILGFVFTL | YES | NO | NO |
| 19221 | TRAV12-2 | TRAJ42 | CALSPGGGSQGNLIF | TRBV21-1 | TRBJ2-1 | CASSRGGLSYNEQFF | GILGFVFTL | YES | NO | YES |
| 19222 | TRAV19 | TRAJ17 | CALSKGAAGNKLTF | TRBV9 | TRBJ1-2 | CASSADPSQGDGNYGYTF | KLGGALQAK | YES | NO | NO |
| 19223 | TRAV5 | TRAJ42 | CAEEGGGGSQGNLIF | TRBV19 | TRBJ1-1 | CASSTRSGTEAFF | GILGFVFTL | YES | NO | YES |
| 19224 | TRAV21 | TRAJ4 | CAVSLSGGYNKLIF | TRBV6-6 | TRBJ2-3 | CASSPAGGTDTQYF | KLGGALQAK | NO | NO | NO |
| 19226 | TRAV27 | TRAJ42 | CAVSGSQGNLIF | TRBV19 | TRBJ2-7 | CASSIRSSYEQYF | GILGFVFTL | YES | NO | YES |
| 19227 | TRAV12-1 | TRAJ39 | CGAAGNMLTF | TRBV11-1 | TRBJ1-1 | CASSPEDRVGDNTEAFF | KLGGALQAK | YES | NO | NO |
| 19228 | TRAV13-1 | TRAJ50 | CAGAETSYDKVIF | TRBV2 | TRBJ2-2 | CASSTPRQGSNTGELFF | RAKFKQLL | YES | NO | YES |
| 19229 | TRAV16 | TRAJ43 | CALKNNDMRF | TRBV19 | TRBJ2-3 | CASSWRSTDTQYF | GILGFVFTL | NO | NO | NO |
| 19230 | TRAV38-1 | TRAJ52 | CAFMKGAGGTSYGKLTF | TRBV19 | TRBJ1-2 | CASSIGSYGYTF | GILGFVFTL | YES | NO | NO |
| 19231 | TRAV29DV5 | TRAJ52 | CAASSGGTSYGKLTF | TRBV3-1 | TRBJ2-5 | CASSPLWVAGGRETQYF | IVTDFSVIK | YES | NO | NO |
| 19232 | TRAV12-1 | TRAJ39 | CGAAGNMLTF | TRBV6-3 | TRBJ2-7 | CASGMTGLTSEQYF | AVFDRKSDAK | YES | NO | NO |
| 19234 | TRAV13-1 | TRAJ50 | CAASETSYDKVIF | TRBV12-3 | TRBJ1-2 | CASSIGDRAYGYTF | RAKFKQLL | YES | YES | YES |
| 19235 | TRAV27 | TRAJ42 | CAGAGGGGSQGNLIF | TRBV19 | TRBJ2-7 | CASSIRAAYEQYF | GILGFVFTL | YES | NO | YES |
| 19236 | TRAV8-3 | TRAJ50 | CAVSETSYDKVIF | TRBV18 | TRBJ1-4 | CASSPRDRGGSNEKLFF | RAKFKQLL | YES | NO | YES |
| 19237 | TRAV12-3 | TRAJ53 | CAMSNSGGSNYKLTF | TRBV29-1 | TRBJ2-7 | CSVGSGDYEQYF | RAKFKQLL | YES | NO | YES |

| | | | | | | | | | | |
|---|---|---|---|---|---|---|---|---|---|---|
| 19238 | TRAV38-2DV8 | TRAJ48 | CAYRSPFGGNEKLTF | TRBV20-1 | TRBJ2-3 | CSARARRGELTDTQYF | KLGGALQAK | NO | NO | NO |
| 19239 | TRAV5 | TRAJ40 | CAEPKGGTYKYIF | TRBV20-1 | TRBJ1-1 | CSAWTGTEAFF | GLCTLVAML | YES | YES | YES |
| 19240 | TRAV5 | TRAJ42 | CAEDSGNLIF | TRBV11-2 | TRBJ1-1 | CASSLNRGDPEAFF | GLCTLVAML | YES | NO | NO |
| 19241 | TRAV26-1 | TRAJ43 | CIVREWDNDMRF | TRBV7-8 | TRBJ2-6 | CASSPRTGNSGANVLTF | RAKFKQLL | YES | NO | YES |
| 19242 | TRAV13-2 | TRAJ42 | CAENIGGGSQGNLIF | TRBV19 | TRBJ2-3 | CASSGRAGDTQYF | GILGFVFTL | YES | NO | YES |
| 19243 | TRAV12-3 | TRAJ21 | CAMSENNFNKFYF | TRBV3-1 | TRBJ1-5 | CASSQASQPQHF | KLGGALQAK | YES | NO | NO |
| 19244 | TRAV17 | TRAJ11 | CATDAMSAGYSTLTF | TRBV19 | TRBJ2-7 | CASSIRSSYEQYF | GILGFVFTL | YES | NO | YES |
| 19245 | TRAV1-2 | TRAJ28 | CAGPPSGAGSYQLTF | TRBV20-1 | TRBJ2-5 | CSARDDSGQGFGFLETQYF | GLCTLVAML | YES | YES | YES |
| 19246 | TRAV38-1 | TRAJ52 | CAFMTNAGGTSYGKLTF | TRBV19 | TRBJ1-2 | CASSLGSYGYTF | GILGFVFTL | YES | NO | YES |
| 19247 | TRAV8-1 | TRAJ21 | CAVNEDNFNKFYF | TRBV4-1 | TRBJ2-1 | CASSLGGTNEQFF | RAKFKQLL | YES | NO | YES |
| 19248 | TRAV21 | TRAJ16 | CAVRRF | TRBV29-1 | TRBJ2-3 | CSGNPGTSAATQYF | KLGGALQAK | NO | NO | NO |
| 19249 | TRAV12-3 | TRAJ9 | CARGGGFKTIF | TRBV7-9 | TRBJ2-6 | CASSPVTGGGSGANVLTF | IVTDFSVIK | YES | NO | NO |
| 19250 | TRAV13-2 | TRAJ50 | CAEMETSYDKVIF | TRBV10-1 | TRBJ1-2 | CASSRPVAVIYGYTF | RAKFKQLL | YES | NO | NO |
| 19251 | TRAV14DV4 | TRAJ17 | CAMRAIKAAGNKLTF | TRBV20-1 | TRBJ1-2 | CSARVSPEGGPENYGYTF | KLGGALQAK | YES | NO | NO |
| 19252 | TRAV29DV5 | TRAJ49 | CAASASGGQFYF | TRBV12-4 | TRBJ1-2 | CASSPGTNYGYTF | KLGGALQAK | YES | NO | NO |
| 19253 | TRAV13-1 | TRAJ50 | CAASETSYDKVIF | TRBV7-9 | TRBJ2-1 | CASSQLGRGDNEQFF | RAKFKQLL | YES | NO | YES |
| 19254 | TRAV13-1 | TRAJ11 | CAAKEGYSTLTF | TRBV5-6 | TRBJ1-1 | CASSLRDGSEAFF | RAKFKQLL | YES | NO | NO |
| 19256 | TRAV26-1 | TRAJ43 | CIVRAYNNNDMRF | TRBV19 | TRBJ2-7 | CASSIRSSYEQYF | GILGFVFTL | YES | NO | YES |
| 19257 | TRAV38-1 | TRAJ53 | CAFMKHVSSGGSNYKLTF | TRBV24-1 | TRBJ2-5 | CATSDNRRGIFQETQYF | KLGGALQAK | YES | NO | NO |
| 19258 | TRAV23DV6 | TRAJ27 | CAALEETNAGKSTF | TRBV19 | TRBJ2-7 | CASSIRSSYEQYF | GILGFVFTL | YES | NO | YES |
| 19259 | TRAV12-3 | TRAJ9 | CARGGGFKTIF | TRBV7-9 | TRBJ2-6 | CASSPVTGGGSGANVLTF | GILGFVFTL | YES | NO | NO |
| 19260 | TRAV27 | TRAJ42 | CAGAGSQGNLIF | TRBV19 | TRBJ2-7 | CASSSRSSYEQYF | GILGFVFTL | YES | NO | NO |
| 19261 | TRAV12-1 | TRAJ39 | CGAAGNMLTF | TRBV6-3 | TRBJ2-7 | CASGMTGLTSEQYF | KLGGALQAK | YES | NO | NO |
| 19262 | TRAV12-2 | TRAJ4 | CAVNGAGGYNKLIF | TRBV27 | TRBJ2-3 | CASSFSSGGAISDTQYF | KLGGALQAK | YES | NO | NO |
| 19263 | TRAV21 | TRAJ36 | CAVIQGPKTGANNLFF | TRBV3-1 | TRBJ2-1 | CASSQGTSGFWQFF | KLGGALQAK | YES | NO | NO |
| 19264 | TRAV8-3 | TRAJ15 | CAVNQAGTALIF | TRBV5-6 | TRBJ1-4 | CASSPAEATNEKLFF | KLGGALQAK | YES | NO | NO |
| 19265 | TRAV1-2 | TRAJ33 | CAVLDSNYQLIW | TRBV20-1 | TRBJ2-7 | CSARGLADPYEQYF | KLGGALQAK | NO | NO | NO |
| 19267 | TRAV38-2DV8 | TRAJ43 | CAYRISLNNNDMRF | TRBV19 | TRBJ2-1 | CASSMYGGSANEQFF | GILGFVFTL | YES | NO | YES |
| 19269 | TRAV19 | TRAJ23 | CALSEGVGKLIF | TRBV9 | TRBJ2-3 | CASSVTSSRGGTDTQYF | KLGGALQAK | YES | NO | NO |
| 19270 | TRAV30 | TRAJ31 | CGTEWEARLMF | TRBV19 | TRBJ2-7 | CASSIRSSYEQYF | GILGFVFTL | YES | NO | NO |
| 19271 | TRAV12-3 | TRAJ8 | CAMSSSFQKLVF | TRBV6-3 | TRBJ2-3 | CASSYPPRTGYGTDTQYF | KLGGALQAK | YES | NO | NO |
| 19273 | TRAV38-2DV8 | TRAJ42 | CAYREGGSQGNLIF | TRBV7-9 | TRBJ2-1 | CASSLTQGIYNEQFF | RAKFKQLL | YES | NO | YES |
| 19274 | TRAV8-3 | TRAJ30 | CAVTRDDKIIF | TRBV28 | TRBJ1-5 | CASSTDRAAQPQHF | RAKFKQLL | YES | NO | YES |
| 19275 | TRAV14DV4 | TRAJ40 | CAMRDPSGTYKYIF | TRBV7-9 | TRBJ2-7 | CASSQDPSRTYEQYF | KLGGALQAK | NO | NO | NO |
| 19279 | TRAV27 | TRAJ42 | CAGSGGGSQGNLIF | TRBV19 | TRBJ2-3 | CASSIRSTDTQYF | GILGFVFTL | YES | NO | YES |
| 19280 | TRAV5 | TRAJ42 | CAEGGNGGSQGNLIF | TRBV19 | TRBJ1-1 | CASSGRSNTEAFF | GILGFVFTL | YES | NO | NO |
| 19281 | TRAV13-1 | TRAJ50 | CAASETSYDKVIF | TRBV19 | TRBJ2-7 | CASSIRSSYEQYF | RAKFKQLL | YES | NO | YES |
| 19282 | TRAV17 | TRAJ11 | CATDGRSTLTF | TRBV11-2 | TRBJ2-7 | CASSSGGPFYEQYF | KLGGALQAK | YES | NO | NO |
| 19284 | TRAV12-2 | TRAJ52 | CAVTGGTSYGKLTF | TRBV24-1 | TRBJ2-2 | CATSDPKTGLNTGELFF | KLGGALQAK | YES | NO | NO |
| 19286 | TRAV8-3 | TRAJ21 | CAVGDNFNKFYF | TRBV5-6 | TRBJ1-1 | CASSLRDGSEAFF | AVFDRKSDAK | YES | NO | YES |
| 19288 | TRAV13-1 | TRAJ11 | CAAKEGYSTLTF | TRBV5-4 | TRBJ1-5 | CASSSTGGGEKDQPQHF | RAKFKQLL | YES | NO | NO |
| 19289 | TRAV2 | TRAJ37 | CVPTGKLIF | TRBV20-1 | TRBJ1-2 | CSARDLDGSNYGYTF | KLGGALQAK | NO | NO | NO |
| 19291 | TRAV13-2 | TRAJ53 | CAETHSGGSNYKLTF | TRBV29-1 | TRBJ2-1 | CSVGSGDHGEQFF | GILGFVFTL | YES | NO | NO |
| 19292 | TRAV27 | TRAJ42 | CAGVGGGSQGNLIF | TRBV19 | TRBJ2-7 | CASSIRSSYEQYF | GILGFVFTL | YES | NO | YES |
| 19293 | TRAV3 | TRAJ23 | CAVRDRENQGGKLIF | TRBV7-9 | TRBJ2-7 | CASSTSEGGLFYEQYF | KLGGALQAK | NO | NO | NO |
| 19294 | TRAV26-1 | TRAJ18 | CIAPEPTLGRLYF | TRBV12-5 | TRBJ2-3 | CASGQELAGTDTQYF | KLGGALQAK | YES | NO | NO |
| 19295 | TRAV25 | TRAJ42 | CAGPIDGGSQGNLIF | TRBV19 | TRBJ1-5 | CASSVYSNQPQHF | GILGFVFTL | YES | NO | YES |
| 19296 | TRAV34 | TRAJ40 | CGADSASGTYKYIF | TRBV29-1 | TRBJ2-7 | CSVGSGEDNEQYF | RAKFKQLL | YES | NO | YES |
| 19297 | TRAV19 | TRAJ17 | CALTPGAAGNKLTF | TRBV9 | TRBJ2-6 | CASSVAVGTGSGANVLTF | AVFDRKSDAK | YES | NO | YES |
| 19298 | TRAV13-1 | TRAJ50 | CAASETSYDKVIF | TRBV29-1 | TRBJ2-7 | CSAGQGPYEQYF | RAKFKQLL | NO | NO | NO |
| 19299 | TRAV8-3 | TRAJ21 | CAVGDNFNKFYF | TRBV19 | TRBJ2-7 | CASSIRSSYEQYF | GILGFVFTL | YES | NO | YES |
| 19300 | TRAV38-1 | TRAJ52 | CAFMISAGGTSYGKLTF | TRBV19 | TRBJ1-2 | CASSIGSYGYTF | GILGFVFTL | YES | NO | NO |
| 19302 | TRAV35 | TRAJ28 | CAGQLILWGWELPTHF | TRBV2 | TRBJ2-2 | CASSTPRQGSNTGELFF | RAKFKQLL | NO | NO | NO |
| 19303 | TRAV16 | TRAJ29 | CALKIPLVF | TRBV19 | TRBJ2-7 | CASSIRSAYEQYF | GILGFVFTL | NO | NO | NO |
| 19304 | TRAV13-1 | TRAJ50 | CAASETSYDKVIF | TRBV4-1 | TRBJ1-2 | CASSTSTGTGYGYTF | RAKFKQLL | YES | NO | YES |
| 19305 | TRAV3 | TRAJ45 | CAVAKDLQNSGGGADGLTF | TRBV19 | TRBJ2-7 | CASSARSSTEQYF | IVTDFSVIK | NO | NO | NO |
| 19306 | TRAV13-1 | TRAJ21 | CAAFDFNKFYF | TRBV7-3 | TRBJ2-5 | CASSSRGGQETQYF | RAKFKQLL | YES | NO | YES |
| 19308 | TRAV6 | TRAJ16 | CALASRSDGQKLLF | TRBV20-1 | TRBJ1-4 | CSARDYRTGDEKLFF | AVFDRKSDAK | NO | NO | NO |
| 19309 | TRAV13-1 | TRAJ50 | CAAAETSYDKVIF | TRBV7-9 | TRBJ2-5 | CASSFVDRAETQYF | RAKFKQLL | YES | NO | NO |
| 19310 | TRAV17 | TRAJ42 | CATDGGGGSQGNLIF | TRBV19 | TRBJ2-3 | CASSGRSTDTQYF | GILGFVFTL | YES | NO | NO |
| 19313 | TRAV13-1 | TRAJ11 | CAAKEGYSTLTF | TRBV5-4 | TRBJ1-5 | CASSSTGGGEKDQPQHF | GILGFVFTL | YES | NO | NO |
| 19314 | TRAV13-1 | TRAJ50 | CAASETSYDKVIF | TRBV5-6 | TRBJ1-1 | CASSLRDGSEAFF | RAKFKQLL | YES | NO | YES |
| 19315 | TRAV13-2 | TRAJ45 | CADSGGGADGLTF | TRBV29-1 | TRBJ2-7 | CSAGQGPYEQYF | RAKFKQLL | NO | NO | NO |
| 19317 | TRAV10 | TRAJ50 | CVVSAGLGKTSYDKVIF | TRBV19 | TRBJ2-3 | CASSIFYSSTDTQYF | GILGFVFTL | YES | NO | YES |
| 19318 | TRAV26-2 | TRAJ43 | CILSESNNNDMRF | TRBV4-1 | TRBJ2-1 | CASSQVTSGAQFF | RAKFKQLL | YES | NO | NO |
| 19319 | TRAV13-1 | TRAJ53 | CAAFNSGGSNYKLTF | TRBV29-1 | TRBJ1-1 | CSVGQGMNTEAFF | RAKFKQLL | YES | NO | YES |
| 19320 | TRAV13-1 | TRAJ11 | CAASRGYSTLTF | TRBV5-4 | TRBJ2-1 | CASSLGGGNNEQFF | KLGGALQAK | YES | NO | NO |
| 19321 | TRAV29DV5 | TRAJ41 | CAASGTSNSGYALNF | TRBV19 | TRBJ1-5 | CASSTGQPQHF | GILGFVFTL | YES | NO | YES |
| 19322 | TRAV8-3 | TRAJ28 | CAVPEAGSYQLTF | TRBV9 | TRBJ1-2 | CASSVGGDYGYTF | KLGGALQAK | YES | NO | NO |
| 19323 | TRAV30 | TRAJ28 | CGTSGAGSYQLTF | TRBV19 | TRBJ2-3 | CASSQRSTDTQYF | GILGFVFTL | YES | YES | YES |
| 19324 | TRAV27 | TRAJ42 | CAGGGSQGNLIF | TRBV6-7 | TRBJ2-1 | CASSSSSRTSNNEQFF | GILGFVFTL | NO | NO | NO |
| 19325 | TRAV10 | TRAJ27 | CVGYTNAGKSTF | TRBV5-5 | TRBJ2-1 | CASSGPGGVDEQFF | RAKFKQLL | YES | NO | YES |
| 19326 | TRAV13-1 | TRAJ50 | CAASETSYDKVIF | TRBV20-1 | TRBJ2-7 | CSASPGDYEQYF | RAKFKQLL | YES | NO | YES |
| 19327 | TRAV19 | TRAJ26 | CALIRGGGQNFVF | TRBV5-4 | TRBJ1-2 | CASSLGTANYGYTF | KLGGALQAK | YES | NO | NO |
| 19328 | TRAV26-1 | TRAJ52 | CIVRVSGAGGTSYGKLTF | TRBV19 | TRBJ2-5 | CASSGRSGETQYF | GILGFVFTL | YES | NO | YES |
| 19329 | TRAV1-2 | TRAJ33 | CAVRDSNYQLIW | TRBV20-1 | TRBJ2-7 | CSARDGRREYEQYF | KLGGALQAK | NO | NO | NO |
| 19331 | TRAV29DV5 | TRAJ38 | CAASRNAGNNRKLIW | TRBV10-1 | TRBJ2-7 | CASSRDTSSYEQYF | KLGGALQAK | YES | NO | NO |
| 19335 | TRAV5 | TRAJ43 | CAERASNNDMRF | TRBV7-2 | TRBJ2-2 | CASSPPGLVGGELFF | GLCTLVAML | NO | NO | NO |
| 19336 | TRAV12-2 | TRAJ42 | CAAPGGGGSQGNLIF | TRBV11-2 | TRBJ1-2 | CASSFPPRLYANYGYTF | GILGFVFTL | YES | NO | YES |
| 19337 | TRAV41 | TRAJ48 | CAVKENEKLTF | TRBV19 | TRBJ2-7 | CASSIRSAYEQYF | GILGFVFTL | YES | NO | YES |
| 19338 | TRAV13-1 | TRAJ11 | CAAKEGYSTLTF | TRBV5-4 | TRBJ1-5 | CASSSTGGGEKDQPQHF | IVTDFSVIK | YES | NO | NO |
| 19339 | TRAV12-3 | TRAJ39 | CAIGPGNMLTF | TRBV29-1 | TRBJ1-2 | CSVDLEANYGYTF | KLGGALQAK | YES | NO | NO |
| 19340 | TRAV35 | TRAJ42 | CAGQSGRVGGSQGNLIF | TRBV6-3 | TRBJ2-1 | CASSYGAGGYNEQFF | KLGGALQAK | YES | NO | NO |
| 19341 | TRAV21 | TRAJ21 | CAVRPGYNFNKFYF | TRBV10-2 | TRBJ2-1 | CASGDRGYNEQFF | KLGGALQAK | YES | NO | NO |
| 19342 | TRAV1-1 | TRAJ24 | CAPKTDSWGKLQF | TRBV27 | TRBJ2-3 | CATMEPGQGPDTQYF | KLGGALQAK | YES | NO | NO |
| 19343 | TRAV14DV4 | TRAJ5 | CAMRDPQGGRRALTF | TRBV7-9 | TRBJ2-1 | CASTGTSGGPTLRDEQFF | KLGGALQAK | YES | NO | NO |
| 19344 | TRAV24 | TRAJ52 | CAFIGDAGGTSYGKLTF | TRBV4-3 | TRBJ1-4 | CASSQDHRMGGHEKLFF | KLGGALQAK | YES | NO | NO |
| 19345 | TRAV8-2 | TRAJ22 | CVGRKGSARQLTF | TRBV20-1 | TRBJ2-7 | CSARDLEVLSYEQYF | KLGGALQAK | YES | NO | NO |
| 19346 | TRAV21 | TRAJ43 | CAVWYMRF | TRBV6-3 | TRBJ2-3 | CASIPAGGPGDTQYF | KLGGALQAK | NO | NO | NO |
| 19347 | TRAV3 | TRAJ37 | CAVRDGGMGSNTGKLIF | TRBV12-4 | TRBJ2-3 | CASSPMDTQYF | KLGGALQAK | NO | NO | NO |
| 19348 | TRAV19 | TRAJ29 | CALSEAFSGNTPLVF | TRBV19 | TRBJ1-1 | CASSMSGPGTEAFF | KLGGALQAK | YES | NO | NO |
| 19349 | TRAV8-4 | TRAJ20 | CAVIQINDYKLSF | TRBV7-8 | TRBJ1-2 | CASSLGQDGYTF | KLGGALQAK | YES | NO | NO |
| 19350 | TRAV5 | TRAJ3 | CAESGYSSASKIIF | TRBV3-1 | TRBJ1-1 | CASSQEGRETEAFF | KLGGALQAK | YES | NO | NO |
| 19351 | TRAV14DV4 | TRAJ4 | CAMRGPISGGYNKLIF | TRBV3-1 | TRBJ1-2 | CASSQGRDRHGYTF | KLGGALQAK | YES | NO | NO |
| 19352 | TRAV12-3 | TRAJ52 | CAIQPDYPLYAGGTSYGKLTF | TRBV9 | TRBJ1-1 | CASSVVGNTEAFF | KLGGALQAK | NO | NO | NO |
| 19353 | TRAV17 | TRAJ36 | CAQTGANNLFF | TRBV11-2 | TRBJ1-1 | CASSLFQGSGTEAFF | KLGGALQAK | YES | NO | NO |
| 19355 | TRAV21 | TRAJ36 | CAVYPGANNLFF | TRBV5-4 | TRBJ1-4 | CASSLDQQGVGEKLFF | KLGGALQAK | YES | NO | NO |
| 19356 | TRAV12-3 | TRAJ34 | CAMSRLEGDKLIF | TRBV4-3 | TRBJ2-3 | CASSQEGGFTDTQYF | KLGGALQAK | YES | NO | NO |
| 19357 | TRAV12-2 | TRAJ31 | CAVNRVRDARLMF | TRBV4-1 | TRBJ1-1 | CASRVGNTEAFF | KLGGALQAK | YES | NO | NO |
| 19358 | TRAV19 | TRAJ40 | CALSGAPSGTYKYIF | TRBV5-4 | TRBJ1-2 | CASSLPSSNYGYTF | KLGGALQAK | YES | NO | NO |
| 19359 | TRAV10 | TRAJ4 | CVVRSGGYNKLIF | TRBV7-9 | TRBJ2-1 | CASSNSLPSGVSYNEQFF | KLGGALQAK | YES | NO | NO |
| 19360 | TRAV27 | TRAJ44 | CAGADGTASKLTF | TRBV27 | TRBJ1-2 | CASSLSGTWGEGYTF | KLGGALQAK | YES | NO | NO |
| 19361 | TRAV12-3 | TRAJ39 | CAIGPGNMLTF | TRBV29-1 | TRBJ1-2 | CSVDLEANYGYTF | AVFDRKSDAK | YES | NO | NO |
| 19362 | TRAV17 | TRAJ53 | CATENSGGSNYKLTF | TRBV19 | TRBJ1-2 | CASSARTGVGYGYTF | KLGGALQAK | YES | NO | NO |

| | | | | | | | | | | |
|---|---|---|---|---|---|---|---|---|---|---|
| 19363 | TRAV17 | TRAJ40 | CATAPSGTYKYIF | TRBV9 | TRBJ1-1 | CASSVDTGFTEAFF | KLGGALQAK | YES | NO | NO |
| 19364 | TRAV21 | TRAJ45 | CAVLGGGADGLTF | TRBV29-1 | TRBJ1-1 | CSFRIGGTEAFF | KLGGALQAK | NO | NO | NO |
| 19365 | TRAV38-2DV8 | TRAJ52 | CAYRRGRNAGGTSYGKLTF | TRBV5-6 | TRBJ2-3 | CASSRDRGEDTQYF | KLGGALQAK | YES | NO | NO |
| 19366 | TRAV14DV4 | TRAJ37 | CAMRGPYSSNTGKLIF | TRBV30 | TRBJ1-2 | CAWGSGLGYGYTF | KLGGALQAK | NO | NO | NO |
| 19367 | TRAV29DV5 | TRAJ36 | CAASSSTGANNLFF | TRBV20-1 | TRBJ2-3 | CSATSNGERTDTQYF | KLGGALQAK | NO | NO | NO |
| 19368 | TRAV5 | TRAJ3 | CAESGYSSASKIIF | TRBV3-1 | TRBJ1-1 | CASSQEGRETEAFF | AVFDRKSDAK | YES | NO | NO |
| 19369 | TRAV19 | TRAJ5 | CALSEALGGRRALTF | TRBV6-5 | TRBJ1-5 | CASRSKTGSKQPQHF | KLGGALQAK | YES | NO | NO |
| 19370 | TRAV29DV5 | TRAJ33 | CAASAVSNYQLIW | TRBV12-3 | TRBJ2-3 | CASSPTGNWNTDTQYF | KLGGALQAK | YES | NO | NO |
| 19371 | TRAV19 | TRAJ45 | CALSEAMGFSGGGADGLTF | TRBV11-2 | TRBJ2-7 | CASSVGRLSAYEQYV | KLGGALQAK | NO | NO | NO |
| 19373 | TRAV26-1 | TRAJ54 | CIVRGADQGAQKLVF | TRBV7-9 | TRBJ2-7 | CASKDGISYEQYF | KLGGALQAK | YES | NO | NO |
| 19374 | TRAV19 | TRAJ22 | CALDFSSGSARQLTF | TRBV14 | TRBJ1-4 | CASSQGPGTVHEKLFF | KLGGALQAK | NO | NO | NO |
| 19375 | TRAV8-4 | TRAJ45 | CAVSYSGGGADGLTF | TRBV27 | TRBJ1-5 | CASSWTGGMWPQHF | KLGGALQAK | YES | NO | NO |
| 19376 | TRAV8-1 | TRAJ22 | CAVRSSGSARQLTF | TRBV7-2 | TRBJ2-7 | CASSLEPSYEQYF | KLGGALQAK | YES | NO | NO |
| 19377 | TRAV26-2 | TRAJ48 | CILRGEFGNEKLTF | TRBV27 | TRBJ1-6 | CASRRTGQSYNSPLHF | KLGGALQAK | YES | NO | NO |
| 19378 | TRAV8-4 | TRAJ20 | CAVIQINDYKLSF | TRBV7-8 | TRBJ1-2 | CASSLGQDGYTF | RLRAEAQVK | YES | NO | NO |
| 19379 | TRAV21 | TRAJ12 | CAVLMDSSYKLIF | TRBV27 | TRBJ2-1 | CASLKRGGSYNEQFF | KLGGALQAK | YES | NO | NO |
| 19380 | TRAV27 | TRAJ26 | CAGAGYWGQNFVF | TRBV14 | TRBJ1-2 | CASSQVPGQGPNYGYTF | KLGGALQAK | NO | NO | NO |
| 19382 | TRAV27 | TRAJ44 | CAGADGTASKLTF | TRBV27 | TRBJ1-2 | CASSLSGTWGEGYTF | AVFDRKSDAK | YES | NO | NO |
| 19383 | TRAV8-4 | TRAJ20 | CAVIPINDYKLSF | TRBV7-8 | TRBJ1-2 | CASSLGQAYGYTF | KLGGALQAK | YES | NO | NO |
| 19384 | TRAV16 | TRAJ39 | CALSRNAGNMLTF | TRBV12-3 | TRBJ2-3 | CASSWGGDTQYF | KLGGALQAK | NO | NO | NO |
| 19385 | TRAV14DV4 | TRAJ4 | CAMRPPFRGGYNKLIF | TRBV5-6 | TRBJ1-5 | CASSSRDSQNQPQHF | KLGGALQAK | YES | NO | NO |
| 19386 | TRAV38-1 | TRAJ31 | CAFGFNNARLMF | TRBV12-5 | TRBJ2-4 | CASGLVEGDSKNIQYF | KLGGALQAK | YES | NO | NO |
| 19387 | TRAV3 | TRAJ37 | CAVRDGGMGSNTGKLIF | TRBV12-4 | TRBJ2-3 | CASSPMDTQYF | AVFDRKSDAK | NO | NO | NO |
| 19388 | TRAV19 | TRAJ37 | CALSEFSSNTGKLIF | TRBV7-9 | TRBJ2-4 | CASSPSRDREFLYIQYF | KLGGALQAK | YES | NO | NO |
| 19389 | TRAV14DV4 | TRAJ23 | CAMREGWSQGGKLIF | TRBV19 | TRBJ1-2 | CASREGQAYGYTF | KLGGALQAK | YES | NO | NO |
| 19390 | TRAV8-4 | TRAJ20 | CAVIQINDYKLSF | TRBV7-8 | TRBJ1-2 | CASSLGQDGYTF | AVFDRKSDAK | YES | NO | NO |
| 19391 | TRAV19 | TRAJ30 | CALSEADDDKIIF | TRBV24-1 | TRBJ2-6 | CATSGAAYSGANVLTF | KLGGALQAK | YES | NO | NO |
| 19392 | TRAV8-6 | TRAJ57 | CAVSYQGGSEKLVF | TRBV11-3 | TRBJ2-5 | CASSQTGLVLGGQETQYF | KLGGALQAK | YES | NO | NO |
| 19393 | TRAV13-2 | TRAJ32 | CAESGATNKLIF | TRBV20-1 | TRBJ1-1 | CSAWDRLNTEAFF | KLGGALQAK | YES | NO | NO |
| 19394 | TRAV38-2DV8 | TRAJ34 | CAYRTSASNTDKLIF | TRBV19 | TRBJ2-2 | CASSISYTGELFF | KLGGALQAK | YES | NO | NO |
| 19395 | TRAV35 | TRAJ20 | CAGHDGNDYKLSF | TRBV7-8 | TRBJ1-2 | CASSLGYGYTF | KLGGALQAK | YES | NO | NO |
| 19396 | TRAV29DV5 | TRAJ37 | CAASALGNTGKLIF | TRBV4-1 | TRBJ1-1 | CASSPGAGDTEAFF | KLGGALQAK | YES | NO | NO |
| 19398 | TRAV14DV4 | TRAJ22 | CAMRRPISSGSARQLTF | TRBV18 | TRBJ2-5 | CASSPPGGQETQYF | KLGGALQAK | YES | NO | NO |
| 19399 | TRAV12-3 | TRAJ39 | CAIGPGNMLTF | TRBV29-1 | TRBJ1-2 | CSVDLEANYGYTF | RLRAEAQVK | YES | NO | NO |
| 19400 | TRAV25 | TRAJ32 | CAGNDYGGATNKLIF | TRBV27 | TRBJ1-4 | CASSLYAGASEKLFF | KLGGALQAK | YES | NO | NO |
| 19401 | TRAV12-3 | TRAJ39 | CAIGPGNMLTF | TRBV7-2 | TRBJ2-7 | CASSLEPSYEQYF | KLGGALQAK | YES | NO | NO |
| 19402 | TRAV21 | TRAJ49 | CAVSGTGNQFYF | TRBV6-6 | TRBJ1-2 | CASSRLGDSYGYTF | KLGGALQAK | NO | NO | NO |
| 19403 | TRAV13-2 | TRAJ17 | CAESLKGGQTAGNKLTF | TRBV3-1 | TRBJ2-2 | CASSQVFRGPPGELFF | KLGGALQAK | YES | NO | NO |
| 19404 | TRAV26-2 | TRAJ58 | CILGPSETSGSRLTF | TRBV7-3 | TRBJ1-1 | CASSLNPVNTEAFF | KLGGALQAK | NO | NO | NO |
| 19405 | TRAV19 | TRAJ30 | CALSEADDDKIIF | TRBV24-1 | TRBJ2-6 | CATSGAAYSGANVLTF | AVFDRKSDAK | YES | NO | NO |
| 19406 | TRAV30 | TRAJ30 | CGTETFRDDKIIF | TRBV6-6 | TRBJ1-2 | CASSYSNEGNGYTF | KLGGALQAK | NO | NO | NO |
| 19407 | TRAV8-6 | TRAJ4 | CAVSASGGYNKLIF | TRBV19 | TRBJ2-1 | CAGGWGDEQFF | KLGGALQAK | YES | NO | NO |
| 19409 | TRAV9-2 | TRAJ6 | CALTVEGGSYIPTF | TRBV15 | TRBJ1-5 | CATSRVVGVDQPQHF | KLGGALQAK | YES | NO | NO |
| 19410 | TRAV17 | TRAJ26 | CVSDYGQNFVF | TRBV11-1 | TRBJ1-6 | CASSSFDGLNSPLHF | KLGGALQAK | YES | NO | NO |
| 19412 | TRAV12-2 | TRAJ20 | CAVNIVNDYKLSF | TRBV7-8 | TRBJ1-2 | CASSLGTANGYTF | KLGGALQAK | YES | NO | NO |
| 19413 | TRAV14DV4 | TRAJ18 | CAMRGPRGSTLGRLYF | TRBV20-1 | TRBJ2-5 | CSARDRAGKETQYF | KLGGALQAK | YES | NO | NO |
| 19415 | TRAV3 | TRAJ17 | CAVRDILKAAGNKLTF | TRBV4-3 | TRBJ1-2 | CASSQTLGGSYGYTF | KLGGALQAK | YES | NO | NO |
| 19417 | TRAV19 | TRAJ56 | CALRVTGANSKLTF | TRBV20-1 | TRBJ2-1 | CSARDIHGQGTYNEQFF | KLGGALQAK | NO | NO | NO |
| 19418 | TRAV26-2 | TRAJ49 | CPDFPTGNQFYF | TRBV10-2 | TRBJ2-2 | CASSEWSGNTGELFF | KLGGALQAK | YES | NO | NO |
| 19419 | TRAV13-2 | TRAJ49 | CAEIRGRDQFYF | TRBV2 | TRBJ1-1 | CASSDRMNTEAFF | AVFDRKSDAK | YES | NO | NO |
| 19420 | TRAV5 | TRAJ27 | CAVLTNAGKSTF | TRBV27 | TRBJ1-1 | CATRLNTEAFF | KLGGALQAK | YES | NO | NO |
| 19421 | TRAV13-1 | TRAJ10 | CAVLPGGGNKLTF | TRBV6-5 | TRBJ2-7 | CASSYSYEQYF | KLGGALQAK | YES | NO | NO |
| 19422 | TRAV12-3 | TRAJ39 | CAIGPGNMLTF | TRBV6-3 | TRBJ2-1 | CASSYGAGGYNEQFF | KLGGALQAK | YES | NO | NO |
| 19424 | TRAV16 | TRAJ29 | CALYNSGNTPLVF | TRBV19 | TRBJ1-2 | CASSILWTGRGYTF | KLGGALQAK | NO | NO | NO |
| 19425 | TRAV8-3 | TRAJ45 | CAVGASGGGADGLTF | TRBV4-2 | TRBJ2-2 | CASSRGGSGNTGELFF | KLGGALQAK | YES | NO | NO |
| 19426 | TRAV22 | TRAJ12 | CAVEDSSYKLIF | TRBV27 | TRBJ2-1 | CASSPTSGSLFTNEQFF | KLGGALQAK | YES | NO | NO |
| 19427 | TRAV8-2 | TRAJ43 | CVVSDSNDMRF | TRBV27 | TRBJ2-7 | CASGNSPYEQYF | KLGGALQAK | YES | NO | NO |
| 19428 | TRAV26-2 | TRAJ37 | CILRDSSNTGKLIF | TRBV9 | TRBJ1-1 | CASSEWGSNTEAFF | KLGGALQAK | NO | NO | NO |
| 19429 | TRAV5 | TRAJ9 | CAESTLDTGGFKTIF | TRBV19 | TRBJ2-1 | CASSIGGSSYNEQFF | KLGGALQAK | YES | NO | NO |
| 19430 | TRAV26-2 | TRAJ15 | CILSGQAGTALIF | TRBV12-4 | TRBJ2-1 | CASSLKRYEQFF | KLGGALQAK | YES | NO | NO |
| 19431 | TRAV29DV5 | TRAJ58 | CAASASSSRLTF | TRBV19 | TRBJ1-1 | CASSKRGPNTEAFF | KLGGALQAK | YES | NO | NO |
| 19432 | TRAV8-1 | TRAJ45 | CAVNGYSGGGADGLTF | TRBV6-5 | TRBJ2-3 | CASSYATGDADTQYF | KLGGALQAK | NO | NO | NO |
| 19433 | TRAV13-1 | TRAJ42 | CAAVLYGGSQGNLIF | TRBV25-1 | TRBJ1-2 | CASRTGNYGYTF | KLGGALQAK | YES | NO | NO |
| 19436 | TRAV9-2 | TRAJ44 | CALRTYGGTASKLTF | TRBV10-2 | TRBJ2-7 | CASSEPQTSNEQYF | KLGGALQAK | YES | NO | NO |
| 19437 | TRAV1-2 | TRAJ33 | CAVRDSNYQLIW | TRBV11-3 | TRBJ2-7 | CASSETSGPSYEQYF | KLGGALQAK | YES | NO | NO |
| 19439 | TRAV12-3 | TRAJ39 | CAIGPGNMLTF | TRBV20-1 | TRBJ2-7 | CSARDLEVLSYEQYF | KLGGALQAK | YES | NO | NO |
| 19440 | TRAV10 | TRAJ4 | CVVRAGGYNKLIF | TRBV7-9 | TRBJ2-7 | CASSDSLGDRGSYEQYF | KLGGALQAK | YES | NO | NO |
| 19441 | TRAV19 | TRAJ22 | CALDFSSGSARQLTF | TRBV14 | TRBJ1-4 | CASSQGPGTVHEKLFF | AVFDRKSDAK | NO | NO | NO |
| 19442 | TRAV22 | TRAJ52 | CAVVGGTSYGKLTF | TRBV12-5 | TRBJ2-7 | CASGNRVSQYF | KLGGALQAK | YES | NO | NO |
| 19443 | TRAV14DV4 | TRAJ5 | CAMRDPQGGRRALTF | TRBV7-9 | TRBJ2-1 | CASTGTSGGPTLRDEQFF | AVFDRKSDAK | YES | NO | NO |
| 19444 | TRAV14DV4 | TRAJ22 | CAMRRPISSGSARQLTF | TRBV20-1 | TRBJ1-4 | CSAPTGTNEKLFF | KLGGALQAK | YES | NO | NO |
| 19445 | TRAV13-2 | TRAJ49 | CAEIRGRDQFYF | TRBV2 | TRBJ1-1 | CASSDRMNTEAFF | KLGGALQAK | YES | NO | NO |
| 19446 | TRAV14DV4 | TRAJ42 | CAMREWLRNYGGSQGNLIF | TRBV19 | TRBJ1-5 | CASSIAWQPQHF | KLGGALQAK | YES | NO | NO |
| 19447 | TRAV21 | TRAJ5 | CAVTPSRAGRRALTF | TRBV4-1 | TRBJ1-3 | CASSQEAGNTIYF | KLGGALQAK | YES | NO | NO |
| 19448 | TRAV8-3 | TRAJ49 | CAVGAFRQFYF | TRBV19 | TRBJ2-1 | CASGAGGLNEQFF | KLGGALQAK | YES | NO | NO |
| 19449 | TRAV5 | TRAJ16 | CAESSVSDGQKLLF | TRBV3-1 | TRBJ1-5 | CASSLTDRVGQPQHF | KLGGALQAK | NO | NO | NO |
| 19450 | TRAV10 | TRAJ36 | CVVTQTGANNLFF | TRBV9 | TRBJ1-6 | CASSVGSNPSSYNSPLHF | KLGGALQAK | YES | NO | NO |
| 19452 | TRAV38-2DV8 | TRAJ54 | CAYRRWGAQKLVF | TRBV2 | TRBJ2-7 | CASKTGTLRTGPYEQYF | KLGGALQAK | YES | NO | NO |
| 19453 | TRAV38-2DV8 | TRAJ42 | CAYRSAYGYGGSQGNLIF | TRBV19 | TRBJ2-5 | CASSIGWETQYF | KLGGALQAK | YES | NO | NO |
| 19455 | TRAV38-2DV8 | TRAJ42 | CAYRILYGGSQGNLIF | TRBV6-5 | TRBJ2-7 | CASSPGLAGGLHPYEQYV | KLGGALQAK | NO | NO | NO |
| 19456 | TRAV29DV5 | TRAJ53 | CAASSNSGGSNYKLTF | TRBV11-2 | TRBJ2-7 | CASSLVLAGGEQYF | KLGGALQAK | YES | NO | NO |
| 19457 | TRAV12-1 | TRAJ4 | CVVHLPPYNKLIF | TRBV11-3 | TRBJ1-1 | CASSFESGGRREAFF | KLGGALQAK | YES | NO | NO |
| 19459 | TRAV12-3 | TRAJ39 | CAIGPGNMLTF | TRBV4-3 | TRBJ1-4 | CASSQDHRMGGHEKLFF | KLGGALQAK | YES | NO | NO |
| 19460 | TRAV24 | TRAJ10 | CAFYILTGGGNKLTF | TRBV19 | TRBJ2-1 | CASSIVGRTGFSYNEQFF | KLGGALQAK | YES | NO | NO |
| 19461 | TRAV13-2 | TRAJ49 | CAEIRGRDQFYF | TRBV29-1 | TRBJ1-2 | CSVDLEANYGYTF | KLGGALQAK | YES | NO | NO |
| 19462 | TRAV27 | TRAJ44 | CAGGSGTASKLTF | TRBV27 | TRBJ2-3 | CASMSGGSYGDTQYF | KLGGALQAK | YES | NO | NO |
| 19463 | TRAV24 | TRAJ40 | CAFPSGTYKYIF | TRBV10-1 | TRBJ1-4 | CASSEFGRTNEKLFF | KLGGALQAK | YES | NO | NO |
| 19464 | TRAV39 | TRAJ31 | CAVDRLGRLMF | TRBV11-2 | TRBJ2-1 | CASSFEDMGGEQFF | KLGGALQAK | YES | NO | NO |
| 19465 | TRAV12-3 | TRAJ9 | CAMTFDAGGFKTIF | TRBV6-5 | TRBJ1-3 | CASSYSNLEGEENTIYF | KLGGALQAK | YES | NO | NO |
| 19466 | TRAV14DV4 | TRAJ22 | CAMRRPISSGSARQLTF | TRBV5-6 | TRBJ2-5 | CASSLKGGPTETQYF | KLGGALQAK | YES | NO | NO |
| 19468 | TRAV17 | TRAJ23 | CATGPYNQGGKLIF | TRBV20-1 | TRBJ2-1 | CSARDIAGGTYNEQFF | KLGGALQAK | NO | NO | NO |
| 19469 | TRAV13-1 | TRAJ11 | CAARGGYSTLTF | TRBV28 | TRBJ2-3 | CASSLRTSGSFTDTQYF | KLGGALQAK | YES | NO | NO |
| 19470 | TRAV24 | TRAJ48 | CASGNFGNEKLTF | TRBV27 | TRBJ1-2 | CASSPGGGVYGYTF | KLGGALQAK | YES | NO | NO |
| 19471 | TRAV17 | TRAJ23 | CATAQYNQGGKLIF | TRBV20-1 | TRBJ2-1 | CSARDALAGTYNEQFF | KLGGALQAK | NO | NO | NO |
| 19474 | TRAV12-3 | TRAJ39 | CAIGPGNMLTF | TRBV7-9 | TRBJ2-7 | CASKDGISYEQYF | KLGGALQAK | YES | NO | NO |
| 19476 | TRAV41 | TRAJ22 | CAVSWAGSARQLTF | TRBV7-6 | TRBJ1-2 | CASSQGGYGYTF | KLGGALQAK | YES | NO | NO |
| 19477 | TRAV6 | TRAJ21 | CARHFNKFYF | TRBV7-9 | TRBJ1-1 | CASSPIQDTEAFF | KLGGALQAK | YES | NO | NO |
| 19478 | TRAV3 | TRAJ13 | CAVRDRTF | TRBV4-1 | TRBJ2-5 | CASSQGSQGAQETQYF | KLGGALQAK | NO | NO | NO |
| 19479 | TRAV14DV4 | TRAJ45 | CAMSDSGGGADGLTF | TRBV7-7 | TRBJ2-5 | CASSWGGASGETQYF | KLGGALQAK | YES | NO | NO |
| 19480 | TRAV14DV4 | TRAJ6 | CAMRGPGGSYIPTF | TRBV12-4 | TRBJ1-1 | CASSPQKGIGTEAFF | KLGGALQAK | YES | NO | NO |
| 19481 | TRAV3 | TRAJ43 | CAVWKGRTDMRF | TRBV5-6 | TRBJ2-3 | CASRQILAANTDTQYF | KLGGALQAK | NO | NO | NO |
| 19483 | TRAV1-1 | TRAJ31 | CASNNNARLMF | TRBV7-9 | TRBJ2-2 | CASTGTSSVSGELFF | KLGGALQAK | YES | NO | NO |
| 19486 | TRAV13-1 | TRAJ41 | CAASPGDSGYALNF | TRBV3-1 | TRBJ2-1 | CASSQDLGGVSYNEQFF | KLGGALQAK | YES | NO | NO |
| 19487 | TRAV1-2 | TRAJ39 | CALNNAGNMLTF | TRBV10-3 | TRBJ2-7 | CAISPGQGESYEQYF | KLGGALQAK | YES | NO | NO |

| | | | | | | | | | | |
|---|---|---|---|---|---|---|---|---|---|---|
| 19489 | TRAV8-4 | TRAJ20 | CAVIQINDYKLSF | TRBV7-2 | TRBJ2-7 | CASSLEPSYEQYF | KLGGALQAK | YES | NO | NO |
| 19492 | TRAV27 | TRAJ34 | CAEPSSYNTDKLIF | TRBV7-9 | TRBJ1-1 | CASSPTSGNTEAFF | KLGGALQAK | YES | NO | NO |
| 19493 | TRAV13-1 | TRAJ49 | CAASWGTGNQFYF | TRBV12-4 | TRBJ2-4 | CASSAGGDIQYF | KLGGALQAK | YES | NO | NO |
| 19494 | TRAV38-1 | TRAJ57 | CAFMTGEQGGSEKLVF | TRBV6-5 | TRBJ2-3 | CASSLSGSTDTQYF | KLGGALQAK | NO | NO | NO |
| 19499 | TRAV8-2 | TRAJ6 | CVVTSFPSGGSYIPTF | TRBV12-4 | TRBJ2-7 | CASSLGTSGGVTYEQYF | KLGGALQAK | YES | NO | NO |
| 19500 | TRAV14DV4 | TRAJ20 | CAMSDLSNDYKLSF | TRBV20-1 | TRBJ2-3 | CSARQPSGRSSLSTDTQYF | KLGGALQAK | YES | NO | NO |
| 19501 | TRAV21 | TRAJ43 | CAVWYMRF | TRBV19 | TRBJ1-2 | CASTPSGDGYTF | KLGGALQAK | NO | NO | NO |
| 19504 | TRAV38-2DV8 | TRAJ34 | CAYIYNTDKLIF | TRBV19 | TRBJ1-6 | CASSITGGLASPLHF | KLGGALQAK | YES | NO | NO |
| 19506 | TRAV12-3 | TRAJ39 | CAIGPGNMLTF | TRBV7-9 | TRBJ2-7 | CASSDSLGDRGSYEQYF | KLGGALQAK | YES | NO | NO |
| 19507 | TRAV17 | TRAJ23 | CATANYNQGGKLIF | TRBV20-1 | TRBJ2-1 | CSARDWASETYNEQFF | KLGGALQAK | NO | NO | NO |
| 19509 | TRAV35 | TRAJ42 | CAGQSGRVGGSQGNLIF | TRBV6-3 | TRBJ2-1 | CASSYGAGGYNEQFF | AVFDRKSDAK | YES | NO | NO |
| 19510 | TRAV13-1 | TRAJ42 | CAAGKSQGNLIF | TRBV5-4 | TRBJ1-2 | CASSLPSSNYGYTF | KLGGALQAK | YES | NO | NO |
| 19512 | TRAV8-3 | TRAJ39 | CAVGDGNAGNMLTF | TRBV6-1 | TRBJ1-1 | CASSTLRVGGRTEAFF | KLGGALQAK | NO | NO | NO |
| 19513 | TRAV27 | TRAJ44 | CAGASGTASKLTF | TRBV27 | TRBJ2-5 | CASSVDGTYGETQYF | KLGGALQAK | YES | NO | NO |
| 19514 | TRAV29DV5 | TRAJ28 | CAARIAYSGAGSYQLTF | TRBV5-5 | TRBJ1-2 | CASSGDSLNYGYTF | KLGGALQAK | YES | NO | NO |
| 19516 | TRAV10 | TRAJ48 | CVVSLPRFGNEKLTF | TRBV18 | TRBJ2-5 | CASSPNTGMETQYF | KLGGALQAK | YES | NO | NO |
| 19517 | TRAV14DV4 | TRAJ22 | CAMRRPISSGSARQLTF | TRBV15 | TRBJ1-4 | CATVNGPQEKLFF | KLGGALQAK | YES | NO | NO |
| 19518 | TRAV19 | TRAJ11 | CALSSGYSTLTF | TRBV19 | TRBJ2-2 | CASTRNAFYSLGTGELFF | KLGGALQAK | YES | NO | NO |
| 19521 | TRAV17 | TRAJ43 | CAGNNNDMRF | TRBV6-3 | TRBJ1-4 | CASSYSYRVELFF | KLGGALQAK | YES | NO | NO |
| 19522 | TRAV38-2DV8 | TRAJ54 | CAYSFQGAQKLVF | TRBV7-9 | TRBJ1-5 | CASSYYGRTVNQPQHF | KLGGALQAK | YES | NO | NO |
| 19523 | TRAV9-2 | TRAJ11 | CALSSHSGYSTLTF | TRBV27 | TRBJ2-1 | CASSQTDNEQFF | KLGGALQAK | YES | NO | NO |
| 19524 | TRAV21 | TRAJ47 | CAVSMEYGNKLVF | TRBV24-1 | TRBJ1-5 | CATSDGAGRASNQPQHF | KLGGALQAK | YES | NO | NO |
| 19525 | TRAV16 | TRAJ39 | CNAGNMLTF | TRBV19 | TRBJ2-3 | CASSSGAGGTDTQYF | KLGGALQAK | NO | NO | NO |
| 19526 | TRAV12-3 | TRAJ35 | CAMKR | TRBV3-1 | TRBJ2-2 | CASSQVFRGPPGELFF | AVFDRKSDAK | NO | NO | NO |
| 19529 | TRAV27 | TRAJ26 | CAGAGSGYGQNFVF | TRBV7-9 | TRBJ2-1 | CASSPRRASSYNEQFF | KLGGALQAK | YES | NO | NO |
| 19532 | TRAV12-2 | TRAJ45 | CAVNAYSGGGADGLTF | TRBV10-3 | TRBJ2-2 | CAISEGASGFTGELFF | KLGGALQAK | YES | NO | NO |
| 19533 | TRAV12-3 | TRAJ39 | CAIGPGNMLTF | TRBV12-4 | TRBJ2-3 | CASSPMDTQYF | KLGGALQAK | YES | NO | NO |
| 19534 | TRAV14DV4 | TRAJ22 | CAMRRPISSGSARQLTF | TRBV29-1 | TRBJ1-2 | CSVDLEANYGYTF | KLGGALQAK | YES | NO | NO |
| 19535 | TRAV29DV5 | TRAJ21 | CAASEYNFNKFYF | TRBV9 | TRBJ1-2 | CASSLGGYGYTF | KLGGALQAK | YES | NO | NO |
| 19536 | TRAV12-2 | TRAJ9 | CAGINTGGFKTIF | TRBV19 | TRBJ2-3 | CASSIDRETSTDTQYF | KLGGALQAK | YES | NO | NO |
| 19537 | TRAV10 | TRAJ4 | CVVRSGGYNKLIF | TRBV29-1 | TRBJ1-2 | CSVDLEANYGYTF | KLGGALQAK | YES | NO | NO |
| 19538 | TRAV8-1 | TRAJ24 | CAVRPPTDSWGKFQF | TRBV7-7 | TRBJ2-5 | CASSSPRDWLYQETQYF | KLGGALQAK | YES | NO | NO |
| 19540 | TRAV1-1 | TRAJ5 | CAVDGDTGRRALTF | TRBV20-1 | TRBJ2-3 | CSARGVTSGSLDTQYF | KLGGALQAK | NO | NO | NO |
| 19543 | TRAV12-2 | TRAJ43 | CAVLLNDMRF | TRBV4-3 | TRBJ2-1 | CASSPGQGHNEQFF | KLGGALQAK | YES | NO | NO |
| 19544 | TRAV1-2 | TRAJ34 | CAVTQSYNTDKLIF | TRBV12-4 | TRBJ2-7 | CASSPRMSPYEQYF | KLGGALQAK | YES | NO | NO |
| 19545 | TRAV21 | TRAJ50 | CAVDLMKTSYDKVIF | TRBV24-1 | TRBJ2-2 | CATSDRLAGGELFF | IVTDFSVIK | YES | NO | YES |
| 19547 | TRAV12-2 | TRAJ4 | CAVLGFSGGYNKLIF | TRBV9 | TRBJ2-1 | CASSVGAGVGNEQFF | KLGGALQAK | YES | NO | NO |
| 19548 | TRAV21 | TRAJ50 | CAVDLMKTSYDKVIF | TRBV20-1 | TRBJ2-7 | CSAMTSGSSYEQYF | IVTDFSVIK | NO | NO | NO |
| 19549 | TRAV21 | TRAJ50 | CAVDLMKTSYDKVIF | TRBV15 | TRBJ2-7 | CATSPGGGRDLYEQYF | IVTDFSVIK | YES | YES | YES |
