## Supplementary Table 2 for "TCRfp: a new fingerprint-based approach for TCR repertoire analysis"

| Genetic Algorithm Parameters | | Main parameters | | | | | Parameters of the Initial Generation | | | | Reproduction Parameters | | | | | | |
|---|---|---|---|---|---|---|---|---|---|---|---|---|---|---|---|---|---|
| Algorithm | | Number of centroids | Dimensions | Initial size | Offspring size | Parent range | Range weighted C | Range weighted P | Range X,Y,Z coordinates | Range C,P values | Mutation number | Mutation range weighted C | Mutation range weighted P | Mutation range X,Y,Z coordinates | Mutation range C,P values | Crossover type | Crossover point |
| Original scoring | TCRfp HS | 6 | 5 | 400 | 200 | 200 | 0/50 | 0/20 | -30/30 | -1/1 | 1/2 | -2/2 | -2/2 | -2/2 | -2/2 | Multiple centroid whole | Random number of centroids swap |
| | TCRfp MaxD | 6 | 5 | 400 | 200 | 200 | 30/50 | 0/5 | -30/30 | -1/1 | 2/4 | -1/1 | -1/1 | -1/1 | -1/1 | Multiple centroid whole | Random number of centroids with swap of the entire centroids |
| | Strict centroid movement | 6 | 5 | 400 | 200 | 200 | 0/100 | 0/50 | Inital close to the tip of the loop position with a range of -1/1 | -30/30 | 1/2 | -2/2 | -2/2 | -2/2 | -2/2 | Multiple centroid whole | Multiple whole centroid swap |
| | New centoid positions | 6 | 5 | 400 | 200 | 200 | 0/100 | 0/50 | 3 centroids starting from CDR3alpha 3 centroids starting from CDR3beta | -30/30 | 1/2 | -5/5 | -5/5 | -10/10 | -5/5 | Multiple centroid whole | Multiple whole centroid swap |
| | TCRfp HS looplength | 6 | 5 | 400 | 200 | 200 | 0/100 | 0/50 | Inital close to the tip - exact position | -30/30 | 1/2 | -2/2 | -2/2 | -2/2 | -2/2 | Multiple centroid whole - box limitation | Multiple whole centroid swap - Mutations can't go further a 3D box |
| | TCRfp MaxD looplength | 6 | 5 | 400 | 200 | 200 | 0/100 | 0/50 | Inital close to the tip - exact position | -30/30 | 2/4 | -1/1 | -1/1 | -1/1 | -1/1 | Multiple centroid whole | Random number of centroids with swap of the entire centroids |
| | Restricted space | 6 | 5 | 400 | 200 | 200 | 0/100 | 0/50 | Inital close to the tip - exact position | -30/30 | 1/2 | -2/2 | -2/2 | -2/2 | -2/2 | Multiple centroid whole - box limitation | Multiple whole centroid swap - Mutations can't go further a 3D box |
| | TCRfp MaxD TOL | 6 | 5 | 400 | 200 | 200 | 0/100 | 0/50 | Inital close to the tip - exact position | -30/30 | 1/2 | -10/10 | -10/10 | -20/20 | -10/10 | Single centroid whole | Single random centroid swap |
| | TCRfp MaxD TOL 2.0 | 6 | 5 | 400 | 200 | 200 | 0/100 | 0/50 | Inital close to the tip - exact position | -1/1 | 1/2 | -10/10 | -10/10 | -20/20 | -10/10 | Multiple centroid break | Random number of centroids swap with random break inside the centroid |
| New scoring | Rank 2 | 6 | 5 | 400 | 200 | 200 | 0/50 | 0/50 | -30/30 | -10/10 | 2/4 | -2/2 | -2/2 | -2/2 | -2/2 | Multiple centroid break | Random number of centroids swap with random break inside the centroid |
| | Rank 5 | 6 | 5 | 400 | 200 | 200 | 0/50 | 0/50 | -30/30 | -10/10 | 2/4 | -2/2 | -2/2 | -2/2 | -2/2 | Multiple centroid break | Random number of centroids swap with random break inside the centroid |
| | Rank 5 relaxed | 6 | 5 | 400 | 200 | 200 | 0/50 | 0/50 | -30/30 | -10/10 | 2/4 | -2/2 | -2/2 | -2/2 | -2/2 | Multiple centroid break | Random number of centroids swap with random break inside the centroid |
