## Supplementary Table 3 for "TCRfp: a new fingerprint-based approach for TCR repertoire analysis"

| Algorithm | | Heuristic search | Initial generation | Training data set | Maximum PI score | Number of runs | Mean final score | Top 5 centroids | | | |
|---|---|---|---|---|---|---|---|---|---|---|---|
| | | | | | | | | Score | C | P | Centroids |
| Original scoring | TCRfp HS | HS | Random | TCRfp HS | 36,5 | 50 | 31,4 | 36,5 | 43 | 5 | 25.816 -7.425 7.769 -1.494 -2.267 1.444 -3.827 3.993 -2.07 -1.145 5.886 23.593 17.028 0.222 -0.479 -23.905 -24.316 -31.71 -2.743 1.113 13.805 -4.239 -29.239 1.191 0.487 -10.432 -4.425 25.446 -2.079 -2.242 |
| | | | | | | | | 34,5 | 37 | 1 | 27.147 -12.329 8.209 -5.032 0.519 6.103 -17.662 -19.221 0.647 0.201 -4.875 1.829 10.115 0.685 0.164 -1.594 -8.229 1.465 -0.405 -1.499 6.838 25.258 15.477 1.289 -0.053 -21.546 -23.554 -29.691 -1.42 3.258 |
| | | | | | | | | 34,3 | 39 | 2 | 9.745 33.625 24.746 -2.123 -1.135 -3.626 17.478 8.719 0.158 1.509 -1.153 -12.066 0.388 -2.482 4.535 -0.468 -4.796 4.5 -0.483 -0.114 -14.789 -6.464 28.624 2.126 2.302 23.685 -9.887 10.863 4.393 -1.084 |
| | | | | | | | | 34,3 | 25 | 1 | 5.74 -17.676 -14.362 -4.474 2.781 -15.863 -7.508 29.805 0.609 0.255 6.839 13.771 -12.089 0.652 1.841 -13.782 -20.527 -19.755 0.774 -3.233 23.584 -5.792 6.108 1.684 -1.03 -16.583 8.482 6.722 -2.955 -1.143 |
| | | | | | | | | 34 | 45 | 10 | 28.317 -11.067 6.225 0.929 0.864 -26.499 -6.132 20.041 -6.147 2.405 -21.264 -15.886 -27.6 1.994 1.286 -3.901 17.982 7.743 -0.48 0.87 5.857 -27.304 -21.059 -4.295 0.018 -13.899 -2.778 27.847 1.942 -1.07 |
| | TCRfp MaxD | MaxD | Random | TCRfp MaxD | 0,63 | 10 | 0,63 | 0,63 | 22 | 1 | 26.996 20.134 -24.869 -0.488 0.065 28.082 -3.472 -11.469 -0.804 -0.36 -21.227 5.444 -8.432 1.311 0.309 24.384 22.306 -2.101 -1.135 -1.021 -25.161 1.728 28.386 0.374 0.085 32.408 7.69 -18.735 -0.122 -0.262 |
| | | | | | | | | 0,63 | 18 | 2 | 29.359 -2.361 -16.318 0.281 -0.85 24.521 14.442 -20.457 0.789 0.513 22.158 21.104 -0.73 -0.885 0.002 27.768 10.775 -9.76 -1.443 -0.052 -21.898 5.952 -9.157 1.381 0.172 -24.141 1.57 27.384 -0.551 -0.461 |
| | | | | | | | | 0,63 | 26 | 0 | -23.859 3.559 27.799 0.647 0.306 27.662 -2.398 -10.598 -0.125 -0.646 -21.196 5.836 -8.776 1.96 -0.196 27.341 24.754 -3.281 -0.337 -0.01 29.477 6.587 -13.63 -0.546 -0.197 29.389 18.926 -25.72 1.802 -0.823 |
| | | | | | | | | 0,63 | 38 | 4 | 31.772 21.833 -28.025 0.622 0.9 -25.995 4.63 32.137 -0.723 -0.291 -22.016 8.95 -7.065 2.684 0.575 30.221 6.175 -15.18 0.314 -0.156 29.873 27.606 -2.809 -1.928 0.277 29.809 -4.297 -15.31 -1.285 -0.577 |
| | | | | | | | | 0,63 | 21 | 0 | 28.31 2.776 -19.898 -0.545 0.398 26.351 -2.069 -8.854 -0.149 1.112 -24.598 0.854 27.434 -0.058 0.822 25.05 19.577 -23.133 -0.069 -0.025 25.735 26.376 0.878 -0.369 -0.088 27.641 19.305 -7.316 -1.065 -1.094 |
| | Strict centroid movement | HS | TOL | TCRfp HS | 34,9 | 60 | 30,03 | 34,9 | 60 | 8 | 5.924 13.697 -12.257 -0.652 -11.692 18.449 12.218 4.02 -20.802 -29.503 0.802 15.505 -4.379 -3.95 1.496 -7.883 10.953 11.972 -12.817 -11.742 -14.665 17.454 1.419 6.234 -31.045 2.738 13.549 5.731 -25.308 -11.222 |
| | | | | | | | | 34,5 | 61 | 7 | 9.042 15.165 -12.019 4.282 -12.164 16.614 10.643 -2.062 23.464 14.151 -0.469 17.215 -7.292 -23.801 6.316 -5.945 11.516 12.094 -12.892 -10.053 -11.468 21.104 3.351 -8.847 -20.749 0.458 11.501 9.29 -22.826 -0.974 |
| | | | | | | | | 34 | 51 | 5 | 6.583 12.072 -11.394 0.214 -11.127 14.133 16.261 0.788 30.665 -26.671 0.874 17.27 -5.895 -24.729 12.554 -4.896 7.198 7.123 -9.401 -6.932 -10.417 11.708 1.853 -4.308 -30.405 0.2 14.692 6.809 -24.956 -13.927 |
| | | | | | | | | 33,9 | 52 | 5 | 6.465 12.492 -11.55 0.007 -11.413 17.81 19.292 -3.375 29.247 -21.726 0.299 16.243 -5.921 -23.075 10.182 -6.333 9.841 10.314 -13.115 -17.412 -9.587 17.467 5.03 -7.339 -21.87 0.192 13.712 7.853 -22.319 7.256 |
| | | | | | | | | 33,8 | 53 | 7 | .817 14.22 -9.502 1.378 -9.917 14.295 15.744 2.122 23.076 -19.54 1.786 15.143 -6.051 -27.509 11.011 -3.988 9.156 10.104 -12.813 -6.117 -12.688 12.495 3.333 -7.92 -25.982 0.055 15.333 5.725 -25.862 -12.747 |
| | New centoid positions | HS | TOL | TCRfp HS | 34,7 | 80 | 30,4 | 34,7 | 50 | 7 | -1.393 15.508 -5.911 -10.189 0.115 8.144 10.102 -9.3 0.569 -17.842 0.177 16.574 -5.801 -32.989 17.372 4.49 3.421 10.492 -23.308 -20.277 -17.089 4.746 10.103 -6.488 3.072 -4.396 9.661 12.074 -14.21 -11.184 |
| | | | | | | | | 34,6 | 63 | 9 | -9.036 12.579 -5.023 -5.291 -26.45 -12.252 2.077 5.64 -10.204 1.911 -2.768 16.446 -7.216 -10.727 -2.776 -5.657 8.959 12.035 -12.896 -11.509 1.992 2.338 8.137 -18.397 -21.732 5.51 12.361 7.52 -27.045 -6.558 |
| | | | | | | | | 34,6 | 60 | 8 | -1.934 17.999 -4.384 -1.458 -30.696 4.504 12.295 -3.506 -2.206 -0.755 1.117 14.632 -7.778 -3.296 21.013 -3.279 8.072 7.949 -20.922 -22.28 -3.949 10.866 11.068 -24.496 -26.15 -6.278 11.38 10.973 -11.775 -12.563 |
| | | | | | | | | 34,5 | 59 | 8 | 5.444 15.773 -2.98 -28.777 2.864 -9.835 18.79 -4.985 -7.594 -20.209 -2.428 17.175 -6.14 -12.506 2.861 2.944 8.657 7.76 -14.57 -12.754 -5.782 6.87 19.458 -16.195 -17.839 -21.281 2.418 9.57 -5.191 12.546 |
| | | | | | | | | 33,8 | 54 | 9 | 3.129 15.73 -6.656 -28.585 2.115 -1.816 16.048 -5.978 -11.796 1.473 -11.271 16.848 -5.574 -3.622 -28.349 -22.604 1.201 10.783 -10.275 10.163 7.848 10.845 16.236 27.758 -28.389 -0.615 4.568 13.697 -10.294 -24.671 |
| | TCRfp HS looplength | HS | TOL | TCRfp HS (whole looplength) | 33,6 | 30 | 28,28 | 33,6 | 54 | 4 | 9.042 15.499 -11.573 7.188 -12.765 13.272 19.498 -0.008 26.711 -30.623 -0.139 18.066 -8.221 -24.665 13.903 -6.333 11.833 11.126 -13.336 -9.608 -13.767 11.013 1.368 -3.976 -27.133 0.541 15.597 6.437 -22.149 -13.985 |
| | | | | | | | | 32,6 | 53 | 15 | 10.396 6.245 -7.913 -31.389 -10.143 18.812 16.305 2.95 -22.761 -18.906 6.178 19.833 -0.407 -14.517 10.077 -7.007 12.522 9.768 -2.859 -22.557 -9.902 13.645 6.349 -13.258 -9.61 -0.021 15.717 4.675 6.294 -17.328 |
| | | | | | | | | 32,2 | 59 | 15 | 9.613 15.165 -11.843 3.537 8.214 18.092 10.297 -1.087 -27.11 -25.863 -0.984 16.243 -6.489 -21.078 -3.392 -6.295 8.376 11.367 -10.111 -12.096 -13.761 13.759 4.689 -25.814 27.003 -2.148 13.666 9.928 25.595 -15.089 |
| | | | | | | | | 31,9 | 52 | 1 | 7.934 13.061 -9.562 -0.908 -9.72 16.397 14.95 -6.007 26.222 -21.866 0.958 16.243 -6.489 -24.053 22.132 -6.333 11.975 10.314 -12.851 -1.69 -12.056 12.7 2.839 -0.937 20.891 1.202 16.766 4.737 -20.685 12.434 |
| | | | | | | | | 31,7 | 62 | 11 | 6.088 8.074 -8.118 -27.509 -0.631 7.275 14.997 -0.112 19.629 19.876 2.022 21.664 3.452 -14.79 8.85 -13.26 11.655 10.52 -14.337 -10.79 -19.716 19.256 -2.878 -7.391 21.916 -6.007 23.724 5.96 20.271 7.927 |
| | TCRfp MaxD looplength | MaxD | TOL | TCRfp MaxD (whole looplength) | 0,67 | 10 | 53 | 0,67 | 97 | 52 | 9.743 14.408 -9.468 -25.666 -5.661 17.097 13.976 -0.896 -25.687 -11.983 -2.087 16.358 -6.379 -28.669 -9.117 -4.905 11.164 11.133 -29.152 -8.005 -10.647 13.702 4.493 -24.875 -9.521 -0.935 15.542 4.845 -27.574 -6.956 |
| | | | | | | | | 0,67 | 95 | 50 | 8.959 16.068 -7.357 -28.91 -2.117 16.397 14.235 -0.825 -26.97 -7.217 0.731 16.243 -6.88 -28.818 -7.587 -6.967 10.904 10.246 -29.039 -5.873 -12.706 15.382 3.585 -26.918 -7.015 1.618 16.04 4.695 -27.549 -8.828 |
| | | | | | | | | 0,66 | 94 | 49 | 9.995 15.424 -9.314 -27.593 -5.992 16.397 14.95 0.006 -28.805 -8.265 1.372 18.222 -6.157 -28.984 -2.935 -5.617 10.92 8.913 -26.723 -7.197 -12.687 15.566 5.48 -20.654 -11.148 0.292 15.597 4.877 -26.475 -8.417 |
| | | | | | | | | 0,66 | 99 | 47 | 8.538 15.768 -9.843 -26.55 -8.569 13.423 14.95 -1.087 -29.096 -4.779 -3.035 16.972 -7.139 -24.445 -9.352 -5.862 12.536 11.695 -27.76 -4.903 -12.903 16.374 5.846 27.014 -25.027 0.879 15.597 6.18 -27.387 -5.457 |
| | | | | | | | | 0,66 | 86 | 56 | 10.082 15.72 -9.545 -24.558 -11.89 15.843 16.379 -1.906 -24.98 -5.963 0.273 16.434 -6.446 -29.371 -7.937 -5.917 11.679 10.265 -18.611 -15.686 -12.269 16.745 5.266 -24.057 -9.276 0.929 16.214 4.292 -25.011 -12.341 |
| | Restricted space | HS | TOL | TCRfp HS | 34,5 | 40 | 29,6 | 34,5 | 50 | 4 | 5.521 13.41 -13.614 1.654 -14.926 18.156 16.399 1.618 29.19 -23.48 -0.062 16.902 -5.464 -27.201 17.416 -3.942 11.302 10.314 -12.383 -8.62 -11.014 8.368 4.712 -8.839 -10.441 -0.389 13.359 6.685 -19.441 6.774 |
| | | | | | | | | 33,7 | 54 | 5 | 10.225 12.645 -11.672 3.192 14.414 15.255 19.03 -0.298 20.965 -21.579 -0.979 20.514 -4.488 -28.495 15.908 -6.333 11.787 11.778 -12.935 -13.701 -8.4 8.325 3.09 -4.916 -16.765 -0.019 15.597 6.958 -29.115 -10.047 |
| | | | | | | | | 33,4 | 59 | 7 | 6.472 12.897 -12.81 0.109 -12.878 16.52 16.824 1.839 26.081 -20.459 2.491 16.243 -7.216 -25.263 9.562 -4.467 8.267 10.314 -9.16 -11.575 -11.181 10.676 5.611 -4.282 -13.289 -0.399 13.358 9.611 -27.856 -1.108 |
| | | | | | | | | 33,4 | 65 | 15 | 9.042 1.187 15.898 -56.169 -52.402 16.397 3.349 13.045 -22.378 -20.905 1.336 14.948 -6.489 -22.155 -2.393 -6.333 0.992 10.314 -7.697 -11.004 -17.245 15.382 -8.296 -8.333 -16.392 0.579 9.973 8.216 -20.539 -5.162 |
| | | | | | | | | 32,2 | 59 | 9 | 4.798 13.834 -13.383 -8.329 11.453 10.542 12.742 0.429 -20.107 -36.25 -0.947 17.408 -5.925 -25.617 7.972 -7.67 5.288 9.428 -11.31 -16.193 -8.038 17.365 5.379 -24.823 20.644 -0.876 16.326 8.328 19.224 2.877 |
| | TCRfp MaxD TOL | MaxD | TOL | TCRfp MaxD | 0,7 | 50 | 0,62 | 0,7 | 118 | 57 | 8.226 15.928 -14.022 -41.381 2.565 23.248 23.223 -0.932 -46.152 -1.66 -0.984 6.239 -4.745 -38.158 -4.196 -6.333 11.843 17.087 -40.971 -4.196 -15.457 10.465 6.94 -33.4 -7.626 0.192 19.555 -8.487 -35.31 -3.445 |
| | | | | | | | | 0,69 | 106 | 61 | 15.868 6.689 -1.236 -37.679 -7.139 23.729 14.95 -17.367 -36.131 -10.036 4.321 26.382 3.51 -34.218 -6.091 -12.097 3.278 8.54 -34.813 -9.846 -7.883 15.382 -1.126 -30.766 -6.76 6.635 15.597 -0.871 -44.903 -0.037 |
| | | | | | | | | 0,69 | 107 | 56 | 11.157 11.023 7.334 -34.873 -6.403 15.773 14.95 -7.985 -32.895 -6.524 -0.984 16.905 -6.387 -27.951 -6.708 -14.045 18.153 22.834 30.874 -27.852 -8.804 15.382 12.12 -44.543 4.416 0.192 8.698 5.934 -31.503 -7.07 |
| | | | | | | | | 0,69 | 98 | 61 | 22.377 15.165 -9.314 -35.301 -6.979 22.473 14.95 -0.932 23.805 -25.253 5.813 11.31 -6.489 -37.694 -6.011 -6.333 0.927 12.042 -25.916 -12.427 -10.438 24.657 4.689 -71.575 4.57 8.478 15.597 3.75 -29.968 -9.162 |
| | | | | | | | | 0,68 | 96 | 57 | 3.948 15.364 -6.54 -26.564 -7.15 39.504 14.95 -1.689 25.068 -24.191 -0.984 16.243 -11.798 -29.446 -9.526 -6.333 10.35 12.306 -26.914 -13.617 -6.408 21.069 4.689 -36.5 -6.154 12.143 15.597 0.671 -28.062 -11.727 |
| | TCRfp MaxD TOL 2.0 | MaxD | TOL | TCRfp MaxD | 0,63 | 30 | 0,61 | 0.63 | 2 | 1 | 25.029 14.692 -16.505 0.295 -0.241 28.579 5.647 -13.195 -0.782 -0.522 -0.984 12.382 -15.828 -0.528 0.001 -19.551 10.067 25.455 -0.127 0.523 -22.267 11.303 -4.435 0.55 -0.74 17.543 14.775 -0.164 0.537 -0.076 |
| | | | | | | | | 0.62 | 3 | 2 | 20.529 17.549 -18.423 -0.505 -0.313 19.595 14.95 -0.932 -0.524 0.482 -1.007 24.126 -19.873 1.384 -0.228 -11.941 11.516 16.675 0.403 0.904 -24.462 15.382 -0.781 -0.243 -0.15 24.214 22.84 -1.666 -0.182 0.213 |
| | | | | | | | | 0.62 | 2 | 8 | 23.705 16.09 -17.28 -0.076 -0.653 17.351 15.305 -1.173 0.109 0.873 -0.692 16.66 -14.765 -0.139 0.086 -16.294 3.262 17.596 -1.495 0.932 -29.235 10.73 -6.079 -0.642 0.811 16.946 13.91 0.322 -0.097 -0.883 |
| | | | | | | | | 0.62 | 5 | 5 | 31.29 7.122 -18.456 -0.763 0.979 18.735 14.95 -0.932 -0.089 0.676 -0.807 16.243 -15.973 0.835 0.659 -14.996 6.045 15.757 0.973 -0.891 -17.893 19.667 29.476 -0.69 -3.528 15.357 12.918 4.845 0.434 0.282 |
| | | | | | | | | 0.62 | 5 | 3 | 23.937 21.919 -24.619 -0.147 -0.821 17.719 17.184 -0.973 -0.287 0.304 -0.508 16.902 -16.524 0.991 -3.759 -13.829 8.147 18.138 -0.783 2.632 -22.955 16.197 -1.174 -0.02 -0.517 16.691 20.71 10.247 0.835 -0.901 |
| New scoring | Rank 2 | HS | Random | TCRfp HS | 30,8 | 10 | 29,44 | 30,8 | 50 | 10 | 9.039 -30.388 -26.323 -0.215 5.405 8.762 12.918 -19.161 -6.232 4.021 -15.415 -21.011 -23.123 -4.448 -2.517 28.459 -17.846 10.42 -5.898 4.004 -0.765 -0.318 7.995 -9.238 4.877 -3.004 -3.342 19.009 -10.619 -10.552 |
| | | | | | | | | 30,5 | 49 | 8 | -8.25 -7.929 10.925 0.642 -6.34 22.51 25.718 -28.171 -10.238 4.091 -20.015 -25.715 -27.713 -5.997 -4.243 27.234 -15.117 10.582 -10.543 3.563 -4.42 -2.854 26.401 -1.085 9.696 8.32 -16.77 -21.068 -4.116 9.792 |
| | | | | | | | | 30,1 | 29 | 1 | 26.646 -9.37 7.55 -3.944 -1.732 8.82 -28.657 -22.399 0.682 -11.755 -1.949 -5.893 26.57 -7.761 8.234 7.578 8.722 10.763 9.951 -0.41 -2.556 4.197 14.005 -1.027 3.776 -18.927 -27.645 -25.863 -3.145 0.562 |
| | | | | | | | | 30 | 48 | 13 | -8.343 -21.567 -16.8 8.412 -6.928 6.727 15.264 -19.192 -4.703 0.891 -2.41 -3.824 19.85 -8.886 -5.48 27.515 -14.578 6.929 -5.514 7.424 10.747 -28.178 -23.19 1.285 -5.703 -19.891 -14.192 11.887 -3.368 9.969 |
| | | | | | | | | 29,5 | 36 | 14 | -16.029 -3.904 16.879 -8.235 5.118 0.128 0.788 28.198 -8.108 -7.473 -11.583 -23.384 -17.924 -5.923 -5.439 6.25 -13.039 -18.299 -5.927 -6.535 2.479 19.424 16.987 6.777 -9.486 26.986 -5.568 3.156 8.123 8.011 |
| | Rank 5 | HS | Random | TCRfp HS | 28,8 | 10 | 27,17 | 28,8 | 52 | 12 | -13.622 -15.172 -23.367 -0.684 -1.808 -3.503 -8.034 27.098 -3.371 -0.753 -16.556 -7.633 14.649 -1.952 5.519 1.456 -15.26 16.543 -13.685 -13.041 10.58 -33.75 -27.085 -0.629 -6.327 31.334 -18.059 10.928 -13.303 0.315 |
| | | | | | | | | 28,3 | 52 | 12 | -30.342 -22.93 16.205 -6.922 2.414 -18.76 -23.137 -29.727 -4.548 4.002 10.675 19.866 -12.755 1.494 4.93 2.401 -9.323 -15.691 -1.468 10.965 28.43 -15.758 8.547 -3.678 8.994 -3.682 -5.114 24.39 -3.946 -1.734 |
| | | | | | | | | 28,3 | 47 | 3 | -1.754 -4.86 7.945 -1.902 4.613 12.167 9.068 -26.007 -4.114 -6.262 30.55 -15.195 7.752 0.526 0.047 -15.232 -18.314 -20.991 1.796 2.329 -2.006 -3.495 26.762 -11.489 -2.89 7.969 -15.96 -20.056 10.745 11.806 |
| | | | | | | | | 28,1 | 48 | 1 | -14.873 -16.928 -20.345 2.171 0.187 -5.387 -1.743 24.645 -0.612 -8.858 31.405 -16.231 9.59 -11.841 -6.543 -0.367 2.689 3.709 -11.139 -2.408 -1.722 -3.579 0.519 -1.711 2.637 15.871 -23.362 -12.786 -6.547 6.455 |
| | | | | | | | | 27,9 | 53 | 12 | 21.222 28.218 -27.937 -4.813 -4.601 -2.778 0.858 19.49 -12.775 -7.073 -16.126 -20.42 -23.248 3.145 -4.73 6.077 -19.437 -20.596 5.249 0.49 -6.297 -18.524 13.177 -5.776 -3.552 31.806 -18.527 9.889 -9.441 10.208 |
| | Rank 5 relaxed | HS | Random | TCRfp HS | 28,7 | 10 | 26,72 | 28,7 | 10 | 13 | -13.486 19.308 25.414 6.867 -7.782 -18.21 -17.699 -25.011 -8.889 -7.504 26.389 -14.222 11.844 2.043 2.293 -6.909 3.485 22.131 -9.443 -1.724 19.302 -26.726 -28.448 -2.936 -3.541 15.295 19.611 -10.154 3.259 -5.317 |
| | | | | | | | | 28,5 | 40 | 6 | 24.302 26.439 -31.274 -9.455 -7.88 28.873 -14.932 10.356 -9.174 0.074 3.835 -18.649 -13.233 -3.704 4.921 -0.359 -2.613 21.625 -3.012 3.237 -11.35 -11.44 -17.448 -4.913 0.254 -1.895 -6.84 1.917 -4.231 -3.625 |
| | | | | | | | | 28,1 | 41 | 2 | 24.608 -10.031 7.668 0.118 2.059 -2.731 -2.823 9.428 -4.435 -0.416 4.48 -13.524 31.853 -8.473 4.845 -13.673 -8.895 -28.007 -6.987 -7.022 7.831 -25.181 -25.258 3.487 4.154 22.897 21.574 -21.866 3.193 -6.723 |
| | | | | | | | | 26,6 | 50 | 3 | -2.975 -13.478 -16.891 6.365 -2.143 -1.153 24.211 21.717 -10.13 -10.61 -1.728 -4.669 -19.28 -3.687 8.105 6.814 -9.568 -25.297 -3.962 -7.133 -26.516 -8.037 26.292 -8.569 3.158 26.92 -9.623 9.111 1.503 8.783 |
| | | | | | | | | 26,6 | 49 | 5 | 18.125 -3.308 -30.282 6.282 1.84 -17.276 -11.529 30.147 -6.985 -5.931 0.968 27.55 25.594 -11.105 4.458 28.95 -13.41 10.812 -4.342 -8.427 6.187 -19.897 -18.833 3.173 0.942 -15.322 -10.11 -17.765 -1.495 8.606 |
